# A moderate water deficit induces profound changes in the proteome of developing maize ovaries

**DOI:** 10.1101/2024.07.12.603247

**Authors:** Mariamawit Ashenafi, Thierry Balliau, Mélisande Blein-Nicolas, Olivier Turc, Michel Zivy, Elodie Marchadier

## Abstract

Water deficit is a major cause of yield loss for maize (*Zea mays*), leading to ovary abortion when applied at flowering time. To help understand the mechanisms involved in this phenomenon, the proteome response to water deficit has been analysed in developing ovaries at silk emergence stage and five days later. Differential analysis, abundance pattern clustering and coexpressions networks were performed in order to draw a general picture of the proteome changes all along ovary development and under the effect of water deficit. The results show that, even mild, water deficit has a major impact on ovary proteome, but that this impact is very different from a response to a stress. A part of the changes can be related to a slowdown of ovary development, while another part cannot. In particular, ovaries submitted to water deficit show an increase in proteins involved in protein biosynthesis and in vesicle transport together with a decrease in proteins involved in amino acid metabolism and proteolysis. According to the functions of increased proteins, the changes may be linked to auxin, brassinosteroids and jasmonate signalling, but not abscisic acid.

## 1. Introduction

Maize (*Zea mays L.*) is one of the most important staple crops and is expected to overtake wheat as the most widely grown crop by 2030 [1]. Maize is mainly cultivated as a direct or indirect feed source through grain and by-products such as stover and silage, playing a crucial role in maintaining the food security of humans and livestock [2]. Maize is also used as a source for bioethanol production providing an alternative to fossil energy [2].

Initially domesticated in Mexico, several millennia allowed maize adaptation from tropical to temperate environments. Maize varieties are now widely cultivated throughout the world including in northern countries but remain particularly susceptible to water stress. Multiple climate models agree on the increase of frequency and severity of drought periods in the near future [3]. Simulations of climate scenarios combining hotter and drier conditions predict higher maize yield losses than climate warming only [4].

In response to drought during vegetative growth, maize plants slow down their development and limit the growth of aerial parts [5]. Cell division and expansion are reduced [6] and stomatal closure diminishes C0_2_ uptake [7] and slows down photosynthetic activity[5]. On the contrary, roots growth is favoured [8].

The reproductive phase is the critical period for maize vulnerability to water stress. A better understanding of the ovary response to water limitation is essential for developing solutions to mitigate its adverse effects.

In maize, only a fraction of ovaries give rise to mature kernels and water stress inhibits ovary growth and decreases kernel number per ear [9]. Ovary growth inhibition is not related to the loss of turgor classically observed in other water-stressed tissues [10]. Water deficit has been shown to delay silking by affecting silk elongation rate in the same manner as leaf growth [11]. Silking delay consequently increases the critical anthesis to silking interval (ASI) resulting in asynchrony between pollen and silk maturations and losses in yield that can reach more than 60% in drought-sensitive maize lines [12]. Drought-induced changes in ASI are related to silk expansion consisting of cell division and cell expansion, both being reduced as a consequence of water deficit. [13]. This growth impairment can lead to ovary abortion. Water stress induced ovary abortion has a major impact on maize productivity. It depends on ovary developmental stage [14] and silk growth rate [15]. The higher sensitivity is particularly observed in ear apical position [5]. As the meristematic region is at the tip of the ear, the ear apical region contains the youngest ovaries.

Due to its importance in the response to environmental stresses such as water limitation, changes in abscisic acid (ABA) concentrations of drought-stressed ovaries were investigated in several studies. This led to different results depending on the tissues: ABA concentration per ovary fresh weight or by kernel increases [16], it decreases when measured by floret [17] and it is stable when measured on an ovary dry-weight basis [18].

Several biochemical studies relating changes in carbohydrate metabolism and transport to ovary abortion have been carried out in the ovaries of plants submitted to a strong water deficit. In these conditions, carbon metabolism is strongly disrupted in ovaries, and soluble sugar and starch contents are particularly affected [5]. Inhibition of ovary growth is associated with an increase in sucrose concentration [10], a decrease in reducing sugars such as glucose and fructose, and a depletion of starch [19]. Invertase activity (hydrolysis of sucrose to glucose and fructose) is inhibited, and starch downstream product is also decreased [19,20]. Sucrose arrives at the ear via phloem transport and ovary growth depends on this continuous C stream from the parent plant. In drought conditions, carbohydrate and sugar delivery into the ovary is inhibited due to photosynthesis inhibition [20,21]. This can be briefly compensated by the conversion of ovary starch to glucose while starch reserves are sufficient [22]. However, in the absence of a starch pool and photosynthesis products arrival, ovary development is aborted [20]. Consistently with the requirement of sucrose for ovary development [23], sucrose supply allows to restore the development of ovaries [19,24] despite invertase activity is only partially recovered [20].

More recent studies [15,25] performed under moderate water deficit investigated its effect on ovaries on a more precise time scale. They showed that ovary abortions are indeed due to their developmental slowdown and a reduction of expansive growth, impairing their silk emergence and thus their pollination. Changes in carbon metabolism occur after this reduction of growth, and can rather be considered as consequences than causes of ovary abortions. This was confirmed by transcriptome studies [15]. Under stronger water stress, a down-regulation of genes coding for sucrose processing enzymes and then the up-regulation of some genes involved in senescence [26] has been described around pollination.

Linking gene expression to metabolism disruption is a difficult task in such a complex and finely-tuned process relying on molecular changes according to space and time. RNA expression studies offer the advantage of accessing fast and finely tuned molecular responses while metabolite contents reflect the composition of the tissues, resulting from multiple upstream regulations and molecular transport. As intermediate molecules, proteins open interesting perspectives to better understand the molecular processes, providing information regarding post-transcriptional regulation events and abundances of enzymes, the active molecules catalyzing metabolic reactions. To go further in the understanding of the effect of water stress on ovary development, this study aims to better characterize their proteome under moderate water deficit.

## 2. Materials and Methods

### 2.1. Experimental design

Plants of the reference line B73 were grown in a greenhouse and watered up to tassel emergence. Irrigation was then stopped for half the plants until they reached a water potential of ca. -0.4 to -0.6 MPa. All plants were then transferred to a growth chamber, where water deficit (WD) or well-watered (WW) conditions were maintained. Starting from silk emergence, ears were daily manually pollinated. Ovary samples were collected at two stages: at the emergence of silks (SE) and 5 days after silk emergence (SE+5d). At each stage, ovaries were collected from five zones of the ear (numbered 1 to 5 from bottom to top, ie. from the oldest to the youngest ovaries), except for zone 5 of SE+5d samples of WD plants because of ovary abortion at the top of the ear. Three biological replicates were taken for each condition*day*zone combination.

### 2.2. Proteomics

Ovary protein extraction and digestion were performed as described in [27]. Mass spectrometry analysis (MS) was performed by using an LTQ-Orbitrap connected to an Ultimate 3000 LC system (Thermo, Waltham, MA, USA) as described in [28]. Protein identification and quantification were performed with i2MassChroq http://pappso.inrae.fr/bioinfo/i2masschroq/, that uses X!Tandem [29] and XtandemPipeline [30] for protein identification and MassChroQ for quantification [31]. The Genome assembly B73 RefGen_v4 database was searched https://ftp.ensemblgenomes.ebi.ac.uk/pub/plants/release-50/fasta/zea_mays/pep/Zea_mays.B73_RefGen_v4.pep.all.fa.gz. Proteins were identified with at least two peptides with an E-value < 0.05 and a protein E-value < 0.0001. The false discovery rate for peptide and protein identification was respectively 0.27% and 0.42%. Peptide quantification was performed by integration of extracted ion current (XIC). Normalization was performed as described [32]. Protein relative quantities were computed by summing the XIC value of specific and reproducible peptides (missing values < 10%), that showed a correlation to each other >0.5. Finally, a quantitative relative abundance was obtained for a total of 800 proteins.

### 2.3. Principal component analysis

Principal component analysis was performed on the whole dataset, with protein mean abundances being considered as variables and the different condition x day x zone combinations as individuals. The prcomp function of the R package [33] was used.

### 2.4. Clustering method

The 800 quantified proteins were distributed in eight clusters by using the K-means method on protein means computed for each condition*day*zone combination (kmeans function of the stats package). Mapman annotations of proteins were collected by using Mercator 4 [34]. The independence between Mapman annotations and cluster assignments was tested using the independence Chi-squared test.

### 2.5. Differential analysis

The following model has been applied to the abundance variable of each of the quantified proteins, with *Y_ijkl_* representing the observed abundances, *µ* the mean abundance of the protein, *α_i_* the condition *i* effect, *β_j_* the zone *j* effect, *δ_k_* the day *k* effect, *γ_ij_*, *θ_ik_* and *ω_jk_* pairwise interactions of the effects and *ɛ_ijkl_* residuals associated to the sample *l*.

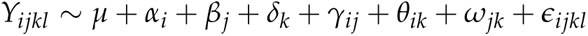

Anova type III has been applied to this model (Anova function of the car R package) and pvalues were corrected using the Benjamini-Hochberg multiple test correction procedure [35]. A significant effect was considered when the adjusted pvalue was lower than 0.05.

### 2.6. Network-based analyses

#### 2.6.1. Identification of protein-protein coexpressions

For each combination of date, zone, and condition, protein abundances of the three replicates were averaged. Then, Pearson correlation coefficients were calculated for each pair of proteins for control and stress conditions independently, leading to a correlation matrix per condition. Pvalues were adjusted using the Benjamini-Hochberg multiple test correction and adjusted pvalues lower than 0.05 were kept as significant.

#### 2.6.2. Analysis of WW network

The density of the WW network was calculated using 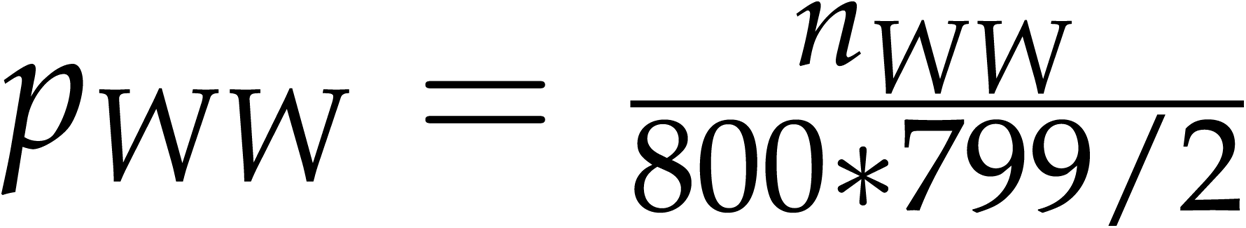 where refers to the number of co-expressed proteins in WW. Then, for each pair of Mapman terms, the proportion of expected protein-protein connections was calculated under the assumption that there were no preferential connections between Mapman terms. These theoretical proportions were compared to observed numbers of connections between pairs of Mapman terms. Pairs of Mapman terms whose proteins were significantly more or less connected together were identified using a hypergeometric test.

#### 2.6.3. Comparison of WW and WD networks

At the general scale, comparisons between WW and WD networks were performed using estimations of the probabilities to observe an edge in each of the networks 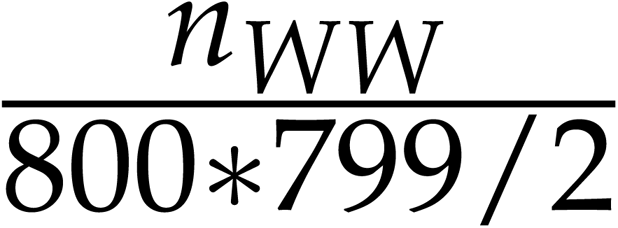 and 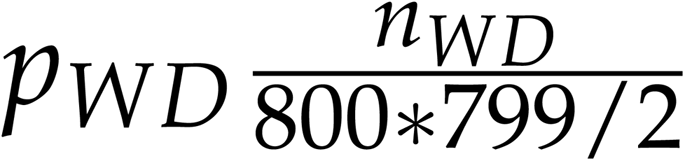 where *n_WW_* and *n_WD_* respectively refer to the number coexpressed proteins in WW and WD networks. Under the assumption of independence between both of the networks, the probability for an edge to be observed in WW and WD networks is equal to *p_WW_ ∗ p_WD_* At a more specific scale, for each pair of Mapman terms, we identified the number of connections involving both of these terms and the number of other connections in WW and in WD. Significant changes of connectivity between WW and WD networks were identified using Chi2 tests of independence and Benjamini-Hochberg [35] adjusted pvalues lower than 0.05 were considered as significant.

### 2.7. KEGG pathway mapping

KEGG orthology (KO) compounds identifiers corresponding to lists of differentially expressed proteins were obtained from Phytozome website (https://data.jgi.doe.gov/refine-download/phytozome?q=zea&expanded=Phytozome-493). KEGG pathways involving the KO compounds were obtained using KEGG API links (https://rest.kegg.jp/link/ko/ pathway and https://rest.kegg.jp/list/pathway). Tables with two columns were created. In the first column, KO compounds corresponding to each of the proteins were stored. In the second column, numerical values corresponding to the over (+1) and under (-1) expression of each protein were indicated. These informations were mapped on selected pathways using KEGG procedure available at https://www.genome.jp/kegg/mapper/.

## 3. Results

To analyze the proteome response of developing ovaries to a moderate water deficit, plants were subjected to water deficit from the tassel emergence stage. Ovaries were then collected from five zones of the ear on the day of silking and five days later. As the meristematic zone is at the top of the ear, the ear zones showed an age gradient, with the lowest zones representing the oldest ovaries and the highest the youngest. Ovaries from well-watered plants were collected in the same way.

### 3.1. Main factors explaining changes in protein abundances

Proteome analysis using mass spectrometry allowed the quantification of 800 proteins (**Appendix A2**). To assess the effects of the condition, day, and zone factors on the proteome, a principal component analysis (PCA) was conducted (Figure 1). Principal components (PC) 1 and 2 represent 26.8% and 22.1% of the total variation, respectively. The PCA reveals a strong condition effect: the WD and WW samples were distinctly separated along the PC1 axis. A pronounced effect of the day is also evident, as SE+5d samples are separated from SE samples within each condition along the second diagonal of the PC1/PC2 plane. PCA also allows to distinguish the zones of the ear mainly at SE, where samples are ordered based on the zone number, with zones 5 (youngest ovaries) and 4 distant from the group formed by zones 1, 2, and 3 along the first diagonal, both for WD and WW conditions.

**Figure 1.**
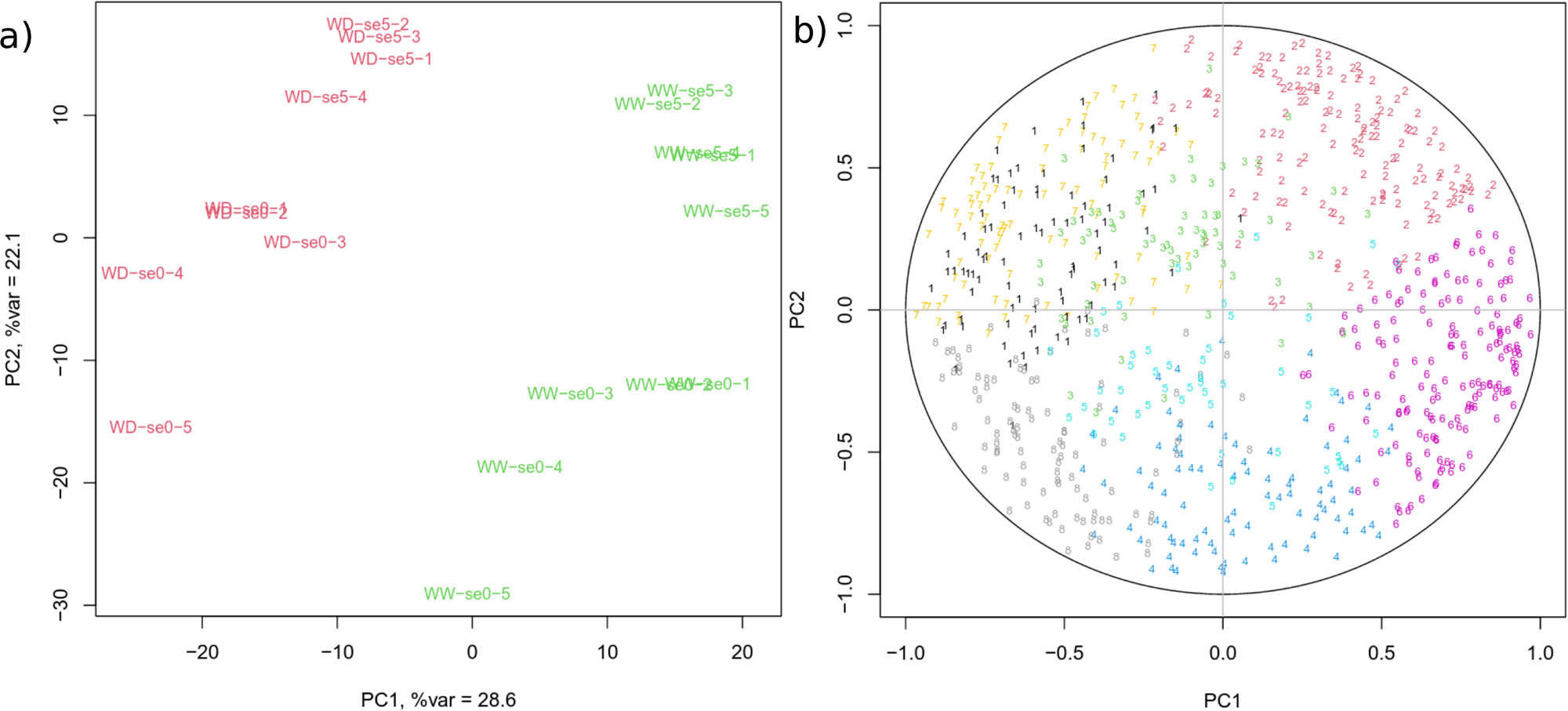
Principal component analysis of ovary proteins. a: Sample are named according to the concatenation of treatment, day and zone. se0 and se5 stand respectively for SE and SE+5d. Red: WD, green: WW. b: Correlation circle. Proteins are represented with the number of the cluster they belong to.

### 3.2. Main functions affected by water deficit and development

To better characterize the patterns of protein abundance variations under the influence of WD and ovary development, a clustering analysis was conducted. Eight protein clusters were identified, which patterns are represented in Figure 2 (**Appendix A3**). Their representation in the PCA is presented in Figure 1.

**Figure 2.**
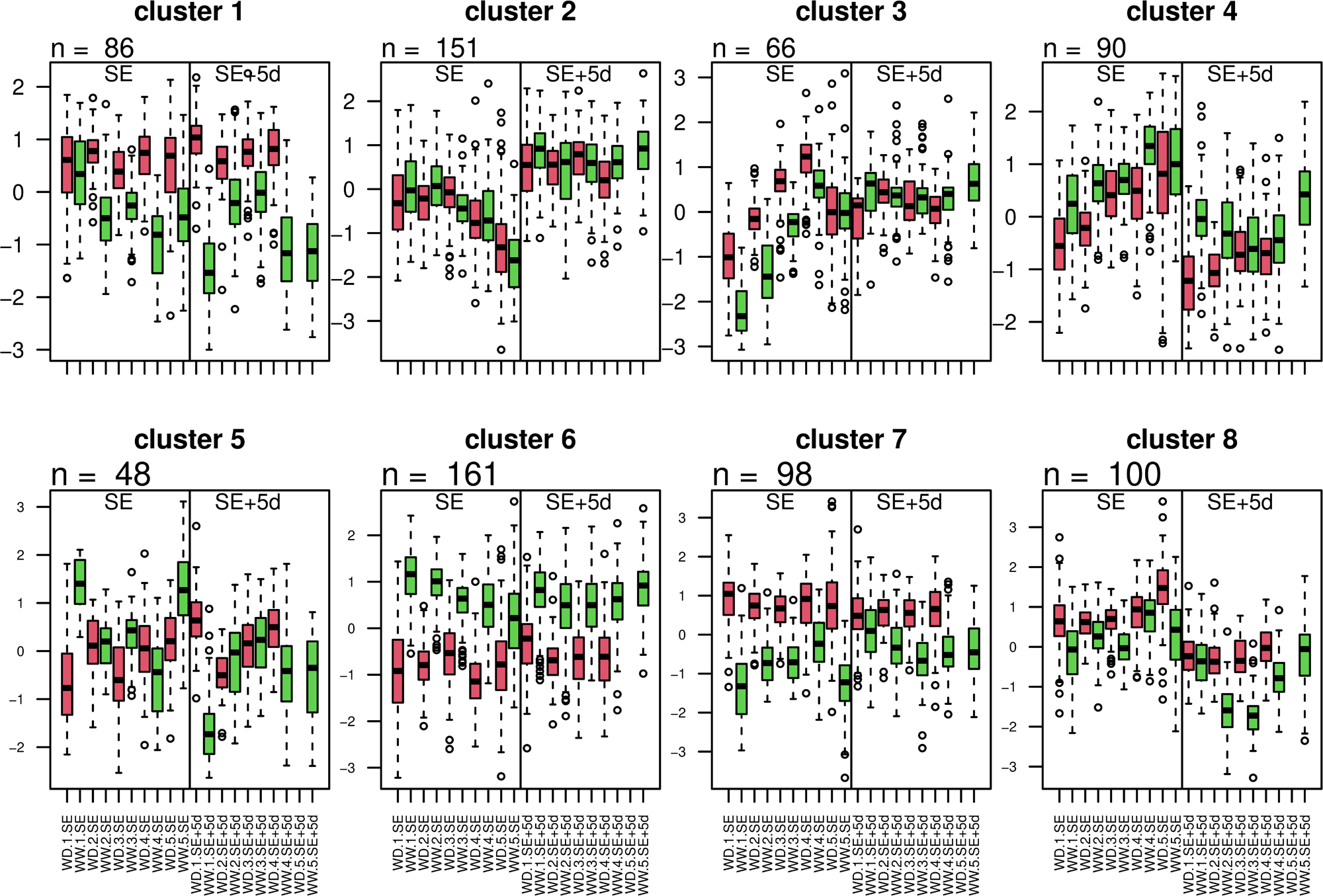
Cluster profiles. Each graph represents the expression profile of the proteins making up a cluster in the different treatment*day*zone combinations. red: WD, green: WW

The expression profiles of proteins belonging to each of these clusters led to group them into four broad categories, according to their expression pattern: clusters exhibiting (i) a developmental effect (ii) a primary condition effect, (iii) a secondary condition effect, and (iv) no effect.

(i) developmental effect

Cluster 2 shows a strong developmental effect, with protein quantity decreasing from the base to the top of the ear at SE. This gradient aligns with an increase in protein quantity during development, as the younger zones are higher up. For these proteins, the quantity would stabilize after this initial increase, as evidenced by stability at SE+5d.

(ii) primary WD effect

We grouped in this category the clusters 1, 6, and 7 for which median protein values in all day*zone combinations are higher in one condition compared to the other. In the PCA, proteins of these clusters contribute maximally to the separation between conditions. Clusters 1 (86 proteins) and 7 (98 proteins) correspond to proteins whose abundances increase under WD. Cluster 6 (161 proteins) corresponds to proteins whose abundances decrease under WD. In summary, clusters 1, 6, and 7 contain proteins for which the response to WD is independent of the ovary developmental stage.

(iii) secondary WD effect

We classified in this category the clusters 3, 4, and 8 where developmental effects (day and zone) dominate, but where a WD effect is visible. In cluster 3 (66 proteins), protein abundance increases from the bottom to the top of the ear at SE, while it is stable along the ear at SE+5d. Cluster 4 (90 proteins) is mostly characterized by a greater abundance in SE than in SE+5d, while proteins are less abundant in the WD than in the WW condition in the lower part of the ear at both stages. Proteins of cluster 8 are also more abundant in SE than in SE+5d, but more abundant in the WD than in the WW condition at both stages.

(iv) no developmental or treatment effect

Cluster 5 exhibits no variation based on any of the three factors. The cluster is located at the center of the PCA correlation circle, indicating that it does not contribute to the observed separation between samples.

Functional categories distribution varies depending on the clusters (Chi-squared test pvalue *<* 2.2 *∗* 10*^−^*^16^). The distribution among the four cluster categories of the major functions (Mapman level 1 categories) represented by more than 20 proteins and of proteins involved in cell division is shown in Figure 3 and described hereafter.

**Figure 3.**
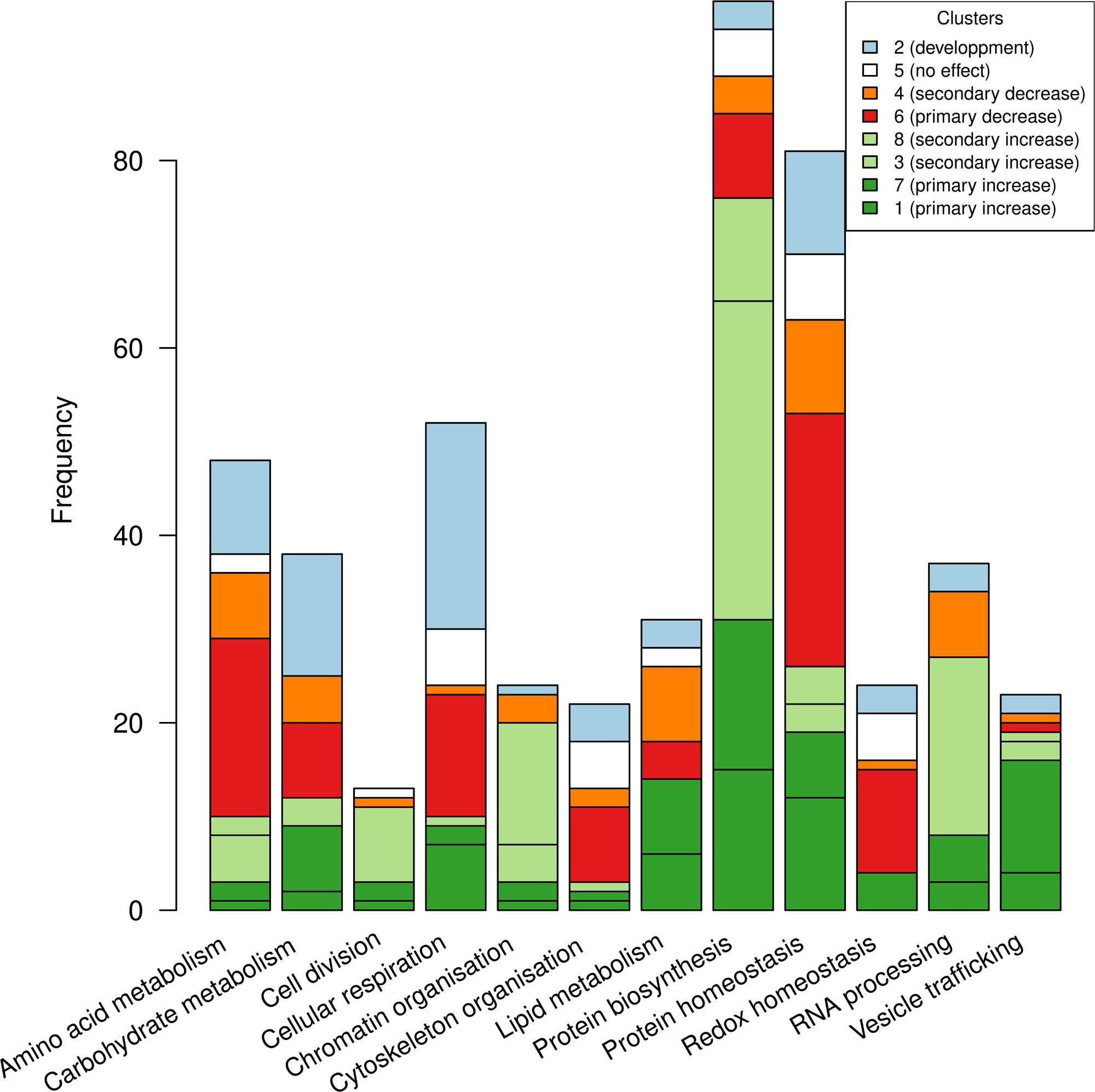
Distribution of functional categories among clusters. Red-scaled colors refers to proteins whose abundances decrease in WD, green-scaled colors refers to proteins whose abundances increase in WD, blue refers to proteins whose abundances change depending on ovary development. Only functional categories represented by more than 20 proteins and the “Cell division” category are shown.

#### Protein Biosynthesis (97 proteins, of which 63 are ribosomal proteins)

Respectively 31 and 45 of them showed a primary or secondary increase in response to WD, while only 13 proteins decreased. Overall, proteins involved in protein synthesis are increased under WD.

#### Protein Homeostasis (81 proteins)

Thirty-five of these proteins are in the ubiquitin-proteasome system category, with 14 being components of the 26S proteasome. Respectively 27 (including 20 from the ubiquitin-proteasome system) and 13 proteins were primarily or secondarily decreased by WD. By contrast, proteins involved in quality control including chaperones and other proteins involved in proteolysis did not show a consistent response to WD.

#### Cellular respiration (52 proteins)

The predominant cluster (22 proteins) is cluster 2, primarily showing a developmental effect. No pronounced trend for response to WD is observed.

#### Amino acid metabolism (48 proteins)

Twenty-six proteins show either a primary or a secondary decrease in abundance in response to WD, while 10 presented an increased abundance. Cluster 2, showing only developmental effects, contains 10 other proteins. Proteins involved in amino acid degradation do not exhibit a specific pattern compared to proteins involved in their synthesis.

#### Carbohydrate metabolism (38 proteins)

The predominant cluster is cluster 2 (13 proteins), indicating a predominant developmental effect with an increase over time. The other proteins do not show a consistent trend. The metabolisms of sucrose and starch react differently: out of the 7 proteins involved in starch metabolism, 5 show a primary increase in response to WD. The effect of WD on proteins involved in sucrose metabolism is more heterogeneous: out of 11 proteins, 4 exhibit a primary or secondary decrease, and 2 show an increase.

#### RNA processing (37 proteins)

RNA processing mainly includes proteins involved in RNA homeostasis (13 proteins) and pre-RNA splicing (12 proteins). Both categories show similar response patterns to WD. Most of them (19 proteins) show a secondary increase in response to WD. Eight other proteins show a primary increase, while a total of 7 proteins show a decrease.

#### Lipid metabolism (31 proteins)

Most of them are involved in fatty acid metabolism (21 proteins). There is no consensus response to WD.

#### Redox homeostasis (24 proteins)

Respectively 11 and 1 proteins showed a primary or secondary decrease in response to WD, while only 4 showed an increase.

#### Chromatin organization (24 proteins)

In total, 20 proteins show a primary or secondary increase in response to WD, while only 3 decreased.

#### Vesicle trafficking (23 proteins)

The majority of these proteins are primarily increased by stress (16 proteins). Three other proteins are secondarily increased, while only two proteins are decreased.

#### Cytoskeleton organization (22 proteins)

These proteins are predominantly decreased in WD (10 proteins). Only 3 proteins are increased.

#### Cell wall organization (20 proteins)

No predominant pattern for these proteins. Eight proteins increase in response to WD and 5 decrease.

#### Cell division (13 proteins)

Eight of the 13 proteins involved in cell division are in cluster 8, where they show a strong ascending gradient at SE and low values in SE+5d. This aligns with the fact that the meristematic region is at the top of the spike, and may indicate that cell division is less active in SE+5d, where cells are more aged.

### 3.3. Pathways affected by water deficit and development: differential analysis

A differential analysis was performed considering developmental (zone and day), condition, and interaction effects (**Appendix A4** and Table 1). Boxplots representing the expression patterns are represented **Appendix A1** and mapping of proteins on KEGG pathways are available on **Appendix A5 and A8**.

**Table 1.**
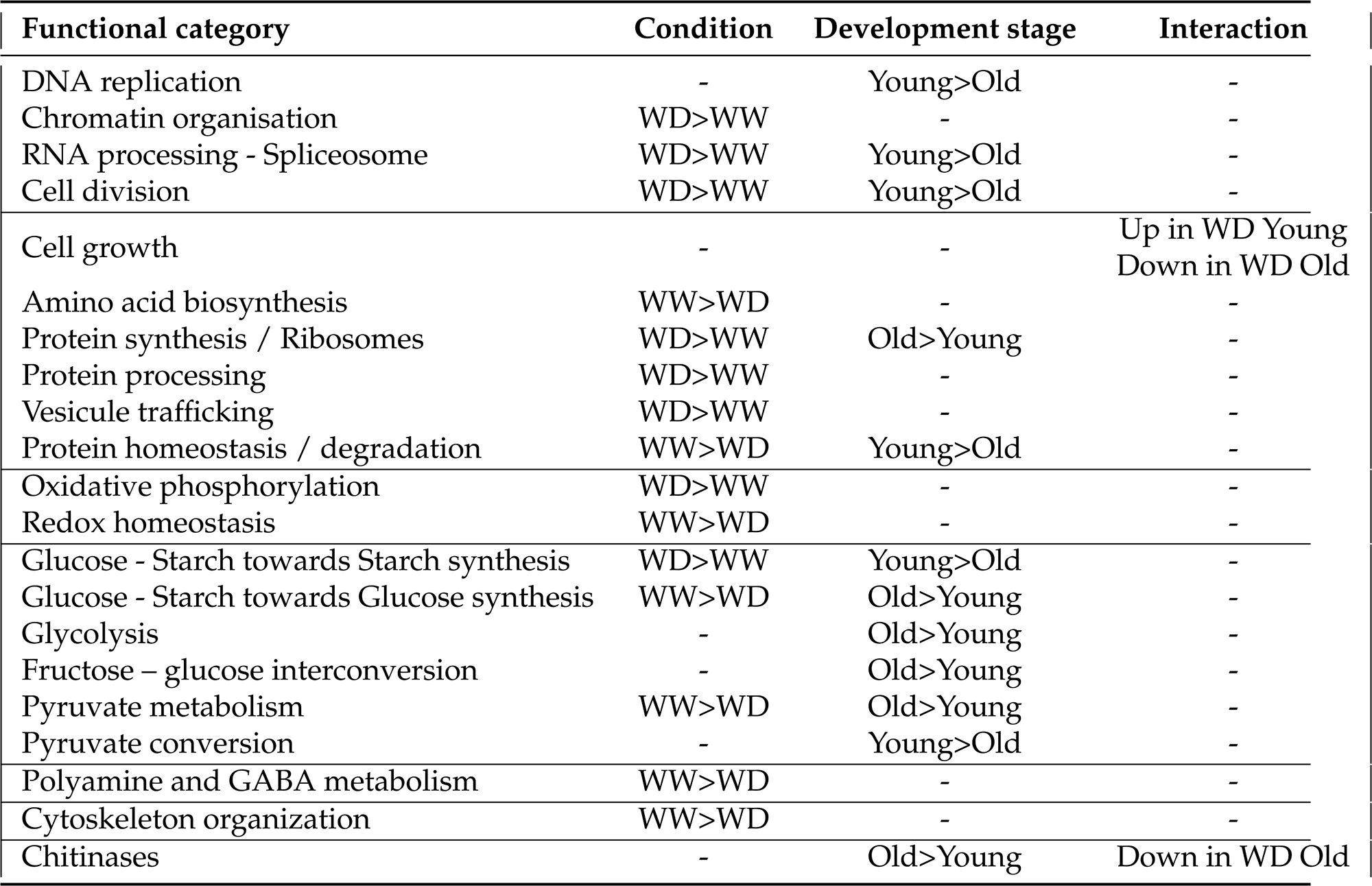
Main functions whose proteins abundance is impacted by developmental and conditions effects.

#### 3.3.1. Developmental effects

Differences observed between zones (162 proteins impacted) are informative regarding the short-term ovary development (from the oldest ovaries in zone 1 to younger ones in zone 5) while those observed between days (327 proteins impacted) refer to a longer time scale (5 days).

The zone and day factors both affect proteins involved in DNA replication, which are more expressed in young ovaries (i.e. ovaries at the SE stage or in the highest zones). Youngest ovaries of higher zones also express more proteins related to the spliceosome, and proteins involved in protein degradation outside the endoplasmic reticulum (ER) are more abundant at the SE stage. Regarding metabolic pathways, enzymes involving pyruvate as a substrate or a product are more abundant in younger ovaries than in older (SE vs SE+5d), such as the ones allowing pyruvate interconversion from phosphoenolpyruvate, acetyl-CoA or malate (Zm00001d052494, Zm00001d001831, Zm00001d023379 and Zm00001d018961 for [EC:2.7.1.40]; Zm00001d006107, Zm00001d000227, Zm00001d052289 and Zm00001d004473 for [EC:1.2.4.1]; Zm00001d037961 for [EC:1.1.1.40]).

Proteins of starch and glucose metabolism acting toward starch synthesis (Zm00001d0038 for [EC:2.4.1.18] and Zm00001d019479 for [EC:2.4.1.242]) are more abundant in young than in older ovaries (higher zones/lower zones).

Globally, glycolysis proteins are more abundant in oldest ovaries (lower zones and SE+5d), proteins of starch and glucose metabolism acting toward glucose synthesis are more abundant in lower zones (like *β*-glucosidases Zm00001d023994, Zm00001d034015 and Zm00001d033649 for [EC:3.2.1.21]) and proteins involved in citrate cycle and fructoseglucose interconversion are more abundant in SE+5d than in SE ovaries.

Proteins of the ER, that are involved in protein processing and ribosome components, are in higher abundance at the SE+5d stage than at the SE stage, indicating that translation is more active in older ovaries.

Numerous proteins involved in the citrate cycle, pyruvate metabolism, and carbon fixation see their abundances impacted depending on the zone of the ear. However, no consistent pattern is observed between proteins of a common pathway. Amino acid metabolism also appears strongly impacted by the day but the direction of the higher abundance varies a lot depending on the reactions which does not allow conclude to a consistent pattern.

The differential analysis identified binary interaction effects for some of the proteins. The interaction day*zone allows to identify 52 proteins, suggesting that the different zones are developing differently over time. Among these proteins, we identified three chitinases (Zm00001d027524, Zm00001d025753, and Zm00001d026415), one of them being involved in the stress response of maize kernels [36]. Two proteins involved in *α*-linolenic acid metabolism were also identified (Zm00001d015852 and Zm00001d029594). This pathway is known to mediate drought response in maize and leads to the biosynthesis of jasmonic acid [37]. In addition, numerous proteins taking part in the ribosomes are identified with a significant day*zone interaction.

#### 3.3.2. Major effect of water deficit

The differential analysis identified 481 proteins whose abundances were significantly impacted by water deficit. Several of them belong to the same pathways and will allow to better understand the metabolic changes induced by WD.

The proteins showing an increase in WD are mostly involved in chromatin organisation, RNA processing, spliceosome, protein processing, and vesicle trafficking, notably coatomer machinery proteins (such as Zm00001d004466, Zm00001d007075, Zm00001d008218, Zm00001d016402, Zm00001d028438, and Zm00001d033734) in WD condition, while proteins decreased in WD are mostly involved in amino acid metabolism, protein homeostasis, carbohydrates metabolism, polyamine metabolism, redox homeostasis, and cytoskeleton organization. Notably, three proteins involved in proline synthesis are decreased (Zm00001d010056, Zm00001d038689, and Zm00001d042117) and one involved in its catabolism (Zm00001d038841) is increased in WD. Additionally, three proteins involved in GABA metabolism are significantly impacted by WD: a *γ*-hydroxybutyrate dehydrogenase and glyoxylate reductase (Zm00001d050810) is decreased while a GABA pyruvate transaminase (Zm00001d002326) and a glutamate decarboxylase (Zm00001d031749) are increased. Among the proteins involved in RNA processing that are increased in WD, there is also a Tudor-SN (Zm00001d050606), a protein involved in the formation of stress granules [38,39]. Among proteins related to chromatin organization, two members of the argonaute family (Zm00001d039214 and Zm00001d040429) as well as a component of of SUVH-DNAJ methylation reader complex (Zm00001d038541) increase in WD.

Protein homeostasis refers mainly to proteasome components whose core components’ expression decreases in WD.

Concerning redox homeostasis, proteins involved in both ascorbate-based (Zm00001d011 Zm00001d024633, Zm00001d032950 and Zm00001d035595) and thiol-based (Zm00001d011581 Zm00001d017823, Zm00001d040341 and Zm00001d046682) homeostasis show a decrease in WD. Therefore, it cannot be concluded that the proteome exhibits a response to oxidative stress. There are few exceptions: the proteins showing a major increase are two 9-lipoxygenases (Zm00001d013493 and Zm00001d042540) that may be involved in brassinosteroid signaling [40].

The effect of WD on carbohydrate metabolism appears complex. A lot of enzymes of carbon metabolism, and especially pyruvate metabolism and citrate cycle, are down-expressed except for a few key enzymes. For instance, enzymes able to interconvert oxaloacetate, citrate, and isocitrate (Zm00001d048358 and Zm00001d009127) are more expressed in WD. Among two enzymes able to convert oxaloacetate to citrate, one is down-expressed (Zm00001d053684, [EC:2.3.3.1]) while another one (Zm00001d009127, [EC:2.3.3.8]) is up in WD, and particularly at SE. Enzymes converting oxaloacetate to phosphoenolpyruvate such as phosphoenolpyruvate carboxylase PEPC (Zm00001d012702, Zm00001d016166, Zm00001d020057 [EC:4.1.1.31]) are also higher expressed in WD. In addition, the enzyme catalyzing phosphoenolpyruvate to aromatic amino acid precursors (Zm00001d006900) is up in WD at SE while the abundances of numerous enzymes involved in amino acid biosynthesis are down WD. Over all the pathways, the most striking pattern is observed for starch and glucose metabolism where proteins catalyzing reactions leading to glucose are down while the ones leading to starch are up in WD. Abundances of proteins involved in oxidative phosphorylation are also higher in WD, especially components of F-type mitochondrial ATP synthase and their related enzymes. On the contrary, a few subunits of V-type ATP synthase (located at other endomembrane compartments [41] are less expressed in WD.

Enzymes playing roles in fatty acid synthesis and degradation and proteins related to cell wall organization have also significant variations of their abundances between WD and WW conditions. However, variations of proteins from the same pathway do not exhibit consistent patterns.

#### 3.3.3. Minor effect of water deficit

The effect of the condition*day interaction is significant for 21 proteins among which 6 are in common with the day*zone interaction. Three of them are enzymes of the starch and glucose metabolism and have glucose and starch as direct products or substrates. Two proteins exhibiting differential abundance are expansins (Zm00001d045861 and Zm00001d034663), a protein family involved in cell wall loosening and to mitigate the effect of drought on maize yield [42]. Their expression profiles reveal their higher abundance in WD condition at SE and their lower abundance in WD at SE+5d (**Appendix A1**).

Only two proteins were showed a For the zone*condition interaction, suggesting that the effect of the condition is not highly different depending on the zone: Zm00001d027524, the previously mentioned chitinase [36] and Zm00001d048021, a protein involved in *α*-linolenic acid metabolism ([EC:4.2.1.92]).

### 3.4. Changes between co-expression networks

To better understand how WD influenced protein expression and to what extent new regulatory networks were set up, we compared the correlation of expression networks performed independently in WW and WD conditions (**Appendix A6 and A7**). Within each condition, the only biological factors that vary are ear zone and sampling date, the combination of which can be used as a proxy for developmental stage. Thus, we compared the two co-expression networks obtained in each condition to better understand how WD affected the coordinated processes required for ear development.

Based on a significant correlation (BH adjusted pvalues < 0.05) the WW and WD protein coexpression networks included 7824 and 2768 edges respectively (Table 2). The number of common edges between both of the networks is much higher than expected by chance (624 in common versus 68 expected under the independence assumption of the networks - pvalue *<* 2.2 *∗* 10*^−^*^16^), showing common protein expression coordination through the ear developmental process in both of the watering conditions.

**Table 2.**
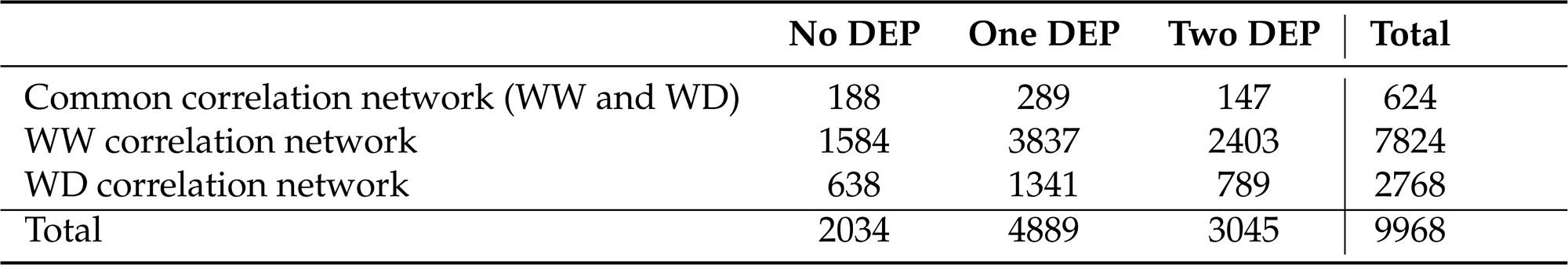
Number of correlations of protein abundances involving 0, 1 or 2 differentially expressed proteins (in columns) per network (in lines).

Table 2 also indicates that nearly 70% of the edges identified in the protein-protein coexpression networks involve at least one protein that was not identified by the differential expression analysis.

To analyze the biological significance of these networks, we used level 1 of the Mapman classification to group proteins involved in the same major process. Although each of these major processes can be made up of different pathways and can include different numbers of proteins, some general trends emerge.

#### 3.4.1. WW network

In a general manner, the network related to the WW condition (Figure 4) is rich in intra-process connections and low in inter-processes connections. As shown in Figure 4, a majority of the Mapman processes have significantly more intra-process correlations than expected by chance. There are two exceptions: “Protein homeostasis” and “Cellular respiration” show less intra-process correlation than expected.

**Figure 4.**
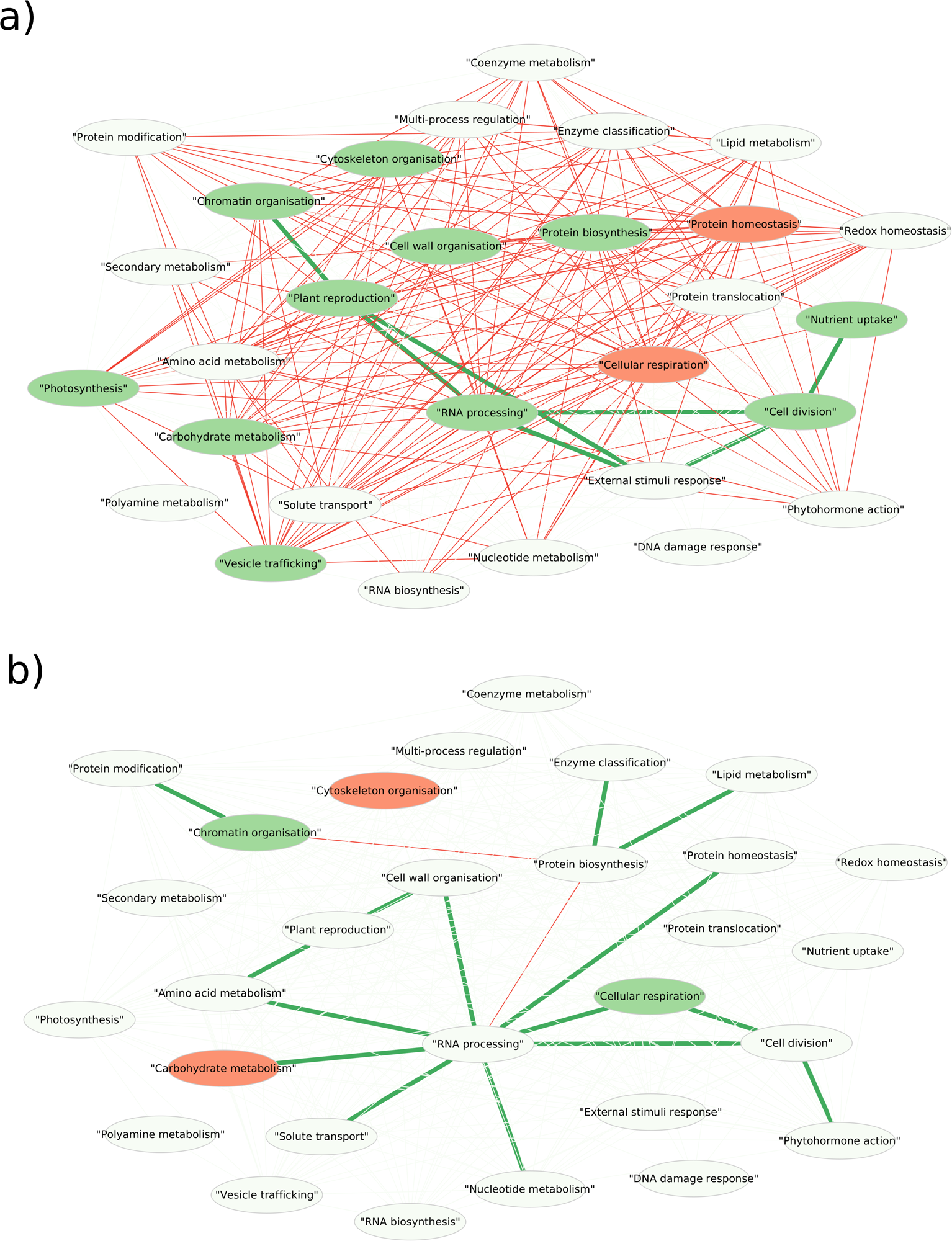
WW network and differences between WW and WD networks. a) WW network: Mapman processes are represented by nodes. A green node represents a process for which the number of pairs of proteins coexpressed is significantly higher than expected by chance while a red node represents a process for which the number of pairs of proteins coexpressed is significantly lower than expected by chance. Similarly, green edges connect processes between which many pairs of proteins are coexpressed and red edges connect processes between which only a few pairs of proteins are coexpressed. b) WD vs WW network: Mapman processes are represented by nodes. Only pairs of Mapman terms whose proportions of connections differ between WD and WW were identified on this WD network. A green node represents a process for which the number of pairs of proteins coexpressed is significantly higher than in the WW network while a red node represents a process for which the number of pairs of proteins coexpressed is significantly lower than in the WW network. Similarly, green edges connect processes between which many pairs of proteins are coexpressed compared to WW network and red edges connect processes between which only a few pairs of proteins are coexpressed compared to WW network.

External stimuli response, nutrient uptake, cell division, plant reproduction, and transcription-related processes are identified as coordinated processes. The absence of connexion with other processes for “Vesicle trafficking”, “Protein biosynthesis”, “Carbohydrate metabolism”, “Photosynthesis”, “Cytoskeleton” and “Cell wall organisation” suggest that these processes independently participate in ovary development in the WW condition.

#### 3.4.2. Effect of WD on the network

The network related to the WD condition was compared to the control network and significant changes in intra and inter-processes connectivities were highlighted (Figure 4). Compared to the WW network, strong losses of intra-process correlations are observed for almost all processes, especially for “Carbohydrate metabolism” and “Cytoskeleton organisation”. Only “Cellular respiration” shows a significant excess of intra-process correlations, while they were significantly fewer in the WW condition.

This global loss of intra-process connections is to the benefit of expression correlations linking proteins of different processes. Three groups of connected processes appear to be independent from each other: (i) a large group of 11 interconnected processes, in which “RNA processing”, occupies a central place since it is directly connected to 8 of the others. Among the latter, “Cell division”, “Phytohormones action” and “Cellular respiration” form a subgroup of coordinated processes. (ii) “Protein biosynthesis”, “Lipid metabolism”, and “Enzymes” and (iii) “Protein modification” and “Chromatin organisation”.

In the WD condition, the plant reproduction process (including 3 proteins involved in floral induction out of 4) appears coordinated with cell wall organisation and amino acid metabolism processes rather than with transcription-related processes as observed in the WW condition.

#### 3.4.3. Proteins involved in the WW/WD network changes

The previous analysis showed that there were profound changes in the co-expression networks between the WD and WW conditions. Some changes were due to loss of connections and some to gains. To better understand the changes between the two conditions, we analysed these gains and losses at the level of the proteins themselves.

Figure 5 reveals that multiple connections were lost due to correlations involving “hub” proteins whose abundances were correlated with multiple other proteins in the WW condition. It is notably the case for the “RNA processing”, “Lipid metabolism”, and “Chromatin organisation” processes. On the contrary, WD-specific correlations are more widespread among proteins of each of the processes (Figures 4 and 5 and **Appendix A9**).

**Figure 5.**
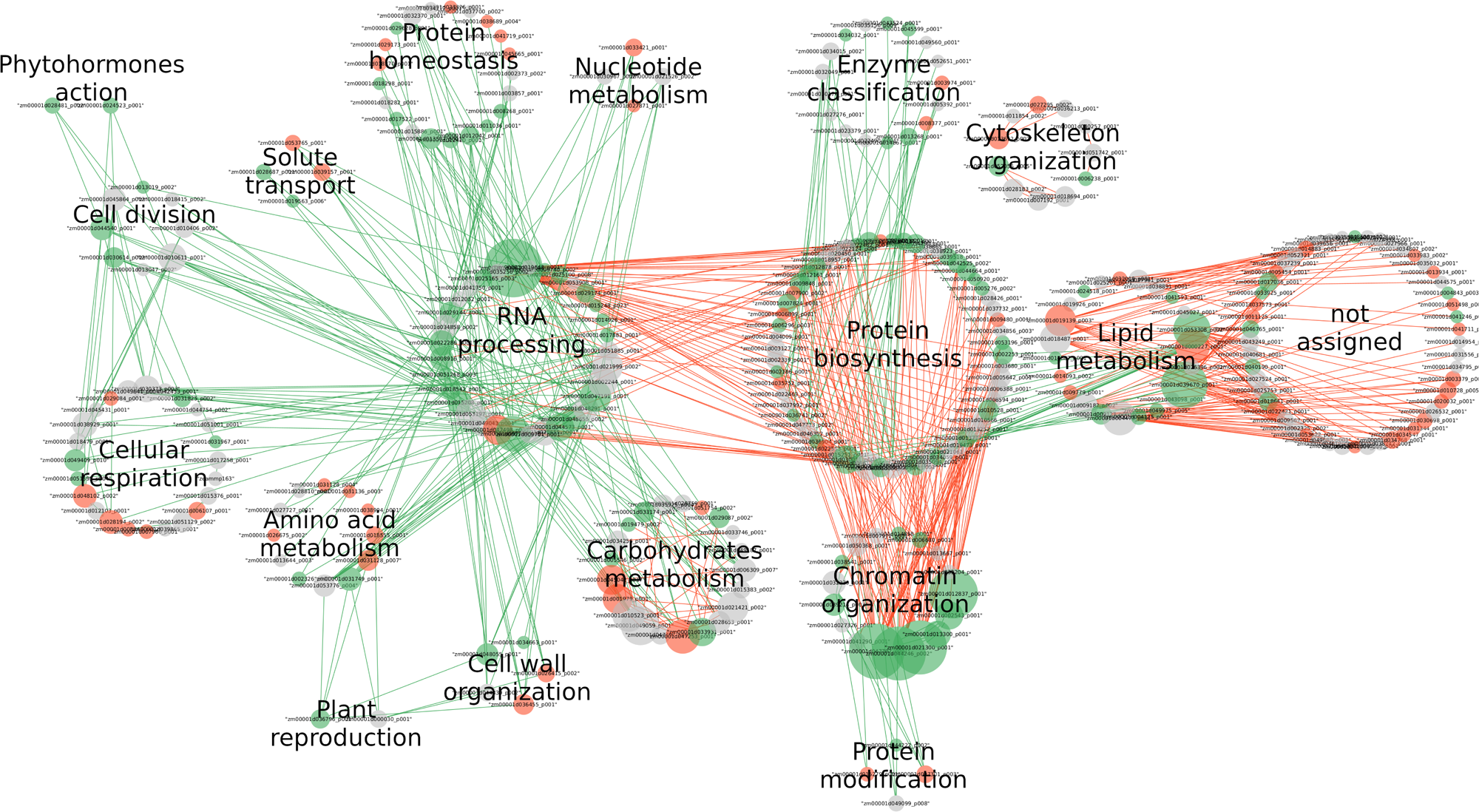
“Water deficit” protein-protein network. Nodes represent proteins involved in changes of inter and intra Mapman processes connections. Node colors refer to the effect of WD on a protein expression (green: highly expressed in WD, red: less expressed in WD, white: no differentially expressed in WD) and node sizes refer to node degrees. Green edges represent correlations of protein abundances that are not observed in WW but are observed in WD while red edges represent correlation of protein abundances that are observed in Ww but lost in WD.

The most striking case is for protein Zm00001d006725, a translational terminator from the “RNA processing” process, that loses 31 connections to proteins of the “Protein biosynthesis” process in the WD condition. Histone proteins from “Chromatin organisation” also lose correlations of expression with proteins involved in the “Protein biosynthesis” process in the WD condition.

Several proteins involved in *α*-linoleic acid are co-expressed with diverse (“not assigned”) proteins in WW and lose these connections in the WD condition, except for “Zm00001d049059”, an alcohol dehydrogenase that loose coexpression with proteins associated to carbohydrate degradation (Zm00001d034256, Zm00001d035925 and Zm00001d049187) and sucrose synthesis (Zm00001d010523) to the benefit of co-expression with several proteins involved in “RNA processing”. A similar change of co-expression partners is observed for the sucrose synthase Zm00001d047253 protein that lost co-expression with other proteins involved in sucrose metabolism (such as Zm00001d045042, Zm00001d029087, and Zm00001d010523) observed in the WW condition and gained several connections with “RNA processing” proteins in the WD condition.

## 4. DISCUSSION

This study investigated the proteome response of developing ovaries to a moderate water deficit, that is sufficient to induce ovary abortion. The effect of water deficit was analyzed in the proteome of ovaries collected along the ear at the stage of silk emergence and five days later. In addition to the differential expression analysis, a clustering procedure allowed us to group together proteins with similar expression profiles and protein abundance correlation networks allowing to study the effect of water deficit on the functional coordination of developing ovaries.

The different approaches used in this study (PCA, ANOVA, cluster analysis, coexpression networks) showed that the effect of water deficit differs according to the stage of ovary development. As a first step, we will analyse the effect of development on ovary proteomes independently of water deficit.

### 4.1. Functions involved in ovary development in the WW condition

Our experimental design allowed to compare ovary developmental stages based on two factors: the zone of the ear allowing the study of early and short-term developmental processes, and the day of the sampling allowing to compare longer developmental time intervals. Results of the PCA reveal that samples of the different zones collected at silking emergence are more distinct than those collected five days later, suggesting that the gradient of ovary development along the ear is larger at the silking emergence stage than five days later: more differentiation takes place during early stages of ovary development.

Previous studies investigating maize ovary development highlight the importance of carbon stream from phloem and sucrose conversion to glucose, fructose, and starch in this process [5,10,19]. In our study, the clustering and differential analysis approaches show that starch synthesis, as well as DNA replication, and transcription-related functions, are enriched in younger ovaries. Proteins involved in cell division are at a higher level in SE than in SE+5d and more than half of them show a strong ascending gradient in SE in agreement with the fact that cell division is more active in young cells. Protein degradation function is also highly expressed at silk emergence. Enzymes allowing the inter-conversion of metabolic hubs such as pyruvate, phosphoenolpyruvate, and acetyl-CoA are also more abundant at the silk emergence stage.

On the contrary, translation is more active in older cells, where ribosomal proteins are more abundant. Carbohydrate metabolism pathways with glycolysis and citrate cycle enzymes are also upregulated in older ovaries. Precursors of jasmonic acid, involved in plant development have also been identified as impacted during ovary development. Interestingly, the differential analysis identified an interaction between the day and zone effects for three chitinases whose expressions were higher in older ovaries (SE+5d), which could correspond to a defense process against fungi that is settled over ovaries development.

To summarize, the following changes were shown to occur during ovary development:

- at the early stages of ovary development, DNA replication, transcription, and protein degradation are active processes while translation is down. Starch and metabolic hub synthesis is also favored
- maturation of ovaries coincides with activation of glucose metabolism, starch to glucose conversion, and glucose catabolism through glycolysis are activated
- mature ovaries exhibit active translation-related functions.

### 4.2. Response to water deficit

Although plants were submitted to a moderate water deficit, our results show that it was the primary cause of protein variation: i) the PCA clearly separated WD from WW samples along the first principal component; ii) the highest number of significant differences identified by ANOVA was for the WD effect, iii) the majority of the clusters showed an effect of water deficit with or without interaction with a developmental effect. In addition, a large difference between the WW and WD co-expression networks was observed.

Independently of the developmental stage, in the WD condition, ovaries are characterized by the higher abundance of starch synthesis proteins and enzymes catalyzing metabolic hubs inter-conversions and by the lower abundance of proteins involved in carbon metabolism. This metabolic pattern is similar to that observed in young ovaries in the WW condition, suggesting that ovaries under WD behave like young ovaries and slow down their carbohydrate metabolism. WD is also associated with the increase of proteins involved in cell division, chromatin organization, and RNA processing.

Consistently, the WD network highlights a loss of coordination between proteins involved in carbohydrate metabolism but also new connections between pathways in WD while the ones observed in the WW condition are still present, suggesting more coordination mechanisms between pathways in the WD than in the WW condition. The inter-pathway coordination could to rely on RNA processing proteins.

This indicates that these proteins have a dominant regulatory role on different pathways under WD, whereas their role would be less important under the WW condition. Interestingly, some of them are indeed involved in stress response. Notably, a Tudor-SN protein known to be induced by stress is primarily increased in the WD condition. It plays a role in sequestering mRNA in stress granules to allow the translation of stress-specific mRNA [38,39]. Enzymes allowing the synthesis and degradation metabolic hubs such as pyruvate, citrate, oxaloacetate, and phosphoenolpyruvate are abundant in the WD condition and could also explain the coordination of pathways. The analysis of protein coexpression networks identified alcohol dehydrogenase and sucrose synthase [43] as potential orchestrators of changes of correlation of expression.

Protein biosynthesis and processing pathways as well as vesicular trafficking are also activated in WD while there is a decrease in most enzymes of the aminoacid metabolism and proteins involved in protein degradation (mostly proteasome subunits). In this aspect, WD ovaries seem to follow their maturation process by maintaining their protein pool through an activation of translation and protein processing (ER) and a limitation of protein degradation. These processes do not appear to rely on amino acid biosynthesis. We can speculate that the increase in proteins involved in vesicular trafficking enables the remobilization of intra- or extracellular protein matter, to be used as amino acid supply. In connection with this increase in vesicle trafficking proteins, two phospholipases D are induced in WD. These proteins play a role in stress signaling and vesicle trafficking [44].

Only two proteins exhibiting a strong condition*zone interaction effect were identified while a significant effect of the day*condition interaction has been observed on 21 proteins. This suggests that WD poorly interferes with short-term ovary developmental program but it does so at a larger developmental time scale. As recently shown on leaves [45], molecular response to water deficit at least partially depends on the age of the tissue. Indeed, as illustrated by cluster 3, some protein abundances are impacted by WD specifically at the silk emergence date. Among them, two expansins (proteins involved in cell expansion) appear induced by WD at silk emergence while their abundances decrease in WD at the later stages of ovary development.

The moderate deficit applied in this study does not induce a typical stress response in the maize ovary proteome. There is no increase in proteins involved in the synthesis of osmoprotectants such as proline (pyrroline-5-carboxylate reductase) or betaine-glycine (betaine-aldehyde dehydrogenase), no general increase in chaperone proteins or dehydrins, and no increase in proteins involved in defense against protein oxidative stress, either ascorbate or thiol-based in WD. Indeed, despite WD globally promote oxidative phosphorylation producing ROS, redox homeostasis is downregulated.

Although there is no typical stress response, the ovary proteome is strongly affected by water deficit. Profound changes in the proteome are observed as soon as at the stage of silk emergence.

Several hormone signaling genes have been detected as upregulated in response to abiotic stress in ovaries under heat stress [46]). Jasmonic acid and abscisic acid signaling are induced upon drought in maize ears [47]. Auxins are involved in growth but also in in seed development [48,49] and response to drought stress [50]. Brassinosteroids interact with auxins to promote cell expansion [51] and play a role in drought stress and senescence [52]. In the present study, the observed changes do not appear to be linked to ABA, as the abundance of proteins involved in its synthesis decrease upon WD. By contrast, other proteins involved in hormonal regulation increase upon WD, such as a topless-related2 protein (TPL/TPR), with relevance to auxins. This protein, as well as nana2, another protein increased upon WD, can also participate in brassinosteroids metabolism [53,54]. Two 9-lipoxygenases involved in brassinosteroid signaling [40] are also increased in WD. An acyl-CoA-binding protein (ACBP6), a protein involved in jasmonic acid synthesis, is increased in WD. Altogether, our results indicate that the changes induced by water deficit are more related to auxins, brassinosteroids, and jasmonate than to abscisic acid signaling. Several responses to WD are typical of young ovaries. Notably, the abundance of proteins involved in transcription and glucose/starch balance are similar to that of young ovaries, suggesting that upon WD the ovaries slow down their development and that some proteins stay at the level that they reached when they were young. Consistently with our results, time course transcriptome analysis of ears under stronger water stress demonstrated an initial downregulation of cell division (DNA replication and cell cycle) related genes followed by a downregulation of genes related to glucose metabolic process, glycolysis, and tricarboxylic acid cycle [47]. Our results also support Oury et al. conclusions [15] showing that under water deficit, the carbon metabolism down-regulation does not participate in the primary response of ears and that expansive growth is firstly impacted by WD. However, similarly to older ovaries, translation is active in WD but the amino acid pool could be provided by alternative ways than amino acid synthesis. Such changes observed in protein synthesis in response to WD have not been described before.

## 5. Conclusion

Several conclusions can be drawn from this study. The proteome of ovaries changes during their development, the proteins involved in cell division and expansion being more abundant in the young than in the older ovaries. A part of the changes induced by mild water deficit is a slowdown of the development: in WD condition the abundance of several proteins looks like their level in younger ovaries. But other changes cannot be related to a developmental slowdown: for example, proteins involved in protein biosynthesis are increased upon water deficit. At the same time, proteins involved in amino acid metabolism and proteolysis decrease, which raises questions about the aminoacid availability for translation. There is a concomitant increase of proteins involved in vesicle traffic. The coexpression networks in WW and WD are very different, confirming that changes are not only related to developmental slowdown. These changes can be directly or indirectly regulated by several of the proteins involved in RNA processing which are increased in WD. Even mild, water deficit has a major and rapid impact on ovary proteome, but its response differs from a response to stress: the proteome does not show the standard symptoms of response to water deficit, like an increased abundance of dehydrins, chaperones or proteins involved in redox homeostasis. The changes may be linked to hormone signaling: several proteins involved in auxin, brassinosteroids, and jasmonate, but not ABA, were increased upon water deficit.

## Author Contributions

MZ and OT designed the experiment. TB and MA performed the experiments. TB, MA, EM and MZ analyzed the results. MBN, EM and MZ discussed the results and wrote the manuscript.

## Data Availability Statement

The mass spectrometry proteomics data have been deposited to the ProteomeXchange Consortium via the PRIDE [55] partner repository with the dataset identifier PXD053569. At the moment only reviewers have access by logging in to the PRIDE website using the following account details: Username: Password: tFL7cxQDcfjO

## Abbreviations

The following abbreviations are used in this manuscript:

WD: Water deficit
WW: Well watered
PCA: Principal Component Analysis
DEP: Differentially Expressed Protein
SE: Silk Emergence

## Appendix A

*Appendix A.1 A1_Boxplot-all-proteins.pdf*

Boxplots representing the abundances of each of the proteins according to the day, zone and condition of the sample.

*Appendix A.2 A2_TableS1.csv*

List of the 800 proteins of the dataset and their Mapman annotations

*Appendix A.3 A3_TableS2_clusters_mapman.csv*

- Observed numbers of protein per cluster and per Mapman term
- Expected numbers of protein per cluster and per Mapman term under the independence hypothesis
- Chi2 components calculated for each couple of cluster / Mapman term
- Number of significantly differentially expressed proteins per couple of cluster / Mapman term

*Appendix A.4 A4_TableS3_differential_analysis.csv*

Pvalues and adjusted pvalues for each of the tested effects and calculated from the differential analysis.

*Appendix A.5 A5_KEGGpathways.pdf*

Map corresponding to several KEGG pathways encompassing differentially expressed proteins. The zone and days maps represent proteins expressed in the younger stages in yellow and proteins expressed in older stages in brown. The conditions maps represent proteins induced in WD in green and proteins induced in WW in red.

*Appendix A.6 A6_TableS4_Cor_SignifWW.csv*

Matrix representing pairs of coexpressed proteins in WW condition.

*Appendix A.7 A7_TableS5_Cor_SignifWD.csv*

Matrix representing pairs of coexpressed proteins in WD condition.

*Appendix A.8 A8_TableS6_KEGG-KO-protnames.csv*

Correspondance table between protein names, KO compounds and KEGG pathways.

*Appendix A.9 A9_TableS7_networkProtHubs.csv*

List of high degree nodes whose connectivities changed between WW and WD deficit condition.

## Disclaimer/Publisher’s Note

The statements, opinions and data contained in all publications are solely those of the individual author(s) and contributor(s) and not of MDPI and/or the editor(s). MDPI and/or the editor(s) disclaim responsibility for any injury to people or property resulting from any ideas, methods, instructions or products referred to in the content.

