## Appendix A1 for "A moderate water deficit induces profound changes in the proteome of developing maize ovaries"

### Individual plots of all quantified proteins

Red : WD
Green : WW

x-axis labels : treatment.zone.day

zeammp036

**Cellular respiration.oxidative phosphorylation**
**description:nad9**

component *(NQO5/ND9) of NADH dehydrogenase electron output (module Q)

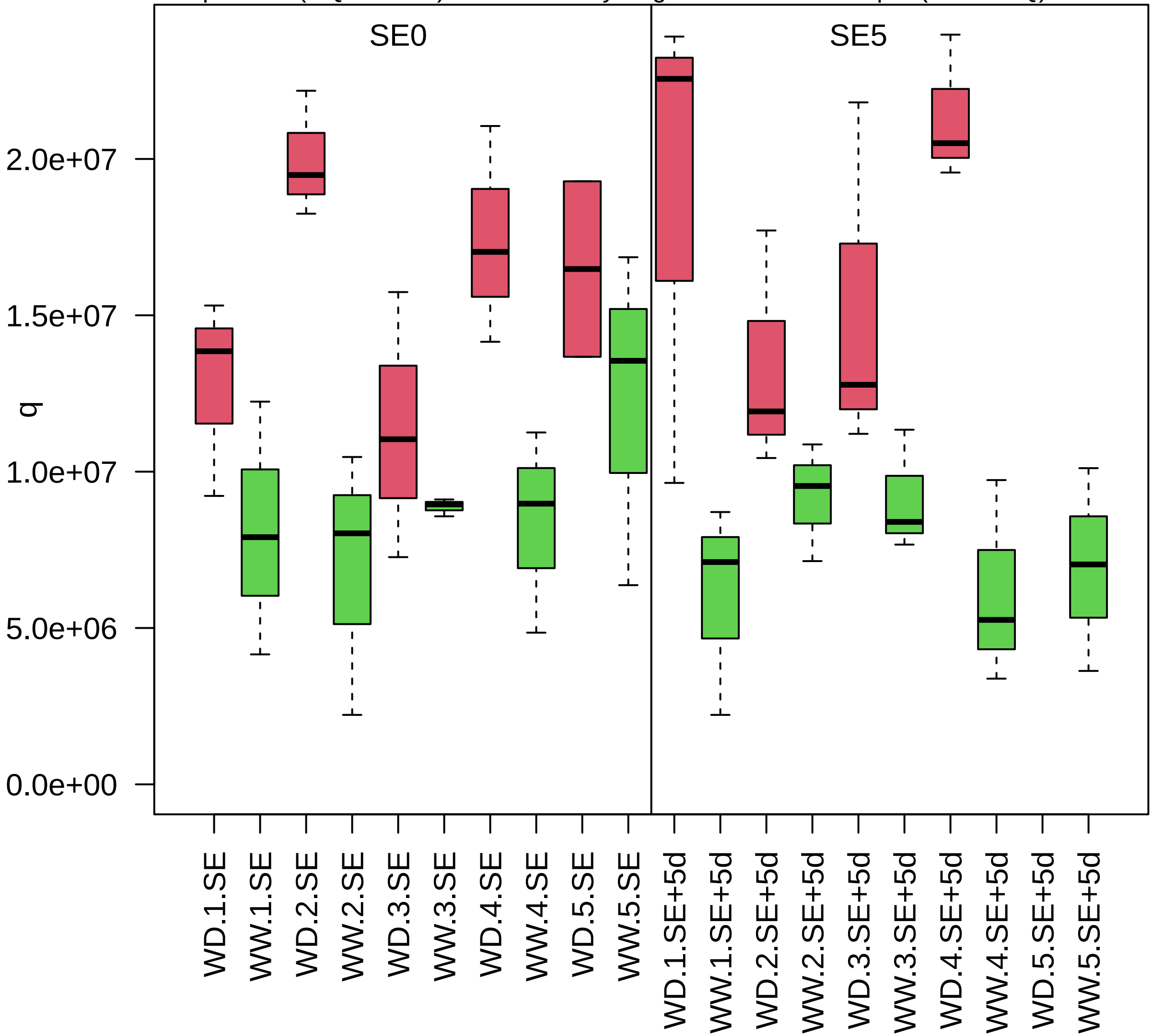

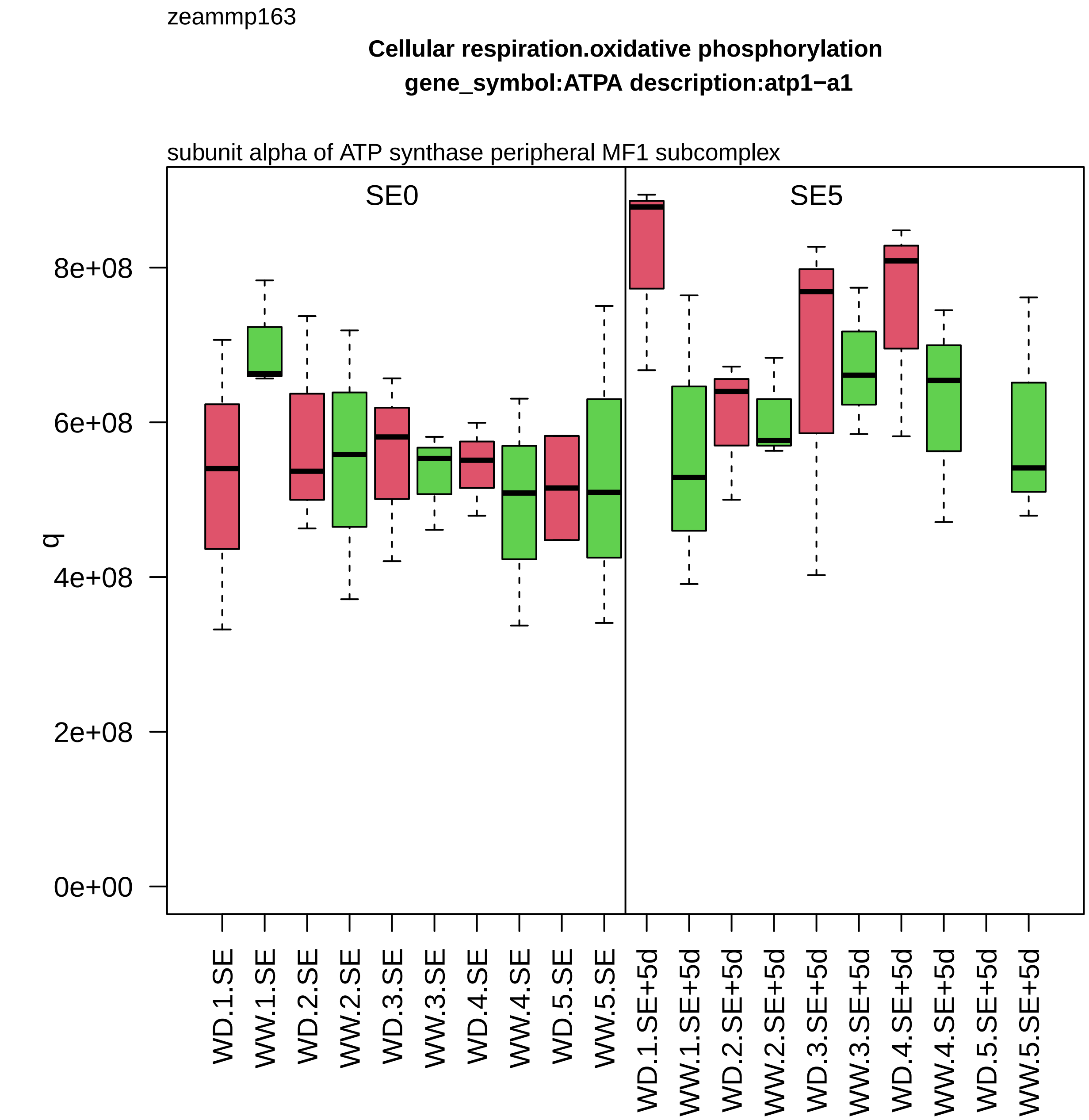
zeammp163
Cellular respiration.oxidative phosphorylation
gene_symbol:ATPA description:atp1−a1
subunit alpha of ATP synthase peripheral MF1 subcomplex
SE0
SE5
q
0e+00
2e+08
4e+08
6e+08
8e+08
WD.1.SE
WW.1.SE
WD.2.SE
WW.2.SE
WD.3.SE
WW.3.SE
WD.4.SE
WW.4.SE
WD.5.SE
WW.5.SE
WD.1.SE+5d
WW.1.SE+5d
WD.2.SE+5d
WW.2.SE+5d
WD.3.SE+5d
WW.3.SE+5d
WD.4.SE+5d
WW.4.SE+5d
WD.5.SE+5d
WW.5.SE+5d

zm00001d000001_p002

**Secondary metabolism.phenolics**

**gene_symbol:polyphenol oxidase1 description: polyphenol oxidase1**

aureusidin synthase

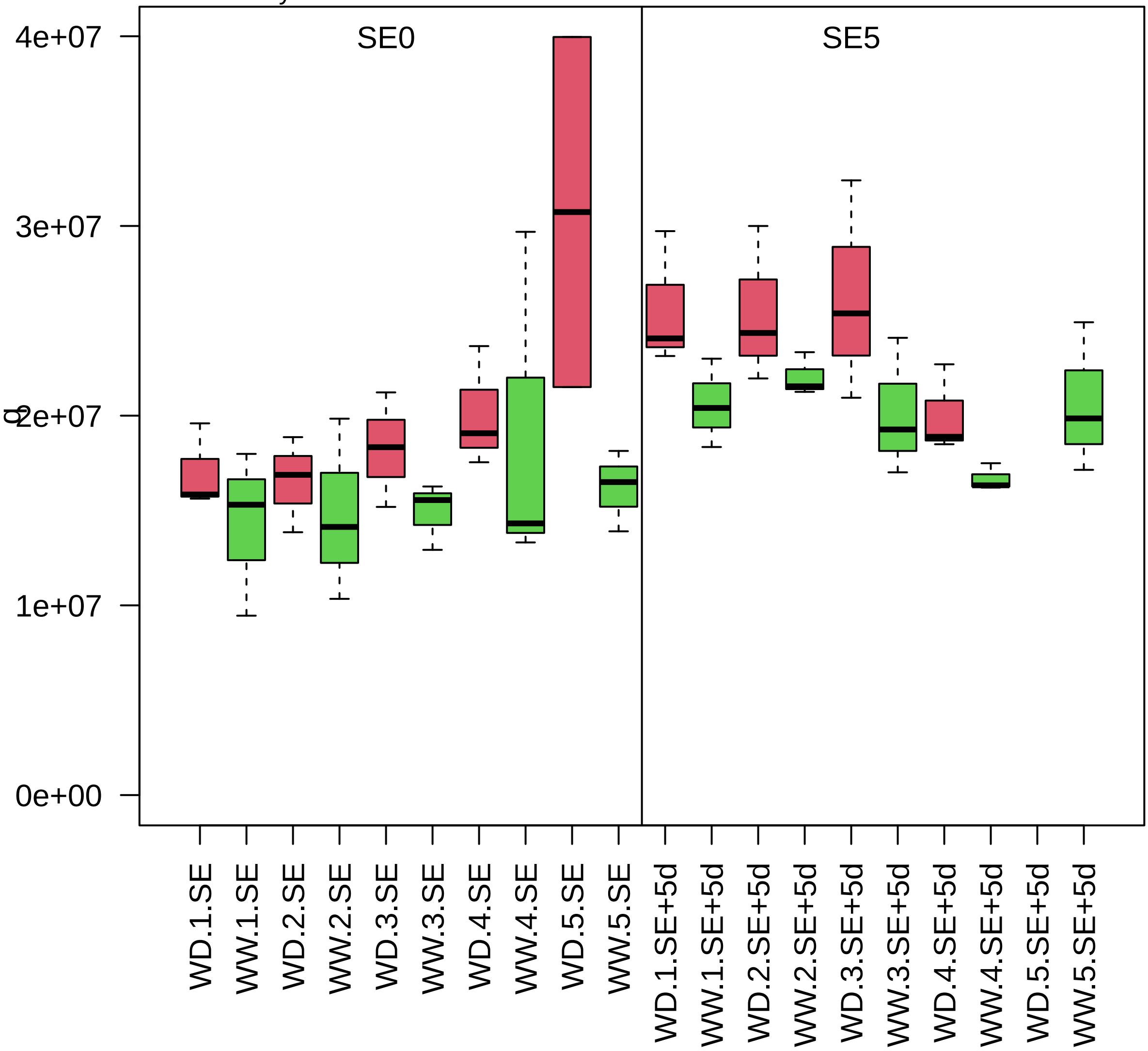

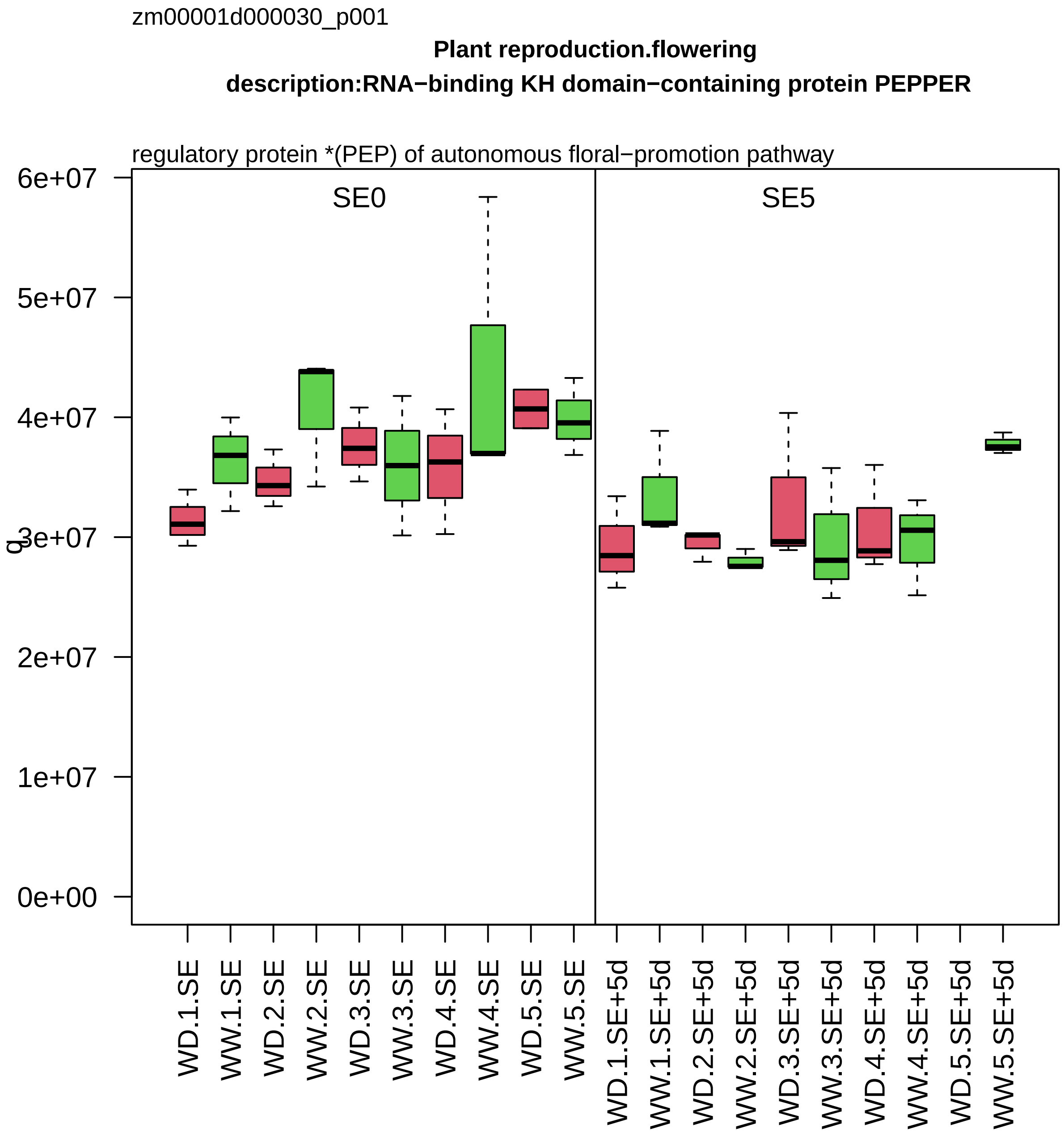
zm00001d000030_p001
Plant reproduction.flowering
description:RNA−binding KH domain−containing protein PEPPER
regulatory protein *(PEP) of autonomous floral−promotion pathway
SE0
SE5
q
6e+07
5e+07
4e+07
3e+07
2e+07
1e+07
0e+00
WD.1.SE
WW.1.SE
WD.2.SE
WW.2.SE
WD.3.SE
WW.3.SE
WD.4.SE
WW.4.SE
WD.5.SE
WW.5.SE
WD.1.SE+5d
WW.1.SE+5d
WD.2.SE+5d
WW.2.SE+5d
WD.3.SE+5d
WW.3.SE+5d
WD.4.SE+5d
WW.4.SE+5d
WD.5.SE+5d
WW.5.SE+5d

zm00001d000227_p001

**Lipid metabolism.fatty acid metabolism**

**description:Pyruvate dehydrogenase E1 component subunit beta**

subunit beta of E1 subcomplex of pyruvate dehydrogenase complex

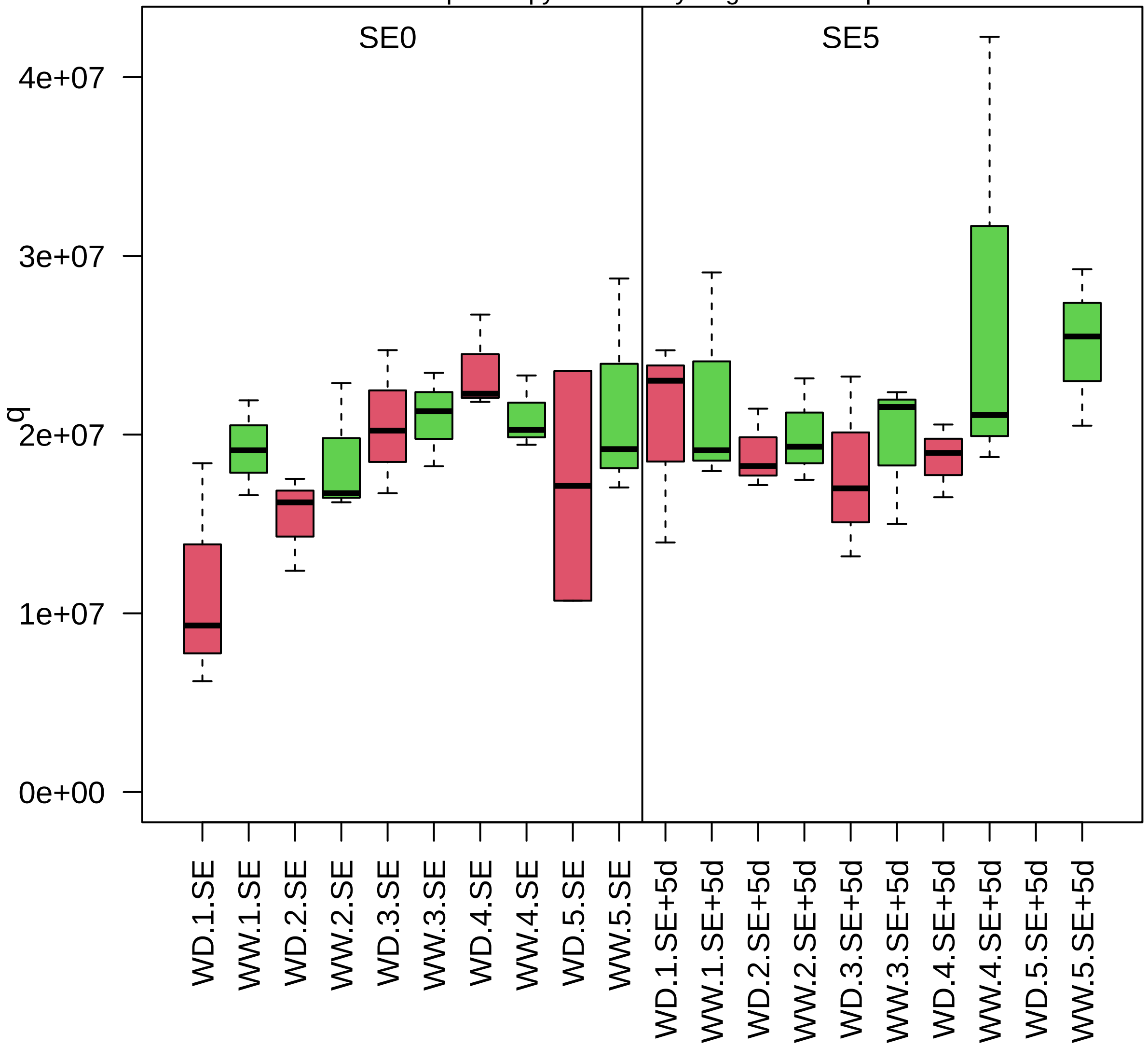

zm00001d001771_p001

**Protein homeostasis.proteolysis**

**description:Eukaryotic aspartyl protease family protein**

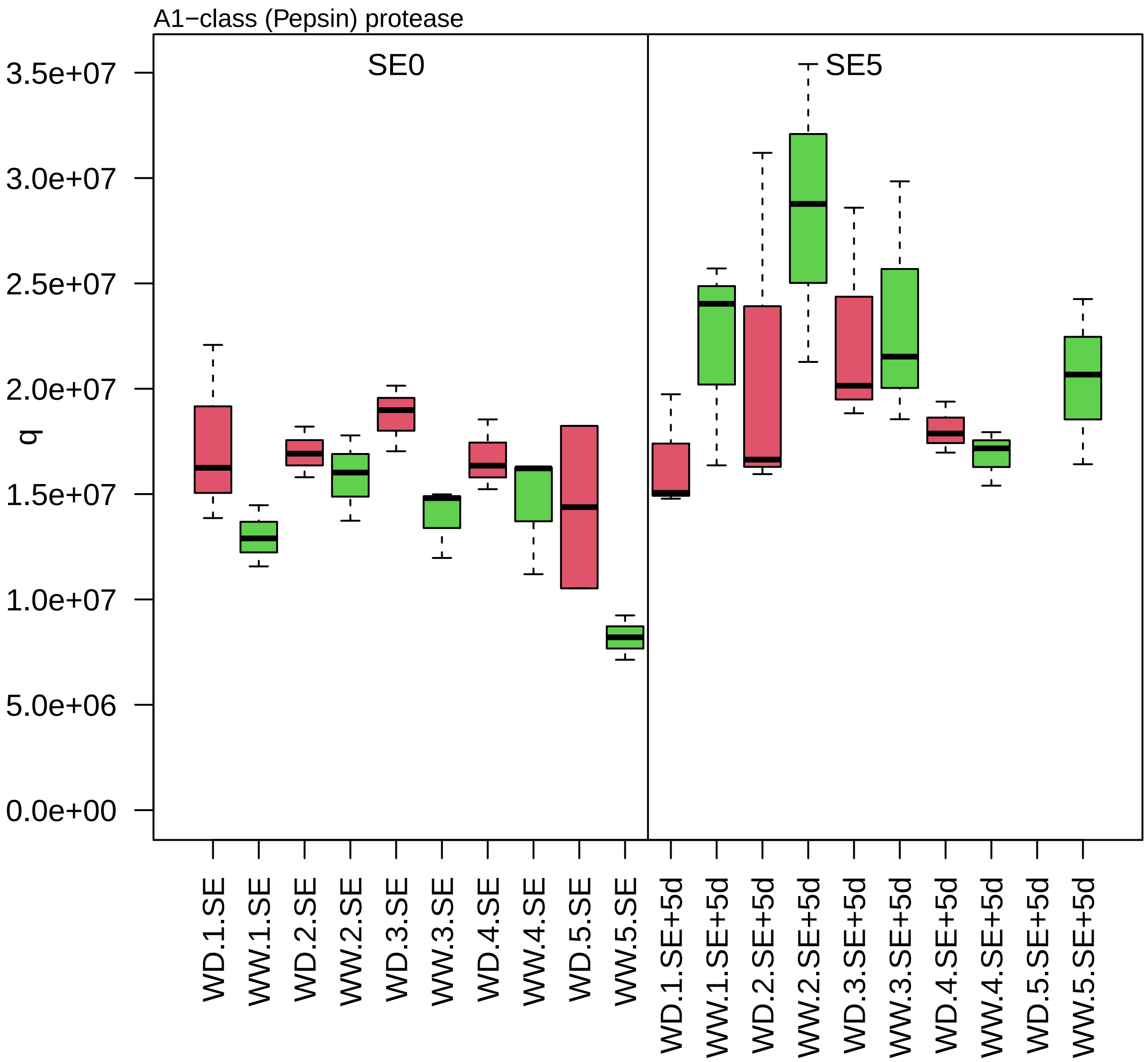

zm00001d001831_p001

**Cellular respiration.glycolysis**
**description:Pyruvate kinase**

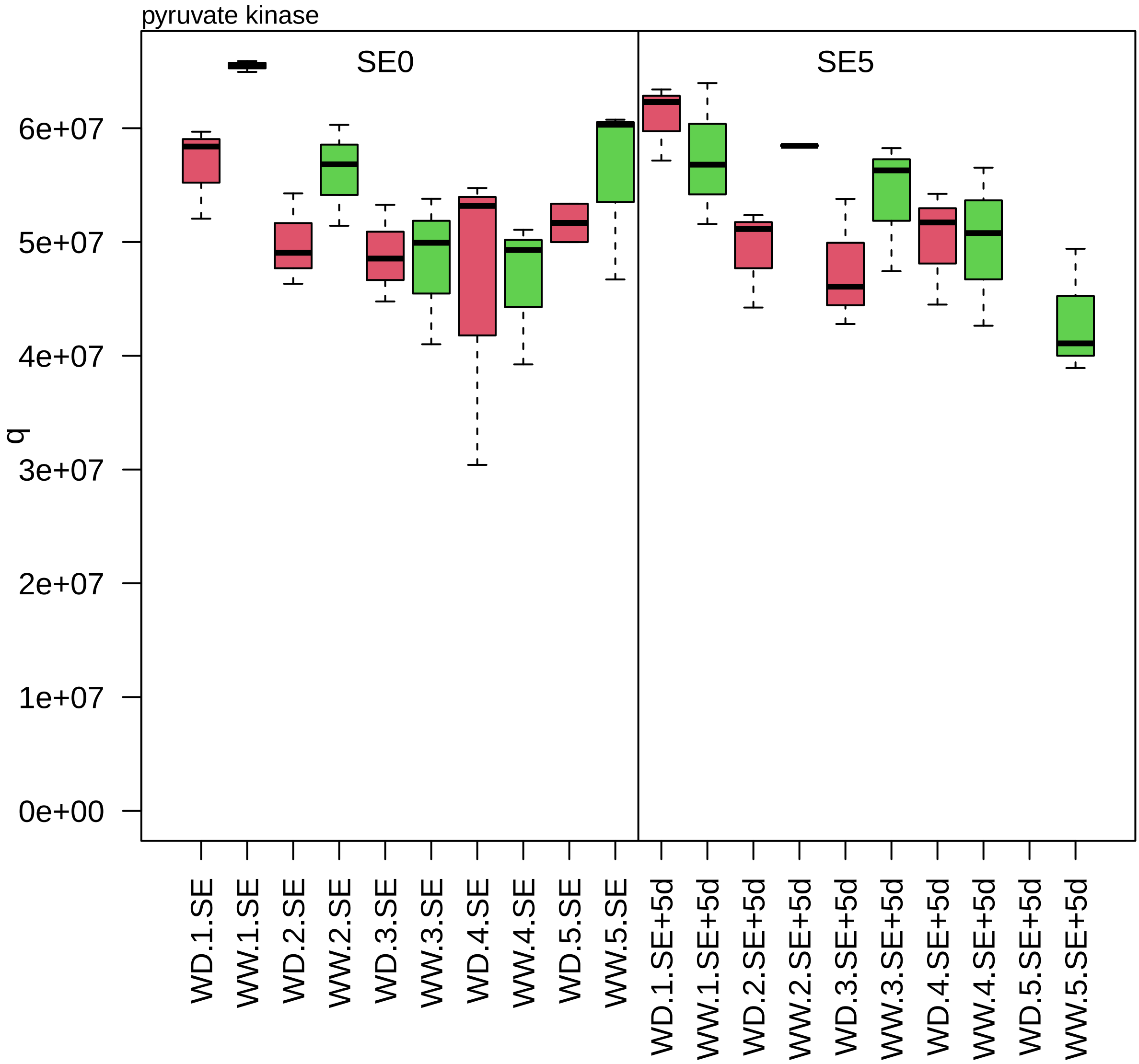

zm00001d001913_p001

**Protein homeostasis.ubiquitin−proteasome system**

**gene_symbol:ubiquitin conjugating enzyme4 description:Ubiquitin−conjugating enzyme E2 11**

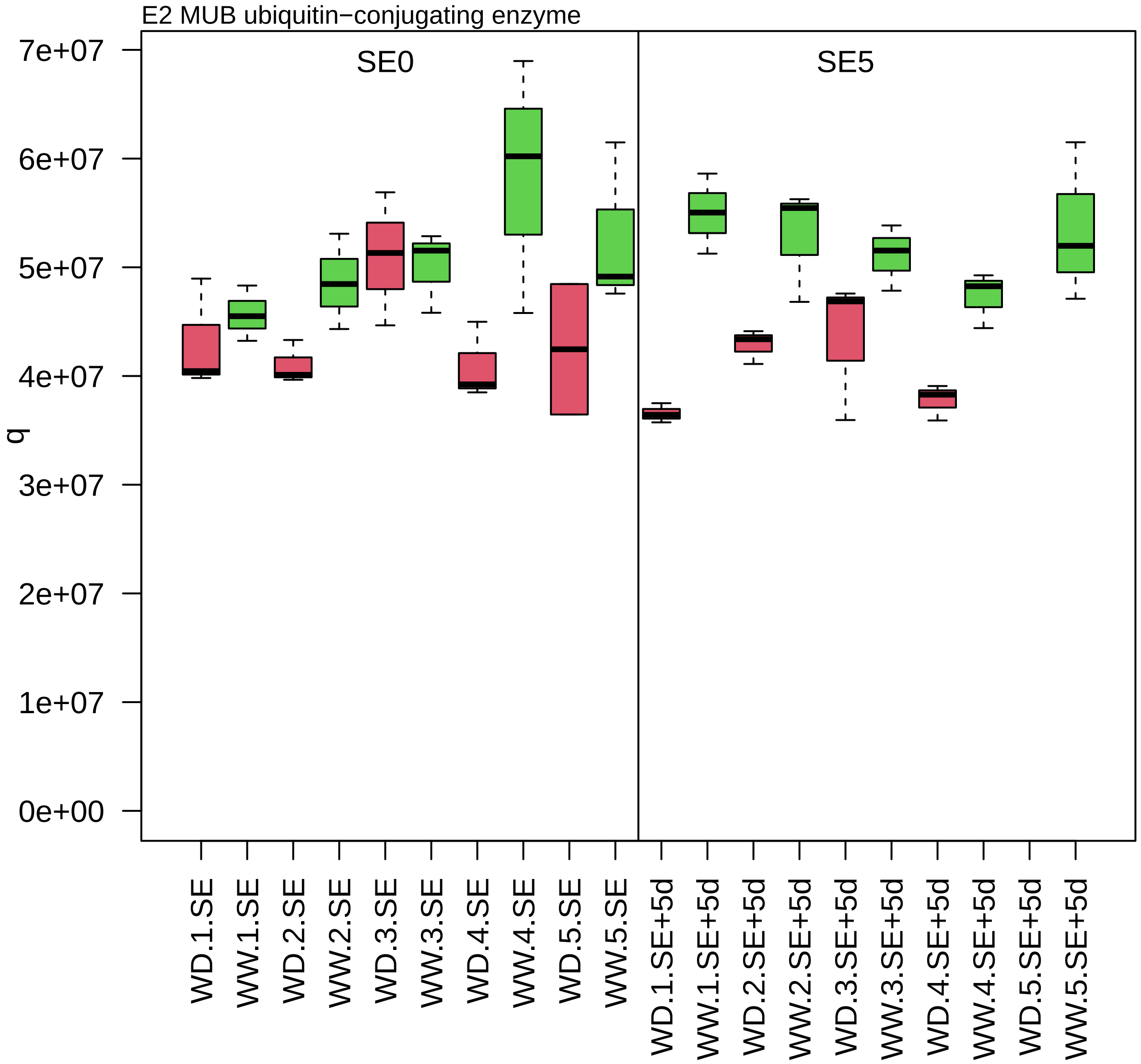

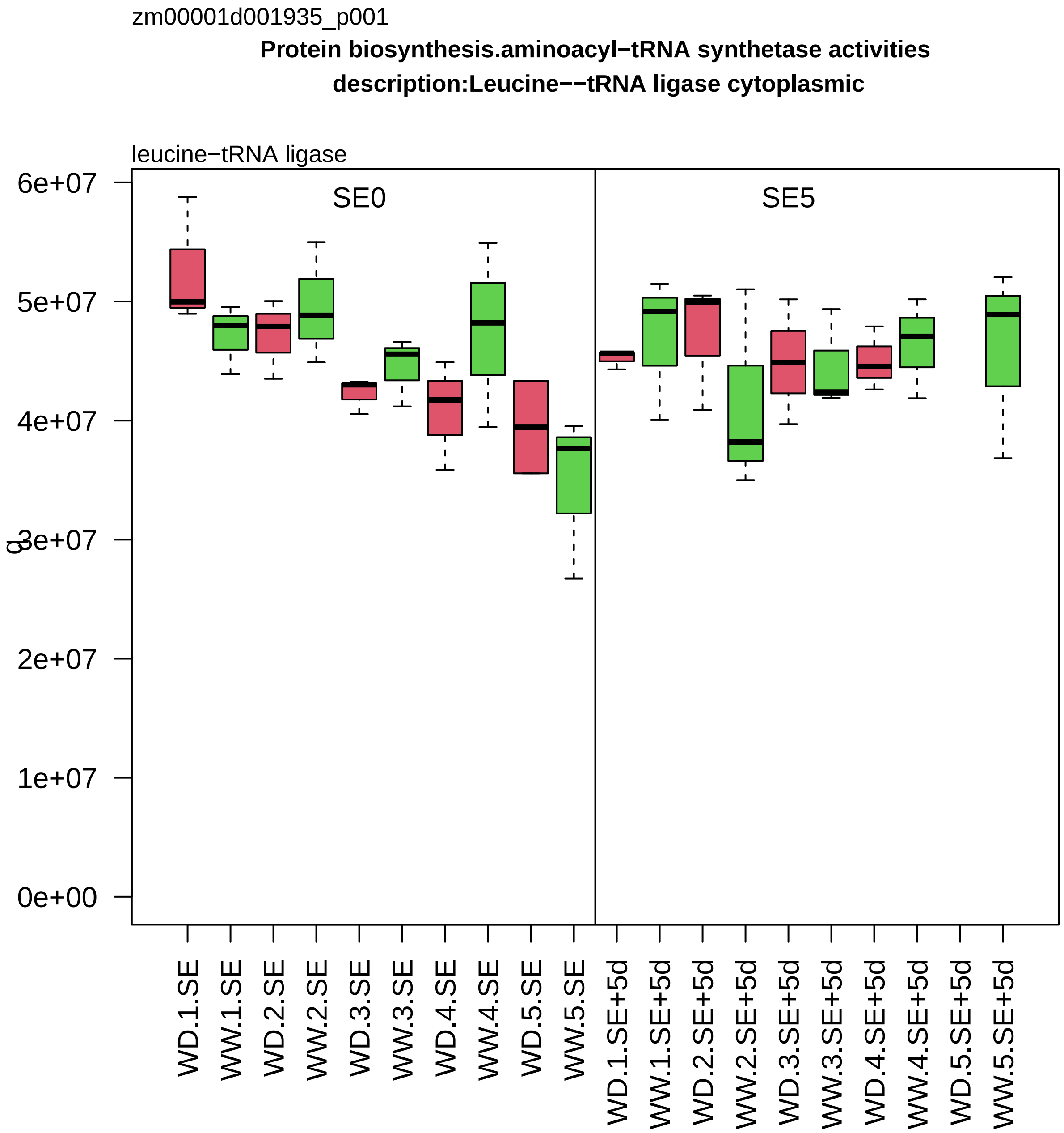
zm00001d001935_p001
Protein biosynthesis.aminoacyl−tRNA synthetase activities
description:Leucine−−tRNA ligase cytoplasmic
leucine−tRNA ligase
SE0
SE5
0e+00
1e+07
2e+07
3e+07
4e+07
5e+07
6e+07
q
WD.1.SE
WW.1.SE
WD.2.SE
WW.2.SE
WD.3.SE
WW.3.SE
WD.4.SE
WW.4.SE
WD.5.SE
WW.5.SE
WD.1.SE+5d
WW.1.SE+5d
WD.2.SE+5d
WW.2.SE+5d
WD.3.SE+5d
WW.3.SE+5d
WD.4.SE+5d
WW.4.SE+5d
WD.5.SE+5d
WW.5.SE+5d

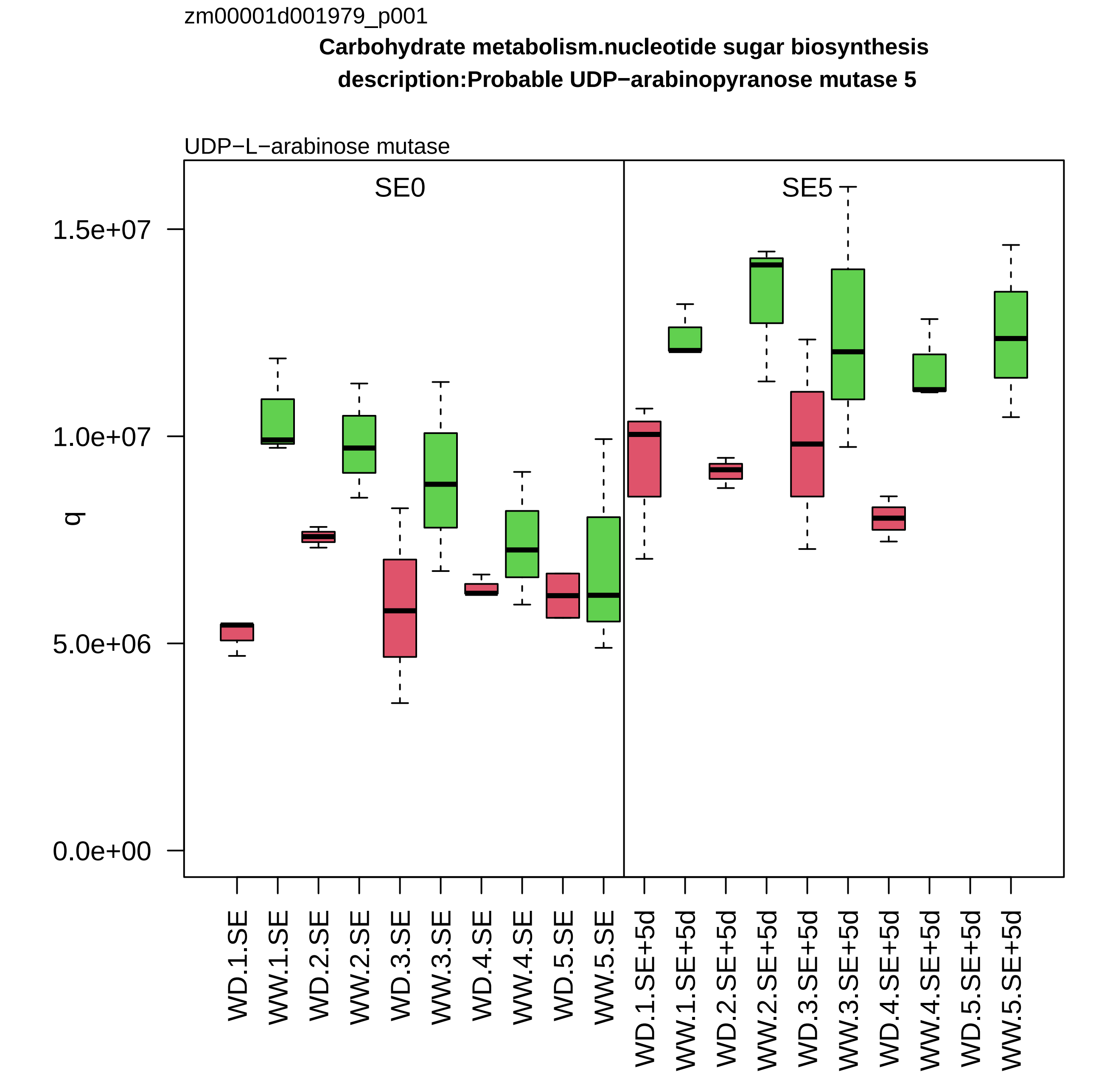
zm00001d001979_p001
Carbohydrate metabolism.nucleotide sugar biosynthesis
description:Probable UDP−arabinopyranose mutase 5
UDP−L−arabinose mutase
SE0
SE5
q
1.5e+07
1.0e+07
5.0e+06
0.0e+00
WD.1.SE
WW.1.SE
WD.2.SE
WW.2.SE
WD.3.SE
WW.3.SE
WD.4.SE
WW.4.SE
WD.5.SE
WW.5.SE
WD.1.SE+5d
WW.1.SE+5d
WD.2.SE+5d
WW.2.SE+5d
WD.3.SE+5d
WW.3.SE+5d
WD.4.SE+5d
WW.4.SE+5d
WD.5.SE+5d
WW.5.SE+5d

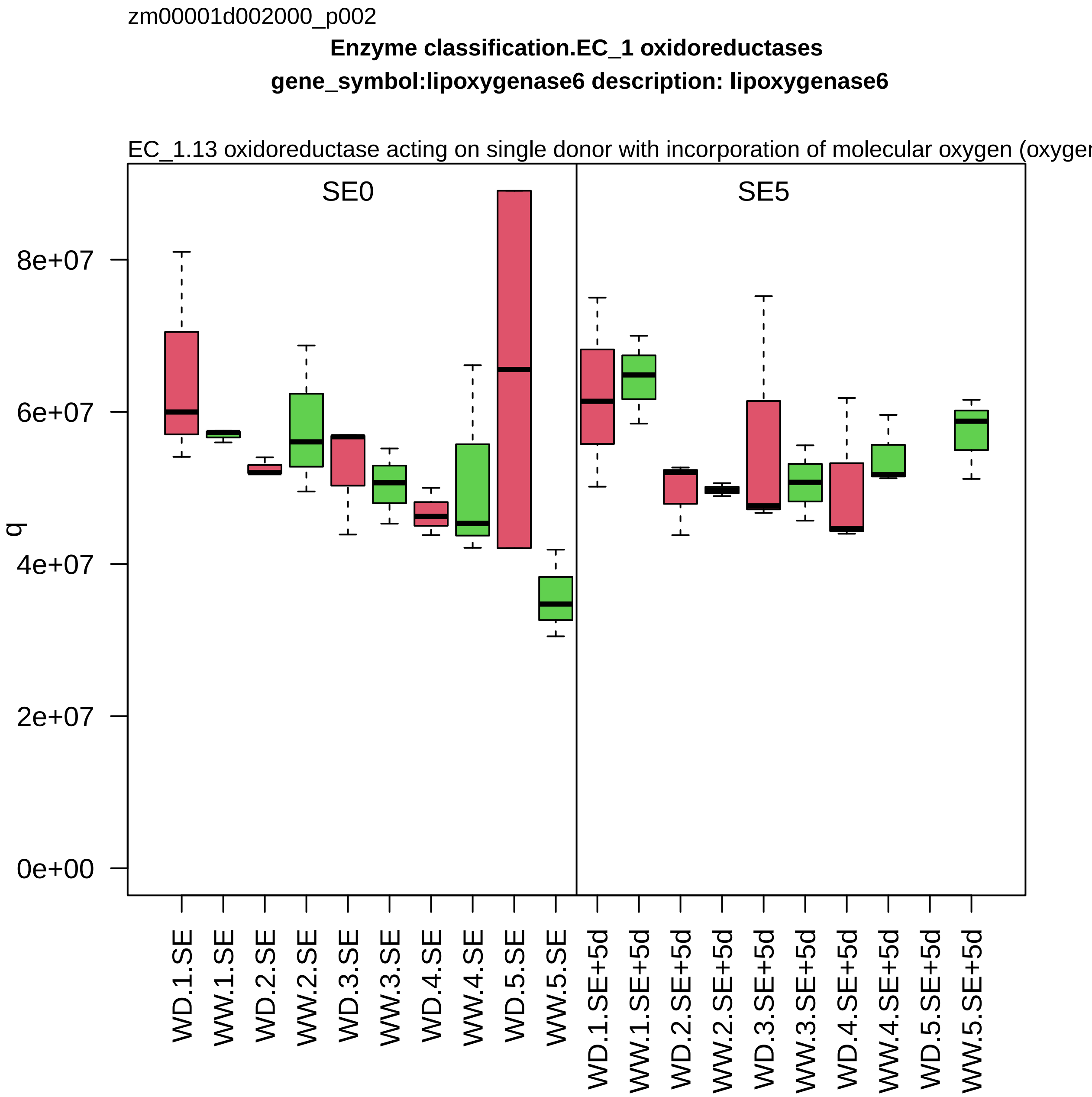
zm00001d002000_p002
Enzyme classification.EC_1 oxidoreductases
gene_symbol:lipoxygenase6 description: lipoxygenase6
EC_1.13 oxidoreductase acting on single donor with incorporation of molecular oxygen (oxygen
SE0
SE5
q
8e+07
6e+07
4e+07
2e+07
0e+00
WD.1.SE
WW.1.SE
WD.2.SE
WW.2.SE
WD.3.SE
WW.3.SE
WD.4.SE
WW.4.SE
WD.5.SE
WW.5.SE
WD.1.SE+5d
WW.1.SE+5d
WD.2.SE+5d
WW.2.SE+5d
WD.3.SE+5d
WW.3.SE+5d
WD.4.SE+5d
WW.4.SE+5d
WD.5.SE+5d
WW.5.SE+5d

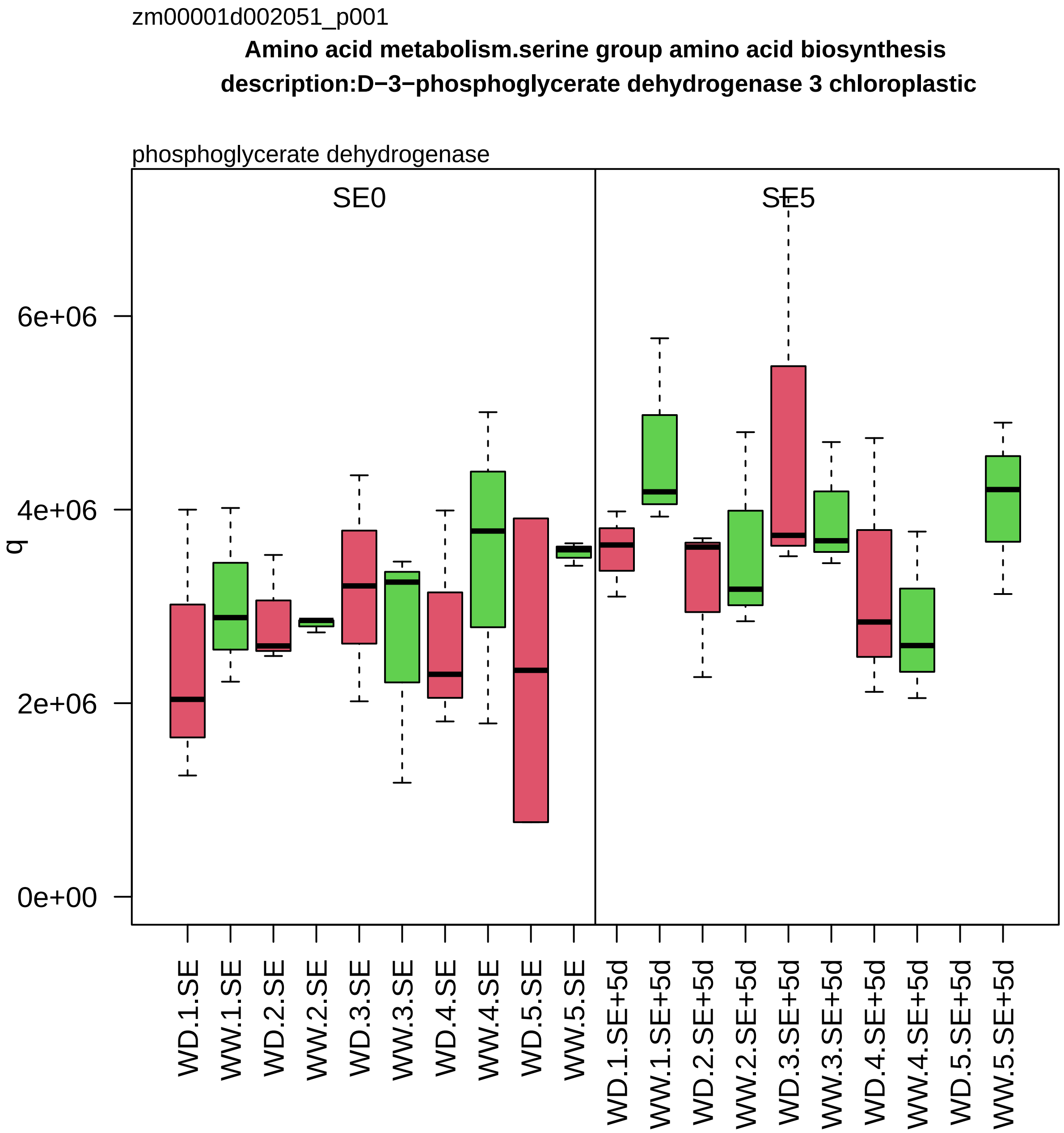
zm00001d002051_p001
Amino acid metabolism.serine group amino acid biosynthesis
description:D−3−phosphoglycerate dehydrogenase 3 chloroplastic
phosphoglycerate dehydrogenase
SE0
SE5
q
0e+00
2e+06
4e+06
6e+06
WD.1.SE
WW.1.SE
WD.2.SE
WW.2.SE
WD.3.SE
WW.3.SE
WD.4.SE
WW.4.SE
WD.5.SE
WW.5.SE
WD.1.SE+5d
WW.1.SE+5d
WD.2.SE+5d
WW.2.SE+5d
WD.3.SE+5d
WW.3.SE+5d
WD.4.SE+5d
WW.4.SE+5d
WD.5.SE+5d
WW.5.SE+5d

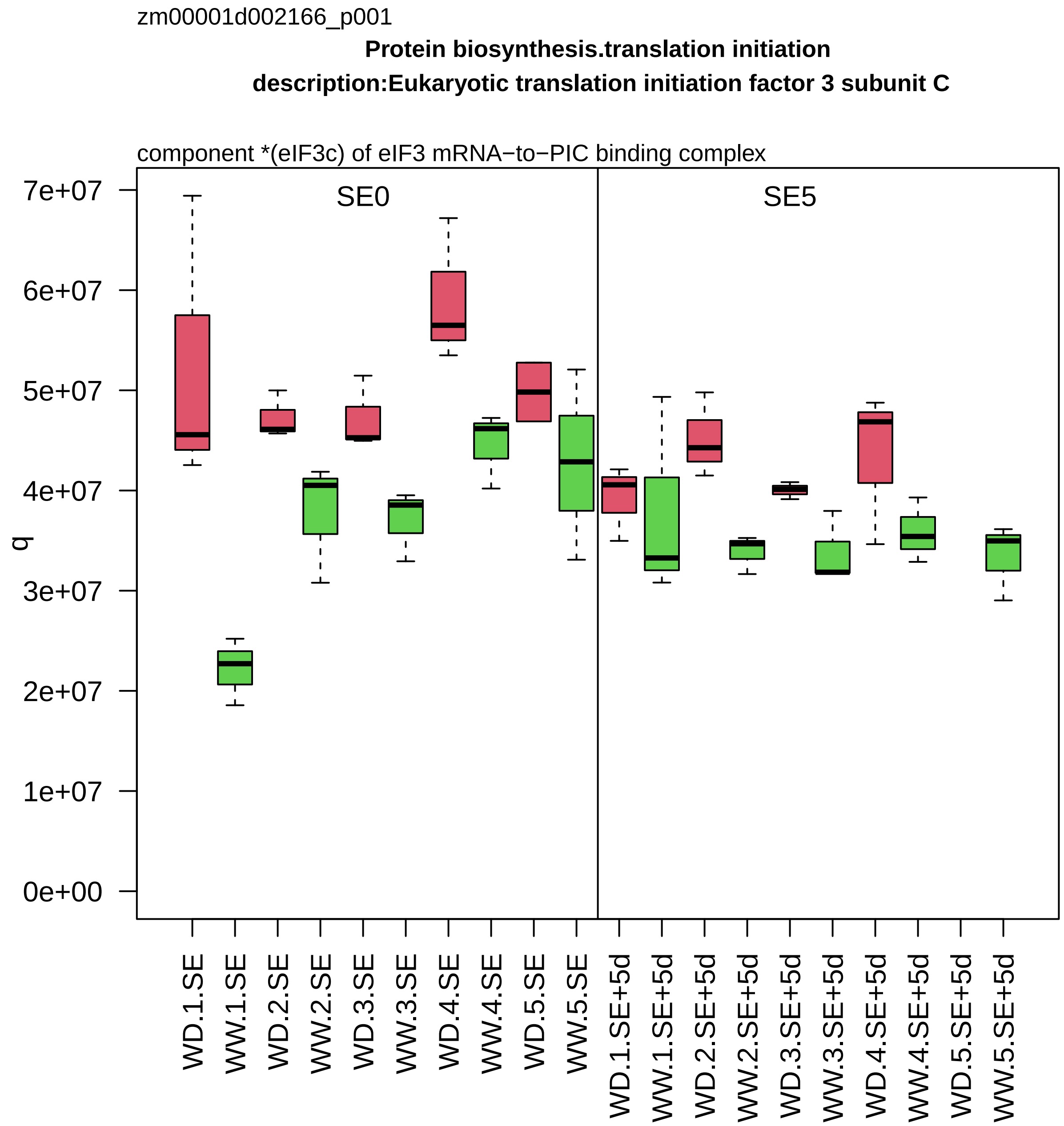

zm00001d002166_p001
Protein biosynthesis.translation initiation
description:Eukaryotic translation initiation factor 3 subunit C
component *(eIF3c) of eIF3 mRNA−to−PIC binding complex
SE0
SE5
q
7e+07
6e+07
5e+07
4e+07
3e+07
2e+07
1e+07
0e+00
WD.1.SE
WW.1.SE
WD.2.SE
WW.2.SE
WD.3.SE
WW.3.SE
WD.4.SE
WW.4.SE
WD.5.SE
WW.5.SE
WD.1.SE+5d
WW.1.SE+5d
WD.2.SE+5d
WW.2.SE+5d
WD.3.SE+5d
WW.3.SE+5d
WD.4.SE+5d
WW.4.SE+5d
WD.5.SE+5d
WW.5.SE+5d

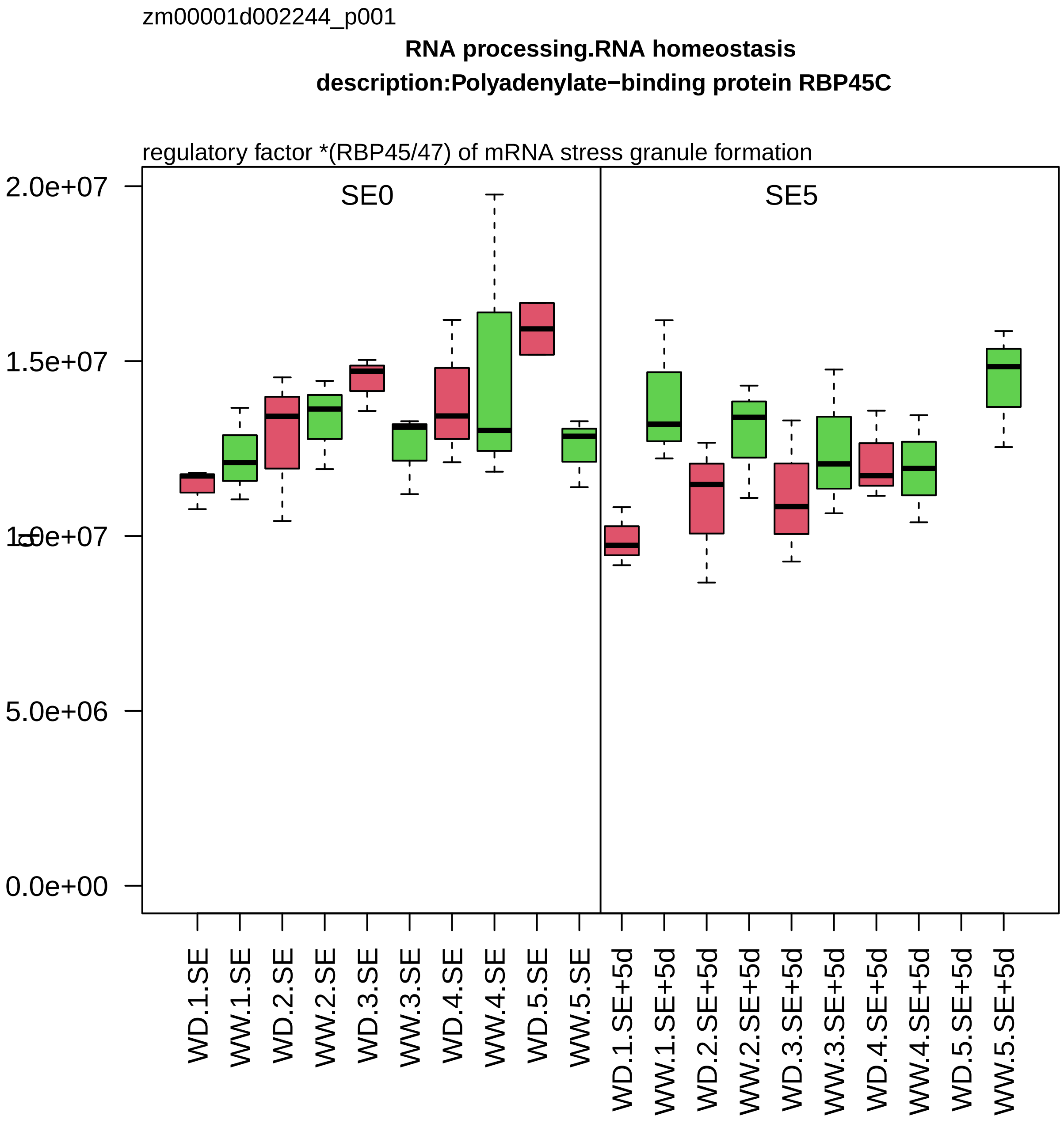
zm00001d002244_p001
RNA processing.RNA homeostasis
description:Polyadenylate−binding protein RBP45C
regulatory factor *(RBP45/47) of mRNA stress granule formation
SE0
SE5
2.0e+07
1.5e+07
1.0e+07
5.0e+06
0.0e+00
WD.1.SE
WW.1.SE
WD.2.SE
WW.2.SE
WD.3.SE
WW.3.SE
WD.4.SE
WW.4.SE
WD.5.SE
WW.5.SE
WD.1.SE+5d
WW.1.SE+5d
WD.2.SE+5d
WW.2.SE+5d
WD.3.SE+5d
WW.3.SE+5d
WD.4.SE+5d
WW.4.SE+5d
WD.5.SE+5d
WW.5.SE+5d

zm00001d002253_p001

**Protein biosynthesis.ribosome biogenesis**
**description:60S ribosomal protein L27**

component *(eL27) of large ribosomal−subunit (LSU) proteome

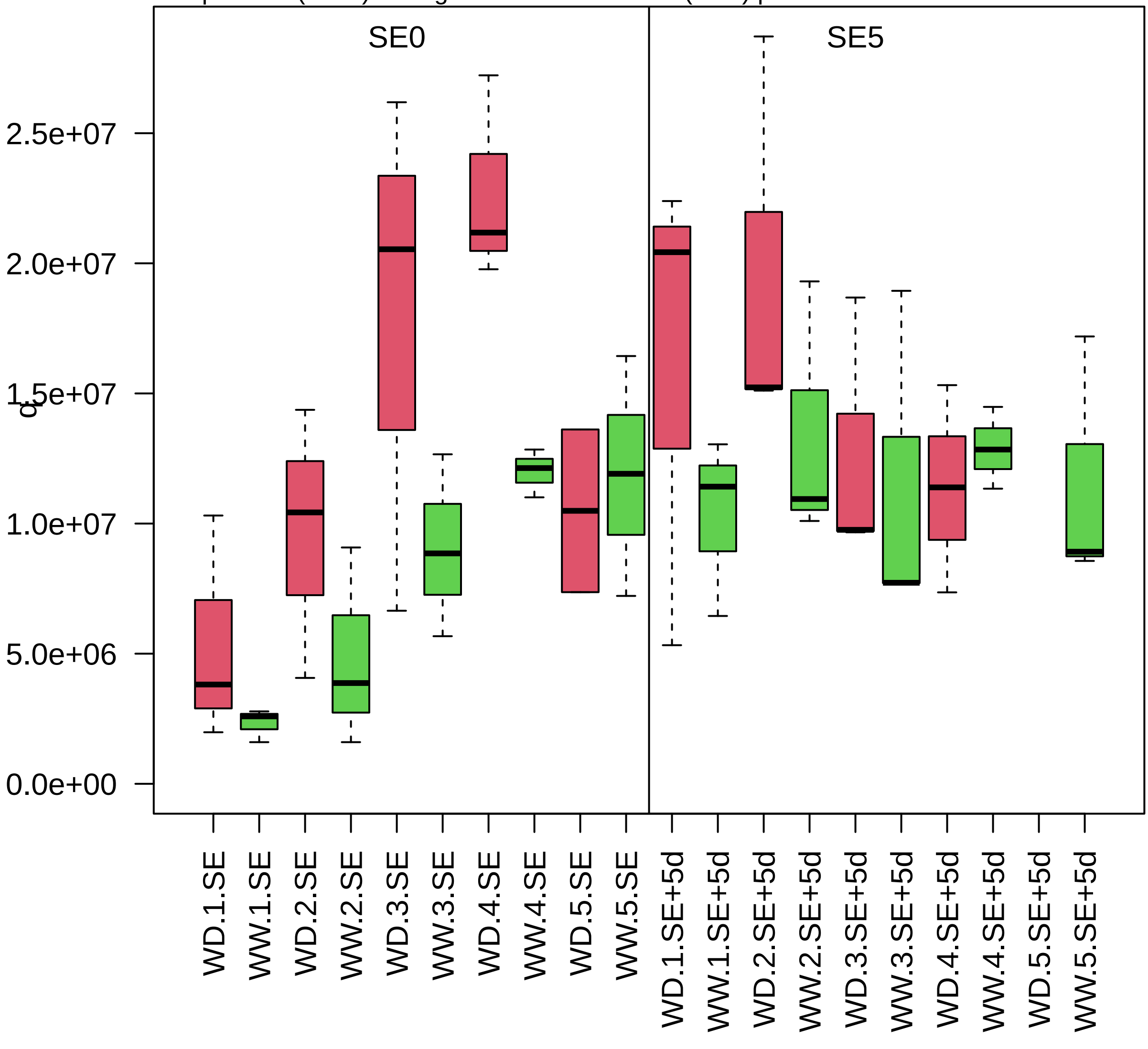

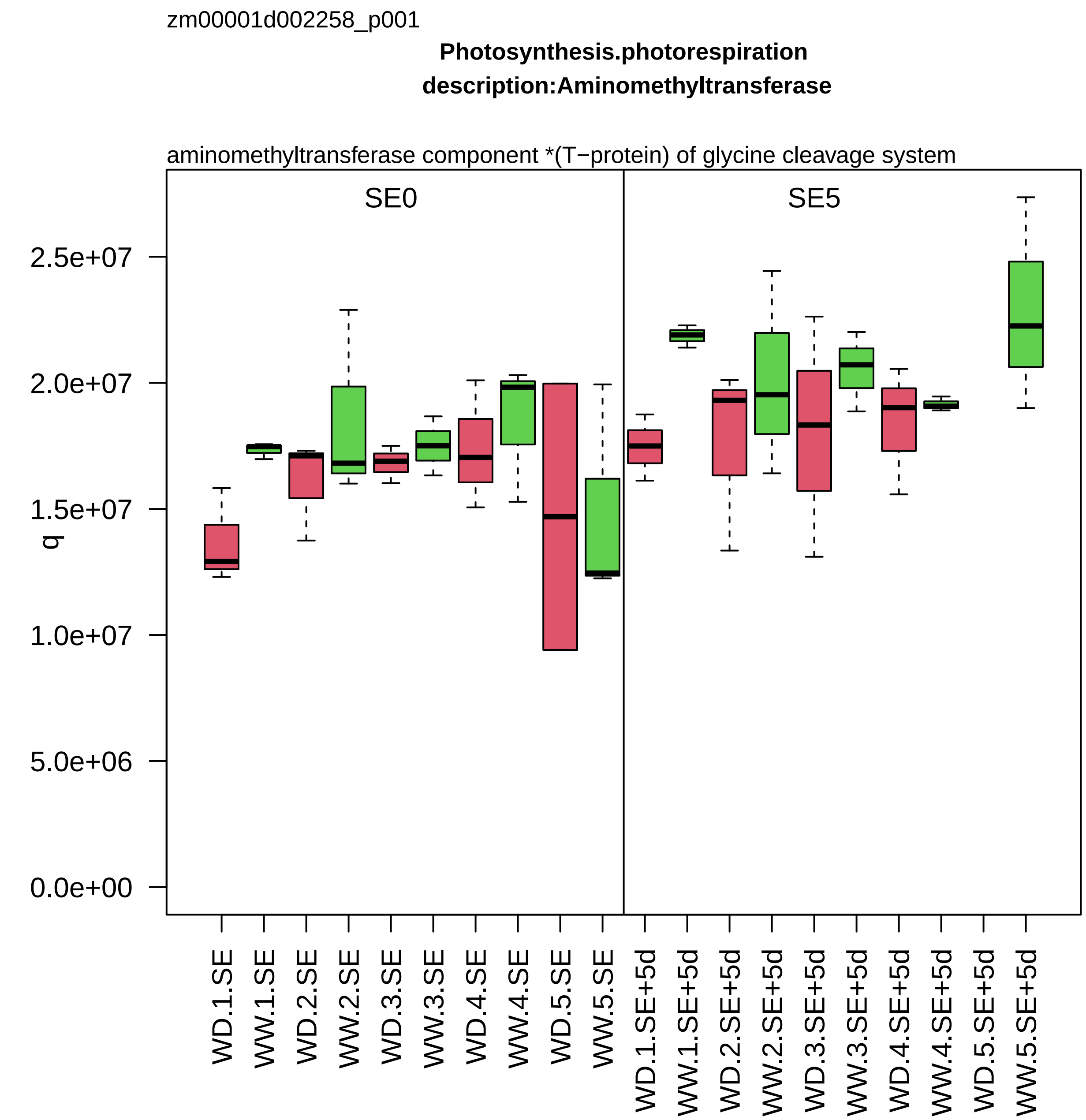

zm00001d002258_p001
Photosynthesis.photorespiration
description:Aminomethyltransferase
aminomethyltransferase component *(T−protein) of glycine cleavage system
SE0
SE5
q
2.5e+07
2.0e+07
1.5e+07
1.0e+07
5.0e+06
0.0e+00
WD.1.SE
WW.1.SE
WD.2.SE
WW.2.SE
WD.3.SE
WW.3.SE
WD.4.SE
WW.4.SE
WD.5.SE
WW.5.SE
WD.1.SE+5d
WW.1.SE+5d
WD.2.SE+5d
WW.2.SE+5d
WD.3.SE+5d
WW.3.SE+5d
WD.4.SE+5d
WW.4.SE+5d
WD.5.SE+5d
WW.5.SE+5d

zm00001d002325_p002

**not assigned.not annotated**

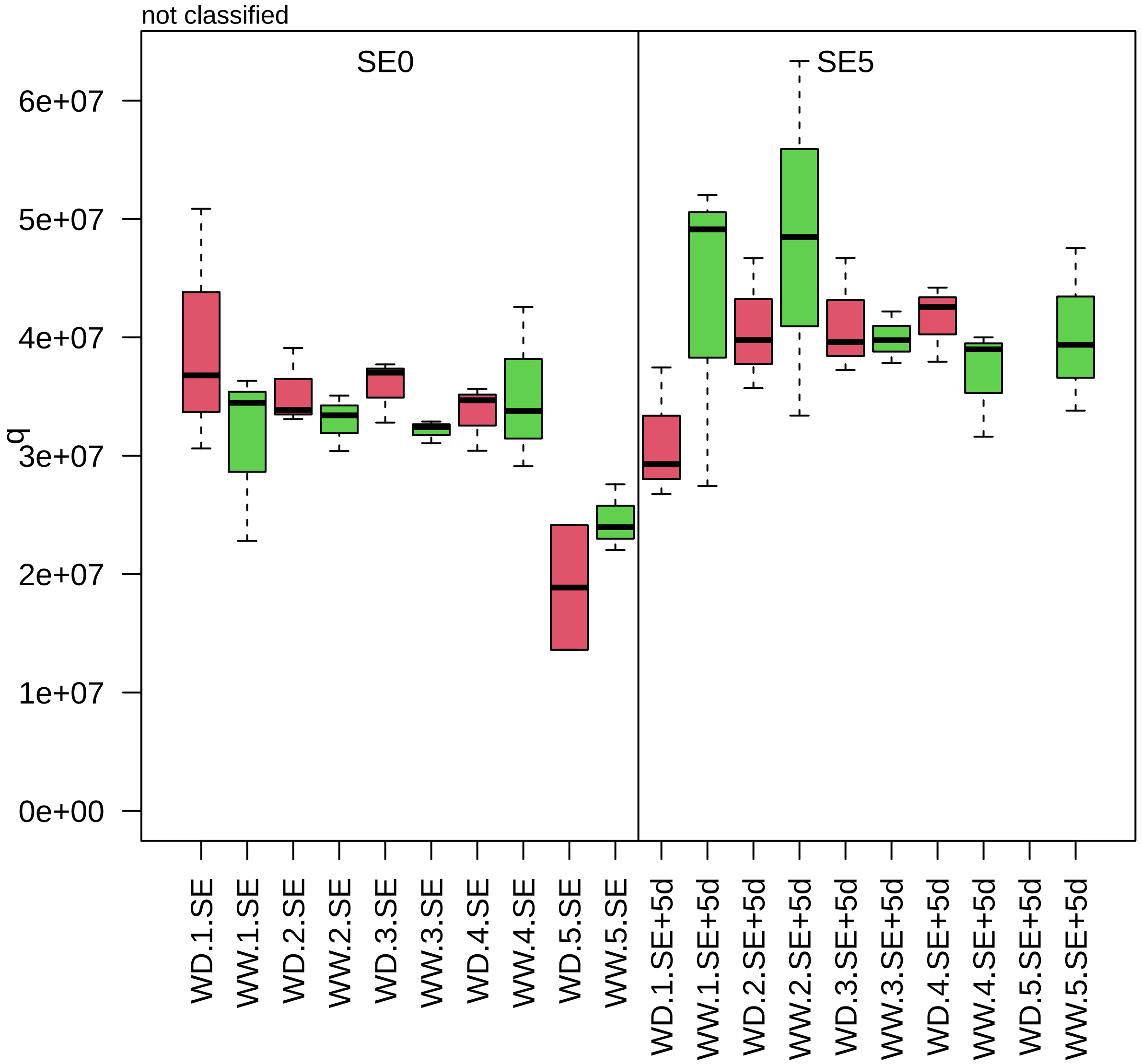

zm00001d002326_p001

**Amino acid metabolism.amino acid degradation**

**gene_symbol:adenosylmethionine aminotransferase1 description: adenosylmethionine aminotransf**

GABA pyruvate transaminase

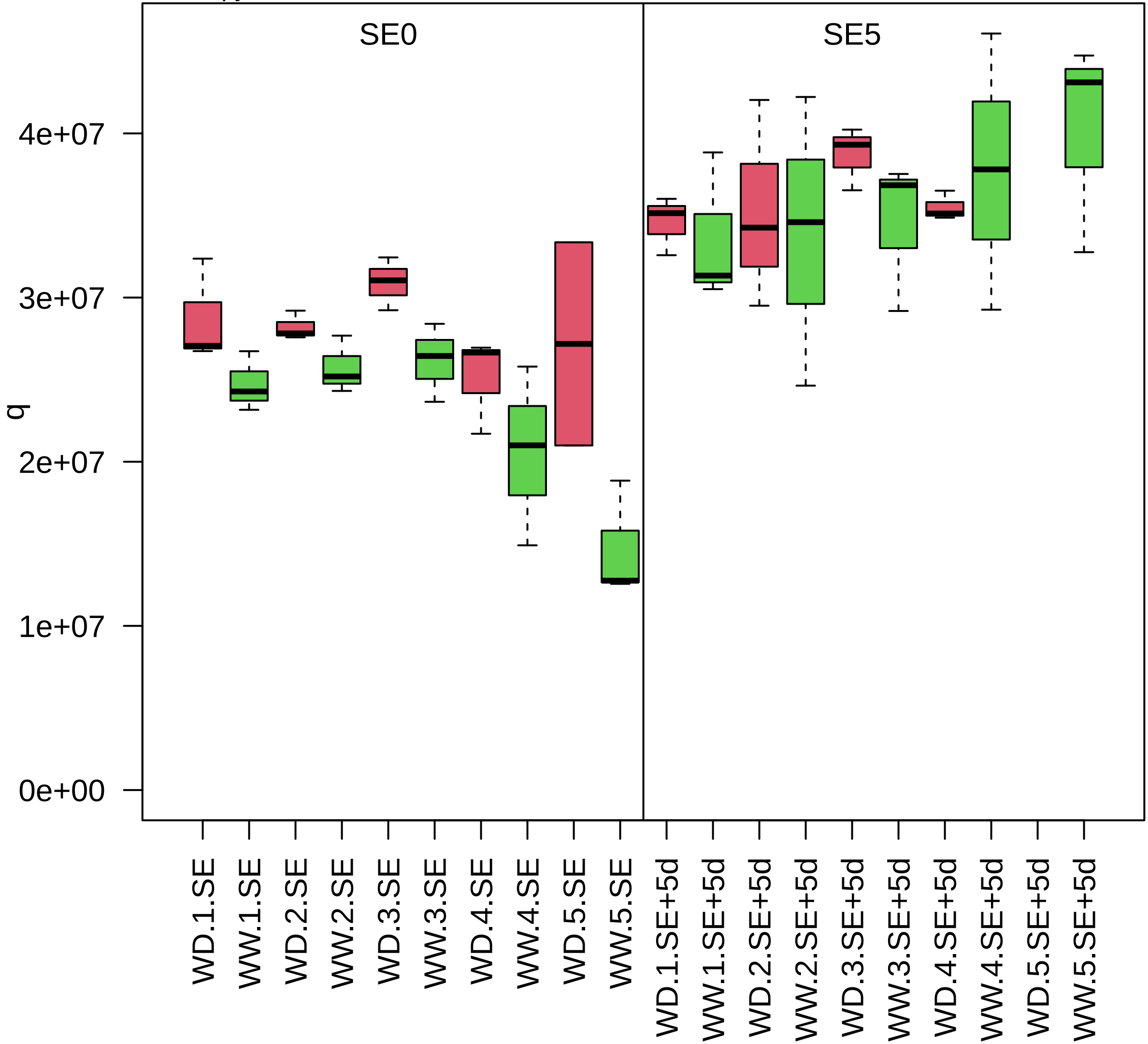

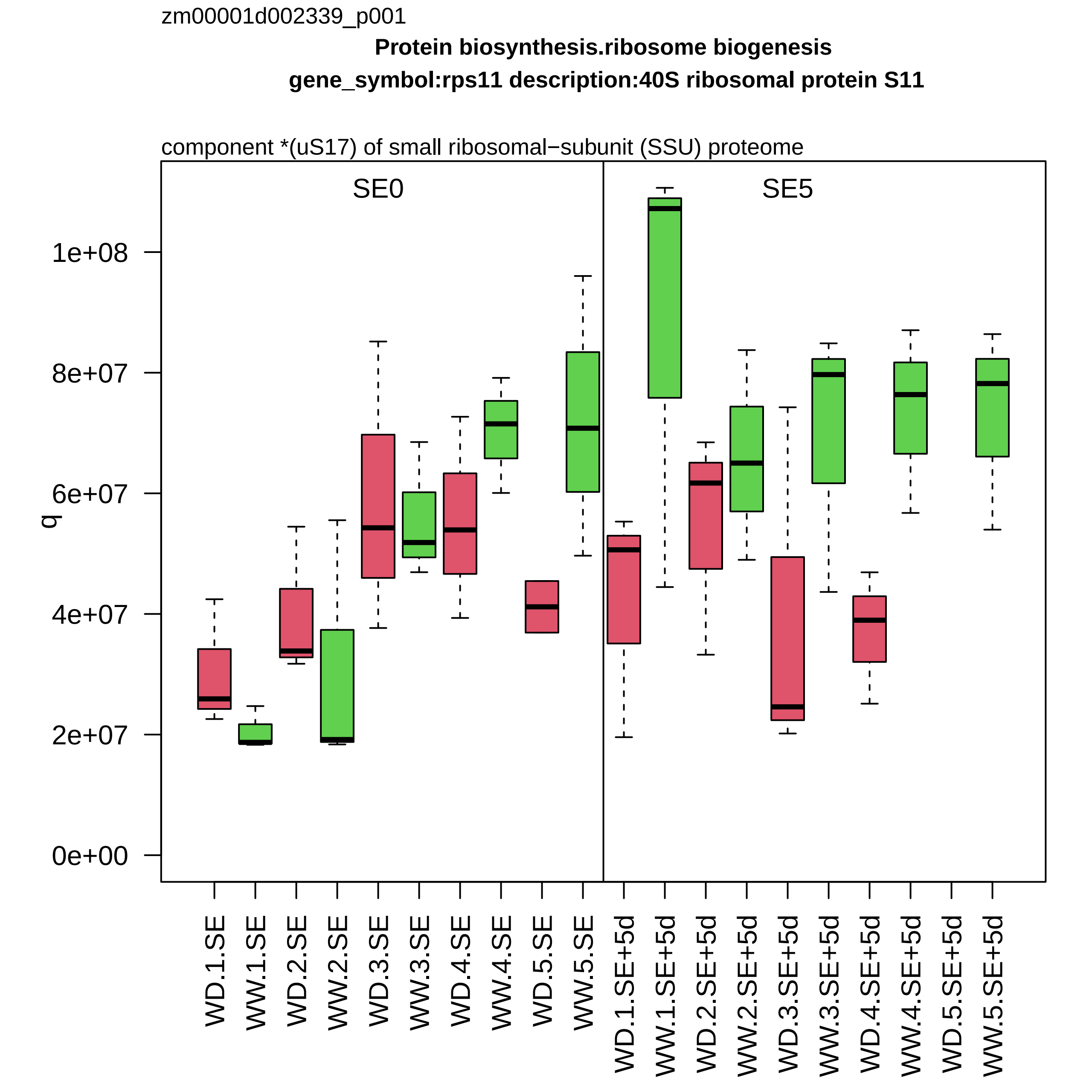

zm00001d002339_p001
Protein biosynthesis.ribosome biogenesis
gene_symbol:rps11 description:40S ribosomal protein S11
component *(uS17) of small ribosomal−subunit (SSU) proteome
SE0
SE5
q
0e+00
2e+07
4e+07
6e+07
8e+07
1e+08
WD.1.SE
WW.1.SE
WD.2.SE
WW.2.SE
WD.3.SE
WW.3.SE
WD.4.SE
WW.4.SE
WD.5.SE
WW.5.SE
WD.1.SE+5d
WW.1.SE+5d
WD.2.SE+5d
WW.2.SE+5d
WD.3.SE+5d
WW.3.SE+5d
WD.4.SE+5d
WW.4.SE+5d
WD.5.SE+5d
WW.5.SE+5d

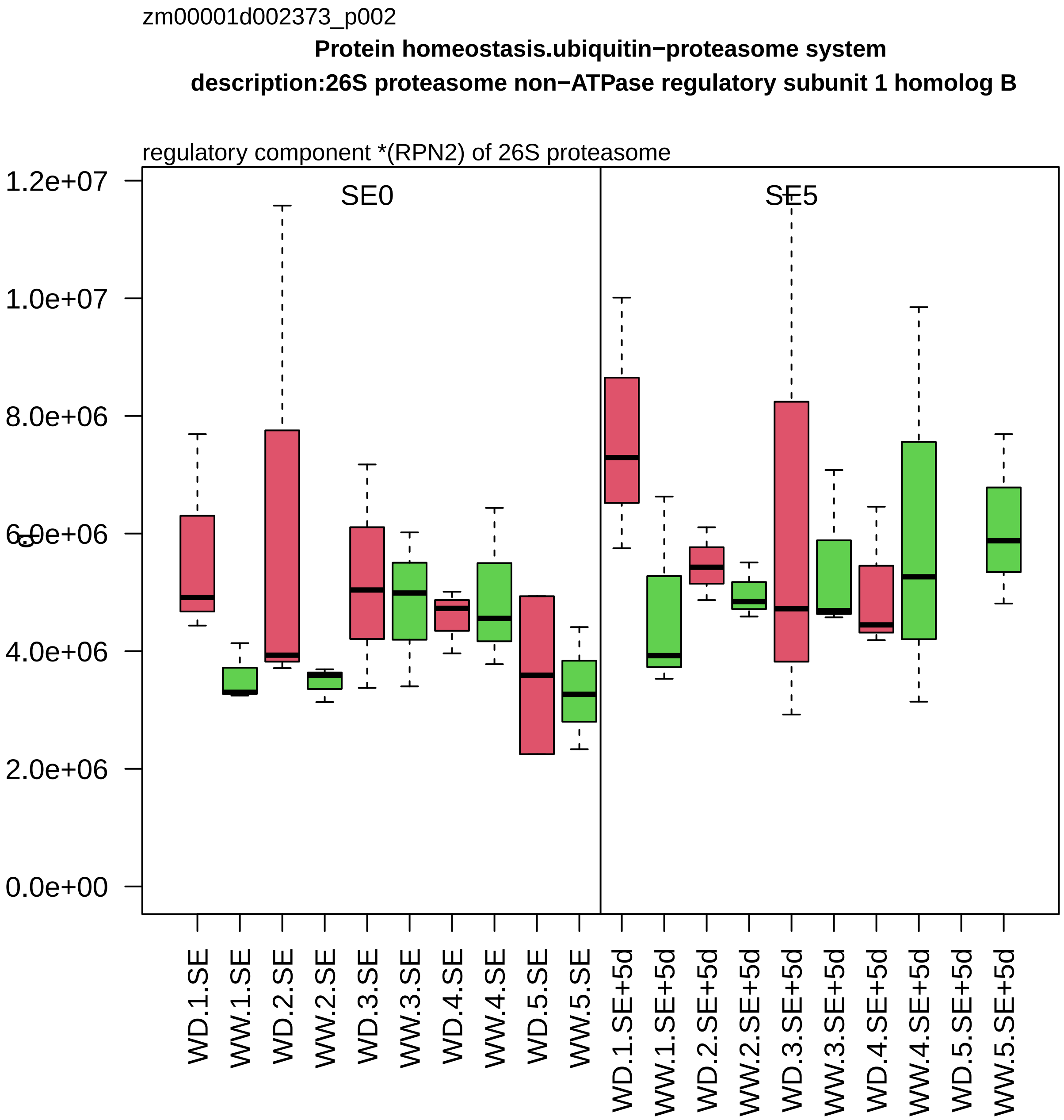

zm00001d002373_p002
Protein homeostasis.ubiquitin−proteasome system
description:26S proteasome non−ATPase regulatory subunit 1 homolog B
regulatory component *(RPN2) of 26S proteasome
SE0
SE5
1.2e+07
1.0e+07
8.0e+06
6.0e+06
4.0e+06
2.0e+06
0.0e+00
WD.1.SE
WW.1.SE
WD.2.SE
WW.2.SE
WD.3.SE
WW.3.SE
WD.4.SE
WW.4.SE
WD.5.SE
WW.5.SE
WD.1.SE+5d
WW.1.SE+5d
WD.2.SE+5d
WW.2.SE+5d
WD.3.SE+5d
WW.3.SE+5d
WD.4.SE+5d
WW.4.SE+5d
WD.5.SE+5d
WW.5.SE+5d

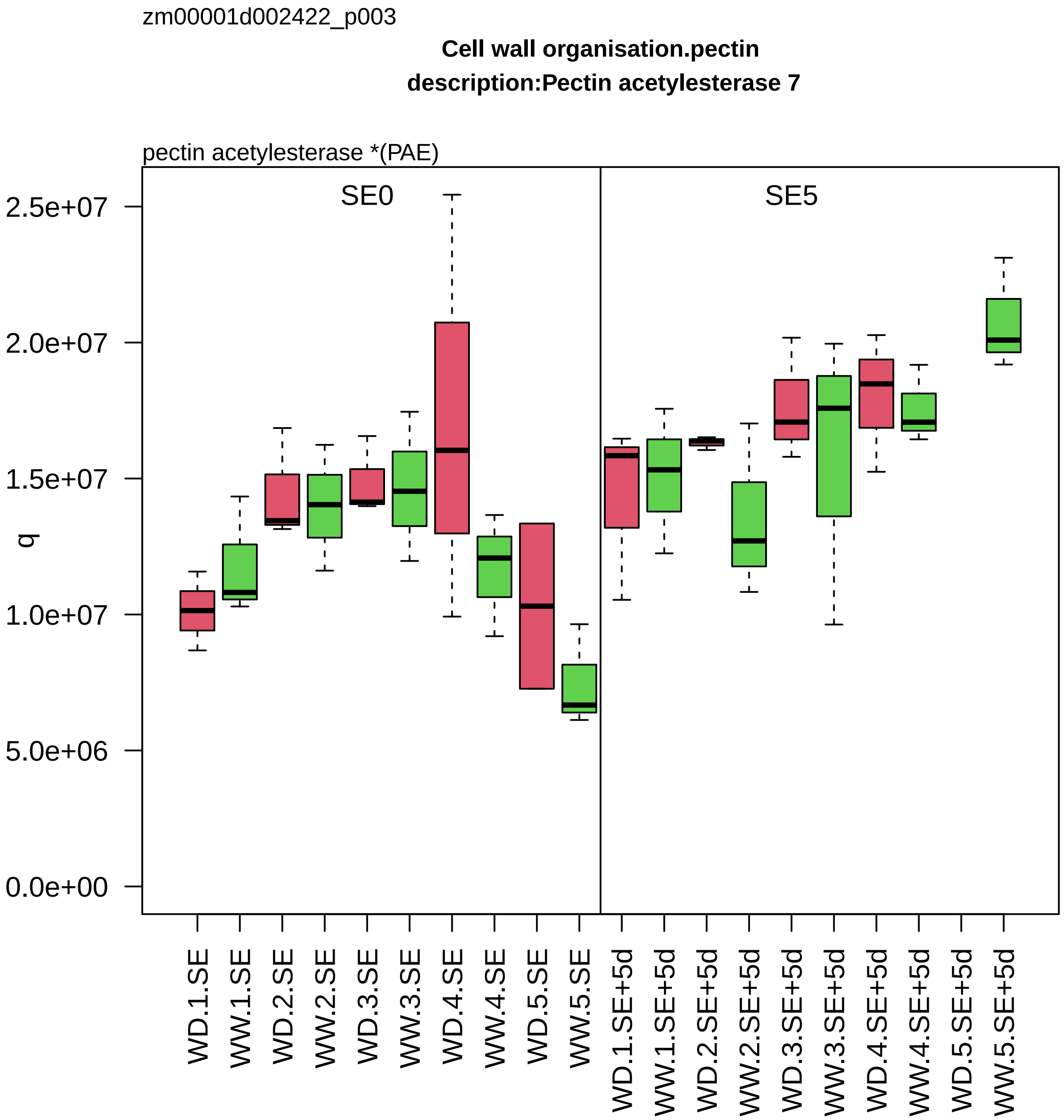
zm00001d002422_p003
Cell wall organisation.pectin
description:Pectin acetylesterase 7
pectin acetylesterase *(PAE)
SE0
SE5
q
2.5e+07
2.0e+07
1.5e+07
1.0e+07
5.0e+06
0.0e+00
WD.1.SE
WW.1.SE
WD.2.SE
WW.2.SE
WD.3.SE
WW.3.SE
WD.4.SE
WW.4.SE
WD.5.SE
WW.5.SE
WD.1.SE+5d
WW.1.SE+5d
WD.2.SE+5d
WW.2.SE+5d
WD.3.SE+5d
WW.3.SE+5d
WD.4.SE+5d
WW.4.SE+5d
WD.5.SE+5d
WW.5.SE+5d

zm00001d002450_p001

**Protein biosynthesis.ribosome biogenesis**
**description:60S ribosomal protein L12−3**

component *(uL11) of large ribosomal−subunit (LSU) proteome

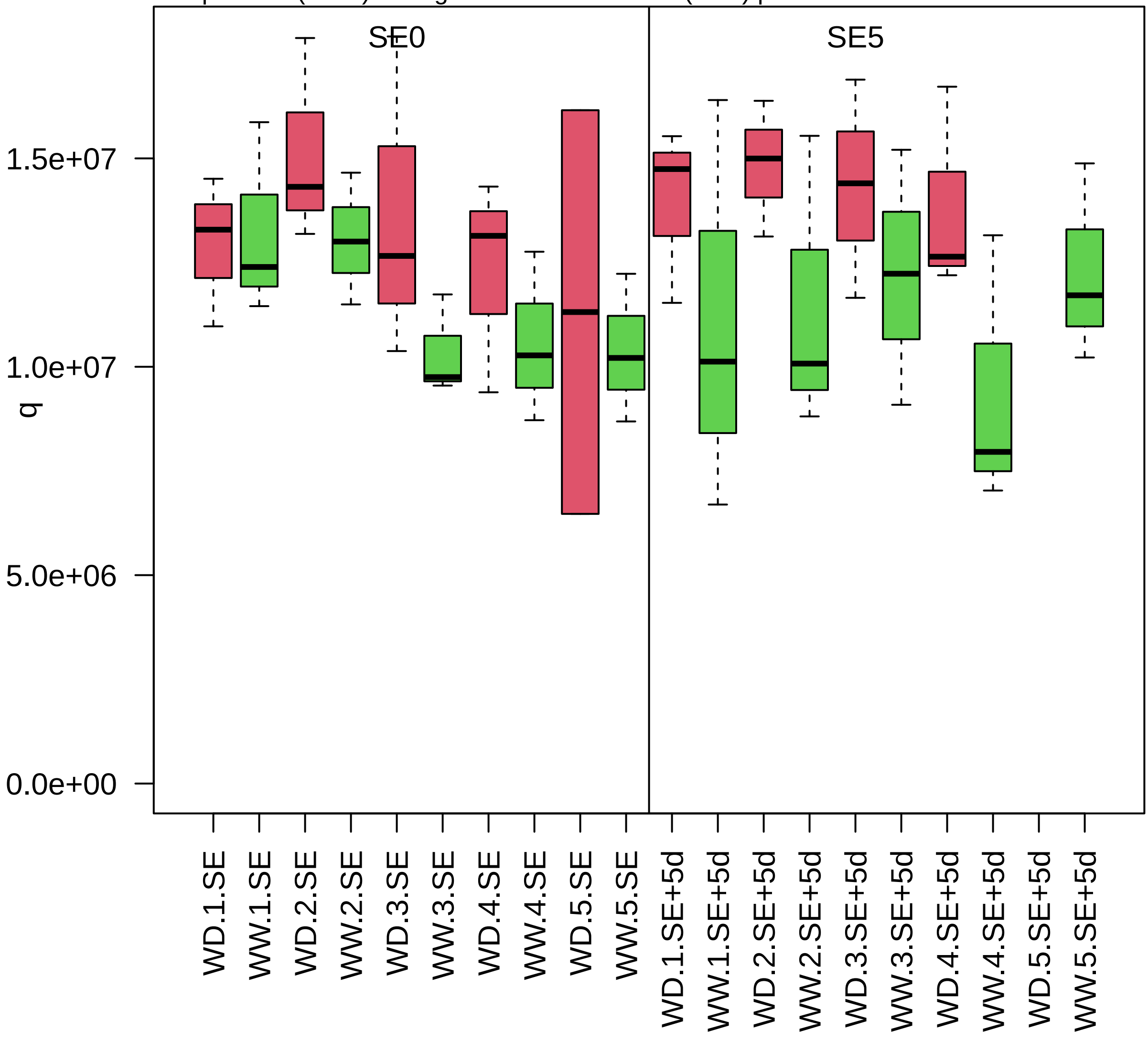

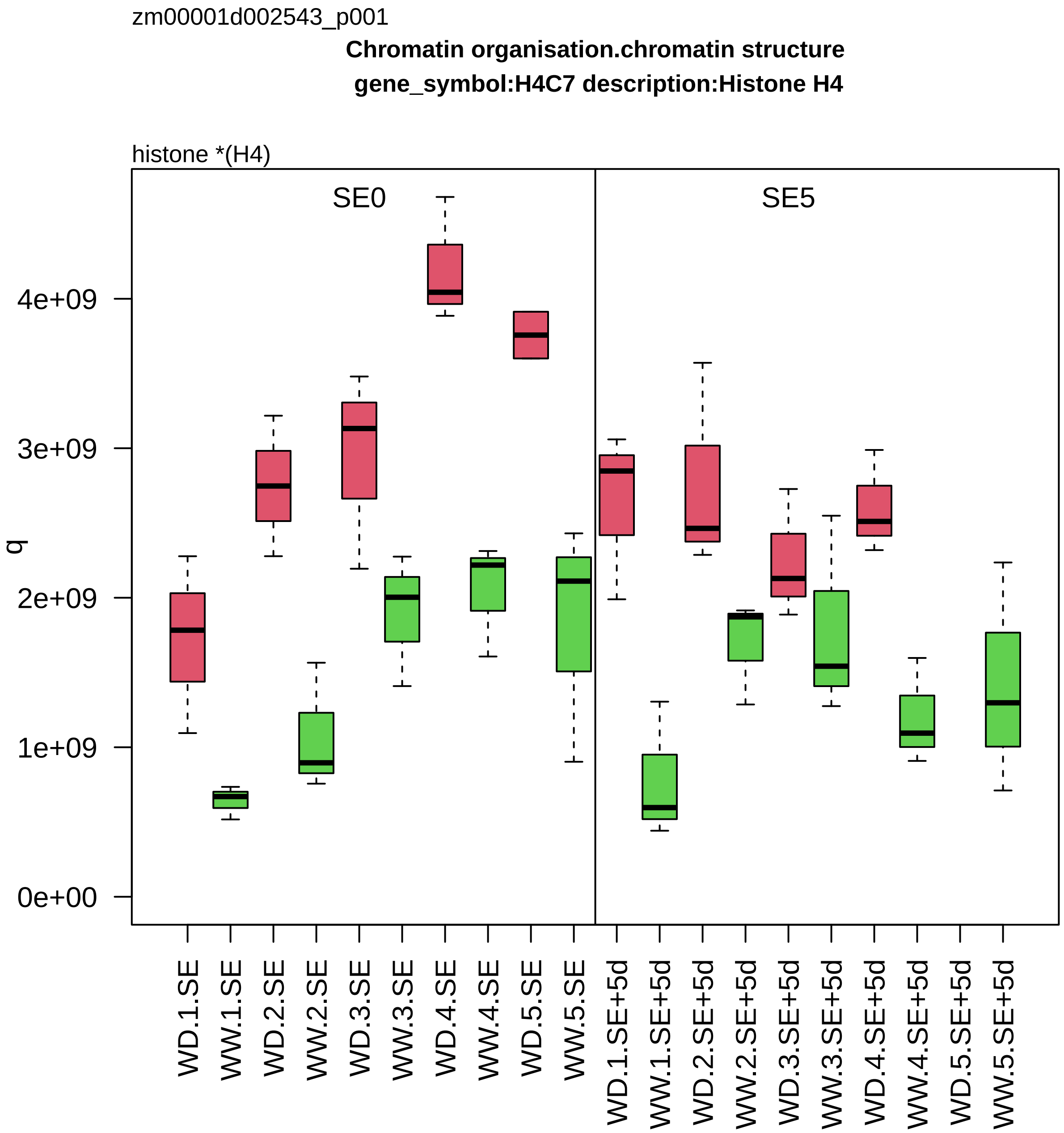
zm00001d002543_p001
Chromatin organisation.chromatin structure
gene_symbol:H4C7 description:Histone H4
histone *(H4)
SE0
SE5
q
0e+00
1e+09
2e+09
3e+09
4e+09
WD.1.SE
WW.1.SE
WD.2.SE
WW.2.SE
WD.3.SE
WW.3.SE
WD.4.SE
WW.4.SE
WD.5.SE
WW.5.SE
WD.1.SE+5d
WW.1.SE+5d
WD.2.SE+5d
WW.2.SE+5d
WD.3.SE+5d
WW.3.SE+5d
WD.4.SE+5d
WW.4.SE+5d
WD.5.SE+5d
WW.5.SE+5d

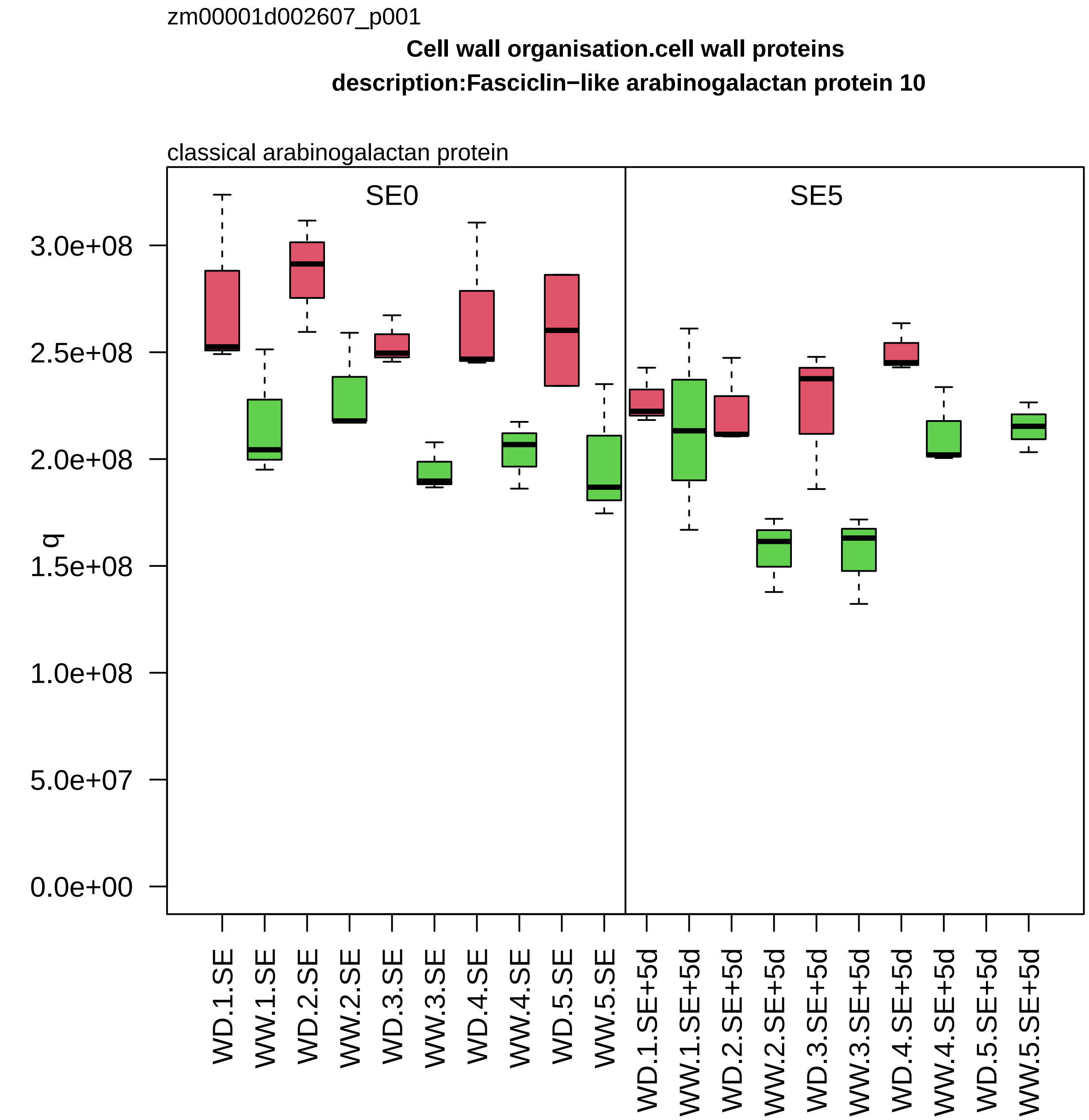
zm00001d002607_p001
Cell wall organisation.cell wall proteins
description:Fasciclin−like arabinogalactan protein 10
classical arabinogalactan protein
SE0
SE5
q
3.0e+08
2.5e+08
2.0e+08
1.5e+08
1.0e+08
5.0e+07
0.0e+00
WD.1.SE
WW.1.SE
WD.2.SE
WW.2.SE
WD.3.SE
WW.3.SE
WD.4.SE
WW.4.SE
WD.5.SE
WW.5.SE
WD.1.SE+5d
WW.1.SE+5d
WD.2.SE+5d
WW.2.SE+5d
WD.3.SE+5d
WW.3.SE+5d
WD.4.SE+5d
WW.4.SE+5d
WD.5.SE+5d
WW.5.SE+5d

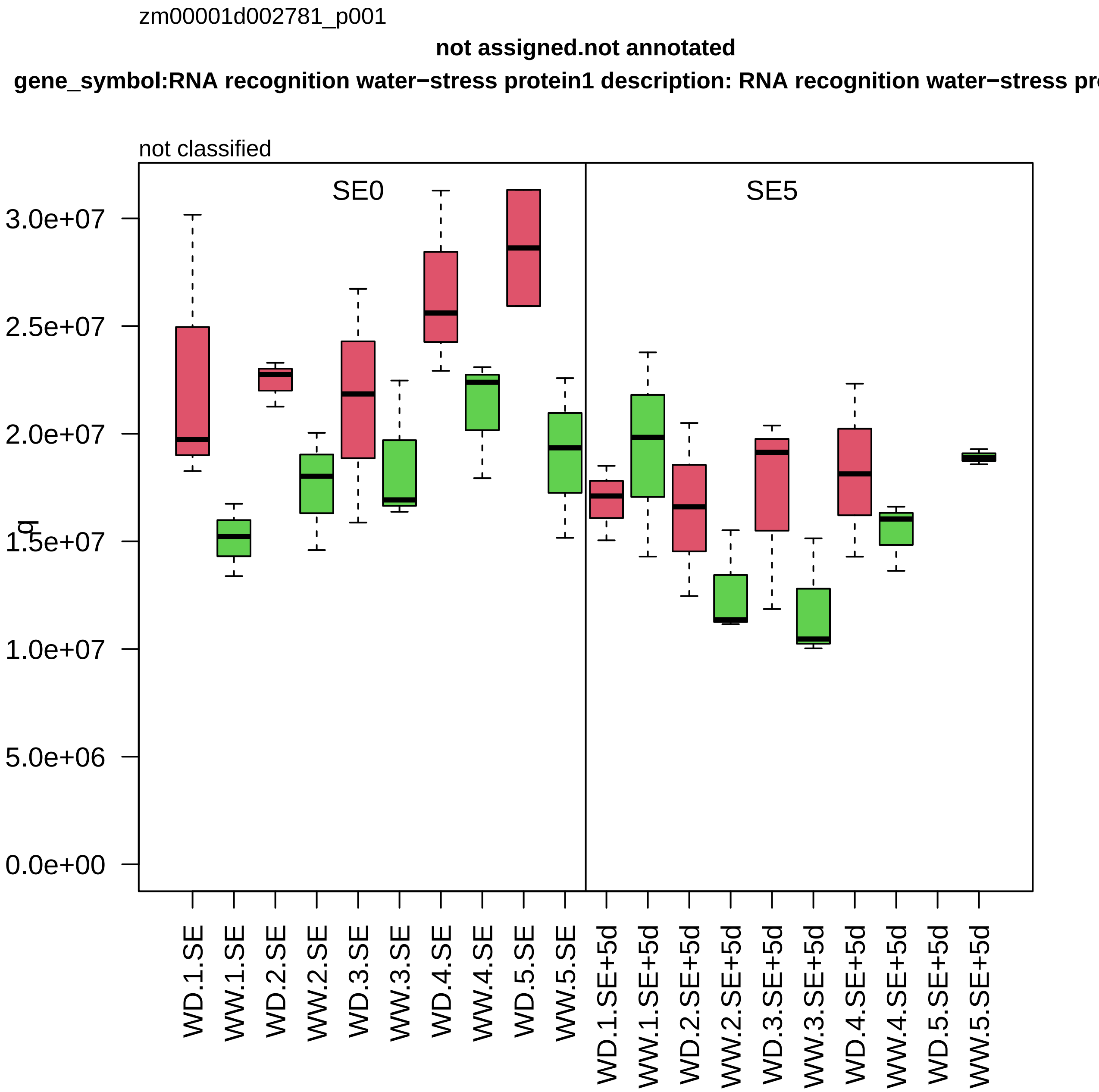

zm00001d002781_p001
not assigned.not annotated
gene_symbol:RNA recognition water−stress protein1 description: RNA recognition water−stress pr
not classified
SE0
SE5
q
3.0e+07
2.5e+07
2.0e+07
1.5e+07
1.0e+07
5.0e+06
0.0e+00
WD.1.SE
WW.1.SE
WD.2.SE
WW.2.SE
WD.3.SE
WW.3.SE
WD.4.SE
WW.4.SE
WD.5.SE
WW.5.SE
WD.1.SE+5d
WW.1.SE+5d
WD.2.SE+5d
WW.2.SE+5d
WD.3.SE+5d
WW.3.SE+5d
WD.4.SE+5d
WW.4.SE+5d
WD.5.SE+5d
WW.5.SE+5d

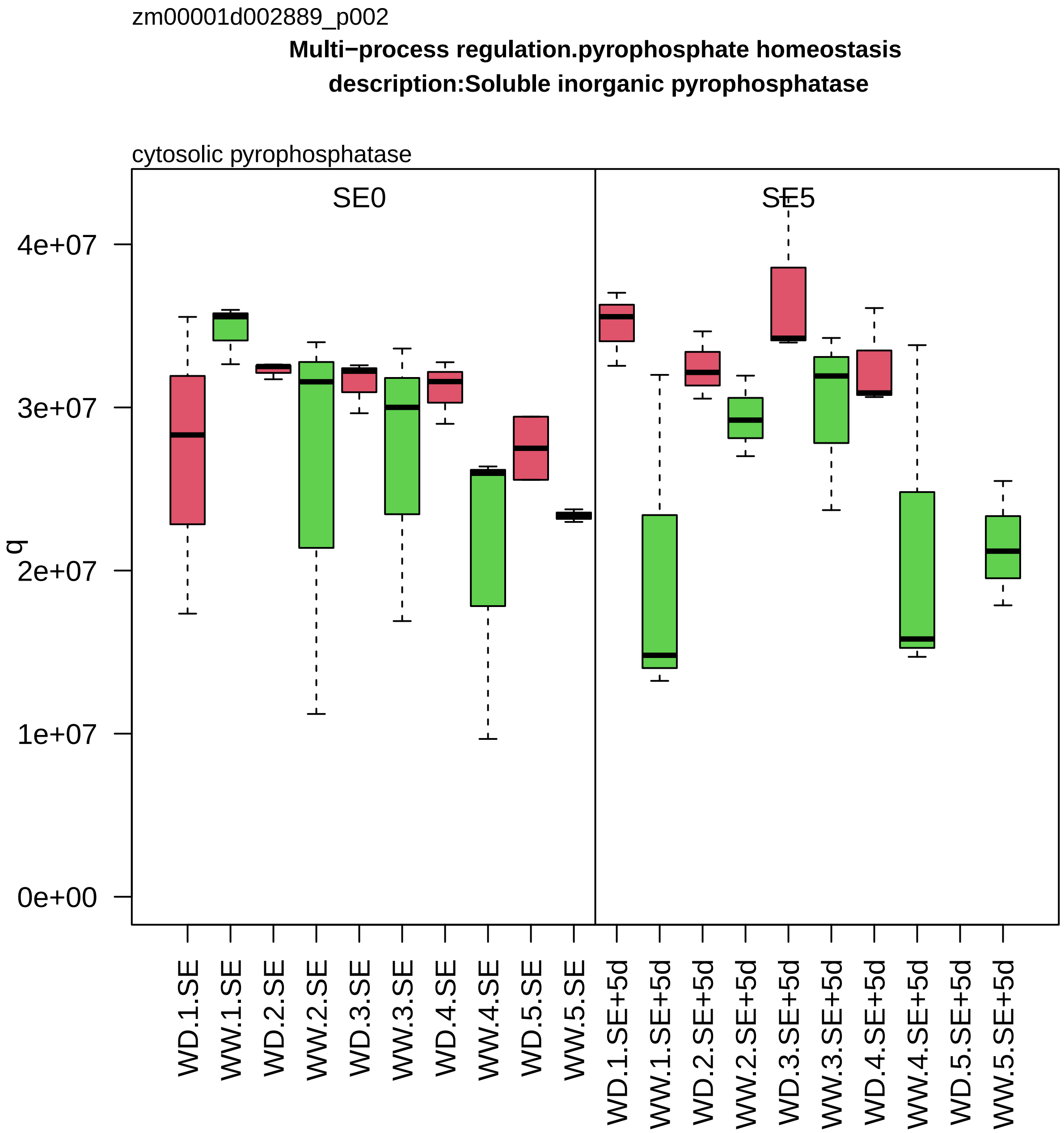

zm00001d002889_p002
Multi−process regulation.pyrophosphate homeostasis
description:Soluble inorganic pyrophosphatase
cytosolic pyrophosphatase
SE0
SE5
q
4e+07
3e+07
2e+07
1e+07
0e+00
WD.1.SE
WW.1.SE
WD.2.SE
WW.2.SE
WD.3.SE
WW.3.SE
WD.4.SE
WW.4.SE
WD.5.SE
WW.5.SE
WD.1.SE+5d
WW.1.SE+5d
WD.2.SE+5d
WW.2.SE+5d
WD.3.SE+5d
WW.3.SE+5d
WD.4.SE+5d
WW.4.SE+5d
WD.5.SE+5d
WW.5.SE+5d

zm00001d003081_p003

**Protein modification.S−glutathionylation**

**gene_symbol:glutaredoxin homolog1 description: glutaredoxin homolog1**

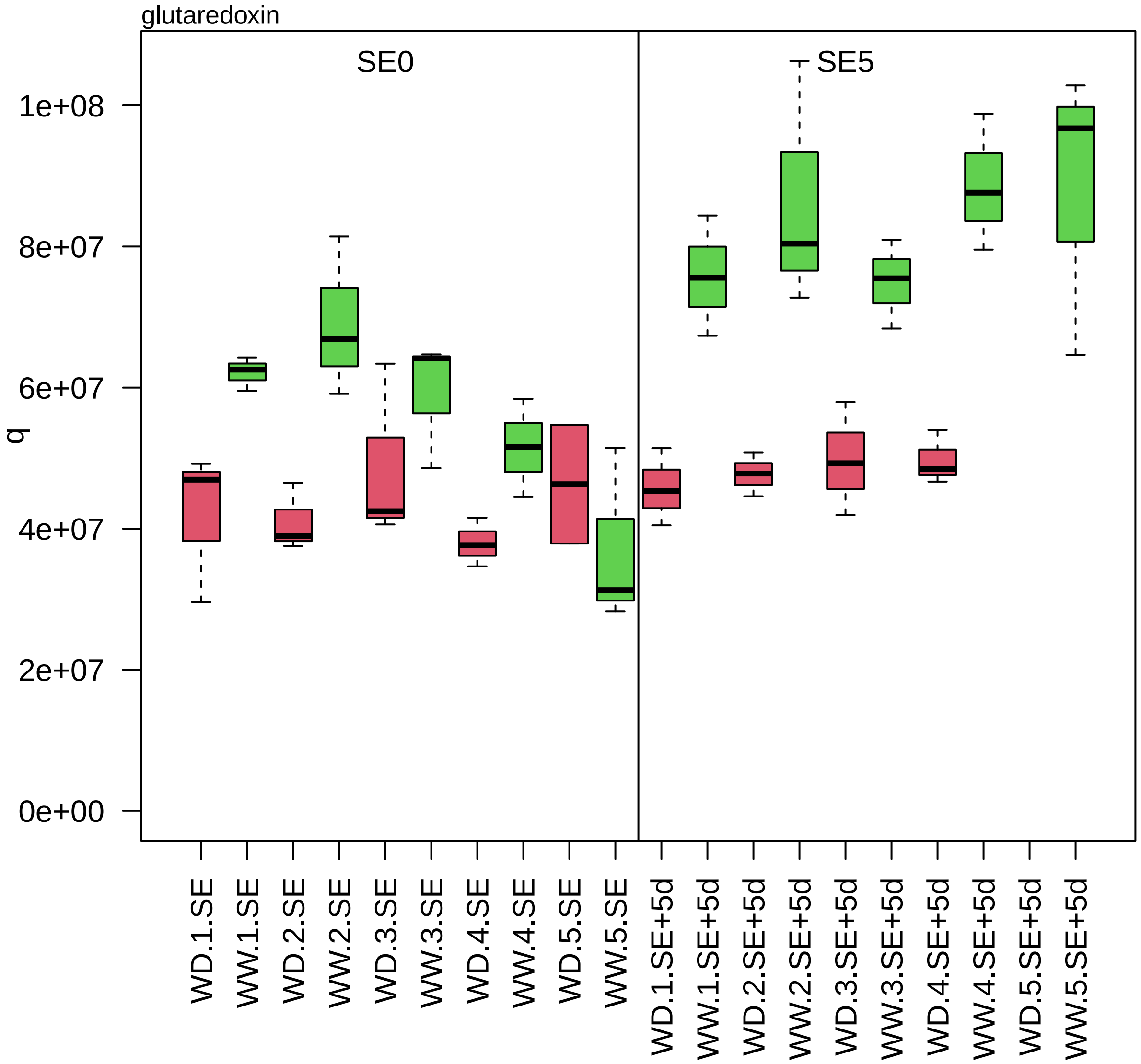

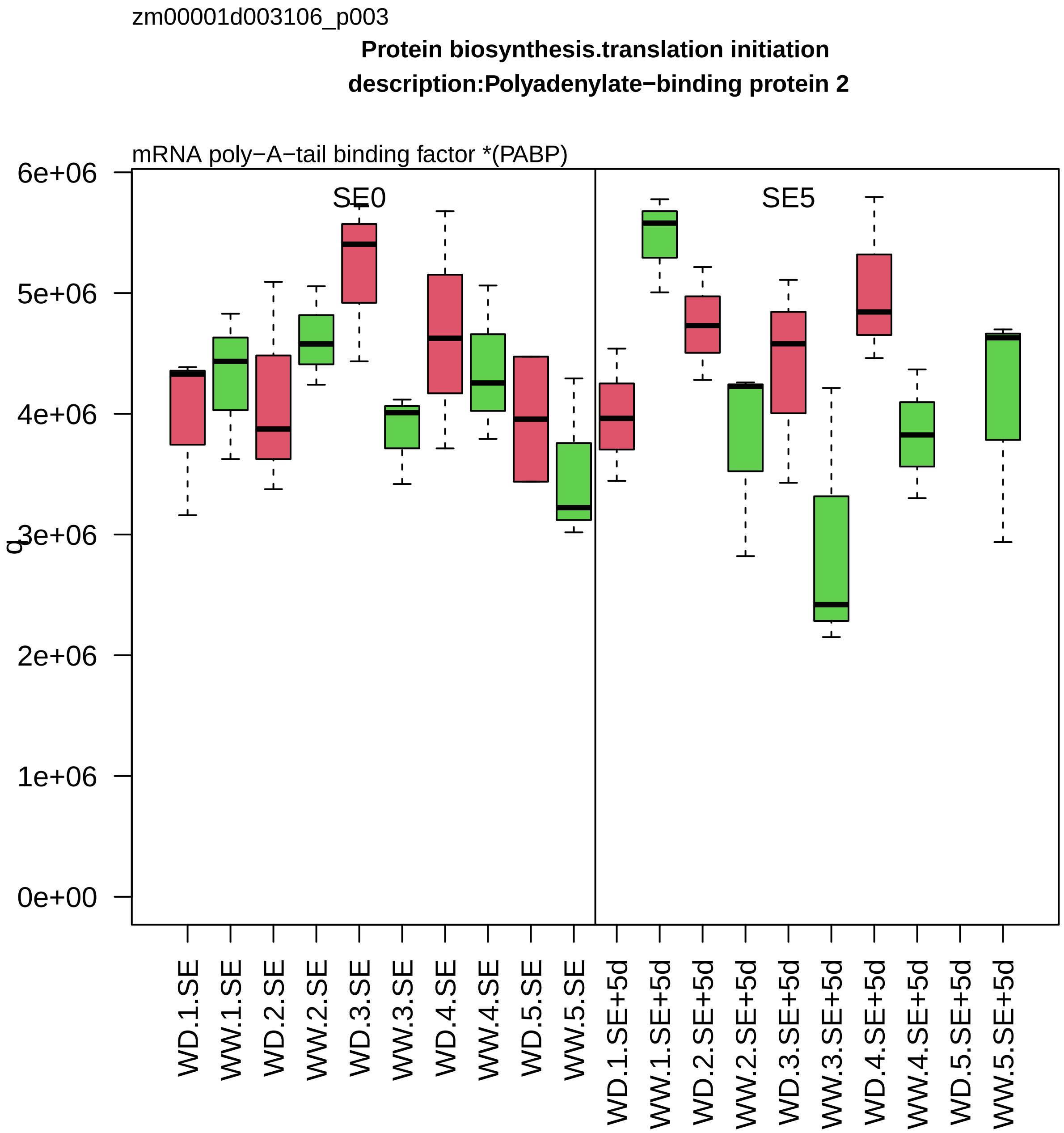
zm00001d003106_p003
Protein biosynthesis.translation initiation
description:Polyadenylate−binding protein 2
mRNA poly−A−tail binding factor *(PABP)
SE0
SE5
6e+06
5e+06
4e+06
3e+06
2e+06
1e+06
0e+00
q
WD.1.SE
WW.1.SE
WD.2.SE
WW.2.SE
WD.3.SE
WW.3.SE
WD.4.SE
WW.4.SE
WD.5.SE
WW.5.SE
WD.1.SE+5d
WW.1.SE+5d
WD.2.SE+5d
WW.2.SE+5d
WD.3.SE+5d
WW.3.SE+5d
WD.4.SE+5d
WW.4.SE+5d
WD.5.SE+5d
WW.5.SE+5d

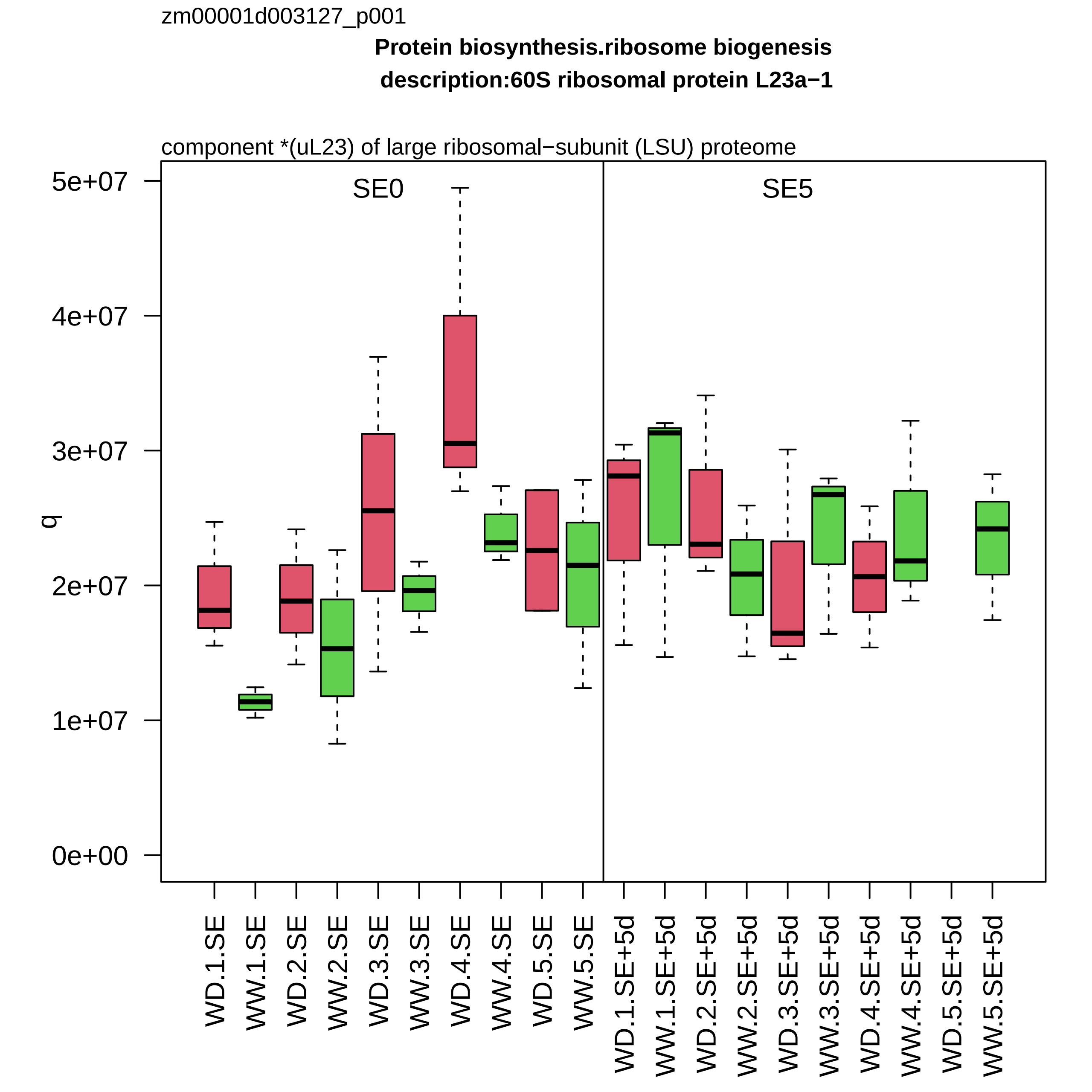

zm00001d003127_p001
Protein biosynthesis.ribosome biogenesis
description:60S ribosomal protein L23a−1
component *(uL23) of large ribosomal−subunit (LSU) proteome
SE0
SE5
q
5e+07
4e+07
3e+07
2e+07
1e+07
0e+00
WD.1.SE
WW.1.SE
WD.2.SE
WW.2.SE
WD.3.SE
WW.3.SE
WD.4.SE
WW.4.SE
WD.5.SE
WW.5.SE
WD.1.SE+5d
WW.1.SE+5d
WD.2.SE+5d
WW.2.SE+5d
WD.3.SE+5d
WW.3.SE+5d
WD.4.SE+5d
WW.4.SE+5d
WD.5.SE+5d
WW.5.SE+5d

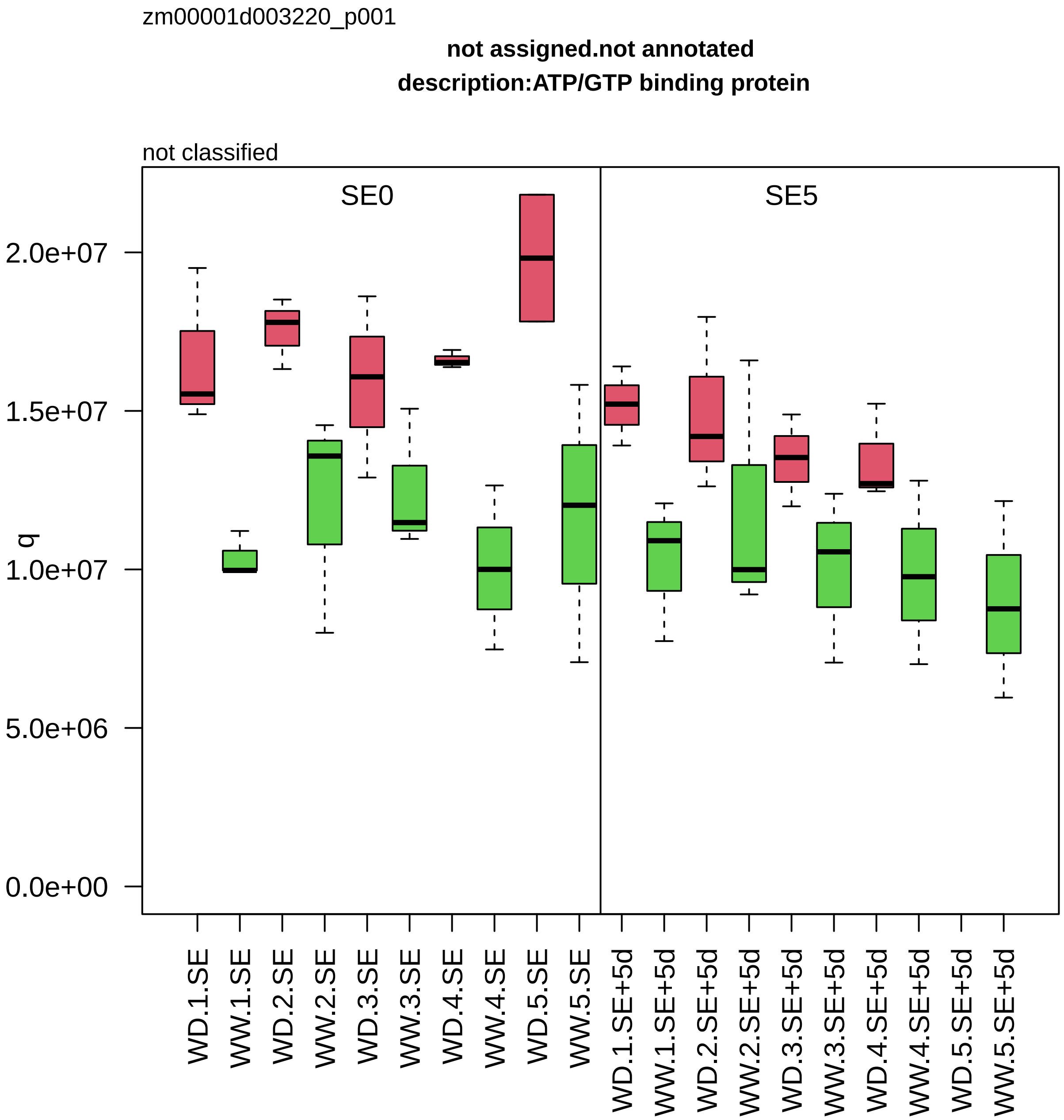

zm00001d003220_p001
not assigned.not annotated
description:ATP/GTP binding protein
not classified
SE0
SE5
q
2.0e+07
1.5e+07
1.0e+07
5.0e+06
0.0e+00
WD.1.SE
WW.1.SE
WD.2.SE
WW.2.SE
WD.3.SE
WW.3.SE
WD.4.SE
WW.4.SE
WD.5.SE
WW.5.SE
WD.1.SE+5d
WW.1.SE+5d
WD.2.SE+5d
WW.2.SE+5d
WD.3.SE+5d
WW.3.SE+5d
WD.4.SE+5d
WW.4.SE+5d
WD.5.SE+5d
WW.5.SE+5d

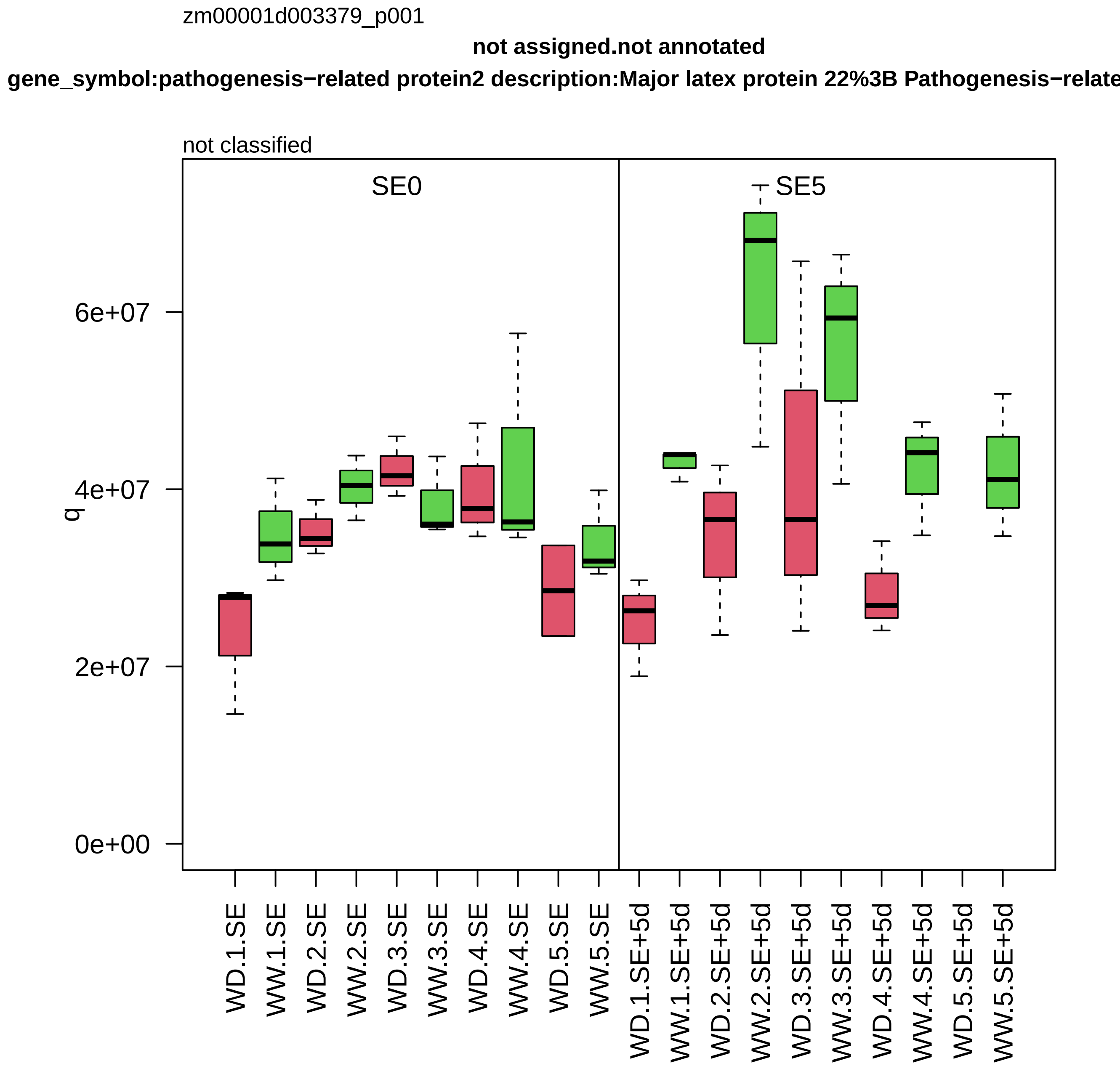
zm00001d003379_p001
not assigned.not annotated
gene_symbol:pathogenesis−related protein2 description:Major latex protein 22%3B Pathogenesis−relate
not classified
SE0
SE5
6e+07
4e+07
2e+07
0e+00
q
WD.1.SE
WW.1.SE
WD.2.SE
WW.2.SE
WD.3.SE
WW.3.SE
WD.4.SE
WW.4.SE
WD.5.SE
WW.5.SE
WD.1.SE+5d
WW.1.SE+5d
WD.2.SE+5d
WW.2.SE+5d
WD.3.SE+5d
WW.3.SE+5d
WD.4.SE+5d
WW.4.SE+5d
WD.5.SE+5d
WW.5.SE+5d

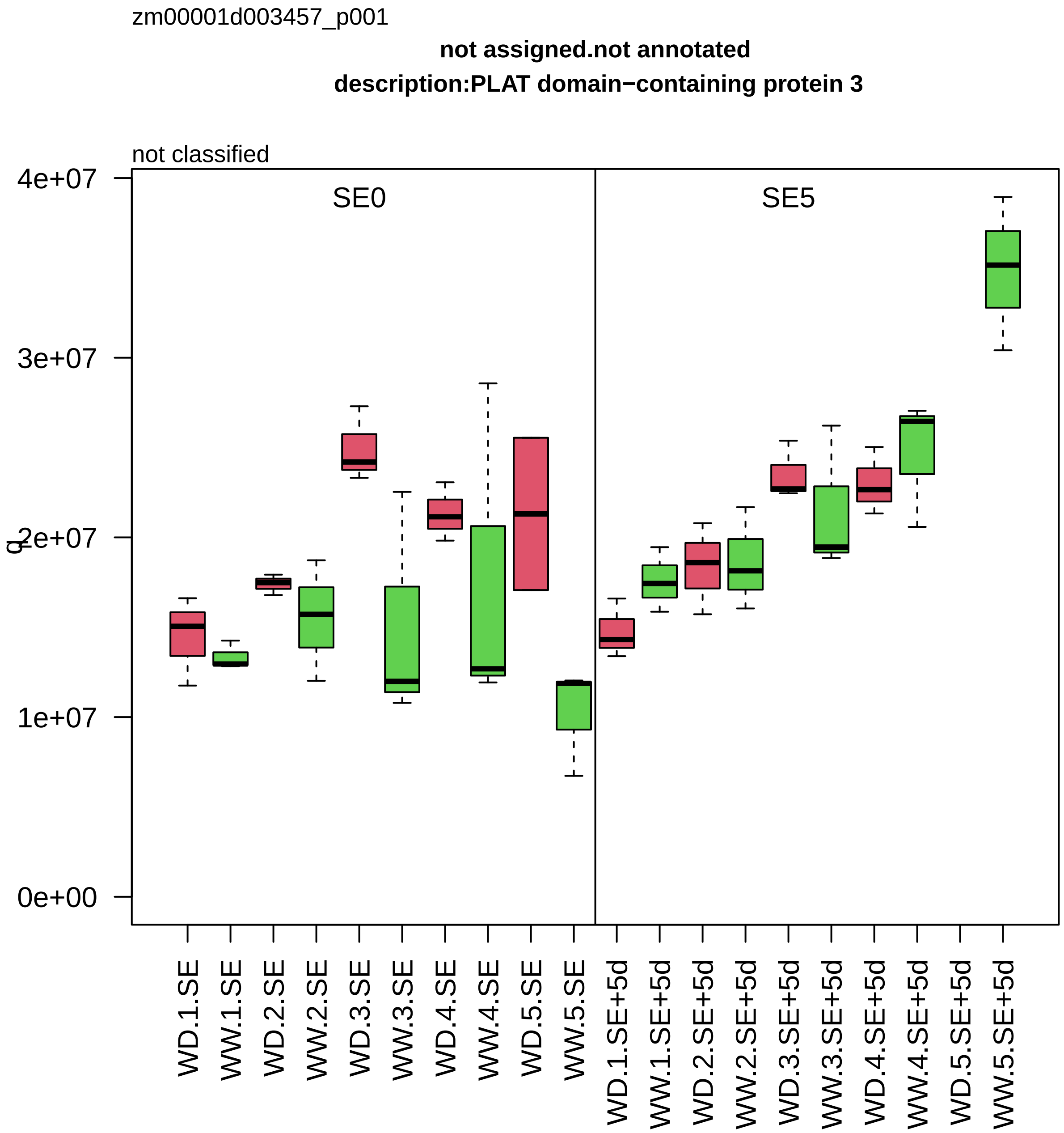
zm00001d003457_p001
not assigned.not annotated
description:PLAT domain−containing protein 3
not classified
SE0
SE5
4e+07
3e+07
2e+07
1e+07
0e+00
q
WD.1.SE
WW.1.SE
WD.2.SE
WW.2.SE
WD.3.SE
WW.3.SE
WD.4.SE
WW.4.SE
WD.5.SE
WW.5.SE
WD.1.SE+5d
WW.1.SE+5d
WD.2.SE+5d
WW.2.SE+5d
WD.3.SE+5d
WW.3.SE+5d
WD.4.SE+5d
WW.4.SE+5d
WD.5.SE+5d
WW.5.SE+5d

zm00001d003632_p002

**External stimuli response.light**

**gene_symbol:regulator of chromosome condensation2 description: regulator of chromosome conden**

photoreceptor *(UVR8)

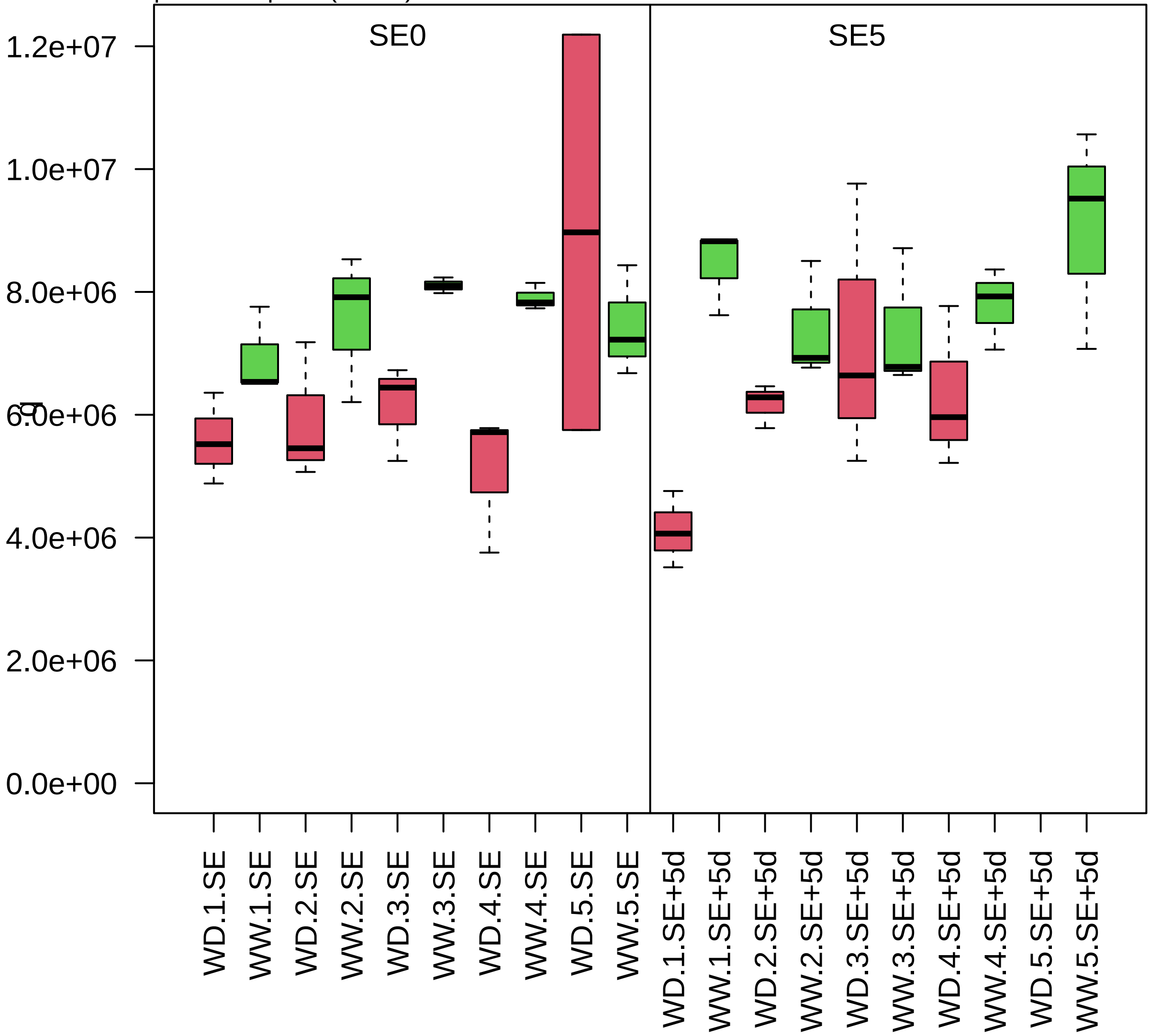

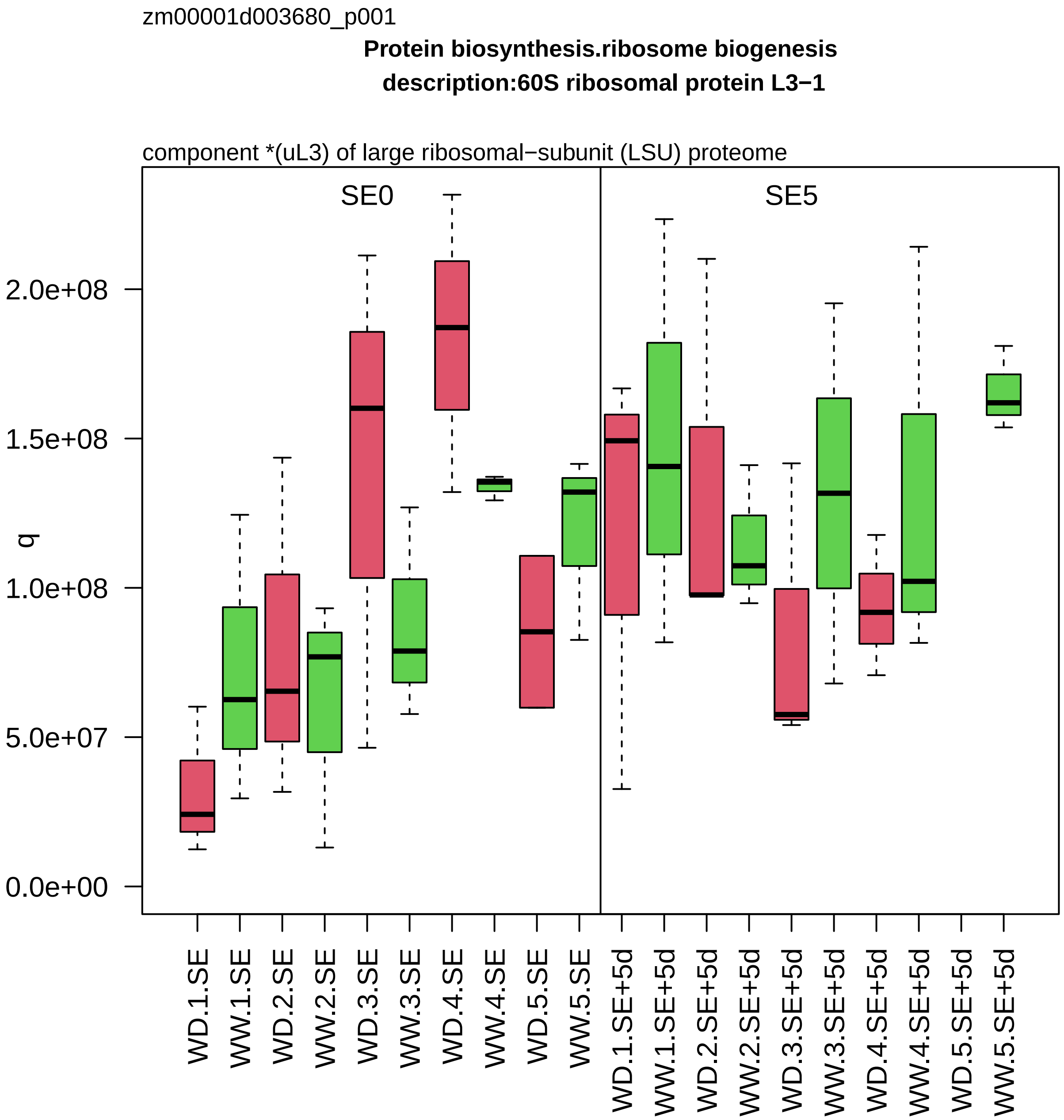

zm00001d003680_p001
Protein biosynthesis.ribosome biogenesis
description:60S ribosomal protein L3−1
component *(uL3) of large ribosomal−subunit (LSU) proteome
SE0
SE5
q
0.0e+00
5.0e+07
1.0e+08
1.5e+08
2.0e+08
WD.1.SE
WW.1.SE
WD.2.SE
WW.2.SE
WD.3.SE
WW.3.SE
WD.4.SE
WW.4.SE
WD.5.SE
WW.5.SE
WD.1.SE+5d
WW.1.SE+5d
WD.2.SE+5d
WW.2.SE+5d
WD.3.SE+5d
WW.3.SE+5d
WD.4.SE+5d
WW.4.SE+5d
WD.5.SE+5d
WW.5.SE+5d

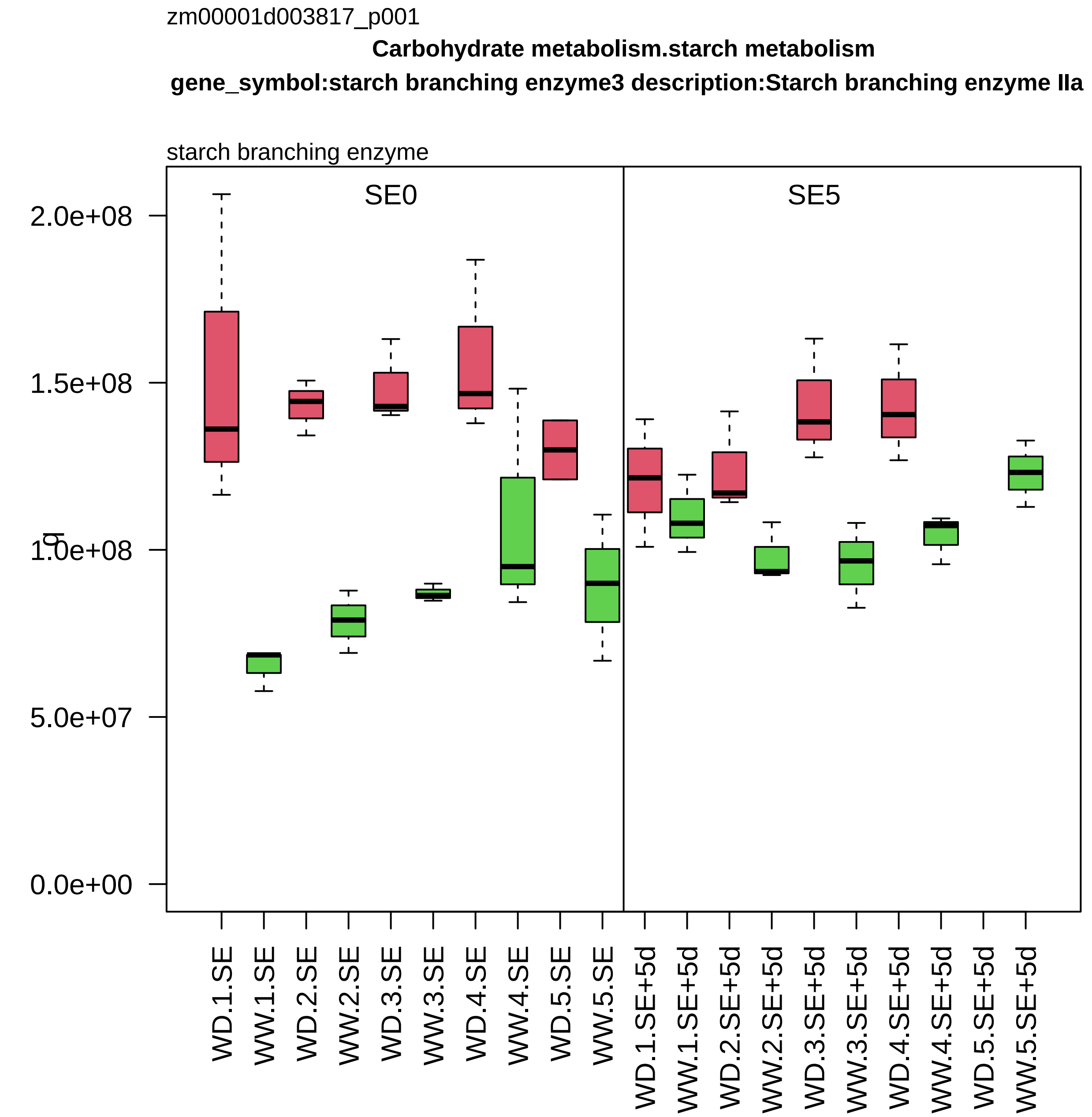
zm00001d003817_p001
Carbohydrate metabolism.starch metabolism
gene_symbol:starch branching enzyme3 description:Starch branching enzyme IIa
starch branching enzyme
SE0
SE5
2.0e+08
1.5e+08
1.0e+08
5.0e+07
0.0e+00
WD.1.SE
WW.1.SE
WD.2.SE
WW.2.SE
WD.3.SE
WW.3.SE
WD.4.SE
WW.4.SE
WD.5.SE
WW.5.SE
WD.1.SE+5d
WW.1.SE+5d
WD.2.SE+5d
WW.2.SE+5d
WD.3.SE+5d
WW.3.SE+5d
WD.4.SE+5d
WW.4.SE+5d
WD.5.SE+5d
WW.5.SE+5d

zm00001d003857_p001

**Protein homeostasis.protein quality control**

**gene_symbol:calnexin homolog1 description: calnexin homolog1**

membrane−anchored lectin chaperone *(CNX)

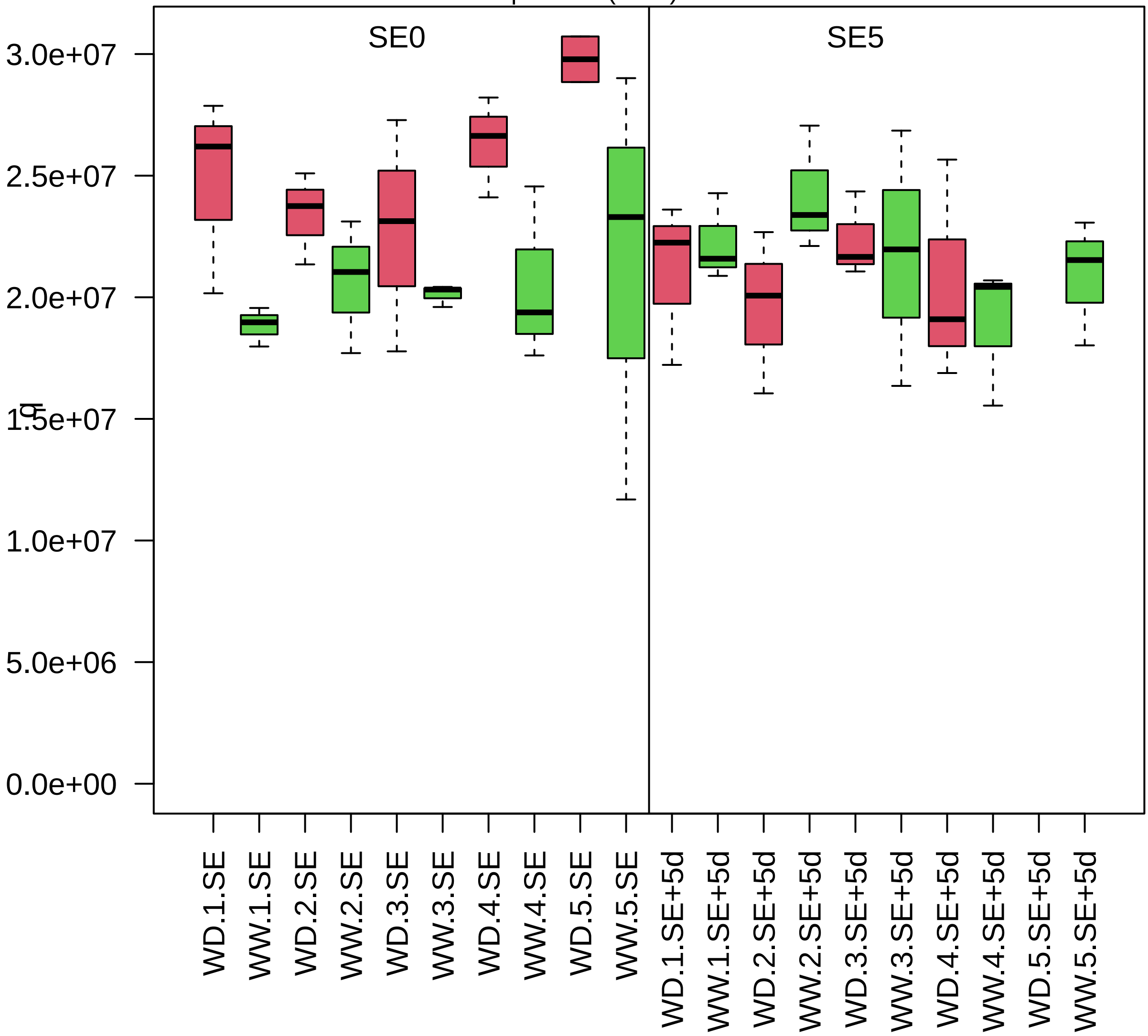

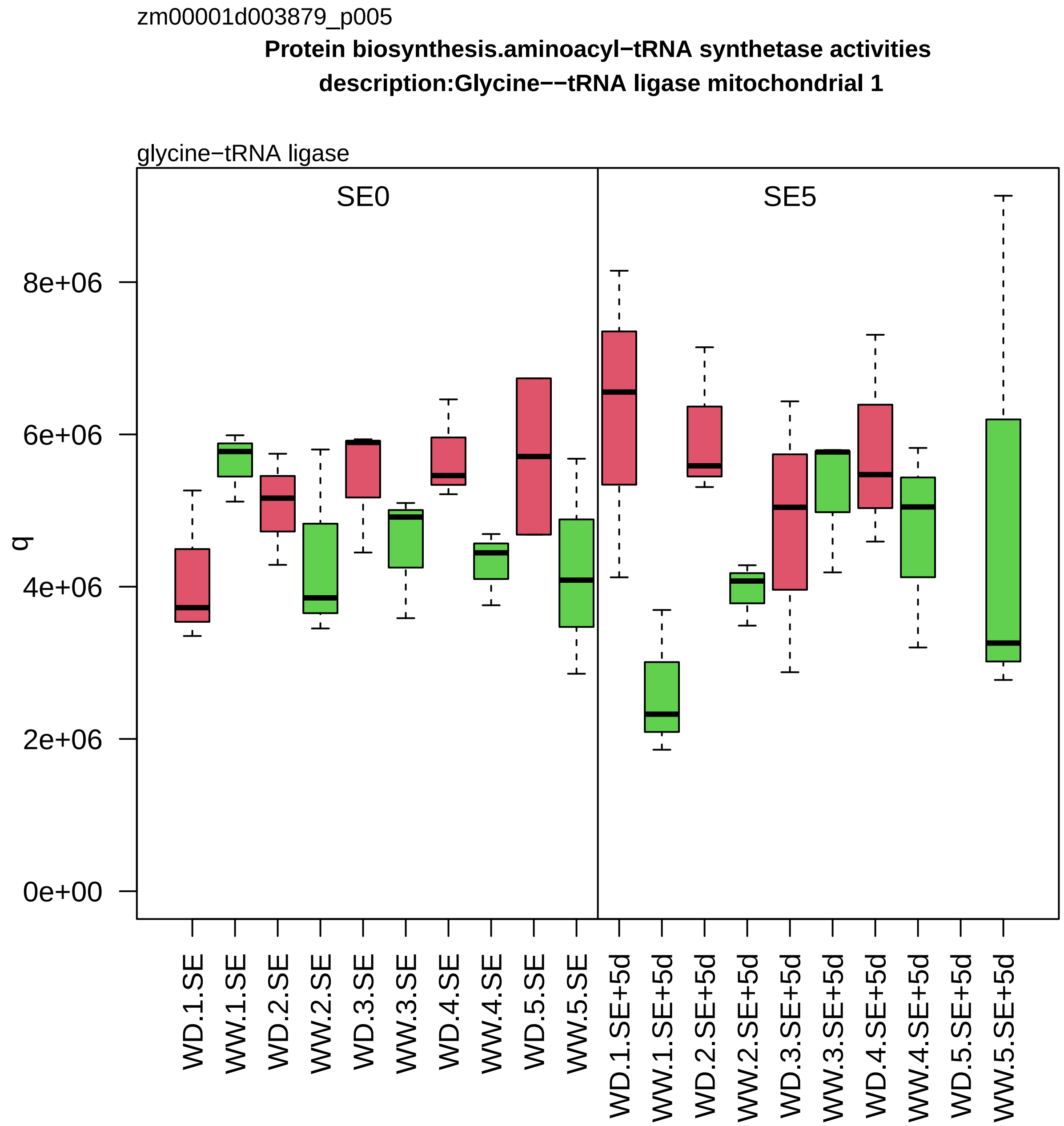
zm00001d003879_p005
Protein biosynthesis.aminoacyl−tRNA synthetase activities
description:Glycine−−tRNA ligase mitochondrial 1
glycine−tRNA ligase
SE0
SE5
q
0e+00
2e+06
4e+06
6e+06
8e+06
WD.1.SE
WW.1.SE
WD.2.SE
WW.2.SE
WD.3.SE
WW.3.SE
WD.4.SE
WW.4.SE
WD.5.SE
WW.5.SE
WD.1.SE+5d
WW.1.SE+5d
WD.2.SE+5d
WW.2.SE+5d
WD.3.SE+5d
WW.3.SE+5d
WD.4.SE+5d
WW.4.SE+5d
WD.5.SE+5d
WW.5.SE+5d

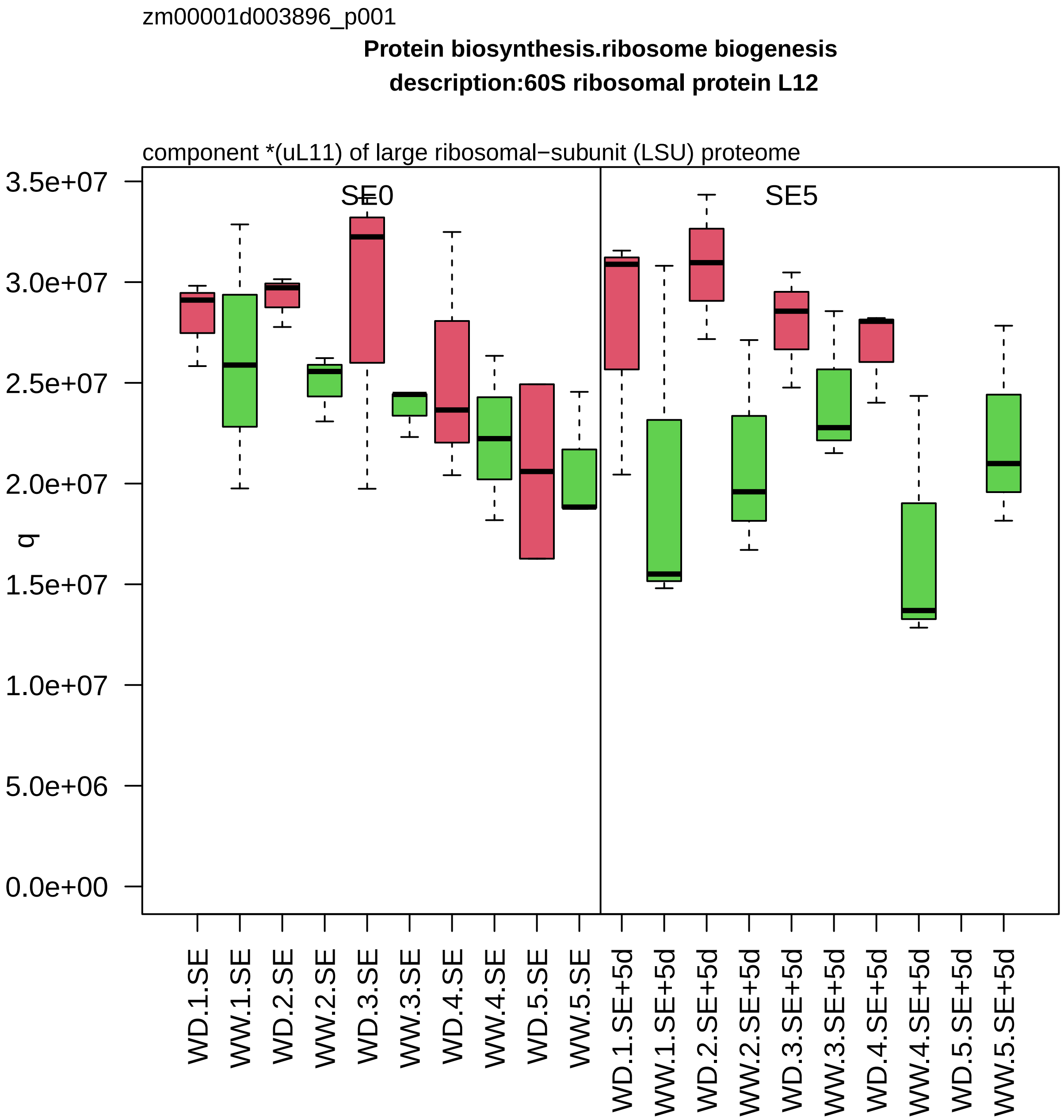
zm00001d003896_p001
Protein biosynthesis.ribosome biogenesis
description:60S ribosomal protein L12
component *(uL11) of large ribosomal−subunit (LSU) proteome
SE0
SE5
q
3.5e+07
3.0e+07
2.5e+07
2.0e+07
1.5e+07
1.0e+07
5.0e+06
0.0e+00
WD.1.SE
WW.1.SE
WD.2.SE
WW.2.SE
WD.3.SE
WW.3.SE
WD.4.SE
WW.4.SE
WD.5.SE
WW.5.SE
WD.1.SE+5d
WW.1.SE+5d
WD.2.SE+5d
WW.2.SE+5d
WD.3.SE+5d
WW.3.SE+5d
WD.4.SE+5d
WW.4.SE+5d
WD.5.SE+5d
WW.5.SE+5d

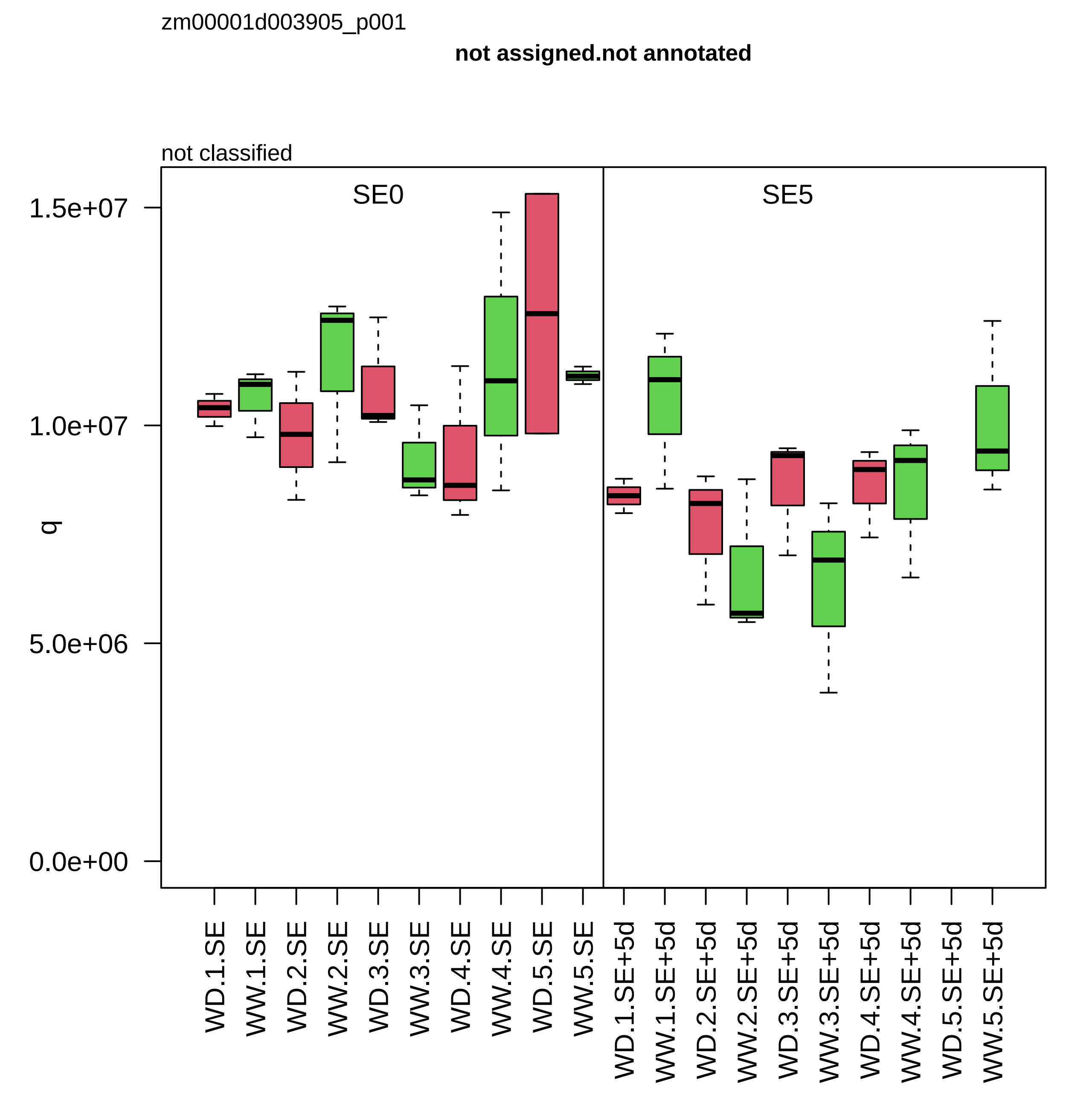
zm00001d003905_p001
not assigned.not annotated
not classified
SE0
SE5
q
1.5e+07
1.0e+07
5.0e+06
0.0e+00
WD.1.SE
WW.1.SE
WD.2.SE
WW.2.SE
WD.3.SE
WW.3.SE
WD.4.SE
WW.4.SE
WD.5.SE
WW.5.SE
WD.1.SE+5d
WW.1.SE+5d
WD.2.SE+5d
WW.2.SE+5d
WD.3.SE+5d
WW.3.SE+5d
WD.4.SE+5d
WW.4.SE+5d
WD.5.SE+5d
WW.5.SE+5d

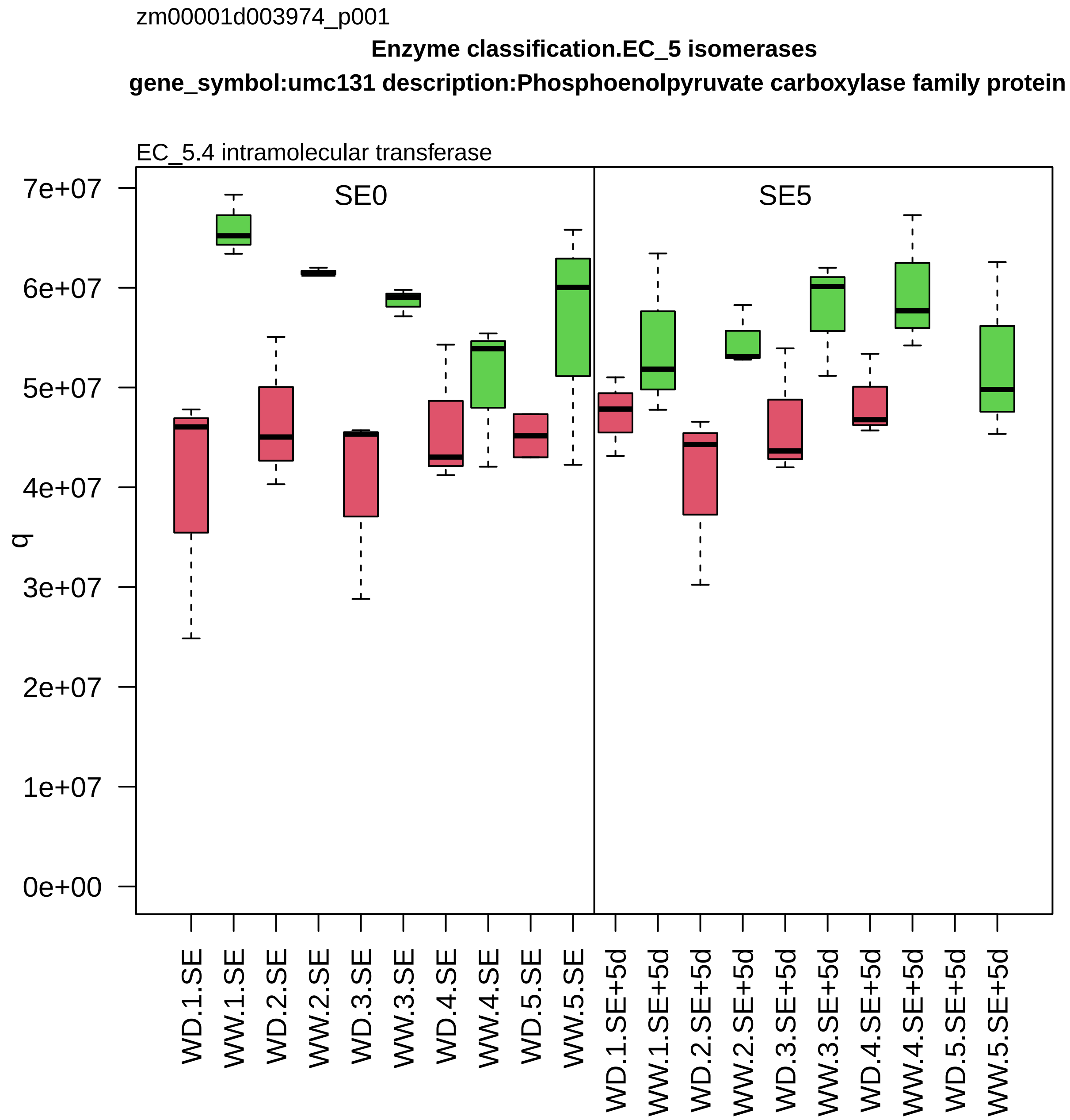

zm00001d003974_p001
Enzyme classification.EC_5 isomerases
gene_symbol:umc131 description:Phosphoenolpyruvate carboxylase family protein
EC_5.4 intramolecular transferase
SE0
SE5
q
0e+00
1e+07
2e+07
3e+07
4e+07
5e+07
6e+07
7e+07
WD.1.SE
WW.1.SE
WD.2.SE
WW.2.SE
WD.3.SE
WW.3.SE
WD.4.SE
WW.4.SE
WD.5.SE
WW.5.SE
WD.1.SE+5d
WW.1.SE+5d
WD.2.SE+5d
WW.2.SE+5d
WD.3.SE+5d
WW.3.SE+5d
WD.4.SE+5d
WW.4.SE+5d
WD.5.SE+5d
WW.5.SE+5d

zm00001d004009_p001

**Protein biosynthesis.ribosome biogenesis**
**description:60S ribosomal protein L37a−2**

component *(eL43) of large ribosomal−subunit (LSU) proteome

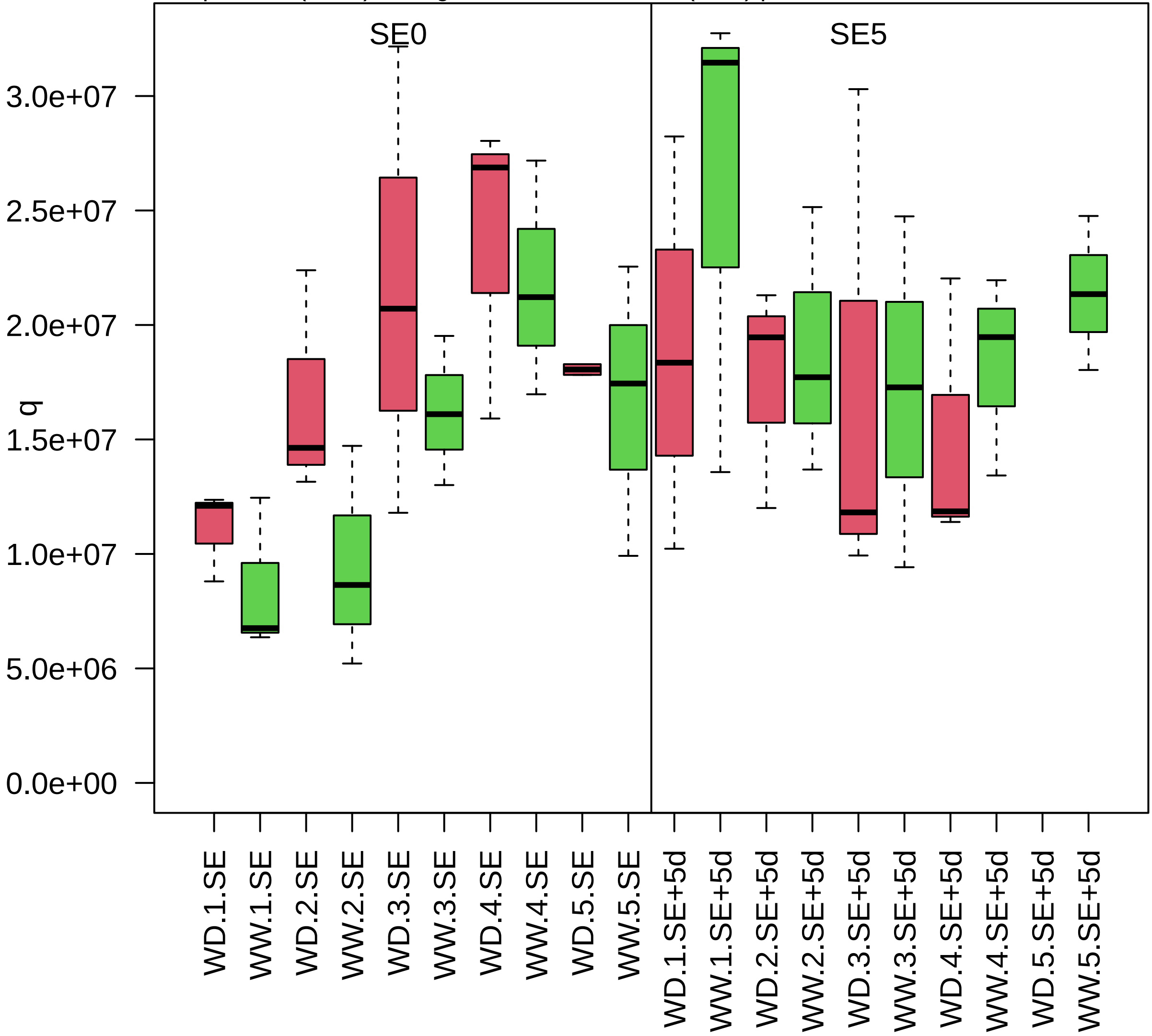

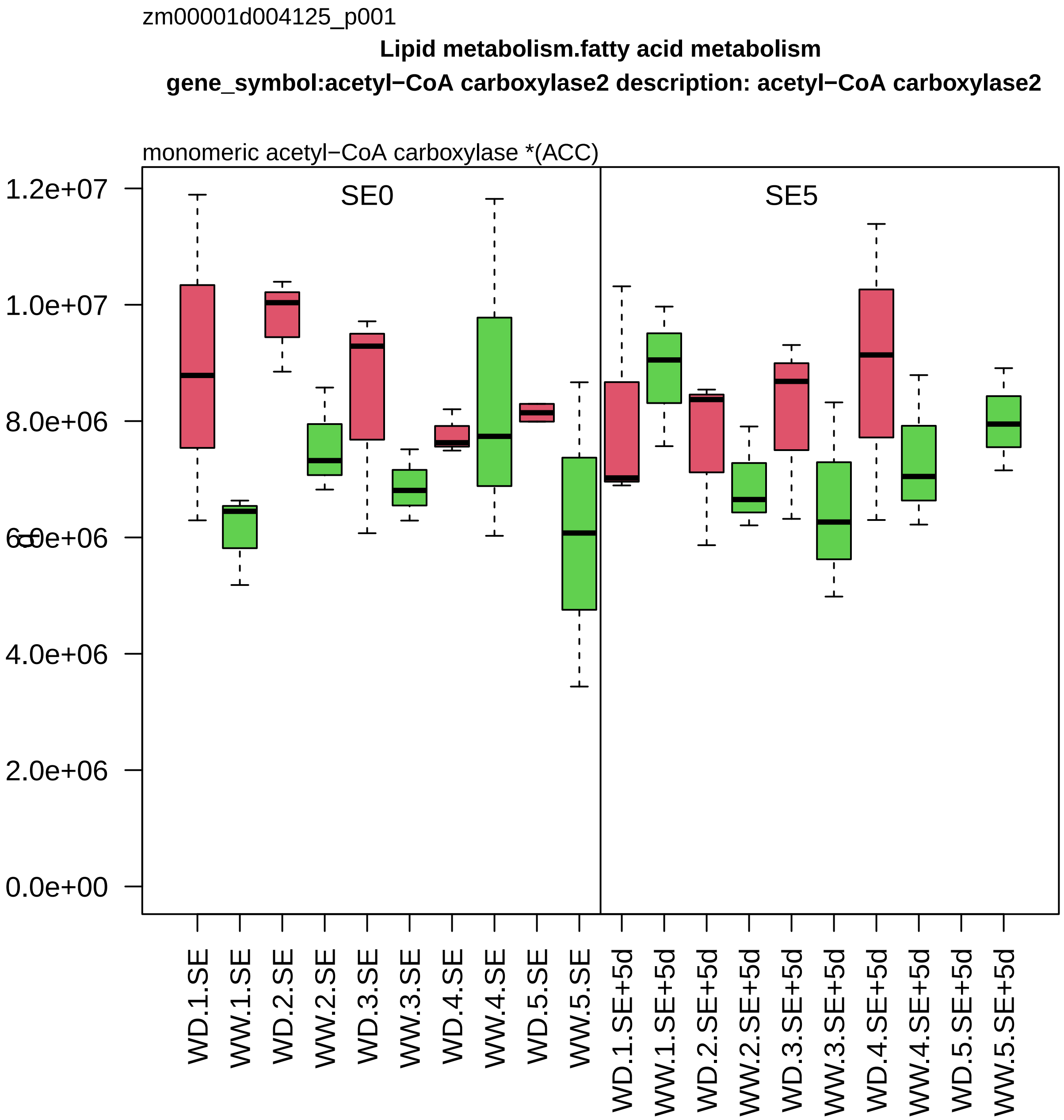
zm00001d004125_p001
Lipid metabolism.fatty acid metabolism
gene_symbol:acetyl−CoA carboxylase2 description: acetyl−CoA carboxylase2
monomeric acetyl−CoA carboxylase *(ACC)
SE0
SE5
1.2e+07
1.0e+07
8.0e+06
6.0e+06
4.0e+06
2.0e+06
0.0e+00
WD.1.SE
WW.1.SE
WD.2.SE
WW.2.SE
WD.3.SE
WW.3.SE
WD.4.SE
WW.4.SE
WD.5.SE
WW.5.SE
WD.1.SE+5d
WW.1.SE+5d
WD.2.SE+5d
WW.2.SE+5d
WD.3.SE+5d
WW.3.SE+5d
WD.4.SE+5d
WW.4.SE+5d
WD.5.SE+5d
WW.5.SE+5d

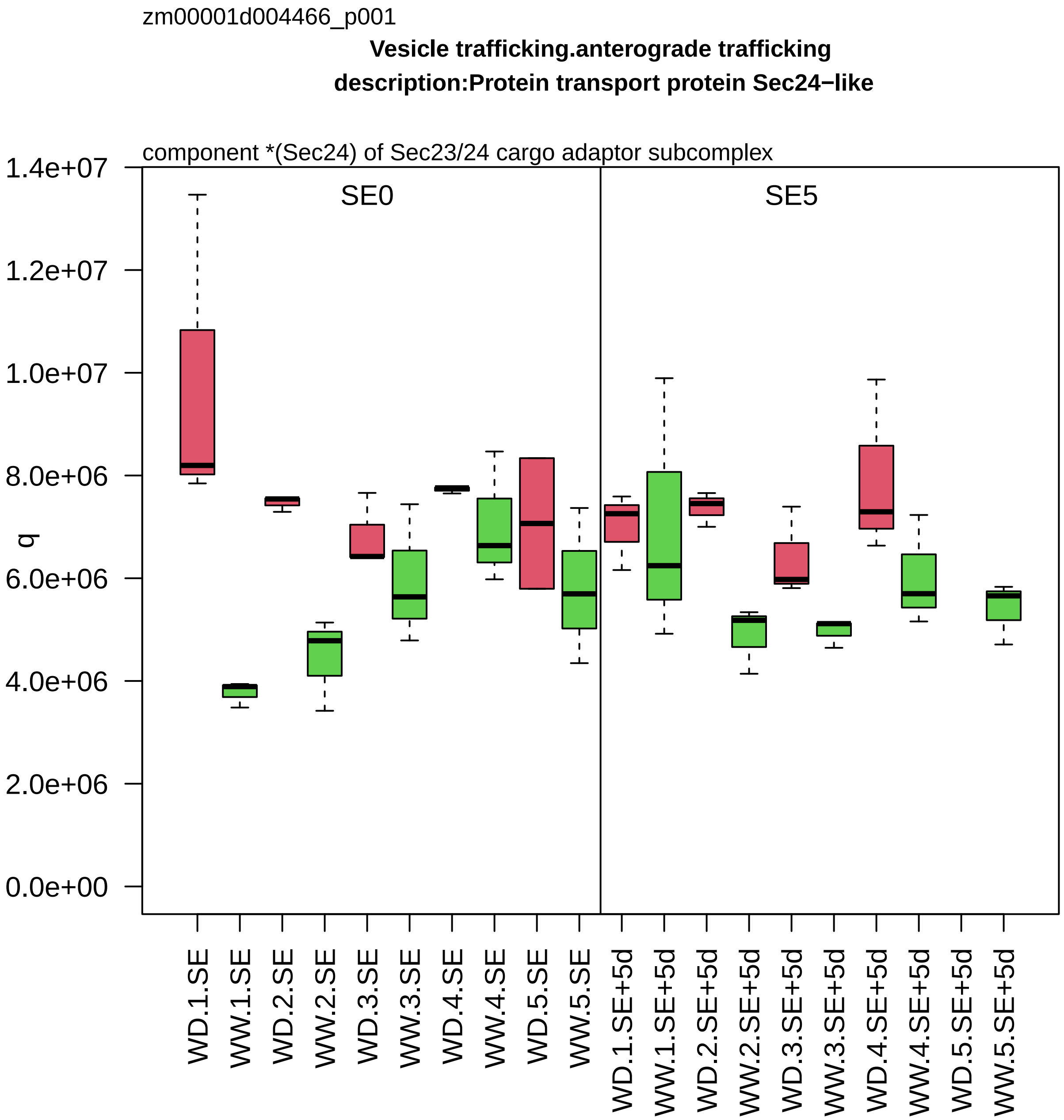

zm00001d004466_p001
Vesicle trafficking.anterograde trafficking
description:Protein transport protein Sec24−like
component *(Sec24) of Sec23/24 cargo adaptor subcomplex
SE0
SE5
q
0.0e+00
2.0e+06
4.0e+06
6.0e+06
8.0e+06
1.0e+07
1.2e+07
1.4e+07
WD.1.SE
WW.1.SE
WD.2.SE
WW.2.SE
WD.3.SE
WW.3.SE
WD.4.SE
WW.4.SE
WD.5.SE
WW.5.SE
WD.1.SE+5d
WW.1.SE+5d
WD.2.SE+5d
WW.2.SE+5d
WD.3.SE+5d
WW.3.SE+5d
WD.4.SE+5d
WW.4.SE+5d
WD.5.SE+5d
WW.5.SE+5d

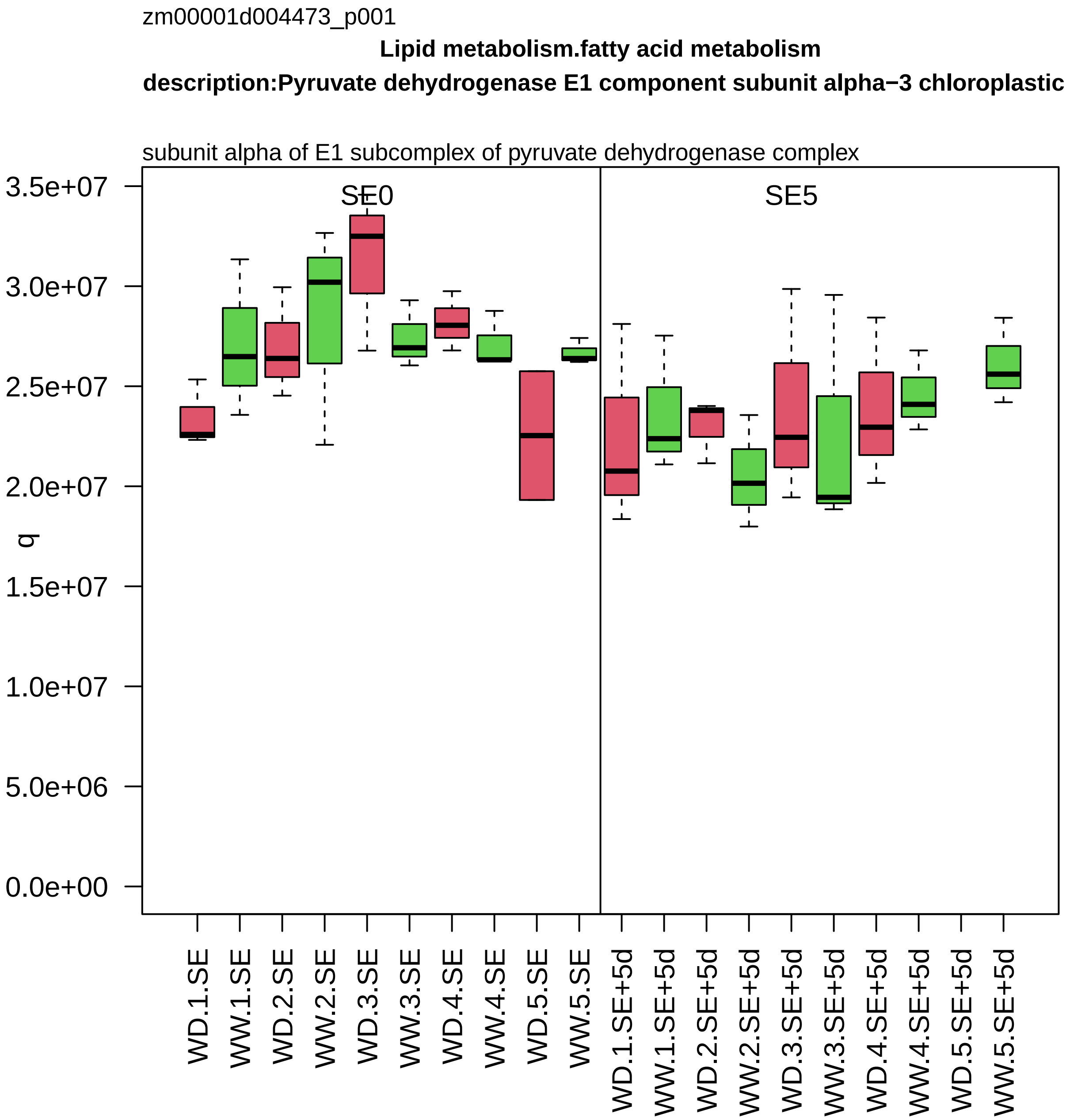
zm00001d004473_p001
Lipid metabolism.fatty acid metabolism
description:Pyruvate dehydrogenase E1 component subunit alpha−3 chloroplastic
subunit alpha of E1 subcomplex of pyruvate dehydrogenase complex
SE0
SE5
q
3.5e+07
3.0e+07
2.5e+07
2.0e+07
1.5e+07
1.0e+07
5.0e+06
0.0e+00
WD.1.SE
WW.1.SE
WD.2.SE
WW.2.SE
WD.3.SE
WW.3.SE
WD.4.SE
WW.4.SE
WD.5.SE
WW.5.SE
WD.1.SE+5d
WW.1.SE+5d
WD.2.SE+5d
WW.2.SE+5d
WD.3.SE+5d
WW.3.SE+5d
WD.4.SE+5d
WW.4.SE+5d
WD.5.SE+5d
WW.5.SE+5d

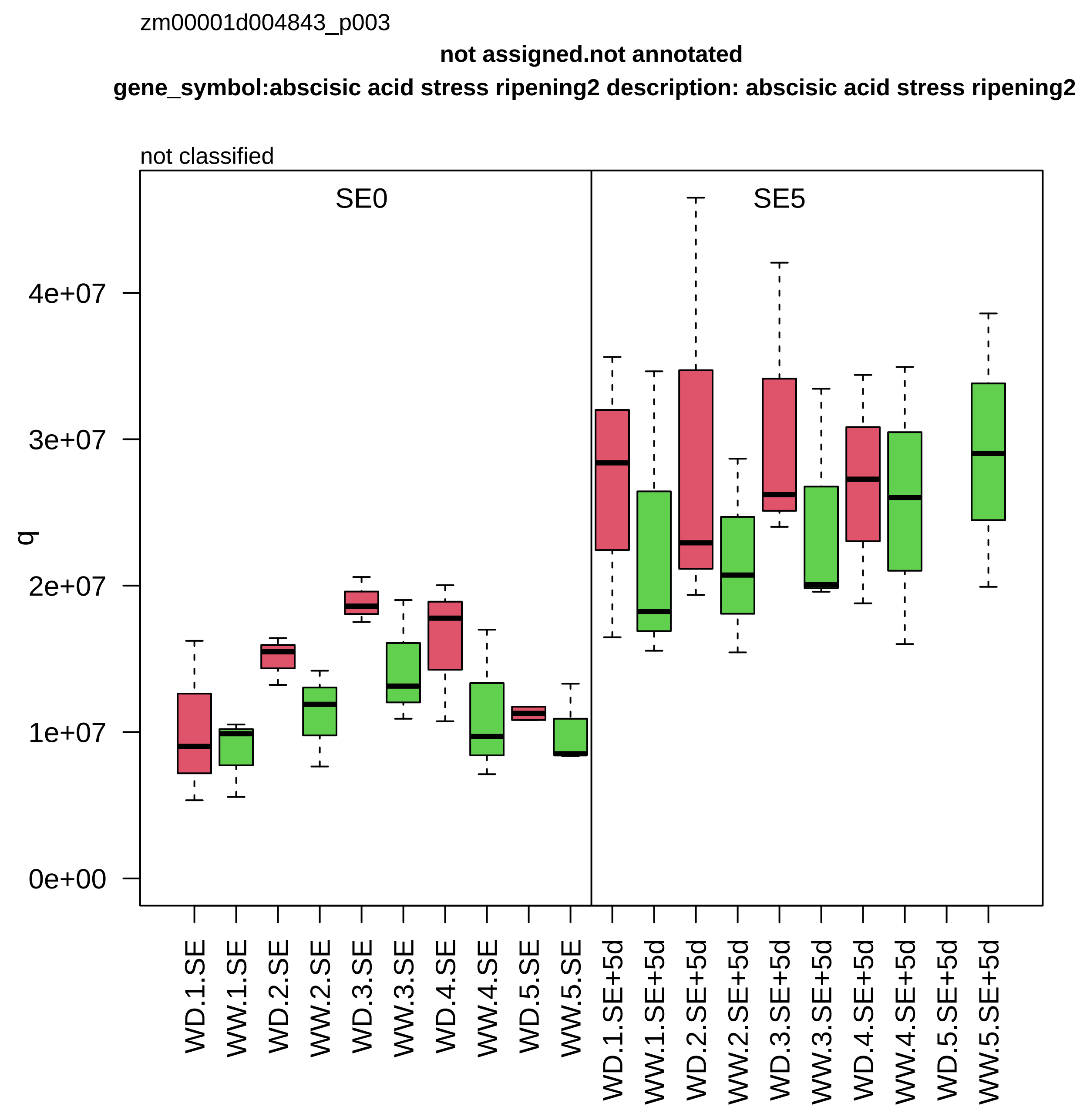
zm00001d004843_p003
not assigned.not annotated
gene_symbol:abscisic acid stress ripening2 description: abscisic acid stress ripening2
not classified
SE0
SE5
q
4e+07
3e+07
2e+07
1e+07
0e+00
WD.1.SE
WW.1.SE
WD.2.SE
WW.2.SE
WD.3.SE
WW.3.SE
WD.4.SE
WW.4.SE
WD.5.SE
WW.5.SE
WD.1.SE+5d
WW.1.SE+5d
WD.2.SE+5d
WW.2.SE+5d
WD.3.SE+5d
WW.3.SE+5d
WD.4.SE+5d
WW.4.SE+5d
WD.5.SE+5d
WW.5.SE+5d

zm00001d004960_p001

**Amino acid metabolism.pyruvate group amino acid biosynthesis description:2−isopropylmalate synthase 1 chloroplastic**

2−isopropylmalate synthase *(IPMS)

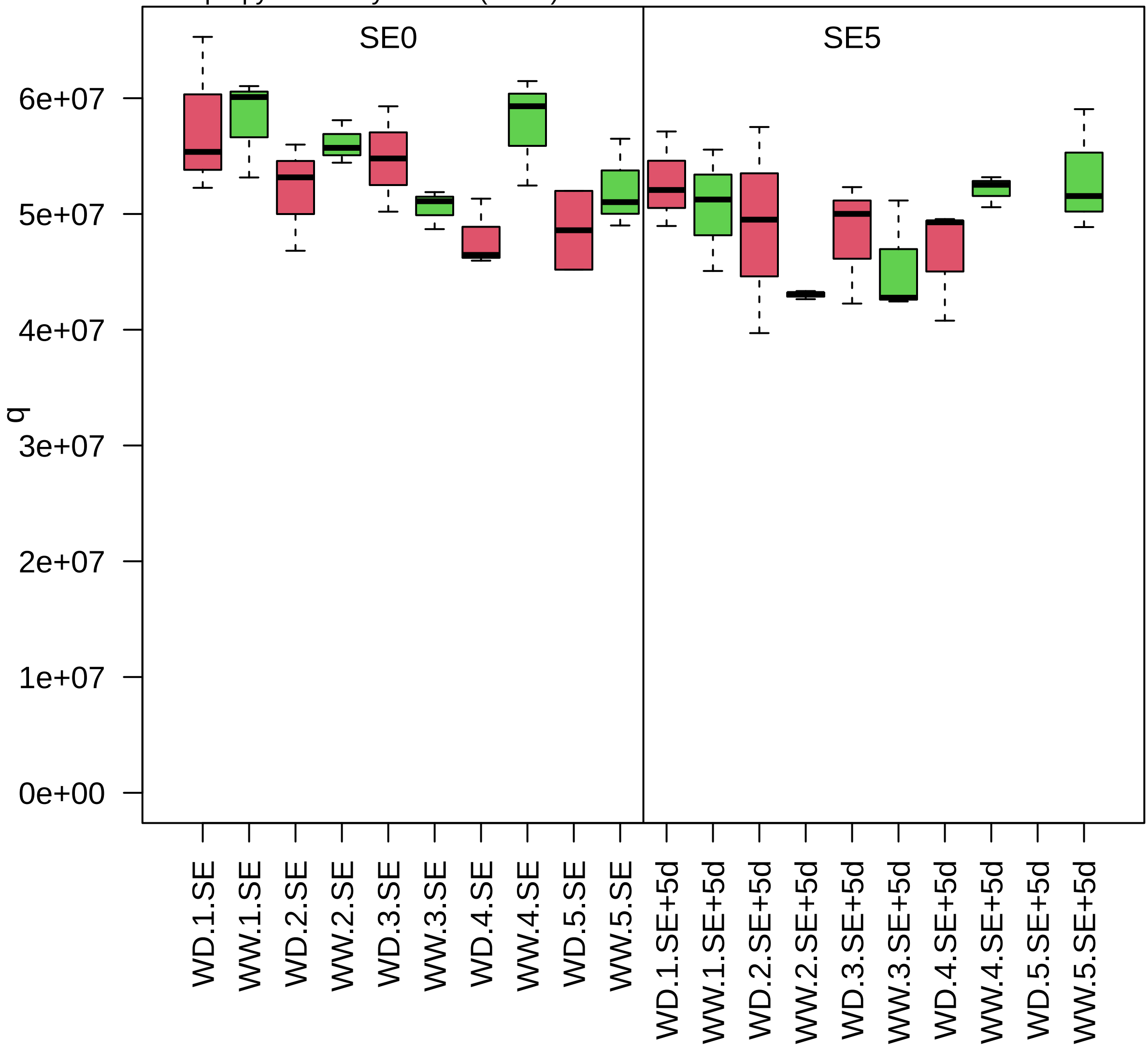

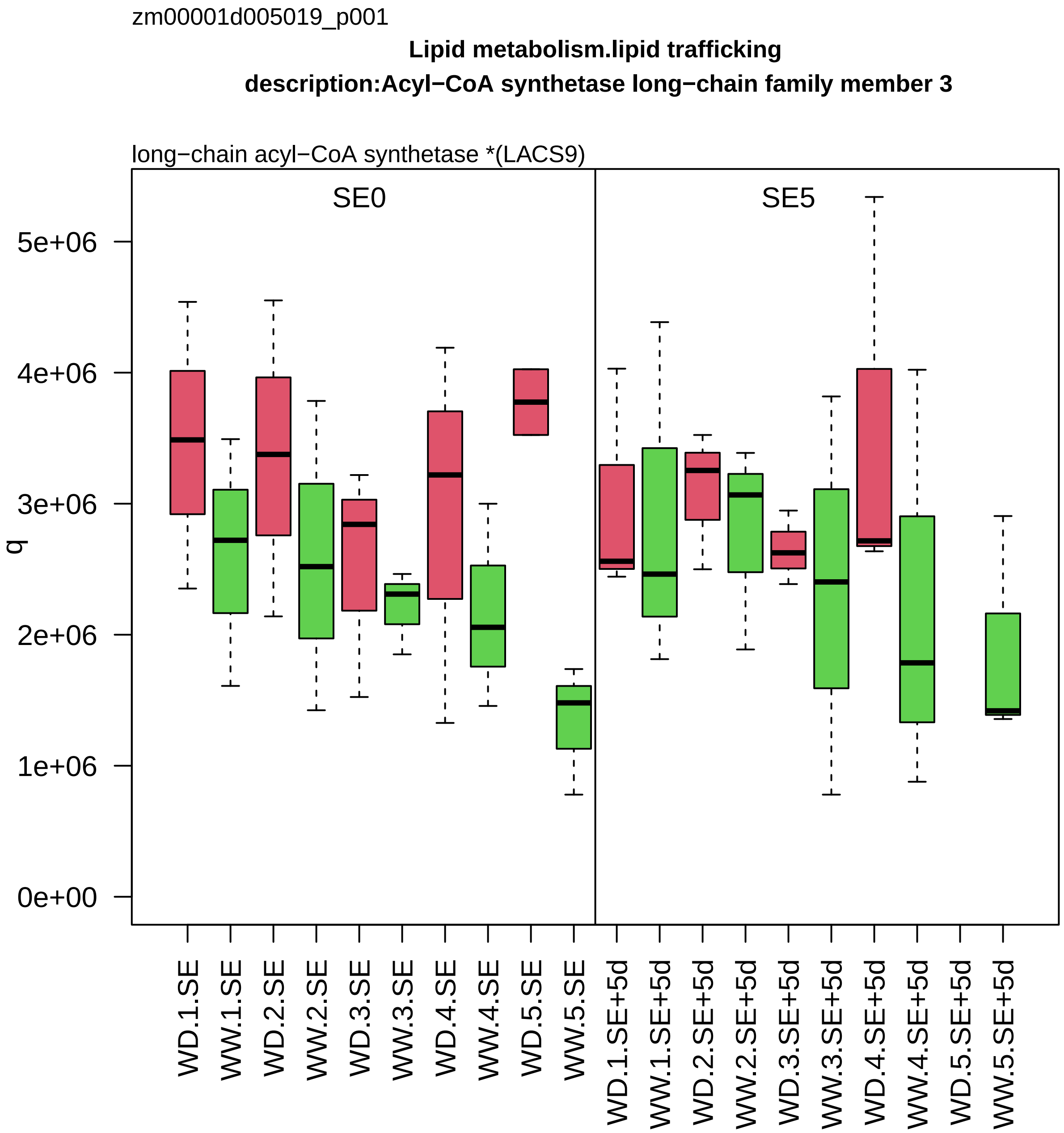
zm00001d005019_p001
Lipid metabolism.lipid trafficking
description:Acyl−CoA synthetase long−chain family member 3
long−chain acyl−CoA synthetase *(LACS9)
SE0
SE5
q
0e+00
1e+06
2e+06
3e+06
4e+06
5e+06
WD.1.SE
WW.1.SE
WD.2.SE
WW.2.SE
WD.3.SE
WW.3.SE
WD.4.SE
WW.4.SE
WD.5.SE
WW.5.SE
WD.1.SE+5d
WW.1.SE+5d
WD.2.SE+5d
WW.2.SE+5d
WD.3.SE+5d
WW.3.SE+5d
WD.4.SE+5d
WW.4.SE+5d
WD.5.SE+5d
WW.5.SE+5d

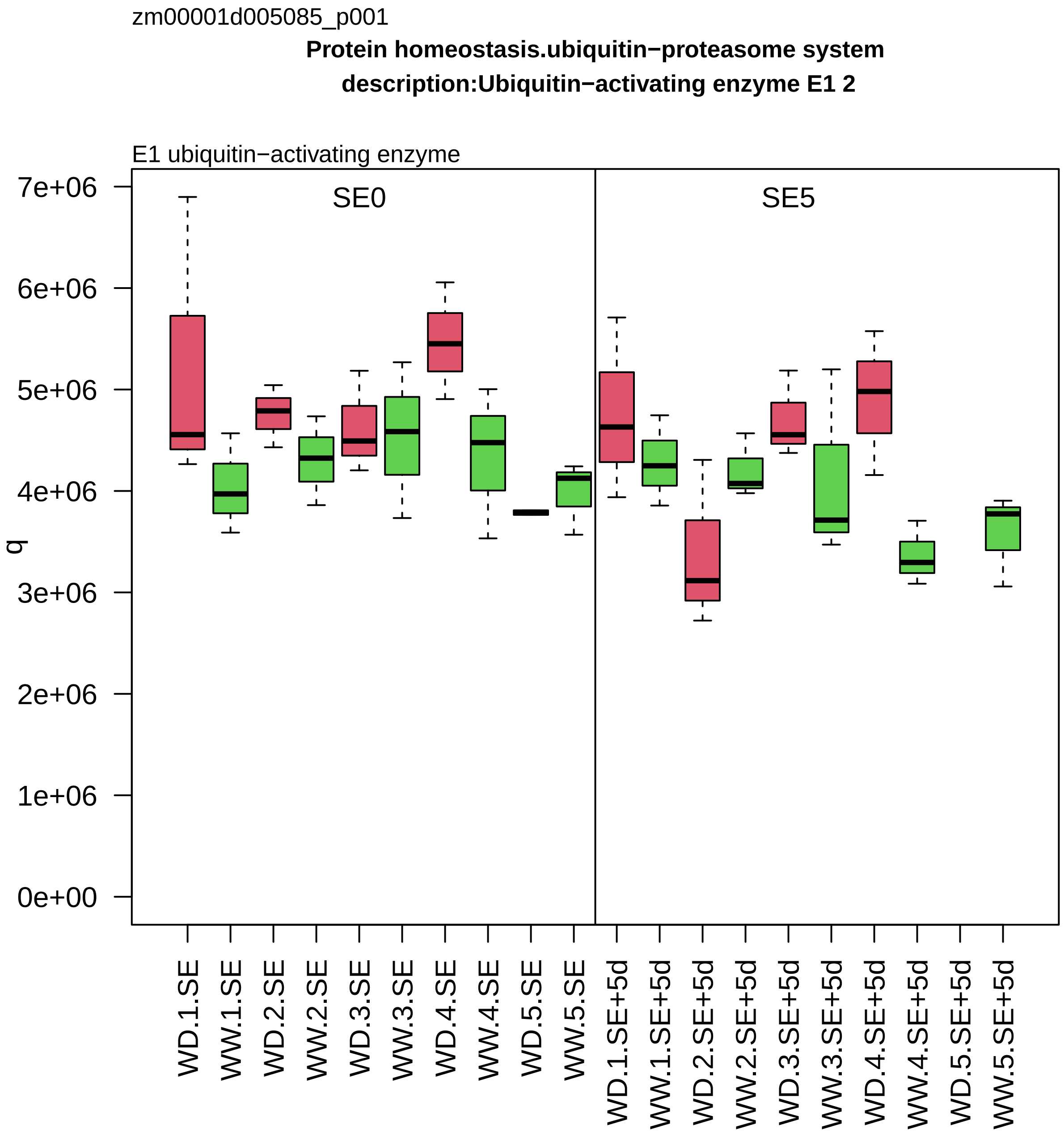

zm00001d005085_p001
Protein homeostasis.ubiquitin−proteasome system
description:Ubiquitin−activating enzyme E1 2
E1 ubiquitin−activating enzyme
SE0
SE5
q
7e+06
6e+06
5e+06
4e+06
3e+06
2e+06
1e+06
0e+00
WD.1.SE
WW.1.SE
WD.2.SE
WW.2.SE
WD.3.SE
WW.3.SE
WD.4.SE
WW.4.SE
WD.5.SE
WW.5.SE
WD.1.SE+5d
WW.1.SE+5d
WD.2.SE+5d
WW.2.SE+5d
WD.3.SE+5d
WW.3.SE+5d
WD.4.SE+5d
WW.4.SE+5d
WD.5.SE+5d
WW.5.SE+5d

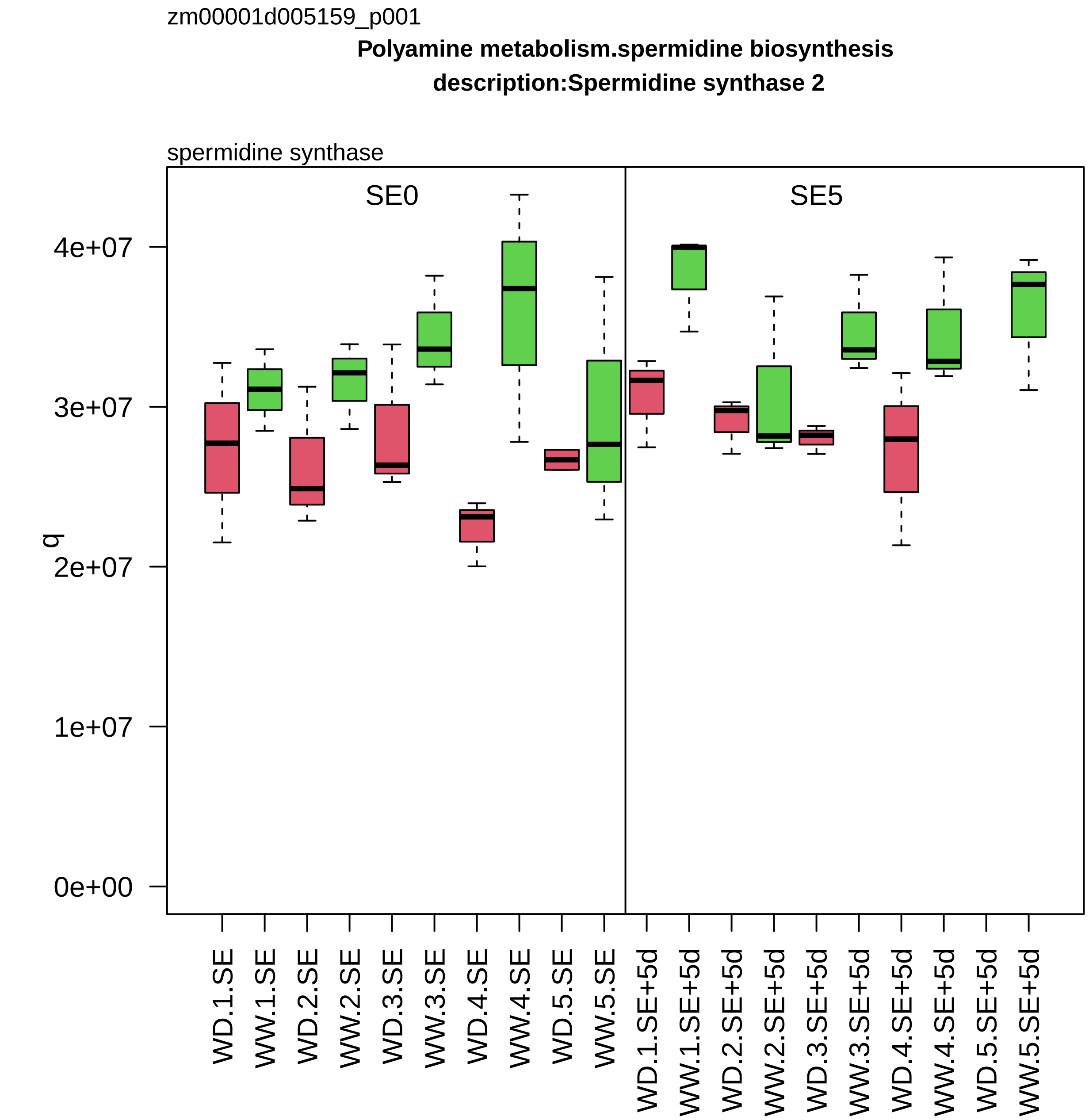
zm00001d005159_p001
Polyamine metabolism.spermidine biosynthesis
description:Spermidine synthase 2
spermidine synthase
SE0
SE5
q
0e+00
1e+07
2e+07
3e+07
4e+07
WD.1.SE
WW.1.SE
WD.2.SE
WW.2.SE
WD.3.SE
WW.3.SE
WD.4.SE
WW.4.SE
WD.5.SE
WW.5.SE
WD.1.SE+5d
WW.1.SE+5d
WD.2.SE+5d
WW.2.SE+5d
WD.3.SE+5d
WW.3.SE+5d
WD.4.SE+5d
WW.4.SE+5d
WD.5.SE+5d
WW.5.SE+5d

zm00001d005231_p001

**Vesicle trafficking.retrograde trafficking**
**description:ADP−ribosylation factor A1F**

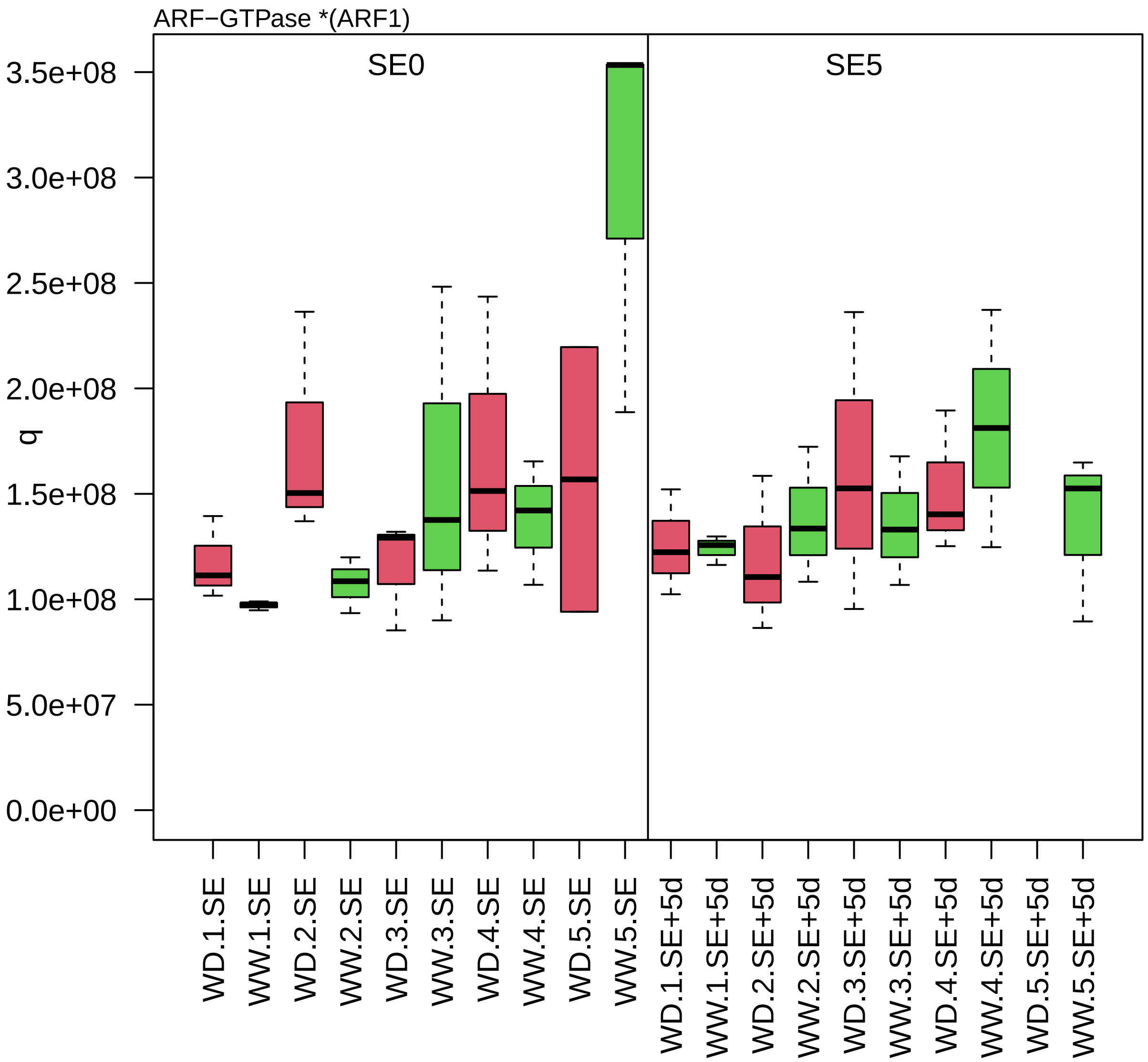

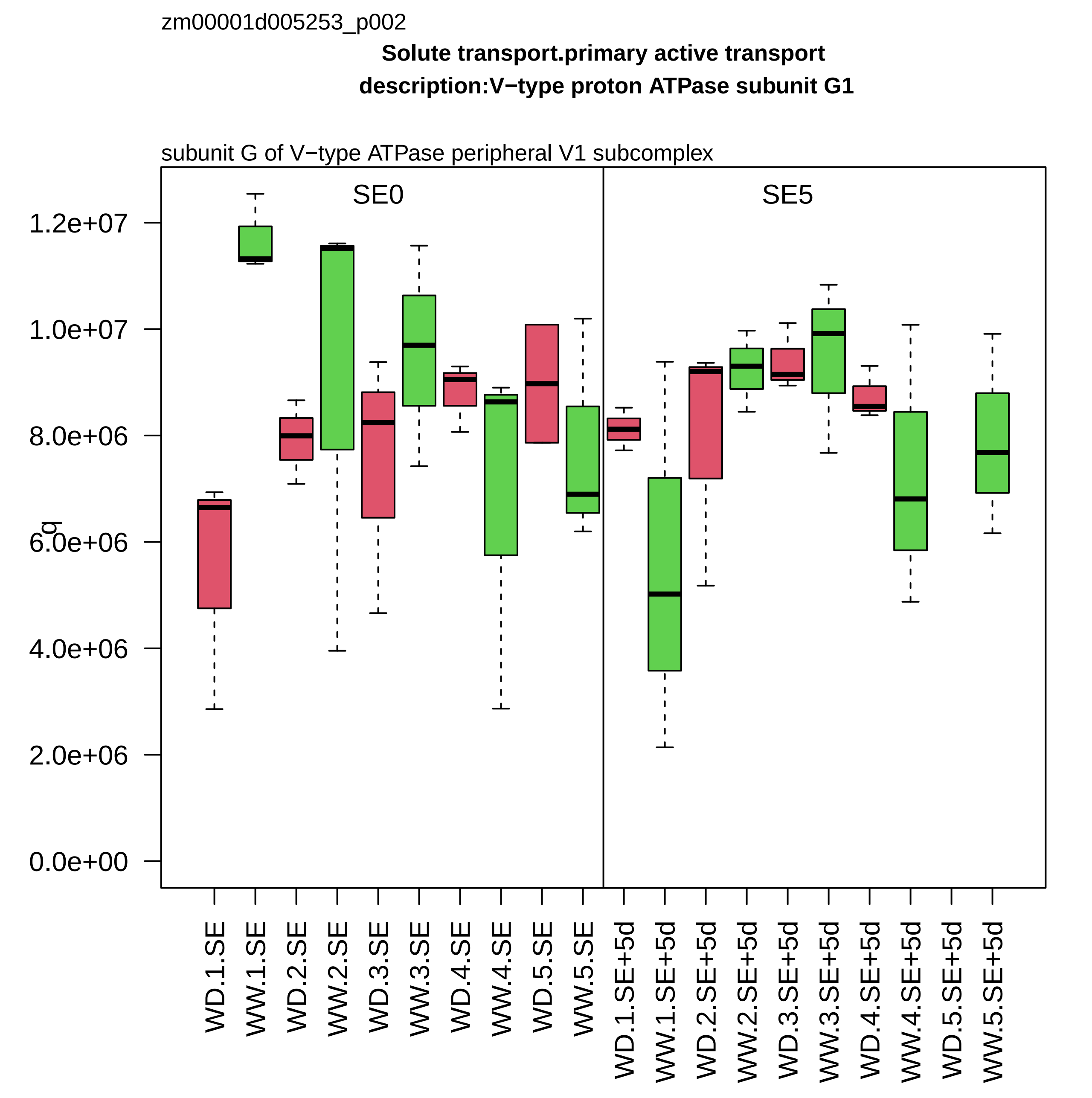

zm00001d005253_p002
Solute transport.primary active transport
description:V−type proton ATPase subunit G1
subunit G of V−type ATPase peripheral V1 subcomplex
SE0
SE5
q
1.2e+07
1.0e+07
8.0e+06
6.0e+06
4.0e+06
2.0e+06
0.0e+00
WD.1.SE
WW.1.SE
WD.2.SE
WW.2.SE
WD.3.SE
WW.3.SE
WD.4.SE
WW.4.SE
WD.5.SE
WW.5.SE
WD.1.SE+5d
WW.1.SE+5d
WD.2.SE+5d
WW.2.SE+5d
WD.3.SE+5d
WW.3.SE+5d
WD.4.SE+5d
WW.4.SE+5d
WD.5.SE+5d
WW.5.SE+5d

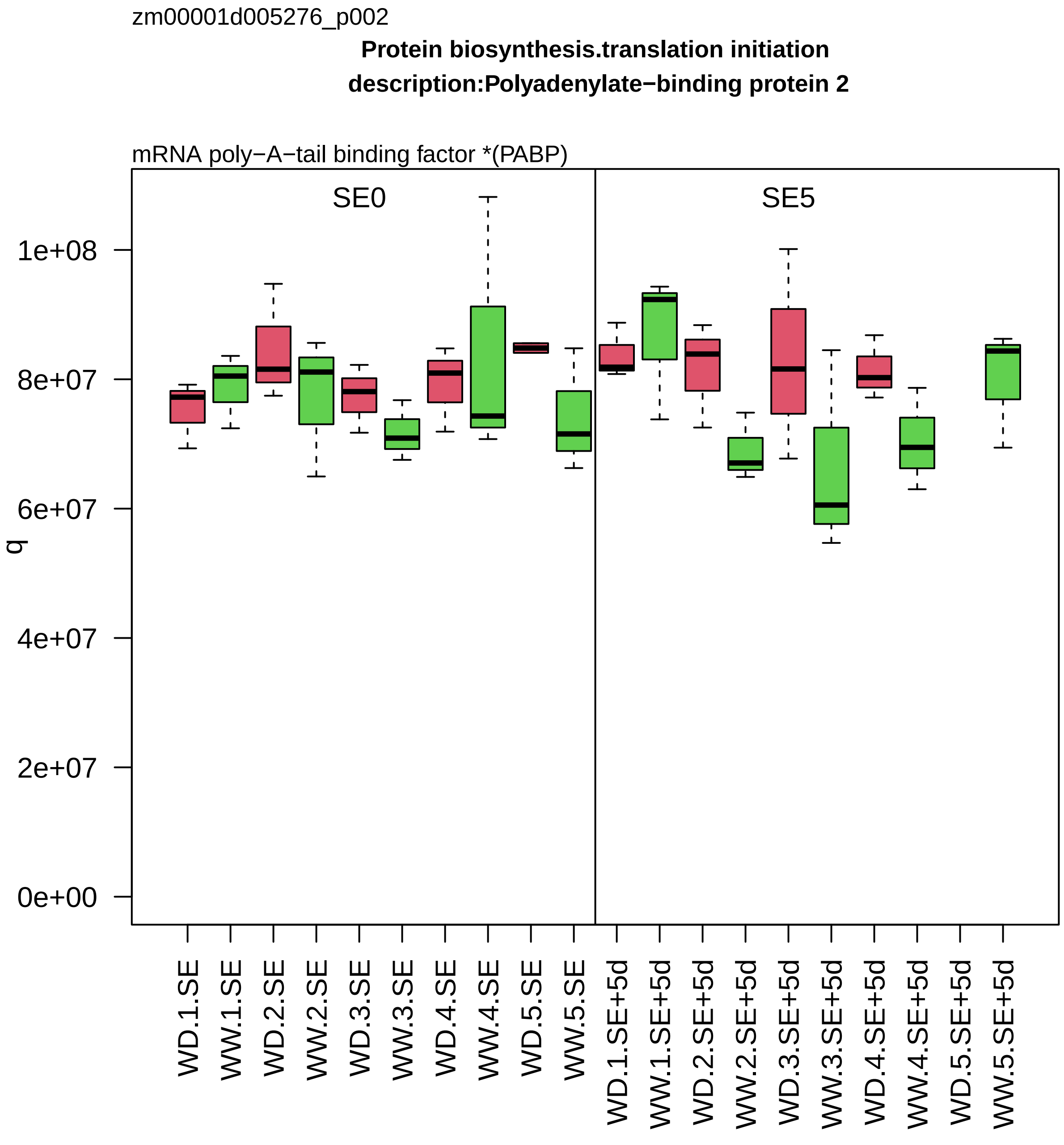
zm00001d005276_p002
Protein biosynthesis.translation initiation
description:Polyadenylate−binding protein 2
mRNA poly−A−tail binding factor *(PABP)
SE0
SE5
q
1e+08
8e+07
6e+07
4e+07
2e+07
0e+00
WD.1.SE
WW.1.SE
WD.2.SE
WW.2.SE
WD.3.SE
WW.3.SE
WD.4.SE
WW.4.SE
WD.5.SE
WW.5.SE
WD.1.SE+5d
WW.1.SE+5d
WD.2.SE+5d
WW.2.SE+5d
WD.3.SE+5d
WW.3.SE+5d
WD.4.SE+5d
WW.4.SE+5d
WD.5.SE+5d
WW.5.SE+5d

zm00001d005334_p001

**Protein biosynthesis.ribosome biogenesis**
**description:Ubiquitin−60S ribosomal protein L40−1**

component *(eL40) of large ribosomal−subunit (LSU) proteome

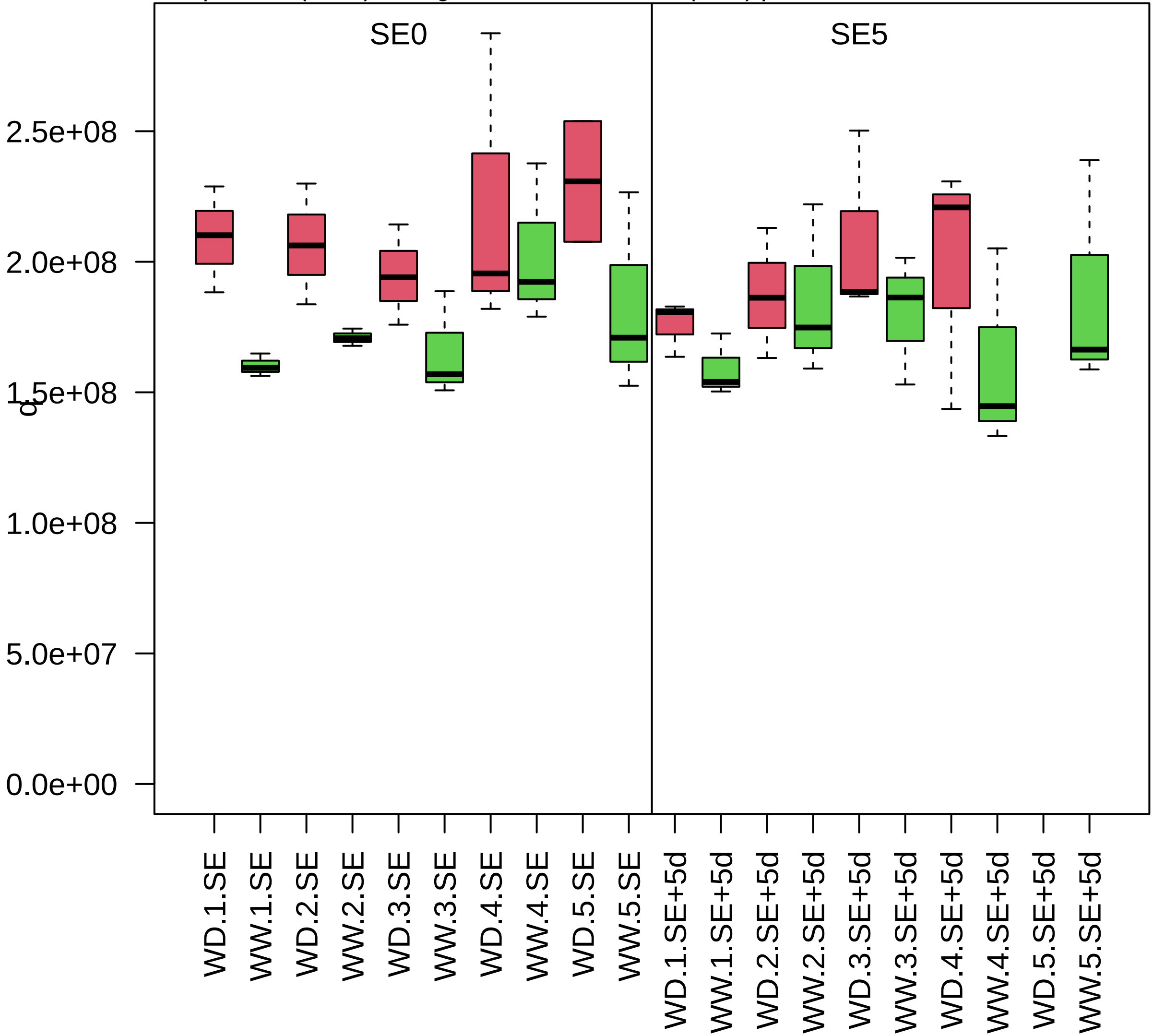

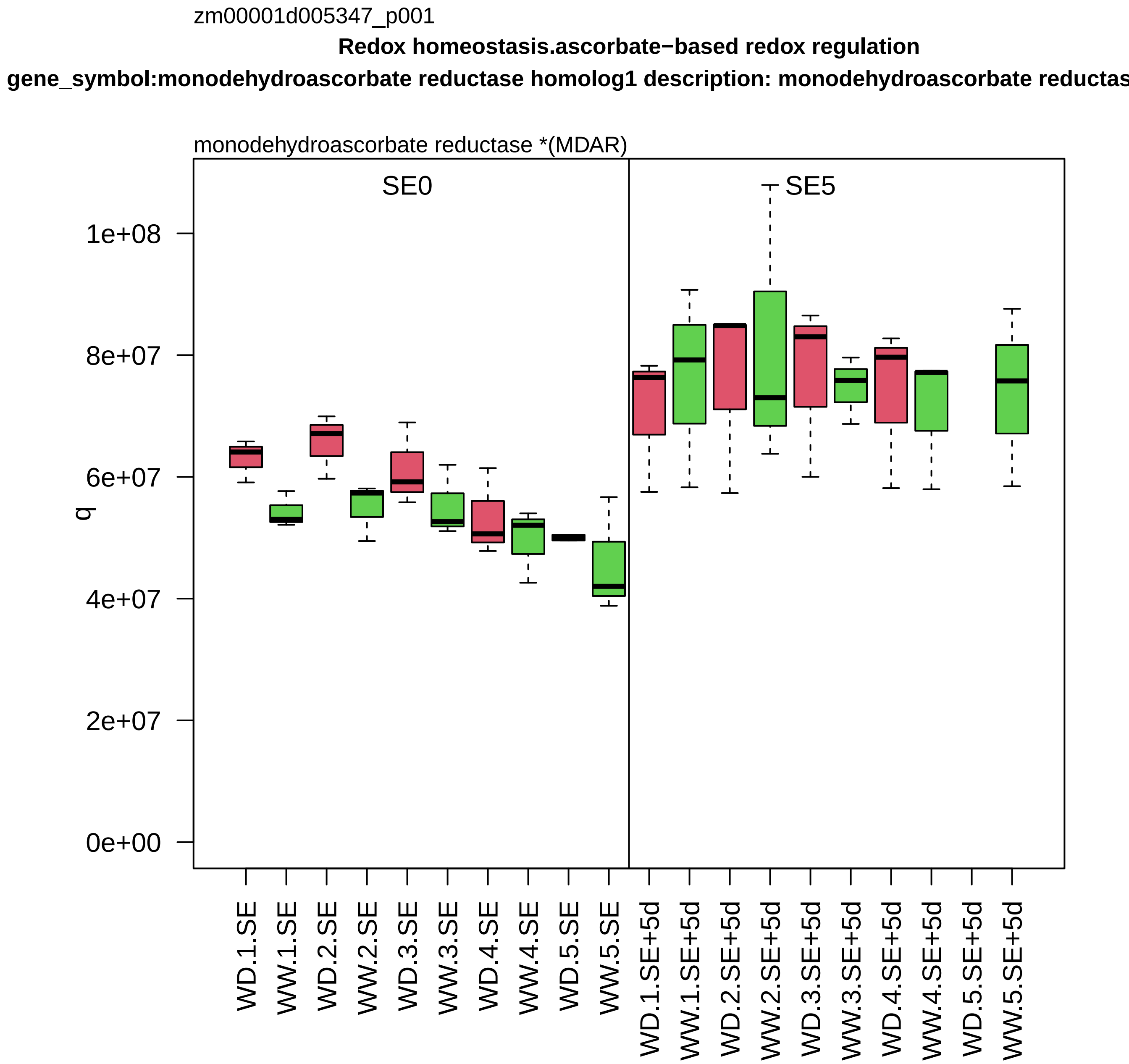
zm00001d005347_p001
Redox homeostasis.ascorbate−based redox regulation
gene_symbol:monodehydroascorbate reductase homolog1 description: monodehydroascorbate reductas
monodehydroascorbate reductase *(MDAR)
SE0
SE5
q
1e+08
8e+07
6e+07
4e+07
2e+07
0e+00
WD.1.SE
WW.1.SE
WD.2.SE
WW.2.SE
WD.3.SE
WW.3.SE
WD.4.SE
WW.4.SE
WD.5.SE
WW.5.SE
WD.1.SE+5d
WW.1.SE+5d
WD.2.SE+5d
WW.2.SE+5d
WD.3.SE+5d
WW.3.SE+5d
WD.4.SE+5d
WW.4.SE+5d
WD.5.SE+5d
WW.5.SE+5d

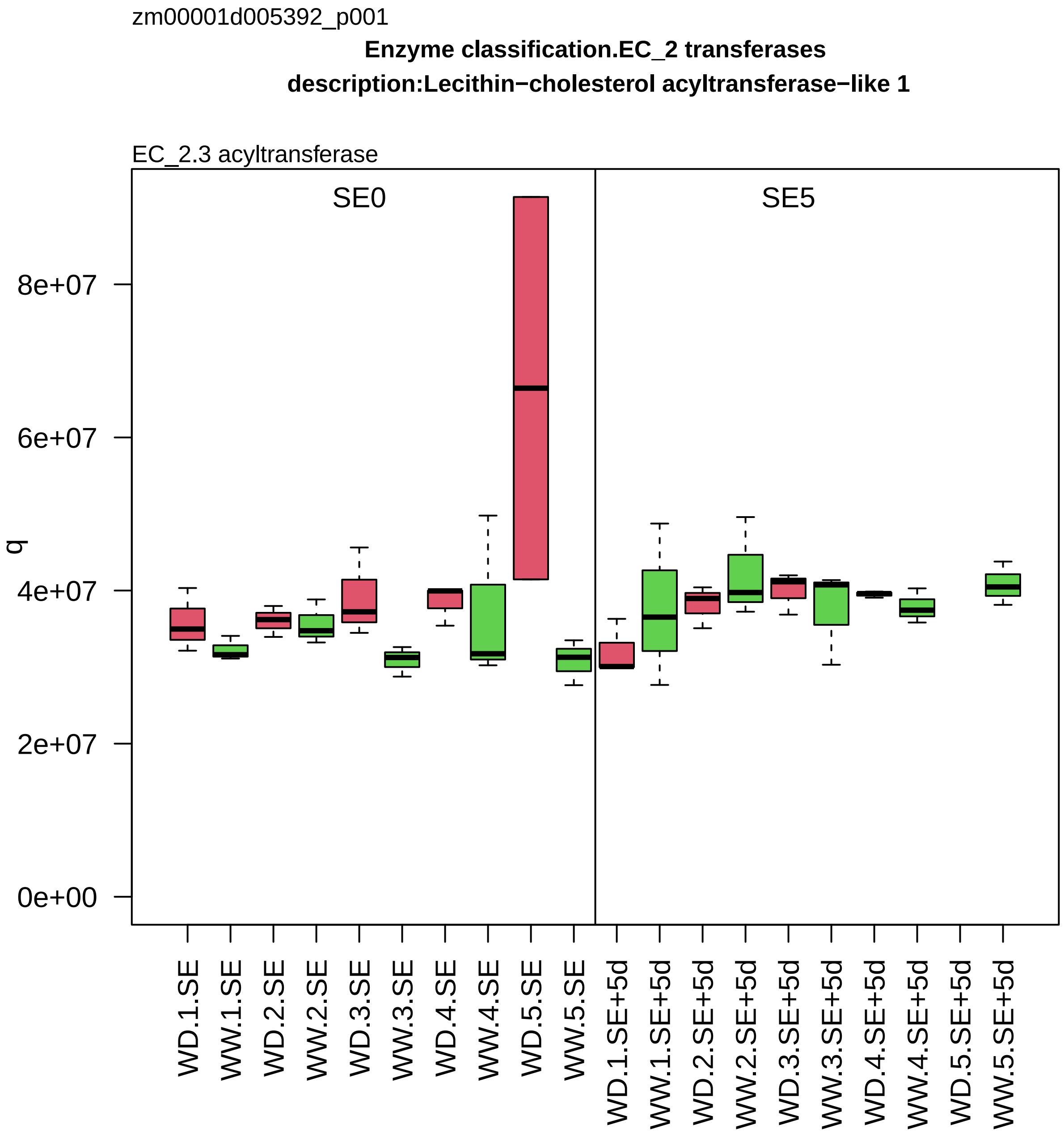

zm00001d005392_p001
Enzyme classification.EC_2 transferases
description:Lecithin−cholesterol acyltransferase−like 1
EC_2.3 acyltransferase
SE0
SE5
q
0e+00
2e+07
4e+07
6e+07
8e+07
WD.1.SE
WW.1.SE
WD.2.SE
WW.2.SE
WD.3.SE
WW.3.SE
WD.4.SE
WW.4.SE
WD.5.SE
WW.5.SE
WD.1.SE+5d
WW.1.SE+5d
WD.2.SE+5d
WW.2.SE+5d
WD.3.SE+5d
WW.3.SE+5d
WD.4.SE+5d
WW.4.SE+5d
WD.5.SE+5d
WW.5.SE+5d

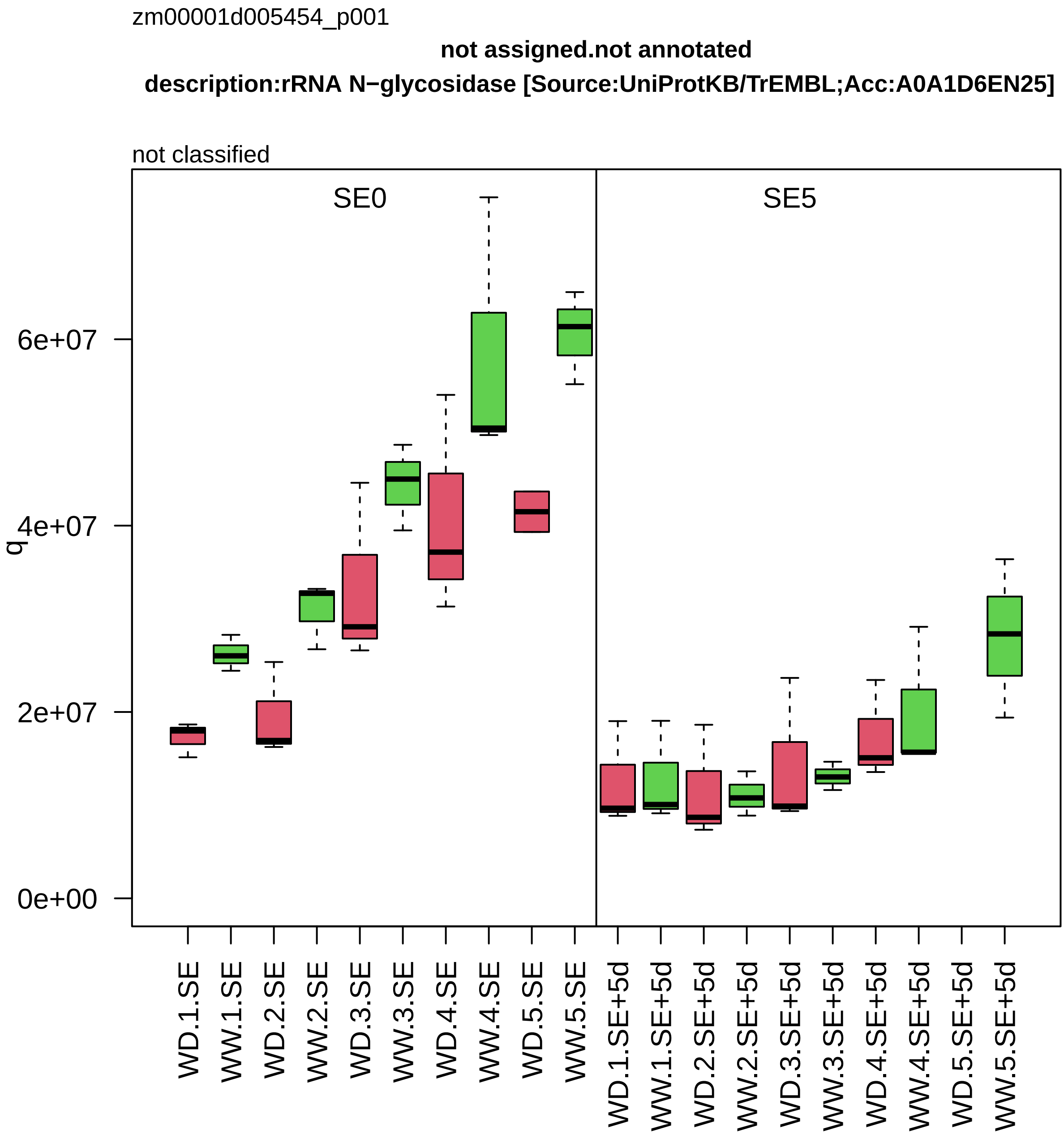
zm00001d005454_p001
not assigned.not annotated
description:rRNA N−glycosidase [Source:UniProtKB/TrEMBL;Acc:A0A1D6EN25]
not classified
SE0
SE5
q
0e+00
2e+07
4e+07
6e+07
WD.1.SE
WW.1.SE
WD.2.SE
WW.2.SE
WD.3.SE
WW.3.SE
WD.4.SE
WW.4.SE
WD.5.SE
WW.5.SE
WD.1.SE+5d
WW.1.SE+5d
WD.2.SE+5d
WW.2.SE+5d
WD.3.SE+5d
WW.3.SE+5d
WD.4.SE+5d
WW.4.SE+5d
WD.5.SE+5d
WW.5.SE+5d

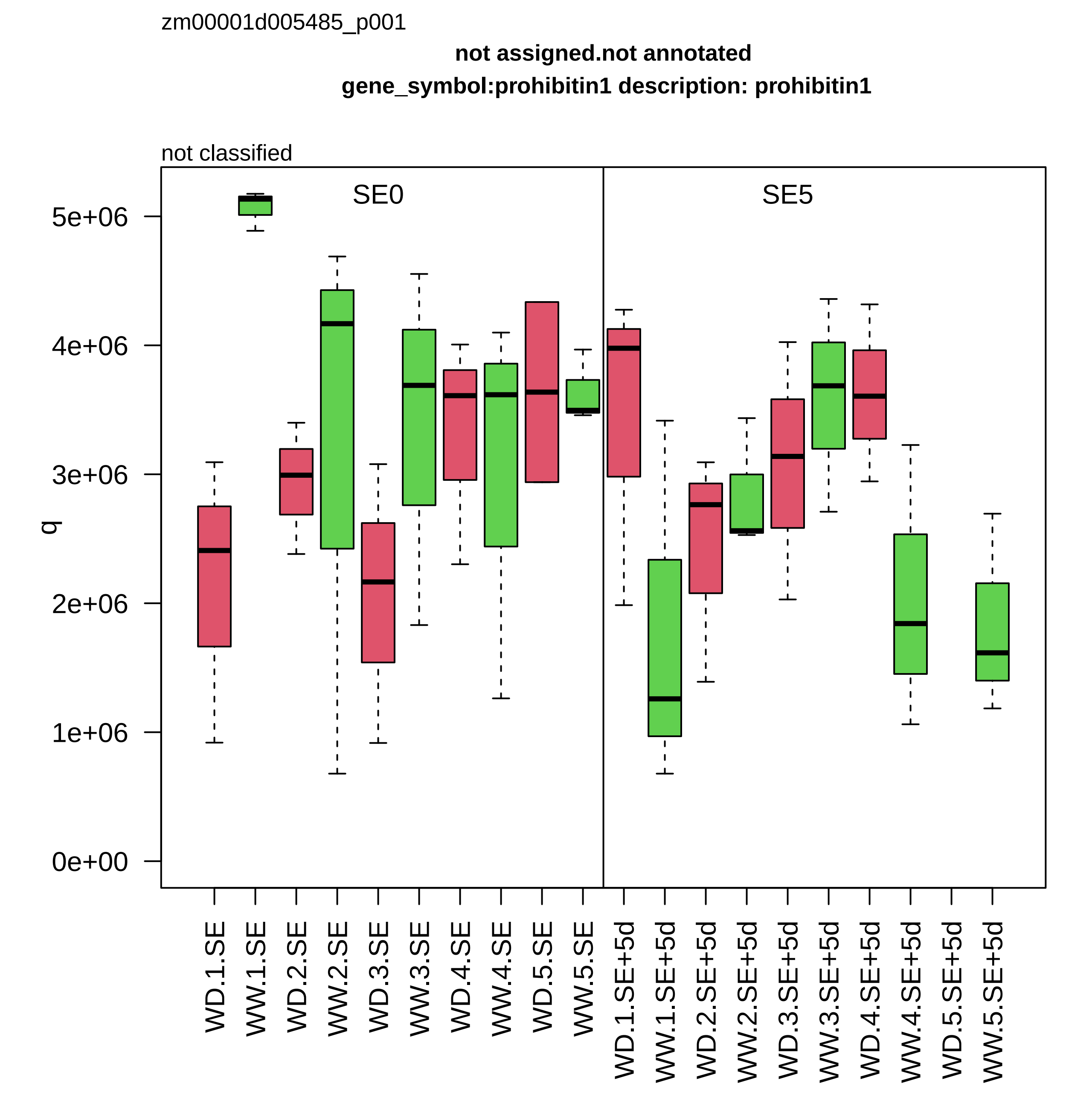

zm00001d005485_p001
not assigned.not annotated
gene_symbol:prohibitin1 description: prohibitin1
not classified
SE0
SE5
q
5e+06
4e+06
3e+06
2e+06
1e+06
0e+00
WD.1.SE
WW.1.SE
WD.2.SE
WW.2.SE
WD.3.SE
WW.3.SE
WD.4.SE
WW.4.SE
WD.5.SE
WW.5.SE
WD.1.SE+5d
WW.1.SE+5d
WD.2.SE+5d
WW.2.SE+5d
WD.3.SE+5d
WW.3.SE+5d
WD.4.SE+5d
WW.4.SE+5d
WD.5.SE+5d
WW.5.SE+5d

zm00001d005546_p002

**Carbohydrate metabolism.starch metabolism**

**ol:ADP glucose pyrophosphorylase small subunit embryo 1 description: ADP glucose pyrophosphorylase**

small subunit *(APS) of ADP−glucose pyrophosphorylase

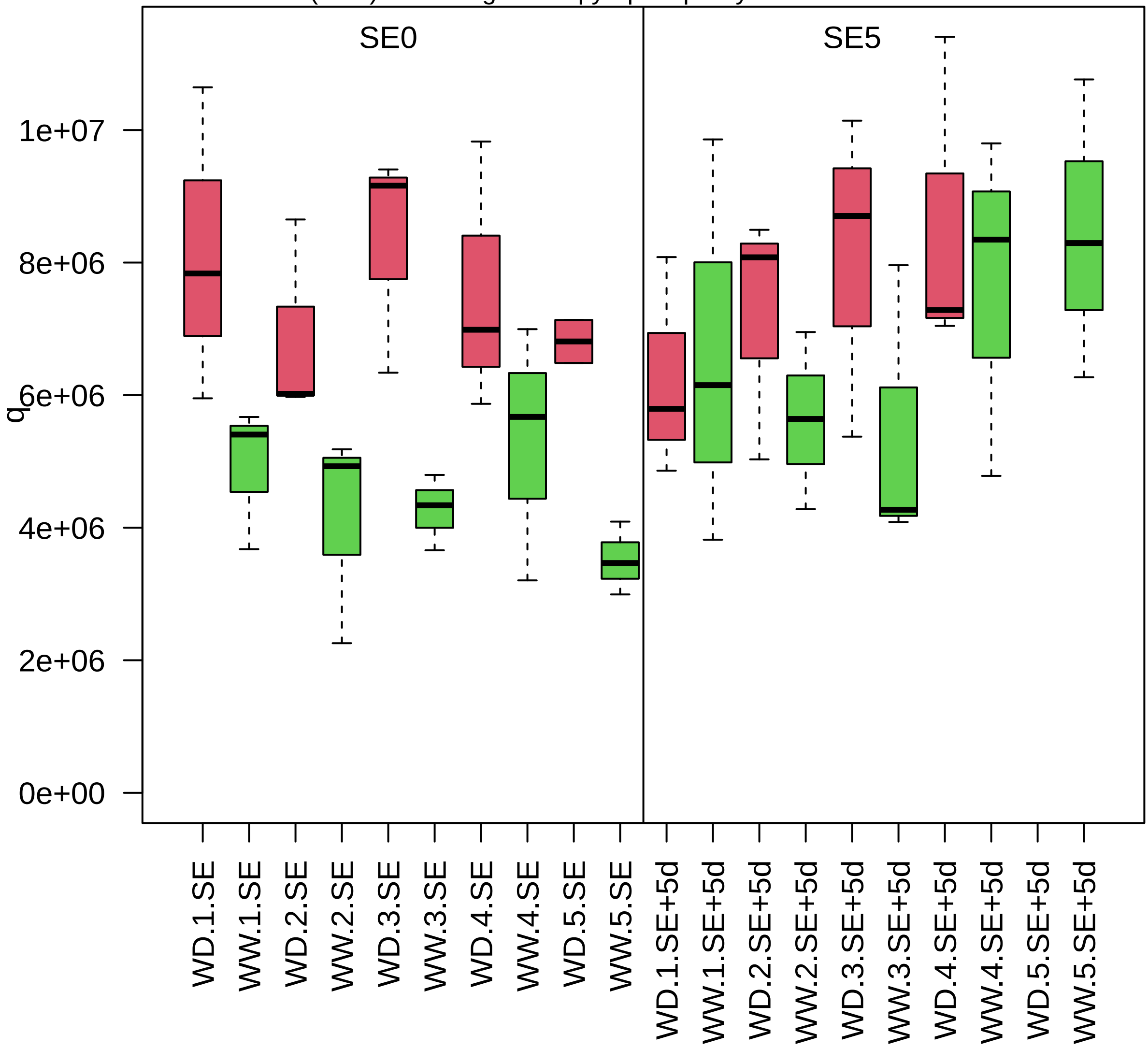

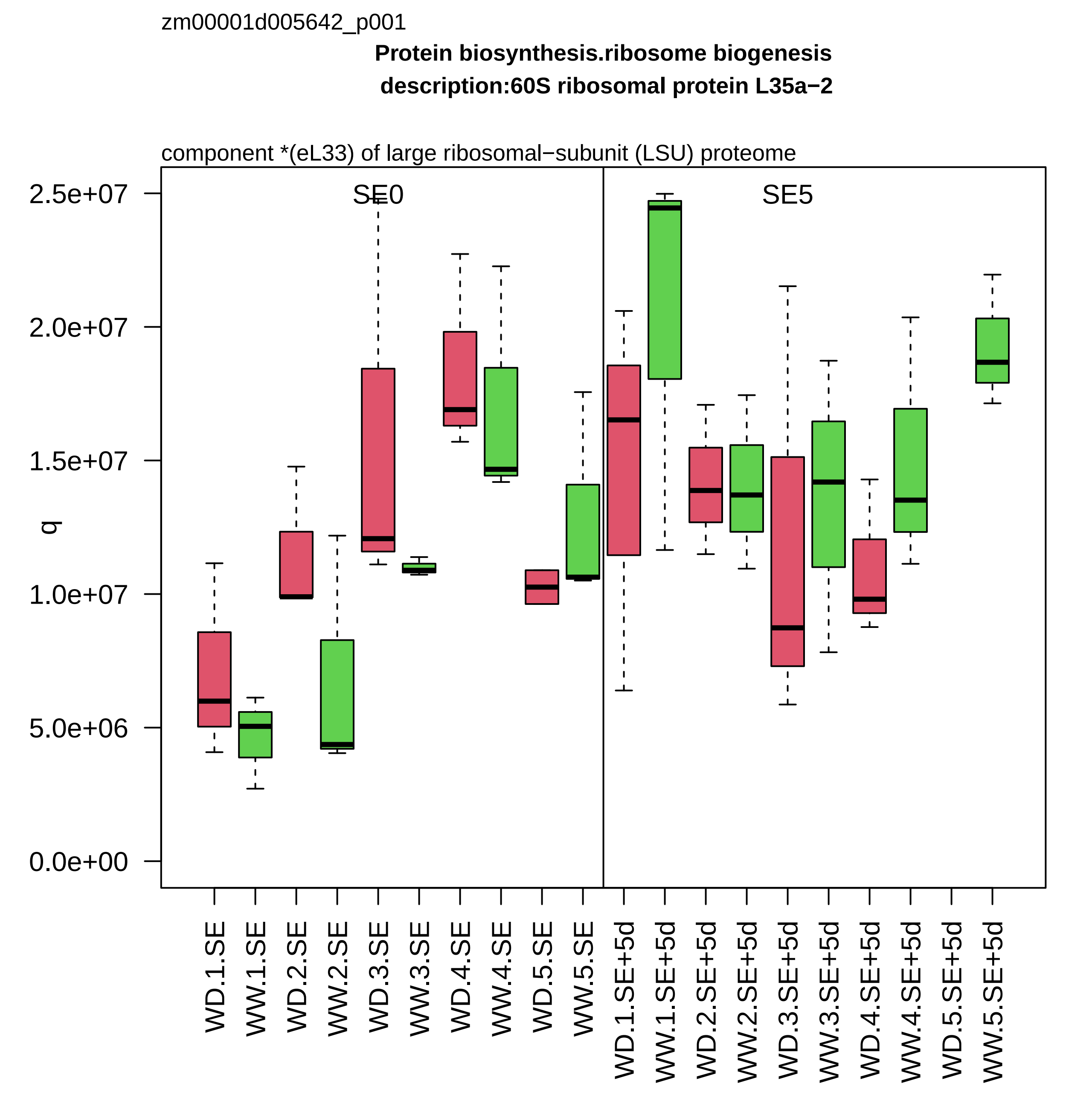

zm00001d005642_p001
Protein biosynthesis.ribosome biogenesis
description:60S ribosomal protein L35a−2
component *(eL33) of large ribosomal−subunit (LSU) proteome
SE0
SE5
q
2.5e+07
2.0e+07
1.5e+07
1.0e+07
5.0e+06
0.0e+00
WD.1.SE
WW.1.SE
WD.2.SE
WW.2.SE
WD.3.SE
WW.3.SE
WD.4.SE
WW.4.SE
WD.5.SE
WW.5.SE
WD.1.SE+5d
WW.1.SE+5d
WD.2.SE+5d
WW.2.SE+5d
WD.3.SE+5d
WW.3.SE+5d
WD.4.SE+5d
WW.4.SE+5d
WD.5.SE+5d
WW.5.SE+5d

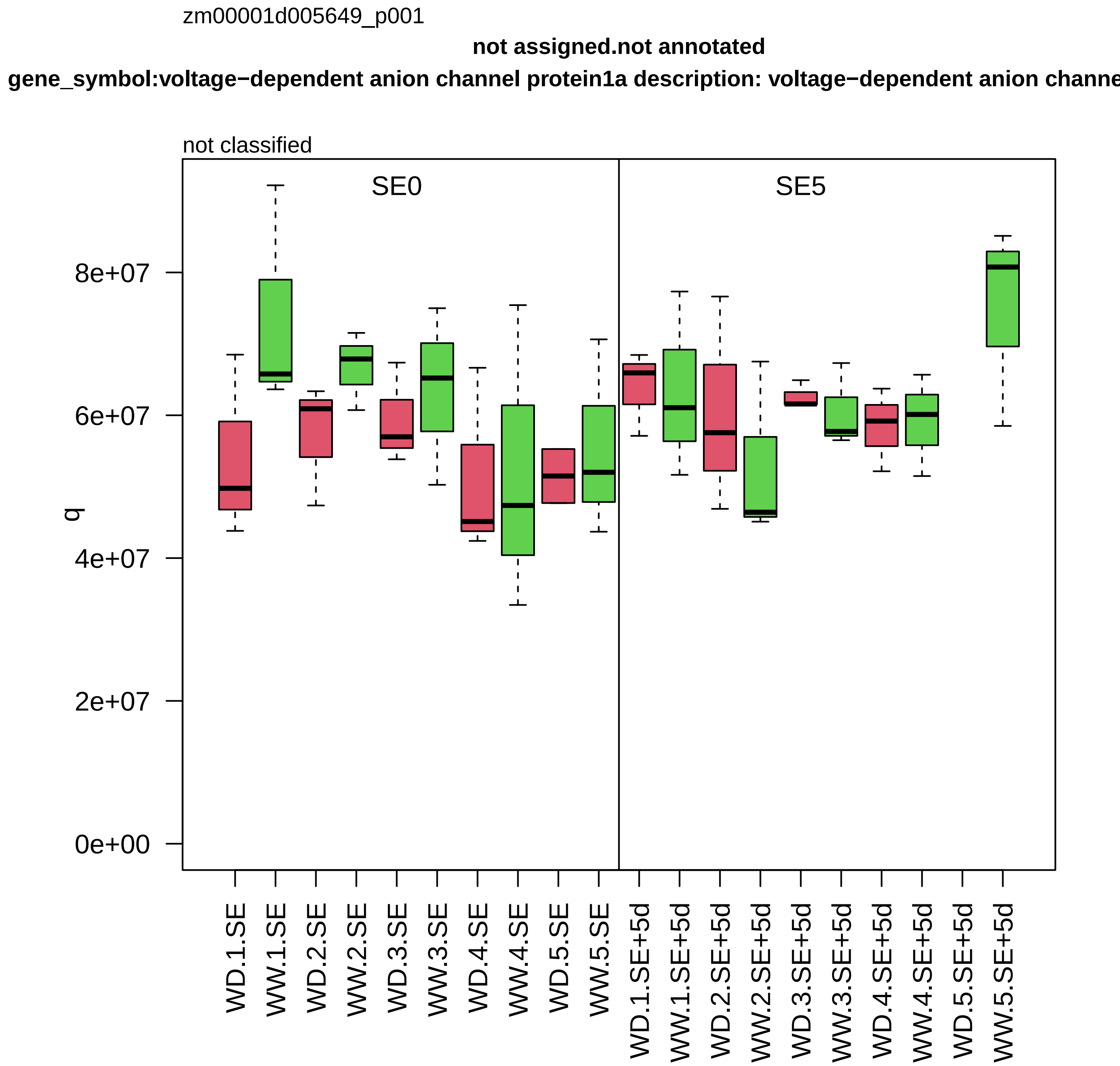

zm00001d005649_p001
not assigned.not annotated
gene_symbol:voltage−dependent anion channel protein1a description: voltage−dependent anion channe
not classified
SE0
SE5
q
8e+07
6e+07
4e+07
2e+07
0e+00
WD.1.SE
WW.1.SE
WD.2.SE
WW.2.SE
WD.3.SE
WW.3.SE
WD.4.SE
WW.4.SE
WD.5.SE
WW.5.SE
WD.1.SE+5d
WW.1.SE+5d
WD.2.SE+5d
WW.2.SE+5d
WD.3.SE+5d
WW.3.SE+5d
WD.4.SE+5d
WW.4.SE+5d
WD.5.SE+5d
WW.5.SE+5d

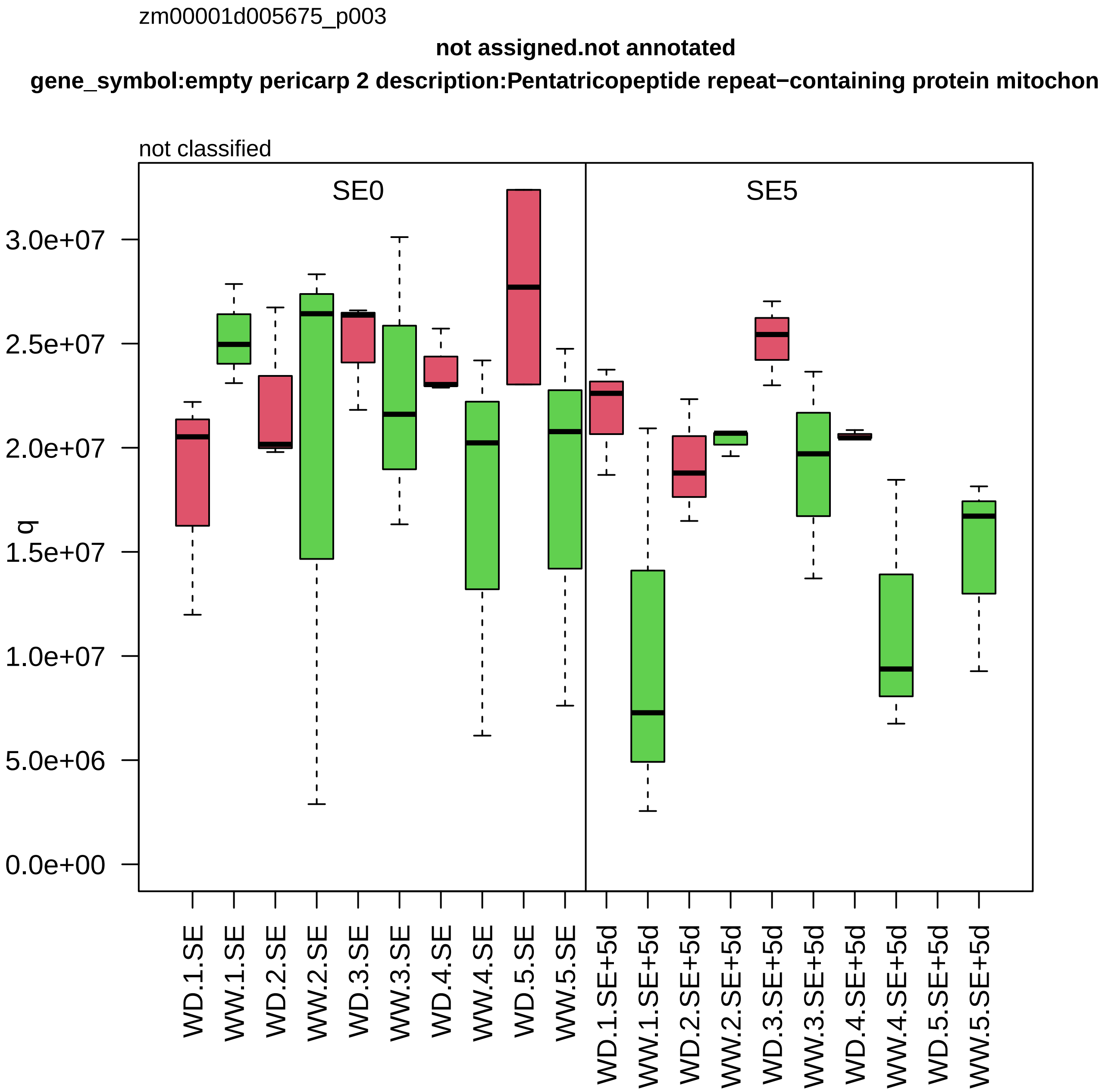

zm00001d005675_p003
not assigned.not annotated
gene_symbol:empty pericarp 2 description:Pentatricopeptide repeat−containing protein mitochon
not classified
SE0
SE5
q
3.0e+07
2.5e+07
2.0e+07
1.5e+07
1.0e+07
5.0e+06
0.0e+00
WD.1.SE
WW.1.SE
WD.2.SE
WW.2.SE
WD.3.SE
WW.3.SE
WD.4.SE
WW.4.SE
WD.5.SE
WW.5.SE
WD.1.SE+5d
WW.1.SE+5d
WD.2.SE+5d
WW.2.SE+5d
WD.3.SE+5d
WW.3.SE+5d
WD.4.SE+5d
WW.4.SE+5d
WD.5.SE+5d
WW.5.SE+5d

zm00001d005818_p001

**Amino acid metabolism.amino acid degradation**

**description:Aldehyde dehydrogenase family 7 member B4**

piperideine−6−carboxylate dehydrogenase

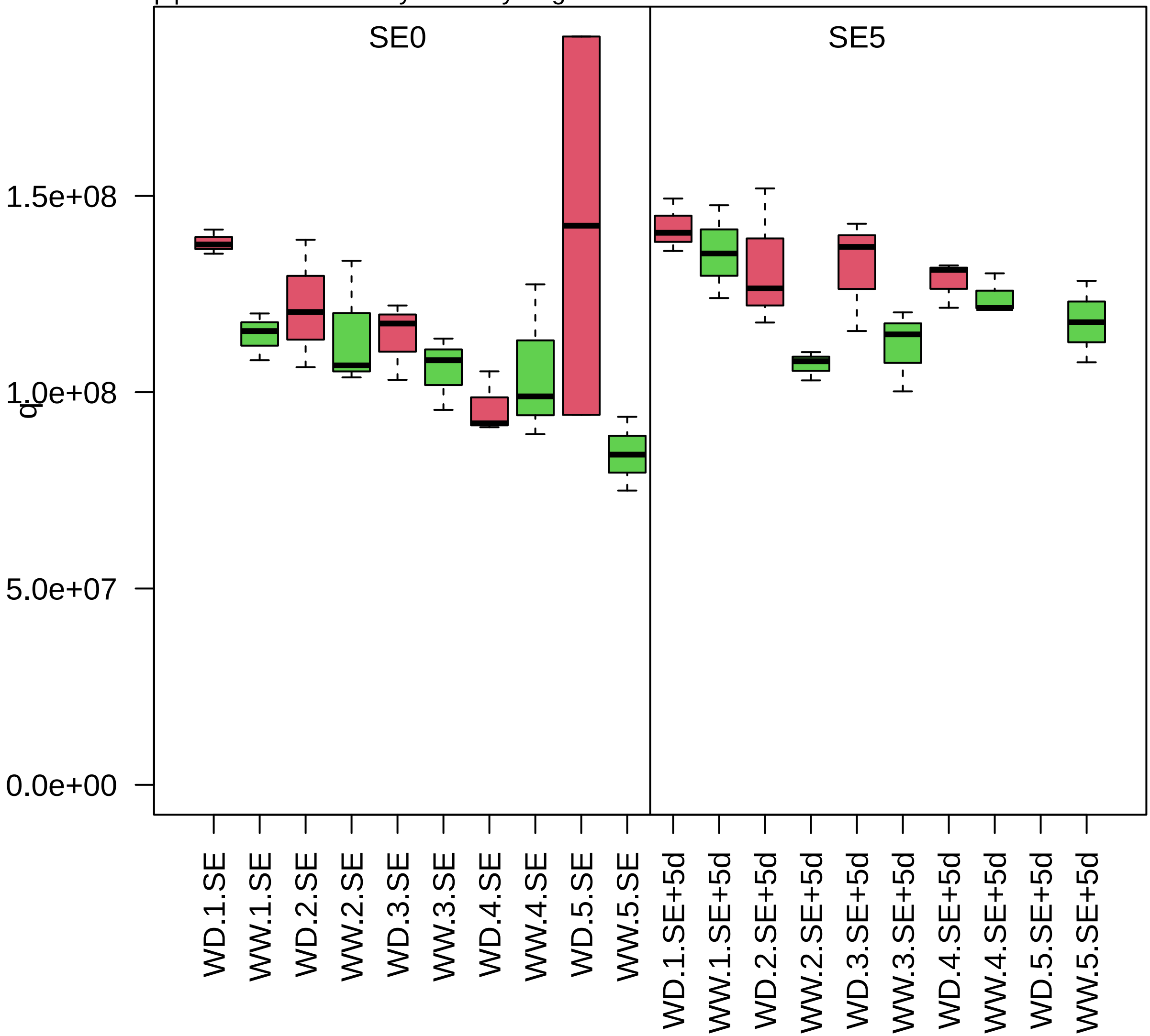

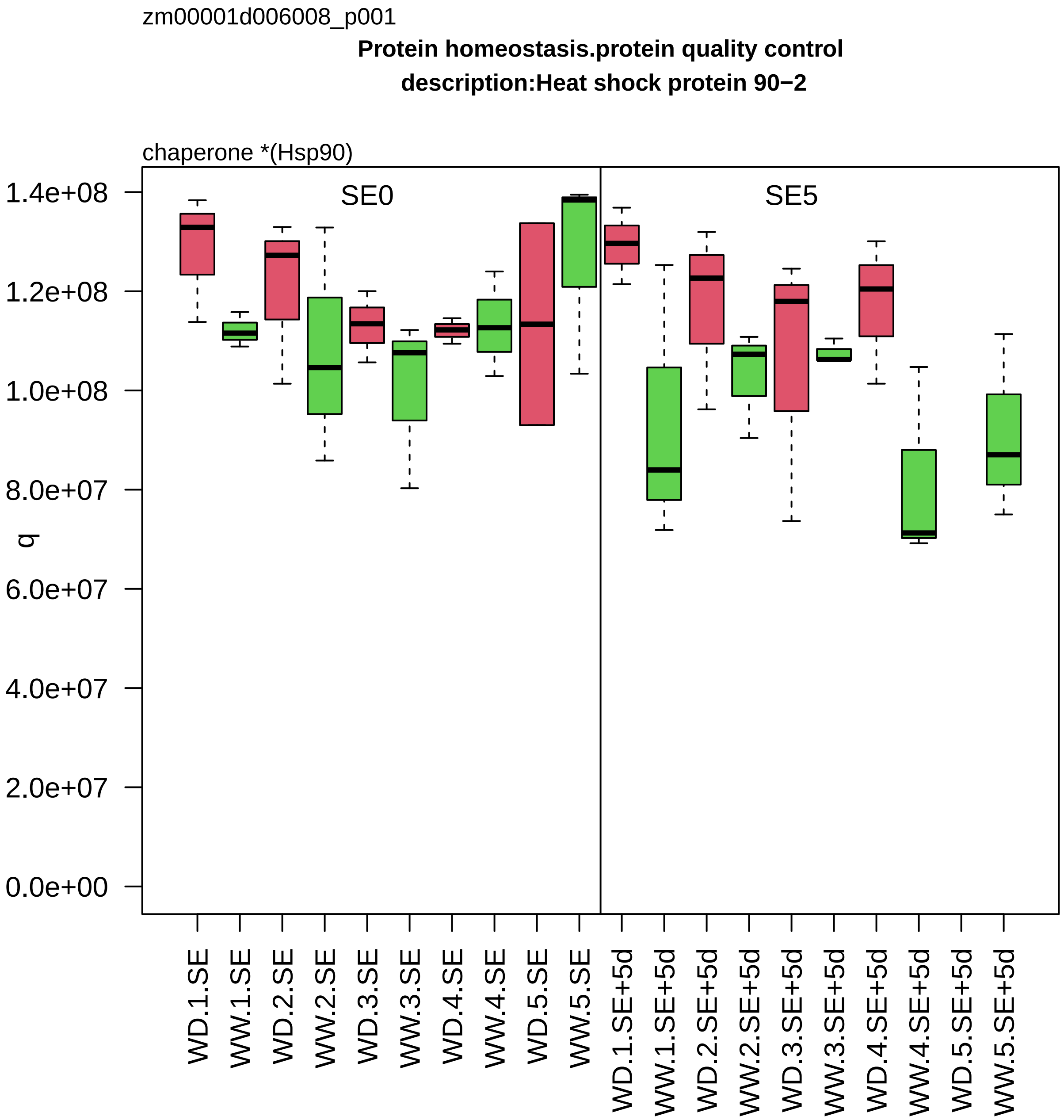

zm00001d006008_p001
Protein homeostasis.protein quality control
description:Heat shock protein 90−2
chaperone *(Hsp90)
SE0
SE5
q
1.4e+08
1.2e+08
1.0e+08
8.0e+07
6.0e+07
4.0e+07
2.0e+07
0.0e+00
WD.1.SE
WW.1.SE
WD.2.SE
WW.2.SE
WD.3.SE
WW.3.SE
WD.4.SE
WW.4.SE
WD.5.SE
WW.5.SE
WD.1.SE+5d
WW.1.SE+5d
WD.2.SE+5d
WW.2.SE+5d
WD.3.SE+5d
WW.3.SE+5d
WD.4.SE+5d
WW.4.SE+5d
WD.5.SE+5d
WW.5.SE+5d

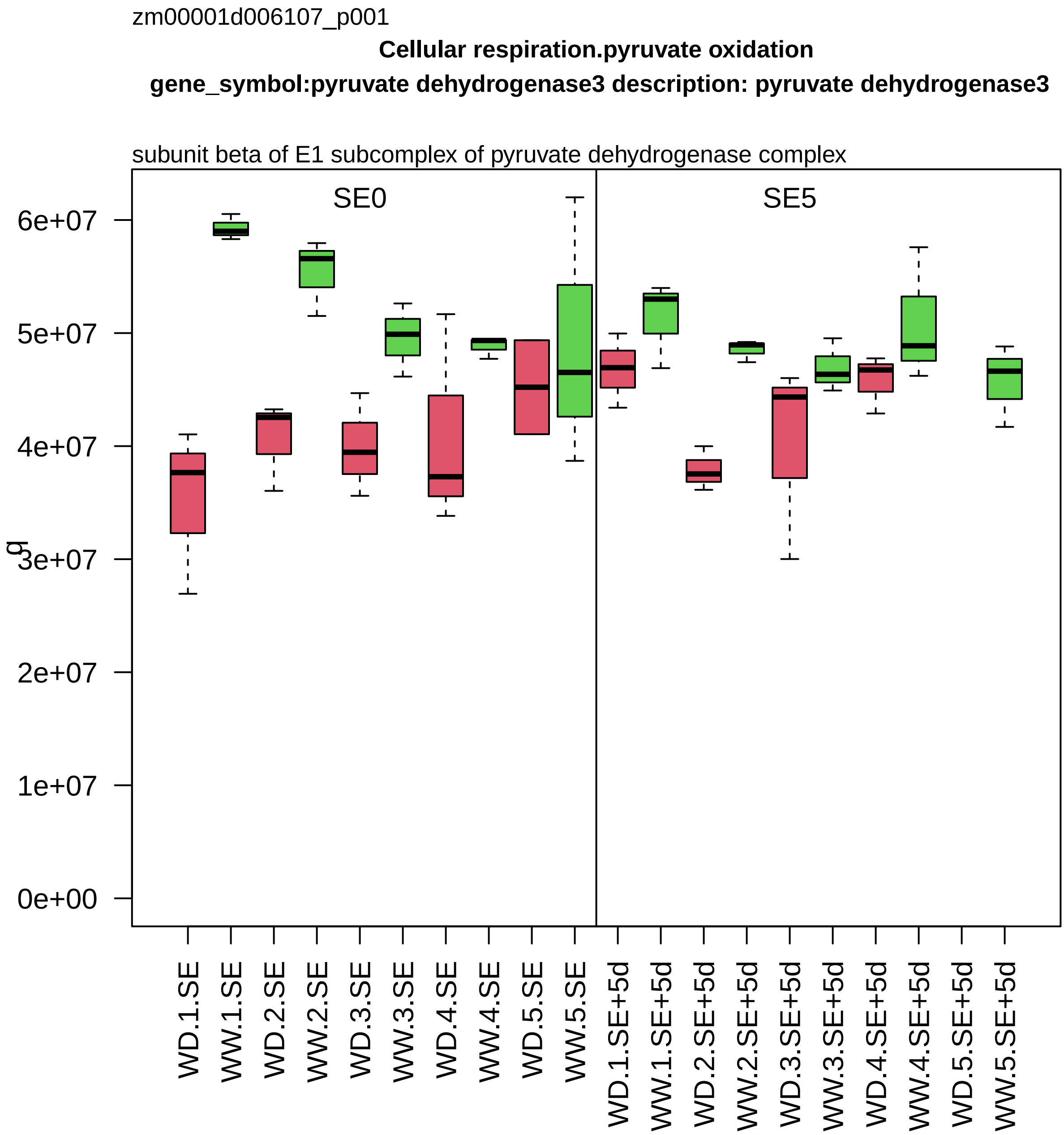
zm00001d006107_p001
Cellular respiration.pyruvate oxidation
gene_symbol:pyruvate dehydrogenase3 description: pyruvate dehydrogenase3
subunit beta of E1 subcomplex of pyruvate dehydrogenase complex
SE0
SE5
g
6e+07
5e+07
4e+07
3e+07
2e+07
1e+07
0e+00
WD.1.SE
WW.1.SE
WD.2.SE
WW.2.SE
WD.3.SE
WW.3.SE
WD.4.SE
WW.4.SE
WD.5.SE
WW.5.SE
WD.1.SE+5d
WW.1.SE+5d
WD.2.SE+5d
WW.2.SE+5d
WD.3.SE+5d
WW.3.SE+5d
WD.4.SE+5d
WW.4.SE+5d
WD.5.SE+5d
WW.5.SE+5d

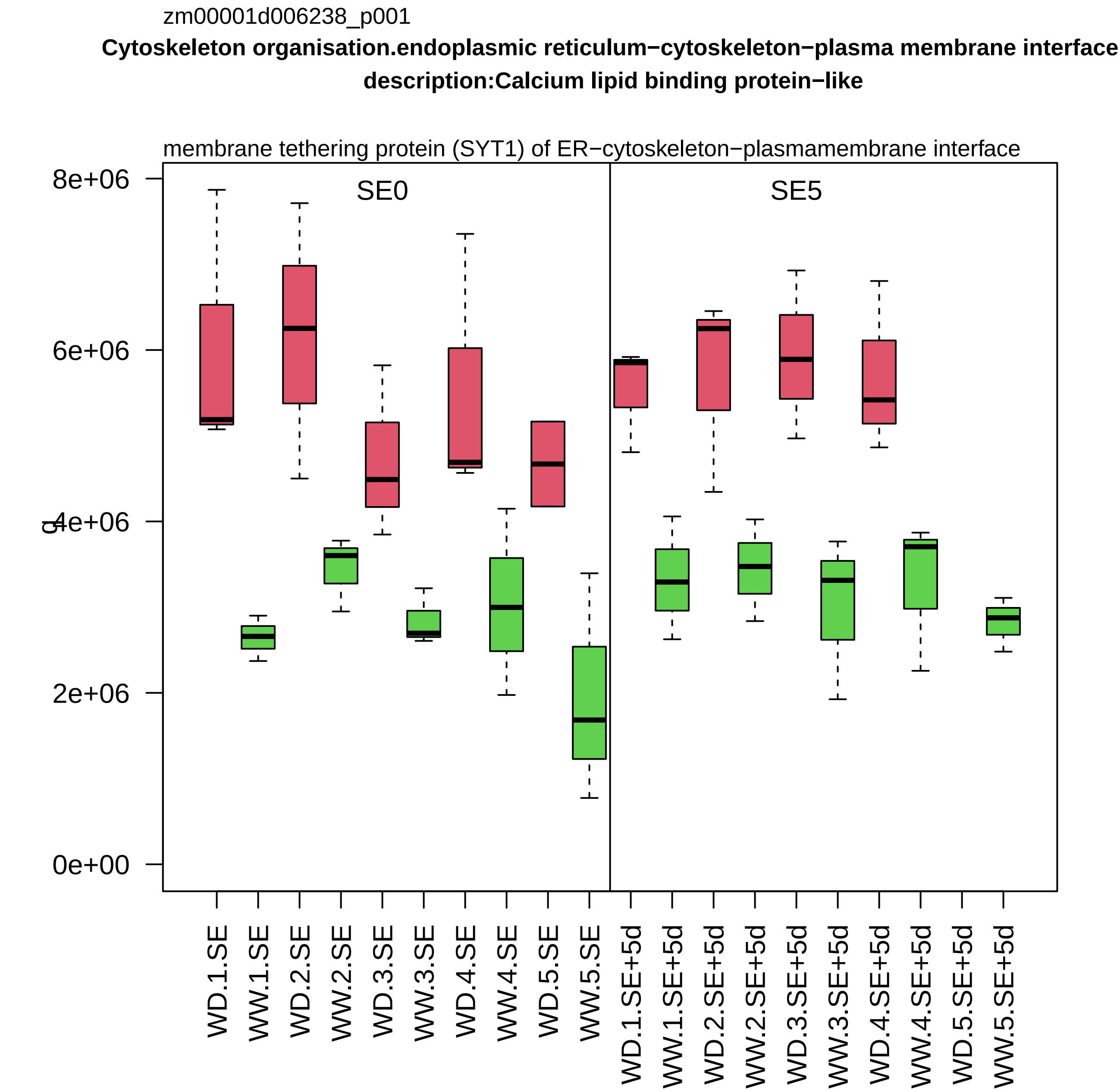

zm00001d006238_p001
Cytoskeleton organisation.endoplasmic reticulum−cytoskeleton−plasma membrane interface
description:Calcium lipid binding protein−like
membrane tethering protein (SYT1) of ER−cytoskeleton−plasmamembrane interface
SE0
SE5
8e+06
6e+06
4e+06
2e+06
0e+00
WD.1.SE
WW.1.SE
WD.2.SE
WW.2.SE
WD.3.SE
WW.3.SE
WD.4.SE
WW.4.SE
WD.5.SE
WW.5.SE
WD.1.SE+5d
WW.1.SE+5d
WD.2.SE+5d
WW.2.SE+5d
WD.3.SE+5d
WW.3.SE+5d
WD.4.SE+5d
WW.4.SE+5d
WD.5.SE+5d
WW.5.SE+5d

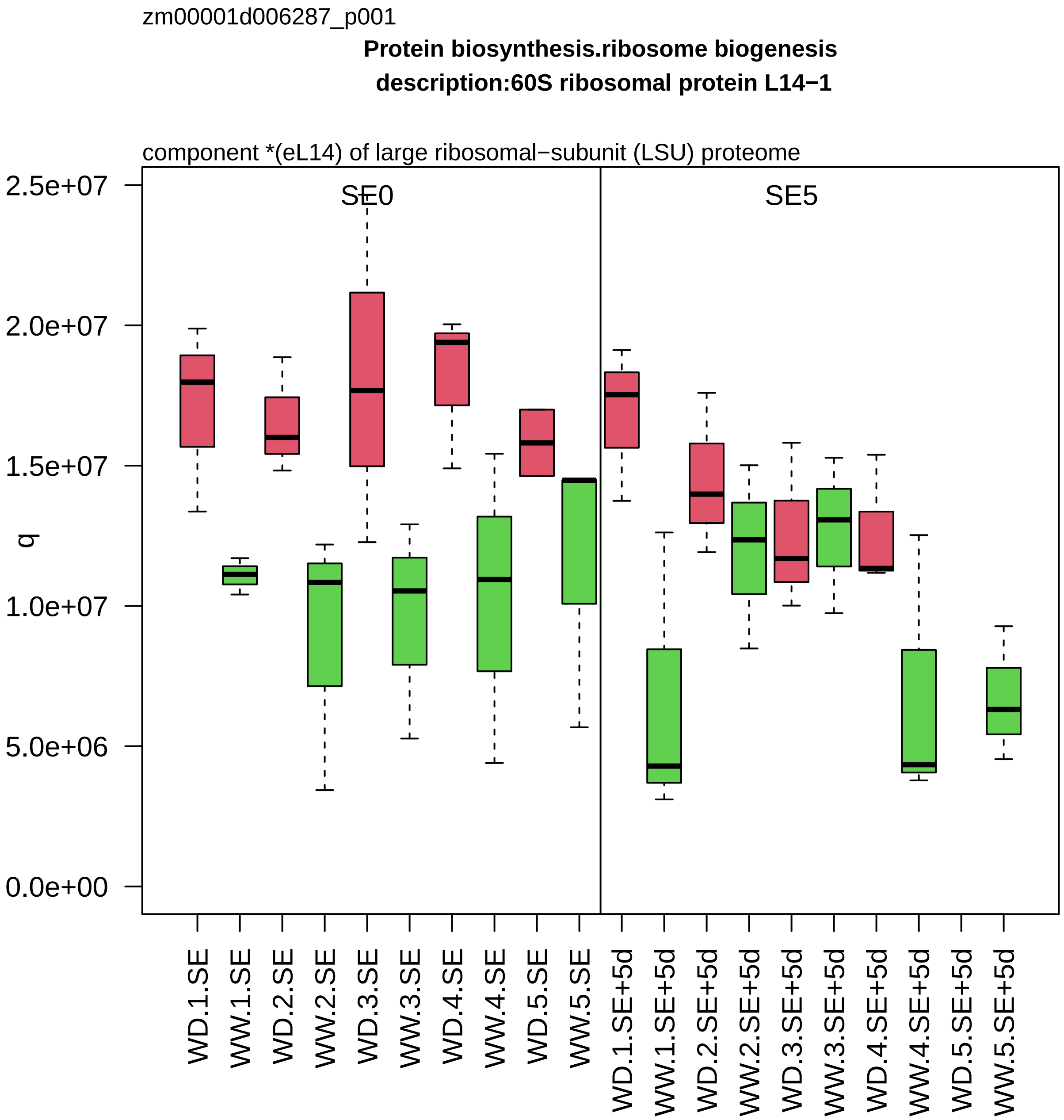
zm00001d006287_p001
Protein biosynthesis.ribosome biogenesis
description:60S ribosomal protein L14−1
component *(eL14) of large ribosomal−subunit (LSU) proteome
SE0
SE5
q
2.5e+07
2.0e+07
1.5e+07
1.0e+07
5.0e+06
0.0e+00
WD.1.SE
WW.1.SE
WD.2.SE
WW.2.SE
WD.3.SE
WW.3.SE
WD.4.SE
WW.4.SE
WD.5.SE
WW.5.SE
WD.1.SE+5d
WW.1.SE+5d
WD.2.SE+5d
WW.2.SE+5d
WD.3.SE+5d
WW.3.SE+5d
WD.4.SE+5d
WW.4.SE+5d
WD.5.SE+5d
WW.5.SE+5d

zm00001d006296_p003

**Protein biosynthesis.aminoacyl−tRNA synthetase activities**
**description:Valine−−tRNA ligase mitochondrial 1**

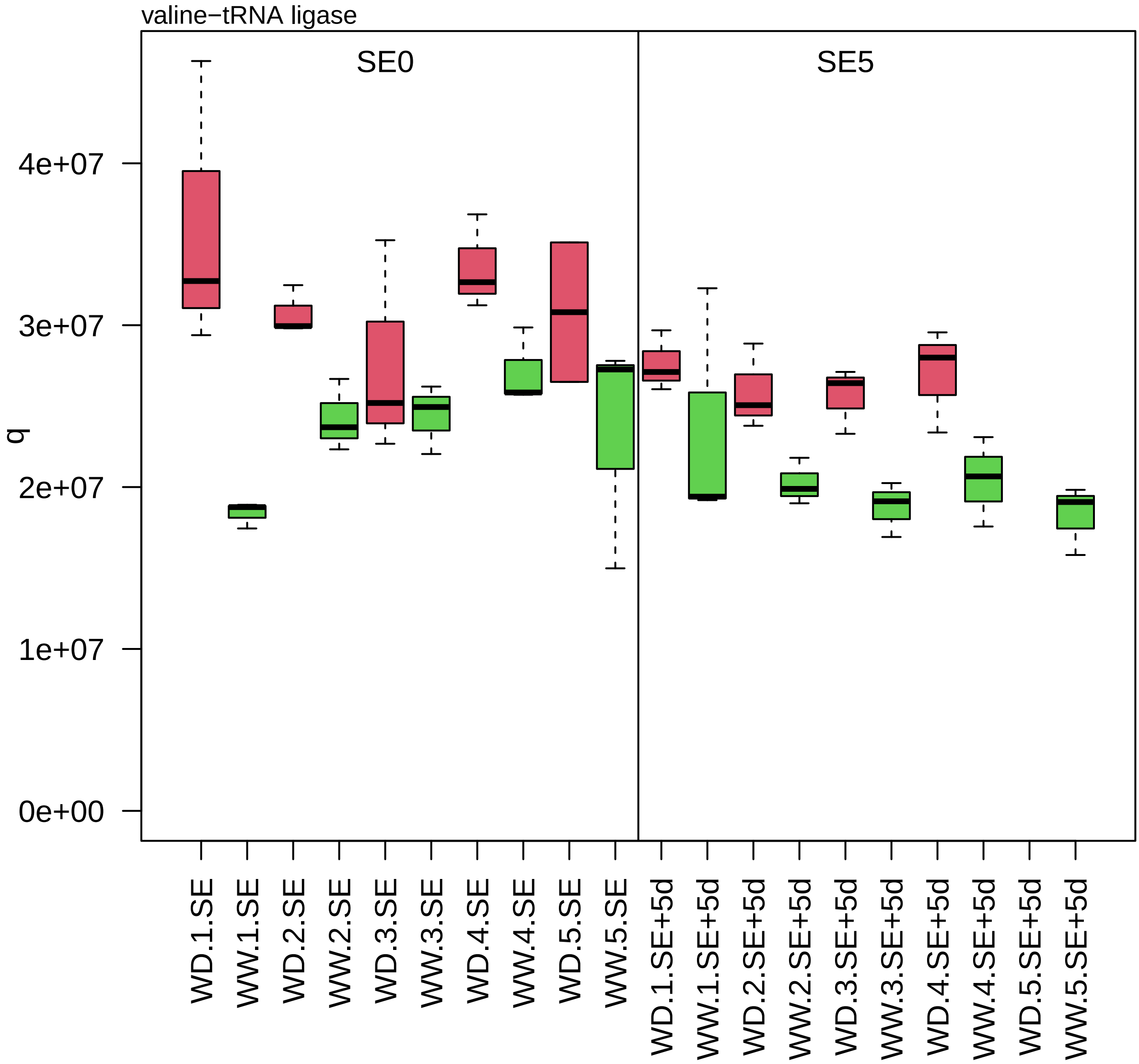

zm00001d006309_p007

**Carbohydrate metabolism.sucrose metabolism**

**gene_symbol:UDP−glucose pyrophosphorylase1 description:UTP−−glucose−1−phosphate uridylyltran**

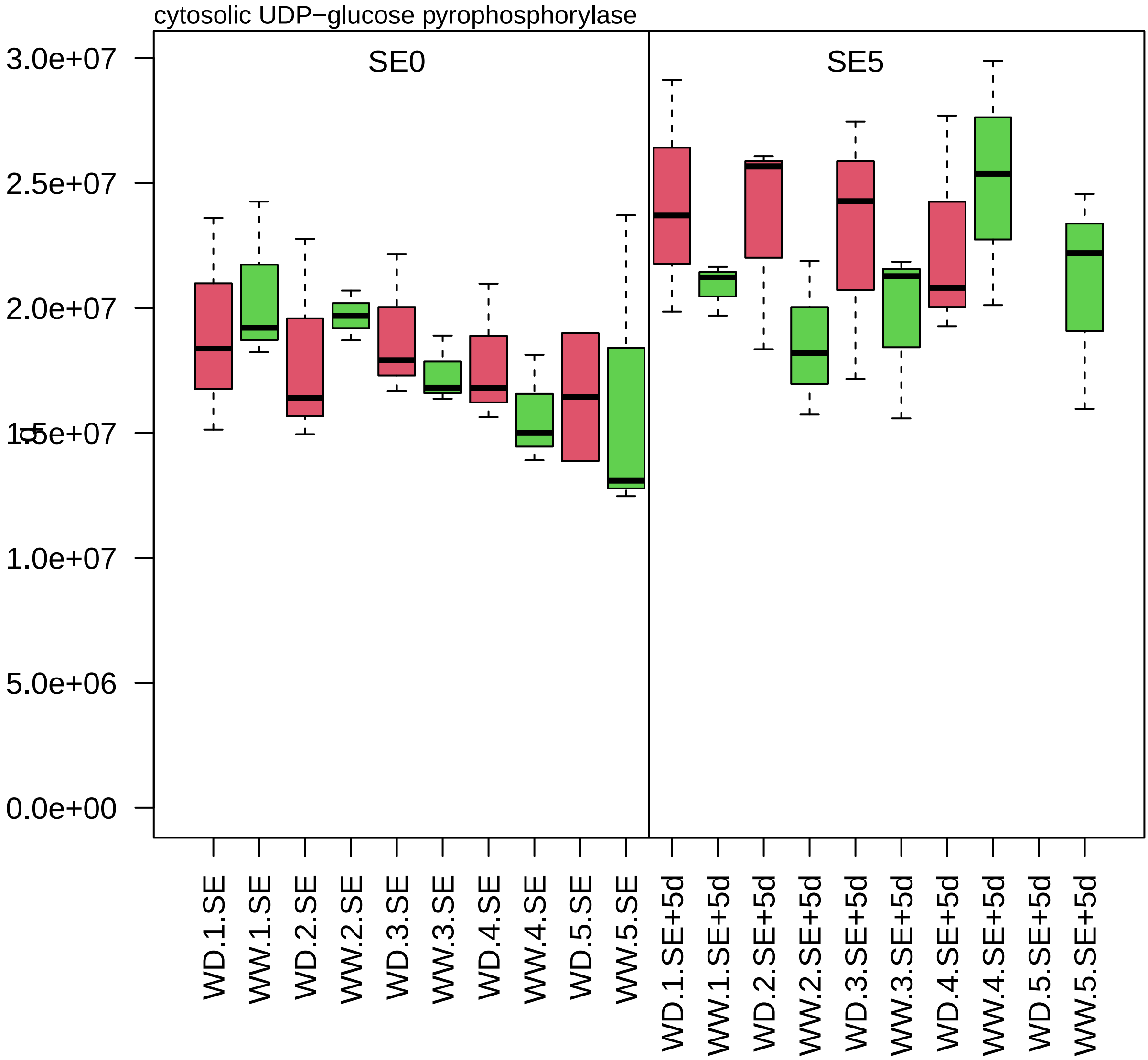

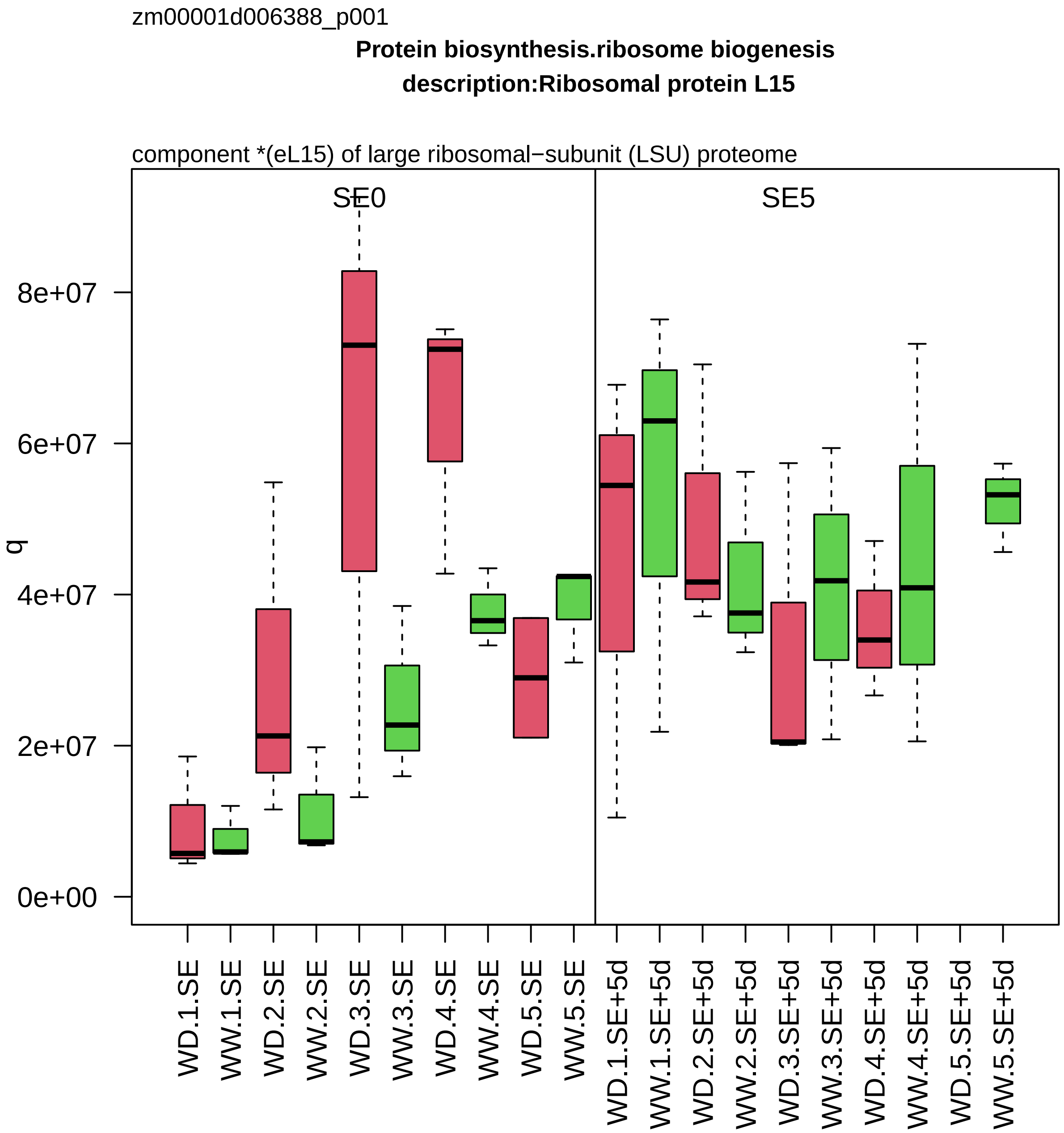

zm00001d006388_p001
Protein biosynthesis.ribosome biogenesis
description:Ribosomal protein L15
component *(eL15) of large ribosomal−subunit (LSU) proteome
SE0
SE5
q
8e+07
6e+07
4e+07
2e+07
0e+00
WD.1.SE
WW.1.SE
WD.2.SE
WW.2.SE
WD.3.SE
WW.3.SE
WD.4.SE
WW.4.SE
WD.5.SE
WW.5.SE
WD.1.SE+5d
WW.1.SE+5d
WD.2.SE+5d
WW.2.SE+5d
WD.3.SE+5d
WW.3.SE+5d
WD.4.SE+5d
WW.4.SE+5d
WD.5.SE+5d
WW.5.SE+5d

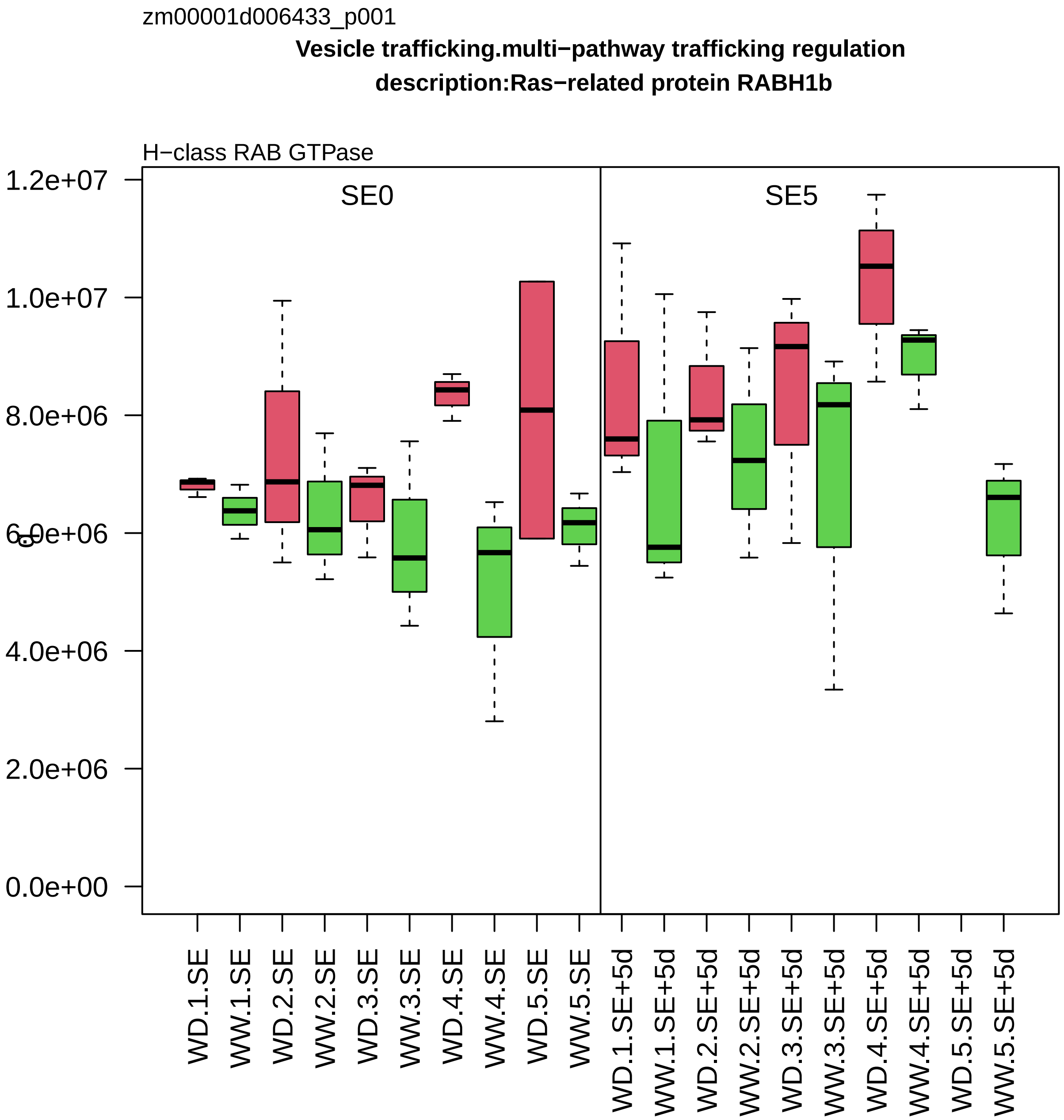

zm00001d006433_p001
Vesicle trafficking.multi−pathway trafficking regulation
description:Ras−related protein RABH1b
H−class RAB GTPase
SE0
SE5
1.2e+07
1.0e+07
8.0e+06
6.0e+06
4.0e+06
2.0e+06
0.0e+00
WD.1.SE
WW.1.SE
WD.2.SE
WW.2.SE
WD.3.SE
WW.3.SE
WD.4.SE
WW.4.SE
WD.5.SE
WW.5.SE
WD.1.SE+5d
WW.1.SE+5d
WD.2.SE+5d
WW.2.SE+5d
WD.3.SE+5d
WW.3.SE+5d
WD.4.SE+5d
WW.4.SE+5d
WD.5.SE+5d
WW.5.SE+5d

zm00001d006439_p001

**Cellular respiration.oxidative phosphorylation**

**gene_symbol:ATP synthase3 description: ATP synthase3**

subunit delta_ of ATP synthase peripheral MF1 subcomplex

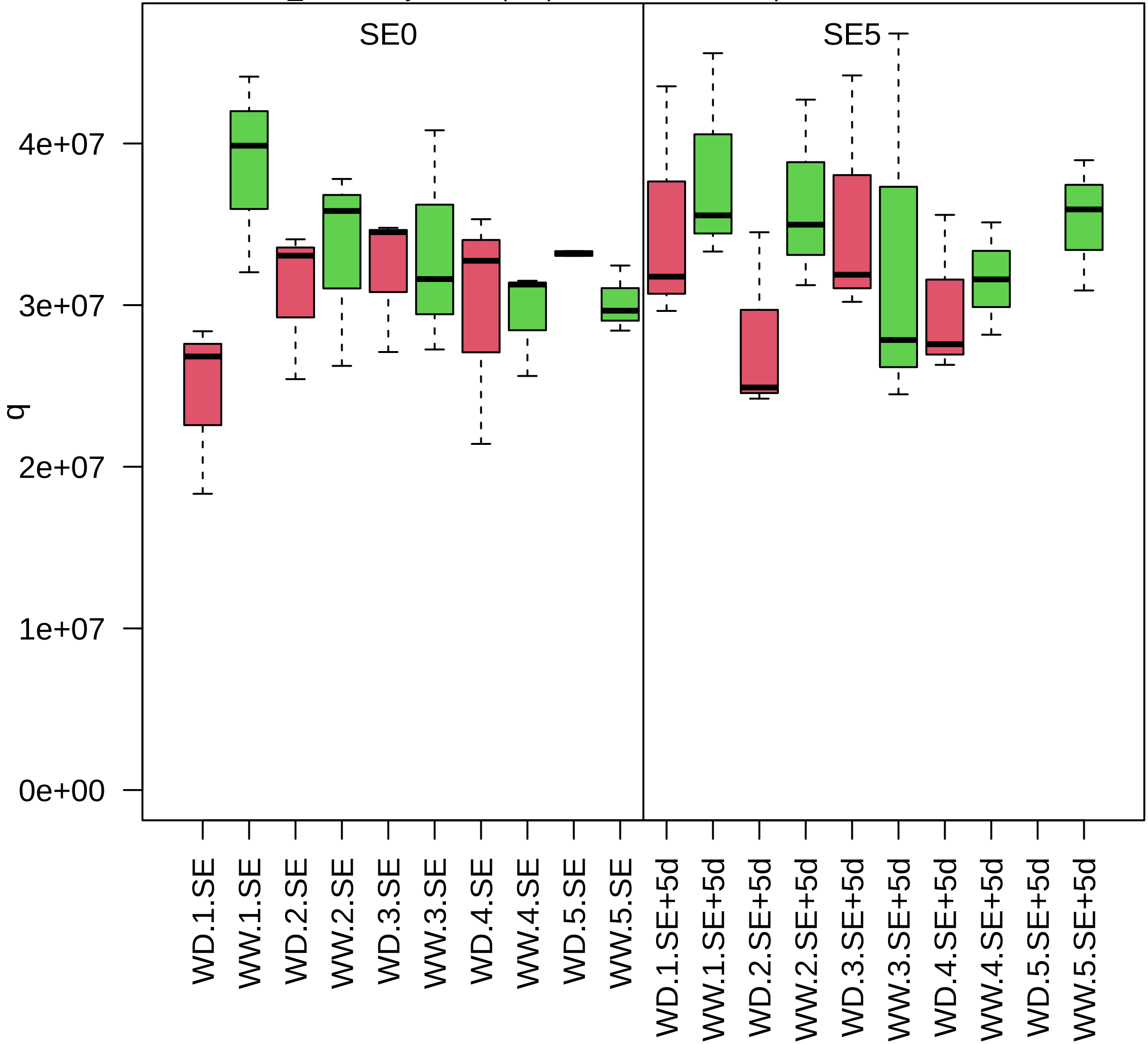

zm00001d006613_p001

**Protein biosynthesis.translation initiation**

**description:eukaryotic translation initiation factor 2 gamma subunit**

component *(eIF2−gamma) of eIF2 Met−tRNA binding factor complex

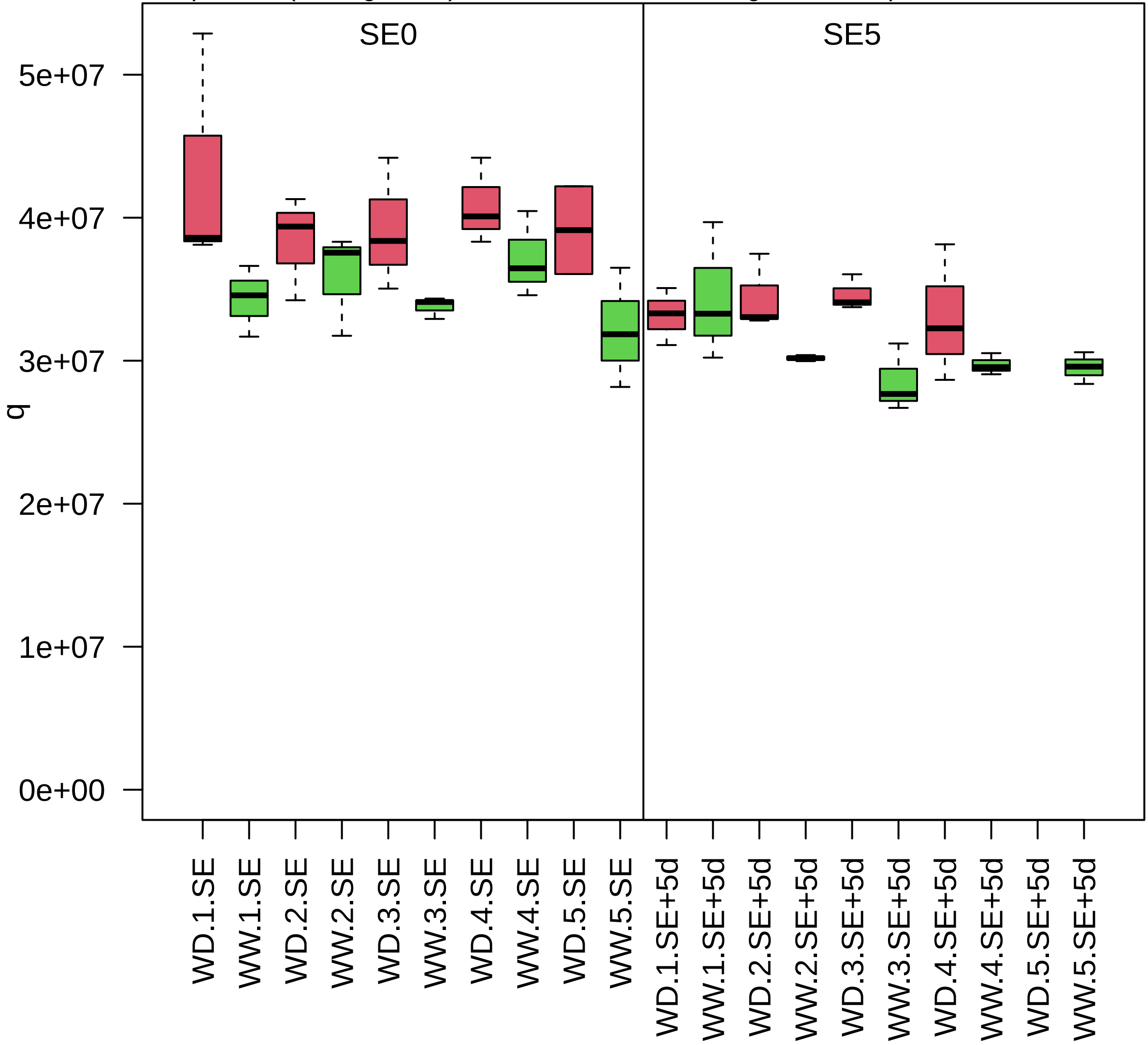

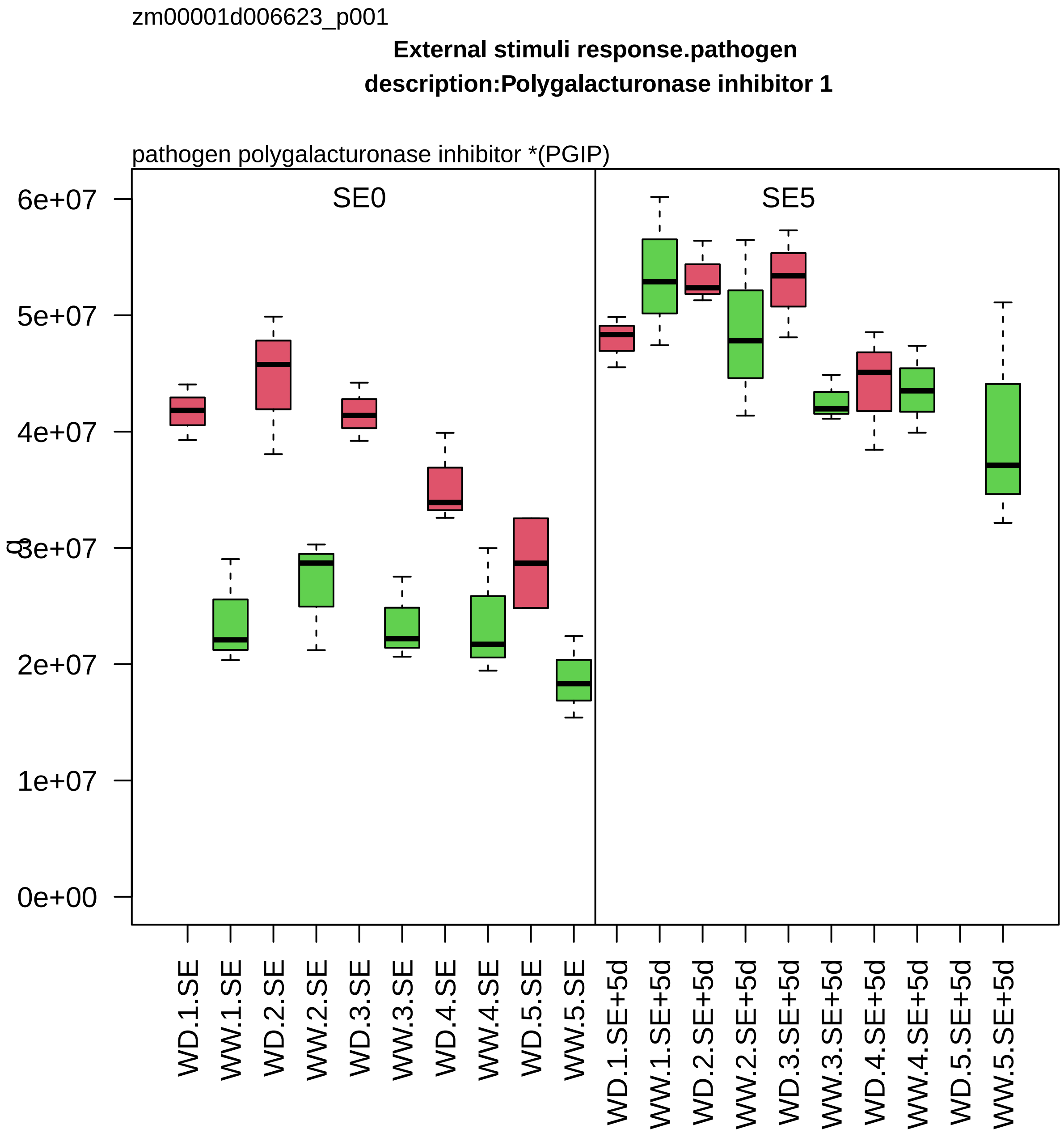
zm00001d006623_p001
External stimuli response.pathogen
description:Polygalacturonase inhibitor 1
pathogen polygalacturonase inhibitor *(PGIP)
SE0
SE5
6e+07
5e+07
4e+07
3e+07
2e+07
1e+07
0e+00
WD.1.SE
WW.1.SE
WD.2.SE
WW.2.SE
WD.3.SE
WW.3.SE
WD.4.SE
WW.4.SE
WD.5.SE
WW.5.SE
WD.1.SE+5d
WW.1.SE+5d
WD.2.SE+5d
WW.2.SE+5d
WD.3.SE+5d
WW.3.SE+5d
WD.4.SE+5d
WW.4.SE+5d
WD.5.SE+5d
WW.5.SE+5d

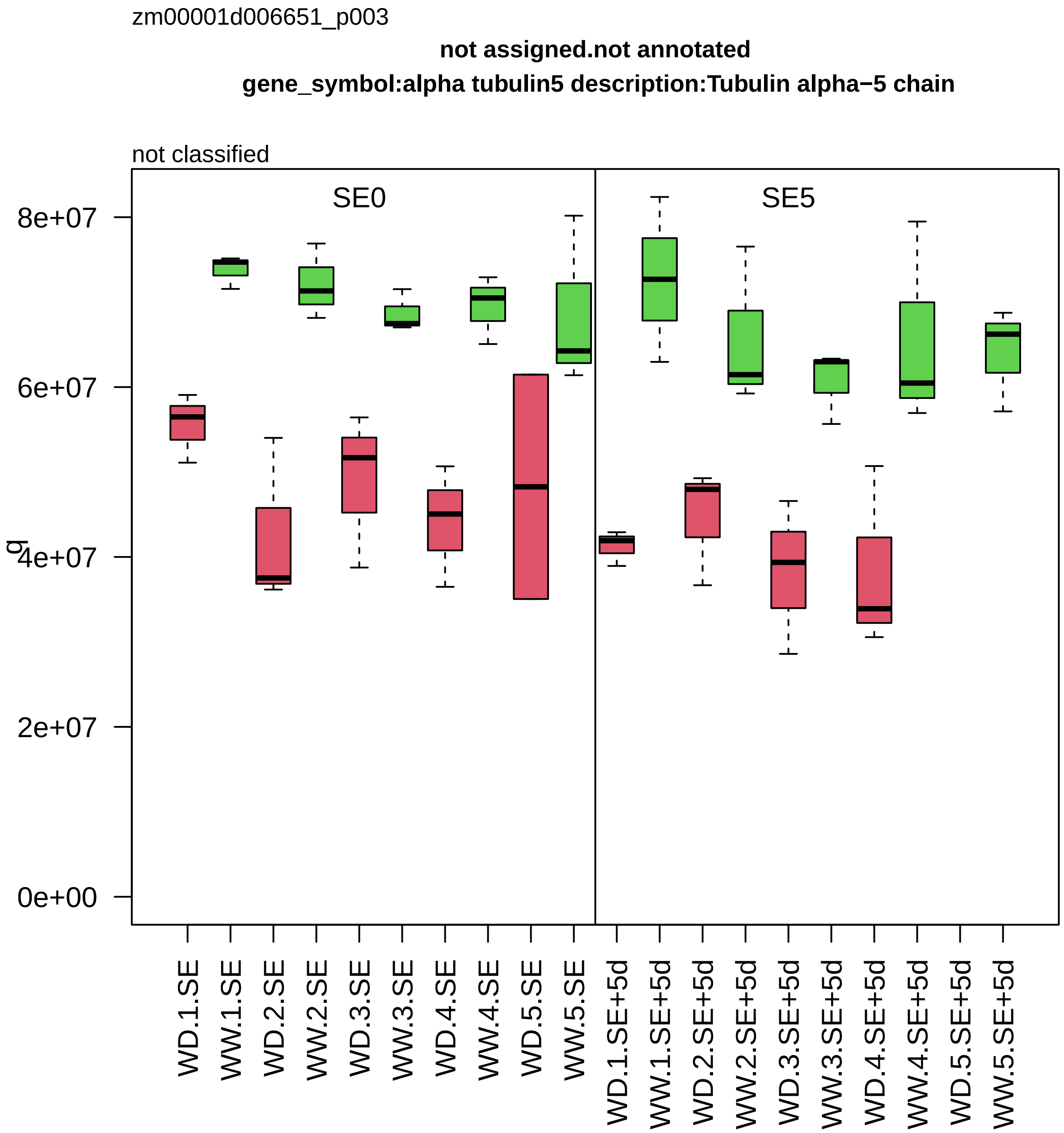

zm00001d006651_p003
not assigned.not annotated
gene_symbol:alpha tubulin5 description:Tubulin alpha−5 chain
not classified
SE0
SE5
q
0e+00
2e+07
4e+07
6e+07
8e+07
WD.1.SE
WW.1.SE
WD.2.SE
WW.2.SE
WD.3.SE
WW.3.SE
WD.4.SE
WW.4.SE
WD.5.SE
WW.5.SE
WD.1.SE+5d
WW.1.SE+5d
WD.2.SE+5d
WW.2.SE+5d
WD.3.SE+5d
WW.3.SE+5d
WD.4.SE+5d
WW.4.SE+5d
WD.5.SE+5d
WW.5.SE+5d

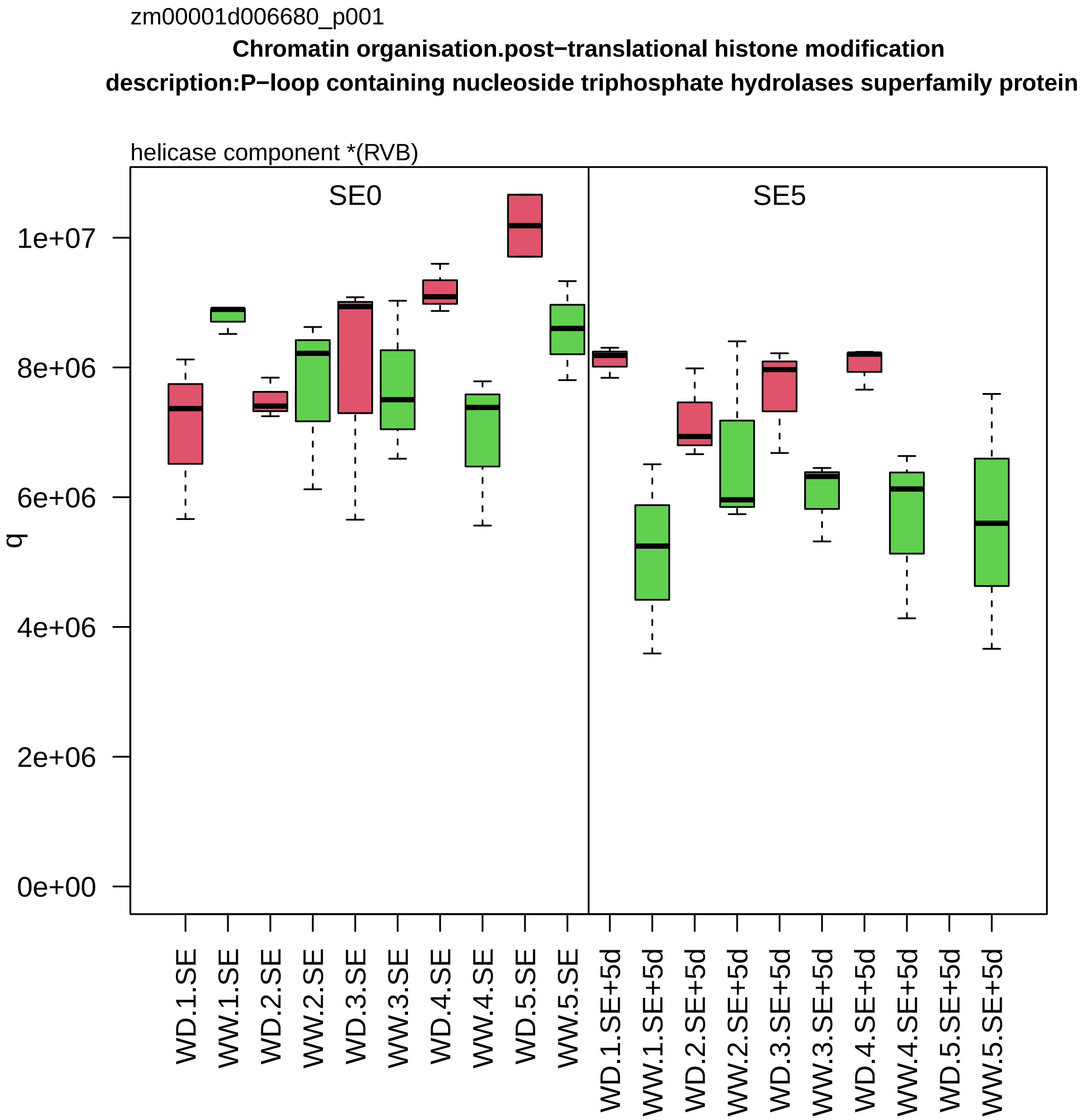

zm00001d006680_p001
Chromatin organisation.post−translational histone modification
description:P−loop containing nucleoside triphosphate hydrolases superfamily protein
helicase component *(RVB)
SE0
SE5
q
1e+07
8e+06
6e+06
4e+06
2e+06
0e+00
WD.1.SE
WW.1.SE
WD.2.SE
WW.2.SE
WD.3.SE
WW.3.SE
WD.4.SE
WW.4.SE
WD.5.SE
WW.5.SE
WD.1.SE+5d
WW.1.SE+5d
WD.2.SE+5d
WW.2.SE+5d
WD.3.SE+5d
WW.3.SE+5d
WD.4.SE+5d
WW.4.SE+5d
WD.5.SE+5d
WW.5.SE+5d

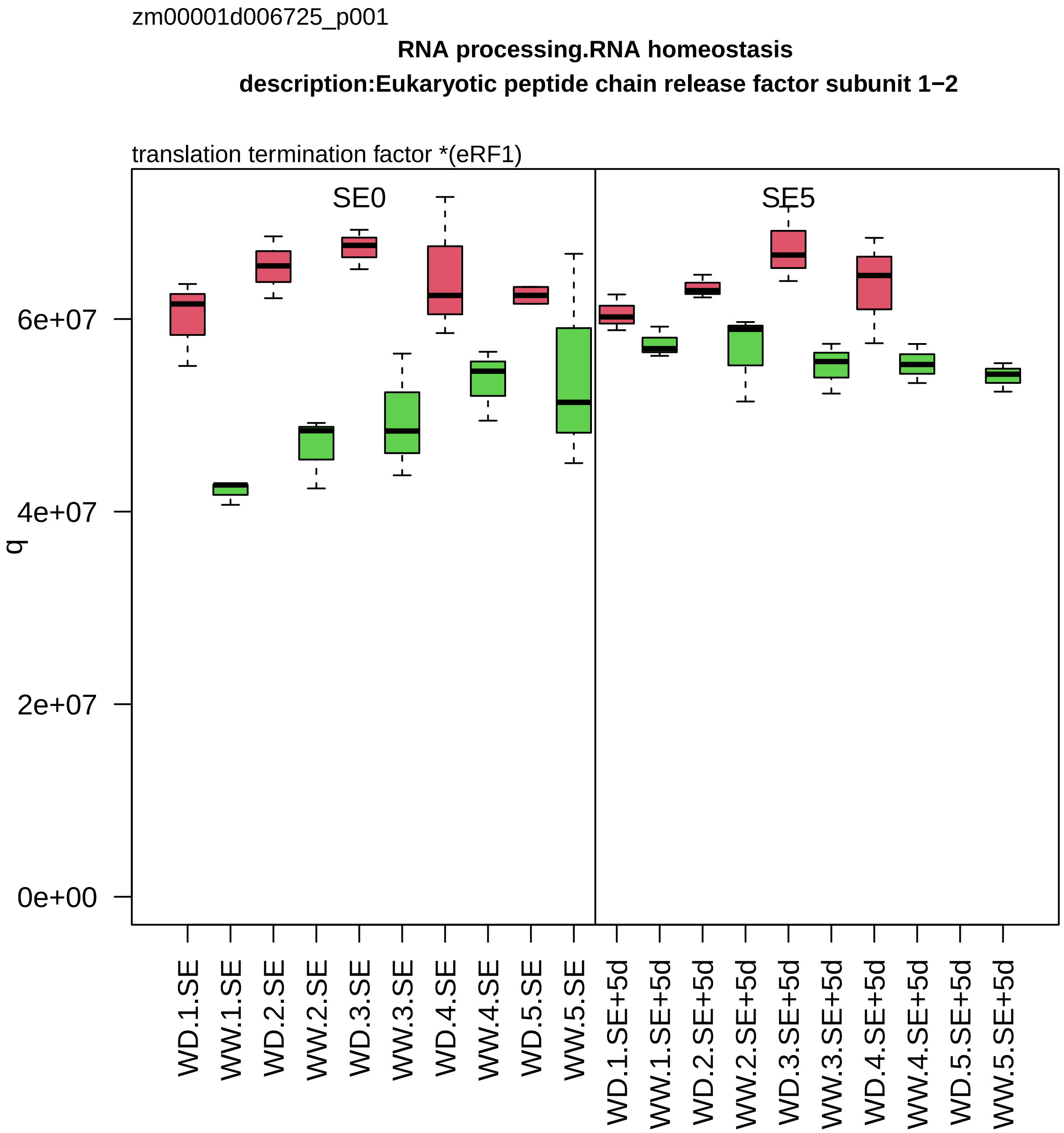

zm00001d006725_p001
RNA processing.RNA homeostasis
description:Eukaryotic peptide chain release factor subunit 1−2
translation termination factor *(eRF1)
SE0
SE5
q
0e+00
2e+07
4e+07
6e+07
WD.1.SE
WW.1.SE
WD.2.SE
WW.2.SE
WD.3.SE
WW.3.SE
WD.4.SE
WW.4.SE
WD.5.SE
WW.5.SE
WD.1.SE+5d
WW.1.SE+5d
WD.2.SE+5d
WW.2.SE+5d
WD.3.SE+5d
WW.3.SE+5d
WD.4.SE+5d
WW.4.SE+5d
WD.5.SE+5d
WW.5.SE+5d

zm00001d006760_p001

**Protein biosynthesis.translation elongation**

**gene_symbol:elongation initiation factor5A description:Eukaryotic translation initiation factor 5**

poly−P/G elongation factor *(eEF5/eIF5A))

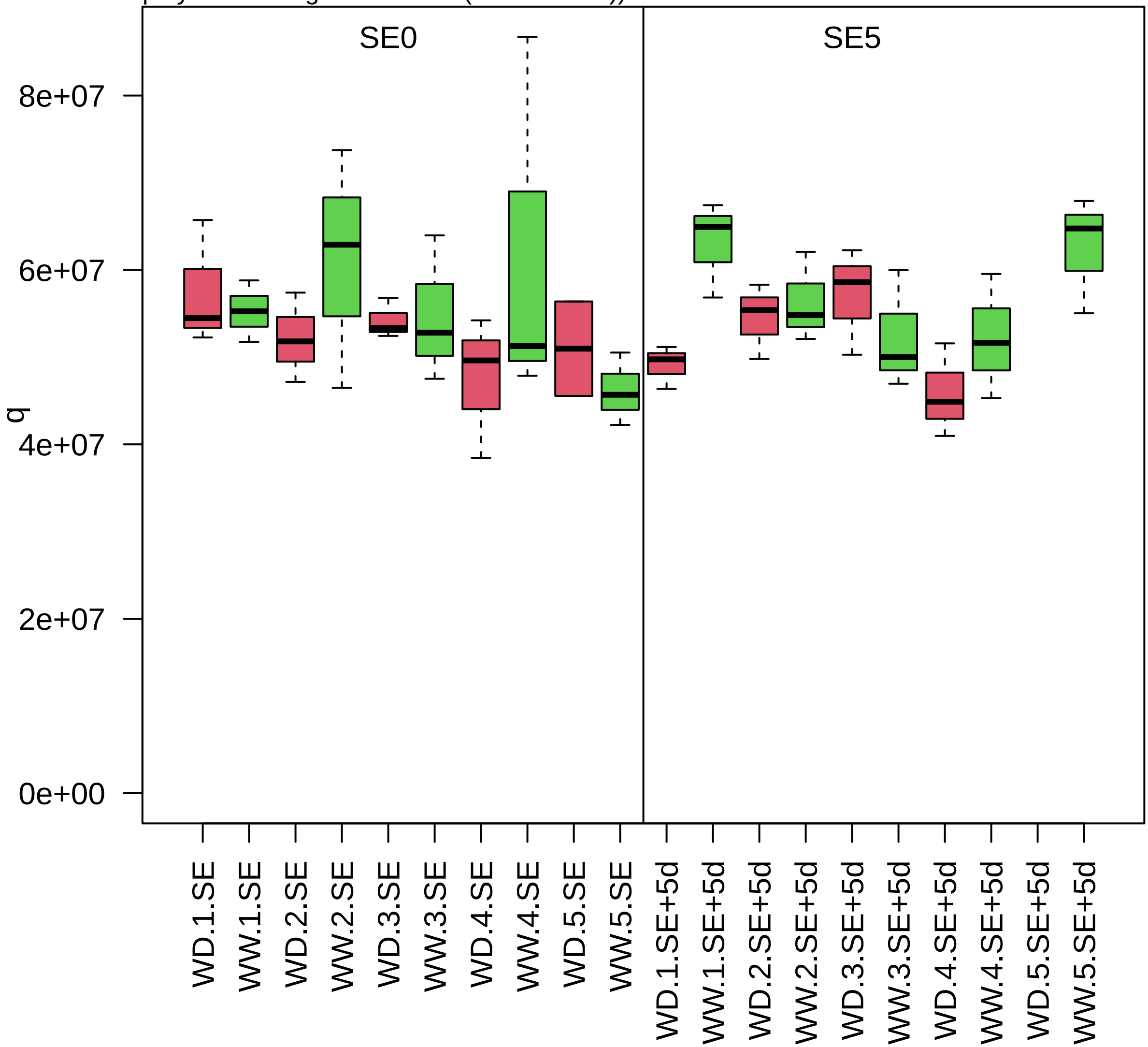

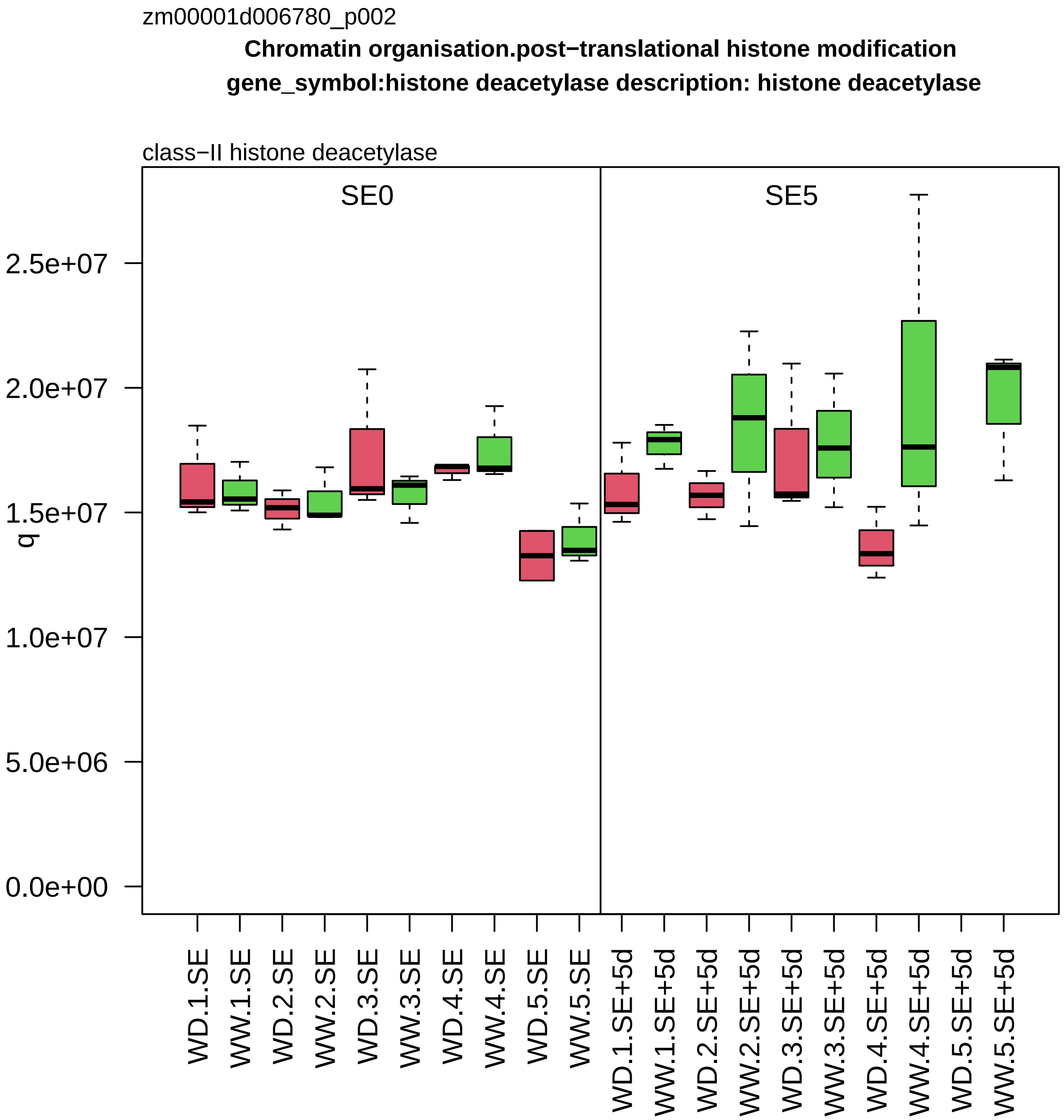

zm00001d006780_p002
Chromatin organisation.post−translational histone modification
gene_symbol:histone deacetylase description: histone deacetylase
class−II histone deacetylase
SE0
SE5
q
2.5e+07
2.0e+07
1.5e+07
1.0e+07
5.0e+06
0.0e+00
WD.1.SE
WW.1.SE
WD.2.SE
WW.2.SE
WD.3.SE
WW.3.SE
WD.4.SE
WW.4.SE
WD.5.SE
WW.5.SE
WD.1.SE+5d
WW.1.SE+5d
WD.2.SE+5d
WW.2.SE+5d
WD.3.SE+5d
WW.3.SE+5d
WD.4.SE+5d
WW.4.SE+5d
WD.5.SE+5d
WW.5.SE+5d

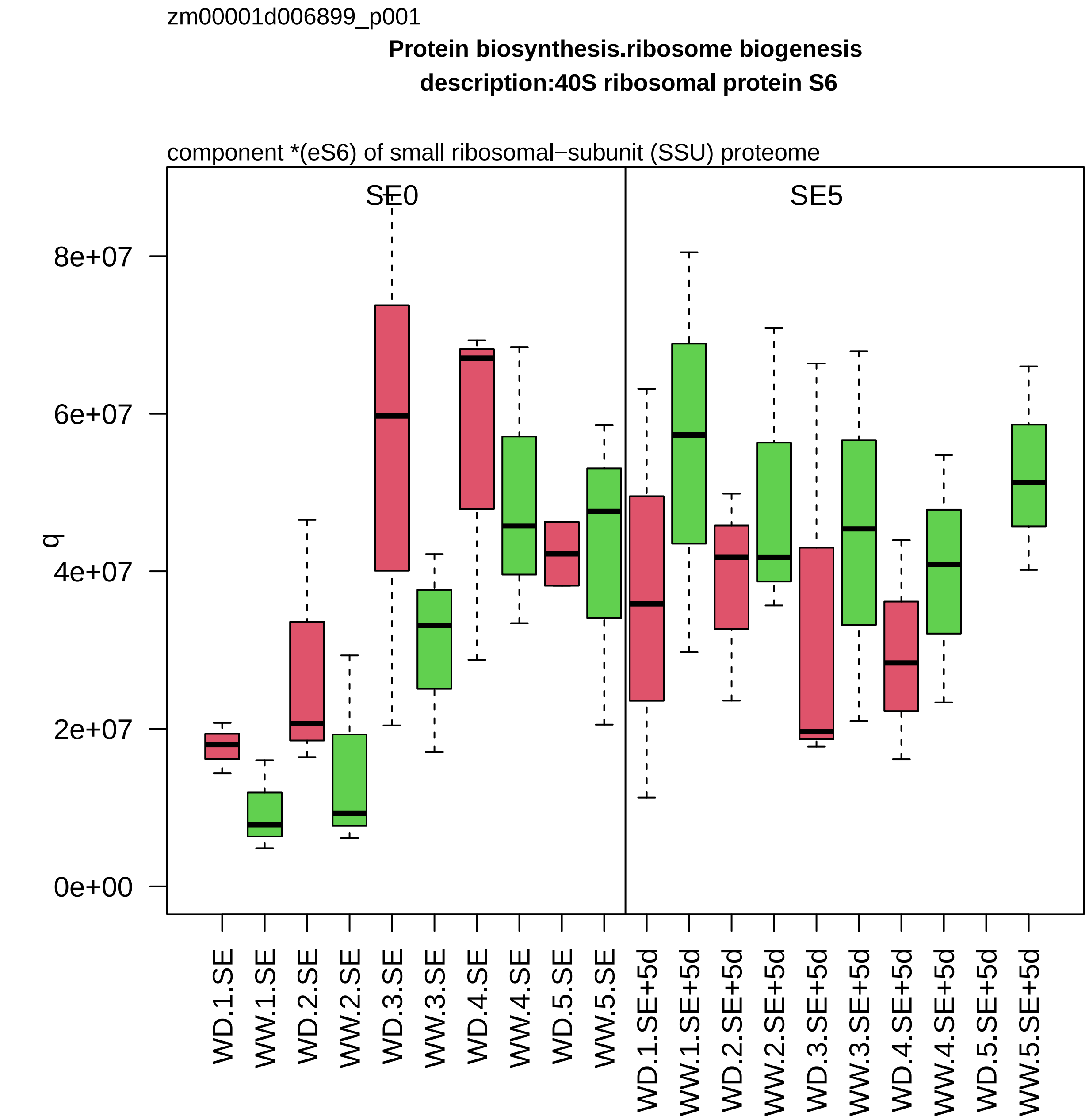

zm00001d006899_p001
Protein biosynthesis.ribosome biogenesis
description:40S ribosomal protein S6
component *(eS6) of small ribosomal−subunit (SSU) proteome
SE0
SE5
q
8e+07
6e+07
4e+07
2e+07
0e+00
WD.1.SE
WW.1.SE
WD.2.SE
WW.2.SE
WD.3.SE
WW.3.SE
WD.4.SE
WW.4.SE
WD.5.SE
WW.5.SE
WD.1.SE+5d
WW.1.SE+5d
WD.2.SE+5d
WW.2.SE+5d
WD.3.SE+5d
WW.3.SE+5d
WD.4.SE+5d
WW.4.SE+5d
WD.5.SE+5d
WW.5.SE+5d

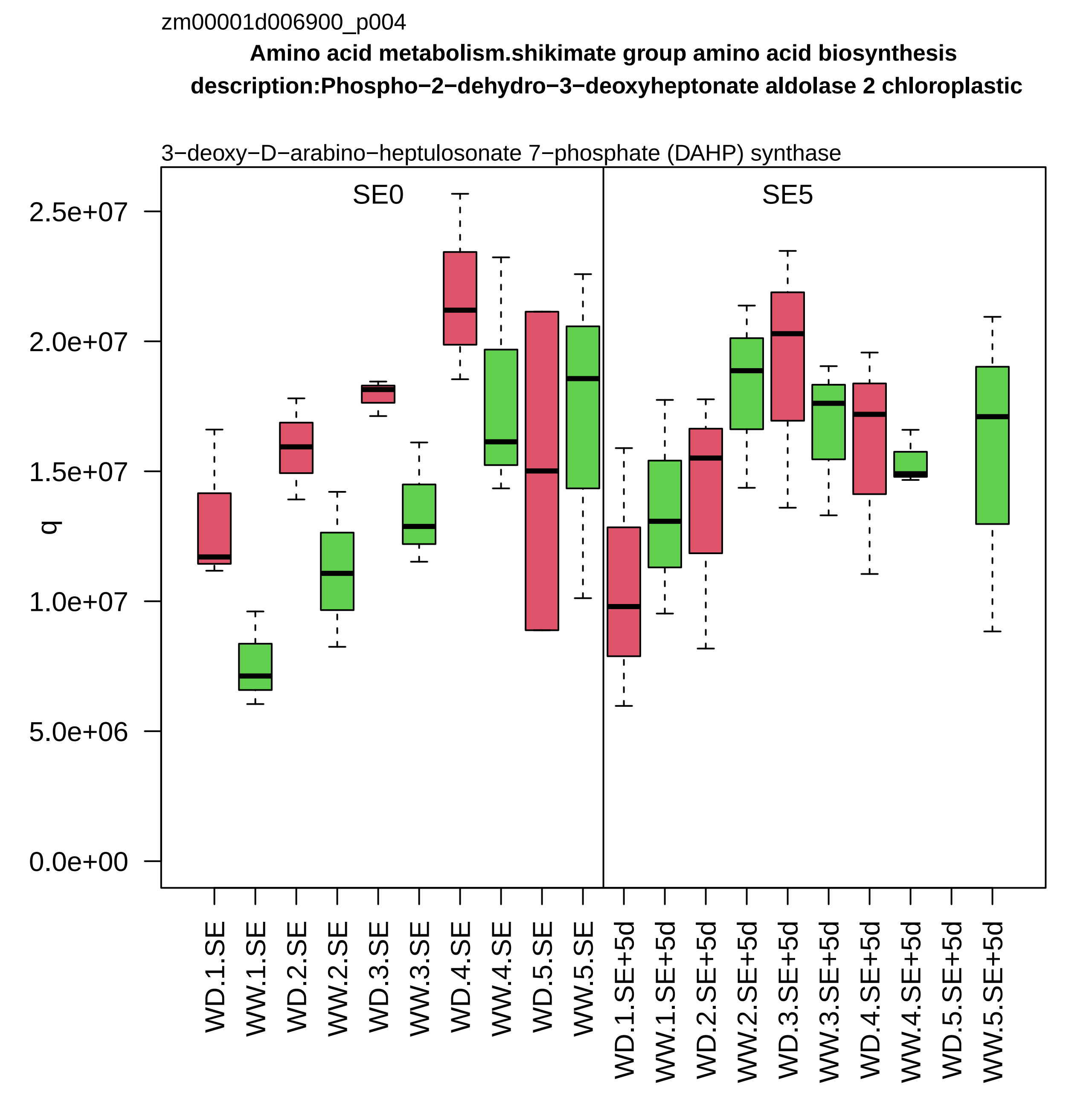
zm00001d006900_p004
Amino acid metabolism.shikimate group amino acid biosynthesis
description:Phospho−2−dehydro−3−deoxyheptonate aldolase 2 chloroplastic
3−deoxy−D−arabino−heptulosonate 7−phosphate (DAHP) synthase
SE0
SE5
q
2.5e+07
2.0e+07
1.5e+07
1.0e+07
5.0e+06
0.0e+00
WD.1.SE
WW.1.SE
WD.2.SE
WW.2.SE
WD.3.SE
WW.3.SE
WD.4.SE
WW.4.SE
WD.5.SE
WW.5.SE
WD.1.SE+5d
WW.1.SE+5d
WD.2.SE+5d
WW.2.SE+5d
WD.3.SE+5d
WW.3.SE+5d
WD.4.SE+5d
WW.4.SE+5d
WD.5.SE+5d
WW.5.SE+5d

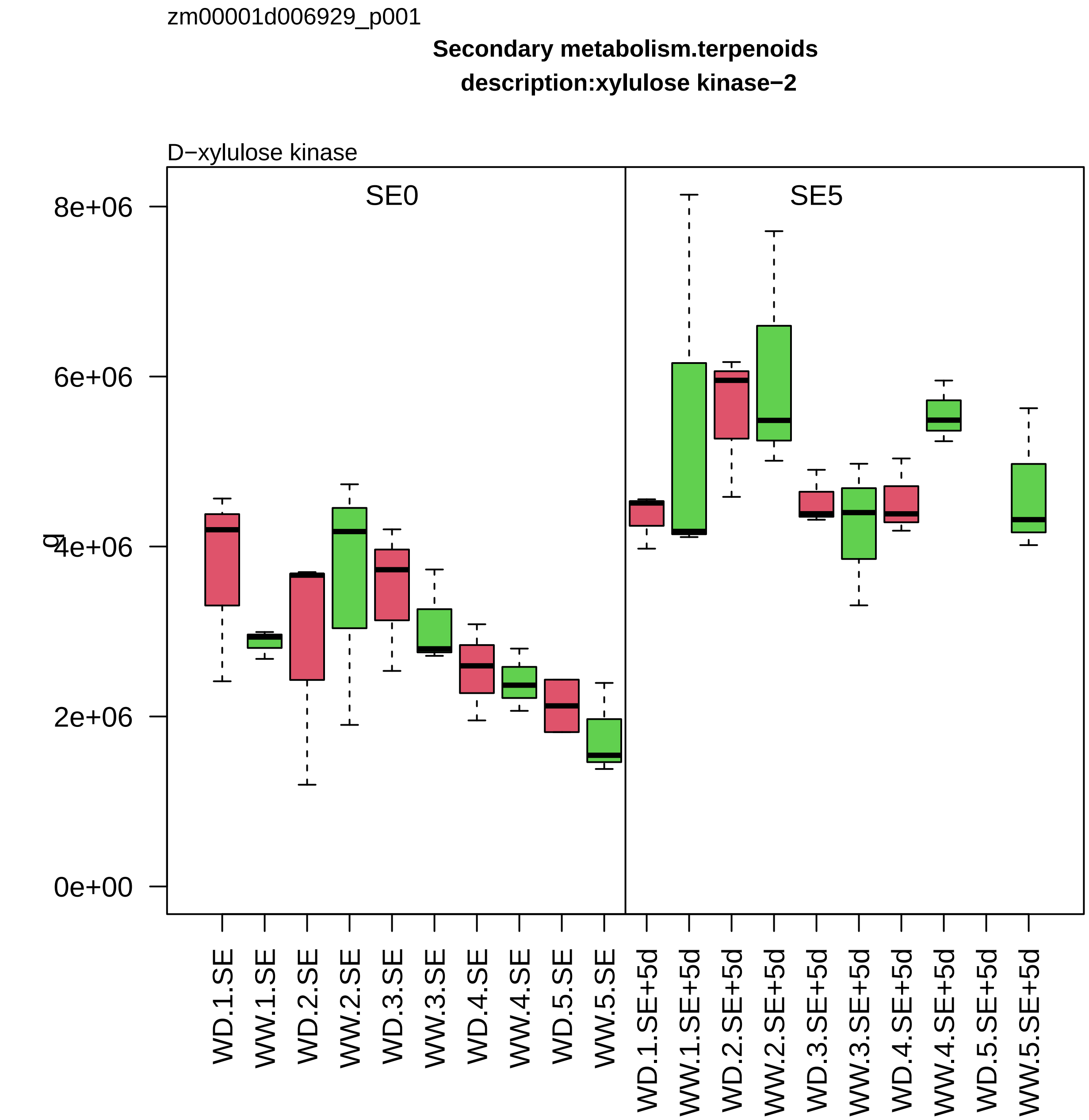
zm00001d006929_p001
Secondary metabolism.terpenoids
description:xylulose kinase−2
D−xylulose kinase
SE0
SE5
8e+06
6e+06
4e+06
2e+06
0e+00
WD.1.SE
WW.1.SE
WD.2.SE
WW.2.SE
WD.3.SE
WW.3.SE
WD.4.SE
WW.4.SE
WD.5.SE
WW.5.SE
WD.1.SE+5d
WW.1.SE+5d
WD.2.SE+5d
WW.2.SE+5d
WD.3.SE+5d
WW.3.SE+5d
WD.4.SE+5d
WW.4.SE+5d
WD.5.SE+5d
WW.5.SE+5d

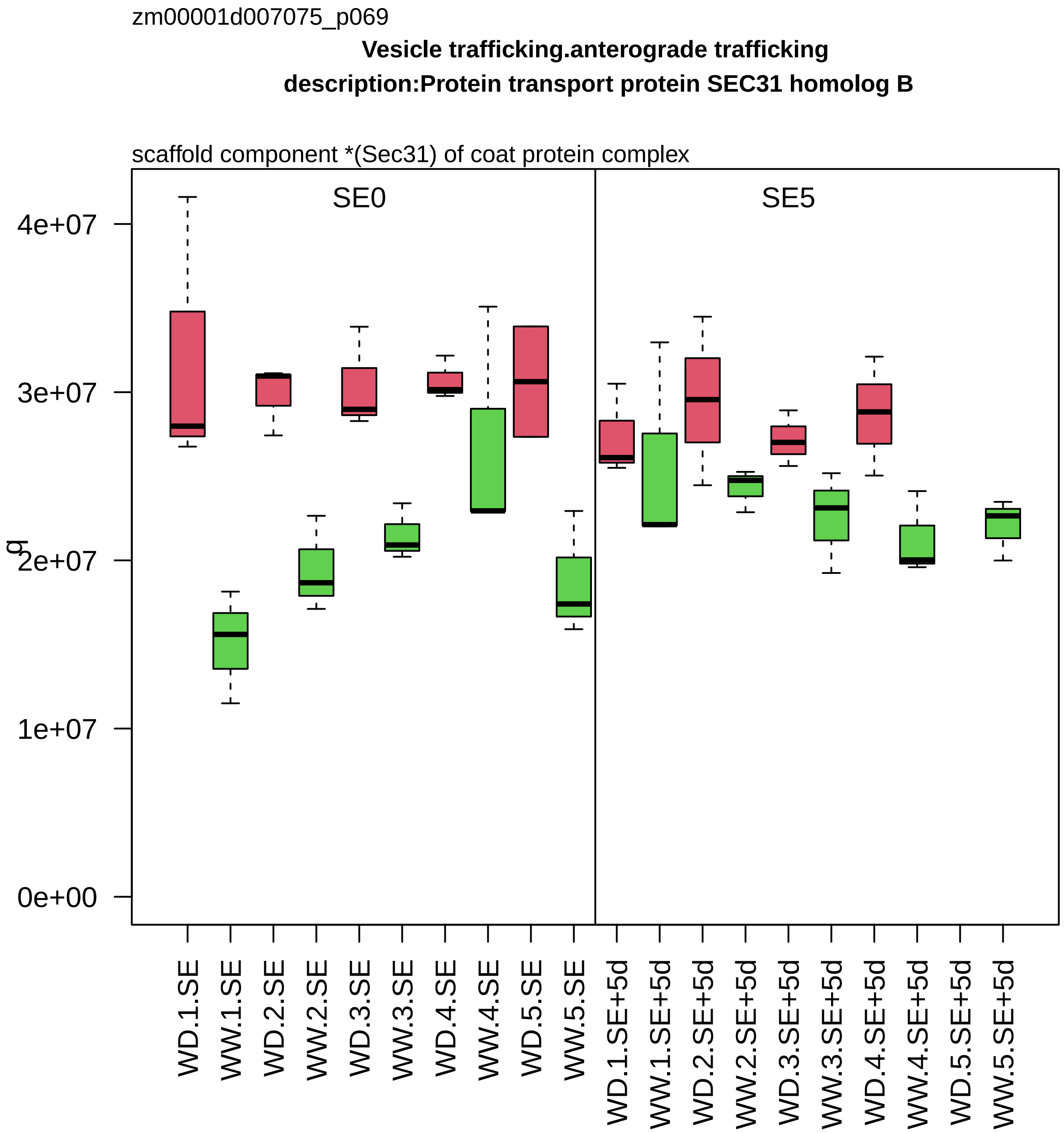

zm00001d007075_p069
Vesicle trafficking.anterograde trafficking
description:Protein transport protein SEC31 homolog B
scaffold component *(Sec31) of coat protein complex
SE0
SE5
q
4e+07
3e+07
2e+07
1e+07
0e+00
WD.1.SE
WW.1.SE
WD.2.SE
WW.2.SE
WD.3.SE
WW.3.SE
WD.4.SE
WW.4.SE
WD.5.SE
WW.5.SE
WD.1.SE+5d
WW.1.SE+5d
WD.2.SE+5d
WW.2.SE+5d
WD.3.SE+5d
WW.3.SE+5d
WD.4.SE+5d
WW.4.SE+5d
WD.5.SE+5d
WW.5.SE+5d

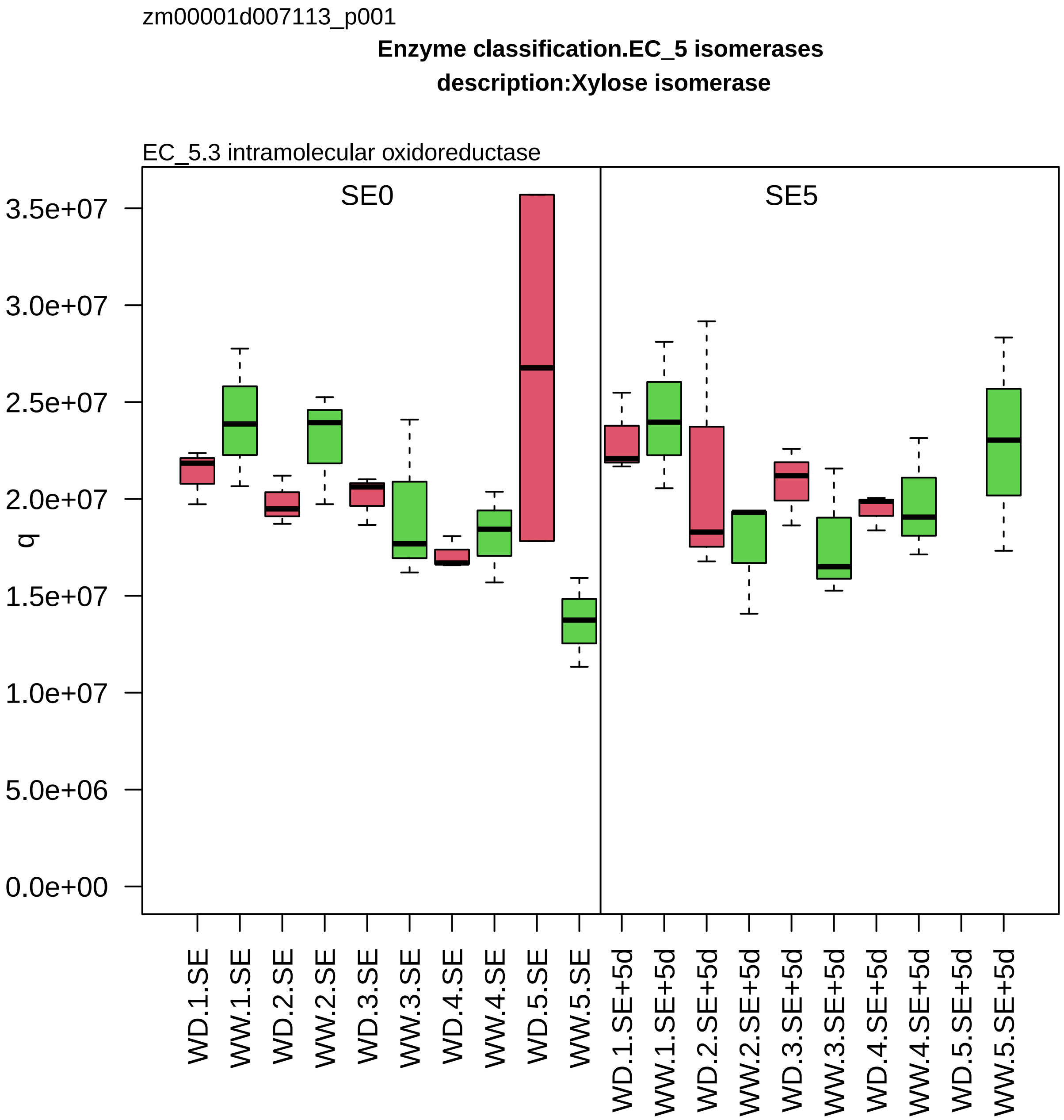

zm00001d007113_p001
Enzyme classification.EC_5 isomerases
description:Xylose isomerase
EC_5.3 intramolecular oxidoreductase
SE0
SE5
q
3.5e+07
3.0e+07
2.5e+07
2.0e+07
1.5e+07
1.0e+07
5.0e+06
0.0e+00
WD.1.SE
WW.1.SE
WD.2.SE
WW.2.SE
WD.3.SE
WW.3.SE
WD.4.SE
WW.4.SE
WD.5.SE
WW.5.SE
WD.1.SE+5d
WW.1.SE+5d
WD.2.SE+5d
WW.2.SE+5d
WD.3.SE+5d
WW.3.SE+5d
WD.4.SE+5d
WW.4.SE+5d
WD.5.SE+5d
WW.5.SE+5d

zm00001d007192_p001

**Cytoskeleton organisation.actin and tubulin folding**
**description:T−complex protein 1 subunit zeta**

subunit zeta *(CCT6) of CCT chaperonin folding complex

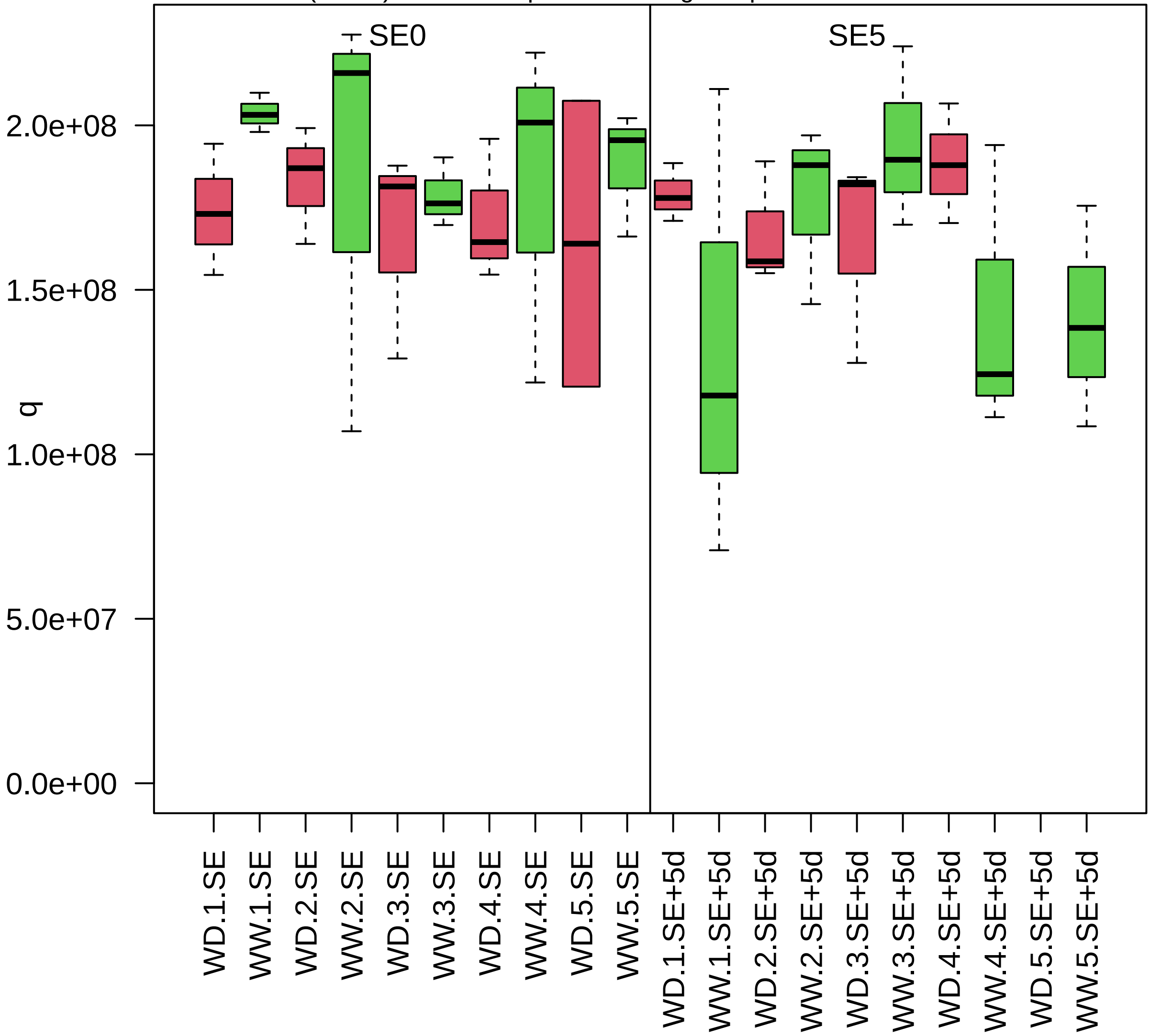

zm00001d007215_p003

**Protein homeostasis.ubiquitin−proteasome system**

**description:26S proteasome regulatory subunit 4 homolog A**

regulatory component *(RPT2) of 26S proteasome

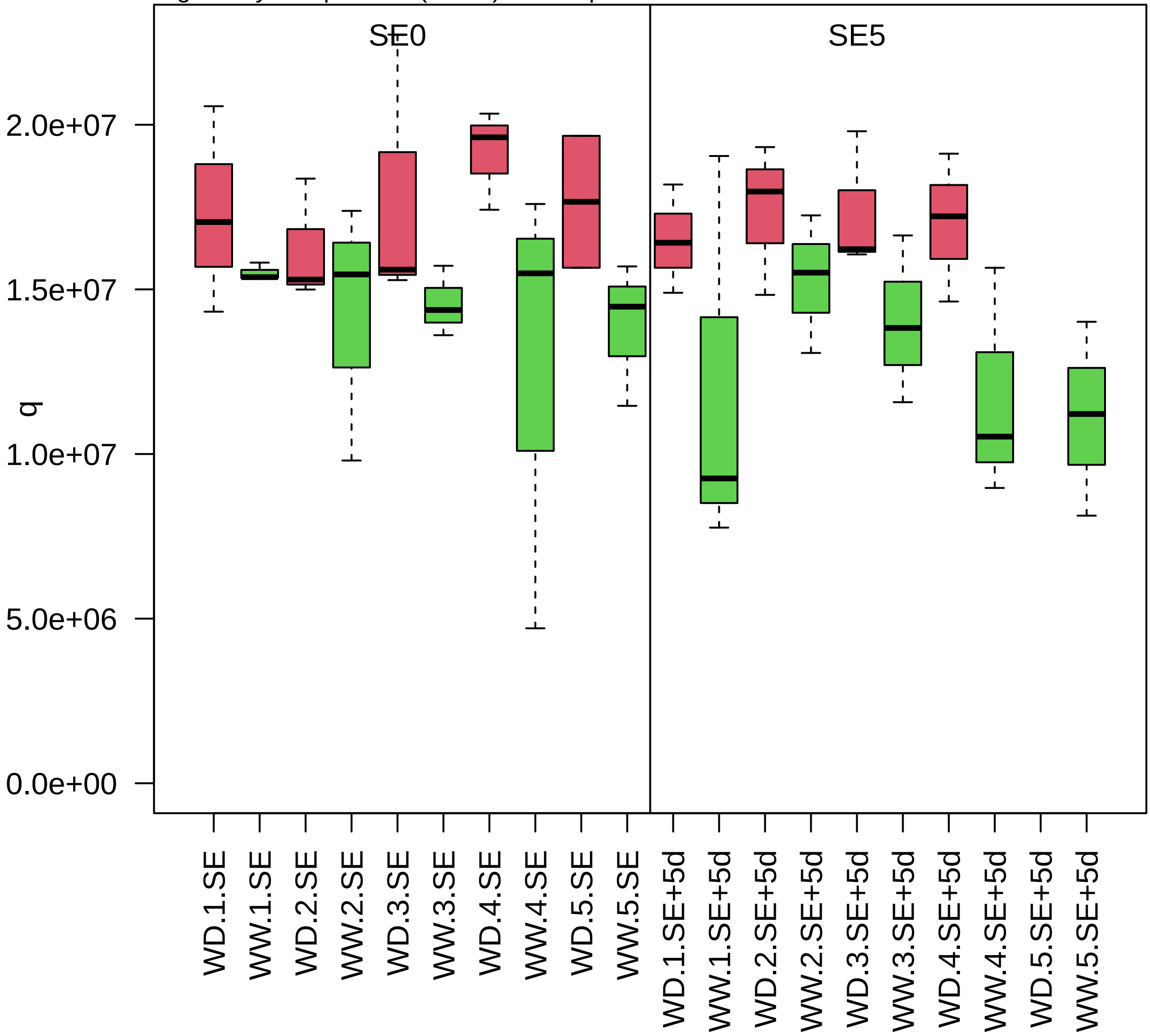

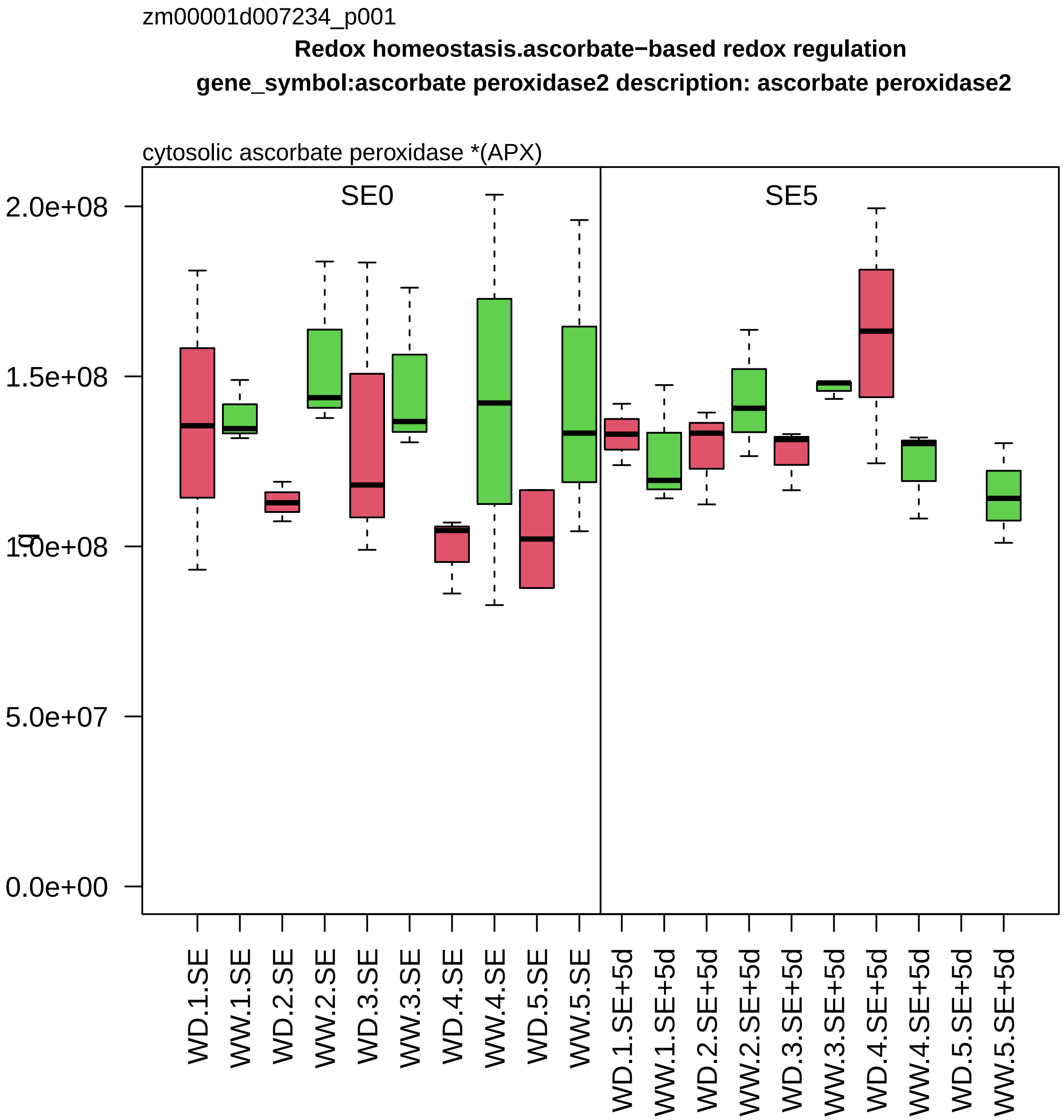
zm00001d007234_p001
Redox homeostasis.ascorbate−based redox regulation
gene_symbol:ascorbate peroxidase2 description: ascorbate peroxidase2
cytosolic ascorbate peroxidase *(APX)
SE0
SE5
2.0e+08
1.5e+08
1.0e+08
5.0e+07
0.0e+00
WD.1.SE
WW.1.SE
WD.2.SE
WW.2.SE
WD.3.SE
WW.3.SE
WD.4.SE
WW.4.SE
WD.5.SE
WW.5.SE
WD.1.SE+5d
WW.1.SE+5d
WD.2.SE+5d
WW.2.SE+5d
WD.3.SE+5d
WW.3.SE+5d
WD.4.SE+5d
WW.4.SE+5d
WD.5.SE+5d
WW.5.SE+5d

zm00001d007301_p003

**Protein modification.cysteine disulfide formation**

**gene_symbol:protein disulfide isomerase2 description: protein disulfide isomerase2**

protein disulfide isomerase *(PDI−L)

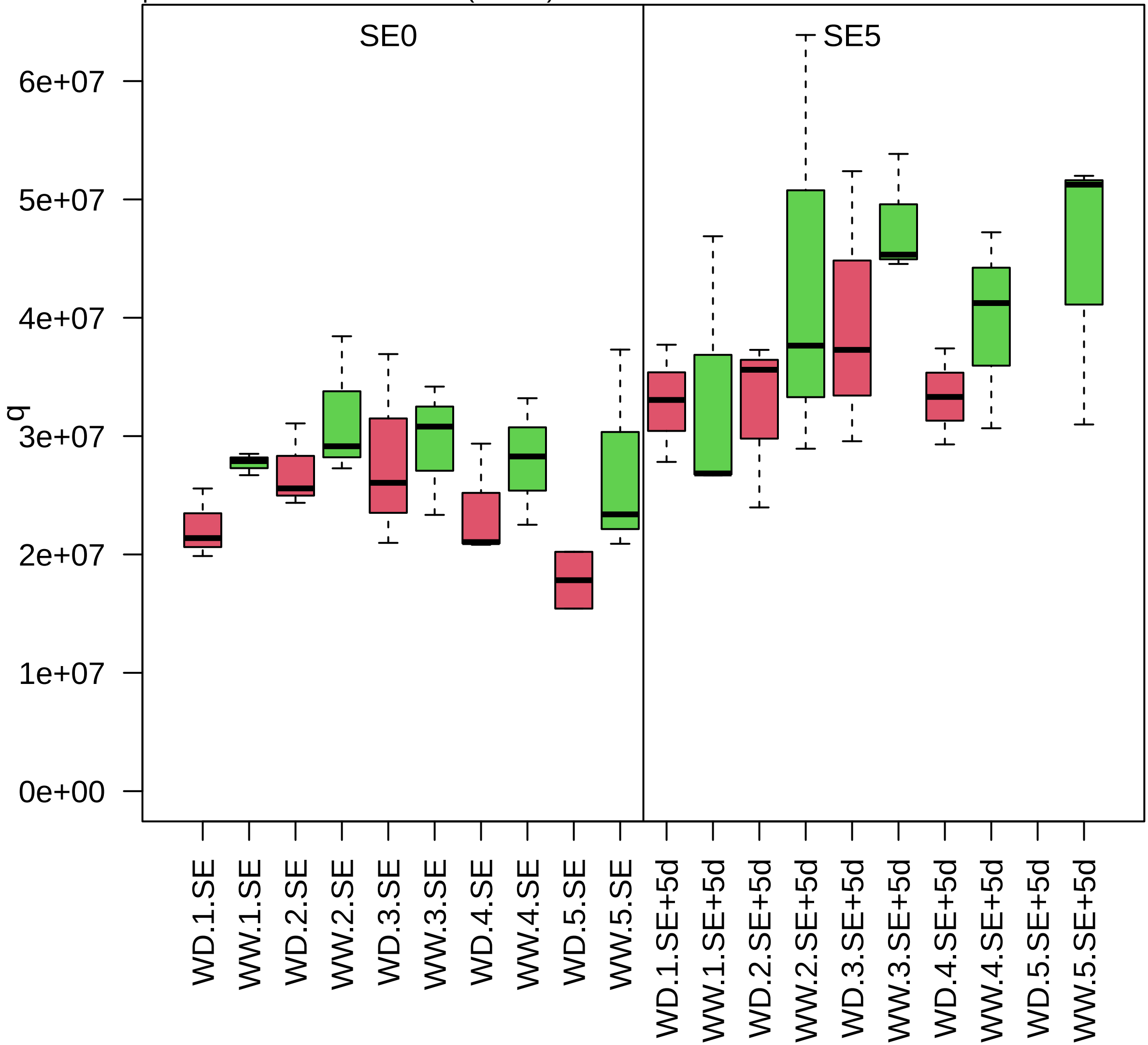

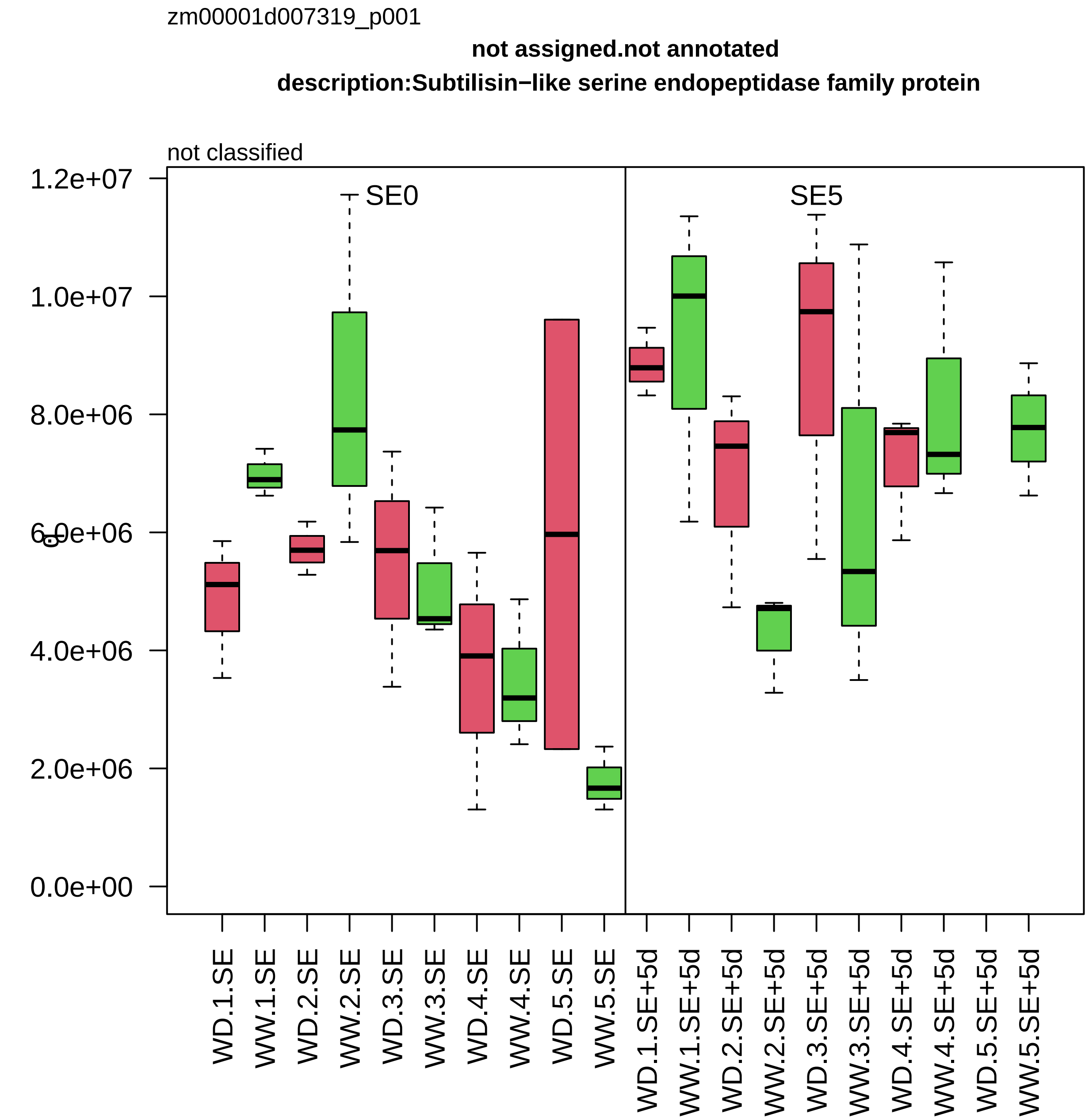

zm00001d007319_p001
not assigned.not annotated
description:Subtilisin−like serine endopeptidase family protein
not classified
SE0
SE5
1.2e+07
1.0e+07
8.0e+06
6.0e+06
4.0e+06
2.0e+06
0.0e+00
WD.1.SE
WW.1.SE
WD.2.SE
WW.2.SE
WD.3.SE
WW.3.SE
WD.4.SE
WW.4.SE
WD.5.SE
WW.5.SE
WD.1.SE+5d
WW.1.SE+5d
WD.2.SE+5d
WW.2.SE+5d
WD.3.SE+5d
WW.3.SE+5d
WD.4.SE+5d
WW.4.SE+5d
WD.5.SE+5d
WW.5.SE+5d

zm00001d007369_p002

**Protein biosynthesis.aminoacyl−tRNA synthetase activities**
**description:Serine−−tRNA ligase cytoplasmic**

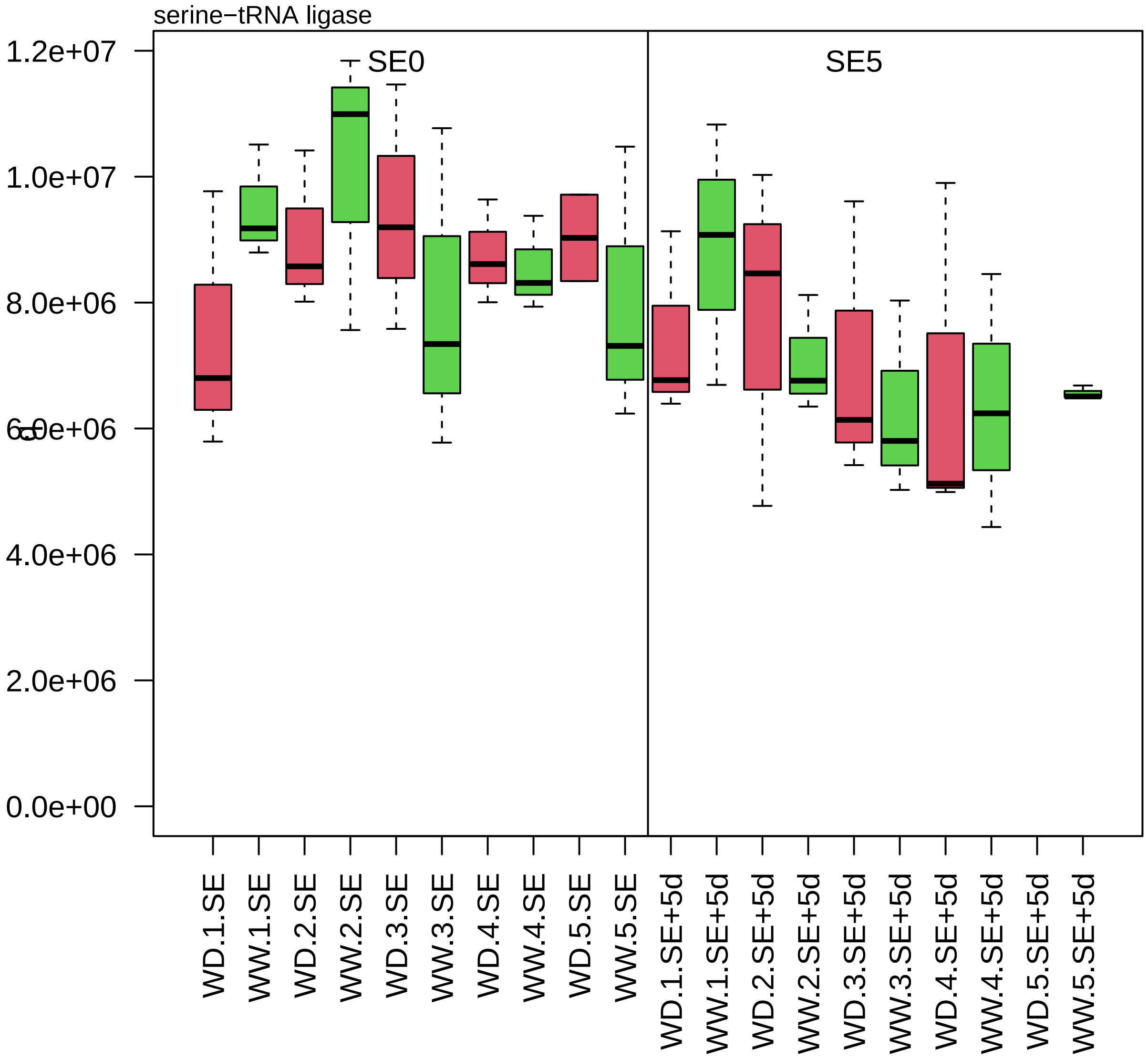

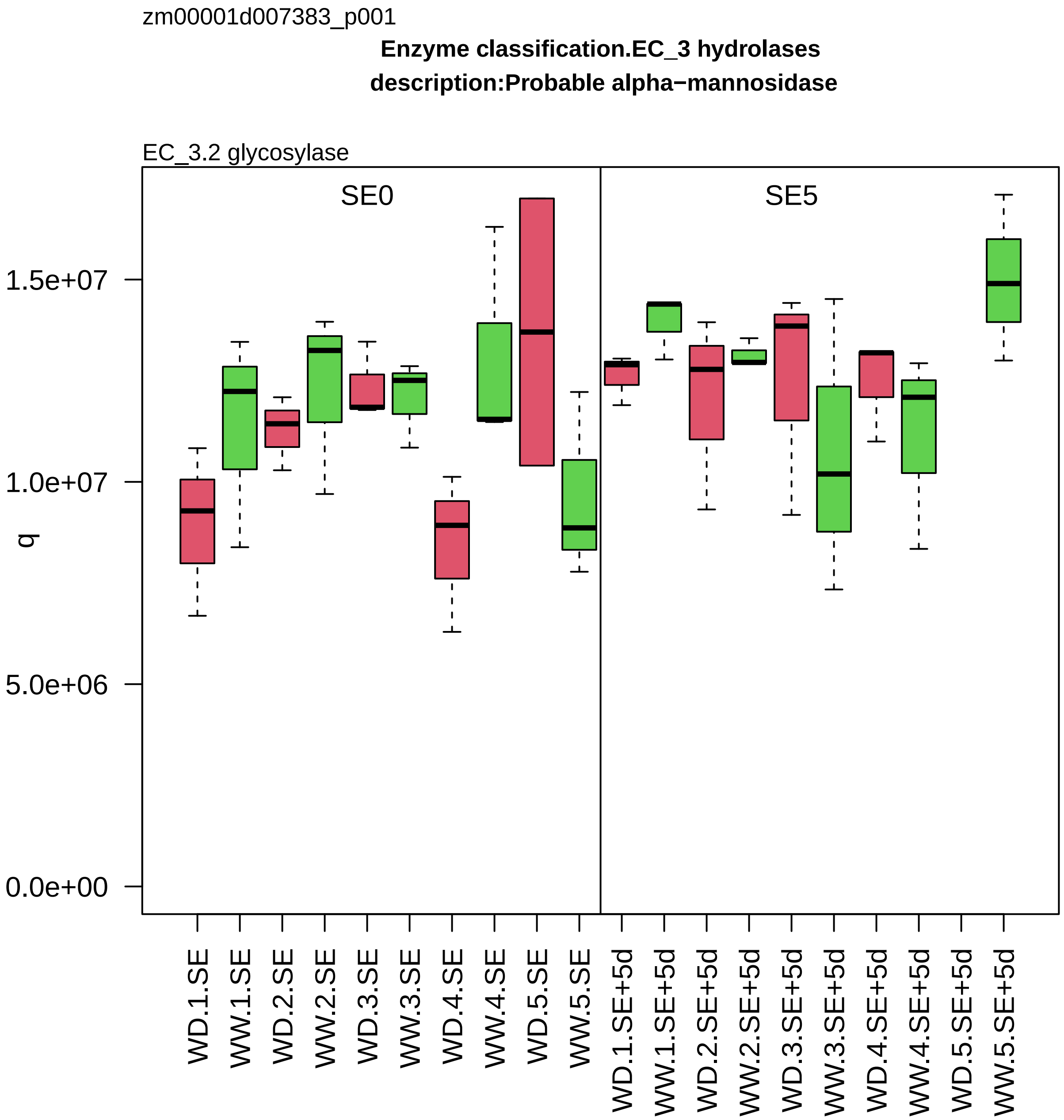

zm00001d007383_p001
Enzyme classification.EC_3 hydrolases
description:Probable alpha−mannosidase
EC_3.2 glycosylase
SE0
SE5
q
1.5e+07
1.0e+07
5.0e+06
0.0e+00
WD.1.SE
WW.1.SE
WD.2.SE
WW.2.SE
WD.3.SE
WW.3.SE
WD.4.SE
WW.4.SE
WD.5.SE
WW.5.SE
WD.1.SE+5d
WW.1.SE+5d
WD.2.SE+5d
WW.2.SE+5d
WD.3.SE+5d
WW.3.SE+5d
WD.4.SE+5d
WW.4.SE+5d
WD.5.SE+5d
WW.5.SE+5d

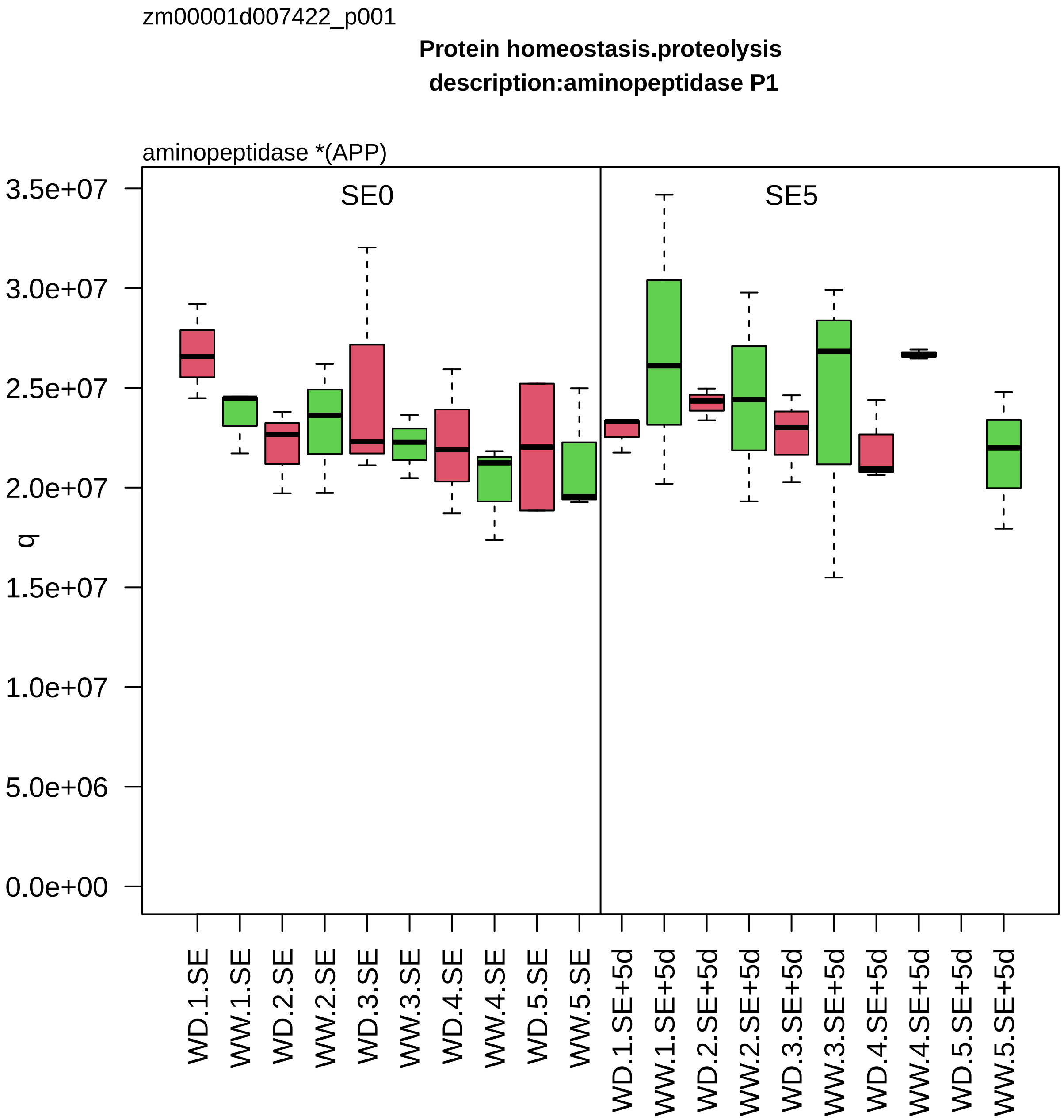

zm00001d007422_p001
Protein homeostasis.proteolysis
description:aminopeptidase P1
aminopeptidase *(APP)
SE0
SE5
q
3.5e+07
3.0e+07
2.5e+07
2.0e+07
1.5e+07
1.0e+07
5.0e+06
0.0e+00
WD.1.SE
WW.1.SE
WD.2.SE
WW.2.SE
WD.3.SE
WW.3.SE
WD.4.SE
WW.4.SE
WD.5.SE
WW.5.SE
WD.1.SE+5d
WW.1.SE+5d
WD.2.SE+5d
WW.2.SE+5d
WD.3.SE+5d
WW.3.SE+5d
WD.4.SE+5d
WW.4.SE+5d
WD.5.SE+5d
WW.5.SE+5d

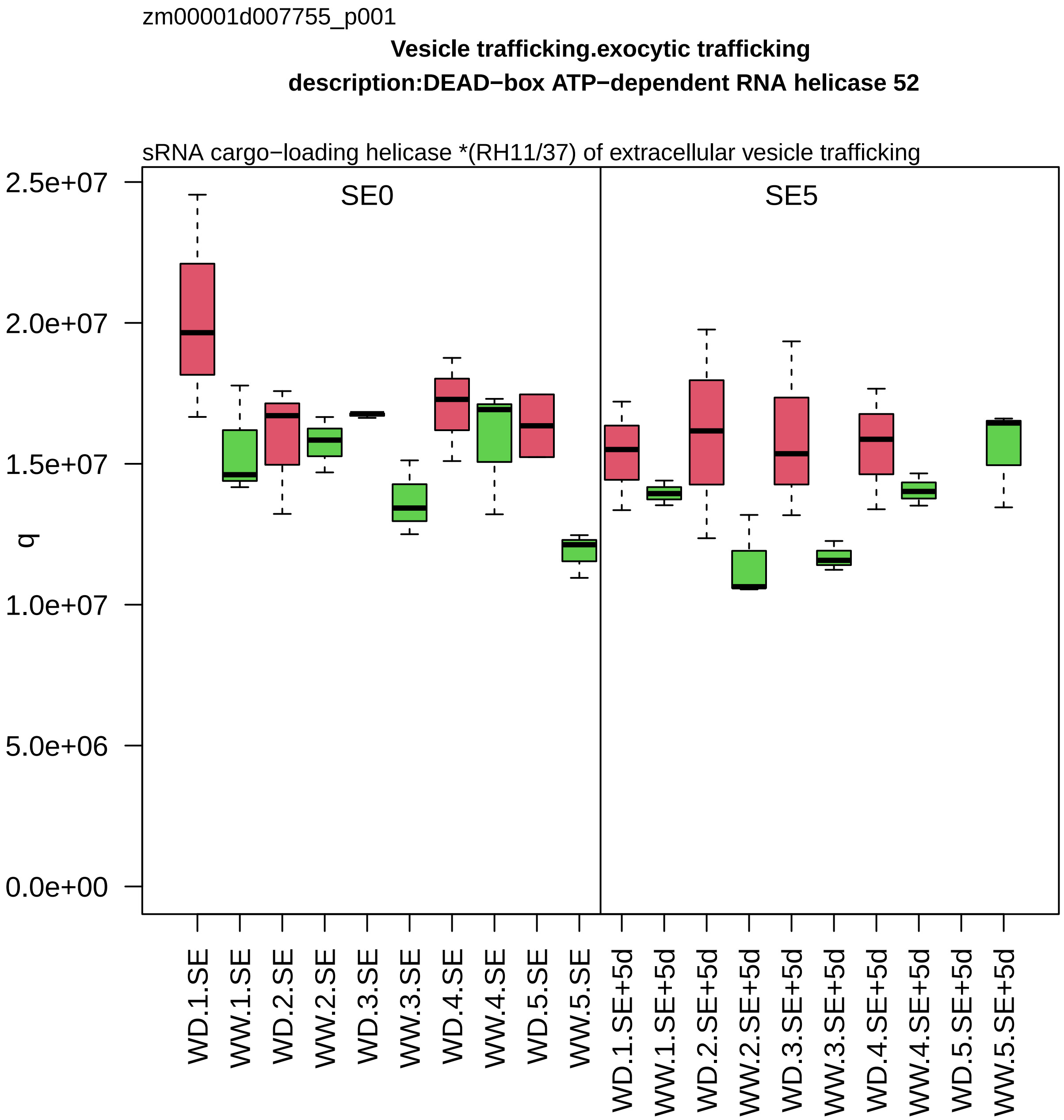

zm00001d007755_p001
Vesicle trafficking.exocytic trafficking
description:DEAD−box ATP−dependent RNA helicase 52
sRNA cargo−loading helicase *(RH11/37) of extracellular vesicle trafficking
SE0
SE5
q
2.5e+07
2.0e+07
1.5e+07
1.0e+07
5.0e+06
0.0e+00
WD.1.SE
WW.1.SE
WD.2.SE
WW.2.SE
WD.3.SE
WW.3.SE
WD.4.SE
WW.4.SE
WD.5.SE
WW.5.SE
WD.1.SE+5d
WW.1.SE+5d
WD.2.SE+5d
WW.2.SE+5d
WD.3.SE+5d
WW.3.SE+5d
WD.4.SE+5d
WW.4.SE+5d
WD.5.SE+5d
WW.5.SE+5d

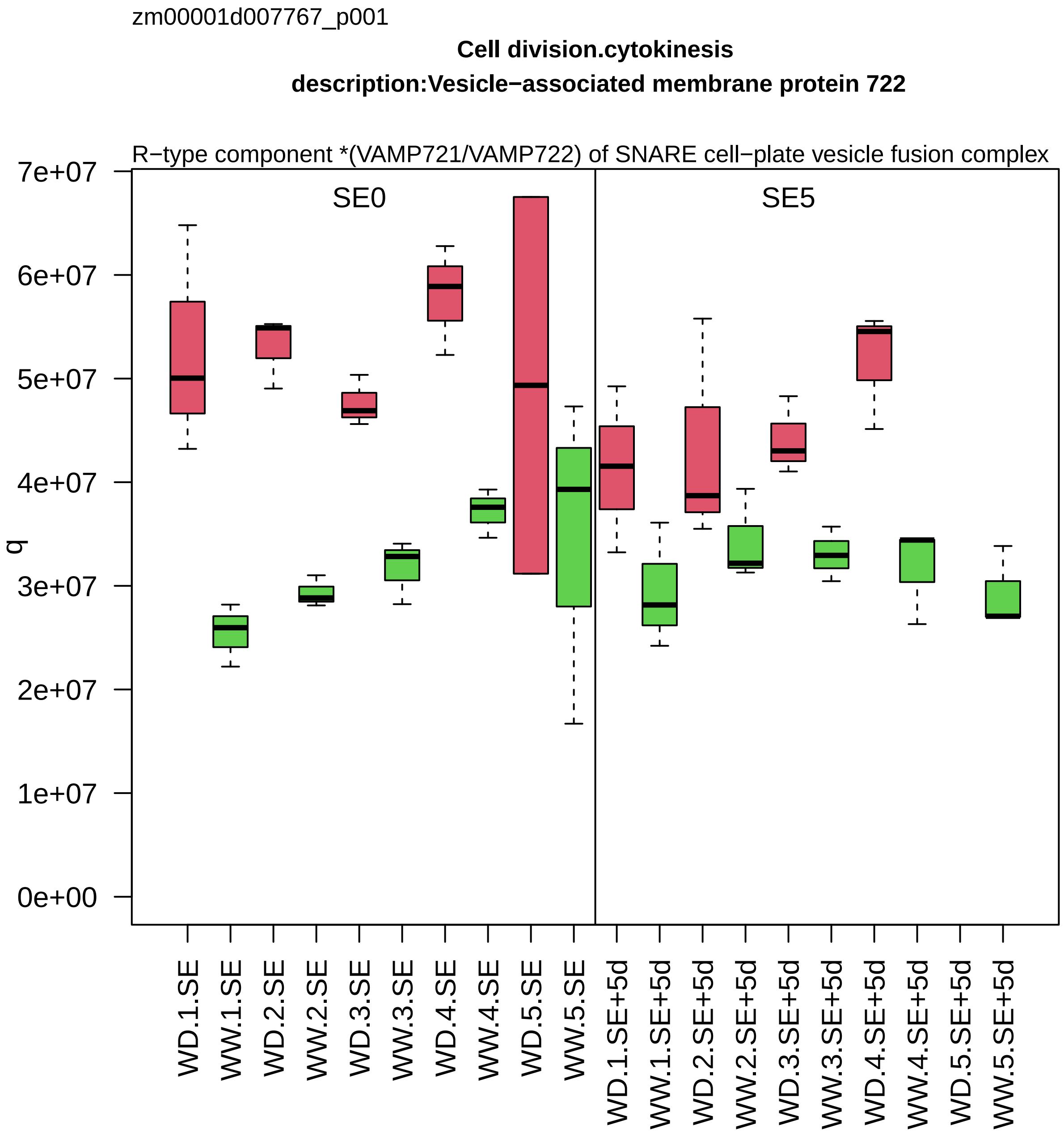
zm00001d007767_p001
Cell division.cytokinesis
description:Vesicle−associated membrane protein 722
R−type component *(VAMP721/VAMP722) of SNARE cell−plate vesicle fusion complex
SE0
SE5
q
7e+07
6e+07
5e+07
4e+07
3e+07
2e+07
1e+07
0e+00
WD.1.SE
WW.1.SE
WD.2.SE
WW.2.SE
WD.3.SE
WW.3.SE
WD.4.SE
WW.4.SE
WD.5.SE
WW.5.SE
WD.1.SE+5d
WW.1.SE+5d
WD.2.SE+5d
WW.2.SE+5d
WD.3.SE+5d
WW.3.SE+5d
WD.4.SE+5d
WW.4.SE+5d
WD.5.SE+5d
WW.5.SE+5d

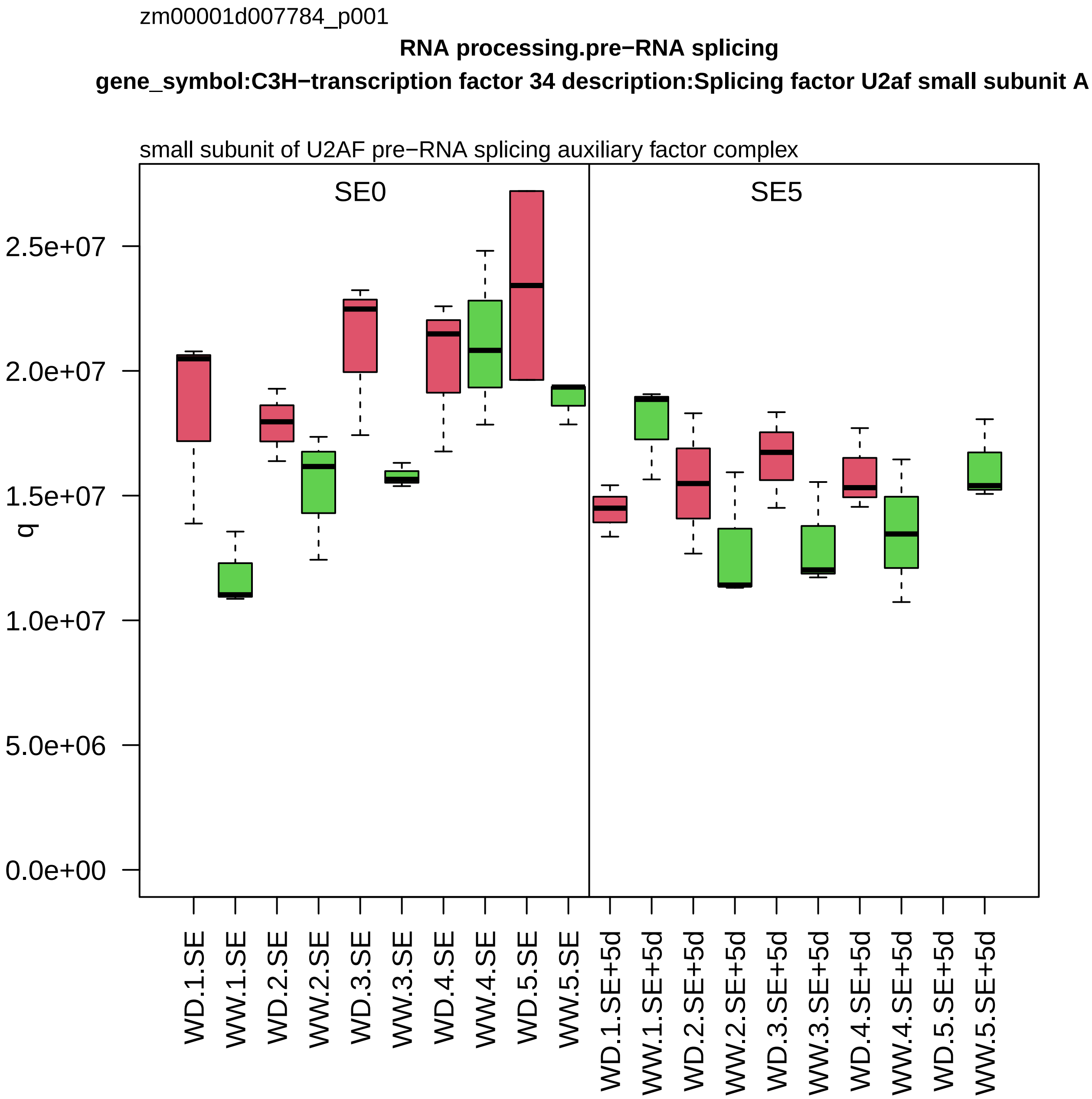

zm00001d007784_p001
RNA processing.pre−RNA splicing
gene_symbol:C3H−transcription factor 34 description:Splicing factor U2af small subunit A
small subunit of U2AF pre−RNA splicing auxiliary factor complex
SE0
SE5
q
0.0e+00
5.0e+06
1.0e+07
1.5e+07
2.0e+07
2.5e+07
WD.1.SE
WW.1.SE
WD.2.SE
WW.2.SE
WD.3.SE
WW.3.SE
WD.4.SE
WW.4.SE
WD.5.SE
WW.5.SE
WD.1.SE+5d
WW.1.SE+5d
WD.2.SE+5d
WW.2.SE+5d
WD.3.SE+5d
WW.3.SE+5d
WD.4.SE+5d
WW.4.SE+5d
WD.5.SE+5d
WW.5.SE+5d

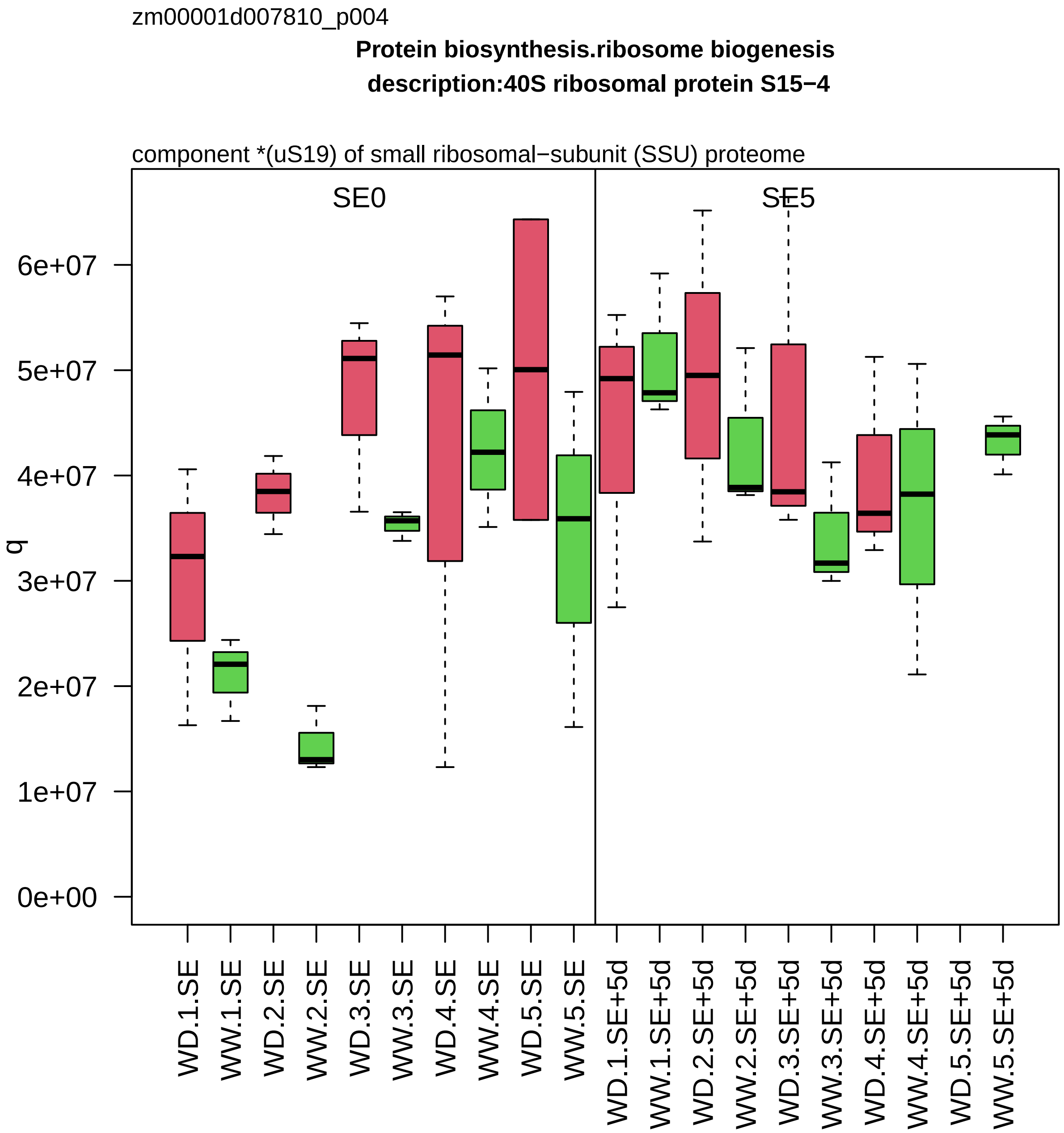
zm00001d007810_p004
Protein biosynthesis.ribosome biogenesis
description:40S ribosomal protein S15−4
component *(uS19) of small ribosomal−subunit (SSU) proteome
SE0
SE5
q
0e+00
1e+07
2e+07
3e+07
4e+07
5e+07
6e+07
WD.1.SE
WW.1.SE
WD.2.SE
WW.2.SE
WD.3.SE
WW.3.SE
WD.4.SE
WW.4.SE
WD.5.SE
WW.5.SE
WD.1.SE+5d
WW.1.SE+5d
WD.2.SE+5d
WW.2.SE+5d
WD.3.SE+5d
WW.3.SE+5d
WD.4.SE+5d
WW.4.SE+5d
WD.5.SE+5d
WW.5.SE+5d

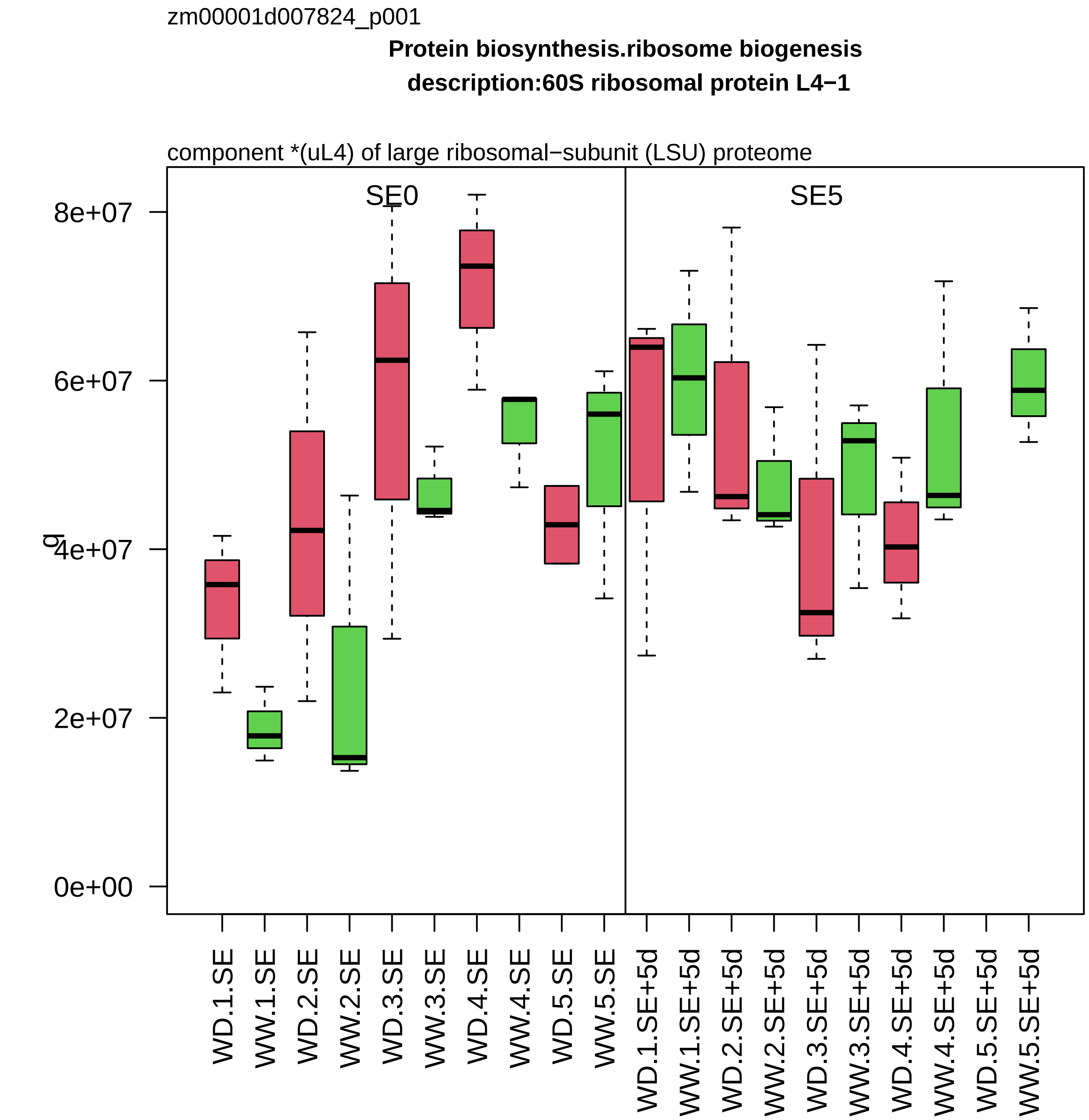
zm00001d007824_p001
Protein biosynthesis.ribosome biogenesis
description:60S ribosomal protein L4−1
component *(uL4) of large ribosomal−subunit (LSU) proteome
SE0
SE5
8e+07
6e+07
4e+07
2e+07
0e+00
q
WD.1.SE
WW.1.SE
WD.2.SE
WW.2.SE
WD.3.SE
WW.3.SE
WD.4.SE
WW.4.SE
WD.5.SE
WW.5.SE
WD.1.SE+5d
WW.1.SE+5d
WD.2.SE+5d
WW.2.SE+5d
WD.3.SE+5d
WW.3.SE+5d
WD.4.SE+5d
WW.4.SE+5d
WD.5.SE+5d
WW.5.SE+5d

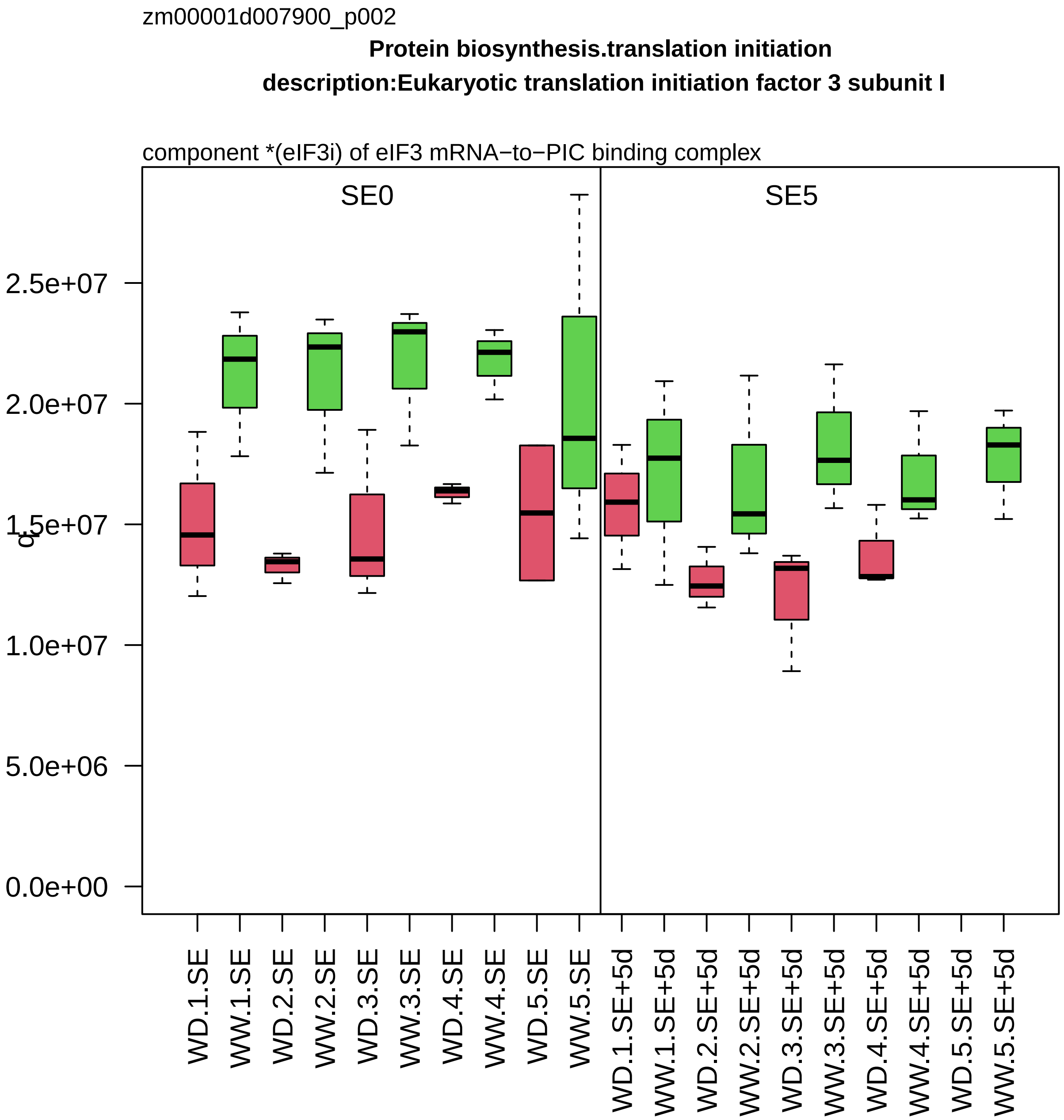

zm00001d007900_p002
Protein biosynthesis.translation initiation
description:Eukaryotic translation initiation factor 3 subunit I
component *(eIF3i) of eIF3 mRNA−to−PIC binding complex
SE0
SE5
2.5e+07
2.0e+07
1.5e+07
1.0e+07
5.0e+06
0.0e+00
WD.1.SE
WW.1.SE
WD.2.SE
WW.2.SE
WD.3.SE
WW.3.SE
WD.4.SE
WW.4.SE
WD.5.SE
WW.5.SE
WD.1.SE+5d
WW.1.SE+5d
WD.2.SE+5d
WW.2.SE+5d
WD.3.SE+5d
WW.3.SE+5d
WD.4.SE+5d
WW.4.SE+5d
WD.5.SE+5d
WW.5.SE+5d

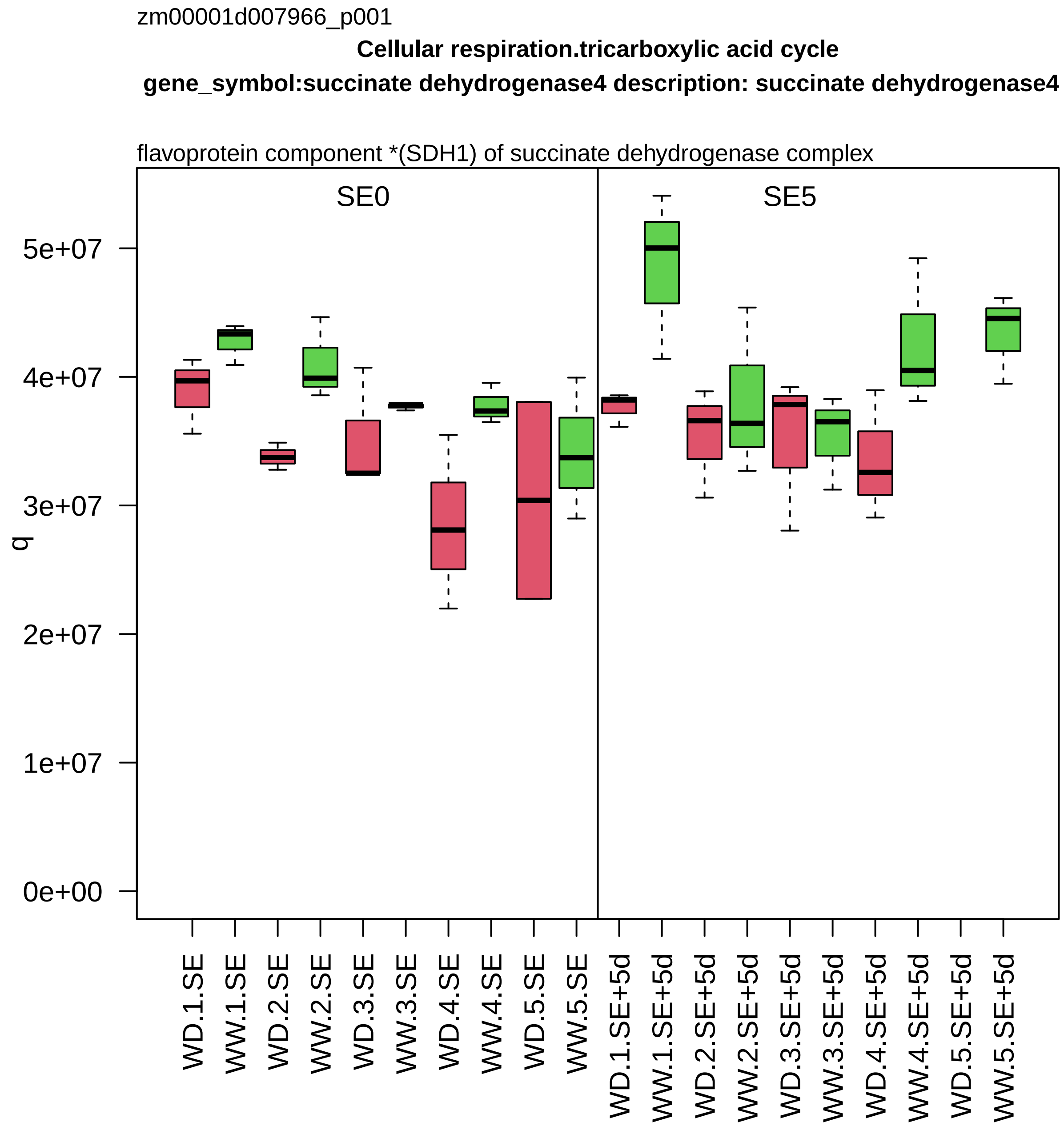
zm00001d007966_p001
Cellular respiration.tricarboxylic acid cycle
gene_symbol:succinate dehydrogenase4 description: succinate dehydrogenase4
flavoprotein component *(SDH1) of succinate dehydrogenase complex
SE0
SE5
q
0e+00
1e+07
2e+07
3e+07
4e+07
5e+07
WD.1.SE
WW.1.SE
WD.2.SE
WW.2.SE
WD.3.SE
WW.3.SE
WD.4.SE
WW.4.SE
WD.5.SE
WW.5.SE
WD.1.SE+5d
WW.1.SE+5d
WD.2.SE+5d
WW.2.SE+5d
WD.3.SE+5d
WW.3.SE+5d
WD.4.SE+5d
WW.4.SE+5d
WD.5.SE+5d
WW.5.SE+5d

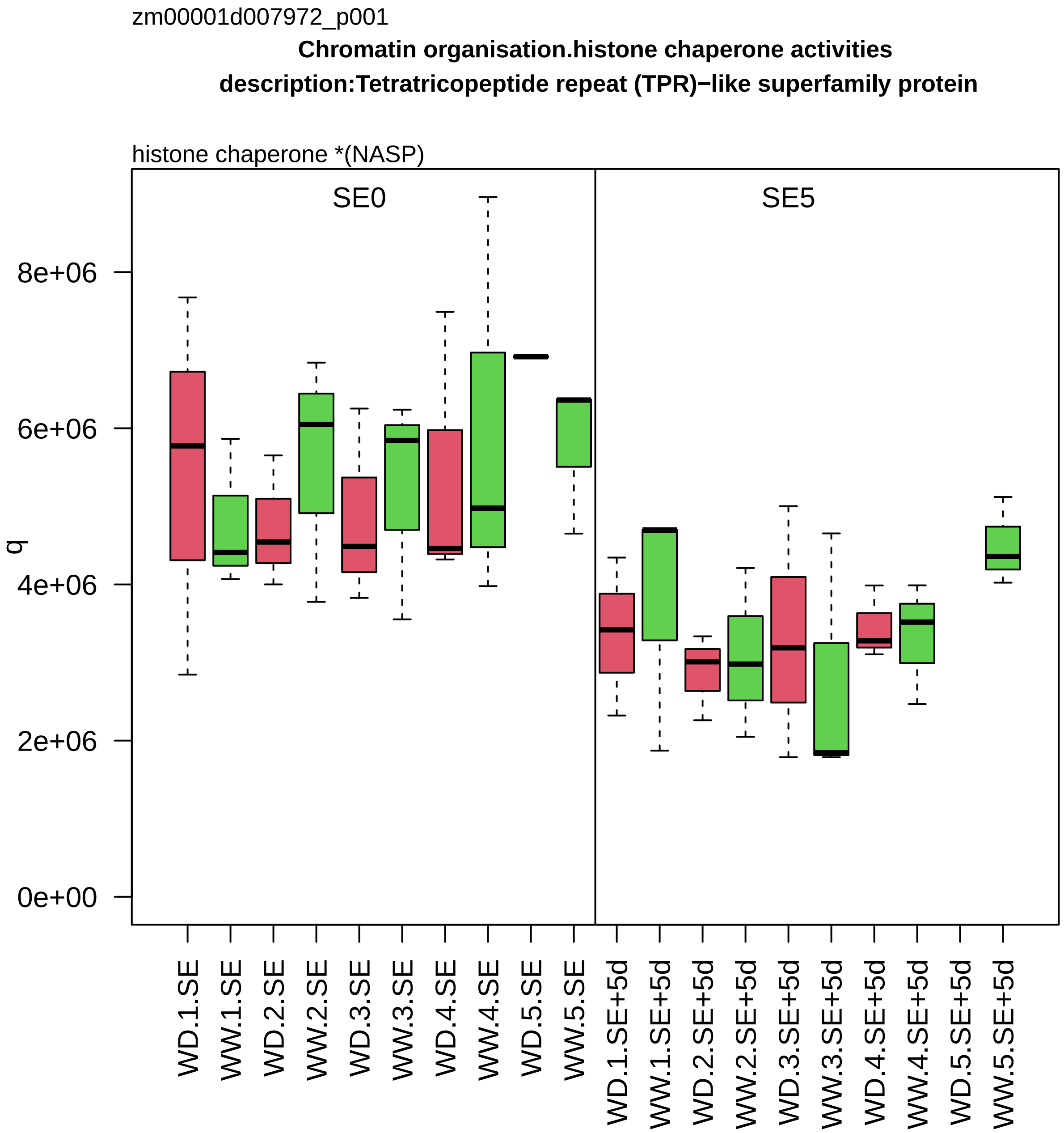
zm00001d007972_p001
Chromatin organisation.histone chaperone activities
description:Tetratricopeptide repeat (TPR)−like superfamily protein
histone chaperone *(NASP)
SE0
SE5
q
0e+00
2e+06
4e+06
6e+06
8e+06
WD.1.SE
WW.1.SE
WD.2.SE
WW.2.SE
WD.3.SE
WW.3.SE
WD.4.SE
WW.4.SE
WD.5.SE
WW.5.SE
WD.1.SE+5d
WW.1.SE+5d
WD.2.SE+5d
WW.2.SE+5d
WD.3.SE+5d
WW.3.SE+5d
WD.4.SE+5d
WW.4.SE+5d
WD.5.SE+5d
WW.5.SE+5d

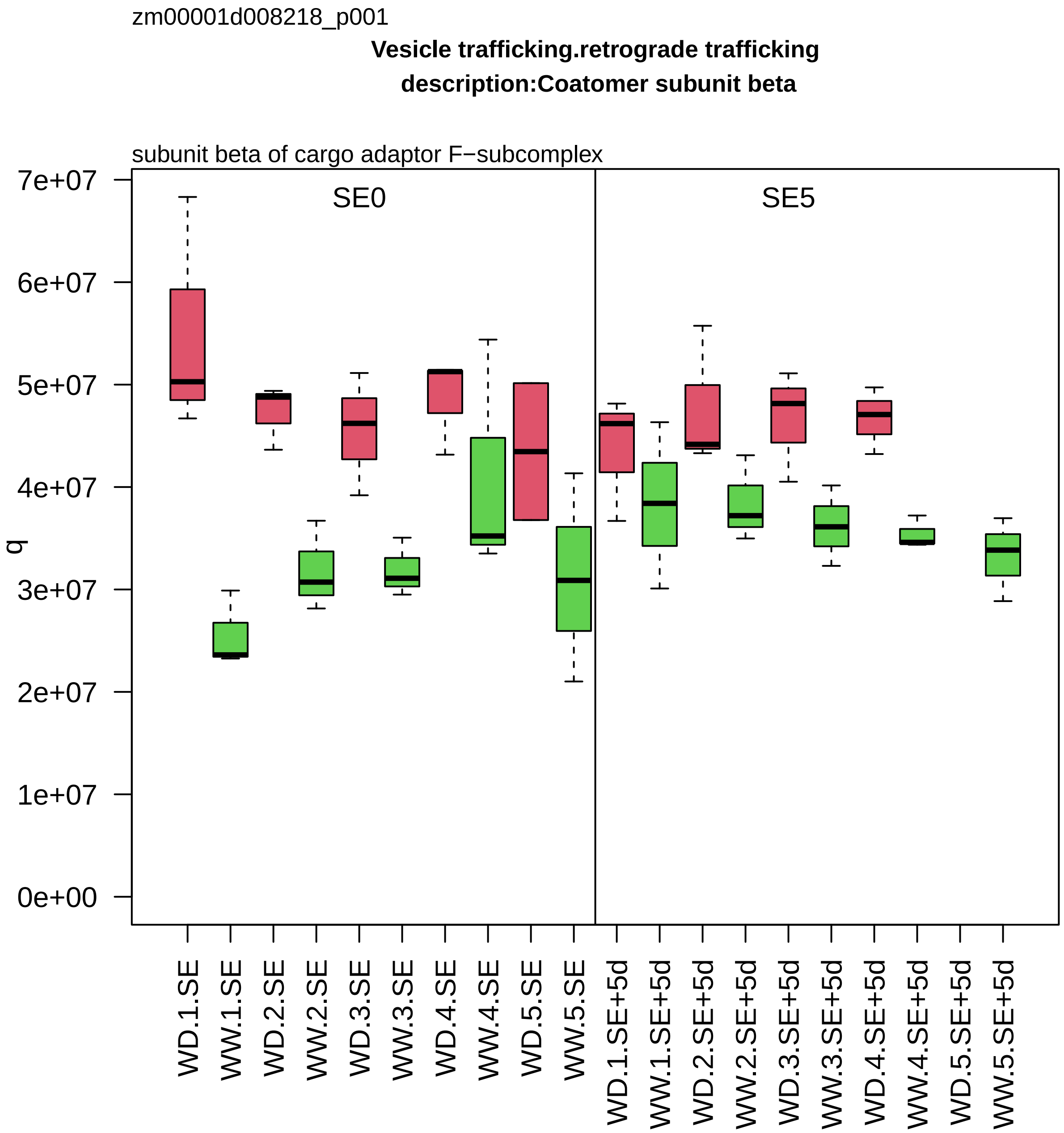
zm00001d008218_p001
Vesicle trafficking.retrograde trafficking
description:Coatomer subunit beta
subunit beta of cargo adaptor F−subcomplex
SE0
SE5
q
7e+07
6e+07
5e+07
4e+07
3e+07
2e+07
1e+07
0e+00
WD.1.SE
WW.1.SE
WD.2.SE
WW.2.SE
WD.3.SE
WW.3.SE
WD.4.SE
WW.4.SE
WD.5.SE
WW.5.SE
WD.1.SE+5d
WW.1.SE+5d
WD.2.SE+5d
WW.2.SE+5d
WD.3.SE+5d
WW.3.SE+5d
WD.4.SE+5d
WW.4.SE+5d
WD.5.SE+5d
WW.5.SE+5d

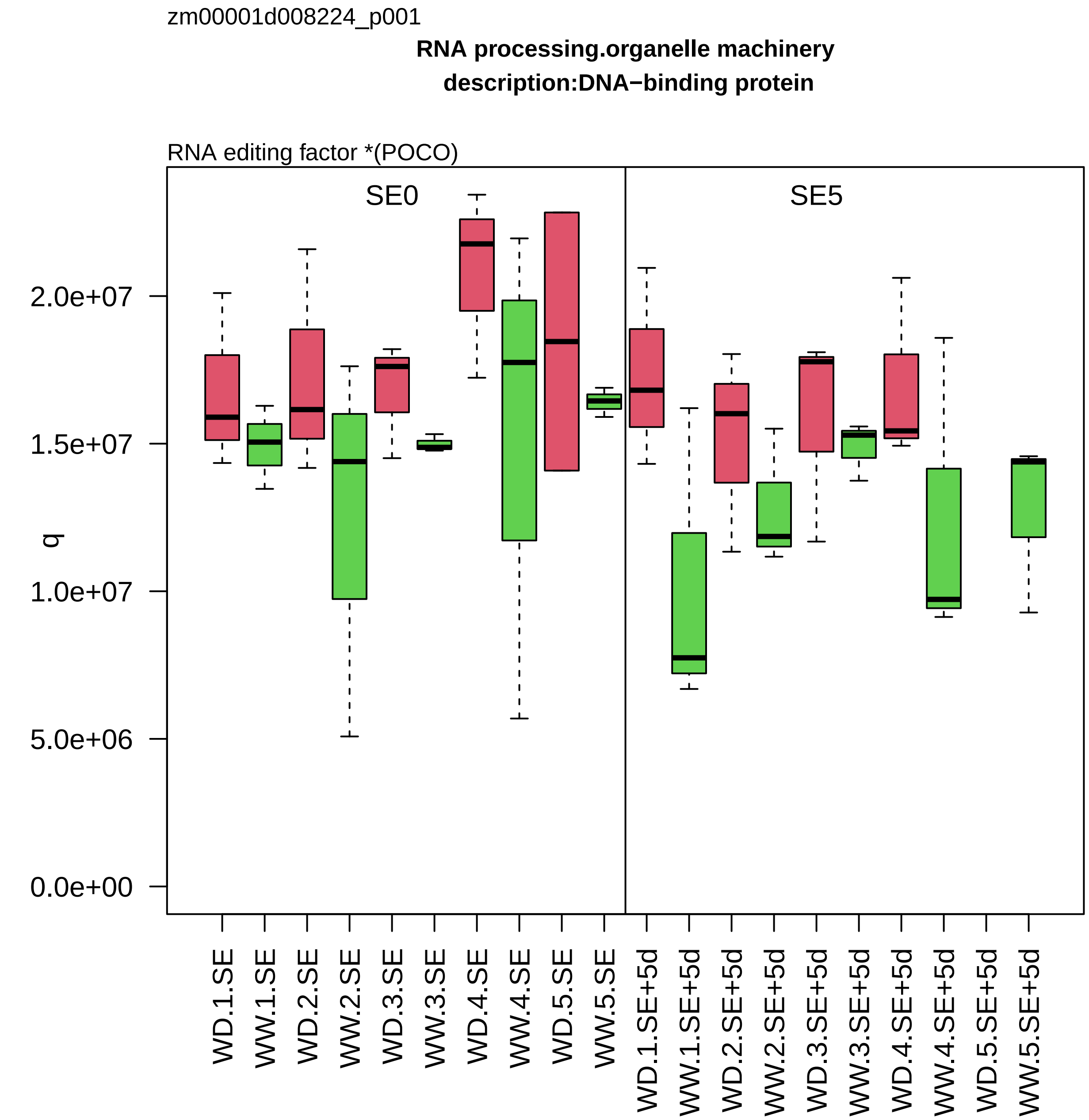

zm00001d008224_p001
RNA processing.organelle machinery
description:DNA−binding protein
RNA editing factor *(POCO)
SE0
SE5
q
2.0e+07
1.5e+07
1.0e+07
5.0e+06
0.0e+00
WD.1.SE
WW.1.SE
WD.2.SE
WW.2.SE
WD.3.SE
WW.3.SE
WD.4.SE
WW.4.SE
WD.5.SE
WW.5.SE
WD.1.SE+5d
WW.1.SE+5d
WD.2.SE+5d
WW.2.SE+5d
WD.3.SE+5d
WW.3.SE+5d
WD.4.SE+5d
WW.4.SE+5d
WD.5.SE+5d
WW.5.SE+5d

zm00001d008242_p001

**Protein translocation.mitochondrion**

**description:Mitochondrial import receptor subunit TOM40−1**

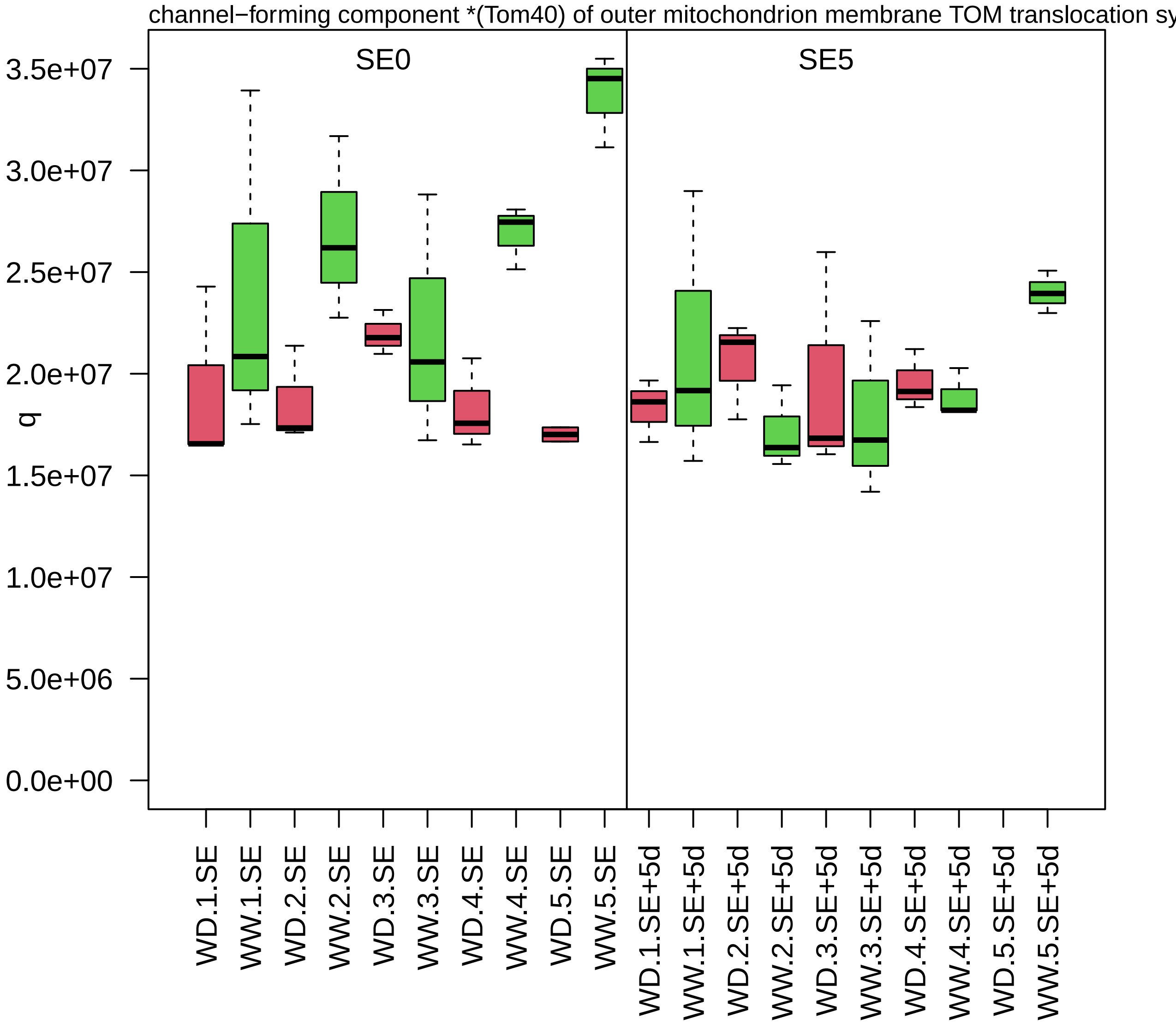

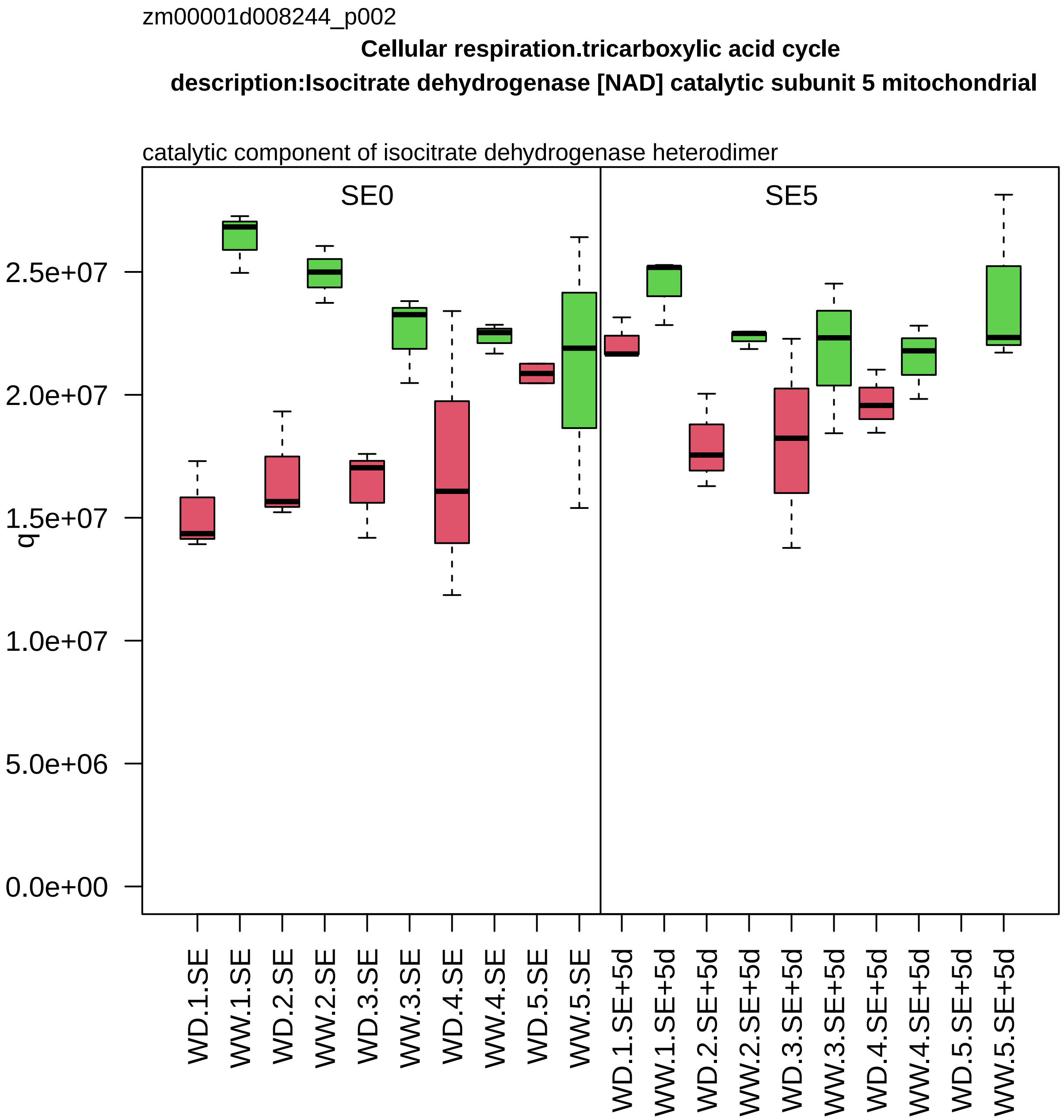

zm00001d008244_p002
Cellular respiration.tricarboxylic acid cycle
description:Isocitrate dehydrogenase [NAD] catalytic subunit 5 mitochondrial
catalytic component of isocitrate dehydrogenase heterodimer
SE0
SE5
q
2.5e+07
2.0e+07
1.5e+07
1.0e+07
5.0e+06
0.0e+00
WD.1.SE
WW.1.SE
WD.2.SE
WW.2.SE
WD.3.SE
WW.3.SE
WD.4.SE
WW.4.SE
WD.5.SE
WW.5.SE
WD.1.SE+5d
WW.1.SE+5d
WD.2.SE+5d
WW.2.SE+5d
WD.3.SE+5d
WW.3.SE+5d
WD.4.SE+5d
WW.4.SE+5d
WD.5.SE+5d
WW.5.SE+5d

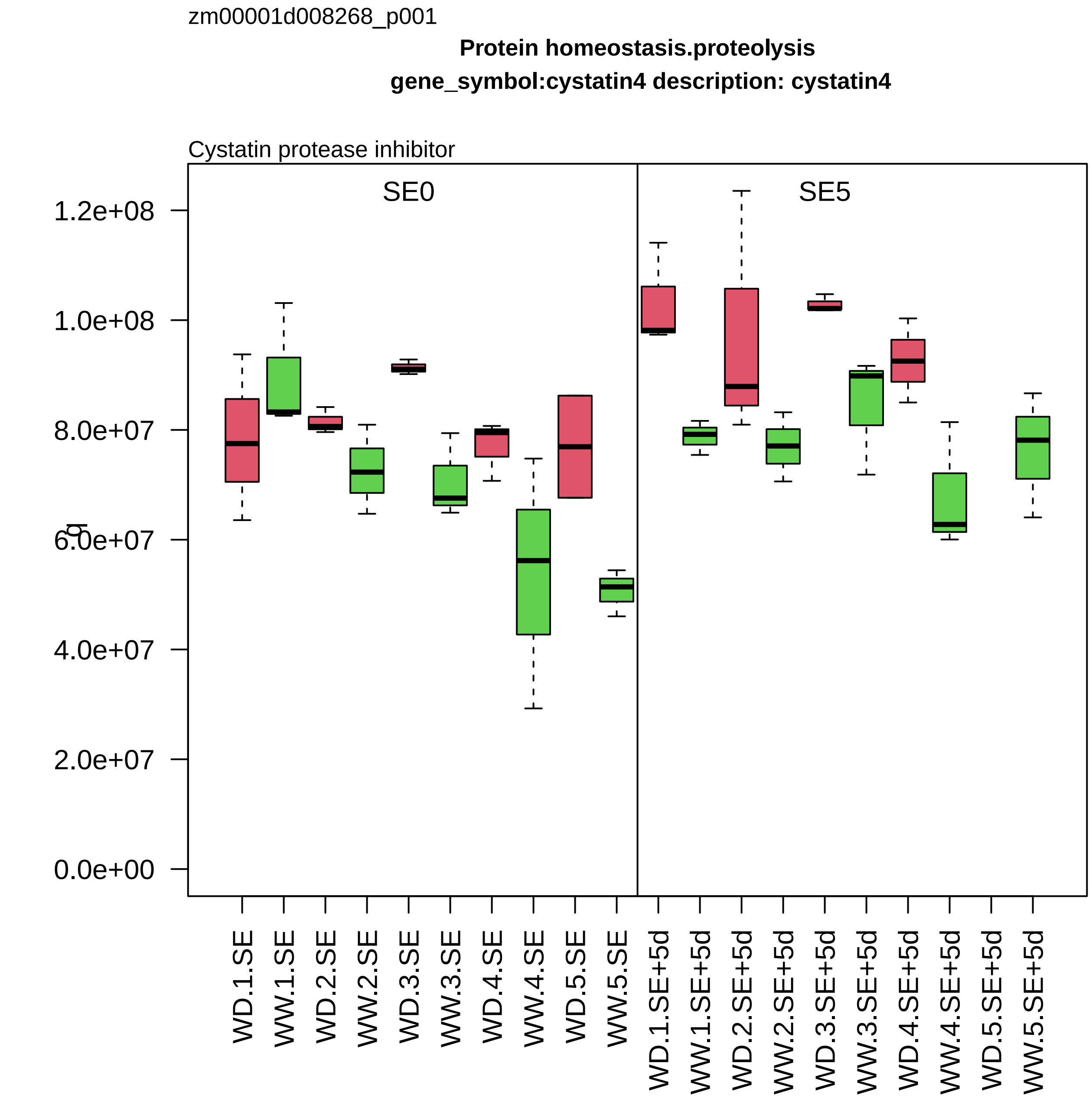
zm00001d008268_p001
Protein homeostasis.proteolysis
gene_symbol:cystatin4 description: cystatin4
Cystatin protease inhibitor
SE0
SE5
1.2e+08
1.0e+08
8.0e+07
6.0e+07
4.0e+07
2.0e+07
0.0e+00
WD.1.SE
WW.1.SE
WD.2.SE
WW.2.SE
WD.3.SE
WW.3.SE
WD.4.SE
WW.4.SE
WD.5.SE
WW.5.SE
WD.1.SE+5d
WW.1.SE+5d
WD.2.SE+5d
WW.2.SE+5d
WD.3.SE+5d
WW.3.SE+5d
WD.4.SE+5d
WW.4.SE+5d
WD.5.SE+5d
WW.5.SE+5d

zm00001d008284_p004

**Protein homeostasis.ubiquitin−proteasome system**

**description:26S proteasome non−ATPase regulatory subunit 14 homolog**

regulatory component *(RPN11) of 26S proteasome

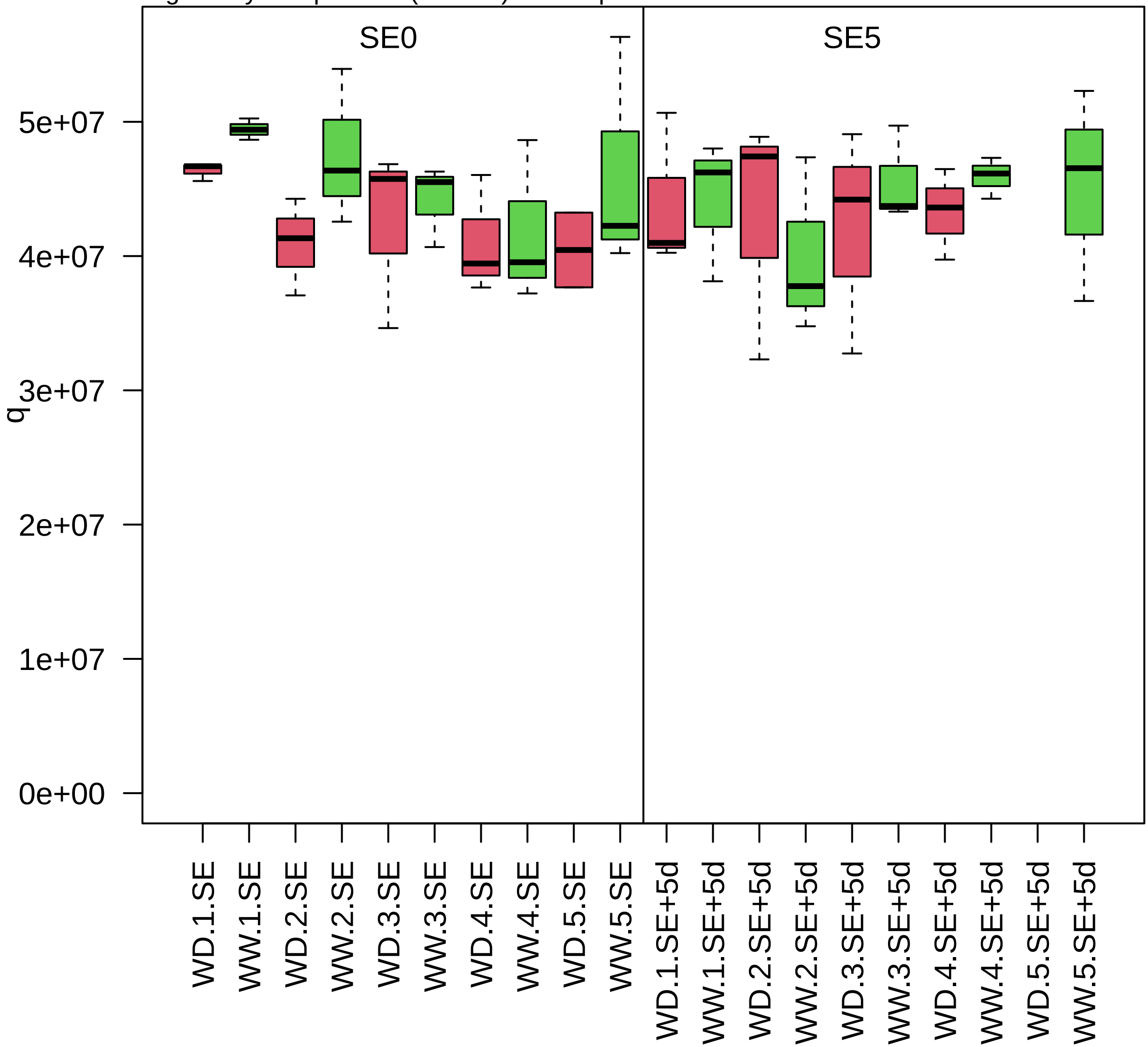

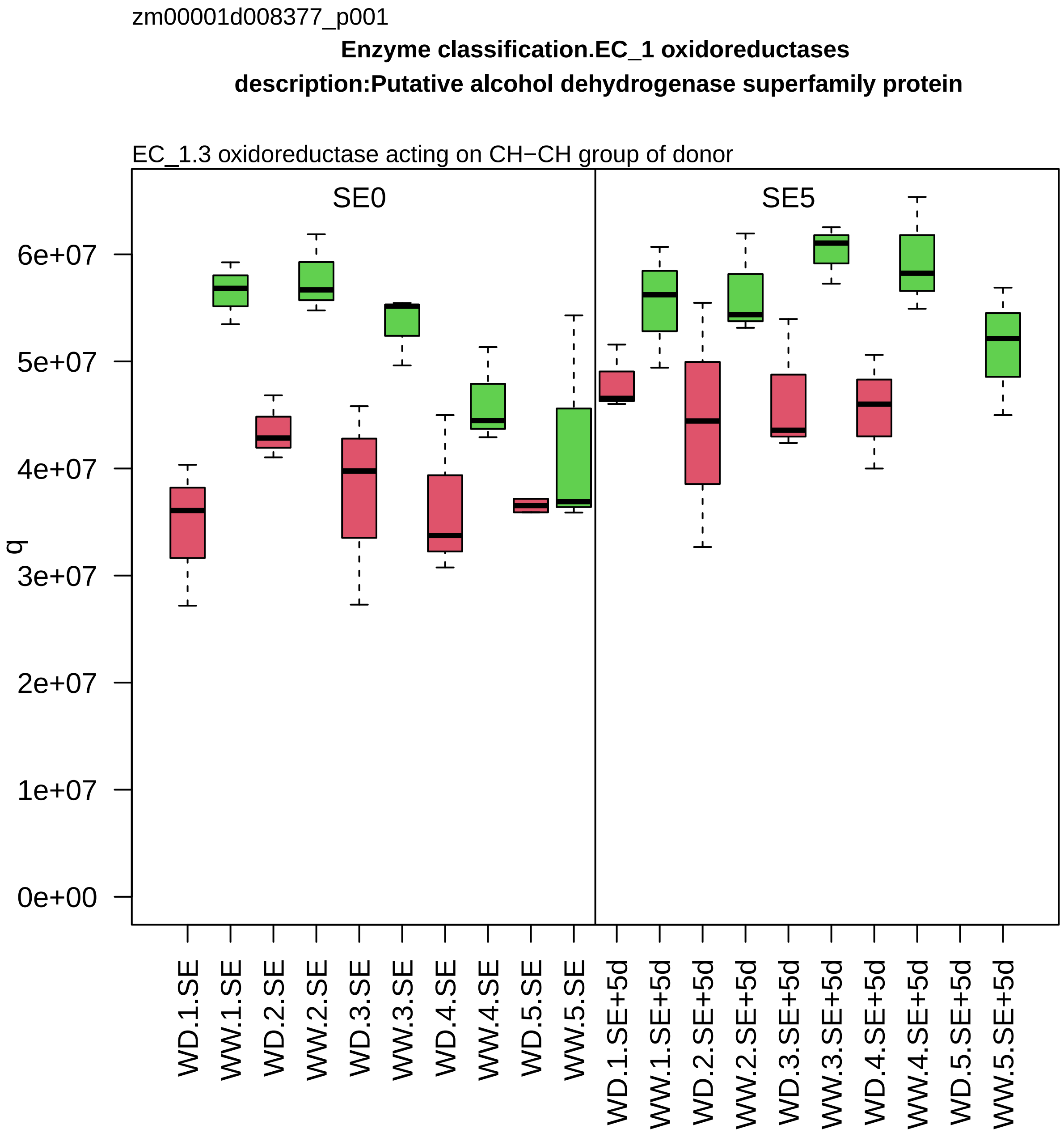

zm00001d008377_p001
Enzyme classification.EC_1 oxidoreductases
description:Putative alcohol dehydrogenase superfamily protein
EC_1.3 oxidoreductase acting on CH−CH group of donor
SE0
SE5
q
0e+00
1e+07
2e+07
3e+07
4e+07
5e+07
6e+07
WD.1.SE
WW.1.SE
WD.2.SE
WW.2.SE
WD.3.SE
WW.3.SE
WD.4.SE
WW.4.SE
WD.5.SE
WW.5.SE
WD.1.SE+5d
WW.1.SE+5d
WD.2.SE+5d
WW.2.SE+5d
WD.3.SE+5d
WW.3.SE+5d
WD.4.SE+5d
WW.4.SE+5d
WD.5.SE+5d
WW.5.SE+5d

zm00001d008387_p001

**Protein biosynthesis.ribosome biogenesis**

**gene_symbol:ribosomal protein S5 description: ribosomal protein S5**

component *(uS7) of small ribosomal−subunit (SSU) proteome

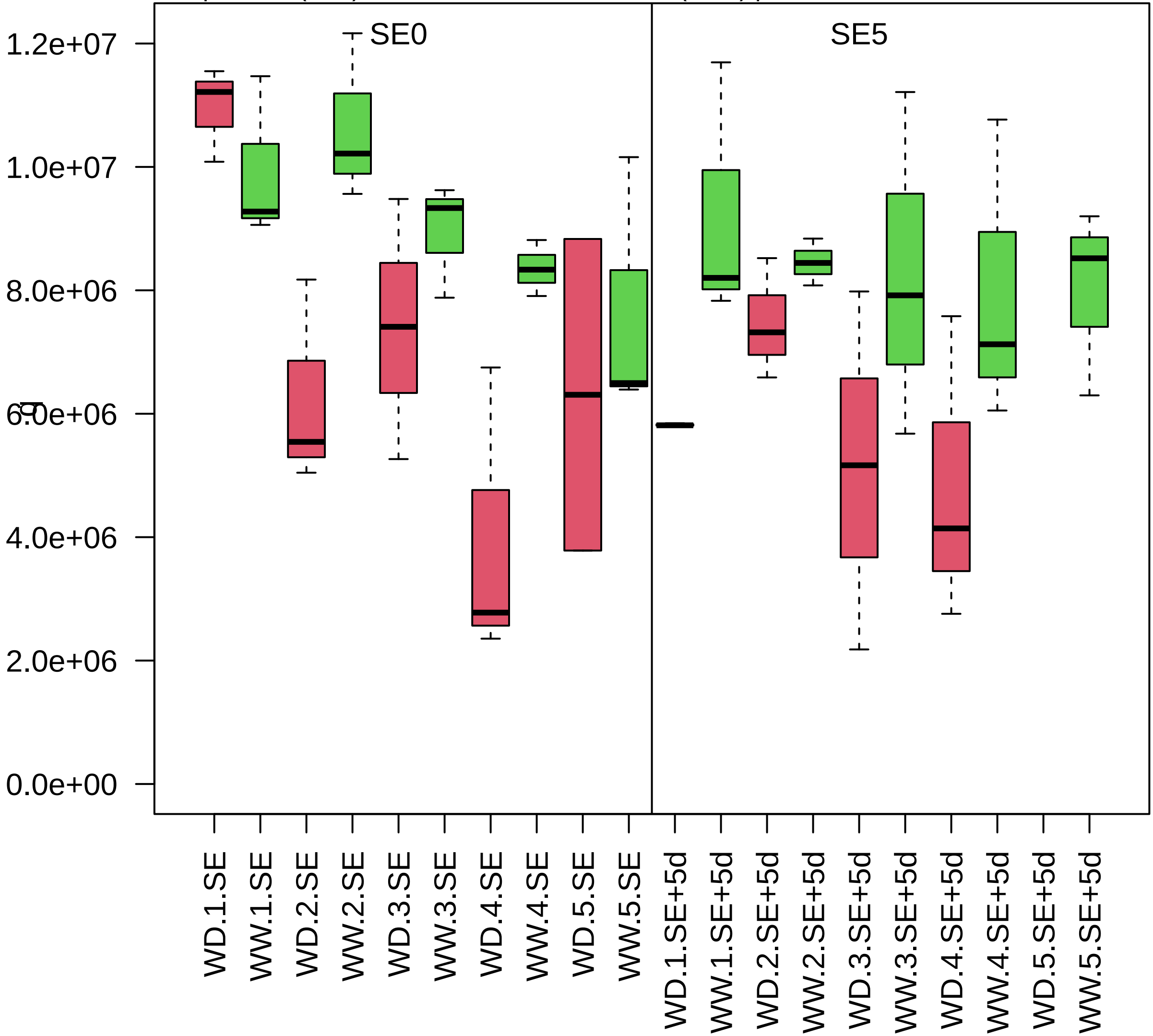

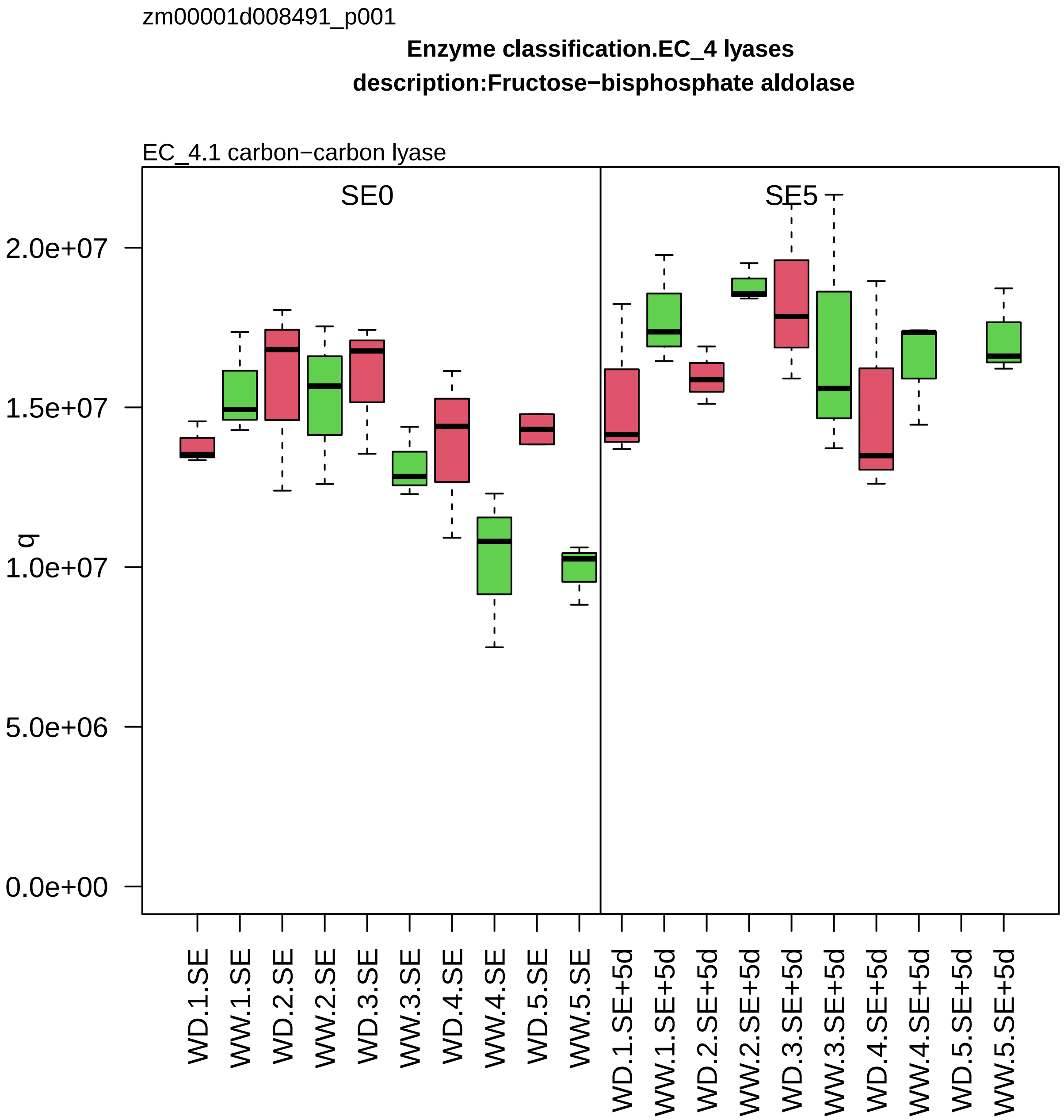
zm00001d008491_p001
Enzyme classification.EC_4 lyases
description:Fructose−bisphosphate aldolase
EC_4.1 carbon−carbon lyase
SE0
SE5
q
2.0e+07
1.5e+07
1.0e+07
5.0e+06
0.0e+00
WD.1.SE
WW.1.SE
WD.2.SE
WW.2.SE
WD.3.SE
WW.3.SE
WD.4.SE
WW.4.SE
WD.5.SE
WW.5.SE
WD.1.SE+5d
WW.1.SE+5d
WD.2.SE+5d
WW.2.SE+5d
WD.3.SE+5d
WW.3.SE+5d
WD.4.SE+5d
WW.4.SE+5d
WD.5.SE+5d
WW.5.SE+5d

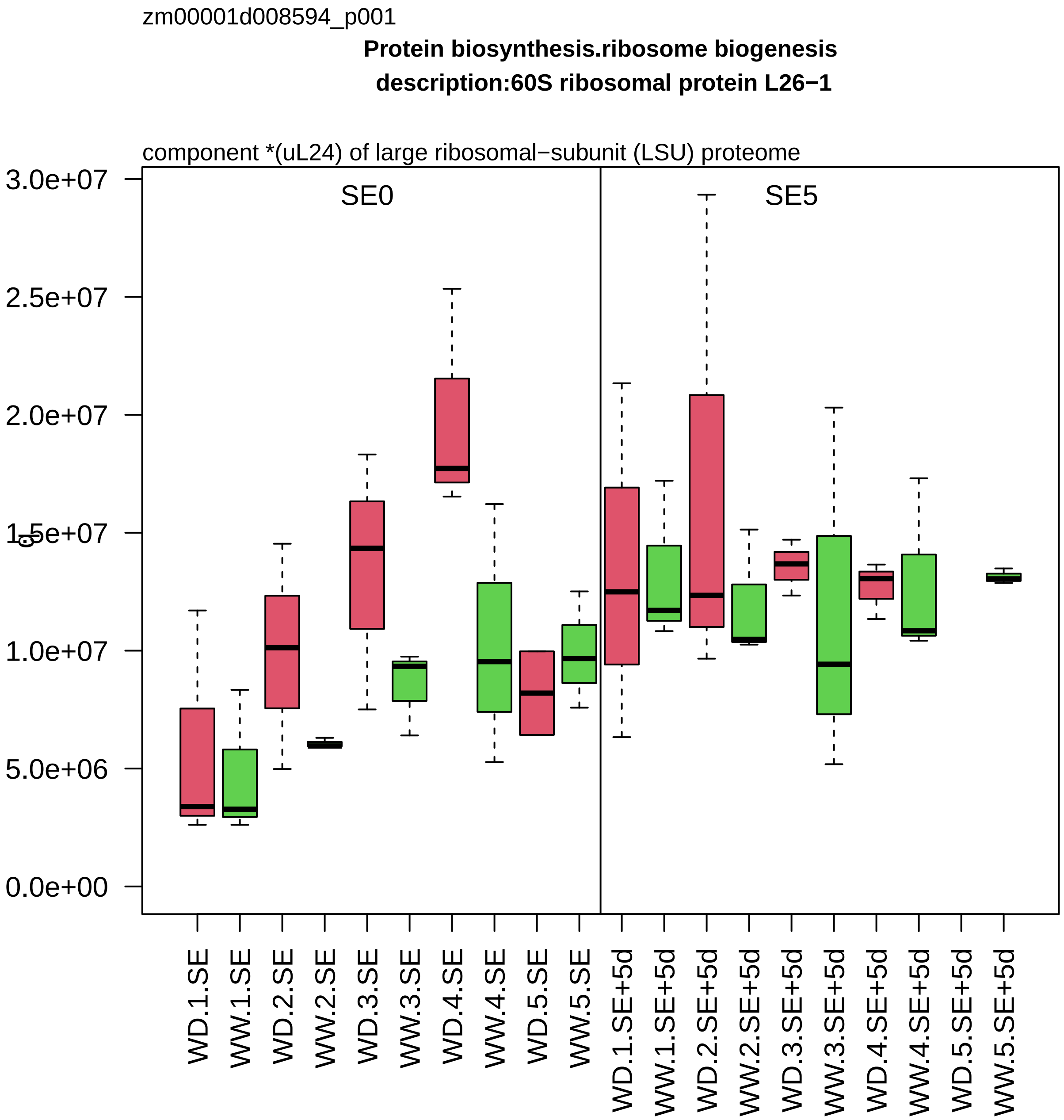

zm00001d008594_p001
Protein biosynthesis.ribosome biogenesis
description:60S ribosomal protein L26−1
component *(uL24) of large ribosomal−subunit (LSU) proteome
SE0
SE5
3.0e+07
2.5e+07
2.0e+07
1.5e+07
1.0e+07
5.0e+06
0.0e+00
WD.1.SE
WW.1.SE
WD.2.SE
WW.2.SE
WD.3.SE
WW.3.SE
WD.4.SE
WW.4.SE
WD.5.SE
WW.5.SE
WD.1.SE+5d
WW.1.SE+5d
WD.2.SE+5d
WW.2.SE+5d
WD.3.SE+5d
WW.3.SE+5d
WD.4.SE+5d
WW.4.SE+5d
WD.5.SE+5d
WW.5.SE+5d

zm00001d008619_p001

**Cellular respiration.glycolysis**

**gene_symbol:triose phosphate isomerase3 description: triose phosphate isomerase3**

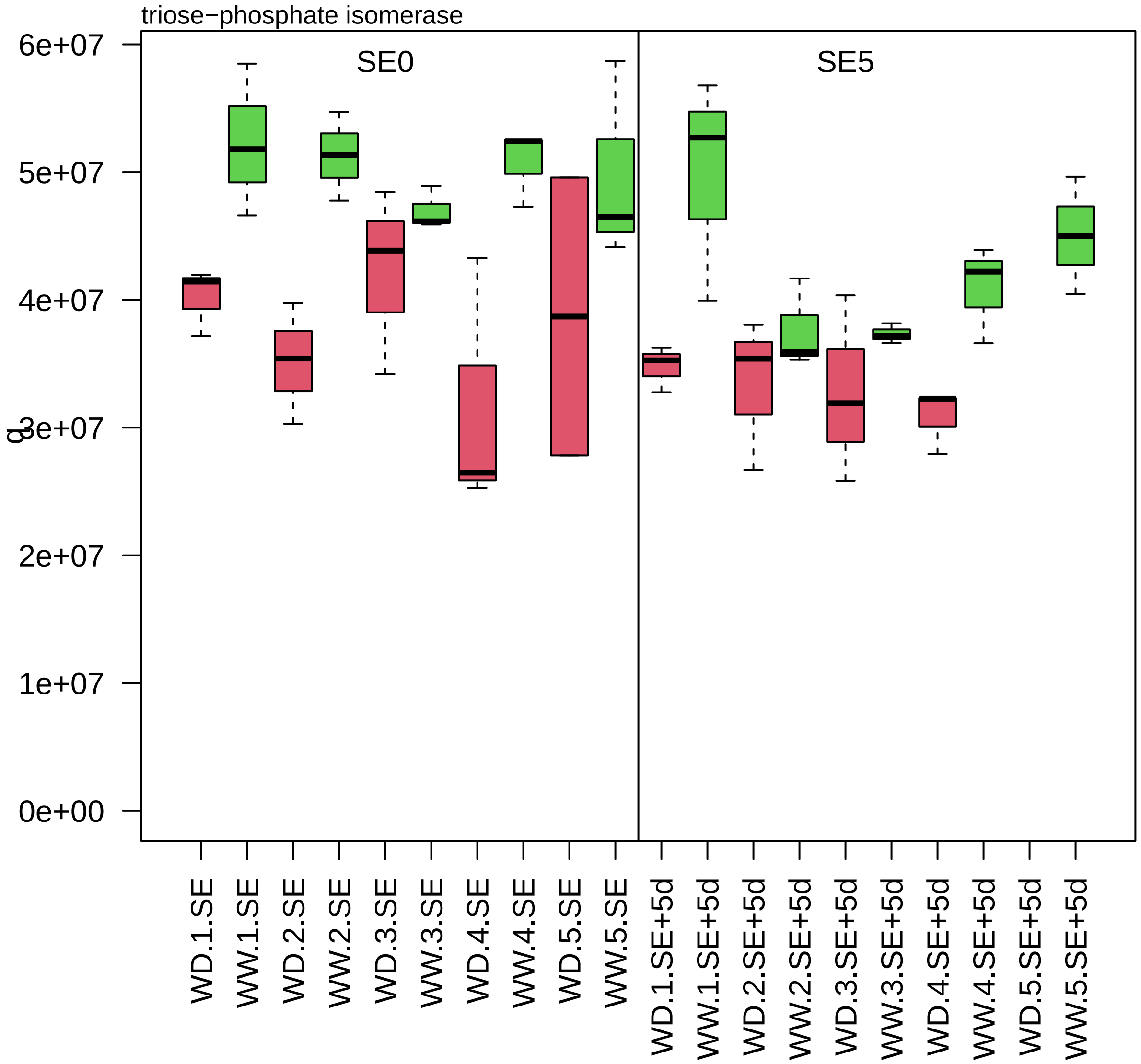

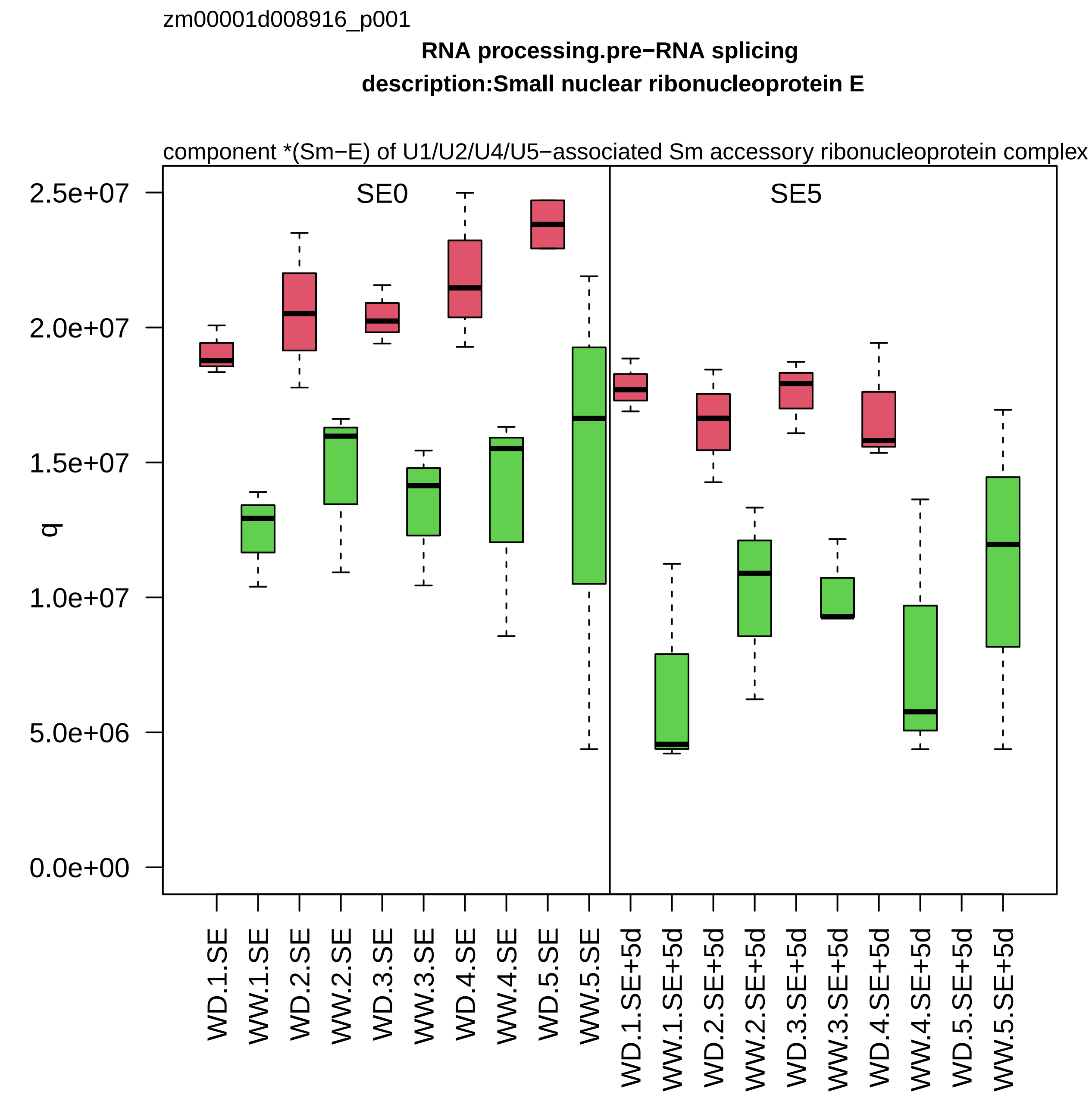

zm00001d008916_p001
RNA processing.pre−RNA splicing
description:Small nuclear ribonucleoprotein E
component *(Sm−E) of U1/U2/U4/U5−associated Sm accessory ribonucleoprotein complex
SE0
SE5
q
2.5e+07
2.0e+07
1.5e+07
1.0e+07
5.0e+06
0.0e+00
WD.1.SE
WW.1.SE
WD.2.SE
WW.2.SE
WD.3.SE
WW.3.SE
WD.4.SE
WW.4.SE
WD.5.SE
WW.5.SE
WD.1.SE+5d
WW.1.SE+5d
WD.2.SE+5d
WW.2.SE+5d
WD.3.SE+5d
WW.3.SE+5d
WD.4.SE+5d
WW.4.SE+5d
WD.5.SE+5d
WW.5.SE+5d

zm00001d009000_p001

**Protein biosynthesis.ribosome biogenesis**
**description:40S ribosomal protein S15a−1**

component *(uS8) of small ribosomal−subunit (SSU) proteome

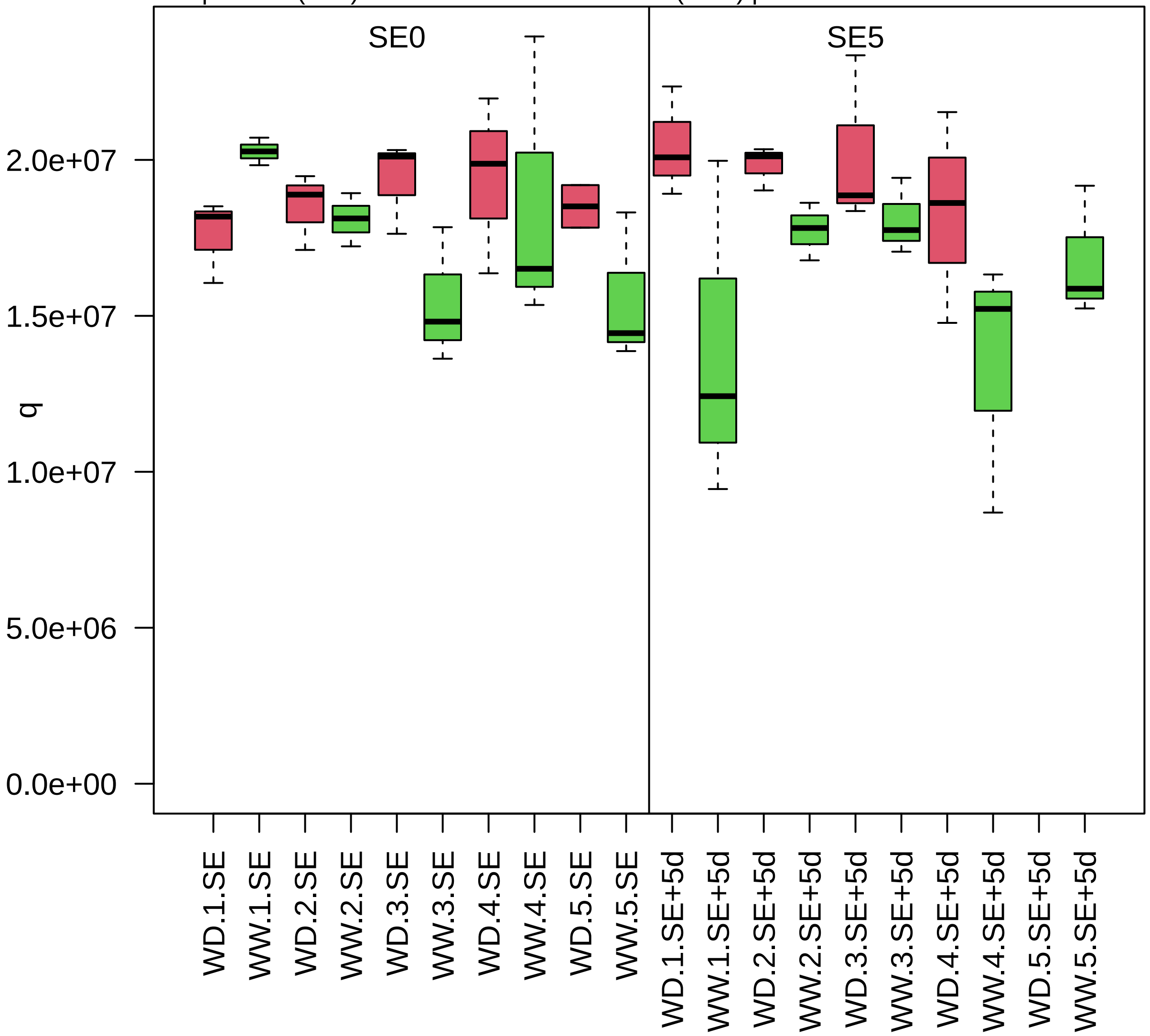

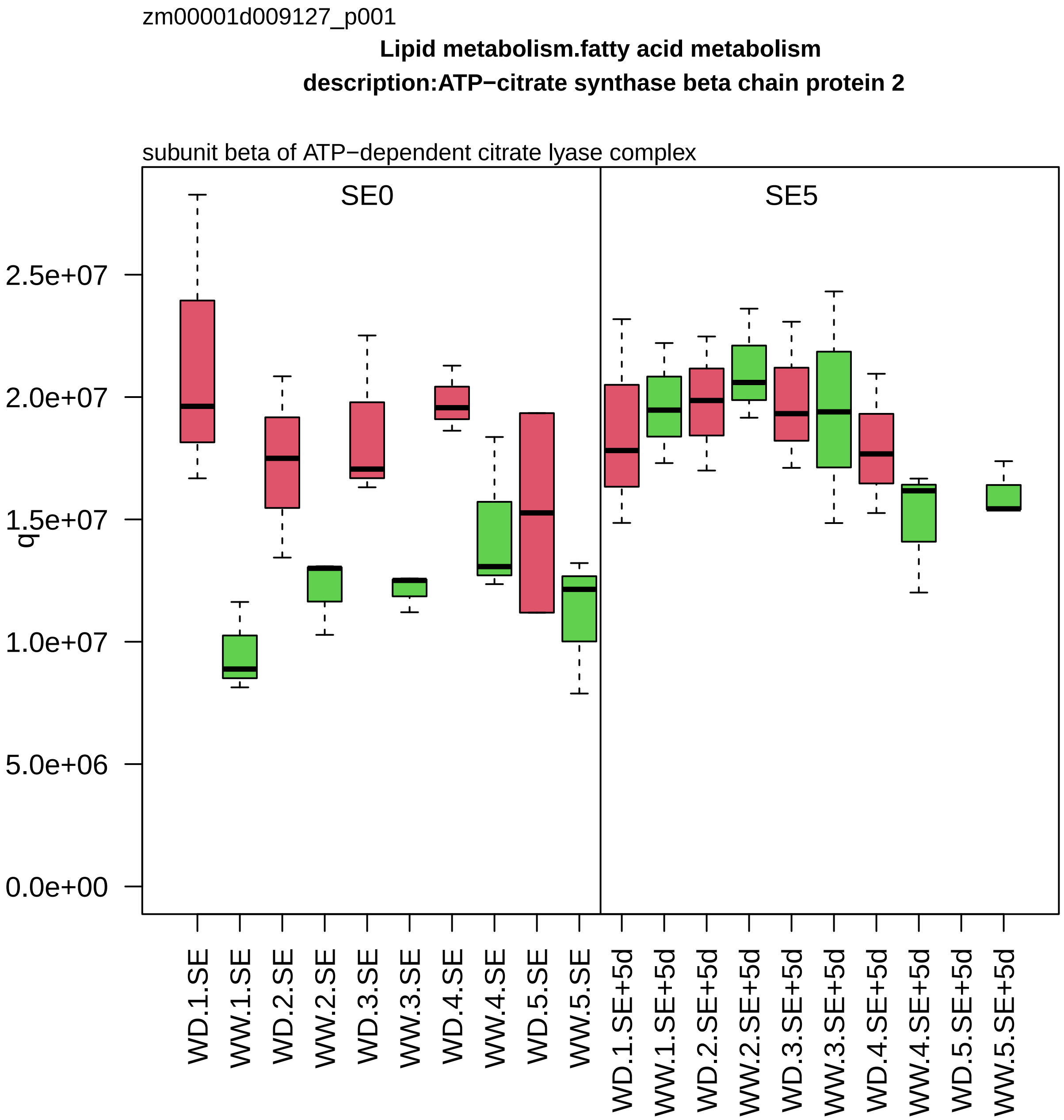

zm00001d009127_p001
Lipid metabolism.fatty acid metabolism
description:ATP−citrate synthase beta chain protein 2
subunit beta of ATP−dependent citrate lyase complex
SE0
SE5
q
2.5e+07
2.0e+07
1.5e+07
1.0e+07
5.0e+06
0.0e+00
WD.1.SE
WW.1.SE
WD.2.SE
WW.2.SE
WD.3.SE
WW.3.SE
WD.4.SE
WW.4.SE
WD.5.SE
WW.5.SE
WD.1.SE+5d
WW.1.SE+5d
WD.2.SE+5d
WW.2.SE+5d
WD.3.SE+5d
WW.3.SE+5d
WD.4.SE+5d
WW.4.SE+5d
WD.5.SE+5d
WW.5.SE+5d

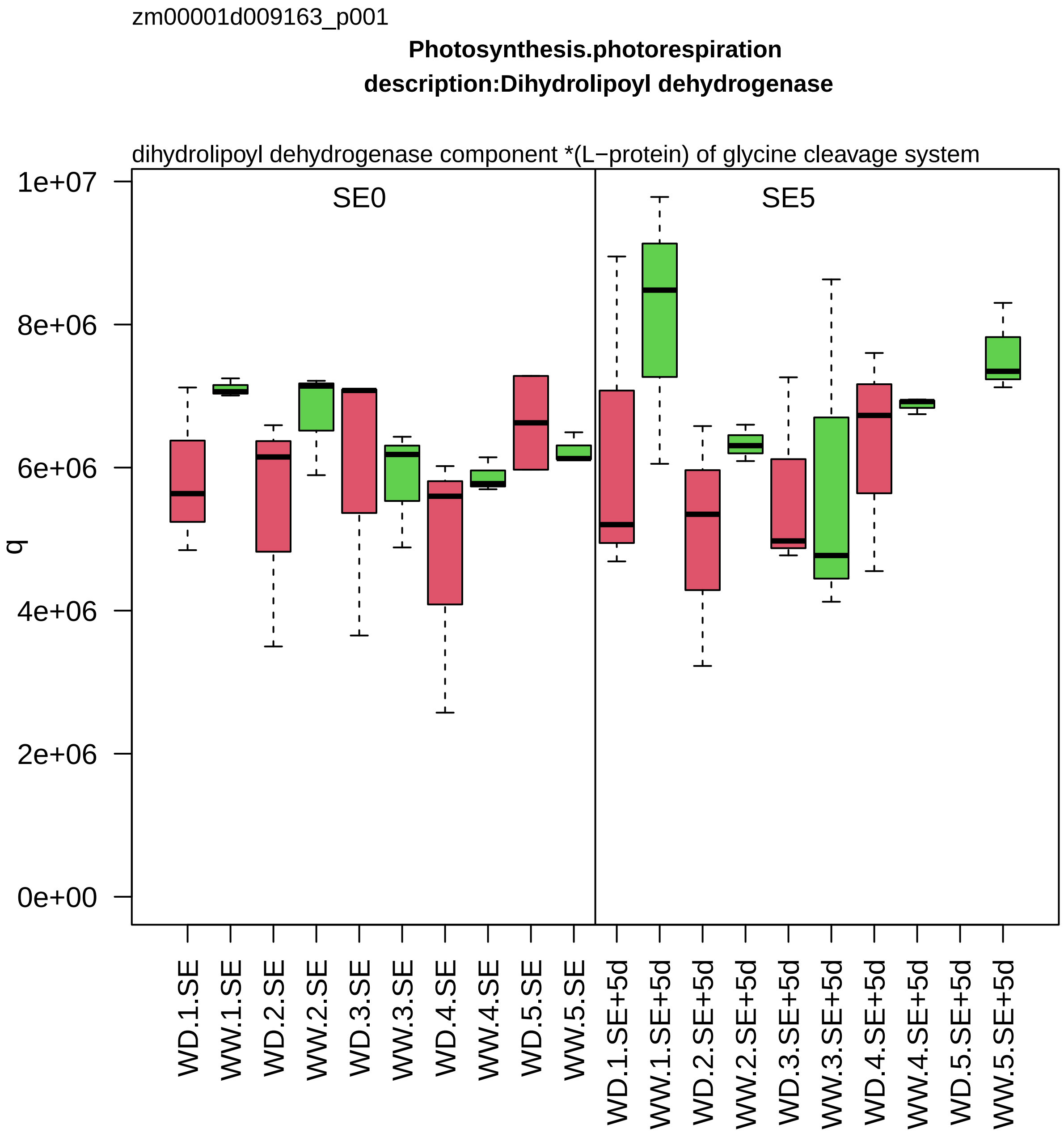

zm00001d009163_p001
Photosynthesis.photorespiration
description:Dihydrolipoyl dehydrogenase
dihydrolipoyl dehydrogenase component *(L−protein) of glycine cleavage system
SE0
SE5
q
1e+07
8e+06
6e+06
4e+06
2e+06
0e+00
WD.1.SE
WW.1.SE
WD.2.SE
WW.2.SE
WD.3.SE
WW.3.SE
WD.4.SE
WW.4.SE
WD.5.SE
WW.5.SE
WD.1.SE+5d
WW.1.SE+5d
WD.2.SE+5d
WW.2.SE+5d
WD.3.SE+5d
WW.3.SE+5d
WD.4.SE+5d
WW.4.SE+5d
WD.5.SE+5d
WW.5.SE+5d

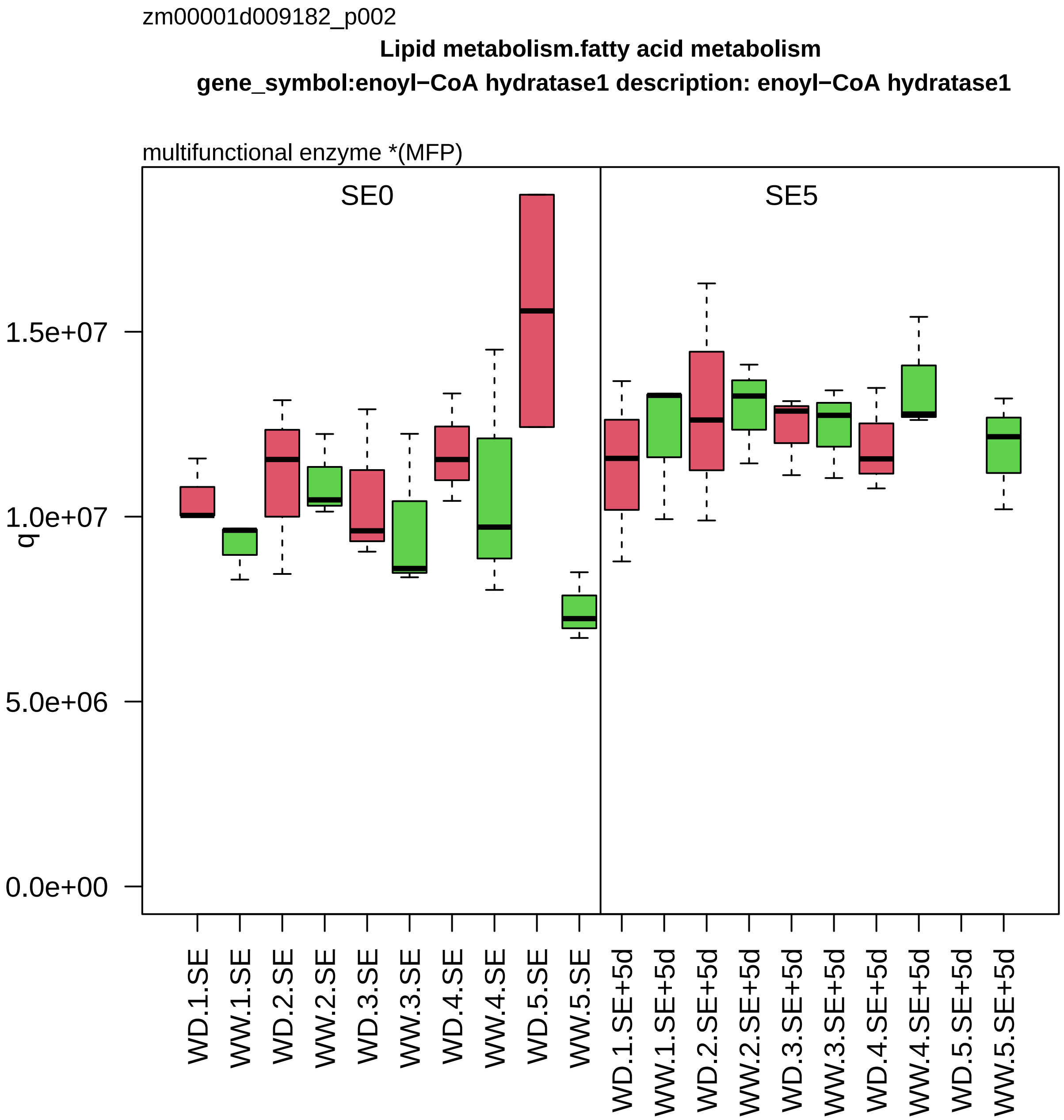
zm00001d009182_p002
Lipid metabolism.fatty acid metabolism
gene_symbol:enoyl−CoA hydratase1 description: enoyl−CoA hydratase1
multifunctional enzyme *(MFP)
SE0
SE5
q
1.5e+07
1.0e+07
5.0e+06
0.0e+00
WD.1.SE
WW.1.SE
WD.2.SE
WW.2.SE
WD.3.SE
WW.3.SE
WD.4.SE
WW.4.SE
WD.5.SE
WW.5.SE
WD.1.SE+5d
WW.1.SE+5d
WD.2.SE+5d
WW.2.SE+5d
WD.3.SE+5d
WW.3.SE+5d
WD.4.SE+5d
WW.4.SE+5d
WD.5.SE+5d
WW.5.SE+5d

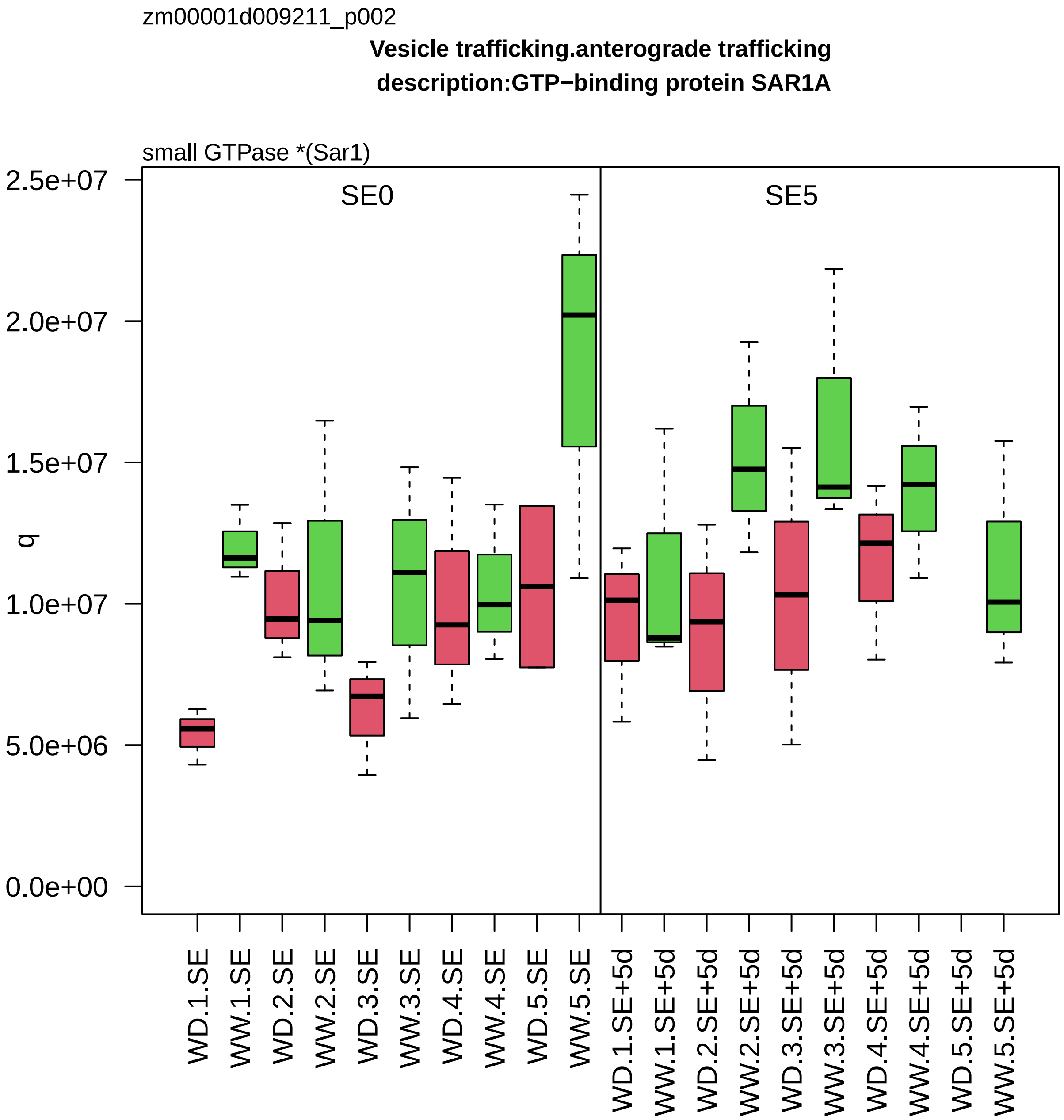

zm00001d009211_p002
Vesicle trafficking.anterograde trafficking
description:GTP−binding protein SAR1A
small GTPase *(Sar1)
q
2.5e+07
2.0e+07
1.5e+07
1.0e+07
5.0e+06
0.0e+00
SE0
SE5
WD.1.SE
WW.1.SE
WD.2.SE
WW.2.SE
WD.3.SE
WW.3.SE
WD.4.SE
WW.4.SE
WD.5.SE
WW.5.SE
WD.1.SE+5d
WW.1.SE+5d
WD.2.SE+5d
WW.2.SE+5d
WD.3.SE+5d
WW.3.SE+5d
WD.4.SE+5d
WW.4.SE+5d
WD.5.SE+5d
WW.5.SE+5d

zm00001d009271_p001

**Protein biosynthesis.aminoacyl−tRNA synthetase activities**
**description:Asparaginyl−tRNA synthetase%2C cytoplasmic 3**

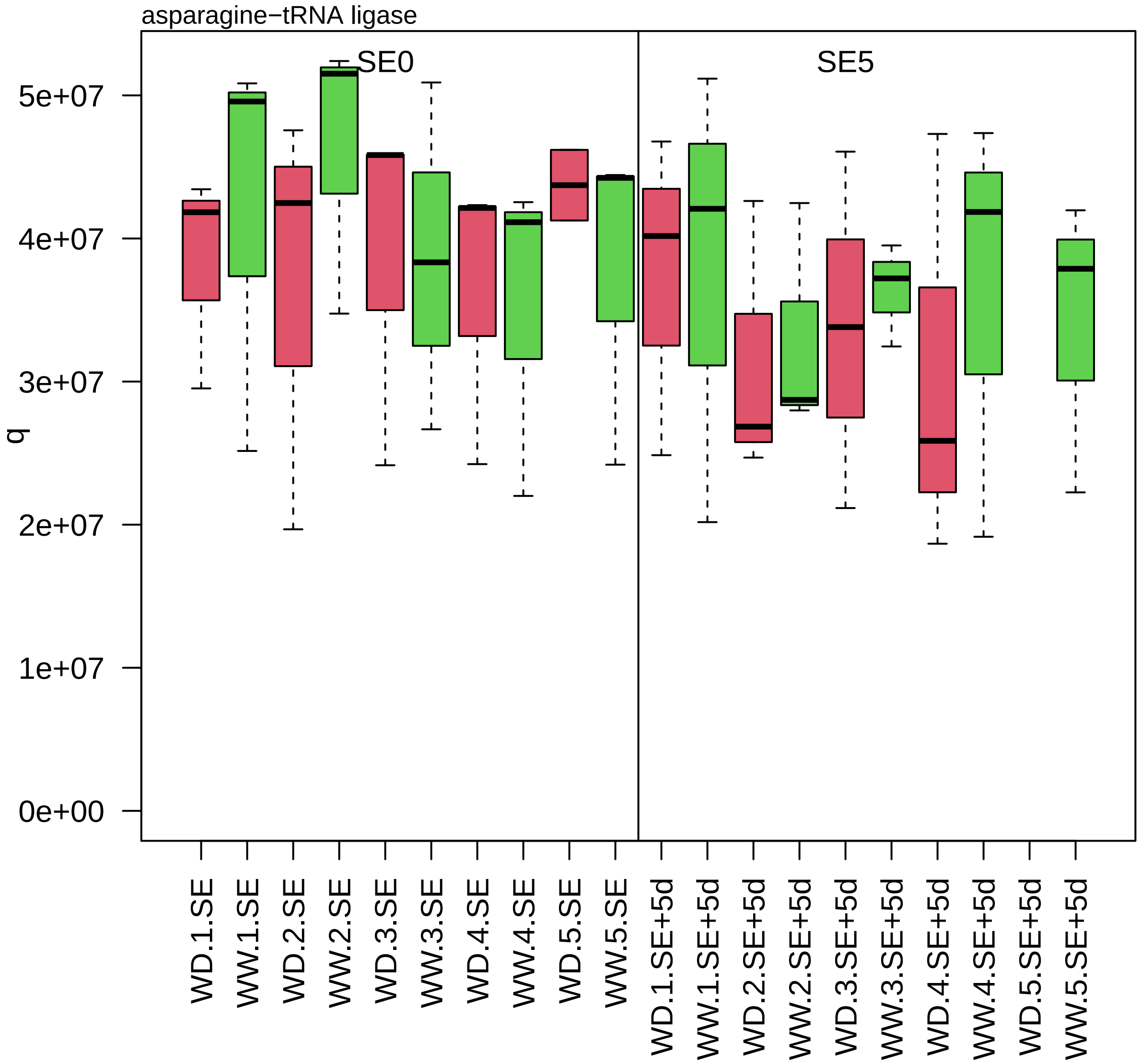

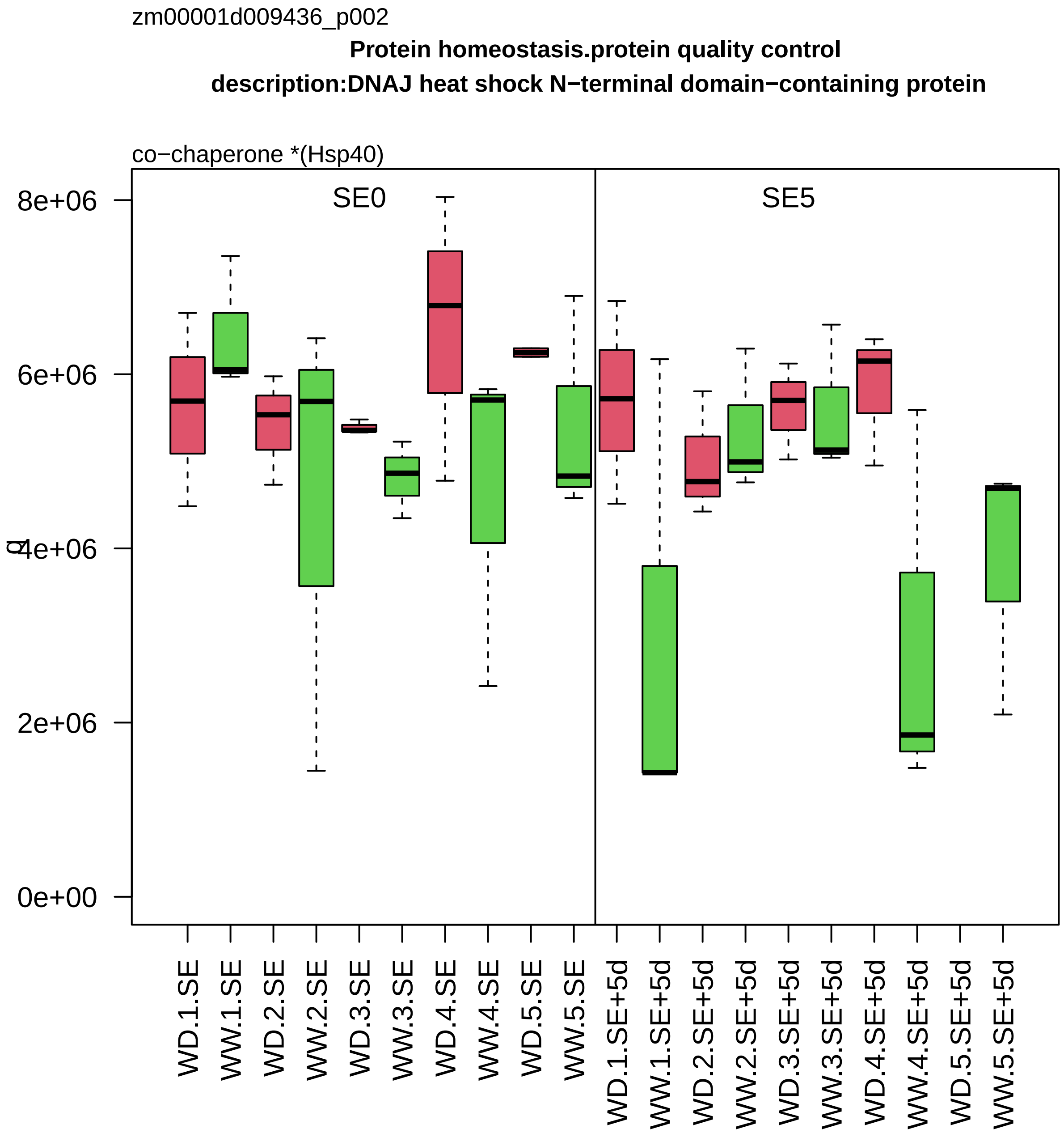

zm00001d009436_p002
Protein homeostasis.protein quality control
description:DNAJ heat shock N−terminal domain−containing protein
co−chaperone *(Hsp40)
SE0
SE5
8e+06
6e+06
4e+06
2e+06
0e+00
WD.1.SE
WW.1.SE
WD.2.SE
WW.2.SE
WD.3.SE
WW.3.SE
WD.4.SE
WW.4.SE
WD.5.SE
WW.5.SE
WD.1.SE+5d
WW.1.SE+5d
WD.2.SE+5d
WW.2.SE+5d
WD.3.SE+5d
WW.3.SE+5d
WD.4.SE+5d
WW.4.SE+5d
WD.5.SE+5d
WW.5.SE+5d

zm00001d009480_p001

**Protein biosynthesis.ribosome biogenesis**

**gene_symbol:receptor for activated C kinase2 description:Receptor for activated C kinase 1A**

component *(RACK1) of small ribosomal−subunit (SSU) proteome

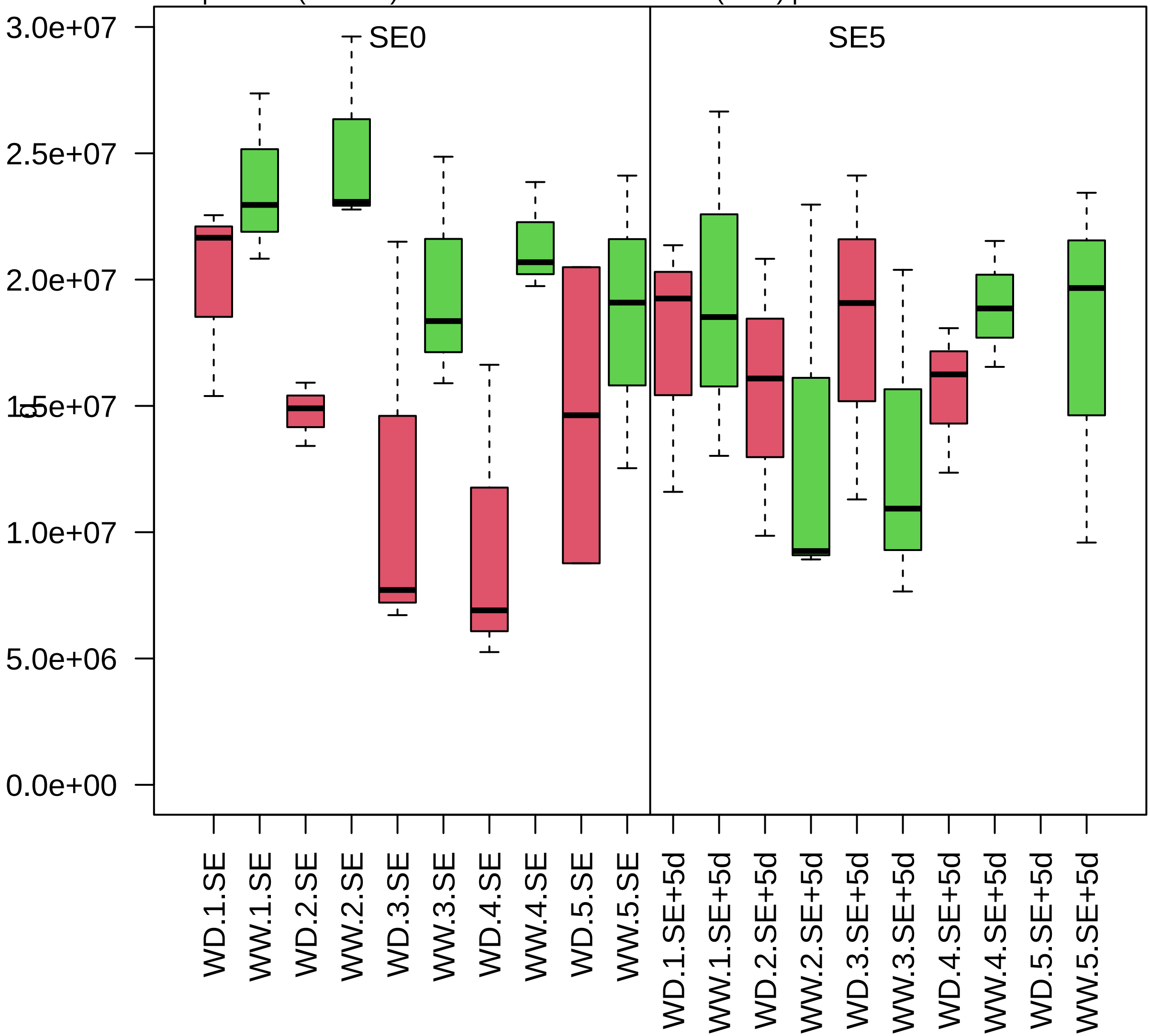

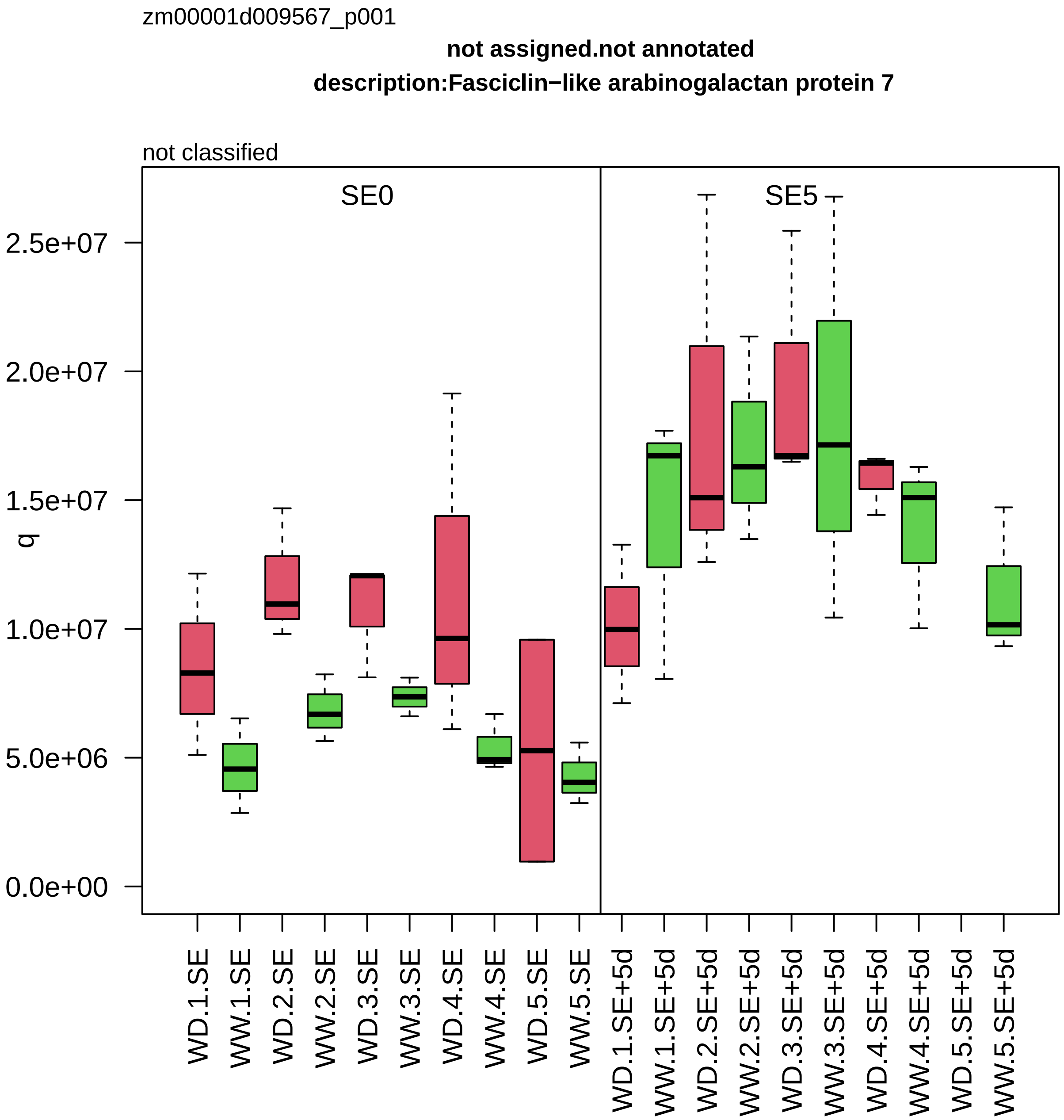

zm00001d009567_p001
not assigned.not annotated
description:Fasciclin−like arabinogalactan protein 7
not classified
SE0
SE5
q
2.5e+07
2.0e+07
1.5e+07
1.0e+07
5.0e+06
0.0e+00
WD.1.SE
WW.1.SE
WD.2.SE
WW.2.SE
WD.3.SE
WW.3.SE
WD.4.SE
WW.4.SE
WD.5.SE
WW.5.SE
WD.1.SE+5d
WW.1.SE+5d
WD.2.SE+5d
WW.2.SE+5d
WD.3.SE+5d
WW.3.SE+5d
WD.4.SE+5d
WW.4.SE+5d
WD.5.SE+5d
WW.5.SE+5d

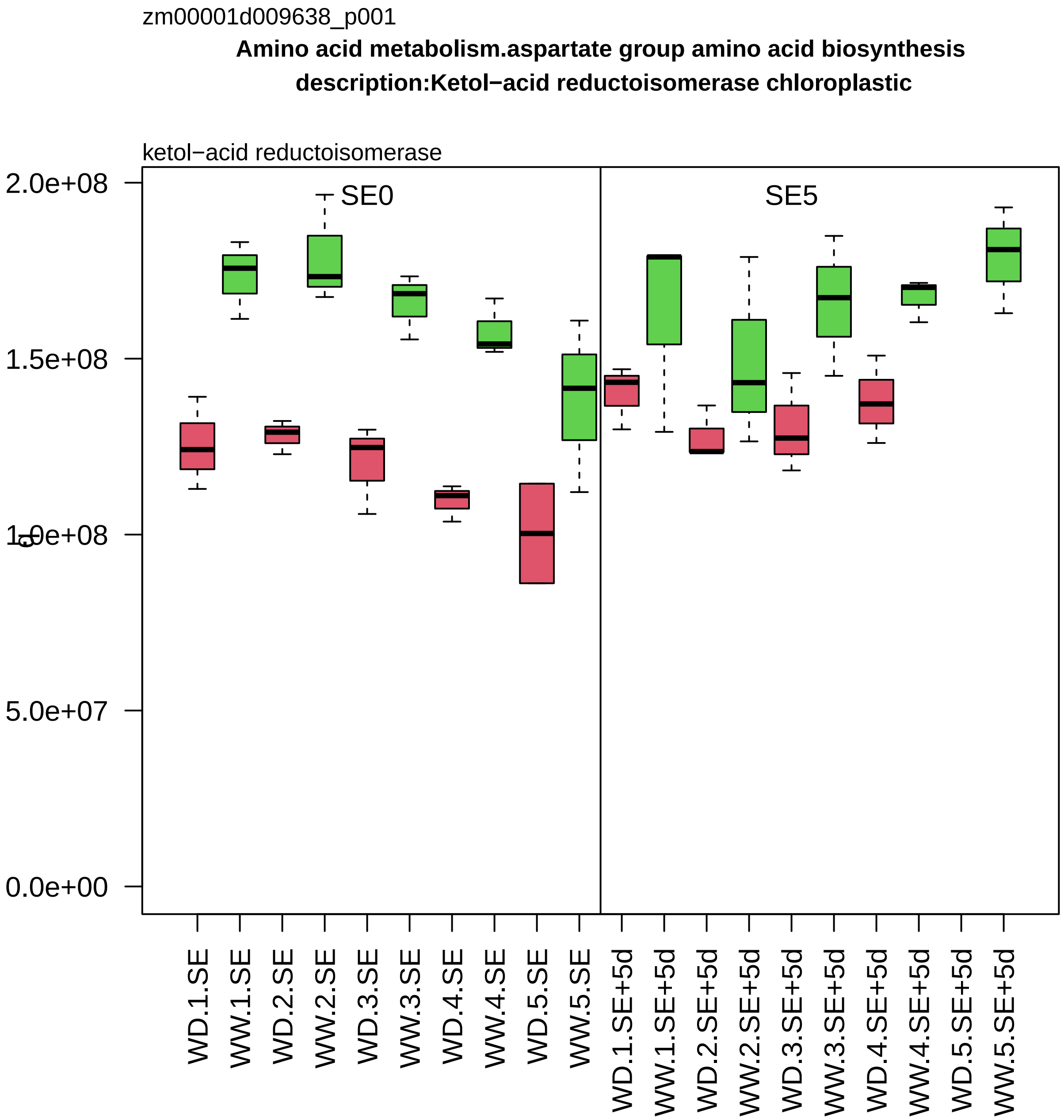
zm00001d009638_p001
Amino acid metabolism.aspartate group amino acid biosynthesis
description:Ketol−acid reductoisomerase chloroplastic
ketol−acid reductoisomerase
SE0
SE5
2.0e+08
1.5e+08
1.0e+08
5.0e+07
0.0e+00
WD.1.SE
WW.1.SE
WD.2.SE
WW.2.SE
WD.3.SE
WW.3.SE
WD.4.SE
WW.4.SE
WD.5.SE
WW.5.SE
WD.1.SE+5d
WW.1.SE+5d
WD.2.SE+5d
WW.2.SE+5d
WD.3.SE+5d
WW.3.SE+5d
WD.4.SE+5d
WW.4.SE+5d
WD.5.SE+5d
WW.5.SE+5d

zm00001d009740_p002

**RNA processing.RNA homeostasis**

**description:Plasminogen activator inhibitor 1 RNA−binding protein**

regulatory protein *(RGG) of mRNA quality control

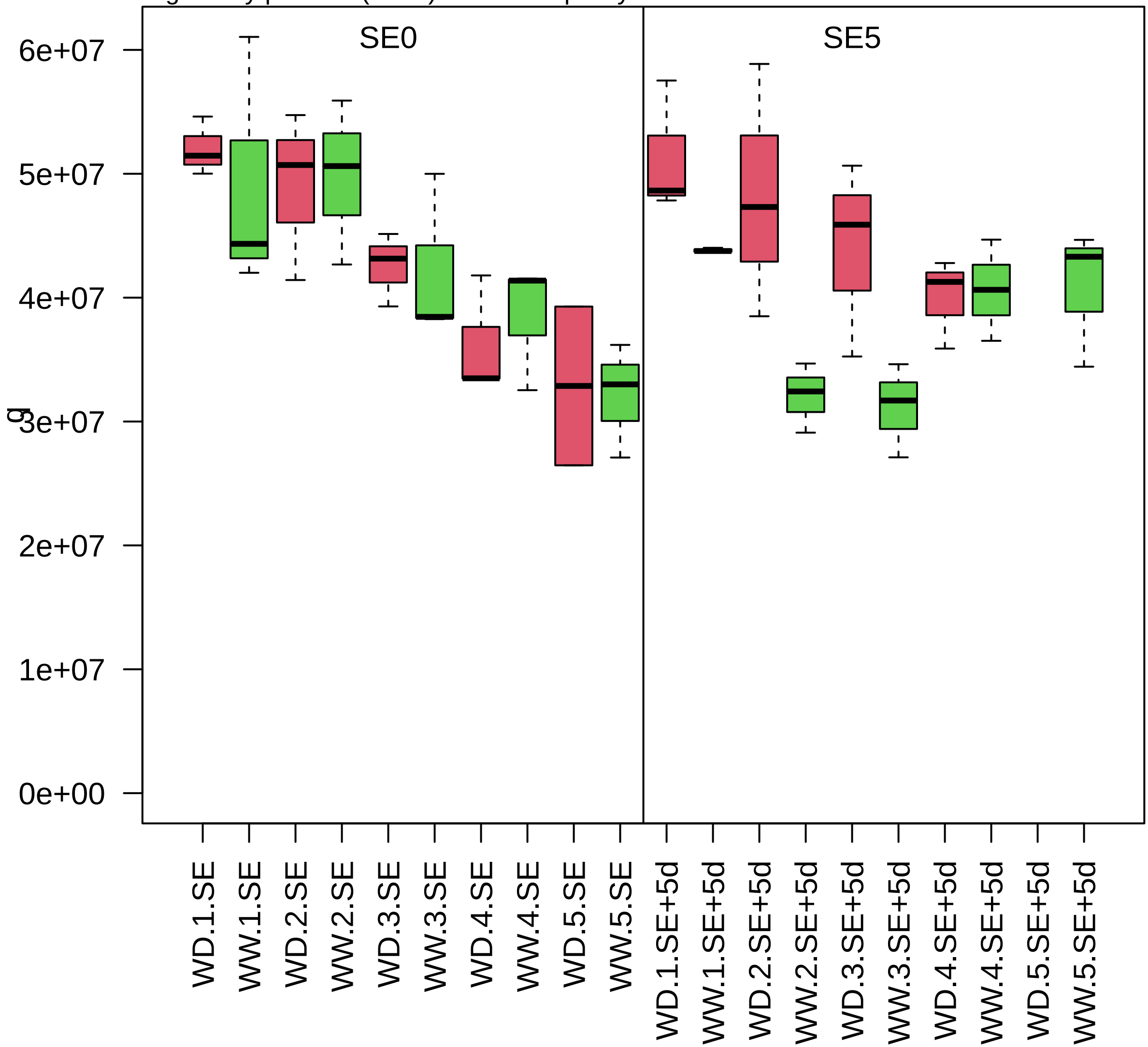

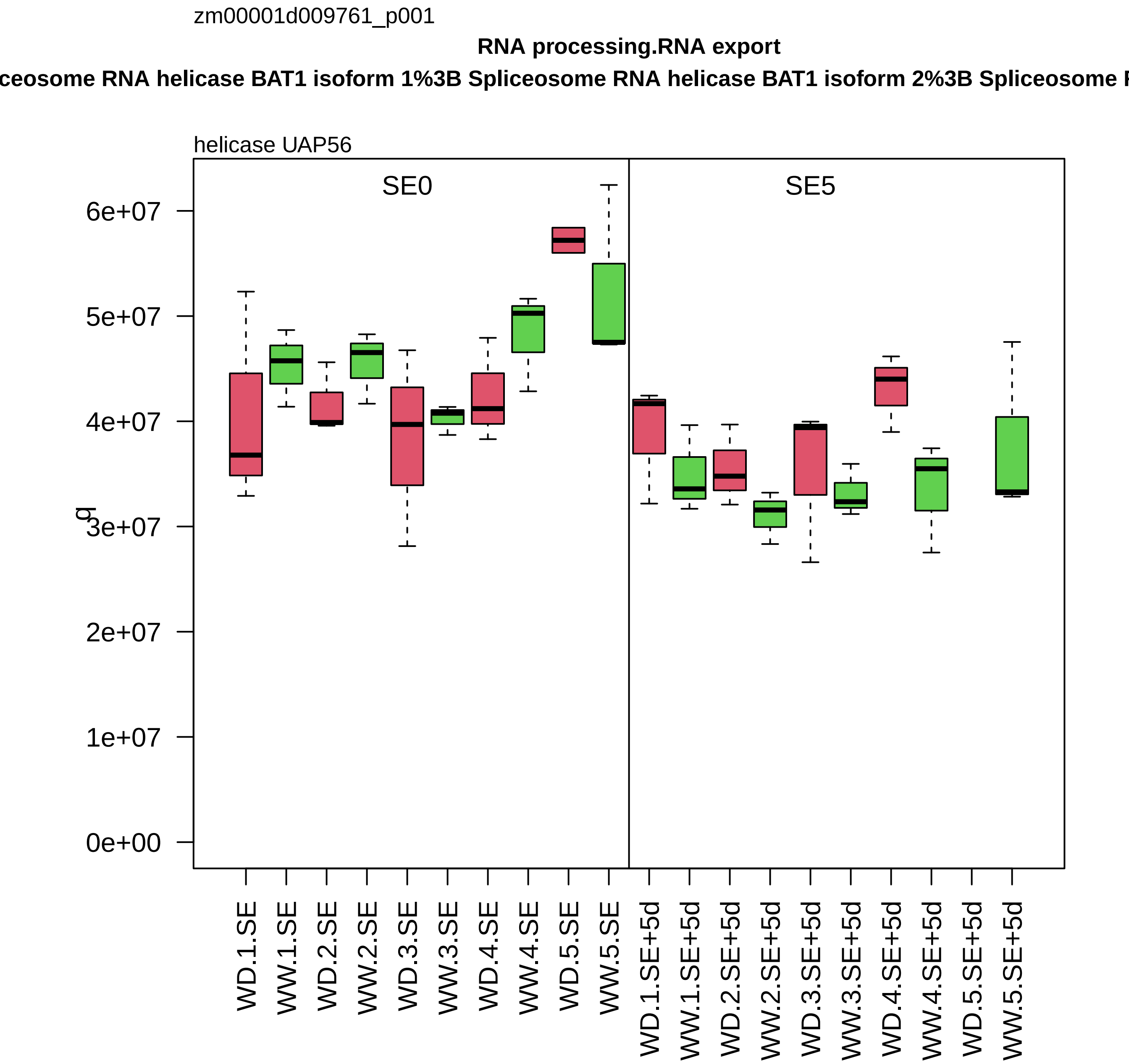

zm00001d009761_p001
RNA processing.RNA export
ceosome RNA helicase BAT1 isoform 1%3B Spliceosome RNA helicase BAT1 isoform 2%3B Spliceosome R
helicase UAP56
SE0
SE5
6e+07
5e+07
4e+07
3e+07
2e+07
1e+07
0e+00
q
WD.1.SE
WW.1.SE
WD.2.SE
WW.2.SE
WD.3.SE
WW.3.SE
WD.4.SE
WW.4.SE
WD.5.SE
WW.5.SE
WD.1.SE+5d
WW.1.SE+5d
WD.2.SE+5d
WW.2.SE+5d
WD.3.SE+5d
WW.3.SE+5d
WD.4.SE+5d
WW.4.SE+5d
WD.5.SE+5d
WW.5.SE+5d

zm00001d009779_p001

**Lipid metabolism.fatty acid metabolism**
**description:dihydrolipoyl dehydrogenases**

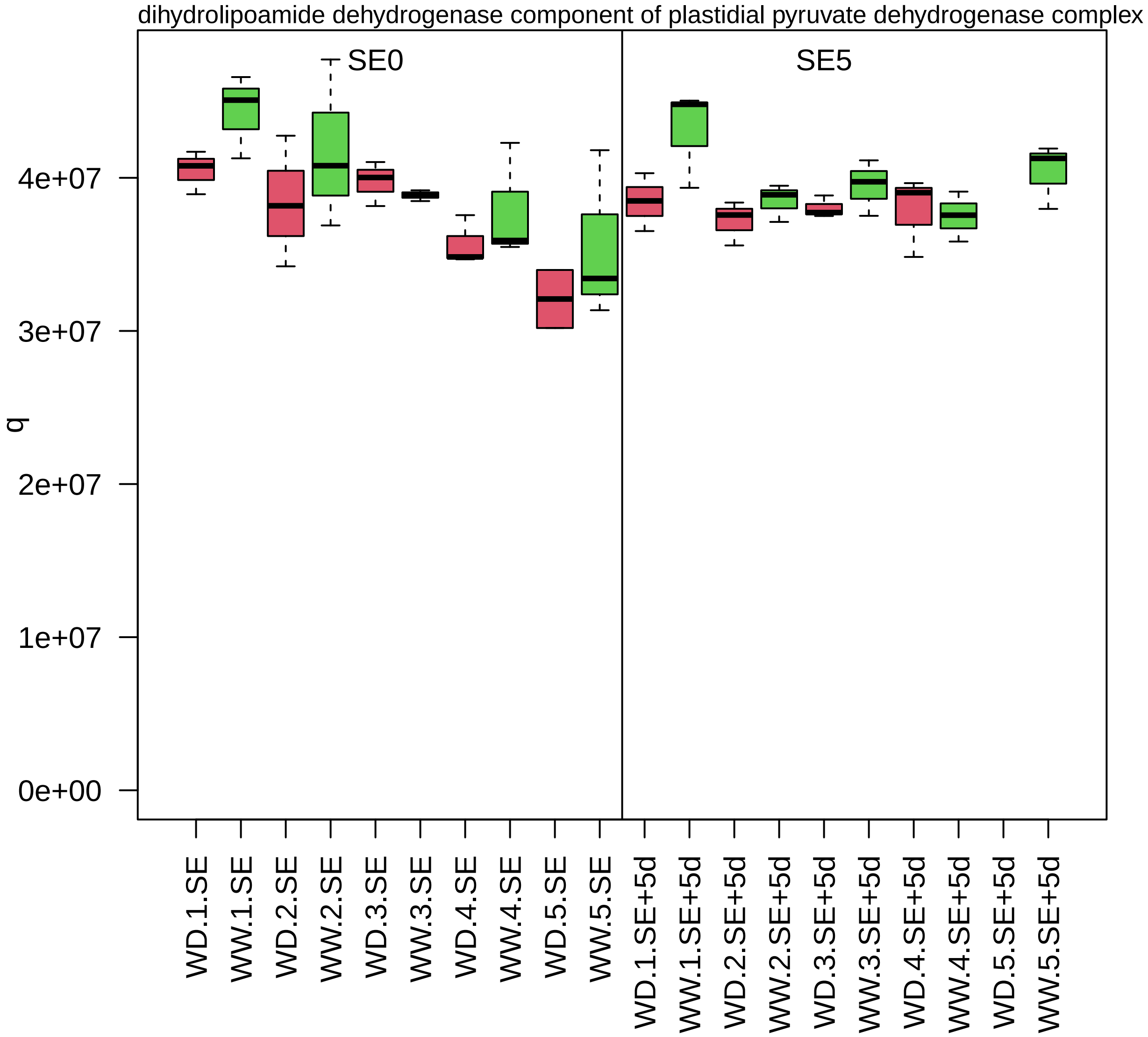

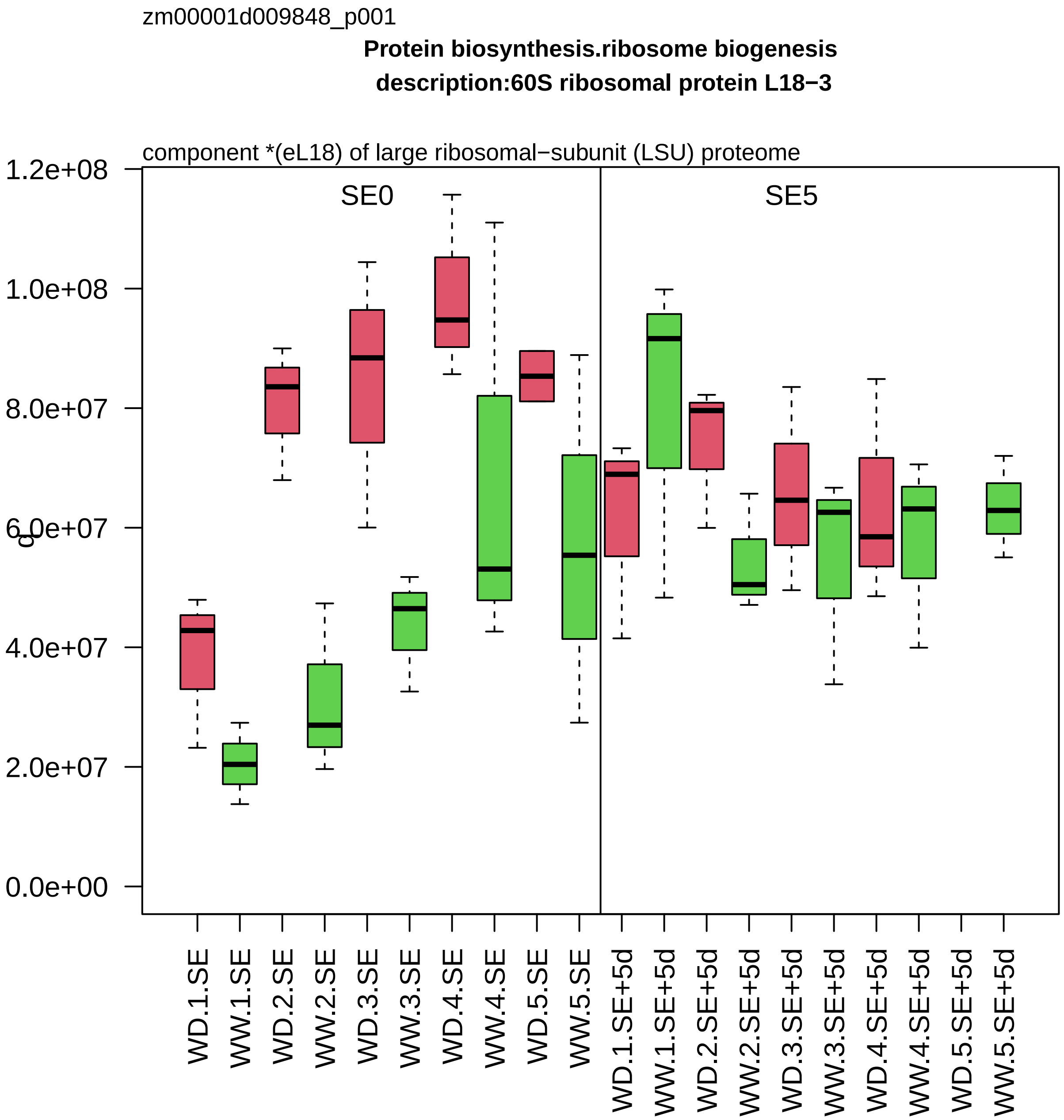
zm00001d009848_p001
Protein biosynthesis.ribosome biogenesis
description:60S ribosomal protein L18−3
component *(eL18) of large ribosomal−subunit (LSU) proteome
SE0
SE5
c
0.0e+00
2.0e+07
4.0e+07
6.0e+07
8.0e+07
1.0e+08
1.2e+08
WD.1.SE
WW.1.SE
WD.2.SE
WW.2.SE
WD.3.SE
WW.3.SE
WD.4.SE
WW.4.SE
WD.5.SE
WW.5.SE
WD.1.SE+5d
WW.1.SE+5d
WD.2.SE+5d
WW.2.SE+5d
WD.3.SE+5d
WW.3.SE+5d
WD.4.SE+5d
WW.4.SE+5d
WD.5.SE+5d
WW.5.SE+5d

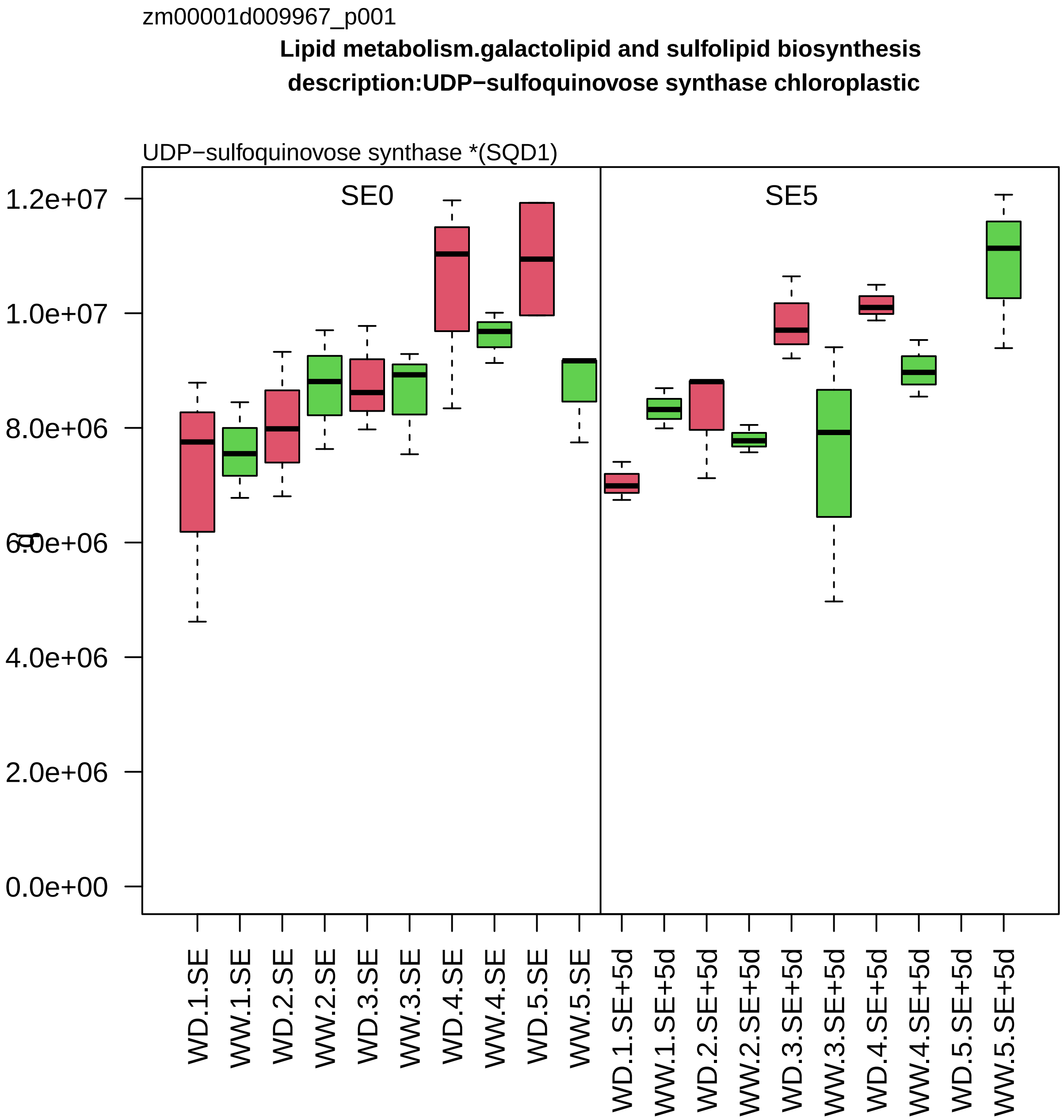
zm00001d009967_p001
Lipid metabolism.galactolipid and sulfolipid biosynthesis
description:UDP−sulfoquinovose synthase chloroplastic
UDP−sulfoquinovose synthase *(SQD1)
SE0
SE5
1.2e+07
1.0e+07
8.0e+06
6.0e+06
4.0e+06
2.0e+06
0.0e+00
WD.1.SE
WW.1.SE
WD.2.SE
WW.2.SE
WD.3.SE
WW.3.SE
WD.4.SE
WW.4.SE
WD.5.SE
WW.5.SE
WD.1.SE+5d
WW.1.SE+5d
WD.2.SE+5d
WW.2.SE+5d
WD.3.SE+5d
WW.3.SE+5d
WD.4.SE+5d
WW.4.SE+5d
WD.5.SE+5d
WW.5.SE+5d

zm00001d010056_p001

**Amino acid metabolism.glutamate group amino acid biosynthesis**

**gene_symbol:proline responding1 description: proline responding1**

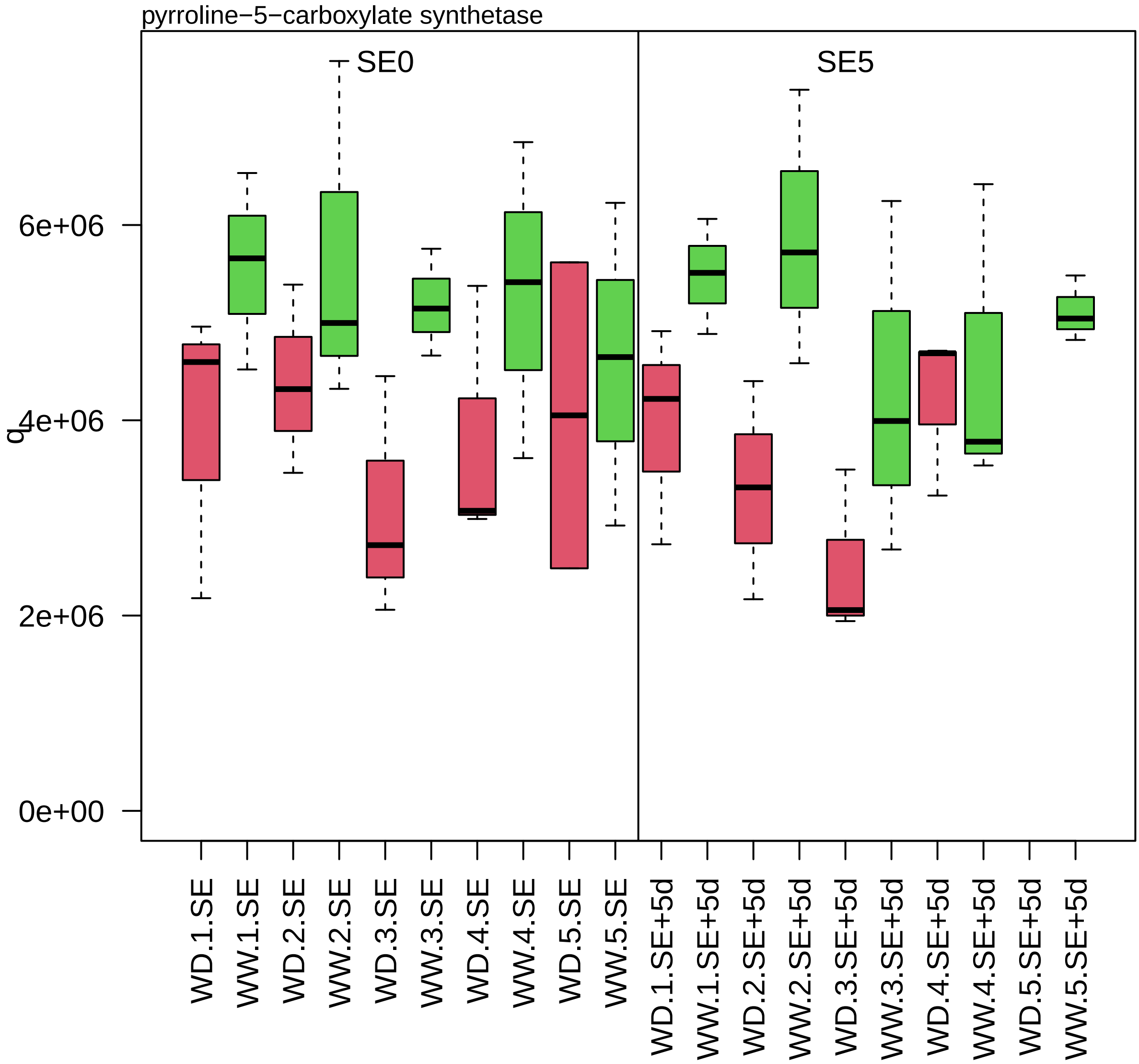

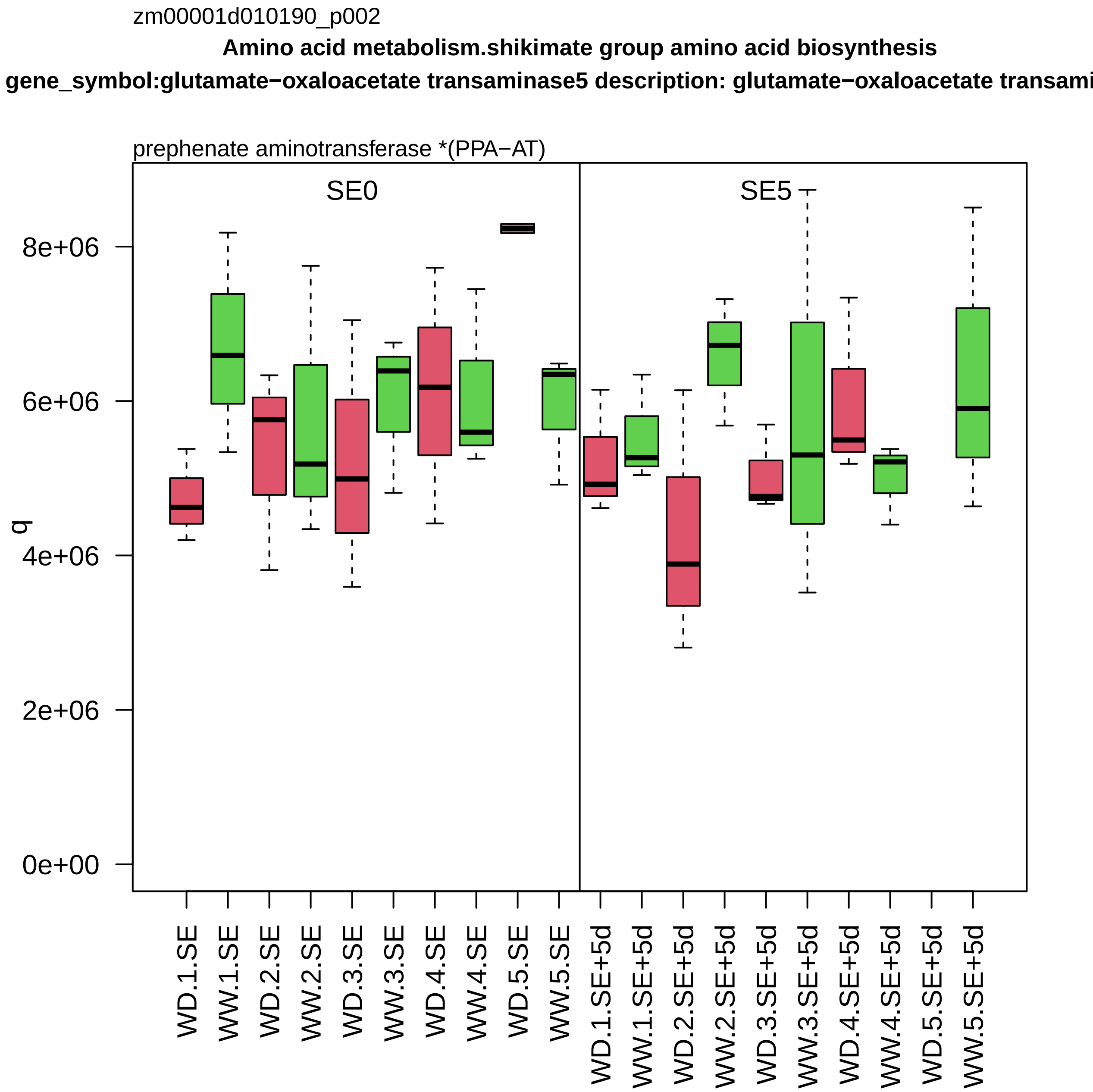
zm00001d010190_p002
Amino acid metabolism.shikimate group amino acid biosynthesis
gene_symbol:glutamate−oxaloacetate transaminase5 description: glutamate−oxaloacetate transami
prephenate aminotransferase *(PPA−AT)
SE0
SE5
q
8e+06
6e+06
4e+06
2e+06
0e+00
WD.1.SE
WW.1.SE
WD.2.SE
WW.2.SE
WD.3.SE
WW.3.SE
WD.4.SE
WW.4.SE
WD.5.SE
WW.5.SE
WD.1.SE+5d
WW.1.SE+5d
WD.2.SE+5d
WW.2.SE+5d
WD.3.SE+5d
WW.3.SE+5d
WD.4.SE+5d
WW.4.SE+5d
WD.5.SE+5d
WW.5.SE+5d

zm00001d010406_p002

**Cell division.DNA replication**

**gene_symbol:minichromosome maintenance6 description:Minichromosome maintenance prote**

component *(MCM6) of MCM replicative DNA helicase complex

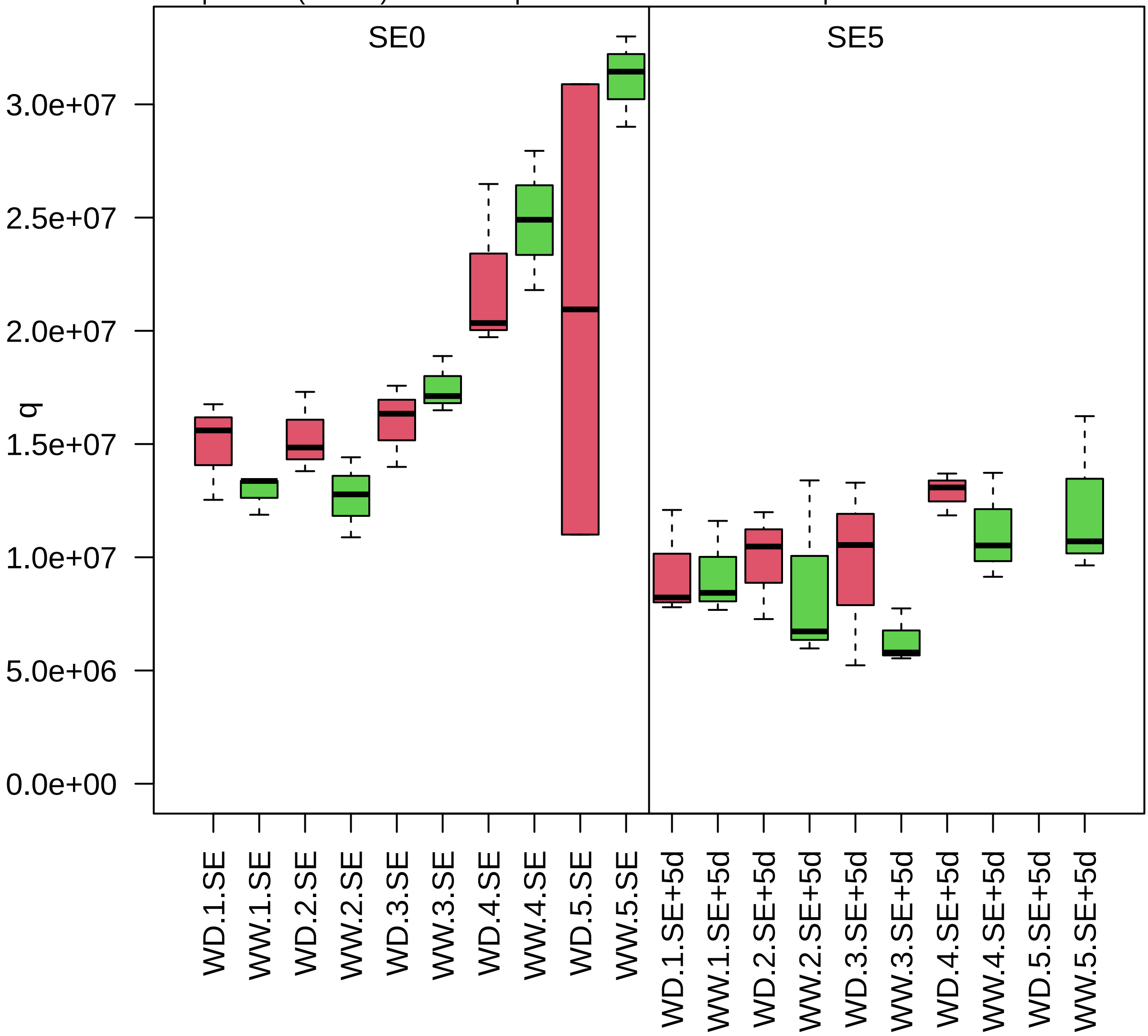

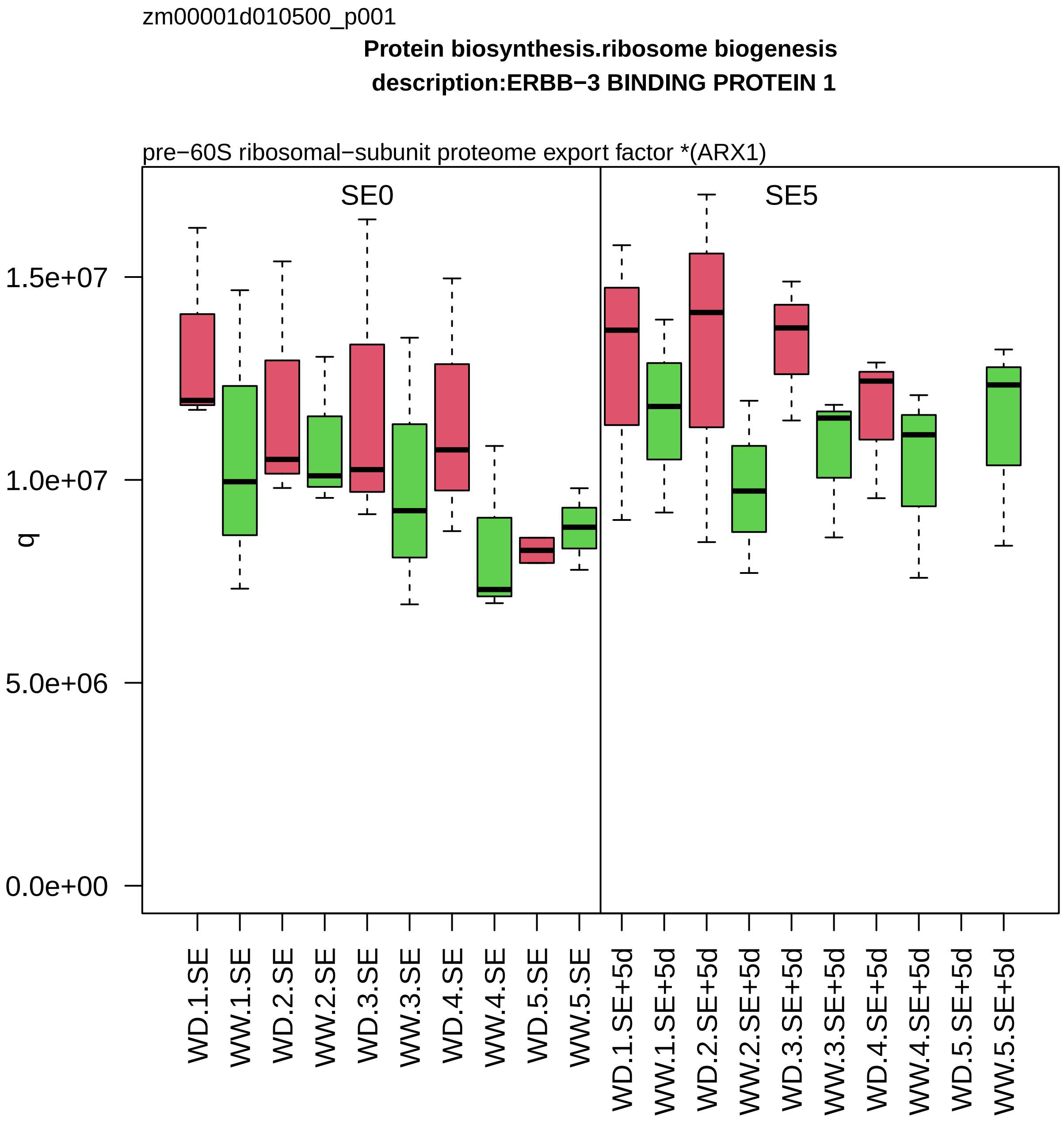

zm00001d010500_p001
Protein biosynthesis.ribosome biogenesis
description:ERBB−3 BINDING PROTEIN 1
pre−60S ribosomal−subunit proteome export factor *(ARX1)
SE0
SE5
q
1.5e+07
1.0e+07
5.0e+06
0.0e+00
WD.1.SE
WW.1.SE
WD.2.SE
WW.2.SE
WD.3.SE
WW.3.SE
WD.4.SE
WW.4.SE
WD.5.SE
WW.5.SE
WD.1.SE+5d
WW.1.SE+5d
WD.2.SE+5d
WW.2.SE+5d
WD.3.SE+5d
WW.3.SE+5d
WD.4.SE+5d
WW.4.SE+5d
WD.5.SE+5d
WW.5.SE+5d

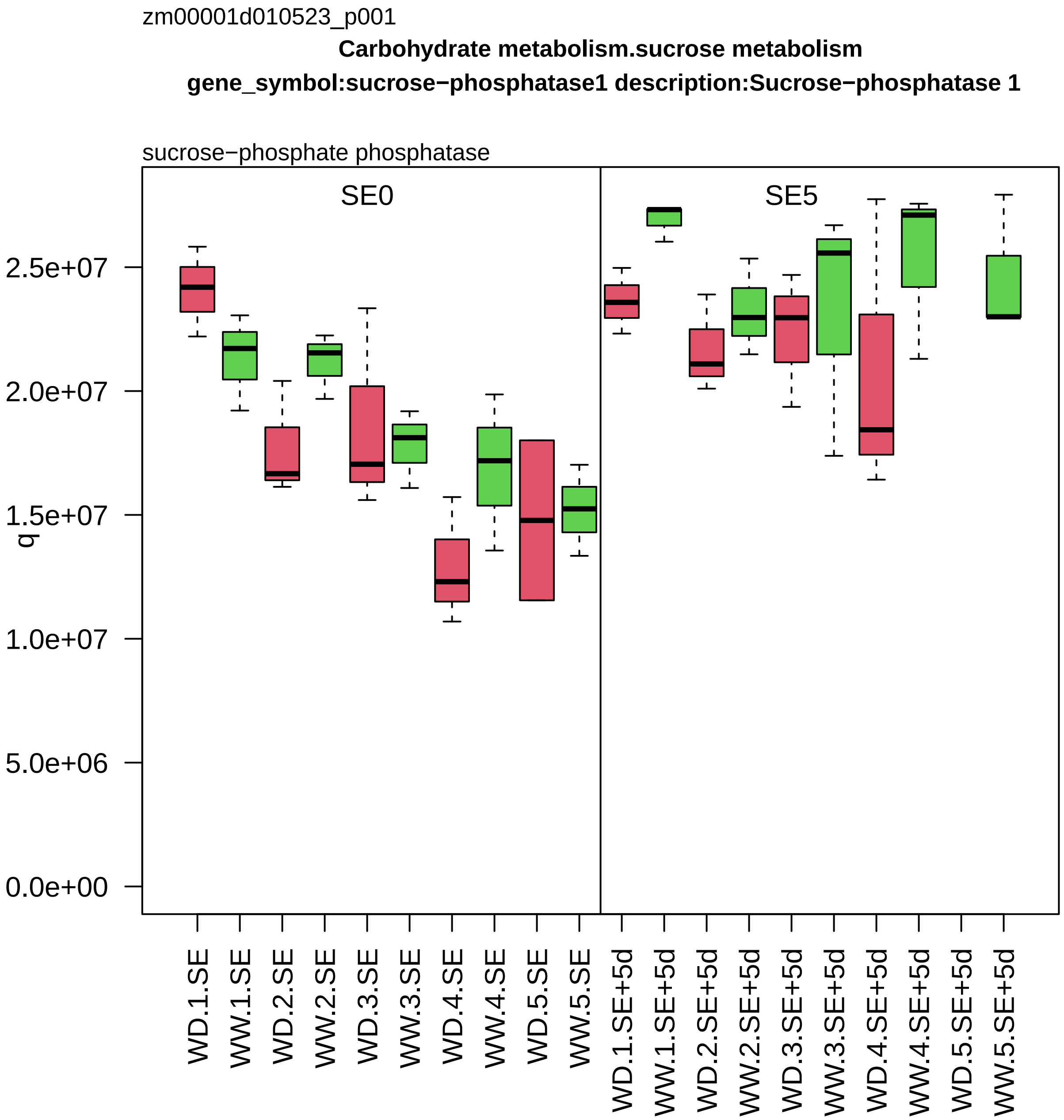

zm00001d010523_p001
Carbohydrate metabolism.sucrose metabolism
gene_symbol:sucrose−phosphatase1 description:Sucrose−phosphatase 1
sucrose−phosphate phosphatase
SE0
SE5
q
2.5e+07
2.0e+07
1.5e+07
1.0e+07
5.0e+06
0.0e+00
WD.1.SE
WW.1.SE
WD.2.SE
WW.2.SE
WD.3.SE
WW.3.SE
WD.4.SE
WW.4.SE
WD.5.SE
WW.5.SE
WD.1.SE+5d
WW.1.SE+5d
WD.2.SE+5d
WW.2.SE+5d
WD.3.SE+5d
WW.3.SE+5d
WD.4.SE+5d
WW.4.SE+5d
WD.5.SE+5d
WW.5.SE+5d

zm00001d010528_p001

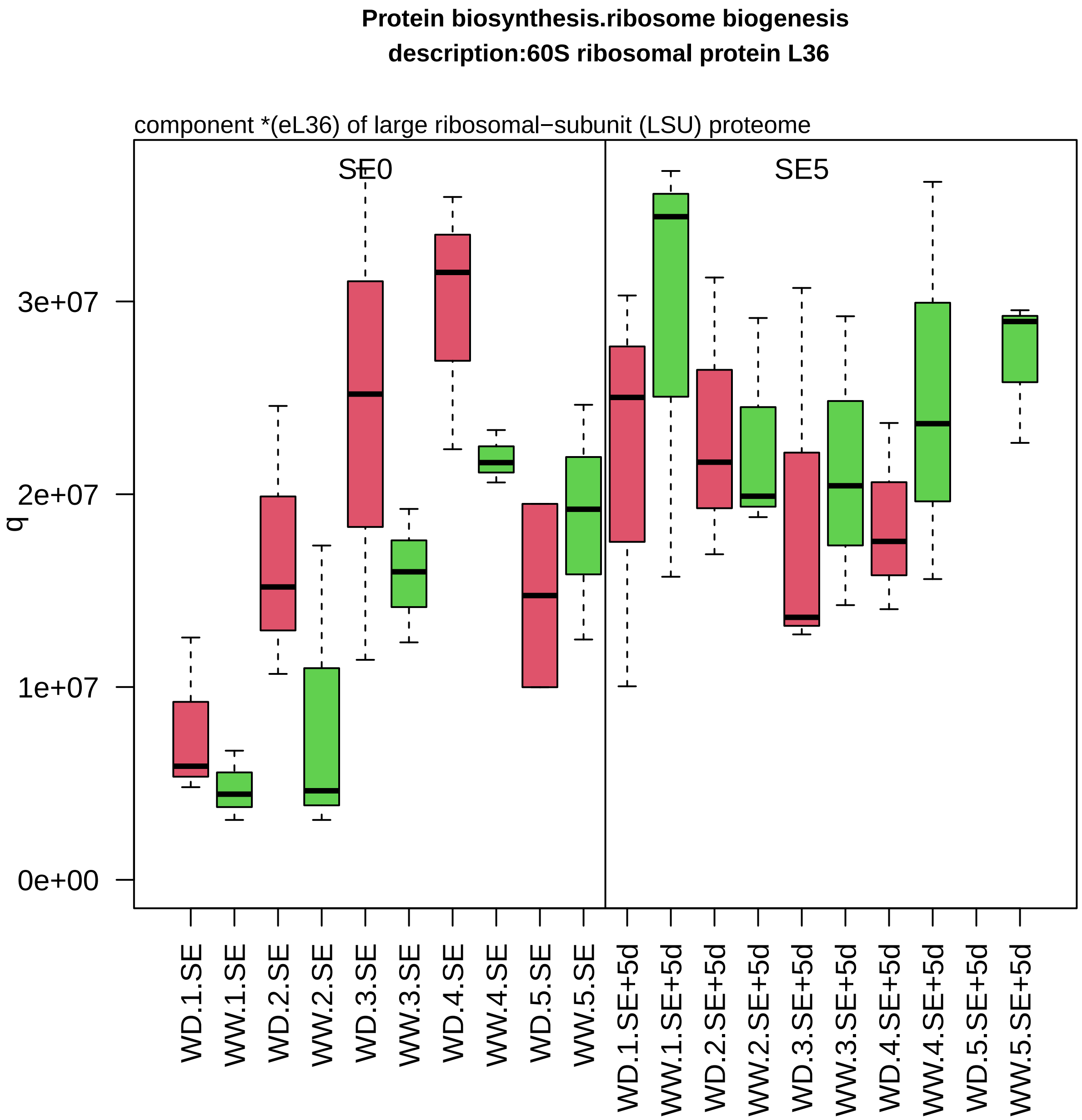

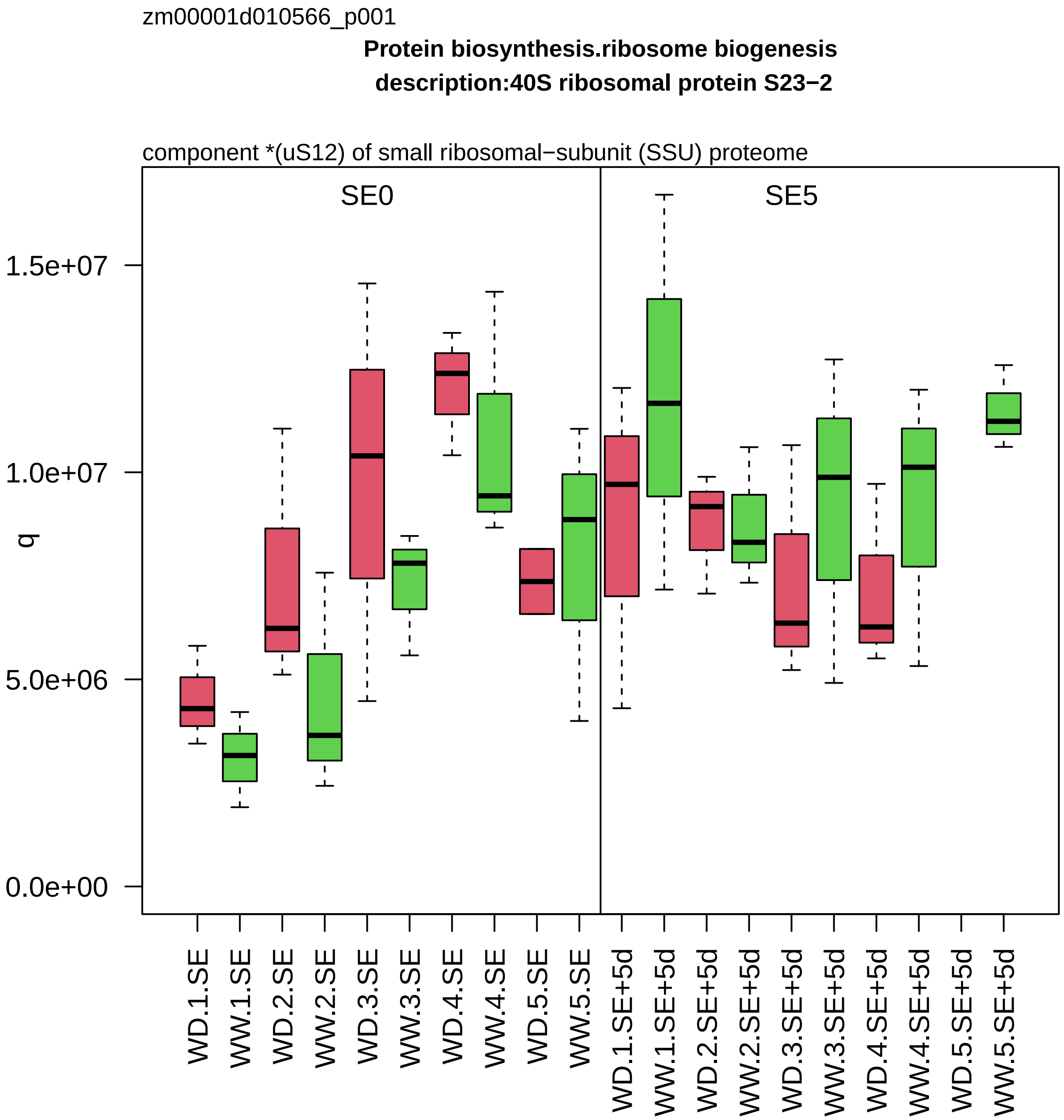

zm00001d010566_p001
Protein biosynthesis.ribosome biogenesis
description:40S ribosomal protein S23−2
component *(uS12) of small ribosomal−subunit (SSU) proteome
SE0
SE5
q
1.5e+07
1.0e+07
5.0e+06
0.0e+00
WD.1.SE
WW.1.SE
WD.2.SE
WW.2.SE
WD.3.SE
WW.3.SE
WD.4.SE
WW.4.SE
WD.5.SE
WW.5.SE
WD.1.SE+5d
WW.1.SE+5d
WD.2.SE+5d
WW.2.SE+5d
WD.3.SE+5d
WW.3.SE+5d
WD.4.SE+5d
WW.4.SE+5d
WD.5.SE+5d
WW.5.SE+5d

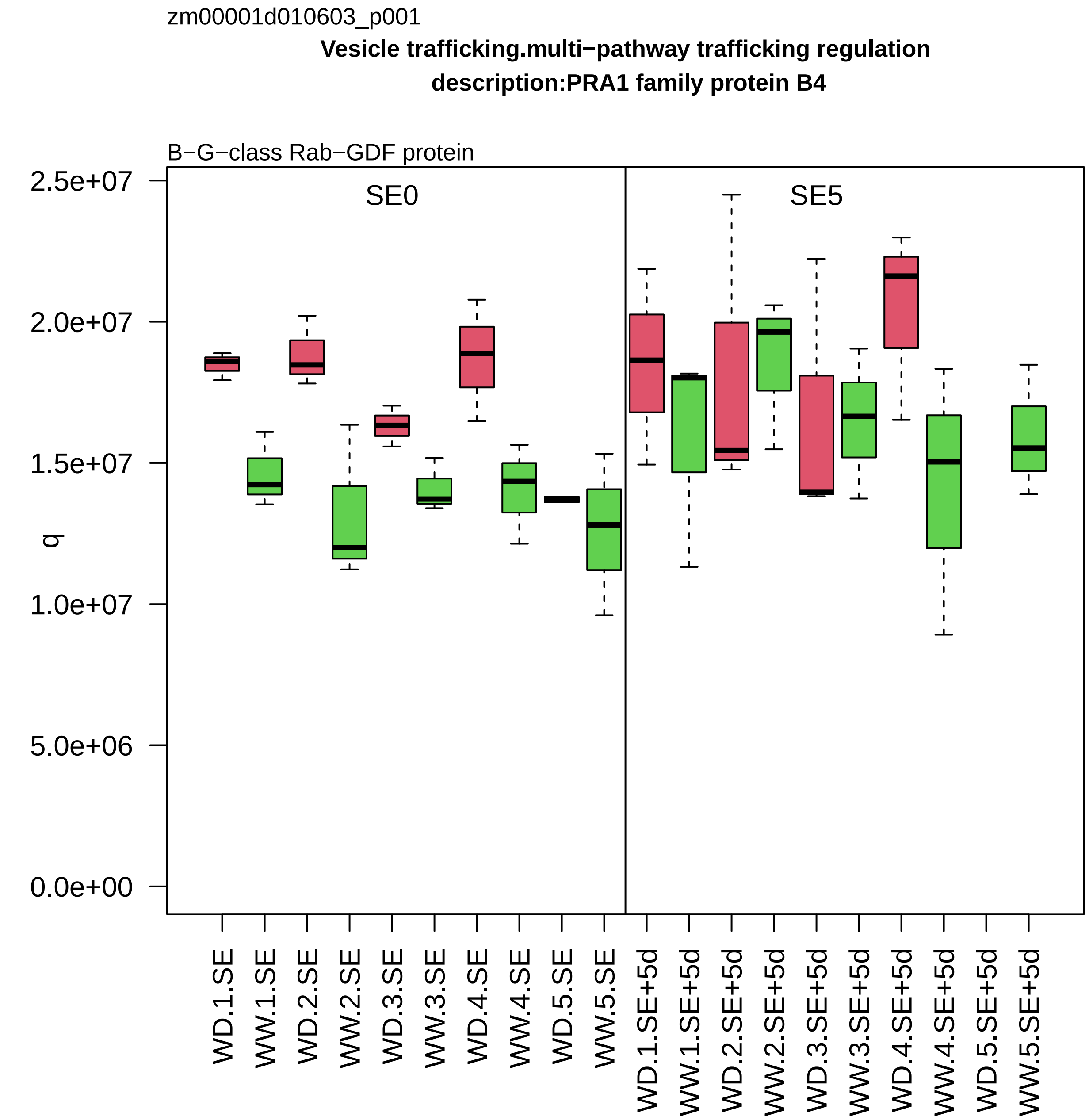
zm00001d010603_p001
Vesicle trafficking.multi−pathway trafficking regulation
description:PRA1 family protein B4
B−G−class Rab−GDF protein
SE0
SE5
q
2.5e+07
2.0e+07
1.5e+07
1.0e+07
5.0e+06
0.0e+00
WD.1.SE
WW.1.SE
WD.2.SE
WW.2.SE
WD.3.SE
WW.3.SE
WD.4.SE
WW.4.SE
WD.5.SE
WW.5.SE
WD.1.SE+5d
WW.1.SE+5d
WD.2.SE+5d
WW.2.SE+5d
WD.3.SE+5d
WW.3.SE+5d
WD.4.SE+5d
WW.4.SE+5d
WD.5.SE+5d
WW.5.SE+5d

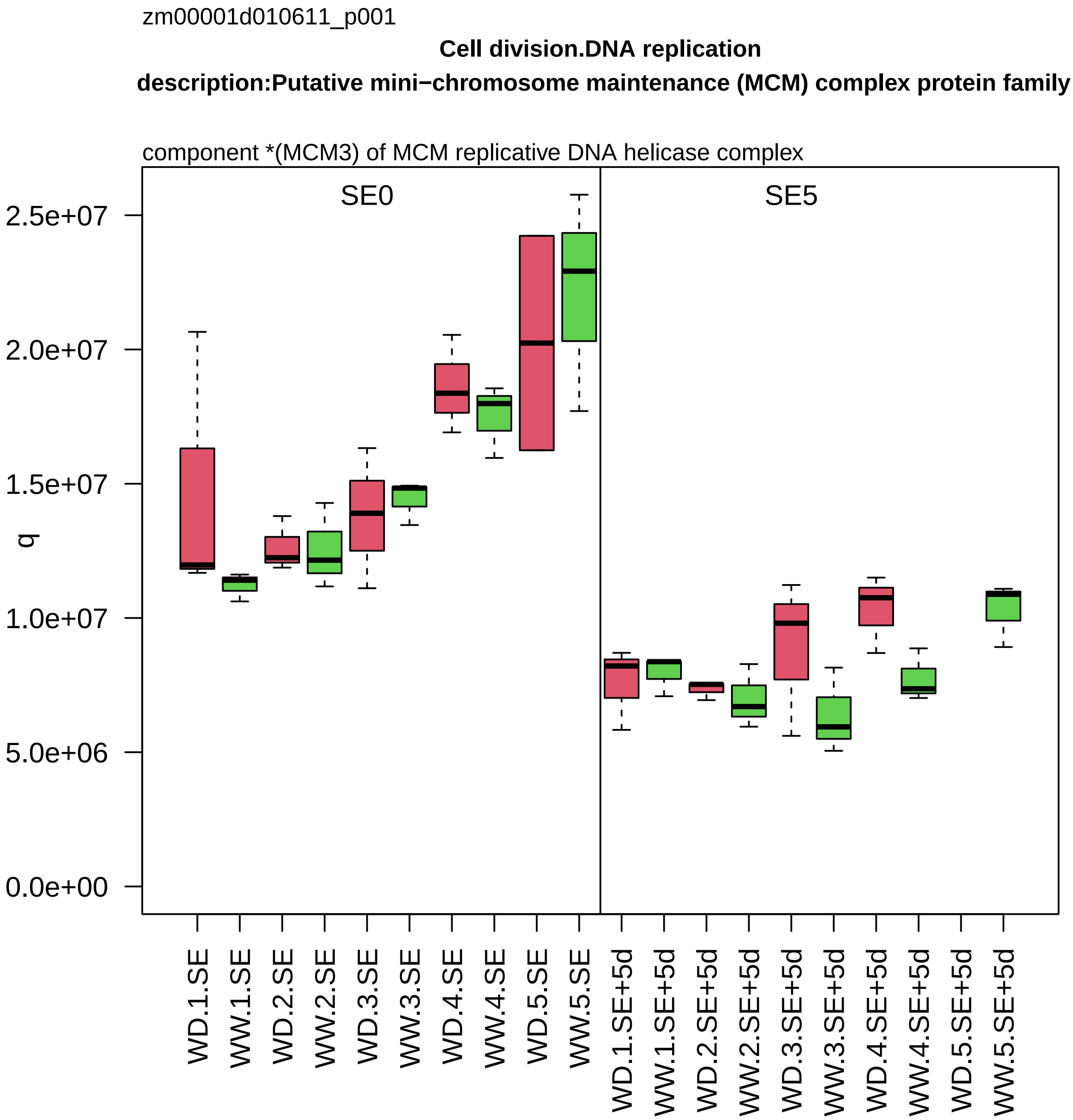

zm00001d010611_p001
Cell division.DNA replication
description:Putative mini−chromosome maintenance (MCM) complex protein family
component *(MCM3) of MCM replicative DNA helicase complex
SE0
SE5
q
2.5e+07
2.0e+07
1.5e+07
1.0e+07
5.0e+06
0.0e+00
WD.1.SE
WW.1.SE
WD.2.SE
WW.2.SE
WD.3.SE
WW.3.SE
WD.4.SE
WW.4.SE
WD.5.SE
WW.5.SE
WD.1.SE+5d
WW.1.SE+5d
WD.2.SE+5d
WW.2.SE+5d
WD.3.SE+5d
WW.3.SE+5d
WD.4.SE+5d
WW.4.SE+5d
WD.5.SE+5d
WW.5.SE+5d

zm00001d010728_p005

**not assigned.not annotated**

**description:PLASMODESMATA CALLOSE−BINDING PROTEIN 2**

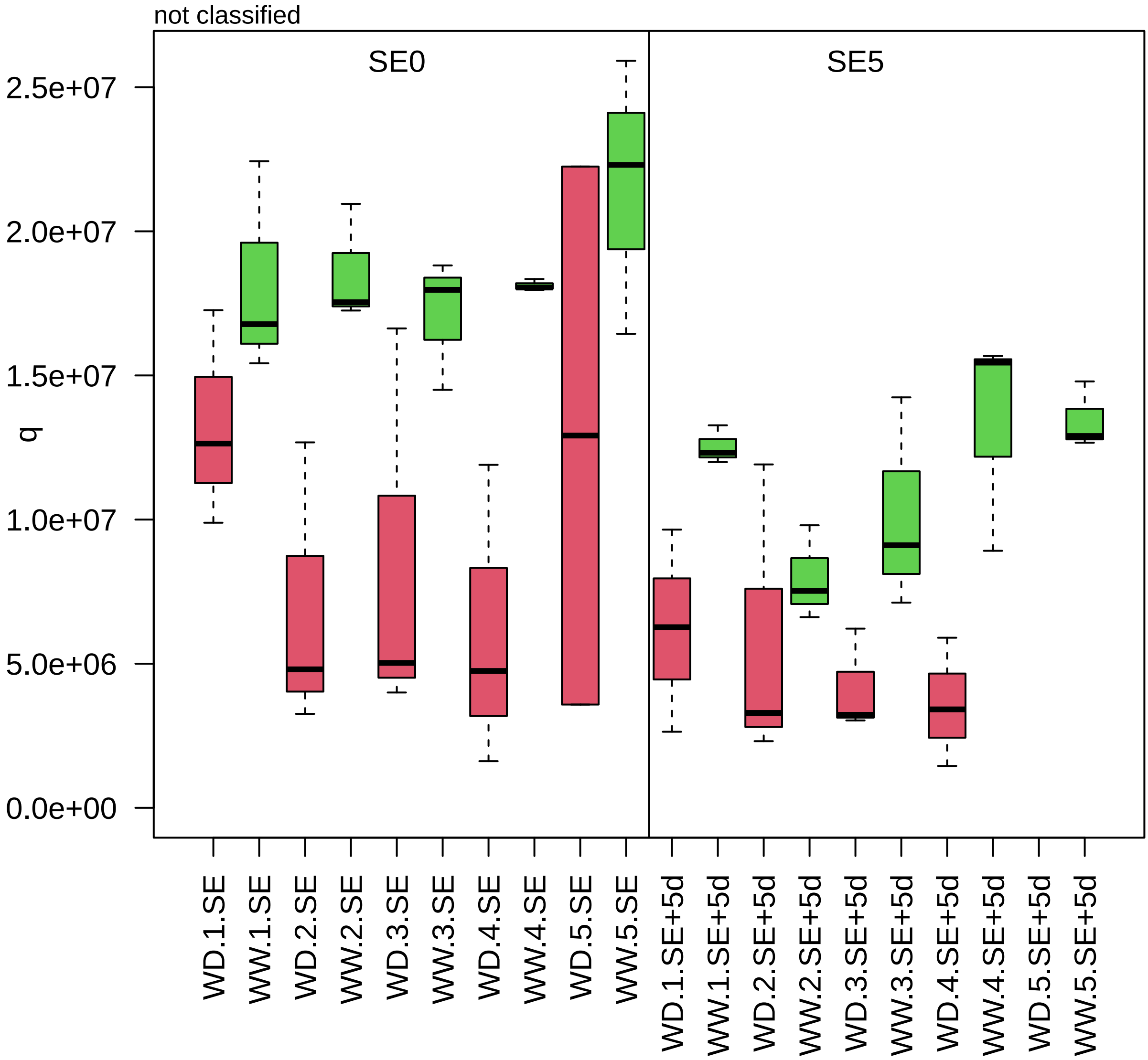

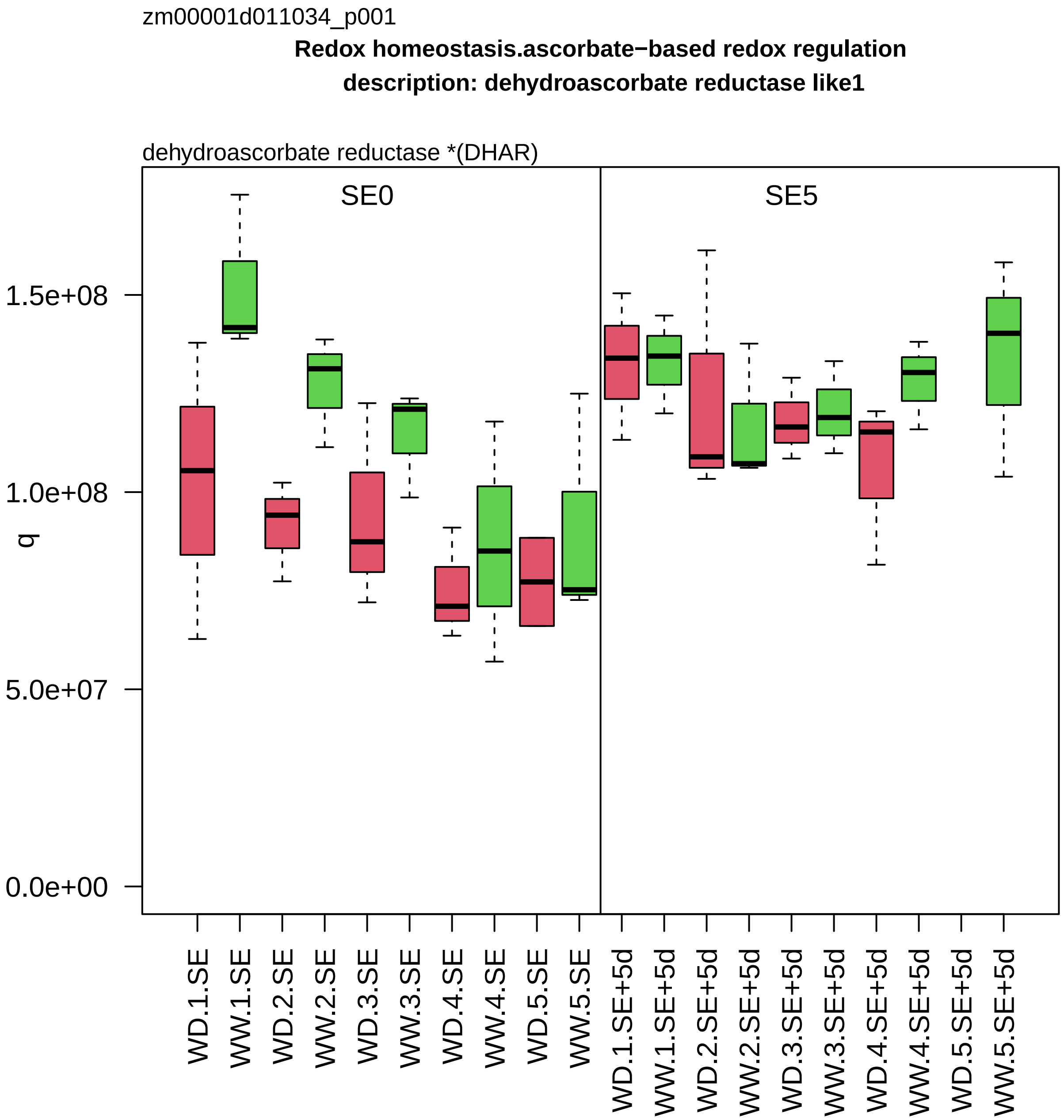
zm00001d011034_p001
Redox homeostasis.ascorbate−based redox regulation
description: dehydroascorbate reductase like1
dehydroascorbate reductase *(DHAR)
SE0
SE5
q
1.5e+08
1.0e+08
5.0e+07
0.0e+00
WD.1.SE
WW.1.SE
WD.2.SE
WW.2.SE
WD.3.SE
WW.3.SE
WD.4.SE
WW.4.SE
WD.5.SE
WW.5.SE
WD.1.SE+5d
WW.1.SE+5d
WD.2.SE+5d
WW.2.SE+5d
WD.3.SE+5d
WW.3.SE+5d
WD.4.SE+5d
WW.4.SE+5d
WD.5.SE+5d
WW.5.SE+5d

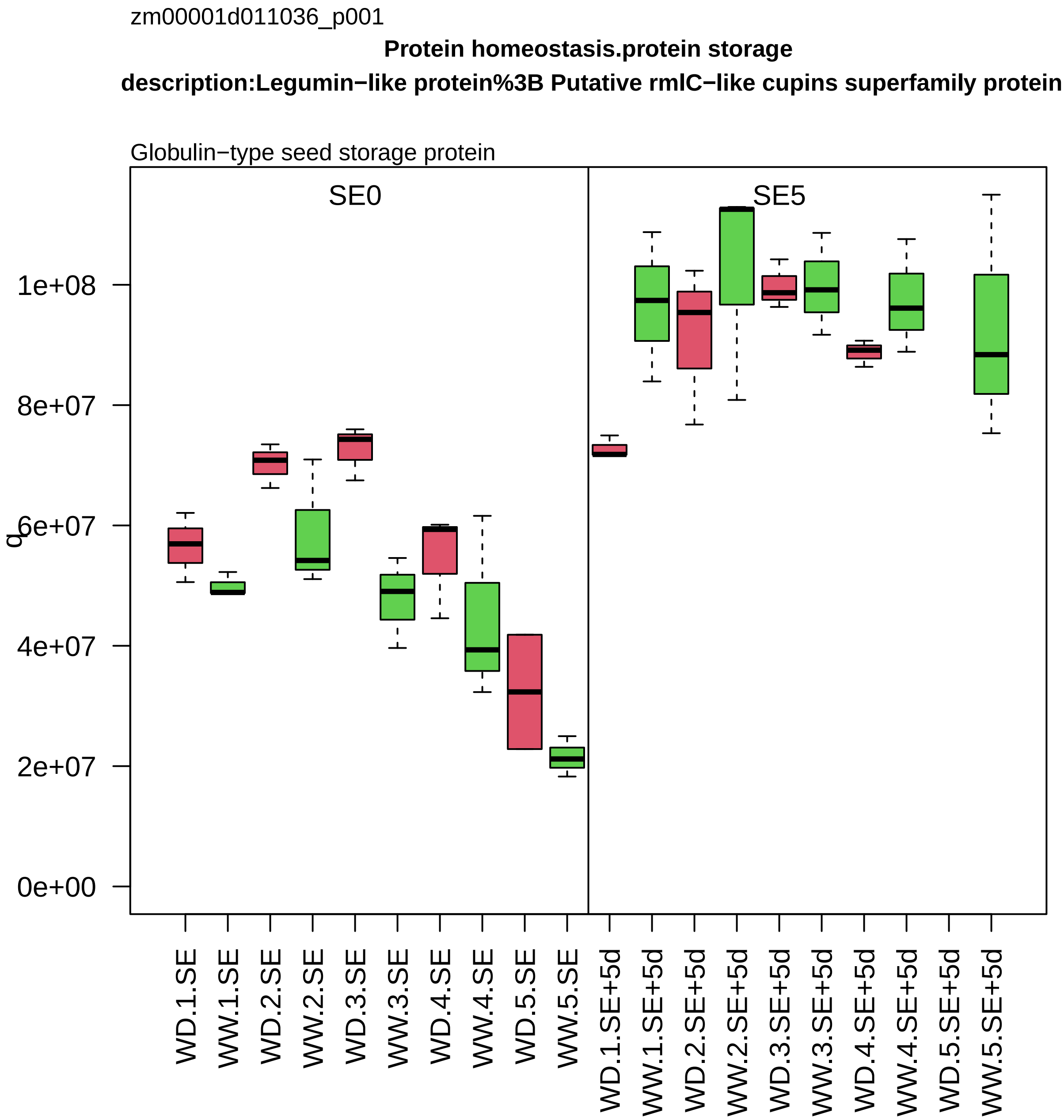
zm00001d011036_p001
Protein homeostasis.protein storage
description:Legumin−like protein%3B Putative rmlC−like cupins superfamily protein
Globulin−type seed storage protein
SE0
SE5
q
0e+00
2e+07
4e+07
6e+07
8e+07
1e+08
WD.1.SE
WW.1.SE
WD.2.SE
WW.2.SE
WD.3.SE
WW.3.SE
WD.4.SE
WW.4.SE
WD.5.SE
WW.5.SE
WD.1.SE+5d
WW.1.SE+5d
WD.2.SE+5d
WW.2.SE+5d
WD.3.SE+5d
WW.3.SE+5d
WD.4.SE+5d
WW.4.SE+5d
WD.5.SE+5d
WW.5.SE+5d

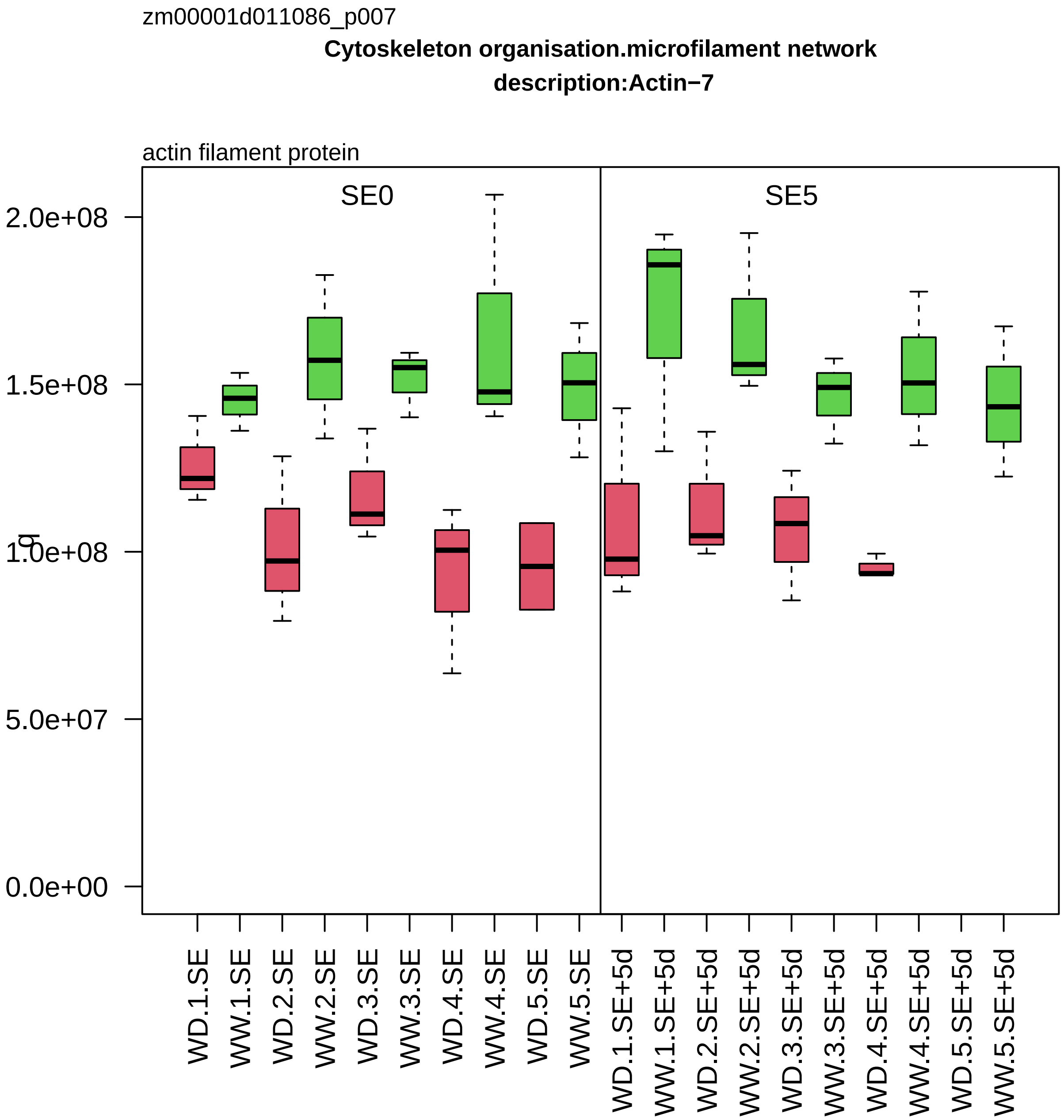

zm00001d011086_p007
Cytoskeleton organisation.microfilament network
description:Actin−7
actin filament protein
SE0
SE5
2.0e+08
1.5e+08
1.0e+08
5.0e+07
0.0e+00
WD.1.SE
WW.1.SE
WD.2.SE
WW.2.SE
WD.3.SE
WW.3.SE
WD.4.SE
WW.4.SE
WD.5.SE
WW.5.SE
WD.1.SE+5d
WW.1.SE+5d
WD.2.SE+5d
WW.2.SE+5d
WD.3.SE+5d
WW.3.SE+5d
WD.4.SE+5d
WW.4.SE+5d
WD.5.SE+5d
WW.5.SE+5d

zm00001d011099_p002

**Amino acid metabolism.aspartate group amino acid biosynthesis**
**description:Methionine S−methyltransferase**

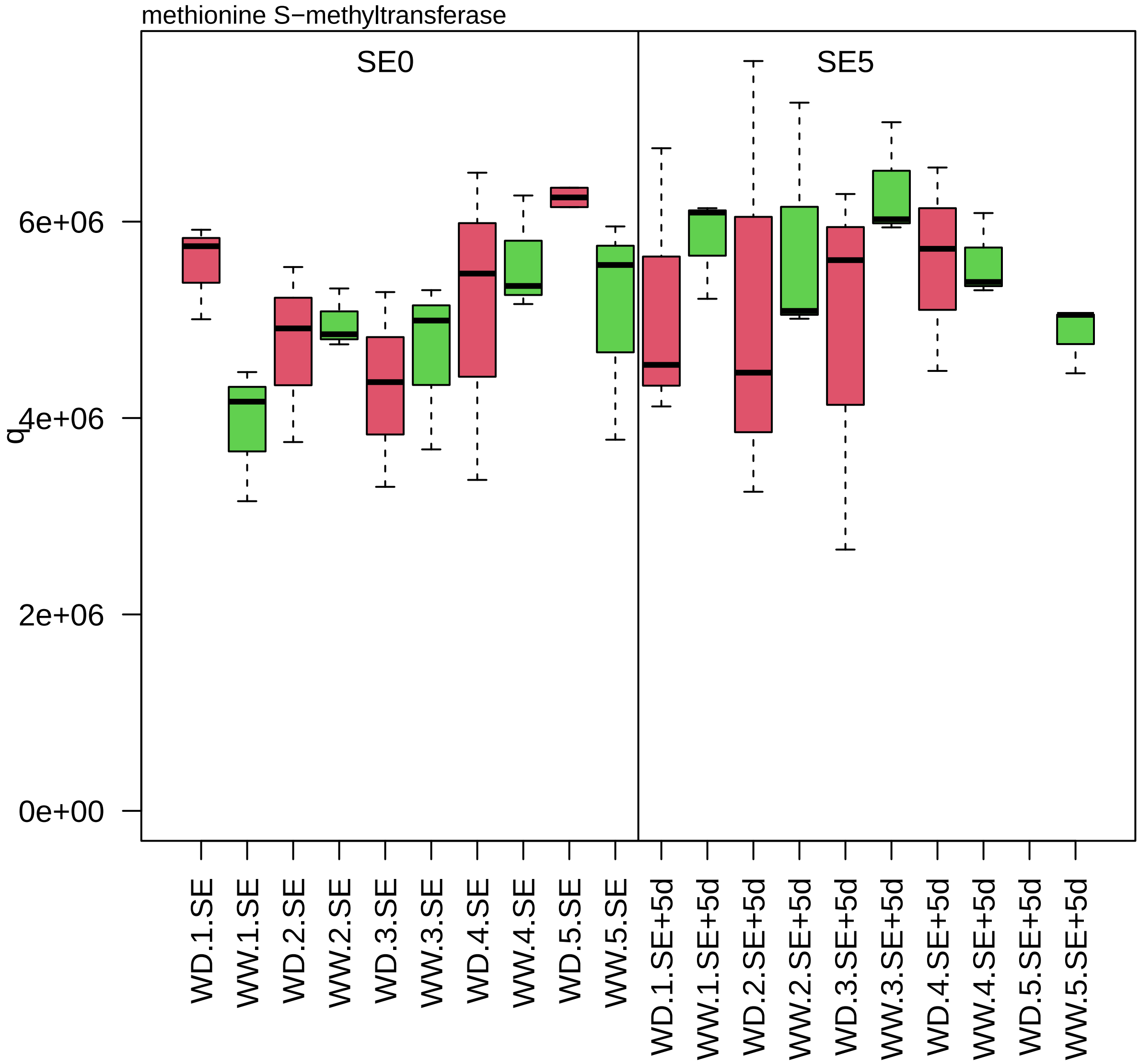

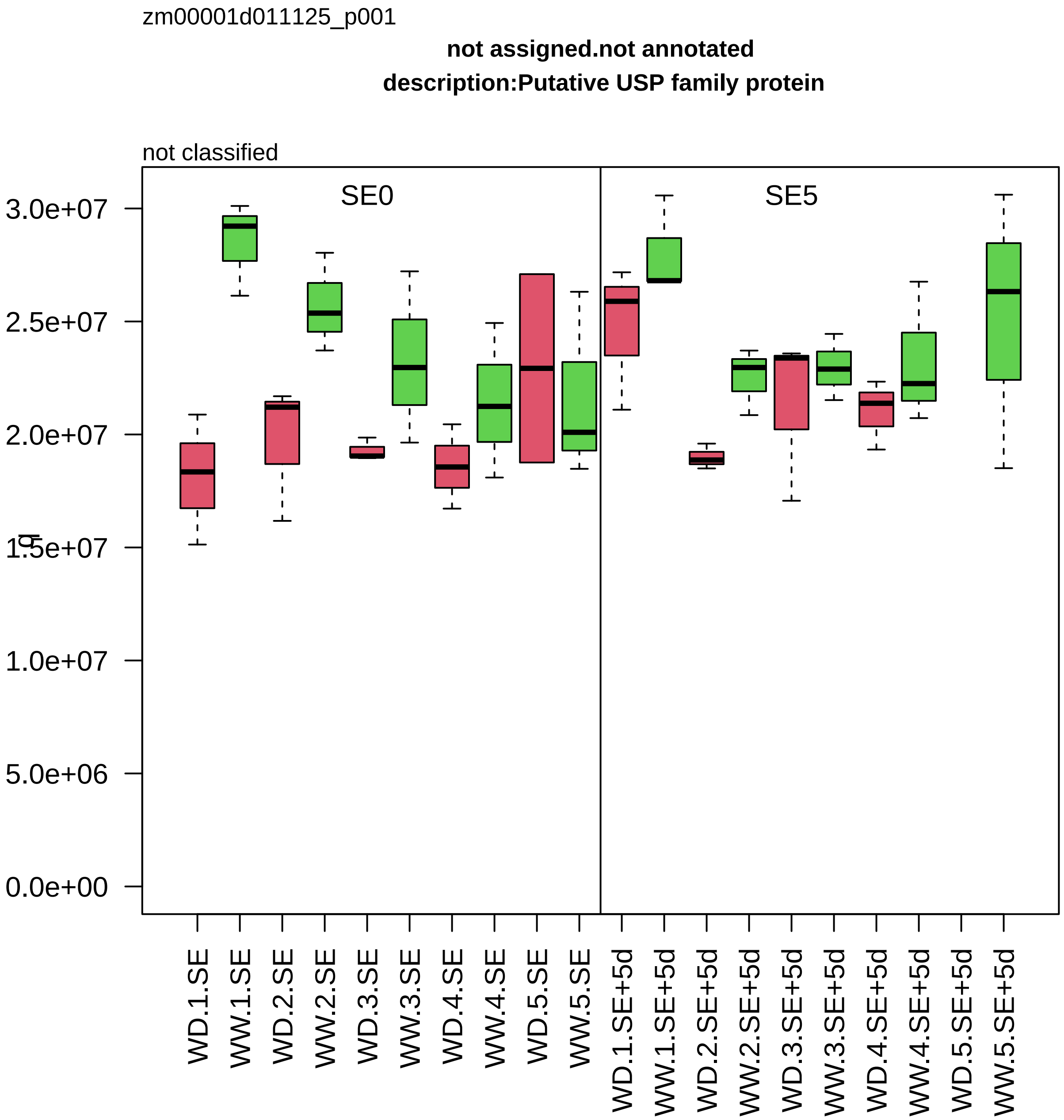

zm00001d011125_p001
not assigned.not annotated
description:Putative USP family protein
not classified
SE0
SE5
3.0e+07
2.5e+07
2.0e+07
1.5e+07
1.0e+07
5.0e+06
0.0e+00
WD.1.SE
WW.1.SE
WD.2.SE
WW.2.SE
WD.3.SE
WW.3.SE
WD.4.SE
WW.4.SE
WD.5.SE
WW.5.SE
WD.1.SE+5d
WW.1.SE+5d
WD.2.SE+5d
WW.2.SE+5d
WD.3.SE+5d
WW.3.SE+5d
WD.4.SE+5d
WW.4.SE+5d
WD.5.SE+5d
WW.5.SE+5d

zm00001d011139_p004

**Chromatin organisation.post−translational histone modification**

**gene_symbol:histone deacetylase102 description: histone deacetylase102**

HD2−type histone deacetylase *(HD2/HDT)

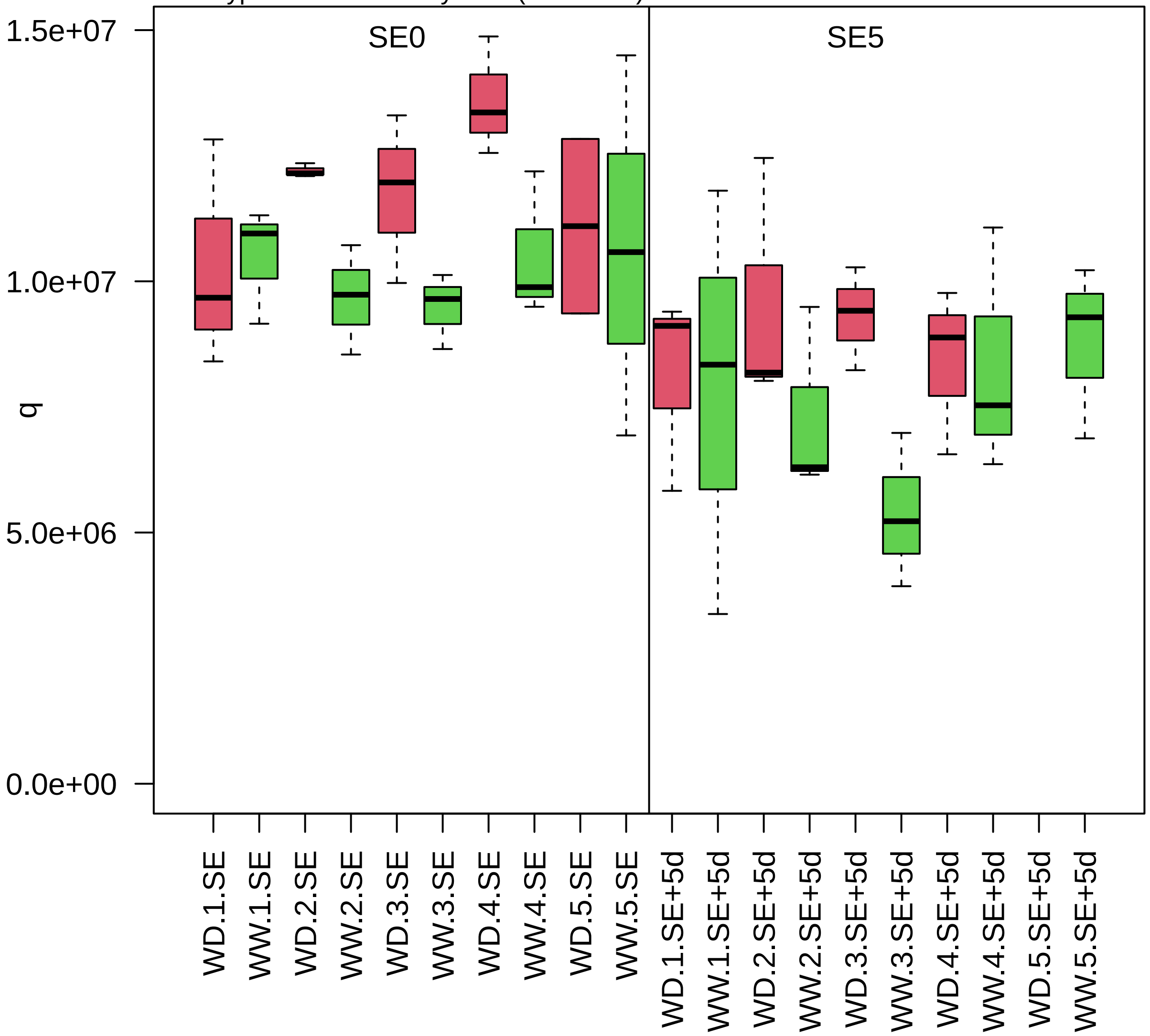

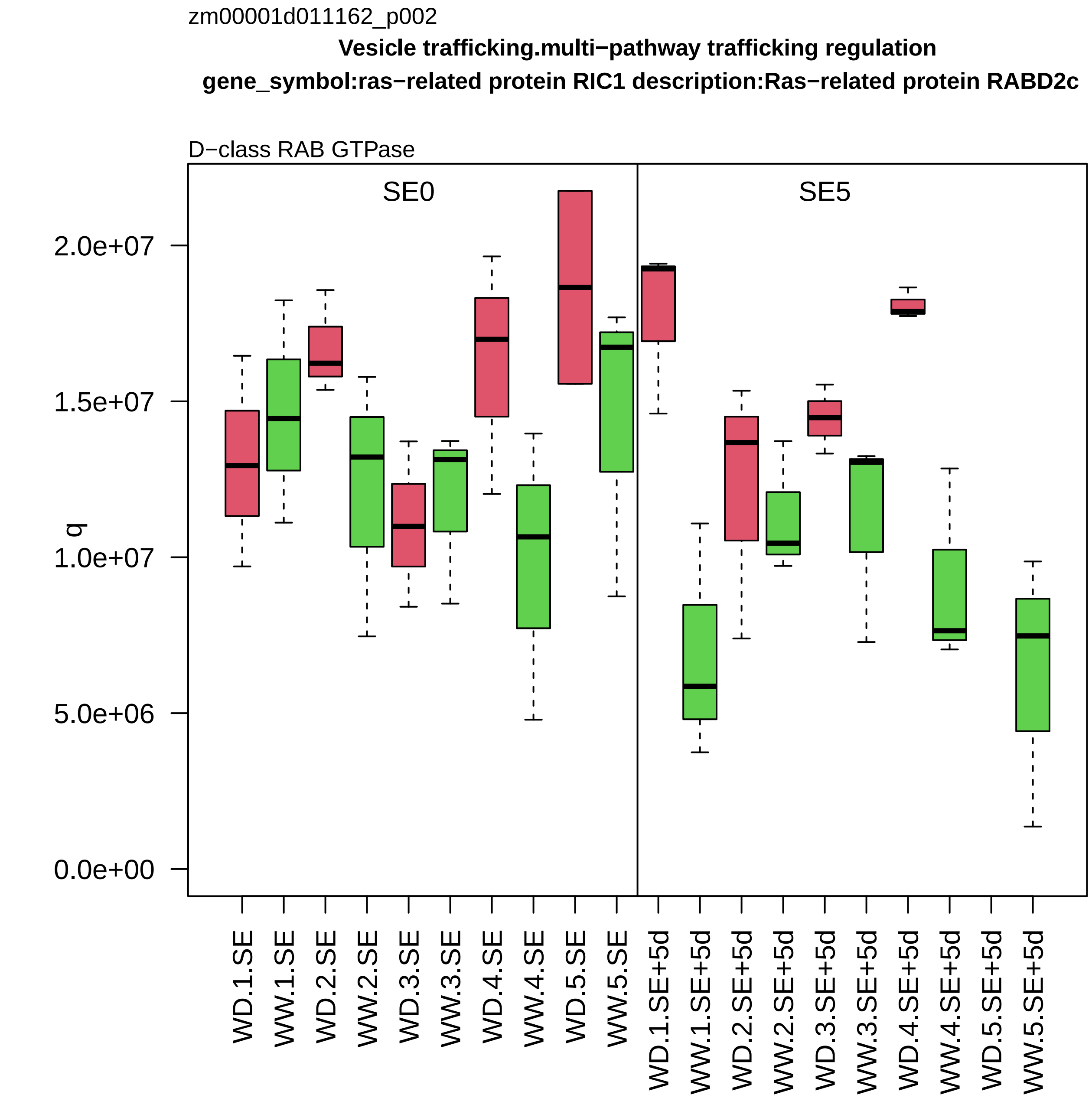

zm00001d011162_p002
Vesicle trafficking.multi−pathway trafficking regulation
gene_symbol:ras−related protein RIC1 description:Ras−related protein RABD2c
D−class RAB GTPase
SE0
SE5
q
0.0e+00
5.0e+06
1.0e+07
1.5e+07
2.0e+07
WD.1.SE
WW.1.SE
WD.2.SE
WW.2.SE
WD.3.SE
WW.3.SE
WD.4.SE
WW.4.SE
WD.5.SE
WW.5.SE
WD.1.SE+5d
WW.1.SE+5d
WD.2.SE+5d
WW.2.SE+5d
WD.3.SE+5d
WW.3.SE+5d
WD.4.SE+5d
WW.4.SE+5d
WD.5.SE+5d
WW.5.SE+5d

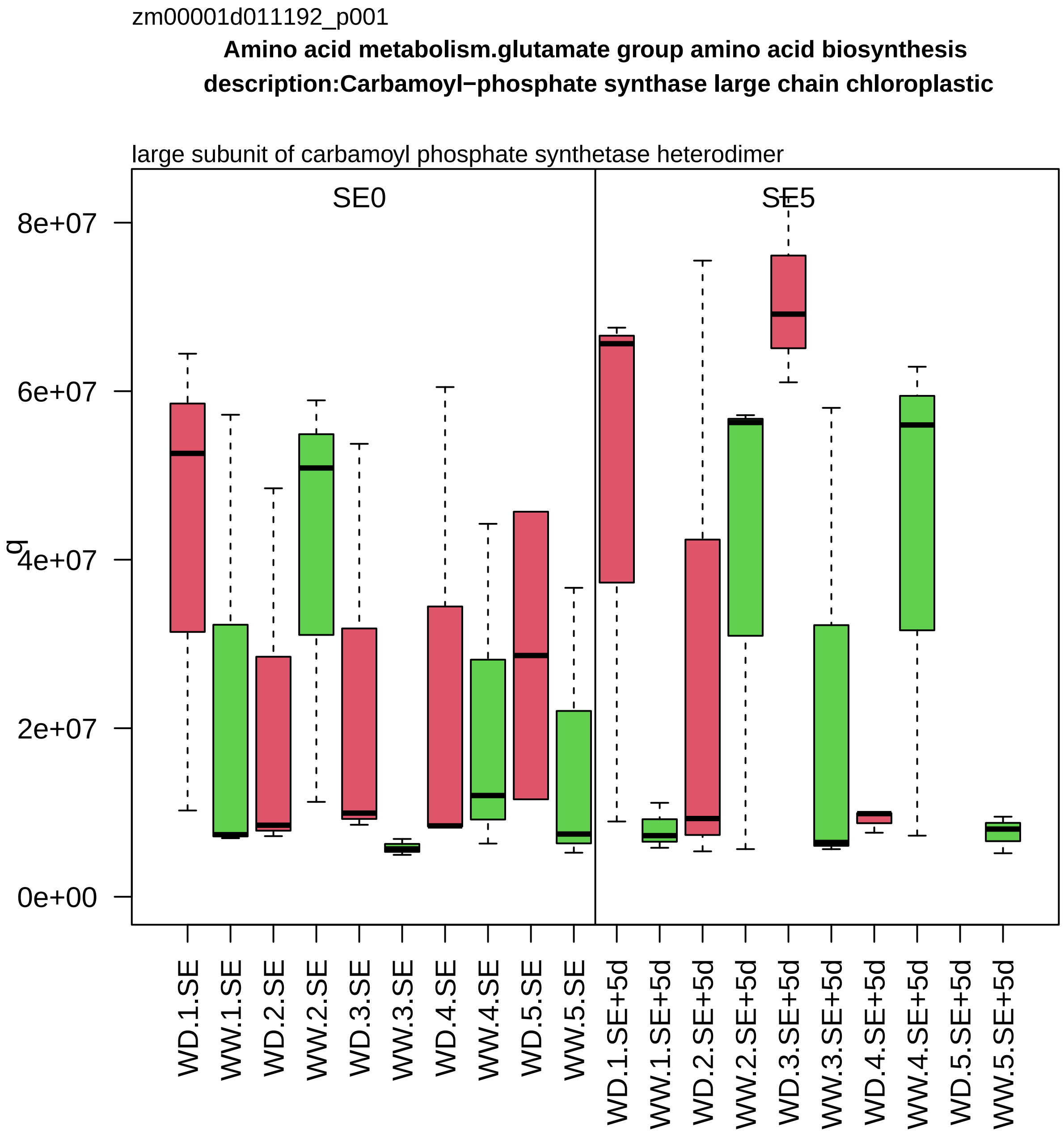
zm00001d011192_p001
Amino acid metabolism.glutamate group amino acid biosynthesis
description:Carbamoyl−phosphate synthase large chain chloroplastic
large subunit of carbamoyl phosphate synthetase heterodimer
SE0
SE5
q
0e+00
2e+07
4e+07
6e+07
8e+07
WD.1.SE
WW.1.SE
WD.2.SE
WW.2.SE
WD.3.SE
WW.3.SE
WD.4.SE
WW.4.SE
WD.5.SE
WW.5.SE
WD.1.SE+5d
WW.1.SE+5d
WD.2.SE+5d
WW.2.SE+5d
WD.3.SE+5d
WW.3.SE+5d
WD.4.SE+5d
WW.4.SE+5d
WD.5.SE+5d
WW.5.SE+5d

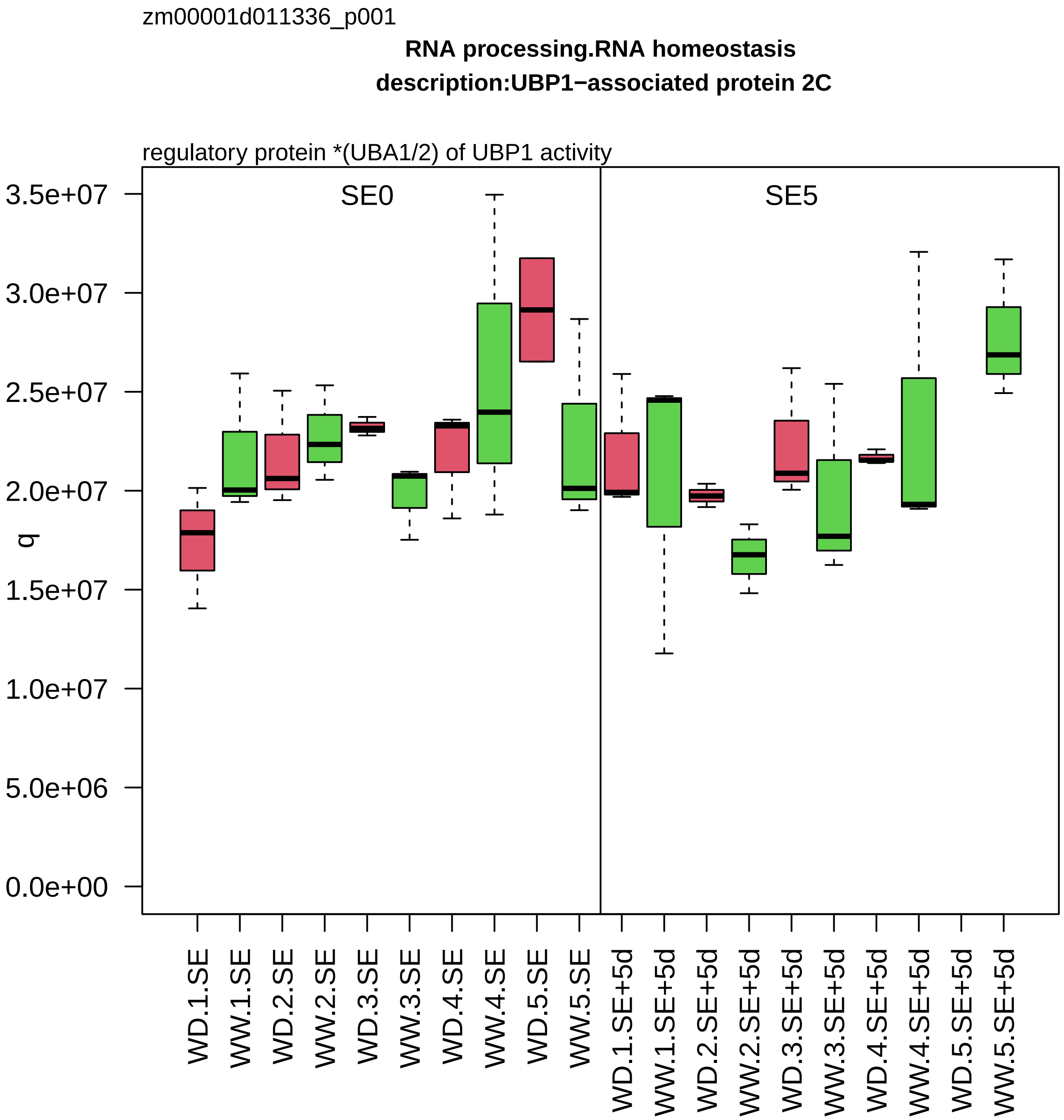

zm00001d011336_p001
RNA processing.RNA homeostasis
description:UBP1−associated protein 2C
regulatory protein *(UBA1/2) of UBP1 activity
SE0
SE5
q
3.5e+07
3.0e+07
2.5e+07
2.0e+07
1.5e+07
1.0e+07
5.0e+06
0.0e+00
WD.1.SE
WW.1.SE
WD.2.SE
WW.2.SE
WD.3.SE
WW.3.SE
WD.4.SE
WW.4.SE
WD.5.SE
WW.5.SE
WD.1.SE+5d
WW.1.SE+5d
WD.2.SE+5d
WW.2.SE+5d
WD.3.SE+5d
WW.3.SE+5d
WD.4.SE+5d
WW.4.SE+5d
WD.5.SE+5d
WW.5.SE+5d

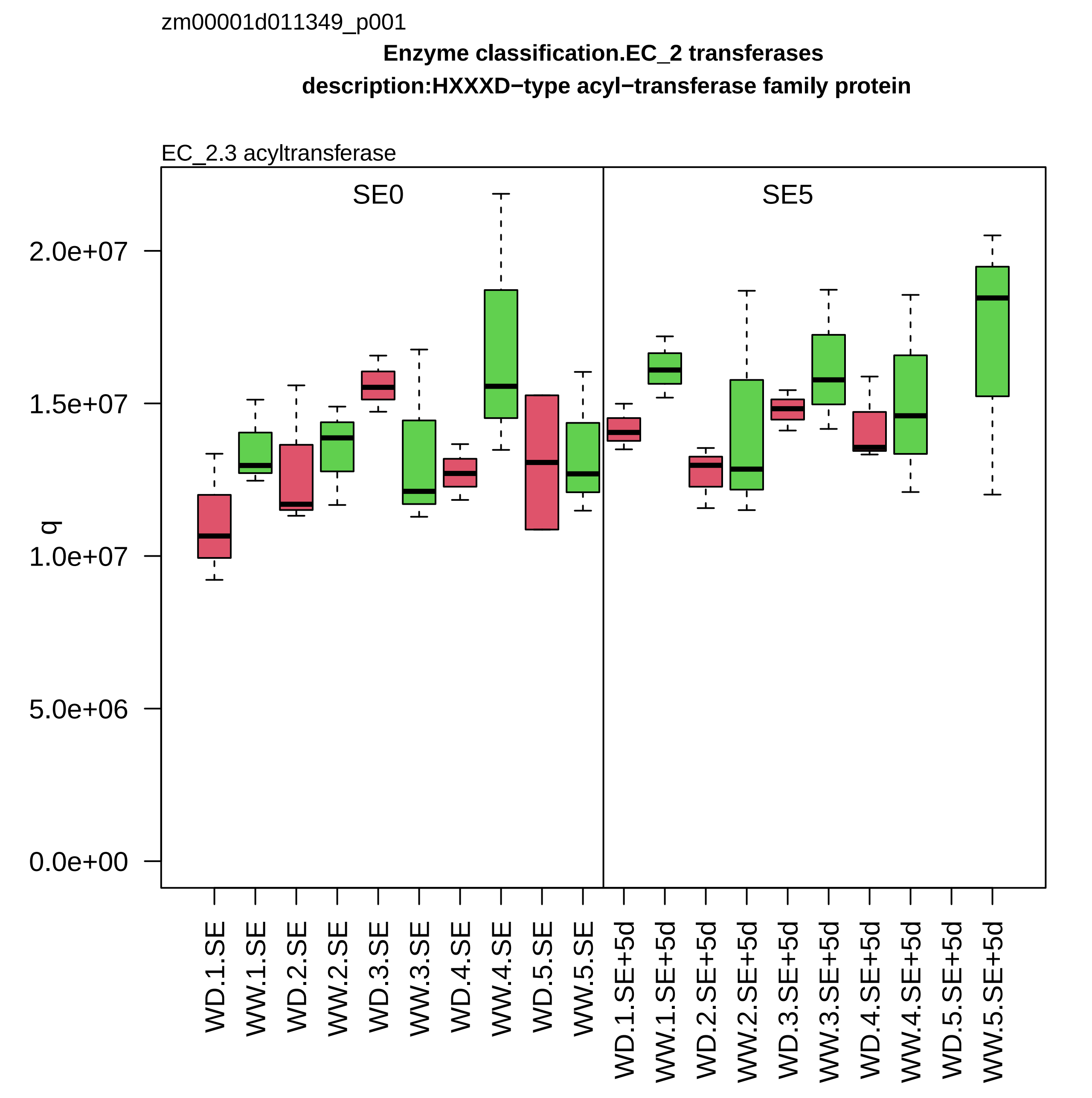
zm00001d011349_p001
Enzyme classification.EC_2 transferases
description:HXXXD−type acyl−transferase family protein
EC_2.3 acyltransferase
SE0
SE5
q
0.0e+00
5.0e+06
1.0e+07
1.5e+07
2.0e+07
WD.1.SE
WW.1.SE
WD.2.SE
WW.2.SE
WD.3.SE
WW.3.SE
WD.4.SE
WW.4.SE
WD.5.SE
WW.5.SE
WD.1.SE+5d
WW.1.SE+5d
WD.2.SE+5d
WW.2.SE+5d
WD.3.SE+5d
WW.3.SE+5d
WD.4.SE+5d
WW.4.SE+5d
WD.5.SE+5d
WW.5.SE+5d

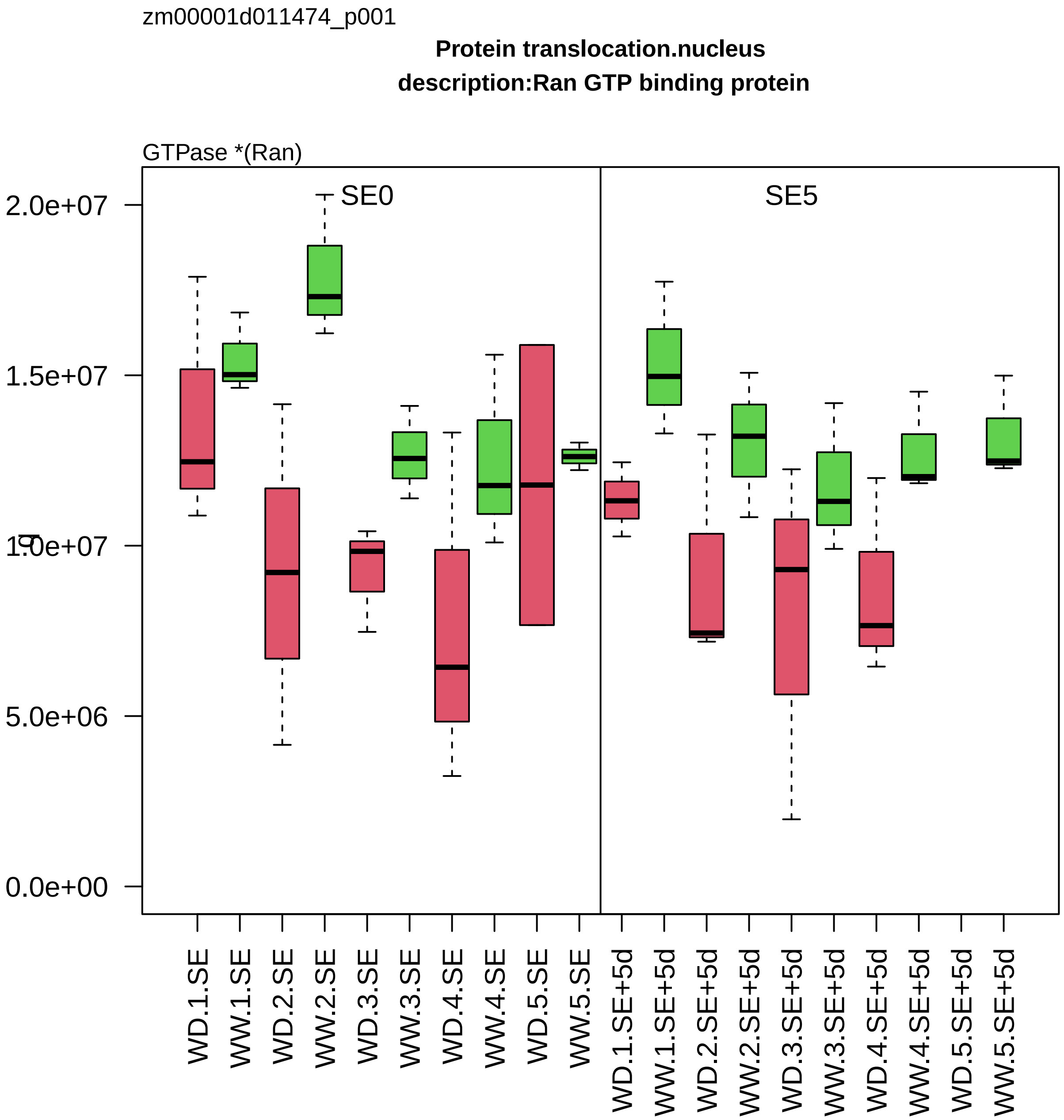
zm00001d011474_p001
Protein translocation.nucleus
description:Ran GTP binding protein
GTPase *(Ran)
SE0
SE5
2.0e+07
1.5e+07
1.0e+07
5.0e+06
0.0e+00
WD.1.SE
WW.1.SE
WD.2.SE
WW.2.SE
WD.3.SE
WW.3.SE
WD.4.SE
WW.4.SE
WD.5.SE
WW.5.SE
WD.1.SE+5d
WW.1.SE+5d
WD.2.SE+5d
WW.2.SE+5d
WD.3.SE+5d
WW.3.SE+5d
WD.4.SE+5d
WW.4.SE+5d
WD.5.SE+5d
WW.5.SE+5d

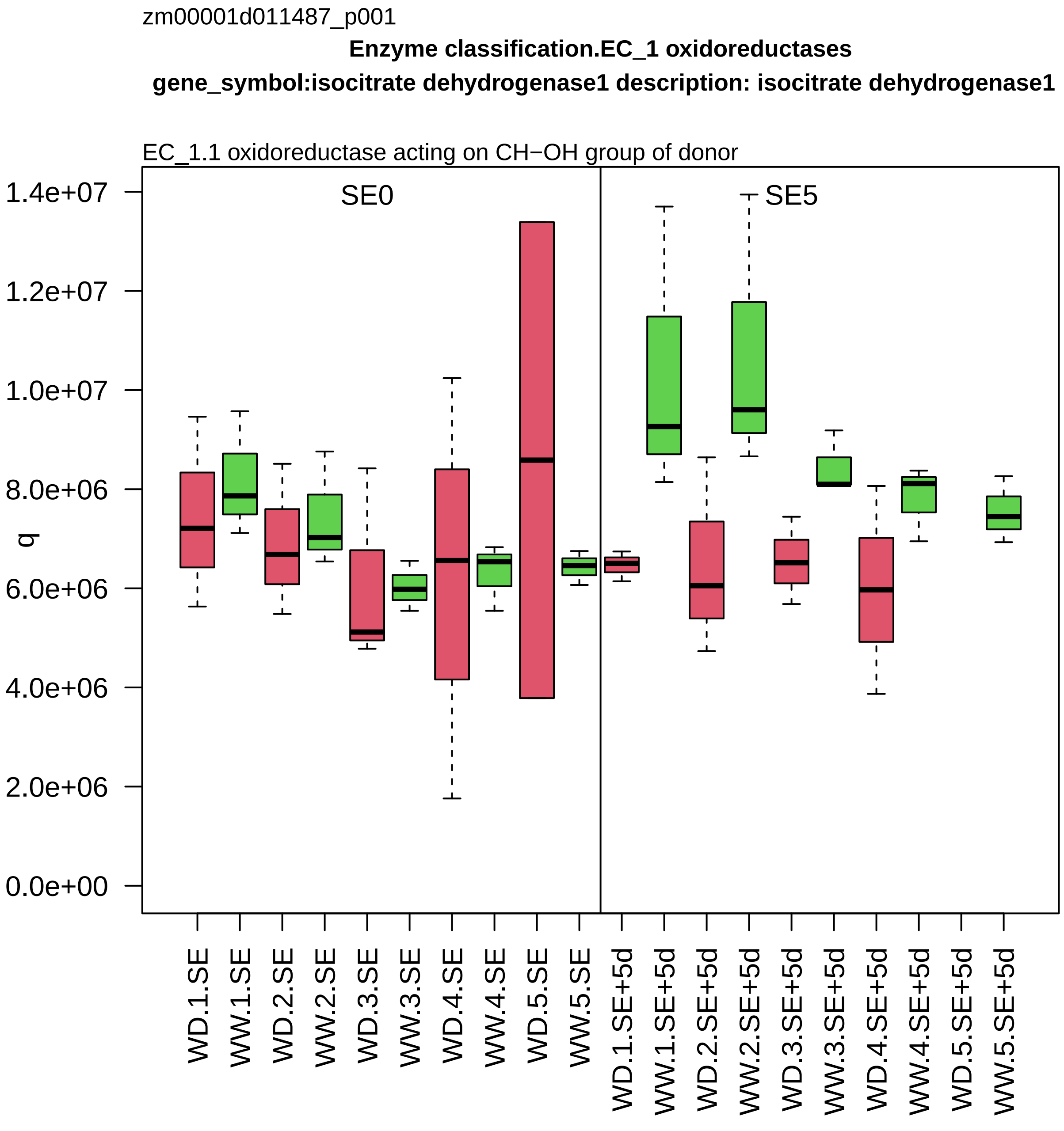

zm00001d011487_p001
Enzyme classification.EC_1 oxidoreductases
gene_symbol:isocitrate dehydrogenase1 description: isocitrate dehydrogenase1
EC_1.1 oxidoreductase acting on CH−OH group of donor
SE0
SE5
q
0.0e+00
2.0e+06
4.0e+06
6.0e+06
8.0e+06
1.0e+07
1.2e+07
1.4e+07
WD.1.SE
WW.1.SE
WD.2.SE
WW.2.SE
WD.3.SE
WW.3.SE
WD.4.SE
WW.4.SE
WD.5.SE
WW.5.SE
WD.1.SE+5d
WW.1.SE+5d
WD.2.SE+5d
WW.2.SE+5d
WD.3.SE+5d
WW.3.SE+5d
WD.4.SE+5d
WW.4.SE+5d
WD.5.SE+5d
WW.5.SE+5d

zm00001d011581_p001

**Redox homeostasis.thiol−based redox regulation**
**description:Peroxiredoxin−2B**

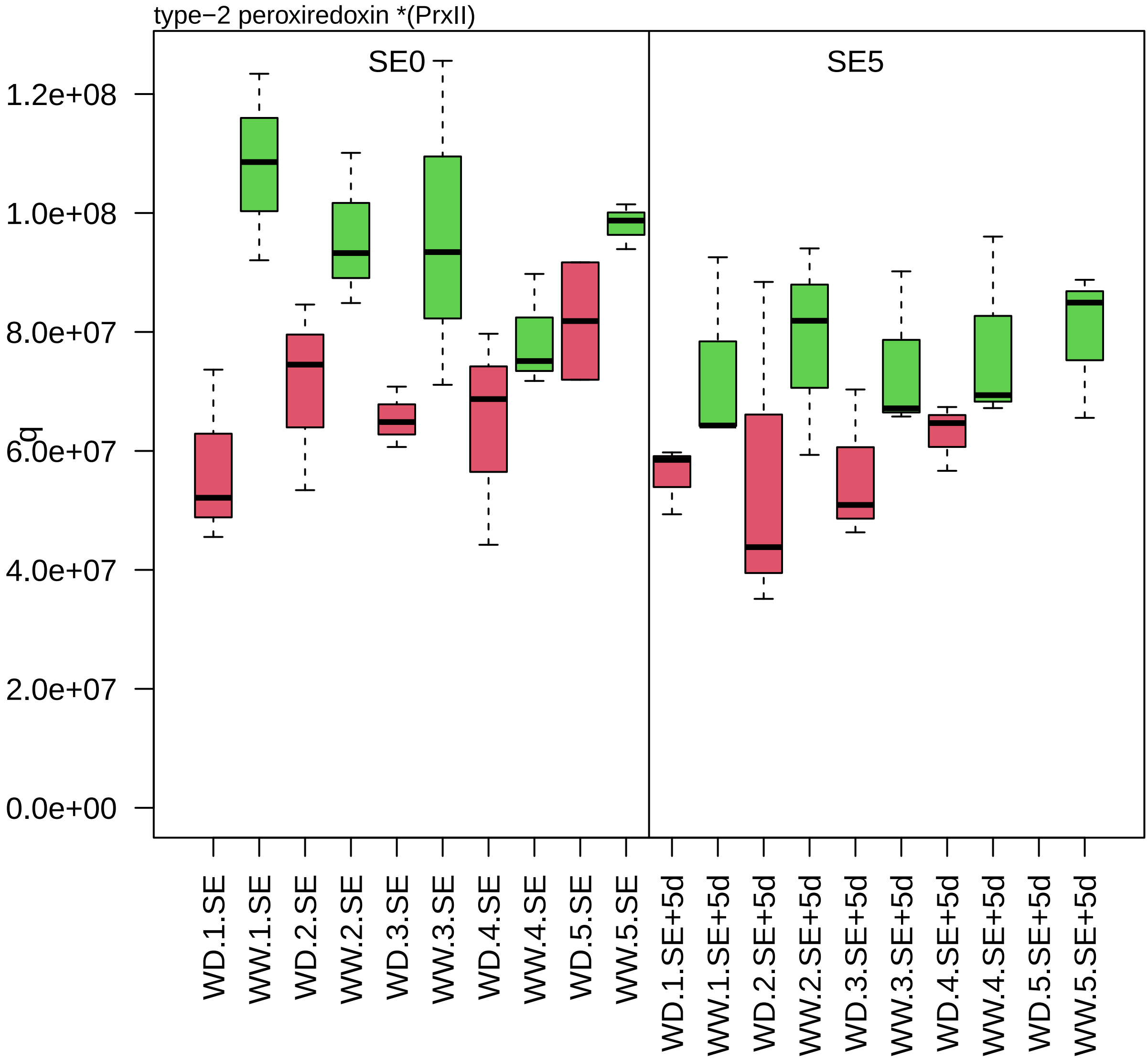

zm00001d011705_p001

**Chromatin organisation.histone chaperone activities**

**description:WD−40 repeat−containing protein MSI4**

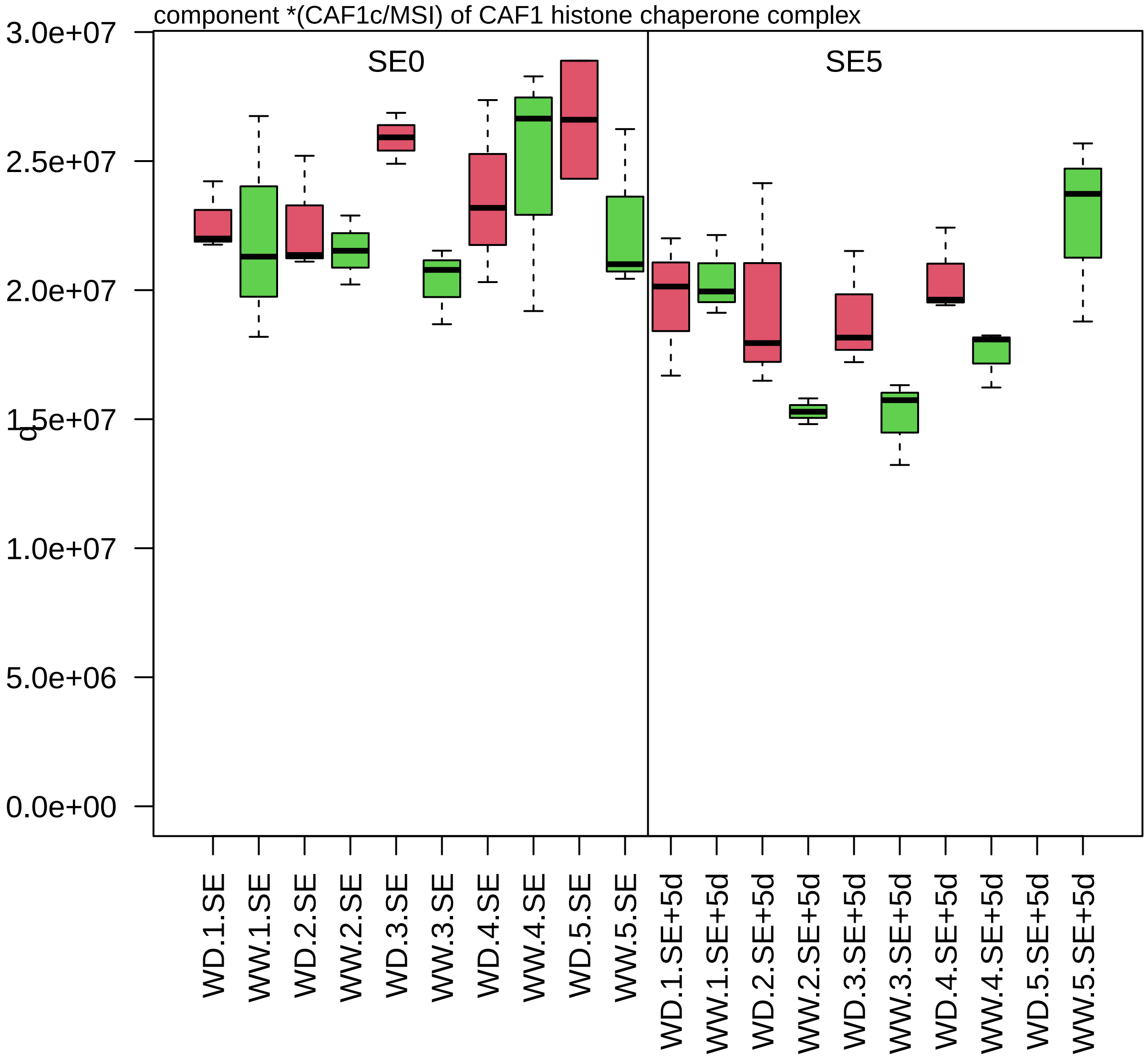

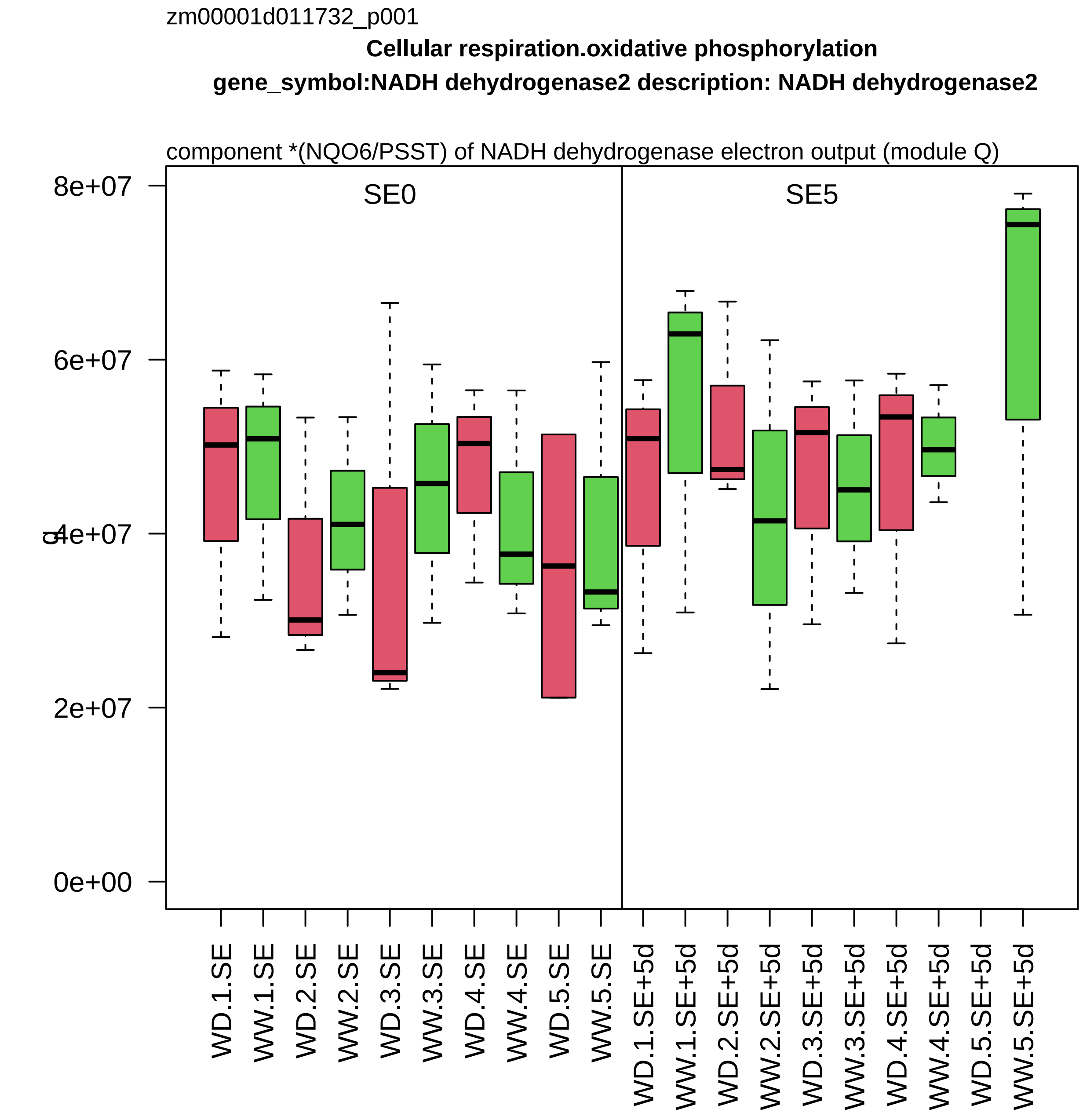
zm00001d011732_p001
Cellular respiration.oxidative phosphorylation
gene_symbol:NADH dehydrogenase2 description: NADH dehydrogenase2
component *(NQO6/PSST) of NADH dehydrogenase electron output (module Q)
SE0
SE5
8e+07
6e+07
4e+07
2e+07
0e+00
WD.1.SE
WW.1.SE
WD.2.SE
WW.2.SE
WD.3.SE
WW.3.SE
WD.4.SE
WW.4.SE
WD.5.SE
WW.5.SE
WD.1.SE+5d
WW.1.SE+5d
WD.2.SE+5d
WW.2.SE+5d
WD.3.SE+5d
WW.3.SE+5d
WD.4.SE+5d
WW.4.SE+5d
WD.5.SE+5d
WW.5.SE+5d

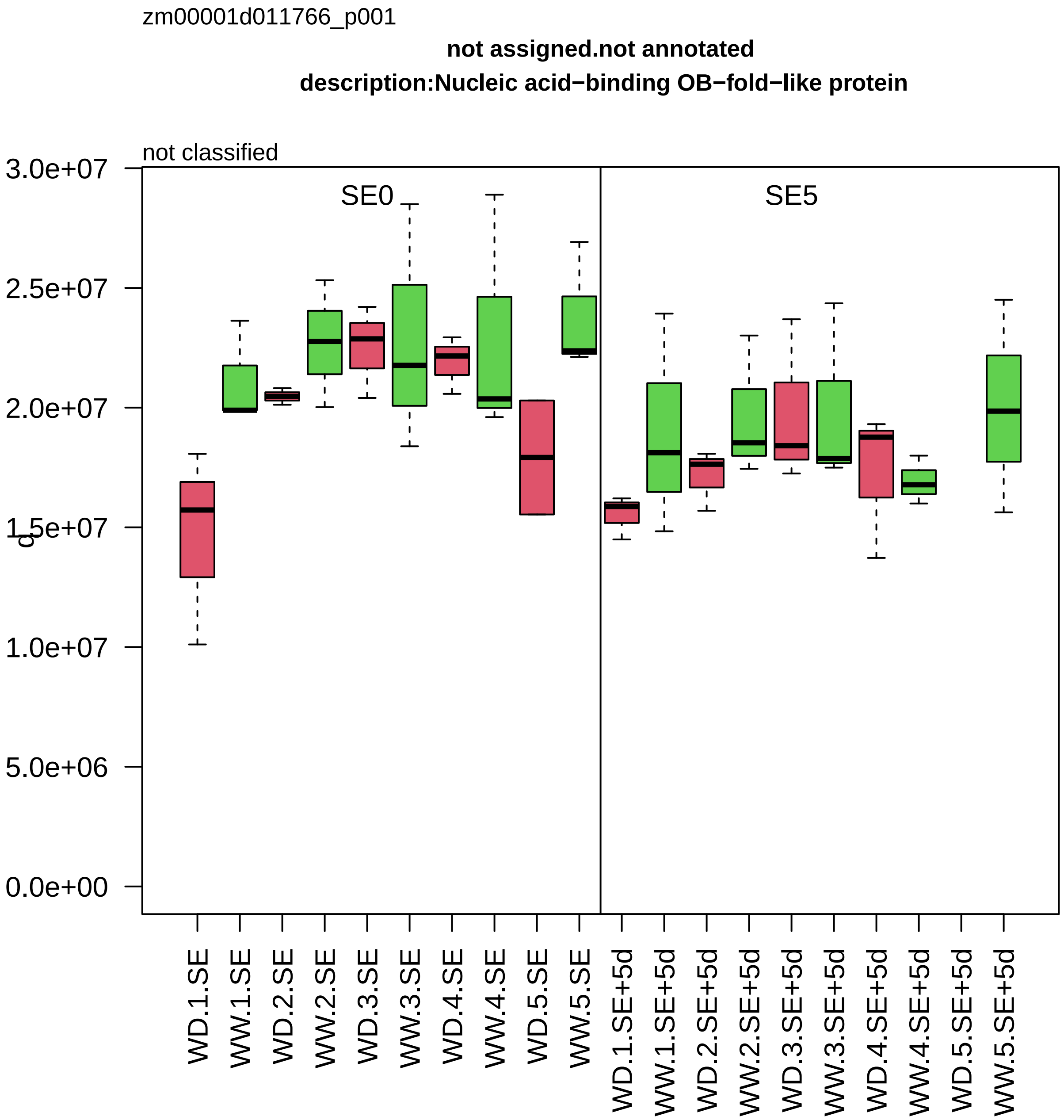

zm00001d011766_p001
not assigned.not annotated
description:Nucleic acid−binding OB−fold−like protein
not classified
SE0
SE5
3.0e+07
2.5e+07
2.0e+07
1.5e+07
1.0e+07
5.0e+06
0.0e+00
c
WD.1.SE
WW.1.SE
WD.2.SE
WW.2.SE
WD.3.SE
WW.3.SE
WD.4.SE
WW.4.SE
WD.5.SE
WW.5.SE
WD.1.SE+5d
WW.1.SE+5d
WD.2.SE+5d
WW.2.SE+5d
WD.3.SE+5d
WW.3.SE+5d
WD.4.SE+5d
WW.4.SE+5d
WD.5.SE+5d
WW.5.SE+5d

zm00001d011816_p001

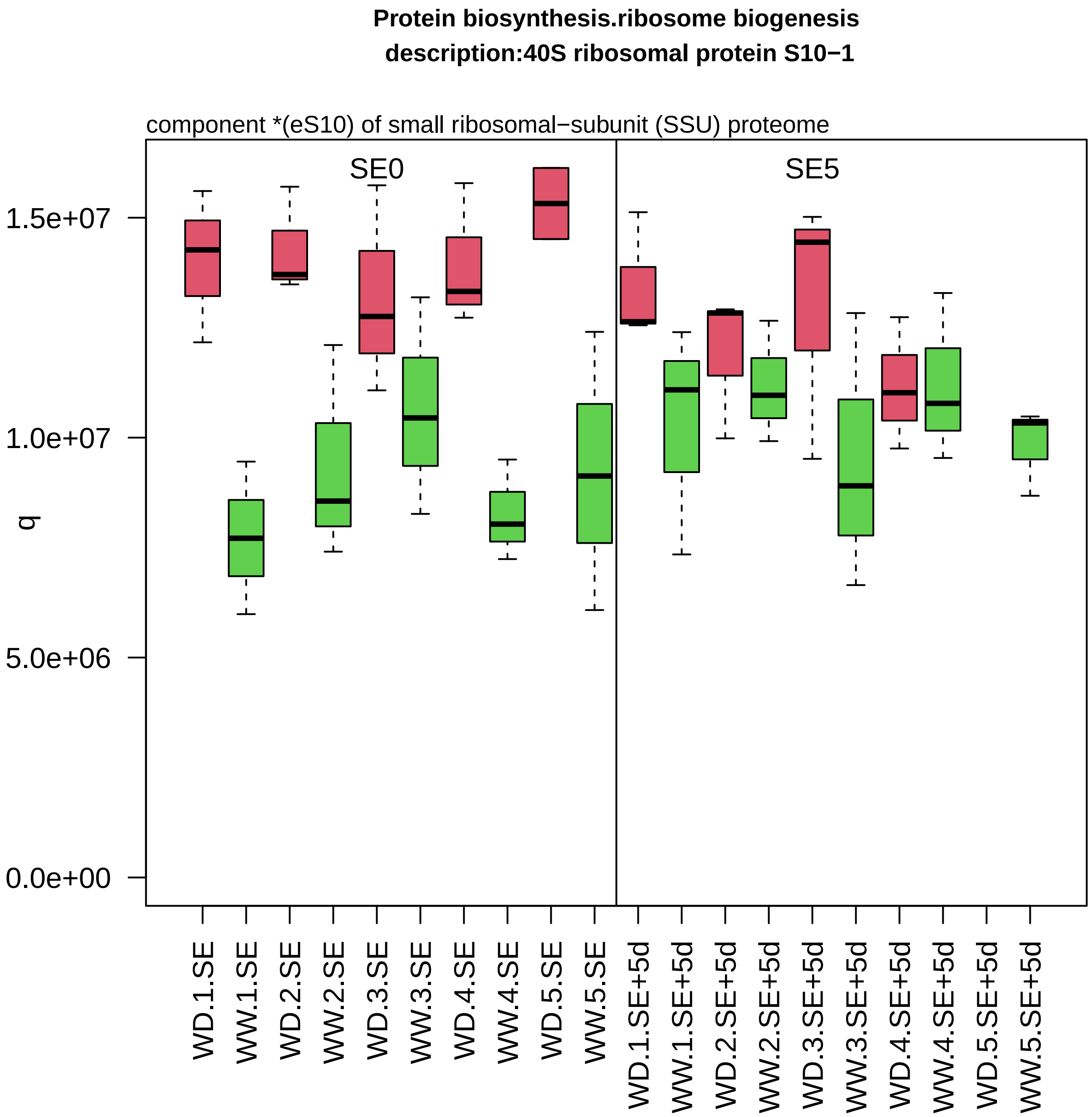

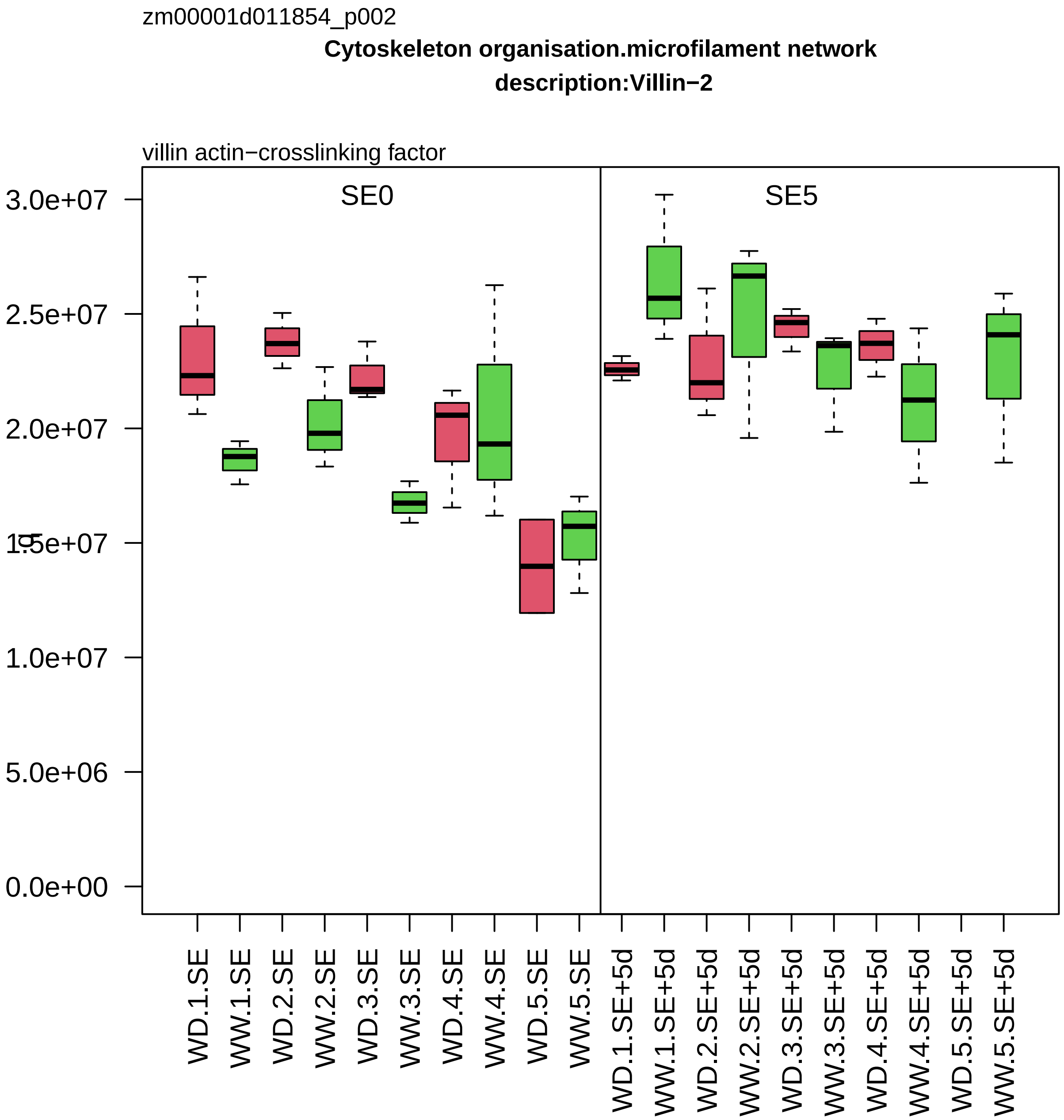

zm00001d011854_p002
Cytoskeleton organisation.microfilament network
description:Villin−2
villin actin−crosslinking factor
SE0
SE5
3.0e+07
2.5e+07
2.0e+07
1.5e+07
1.0e+07
5.0e+06
0.0e+00
WD.1.SE
WW.1.SE
WD.2.SE
WW.2.SE
WD.3.SE
WW.3.SE
WD.4.SE
WW.4.SE
WD.5.SE
WW.5.SE
WD.1.SE+5d
WW.1.SE+5d
WD.2.SE+5d
WW.2.SE+5d
WD.3.SE+5d
WW.3.SE+5d
WD.4.SE+5d
WW.4.SE+5d
WD.5.SE+5d
WW.5.SE+5d

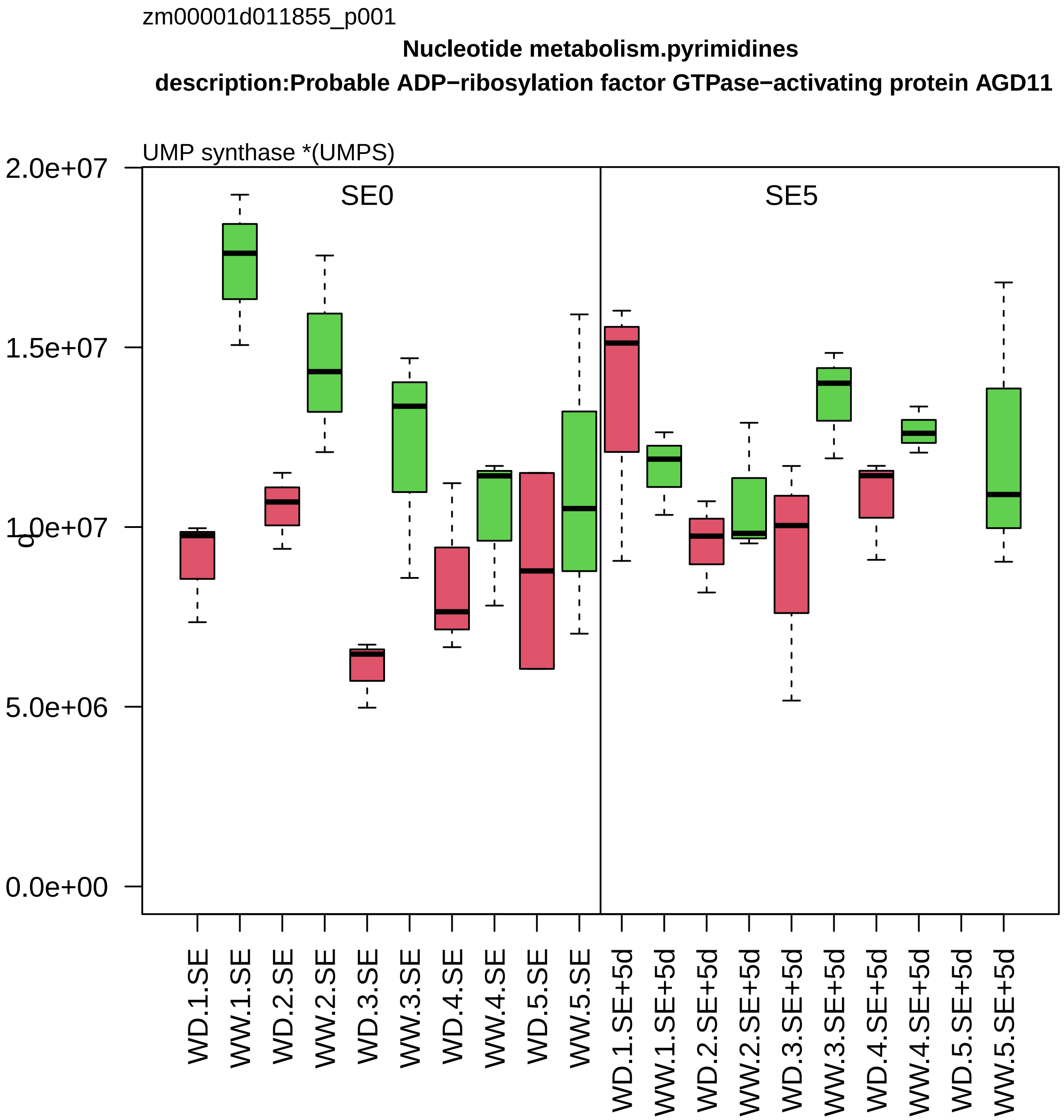

zm00001d011855_p001
Nucleotide metabolism.pyrimidines
description:Probable ADP−ribosylation factor GTPase−activating protein AGD11
UMP synthase *(UMPS)
SE0
SE5
2.0e+07
1.5e+07
1.0e+07
5.0e+06
0.0e+00
q
WD.1.SE
WW.1.SE
WD.2.SE
WW.2.SE
WD.3.SE
WW.3.SE
WD.4.SE
WW.4.SE
WD.5.SE
WW.5.SE
WD.1.SE+5d
WW.1.SE+5d
WD.2.SE+5d
WW.2.SE+5d
WD.3.SE+5d
WW.3.SE+5d
WD.4.SE+5d
WW.4.SE+5d
WD.5.SE+5d
WW.5.SE+5d

zm00001d011891_p001

**External stimuli response.toxic compounds**
**description:S−formylglutathione hydrolase**

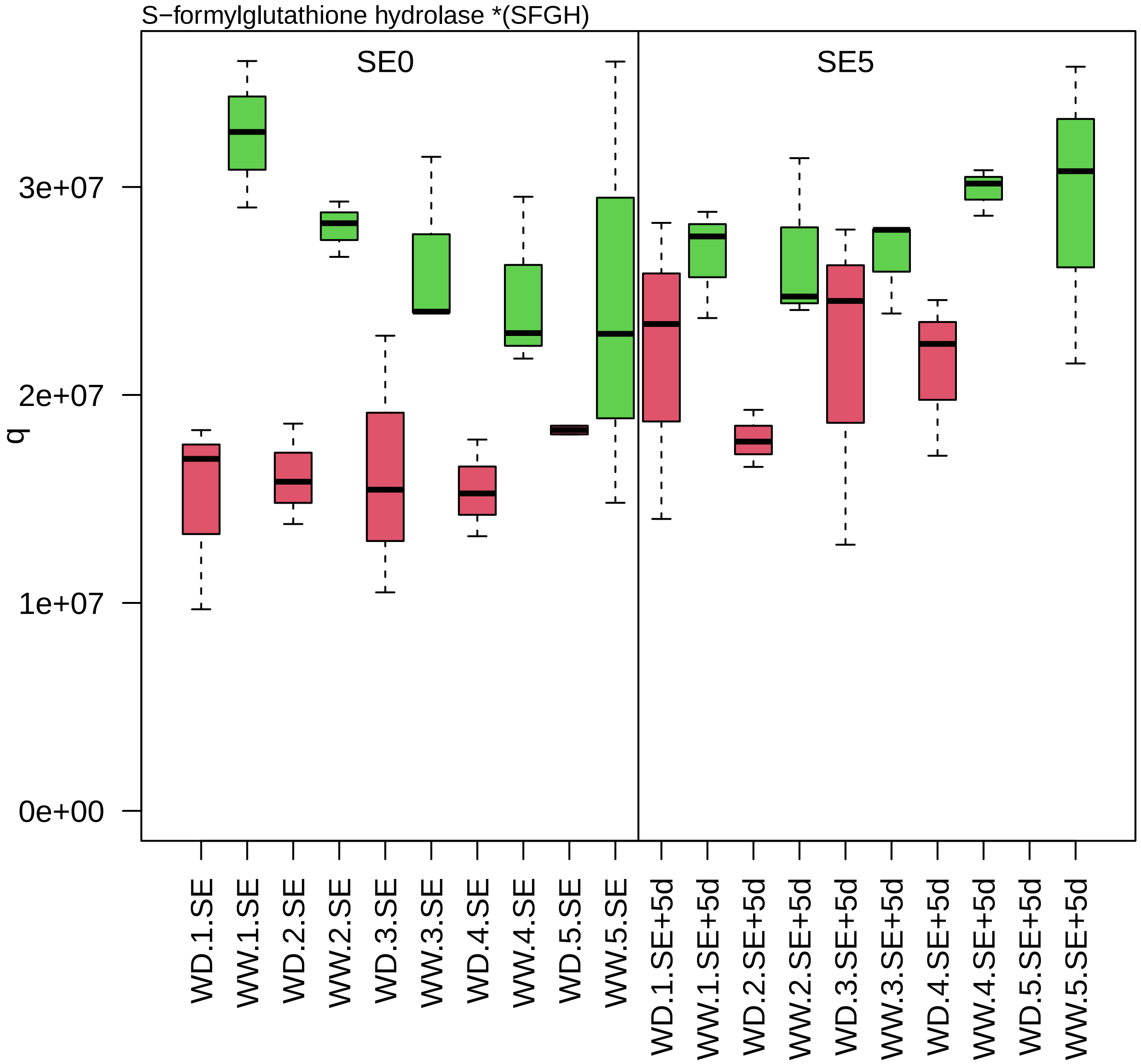

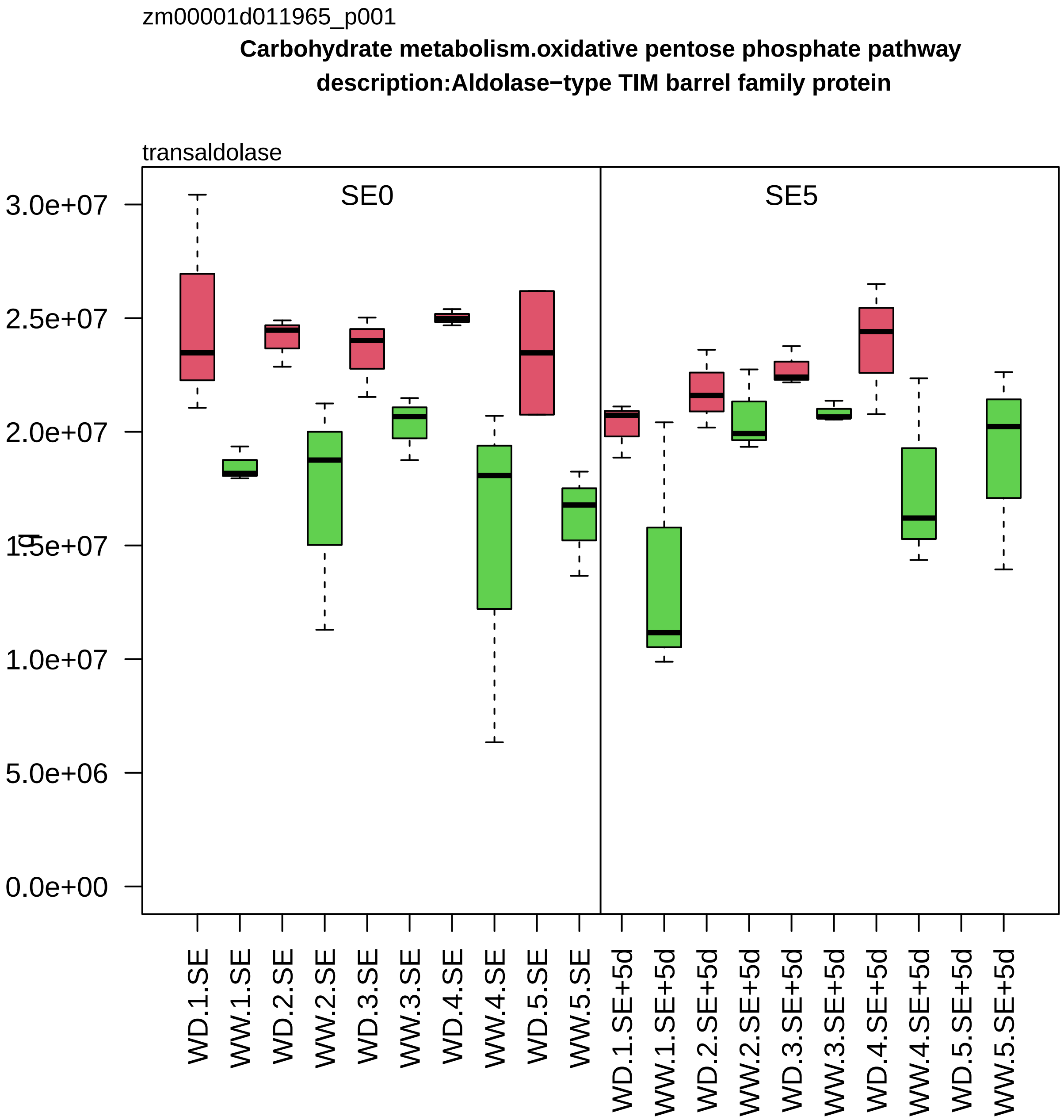

zm00001d011965_p001
Carbohydrate metabolism.oxidative pentose phosphate pathway
description:Aldolase−type TIM barrel family protein
transaldolase
SE0
SE5
3.0e+07
2.5e+07
2.0e+07
1.5e+07
1.0e+07
5.0e+06
0.0e+00
WD.1.SE
WW.1.SE
WD.2.SE
WW.2.SE
WD.3.SE
WW.3.SE
WD.4.SE
WW.4.SE
WD.5.SE
WW.5.SE
WD.1.SE+5d
WW.1.SE+5d
WD.2.SE+5d
WW.2.SE+5d
WD.3.SE+5d
WW.3.SE+5d
WD.4.SE+5d
WW.4.SE+5d
WD.5.SE+5d
WW.5.SE+5d

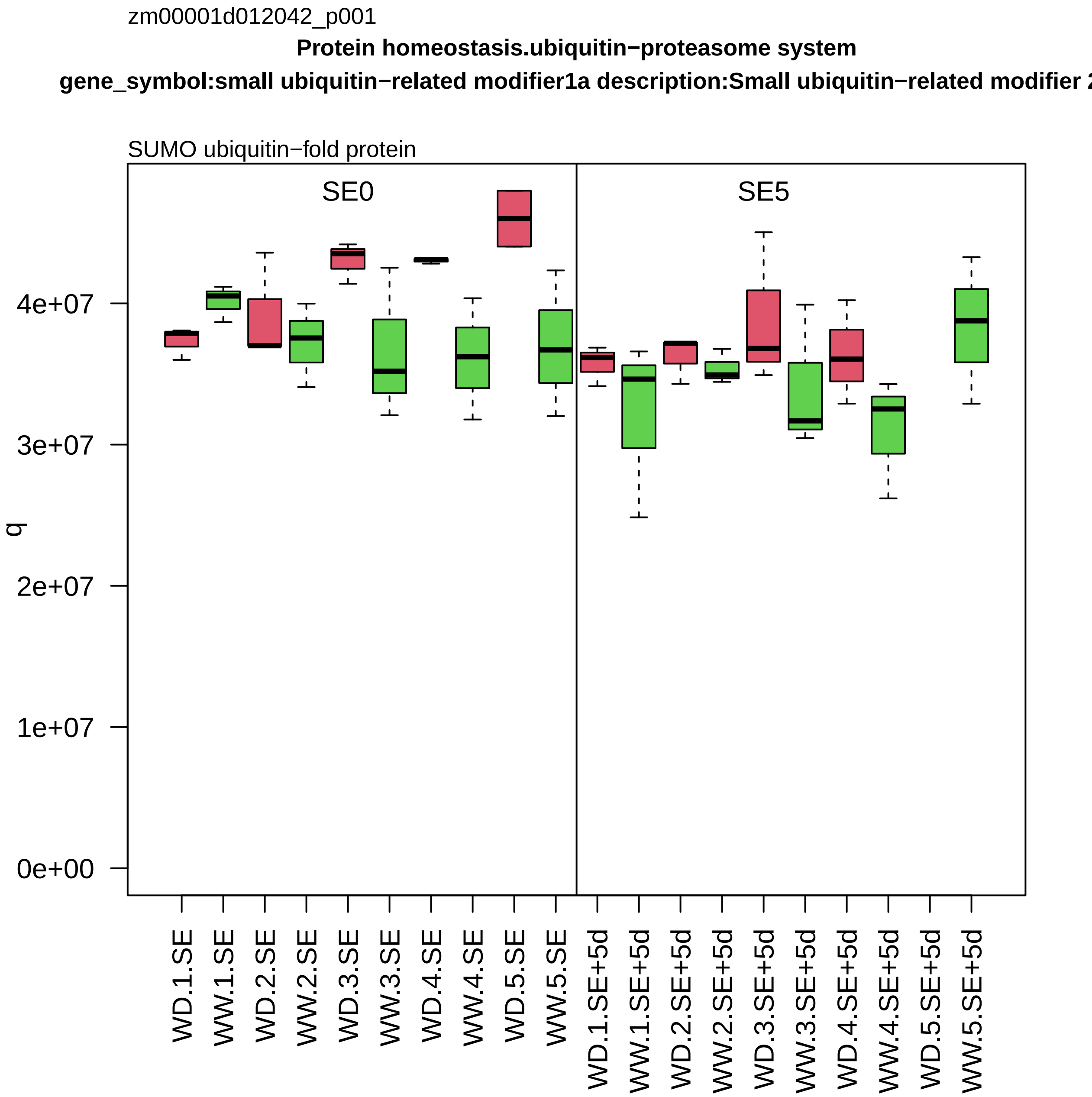
zm00001d012042_p001
Protein homeostasis.ubiquitin−proteasome system
gene_symbol:small ubiquitin−related modifier1a description:Small ubiquitin−related modifier 2
SUMO ubiquitin−fold protein
SE0
SE5
q
0e+00
1e+07
2e+07
3e+07
4e+07
WD.1.SE
WW.1.SE
WD.2.SE
WW.2.SE
WD.3.SE
WW.3.SE
WD.4.SE
WW.4.SE
WD.5.SE
WW.5.SE
WD.1.SE+5d
WW.1.SE+5d
WD.2.SE+5d
WW.2.SE+5d
WD.3.SE+5d
WW.3.SE+5d
WD.4.SE+5d
WW.4.SE+5d
WD.5.SE+5d
WW.5.SE+5d

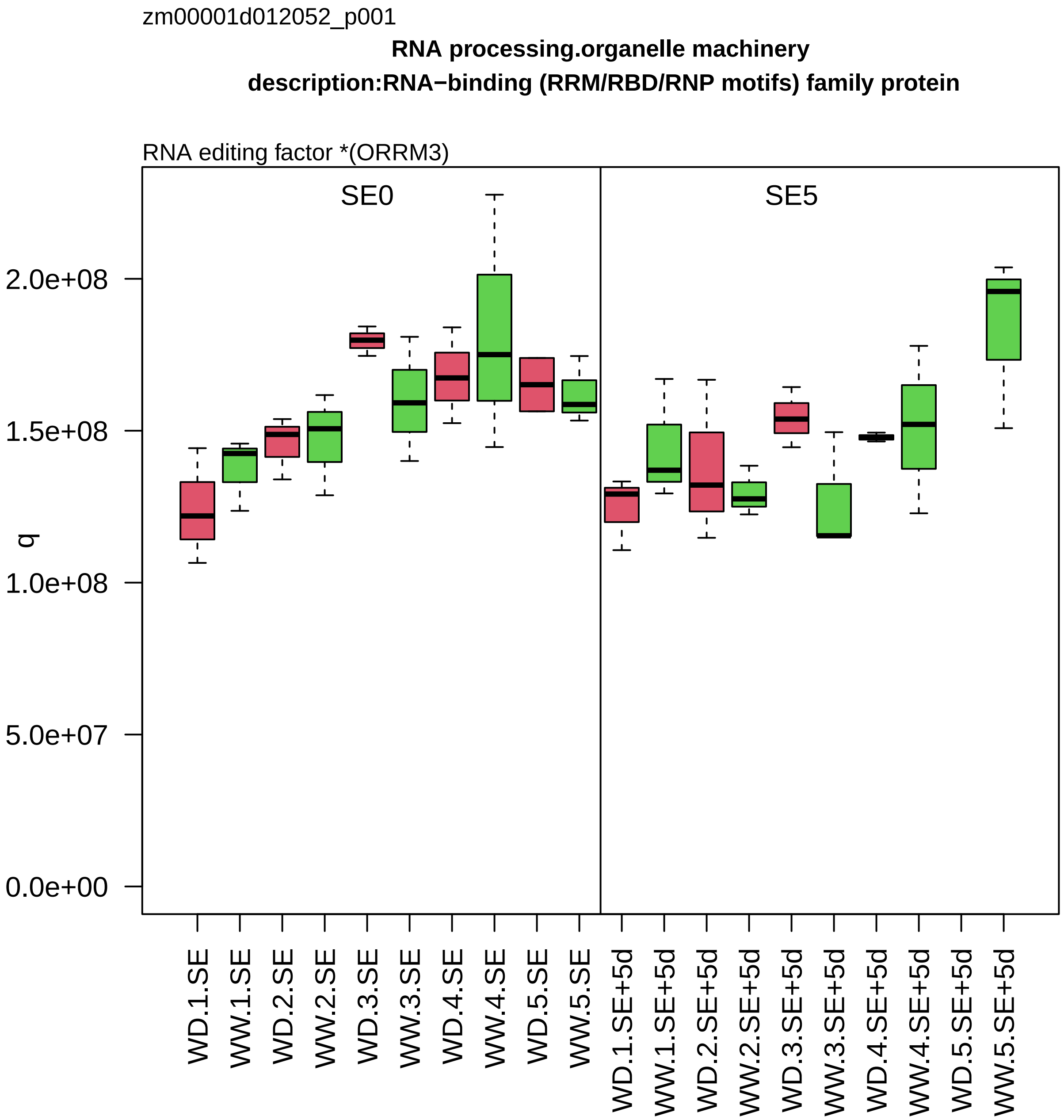

zm00001d012052_p001
RNA processing.organelle machinery
description:RNA−binding (RRM/RBD/RNP motifs) family protein
RNA editing factor *(ORRM3)
SE0
SE5
q
0.0e+00
5.0e+07
1.0e+08
1.5e+08
2.0e+08
WD.1.SE
WW.1.SE
WD.2.SE
WW.2.SE
WD.3.SE
WW.3.SE
WD.4.SE
WW.4.SE
WD.5.SE
WW.5.SE
WD.1.SE+5d
WW.1.SE+5d
WD.2.SE+5d
WW.2.SE+5d
WD.3.SE+5d
WW.3.SE+5d
WD.4.SE+5d
WW.4.SE+5d
WD.5.SE+5d
WW.5.SE+5d

zm00001d012103_p001

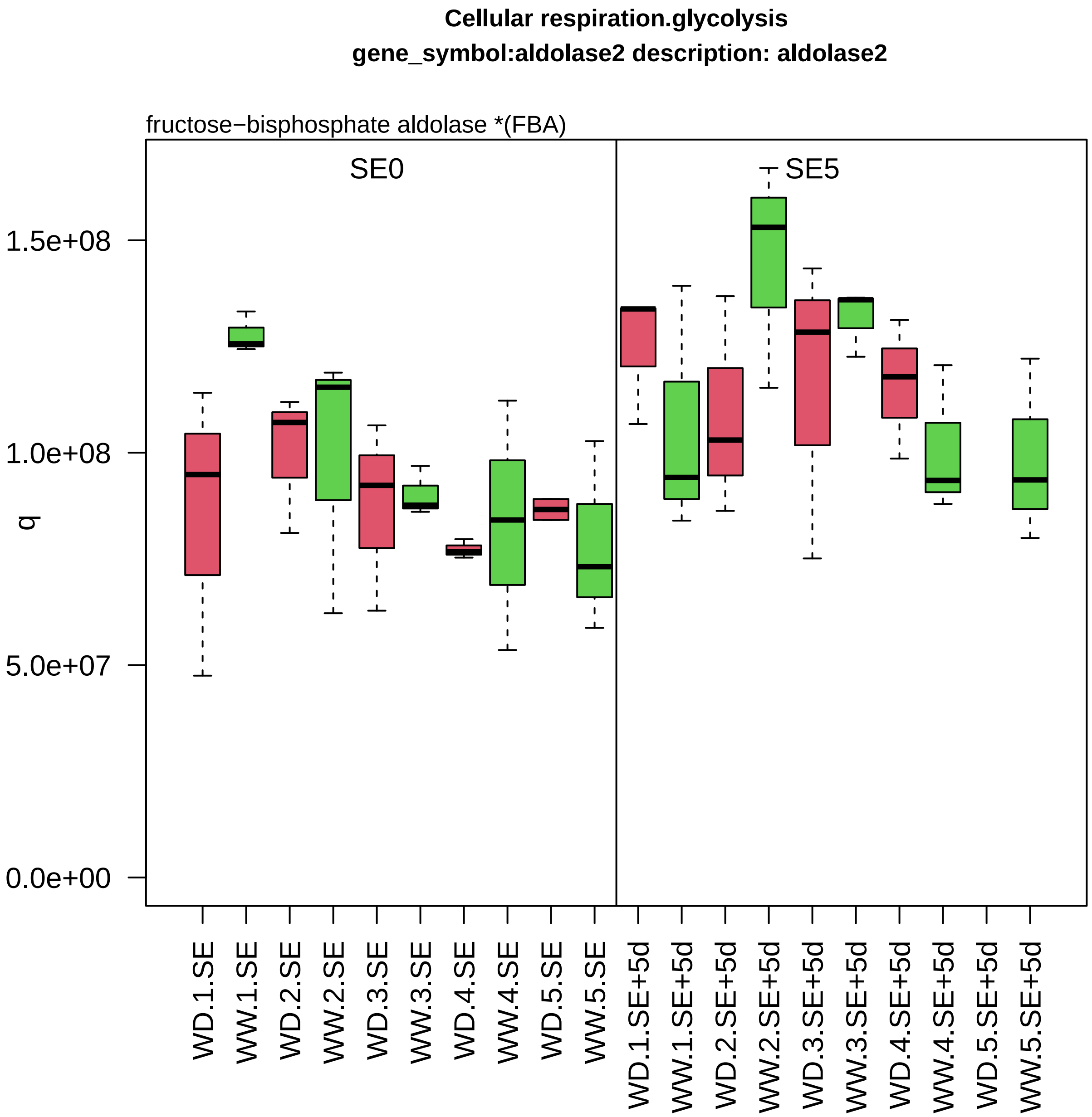

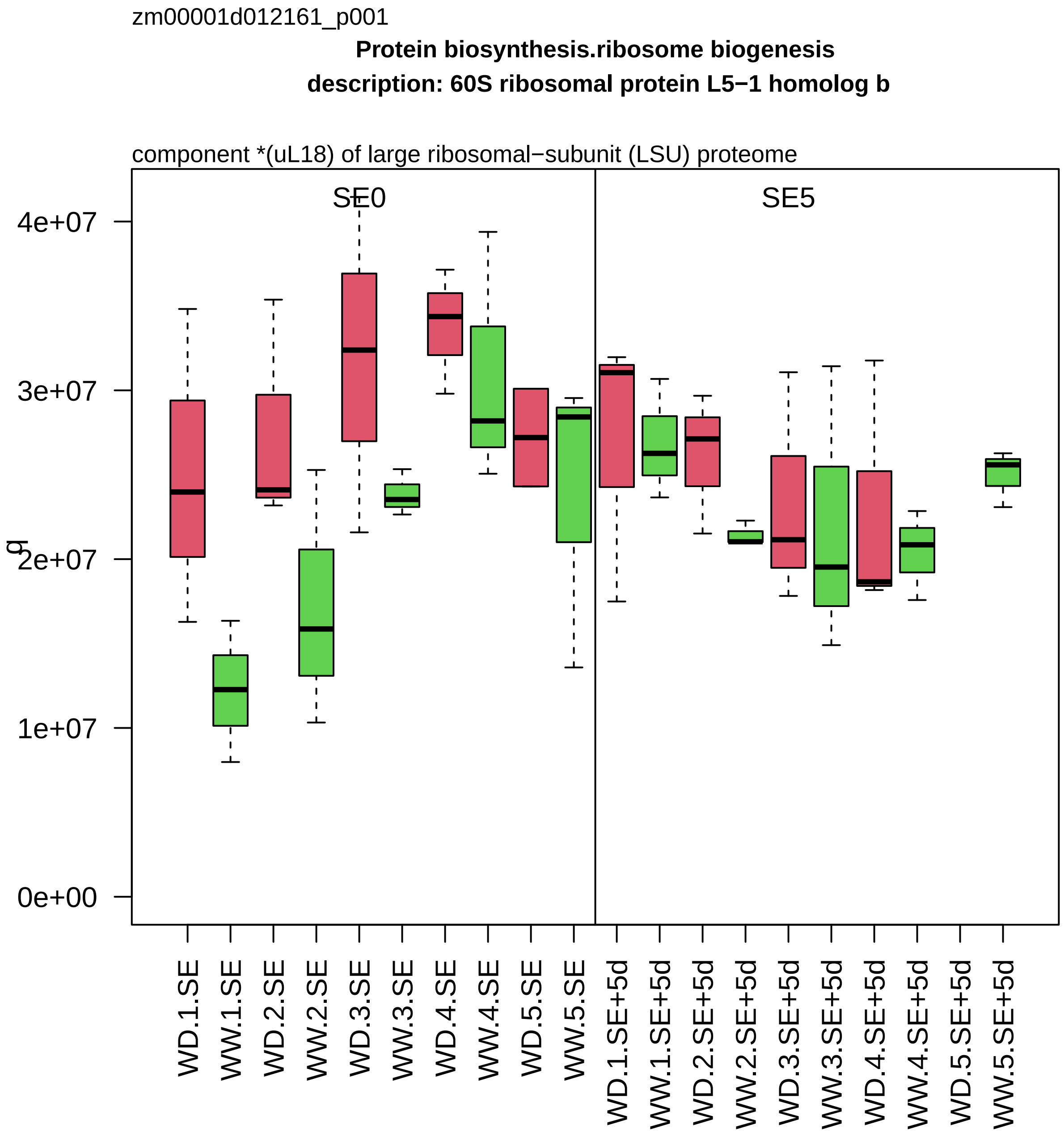
zm00001d012161_p001
Protein biosynthesis.ribosome biogenesis
description: 60S ribosomal protein L5−1 homolog b
component *(uL18) of large ribosomal−subunit (LSU) proteome
SE0
SE5
q
4e+07
3e+07
2e+07
1e+07
0e+00
WD.1.SE
WW.1.SE
WD.2.SE
WW.2.SE
WD.3.SE
WW.3.SE
WD.4.SE
WW.4.SE
WD.5.SE
WW.5.SE
WD.1.SE+5d
WW.1.SE+5d
WD.2.SE+5d
WW.2.SE+5d
WD.3.SE+5d
WW.3.SE+5d
WD.4.SE+5d
WW.4.SE+5d
WD.5.SE+5d
WW.5.SE+5d

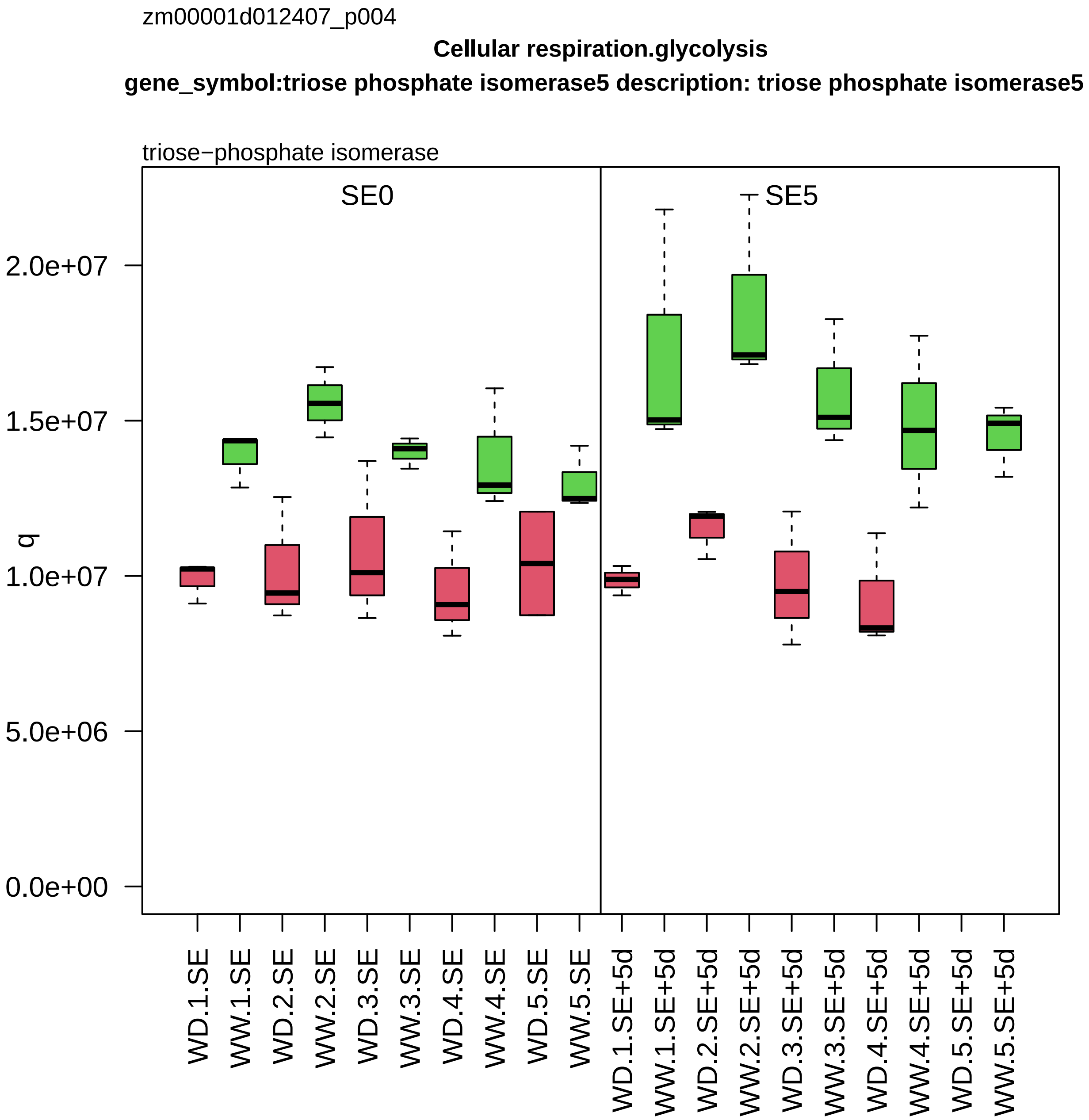
zm00001d012407_p004
Cellular respiration.glycolysis
gene_symbol:triose phosphate isomerase5 description: triose phosphate isomerase5
triose−phosphate isomerase
SE0
SE5
q
0.0e+00
5.0e+06
1.0e+07
1.5e+07
2.0e+07
WD.1.SE
WW.1.SE
WD.2.SE
WW.2.SE
WD.3.SE
WW.3.SE
WD.4.SE
WW.4.SE
WD.5.SE
WW.5.SE
WD.1.SE+5d
WW.1.SE+5d
WD.2.SE+5d
WW.2.SE+5d
WD.3.SE+5d
WW.3.SE+5d
WD.4.SE+5d
WW.4.SE+5d
WD.5.SE+5d
WW.5.SE+5d

zm00001d012420_p001

**Protein homeostasis.protein quality control**

**gene_symbol:heat shock protein1 description:Heat shock 70 kDa protein**

chaperone *(Hsp70)

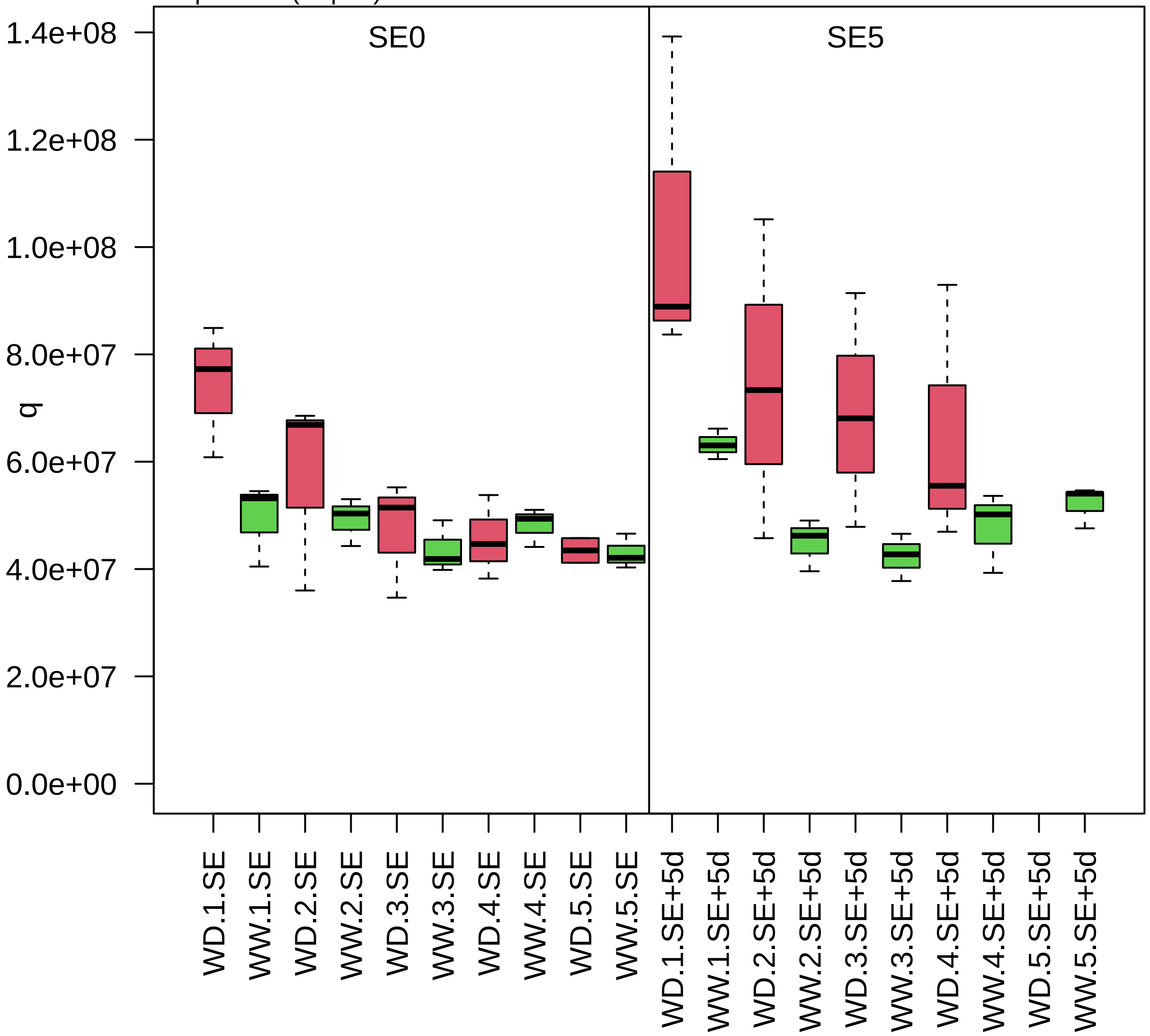

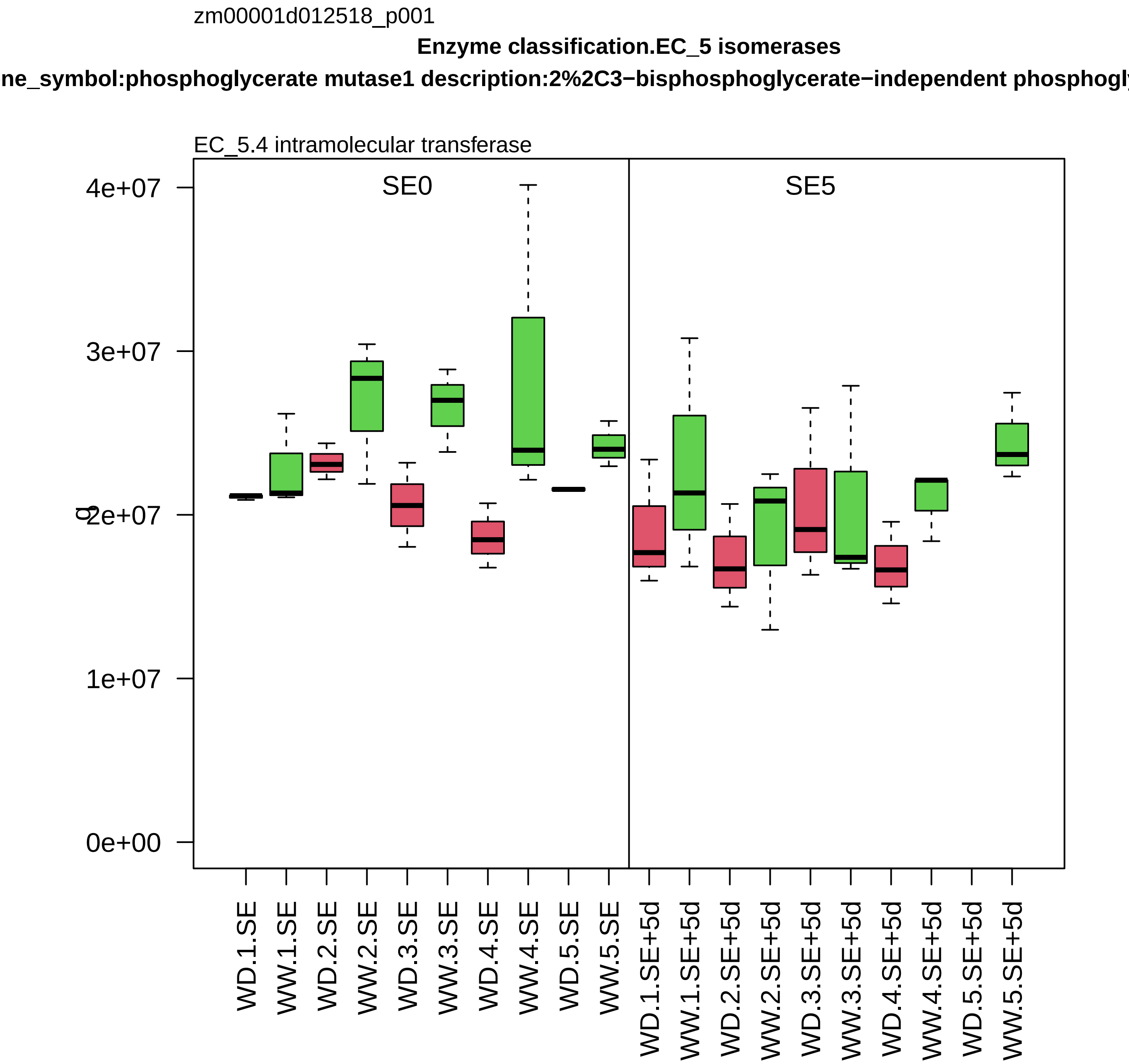
zm00001d012518_p001
Enzyme classification.EC_5 isomerases
ne_symbol:phosphoglycerate mutase1 description:2%2C3−bisphosphoglycerate−independent phosphogl
EC_5.4 intramolecular transferase
SE0
SE5
4e+07
3e+07
2e+07
1e+07
0e+00
WD.1.SE
WW.1.SE
WD.2.SE
WW.2.SE
WD.3.SE
WW.3.SE
WD.4.SE
WW.4.SE
WD.5.SE
WW.5.SE
WD.1.SE+5d
WW.1.SE+5d
WD.2.SE+5d
WW.2.SE+5d
WD.3.SE+5d
WW.3.SE+5d
WD.4.SE+5d
WW.4.SE+5d
WD.5.SE+5d
WW.5.SE+5d

zm00001d012546_p001

**Protein homeostasis.ubiquitin−proteasome system**
**description:Proteasome subunit alpha type−3**

alpha−type−7 component *(PAG) of 26S proteasome

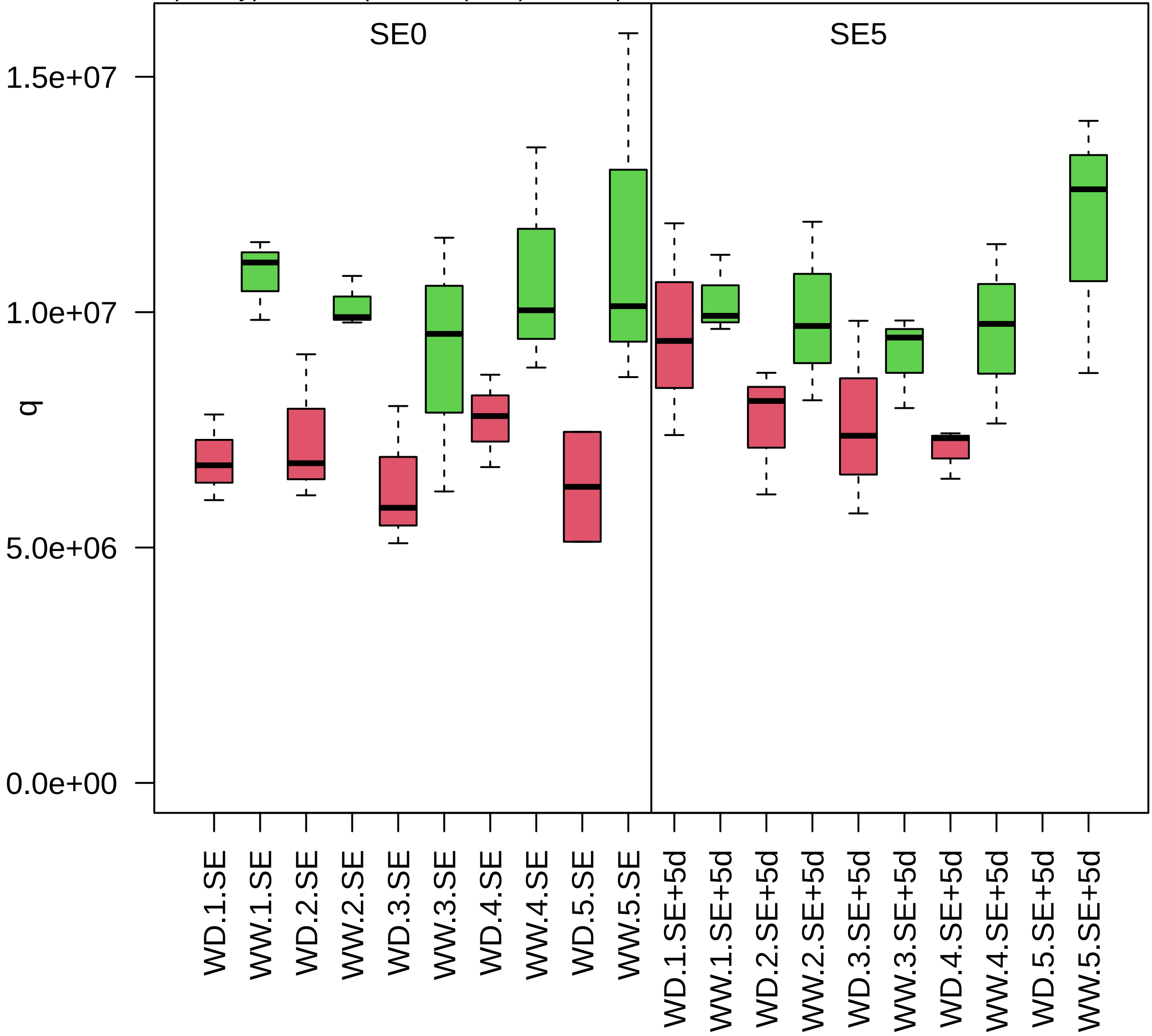

zm00001d012675_p001

**Redox homeostasis.glutathione−based redox regulation**

**gene_symbol:glutathione−S−transferase1 description:Glutathione S−transferase 1**

class phi glutathione S−transferase

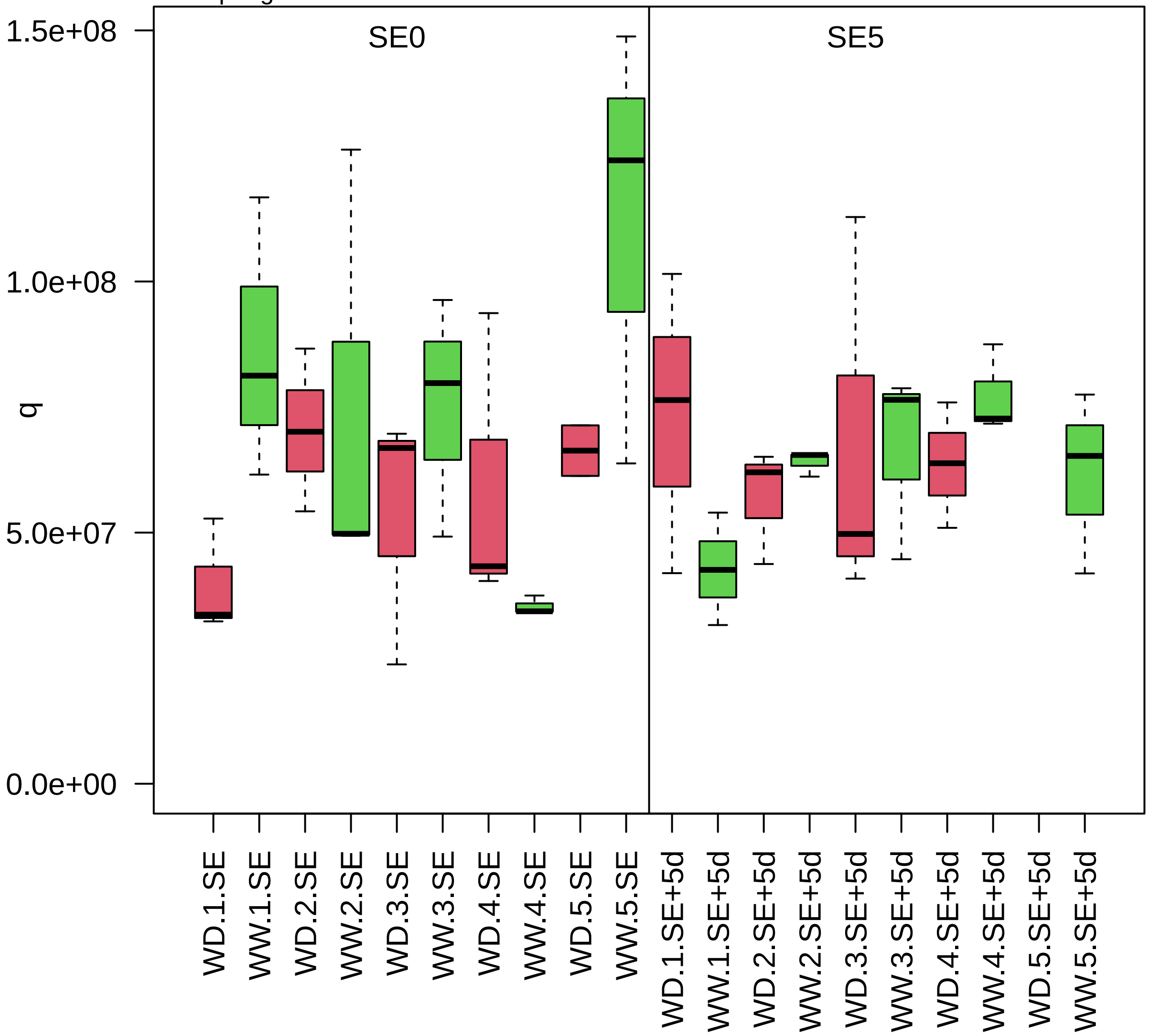

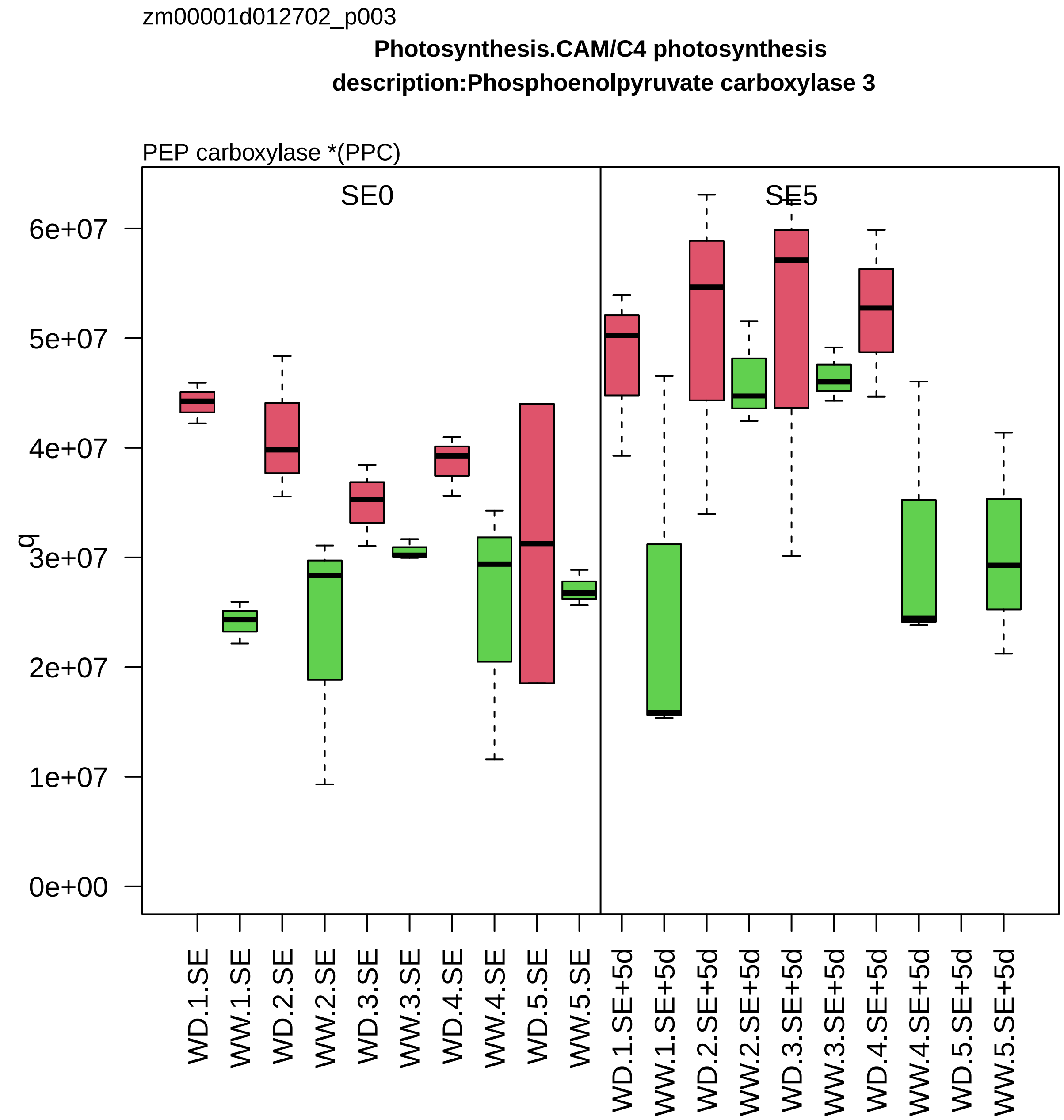

zm00001d012702_p003
Photosynthesis.CAM/C4 photosynthesis
description:Phosphoenolpyruvate carboxylase 3
PEP carboxylase *(PPC)
SE0
SE5
q
6e+07
5e+07
4e+07
3e+07
2e+07
1e+07
0e+00
WD.1.SE
WW.1.SE
WD.2.SE
WW.2.SE
WD.3.SE
WW.3.SE
WD.4.SE
WW.4.SE
WD.5.SE
WW.5.SE
WD.1.SE+5d
WW.1.SE+5d
WD.2.SE+5d
WW.2.SE+5d
WD.3.SE+5d
WW.3.SE+5d
WD.4.SE+5d
WW.4.SE+5d
WD.5.SE+5d
WW.5.SE+5d

zm00001d012837_p001

**Chromatin organisation.chromatin structure**
**description:Histone H2A**

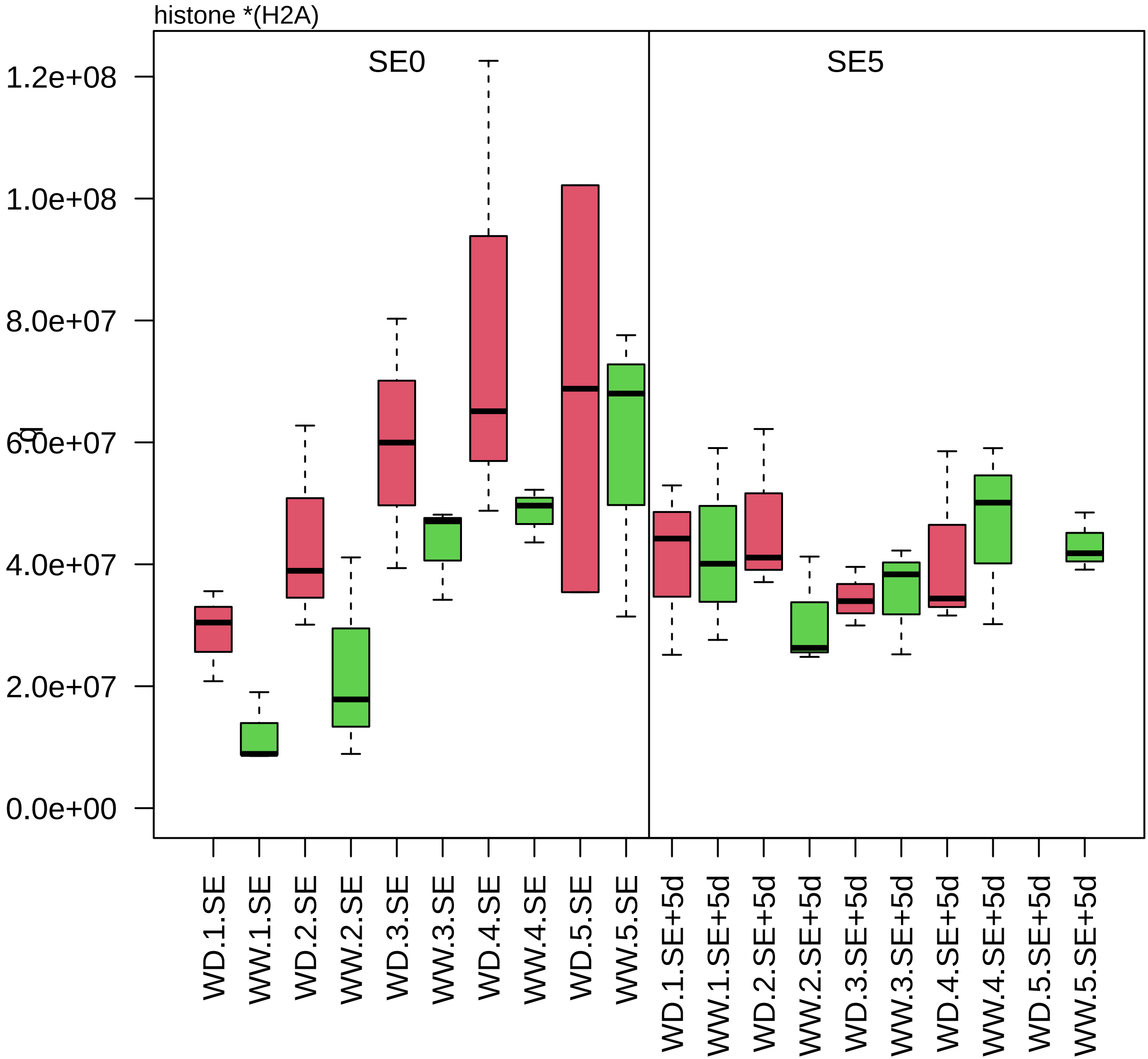

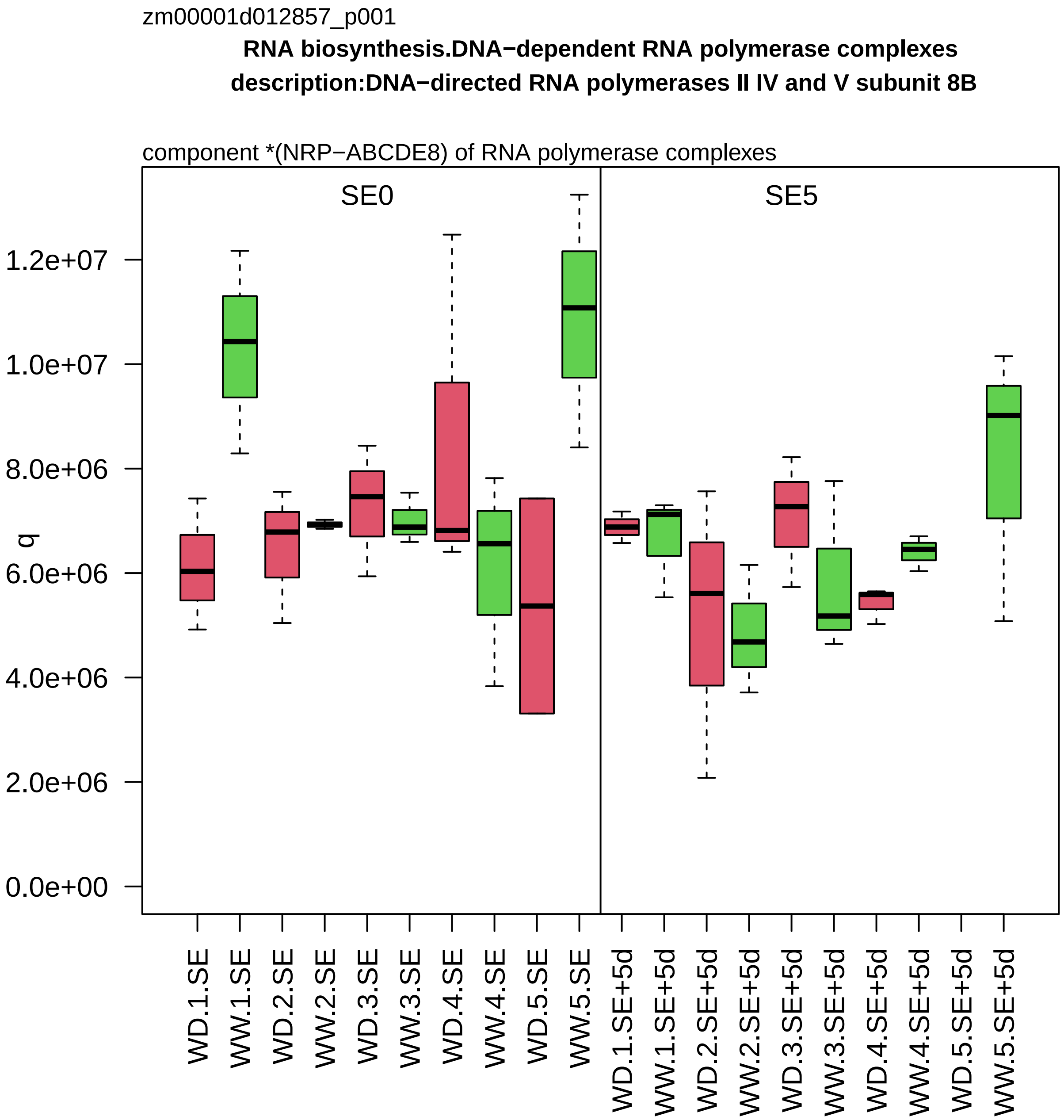

zm00001d012857_p001
RNA biosynthesis.DNA−dependent RNA polymerase complexes
description:DNA−directed RNA polymerases II IV and V subunit 8B
component *(NRP−ABCDE8) of RNA polymerase complexes
SE0
SE5
q
0.0e+00
2.0e+06
4.0e+06
6.0e+06
8.0e+06
1.0e+07
1.2e+07
WD.1.SE
WW.1.SE
WD.2.SE
WW.2.SE
WD.3.SE
WW.3.SE
WD.4.SE
WW.4.SE
WD.5.SE
WW.5.SE
WD.1.SE+5d
WW.1.SE+5d
WD.2.SE+5d
WW.2.SE+5d
WD.3.SE+5d
WW.3.SE+5d
WD.4.SE+5d
WW.4.SE+5d
WD.5.SE+5d
WW.5.SE+5d

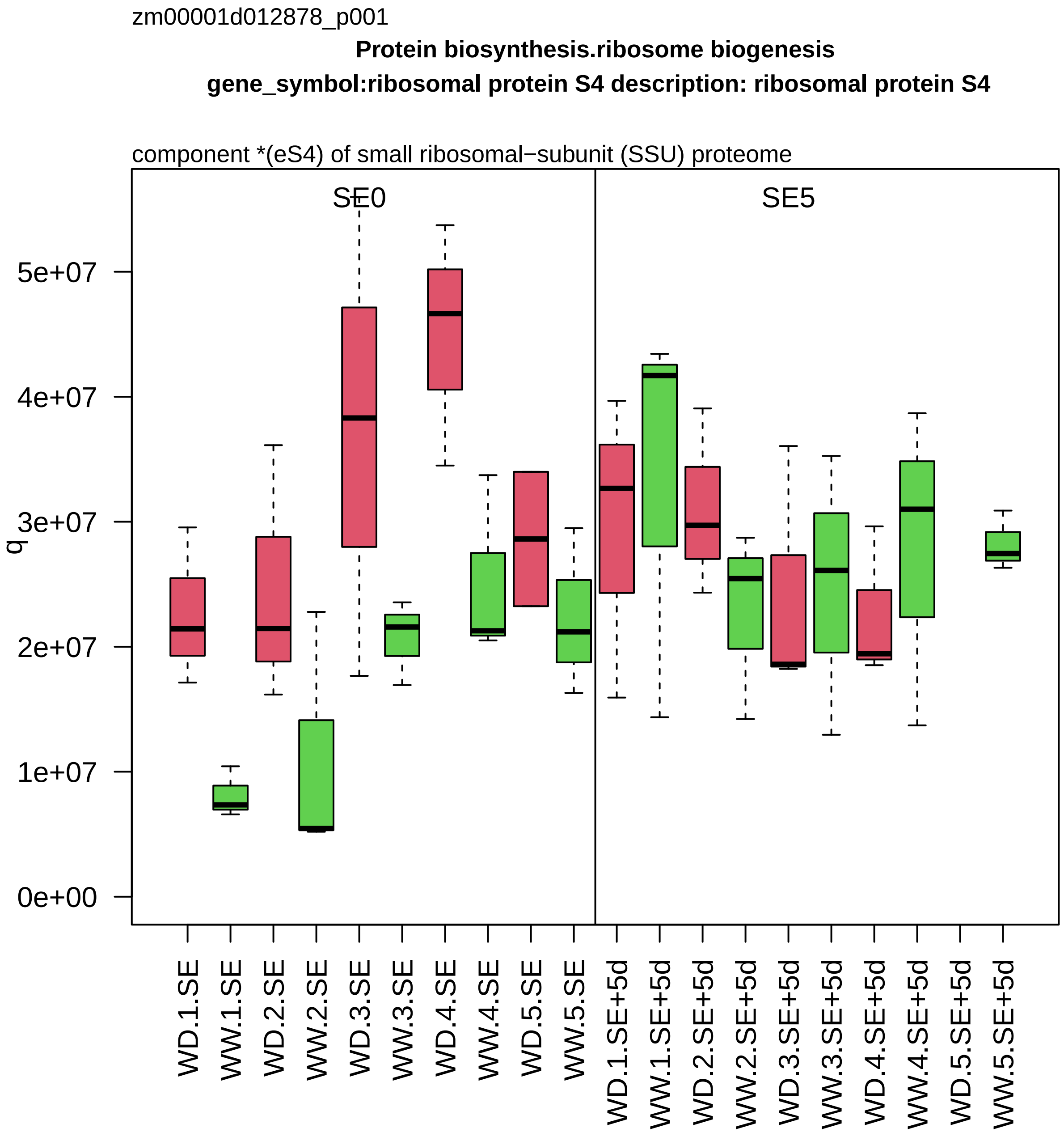

zm00001d012878_p001
Protein biosynthesis.ribosome biogenesis
gene_symbol:ribosomal protein S4 description: ribosomal protein S4
component *(eS4) of small ribosomal−subunit (SSU) proteome
SE0
SE5
q
0e+00
1e+07
2e+07
3e+07
4e+07
5e+07
WD.1.SE
WW.1.SE
WD.2.SE
WW.2.SE
WD.3.SE
WW.3.SE
WD.4.SE
WW.4.SE
WD.5.SE
WW.5.SE
WD.1.SE+5d
WW.1.SE+5d
WD.2.SE+5d
WW.2.SE+5d
WD.3.SE+5d
WW.3.SE+5d
WD.4.SE+5d
WW.4.SE+5d
WD.5.SE+5d
WW.5.SE+5d

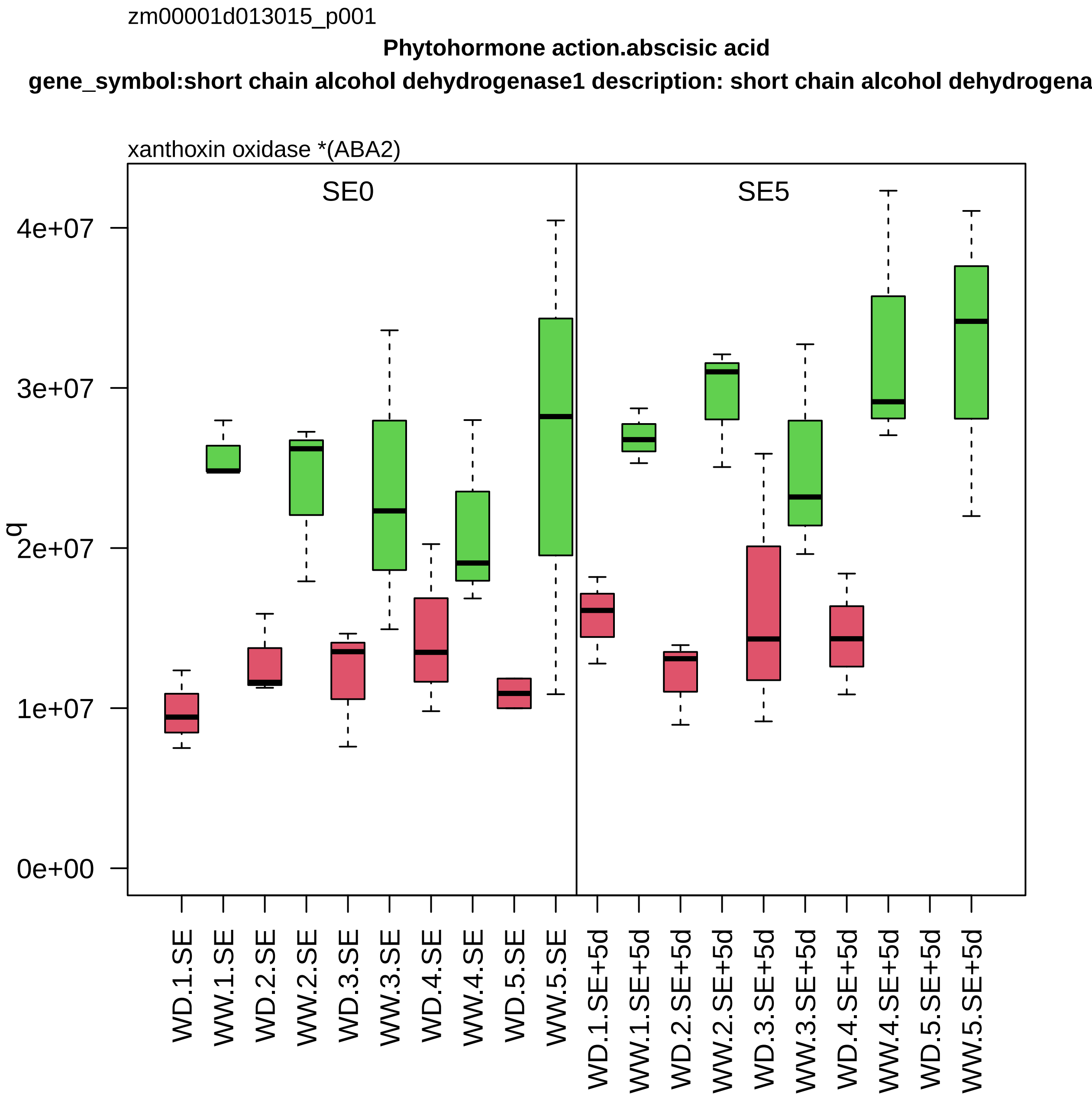
zm00001d013015_p001
Phytohormone action.abscisic acid
gene_symbol:short chain alcohol dehydrogenase1 description: short chain alcohol dehydrogena
xanthoxin oxidase *(ABA2)
SE0
SE5
q
0e+00
1e+07
2e+07
3e+07
4e+07
WD.1.SE
WW.1.SE
WD.2.SE
WW.2.SE
WD.3.SE
WW.3.SE
WD.4.SE
WW.4.SE
WD.5.SE
WW.5.SE
WD.1.SE+5d
WW.1.SE+5d
WD.2.SE+5d
WW.2.SE+5d
WD.3.SE+5d
WW.3.SE+5d
WD.4.SE+5d
WW.4.SE+5d
WD.5.SE+5d
WW.5.SE+5d

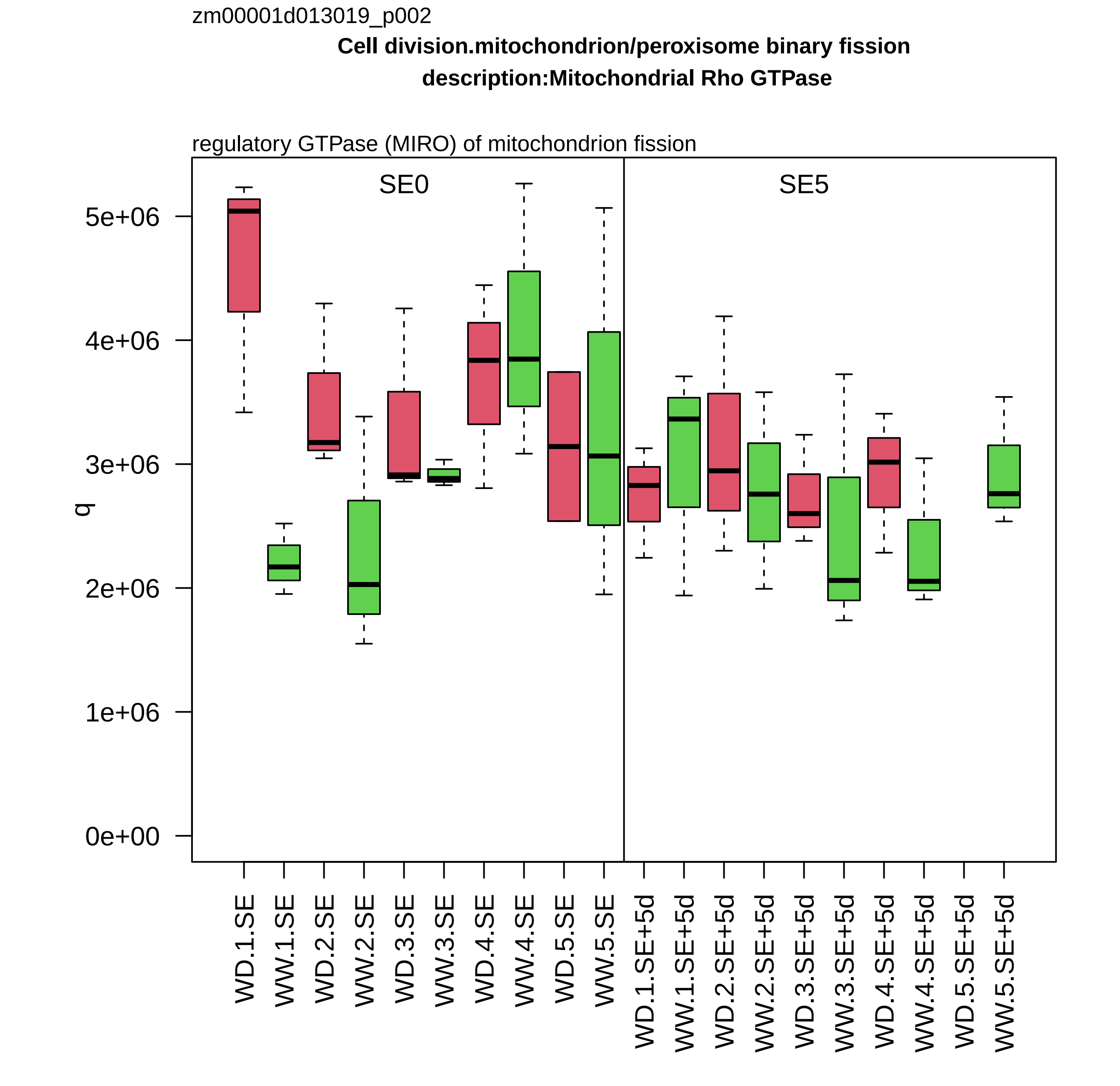

zm00001d013019_p002
Cell division.mitochondrion/peroxisome binary fission
description:Mitochondrial Rho GTPase
regulatory GTPase (MIRO) of mitochondrion fission
SE0
SE5
q
5e+06
4e+06
3e+06
2e+06
1e+06
0e+00
WD.1.SE
WW.1.SE
WD.2.SE
WW.2.SE
WD.3.SE
WW.3.SE
WD.4.SE
WW.4.SE
WD.5.SE
WW.5.SE
WD.1.SE+5d
WW.1.SE+5d
WD.2.SE+5d
WW.2.SE+5d
WD.3.SE+5d
WW.3.SE+5d
WD.4.SE+5d
WW.4.SE+5d
WD.5.SE+5d
WW.5.SE+5d

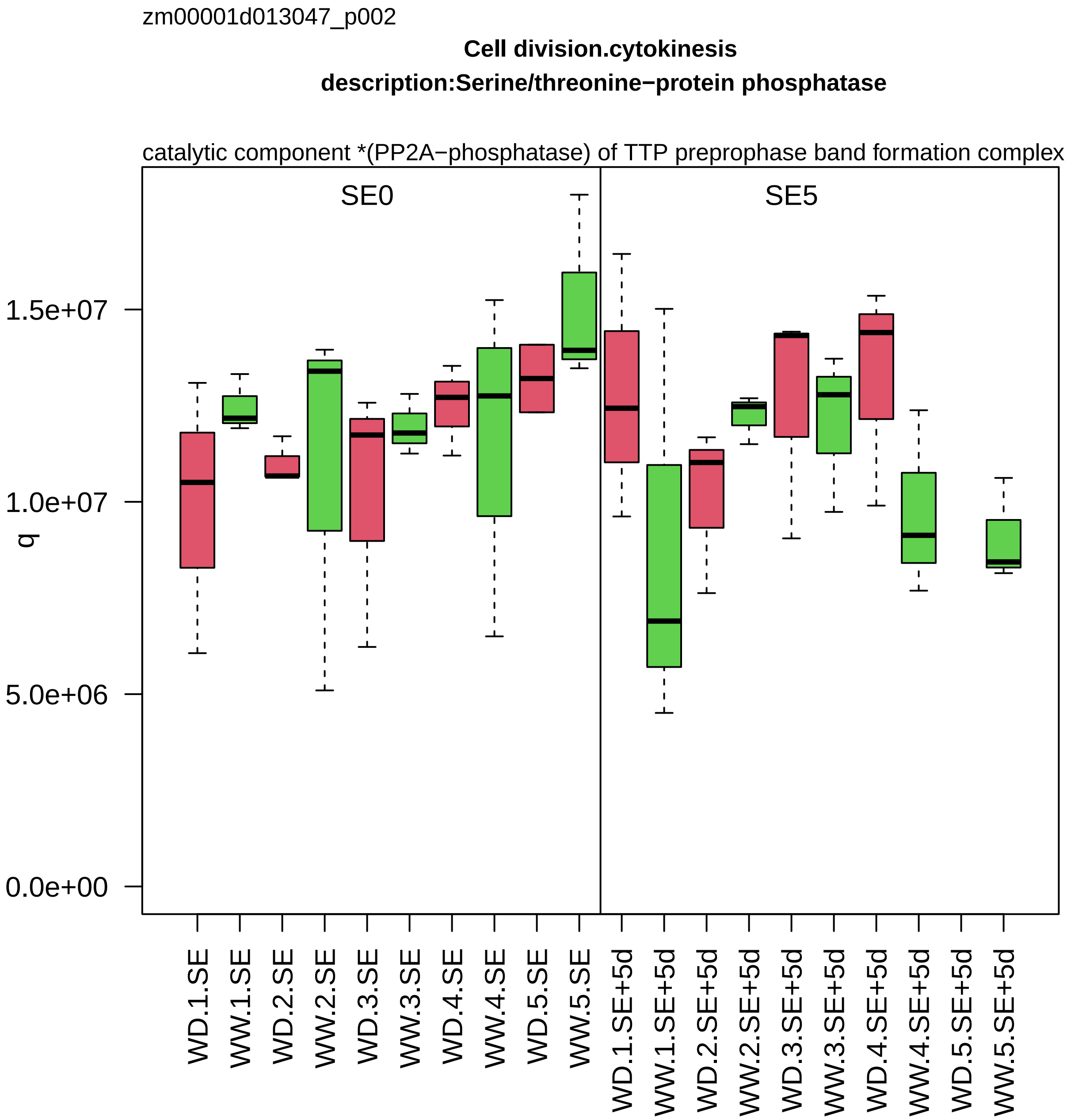
zm00001d013047_p002
Cell division.cytokinesis
description:Serine/threonine−protein phosphatase
catalytic component *(PP2A−phosphatase) of TTP preprophase band formation complex
SE0
SE5
q
1.5e+07
1.0e+07
5.0e+06
0.0e+00
WD.1.SE
WW.1.SE
WD.2.SE
WW.2.SE
WD.3.SE
WW.3.SE
WD.4.SE
WW.4.SE
WD.5.SE
WW.5.SE
WD.1.SE+5d
WW.1.SE+5d
WD.2.SE+5d
WW.2.SE+5d
WD.3.SE+5d
WW.3.SE+5d
WD.4.SE+5d
WW.4.SE+5d
WD.5.SE+5d
WW.5.SE+5d

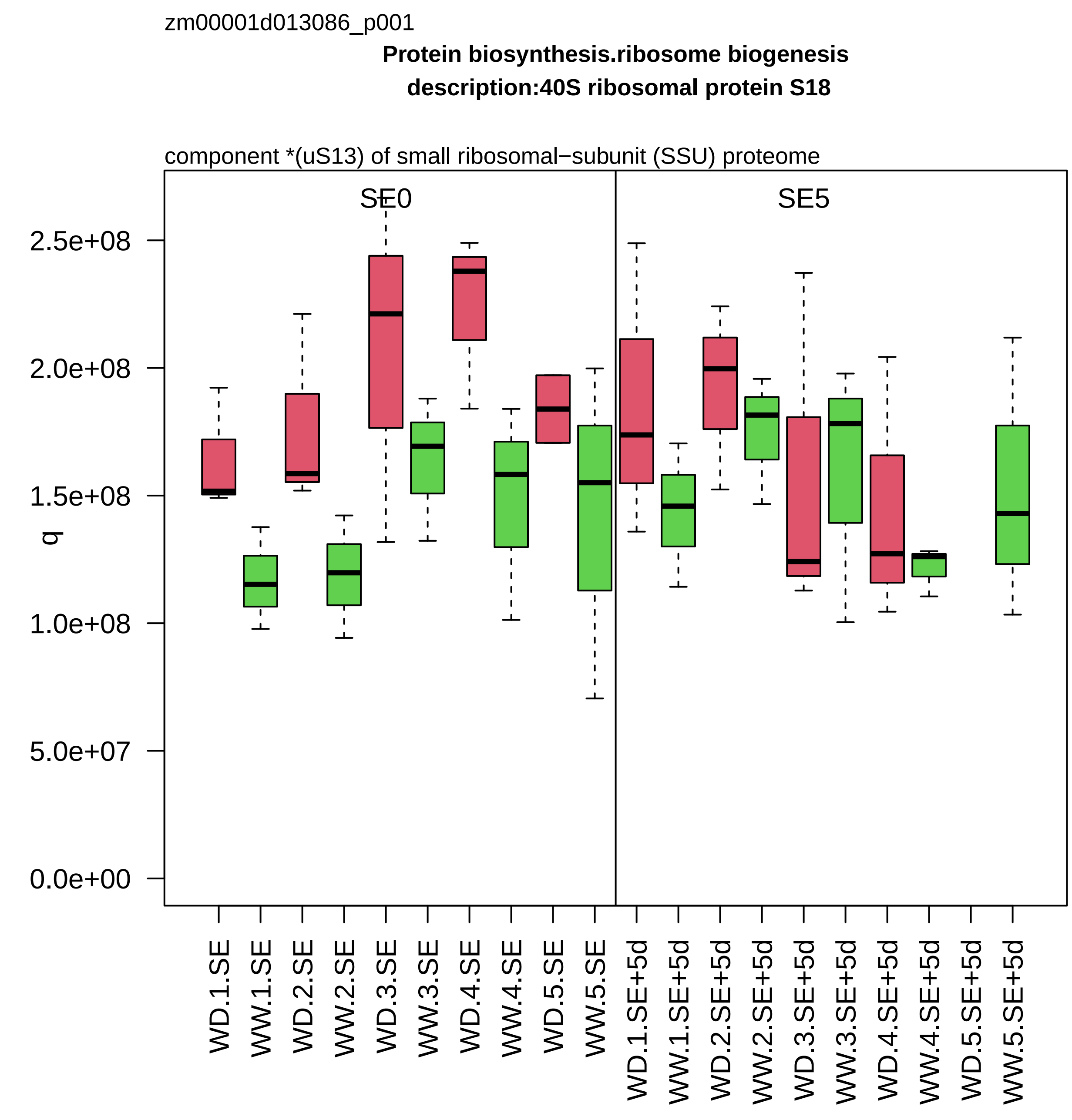

zm00001d013086_p001
Protein biosynthesis.ribosome biogenesis
description:40S ribosomal protein S18
component *(uS13) of small ribosomal−subunit (SSU) proteome
SE0
SE5
q
2.5e+08
2.0e+08
1.5e+08
1.0e+08
5.0e+07
0.0e+00
WD.1.SE
WW.1.SE
WD.2.SE
WW.2.SE
WD.3.SE
WW.3.SE
WD.4.SE
WW.4.SE
WD.5.SE
WW.5.SE
WD.1.SE+5d
WW.1.SE+5d
WD.2.SE+5d
WW.2.SE+5d
WD.3.SE+5d
WW.3.SE+5d
WD.4.SE+5d
WW.4.SE+5d
WD.5.SE+5d
WW.5.SE+5d

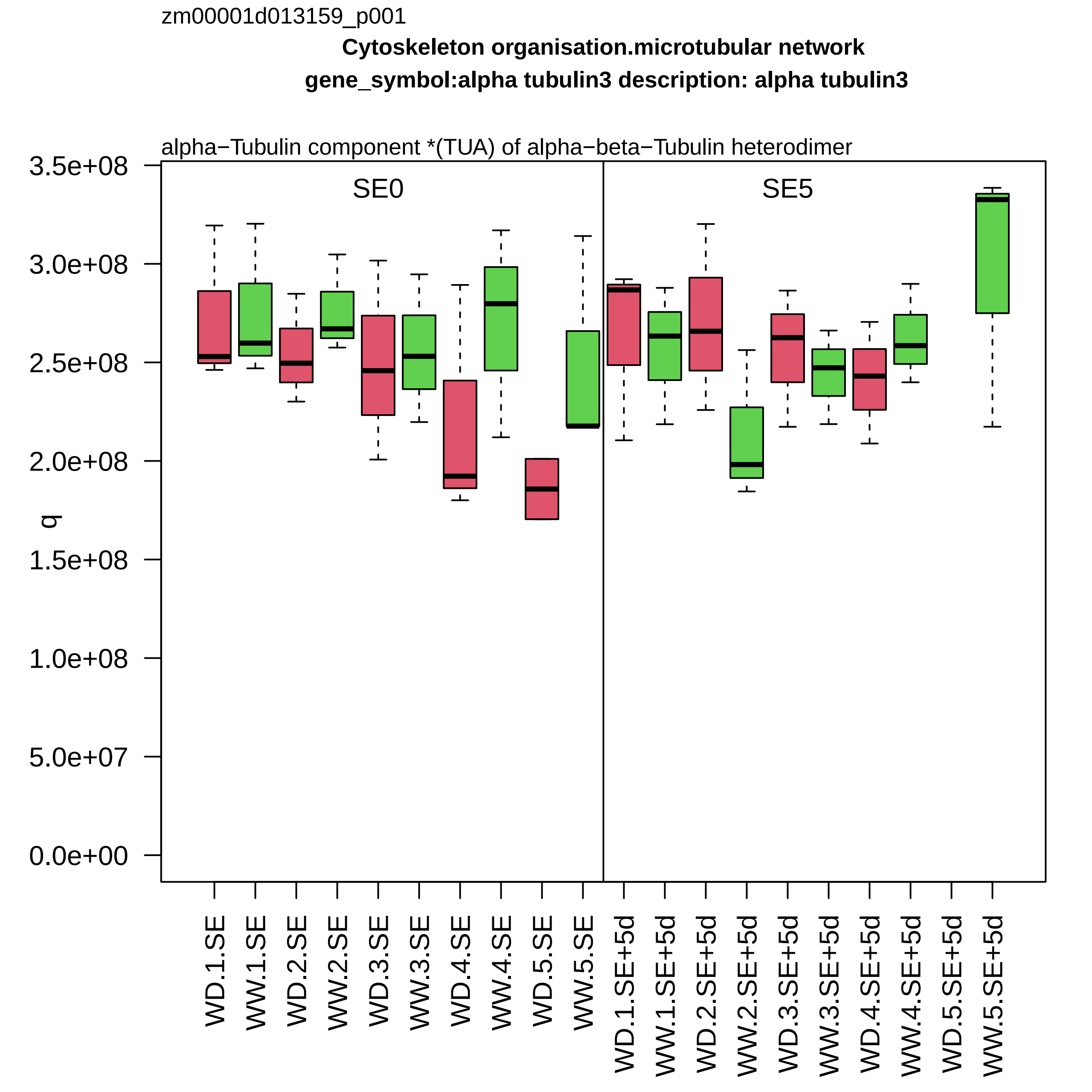

zm00001d013159_p001
Cytoskeleton organisation.microtubular network
gene_symbol:alpha tubulin3 description: alpha tubulin3
alpha−Tubulin component *(TUA) of alpha−beta−Tubulin heterodimer
SE0
SE5
q
3.5e+08
3.0e+08
2.5e+08
2.0e+08
1.5e+08
1.0e+08
5.0e+07
0.0e+00
WD.1.SE
WW.1.SE
WD.2.SE
WW.2.SE
WD.3.SE
WW.3.SE
WD.4.SE
WW.4.SE
WD.5.SE
WW.5.SE
WD.1.SE+5d
WW.1.SE+5d
WD.2.SE+5d
WW.2.SE+5d
WD.3.SE+5d
WW.3.SE+5d
WD.4.SE+5d
WW.4.SE+5d
WD.5.SE+5d
WW.5.SE+5d

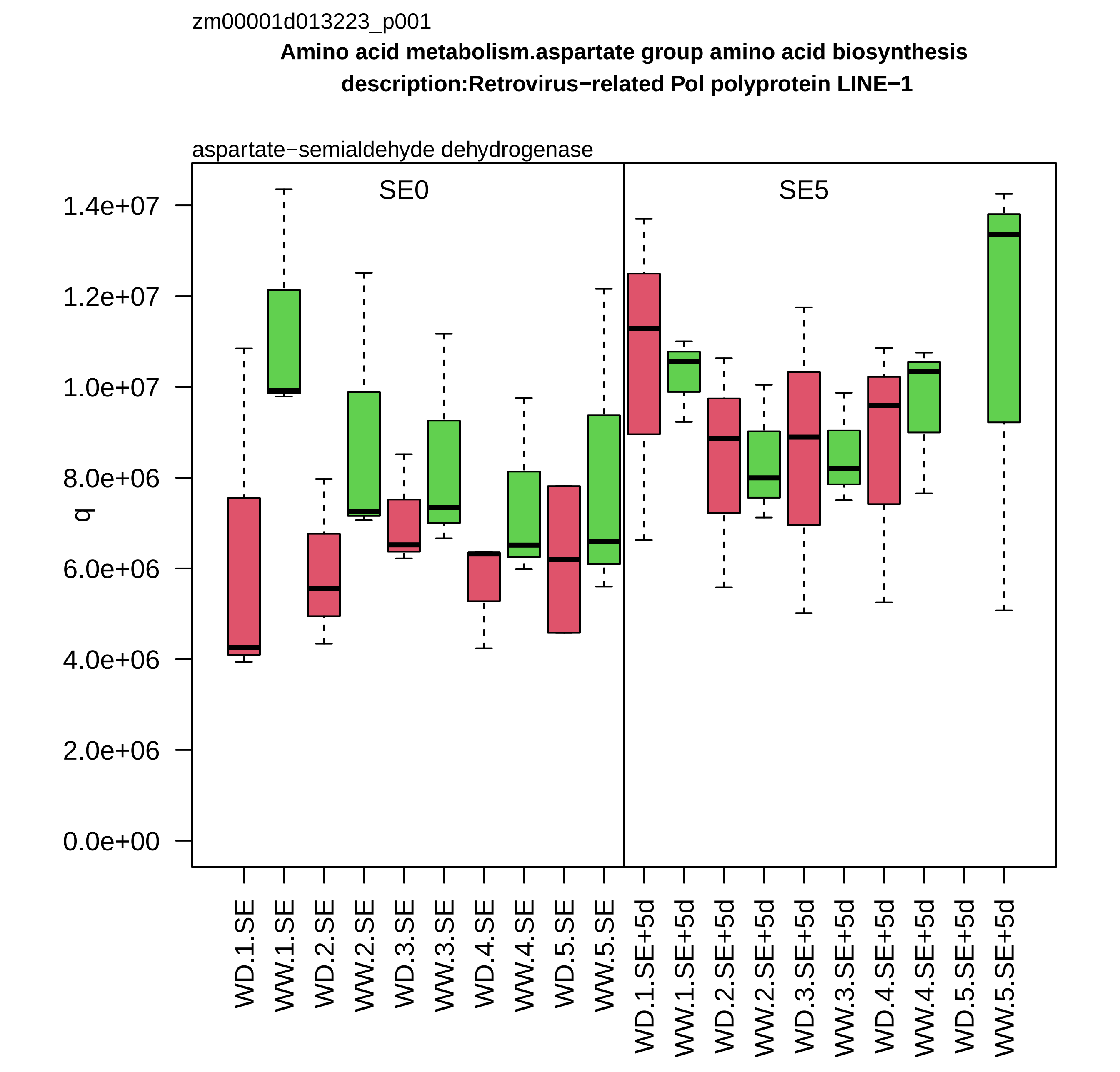
zm00001d013223_p001
Amino acid metabolism.aspartate group amino acid biosynthesis
description:Retrovirus−related Pol polyprotein LINE−1
aspartate−semialdehyde dehydrogenase
SE0
SE5
q
1.4e+07
1.2e+07
1.0e+07
8.0e+06
6.0e+06
4.0e+06
2.0e+06
0.0e+00
WD.1.SE
WW.1.SE
WD.2.SE
WW.2.SE
WD.3.SE
WW.3.SE
WD.4.SE
WW.4.SE
WD.5.SE
WW.5.SE
WD.1.SE+5d
WW.1.SE+5d
WD.2.SE+5d
WW.2.SE+5d
WD.3.SE+5d
WW.3.SE+5d
WD.4.SE+5d
WW.4.SE+5d
WD.5.SE+5d
WW.5.SE+5d

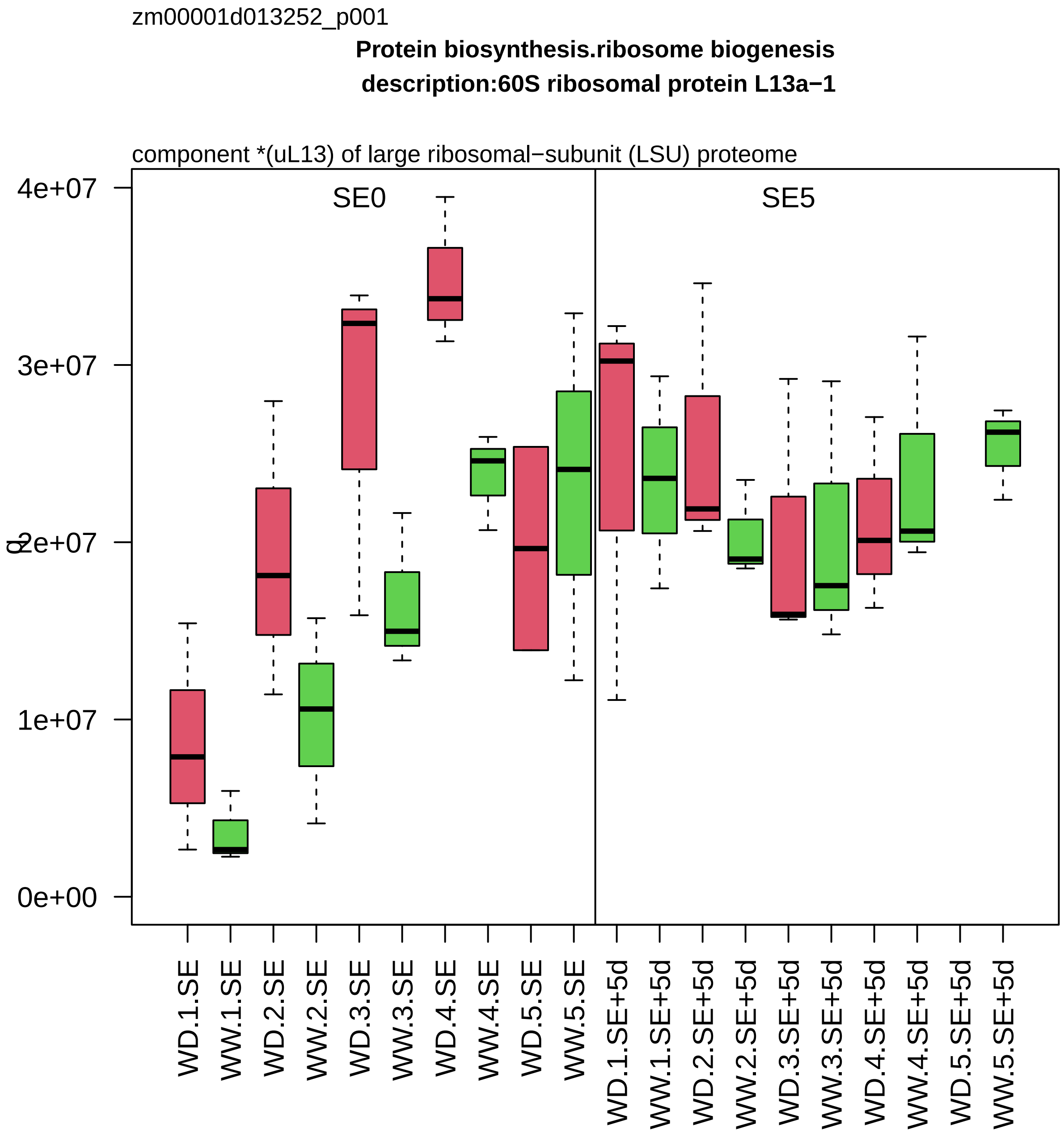
zm00001d013252_p001
Protein biosynthesis.ribosome biogenesis
description:60S ribosomal protein L13a−1
component *(uL13) of large ribosomal−subunit (LSU) proteome
SE0
SE5
0e+00
1e+07
2e+07
3e+07
4e+07
WD.1.SE
WW.1.SE
WD.2.SE
WW.2.SE
WD.3.SE
WW.3.SE
WD.4.SE
WW.4.SE
WD.5.SE
WW.5.SE
WD.1.SE+5d
WW.1.SE+5d
WD.2.SE+5d
WW.2.SE+5d
WD.3.SE+5d
WW.3.SE+5d
WD.4.SE+5d
WW.4.SE+5d
WD.5.SE+5d
WW.5.SE+5d

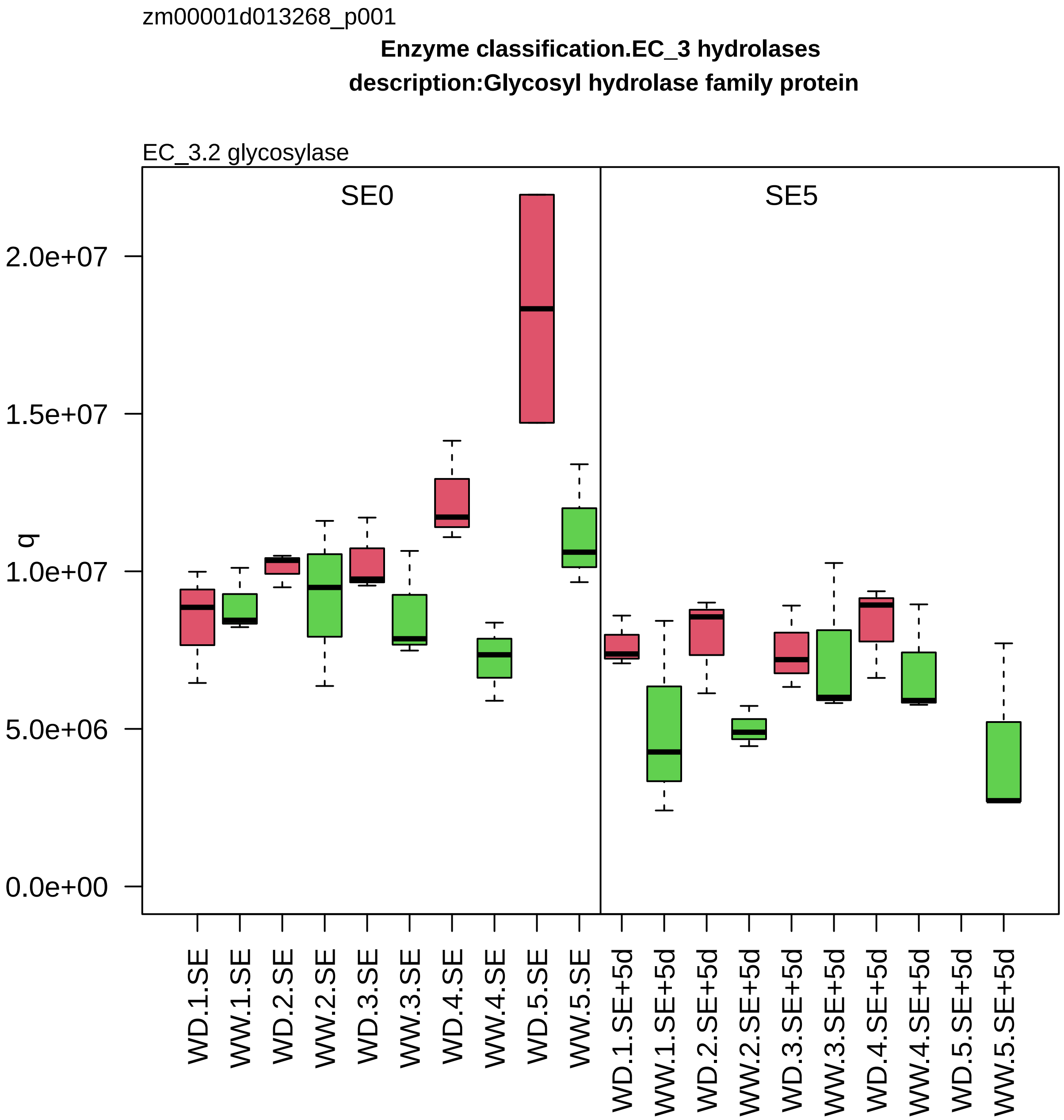
zm00001d013268_p001
Enzyme classification.EC_3 hydrolases
description:Glycosyl hydrolase family protein
EC_3.2 glycosylase
SE0
SE5
q
0.0e+00
5.0e+06
1.0e+07
1.5e+07
2.0e+07
WD.1.SE
WW.1.SE
WD.2.SE
WW.2.SE
WD.3.SE
WW.3.SE
WD.4.SE
WW.4.SE
WD.5.SE
WW.5.SE
WD.1.SE+5d
WW.1.SE+5d
WD.2.SE+5d
WW.2.SE+5d
WD.3.SE+5d
WW.3.SE+5d
WD.4.SE+5d
WW.4.SE+5d
WD.5.SE+5d
WW.5.SE+5d

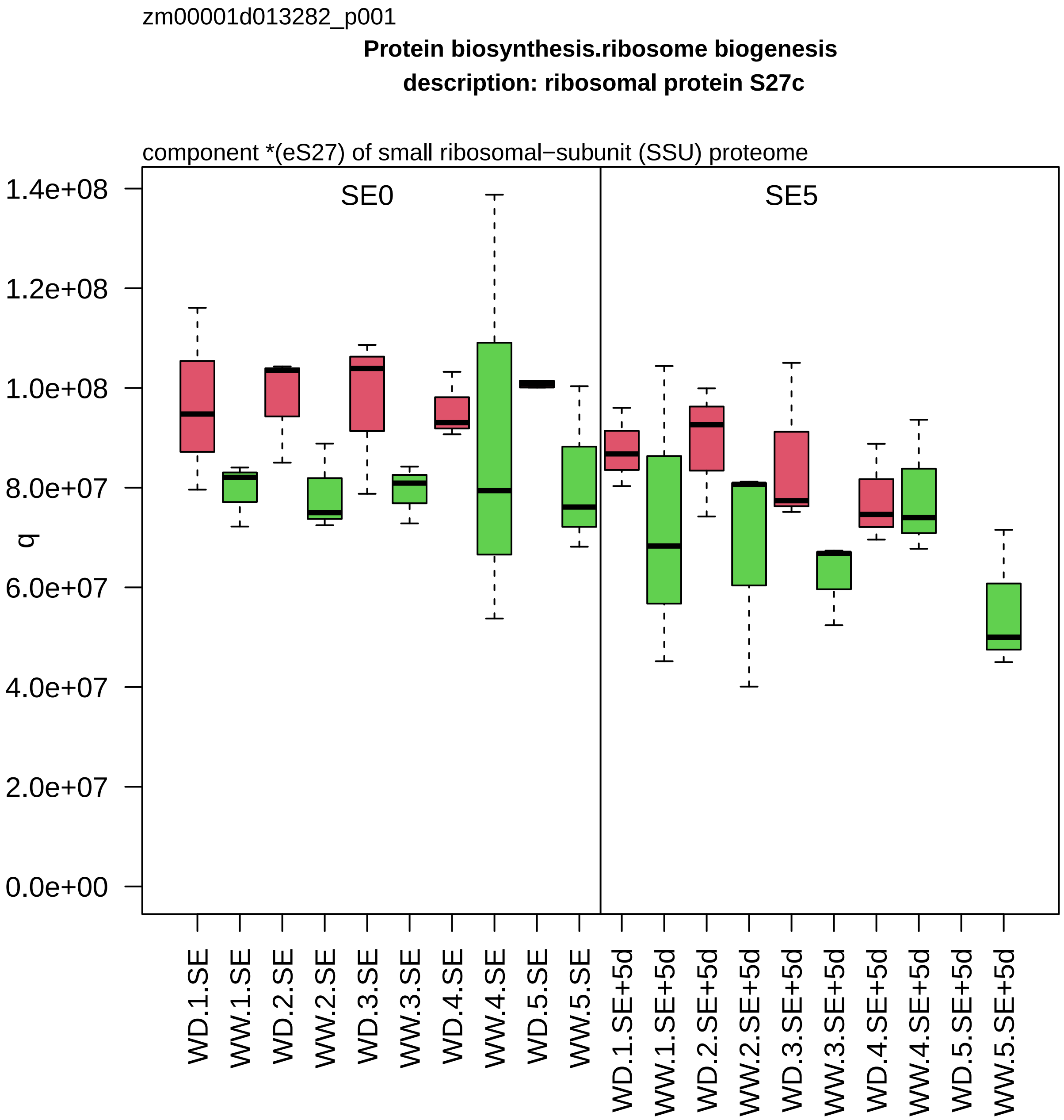

zm00001d013282_p001
Protein biosynthesis.ribosome biogenesis
description: ribosomal protein S27c
component *(eS27) of small ribosomal−subunit (SSU) proteome
SE0
SE5
q
1.4e+08
1.2e+08
1.0e+08
8.0e+07
6.0e+07
4.0e+07
2.0e+07
0.0e+00
WD.1.SE
WW.1.SE
WD.2.SE
WW.2.SE
WD.3.SE
WW.3.SE
WD.4.SE
WW.4.SE
WD.5.SE
WW.5.SE
WD.1.SE+5d
WW.1.SE+5d
WD.2.SE+5d
WW.2.SE+5d
WD.3.SE+5d
WW.3.SE+5d
WD.4.SE+5d
WW.4.SE+5d
WD.5.SE+5d
WW.5.SE+5d

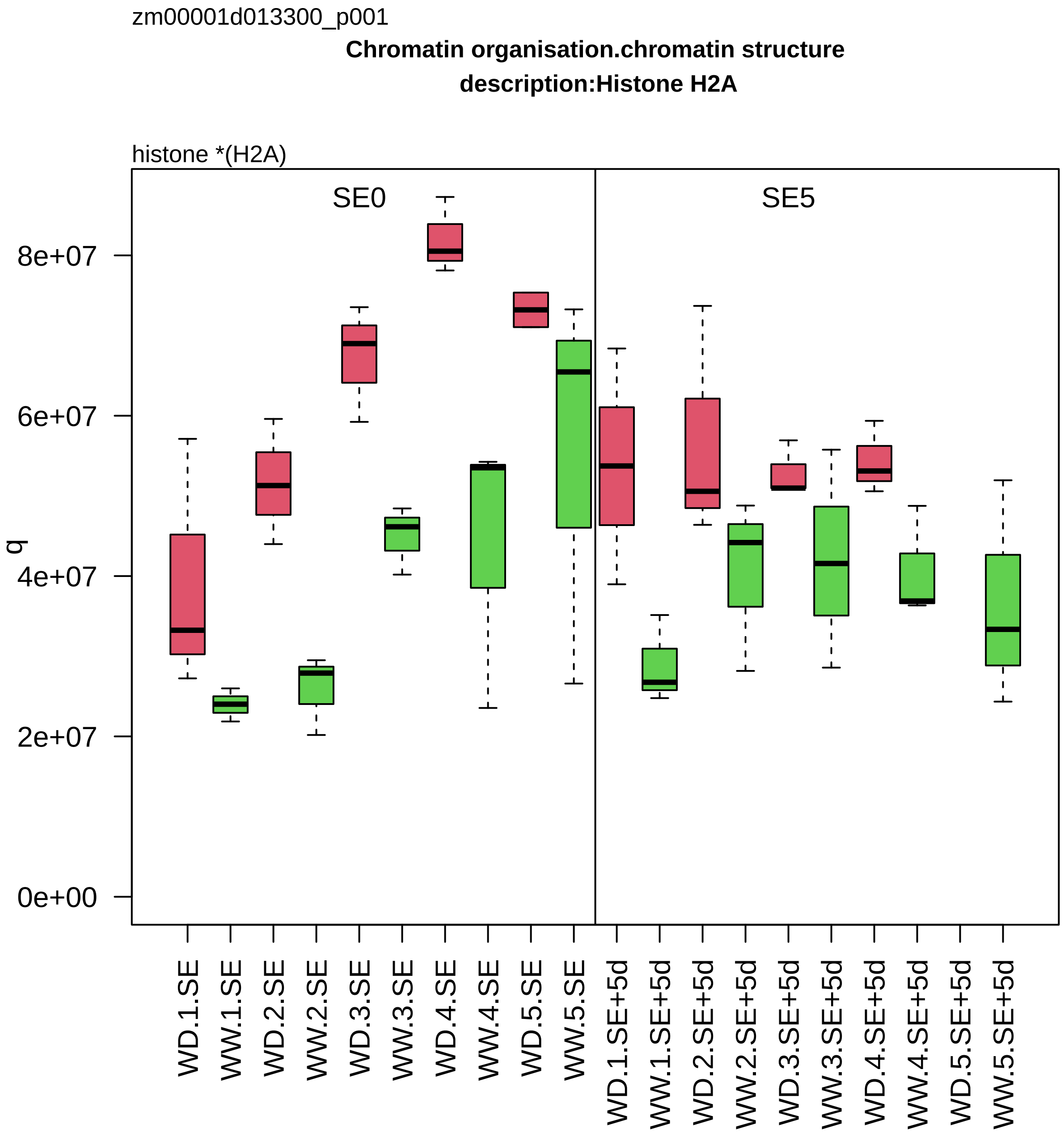

zm00001d013300_p001
Chromatin organisation.chromatin structure
description:Histone H2A
histone *(H2A)
SE0
SE5
q
8e+07
6e+07
4e+07
2e+07
0e+00
WD.1.SE
WW.1.SE
WD.2.SE
WW.2.SE
WD.3.SE
WW.3.SE
WD.4.SE
WW.4.SE
WD.5.SE
WW.5.SE
WD.1.SE+5d
WW.1.SE+5d
WD.2.SE+5d
WW.2.SE+5d
WD.3.SE+5d
WW.3.SE+5d
WD.4.SE+5d
WW.4.SE+5d
WD.5.SE+5d
WW.5.SE+5d

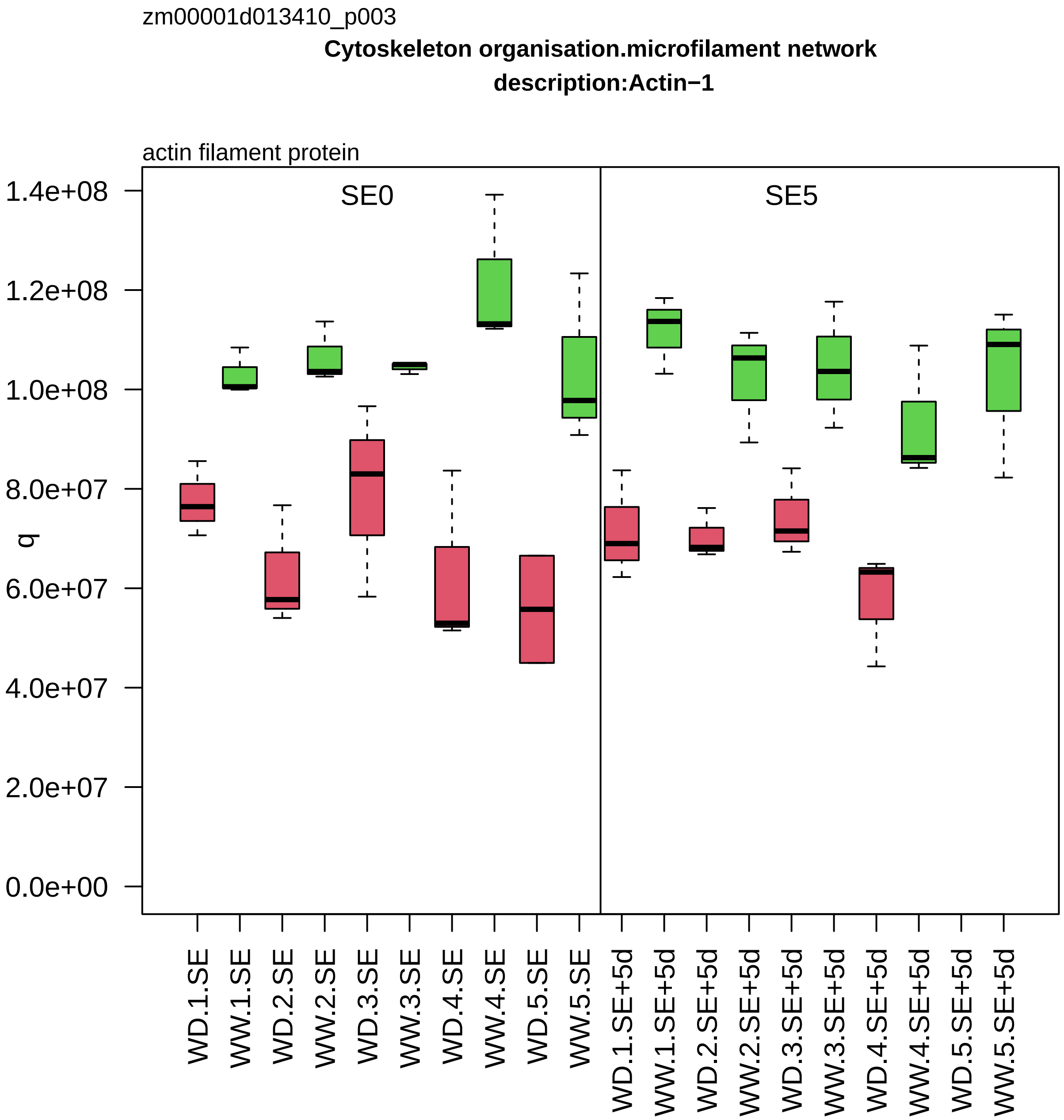

zm00001d013410_p003
Cytoskeleton organisation.microfilament network
description:Actin−1
actin filament protein
SE0
SE5
q
1.4e+08
1.2e+08
1.0e+08
8.0e+07
6.0e+07
4.0e+07
2.0e+07
0.0e+00
WD.1.SE
WW.1.SE
WD.2.SE
WW.2.SE
WD.3.SE
WW.3.SE
WD.4.SE
WW.4.SE
WD.5.SE
WW.5.SE
WD.1.SE+5d
WW.1.SE+5d
WD.2.SE+5d
WW.2.SE+5d
WD.3.SE+5d
WW.3.SE+5d
WD.4.SE+5d
WW.4.SE+5d
WD.5.SE+5d
WW.5.SE+5d

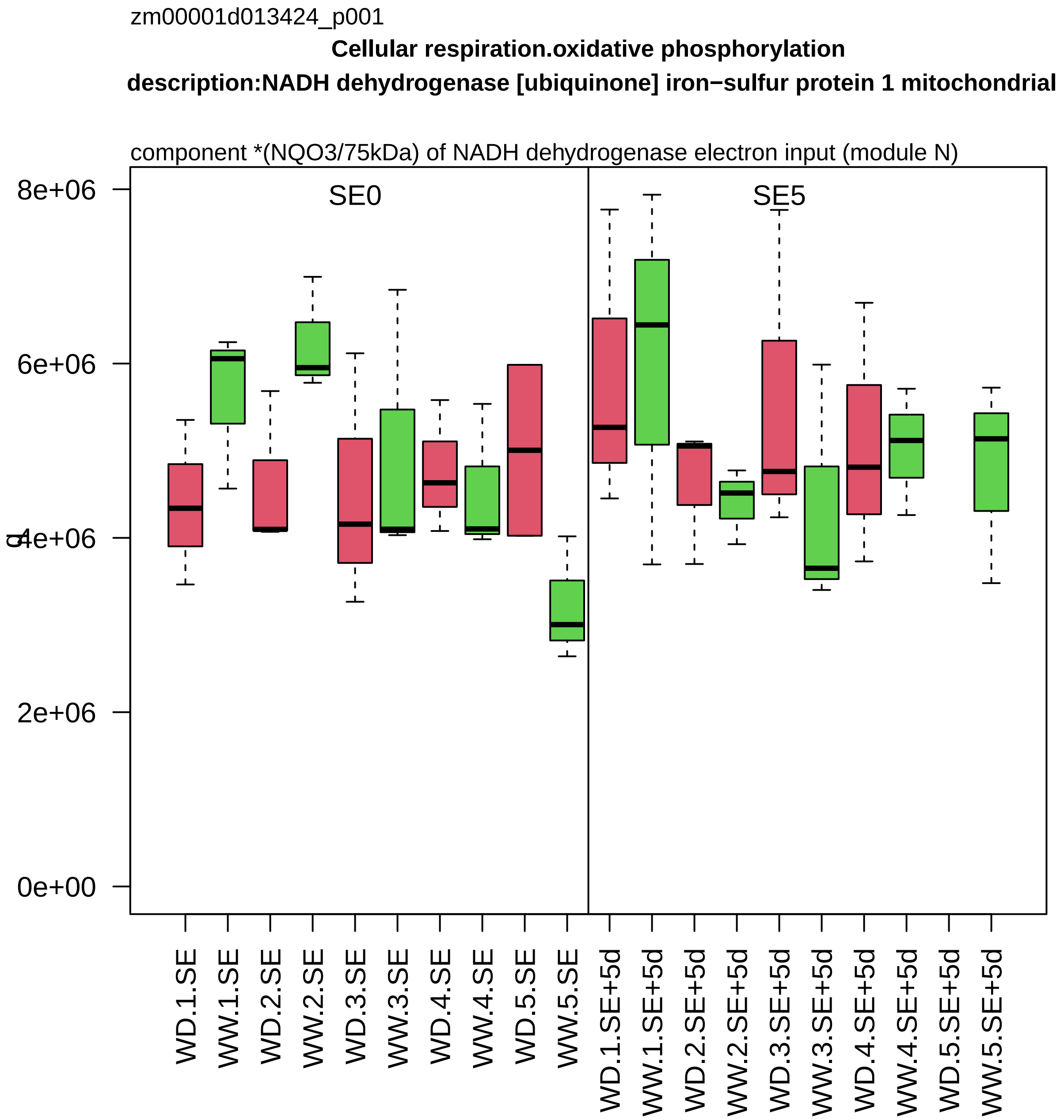
zm00001d013424_p001
Cellular respiration.oxidative phosphorylation
description:NADH dehydrogenase [ubiquinone] iron−sulfur protein 1 mitochondrial
component *(NQO3/75kDa) of NADH dehydrogenase electron input (module N)
SE0
SE5
8e+06
6e+06
4e+06
2e+06
0e+00
WD.1.SE
WW.1.SE
WD.2.SE
WW.2.SE
WD.3.SE
WW.3.SE
WD.4.SE
WW.4.SE
WD.5.SE
WW.5.SE
WD.1.SE+5d
WW.1.SE+5d
WD.2.SE+5d
WW.2.SE+5d
WD.3.SE+5d
WW.3.SE+5d
WD.4.SE+5d
WW.4.SE+5d
WD.5.SE+5d
WW.5.SE+5d

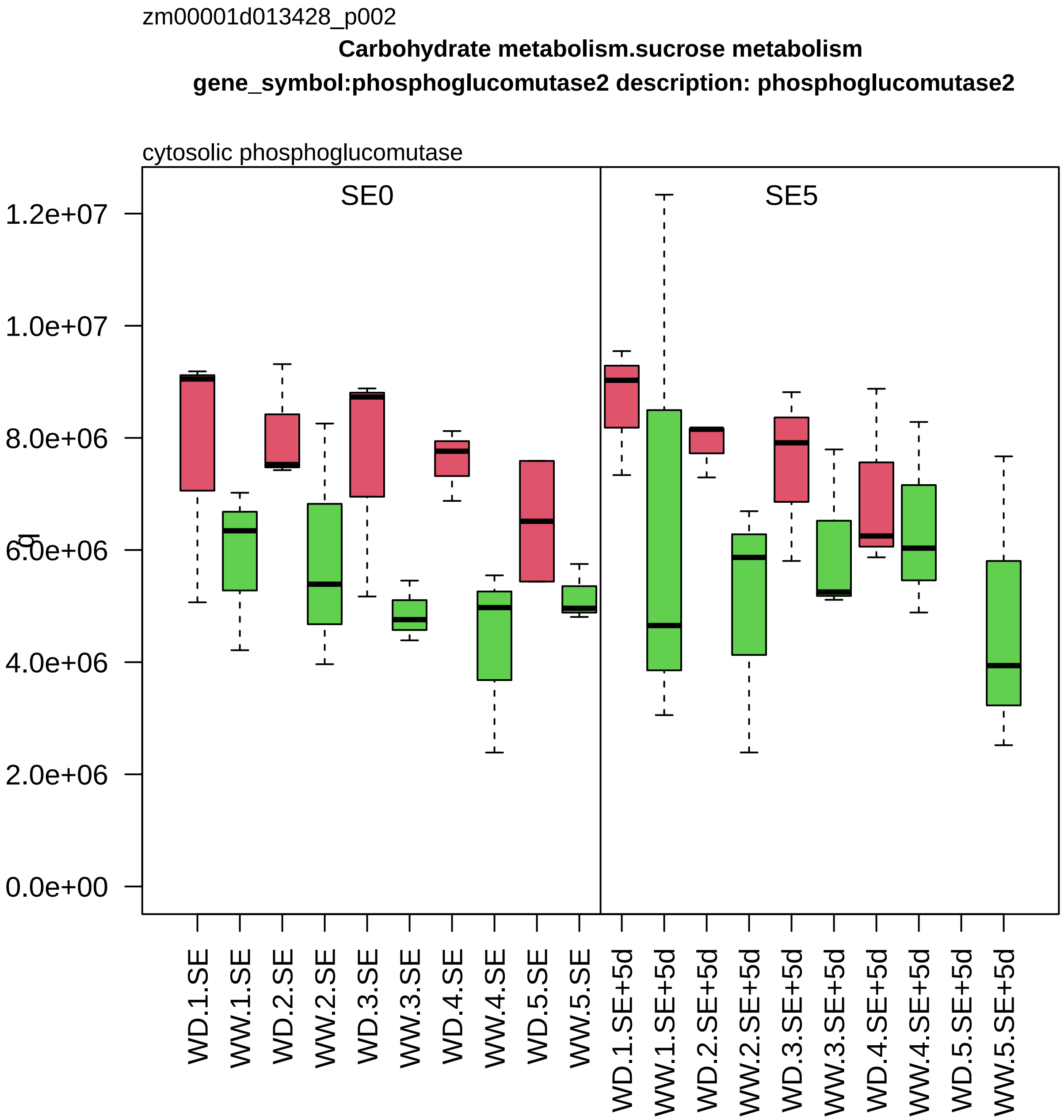

zm00001d013428_p002
Carbohydrate metabolism.sucrose metabolism
gene_symbol:phosphoglucomutase2 description: phosphoglucomutase2
cytosolic phosphoglucomutase
SE0
SE5
1.2e+07
1.0e+07
8.0e+06
6.0e+06
4.0e+06
2.0e+06
0.0e+00
WD.1.SE
WW.1.SE
WD.2.SE
WW.2.SE
WD.3.SE
WW.3.SE
WD.4.SE
WW.4.SE
WD.5.SE
WW.5.SE
WD.1.SE+5d
WW.1.SE+5d
WD.2.SE+5d
WW.2.SE+5d
WD.3.SE+5d
WW.3.SE+5d
WD.4.SE+5d
WW.4.SE+5d
WD.5.SE+5d
WW.5.SE+5d

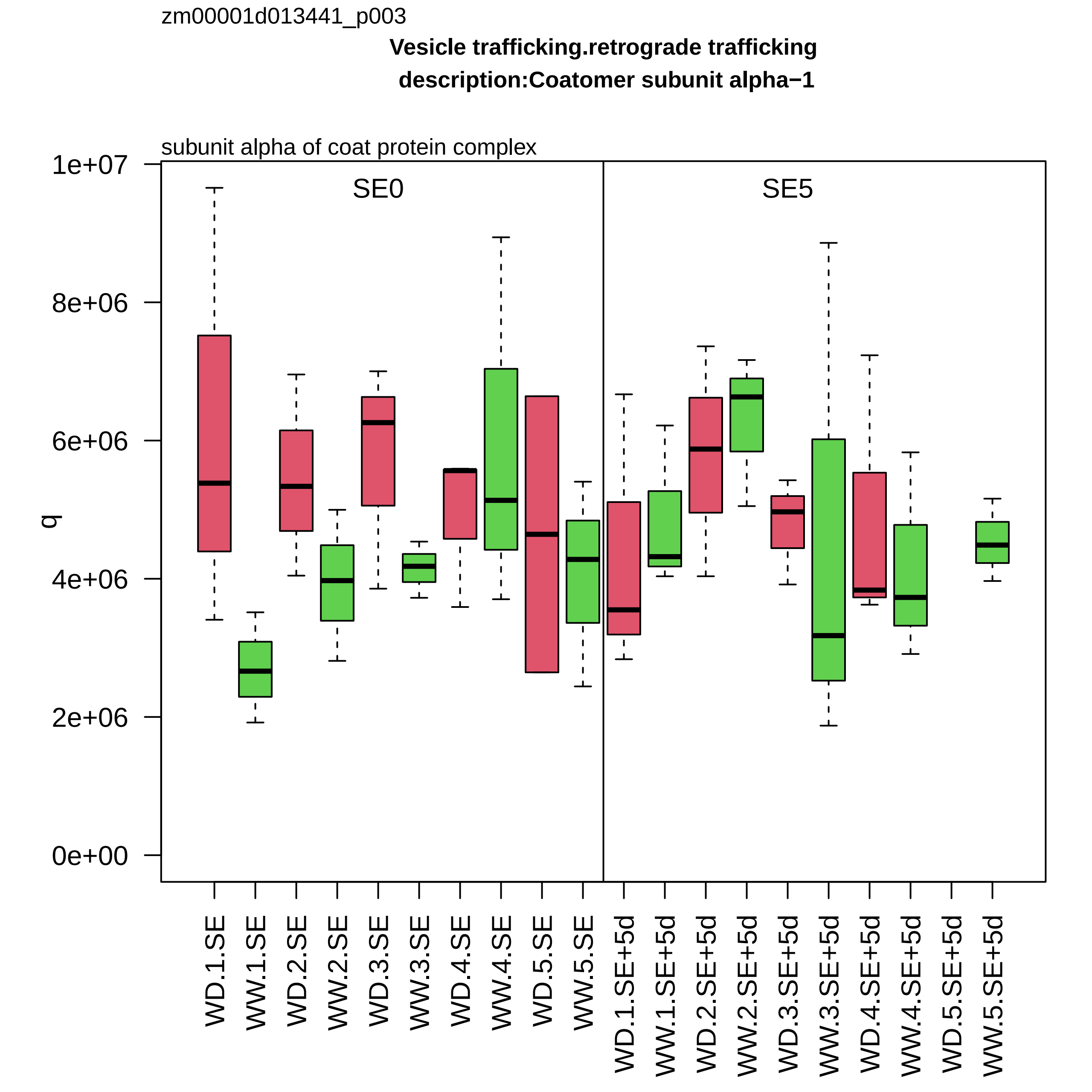
zm00001d013441_p003
Vesicle trafficking.retrograde trafficking
description:Coatomer subunit alpha−1
subunit alpha of coat protein complex
SE0
SE5
q
1e+07
8e+06
6e+06
4e+06
2e+06
0e+00
WD.1.SE
WW.1.SE
WD.2.SE
WW.2.SE
WD.3.SE
WW.3.SE
WD.4.SE
WW.4.SE
WD.5.SE
WW.5.SE
WD.1.SE+5d
WW.1.SE+5d
WD.2.SE+5d
WW.2.SE+5d
WD.3.SE+5d
WW.3.SE+5d
WD.4.SE+5d
WW.4.SE+5d
WD.5.SE+5d
WW.5.SE+5d

zm00001d013493_p001

**Redox homeostasis.reactive electrophilic lipid homeostasis**

**gene_symbol:lipoxygenase5 description: lipoxygenase5**

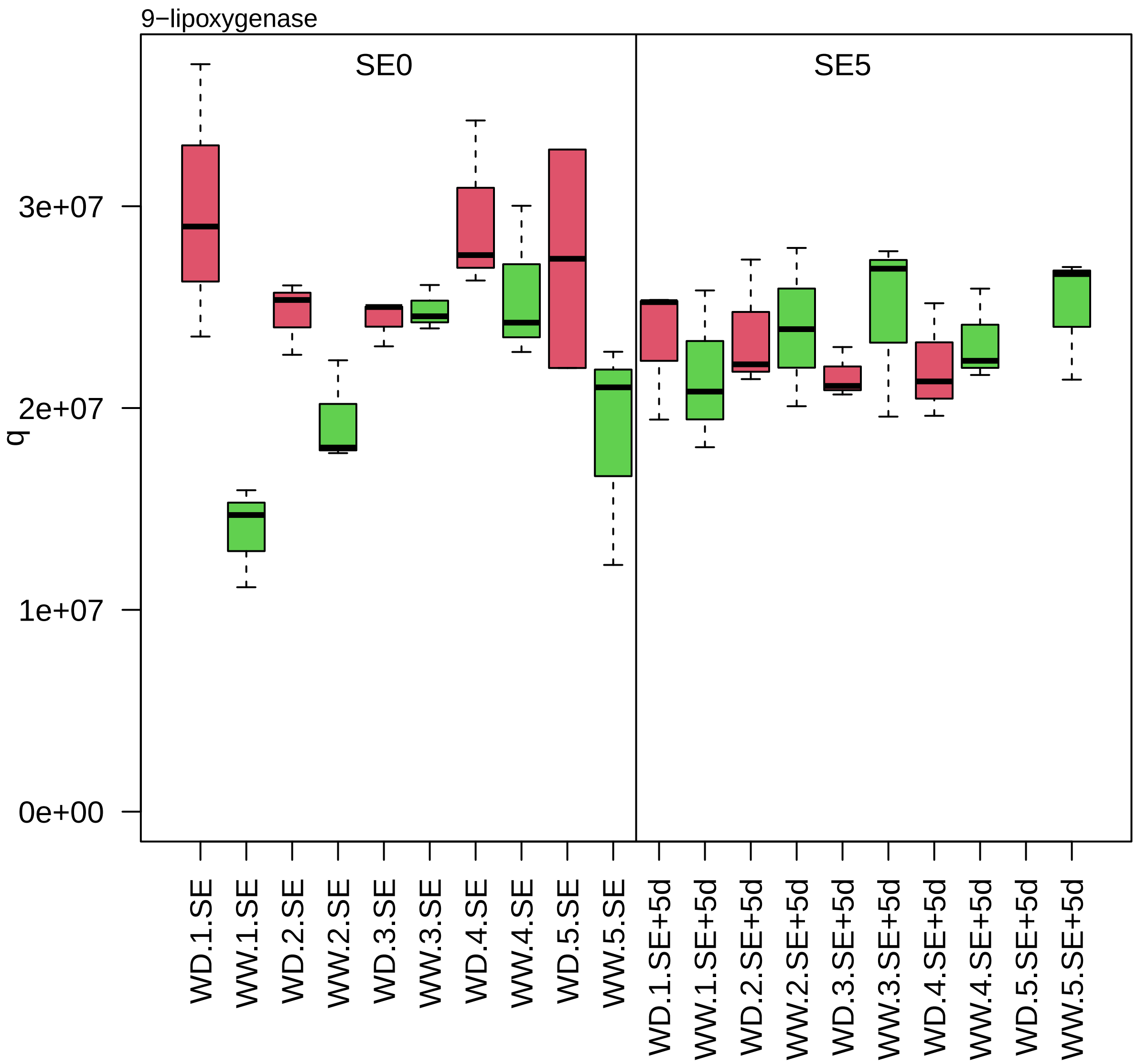

zm00001d013505_p001

**Protein homeostasis.ubiquitin−proteasome system**

**gene_symbol:proteasome component2 description: proteasome component2**

beta−type−4 component *(PBD) of 26S proteasome

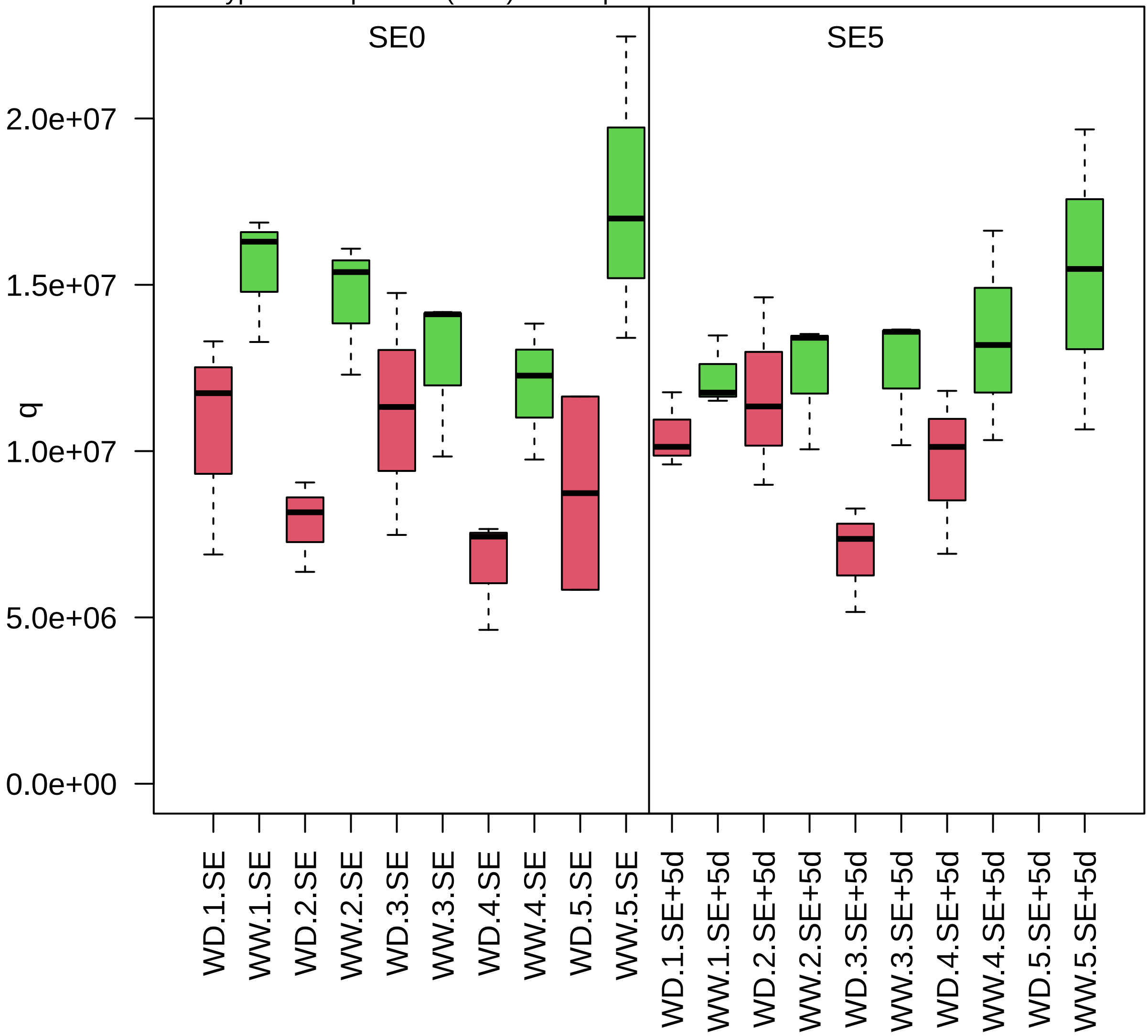

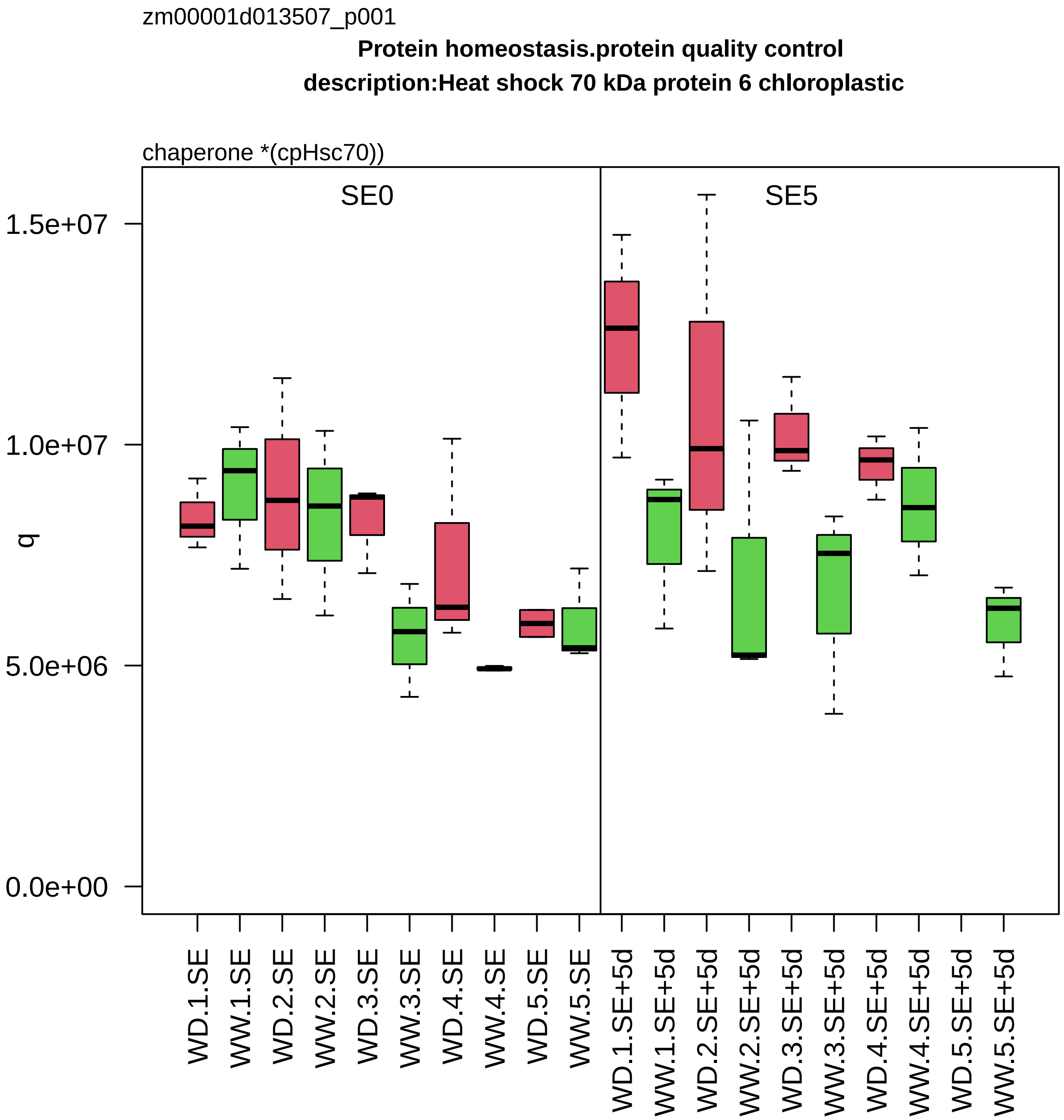
zm00001d013507_p001
Protein homeostasis.protein quality control
description:Heat shock 70 kDa protein 6 chloroplastic
chaperone *(cpHsc70))
SE0
SE5
1.5e+07
1.0e+07
5.0e+06
0.0e+00
q
WD.1.SE
WW.1.SE
WD.2.SE
WW.2.SE
WD.3.SE
WW.3.SE
WD.4.SE
WW.4.SE
WD.5.SE
WW.5.SE
WD.1.SE+5d
WW.1.SE+5d
WD.2.SE+5d
WW.2.SE+5d
WD.3.SE+5d
WW.3.SE+5d
WD.4.SE+5d
WW.4.SE+5d
WD.5.SE+5d
WW.5.SE+5d

zm00001d013568_p001

**not assigned.not annotated**

**gene_symbol:responsive to abscisic acid15 description:Glycine−rich RNA−binding%2C abscisic acid−indu**

not classified

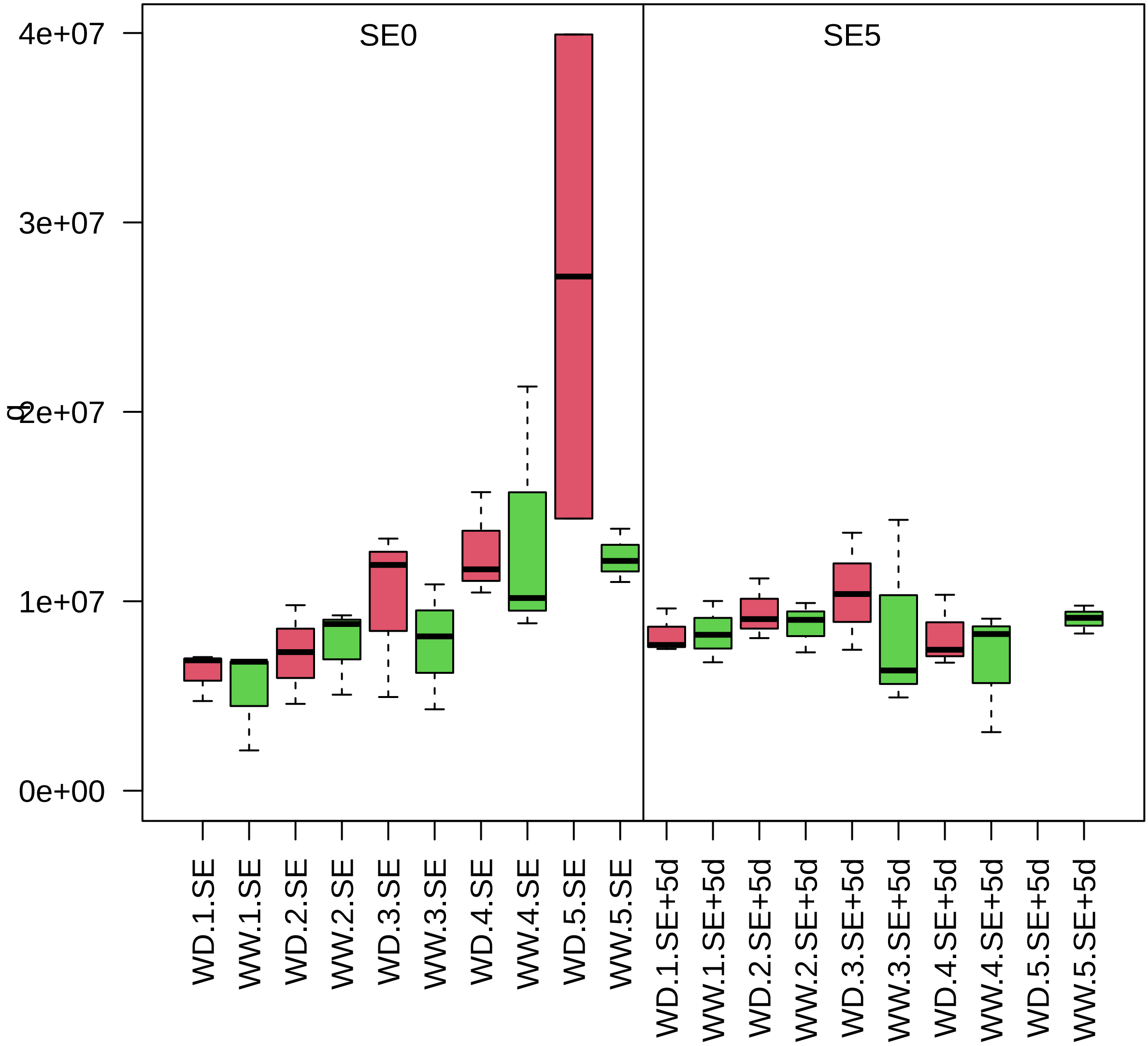

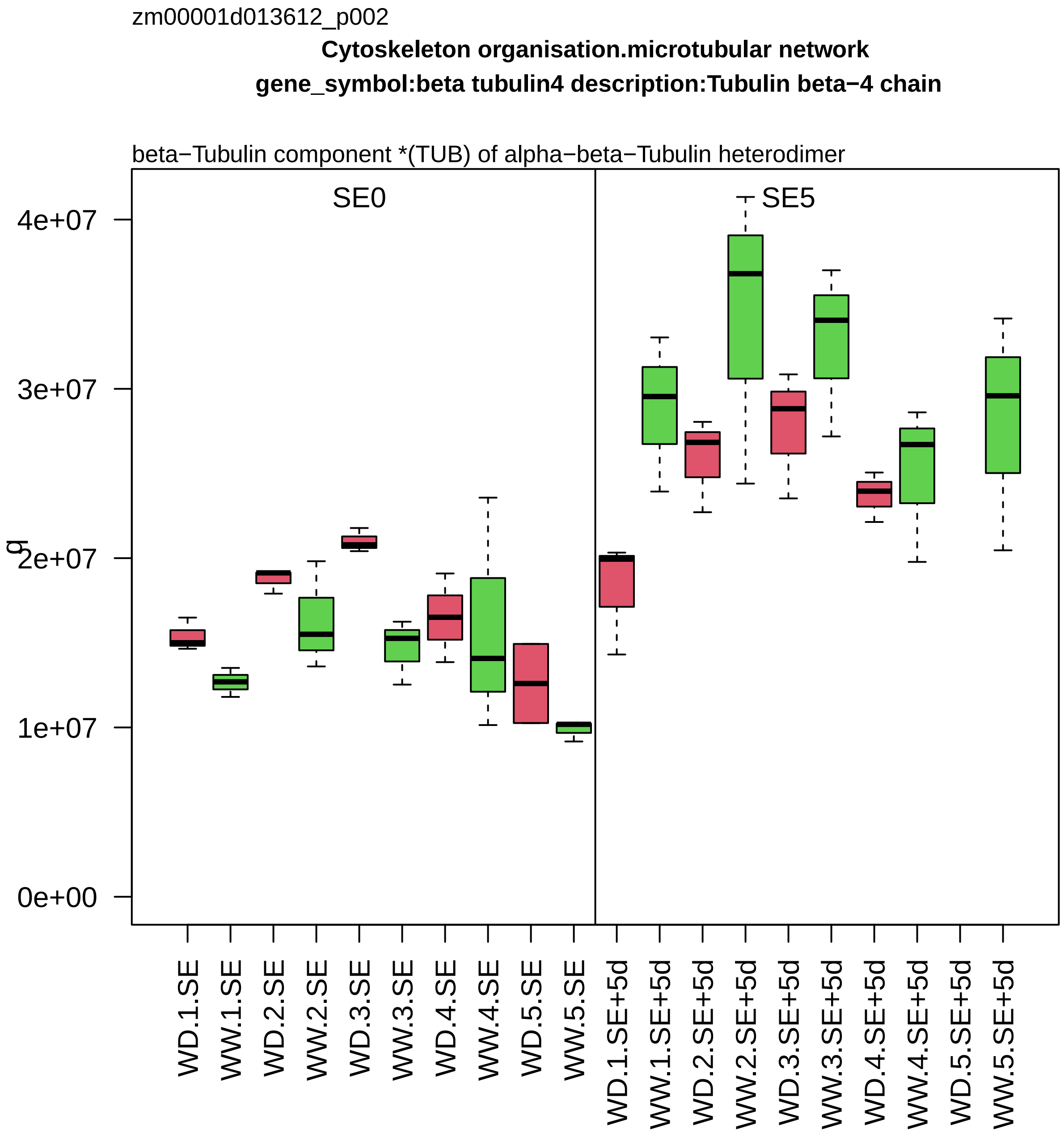
zm00001d013612_p002
Cytoskeleton organisation.microtubular network
gene_symbol:beta tubulin4 description:Tubulin beta−4 chain
beta−Tubulin component *(TUB) of alpha−beta−Tubulin heterodimer
SE0
SE5
q
4e+07
3e+07
2e+07
1e+07
0e+00
WD.1.SE
WW.1.SE
WD.2.SE
WW.2.SE
WD.3.SE
WW.3.SE
WD.4.SE
WW.4.SE
WD.5.SE
WW.5.SE
WD.1.SE+5d
WW.1.SE+5d
WD.2.SE+5d
WW.2.SE+5d
WD.3.SE+5d
WW.3.SE+5d
WD.4.SE+5d
WW.4.SE+5d
WD.5.SE+5d
WW.5.SE+5d

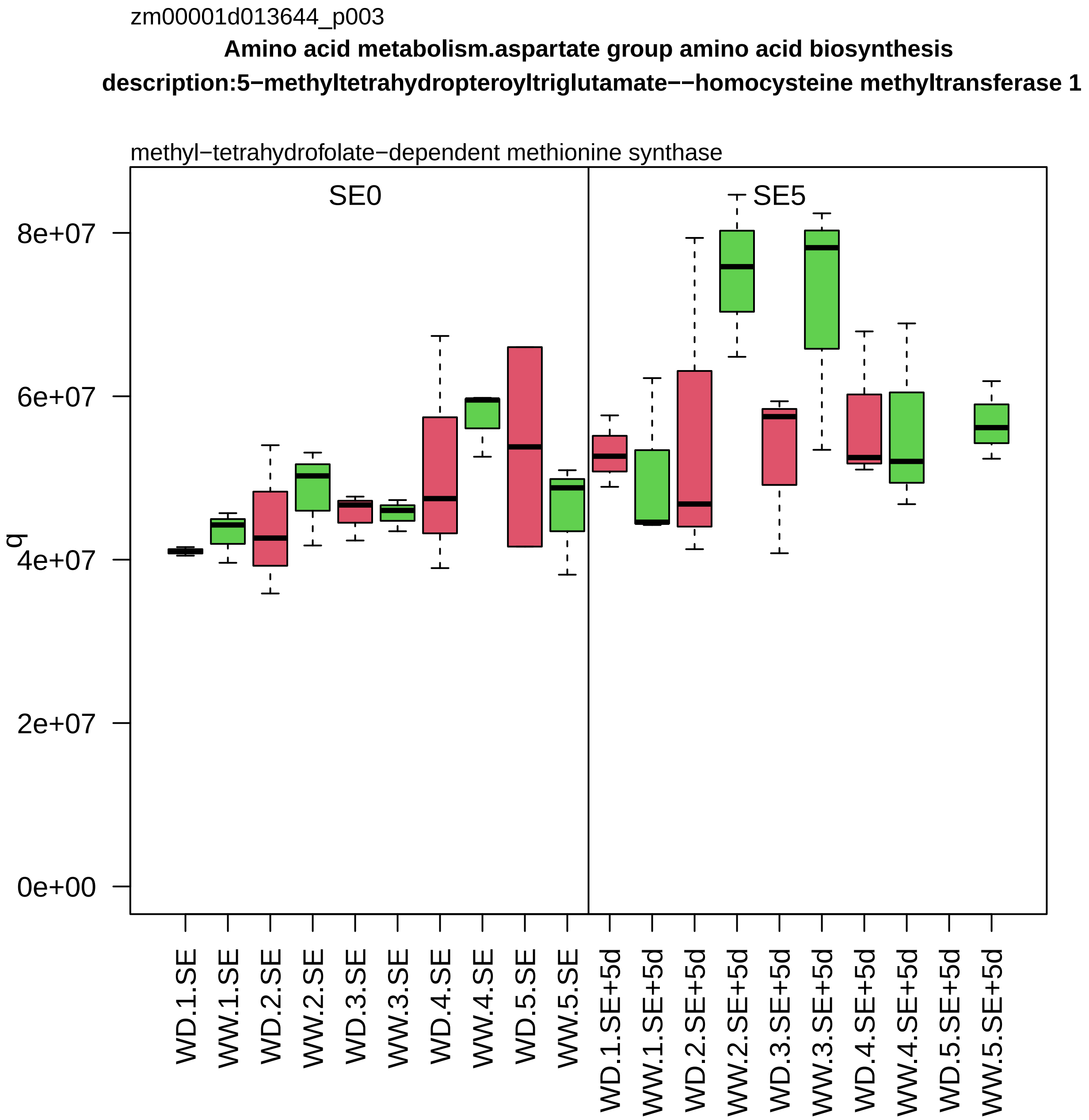

zm00001d013644_p003
Amino acid metabolism.aspartate group amino acid biosynthesis
description:5−methyltetrahydropteroyltriglutamate−−homocysteine methyltransferase 1
methyl−tetrahydrofolate−dependent methionine synthase
SE0
SE5
q
8e+07
6e+07
4e+07
2e+07
0e+00
WD.1.SE
WW.1.SE
WD.2.SE
WW.2.SE
WD.3.SE
WW.3.SE
WD.4.SE
WW.4.SE
WD.5.SE
WW.5.SE
WD.1.SE+5d
WW.1.SE+5d
WD.2.SE+5d
WW.2.SE+5d
WD.3.SE+5d
WW.3.SE+5d
WD.4.SE+5d
WW.4.SE+5d
WD.5.SE+5d
WW.5.SE+5d

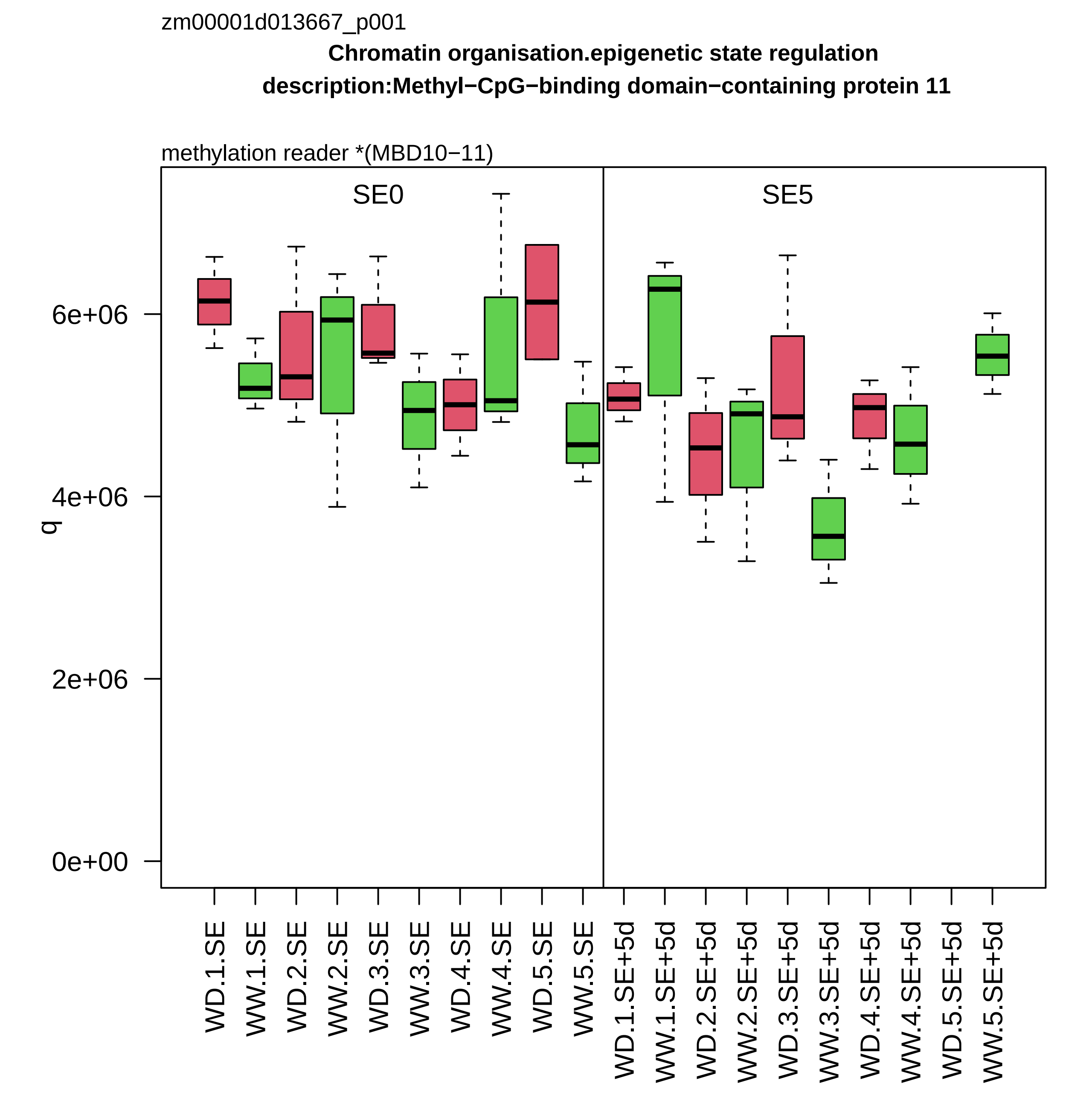

zm00001d013667_p001
Chromatin organisation.epigenetic state regulation
description:Methyl−CpG−binding domain−containing protein 11
methylation reader *(MBD10−11)
SE0
SE5
q
0e+00
2e+06
4e+06
6e+06
WD.1.SE
WW.1.SE
WD.2.SE
WW.2.SE
WD.3.SE
WW.3.SE
WD.4.SE
WW.4.SE
WD.5.SE
WW.5.SE
WD.1.SE+5d
WW.1.SE+5d
WD.2.SE+5d
WW.2.SE+5d
WD.3.SE+5d
WW.3.SE+5d
WD.4.SE+5d
WW.4.SE+5d
WD.5.SE+5d
WW.5.SE+5d

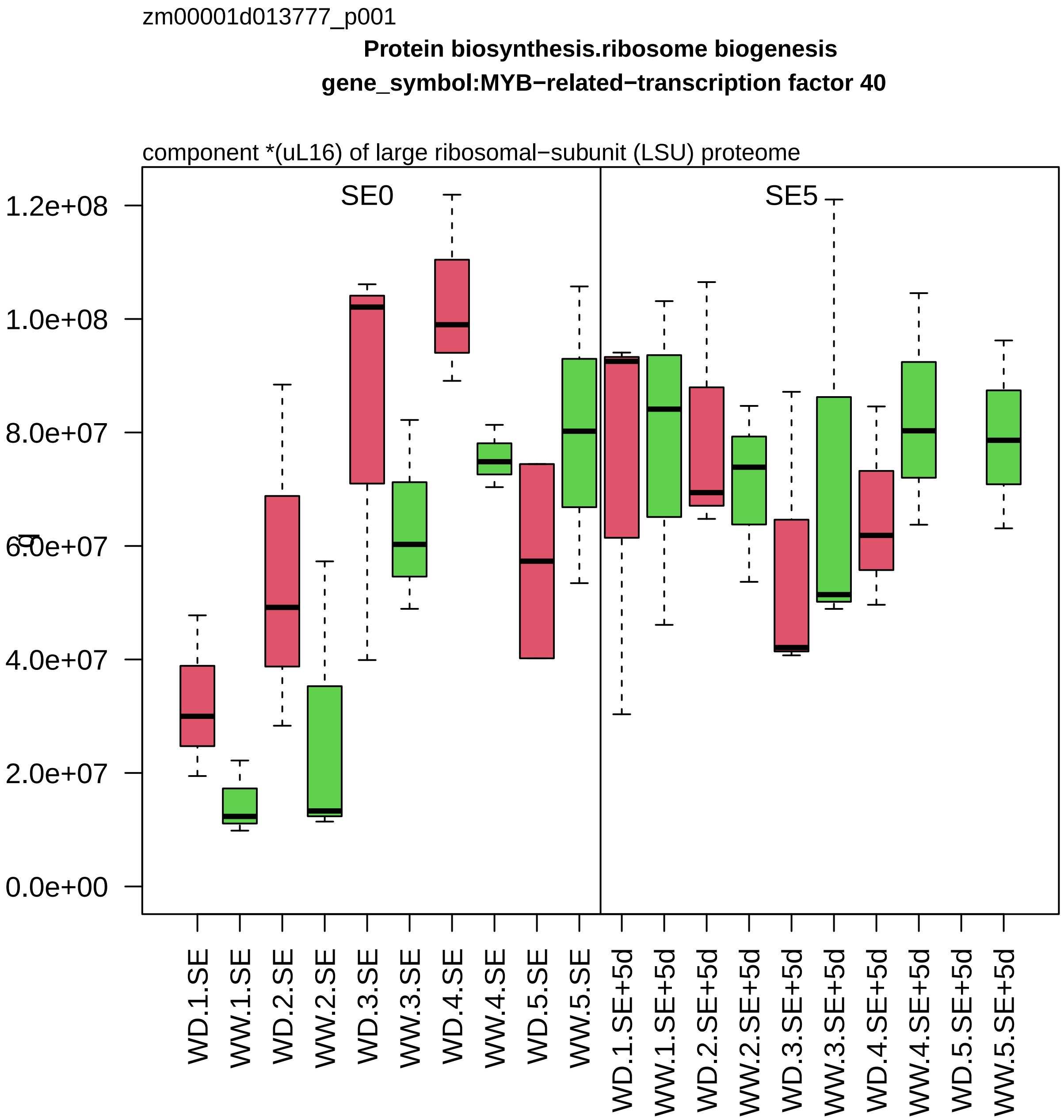

zm00001d013777_p001
Protein biosynthesis.ribosome biogenesis
gene_symbol:MYB−related−transcription factor 40
component *(uL16) of large ribosomal−subunit (LSU) proteome
SE0
SE5
1.2e+08
1.0e+08
8.0e+07
6.0e+07
4.0e+07
2.0e+07
0.0e+00
WD.1.SE
WW.1.SE
WD.2.SE
WW.2.SE
WD.3.SE
WW.3.SE
WD.4.SE
WW.4.SE
WD.5.SE
WW.5.SE
WD.1.SE+5d
WW.1.SE+5d
WD.2.SE+5d
WW.2.SE+5d
WD.3.SE+5d
WW.3.SE+5d
WD.4.SE+5d
WW.4.SE+5d
WD.5.SE+5d
WW.5.SE+5d

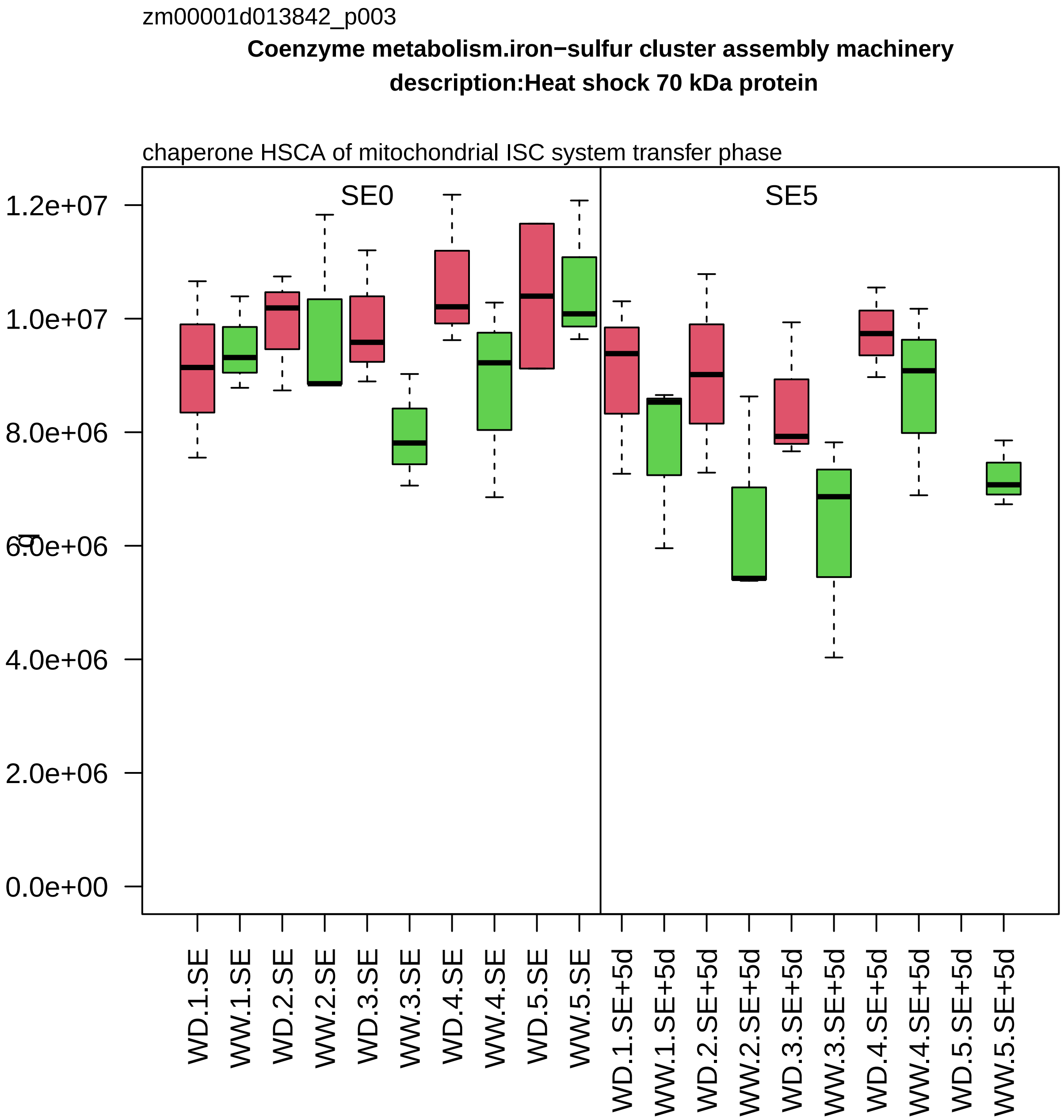
zm00001d013842_p003
Coenzyme metabolism.iron−sulfur cluster assembly machinery
description:Heat shock 70 kDa protein
chaperone HSCA of mitochondrial ISC system transfer phase
SE0
SE5
1.2e+07
1.0e+07
8.0e+06
6.0e+06
4.0e+06
2.0e+06
0.0e+00
WD.1.SE
WW.1.SE
WD.2.SE
WW.2.SE
WD.3.SE
WW.3.SE
WD.4.SE
WW.4.SE
WD.5.SE
WW.5.SE
WD.1.SE+5d
WW.1.SE+5d
WD.2.SE+5d
WW.2.SE+5d
WD.3.SE+5d
WW.3.SE+5d
WD.4.SE+5d
WW.4.SE+5d
WD.5.SE+5d
WW.5.SE+5d

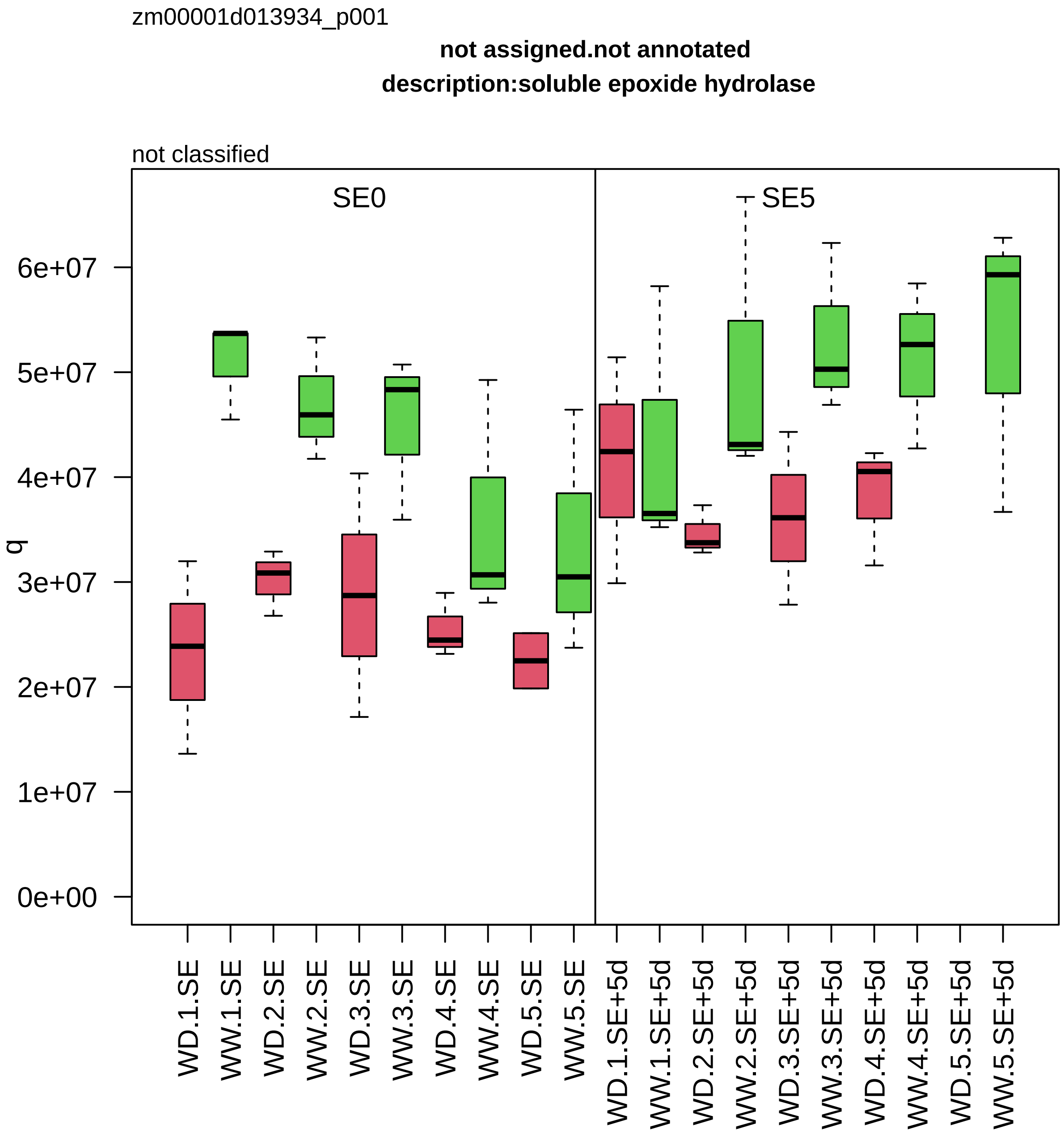

zm00001d013934_p001
not assigned.not annotated
description:soluble epoxide hydrolase
not classified
SE0
SE5
q
0e+00
1e+07
2e+07
3e+07
4e+07
5e+07
6e+07
WD.1.SE
WW.1.SE
WD.2.SE
WW.2.SE
WD.3.SE
WW.3.SE
WD.4.SE
WW.4.SE
WD.5.SE
WW.5.SE
WD.1.SE+5d
WW.1.SE+5d
WD.2.SE+5d
WW.2.SE+5d
WD.3.SE+5d
WW.3.SE+5d
WD.4.SE+5d
WW.4.SE+5d
WD.5.SE+5d
WW.5.SE+5d

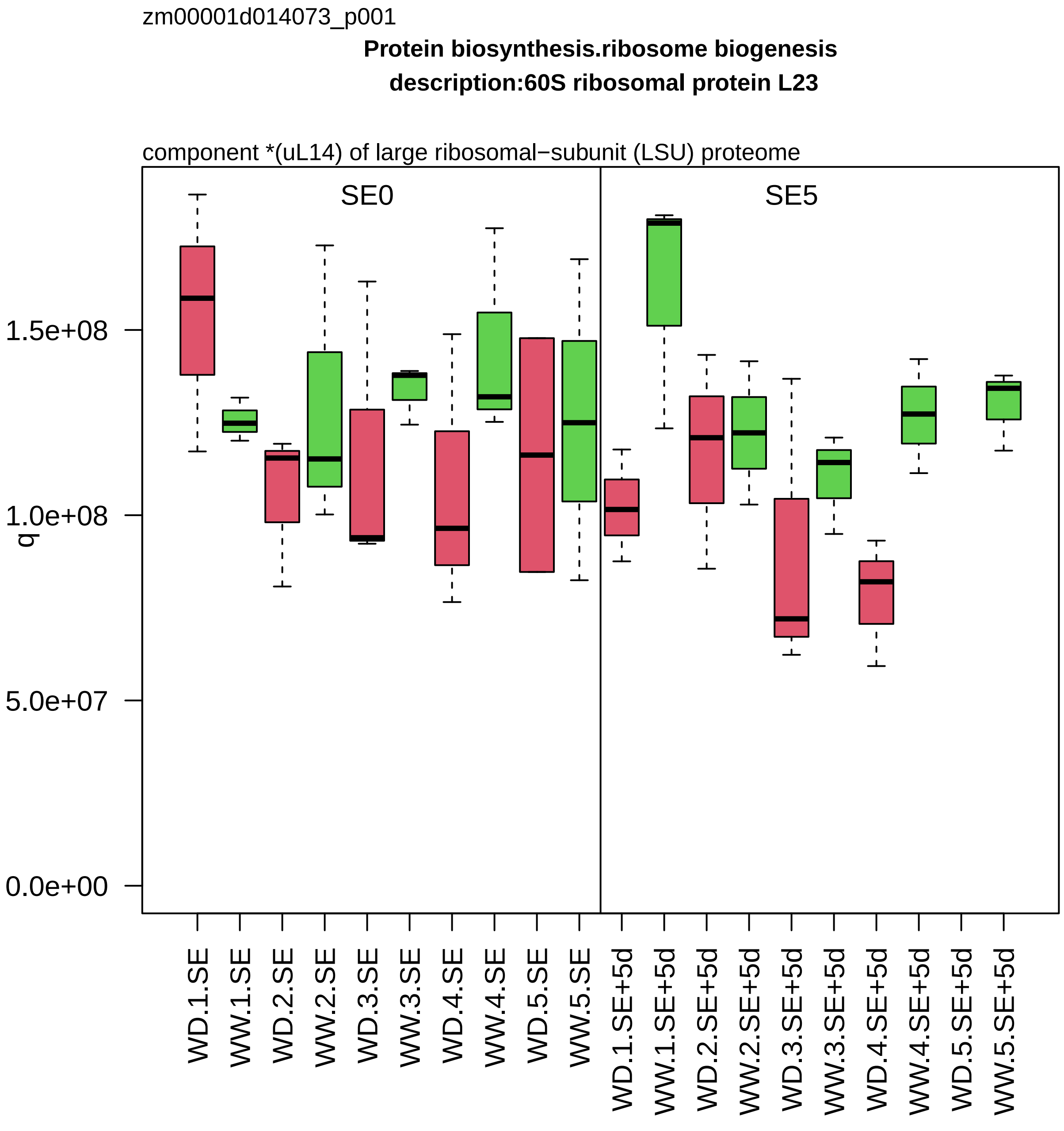
zm00001d014073_p001
Protein biosynthesis.ribosome biogenesis
description:60S ribosomal protein L23
component *(uL14) of large ribosomal−subunit (LSU) proteome
SE0
SE5
q
1.5e+08
1.0e+08
5.0e+07
0.0e+00
WD.1.SE
WW.1.SE
WD.2.SE
WW.2.SE
WD.3.SE
WW.3.SE
WD.4.SE
WW.4.SE
WD.5.SE
WW.5.SE
WD.1.SE+5d
WW.1.SE+5d
WD.2.SE+5d
WW.2.SE+5d
WD.3.SE+5d
WW.3.SE+5d
WD.4.SE+5d
WW.4.SE+5d
WD.5.SE+5d
WW.5.SE+5d

zm00001d014090_p001

**Protein homeostasis.protein quality control**

**gene_symbol:chaperonin 1 description: chaperonin 1**

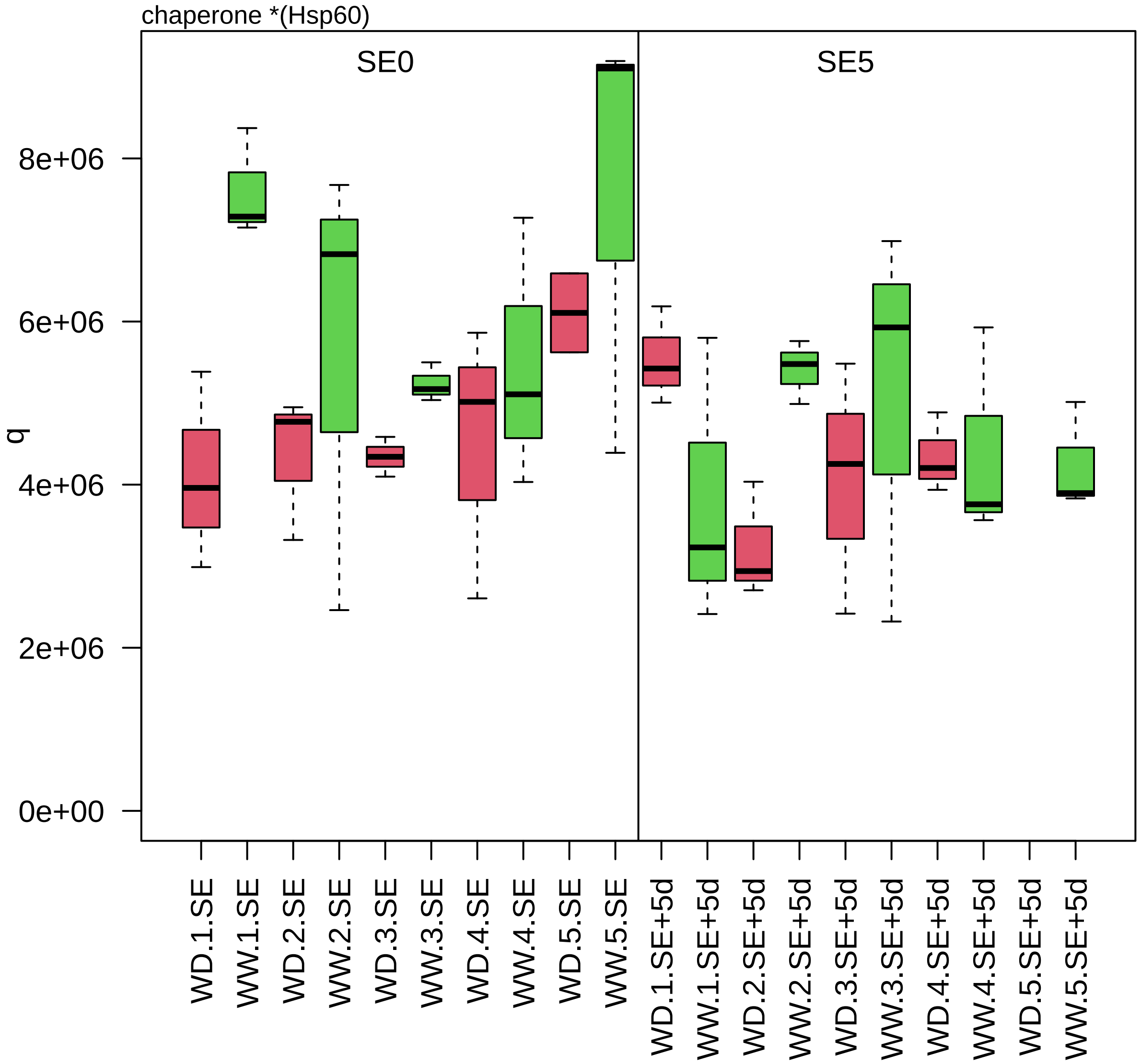

zm00001d014093_p002

**Lipid metabolism.fatty acid metabolism**

**description:3−ketoacyl−CoA thiolase 2 peroxisomal**

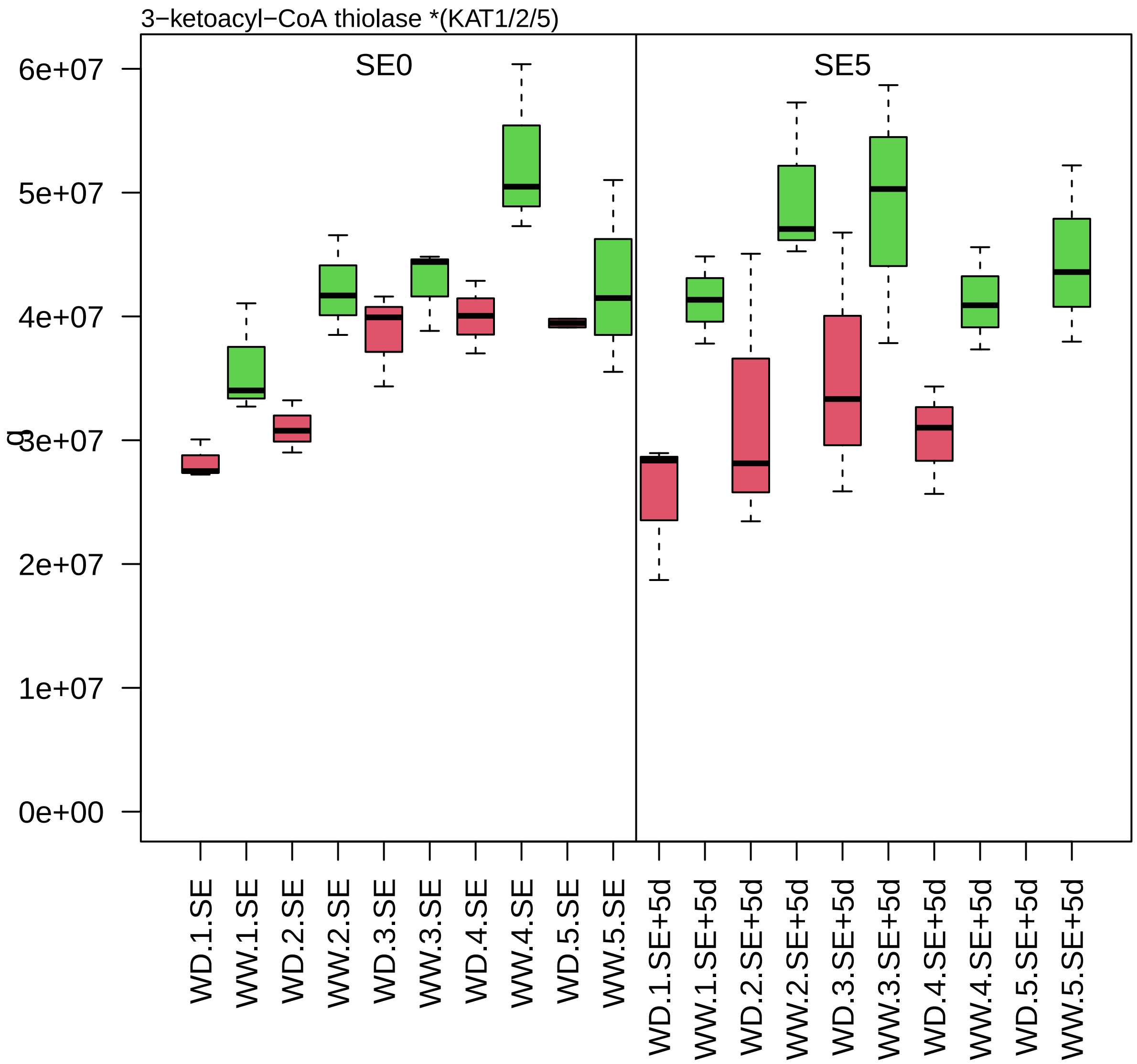

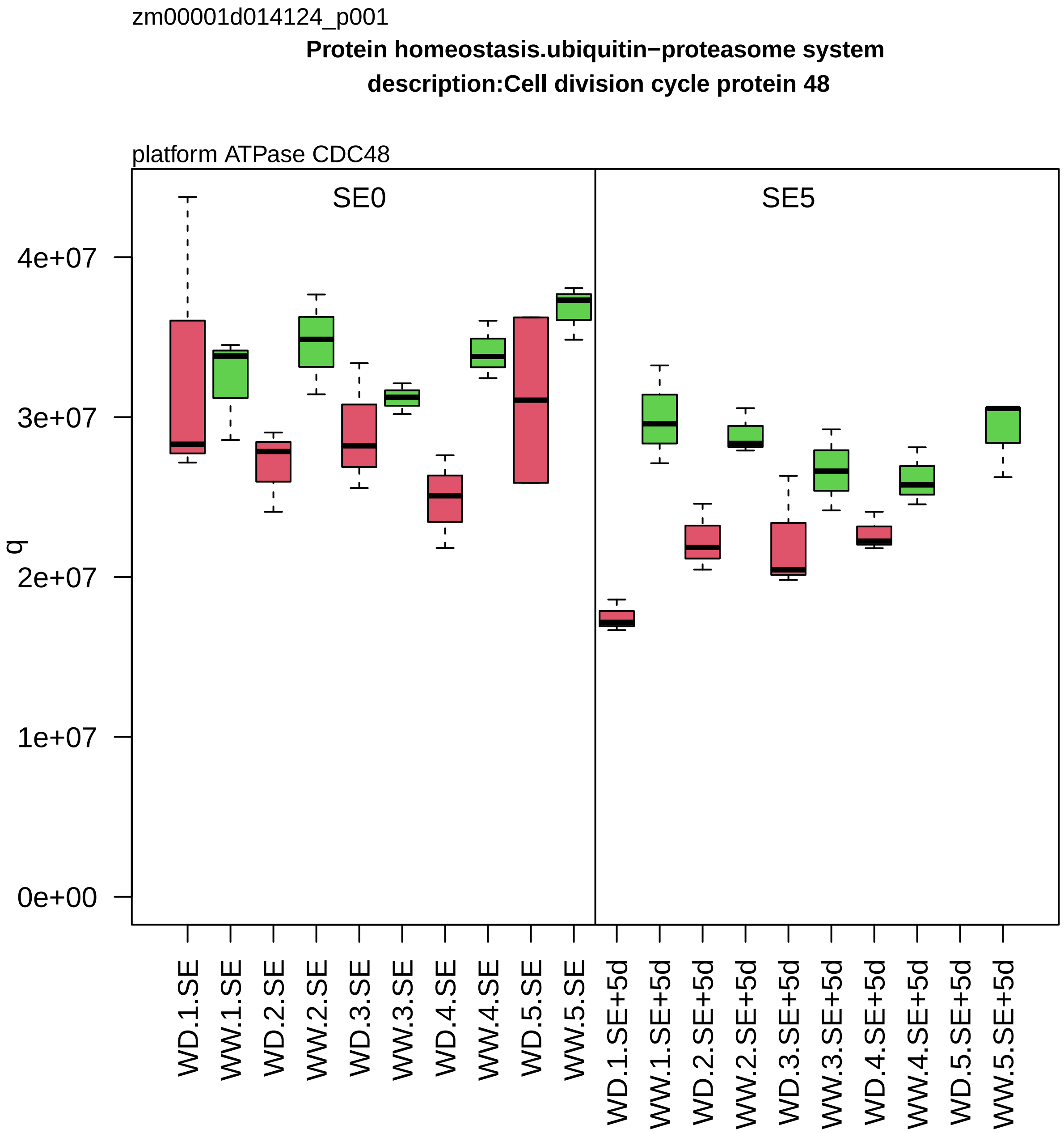
zm00001d014124_p001
Protein homeostasis.ubiquitin−proteasome system
description:Cell division cycle protein 48
platform ATPase CDC48
SE0
SE5
q
4e+07
3e+07
2e+07
1e+07
0e+00
WD.1.SE
WW.1.SE
WD.2.SE
WW.2.SE
WD.3.SE
WW.3.SE
WD.4.SE
WW.4.SE
WD.5.SE
WW.5.SE
WD.1.SE+5d
WW.1.SE+5d
WD.2.SE+5d
WW.2.SE+5d
WD.3.SE+5d
WW.3.SE+5d
WD.4.SE+5d
WW.4.SE+5d
WD.5.SE+5d
WW.5.SE+5d

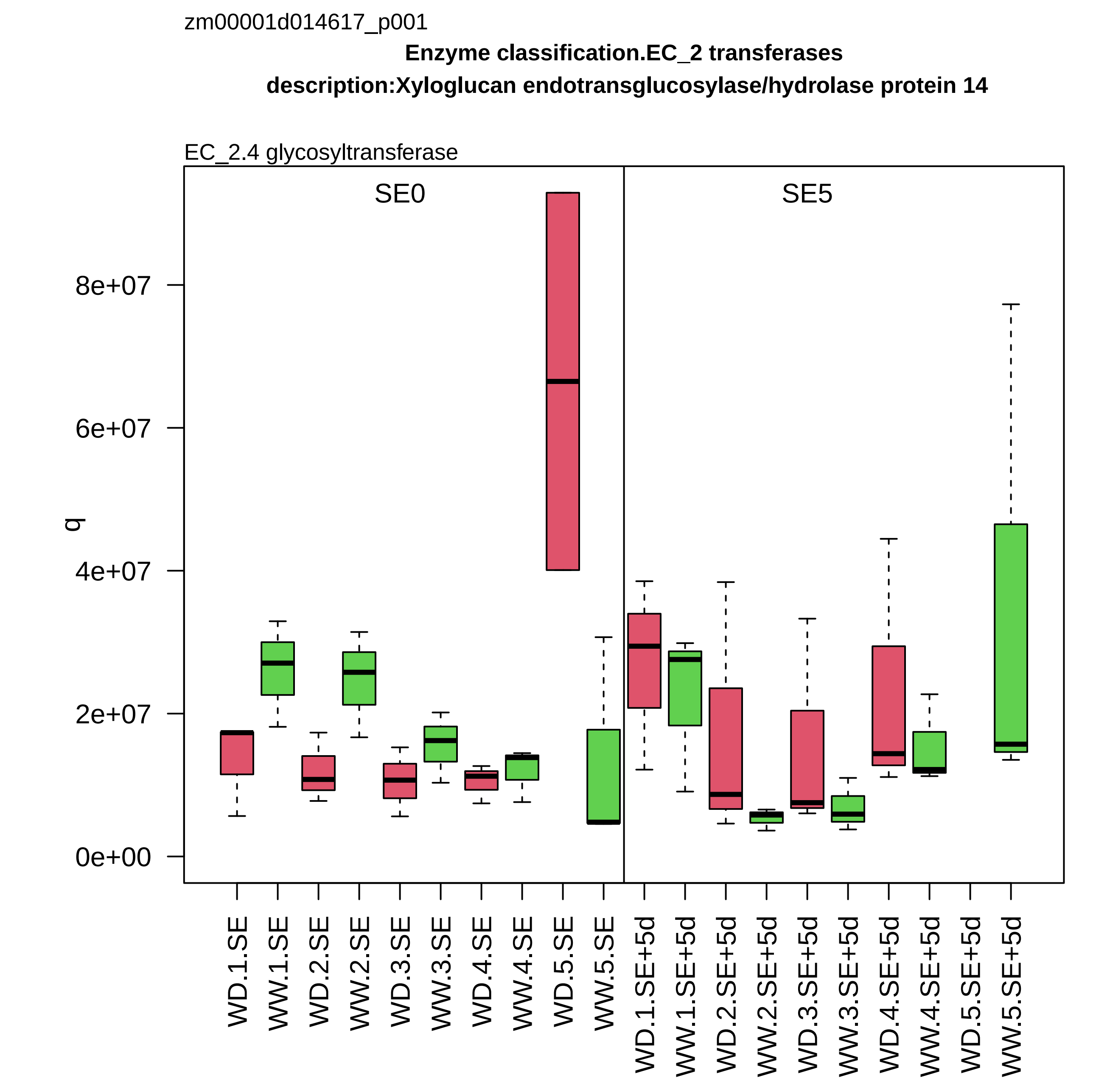

zm00001d014617_p001
Enzyme classification.EC_2 transferases
description:Xyloglucan endotransglucosylase/hydrolase protein 14
EC_2.4 glycosyltransferase
SE0
SE5
q
0e+00
2e+07
4e+07
6e+07
8e+07
WD.1.SE
WW.1.SE
WD.2.SE
WW.2.SE
WD.3.SE
WW.3.SE
WD.4.SE
WW.4.SE
WD.5.SE
WW.5.SE
WD.1.SE+5d
WW.1.SE+5d
WD.2.SE+5d
WW.2.SE+5d
WD.3.SE+5d
WW.3.SE+5d
WD.4.SE+5d
WW.4.SE+5d
WD.5.SE+5d
WW.5.SE+5d

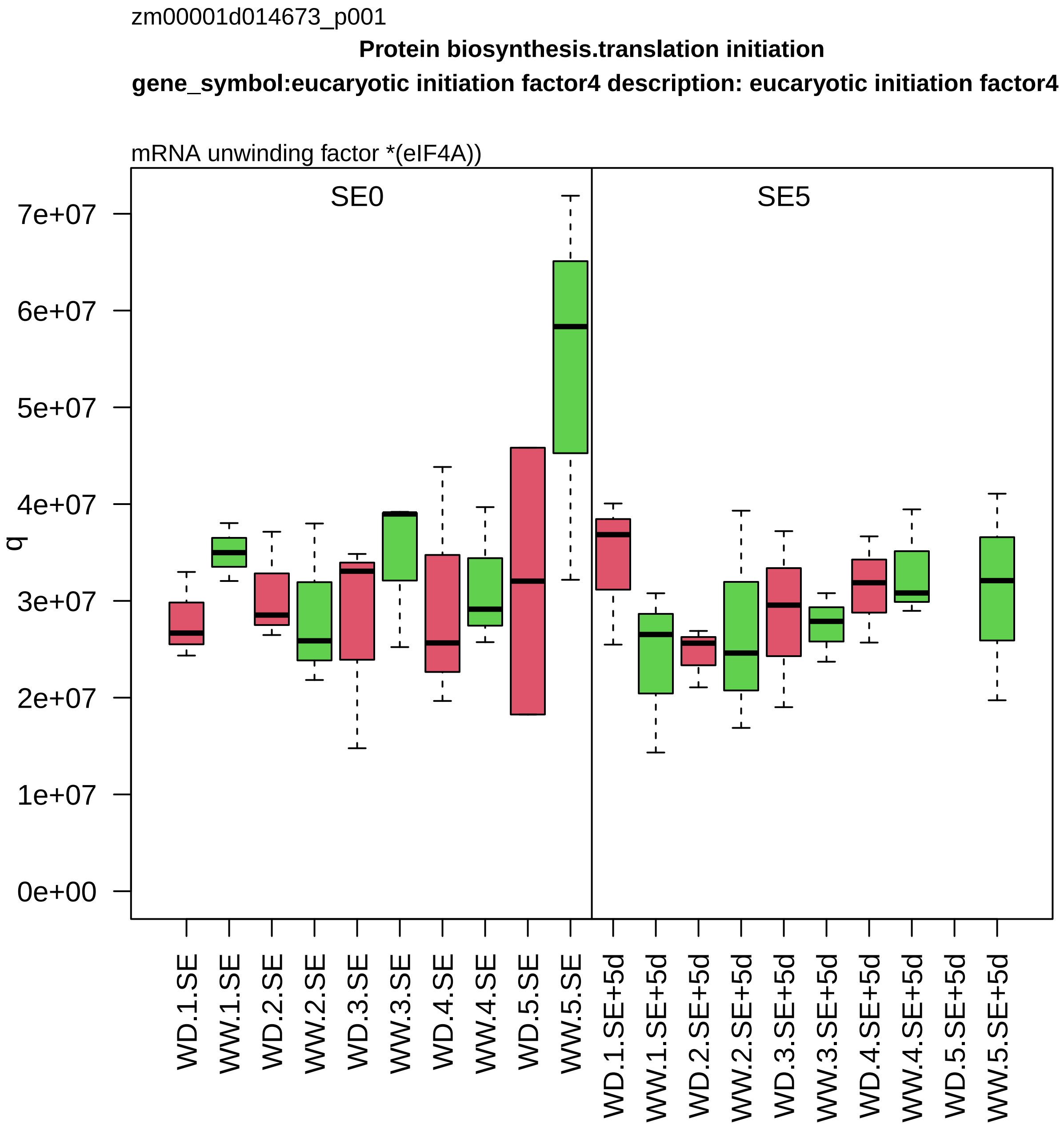

zm00001d014673_p001
Protein biosynthesis.translation initiation
gene_symbol:eucaryotic initiation factor4 description: eucaryotic initiation factor4
mRNA unwinding factor *(eIF4A))
SE0
SE5
q
0e+00
1e+07
2e+07
3e+07
4e+07
5e+07
6e+07
7e+07
WD.1.SE
WW.1.SE
WD.2.SE
WW.2.SE
WD.3.SE
WW.3.SE
WD.4.SE
WW.4.SE
WD.5.SE
WW.5.SE
WD.1.SE+5d
WW.1.SE+5d
WD.2.SE+5d
WW.2.SE+5d
WD.3.SE+5d
WW.3.SE+5d
WD.4.SE+5d
WW.4.SE+5d
WD.5.SE+5d
WW.5.SE+5d

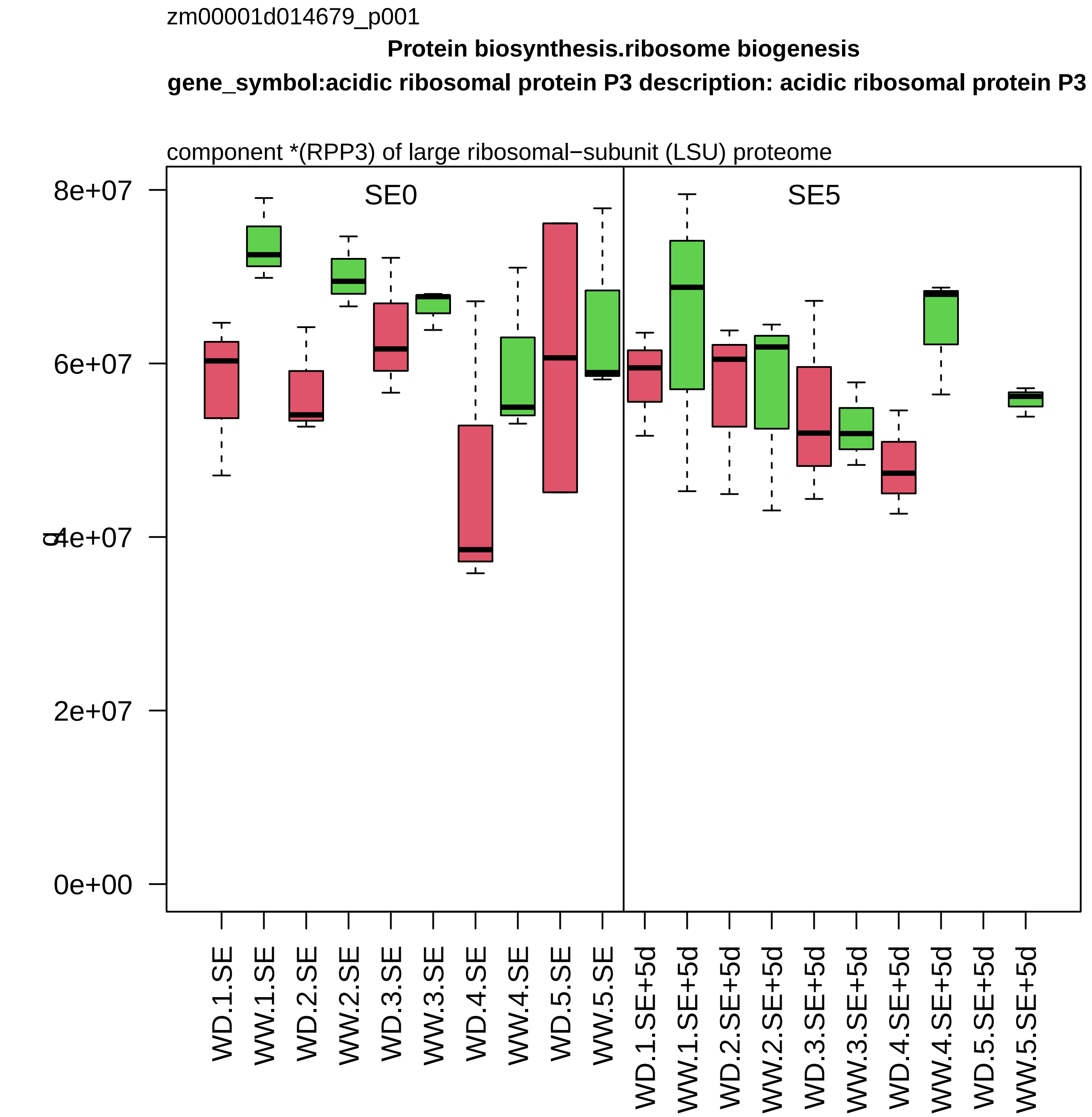

zm00001d014679_p001
Protein biosynthesis.ribosome biogenesis
gene_symbol:acidic ribosomal protein P3 description: acidic ribosomal protein P3
component *(RPP3) of large ribosomal−subunit (LSU) proteome
SE0
SE5
8e+07
6e+07
4e+07
2e+07
0e+00
WD.1.SE
WW.1.SE
WD.2.SE
WW.2.SE
WD.3.SE
WW.3.SE
WD.4.SE
WW.4.SE
WD.5.SE
WW.5.SE
WD.1.SE+5d
WW.1.SE+5d
WD.2.SE+5d
WW.2.SE+5d
WD.3.SE+5d
WW.3.SE+5d
WD.4.SE+5d
WW.4.SE+5d
WD.5.SE+5d
WW.5.SE+5d

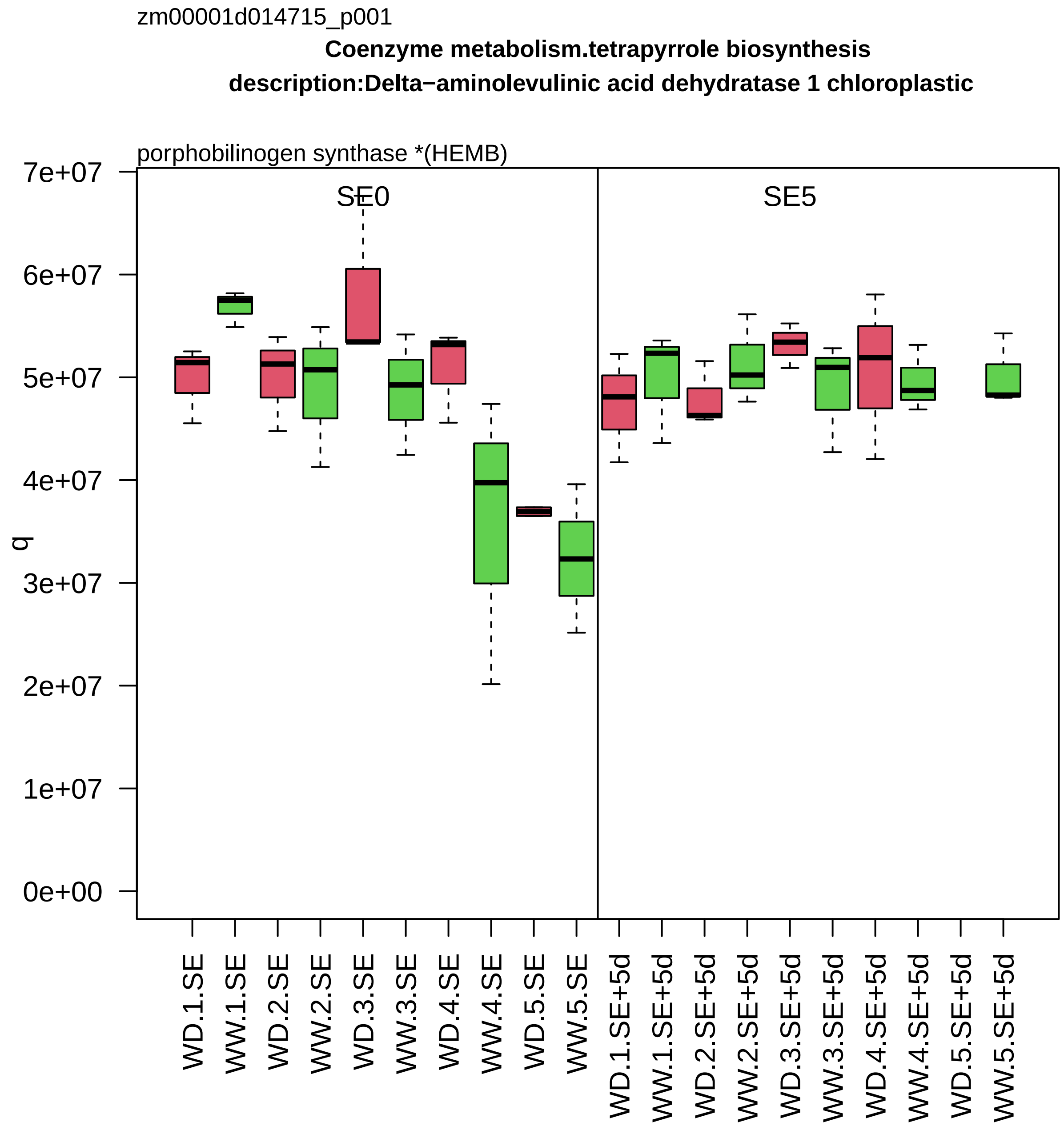

zm00001d014715_p001
Coenzyme metabolism.tetrapyrrole biosynthesis
description:Delta−aminolevulinic acid dehydratase 1 chloroplastic
porphobilinogen synthase *(HEMB)
SE0
SE5
q
7e+07
6e+07
5e+07
4e+07
3e+07
2e+07
1e+07
0e+00
WD.1.SE
WW.1.SE
WD.2.SE
WW.2.SE
WD.3.SE
WW.3.SE
WD.4.SE
WW.4.SE
WD.5.SE
WW.5.SE
WD.1.SE+5d
WW.1.SE+5d
WD.2.SE+5d
WW.2.SE+5d
WD.3.SE+5d
WW.3.SE+5d
WD.4.SE+5d
WW.4.SE+5d
WD.5.SE+5d
WW.5.SE+5d

zm00001d014792_p003

**Protein homeostasis.protein quality control**
**description: shepherd−like1**

chaperone *(Hsp90)

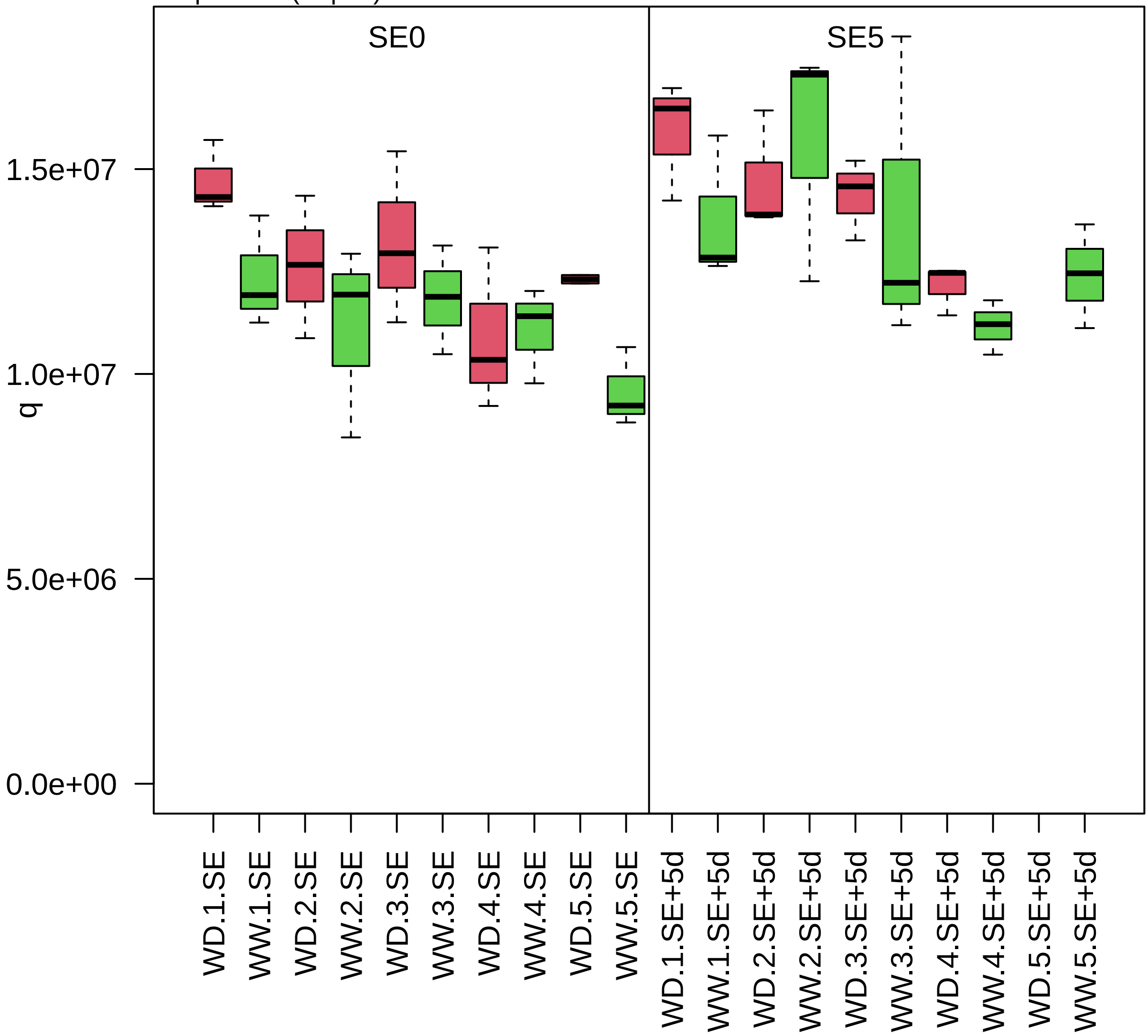

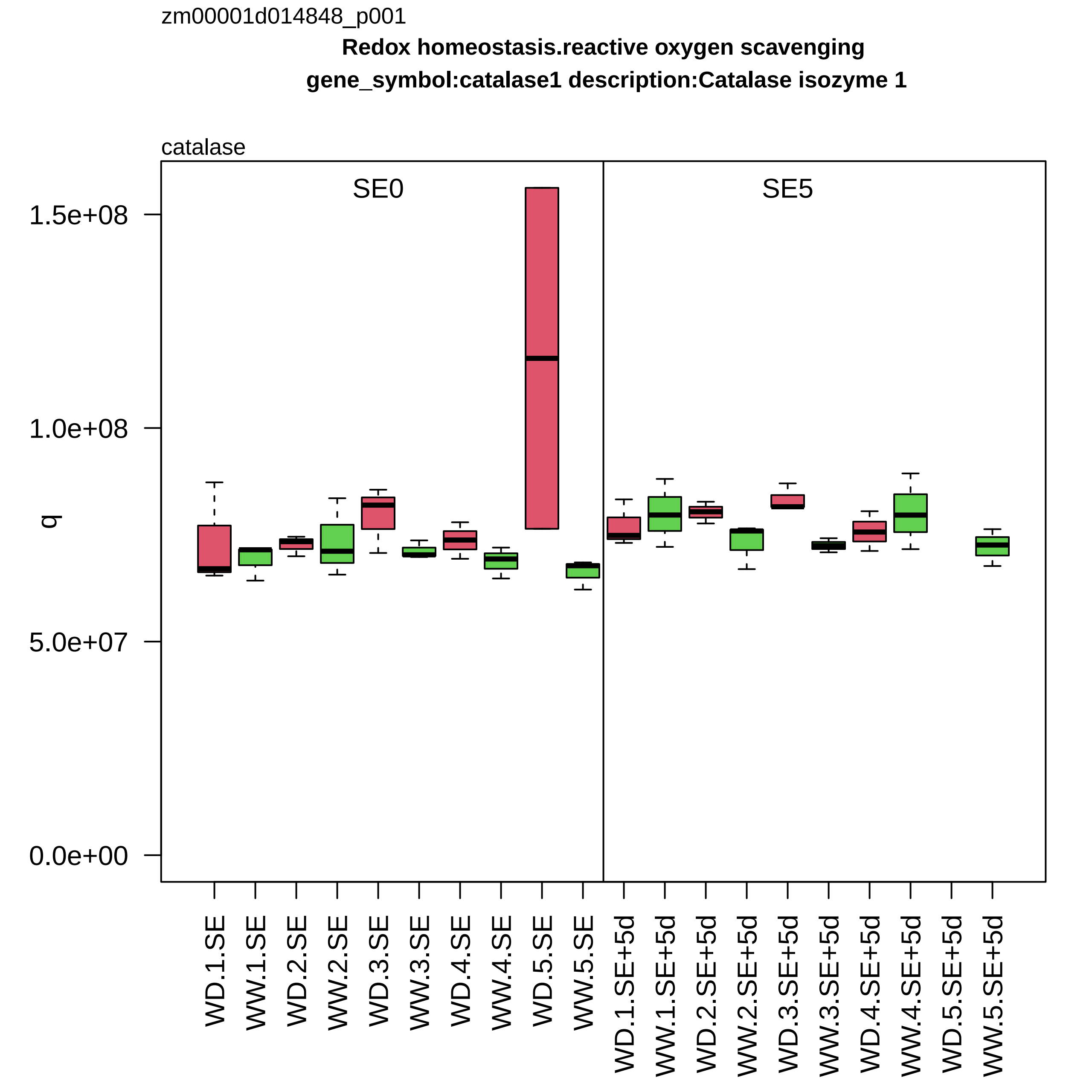

zm00001d014848_p001
Redox homeostasis.reactive oxygen scavenging
gene_symbol:catalase1 description:Catalase isozyme 1
catalase
SE0
SE5
q
1.5e+08
1.0e+08
5.0e+07
0.0e+00
WD.1.SE
WW.1.SE
WD.2.SE
WW.2.SE
WD.3.SE
WW.3.SE
WD.4.SE
WW.4.SE
WD.5.SE
WW.5.SE
WD.1.SE+5d
WW.1.SE+5d
WD.2.SE+5d
WW.2.SE+5d
WD.3.SE+5d
WW.3.SE+5d
WD.4.SE+5d
WW.4.SE+5d
WD.5.SE+5d
WW.5.SE+5d

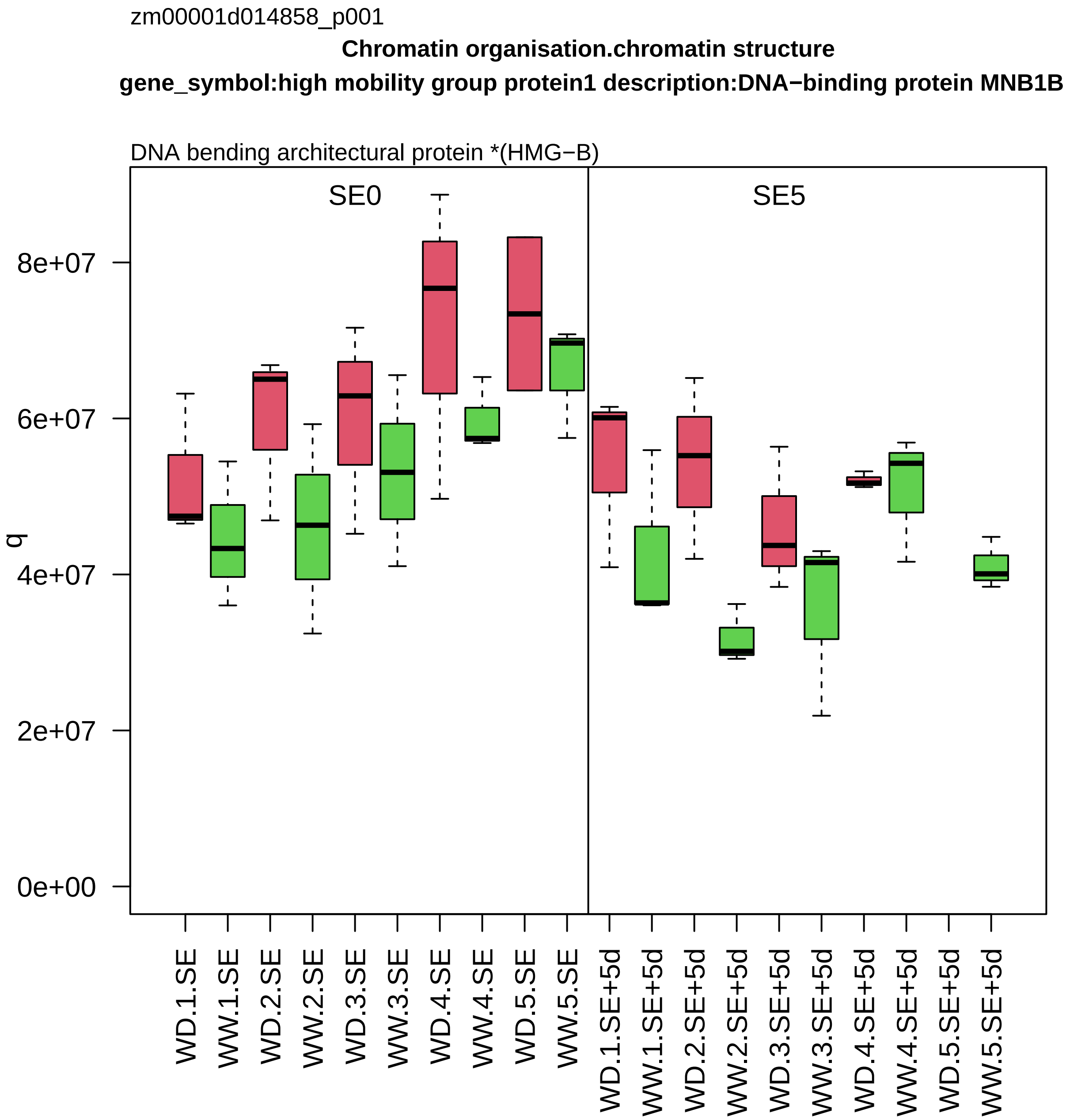

zm00001d014858_p001
Chromatin organisation.chromatin structure
gene_symbol:high mobility group protein1 description:DNA−binding protein MNB1B
DNA bending architectural protein *(HMG−B)
SE0
SE5
q
0e+00
2e+07
4e+07
6e+07
8e+07
WD.1.SE
WW.1.SE
WD.2.SE
WW.2.SE
WD.3.SE
WW.3.SE
WD.4.SE
WW.4.SE
WD.5.SE
WW.5.SE
WD.1.SE+5d
WW.1.SE+5d
WD.2.SE+5d
WW.2.SE+5d
WD.3.SE+5d
WW.3.SE+5d
WD.4.SE+5d
WW.4.SE+5d
WD.5.SE+5d
WW.5.SE+5d

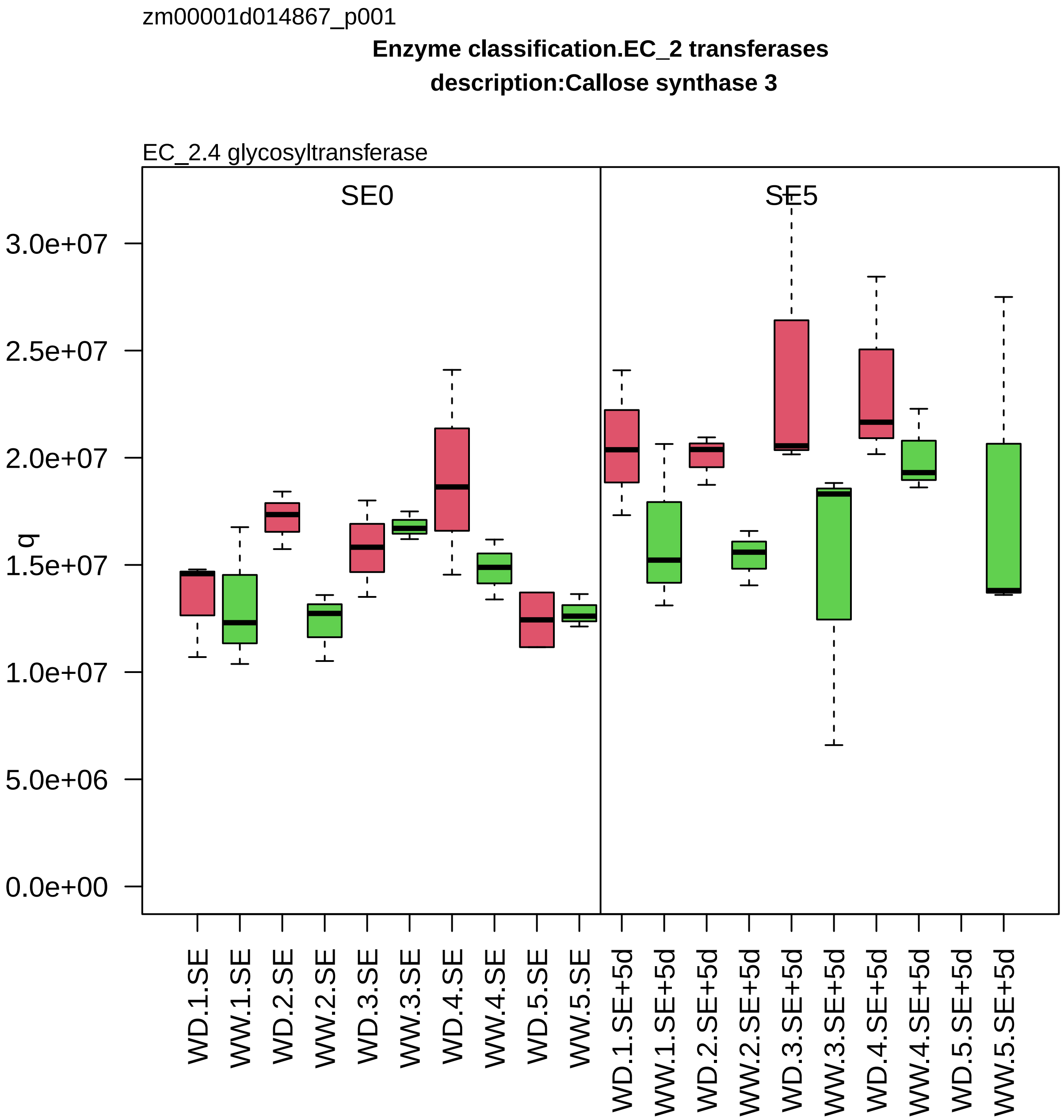

zm00001d014867_p001
Enzyme classification.EC_2 transferases
description:Callose synthase 3
EC_2.4 glycosyltransferase
SE0
SE5
q
3.0e+07
2.5e+07
2.0e+07
1.5e+07
1.0e+07
5.0e+06
0.0e+00
WD.1.SE
WW.1.SE
WD.2.SE
WW.2.SE
WD.3.SE
WW.3.SE
WD.4.SE
WW.4.SE
WD.5.SE
WW.5.SE
WD.1.SE+5d
WW.1.SE+5d
WD.2.SE+5d
WW.2.SE+5d
WD.3.SE+5d
WW.3.SE+5d
WD.4.SE+5d
WW.4.SE+5d
WD.5.SE+5d
WW.5.SE+5d

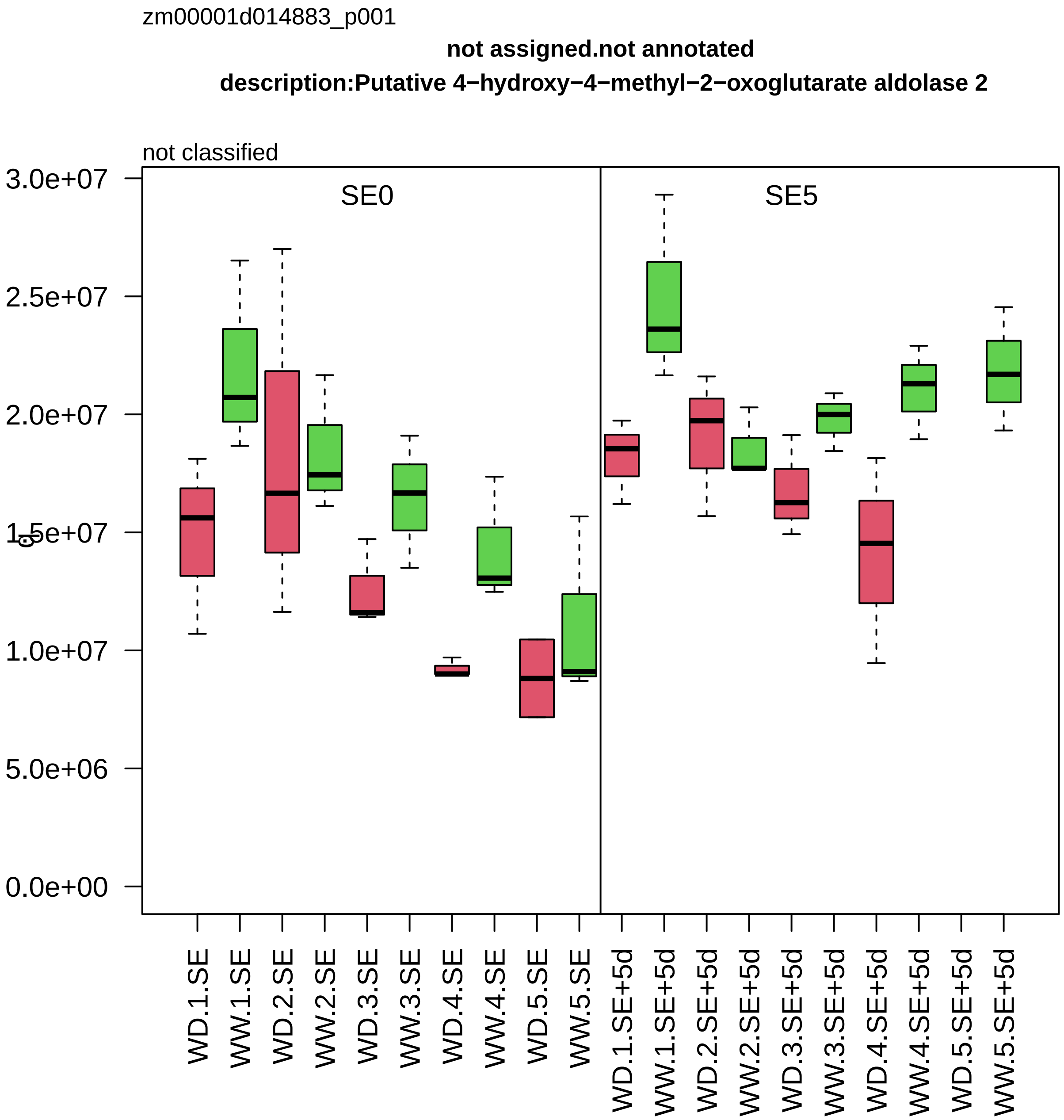
zm00001d014883_p001
not assigned.not annotated
description:Putative 4−hydroxy−4−methyl−2−oxoglutarate aldolase 2
not classified
SE0
SE5
3.0e+07
2.5e+07
2.0e+07
1.5e+07
1.0e+07
5.0e+06
0.0e+00
WD.1.SE
WW.1.SE
WD.2.SE
WW.2.SE
WD.3.SE
WW.3.SE
WD.4.SE
WW.4.SE
WD.5.SE
WW.5.SE
WD.1.SE+5d
WW.1.SE+5d
WD.2.SE+5d
WW.2.SE+5d
WD.3.SE+5d
WW.3.SE+5d
WD.4.SE+5d
WW.4.SE+5d
WD.5.SE+5d
WW.5.SE+5d

zm00001d014887_p001

**Lipid metabolism.phytosterol metabolism**

**gene_symbol:nana plant2 description: nana plant2**

sterol delta24 reductase

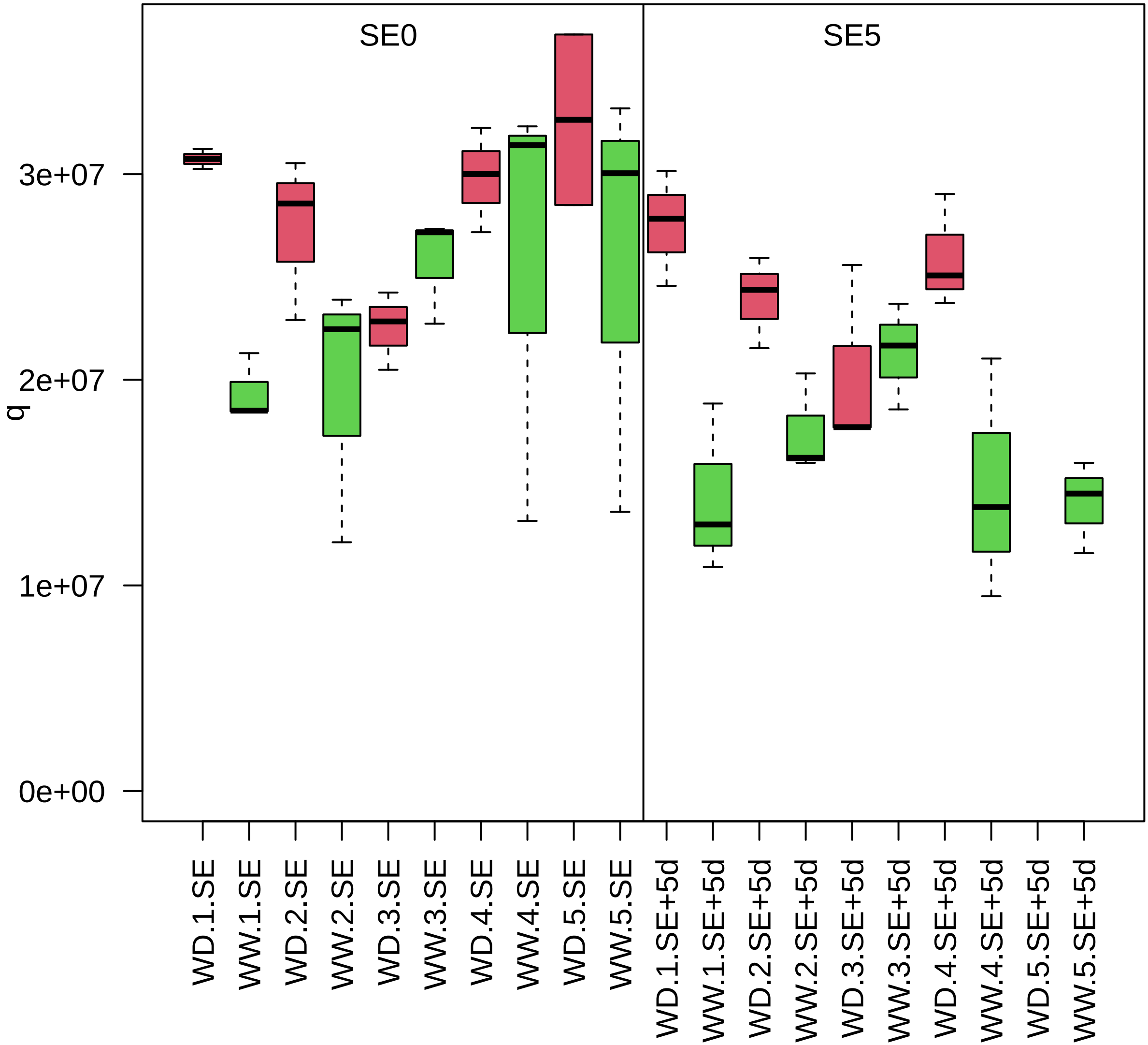

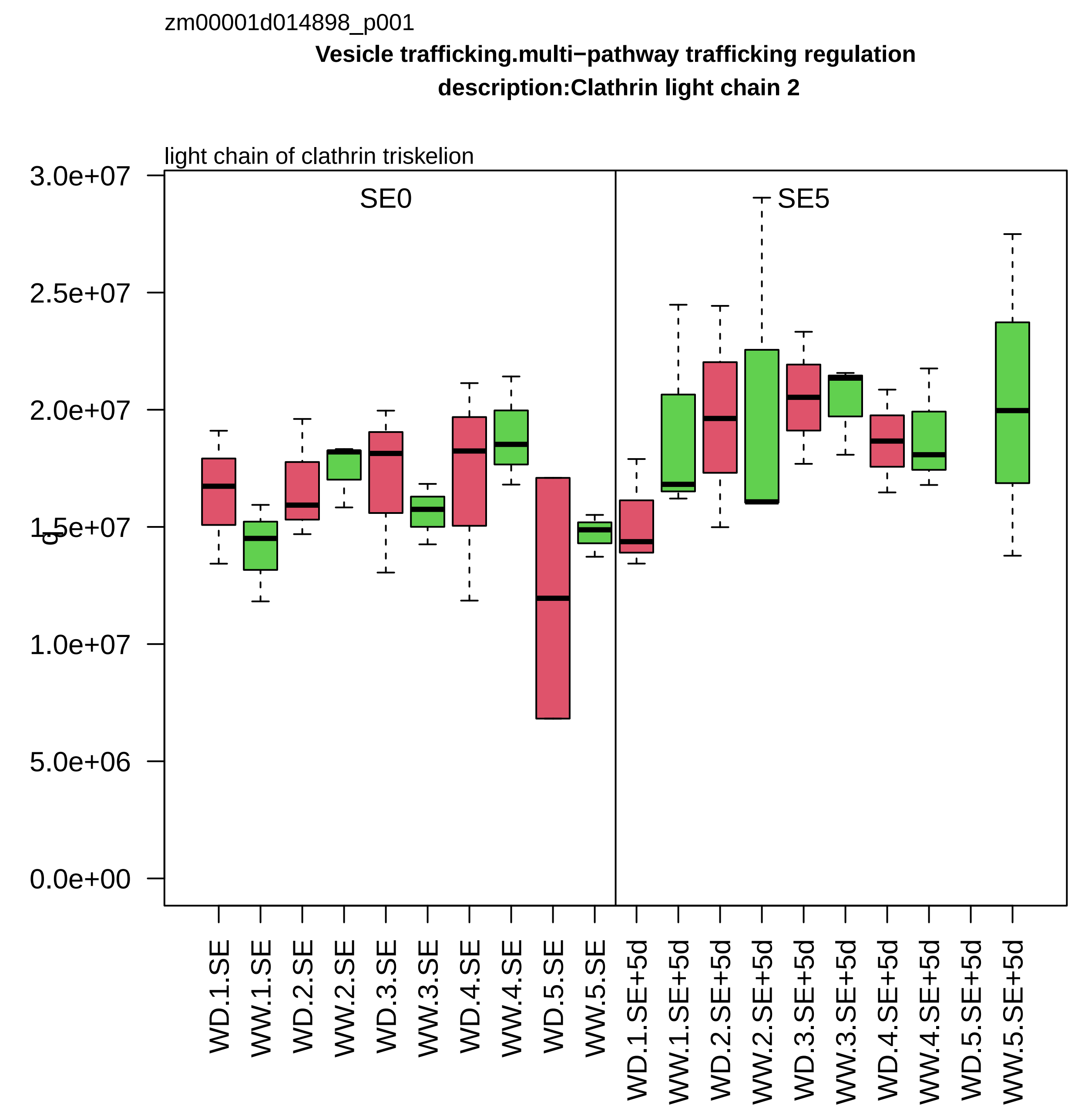
zm00001d014898_p001
Vesicle trafficking.multi−pathway trafficking regulation
description:Clathrin light chain 2
light chain of clathrin triskelion
SE0
SE5
3.0e+07
2.5e+07
2.0e+07
1.5e+07
1.0e+07
5.0e+06
0.0e+00
WD.1.SE
WW.1.SE
WD.2.SE
WW.2.SE
WD.3.SE
WW.3.SE
WD.4.SE
WW.4.SE
WD.5.SE
WW.5.SE
WD.1.SE+5d
WW.1.SE+5d
WD.2.SE+5d
WW.2.SE+5d
WD.3.SE+5d
WW.3.SE+5d
WD.4.SE+5d
WW.4.SE+5d
WD.5.SE+5d
WW.5.SE+5d

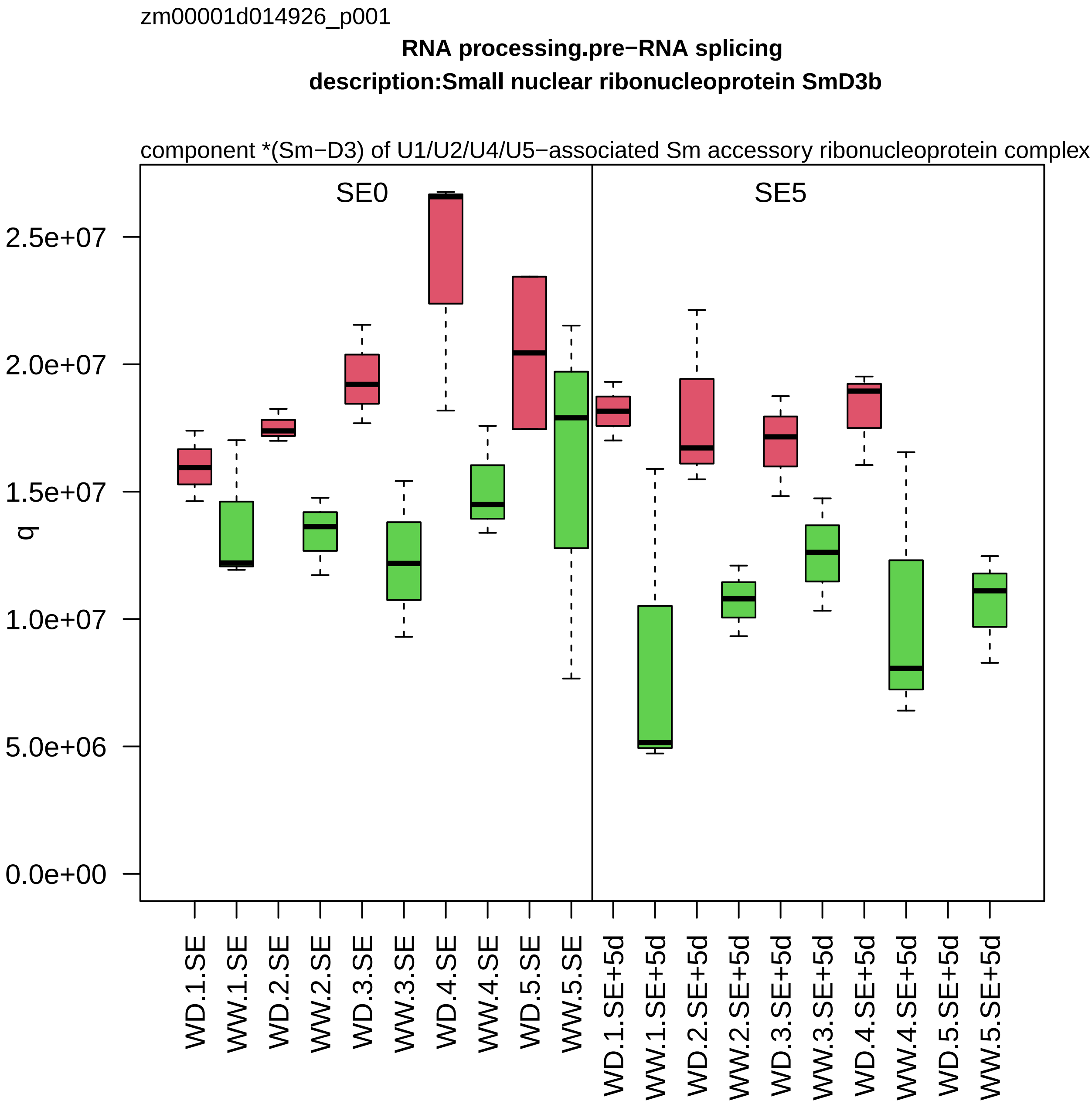

zm00001d014926_p001
RNA processing.pre−RNA splicing
description:Small nuclear ribonucleoprotein SmD3b
component *(Sm−D3) of U1/U2/U4/U5−associated Sm accessory ribonucleoprotein complex
SE0
SE5
q
0.0e+00
5.0e+06
1.0e+07
1.5e+07
2.0e+07
2.5e+07
WD.1.SE
WW.1.SE
WD.2.SE
WW.2.SE
WD.3.SE
WW.3.SE
WD.4.SE
WW.4.SE
WD.5.SE
WW.5.SE
WD.1.SE+5d
WW.1.SE+5d
WD.2.SE+5d
WW.2.SE+5d
WD.3.SE+5d
WW.3.SE+5d
WD.4.SE+5d
WW.4.SE+5d
WD.5.SE+5d
WW.5.SE+5d

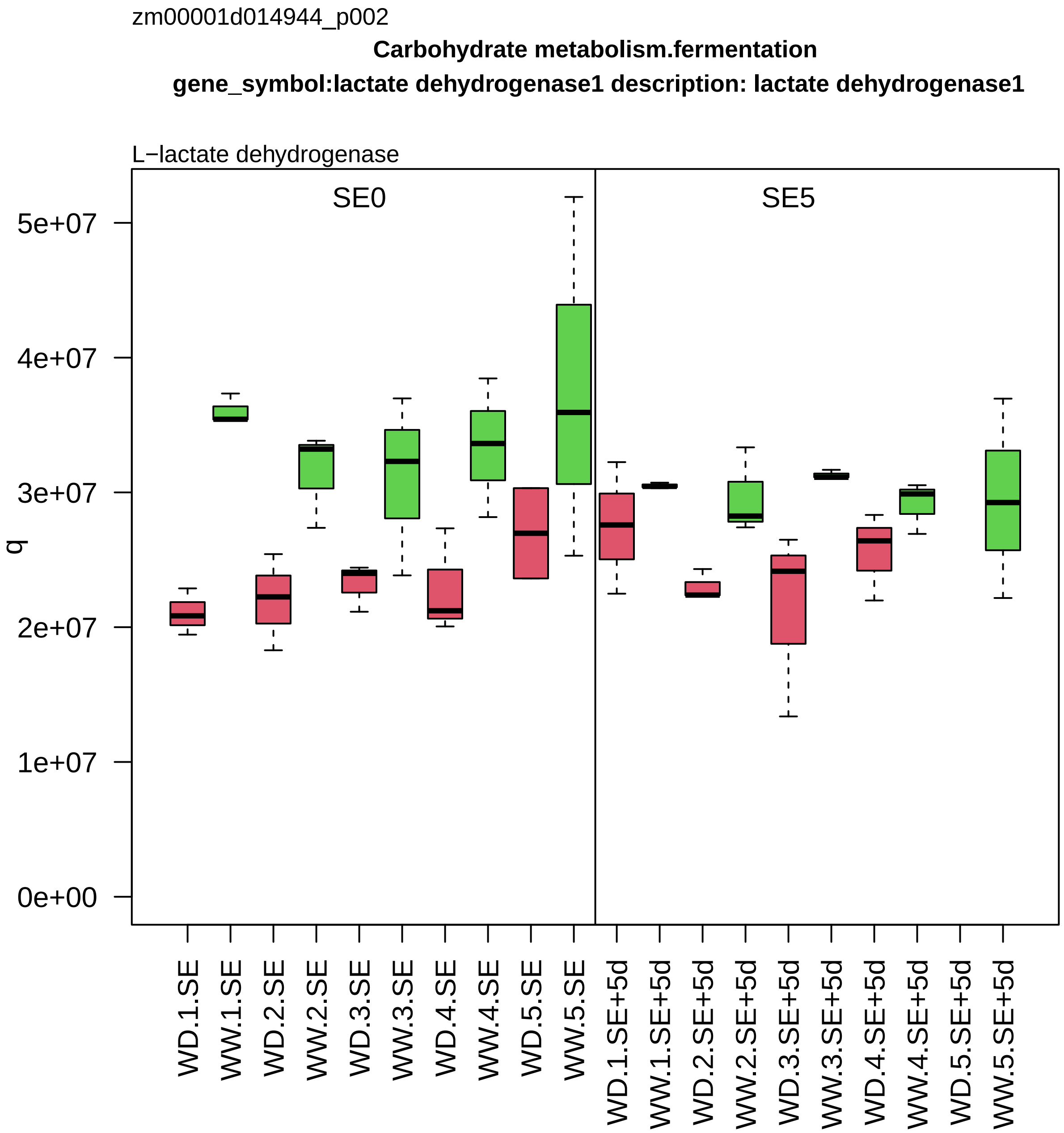
zm00001d014944_p002
Carbohydrate metabolism.fermentation
gene_symbol:lactate dehydrogenase1 description: lactate dehydrogenase1
L−lactate dehydrogenase
SE0
SE5
q
0e+00
1e+07
2e+07
3e+07
4e+07
5e+07
WD.1.SE
WW.1.SE
WD.2.SE
WW.2.SE
WD.3.SE
WW.3.SE
WD.4.SE
WW.4.SE
WD.5.SE
WW.5.SE
WD.1.SE+5d
WW.1.SE+5d
WD.2.SE+5d
WW.2.SE+5d
WD.3.SE+5d
WW.3.SE+5d
WD.4.SE+5d
WW.4.SE+5d
WD.5.SE+5d
WW.5.SE+5d

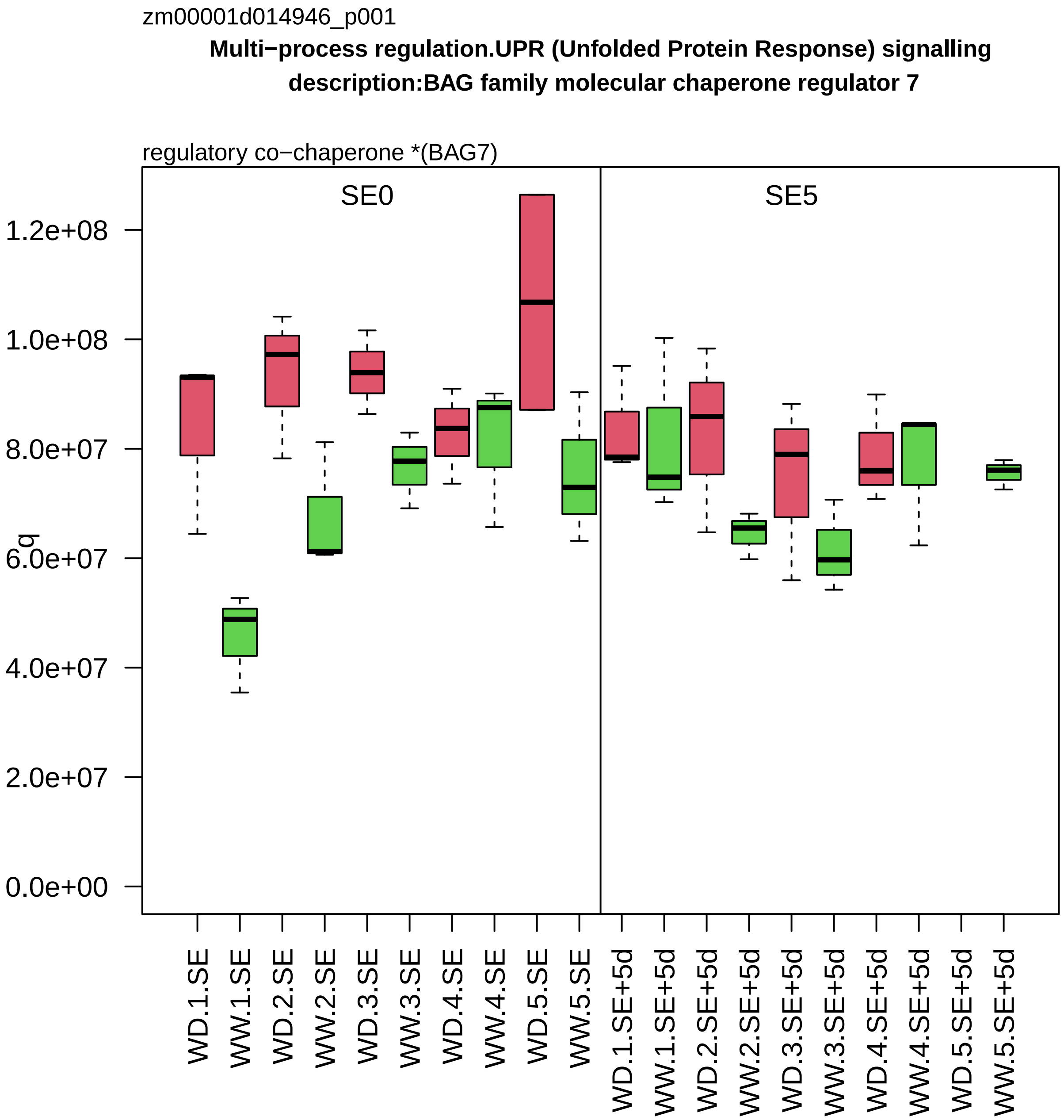
zm00001d014946_p001
Multi−process regulation.UPR (Unfolded Protein Response) signalling
description:BAG family molecular chaperone regulator 7
regulatory co−chaperone *(BAG7)
SE0
SE5
q
1.2e+08
1.0e+08
8.0e+07
6.0e+07
4.0e+07
2.0e+07
0.0e+00
WD.1.SE
WW.1.SE
WD.2.SE
WW.2.SE
WD.3.SE
WW.3.SE
WD.4.SE
WW.4.SE
WD.5.SE
WW.5.SE
WD.1.SE+5d
WW.1.SE+5d
WD.2.SE+5d
WW.2.SE+5d
WD.3.SE+5d
WW.3.SE+5d
WD.4.SE+5d
WW.4.SE+5d
WD.5.SE+5d
WW.5.SE+5d

zm00001d014954_p001

**not assigned.not annotated**

**gene_symbol:translationally controlled tumor1 description: translationally controlled tumor1**

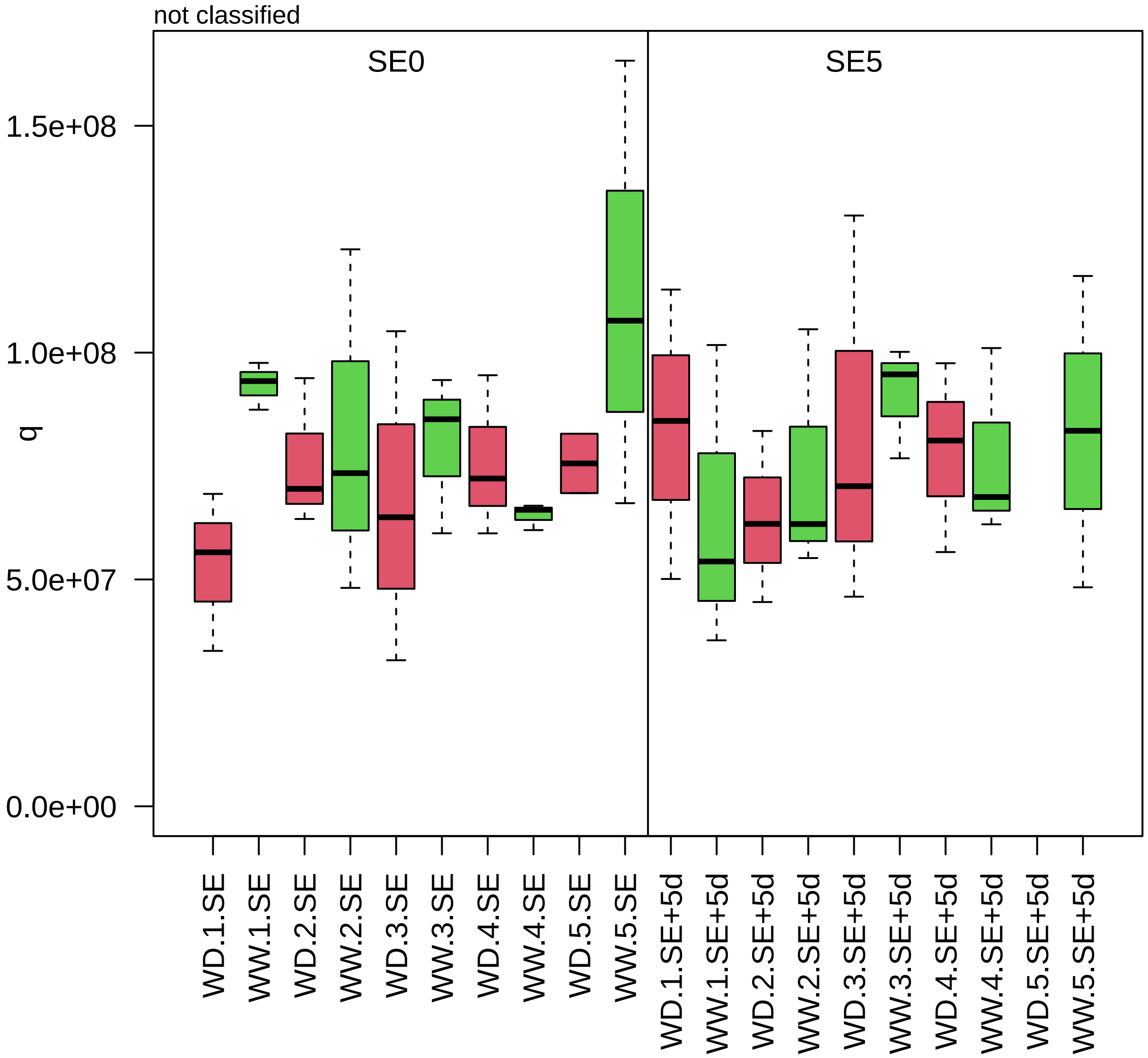

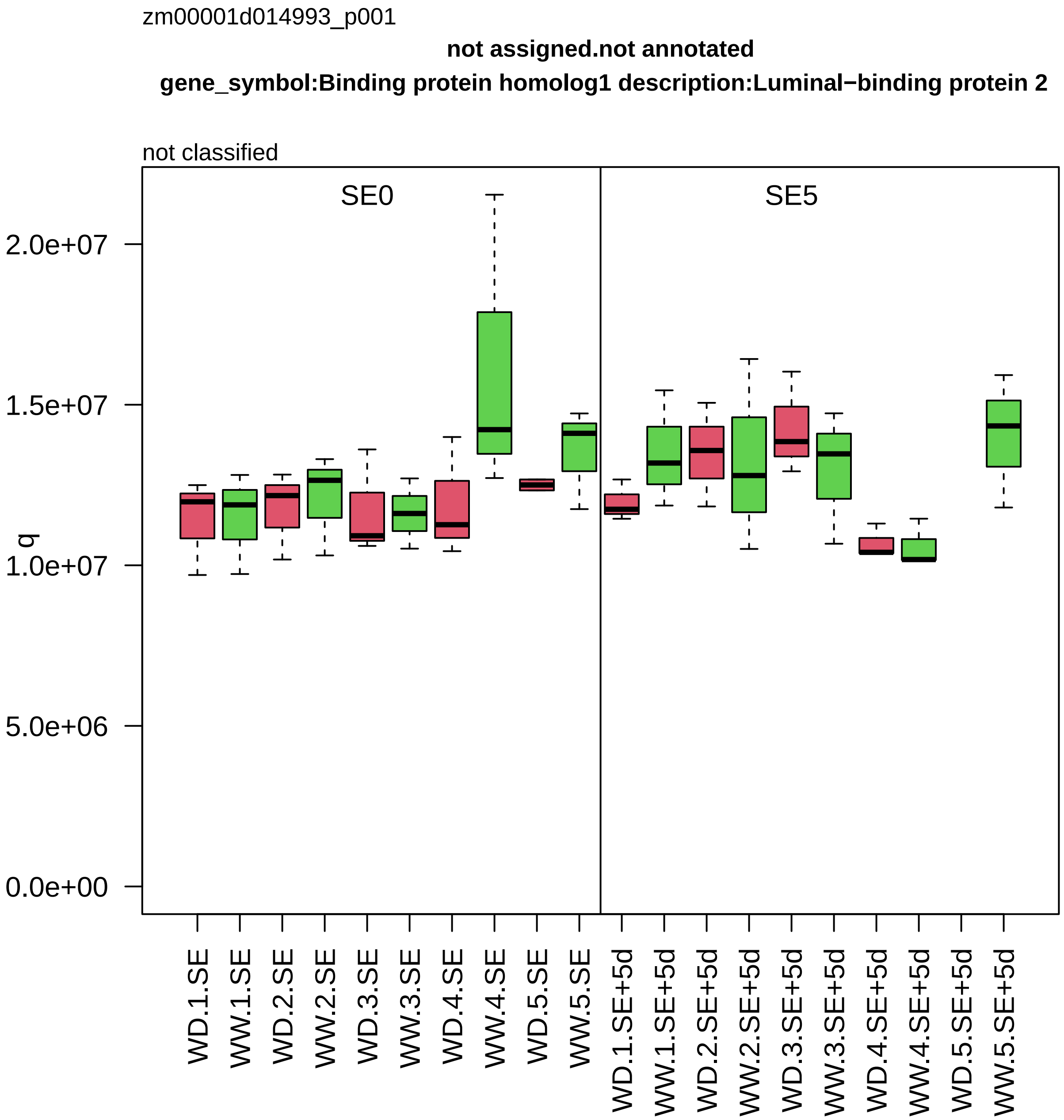

zm00001d014993_p001
not assigned.not annotated
gene_symbol:Binding protein homolog1 description:Luminal−binding protein 2
not classified
SE0
SE5
q
2.0e+07
1.5e+07
1.0e+07
5.0e+06
0.0e+00
WD.1.SE
WW.1.SE
WD.2.SE
WW.2.SE
WD.3.SE
WW.3.SE
WD.4.SE
WW.4.SE
WD.5.SE
WW.5.SE
WD.1.SE+5d
WW.1.SE+5d
WD.2.SE+5d
WW.2.SE+5d
WD.3.SE+5d
WW.3.SE+5d
WD.4.SE+5d
WW.4.SE+5d
WD.5.SE+5d
WW.5.SE+5d

zm00001d015088_p001

**Amino acid metabolism.pyruvate group amino acid biosynthesis description:3−isopropylmalate dehydratase large subunit**

large subunit of isopropylmalate isomerase heterodimer

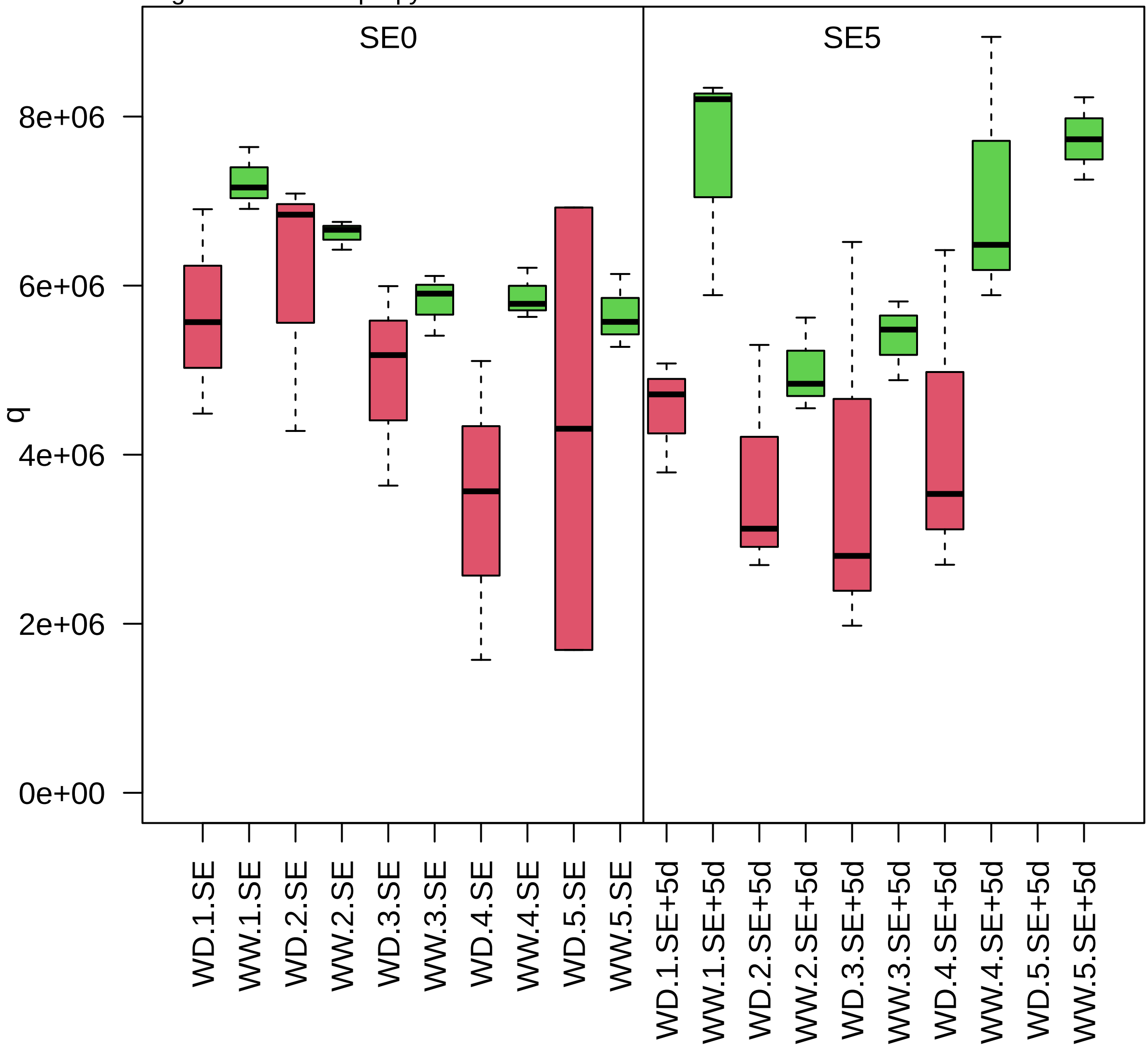

zm00001d015120_p001

**Chromatin organisation.post−translational histone modification**
**description:RuvB−like protein 1**

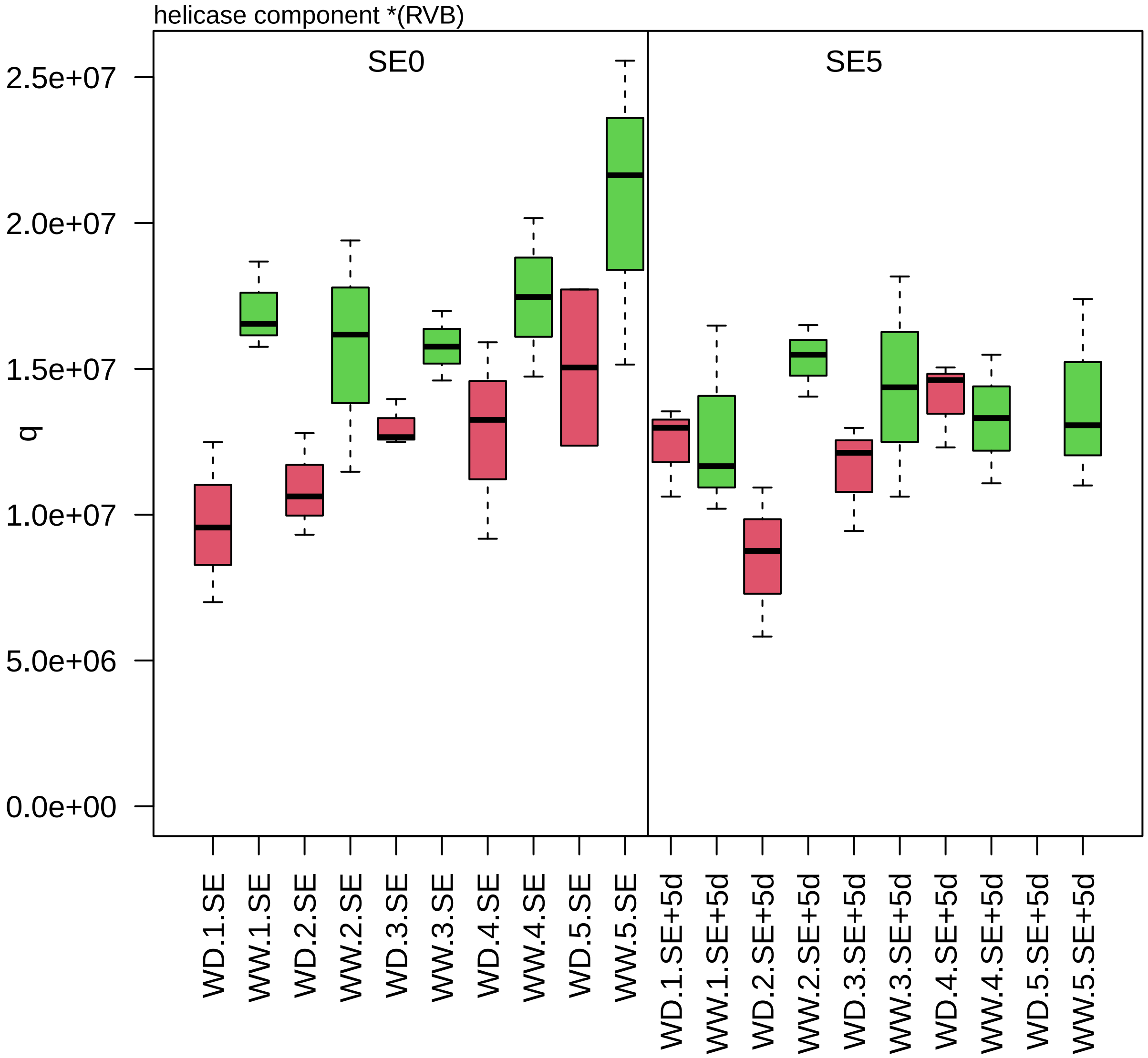

zm00001d015180_p001

**Protein homeostasis.ubiquitin−proteasome system**
**description:Proteasome subunit alpha type−1−B**

alpha−type−6 component *(PAF) of 26S proteasome

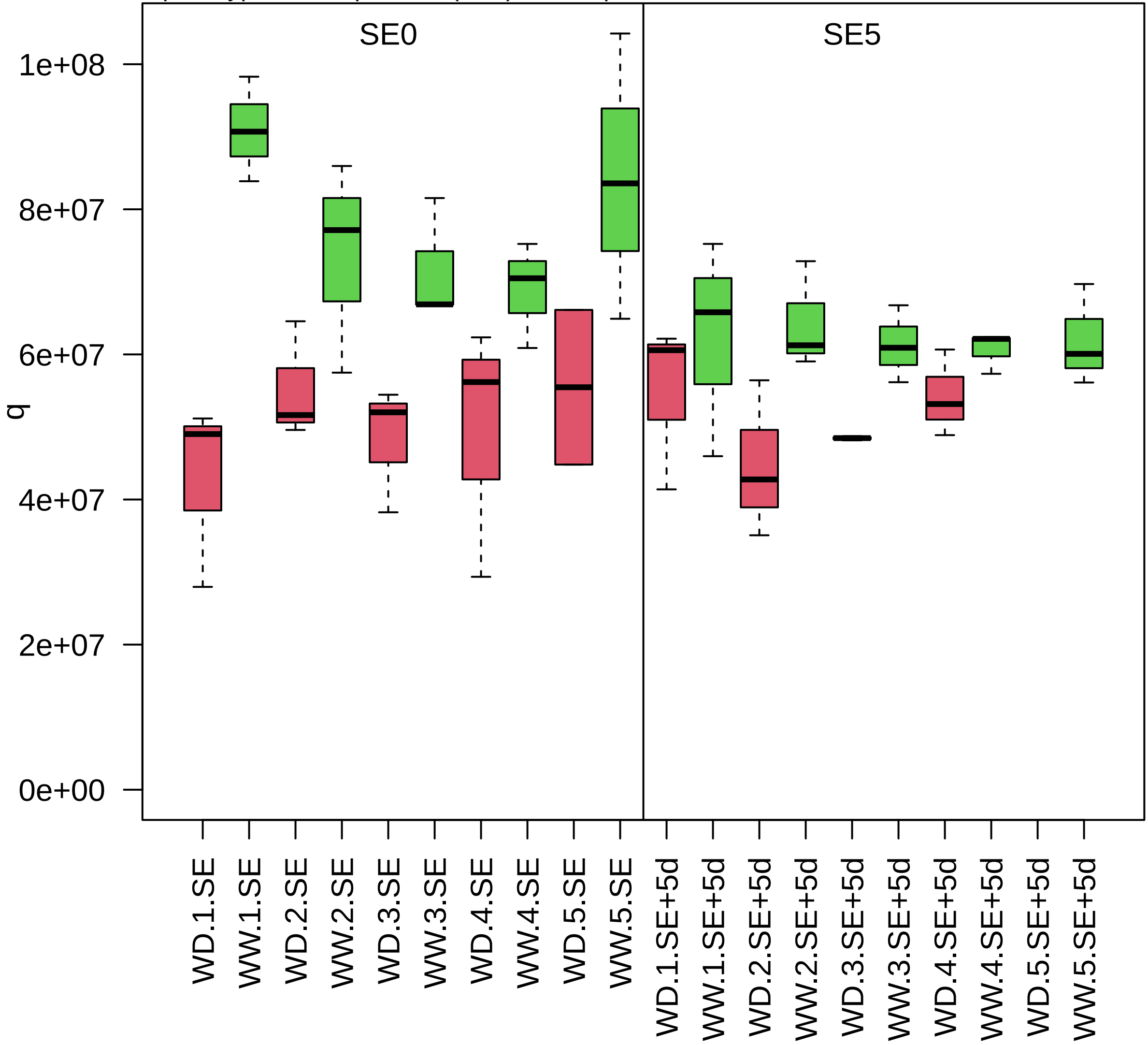

zm00001d015208_p001

**RNA processing.pre−RNA splicing**

**description:Cleavage and polyadenylation specificity factor (CPSF) A subunit protein**

component *(SF3B3) of splicing factor 3B complex

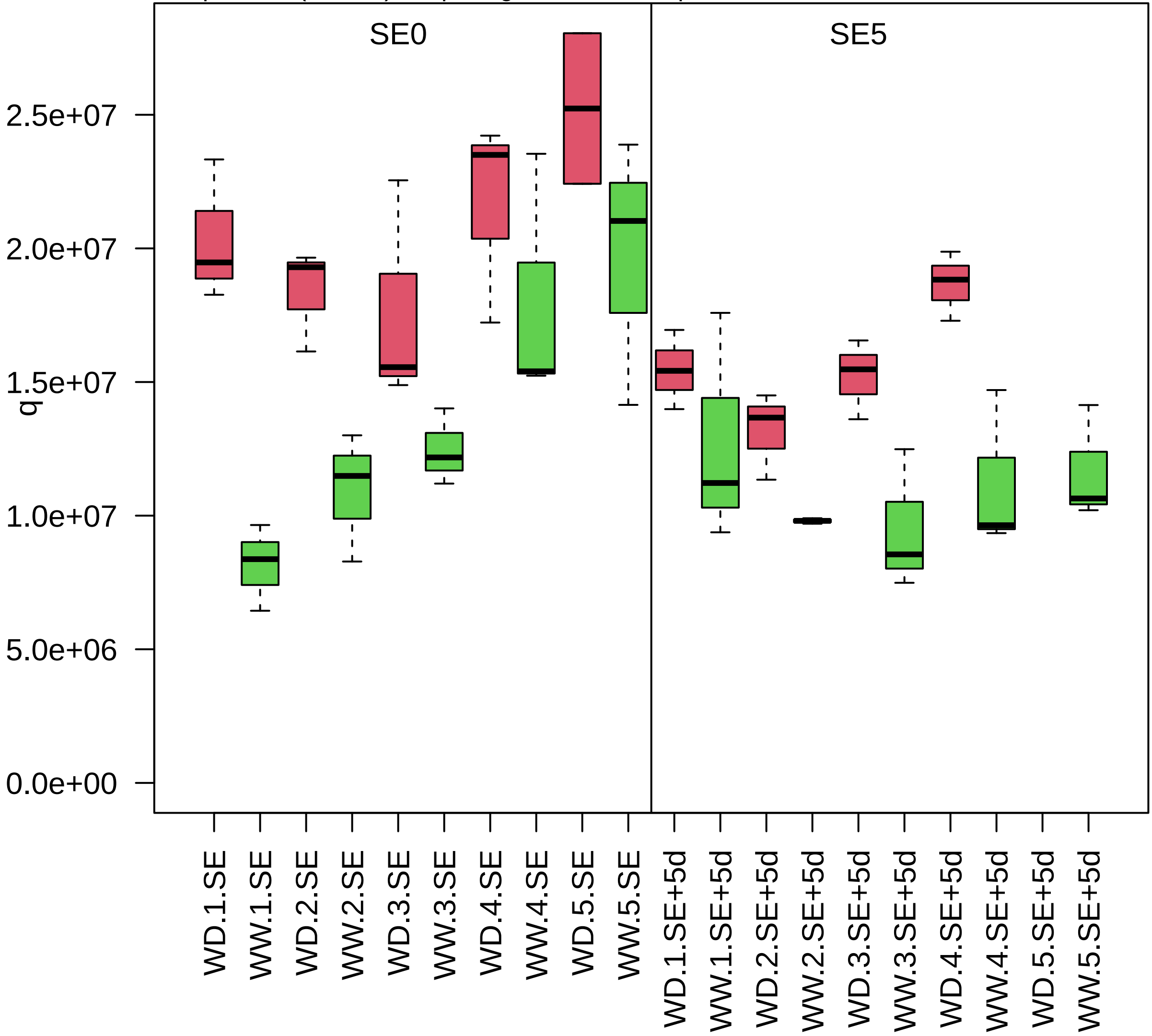

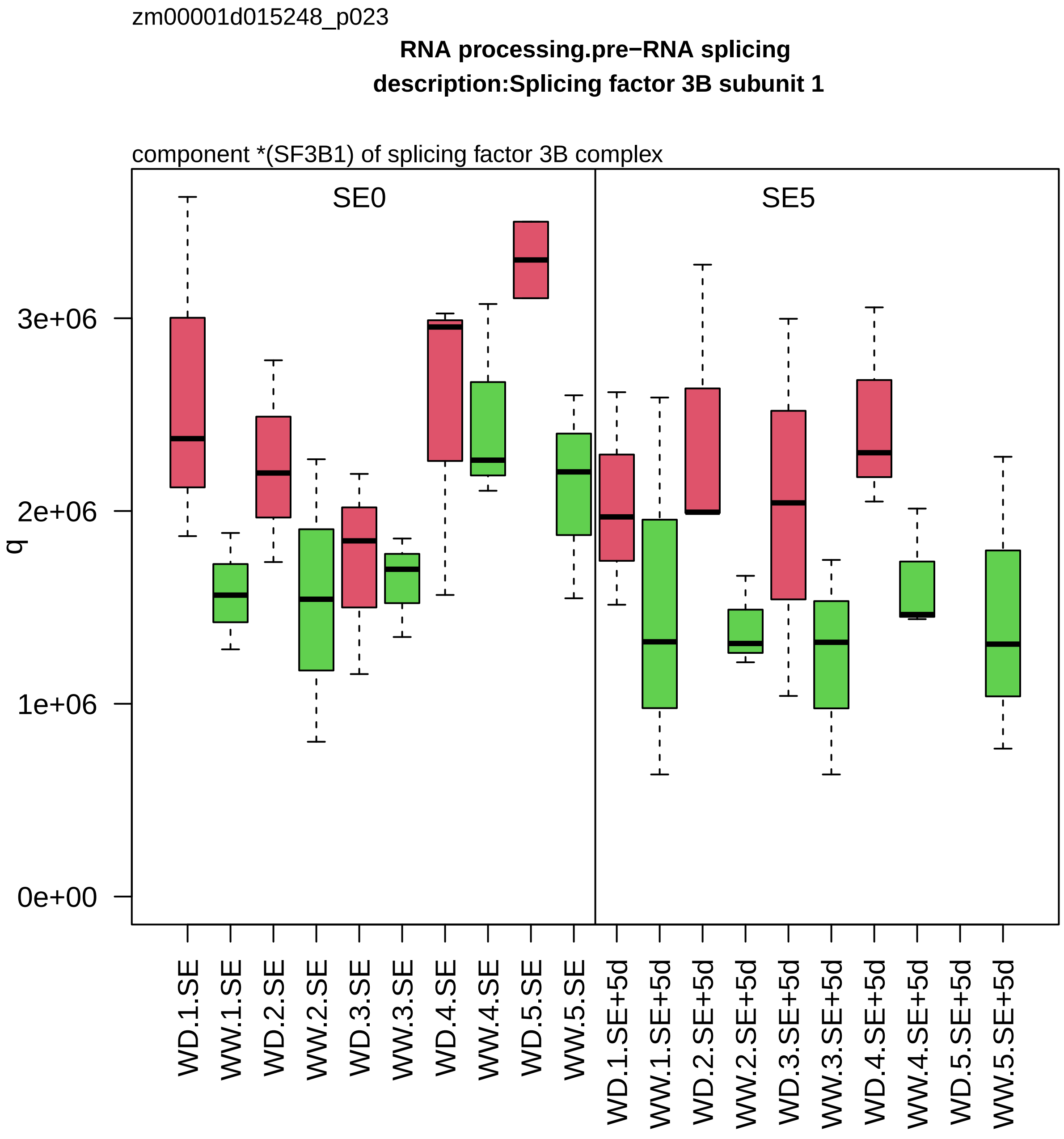

zm00001d015248_p023
RNA processing.pre−RNA splicing
description:Splicing factor 3B subunit 1
component *(SF3B1) of splicing factor 3B complex
SE0
SE5
q
0e+00
1e+06
2e+06
3e+06
WD.1.SE
WW.1.SE
WD.2.SE
WW.2.SE
WD.3.SE
WW.3.SE
WD.4.SE
WW.4.SE
WD.5.SE
WW.5.SE
WD.1.SE+5d
WW.1.SE+5d
WD.2.SE+5d
WW.2.SE+5d
WD.3.SE+5d
WW.3.SE+5d
WD.4.SE+5d
WW.4.SE+5d
WD.5.SE+5d
WW.5.SE+5d

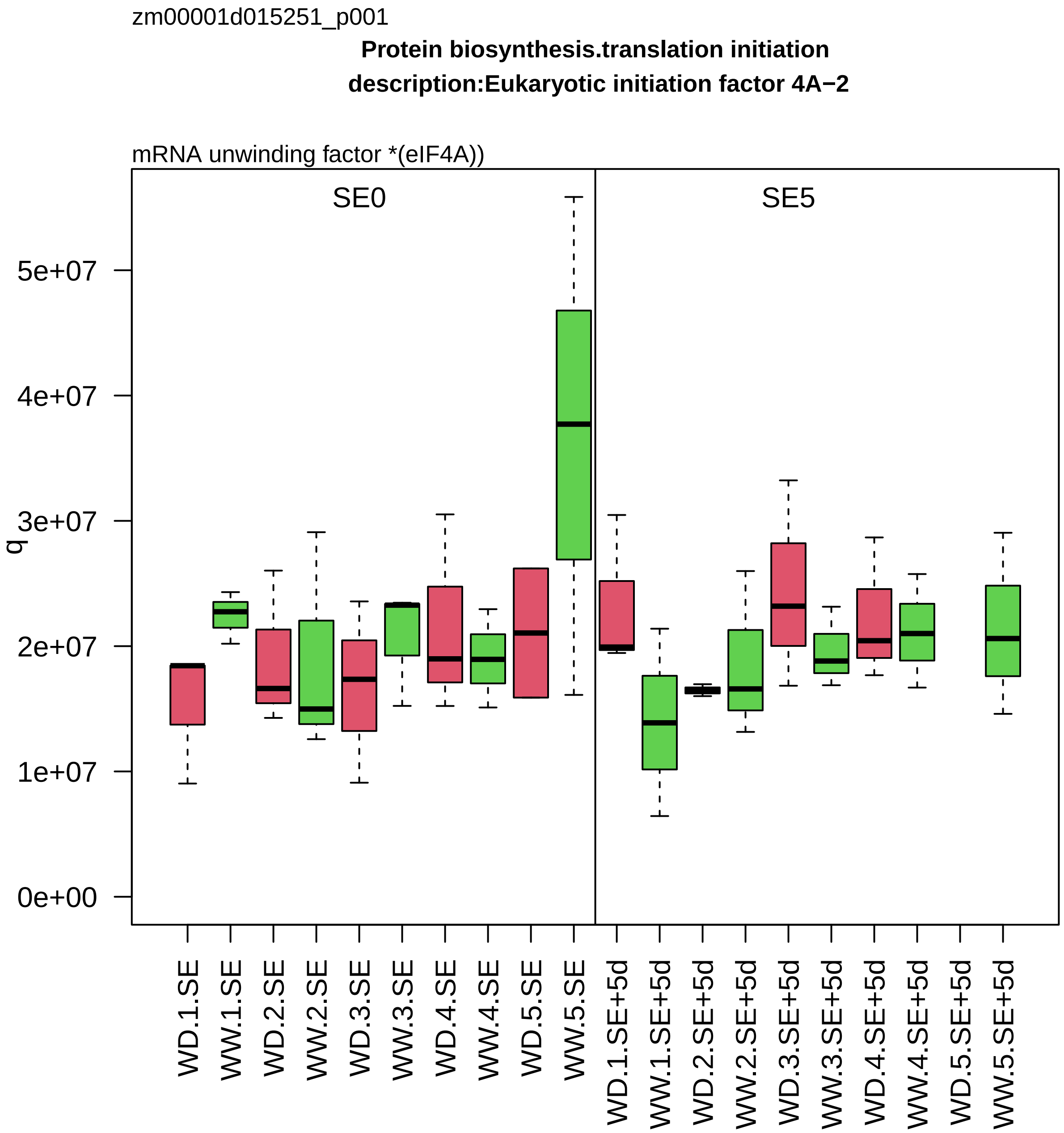

zm00001d015251_p001
Protein biosynthesis.translation initiation
description:Eukaryotic initiation factor 4A−2
mRNA unwinding factor *(eIF4A))
SE0
SE5
q
0e+00
1e+07
2e+07
3e+07
4e+07
5e+07
WD.1.SE
WW.1.SE
WD.2.SE
WW.2.SE
WD.3.SE
WW.3.SE
WD.4.SE
WW.4.SE
WD.5.SE
WW.5.SE
WD.1.SE+5d
WW.1.SE+5d
WD.2.SE+5d
WW.2.SE+5d
WD.3.SE+5d
WW.3.SE+5d
WD.4.SE+5d
WW.4.SE+5d
WD.5.SE+5d
WW.5.SE+5d

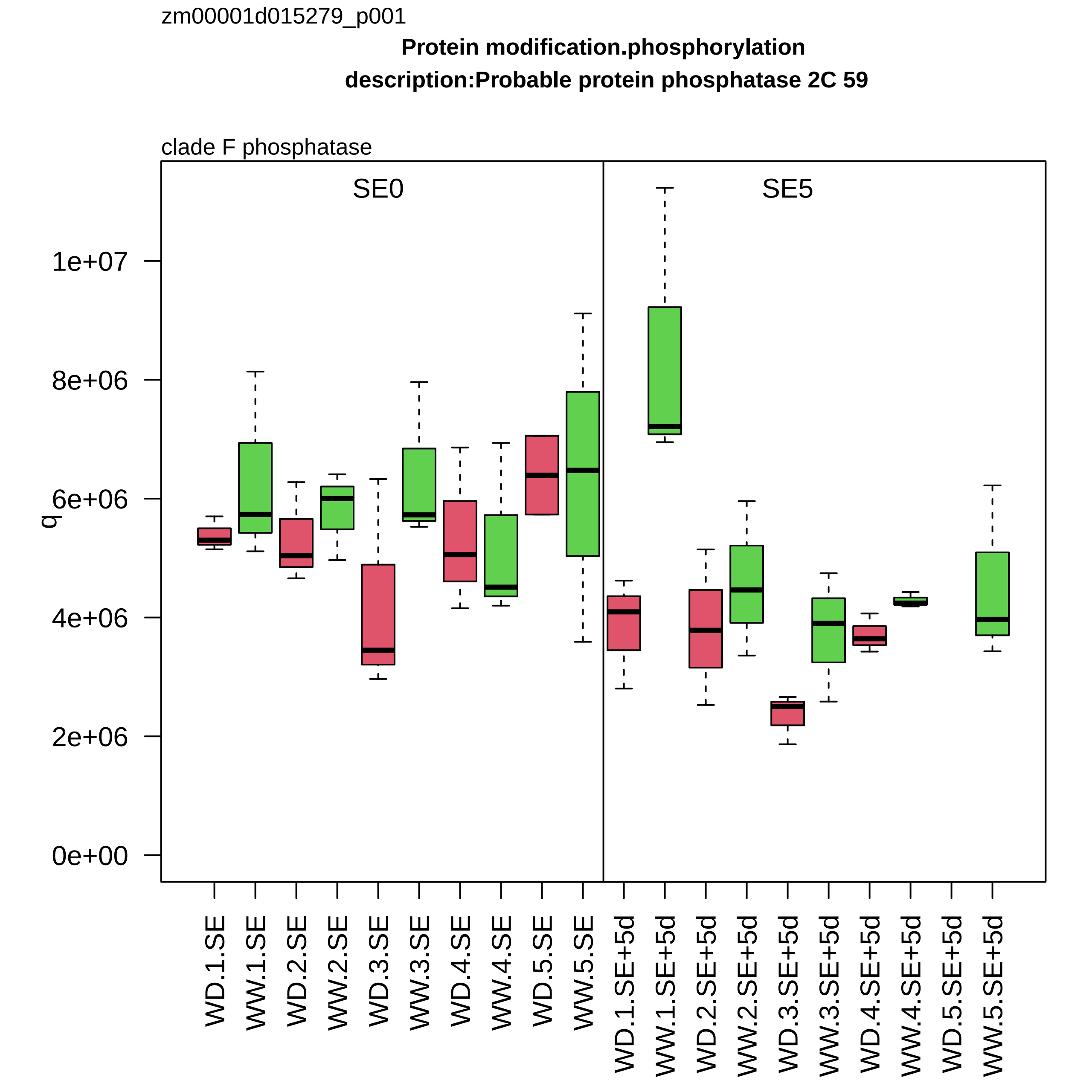

zm00001d015279_p001
Protein modification.phosphorylation
description:Probable protein phosphatase 2C 59
clade F phosphatase
SE0
SE5
q
0e+00
2e+06
4e+06
6e+06
8e+06
1e+07
WD.1.SE
WW.1.SE
WD.2.SE
WW.2.SE
WD.3.SE
WW.3.SE
WD.4.SE
WW.4.SE
WD.5.SE
WW.5.SE
WD.1.SE+5d
WW.1.SE+5d
WD.2.SE+5d
WW.2.SE+5d
WD.3.SE+5d
WW.3.SE+5d
WD.4.SE+5d
WW.4.SE+5d
WD.5.SE+5d
WW.5.SE+5d

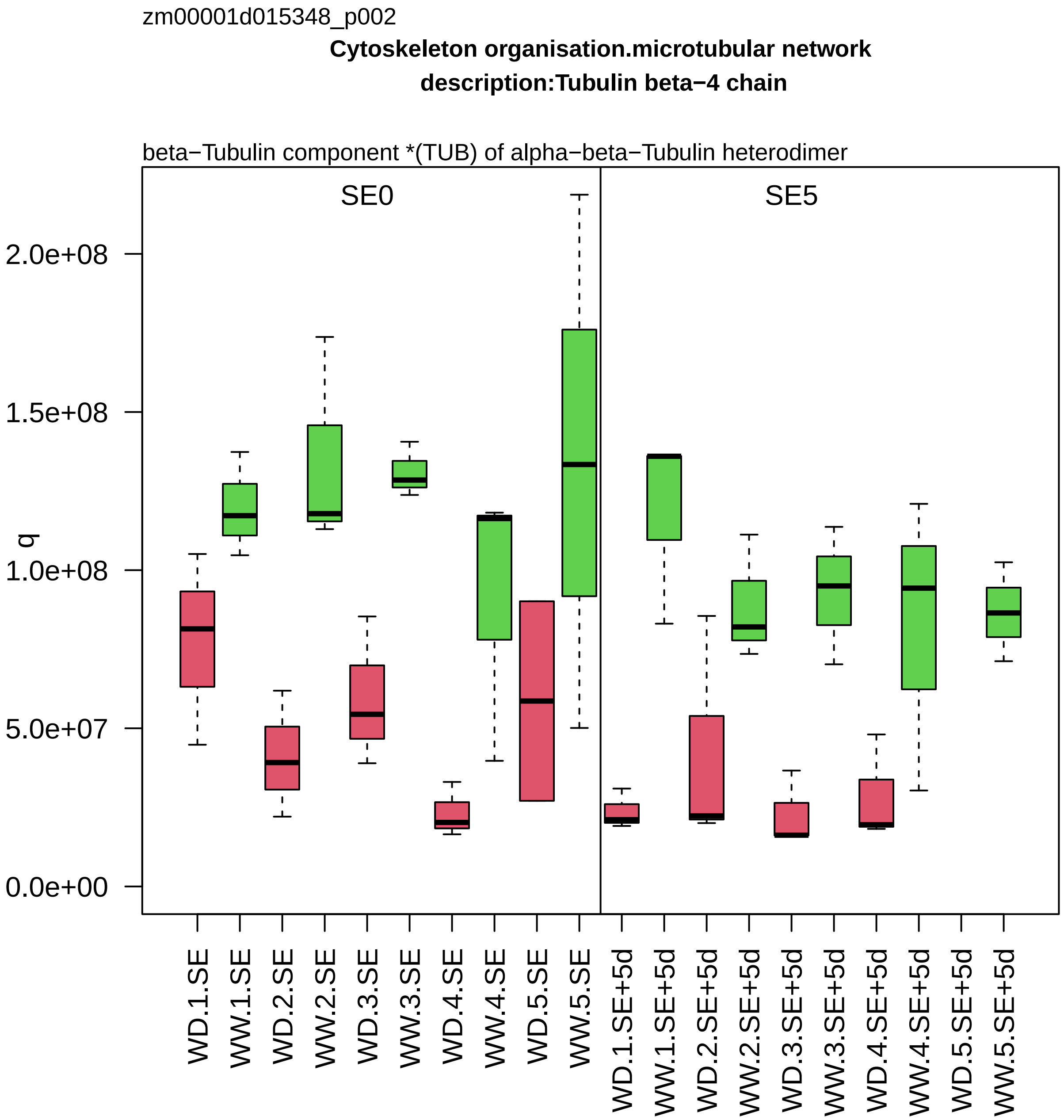
zm00001d015348_p002
Cytoskeleton organisation.microtubular network
description:Tubulin beta−4 chain
beta−Tubulin component *(TUB) of alpha−beta−Tubulin heterodimer
SE0
SE5
q
2.0e+08
1.5e+08
1.0e+08
5.0e+07
0.0e+00
WD.1.SE
WW.1.SE
WD.2.SE
WW.2.SE
WD.3.SE
WW.3.SE
WD.4.SE
WW.4.SE
WD.5.SE
WW.5.SE
WD.1.SE+5d
WW.1.SE+5d
WD.2.SE+5d
WW.2.SE+5d
WD.3.SE+5d
WW.3.SE+5d
WD.4.SE+5d
WW.4.SE+5d
WD.5.SE+5d
WW.5.SE+5d

zm00001d015376_p001

**Cellular respiration.glycolysis**
**description:Phosphoglycerate kinase**

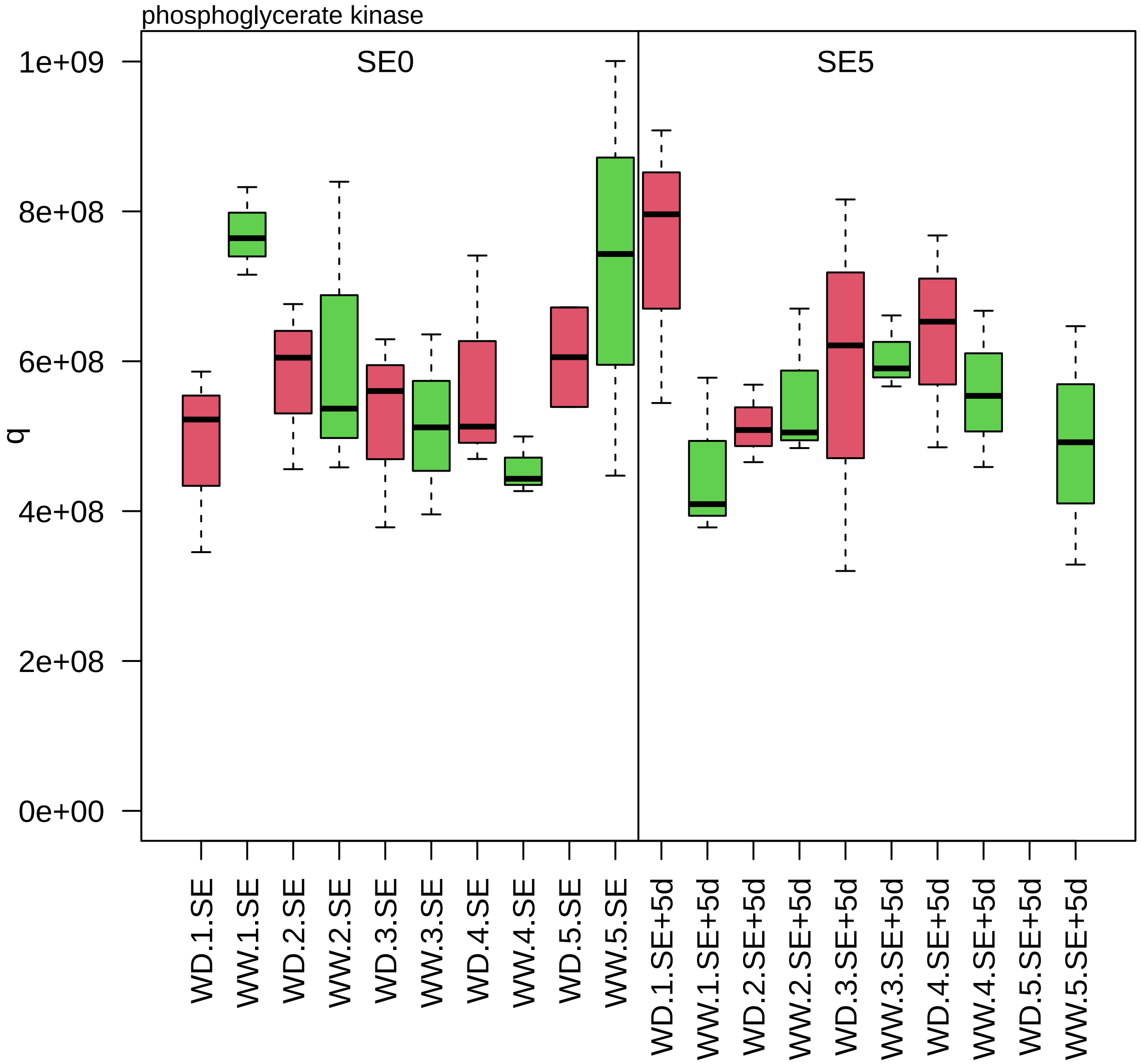

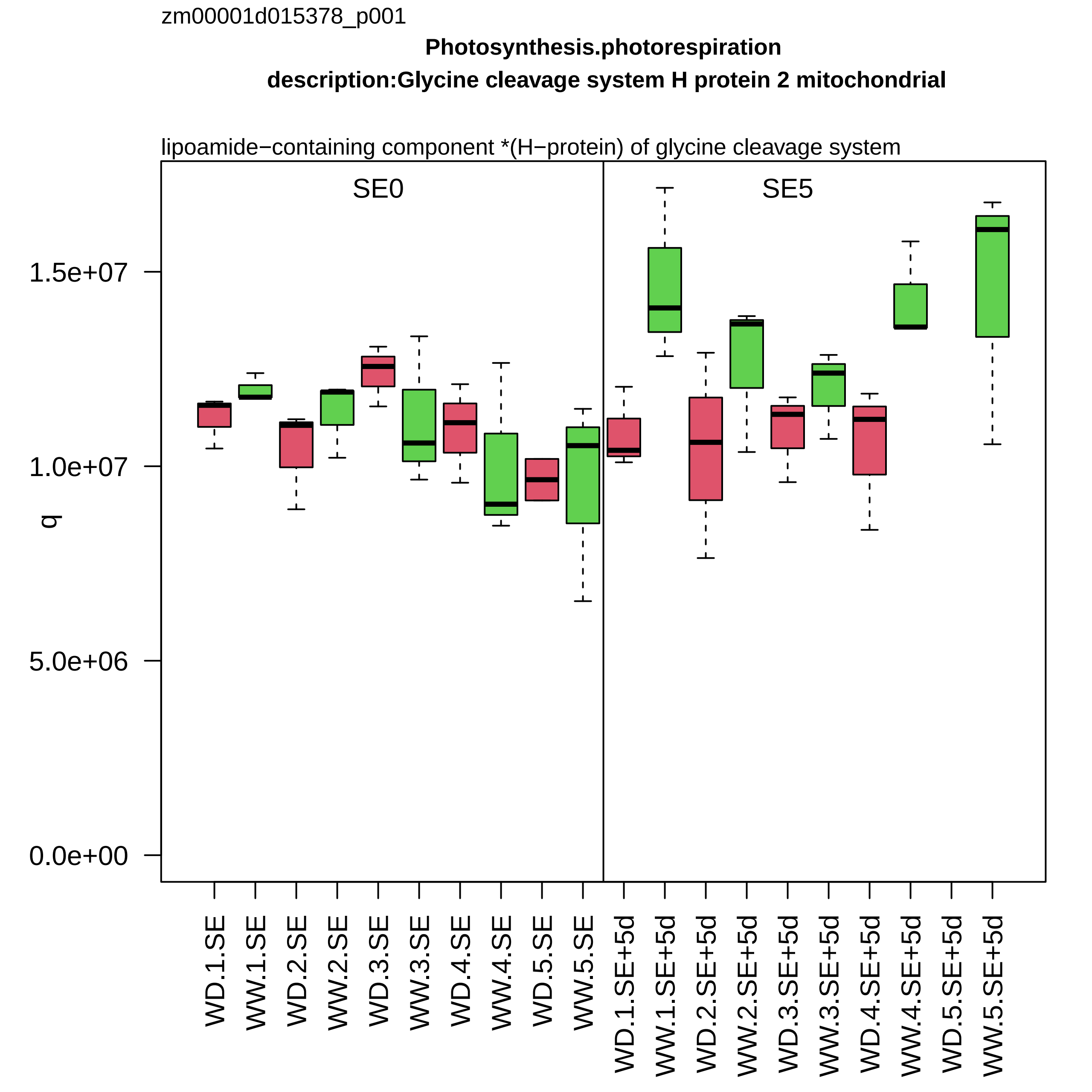

zm00001d015378_p001
Photosynthesis.photorespiration
description:Glycine cleavage system H protein 2 mitochondrial
lipoamide−containing component *(H−protein) of glycine cleavage system
SE0
SE5
q
1.5e+07
1.0e+07
5.0e+06
0.0e+00
WD.1.SE
WW.1.SE
WD.2.SE
WW.2.SE
WD.3.SE
WW.3.SE
WD.4.SE
WW.4.SE
WD.5.SE
WW.5.SE
WD.1.SE+5d
WW.1.SE+5d
WD.2.SE+5d
WW.2.SE+5d
WD.3.SE+5d
WW.3.SE+5d
WD.4.SE+5d
WW.4.SE+5d
WD.5.SE+5d
WW.5.SE+5d

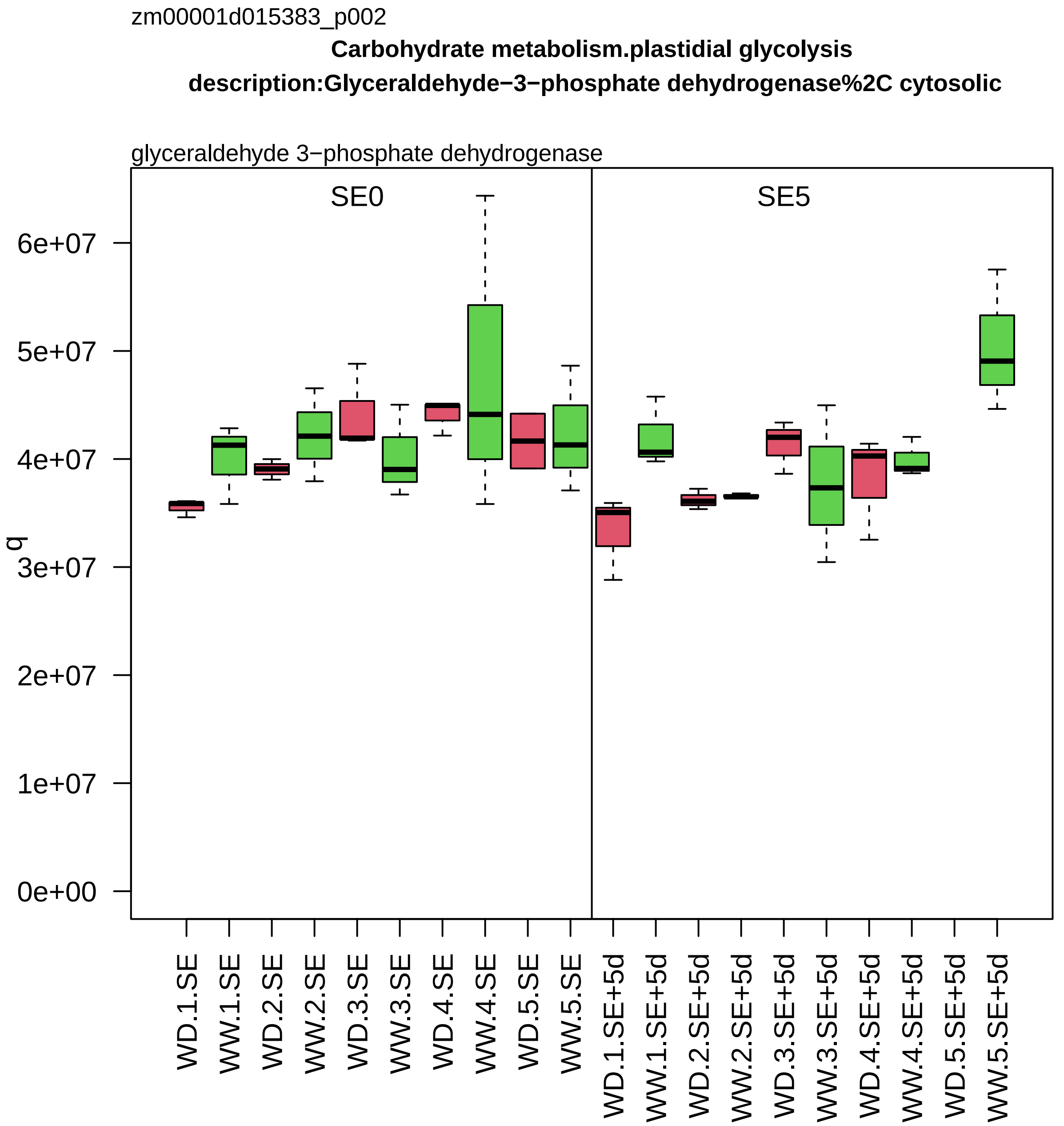
zm00001d015383_p002
Carbohydrate metabolism.plastidial glycolysis
description:Glyceraldehyde−3−phosphate dehydrogenase%2C cytosolic
glyceraldehyde 3−phosphate dehydrogenase
SE0
SE5
q
0e+00
1e+07
2e+07
3e+07
4e+07
5e+07
6e+07
WD.1.SE
WW.1.SE
WD.2.SE
WW.2.SE
WD.3.SE
WW.3.SE
WD.4.SE
WW.4.SE
WD.5.SE
WW.5.SE
WD.1.SE+5d
WW.1.SE+5d
WD.2.SE+5d
WW.2.SE+5d
WD.3.SE+5d
WW.3.SE+5d
WD.4.SE+5d
WW.4.SE+5d
WD.5.SE+5d
WW.5.SE+5d

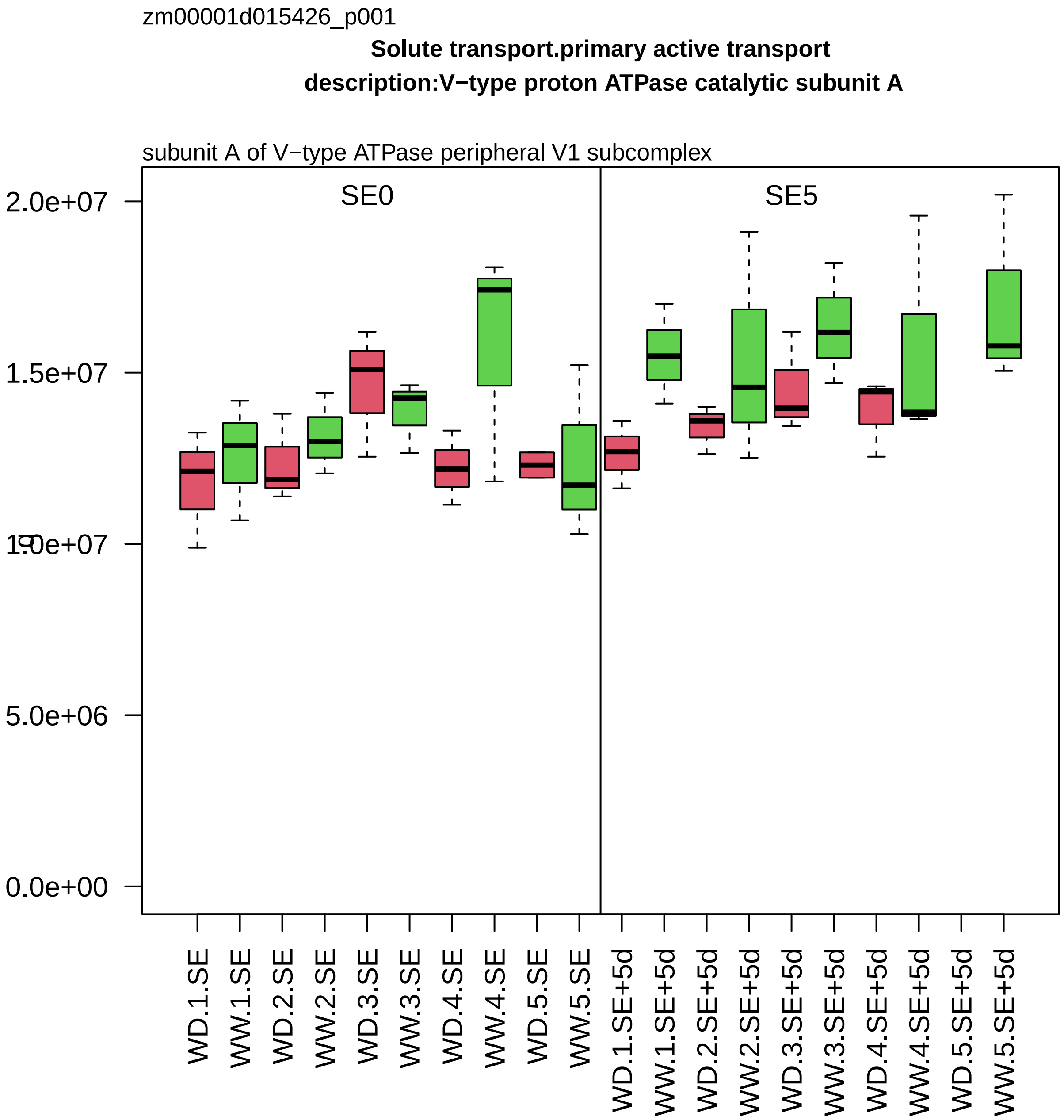

zm00001d015426_p001
Solute transport.primary active transport
description:V−type proton ATPase catalytic subunit A
subunit A of V−type ATPase peripheral V1 subcomplex
SE0
SE5
2.0e+07
1.5e+07
1.0e+07
5.0e+06
0.0e+00
WD.1.SE
WW.1.SE
WD.2.SE
WW.2.SE
WD.3.SE
WW.3.SE
WD.4.SE
WW.4.SE
WD.5.SE
WW.5.SE
WD.1.SE+5d
WW.1.SE+5d
WD.2.SE+5d
WW.2.SE+5d
WD.3.SE+5d
WW.3.SE+5d
WD.4.SE+5d
WW.4.SE+5d
WD.5.SE+5d
WW.5.SE+5d

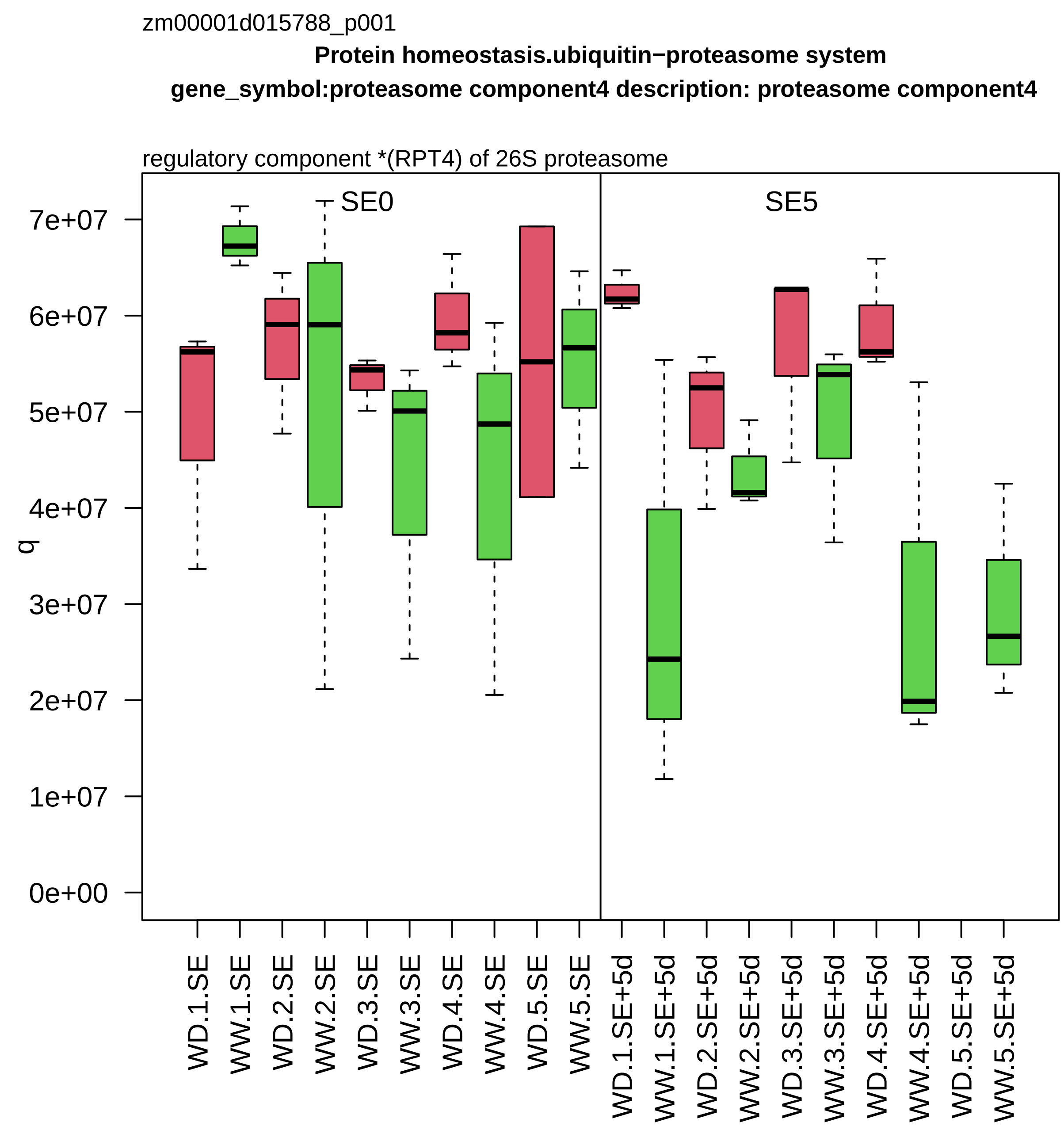

zm00001d015788_p001
Protein homeostasis.ubiquitin−proteasome system
gene_symbol:proteasome component4 description: proteasome component4
regulatory component *(RPT4) of 26S proteasome
SE0
SE5
q
7e+07
6e+07
5e+07
4e+07
3e+07
2e+07
1e+07
0e+00
WD.1.SE
WW.1.SE
WD.2.SE
WW.2.SE
WD.3.SE
WW.3.SE
WD.4.SE
WW.4.SE
WD.5.SE
WW.5.SE
WD.1.SE+5d
WW.1.SE+5d
WD.2.SE+5d
WW.2.SE+5d
WD.3.SE+5d
WW.3.SE+5d
WD.4.SE+5d
WW.4.SE+5d
WD.5.SE+5d
WW.5.SE+5d

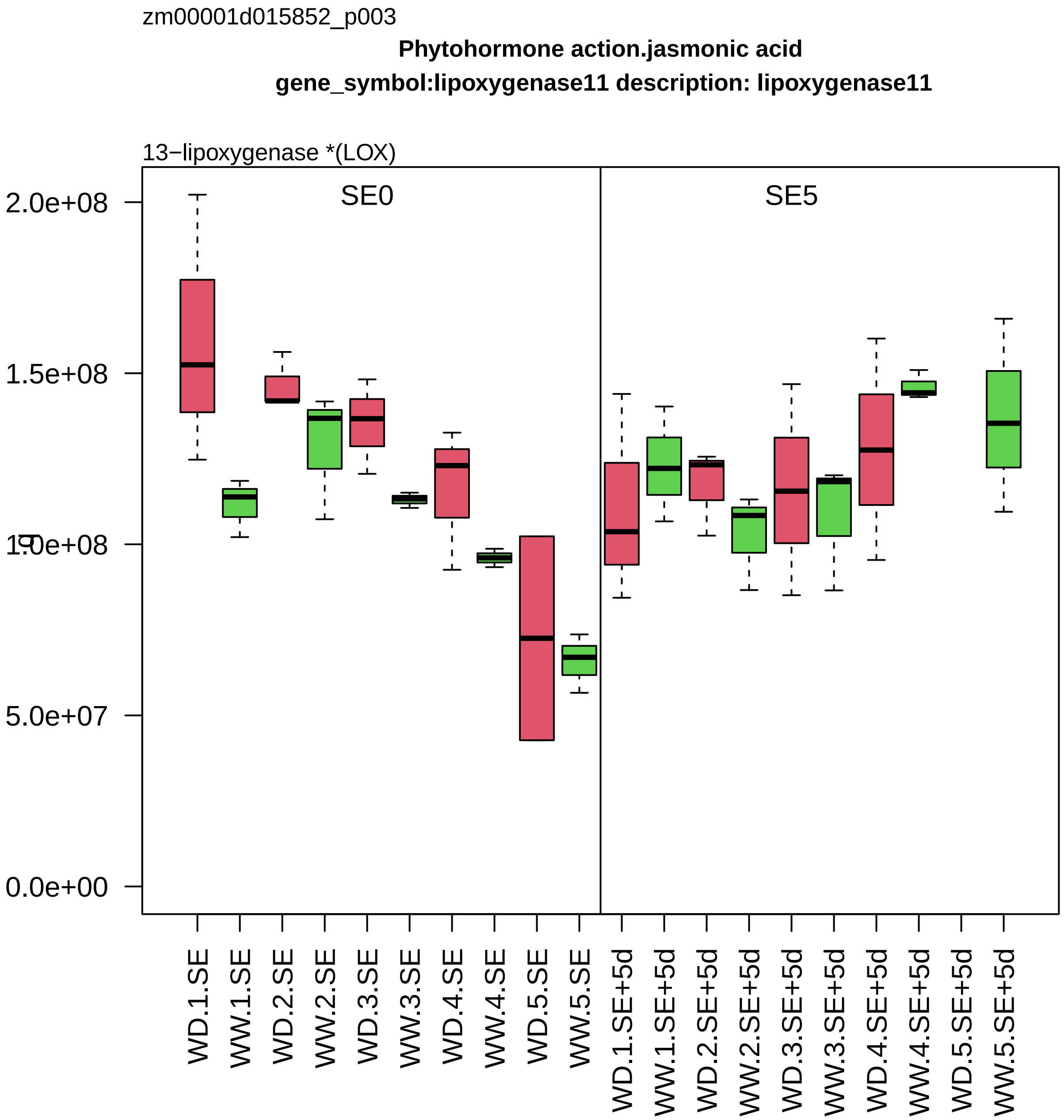

zm00001d015852_p003
Phytohormone action.jasmonic acid
gene_symbol:lipoxygenase11 description: lipoxygenase11
13−lipoxygenase *(LOX)
SE0
SE5
2.0e+08
1.5e+08
1.0e+08
5.0e+07
0.0e+00
WD.1.SE
WW.1.SE
WD.2.SE
WW.2.SE
WD.3.SE
WW.3.SE
WD.4.SE
WW.4.SE
WD.5.SE
WW.5.SE
WD.1.SE+5d
WW.1.SE+5d
WD.2.SE+5d
WW.2.SE+5d
WD.3.SE+5d
WW.3.SE+5d
WD.4.SE+5d
WW.4.SE+5d
WD.5.SE+5d
WW.5.SE+5d

zm00001d015886_p001

**Protein homeostasis.ubiquitin−proteasome system**
**description:26S protease regulatory subunit 6B homolog**

regulatory component *(RPT3) of 26S proteasome

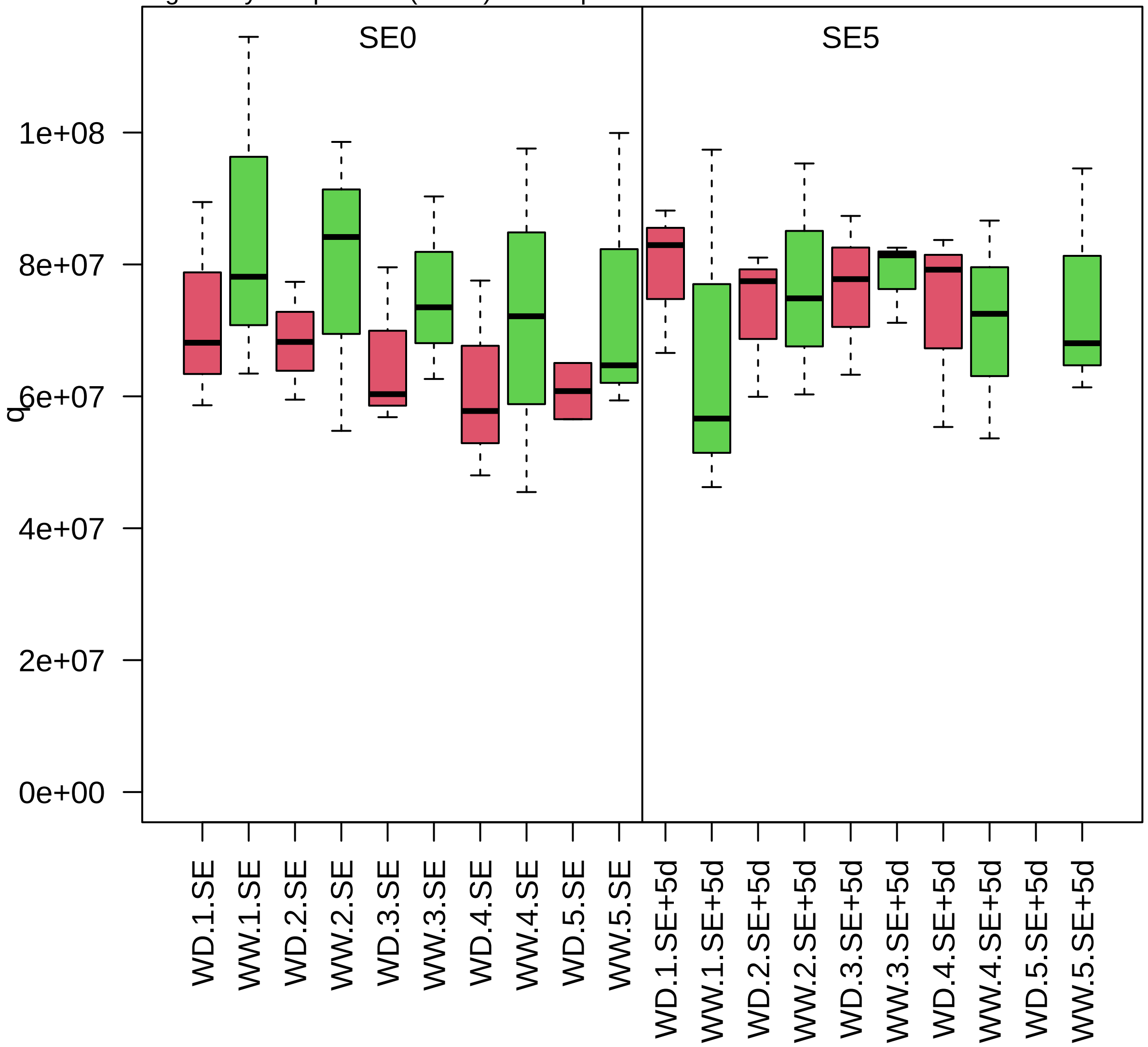

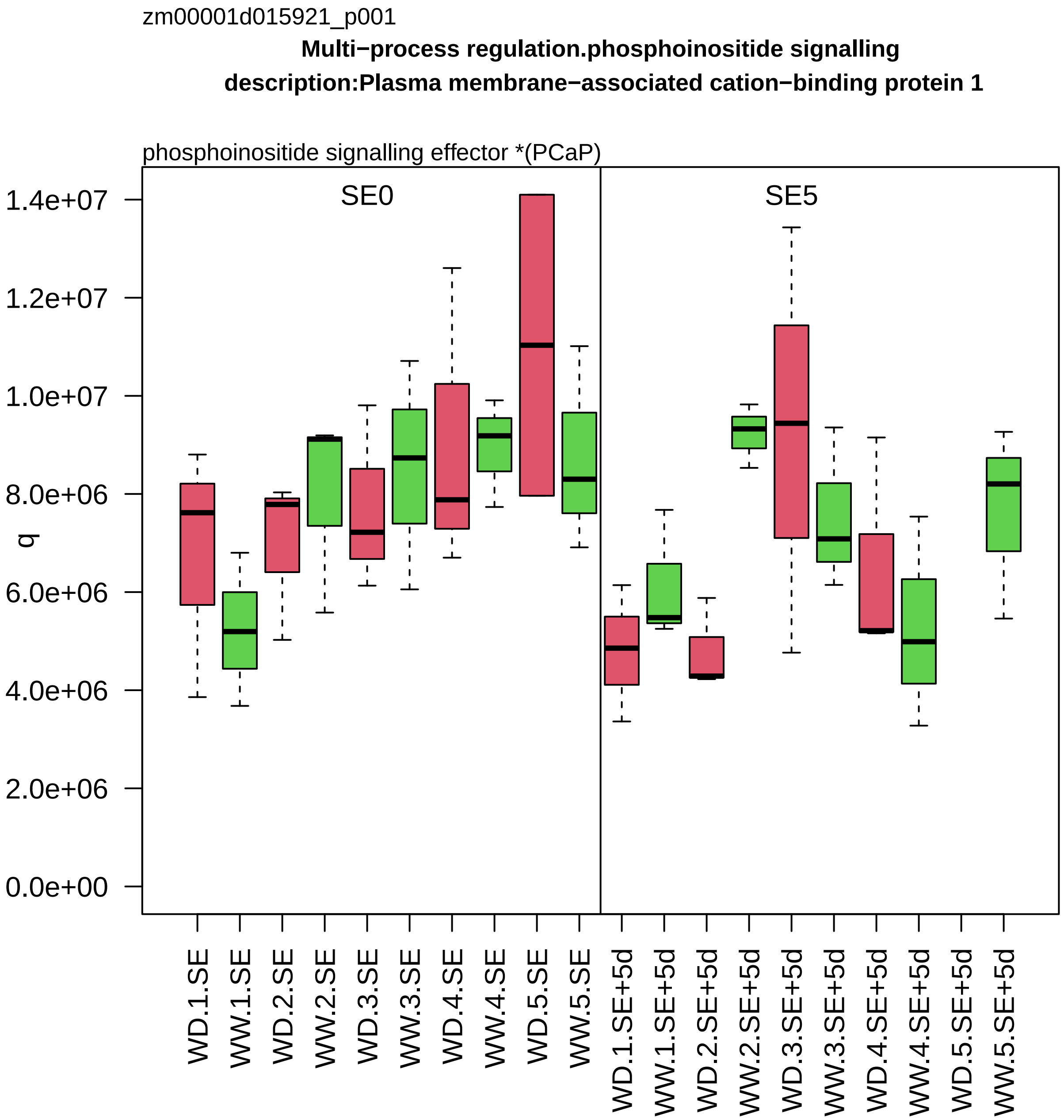

zm00001d015921_p001
Multi−process regulation.phosphoinositide signalling
description:Plasma membrane−associated cation−binding protein 1
phosphoinositide signalling effector *(PCaP)
SE0
SE5
q
1.4e+07
1.2e+07
1.0e+07
8.0e+06
6.0e+06
4.0e+06
2.0e+06
0.0e+00
WD.1.SE
WW.1.SE
WD.2.SE
WW.2.SE
WD.3.SE
WW.3.SE
WD.4.SE
WW.4.SE
WD.5.SE
WW.5.SE
WD.1.SE+5d
WW.1.SE+5d
WD.2.SE+5d
WW.2.SE+5d
WD.3.SE+5d
WW.3.SE+5d
WD.4.SE+5d
WW.4.SE+5d
WD.5.SE+5d
WW.5.SE+5d

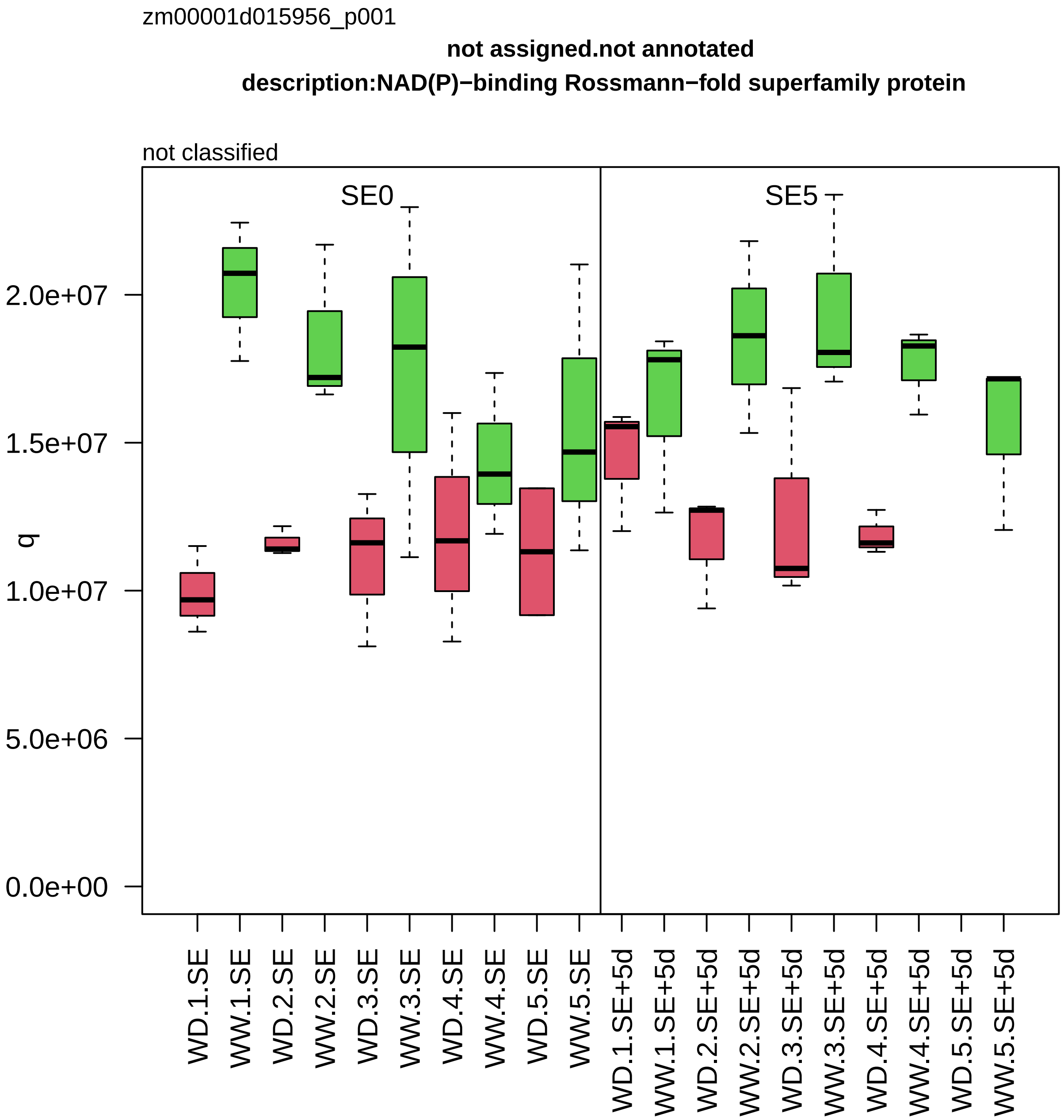

zm00001d015956_p001
not assigned.not annotated
description:NAD(P)−binding Rossmann−fold superfamily protein
not classified
SE0
SE5
q
0.0e+00
5.0e+06
1.0e+07
1.5e+07
2.0e+07
WD.1.SE
WW.1.SE
WD.2.SE
WW.2.SE
WD.3.SE
WW.3.SE
WD.4.SE
WW.4.SE
WD.5.SE
WW.5.SE
WD.1.SE+5d
WW.1.SE+5d
WD.2.SE+5d
WW.2.SE+5d
WD.3.SE+5d
WW.3.SE+5d
WD.4.SE+5d
WW.4.SE+5d
WD.5.SE+5d
WW.5.SE+5d

zm00001d016166_p001

**Photosynthesis.CAM/C4 photosynthesis**

**gene_symbol:phosphoenolpyruvate carboxylase2 description: phosphoenolpyruvate carboxylas**

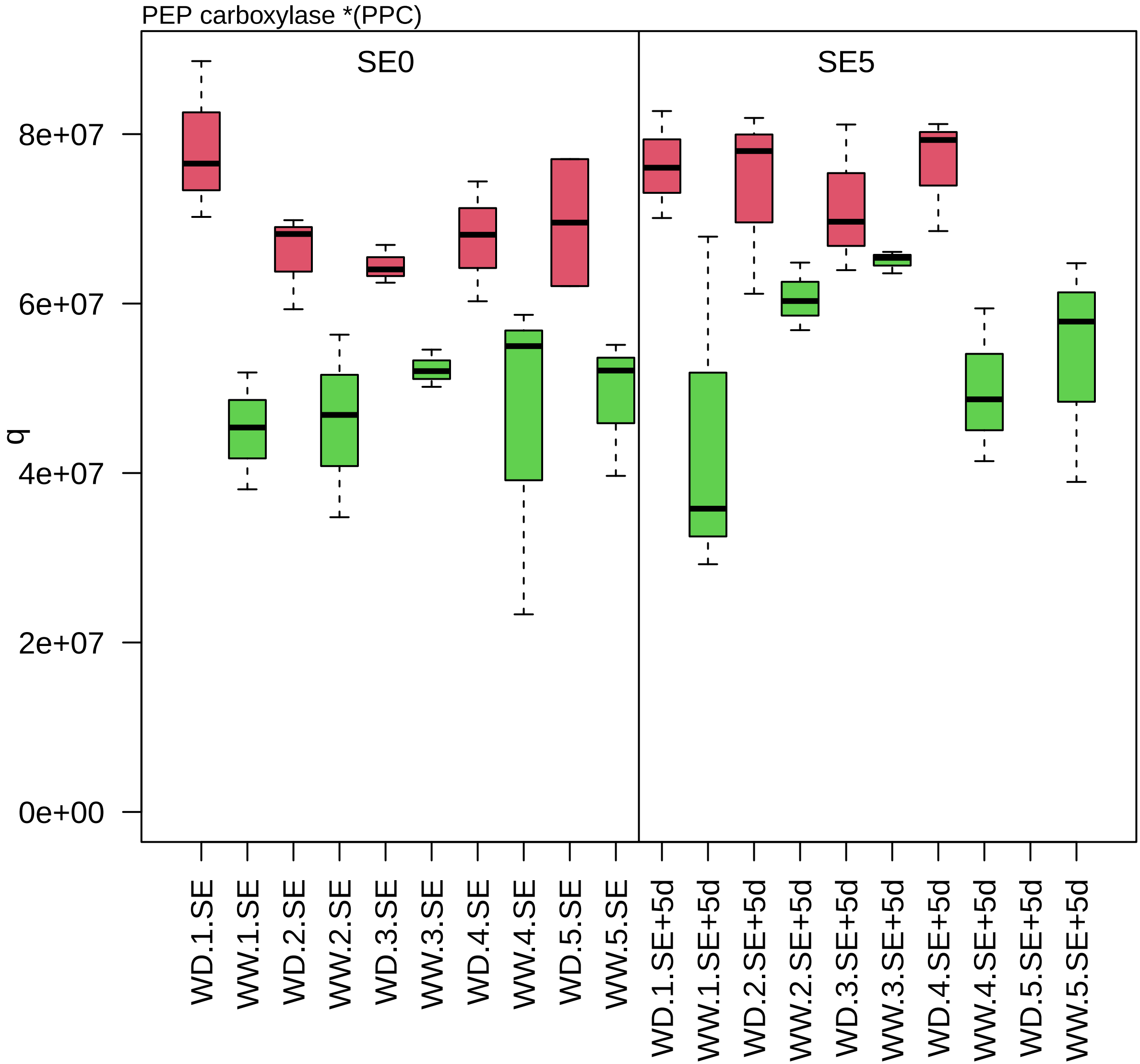

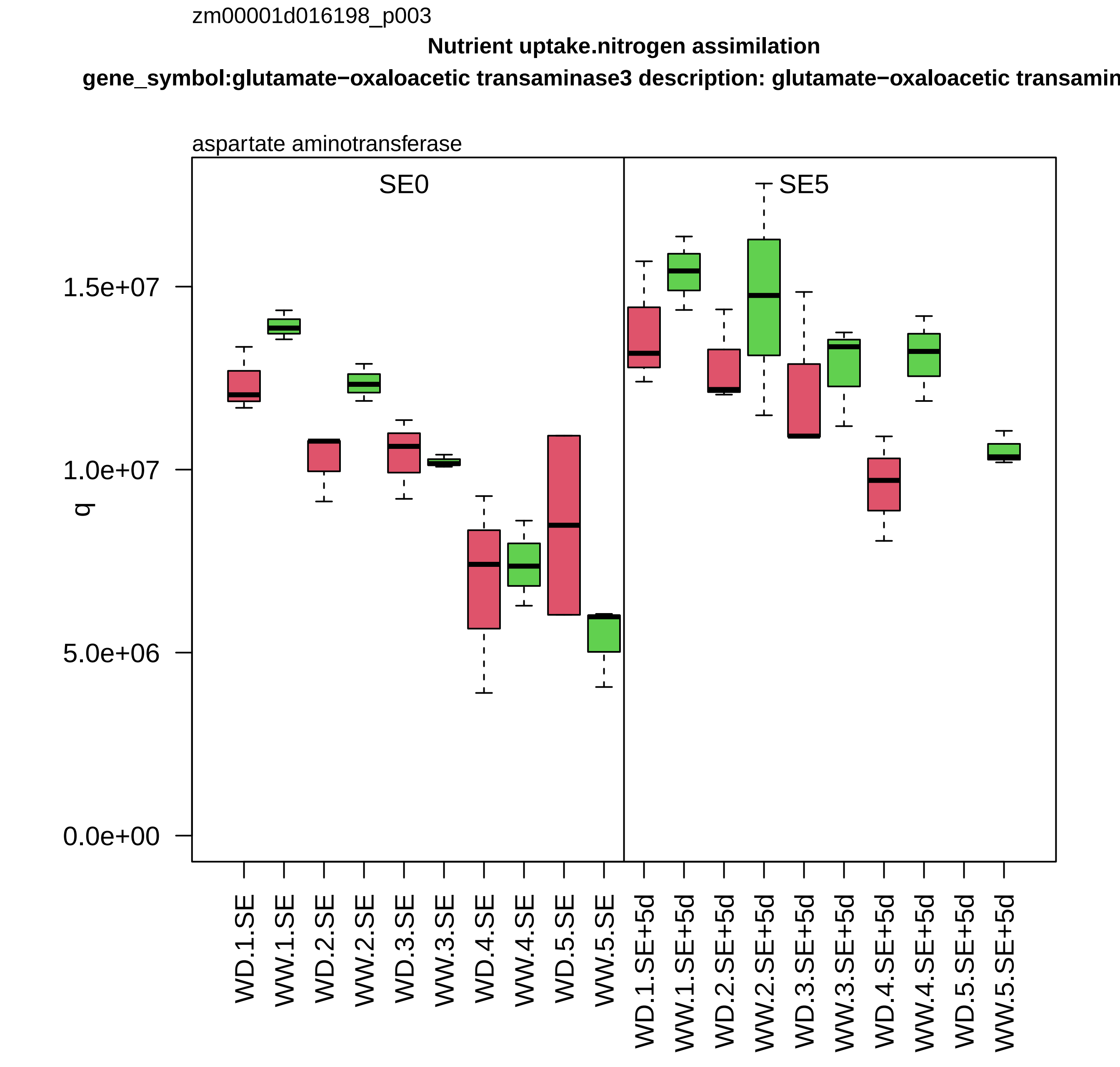
zm00001d016198_p003
Nutrient uptake.nitrogen assimilation
gene_symbol:glutamate−oxaloacetic transaminase3 description: glutamate−oxaloacetic transamin
aspartate aminotransferase
SE0
SE5
q
1.5e+07
1.0e+07
5.0e+06
0.0e+00
WD.1.SE
WW.1.SE
WD.2.SE
WW.2.SE
WD.3.SE
WW.3.SE
WD.4.SE
WW.4.SE
WD.5.SE
WW.5.SE
WD.1.SE+5d
WW.1.SE+5d
WD.2.SE+5d
WW.2.SE+5d
WD.3.SE+5d
WW.3.SE+5d
WD.4.SE+5d
WW.4.SE+5d
WD.5.SE+5d
WW.5.SE+5d

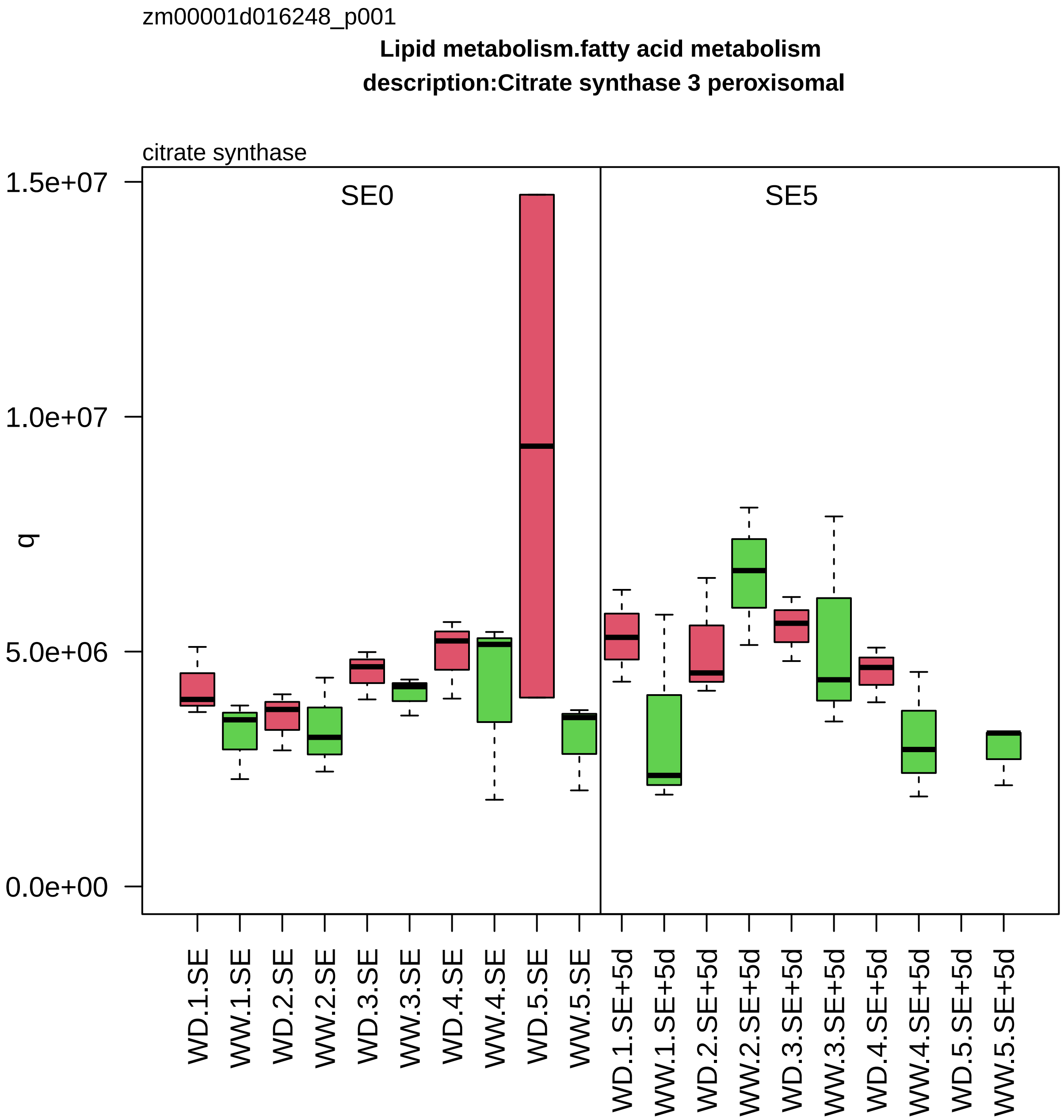

zm00001d016248_p001
Lipid metabolism.fatty acid metabolism
description:Citrate synthase 3 peroxisomal
citrate synthase
SE0
SE5
q
1.5e+07
1.0e+07
5.0e+06
0.0e+00
WD.1.SE
WW.1.SE
WD.2.SE
WW.2.SE
WD.3.SE
WW.3.SE
WD.4.SE
WW.4.SE
WD.5.SE
WW.5.SE
WD.1.SE+5d
WW.1.SE+5d
WD.2.SE+5d
WW.2.SE+5d
WD.3.SE+5d
WW.3.SE+5d
WD.4.SE+5d
WW.4.SE+5d
WD.5.SE+5d
WW.5.SE+5d

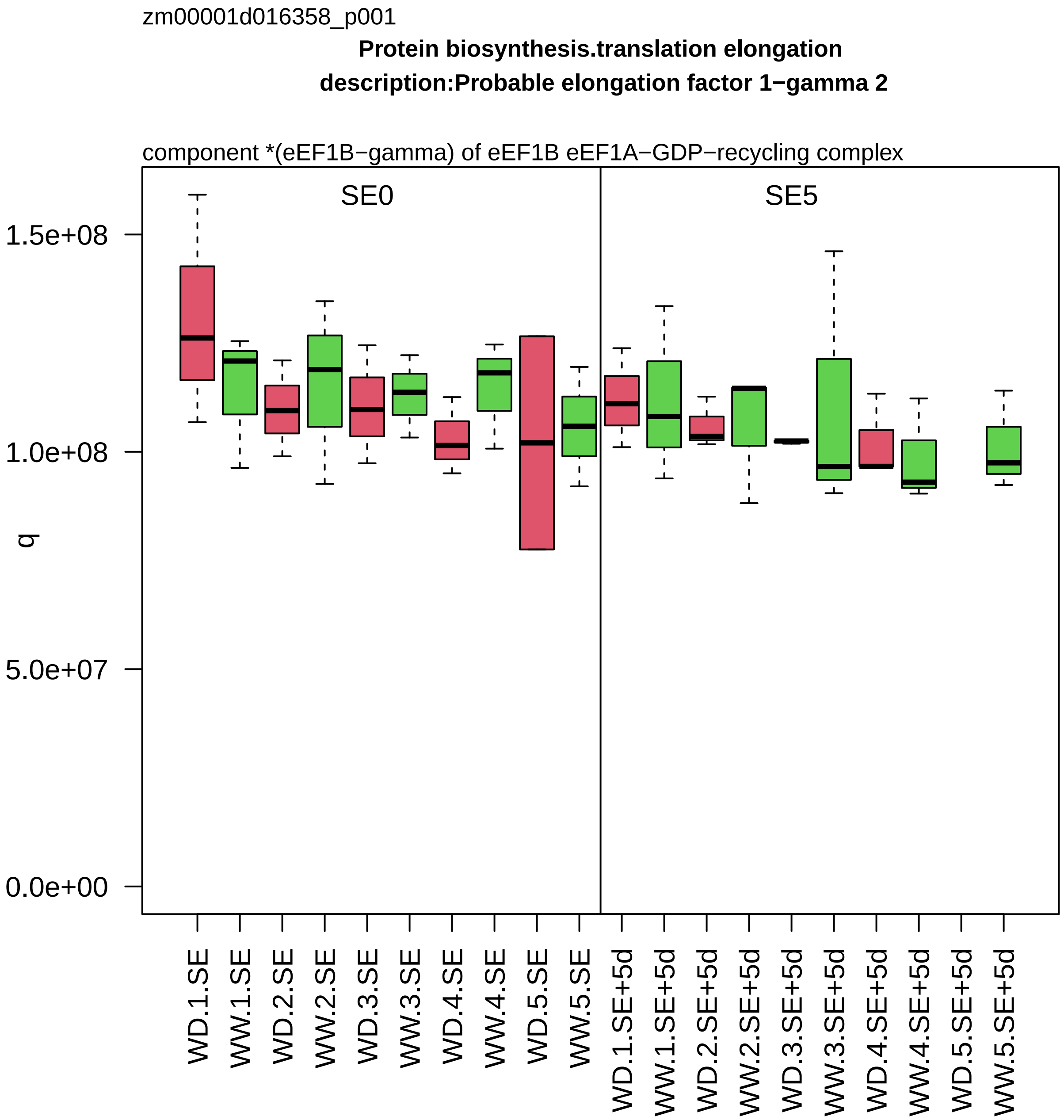
zm00001d016358_p001
Protein biosynthesis.translation elongation
description:Probable elongation factor 1−gamma 2
component *(eEF1B−gamma) of eEF1B eEF1A−GDP−recycling complex
SE0
SE5
q
1.5e+08
1.0e+08
5.0e+07
0.0e+00
WD.1.SE
WW.1.SE
WD.2.SE
WW.2.SE
WD.3.SE
WW.3.SE
WD.4.SE
WW.4.SE
WD.5.SE
WW.5.SE
WD.1.SE+5d
WW.1.SE+5d
WD.2.SE+5d
WW.2.SE+5d
WD.3.SE+5d
WW.3.SE+5d
WD.4.SE+5d
WW.4.SE+5d
WD.5.SE+5d
WW.5.SE+5d

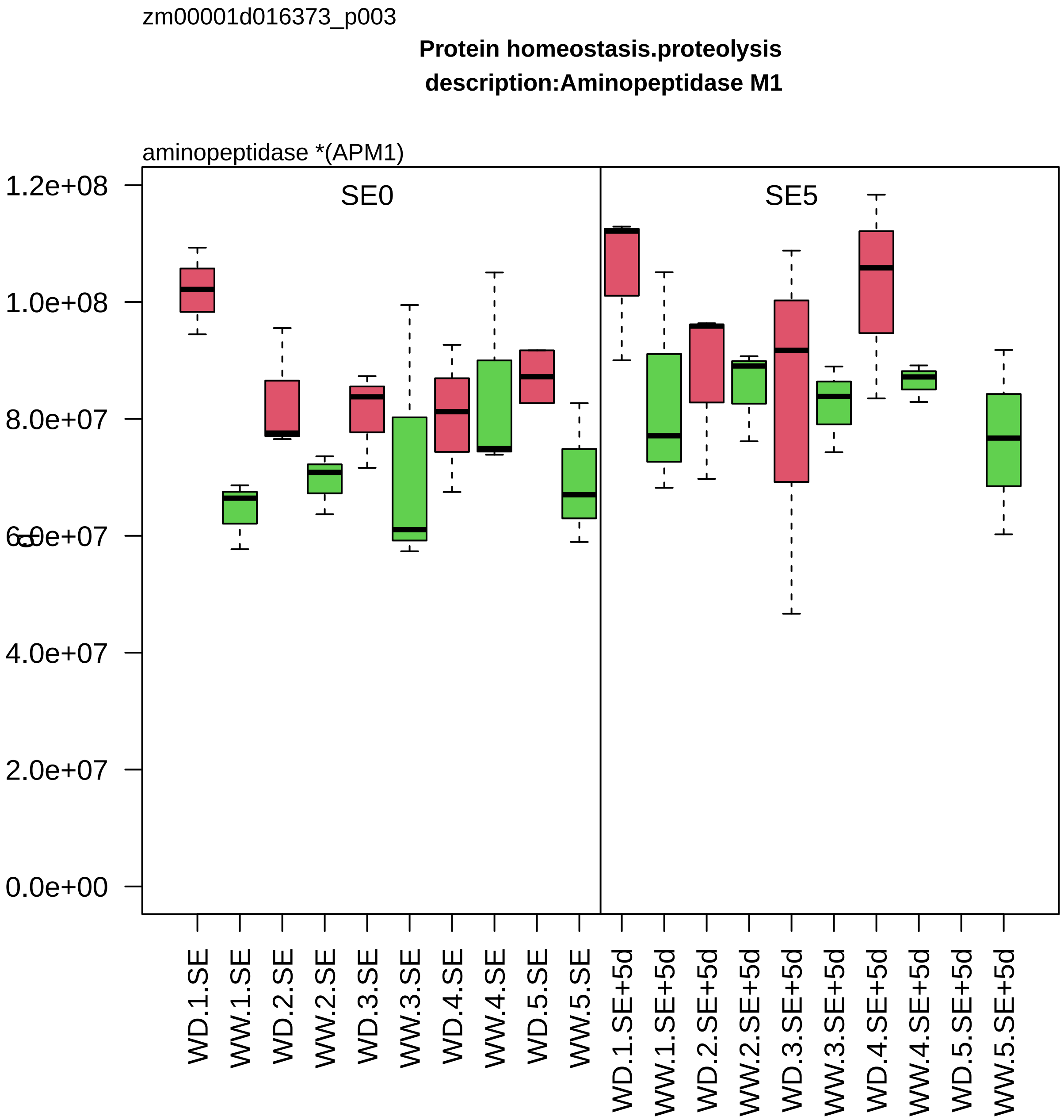
zm00001d016373_p003
Protein homeostasis.proteolysis
description:Aminopeptidase M1
aminopeptidase *(APM1)
SE0
SE5
1.2e+08
1.0e+08
8.0e+07
6.0e+07
4.0e+07
2.0e+07
0.0e+00
WD.1.SE
WW.1.SE
WD.2.SE
WW.2.SE
WD.3.SE
WW.3.SE
WD.4.SE
WW.4.SE
WD.5.SE
WW.5.SE
WD.1.SE+5d
WW.1.SE+5d
WD.2.SE+5d
WW.2.SE+5d
WD.3.SE+5d
WW.3.SE+5d
WD.4.SE+5d
WW.4.SE+5d
WD.5.SE+5d
WW.5.SE+5d

zm00001d016402_p001

**Vesicle trafficking.retrograde trafficking**
**description:Coatomer subunit beta'−3**

subunit beta2 of coat protein complex

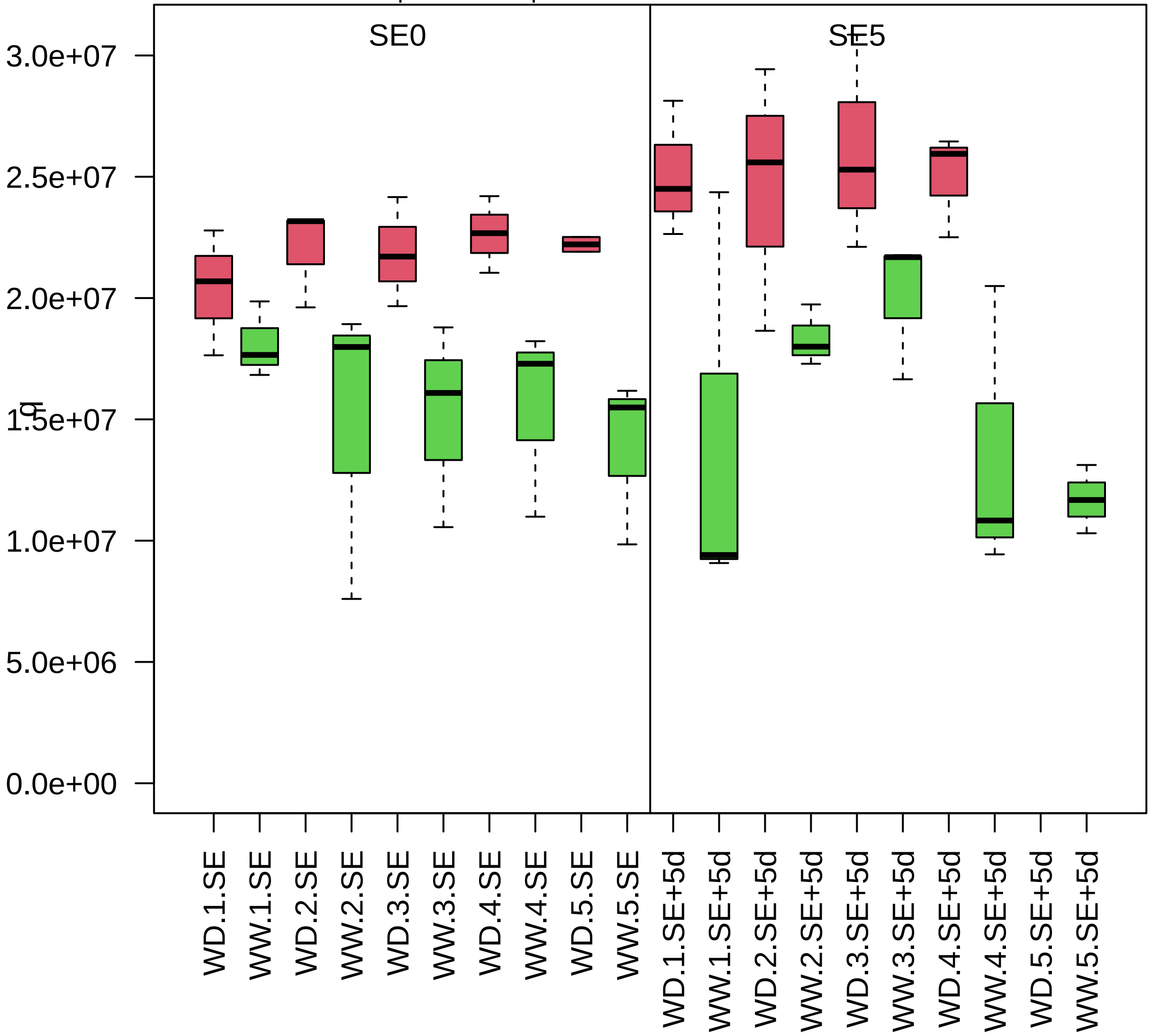

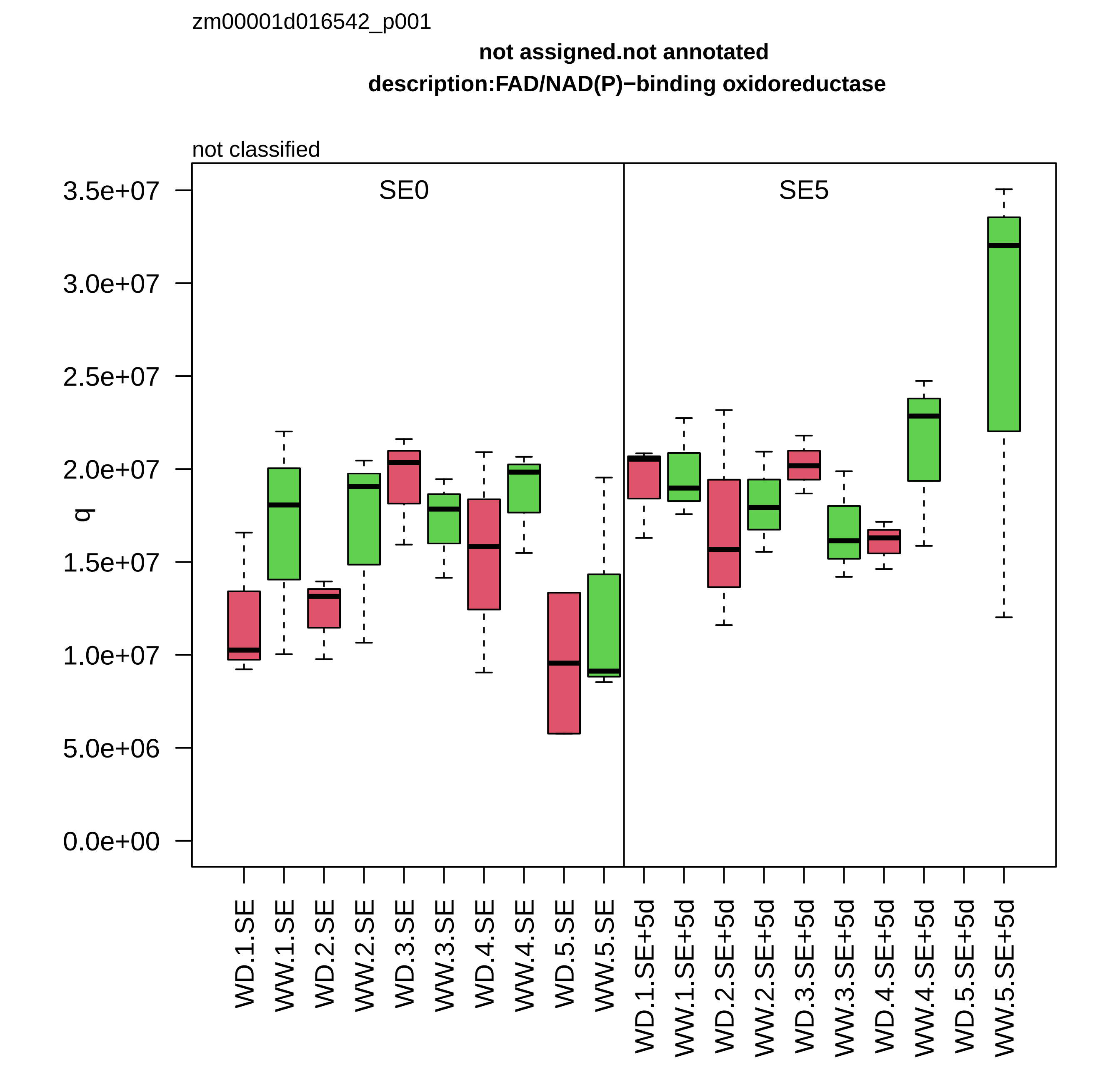
zm00001d016542_p001
not assigned.not annotated
description:FAD/NAD(P)−binding oxidoreductase
not classified
SE0
SE5
q
3.5e+07
3.0e+07
2.5e+07
2.0e+07
1.5e+07
1.0e+07
5.0e+06
0.0e+00
WD.1.SE
WW.1.SE
WD.2.SE
WW.2.SE
WD.3.SE
WW.3.SE
WD.4.SE
WW.4.SE
WD.5.SE
WW.5.SE
WD.1.SE+5d
WW.1.SE+5d
WD.2.SE+5d
WW.2.SE+5d
WD.3.SE+5d
WW.3.SE+5d
WD.4.SE+5d
WW.4.SE+5d
WD.5.SE+5d
WW.5.SE+5d

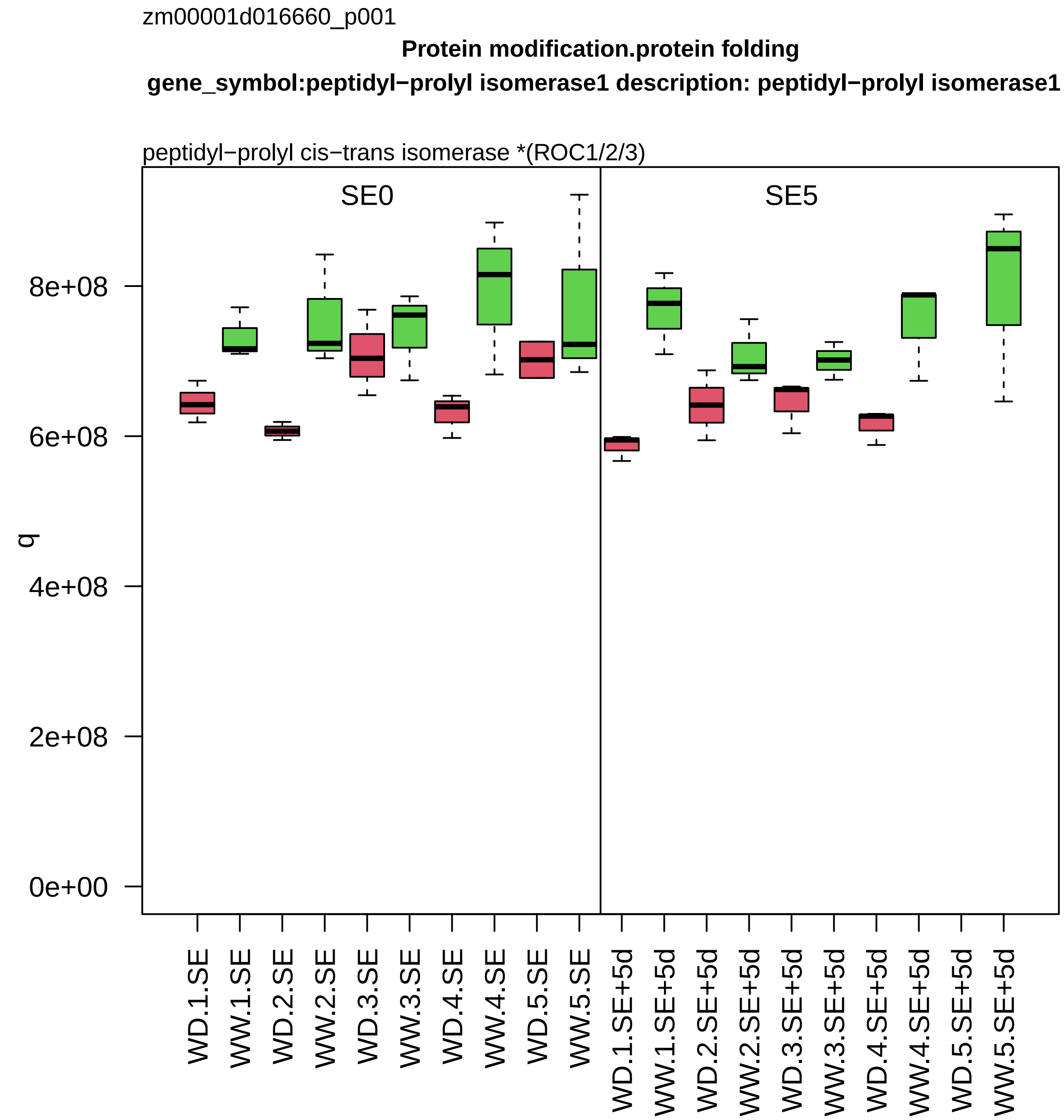

zm00001d016660_p001
Protein modification.protein folding
gene_symbol:peptidyl−prolyl isomerase1 description: peptidyl−prolyl isomerase1
peptidyl−prolyl cis−trans isomerase *(ROC1/2/3)
SE0
SE5
q
0e+00
2e+08
4e+08
6e+08
8e+08
WD.1.SE
WW.1.SE
WD.2.SE
WW.2.SE
WD.3.SE
WW.3.SE
WD.4.SE
WW.4.SE
WD.5.SE
WW.5.SE
WD.1.SE+5d
WW.1.SE+5d
WD.2.SE+5d
WW.2.SE+5d
WD.3.SE+5d
WW.3.SE+5d
WD.4.SE+5d
WW.4.SE+5d
WD.5.SE+5d
WW.5.SE+5d

zm00001d016691_p001

**Nutrient uptake.transition metal homeostasis**
**description:Copper transport protein CCH**

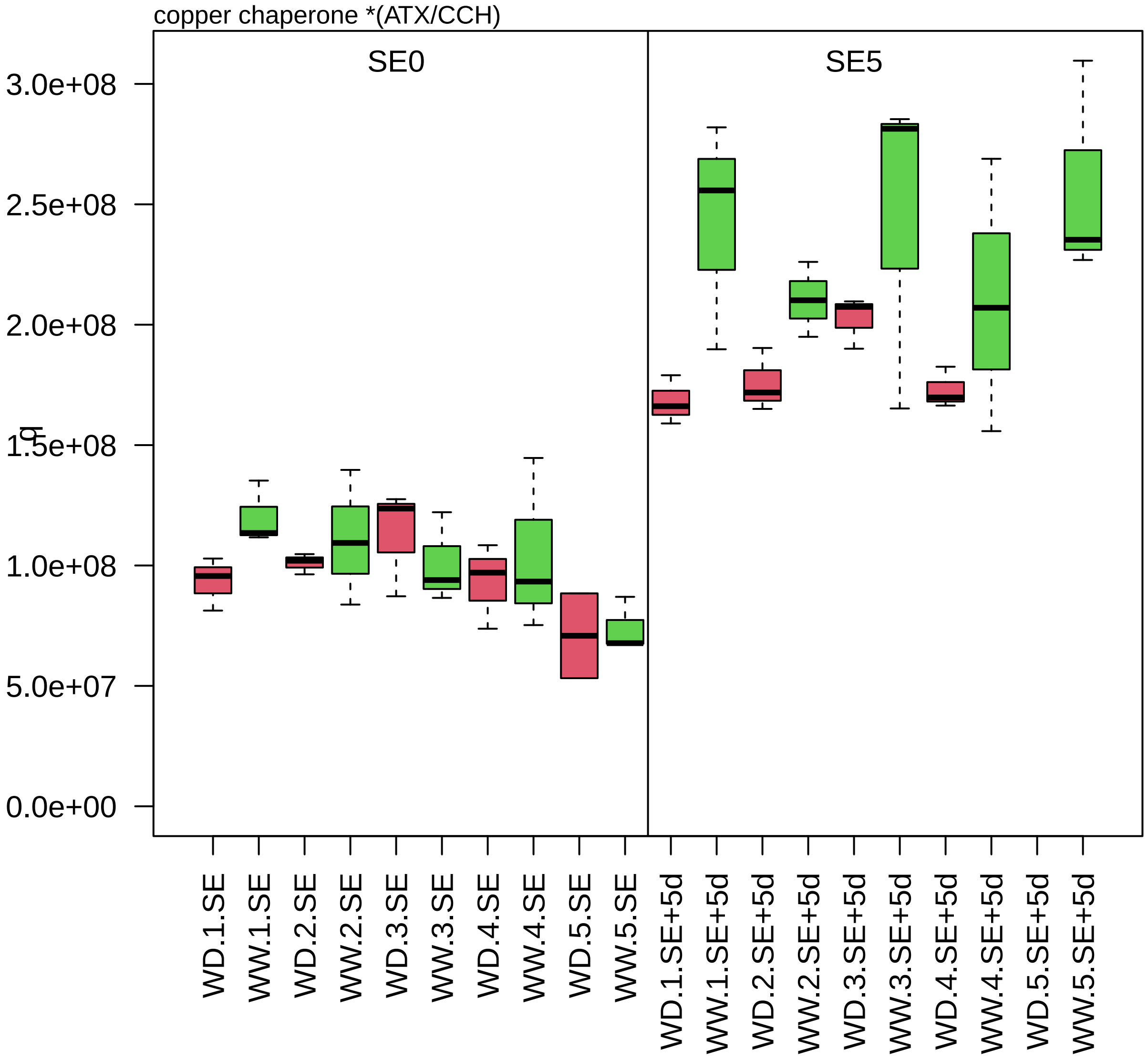

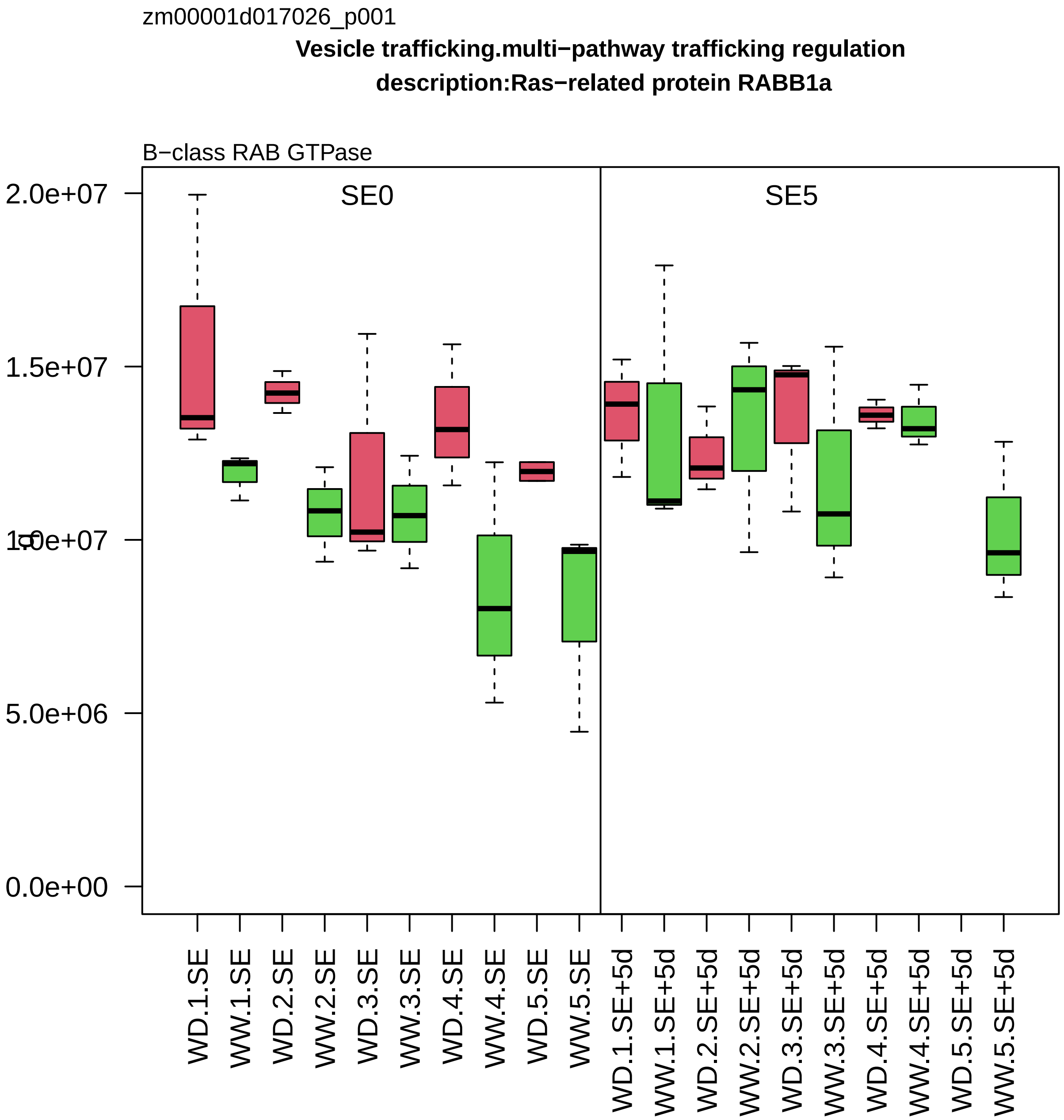

zm00001d017026_p001
Vesicle trafficking.multi−pathway trafficking regulation
description:Ras−related protein RABB1a
B−class RAB GTPase
SE0
SE5
2.0e+07
1.5e+07
1.0e+07
5.0e+06
0.0e+00
WD.1.SE
WW.1.SE
WD.2.SE
WW.2.SE
WD.3.SE
WW.3.SE
WD.4.SE
WW.4.SE
WD.5.SE
WW.5.SE
WD.1.SE+5d
WW.1.SE+5d
WD.2.SE+5d
WW.2.SE+5d
WD.3.SE+5d
WW.3.SE+5d
WD.4.SE+5d
WW.4.SE+5d
WD.5.SE+5d
WW.5.SE+5d

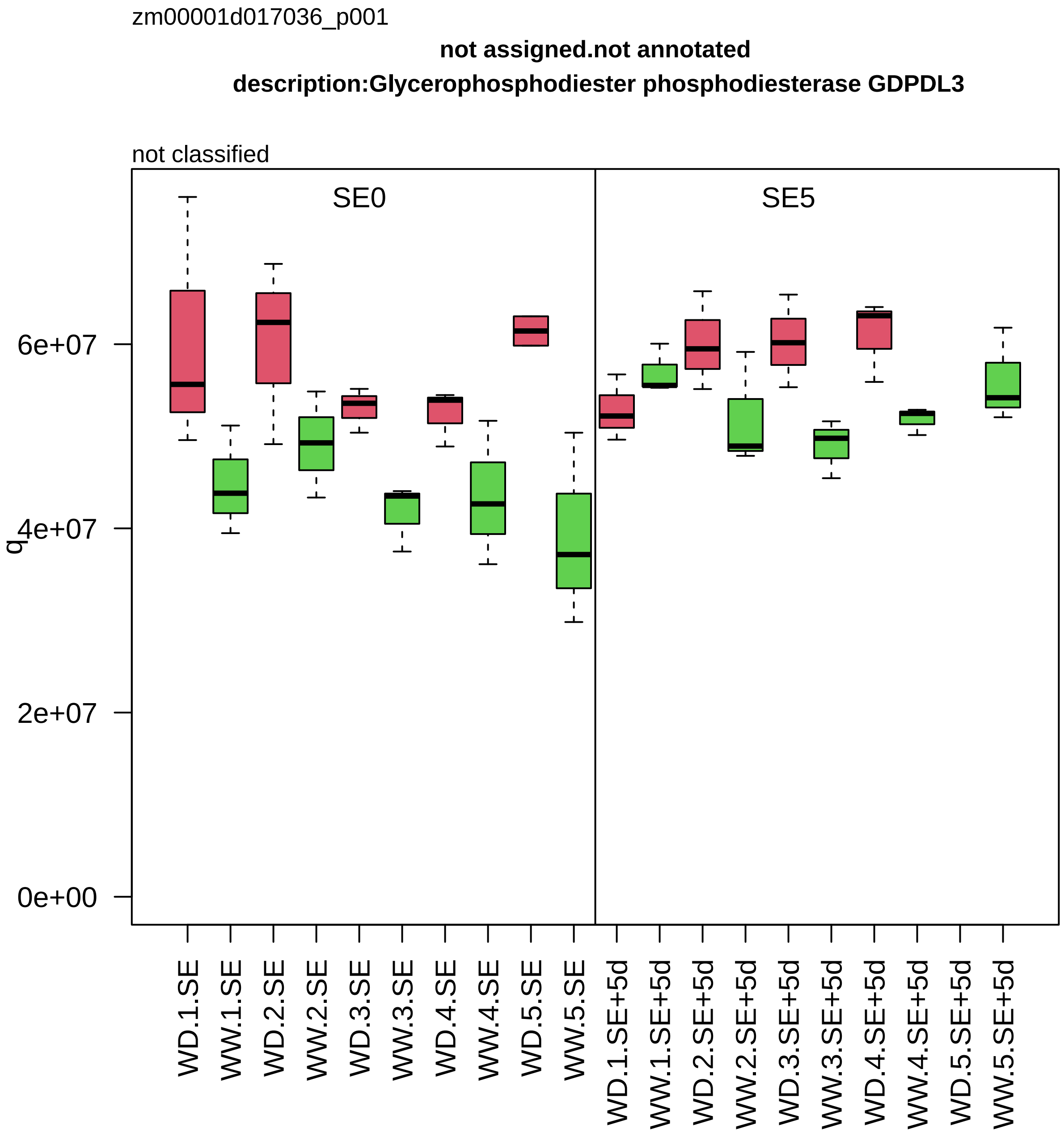

zm00001d017036_p001
not assigned.not annotated
description:Glycerophosphodiester phosphodiesterase GDPDL3
not classified
SE0
SE5
q
0e+00
2e+07
4e+07
6e+07
WD.1.SE
WW.1.SE
WD.2.SE
WW.2.SE
WD.3.SE
WW.3.SE
WD.4.SE
WW.4.SE
WD.5.SE
WW.5.SE
WD.1.SE+5d
WW.1.SE+5d
WD.2.SE+5d
WW.2.SE+5d
WD.3.SE+5d
WW.3.SE+5d
WD.4.SE+5d
WW.4.SE+5d
WD.5.SE+5d
WW.5.SE+5d

zm00001d017047_p001

**Protein biosynthesis.ribosome biogenesis**

**description:60S ribosomal protein L6**

component *(eL6) of large ribosomal−subunit (LSU) proteome

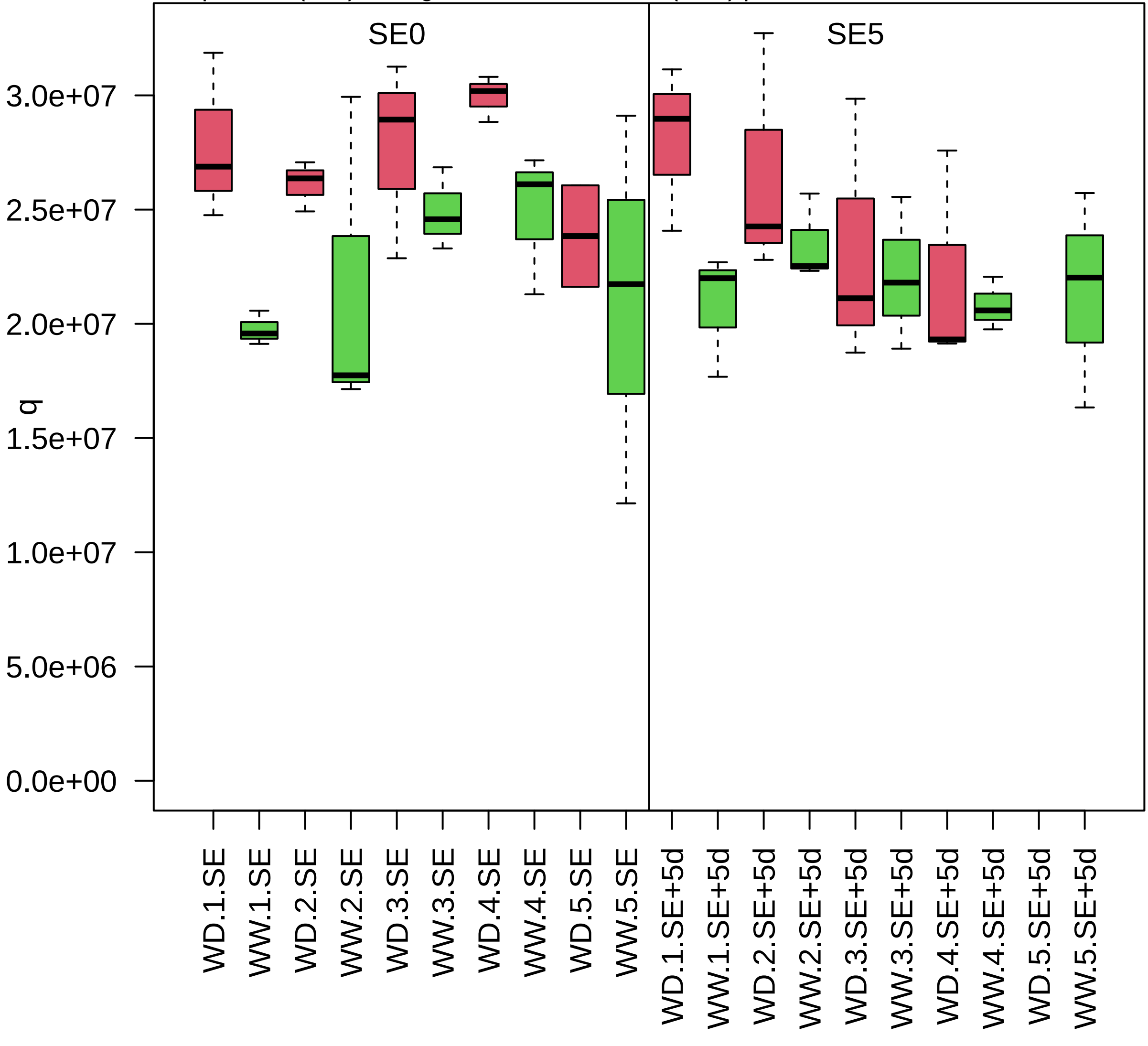

zm00001d017239_p001

**Protein translocation.nucleus**
**description:RAN GTPase−activating protein 2**

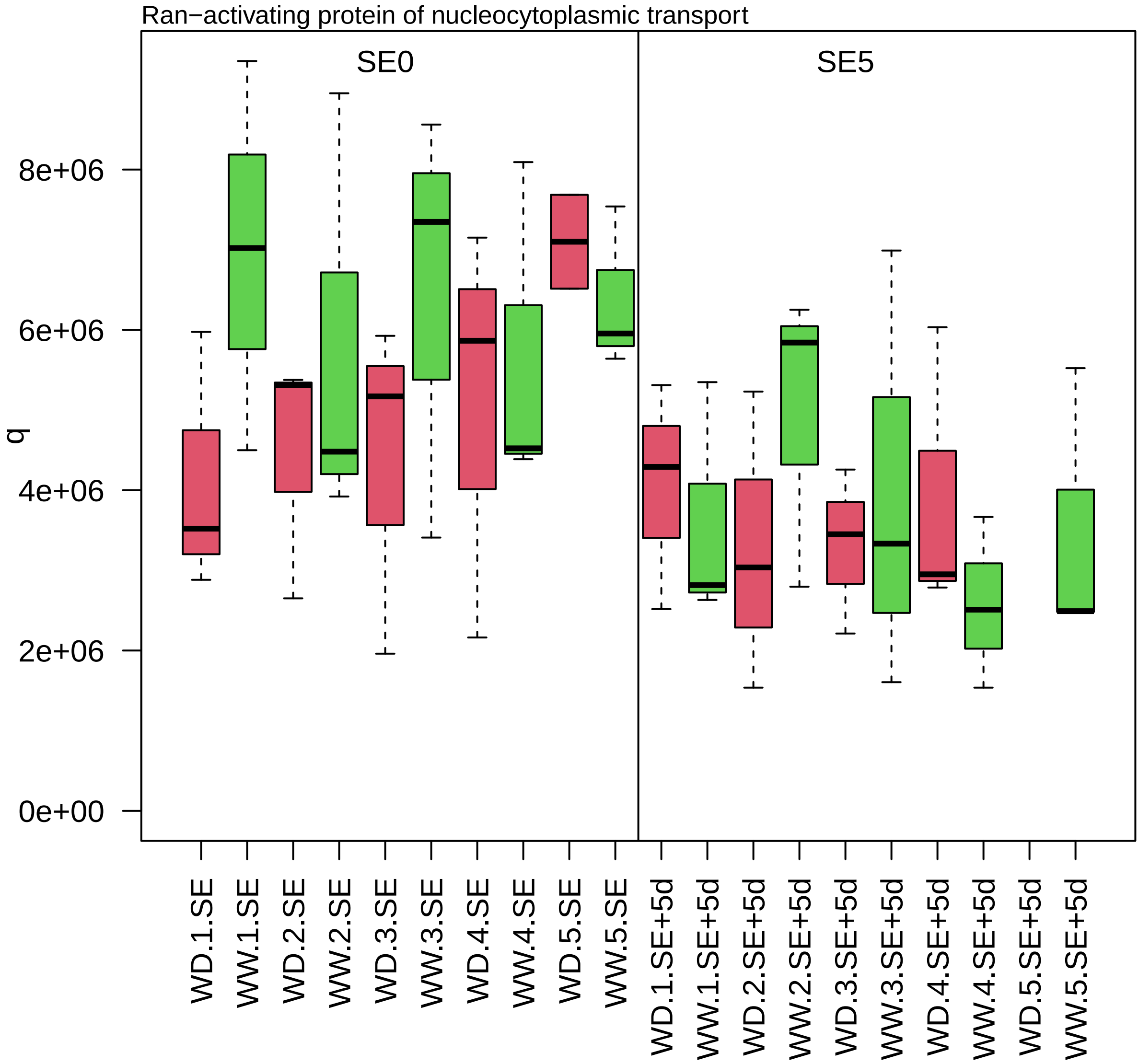

zm00001d017258_p001

**Cellular respiration.tricarboxylic acid cycle**
**description:Succinate−−CoA ligase [ADP−forming] subunit beta mitochondrial**

subunit beta of succinyl−CoA ligase heterodimer

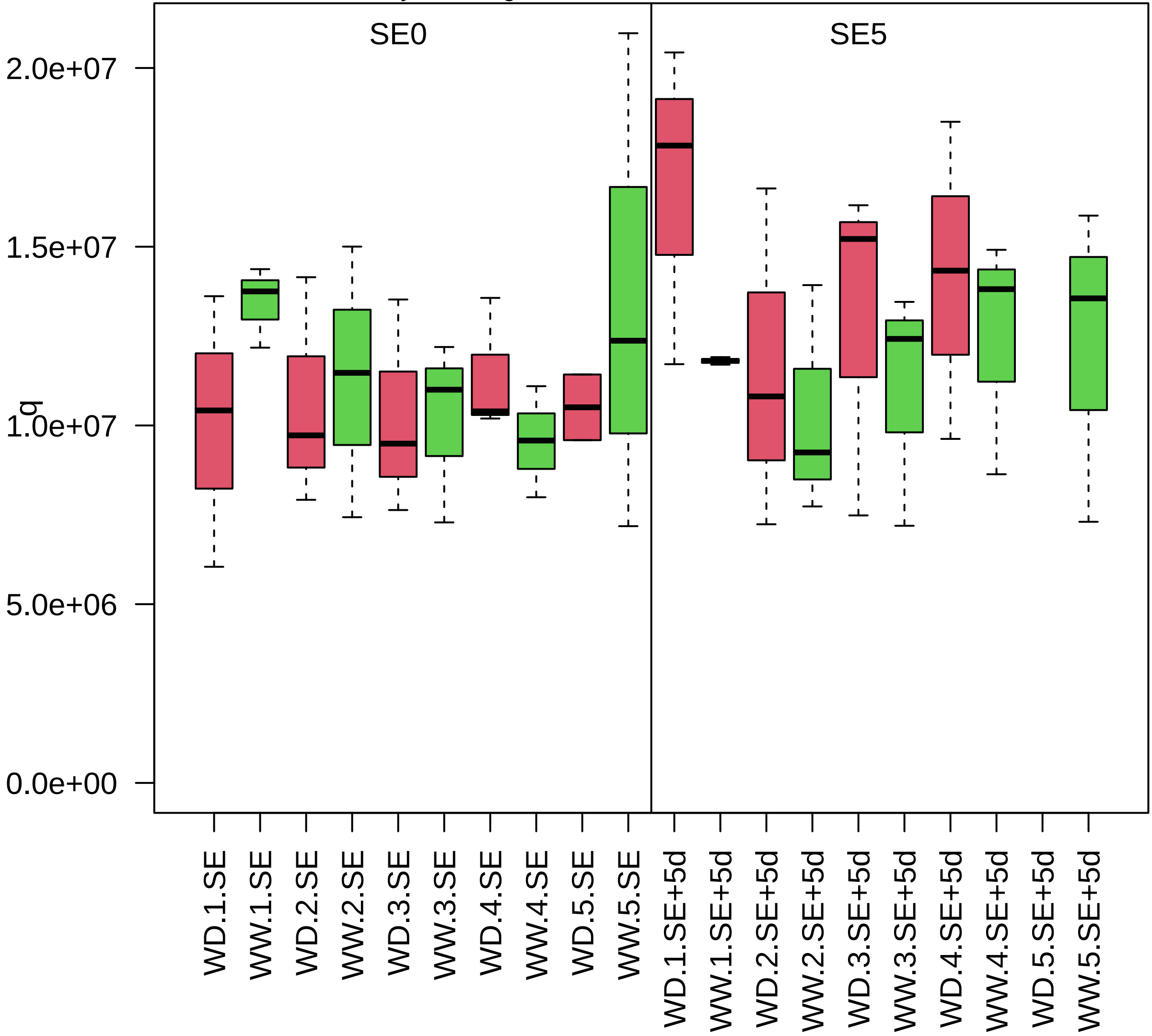

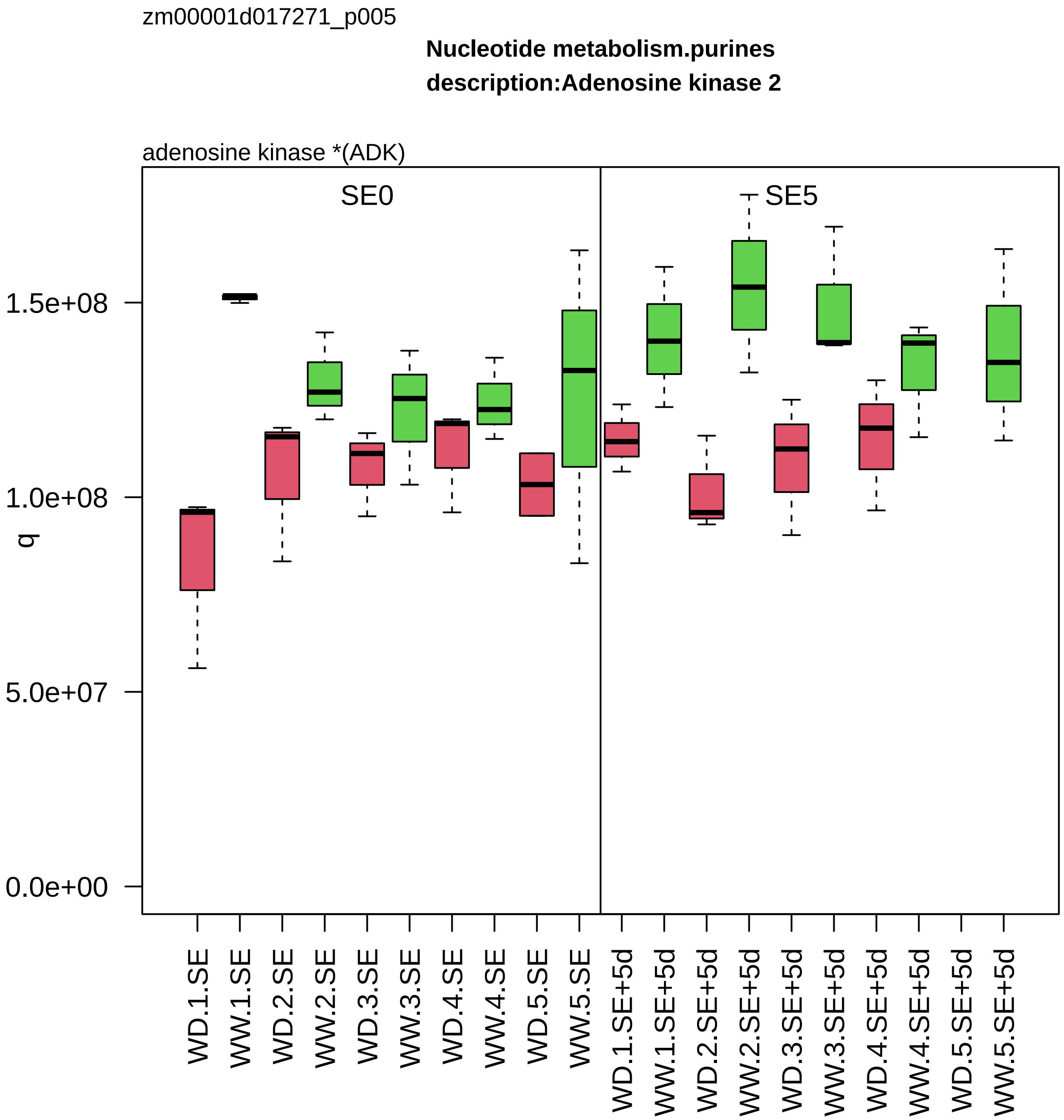
zm00001d017271_p005
Nucleotide metabolism.purines
description:Adenosine kinase 2
adenosine kinase *(ADK)
SE0
SE5
q
1.5e+08
1.0e+08
5.0e+07
0.0e+00
WD.1.SE
WW.1.SE
WD.2.SE
WW.2.SE
WD.3.SE
WW.3.SE
WD.4.SE
WW.4.SE
WD.5.SE
WW.5.SE
WD.1.SE+5d
WW.1.SE+5d
WD.2.SE+5d
WW.2.SE+5d
WD.3.SE+5d
WW.3.SE+5d
WD.4.SE+5d
WW.4.SE+5d
WD.5.SE+5d
WW.5.SE+5d

zm00001d017336_p001

**Protein homeostasis.ubiquitin−proteasome system**
**description:Proteasome subunit alpha type−2−B**

alpha−type−2 component *(PAB) of 26S proteasome

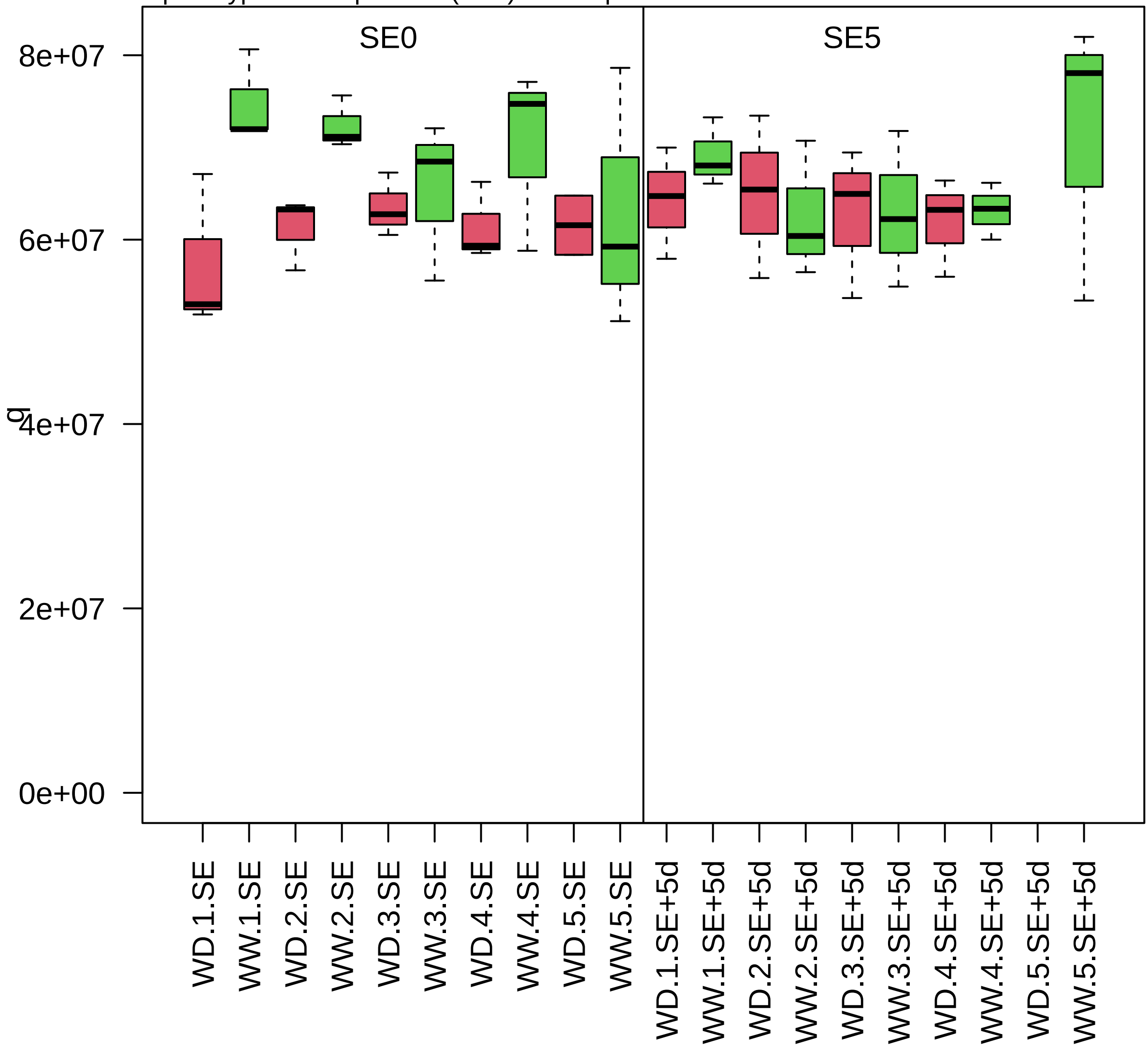

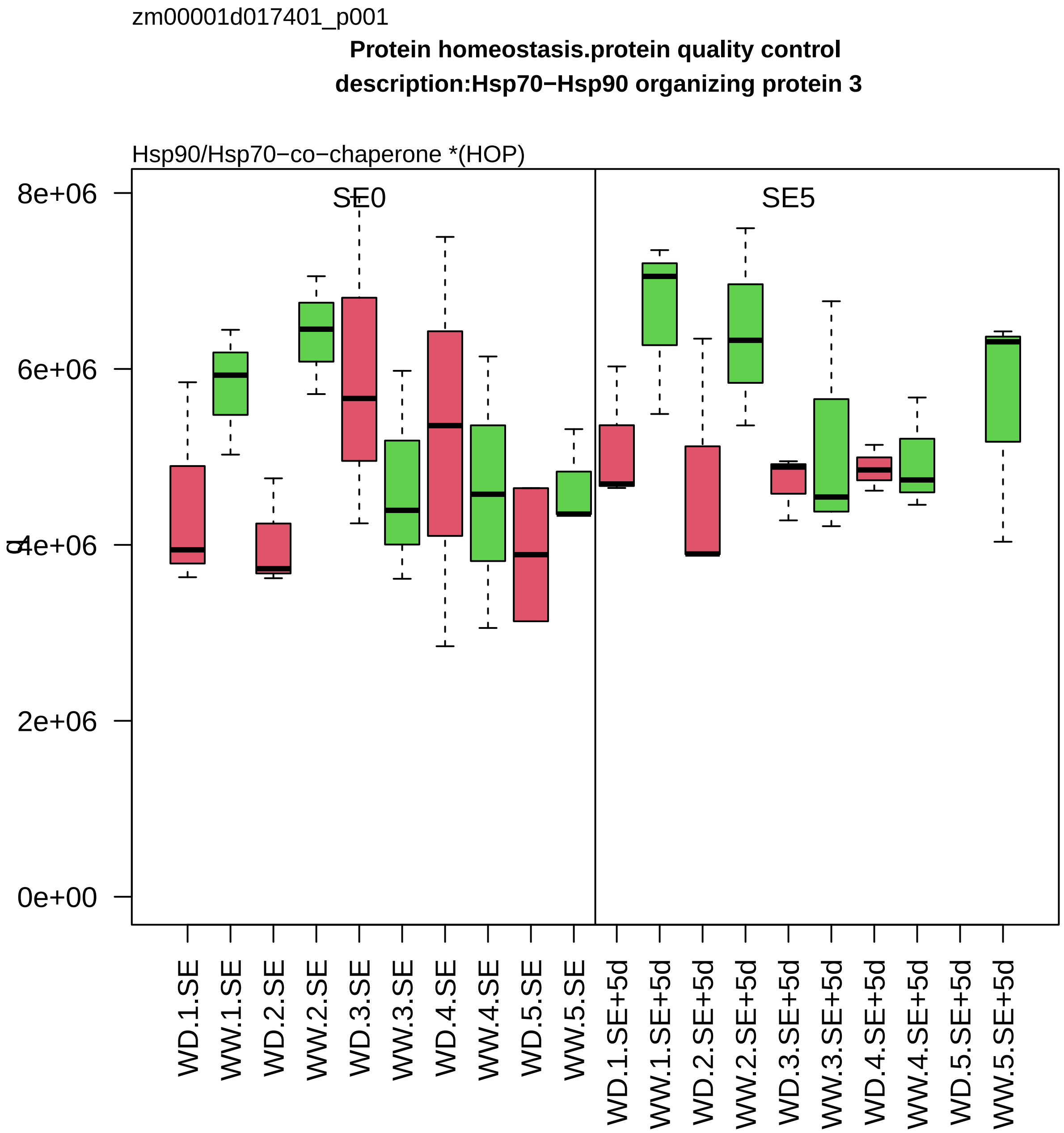

zm00001d017401_p001
Protein homeostasis.protein quality control
description:Hsp70−Hsp90 organizing protein 3
Hsp90/Hsp70−co−chaperone *(HOP)
SE0
SE5
8e+06
6e+06
4e+06
2e+06
0e+00
WD.1.SE
WW.1.SE
WD.2.SE
WW.2.SE
WD.3.SE
WW.3.SE
WD.4.SE
WW.4.SE
WD.5.SE
WW.5.SE
WD.1.SE+5d
WW.1.SE+5d
WD.2.SE+5d
WW.2.SE+5d
WD.3.SE+5d
WW.3.SE+5d
WD.4.SE+5d
WW.4.SE+5d
WD.5.SE+5d
WW.5.SE+5d

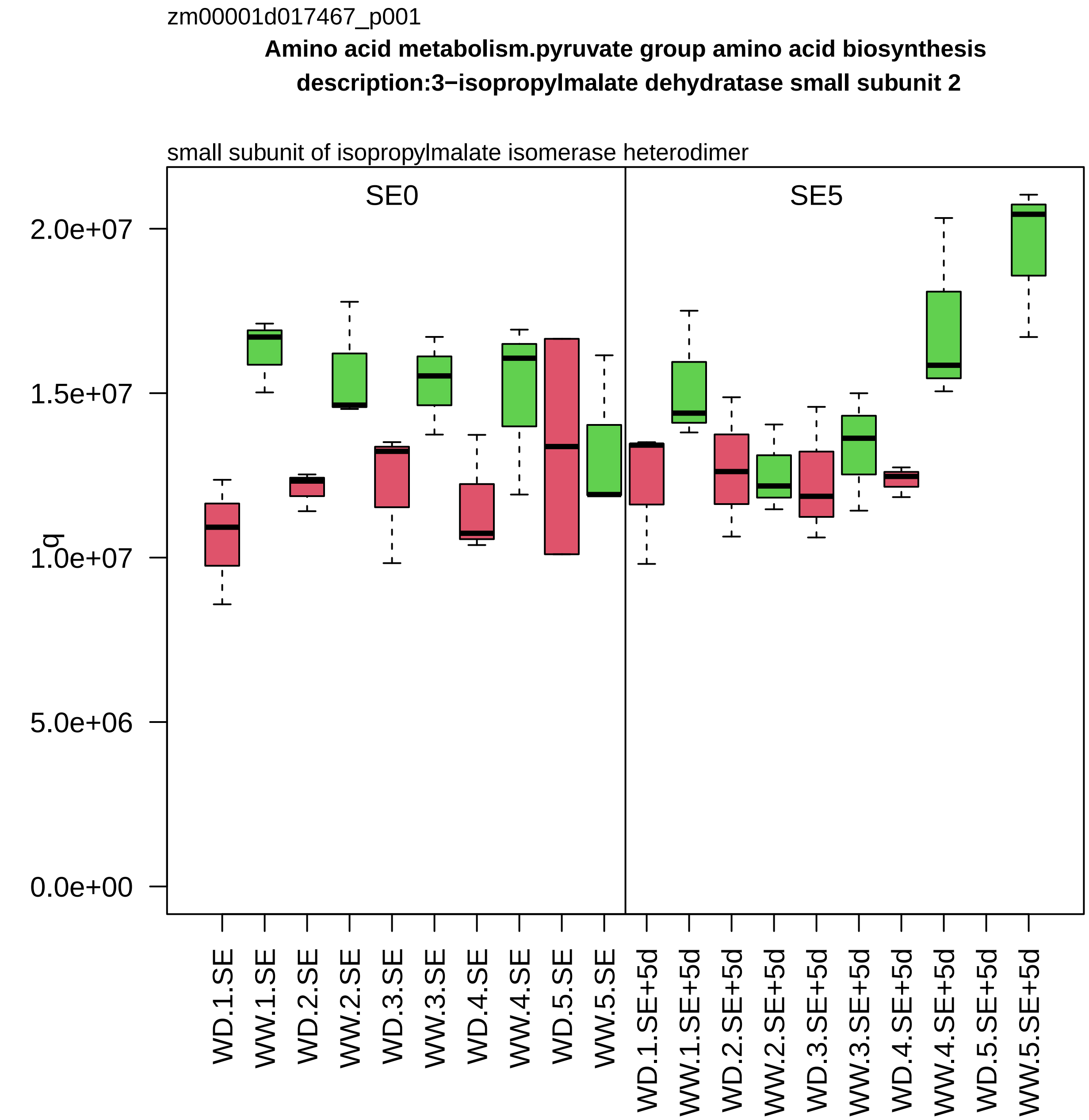

zm00001d017467_p001
Amino acid metabolism.pyruvate group amino acid biosynthesis
description:3−isopropylmalate dehydratase small subunit 2
small subunit of isopropylmalate isomerase heterodimer
SE0
SE5
q
2.0e+07
1.5e+07
1.0e+07
5.0e+06
0.0e+00
WD.1.SE
WW.1.SE
WD.2.SE
WW.2.SE
WD.3.SE
WW.3.SE
WD.4.SE
WW.4.SE
WD.5.SE
WW.5.SE
WD.1.SE+5d
WW.1.SE+5d
WD.2.SE+5d
WW.2.SE+5d
WD.3.SE+5d
WW.3.SE+5d
WD.4.SE+5d
WW.4.SE+5d
WD.5.SE+5d
WW.5.SE+5d

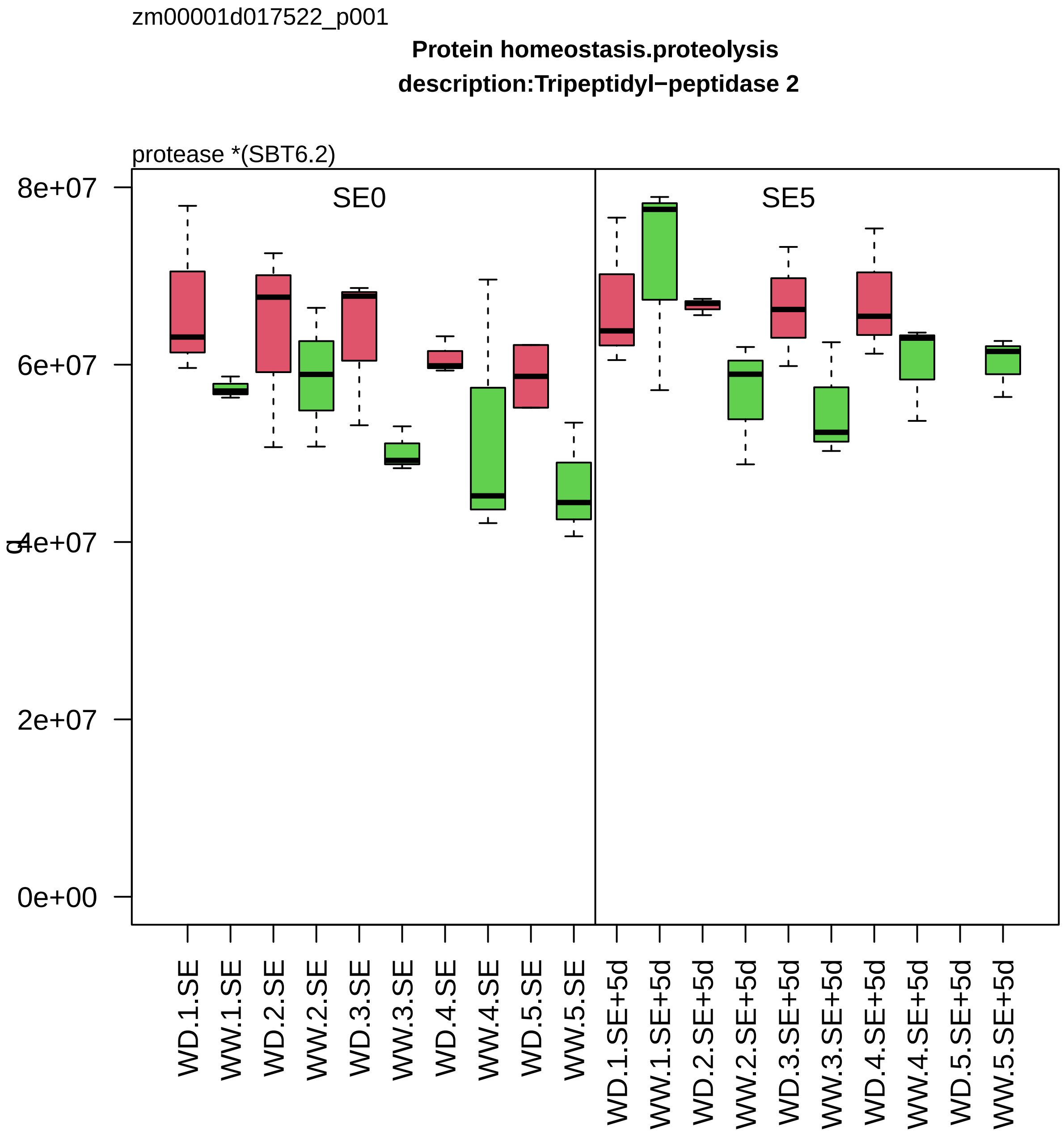

zm00001d017522_p001
Protein homeostasis.proteolysis
description:Tripeptidyl−peptidase 2
protease *(SBT6.2)
SE0
SE5
8e+07
6e+07
4e+07
2e+07
0e+00
WD.1.SE
WW.1.SE
WD.2.SE
WW.2.SE
WD.3.SE
WW.3.SE
WD.4.SE
WW.4.SE
WD.5.SE
WW.5.SE
WD.1.SE+5d
WW.1.SE+5d
WD.2.SE+5d
WW.2.SE+5d
WD.3.SE+5d
WW.3.SE+5d
WD.4.SE+5d
WW.4.SE+5d
WD.5.SE+5d
WW.5.SE+5d

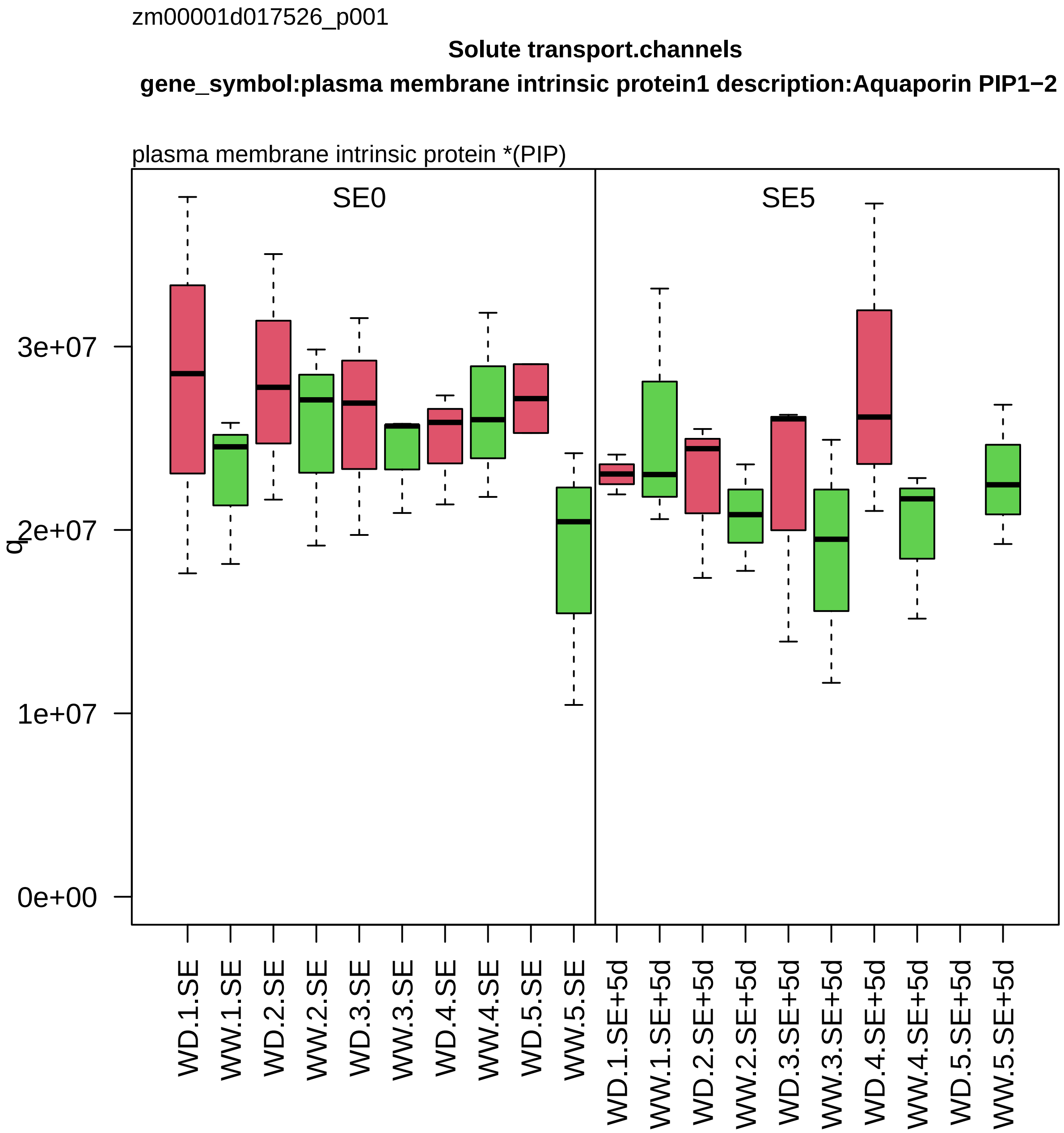

zm00001d017526_p001
Solute transport.channels
gene_symbol:plasma membrane intrinsic protein1 description:Aquaporin PIP1−2
plasma membrane intrinsic protein *(PIP)
SE0
SE5
q
3e+07
2e+07
1e+07
0e+00
WD.1.SE
WW.1.SE
WD.2.SE
WW.2.SE
WD.3.SE
WW.3.SE
WD.4.SE
WW.4.SE
WD.5.SE
WW.5.SE
WD.1.SE+5d
WW.1.SE+5d
WD.2.SE+5d
WW.2.SE+5d
WD.3.SE+5d
WW.3.SE+5d
WD.4.SE+5d
WW.4.SE+5d
WD.5.SE+5d
WW.5.SE+5d

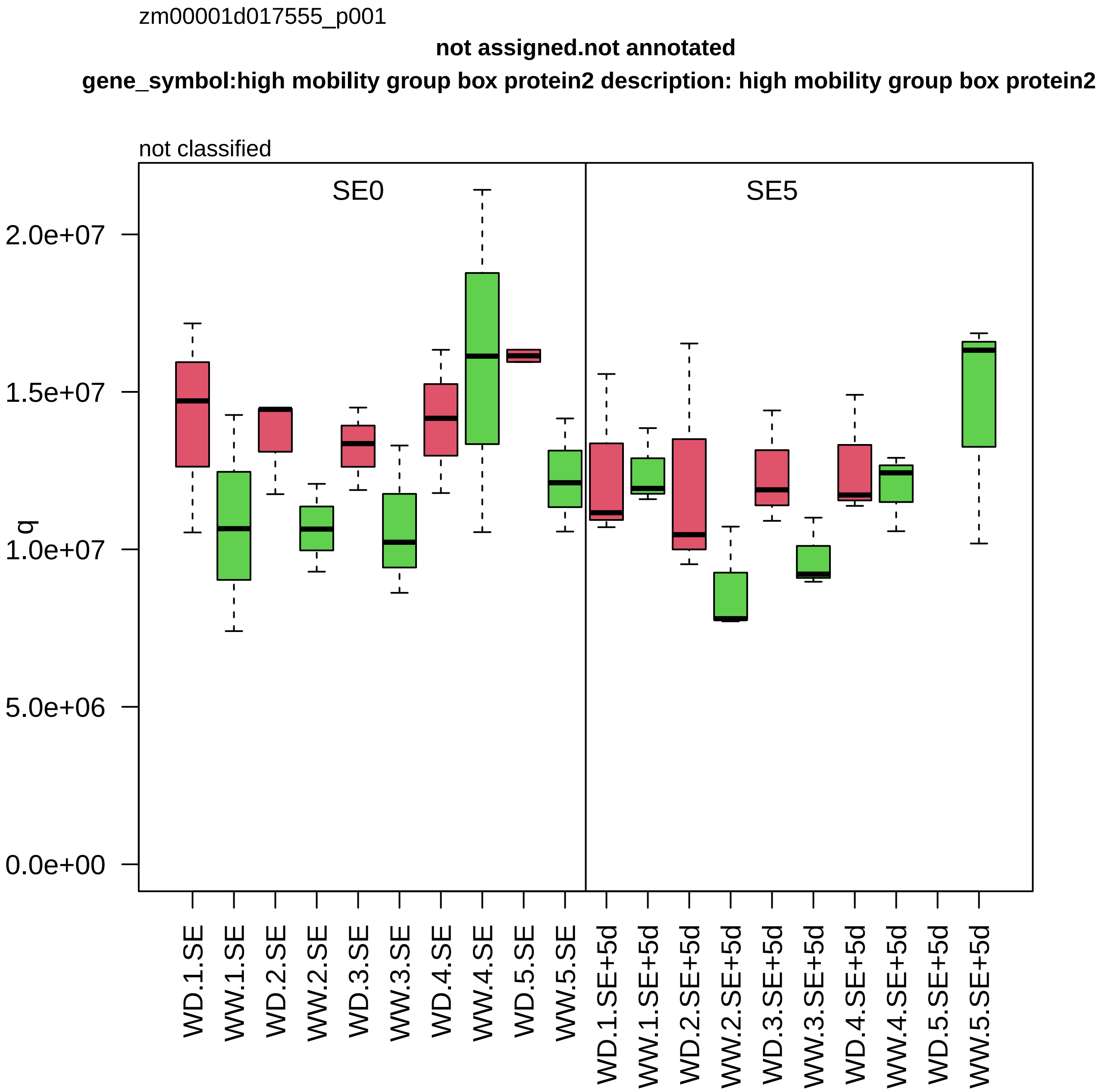
zm00001d017555_p001
not assigned.not annotated
gene_symbol:high mobility group box protein2 description: high mobility group box protein2
not classified
SE0
SE5
q
2.0e+07
1.5e+07
1.0e+07
5.0e+06
0.0e+00
WD.1.SE
WW.1.SE
WD.2.SE
WW.2.SE
WD.3.SE
WW.3.SE
WD.4.SE
WW.4.SE
WD.5.SE
WW.5.SE
WD.1.SE+5d
WW.1.SE+5d
WD.2.SE+5d
WW.2.SE+5d
WD.3.SE+5d
WW.3.SE+5d
WD.4.SE+5d
WW.4.SE+5d
WD.5.SE+5d
WW.5.SE+5d

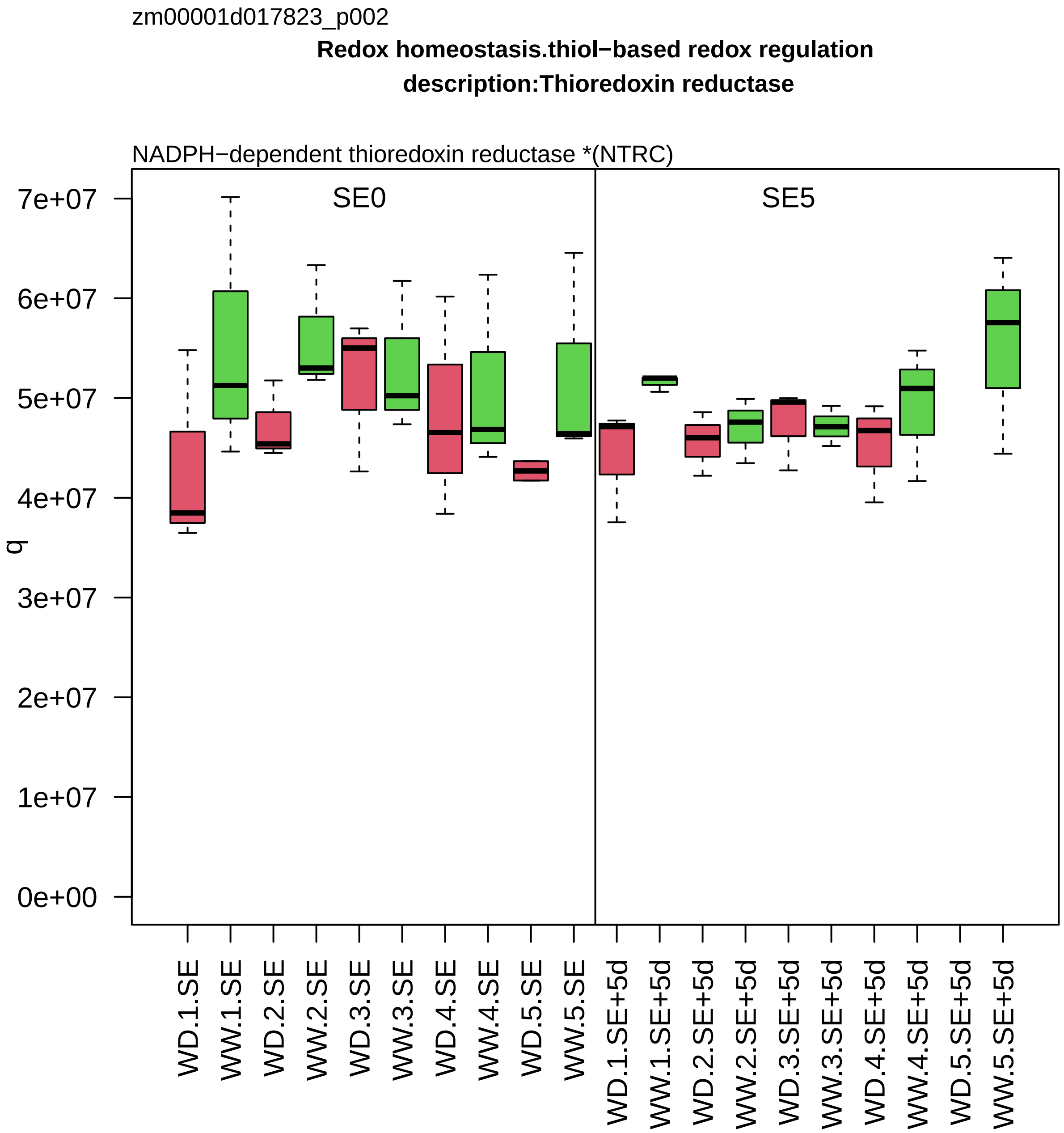

zm00001d017823_p002
Redox homeostasis.thiol−based redox regulation
description:Thioredoxin reductase
NADPH−dependent thioredoxin reductase *(NTRC)
SE0
SE5
q
7e+07
6e+07
5e+07
4e+07
3e+07
2e+07
1e+07
0e+00
WD.1.SE
WW.1.SE
WD.2.SE
WW.2.SE
WD.3.SE
WW.3.SE
WD.4.SE
WW.4.SE
WD.5.SE
WW.5.SE
WD.1.SE+5d
WW.1.SE+5d
WD.2.SE+5d
WW.2.SE+5d
WD.3.SE+5d
WW.3.SE+5d
WD.4.SE+5d
WW.4.SE+5d
WD.5.SE+5d
WW.5.SE+5d

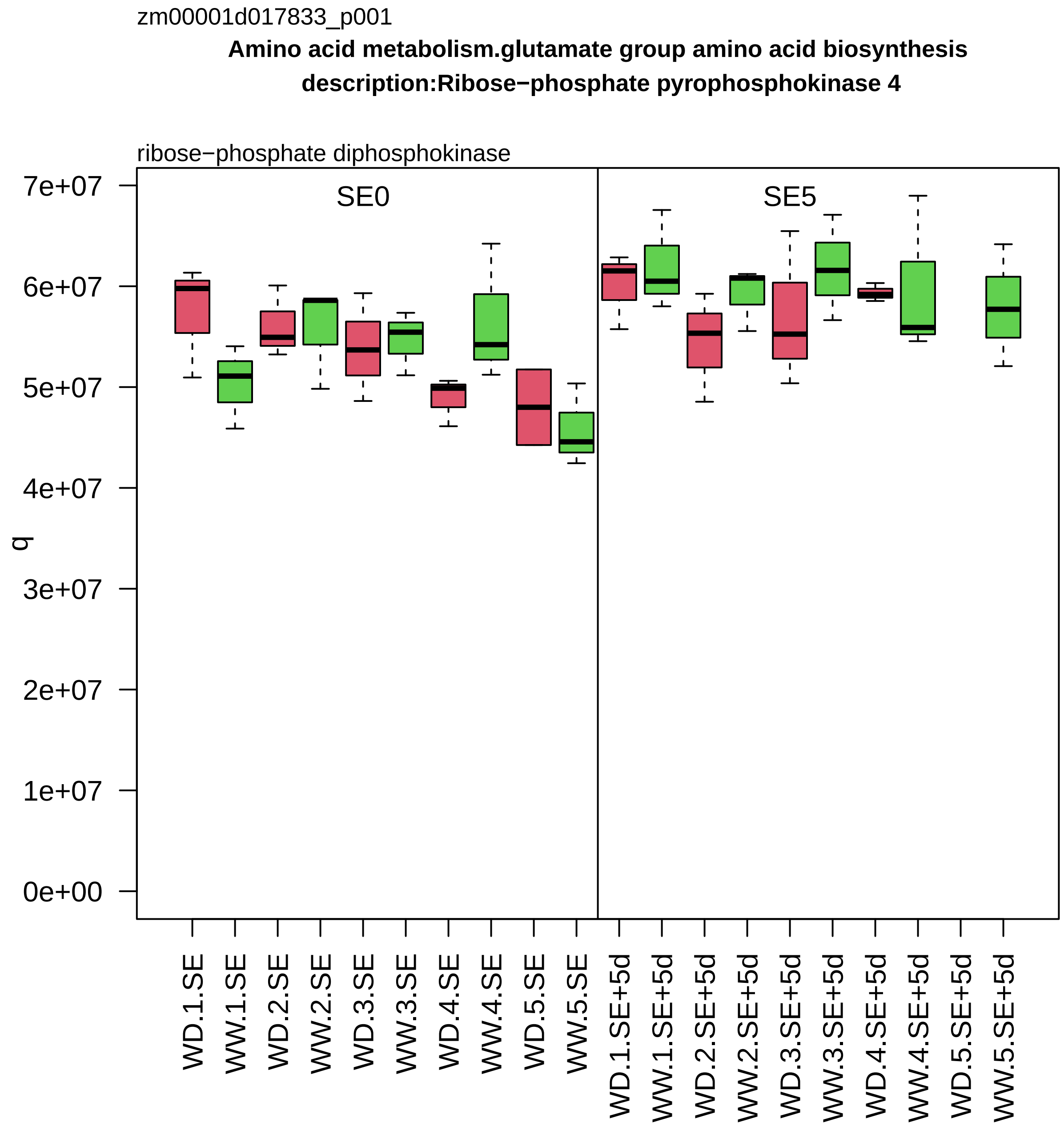
zm00001d017833_p001
Amino acid metabolism.glutamate group amino acid biosynthesis
description:Ribose−phosphate pyrophosphokinase 4
ribose−phosphate diphosphokinase
SE0
SE5
q
7e+07
6e+07
5e+07
4e+07
3e+07
2e+07
1e+07
0e+00
WD.1.SE
WW.1.SE
WD.2.SE
WW.2.SE
WD.3.SE
WW.3.SE
WD.4.SE
WW.4.SE
WD.5.SE
WW.5.SE
WD.1.SE+5d
WW.1.SE+5d
WD.2.SE+5d
WW.2.SE+5d
WD.3.SE+5d
WW.3.SE+5d
WD.4.SE+5d
WW.4.SE+5d
WD.5.SE+5d
WW.5.SE+5d

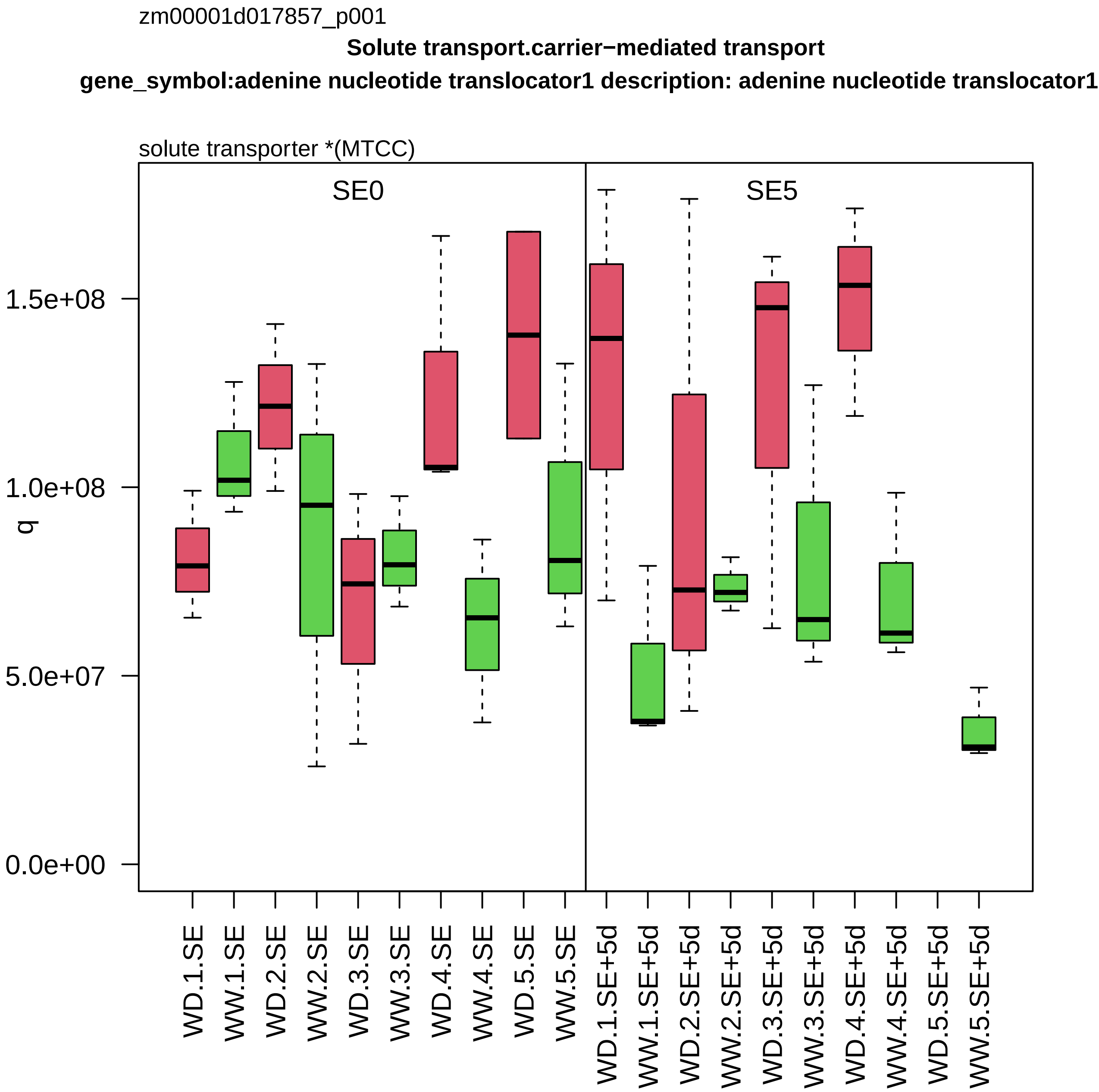
zm00001d017857_p001
Solute transport.carrier−mediated transport
gene_symbol:adenine nucleotide translocator1 description: adenine nucleotide translocator1
solute transporter *(MTCC)
SE0
SE5
q
1.5e+08
1.0e+08
5.0e+07
0.0e+00
WD.1.SE
WW.1.SE
WD.2.SE
WW.2.SE
WD.3.SE
WW.3.SE
WD.4.SE
WW.4.SE
WD.5.SE
WW.5.SE
WD.1.SE+5d
WW.1.SE+5d
WD.2.SE+5d
WW.2.SE+5d
WD.3.SE+5d
WW.3.SE+5d
WD.4.SE+5d
WW.4.SE+5d
WD.5.SE+5d
WW.5.SE+5d

zm00001d017883_p001

**RNA processing.RNA homeostasis**

**description:Enhancer of mRNA−decapping protein 4**

scaffold component *(VCS) of mRNA decapping complex

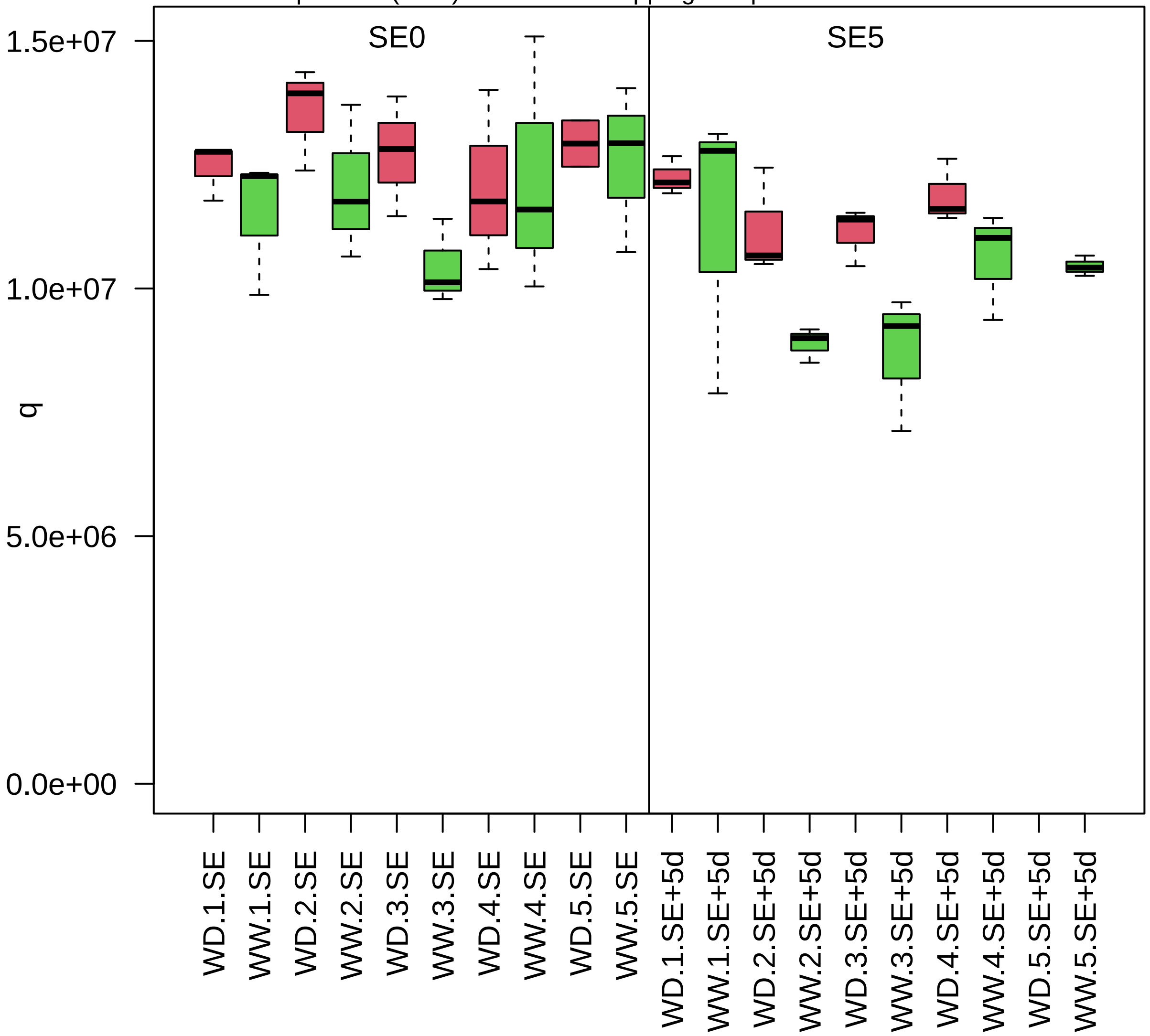

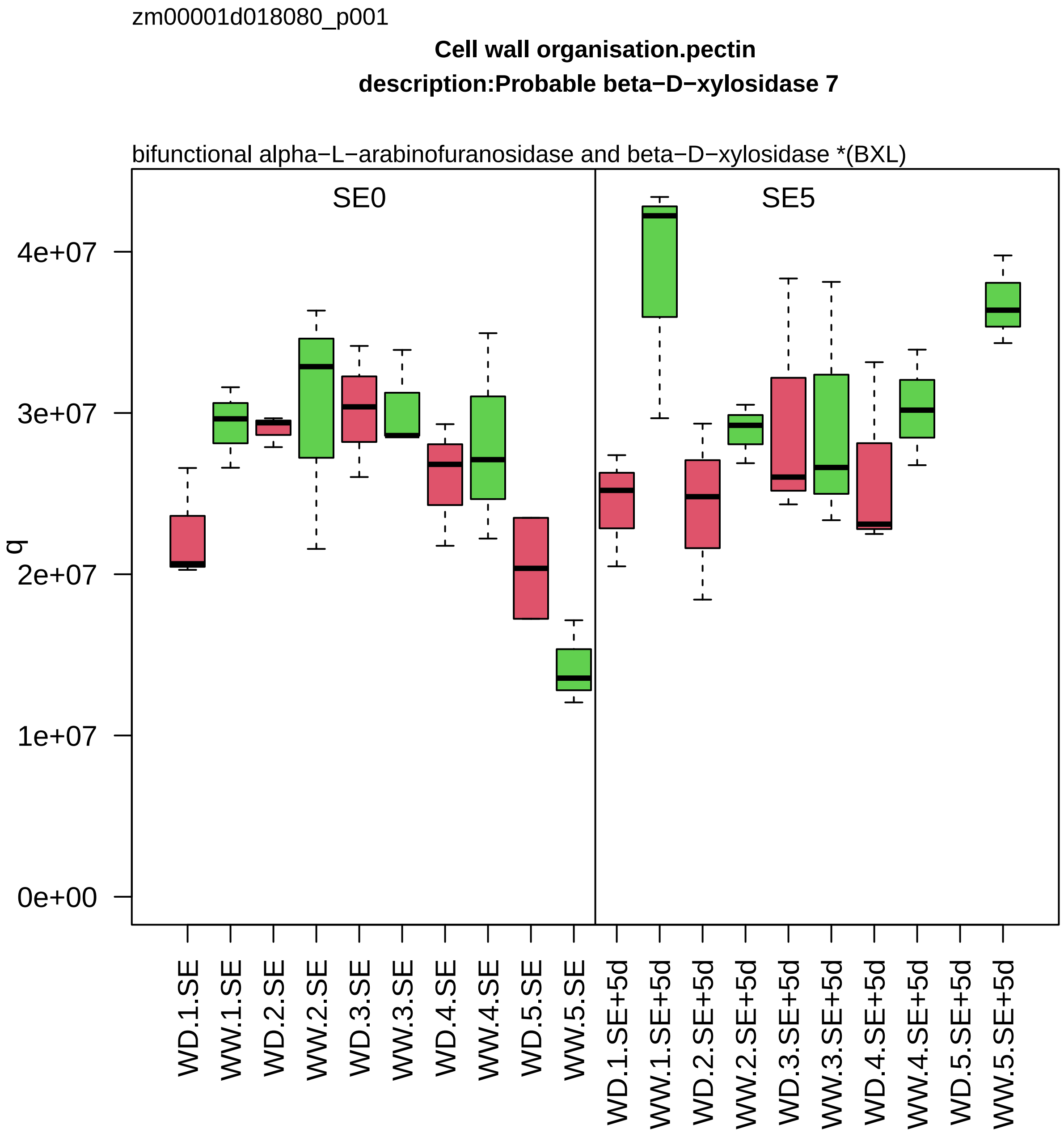
zm00001d018080_p001
Cell wall organisation.pectin
description:Probable beta−D−xylosidase 7
bifunctional alpha−L−arabinofuranosidase and beta−D−xylosidase *(BXL)
SE0
SE5
q
4e+07
3e+07
2e+07
1e+07
0e+00
WD.1.SE
WW.1.SE
WD.2.SE
WW.2.SE
WD.3.SE
WW.3.SE
WD.4.SE
WW.4.SE
WD.5.SE
WW.5.SE
WD.1.SE+5d
WW.1.SE+5d
WD.2.SE+5d
WW.2.SE+5d
WD.3.SE+5d
WW.3.SE+5d
WD.4.SE+5d
WW.4.SE+5d
WD.5.SE+5d
WW.5.SE+5d

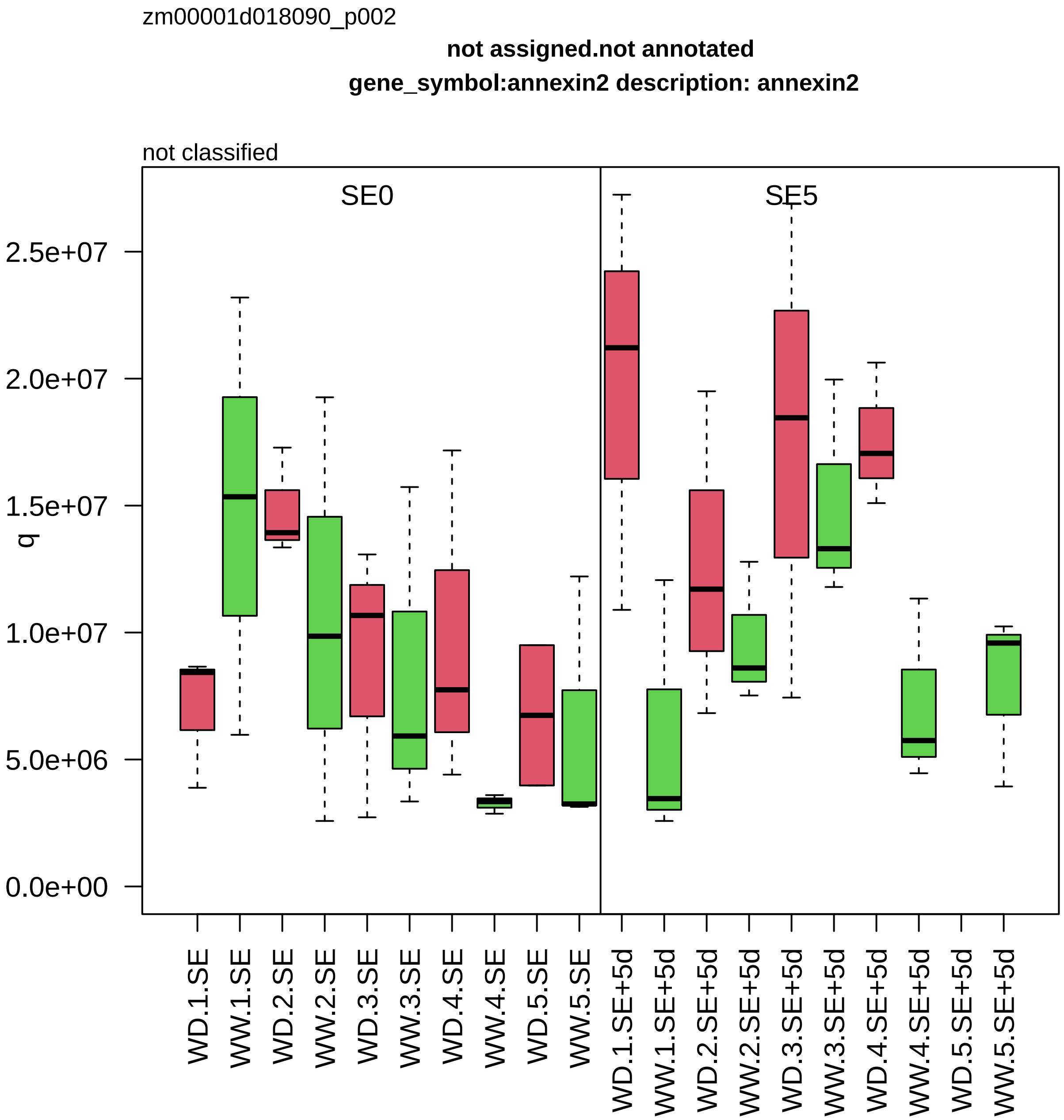

zm00001d018090_p002
not assigned.not annotated
gene_symbol:annexin2 description: annexin2
not classified
SE0
SE5
q
2.5e+07
2.0e+07
1.5e+07
1.0e+07
5.0e+06
0.0e+00
WD.1.SE
WW.1.SE
WD.2.SE
WW.2.SE
WD.3.SE
WW.3.SE
WD.4.SE
WW.4.SE
WD.5.SE
WW.5.SE
WD.1.SE+5d
WW.1.SE+5d
WD.2.SE+5d
WW.2.SE+5d
WD.3.SE+5d
WW.3.SE+5d
WD.4.SE+5d
WW.4.SE+5d
WD.5.SE+5d
WW.5.SE+5d

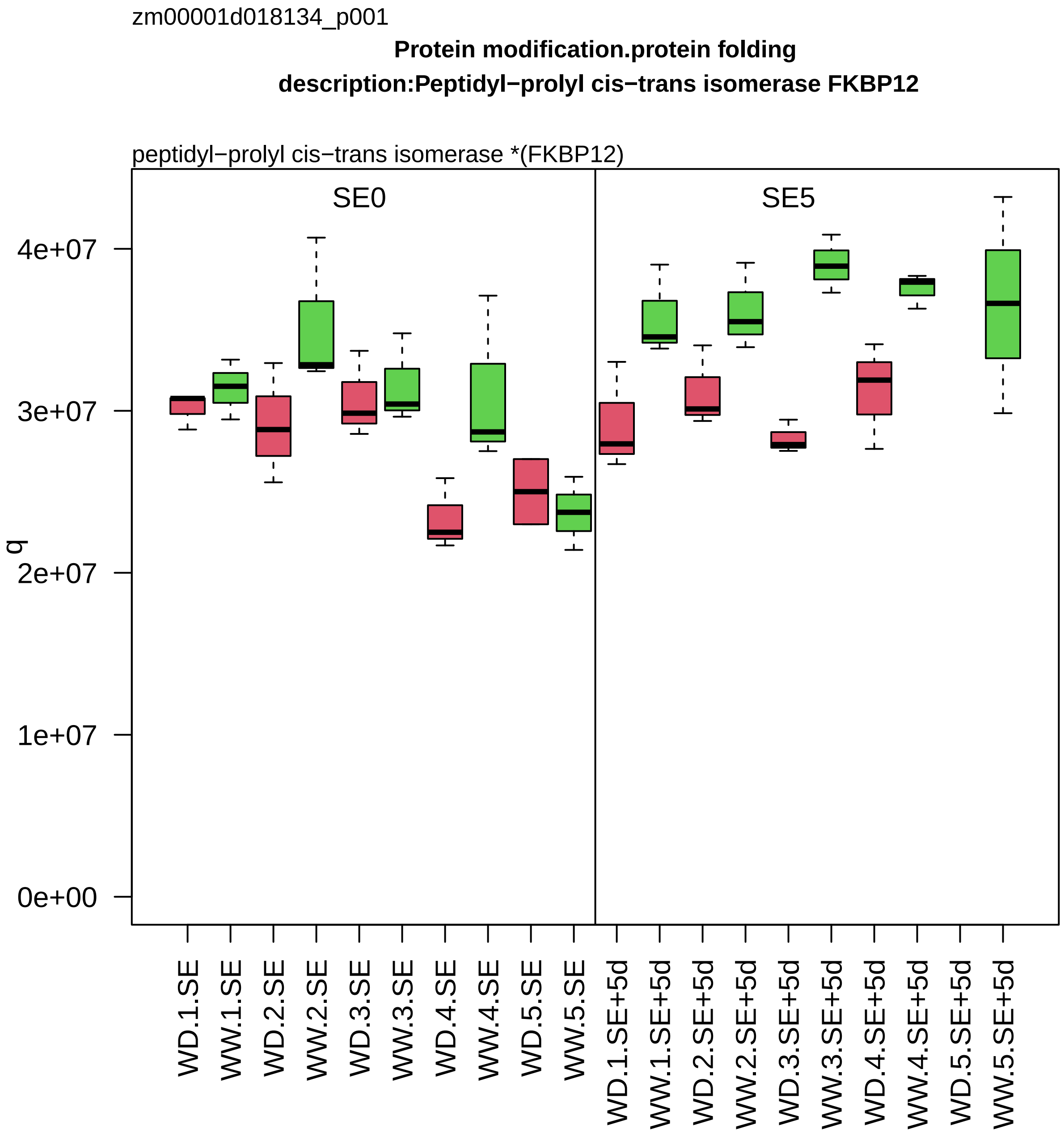
zm00001d018134_p001
Protein modification.protein folding
description:Peptidyl−prolyl cis−trans isomerase FKBP12
peptidyl−prolyl cis−trans isomerase *(FKBP12)
SE0
SE5
q
0e+00
1e+07
2e+07
3e+07
4e+07
WD.1.SE
WW.1.SE
WD.2.SE
WW.2.SE
WD.3.SE
WW.3.SE
WD.4.SE
WW.4.SE
WD.5.SE
WW.5.SE
WD.1.SE+5d
WW.1.SE+5d
WD.2.SE+5d
WW.2.SE+5d
WD.3.SE+5d
WW.3.SE+5d
WD.4.SE+5d
WW.4.SE+5d
WD.5.SE+5d
WW.5.SE+5d

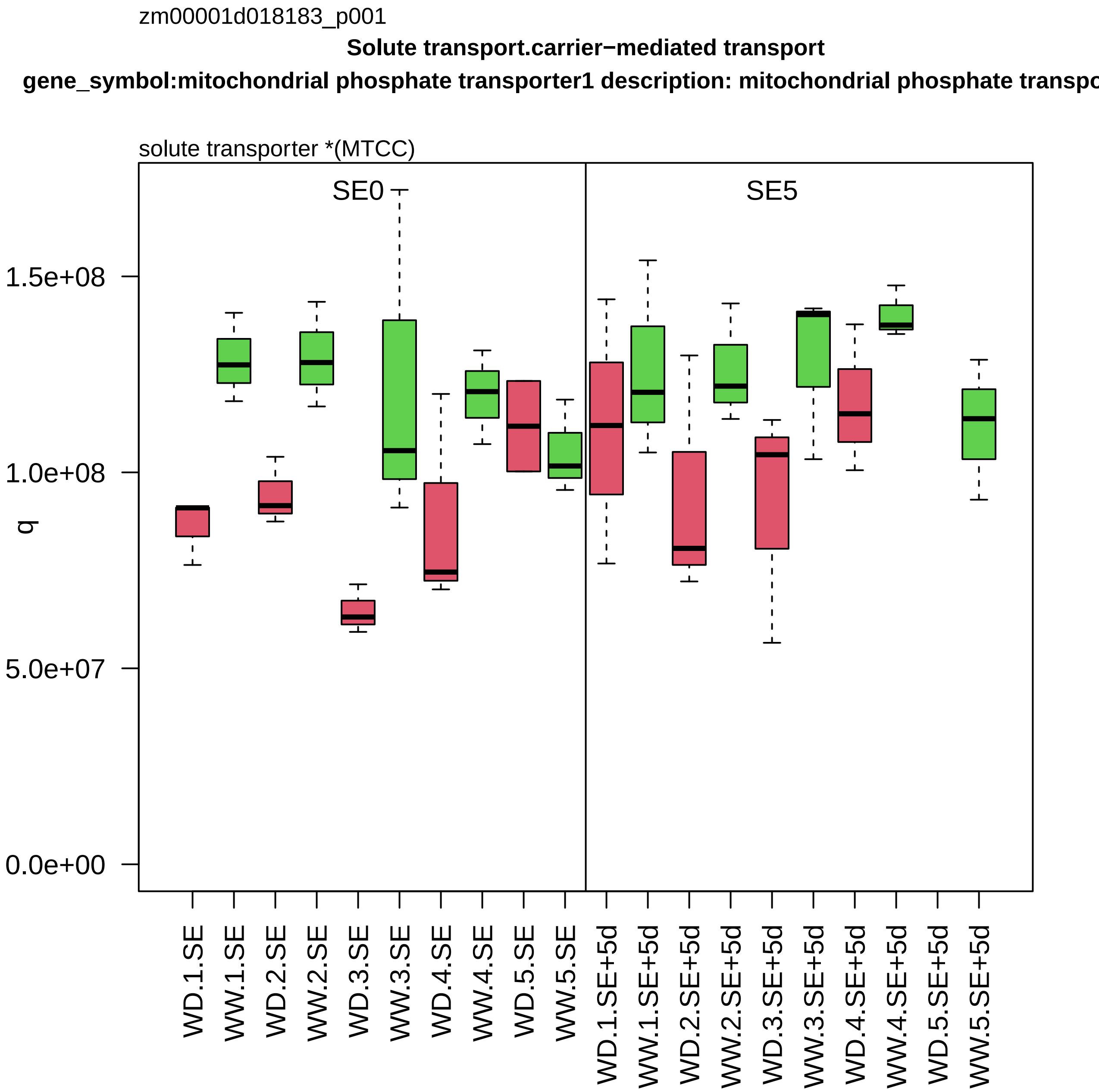
zm00001d018183_p001
Solute transport.carrier−mediated transport
gene_symbol:mitochondrial phosphate transporter1 description: mitochondrial phosphate transpo
solute transporter *(MTCC)
SE0
SE5
q
1.5e+08
1.0e+08
5.0e+07
0.0e+00
WD.1.SE
WW.1.SE
WD.2.SE
WW.2.SE
WD.3.SE
WW.3.SE
WD.4.SE
WW.4.SE
WD.5.SE
WW.5.SE
WD.1.SE+5d
WW.1.SE+5d
WD.2.SE+5d
WW.2.SE+5d
WD.3.SE+5d
WW.3.SE+5d
WD.4.SE+5d
WW.4.SE+5d
WD.5.SE+5d
WW.5.SE+5d

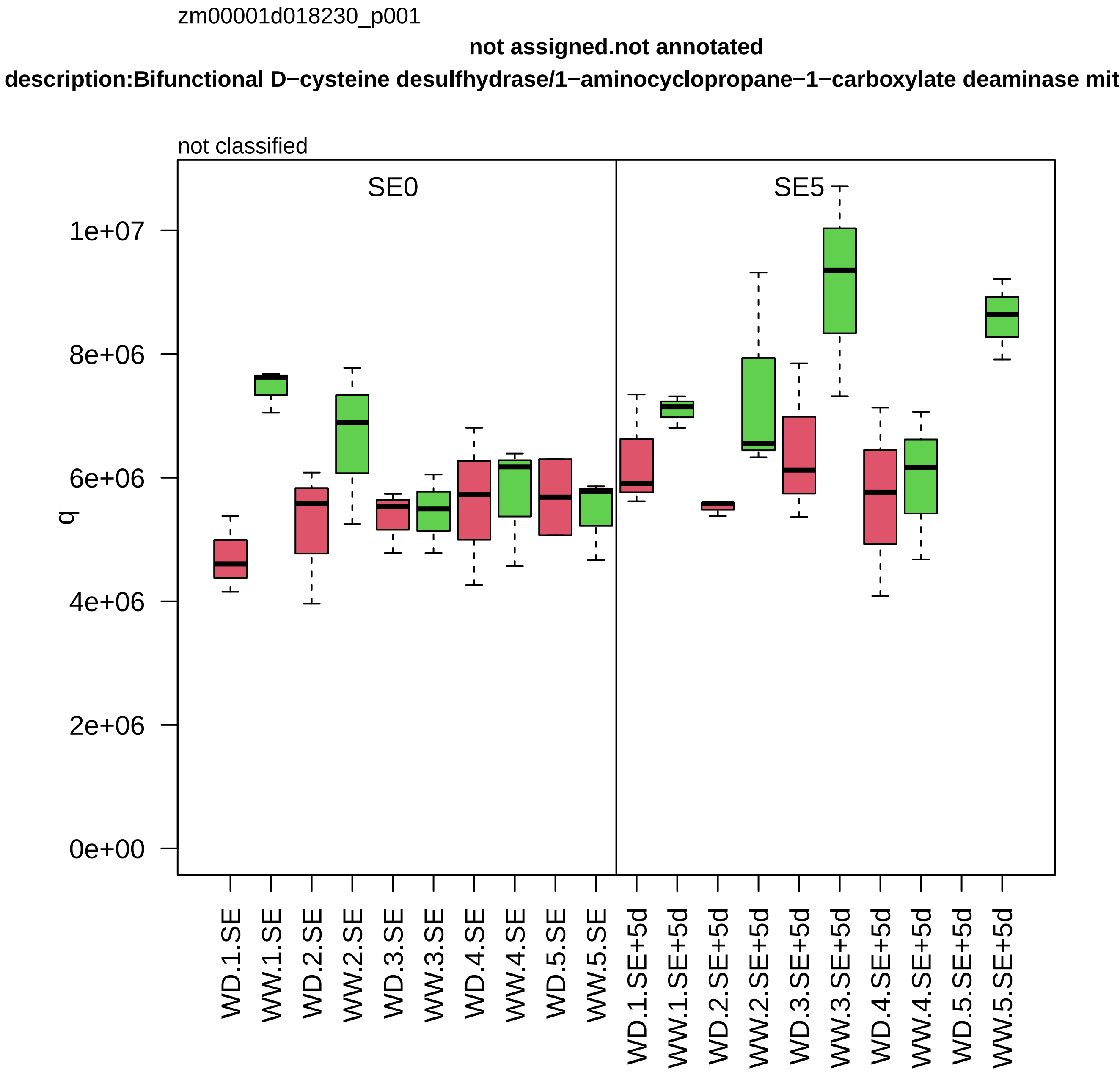

zm00001d018230_p001
not assigned.not annotated
description:Bifunctional D−cysteine desulfhydrase/1−aminocyclopropane−1−carboxylate deaminase mit
not classified
SE0
SE5
q
1e+07
8e+06
6e+06
4e+06
2e+06
0e+00
WD.1.SE
WW.1.SE
WD.2.SE
WW.2.SE
WD.3.SE
WW.3.SE
WD.4.SE
WW.4.SE
WD.5.SE
WW.5.SE
WD.1.SE+5d
WW.1.SE+5d
WD.2.SE+5d
WW.2.SE+5d
WD.3.SE+5d
WW.3.SE+5d
WD.4.SE+5d
WW.4.SE+5d
WD.5.SE+5d
WW.5.SE+5d

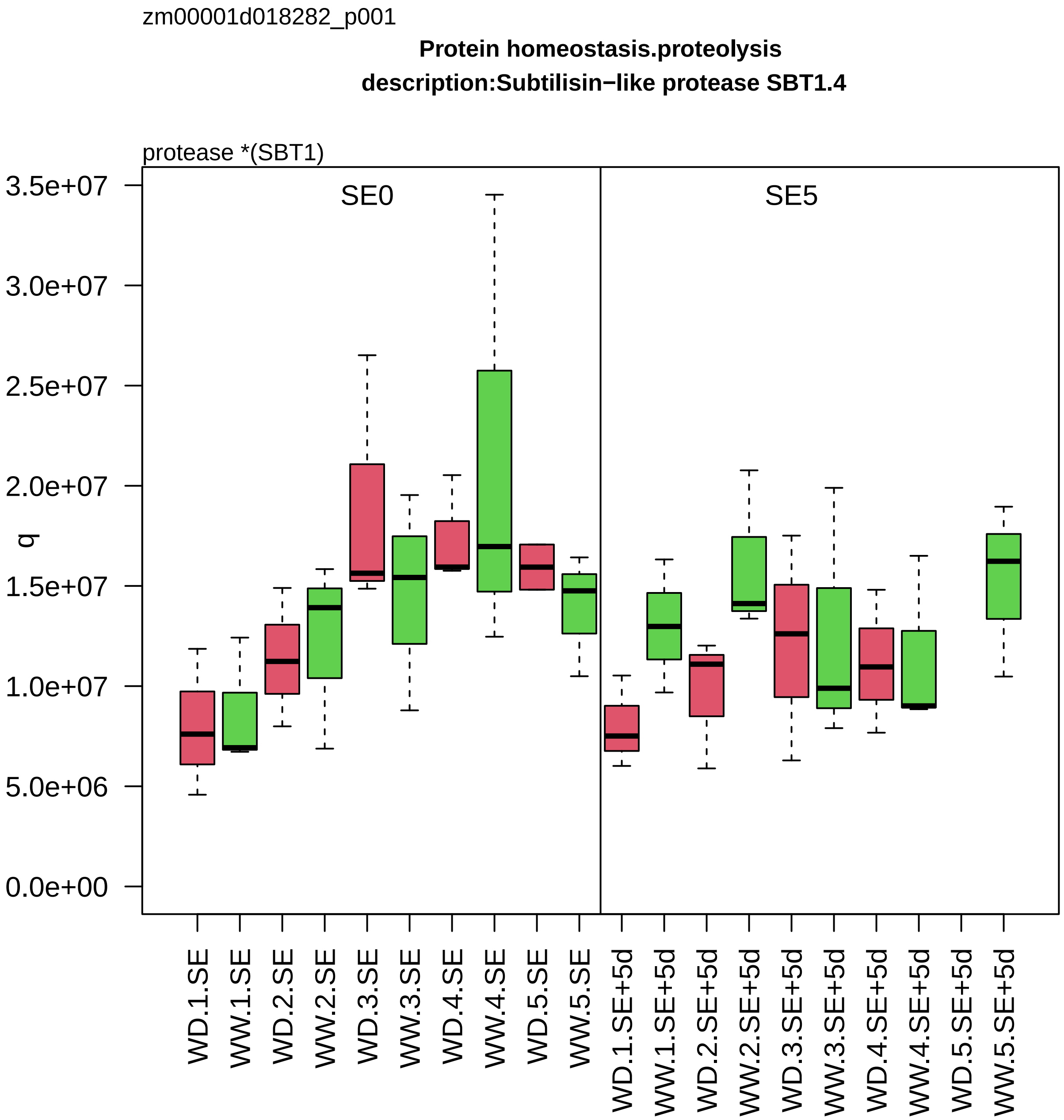

zm00001d018282_p001
Protein homeostasis.proteolysis
description:Subtilisin−like protease SBT1.4
protease *(SBT1)
q
SE0
SE5
3.5e+07
3.0e+07
2.5e+07
2.0e+07
1.5e+07
1.0e+07
5.0e+06
0.0e+00
WD.1.SE
WW.1.SE
WD.2.SE
WW.2.SE
WD.3.SE
WW.3.SE
WD.4.SE
WW.4.SE
WD.5.SE
WW.5.SE
WD.1.SE+5d
WW.1.SE+5d
WD.2.SE+5d
WW.2.SE+5d
WD.3.SE+5d
WW.3.SE+5d
WD.4.SE+5d
WW.4.SE+5d
WD.5.SE+5d
WW.5.SE+5d

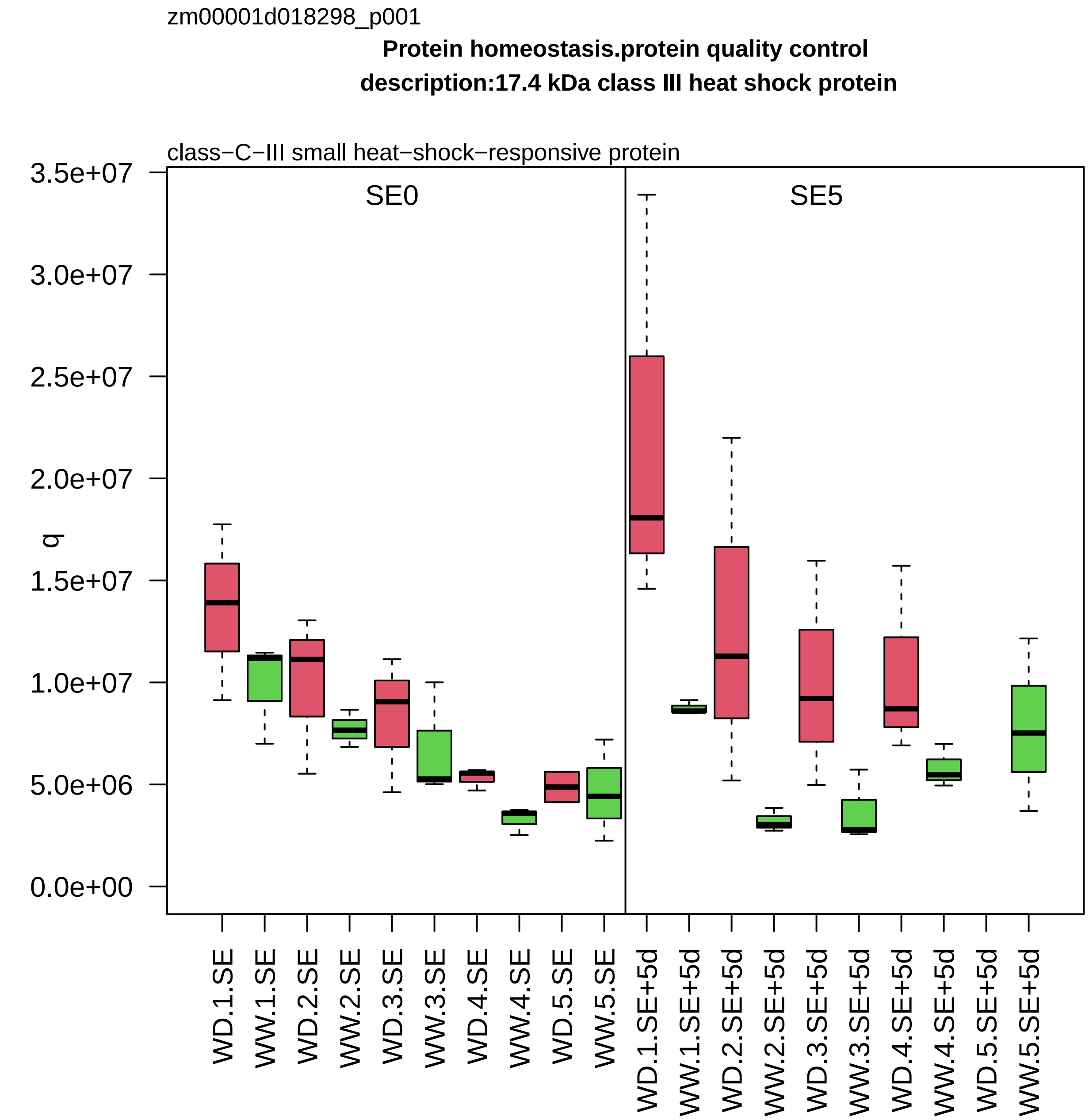
zm00001d018298_p001
Protein homeostasis.protein quality control
description:17.4 kDa class III heat shock protein
class−C−III small heat−shock−responsive protein
SE0
SE5
q
3.5e+07
3.0e+07
2.5e+07
2.0e+07
1.5e+07
1.0e+07
5.0e+06
0.0e+00
WD.1.SE
WW.1.SE
WD.2.SE
WW.2.SE
WD.3.SE
WW.3.SE
WD.4.SE
WW.4.SE
WD.5.SE
WW.5.SE
WD.1.SE+5d
WW.1.SE+5d
WD.2.SE+5d
WW.2.SE+5d
WD.3.SE+5d
WW.3.SE+5d
WD.4.SE+5d
WW.4.SE+5d
WD.5.SE+5d
WW.5.SE+5d

zm00001d018309_p003

**Protein homeostasis.ubiquitin−proteasome system**

**gene_symbol:MSS1 homolog description: MSS1 homolog**

regulatory component *(RPT1) of 26S proteasome

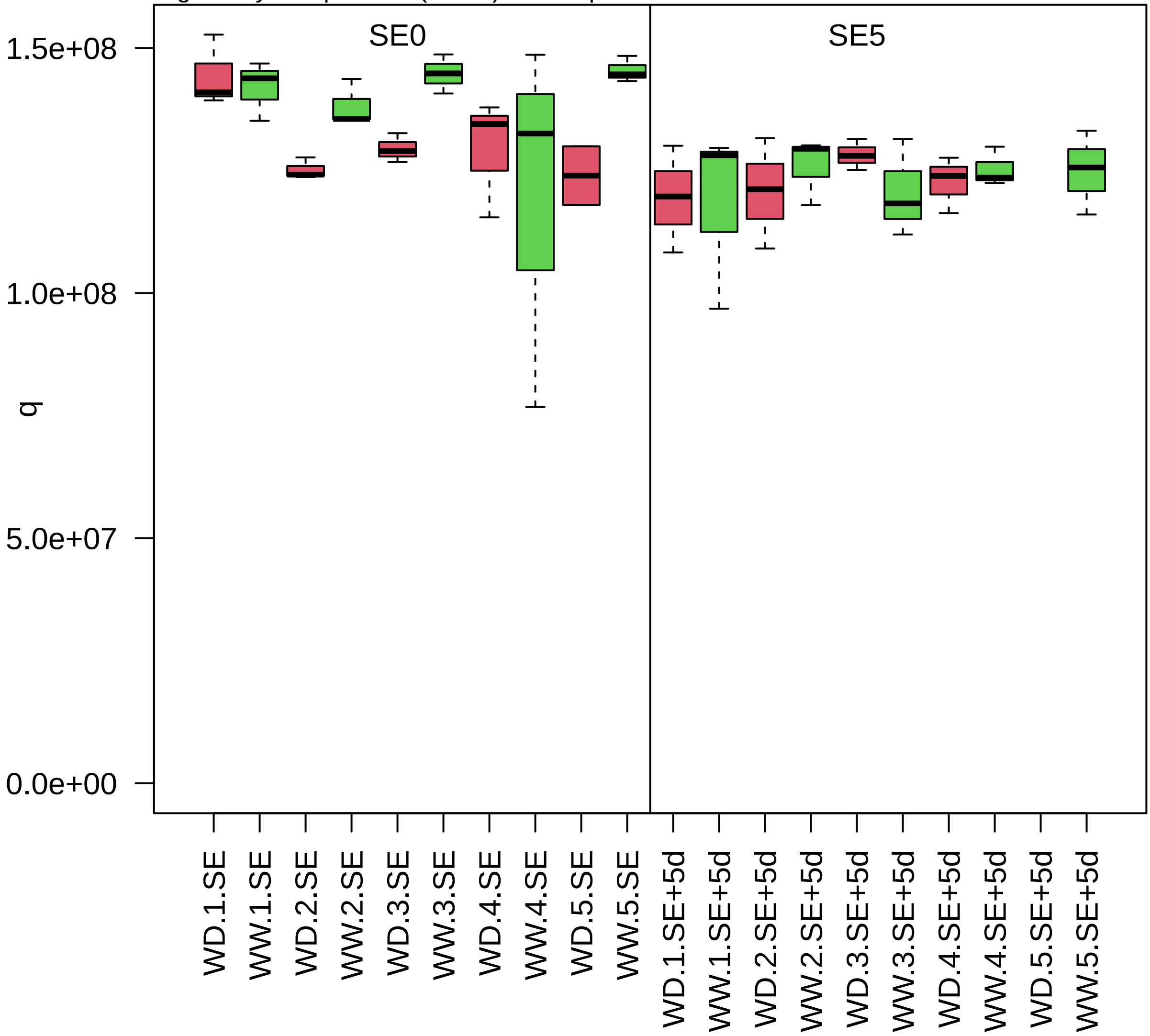

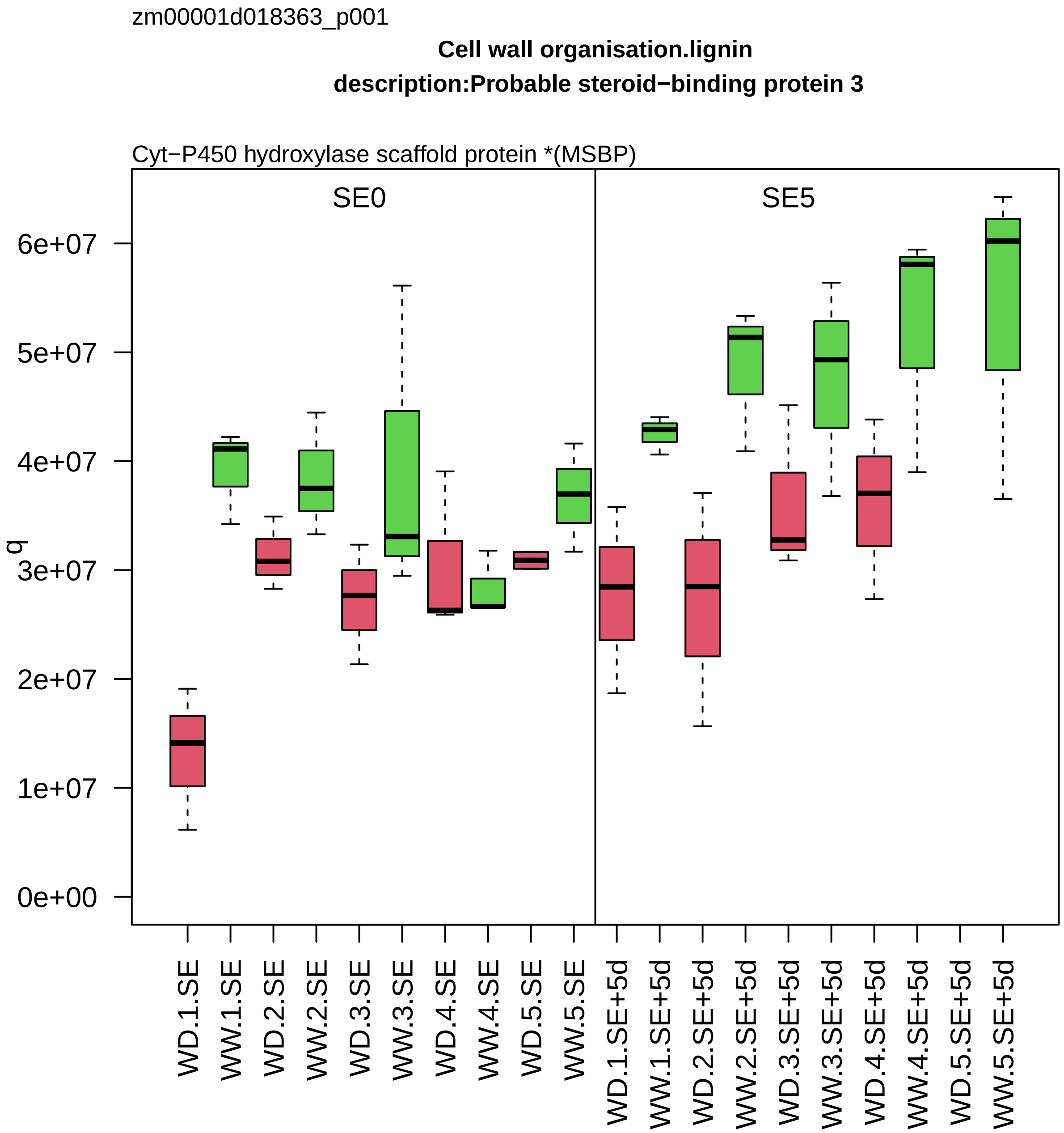
zm00001d018363_p001
Cell wall organisation.lignin
description:Probable steroid−binding protein 3
Cyt−P450 hydroxylase scaffold protein *(MSBP)
SE0
SE5
q
0e+00
1e+07
2e+07
3e+07
4e+07
5e+07
6e+07
WD.1.SE
WW.1.SE
WD.2.SE
WW.2.SE
WD.3.SE
WW.3.SE
WD.4.SE
WW.4.SE
WD.5.SE
WW.5.SE
WD.1.SE+5d
WW.1.SE+5d
WD.2.SE+5d
WW.2.SE+5d
WD.3.SE+5d
WW.3.SE+5d
WD.4.SE+5d
WW.4.SE+5d
WD.5.SE+5d
WW.5.SE+5d

zm00001d018370_p001

**Protein homeostasis.proteolysis**

**gene_symbol:pco101682(581) description:Leucine aminopeptidase 2 chloroplastic**

M17−class leucyl aminopeptidase *(LAP)

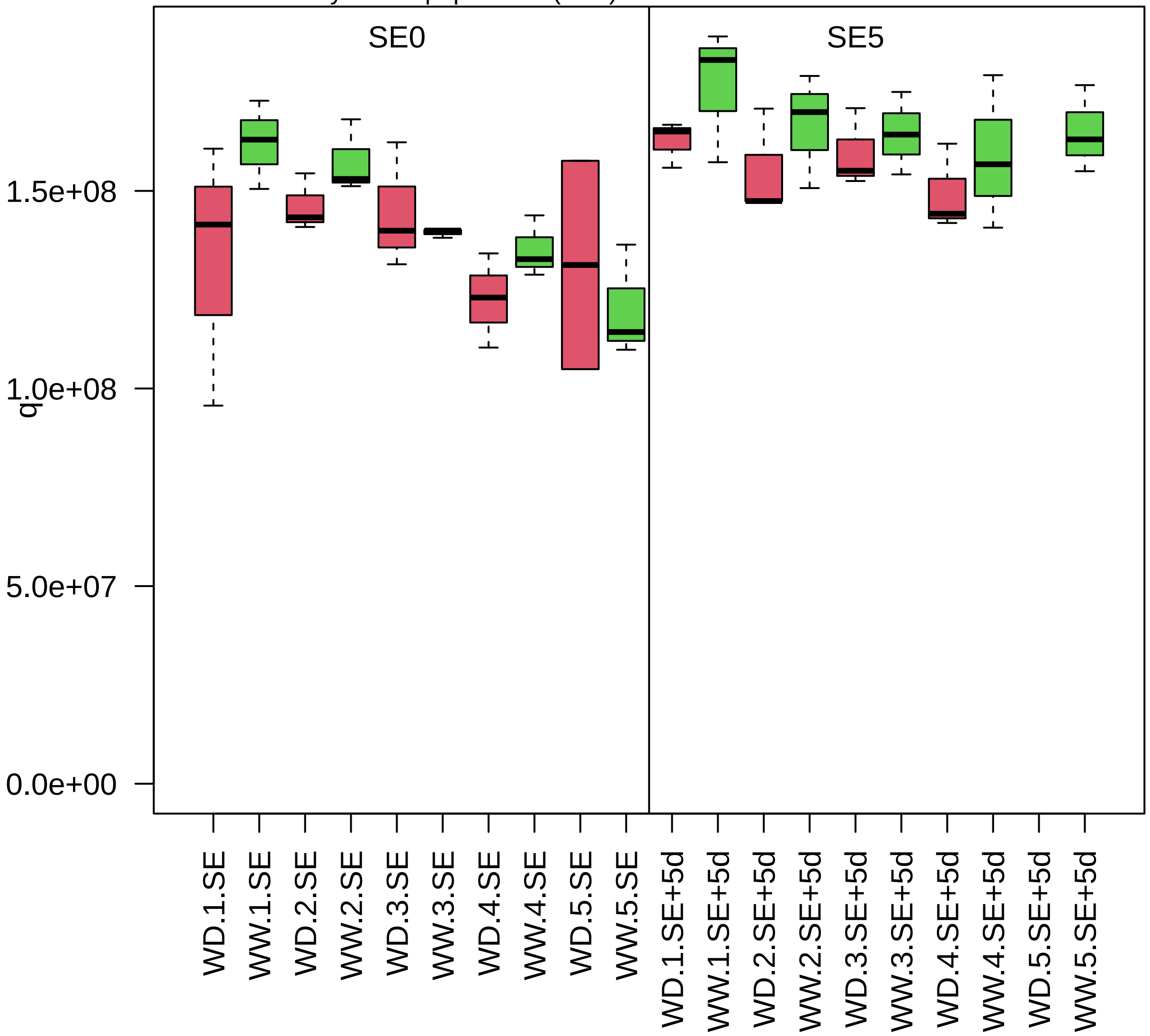

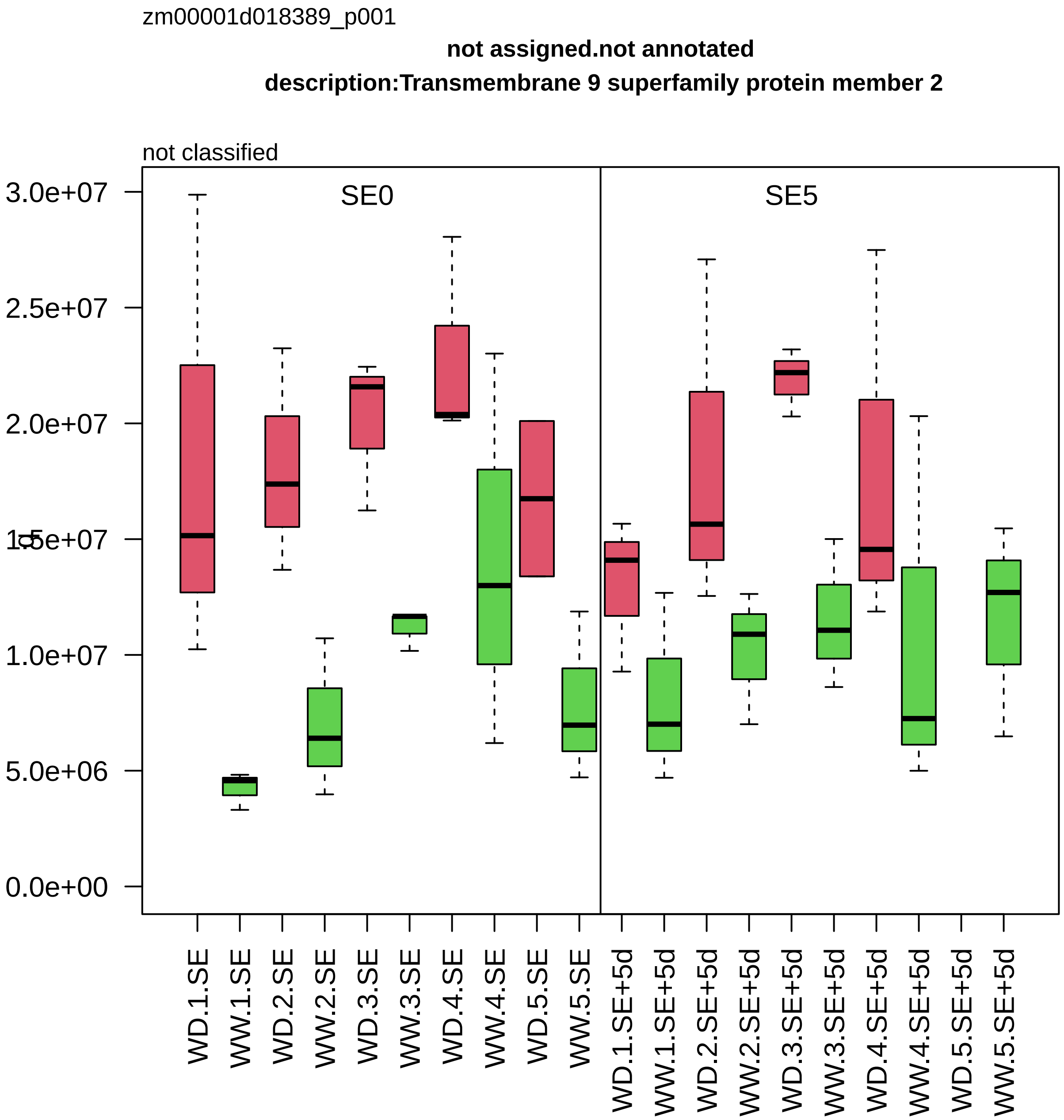

zm00001d018389_p001
not assigned.not annotated
description:Transmembrane 9 superfamily protein member 2
not classified
SE0
SE5
3.0e+07
2.5e+07
2.0e+07
1.5e+07
1.0e+07
5.0e+06
0.0e+00
WD.1.SE
WW.1.SE
WD.2.SE
WW.2.SE
WD.3.SE
WW.3.SE
WD.4.SE
WW.4.SE
WD.5.SE
WW.5.SE
WD.1.SE+5d
WW.1.SE+5d
WD.2.SE+5d
WW.2.SE+5d
WD.3.SE+5d
WW.3.SE+5d
WD.4.SE+5d
WW.4.SE+5d
WD.5.SE+5d
WW.5.SE+5d

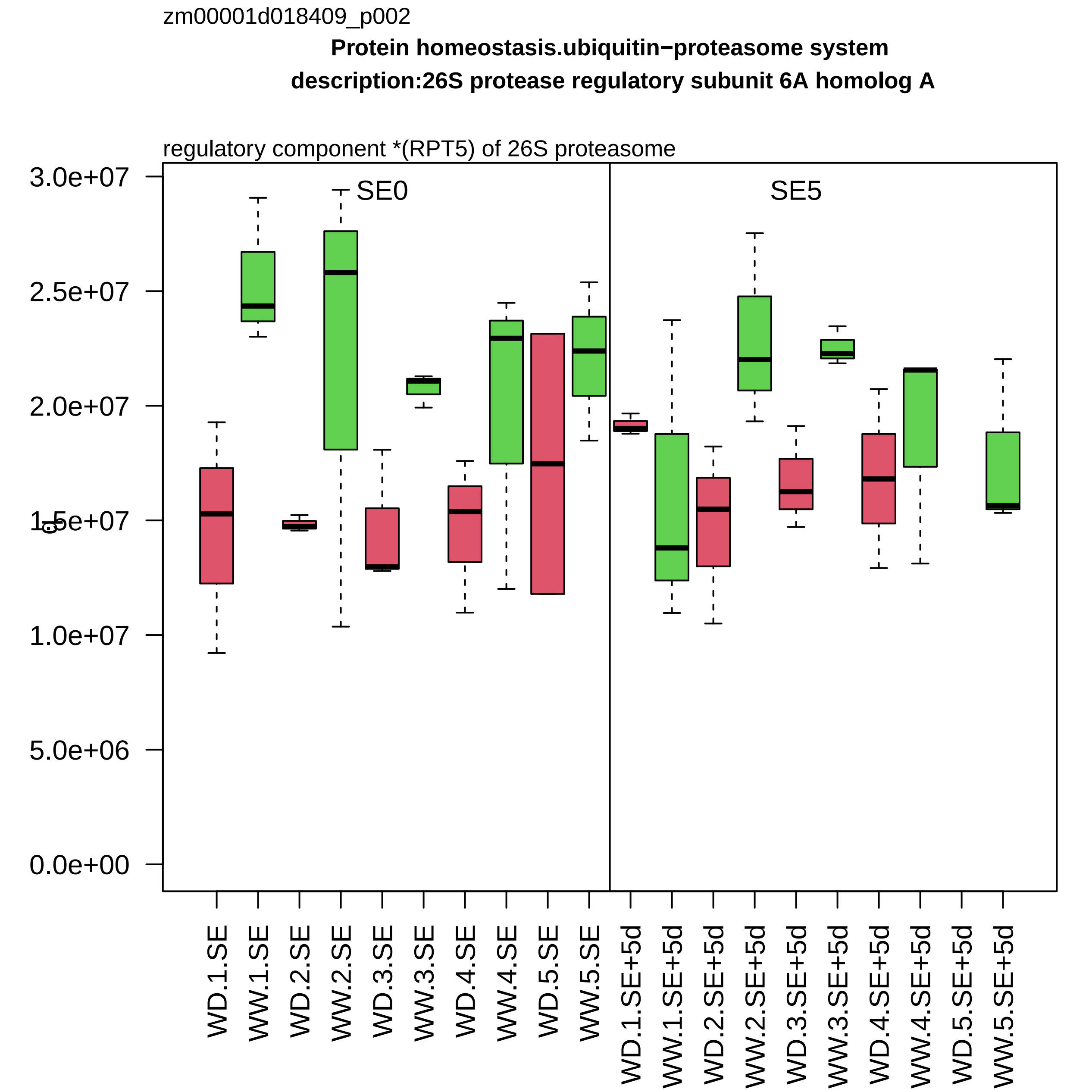

zm00001d018409_p002
Protein homeostasis.ubiquitin−proteasome system
description:26S protease regulatory subunit 6A homolog A
regulatory component *(RPT5) of 26S proteasome
SE0
SE5
3.0e+07
2.5e+07
2.0e+07
1.5e+07
1.0e+07
5.0e+06
0.0e+00
WD.1.SE
WW.1.SE
WD.2.SE
WW.2.SE
WD.3.SE
WW.3.SE
WD.4.SE
WW.4.SE
WD.5.SE
WW.5.SE
WD.1.SE+5d
WW.1.SE+5d
WD.2.SE+5d
WW.2.SE+5d
WD.3.SE+5d
WW.3.SE+5d
WD.4.SE+5d
WW.4.SE+5d
WD.5.SE+5d
WW.5.SE+5d

zm00001d018415_p002

**Cell division.DNA replication**

**gene_symbol:proliferating cell nuclear antigen1 description:Proliferating cell nuclear antigen**

sliding clamp protein *(PCNA)

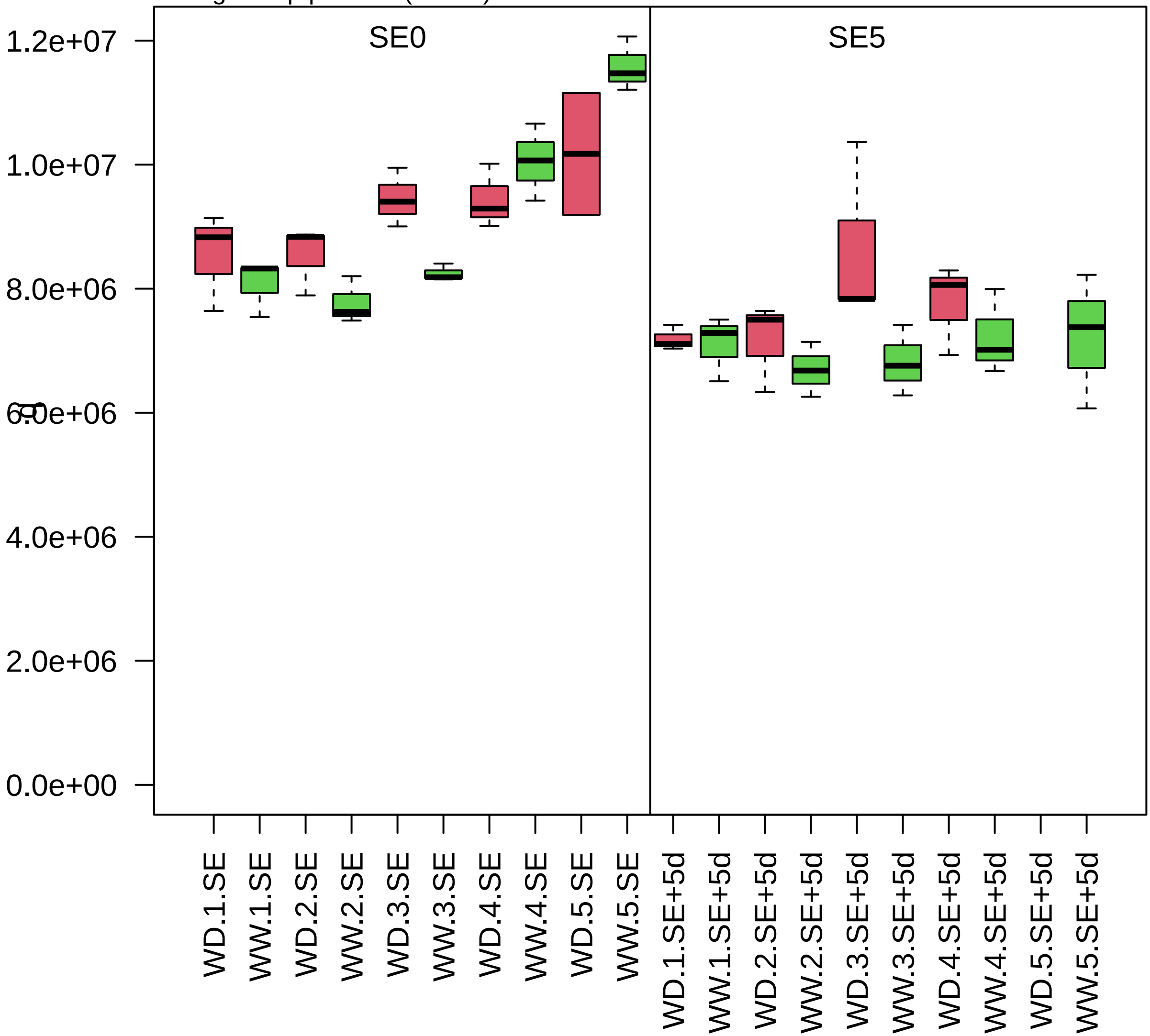

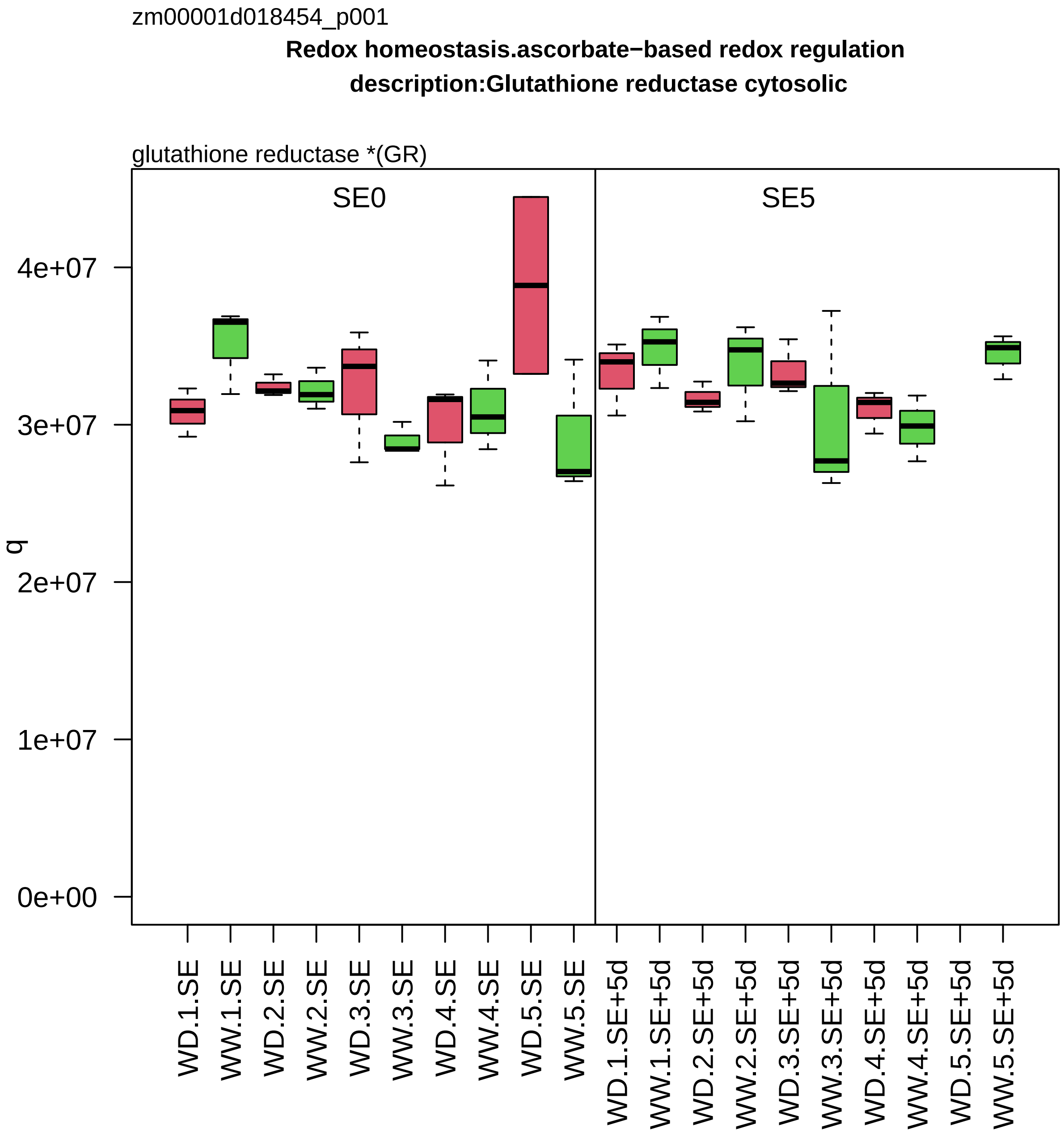
zm00001d018454_p001
Redox homeostasis.ascorbate−based redox regulation
description:Glutathione reductase cytosolic
glutathione reductase *(GR)
SE0
SE5
q
0e+00
1e+07
2e+07
3e+07
4e+07
WD.1.SE
WW.1.SE
WD.2.SE
WW.2.SE
WD.3.SE
WW.3.SE
WD.4.SE
WW.4.SE
WD.5.SE
WW.5.SE
WD.1.SE+5d
WW.1.SE+5d
WD.2.SE+5d
WW.2.SE+5d
WD.3.SE+5d
WW.3.SE+5d
WD.4.SE+5d
WW.4.SE+5d
WD.5.SE+5d
WW.5.SE+5d

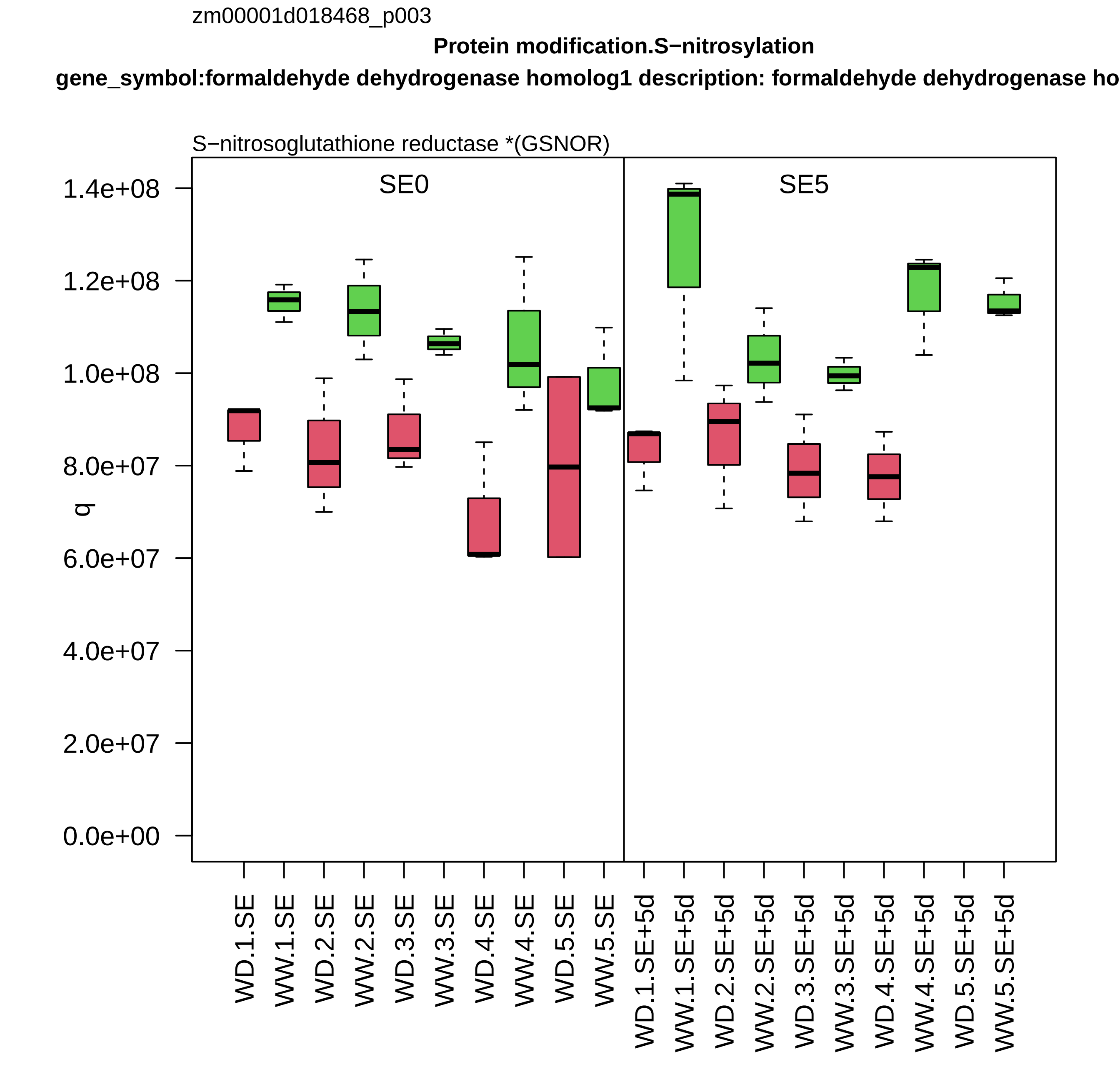

zm00001d018468_p003
Protein modification.S−nitrosylation
gene_symbol:formaldehyde dehydrogenase homolog1 description: formaldehyde dehydrogenase ho
S−nitrosoglutathione reductase *(GSNOR)
SE0
SE5
q
1.4e+08
1.2e+08
1.0e+08
8.0e+07
6.0e+07
4.0e+07
2.0e+07
0.0e+00
WD.1.SE
WW.1.SE
WD.2.SE
WW.2.SE
WD.3.SE
WW.3.SE
WD.4.SE
WW.4.SE
WD.5.SE
WW.5.SE
WD.1.SE+5d
WW.1.SE+5d
WD.2.SE+5d
WW.2.SE+5d
WD.3.SE+5d
WW.3.SE+5d
WD.4.SE+5d
WW.4.SE+5d
WD.5.SE+5d
WW.5.SE+5d

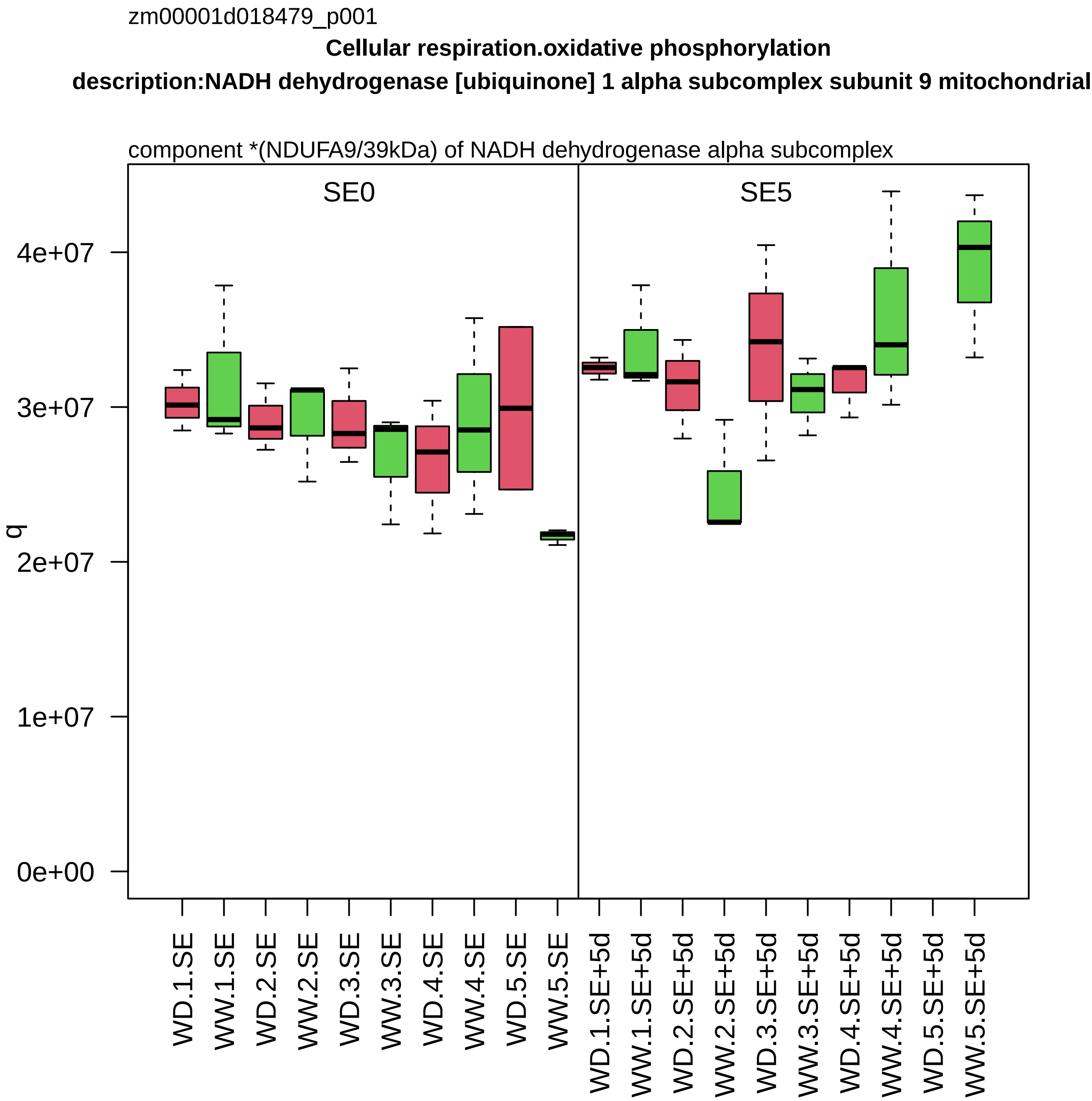

zm00001d018479_p001
Cellular respiration.oxidative phosphorylation
description:NADH dehydrogenase [ubiquinone] 1 alpha subcomplex subunit 9 mitochondrial
component *(NDUFA9/39kDa) of NADH dehydrogenase alpha subcomplex
SE0
SE5
q
0e+00
1e+07
2e+07
3e+07
4e+07
WD.1.SE
WW.1.SE
WD.2.SE
WW.2.SE
WD.3.SE
WW.3.SE
WD.4.SE
WW.4.SE
WD.5.SE
WW.5.SE
WD.1.SE+5d
WW.1.SE+5d
WD.2.SE+5d
WW.2.SE+5d
WD.3.SE+5d
WW.3.SE+5d
WD.4.SE+5d
WW.4.SE+5d
WD.5.SE+5d
WW.5.SE+5d

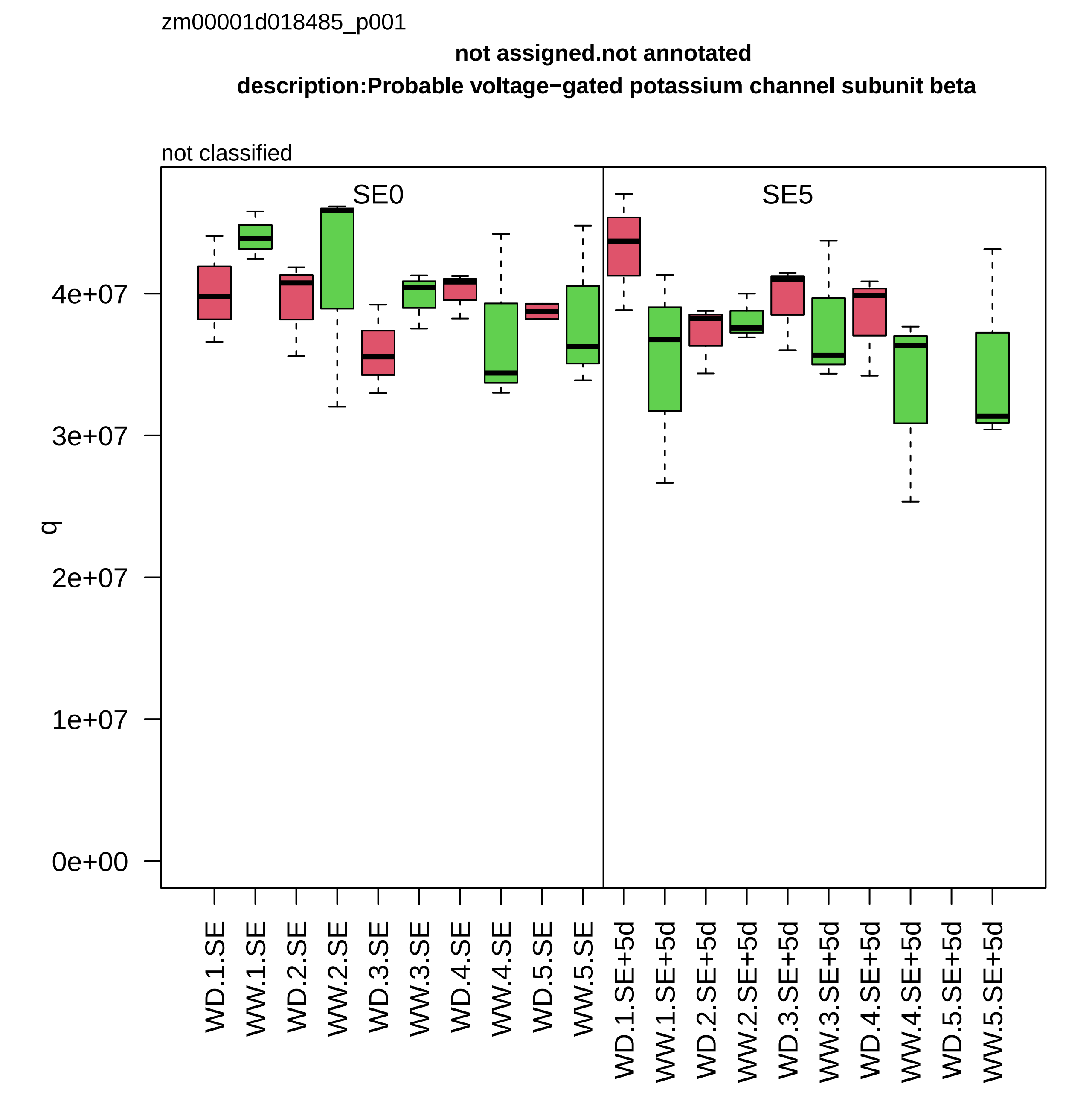
zm00001d018485_p001
not assigned.not annotated
description:Probable voltage−gated potassium channel subunit beta
not classified
SE0
SE5
q
0e+00
1e+07
2e+07
3e+07
4e+07
WD.1.SE
WW.1.SE
WD.2.SE
WW.2.SE
WD.3.SE
WW.3.SE
WD.4.SE
WW.4.SE
WD.5.SE
WW.5.SE
WD.1.SE+5d
WW.1.SE+5d
WD.2.SE+5d
WW.2.SE+5d
WD.3.SE+5d
WW.3.SE+5d
WD.4.SE+5d
WW.4.SE+5d
WD.5.SE+5d
WW.5.SE+5d

zm00001d018487_p001

**Lipid metabolism.fatty acid metabolism**

**description:3−ketoacyl−CoA thiolase 2 peroxisomal**

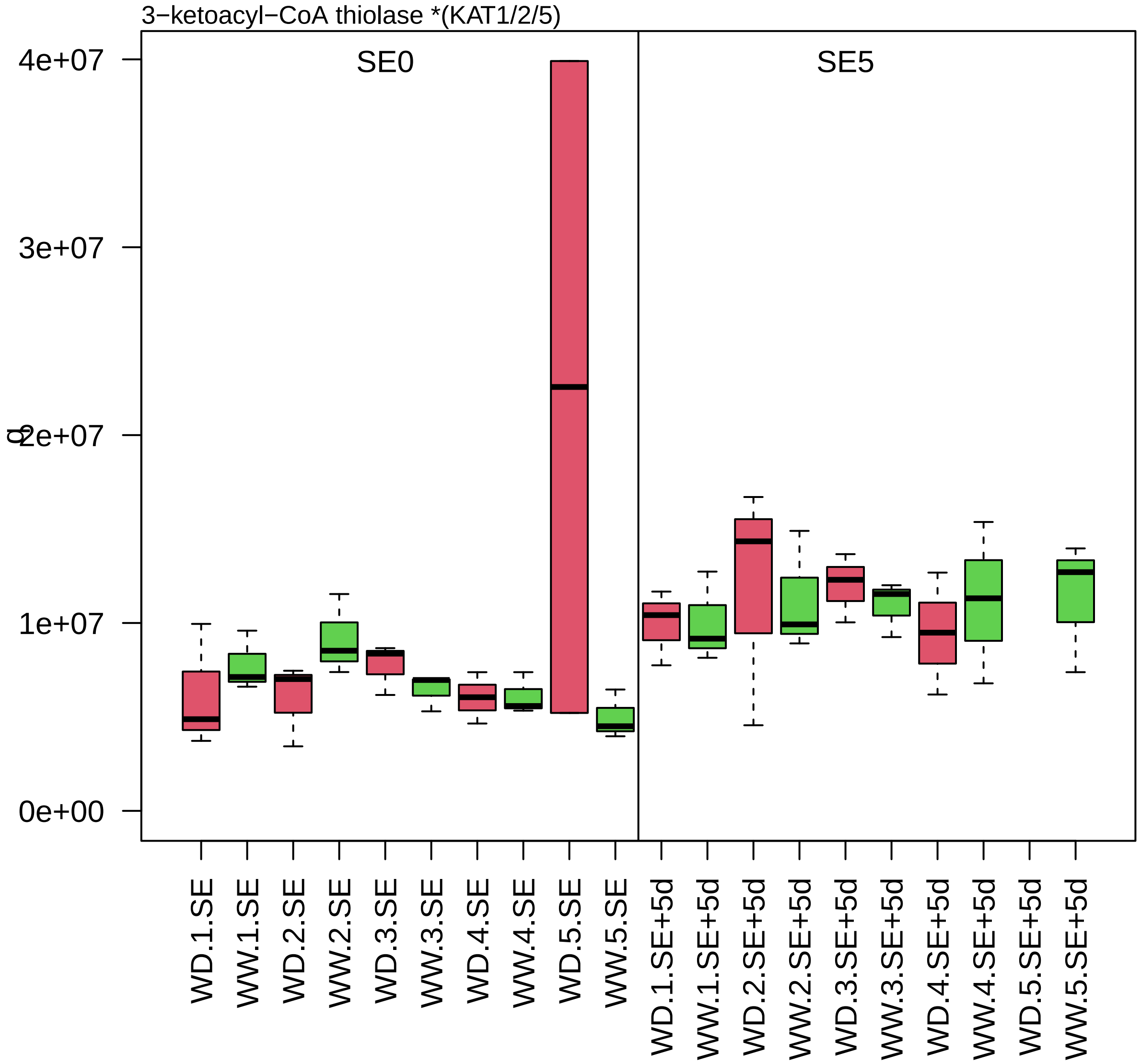

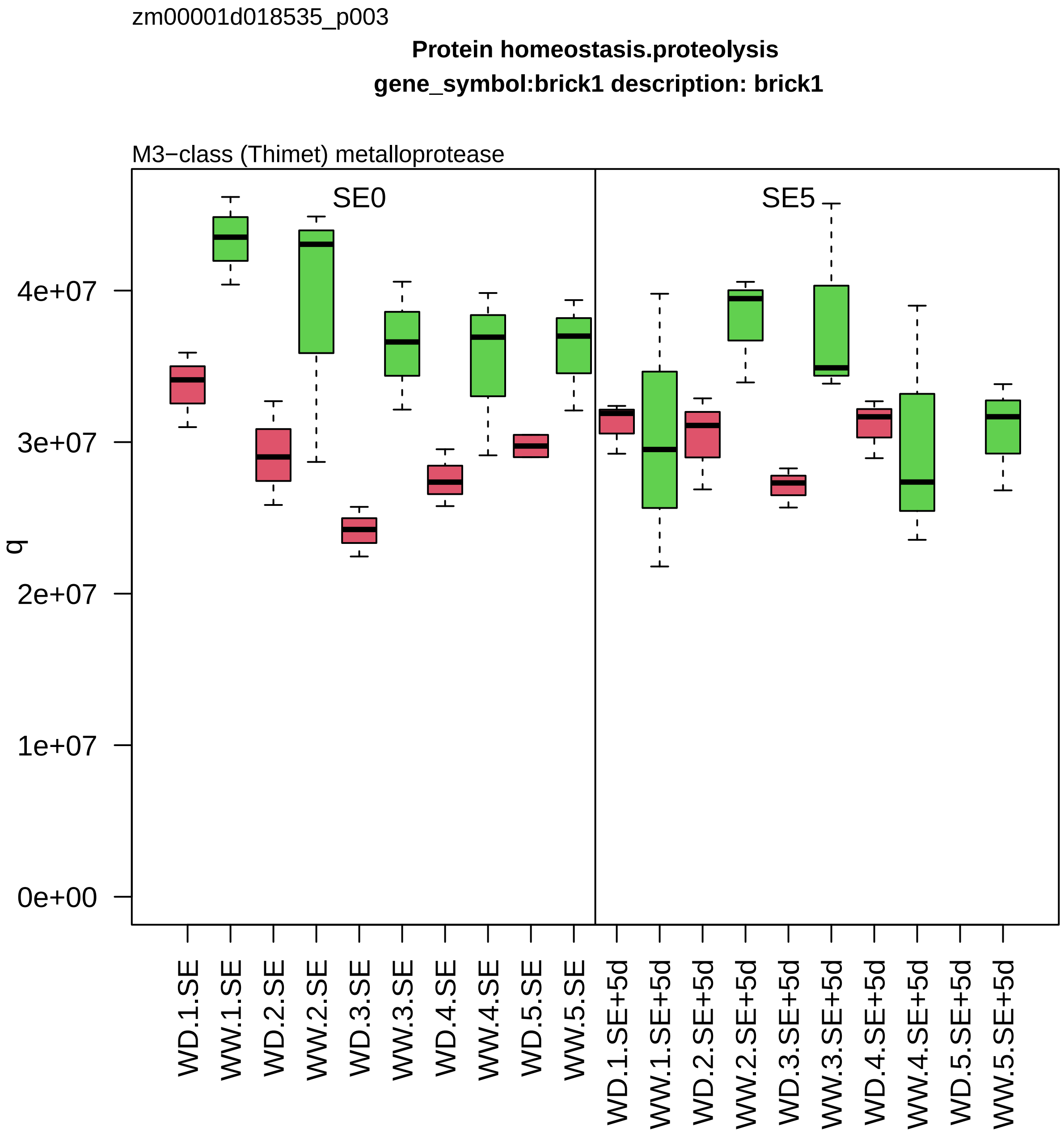

zm00001d018535_p003
Protein homeostasis.proteolysis
gene_symbol:brick1 description: brick1
M3−class (Thimet) metalloprotease
SE0
SE5
q
0e+00
1e+07
2e+07
3e+07
4e+07
WD.1.SE
WW.1.SE
WD.2.SE
WW.2.SE
WD.3.SE
WW.3.SE
WD.4.SE
WW.4.SE
WD.5.SE
WW.5.SE
WD.1.SE+5d
WW.1.SE+5d
WD.2.SE+5d
WW.2.SE+5d
WD.3.SE+5d
WW.3.SE+5d
WD.4.SE+5d
WW.4.SE+5d
WD.5.SE+5d
WW.5.SE+5d

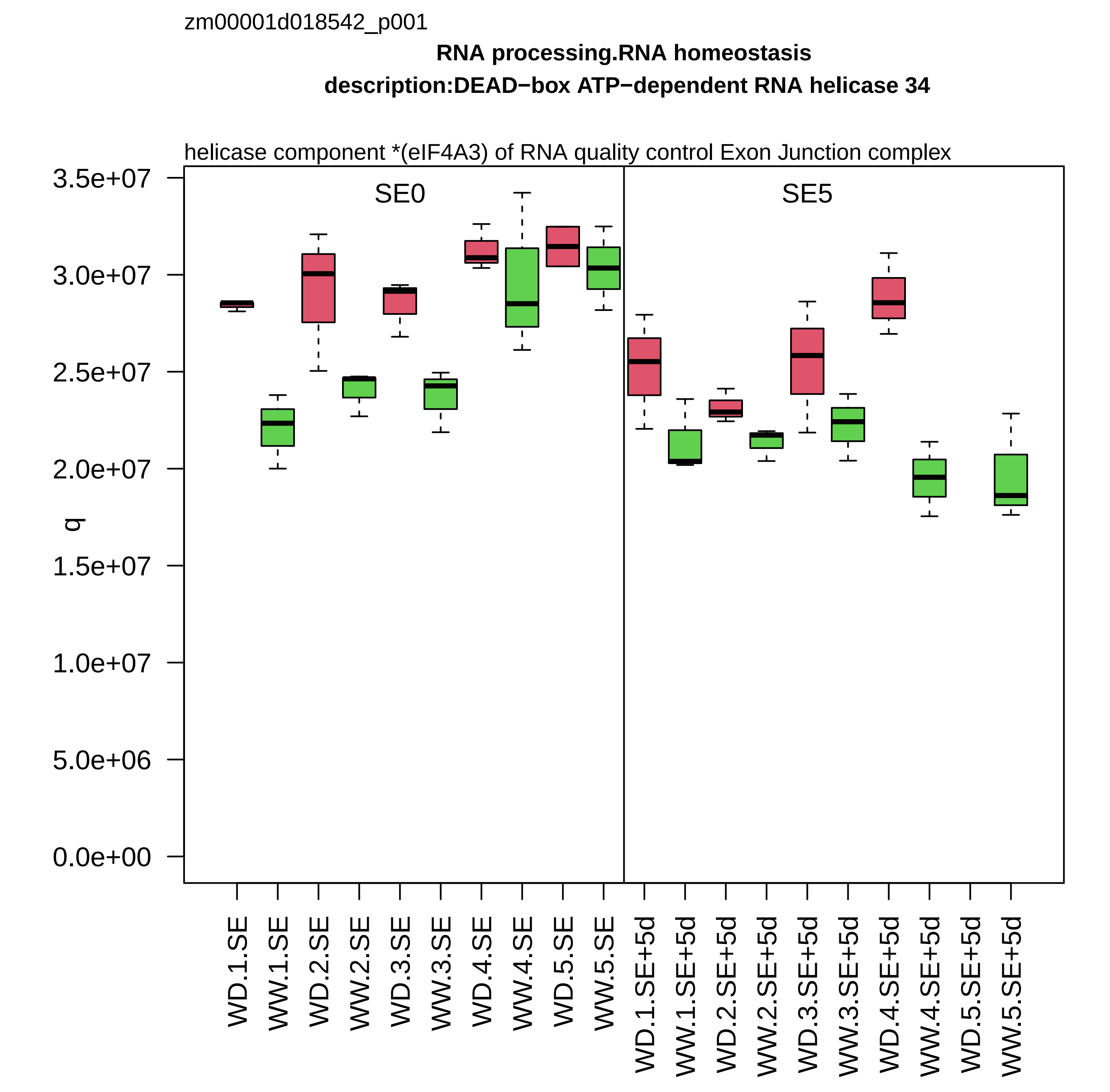

zm00001d018542_p001
RNA processing.RNA homeostasis
description:DEAD−box ATP−dependent RNA helicase 34
helicase component *(eIF4A3) of RNA quality control Exon Junction complex
SE0
SE5
q
3.5e+07
3.0e+07
2.5e+07
2.0e+07
1.5e+07
1.0e+07
5.0e+06
0.0e+00
WD.1.SE
WW.1.SE
WD.2.SE
WW.2.SE
WD.3.SE
WW.3.SE
WD.4.SE
WW.4.SE
WD.5.SE
WW.5.SE
WD.1.SE+5d
WW.1.SE+5d
WD.2.SE+5d
WW.2.SE+5d
WD.3.SE+5d
WW.3.SE+5d
WD.4.SE+5d
WW.4.SE+5d
WD.5.SE+5d
WW.5.SE+5d

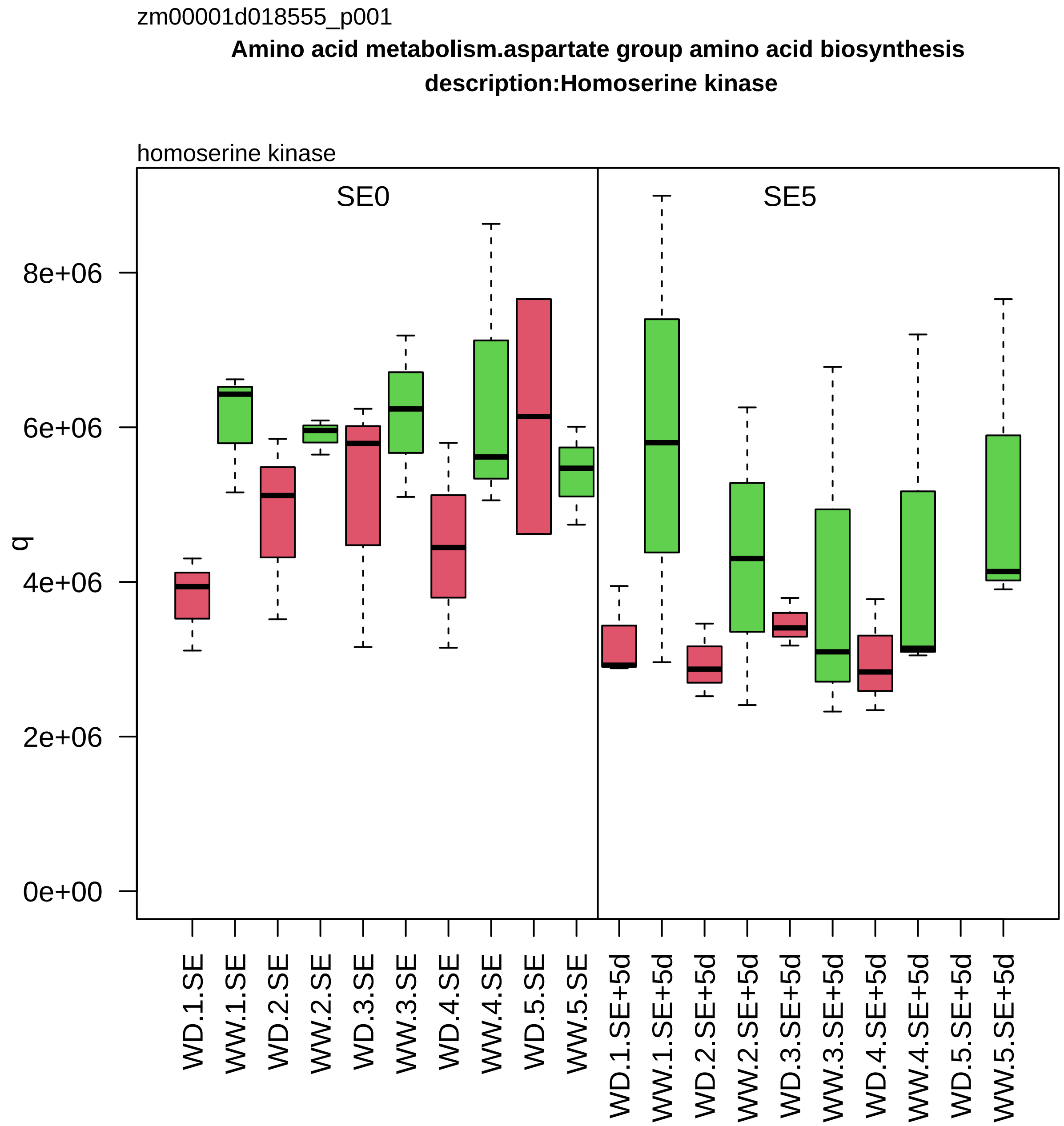
zm00001d018555_p001
Amino acid metabolism.aspartate group amino acid biosynthesis
description:Homoserine kinase
homoserine kinase
SE0
SE5
q
0e+00
2e+06
4e+06
6e+06
8e+06
WD.1.SE
WW.1.SE
WD.2.SE
WW.2.SE
WD.3.SE
WW.3.SE
WD.4.SE
WW.4.SE
WD.5.SE
WW.5.SE
WD.1.SE+5d
WW.1.SE+5d
WD.2.SE+5d
WW.2.SE+5d
WD.3.SE+5d
WW.3.SE+5d
WD.4.SE+5d
WW.4.SE+5d
WD.5.SE+5d
WW.5.SE+5d

zm00001d018632_p001

**Coenzyme metabolism.pyridoxal−phosphate homeostasis**

**description:Pyridoxal 5'−phosphate synthase−like subunit PDX1.2**

synthase component of pyridoxal 5−phosphate synthase complex

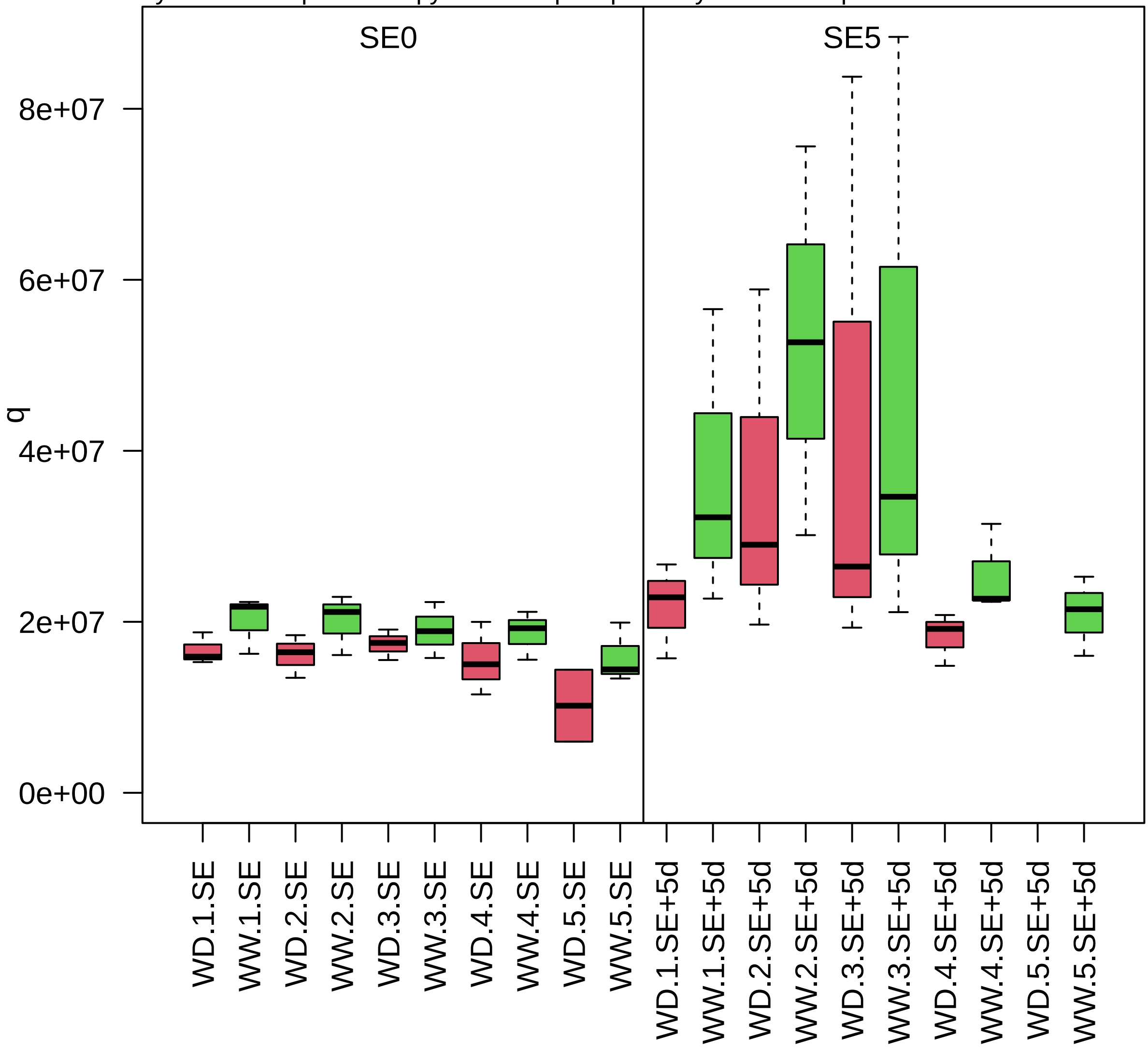

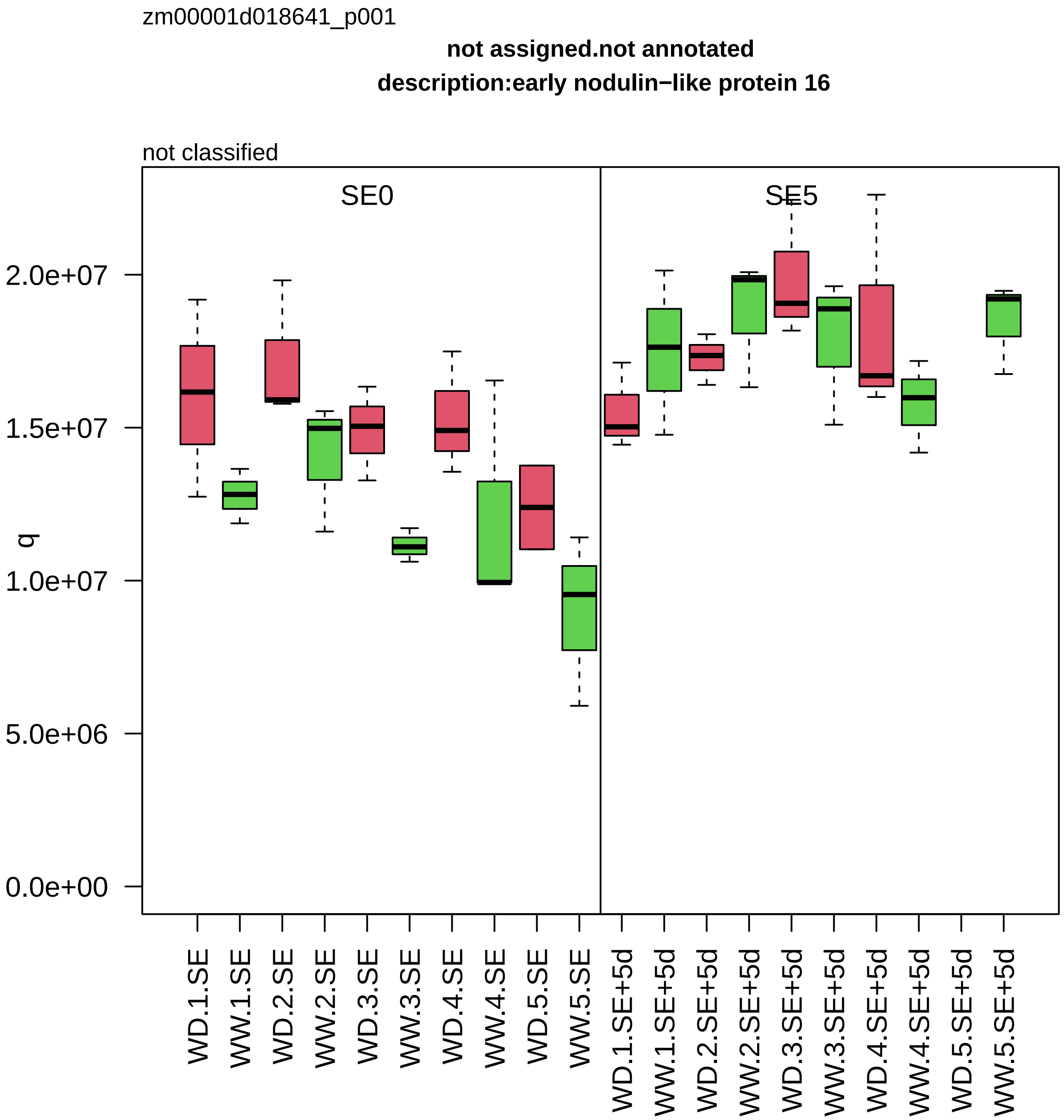

zm00001d018641_p001
not assigned.not annotated
description:early nodulin−like protein 16
not classified
SE0
SE5
q
2.0e+07
1.5e+07
1.0e+07
5.0e+06
0.0e+00
WD.1.SE
WW.1.SE
WD.2.SE
WW.2.SE
WD.3.SE
WW.3.SE
WD.4.SE
WW.4.SE
WD.5.SE
WW.5.SE
WD.1.SE+5d
WW.1.SE+5d
WD.2.SE+5d
WW.2.SE+5d
WD.3.SE+5d
WW.3.SE+5d
WD.4.SE+5d
WW.4.SE+5d
WD.5.SE+5d
WW.5.SE+5d

zm00001d018694_p001

**Cytoskeleton organisation.actin and tubulin folding**
**description:T−complex protein 1 subunit epsilon**

subunit epsilon *(CCT5) of CCT chaperonin folding complex

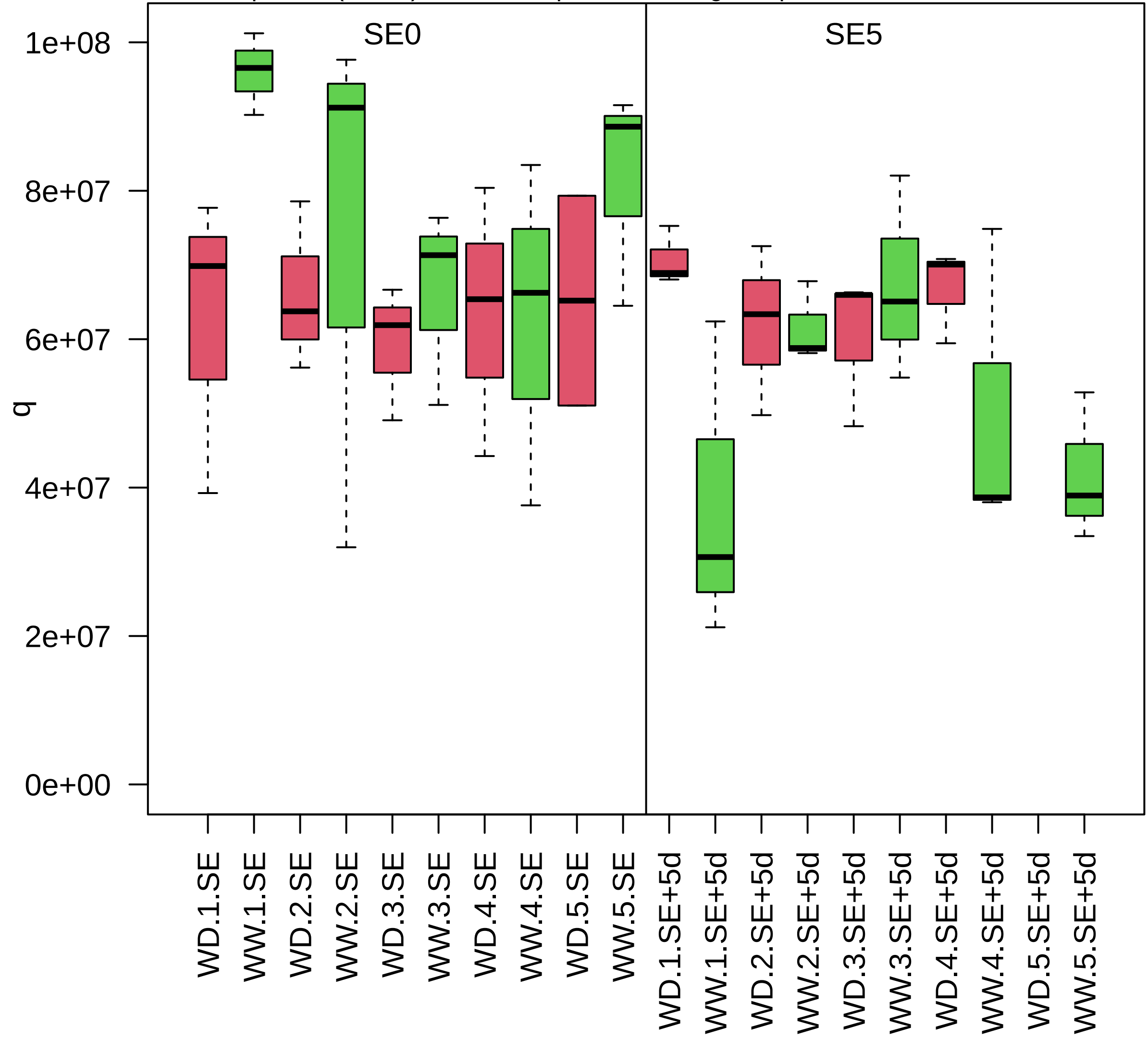

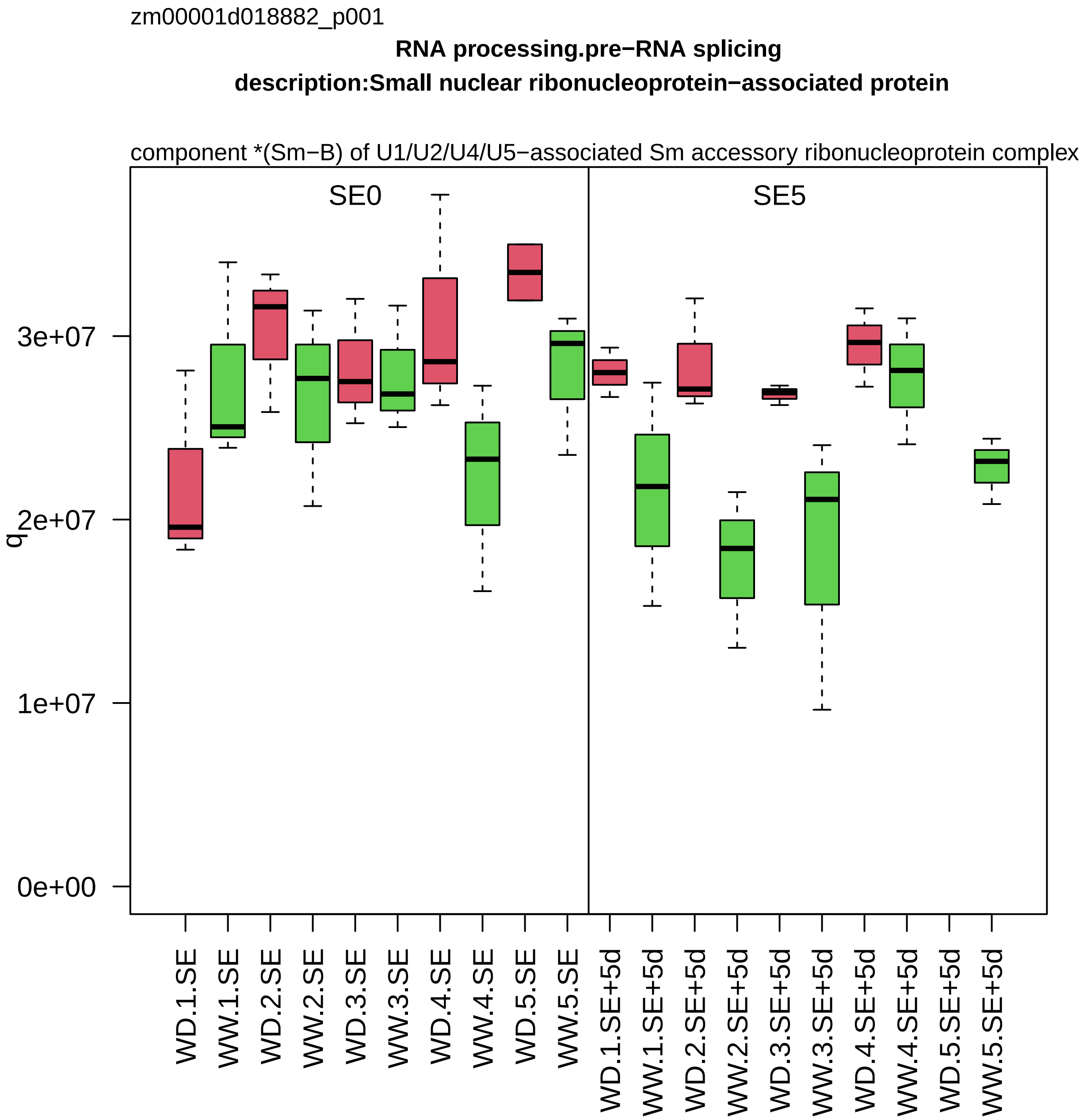
zm00001d018882_p001
RNA processing.pre−RNA splicing
description:Small nuclear ribonucleoprotein−associated protein
component *(Sm−B) of U1/U2/U4/U5−associated Sm accessory ribonucleoprotein complex
SE0
SE5
q
3e+07
2e+07
1e+07
0e+00
WD.1.SE
WW.1.SE
WD.2.SE
WW.2.SE
WD.3.SE
WW.3.SE
WD.4.SE
WW.4.SE
WD.5.SE
WW.5.SE
WD.1.SE+5d
WW.1.SE+5d
WD.2.SE+5d
WW.2.SE+5d
WD.3.SE+5d
WW.3.SE+5d
WD.4.SE+5d
WW.4.SE+5d
WD.5.SE+5d
WW.5.SE+5d

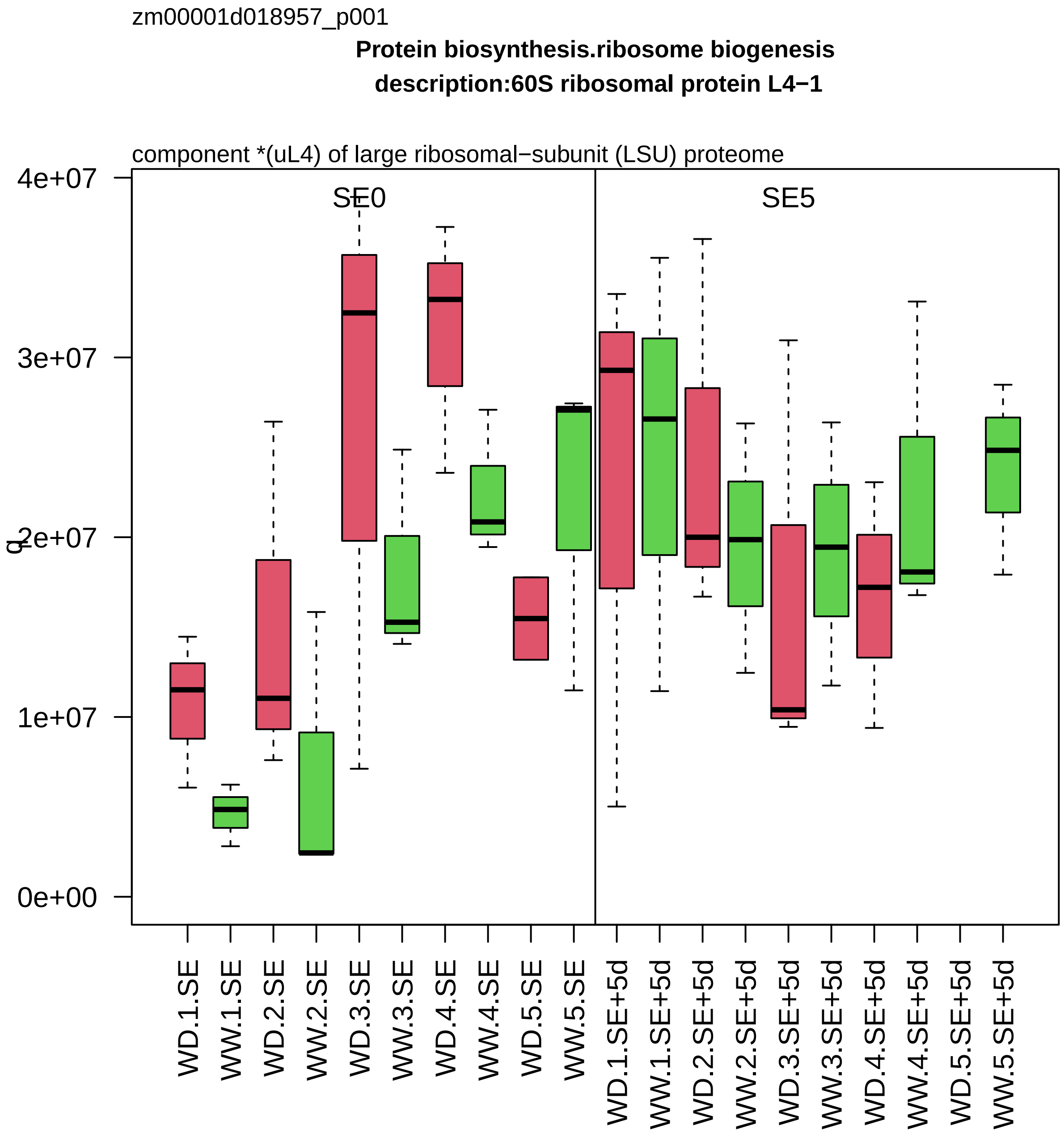

zm00001d018957_p001
Protein biosynthesis.ribosome biogenesis
description:60S ribosomal protein L4−1
component *(uL4) of large ribosomal−subunit (LSU) proteome
SE0
SE5
q
4e+07
3e+07
2e+07
1e+07
0e+00
WD.1.SE
WW.1.SE
WD.2.SE
WW.2.SE
WD.3.SE
WW.3.SE
WD.4.SE
WW.4.SE
WD.5.SE
WW.5.SE
WD.1.SE+5d
WW.1.SE+5d
WD.2.SE+5d
WW.2.SE+5d
WD.3.SE+5d
WW.3.SE+5d
WD.4.SE+5d
WW.4.SE+5d
WD.5.SE+5d
WW.5.SE+5d

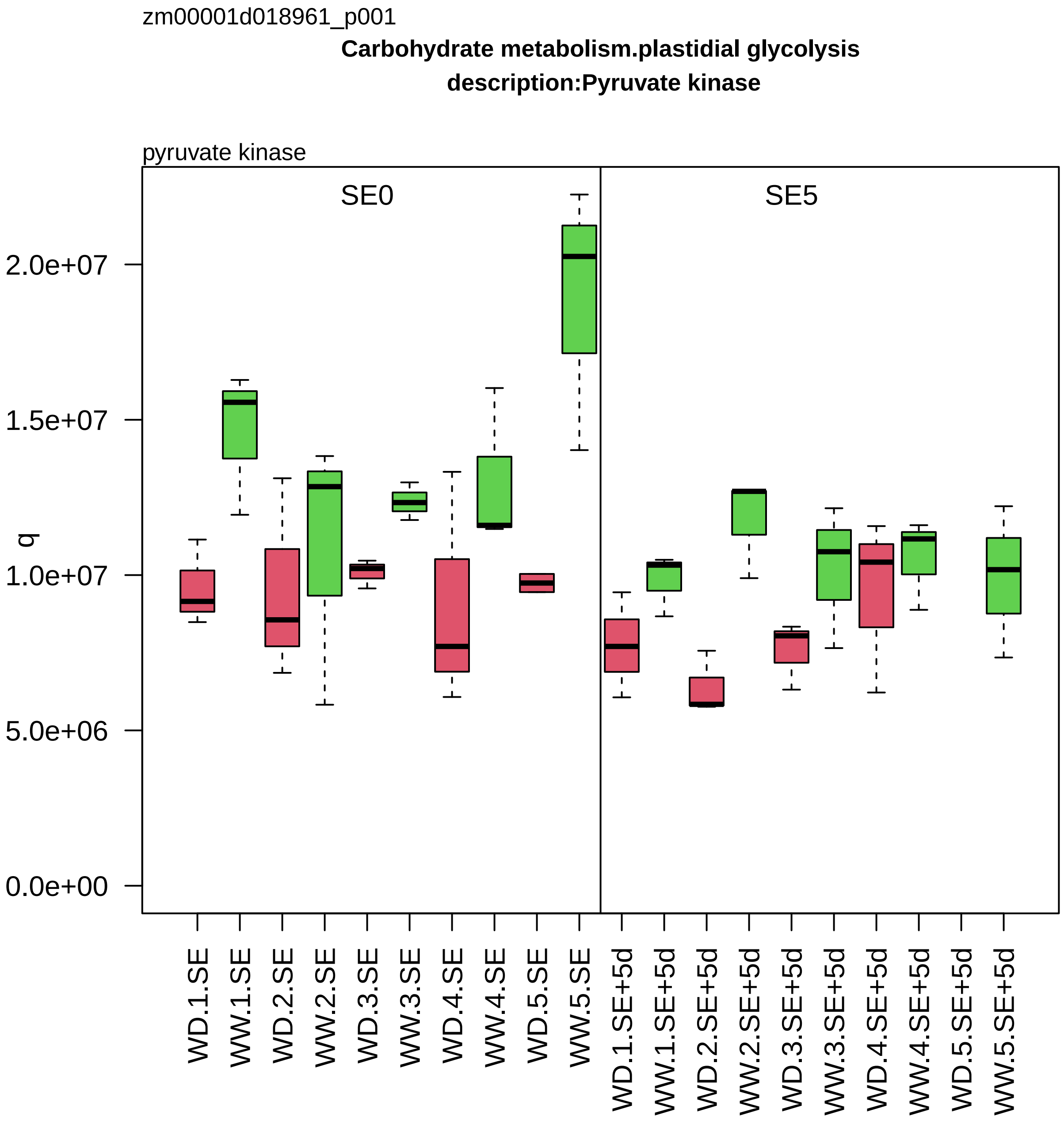
zm00001d018961_p001
Carbohydrate metabolism.plastidial glycolysis
description:Pyruvate kinase
pyruvate kinase
SE0
SE5
q
2.0e+07
1.5e+07
1.0e+07
5.0e+06
0.0e+00
WD.1.SE
WW.1.SE
WD.2.SE
WW.2.SE
WD.3.SE
WW.3.SE
WD.4.SE
WW.4.SE
WD.5.SE
WW.5.SE
WD.1.SE+5d
WW.1.SE+5d
WD.2.SE+5d
WW.2.SE+5d
WD.3.SE+5d
WW.3.SE+5d
WD.4.SE+5d
WW.4.SE+5d
WD.5.SE+5d
WW.5.SE+5d

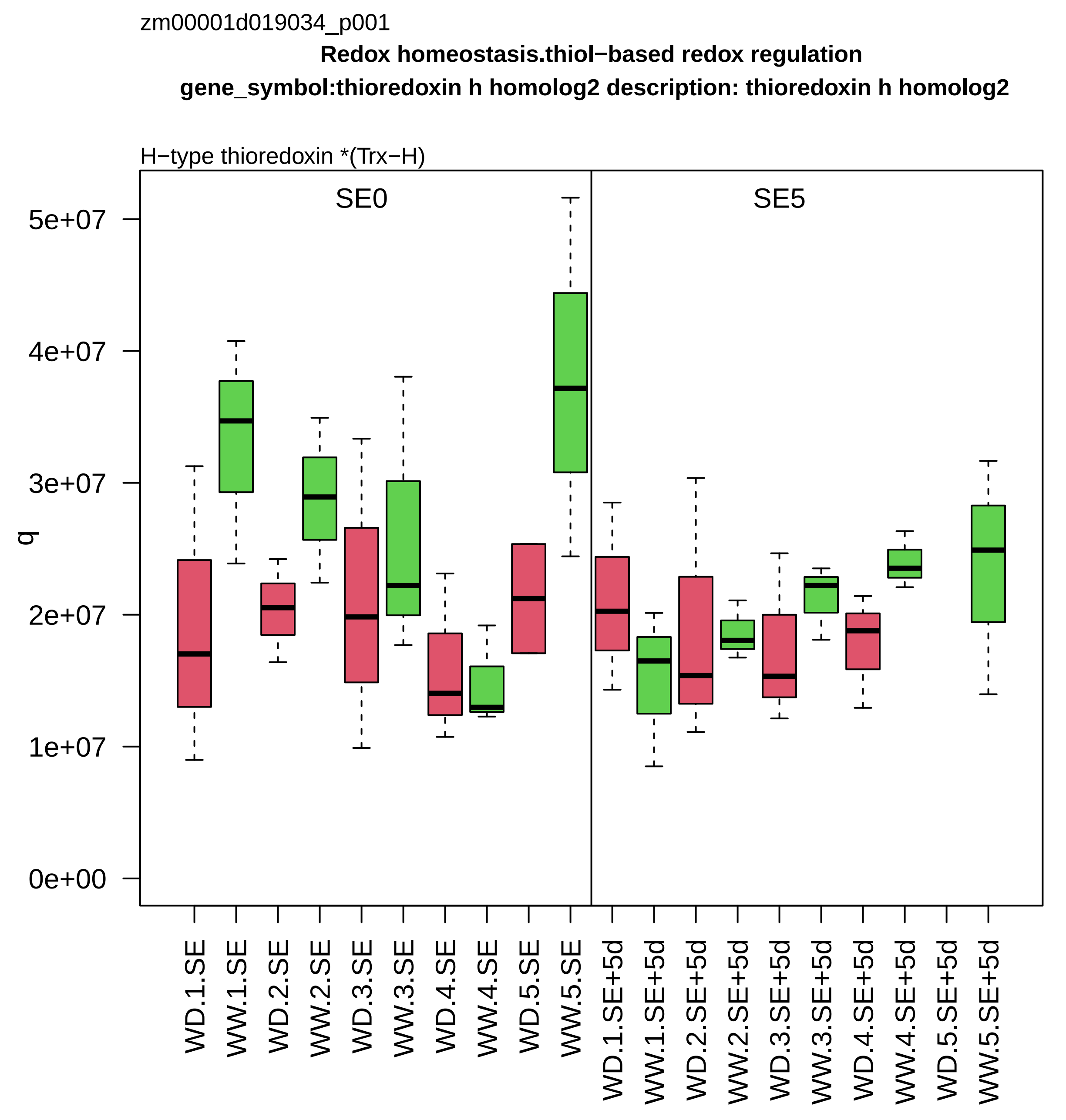
zm00001d019034_p001
Redox homeostasis.thiol−based redox regulation
gene_symbol:thioredoxin h homolog2 description: thioredoxin h homolog2
H−type thioredoxin *(Trx−H)
SE0
SE5
q
0e+00
1e+07
2e+07
3e+07
4e+07
5e+07
WD.1.SE
WW.1.SE
WD.2.SE
WW.2.SE
WD.3.SE
WW.3.SE
WD.4.SE
WW.4.SE
WD.5.SE
WW.5.SE
WD.1.SE+5d
WW.1.SE+5d
WD.2.SE+5d
WW.2.SE+5d
WD.3.SE+5d
WW.3.SE+5d
WD.4.SE+5d
WW.4.SE+5d
WD.5.SE+5d
WW.5.SE+5d

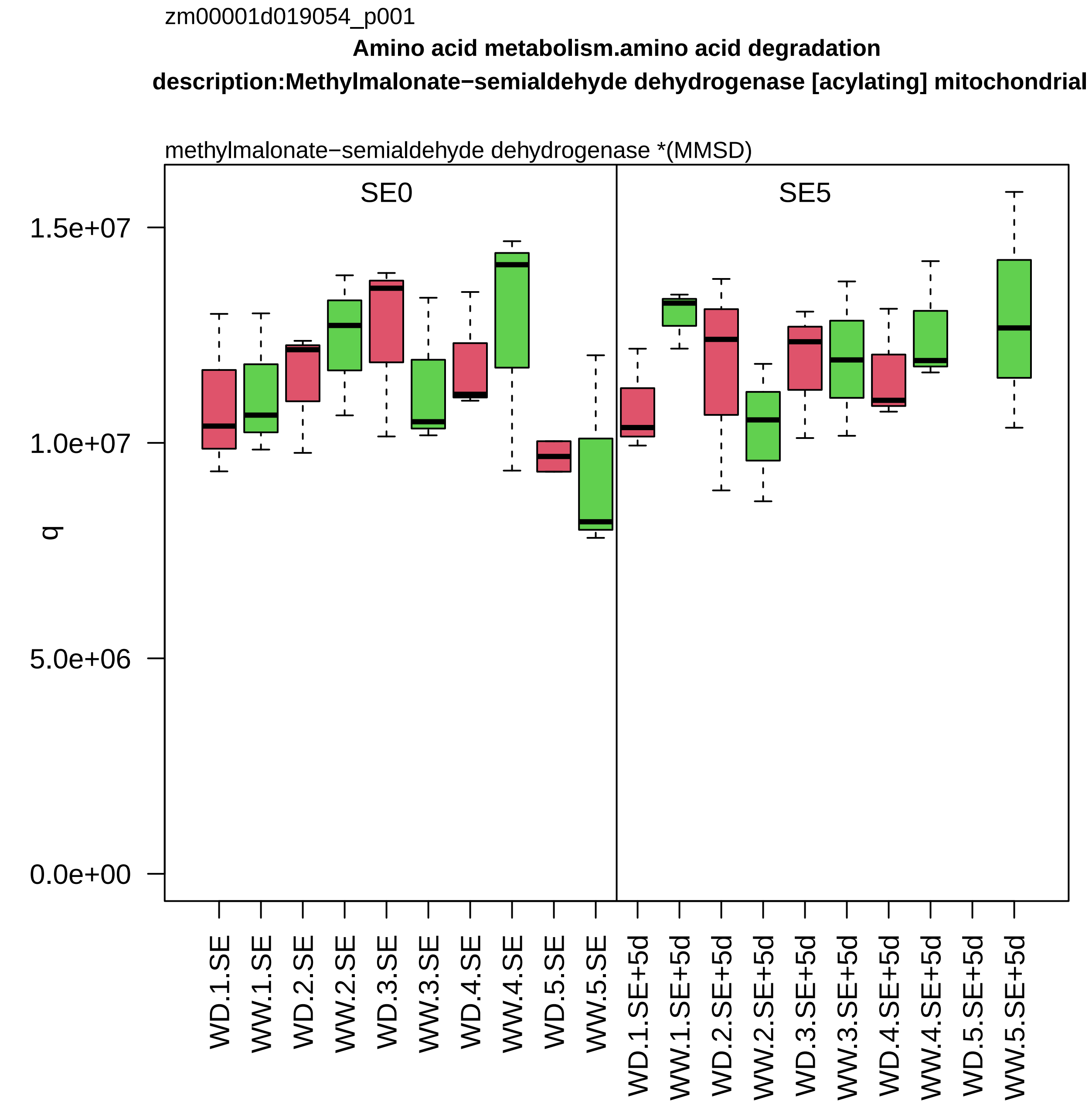
zm00001d019054_p001
Amino acid metabolism.amino acid degradation
description:Methylmalonate−semialdehyde dehydrogenase [acylating] mitochondrial
methylmalonate−semialdehyde dehydrogenase *(MMSD)
SE0
SE5
1.5e+07
1.0e+07
5.0e+06
0.0e+00
q
WD.1.SE
WW.1.SE
WD.2.SE
WW.2.SE
WD.3.SE
WW.3.SE
WD.4.SE
WW.4.SE
WD.5.SE
WW.5.SE
WD.1.SE+5d
WW.1.SE+5d
WD.2.SE+5d
WW.2.SE+5d
WD.3.SE+5d
WW.3.SE+5d
WD.4.SE+5d
WW.4.SE+5d
WD.5.SE+5d
WW.5.SE+5d

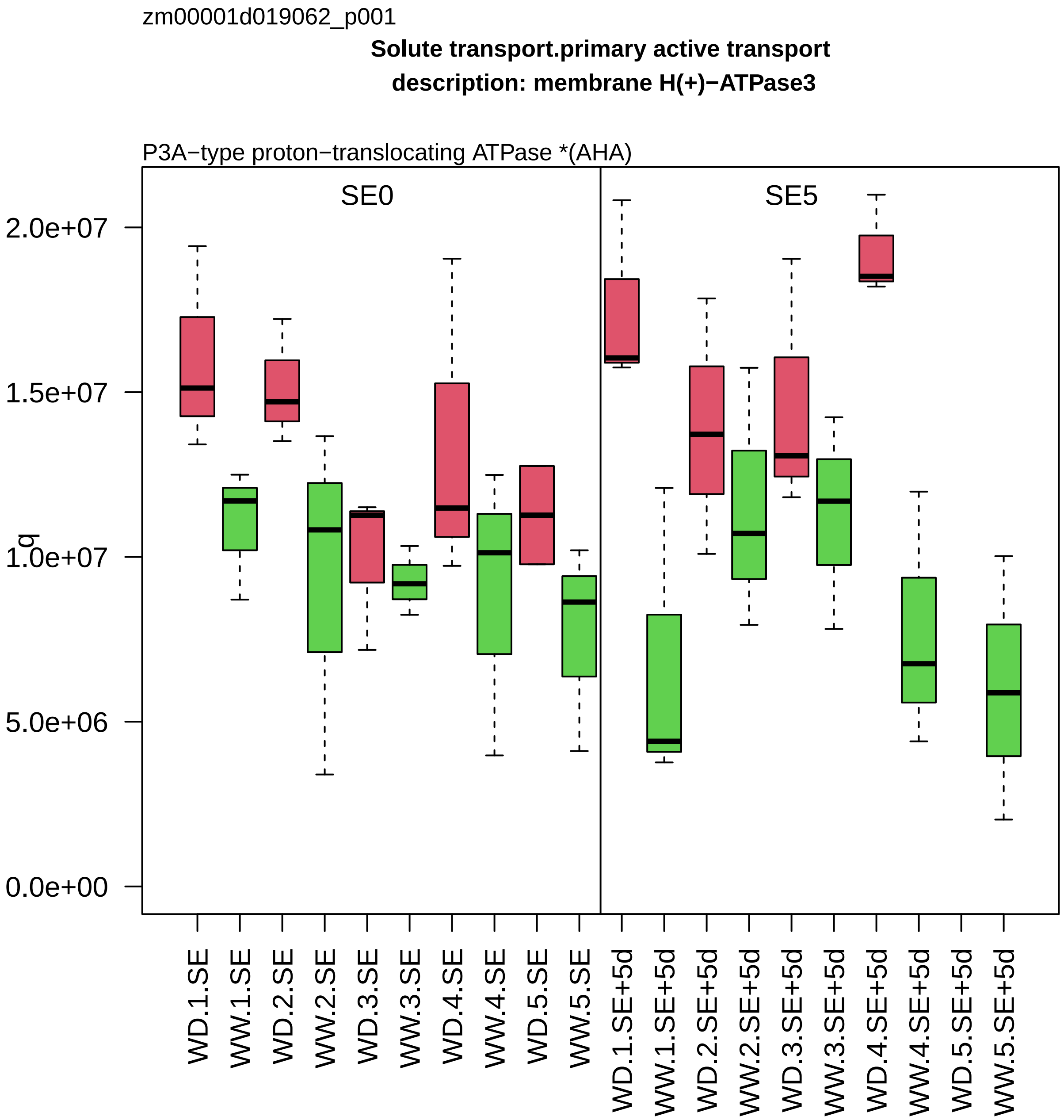
zm00001d019062_p001
Solute transport.primary active transport
description: membrane H(+)−ATPase3
P3A−type proton−translocating ATPase *(AHA)
SE0
SE5
q
2.0e+07
1.5e+07
1.0e+07
5.0e+06
0.0e+00
WD.1.SE
WW.1.SE
WD.2.SE
WW.2.SE
WD.3.SE
WW.3.SE
WD.4.SE
WW.4.SE
WD.5.SE
WW.5.SE
WD.1.SE+5d
WW.1.SE+5d
WD.2.SE+5d
WW.2.SE+5d
WD.3.SE+5d
WW.3.SE+5d
WD.4.SE+5d
WW.4.SE+5d
WD.5.SE+5d
WW.5.SE+5d

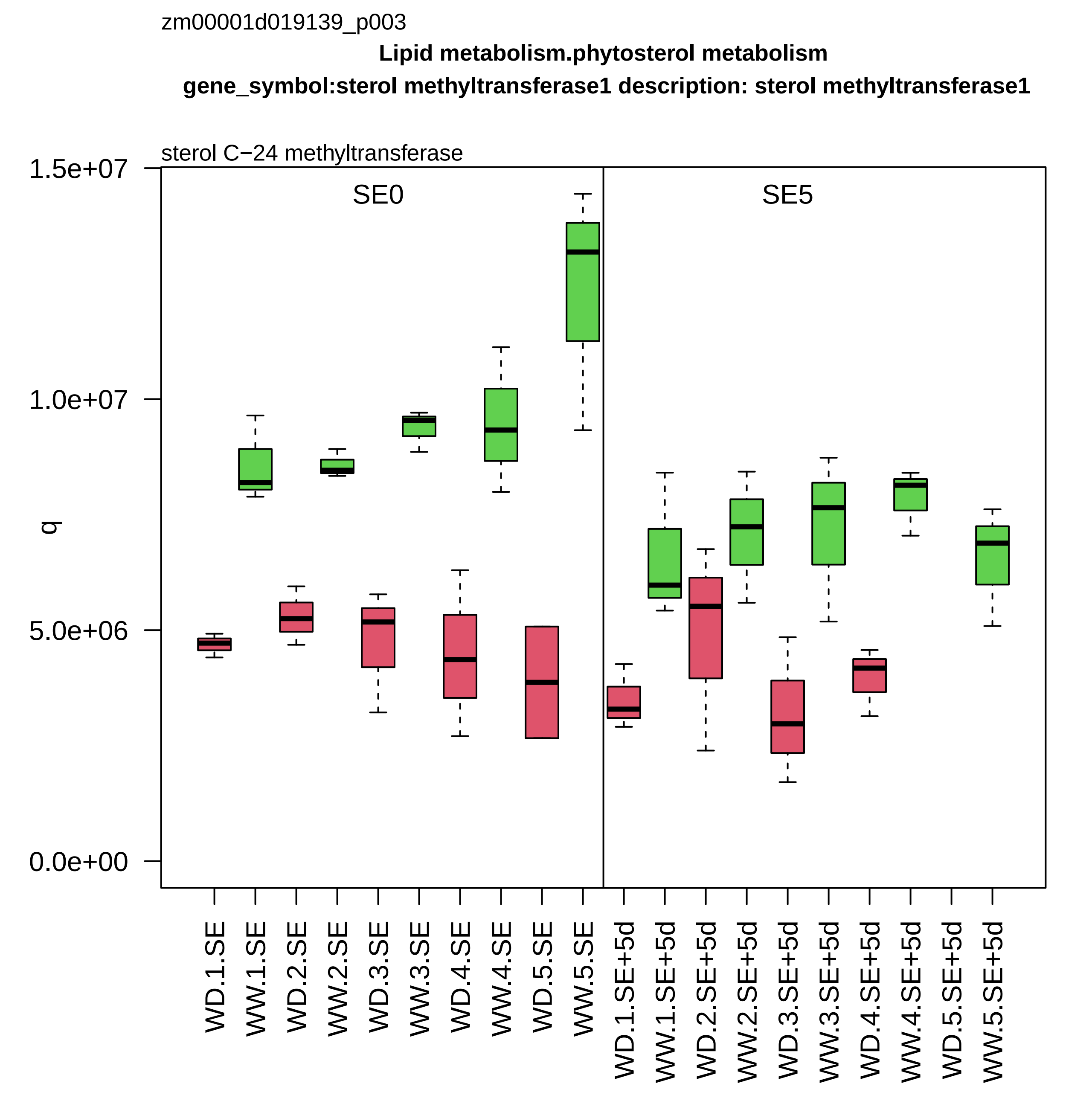

zm00001d019139_p003
Lipid metabolism.phytosterol metabolism
gene_symbol:sterol methyltransferase1 description: sterol methyltransferase1
sterol C−24 methyltransferase
SE0
SE5
q
1.5e+07
1.0e+07
5.0e+06
0.0e+00
WD.1.SE
WW.1.SE
WD.2.SE
WW.2.SE
WD.3.SE
WW.3.SE
WD.4.SE
WW.4.SE
WD.5.SE
WW.5.SE
WD.1.SE+5d
WW.1.SE+5d
WD.2.SE+5d
WW.2.SE+5d
WD.3.SE+5d
WW.3.SE+5d
WD.4.SE+5d
WW.4.SE+5d
WD.5.SE+5d
WW.5.SE+5d

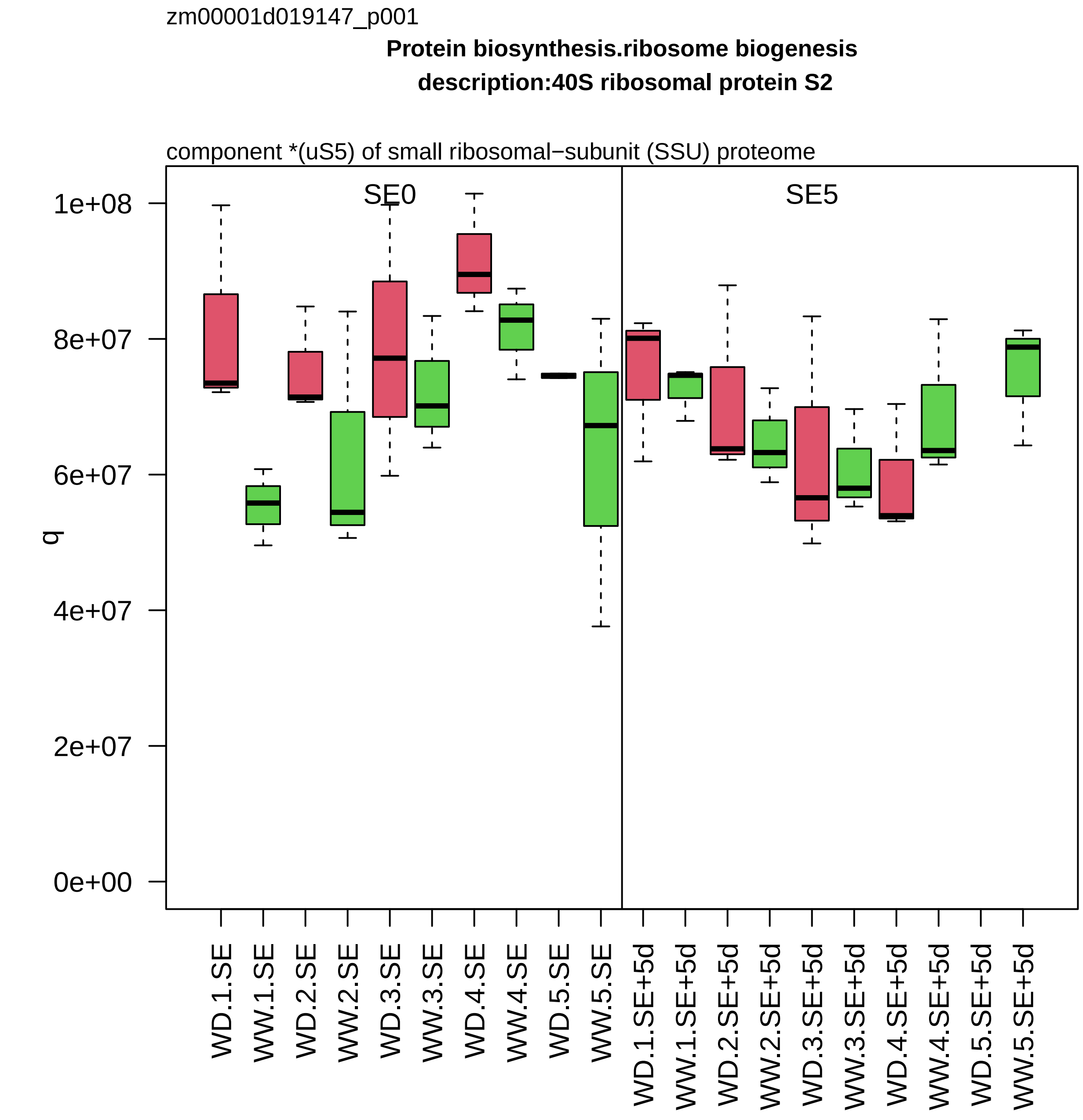

zm00001d019147_p001
Protein biosynthesis.ribosome biogenesis
description:40S ribosomal protein S2
component *(uS5) of small ribosomal−subunit (SSU) proteome
SE0
SE5
q
1e+08
8e+07
6e+07
4e+07
2e+07
0e+00
WD.1.SE
WW.1.SE
WD.2.SE
WW.2.SE
WD.3.SE
WW.3.SE
WD.4.SE
WW.4.SE
WD.5.SE
WW.5.SE
WD.1.SE+5d
WW.1.SE+5d
WD.2.SE+5d
WW.2.SE+5d
WD.3.SE+5d
WW.3.SE+5d
WD.4.SE+5d
WW.4.SE+5d
WD.5.SE+5d
WW.5.SE+5d

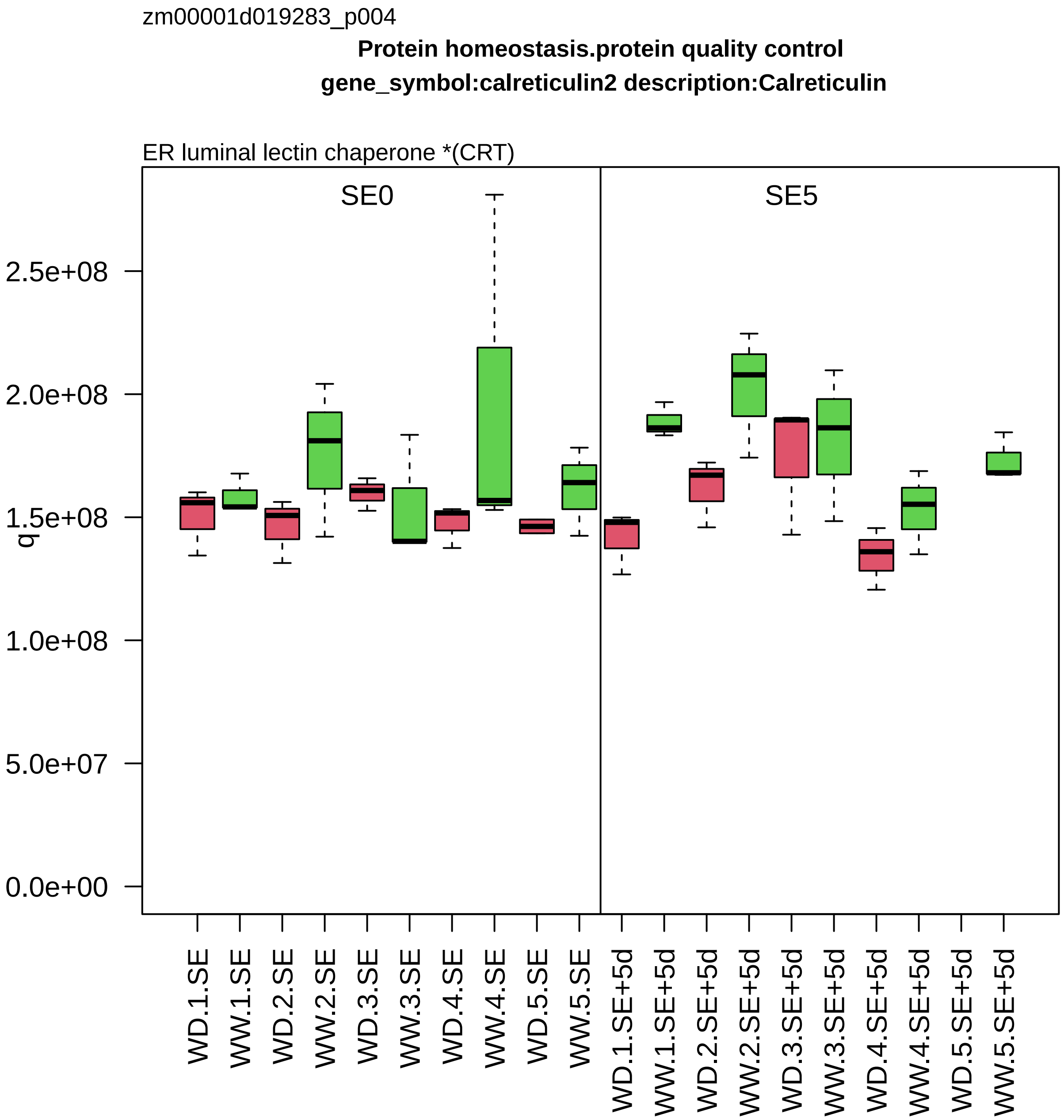
zm00001d019283_p004
Protein homeostasis.protein quality control
gene_symbol:calreticulin2 description:Calreticulin
ER luminal lectin chaperone *(CRT)
SE0
SE5
q
2.5e+08
2.0e+08
1.5e+08
1.0e+08
5.0e+07
0.0e+00
WD.1.SE
WW.1.SE
WD.2.SE
WW.2.SE
WD.3.SE
WW.3.SE
WD.4.SE
WW.4.SE
WD.5.SE
WW.5.SE
WD.1.SE+5d
WW.1.SE+5d
WD.2.SE+5d
WW.2.SE+5d
WD.3.SE+5d
WW.3.SE+5d
WD.4.SE+5d
WW.4.SE+5d
WD.5.SE+5d
WW.5.SE+5d

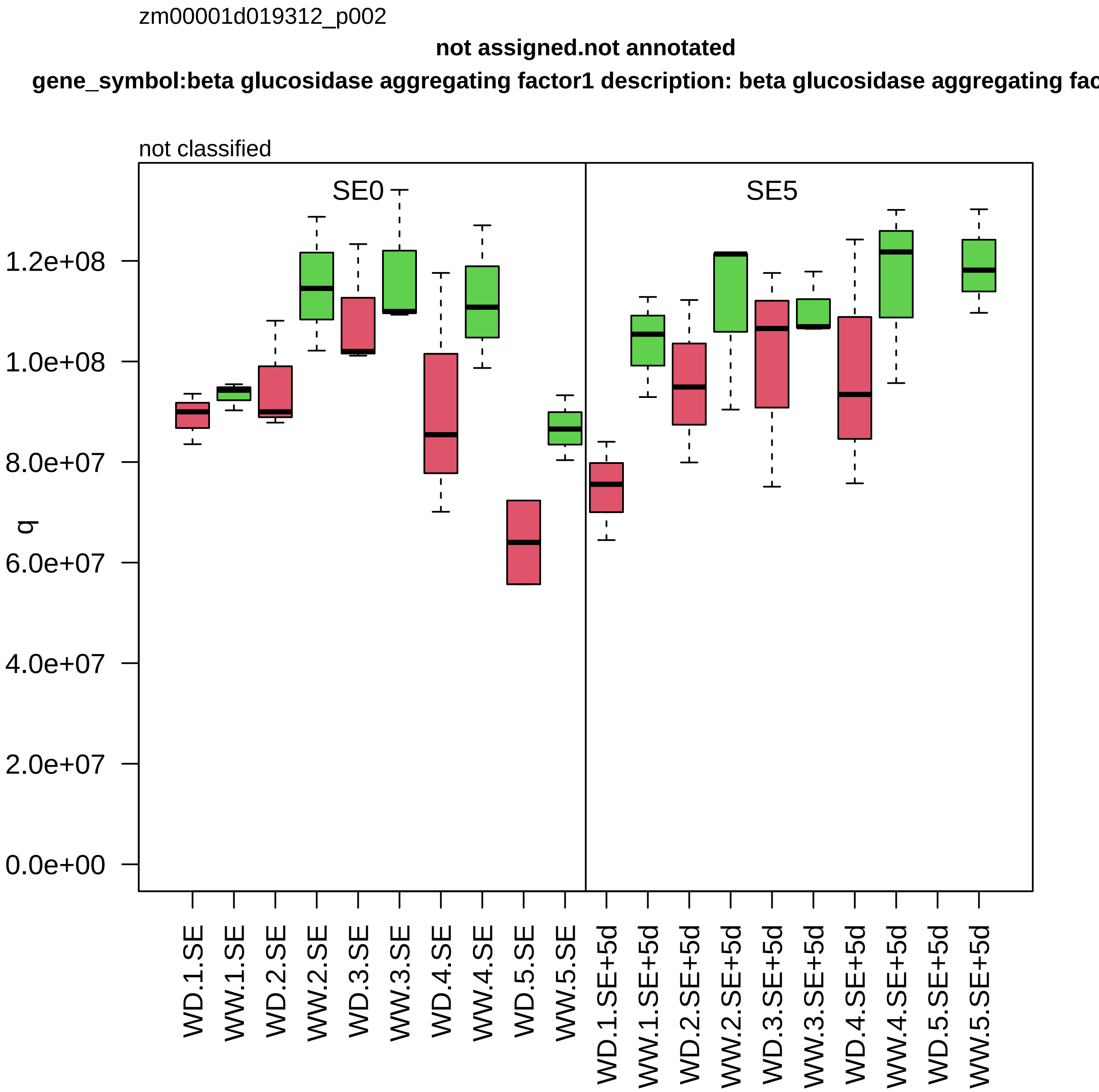

zm00001d019312_p002
not assigned.not annotated
gene_symbol:beta glucosidase aggregating factor1 description: beta glucosidase aggregating fac
not classified
SE0
SE5
q
1.2e+08
1.0e+08
8.0e+07
6.0e+07
4.0e+07
2.0e+07
0.0e+00
WD.1.SE
WW.1.SE
WD.2.SE
WW.2.SE
WD.3.SE
WW.3.SE
WD.4.SE
WW.4.SE
WD.5.SE
WW.5.SE
WD.1.SE+5d
WW.1.SE+5d
WD.2.SE+5d
WW.2.SE+5d
WD.3.SE+5d
WW.3.SE+5d
WD.4.SE+5d
WW.4.SE+5d
WD.5.SE+5d
WW.5.SE+5d

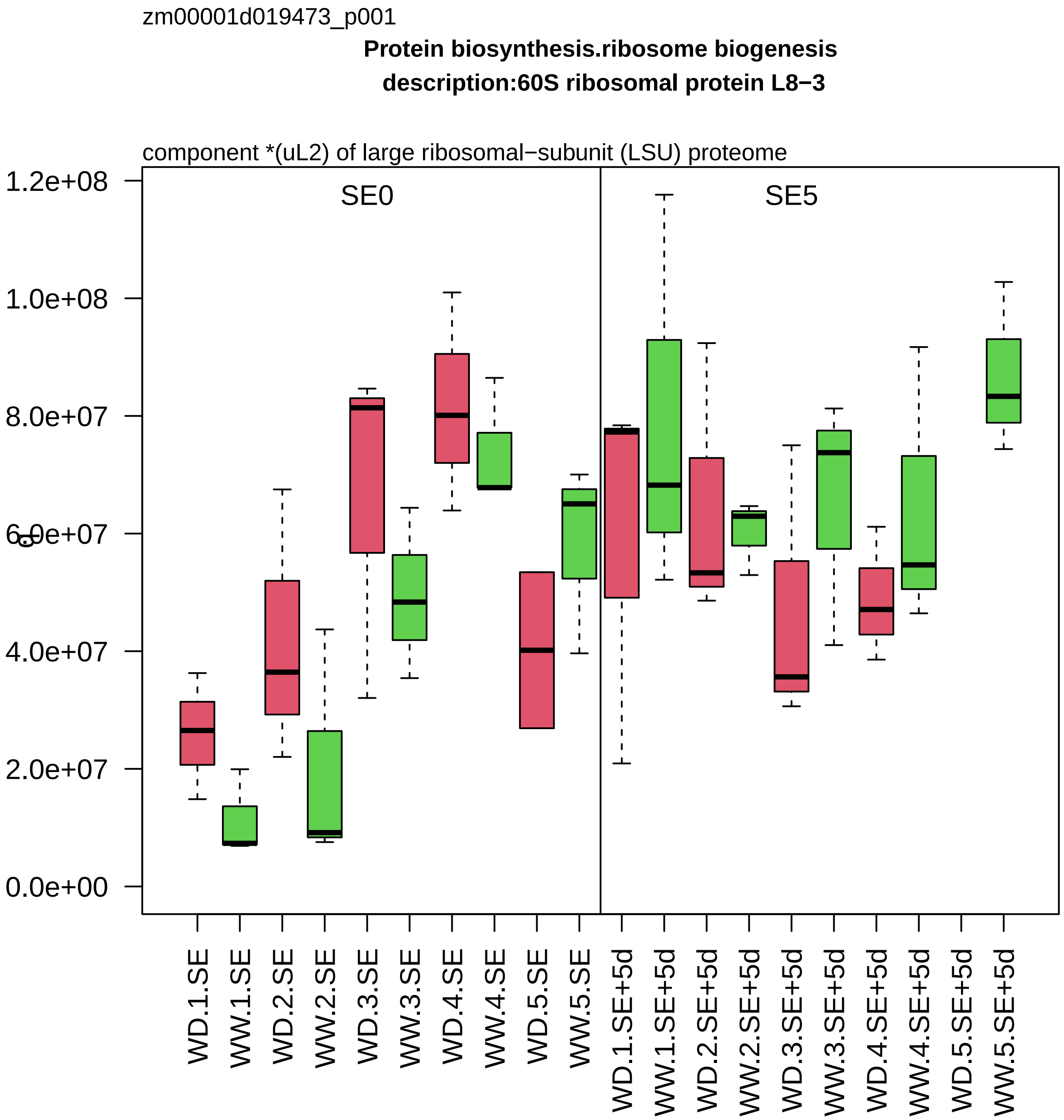

zm00001d019473_p001
Protein biosynthesis.ribosome biogenesis
description:60S ribosomal protein L8−3
component *(uL2) of large ribosomal−subunit (LSU) proteome
SE0
SE5
1.2e+08
1.0e+08
8.0e+07
6.0e+07
4.0e+07
2.0e+07
0.0e+00
WD.1.SE
WW.1.SE
WD.2.SE
WW.2.SE
WD.3.SE
WW.3.SE
WD.4.SE
WW.4.SE
WD.5.SE
WW.5.SE
WD.1.SE+5d
WW.1.SE+5d
WD.2.SE+5d
WW.2.SE+5d
WD.3.SE+5d
WW.3.SE+5d
WD.4.SE+5d
WW.4.SE+5d
WD.5.SE+5d
WW.5.SE+5d

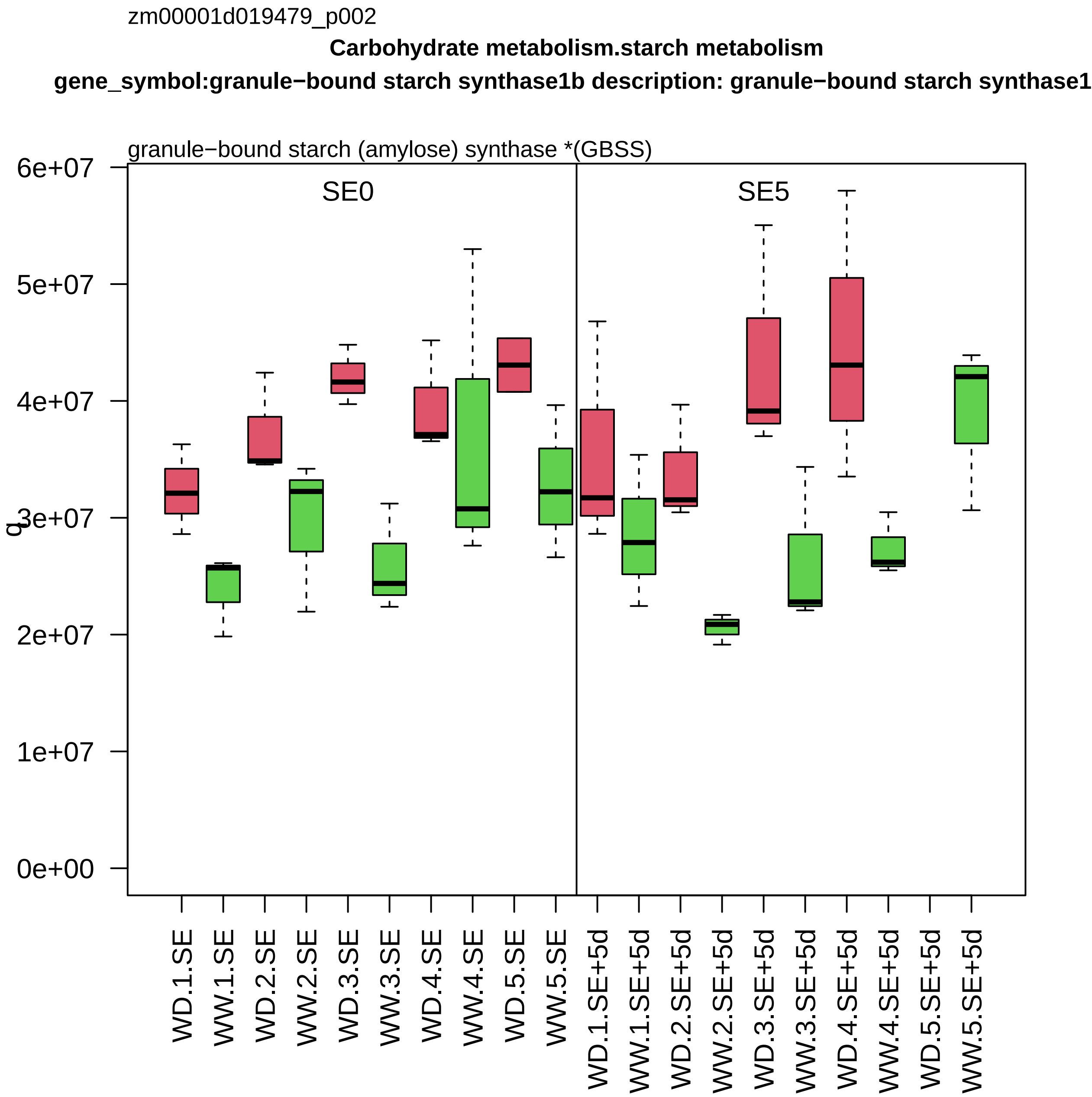
zm00001d019479_p002
Carbohydrate metabolism.starch metabolism
gene_symbol:granule−bound starch synthase1b description: granule−bound starch synthase1
granule−bound starch (amylose) synthase *(GBSS)
SE0
SE5
q
6e+07
5e+07
4e+07
3e+07
2e+07
1e+07
0e+00
WD.1.SE
WW.1.SE
WD.2.SE
WW.2.SE
WD.3.SE
WW.3.SE
WD.4.SE
WW.4.SE
WD.5.SE
WW.5.SE
WD.1.SE+5d
WW.1.SE+5d
WD.2.SE+5d
WW.2.SE+5d
WD.3.SE+5d
WW.3.SE+5d
WD.4.SE+5d
WW.4.SE+5d
WD.5.SE+5d
WW.5.SE+5d

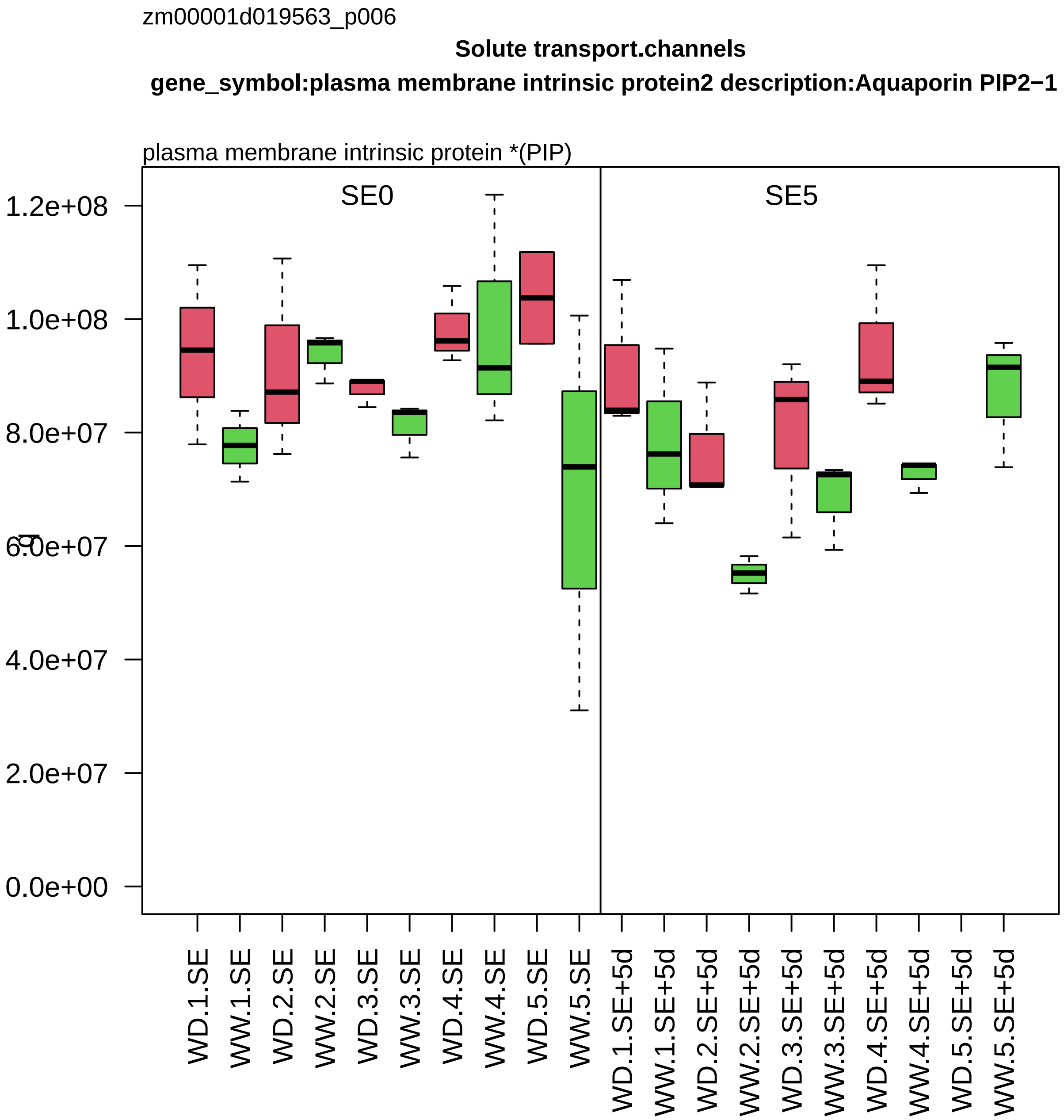

zm00001d019563_p006
Solute transport.channels
gene_symbol:plasma membrane intrinsic protein2 description:Aquaporin PIP2−1
plasma membrane intrinsic protein *(PIP)
SE0
SE5
1.2e+08
1.0e+08
8.0e+07
6.0e+07
4.0e+07
2.0e+07
0.0e+00
WD.1.SE
WW.1.SE
WD.2.SE
WW.2.SE
WD.3.SE
WW.3.SE
WD.4.SE
WW.4.SE
WD.5.SE
WW.5.SE
WD.1.SE+5d
WW.1.SE+5d
WD.2.SE+5d
WW.2.SE+5d
WD.3.SE+5d
WW.3.SE+5d
WD.4.SE+5d
WW.4.SE+5d
WD.5.SE+5d
WW.5.SE+5d

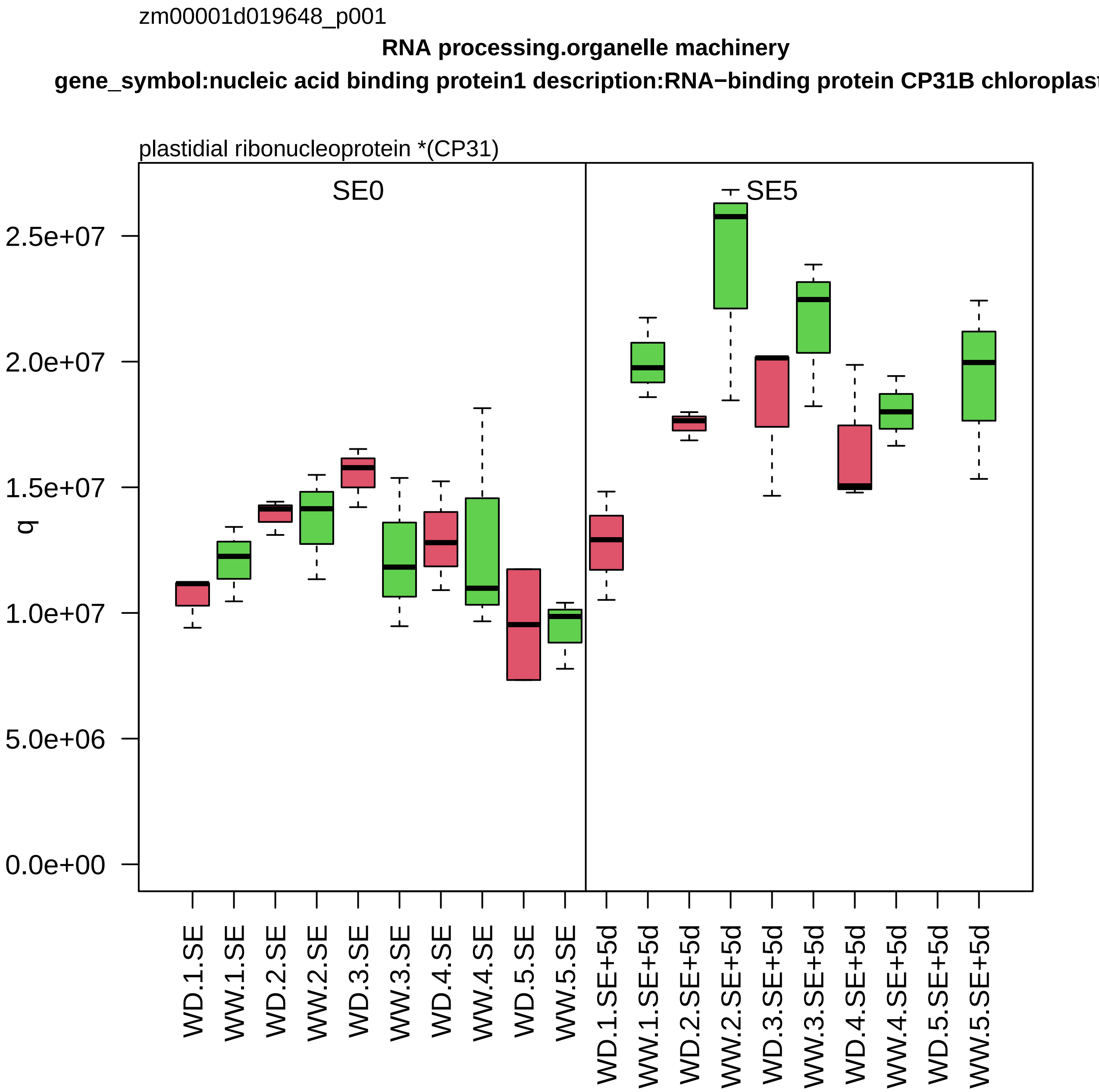
zm00001d019648_p001
RNA processing.organelle machinery
gene_symbol:nucleic acid binding protein1 description:RNA−binding protein CP31B chloroplast
plastidial ribonucleoprotein *(CP31)
SE0
SE5
q
0.0e+00
5.0e+06
1.0e+07
1.5e+07
2.0e+07
2.5e+07
WD.1.SE
WW.1.SE
WD.2.SE
WW.2.SE
WD.3.SE
WW.3.SE
WD.4.SE
WW.4.SE
WD.5.SE
WW.5.SE
WD.1.SE+5d
WW.1.SE+5d
WD.2.SE+5d
WW.2.SE+5d
WD.3.SE+5d
WW.3.SE+5d
WD.4.SE+5d
WW.4.SE+5d
WD.5.SE+5d
WW.5.SE+5d

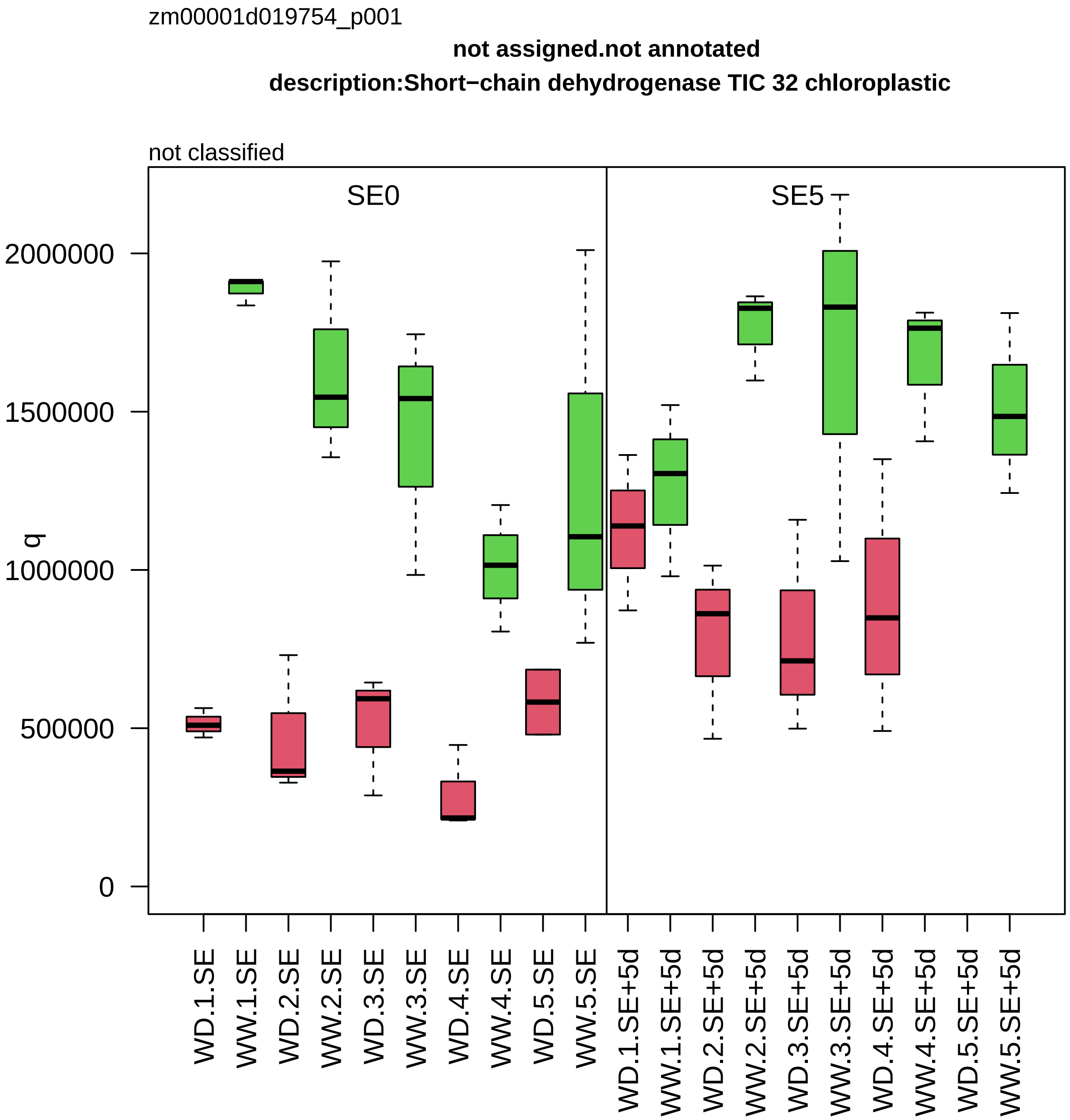

zm00001d019754_p001
not assigned.not annotated
description:Short−chain dehydrogenase TIC 32 chloroplastic
not classified
SE0
SE5
q
2000000
1500000
1000000
500000
0
WD.1.SE
WW.1.SE
WD.2.SE
WW.2.SE
WD.3.SE
WW.3.SE
WD.4.SE
WW.4.SE
WD.5.SE
WW.5.SE
WD.1.SE+5d
WW.1.SE+5d
WD.2.SE+5d
WW.2.SE+5d
WD.3.SE+5d
WW.3.SE+5d
WD.4.SE+5d
WW.4.SE+5d
WD.5.SE+5d
WW.5.SE+5d

zm00001d019926_p001

**Lipid metabolism.fatty acid metabolism**

**description:Enoyl−[acyl−carrier−protein] reductase [NADH] chloroplastic**

enoyl−ACP reductase

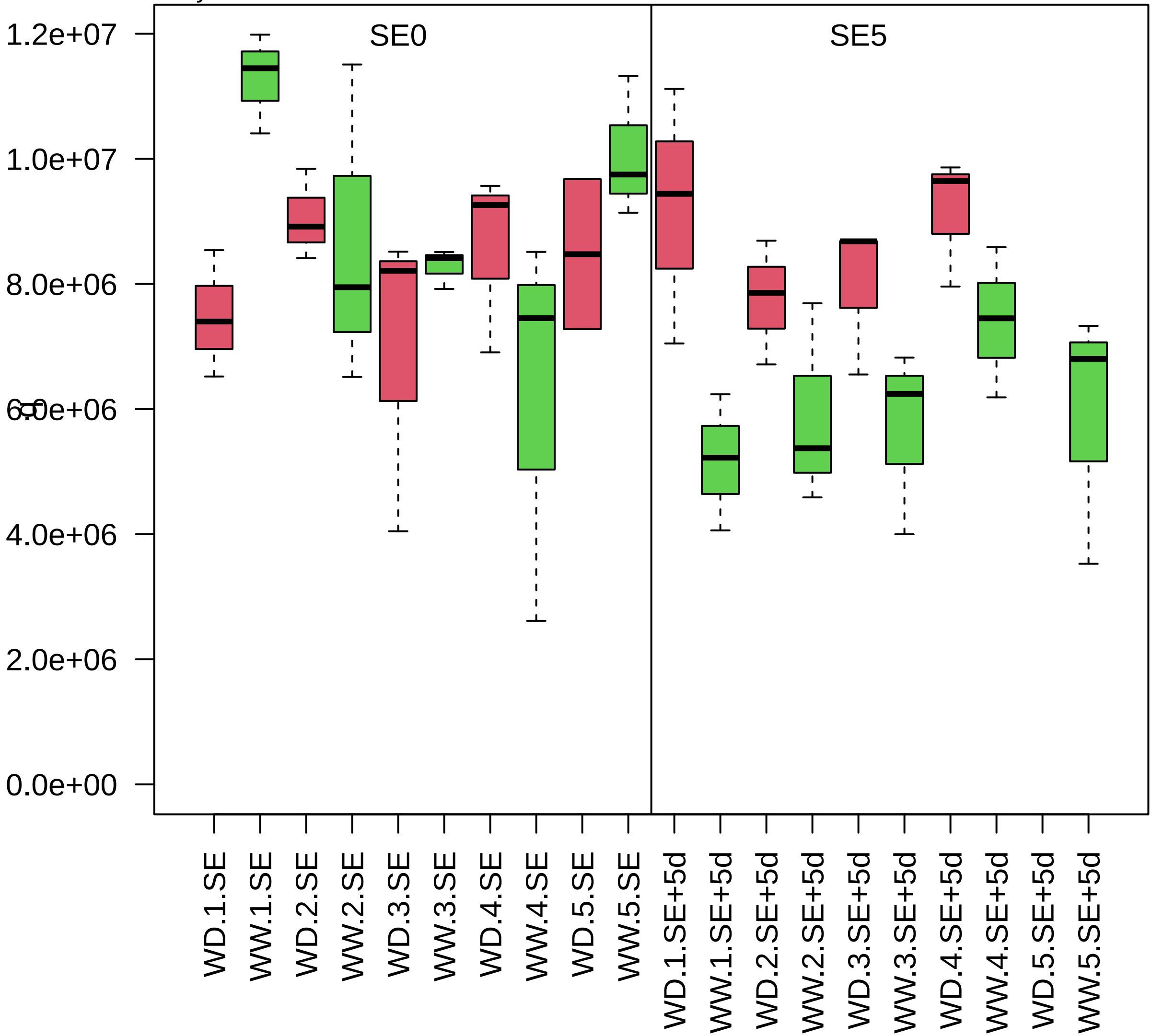

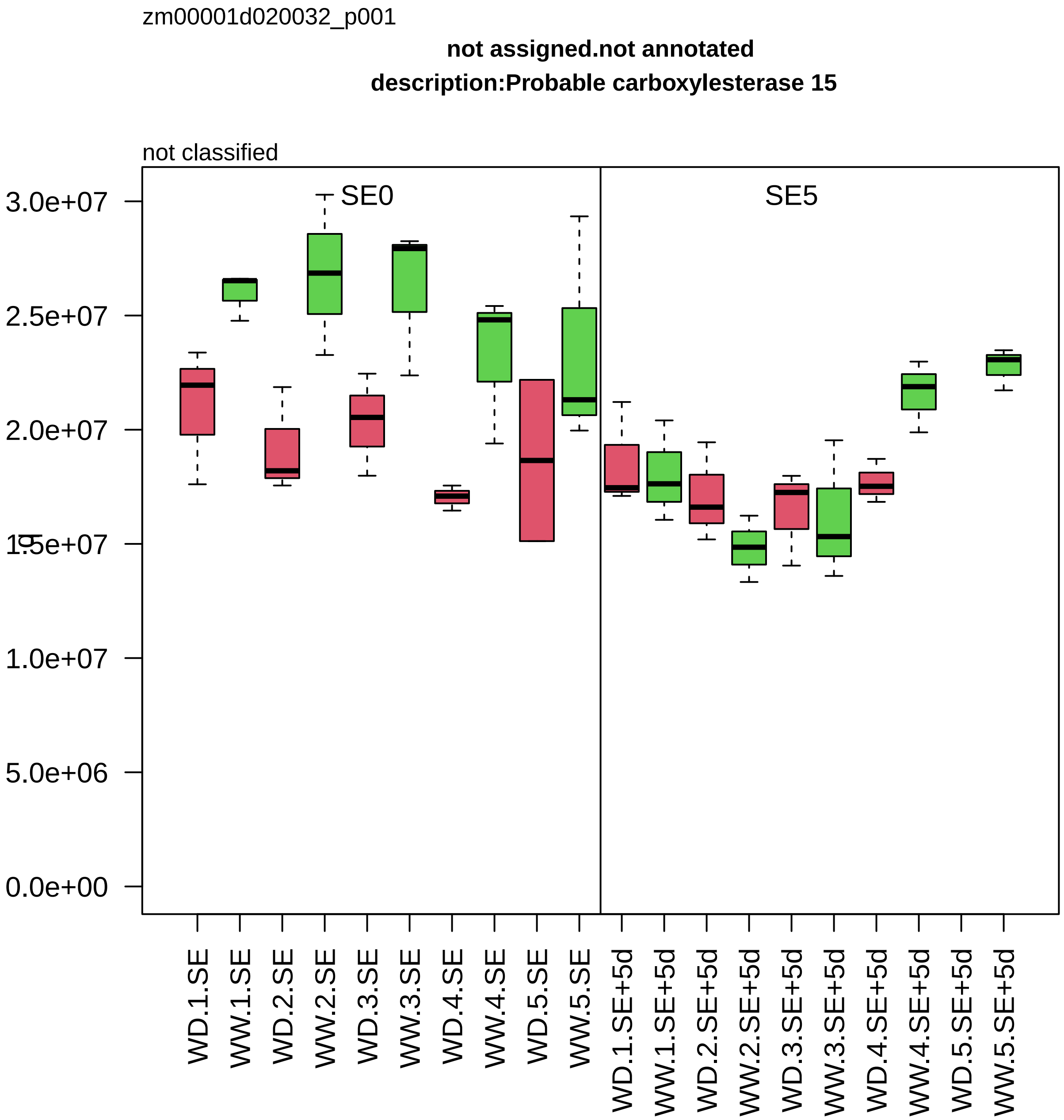

zm00001d020032_p001
not assigned.not annotated
description:Probable carboxylesterase 15
not classified
SE0
SE5
3.0e+07
2.5e+07
2.0e+07
1.5e+07
1.0e+07
5.0e+06
0.0e+00
WD.1.SE
WW.1.SE
WD.2.SE
WW.2.SE
WD.3.SE
WW.3.SE
WD.4.SE
WW.4.SE
WD.5.SE
WW.5.SE
WD.1.SE+5d
WW.1.SE+5d
WD.2.SE+5d
WW.2.SE+5d
WD.3.SE+5d
WW.3.SE+5d
WD.4.SE+5d
WW.4.SE+5d
WD.5.SE+5d
WW.5.SE+5d

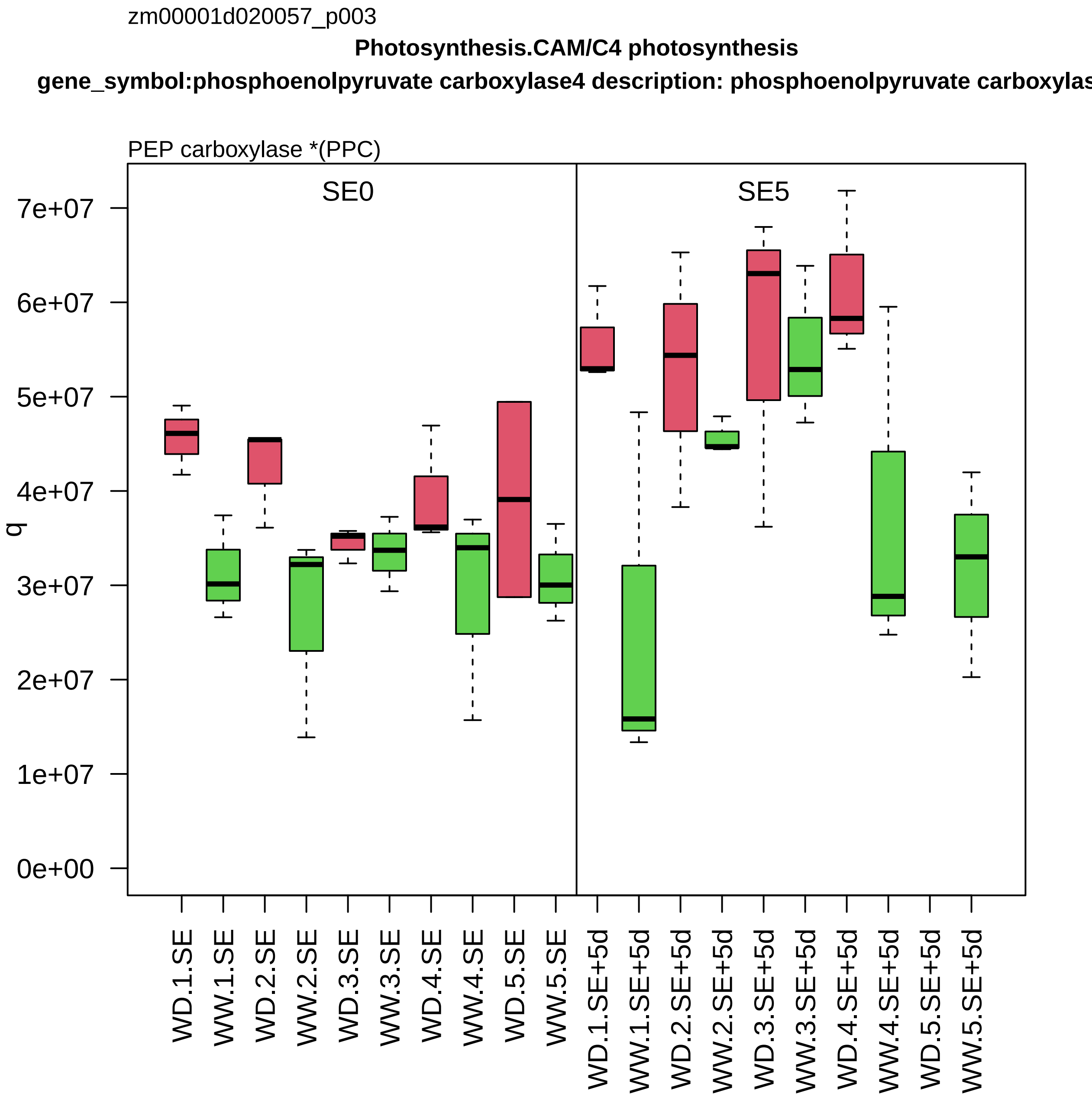
zm00001d020057_p003
Photosynthesis.CAM/C4 photosynthesis
gene_symbol:phosphoenolpyruvate carboxylase4 description: phosphoenolpyruvate carboxylas
PEP carboxylase *(PPC)
SE0
SE5
q
7e+07
6e+07
5e+07
4e+07
3e+07
2e+07
1e+07
0e+00
WD.1.SE
WW.1.SE
WD.2.SE
WW.2.SE
WD.3.SE
WW.3.SE
WD.4.SE
WW.4.SE
WD.5.SE
WW.5.SE
WD.1.SE+5d
WW.1.SE+5d
WD.2.SE+5d
WW.2.SE+5d
WD.3.SE+5d
WW.3.SE+5d
WD.4.SE+5d
WW.4.SE+5d
WD.5.SE+5d
WW.5.SE+5d

zm00001d020219_p002

**Solute transport.porins**

**gene_symbol:voltage−dependent anion channel protein1b description: voltage−dependent anion channe**

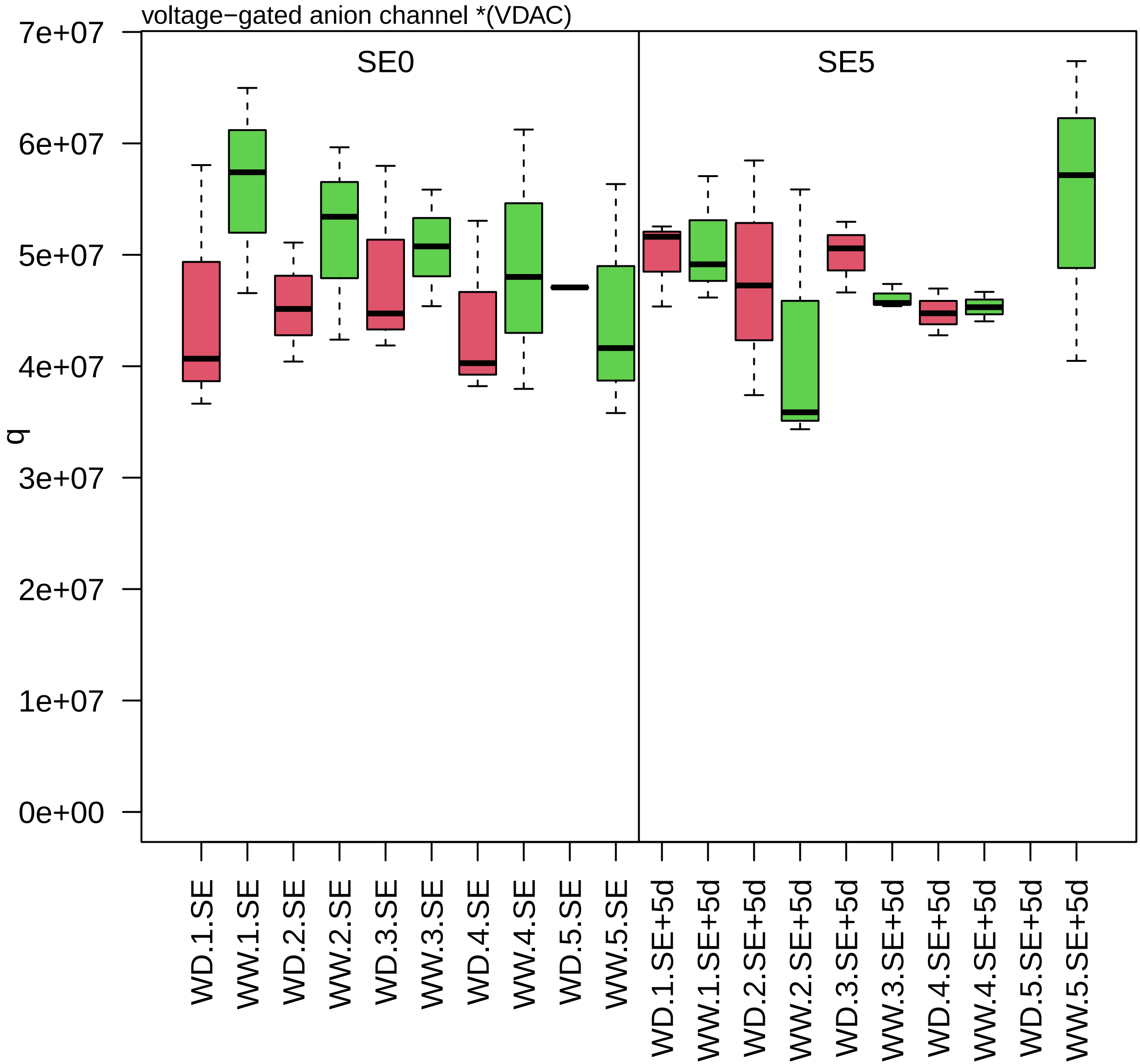

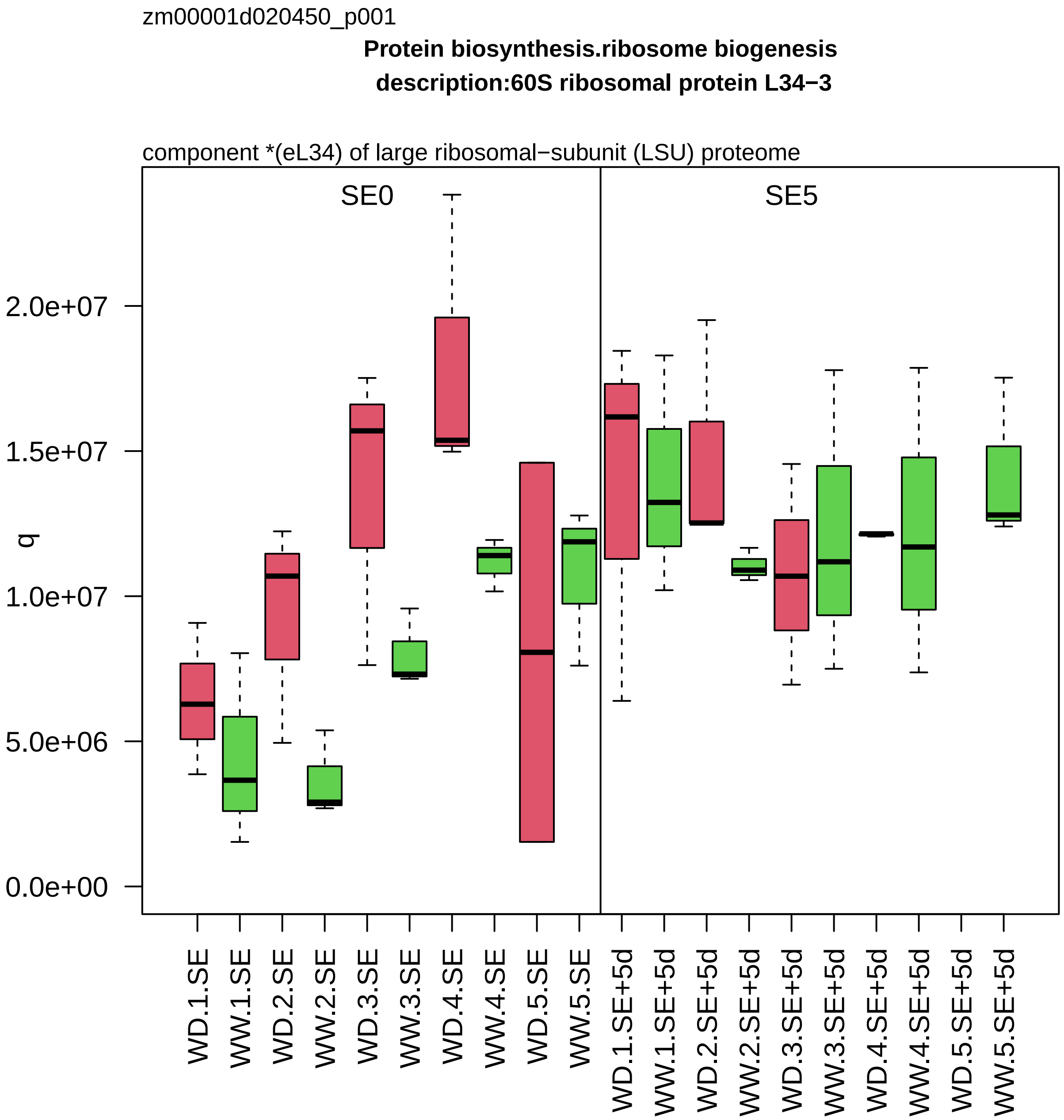
zm00001d020450_p001
Protein biosynthesis.ribosome biogenesis
description:60S ribosomal protein L34−3
component *(eL34) of large ribosomal−subunit (LSU) proteome
SE0
SE5
q
2.0e+07
1.5e+07
1.0e+07
5.0e+06
0.0e+00
WD.1.SE
WW.1.SE
WD.2.SE
WW.2.SE
WD.3.SE
WW.3.SE
WD.4.SE
WW.4.SE
WD.5.SE
WW.5.SE
WD.1.SE+5d
WW.1.SE+5d
WD.2.SE+5d
WW.2.SE+5d
WD.3.SE+5d
WW.3.SE+5d
WD.4.SE+5d
WW.4.SE+5d
WD.5.SE+5d
WW.5.SE+5d

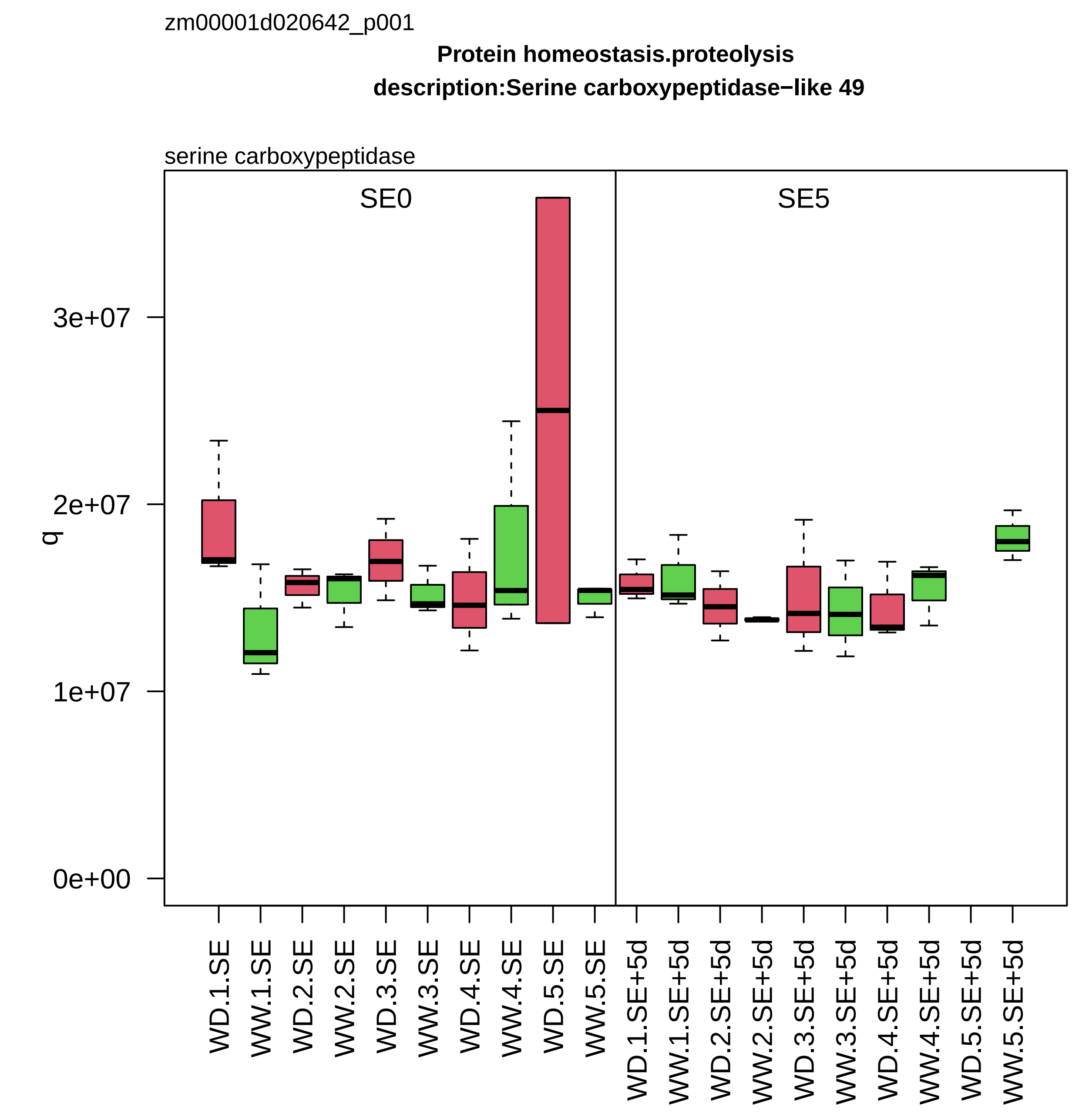
zm00001d020642_p001
Protein homeostasis.proteolysis
description:Serine carboxypeptidase−like 49
serine carboxypeptidase
SE0
SE5
q
3e+07
2e+07
1e+07
0e+00
WD.1.SE
WW.1.SE
WD.2.SE
WW.2.SE
WD.3.SE
WW.3.SE
WD.4.SE
WW.4.SE
WD.5.SE
WW.5.SE
WD.1.SE+5d
WW.1.SE+5d
WD.2.SE+5d
WW.2.SE+5d
WD.3.SE+5d
WW.3.SE+5d
WD.4.SE+5d
WW.4.SE+5d
WD.5.SE+5d
WW.5.SE+5d

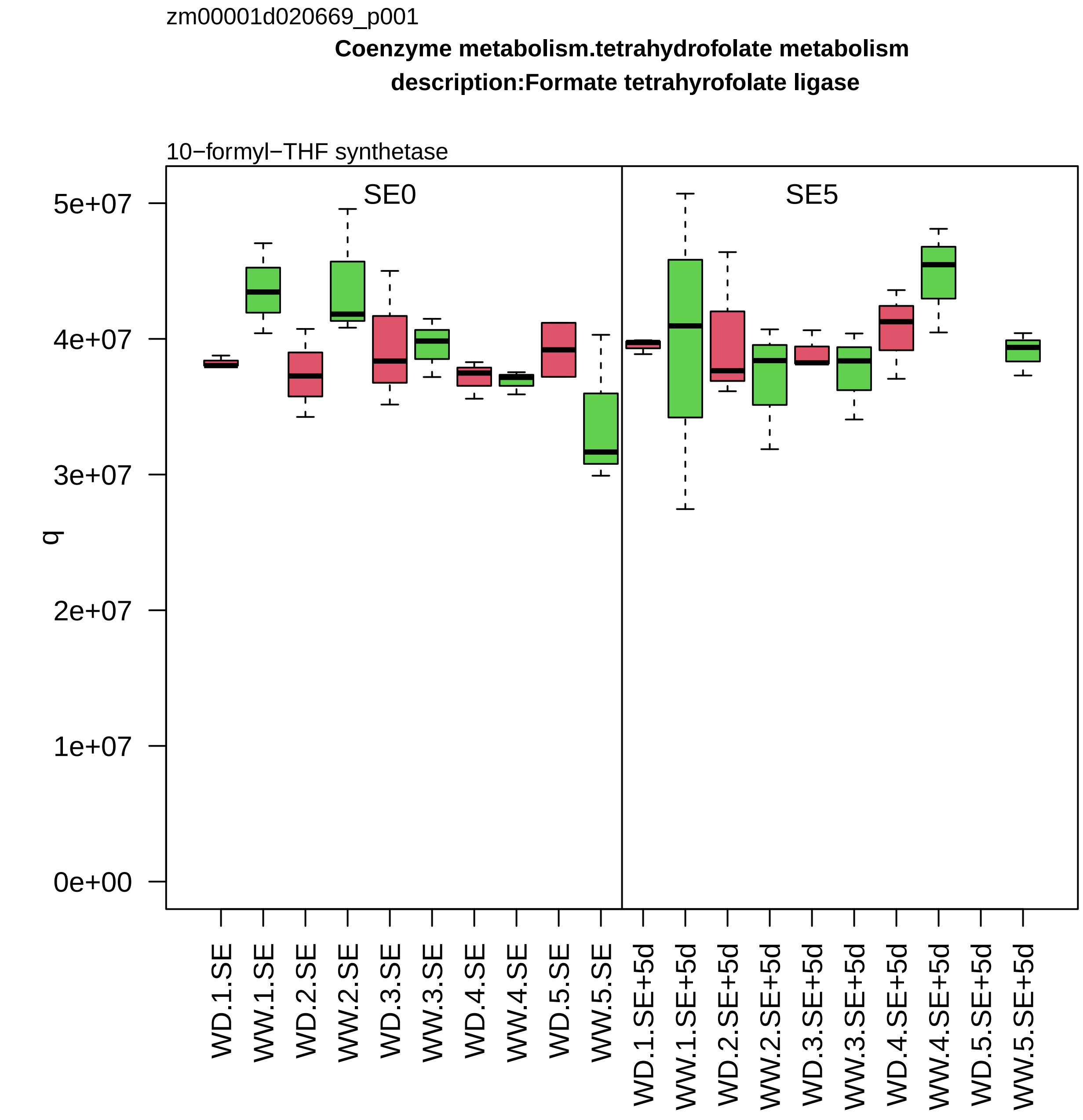
zm00001d020669_p001
Coenzyme metabolism.tetrahydrofolate metabolism
description:Formate tetrahyrofolate ligase
10−formyl−THF synthetase
SE0
SE5
q
0e+00
1e+07
2e+07
3e+07
4e+07
5e+07
WD.1.SE
WW.1.SE
WD.2.SE
WW.2.SE
WD.3.SE
WW.3.SE
WD.4.SE
WW.4.SE
WD.5.SE
WW.5.SE
WD.1.SE+5d
WW.1.SE+5d
WD.2.SE+5d
WW.2.SE+5d
WD.3.SE+5d
WW.3.SE+5d
WD.4.SE+5d
WW.4.SE+5d
WD.5.SE+5d
WW.5.SE+5d

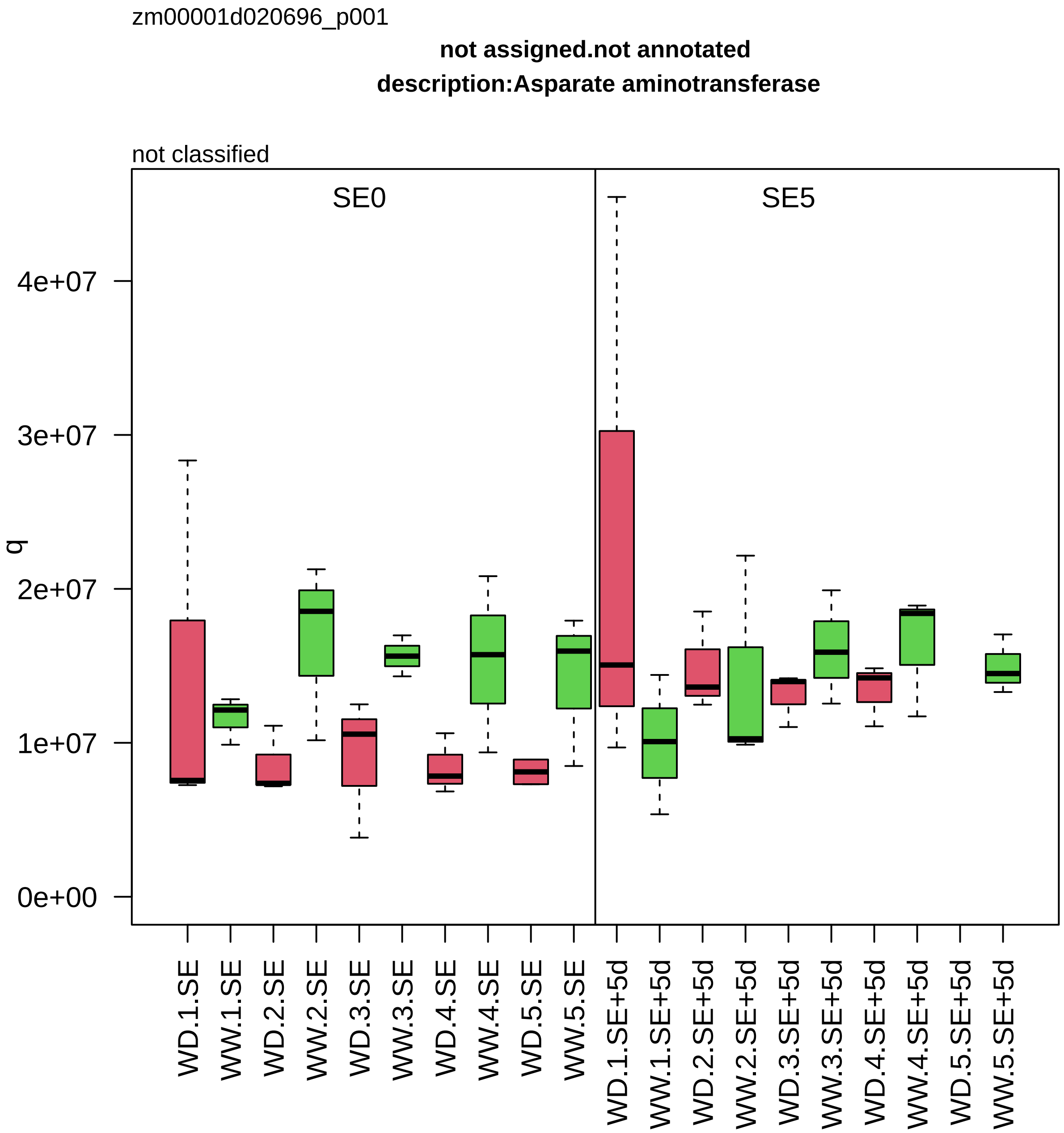

zm00001d020696_p001
not assigned.not annotated
description:Asparate aminotransferase
not classified
SE0
SE5
q
0e+00
1e+07
2e+07
3e+07
4e+07
WD.1.SE
WW.1.SE
WD.2.SE
WW.2.SE
WD.3.SE
WW.3.SE
WD.4.SE
WW.4.SE
WD.5.SE
WW.5.SE
WD.1.SE+5d
WW.1.SE+5d
WD.2.SE+5d
WW.2.SE+5d
WD.3.SE+5d
WW.3.SE+5d
WD.4.SE+5d
WW.4.SE+5d
WD.5.SE+5d
WW.5.SE+5d

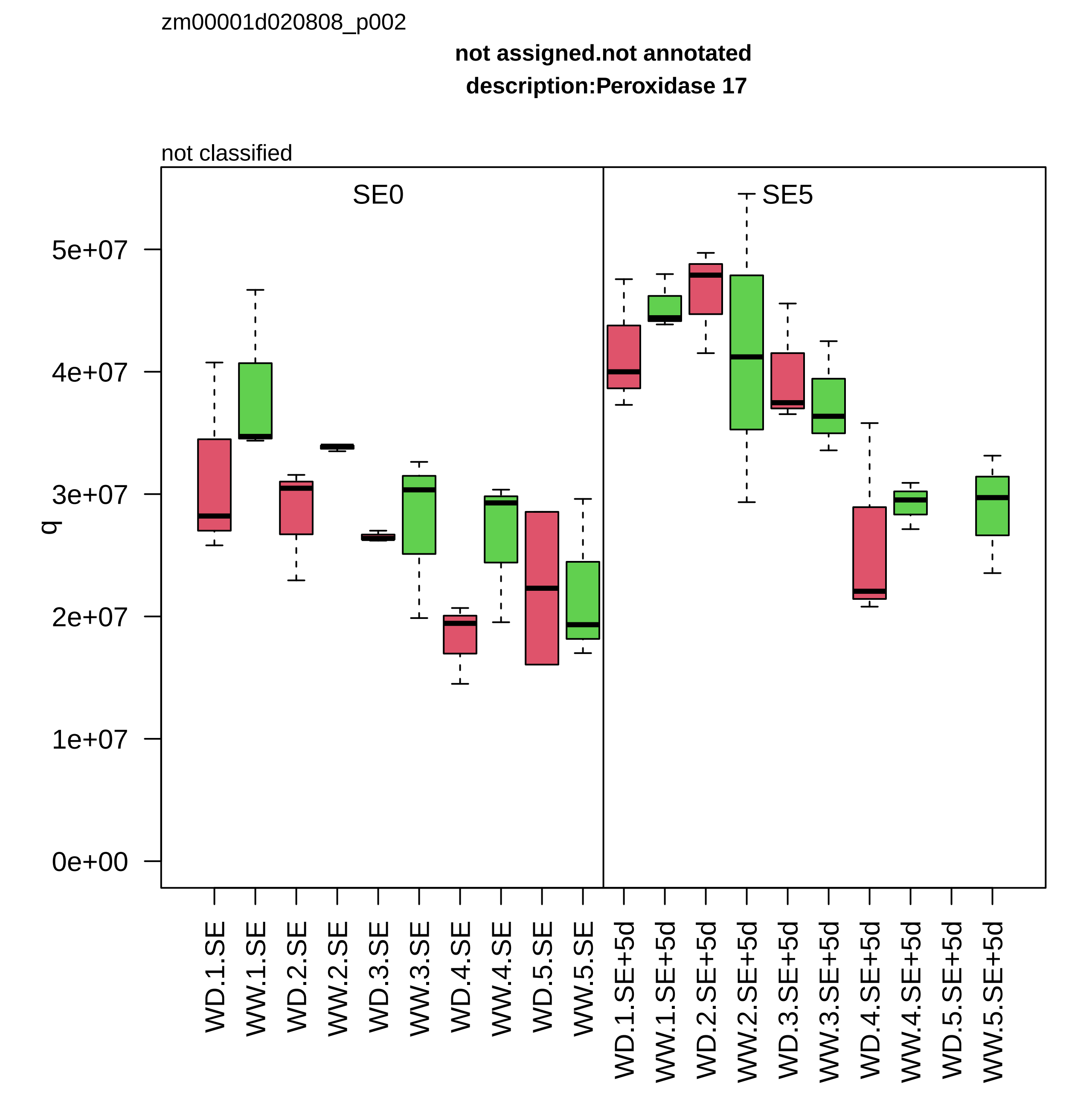
zm00001d020808_p002
not assigned.not annotated
description:Peroxidase 17
not classified
SE0
SE5
q
0e+00
1e+07
2e+07
3e+07
4e+07
5e+07
WD.1.SE
WW.1.SE
WD.2.SE
WW.2.SE
WD.3.SE
WW.3.SE
WD.4.SE
WW.4.SE
WD.5.SE
WW.5.SE
WD.1.SE+5d
WW.1.SE+5d
WD.2.SE+5d
WW.2.SE+5d
WD.3.SE+5d
WW.3.SE+5d
WD.4.SE+5d
WW.4.SE+5d
WD.5.SE+5d
WW.5.SE+5d

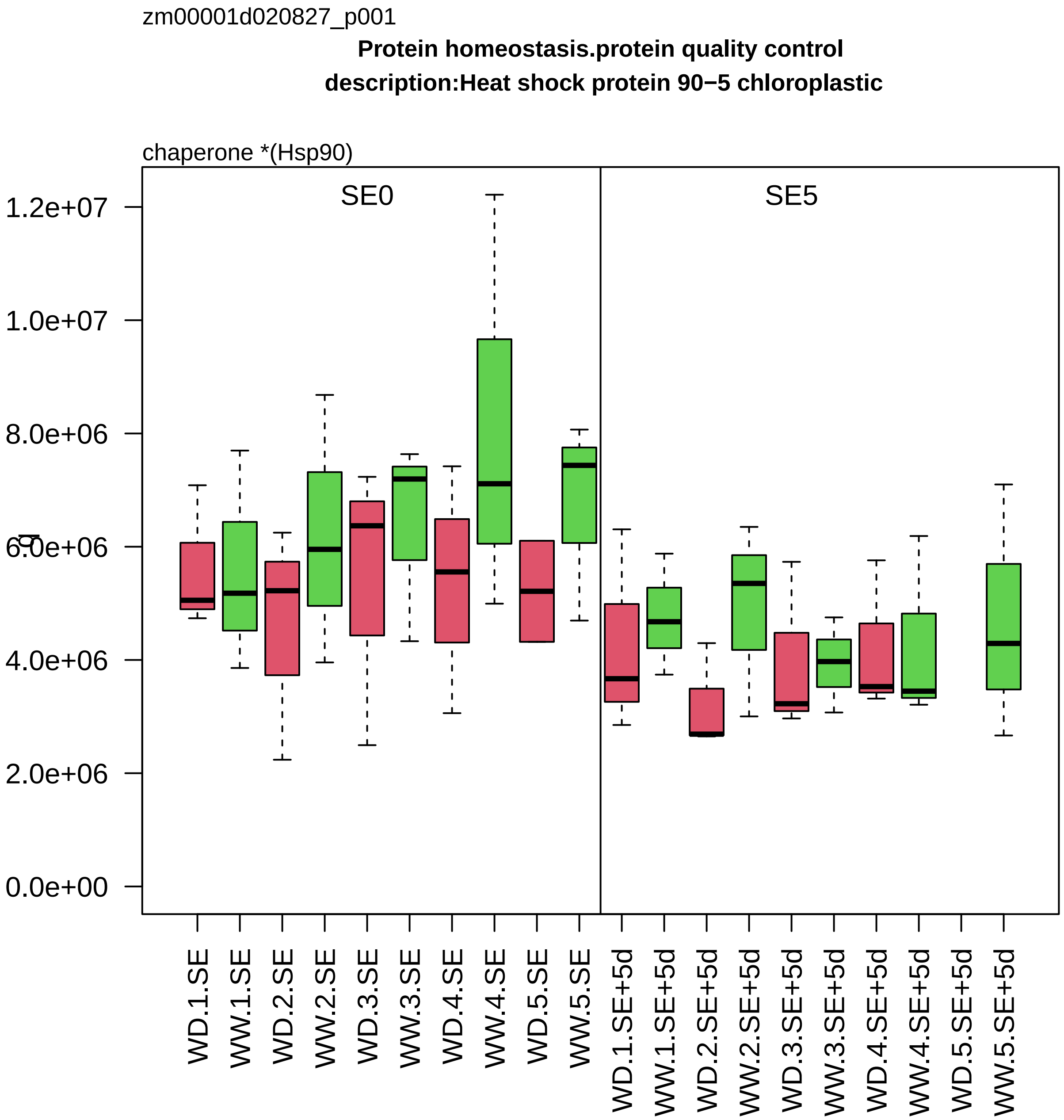

zm00001d020827_p001
Protein homeostasis.protein quality control
description:Heat shock protein 90−5 chloroplastic
chaperone *(Hsp90)
SE0
SE5
1.2e+07
1.0e+07
8.0e+06
6.0e+06
4.0e+06
2.0e+06
0.0e+00
WD.1.SE
WW.1.SE
WD.2.SE
WW.2.SE
WD.3.SE
WW.3.SE
WD.4.SE
WW.4.SE
WD.5.SE
WW.5.SE
WD.1.SE+5d
WW.1.SE+5d
WD.2.SE+5d
WW.2.SE+5d
WD.3.SE+5d
WW.3.SE+5d
WD.4.SE+5d
WW.4.SE+5d
WD.5.SE+5d
WW.5.SE+5d

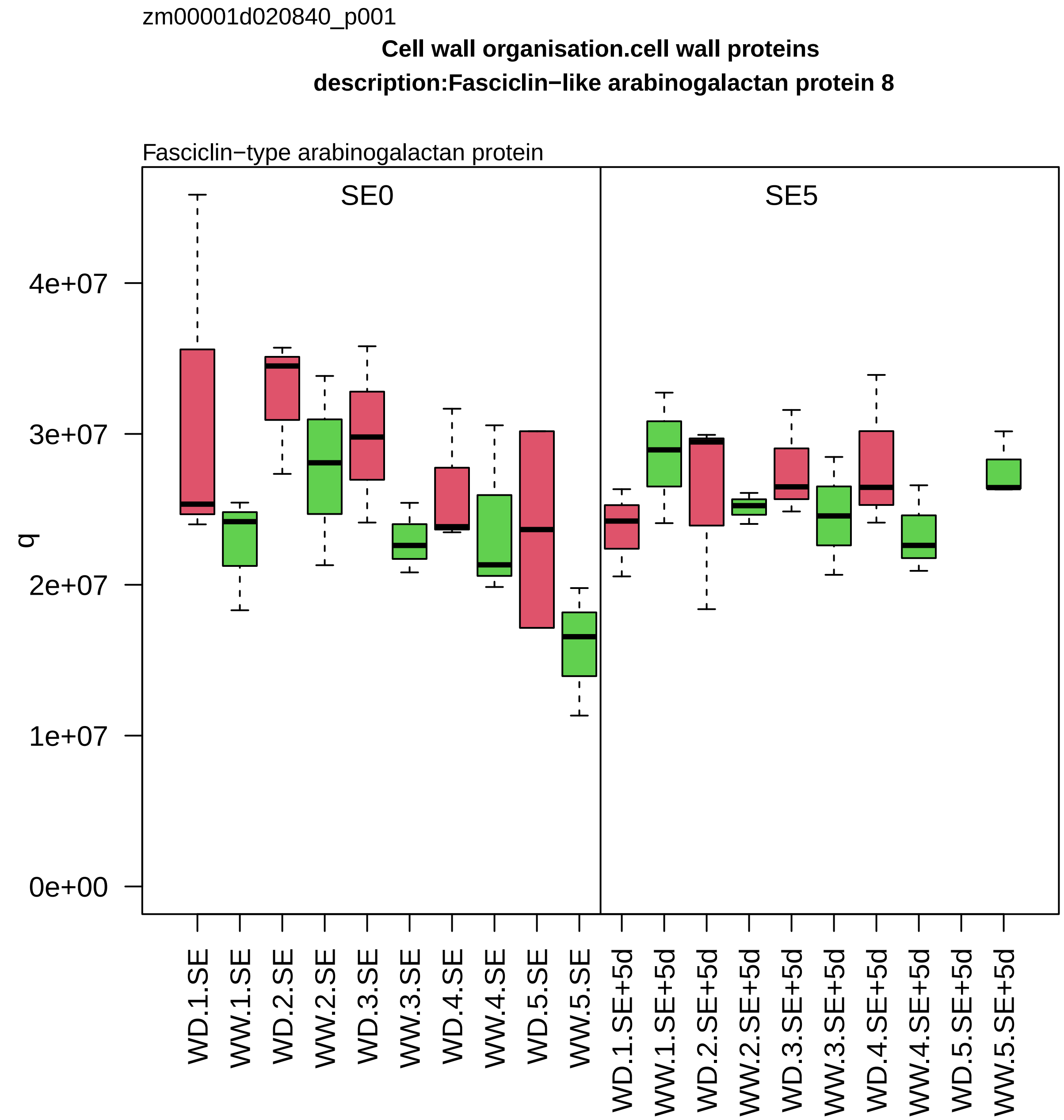
zm00001d020840_p001
Cell wall organisation.cell wall proteins
description:Fasciclin−like arabinogalactan protein 8
Fasciclin−type arabinogalactan protein
SE0
SE5
q
0e+00
1e+07
2e+07
3e+07
4e+07
WD.1.SE
WW.1.SE
WD.2.SE
WW.2.SE
WD.3.SE
WW.3.SE
WD.4.SE
WW.4.SE
WD.5.SE
WW.5.SE
WD.1.SE+5d
WW.1.SE+5d
WD.2.SE+5d
WW.2.SE+5d
WD.3.SE+5d
WW.3.SE+5d
WD.4.SE+5d
WW.4.SE+5d
WD.5.SE+5d
WW.5.SE+5d

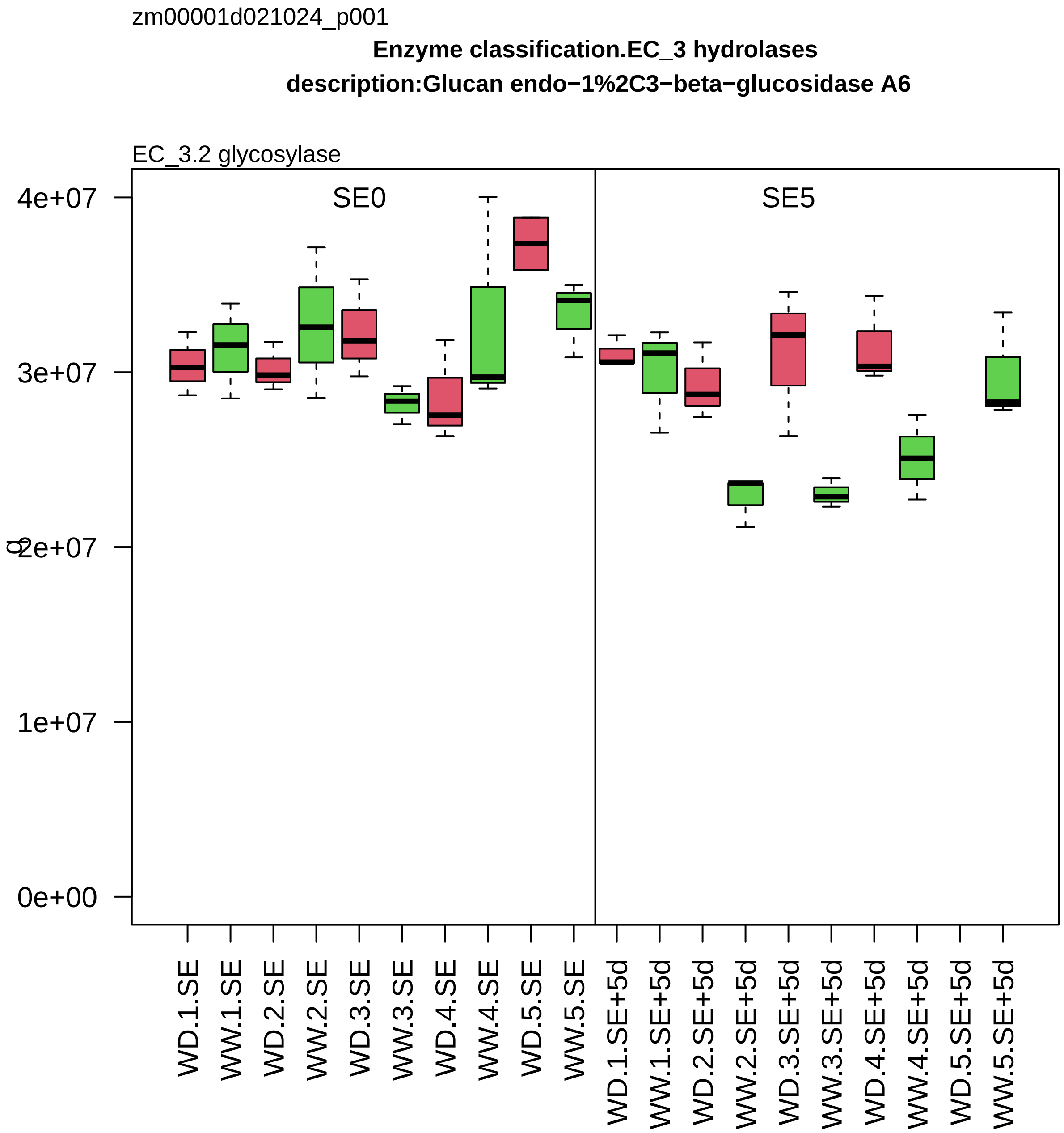
zm00001d021024_p001
Enzyme classification.EC_3 hydrolases
description:Glucan endo−1%2C3−beta−glucosidase A6
EC_3.2 glycosylase
SE0
SE5
4e+07
3e+07
2e+07
1e+07
0e+00
WD.1.SE
WW.1.SE
WD.2.SE
WW.2.SE
WD.3.SE
WW.3.SE
WD.4.SE
WW.4.SE
WD.5.SE
WW.5.SE
WD.1.SE+5d
WW.1.SE+5d
WD.2.SE+5d
WW.2.SE+5d
WD.3.SE+5d
WW.3.SE+5d
WD.4.SE+5d
WW.4.SE+5d
WD.5.SE+5d
WW.5.SE+5d

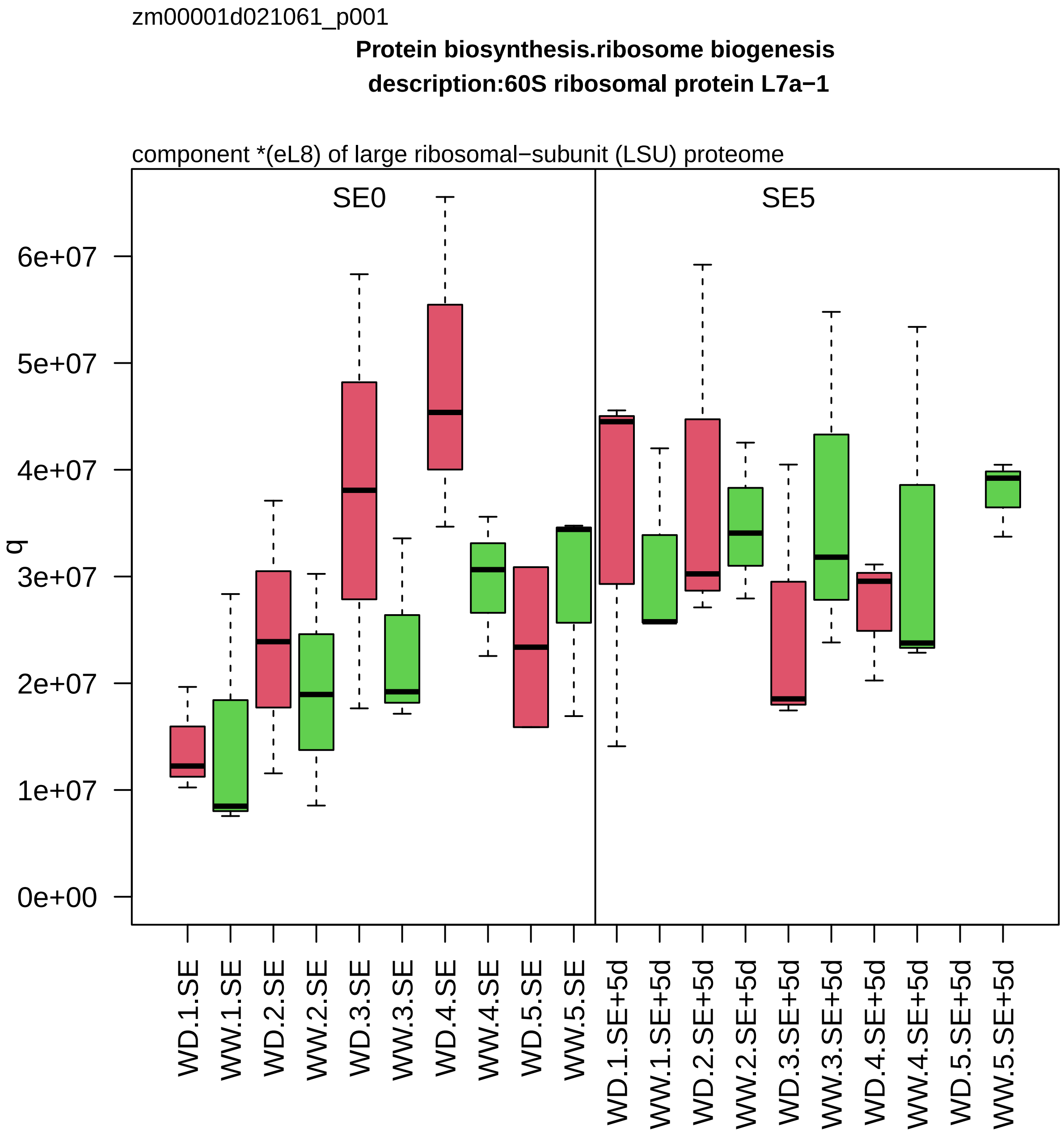
zm00001d021061_p001
Protein biosynthesis.ribosome biogenesis
description:60S ribosomal protein L7a−1
component *(eL8) of large ribosomal−subunit (LSU) proteome
SE0
SE5
q
0e+00
1e+07
2e+07
3e+07
4e+07
5e+07
6e+07
WD.1.SE
WW.1.SE
WD.2.SE
WW.2.SE
WD.3.SE
WW.3.SE
WD.4.SE
WW.4.SE
WD.5.SE
WW.5.SE
WD.1.SE+5d
WW.1.SE+5d
WD.2.SE+5d
WW.2.SE+5d
WD.3.SE+5d
WW.3.SE+5d
WD.4.SE+5d
WW.4.SE+5d
WD.5.SE+5d
WW.5.SE+5d

zm00001d021147_p001

**Protein homeostasis.ubiquitin−proteasome system**
**description:Proteasome subunit beta type−4**

beta−type−7 component *(PBG) of 26S proteasome

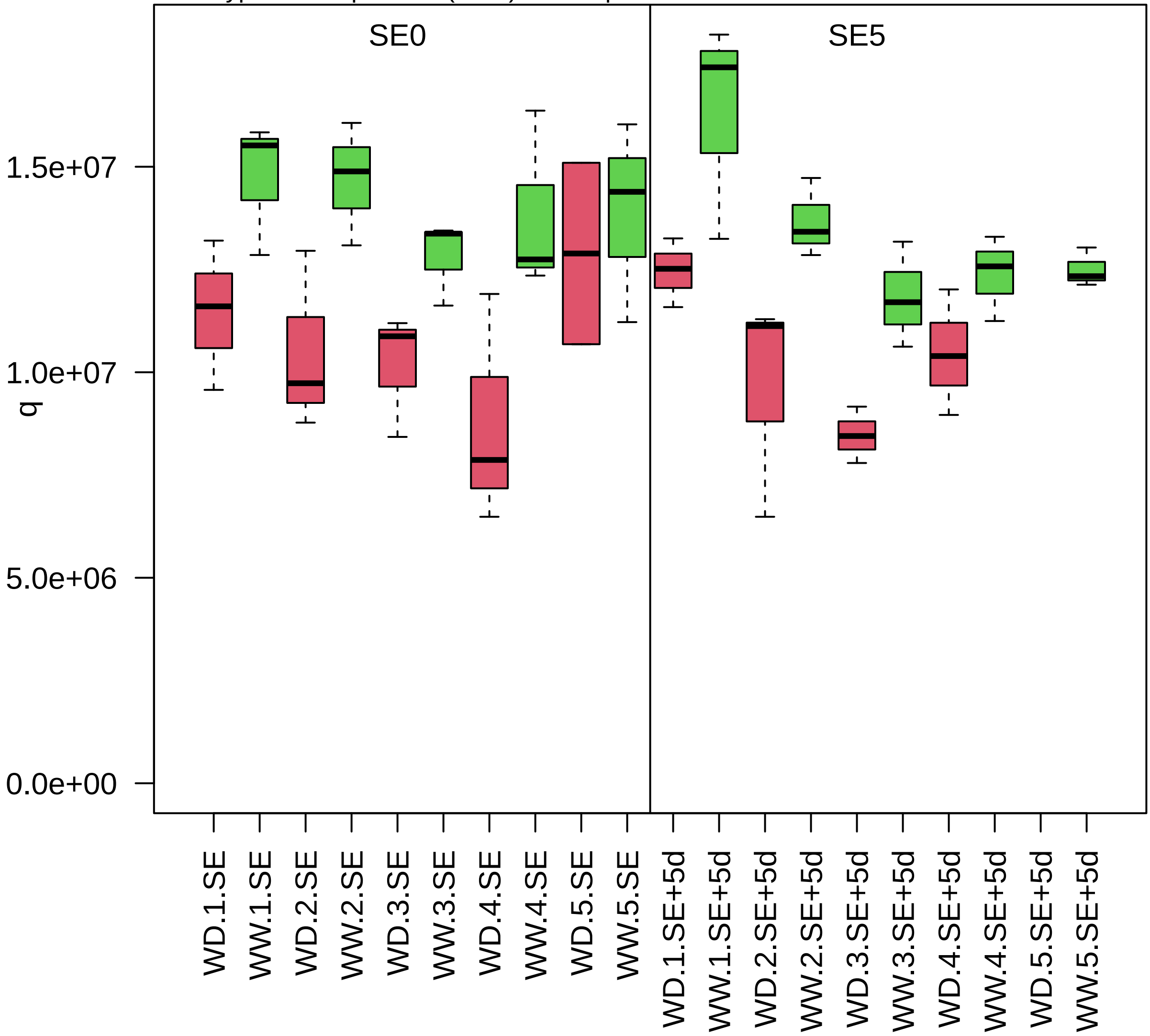

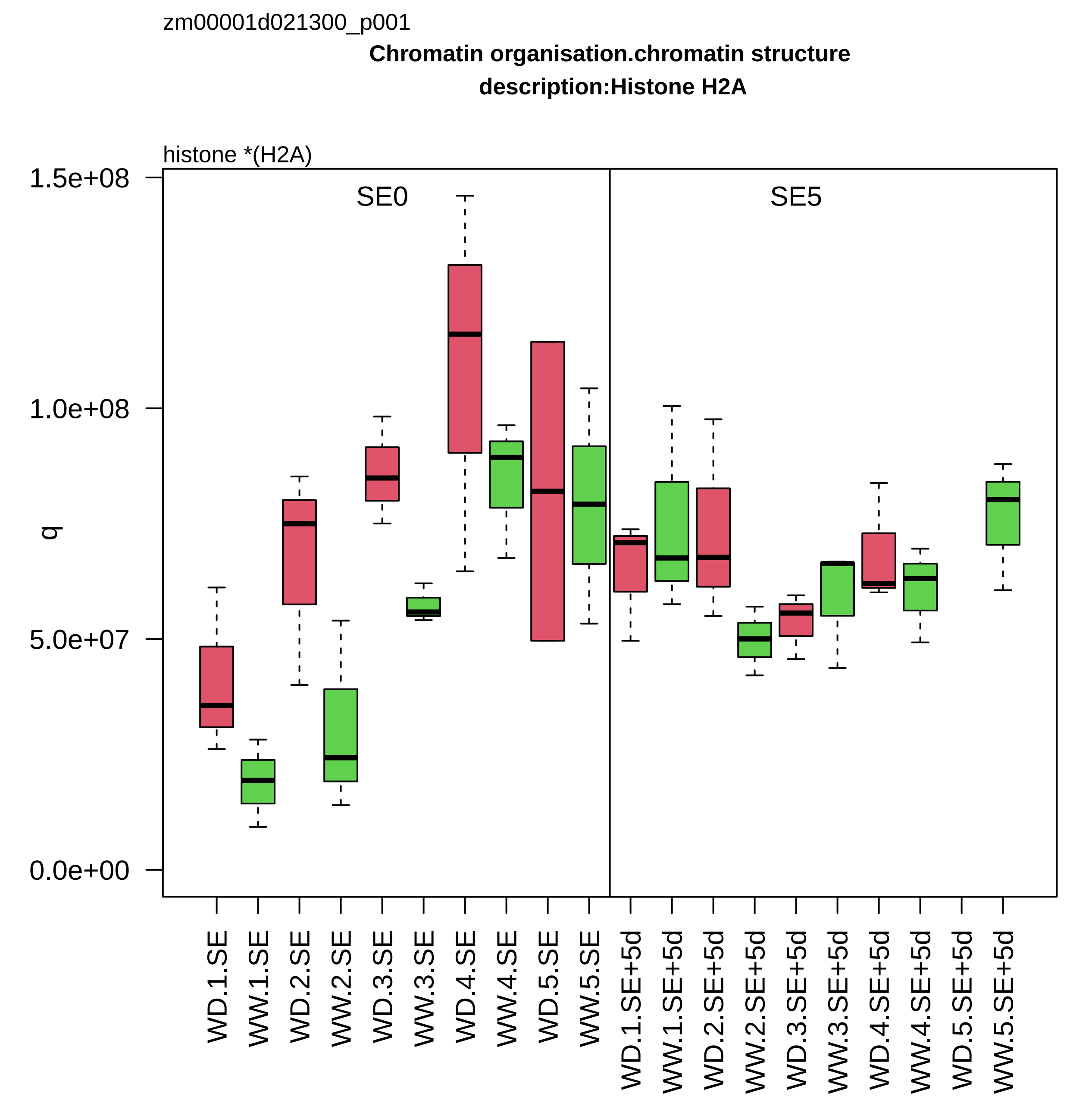

zm00001d021300_p001
Chromatin organisation.chromatin structure
description:Histone H2A
histone *(H2A)
SE0
SE5
q
0.0e+00
5.0e+07
1.0e+08
1.5e+08
WD.1.SE
WW.1.SE
WD.2.SE
WW.2.SE
WD.3.SE
WW.3.SE
WD.4.SE
WW.4.SE
WD.5.SE
WW.5.SE
WD.1.SE+5d
WW.1.SE+5d
WD.2.SE+5d
WW.2.SE+5d
WD.3.SE+5d
WW.3.SE+5d
WD.4.SE+5d
WW.4.SE+5d
WD.5.SE+5d
WW.5.SE+5d

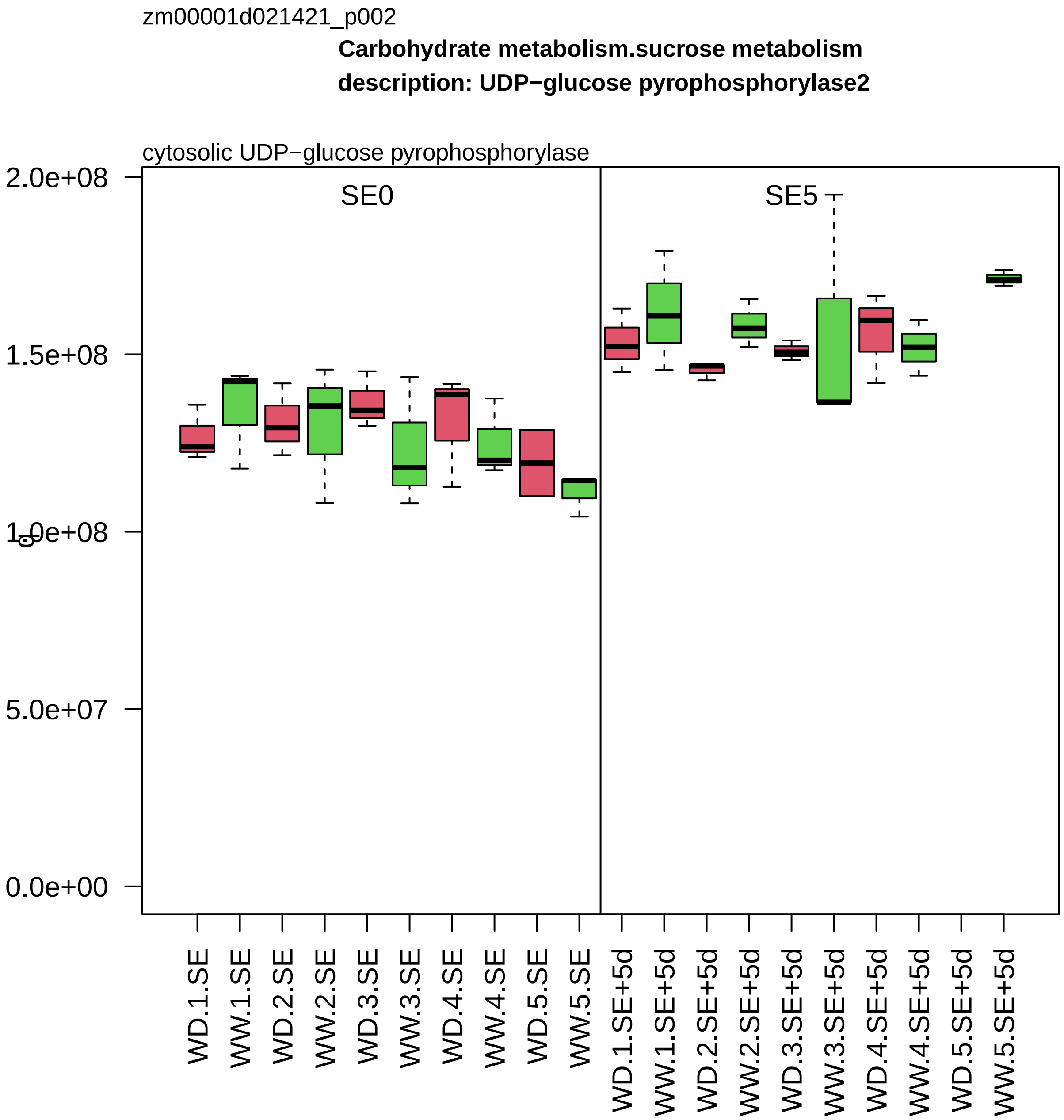

zm00001d021421_p002
Carbohydrate metabolism.sucrose metabolism
description: UDP−glucose pyrophosphorylase2
cytosolic UDP−glucose pyrophosphorylase
SE0
SE5
2.0e+08
1.5e+08
1.0e+08
5.0e+07
0.0e+00
WD.1.SE
WW.1.SE
WD.2.SE
WW.2.SE
WD.3.SE
WW.3.SE
WD.4.SE
WW.4.SE
WD.5.SE
WW.5.SE
WD.1.SE+5d
WW.1.SE+5d
WD.2.SE+5d
WW.2.SE+5d
WD.3.SE+5d
WW.3.SE+5d
WD.4.SE+5d
WW.4.SE+5d
WD.5.SE+5d
WW.5.SE+5d

zm00001d021447_p001

**External stimuli response.pathogen**

**description:LysM domain−containing GPI−anchored protein 2**

chitin receptor protein kinase *(CEBiP)

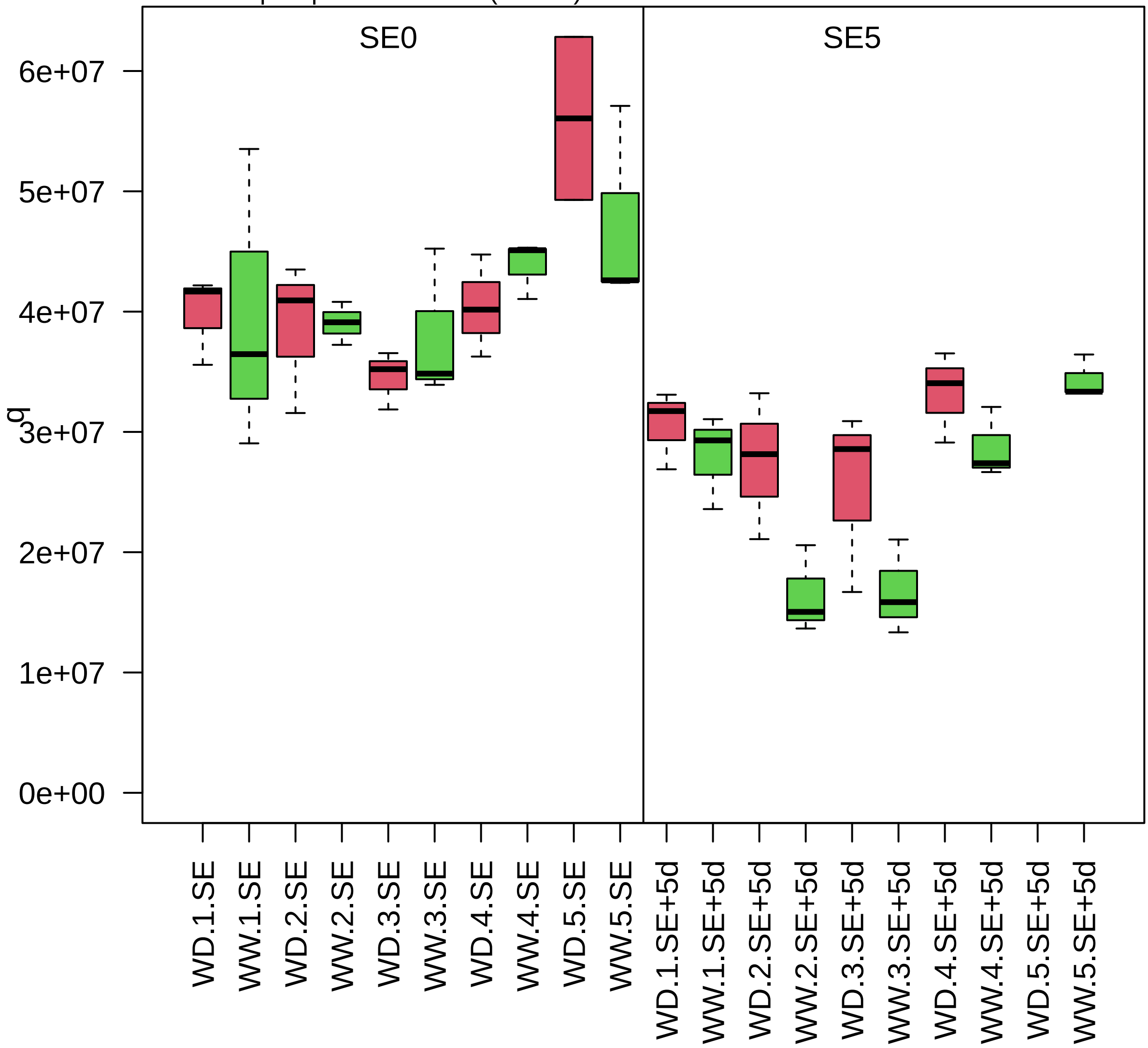

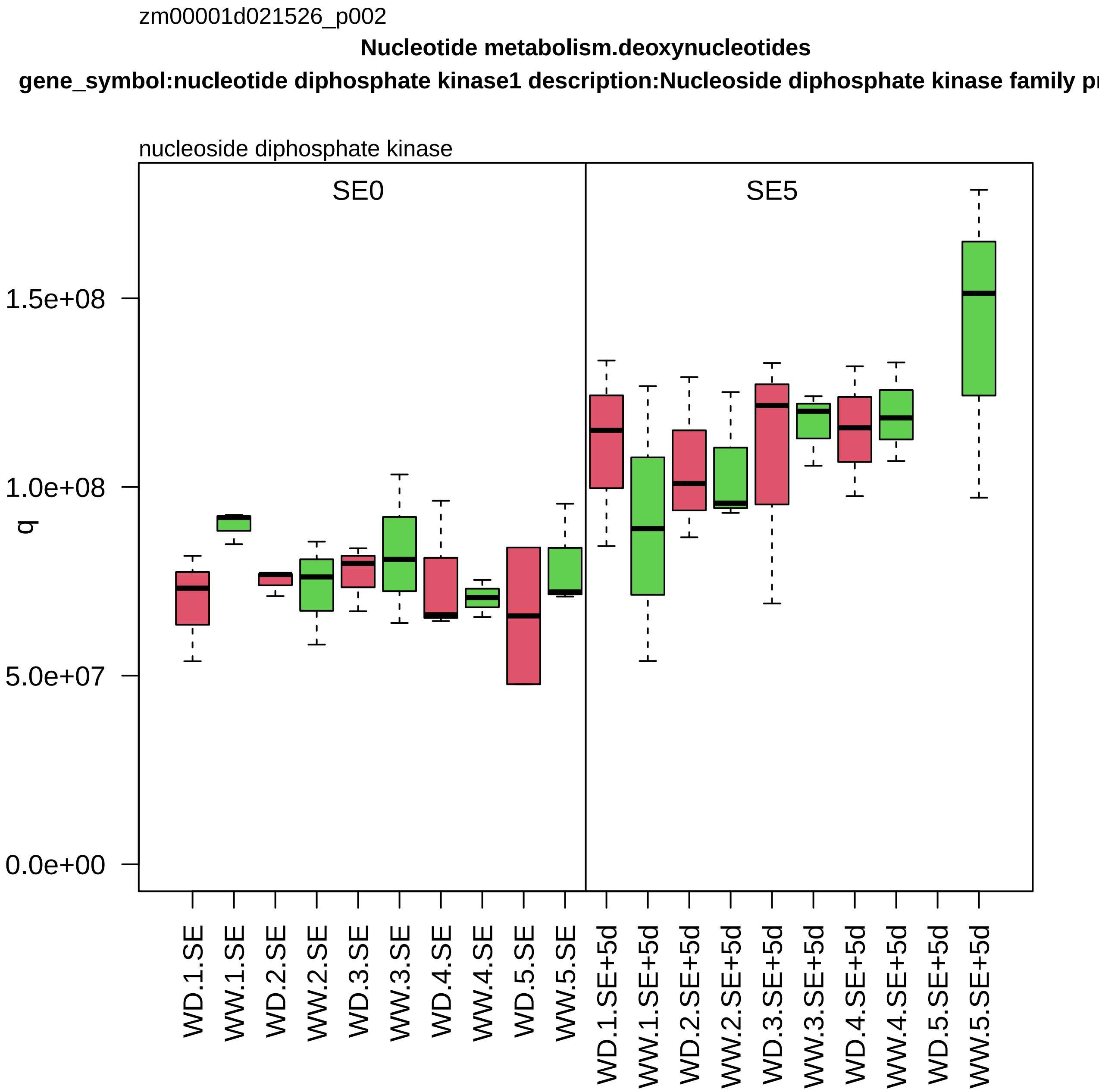
zm00001d021526_p002
Nucleotide metabolism.deoxynucleotides
gene_symbol:nucleotide diphosphate kinase1 description:Nucleoside diphosphate kinase family pr
nucleoside diphosphate kinase
SE0
SE5
q
1.5e+08
1.0e+08
5.0e+07
0.0e+00
WD.1.SE
WW.1.SE
WD.2.SE
WW.2.SE
WD.3.SE
WW.3.SE
WD.4.SE
WW.4.SE
WD.5.SE
WW.5.SE
WD.1.SE+5d
WW.1.SE+5d
WD.2.SE+5d
WW.2.SE+5d
WD.3.SE+5d
WW.3.SE+5d
WD.4.SE+5d
WW.4.SE+5d
WD.5.SE+5d
WW.5.SE+5d

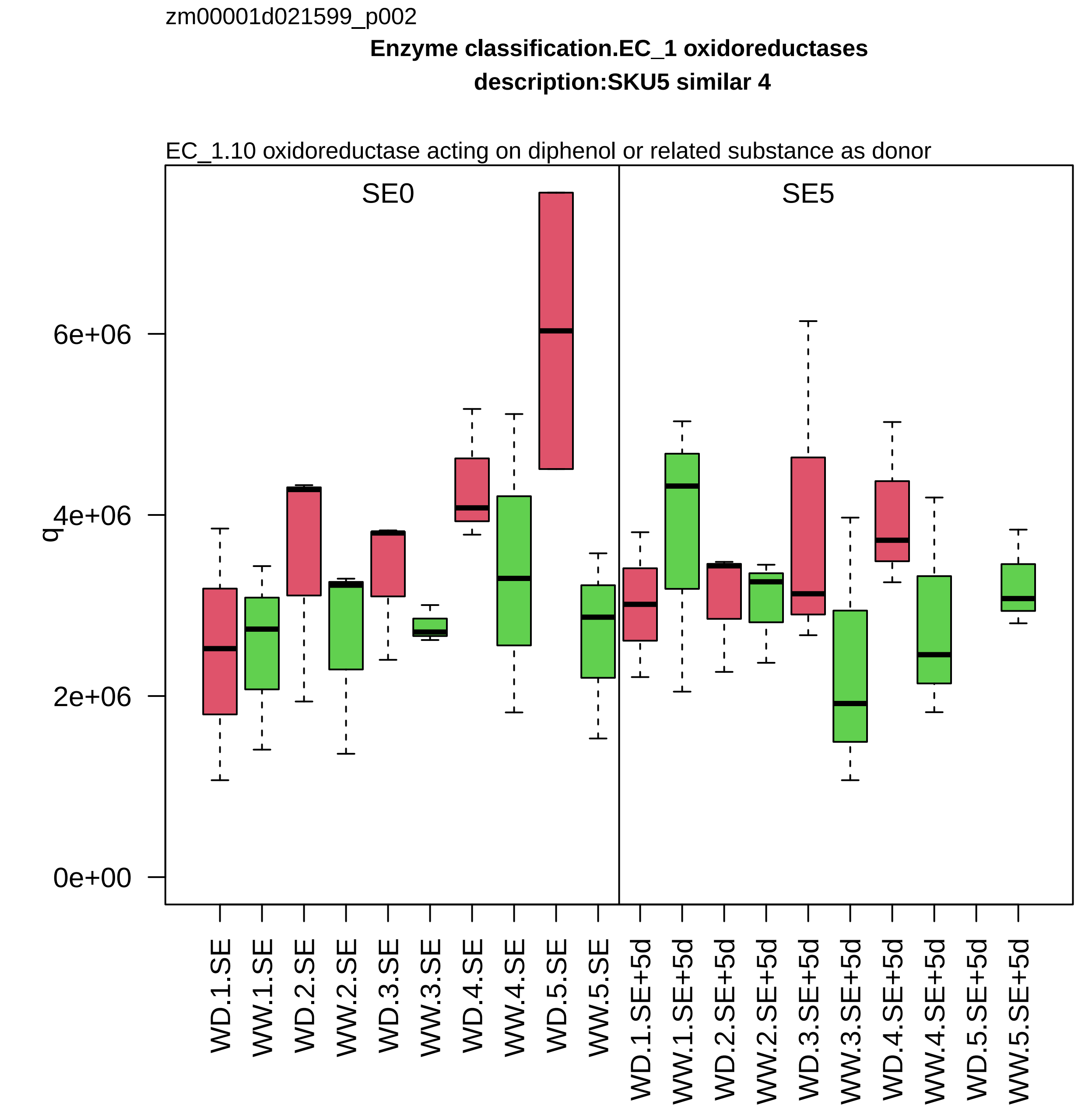

zm00001d021599_p002
Enzyme classification.EC_1 oxidoreductases
description:SKU5 similar 4
EC_1.10 oxidoreductase acting on diphenol or related substance as donor
SE0
SE5
q
0e+00
2e+06
4e+06
6e+06
WD.1.SE
WW.1.SE
WD.2.SE
WW.2.SE
WD.3.SE
WW.3.SE
WD.4.SE
WW.4.SE
WD.5.SE
WW.5.SE
WD.1.SE+5d
WW.1.SE+5d
WD.2.SE+5d
WW.2.SE+5d
WD.3.SE+5d
WW.3.SE+5d
WD.4.SE+5d
WW.4.SE+5d
WD.5.SE+5d
WW.5.SE+5d

zm00001d021626_p002

**Amino acid metabolism.aspartate group amino acid biosynthesis**

**description:Reactive Intermediate Deaminase A chloroplastic**

threonine reactive imine intermediate deaminase *(RidA)

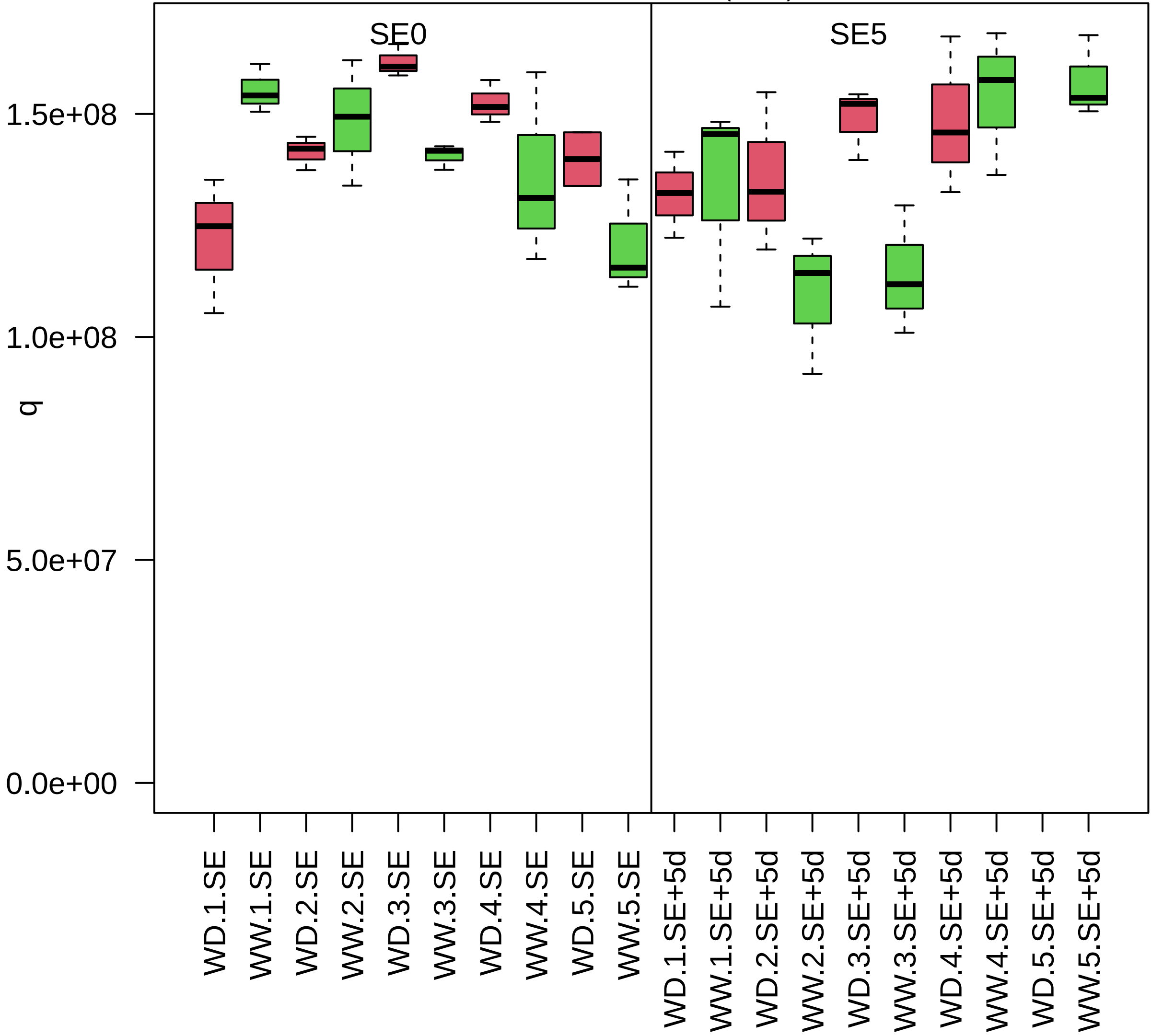

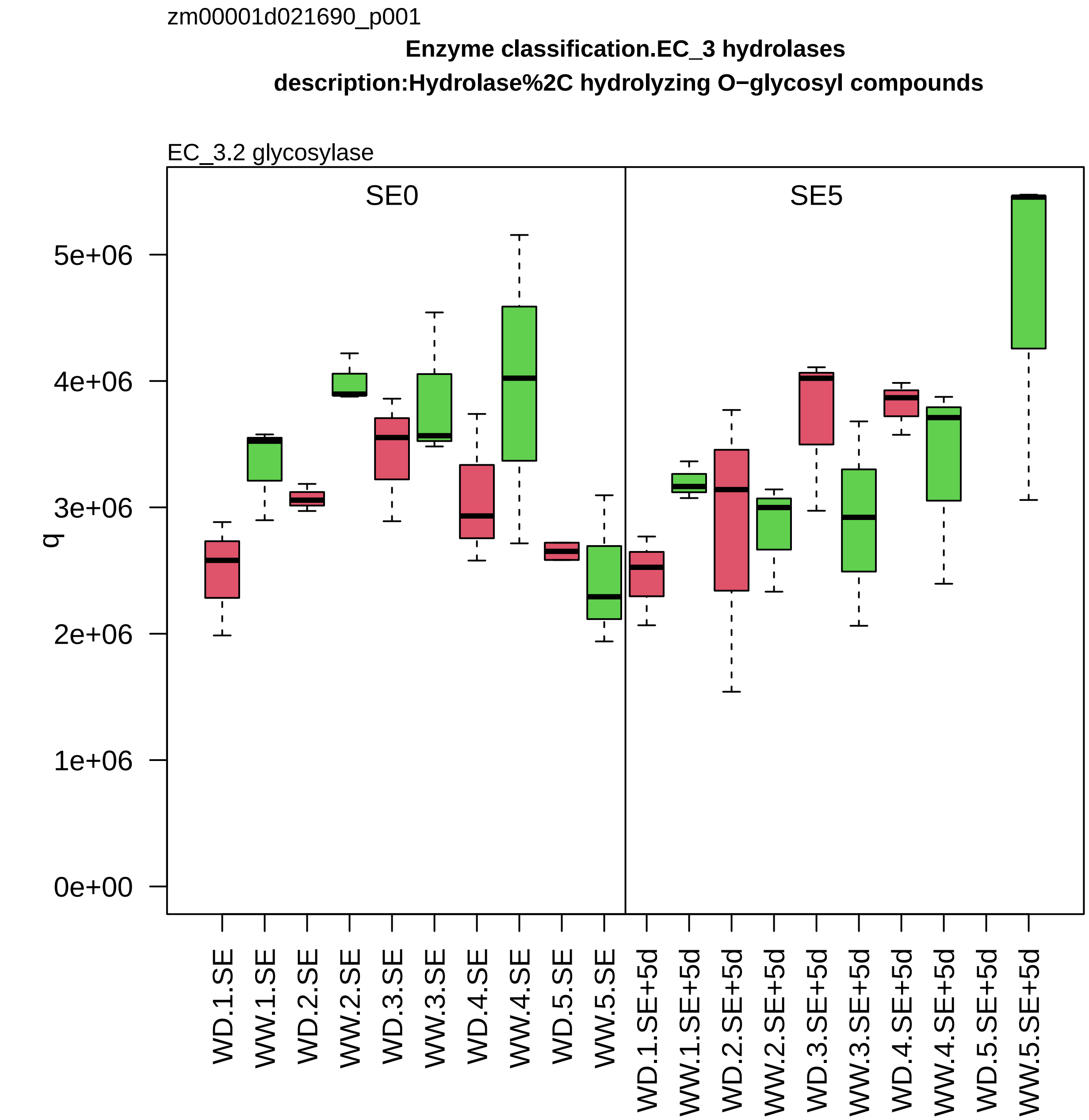
zm00001d021690_p001
Enzyme classification.EC_3 hydrolases
description:Hydrolase%2C hydrolyzing O−glycosyl compounds
EC_3.2 glycosylase
SE0
SE5
q
0e+00
1e+06
2e+06
3e+06
4e+06
5e+06
WD.1.SE
WW.1.SE
WD.2.SE
WW.2.SE
WD.3.SE
WW.3.SE
WD.4.SE
WW.4.SE
WD.5.SE
WW.5.SE
WD.1.SE+5d
WW.1.SE+5d
WD.2.SE+5d
WW.2.SE+5d
WD.3.SE+5d
WW.3.SE+5d
WD.4.SE+5d
WW.4.SE+5d
WD.5.SE+5d
WW.5.SE+5d

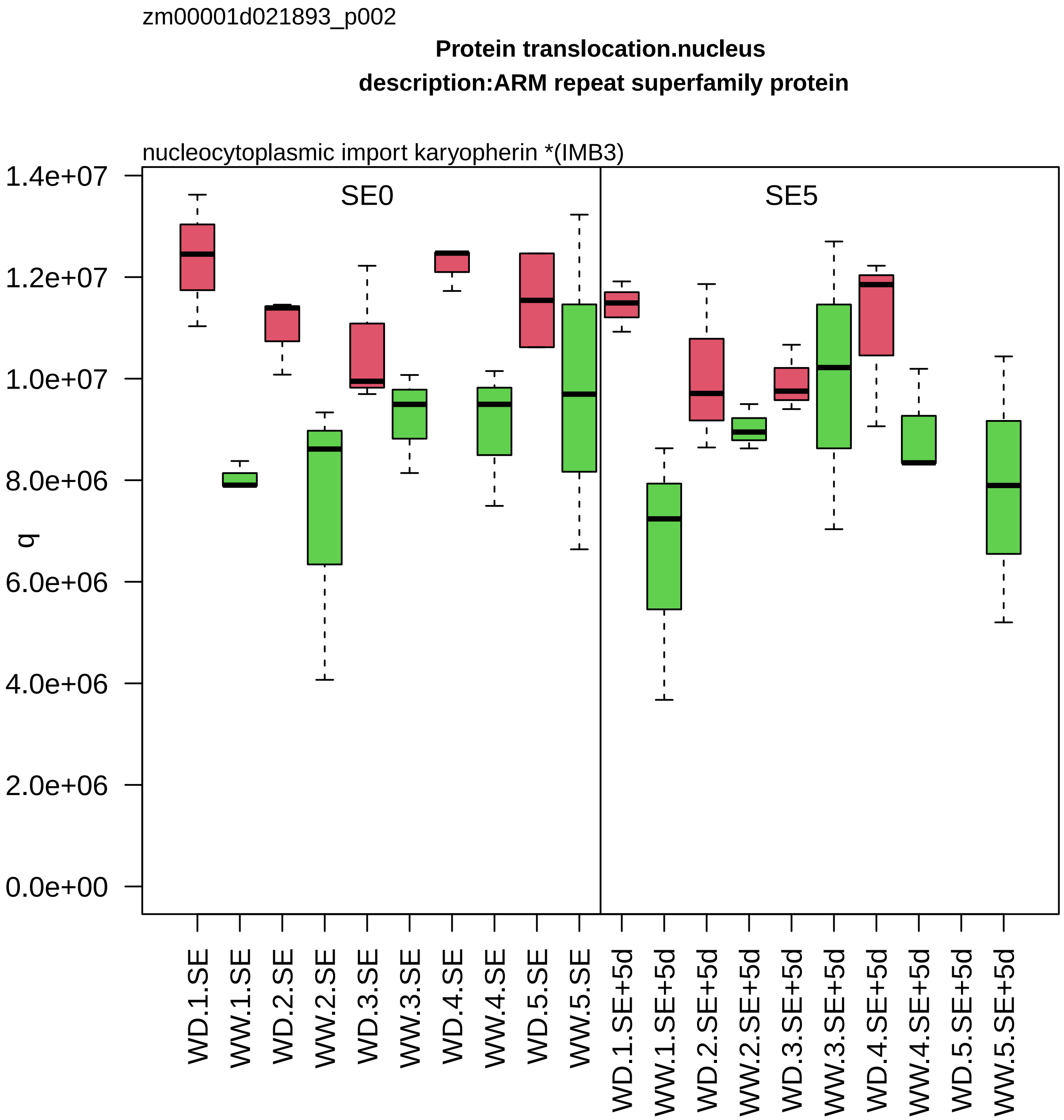
zm00001d021893_p002
Protein translocation.nucleus
description:ARM repeat superfamily protein
nucleocytoplasmic import karyopherin *(IMB3)
SE0
SE5
q
1.4e+07
1.2e+07
1.0e+07
8.0e+06
6.0e+06
4.0e+06
2.0e+06
0.0e+00
WD.1.SE
WW.1.SE
WD.2.SE
WW.2.SE
WD.3.SE
WW.3.SE
WD.4.SE
WW.4.SE
WD.5.SE
WW.5.SE
WD.1.SE+5d
WW.1.SE+5d
WD.2.SE+5d
WW.2.SE+5d
WD.3.SE+5d
WW.3.SE+5d
WD.4.SE+5d
WW.4.SE+5d
WD.5.SE+5d
WW.5.SE+5d

zm00001d021999_p002

**RNA processing.RNA homeostasis**

**description:Nuclear transport factor 2 (NTF2) family protein with RNA binding (RRM−RBD−RNP motifs)**

mRNA endoribonuclease *(G3BP)

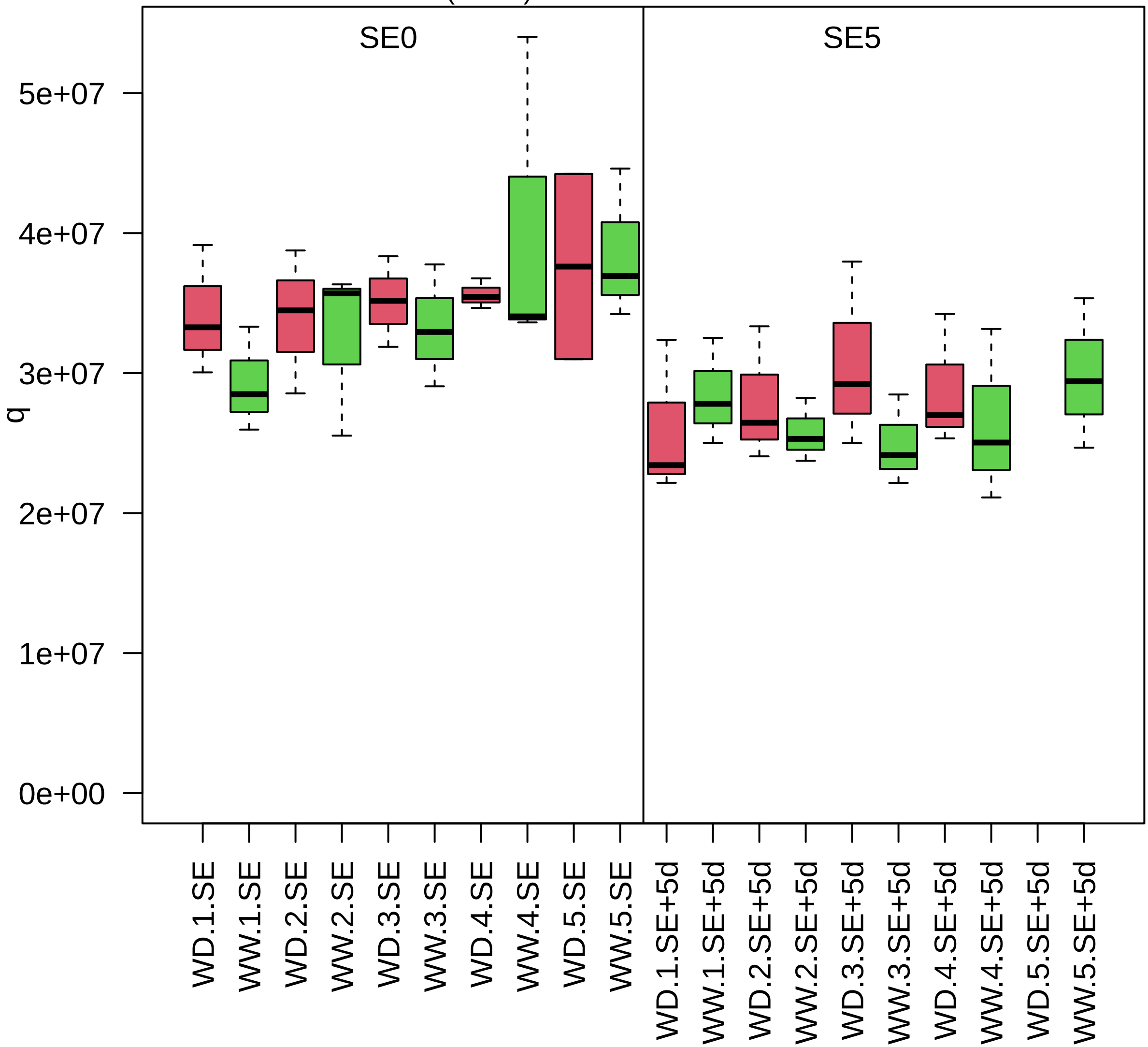

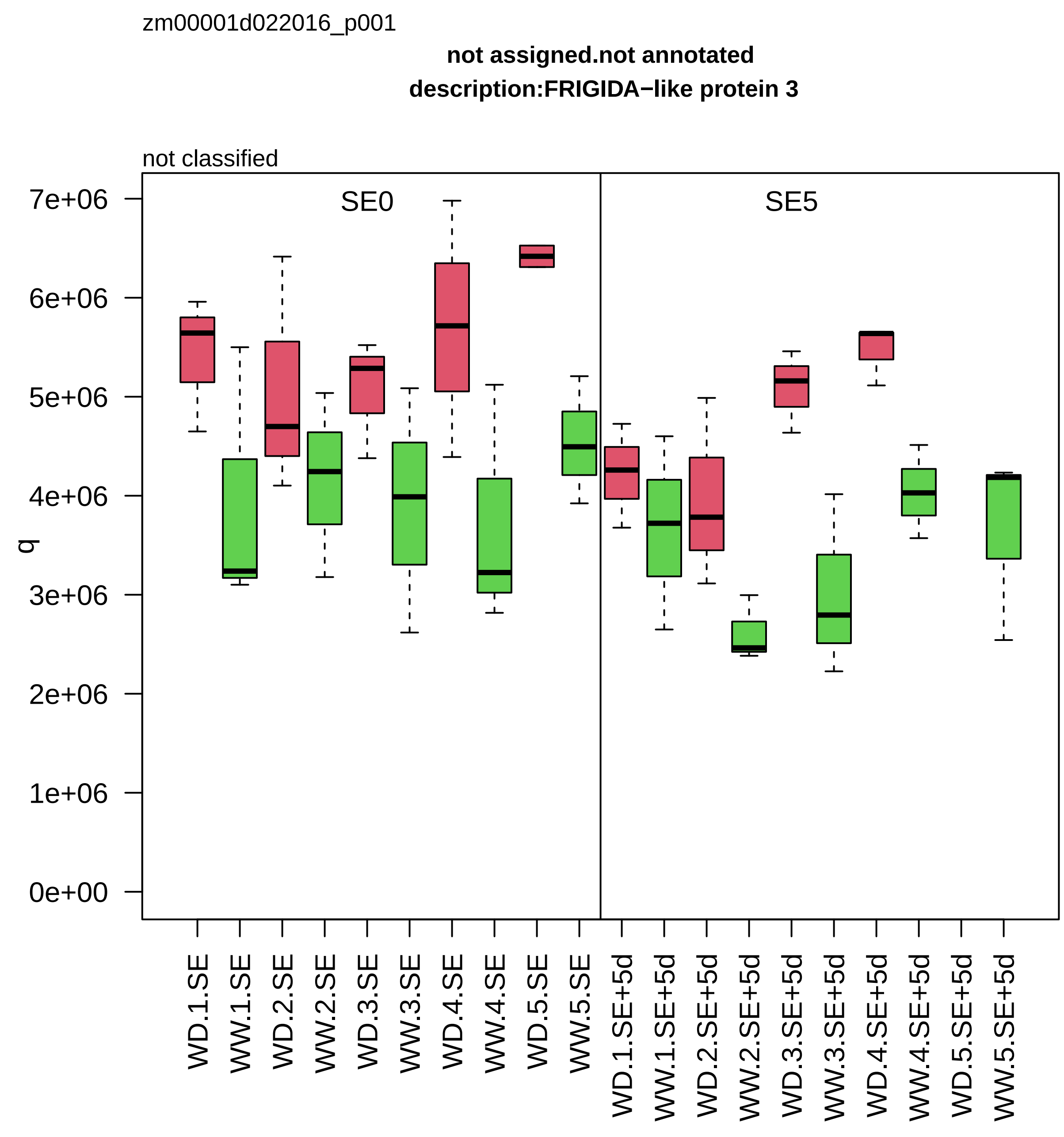
zm00001d022016_p001
not assigned.not annotated
description:FRIGIDA−like protein 3
not classified
SE0
SE5
q
7e+06
6e+06
5e+06
4e+06
3e+06
2e+06
1e+06
0e+00
WD.1.SE
WW.1.SE
WD.2.SE
WW.2.SE
WD.3.SE
WW.3.SE
WD.4.SE
WW.4.SE
WD.5.SE
WW.5.SE
WD.1.SE+5d
WW.1.SE+5d
WD.2.SE+5d
WW.2.SE+5d
WD.3.SE+5d
WW.3.SE+5d
WD.4.SE+5d
WW.4.SE+5d
WD.5.SE+5d
WW.5.SE+5d

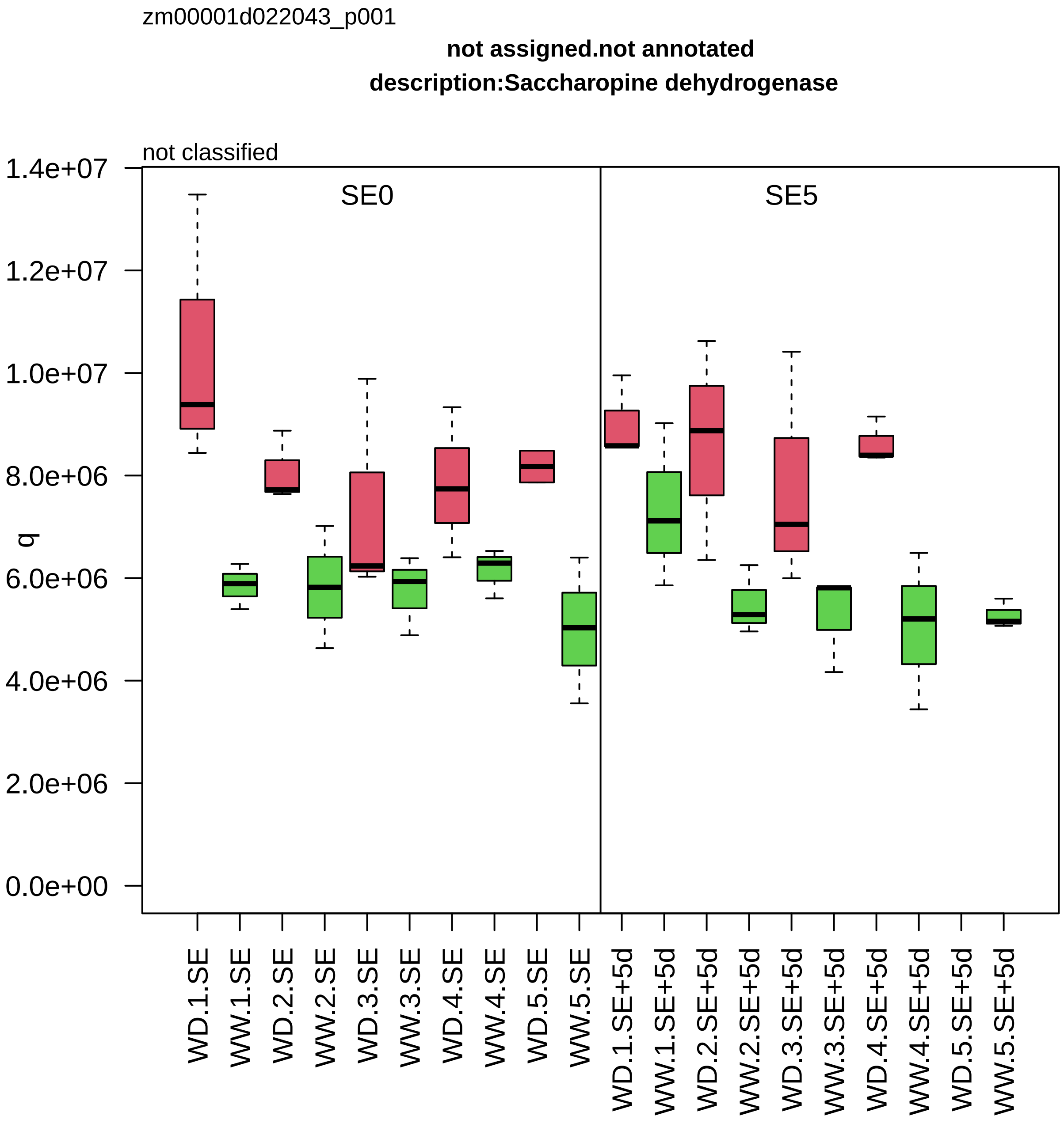

zm00001d022043_p001
not assigned.not annotated
description:Saccharopine dehydrogenase
not classified
SE0
SE5
q
1.4e+07
1.2e+07
1.0e+07
8.0e+06
6.0e+06
4.0e+06
2.0e+06
0.0e+00
WD.1.SE
WW.1.SE
WD.2.SE
WW.2.SE
WD.3.SE
WW.3.SE
WD.4.SE
WW.4.SE
WD.5.SE
WW.5.SE
WD.1.SE+5d
WW.1.SE+5d
WD.2.SE+5d
WW.2.SE+5d
WD.3.SE+5d
WW.3.SE+5d
WD.4.SE+5d
WW.4.SE+5d
WD.5.SE+5d
WW.5.SE+5d

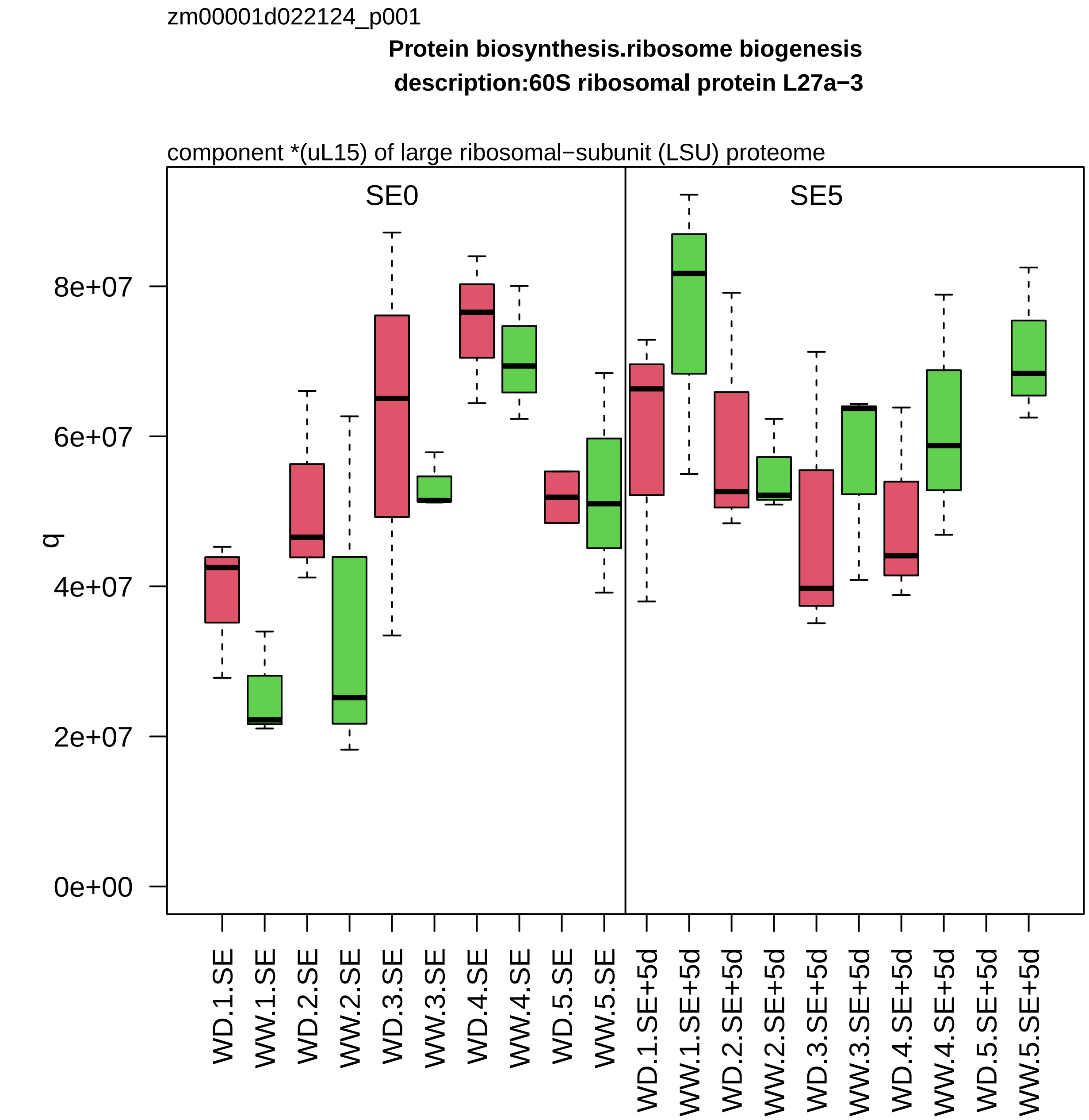

zm00001d022124_p001
Protein biosynthesis.ribosome biogenesis
description:60S ribosomal protein L27a−3
component *(uL15) of large ribosomal−subunit (LSU) proteome
SE0
SE5
q
0e+00
2e+07
4e+07
6e+07
8e+07
WD.1.SE
WW.1.SE
WD.2.SE
WW.2.SE
WD.3.SE
WW.3.SE
WD.4.SE
WW.4.SE
WD.5.SE
WW.5.SE
WD.1.SE+5d
WW.1.SE+5d
WD.2.SE+5d
WW.2.SE+5d
WD.3.SE+5d
WW.3.SE+5d
WD.4.SE+5d
WW.4.SE+5d
WD.5.SE+5d
WW.5.SE+5d

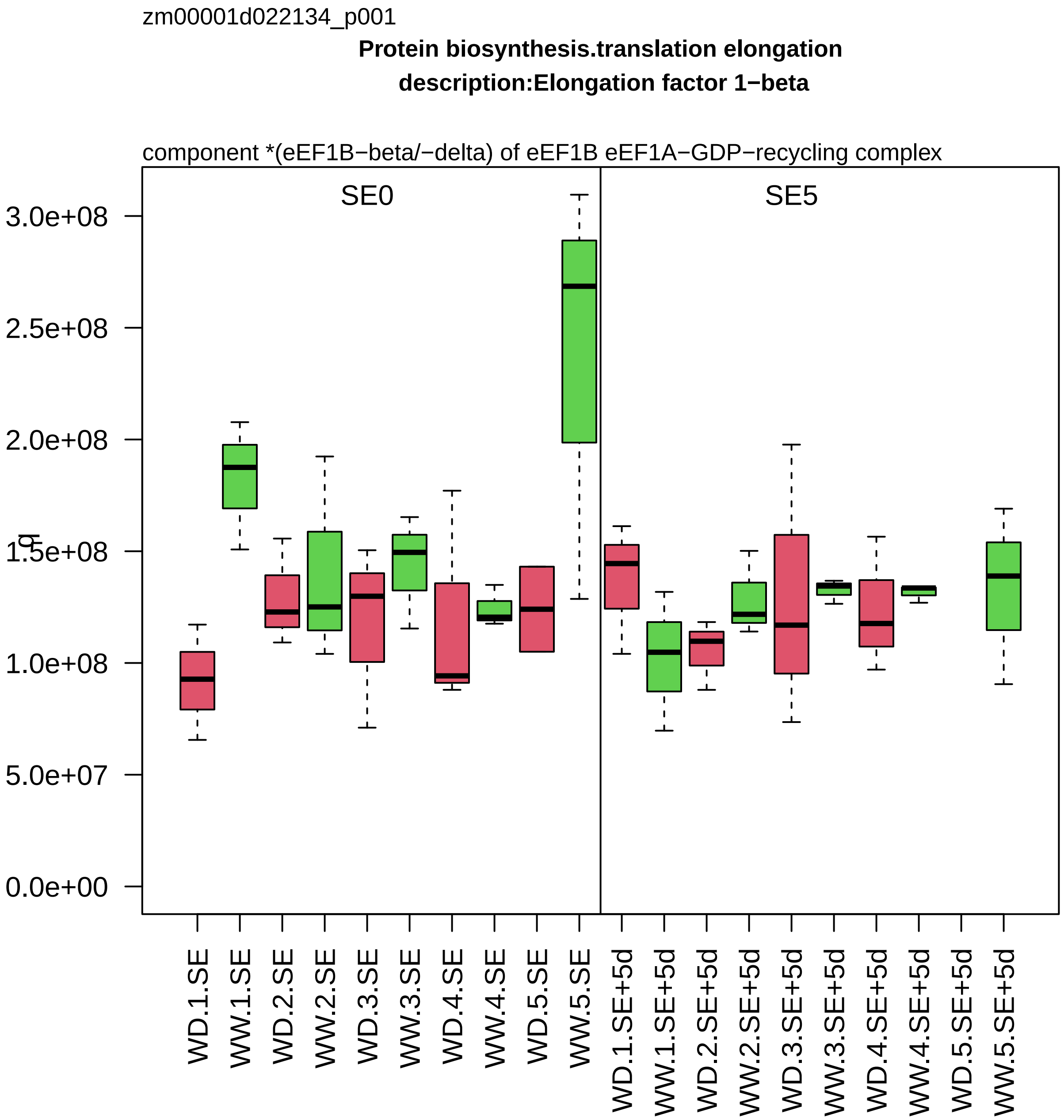
zm00001d022134_p001
Protein biosynthesis.translation elongation
description:Elongation factor 1−beta
component *(eEF1B−beta/−delta) of eEF1B eEF1A−GDP−recycling complex
SE0
SE5
q
3.0e+08
2.5e+08
2.0e+08
1.5e+08
1.0e+08
5.0e+07
0.0e+00
WD.1.SE
WW.1.SE
WD.2.SE
WW.2.SE
WD.3.SE
WW.3.SE
WD.4.SE
WW.4.SE
WD.5.SE
WW.5.SE
WD.1.SE+5d
WW.1.SE+5d
WD.2.SE+5d
WW.2.SE+5d
WD.3.SE+5d
WW.3.SE+5d
WD.4.SE+5d
WW.4.SE+5d
WD.5.SE+5d
WW.5.SE+5d

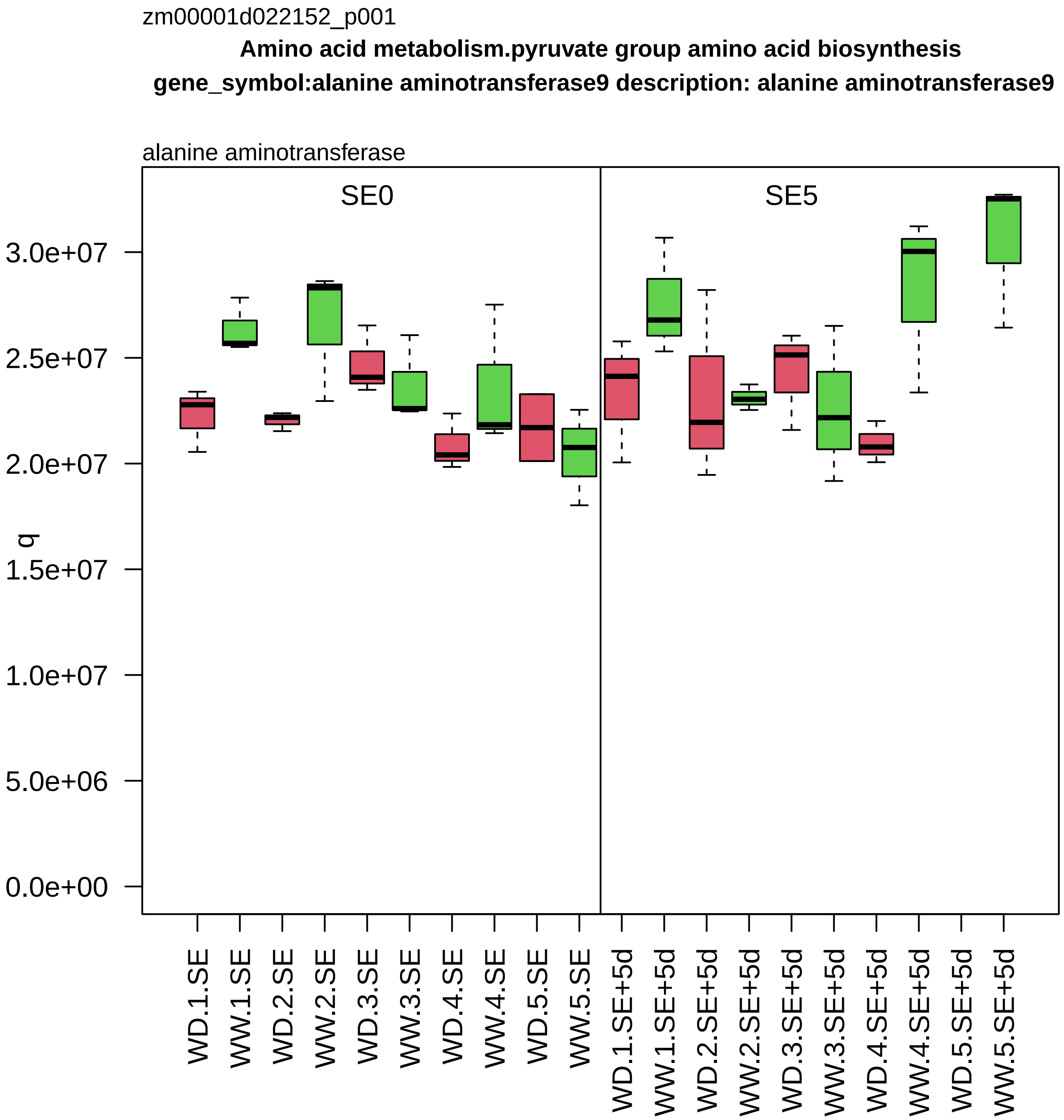

zm00001d022152_p001
Amino acid metabolism.pyruvate group amino acid biosynthesis
gene_symbol:alanine aminotransferase9 description: alanine aminotransferase9
alanine aminotransferase
SE0
SE5
q
3.0e+07
2.5e+07
2.0e+07
1.5e+07
1.0e+07
5.0e+06
0.0e+00
WD.1.SE
WW.1.SE
WD.2.SE
WW.2.SE
WD.3.SE
WW.3.SE
WD.4.SE
WW.4.SE
WD.5.SE
WW.5.SE
WD.1.SE+5d
WW.1.SE+5d
WD.2.SE+5d
WW.2.SE+5d
WD.3.SE+5d
WW.3.SE+5d
WD.4.SE+5d
WW.4.SE+5d
WD.5.SE+5d
WW.5.SE+5d

zm00001d022181_p004

**Amino acid metabolism.shikimate group amino acid biosynthesis**
**description:Phospho−2−dehydro−3−deoxyheptonate aldolase 1**

3−deoxy−D−arabino−heptulosonate 7−phosphate (DAHP) synthase

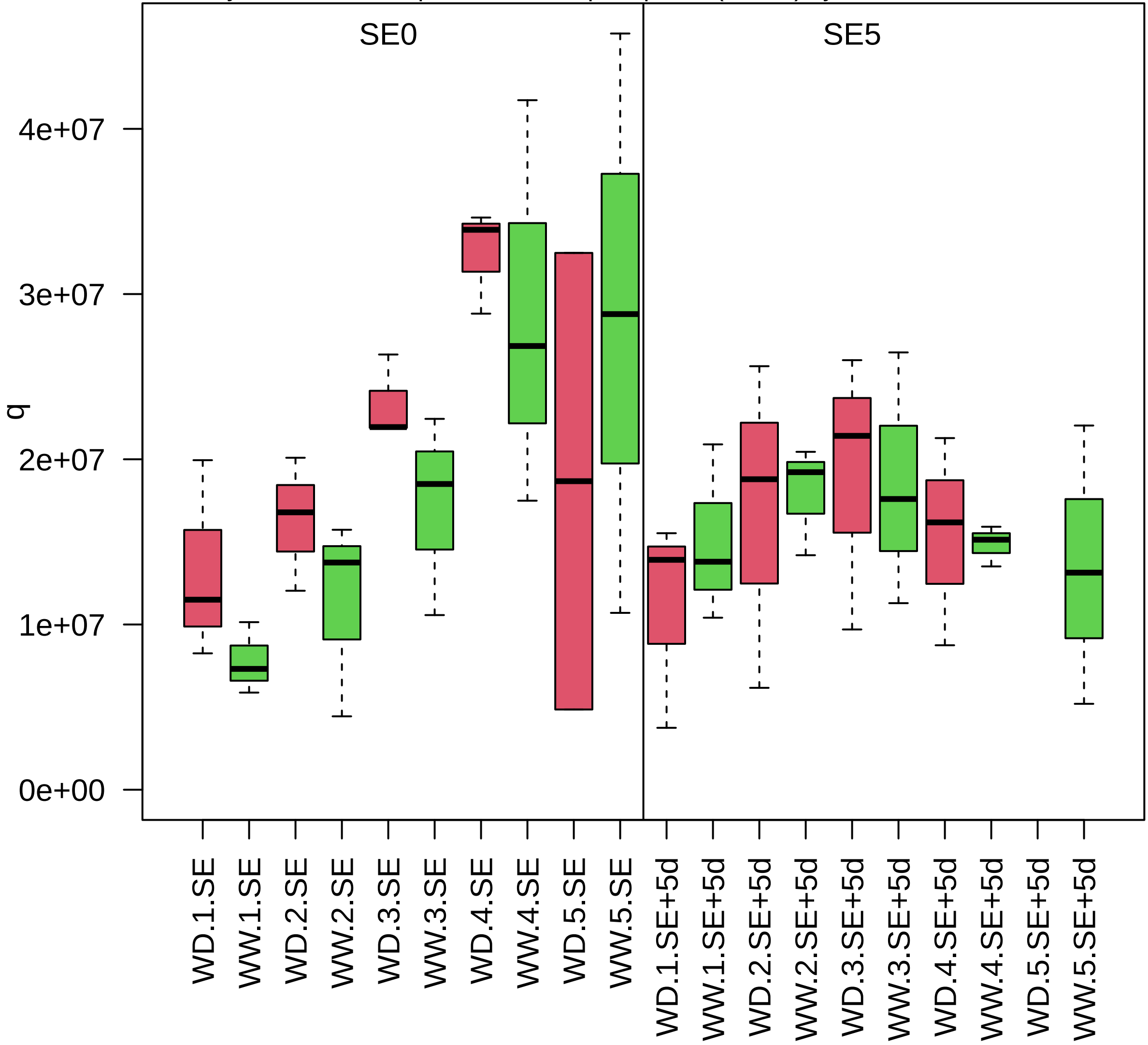

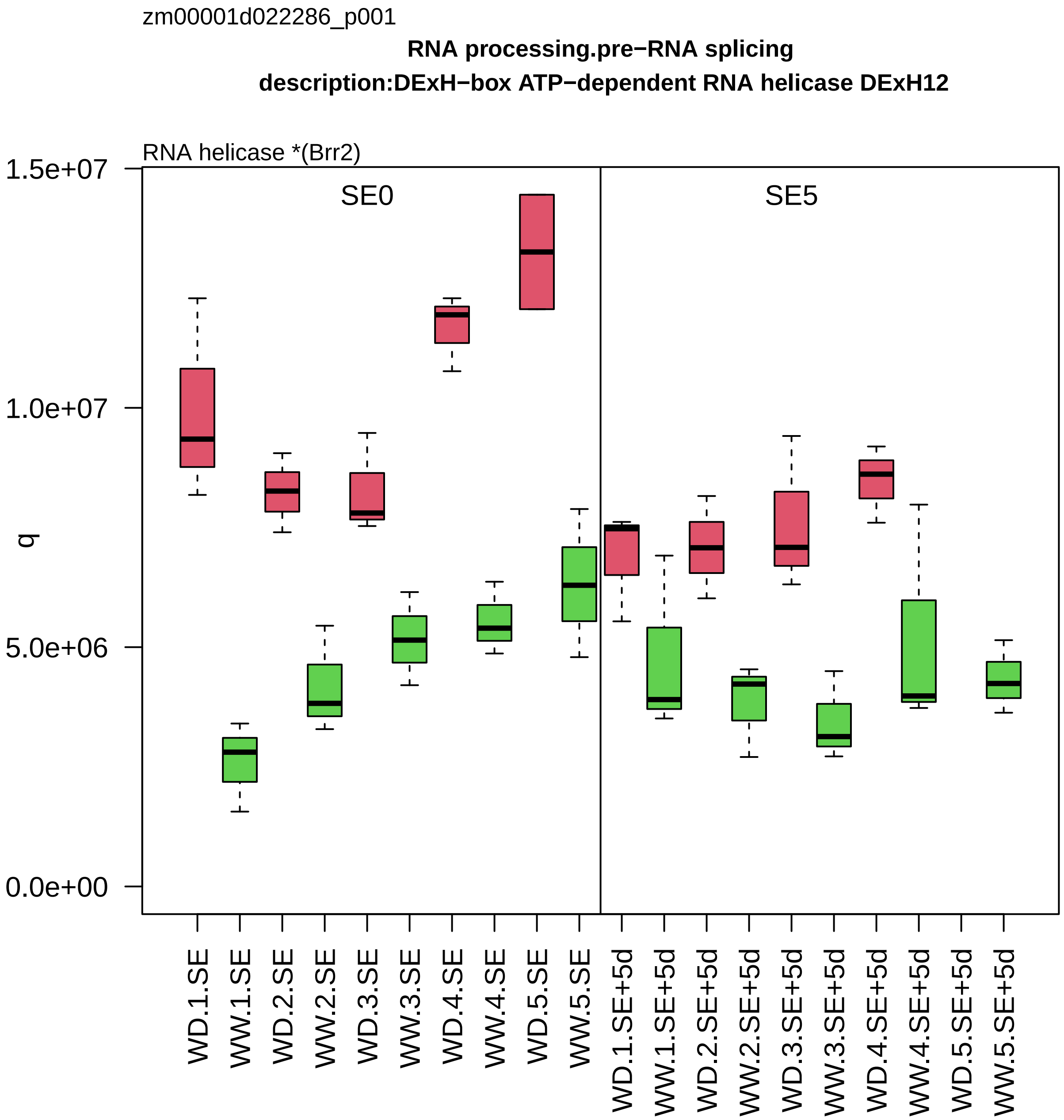

zm00001d022286_p001
RNA processing.pre−RNA splicing
description:DExH−box ATP−dependent RNA helicase DExH12
RNA helicase *(Brr2)
SE0
SE5
q
1.5e+07
1.0e+07
5.0e+06
0.0e+00
WD.1.SE
WW.1.SE
WD.2.SE
WW.2.SE
WD.3.SE
WW.3.SE
WD.4.SE
WW.4.SE
WD.5.SE
WW.5.SE
WD.1.SE+5d
WW.1.SE+5d
WD.2.SE+5d
WW.2.SE+5d
WD.3.SE+5d
WW.3.SE+5d
WD.4.SE+5d
WW.4.SE+5d
WD.5.SE+5d
WW.5.SE+5d

zm00001d022307_p001

**Protein homeostasis.protein quality control**

**description:10 kDa chaperonin**

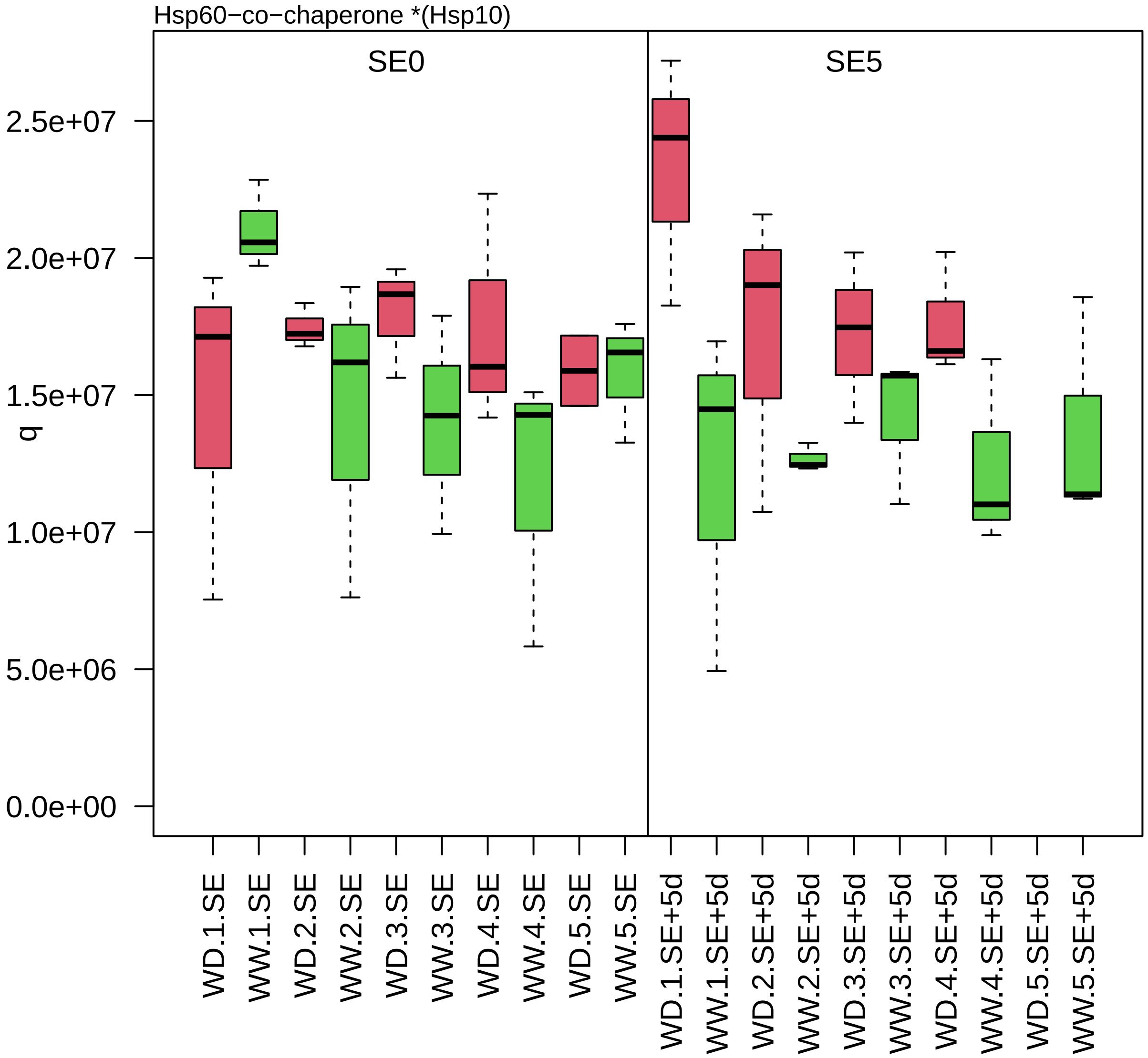

zm00001d022324_p001

**Cellular respiration.oxidative phosphorylation**

**description:Gamma carbonic anhydrase 2 mitochondrial**

component *(CA) of NADH dehydrogenase carbonic anhydrase module

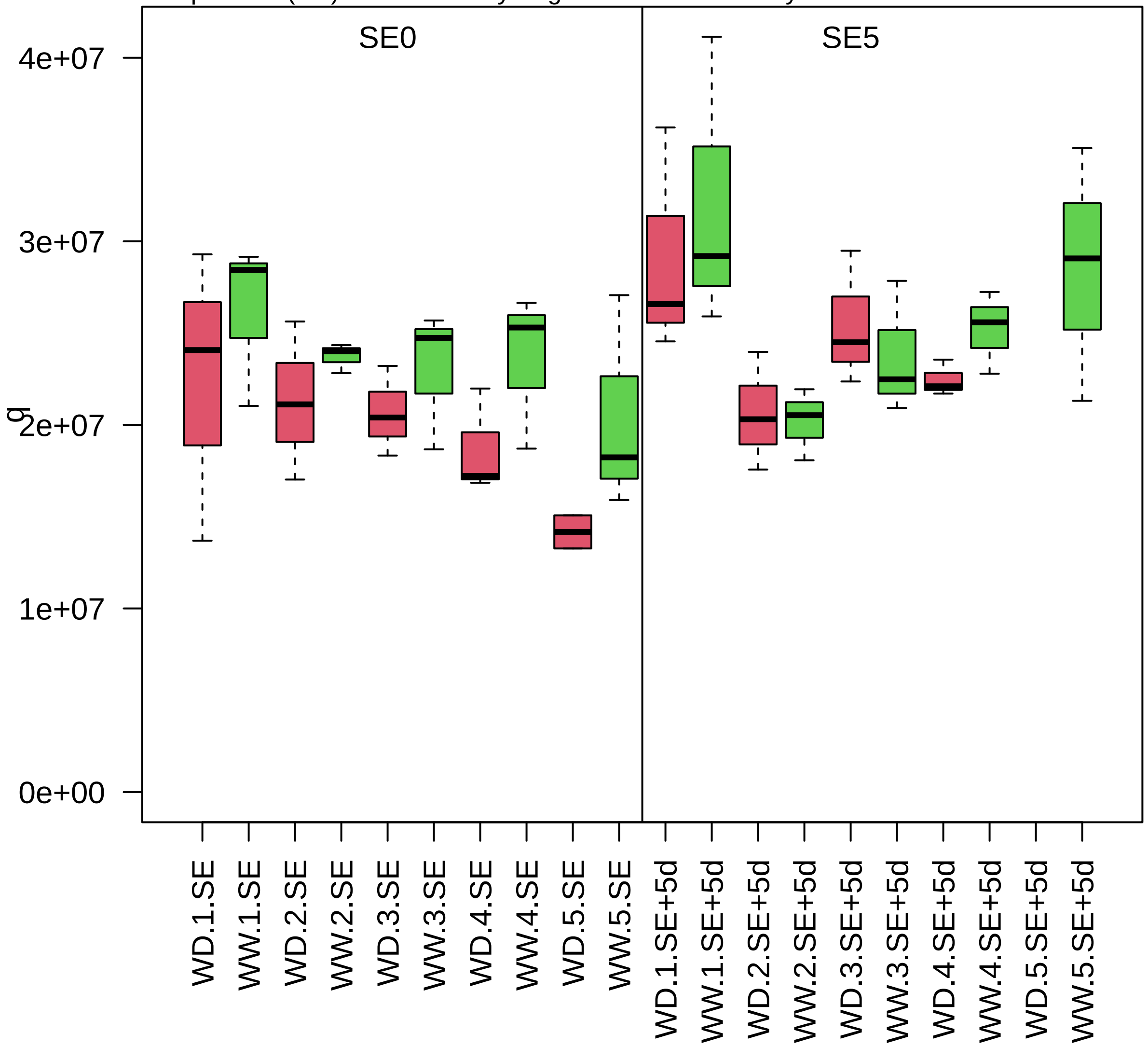

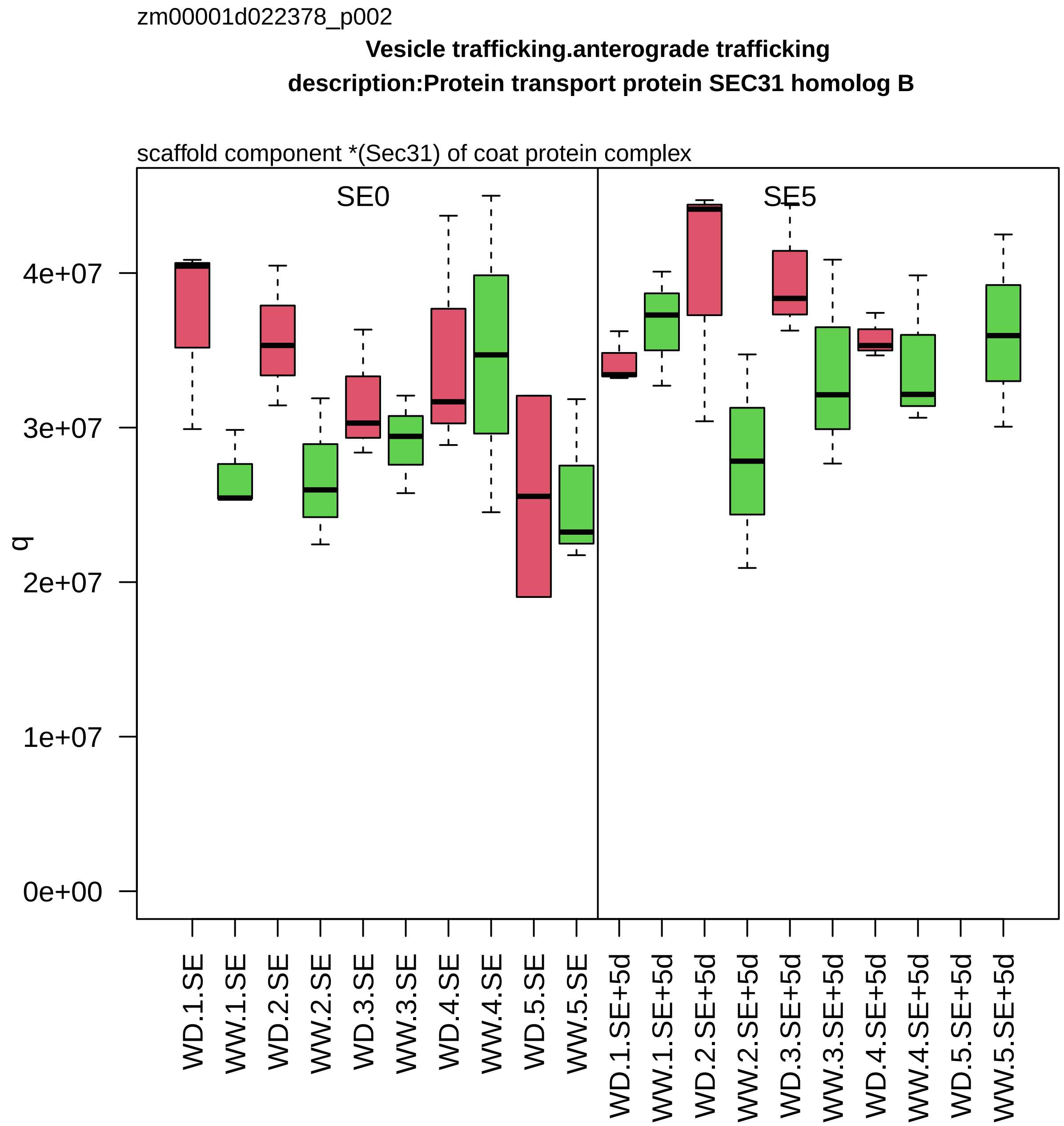
zm00001d022378_p002
Vesicle trafficking.anterograde trafficking
description:Protein transport protein SEC31 homolog B
scaffold component *(Sec31) of coat protein complex
SE0
SE5
q
0e+00
1e+07
2e+07
3e+07
4e+07
WD.1.SE
WW.1.SE
WD.2.SE
WW.2.SE
WD.3.SE
WW.3.SE
WD.4.SE
WW.4.SE
WD.5.SE
WW.5.SE
WD.1.SE+5d
WW.1.SE+5d
WD.2.SE+5d
WW.2.SE+5d
WD.3.SE+5d
WW.3.SE+5d
WD.4.SE+5d
WW.4.SE+5d
WD.5.SE+5d
WW.5.SE+5d

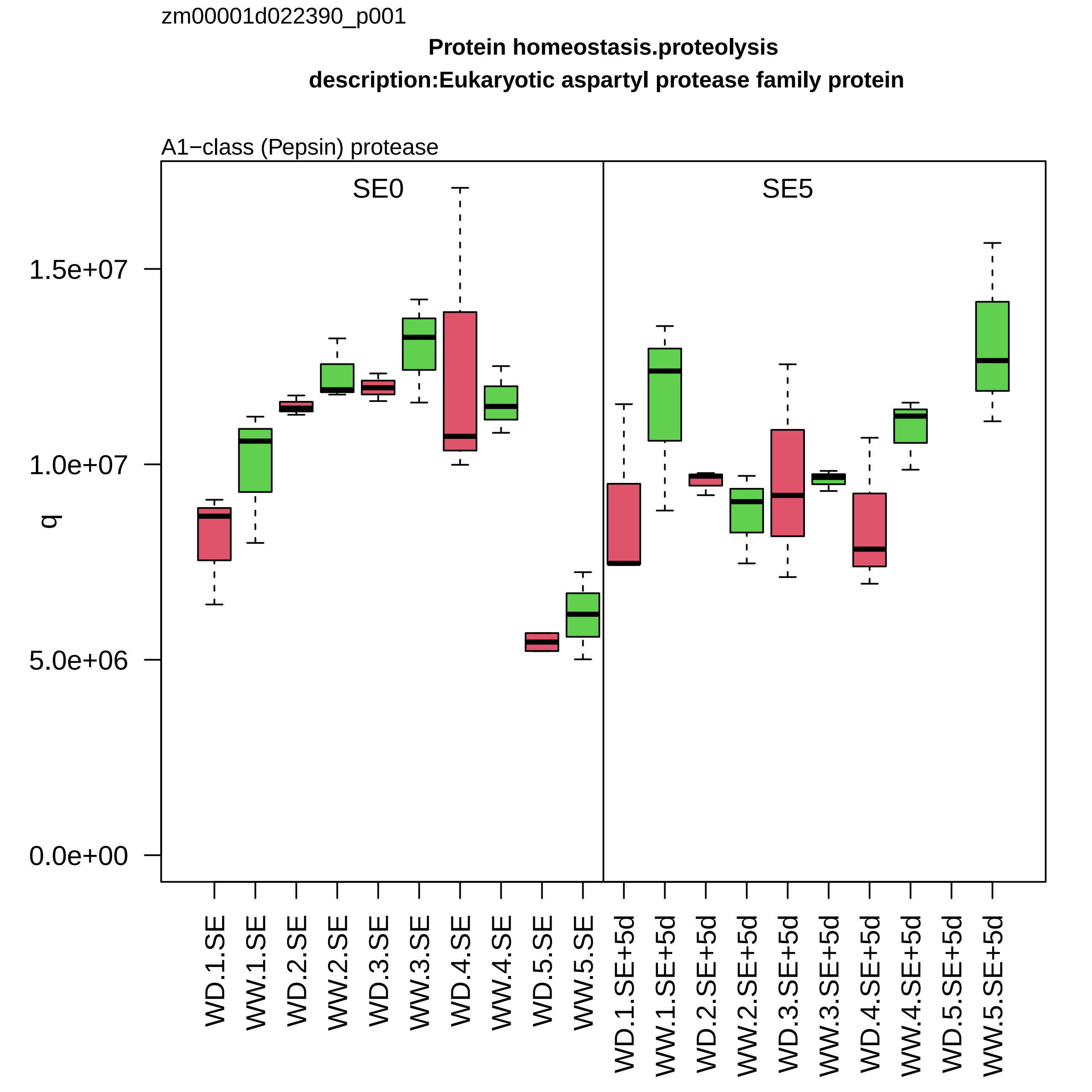

zm00001d022390_p001
Protein homeostasis.proteolysis
description:Eukaryotic aspartyl protease family protein
A1−class (Pepsin) protease
SE0
SE5
q
1.5e+07
1.0e+07
5.0e+06
0.0e+00
WD.1.SE
WW.1.SE
WD.2.SE
WW.2.SE
WD.3.SE
WW.3.SE
WD.4.SE
WW.4.SE
WD.5.SE
WW.5.SE
WD.1.SE+5d
WW.1.SE+5d
WD.2.SE+5d
WW.2.SE+5d
WD.3.SE+5d
WW.3.SE+5d
WD.4.SE+5d
WW.4.SE+5d
WD.5.SE+5d
WW.5.SE+5d

zm00001d022421_p001

**not assigned.not annotated**

**description:hydroxyproline−rich glycoprotein family protein**

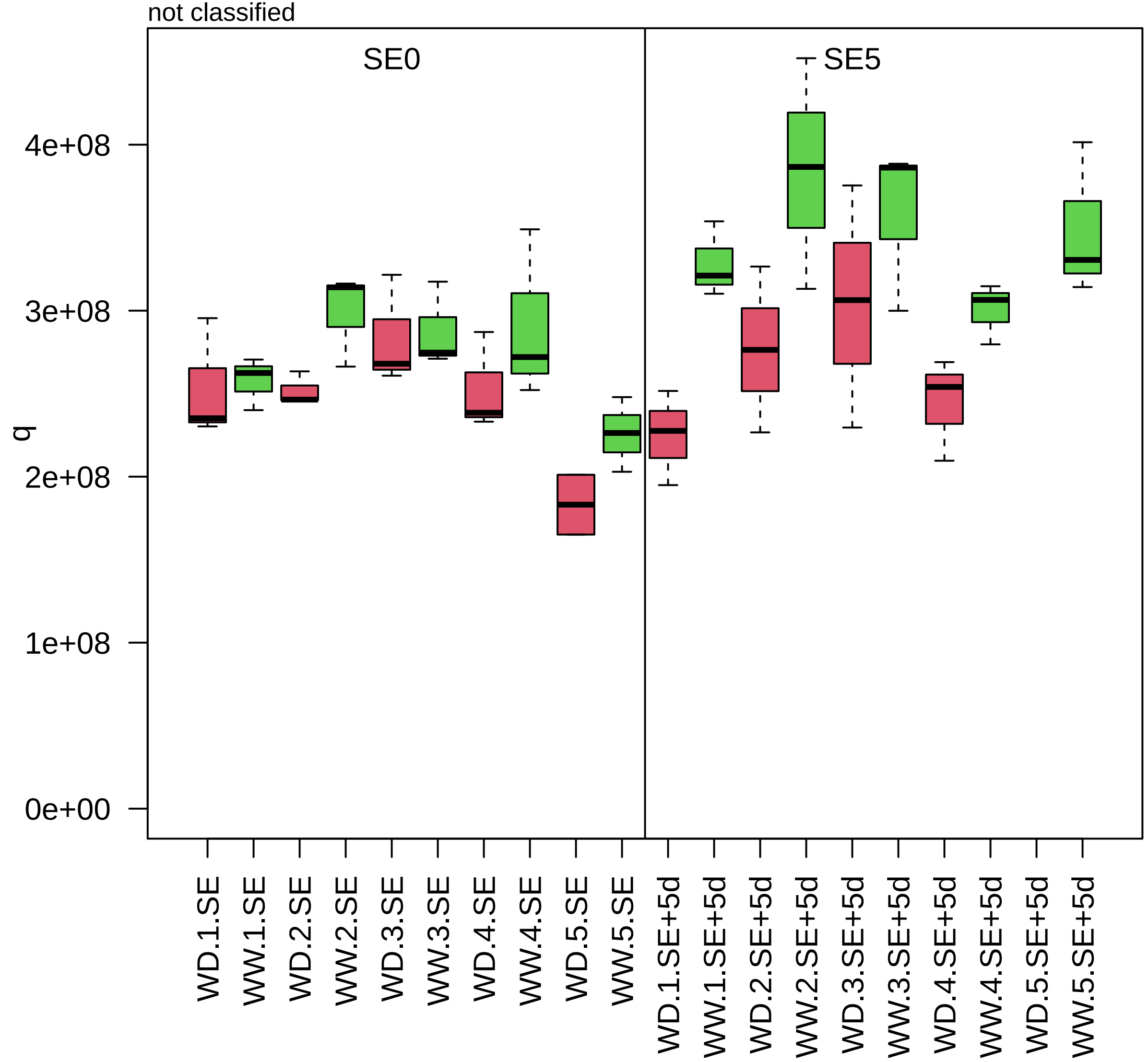

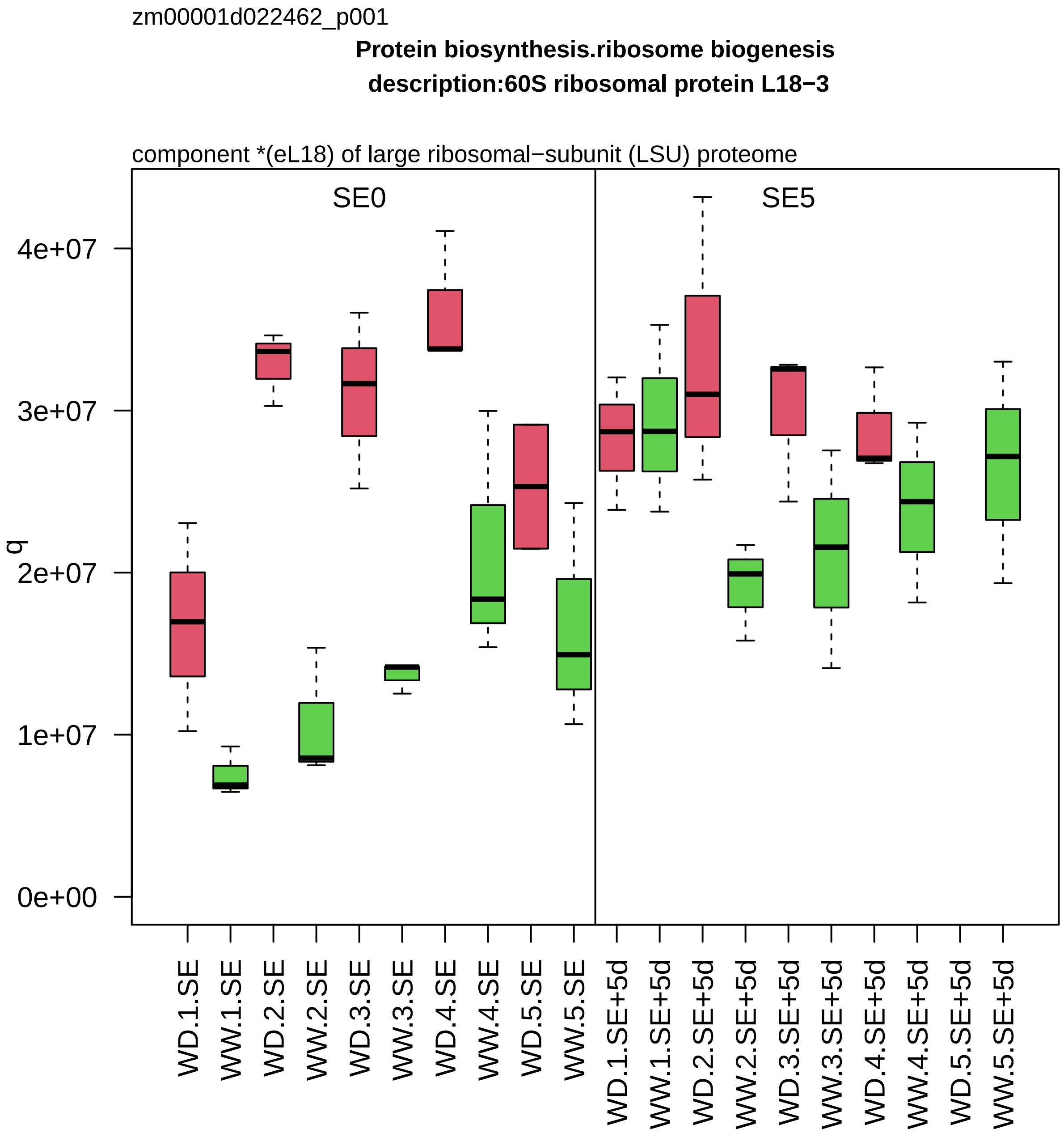

zm00001d022462_p001
Protein biosynthesis.ribosome biogenesis
description:60S ribosomal protein L18−3
component *(eL18) of large ribosomal−subunit (LSU) proteome
SE0
SE5
q
0e+00
1e+07
2e+07
3e+07
4e+07
WD.1.SE
WW.1.SE
WD.2.SE
WW.2.SE
WD.3.SE
WW.3.SE
WD.4.SE
WW.4.SE
WD.5.SE
WW.5.SE
WD.1.SE+5d
WW.1.SE+5d
WD.2.SE+5d
WW.2.SE+5d
WD.3.SE+5d
WW.3.SE+5d
WD.4.SE+5d
WW.4.SE+5d
WD.5.SE+5d
WW.5.SE+5d

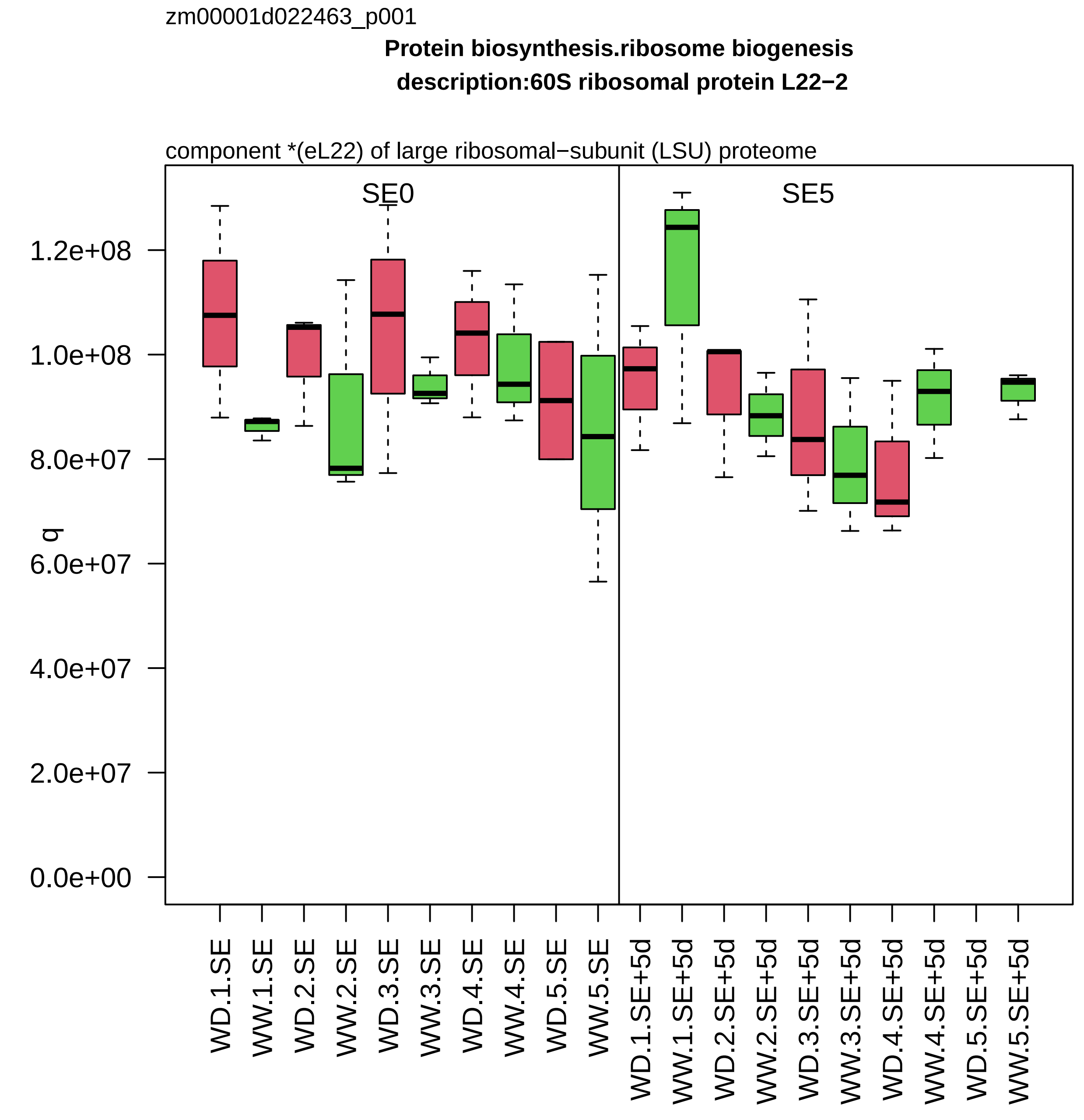

zm00001d022463_p001
Protein biosynthesis.ribosome biogenesis
description:60S ribosomal protein L22−2
component *(eL22) of large ribosomal−subunit (LSU) proteome
SE0
SE5
q
1.2e+08
1.0e+08
8.0e+07
6.0e+07
4.0e+07
2.0e+07
0.0e+00
WD.1.SE
WW.1.SE
WD.2.SE
WW.2.SE
WD.3.SE
WW.3.SE
WD.4.SE
WW.4.SE
WD.5.SE
WW.5.SE
WD.1.SE+5d
WW.1.SE+5d
WD.2.SE+5d
WW.2.SE+5d
WD.3.SE+5d
WW.3.SE+5d
WD.4.SE+5d
WW.4.SE+5d
WD.5.SE+5d
WW.5.SE+5d

zm00001d022490_p001

**Enzyme classification.EC_5 isomerases**
**description:Xylose isomerase**

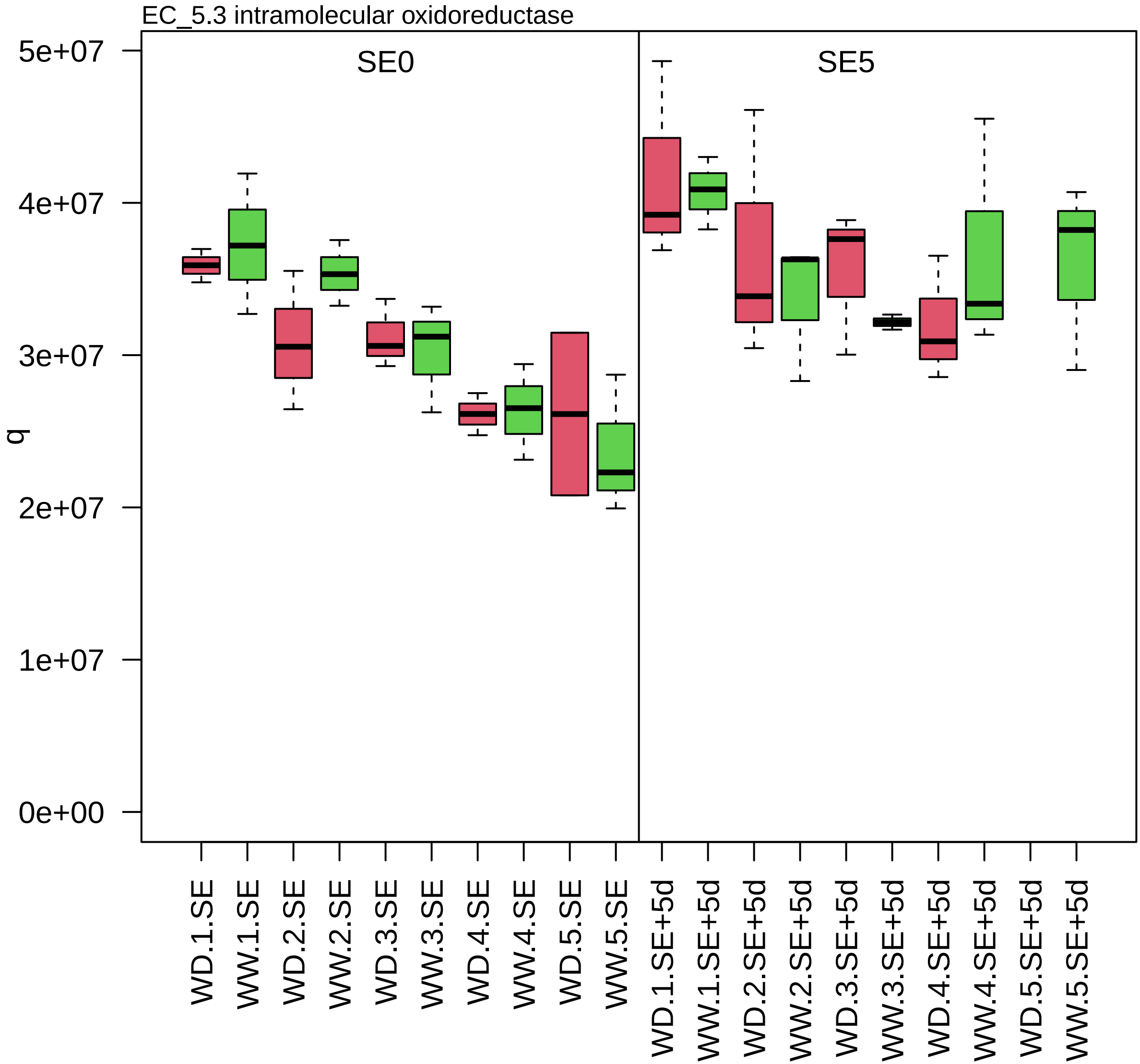

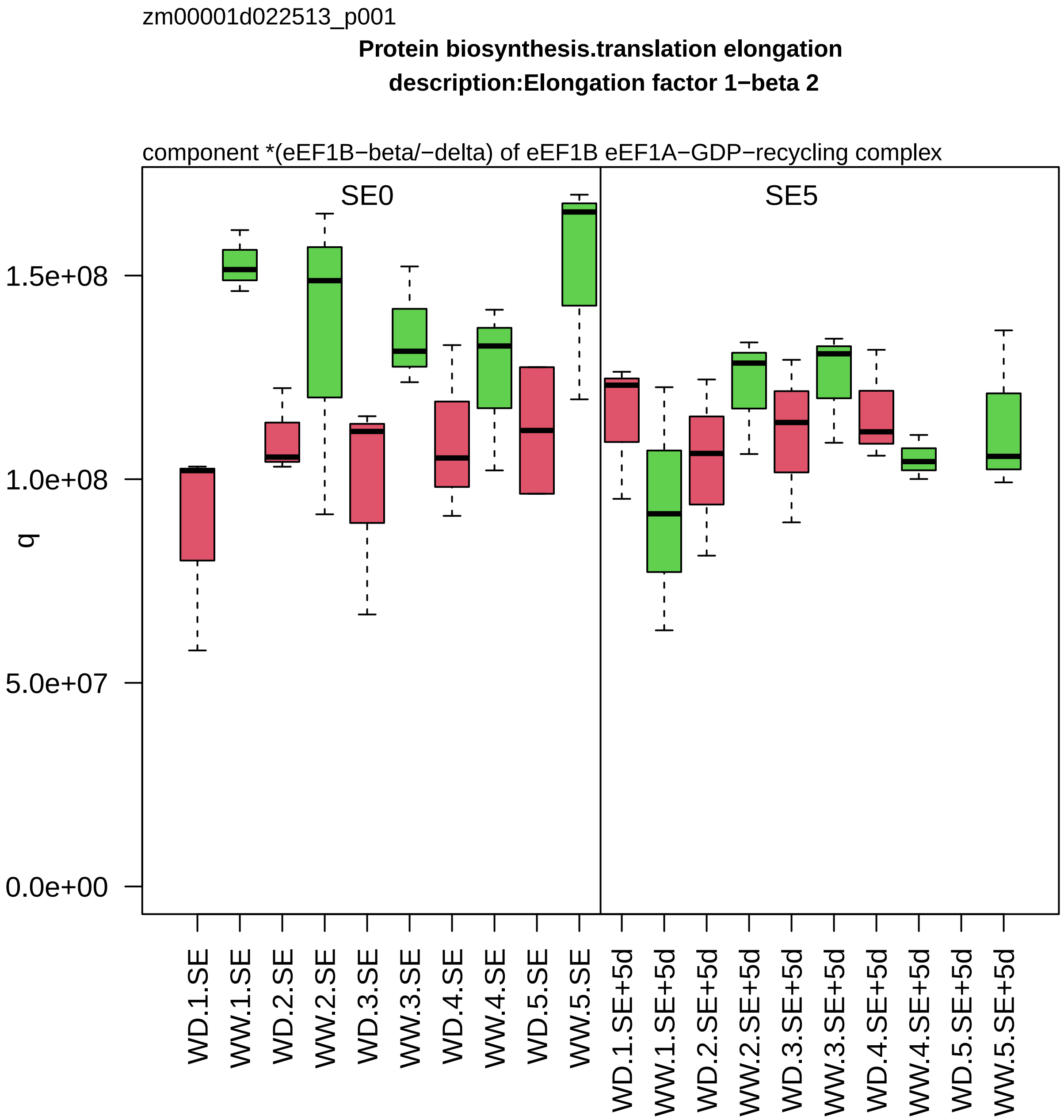
zm00001d022513_p001
Protein biosynthesis.translation elongation
description:Elongation factor 1−beta 2
component *(eEF1B−beta/−delta) of eEF1B eEF1A−GDP−recycling complex
SE0
SE5
q
1.5e+08
1.0e+08
5.0e+07
0.0e+00
WD.1.SE
WW.1.SE
WD.2.SE
WW.2.SE
WD.3.SE
WW.3.SE
WD.4.SE
WW.4.SE
WD.5.SE
WW.5.SE
WD.1.SE+5d
WW.1.SE+5d
WD.2.SE+5d
WW.2.SE+5d
WD.3.SE+5d
WW.3.SE+5d
WD.4.SE+5d
WW.4.SE+5d
WD.5.SE+5d
WW.5.SE+5d

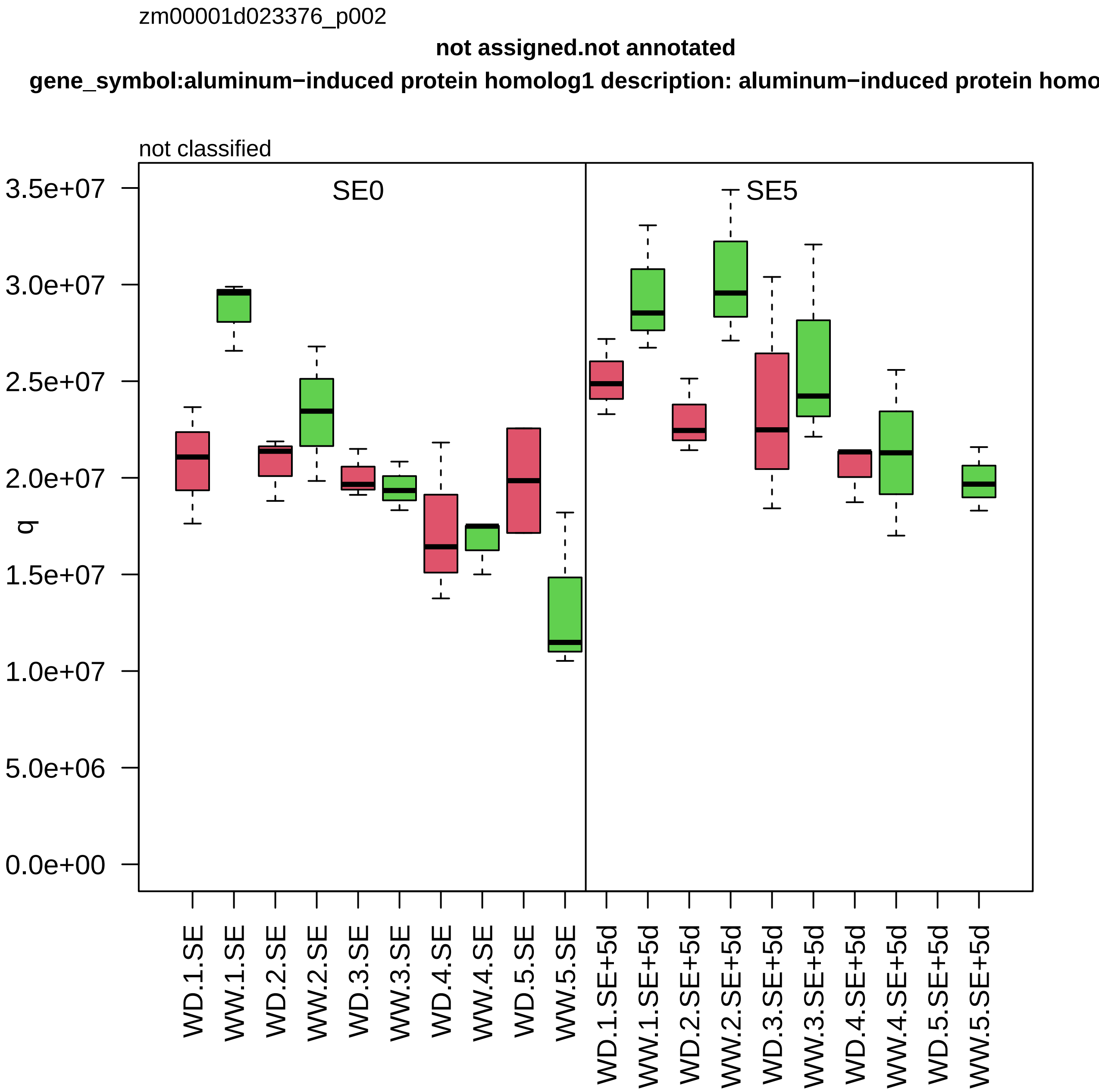

zm00001d023376_p002
not assigned.not annotated
gene_symbol:aluminum−induced protein homolog1 description: aluminum−induced protein homo
not classified
SE0
SE5
q
3.5e+07
3.0e+07
2.5e+07
2.0e+07
1.5e+07
1.0e+07
5.0e+06
0.0e+00
WD.1.SE
WW.1.SE
WD.2.SE
WW.2.SE
WD.3.SE
WW.3.SE
WD.4.SE
WW.4.SE
WD.5.SE
WW.5.SE
WD.1.SE+5d
WW.1.SE+5d
WD.2.SE+5d
WW.2.SE+5d
WD.3.SE+5d
WW.3.SE+5d
WD.4.SE+5d
WW.4.SE+5d
WD.5.SE+5d
WW.5.SE+5d

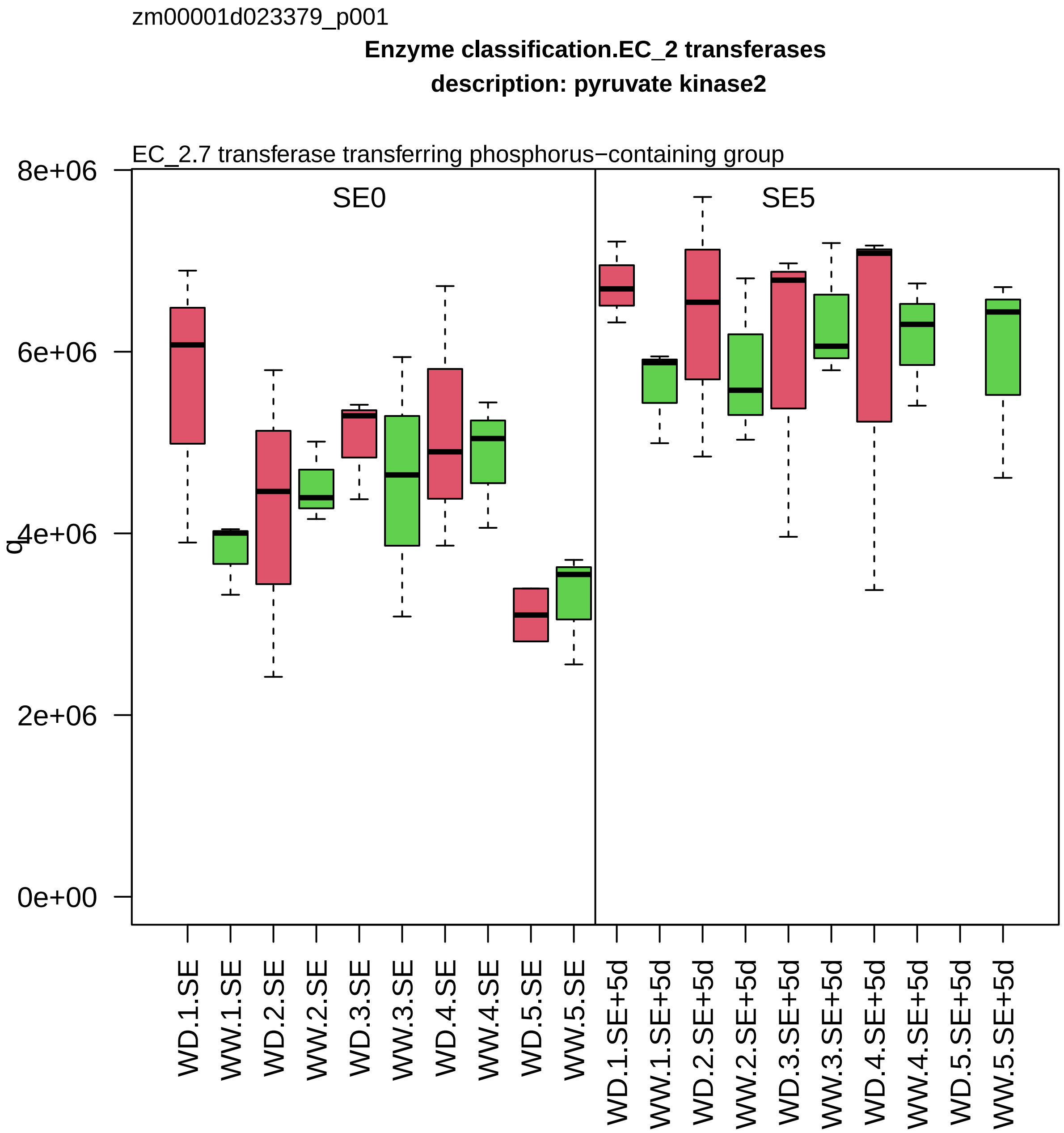
zm00001d023379_p001
Enzyme classification.EC_2 transferases
description: pyruvate kinase2
EC_2.7 transferase transferring phosphorus−containing group
SE0
SE5
8e+06
6e+06
4e+06
2e+06
0e+00
q
WD.1.SE
WW.1.SE
WD.2.SE
WW.2.SE
WD.3.SE
WW.3.SE
WD.4.SE
WW.4.SE
WD.5.SE
WW.5.SE
WD.1.SE+5d
WW.1.SE+5d
WD.2.SE+5d
WW.2.SE+5d
WD.3.SE+5d
WW.3.SE+5d
WD.4.SE+5d
WW.4.SE+5d
WD.5.SE+5d
WW.5.SE+5d

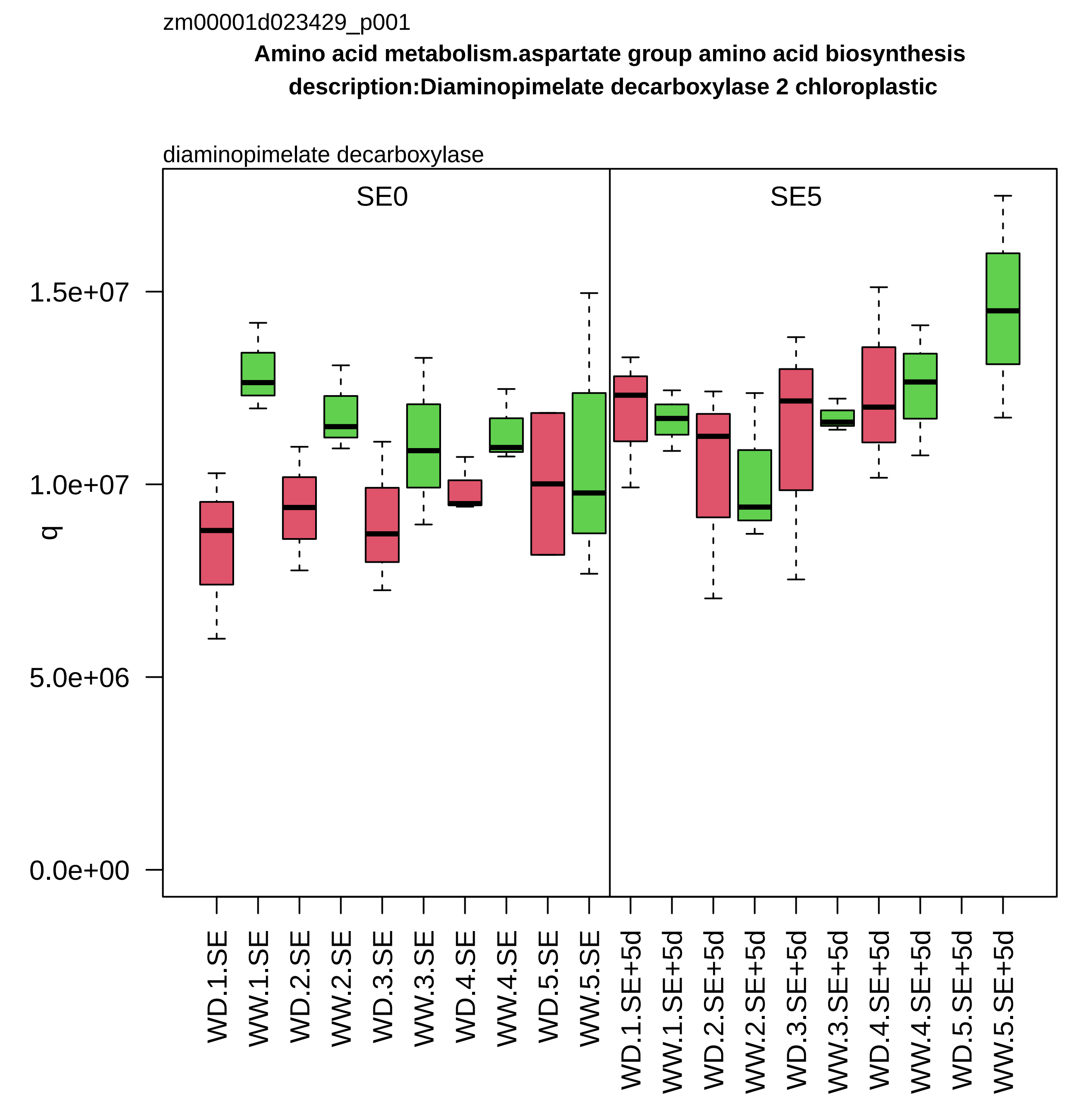
zm00001d023429_p001
Amino acid metabolism.aspartate group amino acid biosynthesis
description:Diaminopimelate decarboxylase 2 chloroplastic
diaminopimelate decarboxylase
SE0
SE5
1.5e+07
1.0e+07
5.0e+06
0.0e+00
q
WD.1.SE
WW.1.SE
WD.2.SE
WW.2.SE
WD.3.SE
WW.3.SE
WD.4.SE
WW.4.SE
WD.5.SE
WW.5.SE
WD.1.SE+5d
WW.1.SE+5d
WD.2.SE+5d
WW.2.SE+5d
WD.3.SE+5d
WW.3.SE+5d
WD.4.SE+5d
WW.4.SE+5d
WD.5.SE+5d
WW.5.SE+5d

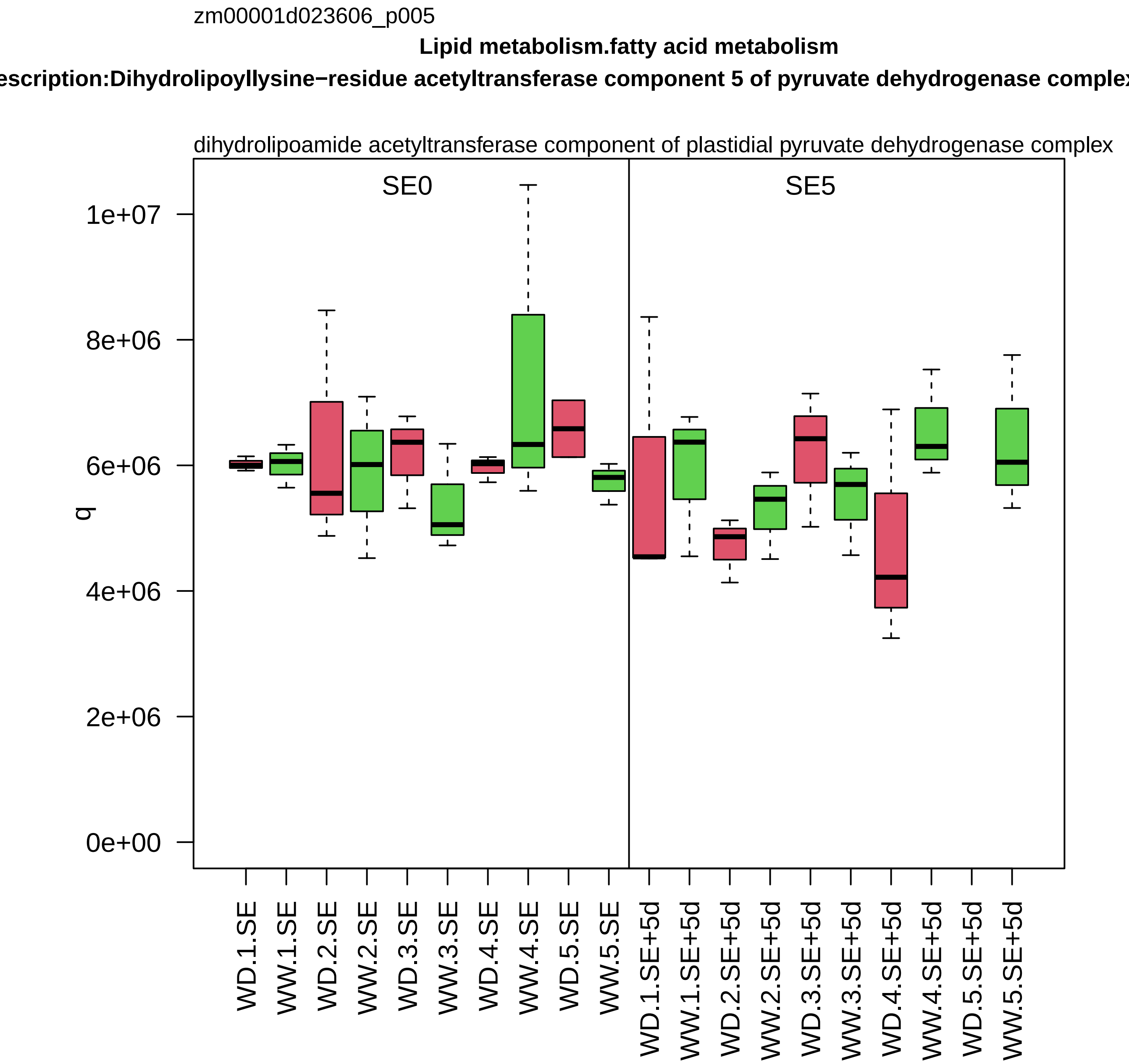
zm00001d023606_p005
Lipid metabolism.fatty acid metabolism
escription:Dihydrolipoyllysine−residue acetyltransferase component 5 of pyruvate dehydrogenase complex
dihydrolipoamide acetyltransferase component of plastidial pyruvate dehydrogenase complex
SE0
SE5
1e+07
8e+06
6e+06
4e+06
2e+06
0e+00
q
WD.1.SE
WW.1.SE
WD.2.SE
WW.2.SE
WD.3.SE
WW.3.SE
WD.4.SE
WW.4.SE
WD.5.SE
WW.5.SE
WD.1.SE+5d
WW.1.SE+5d
WD.2.SE+5d
WW.2.SE+5d
WD.3.SE+5d
WW.3.SE+5d
WD.4.SE+5d
WW.4.SE+5d
WD.5.SE+5d
WW.5.SE+5d

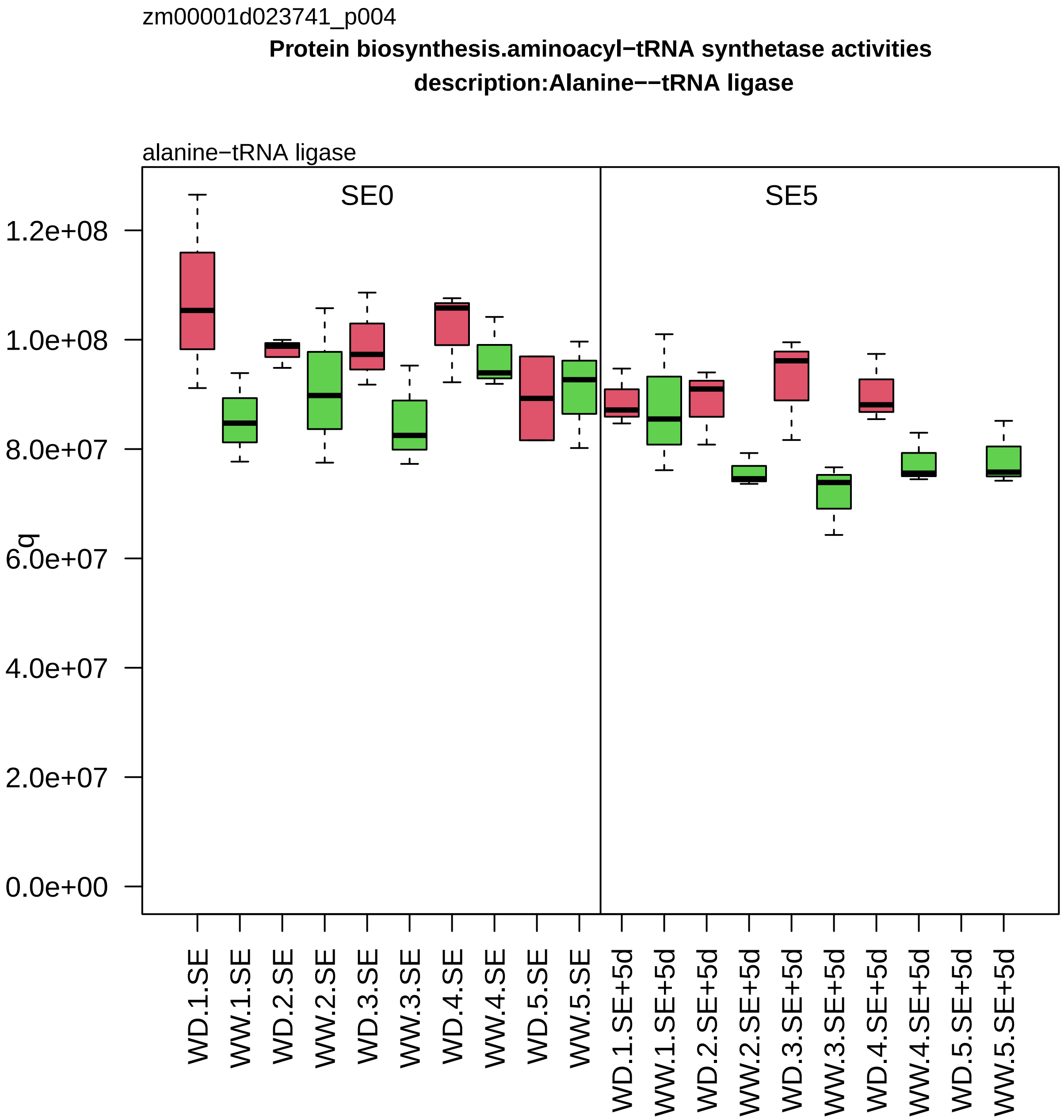
zm00001d023741_p004
Protein biosynthesis.aminoacyl−tRNA synthetase activities
description:Alanine−−tRNA ligase
alanine−tRNA ligase
SE0
SE5
q
1.2e+08
1.0e+08
8.0e+07
6.0e+07
4.0e+07
2.0e+07
0.0e+00
WD.1.SE
WW.1.SE
WD.2.SE
WW.2.SE
WD.3.SE
WW.3.SE
WD.4.SE
WW.4.SE
WD.5.SE
WW.5.SE
WD.1.SE+5d
WW.1.SE+5d
WD.2.SE+5d
WW.2.SE+5d
WD.3.SE+5d
WW.3.SE+5d
WD.4.SE+5d
WW.4.SE+5d
WD.5.SE+5d
WW.5.SE+5d

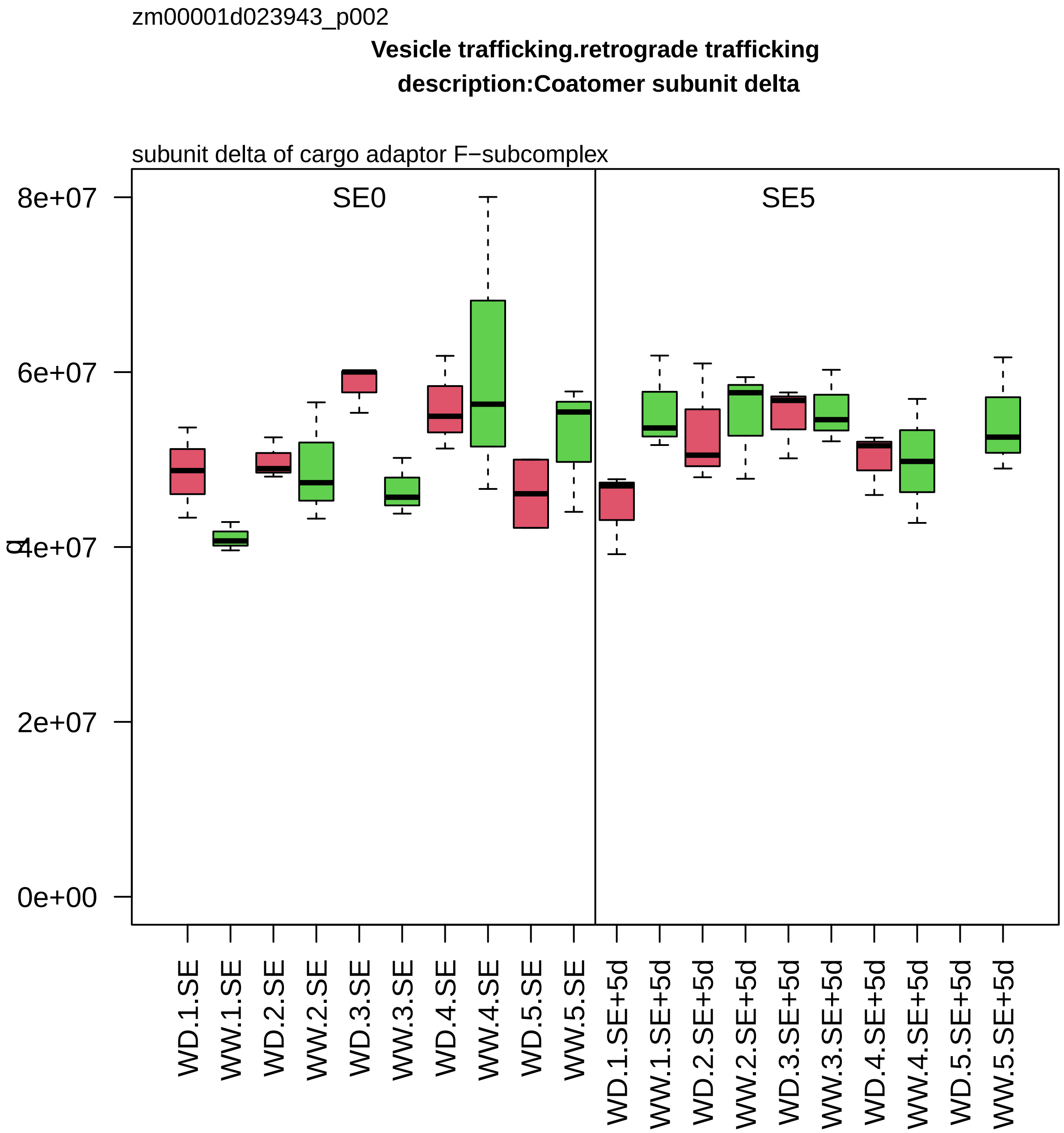

zm00001d023943_p002
Vesicle trafficking.retrograde trafficking
description:Coatomer subunit delta
subunit delta of cargo adaptor F−subcomplex
SE0
SE5
8e+07
6e+07
4e+07
2e+07
0e+00
WD.1.SE
WW.1.SE
WD.2.SE
WW.2.SE
WD.3.SE
WW.3.SE
WD.4.SE
WW.4.SE
WD.5.SE
WW.5.SE
WD.1.SE+5d
WW.1.SE+5d
WD.2.SE+5d
WW.2.SE+5d
WD.3.SE+5d
WW.3.SE+5d
WD.4.SE+5d
WW.4.SE+5d
WD.5.SE+5d
WW.5.SE+5d

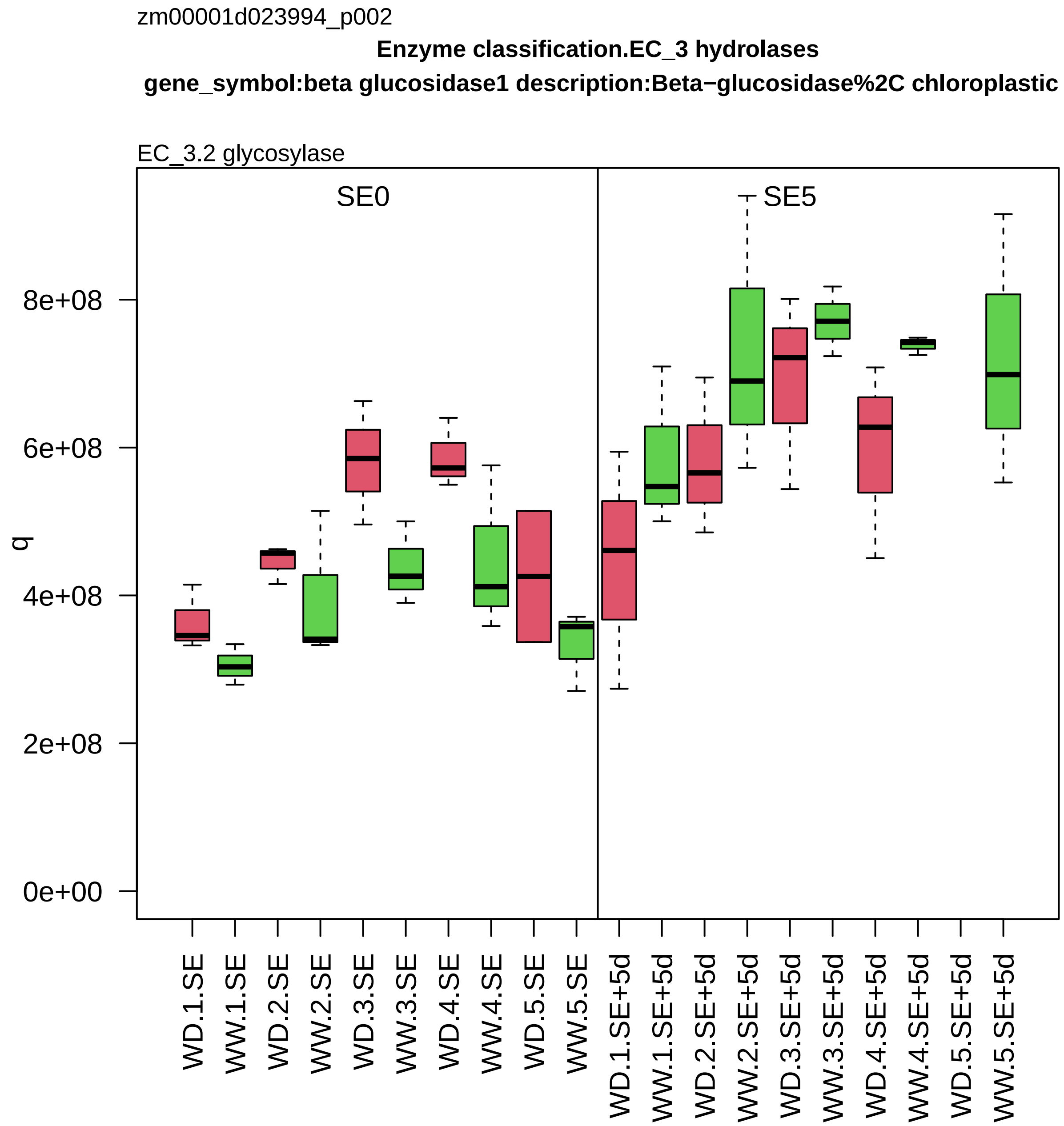
zm00001d023994_p002
Enzyme classification.EC_3 hydrolases
gene_symbol:beta glucosidase1 description:Beta−glucosidase%2C chloroplastic
EC_3.2 glycosylase
SE0
SE5
q
0e+00
2e+08
4e+08
6e+08
8e+08
WD.1.SE
WW.1.SE
WD.2.SE
WW.2.SE
WD.3.SE
WW.3.SE
WD.4.SE
WW.4.SE
WD.5.SE
WW.5.SE
WD.1.SE+5d
WW.1.SE+5d
WD.2.SE+5d
WW.2.SE+5d
WD.3.SE+5d
WW.3.SE+5d
WD.4.SE+5d
WW.4.SE+5d
WD.5.SE+5d
WW.5.SE+5d

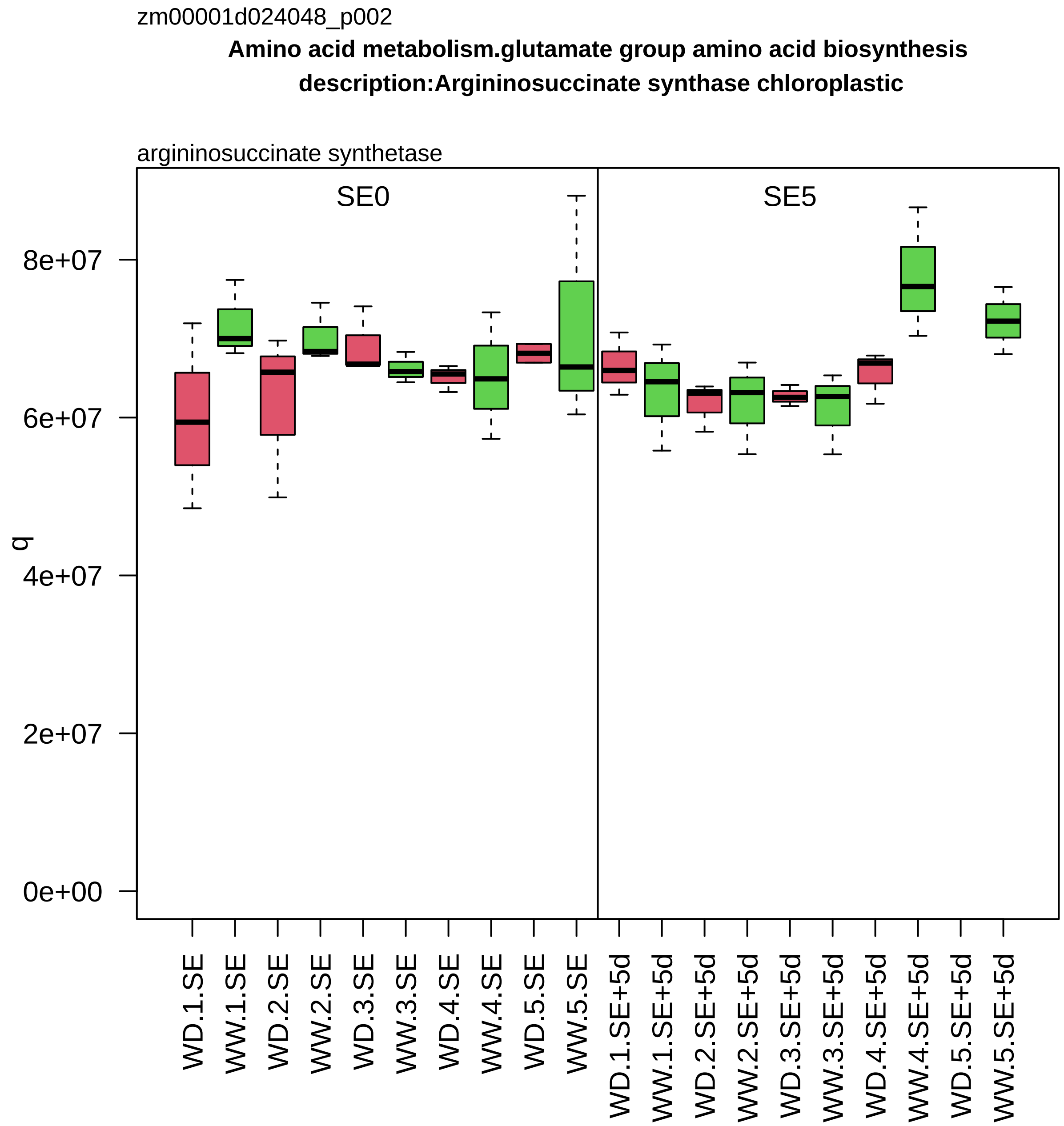
zm00001d024048_p002
Amino acid metabolism.glutamate group amino acid biosynthesis
description:Argininosuccinate synthase chloroplastic
argininosuccinate synthetase
SE0
SE5
q
0e+00
2e+07
4e+07
6e+07
8e+07
WD.1.SE
WW.1.SE
WD.2.SE
WW.2.SE
WD.3.SE
WW.3.SE
WD.4.SE
WW.4.SE
WD.5.SE
WW.5.SE
WD.1.SE+5d
WW.1.SE+5d
WD.2.SE+5d
WW.2.SE+5d
WD.3.SE+5d
WW.3.SE+5d
WD.4.SE+5d
WW.4.SE+5d
WD.5.SE+5d
WW.5.SE+5d

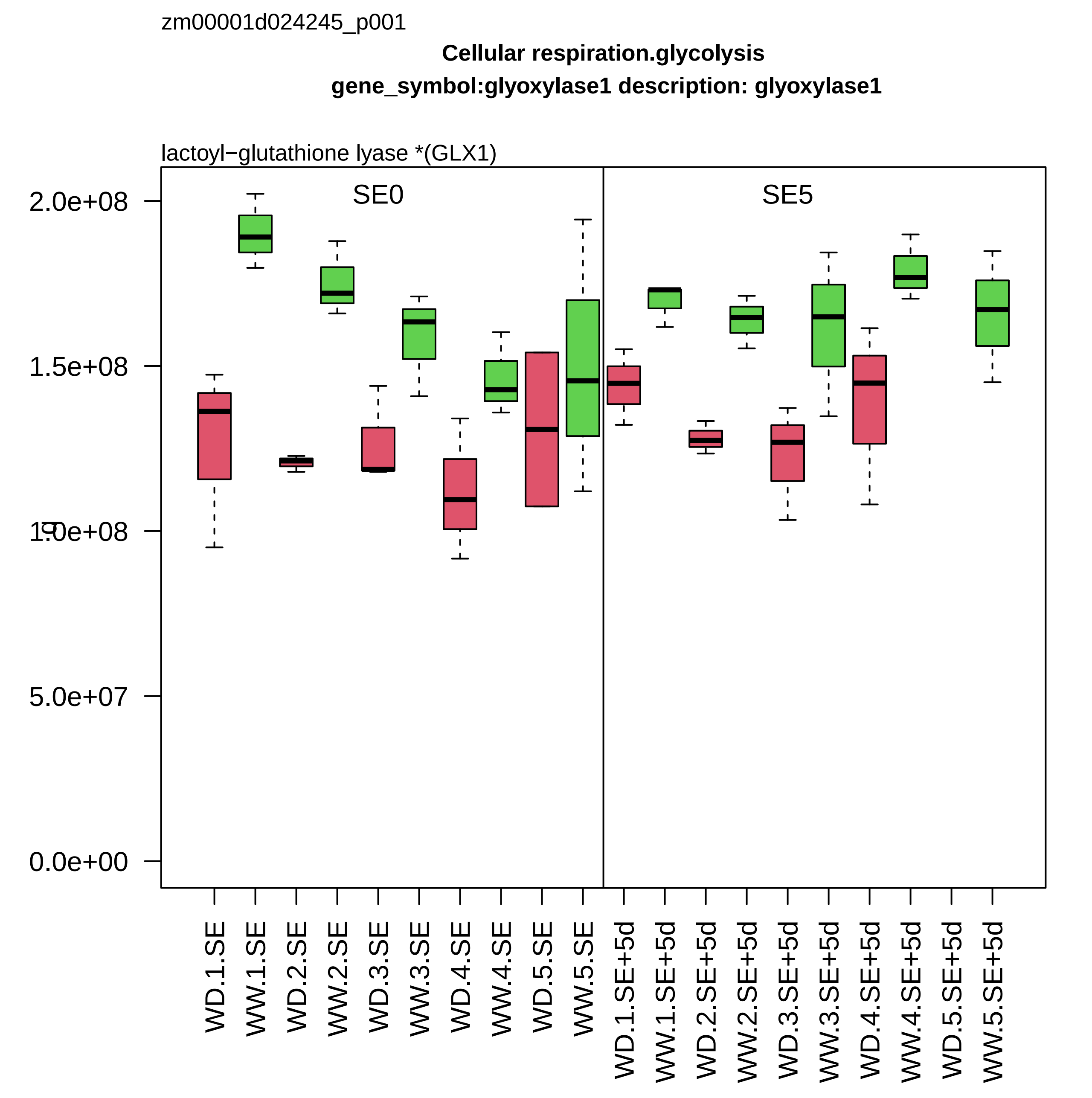
zm00001d024245_p001
Cellular respiration.glycolysis
gene_symbol:glyoxylase1 description: glyoxylase1
lactoyl−glutathione lyase *(GLX1)
SE0
SE5
2.0e+08
1.5e+08
1.0e+08
5.0e+07
0.0e+00
WD.1.SE
WW.1.SE
WD.2.SE
WW.2.SE
WD.3.SE
WW.3.SE
WD.4.SE
WW.4.SE
WD.5.SE
WW.5.SE
WD.1.SE+5d
WW.1.SE+5d
WD.2.SE+5d
WW.2.SE+5d
WD.3.SE+5d
WW.3.SE+5d
WD.4.SE+5d
WW.4.SE+5d
WD.5.SE+5d
WW.5.SE+5d

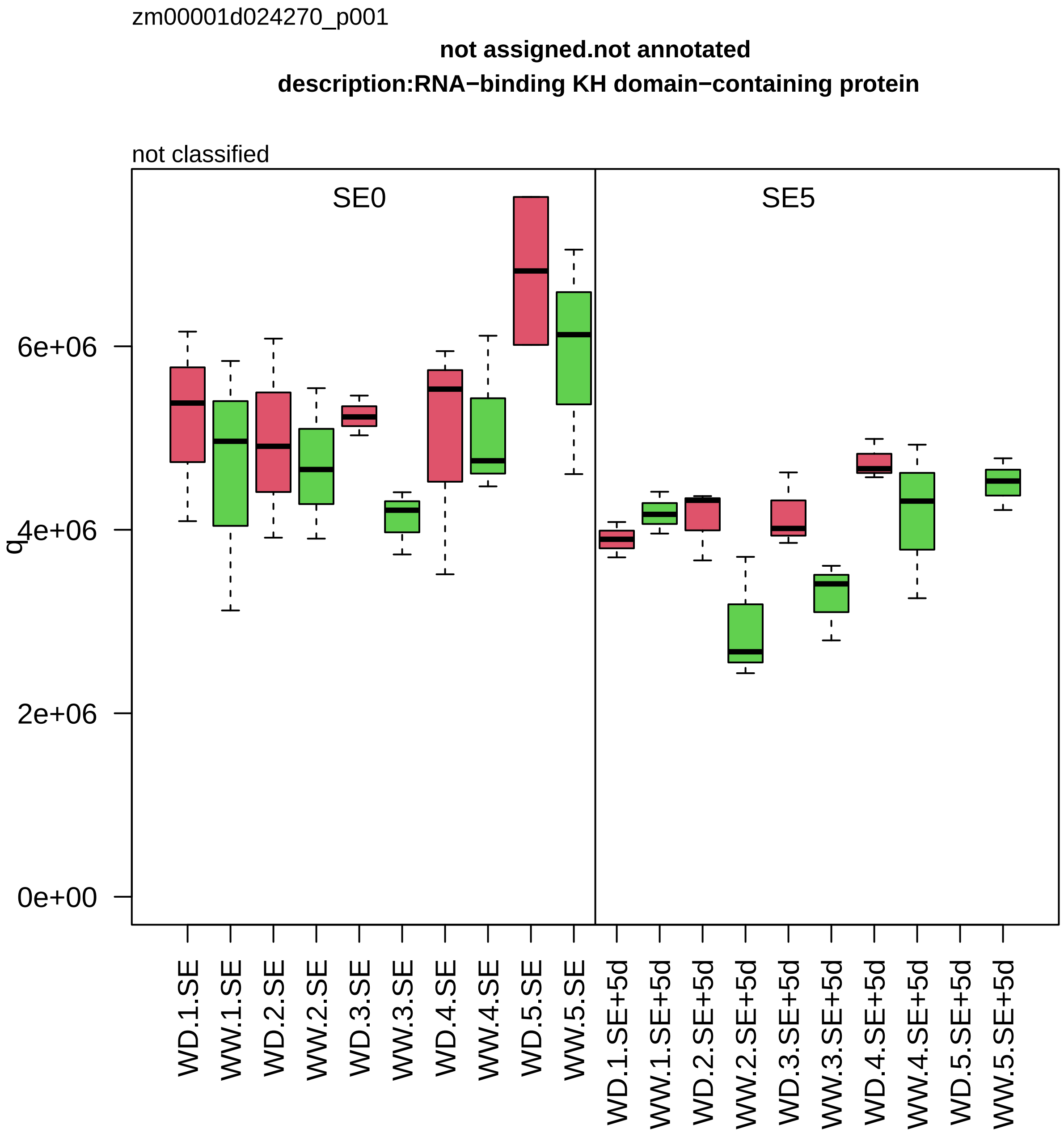

zm00001d024270_p001
not assigned.not annotated
description:RNA−binding KH domain−containing protein
not classified
SE0
SE5
q
0e+00
2e+06
4e+06
6e+06
WD.1.SE
WW.1.SE
WD.2.SE
WW.2.SE
WD.3.SE
WW.3.SE
WD.4.SE
WW.4.SE
WD.5.SE
WW.5.SE
WD.1.SE+5d
WW.1.SE+5d
WD.2.SE+5d
WW.2.SE+5d
WD.3.SE+5d
WW.3.SE+5d
WD.4.SE+5d
WW.4.SE+5d
WD.5.SE+5d
WW.5.SE+5d

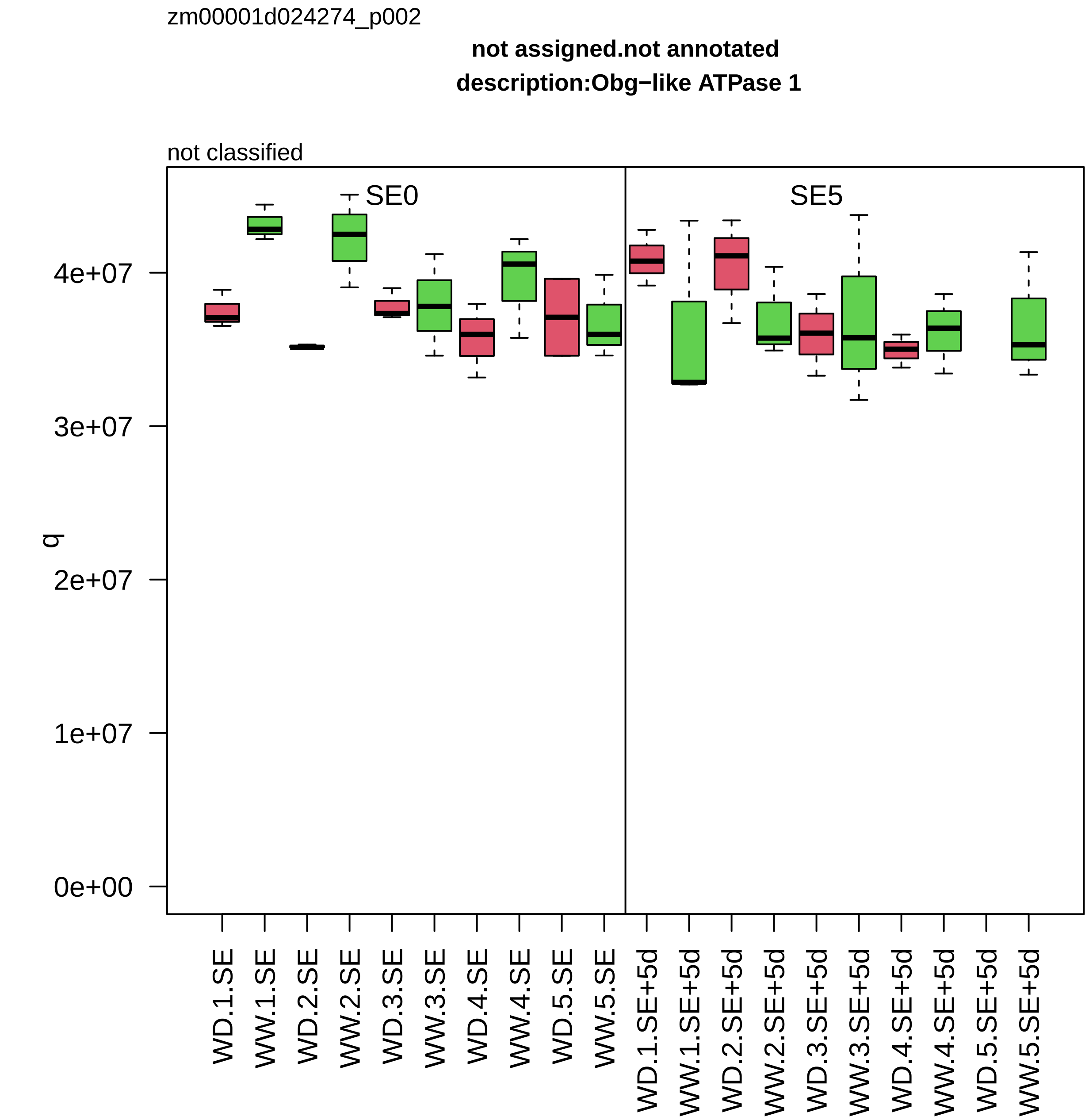
zm00001d024274_p002
not assigned.not annotated
description:Obg−like ATPase 1
not classified
SE0
SE5
q
0e+00
1e+07
2e+07
3e+07
4e+07
WD.1.SE
WW.1.SE
WD.2.SE
WW.2.SE
WD.3.SE
WW.3.SE
WD.4.SE
WW.4.SE
WD.5.SE
WW.5.SE
WD.1.SE+5d
WW.1.SE+5d
WD.2.SE+5d
WW.2.SE+5d
WD.3.SE+5d
WW.3.SE+5d
WD.4.SE+5d
WW.4.SE+5d
WD.5.SE+5d
WW.5.SE+5d

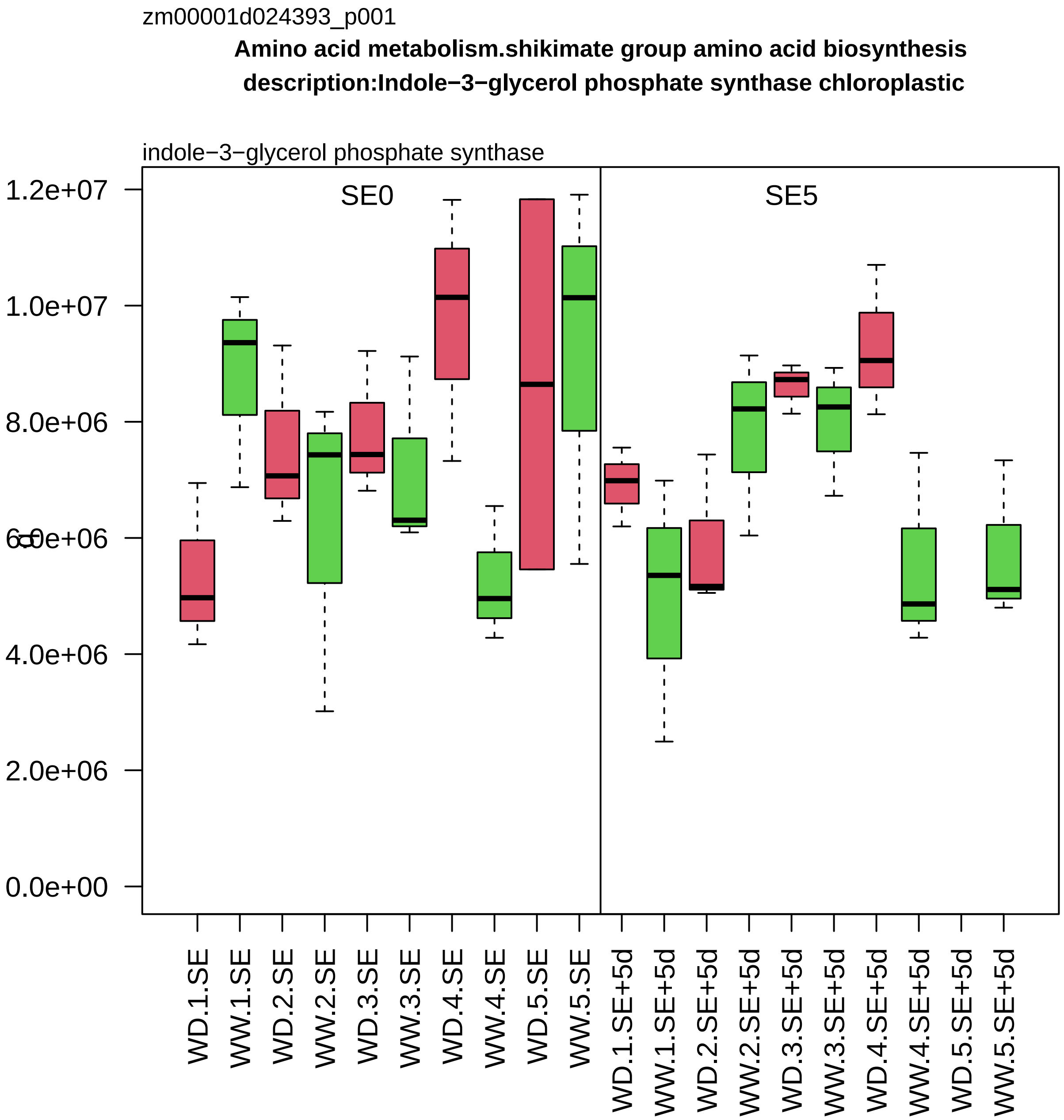
zm00001d024393_p001
Amino acid metabolism.shikimate group amino acid biosynthesis
description:Indole−3−glycerol phosphate synthase chloroplastic
indole−3−glycerol phosphate synthase
SE0
SE5
1.2e+07
1.0e+07
8.0e+06
6.0e+06
4.0e+06
2.0e+06
0.0e+00
WD.1.SE
WW.1.SE
WD.2.SE
WW.2.SE
WD.3.SE
WW.3.SE
WD.4.SE
WW.4.SE
WD.5.SE
WW.5.SE
WD.1.SE+5d
WW.1.SE+5d
WD.2.SE+5d
WW.2.SE+5d
WD.3.SE+5d
WW.3.SE+5d
WD.4.SE+5d
WW.4.SE+5d
WD.5.SE+5d
WW.5.SE+5d

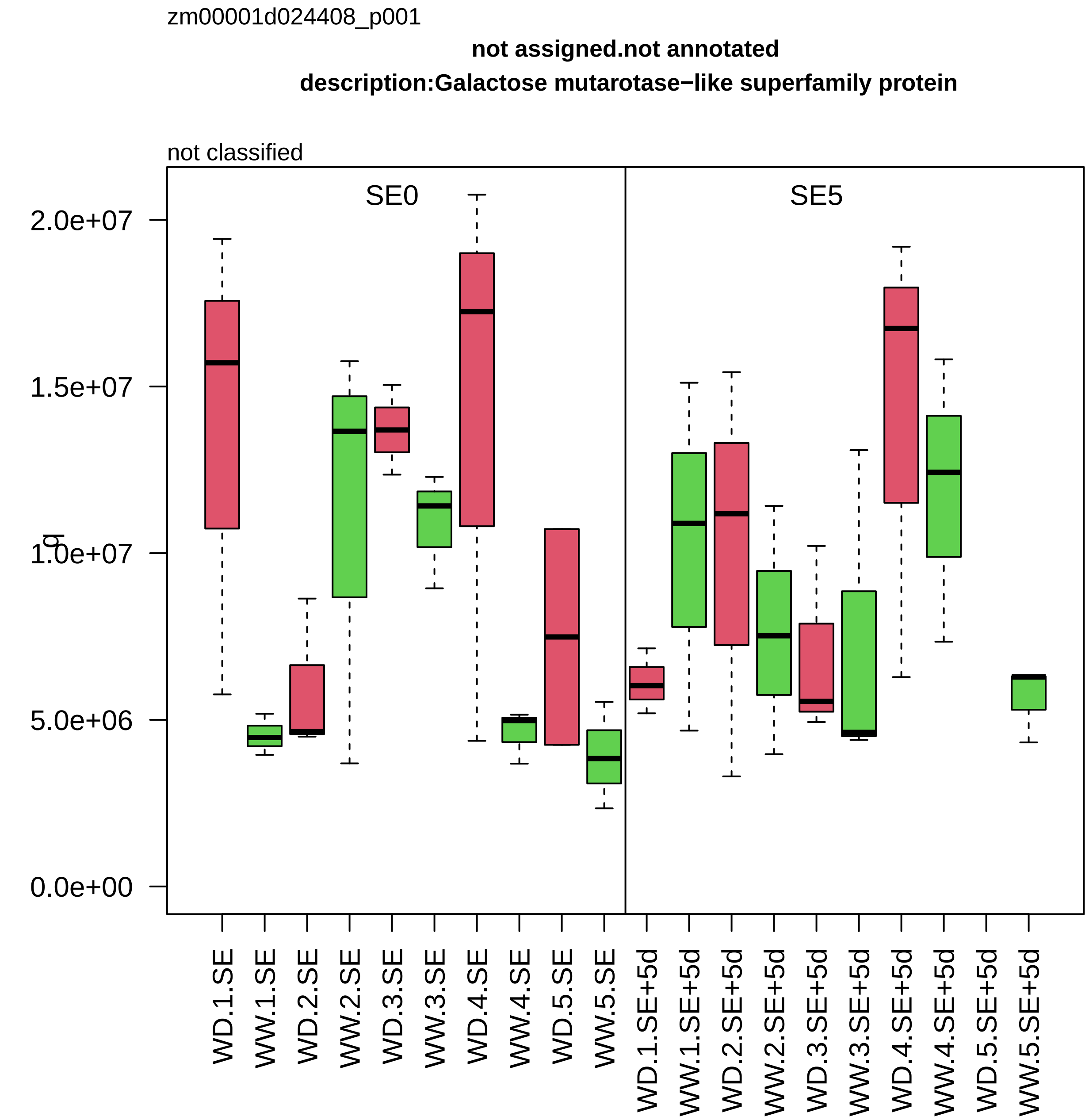

zm00001d024408_p001
not assigned.not annotated
description:Galactose mutarotase−like superfamily protein
not classified
SE0
SE5
q
2.0e+07
1.5e+07
1.0e+07
5.0e+06
0.0e+00
WD.1.SE
WW.1.SE
WD.2.SE
WW.2.SE
WD.3.SE
WW.3.SE
WD.4.SE
WW.4.SE
WD.5.SE
WW.5.SE
WD.1.SE+5d
WW.1.SE+5d
WD.2.SE+5d
WW.2.SE+5d
WD.3.SE+5d
WW.3.SE+5d
WD.4.SE+5d
WW.4.SE+5d
WD.5.SE+5d
WW.5.SE+5d

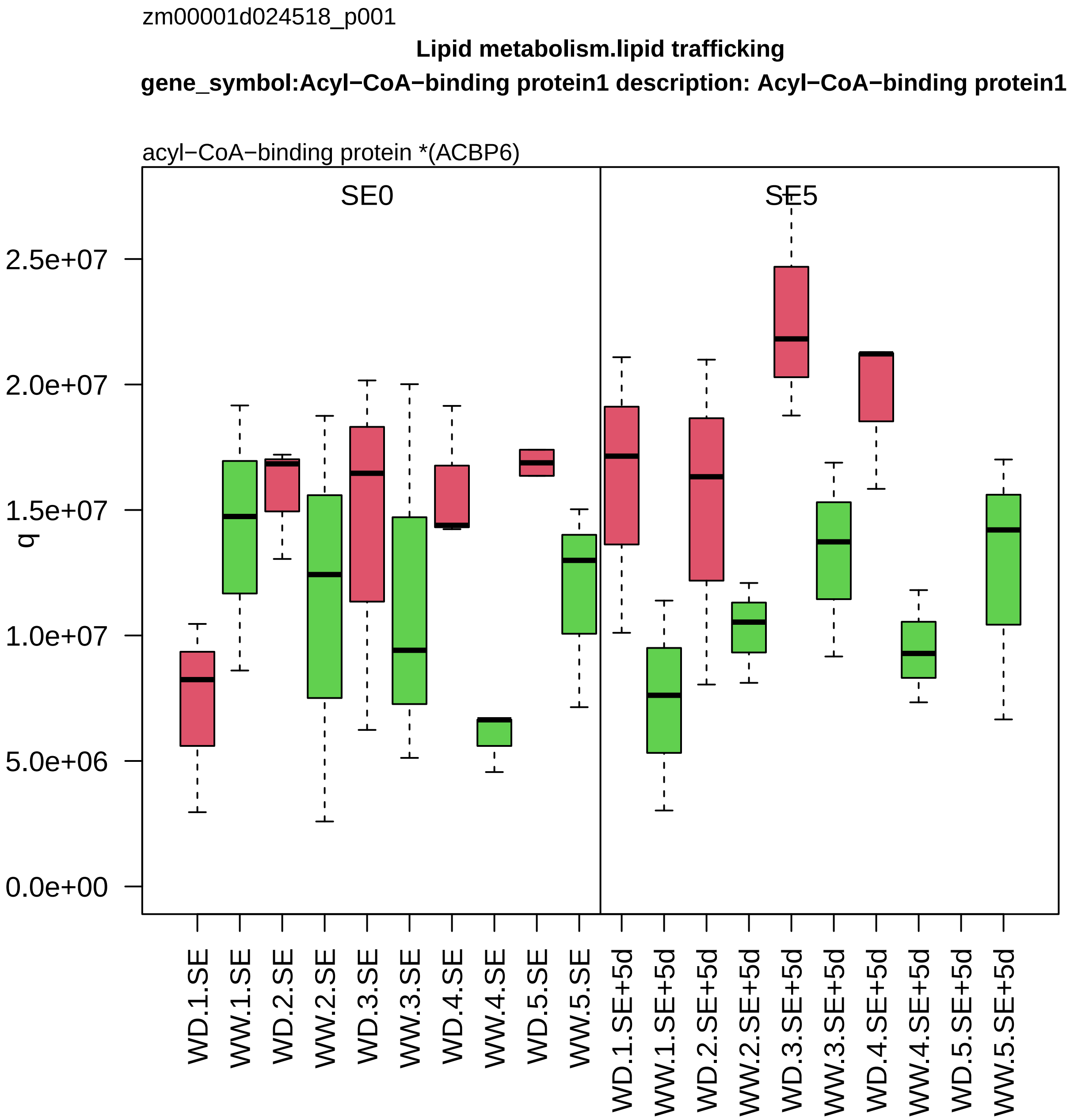
zm00001d024518_p001
Lipid metabolism.lipid trafficking
gene_symbol:Acyl−CoA−binding protein1 description: Acyl−CoA−binding protein1
acyl−CoA−binding protein *(ACBP6)
SE0
SE5
q
0.0e+00
5.0e+06
1.0e+07
1.5e+07
2.0e+07
2.5e+07
WD.1.SE
WW.1.SE
WD.2.SE
WW.2.SE
WD.3.SE
WW.3.SE
WD.4.SE
WW.4.SE
WD.5.SE
WW.5.SE
WD.1.SE+5d
WW.1.SE+5d
WD.2.SE+5d
WW.2.SE+5d
WD.3.SE+5d
WW.3.SE+5d
WD.4.SE+5d
WW.4.SE+5d
WD.5.SE+5d
WW.5.SE+5d

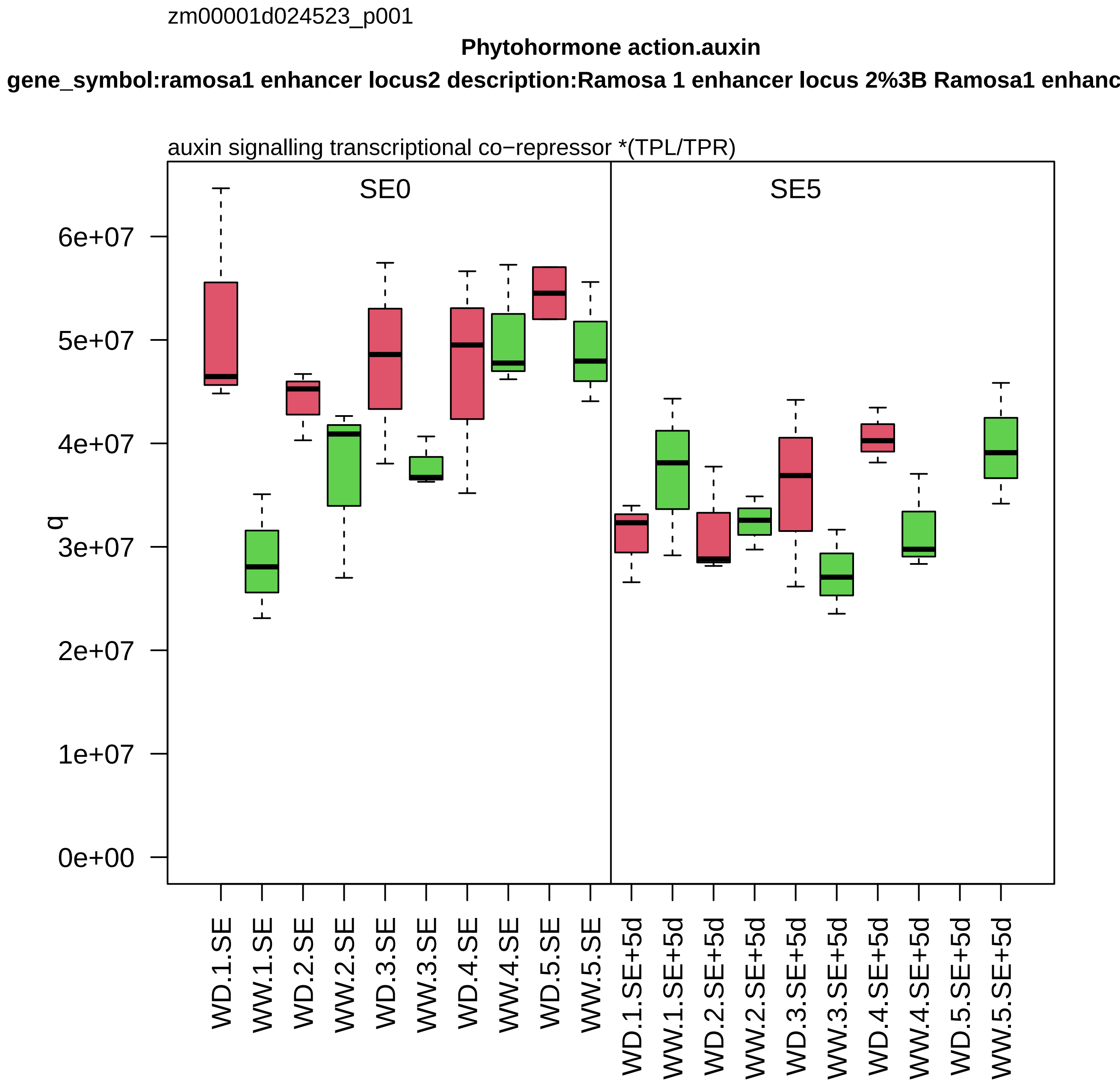
zm00001d024523_p001
Phytohormone action.auxin
gene_symbol:ramosa1 enhancer locus2 description:Ramosa 1 enhancer locus 2%3B Ramosa1 enhanc
auxin signalling transcriptional co−repressor *(TPL/TPR)
SE0
SE5
q
0e+00
1e+07
2e+07
3e+07
4e+07
5e+07
6e+07
WD.1.SE
WW.1.SE
WD.2.SE
WW.2.SE
WD.3.SE
WW.3.SE
WD.4.SE
WW.4.SE
WD.5.SE
WW.5.SE
WD.1.SE+5d
WW.1.SE+5d
WD.2.SE+5d
WW.2.SE+5d
WD.3.SE+5d
WW.3.SE+5d
WD.4.SE+5d
WW.4.SE+5d
WD.5.SE+5d
WW.5.SE+5d

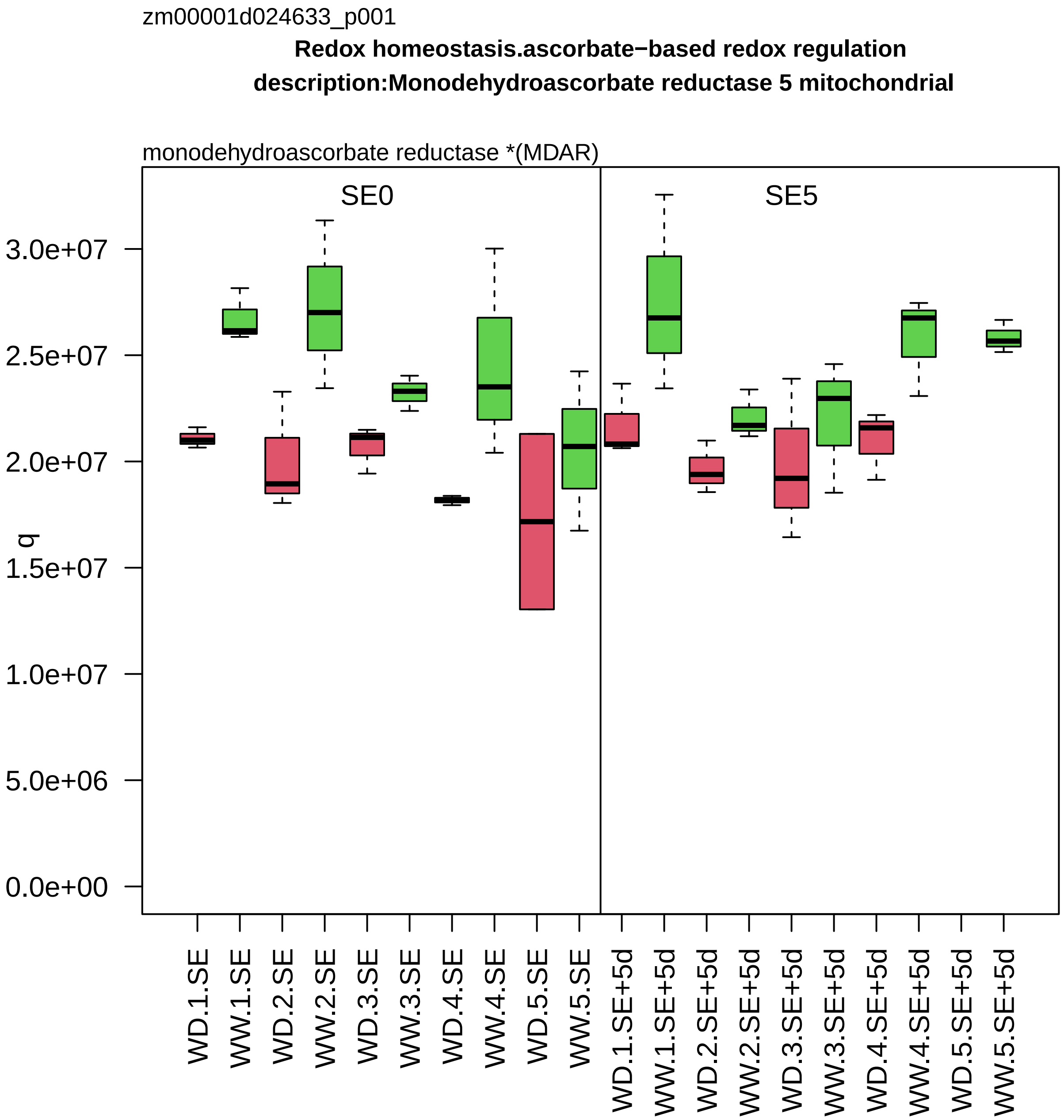

zm00001d024633_p001
Redox homeostasis.ascorbate−based redox regulation
description:Monodehydroascorbate reductase 5 mitochondrial
monodehydroascorbate reductase *(MDAR)
SE0
SE5
q
3.0e+07
2.5e+07
2.0e+07
1.5e+07
1.0e+07
5.0e+06
0.0e+00
WD.1.SE
WW.1.SE
WD.2.SE
WW.2.SE
WD.3.SE
WW.3.SE
WD.4.SE
WW.4.SE
WD.5.SE
WW.5.SE
WD.1.SE+5d
WW.1.SE+5d
WD.2.SE+5d
WW.2.SE+5d
WD.3.SE+5d
WW.3.SE+5d
WD.4.SE+5d
WW.4.SE+5d
WD.5.SE+5d
WW.5.SE+5d

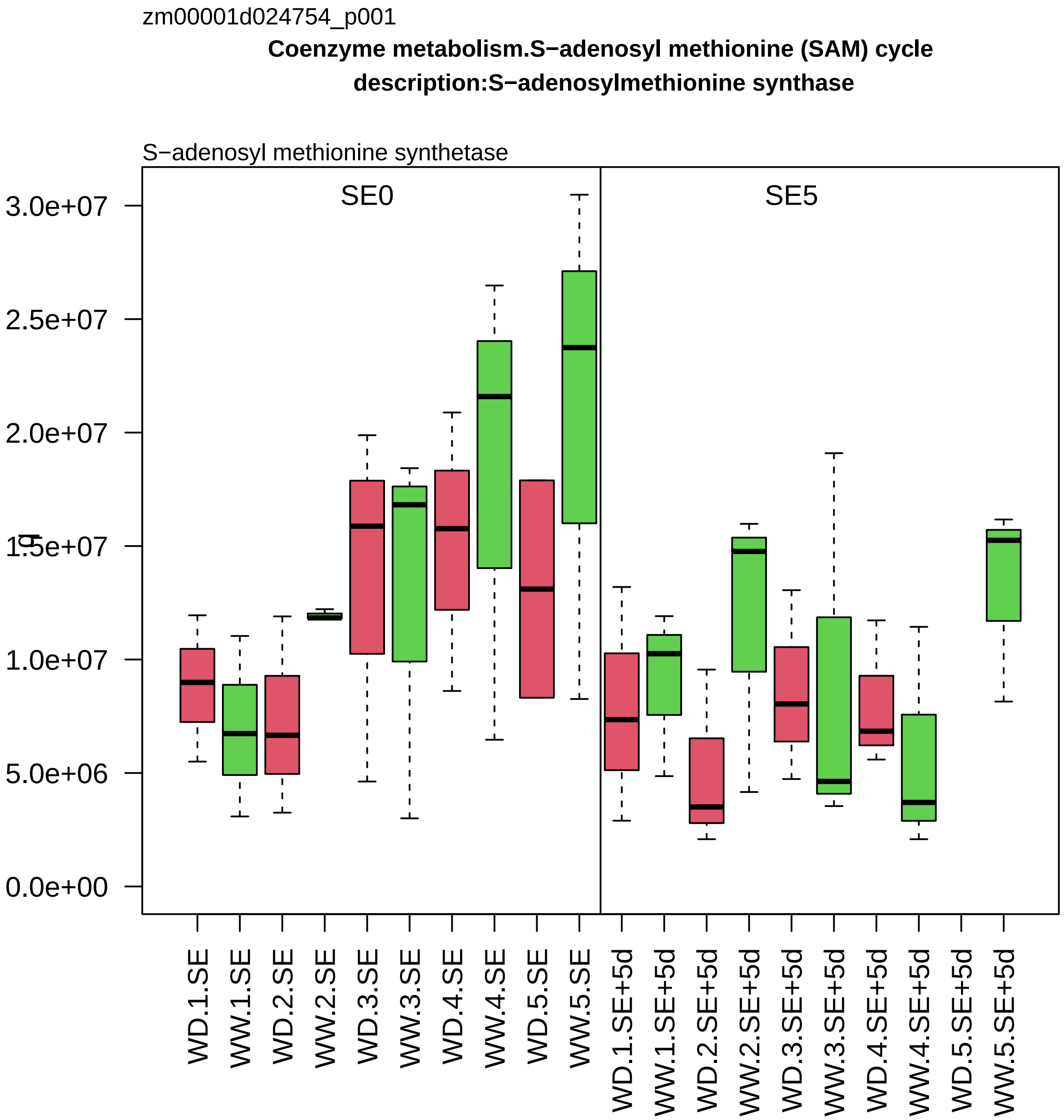

zm00001d024754_p001
Coenzyme metabolism.S−adenosyl methionine (SAM) cycle
description:S−adenosylmethionine synthase
S−adenosyl methionine synthetase
SE0
SE5
3.0e+07
2.5e+07
2.0e+07
1.5e+07
1.0e+07
5.0e+06
0.0e+00
WD.1.SE
WW.1.SE
WD.2.SE
WW.2.SE
WD.3.SE
WW.3.SE
WD.4.SE
WW.4.SE
WD.5.SE
WW.5.SE
WD.1.SE+5d
WW.1.SE+5d
WD.2.SE+5d
WW.2.SE+5d
WD.3.SE+5d
WW.3.SE+5d
WD.4.SE+5d
WW.4.SE+5d
WD.5.SE+5d
WW.5.SE+5d

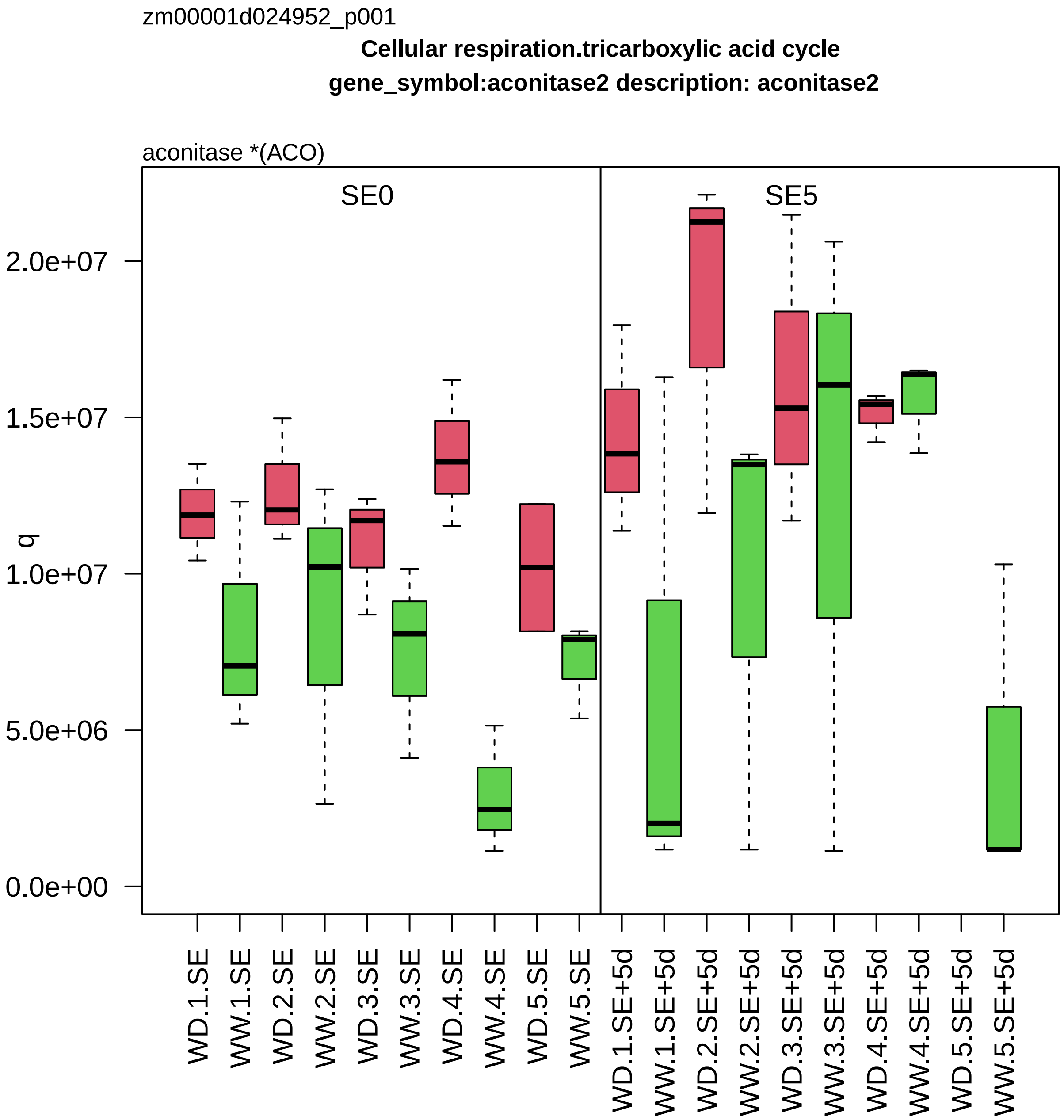
zm00001d024952_p001
Cellular respiration.tricarboxylic acid cycle
gene_symbol:aconitase2 description: aconitase2
aconitase *(ACO)
SE0
SE5
q
0.0e+00
5.0e+06
1.0e+07
1.5e+07
2.0e+07
WD.1.SE
WW.1.SE
WD.2.SE
WW.2.SE
WD.3.SE
WW.3.SE
WD.4.SE
WW.4.SE
WD.5.SE
WW.5.SE
WD.1.SE+5d
WW.1.SE+5d
WD.2.SE+5d
WW.2.SE+5d
WD.3.SE+5d
WW.3.SE+5d
WD.4.SE+5d
WW.4.SE+5d
WD.5.SE+5d
WW.5.SE+5d

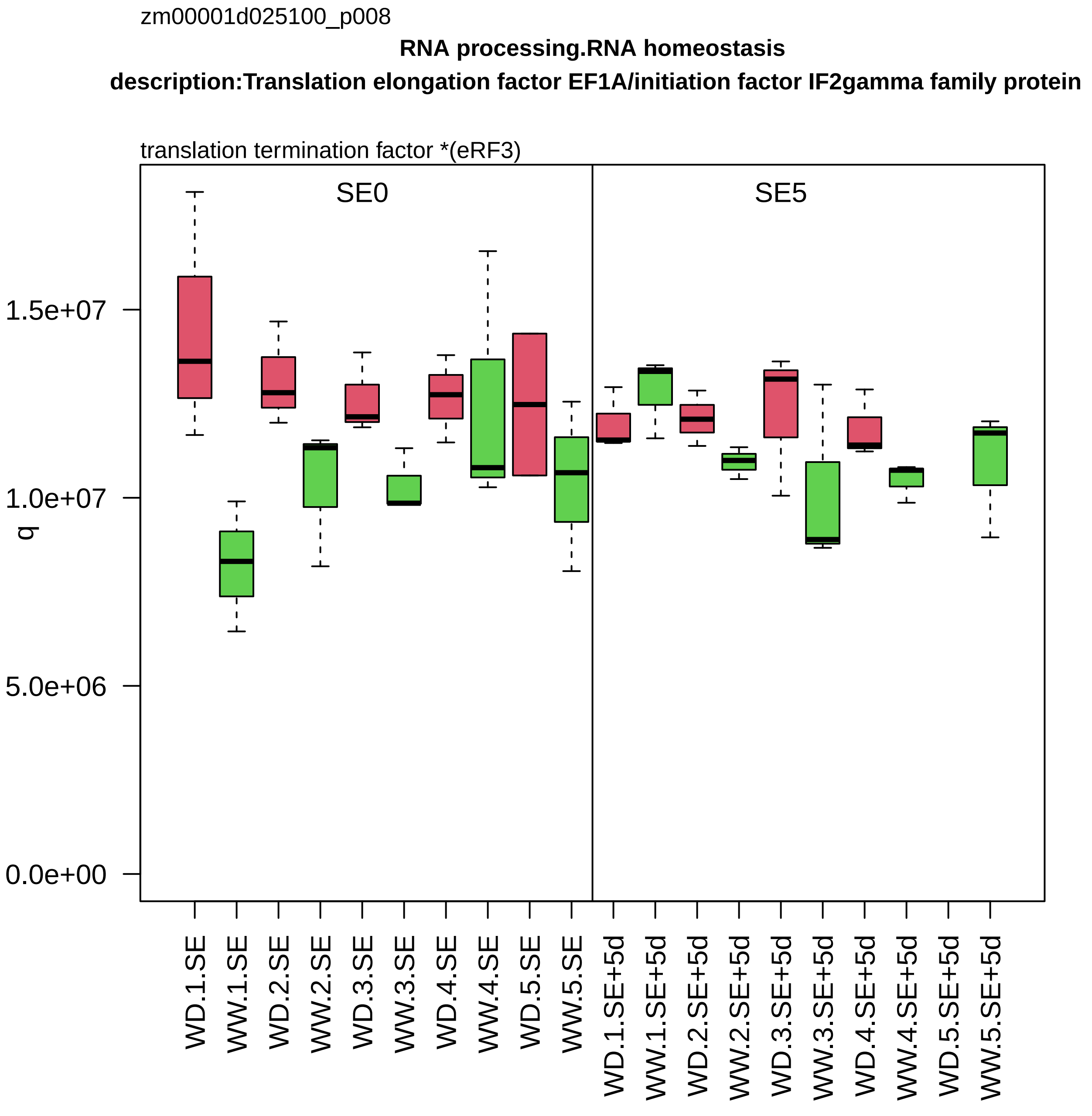

zm00001d025100_p008
RNA processing.RNA homeostasis
description:Translation elongation factor EF1A/initiation factor IF2gamma family protein
translation termination factor *(eRF3)
SE0
SE5
q
1.5e+07
1.0e+07
5.0e+06
0.0e+00
WD.1.SE
WW.1.SE
WD.2.SE
WW.2.SE
WD.3.SE
WW.3.SE
WD.4.SE
WW.4.SE
WD.5.SE
WW.5.SE
WD.1.SE+5d
WW.1.SE+5d
WD.2.SE+5d
WW.2.SE+5d
WD.3.SE+5d
WW.3.SE+5d
WD.4.SE+5d
WW.4.SE+5d
WD.5.SE+5d
WW.5.SE+5d

zm00001d025165_p003

**RNA processing.RNA homeostasis**

**description:Nuclear transport factor 2 (NTF2) family protein with RNA binding (RRM−RBD−RNP motifs)**

mRNA endoribonuclease *(G3BP)

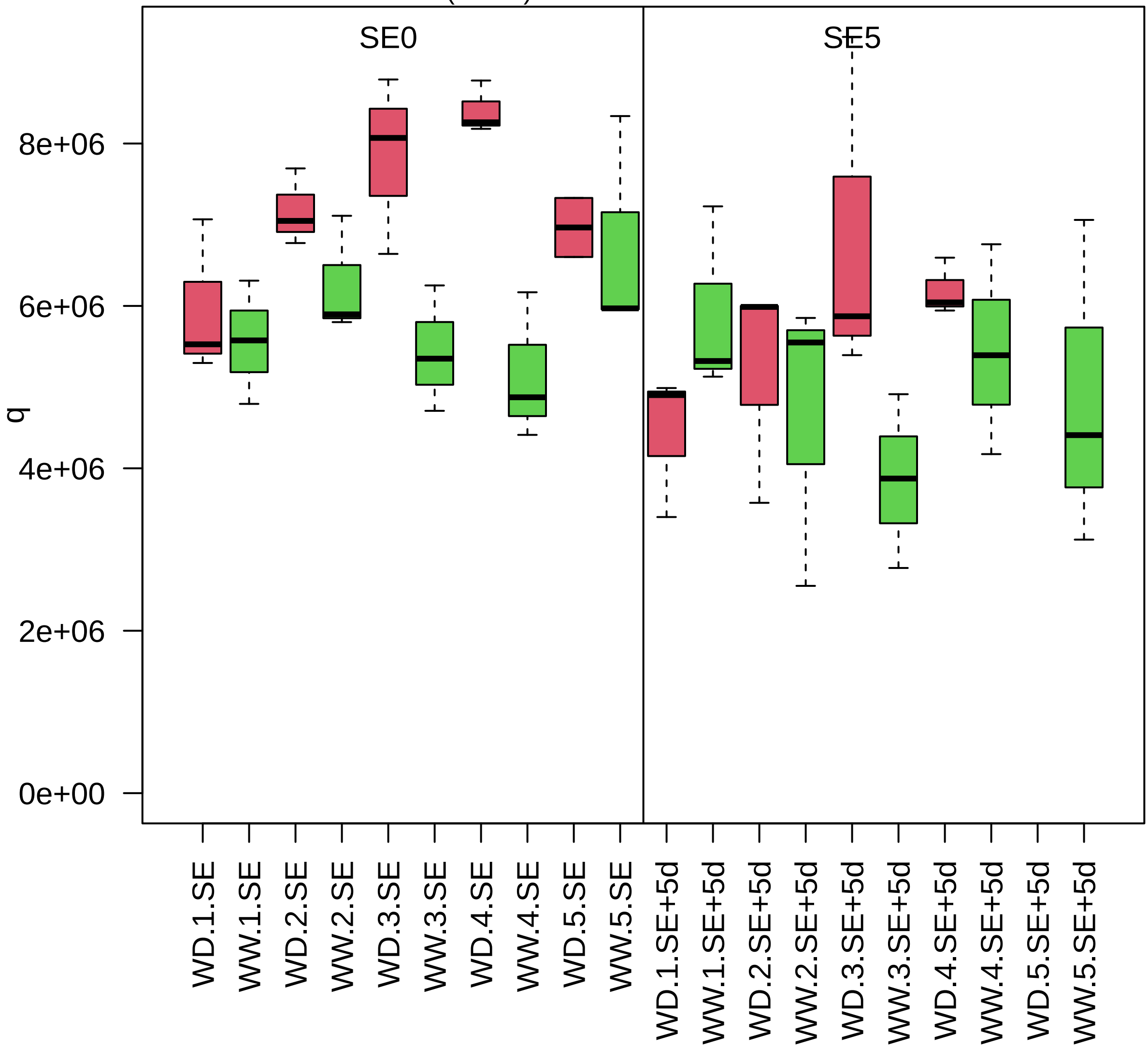

zm00001d025185_p001

**Protein biosynthesis.translation initiation**

**description:Eukaryotic translation initiation factor 3 subunit H**

component *(eIF3h) of eIF3 mRNA−to−PIC binding complex

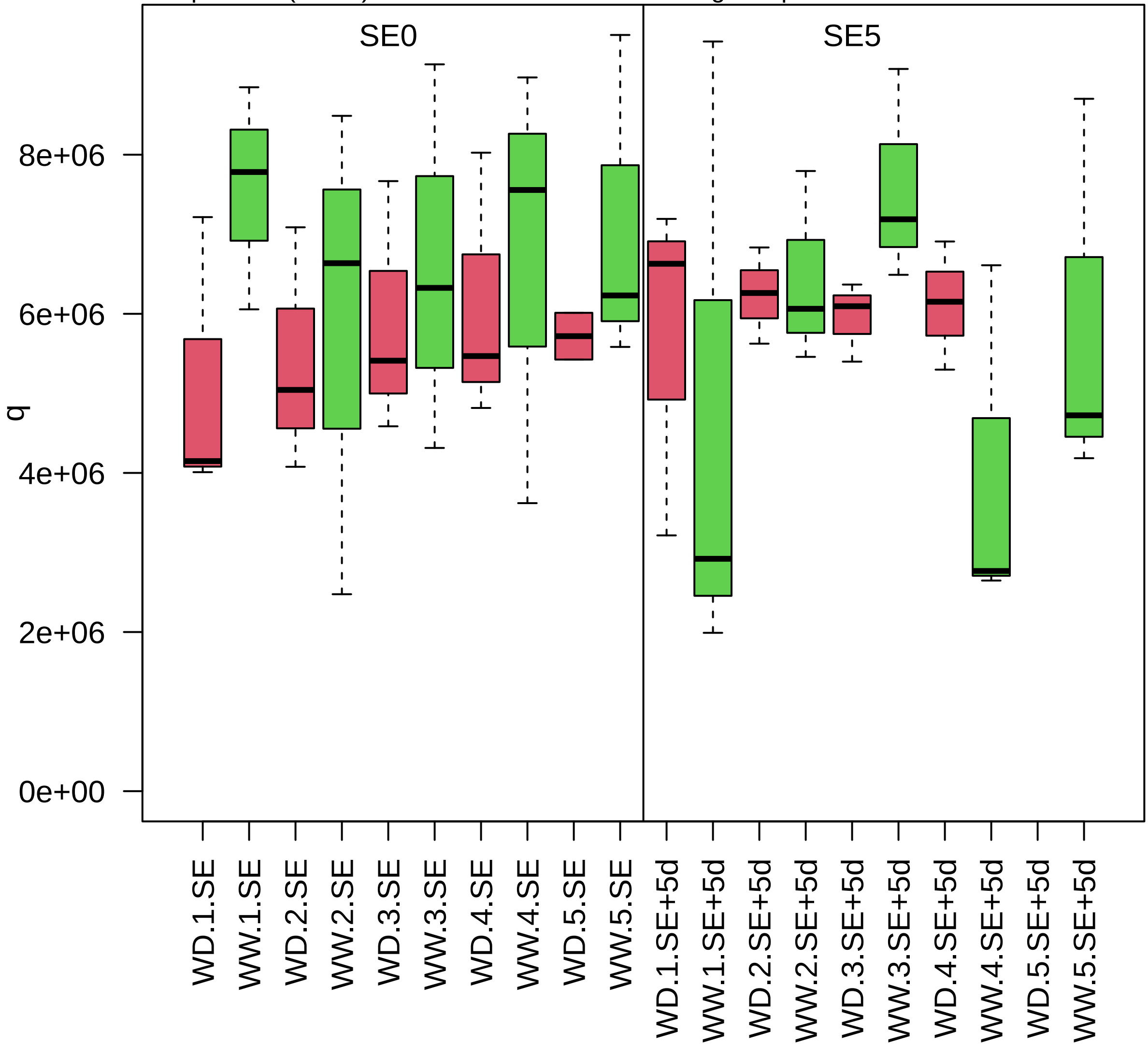

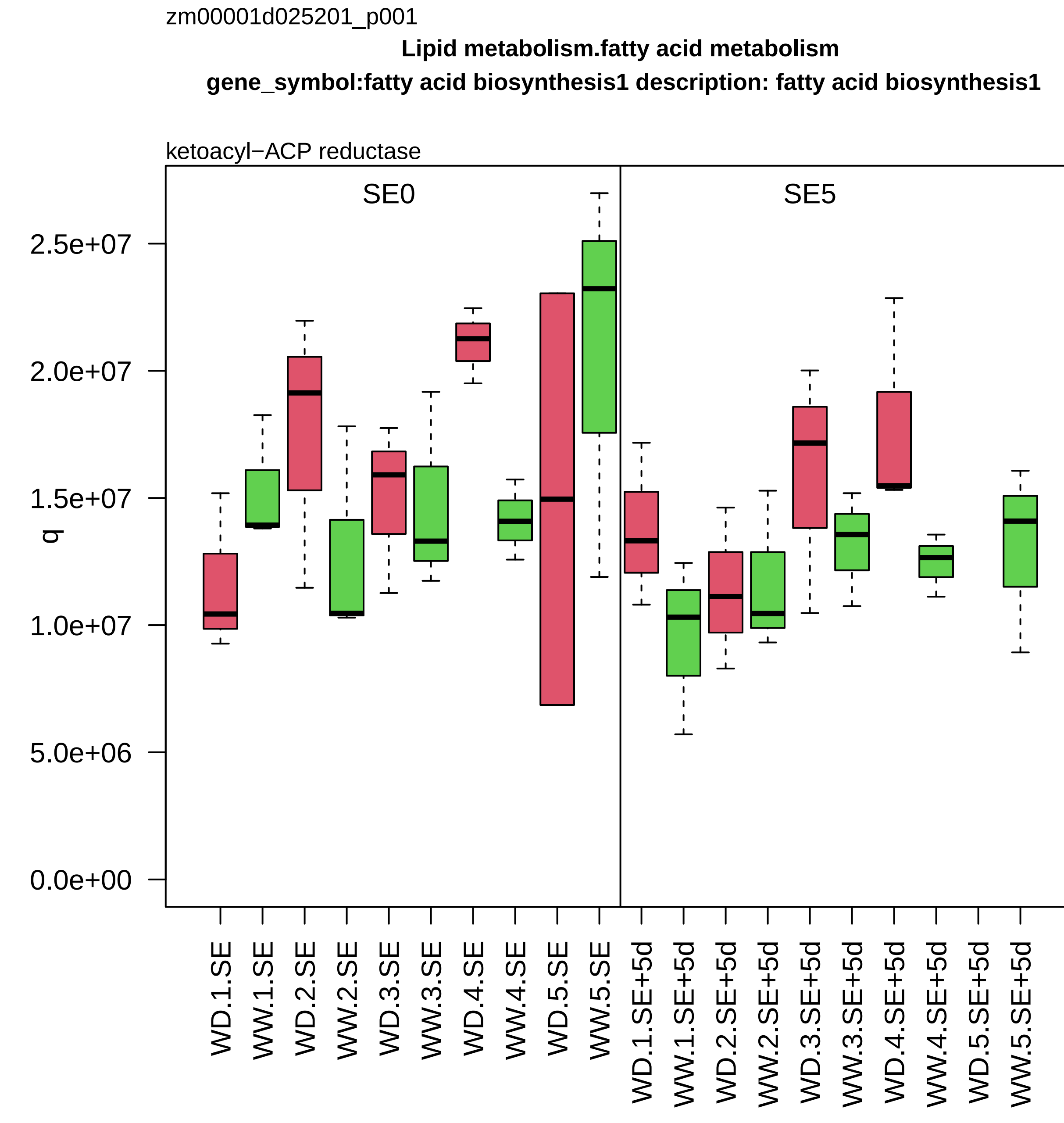

zm00001d025201_p001
Lipid metabolism.fatty acid metabolism
gene_symbol:fatty acid biosynthesis1 description: fatty acid biosynthesis1
ketoacyl−ACP reductase
SE0
SE5
q
2.5e+07
2.0e+07
1.5e+07
1.0e+07
5.0e+06
0.0e+00
WD.1.SE
WW.1.SE
WD.2.SE
WW.2.SE
WD.3.SE
WW.3.SE
WD.4.SE
WW.4.SE
WD.5.SE
WW.5.SE
WD.1.SE+5d
WW.1.SE+5d
WD.2.SE+5d
WW.2.SE+5d
WD.3.SE+5d
WW.3.SE+5d
WD.4.SE+5d
WW.4.SE+5d
WD.5.SE+5d
WW.5.SE+5d

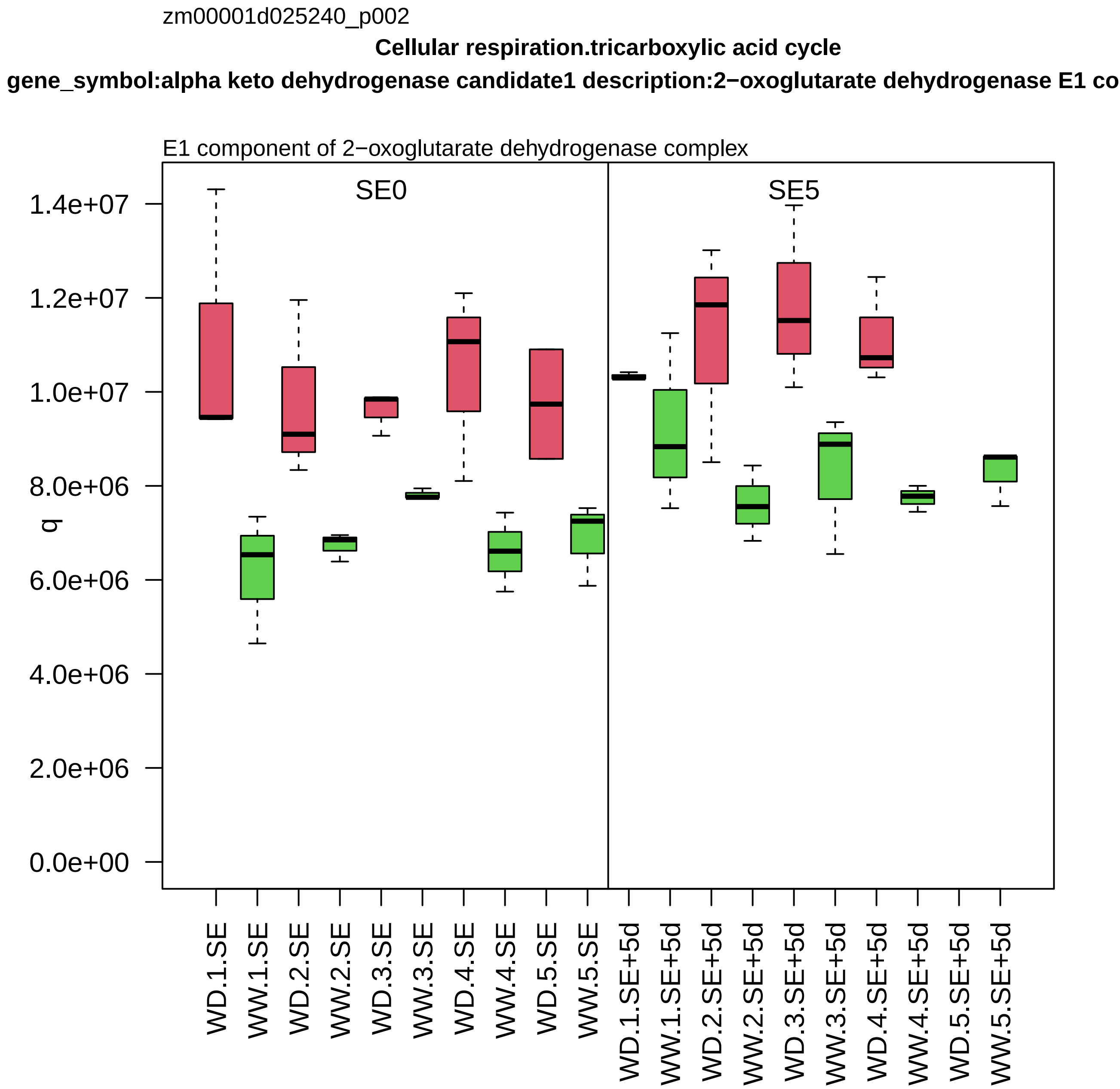
zm00001d025240_p002
Cellular respiration.tricarboxylic acid cycle
gene_symbol:alpha keto dehydrogenase candidate1 description:2−oxoglutarate dehydrogenase E1 co
E1 component of 2−oxoglutarate dehydrogenase complex
SE0
SE5
q
1.4e+07
1.2e+07
1.0e+07
8.0e+06
6.0e+06
4.0e+06
2.0e+06
0.0e+00
WD.1.SE
WW.1.SE
WD.2.SE
WW.2.SE
WD.3.SE
WW.3.SE
WD.4.SE
WW.4.SE
WD.5.SE
WW.5.SE
WD.1.SE+5d
WW.1.SE+5d
WD.2.SE+5d
WW.2.SE+5d
WD.3.SE+5d
WW.3.SE+5d
WD.4.SE+5d
WW.4.SE+5d
WD.5.SE+5d
WW.5.SE+5d

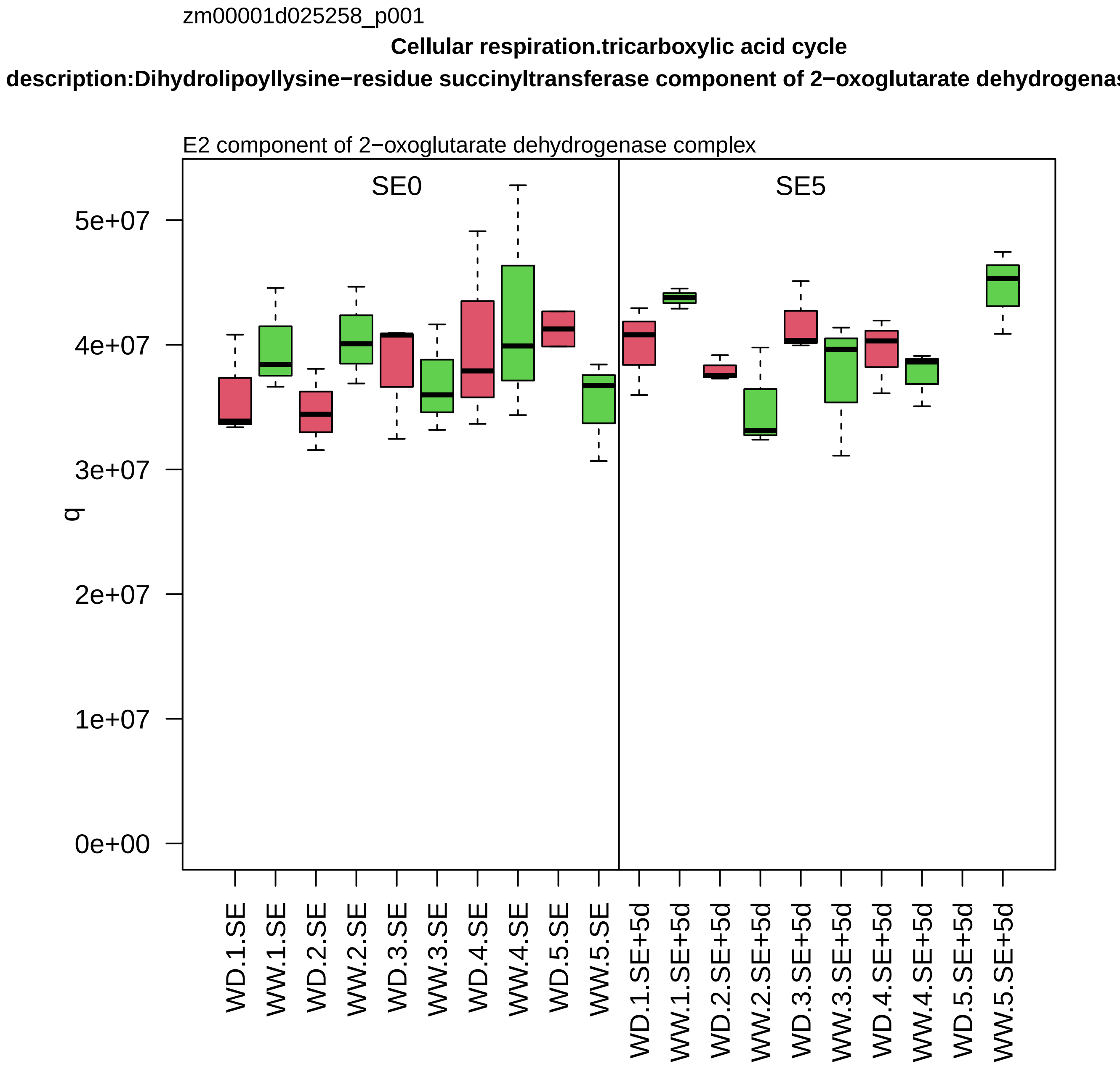
zm00001d025258_p001
Cellular respiration.tricarboxylic acid cycle
description:Dihydrolipoyllysine−residue succinyltransferase component of 2−oxoglutarate dehydrogenas
E2 component of 2−oxoglutarate dehydrogenase complex
SE0
SE5
q
0e+00
1e+07
2e+07
3e+07
4e+07
5e+07
WD.1.SE
WW.1.SE
WD.2.SE
WW.2.SE
WD.3.SE
WW.3.SE
WD.4.SE
WW.4.SE
WD.5.SE
WW.5.SE
WD.1.SE+5d
WW.1.SE+5d
WD.2.SE+5d
WW.2.SE+5d
WD.3.SE+5d
WW.3.SE+5d
WD.4.SE+5d
WW.4.SE+5d
WD.5.SE+5d
WW.5.SE+5d

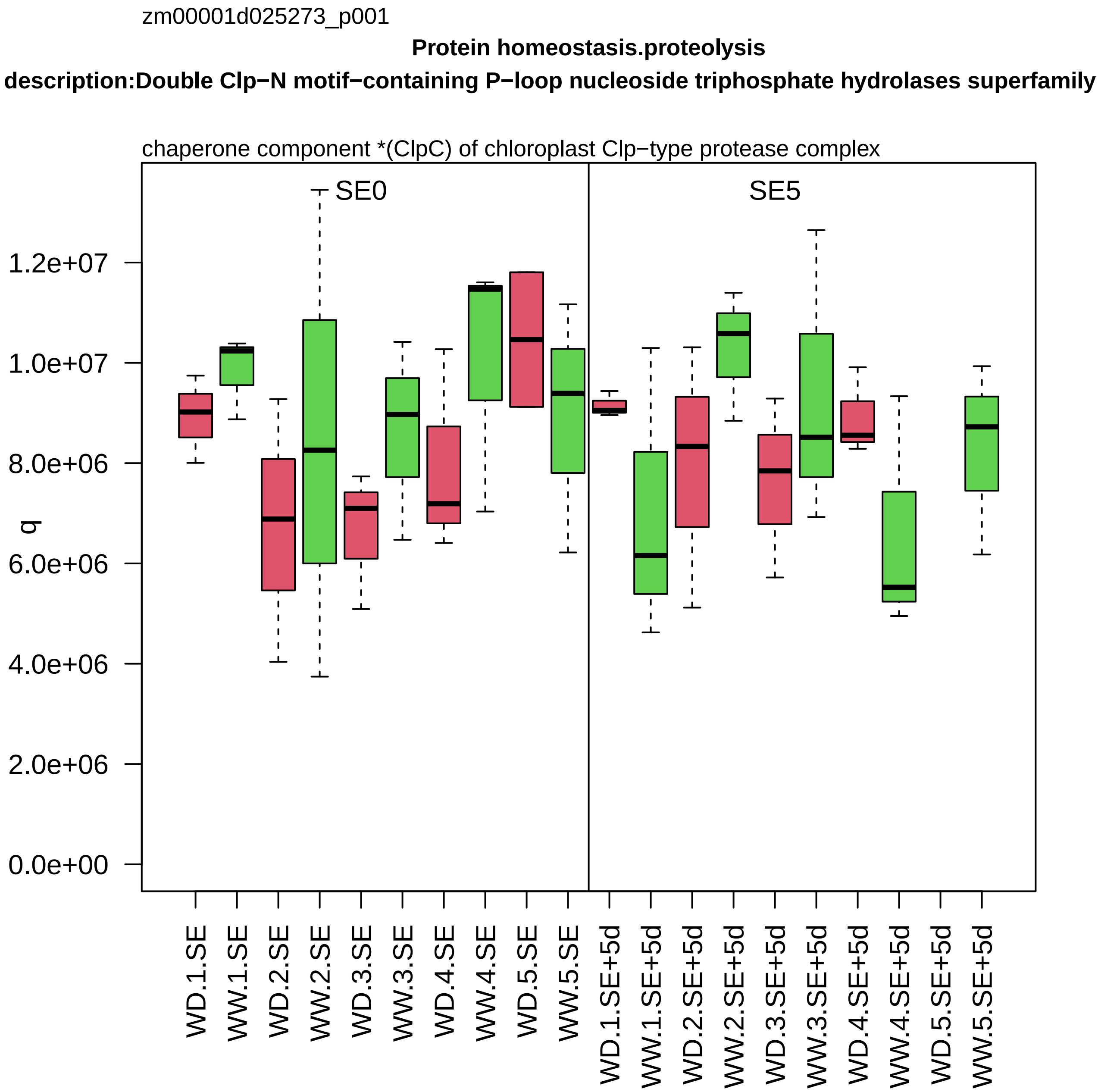

zm00001d025273_p001
Protein homeostasis.proteolysis
description:Double Clp−N motif−containing P−loop nucleoside triphosphate hydrolases superfamily
chaperone component *(ClpC) of chloroplast Clp−type protease complex
SE0
SE5
q
0.0e+00
2.0e+06
4.0e+06
6.0e+06
8.0e+06
1.0e+07
1.2e+07
WD.1.SE
WW.1.SE
WD.2.SE
WW.2.SE
WD.3.SE
WW.3.SE
WD.4.SE
WW.4.SE
WD.5.SE
WW.5.SE
WD.1.SE+5d
WW.1.SE+5d
WD.2.SE+5d
WW.2.SE+5d
WD.3.SE+5d
WW.3.SE+5d
WD.4.SE+5d
WW.4.SE+5d
WD.5.SE+5d
WW.5.SE+5d

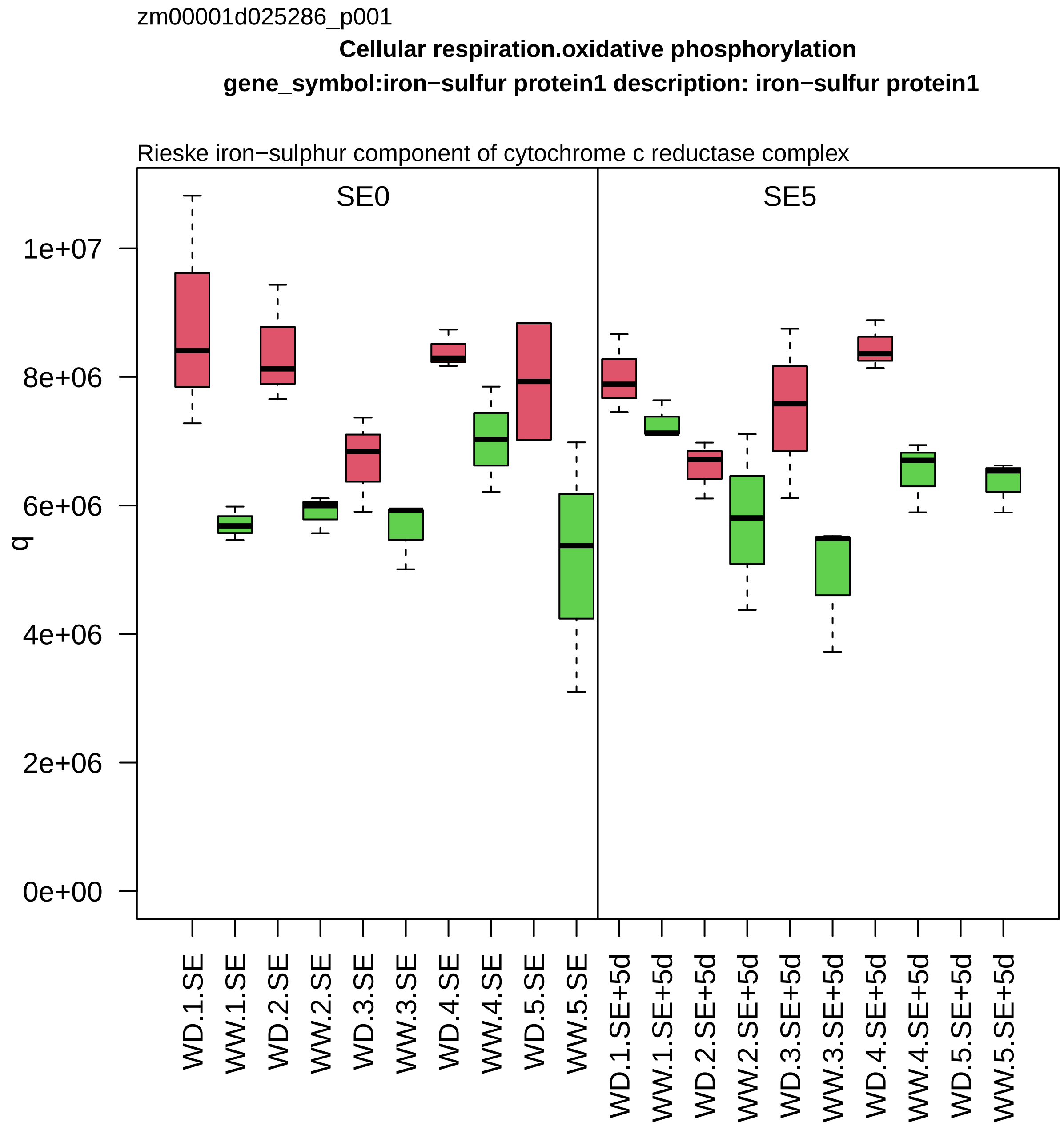
zm00001d025286_p001
Cellular respiration.oxidative phosphorylation
gene_symbol:iron−sulfur protein1 description: iron−sulfur protein1
Rieske iron−sulphur component of cytochrome c reductase complex
SE0
SE5
q
1e+07
8e+06
6e+06
4e+06
2e+06
0e+00
WD.1.SE
WW.1.SE
WD.2.SE
WW.2.SE
WD.3.SE
WW.3.SE
WD.4.SE
WW.4.SE
WD.5.SE
WW.5.SE
WD.1.SE+5d
WW.1.SE+5d
WD.2.SE+5d
WW.2.SE+5d
WD.3.SE+5d
WW.3.SE+5d
WD.4.SE+5d
WW.4.SE+5d
WD.5.SE+5d
WW.5.SE+5d

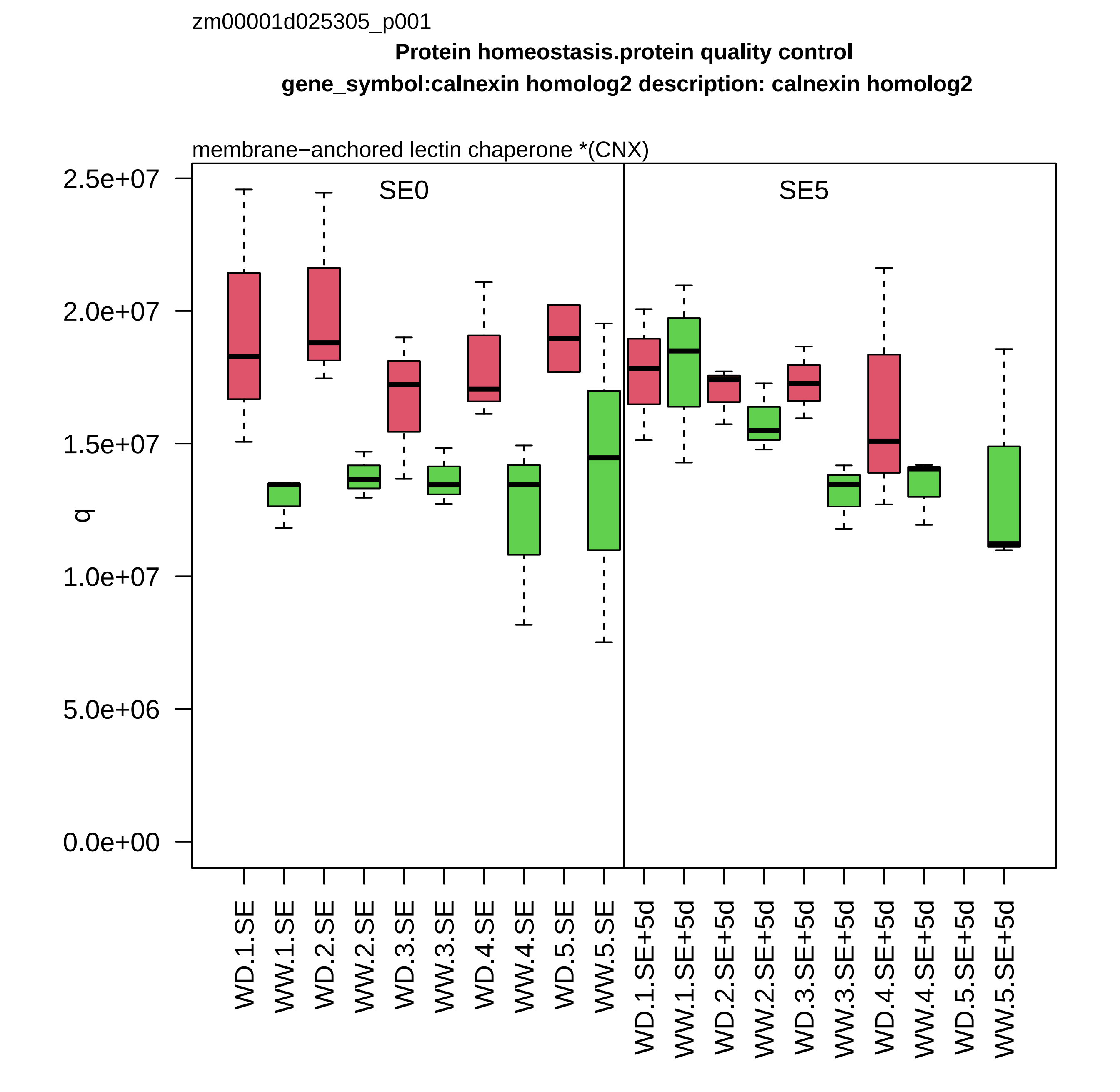

zm00001d025305_p001
Protein homeostasis.protein quality control
gene_symbol:calnexin homolog2 description: calnexin homolog2
membrane−anchored lectin chaperone *(CNX)
SE0
SE5
q
2.5e+07
2.0e+07
1.5e+07
1.0e+07
5.0e+06
0.0e+00
WD.1.SE
WW.1.SE
WD.2.SE
WW.2.SE
WD.3.SE
WW.3.SE
WD.4.SE
WW.4.SE
WD.5.SE
WW.5.SE
WD.1.SE+5d
WW.1.SE+5d
WD.2.SE+5d
WW.2.SE+5d
WD.3.SE+5d
WW.3.SE+5d
WD.4.SE+5d
WW.4.SE+5d
WD.5.SE+5d
WW.5.SE+5d

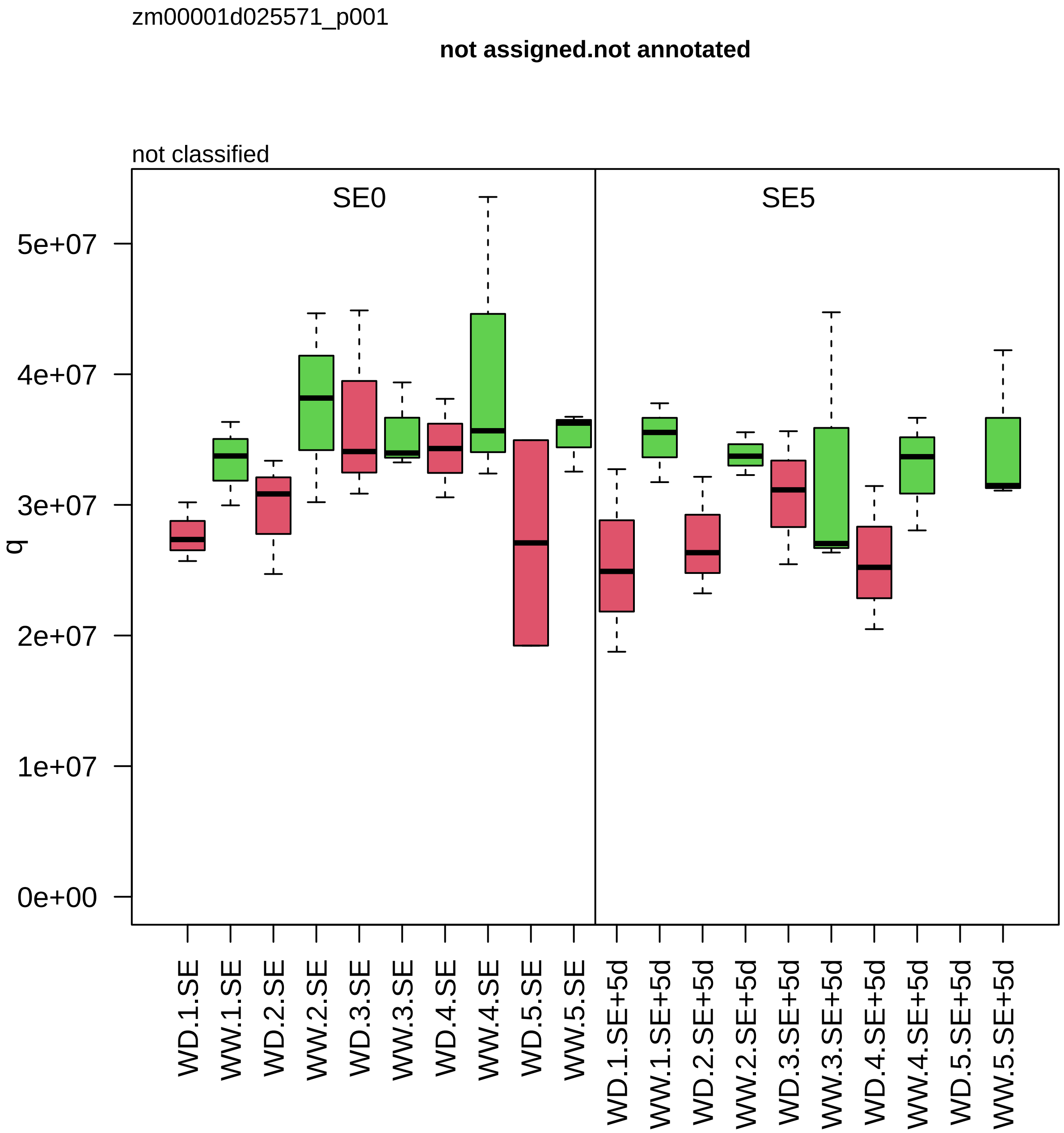
zm00001d025571_p001
not assigned.not annotated
not classified
SE0
SE5
q
0e+00
1e+07
2e+07
3e+07
4e+07
5e+07
WD.1.SE
WW.1.SE
WD.2.SE
WW.2.SE
WD.3.SE
WW.3.SE
WD.4.SE
WW.4.SE
WD.5.SE
WW.5.SE
WD.1.SE+5d
WW.1.SE+5d
WD.2.SE+5d
WW.2.SE+5d
WD.3.SE+5d
WW.3.SE+5d
WD.4.SE+5d
WW.4.SE+5d
WD.5.SE+5d
WW.5.SE+5d

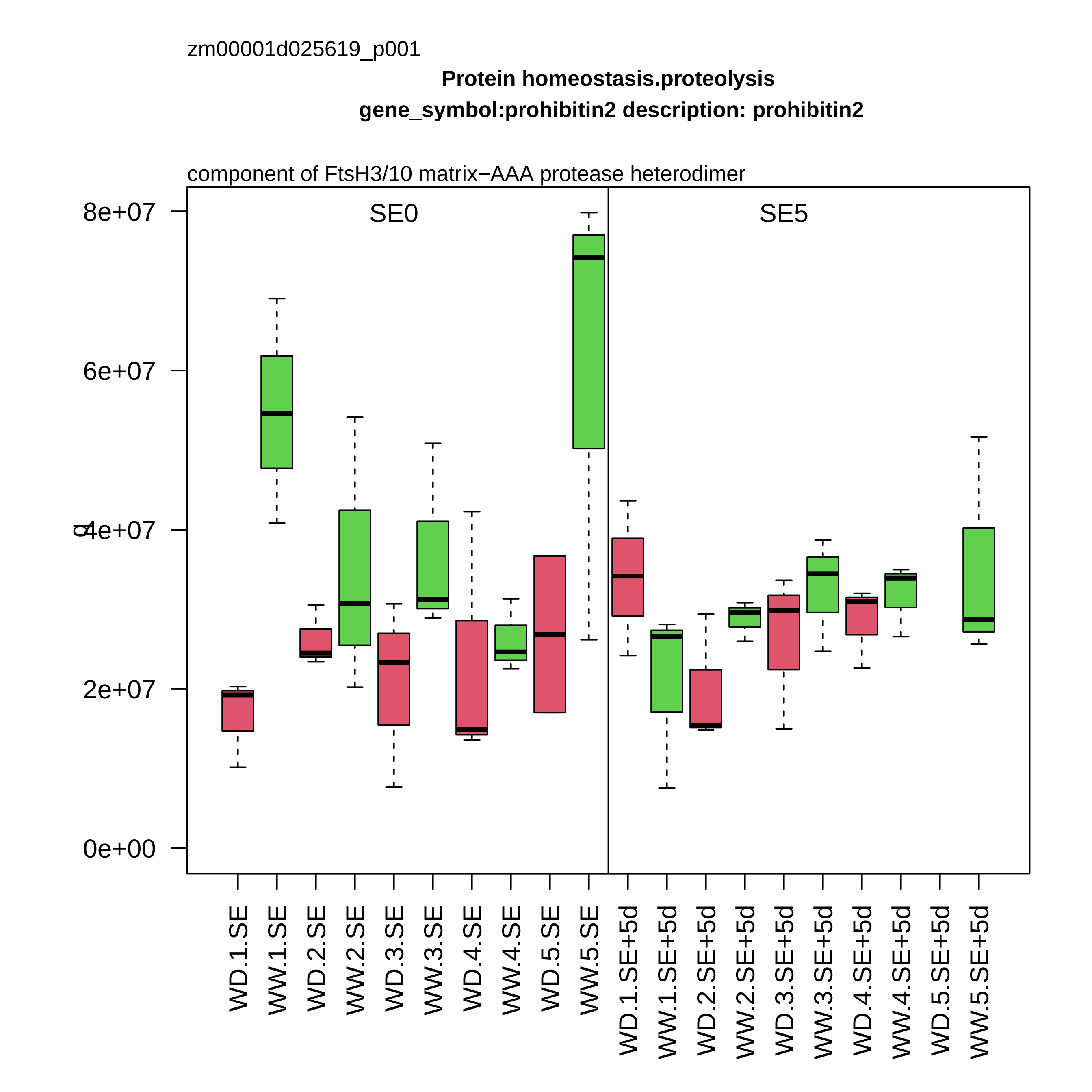

zm00001d025619_p001
Protein homeostasis.proteolysis
gene_symbol:prohibitin2 description: prohibitin2
component of FtsH3/10 matrix−AAA protease heterodimer
SE0
SE5
8e+07
6e+07
4e+07
2e+07
0e+00
WD.1.SE
WW.1.SE
WD.2.SE
WW.2.SE
WD.3.SE
WW.3.SE
WD.4.SE
WW.4.SE
WD.5.SE
WW.5.SE
WD.1.SE+5d
WW.1.SE+5d
WD.2.SE+5d
WW.2.SE+5d
WD.3.SE+5d
WW.3.SE+5d
WD.4.SE+5d
WW.4.SE+5d
WD.5.SE+5d
WW.5.SE+5d

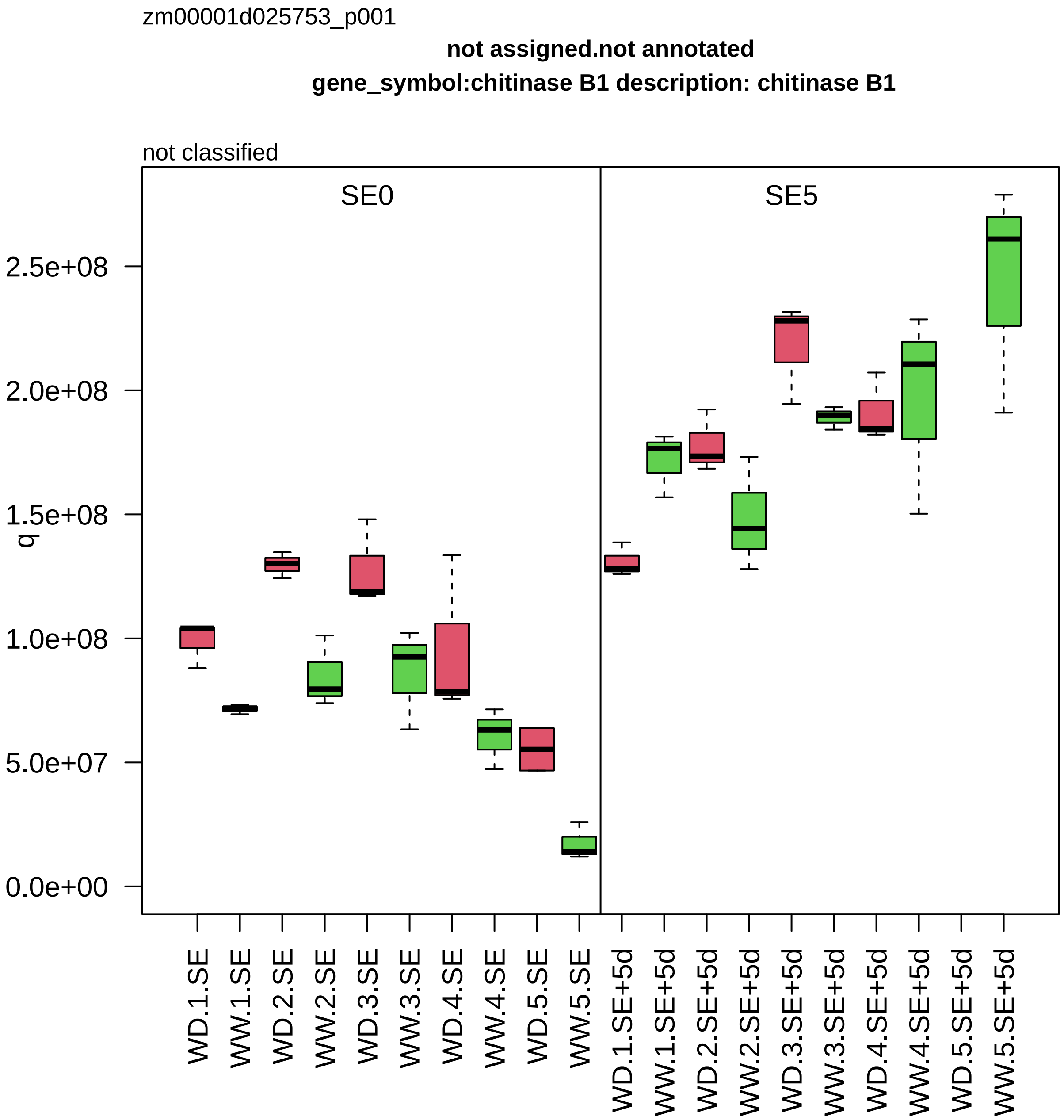
zm00001d025753_p001
not assigned.not annotated
gene_symbol:chitinase B1 description: chitinase B1
not classified
SE0
SE5
q
0.0e+00
5.0e+07
1.0e+08
1.5e+08
2.0e+08
2.5e+08
WD.1.SE
WW.1.SE
WD.2.SE
WW.2.SE
WD.3.SE
WW.3.SE
WD.4.SE
WW.4.SE
WD.5.SE
WW.5.SE
WD.1.SE+5d
WW.1.SE+5d
WD.2.SE+5d
WW.2.SE+5d
WD.3.SE+5d
WW.3.SE+5d
WD.4.SE+5d
WW.4.SE+5d
WD.5.SE+5d
WW.5.SE+5d

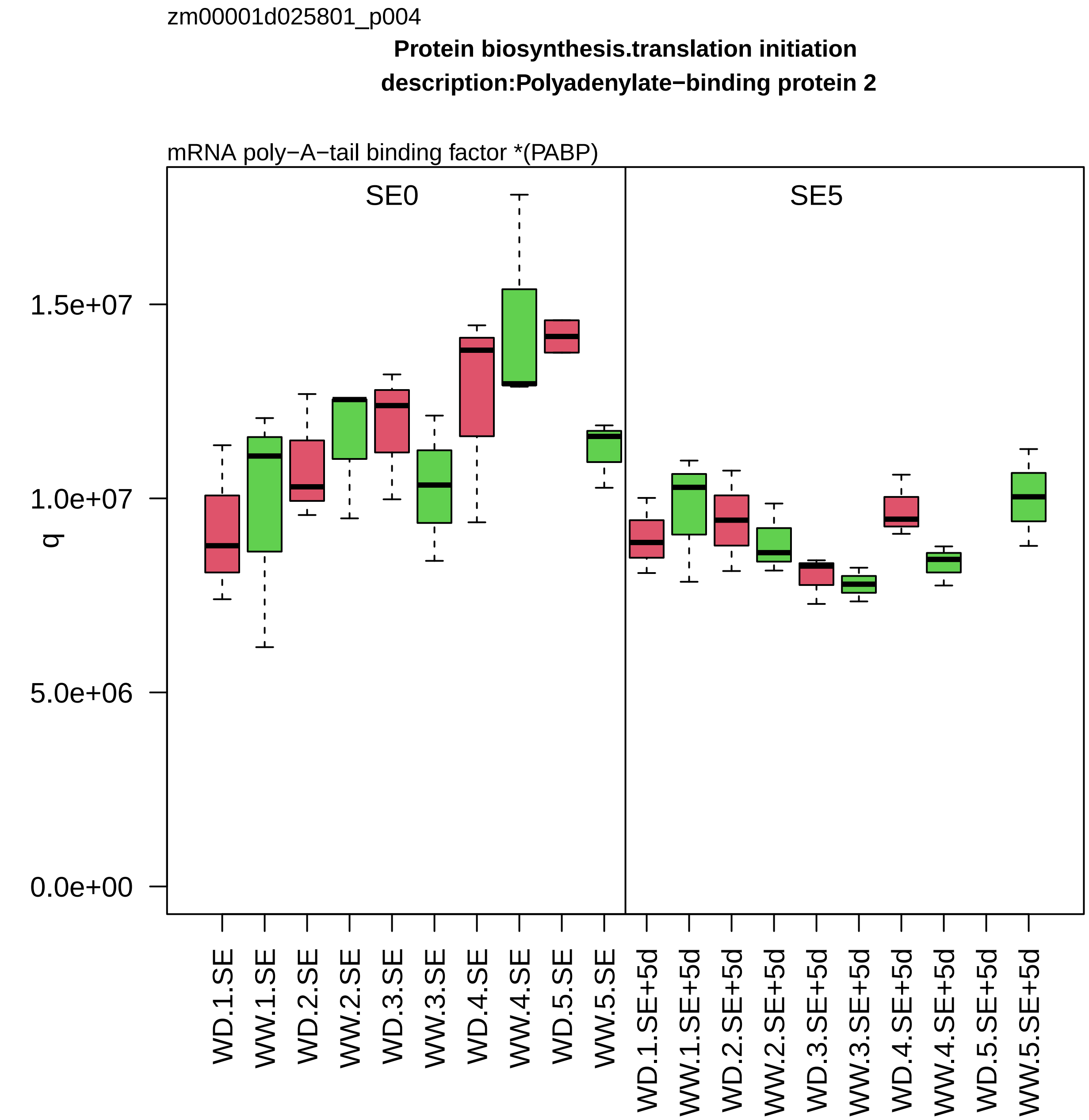
zm00001d025801_p004
Protein biosynthesis.translation initiation
description:Polyadenylate−binding protein 2
mRNA poly−A−tail binding factor *(PABP)
SE0
SE5
q
1.5e+07
1.0e+07
5.0e+06
0.0e+00
WD.1.SE
WW.1.SE
WD.2.SE
WW.2.SE
WD.3.SE
WW.3.SE
WD.4.SE
WW.4.SE
WD.5.SE
WW.5.SE
WD.1.SE+5d
WW.1.SE+5d
WD.2.SE+5d
WW.2.SE+5d
WD.3.SE+5d
WW.3.SE+5d
WD.4.SE+5d
WW.4.SE+5d
WD.5.SE+5d
WW.5.SE+5d

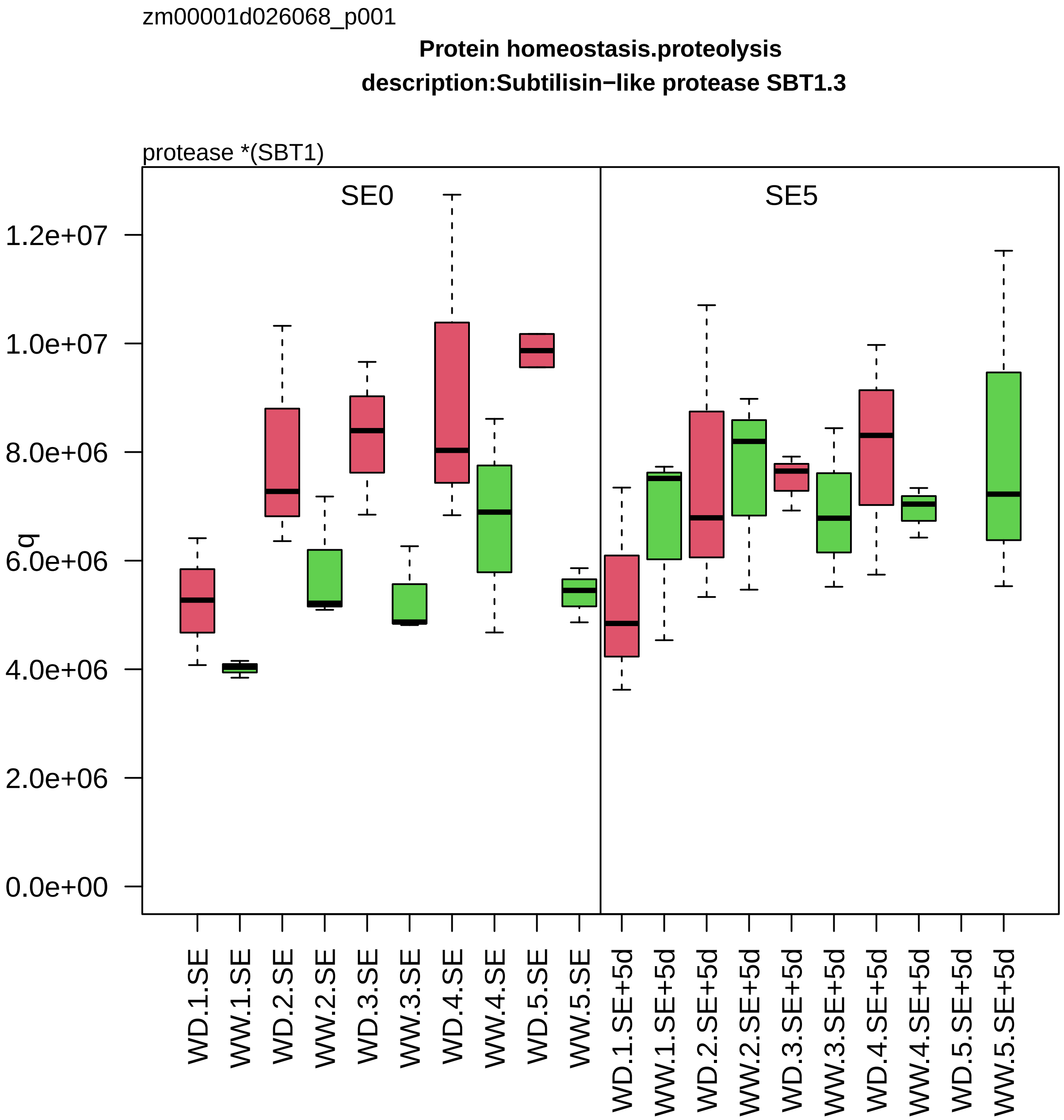
zm00001d026068_p001
Protein homeostasis.proteolysis
description:Subtilisin−like protease SBT1.3
protease *(SBT1)
SE0
SE5
q
1.2e+07
1.0e+07
8.0e+06
6.0e+06
4.0e+06
2.0e+06
0.0e+00
WD.1.SE
WW.1.SE
WD.2.SE
WW.2.SE
WD.3.SE
WW.3.SE
WD.4.SE
WW.4.SE
WD.5.SE
WW.5.SE
WD.1.SE+5d
WW.1.SE+5d
WD.2.SE+5d
WW.2.SE+5d
WD.3.SE+5d
WW.3.SE+5d
WD.4.SE+5d
WW.4.SE+5d
WD.5.SE+5d
WW.5.SE+5d

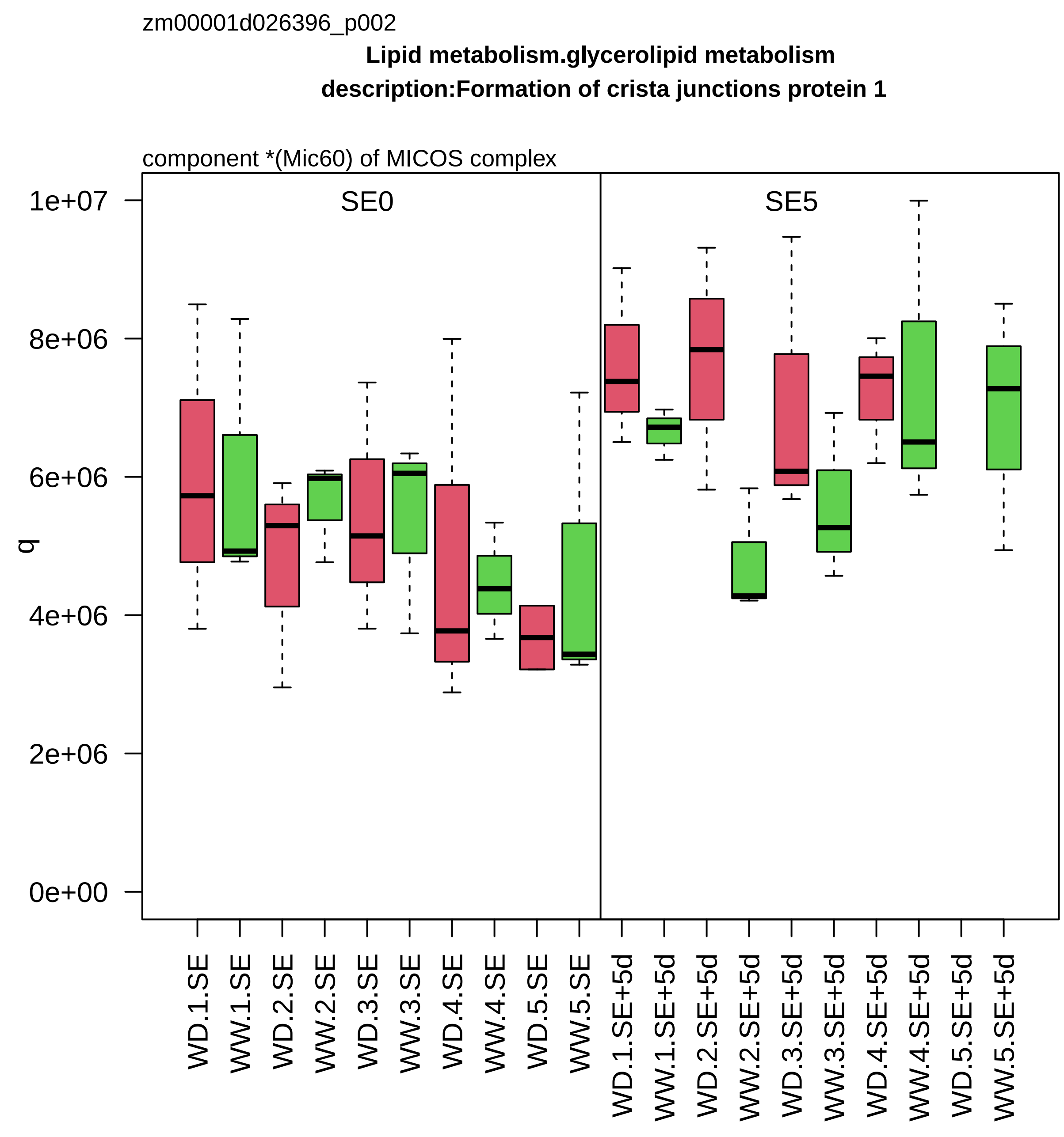

zm00001d026396_p002
Lipid metabolism.glycerolipid metabolism
description:Formation of crista junctions protein 1
component *(Mic60) of MICOS complex
SE0
SE5
q
1e+07
8e+06
6e+06
4e+06
2e+06
0e+00
WD.1.SE
WW.1.SE
WD.2.SE
WW.2.SE
WD.3.SE
WW.3.SE
WD.4.SE
WW.4.SE
WD.5.SE
WW.5.SE
WD.1.SE+5d
WW.1.SE+5d
WD.2.SE+5d
WW.2.SE+5d
WD.3.SE+5d
WW.3.SE+5d
WD.4.SE+5d
WW.4.SE+5d
WD.5.SE+5d
WW.5.SE+5d

zm00001d026415_p002

**Cell wall organisation.pectin**
**description:Beta−D−xylosidase 4**

bifunctional alpha−L−arabinofuranosidase and beta−D−xylosidase *(BXL)

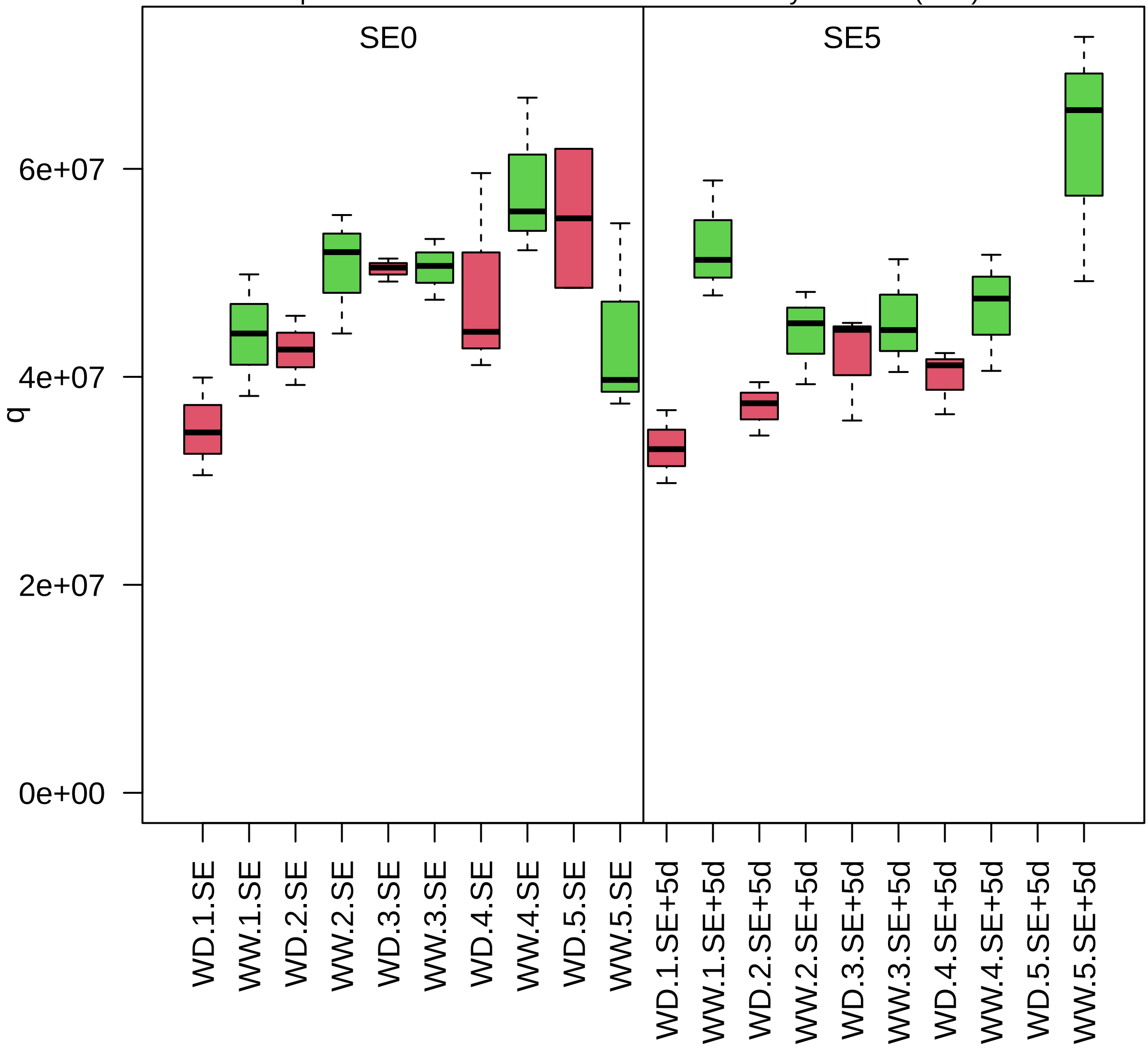

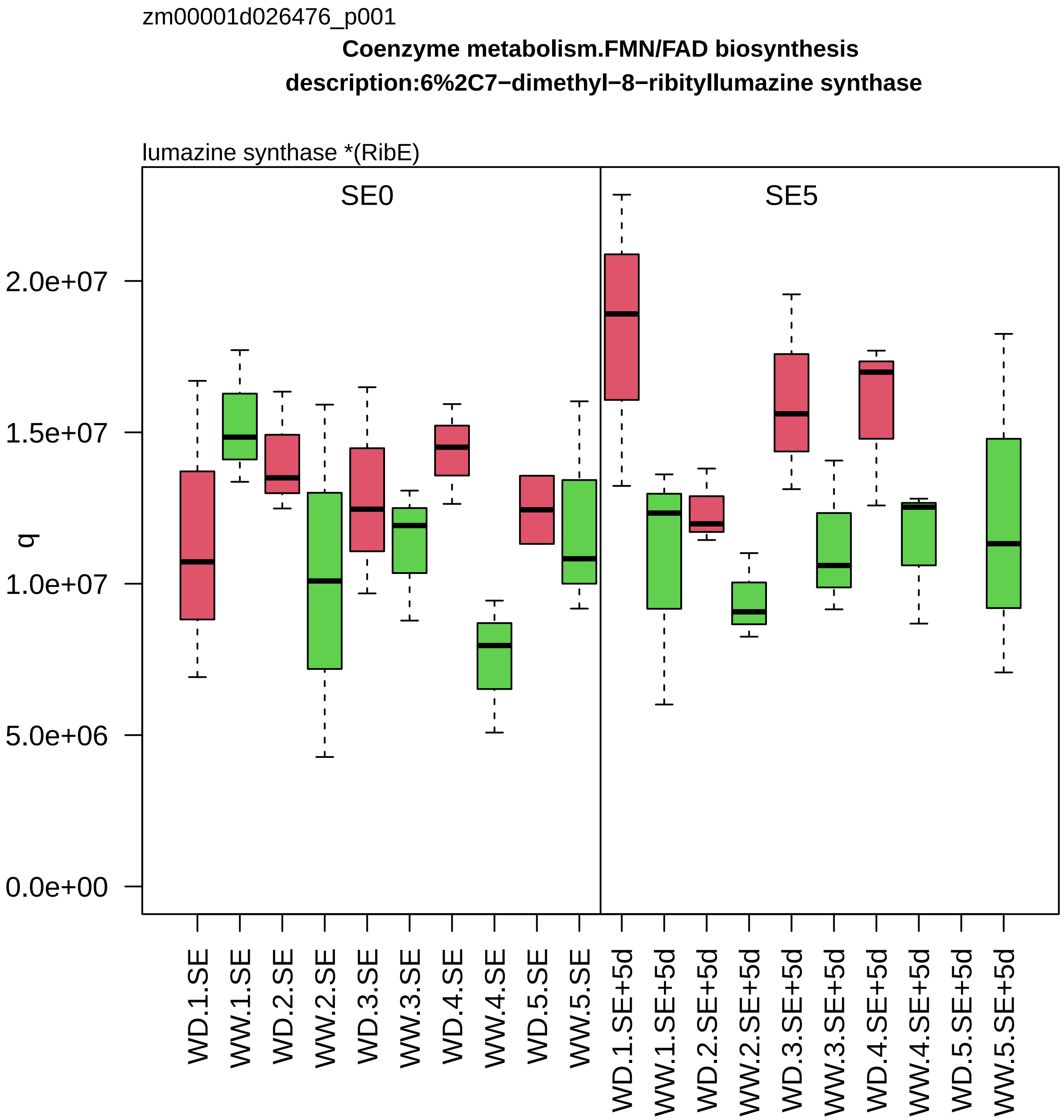

zm00001d026476_p001
Coenzyme metabolism.FMN/FAD biosynthesis
description:6%2C7−dimethyl−8−ribityllumazine synthase
lumazine synthase *(RibE)
SE0
SE5
q
0.0e+00
5.0e+06
1.0e+07
1.5e+07
2.0e+07
WD.1.SE
WW.1.SE
WD.2.SE
WW.2.SE
WD.3.SE
WW.3.SE
WD.4.SE
WW.4.SE
WD.5.SE
WW.5.SE
WD.1.SE+5d
WW.1.SE+5d
WD.2.SE+5d
WW.2.SE+5d
WD.3.SE+5d
WW.3.SE+5d
WD.4.SE+5d
WW.4.SE+5d
WD.5.SE+5d
WW.5.SE+5d

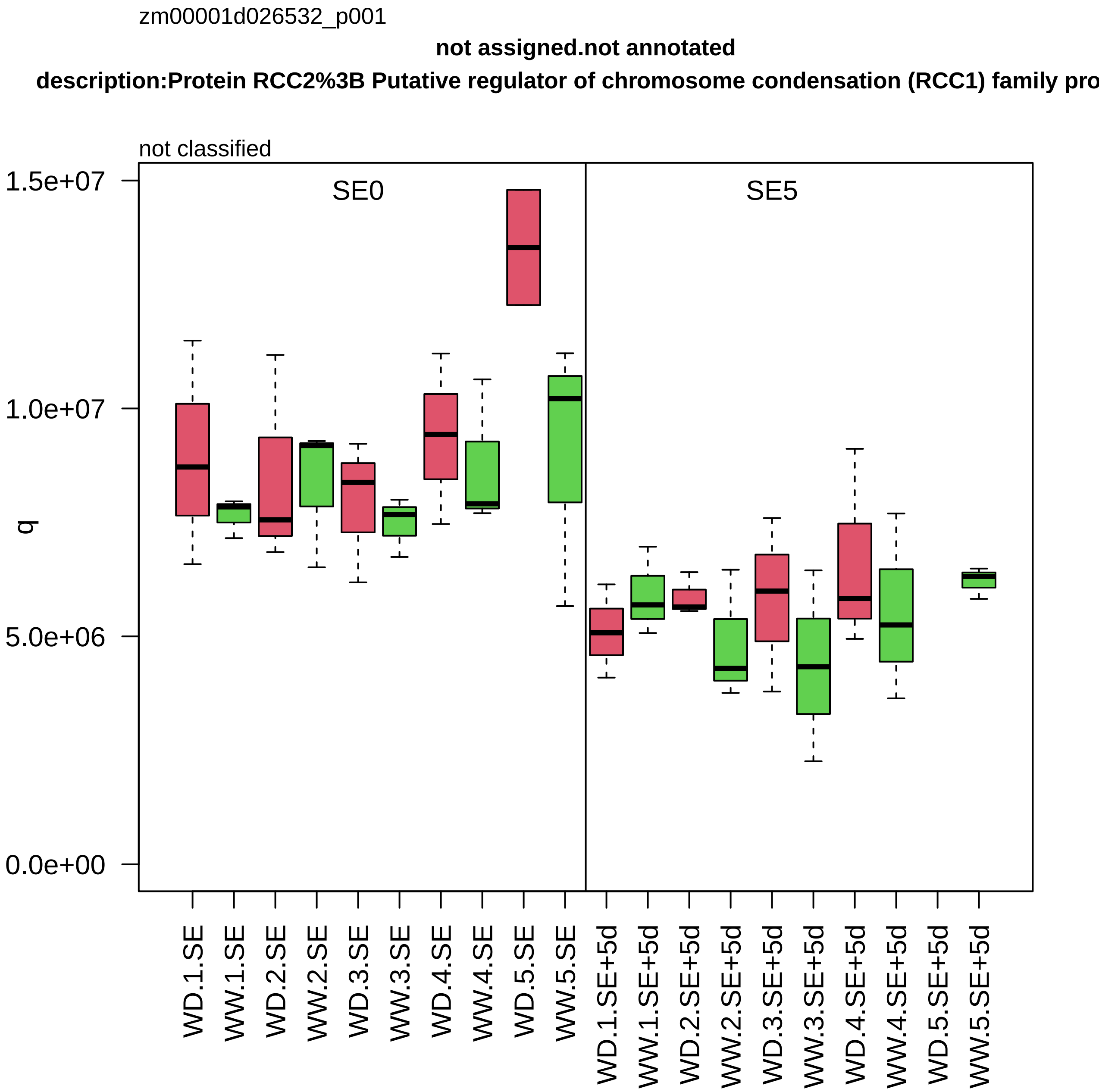
zm00001d026532_p001
not assigned.not annotated
description:Protein RCC2%3B Putative regulator of chromosome condensation (RCC1) family pro
not classified
SE0
SE5
q
1.5e+07
1.0e+07
5.0e+06
0.0e+00
WD.1.SE
WW.1.SE
WD.2.SE
WW.2.SE
WD.3.SE
WW.3.SE
WD.4.SE
WW.4.SE
WD.5.SE
WW.5.SE
WD.1.SE+5d
WW.1.SE+5d
WD.2.SE+5d
WW.2.SE+5d
WD.3.SE+5d
WW.3.SE+5d
WD.4.SE+5d
WW.4.SE+5d
WD.5.SE+5d
WW.5.SE+5d

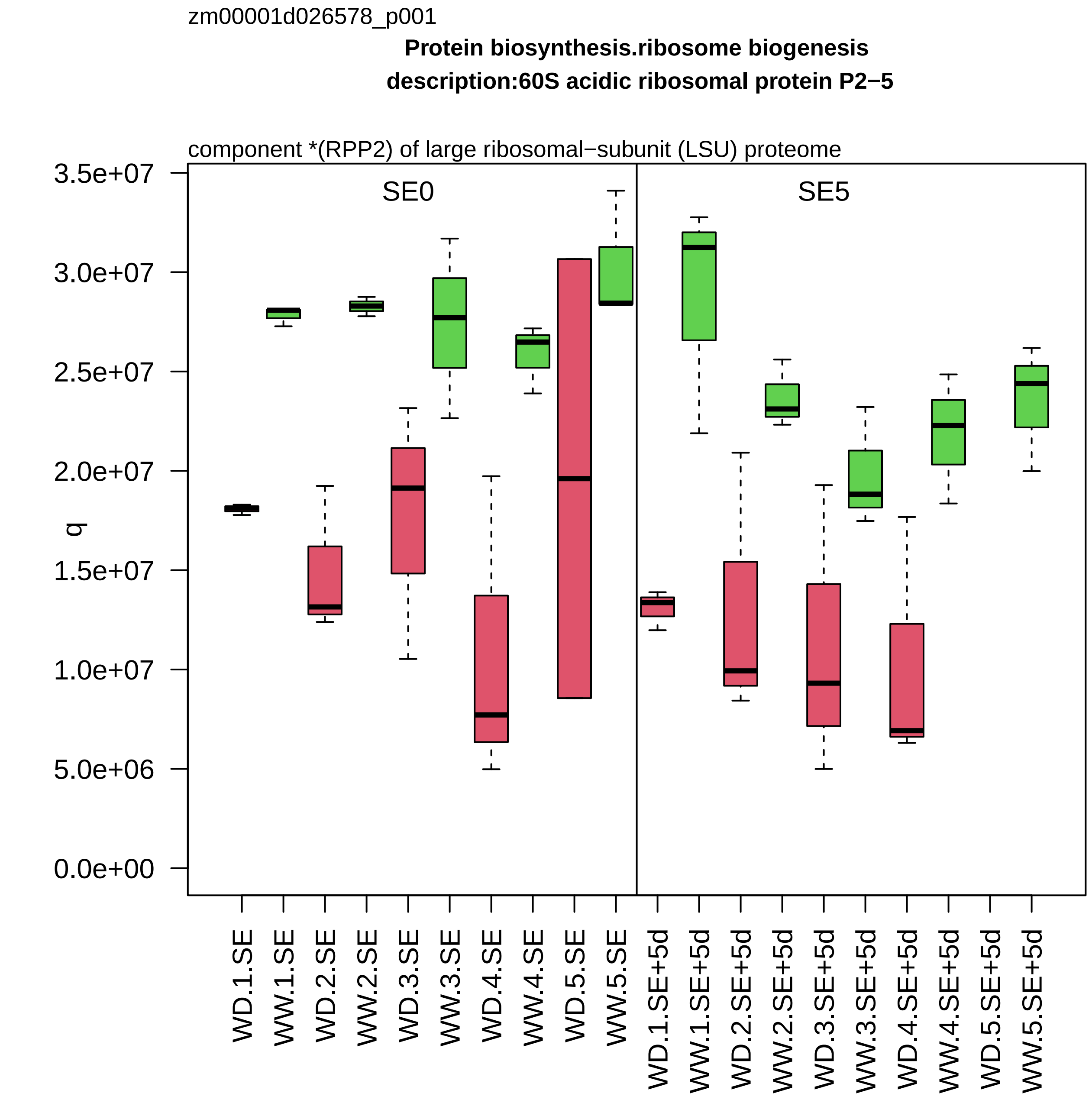

zm00001d026578_p001
Protein biosynthesis.ribosome biogenesis
description:60S acidic ribosomal protein P2−5
component *(RPP2) of large ribosomal−subunit (LSU) proteome
SE0
SE5
q
3.5e+07
3.0e+07
2.5e+07
2.0e+07
1.5e+07
1.0e+07
5.0e+06
0.0e+00
WD.1.SE
WW.1.SE
WD.2.SE
WW.2.SE
WD.3.SE
WW.3.SE
WD.4.SE
WW.4.SE
WD.5.SE
WW.5.SE
WD.1.SE+5d
WW.1.SE+5d
WD.2.SE+5d
WW.2.SE+5d
WD.3.SE+5d
WW.3.SE+5d
WD.4.SE+5d
WW.4.SE+5d
WD.5.SE+5d
WW.5.SE+5d

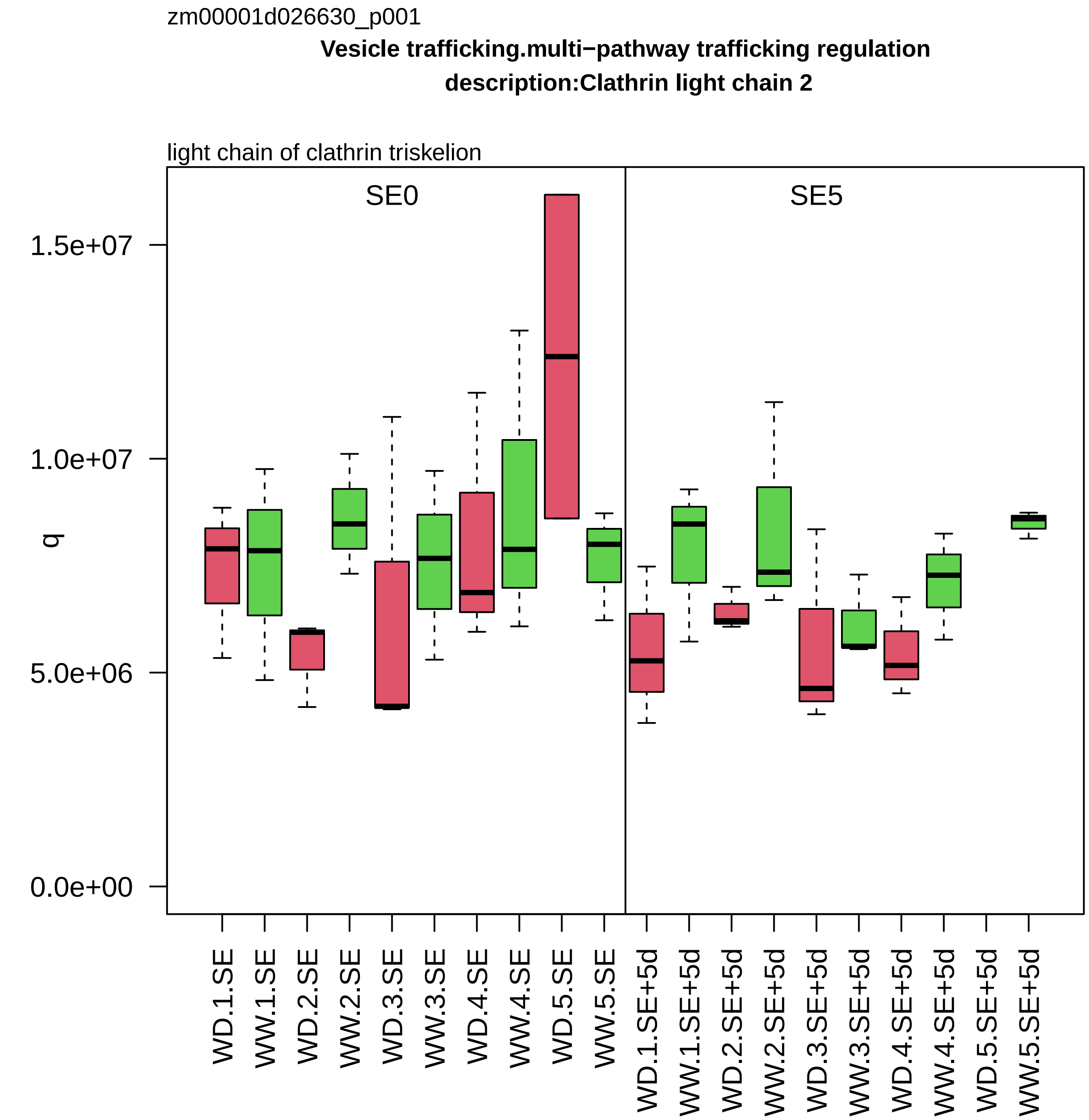
zm00001d026630_p001
Vesicle trafficking.multi−pathway trafficking regulation
description:Clathrin light chain 2
light chain of clathrin triskelion
SE0
SE5
1.5e+07
1.0e+07
5.0e+06
0.0e+00
q
WD.1.SE
WW.1.SE
WD.2.SE
WW.2.SE
WD.3.SE
WW.3.SE
WD.4.SE
WW.4.SE
WD.5.SE
WW.5.SE
WD.1.SE+5d
WW.1.SE+5d
WD.2.SE+5d
WW.2.SE+5d
WD.3.SE+5d
WW.3.SE+5d
WD.4.SE+5d
WW.4.SE+5d
WD.5.SE+5d
WW.5.SE+5d

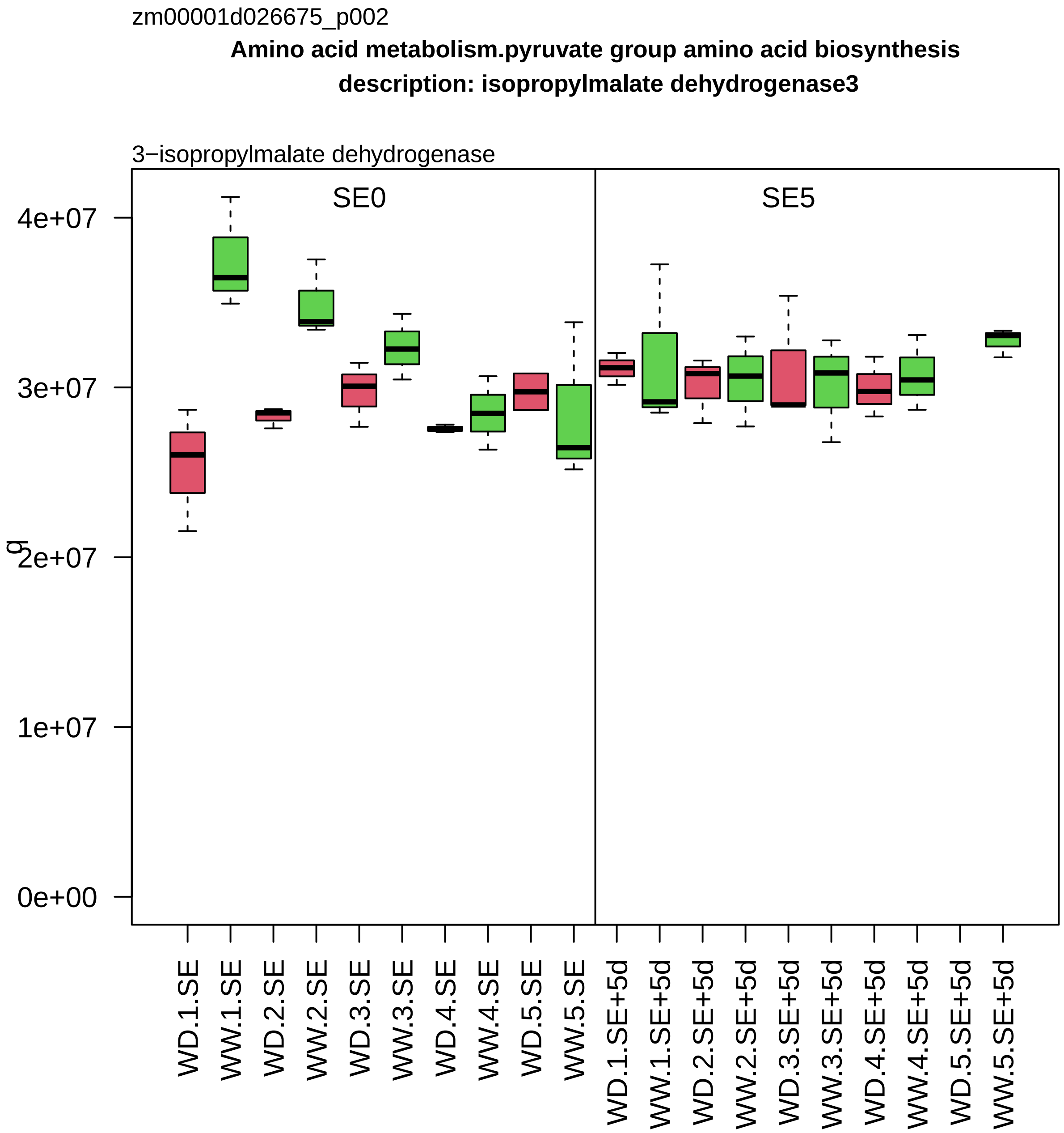

zm00001d026675_p002
Amino acid metabolism.pyruvate group amino acid biosynthesis
description: isopropylmalate dehydrogenase3
3−isopropylmalate dehydrogenase
SE0
SE5
g
0e+00
1e+07
2e+07
3e+07
4e+07
WD.1.SE
WW.1.SE
WD.2.SE
WW.2.SE
WD.3.SE
WW.3.SE
WD.4.SE
WW.4.SE
WD.5.SE
WW.5.SE
WD.1.SE+5d
WW.1.SE+5d
WD.2.SE+5d
WW.2.SE+5d
WD.3.SE+5d
WW.3.SE+5d
WD.4.SE+5d
WW.4.SE+5d
WD.5.SE+5d
WW.5.SE+5d

zm00001d027276_p001

**Enzyme classification.EC_1 oxidoreductases**

**description:GroES−like zinc−binding alcohol dehydrogenase family protein**

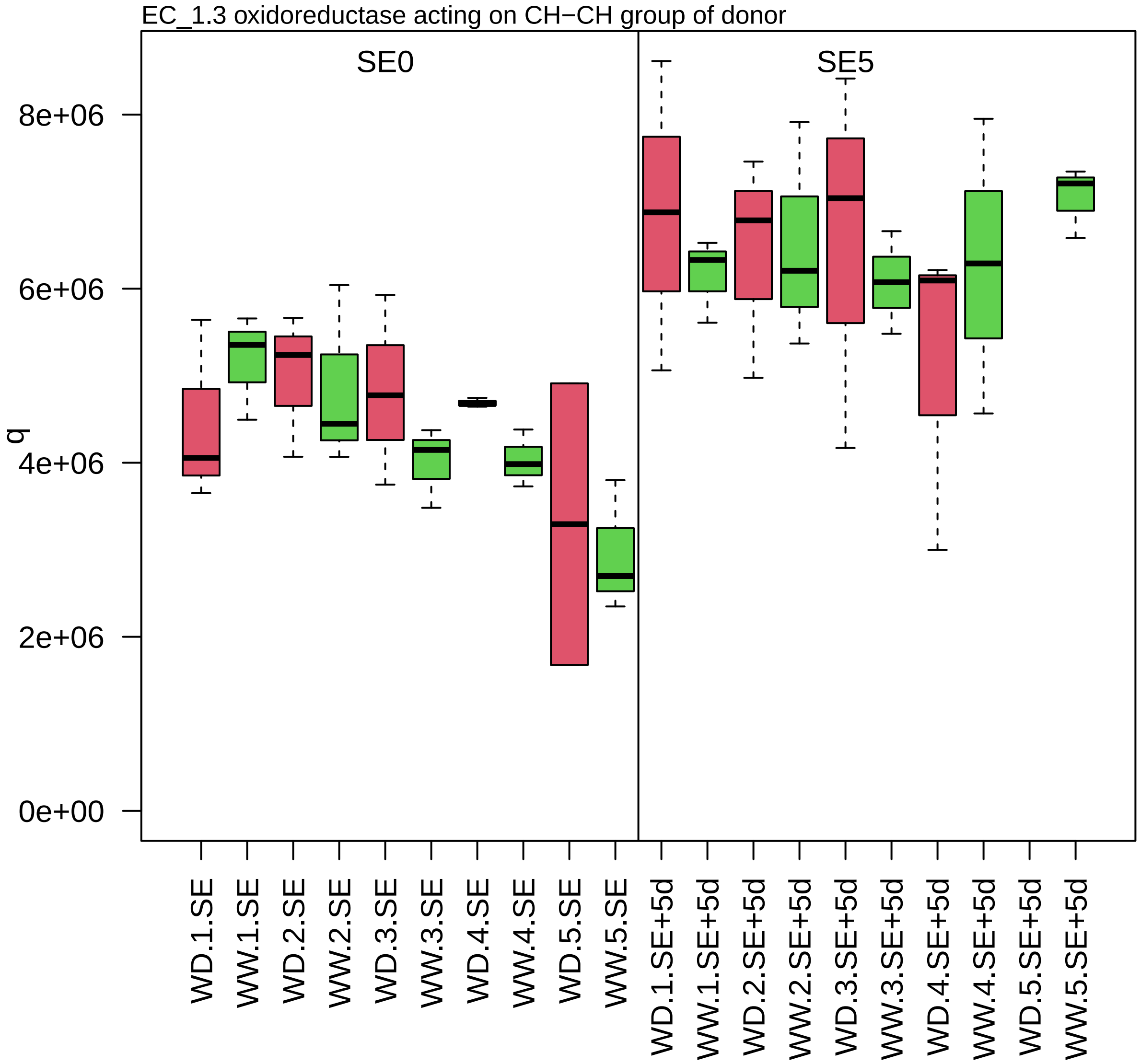

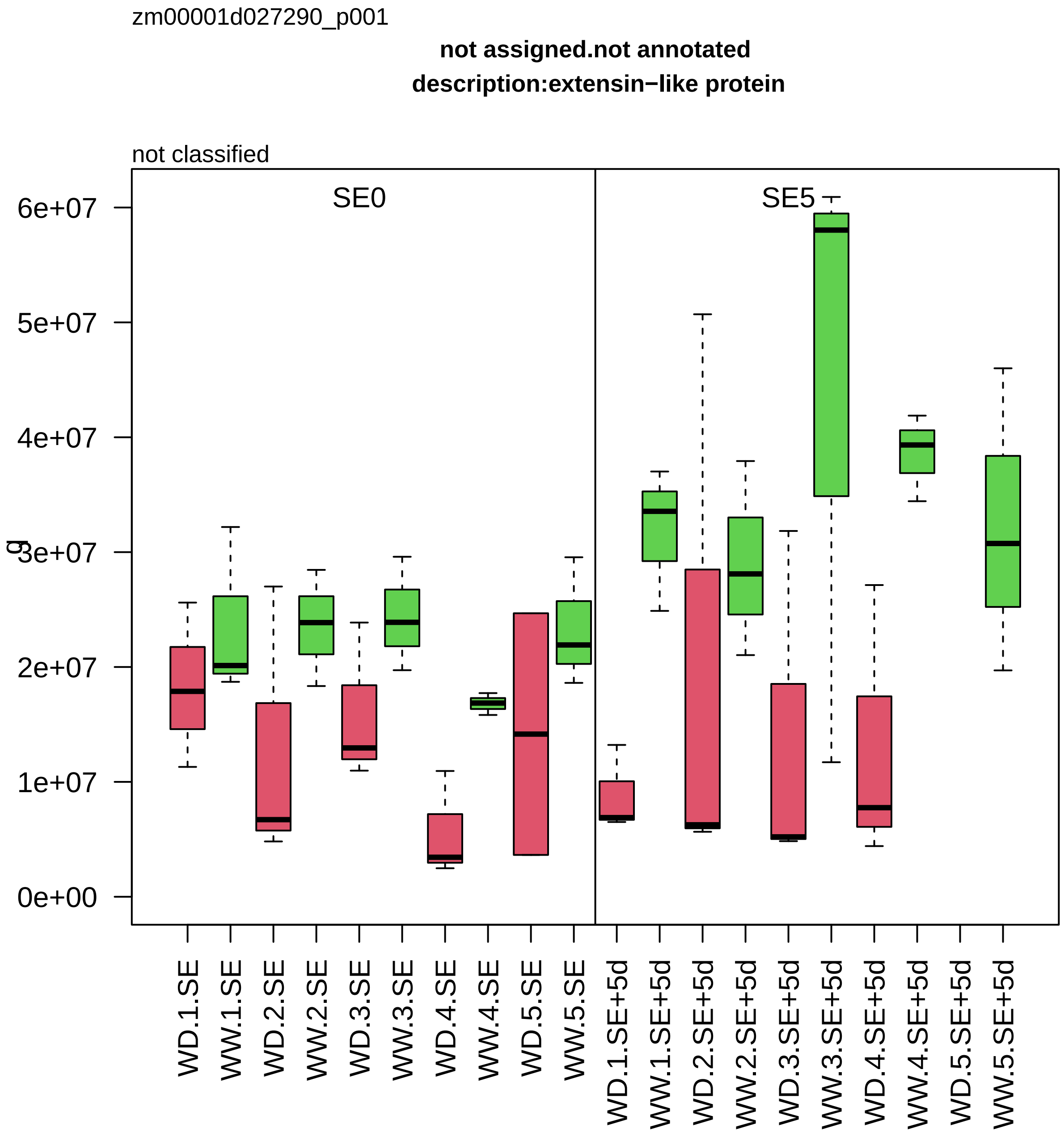
zm00001d027290_p001
not assigned.not annotated
description:extensin−like protein
not classified
SE0
SE5
g
6e+07
5e+07
4e+07
3e+07
2e+07
1e+07
0e+00
WD.1.SE
WW.1.SE
WD.2.SE
WW.2.SE
WD.3.SE
WW.3.SE
WD.4.SE
WW.4.SE
WD.5.SE
WW.5.SE
WD.1.SE+5d
WW.1.SE+5d
WD.2.SE+5d
WW.2.SE+5d
WD.3.SE+5d
WW.3.SE+5d
WD.4.SE+5d
WW.4.SE+5d
WD.5.SE+5d
WW.5.SE+5d

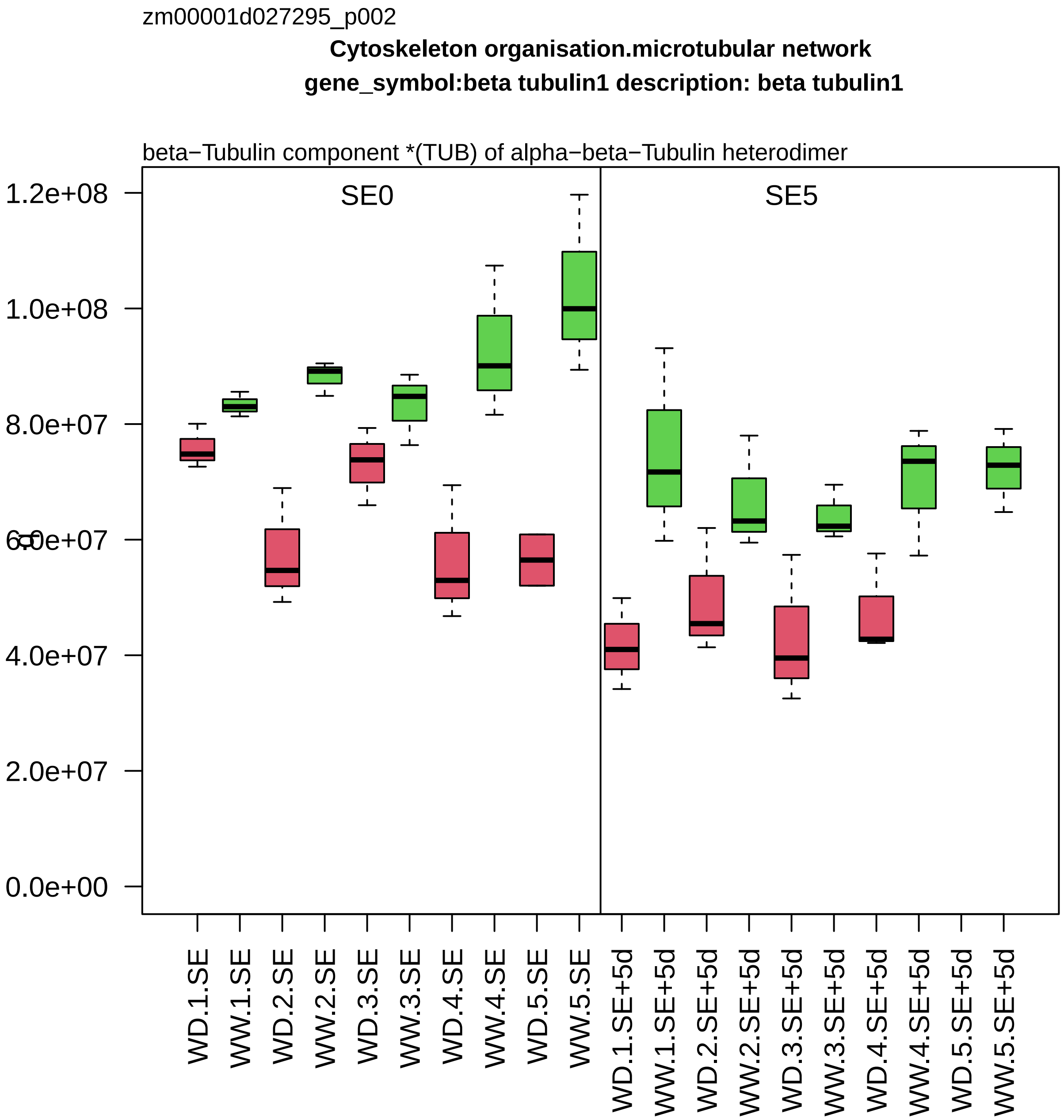

zm00001d027295_p002
Cytoskeleton organisation.microtubular network
gene_symbol:beta tubulin1 description: beta tubulin1
beta−Tubulin component *(TUB) of alpha−beta−Tubulin heterodimer
SE0
SE5
1.2e+08
1.0e+08
8.0e+07
6.0e+07
4.0e+07
2.0e+07
0.0e+00
WD.1.SE
WW.1.SE
WD.2.SE
WW.2.SE
WD.3.SE
WW.3.SE
WD.4.SE
WW.4.SE
WD.5.SE
WW.5.SE
WD.1.SE+5d
WW.1.SE+5d
WD.2.SE+5d
WW.2.SE+5d
WD.3.SE+5d
WW.3.SE+5d
WD.4.SE+5d
WW.4.SE+5d
WD.5.SE+5d
WW.5.SE+5d

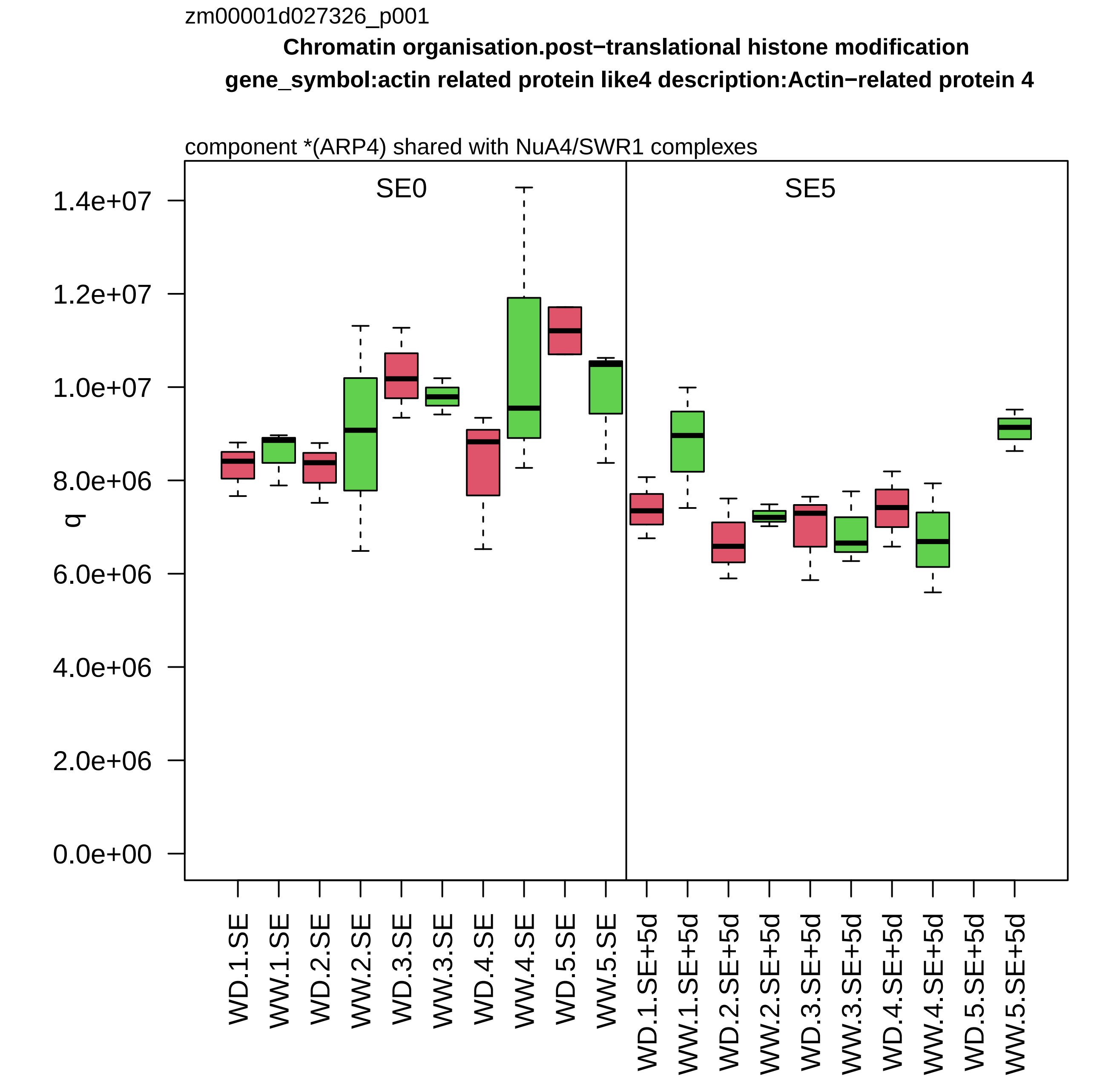

zm00001d027326_p001
Chromatin organisation.post−translational histone modification
gene_symbol:actin related protein like4 description:Actin−related protein 4
component *(ARP4) shared with NuA4/SWR1 complexes
SE0
SE5
1.4e+07
1.2e+07
1.0e+07
8.0e+06
6.0e+06
4.0e+06
2.0e+06
0.0e+00
q
WD.1.SE
WW.1.SE
WD.2.SE
WW.2.SE
WD.3.SE
WW.3.SE
WD.4.SE
WW.4.SE
WD.5.SE
WW.5.SE
WD.1.SE+5d
WW.1.SE+5d
WD.2.SE+5d
WW.2.SE+5d
WD.3.SE+5d
WW.3.SE+5d
WD.4.SE+5d
WW.4.SE+5d
WD.5.SE+5d
WW.5.SE+5d

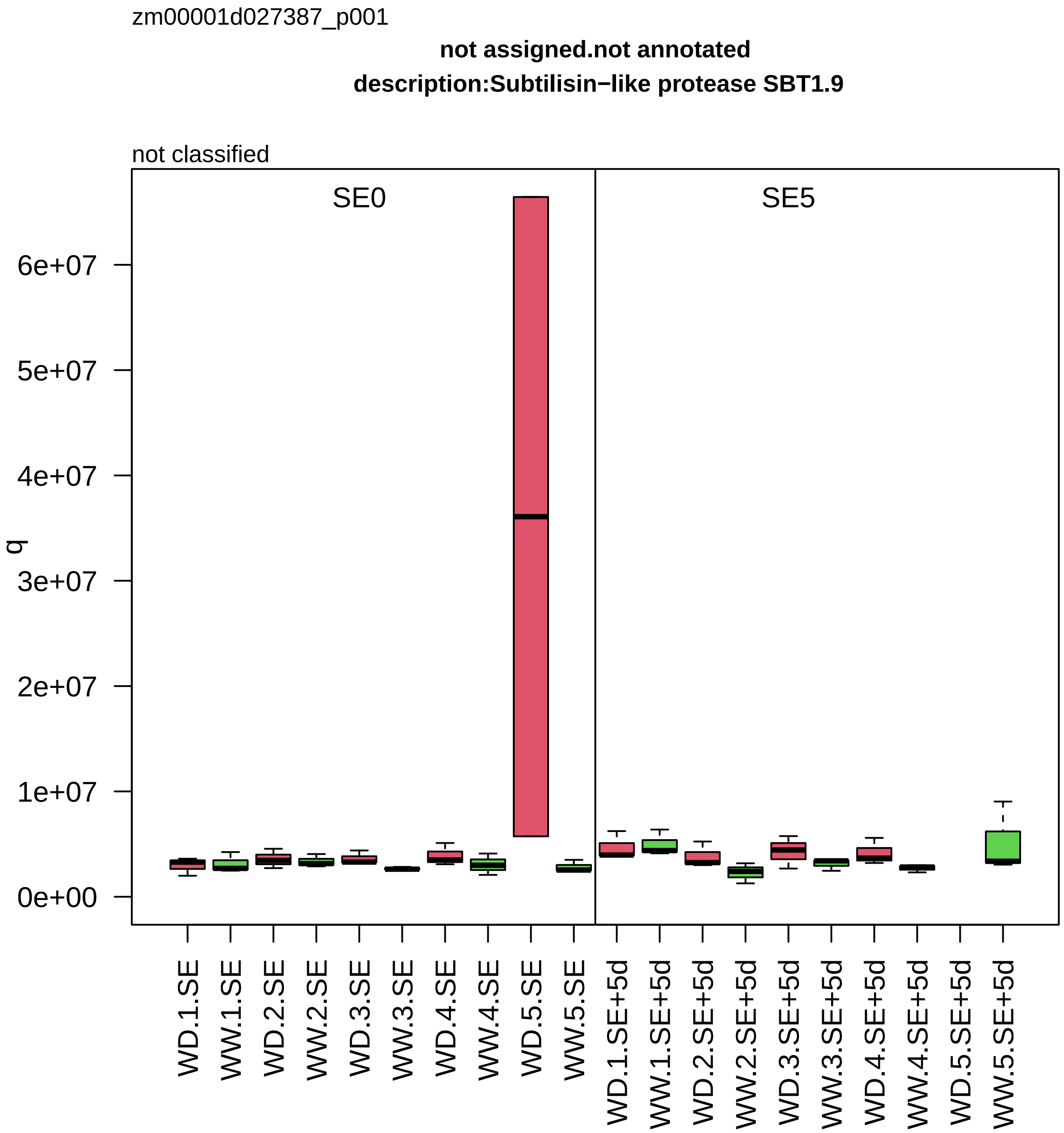

zm00001d027387_p001
not assigned.not annotated
description:Subtilisin−like protease SBT1.9
not classified
SE0
SE5
q
6e+07
5e+07
4e+07
3e+07
2e+07
1e+07
0e+00
WD.1.SE
WW.1.SE
WD.2.SE
WW.2.SE
WD.3.SE
WW.3.SE
WD.4.SE
WW.4.SE
WD.5.SE
WW.5.SE
WD.1.SE+5d
WW.1.SE+5d
WD.2.SE+5d
WW.2.SE+5d
WD.3.SE+5d
WW.3.SE+5d
WD.4.SE+5d
WW.4.SE+5d
WD.5.SE+5d
WW.5.SE+5d

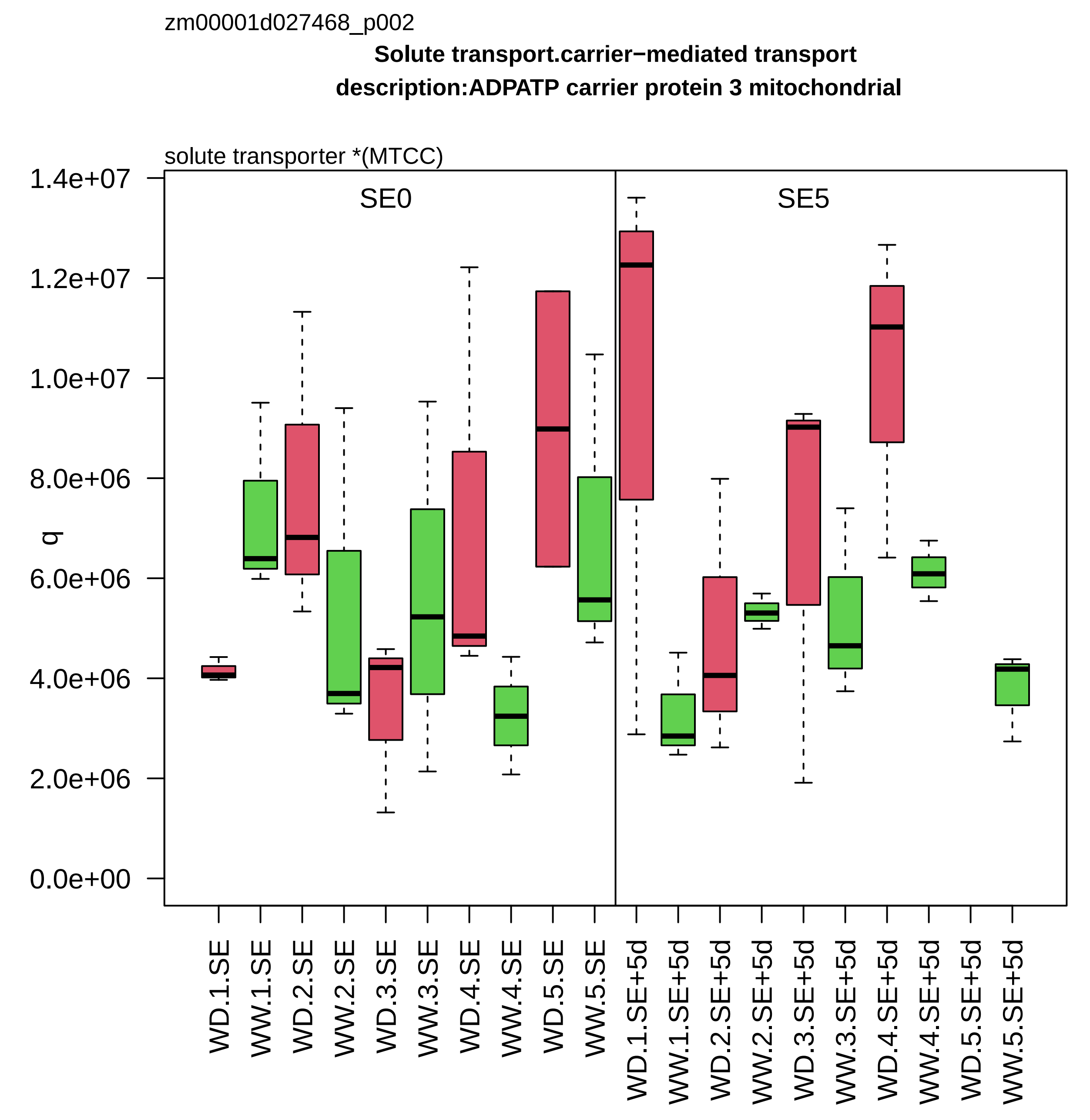
zm00001d027468_p002
Solute transport.carrier−mediated transport
description:ADPATP carrier protein 3 mitochondrial
solute transporter *(MTCC)
SE0
SE5
q
1.4e+07
1.2e+07
1.0e+07
8.0e+06
6.0e+06
4.0e+06
2.0e+06
0.0e+00
WD.1.SE
WW.1.SE
WD.2.SE
WW.2.SE
WD.3.SE
WW.3.SE
WD.4.SE
WW.4.SE
WD.5.SE
WW.5.SE
WD.1.SE+5d
WW.1.SE+5d
WD.2.SE+5d
WW.2.SE+5d
WD.3.SE+5d
WW.3.SE+5d
WD.4.SE+5d
WW.4.SE+5d
WD.5.SE+5d
WW.5.SE+5d

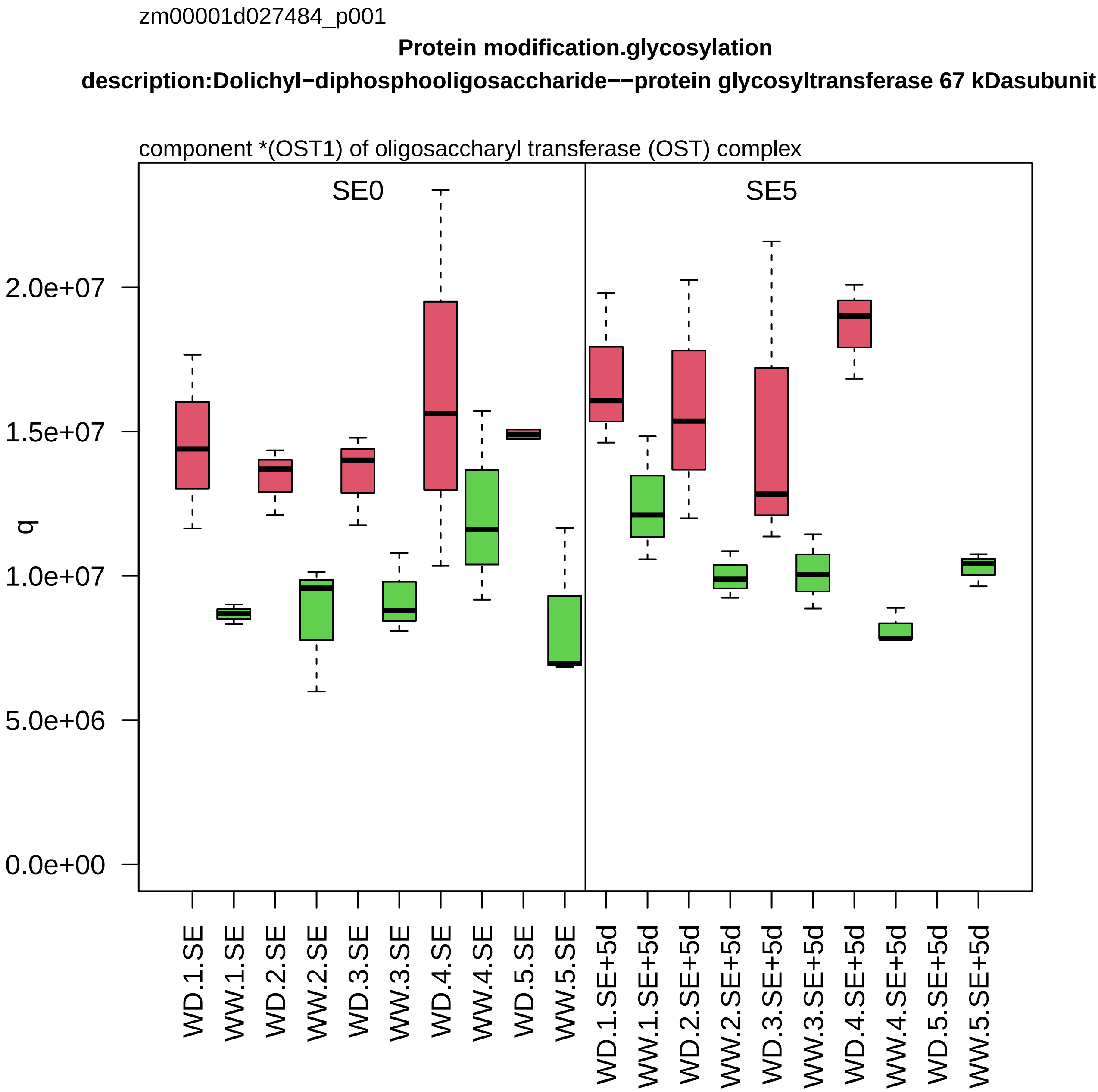
zm00001d027484_p001
Protein modification.glycosylation
description:Dolichyl−diphosphooligosaccharide−−protein glycosyltransferase 67 kDasubunit
component *(OST1) of oligosaccharyl transferase (OST) complex
SE0
SE5
q
0.0e+00
5.0e+06
1.0e+07
1.5e+07
2.0e+07
WD.1.SE
WW.1.SE
WD.2.SE
WW.2.SE
WD.3.SE
WW.3.SE
WD.4.SE
WW.4.SE
WD.5.SE
WW.5.SE
WD.1.SE+5d
WW.1.SE+5d
WD.2.SE+5d
WW.2.SE+5d
WD.3.SE+5d
WW.3.SE+5d
WD.4.SE+5d
WW.4.SE+5d
WD.5.SE+5d
WW.5.SE+5d

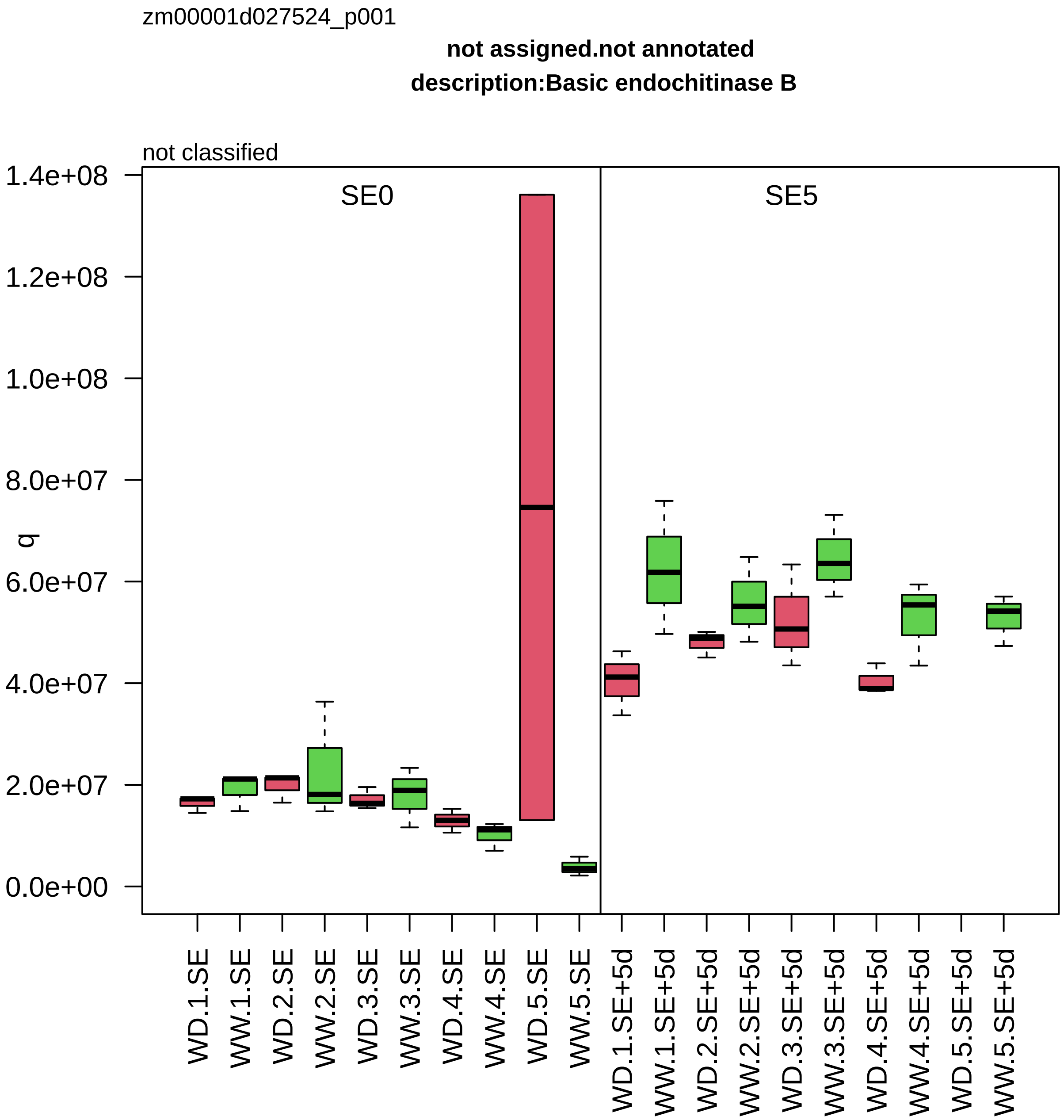

zm00001d027524_p001
not assigned.not annotated
description:Basic endochitinase B
not classified
SE0
SE5
q
1.4e+08
1.2e+08
1.0e+08
8.0e+07
6.0e+07
4.0e+07
2.0e+07
0.0e+00
WD.1.SE
WW.1.SE
WD.2.SE
WW.2.SE
WD.3.SE
WW.3.SE
WD.4.SE
WW.4.SE
WD.5.SE
WW.5.SE
WD.1.SE+5d
WW.1.SE+5d
WD.2.SE+5d
WW.2.SE+5d
WD.3.SE+5d
WW.3.SE+5d
WD.4.SE+5d
WW.4.SE+5d
WD.5.SE+5d
WW.5.SE+5d

zm00001d027540_p001

**Redox homeostasis.glutathione−based redox regulation**

**gene_symbol:glutathione S−transferase12 description: glutathione S−transferase12**

class phi glutathione S−transferase

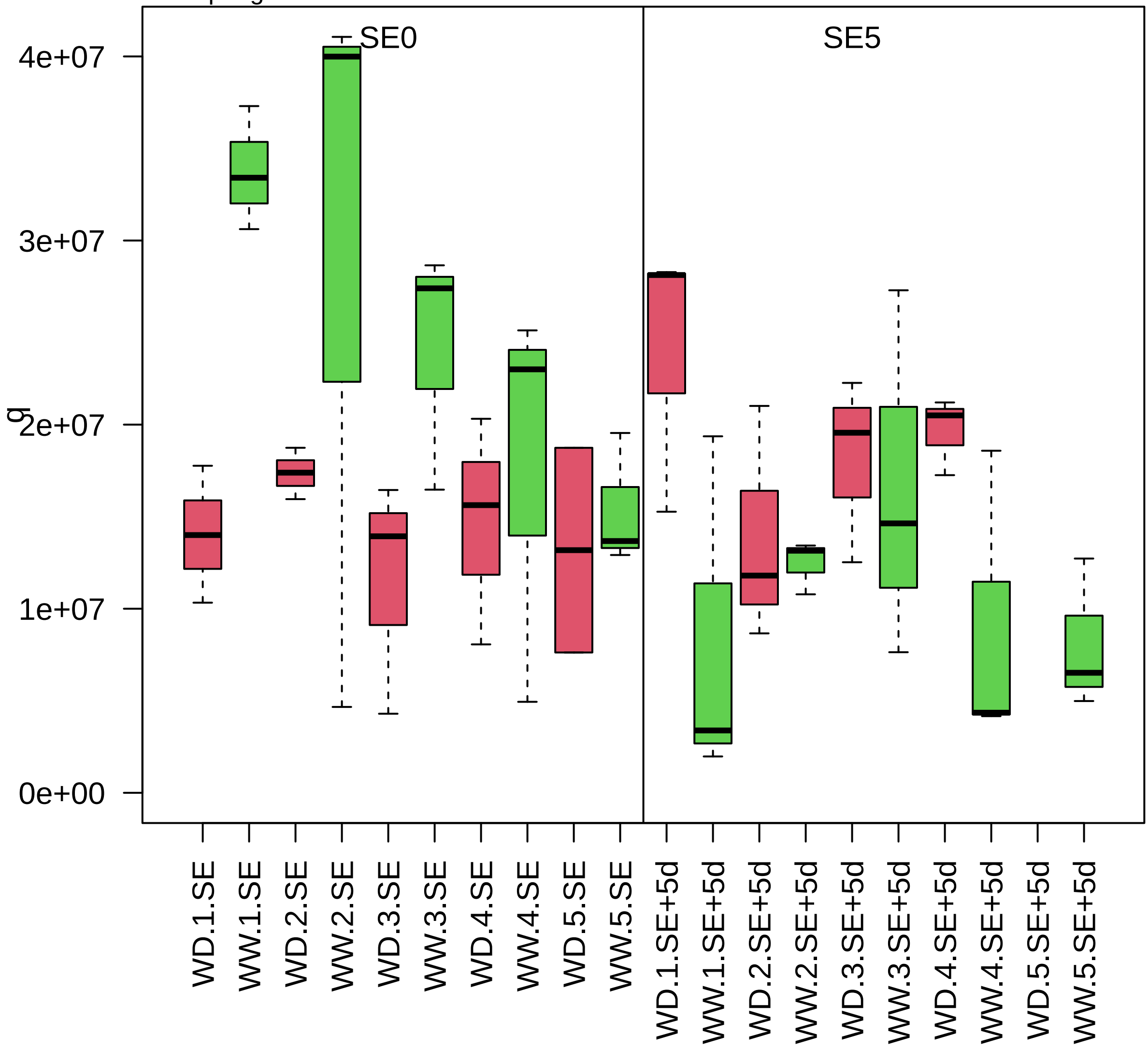

zm00001d027548_p001

**Lipid metabolism.phytosterol metabolism**
**description: sterol methyl transferase2**

sterol C−24 methyltransferase

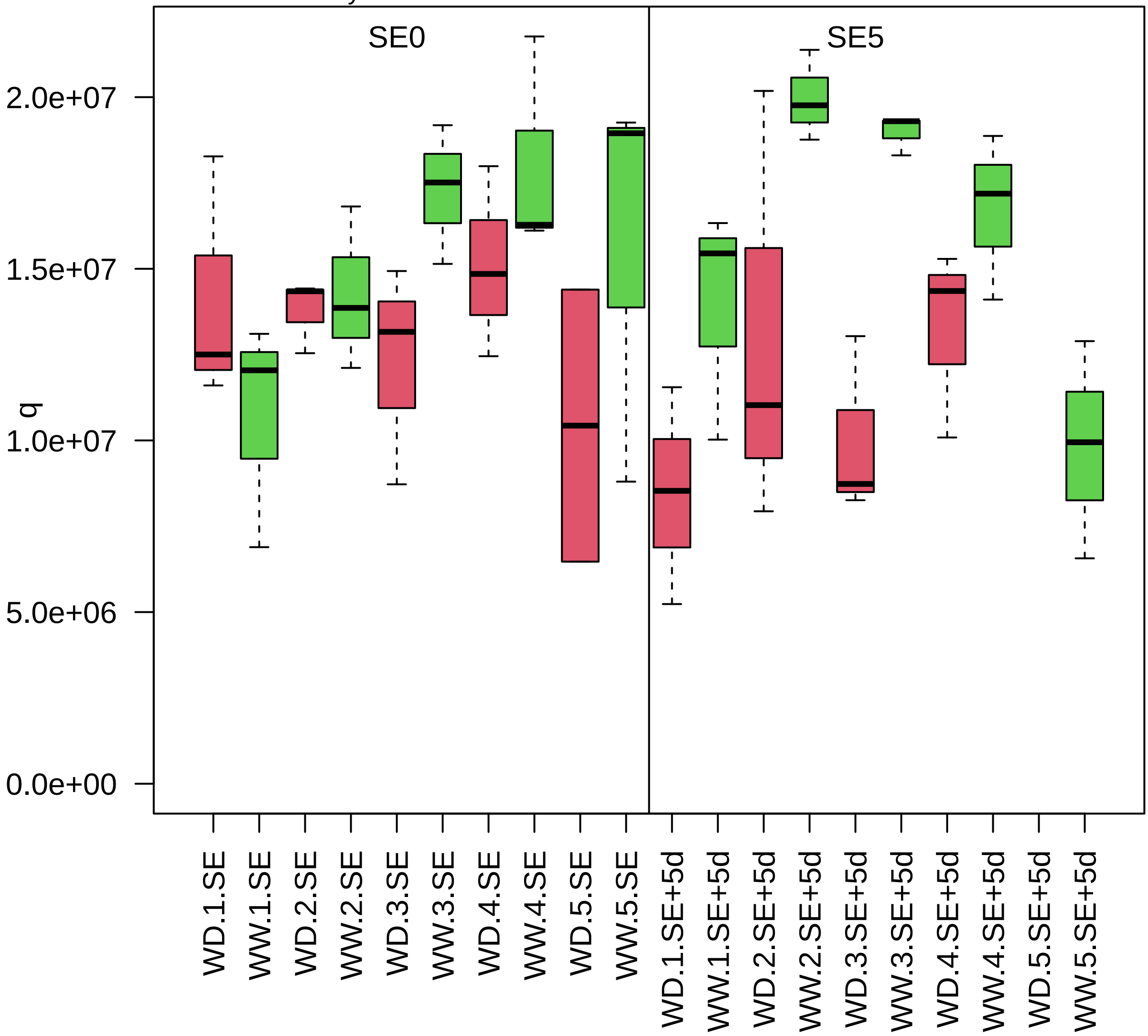

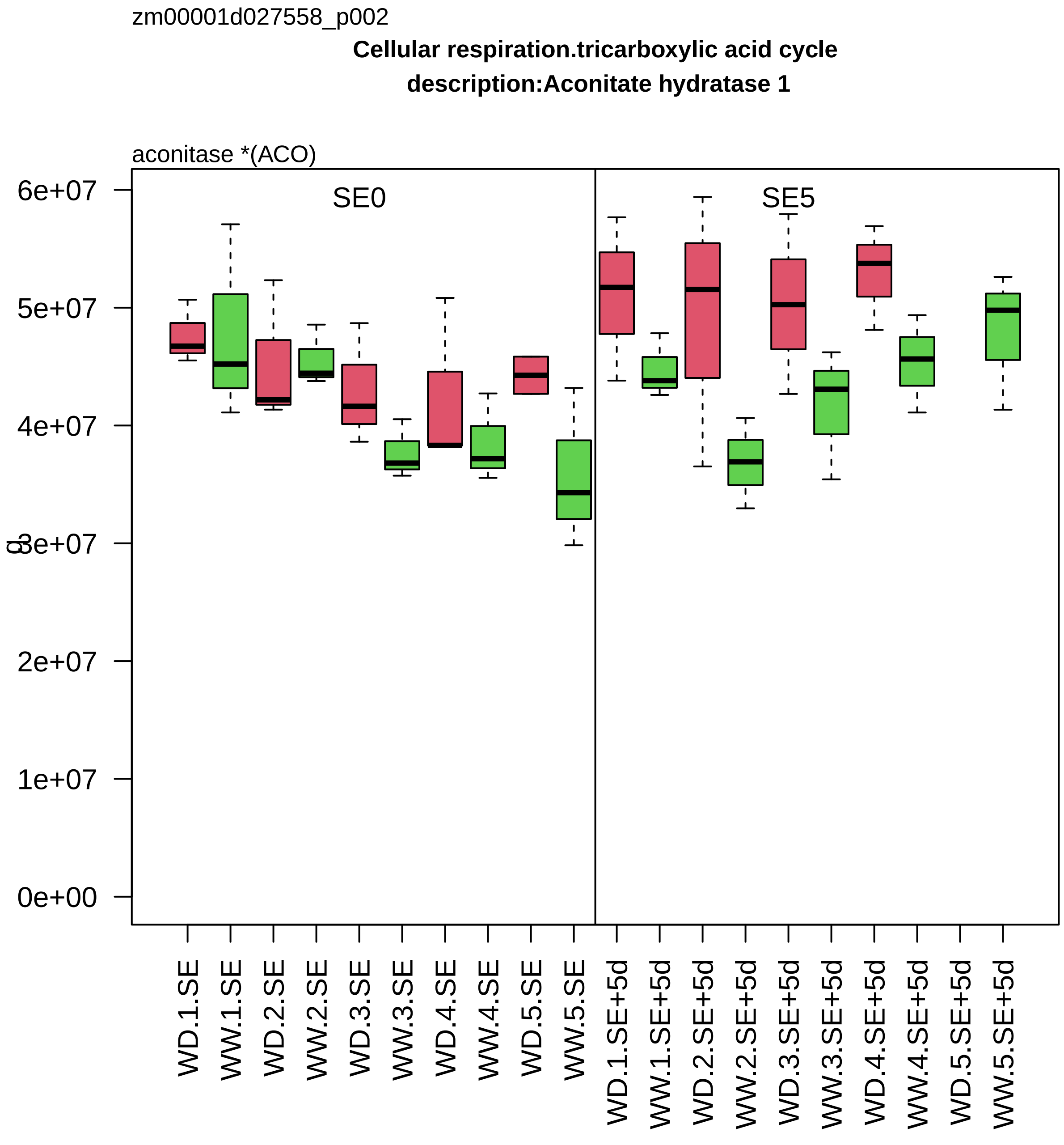

zm00001d027558_p002
Cellular respiration.tricarboxylic acid cycle
description:Aconitate hydratase 1
aconitase *(ACO)
SE0
SE5
6e+07
5e+07
4e+07
3e+07
2e+07
1e+07
0e+00
WD.1.SE
WW.1.SE
WD.2.SE
WW.2.SE
WD.3.SE
WW.3.SE
WD.4.SE
WW.4.SE
WD.5.SE
WW.5.SE
WD.1.SE+5d
WW.1.SE+5d
WD.2.SE+5d
WW.2.SE+5d
WD.3.SE+5d
WW.3.SE+5d
WD.4.SE+5d
WW.4.SE+5d
WD.5.SE+5d
WW.5.SE+5d

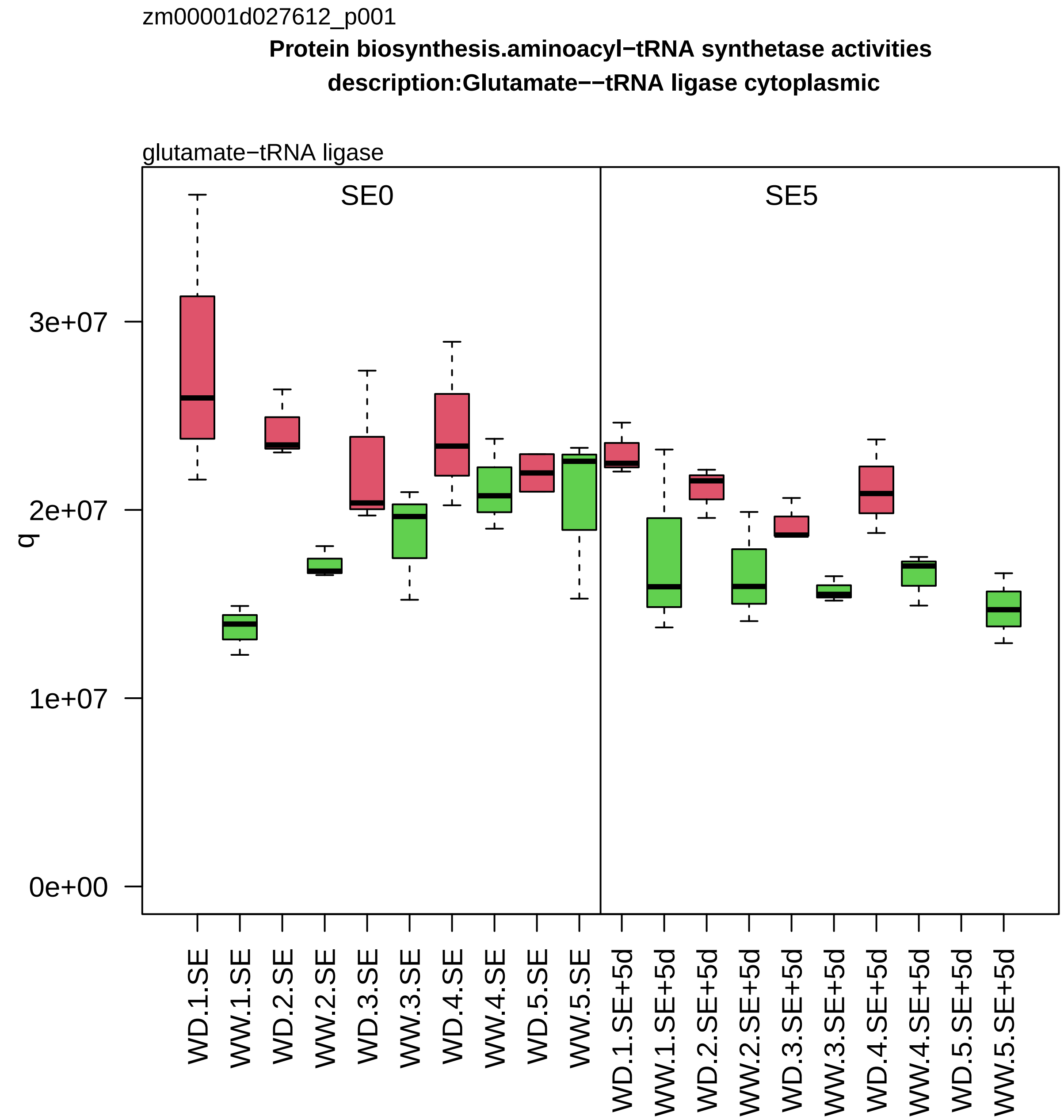
zm00001d027612_p001
Protein biosynthesis.aminoacyl−tRNA synthetase activities
description:Glutamate−−tRNA ligase cytoplasmic
glutamate−tRNA ligase
SE0
SE5
q
0e+00
1e+07
2e+07
3e+07
WD.1.SE
WW.1.SE
WD.2.SE
WW.2.SE
WD.3.SE
WW.3.SE
WD.4.SE
WW.4.SE
WD.5.SE
WW.5.SE
WD.1.SE+5d
WW.1.SE+5d
WD.2.SE+5d
WW.2.SE+5d
WD.3.SE+5d
WW.3.SE+5d
WD.4.SE+5d
WW.4.SE+5d
WD.5.SE+5d
WW.5.SE+5d

zm00001d027680_p001

**not assigned.not annotated**

**description:HR−like lesion−inducing protein−related**

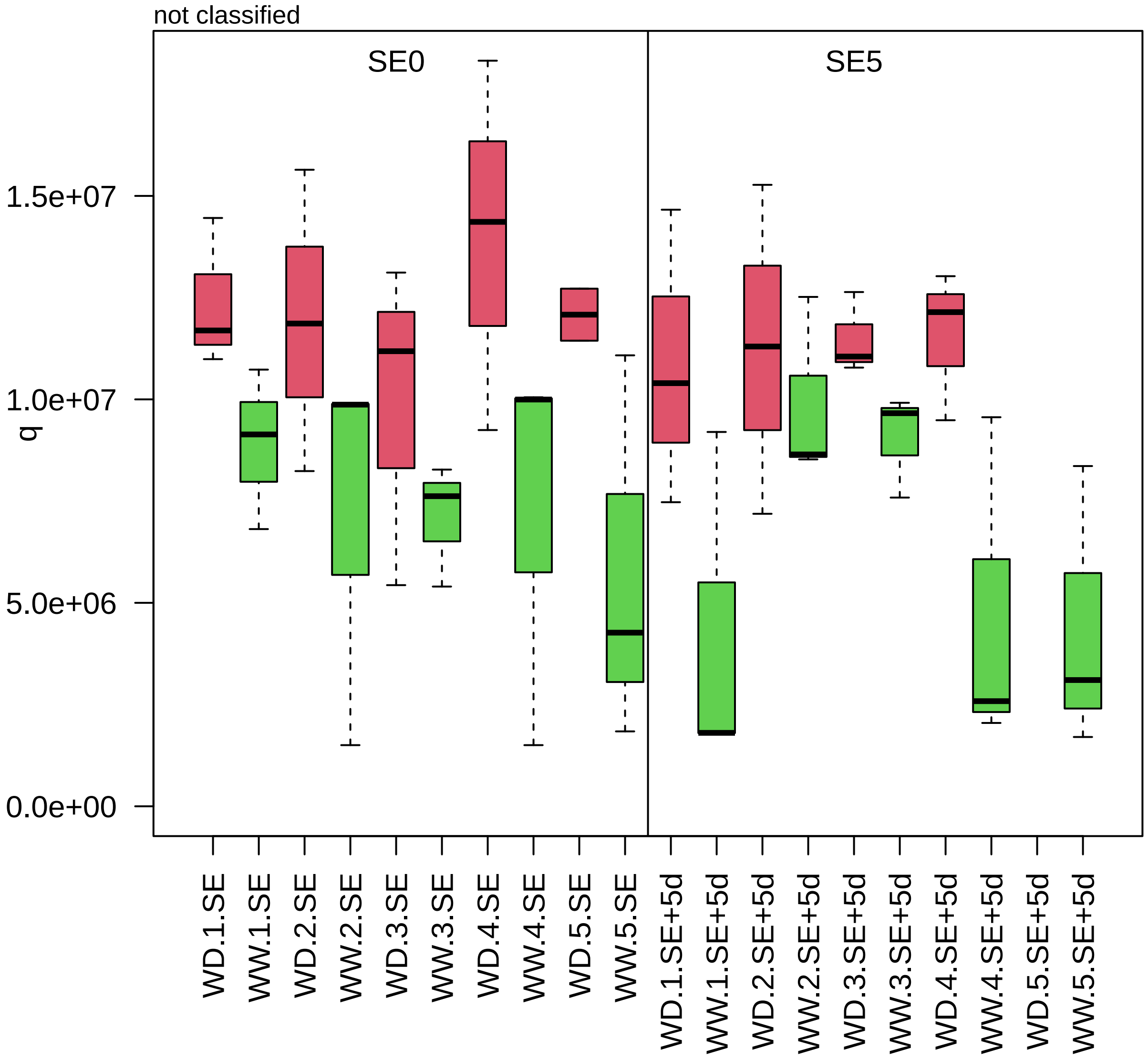

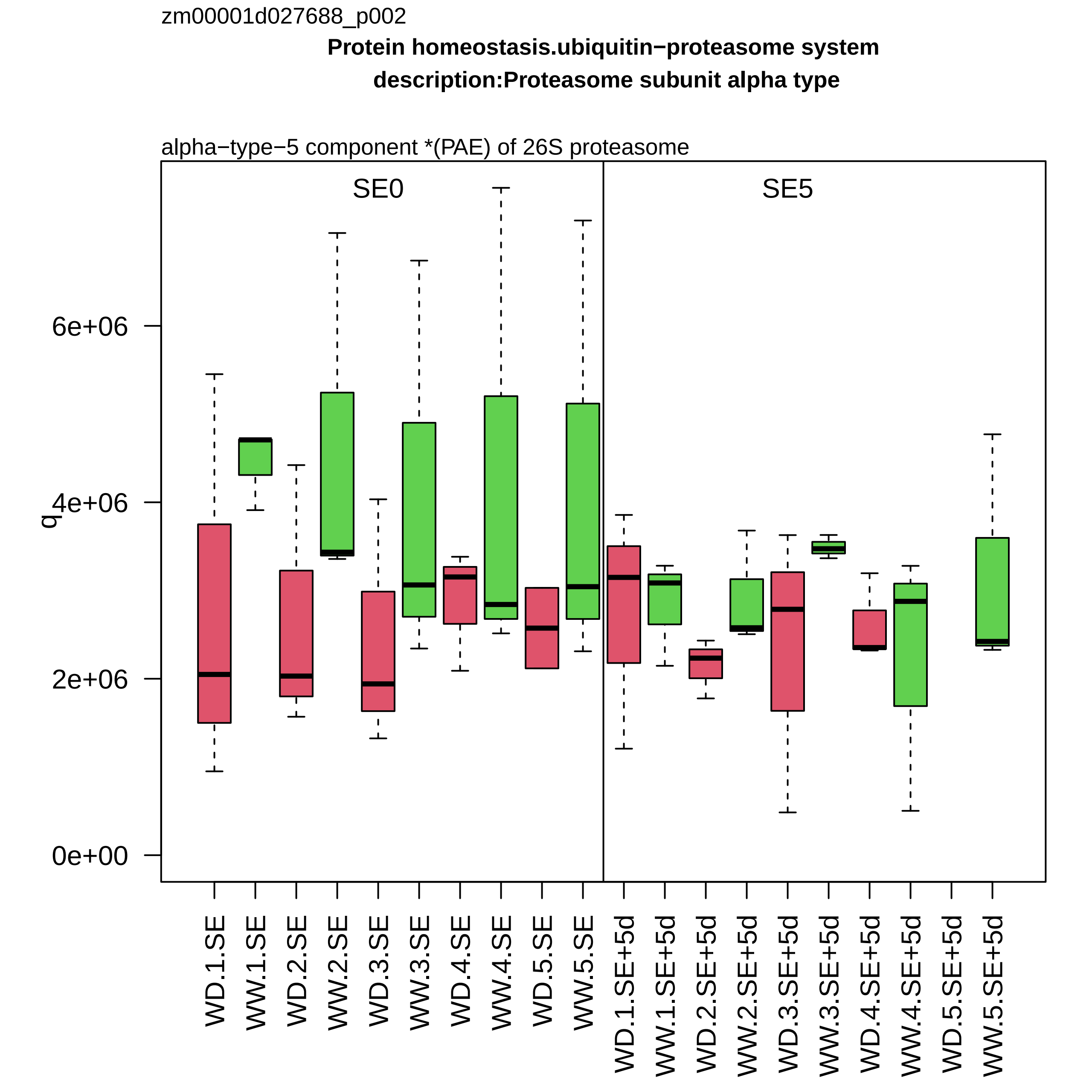
zm00001d027688_p002
Protein homeostasis.ubiquitin−proteasome system
description:Proteasome subunit alpha type
alpha−type−5 component *(PAE) of 26S proteasome
SE0
SE5
q
6e+06
4e+06
2e+06
0e+00
WD.1.SE
WW.1.SE
WD.2.SE
WW.2.SE
WD.3.SE
WW.3.SE
WD.4.SE
WW.4.SE
WD.5.SE
WW.5.SE
WD.1.SE+5d
WW.1.SE+5d
WD.2.SE+5d
WW.2.SE+5d
WD.3.SE+5d
WW.3.SE+5d
WD.4.SE+5d
WW.4.SE+5d
WD.5.SE+5d
WW.5.SE+5d

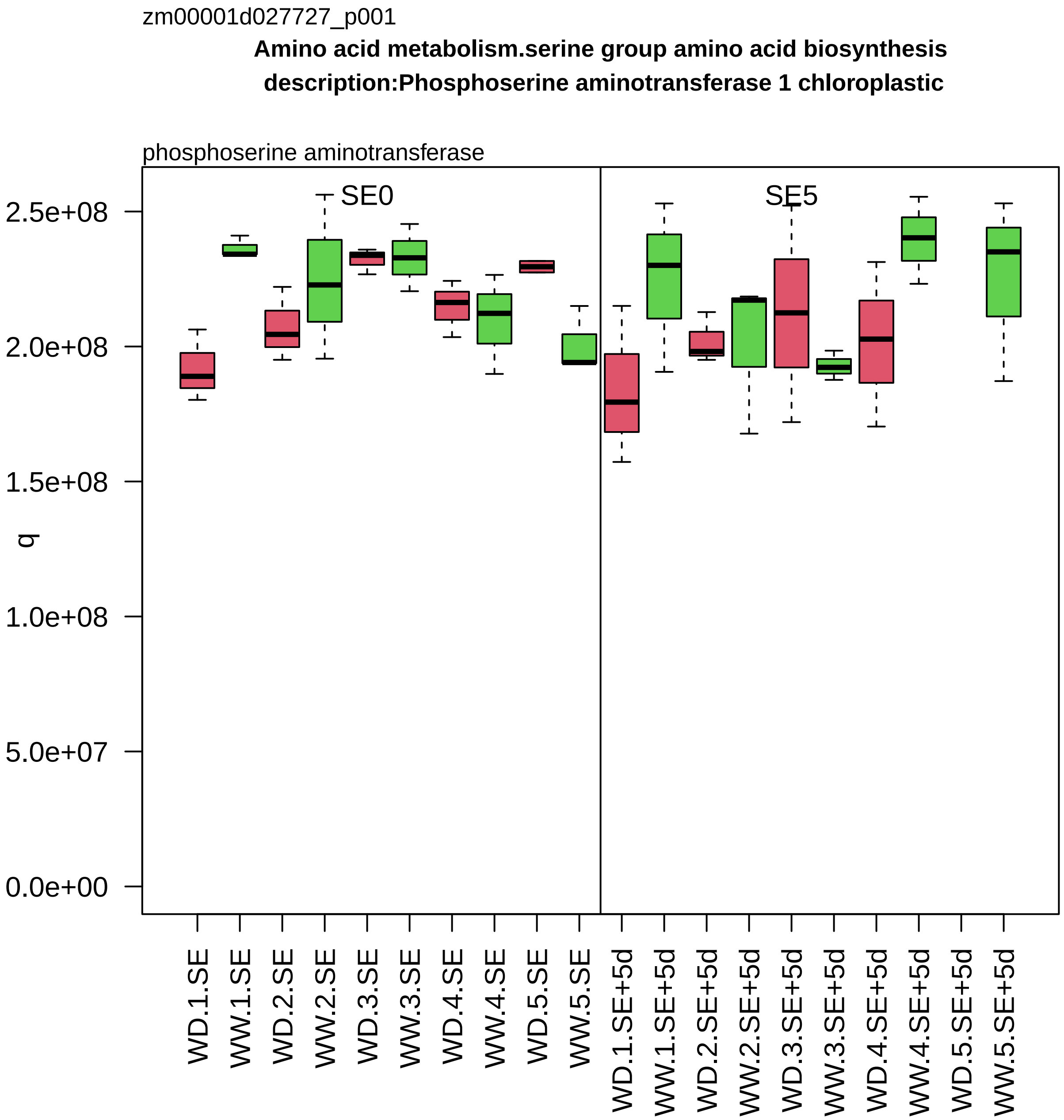
zm00001d027727_p001
Amino acid metabolism.serine group amino acid biosynthesis
description:Phosphoserine aminotransferase 1 chloroplastic
phosphoserine aminotransferase
SE0
SE5
q
2.5e+08
2.0e+08
1.5e+08
1.0e+08
5.0e+07
0.0e+00
WD.1.SE
WW.1.SE
WD.2.SE
WW.2.SE
WD.3.SE
WW.3.SE
WD.4.SE
WW.4.SE
WD.5.SE
WW.5.SE
WD.1.SE+5d
WW.1.SE+5d
WD.2.SE+5d
WW.2.SE+5d
WD.3.SE+5d
WW.3.SE+5d
WD.4.SE+5d
WW.4.SE+5d
WD.5.SE+5d
WW.5.SE+5d

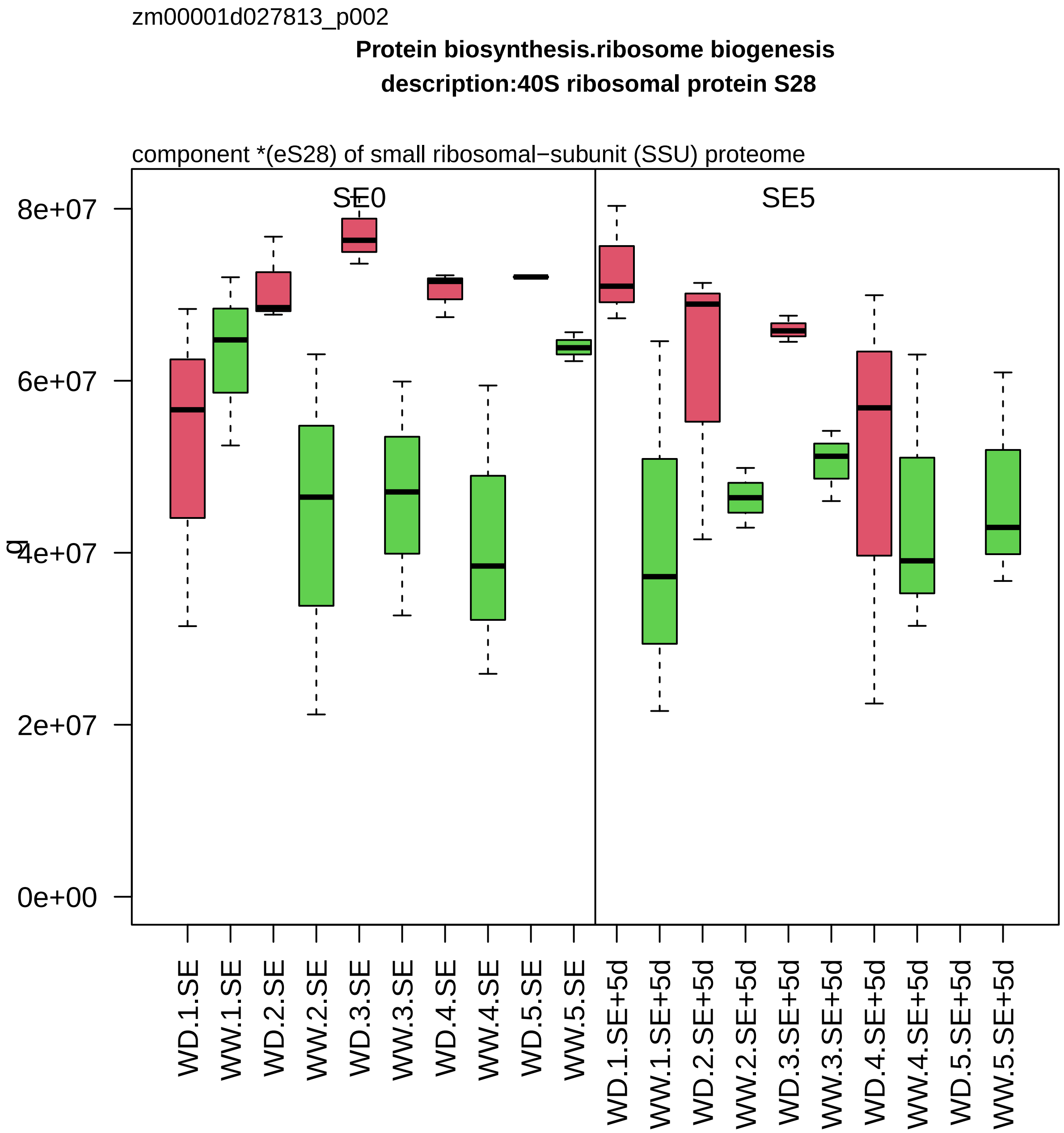
zm00001d027813_p002
Protein biosynthesis.ribosome biogenesis
description:40S ribosomal protein S28
component *(eS28) of small ribosomal−subunit (SSU) proteome
SE0
SE5
g
8e+07
6e+07
4e+07
2e+07
0e+00
WD.1.SE
WW.1.SE
WD.2.SE
WW.2.SE
WD.3.SE
WW.3.SE
WD.4.SE
WW.4.SE
WD.5.SE
WW.5.SE
WD.1.SE+5d
WW.1.SE+5d
WD.2.SE+5d
WW.2.SE+5d
WD.3.SE+5d
WW.3.SE+5d
WD.4.SE+5d
WW.4.SE+5d
WD.5.SE+5d
WW.5.SE+5d

zm00001d027871_p001

**Nucleotide metabolism.purines**

**gene_symbol:adenylosuccinate synthetase description: adenylosuccinate synthetase**

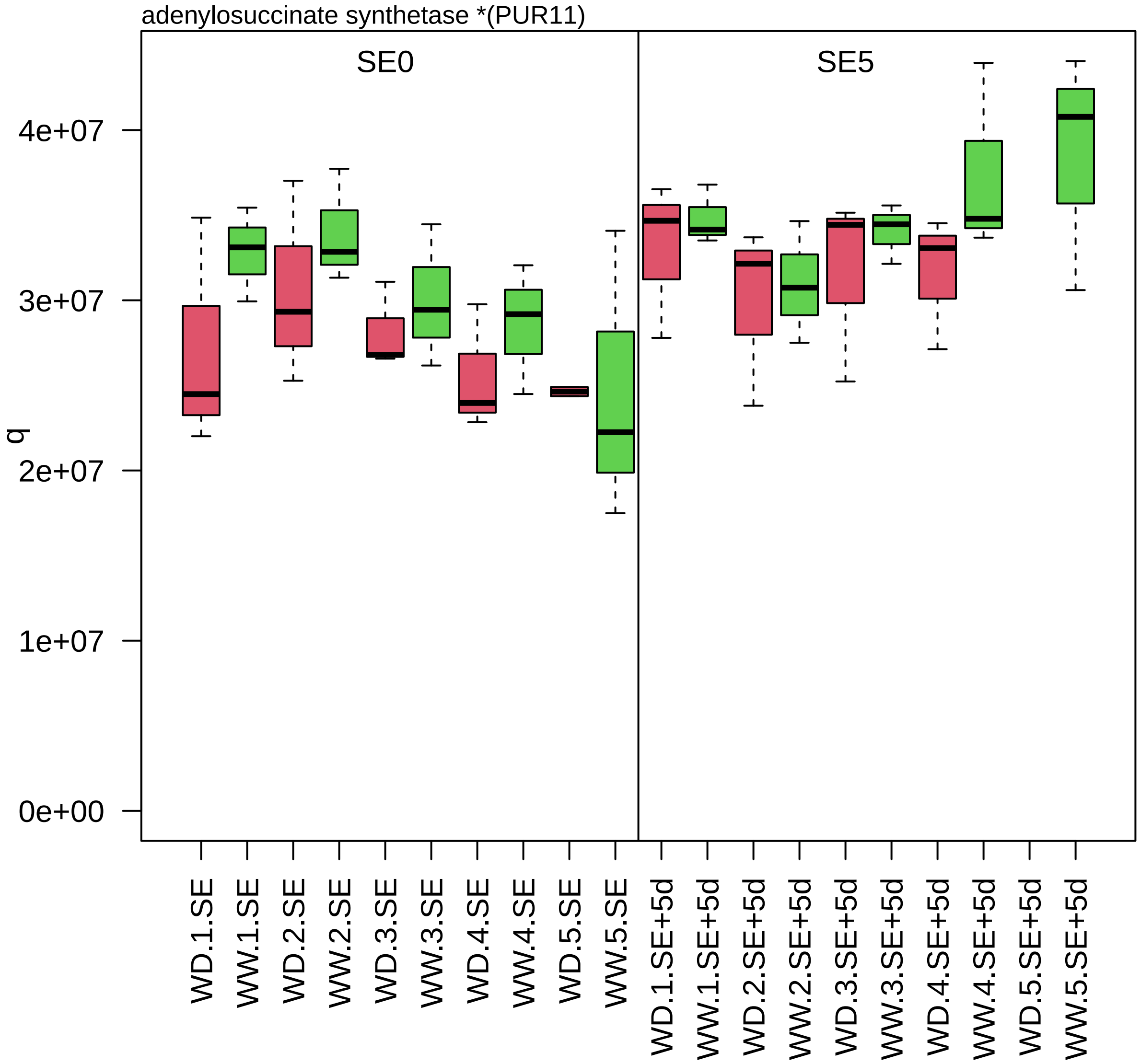

zm00001d027965_p001

**Protein homeostasis.proteolysis**
**description:Aspartyl protease AED3**

A1−class (Pepsin) protease

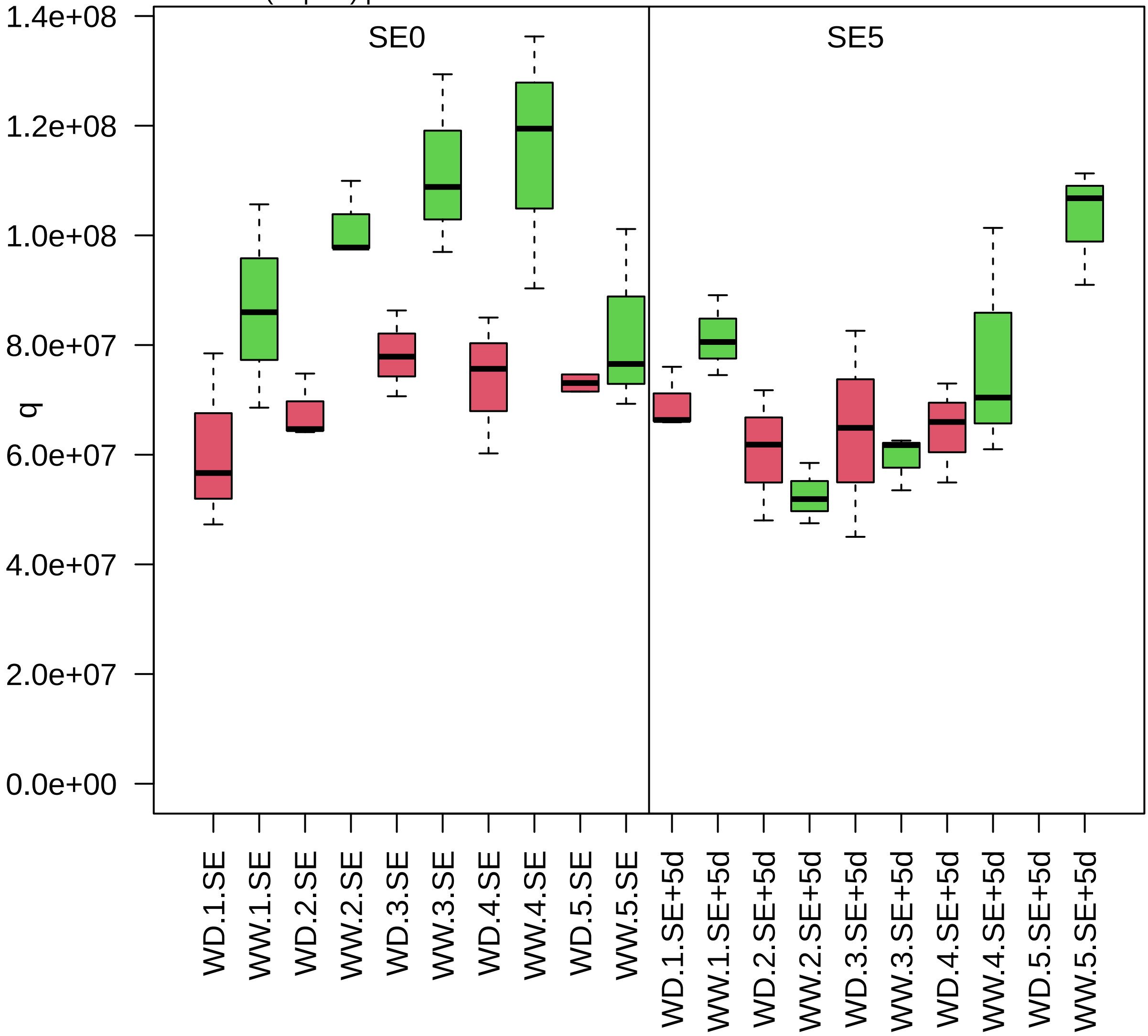

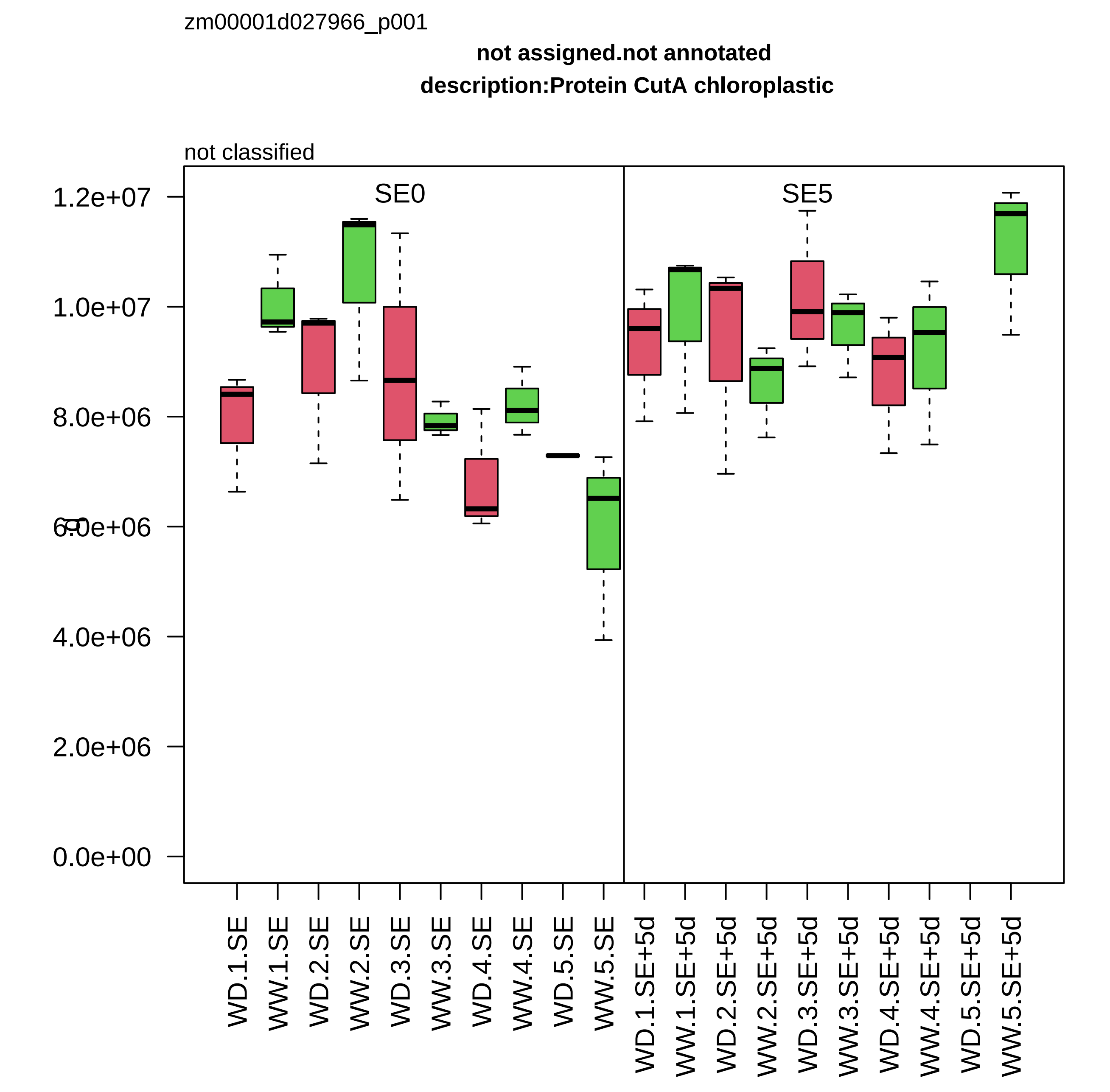

zm00001d027966_p001
not assigned.not annotated
description:Protein CutA chloroplastic
not classified
SE0
SE5
1.2e+07
1.0e+07
8.0e+06
6.0e+06
4.0e+06
2.0e+06
0.0e+00
WD.1.SE
WW.1.SE
WD.2.SE
WW.2.SE
WD.3.SE
WW.3.SE
WD.4.SE
WW.4.SE
WD.5.SE
WW.5.SE
WD.1.SE+5d
WW.1.SE+5d
WD.2.SE+5d
WW.2.SE+5d
WD.3.SE+5d
WW.3.SE+5d
WD.4.SE+5d
WW.4.SE+5d
WD.5.SE+5d
WW.5.SE+5d

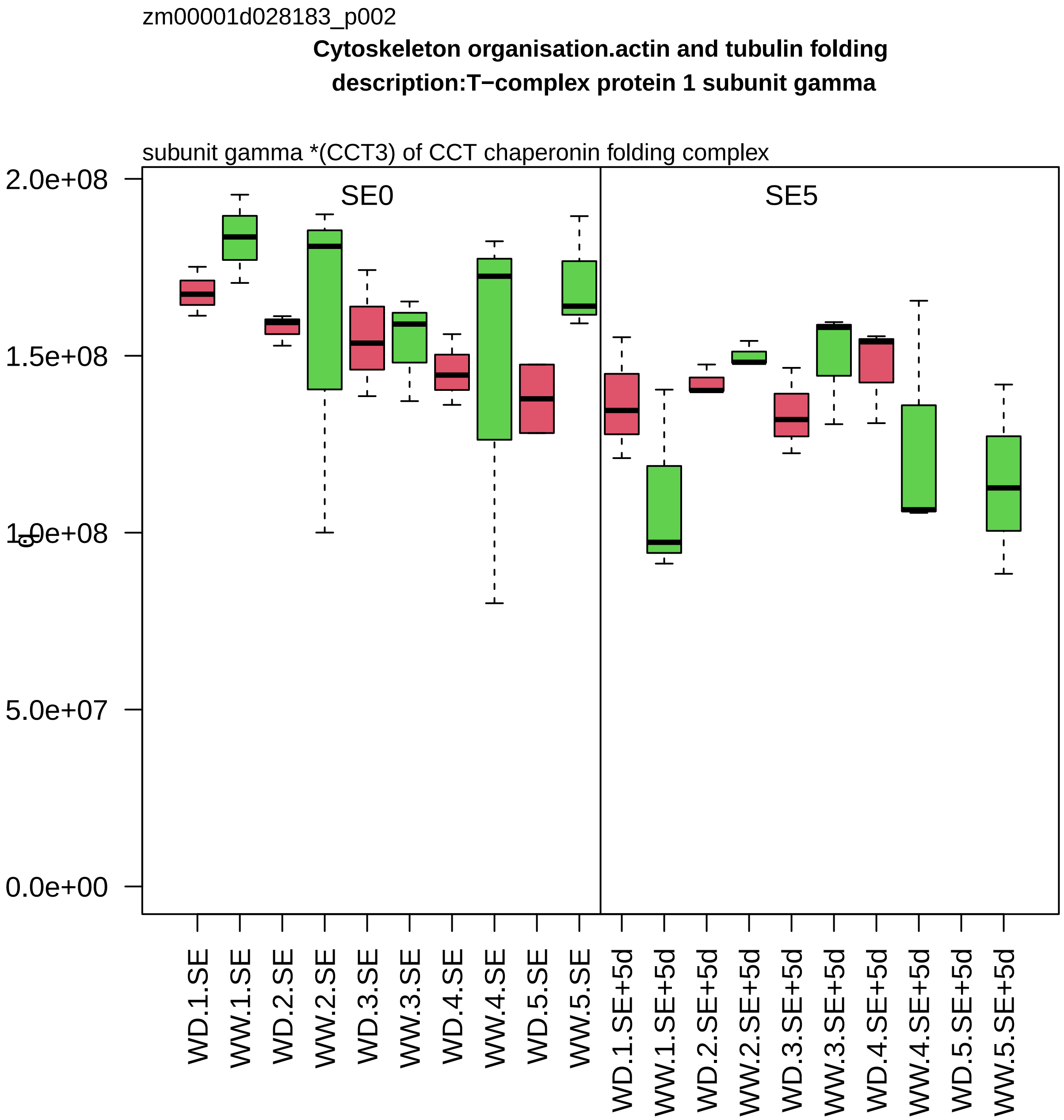
zm00001d028183_p002
Cytoskeleton organisation.actin and tubulin folding
description:T−complex protein 1 subunit gamma
subunit gamma *(CCT3) of CCT chaperonin folding complex
SE0
SE5
2.0e+08
1.5e+08
1.0e+08
5.0e+07
0.0e+00
WD.1.SE
WW.1.SE
WD.2.SE
WW.2.SE
WD.3.SE
WW.3.SE
WD.4.SE
WW.4.SE
WD.5.SE
WW.5.SE
WD.1.SE+5d
WW.1.SE+5d
WD.2.SE+5d
WW.2.SE+5d
WD.3.SE+5d
WW.3.SE+5d
WD.4.SE+5d
WW.4.SE+5d
WD.5.SE+5d
WW.5.SE+5d

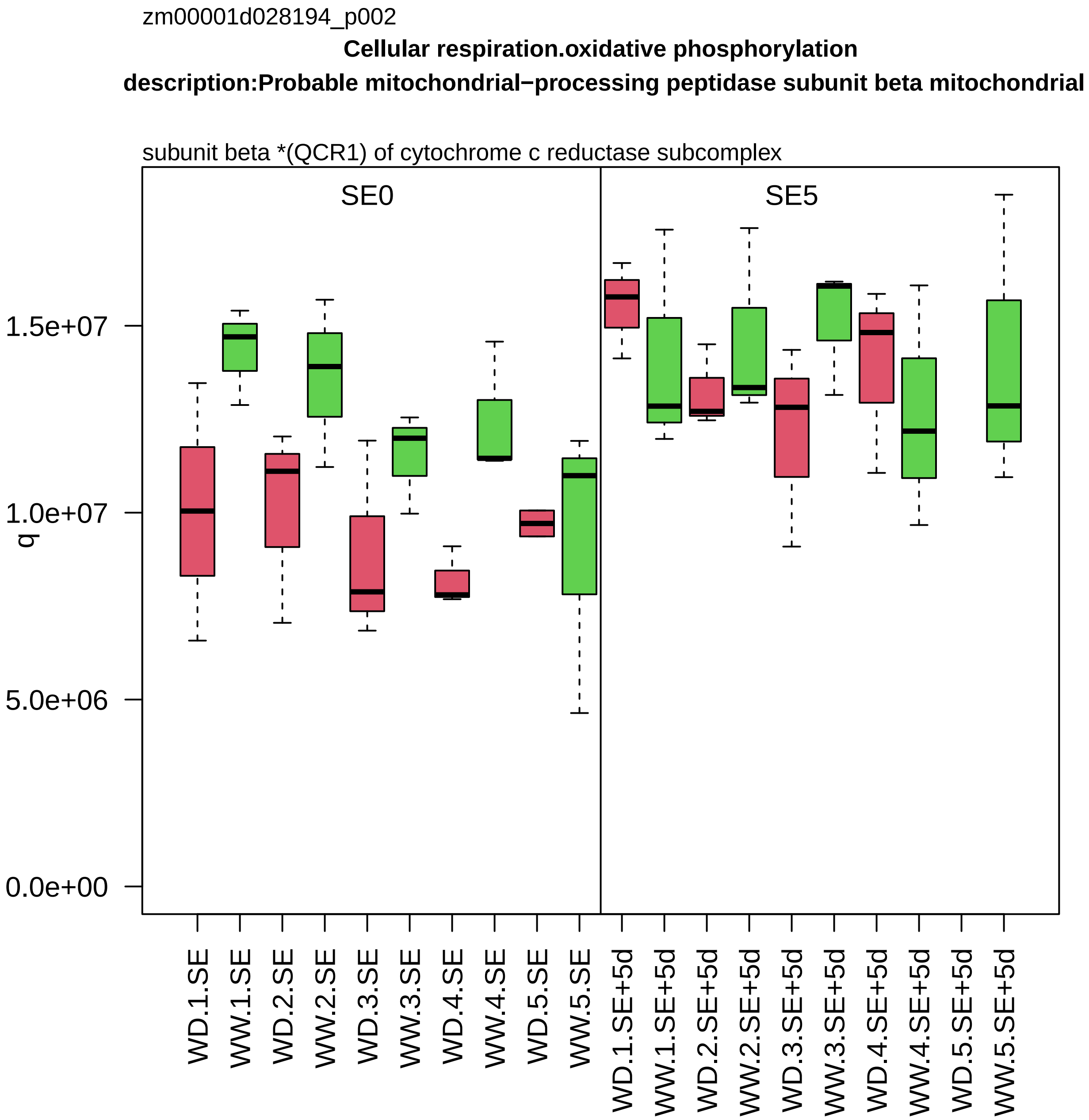

zm00001d028194_p002
Cellular respiration.oxidative phosphorylation
description:Probable mitochondrial−processing peptidase subunit beta mitochondrial
subunit beta *(QCR1) of cytochrome c reductase subcomplex
SE0
SE5
q
1.5e+07
1.0e+07
5.0e+06
0.0e+00
WD.1.SE
WW.1.SE
WD.2.SE
WW.2.SE
WD.3.SE
WW.3.SE
WD.4.SE
WW.4.SE
WD.5.SE
WW.5.SE
WD.1.SE+5d
WW.1.SE+5d
WD.2.SE+5d
WW.2.SE+5d
WD.3.SE+5d
WW.3.SE+5d
WD.4.SE+5d
WW.4.SE+5d
WD.5.SE+5d
WW.5.SE+5d

zm00001d028218_p001

**Protein biosynthesis.aminoacyl−tRNA synthetase activities**

**description:Methionine−−tRNA ligase cytoplasmic**

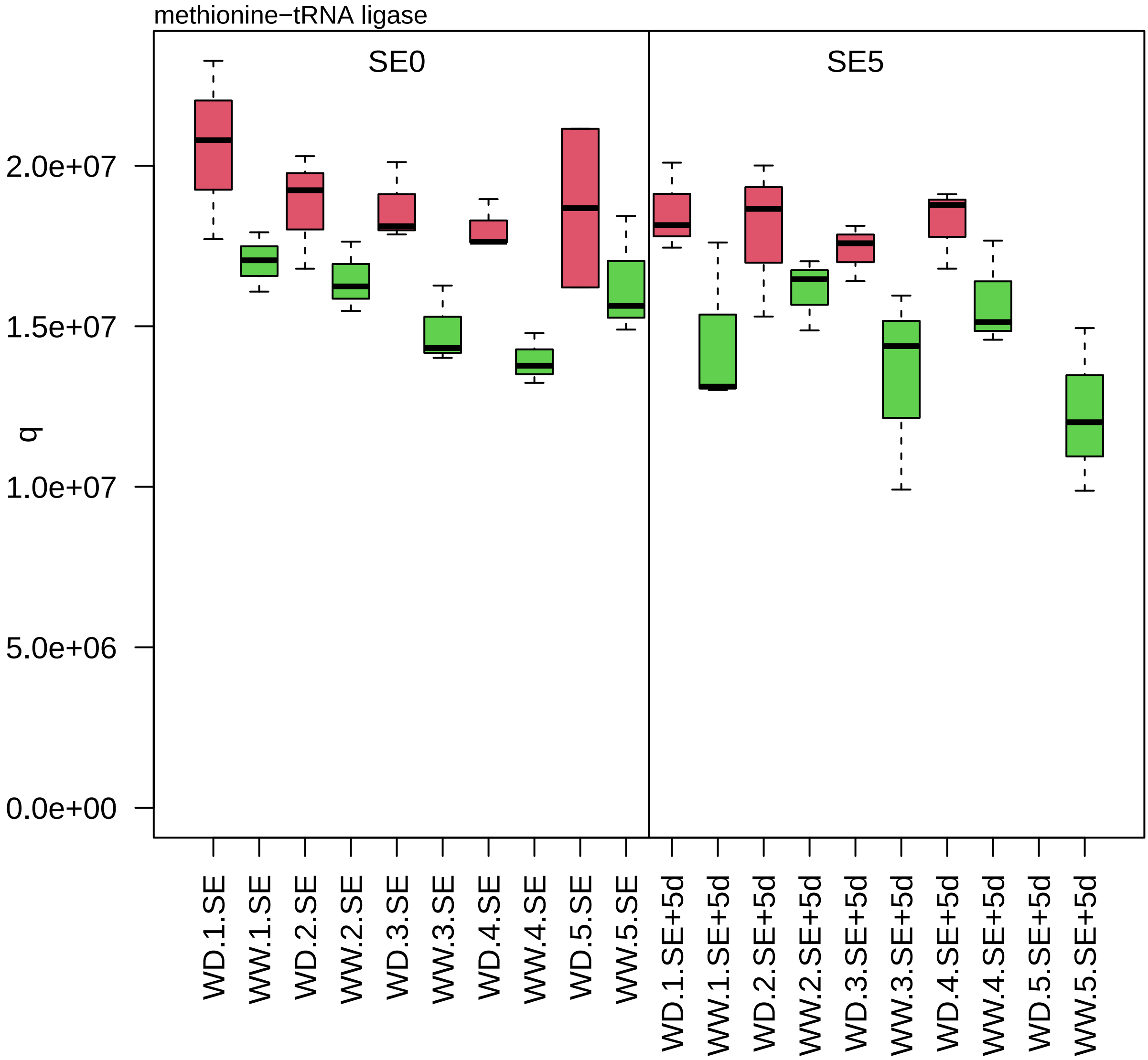

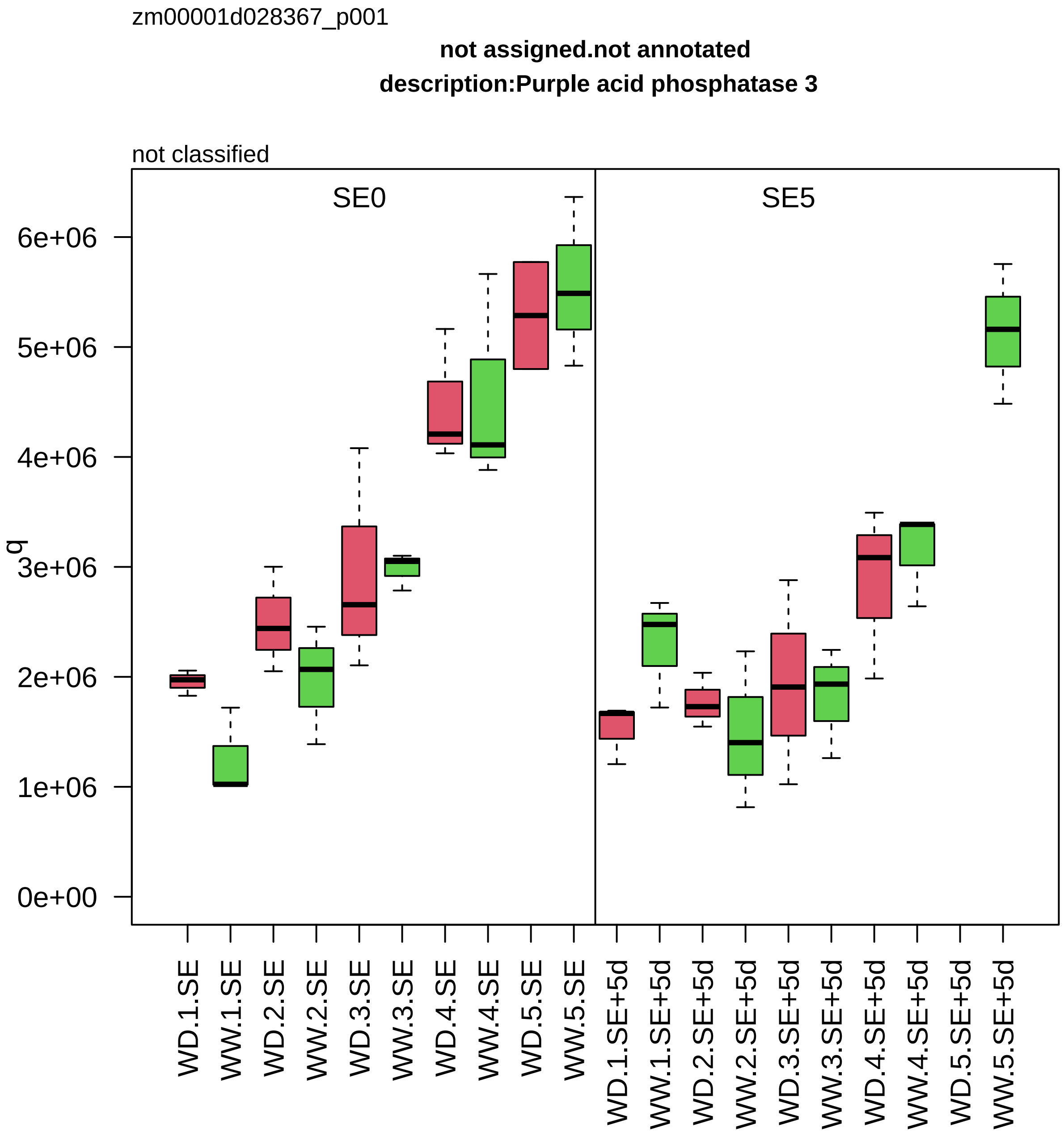

zm00001d028367_p001
not assigned.not annotated
description:Purple acid phosphatase 3
not classified
SE0
SE5
q
0e+00
1e+06
2e+06
3e+06
4e+06
5e+06
6e+06
WD.1.SE
WW.1.SE
WD.2.SE
WW.2.SE
WD.3.SE
WW.3.SE
WD.4.SE
WW.4.SE
WD.5.SE
WW.5.SE
WD.1.SE+5d
WW.1.SE+5d
WD.2.SE+5d
WW.2.SE+5d
WD.3.SE+5d
WW.3.SE+5d
WD.4.SE+5d
WW.4.SE+5d
WD.5.SE+5d
WW.5.SE+5d

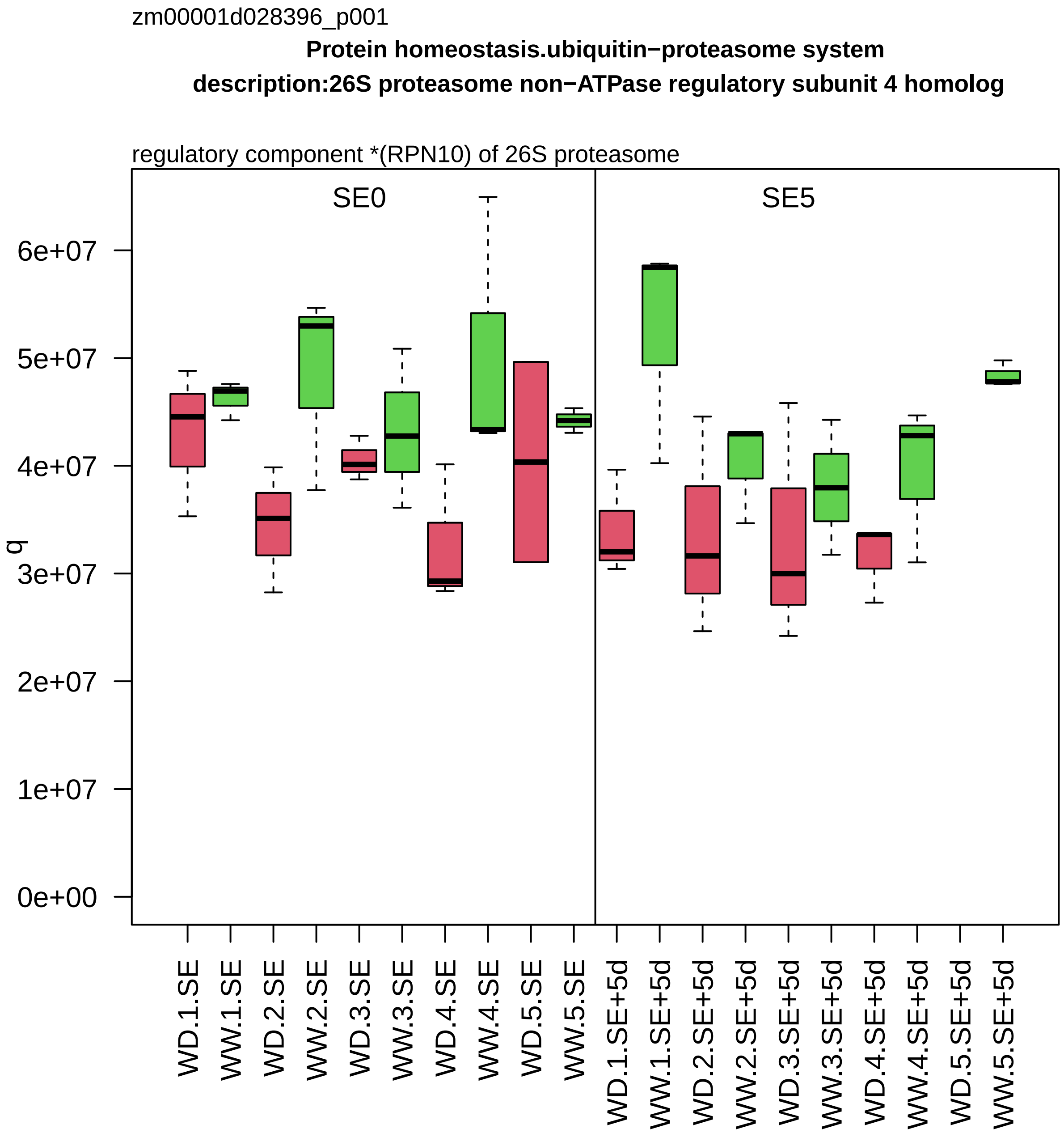

zm00001d028396_p001
Protein homeostasis.ubiquitin−proteasome system
description:26S proteasome non−ATPase regulatory subunit 4 homolog
regulatory component *(RPN10) of 26S proteasome
SE0
SE5
q
6e+07
5e+07
4e+07
3e+07
2e+07
1e+07
0e+00
WD.1.SE
WW.1.SE
WD.2.SE
WW.2.SE
WD.3.SE
WW.3.SE
WD.4.SE
WW.4.SE
WD.5.SE
WW.5.SE
WD.1.SE+5d
WW.1.SE+5d
WD.2.SE+5d
WW.2.SE+5d
WD.3.SE+5d
WW.3.SE+5d
WD.4.SE+5d
WW.4.SE+5d
WD.5.SE+5d
WW.5.SE+5d

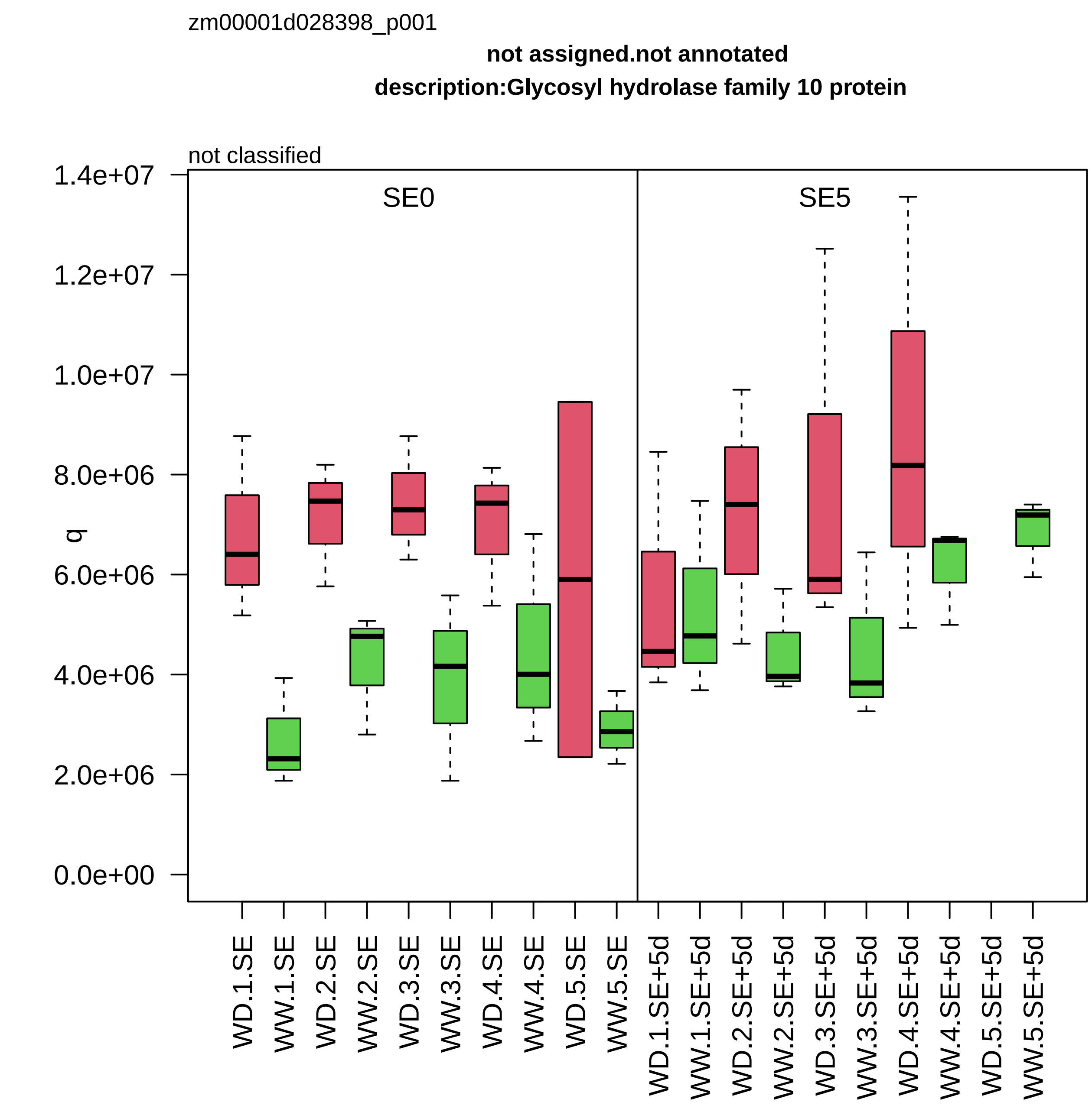
zm00001d028398_p001
not assigned.not annotated
description:Glycosyl hydrolase family 10 protein
not classified
SE0
SE5
q
1.4e+07
1.2e+07
1.0e+07
8.0e+06
6.0e+06
4.0e+06
2.0e+06
0.0e+00
WD.1.SE
WW.1.SE
WD.2.SE
WW.2.SE
WD.3.SE
WW.3.SE
WD.4.SE
WW.4.SE
WD.5.SE
WW.5.SE
WD.1.SE+5d
WW.1.SE+5d
WD.2.SE+5d
WW.2.SE+5d
WD.3.SE+5d
WW.3.SE+5d
WD.4.SE+5d
WW.4.SE+5d
WD.5.SE+5d
WW.5.SE+5d

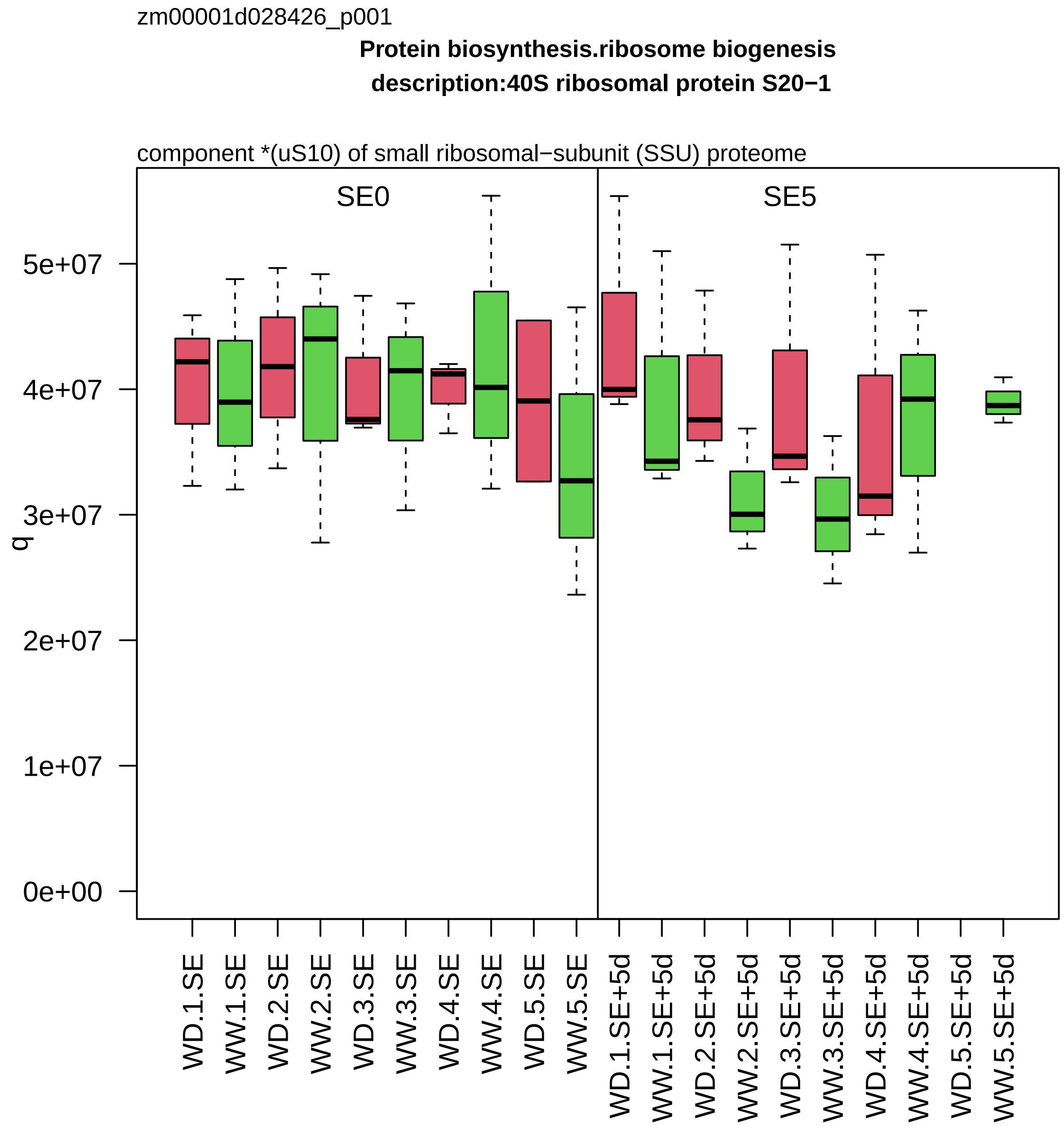

zm00001d028426_p001
Protein biosynthesis.ribosome biogenesis
description:40S ribosomal protein S20−1
component *(uS10) of small ribosomal−subunit (SSU) proteome
SE0
SE5
q
0e+00
1e+07
2e+07
3e+07
4e+07
5e+07
WD.1.SE
WW.1.SE
WD.2.SE
WW.2.SE
WD.3.SE
WW.3.SE
WD.4.SE
WW.4.SE
WD.5.SE
WW.5.SE
WD.1.SE+5d
WW.1.SE+5d
WD.2.SE+5d
WW.2.SE+5d
WD.3.SE+5d
WW.3.SE+5d
WD.4.SE+5d
WW.4.SE+5d
WD.5.SE+5d
WW.5.SE+5d

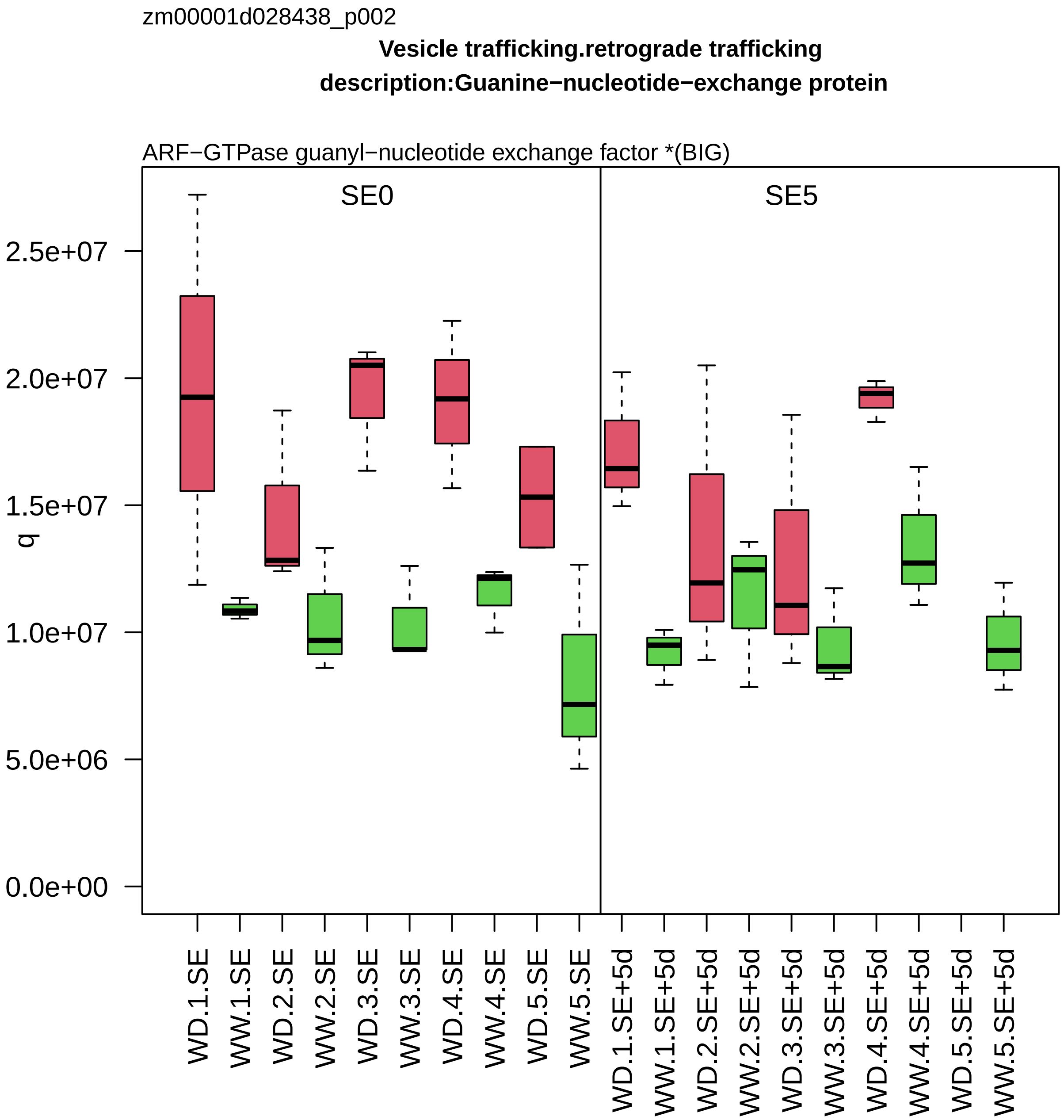
zm00001d028438_p002
Vesicle trafficking.retrograde trafficking
description:Guanine−nucleotide−exchange protein
ARF−GTPase guanyl−nucleotide exchange factor *(BIG)
SE0
SE5
q
0.0e+00
5.0e+06
1.0e+07
1.5e+07
2.0e+07
2.5e+07
WD.1.SE
WW.1.SE
WD.2.SE
WW.2.SE
WD.3.SE
WW.3.SE
WD.4.SE
WW.4.SE
WD.5.SE
WW.5.SE
WD.1.SE+5d
WW.1.SE+5d
WD.2.SE+5d
WW.2.SE+5d
WD.3.SE+5d
WW.3.SE+5d
WD.4.SE+5d
WW.4.SE+5d
WD.5.SE+5d
WW.5.SE+5d

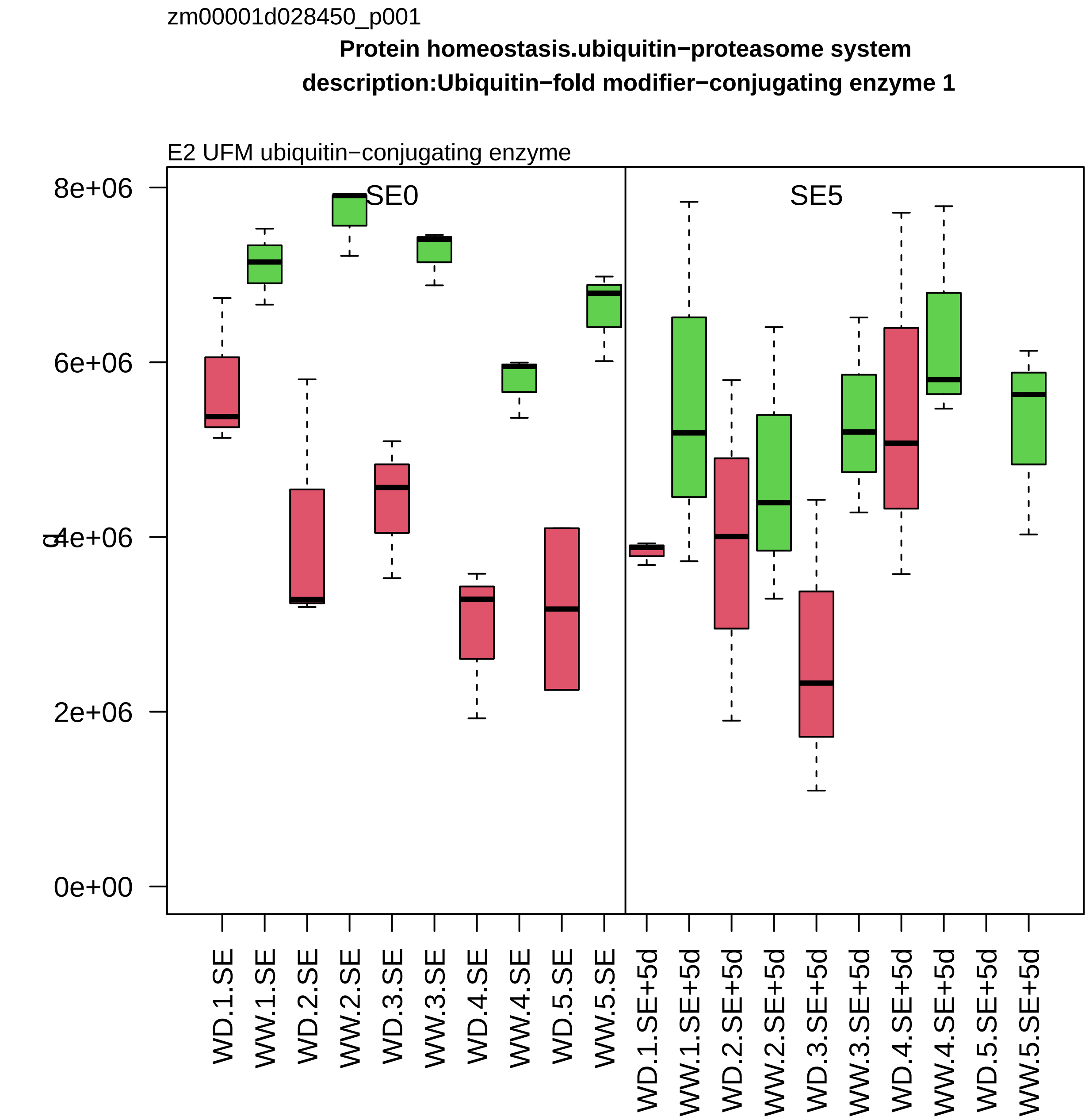
zm00001d028450_p001
Protein homeostasis.ubiquitin−proteasome system
description:Ubiquitin−fold modifier−conjugating enzyme 1
E2 UFM ubiquitin−conjugating enzyme
SE0
SE5
8e+06
6e+06
4e+06
2e+06
0e+00
WD.1.SE
WW.1.SE
WD.2.SE
WW.2.SE
WD.3.SE
WW.3.SE
WD.4.SE
WW.4.SE
WD.5.SE
WW.5.SE
WD.1.SE+5d
WW.1.SE+5d
WD.2.SE+5d
WW.2.SE+5d
WD.3.SE+5d
WW.3.SE+5d
WD.4.SE+5d
WW.4.SE+5d
WD.5.SE+5d
WW.5.SE+5d

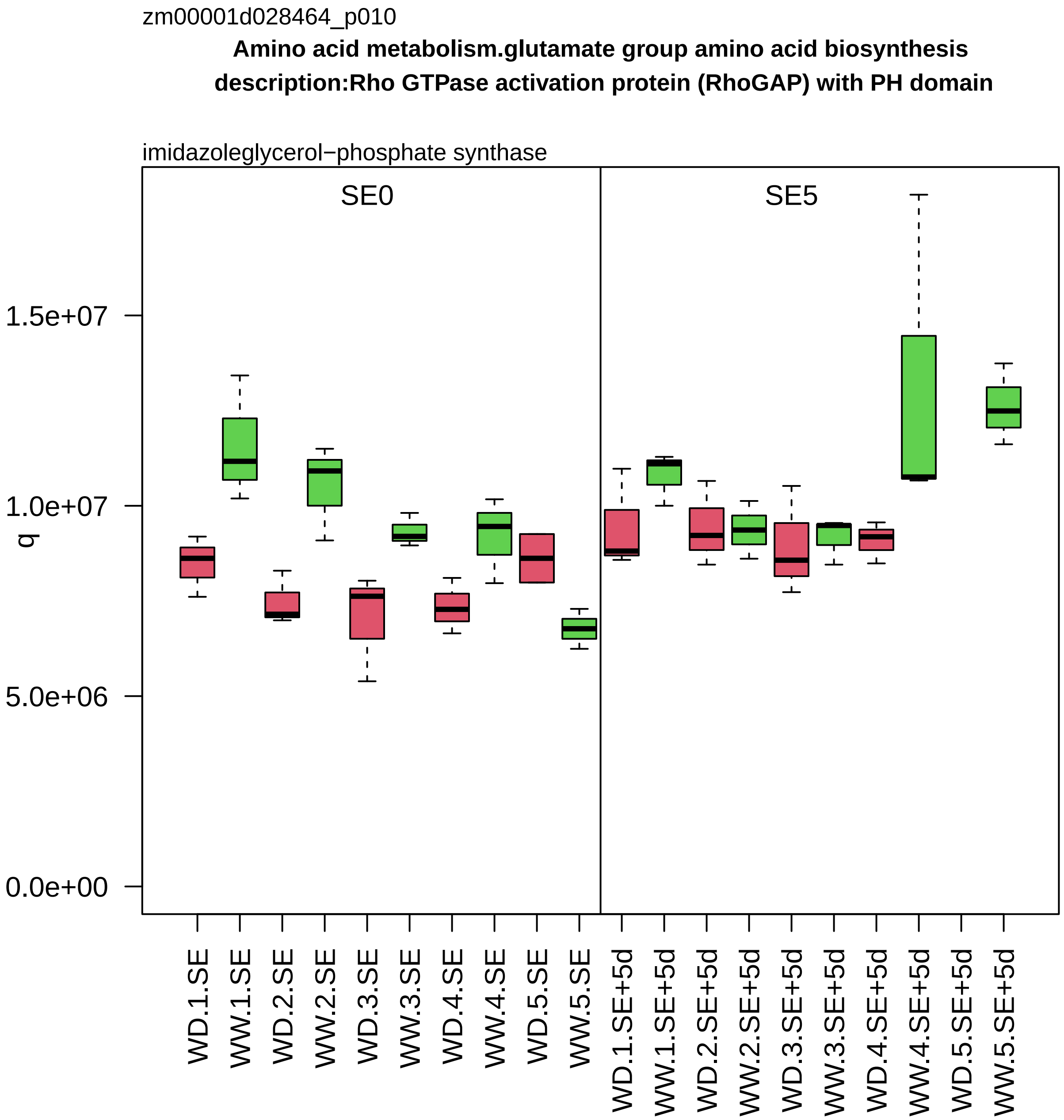
zm00001d028464_p010
Amino acid metabolism.glutamate group amino acid biosynthesis
description:Rho GTPase activation protein (RhoGAP) with PH domain
imidazoleglycerol−phosphate synthase
SE0
SE5
q
1.5e+07
1.0e+07
5.0e+06
0.0e+00
WD.1.SE
WW.1.SE
WD.2.SE
WW.2.SE
WD.3.SE
WW.3.SE
WD.4.SE
WW.4.SE
WD.5.SE
WW.5.SE
WD.1.SE+5d
WW.1.SE+5d
WD.2.SE+5d
WW.2.SE+5d
WD.3.SE+5d
WW.3.SE+5d
WD.4.SE+5d
WW.4.SE+5d
WD.5.SE+5d
WW.5.SE+5d

zm00001d028474_p001

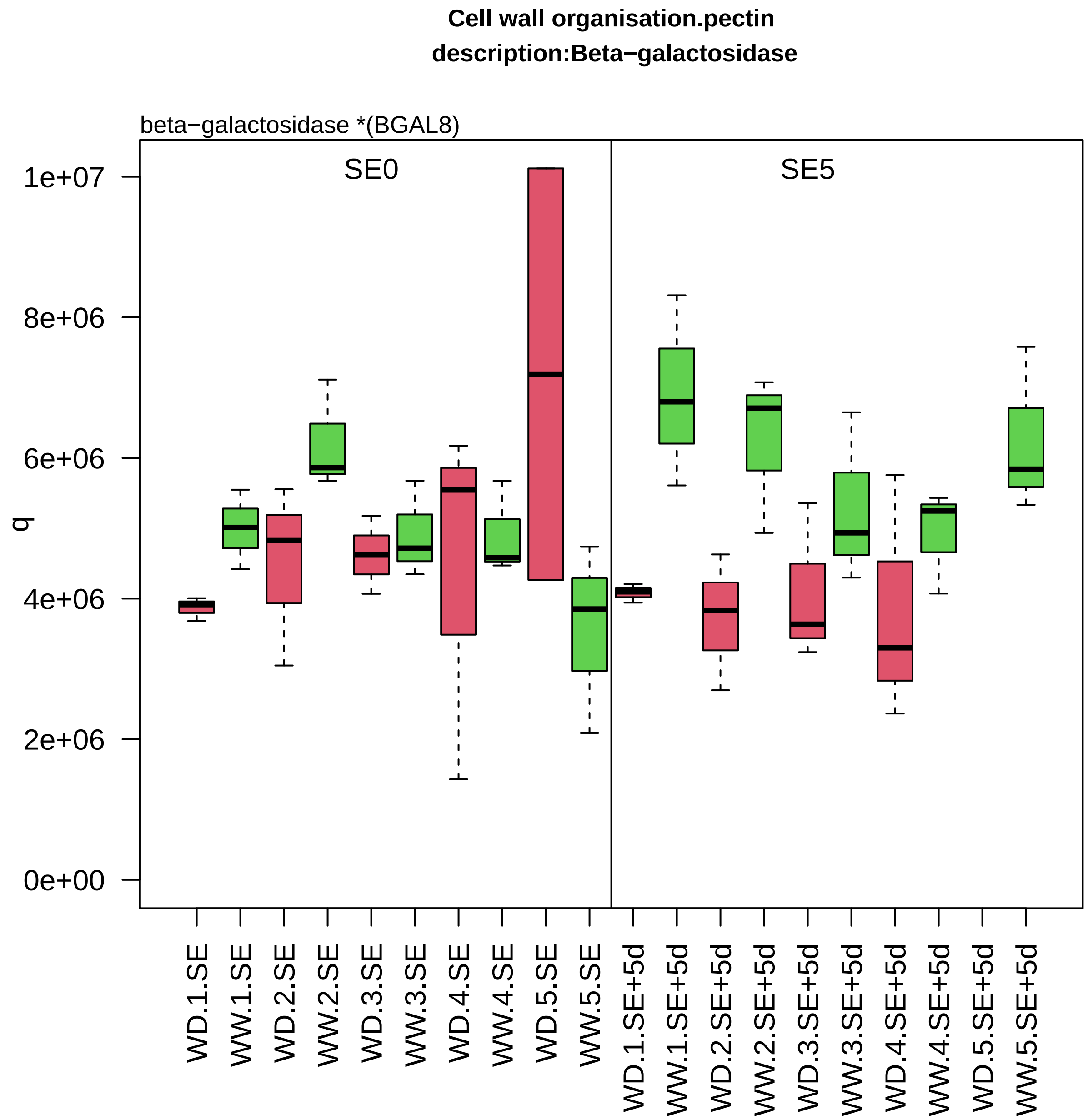

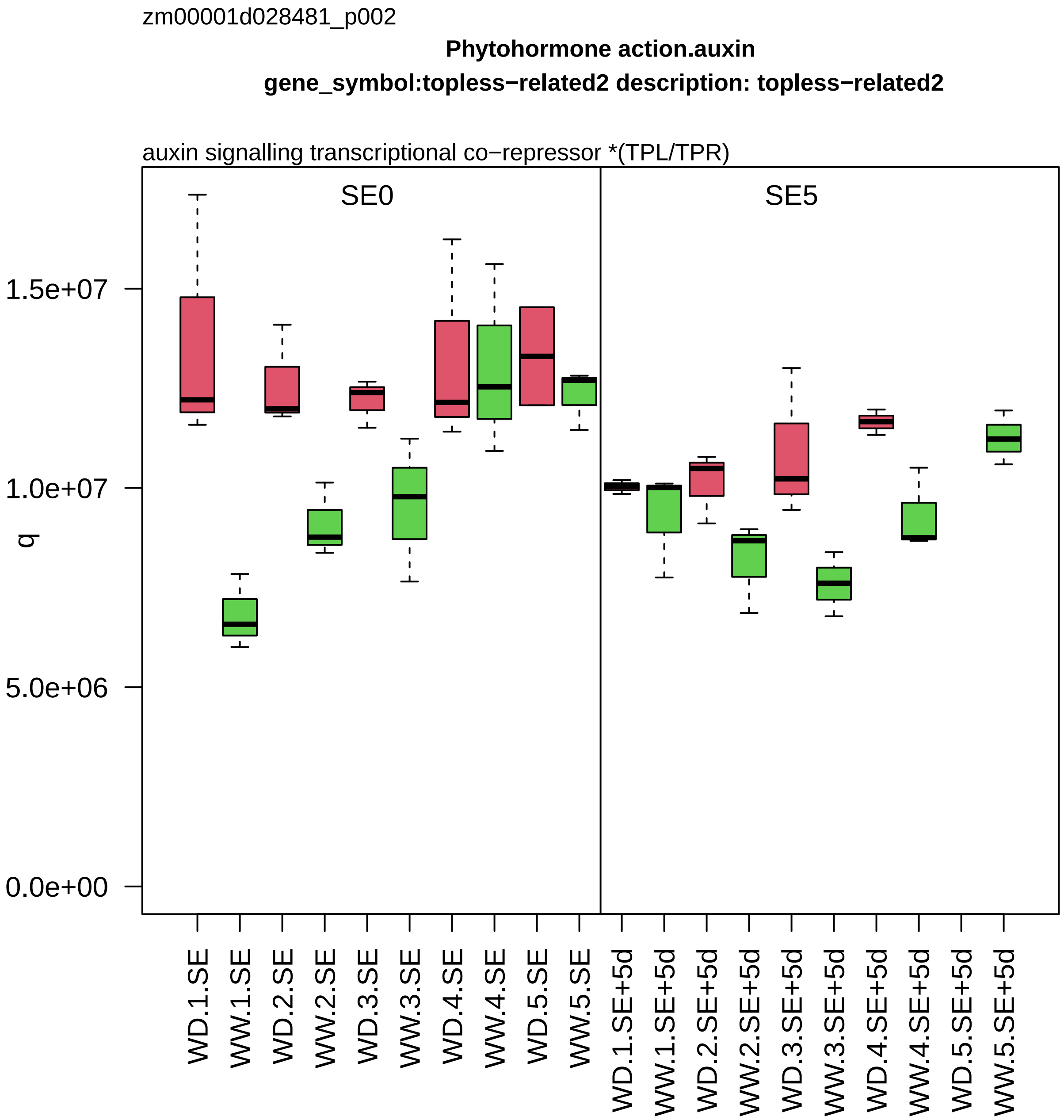
zm00001d028481_p002
Phytohormone action.auxin
gene_symbol:topless−related2 description: topless−related2
auxin signalling transcriptional co−repressor *(TPL/TPR)
SE0
SE5
q
1.5e+07
1.0e+07
5.0e+06
0.0e+00
WD.1.SE
WW.1.SE
WD.2.SE
WW.2.SE
WD.3.SE
WW.3.SE
WD.4.SE
WW.4.SE
WD.5.SE
WW.5.SE
WD.1.SE+5d
WW.1.SE+5d
WD.2.SE+5d
WW.2.SE+5d
WD.3.SE+5d
WW.3.SE+5d
WD.4.SE+5d
WW.4.SE+5d
WD.5.SE+5d
WW.5.SE+5d

zm00001d028653_p001

**Carbohydrate metabolism.nucleotide sugar biosynthesis**

**gene_symbol:dTDP−glucose dehydratase homolog csu219 description: dTDP−glucose dehydratase homo**

UDP−D−glucuronic acid decarboxylase

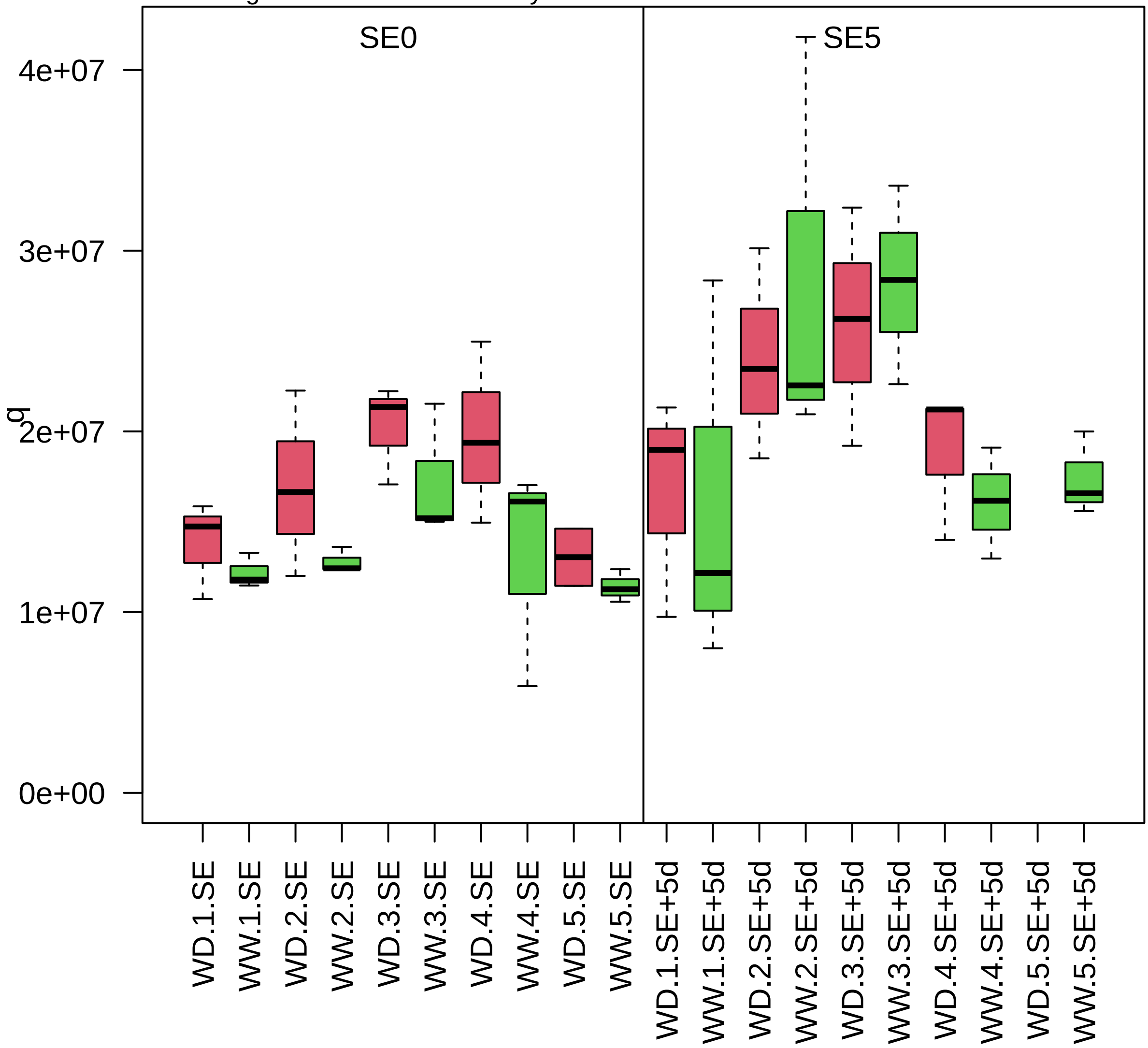

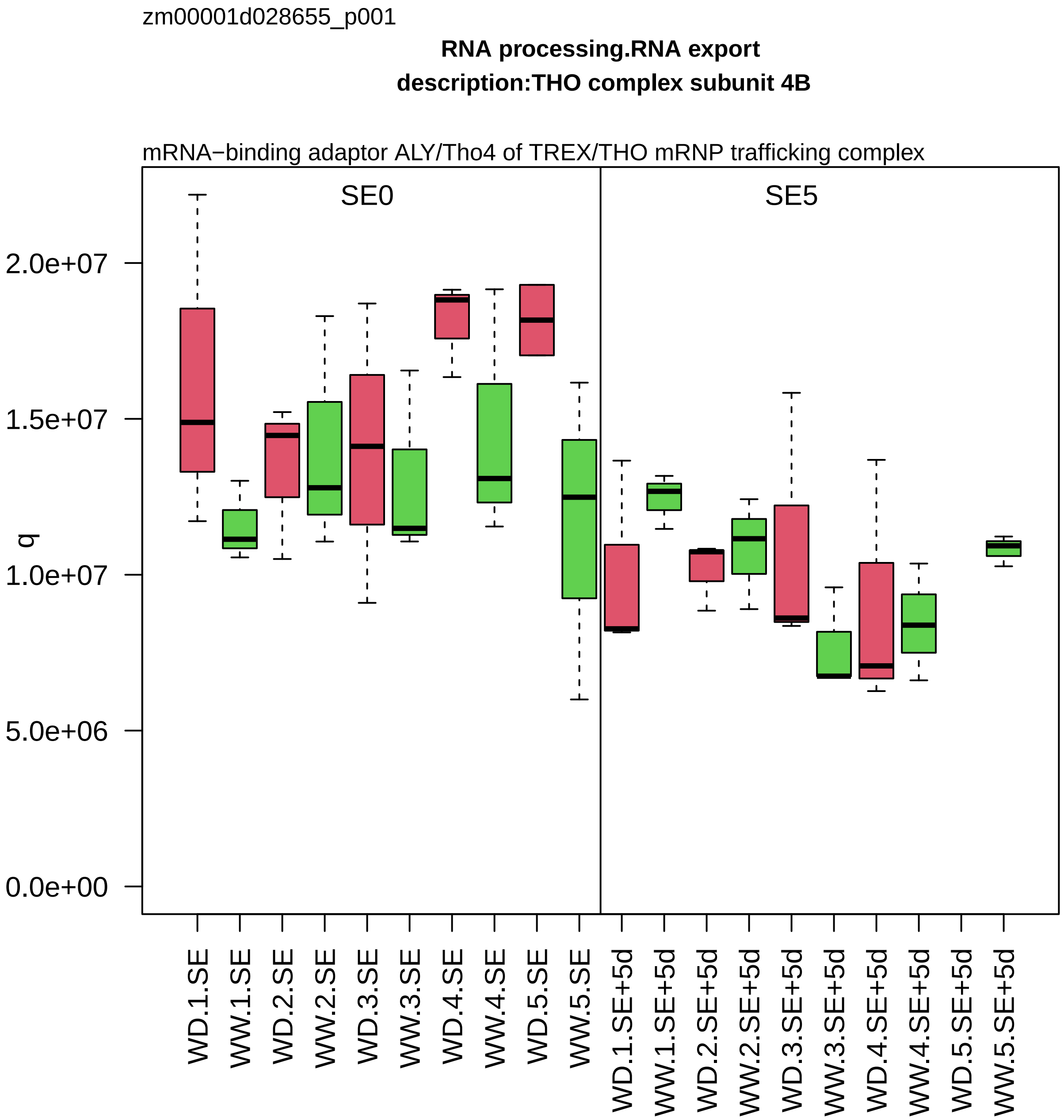
zm00001d028655_p001
RNA processing.RNA export
description:THO complex subunit 4B
mRNA−binding adaptor ALY/Tho4 of TREX/THO mRNP trafficking complex
SE0
SE5
q
2.0e+07
1.5e+07
1.0e+07
5.0e+06
0.0e+00
WD.1.SE
WW.1.SE
WD.2.SE
WW.2.SE
WD.3.SE
WW.3.SE
WD.4.SE
WW.4.SE
WD.5.SE
WW.5.SE
WD.1.SE+5d
WW.1.SE+5d
WD.2.SE+5d
WW.2.SE+5d
WD.3.SE+5d
WW.3.SE+5d
WD.4.SE+5d
WW.4.SE+5d
WD.5.SE+5d
WW.5.SE+5d

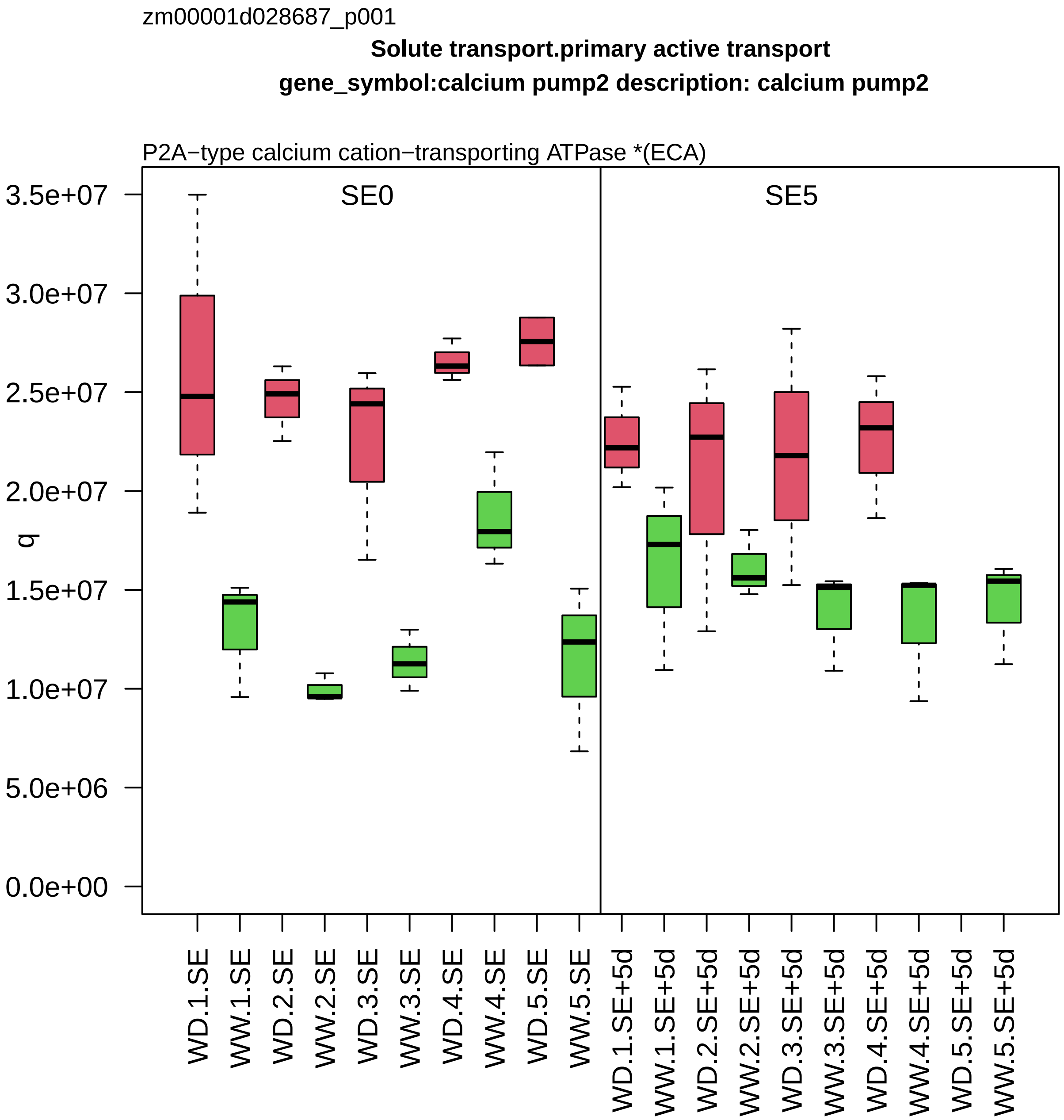
zm00001d028687_p001
Solute transport.primary active transport
gene_symbol:calcium pump2 description: calcium pump2
P2A−type calcium cation−transporting ATPase *(ECA)
SE0
SE5
q
3.5e+07
3.0e+07
2.5e+07
2.0e+07
1.5e+07
1.0e+07
5.0e+06
0.0e+00
WD.1.SE
WW.1.SE
WD.2.SE
WW.2.SE
WD.3.SE
WW.3.SE
WD.4.SE
WW.4.SE
WD.5.SE
WW.5.SE
WD.1.SE+5d
WW.1.SE+5d
WD.2.SE+5d
WW.2.SE+5d
WD.3.SE+5d
WW.3.SE+5d
WD.4.SE+5d
WW.4.SE+5d
WD.5.SE+5d
WW.5.SE+5d

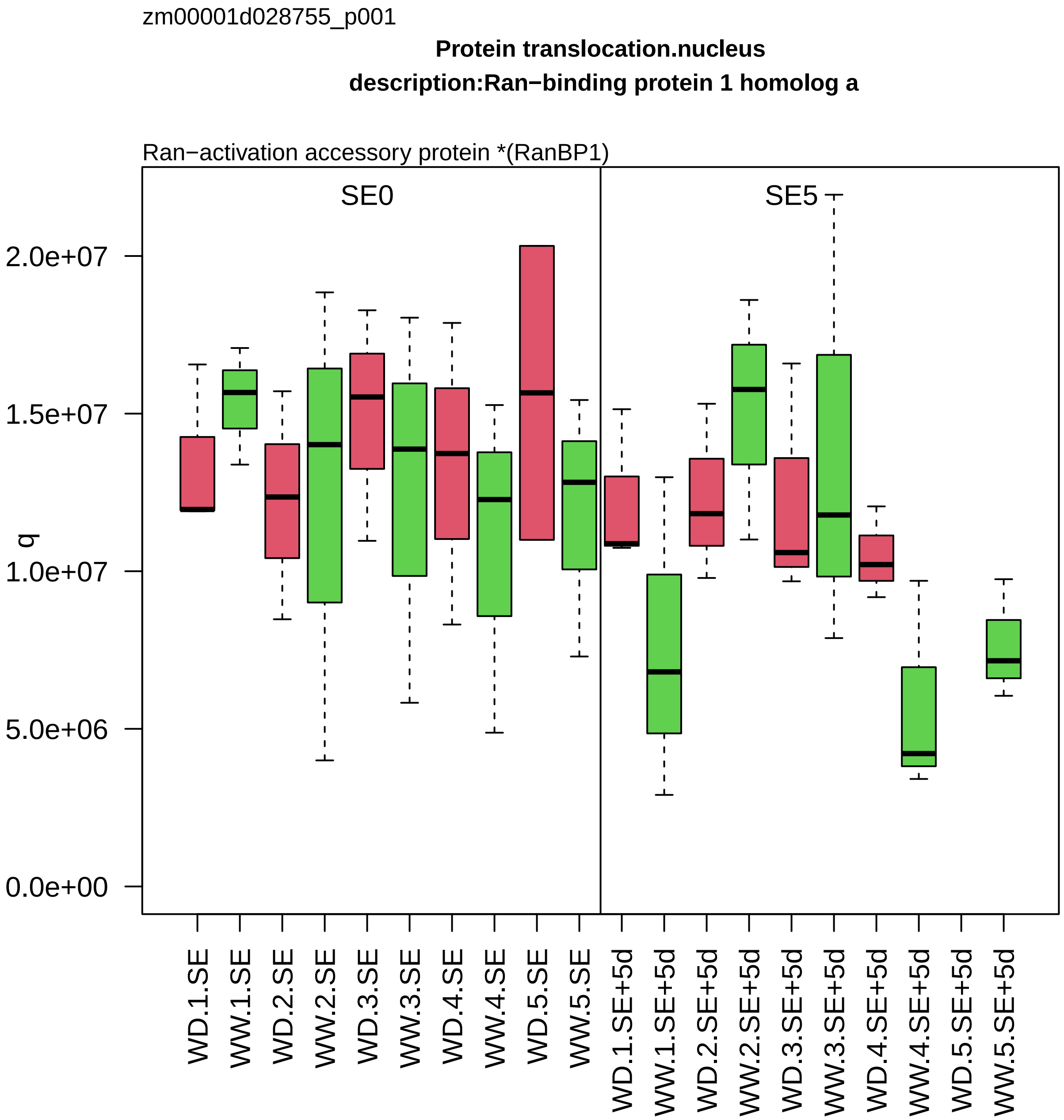
zm00001d028755_p001
Protein translocation.nucleus
description:Ran−binding protein 1 homolog a
Ran−activation accessory protein *(RanBP1)
SE0
SE5
q
2.0e+07
1.5e+07
1.0e+07
5.0e+06
0.0e+00
WD.1.SE
WW.1.SE
WD.2.SE
WW.2.SE
WD.3.SE
WW.3.SE
WD.4.SE
WW.4.SE
WD.5.SE
WW.5.SE
WD.1.SE+5d
WW.1.SE+5d
WD.2.SE+5d
WW.2.SE+5d
WD.3.SE+5d
WW.3.SE+5d
WD.4.SE+5d
WW.4.SE+5d
WD.5.SE+5d
WW.5.SE+5d

zm00001d028759_p001

**Carbohydrate metabolism.fermentation**

**gene_symbol:pyruvate decarboxylase3 description:Pyruvate decarboxylase isozyme 3**

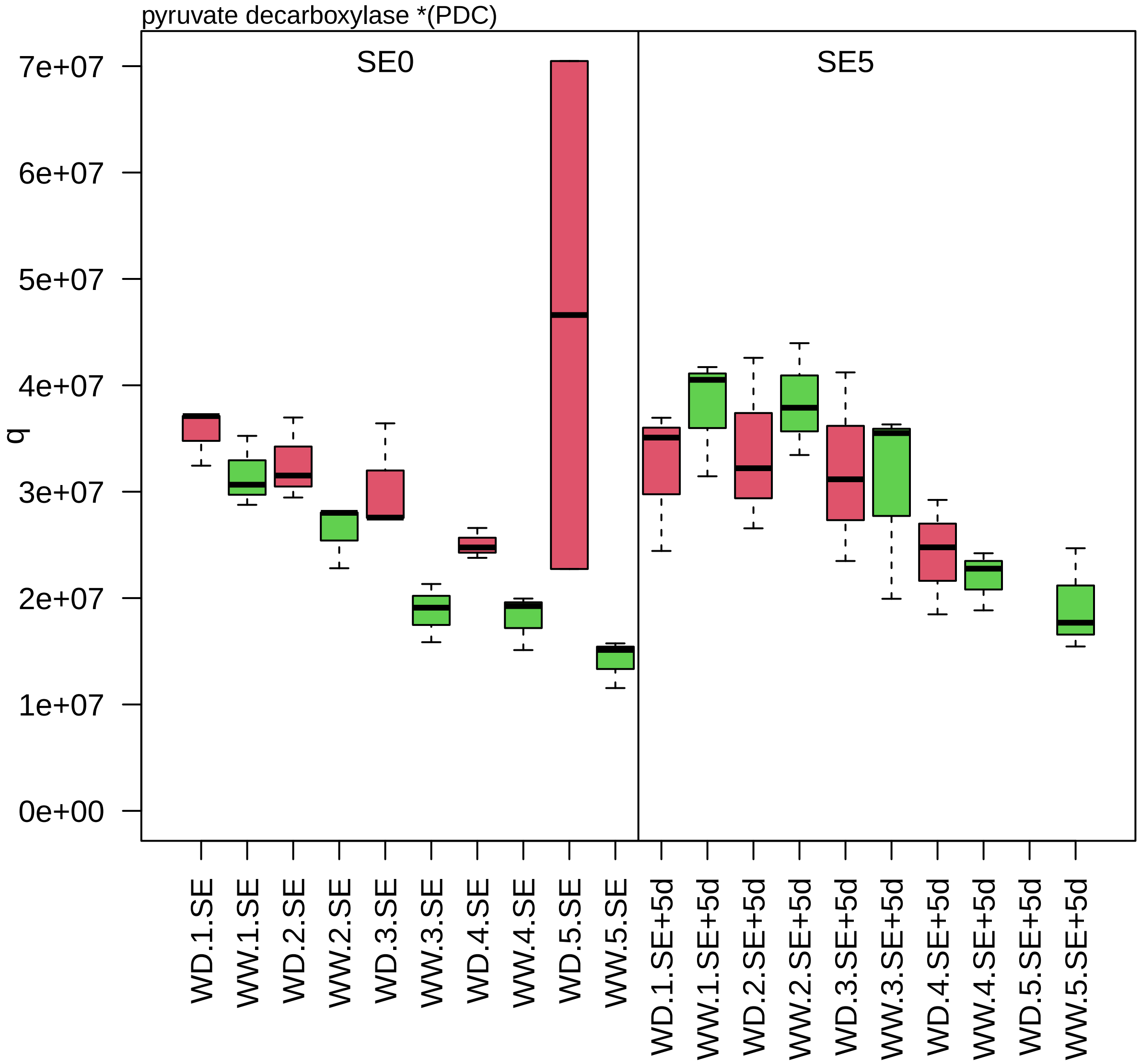

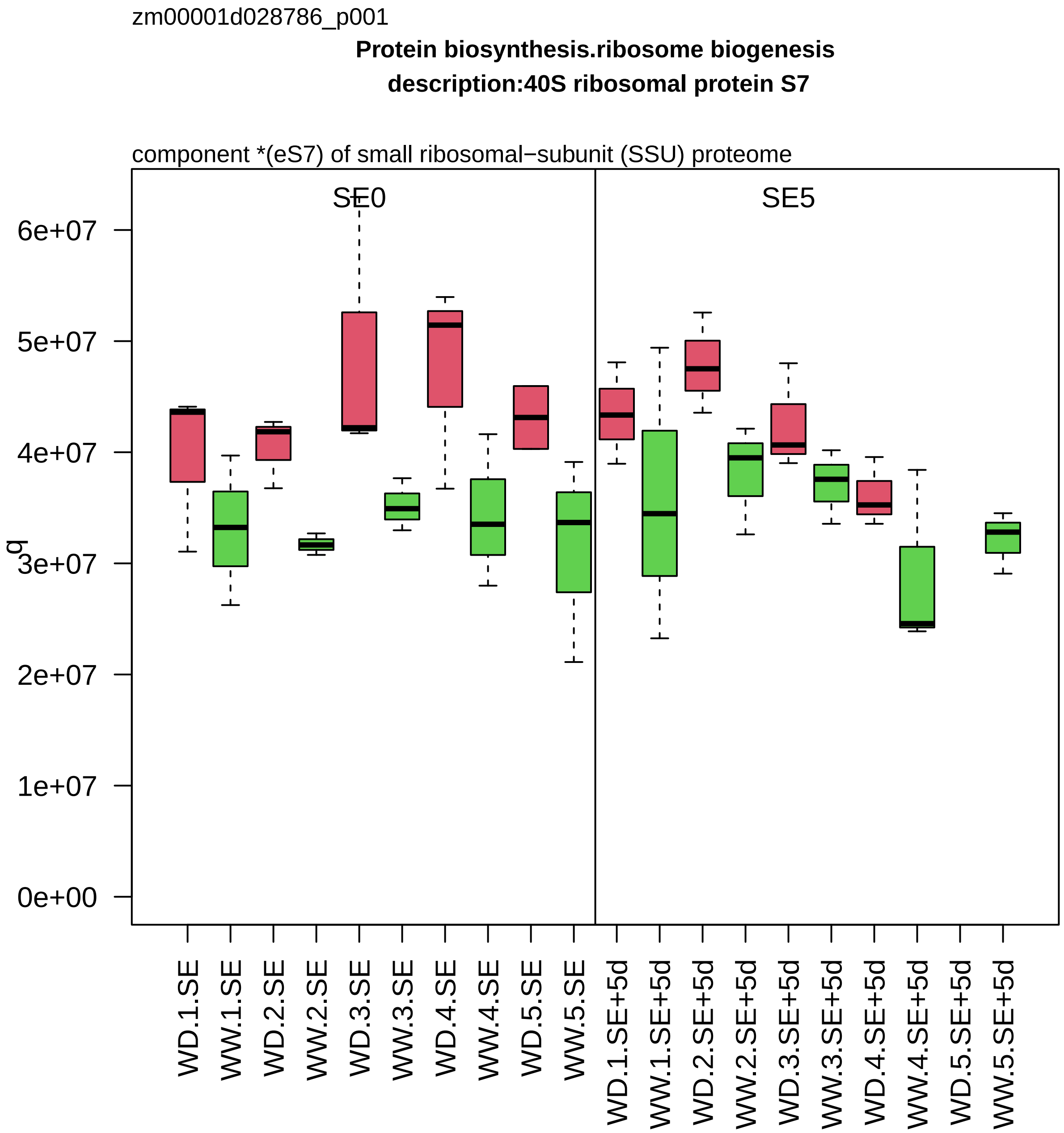
zm00001d028786_p001
Protein biosynthesis.ribosome biogenesis
description:40S ribosomal protein S7
component *(eS7) of small ribosomal−subunit (SSU) proteome
SE0
SE5
q
0e+00
1e+07
2e+07
3e+07
4e+07
5e+07
6e+07
WD.1.SE
WW.1.SE
WD.2.SE
WW.2.SE
WD.3.SE
WW.3.SE
WD.4.SE
WW.4.SE
WD.5.SE
WW.5.SE
WD.1.SE+5d
WW.1.SE+5d
WD.2.SE+5d
WW.2.SE+5d
WD.3.SE+5d
WW.3.SE+5d
WD.4.SE+5d
WW.4.SE+5d
WD.5.SE+5d
WW.5.SE+5d

zm00001d028810_p001

**Amino acid metabolism.aspartate group amino acid biosynthesis**

**description:LL−diaminopimelate aminotransferase chloroplastic**

LL−diaminopimelate aminotransferase

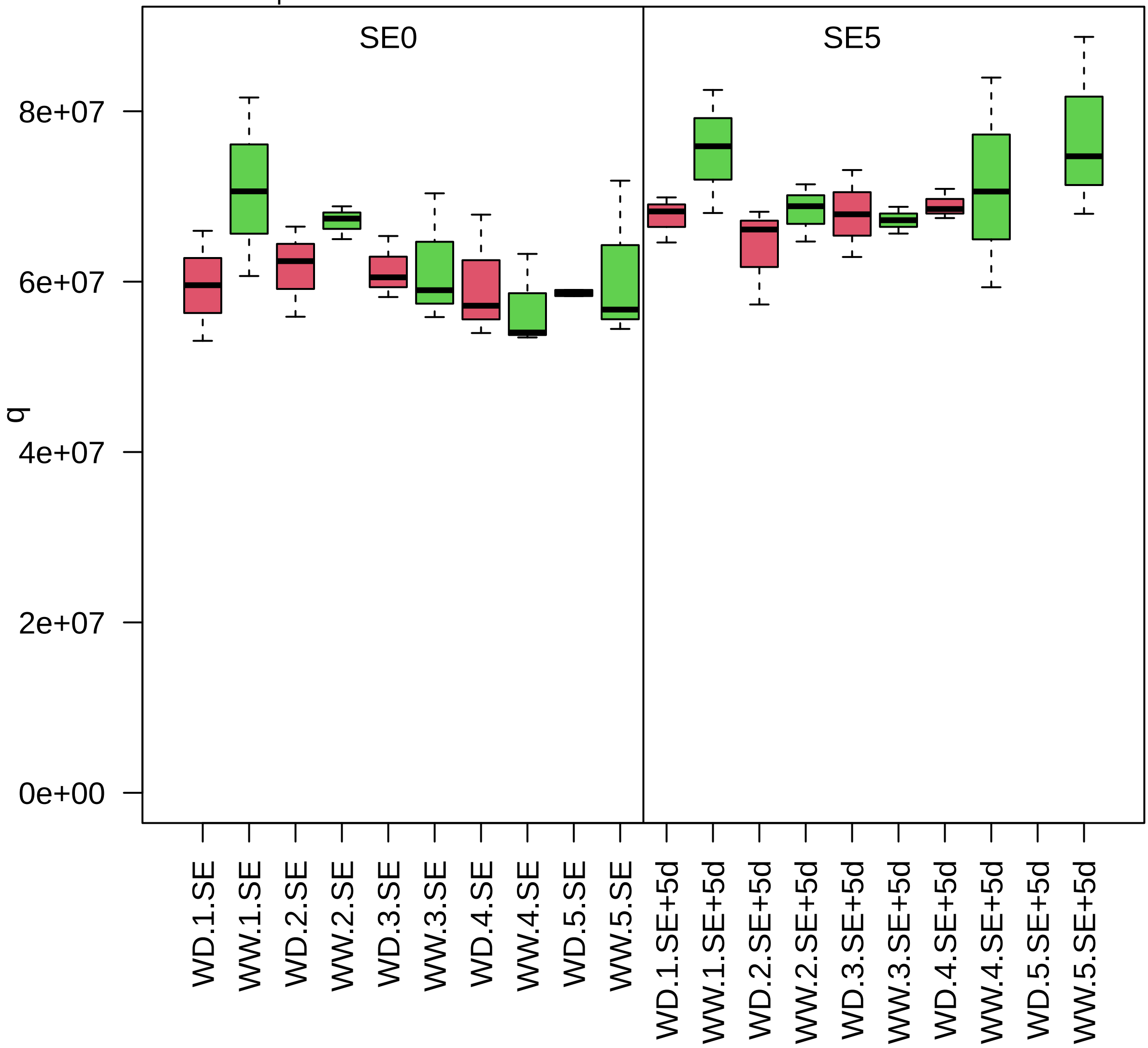

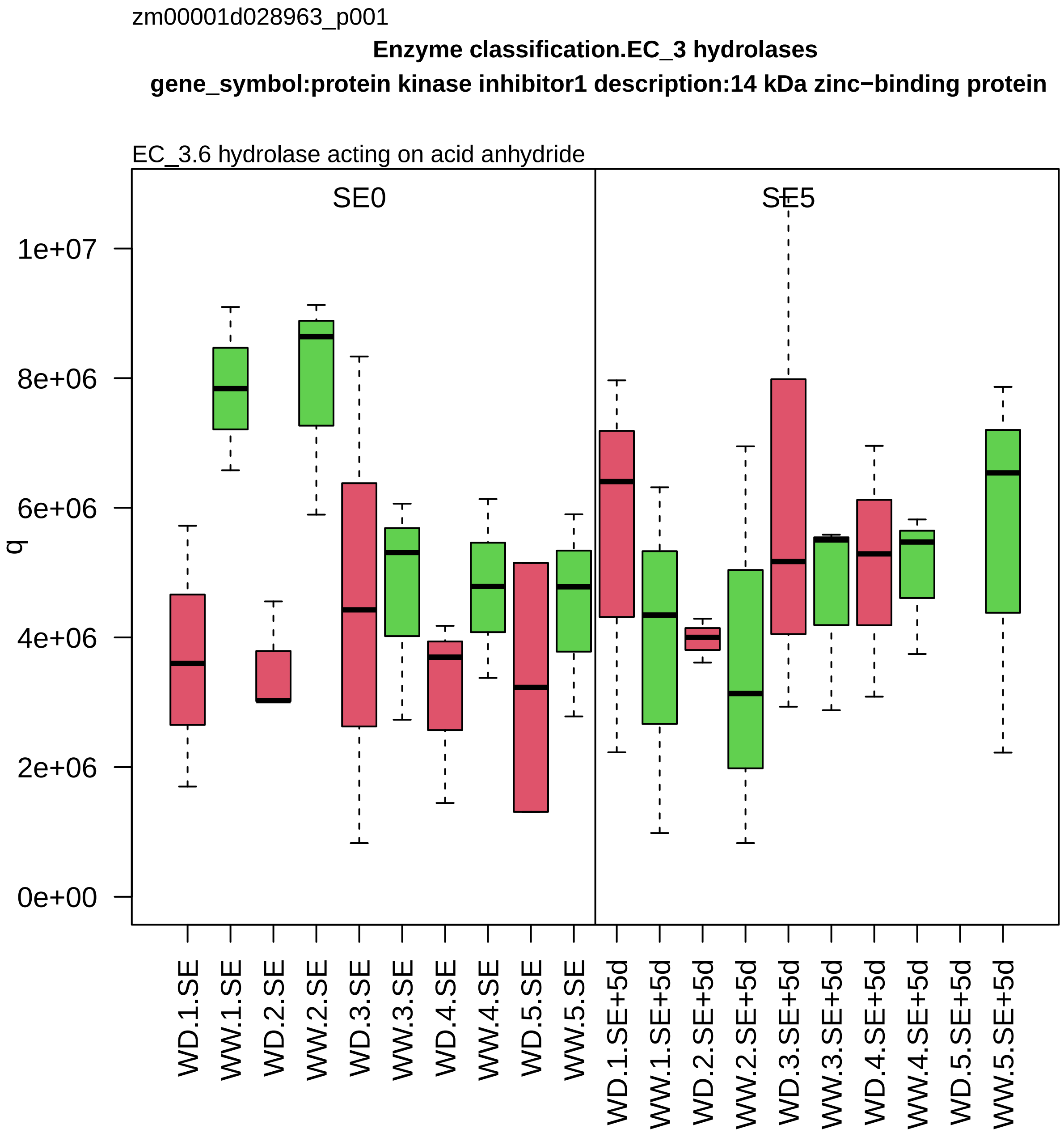

zm00001d028963_p001
Enzyme classification.EC_3 hydrolases
gene_symbol:protein kinase inhibitor1 description:14 kDa zinc−binding protein
EC_3.6 hydrolase acting on acid anhydride
SE0
SE5
q
0e+00
2e+06
4e+06
6e+06
8e+06
1e+07
WD.1.SE
WW.1.SE
WD.2.SE
WW.2.SE
WD.3.SE
WW.3.SE
WD.4.SE
WW.4.SE
WD.5.SE
WW.5.SE
WD.1.SE+5d
WW.1.SE+5d
WD.2.SE+5d
WW.2.SE+5d
WD.3.SE+5d
WW.3.SE+5d
WD.4.SE+5d
WW.4.SE+5d
WD.5.SE+5d
WW.5.SE+5d

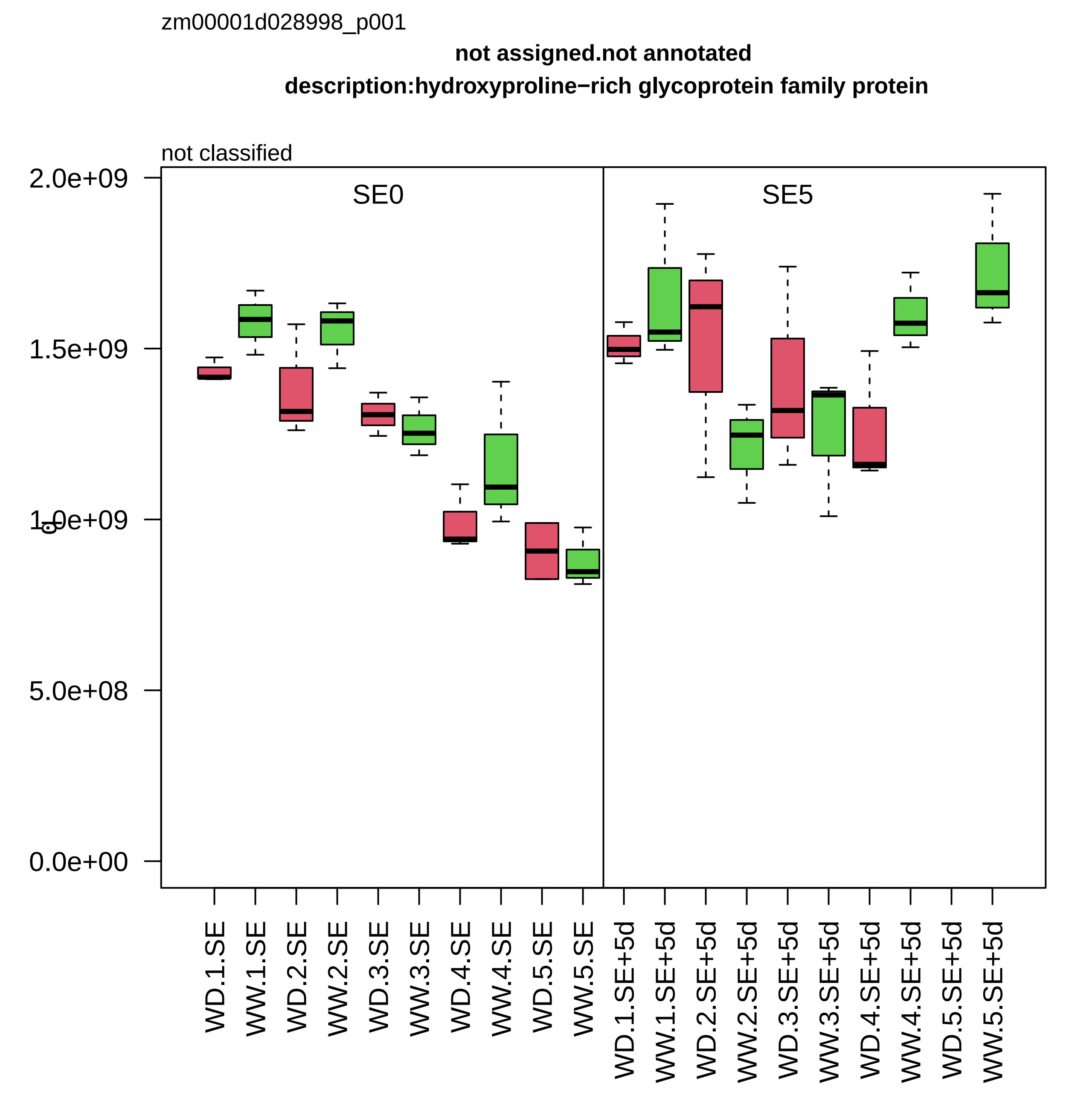

zm00001d028998_p001
not assigned.not annotated
description:hydroxyproline−rich glycoprotein family protein
not classified
SE0
SE5
2.0e+09
1.5e+09
1.0e+09
5.0e+08
0.0e+00
WD.1.SE
WW.1.SE
WD.2.SE
WW.2.SE
WD.3.SE
WW.3.SE
WD.4.SE
WW.4.SE
WD.5.SE
WW.5.SE
WD.1.SE+5d
WW.1.SE+5d
WD.2.SE+5d
WW.2.SE+5d
WD.3.SE+5d
WW.3.SE+5d
WD.4.SE+5d
WW.4.SE+5d
WD.5.SE+5d
WW.5.SE+5d

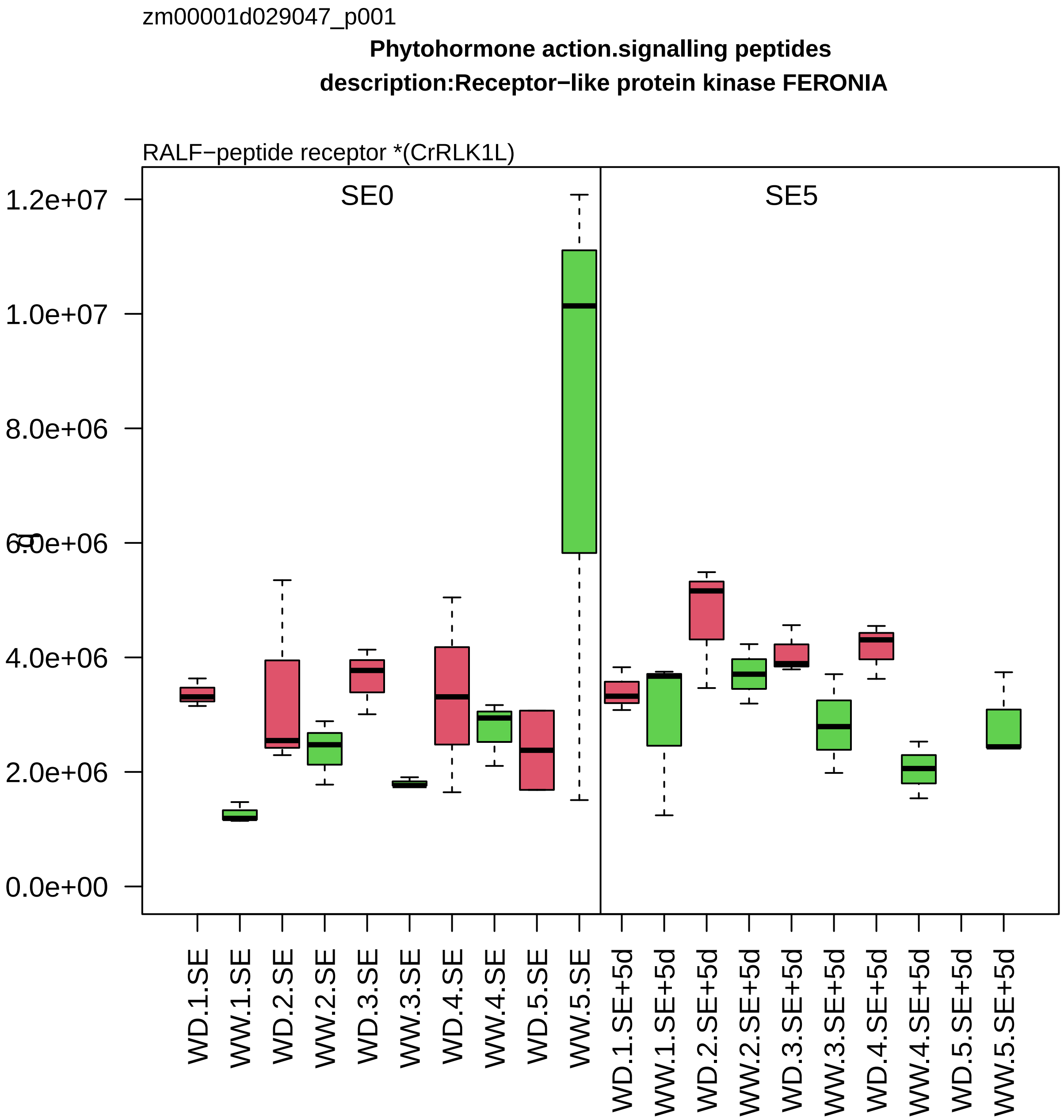

zm00001d029047_p001
Phytohormone action.signalling peptides
description:Receptor−like protein kinase FERONIA
RALF−peptide receptor *(CrRLK1L)
SE0
SE5
1.2e+07
1.0e+07
8.0e+06
6.0e+06
4.0e+06
2.0e+06
0.0e+00
WD.1.SE
WW.1.SE
WD.2.SE
WW.2.SE
WD.3.SE
WW.3.SE
WD.4.SE
WW.4.SE
WD.5.SE
WW.5.SE
WD.1.SE+5d
WW.1.SE+5d
WD.2.SE+5d
WW.2.SE+5d
WD.3.SE+5d
WW.3.SE+5d
WD.4.SE+5d
WW.4.SE+5d
WD.5.SE+5d
WW.5.SE+5d

zm00001d029084_p001

**Cellular respiration.tricarboxylic acid cycle**
**description:Fumarate hydratase 1 mitochondrial**

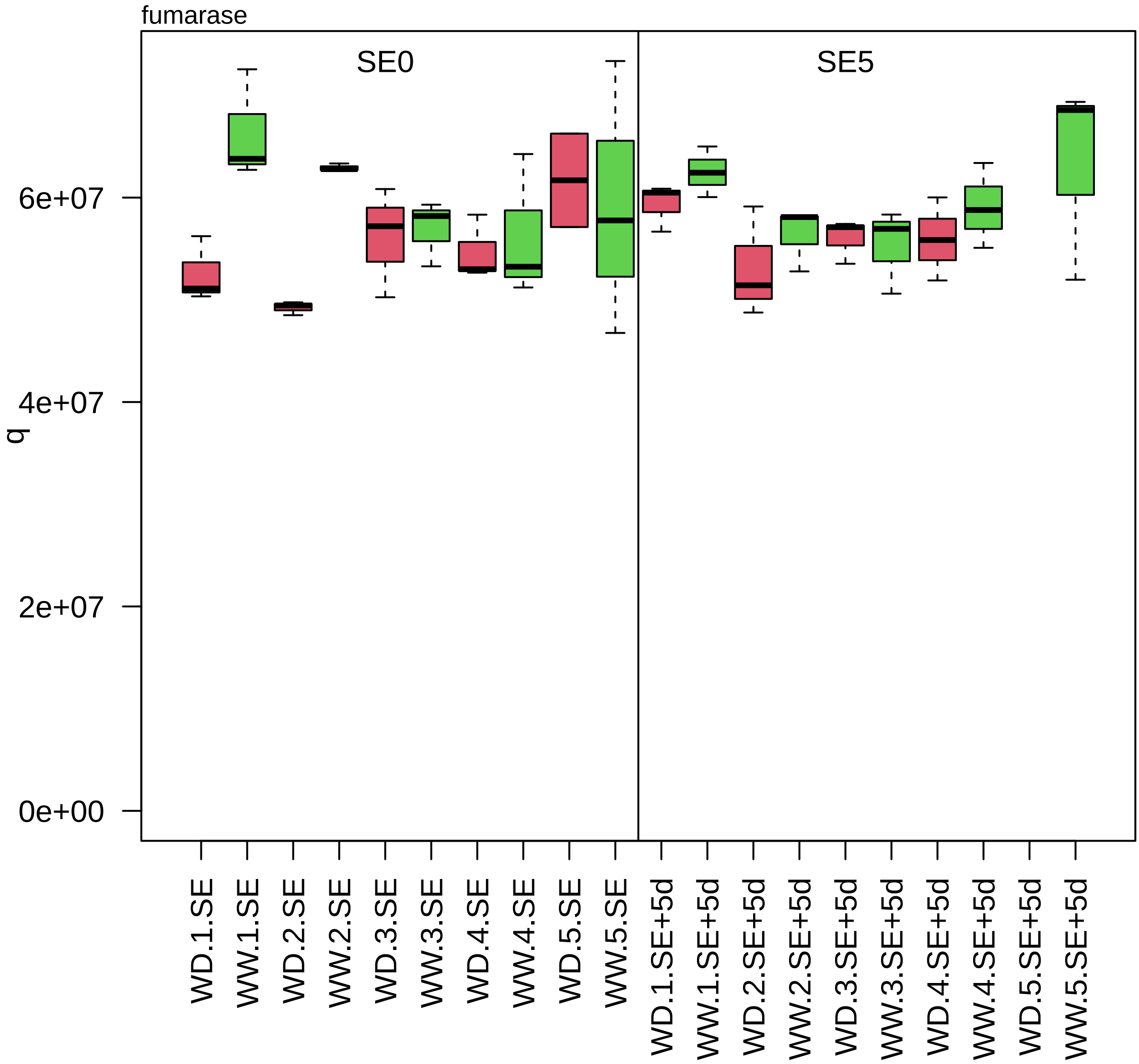

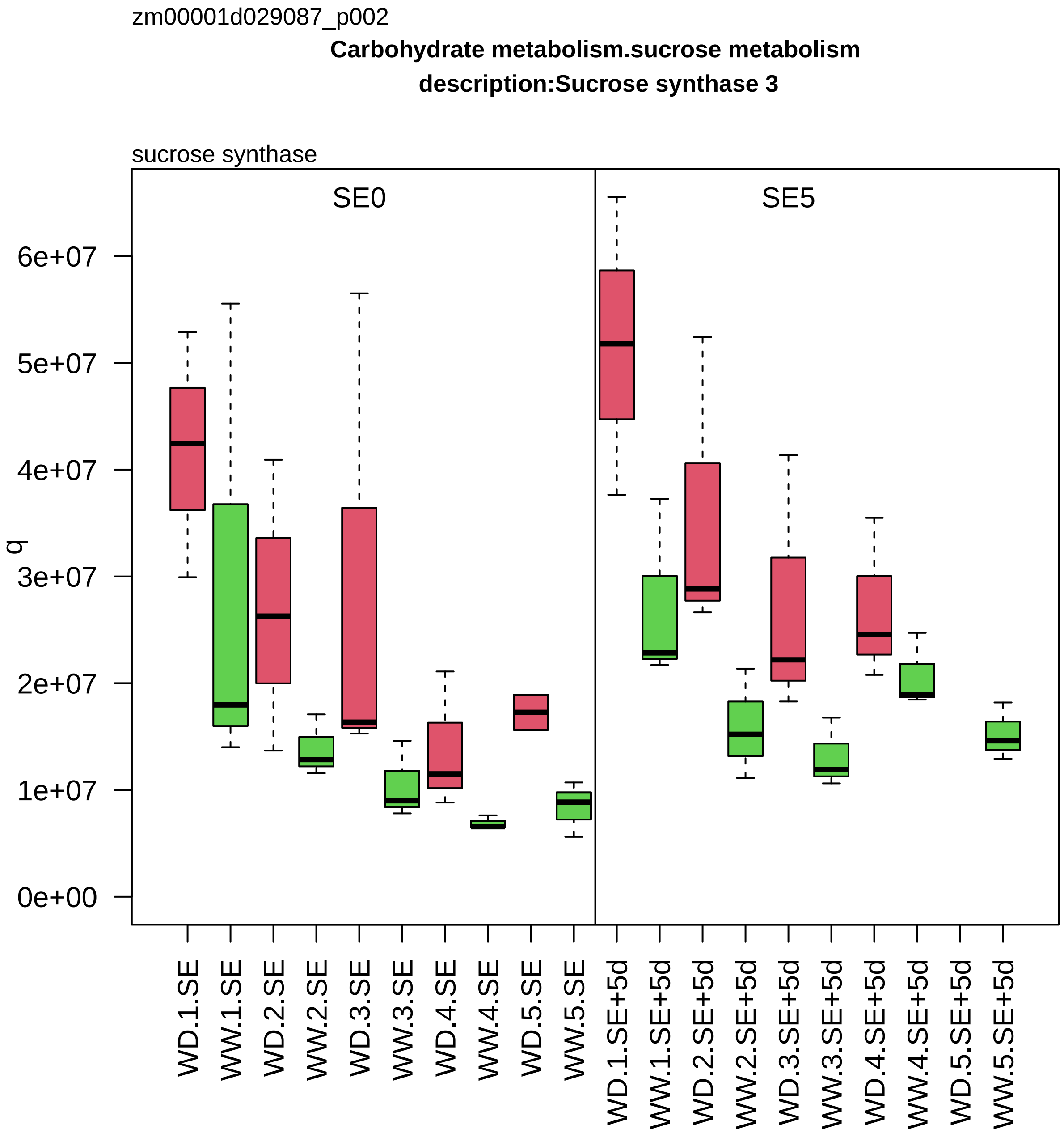

zm00001d029087_p002
Carbohydrate metabolism.sucrose metabolism
description:Sucrose synthase 3
sucrose synthase
SE0
SE5
q
0e+00
1e+07
2e+07
3e+07
4e+07
5e+07
6e+07
WD.1.SE
WW.1.SE
WD.2.SE
WW.2.SE
WD.3.SE
WW.3.SE
WD.4.SE
WW.4.SE
WD.5.SE
WW.5.SE
WD.1.SE+5d
WW.1.SE+5d
WD.2.SE+5d
WW.2.SE+5d
WD.3.SE+5d
WW.3.SE+5d
WD.4.SE+5d
WW.4.SE+5d
WD.5.SE+5d
WW.5.SE+5d

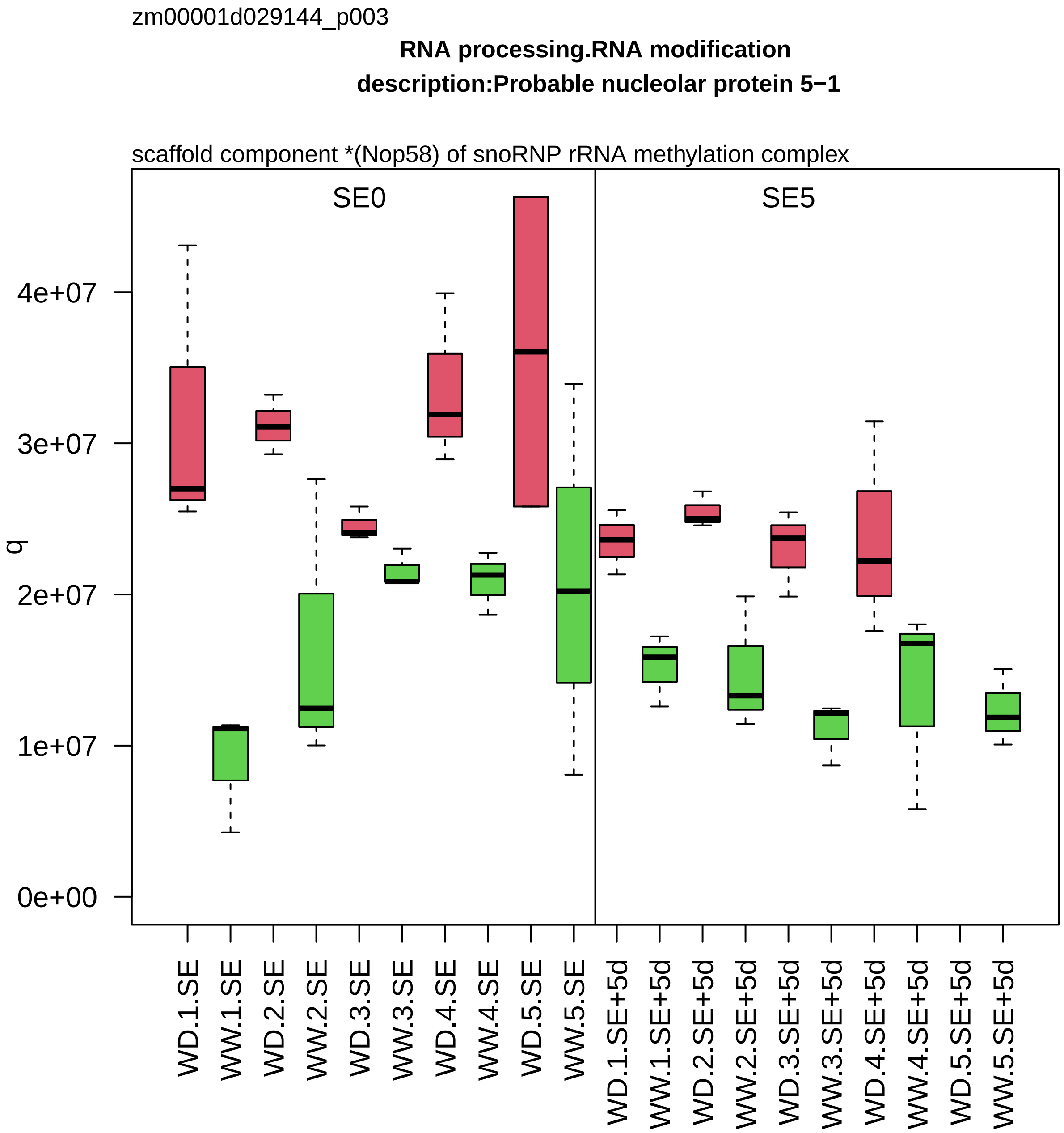
zm00001d029144_p003
RNA processing.RNA modification
description:Probable nucleolar protein 5−1
scaffold component *(Nop58) of snoRNP rRNA methylation complex
SE0
SE5
q
4e+07
3e+07
2e+07
1e+07
0e+00
WD.1.SE
WW.1.SE
WD.2.SE
WW.2.SE
WD.3.SE
WW.3.SE
WD.4.SE
WW.4.SE
WD.5.SE
WW.5.SE
WD.1.SE+5d
WW.1.SE+5d
WD.2.SE+5d
WW.2.SE+5d
WD.3.SE+5d
WW.3.SE+5d
WD.4.SE+5d
WW.4.SE+5d
WD.5.SE+5d
WW.5.SE+5d

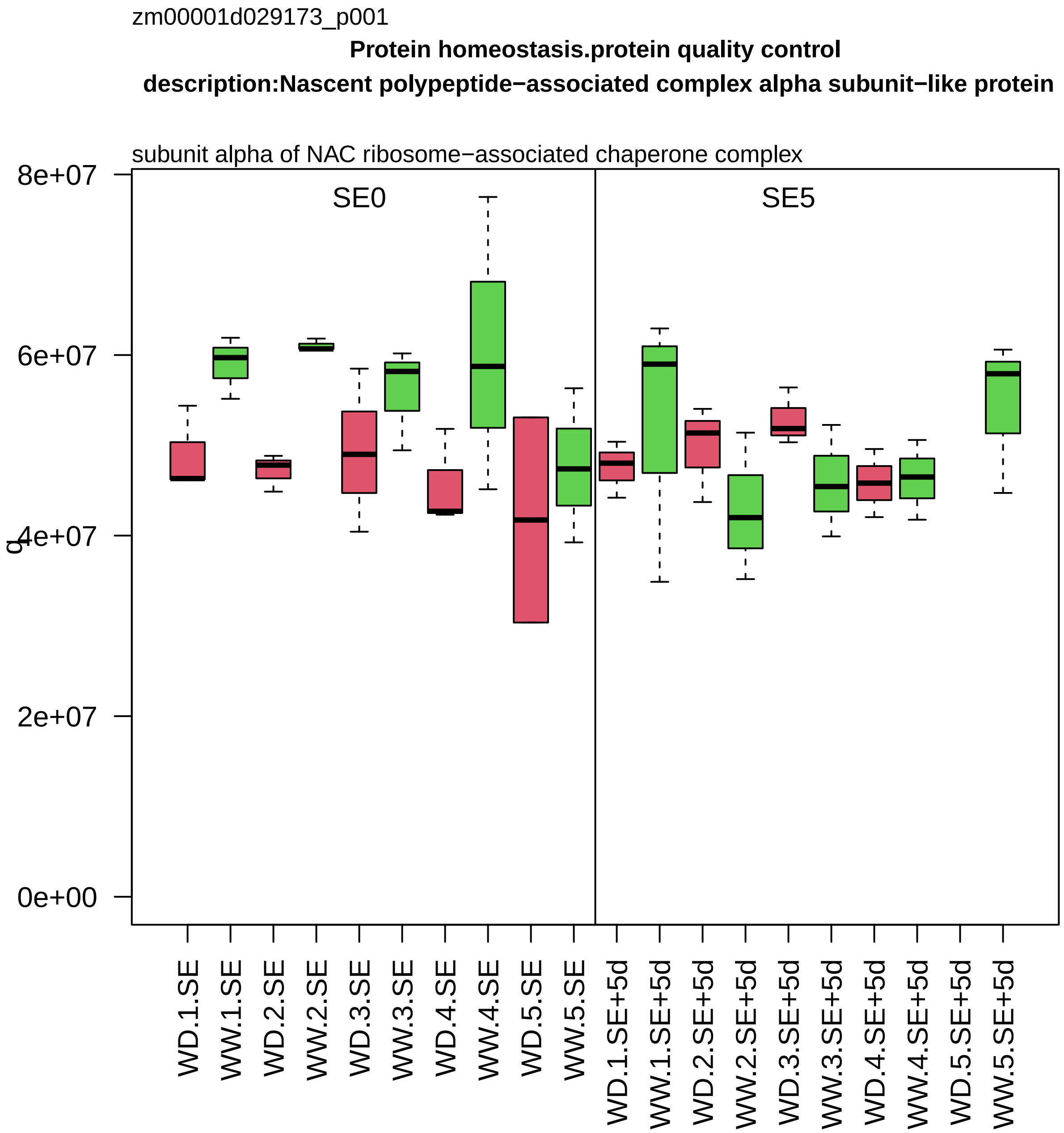

zm00001d029173_p001
Protein homeostasis.protein quality control
description:Nascent polypeptide−associated complex alpha subunit−like protein
subunit alpha of NAC ribosome−associated chaperone complex
SE0
SE5
8e+07
6e+07
4e+07
2e+07
0e+00
q
WD.1.SE
WW.1.SE
WD.2.SE
WW.2.SE
WD.3.SE
WW.3.SE
WD.4.SE
WW.4.SE
WD.5.SE
WW.5.SE
WD.1.SE+5d
WW.1.SE+5d
WD.2.SE+5d
WW.2.SE+5d
WD.3.SE+5d
WW.3.SE+5d
WD.4.SE+5d
WW.4.SE+5d
WD.5.SE+5d
WW.5.SE+5d

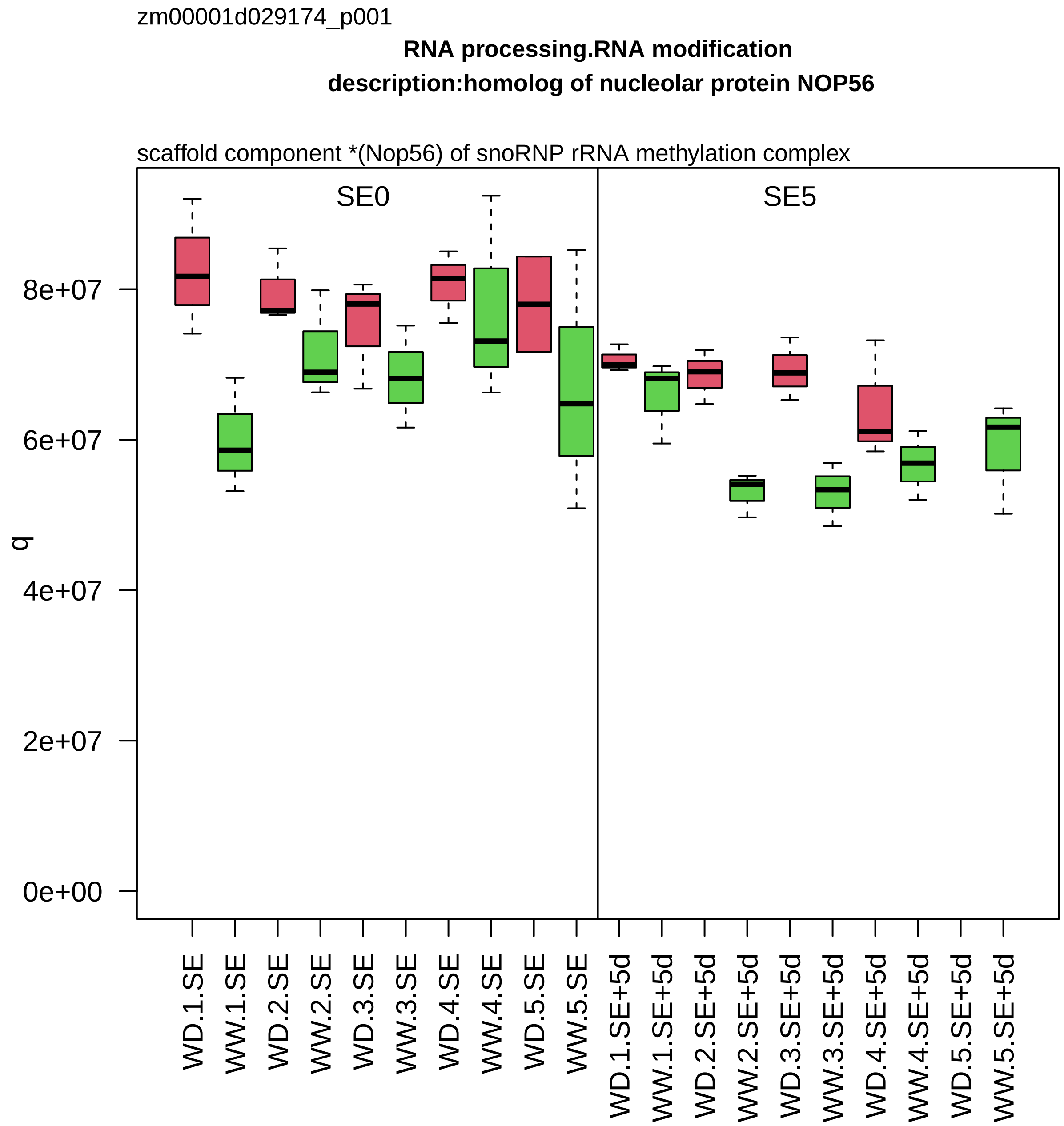

zm00001d029174_p001
RNA processing.RNA modification
description:homolog of nucleolar protein NOP56
scaffold component *(Nop56) of snoRNP rRNA methylation complex
SE0
SE5
q
8e+07
6e+07
4e+07
2e+07
0e+00
WD.1.SE
WW.1.SE
WD.2.SE
WW.2.SE
WD.3.SE
WW.3.SE
WD.4.SE
WW.4.SE
WD.5.SE
WW.5.SE
WD.1.SE+5d
WW.1.SE+5d
WD.2.SE+5d
WW.2.SE+5d
WD.3.SE+5d
WW.3.SE+5d
WD.4.SE+5d
WW.4.SE+5d
WD.5.SE+5d
WW.5.SE+5d

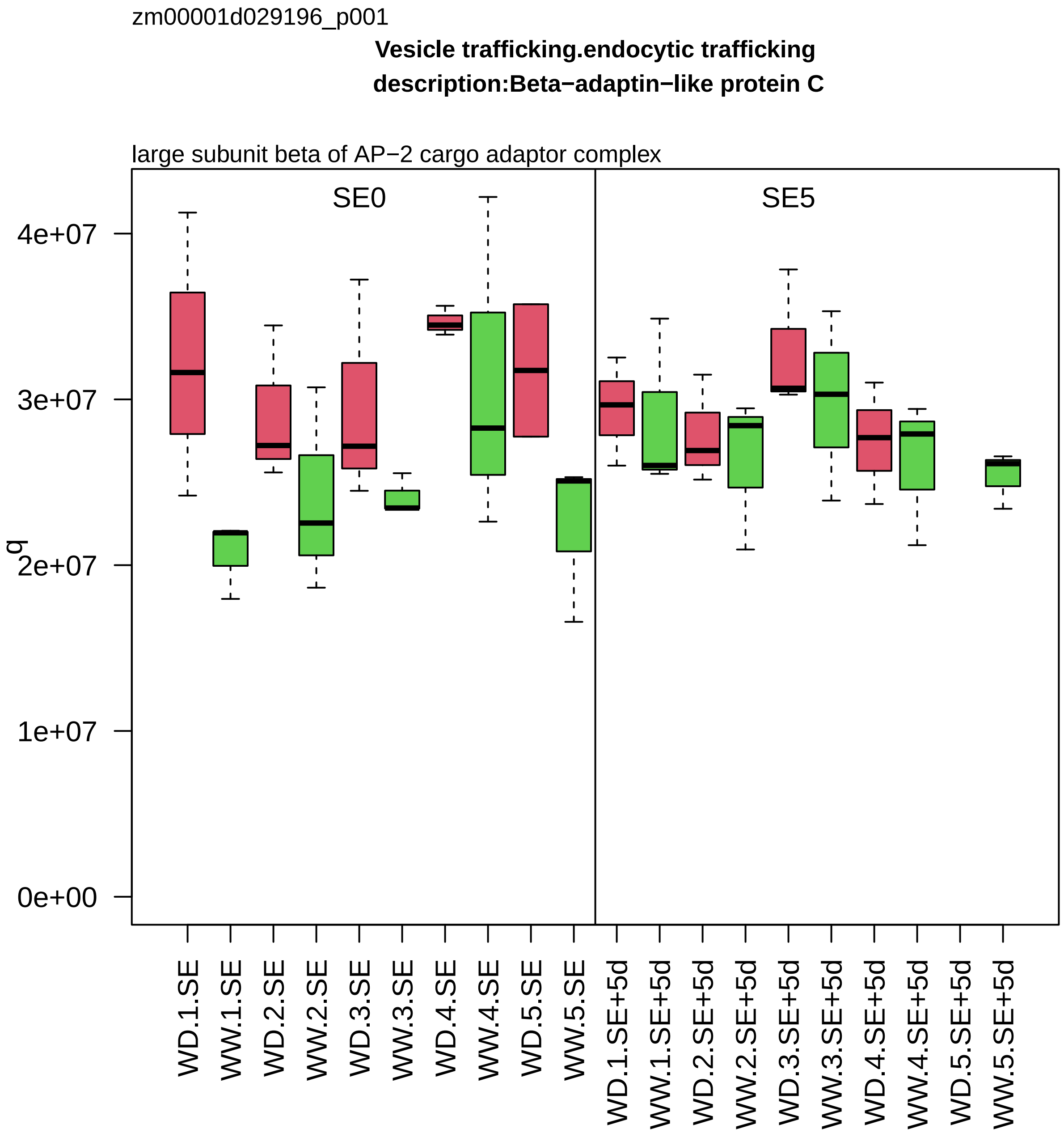

zm00001d029196_p001
Vesicle trafficking.endocytic trafficking
description:Beta−adaptin−like protein C
large subunit beta of AP−2 cargo adaptor complex
SE0
SE5
q
4e+07
3e+07
2e+07
1e+07
0e+00
WD.1.SE
WW.1.SE
WD.2.SE
WW.2.SE
WD.3.SE
WW.3.SE
WD.4.SE
WW.4.SE
WD.5.SE
WW.5.SE
WD.1.SE+5d
WW.1.SE+5d
WD.2.SE+5d
WW.2.SE+5d
WD.3.SE+5d
WW.3.SE+5d
WD.4.SE+5d
WW.4.SE+5d
WD.5.SE+5d
WW.5.SE+5d

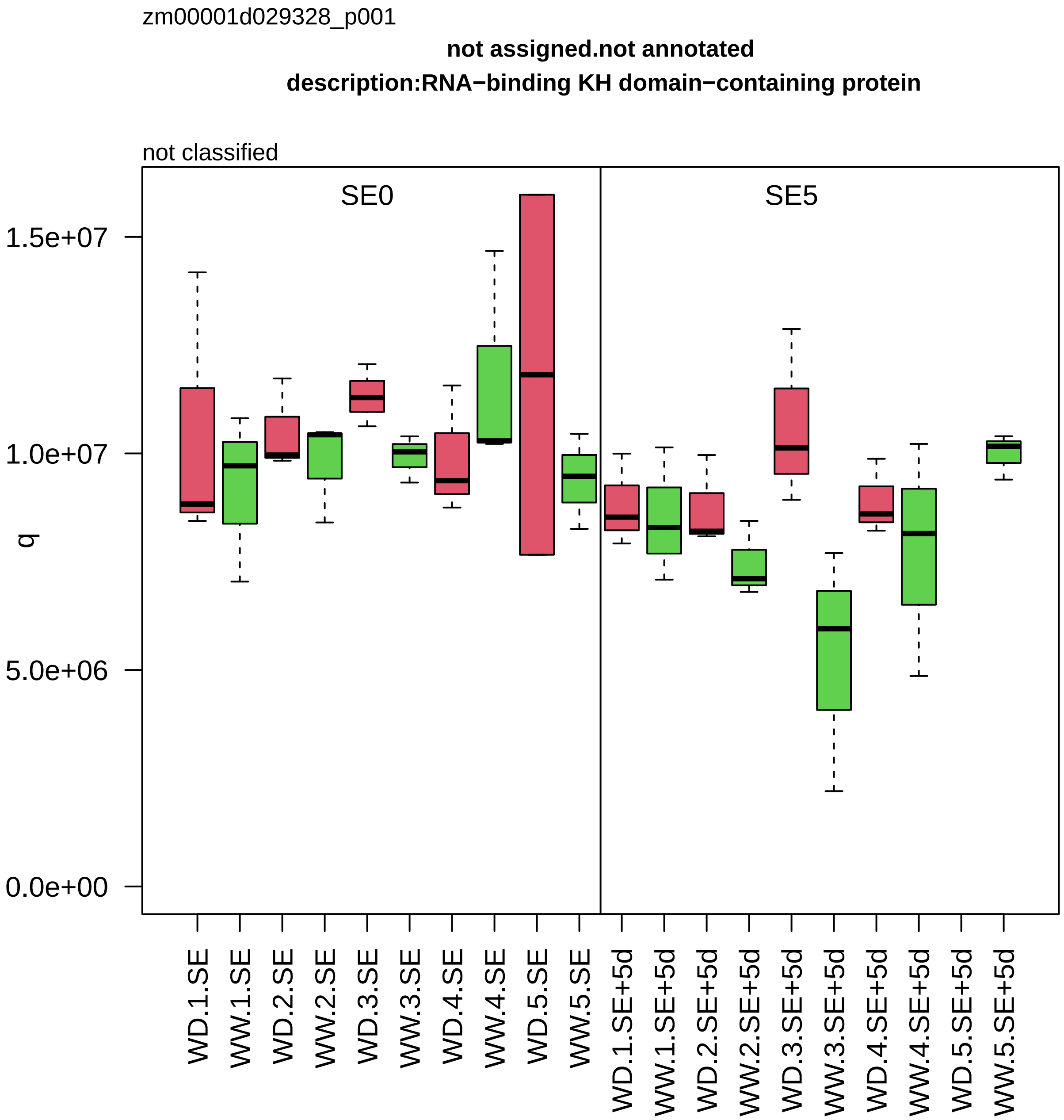

zm00001d029328_p001
not assigned.not annotated
description:RNA−binding KH domain−containing protein
not classified
SE0
SE5
1.5e+07
1.0e+07
5.0e+06
0.0e+00
q
WD.1.SE
WW.1.SE
WD.2.SE
WW.2.SE
WD.3.SE
WW.3.SE
WD.4.SE
WW.4.SE
WD.5.SE
WW.5.SE
WD.1.SE+5d
WW.1.SE+5d
WD.2.SE+5d
WW.2.SE+5d
WD.3.SE+5d
WW.3.SE+5d
WD.4.SE+5d
WW.4.SE+5d
WD.5.SE+5d
WW.5.SE+5d

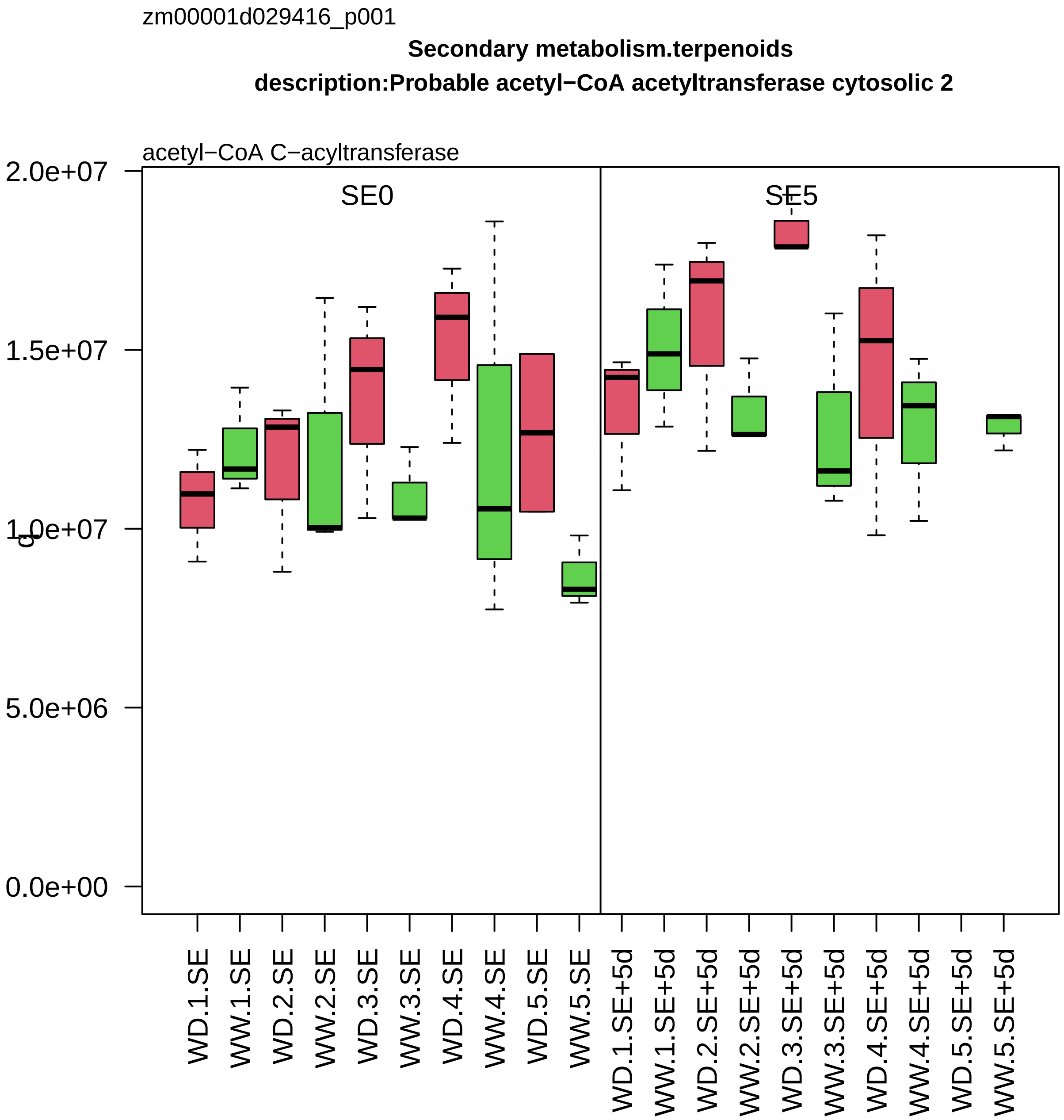

zm00001d029416_p001
Secondary metabolism.terpenoids
description:Probable acetyl−CoA acetyltransferase cytosolic 2
acetyl−CoA C−acyltransferase
SE0
SE5
2.0e+07
1.5e+07
1.0e+07
5.0e+06
0.0e+00
WD.1.SE
WW.1.SE
WD.2.SE
WW.2.SE
WD.3.SE
WW.3.SE
WD.4.SE
WW.4.SE
WD.5.SE
WW.5.SE
WD.1.SE+5d
WW.1.SE+5d
WD.2.SE+5d
WW.2.SE+5d
WD.3.SE+5d
WW.3.SE+5d
WD.4.SE+5d
WW.4.SE+5d
WD.5.SE+5d
WW.5.SE+5d

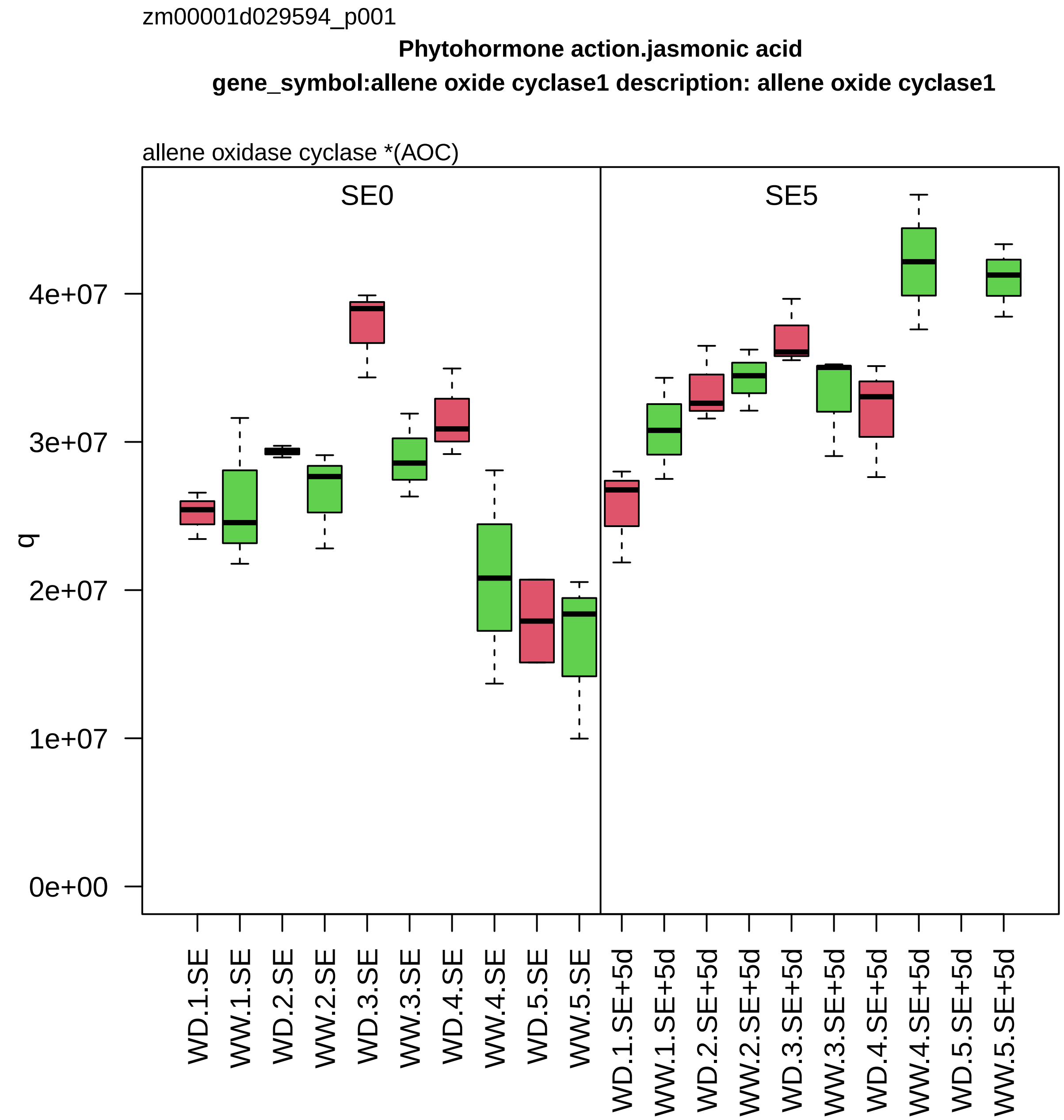

zm00001d029594_p001
Phytohormone action.jasmonic acid
gene_symbol:allene oxide cyclase1 description: allene oxide cyclase1
allene oxidase cyclase *(AOC)
SE0
SE5
q
0e+00
1e+07
2e+07
3e+07
4e+07
WD.1.SE
WW.1.SE
WD.2.SE
WW.2.SE
WD.3.SE
WW.3.SE
WD.4.SE
WW.4.SE
WD.5.SE
WW.5.SE
WD.1.SE+5d
WW.1.SE+5d
WD.2.SE+5d
WW.2.SE+5d
WD.3.SE+5d
WW.3.SE+5d
WD.4.SE+5d
WW.4.SE+5d
WD.5.SE+5d
WW.5.SE+5d

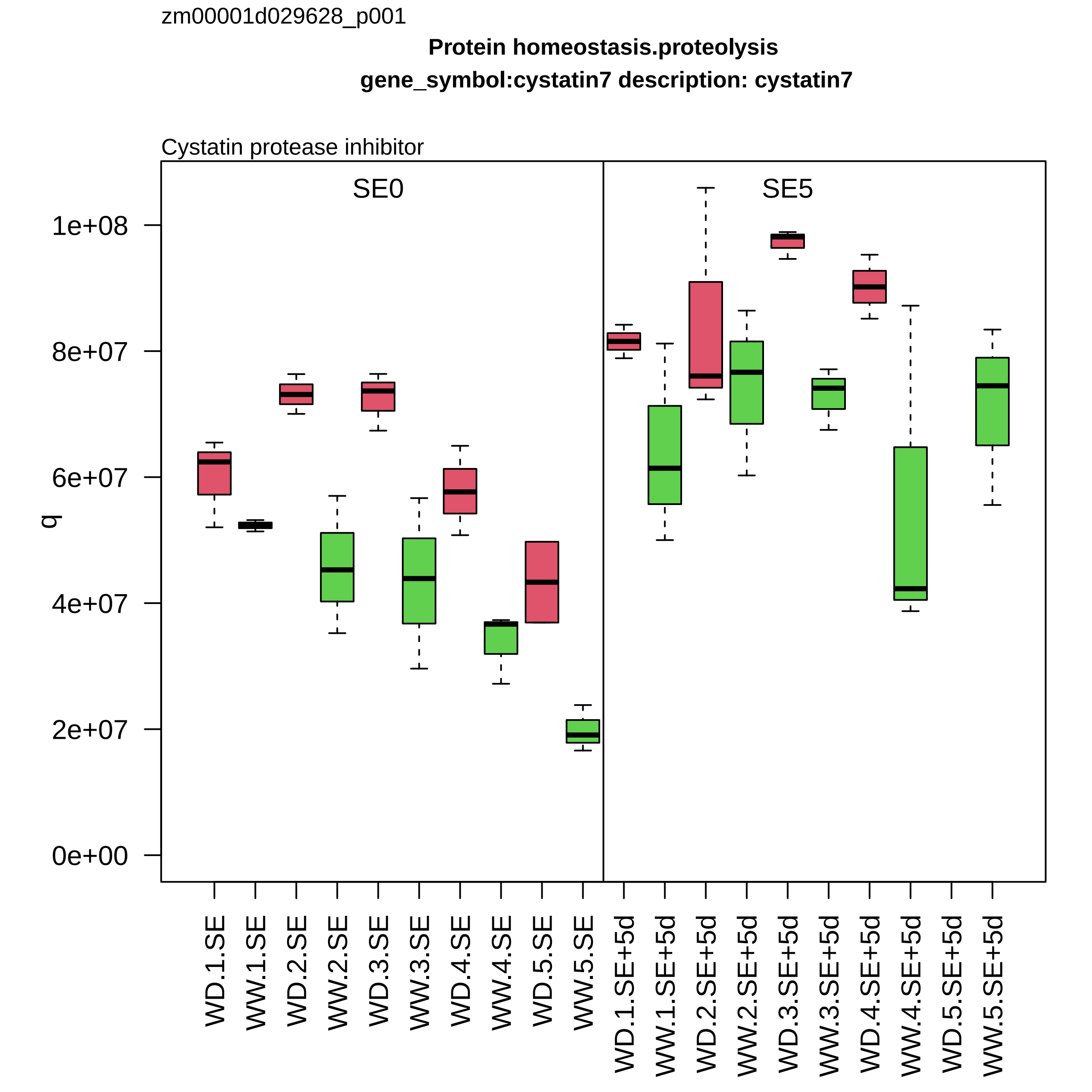
zm00001d029628_p001
Protein homeostasis.proteolysis
gene_symbol:cystatin7 description: cystatin7
Cystatin protease inhibitor
SE0
SE5
q
1e+08
8e+07
6e+07
4e+07
2e+07
0e+00
WD.1.SE
WW.1.SE
WD.2.SE
WW.2.SE
WD.3.SE
WW.3.SE
WD.4.SE
WW.4.SE
WD.5.SE
WW.5.SE
WD.1.SE+5d
WW.1.SE+5d
WD.2.SE+5d
WW.2.SE+5d
WD.3.SE+5d
WW.3.SE+5d
WD.4.SE+5d
WW.4.SE+5d
WD.5.SE+5d
WW.5.SE+5d

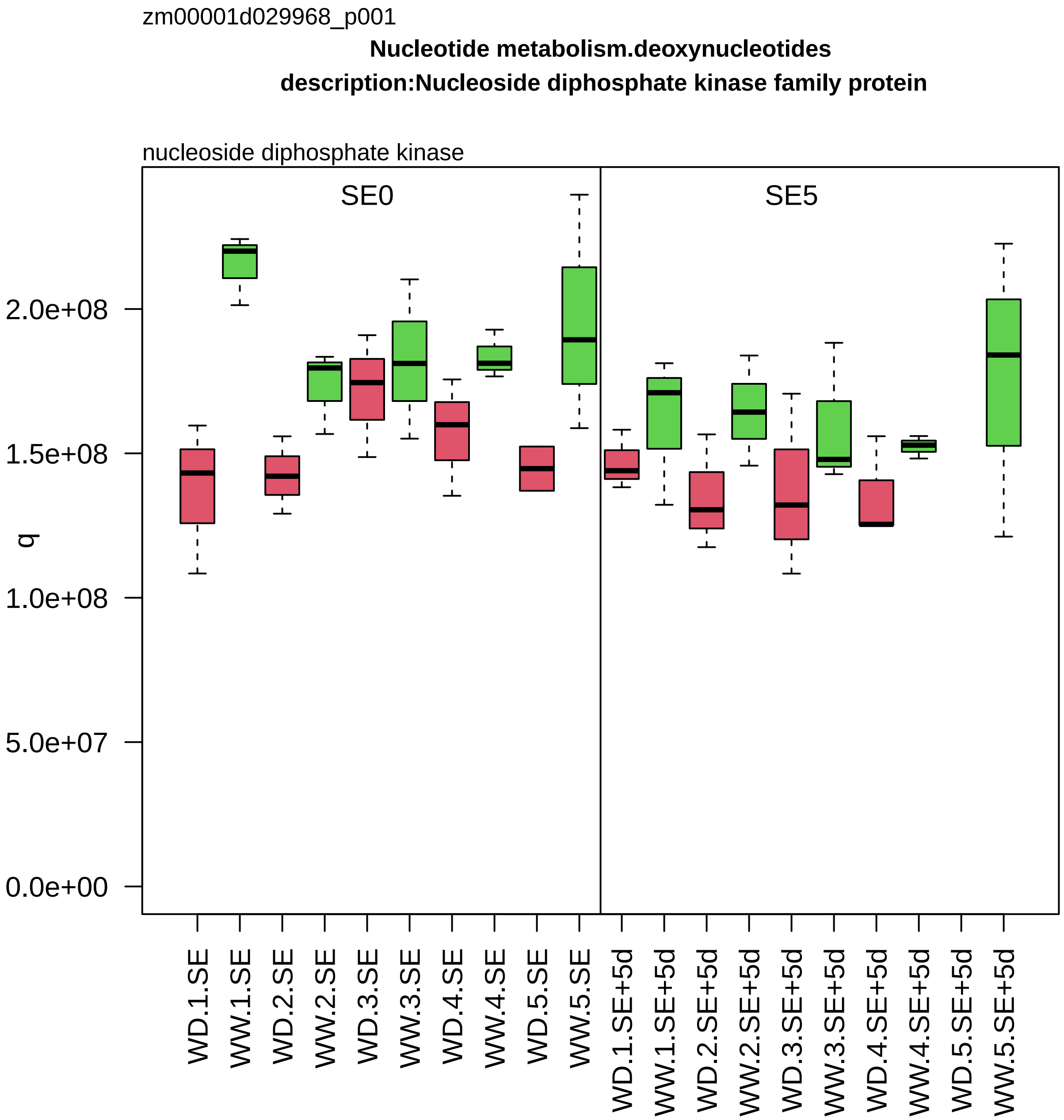

zm00001d029968_p001
Nucleotide metabolism.deoxynucleotides
description:Nucleoside diphosphate kinase family protein
nucleoside diphosphate kinase
SE0
SE5
q
0.0e+00
5.0e+07
1.0e+08
1.5e+08
2.0e+08
WD.1.SE
WW.1.SE
WD.2.SE
WW.2.SE
WD.3.SE
WW.3.SE
WD.4.SE
WW.4.SE
WD.5.SE
WW.5.SE
WD.1.SE+5d
WW.1.SE+5d
WD.2.SE+5d
WW.2.SE+5d
WD.3.SE+5d
WW.3.SE+5d
WD.4.SE+5d
WW.4.SE+5d
WD.5.SE+5d
WW.5.SE+5d

zm00001d030011_p001

**Protein biosynthesis.translation initiation**

**description:Eukaryotic translation initiation factor 3 subunit B**

component *(eIF3b) of eIF3 mRNA−to−PIC binding complex

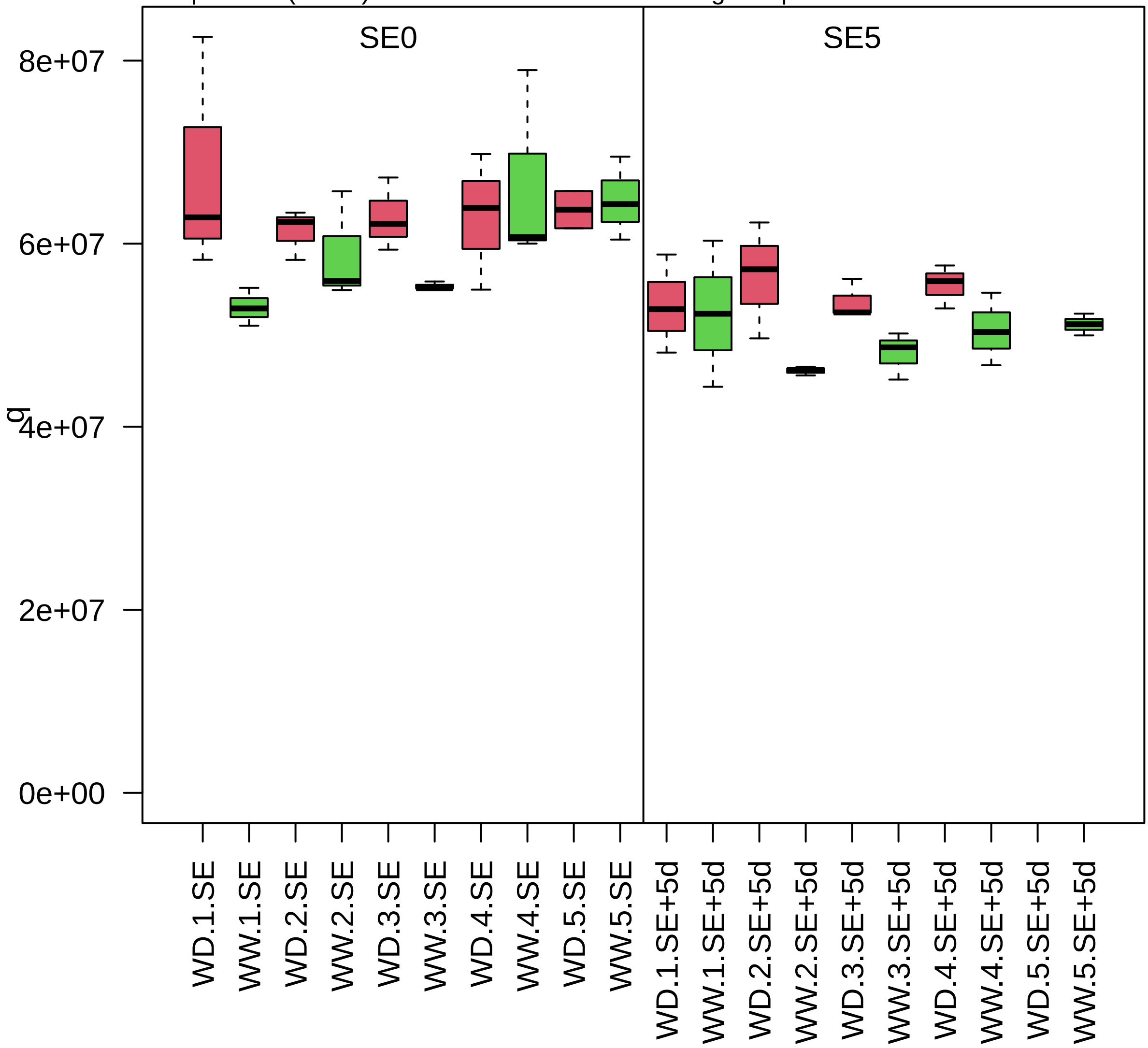

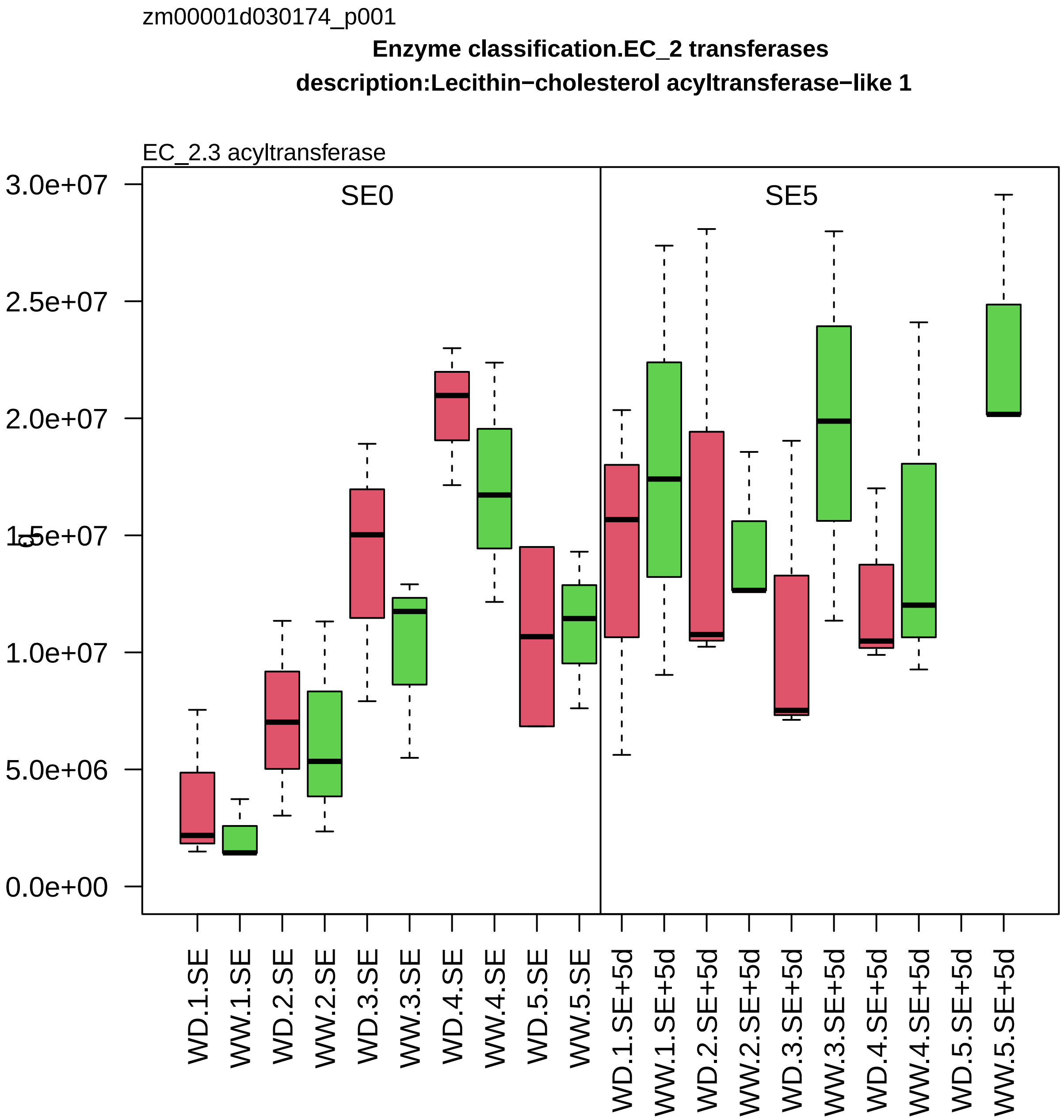

zm00001d030174_p001
Enzyme classification.EC_2 transferases
description:Lecithin−cholesterol acyltransferase−like 1
EC_2.3 acyltransferase
SE0
SE5
3.0e+07
2.5e+07
2.0e+07
1.5e+07
1.0e+07
5.0e+06
0.0e+00
WD.1.SE
WW.1.SE
WD.2.SE
WW.2.SE
WD.3.SE
WW.3.SE
WD.4.SE
WW.4.SE
WD.5.SE
WW.5.SE
WD.1.SE+5d
WW.1.SE+5d
WD.2.SE+5d
WW.2.SE+5d
WD.3.SE+5d
WW.3.SE+5d
WD.4.SE+5d
WW.4.SE+5d
WD.5.SE+5d
WW.5.SE+5d

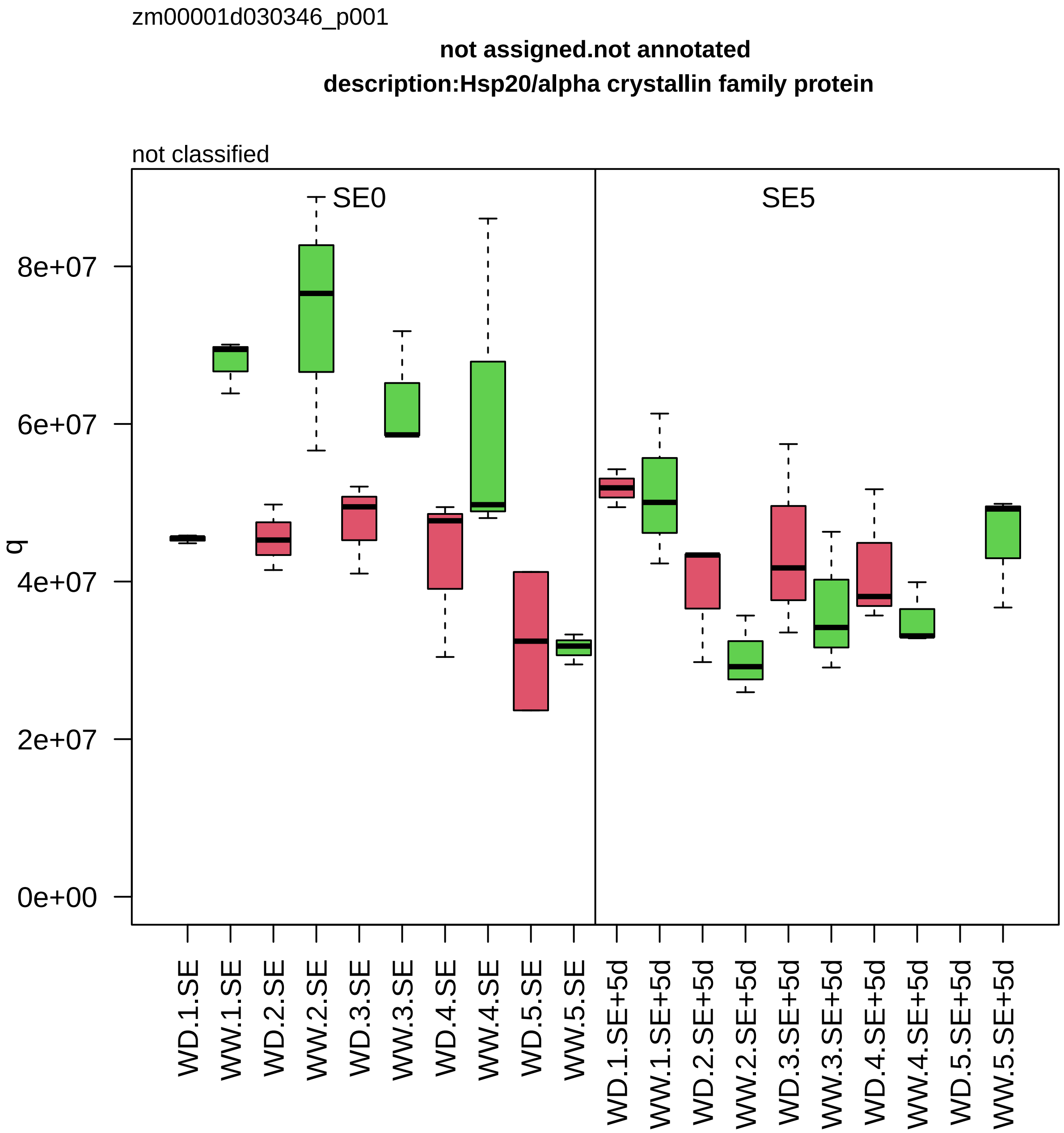

zm00001d030346_p001
not assigned.not annotated
description:Hsp20/alpha crystallin family protein
not classified
SE0
SE5
q
8e+07
6e+07
4e+07
2e+07
0e+00
WD.1.SE
WW.1.SE
WD.2.SE
WW.2.SE
WD.3.SE
WW.3.SE
WD.4.SE
WW.4.SE
WD.5.SE
WW.5.SE
WD.1.SE+5d
WW.1.SE+5d
WD.2.SE+5d
WW.2.SE+5d
WD.3.SE+5d
WW.3.SE+5d
WD.4.SE+5d
WW.4.SE+5d
WD.5.SE+5d
WW.5.SE+5d

zm00001d030373_p003

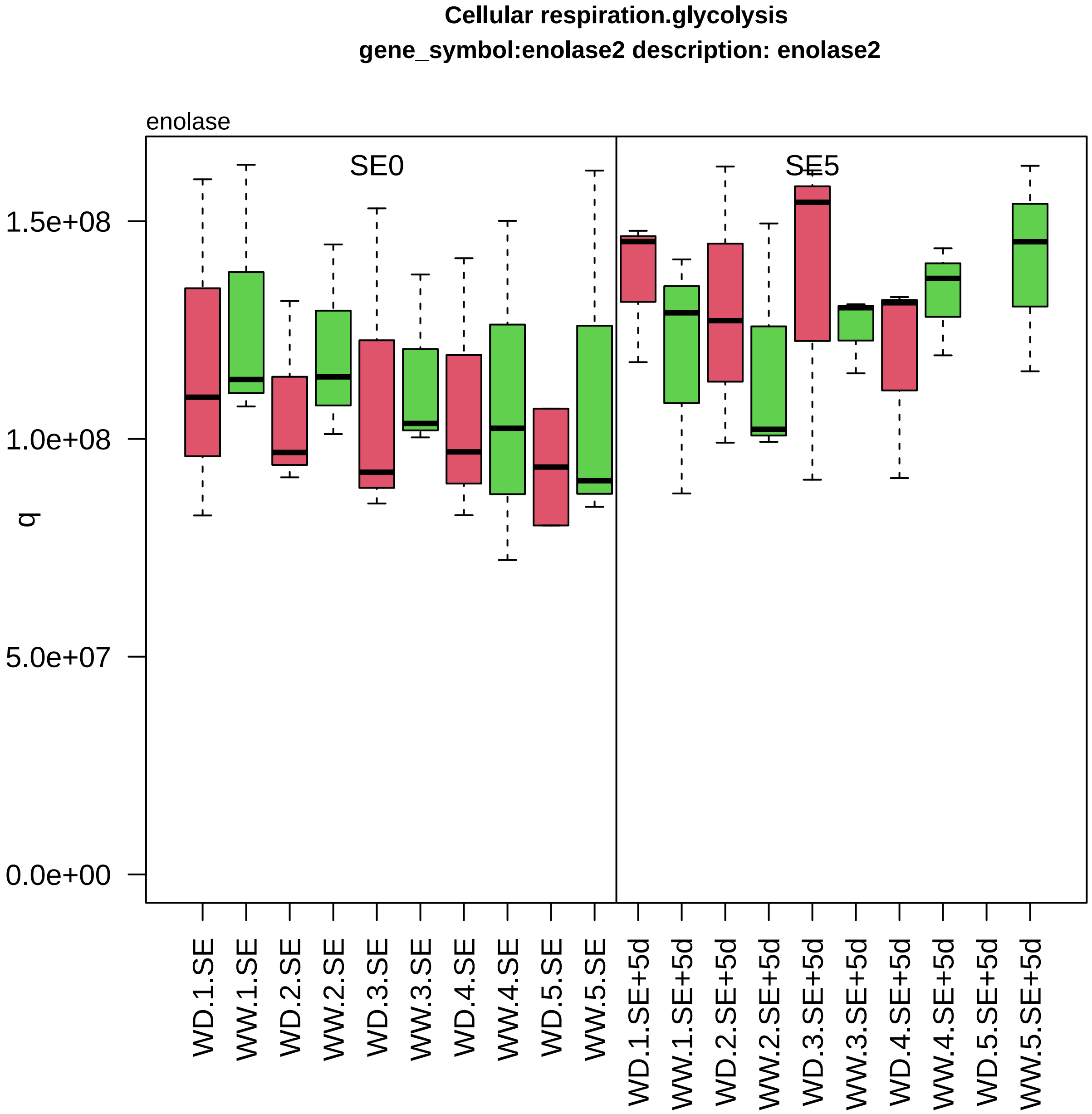

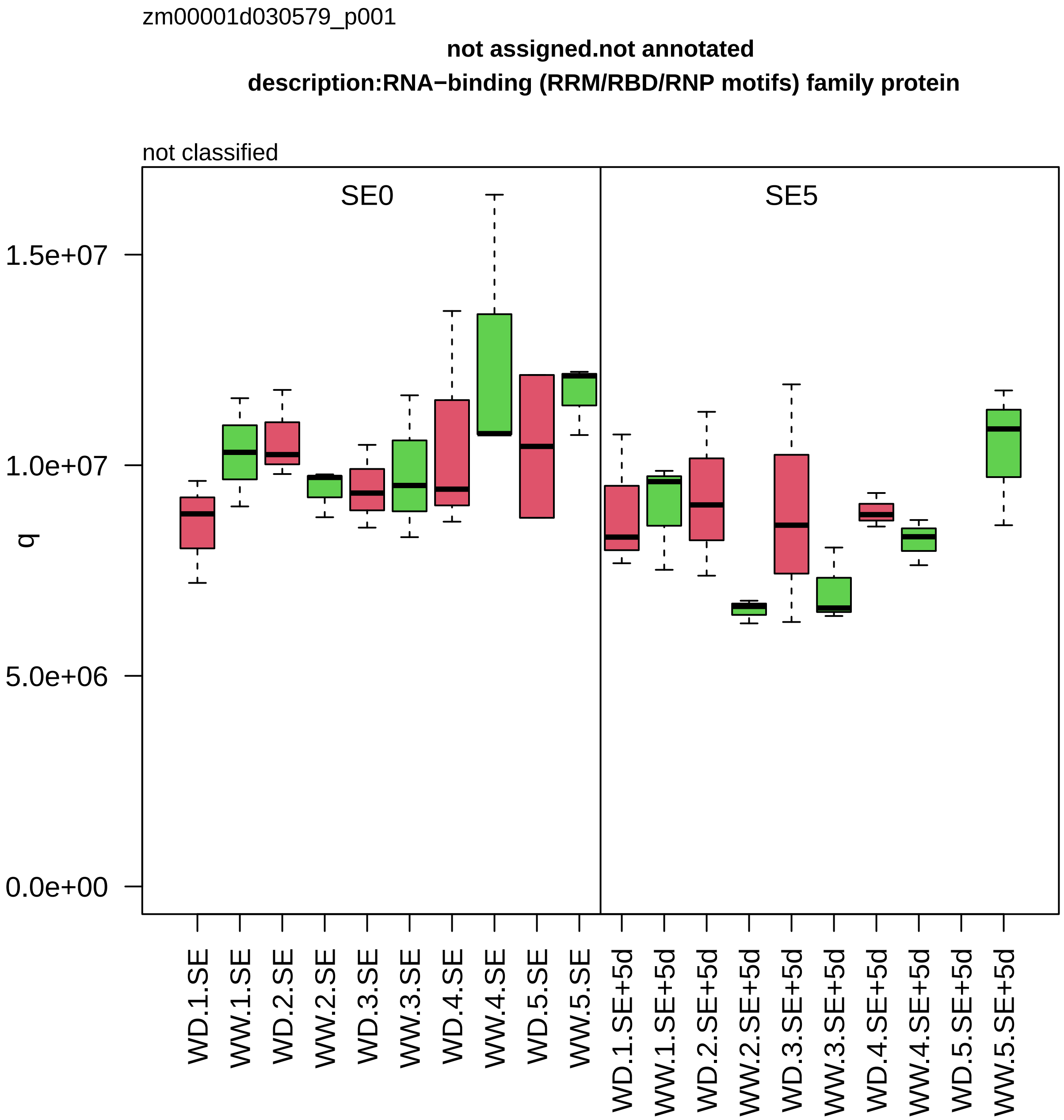

zm00001d030579_p001
not assigned.not annotated
description:RNA−binding (RRM/RBD/RNP motifs) family protein
not classified
SE0
SE5
q
1.5e+07
1.0e+07
5.0e+06
0.0e+00
WD.1.SE
WW.1.SE
WD.2.SE
WW.2.SE
WD.3.SE
WW.3.SE
WD.4.SE
WW.4.SE
WD.5.SE
WW.5.SE
WD.1.SE+5d
WW.1.SE+5d
WD.2.SE+5d
WW.2.SE+5d
WD.3.SE+5d
WW.3.SE+5d
WD.4.SE+5d
WW.4.SE+5d
WD.5.SE+5d
WW.5.SE+5d

zm00001d030614_p002

**Cell division.DNA replication**

**gene_symbol:minichromosome maintenance1 description:DNA replication licensing factor MCM**

component *(MCM7) of MCM replicative DNA helicase complex

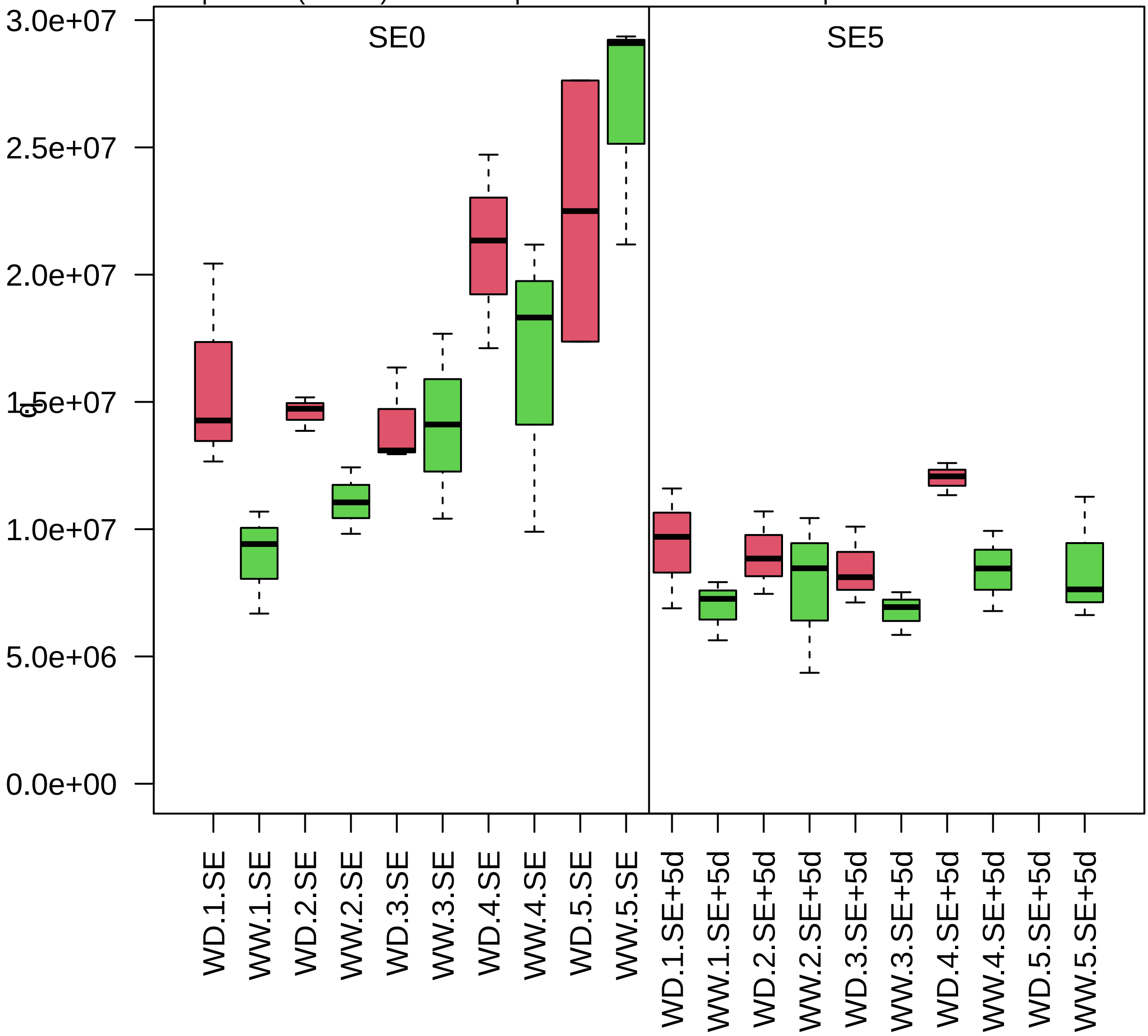

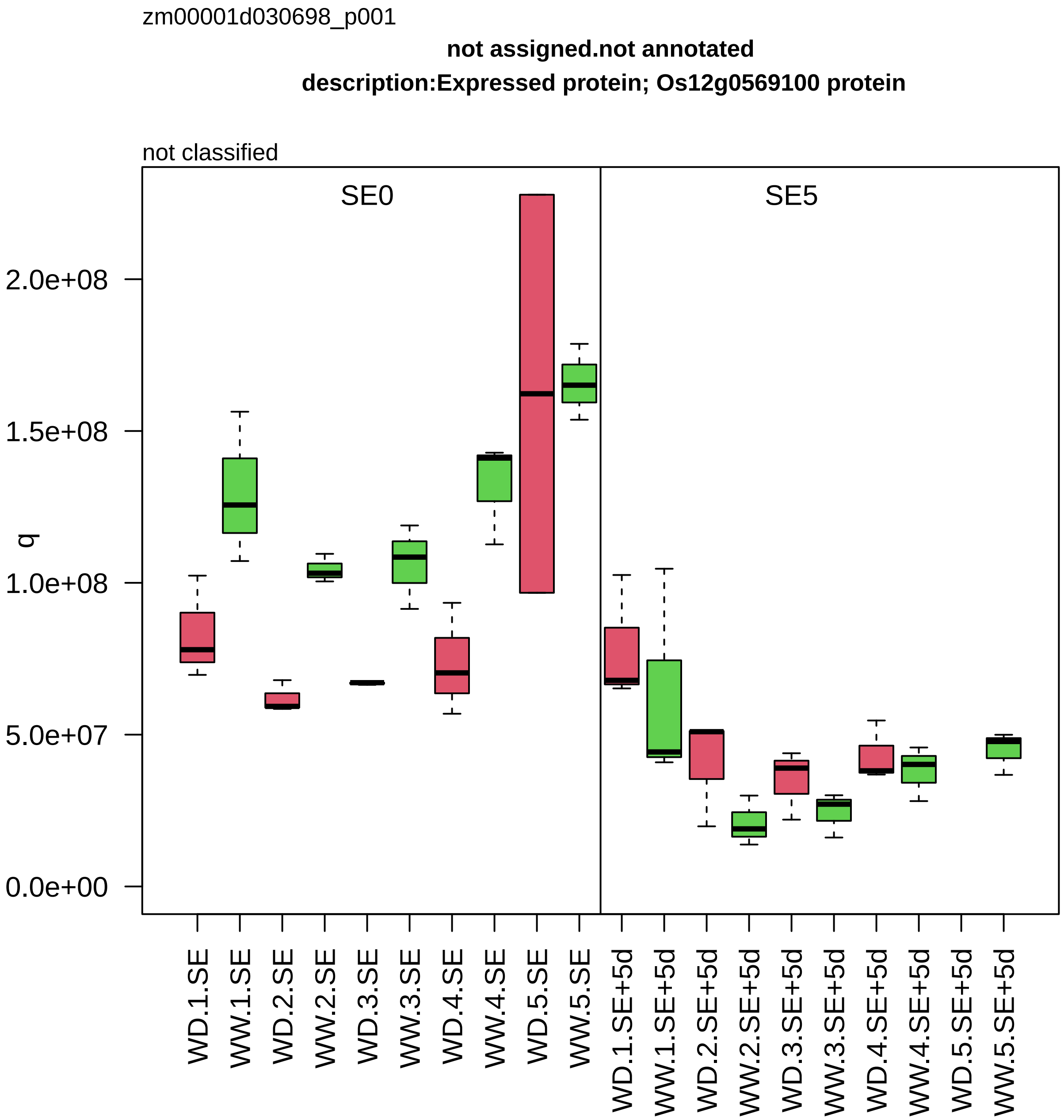

zm00001d030698_p001
not assigned.not annotated
description:Expressed protein; Os12g0569100 protein
not classified
SE0
SE5
q
0.0e+00
5.0e+07
1.0e+08
1.5e+08
2.0e+08
WD.1.SE
WW.1.SE
WD.2.SE
WW.2.SE
WD.3.SE
WW.3.SE
WD.4.SE
WW.4.SE
WD.5.SE
WW.5.SE
WD.1.SE+5d
WW.1.SE+5d
WD.2.SE+5d
WW.2.SE+5d
WD.3.SE+5d
WW.3.SE+5d
WD.4.SE+5d
WW.4.SE+5d
WD.5.SE+5d
WW.5.SE+5d

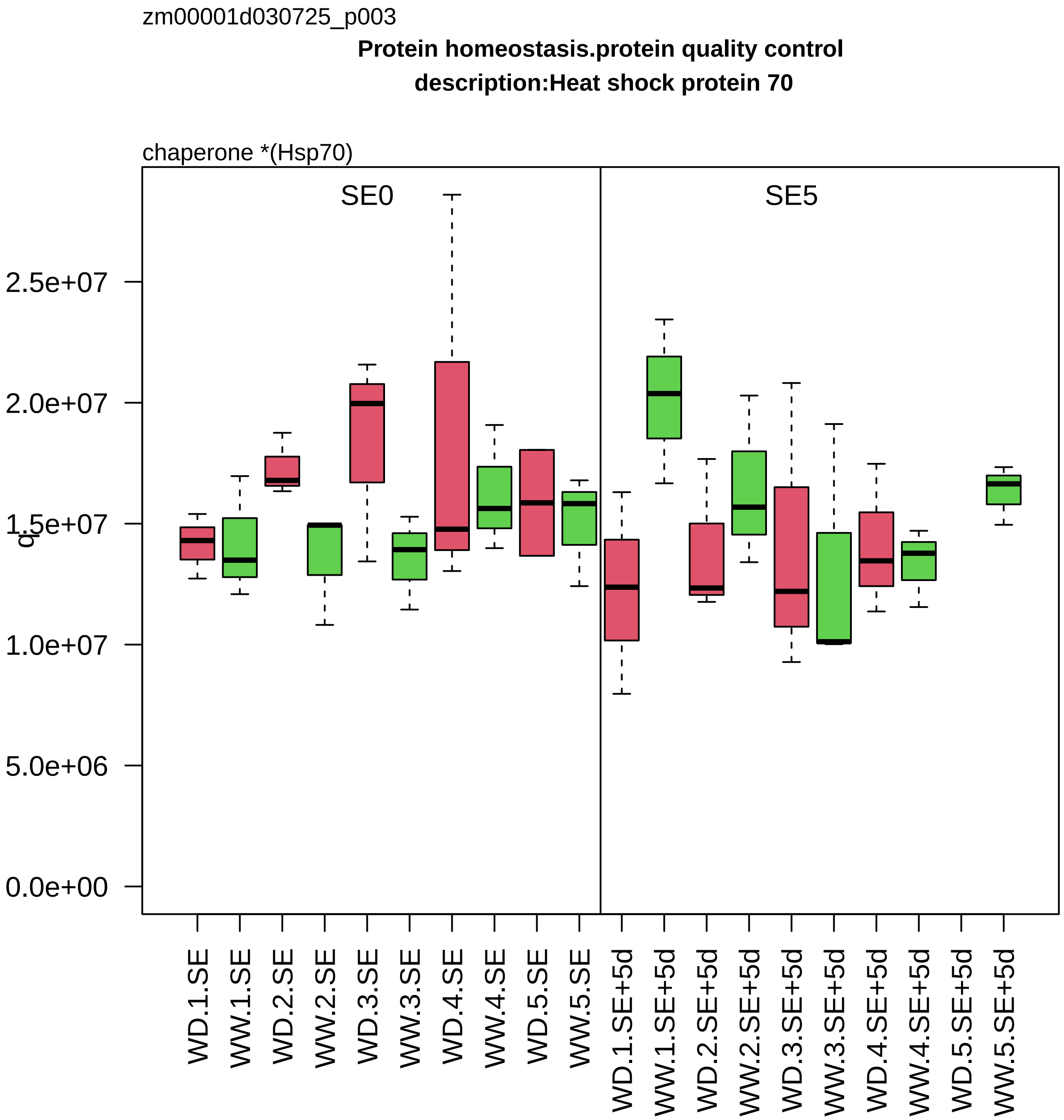
zm00001d030725_p003
Protein homeostasis.protein quality control
description:Heat shock protein 70
chaperone *(Hsp70)
SE0
SE5
2.5e+07
2.0e+07
1.5e+07
1.0e+07
5.0e+06
0.0e+00
WD.1.SE
WW.1.SE
WD.2.SE
WW.2.SE
WD.3.SE
WW.3.SE
WD.4.SE
WW.4.SE
WD.5.SE
WW.5.SE
WD.1.SE+5d
WW.1.SE+5d
WD.2.SE+5d
WW.2.SE+5d
WD.3.SE+5d
WW.3.SE+5d
WD.4.SE+5d
WW.4.SE+5d
WD.5.SE+5d
WW.5.SE+5d

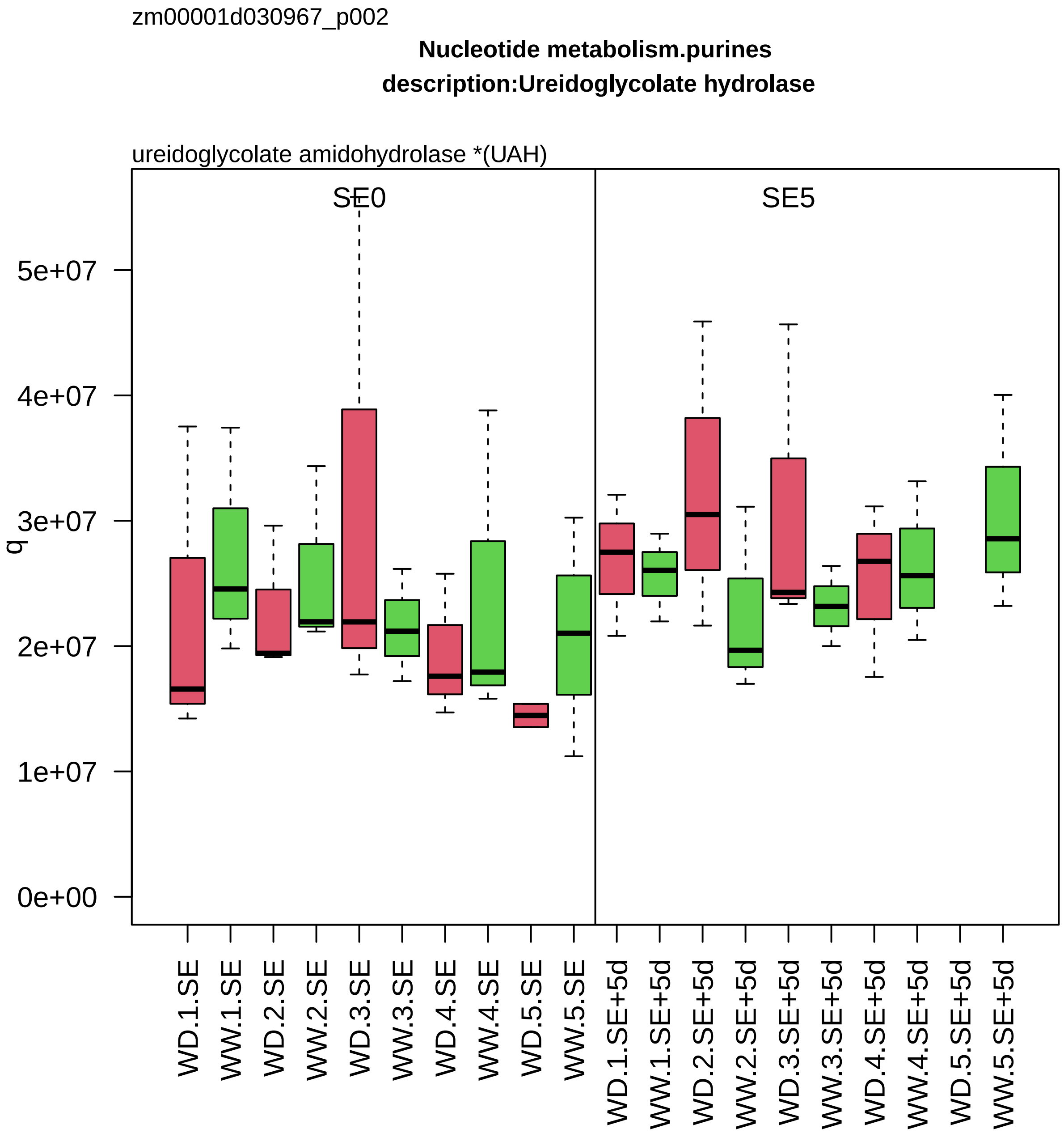

zm00001d030967_p002
Nucleotide metabolism.purines
description:Ureidoglycolate hydrolase
ureidoglycolate amidohydrolase *(UAH)
SE0
SE5
q
0e+00
1e+07
2e+07
3e+07
4e+07
5e+07
WD.1.SE
WW.1.SE
WD.2.SE
WW.2.SE
WD.3.SE
WW.3.SE
WD.4.SE
WW.4.SE
WD.5.SE
WW.5.SE
WD.1.SE+5d
WW.1.SE+5d
WD.2.SE+5d
WW.2.SE+5d
WD.3.SE+5d
WW.3.SE+5d
WD.4.SE+5d
WW.4.SE+5d
WD.5.SE+5d
WW.5.SE+5d

zm00001d030968_p006

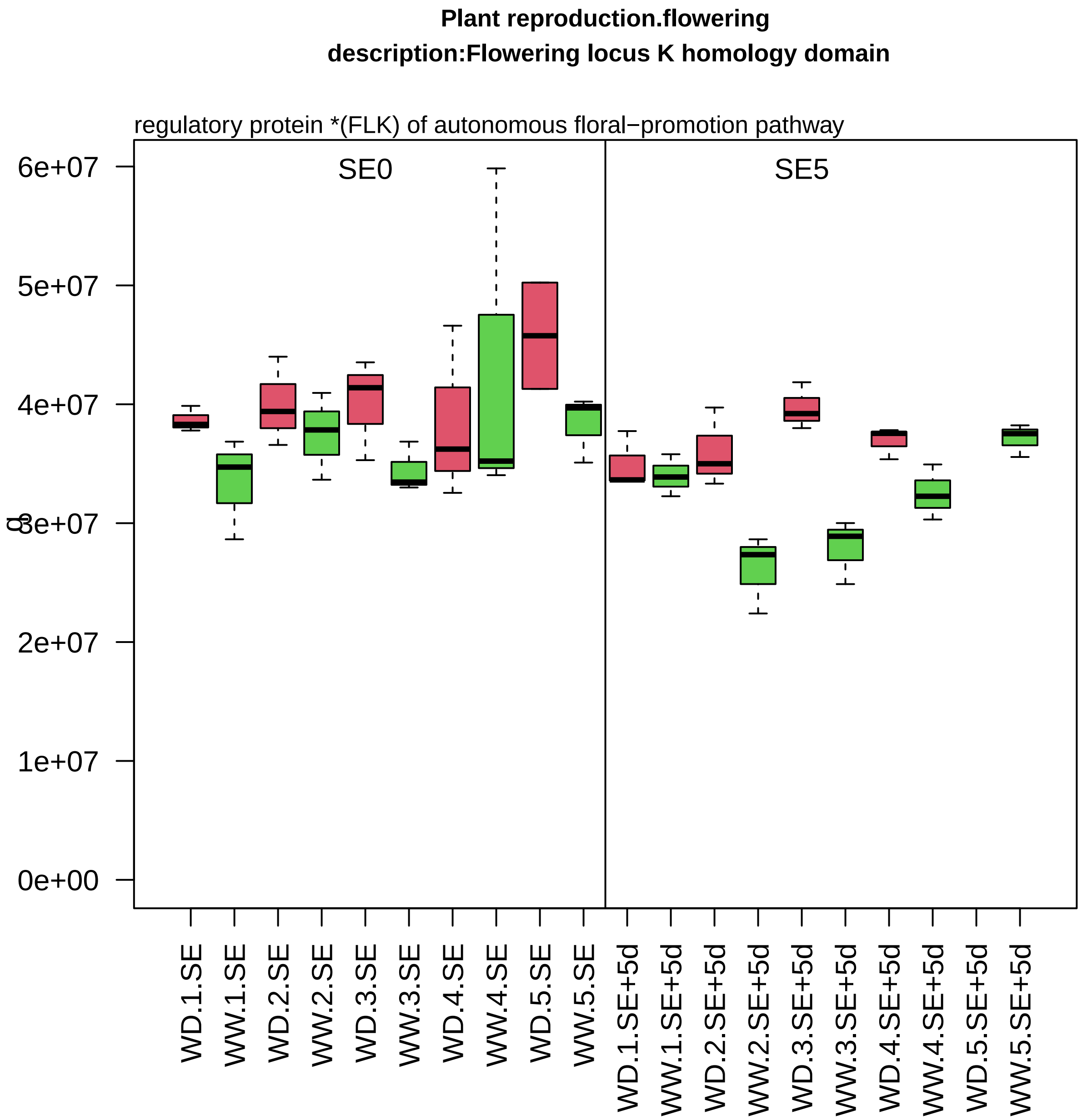

zm00001d031024_p001

**Amino acid metabolism.amino acid degradation**
**description:Sulfurtransferase**

mercaptopyruvate sulfurtransferase

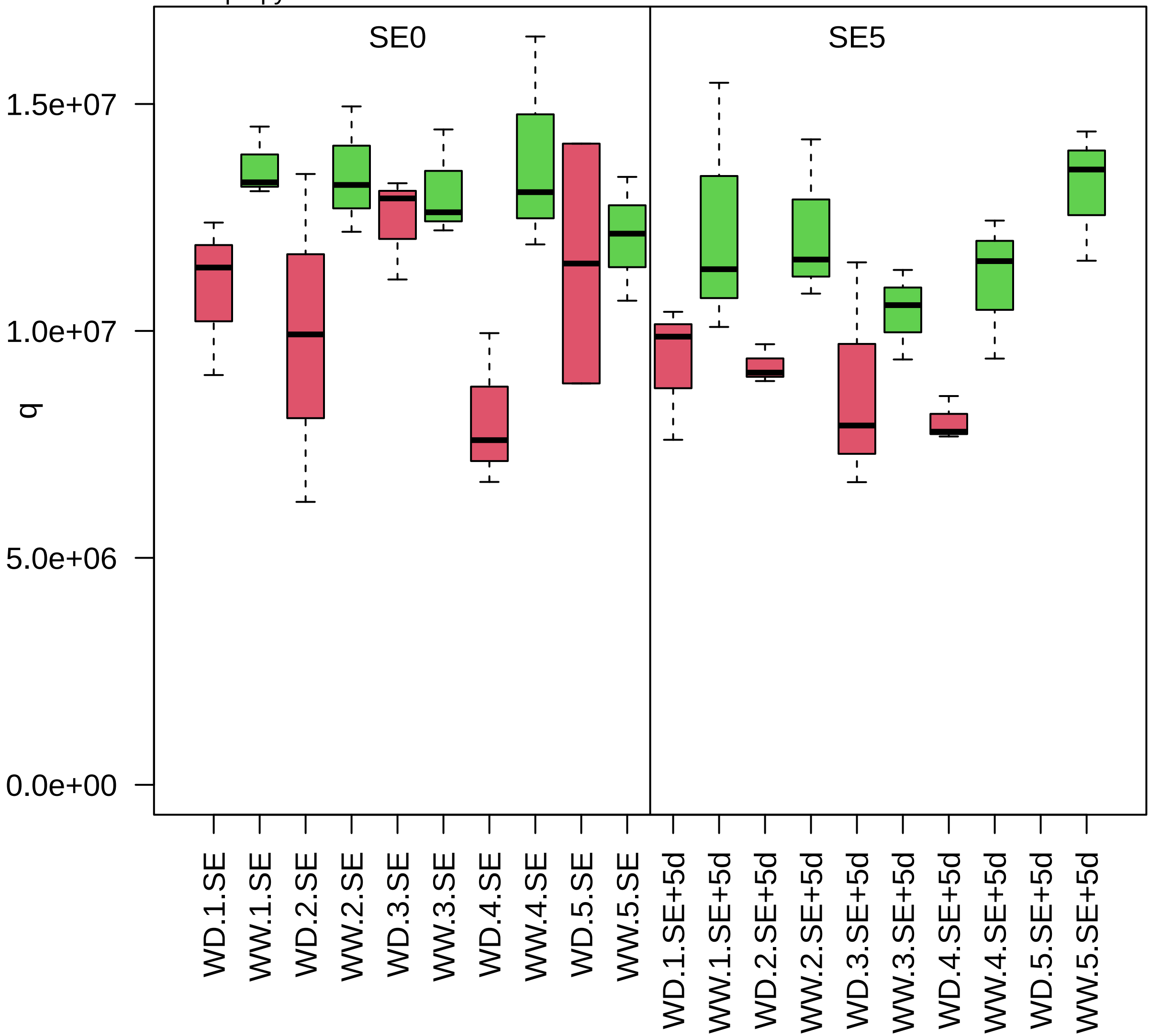

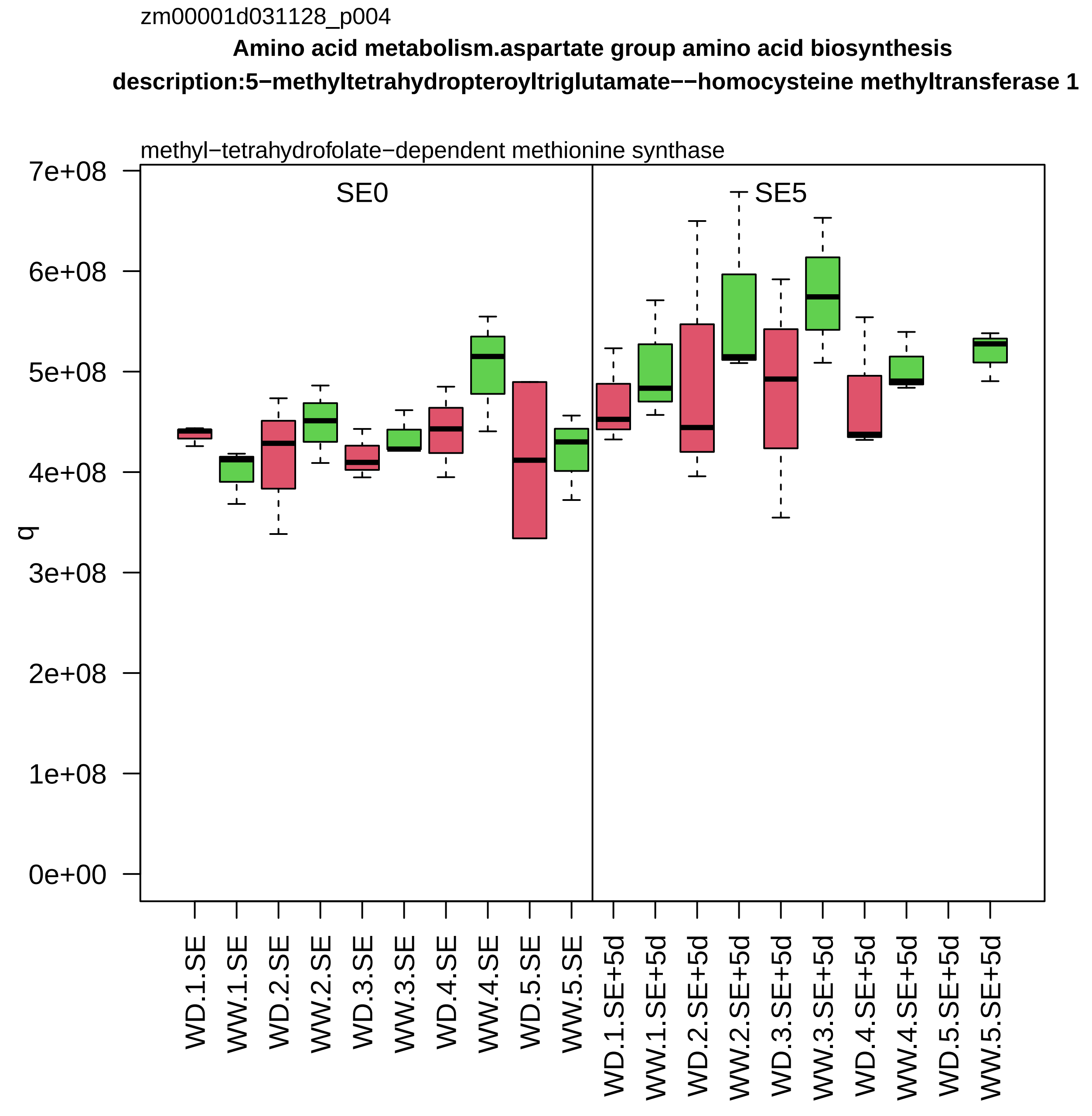
zm00001d031128_p004
Amino acid metabolism.aspartate group amino acid biosynthesis
description:5−methyltetrahydropteroyltriglutamate−−homocysteine methyltransferase 1
methyl−tetrahydrofolate−dependent methionine synthase
SE0
SE5
q
0e+00
1e+08
2e+08
3e+08
4e+08
5e+08
6e+08
7e+08
WD.1.SE
WW.1.SE
WD.2.SE
WW.2.SE
WD.3.SE
WW.3.SE
WD.4.SE
WW.4.SE
WD.5.SE
WW.5.SE
WD.1.SE+5d
WW.1.SE+5d
WD.2.SE+5d
WW.2.SE+5d
WD.3.SE+5d
WW.3.SE+5d
WD.4.SE+5d
WW.4.SE+5d
WD.5.SE+5d
WW.5.SE+5d

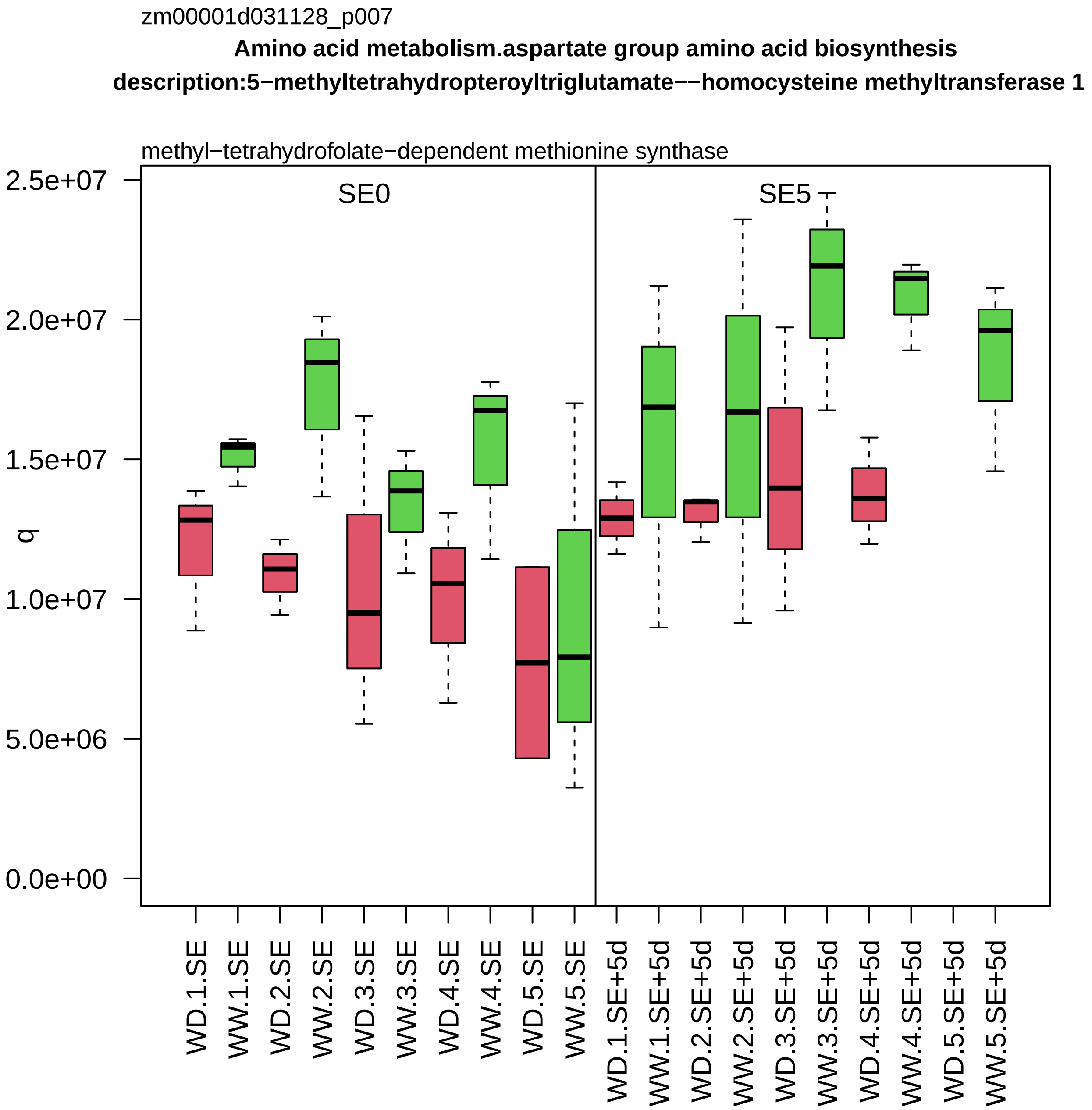

zm00001d031128_p007
Amino acid metabolism.aspartate group amino acid biosynthesis
description:5−methyltetrahydropteroyltriglutamate−−homocysteine methyltransferase 1
methyl−tetrahydrofolate−dependent methionine synthase
SE0
SE5
q
2.5e+07
2.0e+07
1.5e+07
1.0e+07
5.0e+06
0.0e+00
WD.1.SE
WW.1.SE
WD.2.SE
WW.2.SE
WD.3.SE
WW.3.SE
WD.4.SE
WW.4.SE
WD.5.SE
WW.5.SE
WD.1.SE+5d
WW.1.SE+5d
WD.2.SE+5d
WW.2.SE+5d
WD.3.SE+5d
WW.3.SE+5d
WD.4.SE+5d
WW.4.SE+5d
WD.5.SE+5d
WW.5.SE+5d

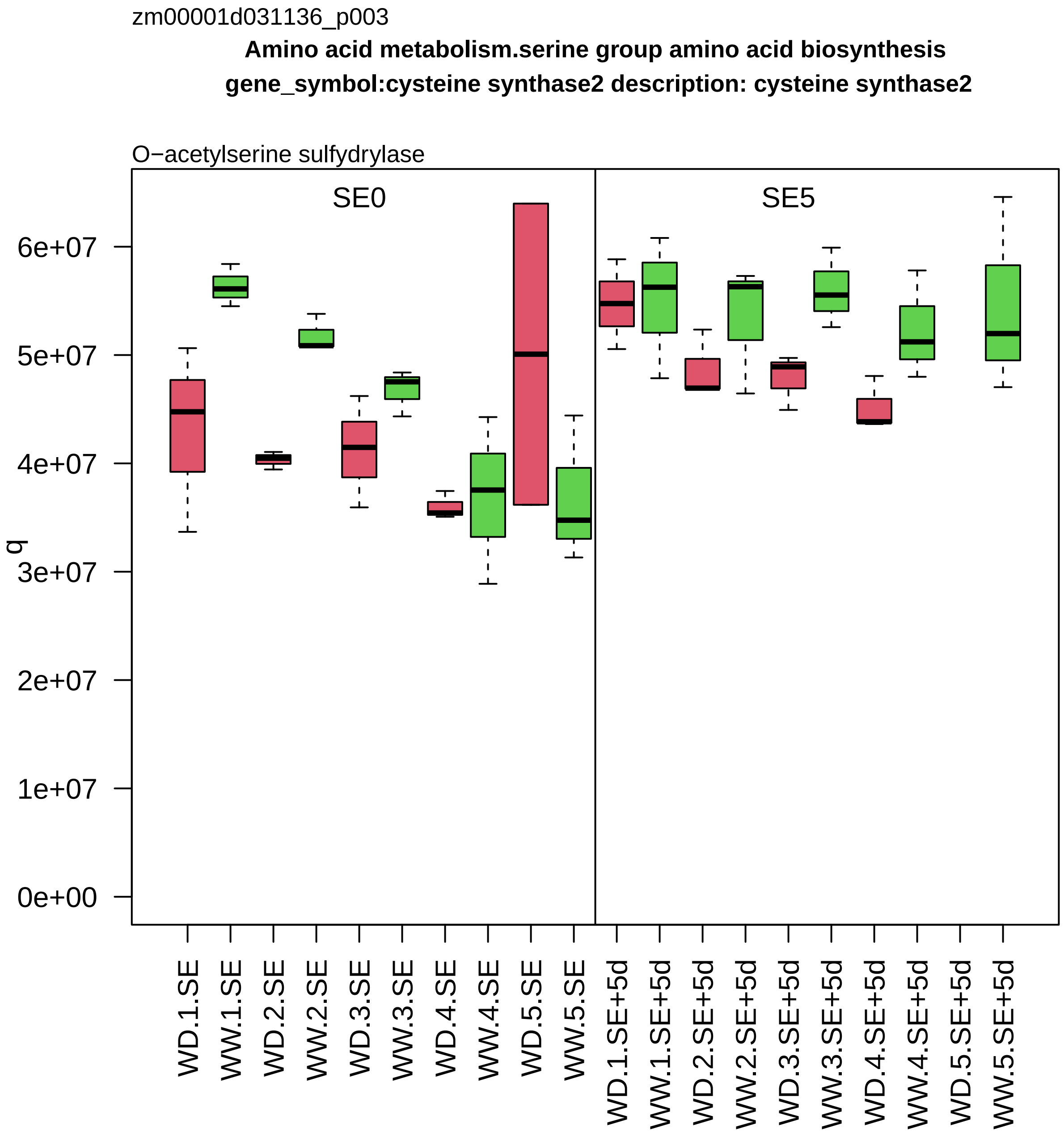

zm00001d031136_p003
Amino acid metabolism.serine group amino acid biosynthesis
gene_symbol:cysteine synthase2 description: cysteine synthase2
O−acetylserine sulfydrylase
SE0
SE5
q
0e+00
1e+07
2e+07
3e+07
4e+07
5e+07
6e+07
WD.1.SE
WW.1.SE
WD.2.SE
WW.2.SE
WD.3.SE
WW.3.SE
WD.4.SE
WW.4.SE
WD.5.SE
WW.5.SE
WD.1.SE+5d
WW.1.SE+5d
WD.2.SE+5d
WW.2.SE+5d
WD.3.SE+5d
WW.3.SE+5d
WD.4.SE+5d
WW.4.SE+5d
WD.5.SE+5d
WW.5.SE+5d

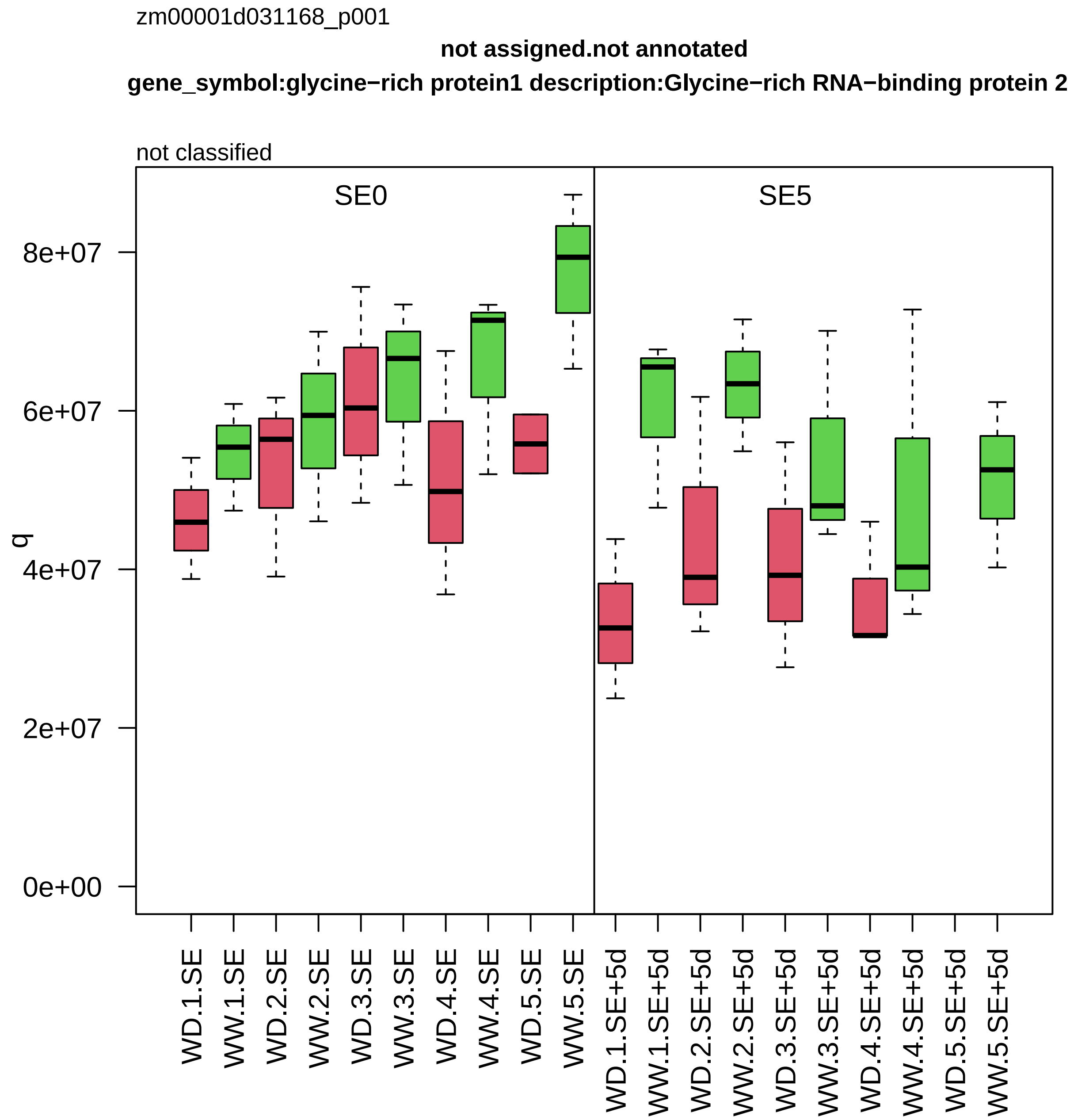

zm00001d031168_p001
not assigned.not annotated
gene_symbol:glycine−rich protein1 description:Glycine−rich RNA−binding protein 2
not classified
SE0
SE5
q
0e+00
2e+07
4e+07
6e+07
8e+07
WD.1.SE
WW.1.SE
WD.2.SE
WW.2.SE
WD.3.SE
WW.3.SE
WD.4.SE
WW.4.SE
WD.5.SE
WW.5.SE
WD.1.SE+5d
WW.1.SE+5d
WD.2.SE+5d
WW.2.SE+5d
WD.3.SE+5d
WW.3.SE+5d
WD.4.SE+5d
WW.4.SE+5d
WD.5.SE+5d
WW.5.SE+5d

zm00001d031209_p001

**Enzyme classification.EC_2 transferases**

**gene_symbol:benzoxazinone synthesis9 description: benzoxazinone synthesis9**

EC_2.4 glycosyltransferase

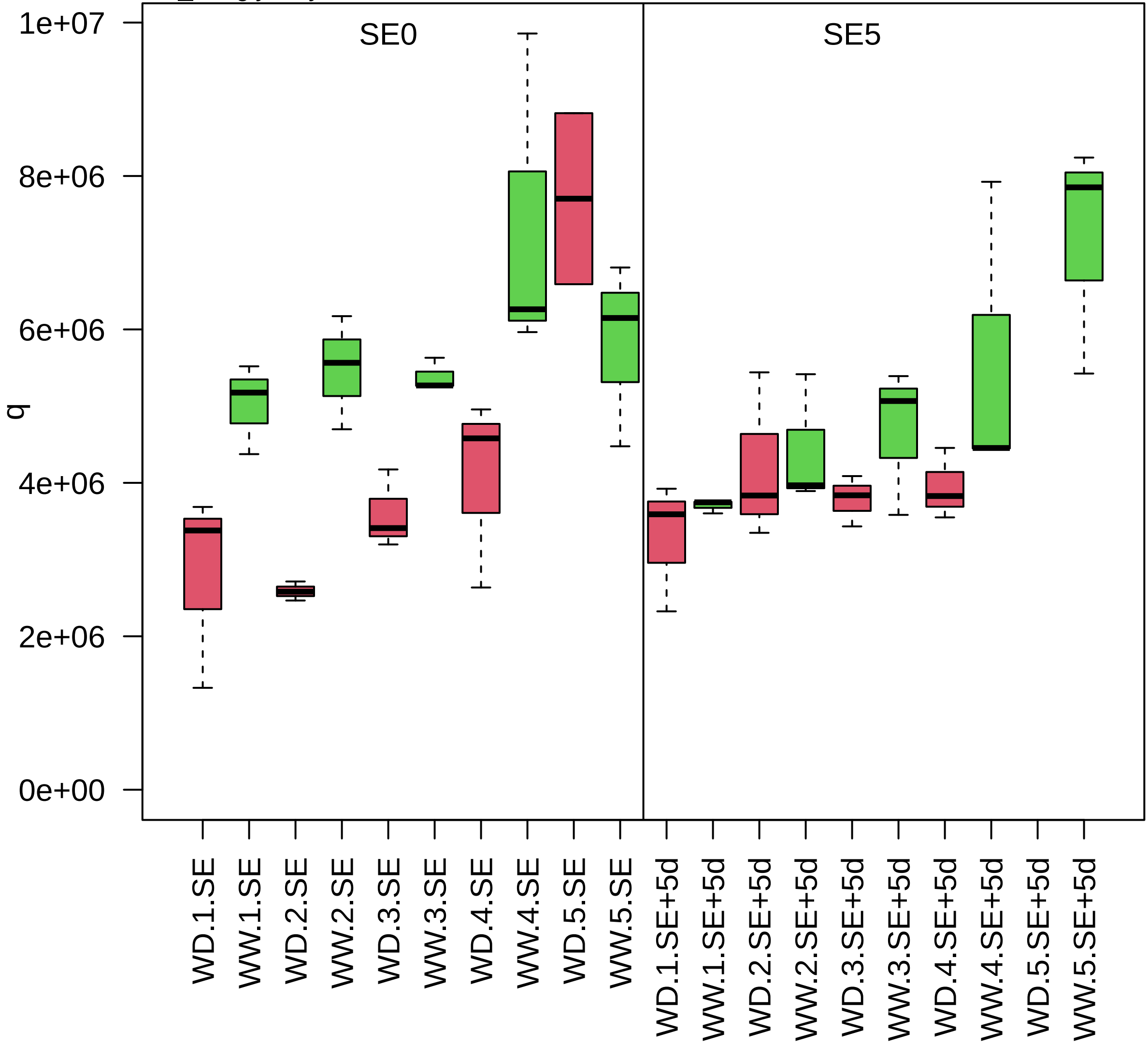

zm00001d031212_p002

**Secondary metabolism.terpenoids**
**description: thiolase1**

acetyl−CoA C−acyltransferase

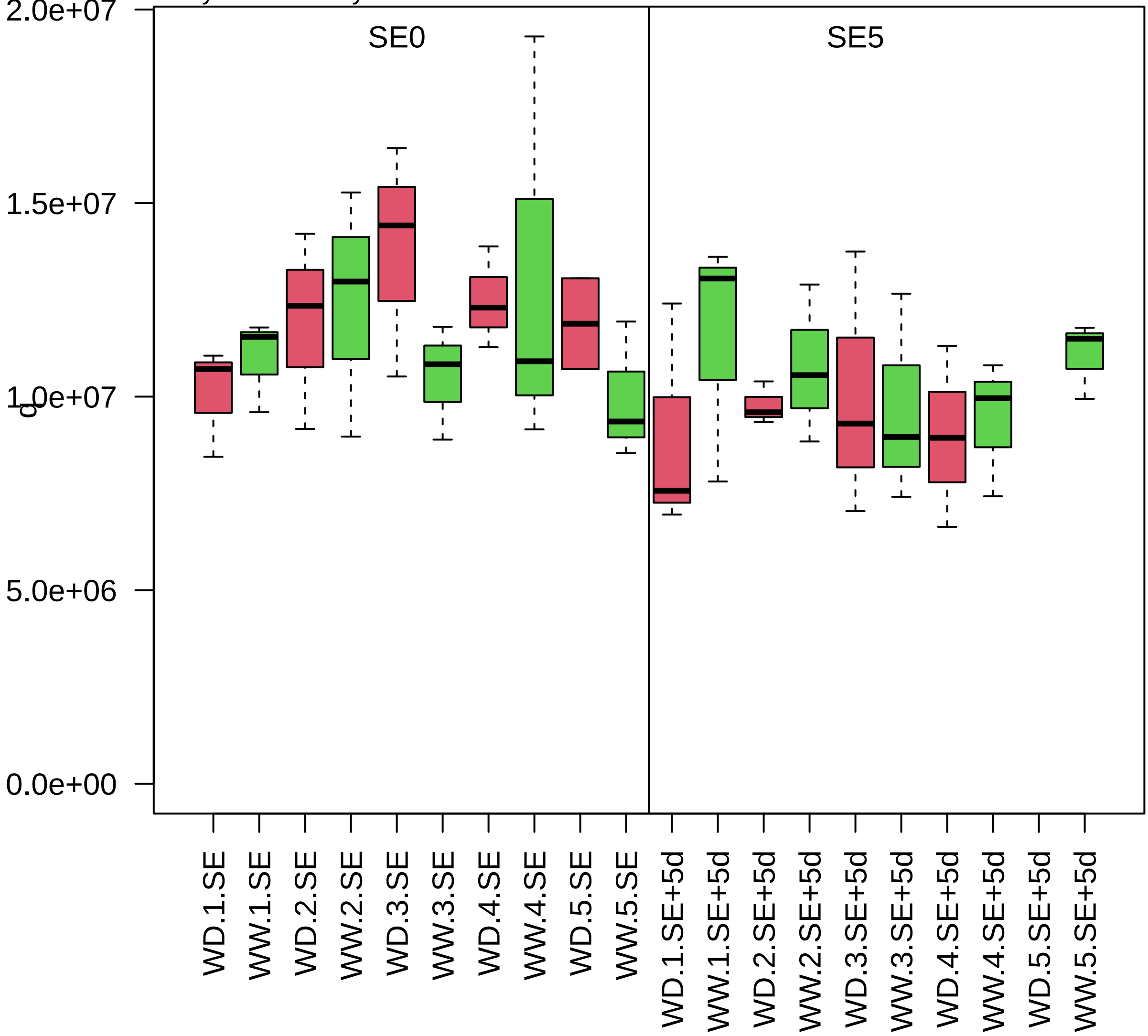

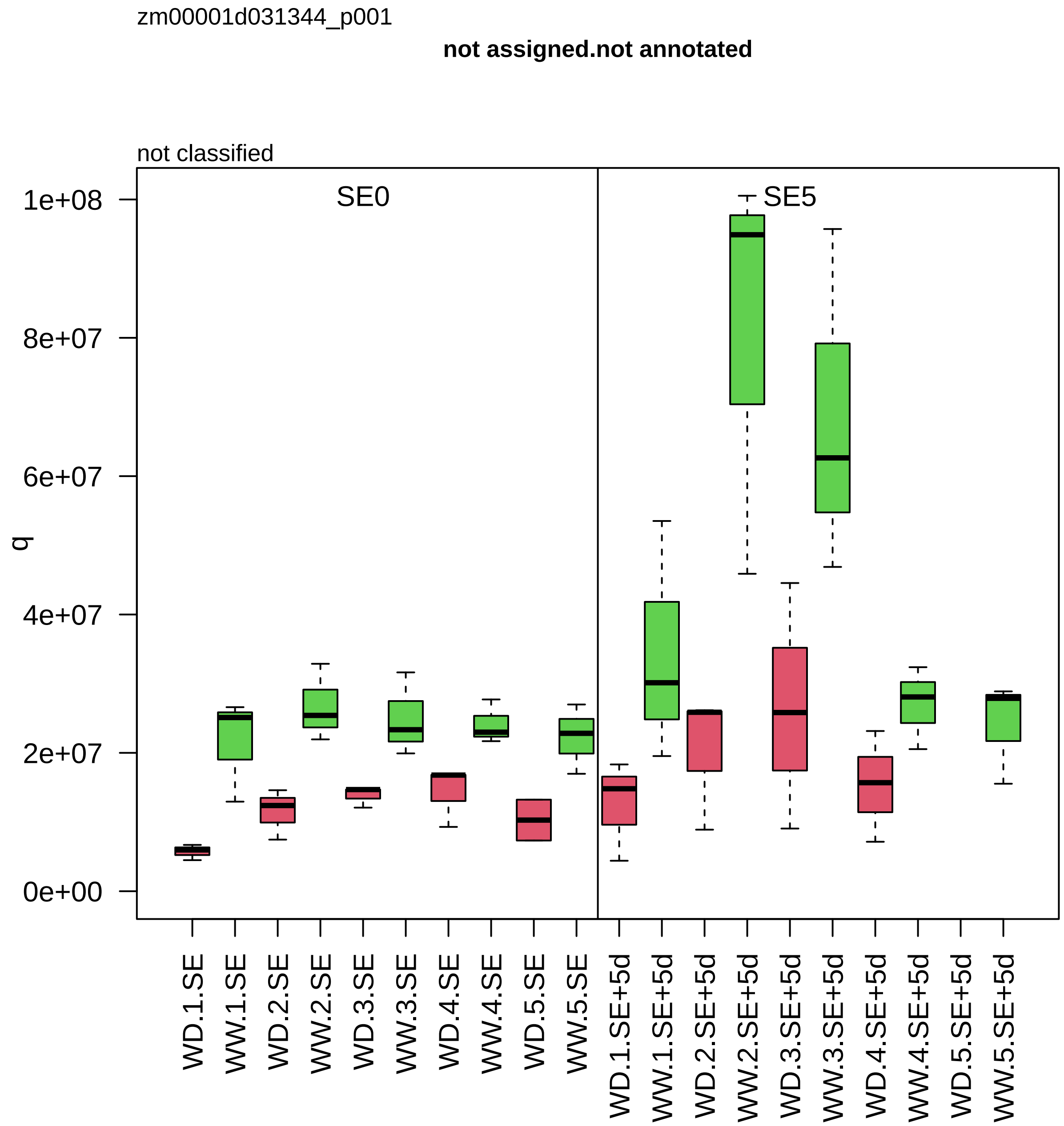
zm00001d031344_p001
not assigned.not annotated
not classified
SE0
SE5
q
1e+08
8e+07
6e+07
4e+07
2e+07
0e+00
WD.1.SE
WW.1.SE
WD.2.SE
WW.2.SE
WD.3.SE
WW.3.SE
WD.4.SE
WW.4.SE
WD.5.SE
WW.5.SE
WD.1.SE+5d
WW.1.SE+5d
WD.2.SE+5d
WW.2.SE+5d
WD.3.SE+5d
WW.3.SE+5d
WD.4.SE+5d
WW.4.SE+5d
WD.5.SE+5d
WW.5.SE+5d

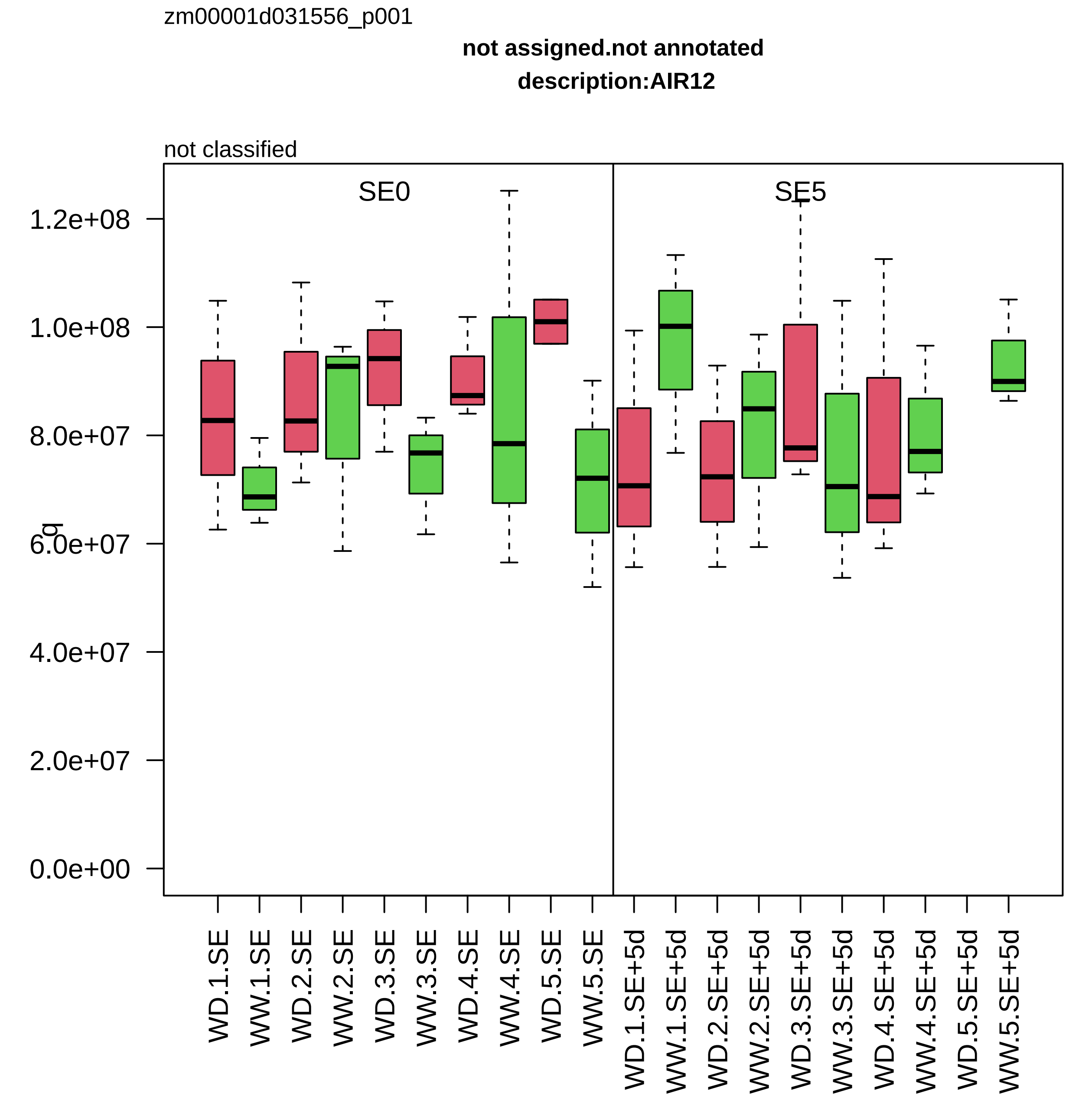
zm00001d031556_p001
not assigned.not annotated
description:AIR12
not classified
SE0
SE5
q
1.2e+08
1.0e+08
8.0e+07
6.0e+07
4.0e+07
2.0e+07
0.0e+00
WD.1.SE
WW.1.SE
WD.2.SE
WW.2.SE
WD.3.SE
WW.3.SE
WD.4.SE
WW.4.SE
WD.5.SE
WW.5.SE
WD.1.SE+5d
WW.1.SE+5d
WD.2.SE+5d
WW.2.SE+5d
WD.3.SE+5d
WW.3.SE+5d
WD.4.SE+5d
WW.4.SE+5d
WD.5.SE+5d
WW.5.SE+5d

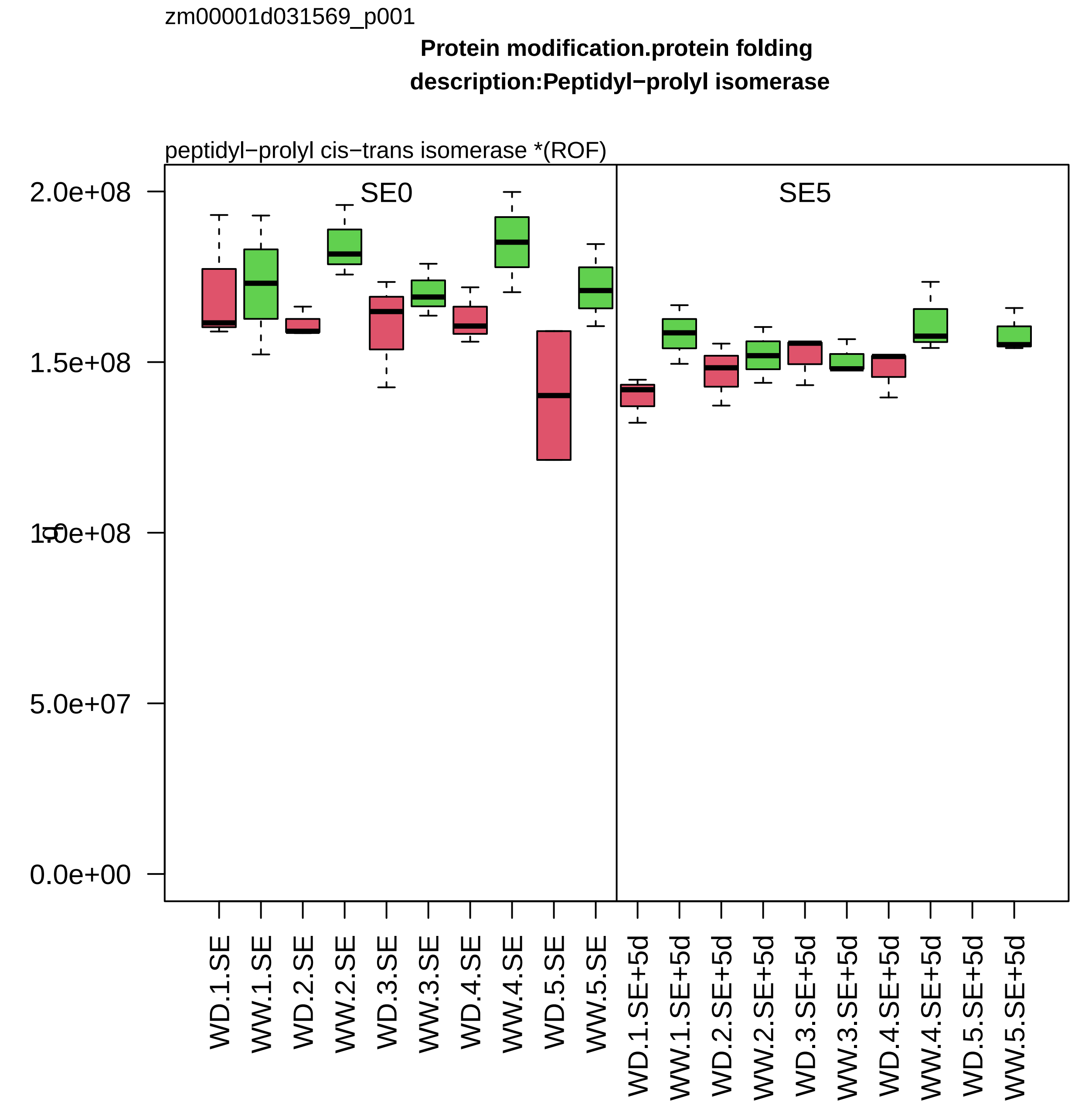

zm00001d031569_p001
Protein modification.protein folding
description:Peptidyl−prolyl isomerase
peptidyl−prolyl cis−trans isomerase *(ROF)
SE0
SE5
2.0e+08
1.5e+08
1.0e+08
5.0e+07
0.0e+00
WD.1.SE
WW.1.SE
WD.2.SE
WW.2.SE
WD.3.SE
WW.3.SE
WD.4.SE
WW.4.SE
WD.5.SE
WW.5.SE
WD.1.SE+5d
WW.1.SE+5d
WD.2.SE+5d
WW.2.SE+5d
WD.3.SE+5d
WW.3.SE+5d
WD.4.SE+5d
WW.4.SE+5d
WD.5.SE+5d
WW.5.SE+5d

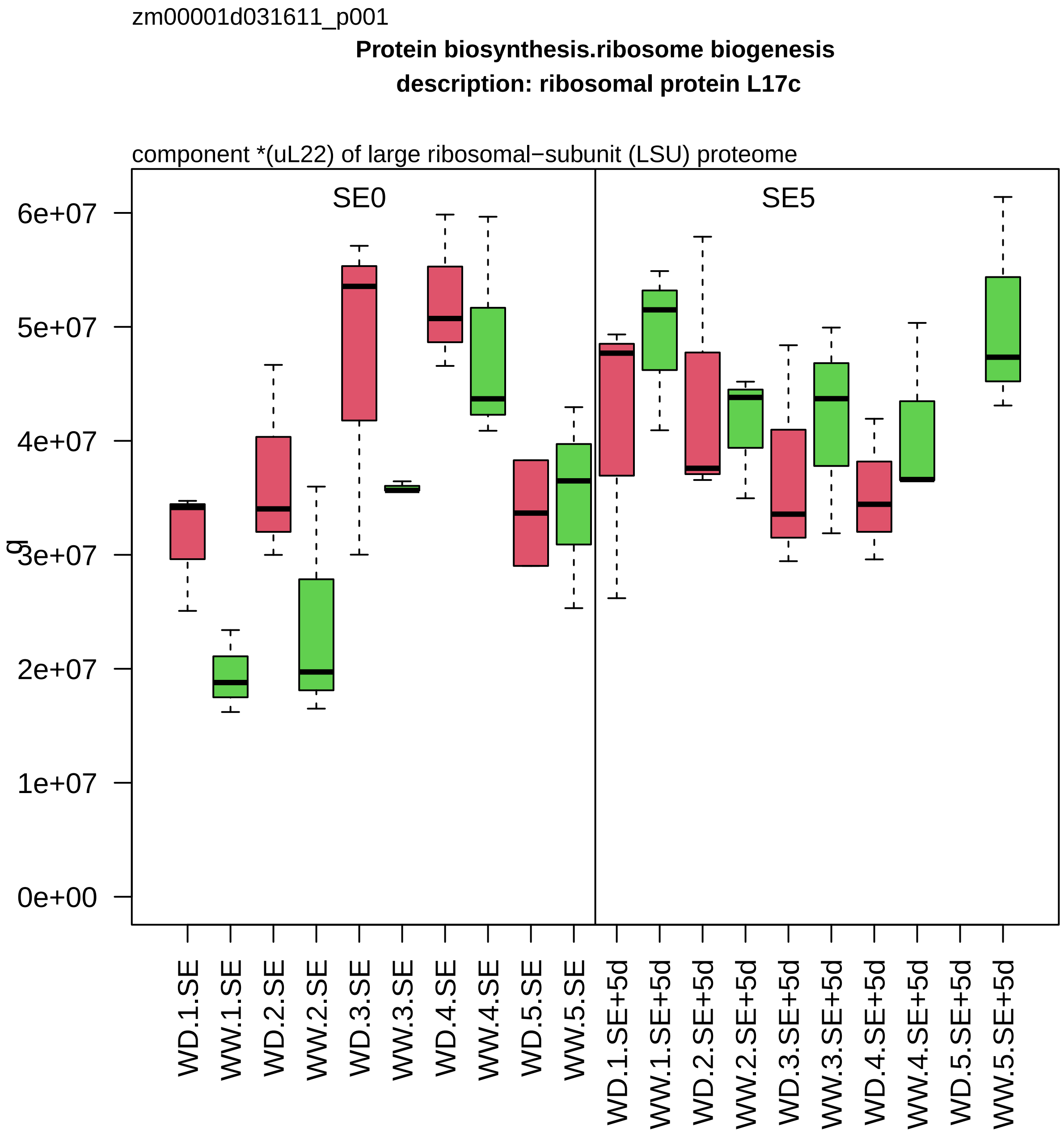
zm00001d031611_p001
Protein biosynthesis.ribosome biogenesis
description: ribosomal protein L17c
component *(uL22) of large ribosomal−subunit (LSU) proteome
SE0
SE5
g
6e+07
5e+07
4e+07
3e+07
2e+07
1e+07
0e+00
WD.1.SE
WW.1.SE
WD.2.SE
WW.2.SE
WD.3.SE
WW.3.SE
WD.4.SE
WW.4.SE
WD.5.SE
WW.5.SE
WD.1.SE+5d
WW.1.SE+5d
WD.2.SE+5d
WW.2.SE+5d
WD.3.SE+5d
WW.3.SE+5d
WD.4.SE+5d
WW.4.SE+5d
WD.5.SE+5d
WW.5.SE+5d

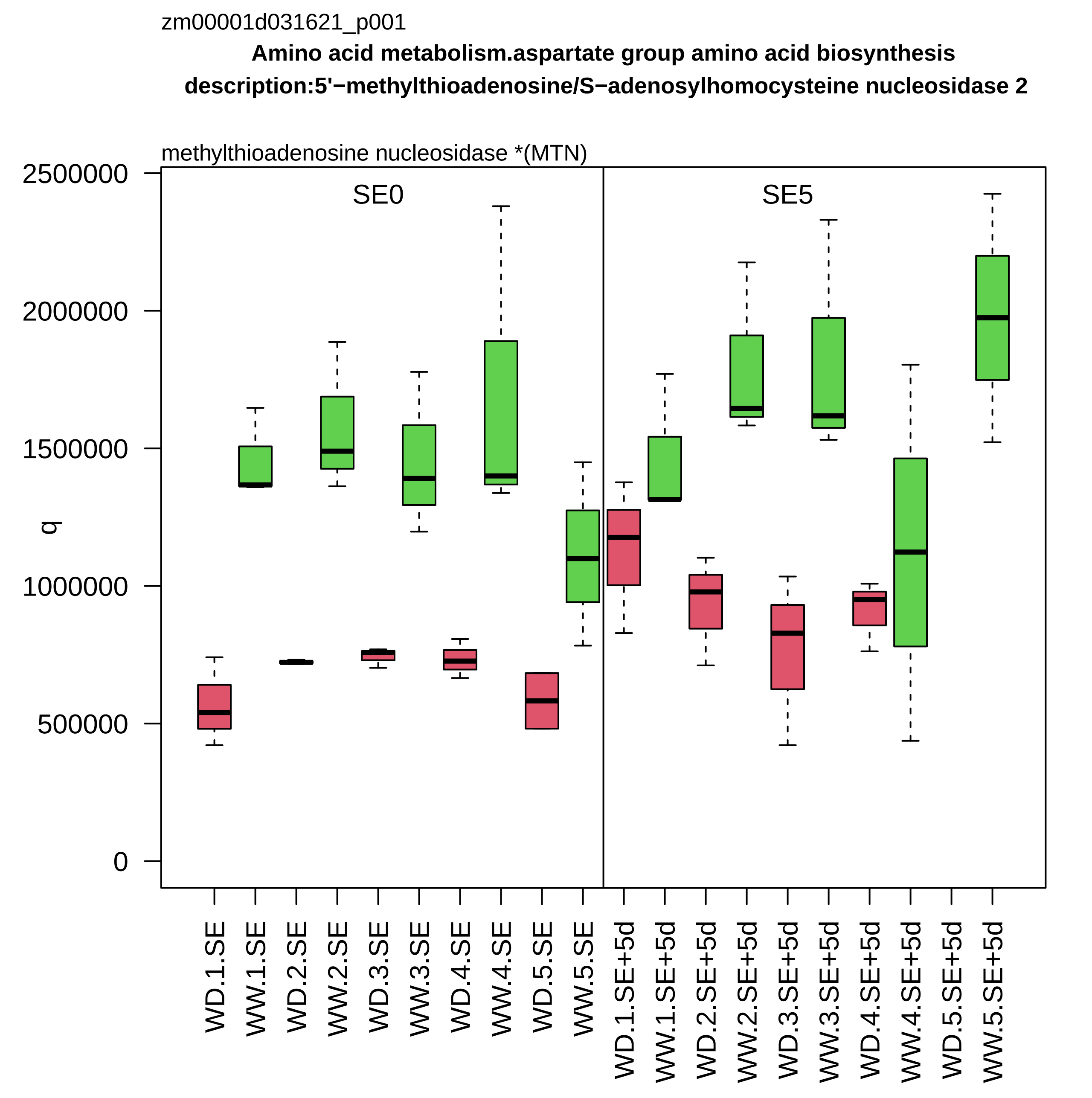

zm00001d031621_p001
Amino acid metabolism.aspartate group amino acid biosynthesis
description:5'−methylthioadenosine/S−adenosylhomocysteine nucleosidase 2
methylthioadenosine nucleosidase *(MTN)
SE0
SE5
q
2500000
2000000
1500000
1000000
500000
0
WD.1.SE
WW.1.SE
WD.2.SE
WW.2.SE
WD.3.SE
WW.3.SE
WD.4.SE
WW.4.SE
WD.5.SE
WW.5.SE
WD.1.SE+5d
WW.1.SE+5d
WD.2.SE+5d
WW.2.SE+5d
WD.3.SE+5d
WW.3.SE+5d
WD.4.SE+5d
WW.4.SE+5d
WD.5.SE+5d
WW.5.SE+5d

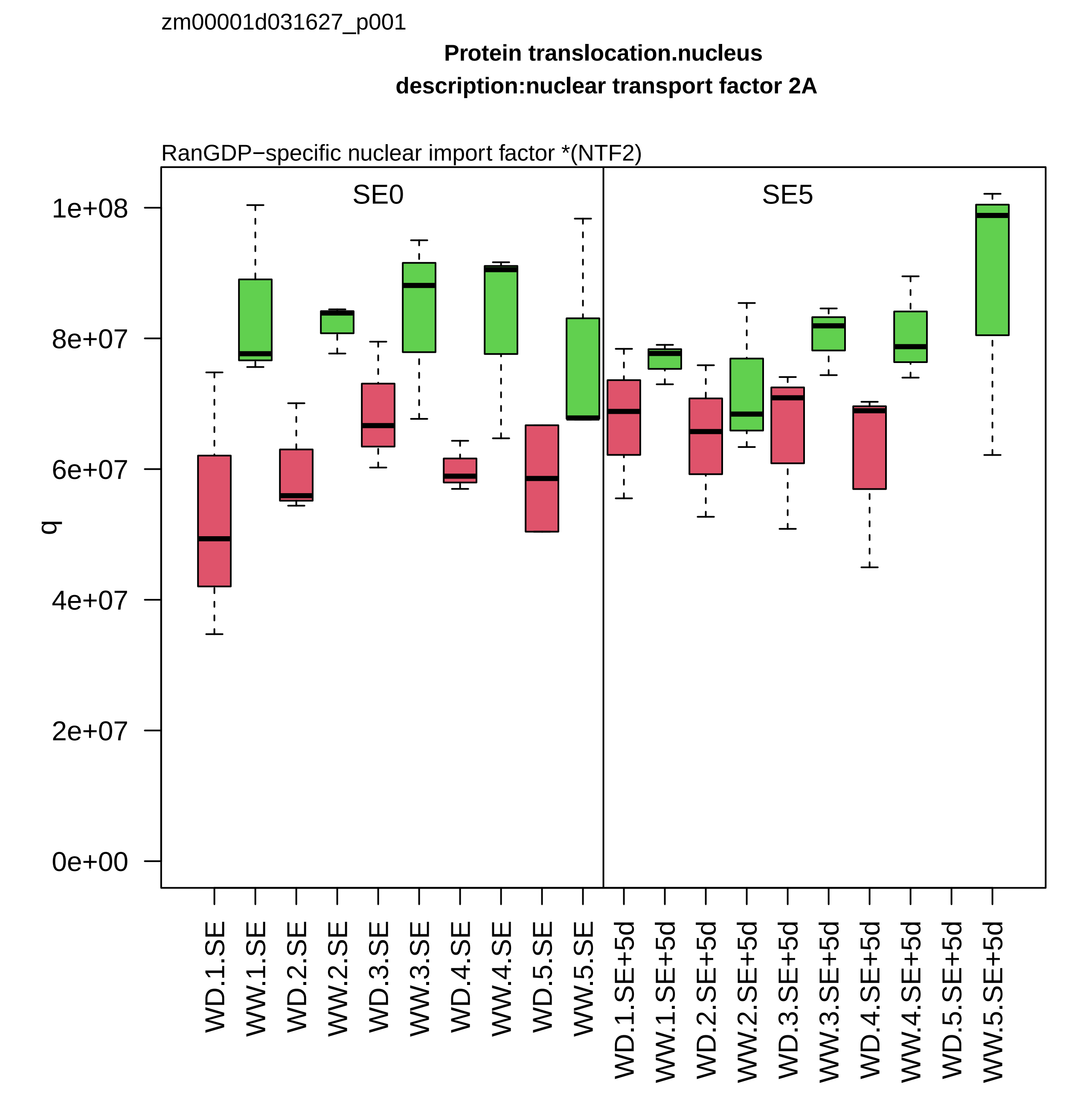
zm00001d031627_p001
Protein translocation.nucleus
description:nuclear transport factor 2A
RanGDP−specific nuclear import factor *(NTF2)
SE0
SE5
q
1e+08
8e+07
6e+07
4e+07
2e+07
0e+00
WD.1.SE
WW.1.SE
WD.2.SE
WW.2.SE
WD.3.SE
WW.3.SE
WD.4.SE
WW.4.SE
WD.5.SE
WW.5.SE
WD.1.SE+5d
WW.1.SE+5d
WD.2.SE+5d
WW.2.SE+5d
WD.3.SE+5d
WW.3.SE+5d
WD.4.SE+5d
WW.4.SE+5d
WD.5.SE+5d
WW.5.SE+5d

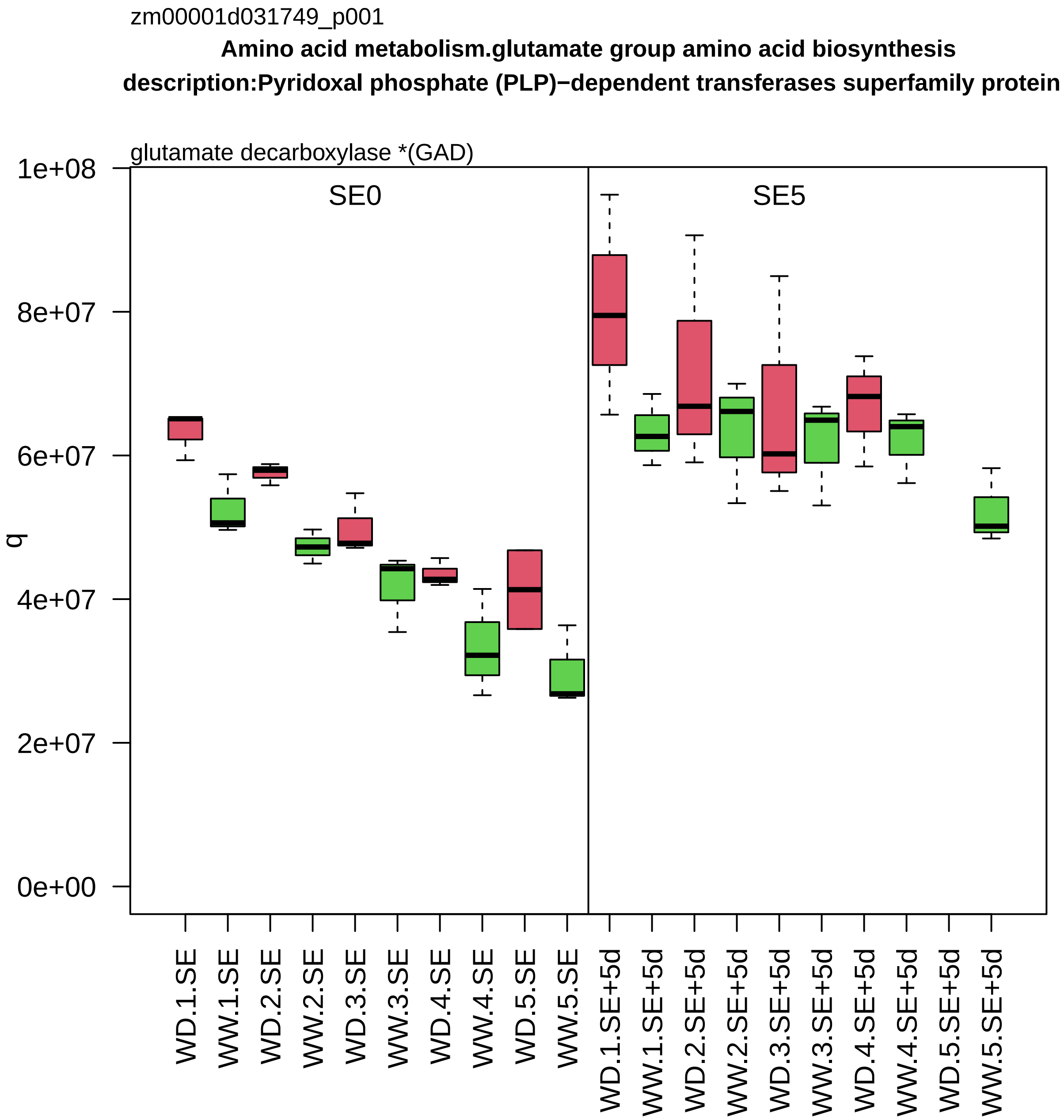
zm00001d031749_p001
Amino acid metabolism.glutamate group amino acid biosynthesis
description:Pyridoxal phosphate (PLP)−dependent transferases superfamily protein
glutamate decarboxylase *(GAD)
SE0
SE5
q
1e+08
8e+07
6e+07
4e+07
2e+07
0e+00
WD.1.SE
WW.1.SE
WD.2.SE
WW.2.SE
WD.3.SE
WW.3.SE
WD.4.SE
WW.4.SE
WD.5.SE
WW.5.SE
WD.1.SE+5d
WW.1.SE+5d
WD.2.SE+5d
WW.2.SE+5d
WD.3.SE+5d
WW.3.SE+5d
WD.4.SE+5d
WW.4.SE+5d
WD.5.SE+5d
WW.5.SE+5d

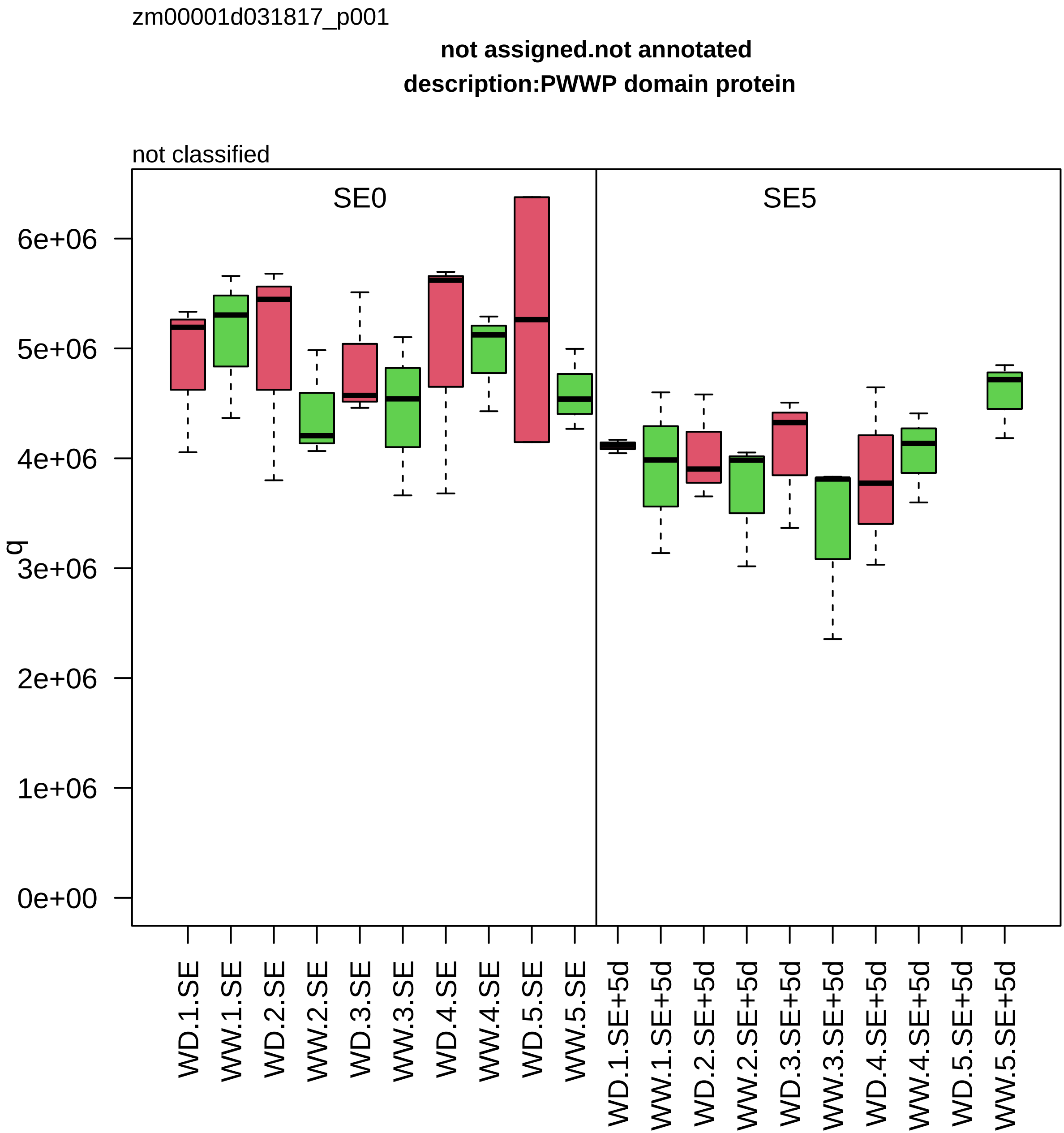

zm00001d031817_p001
not assigned.not annotated
description:PWWP domain protein
not classified
SE0
SE5
q
6e+06
5e+06
4e+06
3e+06
2e+06
1e+06
0e+00
WD.1.SE
WW.1.SE
WD.2.SE
WW.2.SE
WD.3.SE
WW.3.SE
WD.4.SE
WW.4.SE
WD.5.SE
WW.5.SE
WD.1.SE+5d
WW.1.SE+5d
WD.2.SE+5d
WW.2.SE+5d
WD.3.SE+5d
WW.3.SE+5d
WD.4.SE+5d
WW.4.SE+5d
WD.5.SE+5d
WW.5.SE+5d

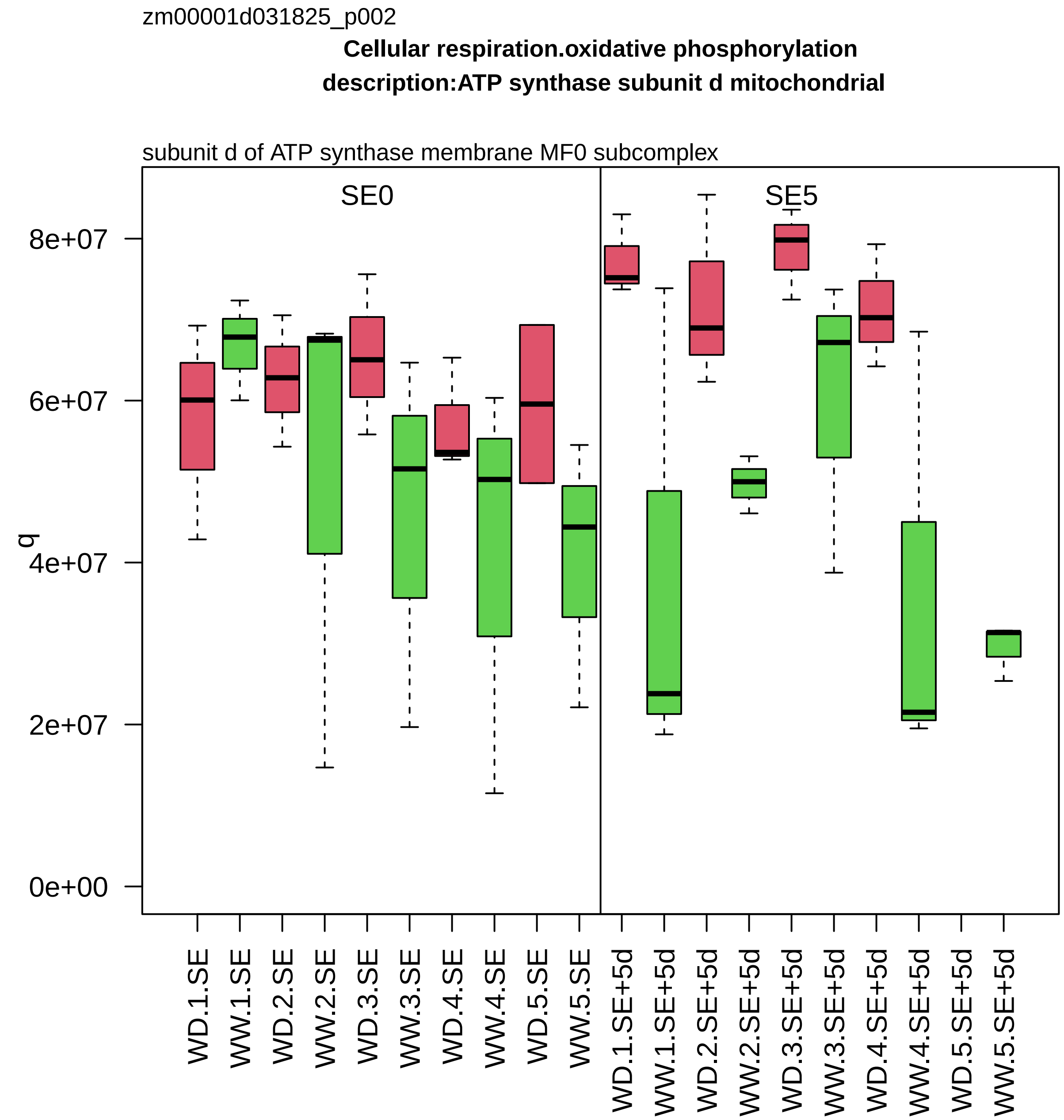
zm00001d031825_p002
Cellular respiration.oxidative phosphorylation
description:ATP synthase subunit d mitochondrial
subunit d of ATP synthase membrane MF0 subcomplex
SE0
SE5
q
8e+07
6e+07
4e+07
2e+07
0e+00
WD.1.SE
WW.1.SE
WD.2.SE
WW.2.SE
WD.3.SE
WW.3.SE
WD.4.SE
WW.4.SE
WD.5.SE
WW.5.SE
WD.1.SE+5d
WW.1.SE+5d
WD.2.SE+5d
WW.2.SE+5d
WD.3.SE+5d
WW.3.SE+5d
WD.4.SE+5d
WW.4.SE+5d
WD.5.SE+5d
WW.5.SE+5d

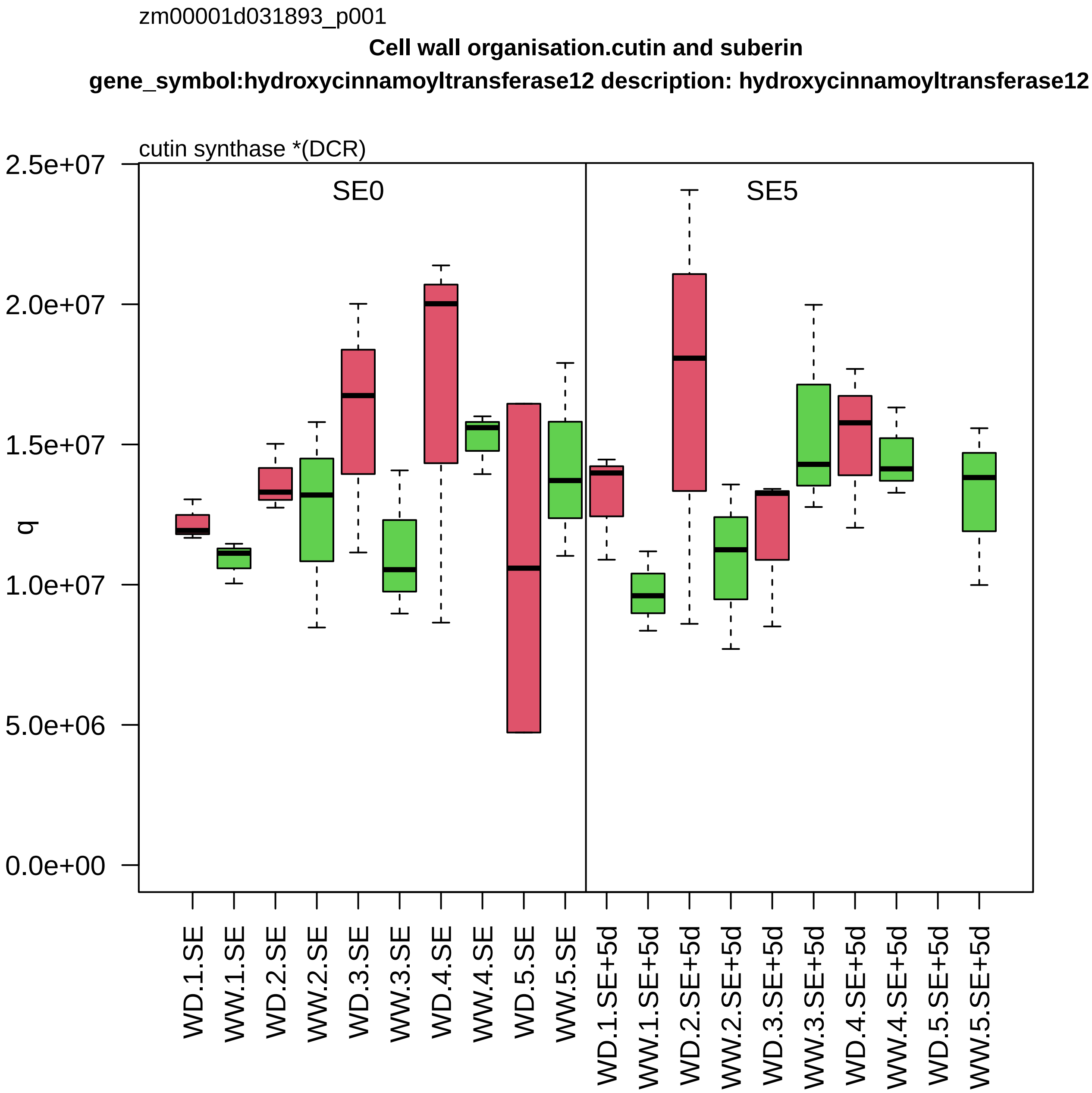
zm00001d031893_p001
Cell wall organisation.cutin and suberin
gene_symbol:hydroxycinnamoyltransferase12 description: hydroxycinnamoyltransferase12
cutin synthase *(DCR)
SE0
SE5
q
2.5e+07
2.0e+07
1.5e+07
1.0e+07
5.0e+06
0.0e+00
WD.1.SE
WW.1.SE
WD.2.SE
WW.2.SE
WD.3.SE
WW.3.SE
WD.4.SE
WW.4.SE
WD.5.SE
WW.5.SE
WD.1.SE+5d
WW.1.SE+5d
WD.2.SE+5d
WW.2.SE+5d
WD.3.SE+5d
WW.3.SE+5d
WD.4.SE+5d
WW.4.SE+5d
WD.5.SE+5d
WW.5.SE+5d

zm00001d031908_p001

**Redox homeostasis.reactive oxygen scavenging**
**description:Superoxide dismutase**

copper/zinc superoxide dismutase *(CSD)

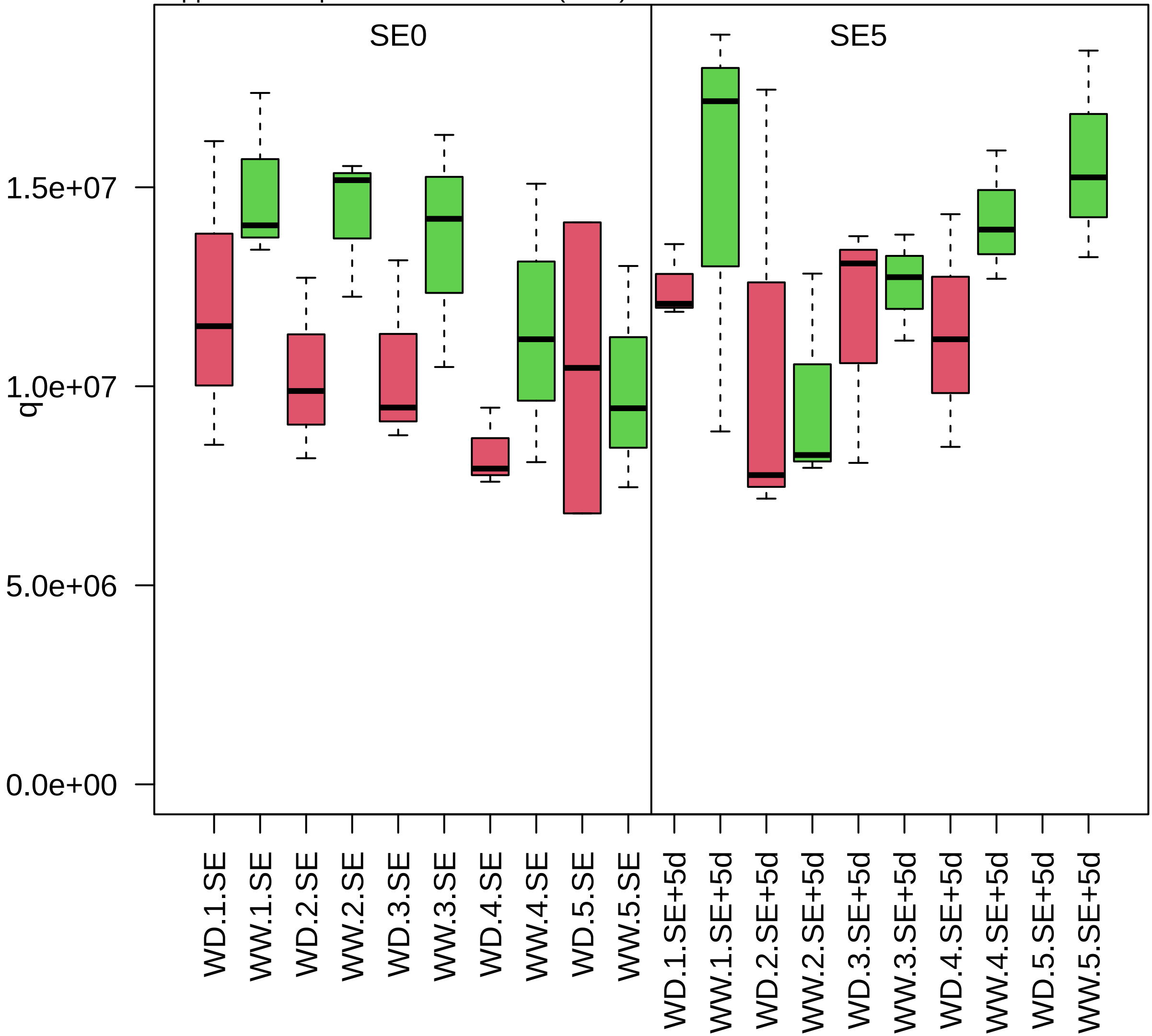

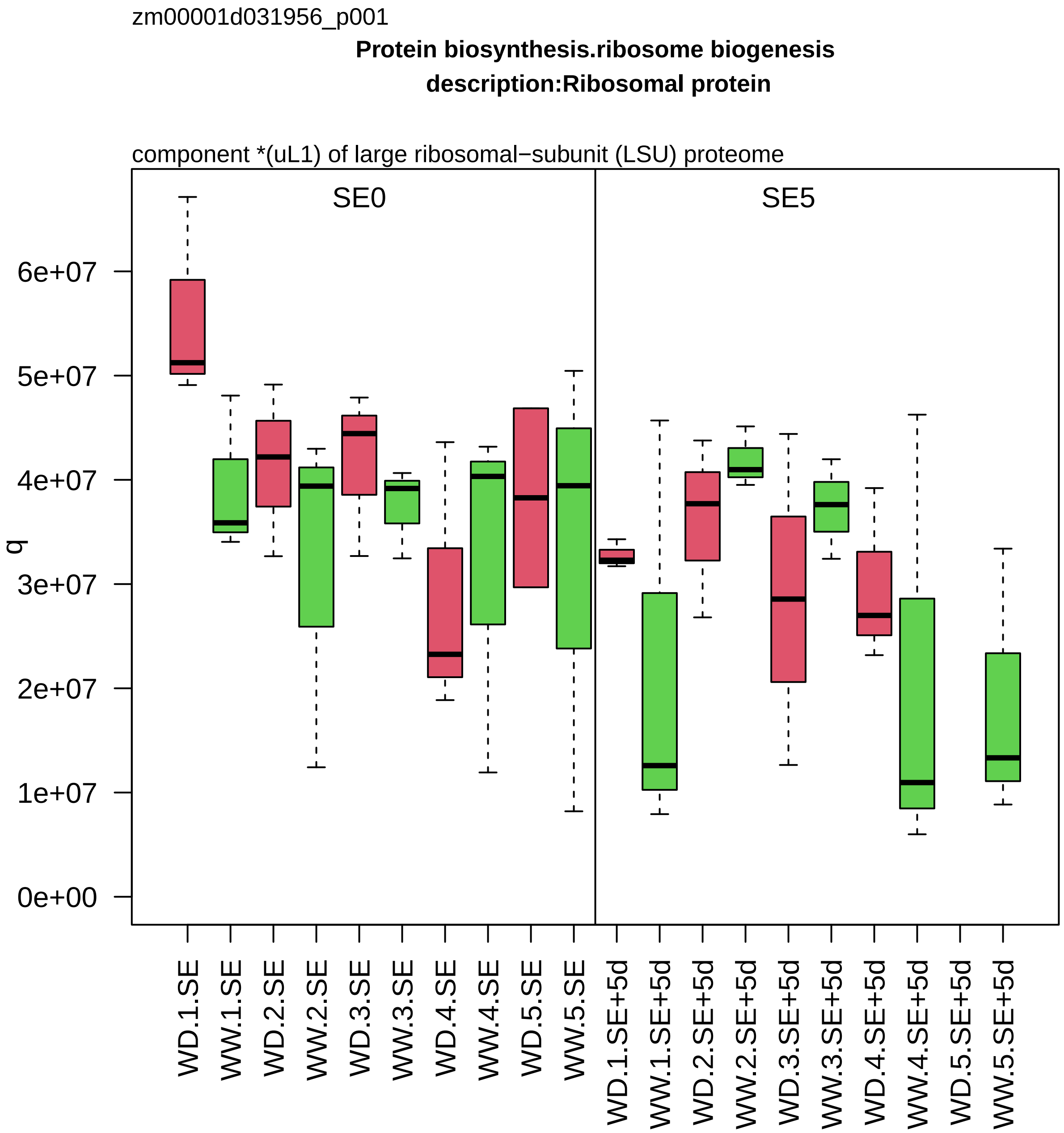
zm00001d031956_p001
Protein biosynthesis.ribosome biogenesis
description:Ribosomal protein
component *(uL1) of large ribosomal−subunit (LSU) proteome
SE0
SE5
q
0e+00
1e+07
2e+07
3e+07
4e+07
5e+07
6e+07
WD.1.SE
WW.1.SE
WD.2.SE
WW.2.SE
WD.3.SE
WW.3.SE
WD.4.SE
WW.4.SE
WD.5.SE
WW.5.SE
WD.1.SE+5d
WW.1.SE+5d
WD.2.SE+5d
WW.2.SE+5d
WD.3.SE+5d
WW.3.SE+5d
WD.4.SE+5d
WW.4.SE+5d
WD.5.SE+5d
WW.5.SE+5d

zm00001d031967_p001

**Cellular respiration.oxidative phosphorylation**

**description:Cytochrome c1 1 heme protein mitochondrial**

apocytochrome c1 component of cytochrome c reductase complex

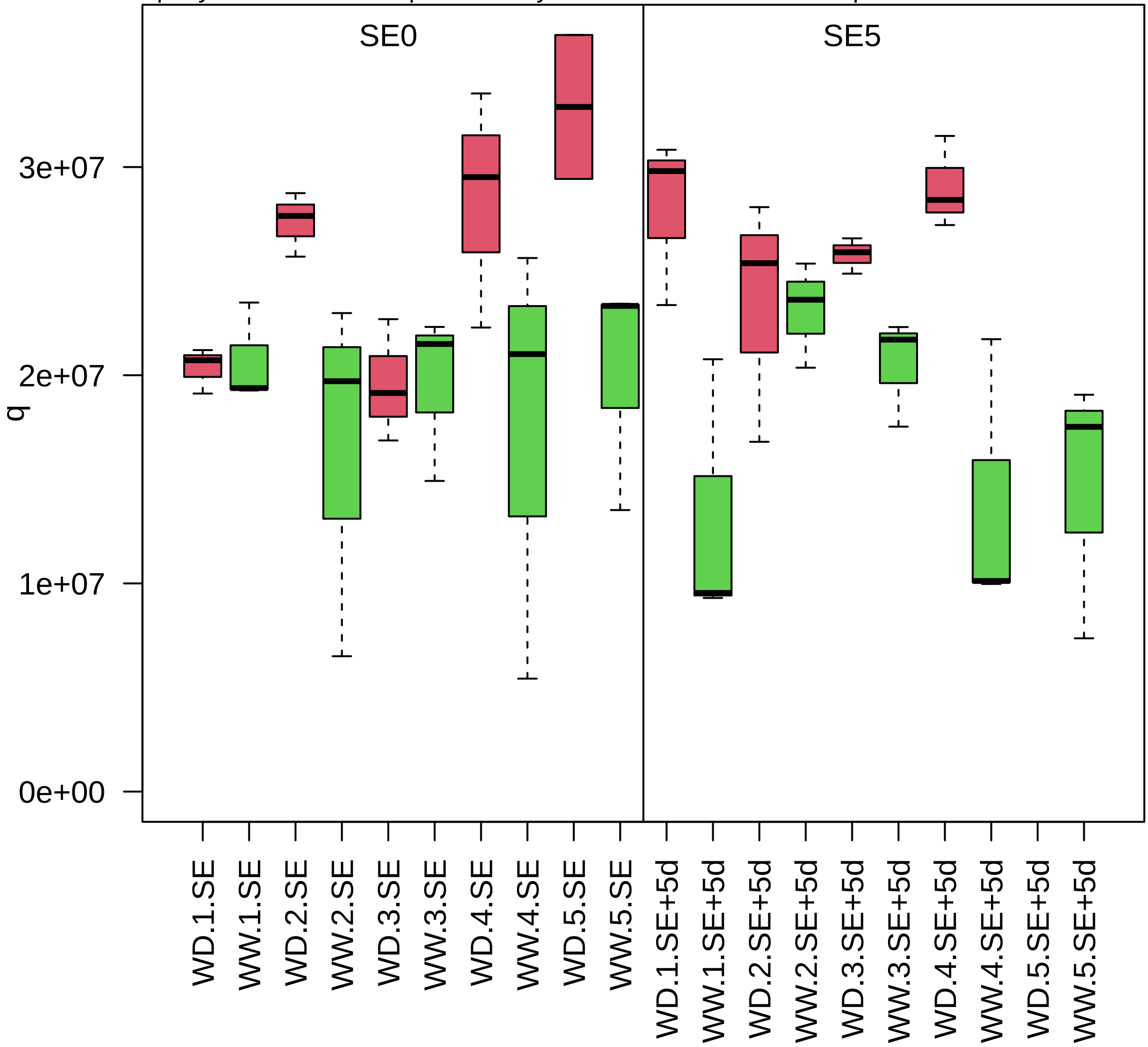

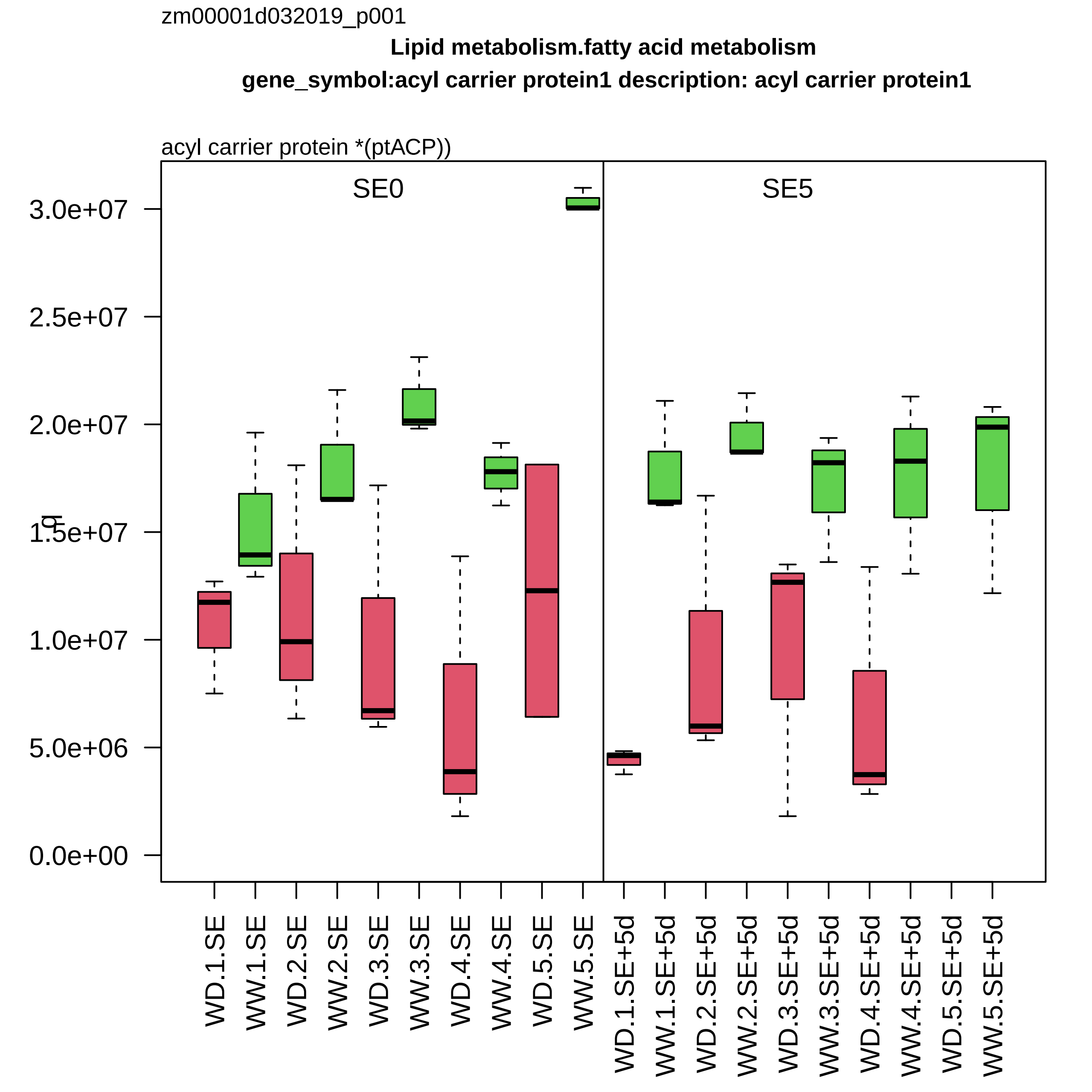

zm00001d032019_p001
Lipid metabolism.fatty acid metabolism
gene_symbol:acyl carrier protein1 description: acyl carrier protein1
acyl carrier protein *(ptACP))
SE0
SE5
3.0e+07
2.5e+07
2.0e+07
1.5e+07
1.0e+07
5.0e+06
0.0e+00
WD.1.SE
WW.1.SE
WD.2.SE
WW.2.SE
WD.3.SE
WW.3.SE
WD.4.SE
WW.4.SE
WD.5.SE
WW.5.SE
WD.1.SE+5d
WW.1.SE+5d
WD.2.SE+5d
WW.2.SE+5d
WD.3.SE+5d
WW.3.SE+5d
WD.4.SE+5d
WW.4.SE+5d
WD.5.SE+5d
WW.5.SE+5d

zm00001d032049_p001

**Enzyme classification.EC_1 oxidoreductases**

**gene_symbol:12−oxo−phytodienoic acid reductase7 description: 12−oxo−phytodienoic acid reduct**

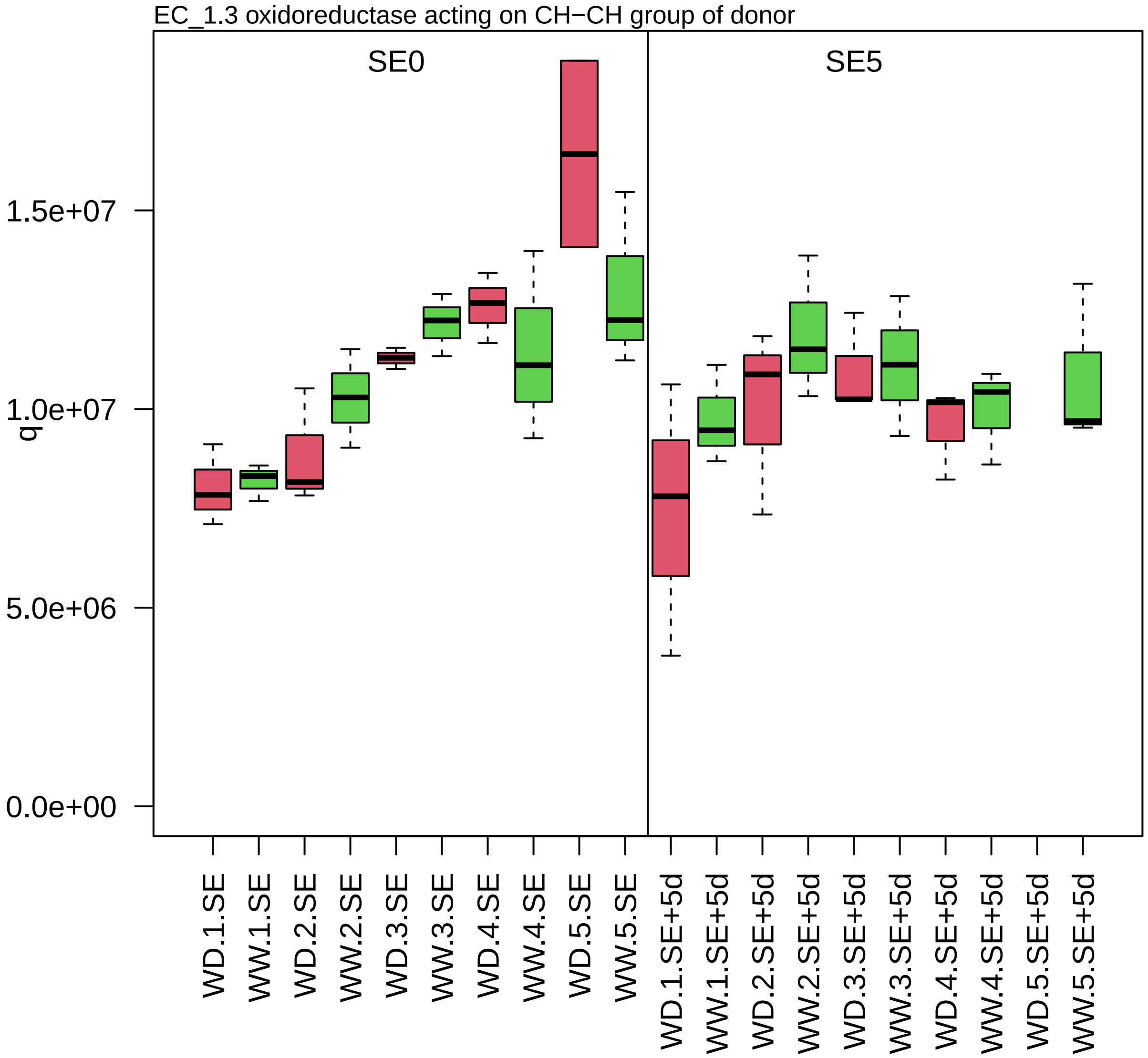

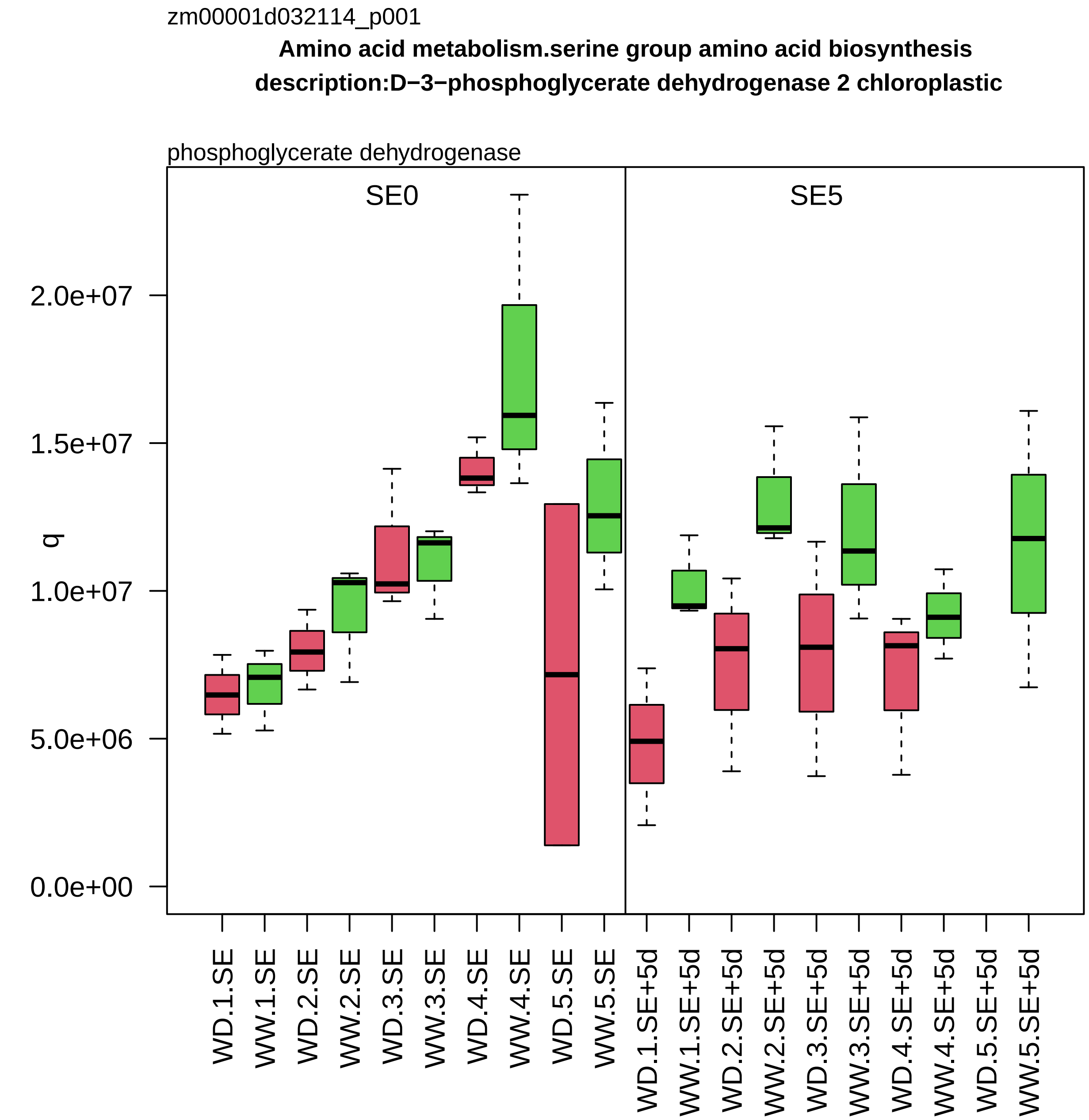
zm00001d032114_p001
Amino acid metabolism.serine group amino acid biosynthesis
description:D−3−phosphoglycerate dehydrogenase 2 chloroplastic
phosphoglycerate dehydrogenase
SE0
SE5
q
0.0e+00
5.0e+06
1.0e+07
1.5e+07
2.0e+07
WD.1.SE
WW.1.SE
WD.2.SE
WW.2.SE
WD.3.SE
WW.3.SE
WD.4.SE
WW.4.SE
WD.5.SE
WW.5.SE
WD.1.SE+5d
WW.1.SE+5d
WD.2.SE+5d
WW.2.SE+5d
WD.3.SE+5d
WW.3.SE+5d
WD.4.SE+5d
WW.4.SE+5d
WD.5.SE+5d
WW.5.SE+5d

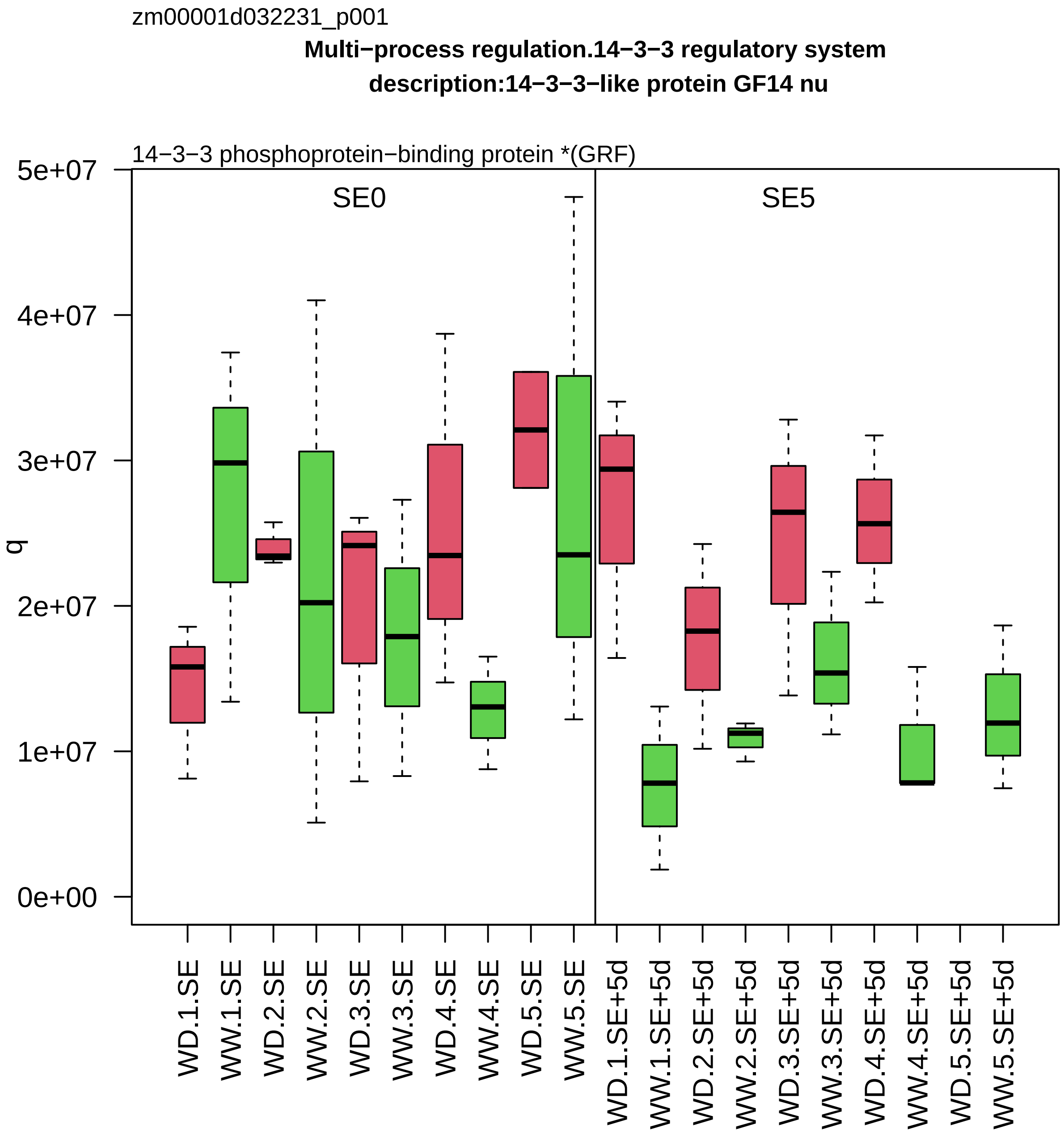
zm00001d032231_p001
Multi−process regulation.14−3−3 regulatory system
description:14−3−3−like protein GF14 nu
14−3−3 phosphoprotein−binding protein *(GRF)
SE0
SE5
q
5e+07
4e+07
3e+07
2e+07
1e+07
0e+00
WD.1.SE
WW.1.SE
WD.2.SE
WW.2.SE
WD.3.SE
WW.3.SE
WD.4.SE
WW.4.SE
WD.5.SE
WW.5.SE
WD.1.SE+5d
WW.1.SE+5d
WD.2.SE+5d
WW.2.SE+5d
WD.3.SE+5d
WW.3.SE+5d
WD.4.SE+5d
WW.4.SE+5d
WD.5.SE+5d
WW.5.SE+5d

zm00001d032239_p001

**Chromatin organisation.chromatin structure**

**gene_symbol:high mobility group family A 102 description: high mobility group family A 102**

chromatin architectural modulator *(HMGA)

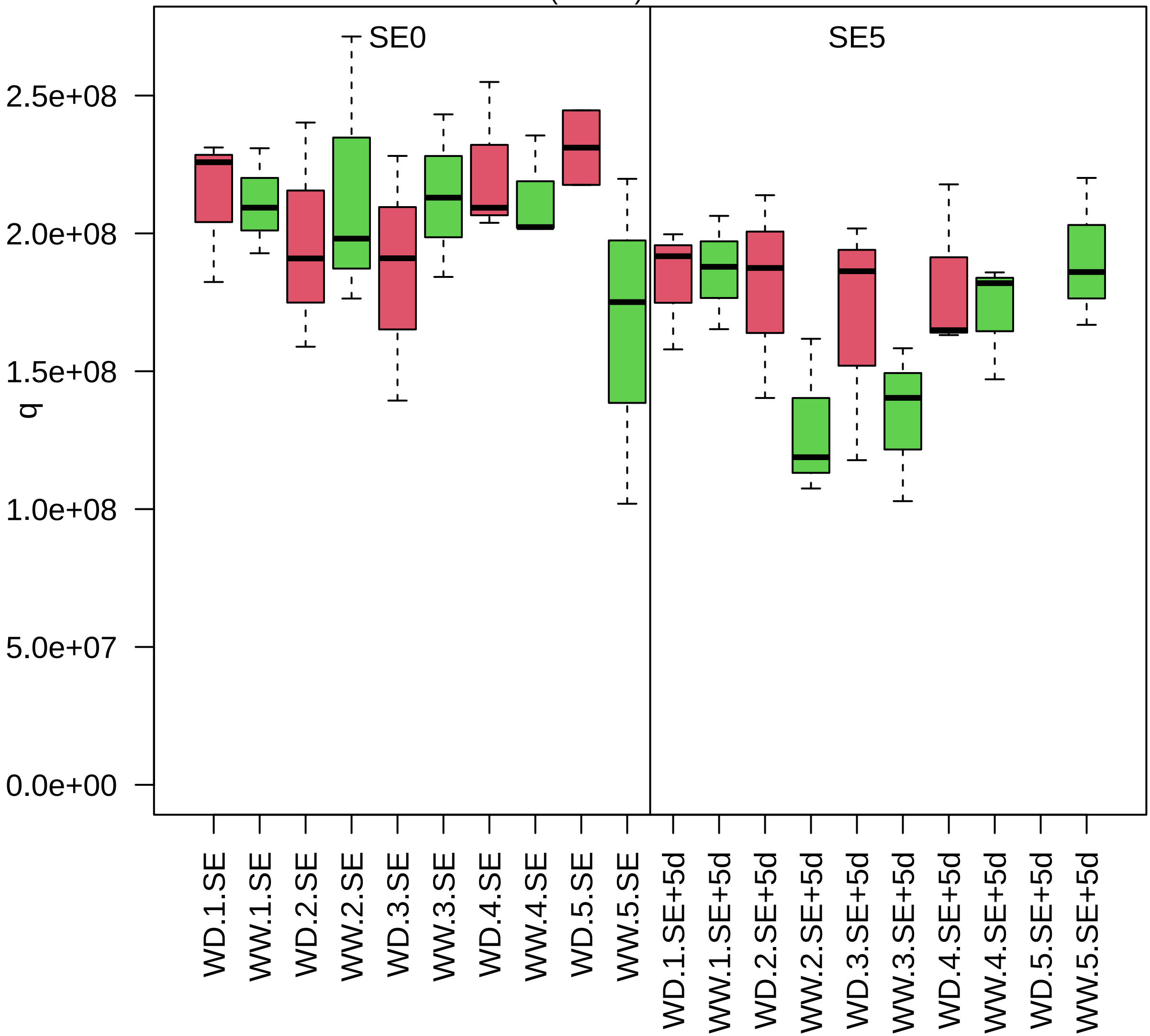

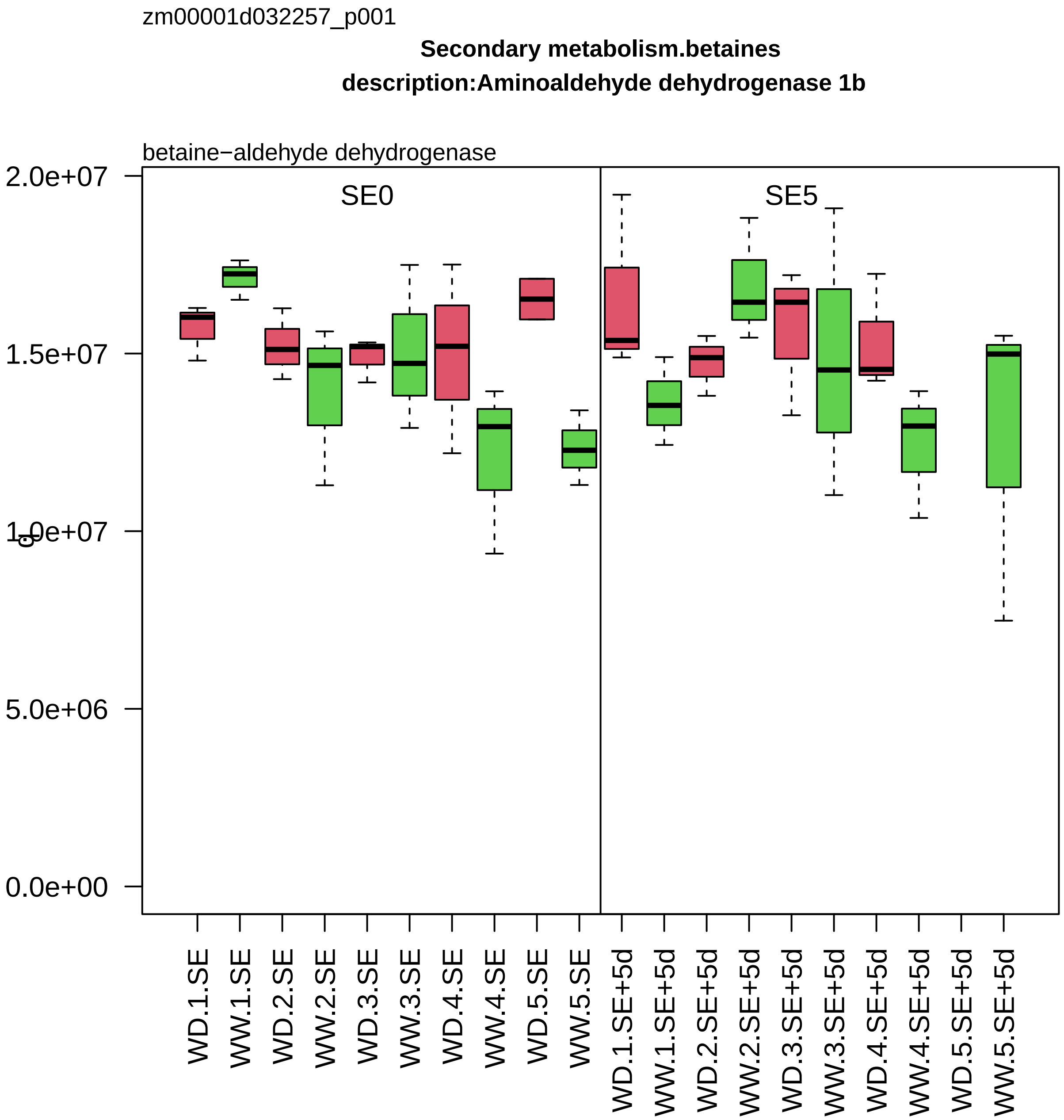
zm00001d032257_p001
Secondary metabolism.betaines
description:Aminoaldehyde dehydrogenase 1b
betaine−aldehyde dehydrogenase
SE0
SE5
2.0e+07
1.5e+07
1.0e+07
5.0e+06
0.0e+00
WD.1.SE
WW.1.SE
WD.2.SE
WW.2.SE
WD.3.SE
WW.3.SE
WD.4.SE
WW.4.SE
WD.5.SE
WW.5.SE
WD.1.SE+5d
WW.1.SE+5d
WD.2.SE+5d
WW.2.SE+5d
WD.3.SE+5d
WW.3.SE+5d
WD.4.SE+5d
WW.4.SE+5d
WD.5.SE+5d
WW.5.SE+5d

zm00001d032282_p012

**Protein homeostasis.proteolysis**
**description:Aminopeptidase M1**

aminopeptidase *(APM1)

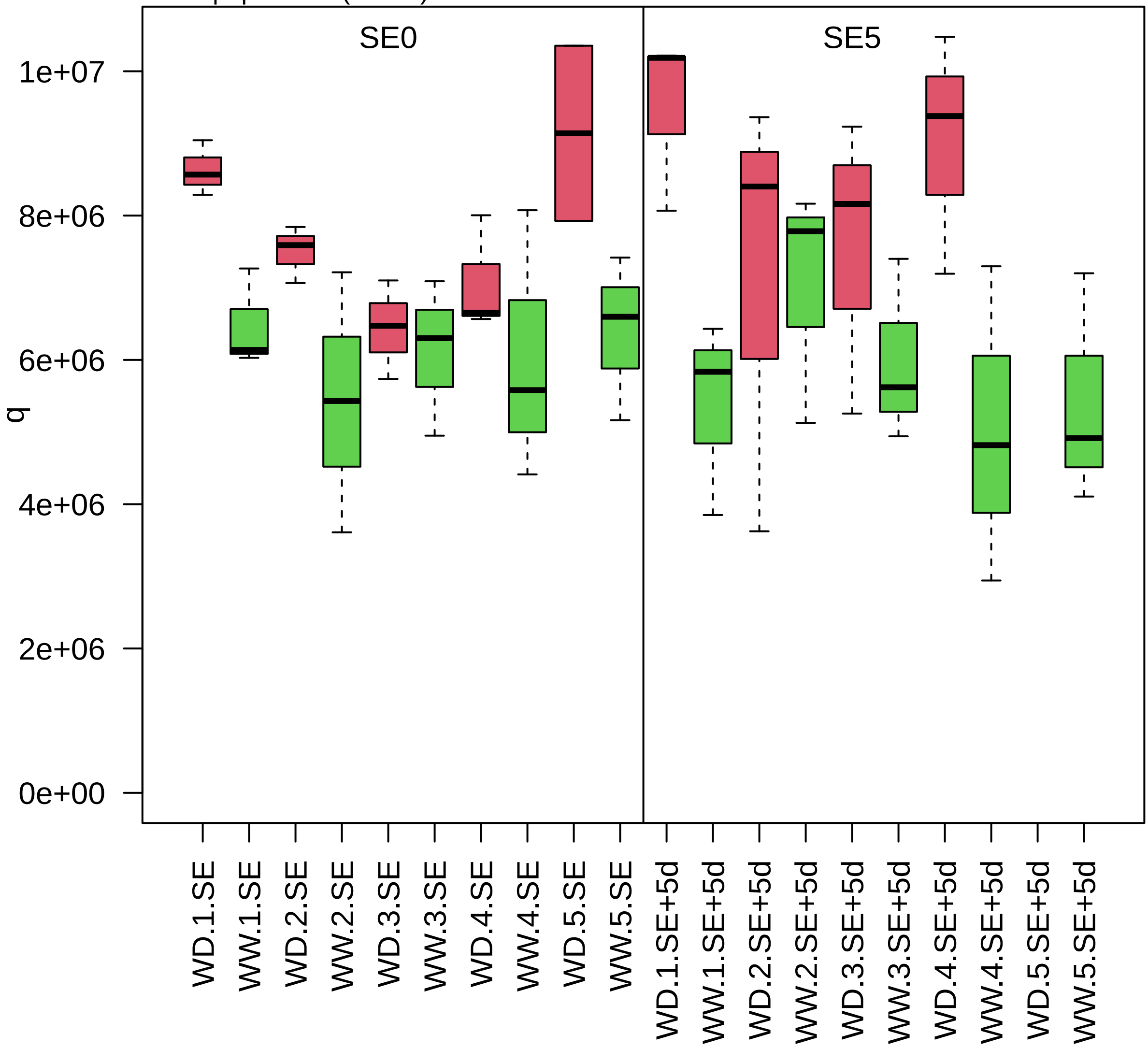

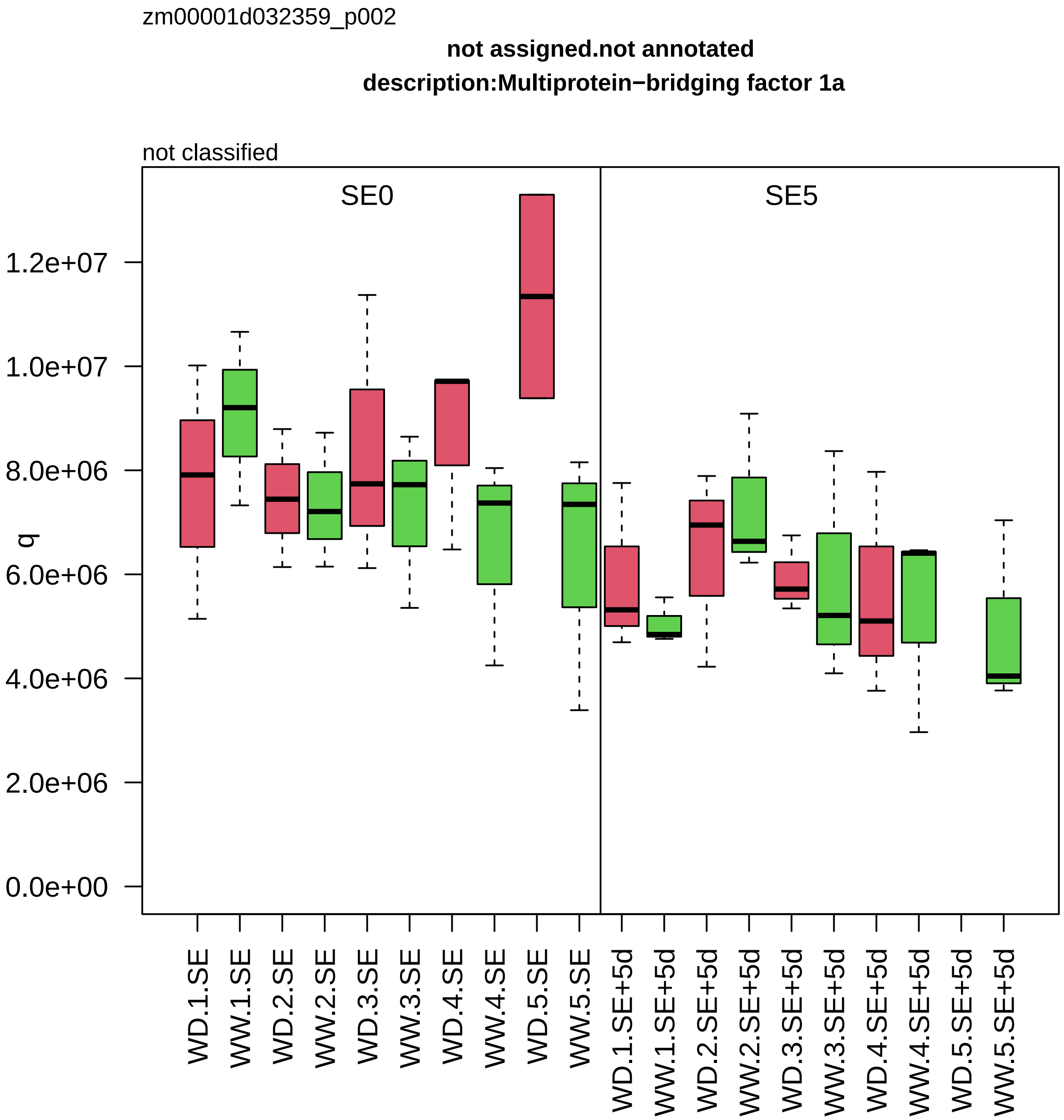
zm00001d032359_p002
not assigned.not annotated
description:Multiprotein−bridging factor 1a
not classified
SE0
SE5
q
1.2e+07
1.0e+07
8.0e+06
6.0e+06
4.0e+06
2.0e+06
0.0e+00
WD.1.SE
WW.1.SE
WD.2.SE
WW.2.SE
WD.3.SE
WW.3.SE
WD.4.SE
WW.4.SE
WD.5.SE
WW.5.SE
WD.1.SE+5d
WW.1.SE+5d
WD.2.SE+5d
WW.2.SE+5d
WD.3.SE+5d
WW.3.SE+5d
WD.4.SE+5d
WW.4.SE+5d
WD.5.SE+5d
WW.5.SE+5d

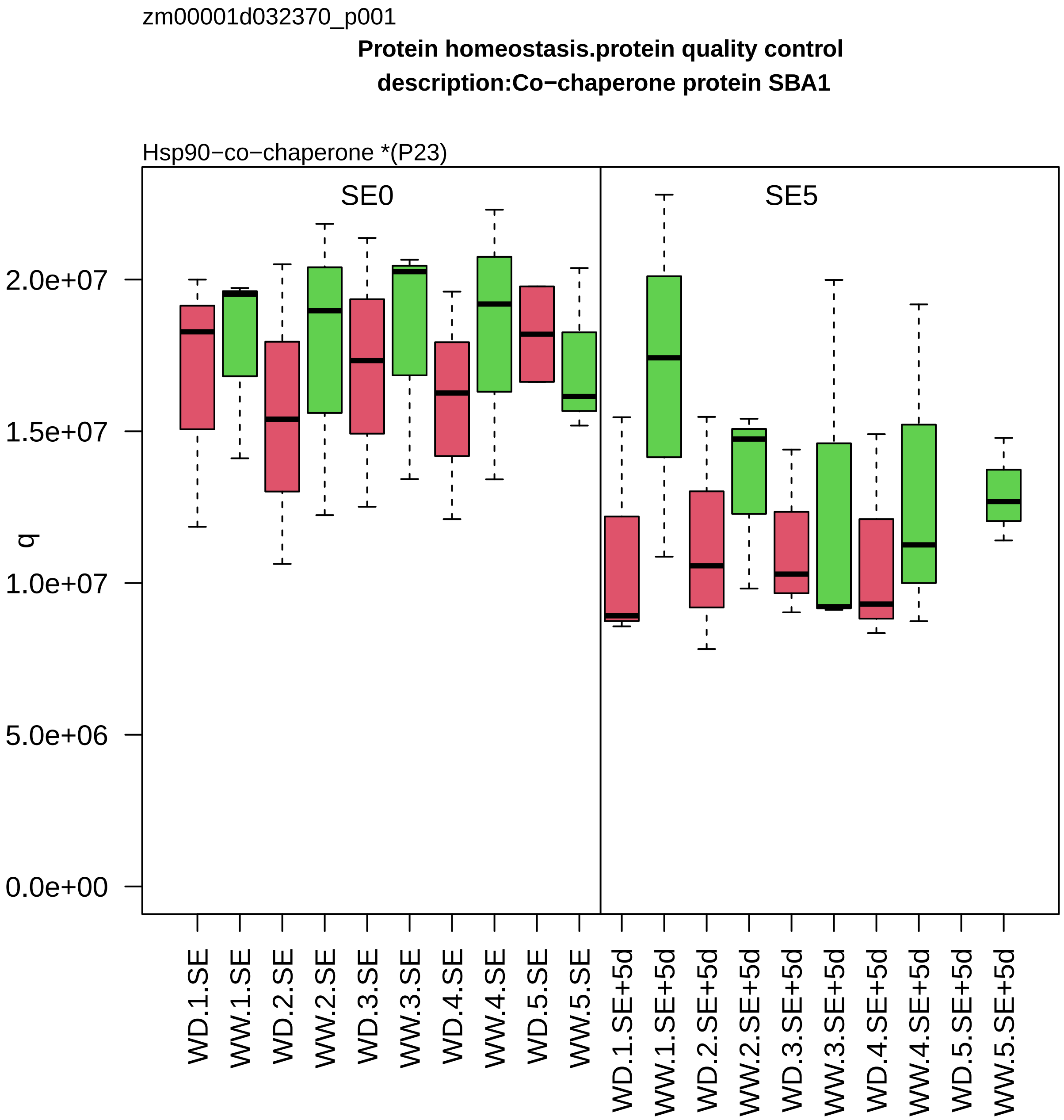
zm00001d032370_p001
Protein homeostasis.protein quality control
description:Co−chaperone protein SBA1
Hsp90−co−chaperone *(P23)
SE0
SE5
q
2.0e+07
1.5e+07
1.0e+07
5.0e+06
0.0e+00
WD.1.SE
WW.1.SE
WD.2.SE
WW.2.SE
WD.3.SE
WW.3.SE
WD.4.SE
WW.4.SE
WD.5.SE
WW.5.SE
WD.1.SE+5d
WW.1.SE+5d
WD.2.SE+5d
WW.2.SE+5d
WD.3.SE+5d
WW.3.SE+5d
WD.4.SE+5d
WW.4.SE+5d
WD.5.SE+5d
WW.5.SE+5d

zm00001d032385_p001

**Carbohydrate metabolism.starch metabolism**

**symbol:ADP glucose pyrophosphorylase small subunit leaf1 description: ADP glucose pyrophosphorylase**

small subunit *(APS) of ADP−glucose pyrophosphorylase

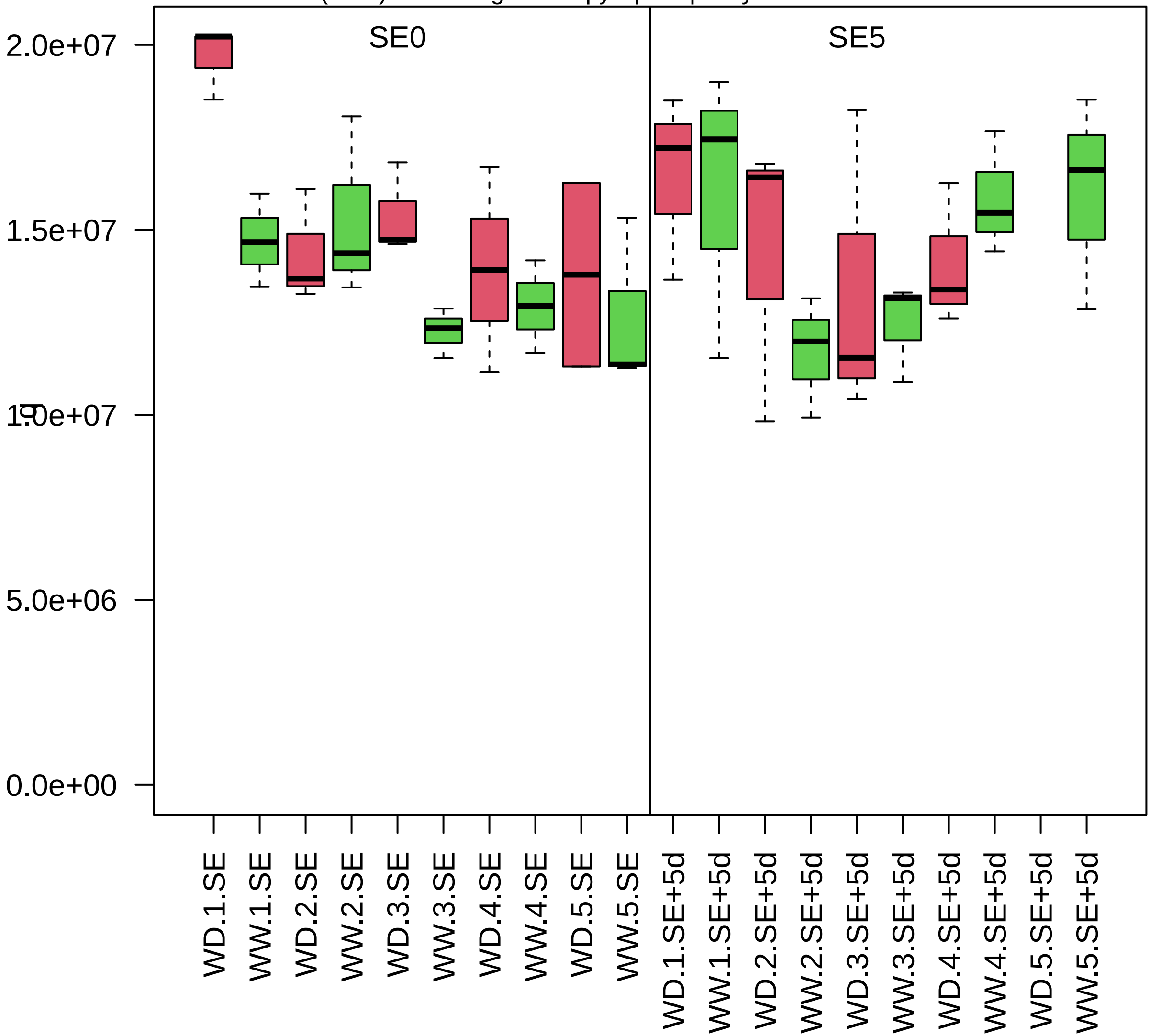

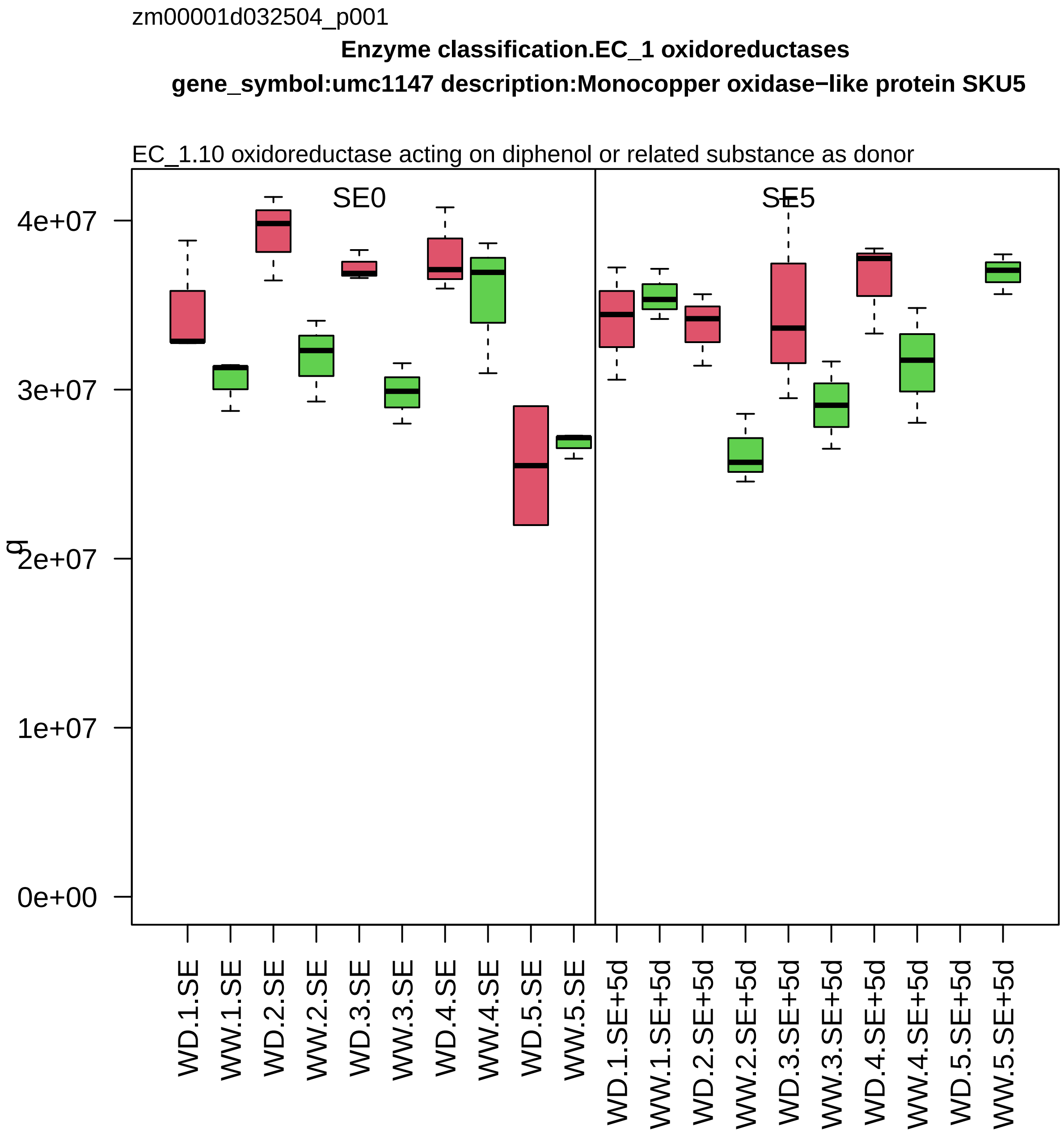

zm00001d032504_p001
Enzyme classification.EC_1 oxidoreductases
gene_symbol:umc1147 description:Monocopper oxidase−like protein SKU5
EC_1.10 oxidoreductase acting on diphenol or related substance as donor
SE0
SE5
g
0e+00
1e+07
2e+07
3e+07
4e+07
WD.1.SE
WW.1.SE
WD.2.SE
WW.2.SE
WD.3.SE
WW.3.SE
WD.4.SE
WW.4.SE
WD.5.SE
WW.5.SE
WD.1.SE+5d
WW.1.SE+5d
WD.2.SE+5d
WW.2.SE+5d
WD.3.SE+5d
WW.3.SE+5d
WD.4.SE+5d
WW.4.SE+5d
WD.5.SE+5d
WW.5.SE+5d

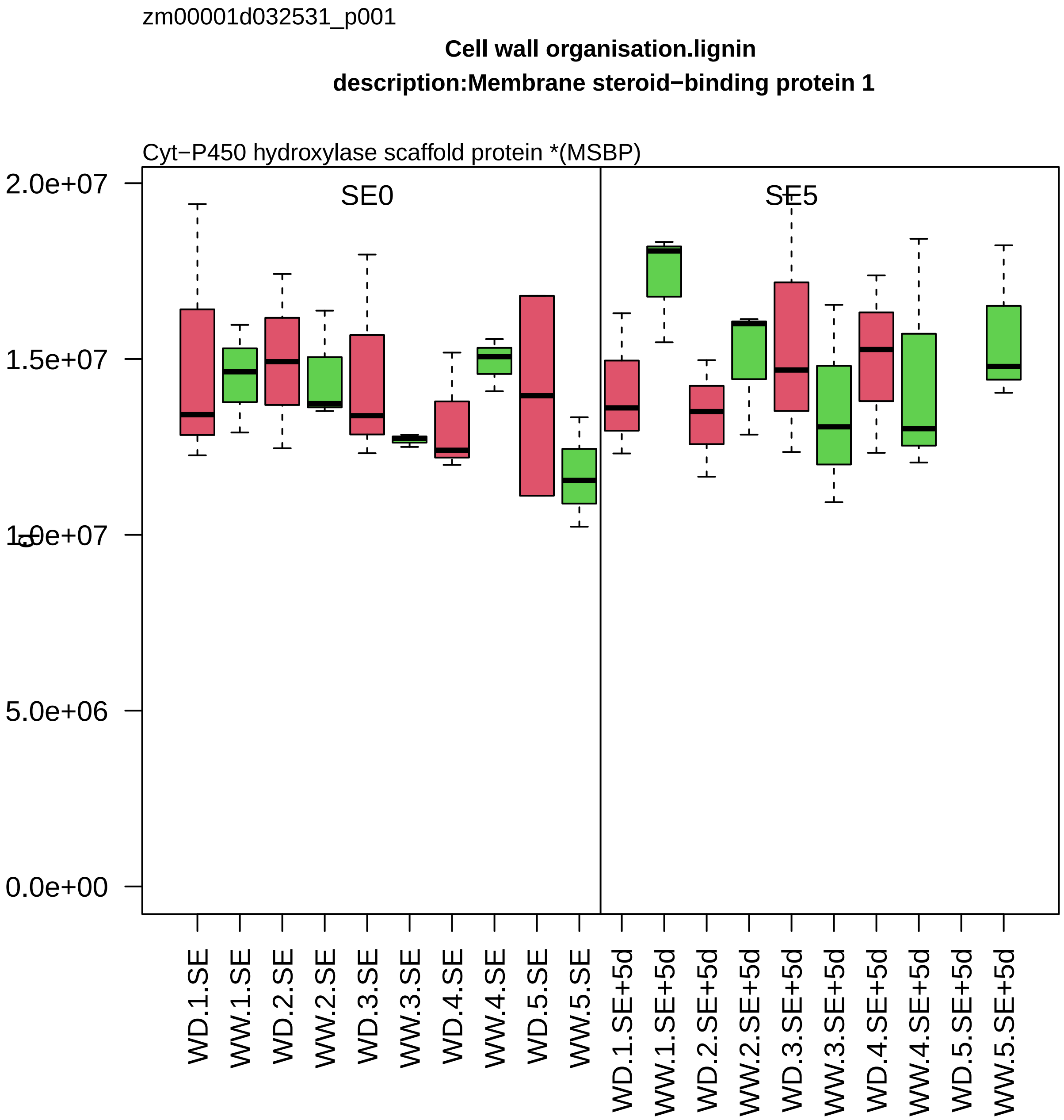
zm00001d032531_p001
Cell wall organisation.lignin
description:Membrane steroid−binding protein 1
Cyt−P450 hydroxylase scaffold protein *(MSBP)
SE0
SE5
2.0e+07
1.5e+07
1.0e+07
5.0e+06
0.0e+00
WD.1.SE
WW.1.SE
WD.2.SE
WW.2.SE
WD.3.SE
WW.3.SE
WD.4.SE
WW.4.SE
WD.5.SE
WW.5.SE
WD.1.SE+5d
WW.1.SE+5d
WD.2.SE+5d
WW.2.SE+5d
WD.3.SE+5d
WW.3.SE+5d
WD.4.SE+5d
WW.4.SE+5d
WD.5.SE+5d
WW.5.SE+5d

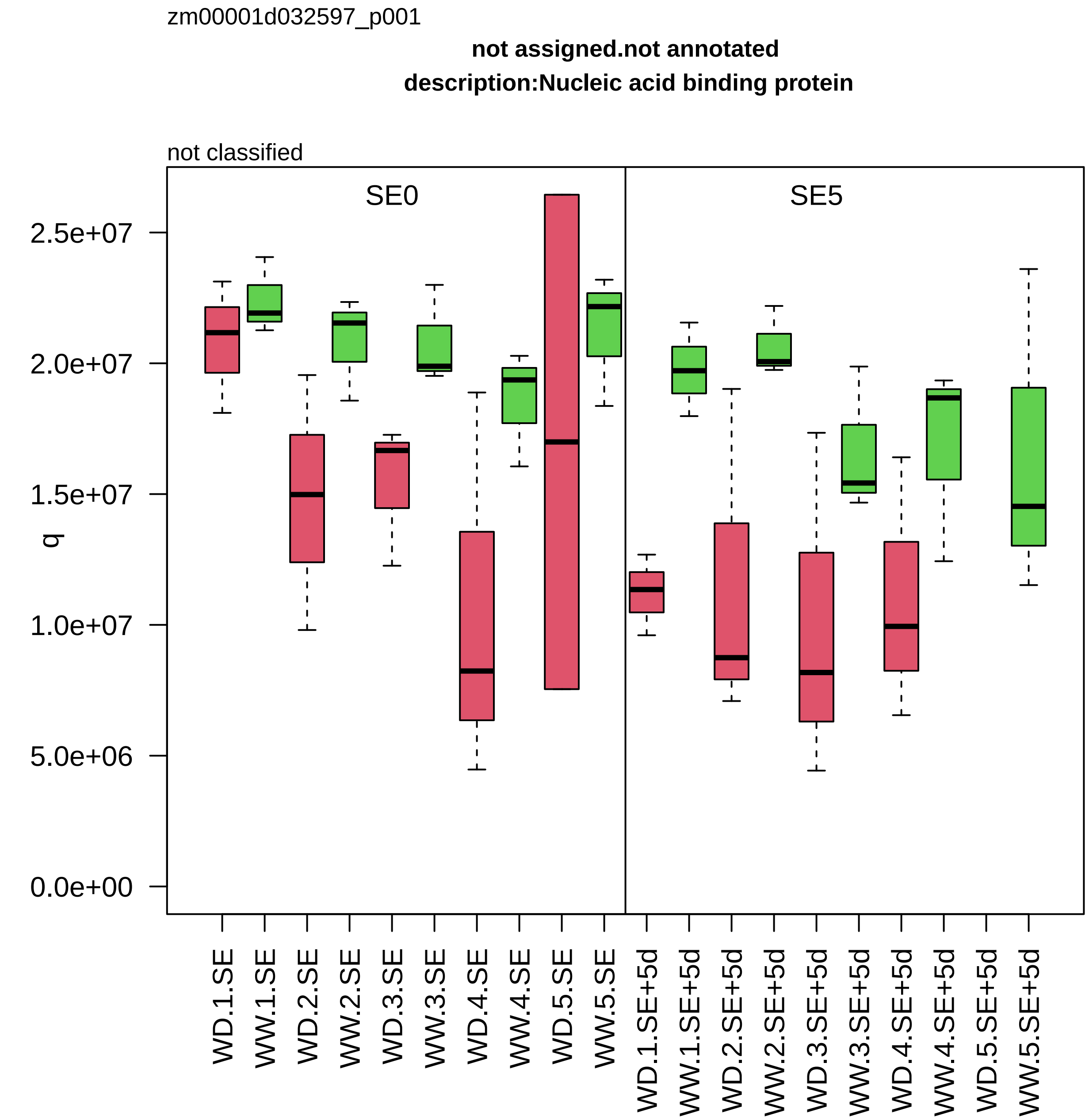

zm00001d032597_p001
not assigned.not annotated
description:Nucleic acid binding protein
not classified
SE0
SE5
q
2.5e+07
2.0e+07
1.5e+07
1.0e+07
5.0e+06
0.0e+00
WD.1.SE
WW.1.SE
WD.2.SE
WW.2.SE
WD.3.SE
WW.3.SE
WD.4.SE
WW.4.SE
WD.5.SE
WW.5.SE
WD.1.SE+5d
WW.1.SE+5d
WD.2.SE+5d
WW.2.SE+5d
WD.3.SE+5d
WW.3.SE+5d
WD.4.SE+5d
WW.4.SE+5d
WD.5.SE+5d
WW.5.SE+5d

zm00001d032695_p001

**Photosynthesis.CAM/C4 photosynthesis**

**gene_symbol:malate dehydrogenase4 description:Malate dehydrogenase%2C cytoplasmic**

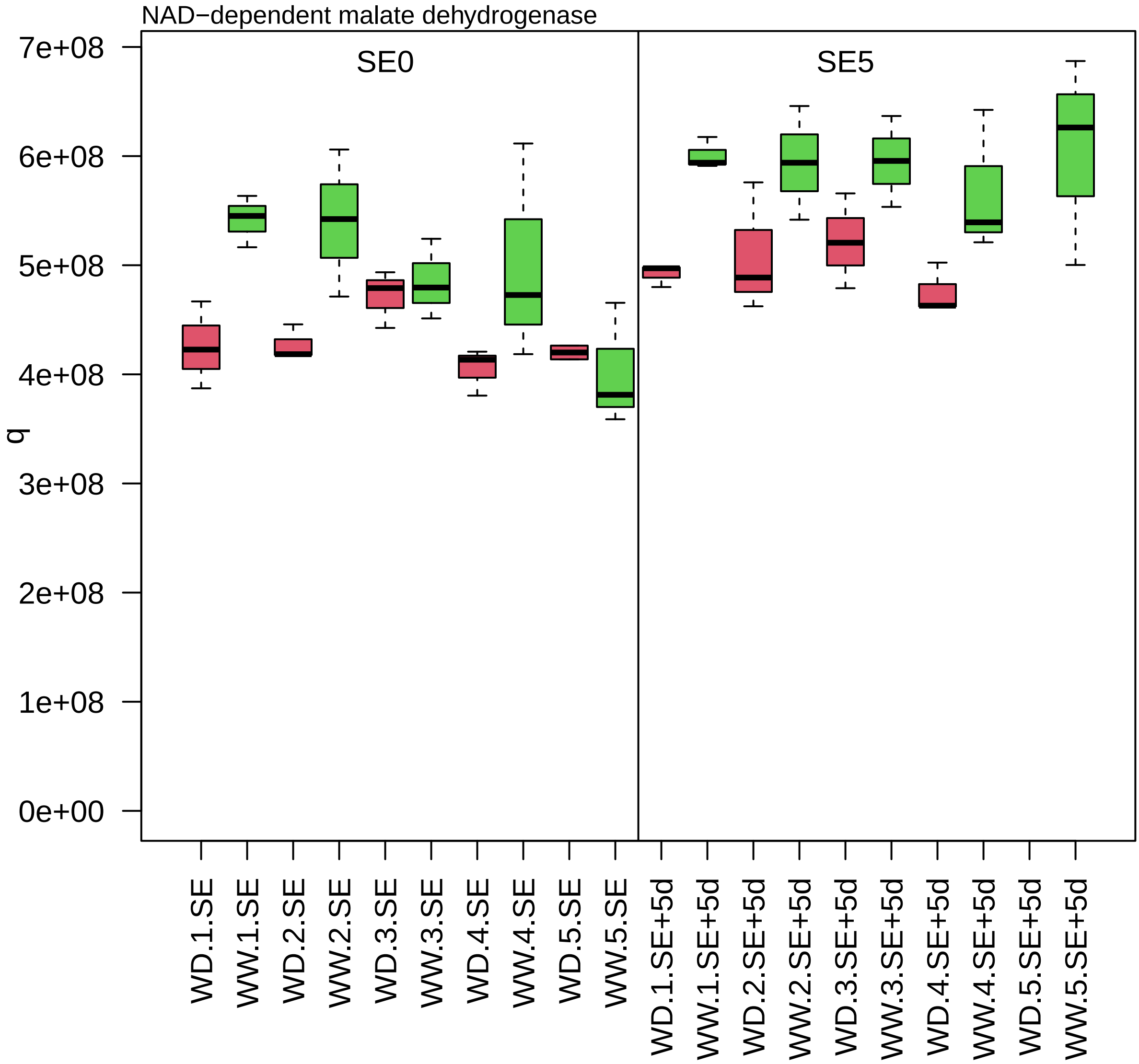

zm00001d032950_p001

**Redox homeostasis.ascorbate−based redox regulation**
**description:GDP−mannose 35−epimerase**

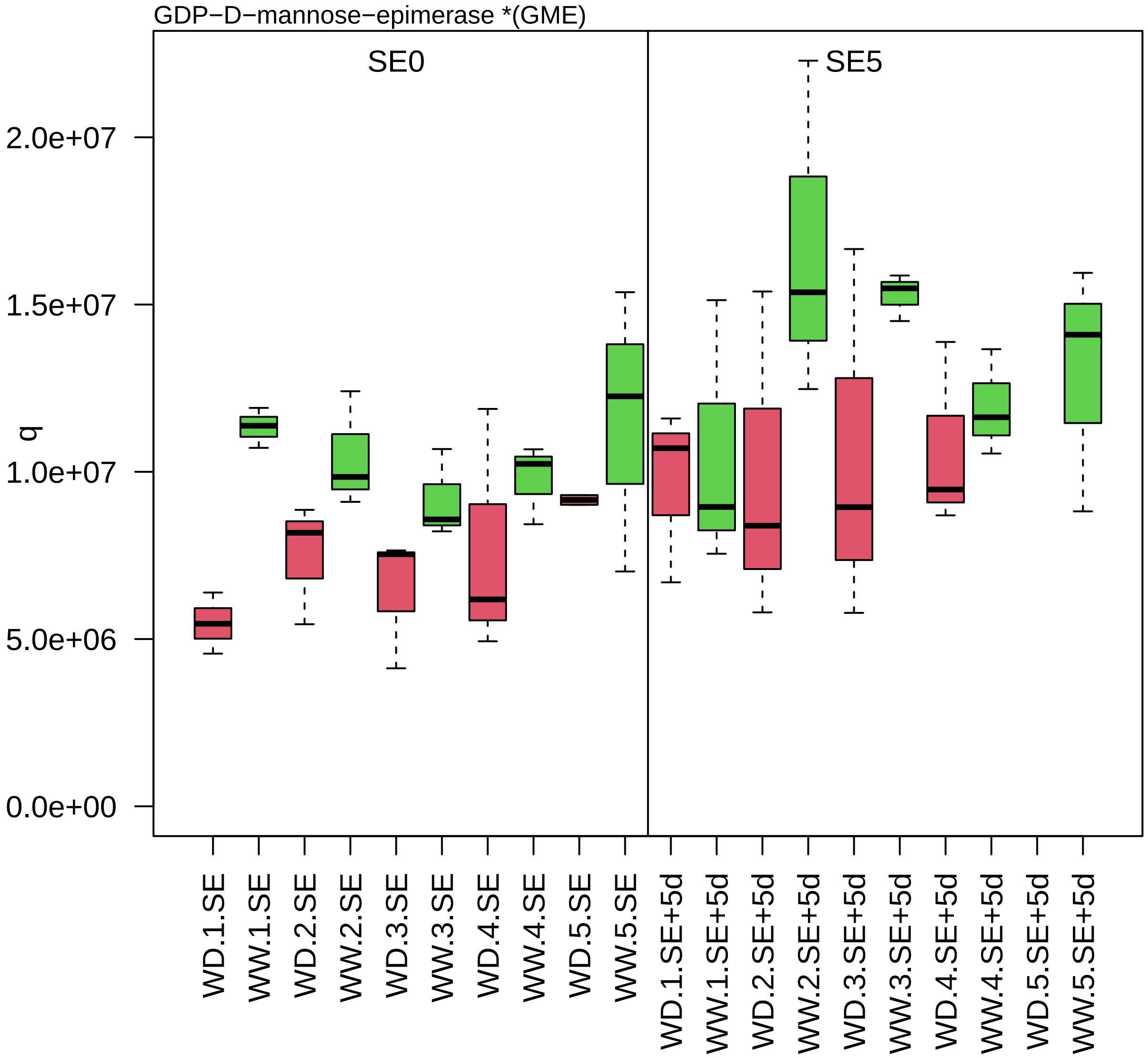

zm00001d032956_p008

**Enzyme classification.EC_3 hydrolases**
**description:Acylamino−acid−releasing enzyme**

EC_3.4 hydrolase acting on peptide bond (peptidase)

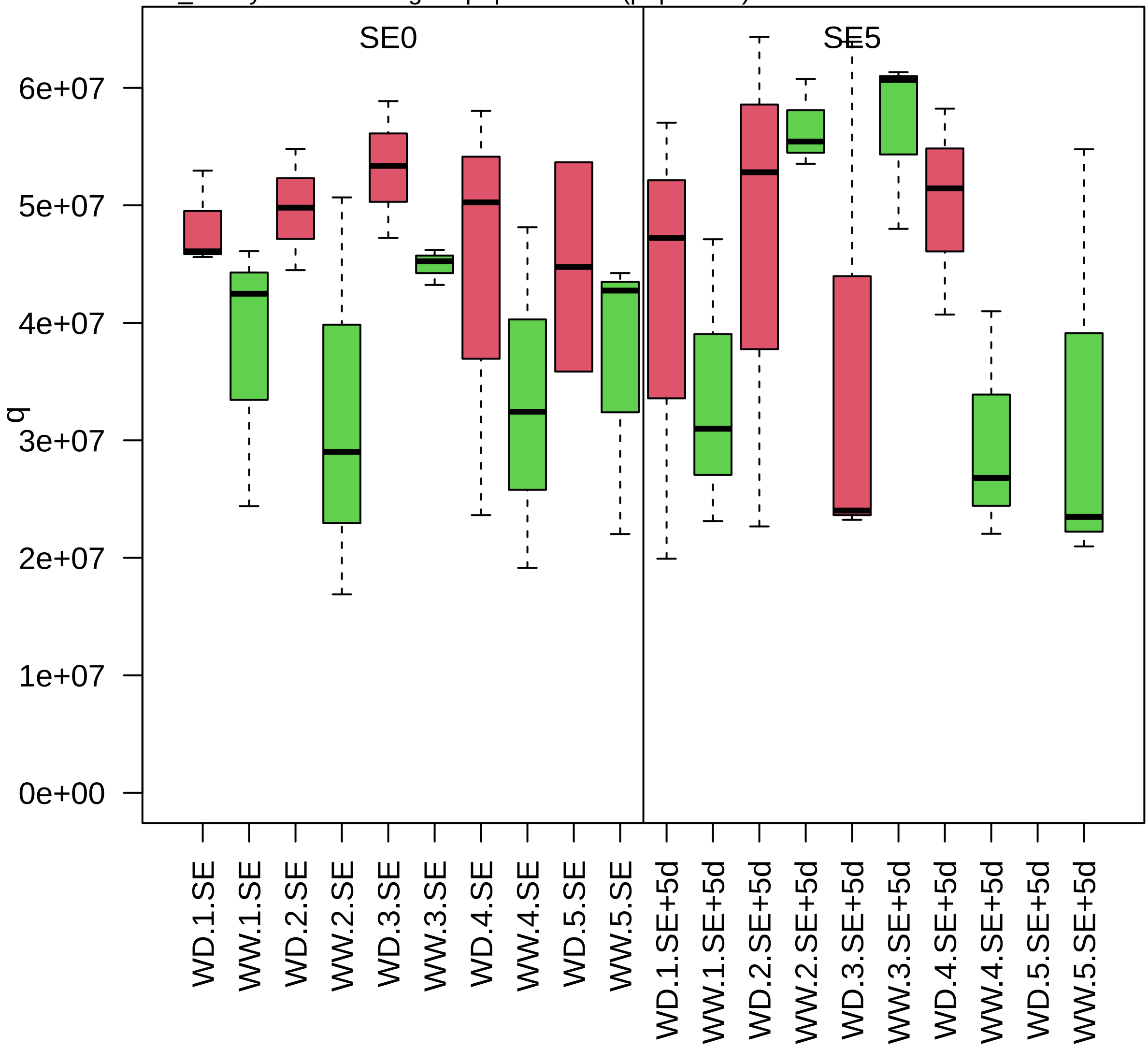

zm00001d033174_p001

**Carbohydrate metabolism.nucleotide sugar biosynthesis**

**gene_symbol:Golgi associated protein homolog description: Golgi associated protein homolog**

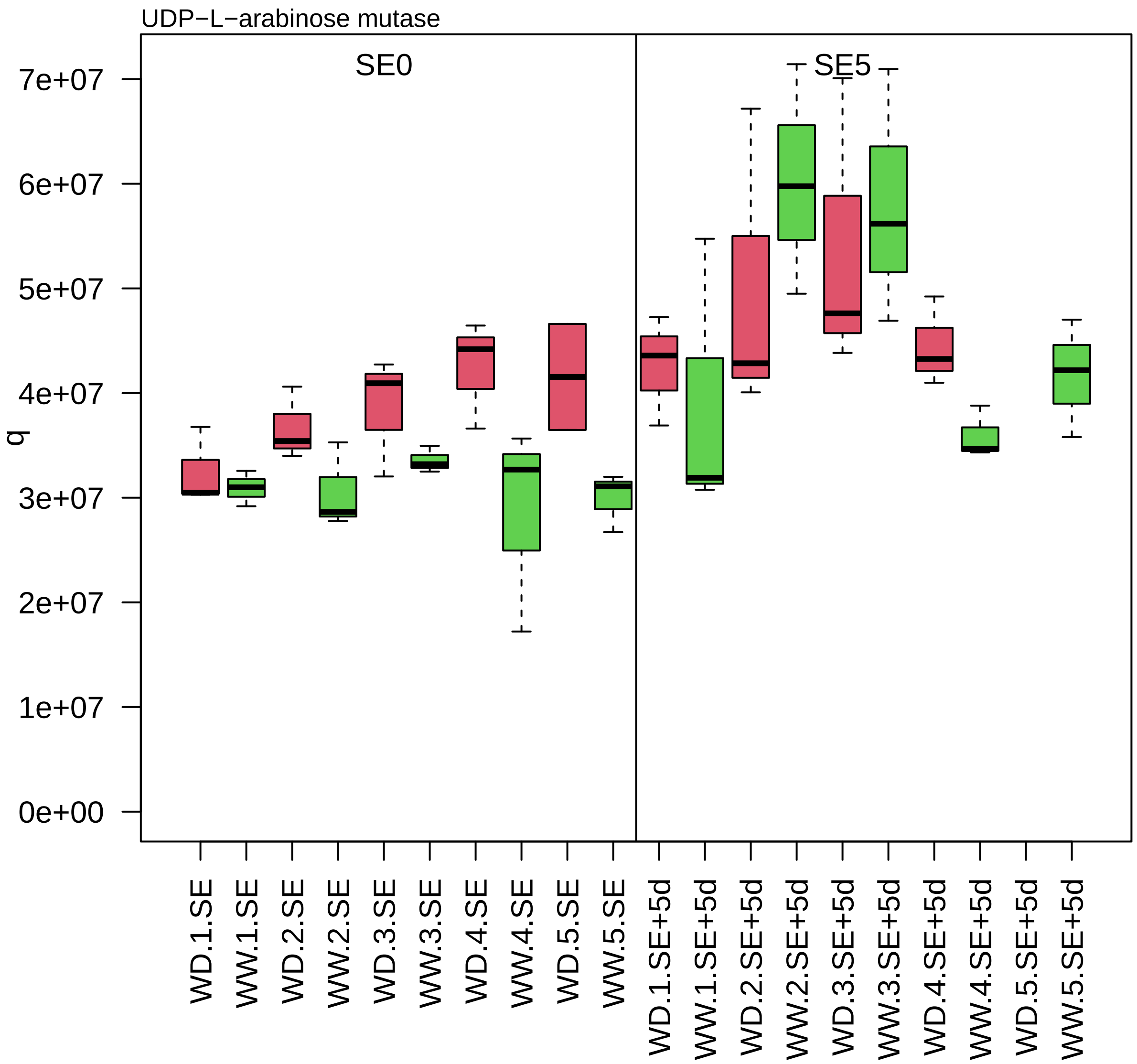

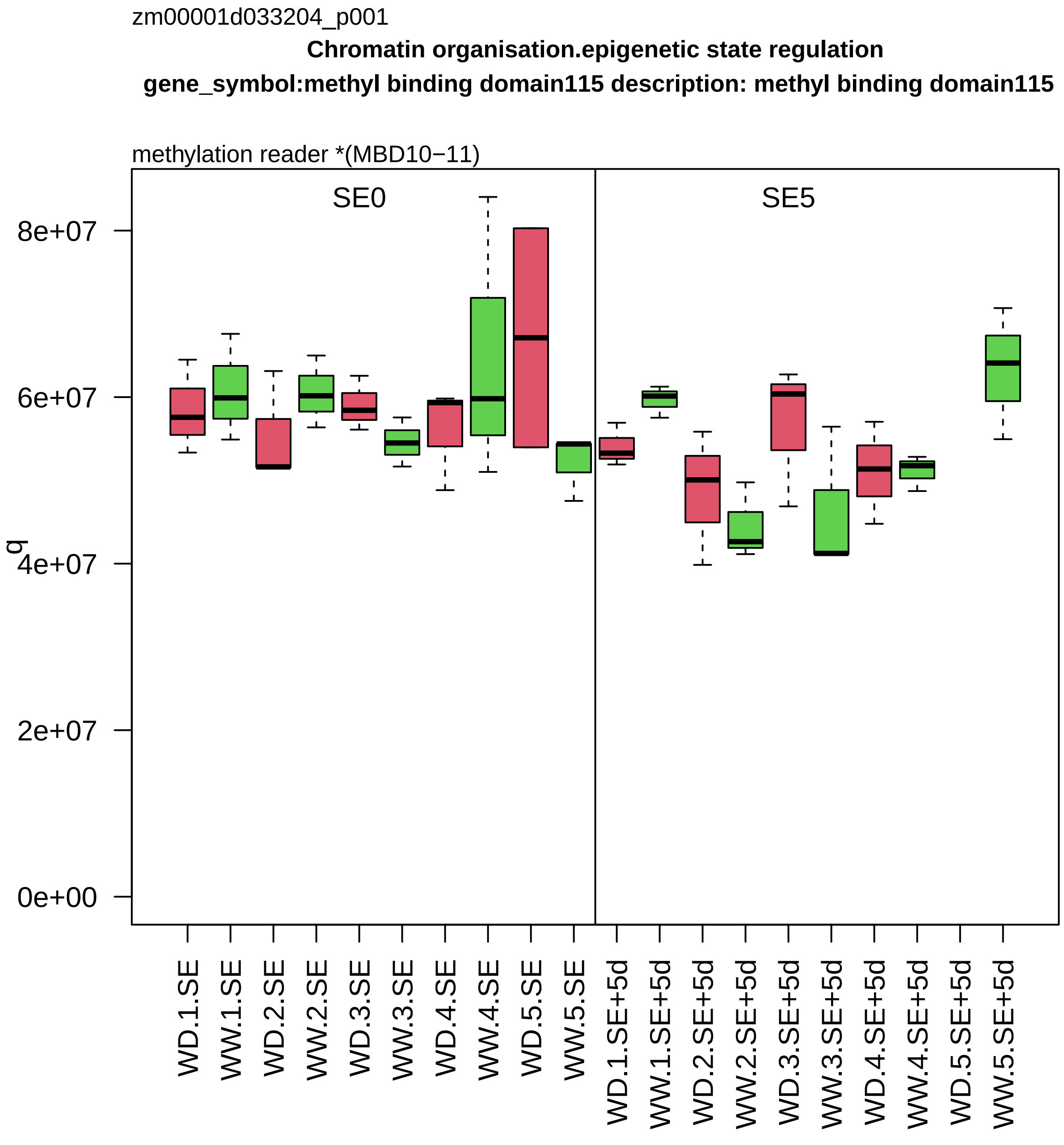
zm00001d033204_p001
Chromatin organisation.epigenetic state regulation
gene_symbol:methyl binding domain115 description: methyl binding domain115
methylation reader *(MBD10−11)
SE0
SE5
q
8e+07
6e+07
4e+07
2e+07
0e+00
WD.1.SE
WW.1.SE
WD.2.SE
WW.2.SE
WD.3.SE
WW.3.SE
WD.4.SE
WW.4.SE
WD.5.SE
WW.5.SE
WD.1.SE+5d
WW.1.SE+5d
WD.2.SE+5d
WW.2.SE+5d
WD.3.SE+5d
WW.3.SE+5d
WD.4.SE+5d
WW.4.SE+5d
WD.5.SE+5d
WW.5.SE+5d

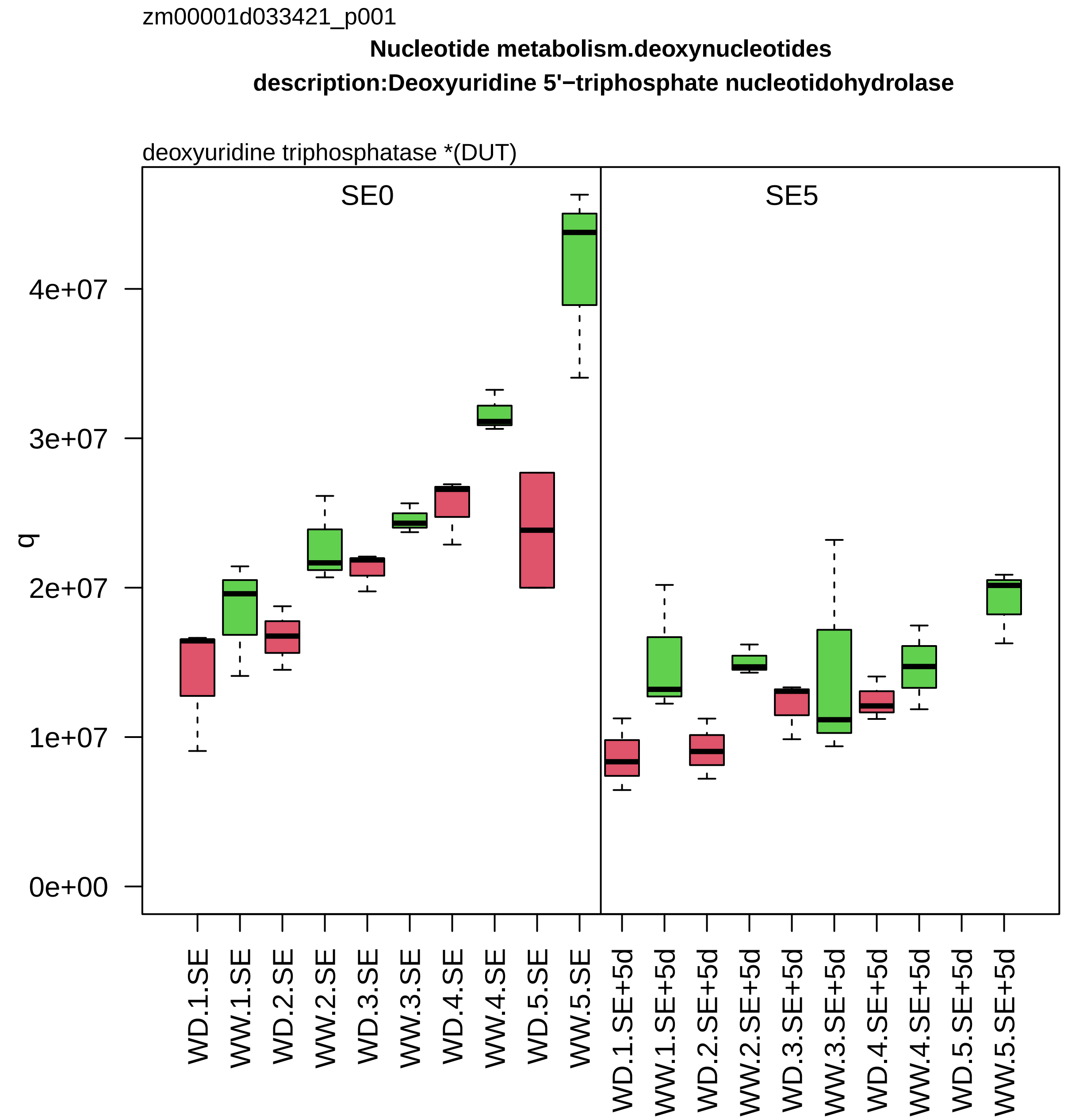
zm00001d033421_p001
Nucleotide metabolism.deoxynucleotides
description:Deoxyuridine 5'−triphosphate nucleotidohydrolase
deoxyuridine triphosphatase *(DUT)
SE0
SE5
q
0e+00
1e+07
2e+07
3e+07
4e+07
WD.1.SE
WW.1.SE
WD.2.SE
WW.2.SE
WD.3.SE
WW.3.SE
WD.4.SE
WW.4.SE
WD.5.SE
WW.5.SE
WD.1.SE+5d
WW.1.SE+5d
WD.2.SE+5d
WW.2.SE+5d
WD.3.SE+5d
WW.3.SE+5d
WD.4.SE+5d
WW.4.SE+5d
WD.5.SE+5d
WW.5.SE+5d

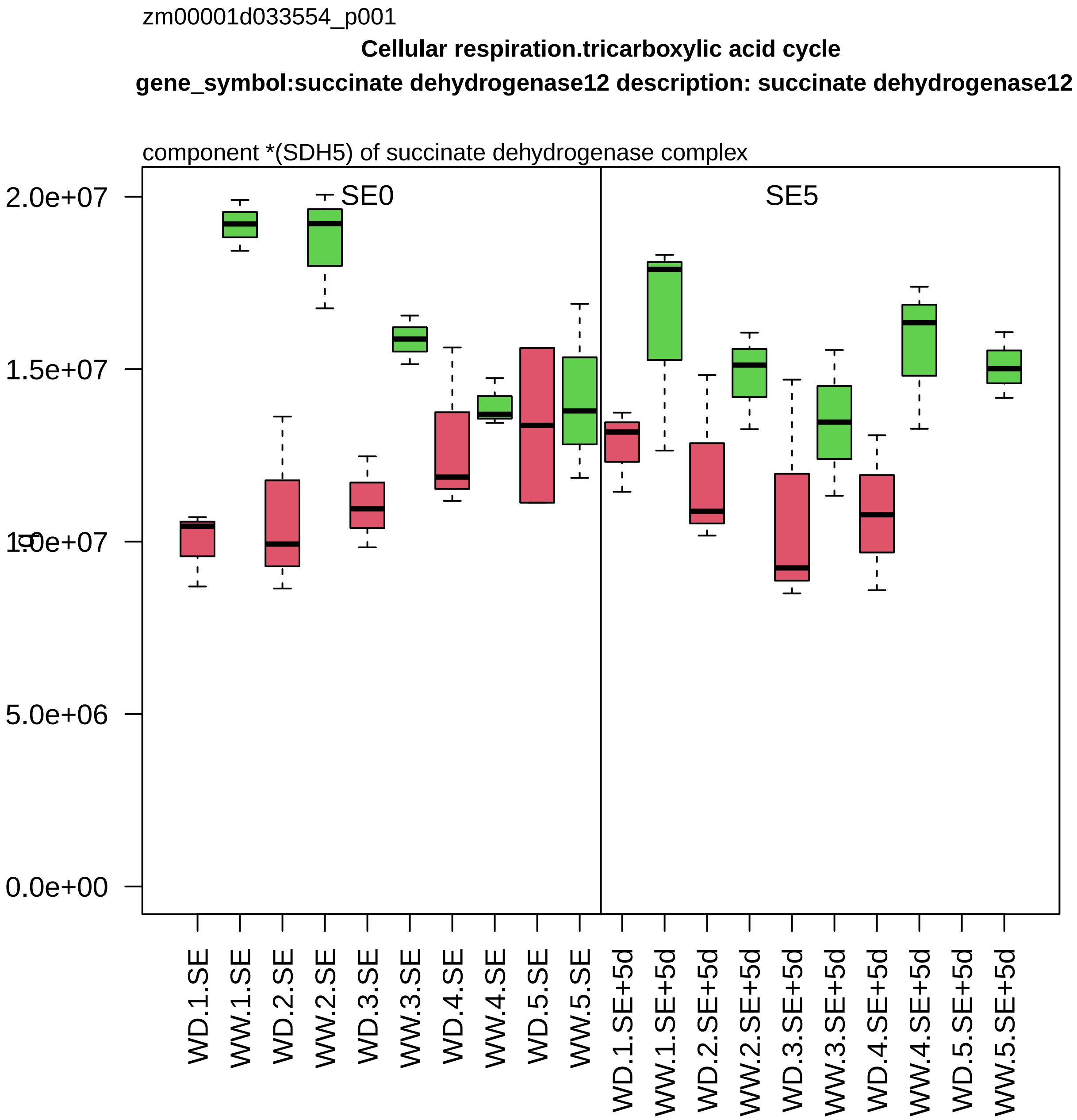

zm00001d033554_p001
Cellular respiration.tricarboxylic acid cycle
gene_symbol:succinate dehydrogenase12 description: succinate dehydrogenase12
component *(SDH5) of succinate dehydrogenase complex
SE0
SE5
2.0e+07
1.5e+07
1.0e+07
5.0e+06
0.0e+00
WD.1.SE
WW.1.SE
WD.2.SE
WW.2.SE
WD.3.SE
WW.3.SE
WD.4.SE
WW.4.SE
WD.5.SE
WW.5.SE
WD.1.SE+5d
WW.1.SE+5d
WD.2.SE+5d
WW.2.SE+5d
WD.3.SE+5d
WW.3.SE+5d
WD.4.SE+5d
WW.4.SE+5d
WD.5.SE+5d
WW.5.SE+5d

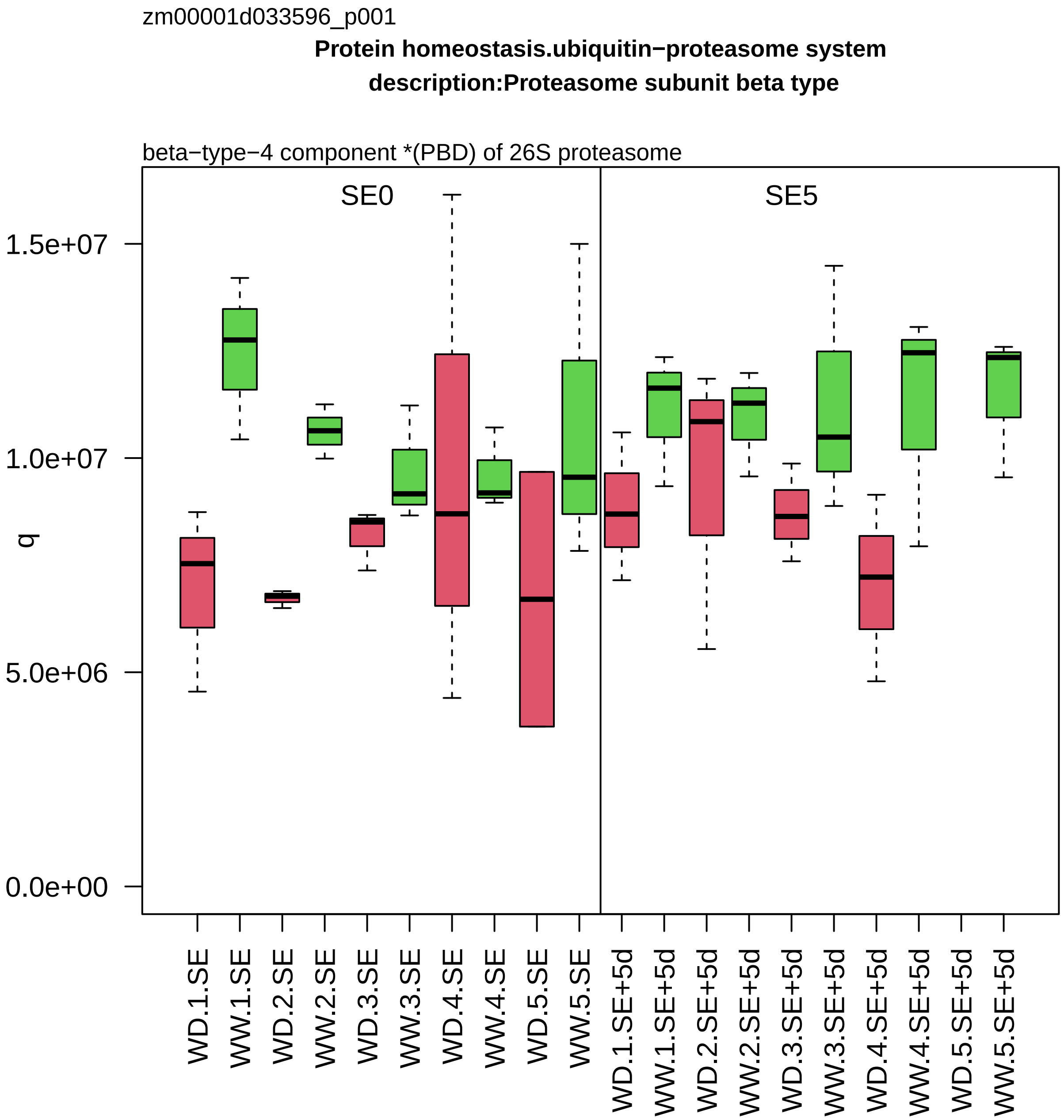

zm00001d033596_p001
Protein homeostasis.ubiquitin−proteasome system
description:Proteasome subunit beta type
beta−type−4 component *(PBD) of 26S proteasome
SE0
SE5
q
1.5e+07
1.0e+07
5.0e+06
0.0e+00
WD.1.SE
WW.1.SE
WD.2.SE
WW.2.SE
WD.3.SE
WW.3.SE
WD.4.SE
WW.4.SE
WD.5.SE
WW.5.SE
WD.1.SE+5d
WW.1.SE+5d
WD.2.SE+5d
WW.2.SE+5d
WD.3.SE+5d
WW.3.SE+5d
WD.4.SE+5d
WW.4.SE+5d
WD.5.SE+5d
WW.5.SE+5d

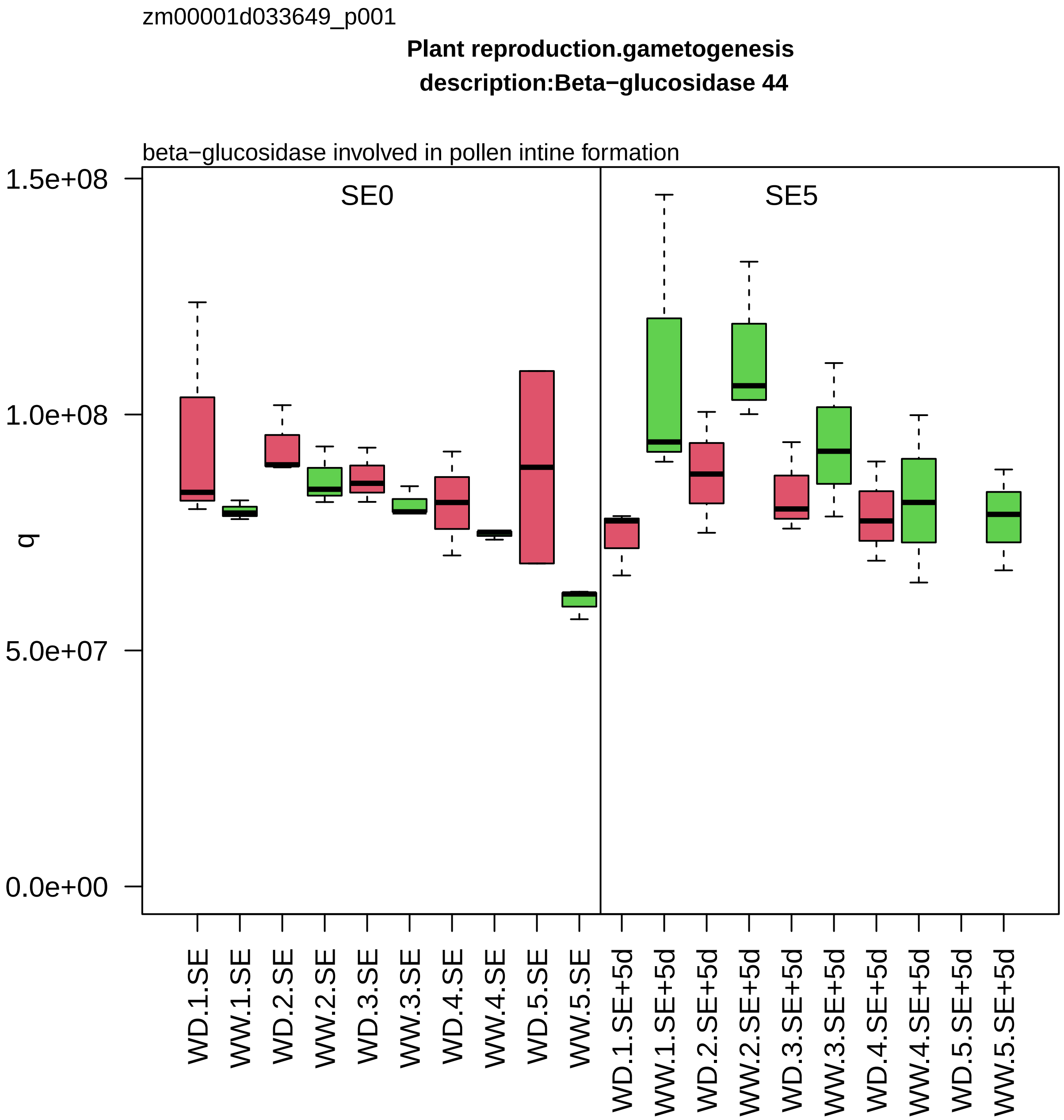

zm00001d033649_p001
Plant reproduction.gametogenesis
description:Beta−glucosidase 44
beta−glucosidase involved in pollen intine formation
SE0
SE5
q
1.5e+08
1.0e+08
5.0e+07
0.0e+00
WD.1.SE
WW.1.SE
WD.2.SE
WW.2.SE
WD.3.SE
WW.3.SE
WD.4.SE
WW.4.SE
WD.5.SE
WW.5.SE
WD.1.SE+5d
WW.1.SE+5d
WD.2.SE+5d
WW.2.SE+5d
WD.3.SE+5d
WW.3.SE+5d
WD.4.SE+5d
WW.4.SE+5d
WD.5.SE+5d
WW.5.SE+5d

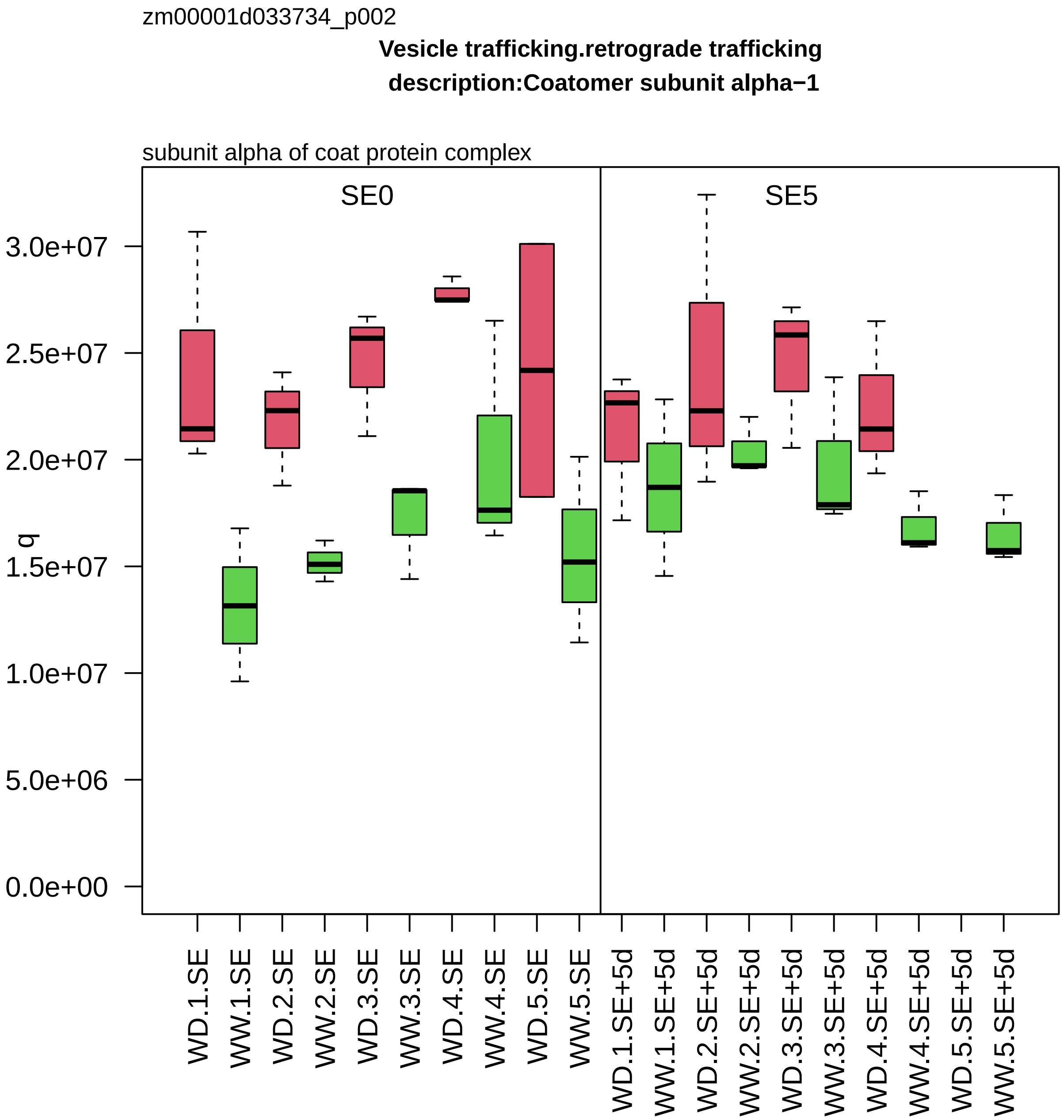

zm00001d033734_p002
Vesicle trafficking.retrograde trafficking
description:Coatomer subunit alpha−1
subunit alpha of coat protein complex
SE0
SE5
q
3.0e+07
2.5e+07
2.0e+07
1.5e+07
1.0e+07
5.0e+06
0.0e+00
WD.1.SE
WW.1.SE
WD.2.SE
WW.2.SE
WD.3.SE
WW.3.SE
WD.4.SE
WW.4.SE
WD.5.SE
WW.5.SE
WD.1.SE+5d
WW.1.SE+5d
WD.2.SE+5d
WW.2.SE+5d
WD.3.SE+5d
WW.3.SE+5d
WD.4.SE+5d
WW.4.SE+5d
WD.5.SE+5d
WW.5.SE+5d

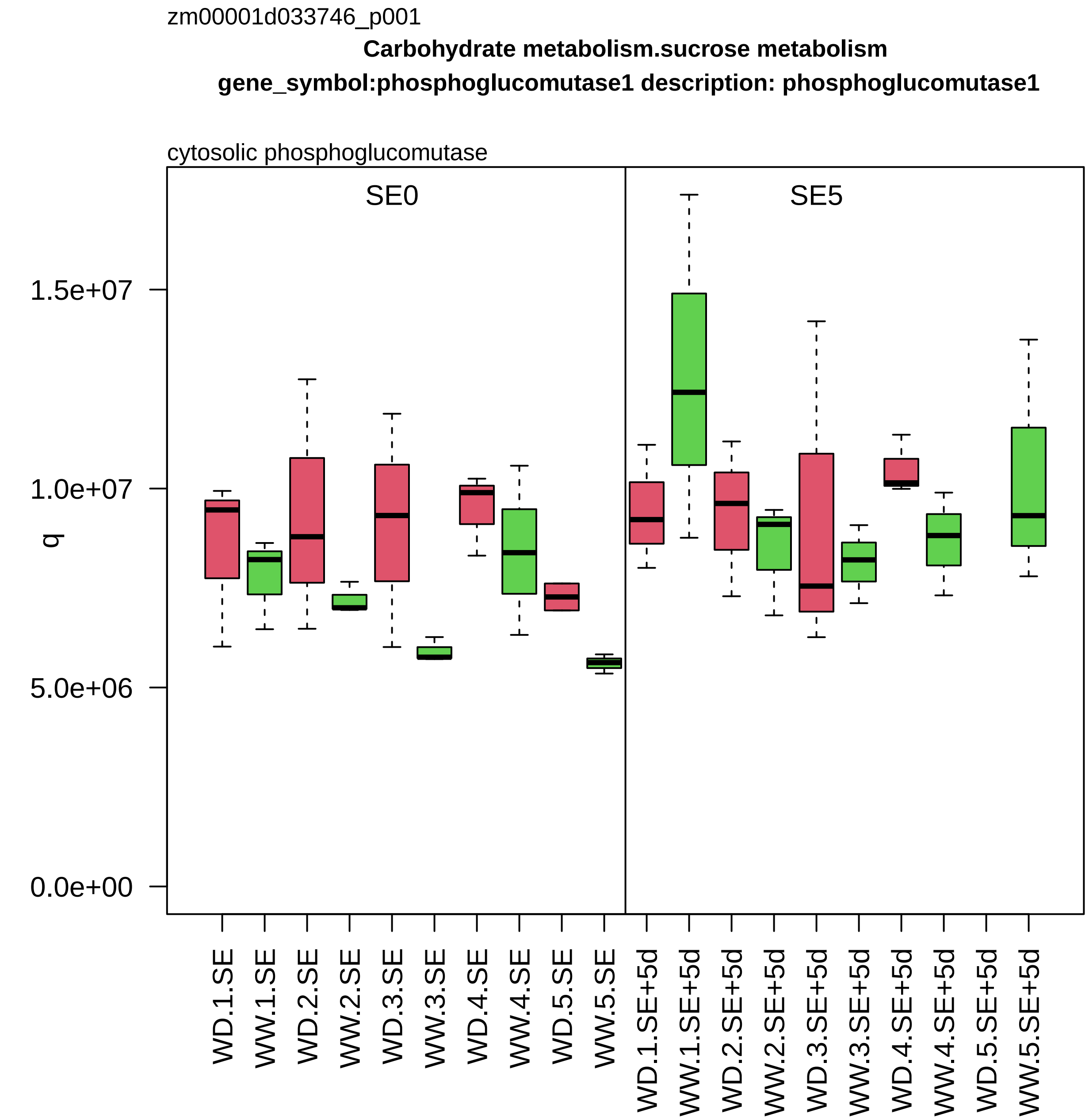
zm00001d033746_p001
Carbohydrate metabolism.sucrose metabolism
gene_symbol:phosphoglucomutase1 description: phosphoglucomutase1
cytosolic phosphoglucomutase
SE0
SE5
q
1.5e+07
1.0e+07
5.0e+06
0.0e+00
WD.1.SE
WW.1.SE
WD.2.SE
WW.2.SE
WD.3.SE
WW.3.SE
WD.4.SE
WW.4.SE
WD.5.SE
WW.5.SE
WD.1.SE+5d
WW.1.SE+5d
WD.2.SE+5d
WW.2.SE+5d
WD.3.SE+5d
WW.3.SE+5d
WD.4.SE+5d
WW.4.SE+5d
WD.5.SE+5d
WW.5.SE+5d

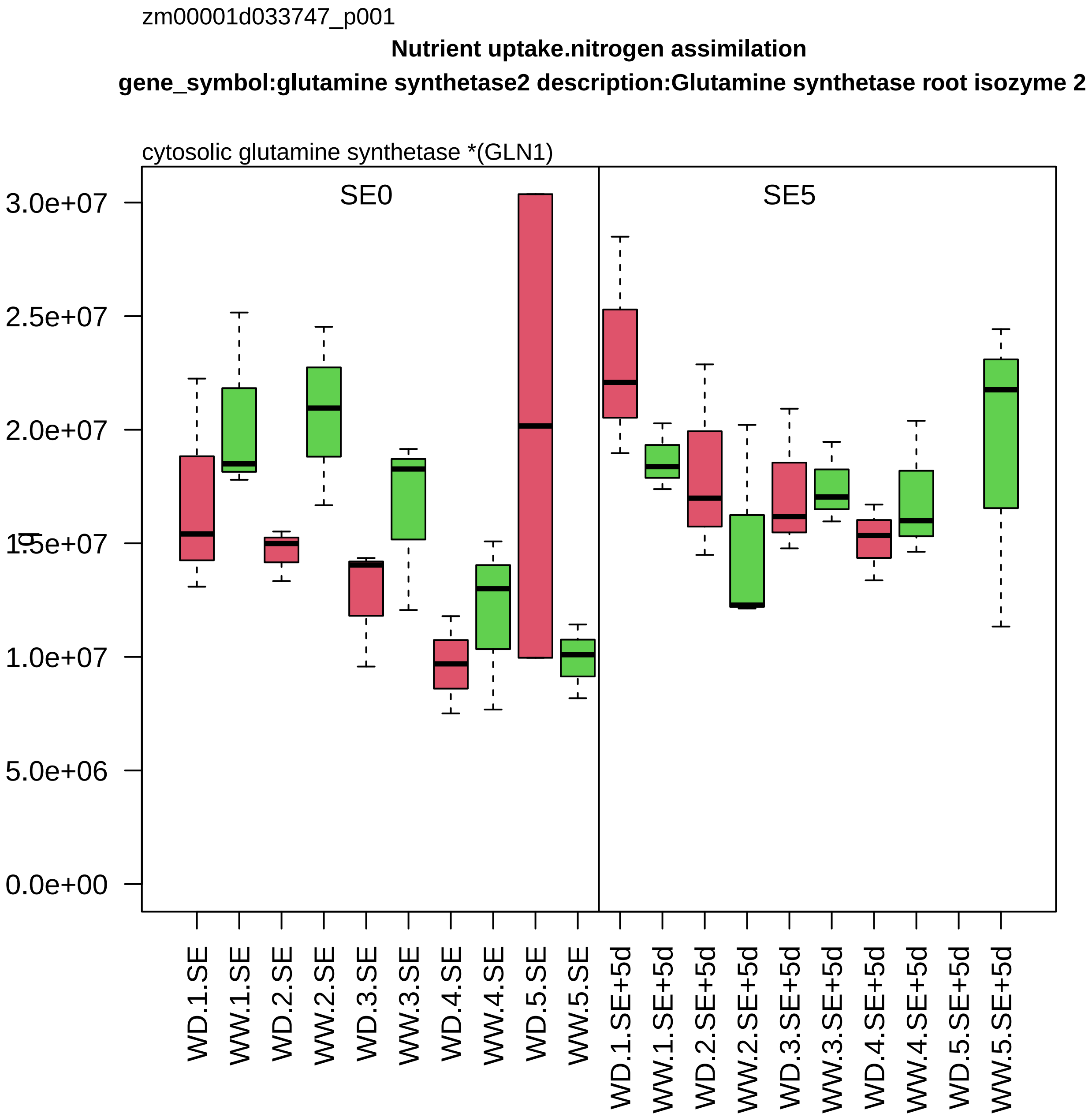

zm00001d033747_p001
Nutrient uptake.nitrogen assimilation
gene_symbol:glutamine synthetase2 description:Glutamine synthetase root isozyme 2
cytosolic glutamine synthetase *(GLN1)
SE0
SE5
3.0e+07
2.5e+07
2.0e+07
1.5e+07
1.0e+07
5.0e+06
0.0e+00
WD.1.SE
WW.1.SE
WD.2.SE
WW.2.SE
WD.3.SE
WW.3.SE
WD.4.SE
WW.4.SE
WD.5.SE
WW.5.SE
WD.1.SE+5d
WW.1.SE+5d
WD.2.SE+5d
WW.2.SE+5d
WD.3.SE+5d
WW.3.SE+5d
WD.4.SE+5d
WW.4.SE+5d
WD.5.SE+5d
WW.5.SE+5d

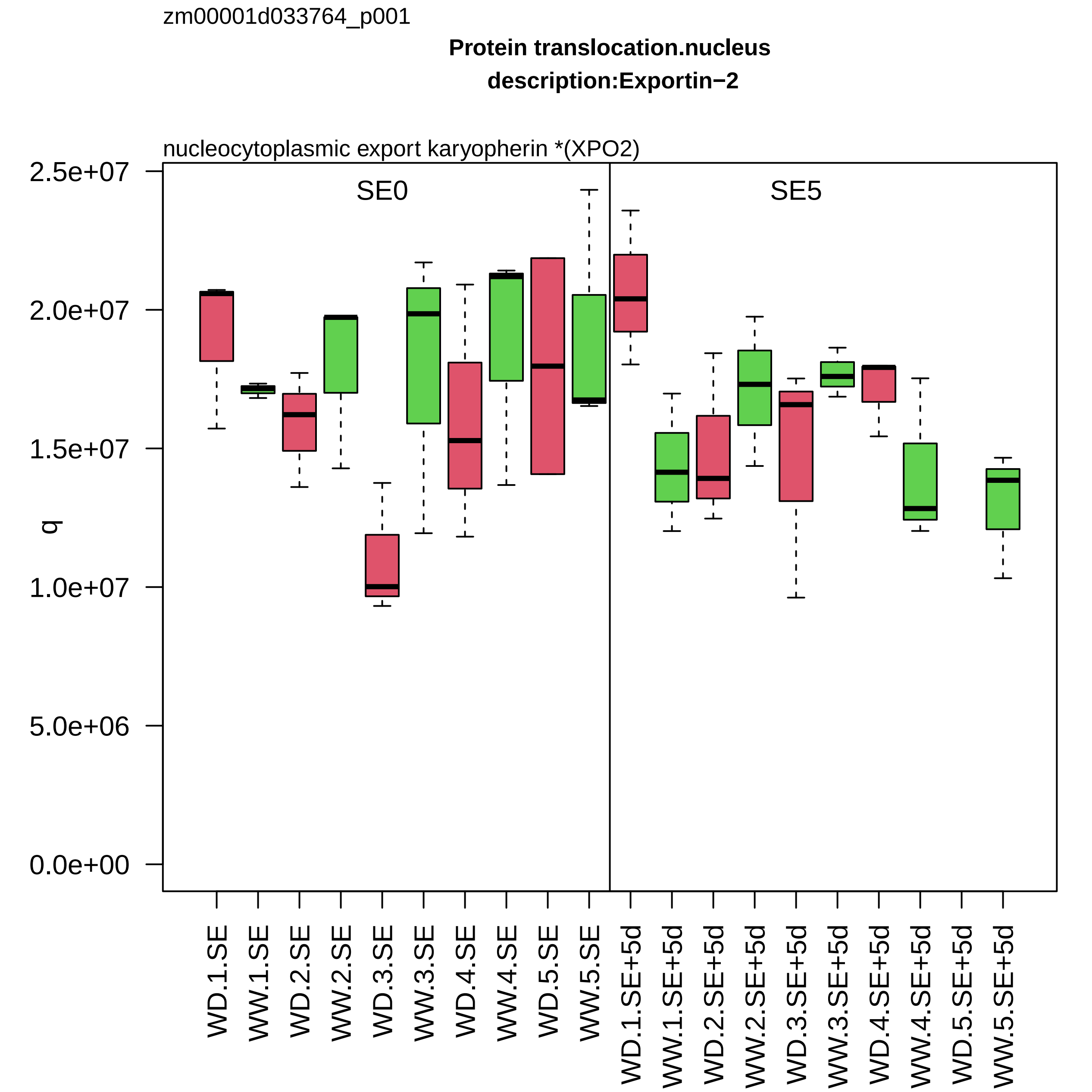

zm00001d033764_p001
Protein translocation.nucleus
description:Exportin−2
nucleocytoplasmic export karyopherin *(XPO2)
SE0
SE5
q
2.5e+07
2.0e+07
1.5e+07
1.0e+07
5.0e+06
0.0e+00
WD.1.SE
WW.1.SE
WD.2.SE
WW.2.SE
WD.3.SE
WW.3.SE
WD.4.SE
WW.4.SE
WD.5.SE
WW.5.SE
WD.1.SE+5d
WW.1.SE+5d
WD.2.SE+5d
WW.2.SE+5d
WD.3.SE+5d
WW.3.SE+5d
WD.4.SE+5d
WW.4.SE+5d
WD.5.SE+5d
WW.5.SE+5d

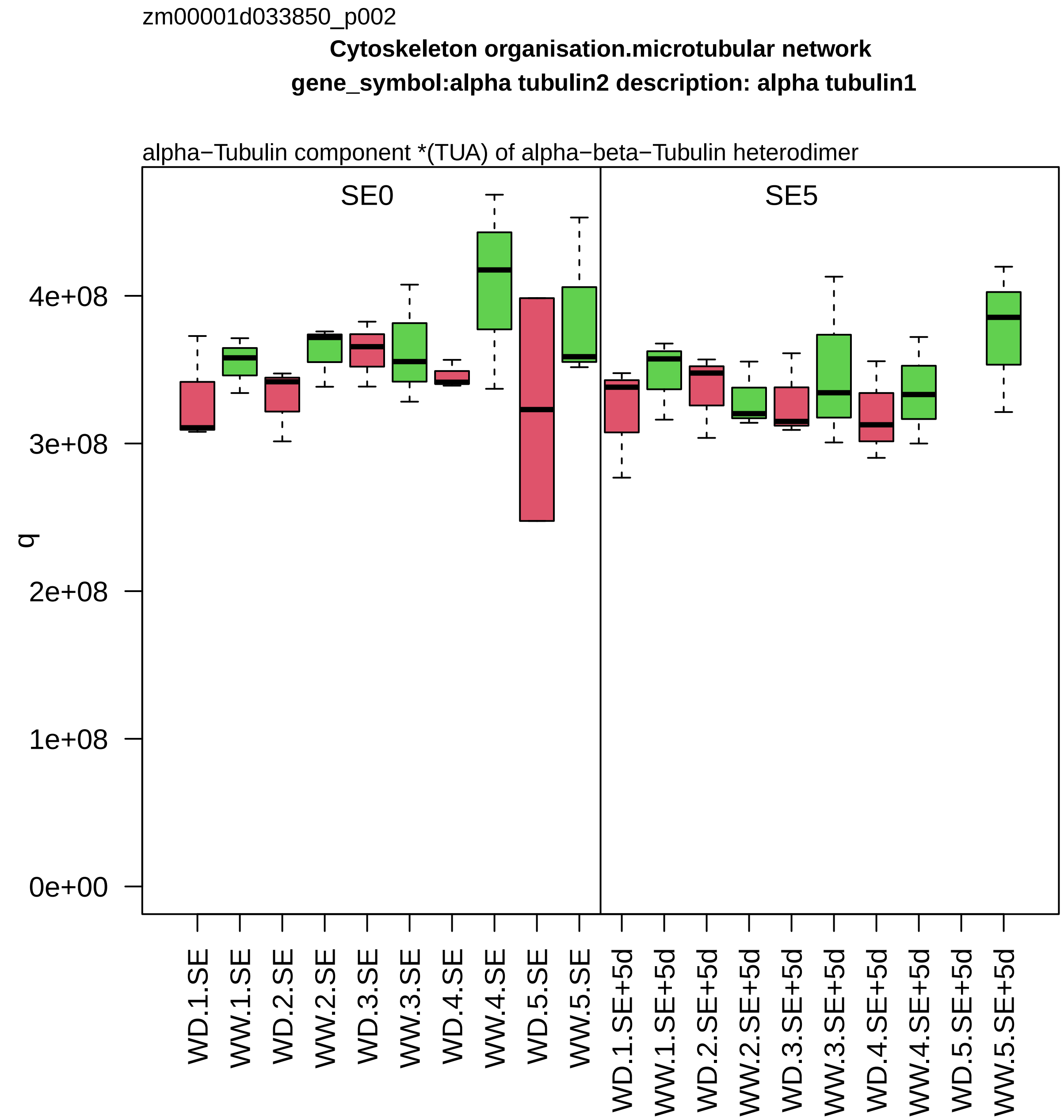

zm00001d033850_p002
Cytoskeleton organisation.microtubular network
gene_symbol:alpha tubulin2 description: alpha tubulin1
alpha−Tubulin component *(TUA) of alpha−beta−Tubulin heterodimer
SE0
SE5
q
4e+08
3e+08
2e+08
1e+08
0e+00
WD.1.SE
WW.1.SE
WD.2.SE
WW.2.SE
WD.3.SE
WW.3.SE
WD.4.SE
WW.4.SE
WD.5.SE
WW.5.SE
WD.1.SE+5d
WW.1.SE+5d
WD.2.SE+5d
WW.2.SE+5d
WD.3.SE+5d
WW.3.SE+5d
WD.4.SE+5d
WW.4.SE+5d
WD.5.SE+5d
WW.5.SE+5d

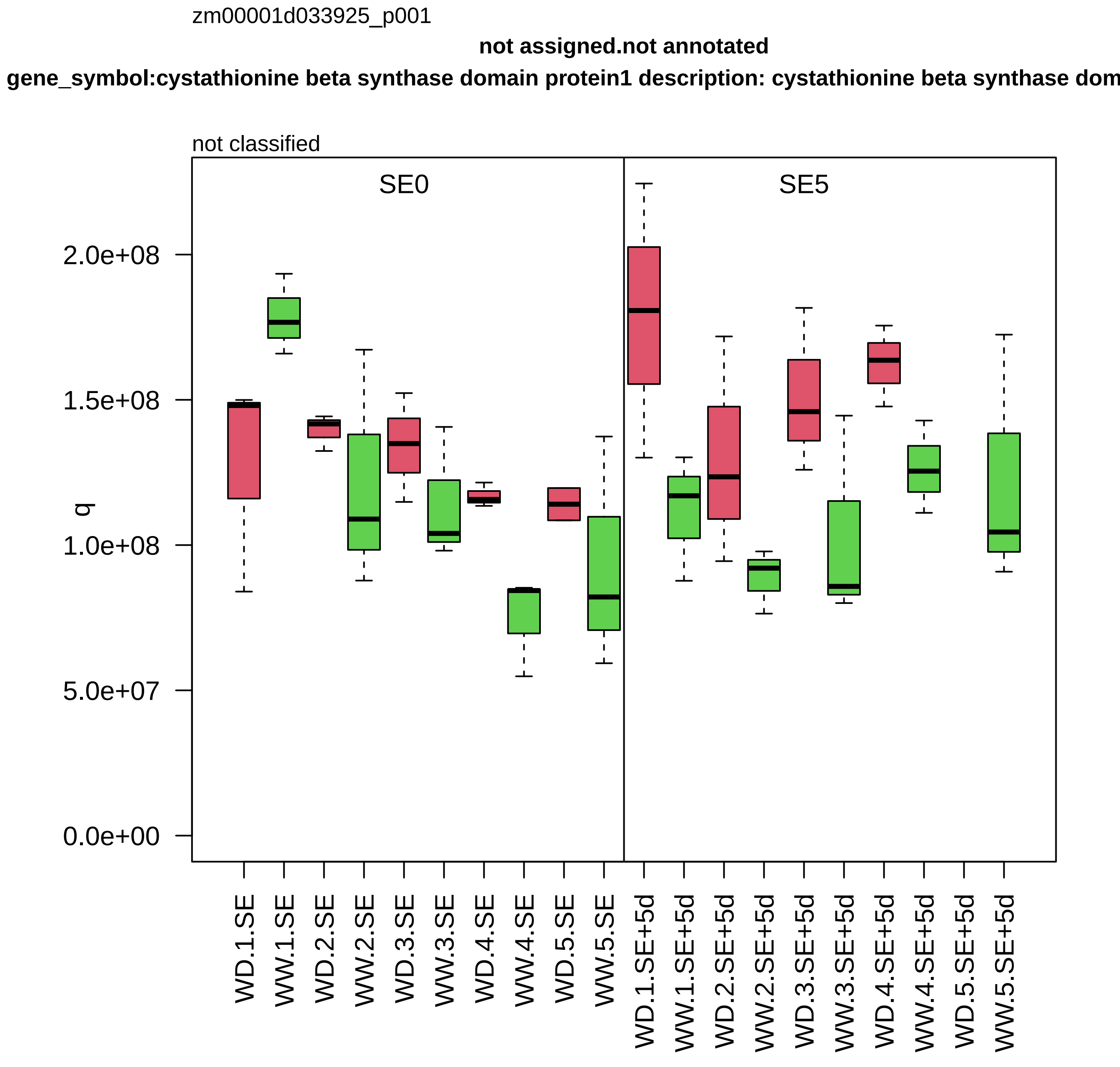
zm00001d033925_p001
not assigned.not annotated
gene_symbol:cystathionine beta synthase domain protein1 description: cystathionine beta synthase dom
not classified
SE0
SE5
q
0.0e+00
5.0e+07
1.0e+08
1.5e+08
2.0e+08
WD.1.SE
WW.1.SE
WD.2.SE
WW.2.SE
WD.3.SE
WW.3.SE
WD.4.SE
WW.4.SE
WD.5.SE
WW.5.SE
WD.1.SE+5d
WW.1.SE+5d
WD.2.SE+5d
WW.2.SE+5d
WD.3.SE+5d
WW.3.SE+5d
WD.4.SE+5d
WW.4.SE+5d
WD.5.SE+5d
WW.5.SE+5d

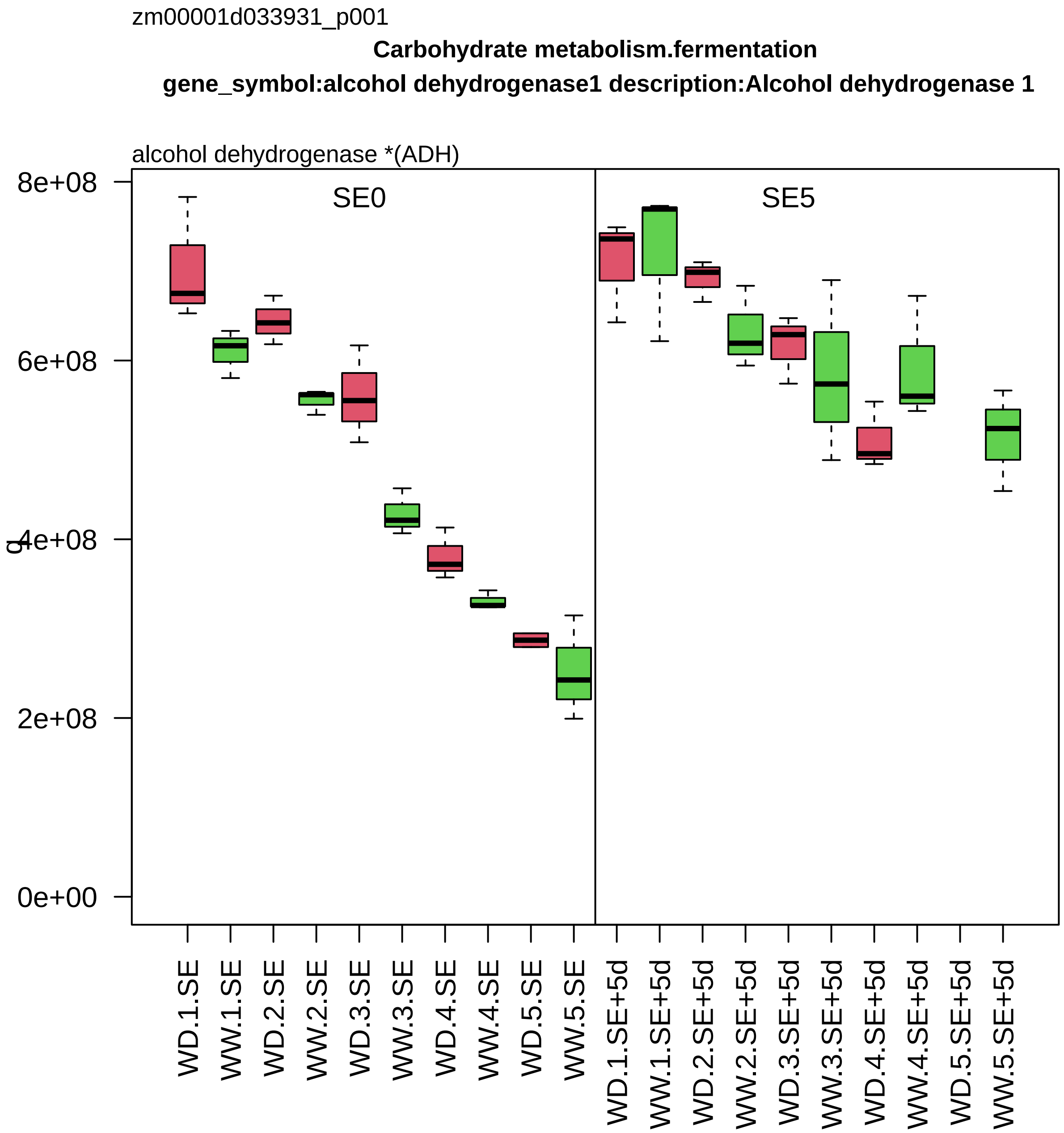
zm00001d033931_p001
Carbohydrate metabolism.fermentation
gene_symbol:alcohol dehydrogenase1 description:Alcohol dehydrogenase 1
alcohol dehydrogenase *(ADH)
SE0
SE5
q
8e+08
6e+08
4e+08
2e+08
0e+00
WD.1.SE
WW.1.SE
WD.2.SE
WW.2.SE
WD.3.SE
WW.3.SE
WD.4.SE
WW.4.SE
WD.5.SE
WW.5.SE
WD.1.SE+5d
WW.1.SE+5d
WD.2.SE+5d
WW.2.SE+5d
WD.3.SE+5d
WW.3.SE+5d
WD.4.SE+5d
WW.4.SE+5d
WD.5.SE+5d
WW.5.SE+5d

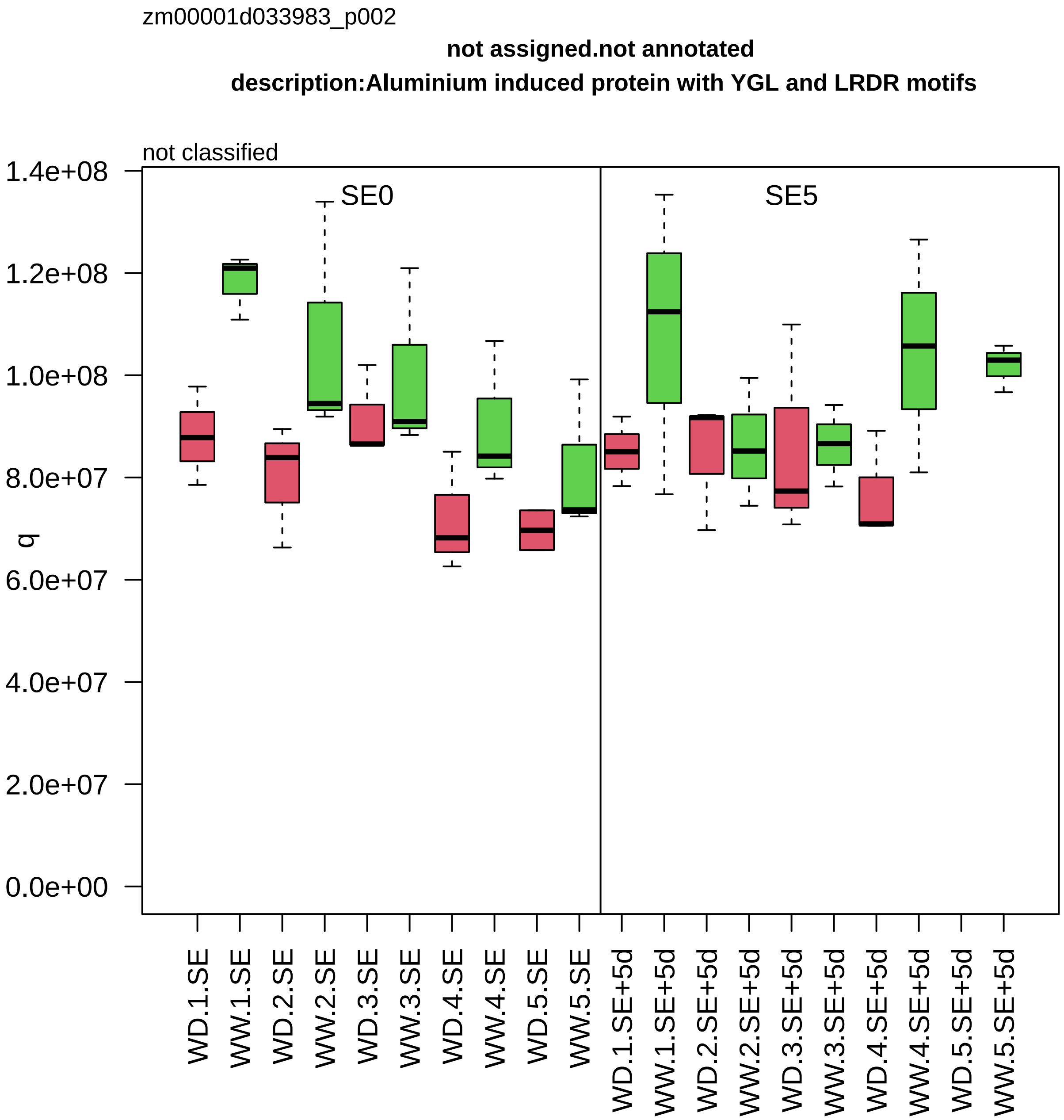

zm00001d033983_p002
not assigned.not annotated
description:Aluminium induced protein with YGL and LRDR motifs
not classified
SE0
SE5
q
1.4e+08
1.2e+08
1.0e+08
8.0e+07
6.0e+07
4.0e+07
2.0e+07
0.0e+00
WD.1.SE
WW.1.SE
WD.2.SE
WW.2.SE
WD.3.SE
WW.3.SE
WD.4.SE
WW.4.SE
WD.5.SE
WW.5.SE
WD.1.SE+5d
WW.1.SE+5d
WD.2.SE+5d
WW.2.SE+5d
WD.3.SE+5d
WW.3.SE+5d
WD.4.SE+5d
WW.4.SE+5d
WD.5.SE+5d
WW.5.SE+5d

zm00001d034015_p002

**Enzyme classification.EC_3 hydrolases**

**gene_symbol:exoglucanase1 description: exoglucanase1**

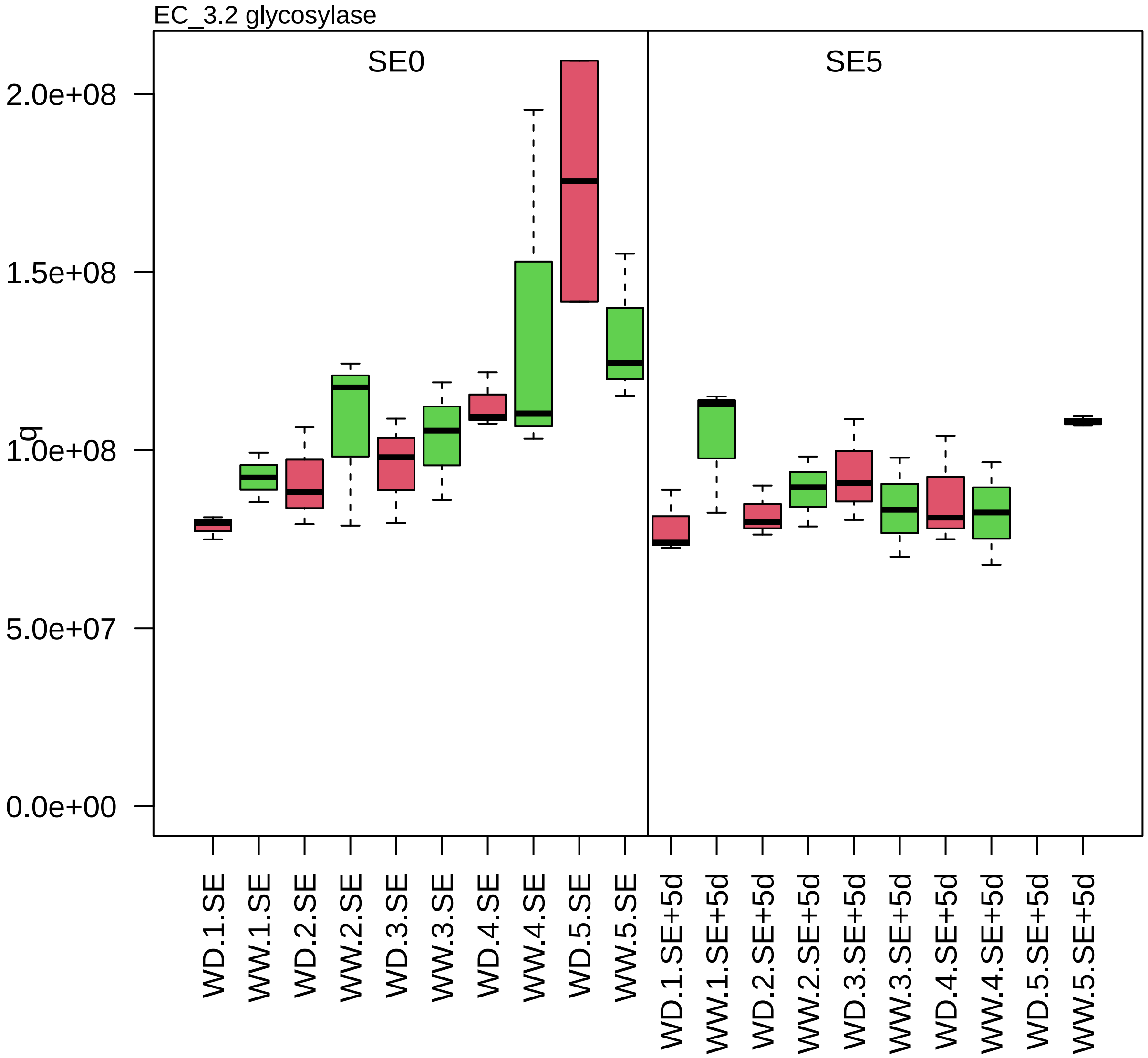

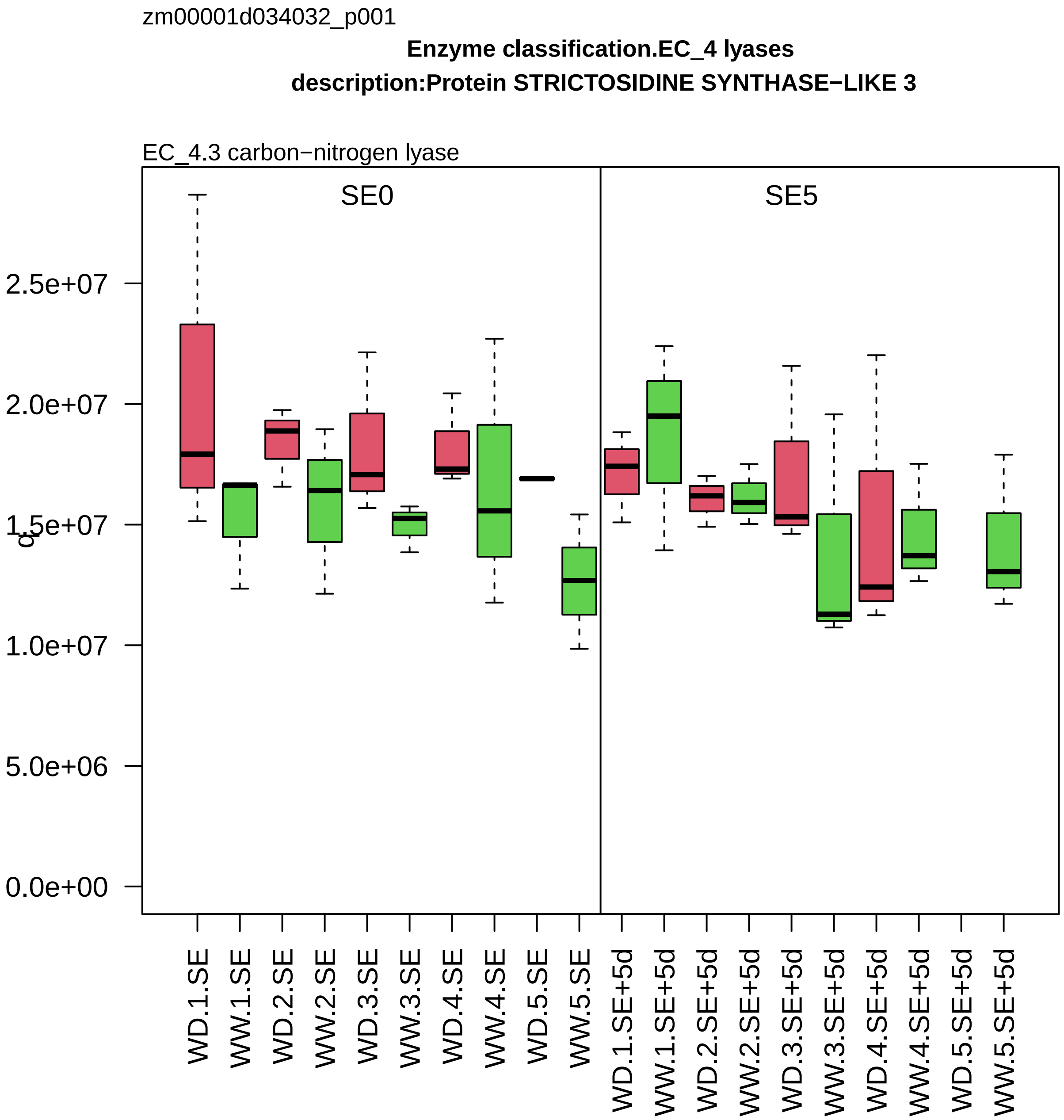
zm00001d034032_p001
Enzyme classification.EC_4 lyases
description:Protein STRICTOSIDINE SYNTHASE−LIKE 3
EC_4.3 carbon−nitrogen lyase
SE0
SE5
2.5e+07
2.0e+07
1.5e+07
1.0e+07
5.0e+06
0.0e+00
WD.1.SE
WW.1.SE
WD.2.SE
WW.2.SE
WD.3.SE
WW.3.SE
WD.4.SE
WW.4.SE
WD.5.SE
WW.5.SE
WD.1.SE+5d
WW.1.SE+5d
WD.2.SE+5d
WW.2.SE+5d
WD.3.SE+5d
WW.3.SE+5d
WD.4.SE+5d
WW.4.SE+5d
WD.5.SE+5d
WW.5.SE+5d

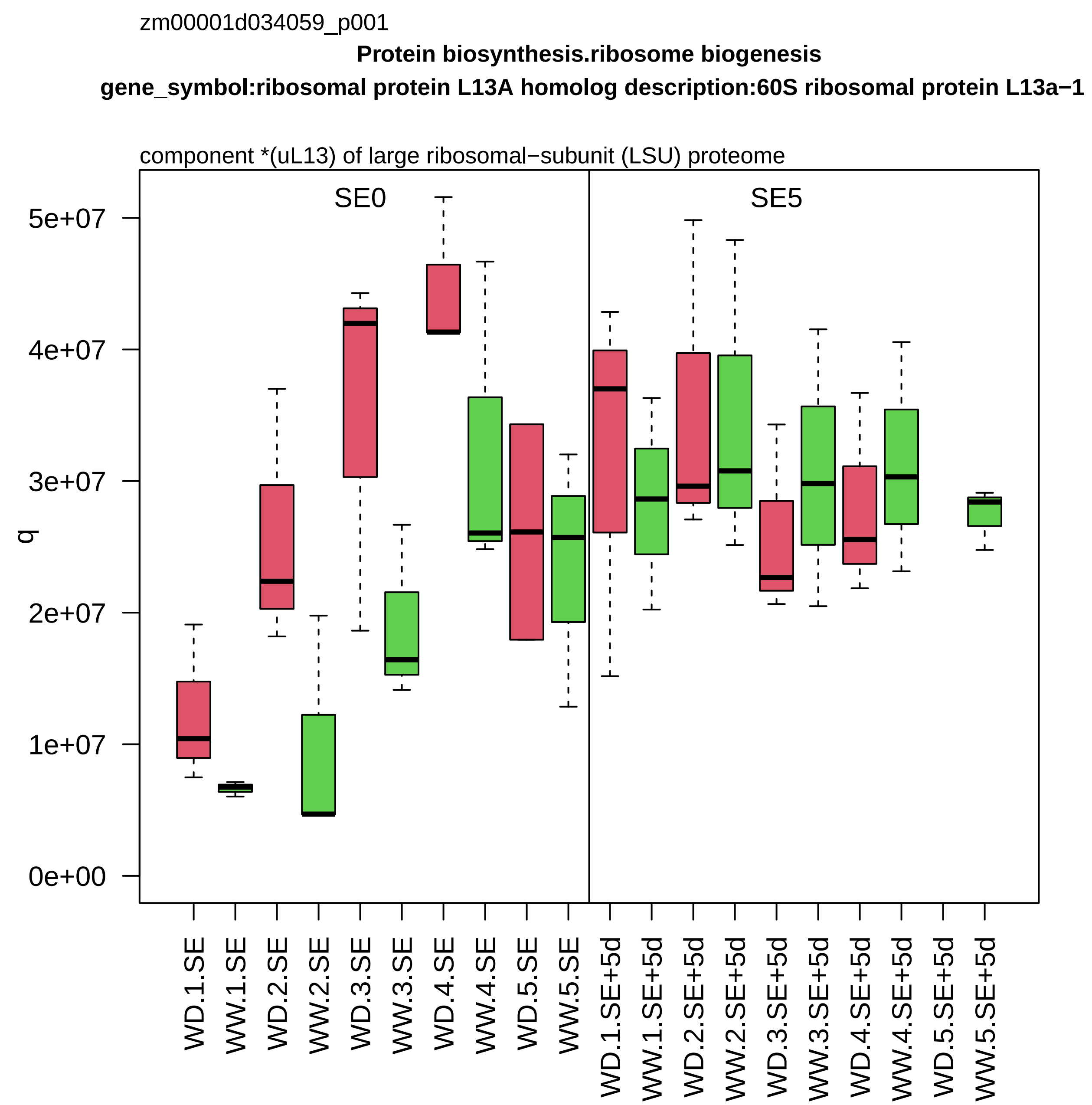
zm00001d034059_p001
Protein biosynthesis.ribosome biogenesis
gene_symbol:ribosomal protein L13A homolog description:60S ribosomal protein L13a−1
component *(uL13) of large ribosomal−subunit (LSU) proteome
SE0
SE5
q
5e+07
4e+07
3e+07
2e+07
1e+07
0e+00
WD.1.SE
WW.1.SE
WD.2.SE
WW.2.SE
WD.3.SE
WW.3.SE
WD.4.SE
WW.4.SE
WD.5.SE
WW.5.SE
WD.1.SE+5d
WW.1.SE+5d
WD.2.SE+5d
WW.2.SE+5d
WD.3.SE+5d
WW.3.SE+5d
WD.4.SE+5d
WW.4.SE+5d
WD.5.SE+5d
WW.5.SE+5d

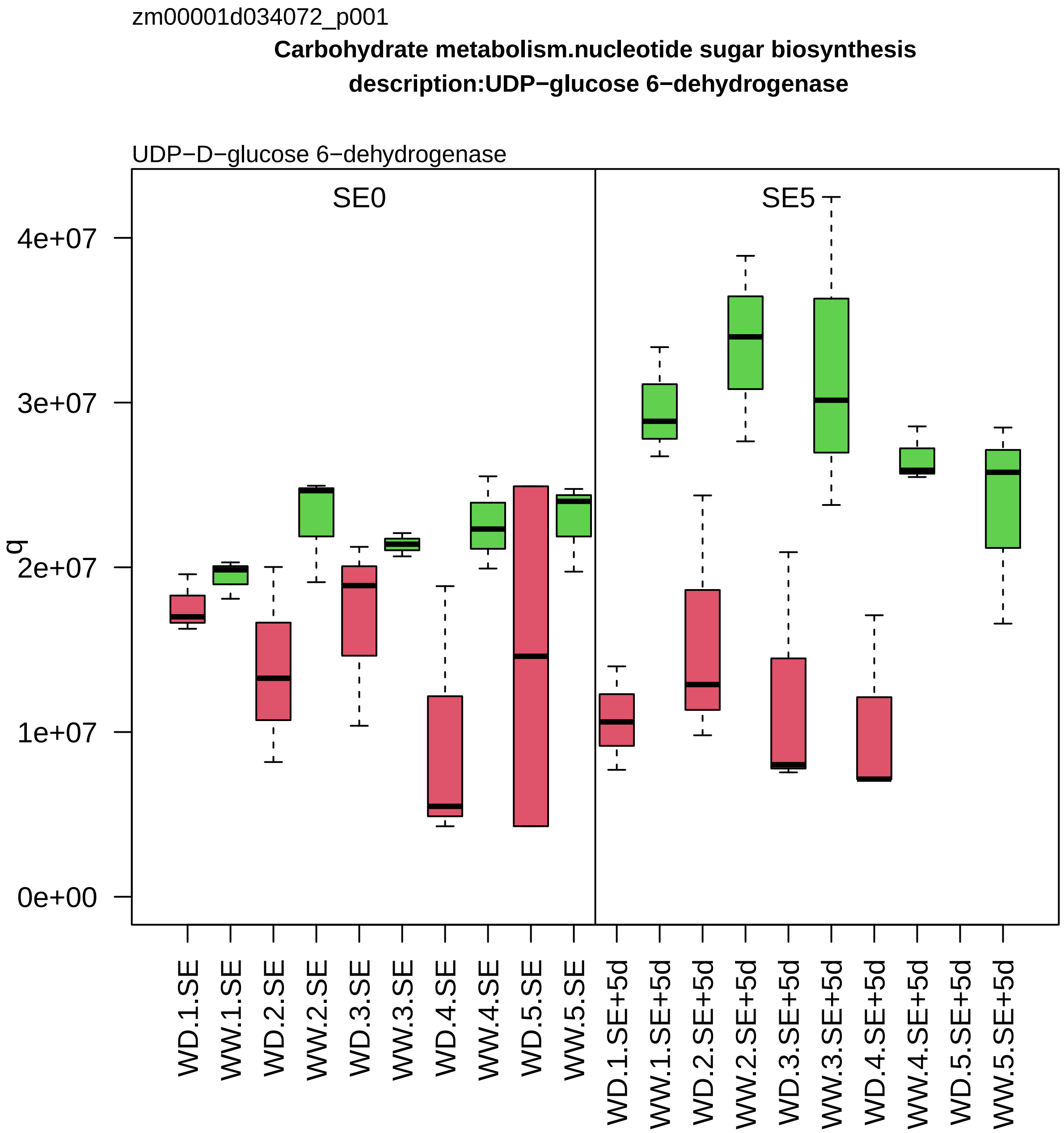
zm00001d034072_p001
Carbohydrate metabolism.nucleotide sugar biosynthesis
description:UDP−glucose 6−dehydrogenase
UDP−D−glucose 6−dehydrogenase
SE0
SE5
q
0e+00
1e+07
2e+07
3e+07
4e+07
WD.1.SE
WW.1.SE
WD.2.SE
WW.2.SE
WD.3.SE
WW.3.SE
WD.4.SE
WW.4.SE
WD.5.SE
WW.5.SE
WD.1.SE+5d
WW.1.SE+5d
WD.2.SE+5d
WW.2.SE+5d
WD.3.SE+5d
WW.3.SE+5d
WD.4.SE+5d
WW.4.SE+5d
WD.5.SE+5d
WW.5.SE+5d

zm00001d034074_p003

**Carbohydrate metabolism.starch metabolism**

**gene_symbol:starch phosphorylase1 description: starch phosphorylase1**

plastidial alpha−glucan phosphorylase *(PHS1)

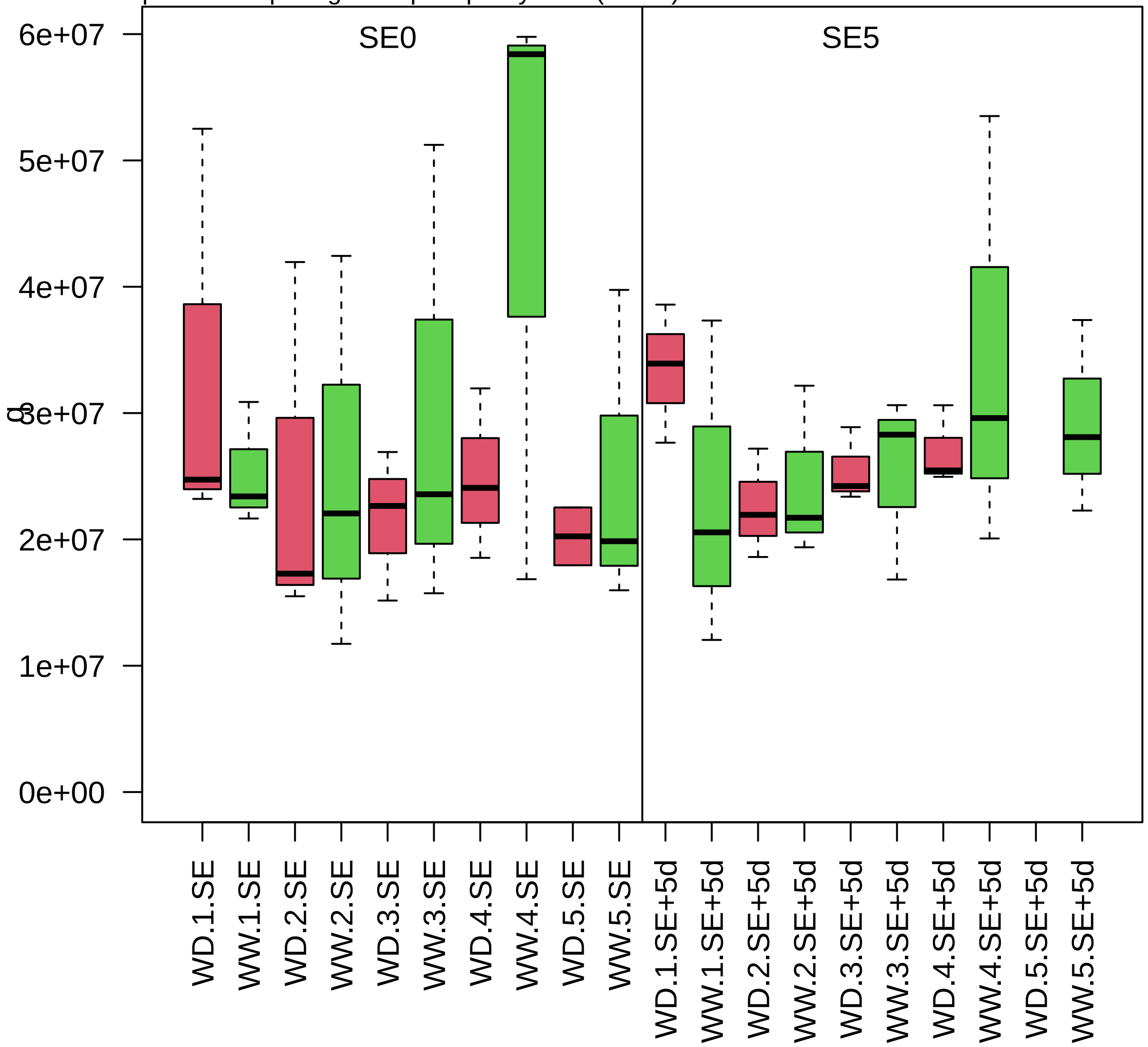

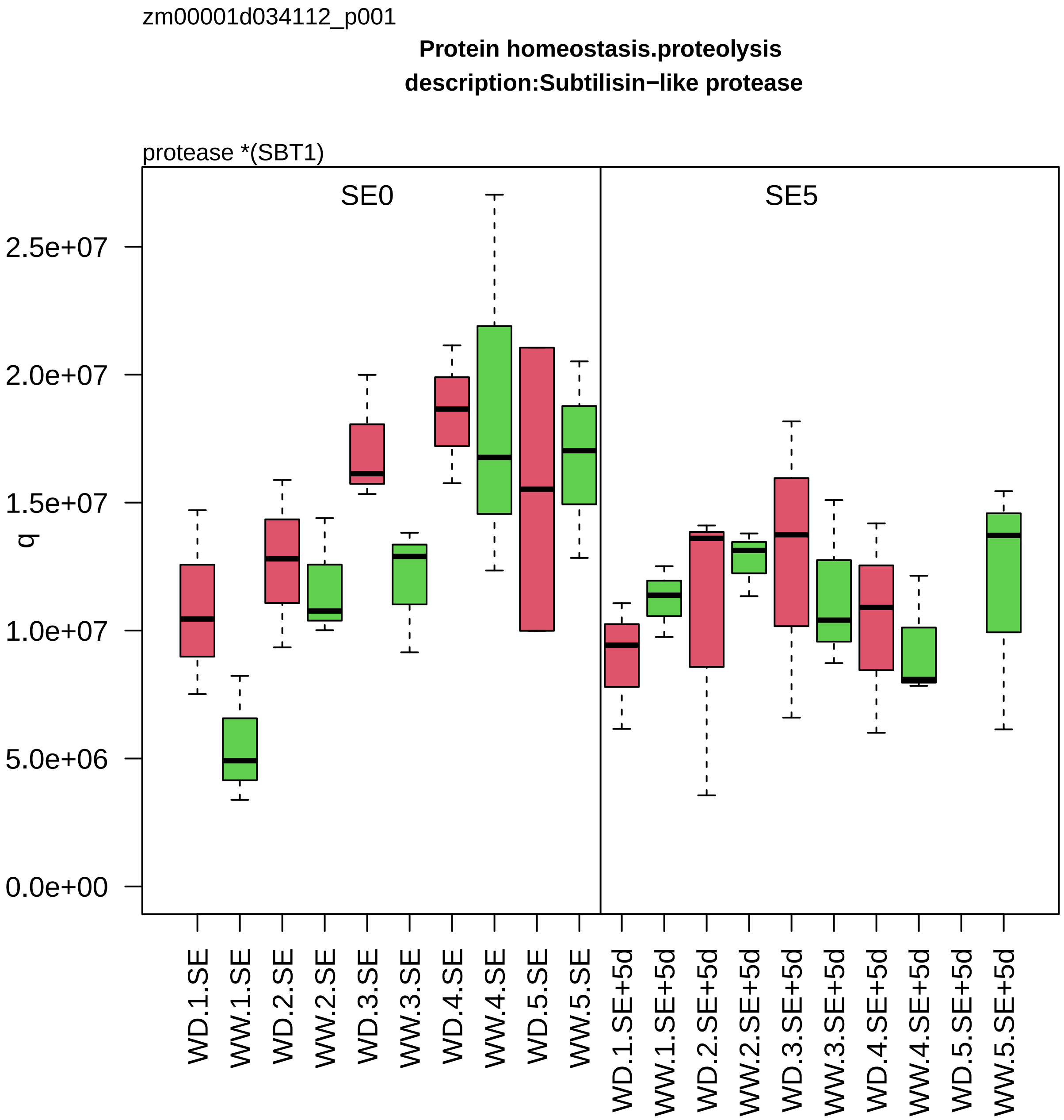

zm00001d034112_p001
Protein homeostasis.proteolysis
description:Subtilisin−like protease
protease *(SBT1)
SE0
SE5
q
2.5e+07
2.0e+07
1.5e+07
1.0e+07
5.0e+06
0.0e+00
WD.1.SE
WW.1.SE
WD.2.SE
WW.2.SE
WD.3.SE
WW.3.SE
WD.4.SE
WW.4.SE
WD.5.SE
WW.5.SE
WD.1.SE+5d
WW.1.SE+5d
WD.2.SE+5d
WW.2.SE+5d
WD.3.SE+5d
WW.3.SE+5d
WD.4.SE+5d
WW.4.SE+5d
WD.5.SE+5d
WW.5.SE+5d

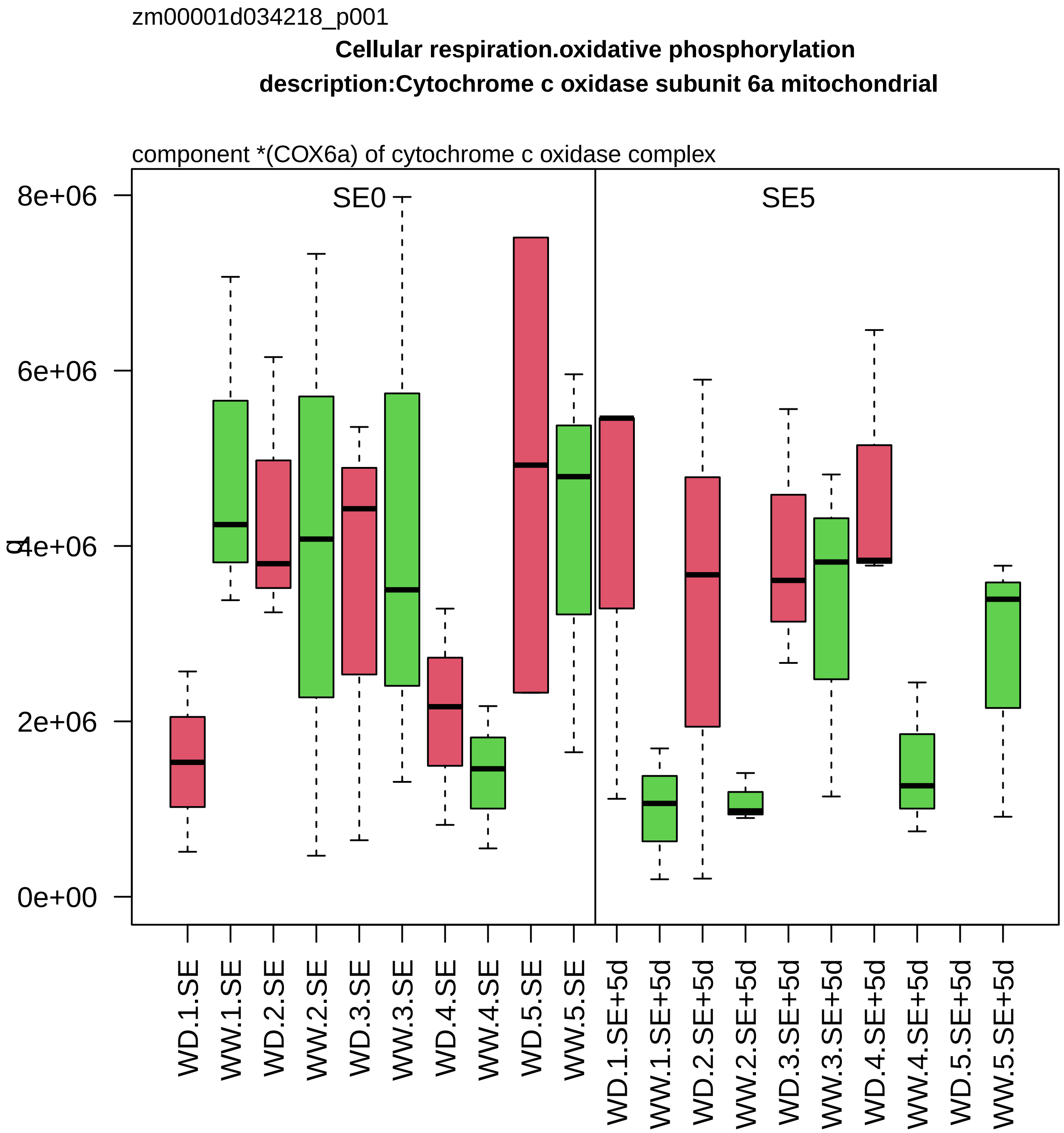

zm00001d034218_p001
Cellular respiration.oxidative phosphorylation
description:Cytochrome c oxidase subunit 6a mitochondrial
component *(COX6a) of cytochrome c oxidase complex
SE0
SE5
8e+06
6e+06
4e+06
2e+06
0e+00
q
WD.1.SE
WW.1.SE
WD.2.SE
WW.2.SE
WD.3.SE
WW.3.SE
WD.4.SE
WW.4.SE
WD.5.SE
WW.5.SE
WD.1.SE+5d
WW.1.SE+5d
WD.2.SE+5d
WW.2.SE+5d
WD.3.SE+5d
WW.3.SE+5d
WD.4.SE+5d
WW.4.SE+5d
WD.5.SE+5d
WW.5.SE+5d

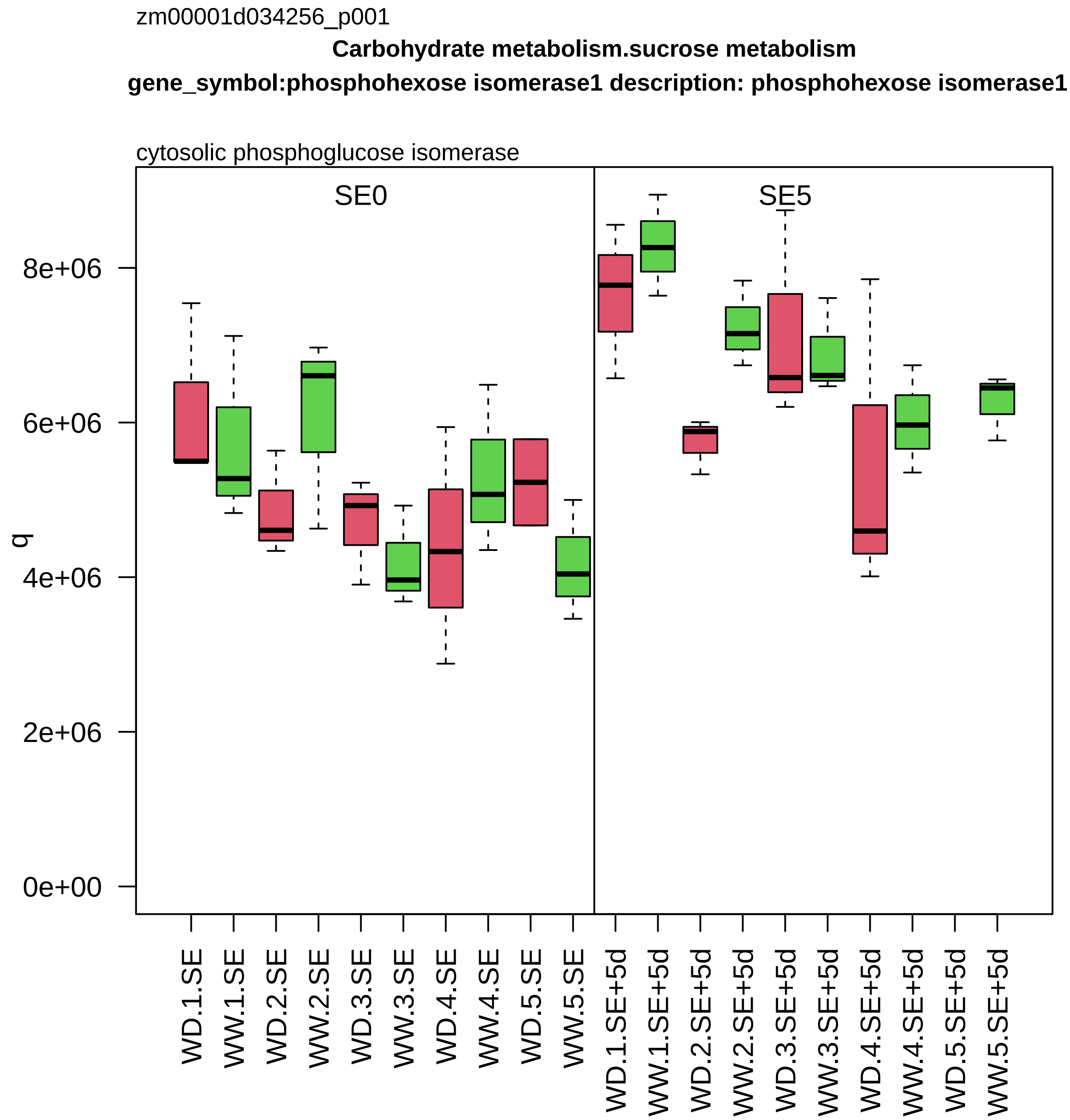

zm00001d034256_p001
Carbohydrate metabolism.sucrose metabolism
gene_symbol:phosphohexose isomerase1 description: phosphohexose isomerase1
cytosolic phosphoglucose isomerase
SE0
SE5
q
0e+00
2e+06
4e+06
6e+06
8e+06
WD.1.SE
WW.1.SE
WD.2.SE
WW.2.SE
WD.3.SE
WW.3.SE
WD.4.SE
WW.4.SE
WD.5.SE
WW.5.SE
WD.1.SE+5d
WW.1.SE+5d
WD.2.SE+5d
WW.2.SE+5d
WD.3.SE+5d
WW.3.SE+5d
WD.4.SE+5d
WW.4.SE+5d
WD.5.SE+5d
WW.5.SE+5d

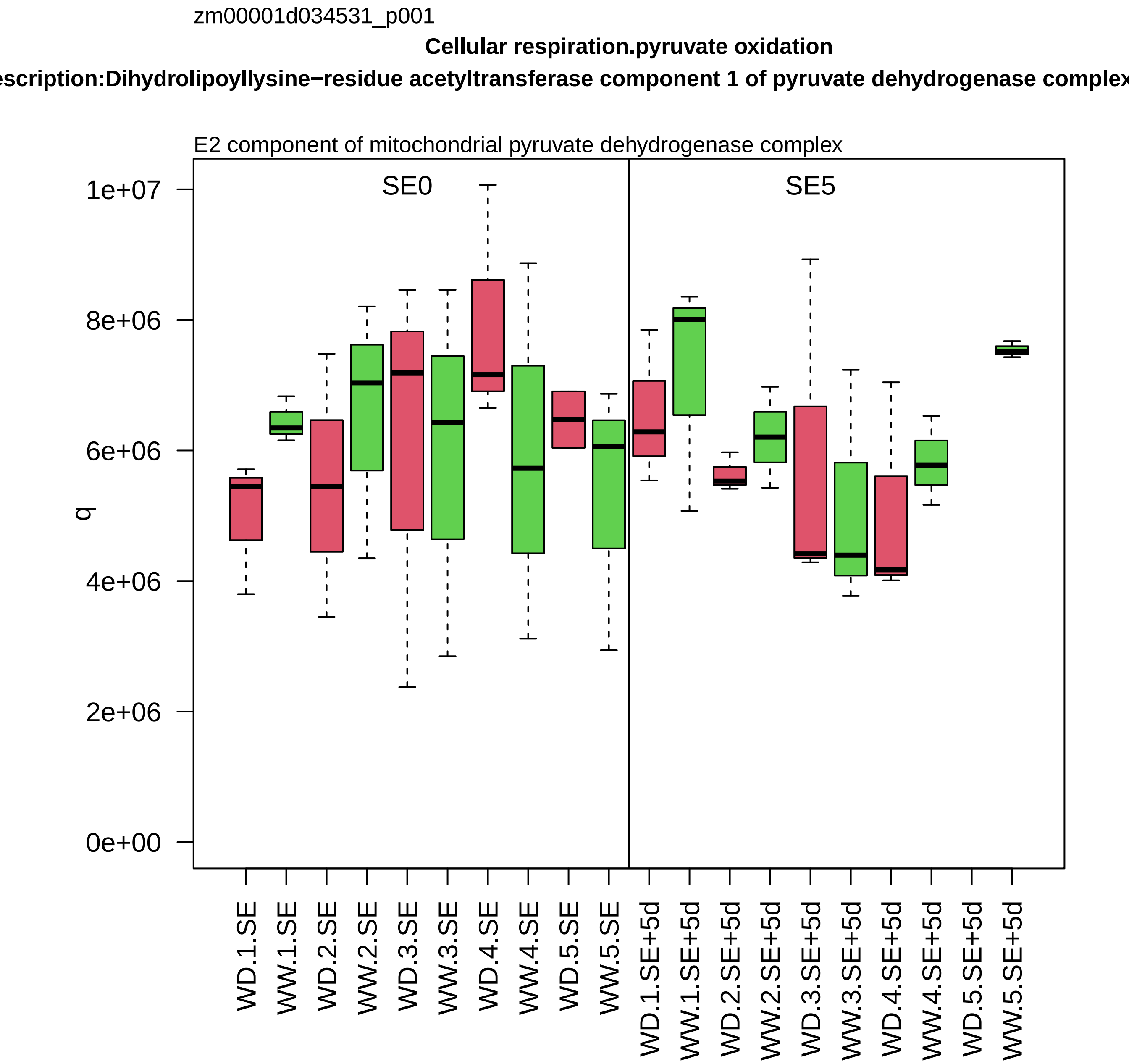
zm00001d034531_p001
Cellular respiration.pyruvate oxidation
escription:Dihydrolipoyllysine−residue acetyltransferase component 1 of pyruvate dehydrogenase complex
E2 component of mitochondrial pyruvate dehydrogenase complex
SE0
SE5
q
0e+00
2e+06
4e+06
6e+06
8e+06
1e+07
WD.1.SE
WW.1.SE
WD.2.SE
WW.2.SE
WD.3.SE
WW.3.SE
WD.4.SE
WW.4.SE
WD.5.SE
WW.5.SE
WD.1.SE+5d
WW.1.SE+5d
WD.2.SE+5d
WW.2.SE+5d
WD.3.SE+5d
WW.3.SE+5d
WD.4.SE+5d
WW.4.SE+5d
WD.5.SE+5d
WW.5.SE+5d

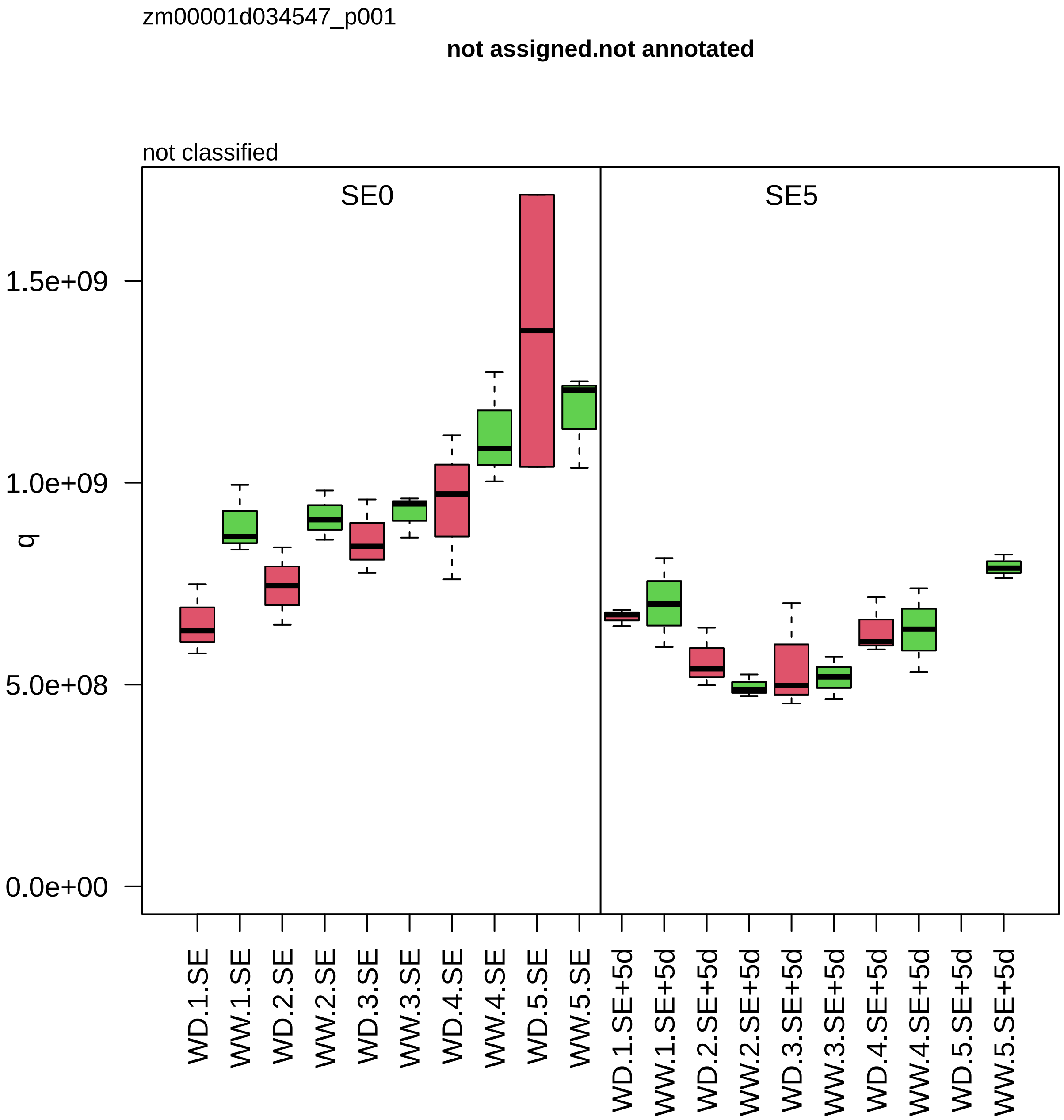

zm00001d034547_p001
not assigned.not annotated
not classified
SE0
SE5
q
1.5e+09
1.0e+09
5.0e+08
0.0e+00
WD.1.SE
WW.1.SE
WD.2.SE
WW.2.SE
WD.3.SE
WW.3.SE
WD.4.SE
WW.4.SE
WD.5.SE
WW.5.SE
WD.1.SE+5d
WW.1.SE+5d
WD.2.SE+5d
WW.2.SE+5d
WD.3.SE+5d
WW.3.SE+5d
WD.4.SE+5d
WW.4.SE+5d
WD.5.SE+5d
WW.5.SE+5d

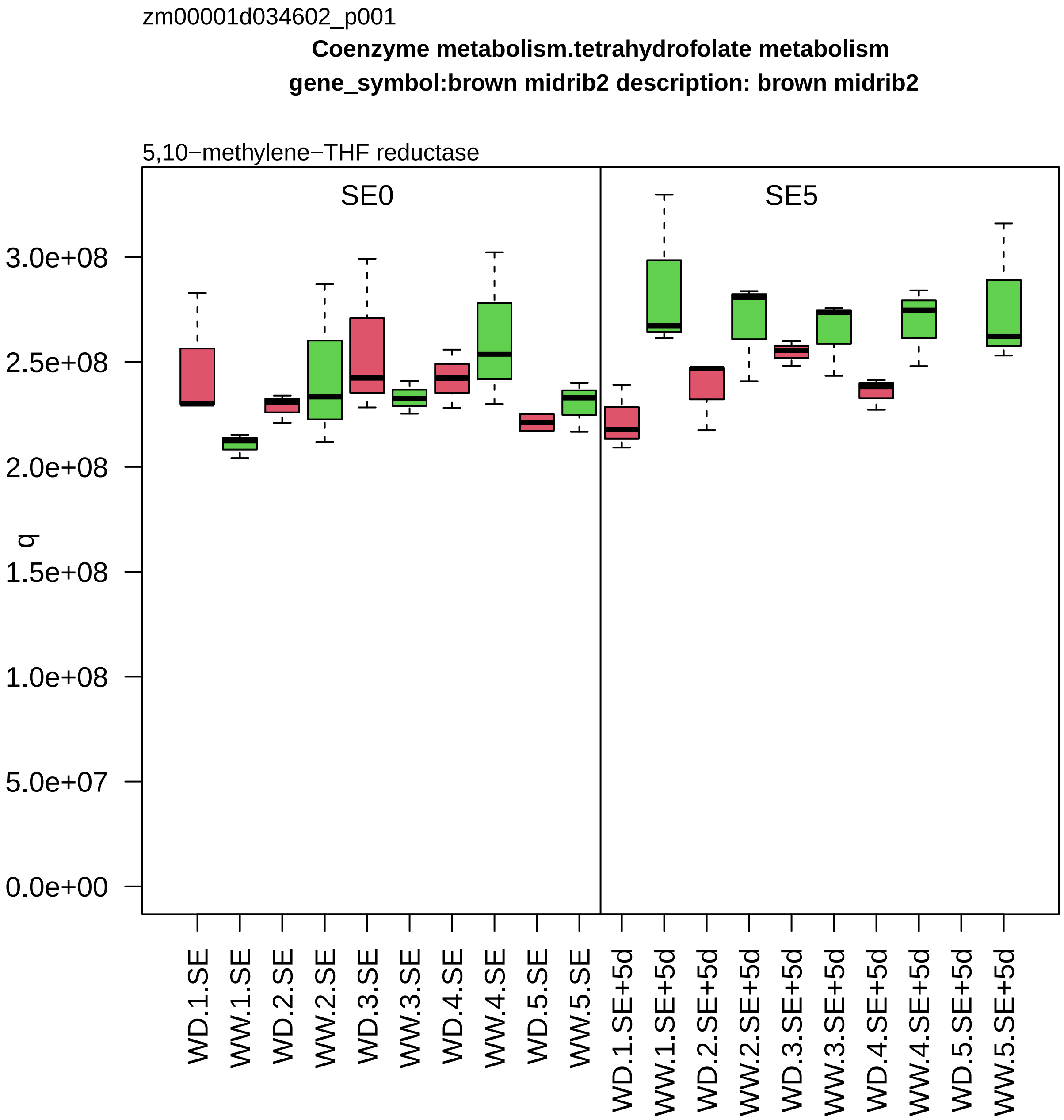

zm00001d034602_p001
Coenzyme metabolism.tetrahydrofolate metabolism
gene_symbol:brown midrib2 description: brown midrib2
5,10−methylene−THF reductase
SE0
SE5
q
3.0e+08
2.5e+08
2.0e+08
1.5e+08
1.0e+08
5.0e+07
0.0e+00
WD.1.SE
WW.1.SE
WD.2.SE
WW.2.SE
WD.3.SE
WW.3.SE
WD.4.SE
WW.4.SE
WD.5.SE
WW.5.SE
WD.1.SE+5d
WW.1.SE+5d
WD.2.SE+5d
WW.2.SE+5d
WD.3.SE+5d
WW.3.SE+5d
WD.4.SE+5d
WW.4.SE+5d
WD.5.SE+5d
WW.5.SE+5d

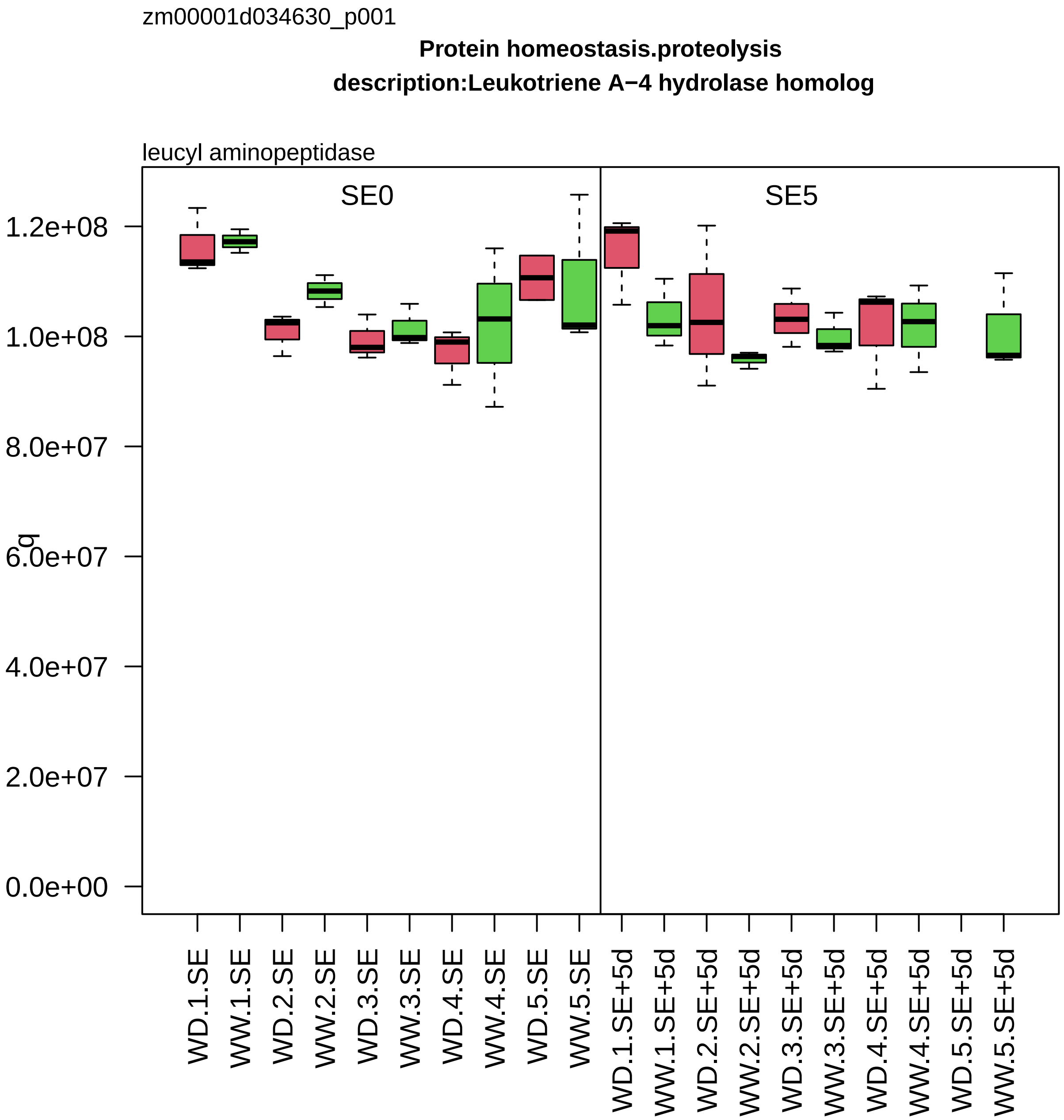

zm00001d034630_p001
Protein homeostasis.proteolysis
description:Leukotriene A−4 hydrolase homolog
leucyl aminopeptidase
SE0
SE5
q
1.2e+08
1.0e+08
8.0e+07
6.0e+07
4.0e+07
2.0e+07
0.0e+00
WD.1.SE
WW.1.SE
WD.2.SE
WW.2.SE
WD.3.SE
WW.3.SE
WD.4.SE
WW.4.SE
WD.5.SE
WW.5.SE
WD.1.SE+5d
WW.1.SE+5d
WD.2.SE+5d
WW.2.SE+5d
WD.3.SE+5d
WW.3.SE+5d
WD.4.SE+5d
WW.4.SE+5d
WD.5.SE+5d
WW.5.SE+5d

zm00001d034635_p001

**Secondary metabolism.phenolics**

**gene_symbol:chalcone flavanone isomerase1 description: chalcone flavanone isomerase1**

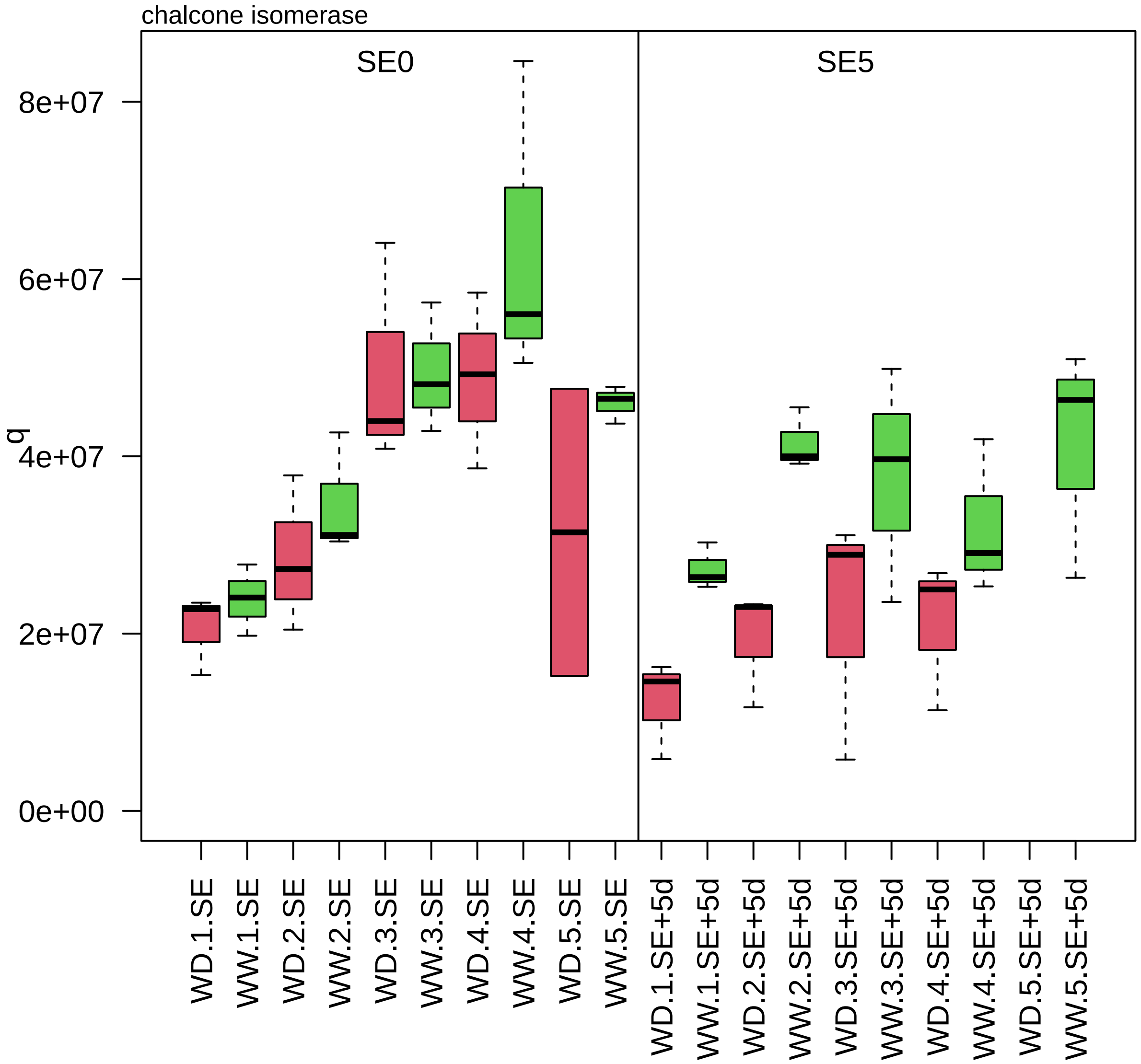

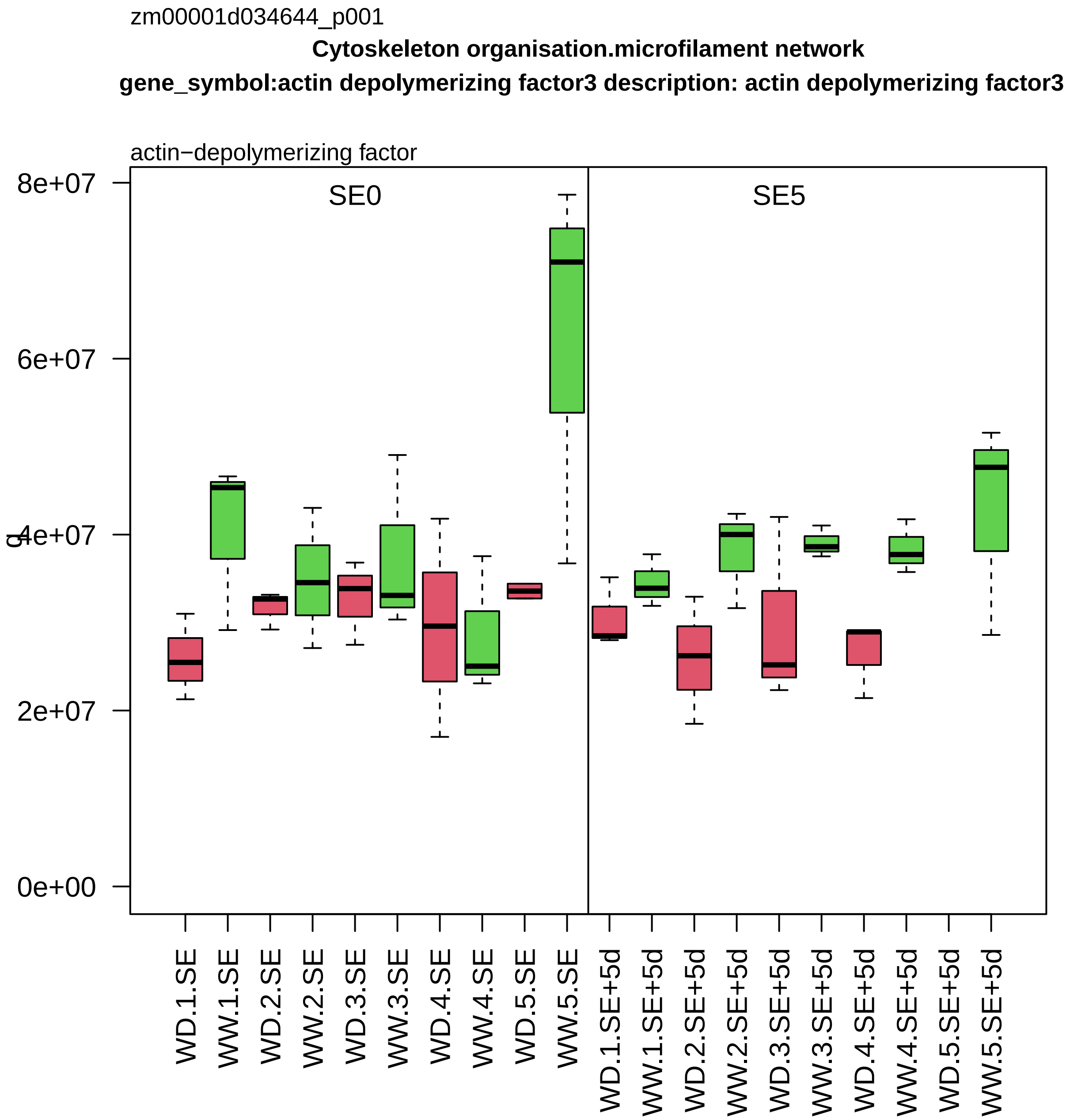
zm00001d034644_p001
Cytoskeleton organisation.microfilament network
gene_symbol:actin depolymerizing factor3 description: actin depolymerizing factor3
actin−depolymerizing factor
SE0
SE5
8e+07
6e+07
4e+07
2e+07
0e+00
WD.1.SE
WW.1.SE
WD.2.SE
WW.2.SE
WD.3.SE
WW.3.SE
WD.4.SE
WW.4.SE
WD.5.SE
WW.5.SE
WD.1.SE+5d
WW.1.SE+5d
WD.2.SE+5d
WW.2.SE+5d
WD.3.SE+5d
WW.3.SE+5d
WD.4.SE+5d
WW.4.SE+5d
WD.5.SE+5d
WW.5.SE+5d

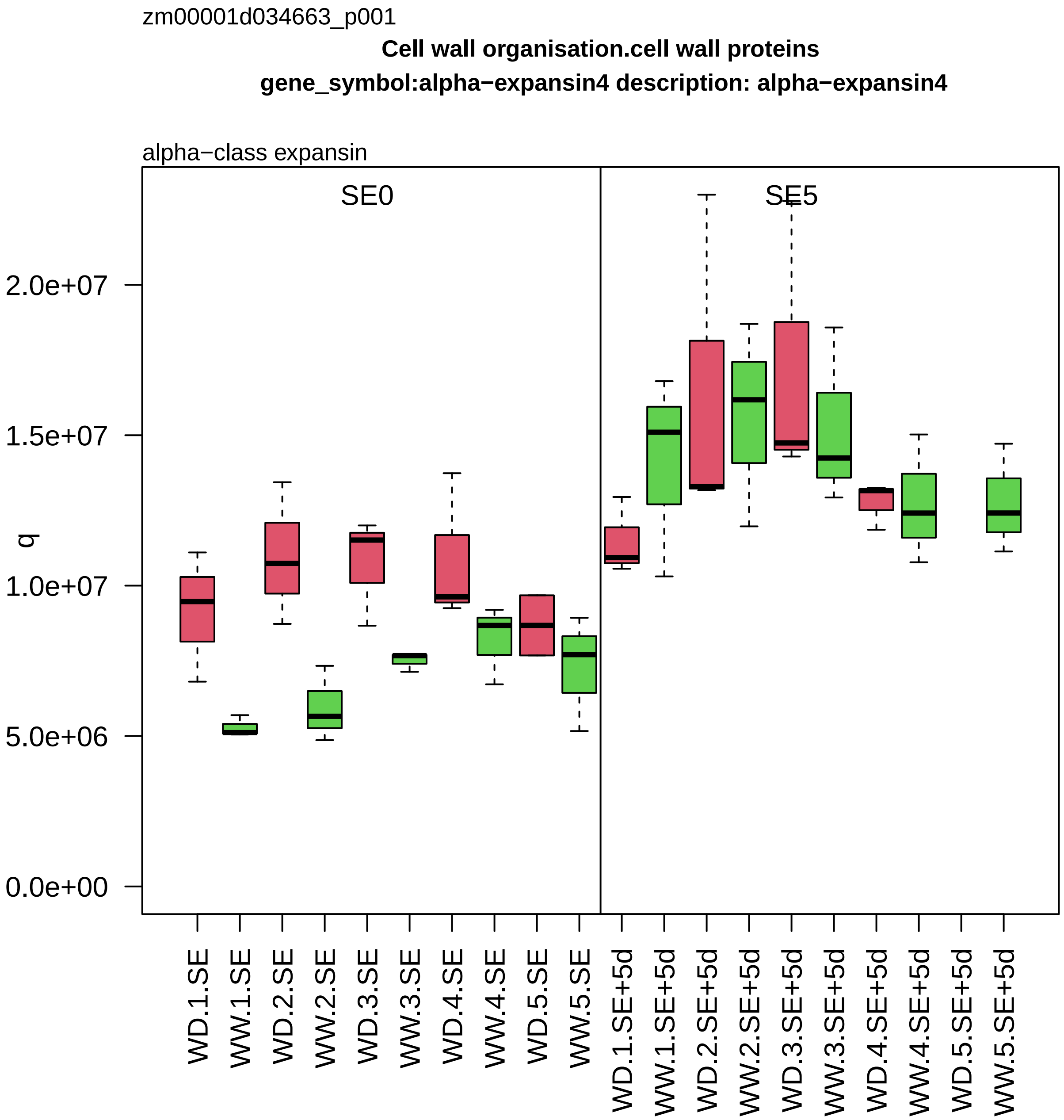
zm00001d034663_p001
Cell wall organisation.cell wall proteins
gene_symbol:alpha−expansin4 description: alpha−expansin4
alpha−class expansin
SE0
SE5
q
2.0e+07
1.5e+07
1.0e+07
5.0e+06
0.0e+00
WD.1.SE
WW.1.SE
WD.2.SE
WW.2.SE
WD.3.SE
WW.3.SE
WD.4.SE
WW.4.SE
WD.5.SE
WW.5.SE
WD.1.SE+5d
WW.1.SE+5d
WD.2.SE+5d
WW.2.SE+5d
WD.3.SE+5d
WW.3.SE+5d
WD.4.SE+5d
WW.4.SE+5d
WD.5.SE+5d
WW.5.SE+5d

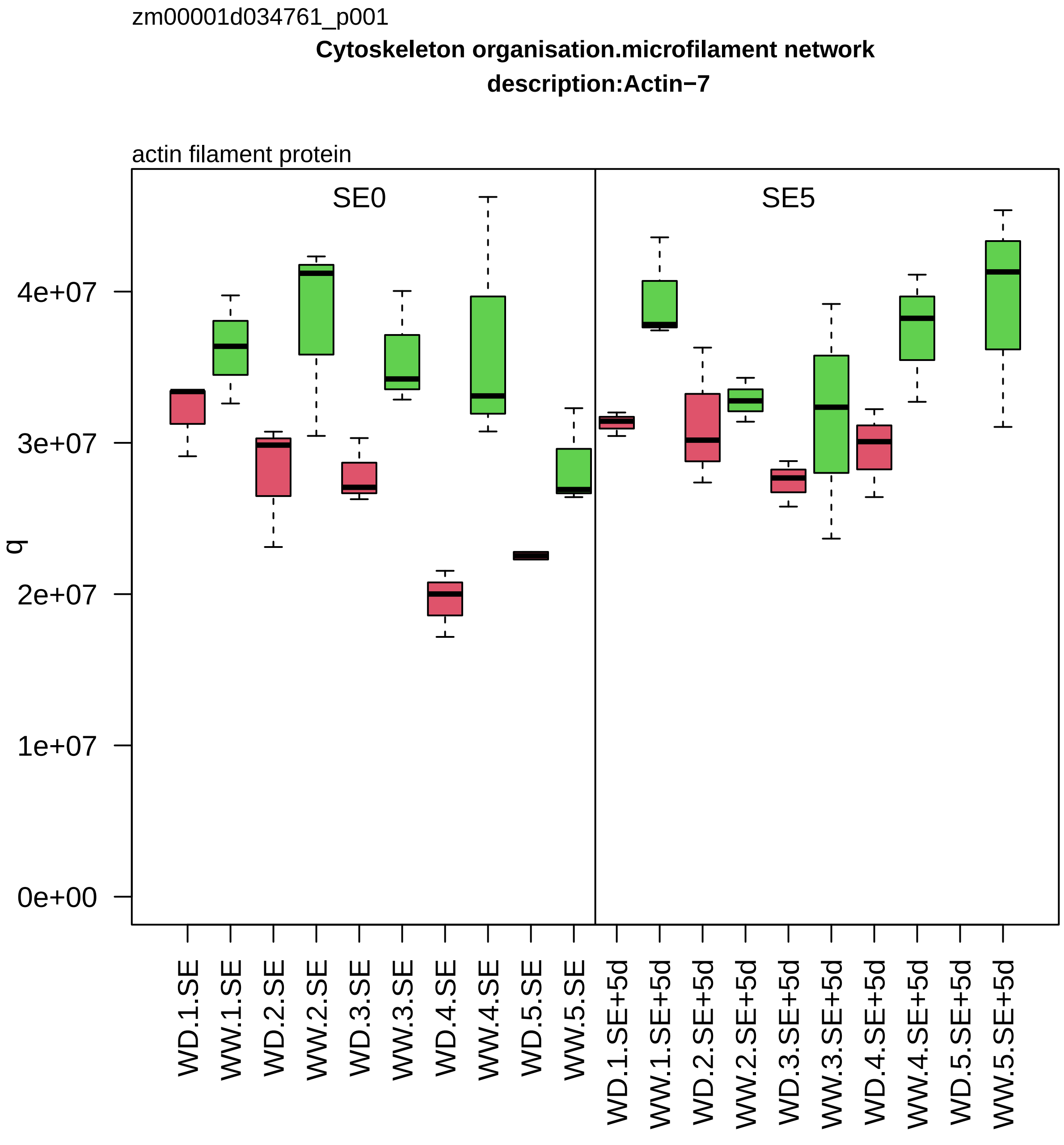
zm00001d034761_p001
Cytoskeleton organisation.microfilament network
description:Actin−7
actin filament protein
SE0
SE5
q
4e+07
3e+07
2e+07
1e+07
0e+00
WD.1.SE
WW.1.SE
WD.2.SE
WW.2.SE
WD.3.SE
WW.3.SE
WD.4.SE
WW.4.SE
WD.5.SE
WW.5.SE
WD.1.SE+5d
WW.1.SE+5d
WD.2.SE+5d
WW.2.SE+5d
WD.3.SE+5d
WW.3.SE+5d
WD.4.SE+5d
WW.4.SE+5d
WD.5.SE+5d
WW.5.SE+5d

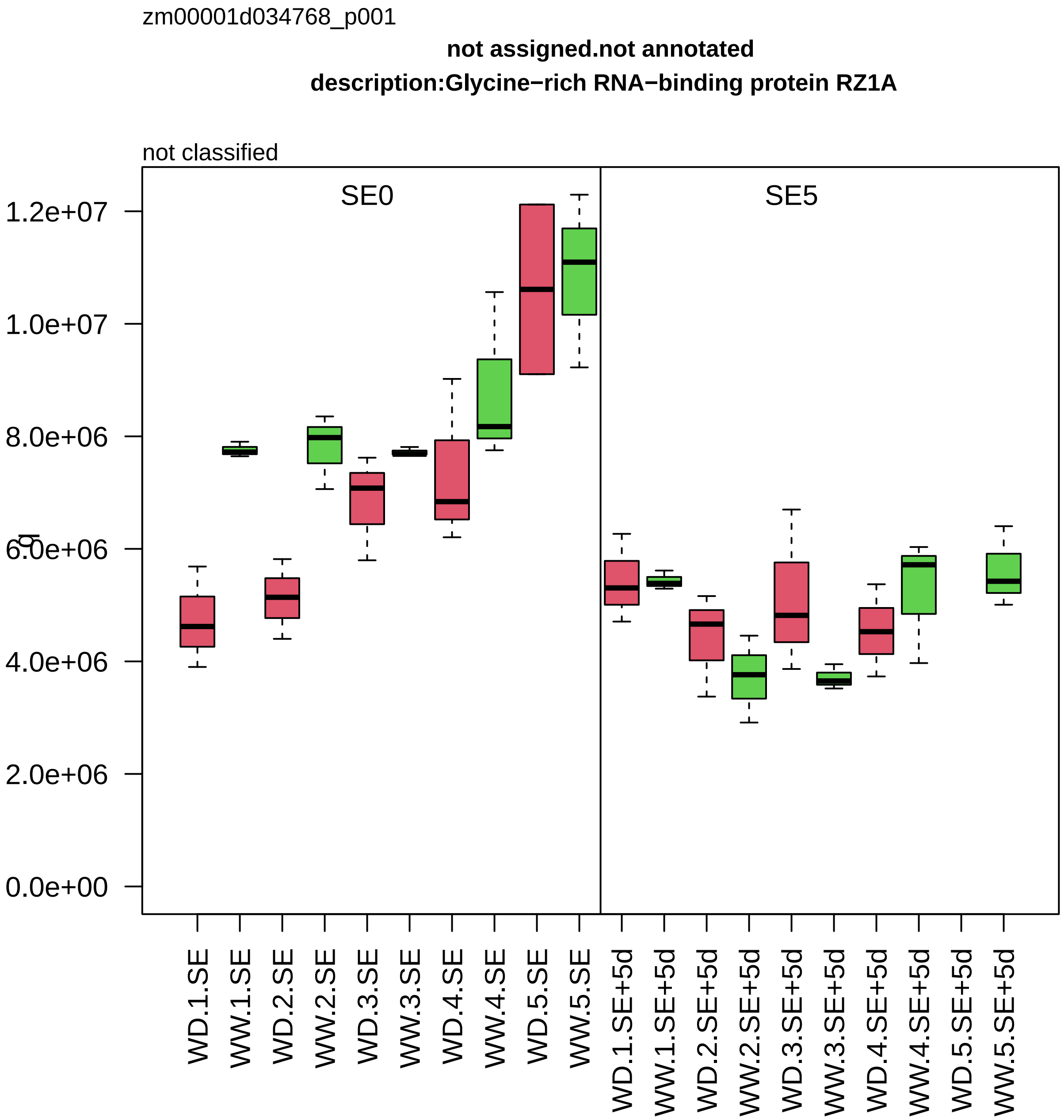
zm00001d034768_p001
not assigned.not annotated
description:Glycine−rich RNA−binding protein RZ1A
not classified
SE0
SE5
0.0e+00
2.0e+06
4.0e+06
6.0e+06
8.0e+06
1.0e+07
1.2e+07
WD.1.SE
WW.1.SE
WD.2.SE
WW.2.SE
WD.3.SE
WW.3.SE
WD.4.SE
WW.4.SE
WD.5.SE
WW.5.SE
WD.1.SE+5d
WW.1.SE+5d
WD.2.SE+5d
WW.2.SE+5d
WD.3.SE+5d
WW.3.SE+5d
WD.4.SE+5d
WW.4.SE+5d
WD.5.SE+5d
WW.5.SE+5d

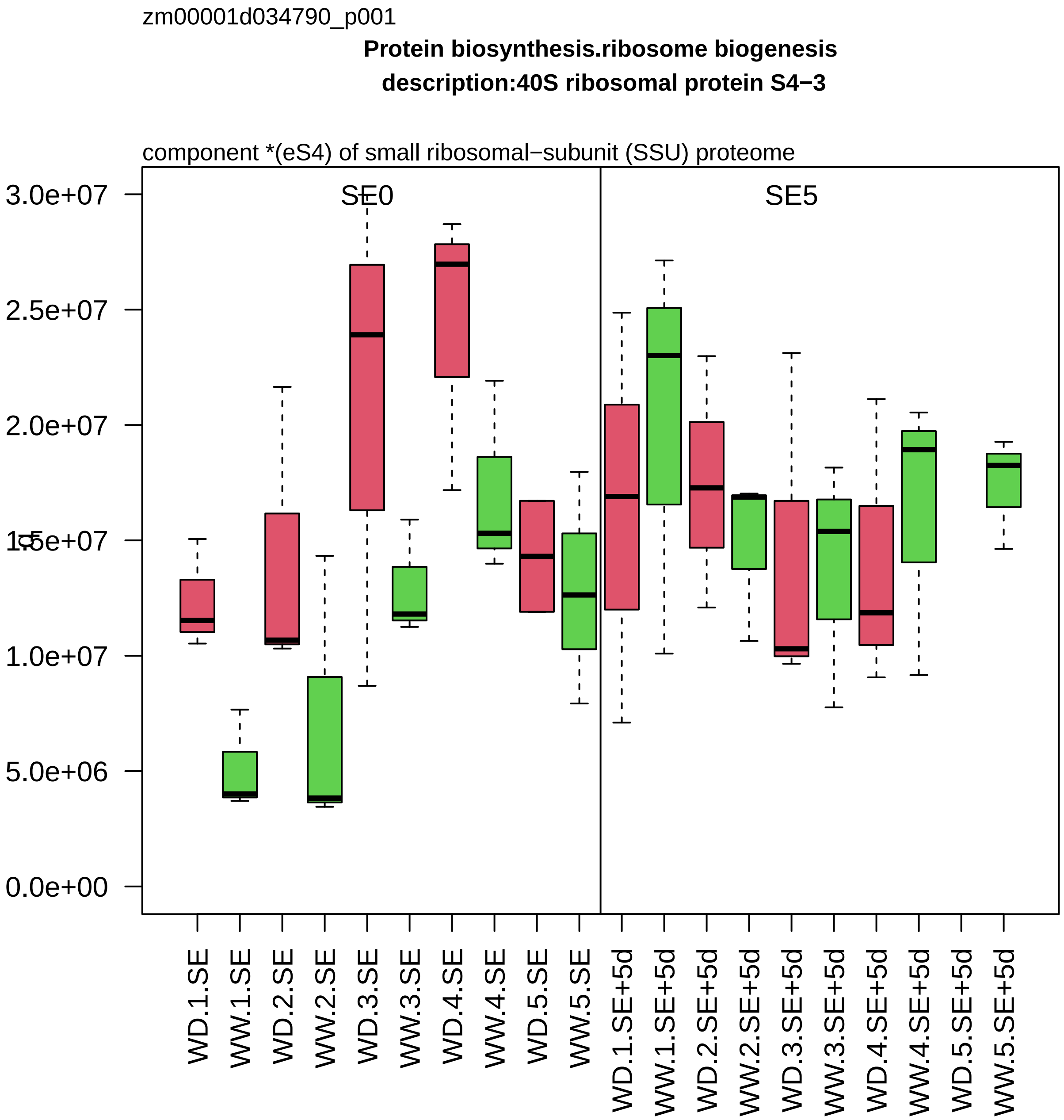

zm00001d034790_p001
Protein biosynthesis.ribosome biogenesis
description:40S ribosomal protein S4−3
component *(eS4) of small ribosomal−subunit (SSU) proteome
SE0
SE5
3.0e+07
2.5e+07
2.0e+07
1.5e+07
1.0e+07
5.0e+06
0.0e+00
WD.1.SE
WW.1.SE
WD.2.SE
WW.2.SE
WD.3.SE
WW.3.SE
WD.4.SE
WW.4.SE
WD.5.SE
WW.5.SE
WD.1.SE+5d
WW.1.SE+5d
WD.2.SE+5d
WW.2.SE+5d
WD.3.SE+5d
WW.3.SE+5d
WD.4.SE+5d
WW.4.SE+5d
WD.5.SE+5d
WW.5.SE+5d

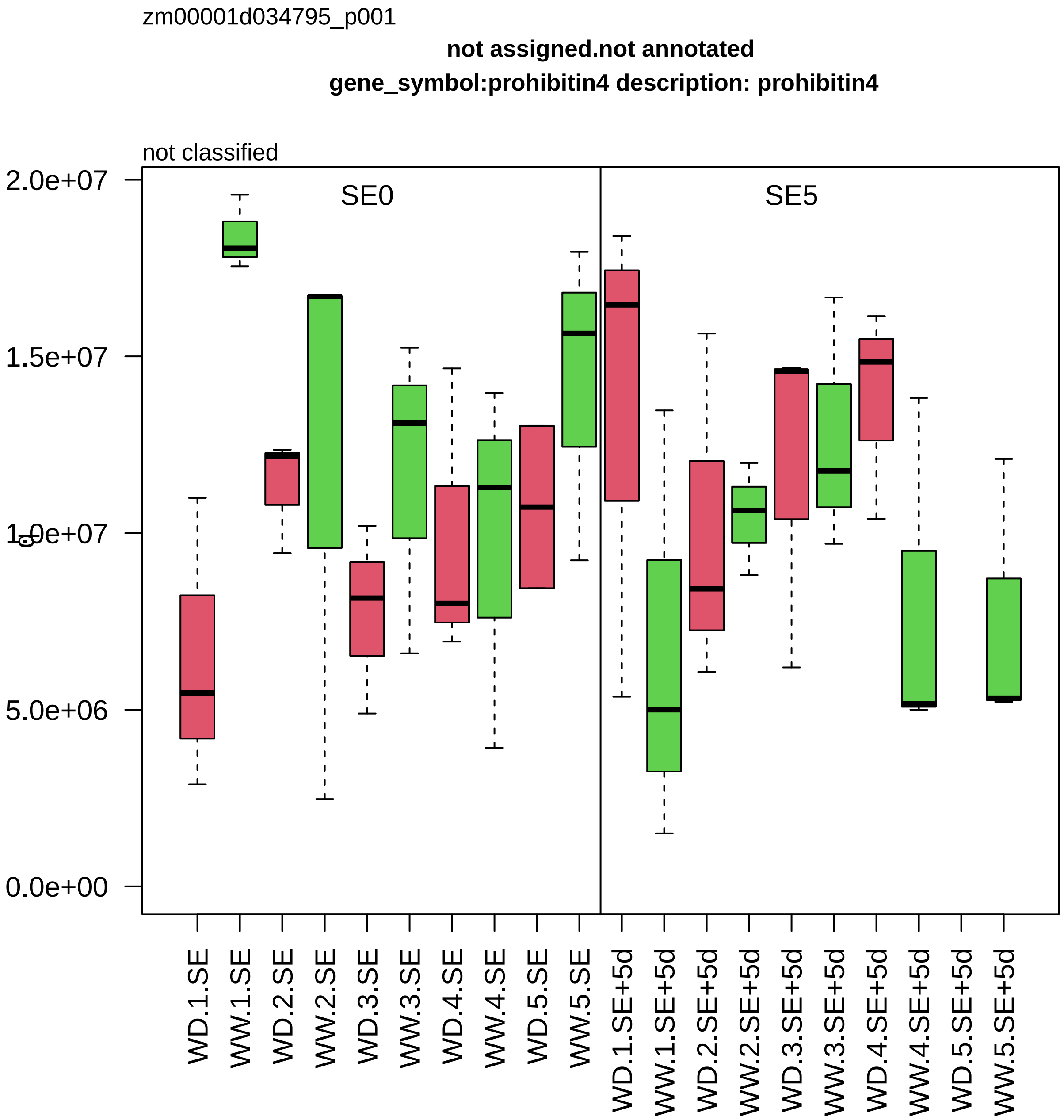
zm00001d034795_p001
not assigned.not annotated
gene_symbol:prohibitin4 description: prohibitin4
not classified
SE0
SE5
2.0e+07
1.5e+07
1.0e+07
5.0e+06
0.0e+00
WD.1.SE
WW.1.SE
WD.2.SE
WW.2.SE
WD.3.SE
WW.3.SE
WD.4.SE
WW.4.SE
WD.5.SE
WW.5.SE
WD.1.SE+5d
WW.1.SE+5d
WD.2.SE+5d
WW.2.SE+5d
WD.3.SE+5d
WW.3.SE+5d
WD.4.SE+5d
WW.4.SE+5d
WD.5.SE+5d
WW.5.SE+5d

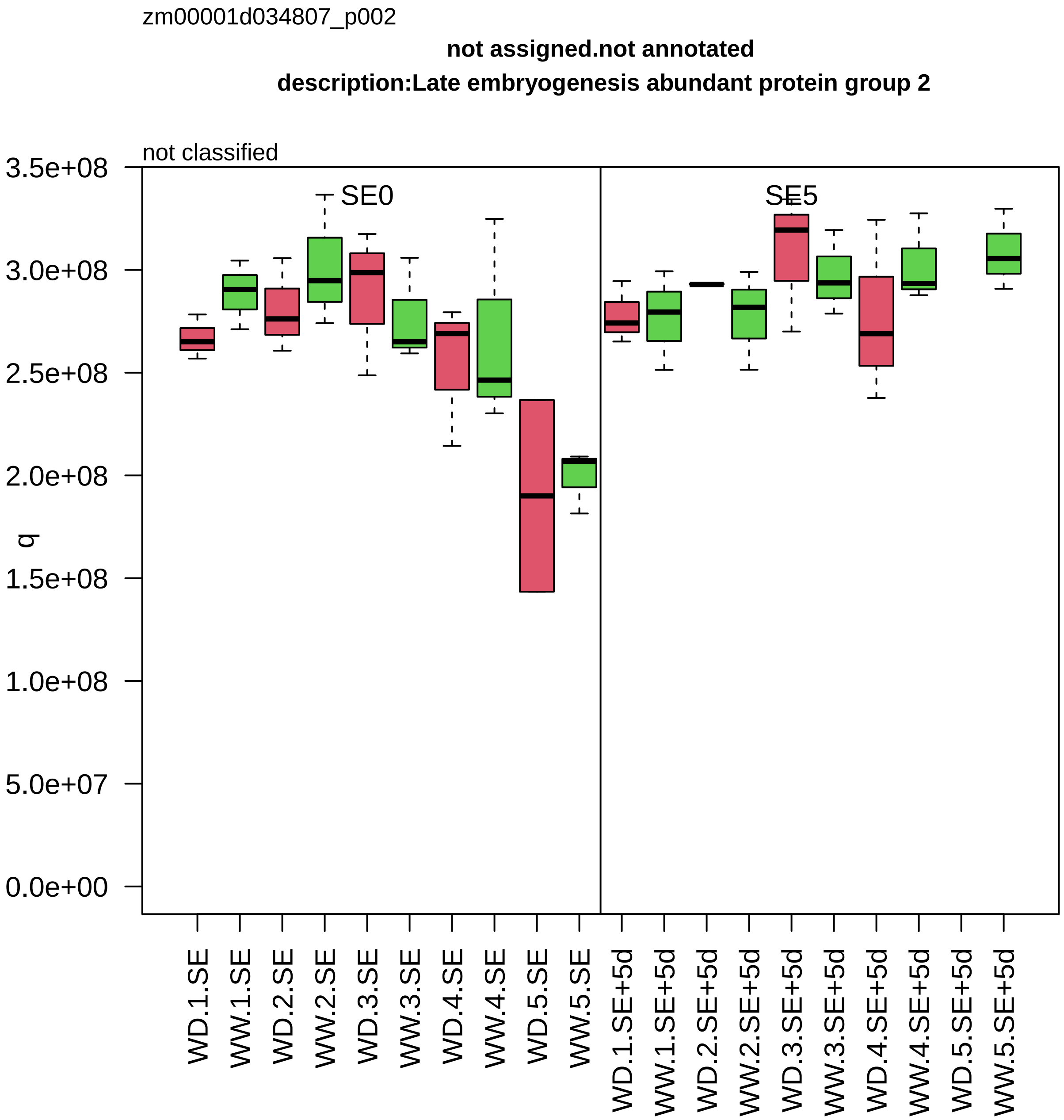

zm00001d034807_p002
not assigned.not annotated
description:Late embryogenesis abundant protein group 2
not classified
SE0
SE5
q
3.5e+08
3.0e+08
2.5e+08
2.0e+08
1.5e+08
1.0e+08
5.0e+07
0.0e+00
WD.1.SE
WW.1.SE
WD.2.SE
WW.2.SE
WD.3.SE
WW.3.SE
WD.4.SE
WW.4.SE
WD.5.SE
WW.5.SE
WD.1.SE+5d
WW.1.SE+5d
WD.2.SE+5d
WW.2.SE+5d
WD.3.SE+5d
WW.3.SE+5d
WD.4.SE+5d
WW.4.SE+5d
WD.5.SE+5d
WW.5.SE+5d

zm00001d034856_p003

**Protein biosynthesis.organelle machinery**

**gene_symbol:elongation factor TU mitochondrial1 description: elongation factor TU mitochondri**

translation elongation factor *(EF−Tu)

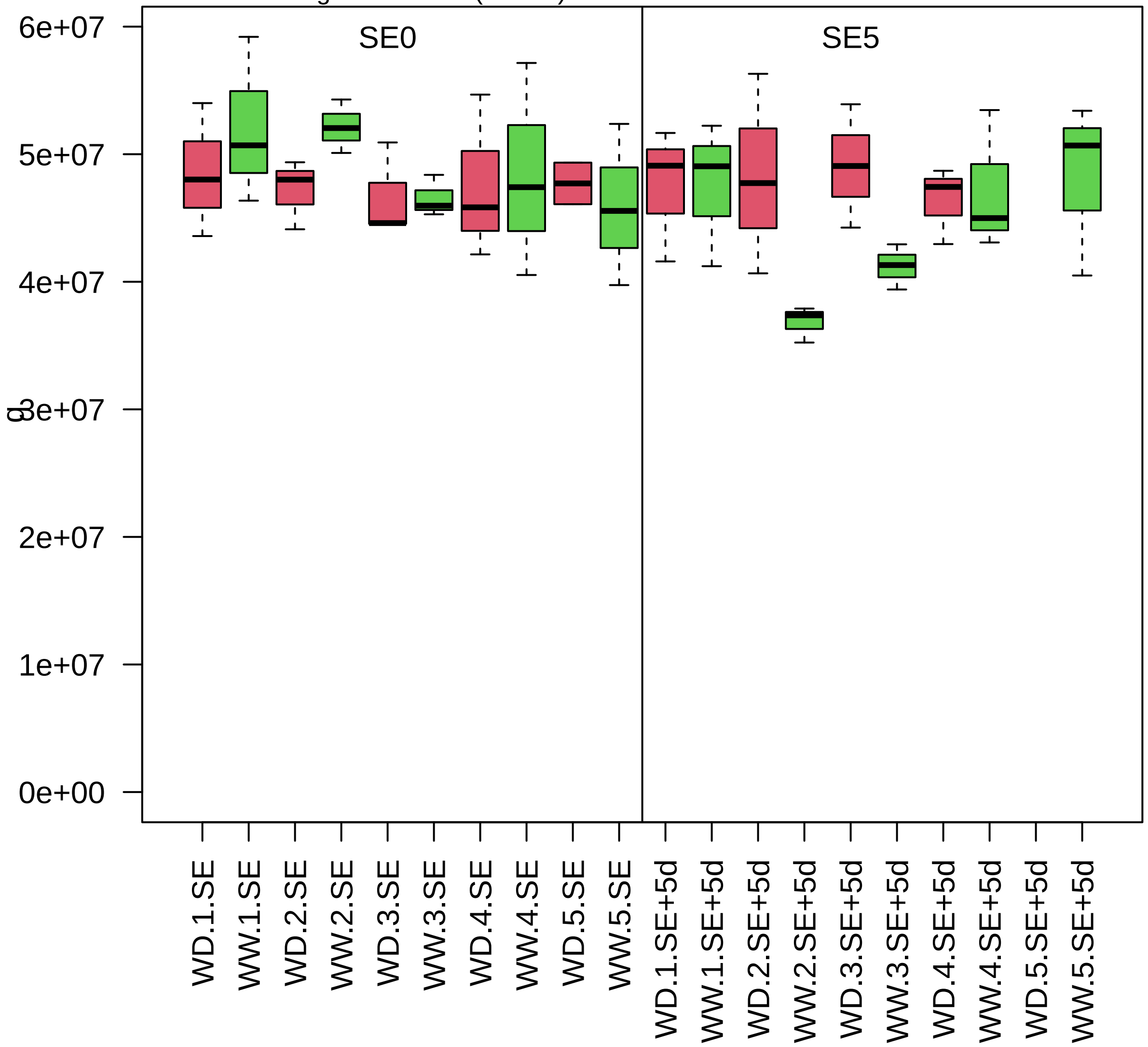

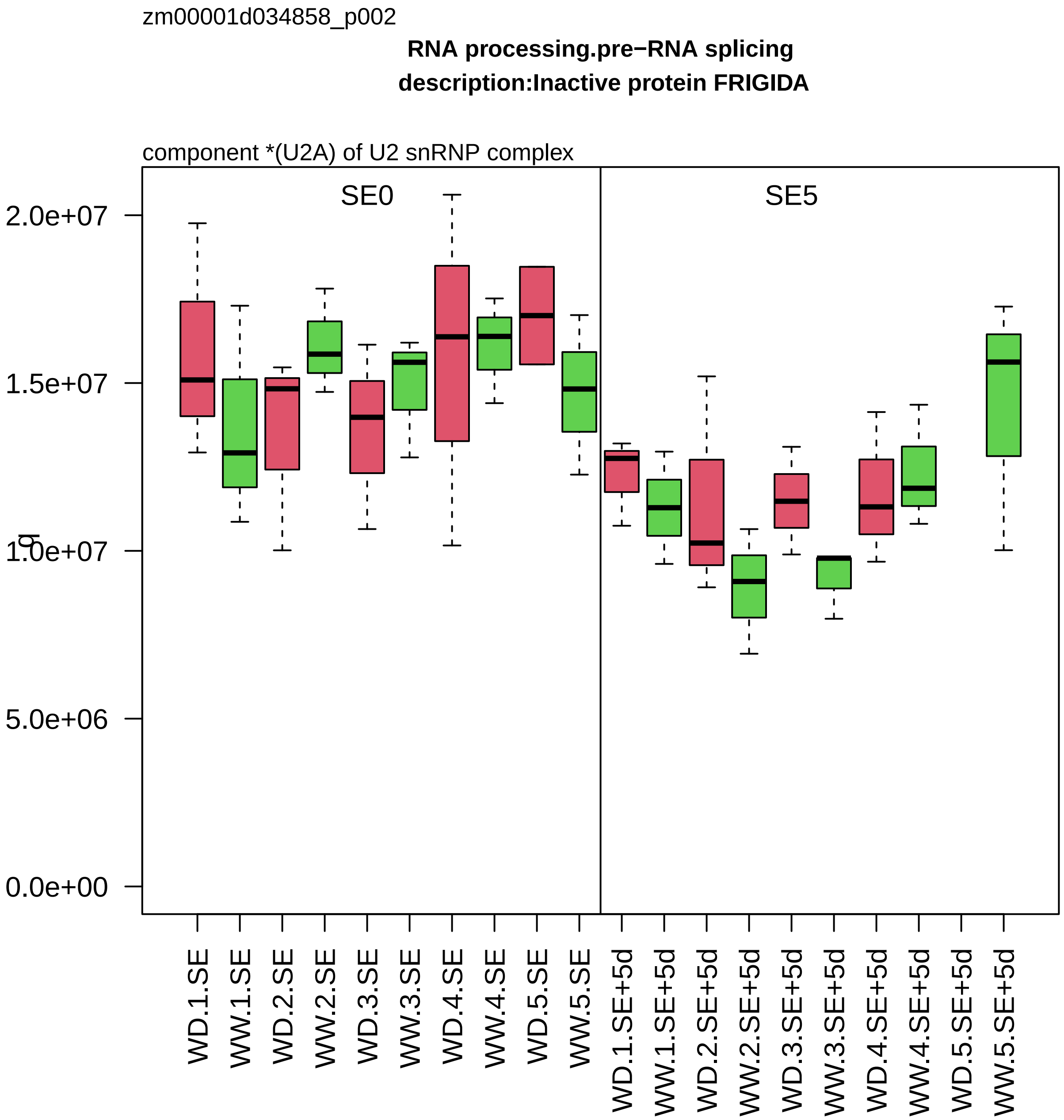

zm00001d034858_p002
RNA processing.pre−RNA splicing
description:Inactive protein FRIGIDA
component *(U2A) of U2 snRNP complex
SE0
SE5
2.0e+07
1.5e+07
1.0e+07
5.0e+06
0.0e+00
WD.1.SE
WW.1.SE
WD.2.SE
WW.2.SE
WD.3.SE
WW.3.SE
WD.4.SE
WW.4.SE
WD.5.SE
WW.5.SE
WD.1.SE+5d
WW.1.SE+5d
WD.2.SE+5d
WW.2.SE+5d
WD.3.SE+5d
WW.3.SE+5d
WD.4.SE+5d
WW.4.SE+5d
WD.5.SE+5d
WW.5.SE+5d

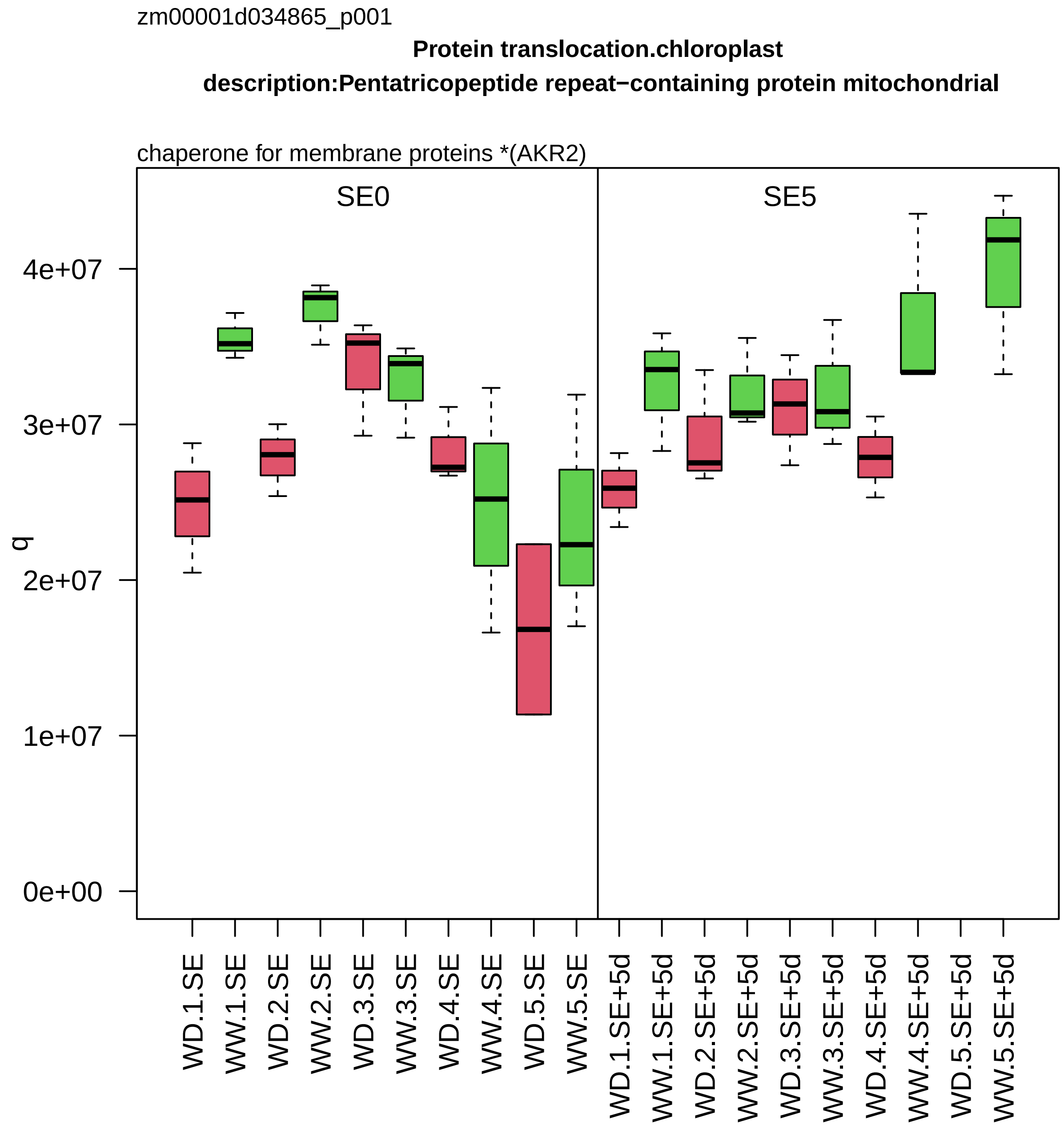
zm00001d034865_p001
Protein translocation.chloroplast
description:Pentatricopeptide repeat−containing protein mitochondrial
chaperone for membrane proteins *(AKR2)
SE0
SE5
q
0e+00
1e+07
2e+07
3e+07
4e+07
WD.1.SE
WW.1.SE
WD.2.SE
WW.2.SE
WD.3.SE
WW.3.SE
WD.4.SE
WW.4.SE
WD.5.SE
WW.5.SE
WD.1.SE+5d
WW.1.SE+5d
WD.2.SE+5d
WW.2.SE+5d
WD.3.SE+5d
WW.3.SE+5d
WD.4.SE+5d
WW.4.SE+5d
WD.5.SE+5d
WW.5.SE+5d

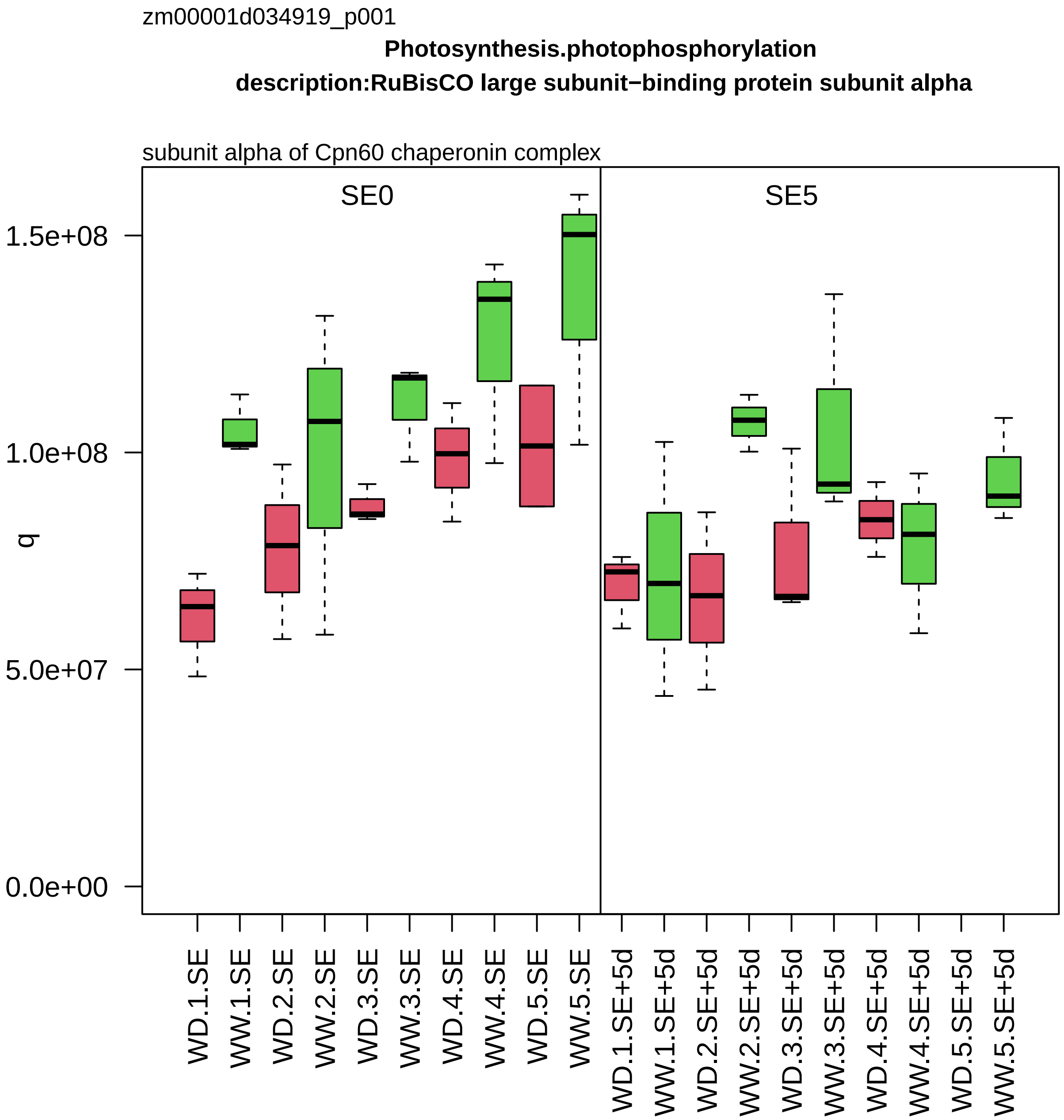

zm00001d034919_p001
Photosynthesis.photophosphorylation
description:RuBisCO large subunit−binding protein subunit alpha
subunit alpha of Cpn60 chaperonin complex
SE0
SE5
q
1.5e+08
1.0e+08
5.0e+07
0.0e+00
WD.1.SE
WW.1.SE
WD.2.SE
WW.2.SE
WD.3.SE
WW.3.SE
WD.4.SE
WW.4.SE
WD.5.SE
WW.5.SE
WD.1.SE+5d
WW.1.SE+5d
WD.2.SE+5d
WW.2.SE+5d
WD.3.SE+5d
WW.3.SE+5d
WD.4.SE+5d
WW.4.SE+5d
WD.5.SE+5d
WW.5.SE+5d

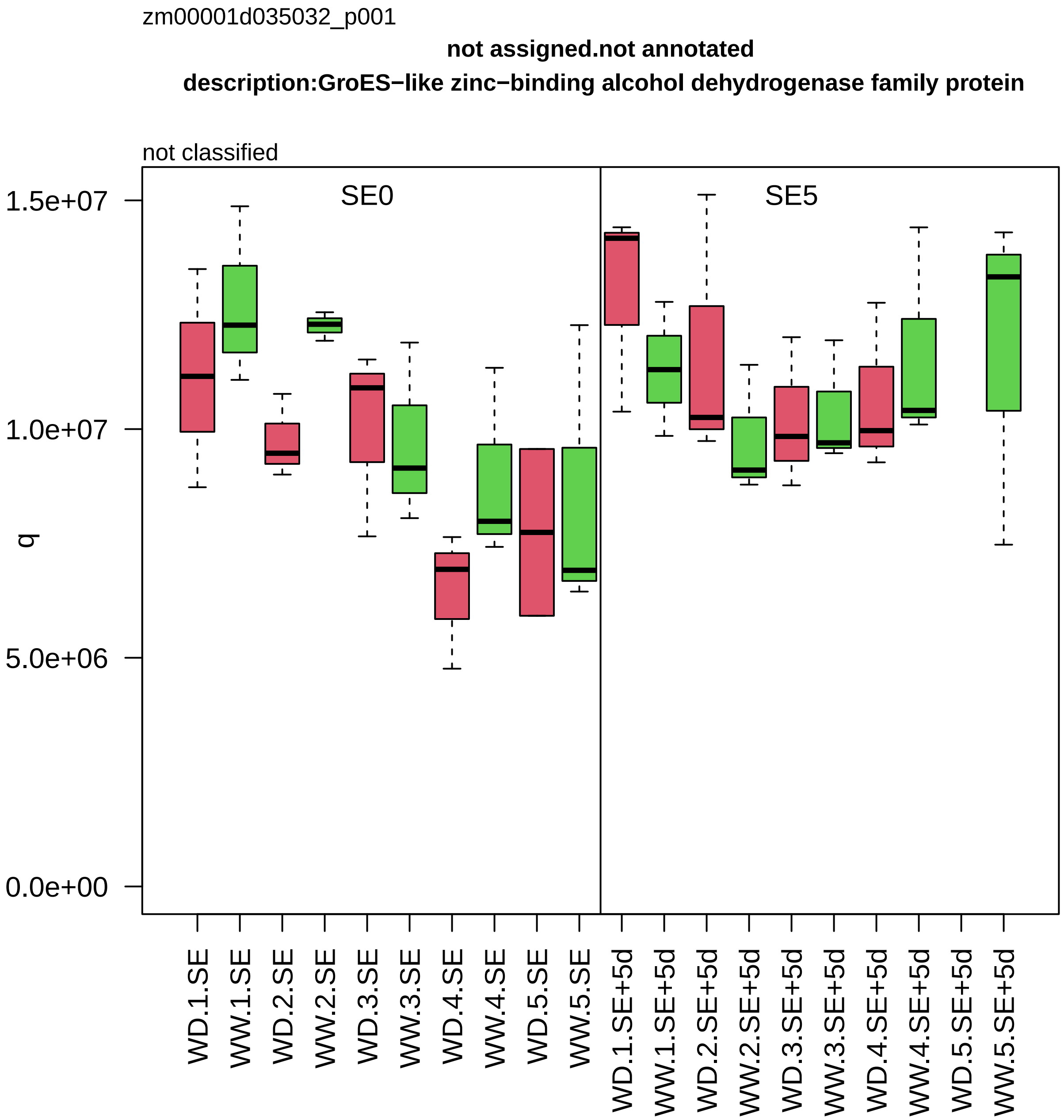
zm00001d035032_p001
not assigned.not annotated
description:GroES−like zinc−binding alcohol dehydrogenase family protein
not classified
SE0
SE5
q
1.5e+07
1.0e+07
5.0e+06
0.0e+00
WD.1.SE
WW.1.SE
WD.2.SE
WW.2.SE
WD.3.SE
WW.3.SE
WD.4.SE
WW.4.SE
WD.5.SE
WW.5.SE
WD.1.SE+5d
WW.1.SE+5d
WD.2.SE+5d
WW.2.SE+5d
WD.3.SE+5d
WW.3.SE+5d
WD.4.SE+5d
WW.4.SE+5d
WD.5.SE+5d
WW.5.SE+5d

zm00001d035037_p001

**Carbohydrate metabolism.sucrose metabolism**
**description:Fructokinase−2**

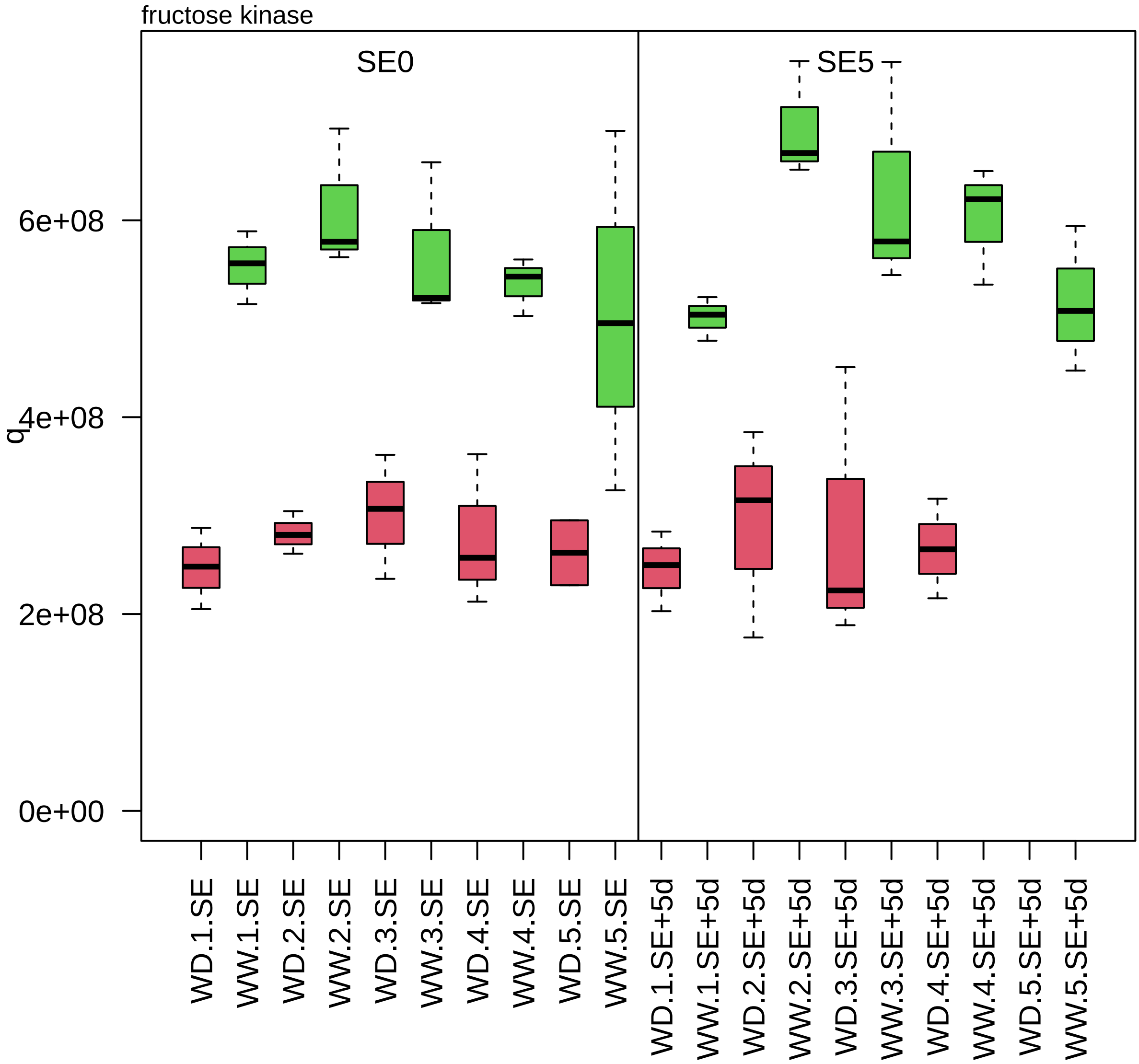

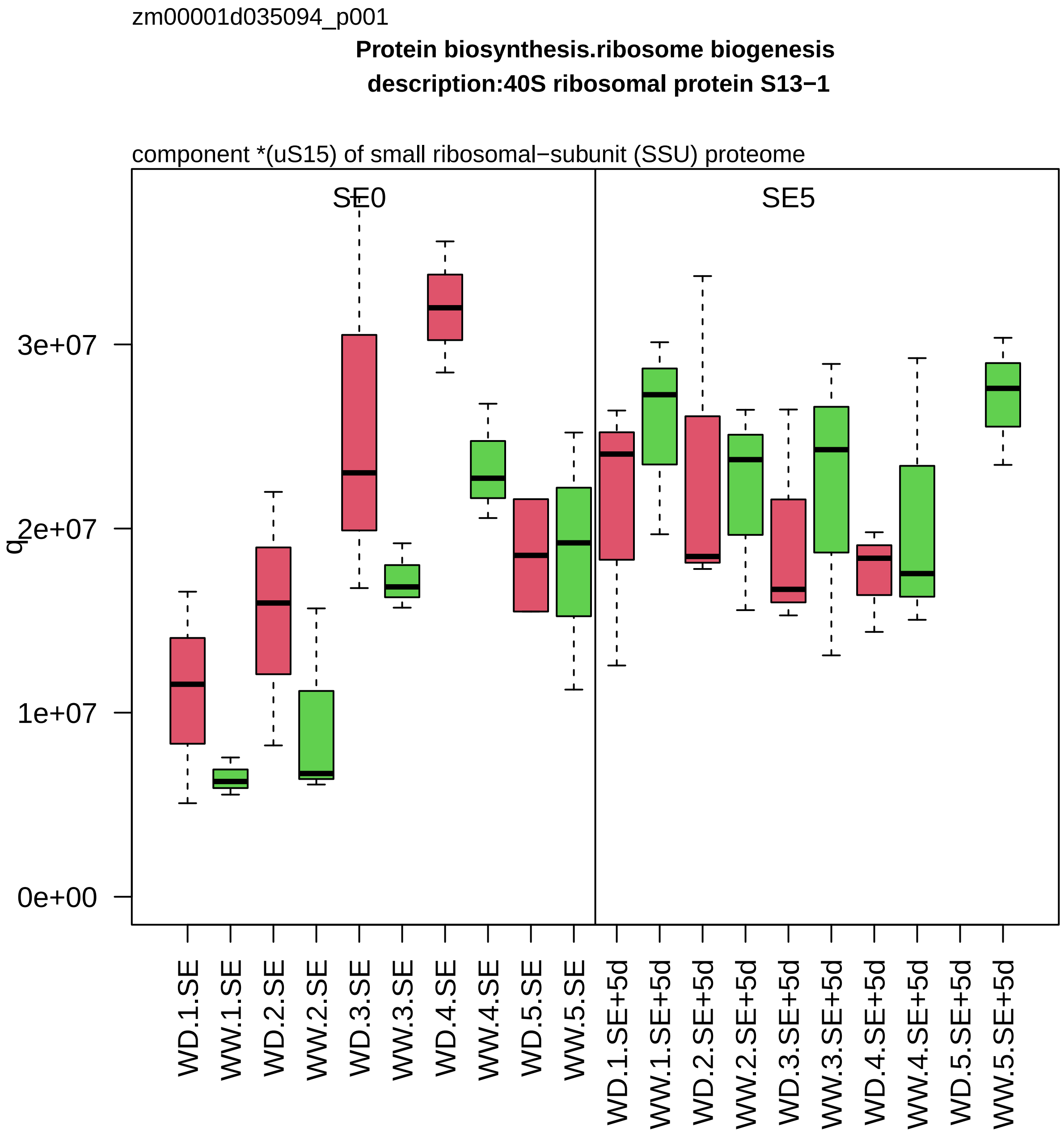

zm00001d035094_p001
Protein biosynthesis.ribosome biogenesis
description:40S ribosomal protein S13−1
component *(uS15) of small ribosomal−subunit (SSU) proteome
SE0
SE5
q
3e+07
2e+07
1e+07
0e+00
WD.1.SE
WW.1.SE
WD.2.SE
WW.2.SE
WD.3.SE
WW.3.SE
WD.4.SE
WW.4.SE
WD.5.SE
WW.5.SE
WD.1.SE+5d
WW.1.SE+5d
WD.2.SE+5d
WW.2.SE+5d
WD.3.SE+5d
WW.3.SE+5d
WD.4.SE+5d
WW.4.SE+5d
WD.5.SE+5d
WW.5.SE+5d

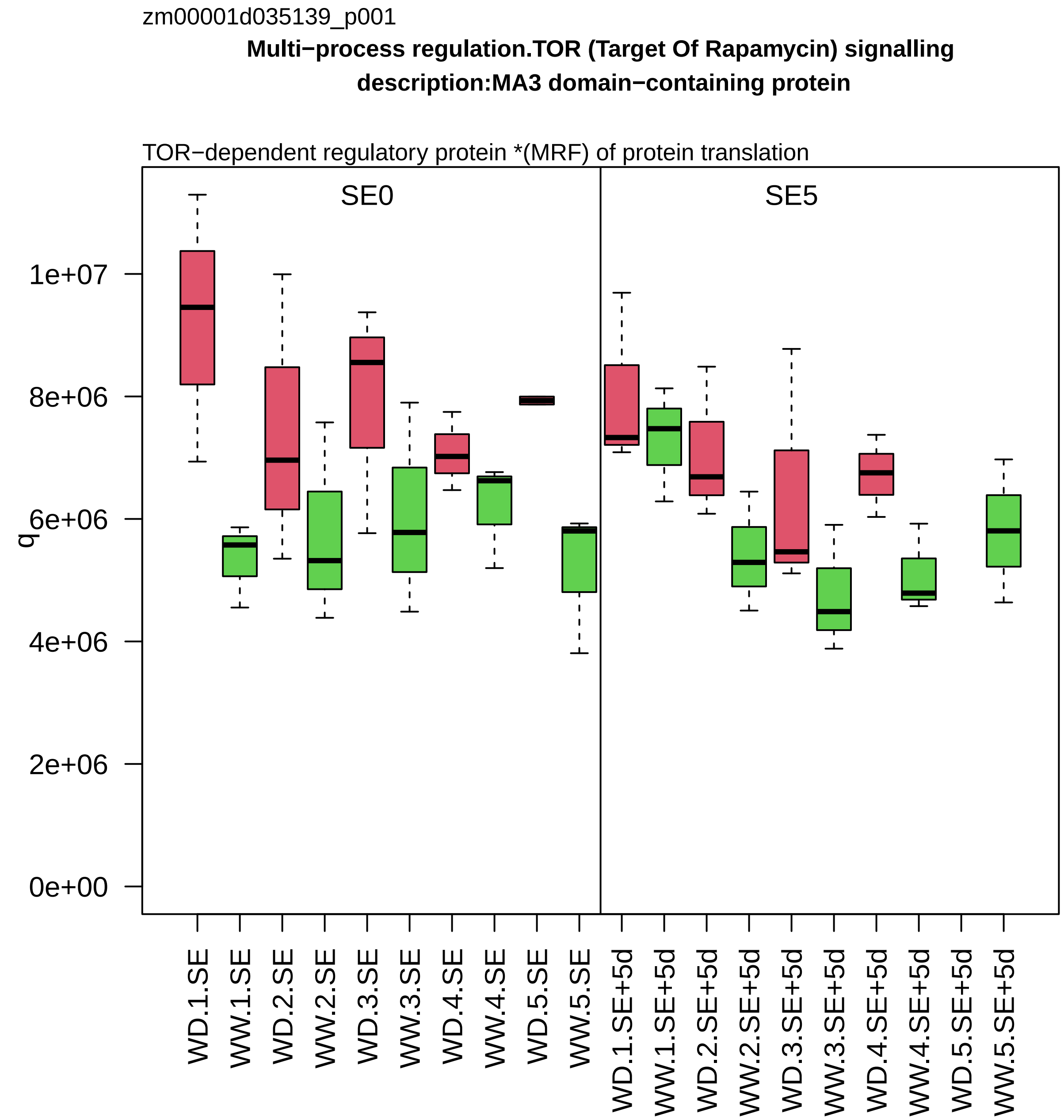

zm00001d035139_p001
Multi−process regulation.TOR (Target Of Rapamycin) signalling
description:MA3 domain−containing protein
TOR−dependent regulatory protein *(MRF) of protein translation
SE0
SE5
q
0e+00
2e+06
4e+06
6e+06
8e+06
1e+07
WD.1.SE
WW.1.SE
WD.2.SE
WW.2.SE
WD.3.SE
WW.3.SE
WD.4.SE
WW.4.SE
WD.5.SE
WW.5.SE
WD.1.SE+5d
WW.1.SE+5d
WD.2.SE+5d
WW.2.SE+5d
WD.3.SE+5d
WW.3.SE+5d
WD.4.SE+5d
WW.4.SE+5d
WD.5.SE+5d
WW.5.SE+5d

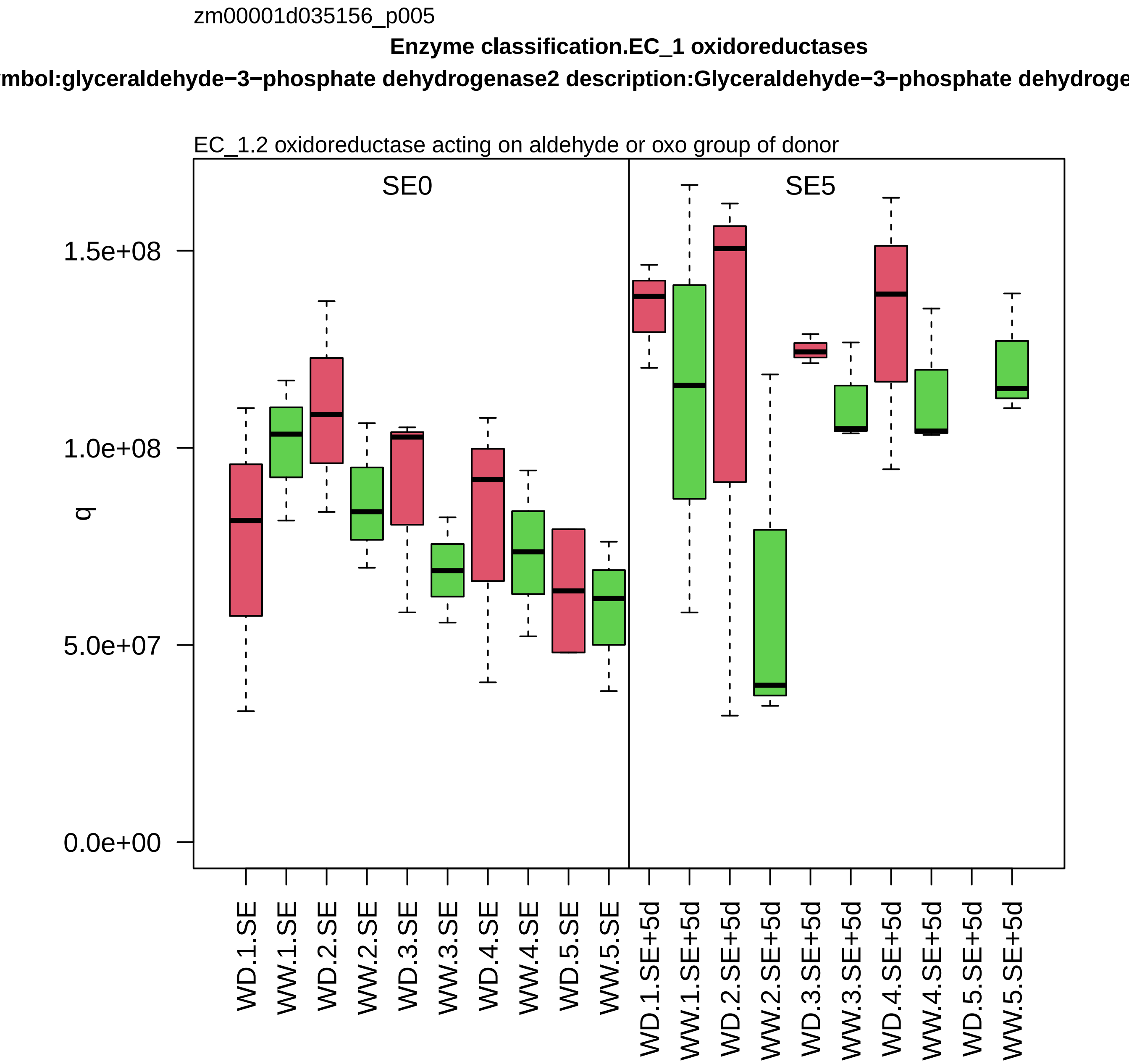

zm00001d035156_p005
Enzyme classification.EC_1 oxidoreductases
ymbol:glyceraldehyde−3−phosphate dehydrogenase2 description:Glyceraldehyde−3−phosphate dehydroge
EC_1.2 oxidoreductase acting on aldehyde or oxo group of donor
SE0
SE5
q
1.5e+08
1.0e+08
5.0e+07
0.0e+00
WD.1.SE
WW.1.SE
WD.2.SE
WW.2.SE
WD.3.SE
WW.3.SE
WD.4.SE
WW.4.SE
WD.5.SE
WW.5.SE
WD.1.SE+5d
WW.1.SE+5d
WD.2.SE+5d
WW.2.SE+5d
WD.3.SE+5d
WW.3.SE+5d
WD.4.SE+5d
WW.4.SE+5d
WD.5.SE+5d
WW.5.SE+5d

zm00001d035156_p009

**Enzyme classification.EC_1 oxidoreductases**

**symbol:glyceraldehyde−3−phosphate dehydrogenase2 description:Glyceraldehyde−3−phosphate dehydrogenase**

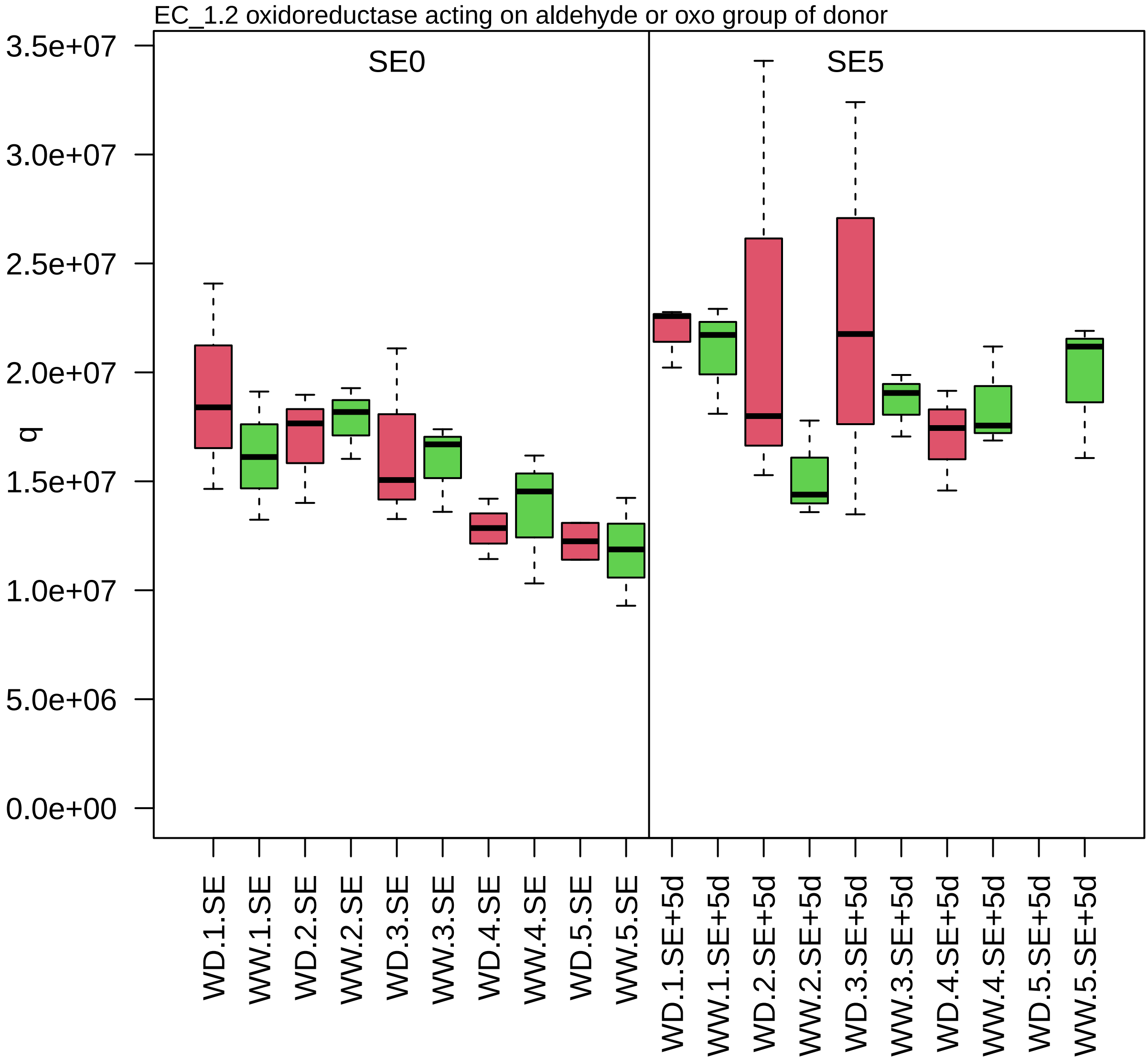

zm00001d035256_p002

**RNA processing.mRNA silencing**

**gene_symbol:ragged seedling1 description:Protein SUPPRESSOR OF GENE SILENCING 3 homo**

stabilization factor *(SGS3) of transacting siRNA pathway

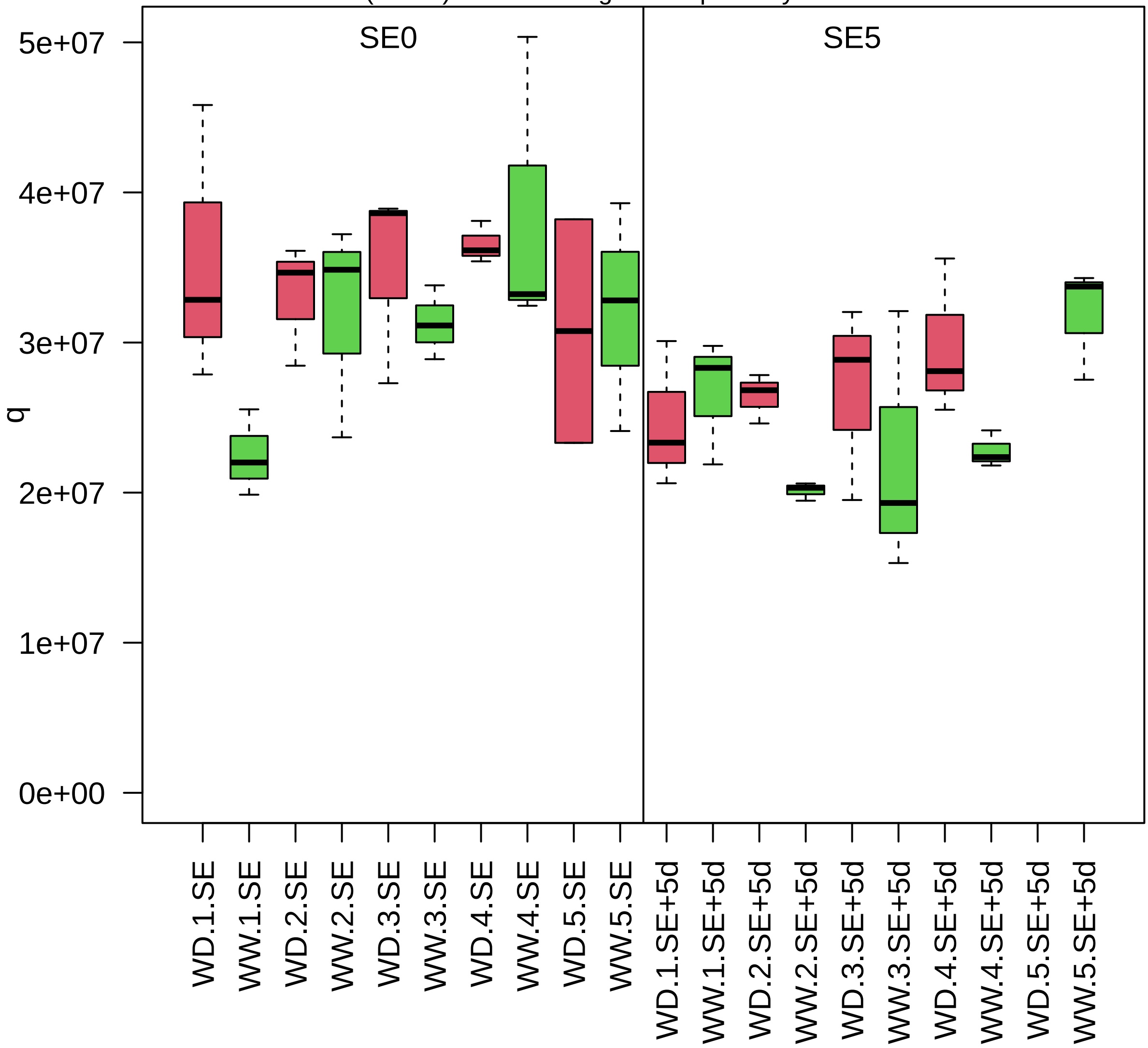

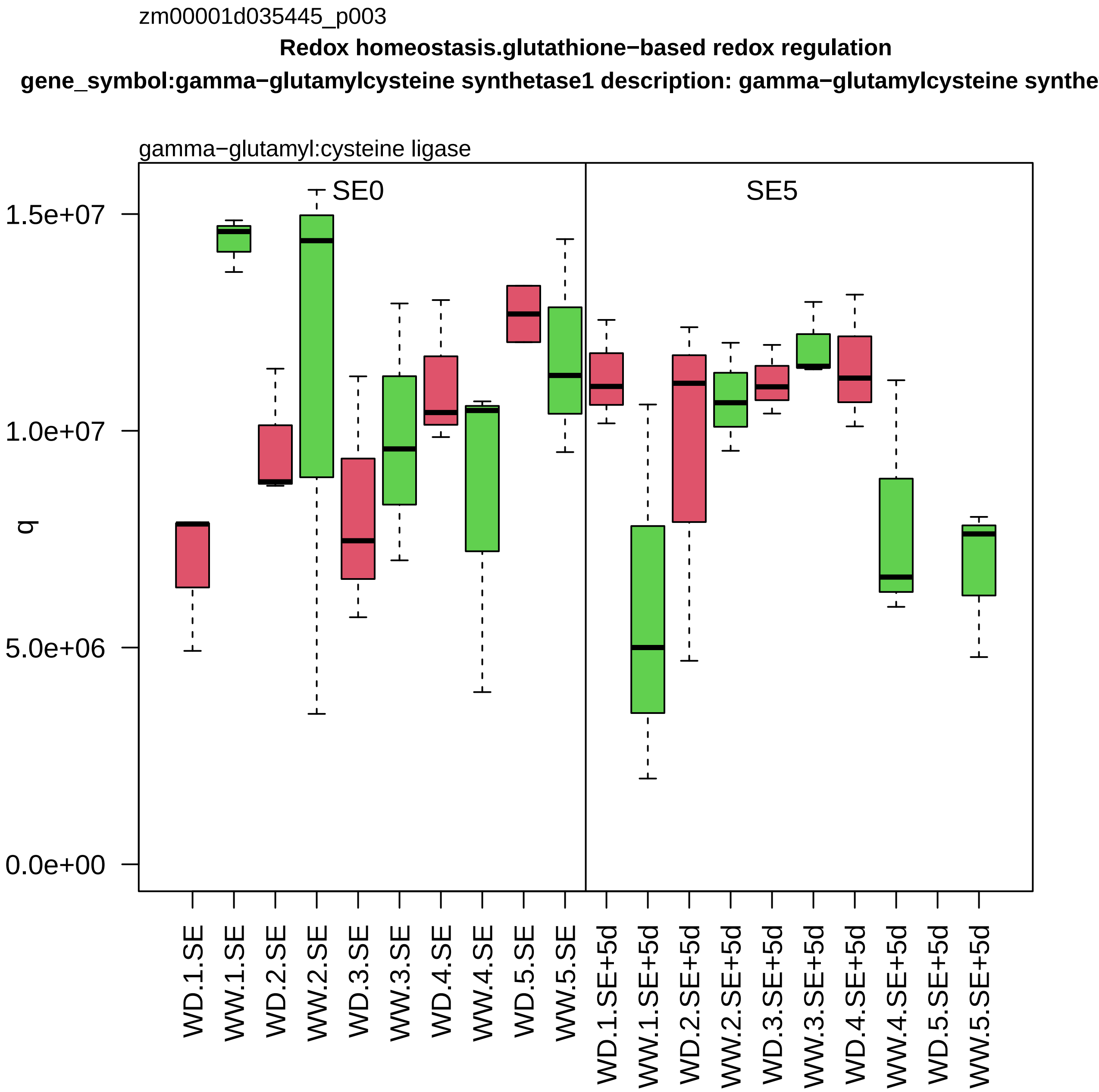

zm00001d035445_p003
Redox homeostasis.glutathione−based redox regulation
gene_symbol:gamma−glutamylcysteine synthetase1 description: gamma−glutamylcysteine synthe
gamma−glutamyl:cysteine ligase
SE0
SE5
q
1.5e+07
1.0e+07
5.0e+06
0.0e+00
WD.1.SE
WW.1.SE
WD.2.SE
WW.2.SE
WD.3.SE
WW.3.SE
WD.4.SE
WW.4.SE
WD.5.SE
WW.5.SE
WD.1.SE+5d
WW.1.SE+5d
WD.2.SE+5d
WW.2.SE+5d
WD.3.SE+5d
WW.3.SE+5d
WD.4.SE+5d
WW.4.SE+5d
WD.5.SE+5d
WW.5.SE+5d

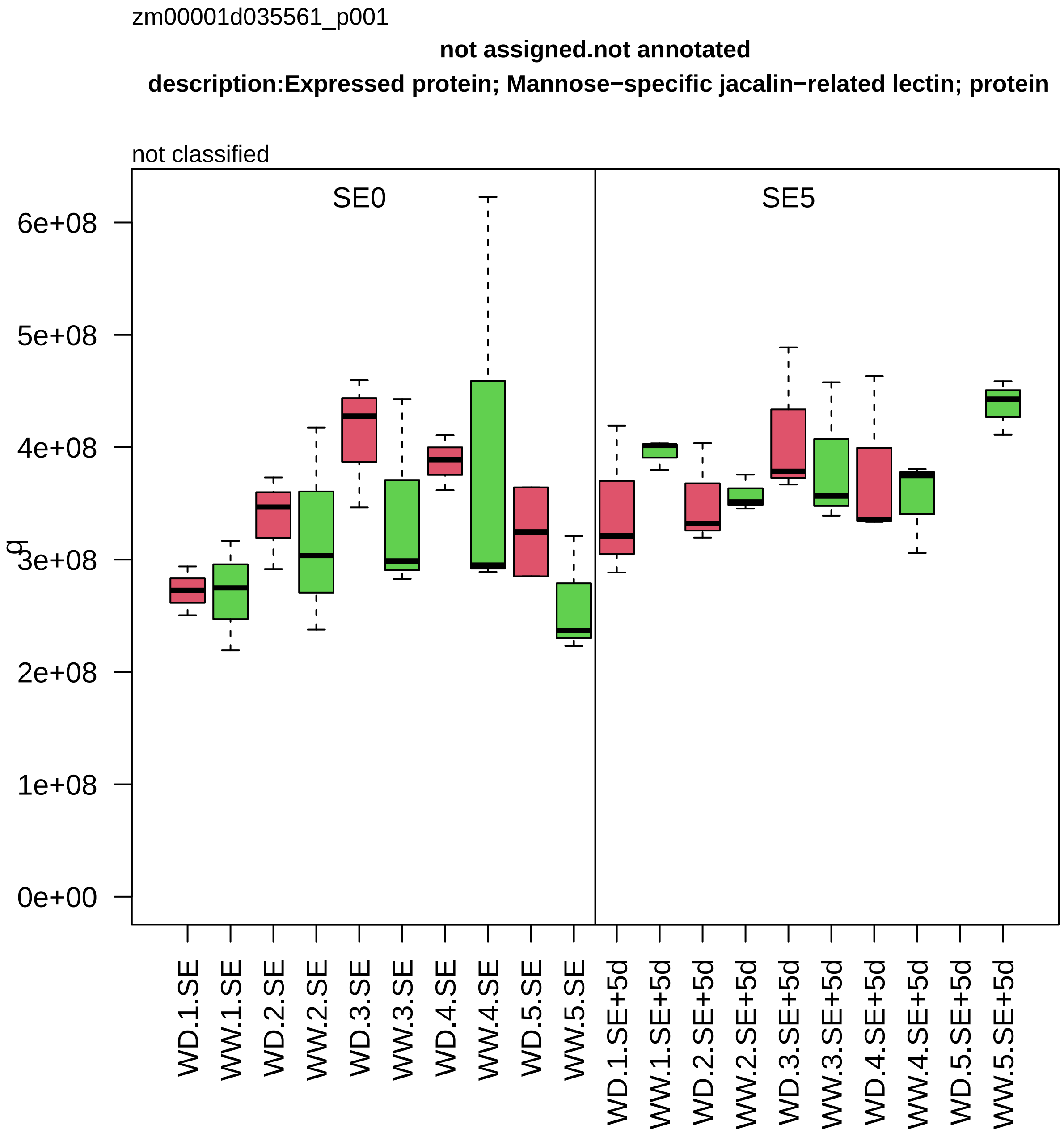

zm00001d035561_p001
not assigned.not annotated
description:Expressed protein; Mannose−specific jacalin−related lectin; protein
not classified
SE0
SE5
6e+08
5e+08
4e+08
3e+08
2e+08
1e+08
0e+00
g
WD.1.SE
WW.1.SE
WD.2.SE
WW.2.SE
WD.3.SE
WW.3.SE
WD.4.SE
WW.4.SE
WD.5.SE
WW.5.SE
WD.1.SE+5d
WW.1.SE+5d
WD.2.SE+5d
WW.2.SE+5d
WD.3.SE+5d
WW.3.SE+5d
WD.4.SE+5d
WW.4.SE+5d
WD.5.SE+5d
WW.5.SE+5d

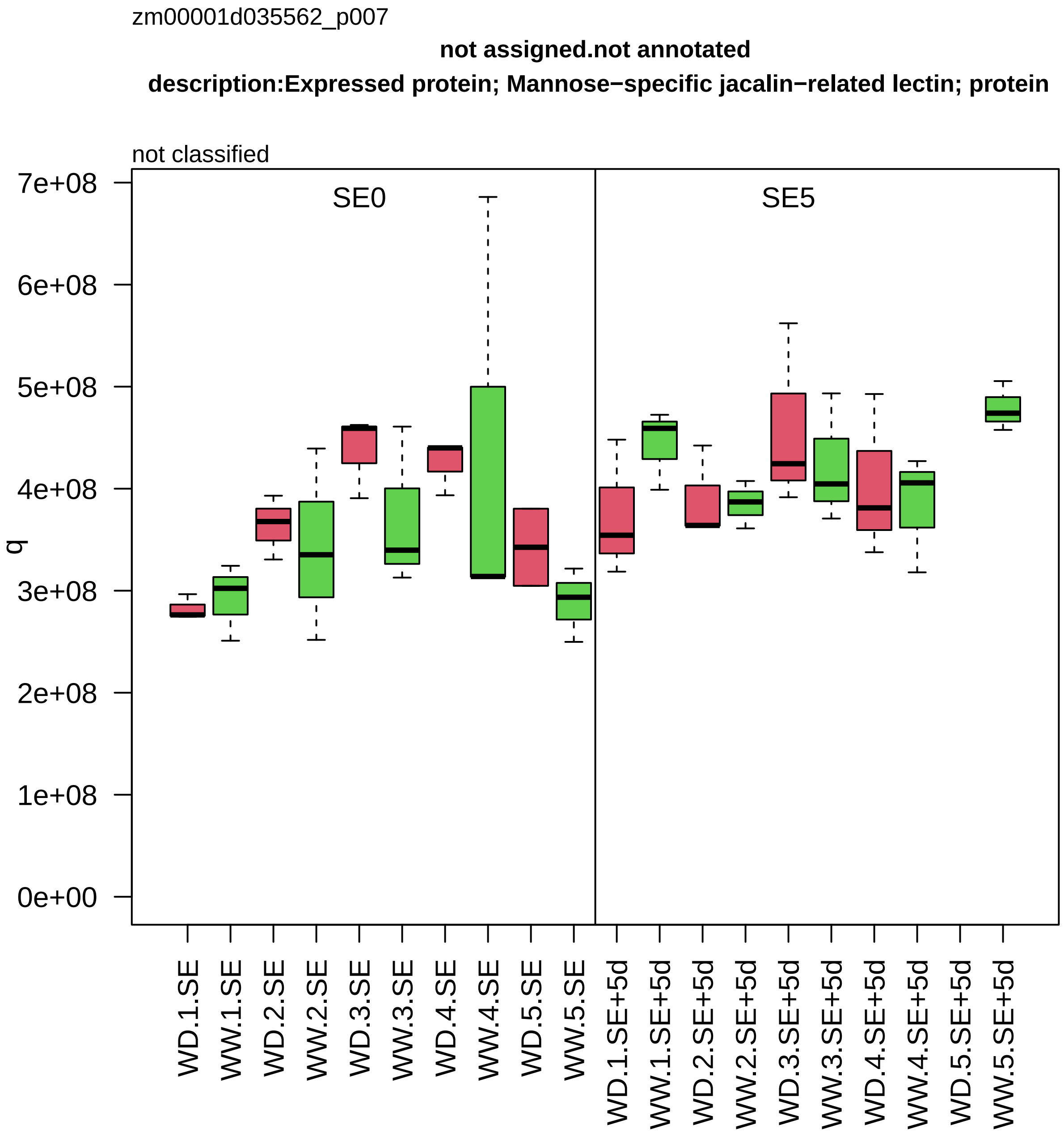

zm00001d035562_p007
not assigned.not annotated
description:Expressed protein; Mannose−specific jacalin−related lectin; protein
not classified
SE0
SE5
q
0e+00
1e+08
2e+08
3e+08
4e+08
5e+08
6e+08
7e+08
WD.1.SE
WW.1.SE
WD.2.SE
WW.2.SE
WD.3.SE
WW.3.SE
WD.4.SE
WW.4.SE
WD.5.SE
WW.5.SE
WD.1.SE+5d
WW.1.SE+5d
WD.2.SE+5d
WW.2.SE+5d
WD.3.SE+5d
WW.3.SE+5d
WD.4.SE+5d
WW.4.SE+5d
WD.5.SE+5d
WW.5.SE+5d

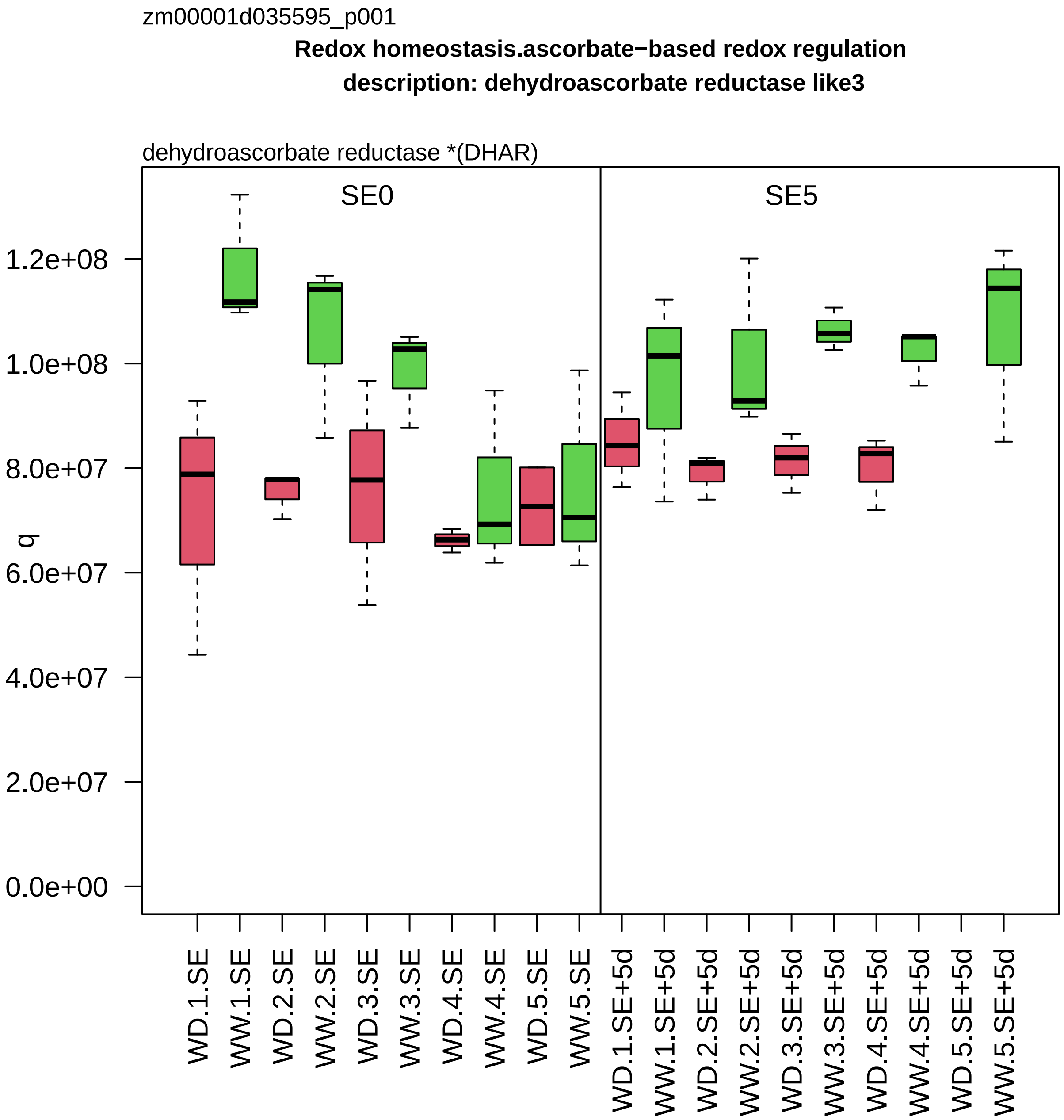

zm00001d035595_p001
Redox homeostasis.ascorbate−based redox regulation
description: dehydroascorbate reductase like3
dehydroascorbate reductase *(DHAR)
SE0
SE5
q
1.2e+08
1.0e+08
8.0e+07
6.0e+07
4.0e+07
2.0e+07
0.0e+00
WD.1.SE
WW.1.SE
WD.2.SE
WW.2.SE
WD.3.SE
WW.3.SE
WD.4.SE
WW.4.SE
WD.5.SE
WW.5.SE
WD.1.SE+5d
WW.1.SE+5d
WD.2.SE+5d
WW.2.SE+5d
WD.3.SE+5d
WW.3.SE+5d
WD.4.SE+5d
WW.4.SE+5d
WD.5.SE+5d
WW.5.SE+5d

zm00001d035597_p001

**Protein homeostasis.protein storage**
**description:RmlC−like cupins superfamily protein**

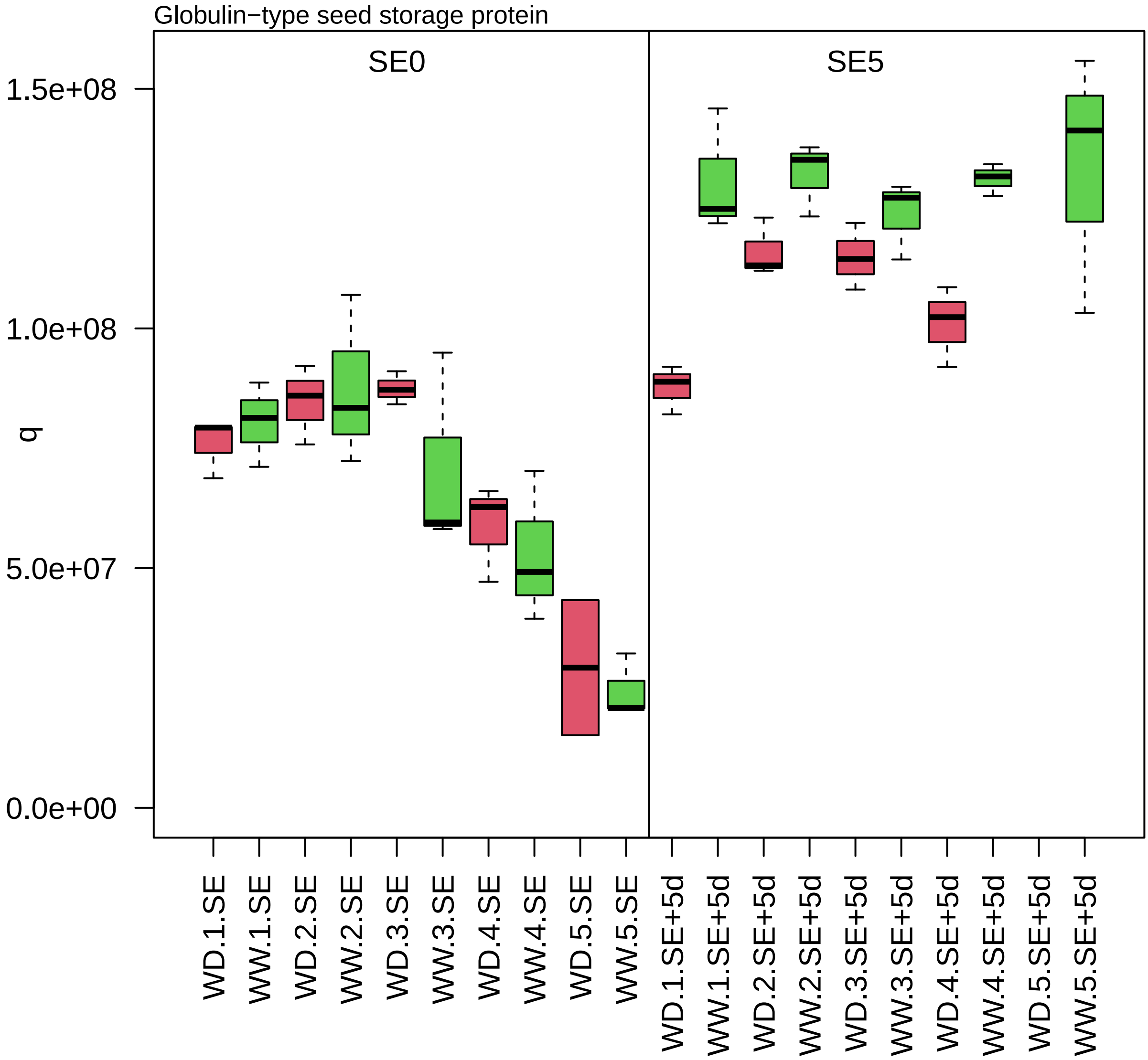

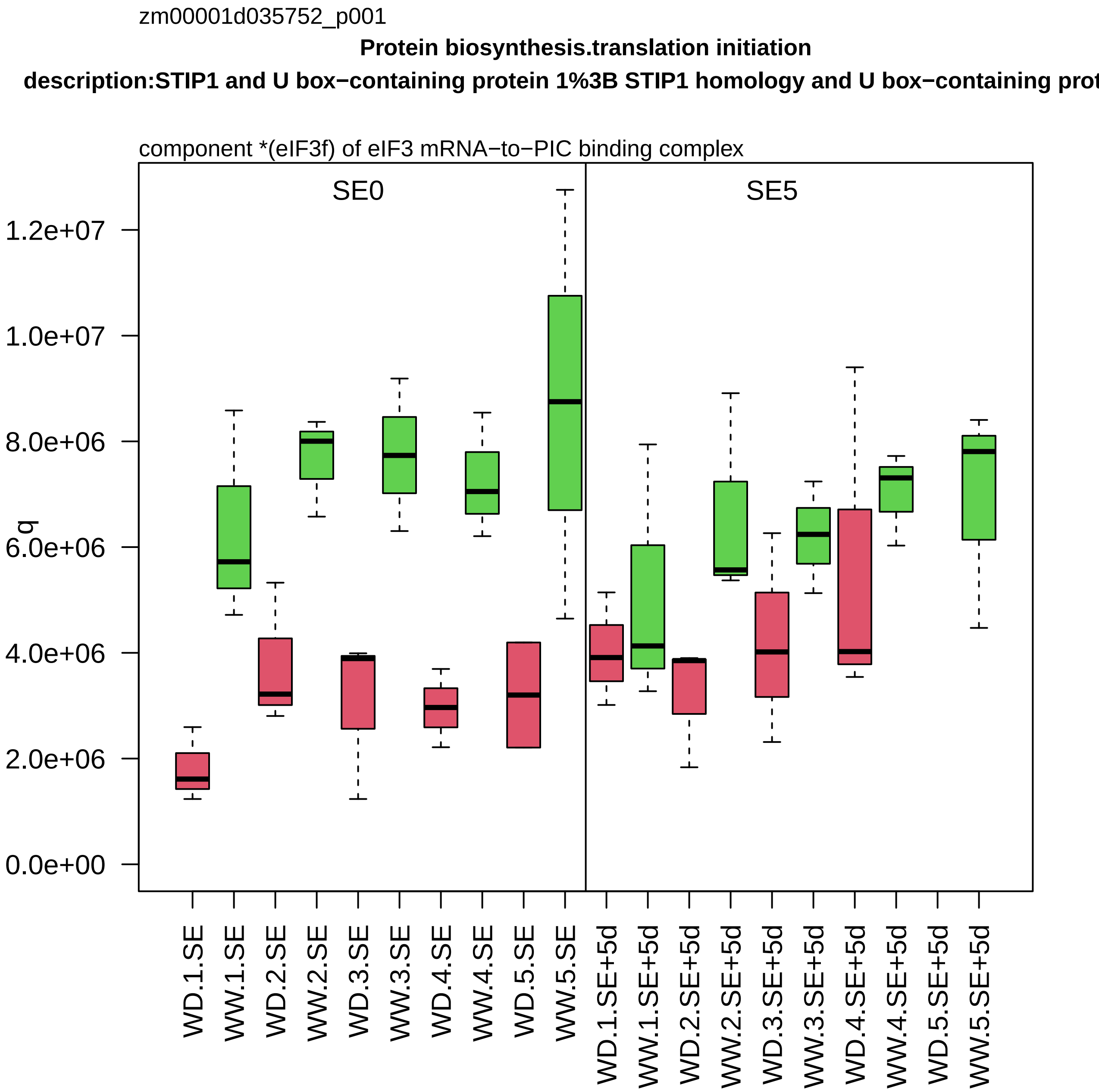

zm00001d035752_p001
Protein biosynthesis.translation initiation
description:STIP1 and U box−containing protein 1%3B STIP1 homology and U box−containing prot
component *(eIF3f) of eIF3 mRNA−to−PIC binding complex
SE0
SE5
q
1.2e+07
1.0e+07
8.0e+06
6.0e+06
4.0e+06
2.0e+06
0.0e+00
WD.1.SE
WW.1.SE
WD.2.SE
WW.2.SE
WD.3.SE
WW.3.SE
WD.4.SE
WW.4.SE
WD.5.SE
WW.5.SE
WD.1.SE+5d
WW.1.SE+5d
WD.2.SE+5d
WW.2.SE+5d
WD.3.SE+5d
WW.3.SE+5d
WD.4.SE+5d
WW.4.SE+5d
WD.5.SE+5d
WW.5.SE+5d

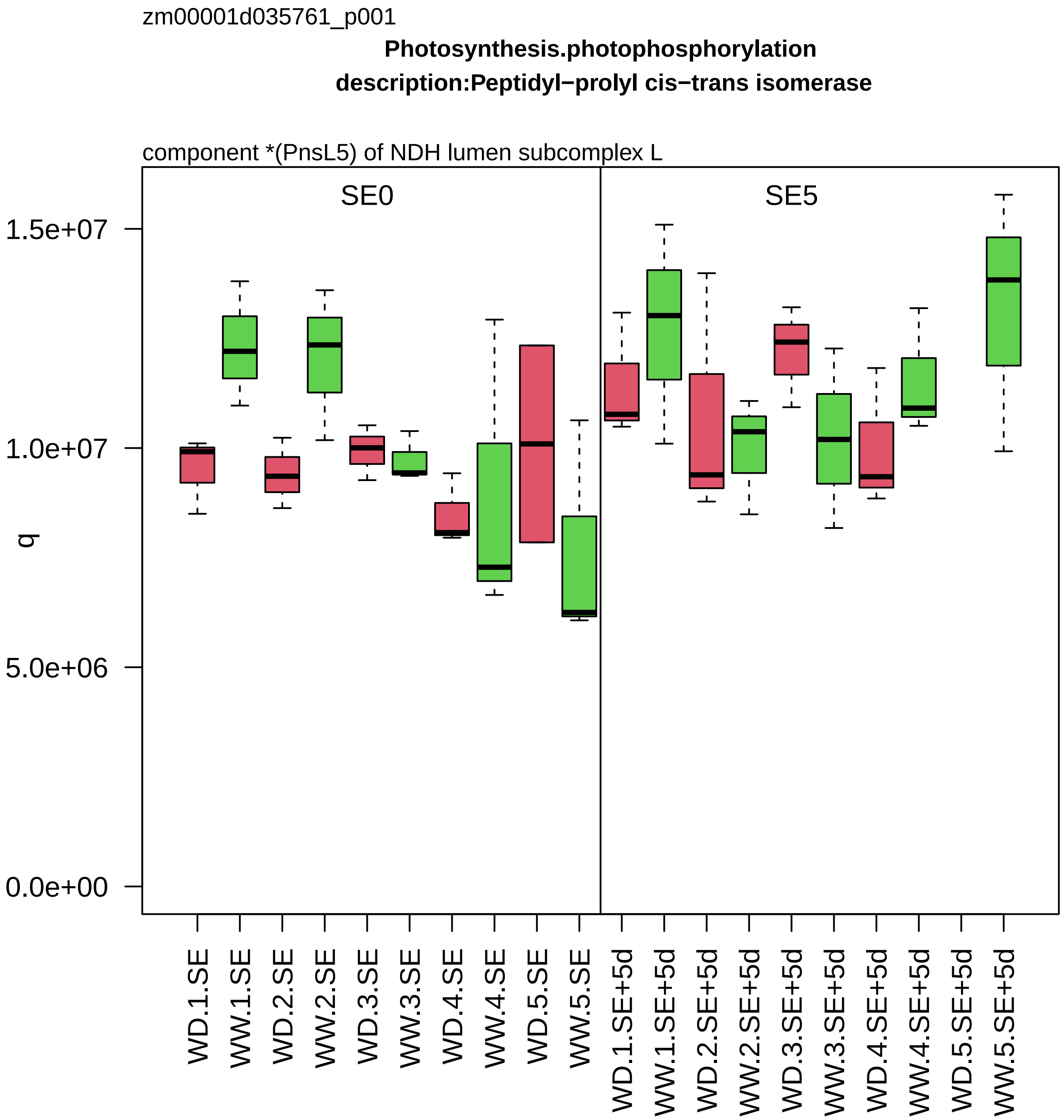

zm00001d035761_p001
Photosynthesis.photophosphorylation
description:Peptidyl−prolyl cis−trans isomerase
component *(PnsL5) of NDH lumen subcomplex L
SE0
SE5
q
1.5e+07
1.0e+07
5.0e+06
0.0e+00
WD.1.SE
WW.1.SE
WD.2.SE
WW.2.SE
WD.3.SE
WW.3.SE
WD.4.SE
WW.4.SE
WD.5.SE
WW.5.SE
WD.1.SE+5d
WW.1.SE+5d
WD.2.SE+5d
WW.2.SE+5d
WD.3.SE+5d
WW.3.SE+5d
WD.4.SE+5d
WW.4.SE+5d
WD.5.SE+5d
WW.5.SE+5d

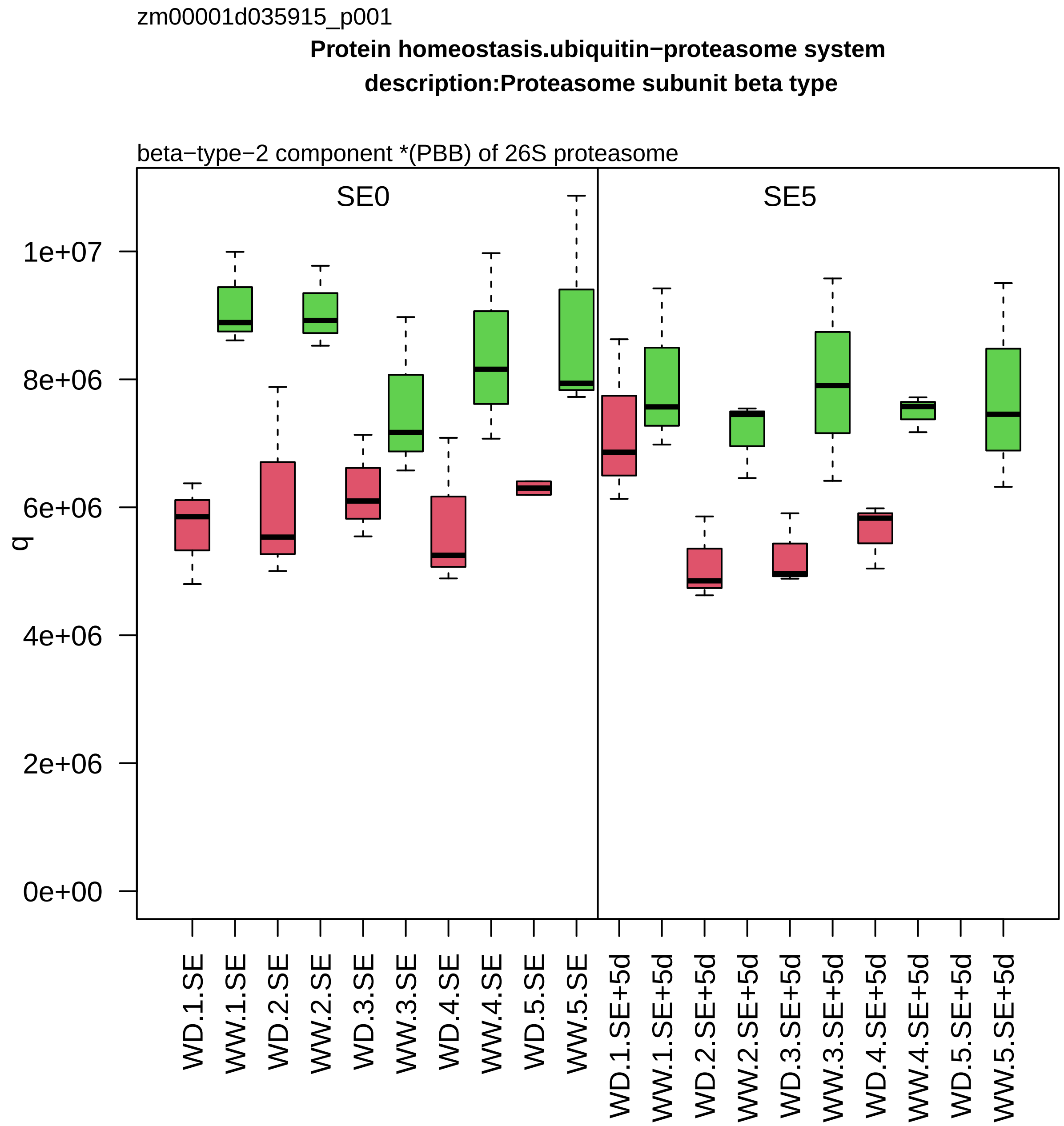
zm00001d035915_p001
Protein homeostasis.ubiquitin−proteasome system
description:Proteasome subunit beta type
beta−type−2 component *(PBB) of 26S proteasome
SE0
SE5
q
0e+00
2e+06
4e+06
6e+06
8e+06
1e+07
WD.1.SE
WW.1.SE
WD.2.SE
WW.2.SE
WD.3.SE
WW.3.SE
WD.4.SE
WW.4.SE
WD.5.SE
WW.5.SE
WD.1.SE+5d
WW.1.SE+5d
WD.2.SE+5d
WW.2.SE+5d
WD.3.SE+5d
WW.3.SE+5d
WD.4.SE+5d
WW.4.SE+5d
WD.5.SE+5d
WW.5.SE+5d

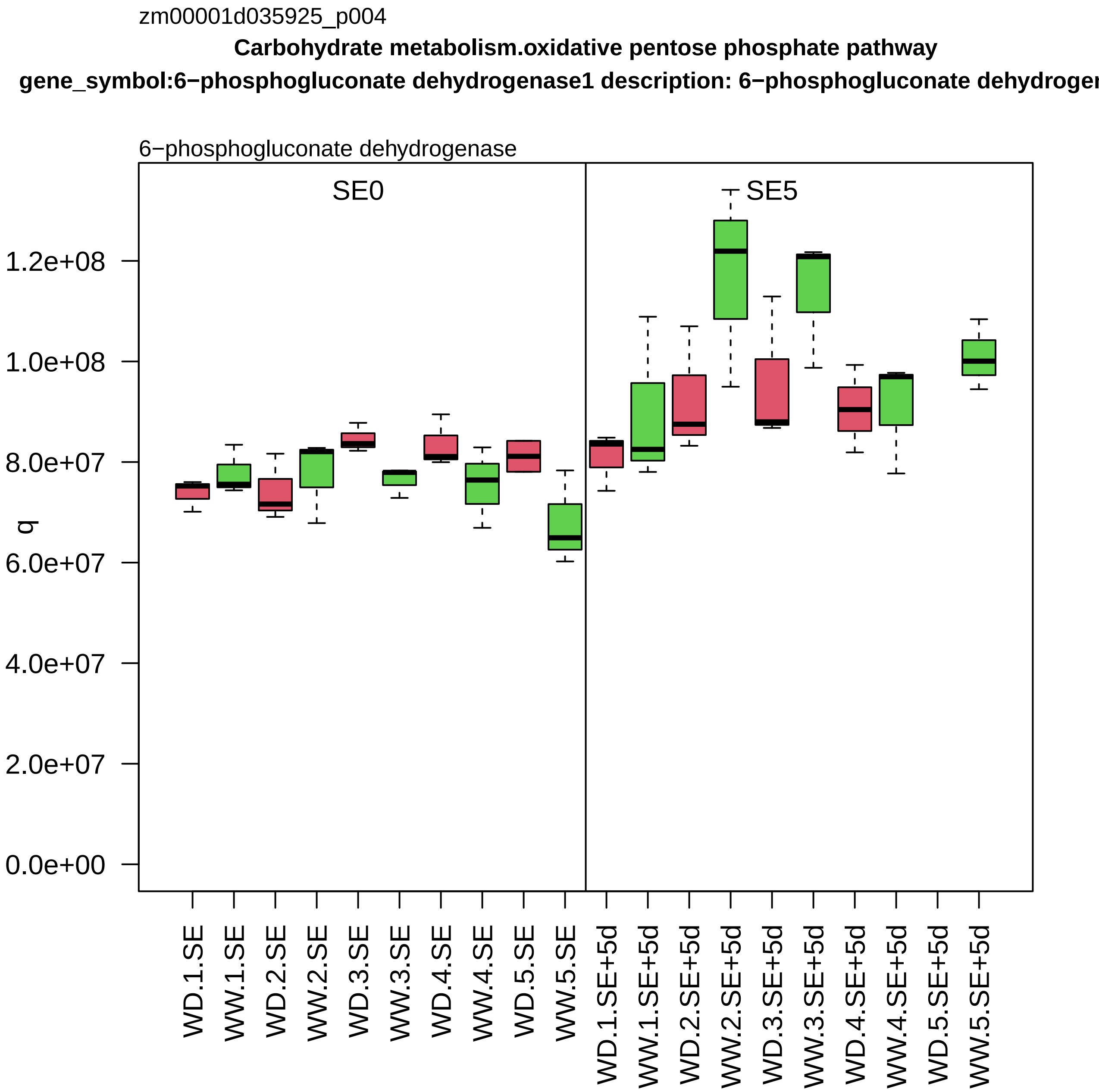

zm00001d035925_p004
Carbohydrate metabolism.oxidative pentose phosphate pathway
gene_symbol:6−phosphogluconate dehydrogenase1 description: 6−phosphogluconate dehydrogen
6−phosphogluconate dehydrogenase
SE0
SE5
q
1.2e+08
1.0e+08
8.0e+07
6.0e+07
4.0e+07
2.0e+07
0.0e+00
WD.1.SE
WW.1.SE
WD.2.SE
WW.2.SE
WD.3.SE
WW.3.SE
WD.4.SE
WW.4.SE
WD.5.SE
WW.5.SE
WD.1.SE+5d
WW.1.SE+5d
WD.2.SE+5d
WW.2.SE+5d
WD.3.SE+5d
WW.3.SE+5d
WD.4.SE+5d
WW.4.SE+5d
WD.5.SE+5d
WW.5.SE+5d

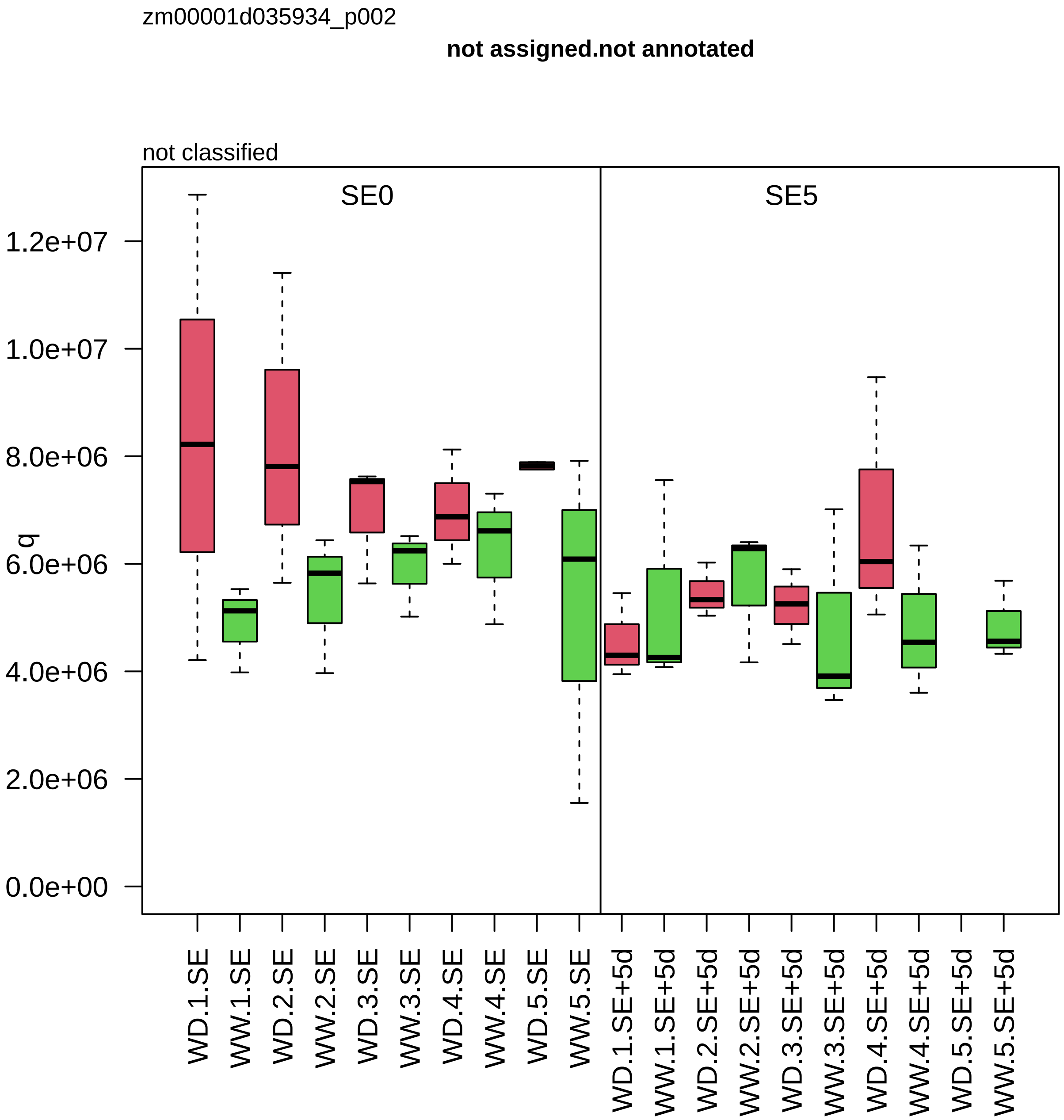

zm00001d035934_p002
not assigned.not annotated
not classified
SE0
SE5
q
1.2e+07
1.0e+07
8.0e+06
6.0e+06
4.0e+06
2.0e+06
0.0e+00
WD.1.SE
WW.1.SE
WD.2.SE
WW.2.SE
WD.3.SE
WW.3.SE
WD.4.SE
WW.4.SE
WD.5.SE
WW.5.SE
WD.1.SE+5d
WW.1.SE+5d
WD.2.SE+5d
WW.2.SE+5d
WD.3.SE+5d
WW.3.SE+5d
WD.4.SE+5d
WW.4.SE+5d
WD.5.SE+5d
WW.5.SE+5d

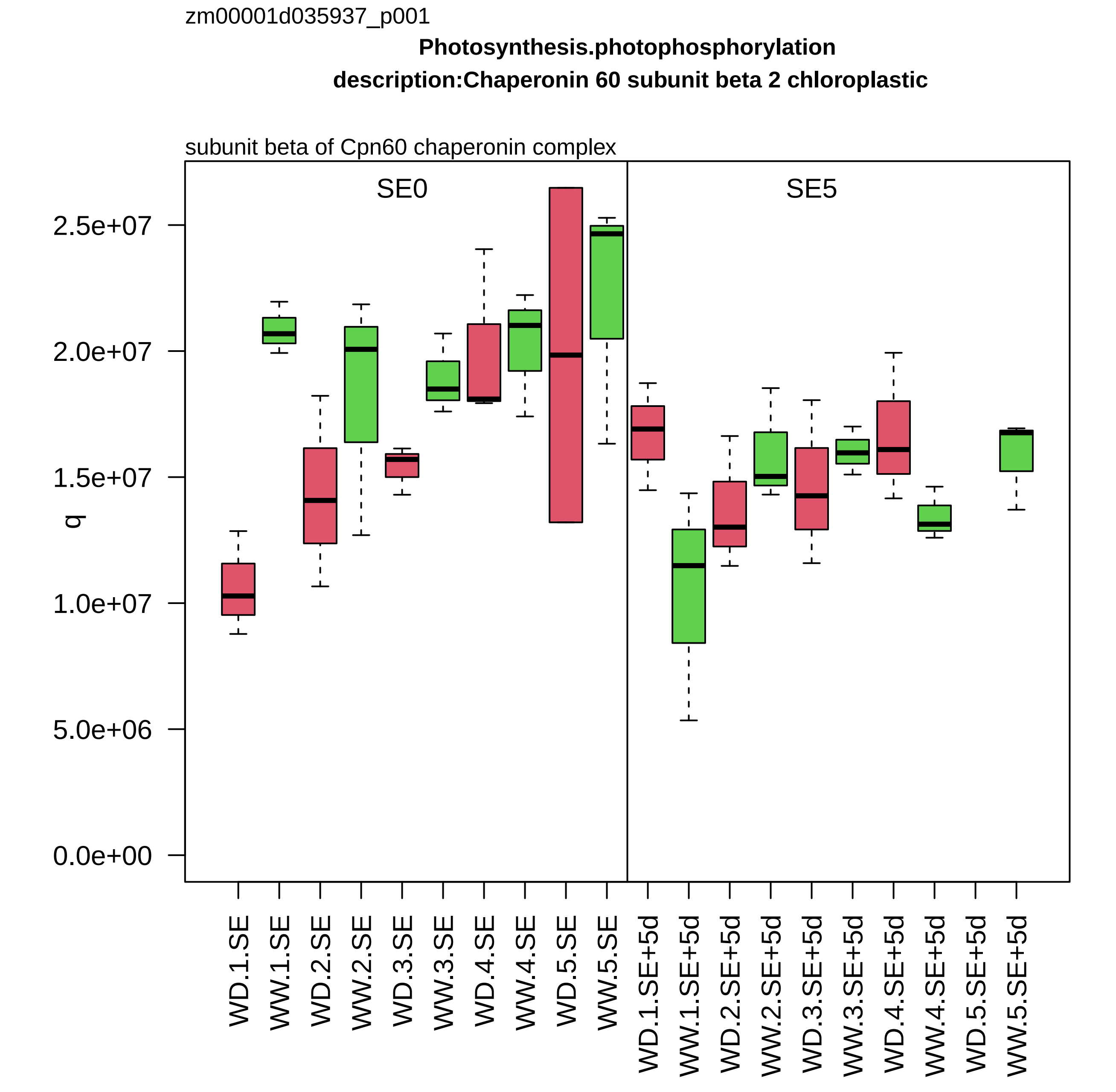
zm00001d035937_p001
Photosynthesis.photophosphorylation
description:Chaperonin 60 subunit beta 2 chloroplastic
subunit beta of Cpn60 chaperonin complex
SE0
SE5
q
2.5e+07
2.0e+07
1.5e+07
1.0e+07
5.0e+06
0.0e+00
WD.1.SE
WW.1.SE
WD.2.SE
WW.2.SE
WD.3.SE
WW.3.SE
WD.4.SE
WW.4.SE
WD.5.SE
WW.5.SE
WD.1.SE+5d
WW.1.SE+5d
WD.2.SE+5d
WW.2.SE+5d
WD.3.SE+5d
WW.3.SE+5d
WD.4.SE+5d
WW.4.SE+5d
WD.5.SE+5d
WW.5.SE+5d

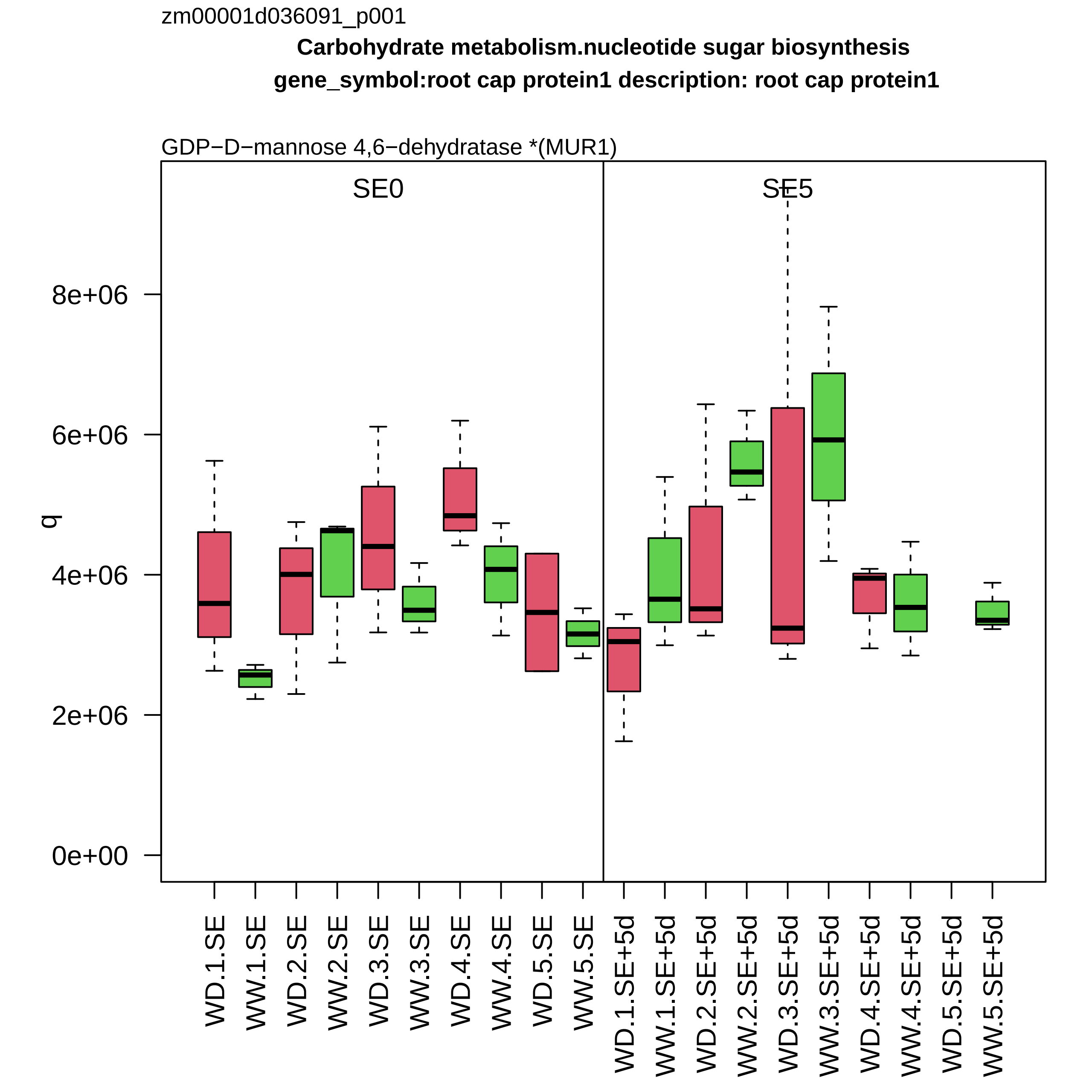

zm00001d036091_p001
Carbohydrate metabolism.nucleotide sugar biosynthesis
gene_symbol:root cap protein1 description: root cap protein1
GDP−D−mannose 4,6−dehydratase *(MUR1)
SE0
SE5
q
0e+00
2e+06
4e+06
6e+06
8e+06
WD.1.SE
WW.1.SE
WD.2.SE
WW.2.SE
WD.3.SE
WW.3.SE
WD.4.SE
WW.4.SE
WD.5.SE
WW.5.SE
WD.1.SE+5d
WW.1.SE+5d
WD.2.SE+5d
WW.2.SE+5d
WD.3.SE+5d
WW.3.SE+5d
WD.4.SE+5d
WW.4.SE+5d
WD.5.SE+5d
WW.5.SE+5d

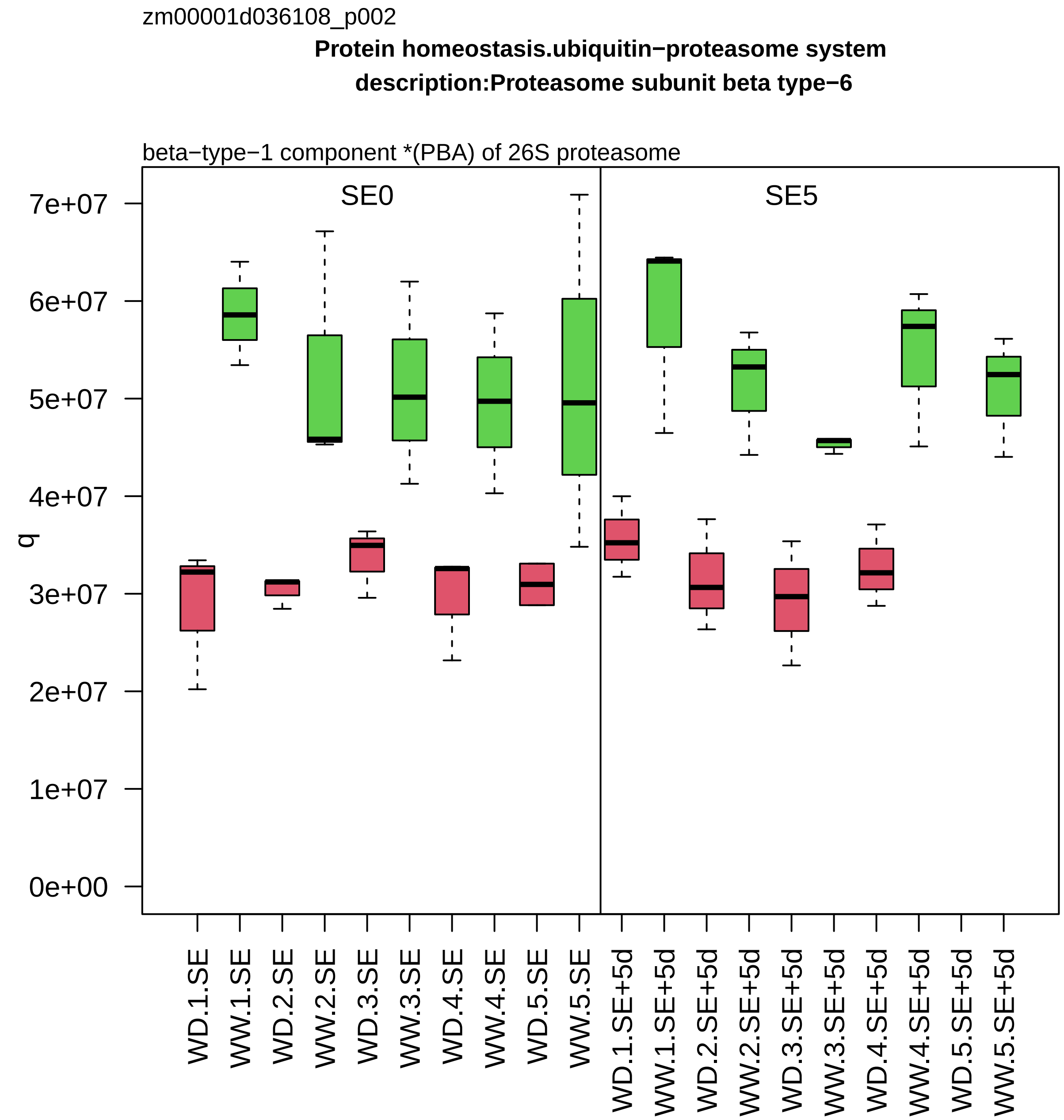

zm00001d036108_p002
Protein homeostasis.ubiquitin−proteasome system
description:Proteasome subunit beta type−6
beta−type−1 component *(PBA) of 26S proteasome
SE0
SE5
q
7e+07
6e+07
5e+07
4e+07
3e+07
2e+07
1e+07
0e+00
WD.1.SE
WW.1.SE
WD.2.SE
WW.2.SE
WD.3.SE
WW.3.SE
WD.4.SE
WW.4.SE
WD.5.SE
WW.5.SE
WD.1.SE+5d
WW.1.SE+5d
WD.2.SE+5d
WW.2.SE+5d
WD.3.SE+5d
WW.3.SE+5d
WD.4.SE+5d
WW.4.SE+5d
WD.5.SE+5d
WW.5.SE+5d

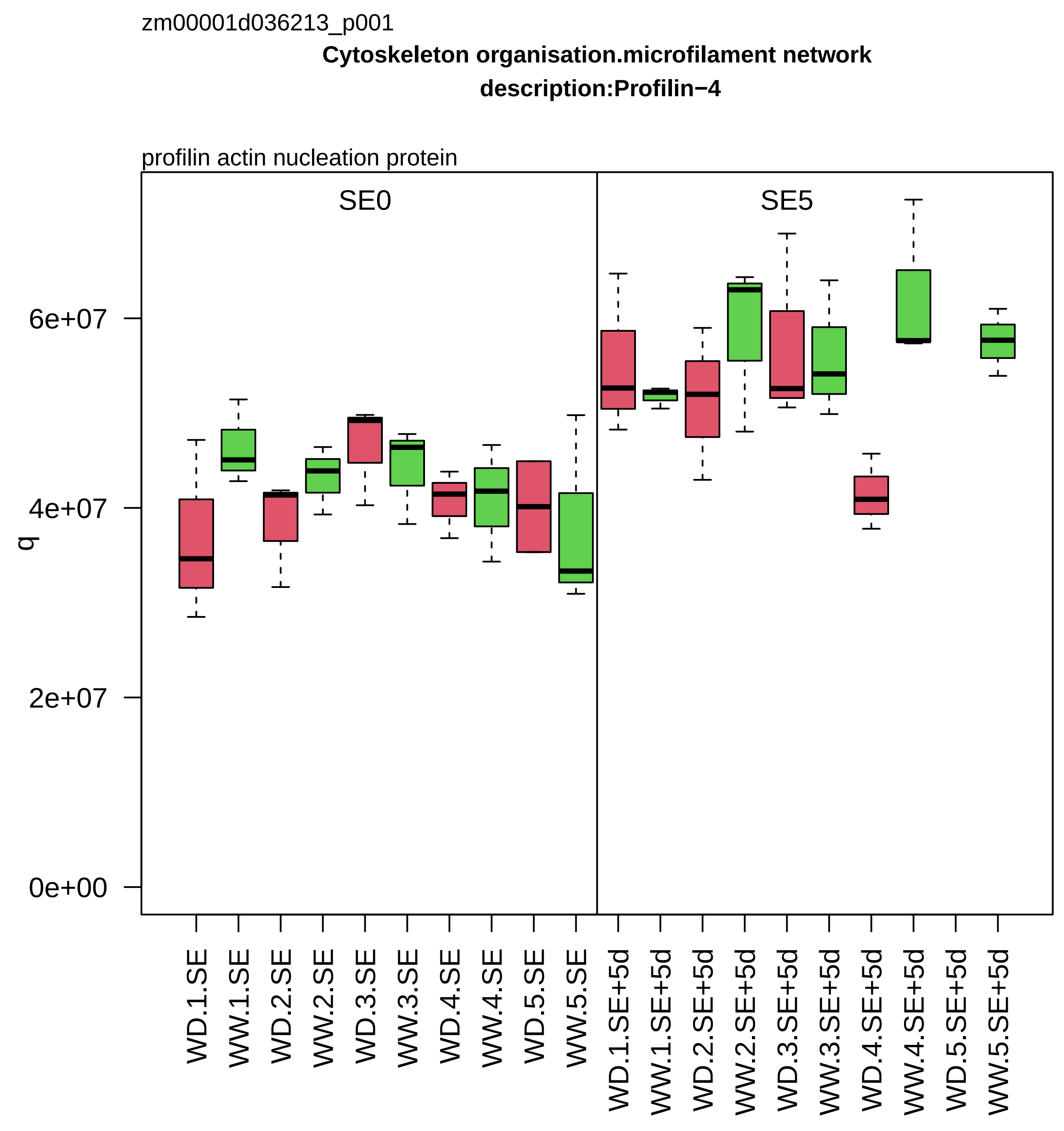
zm00001d036213_p001
Cytoskeleton organisation.microfilament network
description:Profilin−4
profilin actin nucleation protein
SE0
SE5
q
6e+07
4e+07
2e+07
0e+00
WD.1.SE
WW.1.SE
WD.2.SE
WW.2.SE
WD.3.SE
WW.3.SE
WD.4.SE
WW.4.SE
WD.5.SE
WW.5.SE
WD.1.SE+5d
WW.1.SE+5d
WD.2.SE+5d
WW.2.SE+5d
WD.3.SE+5d
WW.3.SE+5d
WD.4.SE+5d
WW.4.SE+5d
WD.5.SE+5d
WW.5.SE+5d

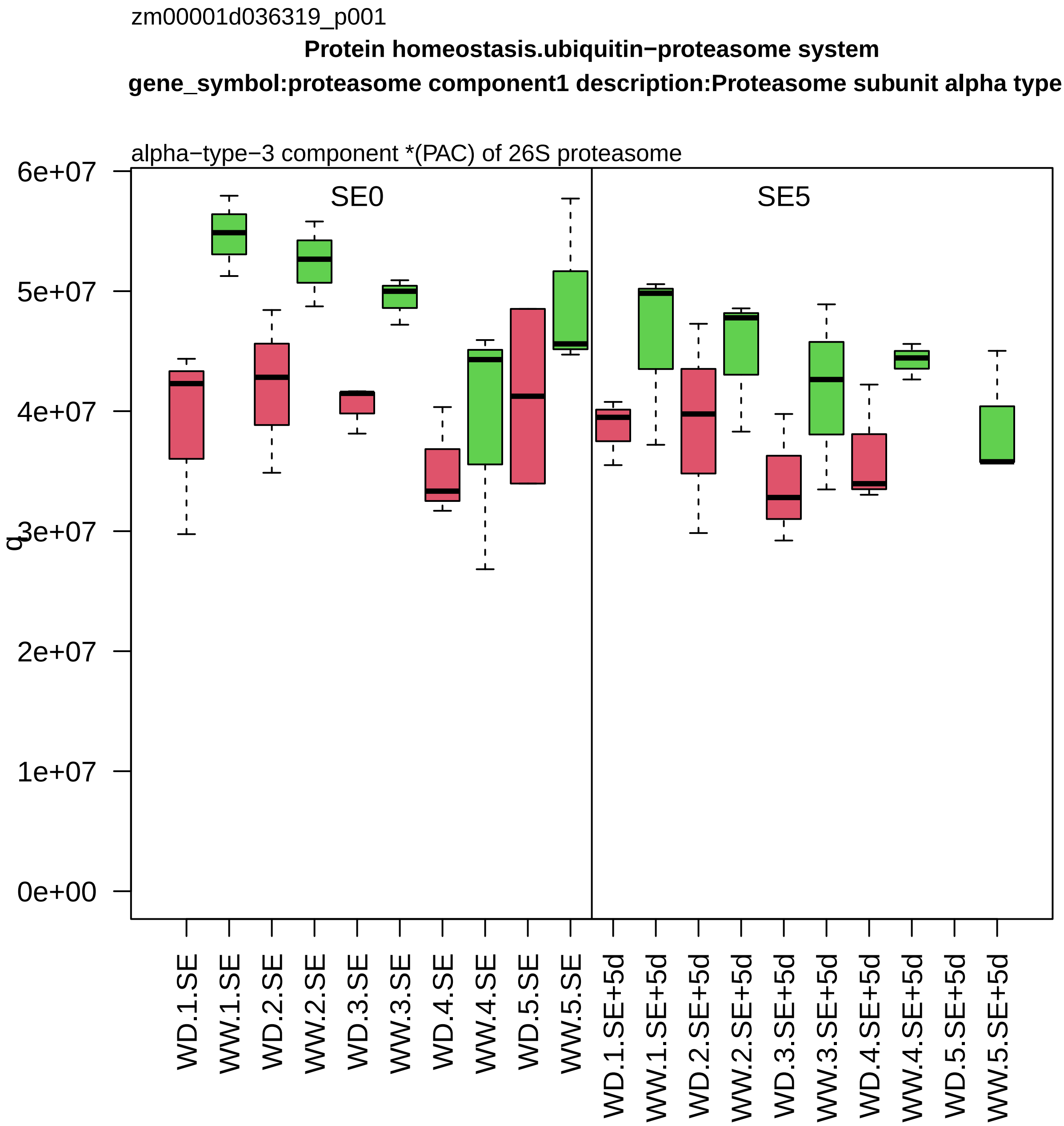
zm00001d036319_p001
Protein homeostasis.ubiquitin−proteasome system
gene_symbol:proteasome component1 description:Proteasome subunit alpha type
alpha−type−3 component *(PAC) of 26S proteasome
SE0
SE5
6e+07
5e+07
4e+07
3e+07
2e+07
1e+07
0e+00
q
WD.1.SE
WW.1.SE
WD.2.SE
WW.2.SE
WD.3.SE
WW.3.SE
WD.4.SE
WW.4.SE
WD.5.SE
WW.5.SE
WD.1.SE+5d
WW.1.SE+5d
WD.2.SE+5d
WW.2.SE+5d
WD.3.SE+5d
WW.3.SE+5d
WD.4.SE+5d
WW.4.SE+5d
WD.5.SE+5d
WW.5.SE+5d

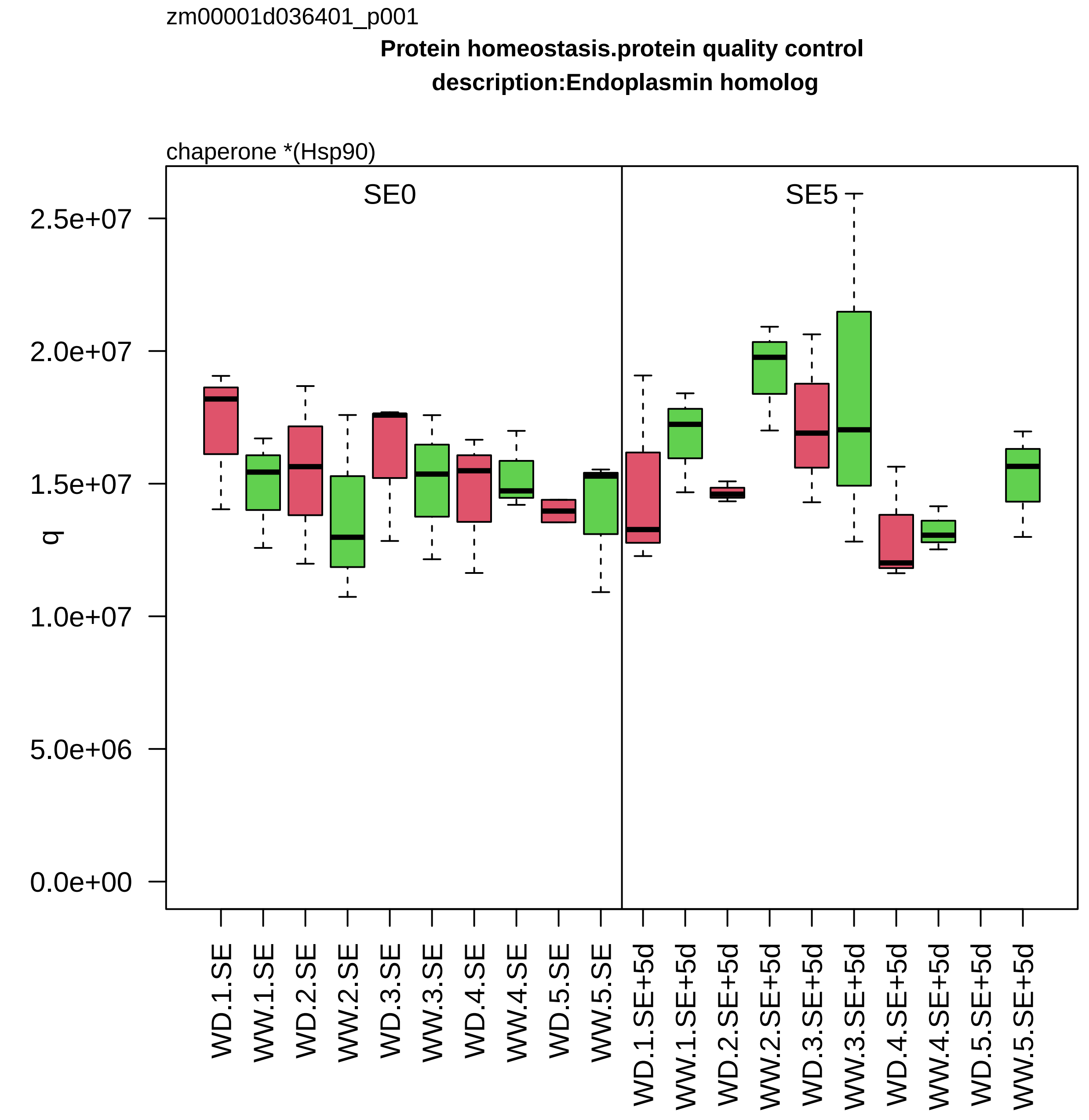
zm00001d036401_p001
Protein homeostasis.protein quality control
description:Endoplasmin homolog
chaperone *(Hsp90)
SE0
SE5
q
2.5e+07
2.0e+07
1.5e+07
1.0e+07
5.0e+06
0.0e+00
WD.1.SE
WW.1.SE
WD.2.SE
WW.2.SE
WD.3.SE
WW.3.SE
WD.4.SE
WW.4.SE
WD.5.SE
WW.5.SE
WD.1.SE+5d
WW.1.SE+5d
WD.2.SE+5d
WW.2.SE+5d
WD.3.SE+5d
WW.3.SE+5d
WD.4.SE+5d
WW.4.SE+5d
WD.5.SE+5d
WW.5.SE+5d

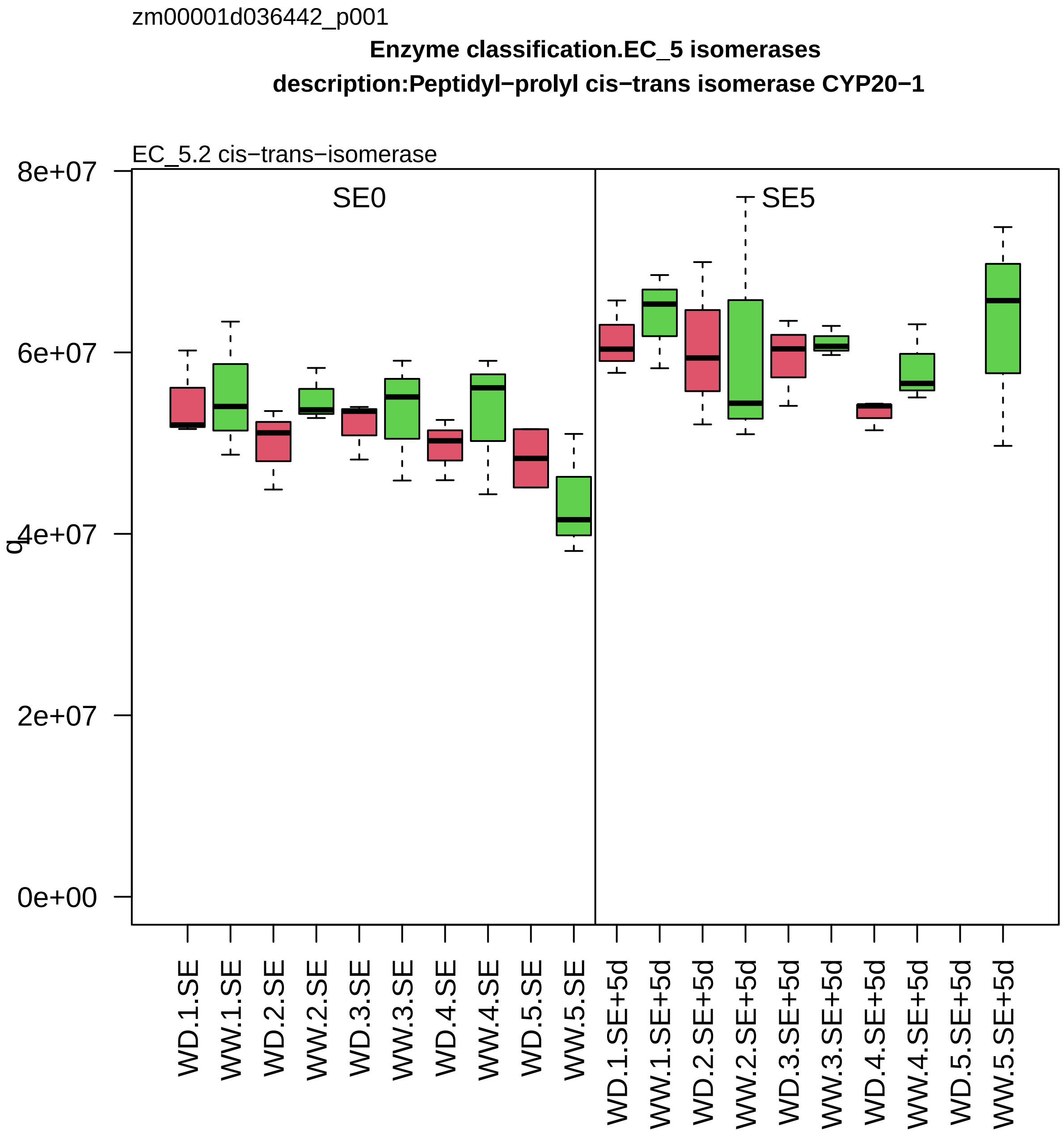
zm00001d036442_p001
Enzyme classification.EC_5 isomerases
description:Peptidyl−prolyl cis−trans isomerase CYP20−1
EC_5.2 cis−trans−isomerase
SE0
SE5
8e+07
6e+07
4e+07
2e+07
0e+00
q
WD.1.SE
WW.1.SE
WD.2.SE
WW.2.SE
WD.3.SE
WW.3.SE
WD.4.SE
WW.4.SE
WD.5.SE
WW.5.SE
WD.1.SE+5d
WW.1.SE+5d
WD.2.SE+5d
WW.2.SE+5d
WD.3.SE+5d
WW.3.SE+5d
WD.4.SE+5d
WW.4.SE+5d
WD.5.SE+5d
WW.5.SE+5d

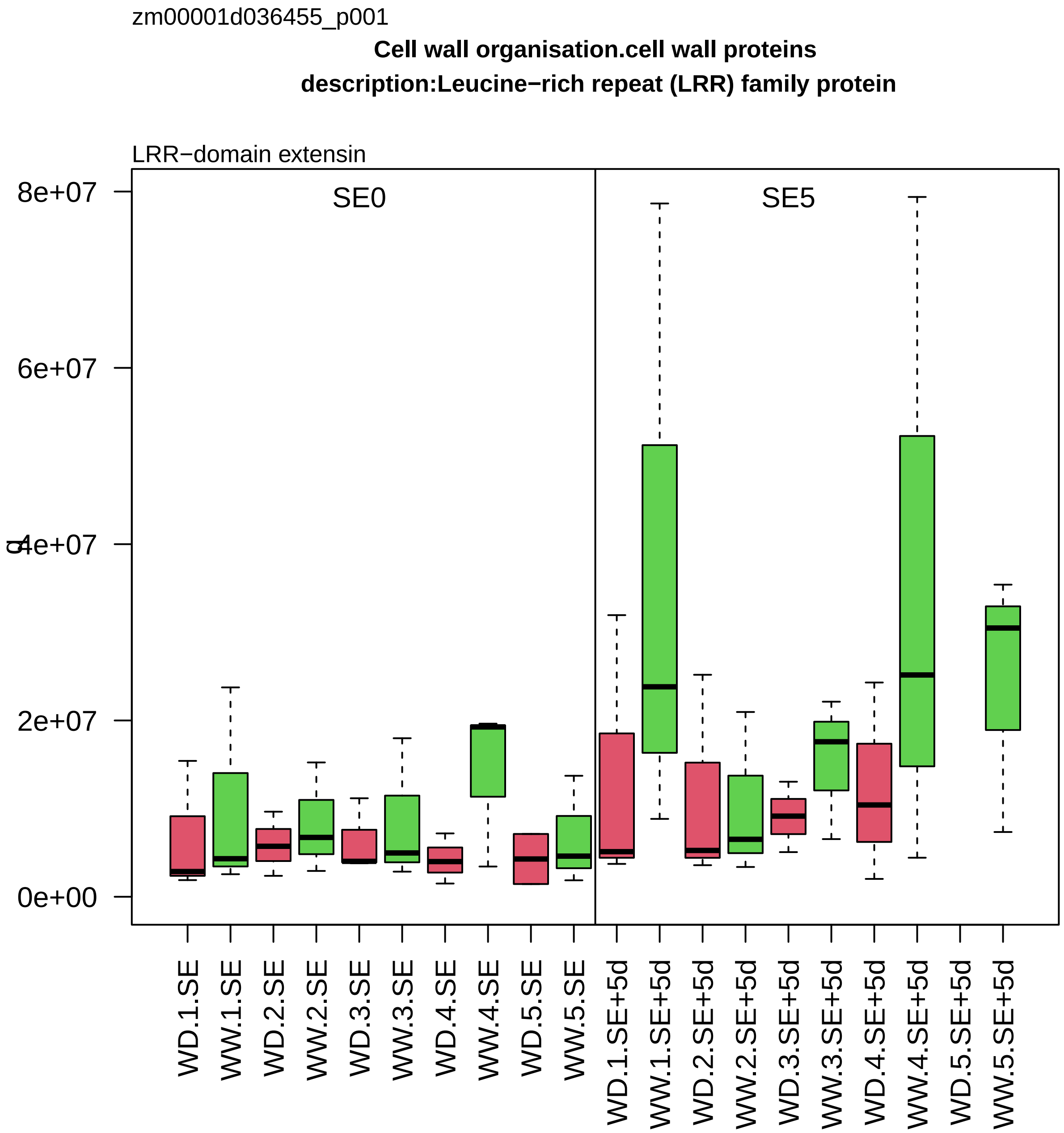

zm00001d036455_p001
Cell wall organisation.cell wall proteins
description:Leucine−rich repeat (LRR) family protein
LRR−domain extensin
SE0
SE5
8e+07
6e+07
4e+07
2e+07
0e+00
WD.1.SE
WW.1.SE
WD.2.SE
WW.2.SE
WD.3.SE
WW.3.SE
WD.4.SE
WW.4.SE
WD.5.SE
WW.5.SE
WD.1.SE+5d
WW.1.SE+5d
WD.2.SE+5d
WW.2.SE+5d
WD.3.SE+5d
WW.3.SE+5d
WD.4.SE+5d
WW.4.SE+5d
WD.5.SE+5d
WW.5.SE+5d

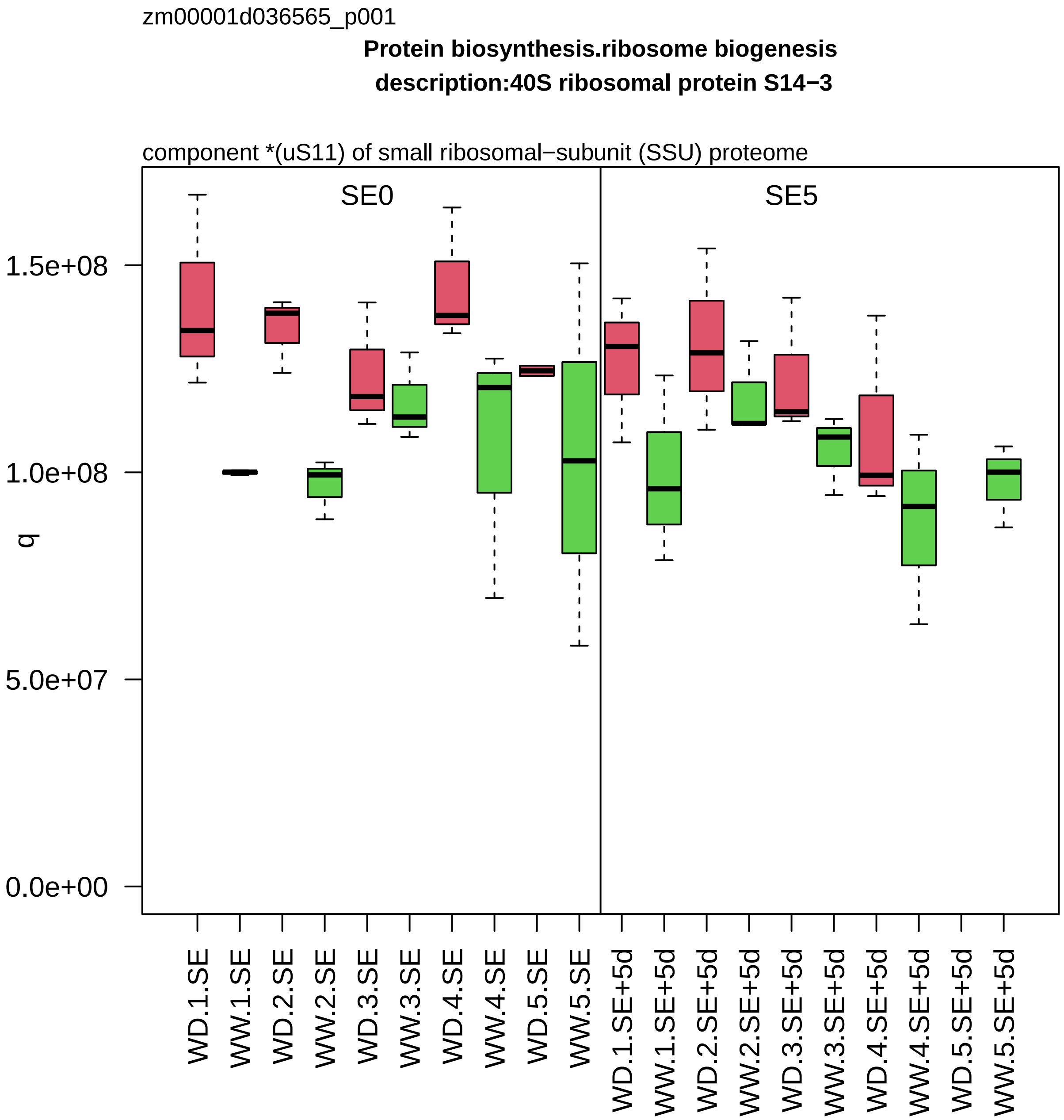

zm00001d036565_p001
Protein biosynthesis.ribosome biogenesis
description:40S ribosomal protein S14−3
component *(uS11) of small ribosomal−subunit (SSU) proteome
SE0
SE5
q
1.5e+08
1.0e+08
5.0e+07
0.0e+00
WD.1.SE
WW.1.SE
WD.2.SE
WW.2.SE
WD.3.SE
WW.3.SE
WD.4.SE
WW.4.SE
WD.5.SE
WW.5.SE
WD.1.SE+5d
WW.1.SE+5d
WD.2.SE+5d
WW.2.SE+5d
WD.3.SE+5d
WW.3.SE+5d
WD.4.SE+5d
WW.4.SE+5d
WD.5.SE+5d
WW.5.SE+5d

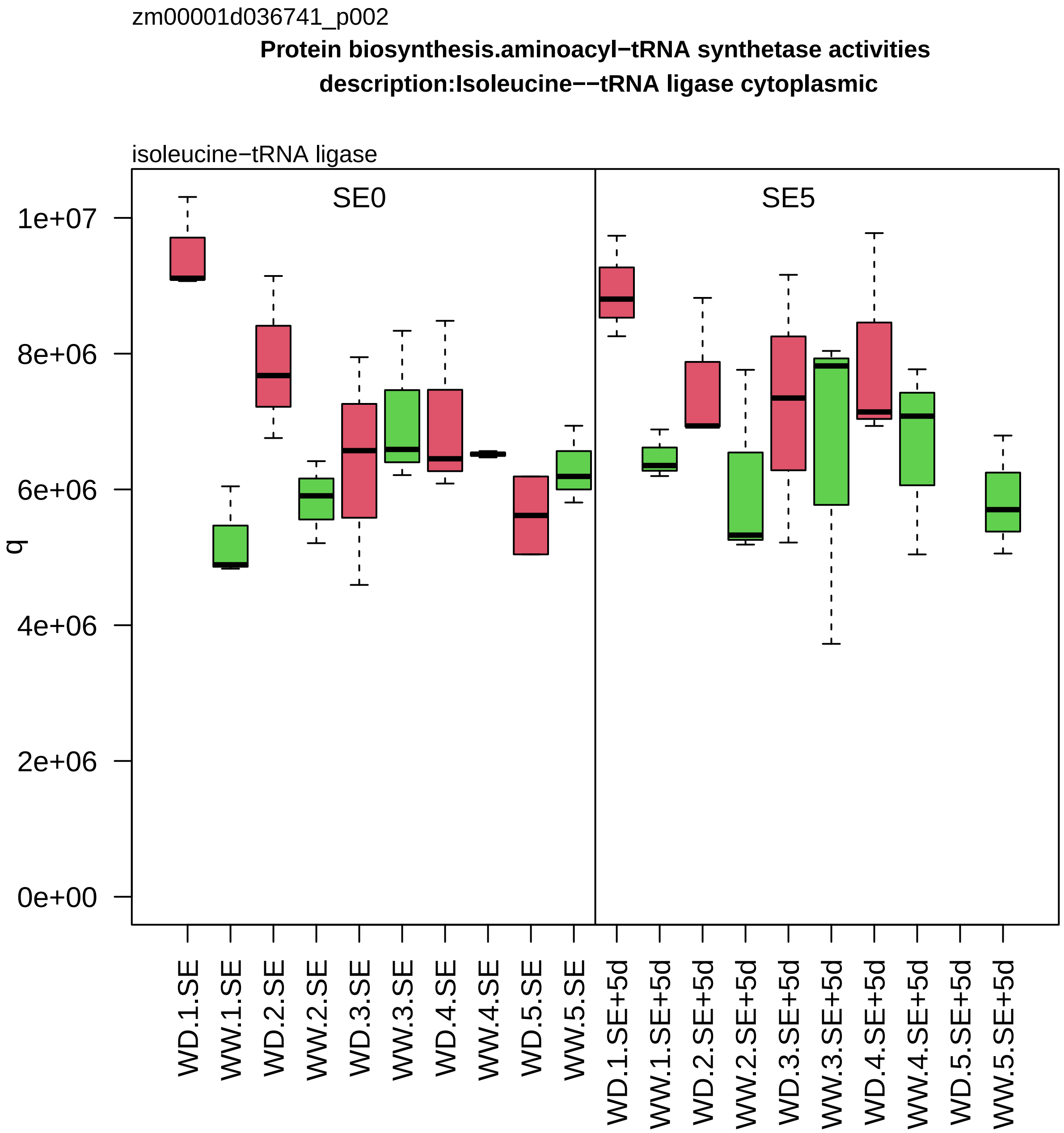
zm00001d036741_p002
Protein biosynthesis.aminoacyl−tRNA synthetase activities
description:Isoleucine−−tRNA ligase cytoplasmic
isoleucine−tRNA ligase
SE0
SE5
q
1e+07
8e+06
6e+06
4e+06
2e+06
0e+00
WD.1.SE
WW.1.SE
WD.2.SE
WW.2.SE
WD.3.SE
WW.3.SE
WD.4.SE
WW.4.SE
WD.5.SE
WW.5.SE
WD.1.SE+5d
WW.1.SE+5d
WD.2.SE+5d
WW.2.SE+5d
WD.3.SE+5d
WW.3.SE+5d
WD.4.SE+5d
WW.4.SE+5d
WD.5.SE+5d
WW.5.SE+5d

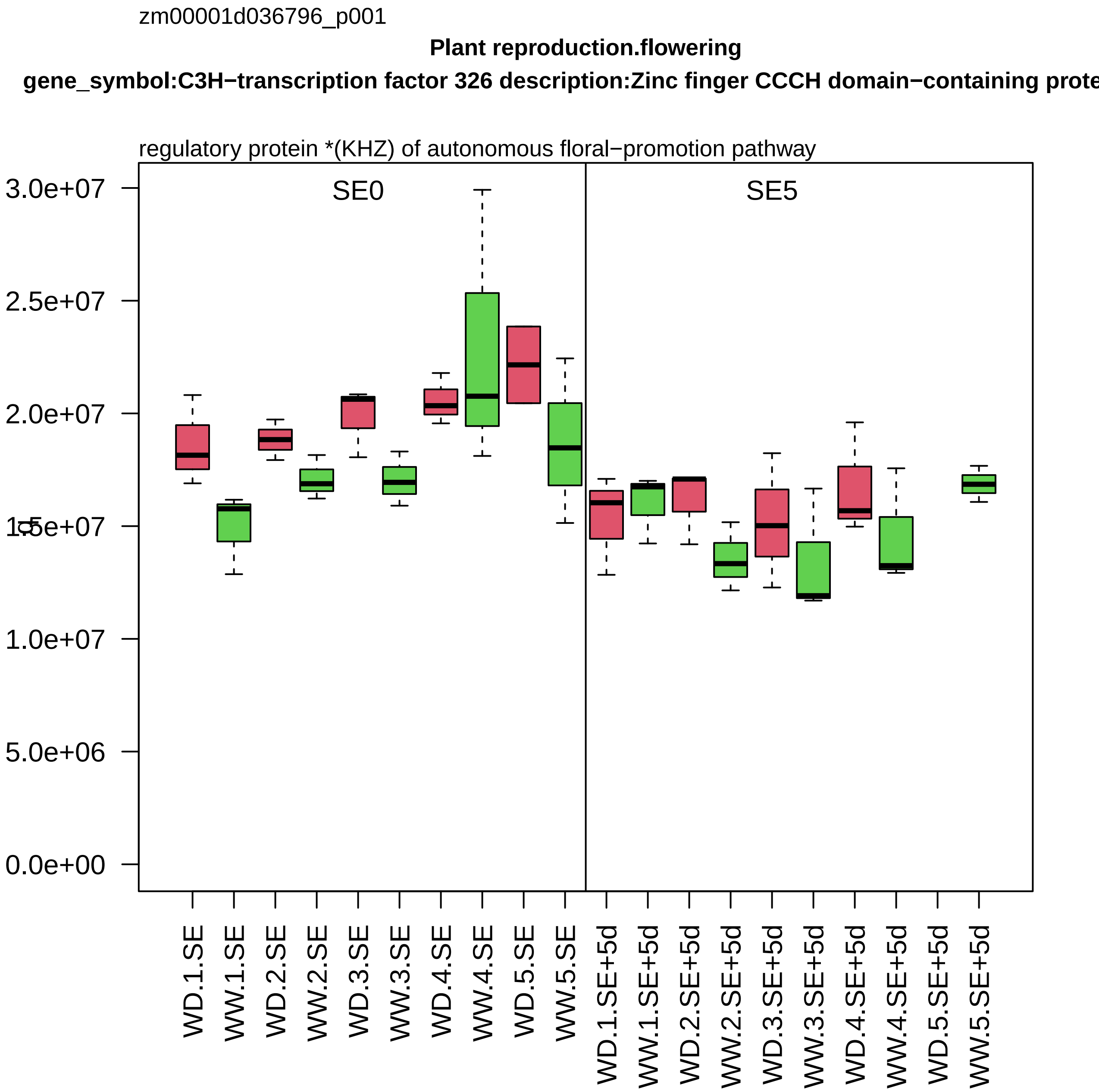

zm00001d036796_p001
Plant reproduction.flowering
gene_symbol:C3H−transcription factor 326 description:Zinc finger CCCH domain−containing prote
regulatory protein *(KHZ) of autonomous floral−promotion pathway
SE0
SE5
3.0e+07
2.5e+07
2.0e+07
1.5e+07
1.0e+07
5.0e+06
0.0e+00
WD.1.SE
WW.1.SE
WD.2.SE
WW.2.SE
WD.3.SE
WW.3.SE
WD.4.SE
WW.4.SE
WD.5.SE
WW.5.SE
WD.1.SE+5d
WW.1.SE+5d
WD.2.SE+5d
WW.2.SE+5d
WD.3.SE+5d
WW.3.SE+5d
WD.4.SE+5d
WW.4.SE+5d
WD.5.SE+5d
WW.5.SE+5d

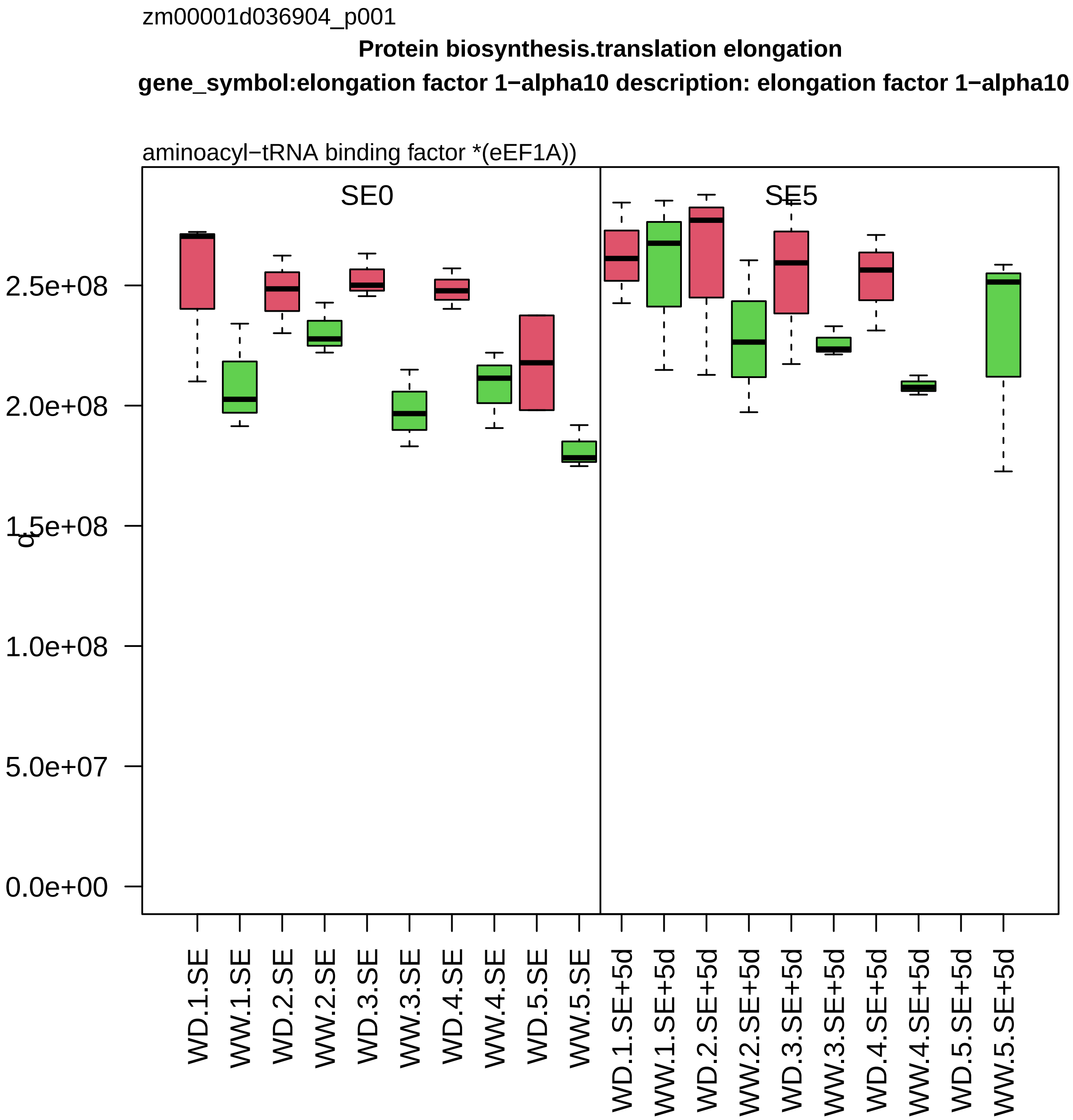

zm00001d036904_p001
Protein biosynthesis.translation elongation
gene_symbol:elongation factor 1−alpha10 description: elongation factor 1−alpha10
aminoacyl−tRNA binding factor *(eEF1A))
SE0
SE5
2.5e+08
2.0e+08
1.5e+08
1.0e+08
5.0e+07
0.0e+00
WD.1.SE
WW.1.SE
WD.2.SE
WW.2.SE
WD.3.SE
WW.3.SE
WD.4.SE
WW.4.SE
WD.5.SE
WW.5.SE
WD.1.SE+5d
WW.1.SE+5d
WD.2.SE+5d
WW.2.SE+5d
WD.3.SE+5d
WW.3.SE+5d
WD.4.SE+5d
WW.4.SE+5d
WD.5.SE+5d
WW.5.SE+5d

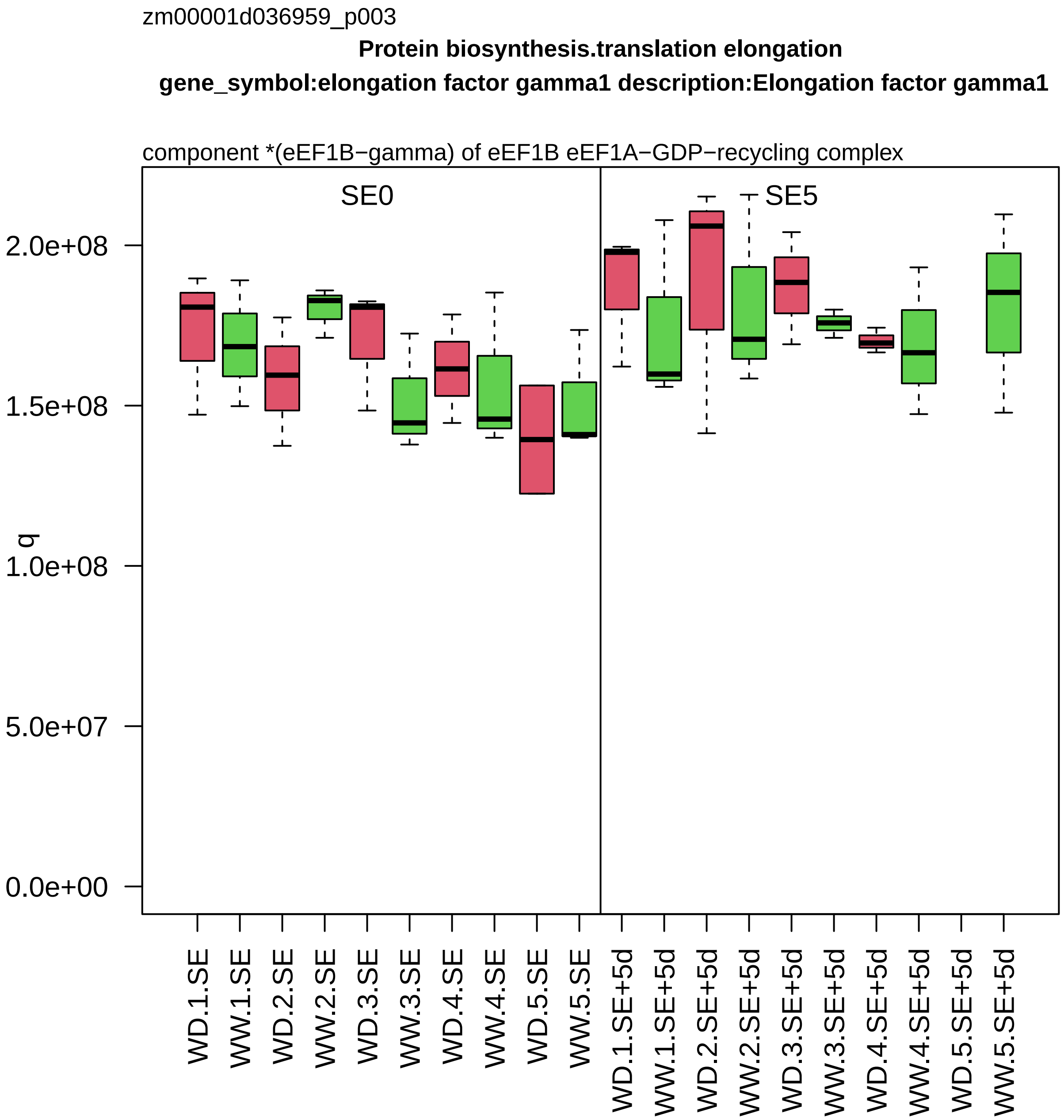

zm00001d036959_p003
Protein biosynthesis.translation elongation
gene_symbol:elongation factor gamma1 description:Elongation factor gamma1
component *(eEF1B−gamma) of eEF1B eEF1A−GDP−recycling complex
SE0
SE5
q
0.0e+00
5.0e+07
1.0e+08
1.5e+08
2.0e+08
WD.1.SE
WW.1.SE
WD.2.SE
WW.2.SE
WD.3.SE
WW.3.SE
WD.4.SE
WW.4.SE
WD.5.SE
WW.5.SE
WD.1.SE+5d
WW.1.SE+5d
WD.2.SE+5d
WW.2.SE+5d
WD.3.SE+5d
WW.3.SE+5d
WD.4.SE+5d
WW.4.SE+5d
WD.5.SE+5d
WW.5.SE+5d

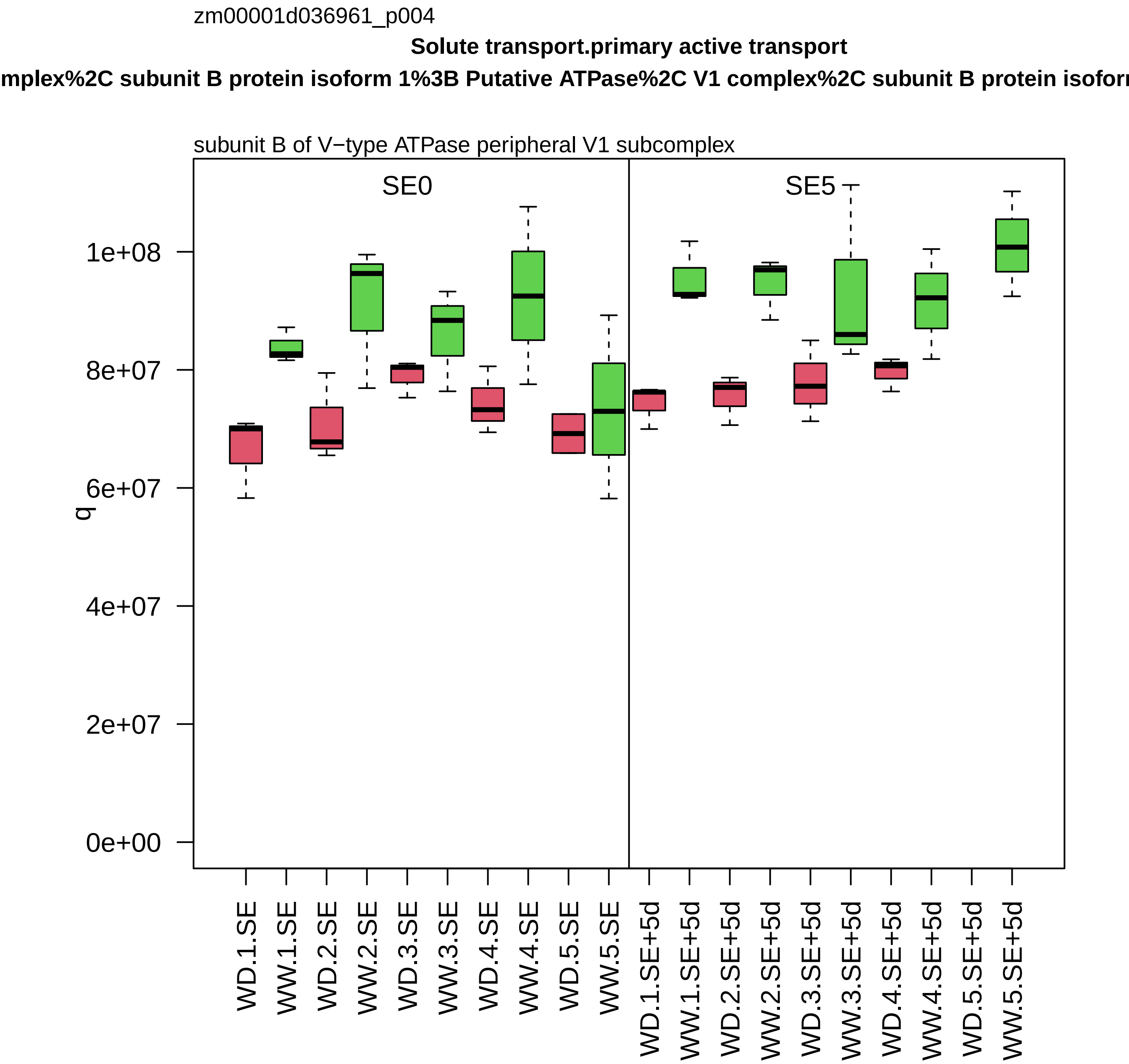

zm00001d036961_p004
Solute transport.primary active transport
mplex%2C subunit B protein isoform 1%3B Putative ATPase%2C V1 complex%2C subunit B protein isofor
subunit B of V−type ATPase peripheral V1 subcomplex
SE0
SE5
q
1e+08
8e+07
6e+07
4e+07
2e+07
0e+00
WD.1.SE
WW.1.SE
WD.2.SE
WW.2.SE
WD.3.SE
WW.3.SE
WD.4.SE
WW.4.SE
WD.5.SE
WW.5.SE
WD.1.SE+5d
WW.1.SE+5d
WD.2.SE+5d
WW.2.SE+5d
WD.3.SE+5d
WW.3.SE+5d
WD.4.SE+5d
WW.4.SE+5d
WD.5.SE+5d
WW.5.SE+5d

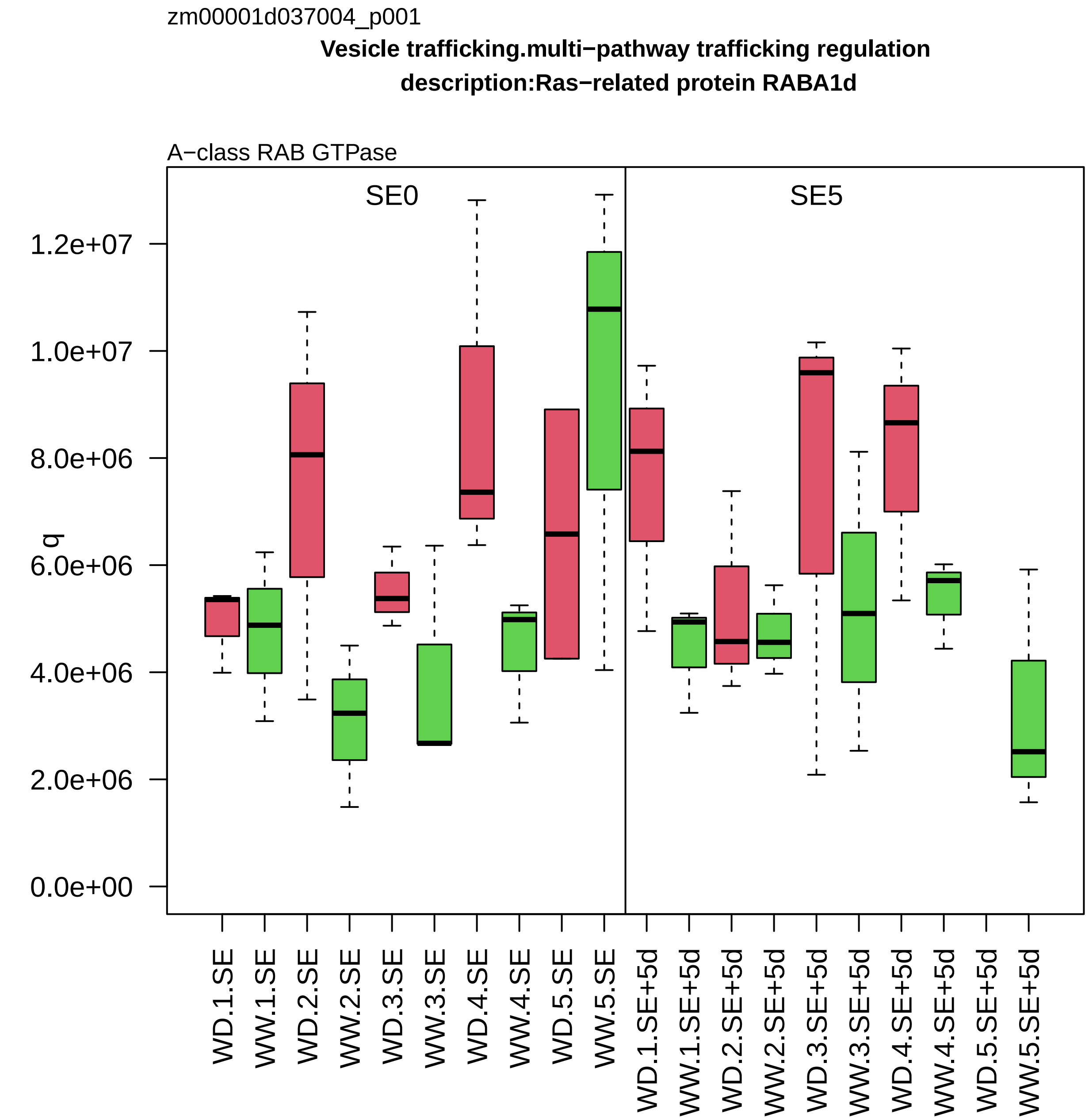

zm00001d037004_p001
Vesicle trafficking.multi−pathway trafficking regulation
description:Ras−related protein RABA1d
A−class RAB GTPase
SE0
SE5
q
0.0e+00
2.0e+06
4.0e+06
6.0e+06
8.0e+06
1.0e+07
1.2e+07
WD.1.SE
WW.1.SE
WD.2.SE
WW.2.SE
WD.3.SE
WW.3.SE
WD.4.SE
WW.4.SE
WD.5.SE
WW.5.SE
WD.1.SE+5d
WW.1.SE+5d
WD.2.SE+5d
WW.2.SE+5d
WD.3.SE+5d
WW.3.SE+5d
WD.4.SE+5d
WW.4.SE+5d
WD.5.SE+5d
WW.5.SE+5d

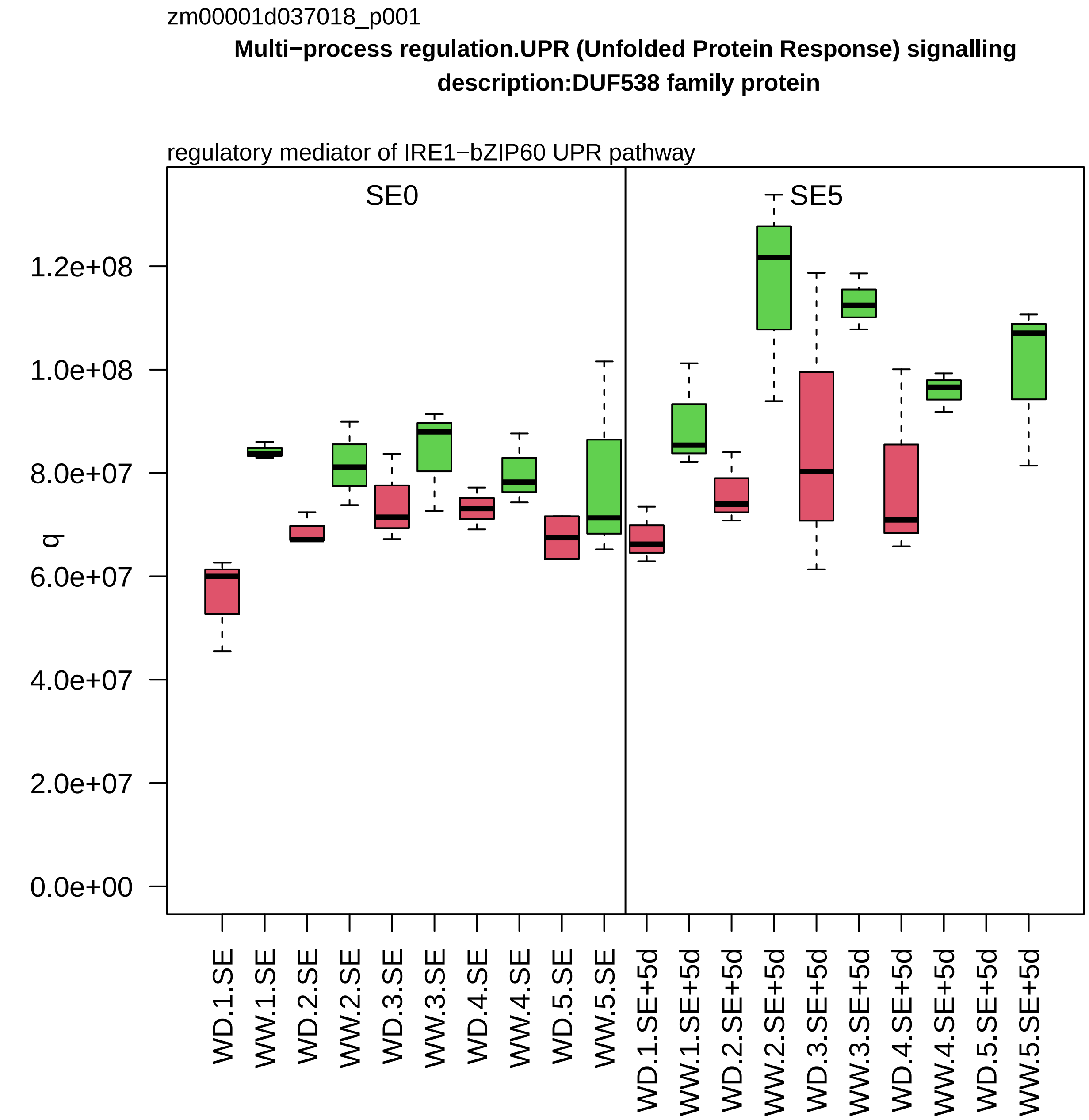

zm00001d037018_p001
Multi−process regulation.UPR (Unfolded Protein Response) signalling
description:DUF538 family protein
regulatory mediator of IRE1−bZIP60 UPR pathway
SE0
SE5
q
0.0e+00
2.0e+07
4.0e+07
6.0e+07
8.0e+07
1.0e+08
1.2e+08
WD.1.SE
WW.1.SE
WD.2.SE
WW.2.SE
WD.3.SE
WW.3.SE
WD.4.SE
WW.4.SE
WD.5.SE
WW.5.SE
WD.1.SE+5d
WW.1.SE+5d
WD.2.SE+5d
WW.2.SE+5d
WD.3.SE+5d
WW.3.SE+5d
WD.4.SE+5d
WW.4.SE+5d
WD.5.SE+5d
WW.5.SE+5d

zm00001d037111_p001

**Protein biosynthesis.ribosome biogenesis**

**description:40S ribosomal protein S17−1**

component *(eS17) of small ribosomal−subunit (SSU) proteome

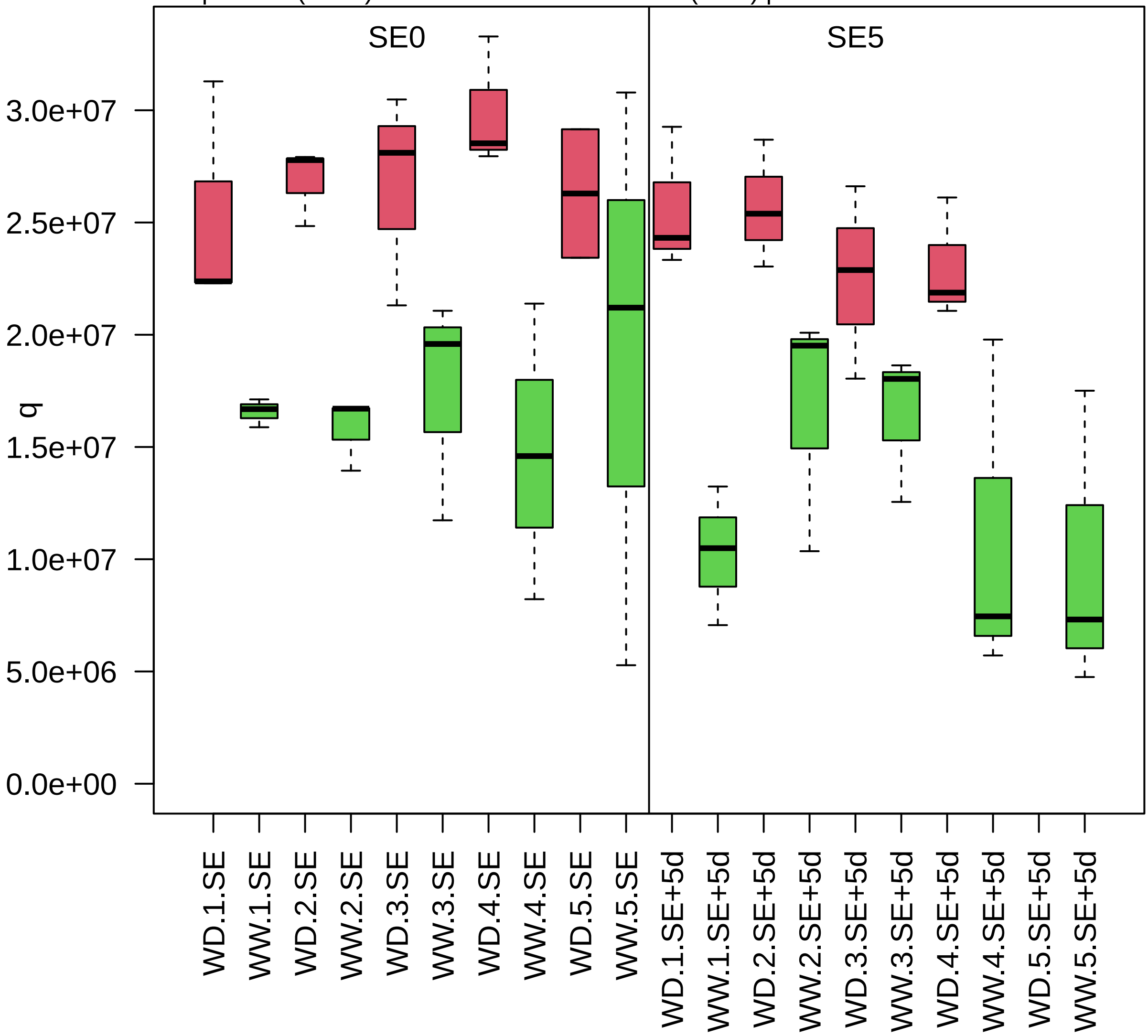

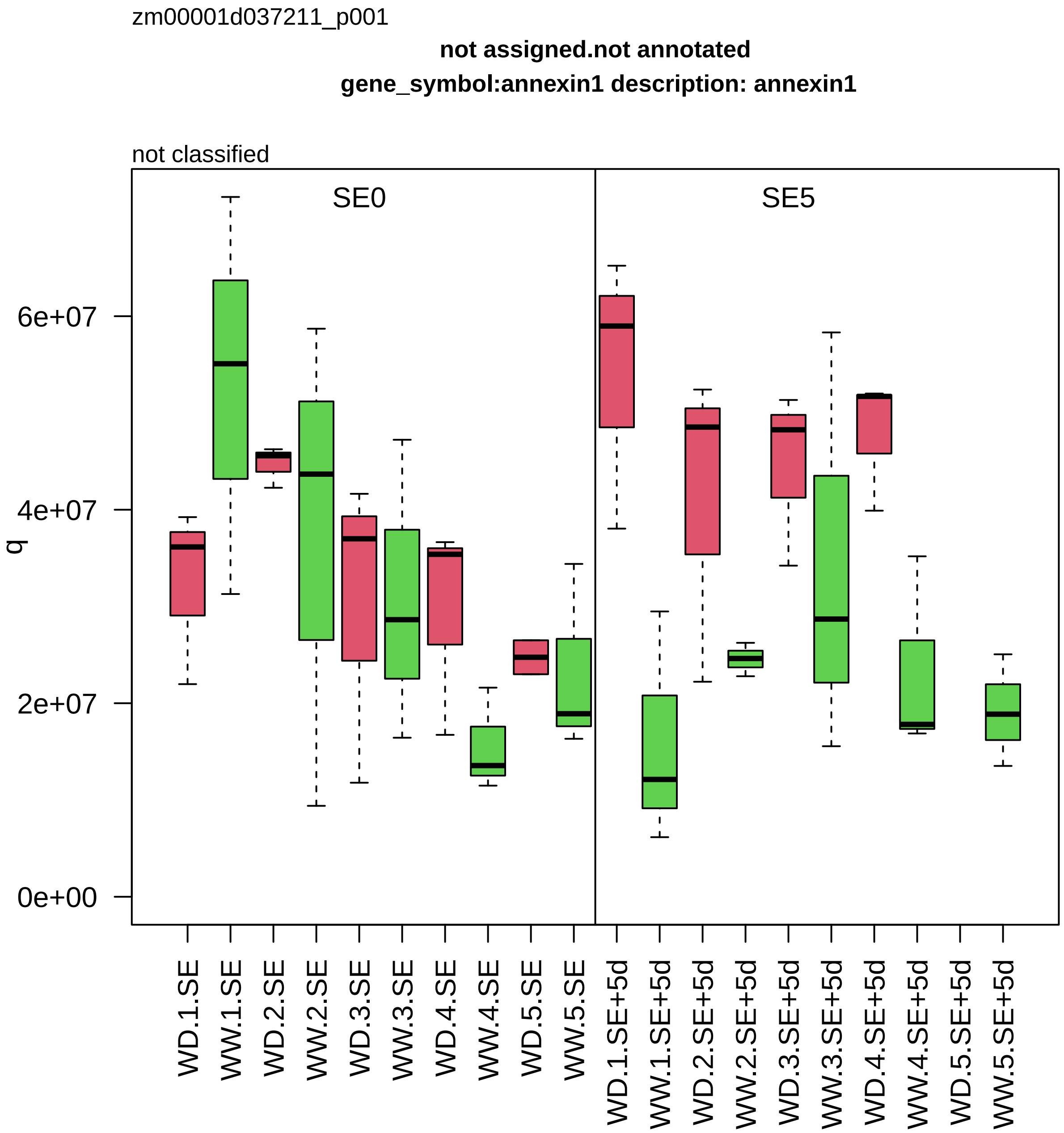
zm00001d037211_p001
not assigned.not annotated
gene_symbol:annexin1 description: annexin1
not classified
SE0
SE5
q
6e+07
4e+07
2e+07
0e+00
WD.1.SE
WW.1.SE
WD.2.SE
WW.2.SE
WD.3.SE
WW.3.SE
WD.4.SE
WW.4.SE
WD.5.SE
WW.5.SE
WD.1.SE+5d
WW.1.SE+5d
WD.2.SE+5d
WW.2.SE+5d
WD.3.SE+5d
WW.3.SE+5d
WD.4.SE+5d
WW.4.SE+5d
WD.5.SE+5d
WW.5.SE+5d

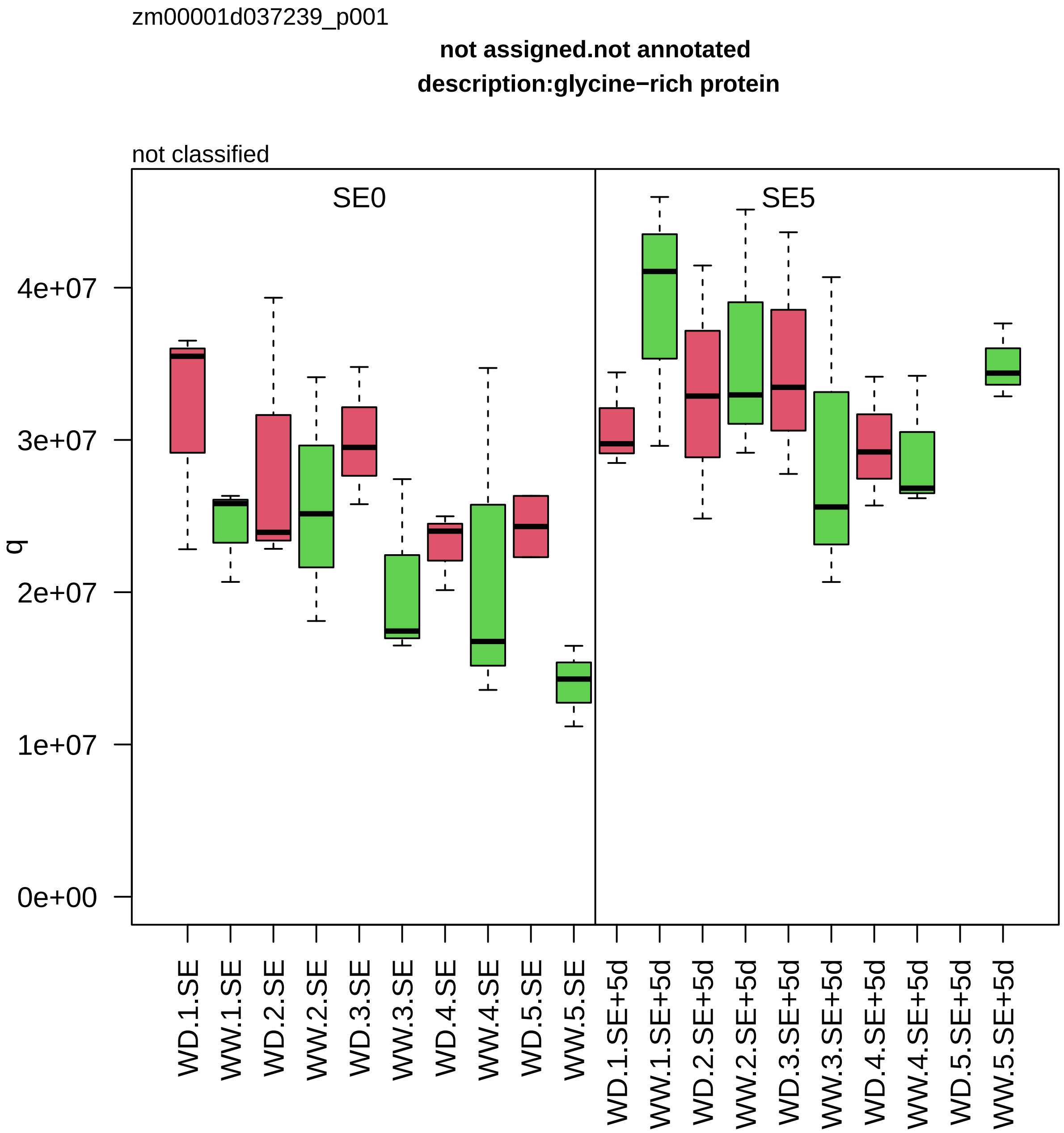

zm00001d037239_p001
not assigned.not annotated
description:glycine−rich protein
not classified
SE0
SE5
q
4e+07
3e+07
2e+07
1e+07
0e+00
WD.1.SE
WW.1.SE
WD.2.SE
WW.2.SE
WD.3.SE
WW.3.SE
WD.4.SE
WW.4.SE
WD.5.SE
WW.5.SE
WD.1.SE+5d
WW.1.SE+5d
WD.2.SE+5d
WW.2.SE+5d
WD.3.SE+5d
WW.3.SE+5d
WD.4.SE+5d
WW.4.SE+5d
WD.5.SE+5d
WW.5.SE+5d

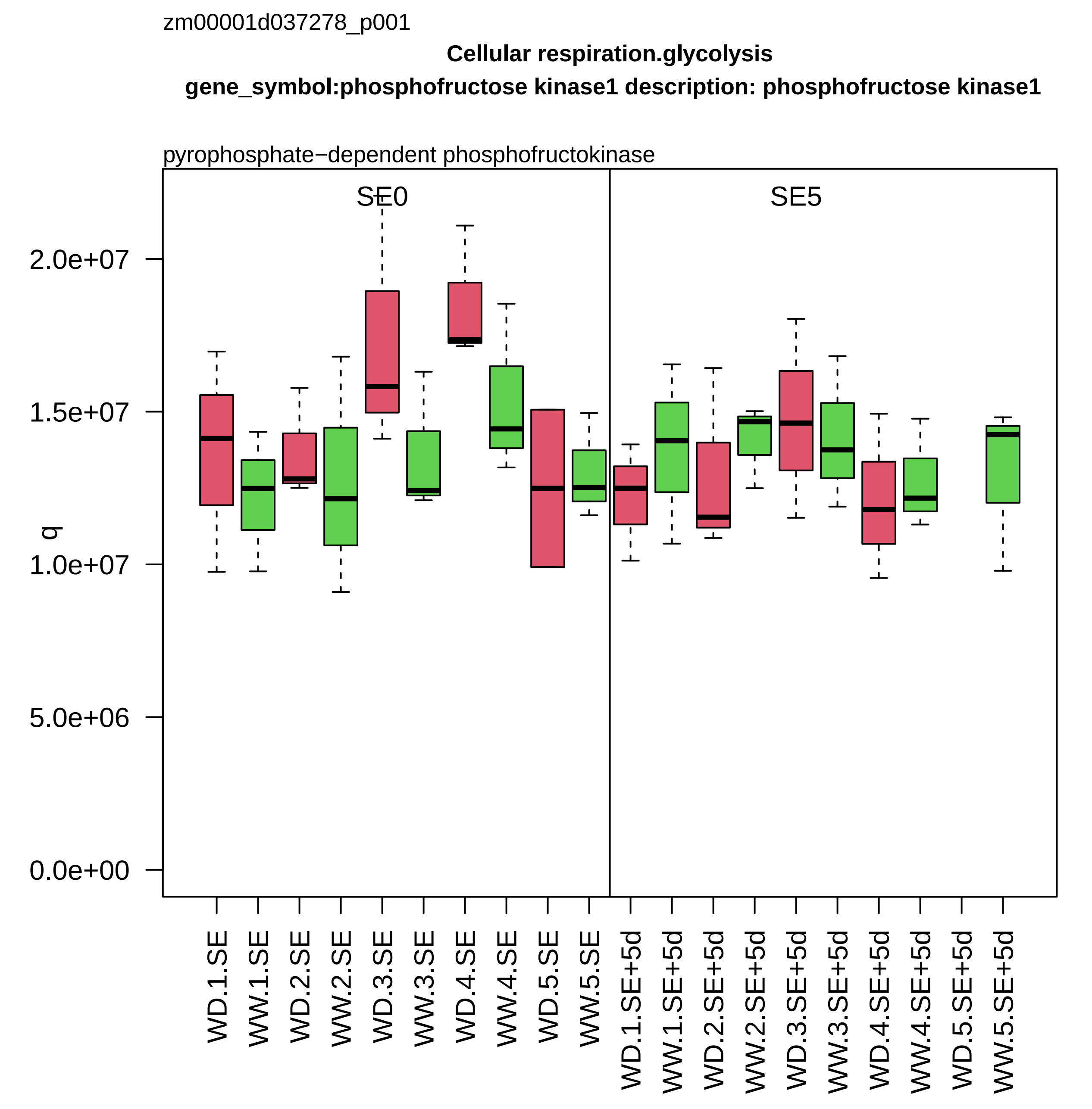

zm00001d037278_p001
Cellular respiration.glycolysis
gene_symbol:phosphofructose kinase1 description: phosphofructose kinase1
pyrophosphate−dependent phosphofructokinase
SE0
SE5
q
0.0e+00
5.0e+06
1.0e+07
1.5e+07
2.0e+07
WD.1.SE
WW.1.SE
WD.2.SE
WW.2.SE
WD.3.SE
WW.3.SE
WD.4.SE
WW.4.SE
WD.5.SE
WW.5.SE
WD.1.SE+5d
WW.1.SE+5d
WD.2.SE+5d
WW.2.SE+5d
WD.3.SE+5d
WW.3.SE+5d
WD.4.SE+5d
WW.4.SE+5d
WD.5.SE+5d
WW.5.SE+5d

zm00001d037326_p001

**Protein homeostasis.ubiquitin−proteasome system**
**description:Ubiquitin receptor RAD23c**

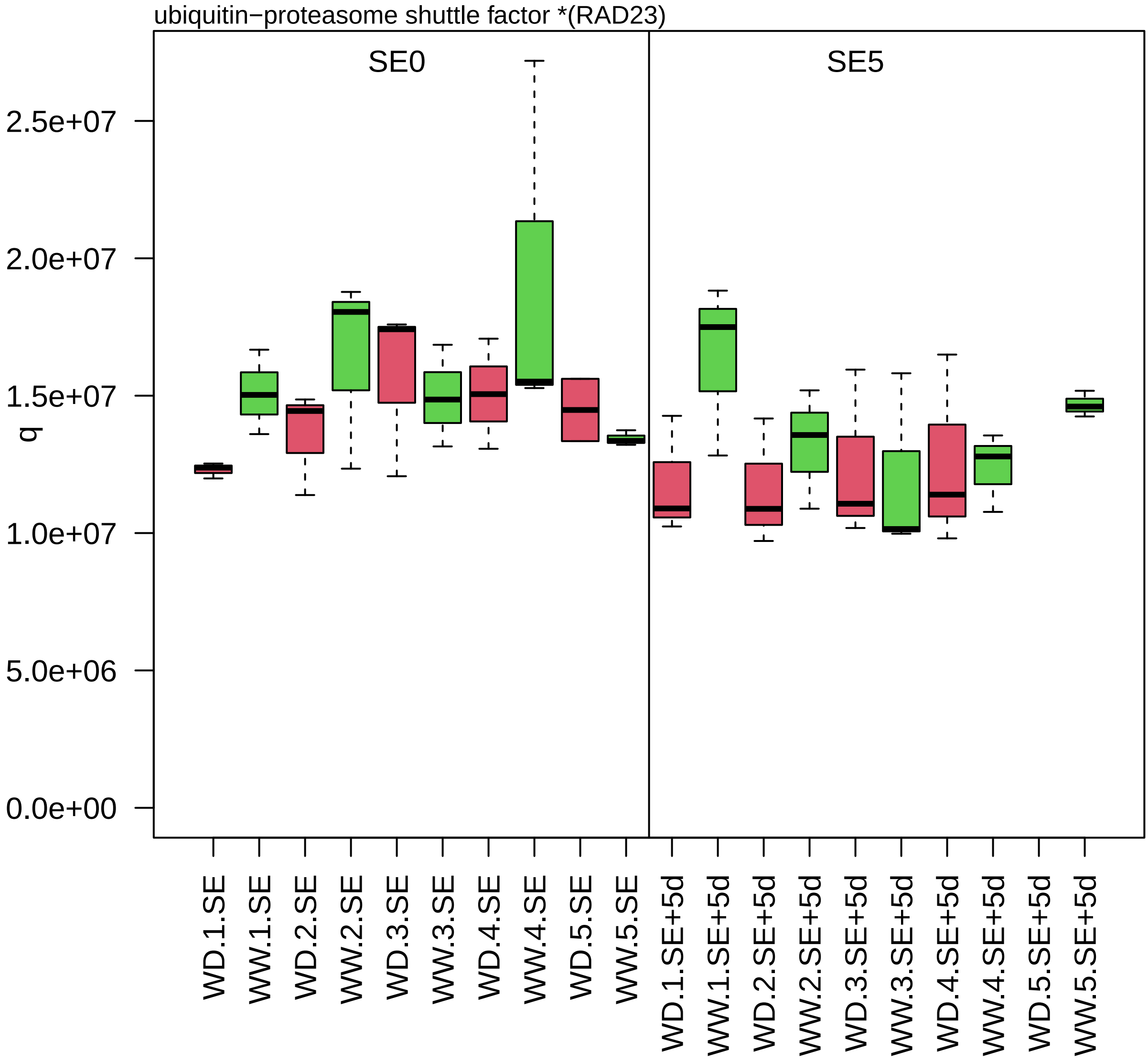

zm00001d037540_p001

**Cell wall organisation.cell wall proteins**

**description:Fasciclin−like arabinogalactan protein 2**

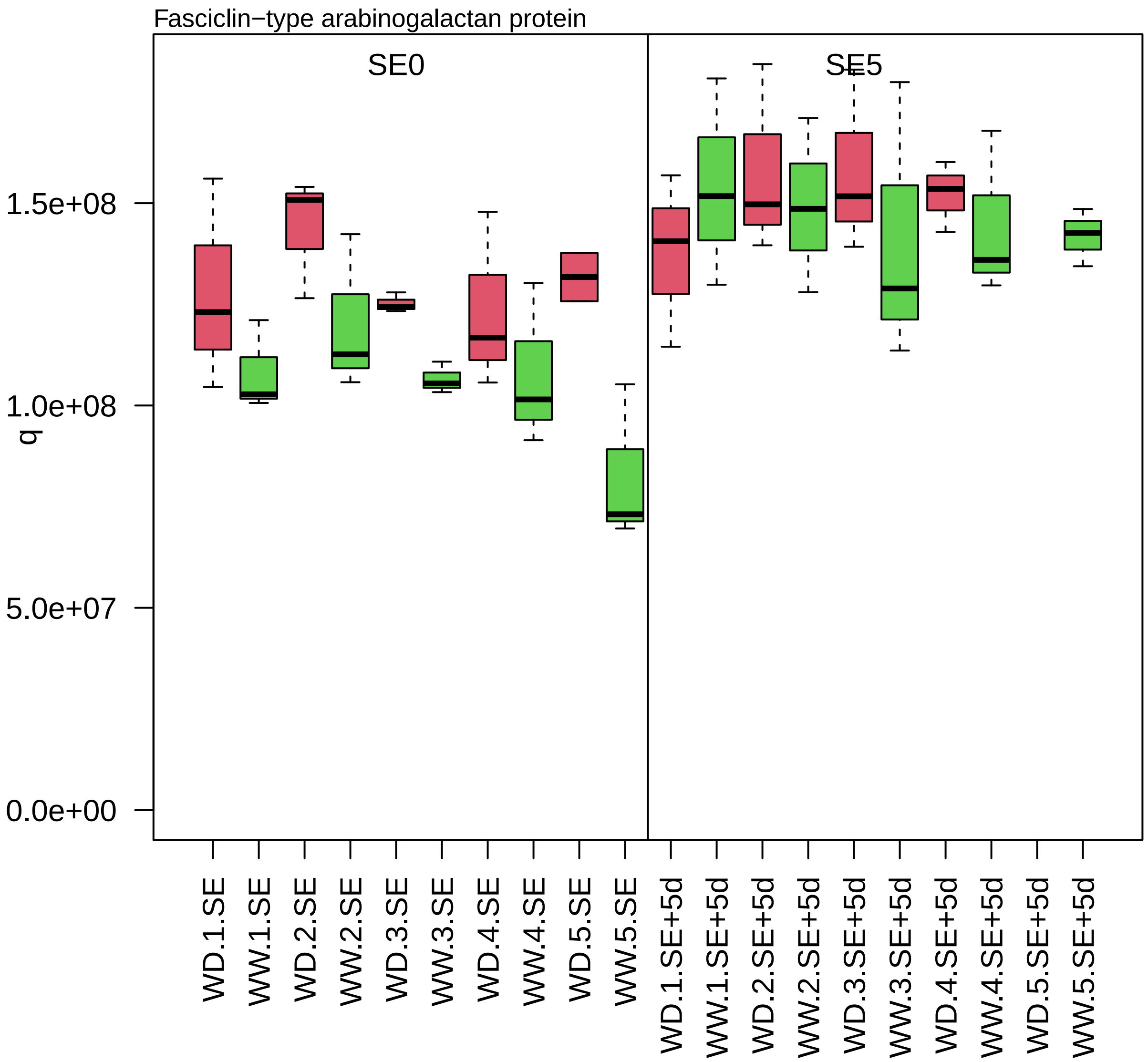

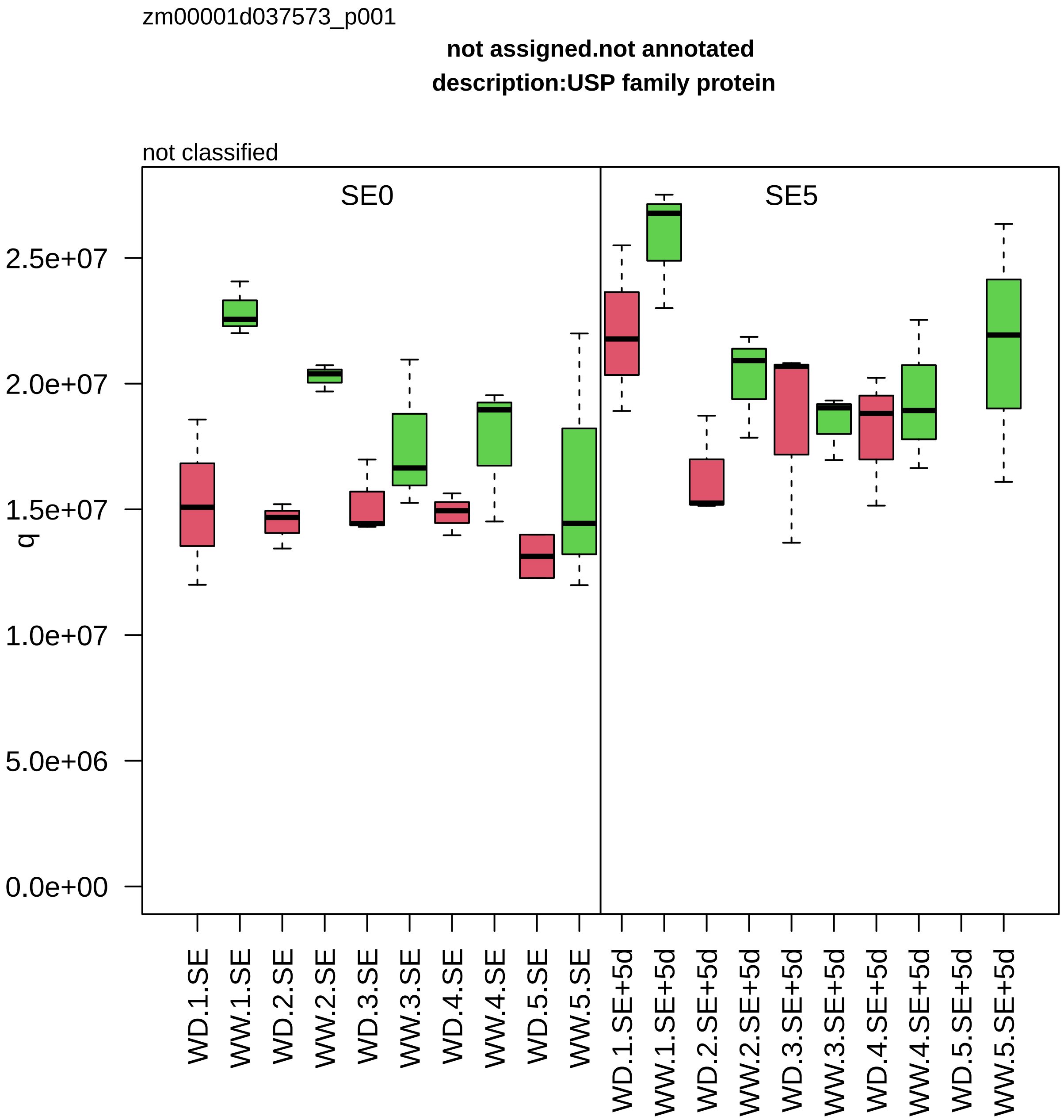

zm00001d037573_p001
not assigned.not annotated
description:USP family protein
not classified
SE0
SE5
q
2.5e+07
2.0e+07
1.5e+07
1.0e+07
5.0e+06
0.0e+00
WD.1.SE
WW.1.SE
WD.2.SE
WW.2.SE
WD.3.SE
WW.3.SE
WD.4.SE
WW.4.SE
WD.5.SE
WW.5.SE
WD.1.SE+5d
WW.1.SE+5d
WD.2.SE+5d
WW.2.SE+5d
WD.3.SE+5d
WW.3.SE+5d
WD.4.SE+5d
WW.4.SE+5d
WD.5.SE+5d
WW.5.SE+5d

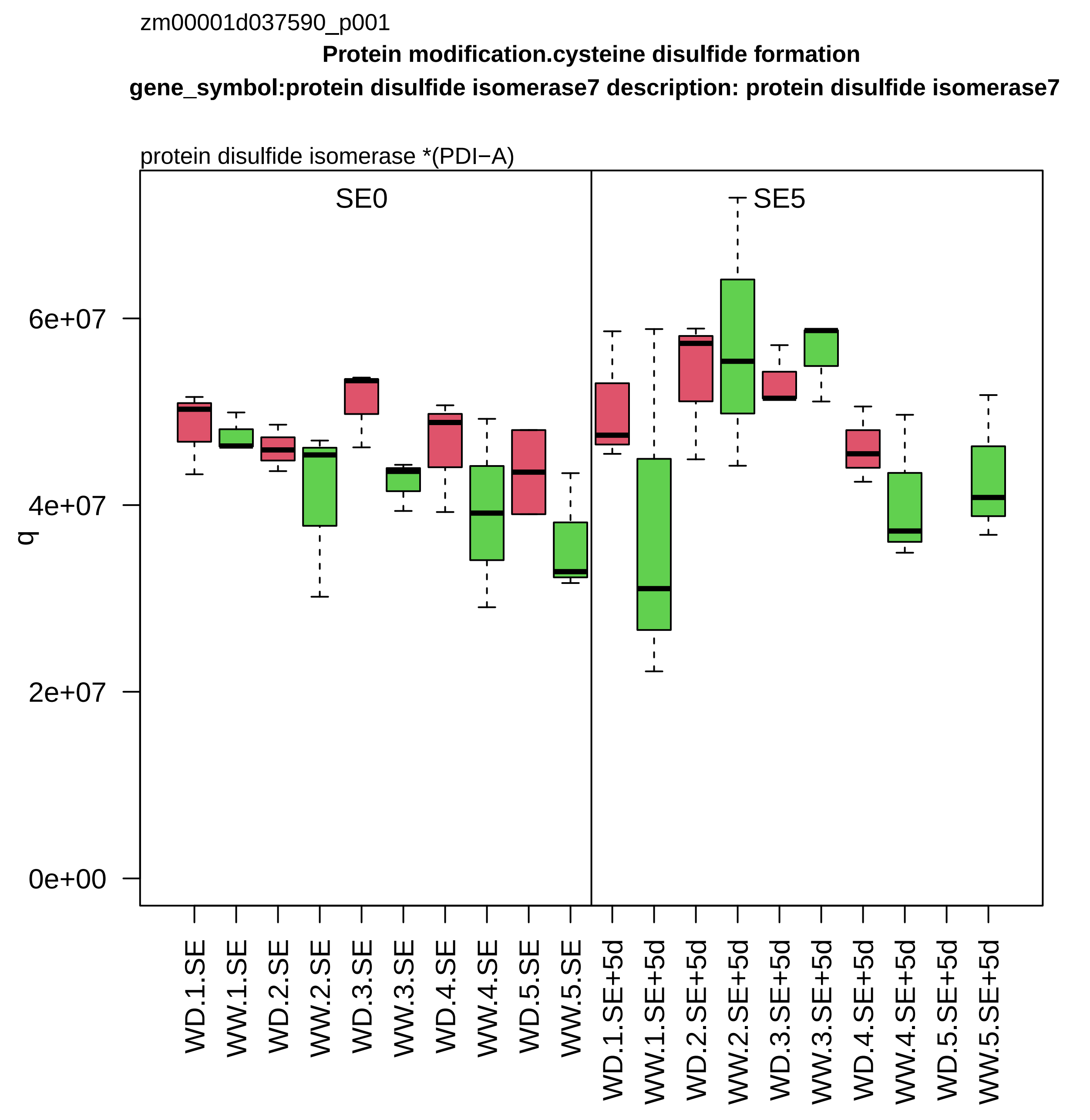

zm00001d037590_p001
Protein modification.cysteine disulfide formation
gene_symbol:protein disulfide isomerase7 description: protein disulfide isomerase7
protein disulfide isomerase *(PDI−A)
SE0
SE5
q
6e+07
4e+07
2e+07
0e+00
WD.1.SE
WW.1.SE
WD.2.SE
WW.2.SE
WD.3.SE
WW.3.SE
WD.4.SE
WW.4.SE
WD.5.SE
WW.5.SE
WD.1.SE+5d
WW.1.SE+5d
WD.2.SE+5d
WW.2.SE+5d
WD.3.SE+5d
WW.3.SE+5d
WD.4.SE+5d
WW.4.SE+5d
WD.5.SE+5d
WW.5.SE+5d

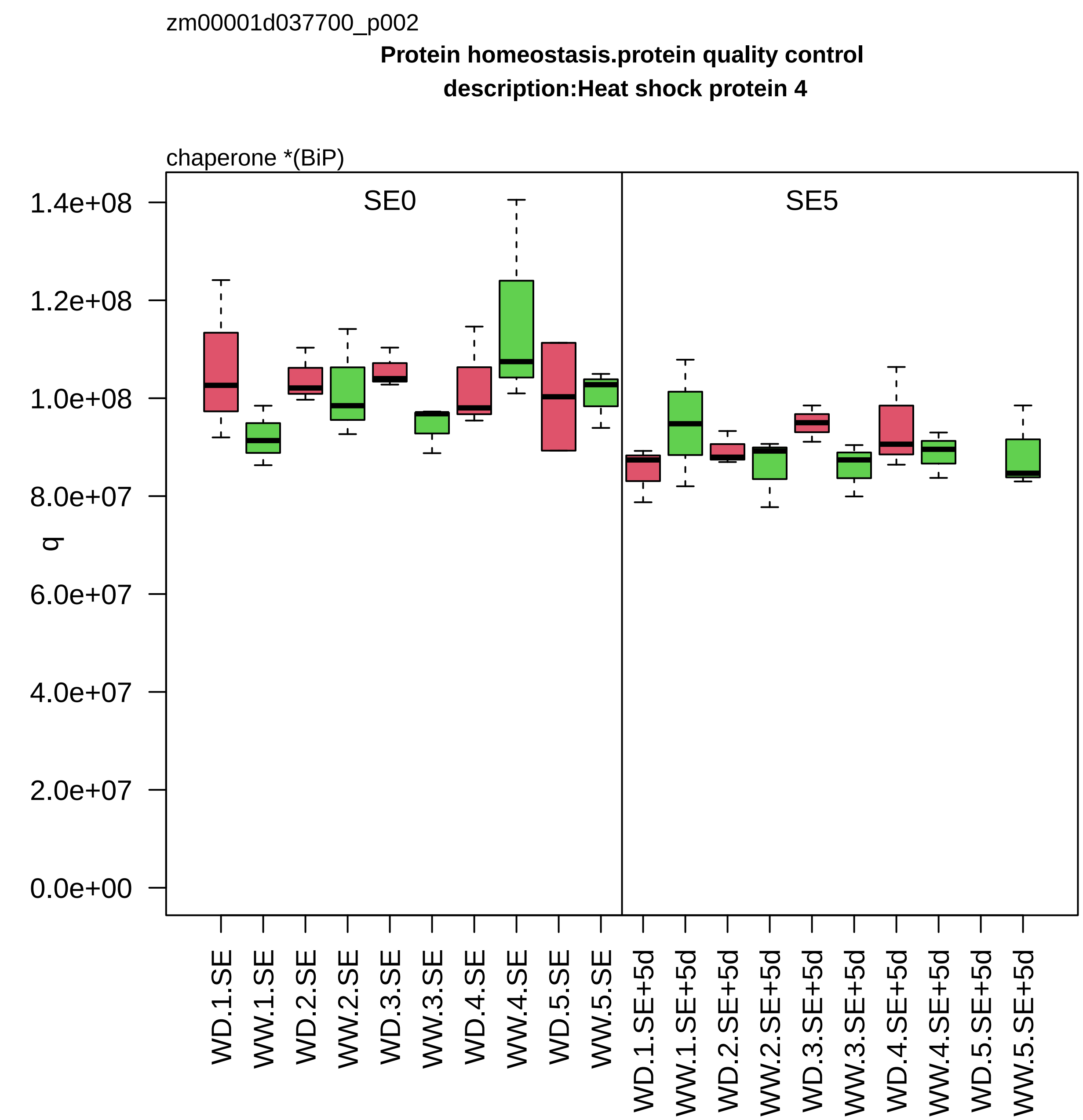

zm00001d037700_p002
Protein homeostasis.protein quality control
description:Heat shock protein 4
chaperone *(BiP)
SE0
SE5
q
1.4e+08
1.2e+08
1.0e+08
8.0e+07
6.0e+07
4.0e+07
2.0e+07
0.0e+00
WD.1.SE
WW.1.SE
WD.2.SE
WW.2.SE
WD.3.SE
WW.3.SE
WD.4.SE
WW.4.SE
WD.5.SE
WW.5.SE
WD.1.SE+5d
WW.1.SE+5d
WD.2.SE+5d
WW.2.SE+5d
WD.3.SE+5d
WW.3.SE+5d
WD.4.SE+5d
WW.4.SE+5d
WD.5.SE+5d
WW.5.SE+5d

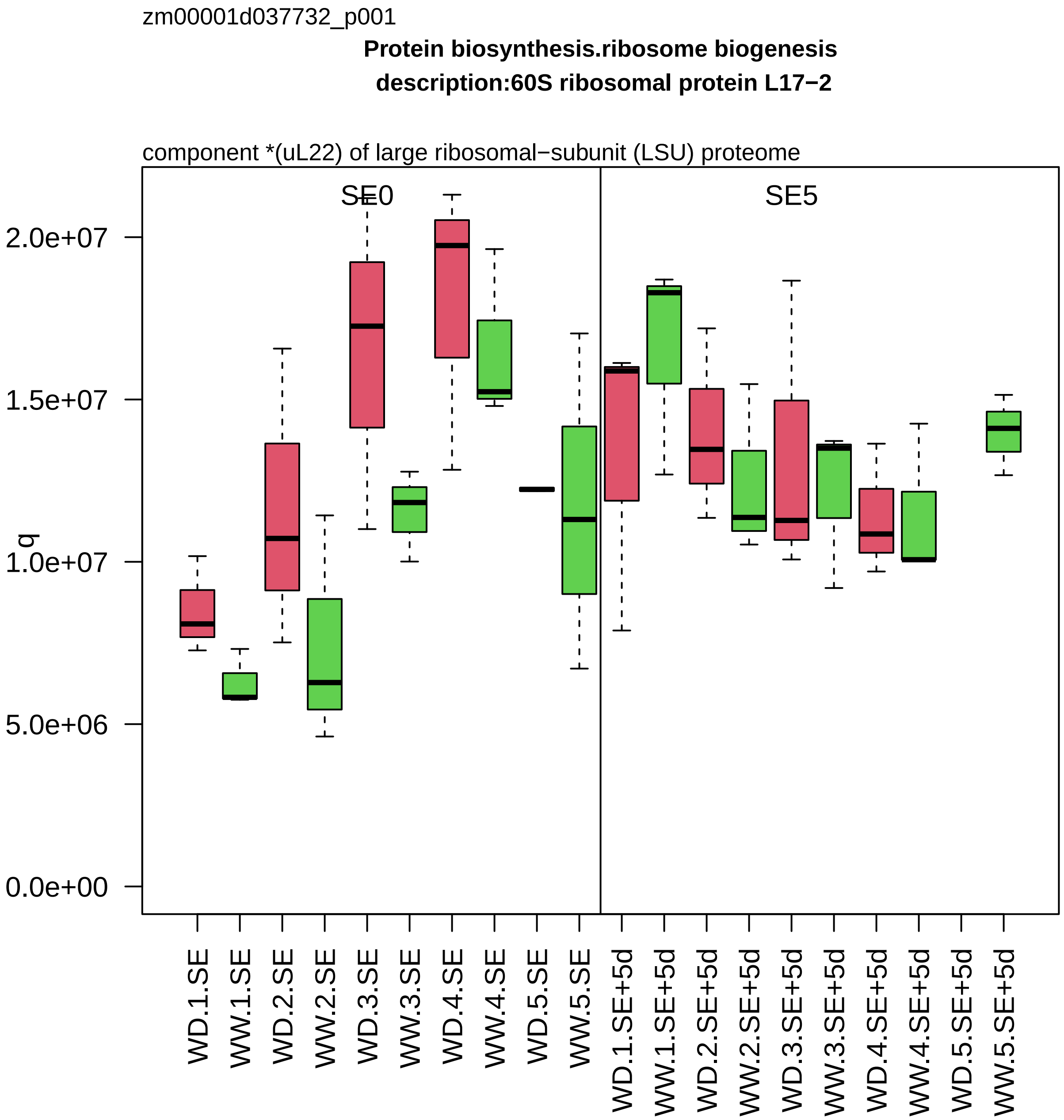
zm00001d037732_p001
Protein biosynthesis.ribosome biogenesis
description:60S ribosomal protein L17−2
component *(uL22) of large ribosomal−subunit (LSU) proteome
SE0
SE5
q
2.0e+07
1.5e+07
1.0e+07
5.0e+06
0.0e+00
WD.1.SE
WW.1.SE
WD.2.SE
WW.2.SE
WD.3.SE
WW.3.SE
WD.4.SE
WW.4.SE
WD.5.SE
WW.5.SE
WD.1.SE+5d
WW.1.SE+5d
WD.2.SE+5d
WW.2.SE+5d
WD.3.SE+5d
WW.3.SE+5d
WD.4.SE+5d
WW.4.SE+5d
WD.5.SE+5d
WW.5.SE+5d

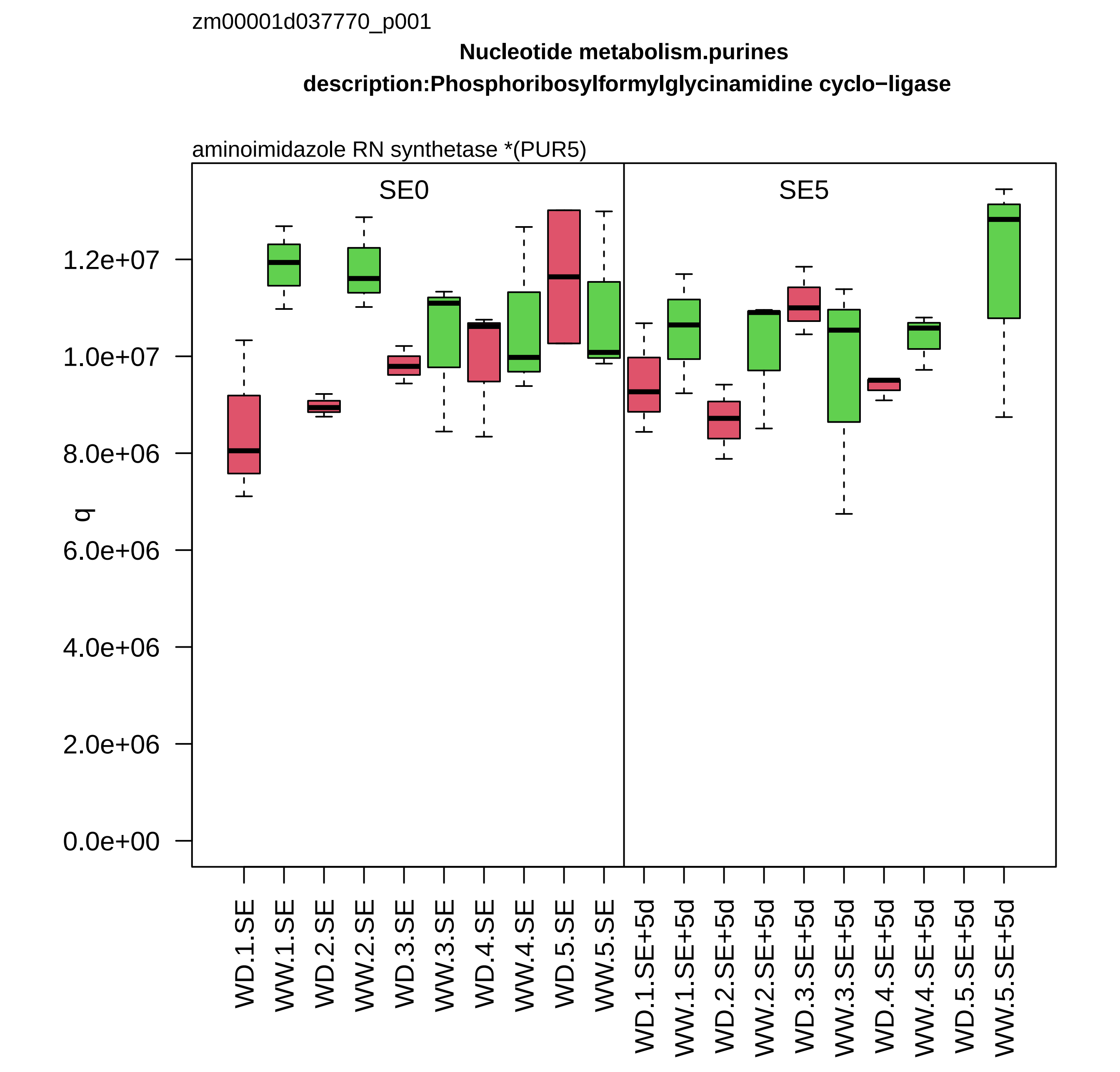

zm00001d037770_p001
Nucleotide metabolism.purines
description:Phosphoribosylformylglycinamidine cyclo−ligase
aminoimidazole RN synthetase *(PUR5)
SE0
SE5
q
1.2e+07
1.0e+07
8.0e+06
6.0e+06
4.0e+06
2.0e+06
0.0e+00
WD.1.SE
WW.1.SE
WD.2.SE
WW.2.SE
WD.3.SE
WW.3.SE
WD.4.SE
WW.4.SE
WD.5.SE
WW.5.SE
WD.1.SE+5d
WW.1.SE+5d
WD.2.SE+5d
WW.2.SE+5d
WD.3.SE+5d
WW.3.SE+5d
WD.4.SE+5d
WW.4.SE+5d
WD.5.SE+5d
WW.5.SE+5d

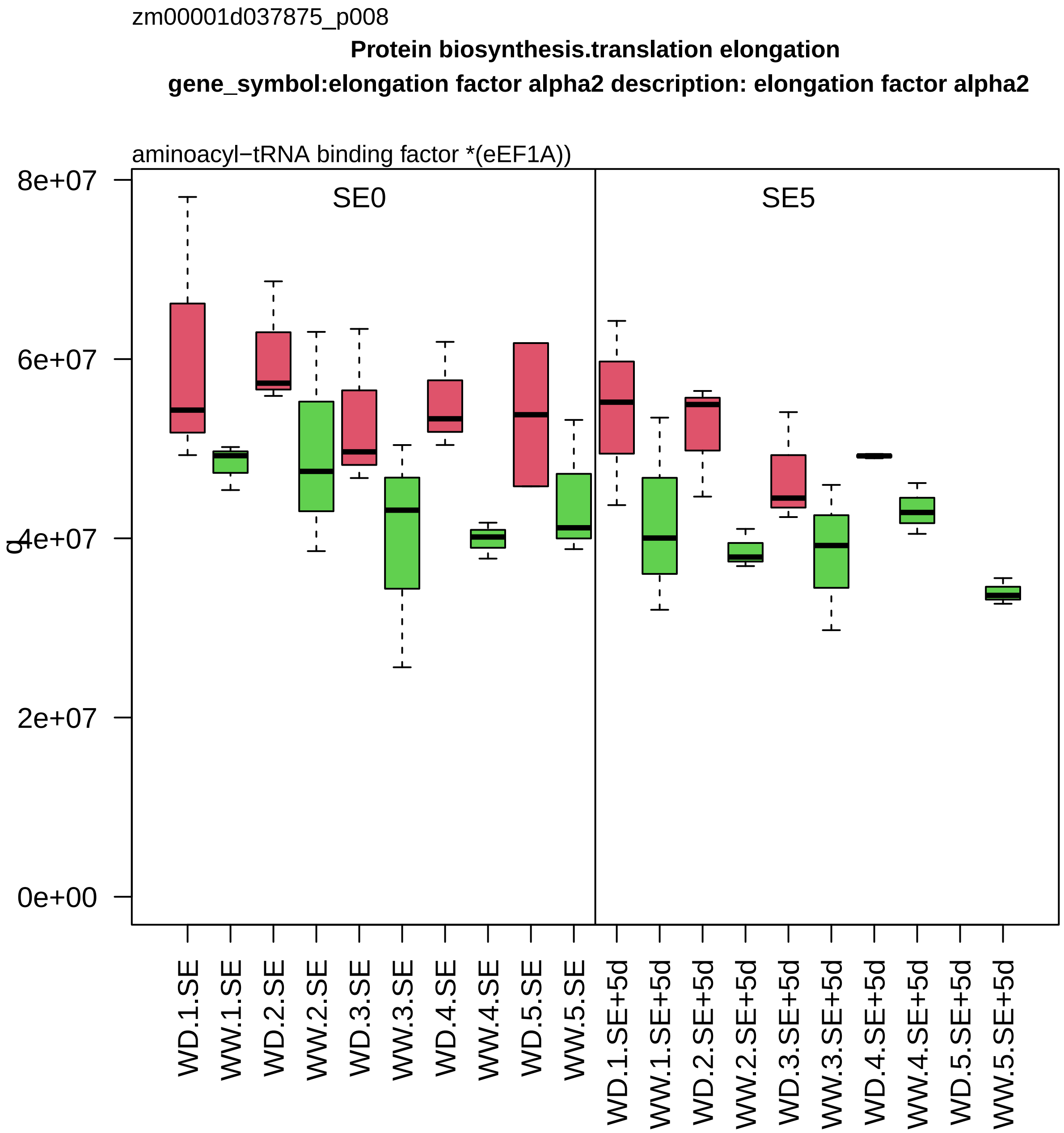

zm00001d037875_p008
Protein biosynthesis.translation elongation
gene_symbol:elongation factor alpha2 description: elongation factor alpha2
aminoacyl−tRNA binding factor *(eEF1A))
SE0
SE5
8e+07
6e+07
4e+07
2e+07
0e+00
q
WD.1.SE
WW.1.SE
WD.2.SE
WW.2.SE
WD.3.SE
WW.3.SE
WD.4.SE
WW.4.SE
WD.5.SE
WW.5.SE
WD.1.SE+5d
WW.1.SE+5d
WD.2.SE+5d
WW.2.SE+5d
WD.3.SE+5d
WW.3.SE+5d
WD.4.SE+5d
WW.4.SE+5d
WD.5.SE+5d
WW.5.SE+5d

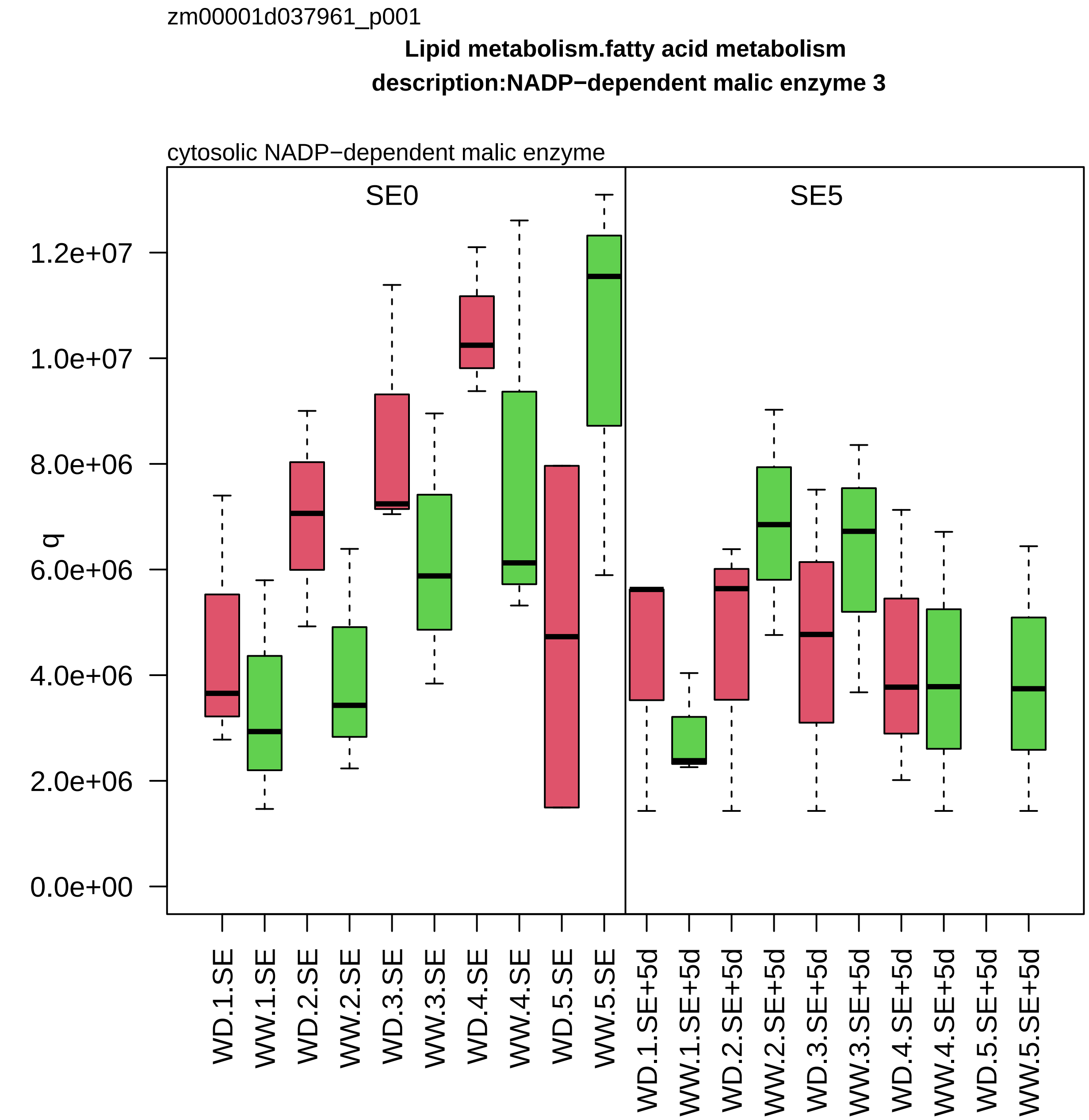

zm00001d037961_p001
Lipid metabolism.fatty acid metabolism
description:NADP−dependent malic enzyme 3
cytosolic NADP−dependent malic enzyme
SE0
SE5
q
1.2e+07
1.0e+07
8.0e+06
6.0e+06
4.0e+06
2.0e+06
0.0e+00
WD.1.SE
WW.1.SE
WD.2.SE
WW.2.SE
WD.3.SE
WW.3.SE
WD.4.SE
WW.4.SE
WD.5.SE
WW.5.SE
WD.1.SE+5d
WW.1.SE+5d
WD.2.SE+5d
WW.2.SE+5d
WD.3.SE+5d
WW.3.SE+5d
WD.4.SE+5d
WW.4.SE+5d
WD.5.SE+5d
WW.5.SE+5d

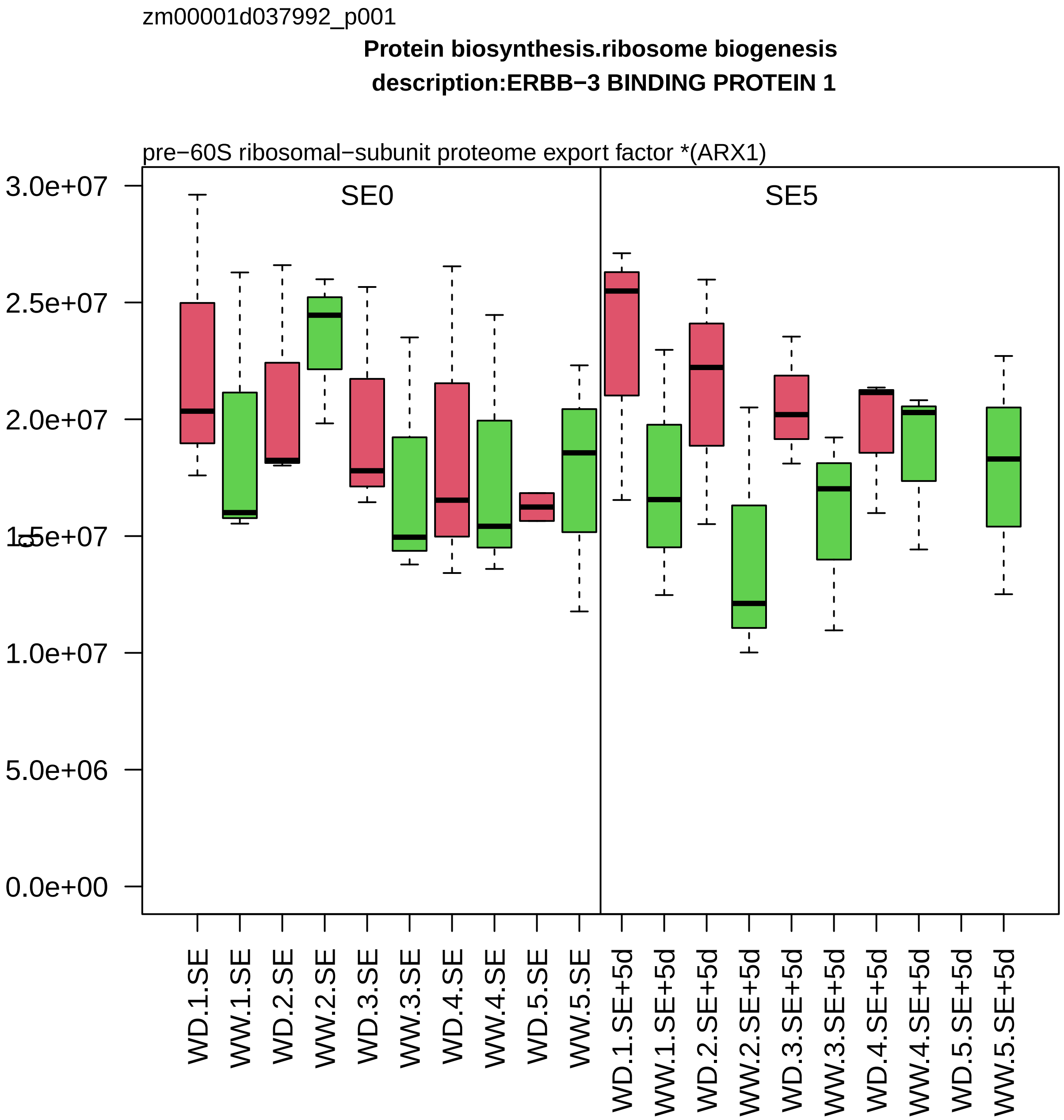
zm00001d037992_p001
Protein biosynthesis.ribosome biogenesis
description:ERBB−3 BINDING PROTEIN 1
pre−60S ribosomal−subunit proteome export factor *(ARX1)
SE0
SE5
3.0e+07
2.5e+07
2.0e+07
1.5e+07
1.0e+07
5.0e+06
0.0e+00
WD.1.SE
WW.1.SE
WD.2.SE
WW.2.SE
WD.3.SE
WW.3.SE
WD.4.SE
WW.4.SE
WD.5.SE
WW.5.SE
WD.1.SE+5d
WW.1.SE+5d
WD.2.SE+5d
WW.2.SE+5d
WD.3.SE+5d
WW.3.SE+5d
WD.4.SE+5d
WW.4.SE+5d
WD.5.SE+5d
WW.5.SE+5d

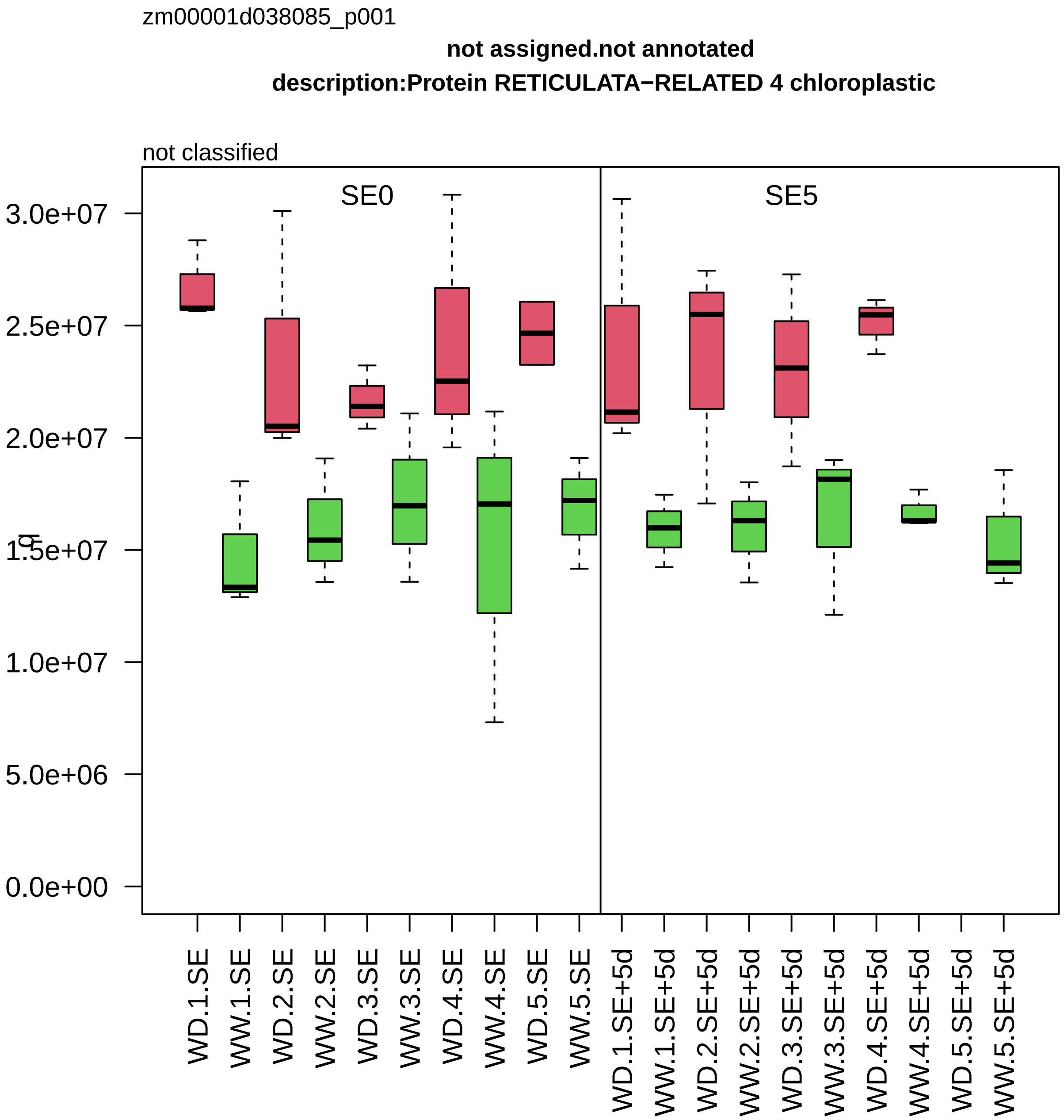
zm00001d038085_p001
not assigned.not annotated
description:Protein RETICULATA−RELATED 4 chloroplastic
not classified
SE0
SE5
0.0e+00
5.0e+06
1.0e+07
1.5e+07
2.0e+07
2.5e+07
3.0e+07
WD.1.SE
WW.1.SE
WD.2.SE
WW.2.SE
WD.3.SE
WW.3.SE
WD.4.SE
WW.4.SE
WD.5.SE
WW.5.SE
WD.1.SE+5d
WW.1.SE+5d
WD.2.SE+5d
WW.2.SE+5d
WD.3.SE+5d
WW.3.SE+5d
WD.4.SE+5d
WW.4.SE+5d
WD.5.SE+5d
WW.5.SE+5d

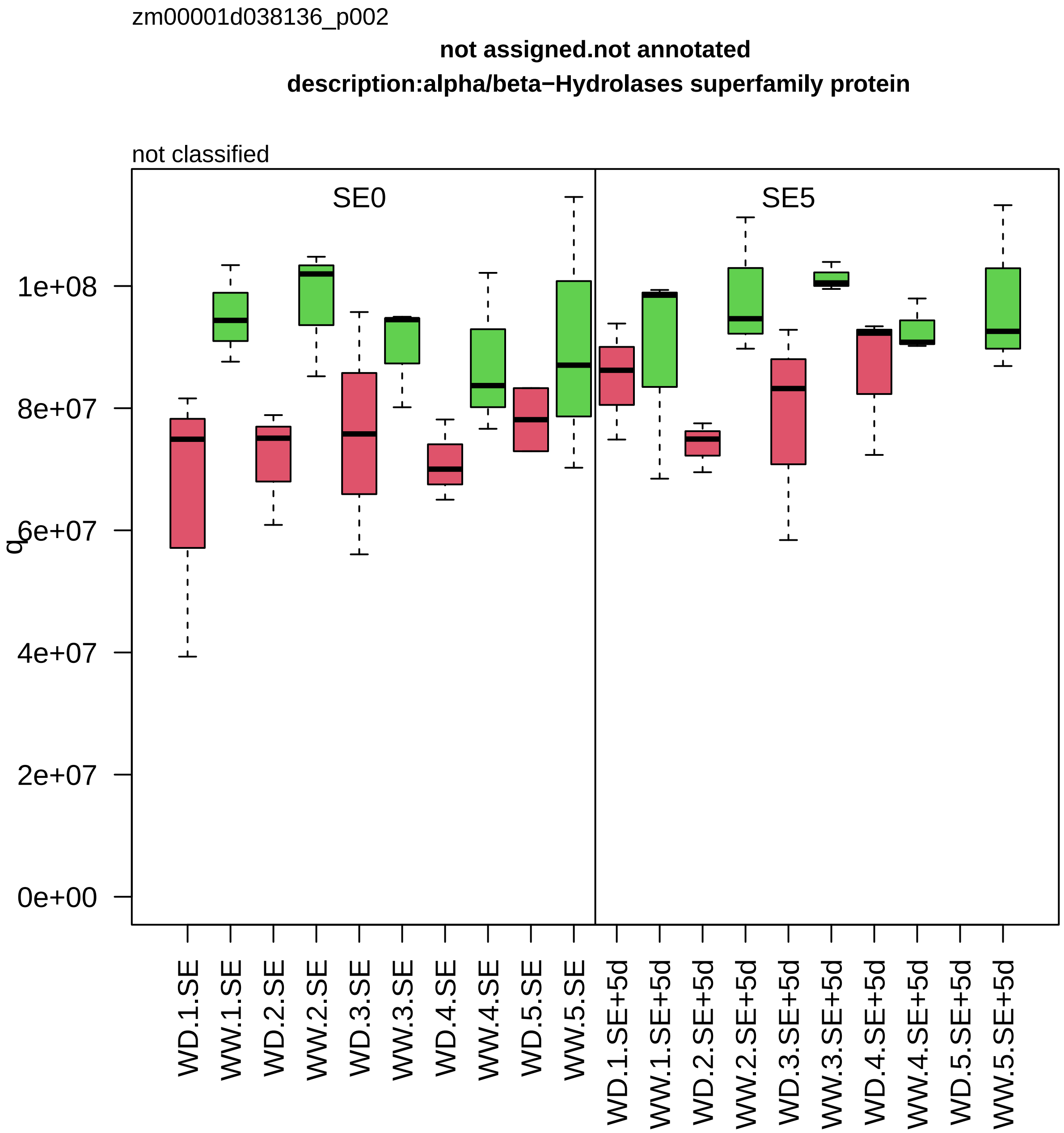

zm00001d038136_p002
not assigned.not annotated
description:alpha/beta−Hydrolases superfamily protein
not classified
SE0
SE5
q
0e+00
2e+07
4e+07
6e+07
8e+07
1e+08
WD.1.SE
WW.1.SE
WD.2.SE
WW.2.SE
WD.3.SE
WW.3.SE
WD.4.SE
WW.4.SE
WD.5.SE
WW.5.SE
WD.1.SE+5d
WW.1.SE+5d
WD.2.SE+5d
WW.2.SE+5d
WD.3.SE+5d
WW.3.SE+5d
WD.4.SE+5d
WW.4.SE+5d
WD.5.SE+5d
WW.5.SE+5d

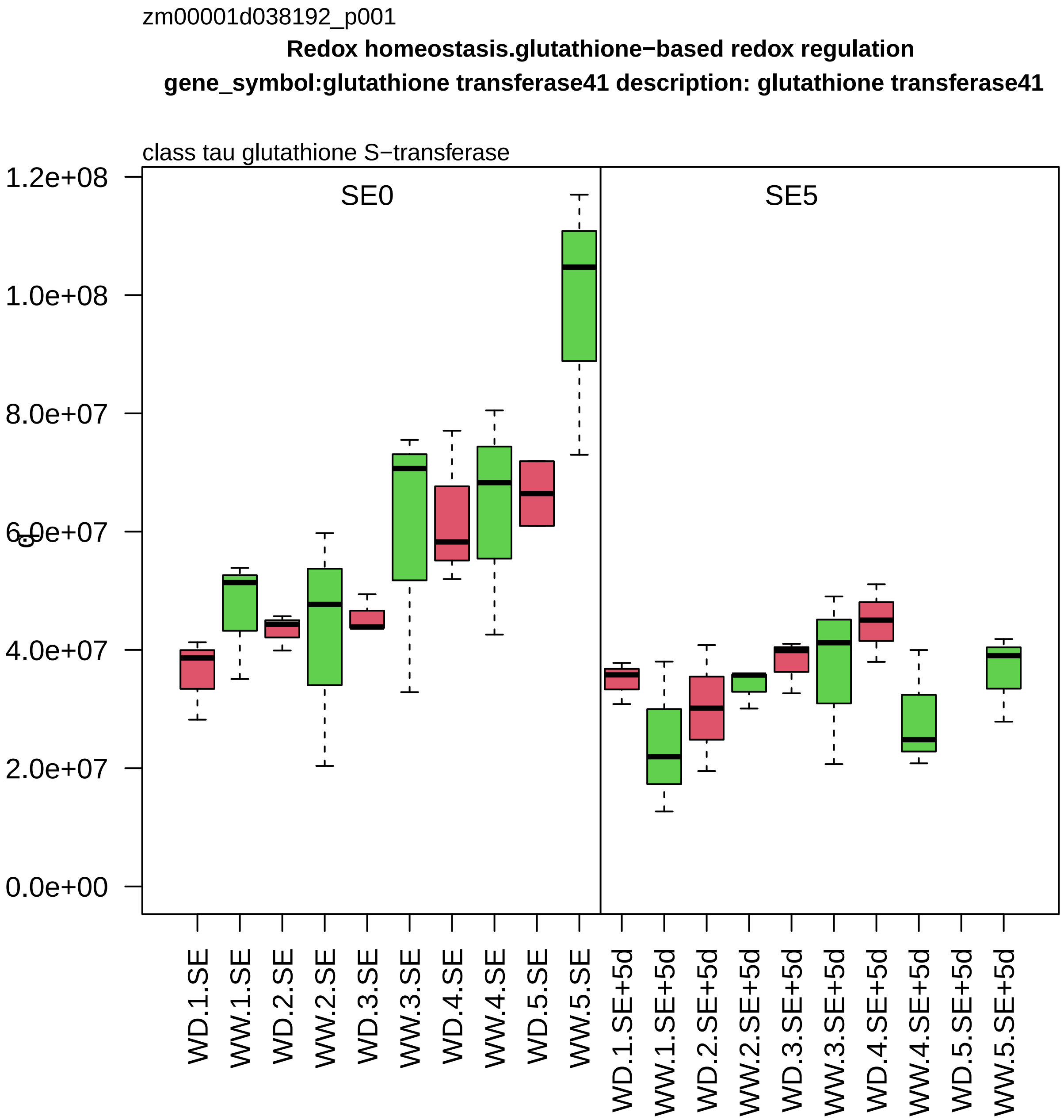

zm00001d038192_p001
Redox homeostasis.glutathione−based redox regulation
gene_symbol:glutathione transferase41 description: glutathione transferase41
class tau glutathione S−transferase
SE0
SE5
1.2e+08
1.0e+08
8.0e+07
6.0e+07
4.0e+07
2.0e+07
0.0e+00
WD.1.SE
WW.1.SE
WD.2.SE
WW.2.SE
WD.3.SE
WW.3.SE
WD.4.SE
WW.4.SE
WD.5.SE
WW.5.SE
WD.1.SE+5d
WW.1.SE+5d
WD.2.SE+5d
WW.2.SE+5d
WD.3.SE+5d
WW.3.SE+5d
WD.4.SE+5d
WW.4.SE+5d
WD.5.SE+5d
WW.5.SE+5d

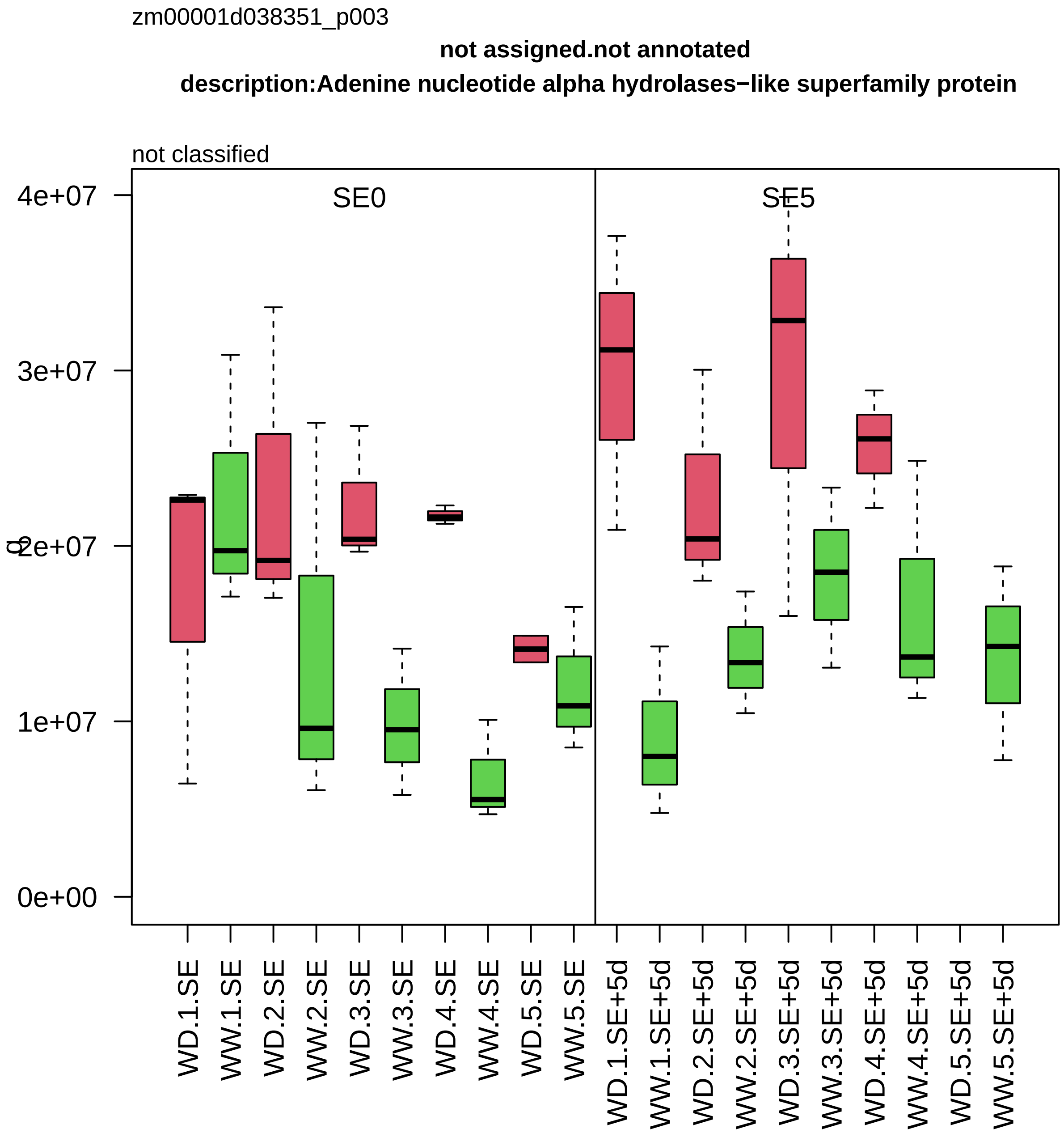
zm00001d038351_p003
not assigned.not annotated
description:Adenine nucleotide alpha hydrolases−like superfamily protein
not classified
SE0
SE5
4e+07
3e+07
2e+07
1e+07
0e+00
WD.1.SE
WW.1.SE
WD.2.SE
WW.2.SE
WD.3.SE
WW.3.SE
WD.4.SE
WW.4.SE
WD.5.SE
WW.5.SE
WD.1.SE+5d
WW.1.SE+5d
WD.2.SE+5d
WW.2.SE+5d
WD.3.SE+5d
WW.3.SE+5d
WD.4.SE+5d
WW.4.SE+5d
WD.5.SE+5d
WW.5.SE+5d

zm00001d038535_p004

**Protein biosynthesis.ribosome biogenesis**
**description:60S ribosomal protein L30−2**

component *(eL30) of large ribosomal−subunit (LSU) proteome

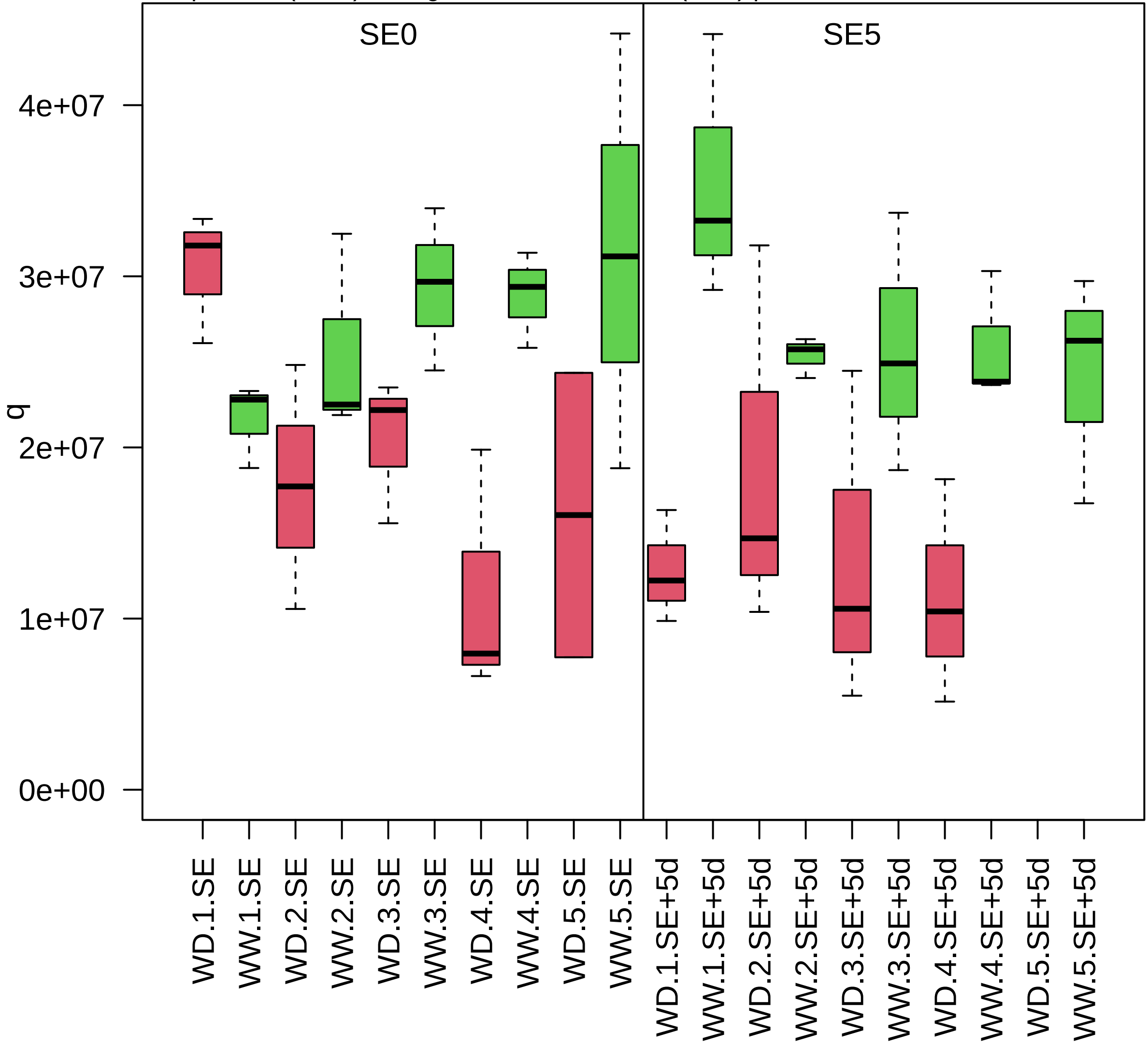

zm00001d038541_p001

**Chromatin organisation.epigenetic state regulation**

**gene_symbol:set domain gene111 description:Histone−lysine N−methyltransferase H3 lysine−9 specifi**

methylated DNA binding component *(SUVH1/3) of SUVH−DNAJ methylation reader complex

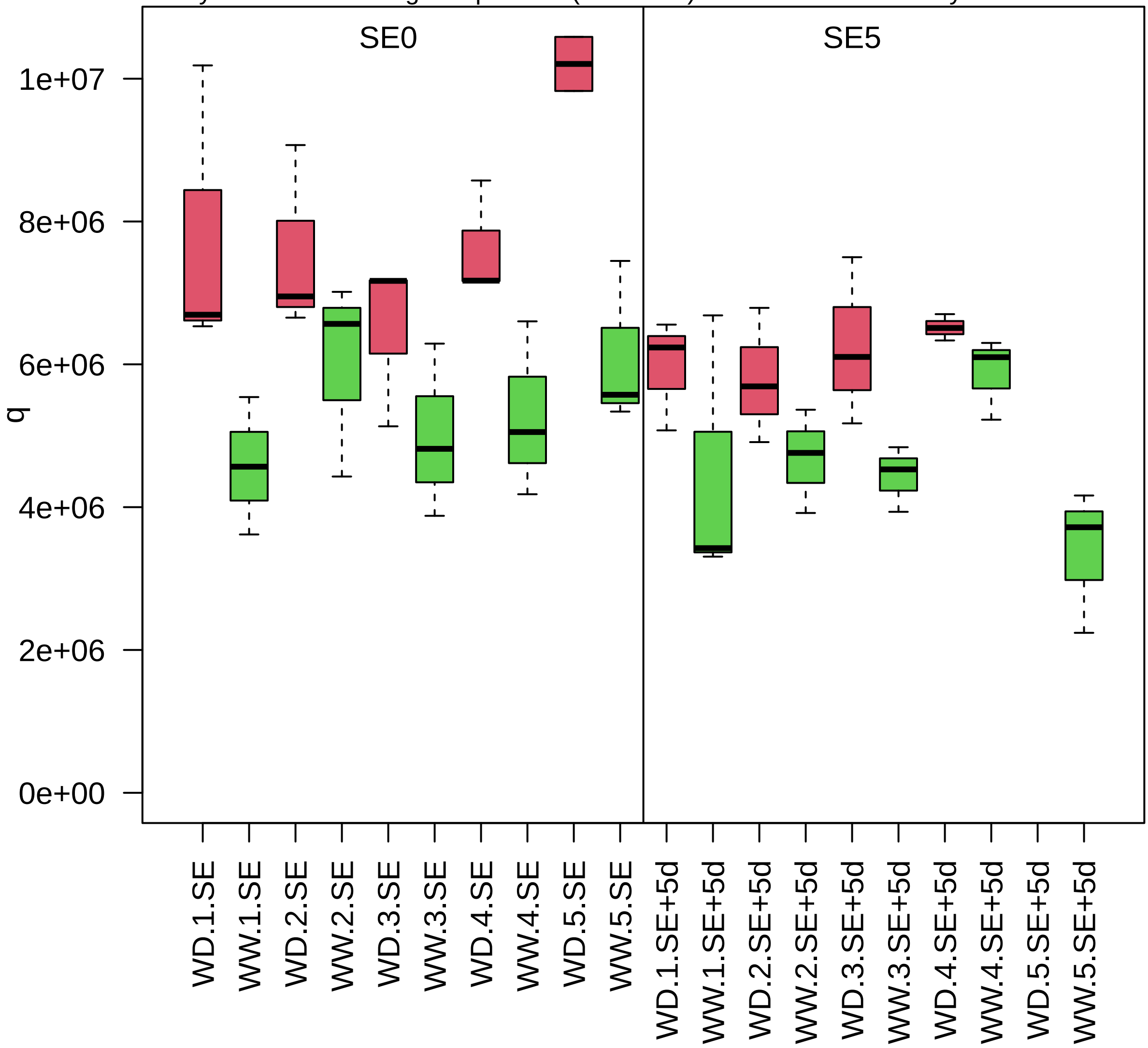

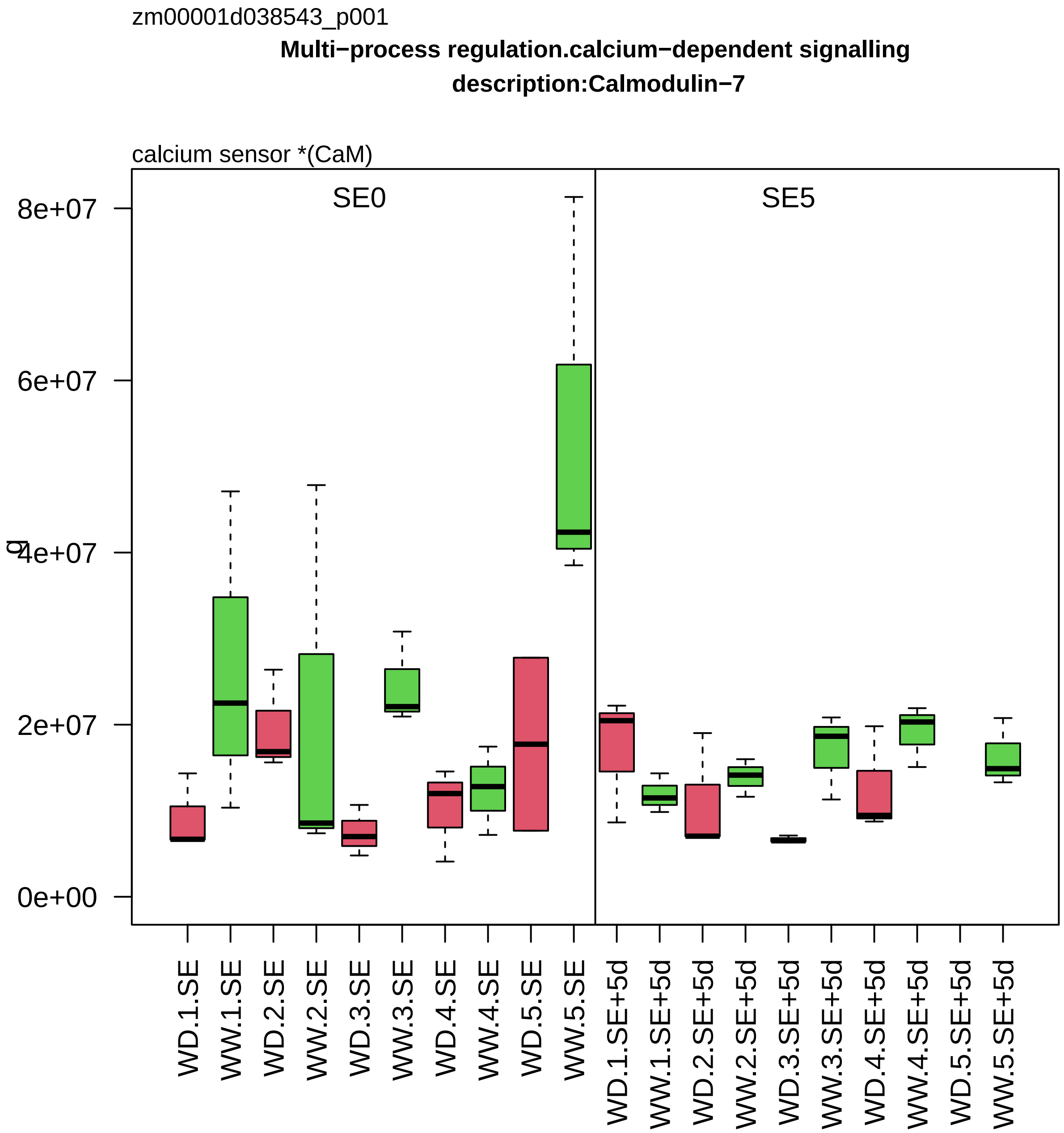
zm00001d038543_p001
Multi−process regulation.calcium−dependent signalling
description:Calmodulin−7
calcium sensor *(CaM)
SE0
SE5
0e+00
2e+07
4e+07
6e+07
8e+07
WD.1.SE
WW.1.SE
WD.2.SE
WW.2.SE
WD.3.SE
WW.3.SE
WD.4.SE
WW.4.SE
WD.5.SE
WW.5.SE
WD.1.SE+5d
WW.1.SE+5d
WD.2.SE+5d
WW.2.SE+5d
WD.3.SE+5d
WW.3.SE+5d
WD.4.SE+5d
WW.4.SE+5d
WD.5.SE+5d
WW.5.SE+5d

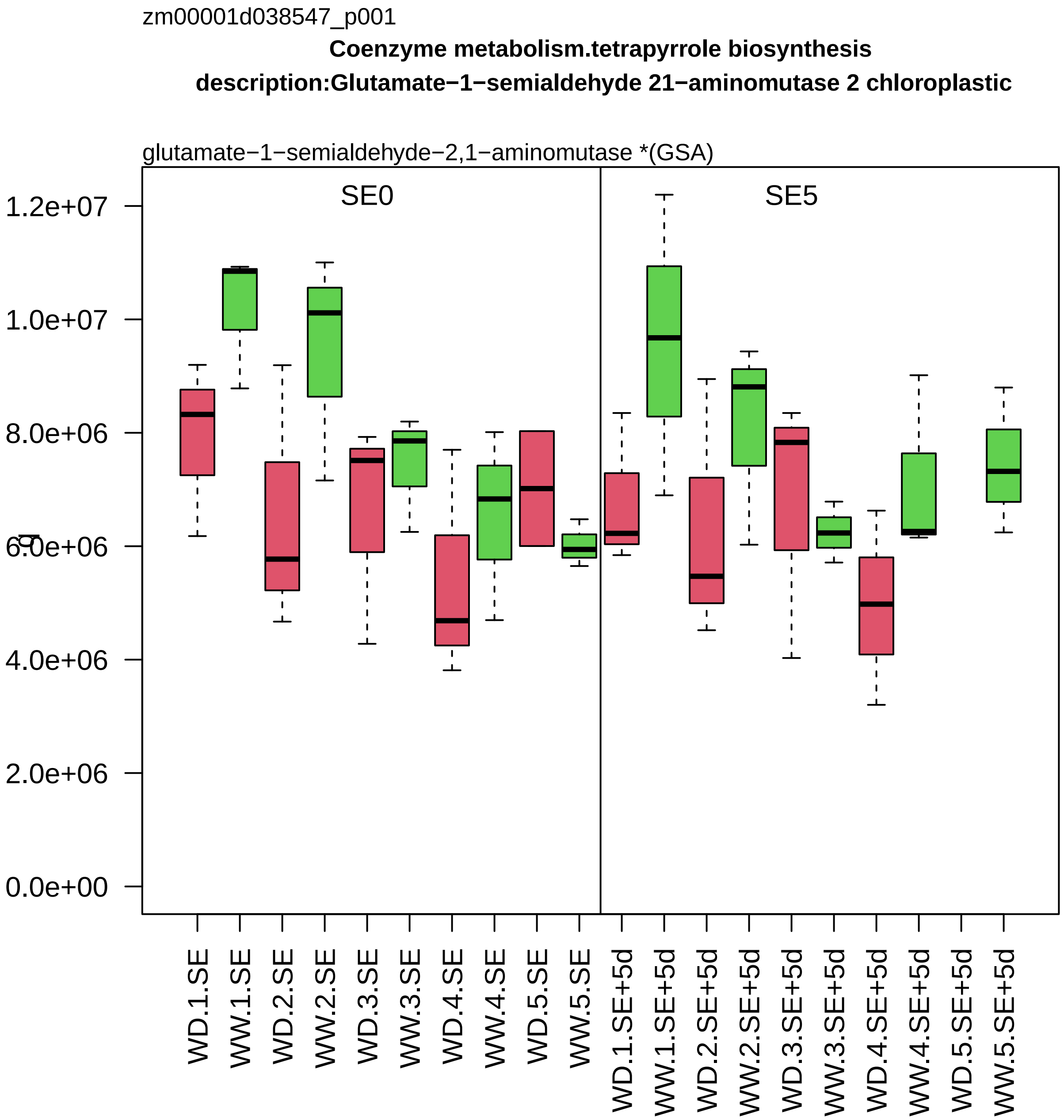
zm00001d038547_p001
Coenzyme metabolism.tetrapyrrole biosynthesis
description:Glutamate−1−semialdehyde 21−aminomutase 2 chloroplastic
glutamate−1−semialdehyde−2,1−aminomutase *(GSA)
SE0
SE5
1.2e+07
1.0e+07
8.0e+06
6.0e+06
4.0e+06
2.0e+06
0.0e+00
WD.1.SE
WW.1.SE
WD.2.SE
WW.2.SE
WD.3.SE
WW.3.SE
WD.4.SE
WW.4.SE
WD.5.SE
WW.5.SE
WD.1.SE+5d
WW.1.SE+5d
WD.2.SE+5d
WW.2.SE+5d
WD.3.SE+5d
WW.3.SE+5d
WD.4.SE+5d
WW.4.SE+5d
WD.5.SE+5d
WW.5.SE+5d

zm00001d038570_p003

**RNA biosynthesis.RNA polymerase II−dependent transcription description:global transcription factor group B1**

component *(SPT6) of SPT6−IWS1 transcription elongation complex

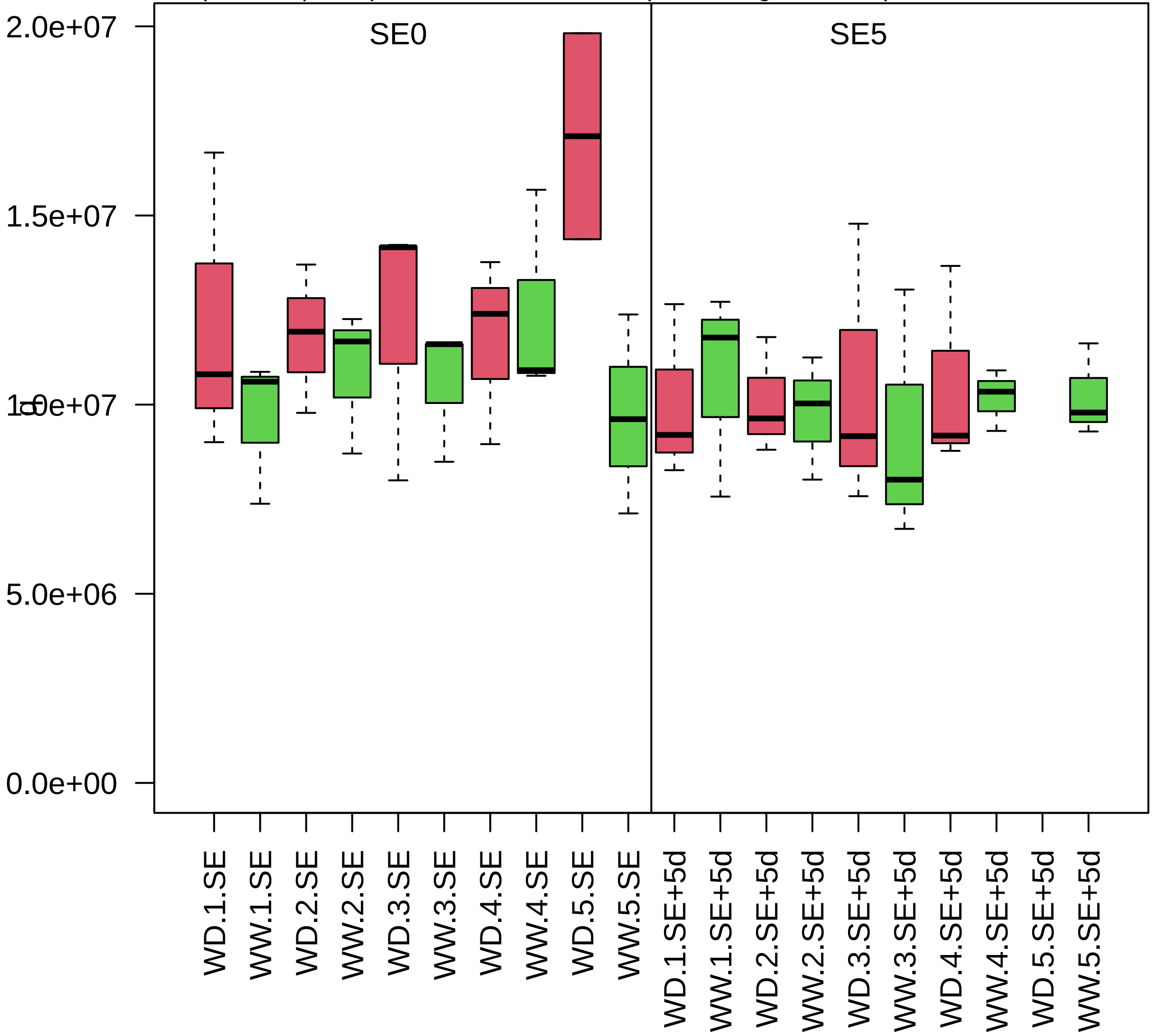

zm00001d038579_p001

**Photosynthesis.calvin cycle**
**description:Phosphoglycerate kinase**

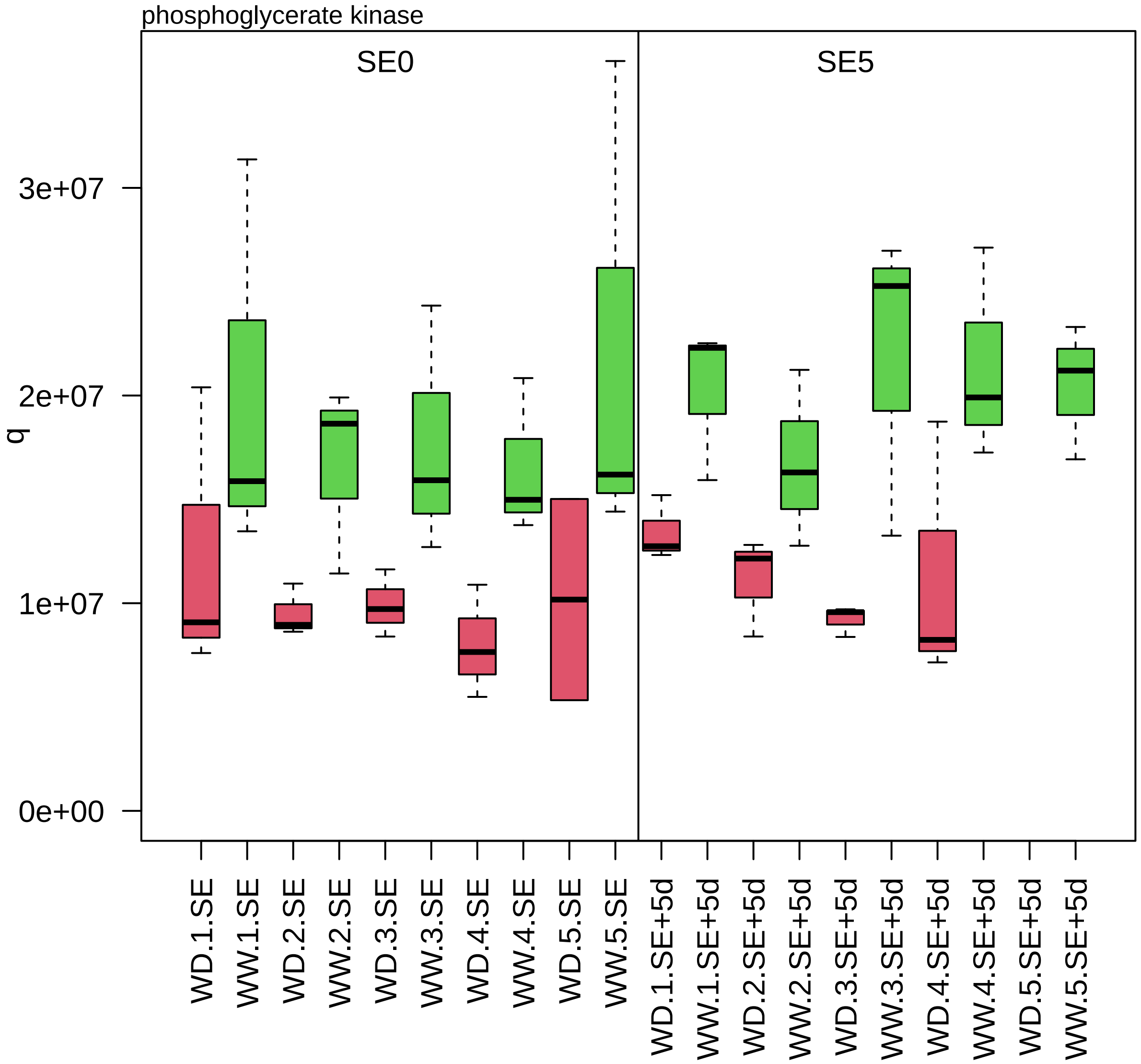

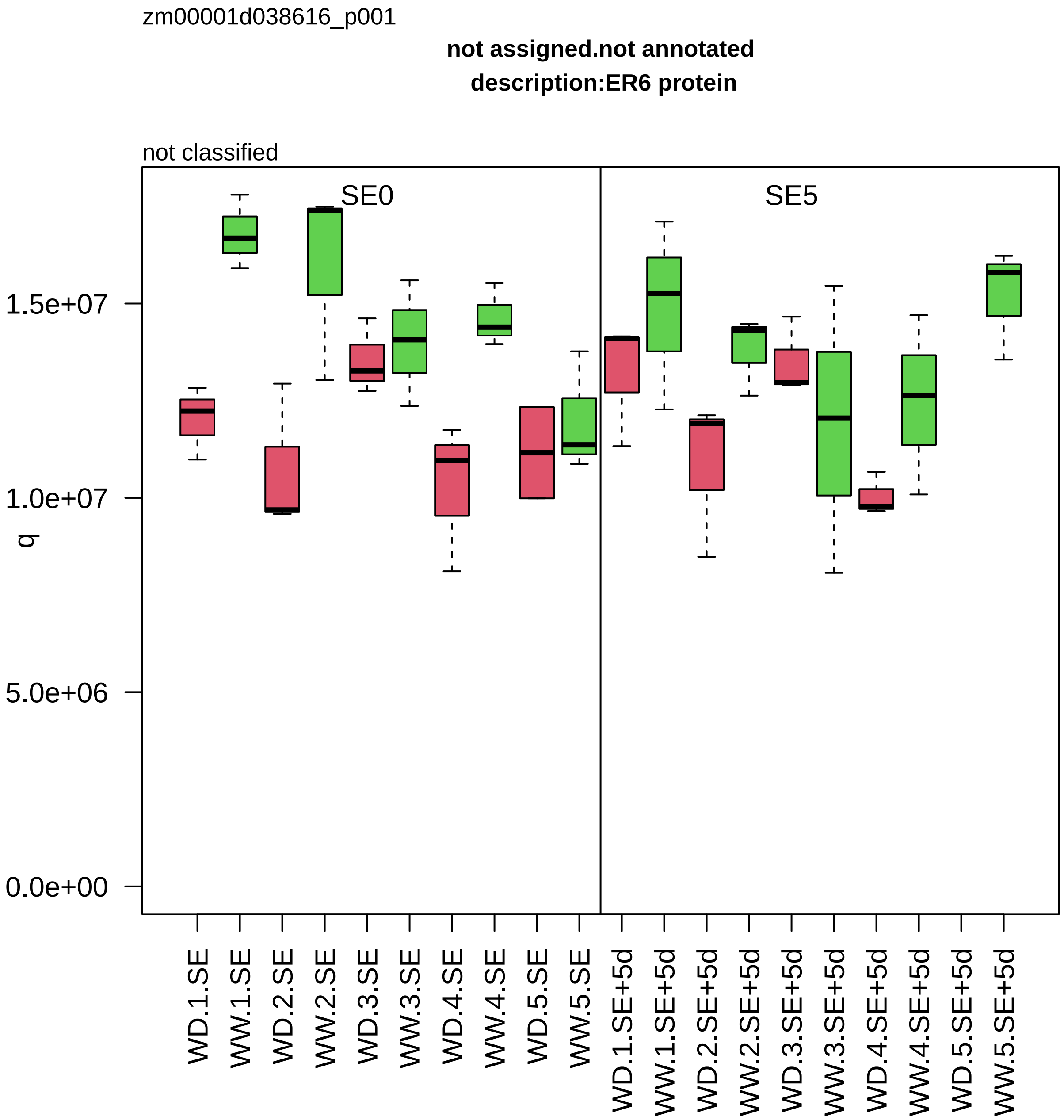
zm00001d038616_p001
not assigned.not annotated
description:ER6 protein
not classified
SE0
SE5
q
1.5e+07
1.0e+07
5.0e+06
0.0e+00
WD.1.SE
WW.1.SE
WD.2.SE
WW.2.SE
WD.3.SE
WW.3.SE
WD.4.SE
WW.4.SE
WD.5.SE
WW.5.SE
WD.1.SE+5d
WW.1.SE+5d
WD.2.SE+5d
WW.2.SE+5d
WD.3.SE+5d
WW.3.SE+5d
WD.4.SE+5d
WW.4.SE+5d
WD.5.SE+5d
WW.5.SE+5d

zm00001d038652_p003

**Redox homeostasis.thiol−based redox regulation**
**description:Thioredoxin H4**

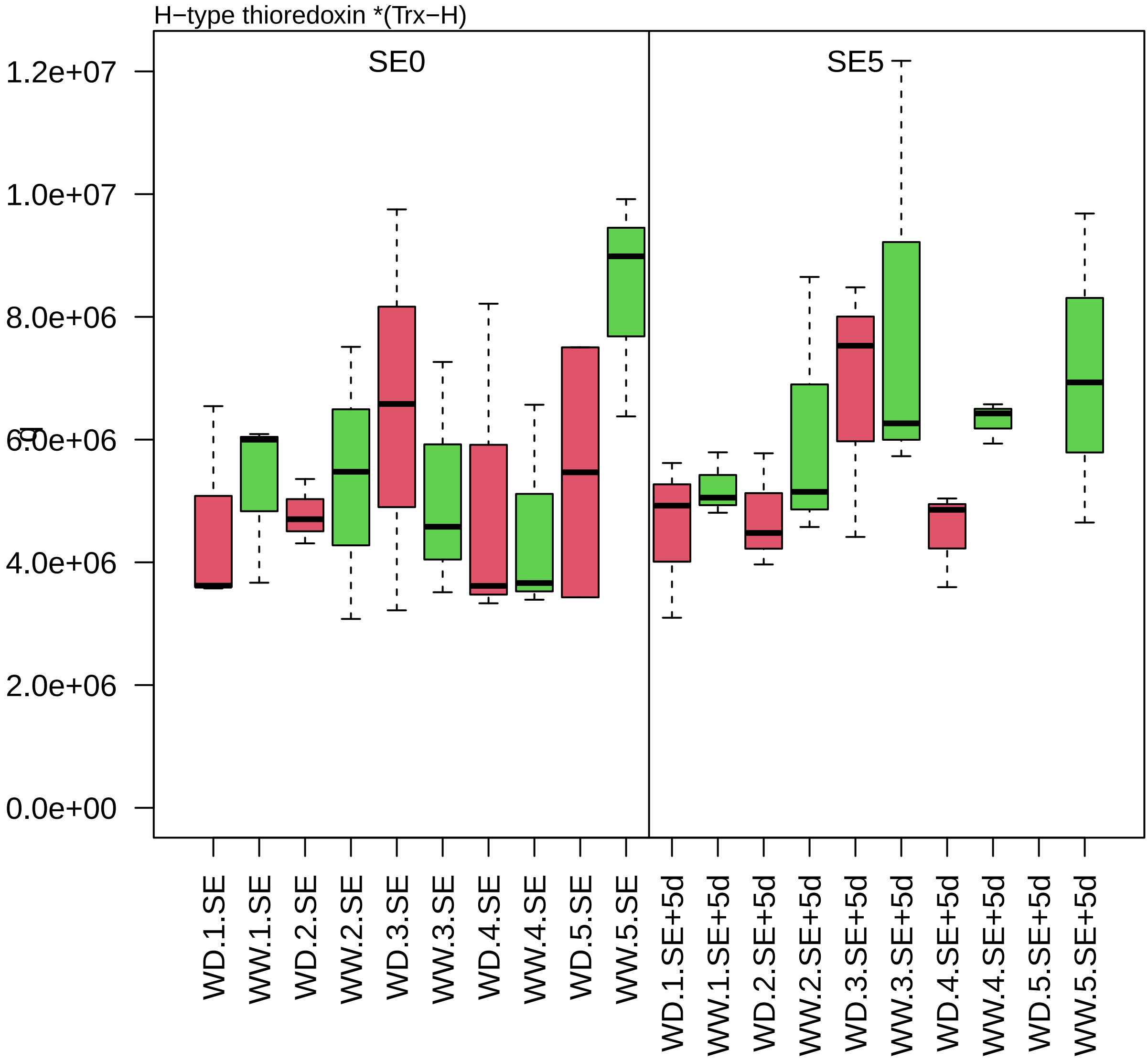

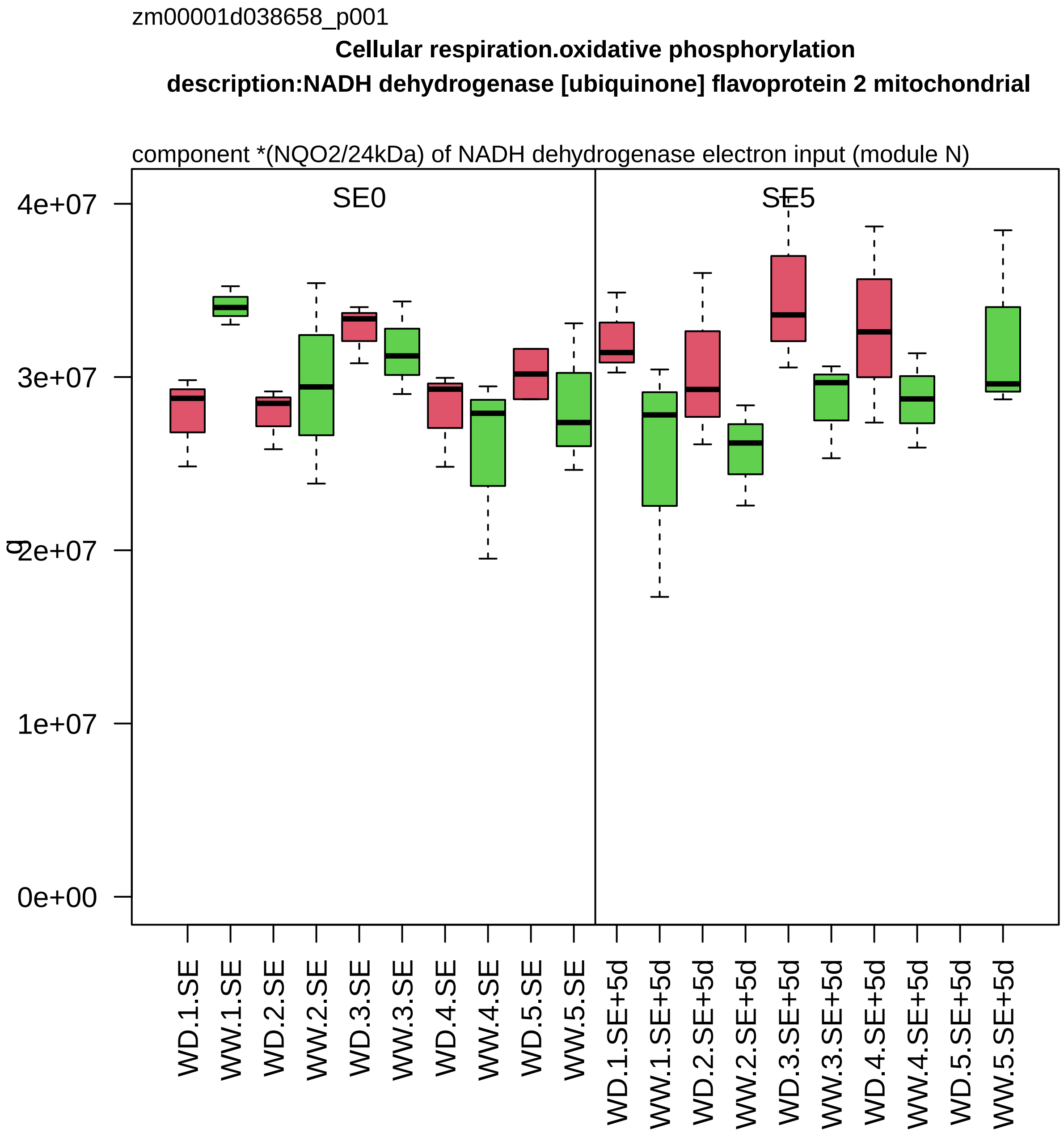

zm00001d038658_p001
Cellular respiration.oxidative phosphorylation
description:NADH dehydrogenase [ubiquinone] flavoprotein 2 mitochondrial
component *(NQO2/24kDa) of NADH dehydrogenase electron input (module N)
SE0
SE5
g
4e+07
3e+07
2e+07
1e+07
0e+00
WD.1.SE
WW.1.SE
WD.2.SE
WW.2.SE
WD.3.SE
WW.3.SE
WD.4.SE
WW.4.SE
WD.5.SE
WW.5.SE
WD.1.SE+5d
WW.1.SE+5d
WD.2.SE+5d
WW.2.SE+5d
WD.3.SE+5d
WW.3.SE+5d
WD.4.SE+5d
WW.4.SE+5d
WD.5.SE+5d
WW.5.SE+5d

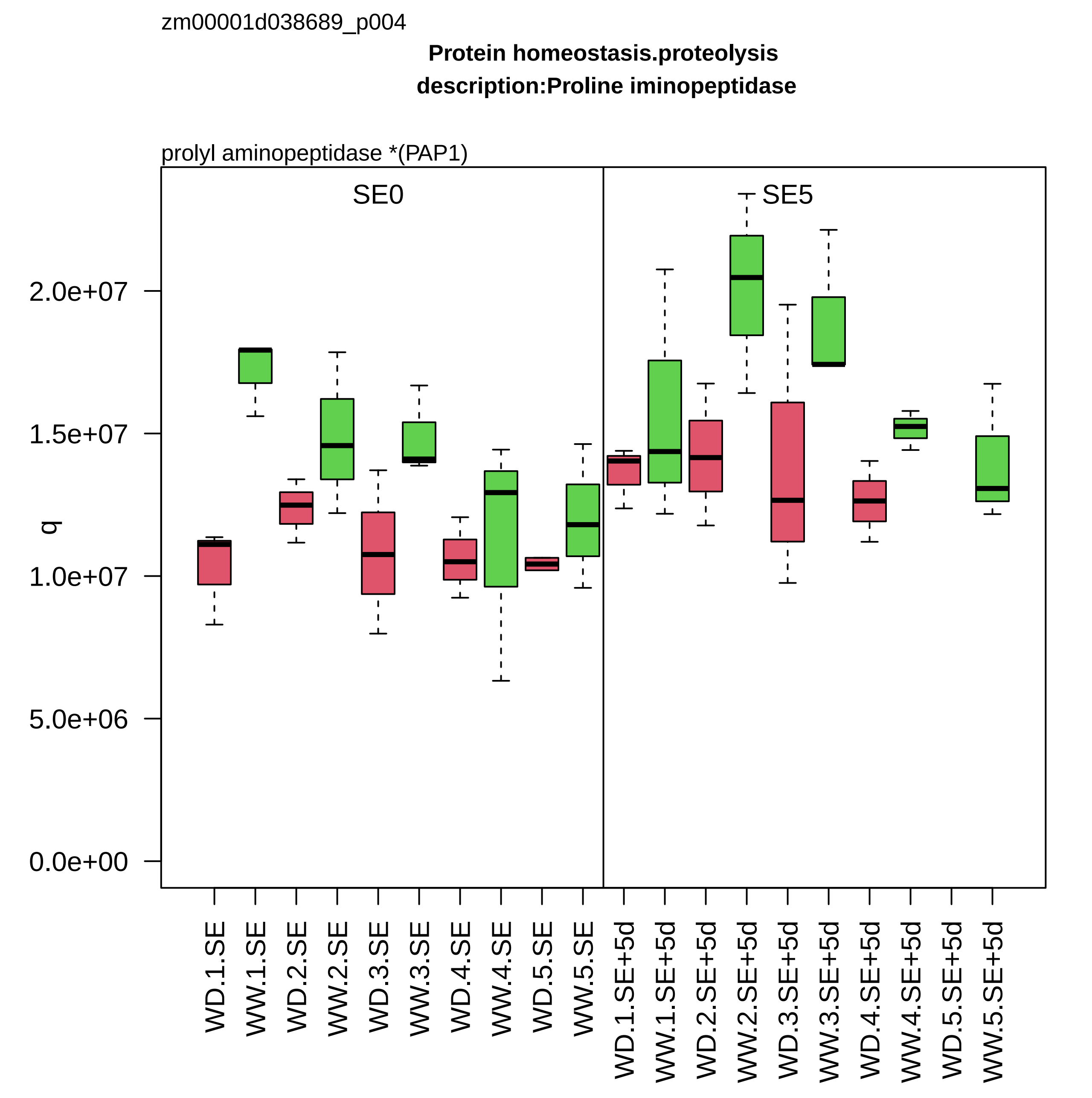

zm00001d038689_p004
Protein homeostasis.proteolysis
description:Proline iminopeptidase
prolyl aminopeptidase *(PAP1)
SE0
SE5
q
2.0e+07
1.5e+07
1.0e+07
5.0e+06
0.0e+00
WD.1.SE
WW.1.SE
WD.2.SE
WW.2.SE
WD.3.SE
WW.3.SE
WD.4.SE
WW.4.SE
WD.5.SE
WW.5.SE
WD.1.SE+5d
WW.1.SE+5d
WD.2.SE+5d
WW.2.SE+5d
WD.3.SE+5d
WW.3.SE+5d
WD.4.SE+5d
WW.4.SE+5d
WD.5.SE+5d
WW.5.SE+5d

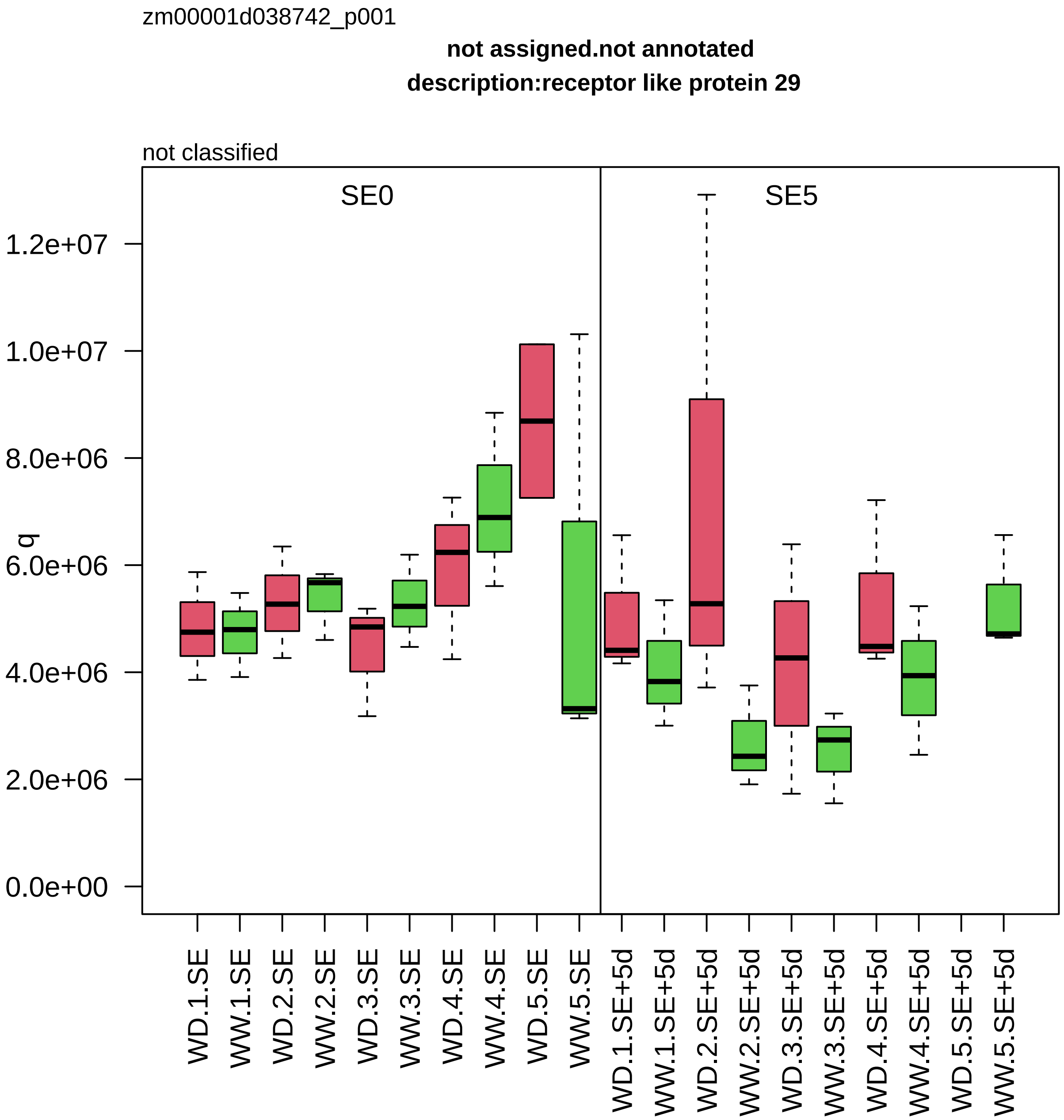
zm00001d038742_p001
not assigned.not annotated
description:receptor like protein 29
not classified
SE0
SE5
q
0.0e+00
2.0e+06
4.0e+06
6.0e+06
8.0e+06
1.0e+07
1.2e+07
WD.1.SE
WW.1.SE
WD.2.SE
WW.2.SE
WD.3.SE
WW.3.SE
WD.4.SE
WW.4.SE
WD.5.SE
WW.5.SE
WD.1.SE+5d
WW.1.SE+5d
WD.2.SE+5d
WW.2.SE+5d
WD.3.SE+5d
WW.3.SE+5d
WD.4.SE+5d
WW.4.SE+5d
WD.5.SE+5d
WW.5.SE+5d

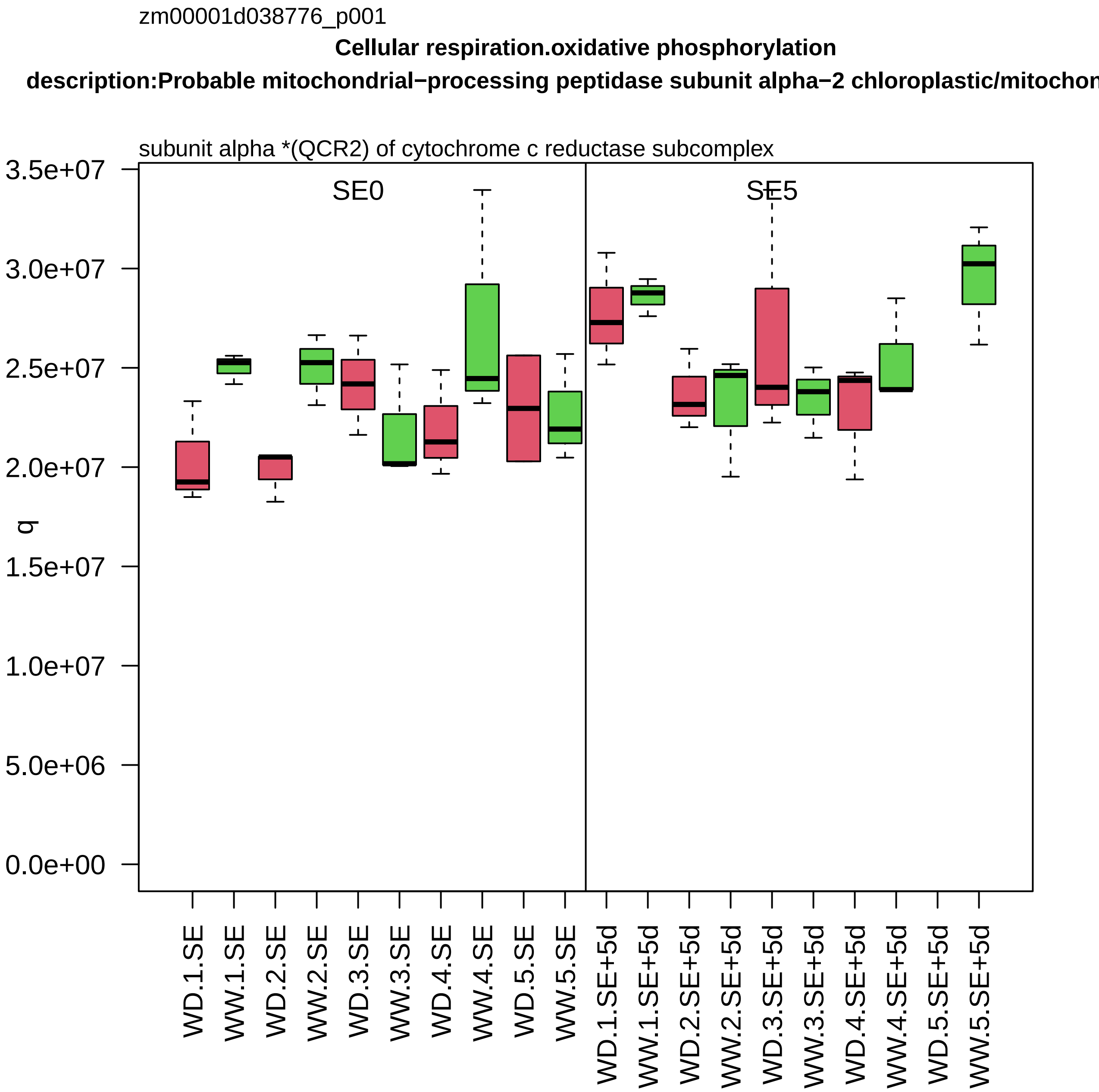

zm00001d038776_p001
Cellular respiration.oxidative phosphorylation
description:Probable mitochondrial−processing peptidase subunit alpha−2 chloroplastic/mitochon
subunit alpha *(QCR2) of cytochrome c reductase subcomplex
SE0
SE5
q
3.5e+07
3.0e+07
2.5e+07
2.0e+07
1.5e+07
1.0e+07
5.0e+06
0.0e+00
WD.1.SE
WW.1.SE
WD.2.SE
WW.2.SE
WD.3.SE
WW.3.SE
WD.4.SE
WW.4.SE
WD.5.SE
WW.5.SE
WD.1.SE+5d
WW.1.SE+5d
WD.2.SE+5d
WW.2.SE+5d
WD.3.SE+5d
WW.3.SE+5d
WD.4.SE+5d
WW.4.SE+5d
WD.5.SE+5d
WW.5.SE+5d

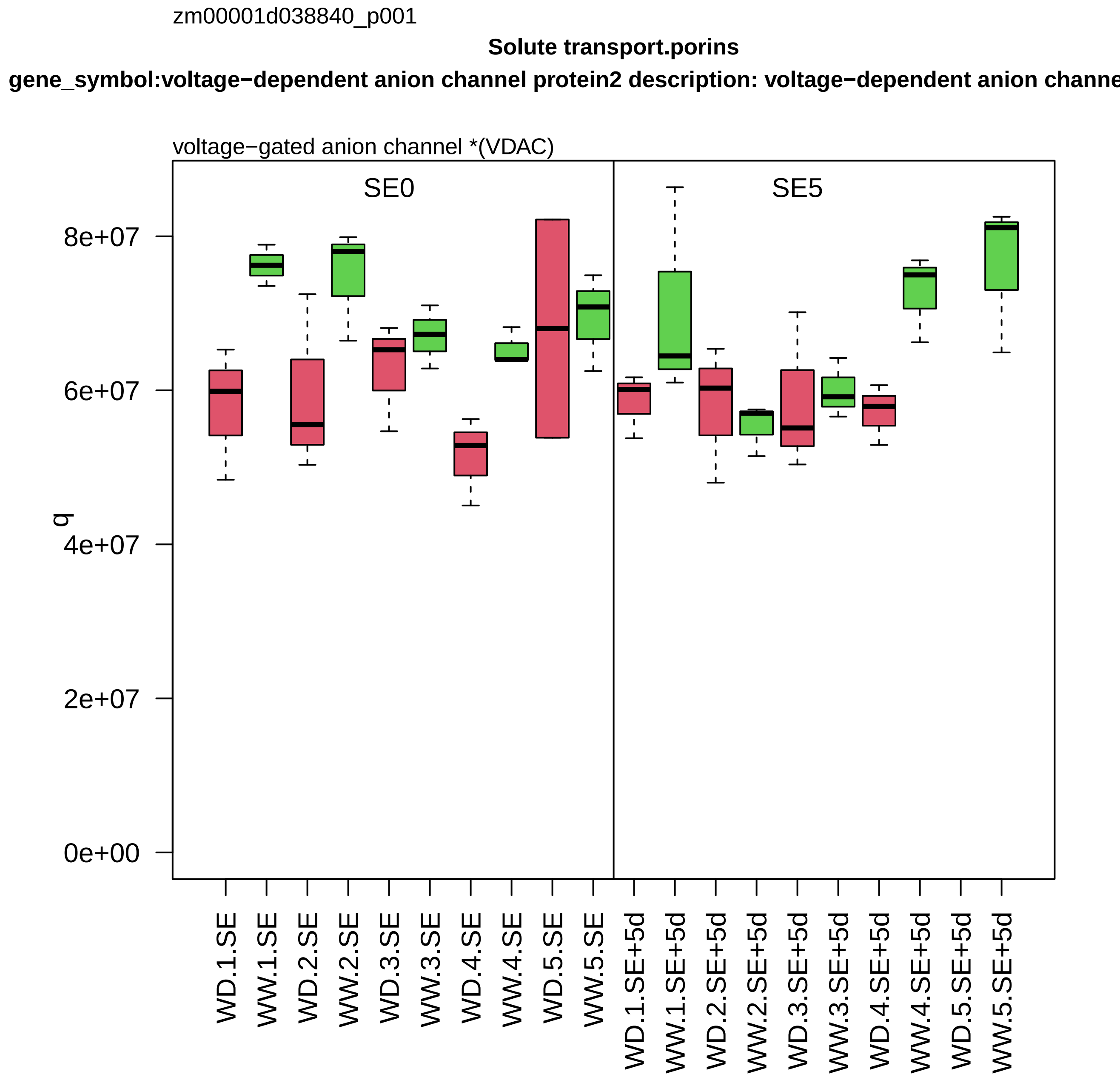

zm00001d038840_p001
Solute transport.porins
gene_symbol:voltage−dependent anion channel protein2 description: voltage−dependent anion channe
voltage−gated anion channel *(VDAC)
SE0
SE5
q
0e+00
2e+07
4e+07
6e+07
8e+07
WD.1.SE
WW.1.SE
WD.2.SE
WW.2.SE
WD.3.SE
WW.3.SE
WD.4.SE
WW.4.SE
WD.5.SE
WW.5.SE
WD.1.SE+5d
WW.1.SE+5d
WD.2.SE+5d
WW.2.SE+5d
WD.3.SE+5d
WW.3.SE+5d
WD.4.SE+5d
WW.4.SE+5d
WD.5.SE+5d
WW.5.SE+5d

zm00001d038841_p010

**Amino acid metabolism.amino acid degradation**

**gene_symbol:putative aldehyde dehydrogenase MIS1 description: putative aldehyde dehydrogenase**

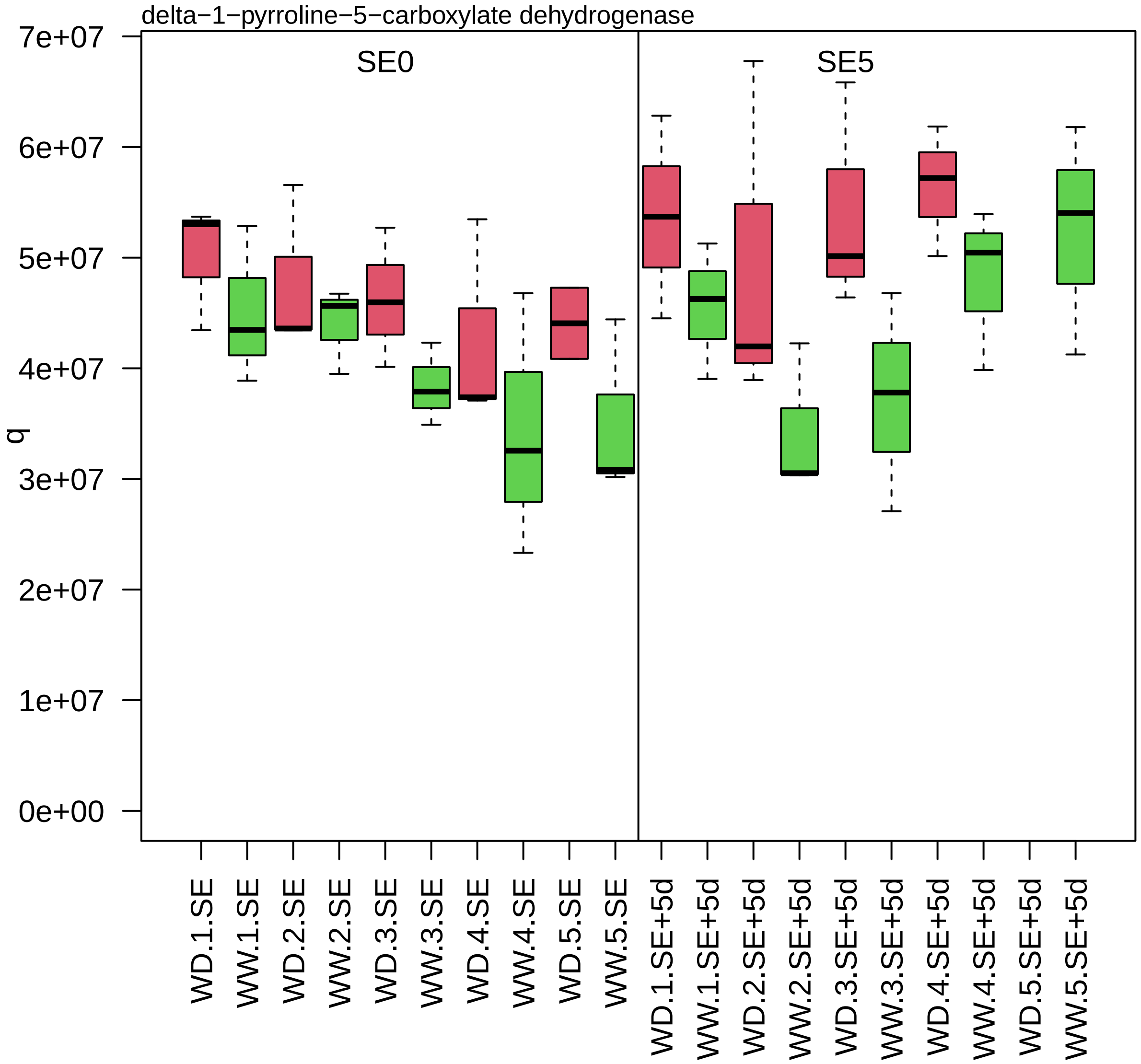

zm00001d038855_p001

**Cell division.DNA replication**

**gene_symbol:flap endonuclease1 description:Flap endonuclease 1**

flap structure−specific endonuclease *(FEN1)

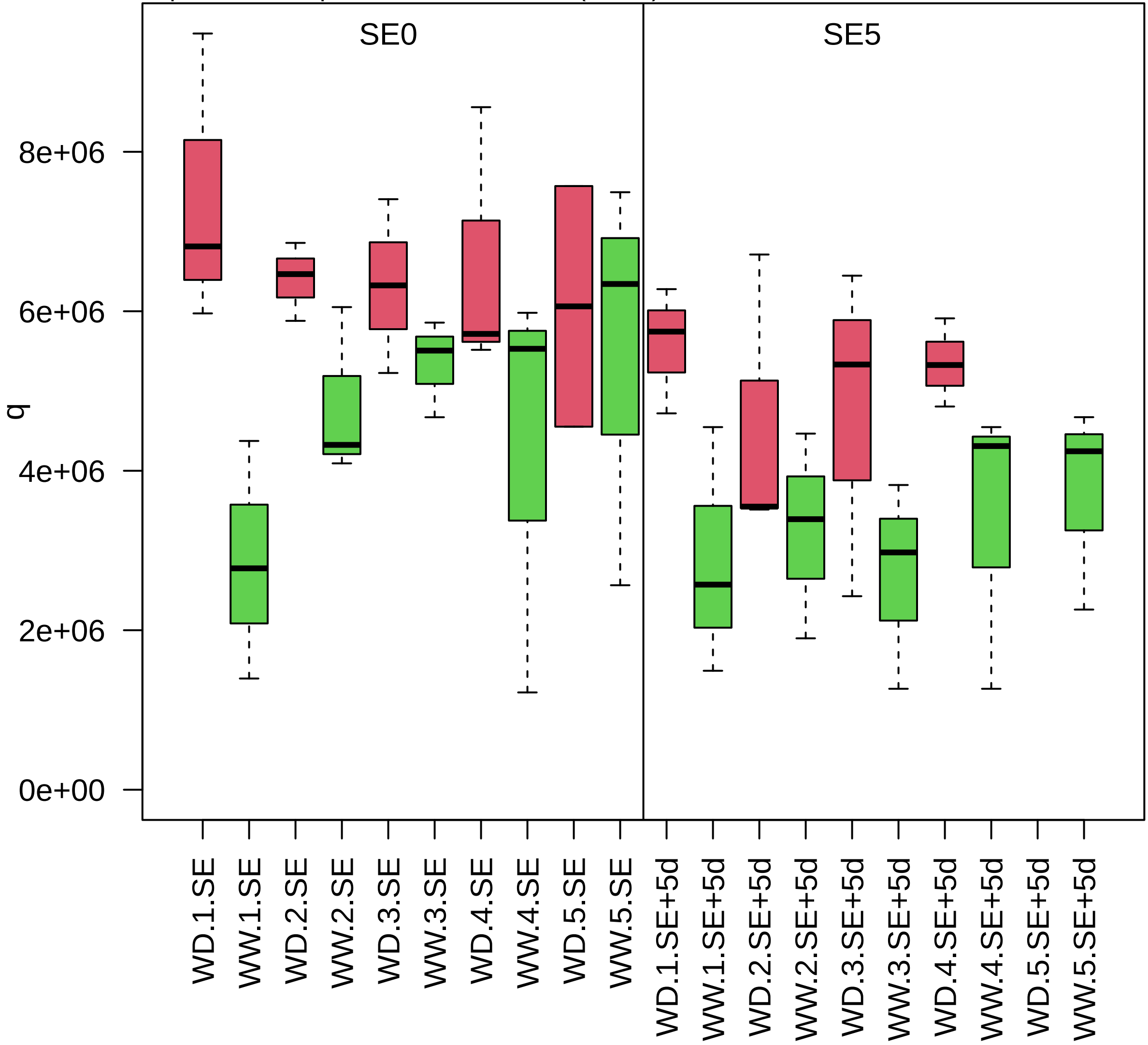

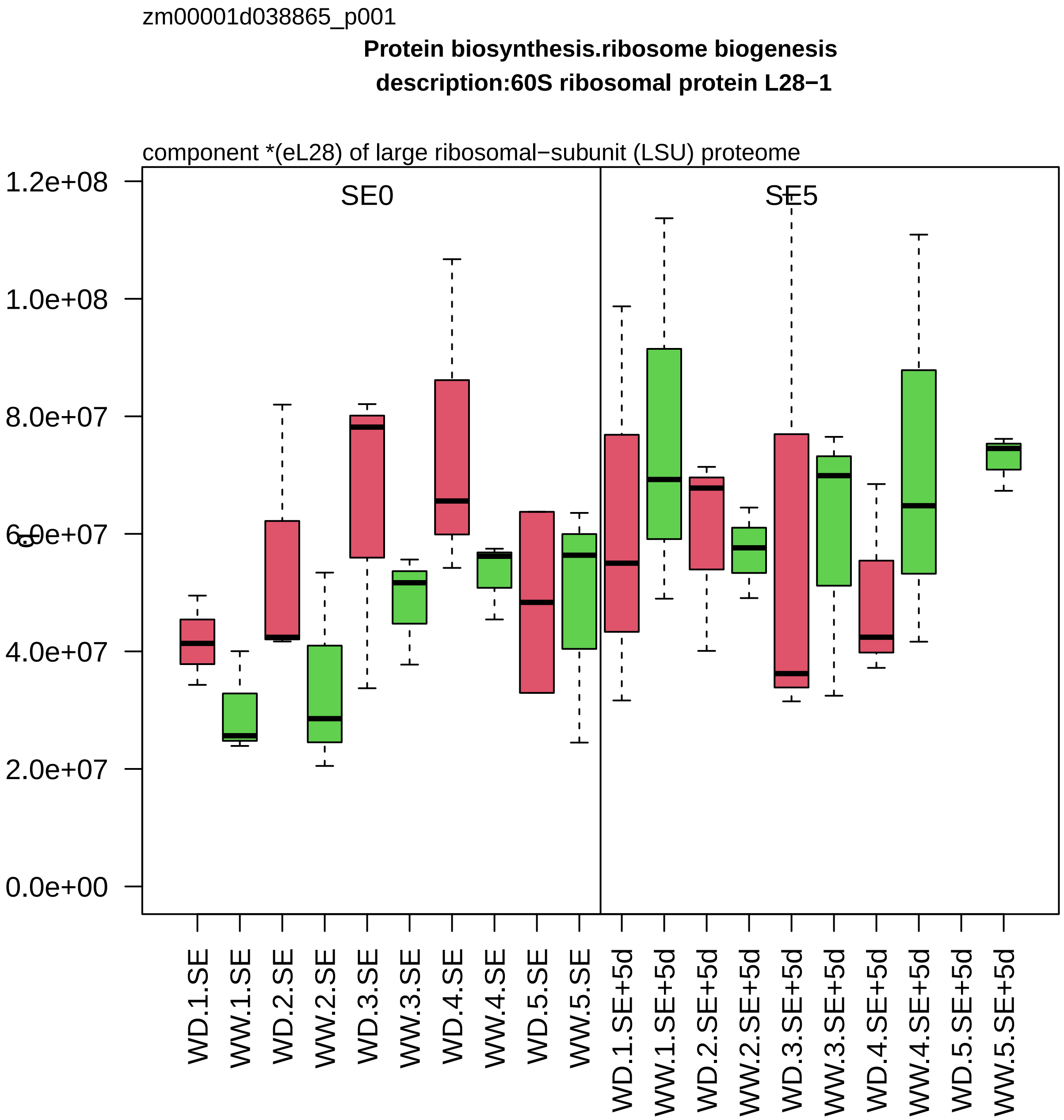
zm00001d038865_p001
Protein biosynthesis.ribosome biogenesis
description:60S ribosomal protein L28−1
component *(eL28) of large ribosomal−subunit (LSU) proteome
SE0
SE5
1.2e+08
1.0e+08
8.0e+07
6.0e+07
4.0e+07
2.0e+07
0.0e+00
WD.1.SE
WW.1.SE
WD.2.SE
WW.2.SE
WD.3.SE
WW.3.SE
WD.4.SE
WW.4.SE
WD.5.SE
WW.5.SE
WD.1.SE+5d
WW.1.SE+5d
WD.2.SE+5d
WW.2.SE+5d
WD.3.SE+5d
WW.3.SE+5d
WD.4.SE+5d
WW.4.SE+5d
WD.5.SE+5d
WW.5.SE+5d

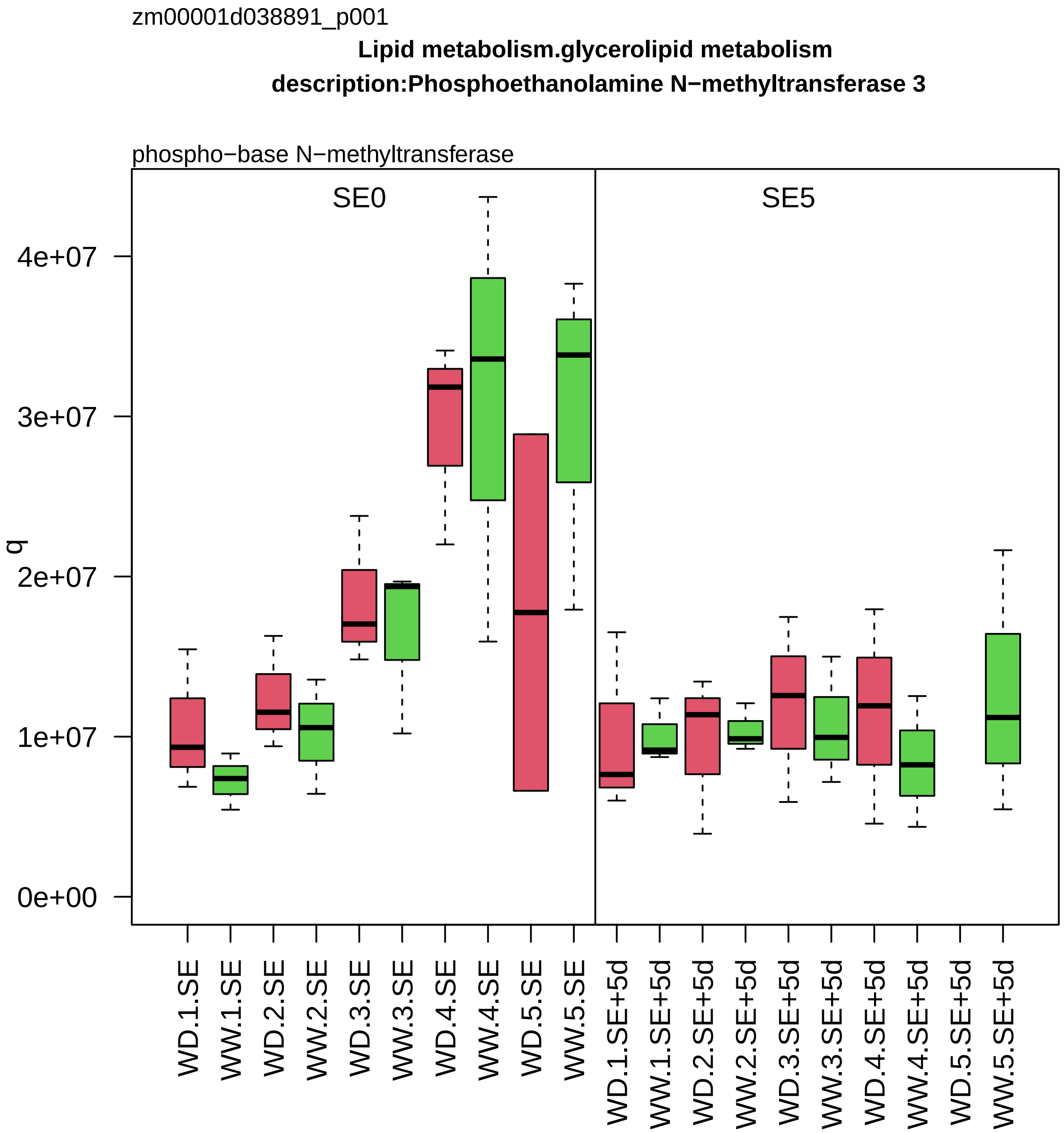

zm00001d038891_p001
Lipid metabolism.glycerolipid metabolism
description:Phosphoethanolamine N−methyltransferase 3
phospho−base N−methyltransferase
SE0
SE5
q
0e+00
1e+07
2e+07
3e+07
4e+07
WD.1.SE
WW.1.SE
WD.2.SE
WW.2.SE
WD.3.SE
WW.3.SE
WD.4.SE
WW.4.SE
WD.5.SE
WW.5.SE
WD.1.SE+5d
WW.1.SE+5d
WD.2.SE+5d
WW.2.SE+5d
WD.3.SE+5d
WW.3.SE+5d
WD.4.SE+5d
WW.4.SE+5d
WD.5.SE+5d
WW.5.SE+5d

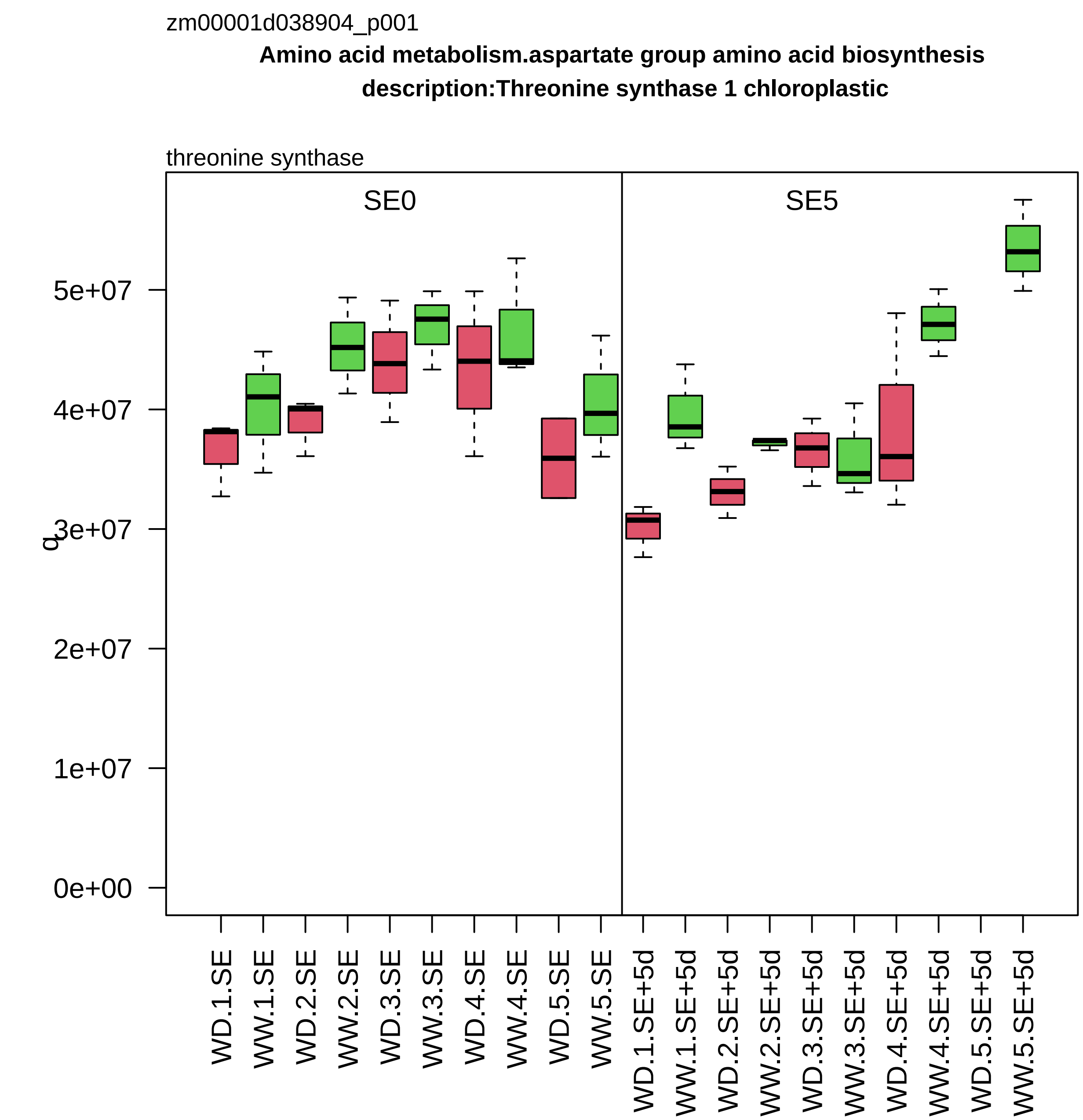

zm00001d038904_p001
Amino acid metabolism.aspartate group amino acid biosynthesis
description:Threonine synthase 1 chloroplastic
threonine synthase
SE0
SE5
q
0e+00
1e+07
2e+07
3e+07
4e+07
5e+07
WD.1.SE
WW.1.SE
WD.2.SE
WW.2.SE
WD.3.SE
WW.3.SE
WD.4.SE
WW.4.SE
WD.5.SE
WW.5.SE
WD.1.SE+5d
WW.1.SE+5d
WD.2.SE+5d
WW.2.SE+5d
WD.3.SE+5d
WW.3.SE+5d
WD.4.SE+5d
WW.4.SE+5d
WD.5.SE+5d
WW.5.SE+5d

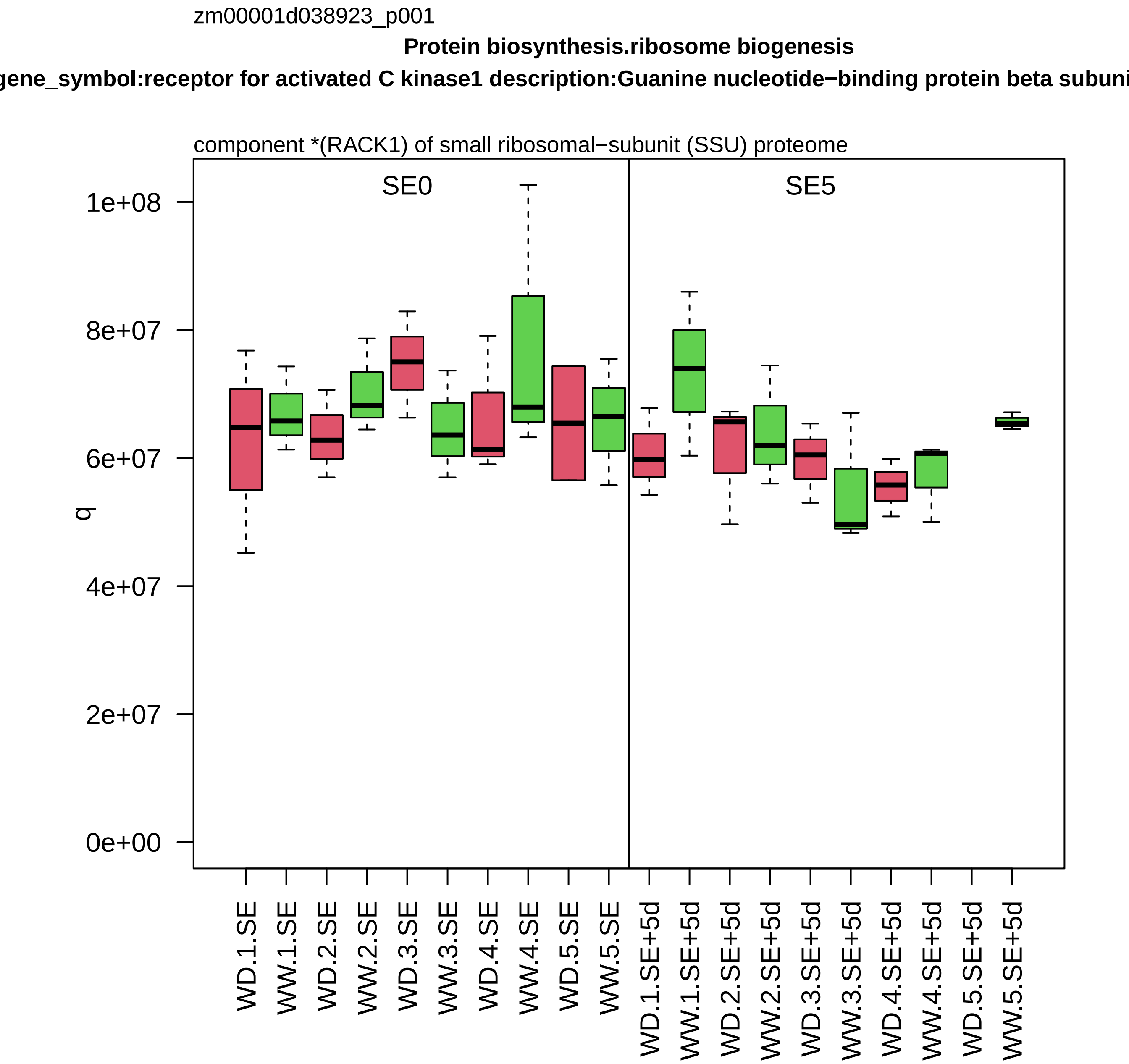

zm00001d038923_p001
Protein biosynthesis.ribosome biogenesis
gene_symbol:receptor for activated C kinase1 description:Guanine nucleotide−binding protein beta subuni
component *(RACK1) of small ribosomal−subunit (SSU) proteome
SE0
SE5
q
0e+00
2e+07
4e+07
6e+07
8e+07
1e+08
WD.1.SE
WW.1.SE
WD.2.SE
WW.2.SE
WD.3.SE
WW.3.SE
WD.4.SE
WW.4.SE
WD.5.SE
WW.5.SE
WD.1.SE+5d
WW.1.SE+5d
WD.2.SE+5d
WW.2.SE+5d
WD.3.SE+5d
WW.3.SE+5d
WD.4.SE+5d
WW.4.SE+5d
WD.5.SE+5d
WW.5.SE+5d

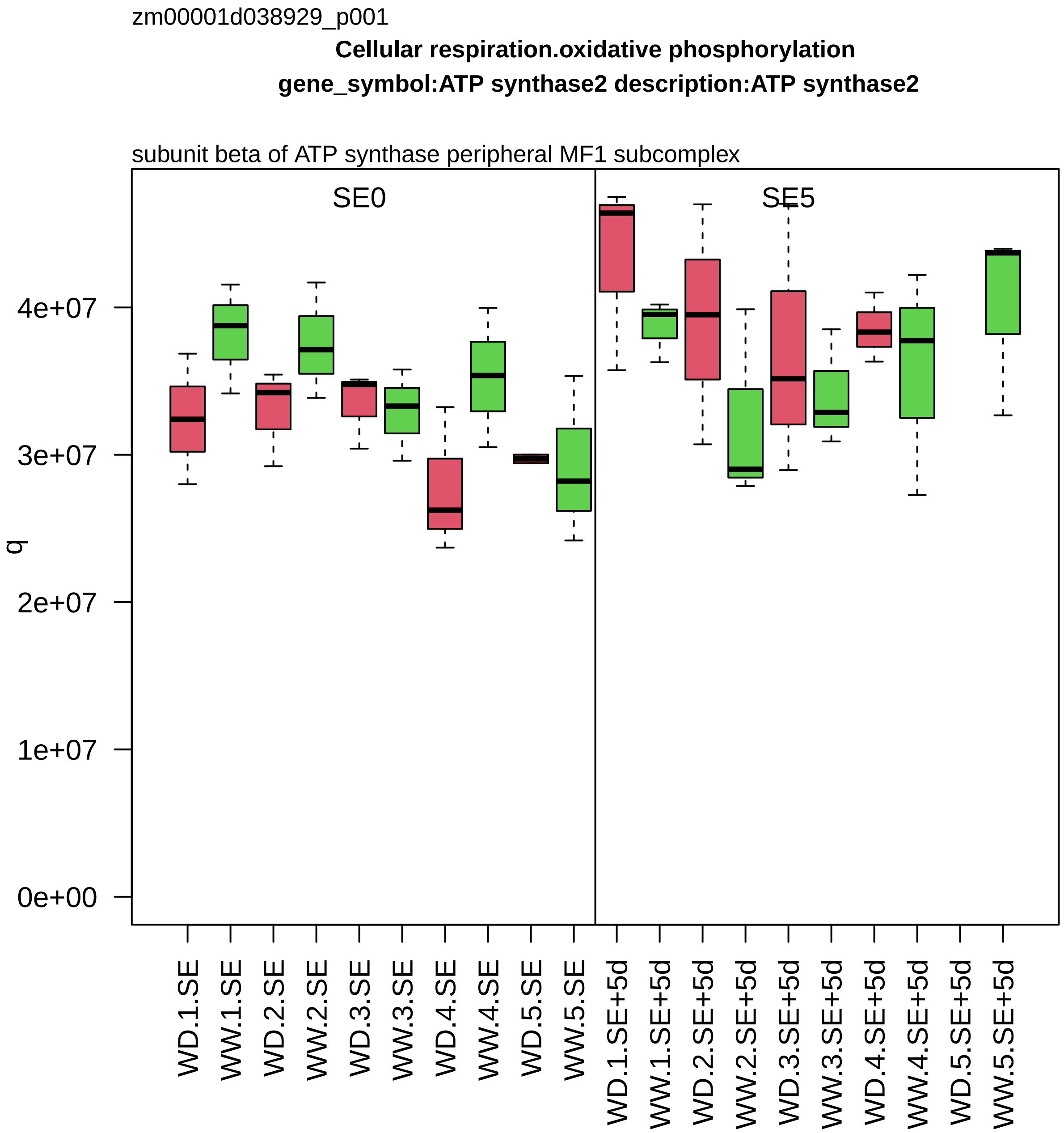
zm00001d038929_p001
Cellular respiration.oxidative phosphorylation
gene_symbol:ATP synthase2 description:ATP synthase2
subunit beta of ATP synthase peripheral MF1 subcomplex
SE0
SE5
q
0e+00
1e+07
2e+07
3e+07
4e+07
WD.1.SE
WW.1.SE
WD.2.SE
WW.2.SE
WD.3.SE
WW.3.SE
WD.4.SE
WW.4.SE
WD.5.SE
WW.5.SE
WD.1.SE+5d
WW.1.SE+5d
WD.2.SE+5d
WW.2.SE+5d
WD.3.SE+5d
WW.3.SE+5d
WD.4.SE+5d
WW.4.SE+5d
WD.5.SE+5d
WW.5.SE+5d

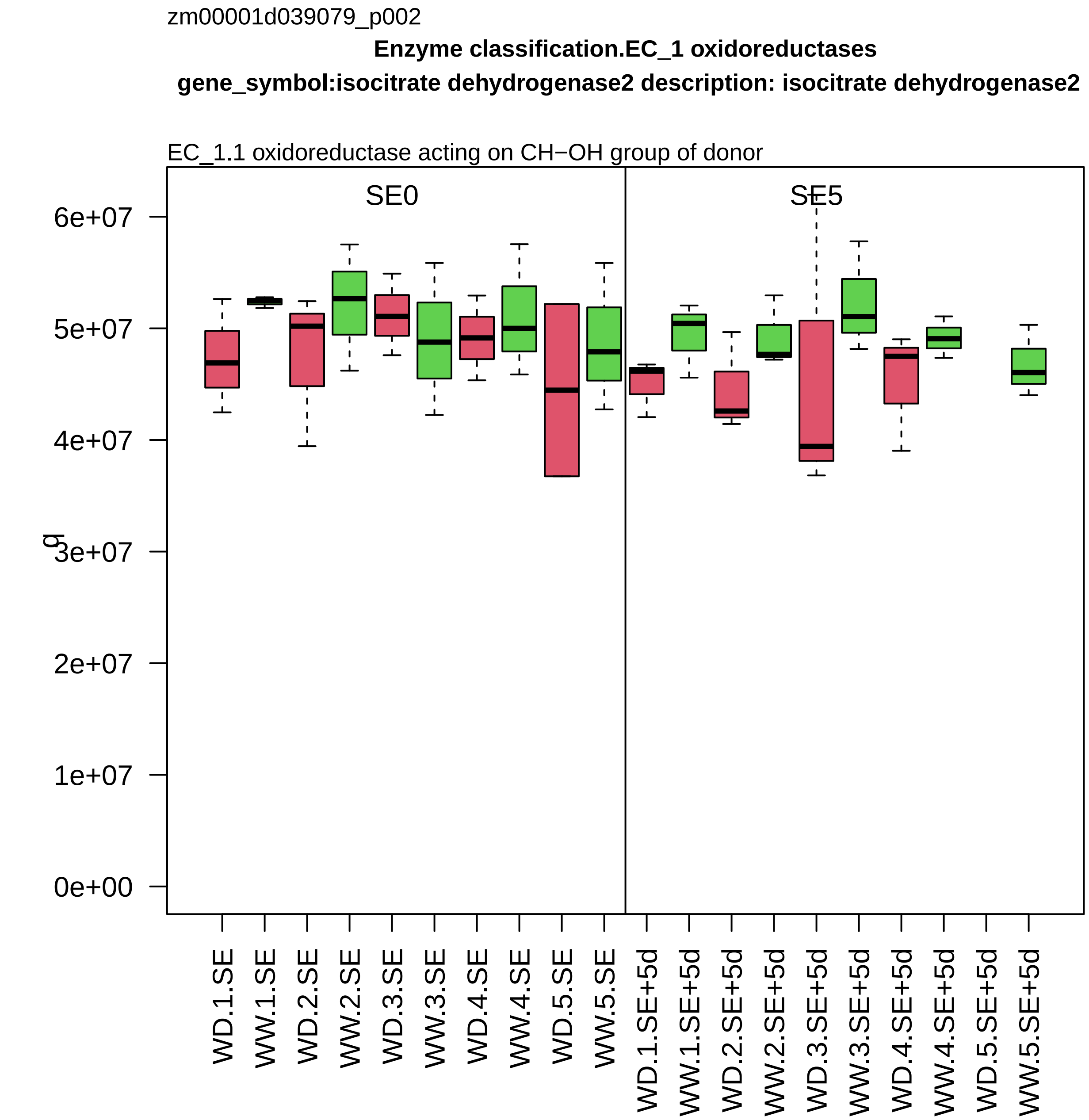

zm00001d039079_p002
Enzyme classification.EC_1 oxidoreductases
gene_symbol:isocitrate dehydrogenase2 description: isocitrate dehydrogenase2
EC_1.1 oxidoreductase acting on CH−OH group of donor
SE0
SE5
g
6e+07
5e+07
4e+07
3e+07
2e+07
1e+07
0e+00
WD.1.SE
WW.1.SE
WD.2.SE
WW.2.SE
WD.3.SE
WW.3.SE
WD.4.SE
WW.4.SE
WD.5.SE
WW.5.SE
WD.1.SE+5d
WW.1.SE+5d
WD.2.SE+5d
WW.2.SE+5d
WD.3.SE+5d
WW.3.SE+5d
WD.4.SE+5d
WW.4.SE+5d
WD.5.SE+5d
WW.5.SE+5d

zm00001d039131_p002

**Carbohydrate metabolism.starch metabolism**

**gene_symbol:ADP glucose pyrophosphorylase2 description: ADP glucose pyrophosphorylase**

large subunit of ADP−glucose pyrophosphorylase

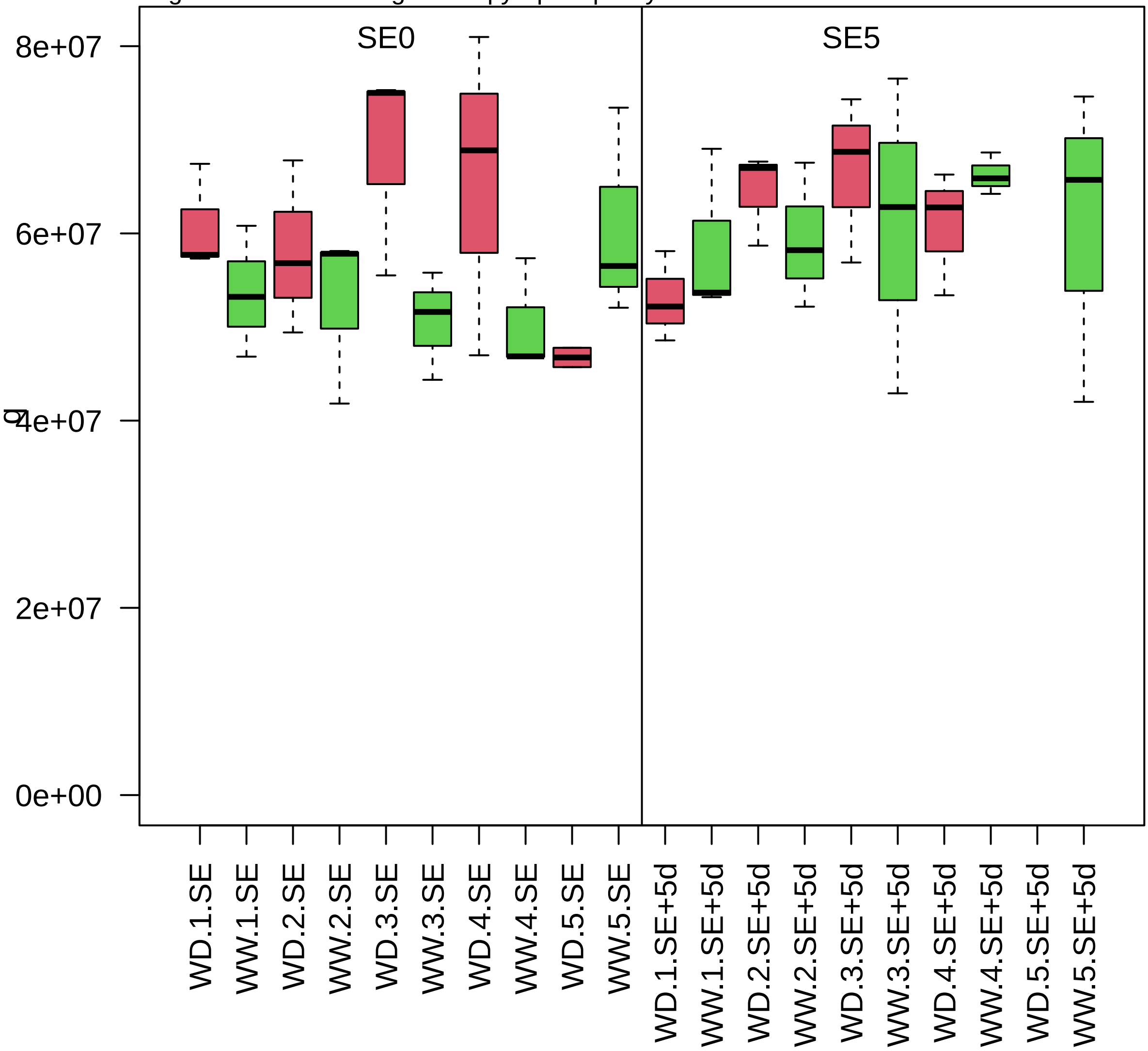

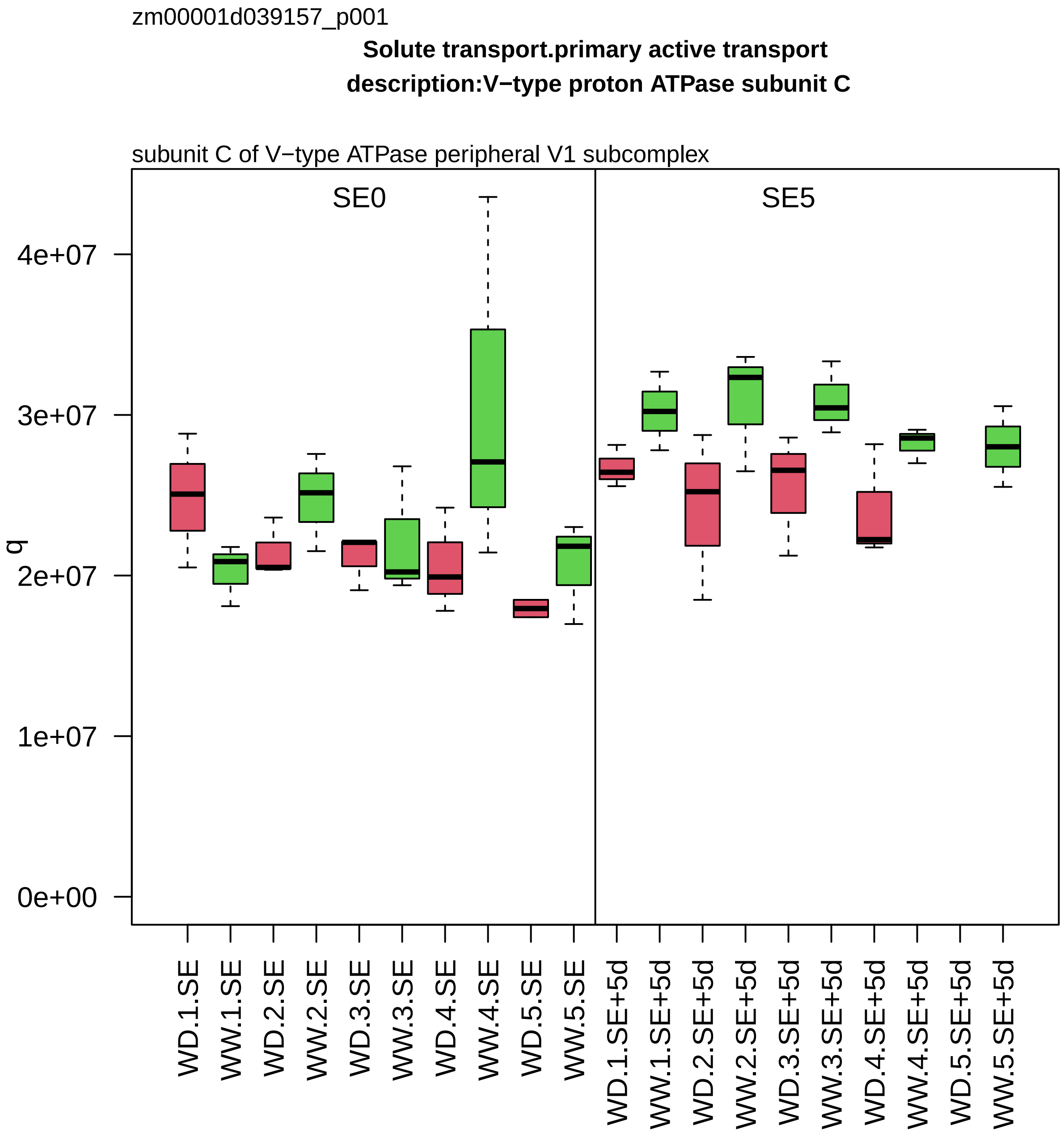

zm00001d039157_p001
Solute transport.primary active transport
description:V−type proton ATPase subunit C
subunit C of V−type ATPase peripheral V1 subcomplex
SE0
SE5
q
4e+07
3e+07
2e+07
1e+07
0e+00
WD.1.SE
WW.1.SE
WD.2.SE
WW.2.SE
WD.3.SE
WW.3.SE
WD.4.SE
WW.4.SE
WD.5.SE
WW.5.SE
WD.1.SE+5d
WW.1.SE+5d
WD.2.SE+5d
WW.2.SE+5d
WD.3.SE+5d
WW.3.SE+5d
WD.4.SE+5d
WW.4.SE+5d
WD.5.SE+5d
WW.5.SE+5d

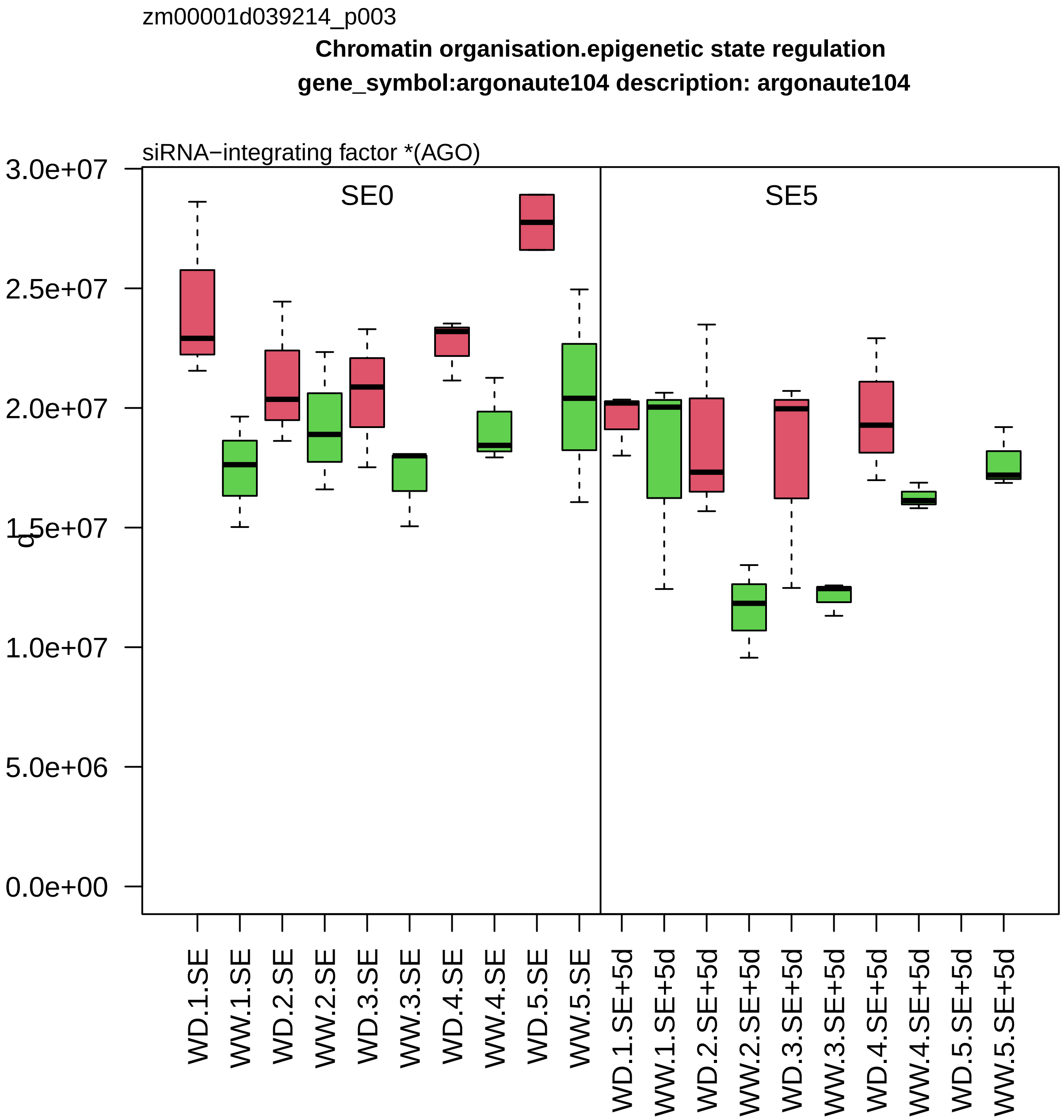

zm00001d039214_p003
Chromatin organisation.epigenetic state regulation
gene_symbol:argonaute104 description: argonaute104
siRNA−integrating factor *(AGO)
SE0
SE5
3.0e+07
2.5e+07
2.0e+07
1.5e+07
1.0e+07
5.0e+06
0.0e+00
WD.1.SE
WW.1.SE
WD.2.SE
WW.2.SE
WD.3.SE
WW.3.SE
WD.4.SE
WW.4.SE
WD.5.SE
WW.5.SE
WD.1.SE+5d
WW.1.SE+5d
WD.2.SE+5d
WW.2.SE+5d
WD.3.SE+5d
WW.3.SE+5d
WD.4.SE+5d
WW.4.SE+5d
WD.5.SE+5d
WW.5.SE+5d

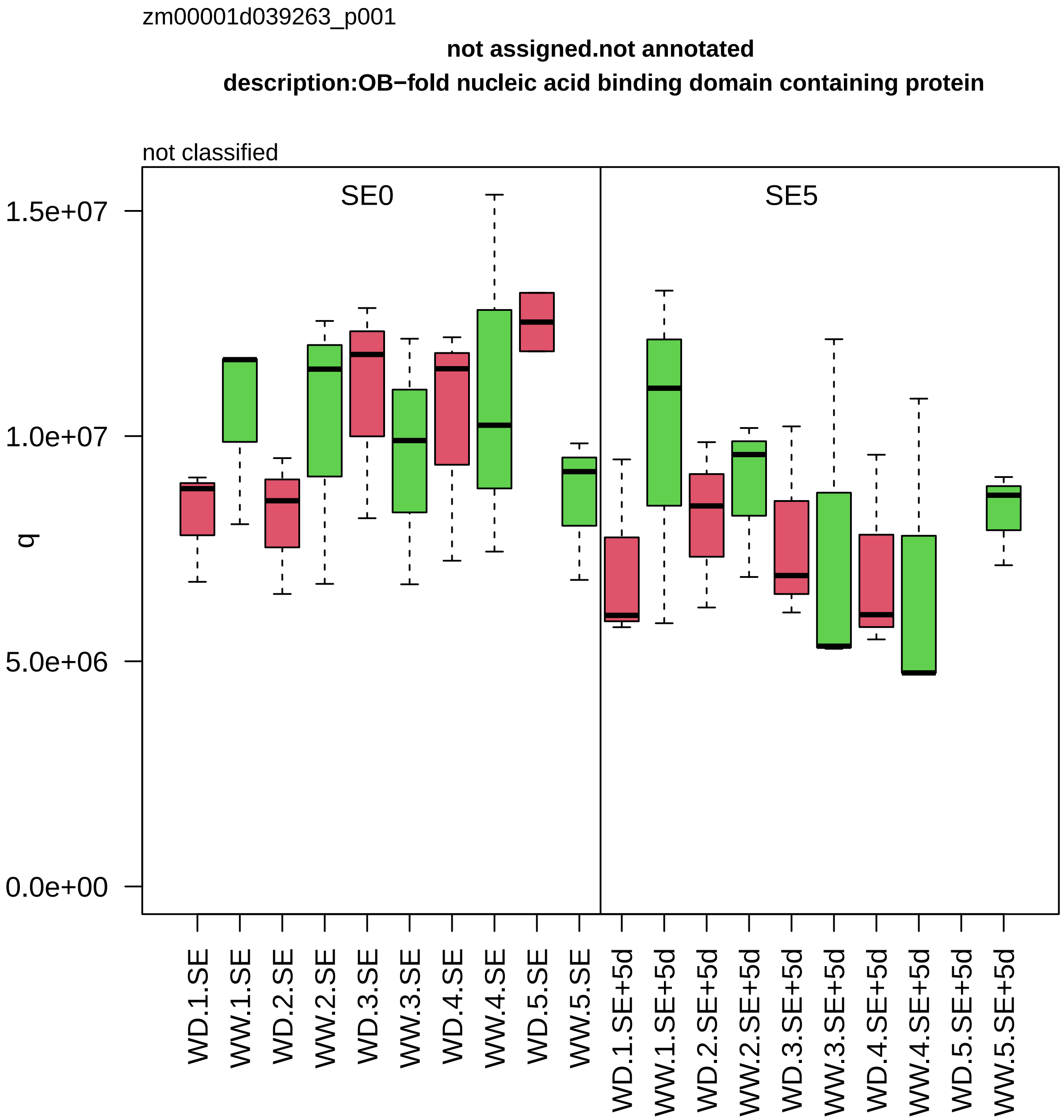

zm00001d039263_p001
not assigned.not annotated
description:OB−fold nucleic acid binding domain containing protein
not classified
SE0
SE5
1.5e+07
1.0e+07
5.0e+06
0.0e+00
q
WD.1.SE
WW.1.SE
WD.2.SE
WW.2.SE
WD.3.SE
WW.3.SE
WD.4.SE
WW.4.SE
WD.5.SE
WW.5.SE
WD.1.SE+5d
WW.1.SE+5d
WD.2.SE+5d
WW.2.SE+5d
WD.3.SE+5d
WW.3.SE+5d
WD.4.SE+5d
WW.4.SE+5d
WD.5.SE+5d
WW.5.SE+5d

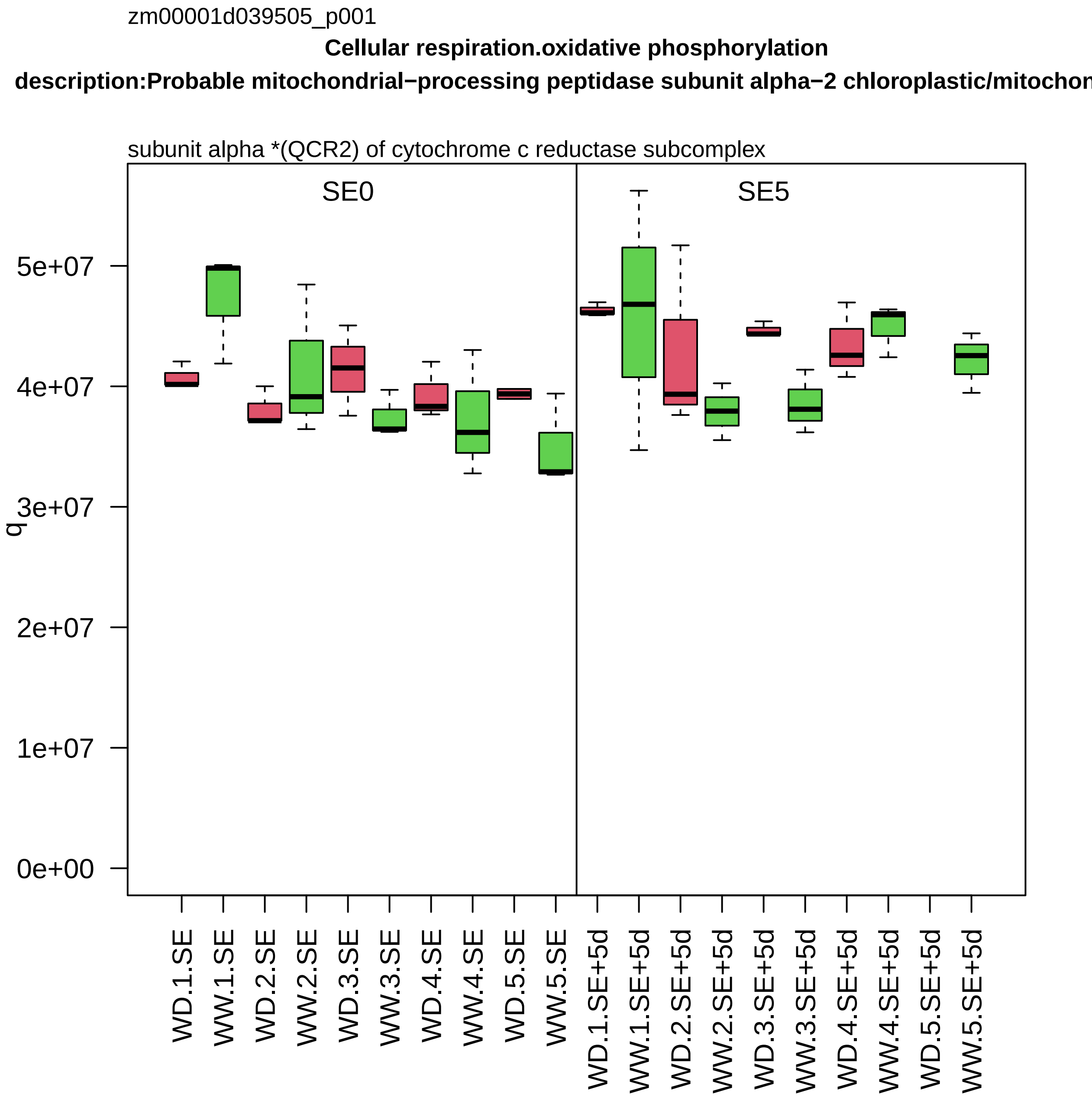

zm00001d039505_p001
Cellular respiration.oxidative phosphorylation
description:Probable mitochondrial−processing peptidase subunit alpha−2 chloroplastic/mitochon
subunit alpha *(QCR2) of cytochrome c reductase subcomplex
SE0
SE5
q
5e+07
4e+07
3e+07
2e+07
1e+07
0e+00
WD.1.SE
WW.1.SE
WD.2.SE
WW.2.SE
WD.3.SE
WW.3.SE
WD.4.SE
WW.4.SE
WD.5.SE
WW.5.SE
WD.1.SE+5d
WW.1.SE+5d
WD.2.SE+5d
WW.2.SE+5d
WD.3.SE+5d
WW.3.SE+5d
WD.4.SE+5d
WW.4.SE+5d
WD.5.SE+5d
WW.5.SE+5d

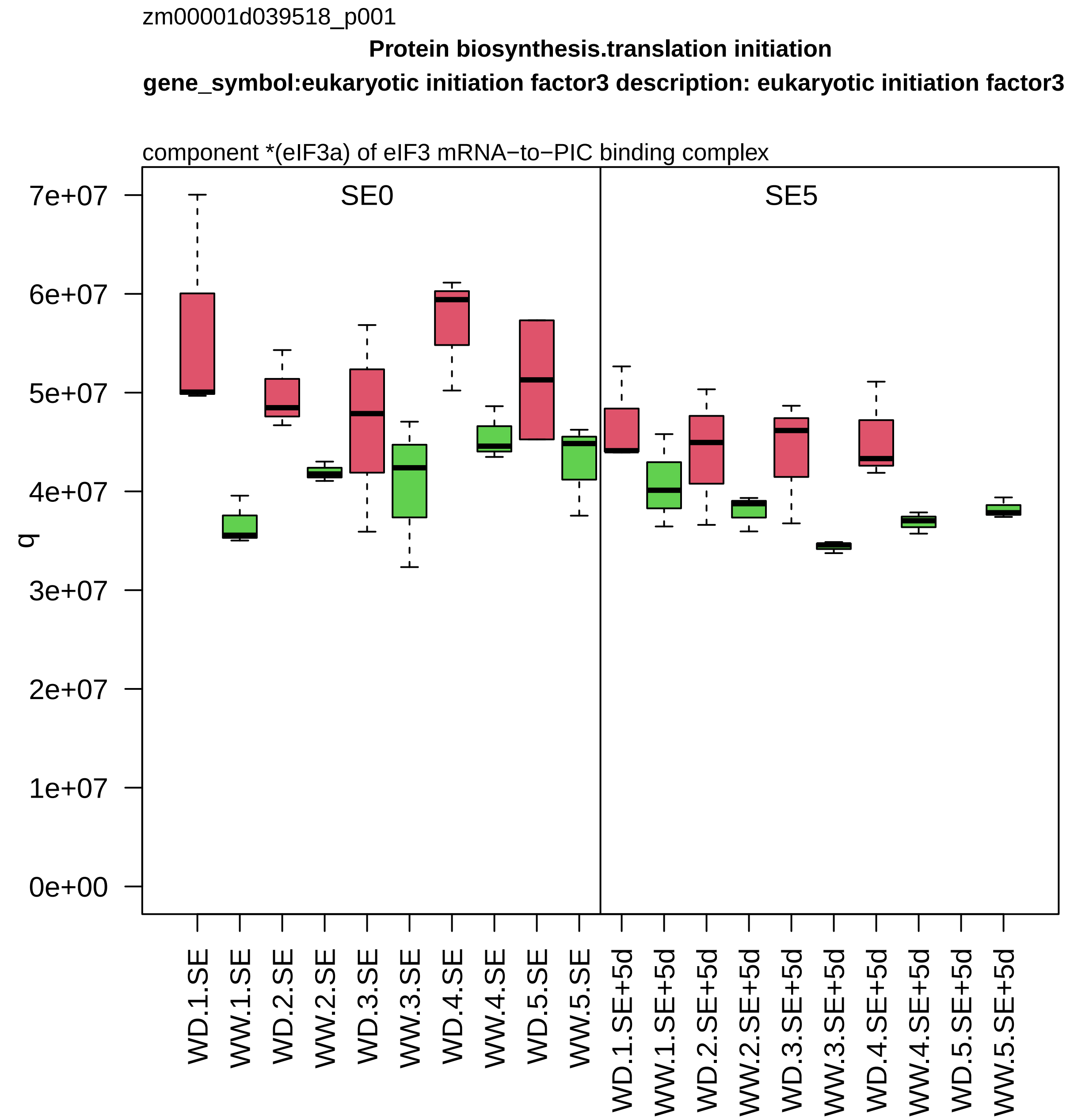
zm00001d039518_p001
Protein biosynthesis.translation initiation
gene_symbol:eukaryotic initiation factor3 description: eukaryotic initiation factor3
component *(eIF3a) of eIF3 mRNA−to−PIC binding complex
SE0
SE5
q
0e+00
1e+07
2e+07
3e+07
4e+07
5e+07
6e+07
7e+07
WD.1.SE
WW.1.SE
WD.2.SE
WW.2.SE
WD.3.SE
WW.3.SE
WD.4.SE
WW.4.SE
WD.5.SE
WW.5.SE
WD.1.SE+5d
WW.1.SE+5d
WD.2.SE+5d
WW.2.SE+5d
WD.3.SE+5d
WW.3.SE+5d
WD.4.SE+5d
WW.4.SE+5d
WD.5.SE+5d
WW.5.SE+5d

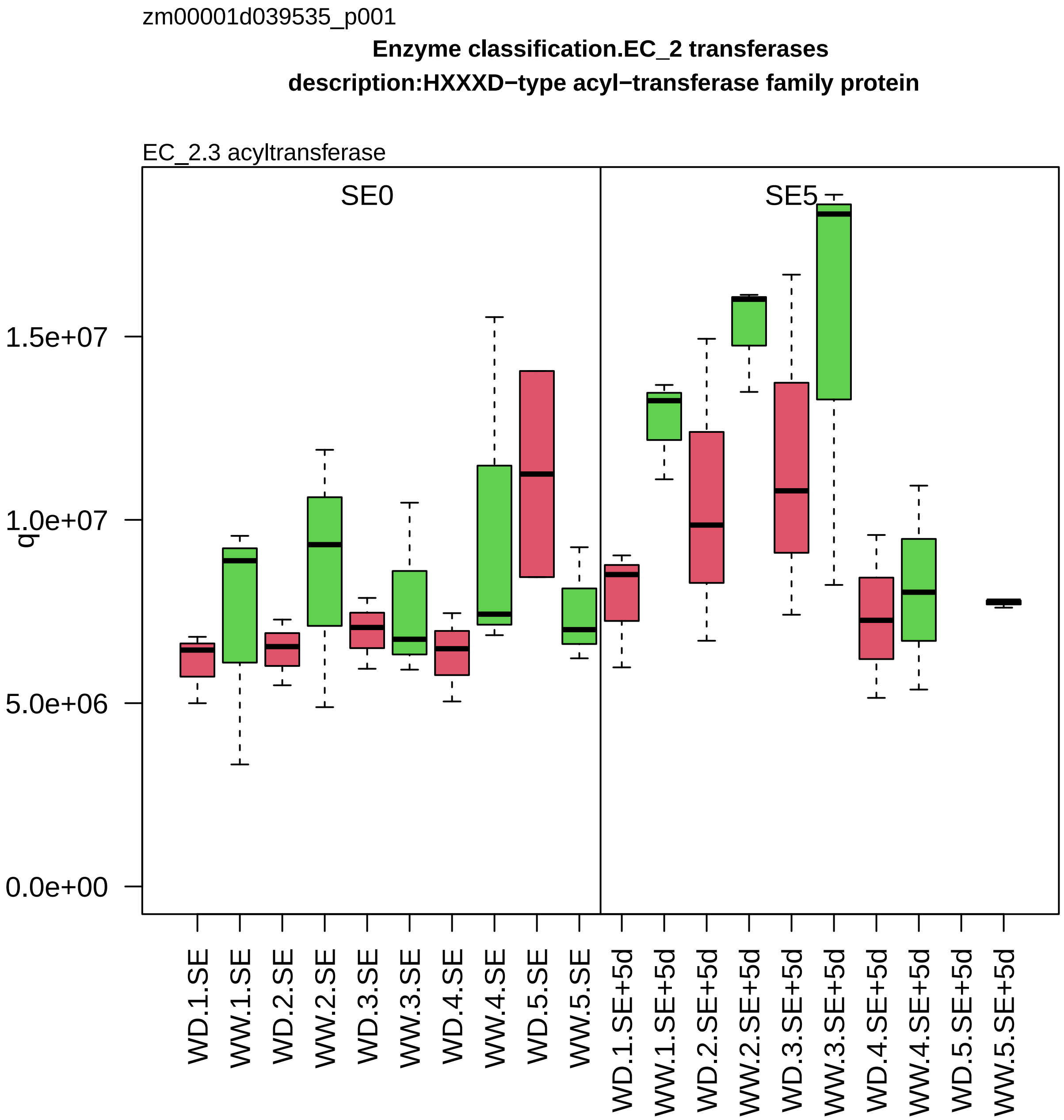
zm00001d039535_p001
Enzyme classification.EC_2 transferases
description:HXXXD−type acyl−transferase family protein
EC_2.3 acyltransferase
SE0
SE5
q
0.0e+00
5.0e+06
1.0e+07
1.5e+07
WD.1.SE
WW.1.SE
WD.2.SE
WW.2.SE
WD.3.SE
WW.3.SE
WD.4.SE
WW.4.SE
WD.5.SE
WW.5.SE
WD.1.SE+5d
WW.1.SE+5d
WD.2.SE+5d
WW.2.SE+5d
WD.3.SE+5d
WW.3.SE+5d
WD.4.SE+5d
WW.4.SE+5d
WD.5.SE+5d
WW.5.SE+5d

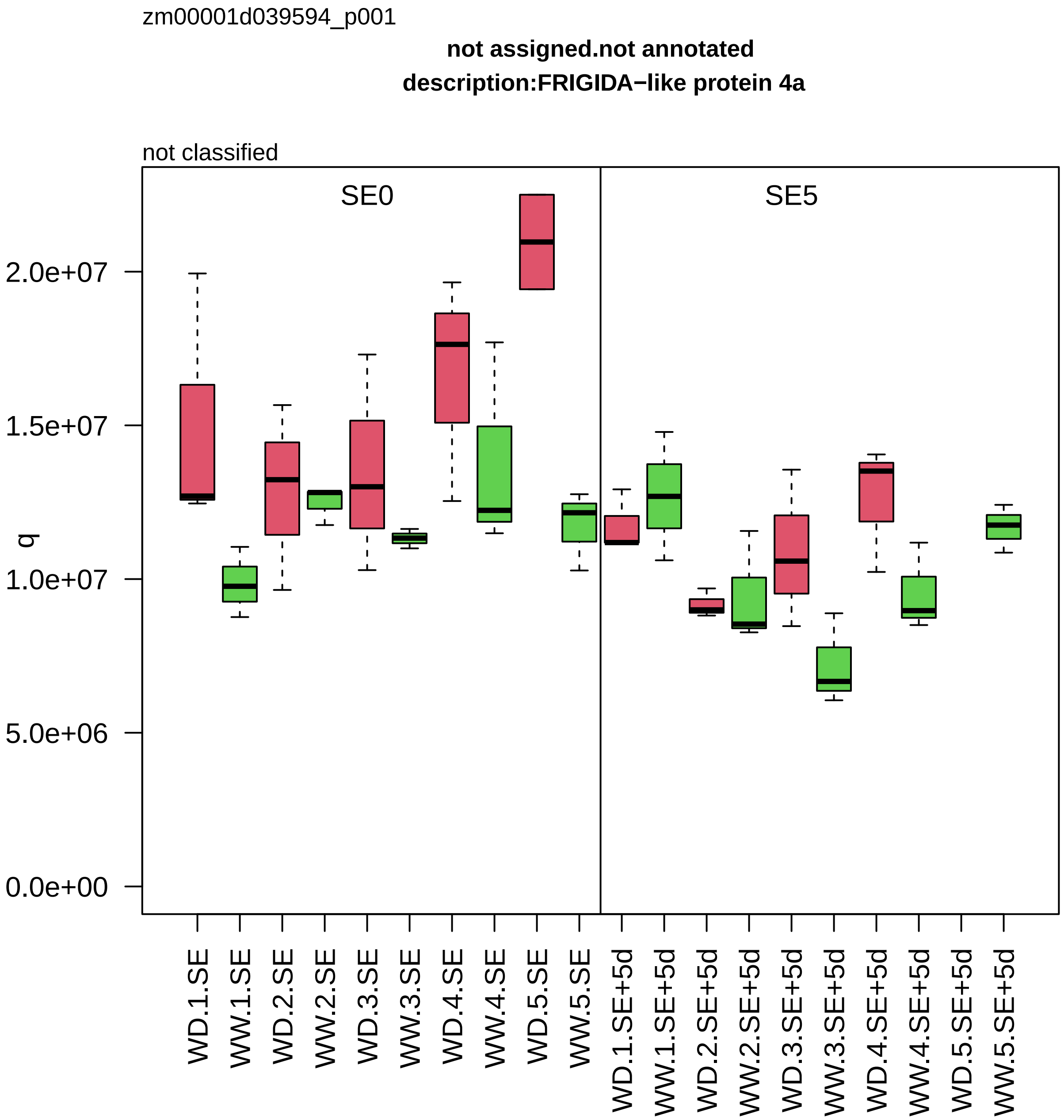

zm00001d039594_p001
not assigned.not annotated
description:FRIGIDA−like protein 4a
not classified
SE0
SE5
q
2.0e+07
1.5e+07
1.0e+07
5.0e+06
0.0e+00
WD.1.SE
WW.1.SE
WD.2.SE
WW.2.SE
WD.3.SE
WW.3.SE
WD.4.SE
WW.4.SE
WD.5.SE
WW.5.SE
WD.1.SE+5d
WW.1.SE+5d
WD.2.SE+5d
WW.2.SE+5d
WD.3.SE+5d
WW.3.SE+5d
WD.4.SE+5d
WW.4.SE+5d
WD.5.SE+5d
WW.5.SE+5d

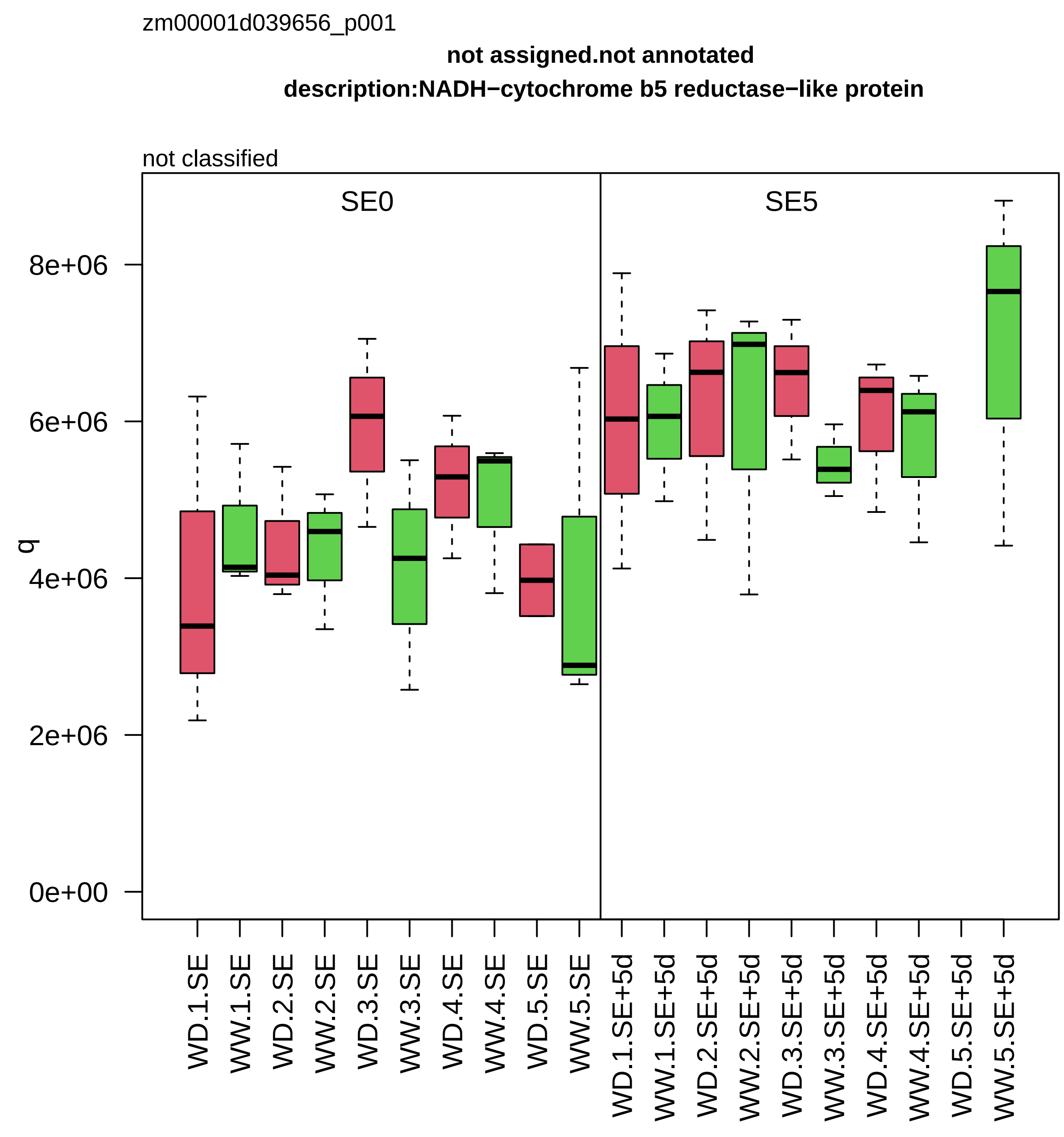

zm00001d039656_p001
not assigned.not annotated
description:NADH−cytochrome b5 reductase−like protein
not classified
SE0
SE5
q
8e+06
6e+06
4e+06
2e+06
0e+00
WD.1.SE
WW.1.SE
WD.2.SE
WW.2.SE
WD.3.SE
WW.3.SE
WD.4.SE
WW.4.SE
WD.5.SE
WW.5.SE
WD.1.SE+5d
WW.1.SE+5d
WD.2.SE+5d
WW.2.SE+5d
WD.3.SE+5d
WW.3.SE+5d
WD.4.SE+5d
WW.4.SE+5d
WD.5.SE+5d
WW.5.SE+5d

zm00001d039663_p001

**RNA biosynthesis.RNA polymerase II−dependent transcription**
**description:Alba DNA/RNA−binding protein**

R−loop reader protein *(ALBA1/2)

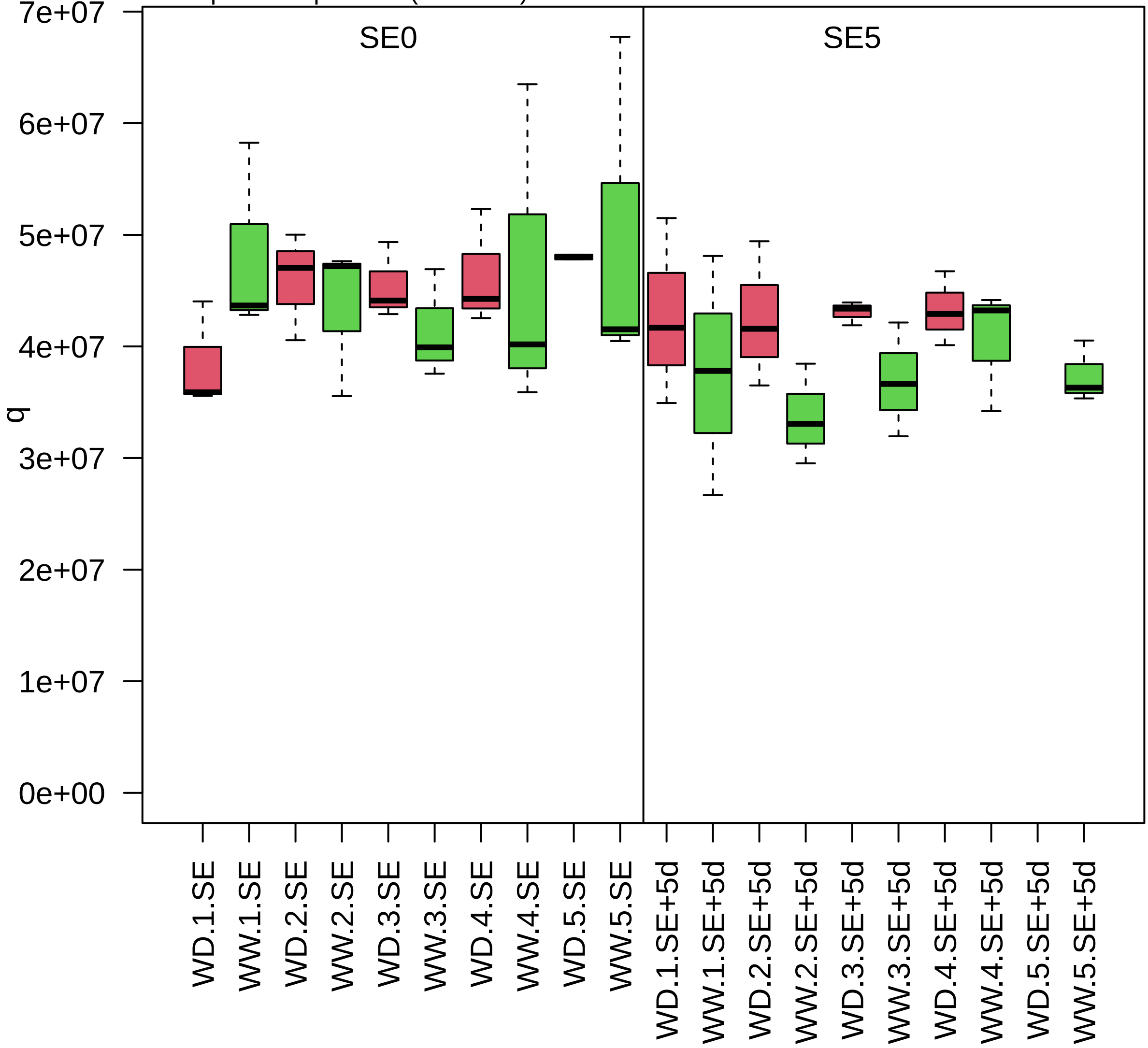

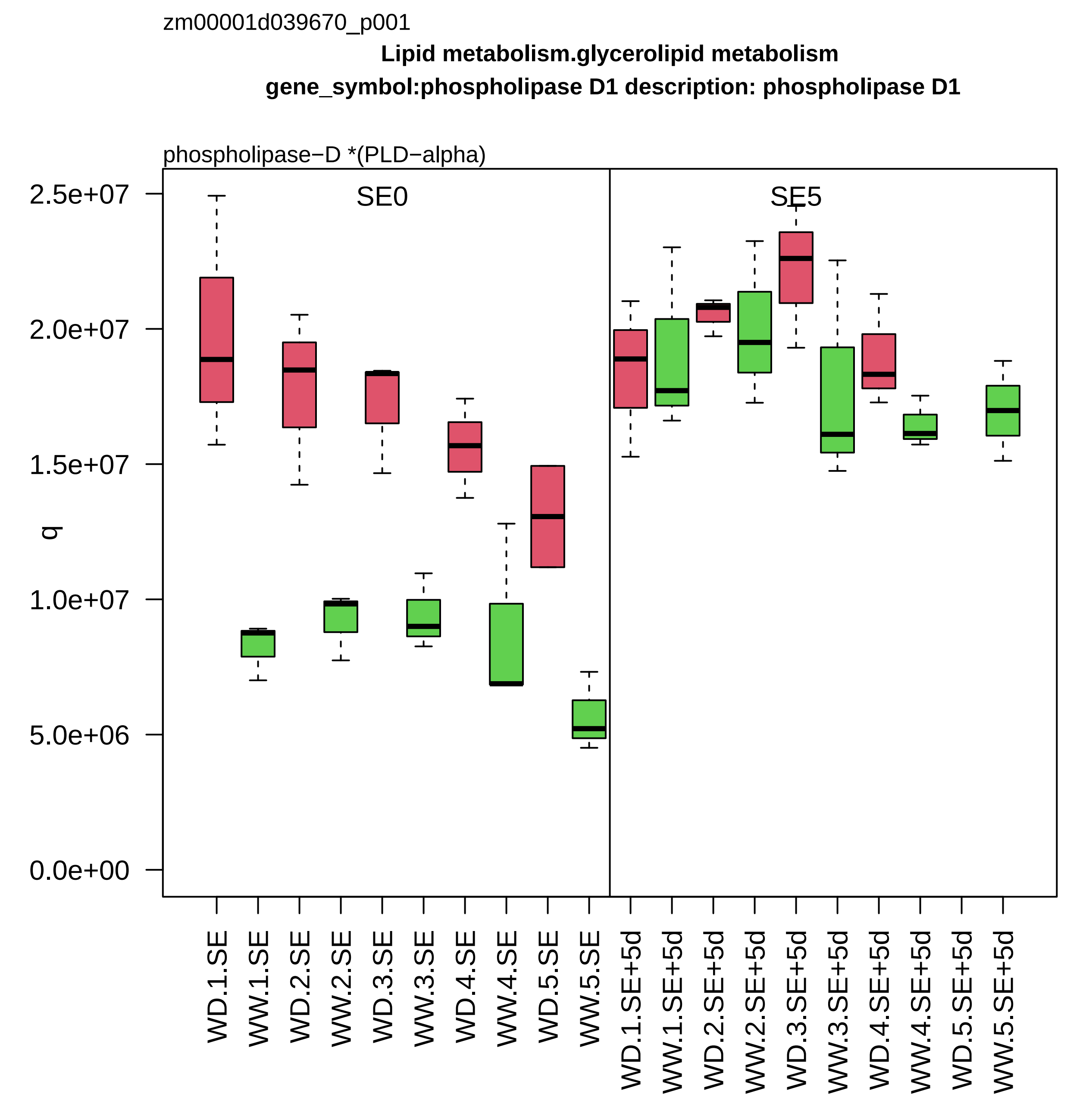

zm00001d039670_p001
Lipid metabolism.glycerolipid metabolism
gene_symbol:phospholipase D1 description: phospholipase D1
phospholipase−D *(PLD−alpha)
SE0
SE5
q
2.5e+07
2.0e+07
1.5e+07
1.0e+07
5.0e+06
0.0e+00
WD.1.SE
WW.1.SE
WD.2.SE
WW.2.SE
WD.3.SE
WW.3.SE
WD.4.SE
WW.4.SE
WD.5.SE
WW.5.SE
WD.1.SE+5d
WW.1.SE+5d
WD.2.SE+5d
WW.2.SE+5d
WD.3.SE+5d
WW.3.SE+5d
WD.4.SE+5d
WW.4.SE+5d
WD.5.SE+5d
WW.5.SE+5d

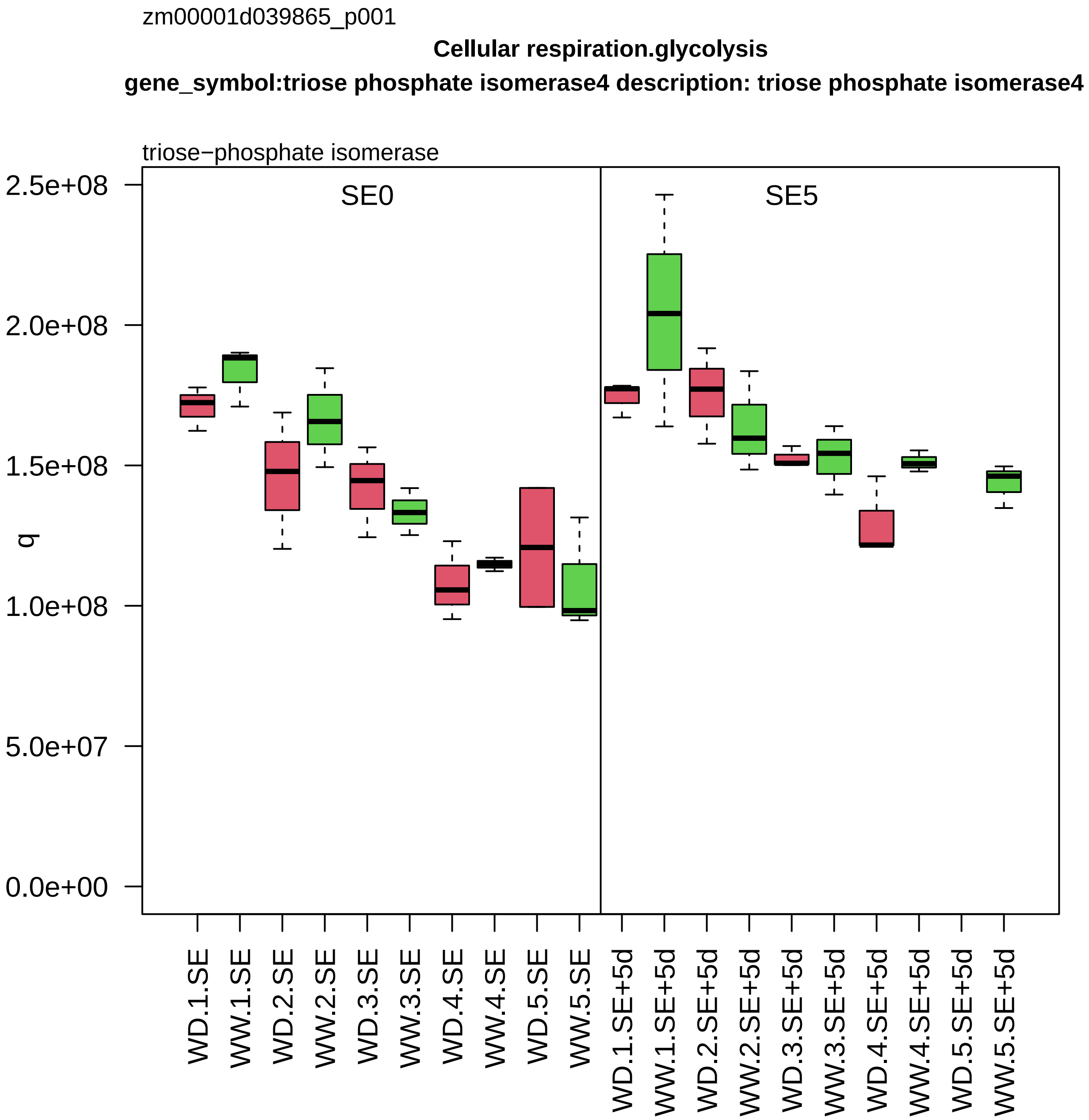
zm00001d039865_p001
Cellular respiration.glycolysis
gene_symbol:triose phosphate isomerase4 description: triose phosphate isomerase4
triose−phosphate isomerase
SE0
SE5
q
2.5e+08
2.0e+08
1.5e+08
1.0e+08
5.0e+07
0.0e+00
WD.1.SE
WW.1.SE
WD.2.SE
WW.2.SE
WD.3.SE
WW.3.SE
WD.4.SE
WW.4.SE
WD.5.SE
WW.5.SE
WD.1.SE+5d
WW.1.SE+5d
WD.2.SE+5d
WW.2.SE+5d
WD.3.SE+5d
WW.3.SE+5d
WD.4.SE+5d
WW.4.SE+5d
WD.5.SE+5d
WW.5.SE+5d

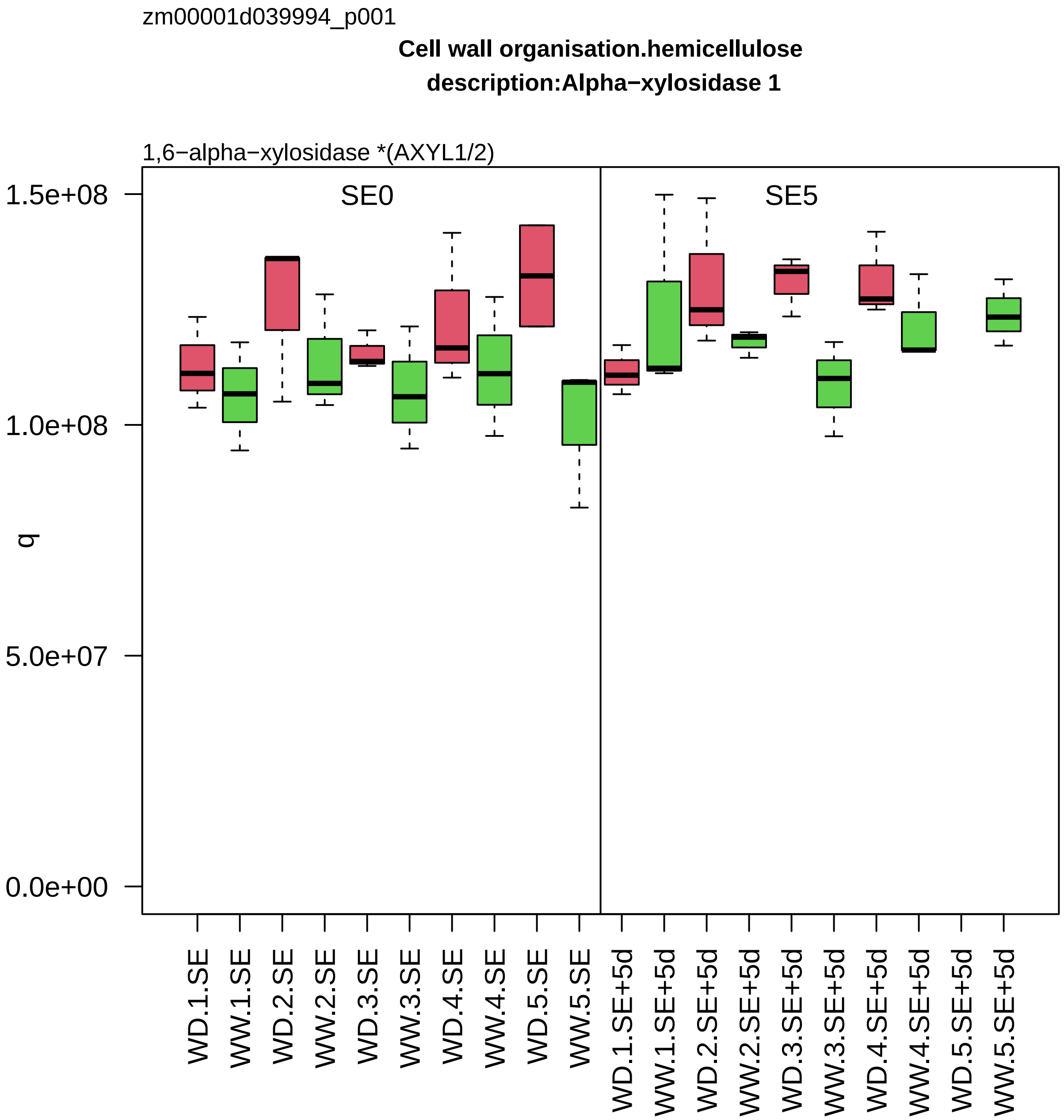

zm00001d039994_p001
Cell wall organisation.hemicellulose
description:Alpha−xylosidase 1
1,6−alpha−xylosidase *(AXYL1/2)
SE0
SE5
1.5e+08
1.0e+08
5.0e+07
0.0e+00
q
WD.1.SE
WW.1.SE
WD.2.SE
WW.2.SE
WD.3.SE
WW.3.SE
WD.4.SE
WW.4.SE
WD.5.SE
WW.5.SE
WD.1.SE+5d
WW.1.SE+5d
WD.2.SE+5d
WW.2.SE+5d
WD.3.SE+5d
WW.3.SE+5d
WD.4.SE+5d
WW.4.SE+5d
WD.5.SE+5d
WW.5.SE+5d

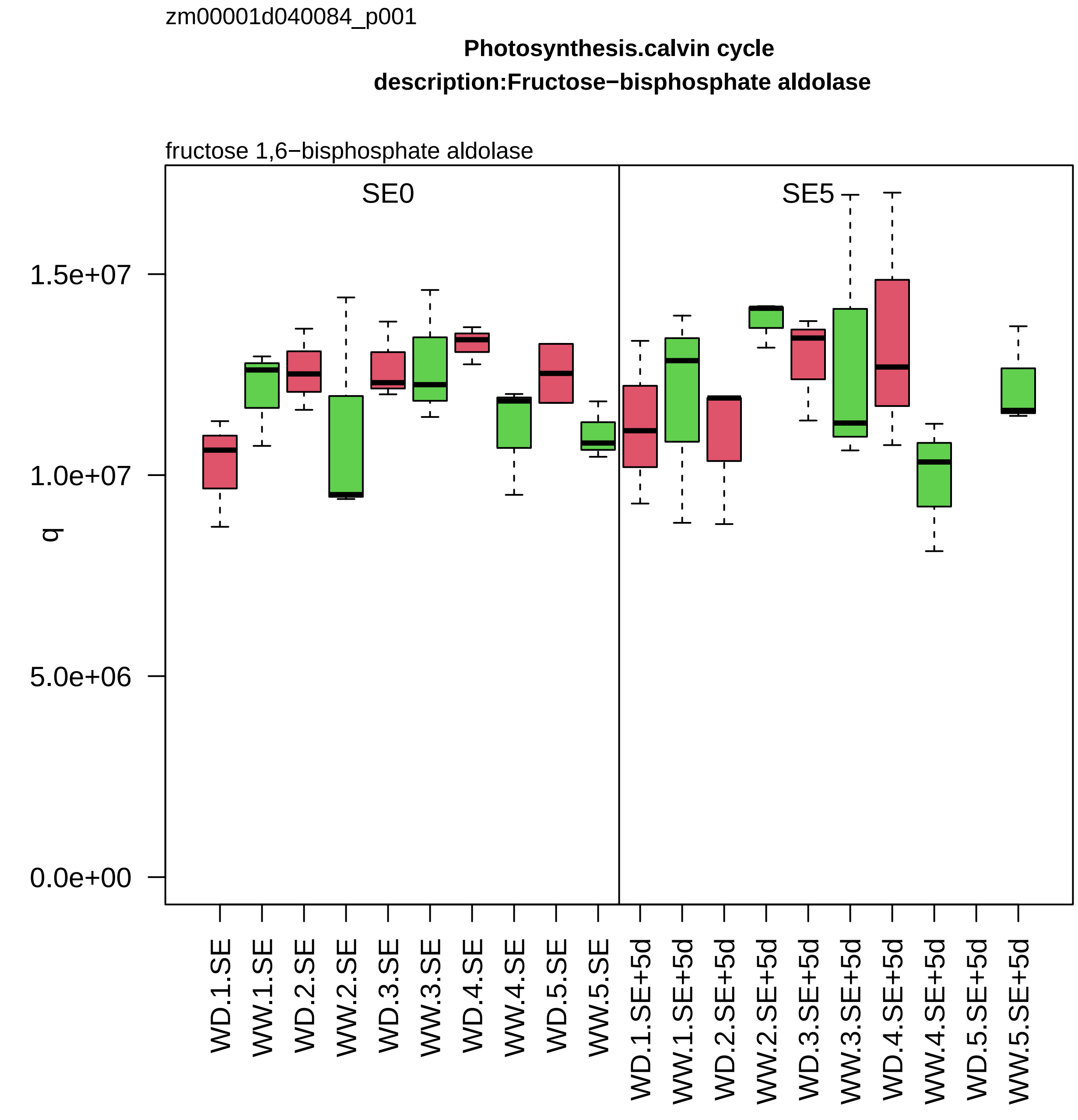

zm00001d040084_p001
Photosynthesis.calvin cycle
description:Fructose−bisphosphate aldolase
fructose 1,6−bisphosphate aldolase
SE0
SE5
q
1.5e+07
1.0e+07
5.0e+06
0.0e+00
WD.1.SE
WW.1.SE
WD.2.SE
WW.2.SE
WD.3.SE
WW.3.SE
WD.4.SE
WW.4.SE
WD.5.SE
WW.5.SE
WD.1.SE+5d
WW.1.SE+5d
WD.2.SE+5d
WW.2.SE+5d
WD.3.SE+5d
WW.3.SE+5d
WD.4.SE+5d
WW.4.SE+5d
WD.5.SE+5d
WW.5.SE+5d

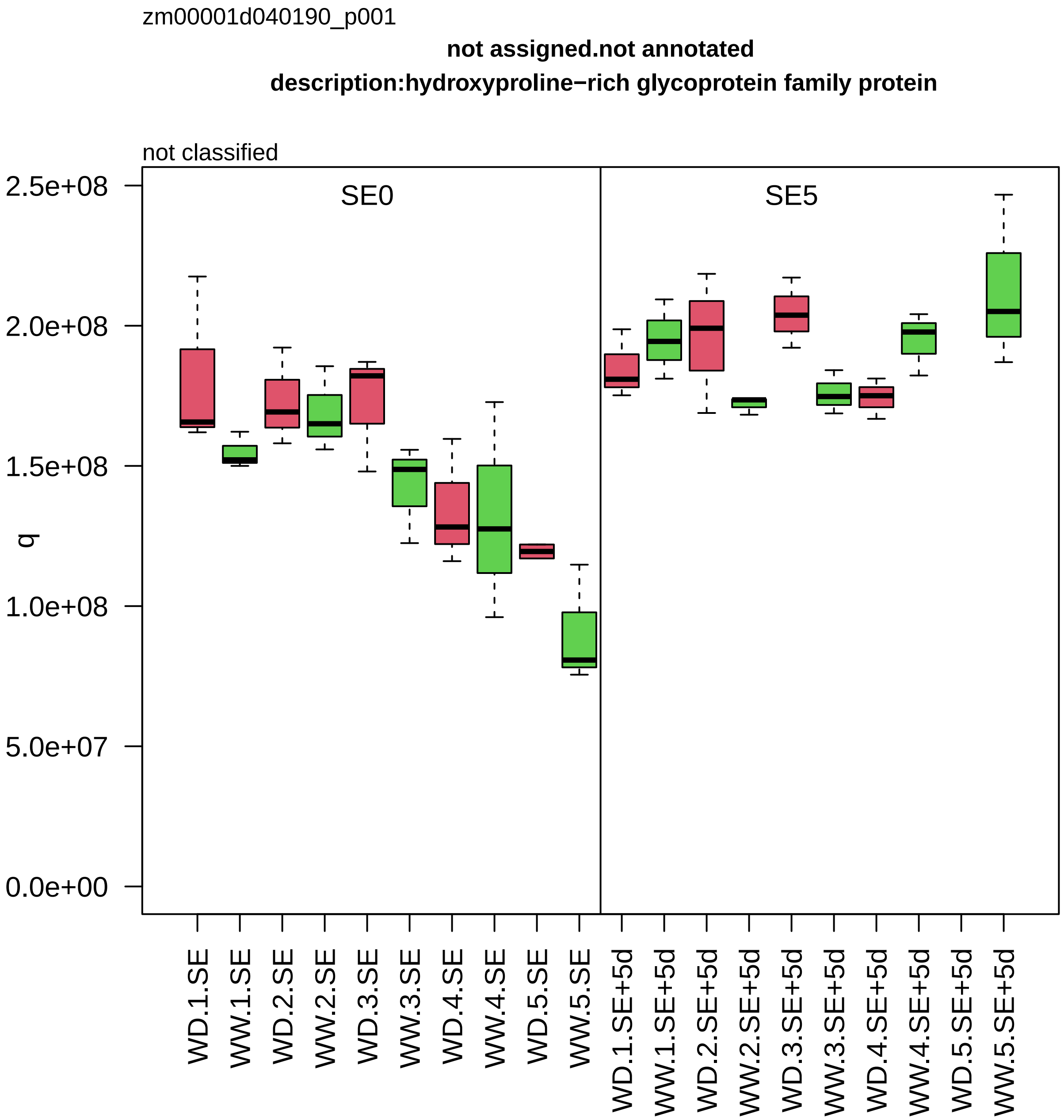

zm00001d040190_p001
not assigned.not annotated
description:hydroxyproline−rich glycoprotein family protein
not classified
SE0
SE5
q
2.5e+08
2.0e+08
1.5e+08
1.0e+08
5.0e+07
0.0e+00
WD.1.SE
WW.1.SE
WD.2.SE
WW.2.SE
WD.3.SE
WW.3.SE
WD.4.SE
WW.4.SE
WD.5.SE
WW.5.SE
WD.1.SE+5d
WW.1.SE+5d
WD.2.SE+5d
WW.2.SE+5d
WD.3.SE+5d
WW.3.SE+5d
WD.4.SE+5d
WW.4.SE+5d
WD.5.SE+5d
WW.5.SE+5d

zm00001d040257_p001

**Cytoskeleton organisation.actin and tubulin folding**
**description:T−complex protein 1 subunit beta**

subunit beta *(CCT2) of CCT chaperonin folding complex

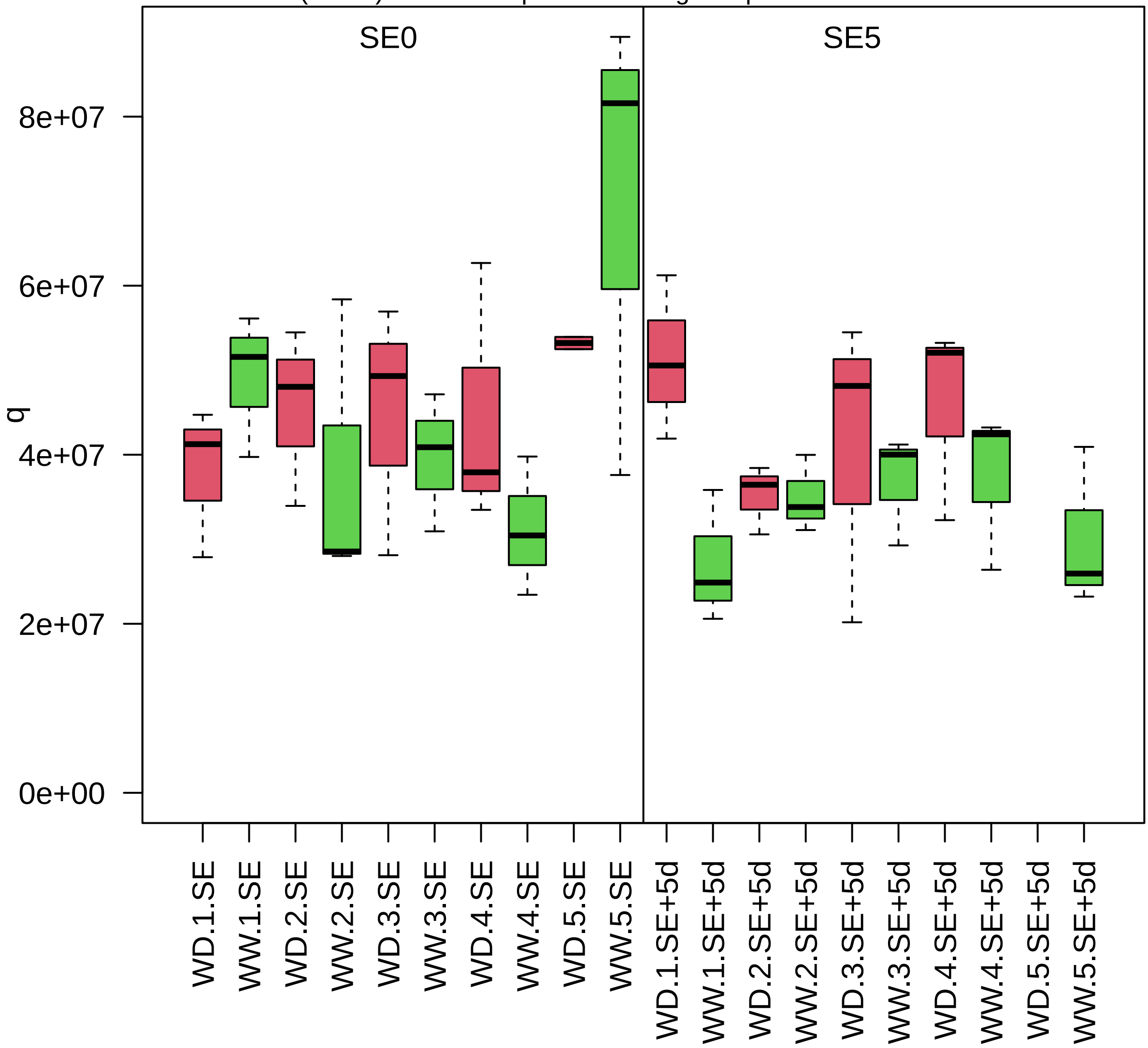

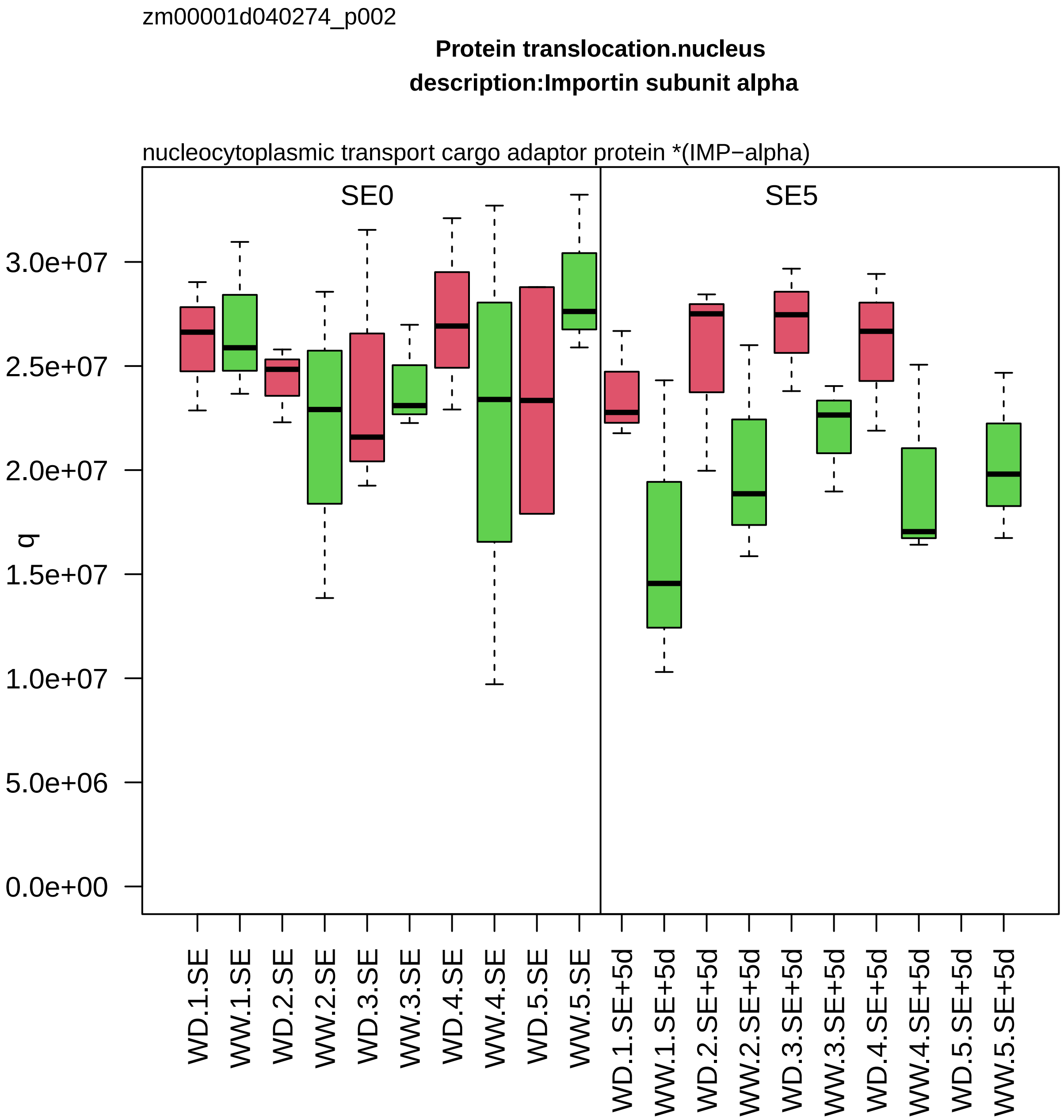

zm00001d040274_p002
Protein translocation.nucleus
description:Importin subunit alpha
nucleocytoplasmic transport cargo adaptor protein *(IMP−alpha)
SE0
SE5
q
3.0e+07
2.5e+07
2.0e+07
1.5e+07
1.0e+07
5.0e+06
0.0e+00
WD.1.SE
WW.1.SE
WD.2.SE
WW.2.SE
WD.3.SE
WW.3.SE
WD.4.SE
WW.4.SE
WD.5.SE
WW.5.SE
WD.1.SE+5d
WW.1.SE+5d
WD.2.SE+5d
WW.2.SE+5d
WD.3.SE+5d
WW.3.SE+5d
WD.4.SE+5d
WW.4.SE+5d
WD.5.SE+5d
WW.5.SE+5d

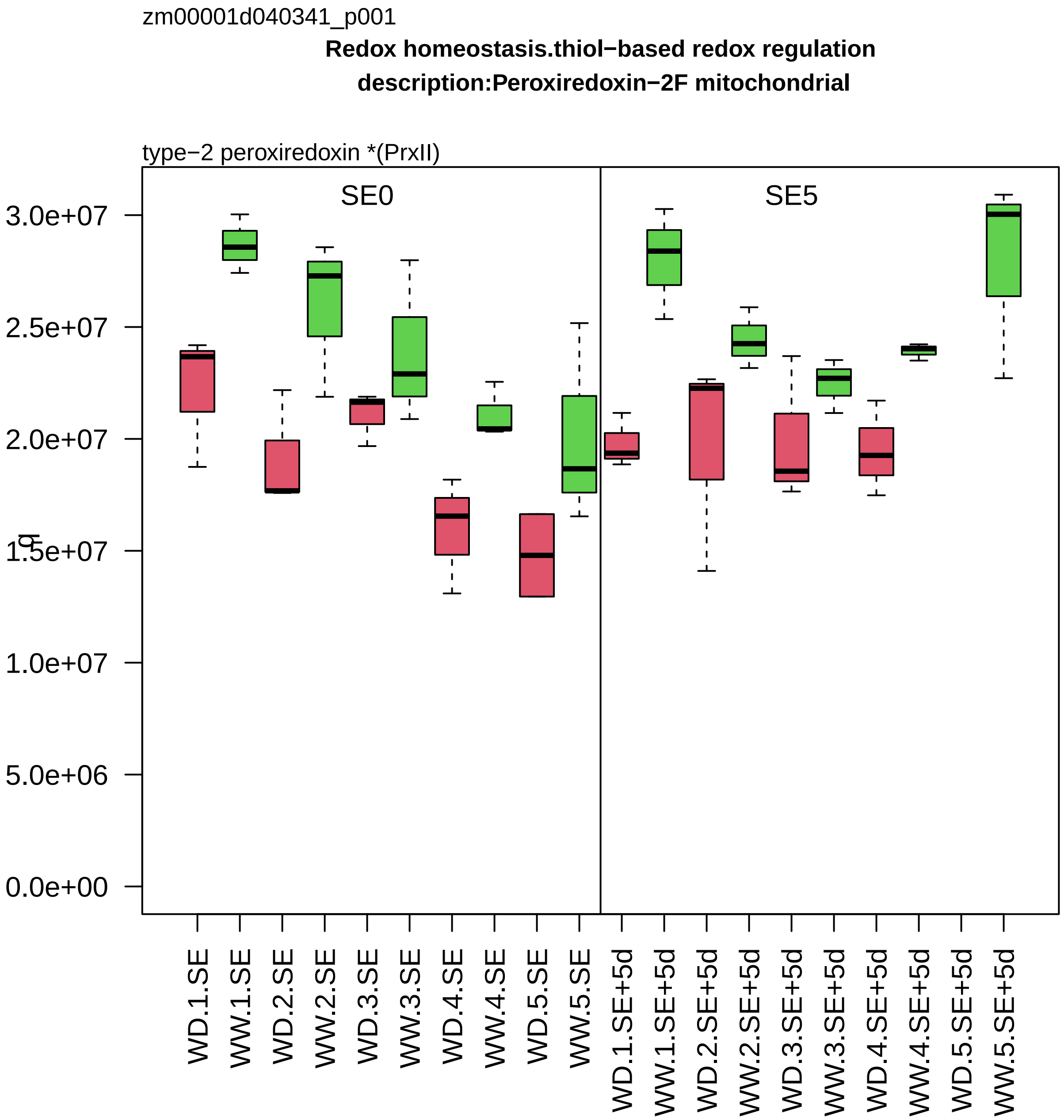

zm00001d040341_p001
Redox homeostasis.thiol−based redox regulation
description:Peroxiredoxin−2F mitochondrial
type−2 peroxiredoxin *(PrxII)
SE0
SE5
3.0e+07
2.5e+07
2.0e+07
1.5e+07
1.0e+07
5.0e+06
0.0e+00
WD.1.SE
WW.1.SE
WD.2.SE
WW.2.SE
WD.3.SE
WW.3.SE
WD.4.SE
WW.4.SE
WD.5.SE
WW.5.SE
WD.1.SE+5d
WW.1.SE+5d
WD.2.SE+5d
WW.2.SE+5d
WD.3.SE+5d
WW.3.SE+5d
WD.4.SE+5d
WW.4.SE+5d
WD.5.SE+5d
WW.5.SE+5d

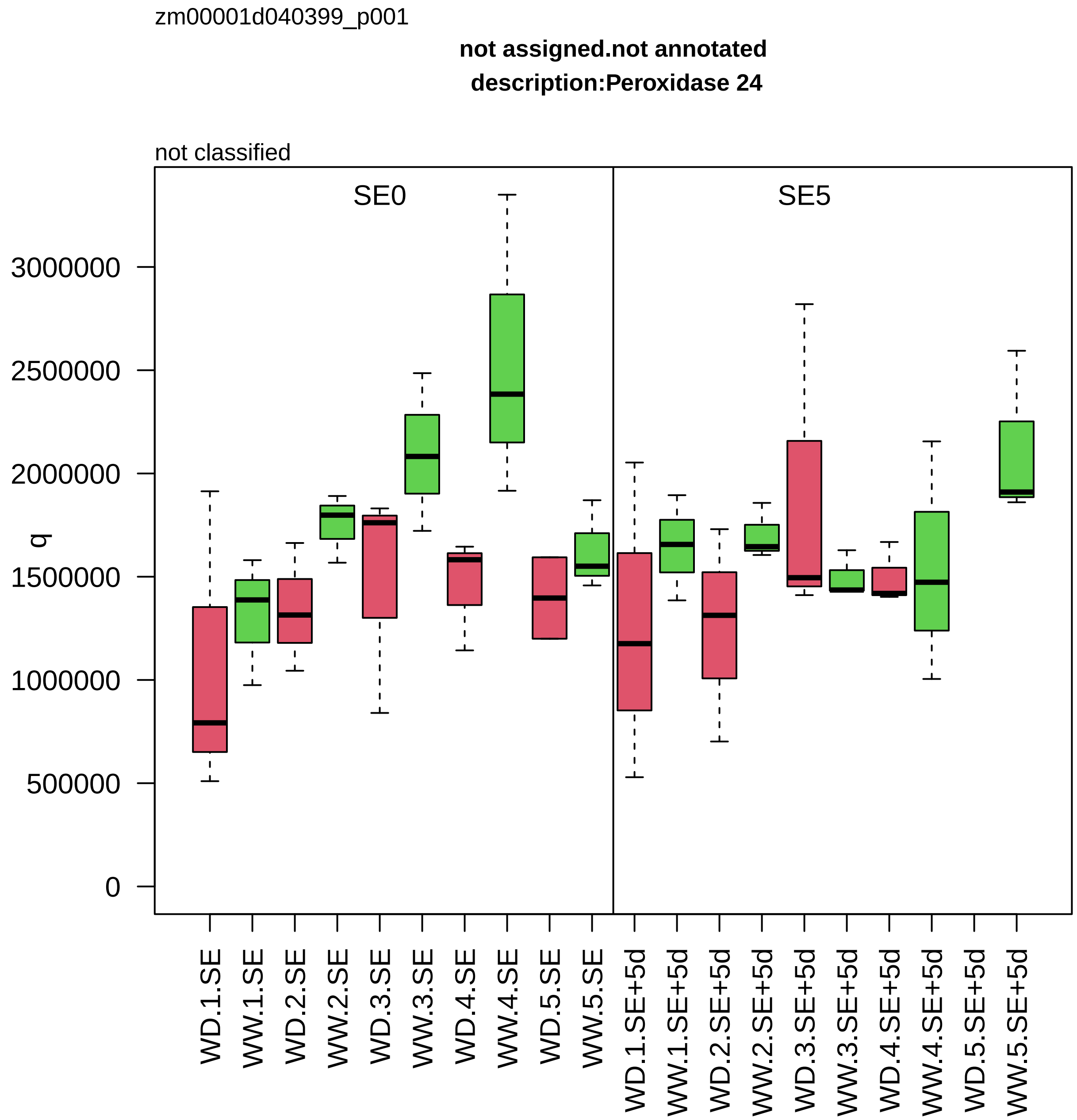

zm00001d040399_p001
not assigned.not annotated
description:Peroxidase 24
not classified
SE0
SE5
q
0
500000
1000000
1500000
2000000
2500000
3000000
WD.1.SE
WW.1.SE
WD.2.SE
WW.2.SE
WD.3.SE
WW.3.SE
WD.4.SE
WW.4.SE
WD.5.SE
WW.5.SE
WD.1.SE+5d
WW.1.SE+5d
WD.2.SE+5d
WW.2.SE+5d
WD.3.SE+5d
WW.3.SE+5d
WD.4.SE+5d
WW.4.SE+5d
WD.5.SE+5d
WW.5.SE+5d

zm00001d040429_p003

**Chromatin organisation.epigenetic state regulation**

**description: argonaute105**

siRNA−integrating factor *(AGO)

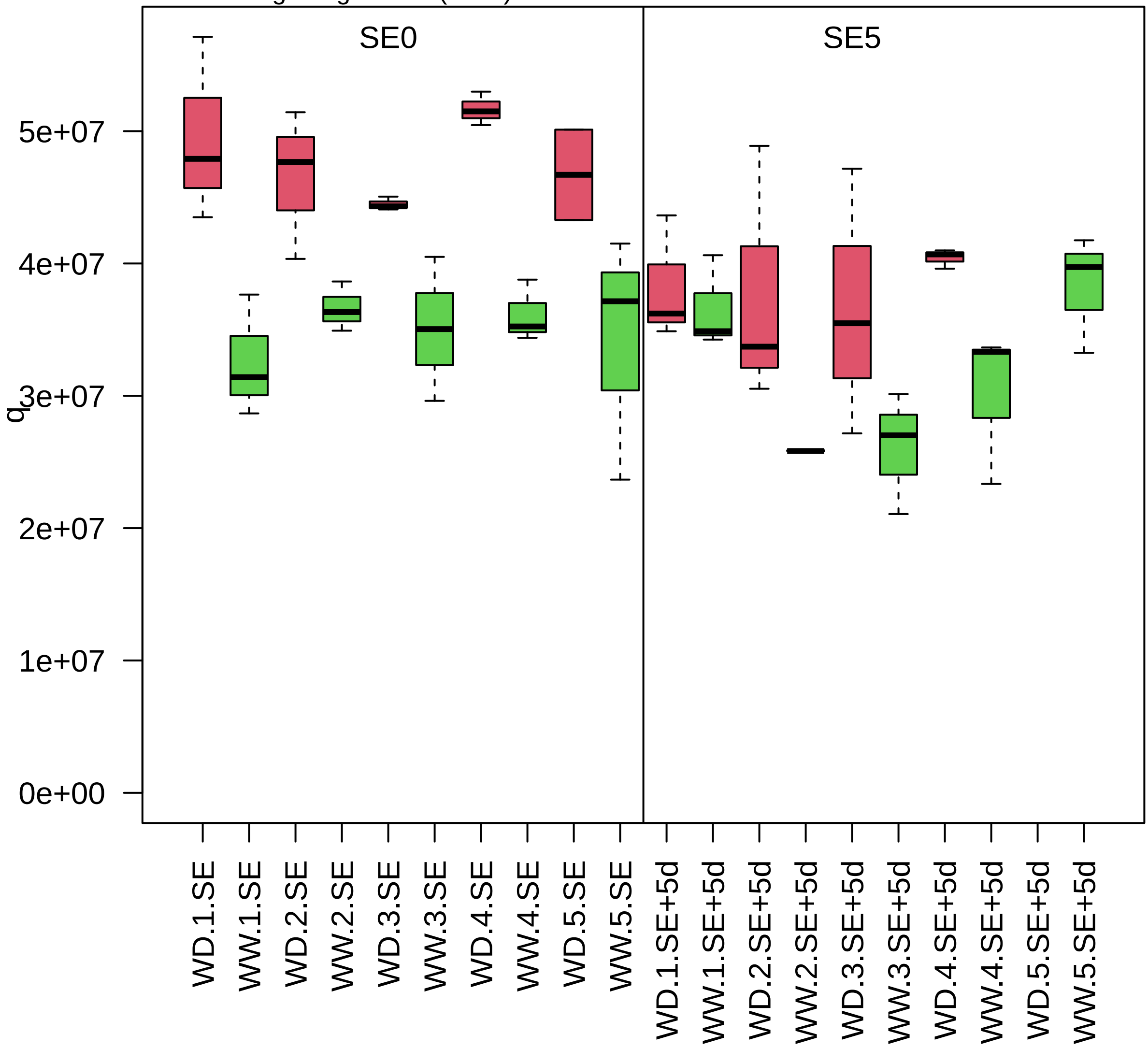

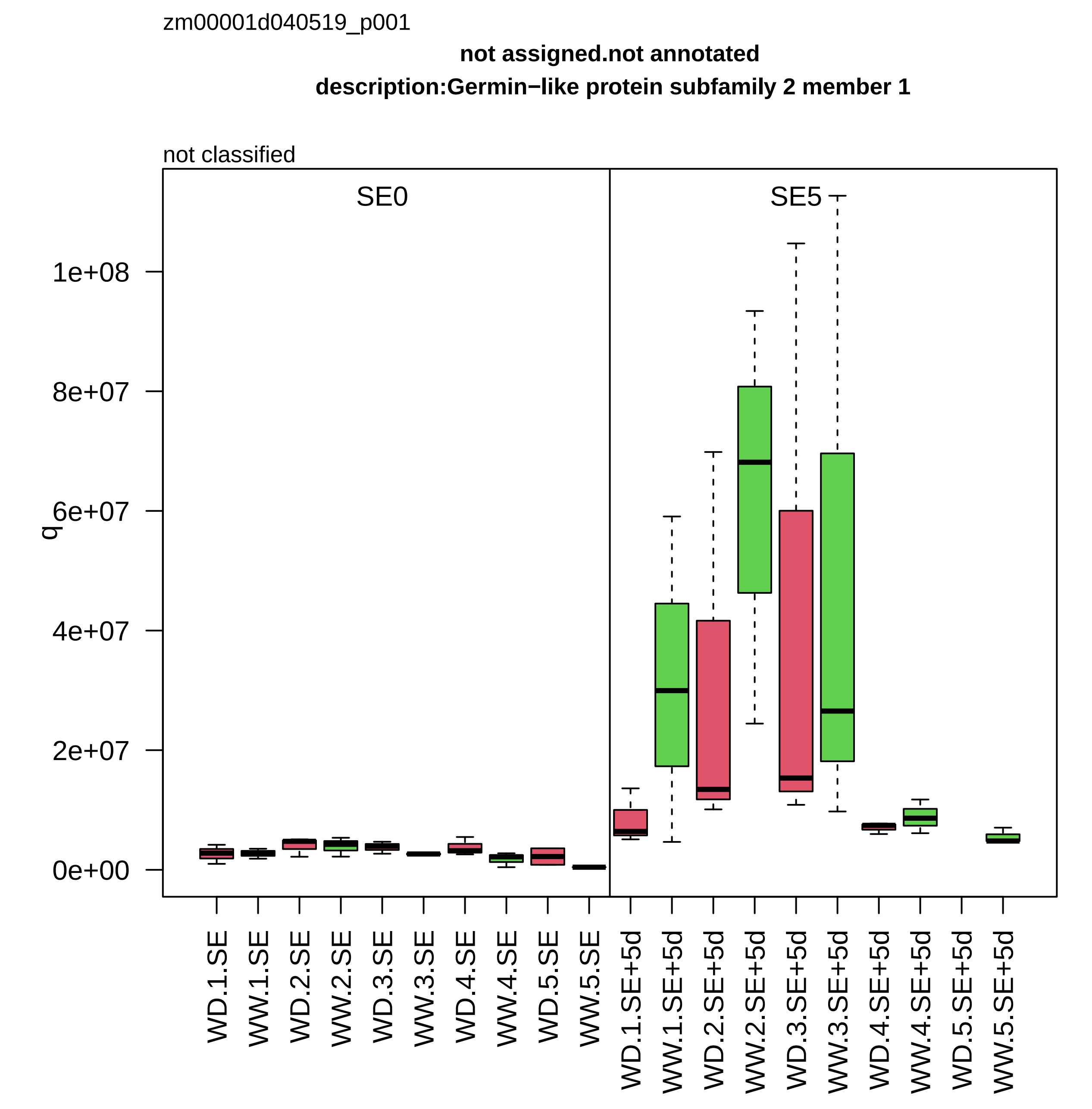

zm00001d040519_p001
not assigned.not annotated
description:Germin−like protein subfamily 2 member 1
not classified
SE0
SE5
q
0e+00
2e+07
4e+07
6e+07
8e+07
1e+08
WD.1.SE
WW.1.SE
WD.2.SE
WW.2.SE
WD.3.SE
WW.3.SE
WD.4.SE
WW.4.SE
WD.5.SE
WW.5.SE
WD.1.SE+5d
WW.1.SE+5d
WD.2.SE+5d
WW.2.SE+5d
WD.3.SE+5d
WW.3.SE+5d
WD.4.SE+5d
WW.4.SE+5d
WD.5.SE+5d
WW.5.SE+5d

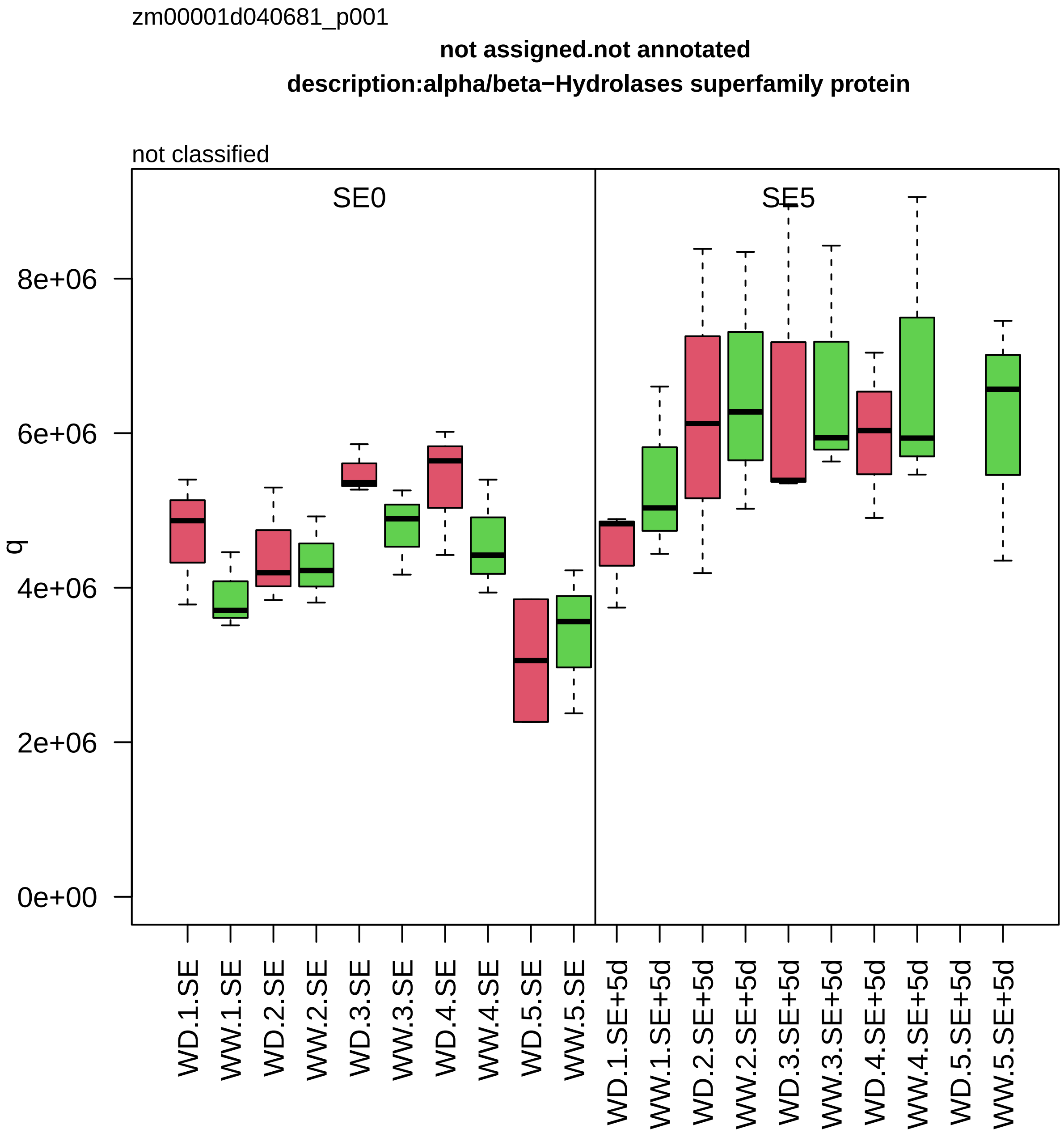

zm00001d040681_p001
not assigned.not annotated
description:alpha/beta−Hydrolases superfamily protein
not classified
SE0
SE5
q
8e+06
6e+06
4e+06
2e+06
0e+00
WD.1.SE
WW.1.SE
WD.2.SE
WW.2.SE
WD.3.SE
WW.3.SE
WD.4.SE
WW.4.SE
WD.5.SE
WW.5.SE
WD.1.SE+5d
WW.1.SE+5d
WD.2.SE+5d
WW.2.SE+5d
WD.3.SE+5d
WW.3.SE+5d
WD.4.SE+5d
WW.4.SE+5d
WD.5.SE+5d
WW.5.SE+5d

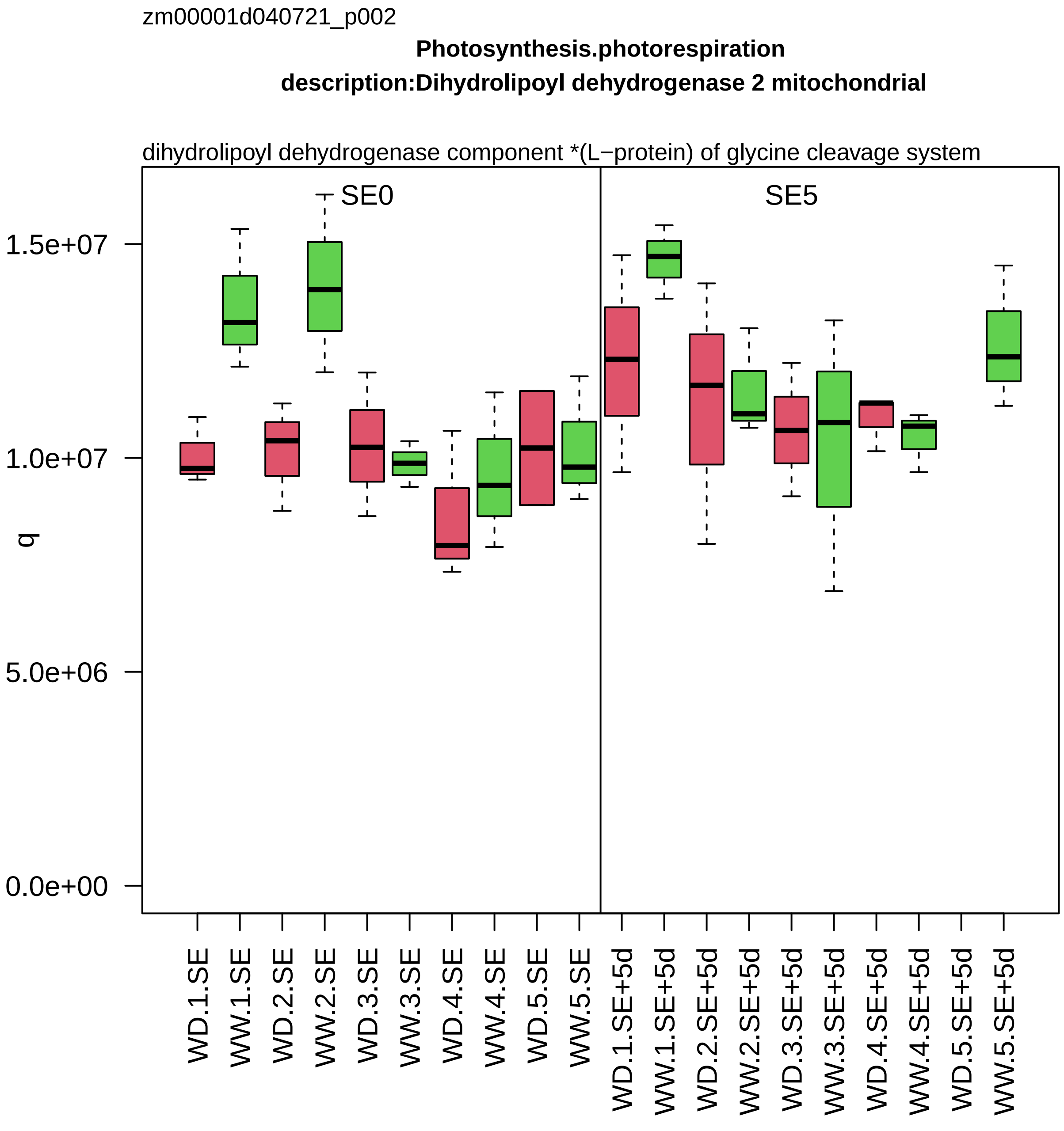
zm00001d040721_p002
Photosynthesis.photorespiration
description:Dihydrolipoyl dehydrogenase 2 mitochondrial
dihydrolipoyl dehydrogenase component *(L−protein) of glycine cleavage system
SE0
SE5
q
1.5e+07
1.0e+07
5.0e+06
0.0e+00
WD.1.SE
WW.1.SE
WD.2.SE
WW.2.SE
WD.3.SE
WW.3.SE
WD.4.SE
WW.4.SE
WD.5.SE
WW.5.SE
WD.1.SE+5d
WW.1.SE+5d
WD.2.SE+5d
WW.2.SE+5d
WD.3.SE+5d
WW.3.SE+5d
WD.4.SE+5d
WW.4.SE+5d
WD.5.SE+5d
WW.5.SE+5d

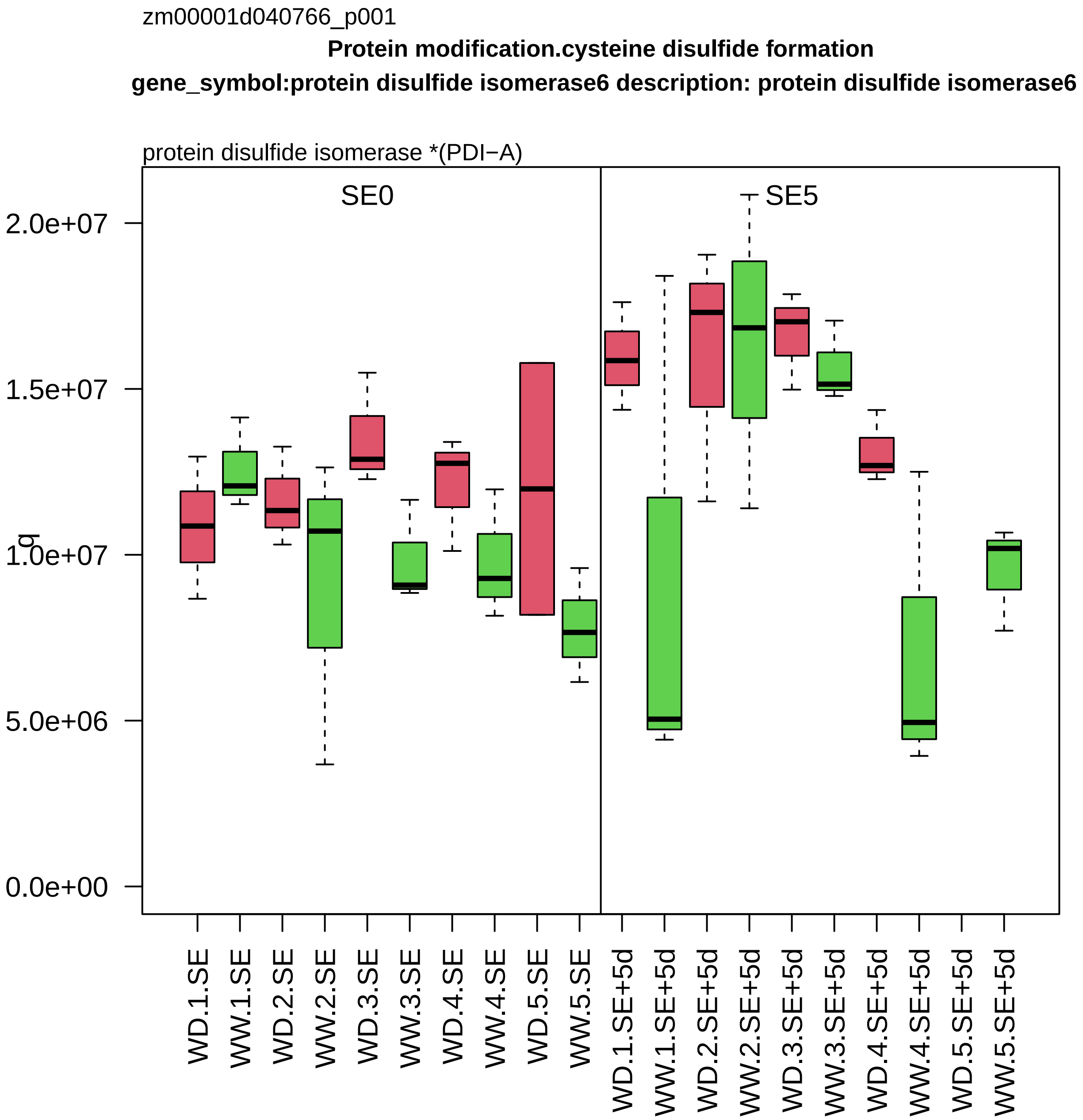
zm00001d040766_p001
Protein modification.cysteine disulfide formation
gene_symbol:protein disulfide isomerase6 description: protein disulfide isomerase6
protein disulfide isomerase *(PDI−A)
SE0
SE5
q
2.0e+07
1.5e+07
1.0e+07
5.0e+06
0.0e+00
WD.1.SE
WW.1.SE
WD.2.SE
WW.2.SE
WD.3.SE
WW.3.SE
WD.4.SE
WW.4.SE
WD.5.SE
WW.5.SE
WD.1.SE+5d
WW.1.SE+5d
WD.2.SE+5d
WW.2.SE+5d
WD.3.SE+5d
WW.3.SE+5d
WD.4.SE+5d
WW.4.SE+5d
WD.5.SE+5d
WW.5.SE+5d

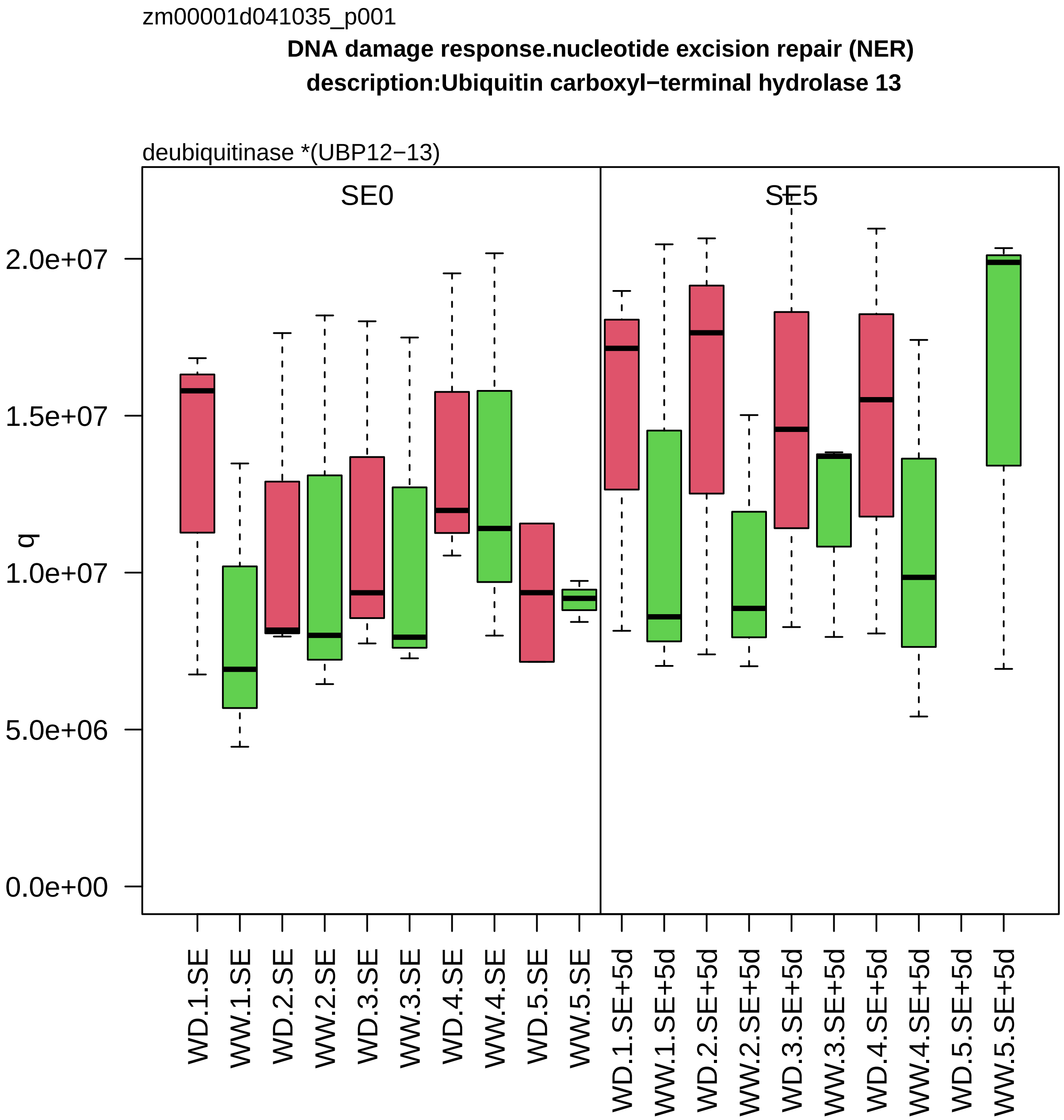

zm00001d041035_p001
DNA damage response.nucleotide excision repair (NER)
description:Ubiquitin carboxyl−terminal hydrolase 13
deubiquitinase *(UBP12−13)
SE0
SE5
q
2.0e+07
1.5e+07
1.0e+07
5.0e+06
0.0e+00
WD.1.SE
WW.1.SE
WD.2.SE
WW.2.SE
WD.3.SE
WW.3.SE
WD.4.SE
WW.4.SE
WD.5.SE
WW.5.SE
WD.1.SE+5d
WW.1.SE+5d
WD.2.SE+5d
WW.2.SE+5d
WD.3.SE+5d
WW.3.SE+5d
WD.4.SE+5d
WW.4.SE+5d
WD.5.SE+5d
WW.5.SE+5d

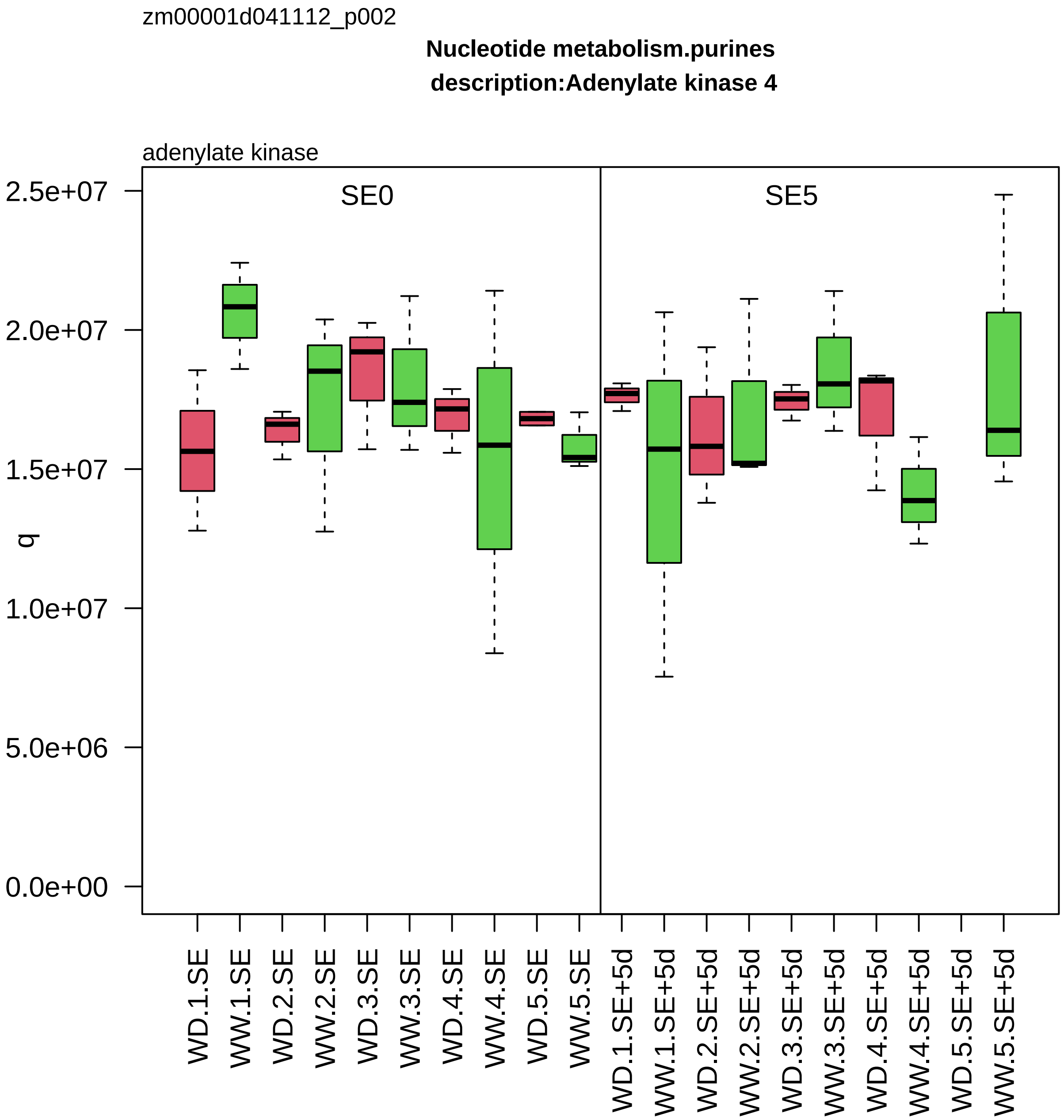

zm00001d041112_p002
Nucleotide metabolism.purines
description:Adenylate kinase 4
adenylate kinase
SE0
SE5
q
2.5e+07
2.0e+07
1.5e+07
1.0e+07
5.0e+06
0.0e+00
WD.1.SE
WW.1.SE
WD.2.SE
WW.2.SE
WD.3.SE
WW.3.SE
WD.4.SE
WW.4.SE
WD.5.SE
WW.5.SE
WD.1.SE+5d
WW.1.SE+5d
WD.2.SE+5d
WW.2.SE+5d
WD.3.SE+5d
WW.3.SE+5d
WD.4.SE+5d
WW.4.SE+5d
WD.5.SE+5d
WW.5.SE+5d

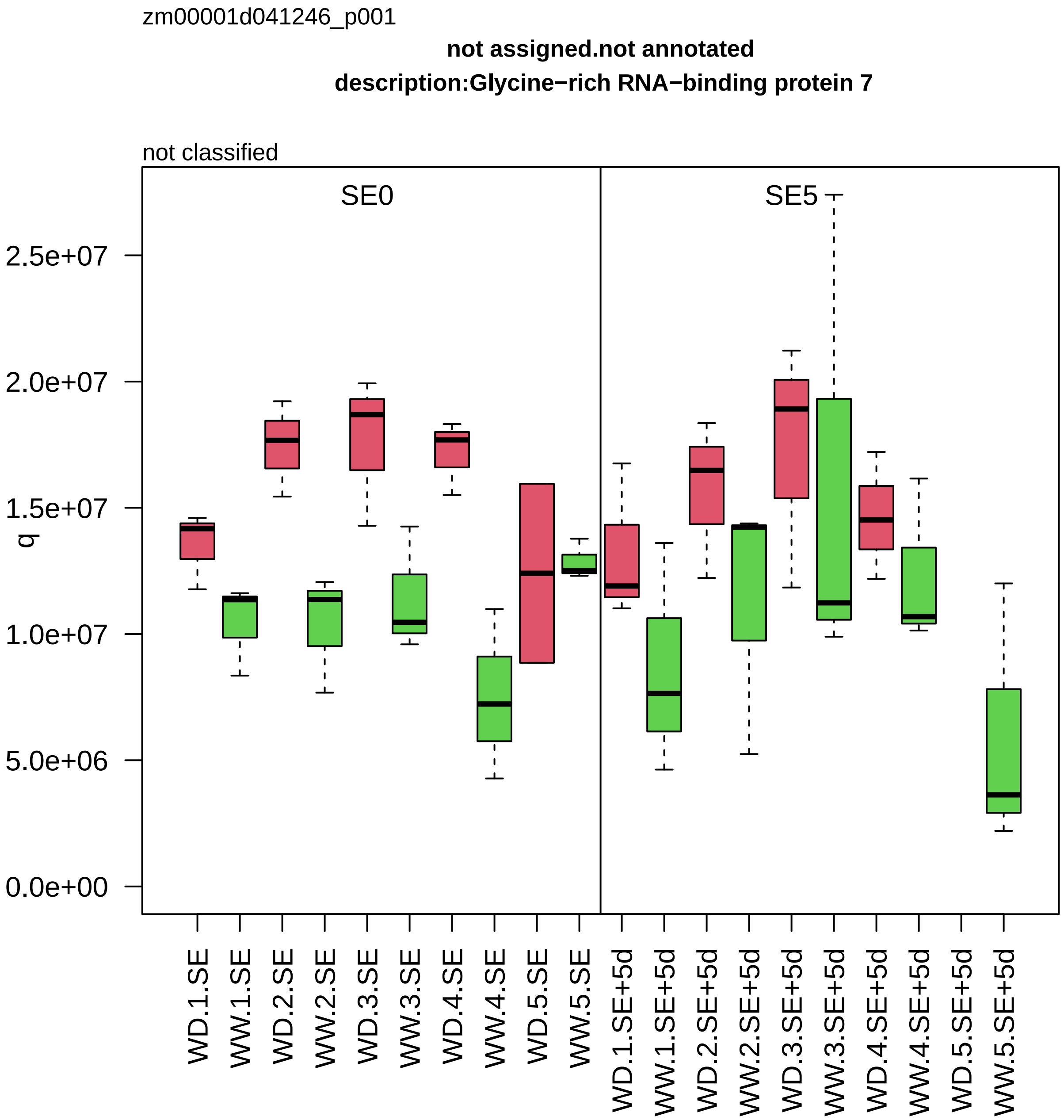
zm00001d041246_p001
not assigned.not annotated
description:Glycine−rich RNA−binding protein 7
not classified
SE0
SE5
q
2.5e+07
2.0e+07
1.5e+07
1.0e+07
5.0e+06
0.0e+00
WD.1.SE
WW.1.SE
WD.2.SE
WW.2.SE
WD.3.SE
WW.3.SE
WD.4.SE
WW.4.SE
WD.5.SE
WW.5.SE
WD.1.SE+5d
WW.1.SE+5d
WD.2.SE+5d
WW.2.SE+5d
WD.3.SE+5d
WW.3.SE+5d
WD.4.SE+5d
WW.4.SE+5d
WD.5.SE+5d
WW.5.SE+5d

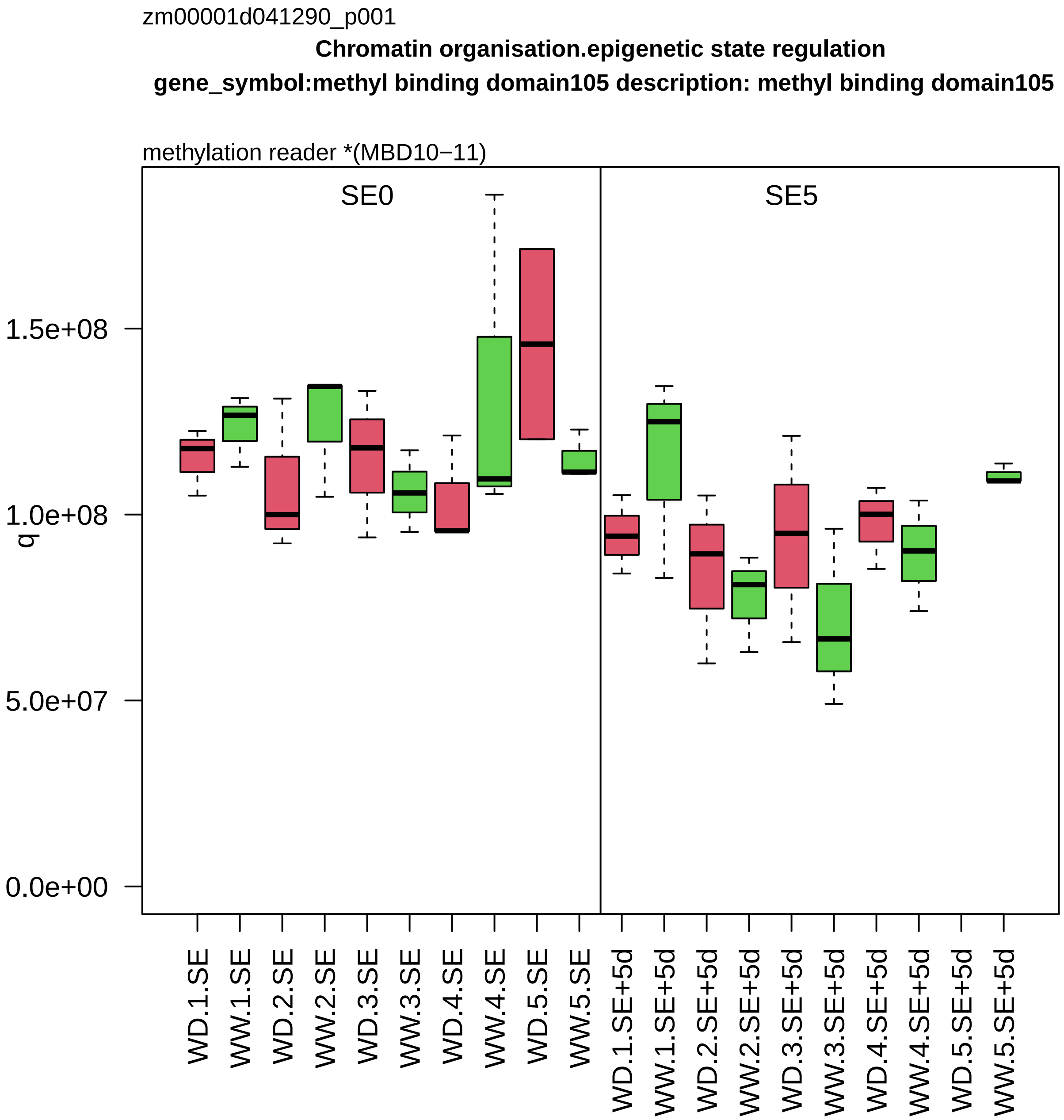

zm00001d041290_p001
Chromatin organisation.epigenetic state regulation
gene_symbol:methyl binding domain105 description: methyl binding domain105
methylation reader *(MBD10−11)
SE0
SE5
q
1.5e+08
1.0e+08
5.0e+07
0.0e+00
WD.1.SE
WW.1.SE
WD.2.SE
WW.2.SE
WD.3.SE
WW.3.SE
WD.4.SE
WW.4.SE
WD.5.SE
WW.5.SE
WD.1.SE+5d
WW.1.SE+5d
WD.2.SE+5d
WW.2.SE+5d
WD.3.SE+5d
WW.3.SE+5d
WD.4.SE+5d
WW.4.SE+5d
WD.5.SE+5d
WW.5.SE+5d

zm00001d041550_p004

**Protein homeostasis.protein quality control**

**gene_symbol:heat shock protein70−4 description:Heat shock 70 kDa protein 3**

chaperone *(Hsp70)

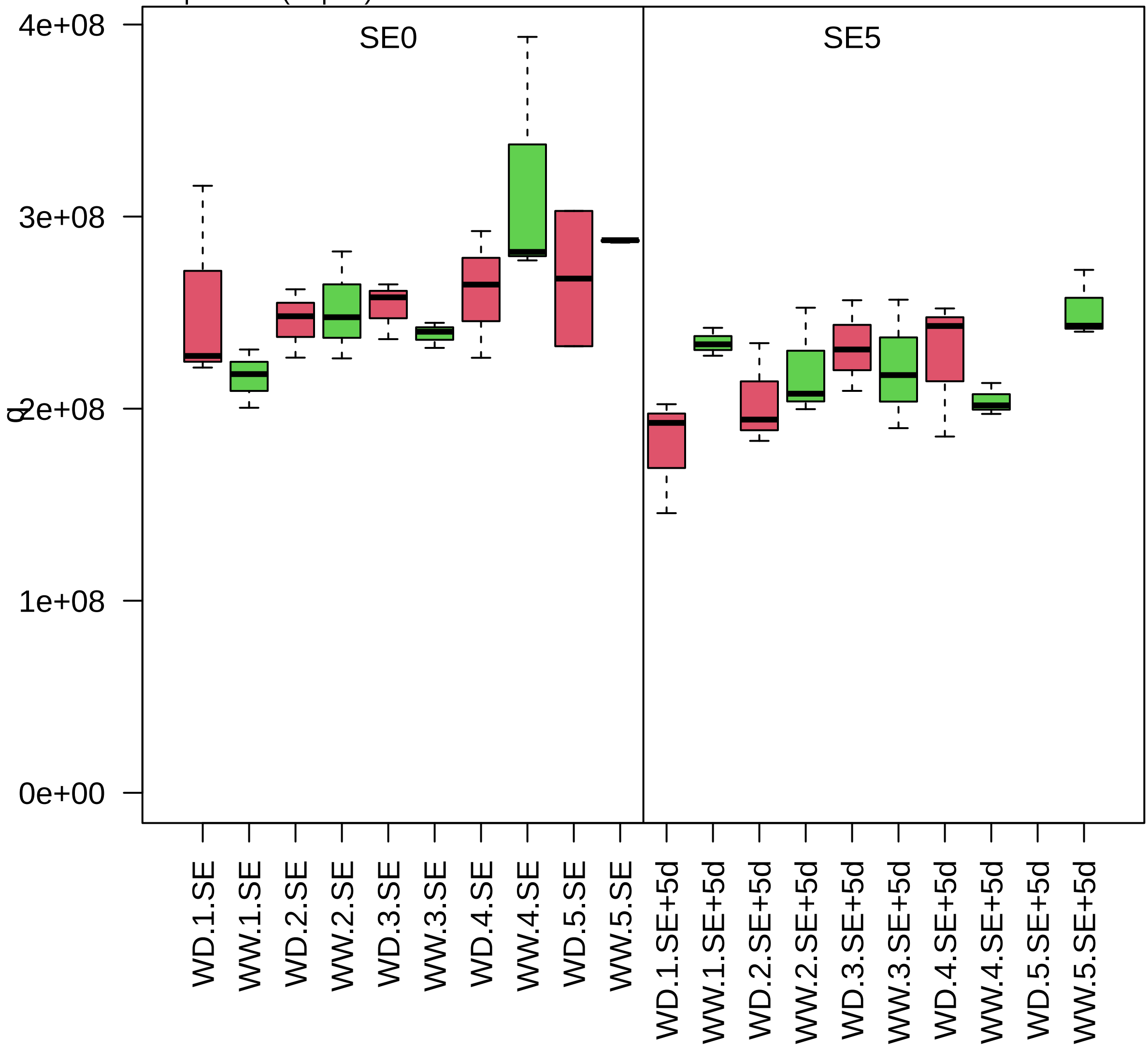

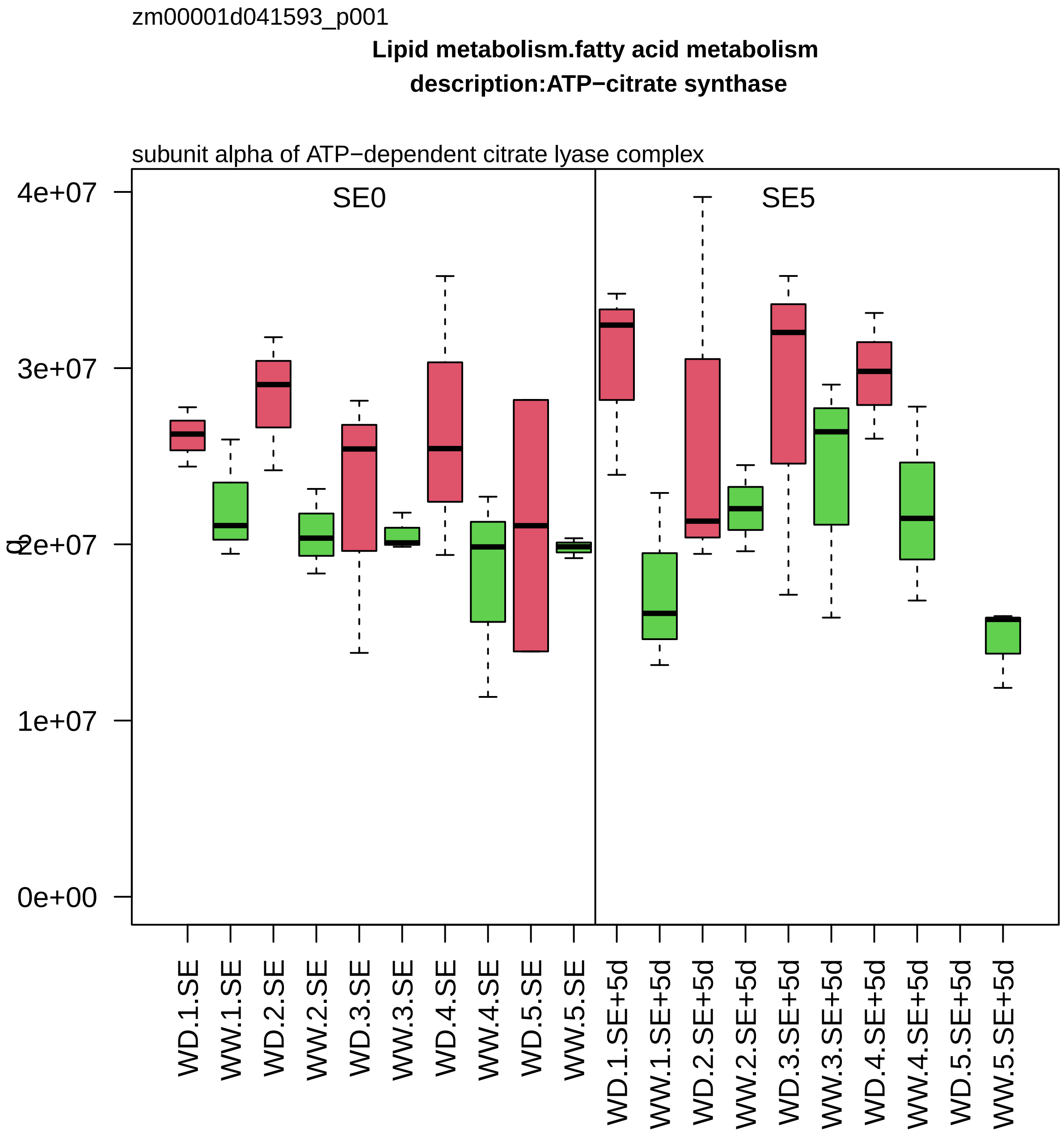

zm00001d041593_p001
Lipid metabolism.fatty acid metabolism
description:ATP−citrate synthase
subunit alpha of ATP−dependent citrate lyase complex
SE0
SE5
4e+07
3e+07
2e+07
1e+07
0e+00
q
WD.1.SE
WW.1.SE
WD.2.SE
WW.2.SE
WD.3.SE
WW.3.SE
WD.4.SE
WW.4.SE
WD.5.SE
WW.5.SE
WD.1.SE+5d
WW.1.SE+5d
WD.2.SE+5d
WW.2.SE+5d
WD.3.SE+5d
WW.3.SE+5d
WD.4.SE+5d
WW.4.SE+5d
WD.5.SE+5d
WW.5.SE+5d

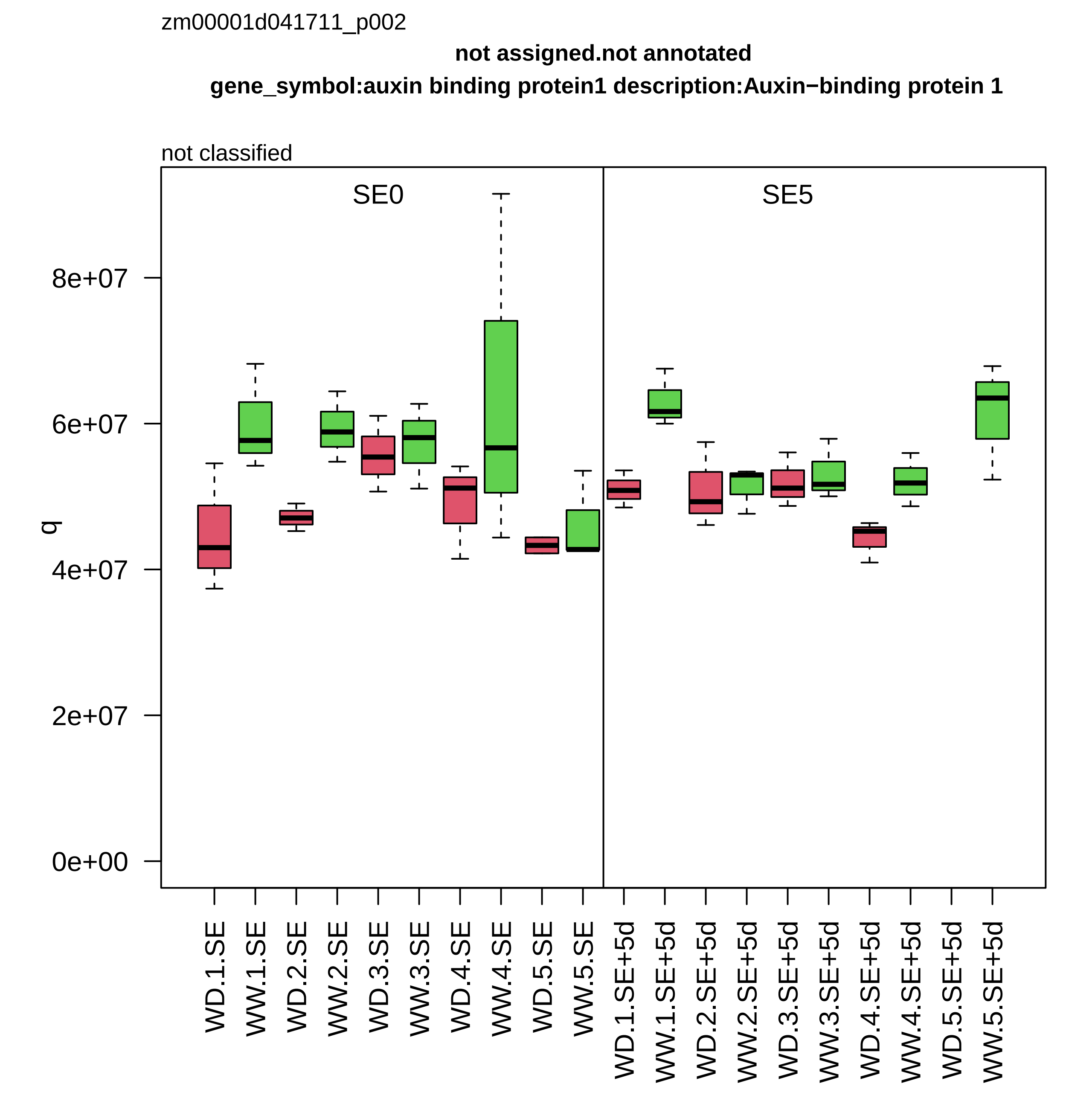

zm00001d041711_p002
not assigned.not annotated
gene_symbol:auxin binding protein1 description:Auxin−binding protein 1
not classified
SE0
SE5
q
8e+07
6e+07
4e+07
2e+07
0e+00
WD.1.SE
WW.1.SE
WD.2.SE
WW.2.SE
WD.3.SE
WW.3.SE
WD.4.SE
WW.4.SE
WD.5.SE
WW.5.SE
WD.1.SE+5d
WW.1.SE+5d
WD.2.SE+5d
WW.2.SE+5d
WD.3.SE+5d
WW.3.SE+5d
WD.4.SE+5d
WW.4.SE+5d
WD.5.SE+5d
WW.5.SE+5d

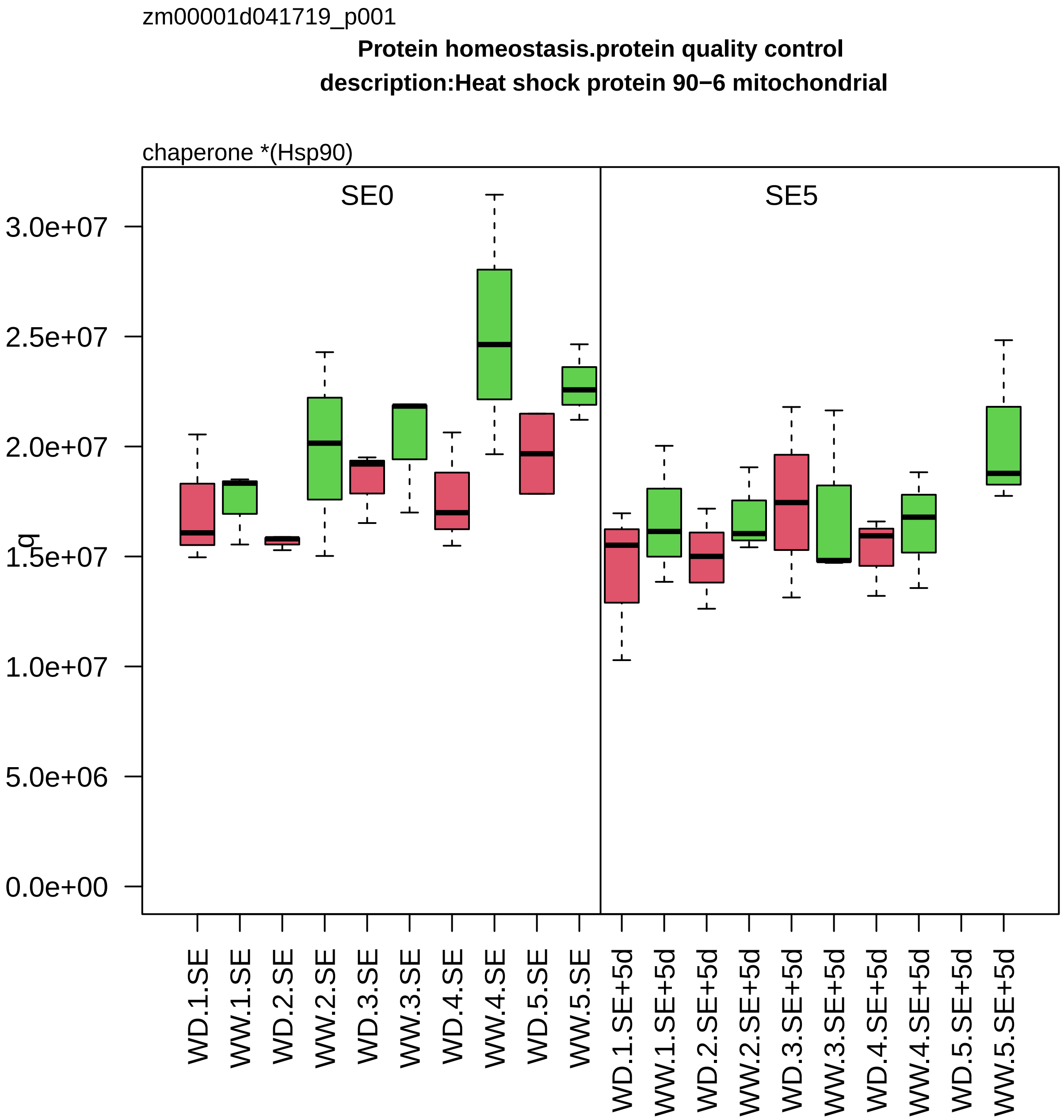

zm00001d041719_p001
Protein homeostasis.protein quality control
description:Heat shock protein 90−6 mitochondrial
chaperone *(Hsp90)
SE0
SE5
q
3.0e+07
2.5e+07
2.0e+07
1.5e+07
1.0e+07
5.0e+06
0.0e+00
WD.1.SE
WW.1.SE
WD.2.SE
WW.2.SE
WD.3.SE
WW.3.SE
WD.4.SE
WW.4.SE
WD.5.SE
WW.5.SE
WD.1.SE+5d
WW.1.SE+5d
WD.2.SE+5d
WW.2.SE+5d
WD.3.SE+5d
WW.3.SE+5d
WD.4.SE+5d
WW.4.SE+5d
WD.5.SE+5d
WW.5.SE+5d

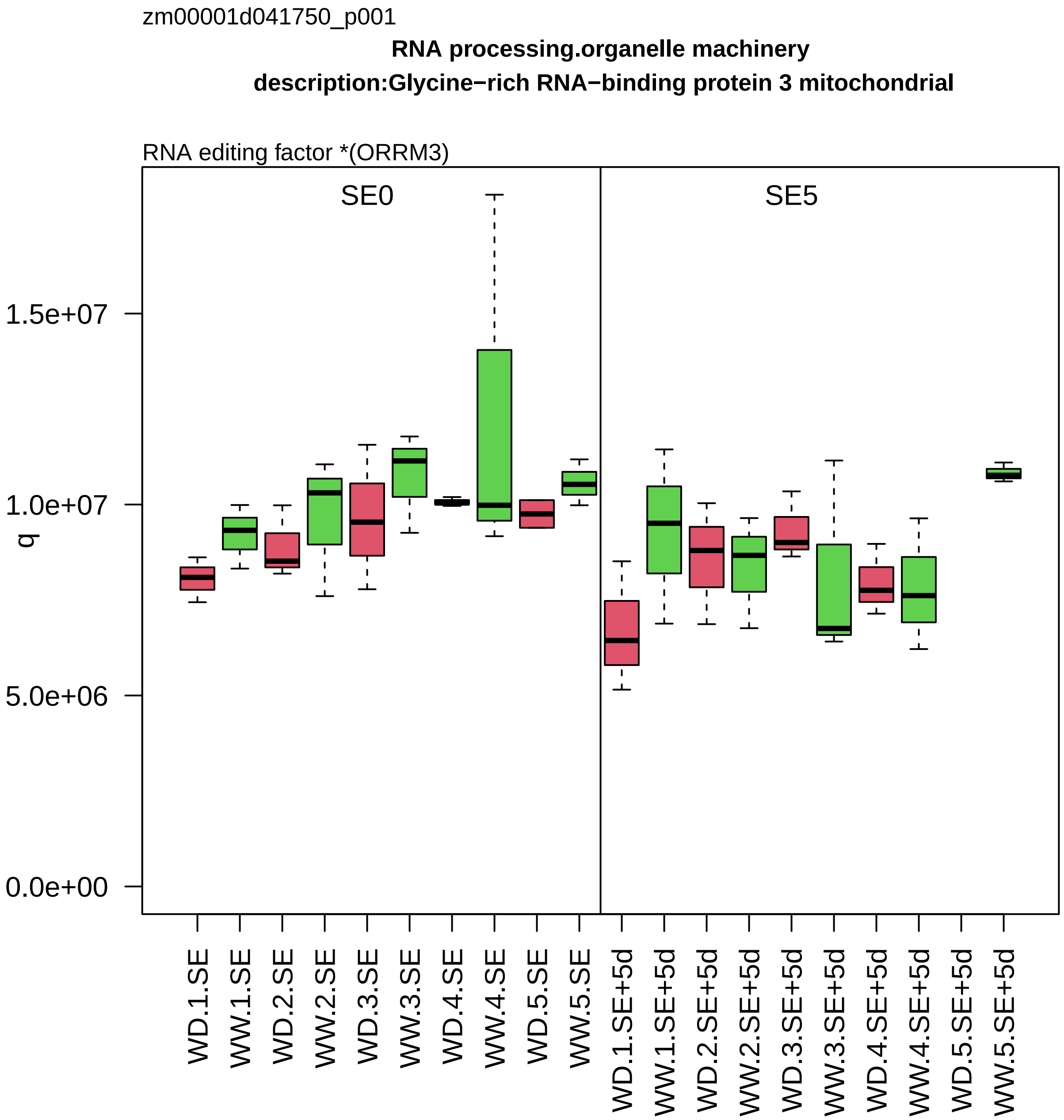

zm00001d041750_p001
RNA processing.organelle machinery
description:Glycine−rich RNA−binding protein 3 mitochondrial
RNA editing factor *(ORRM3)
SE0
SE5
q
1.5e+07
1.0e+07
5.0e+06
0.0e+00
WD.1.SE
WW.1.SE
WD.2.SE
WW.2.SE
WD.3.SE
WW.3.SE
WD.4.SE
WW.4.SE
WD.5.SE
WW.5.SE
WD.1.SE+5d
WW.1.SE+5d
WD.2.SE+5d
WW.2.SE+5d
WD.3.SE+5d
WW.3.SE+5d
WD.4.SE+5d
WW.4.SE+5d
WD.5.SE+5d
WW.5.SE+5d

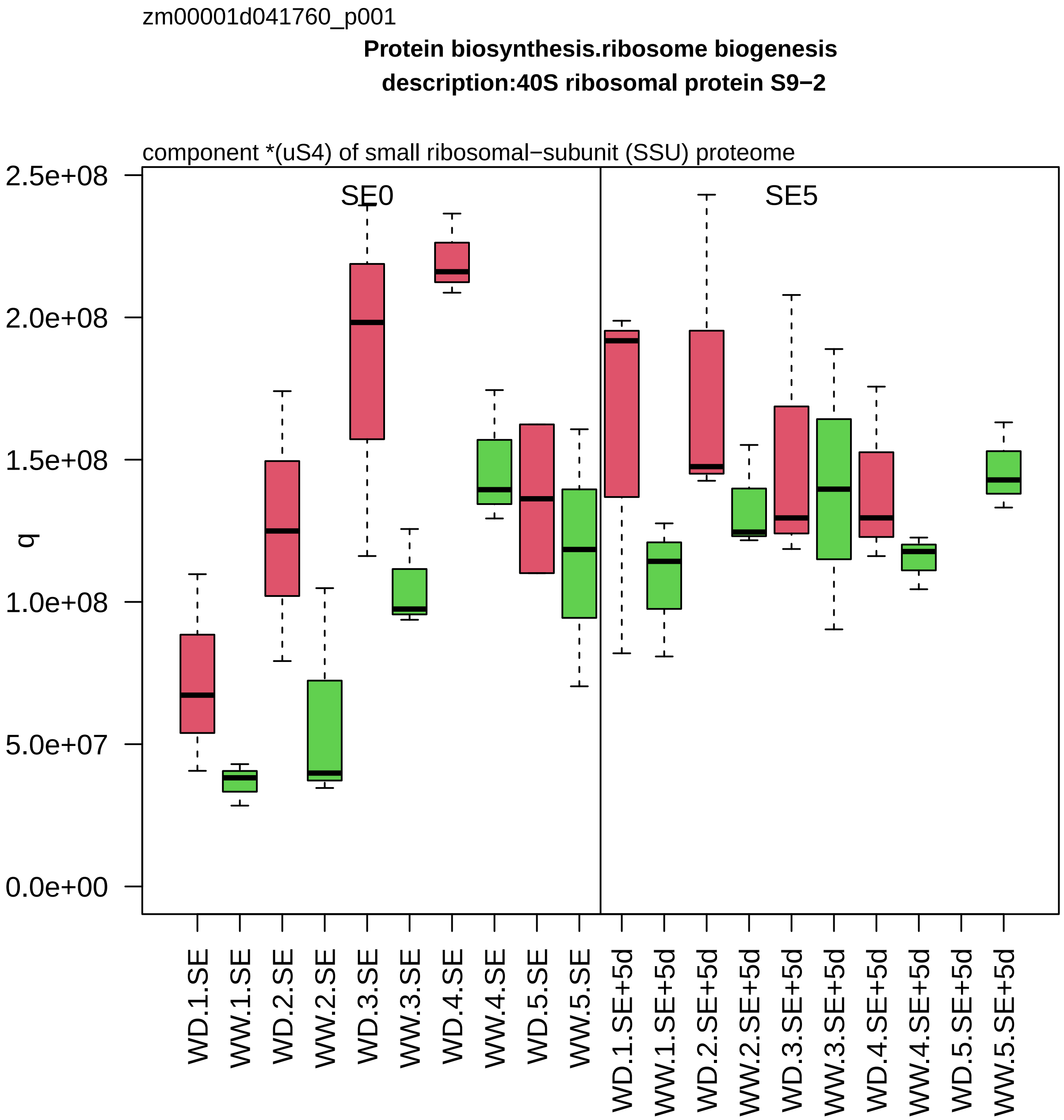

zm00001d041760_p001
Protein biosynthesis.ribosome biogenesis
description:40S ribosomal protein S9−2
component *(uS4) of small ribosomal−subunit (SSU) proteome
SE0
SE5
q
2.5e+08
2.0e+08
1.5e+08
1.0e+08
5.0e+07
0.0e+00
WD.1.SE
WW.1.SE
WD.2.SE
WW.2.SE
WD.3.SE
WW.3.SE
WD.4.SE
WW.4.SE
WD.5.SE
WW.5.SE
WD.1.SE+5d
WW.1.SE+5d
WD.2.SE+5d
WW.2.SE+5d
WD.3.SE+5d
WW.3.SE+5d
WD.4.SE+5d
WW.4.SE+5d
WD.5.SE+5d
WW.5.SE+5d

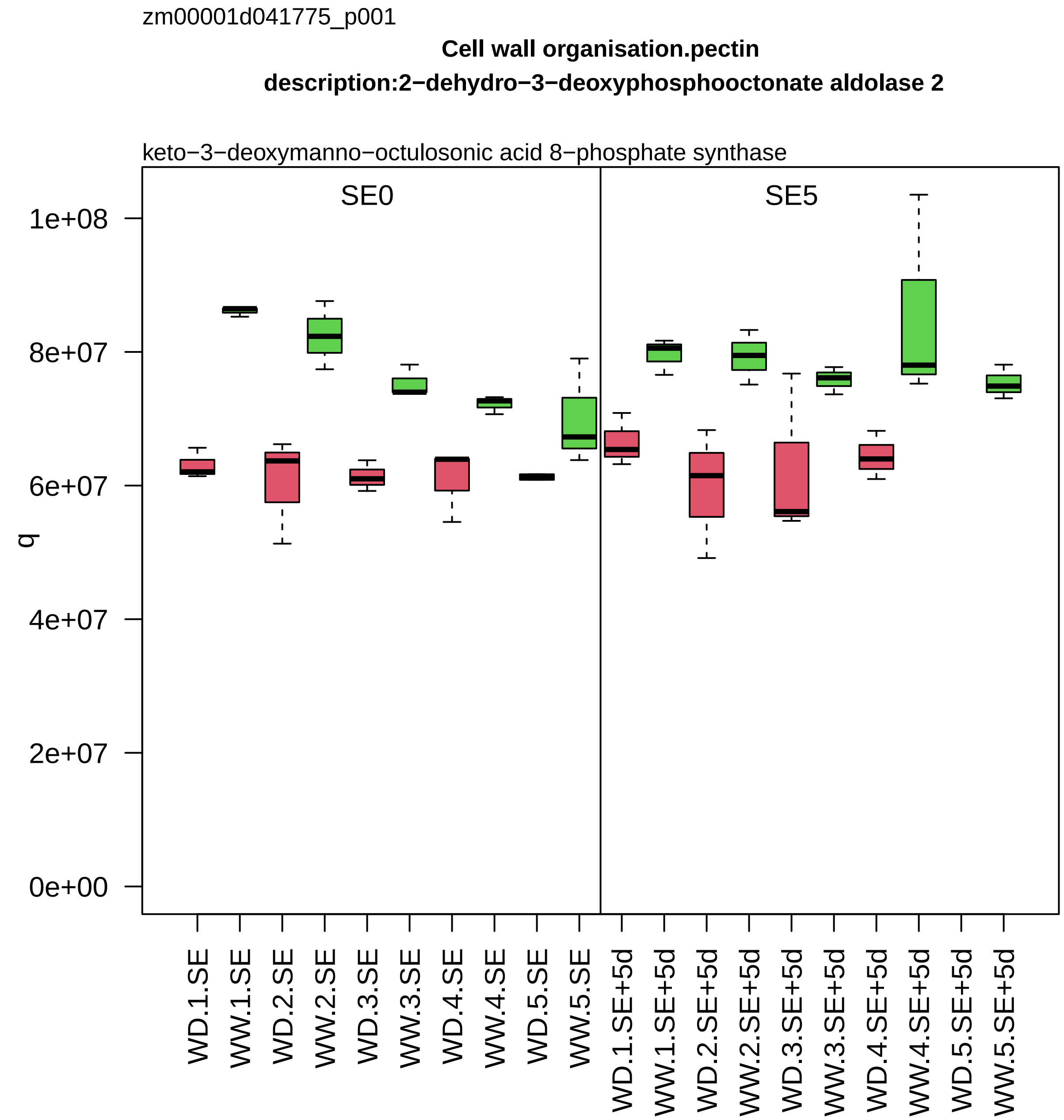
zm00001d041775_p001
Cell wall organisation.pectin
description:2−dehydro−3−deoxyphosphooctonate aldolase 2
keto−3−deoxymanno−octulosonic acid 8−phosphate synthase
SE0
SE5
q
1e+08
8e+07
6e+07
4e+07
2e+07
0e+00
WD.1.SE
WW.1.SE
WD.2.SE
WW.2.SE
WD.3.SE
WW.3.SE
WD.4.SE
WW.4.SE
WD.5.SE
WW.5.SE
WD.1.SE+5d
WW.1.SE+5d
WD.2.SE+5d
WW.2.SE+5d
WD.3.SE+5d
WW.3.SE+5d
WD.4.SE+5d
WW.4.SE+5d
WD.5.SE+5d
WW.5.SE+5d

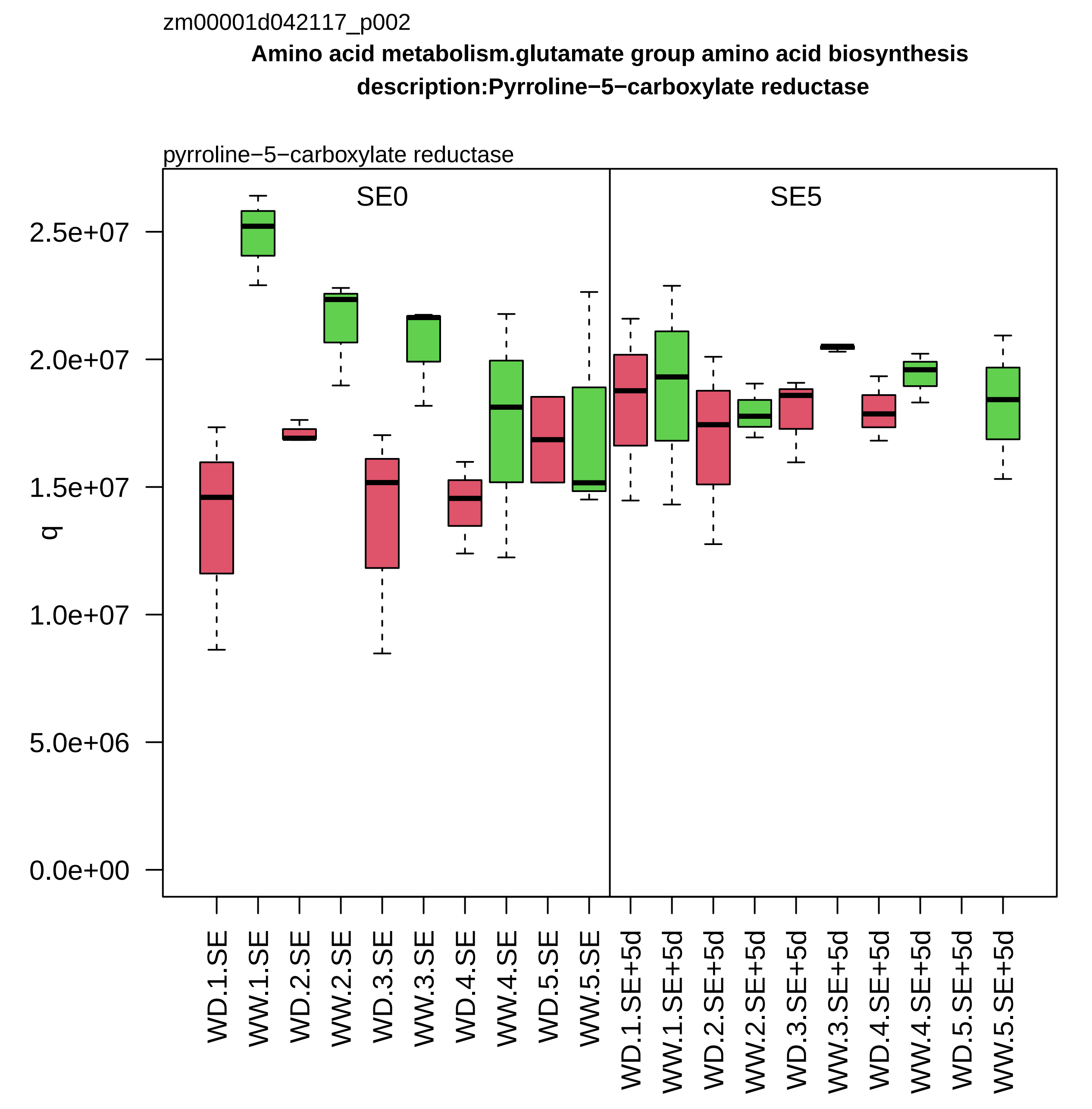

zm00001d042117_p002
Amino acid metabolism.glutamate group amino acid biosynthesis
description:Pyrroline−5−carboxylate reductase
pyrroline−5−carboxylate reductase
SE0
SE5
q
2.5e+07
2.0e+07
1.5e+07
1.0e+07
5.0e+06
0.0e+00
WD.1.SE
WW.1.SE
WD.2.SE
WW.2.SE
WD.3.SE
WW.3.SE
WD.4.SE
WW.4.SE
WD.5.SE
WW.5.SE
WD.1.SE+5d
WW.1.SE+5d
WD.2.SE+5d
WW.2.SE+5d
WD.3.SE+5d
WW.3.SE+5d
WD.4.SE+5d
WW.4.SE+5d
WD.5.SE+5d
WW.5.SE+5d

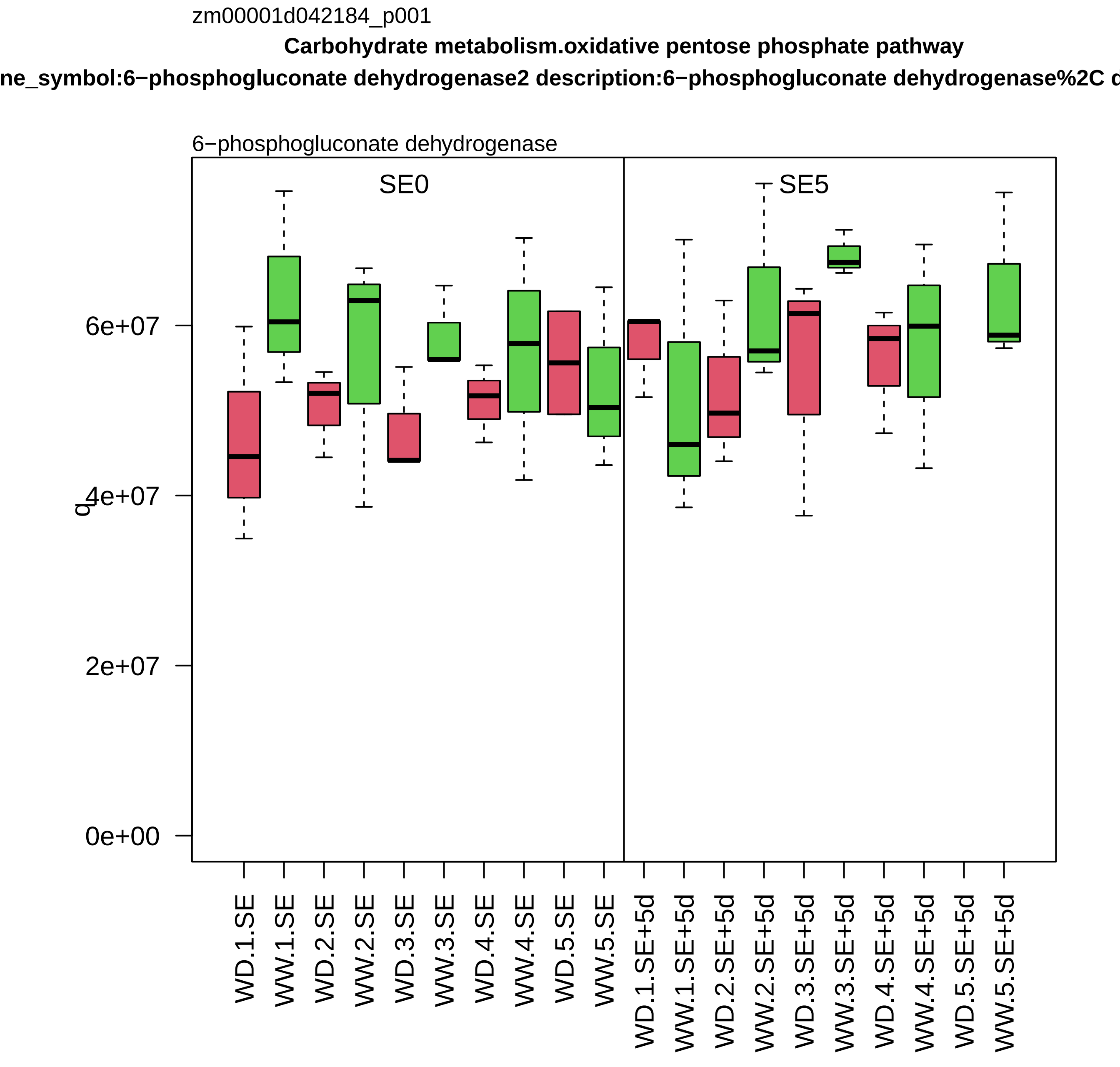
zm00001d042184_p001
Carbohydrate metabolism.oxidative pentose phosphate pathway
ne_symbol:6−phosphogluconate dehydrogenase2 description:6−phosphogluconate dehydrogenase%2C d
6−phosphogluconate dehydrogenase
SE0
SE5
q
0e+00
2e+07
4e+07
6e+07
WD.1.SE
WW.1.SE
WD.2.SE
WW.2.SE
WD.3.SE
WW.3.SE
WD.4.SE
WW.4.SE
WD.5.SE
WW.5.SE
WD.1.SE+5d
WW.1.SE+5d
WD.2.SE+5d
WW.2.SE+5d
WD.3.SE+5d
WW.3.SE+5d
WD.4.SE+5d
WW.4.SE+5d
WD.5.SE+5d
WW.5.SE+5d

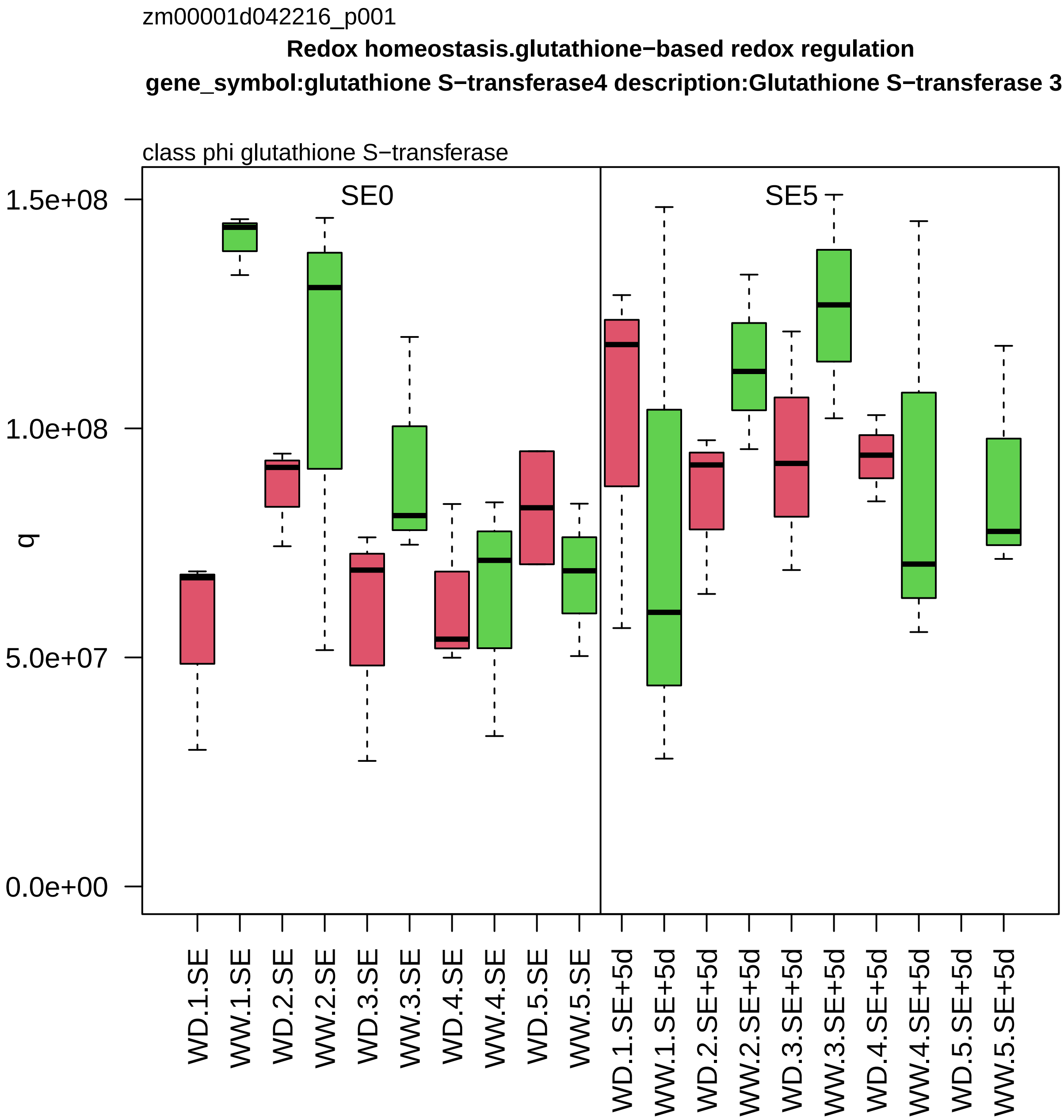

zm00001d042216_p001
Redox homeostasis.glutathione−based redox regulation
gene_symbol:glutathione S−transferase4 description:Glutathione S−transferase 3
class phi glutathione S−transferase
SE0
SE5
q
1.5e+08
1.0e+08
5.0e+07
0.0e+00
WD.1.SE
WW.1.SE
WD.2.SE
WW.2.SE
WD.3.SE
WW.3.SE
WD.4.SE
WW.4.SE
WD.5.SE
WW.5.SE
WD.1.SE+5d
WW.1.SE+5d
WD.2.SE+5d
WW.2.SE+5d
WD.3.SE+5d
WW.3.SE+5d
WD.4.SE+5d
WW.4.SE+5d
WD.5.SE+5d
WW.5.SE+5d

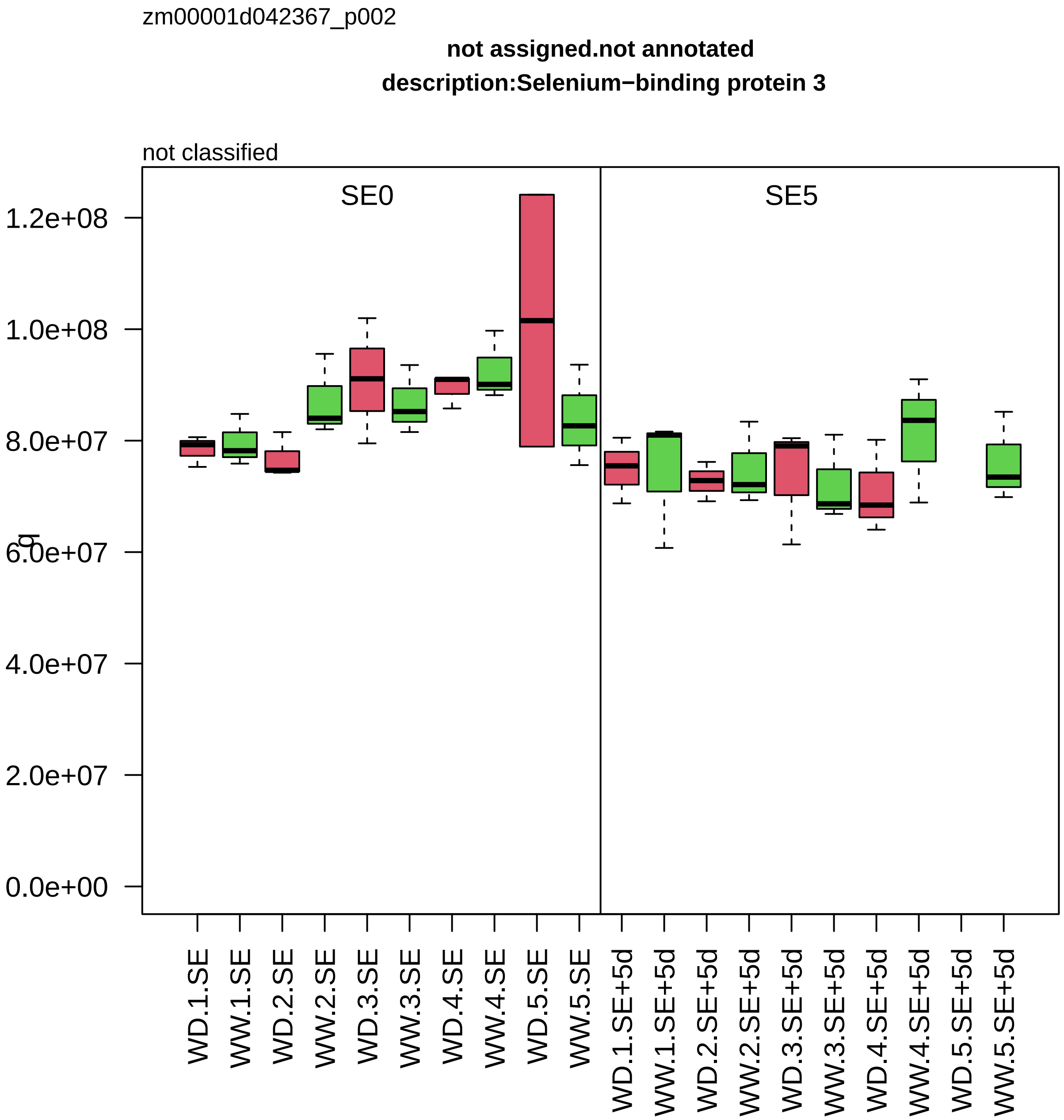

zm00001d042367_p002
not assigned.not annotated
description:Selenium−binding protein 3
not classified
SE0
SE5
1.2e+08
1.0e+08
8.0e+07
6.0e+07
4.0e+07
2.0e+07
0.0e+00
WD.1.SE
WW.1.SE
WD.2.SE
WW.2.SE
WD.3.SE
WW.3.SE
WD.4.SE
WW.4.SE
WD.5.SE
WW.5.SE
WD.1.SE+5d
WW.1.SE+5d
WD.2.SE+5d
WW.2.SE+5d
WD.3.SE+5d
WW.3.SE+5d
WD.4.SE+5d
WW.4.SE+5d
WD.5.SE+5d
WW.5.SE+5d

zm00001d042398_p002

**Protein modification.glycosylation**

**description:Dolichyl−diphosphooligosaccharide−−protein glycosyltransferase subunit 2**

component *(HAP6) of oligosaccharyl transferase (OST) complex

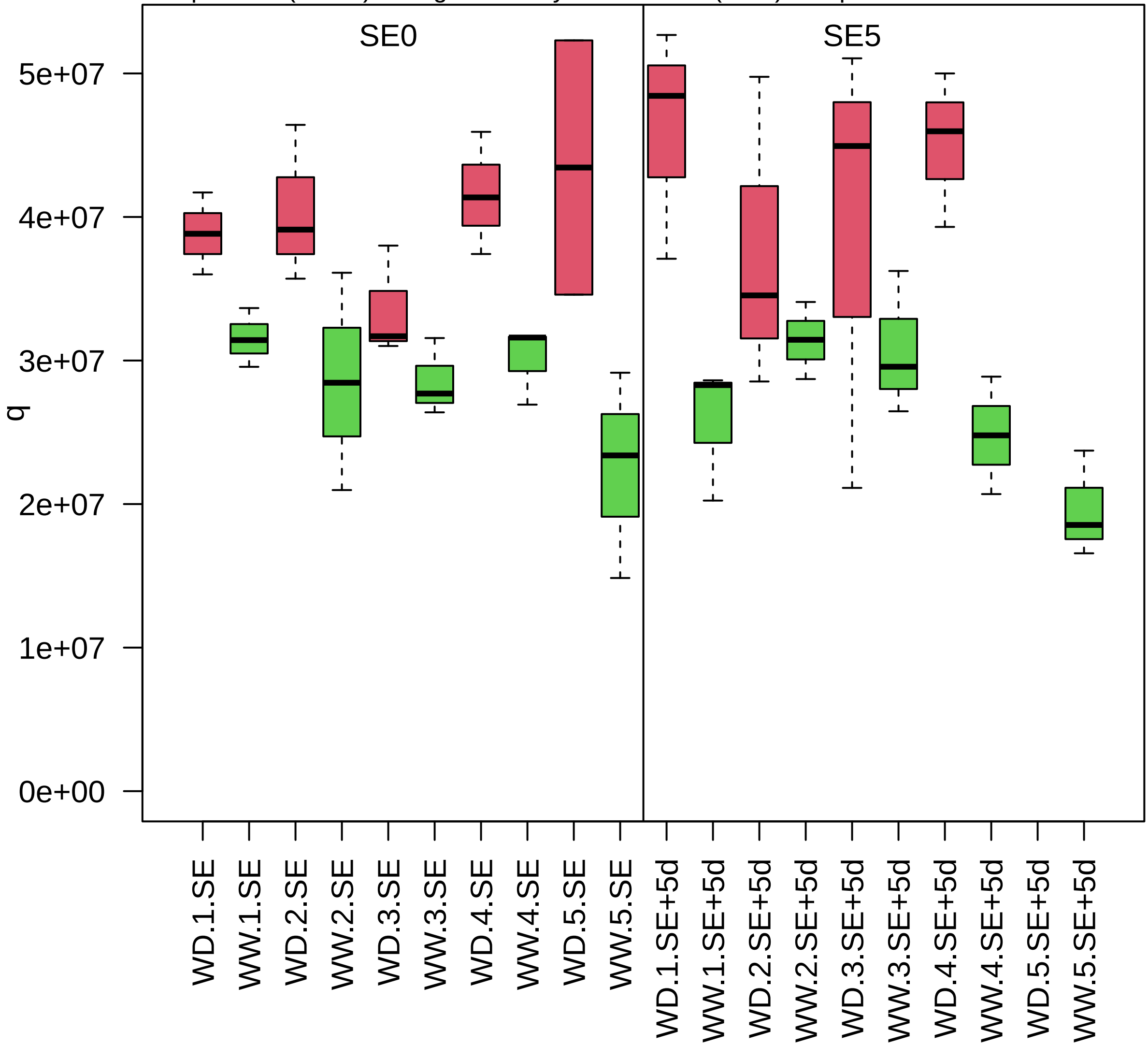

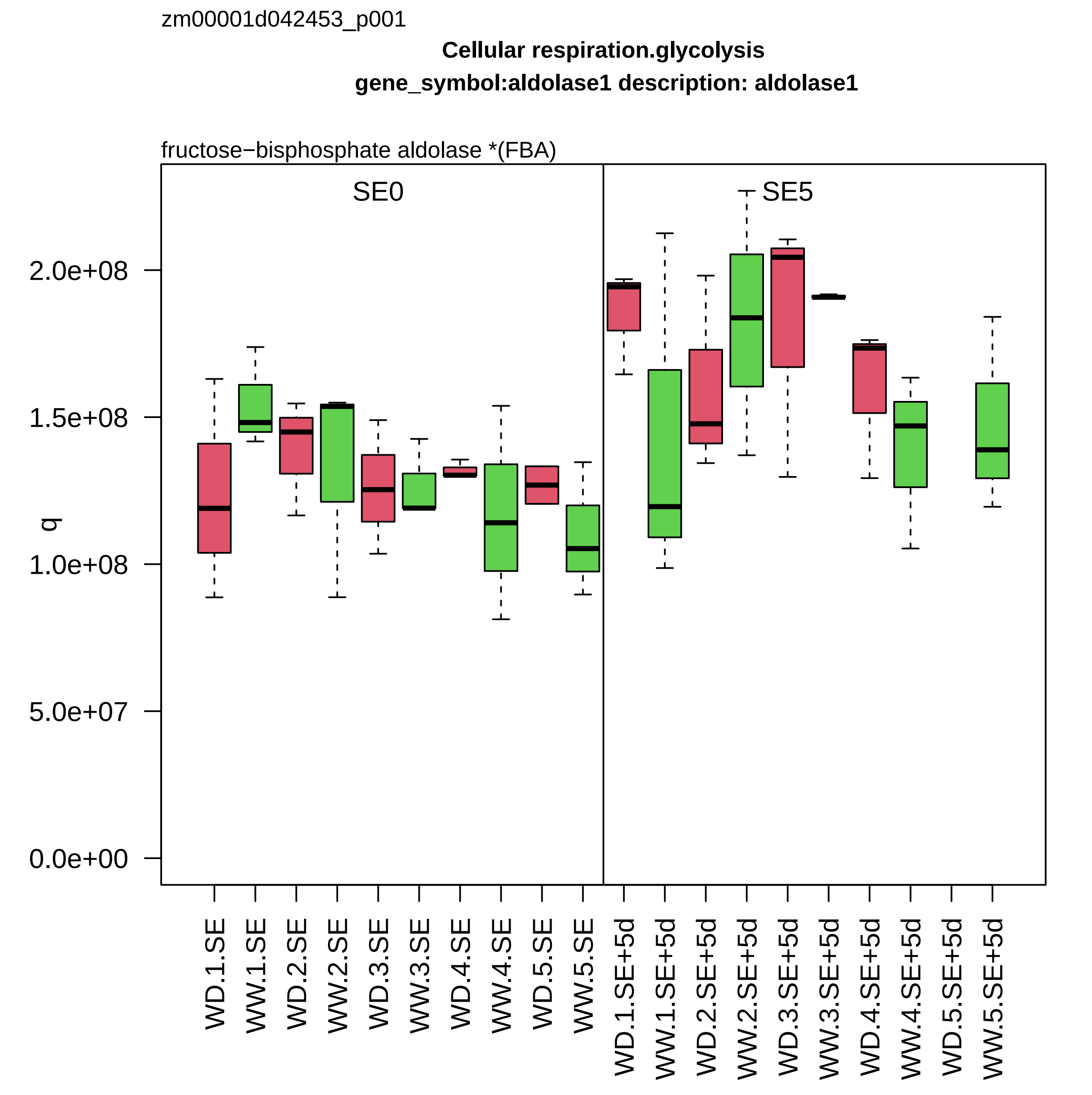
zm00001d042453_p001
Cellular respiration.glycolysis
gene_symbol:aldolase1 description: aldolase1
fructose−bisphosphate aldolase *(FBA)
SE0
SE5
q
2.0e+08
1.5e+08
1.0e+08
5.0e+07
0.0e+00
WD.1.SE
WW.1.SE
WD.2.SE
WW.2.SE
WD.3.SE
WW.3.SE
WD.4.SE
WW.4.SE
WD.5.SE
WW.5.SE
WD.1.SE+5d
WW.1.SE+5d
WD.2.SE+5d
WW.2.SE+5d
WD.3.SE+5d
WW.3.SE+5d
WD.4.SE+5d
WW.4.SE+5d
WD.5.SE+5d
WW.5.SE+5d

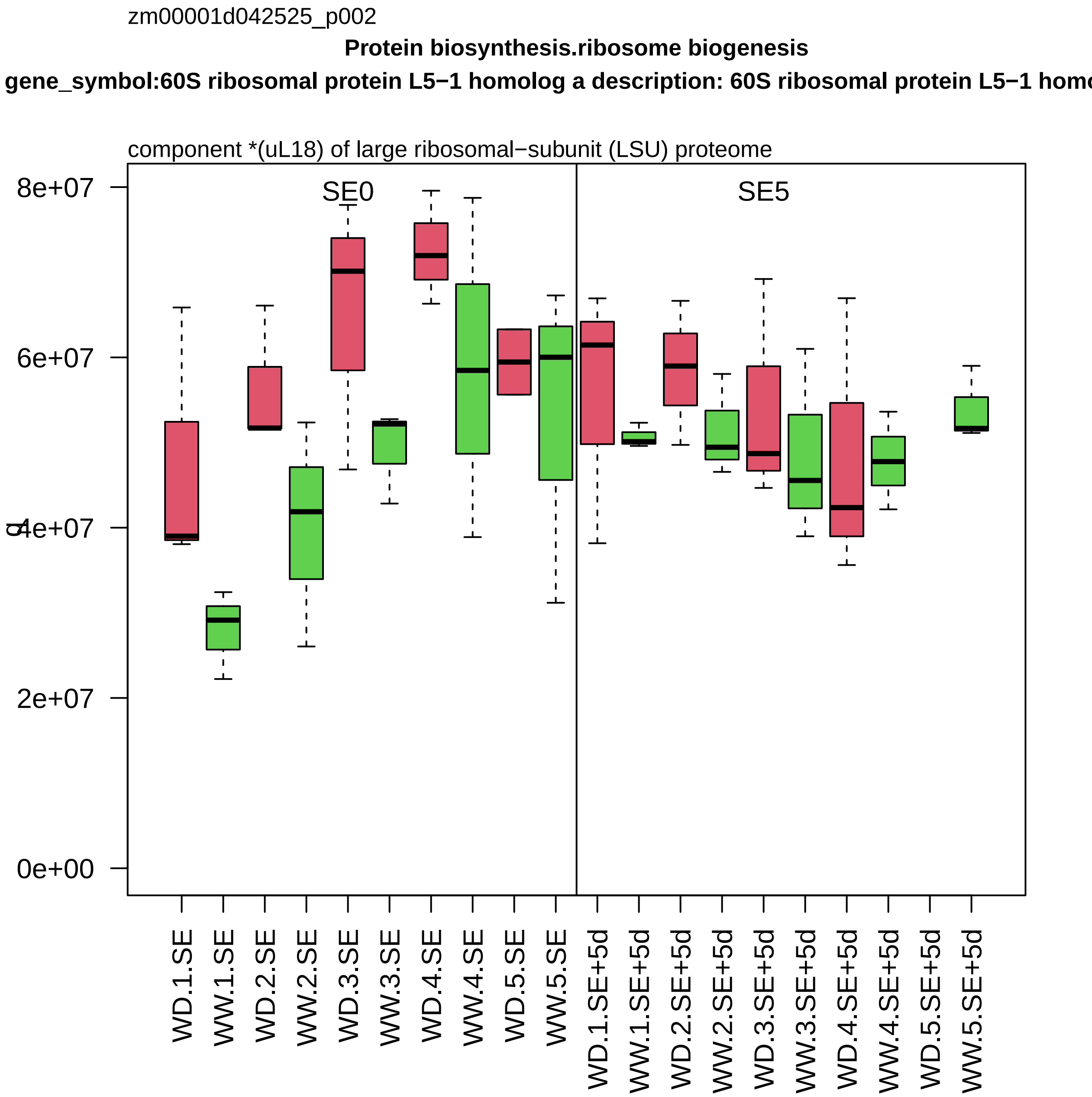
zm00001d042525_p002
Protein biosynthesis.ribosome biogenesis
gene_symbol:60S ribosomal protein L5−1 homolog a description: 60S ribosomal protein L5−1 homo
component *(uL18) of large ribosomal−subunit (LSU) proteome
SE0
SE5
8e+07
6e+07
4e+07
2e+07
0e+00
WD.1.SE
WW.1.SE
WD.2.SE
WW.2.SE
WD.3.SE
WW.3.SE
WD.4.SE
WW.4.SE
WD.5.SE
WW.5.SE
WD.1.SE+5d
WW.1.SE+5d
WD.2.SE+5d
WW.2.SE+5d
WD.3.SE+5d
WW.3.SE+5d
WD.4.SE+5d
WW.4.SE+5d
WD.5.SE+5d
WW.5.SE+5d

zm00001d042536_p001

**Carbohydrate metabolism.sucrose metabolism**

**gene_symbol:fructokinase1 description: fructokinase1**

fructose kinase

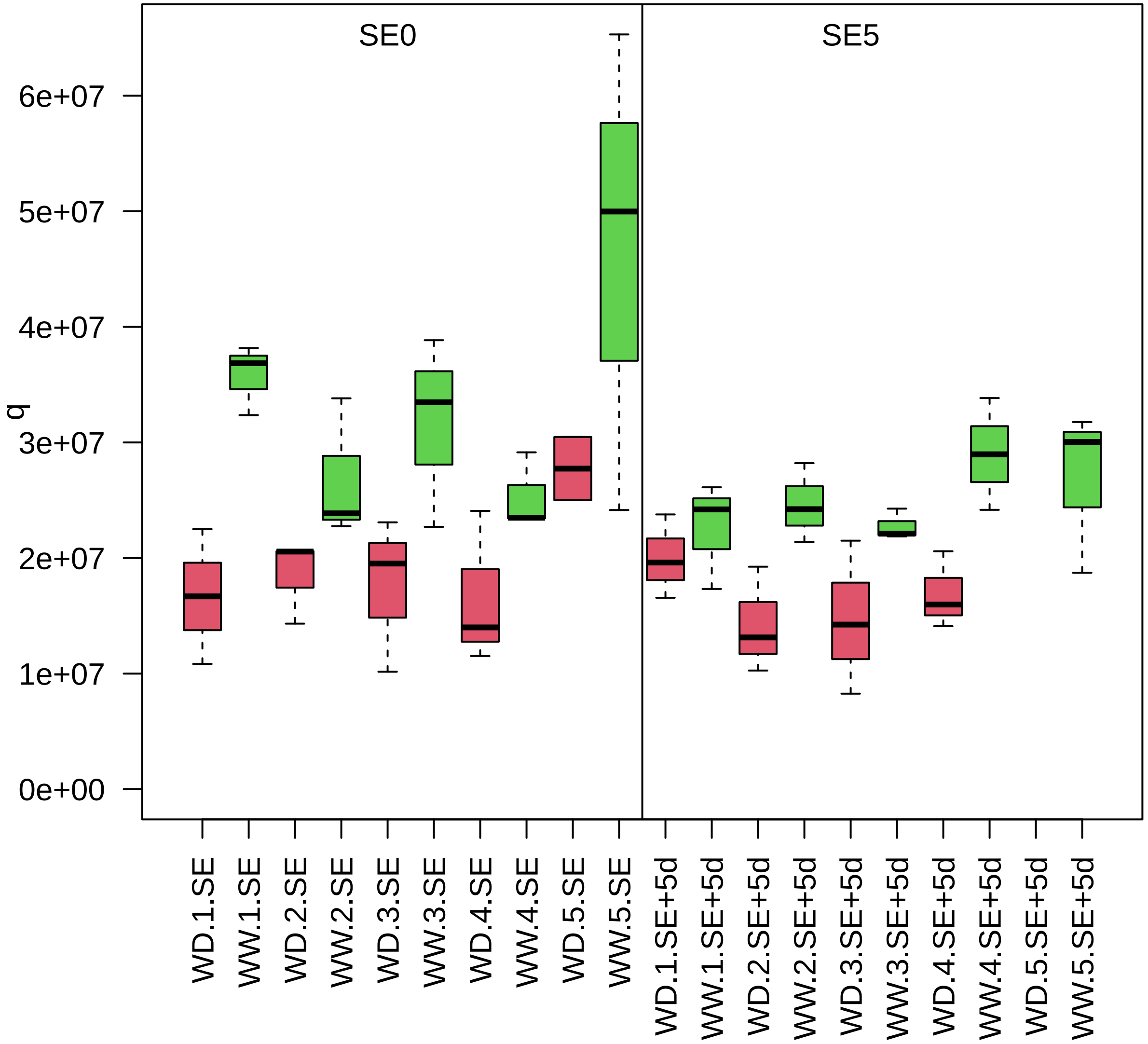

zm00001d042540_p007

**Redox homeostasis.reactive electrophilic lipid homeostasis**
**description:Linoleate 9S−lipoxygenase 1**

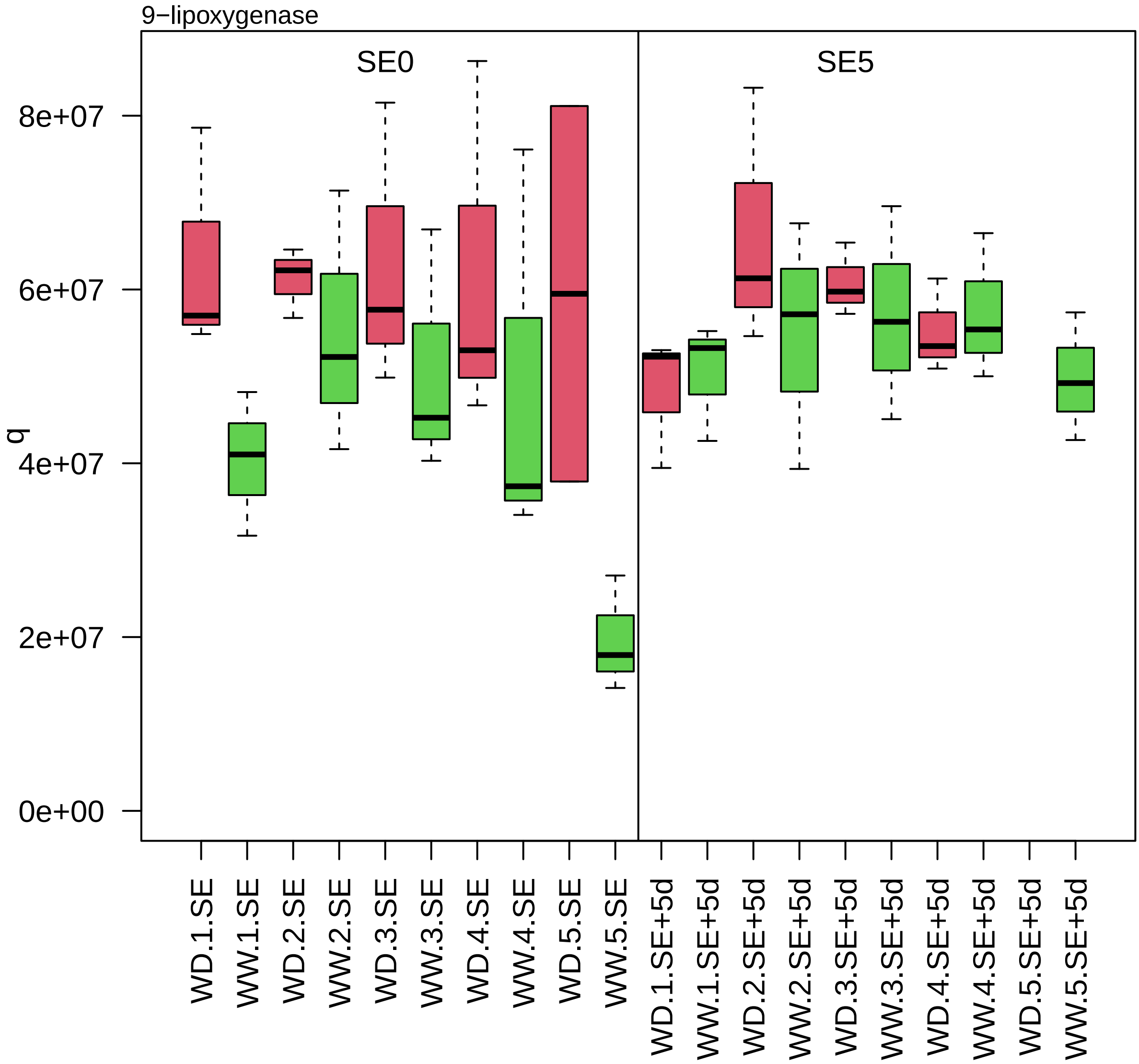

zm00001d042555_p001

**Secondary metabolism.phenolics**

**description:Zinc−binding dehydrogenase family protein**

hydroxyacyl−CoA dehydrogenase *(HCD)

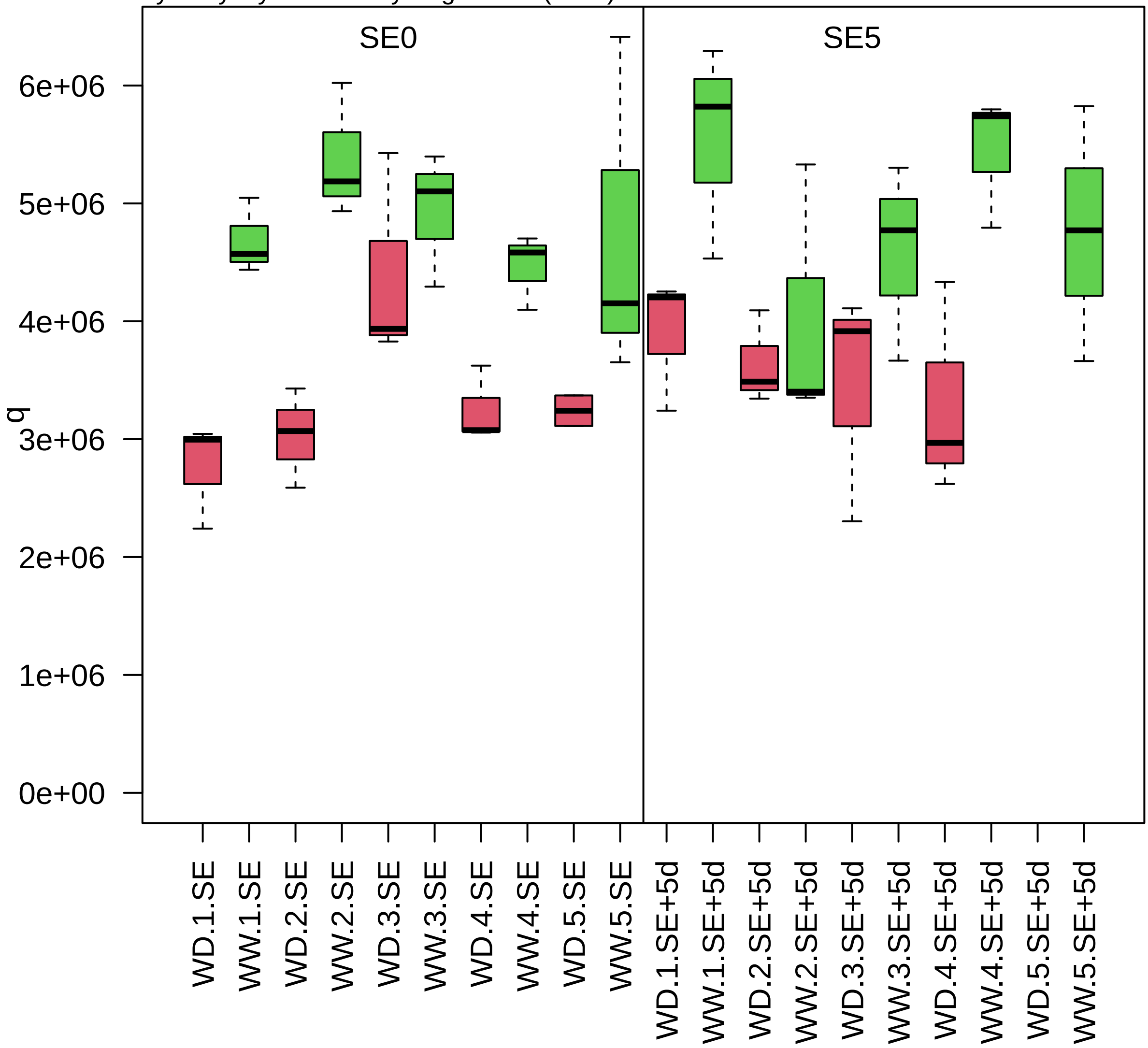

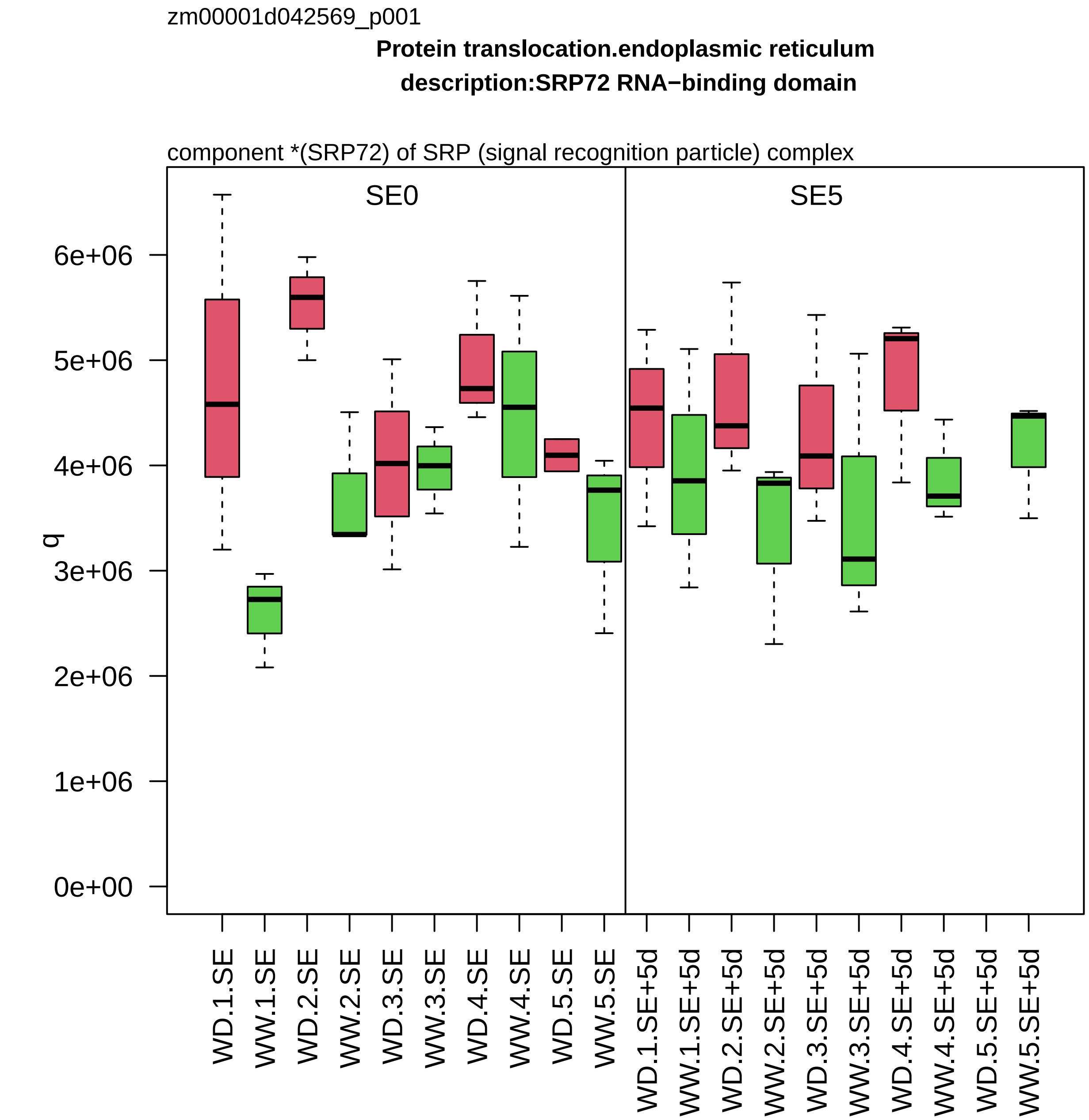
zm00001d042569_p001
Protein translocation.endoplasmic reticulum
description:SRP72 RNA−binding domain
component *(SRP72) of SRP (signal recognition particle) complex
SE0
SE5
q
0e+00
1e+06
2e+06
3e+06
4e+06
5e+06
6e+06
WD.1.SE
WW.1.SE
WD.2.SE
WW.2.SE
WD.3.SE
WW.3.SE
WD.4.SE
WW.4.SE
WD.5.SE
WW.5.SE
WD.1.SE+5d
WW.1.SE+5d
WD.2.SE+5d
WW.2.SE+5d
WD.3.SE+5d
WW.3.SE+5d
WD.4.SE+5d
WW.4.SE+5d
WD.5.SE+5d
WW.5.SE+5d

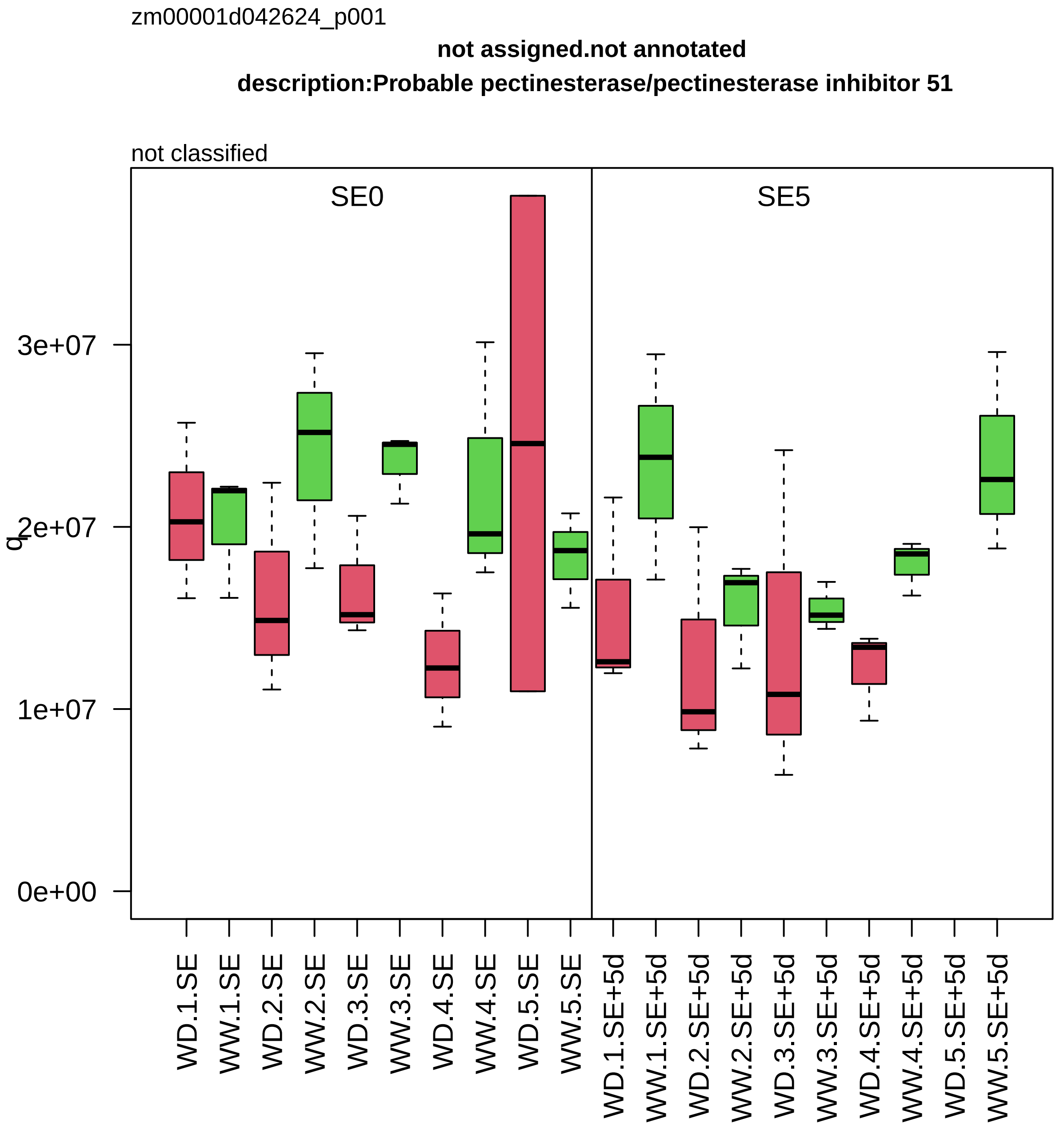
zm00001d042624_p001
not assigned.not annotated
description:Probable pectinesterase/pectinesterase inhibitor 51
not classified
SE0
SE5
q
3e+07
2e+07
1e+07
0e+00
WD.1.SE
WW.1.SE
WD.2.SE
WW.2.SE
WD.3.SE
WW.3.SE
WD.4.SE
WW.4.SE
WD.5.SE
WW.5.SE
WD.1.SE+5d
WW.1.SE+5d
WD.2.SE+5d
WW.2.SE+5d
WD.3.SE+5d
WW.3.SE+5d
WD.4.SE+5d
WW.4.SE+5d
WD.5.SE+5d
WW.5.SE+5d

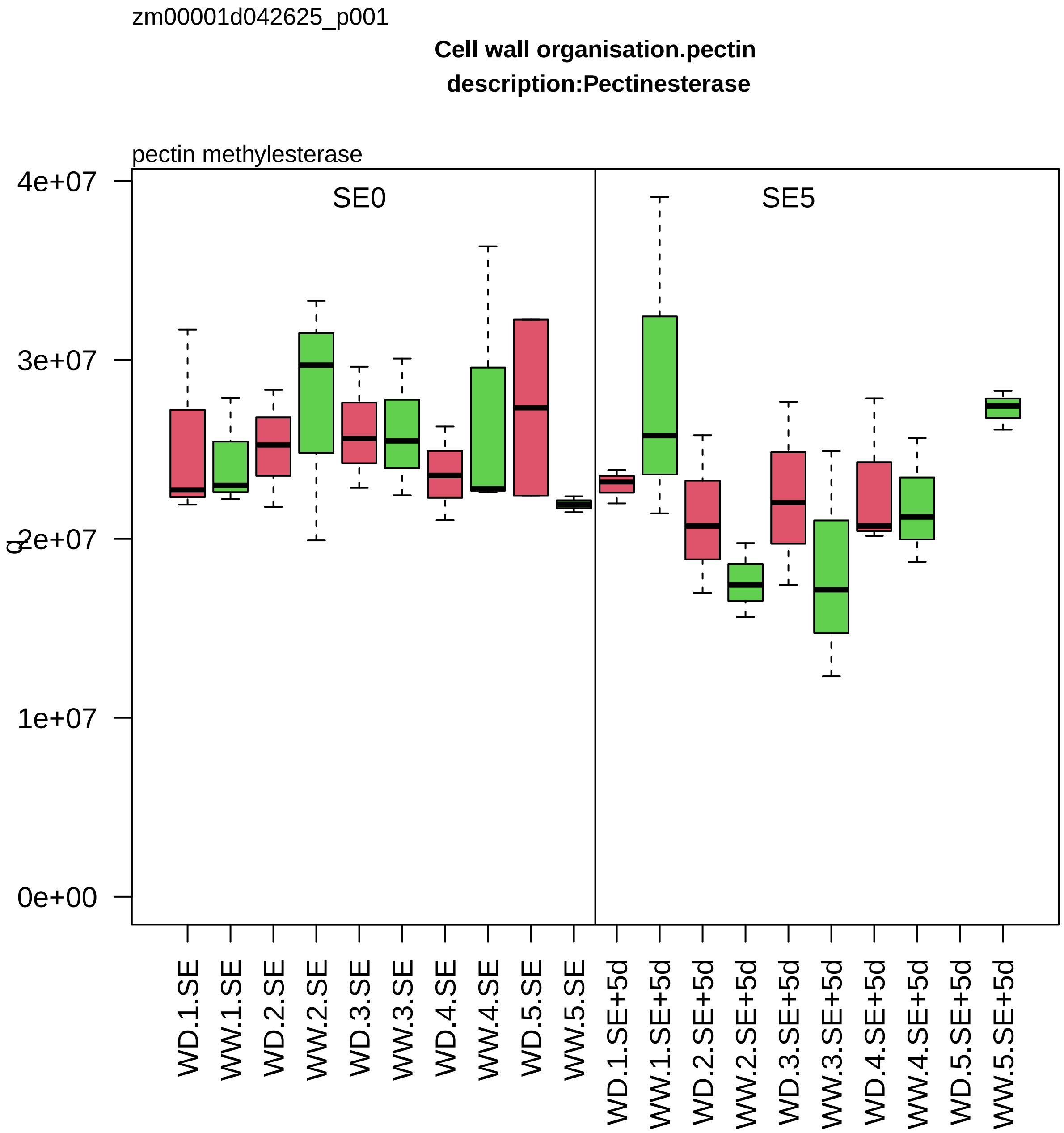
zm00001d042625_p001
Cell wall organisation.pectin
description:Pectinesterase
pectin methylesterase
SE0
SE5
4e+07
3e+07
2e+07
1e+07
0e+00
WD.1.SE
WW.1.SE
WD.2.SE
WW.2.SE
WD.3.SE
WW.3.SE
WD.4.SE
WW.4.SE
WD.5.SE
WW.5.SE
WD.1.SE+5d
WW.1.SE+5d
WD.2.SE+5d
WW.2.SE+5d
WD.3.SE+5d
WW.3.SE+5d
WD.4.SE+5d
WW.4.SE+5d
WD.5.SE+5d
WW.5.SE+5d

zm00001d042654_p003

**Cell wall organisation.hemicellulose**
**description:Beta−galactosidase**

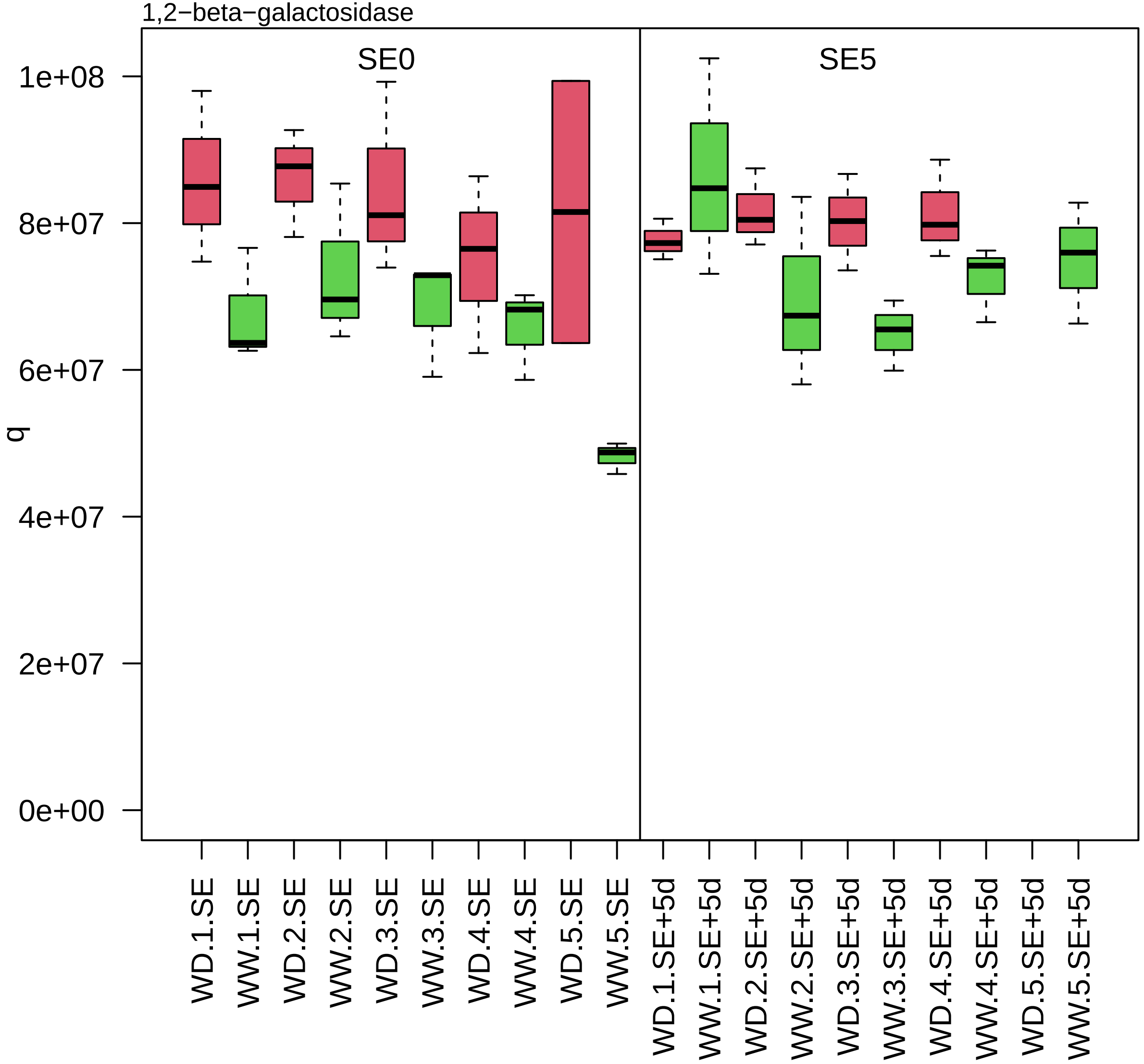

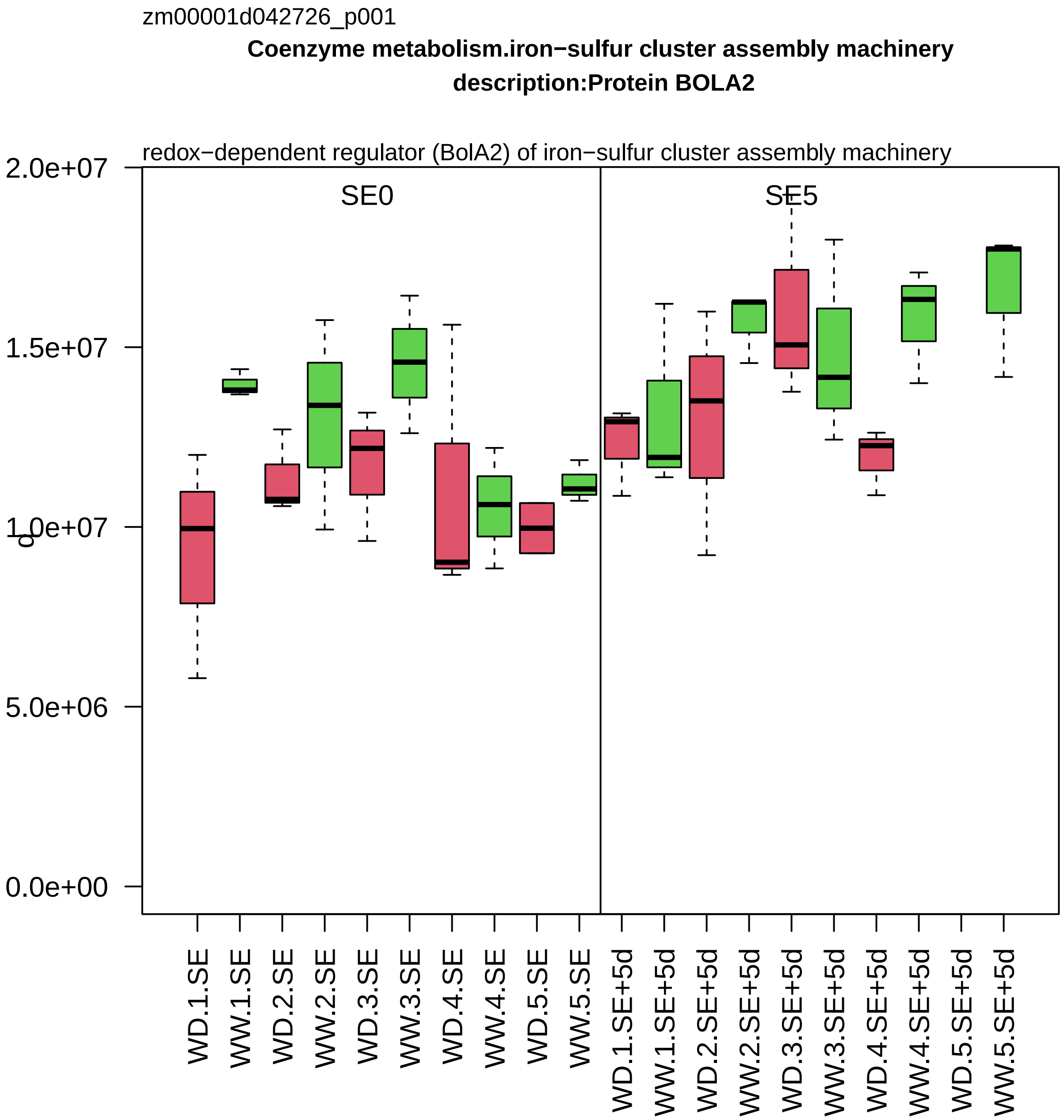

zm00001d042726_p001
Coenzyme metabolism.iron−sulfur cluster assembly machinery
description:Protein BOLA2
redox−dependent regulator (BolA2) of iron−sulfur cluster assembly machinery
SE0
SE5
2.0e+07
1.5e+07
1.0e+07
5.0e+06
0.0e+00
WD.1.SE
WW.1.SE
WD.2.SE
WW.2.SE
WD.3.SE
WW.3.SE
WD.4.SE
WW.4.SE
WD.5.SE
WW.5.SE
WD.1.SE+5d
WW.1.SE+5d
WD.2.SE+5d
WW.2.SE+5d
WD.3.SE+5d
WW.3.SE+5d
WD.4.SE+5d
WW.4.SE+5d
WD.5.SE+5d
WW.5.SE+5d

zm00001d042746_p002

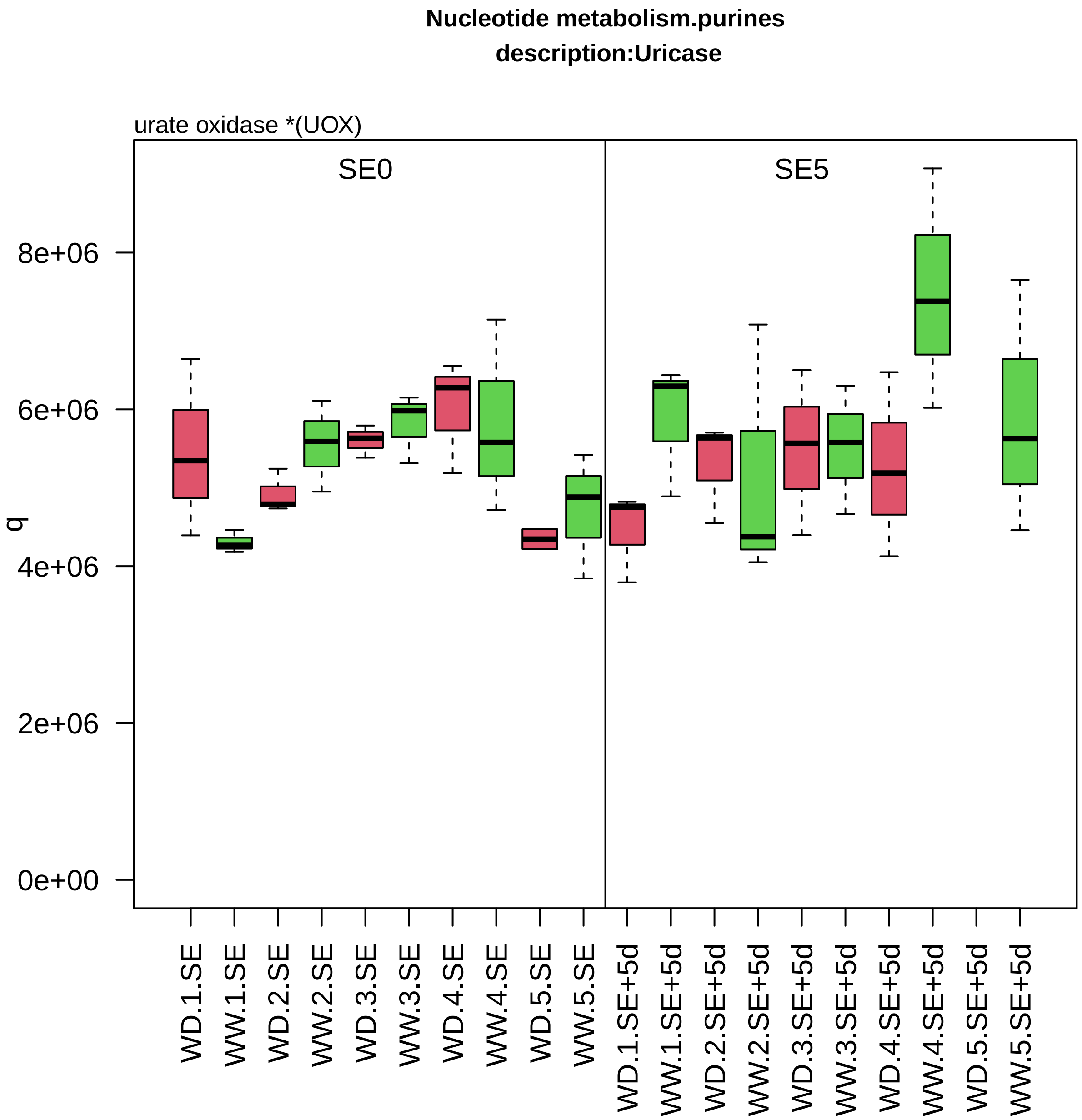

zm00001d042842_p003

**Carbohydrate metabolism.starch metabolism**

**gene_symbol:starch phoshorylase2 description: starch phoshorylase2**

cytosolic alpha−glucan phosphorylase

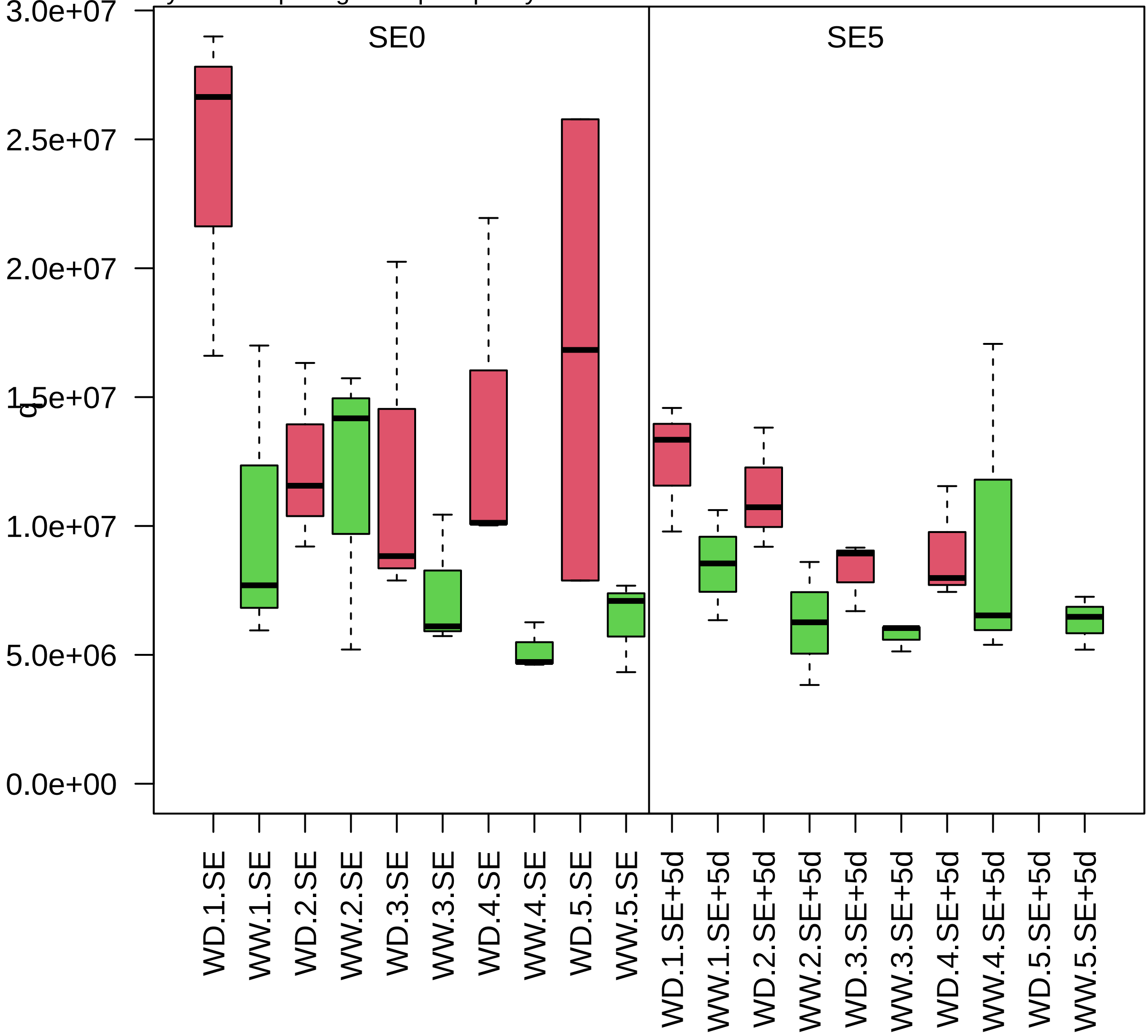

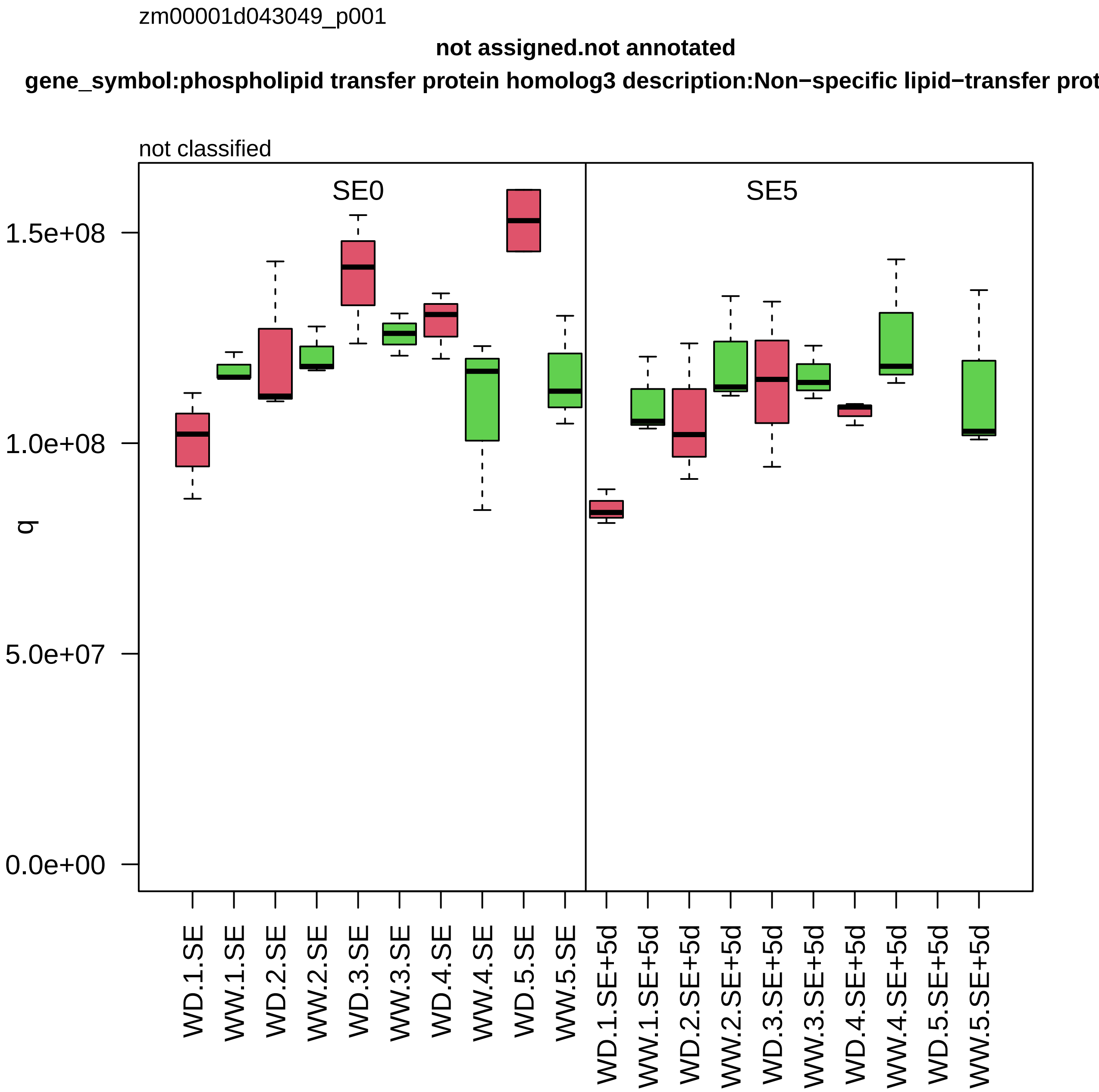
zm00001d043049_p001
not assigned.not annotated
gene_symbol:phospholipid transfer protein homolog3 description:Non−specific lipid−transfer prot
not classified
SE0
SE5
q
1.5e+08
1.0e+08
5.0e+07
0.0e+00
WD.1.SE
WW.1.SE
WD.2.SE
WW.2.SE
WD.3.SE
WW.3.SE
WD.4.SE
WW.4.SE
WD.5.SE
WW.5.SE
WD.1.SE+5d
WW.1.SE+5d
WD.2.SE+5d
WW.2.SE+5d
WD.3.SE+5d
WW.3.SE+5d
WD.4.SE+5d
WW.4.SE+5d
WD.5.SE+5d
WW.5.SE+5d

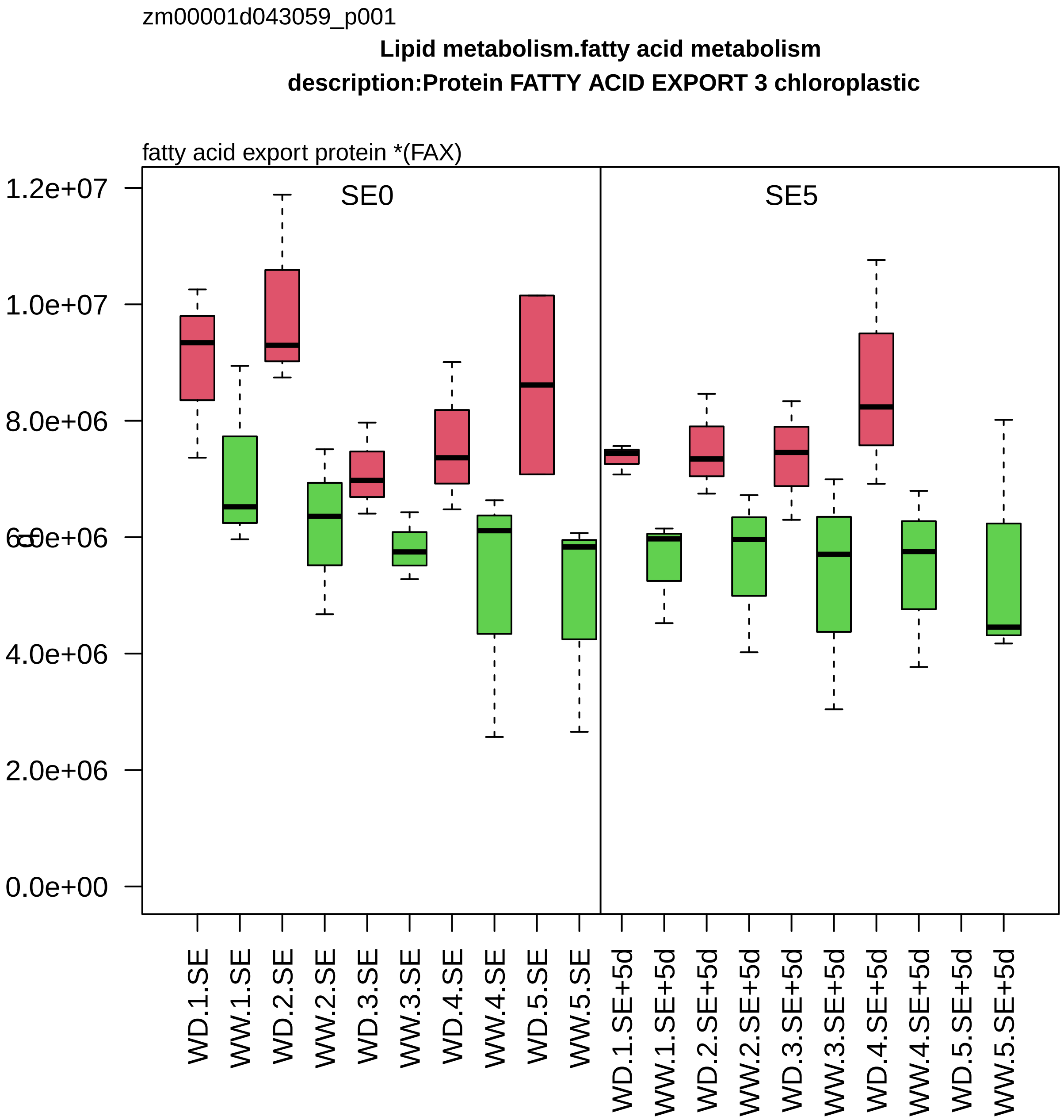
zm00001d043059_p001
Lipid metabolism.fatty acid metabolism
description:Protein FATTY ACID EXPORT 3 chloroplastic
fatty acid export protein *(FAX)
SE0
SE5
1.2e+07
1.0e+07
8.0e+06
6.0e+06
4.0e+06
2.0e+06
0.0e+00
WD.1.SE
WW.1.SE
WD.2.SE
WW.2.SE
WD.3.SE
WW.3.SE
WD.4.SE
WW.4.SE
WD.5.SE
WW.5.SE
WD.1.SE+5d
WW.1.SE+5d
WD.2.SE+5d
WW.2.SE+5d
WD.3.SE+5d
WW.3.SE+5d
WD.4.SE+5d
WW.4.SE+5d
WD.5.SE+5d
WW.5.SE+5d

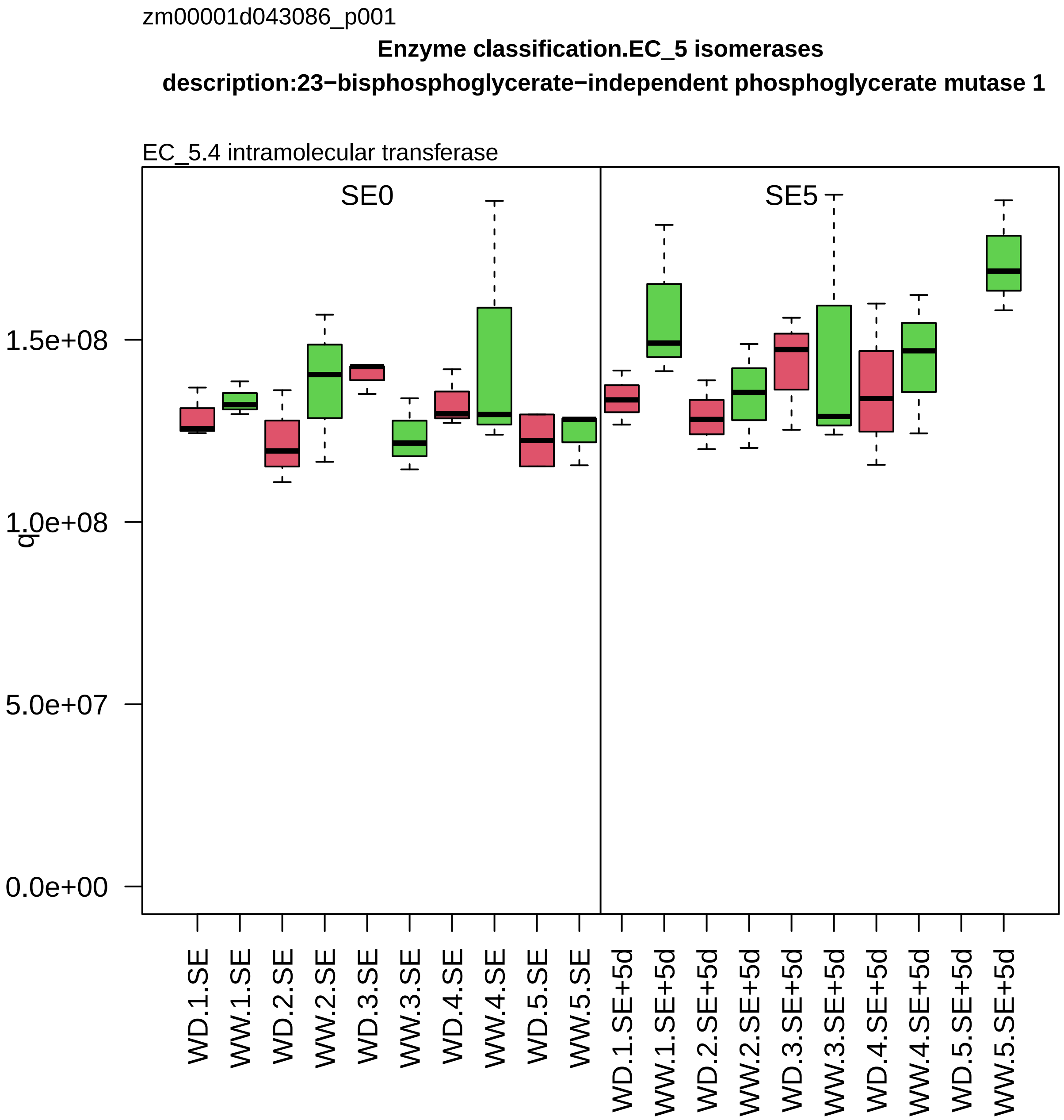

zm00001d043086_p001
Enzyme classification.EC_5 isomerases
description:23−bisphosphoglycerate−independent phosphoglycerate mutase 1
EC_5.4 intramolecular transferase
SE0
SE5
q
0.0e+00
5.0e+07
1.0e+08
1.5e+08
WD.1.SE
WW.1.SE
WD.2.SE
WW.2.SE
WD.3.SE
WW.3.SE
WD.4.SE
WW.4.SE
WD.5.SE
WW.5.SE
WD.1.SE+5d
WW.1.SE+5d
WD.2.SE+5d
WW.2.SE+5d
WD.3.SE+5d
WW.3.SE+5d
WD.4.SE+5d
WW.4.SE+5d
WD.5.SE+5d
WW.5.SE+5d

zm00001d043098_p001

**Lipid metabolism.fatty acid metabolism**

**gene_symbol:ferric−chelate reductase (NADH)1 description:Cytochrome b5 reductase**

NADH:cytochrome−b5 reductase *(CBR1)

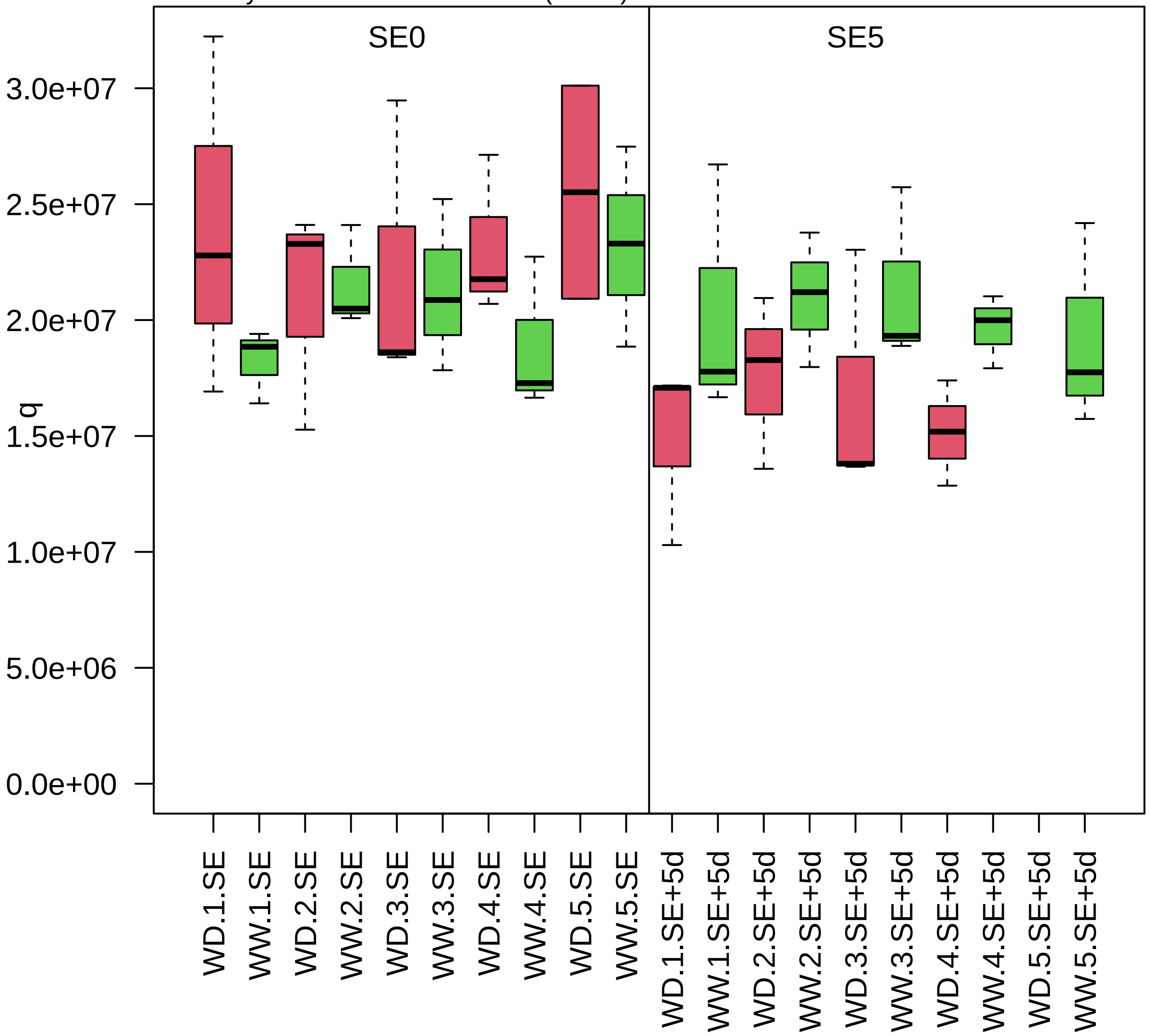

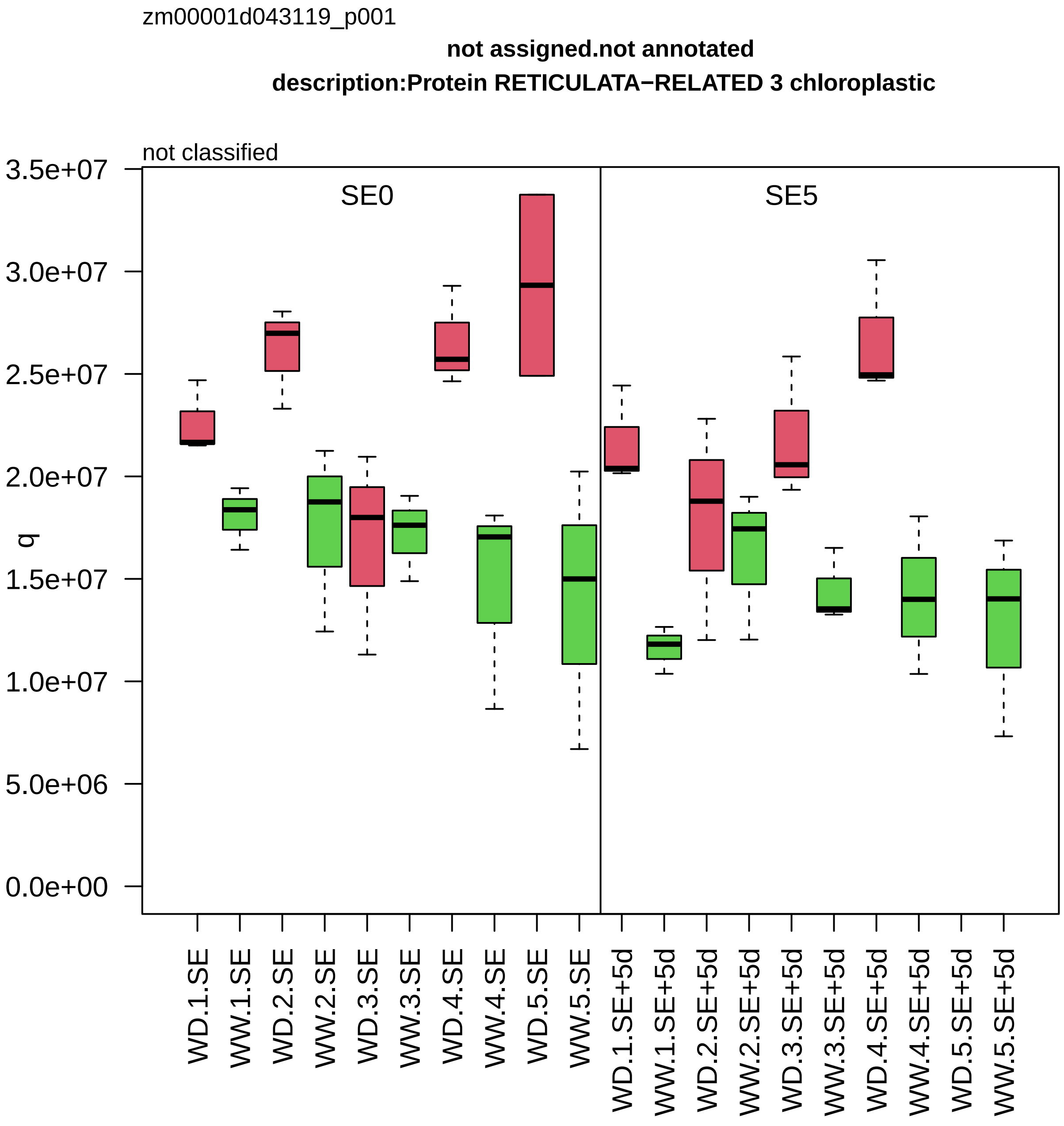

zm00001d043119_p001
not assigned.not annotated
description:Protein RETICULATA−RELATED 3 chloroplastic
not classified
SE0
SE5
q
3.5e+07
3.0e+07
2.5e+07
2.0e+07
1.5e+07
1.0e+07
5.0e+06
0.0e+00
WD.1.SE
WW.1.SE
WD.2.SE
WW.2.SE
WD.3.SE
WW.3.SE
WD.4.SE
WW.4.SE
WD.5.SE
WW.5.SE
WD.1.SE+5d
WW.1.SE+5d
WD.2.SE+5d
WW.2.SE+5d
WD.3.SE+5d
WW.3.SE+5d
WD.4.SE+5d
WW.4.SE+5d
WD.5.SE+5d
WW.5.SE+5d

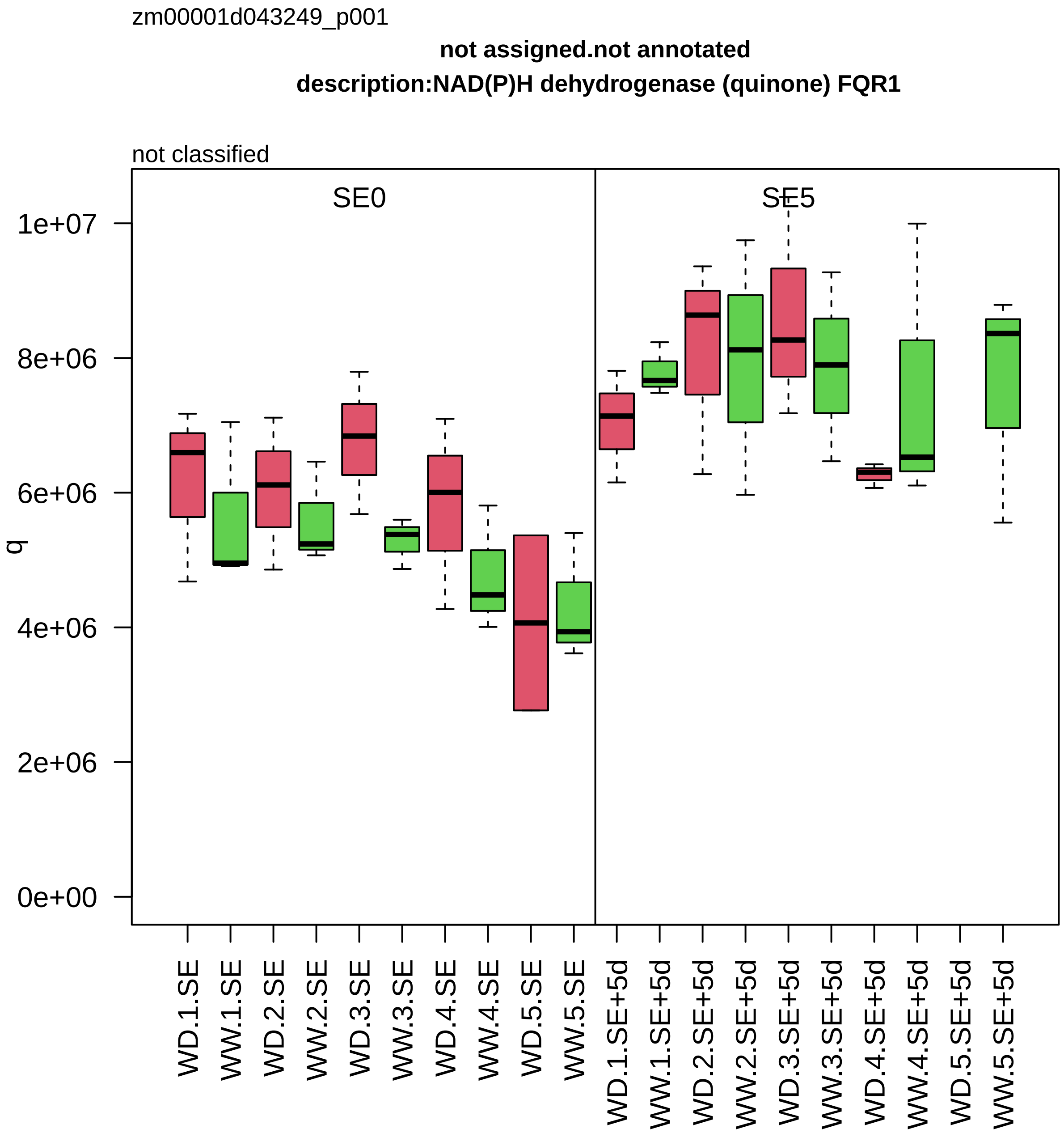

zm00001d043249_p001
not assigned.not annotated
description:NAD(P)H dehydrogenase (quinone) FQR1
not classified
SE0
SE5
q
1e+07
8e+06
6e+06
4e+06
2e+06
0e+00
WD.1.SE
WW.1.SE
WD.2.SE
WW.2.SE
WD.3.SE
WW.3.SE
WD.4.SE
WW.4.SE
WD.5.SE
WW.5.SE
WD.1.SE+5d
WW.1.SE+5d
WD.2.SE+5d
WW.2.SE+5d
WD.3.SE+5d
WW.3.SE+5d
WD.4.SE+5d
WW.4.SE+5d
WD.5.SE+5d
WW.5.SE+5d

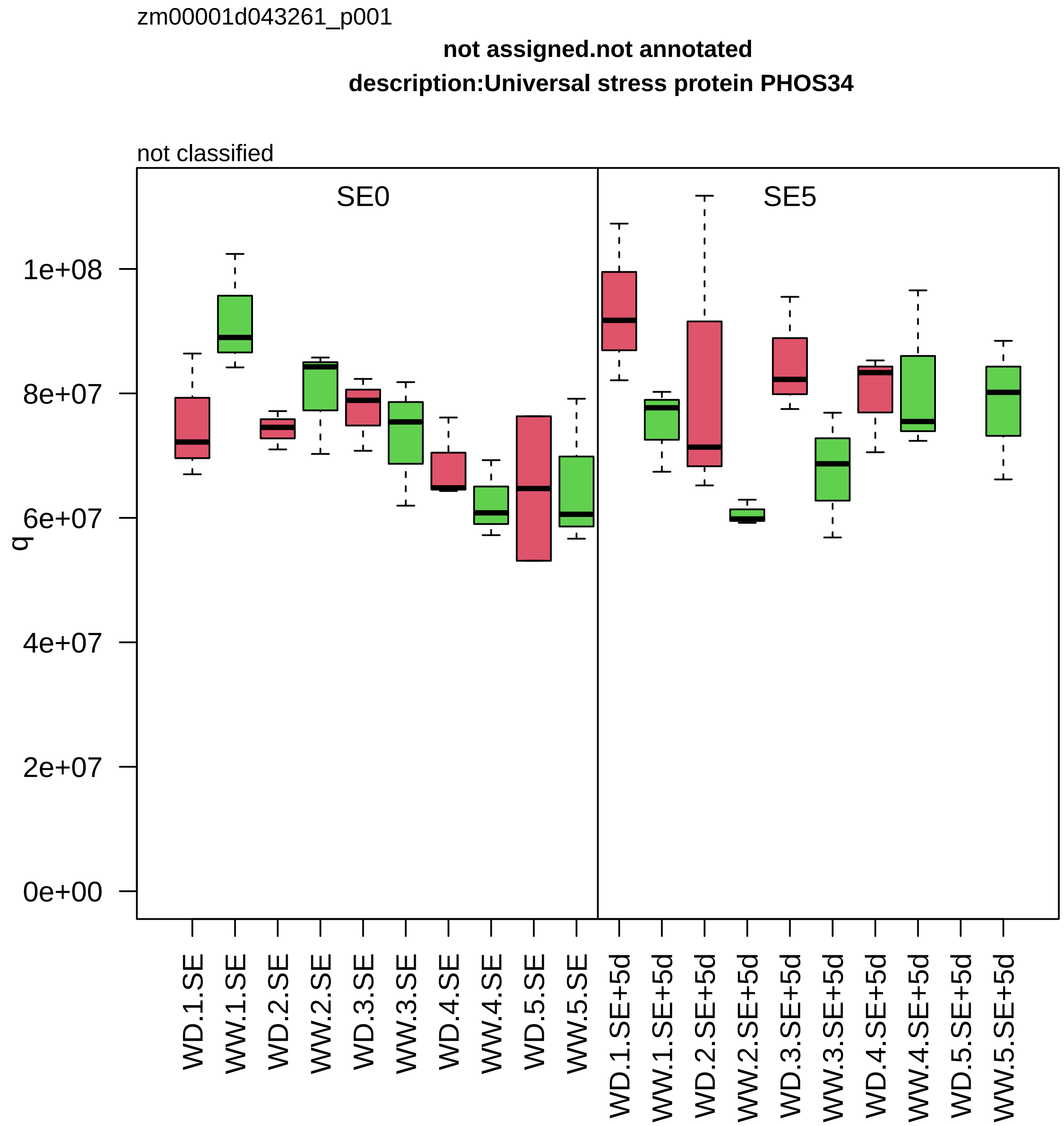

zm00001d043261_p001
not assigned.not annotated
description:Universal stress protein PHOS34
not classified
SE0
SE5
q
1e+08
8e+07
6e+07
4e+07
2e+07
0e+00
WD.1.SE
WW.1.SE
WD.2.SE
WW.2.SE
WD.3.SE
WW.3.SE
WD.4.SE
WW.4.SE
WD.5.SE
WW.5.SE
WD.1.SE+5d
WW.1.SE+5d
WD.2.SE+5d
WW.2.SE+5d
WD.3.SE+5d
WW.3.SE+5d
WD.4.SE+5d
WW.4.SE+5d
WD.5.SE+5d
WW.5.SE+5d

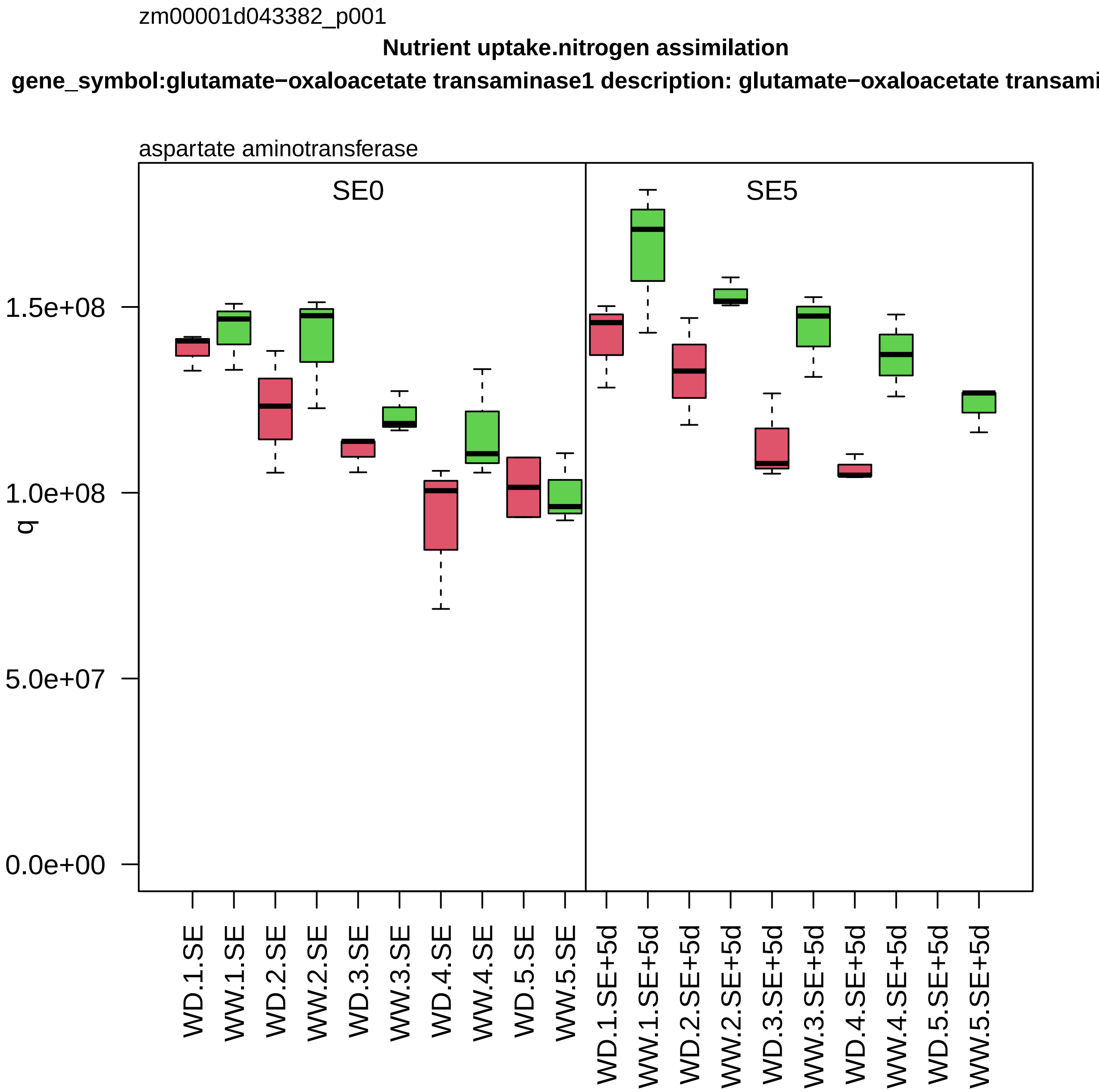

zm00001d043382_p001
Nutrient uptake.nitrogen assimilation
gene_symbol:glutamate−oxaloacetate transaminase1 description: glutamate−oxaloacetate transami
aspartate aminotransferase
SE0
SE5
q
0.0e+00
5.0e+07
1.0e+08
1.5e+08
WD.1.SE
WW.1.SE
WD.2.SE
WW.2.SE
WD.3.SE
WW.3.SE
WD.4.SE
WW.4.SE
WD.5.SE
WW.5.SE
WD.1.SE+5d
WW.1.SE+5d
WD.2.SE+5d
WW.2.SE+5d
WD.3.SE+5d
WW.3.SE+5d
WD.4.SE+5d
WW.4.SE+5d
WD.5.SE+5d
WW.5.SE+5d

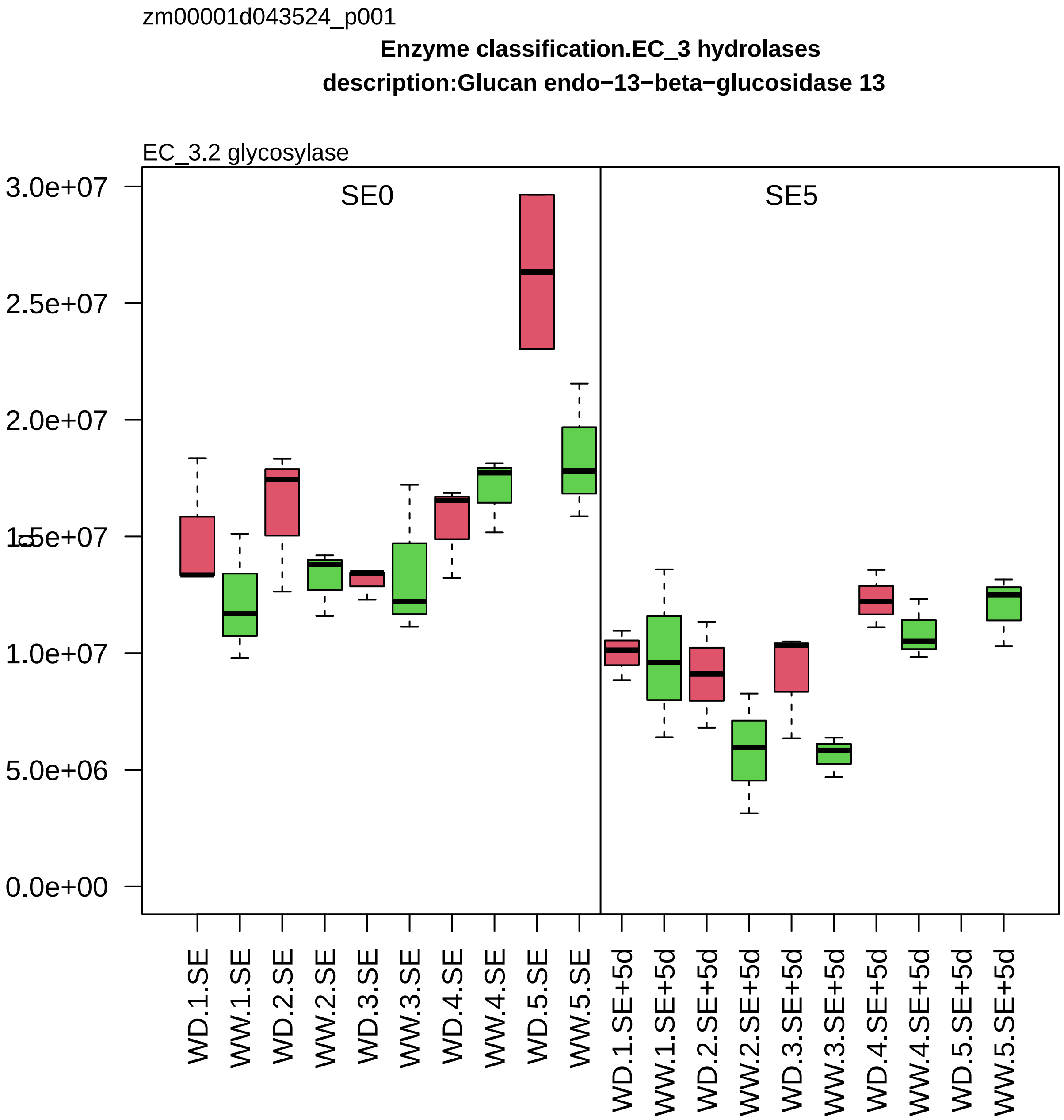
zm00001d043524_p001
Enzyme classification.EC_3 hydrolases
description:Glucan endo−13−beta−glucosidase 13
EC_3.2 glycosylase
SE0
SE5
3.0e+07
2.5e+07
2.0e+07
1.5e+07
1.0e+07
5.0e+06
0.0e+00
WD.1.SE
WW.1.SE
WD.2.SE
WW.2.SE
WD.3.SE
WW.3.SE
WD.4.SE
WW.4.SE
WD.5.SE
WW.5.SE
WD.1.SE+5d
WW.1.SE+5d
WD.2.SE+5d
WW.2.SE+5d
WD.3.SE+5d
WW.3.SE+5d
WD.4.SE+5d
WW.4.SE+5d
WD.5.SE+5d
WW.5.SE+5d

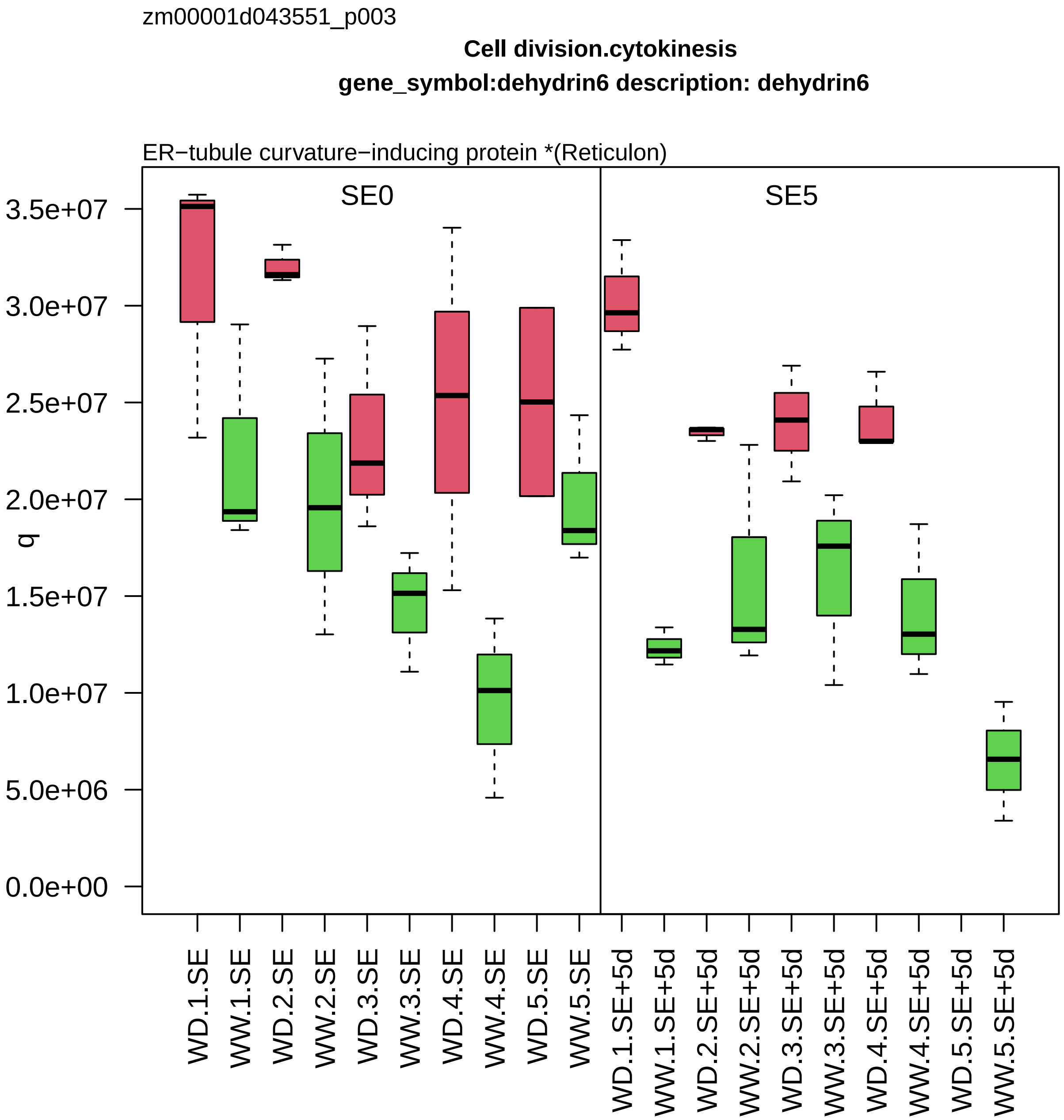

zm00001d043551_p003
Cell division.cytokinesis
gene_symbol:dehydrin6 description: dehydrin6
ER−tubule curvature−inducing protein *(Reticulon)
SE0
SE5
q
3.5e+07
3.0e+07
2.5e+07
2.0e+07
1.5e+07
1.0e+07
5.0e+06
0.0e+00
WD.1.SE
WW.1.SE
WD.2.SE
WW.2.SE
WD.3.SE
WW.3.SE
WD.4.SE
WW.4.SE
WD.5.SE
WW.5.SE
WD.1.SE+5d
WW.1.SE+5d
WD.2.SE+5d
WW.2.SE+5d
WD.3.SE+5d
WW.3.SE+5d
WD.4.SE+5d
WW.4.SE+5d
WD.5.SE+5d
WW.5.SE+5d

zm00001d044035_p001

**Amino acid metabolism.aspartate group amino acid biosynthesis**
**description:Ketol−acid reductoisomerase chloroplastic**

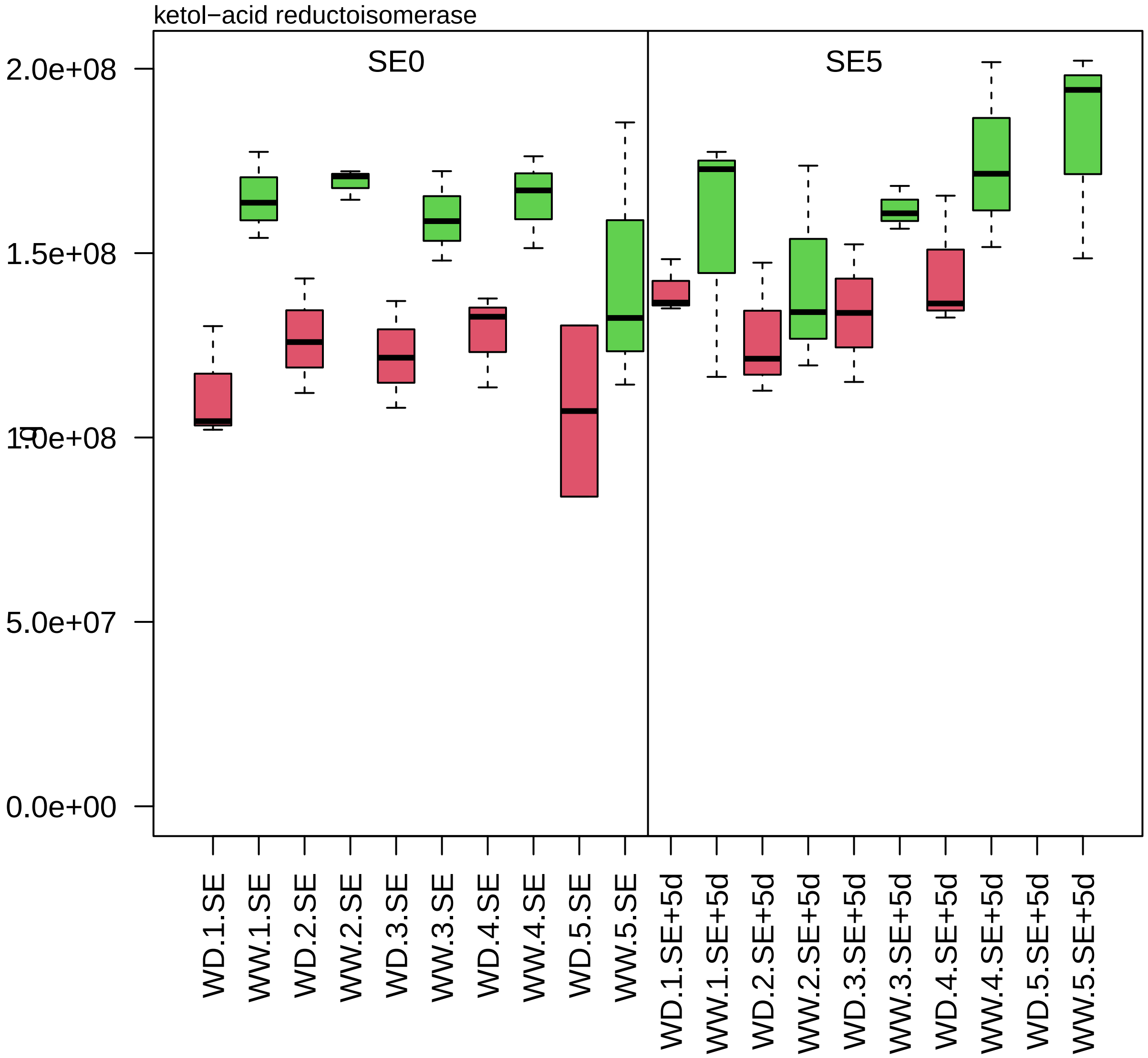

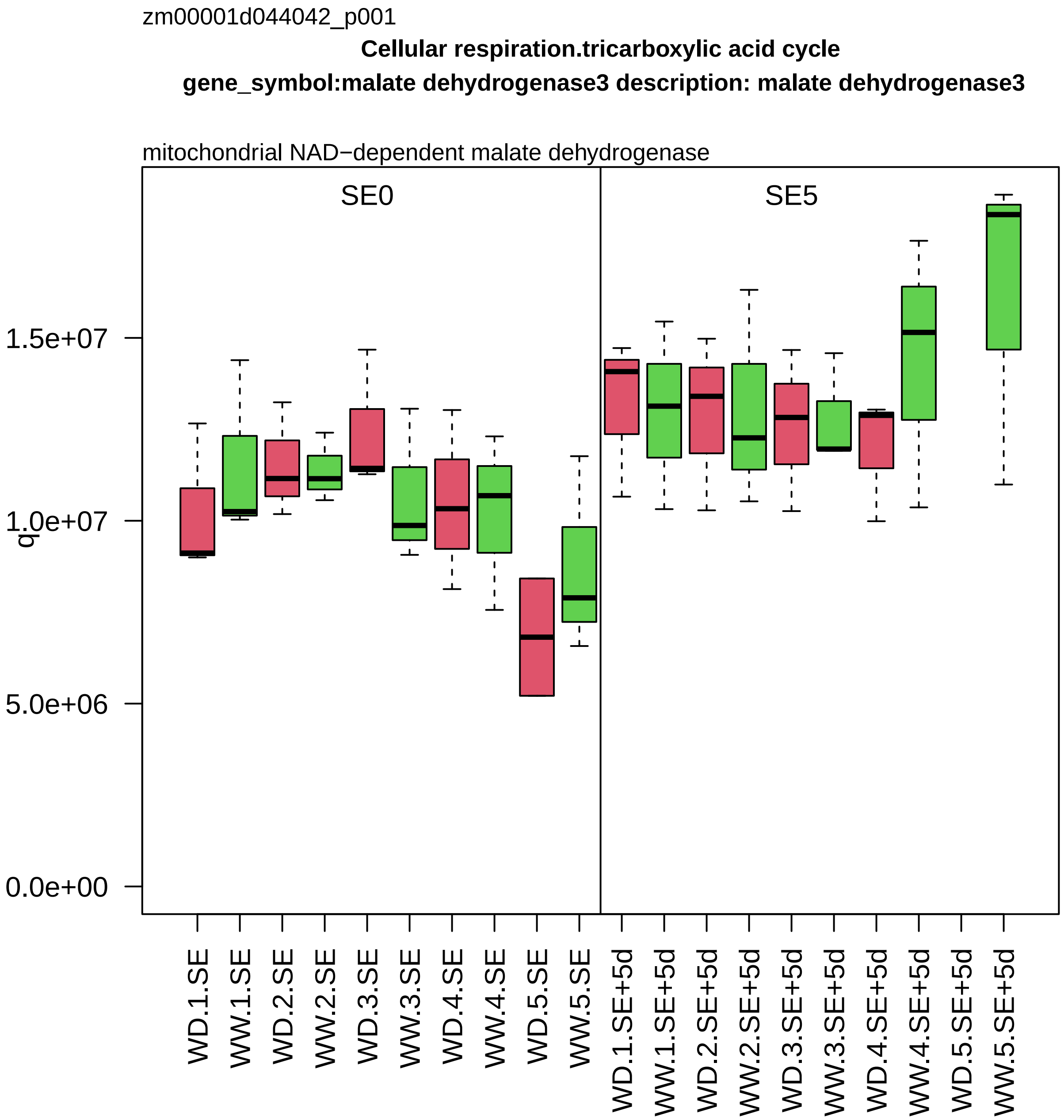
zm00001d044042_p001
Cellular respiration.tricarboxylic acid cycle
gene_symbol:malate dehydrogenase3 description: malate dehydrogenase3
mitochondrial NAD−dependent malate dehydrogenase
SE0
SE5
q
0.0e+00
5.0e+06
1.0e+07
1.5e+07
WD.1.SE
WW.1.SE
WD.2.SE
WW.2.SE
WD.3.SE
WW.3.SE
WD.4.SE
WW.4.SE
WD.5.SE
WW.5.SE
WD.1.SE+5d
WW.1.SE+5d
WD.2.SE+5d
WW.2.SE+5d
WD.3.SE+5d
WW.3.SE+5d
WD.4.SE+5d
WW.4.SE+5d
WD.5.SE+5d
WW.5.SE+5d

zm00001d044212_p001

**Enzyme classification.EC_1 oxidoreductases**
**description:Probable aldo−keto reductase 4**

EC_1.1 oxidoreductase acting on CH−OH group of donor

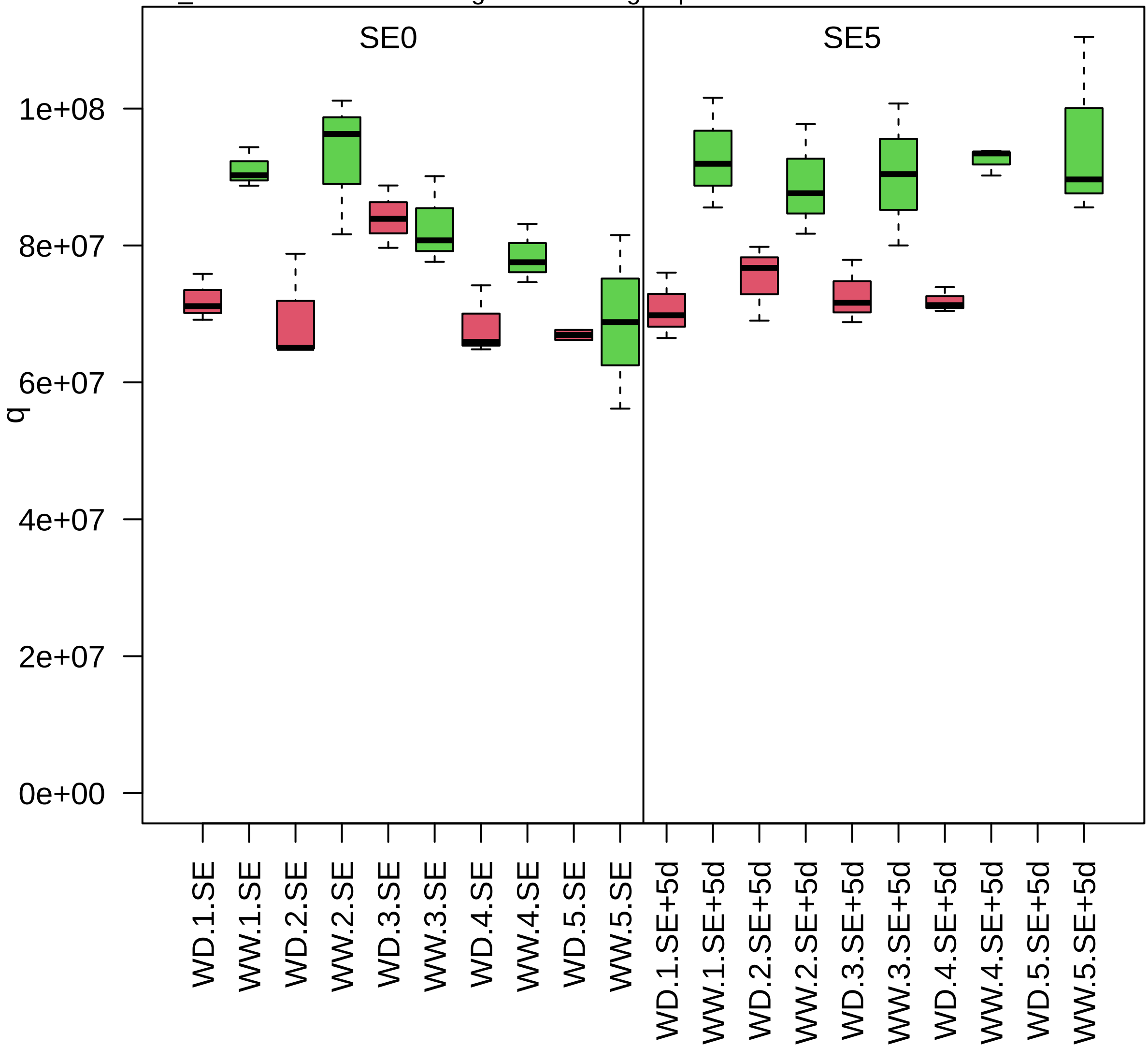

zm00001d044222_p002

**Protein modification.acetylation**
**description:tetratricopeptide repeat (TPR)−containing protein**

auxiliary component *(NAA15) of NatA/NatE N−terminal acetylase complexes

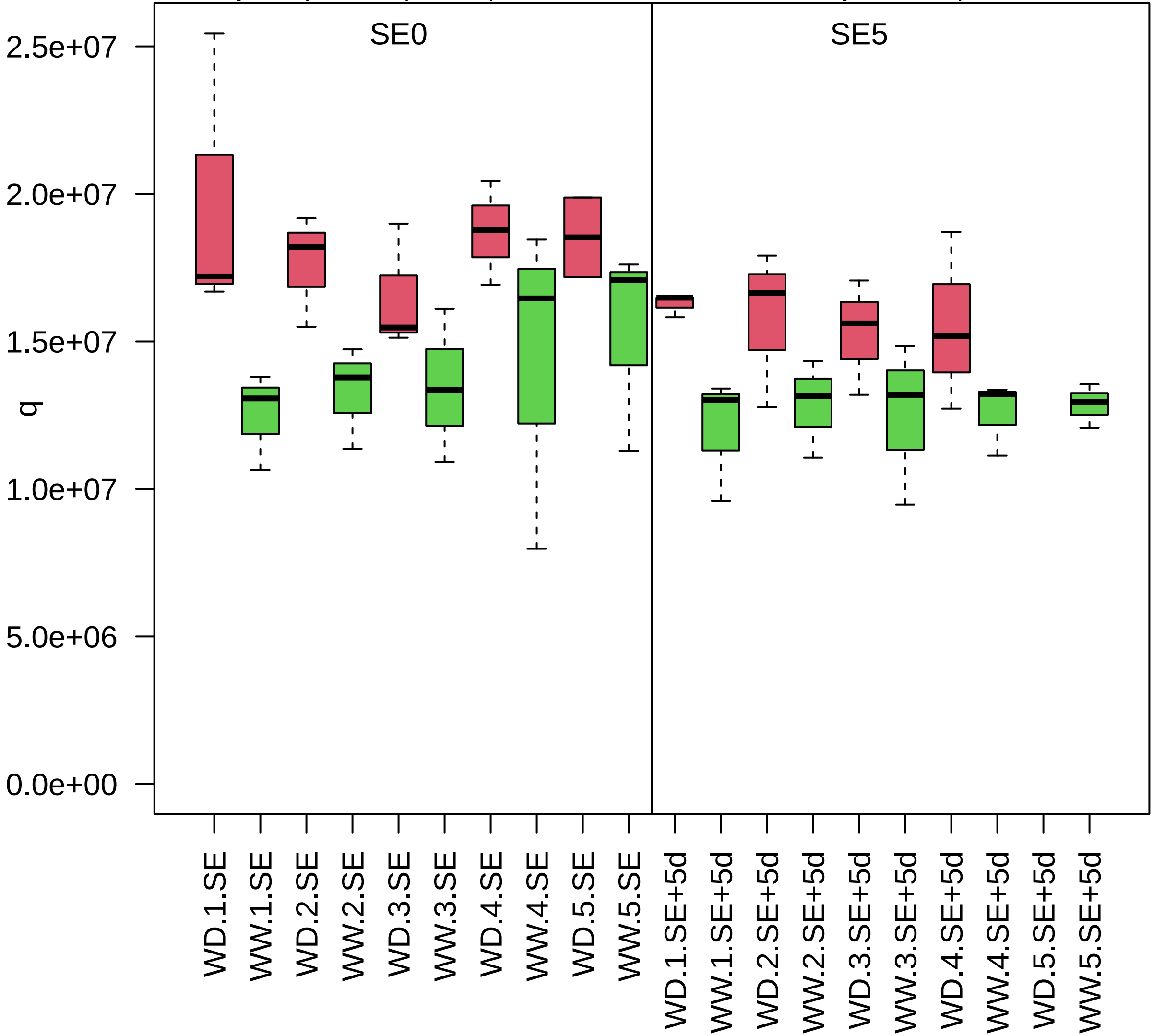

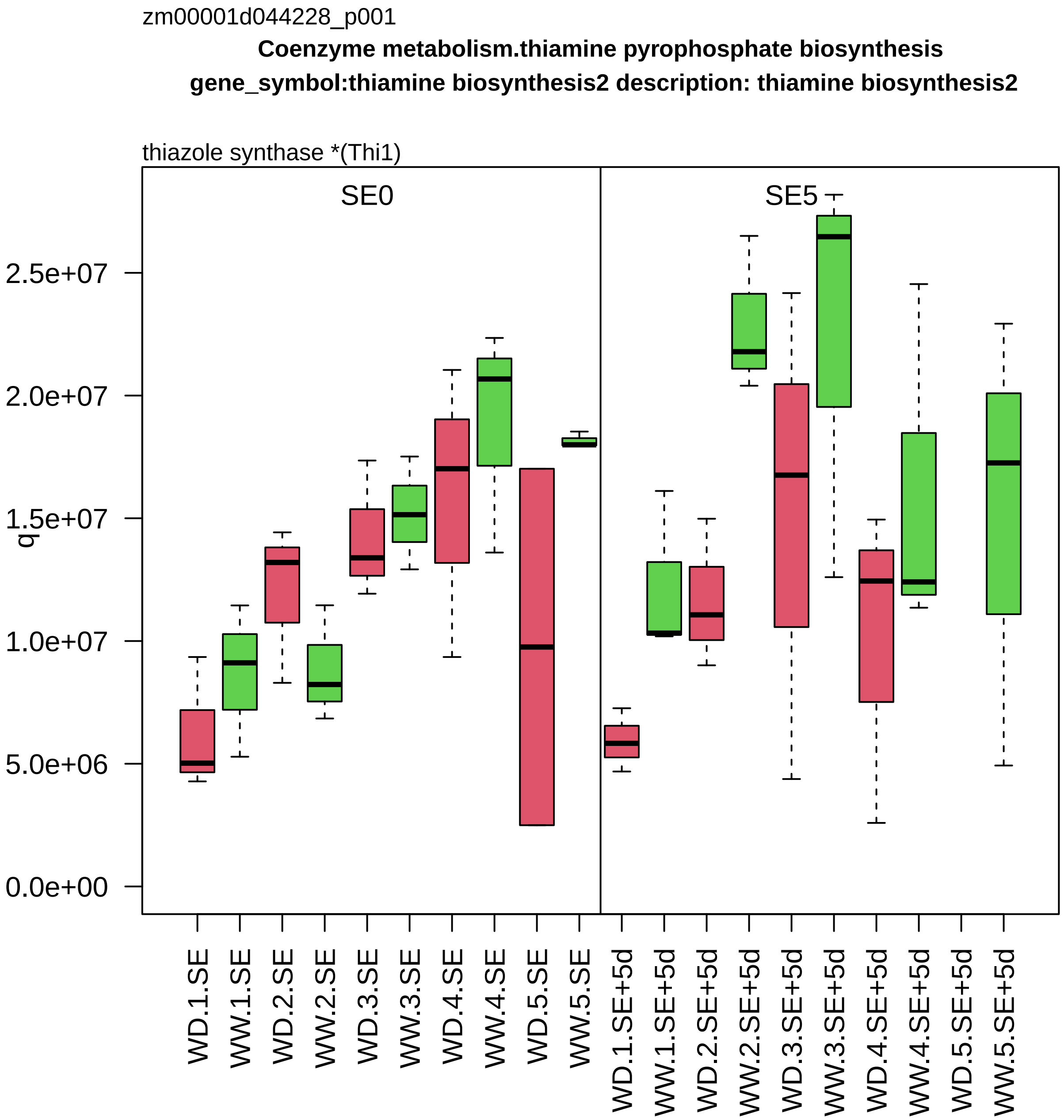
zm00001d044228_p001
Coenzyme metabolism.thiamine pyrophosphate biosynthesis
gene_symbol:thiamine biosynthesis2 description: thiamine biosynthesis2
thiazole synthase *(Thi1)
SE0
SE5
q
2.5e+07
2.0e+07
1.5e+07
1.0e+07
5.0e+06
0.0e+00
WD.1.SE
WW.1.SE
WD.2.SE
WW.2.SE
WD.3.SE
WW.3.SE
WD.4.SE
WW.4.SE
WD.5.SE
WW.5.SE
WD.1.SE+5d
WW.1.SE+5d
WD.2.SE+5d
WW.2.SE+5d
WD.3.SE+5d
WW.3.SE+5d
WD.4.SE+5d
WW.4.SE+5d
WD.5.SE+5d
WW.5.SE+5d

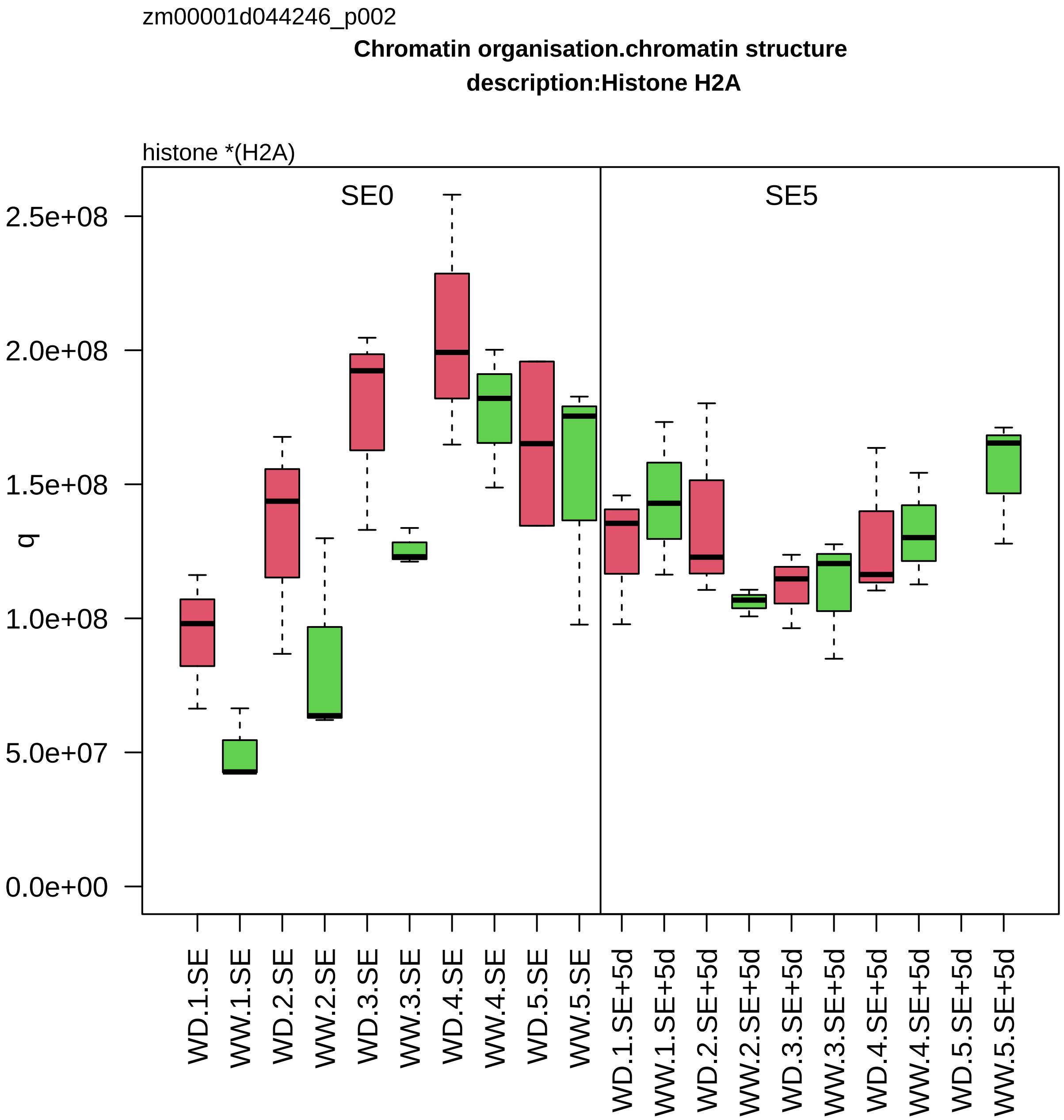
zm00001d044246_p002
Chromatin organisation.chromatin structure
description:Histone H2A
histone *(H2A)
SE0
SE5
q
2.5e+08
2.0e+08
1.5e+08
1.0e+08
5.0e+07
0.0e+00
WD.1.SE
WW.1.SE
WD.2.SE
WW.2.SE
WD.3.SE
WW.3.SE
WD.4.SE
WW.4.SE
WD.5.SE
WW.5.SE
WD.1.SE+5d
WW.1.SE+5d
WD.2.SE+5d
WW.2.SE+5d
WD.3.SE+5d
WW.3.SE+5d
WD.4.SE+5d
WW.4.SE+5d
WD.5.SE+5d
WW.5.SE+5d

zm00001d044271_p001

**Cell division.cytokinesis**

**description:Protein ROOT HAIR DEFECTIVE 3**

ER tubulae formation factor *(RHD3/RL)

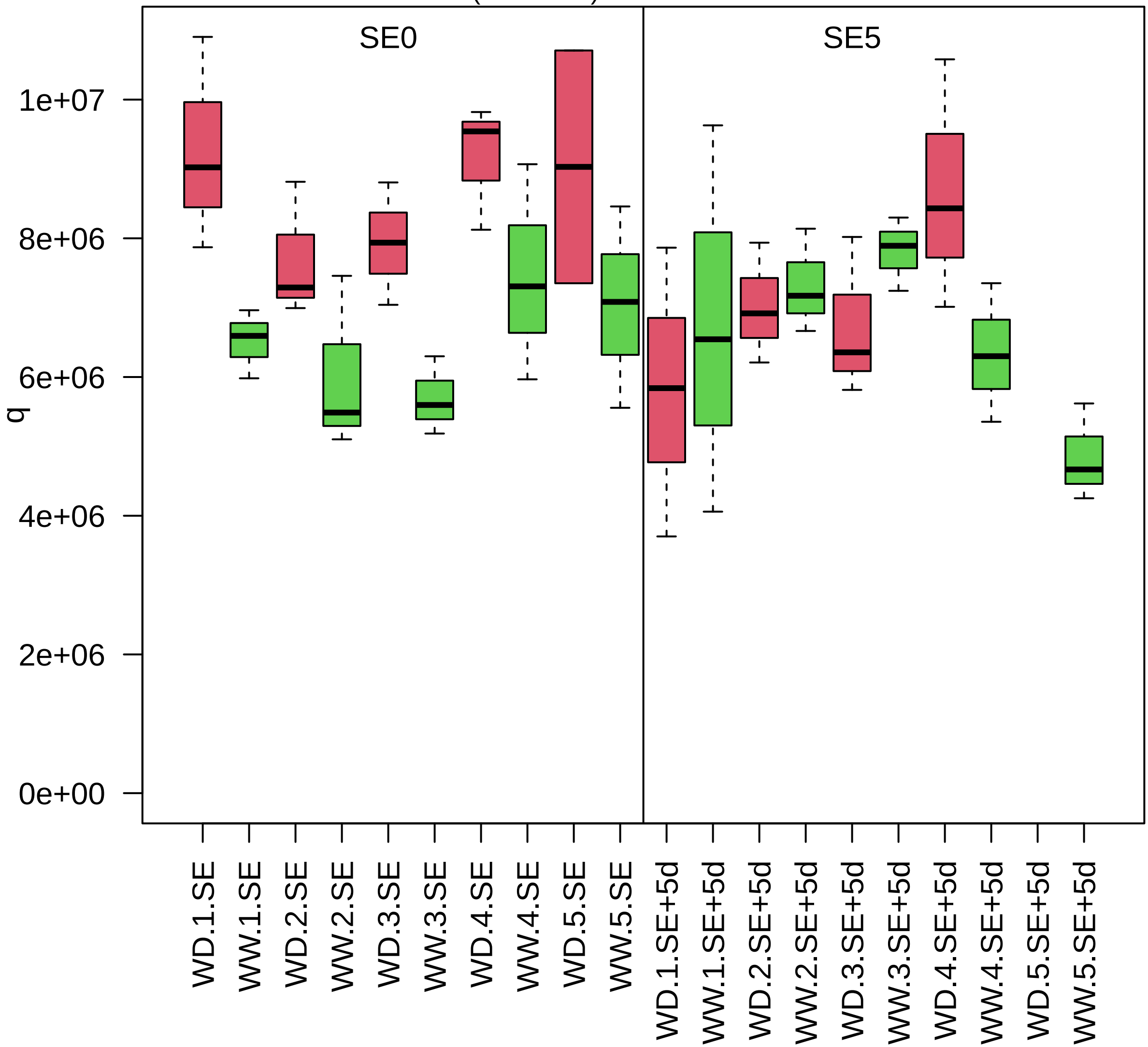

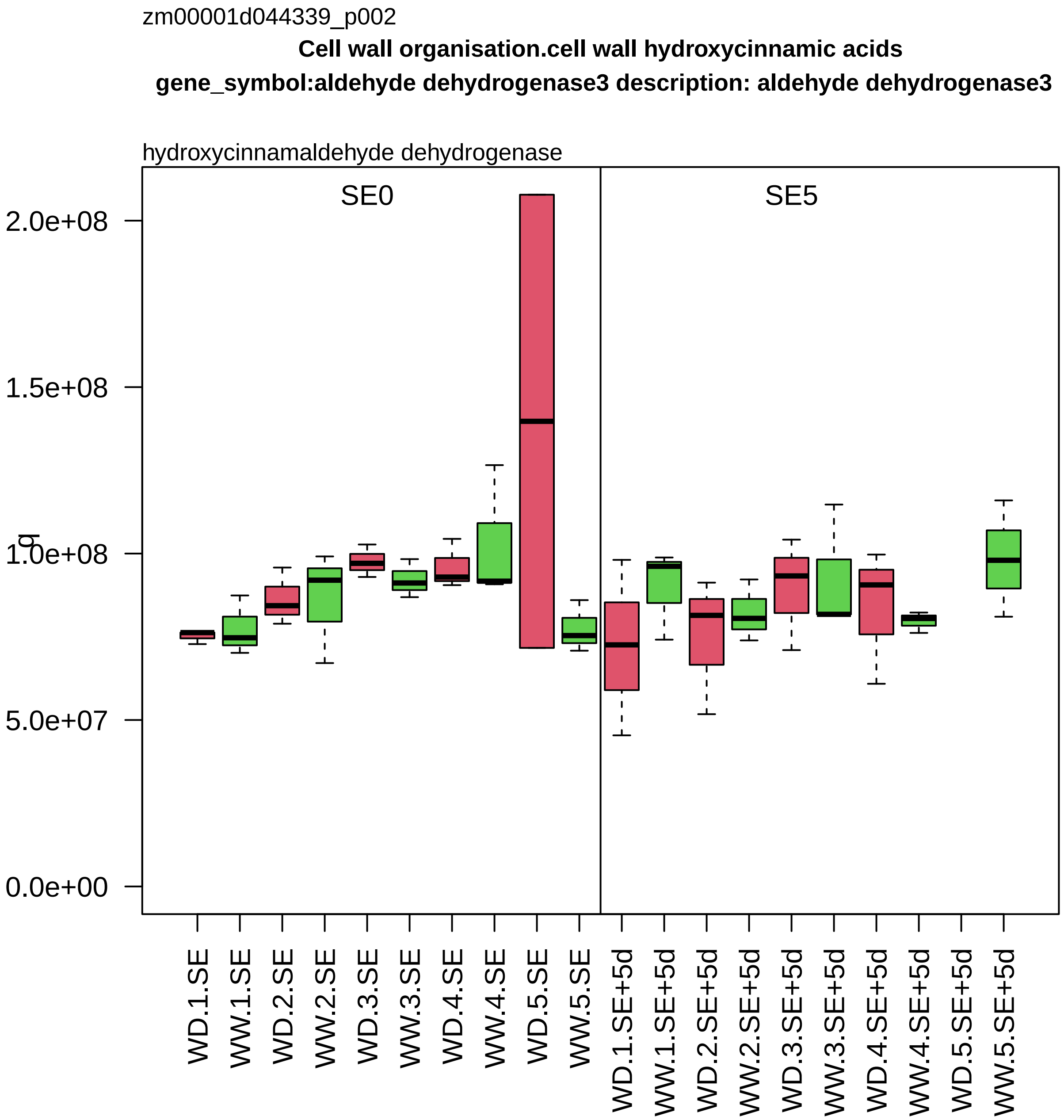

zm00001d044339_p002
Cell wall organisation.cell wall hydroxycinnamic acids
gene_symbol:aldehyde dehydrogenase3 description: aldehyde dehydrogenase3
hydroxycinnamaldehyde dehydrogenase
SE0
SE5
q
2.0e+08
1.5e+08
1.0e+08
5.0e+07
0.0e+00
WD.1.SE
WW.1.SE
WD.2.SE
WW.2.SE
WD.3.SE
WW.3.SE
WD.4.SE
WW.4.SE
WD.5.SE
WW.5.SE
WD.1.SE+5d
WW.1.SE+5d
WD.2.SE+5d
WW.2.SE+5d
WD.3.SE+5d
WW.3.SE+5d
WD.4.SE+5d
WW.4.SE+5d
WD.5.SE+5d
WW.5.SE+5d

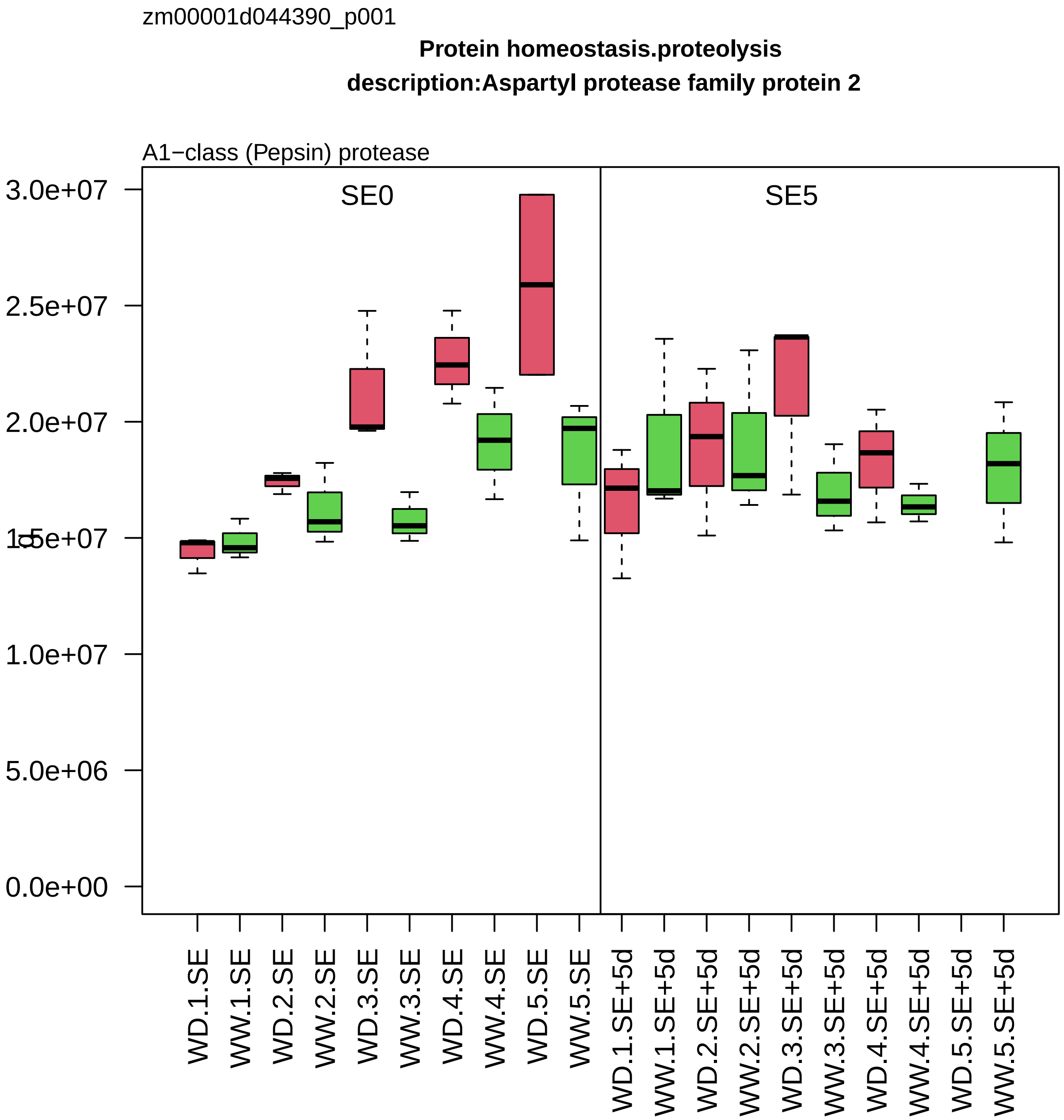

zm00001d044390_p001
Protein homeostasis.proteolysis
description:Aspartyl protease family protein 2
A1−class (Pepsin) protease
SE0
SE5
3.0e+07
2.5e+07
2.0e+07
1.5e+07
1.0e+07
5.0e+06
0.0e+00
WD.1.SE
WW.1.SE
WD.2.SE
WW.2.SE
WD.3.SE
WW.3.SE
WD.4.SE
WW.4.SE
WD.5.SE
WW.5.SE
WD.1.SE+5d
WW.1.SE+5d
WD.2.SE+5d
WW.2.SE+5d
WD.3.SE+5d
WW.3.SE+5d
WD.4.SE+5d
WW.4.SE+5d
WD.5.SE+5d
WW.5.SE+5d

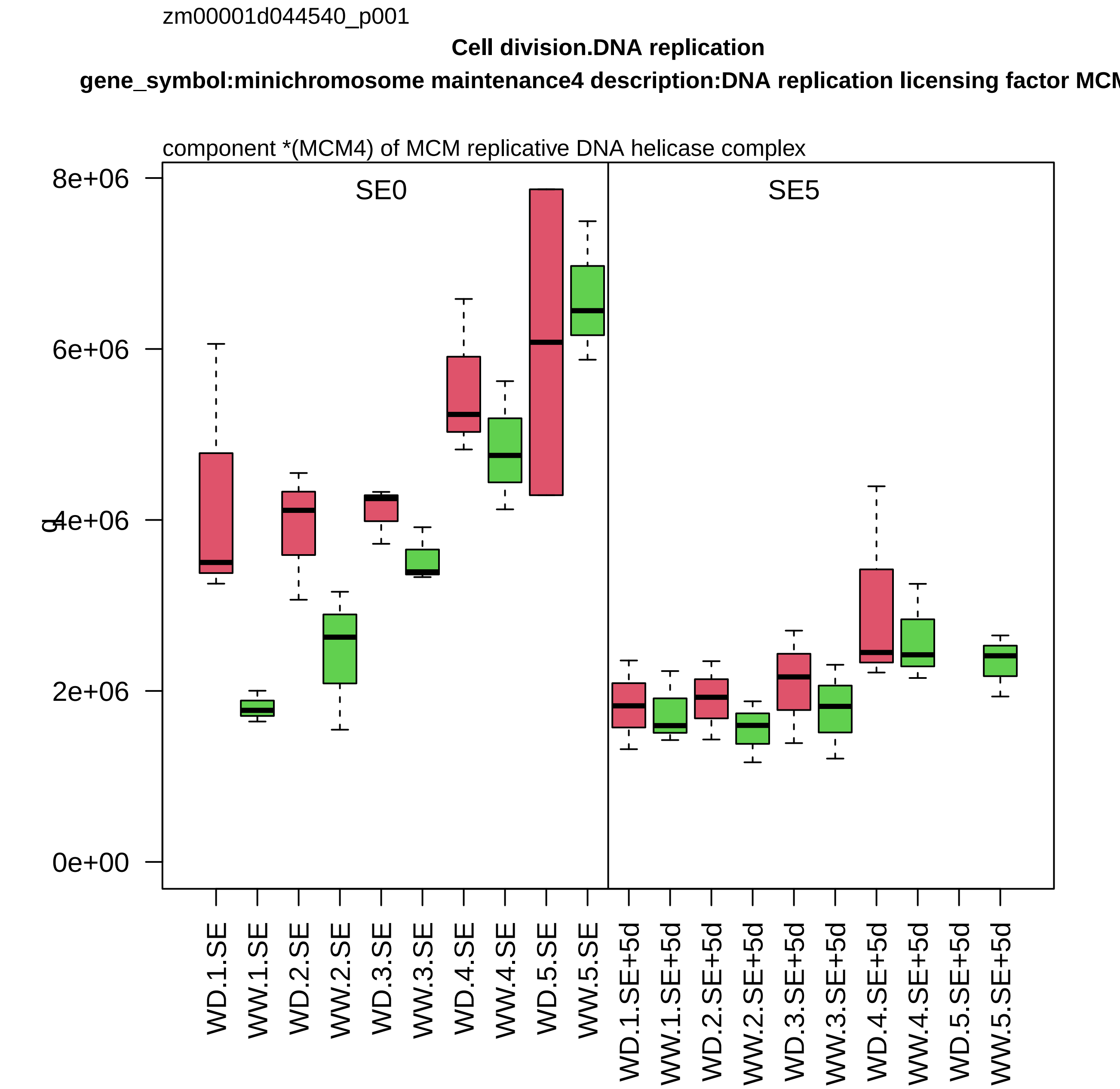

zm00001d044540_p001
Cell division.DNA replication
gene_symbol:minichromosome maintenance4 description:DNA replication licensing factor MCM
component *(MCM4) of MCM replicative DNA helicase complex
SE0
SE5
0e+00
2e+06
4e+06
6e+06
8e+06
q
WD.1.SE
WW.1.SE
WD.2.SE
WW.2.SE
WD.3.SE
WW.3.SE
WD.4.SE
WW.4.SE
WD.5.SE
WW.5.SE
WD.1.SE+5d
WW.1.SE+5d
WD.2.SE+5d
WW.2.SE+5d
WD.3.SE+5d
WW.3.SE+5d
WD.4.SE+5d
WW.4.SE+5d
WD.5.SE+5d
WW.5.SE+5d

zm00001d044573_p001

**RNA processing.pre−RNA splicing**
**description:Pre−mRNA−processing protein 40A**

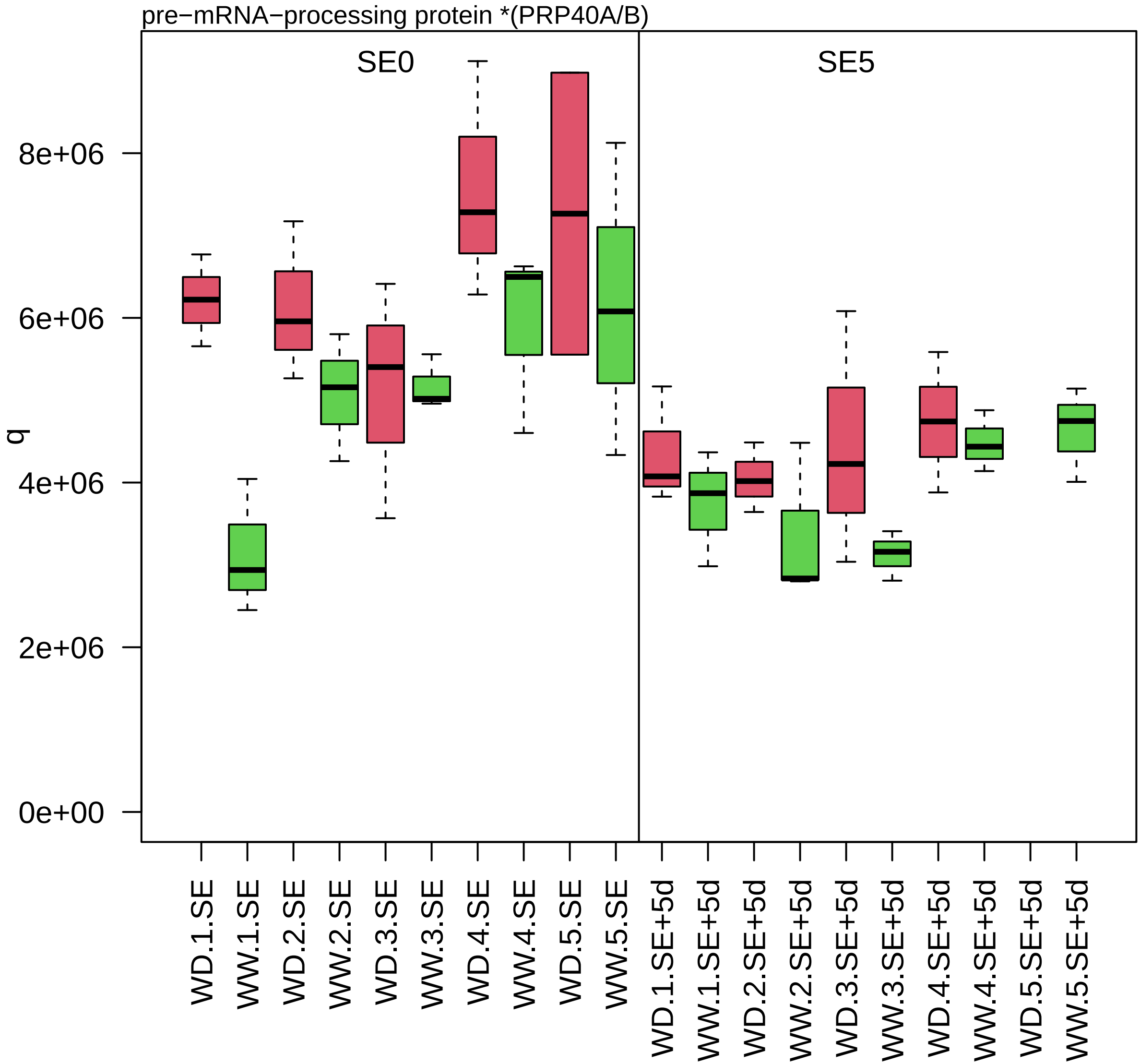

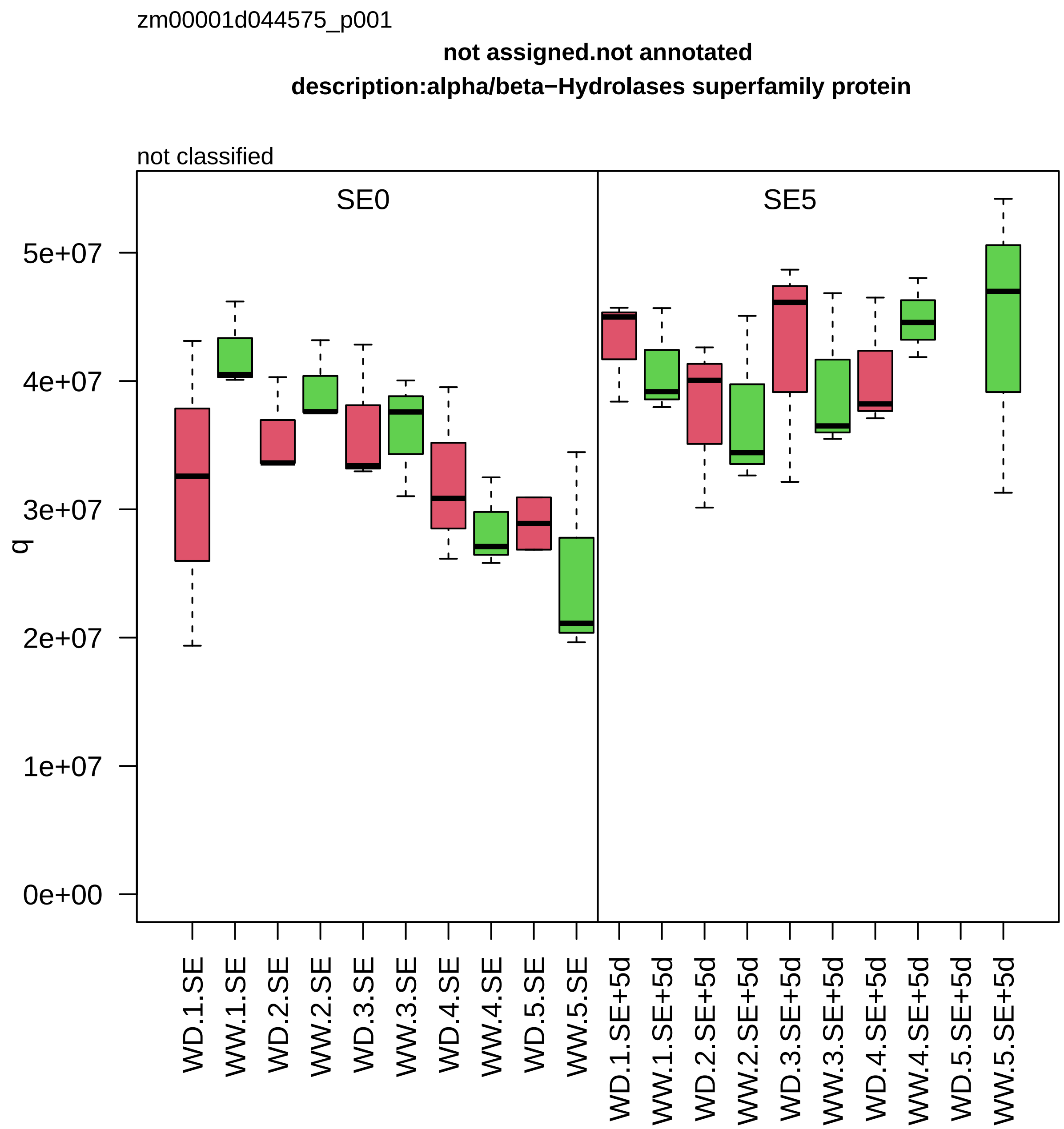
zm00001d044575_p001
not assigned.not annotated
description:alpha/beta−Hydrolases superfamily protein
not classified
SE0
SE5
q
0e+00
1e+07
2e+07
3e+07
4e+07
5e+07
WD.1.SE
WW.1.SE
WD.2.SE
WW.2.SE
WD.3.SE
WW.3.SE
WD.4.SE
WW.4.SE
WD.5.SE
WW.5.SE
WD.1.SE+5d
WW.1.SE+5d
WD.2.SE+5d
WW.2.SE+5d
WD.3.SE+5d
WW.3.SE+5d
WD.4.SE+5d
WW.4.SE+5d
WD.5.SE+5d
WW.5.SE+5d

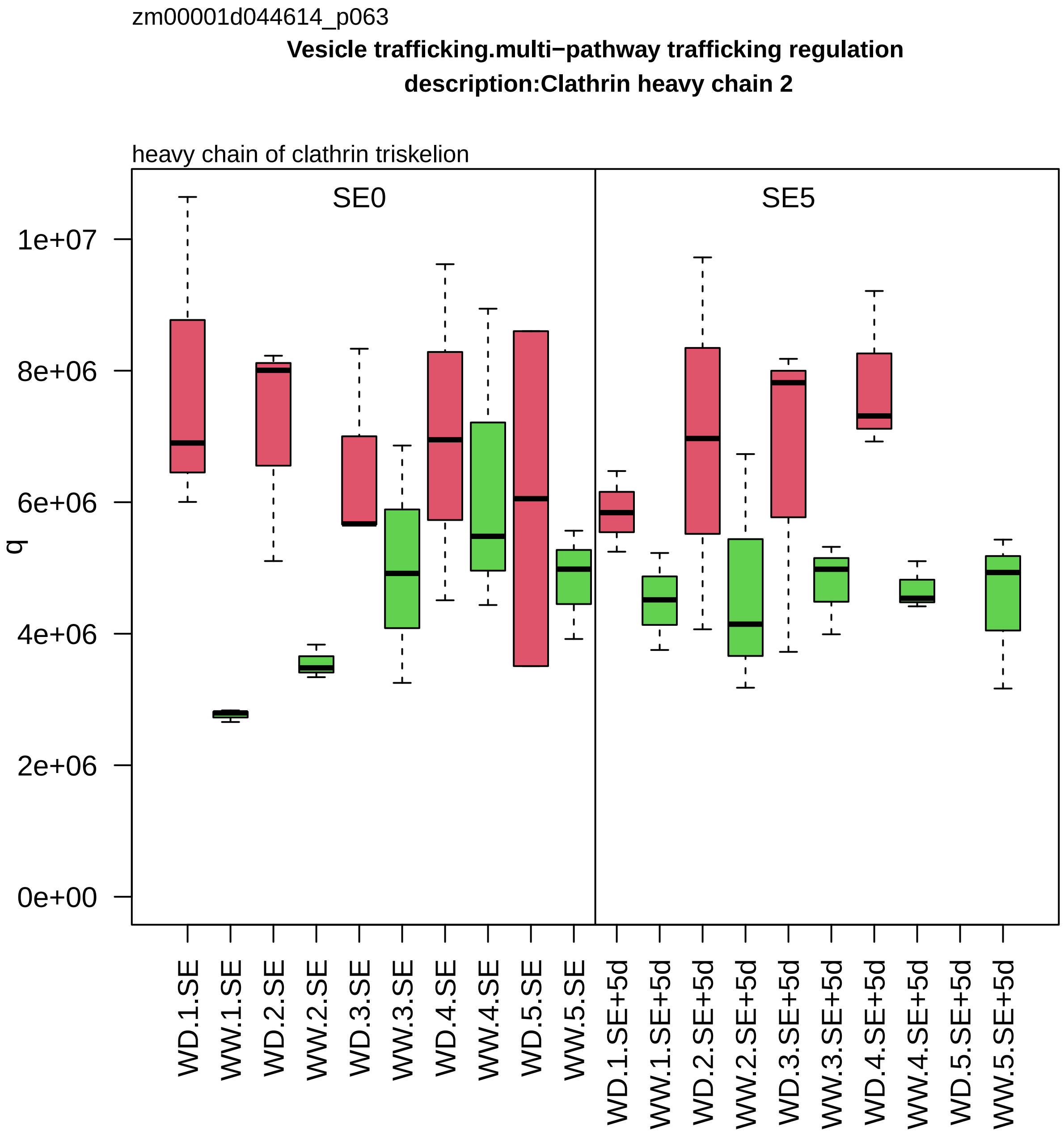
zm00001d044614_p063
Vesicle trafficking.multi−pathway trafficking regulation
description:Clathrin heavy chain 2
heavy chain of clathrin triskelion
SE0
SE5
q
1e+07
8e+06
6e+06
4e+06
2e+06
0e+00
WD.1.SE
WW.1.SE
WD.2.SE
WW.2.SE
WD.3.SE
WW.3.SE
WD.4.SE
WW.4.SE
WD.5.SE
WW.5.SE
WD.1.SE+5d
WW.1.SE+5d
WD.2.SE+5d
WW.2.SE+5d
WD.3.SE+5d
WW.3.SE+5d
WD.4.SE+5d
WW.4.SE+5d
WD.5.SE+5d
WW.5.SE+5d

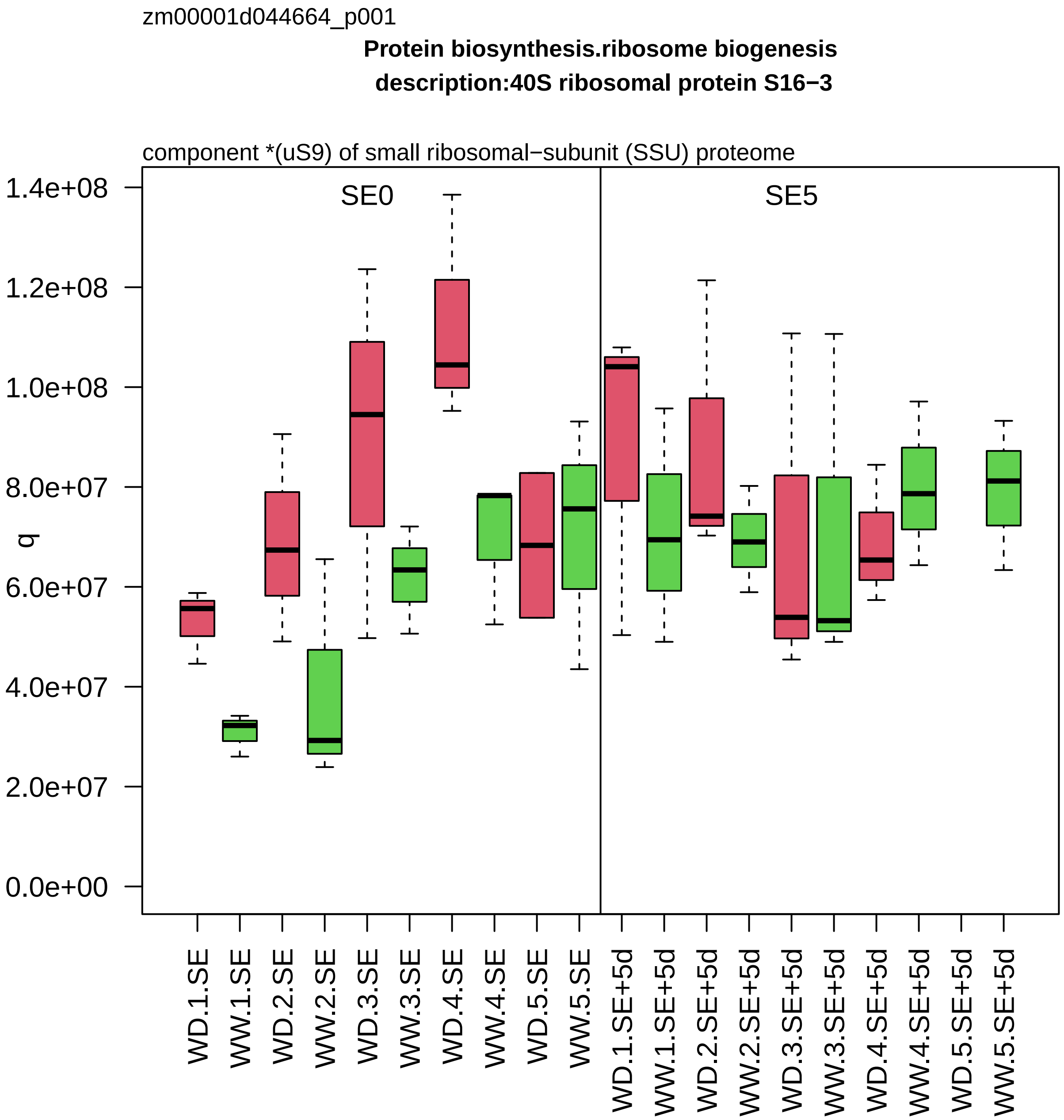
zm00001d044664_p001
Protein biosynthesis.ribosome biogenesis
description:40S ribosomal protein S16−3
component *(uS9) of small ribosomal−subunit (SSU) proteome
SE0
SE5
q
1.4e+08
1.2e+08
1.0e+08
8.0e+07
6.0e+07
4.0e+07
2.0e+07
0.0e+00
WD.1.SE
WW.1.SE
WD.2.SE
WW.2.SE
WD.3.SE
WW.3.SE
WD.4.SE
WW.4.SE
WD.5.SE
WW.5.SE
WD.1.SE+5d
WW.1.SE+5d
WD.2.SE+5d
WW.2.SE+5d
WD.3.SE+5d
WW.3.SE+5d
WD.4.SE+5d
WW.4.SE+5d
WD.5.SE+5d
WW.5.SE+5d

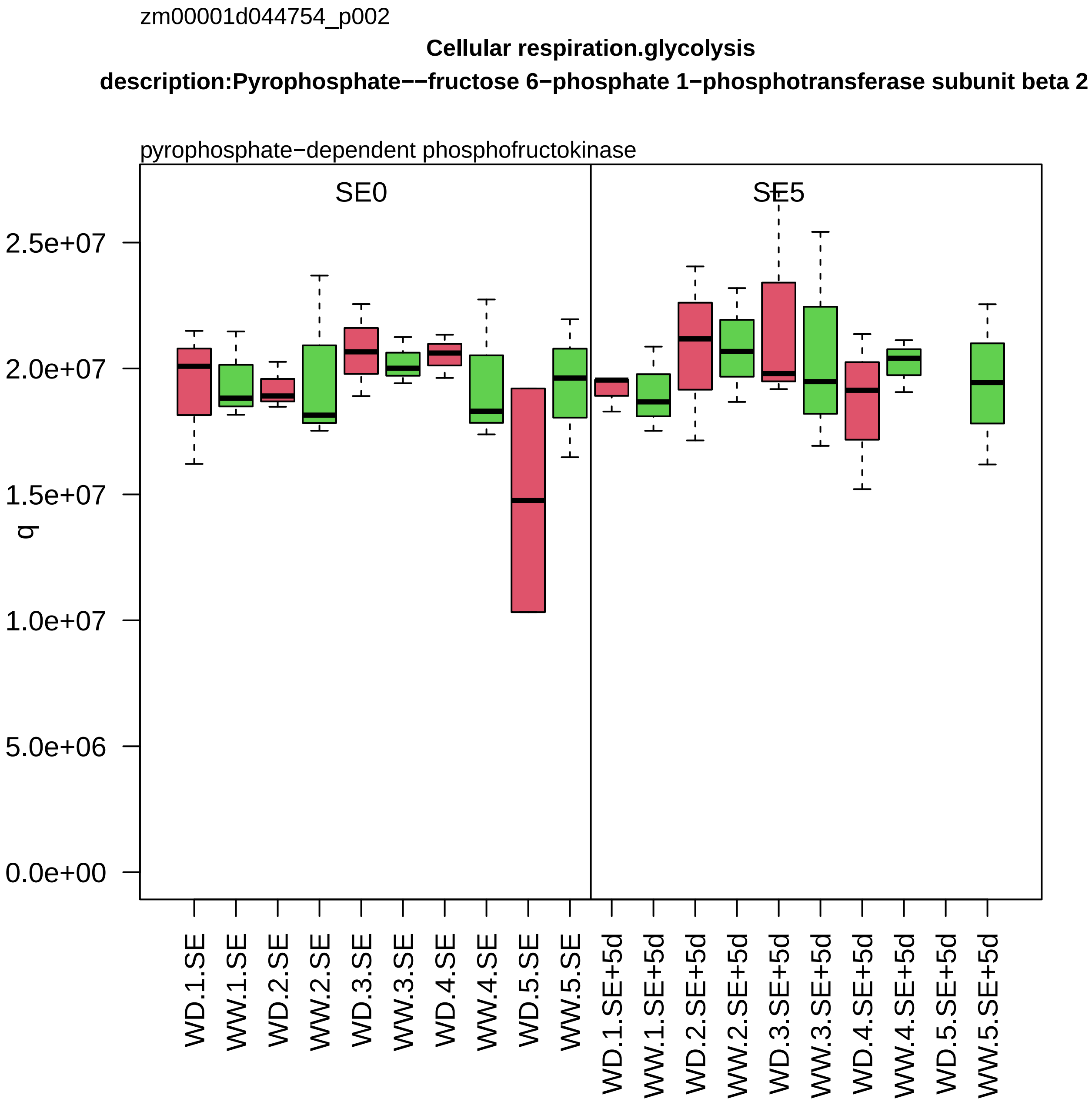

zm00001d044754_p002
Cellular respiration.glycolysis
description:Pyrophosphate−−fructose 6−phosphate 1−phosphotransferase subunit beta 2
pyrophosphate−dependent phosphofructokinase
SE0
SE5
q
2.5e+07
2.0e+07
1.5e+07
1.0e+07
5.0e+06
0.0e+00
WD.1.SE
WW.1.SE
WD.2.SE
WW.2.SE
WD.3.SE
WW.3.SE
WD.4.SE
WW.4.SE
WD.5.SE
WW.5.SE
WD.1.SE+5d
WW.1.SE+5d
WD.2.SE+5d
WW.2.SE+5d
WD.3.SE+5d
WW.3.SE+5d
WD.4.SE+5d
WW.4.SE+5d
WD.5.SE+5d
WW.5.SE+5d

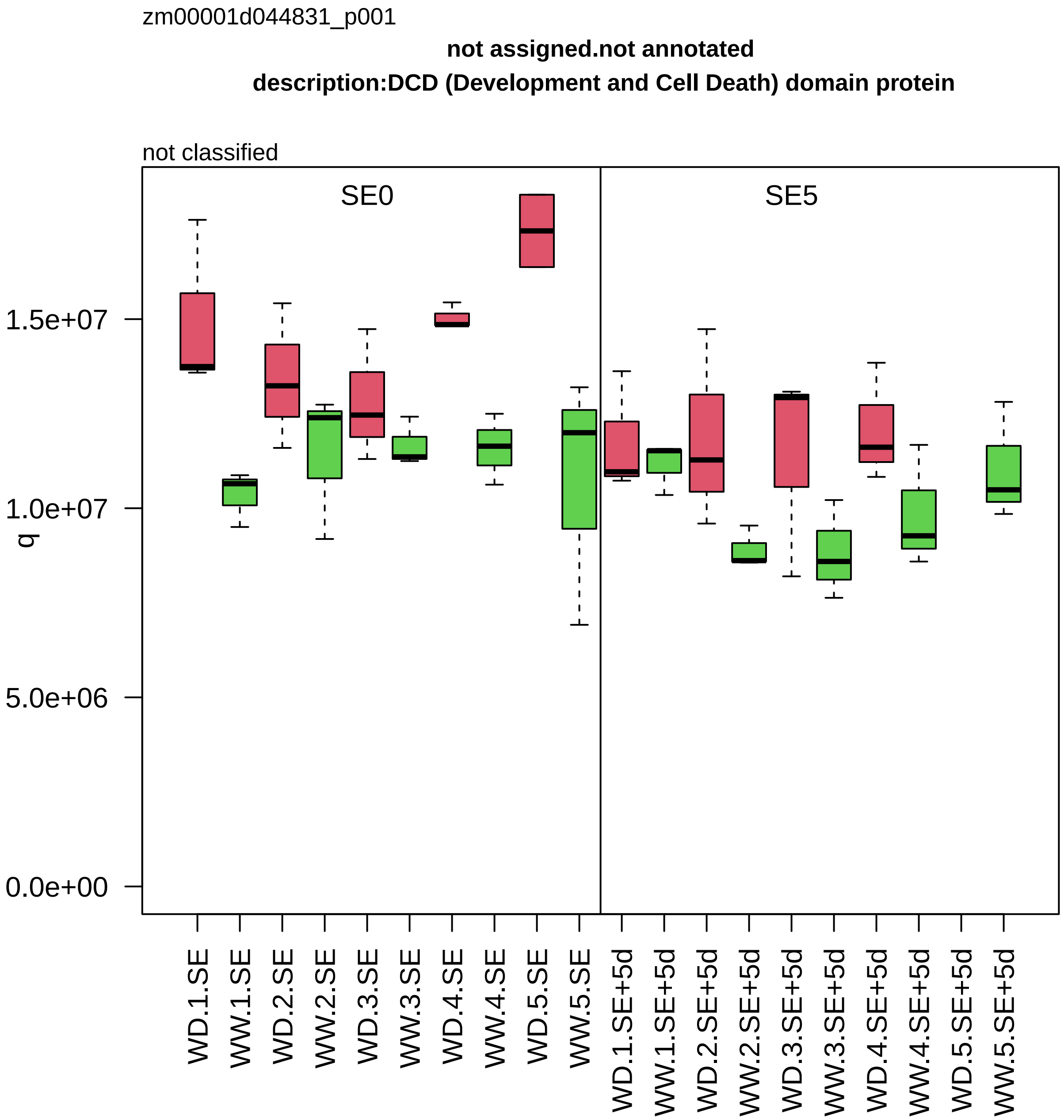
zm00001d044831_p001
not assigned.not annotated
description:DCD (Development and Cell Death) domain protein
not classified
SE0
SE5
q
1.5e+07
1.0e+07
5.0e+06
0.0e+00
WD.1.SE
WW.1.SE
WD.2.SE
WW.2.SE
WD.3.SE
WW.3.SE
WD.4.SE
WW.4.SE
WD.5.SE
WW.5.SE
WD.1.SE+5d
WW.1.SE+5d
WD.2.SE+5d
WW.2.SE+5d
WD.3.SE+5d
WW.3.SE+5d
WD.4.SE+5d
WW.4.SE+5d
WD.5.SE+5d
WW.5.SE+5d

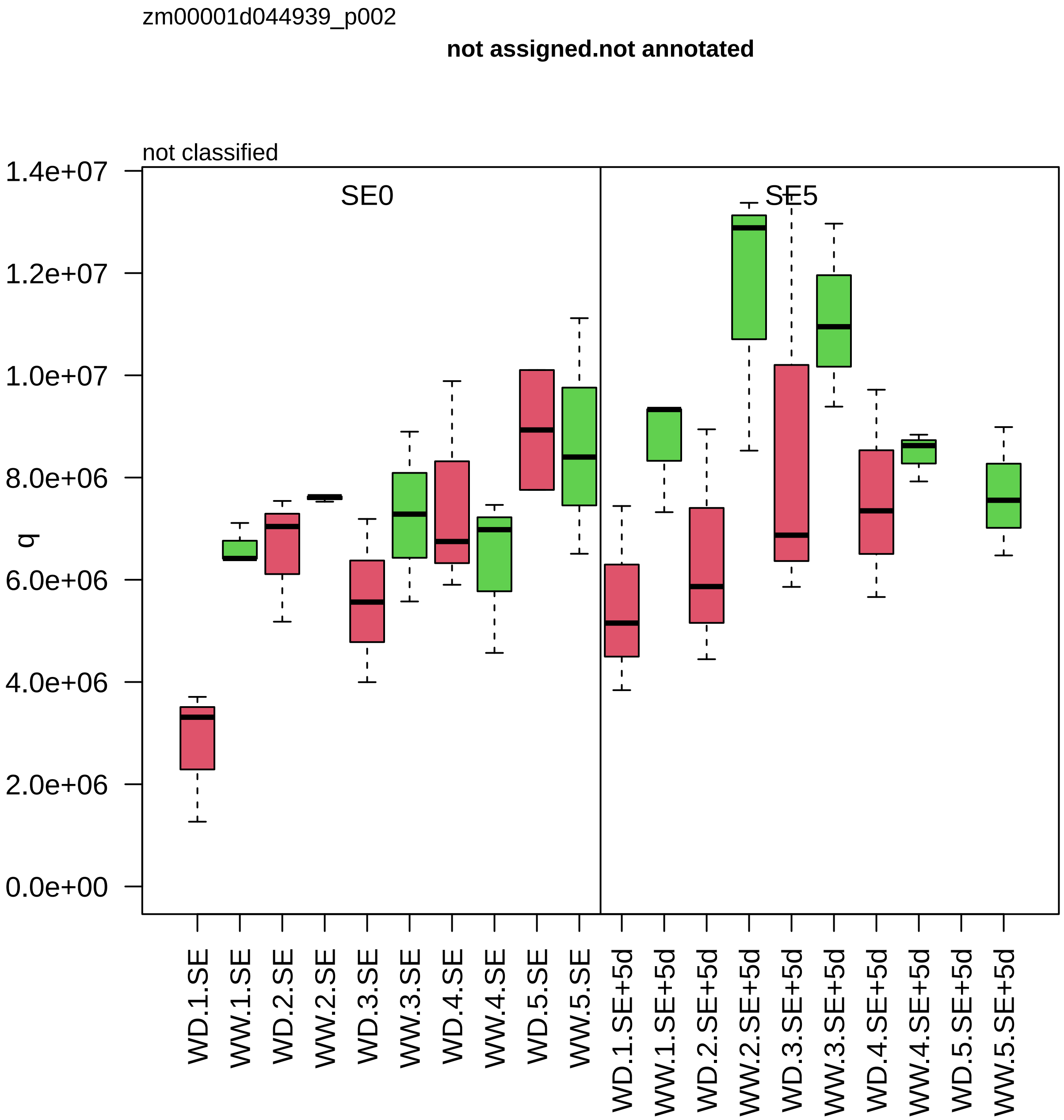
zm00001d044939_p002
not assigned.not annotated
not classified
SE0
SE5
q
1.4e+07
1.2e+07
1.0e+07
8.0e+06
6.0e+06
4.0e+06
2.0e+06
0.0e+00
WD.1.SE
WW.1.SE
WD.2.SE
WW.2.SE
WD.3.SE
WW.3.SE
WD.4.SE
WW.4.SE
WD.5.SE
WW.5.SE
WD.1.SE+5d
WW.1.SE+5d
WD.2.SE+5d
WW.2.SE+5d
WD.3.SE+5d
WW.3.SE+5d
WD.4.SE+5d
WW.4.SE+5d
WD.5.SE+5d
WW.5.SE+5d

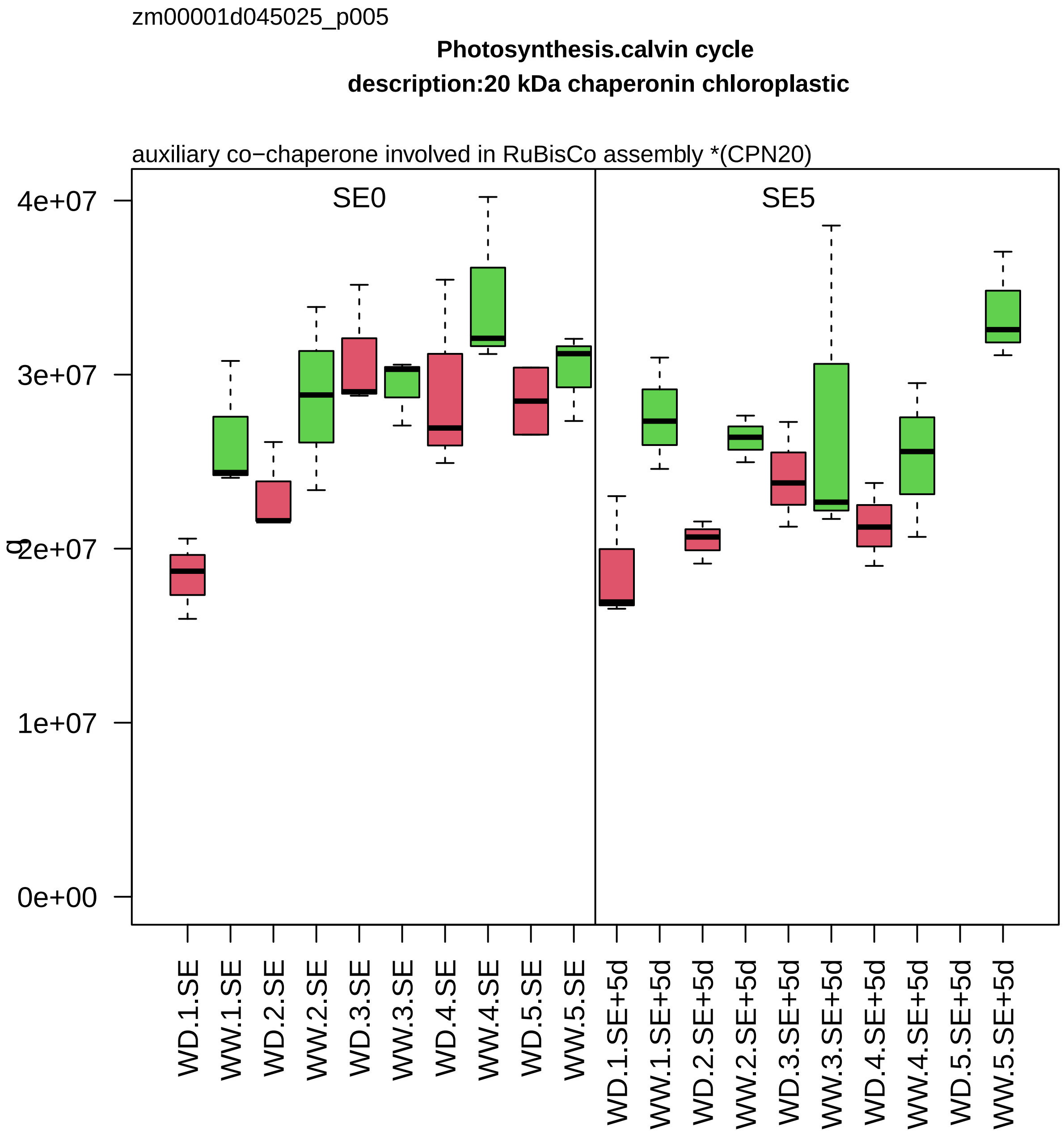

zm00001d045025_p005
Photosynthesis.calvin cycle
description:20 kDa chaperonin chloroplastic
auxiliary co−chaperone involved in RuBisCo assembly *(CPN20)
SE0
SE5
4e+07
3e+07
2e+07
1e+07
0e+00
WD.1.SE
WW.1.SE
WD.2.SE
WW.2.SE
WD.3.SE
WW.3.SE
WD.4.SE
WW.4.SE
WD.5.SE
WW.5.SE
WD.1.SE+5d
WW.1.SE+5d
WD.2.SE+5d
WW.2.SE+5d
WD.3.SE+5d
WW.3.SE+5d
WD.4.SE+5d
WW.4.SE+5d
WD.5.SE+5d
WW.5.SE+5d

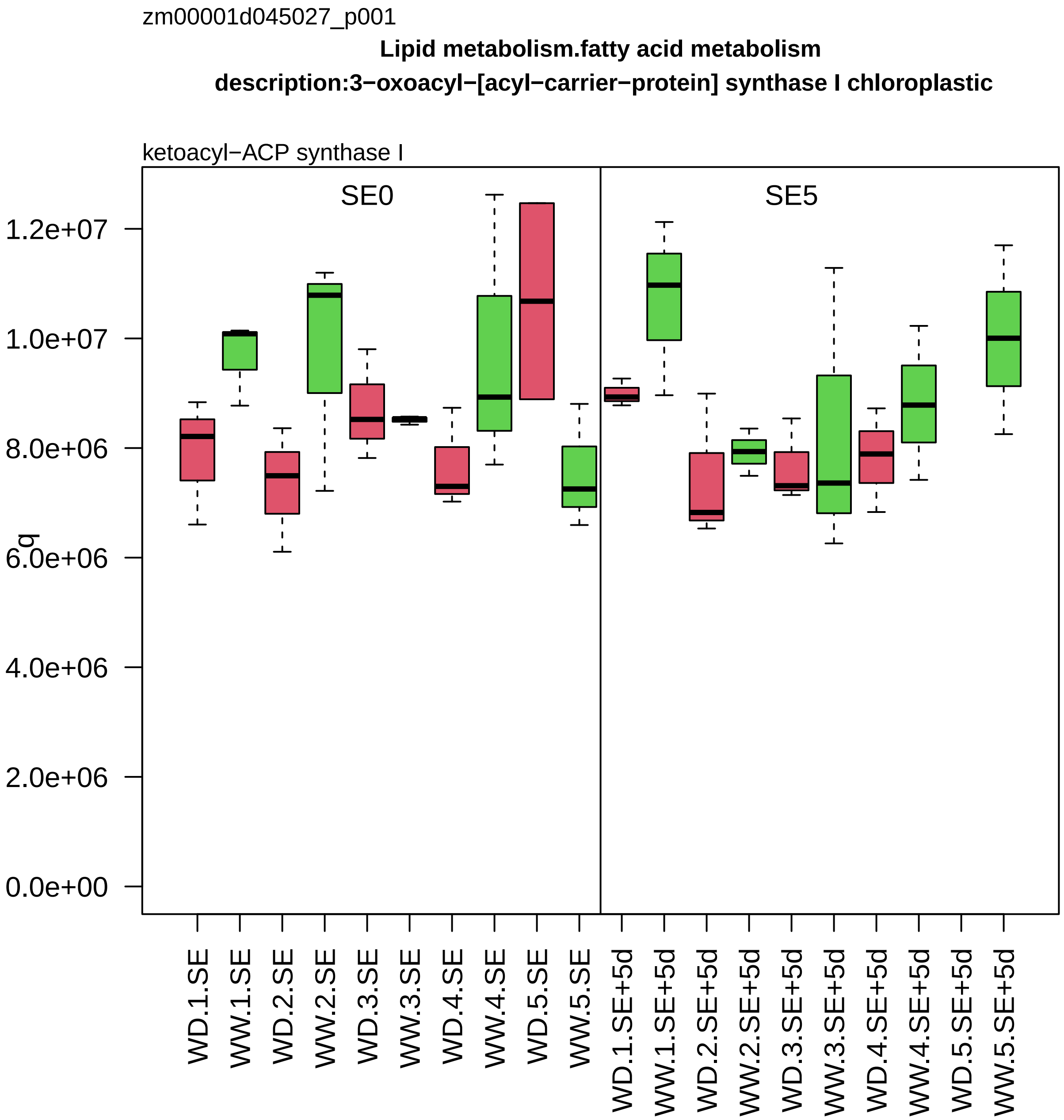
zm00001d045027_p001
Lipid metabolism.fatty acid metabolism
description:3−oxoacyl−[acyl−carrier−protein] synthase I chloroplastic
ketoacyl−ACP synthase I
SE0
SE5
q
1.2e+07
1.0e+07
8.0e+06
6.0e+06
4.0e+06
2.0e+06
0.0e+00
WD.1.SE
WW.1.SE
WD.2.SE
WW.2.SE
WD.3.SE
WW.3.SE
WD.4.SE
WW.4.SE
WD.5.SE
WW.5.SE
WD.1.SE+5d
WW.1.SE+5d
WD.2.SE+5d
WW.2.SE+5d
WD.3.SE+5d
WW.3.SE+5d
WD.4.SE+5d
WW.4.SE+5d
WD.5.SE+5d
WW.5.SE+5d

zm00001d045042_p017

**Carbohydrate metabolism.sucrose metabolism**

**gene_symbol:shrunken1 description:Sucrose synthase 1**

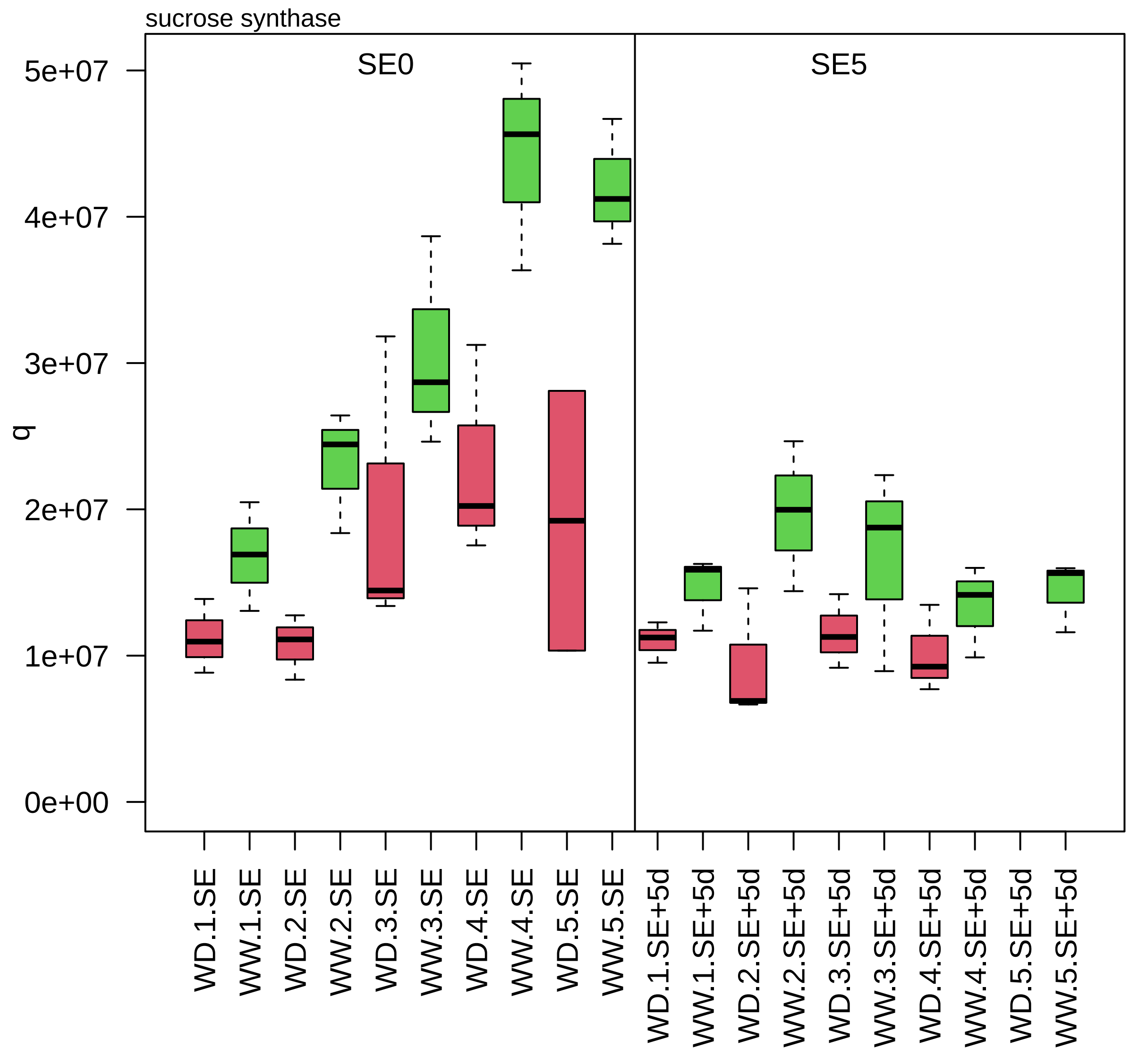

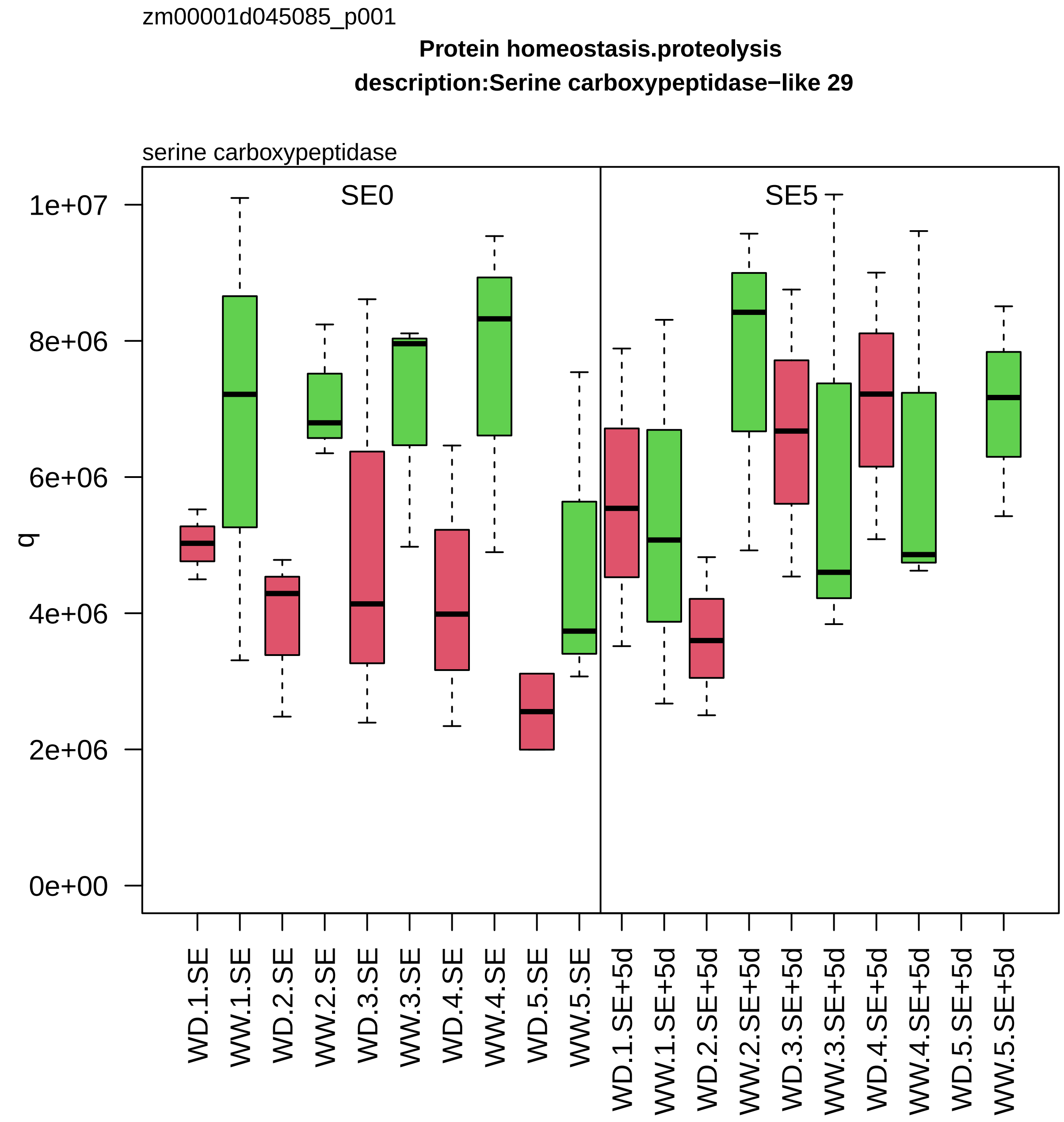
zm00001d045085_p001
Protein homeostasis.proteolysis
description:Serine carboxypeptidase−like 29
serine carboxypeptidase
SE0
SE5
q
1e+07
8e+06
6e+06
4e+06
2e+06
0e+00
WD.1.SE
WW.1.SE
WD.2.SE
WW.2.SE
WD.3.SE
WW.3.SE
WD.4.SE
WW.4.SE
WD.5.SE
WW.5.SE
WD.1.SE+5d
WW.1.SE+5d
WD.2.SE+5d
WW.2.SE+5d
WD.3.SE+5d
WW.3.SE+5d
WD.4.SE+5d
WW.4.SE+5d
WD.5.SE+5d
WW.5.SE+5d

zm00001d045315_p004

**Protein homeostasis.ubiquitin−proteasome system**
**description:Proteasome subunit beta type**

beta−type−5 component *(PBE) of 26S proteasome

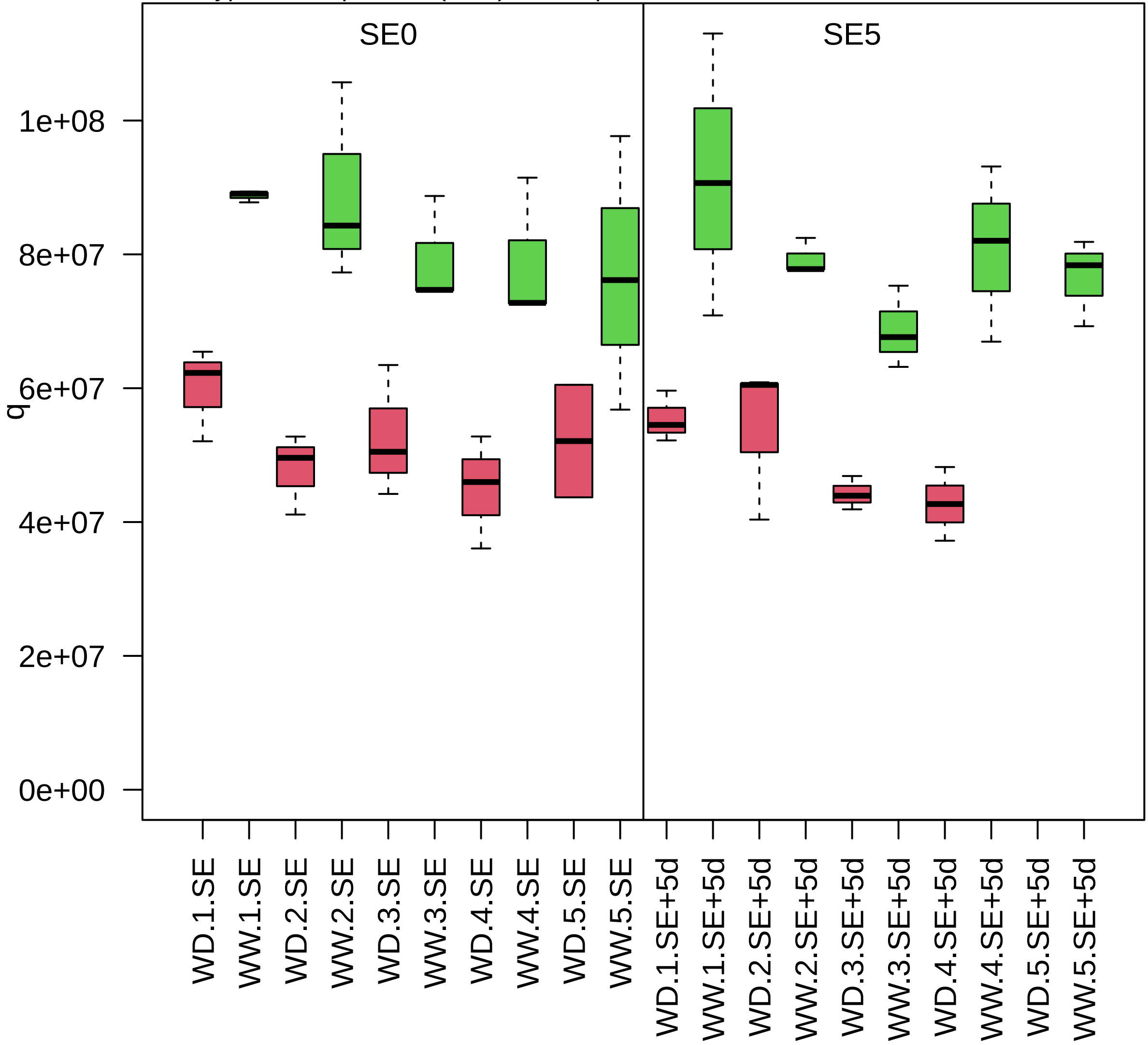

zm00001d045352_p002

**Chromatin organisation.histone chaperone activities**

**description:Nucleosome assembly protein 1;2**

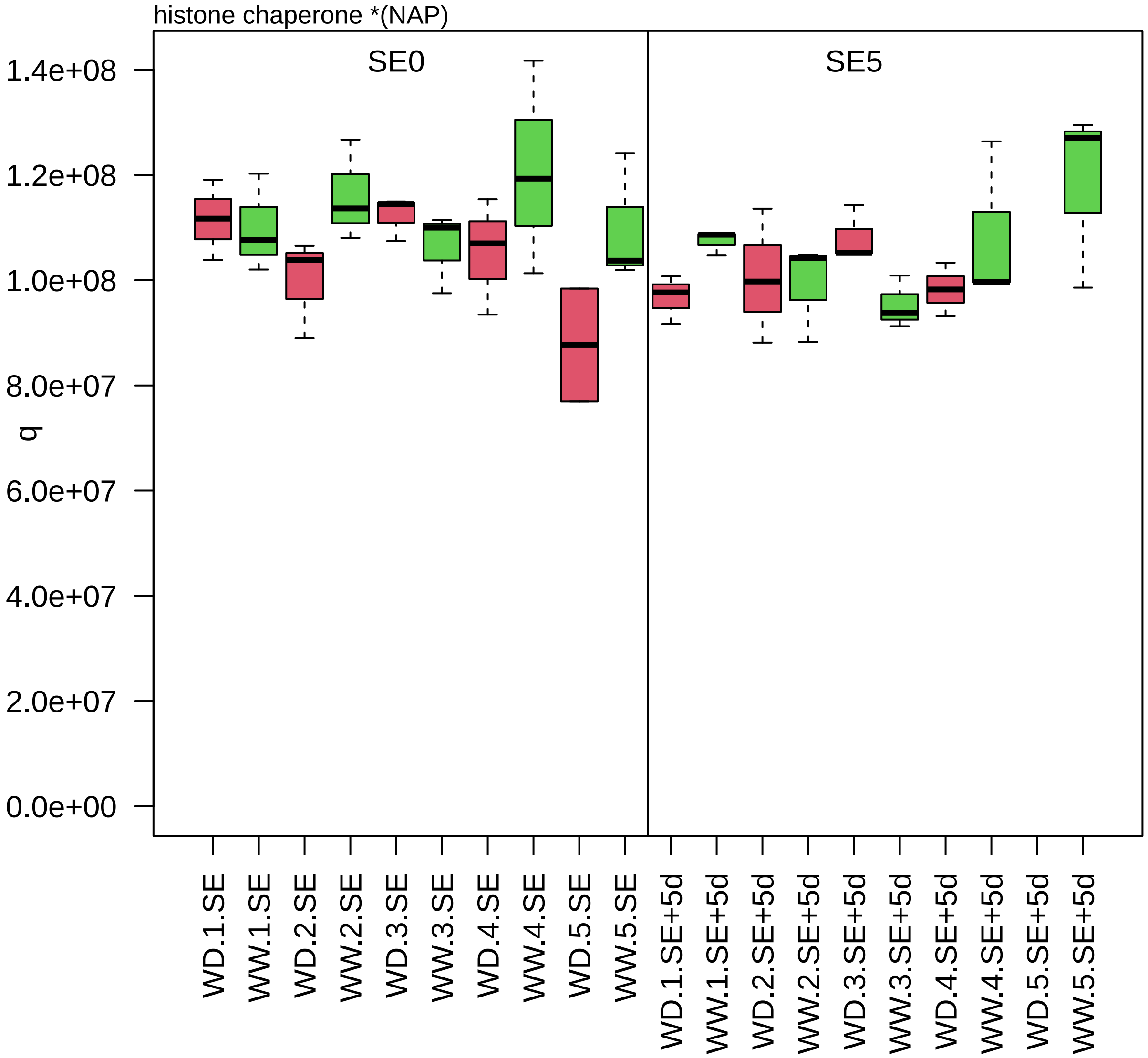

zm00001d045431_p001

**Cellular respiration.glycolysis**
**gene_symbol:enolase1 description:Enolase 1**

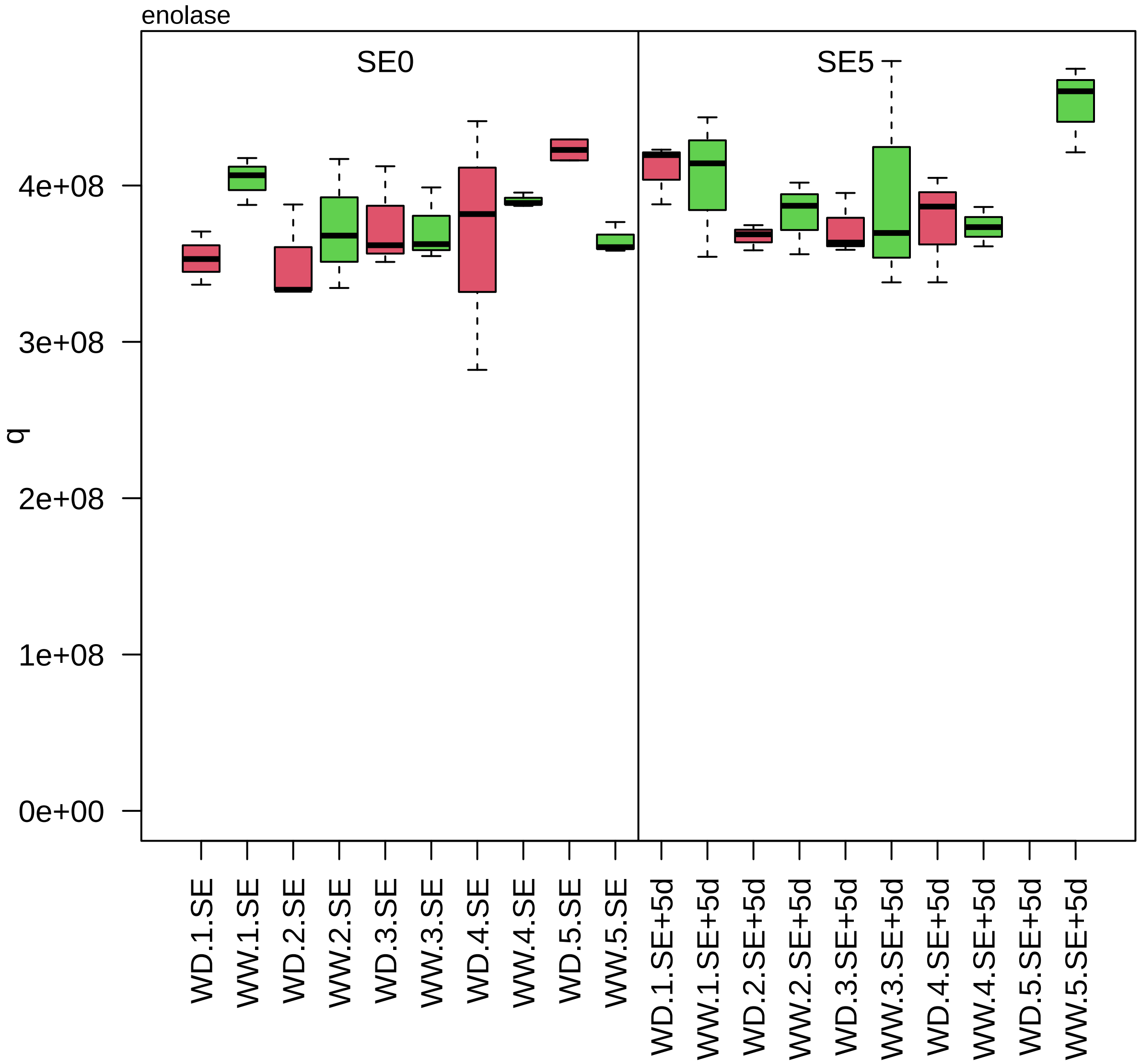

zm00001d045450_p001

**Amino acid metabolism.shikimate group amino acid biosynthesis**
**description: enolpyruvylshikimate phosphate synthase1**

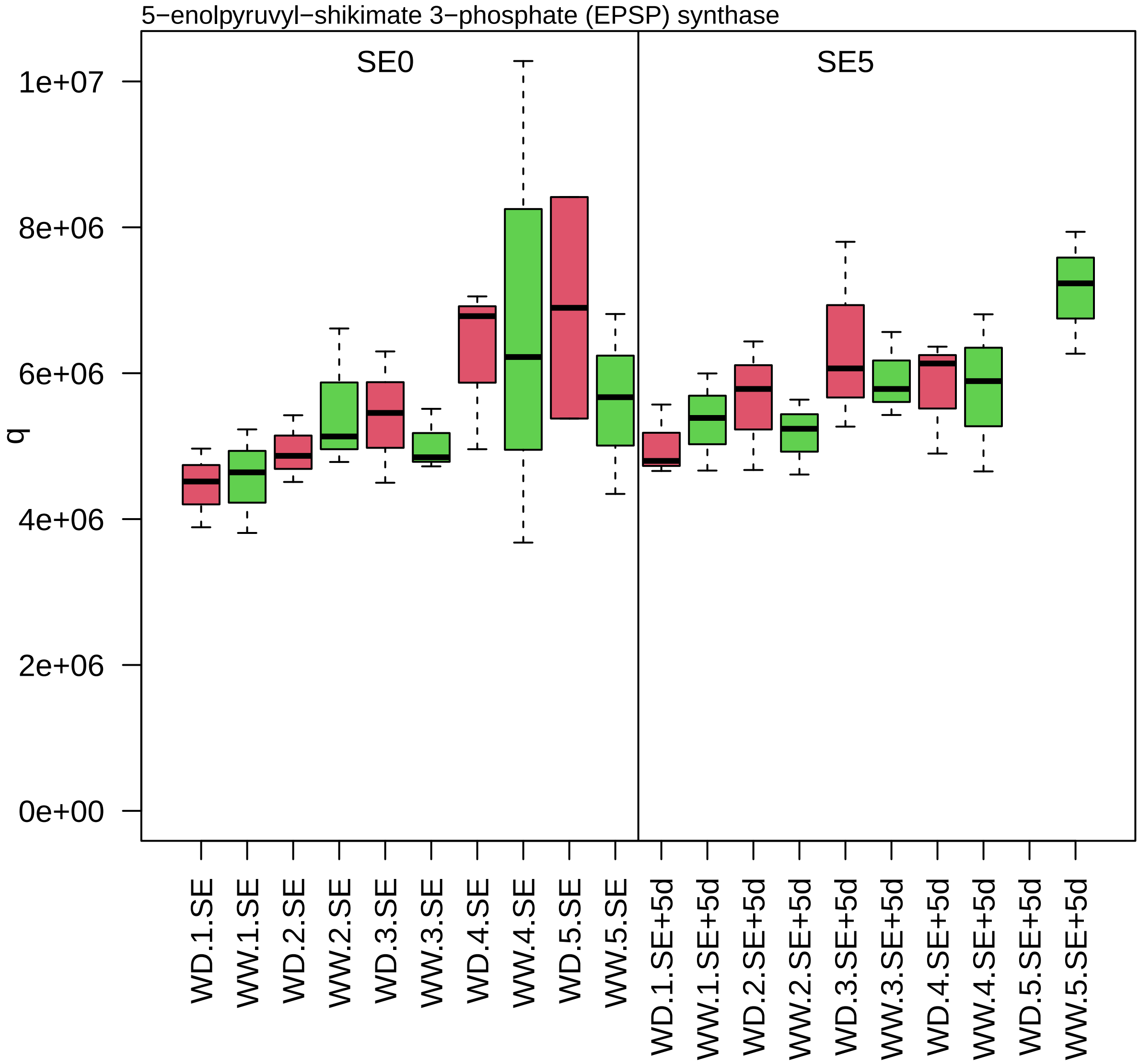

zm00001d045451_p001

**Photosynthesis.calvin cycle**

**gene_symbol:transketolase 1 description: transketolase 1**

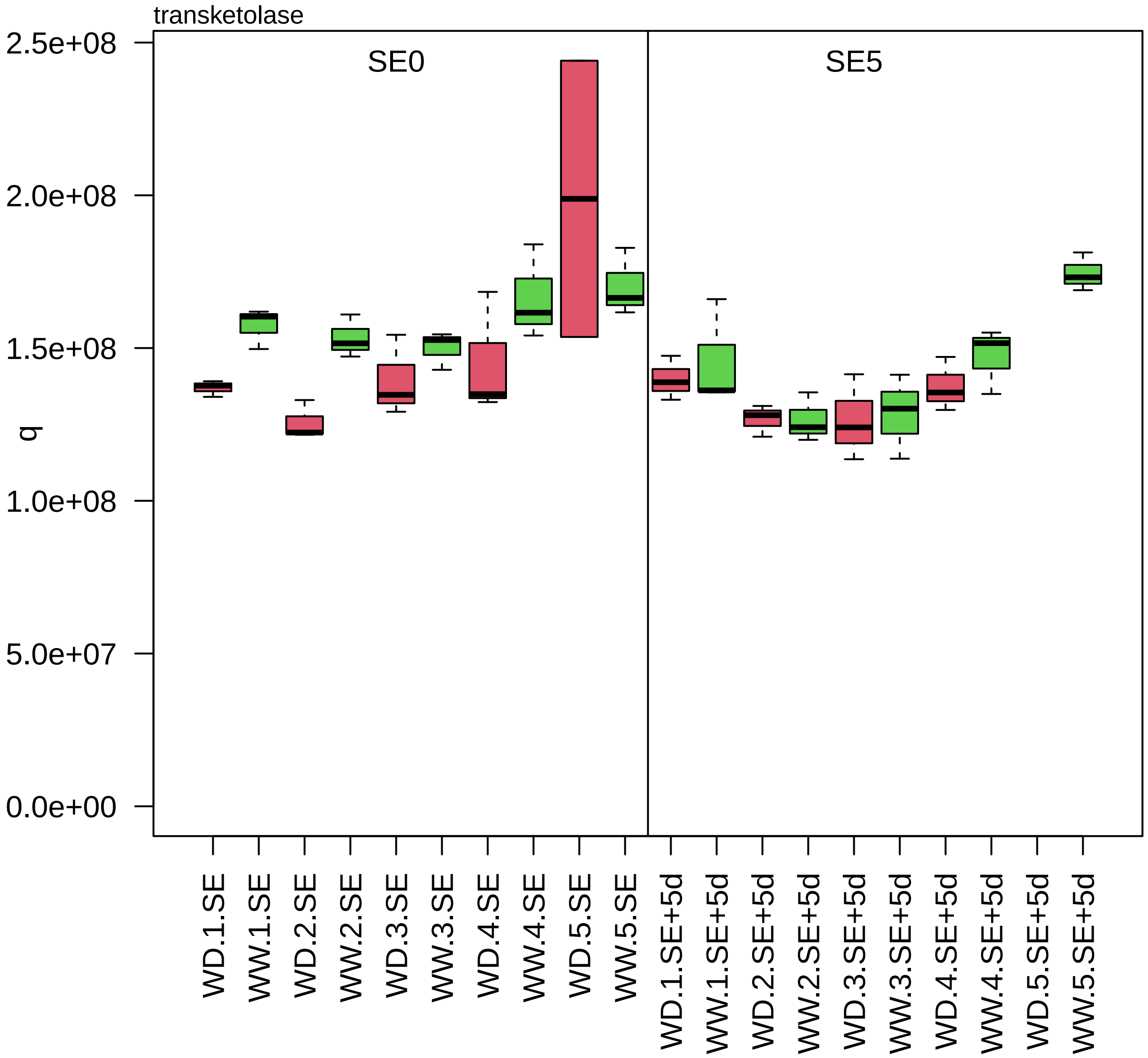

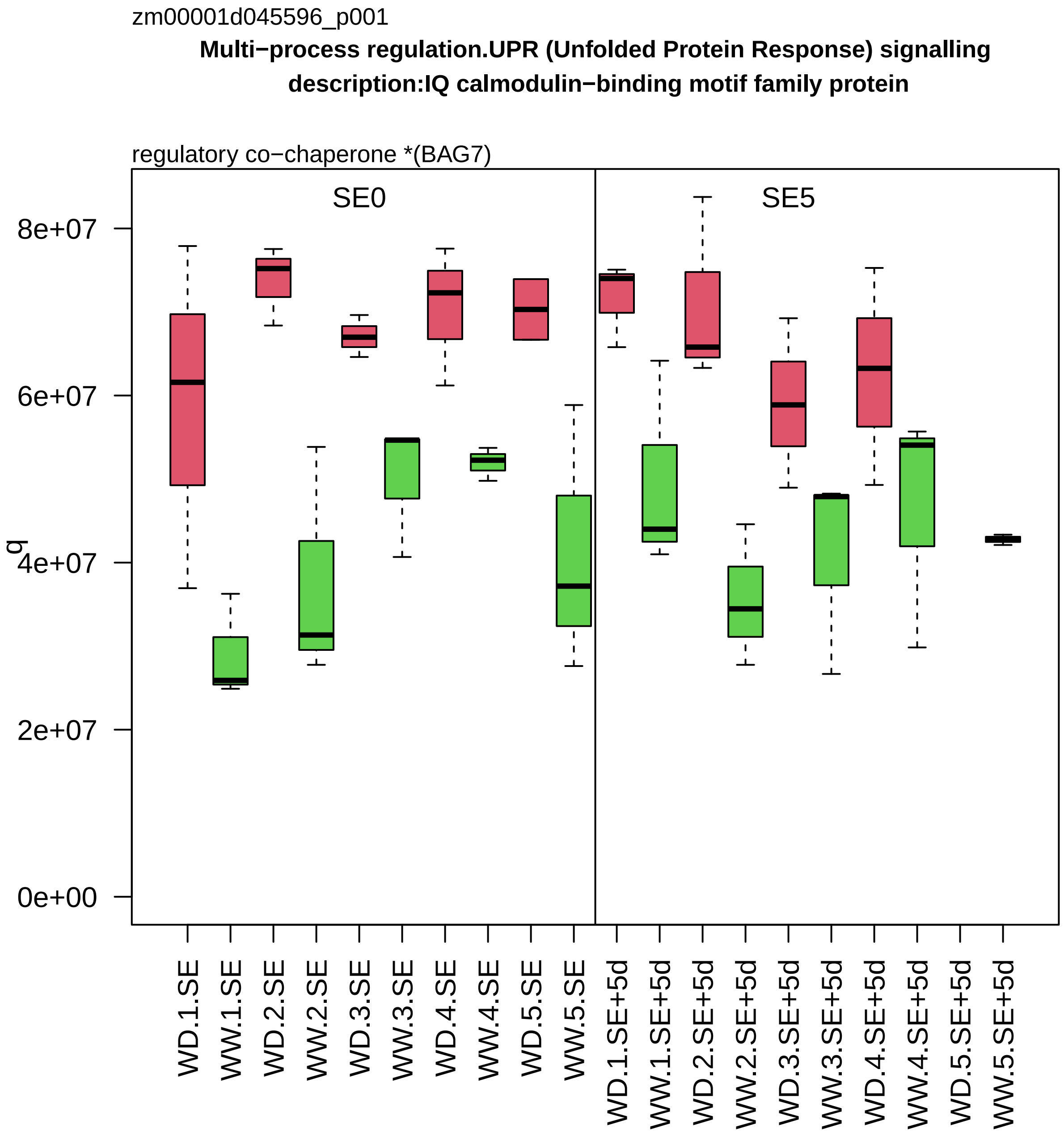
zm00001d045596_p001
Multi−process regulation.UPR (Unfolded Protein Response) signalling
description:IQ calmodulin−binding motif family protein
regulatory co−chaperone *(BAG7)
SE0
SE5
q
8e+07
6e+07
4e+07
2e+07
0e+00
WD.1.SE
WW.1.SE
WD.2.SE
WW.2.SE
WD.3.SE
WW.3.SE
WD.4.SE
WW.4.SE
WD.5.SE
WW.5.SE
WD.1.SE+5d
WW.1.SE+5d
WD.2.SE+5d
WW.2.SE+5d
WD.3.SE+5d
WW.3.SE+5d
WD.4.SE+5d
WW.4.SE+5d
WD.5.SE+5d
WW.5.SE+5d

zm00001d045599_p001

**Enzyme classification.EC_1 oxidoreductases**
**description:Monocopper oxidase−like protein SKU5**

EC_1.10 oxidoreductase acting on diphenol or related substance as donor

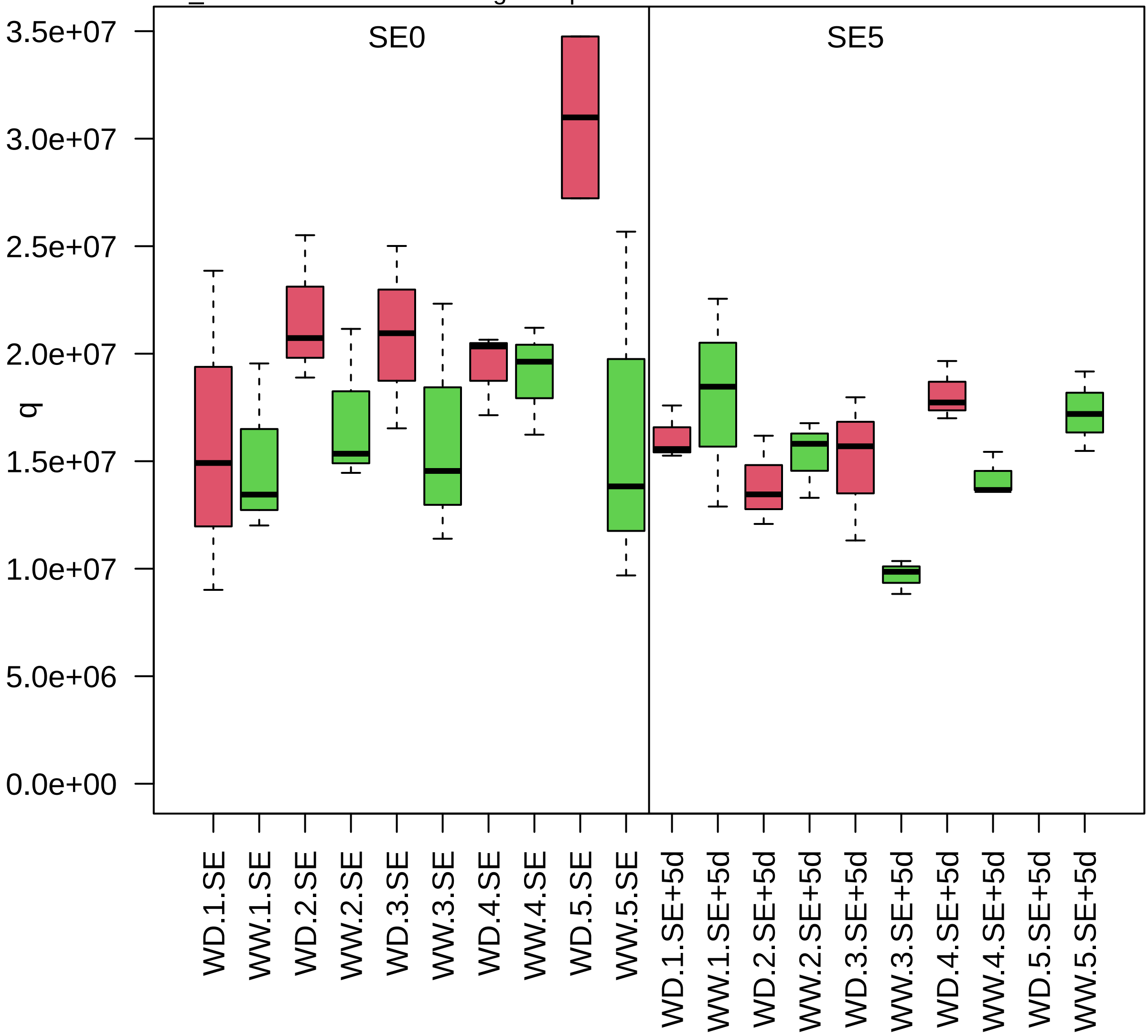

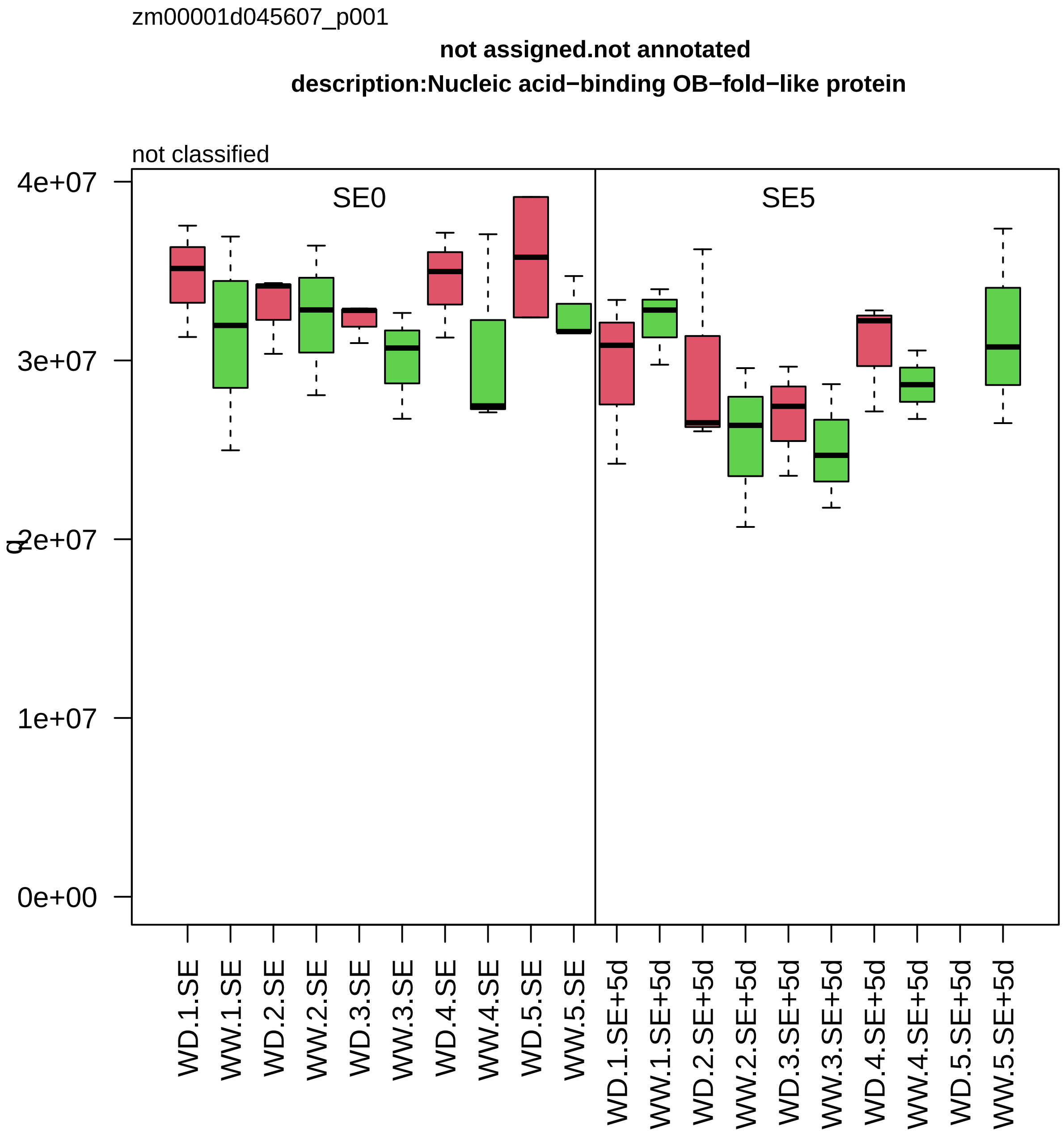

zm00001d045607_p001
not assigned.not annotated
description:Nucleic acid−binding OB−fold−like protein
not classified
SE0
SE5
0e+00
1e+07
2e+07
3e+07
4e+07
WD.1.SE
WW.1.SE
WD.2.SE
WW.2.SE
WD.3.SE
WW.3.SE
WD.4.SE
WW.4.SE
WD.5.SE
WW.5.SE
WD.1.SE+5d
WW.1.SE+5d
WD.2.SE+5d
WW.2.SE+5d
WD.3.SE+5d
WW.3.SE+5d
WD.4.SE+5d
WW.4.SE+5d
WD.5.SE+5d
WW.5.SE+5d

zm00001d045665_p001

**Protein homeostasis.ubiquitin−proteasome system**

**description:Ubiquitin receptor RAD23c**

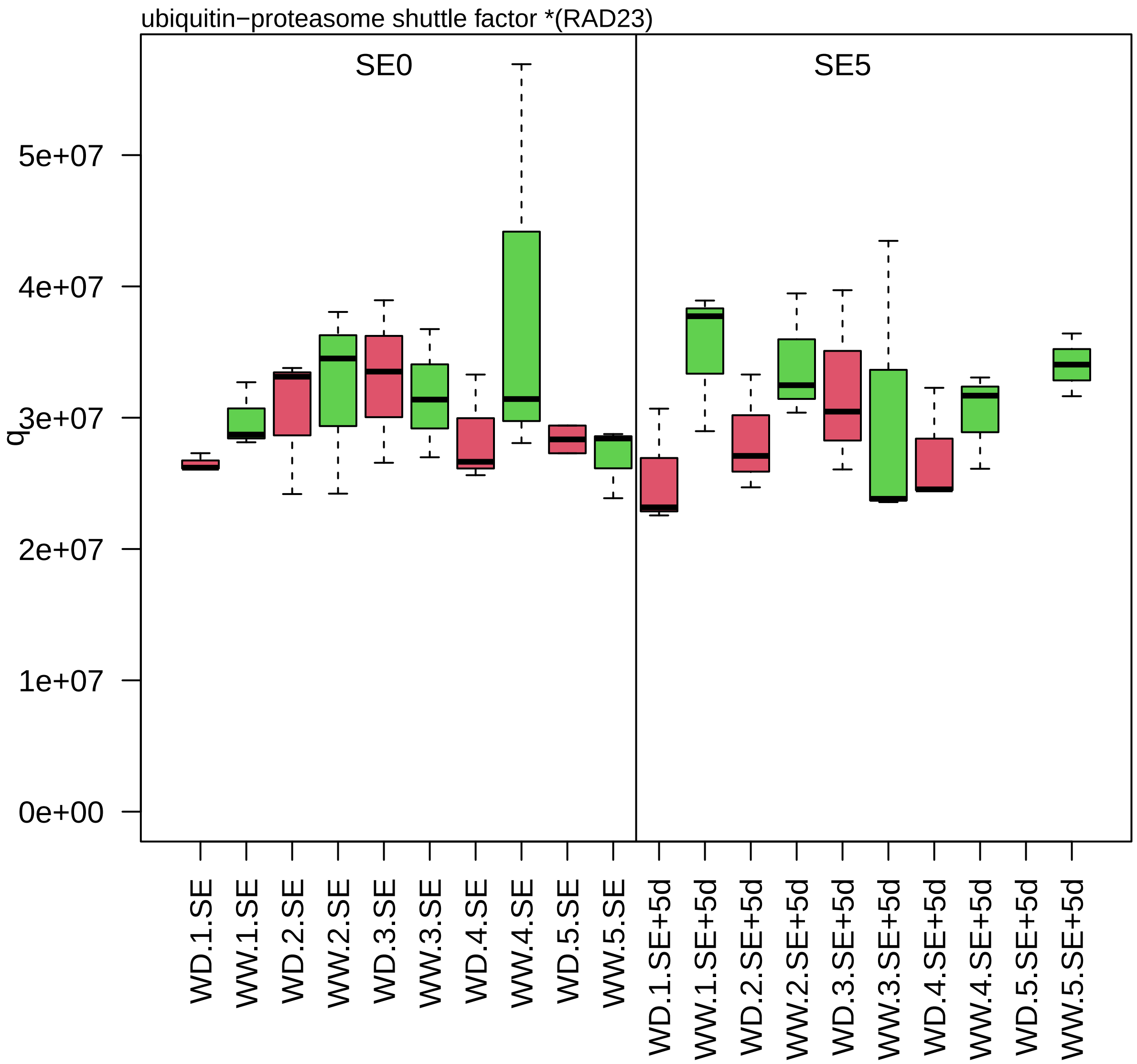

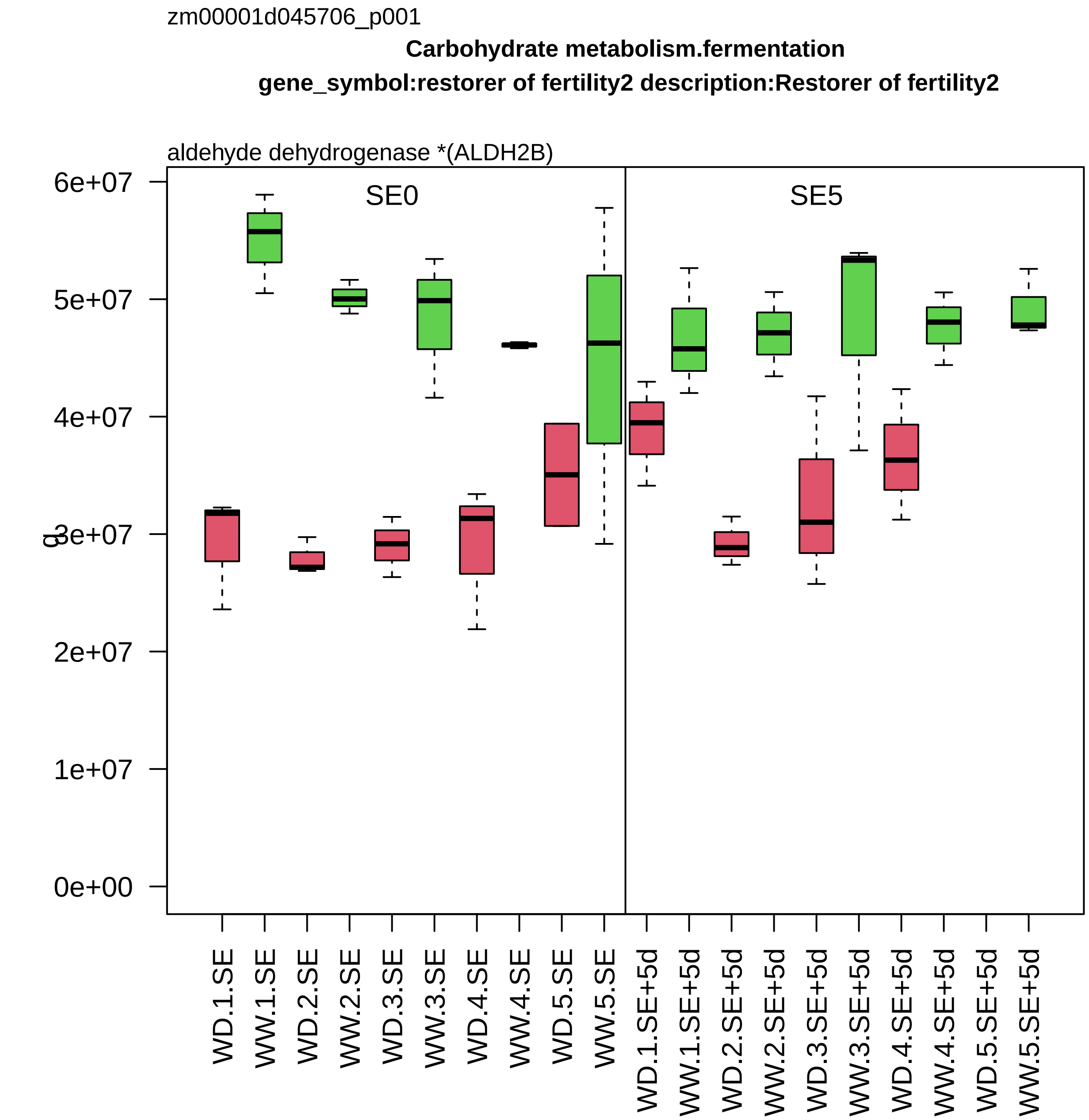

zm00001d045706_p001
Carbohydrate metabolism.fermentation
gene_symbol:restorer of fertility2 description:Restorer of fertility2
aldehyde dehydrogenase *(ALDH2B)
SE0
SE5
0e+00
1e+07
2e+07
3e+07
4e+07
5e+07
6e+07
WD.1.SE
WW.1.SE
WD.2.SE
WW.2.SE
WD.3.SE
WW.3.SE
WD.4.SE
WW.4.SE
WD.5.SE
WW.5.SE
WD.1.SE+5d
WW.1.SE+5d
WD.2.SE+5d
WW.2.SE+5d
WD.3.SE+5d
WW.3.SE+5d
WD.4.SE+5d
WW.4.SE+5d
WD.5.SE+5d
WW.5.SE+5d

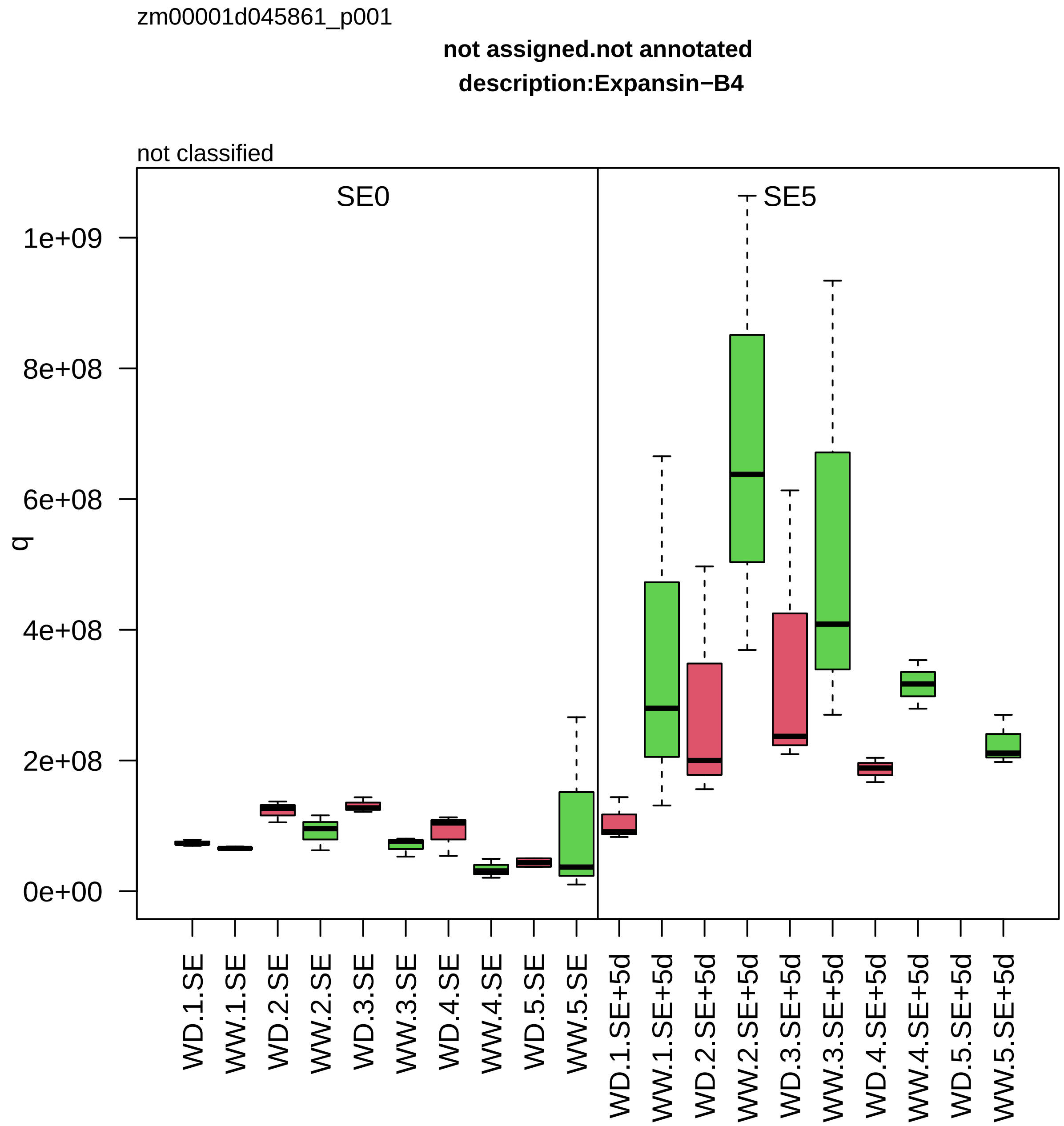

zm00001d045861_p001
not assigned.not annotated
description:Expansin−B4
not classified
SE0
SE5
q
1e+09
8e+08
6e+08
4e+08
2e+08
0e+00
WD.1.SE
WW.1.SE
WD.2.SE
WW.2.SE
WD.3.SE
WW.3.SE
WD.4.SE
WW.4.SE
WD.5.SE
WW.5.SE
WD.1.SE+5d
WW.1.SE+5d
WD.2.SE+5d
WW.2.SE+5d
WD.3.SE+5d
WW.3.SE+5d
WD.4.SE+5d
WW.4.SE+5d
WD.5.SE+5d
WW.5.SE+5d

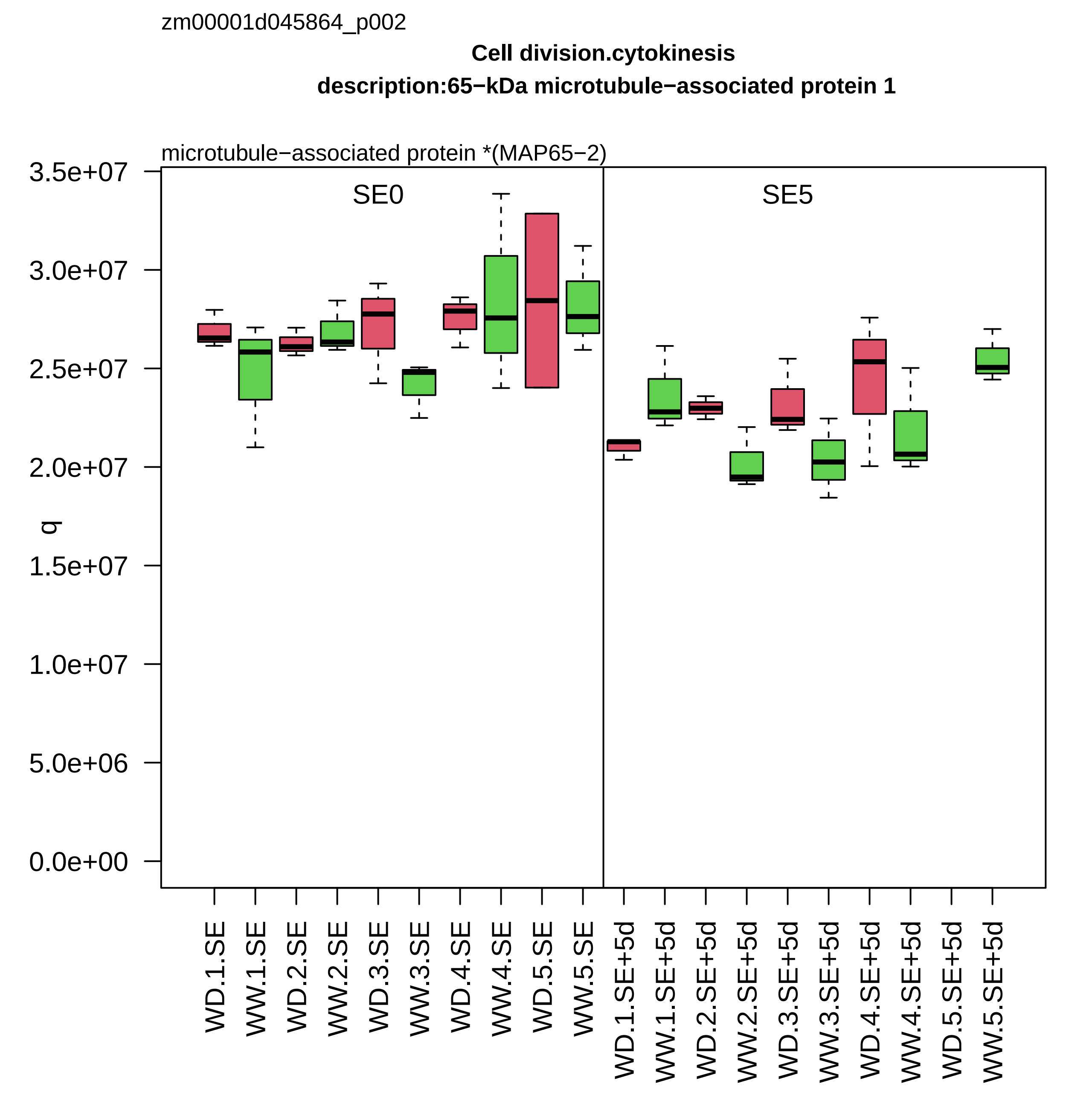

zm00001d045864_p002
Cell division.cytokinesis
description:65−kDa microtubule−associated protein 1
microtubule−associated protein *(MAP65−2)
SE0
SE5
q
3.5e+07
3.0e+07
2.5e+07
2.0e+07
1.5e+07
1.0e+07
5.0e+06
0.0e+00
WD.1.SE
WW.1.SE
WD.2.SE
WW.2.SE
WD.3.SE
WW.3.SE
WD.4.SE
WW.4.SE
WD.5.SE
WW.5.SE
WD.1.SE+5d
WW.1.SE+5d
WD.2.SE+5d
WW.2.SE+5d
WD.3.SE+5d
WW.3.SE+5d
WD.4.SE+5d
WW.4.SE+5d
WD.5.SE+5d
WW.5.SE+5d

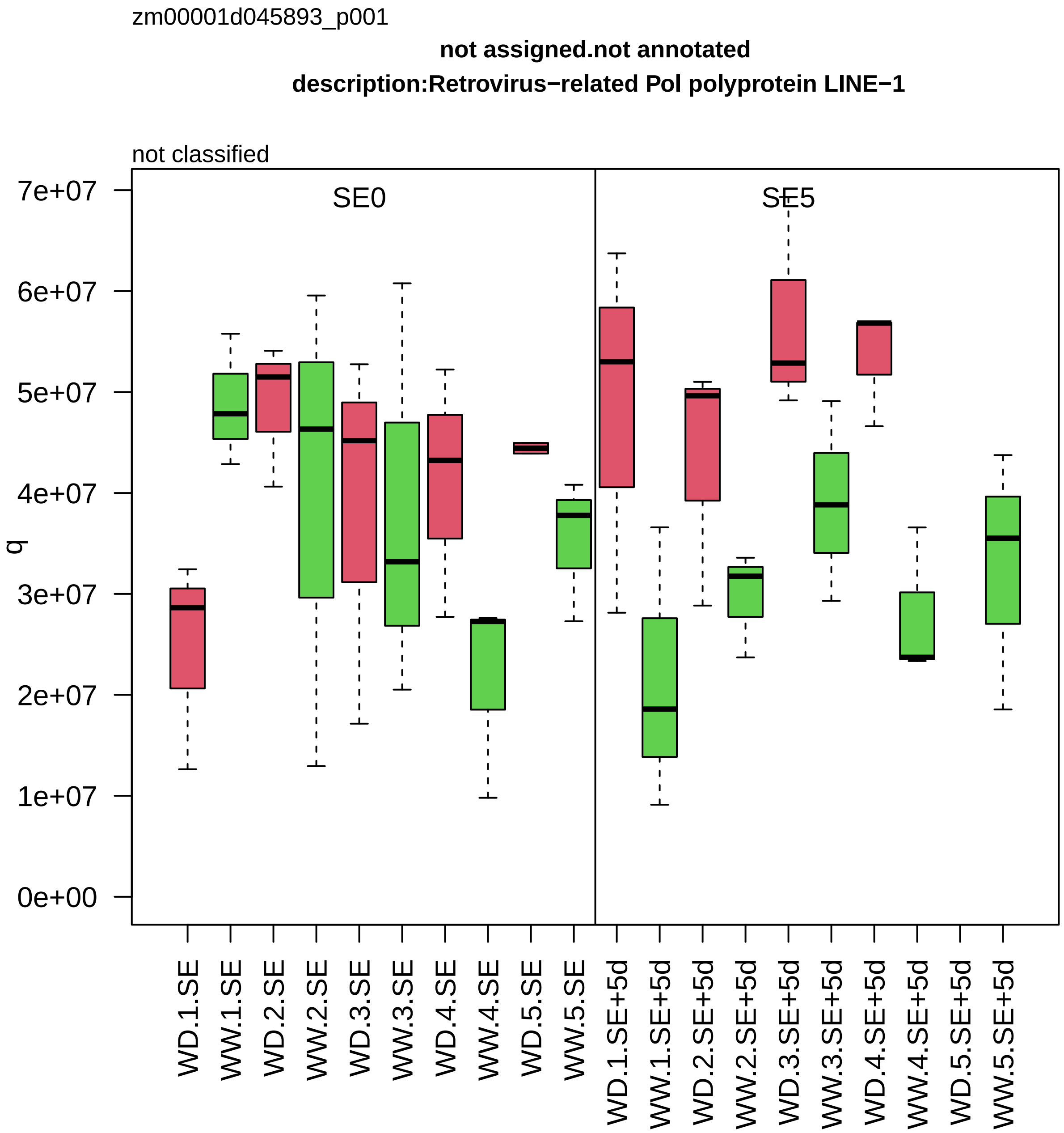

zm00001d045893_p001
not assigned.not annotated
description:Retrovirus−related Pol polyprotein LINE−1
not classified
SE0
SE5
q
0e+00
1e+07
2e+07
3e+07
4e+07
5e+07
6e+07
7e+07
WD.1.SE
WW.1.SE
WD.2.SE
WW.2.SE
WD.3.SE
WW.3.SE
WD.4.SE
WW.4.SE
WD.5.SE
WW.5.SE
WD.1.SE+5d
WW.1.SE+5d
WD.2.SE+5d
WW.2.SE+5d
WD.3.SE+5d
WW.3.SE+5d
WD.4.SE+5d
WW.4.SE+5d
WD.5.SE+5d
WW.5.SE+5d

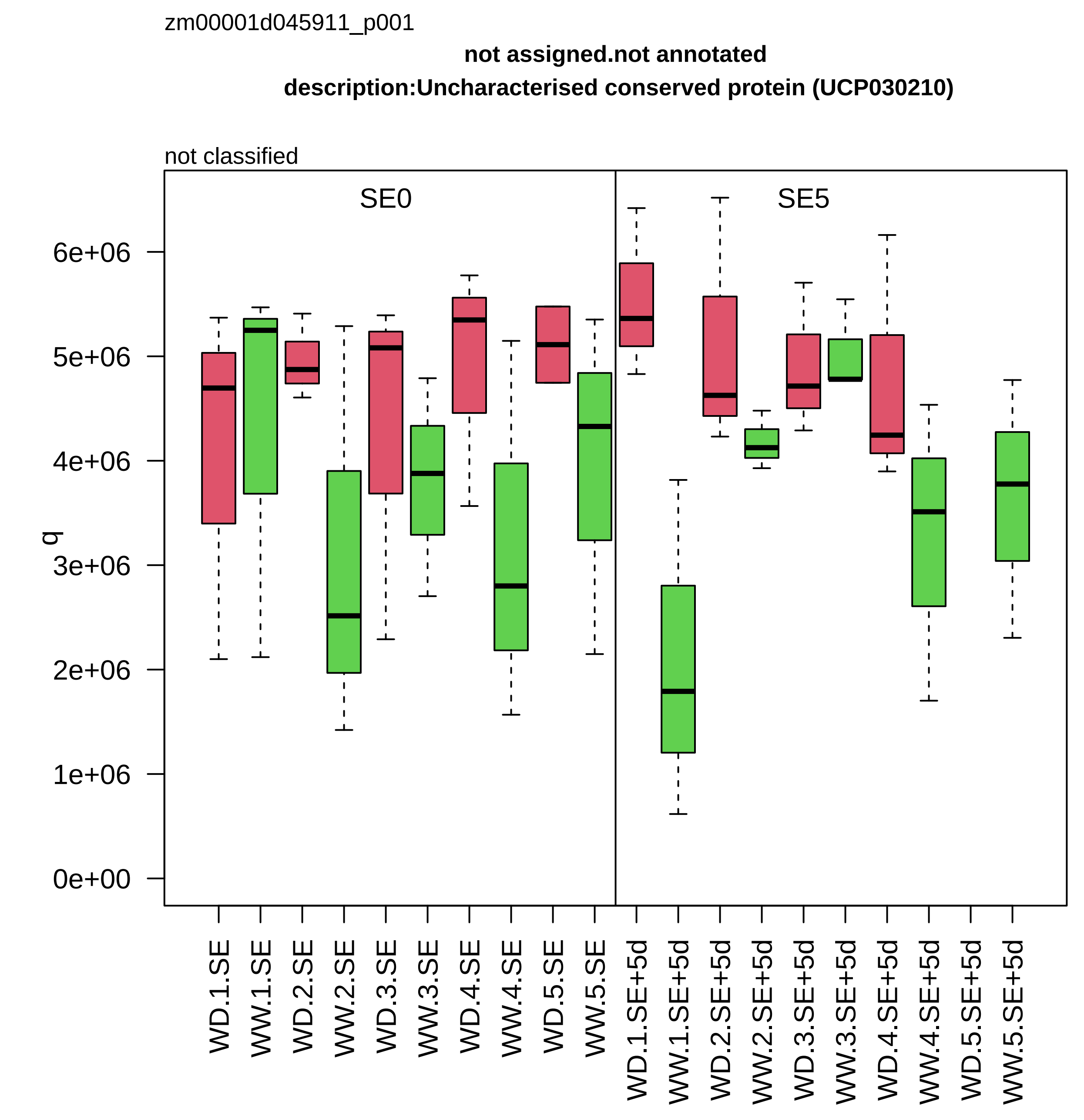

zm00001d045911_p001
not assigned.not annotated
description:Uncharacterised conserved protein (UCP030210)
not classified
SE0
SE5
q
6e+06
5e+06
4e+06
3e+06
2e+06
1e+06
0e+00
WD.1.SE
WW.1.SE
WD.2.SE
WW.2.SE
WD.3.SE
WW.3.SE
WD.4.SE
WW.4.SE
WD.5.SE
WW.5.SE
WD.1.SE+5d
WW.1.SE+5d
WD.2.SE+5d
WW.2.SE+5d
WD.3.SE+5d
WW.3.SE+5d
WD.4.SE+5d
WW.4.SE+5d
WD.5.SE+5d
WW.5.SE+5d

zm00001d045965_p001

**Enzyme classification.EC_1 oxidoreductases**

**description:Formate dehydrogenase chloroplastic/mitochondrial**

EC_1.17 oxidoreductase acting on CH or CH2 group

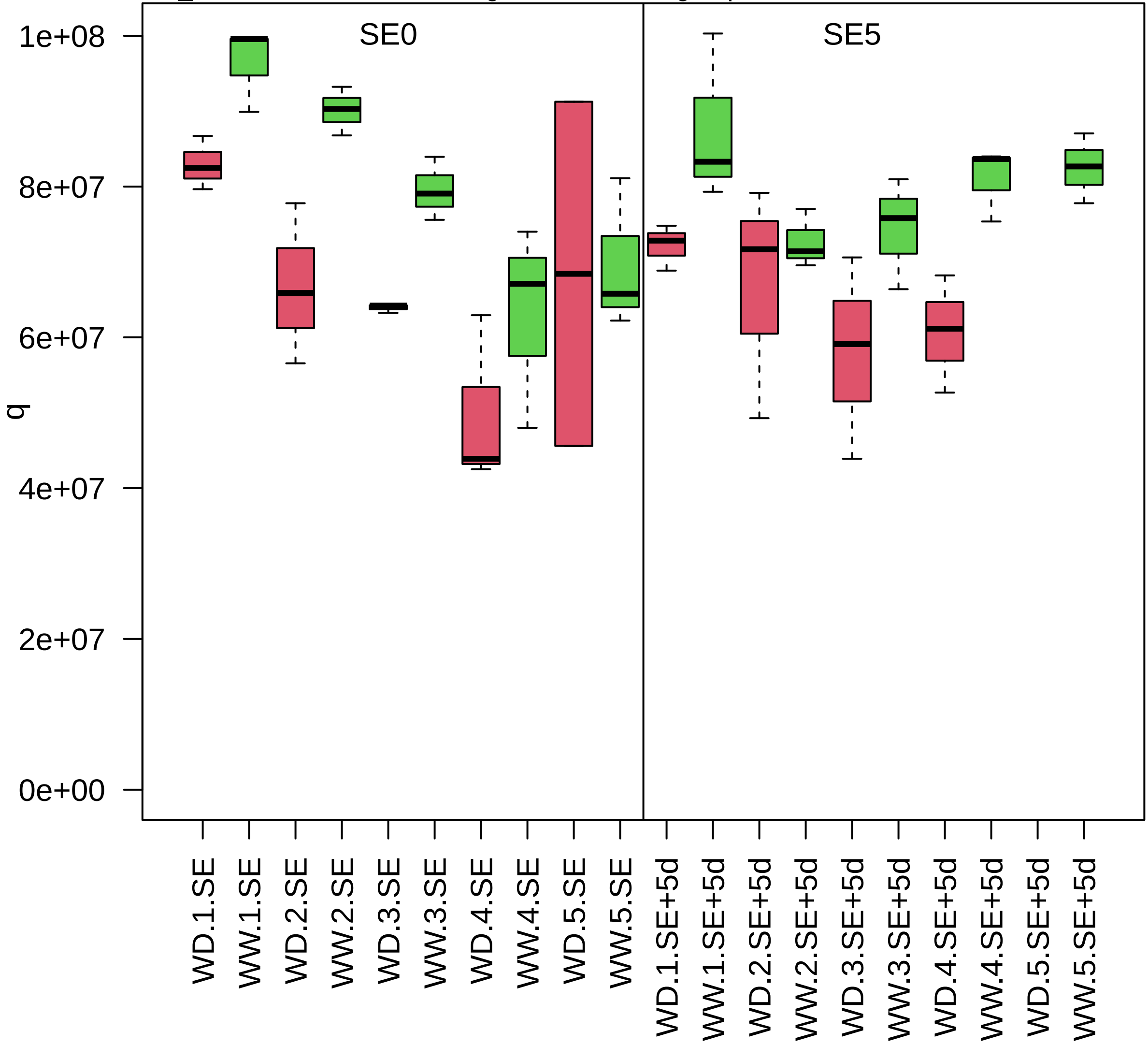

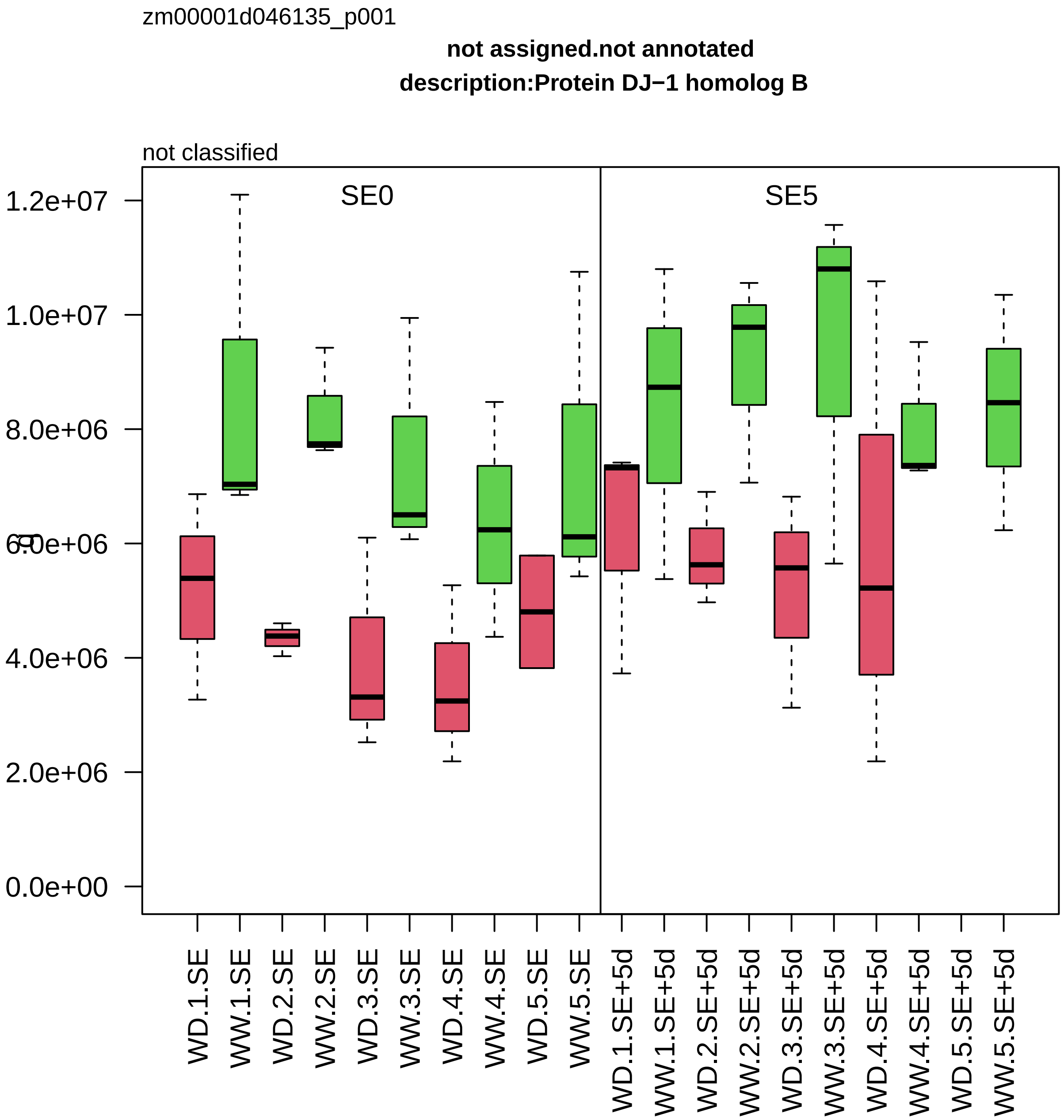

zm00001d046135_p001
not assigned.not annotated
description:Protein DJ−1 homolog B
not classified
SE0
SE5
1.2e+07
1.0e+07
8.0e+06
6.0e+06
4.0e+06
2.0e+06
0.0e+00
WD.1.SE
WW.1.SE
WD.2.SE
WW.2.SE
WD.3.SE
WW.3.SE
WD.4.SE
WW.4.SE
WD.5.SE
WW.5.SE
WD.1.SE+5d
WW.1.SE+5d
WD.2.SE+5d
WW.2.SE+5d
WD.3.SE+5d
WW.3.SE+5d
WD.4.SE+5d
WW.4.SE+5d
WD.5.SE+5d
WW.5.SE+5d

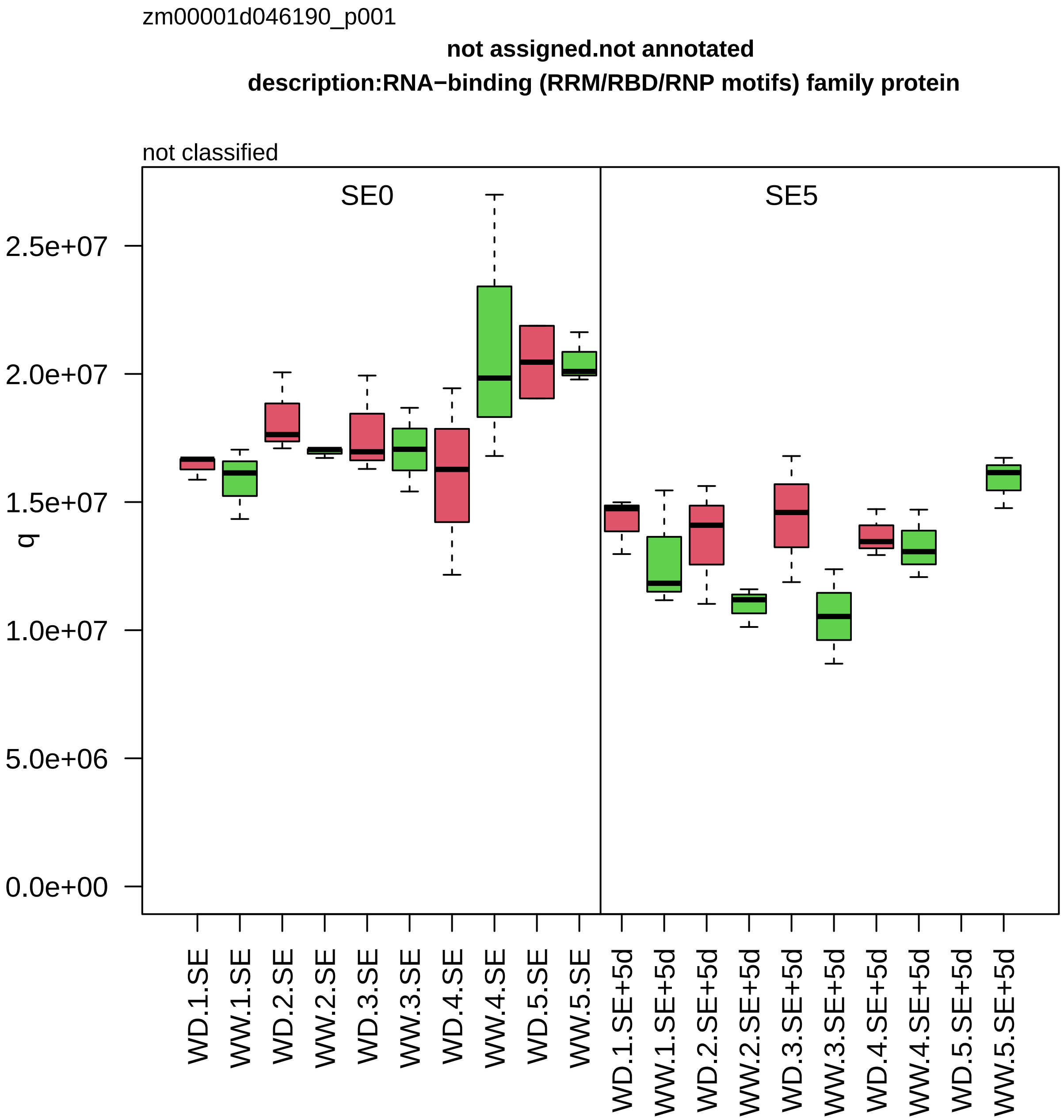

zm00001d046190_p001
not assigned.not annotated
description:RNA−binding (RRM/RBD/RNP motifs) family protein
not classified
SE0
SE5
q
2.5e+07
2.0e+07
1.5e+07
1.0e+07
5.0e+06
0.0e+00
WD.1.SE
WW.1.SE
WD.2.SE
WW.2.SE
WD.3.SE
WW.3.SE
WD.4.SE
WW.4.SE
WD.5.SE
WW.5.SE
WD.1.SE+5d
WW.1.SE+5d
WD.2.SE+5d
WW.2.SE+5d
WD.3.SE+5d
WW.3.SE+5d
WD.4.SE+5d
WW.4.SE+5d
WD.5.SE+5d
WW.5.SE+5d

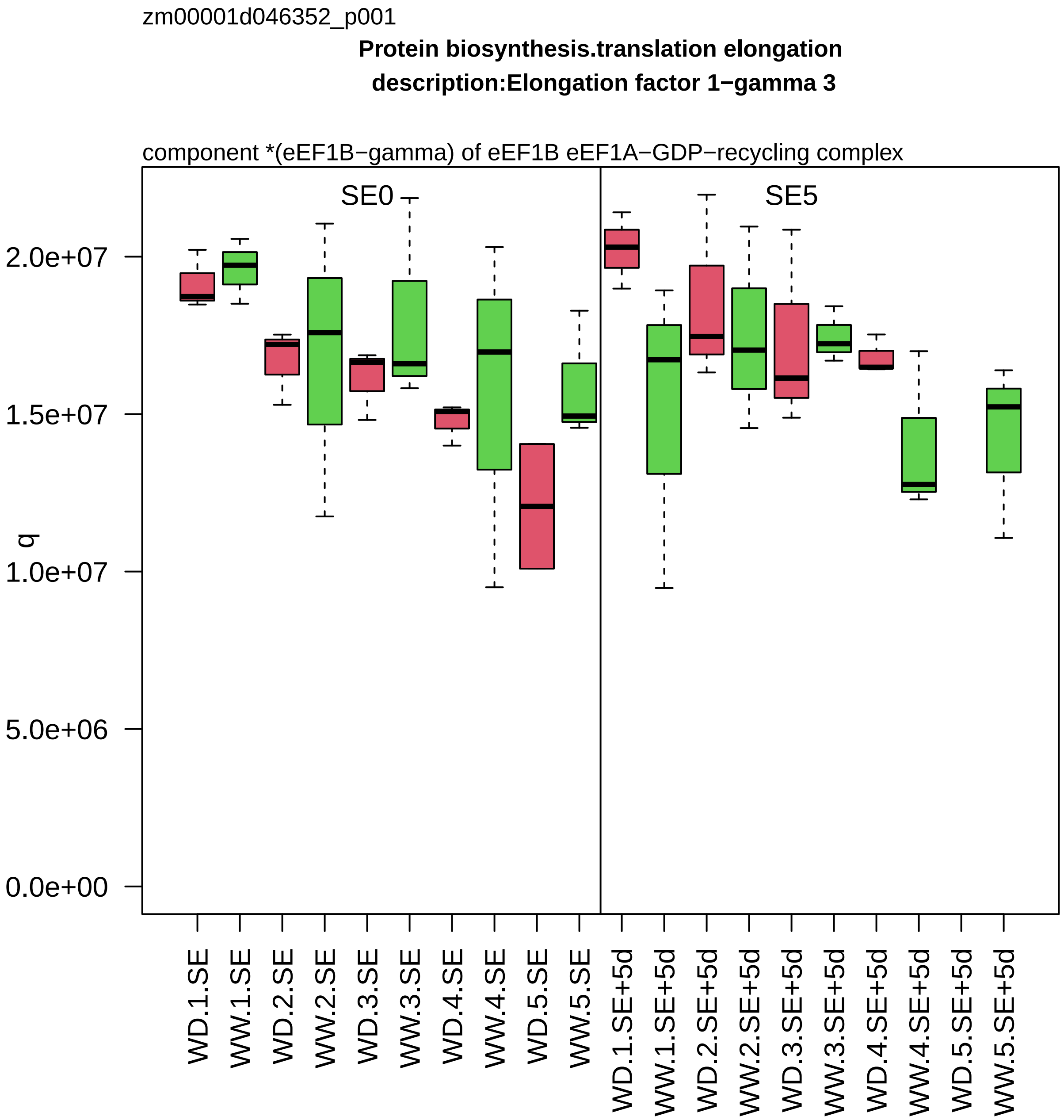

zm00001d046352_p001
Protein biosynthesis.translation elongation
description:Elongation factor 1−gamma 3
component *(eEF1B−gamma) of eEF1B eEF1A−GDP−recycling complex
SE0
SE5
q
2.0e+07
1.5e+07
1.0e+07
5.0e+06
0.0e+00
WD.1.SE
WW.1.SE
WD.2.SE
WW.2.SE
WD.3.SE
WW.3.SE
WD.4.SE
WW.4.SE
WD.5.SE
WW.5.SE
WD.1.SE+5d
WW.1.SE+5d
WD.2.SE+5d
WW.2.SE+5d
WD.3.SE+5d
WW.3.SE+5d
WD.4.SE+5d
WW.4.SE+5d
WD.5.SE+5d
WW.5.SE+5d

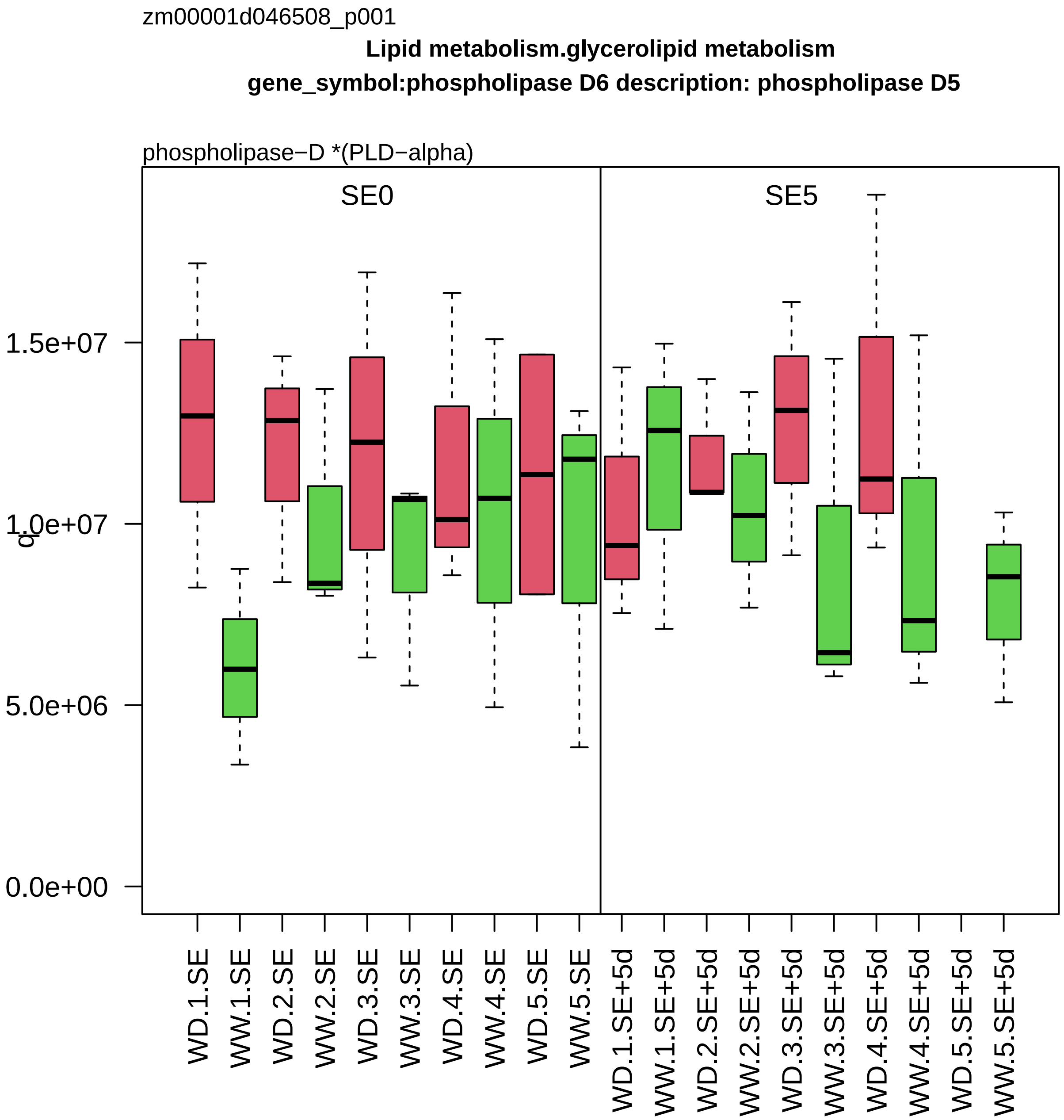
zm00001d046508_p001
Lipid metabolism.glycerolipid metabolism
gene_symbol:phospholipase D6 description: phospholipase D5
phospholipase−D *(PLD−alpha)
SE0
SE5
1.5e+07
1.0e+07
5.0e+06
0.0e+00
q
WD.1.SE
WW.1.SE
WD.2.SE
WW.2.SE
WD.3.SE
WW.3.SE
WD.4.SE
WW.4.SE
WD.5.SE
WW.5.SE
WD.1.SE+5d
WW.1.SE+5d
WD.2.SE+5d
WW.2.SE+5d
WD.3.SE+5d
WW.3.SE+5d
WD.4.SE+5d
WW.4.SE+5d
WD.5.SE+5d
WW.5.SE+5d

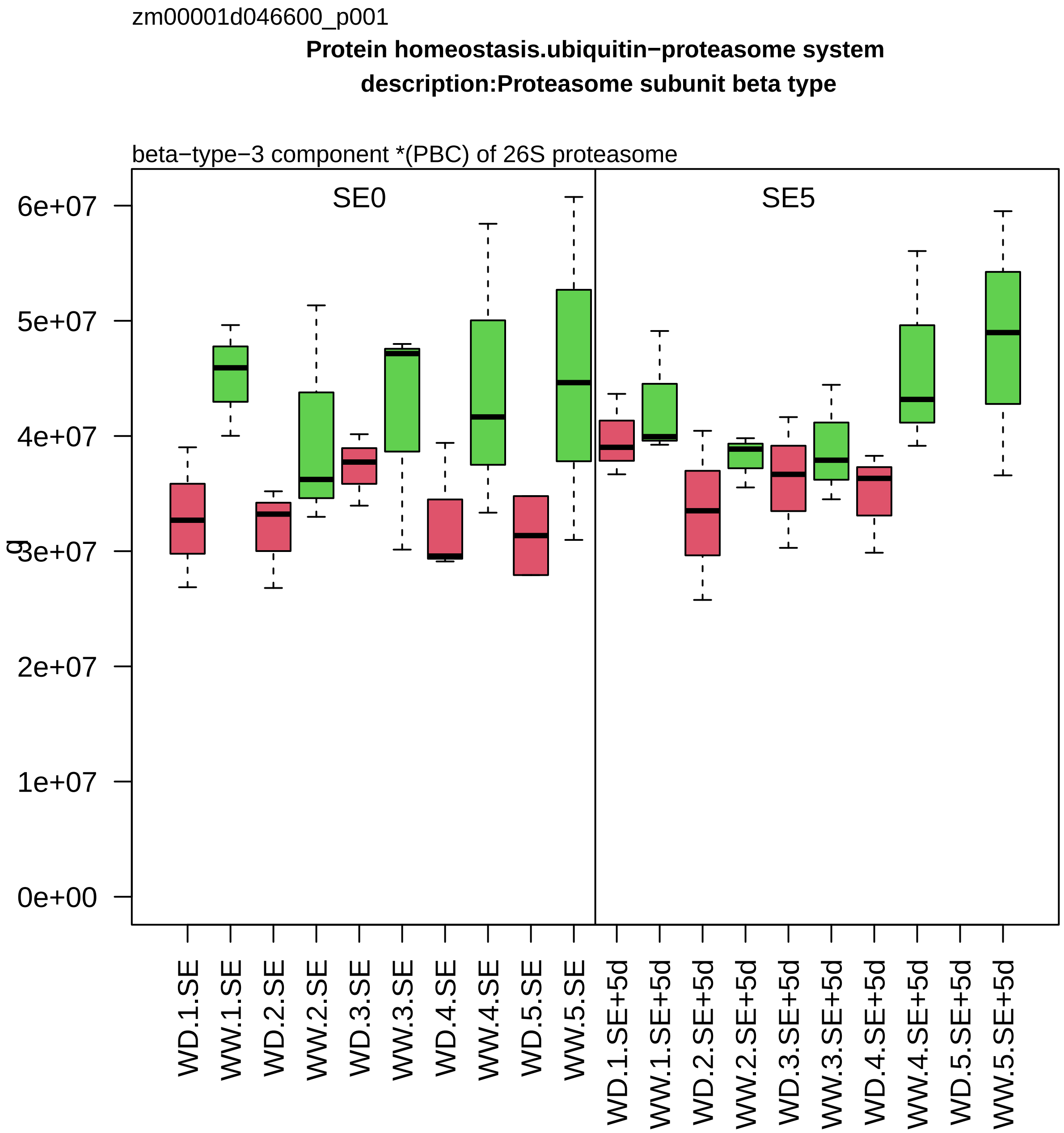
zm00001d046600_p001
Protein homeostasis.ubiquitin−proteasome system
description:Proteasome subunit beta type
beta−type−3 component *(PBC) of 26S proteasome
SE0
SE5
g
6e+07
5e+07
4e+07
3e+07
2e+07
1e+07
0e+00
WD.1.SE
WW.1.SE
WD.2.SE
WW.2.SE
WD.3.SE
WW.3.SE
WD.4.SE
WW.4.SE
WD.5.SE
WW.5.SE
WD.1.SE+5d
WW.1.SE+5d
WD.2.SE+5d
WW.2.SE+5d
WD.3.SE+5d
WW.3.SE+5d
WD.4.SE+5d
WW.4.SE+5d
WD.5.SE+5d
WW.5.SE+5d

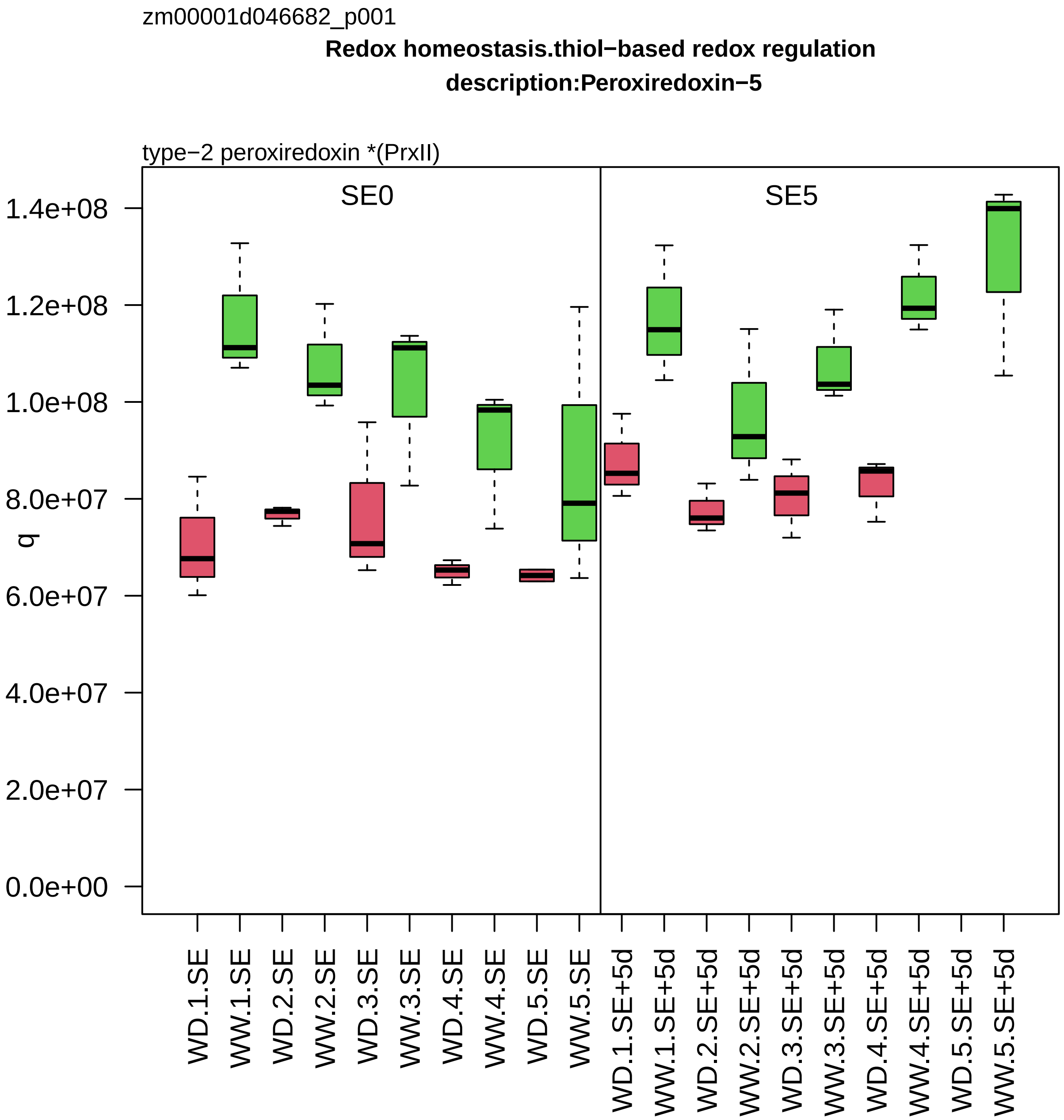
zm00001d046682_p001
Redox homeostasis.thiol−based redox regulation
description:Peroxiredoxin−5
type−2 peroxiredoxin *(PrxII)
SE0
SE5
q
1.4e+08
1.2e+08
1.0e+08
8.0e+07
6.0e+07
4.0e+07
2.0e+07
0.0e+00
WD.1.SE
WW.1.SE
WD.2.SE
WW.2.SE
WD.3.SE
WW.3.SE
WD.4.SE
WW.4.SE
WD.5.SE
WW.5.SE
WD.1.SE+5d
WW.1.SE+5d
WD.2.SE+5d
WW.2.SE+5d
WD.3.SE+5d
WW.3.SE+5d
WD.4.SE+5d
WW.4.SE+5d
WD.5.SE+5d
WW.5.SE+5d

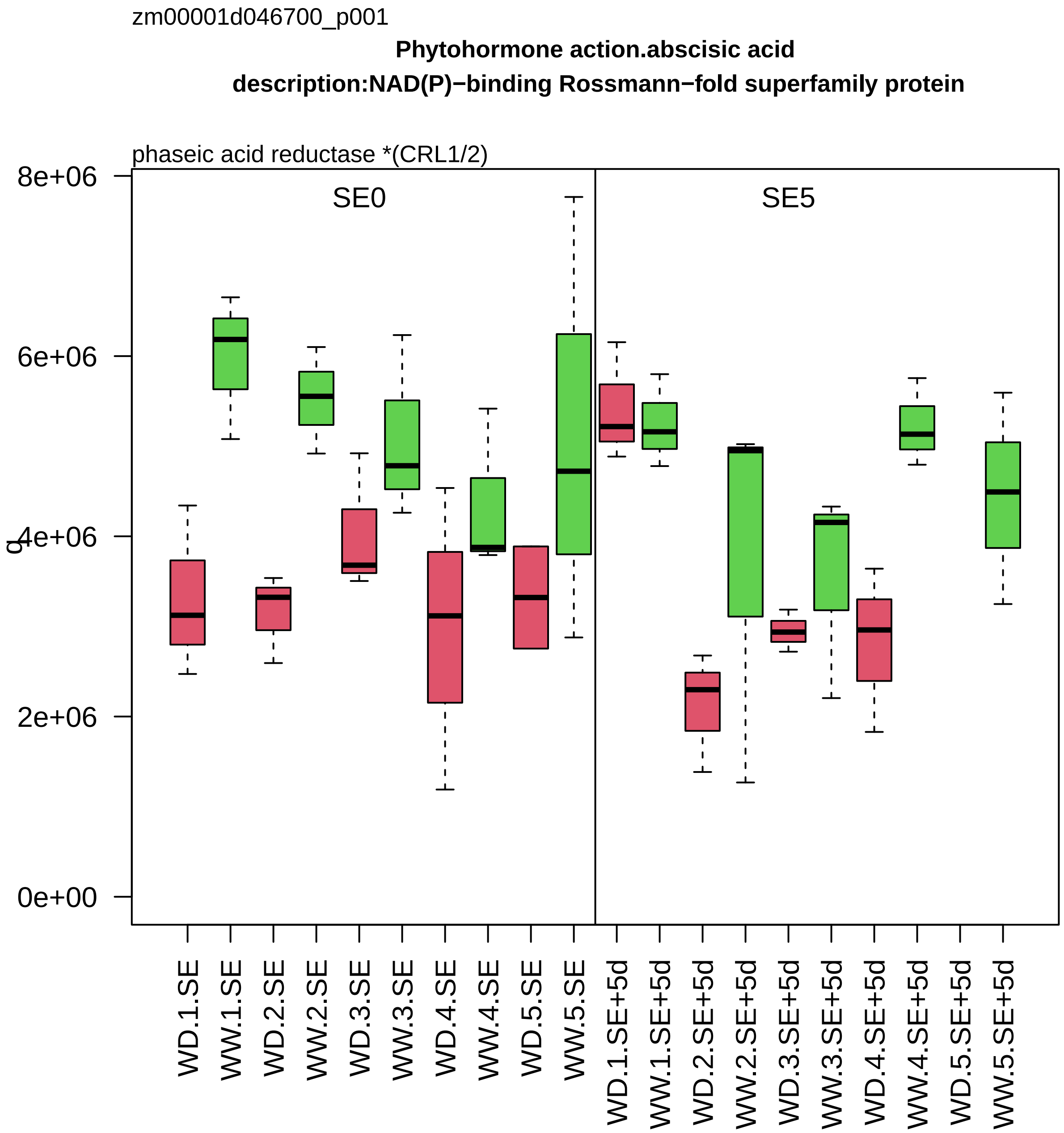
zm00001d046700_p001
Phytohormone action.abscisic acid
description:NAD(P)−binding Rossmann−fold superfamily protein
phaseic acid reductase *(CRL1/2)
SE0
SE5
q
8e+06
6e+06
4e+06
2e+06
0e+00
WD.1.SE
WW.1.SE
WD.2.SE
WW.2.SE
WD.3.SE
WW.3.SE
WD.4.SE
WW.4.SE
WD.5.SE
WW.5.SE
WD.1.SE+5d
WW.1.SE+5d
WD.2.SE+5d
WW.2.SE+5d
WD.3.SE+5d
WW.3.SE+5d
WD.4.SE+5d
WW.4.SE+5d
WD.5.SE+5d
WW.5.SE+5d

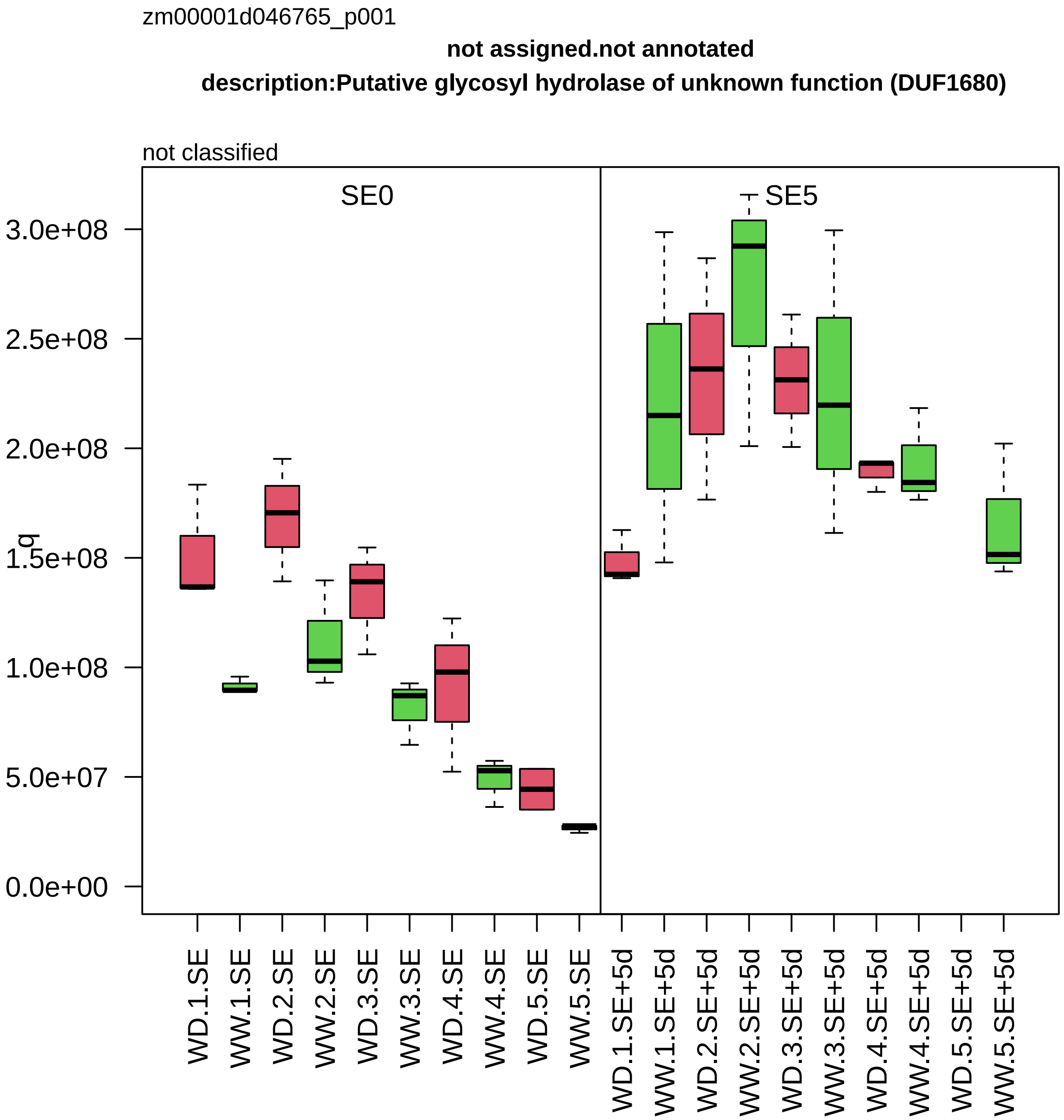

zm00001d046765_p001
not assigned.not annotated
description:Putative glycosyl hydrolase of unknown function (DUF1680)
not classified
SE0
SE5
q
3.0e+08
2.5e+08
2.0e+08
1.5e+08
1.0e+08
5.0e+07
0.0e+00
WD.1.SE
WW.1.SE
WD.2.SE
WW.2.SE
WD.3.SE
WW.3.SE
WD.4.SE
WW.4.SE
WD.5.SE
WW.5.SE
WD.1.SE+5d
WW.1.SE+5d
WD.2.SE+5d
WW.2.SE+5d
WD.3.SE+5d
WW.3.SE+5d
WD.4.SE+5d
WW.4.SE+5d
WD.5.SE+5d
WW.5.SE+5d

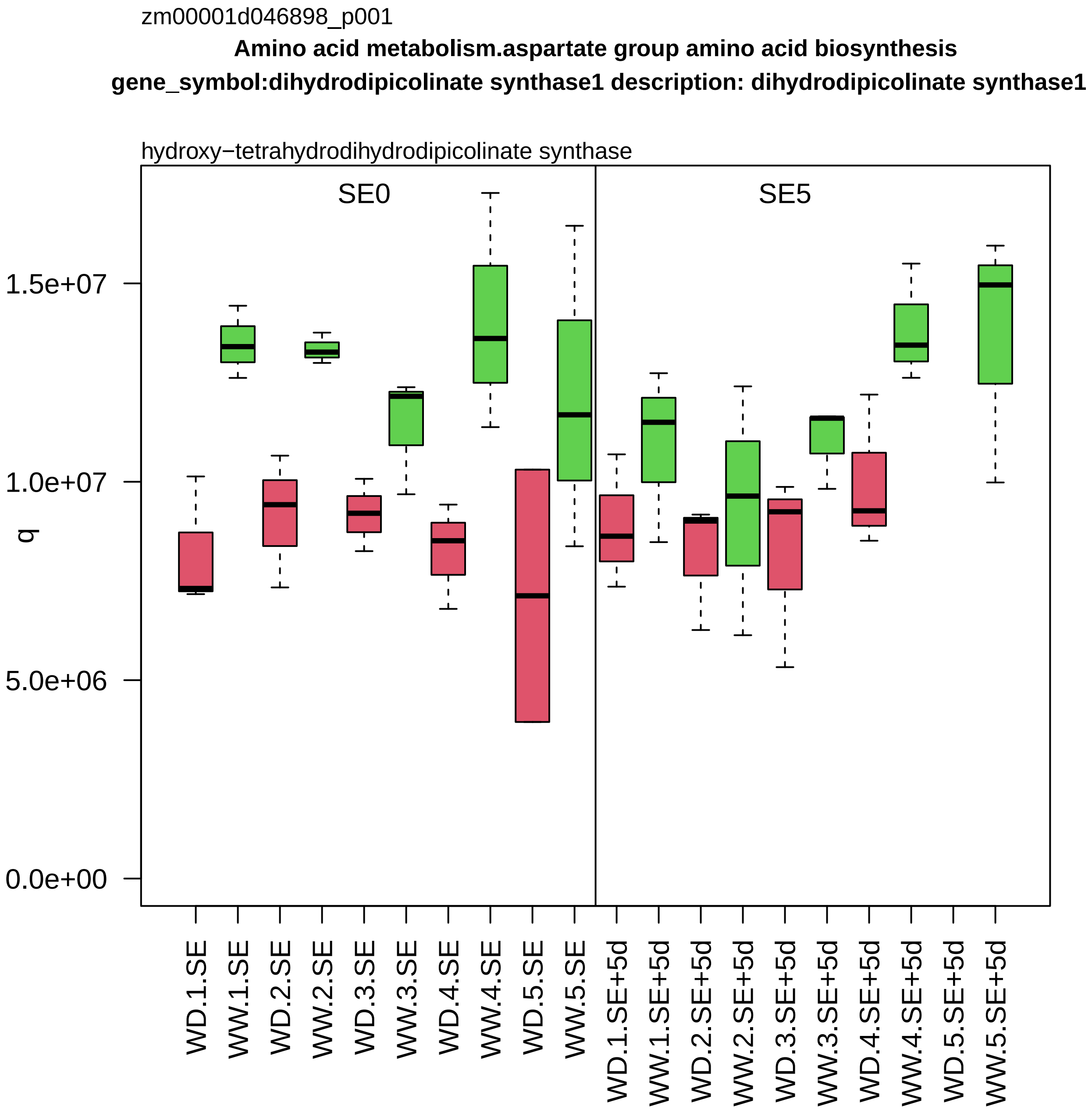
zm00001d046898_p001
Amino acid metabolism.aspartate group amino acid biosynthesis
gene_symbol:dihydrodipicolinate synthase1 description: dihydrodipicolinate synthase1
hydroxy−tetrahydrodihydrodipicolinate synthase
SE0
SE5
q
1.5e+07
1.0e+07
5.0e+06
0.0e+00
WD.1.SE
WW.1.SE
WD.2.SE
WW.2.SE
WD.3.SE
WW.3.SE
WD.4.SE
WW.4.SE
WD.5.SE
WW.5.SE
WD.1.SE+5d
WW.1.SE+5d
WD.2.SE+5d
WW.2.SE+5d
WD.3.SE+5d
WW.3.SE+5d
WD.4.SE+5d
WW.4.SE+5d
WD.5.SE+5d
WW.5.SE+5d

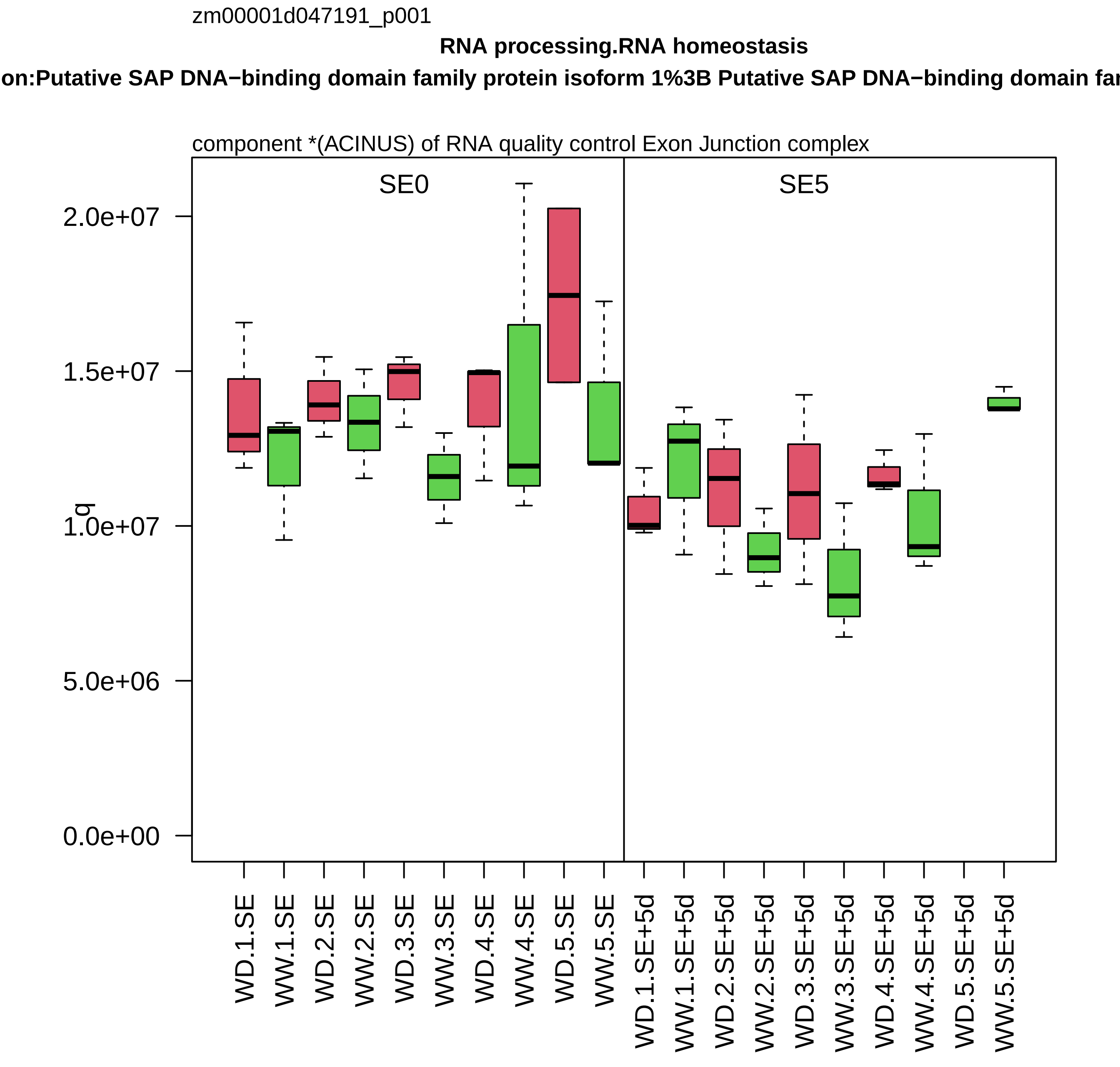

zm00001d047191_p001
RNA processing.RNA homeostasis
on:Putative SAP DNA−binding domain family protein isoform 1%3B Putative SAP DNA−binding domain fa
component *(ACINUS) of RNA quality control Exon Junction complex
SE0
SE5
q
2.0e+07
1.5e+07
1.0e+07
5.0e+06
0.0e+00
WD.1.SE
WW.1.SE
WD.2.SE
WW.2.SE
WD.3.SE
WW.3.SE
WD.4.SE
WW.4.SE
WD.5.SE
WW.5.SE
WD.1.SE+5d
WW.1.SE+5d
WD.2.SE+5d
WW.2.SE+5d
WD.3.SE+5d
WW.3.SE+5d
WD.4.SE+5d
WW.4.SE+5d
WD.5.SE+5d
WW.5.SE+5d

zm00001d047253_p001

**Carbohydrate metabolism.sucrose metabolism**

**gene_symbol:sucrose synthase1 description: sucrose synthase1**

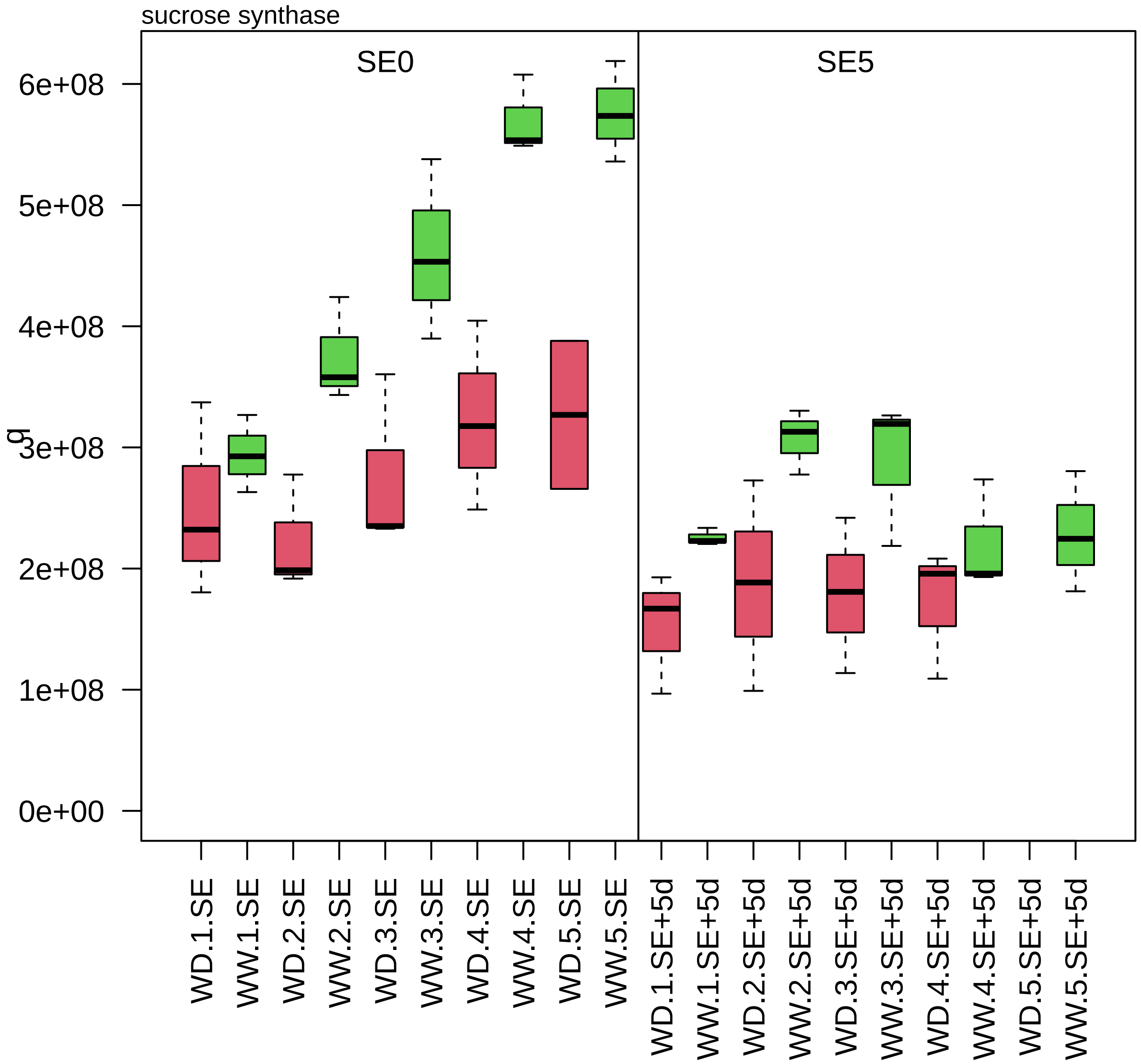

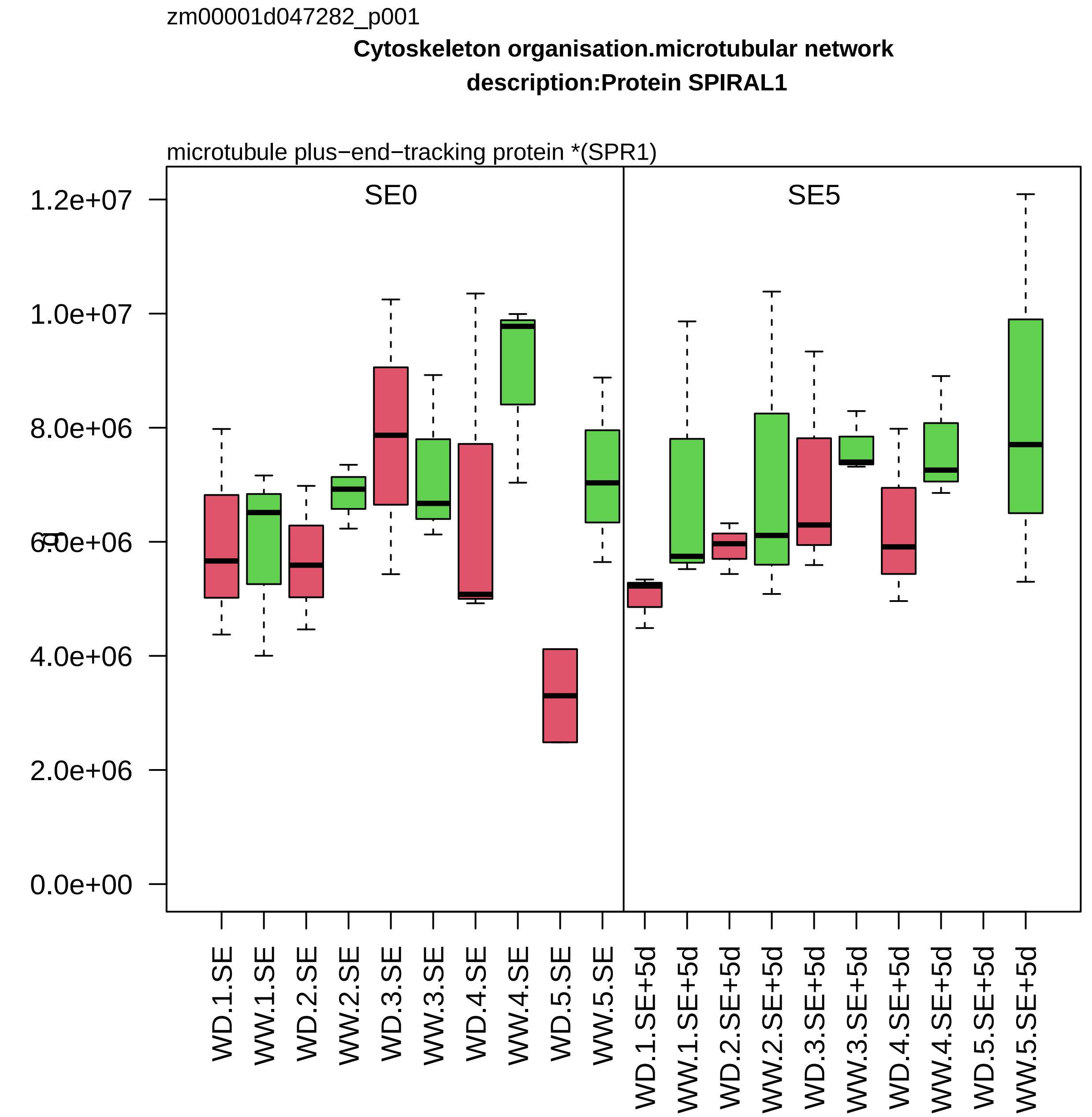
zm00001d047282_p001
Cytoskeleton organisation.microtubular network
description:Protein SPIRAL1
microtubule plus−end−tracking protein *(SPR1)
SE0
SE5
1.2e+07
1.0e+07
8.0e+06
6.0e+06
4.0e+06
2.0e+06
0.0e+00
WD.1.SE
WW.1.SE
WD.2.SE
WW.2.SE
WD.3.SE
WW.3.SE
WD.4.SE
WW.4.SE
WD.5.SE
WW.5.SE
WD.1.SE+5d
WW.1.SE+5d
WD.2.SE+5d
WW.2.SE+5d
WD.3.SE+5d
WW.3.SE+5d
WD.4.SE+5d
WW.4.SE+5d
WD.5.SE+5d
WW.5.SE+5d

zm00001d047498_p001

**RNA processing.pre−RNA splicing**

**description:Serine/arginine−rich−splicing factor SR34**

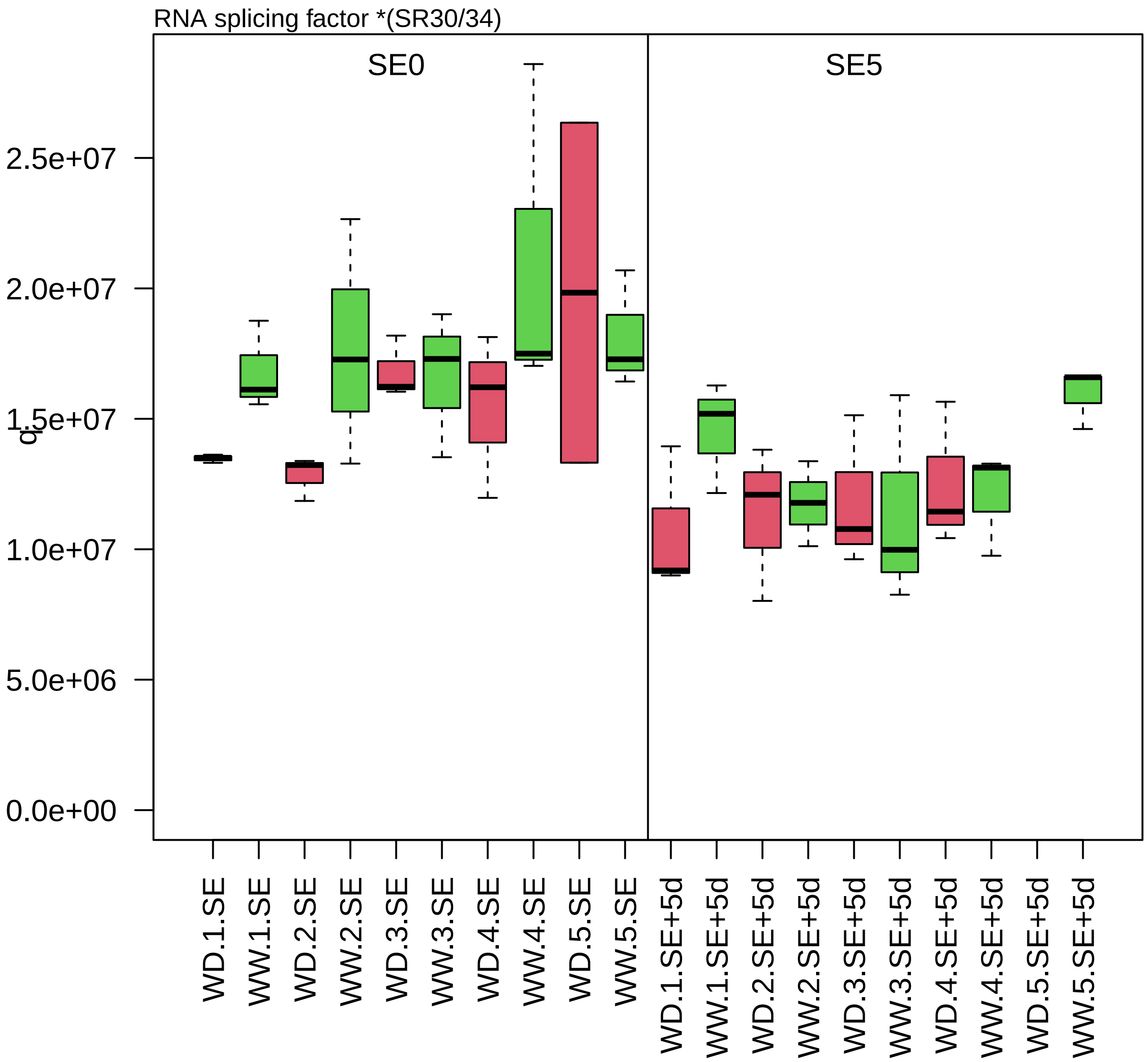

zm00001d047663_p001

**Lipid metabolism.fatty acid metabolism**

**description:Acetate/butyrate−−CoA ligase AAE7 peroxisomal**

peroxisomal acetyl−CoA synthetase *(ACN/BZU)

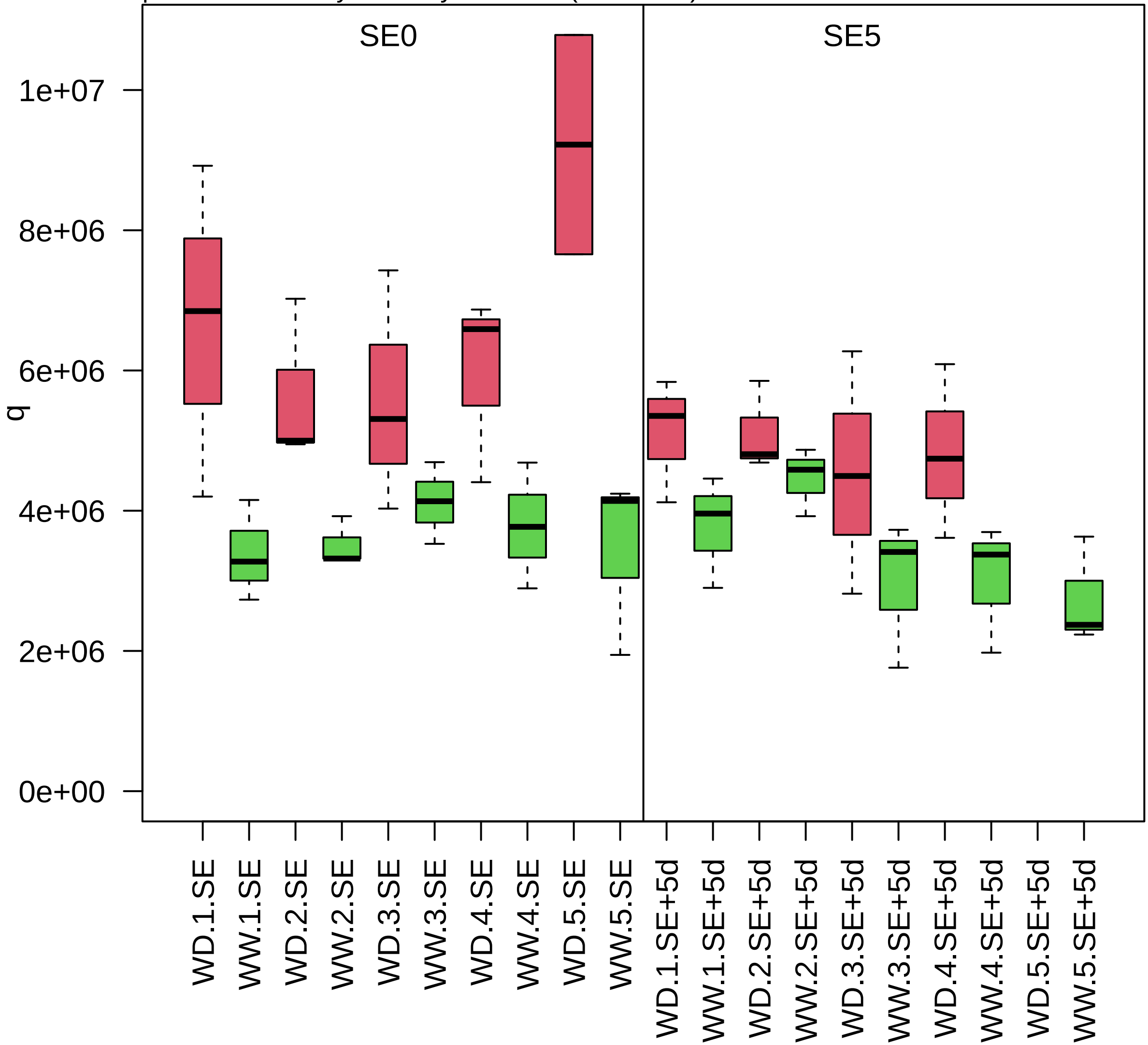

zm00001d047707_p001

**Protein biosynthesis.ribosome biogenesis**

**description:40S ribosomal protein S7−2**

component *(eS7) of small ribosomal−subunit (SSU) proteome

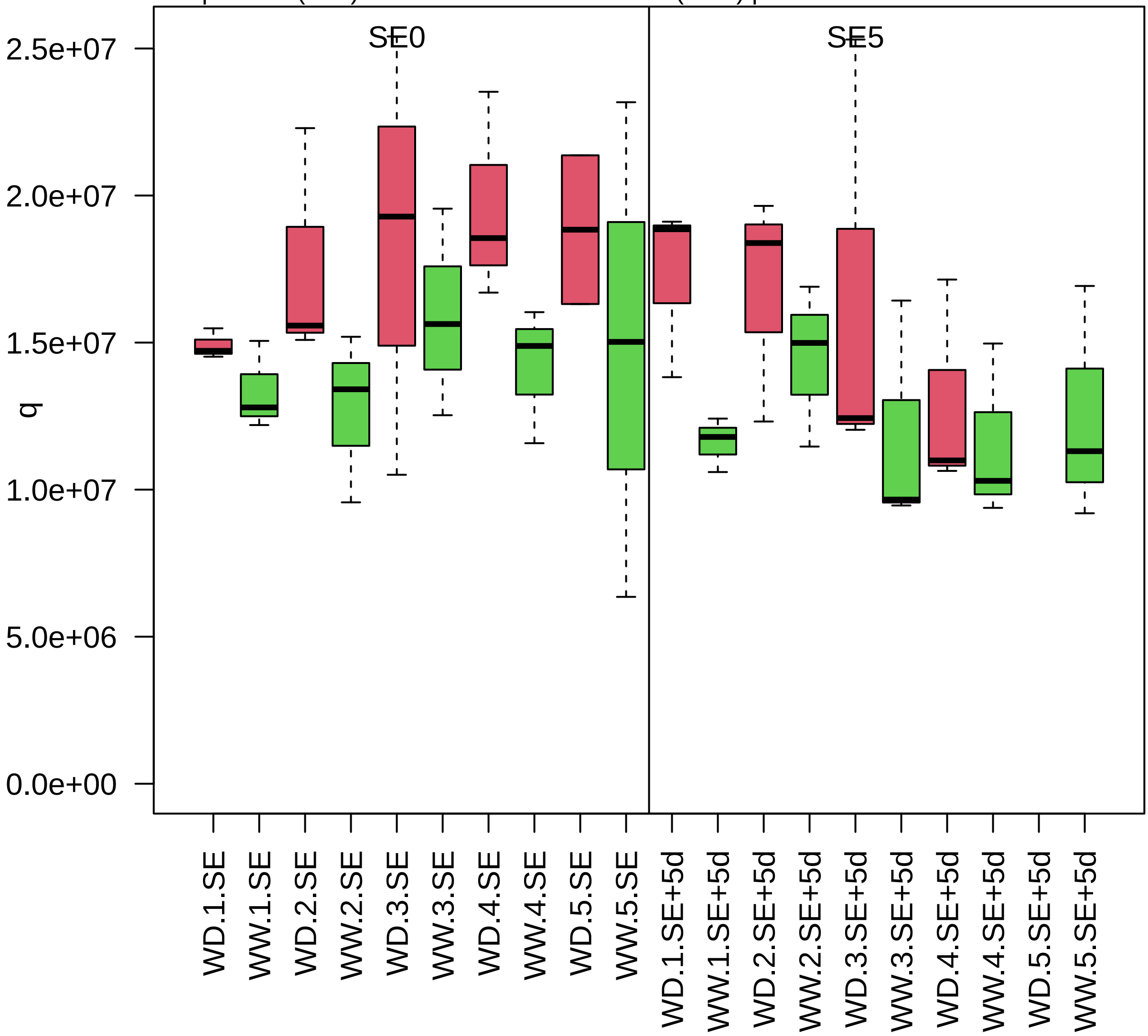

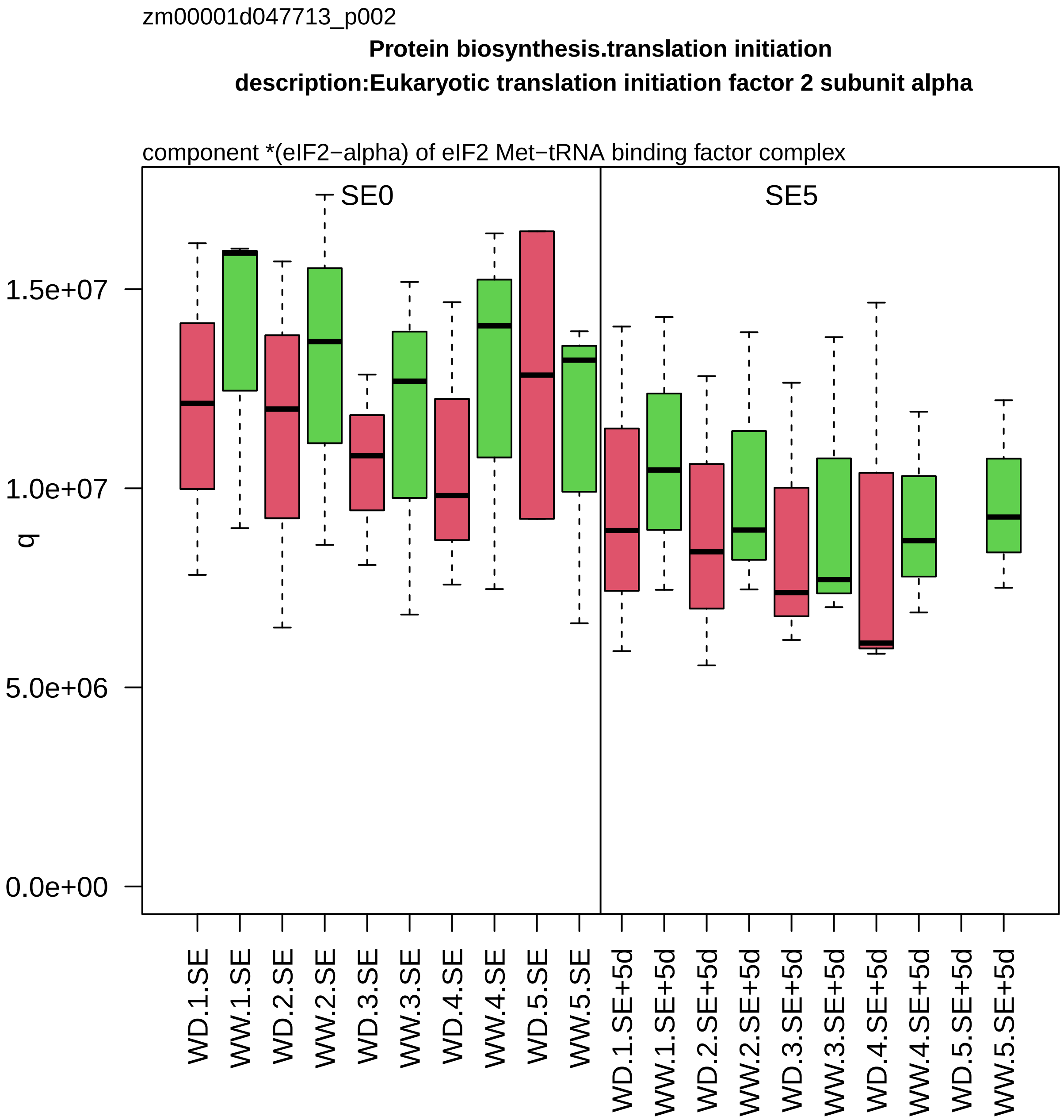

zm00001d047713_p002
Protein biosynthesis.translation initiation
description:Eukaryotic translation initiation factor 2 subunit alpha
component *(eIF2−alpha) of eIF2 Met−tRNA binding factor complex
SE0
SE5
q
1.5e+07
1.0e+07
5.0e+06
0.0e+00
WD.1.SE
WW.1.SE
WD.2.SE
WW.2.SE
WD.3.SE
WW.3.SE
WD.4.SE
WW.4.SE
WD.5.SE
WW.5.SE
WD.1.SE+5d
WW.1.SE+5d
WD.2.SE+5d
WW.2.SE+5d
WD.3.SE+5d
WW.3.SE+5d
WD.4.SE+5d
WW.4.SE+5d
WD.5.SE+5d
WW.5.SE+5d

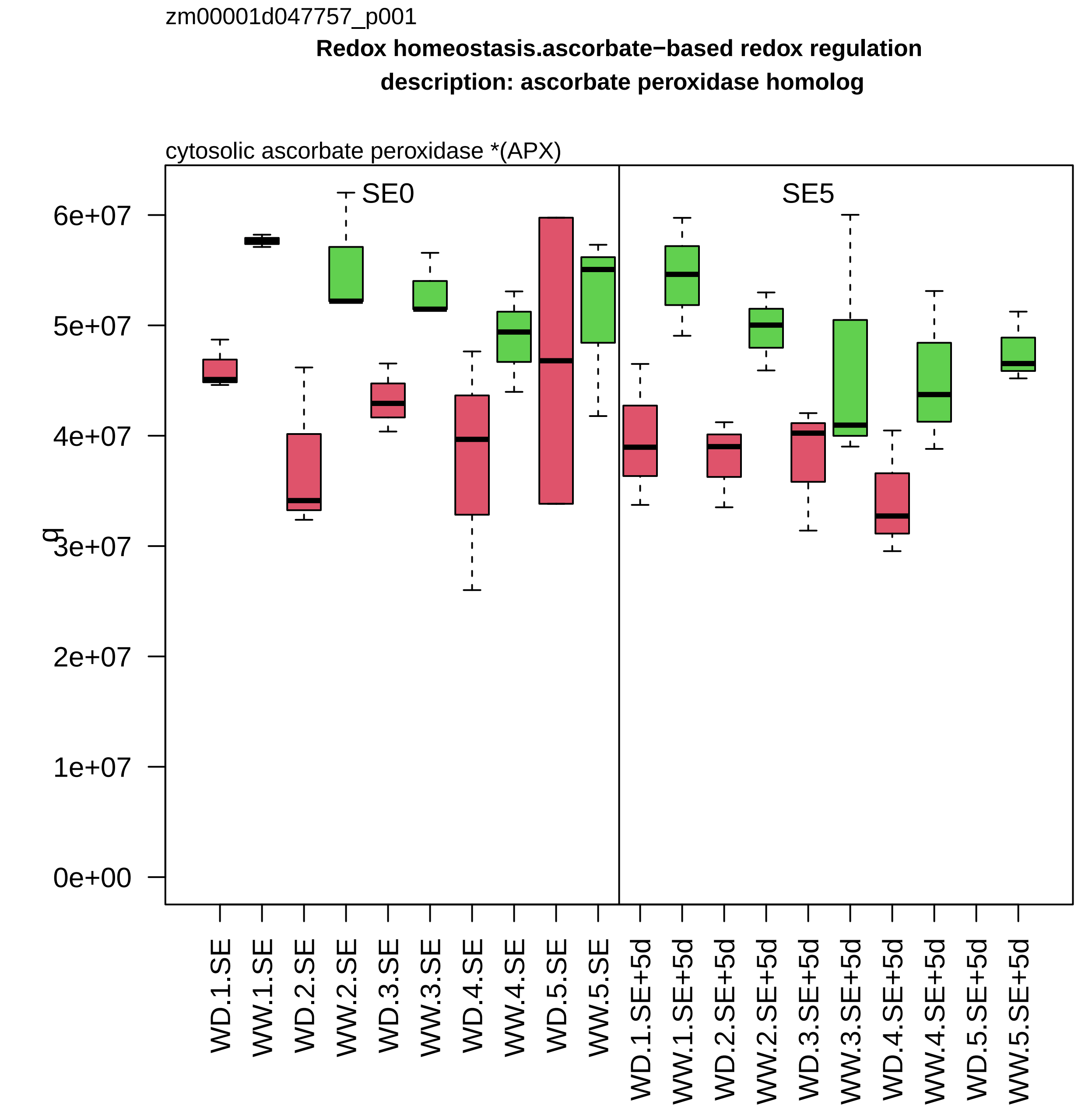
zm00001d047757_p001
Redox homeostasis.ascorbate−based redox regulation
description: ascorbate peroxidase homolog
cytosolic ascorbate peroxidase *(APX)
SE0
SE5
g
0e+00
1e+07
2e+07
3e+07
4e+07
5e+07
6e+07
WD.1.SE
WW.1.SE
WD.2.SE
WW.2.SE
WD.3.SE
WW.3.SE
WD.4.SE
WW.4.SE
WD.5.SE
WW.5.SE
WD.1.SE+5d
WW.1.SE+5d
WD.2.SE+5d
WW.2.SE+5d
WD.3.SE+5d
WW.3.SE+5d
WD.4.SE+5d
WW.4.SE+5d
WD.5.SE+5d
WW.5.SE+5d

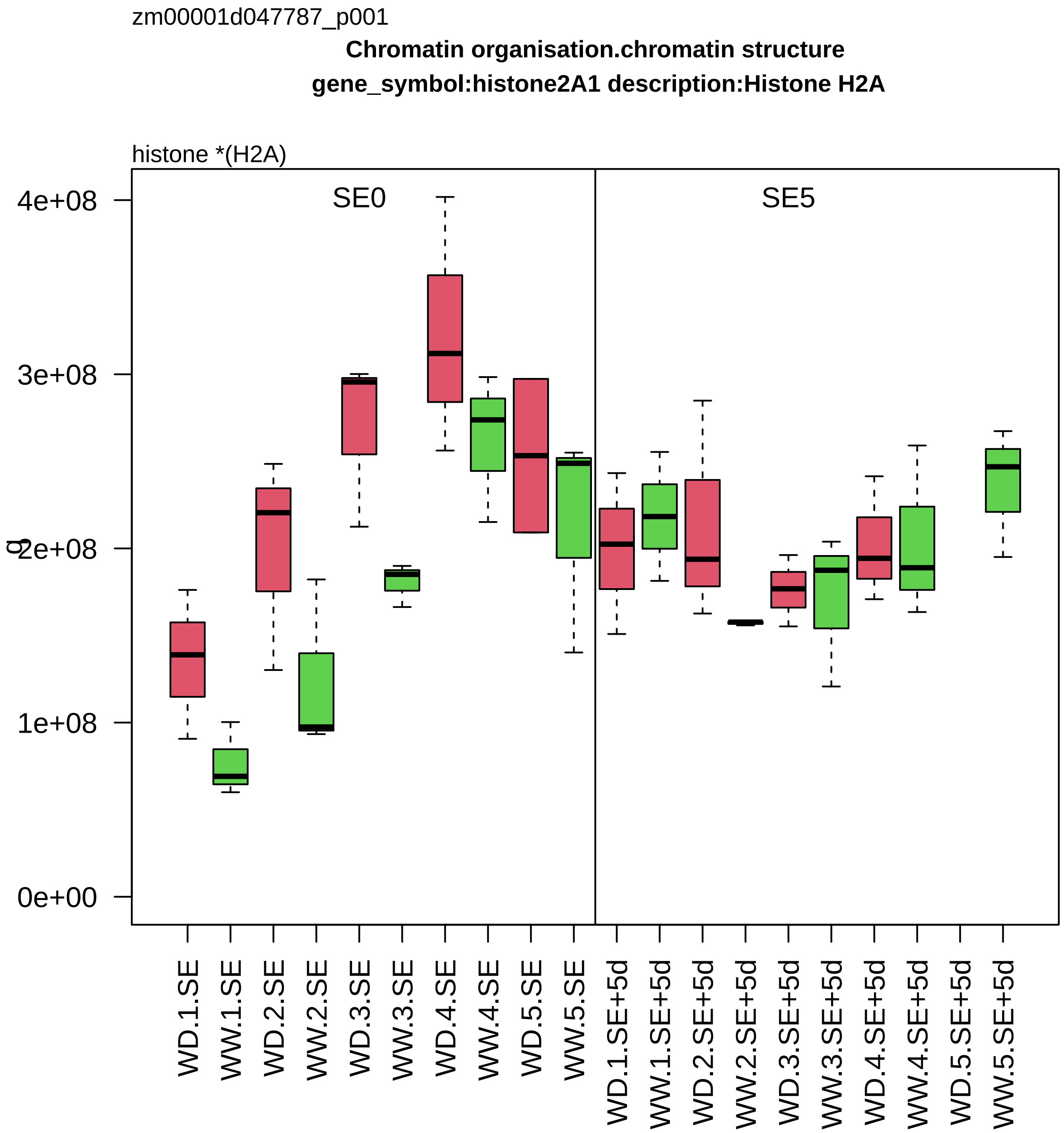

zm00001d047787_p001
Chromatin organisation.chromatin structure
gene_symbol:histone2A1 description:Histone H2A
histone *(H2A)
SE0
SE5
4e+08
3e+08
2e+08
1e+08
0e+00
WD.1.SE
WW.1.SE
WD.2.SE
WW.2.SE
WD.3.SE
WW.3.SE
WD.4.SE
WW.4.SE
WD.5.SE
WW.5.SE
WD.1.SE+5d
WW.1.SE+5d
WD.2.SE+5d
WW.2.SE+5d
WD.3.SE+5d
WW.3.SE+5d
WD.4.SE+5d
WW.4.SE+5d
WD.5.SE+5d
WW.5.SE+5d

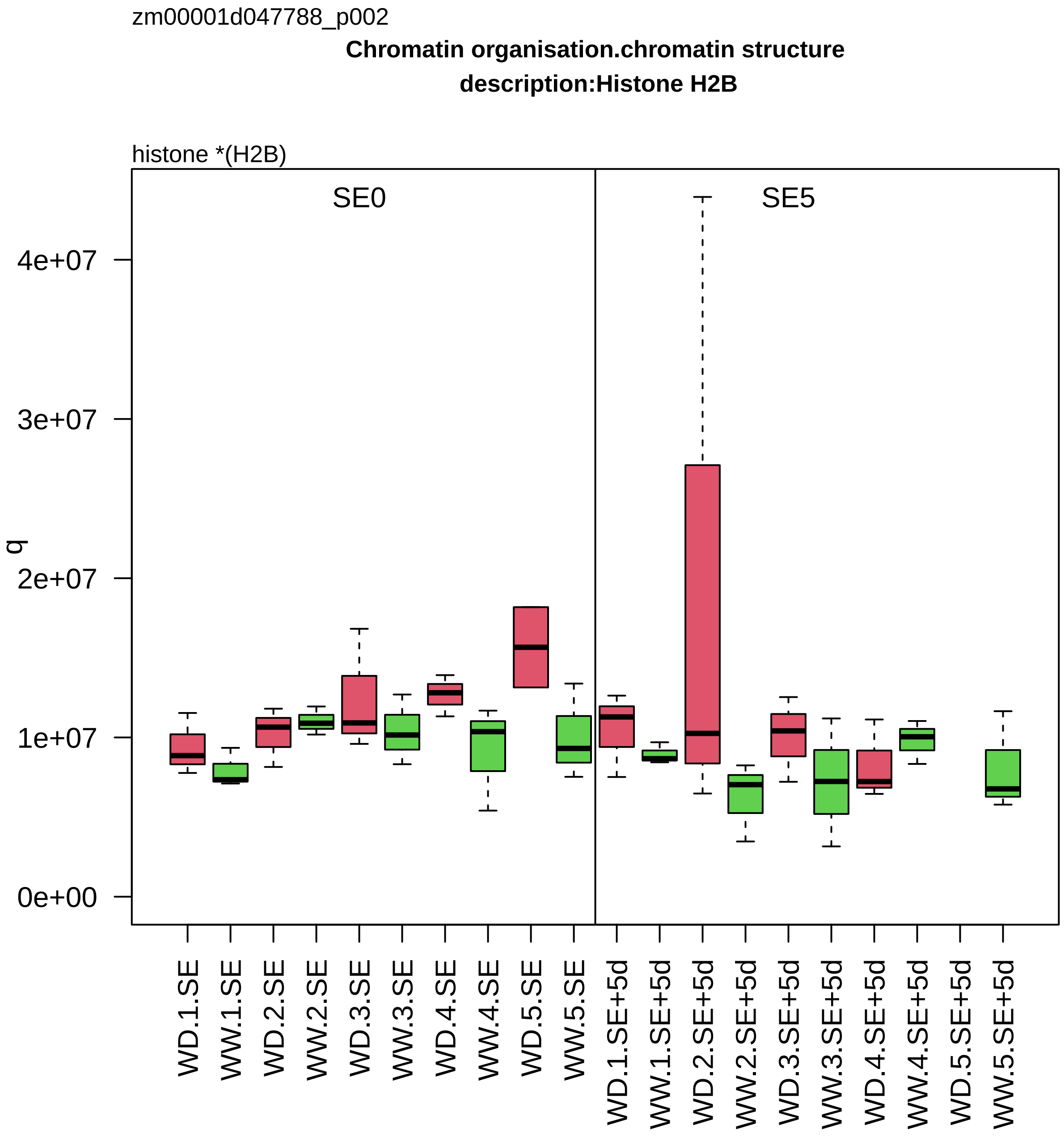
zm00001d047788_p002
Chromatin organisation.chromatin structure
description:Histone H2B
histone *(H2B)
SE0
SE5
q
0e+00
1e+07
2e+07
3e+07
4e+07
WD.1.SE
WW.1.SE
WD.2.SE
WW.2.SE
WD.3.SE
WW.3.SE
WD.4.SE
WW.4.SE
WD.5.SE
WW.5.SE
WD.1.SE+5d
WW.1.SE+5d
WD.2.SE+5d
WW.2.SE+5d
WD.3.SE+5d
WW.3.SE+5d
WD.4.SE+5d
WW.4.SE+5d
WD.5.SE+5d
WW.5.SE+5d

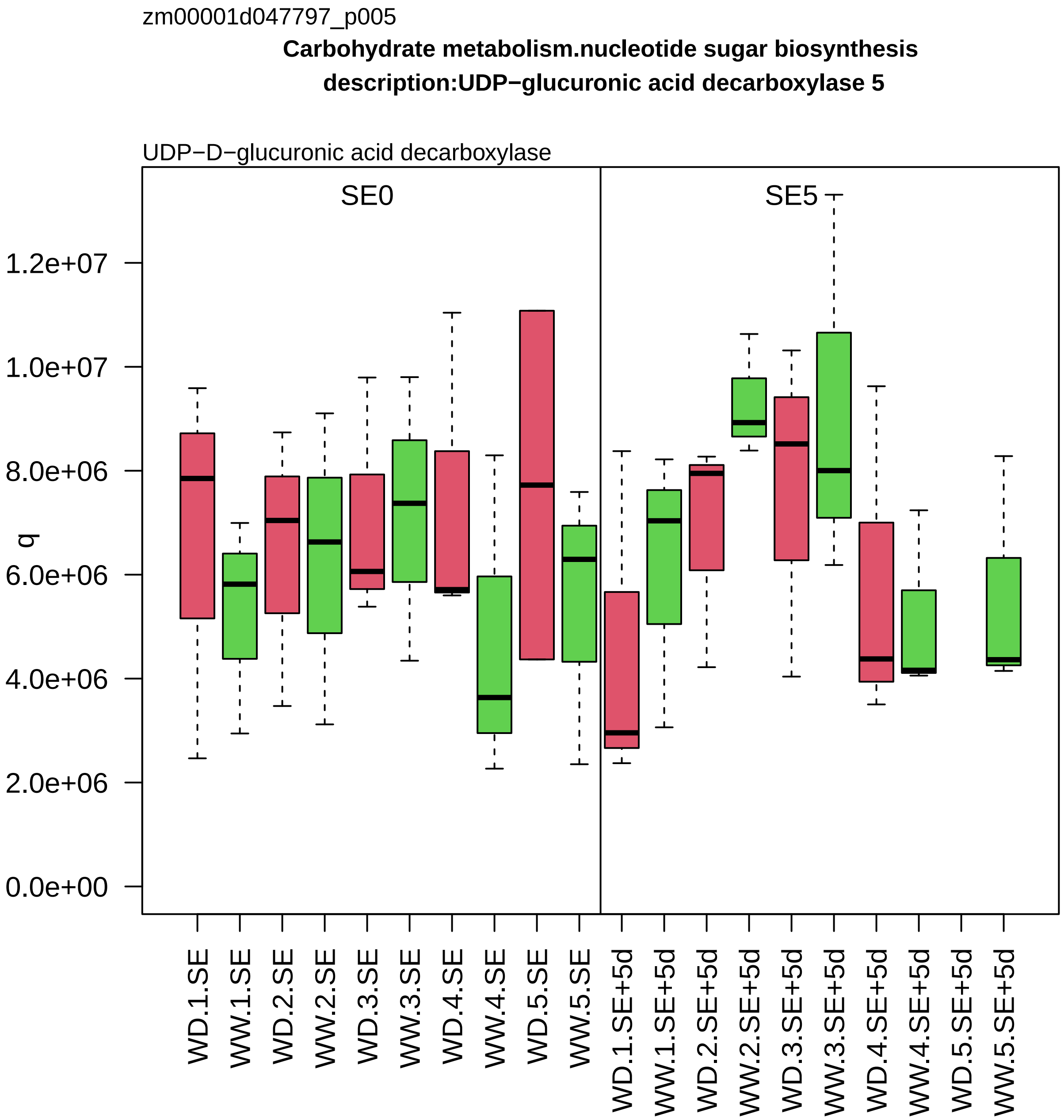

zm00001d047797_p005
Carbohydrate metabolism.nucleotide sugar biosynthesis
description:UDP−glucuronic acid decarboxylase 5
UDP−D−glucuronic acid decarboxylase
SE0
SE5
q
0.0e+00
2.0e+06
4.0e+06
6.0e+06
8.0e+06
1.0e+07
1.2e+07
WD.1.SE
WW.1.SE
WD.2.SE
WW.2.SE
WD.3.SE
WW.3.SE
WD.4.SE
WW.4.SE
WD.5.SE
WW.5.SE
WD.1.SE+5d
WW.1.SE+5d
WD.2.SE+5d
WW.2.SE+5d
WD.3.SE+5d
WW.3.SE+5d
WD.4.SE+5d
WW.4.SE+5d
WD.5.SE+5d
WW.5.SE+5d

zm00001d047893_p003

**Carbohydrate metabolism.gluconeogenesis**

**ene_symbol:phosphoenolpyruvate carboxykinase homolog2 description: phosphoenolpyruvate carboxykir**

phosphoenolpyruvate carboxykinase

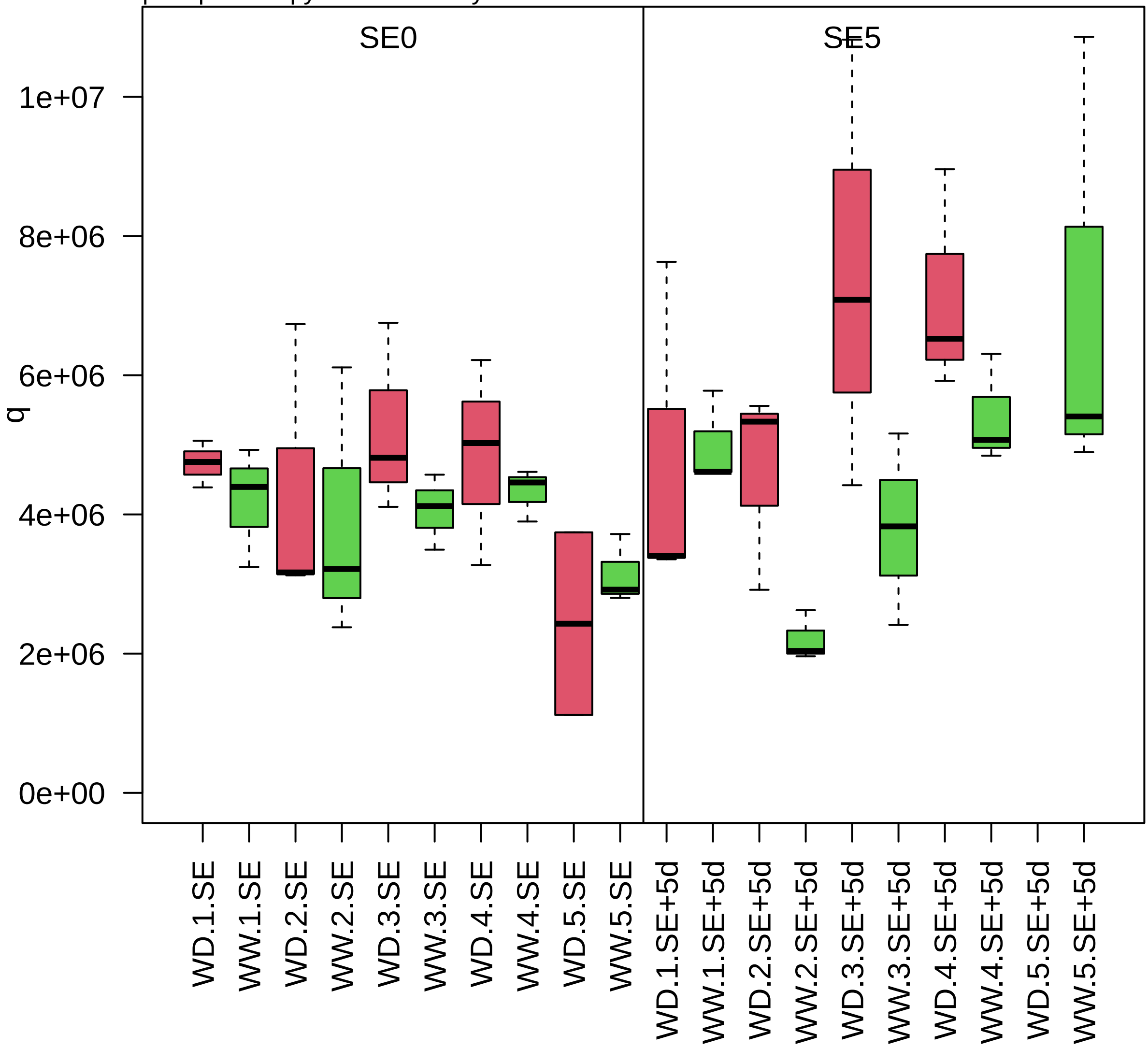

zm00001d047910_p001

**Solute transport.primary active transport**
**description:V−type proton ATPase subunit a3**

subunit a of V−type ATPase membrane V0 subcomplex

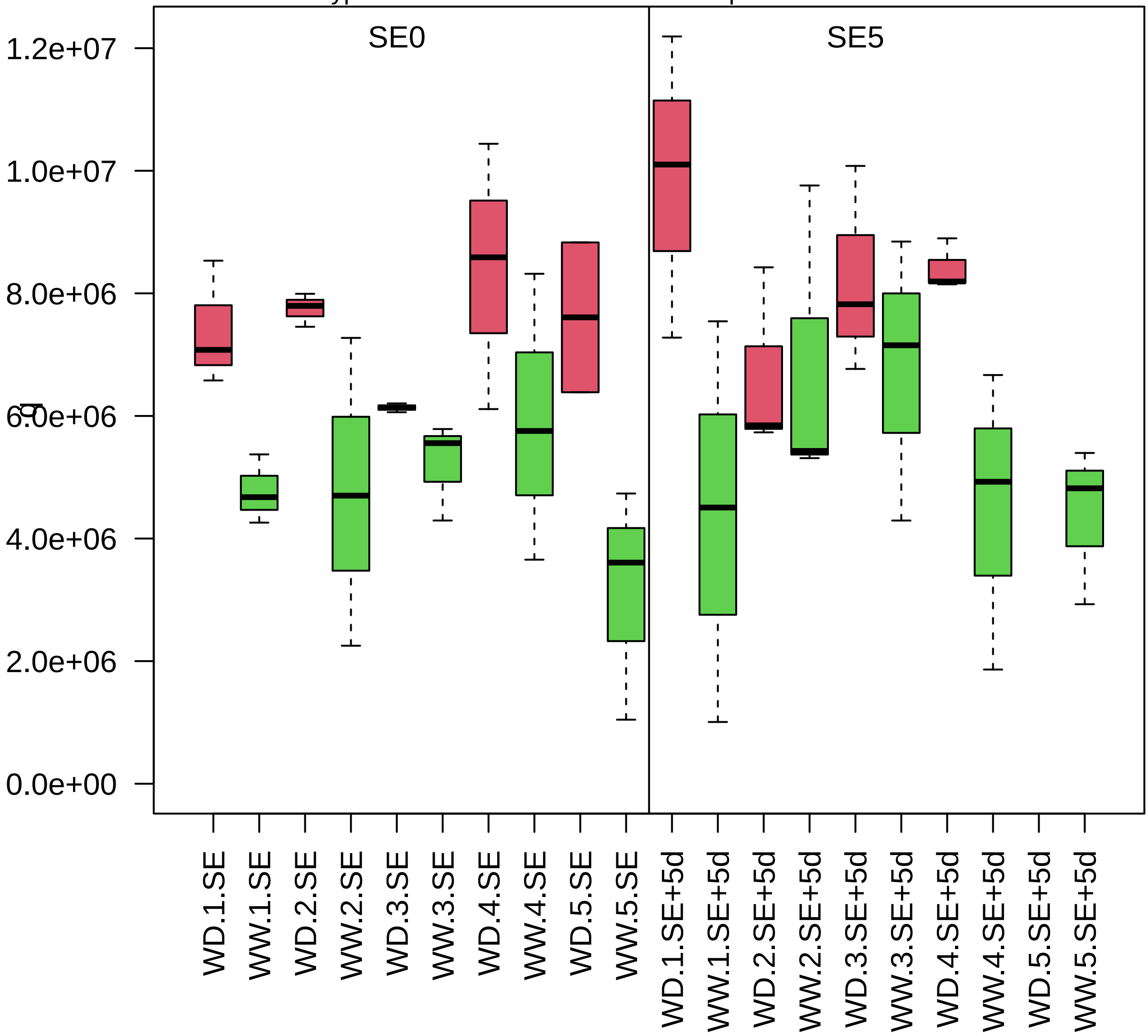

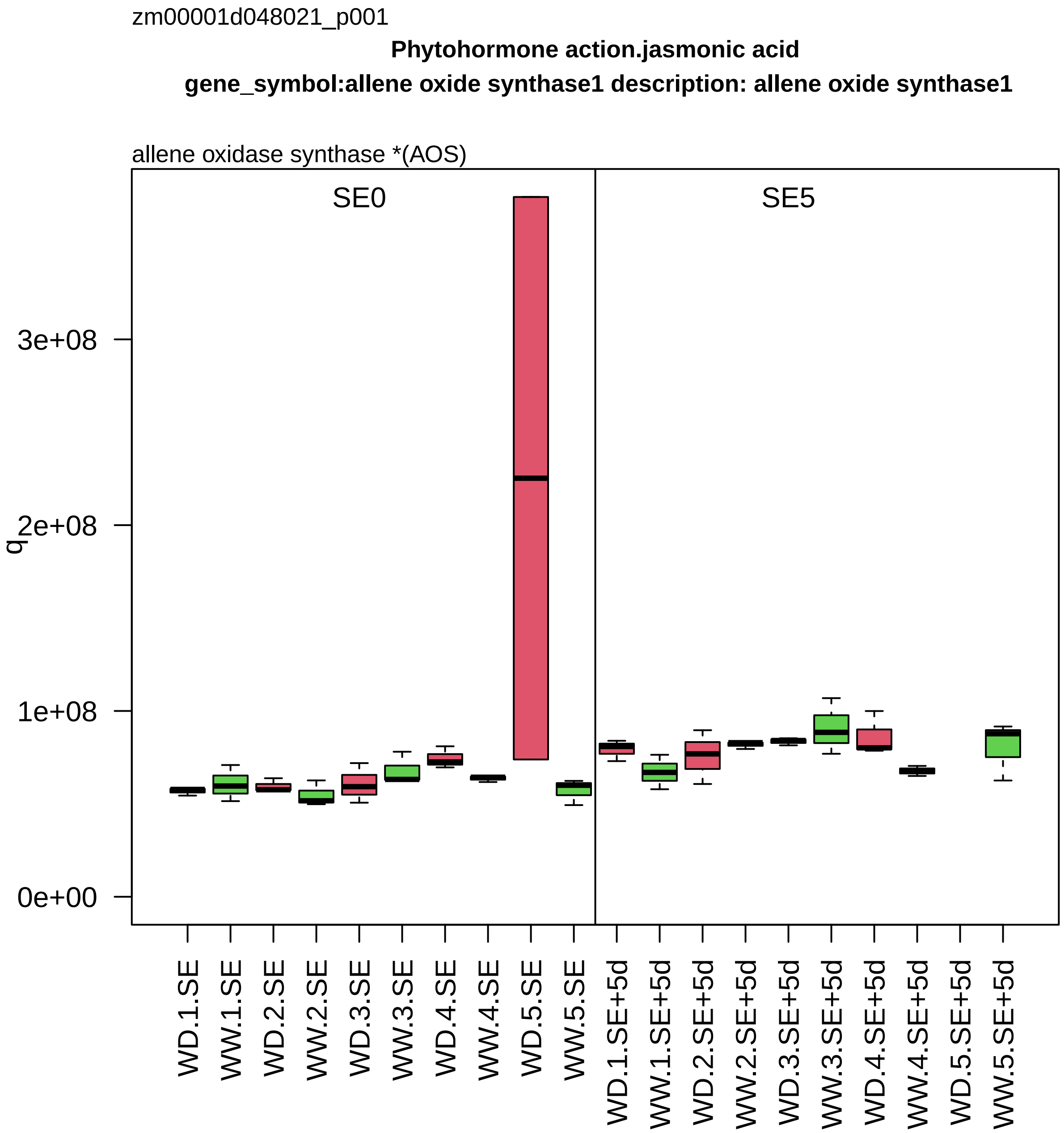
zm00001d048021_p001
Phytohormone action.jasmonic acid
gene_symbol:allene oxide synthase1 description: allene oxide synthase1
allene oxidase synthase *(AOS)
SE0
SE5
q
3e+08
2e+08
1e+08
0e+00
WD.1.SE
WW.1.SE
WD.2.SE
WW.2.SE
WD.3.SE
WW.3.SE
WD.4.SE
WW.4.SE
WD.5.SE
WW.5.SE
WD.1.SE+5d
WW.1.SE+5d
WD.2.SE+5d
WW.2.SE+5d
WD.3.SE+5d
WW.3.SE+5d
WD.4.SE+5d
WW.4.SE+5d
WD.5.SE+5d
WW.5.SE+5d

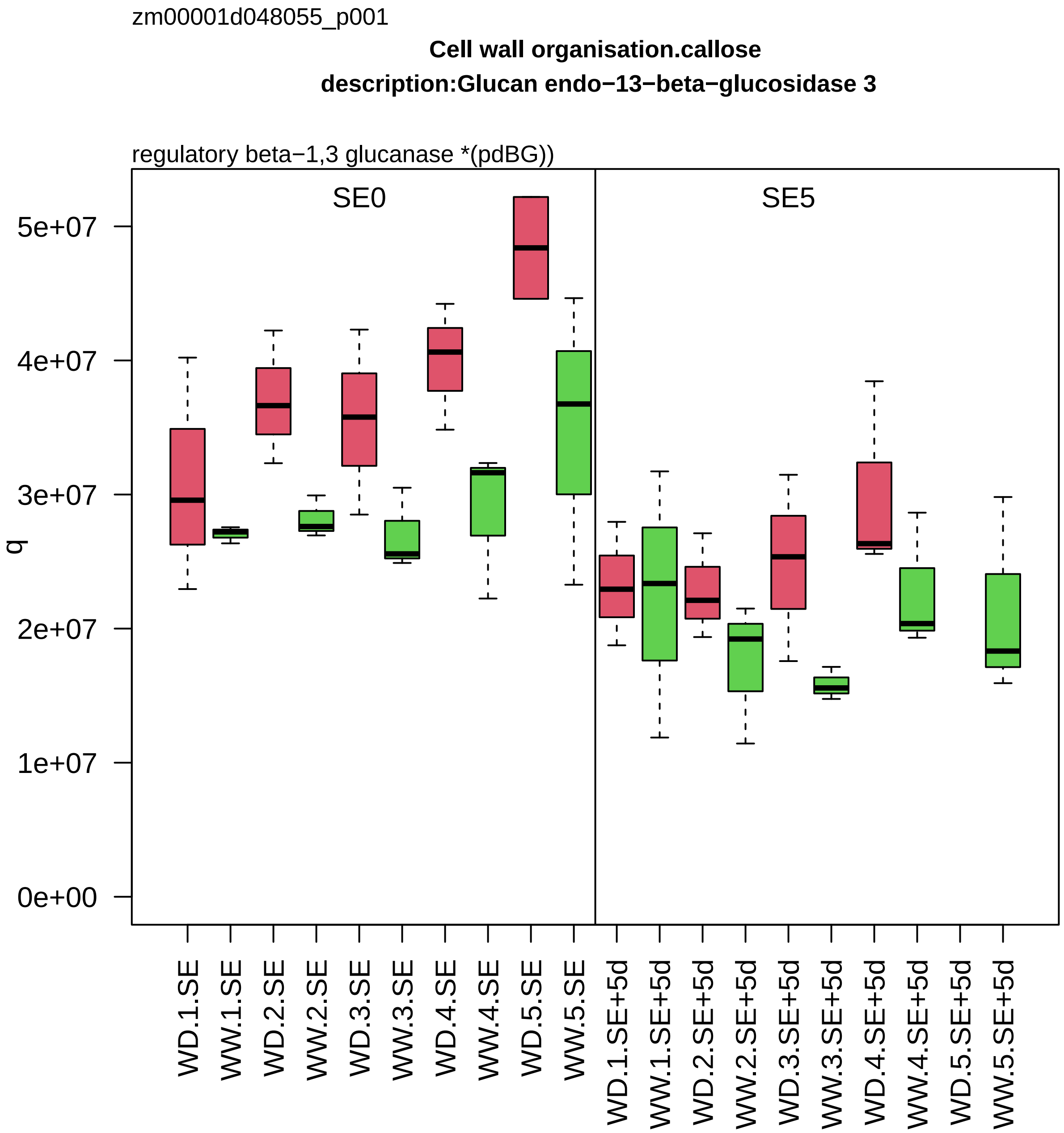
zm00001d048055_p001
Cell wall organisation.callose
description:Glucan endo−13−beta−glucosidase 3
regulatory beta−1,3 glucanase *(pdBG))
SE0
SE5
q
5e+07
4e+07
3e+07
2e+07
1e+07
0e+00
WD.1.SE
WW.1.SE
WD.2.SE
WW.2.SE
WD.3.SE
WW.3.SE
WD.4.SE
WW.4.SE
WD.5.SE
WW.5.SE
WD.1.SE+5d
WW.1.SE+5d
WD.2.SE+5d
WW.2.SE+5d
WD.3.SE+5d
WW.3.SE+5d
WD.4.SE+5d
WW.4.SE+5d
WD.5.SE+5d
WW.5.SE+5d

zm00001d048091_p002

**Cellular respiration.oxidative phosphorylation**
**description:ATP synthase subunit gamma mitochondrial**

subunit gamma of ATP synthase peripheral MF1 subcomplex

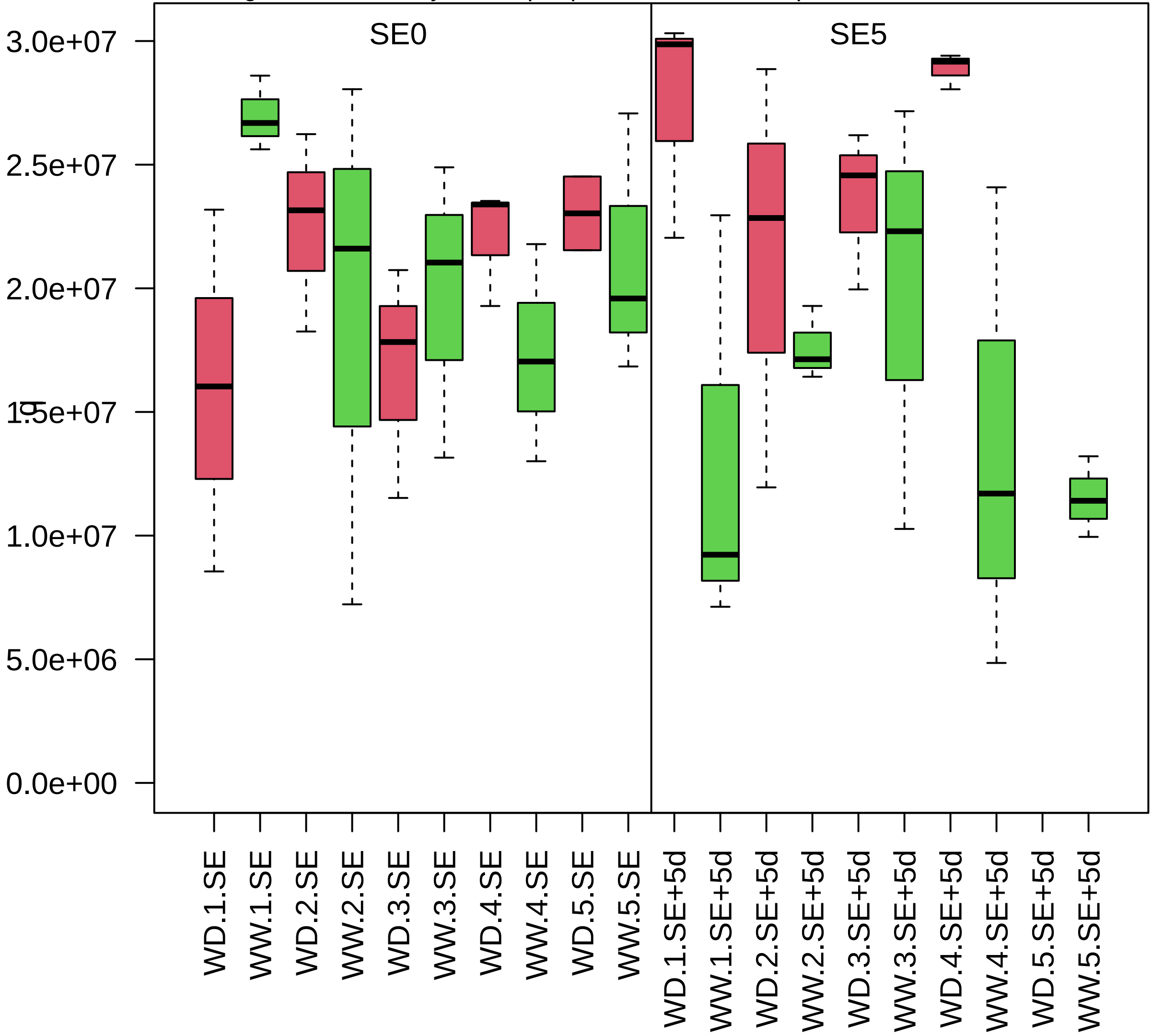

zm00001d048102_p002

**Cellular respiration.oxidative phosphorylation**

**description:Peptidase beta subunit**

subunit beta *(QCR1) of cytochrome c reductase subcomplex

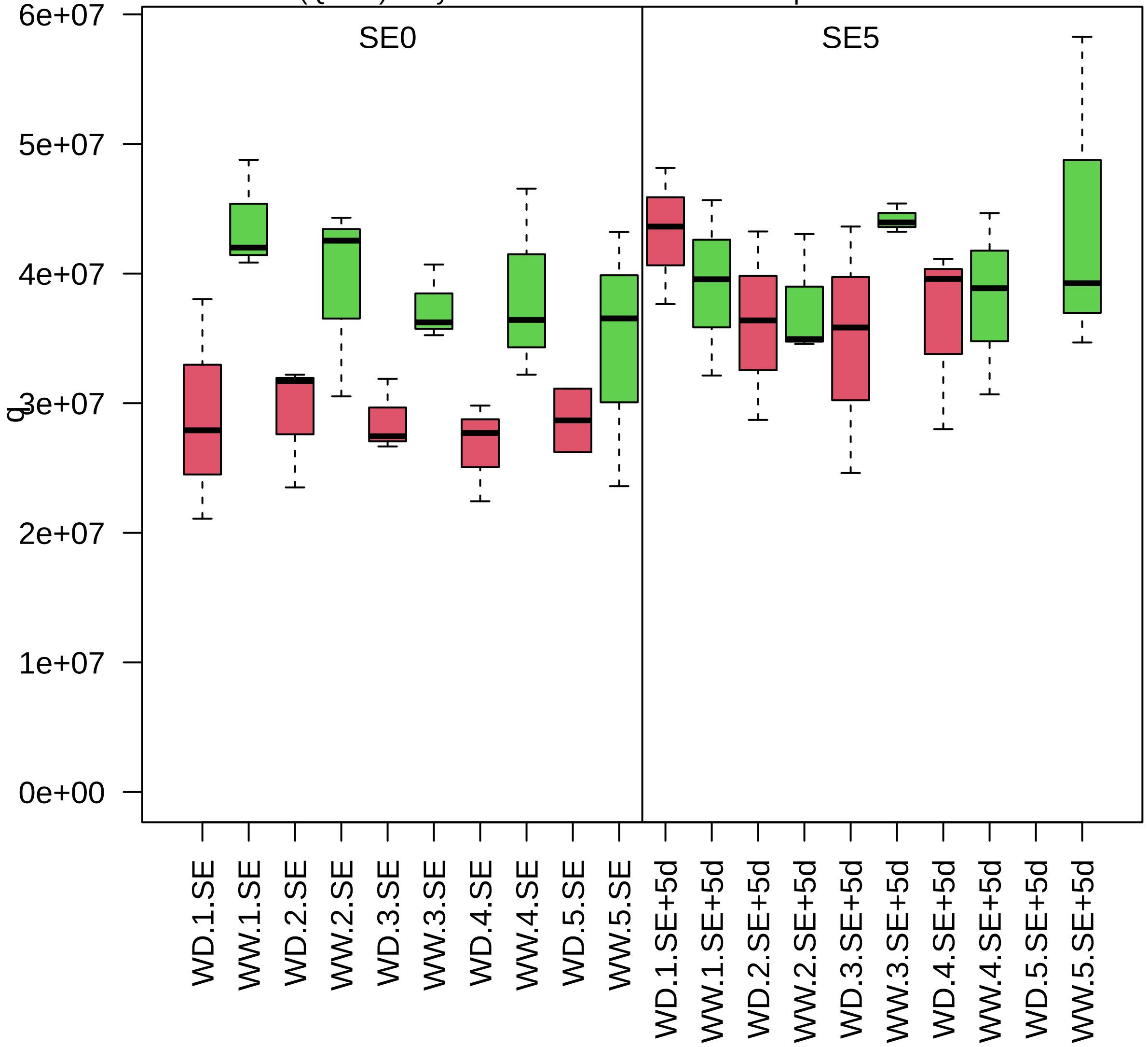

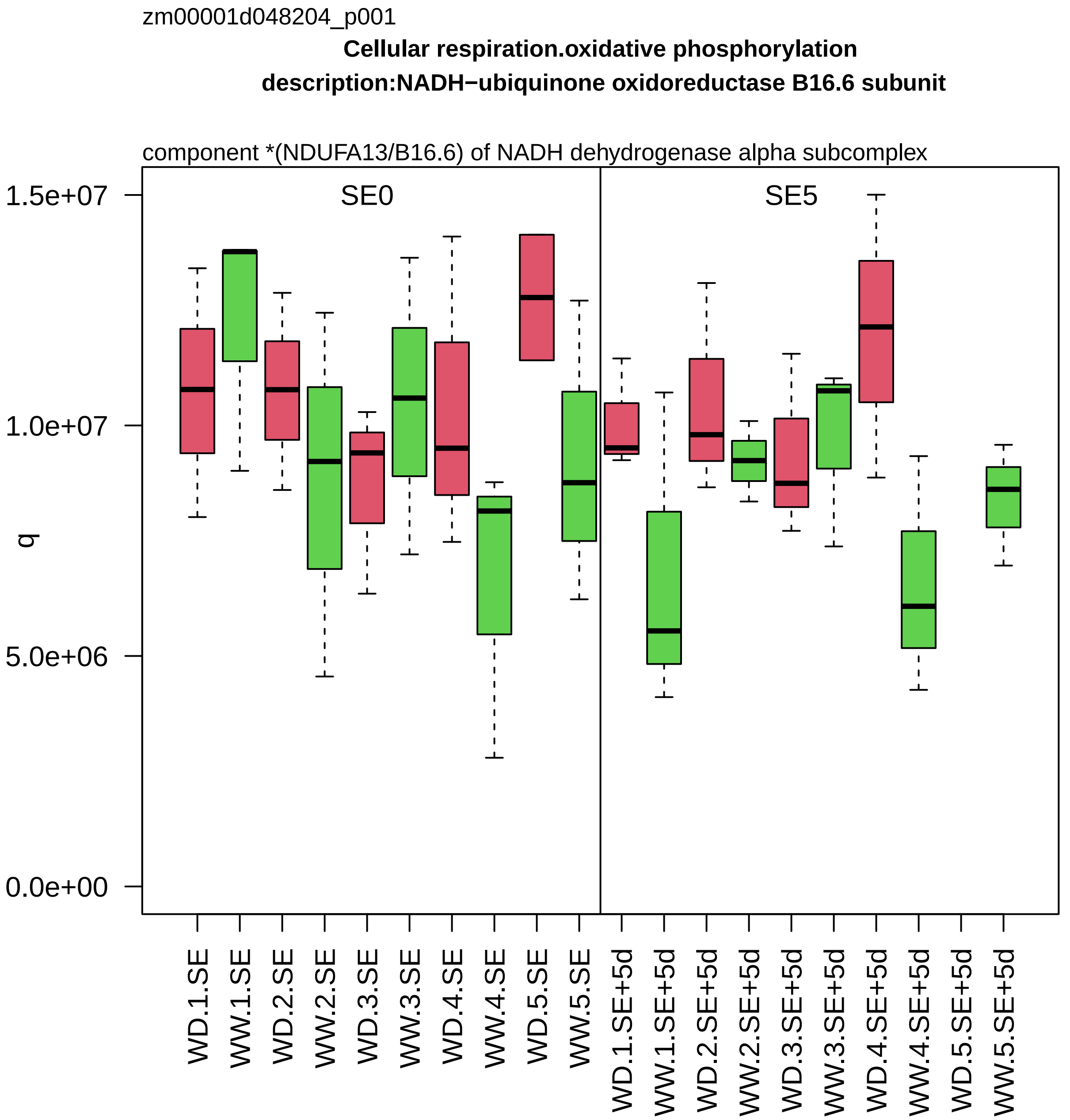

zm00001d048204_p001
Cellular respiration.oxidative phosphorylation
description:NADH−ubiquinone oxidoreductase B16.6 subunit
component *(NDUFA13/B16.6) of NADH dehydrogenase alpha subcomplex
SE0
SE5
q
1.5e+07
1.0e+07
5.0e+06
0.0e+00
WD.1.SE
WW.1.SE
WD.2.SE
WW.2.SE
WD.3.SE
WW.3.SE
WD.4.SE
WW.4.SE
WD.5.SE
WW.5.SE
WD.1.SE+5d
WW.1.SE+5d
WD.2.SE+5d
WW.2.SE+5d
WD.3.SE+5d
WW.3.SE+5d
WD.4.SE+5d
WW.4.SE+5d
WD.5.SE+5d
WW.5.SE+5d

zm00001d048291_p001

**RNA processing.organelle machinery**

**description:Multiple organellar RNA editing factor 8 chloroplastic/mitochondrial**

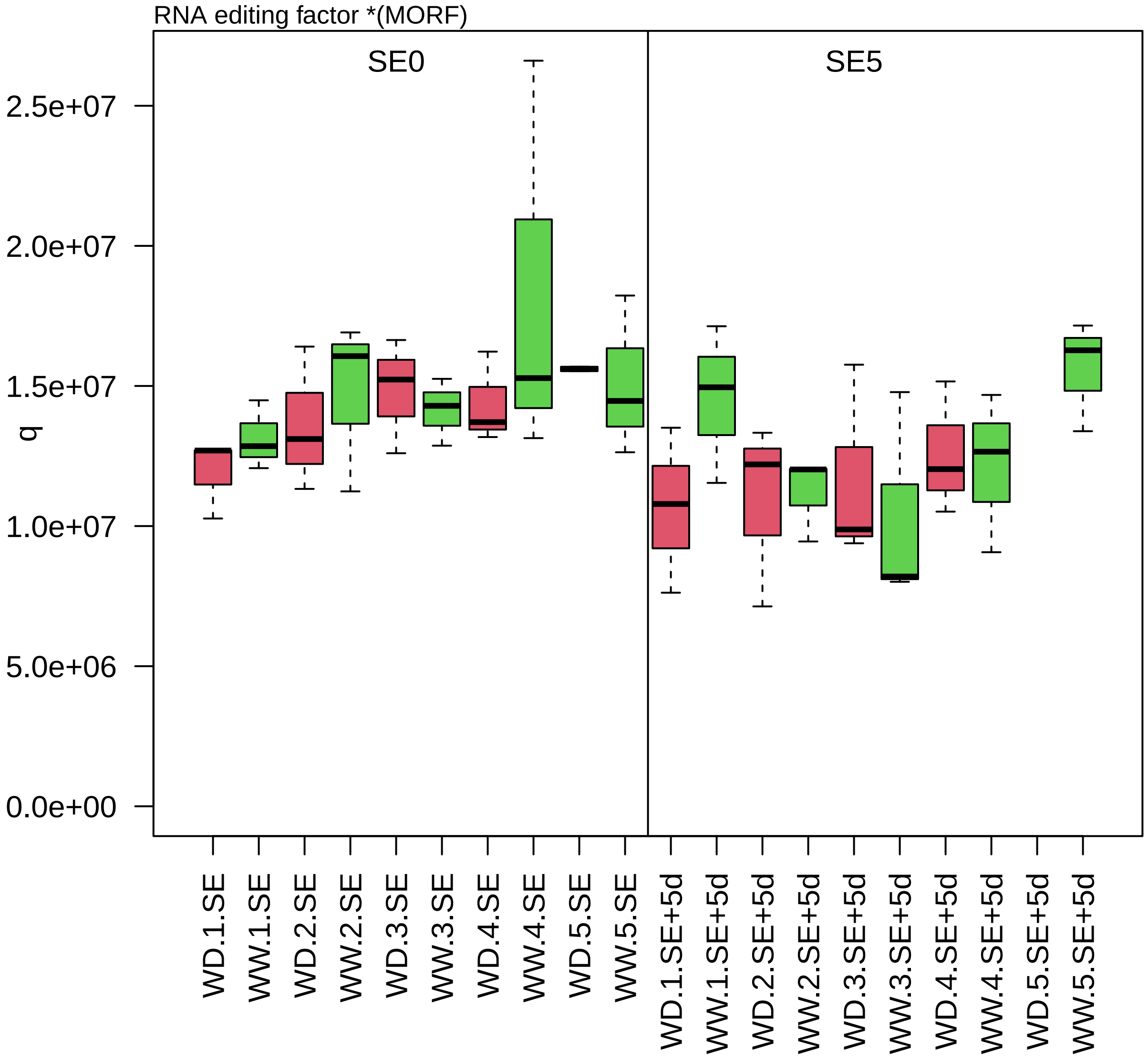

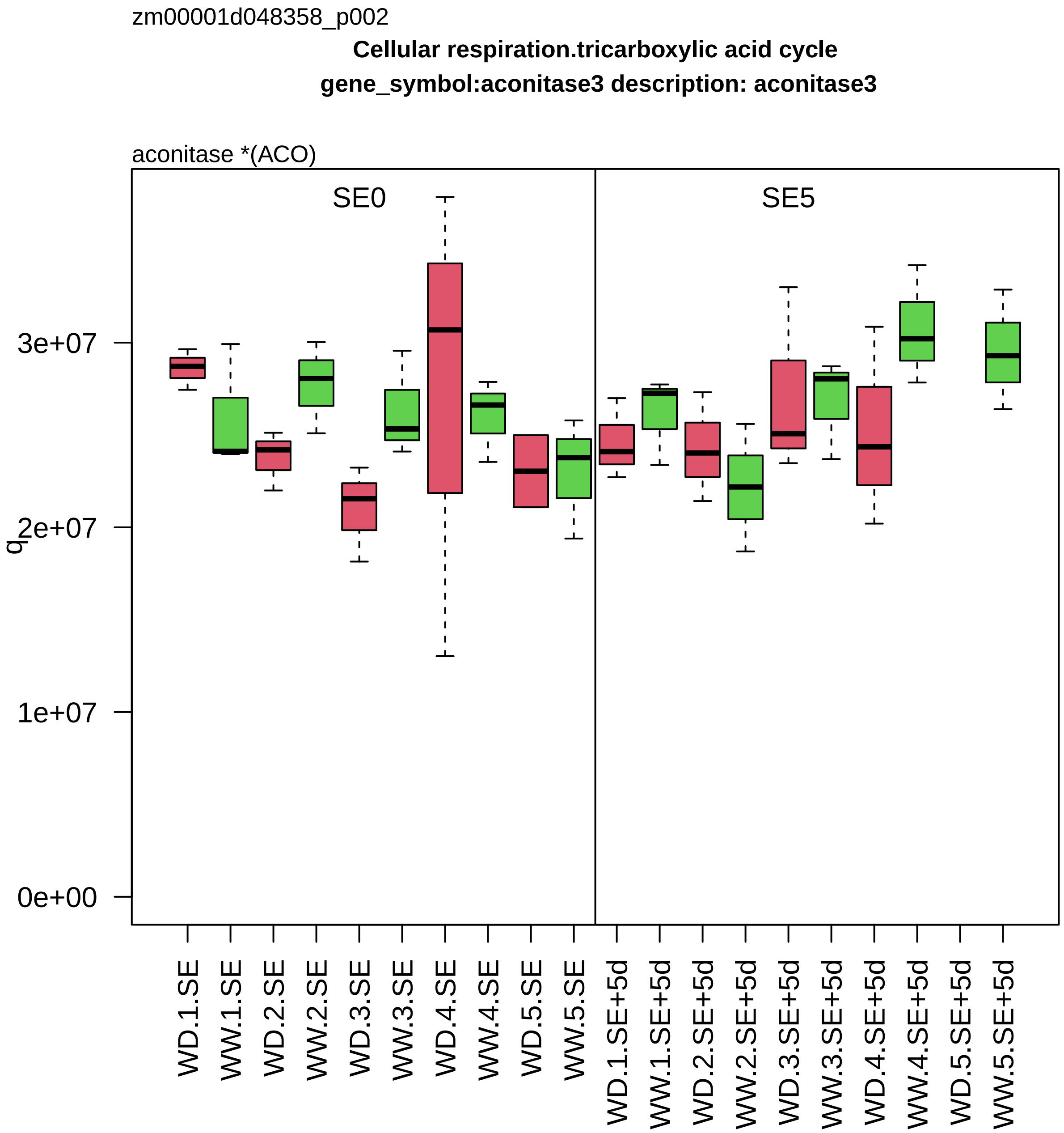
zm00001d048358_p002
Cellular respiration.tricarboxylic acid cycle
gene_symbol:aconitase3 description: aconitase3
aconitase *(ACO)
SE0
SE5
q
3e+07
2e+07
1e+07
0e+00
WD.1.SE
WW.1.SE
WD.2.SE
WW.2.SE
WD.3.SE
WW.3.SE
WD.4.SE
WW.4.SE
WD.5.SE
WW.5.SE
WD.1.SE+5d
WW.1.SE+5d
WD.2.SE+5d
WW.2.SE+5d
WD.3.SE+5d
WW.3.SE+5d
WD.4.SE+5d
WW.4.SE+5d
WD.5.SE+5d
WW.5.SE+5d

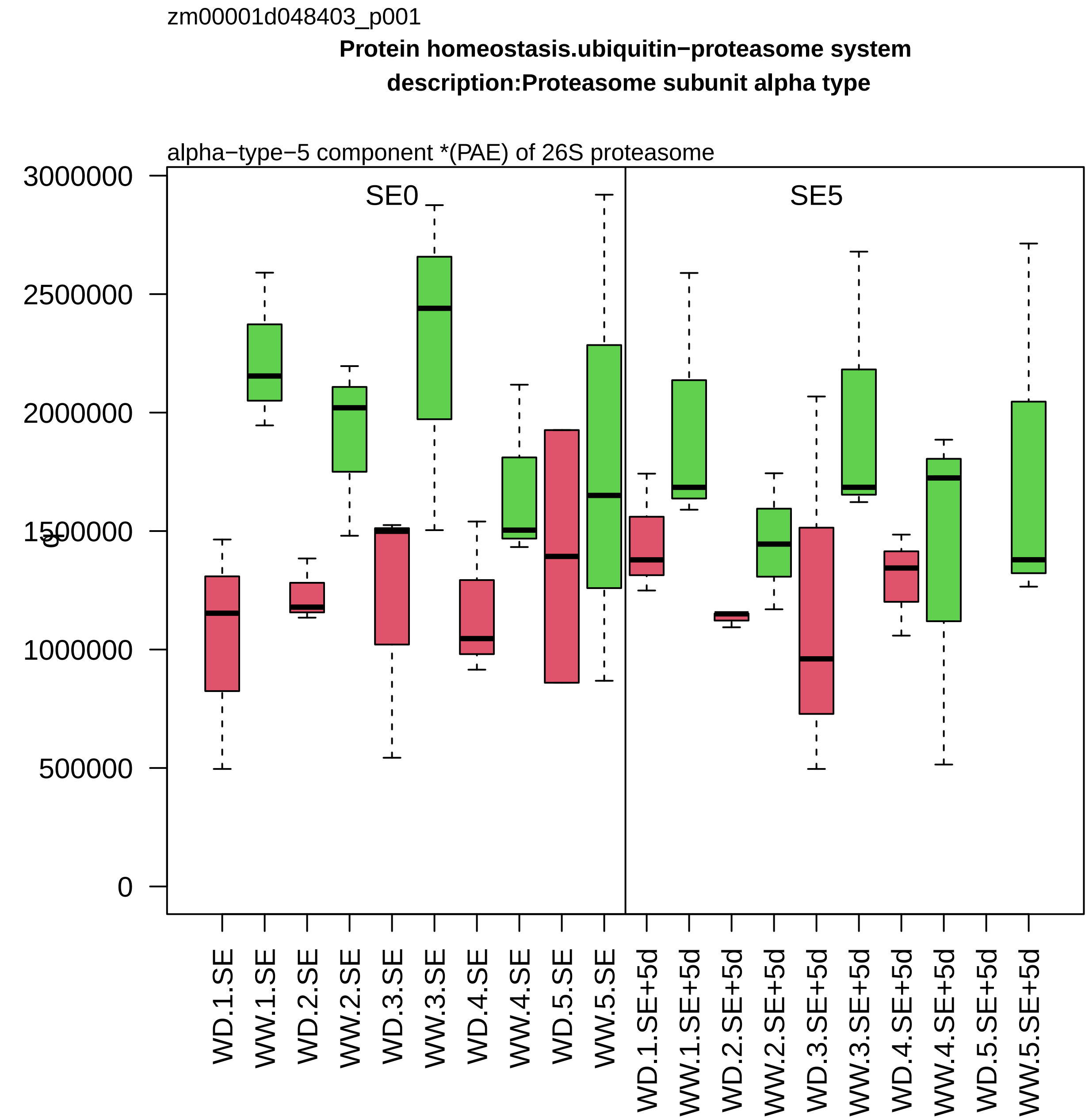

zm00001d048403_p001
Protein homeostasis.ubiquitin−proteasome system
description:Proteasome subunit alpha type
alpha−type−5 component *(PAE) of 26S proteasome
SE0
SE5
3000000
2500000
2000000
1500000
1000000
500000
0
WD.1.SE
WW.1.SE
WD.2.SE
WW.2.SE
WD.3.SE
WW.3.SE
WD.4.SE
WW.4.SE
WD.5.SE
WW.5.SE
WD.1.SE+5d
WW.1.SE+5d
WD.2.SE+5d
WW.2.SE+5d
WD.3.SE+5d
WW.3.SE+5d
WD.4.SE+5d
WW.4.SE+5d
WD.5.SE+5d
WW.5.SE+5d

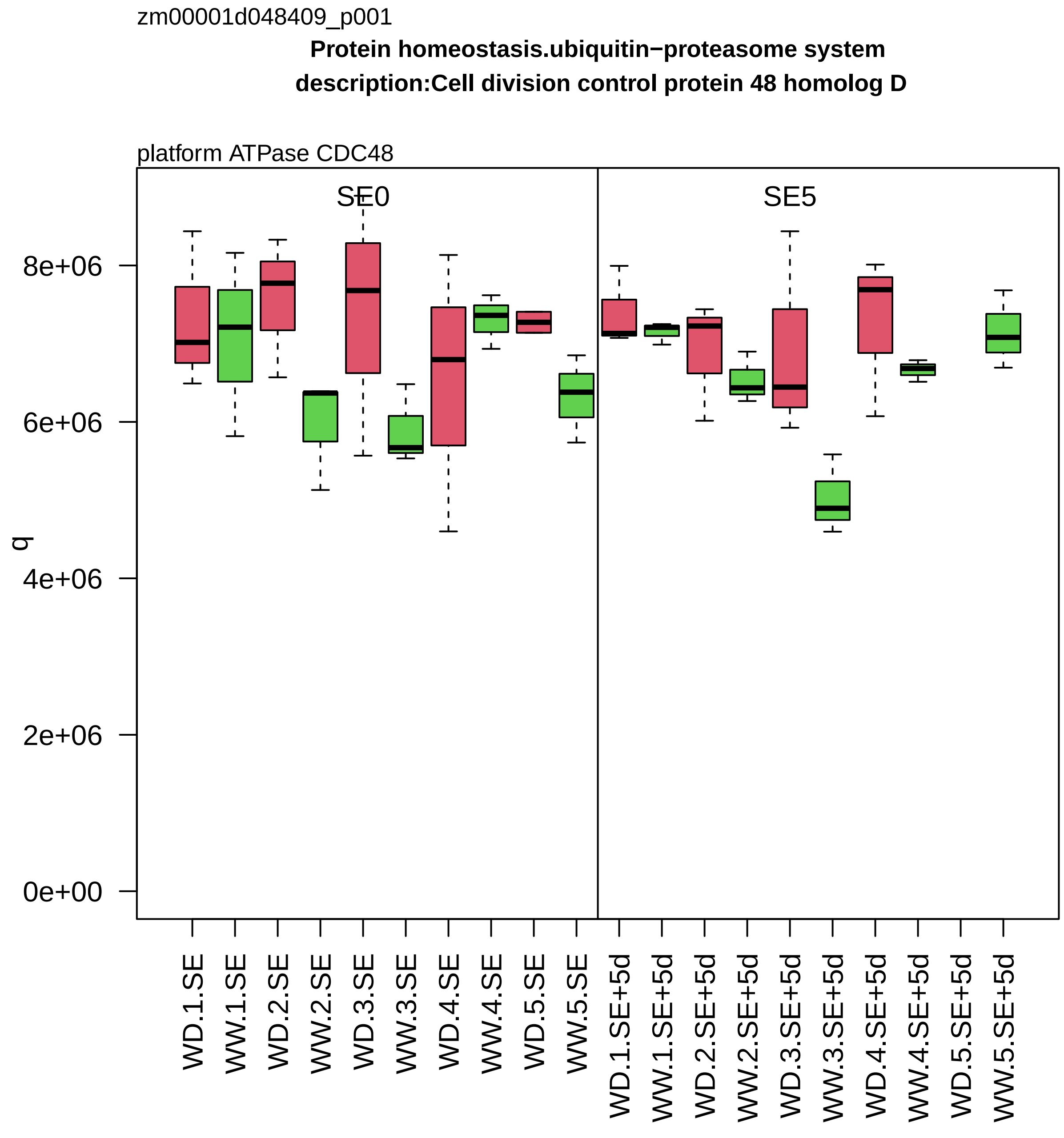
zm00001d048409_p001
Protein homeostasis.ubiquitin−proteasome system
description:Cell division control protein 48 homolog D
platform ATPase CDC48
SE0
SE5
q
0e+00
2e+06
4e+06
6e+06
8e+06
WD.1.SE
WW.1.SE
WD.2.SE
WW.2.SE
WD.3.SE
WW.3.SE
WD.4.SE
WW.4.SE
WD.5.SE
WW.5.SE
WD.1.SE+5d
WW.1.SE+5d
WD.2.SE+5d
WW.2.SE+5d
WD.3.SE+5d
WW.3.SE+5d
WD.4.SE+5d
WW.4.SE+5d
WD.5.SE+5d
WW.5.SE+5d

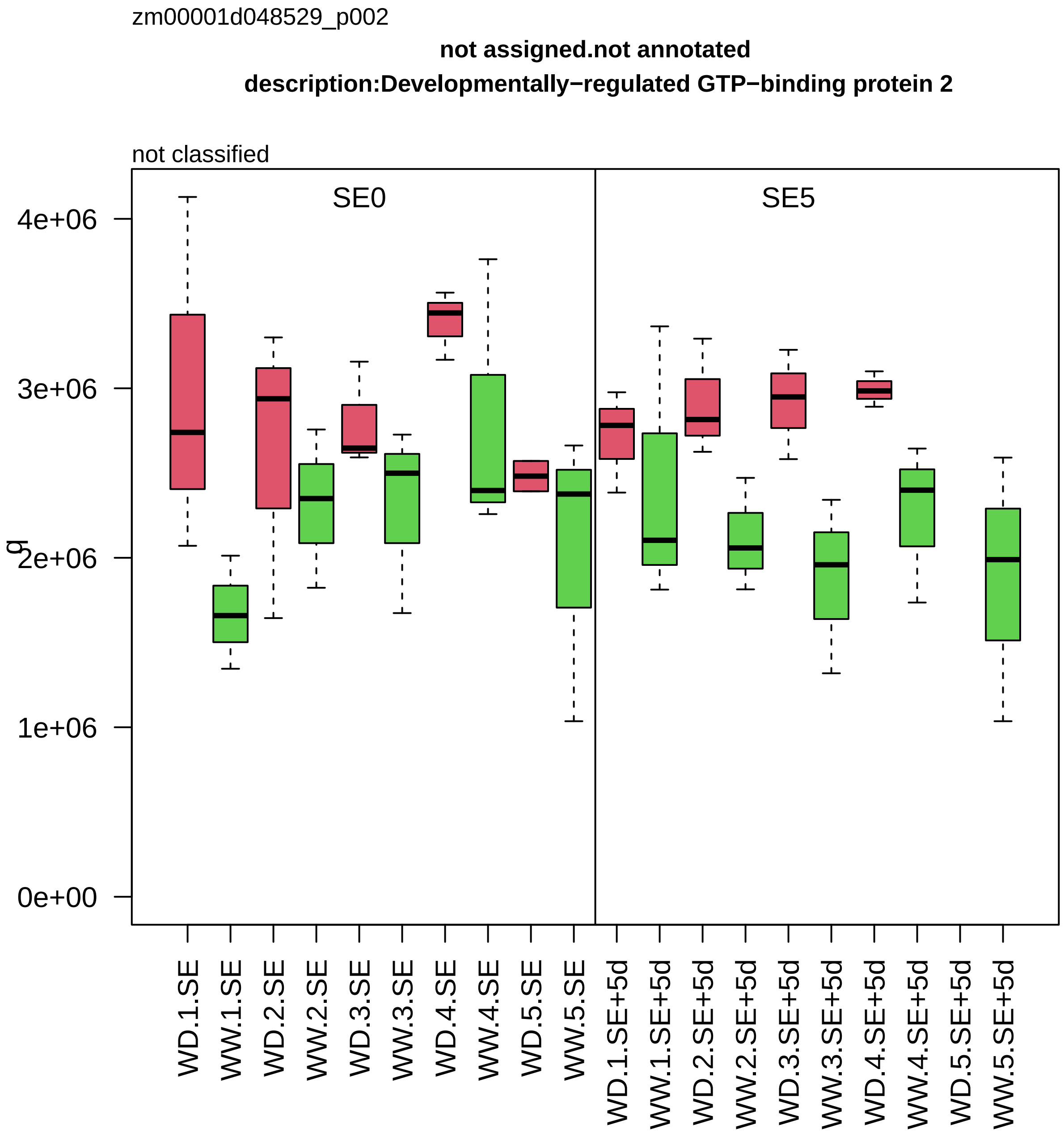

zm00001d048529_p002
not assigned.not annotated
description:Developmentally−regulated GTP−binding protein 2
not classified
SE0
SE5
4e+06
3e+06
2e+06
1e+06
0e+00
q
WD.1.SE
WW.1.SE
WD.2.SE
WW.2.SE
WD.3.SE
WW.3.SE
WD.4.SE
WW.4.SE
WD.5.SE
WW.5.SE
WD.1.SE+5d
WW.1.SE+5d
WD.2.SE+5d
WW.2.SE+5d
WD.3.SE+5d
WW.3.SE+5d
WD.4.SE+5d
WW.4.SE+5d
WD.5.SE+5d
WW.5.SE+5d

zm00001d048634_p001

**Enzyme classification.EC_1 oxidoreductases**

**gene_symbol:benzoxazinone synthesis6 description:DIBOA−glucoside dioxygenase BX6**

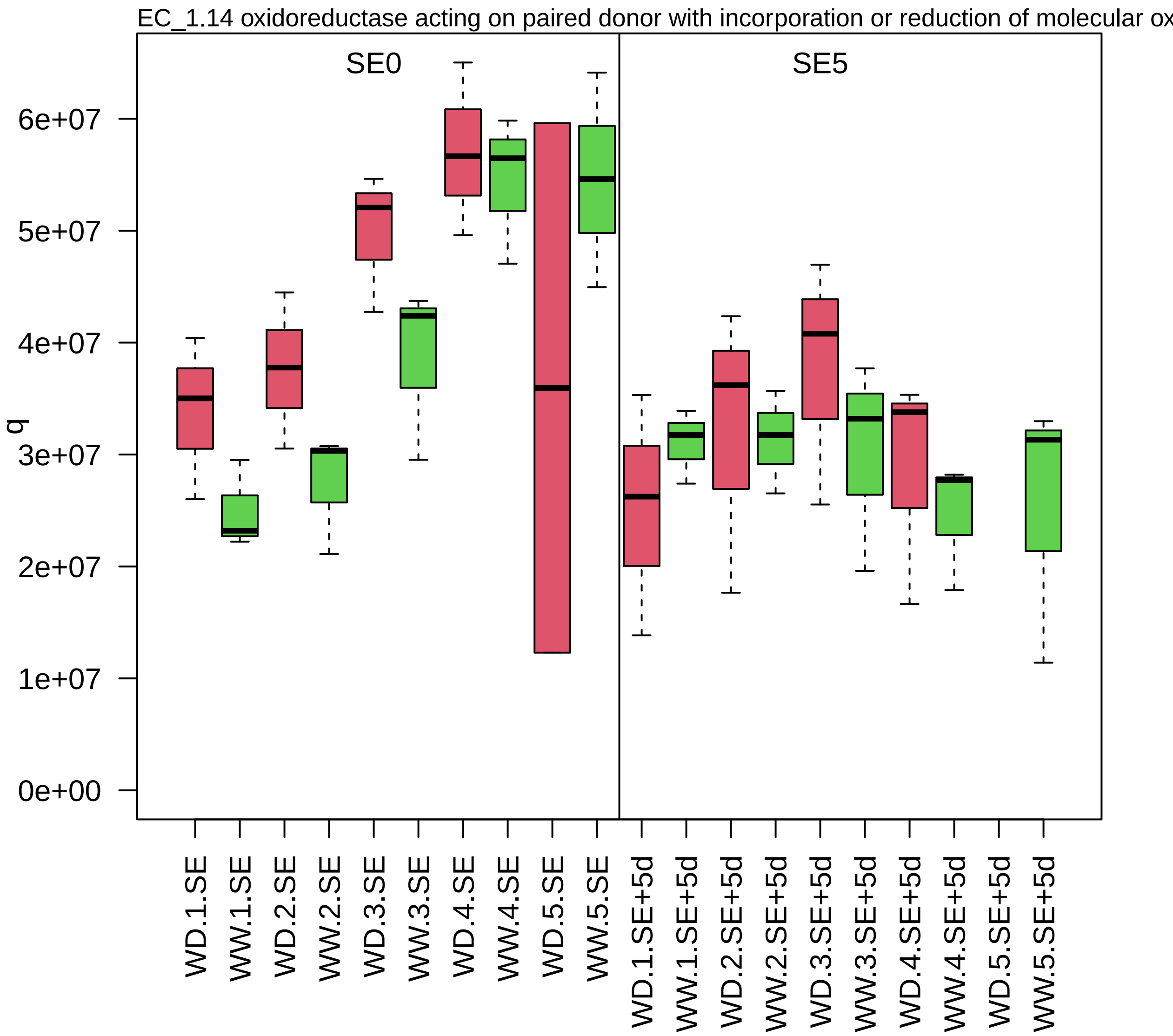

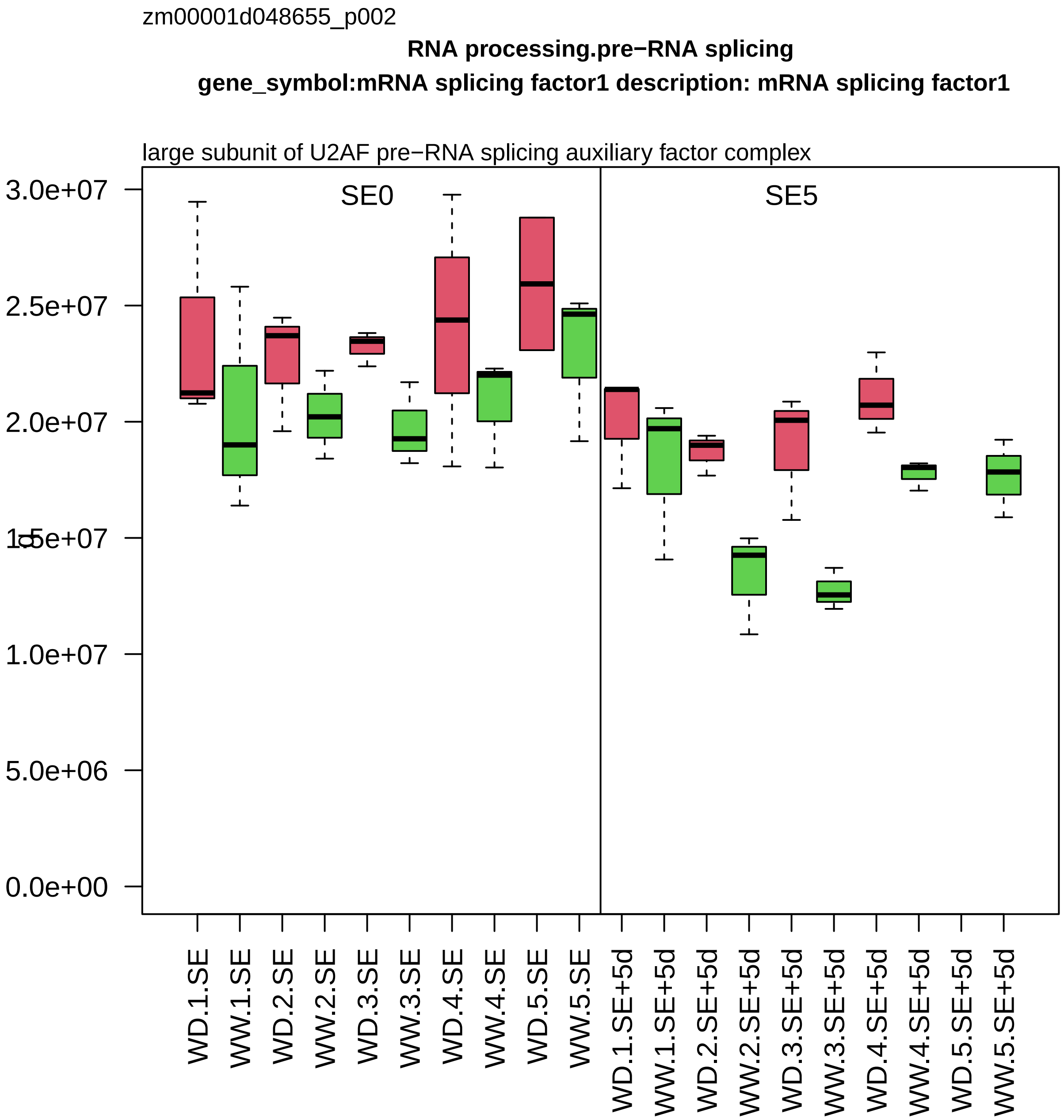
zm00001d048655_p002
RNA processing.pre−RNA splicing
gene_symbol:mRNA splicing factor1 description: mRNA splicing factor1
large subunit of U2AF pre−RNA splicing auxiliary factor complex
SE0
SE5
3.0e+07
2.5e+07
2.0e+07
1.5e+07
1.0e+07
5.0e+06
0.0e+00
WD.1.SE
WW.1.SE
WD.2.SE
WW.2.SE
WD.3.SE
WW.3.SE
WD.4.SE
WW.4.SE
WD.5.SE
WW.5.SE
WD.1.SE+5d
WW.1.SE+5d
WD.2.SE+5d
WW.2.SE+5d
WD.3.SE+5d
WW.3.SE+5d
WD.4.SE+5d
WW.4.SE+5d
WD.5.SE+5d
WW.5.SE+5d

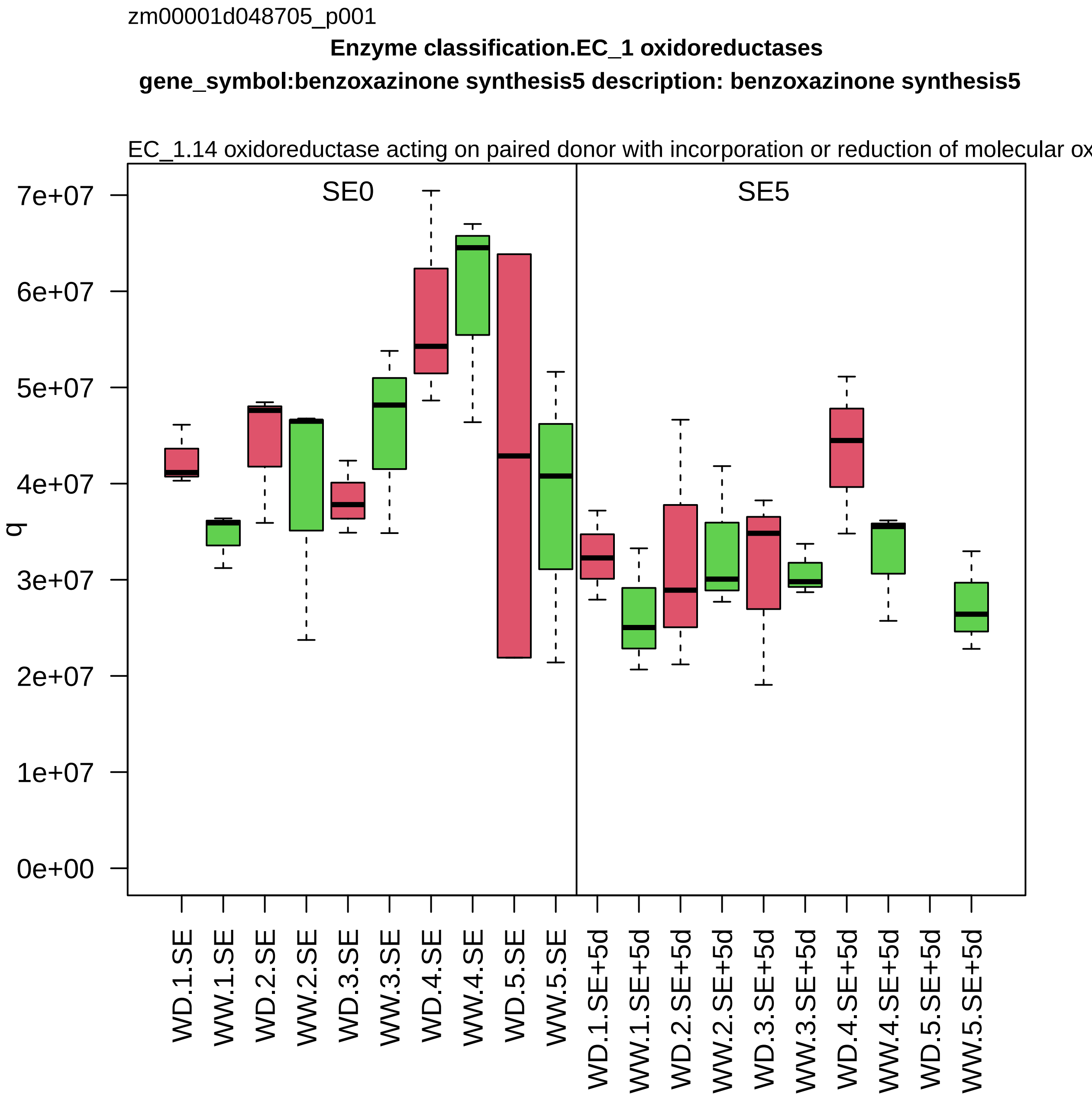
zm00001d048705_p001
Enzyme classification.EC_1 oxidoreductases
gene_symbol:benzoxazinone synthesis5 description: benzoxazinone synthesis5
EC_1.14 oxidoreductase acting on paired donor with incorporation or reduction of molecular ox
SE0
SE5
q
0e+00
1e+07
2e+07
3e+07
4e+07
5e+07
6e+07
7e+07
WD.1.SE
WW.1.SE
WD.2.SE
WW.2.SE
WD.3.SE
WW.3.SE
WD.4.SE
WW.4.SE
WD.5.SE
WW.5.SE
WD.1.SE+5d
WW.1.SE+5d
WD.2.SE+5d
WW.2.SE+5d
WD.3.SE+5d
WW.3.SE+5d
WD.4.SE+5d
WW.4.SE+5d
WD.5.SE+5d
WW.5.SE+5d

zm00001d048707_p001

**Enzyme classification.EC_2 transferases**

**gene_symbol:benzoxazinone synthesis8 description: benzoxazinone synthesis8**

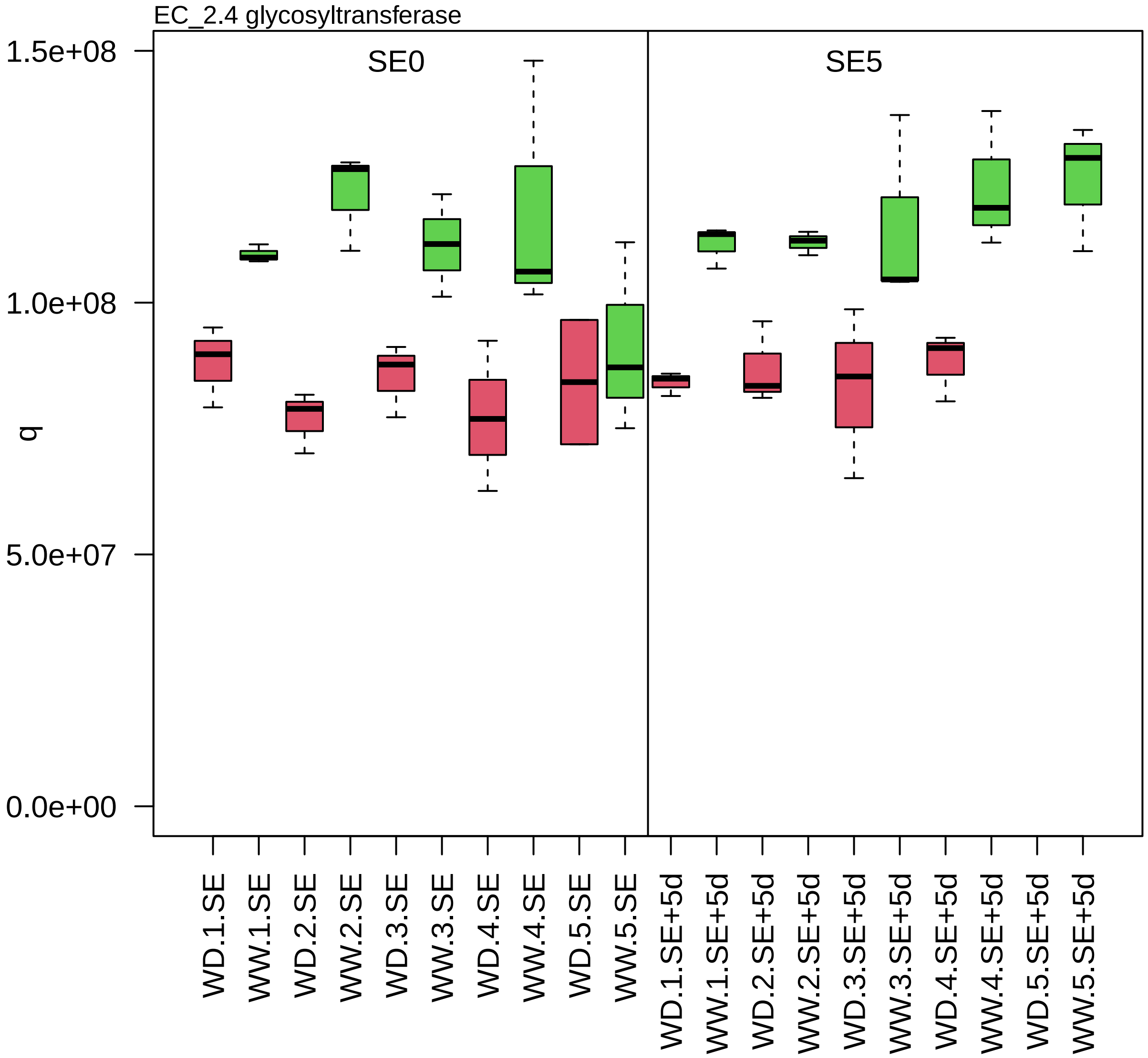

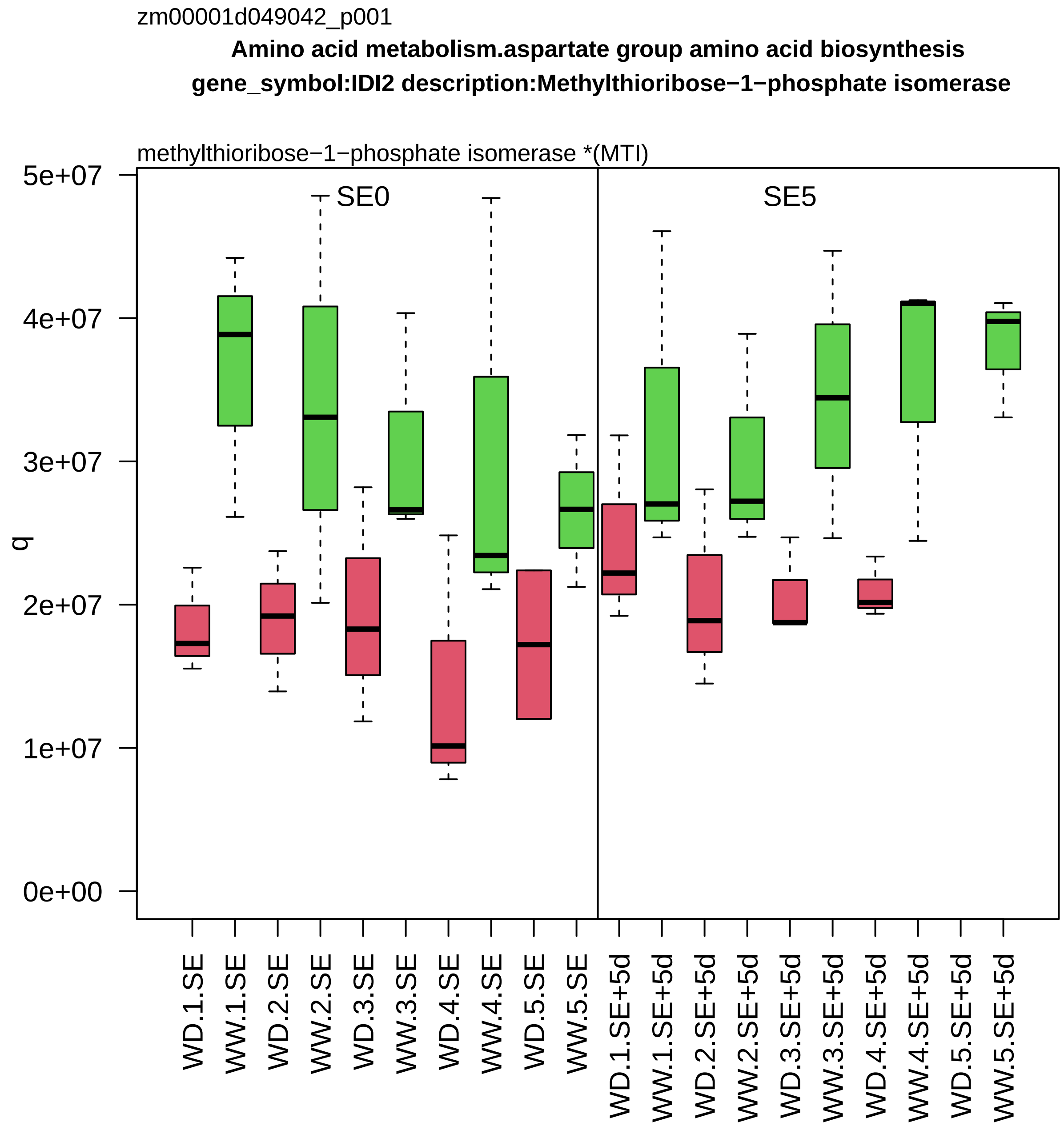
zm00001d049042_p001
Amino acid metabolism.aspartate group amino acid biosynthesis
gene_symbol:IDI2 description:Methylthioribose−1−phosphate isomerase
methylthioribose−1−phosphate isomerase *(MTI)
SE0
SE5
q
5e+07
4e+07
3e+07
2e+07
1e+07
0e+00
WD.1.SE
WW.1.SE
WD.2.SE
WW.2.SE
WD.3.SE
WW.3.SE
WD.4.SE
WW.4.SE
WD.5.SE
WW.5.SE
WD.1.SE+5d
WW.1.SE+5d
WD.2.SE+5d
WW.2.SE+5d
WD.3.SE+5d
WW.3.SE+5d
WD.4.SE+5d
WW.4.SE+5d
WD.5.SE+5d
WW.5.SE+5d

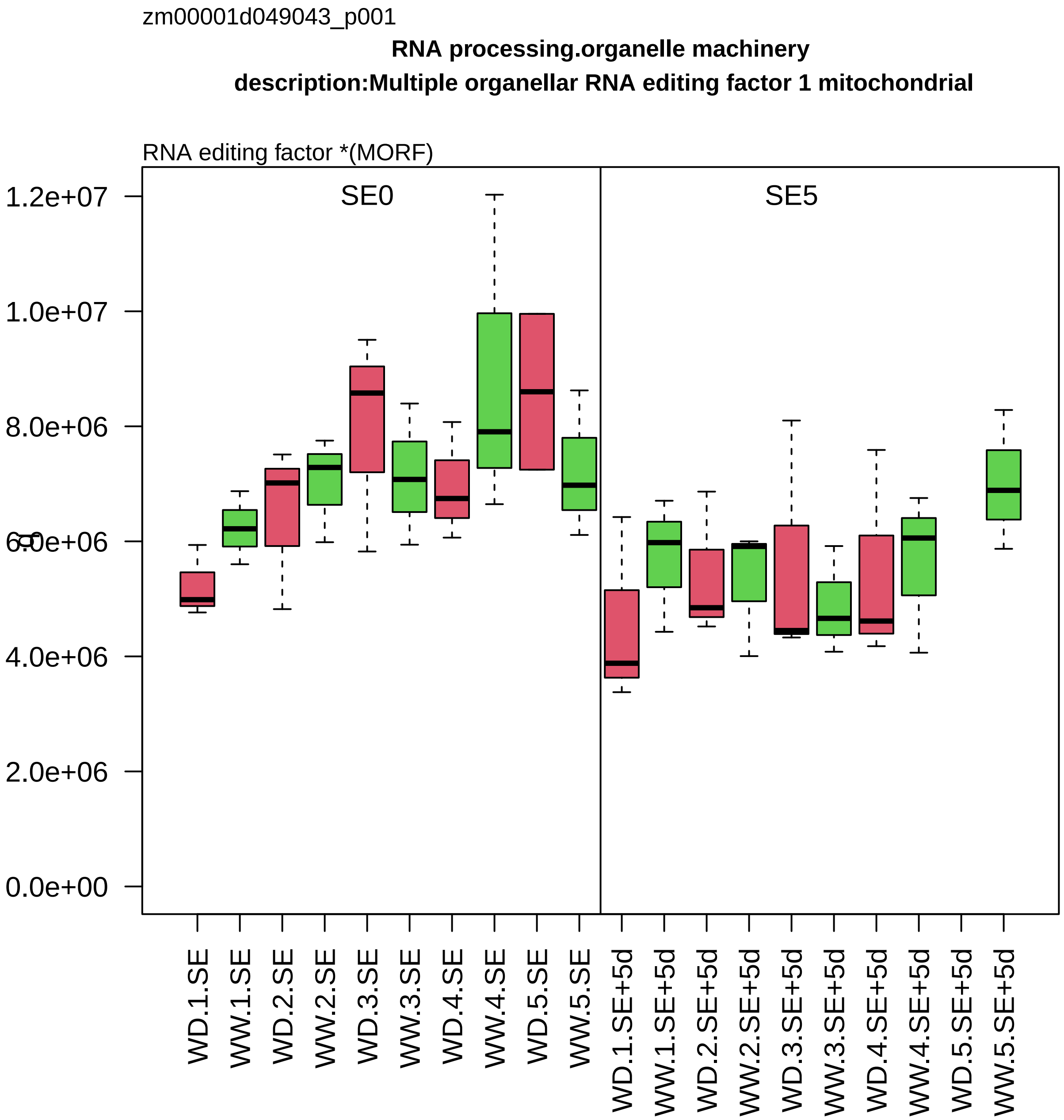
zm00001d049043_p001
RNA processing.organelle machinery
description:Multiple organellar RNA editing factor 1 mitochondrial
RNA editing factor *(MORF)
SE0
SE5
0.0e+00
2.0e+06
4.0e+06
6.0e+06
8.0e+06
1.0e+07
1.2e+07
WD.1.SE
WW.1.SE
WD.2.SE
WW.2.SE
WD.3.SE
WW.3.SE
WD.4.SE
WW.4.SE
WD.5.SE
WW.5.SE
WD.1.SE+5d
WW.1.SE+5d
WD.2.SE+5d
WW.2.SE+5d
WD.3.SE+5d
WW.3.SE+5d
WD.4.SE+5d
WW.4.SE+5d
WD.5.SE+5d
WW.5.SE+5d

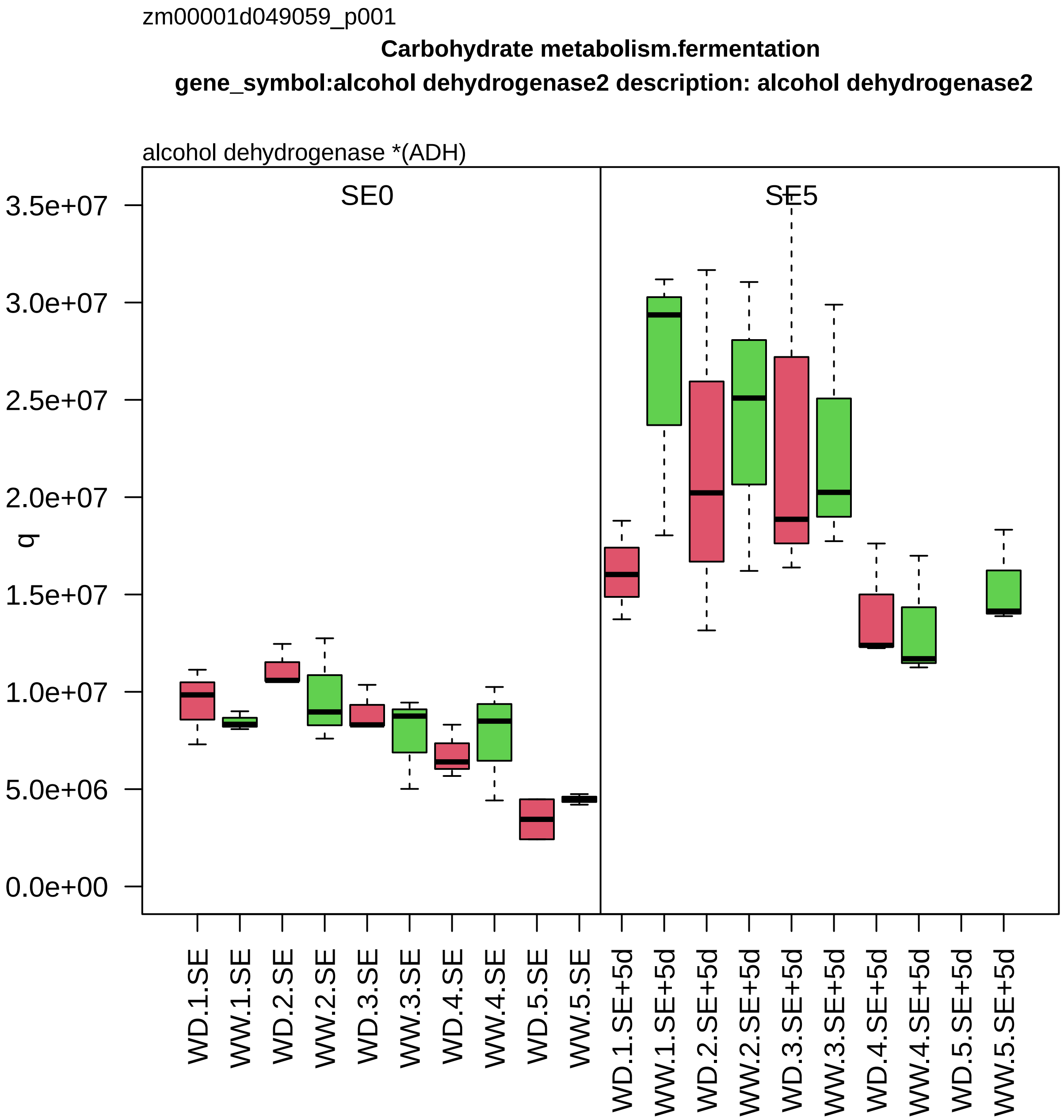
zm00001d049059_p001
Carbohydrate metabolism.fermentation
gene_symbol:alcohol dehydrogenase2 description: alcohol dehydrogenase2
alcohol dehydrogenase *(ADH)
SE0
SE5
q
3.5e+07
3.0e+07
2.5e+07
2.0e+07
1.5e+07
1.0e+07
5.0e+06
0.0e+00
WD.1.SE
WW.1.SE
WD.2.SE
WW.2.SE
WD.3.SE
WW.3.SE
WD.4.SE
WW.4.SE
WD.5.SE
WW.5.SE
WD.1.SE+5d
WW.1.SE+5d
WD.2.SE+5d
WW.2.SE+5d
WD.3.SE+5d
WW.3.SE+5d
WD.4.SE+5d
WW.4.SE+5d
WD.5.SE+5d
WW.5.SE+5d

zm00001d049099_p008

**Protein modification.cysteine disulfide formation**

**gene_symbol:protein disulfide isomerase1 description: protein disulfide isomerase1**

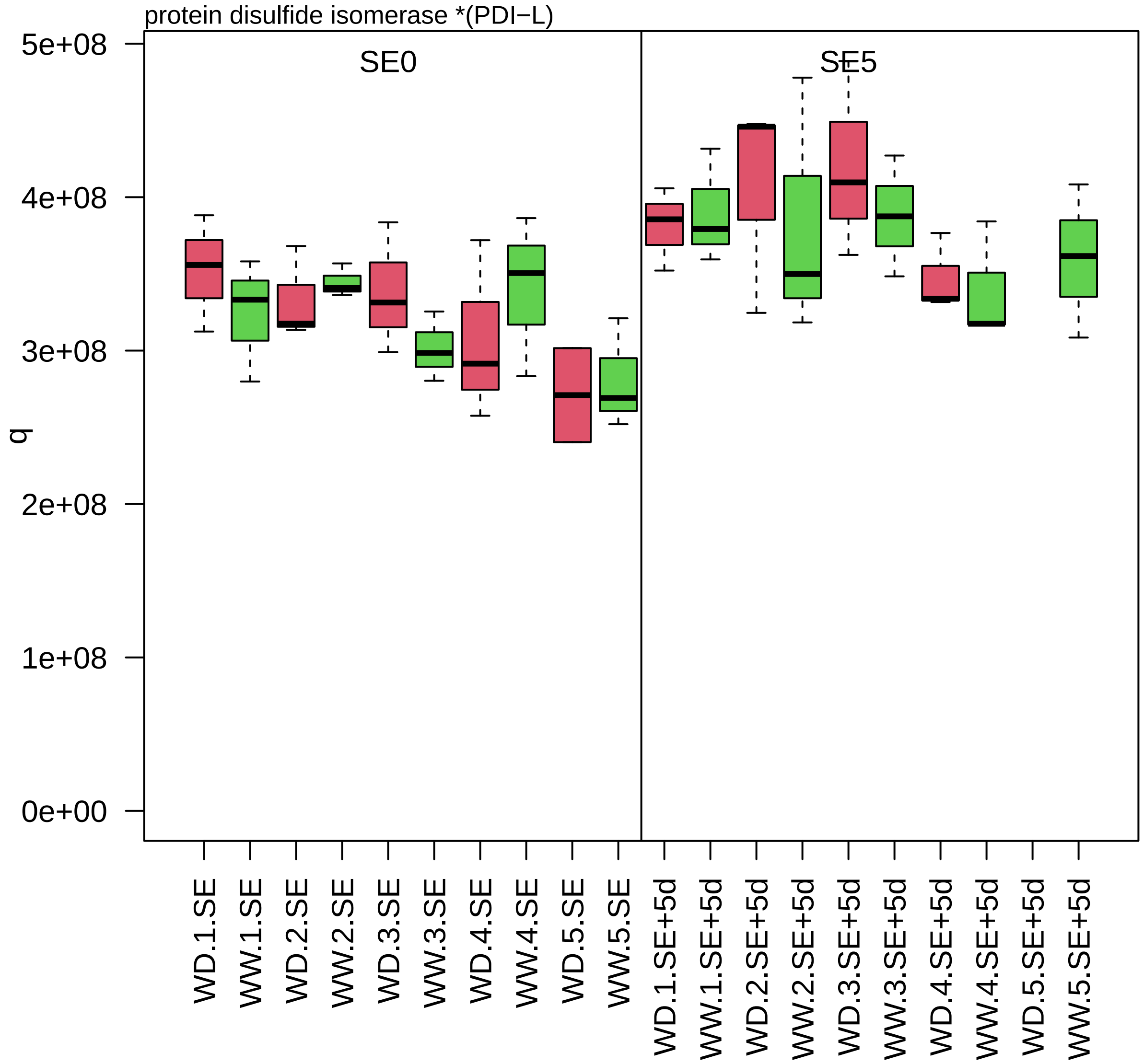

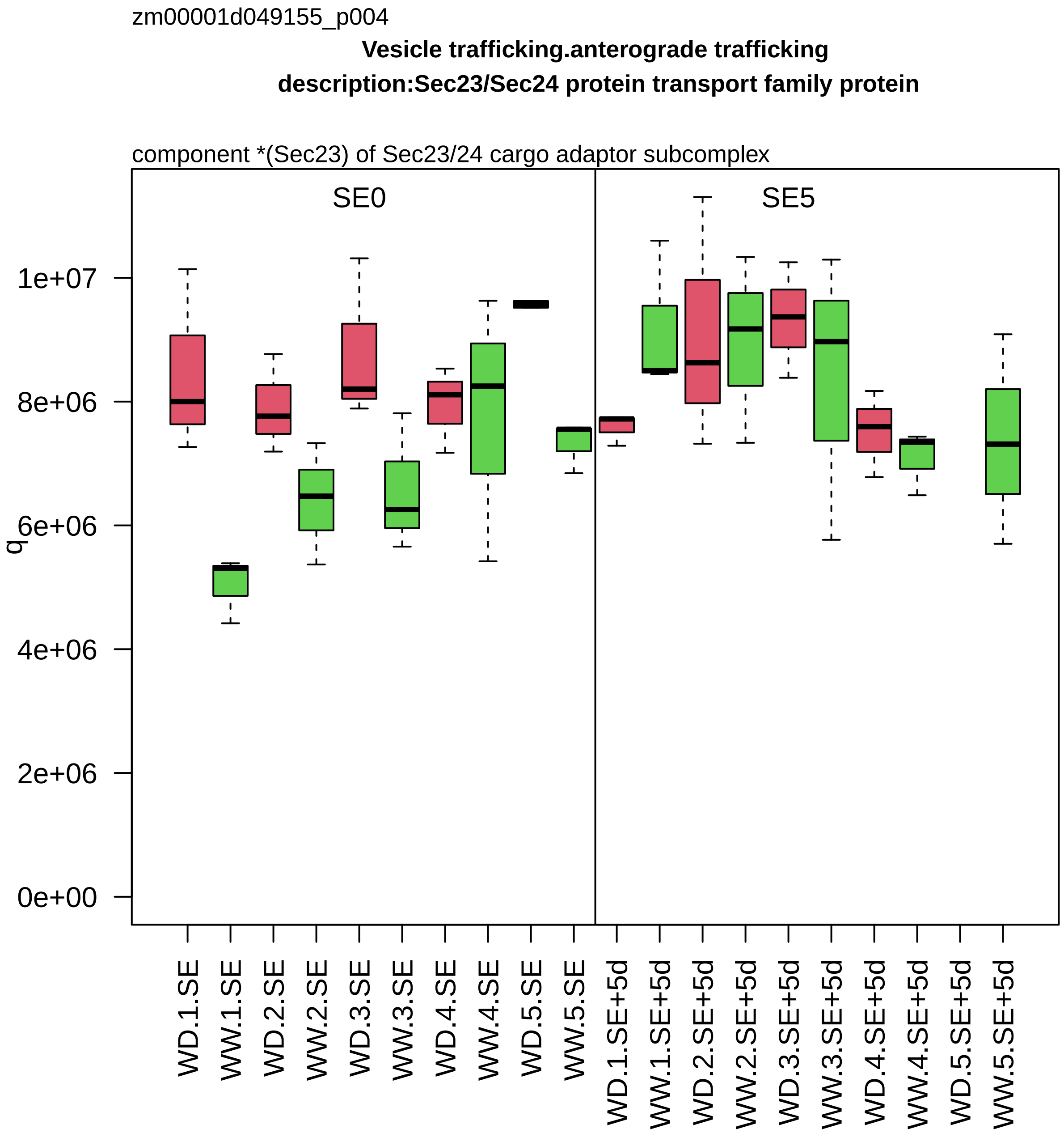
zm00001d049155_p004
Vesicle trafficking.anterograde trafficking
description:Sec23/Sec24 protein transport family protein
component *(Sec23) of Sec23/24 cargo adaptor subcomplex
SE0
SE5
q
0e+00
2e+06
4e+06
6e+06
8e+06
1e+07
WD.1.SE
WW.1.SE
WD.2.SE
WW.2.SE
WD.3.SE
WW.3.SE
WD.4.SE
WW.4.SE
WD.5.SE
WW.5.SE
WD.1.SE+5d
WW.1.SE+5d
WD.2.SE+5d
WW.2.SE+5d
WD.3.SE+5d
WW.3.SE+5d
WD.4.SE+5d
WW.4.SE+5d
WD.5.SE+5d
WW.5.SE+5d

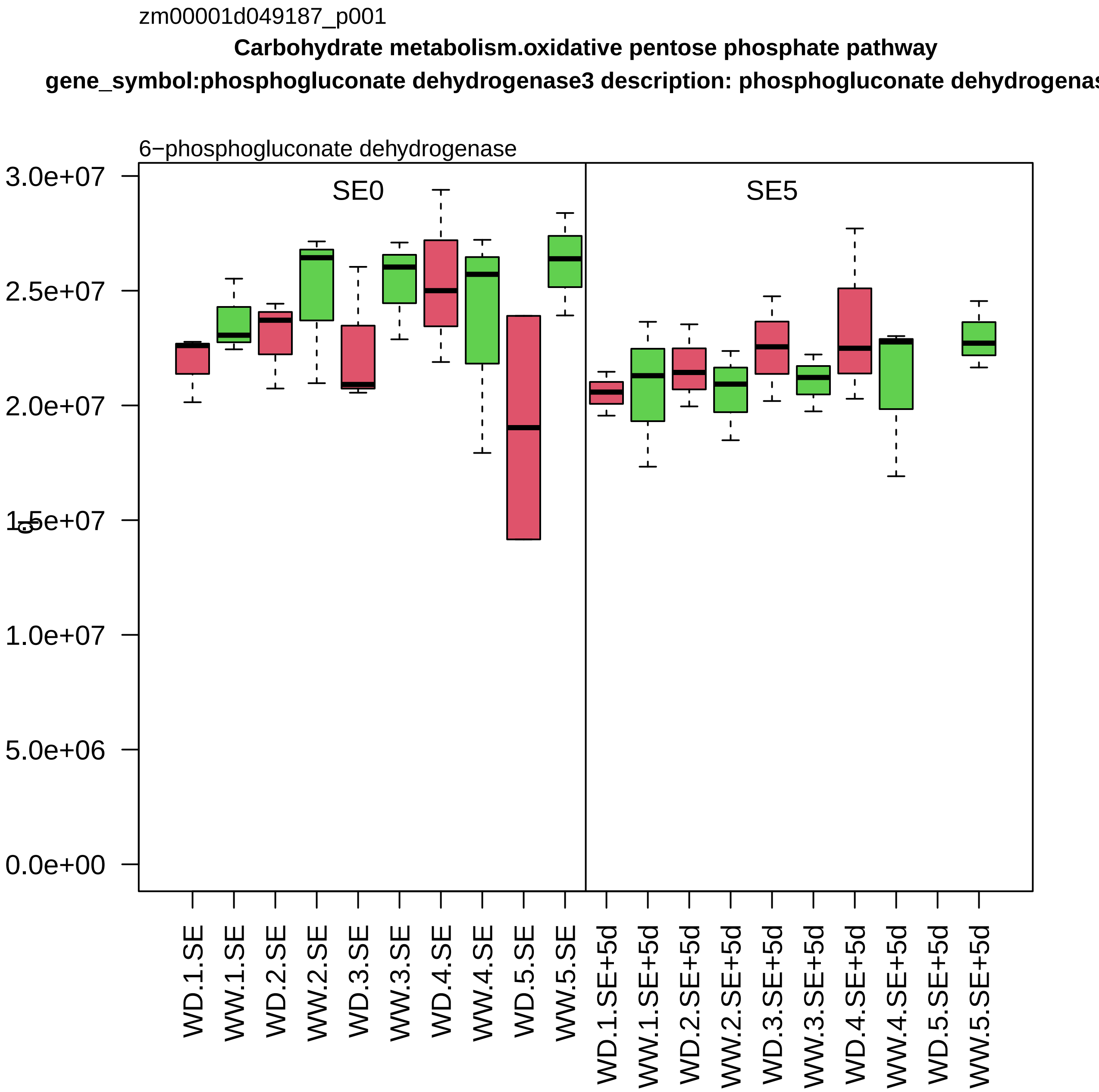

zm00001d049187_p001
Carbohydrate metabolism.oxidative pentose phosphate pathway
gene_symbol:phosphogluconate dehydrogenase3 description: phosphogluconate dehydrogenas
6−phosphogluconate dehydrogenase
SE0
SE5
3.0e+07
2.5e+07
2.0e+07
1.5e+07
1.0e+07
5.0e+06
0.0e+00
WD.1.SE
WW.1.SE
WD.2.SE
WW.2.SE
WD.3.SE
WW.3.SE
WD.4.SE
WW.4.SE
WD.5.SE
WW.5.SE
WD.1.SE+5d
WW.1.SE+5d
WD.2.SE+5d
WW.2.SE+5d
WD.3.SE+5d
WW.3.SE+5d
WD.4.SE+5d
WW.4.SE+5d
WD.5.SE+5d
WW.5.SE+5d

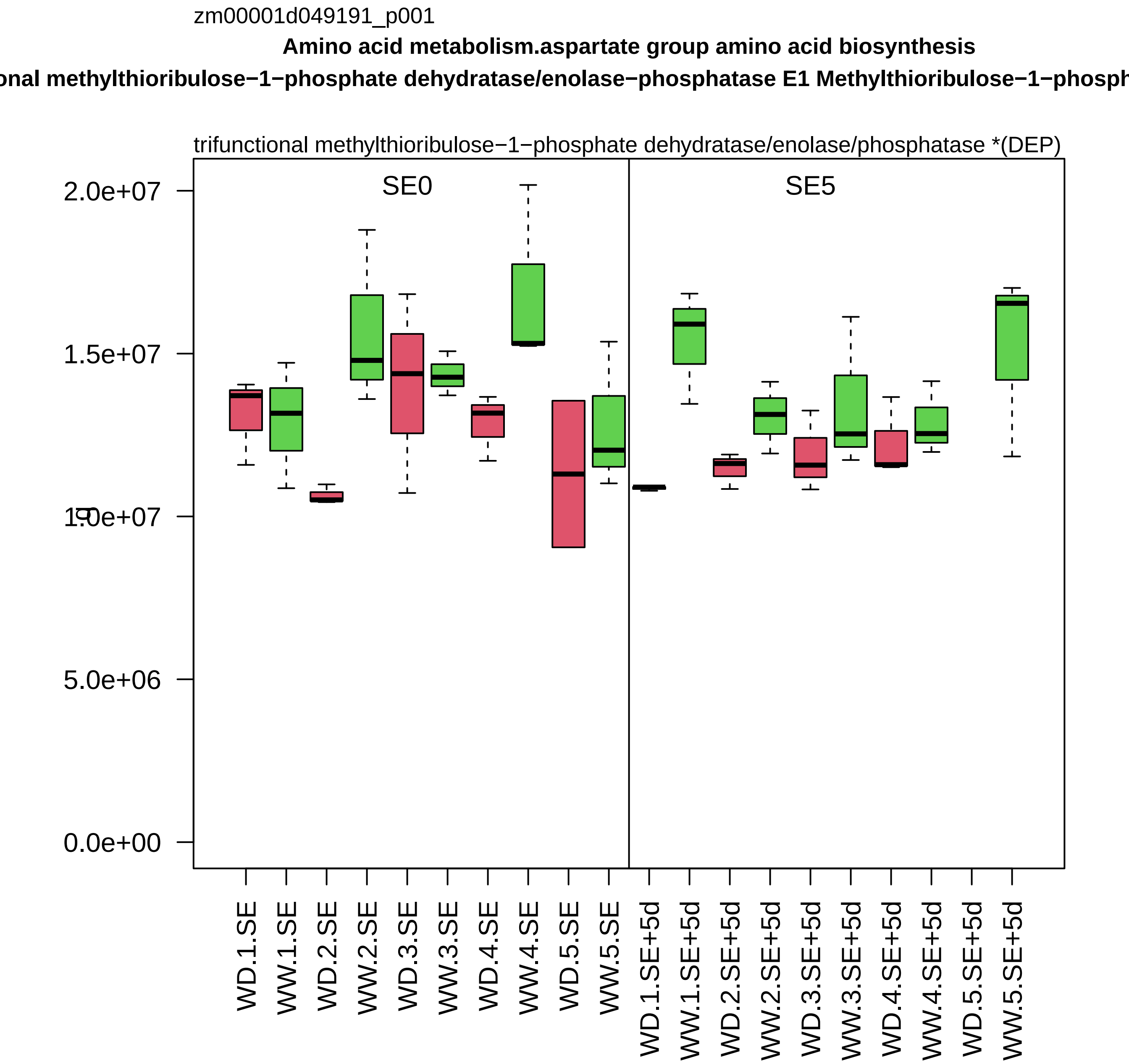

zm00001d049191_p001
Amino acid metabolism.aspartate group amino acid biosynthesis
onal methylthioribulose−1−phosphate dehydratase/enolase−phosphatase E1 Methylthioribulose−1−phosph
trifunctional methylthioribulose−1−phosphate dehydratase/enolase/phosphatase *(DEP)
SE0
SE5
2.0e+07
1.5e+07
1.0e+07
5.0e+06
0.0e+00
WD.1.SE
WW.1.SE
WD.2.SE
WW.2.SE
WD.3.SE
WW.3.SE
WD.4.SE
WW.4.SE
WD.5.SE
WW.5.SE
WD.1.SE+5d
WW.1.SE+5d
WD.2.SE+5d
WW.2.SE+5d
WD.3.SE+5d
WW.3.SE+5d
WD.4.SE+5d
WW.4.SE+5d
WD.5.SE+5d
WW.5.SE+5d

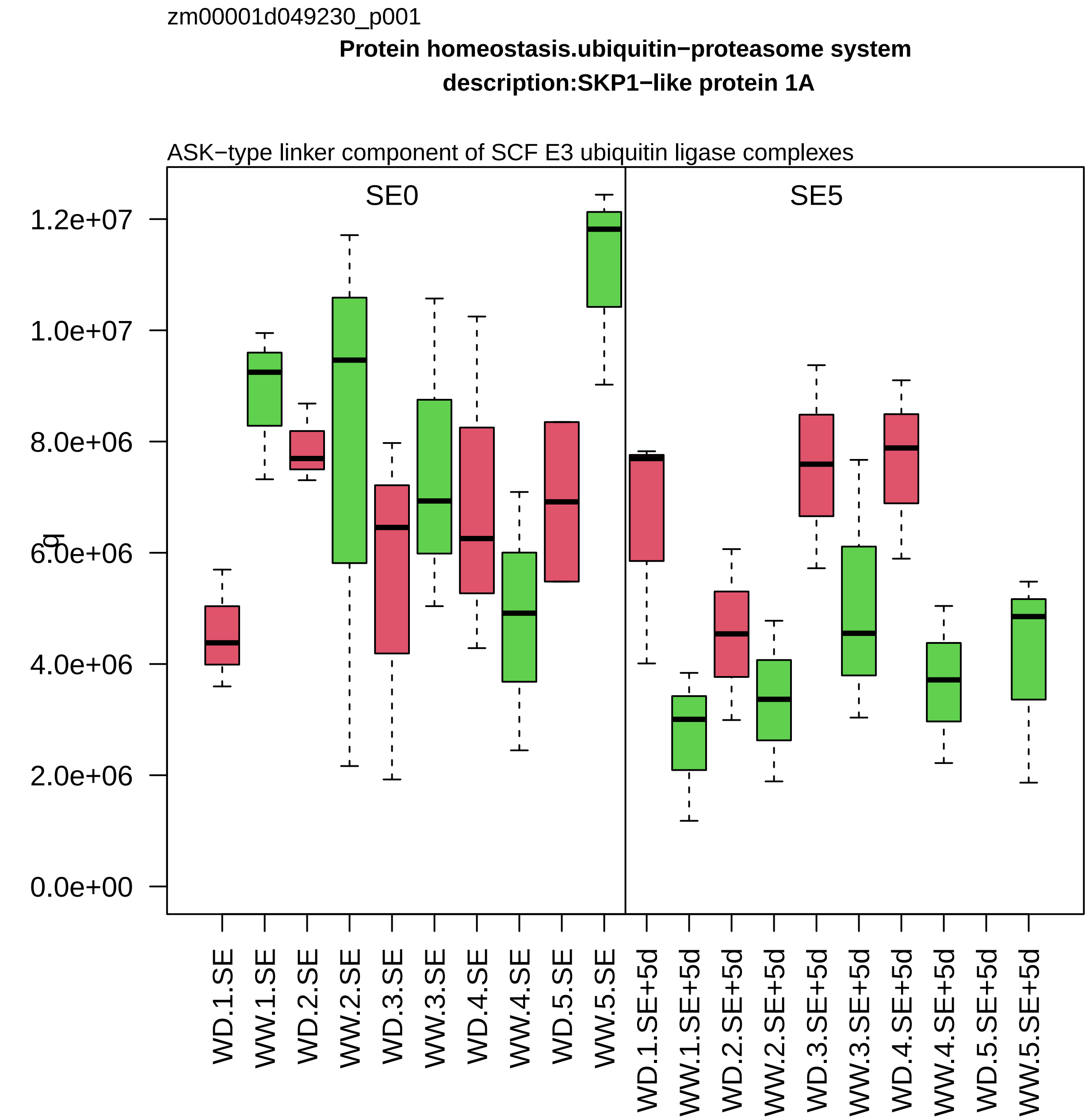

zm00001d049230_p001
Protein homeostasis.ubiquitin−proteasome system
description:SKP1−like protein 1A
ASK−type linker component of SCF E3 ubiquitin ligase complexes
SE0
SE5
1.2e+07
1.0e+07
8.0e+06
6.0e+06
4.0e+06
2.0e+06
0.0e+00
WD.1.SE
WW.1.SE
WD.2.SE
WW.2.SE
WD.3.SE
WW.3.SE
WD.4.SE
WW.4.SE
WD.5.SE
WW.5.SE
WD.1.SE+5d
WW.1.SE+5d
WD.2.SE+5d
WW.2.SE+5d
WD.3.SE+5d
WW.3.SE+5d
WD.4.SE+5d
WW.4.SE+5d
WD.5.SE+5d
WW.5.SE+5d

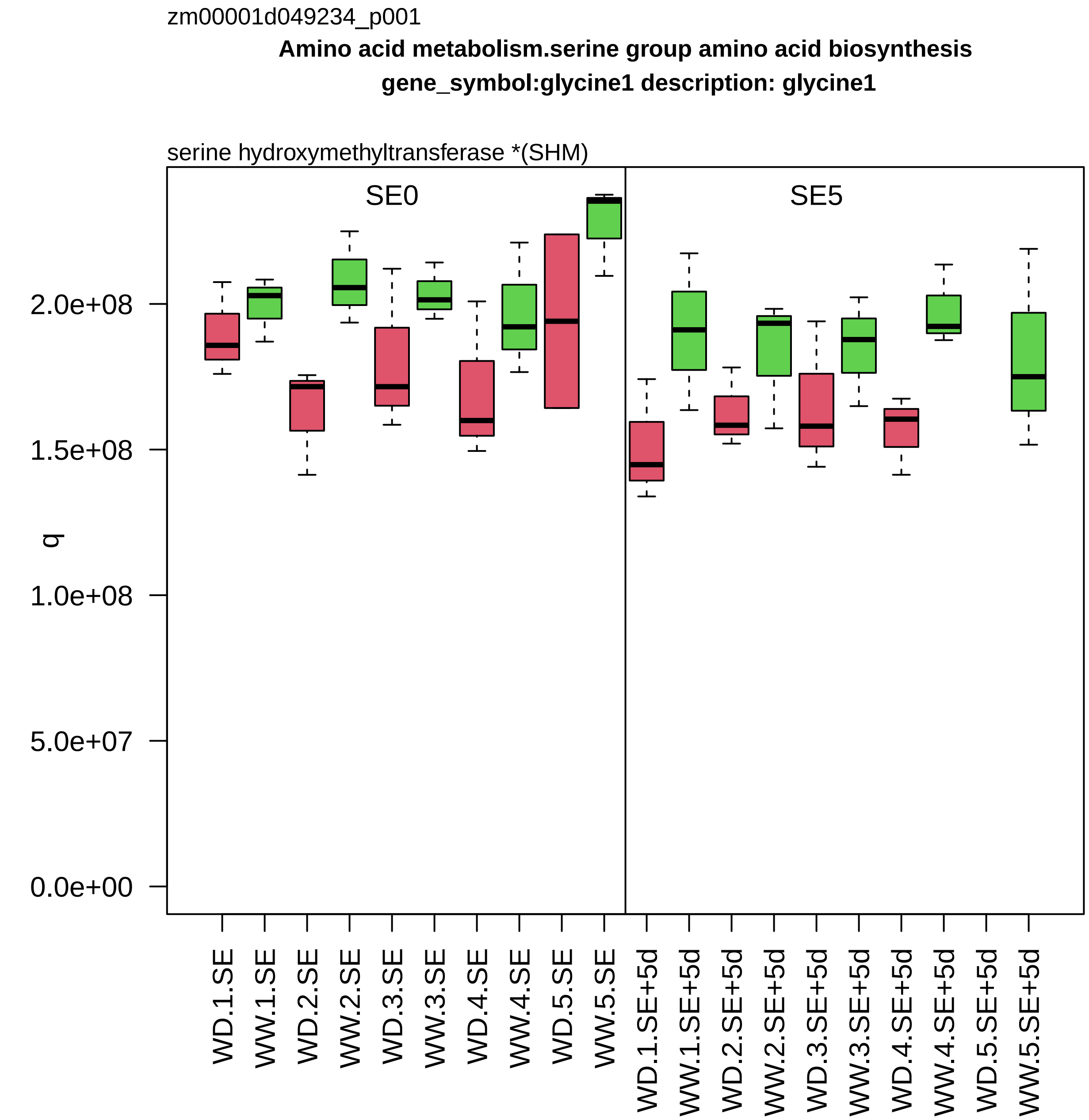

zm00001d049234_p001
Amino acid metabolism.serine group amino acid biosynthesis
gene_symbol:glycine1 description: glycine1
serine hydroxymethyltransferase *(SHM)
SE0
SE5
q
2.0e+08
1.5e+08
1.0e+08
5.0e+07
0.0e+00
WD.1.SE
WW.1.SE
WD.2.SE
WW.2.SE
WD.3.SE
WW.3.SE
WD.4.SE
WW.4.SE
WD.5.SE
WW.5.SE
WD.1.SE+5d
WW.1.SE+5d
WD.2.SE+5d
WW.2.SE+5d
WD.3.SE+5d
WW.3.SE+5d
WD.4.SE+5d
WW.4.SE+5d
WD.5.SE+5d
WW.5.SE+5d

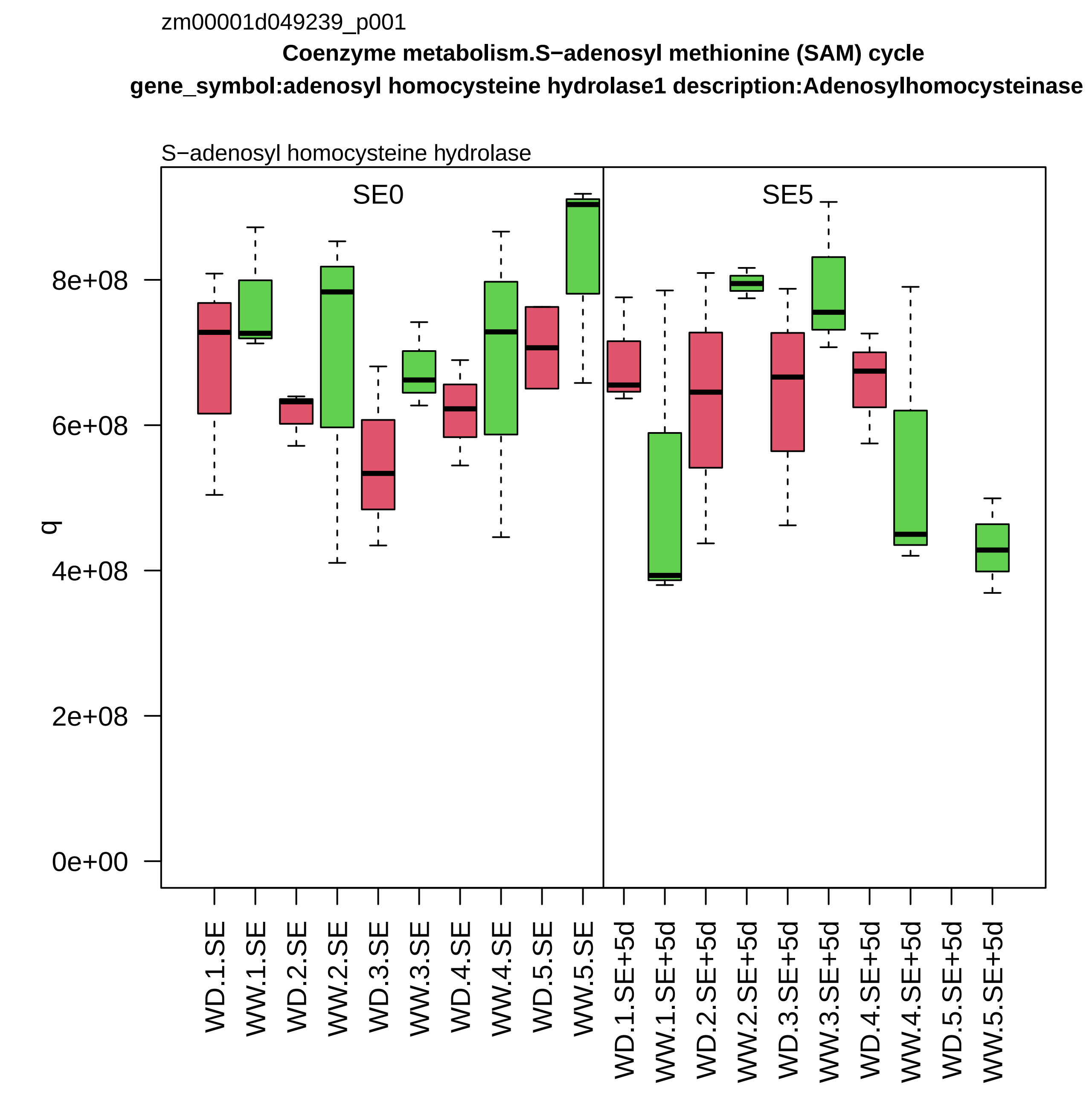
zm00001d049239_p001
Coenzyme metabolism.S−adenosyl methionine (SAM) cycle
gene_symbol:adenosyl homocysteine hydrolase1 description:Adenosylhomocysteinase
S−adenosyl homocysteine hydrolase
SE0
SE5
q
0e+00
2e+08
4e+08
6e+08
8e+08
WD.1.SE
WW.1.SE
WD.2.SE
WW.2.SE
WD.3.SE
WW.3.SE
WD.4.SE
WW.4.SE
WD.5.SE
WW.5.SE
WD.1.SE+5d
WW.1.SE+5d
WD.2.SE+5d
WW.2.SE+5d
WD.3.SE+5d
WW.3.SE+5d
WD.4.SE+5d
WW.4.SE+5d
WD.5.SE+5d
WW.5.SE+5d

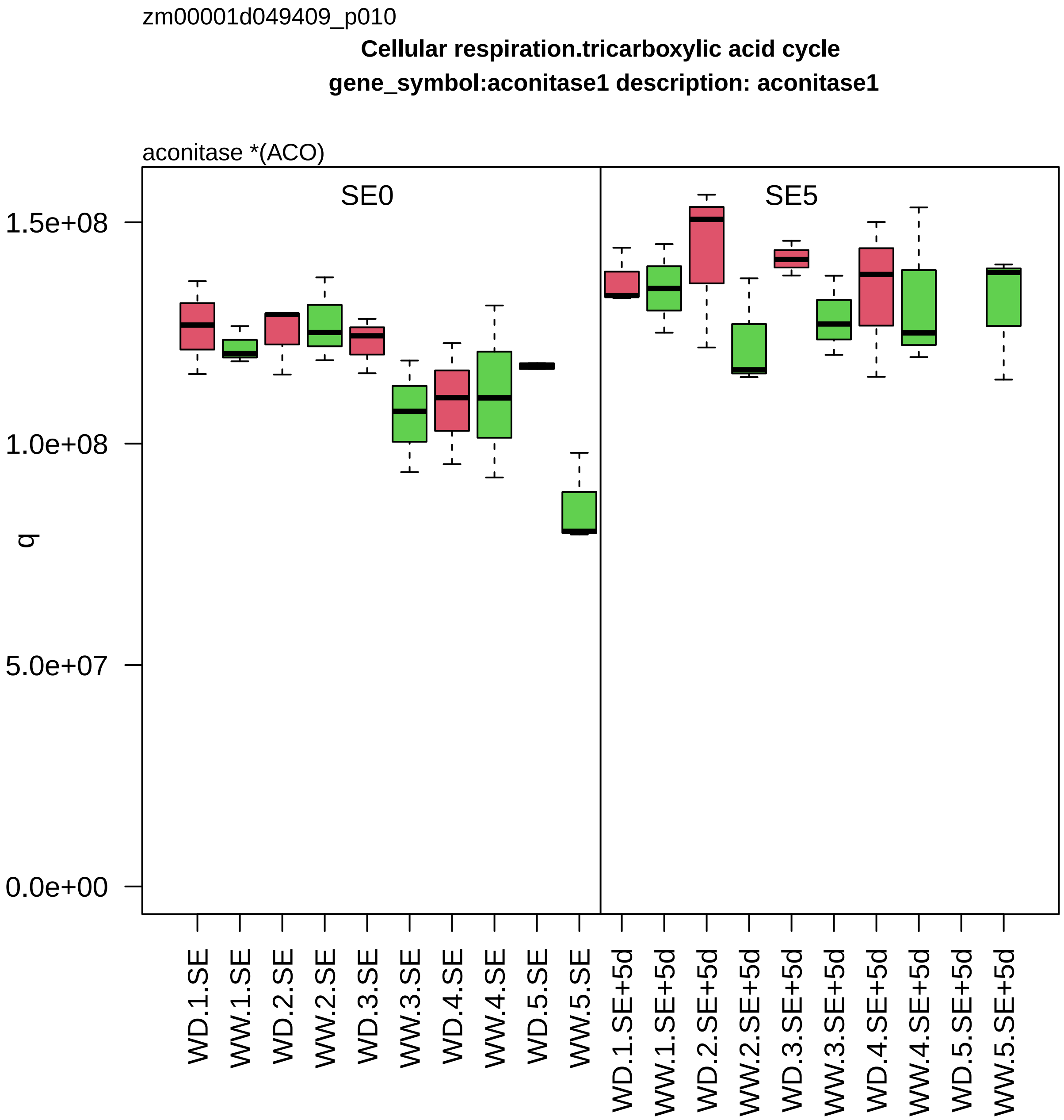

zm00001d049409_p010
Cellular respiration.tricarboxylic acid cycle
gene_symbol:aconitase1 description: aconitase1
aconitase *(ACO)
SE0
SE5
1.5e+08
1.0e+08
5.0e+07
0.0e+00
q
WD.1.SE
WW.1.SE
WD.2.SE
WW.2.SE
WD.3.SE
WW.3.SE
WD.4.SE
WW.4.SE
WD.5.SE
WW.5.SE
WD.1.SE+5d
WW.1.SE+5d
WD.2.SE+5d
WW.2.SE+5d
WD.3.SE+5d
WW.3.SE+5d
WD.4.SE+5d
WW.4.SE+5d
WD.5.SE+5d
WW.5.SE+5d

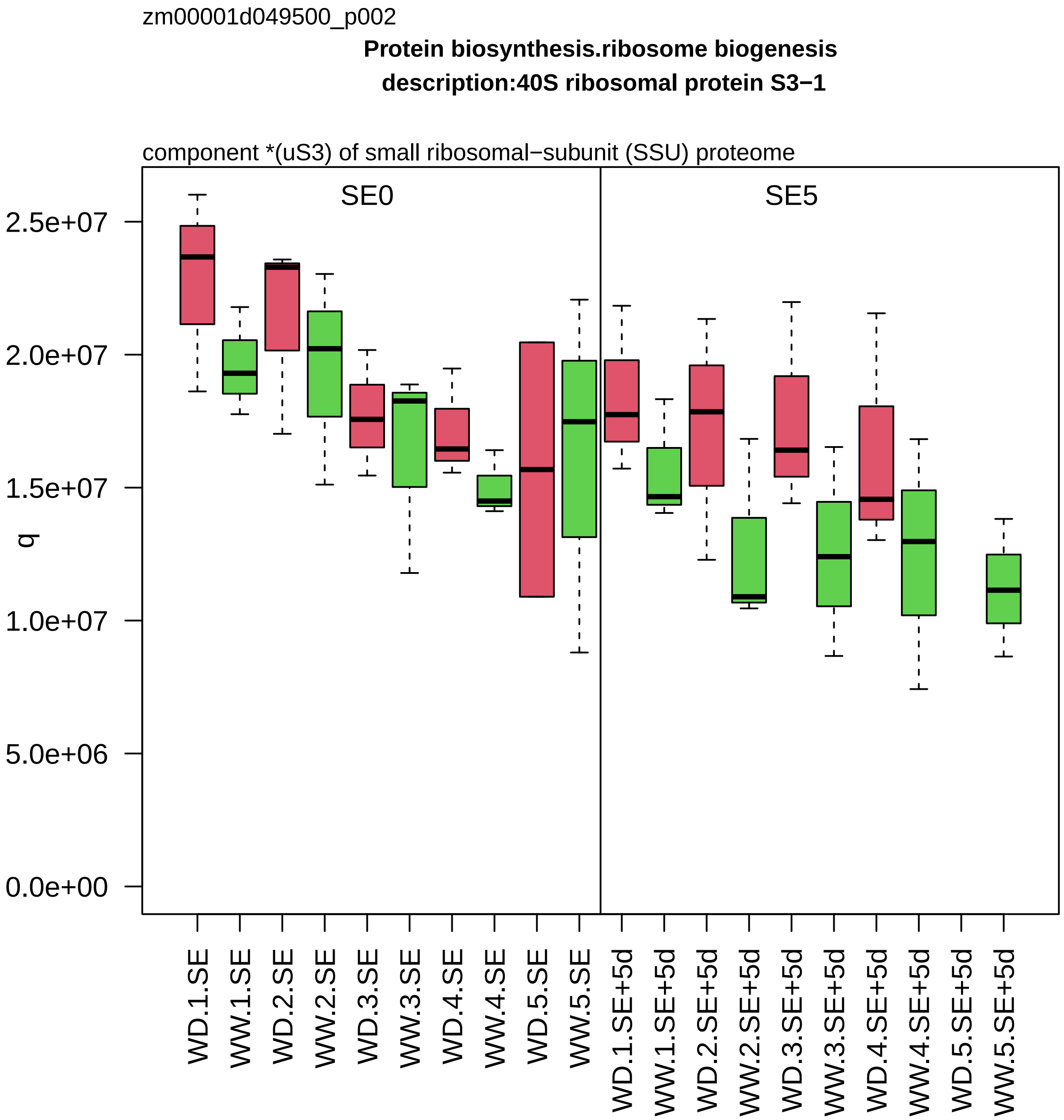

zm00001d049500_p002
Protein biosynthesis.ribosome biogenesis
description:40S ribosomal protein S3−1
component *(uS3) of small ribosomal−subunit (SSU) proteome
SE0
SE5
q
2.5e+07
2.0e+07
1.5e+07
1.0e+07
5.0e+06
0.0e+00
WD.1.SE
WW.1.SE
WD.2.SE
WW.2.SE
WD.3.SE
WW.3.SE
WD.4.SE
WW.4.SE
WD.5.SE
WW.5.SE
WD.1.SE+5d
WW.1.SE+5d
WD.2.SE+5d
WW.2.SE+5d
WD.3.SE+5d
WW.3.SE+5d
WD.4.SE+5d
WW.4.SE+5d
WD.5.SE+5d
WW.5.SE+5d

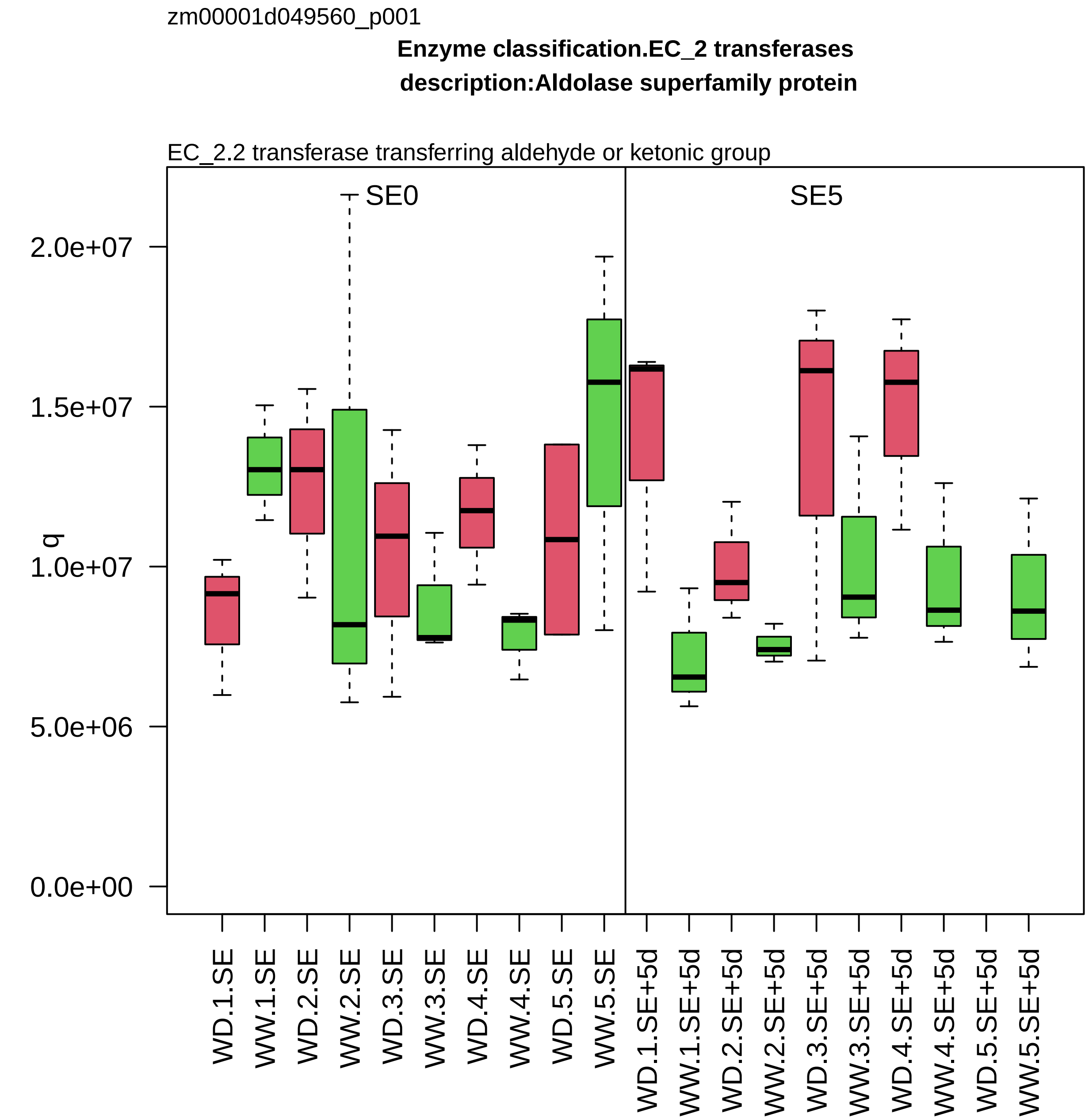

zm00001d049560_p001
Enzyme classification.EC_2 transferases
description:Aldolase superfamily protein
EC_2.2 transferase transferring aldehyde or ketonic group
SE0
SE5
q
2.0e+07
1.5e+07
1.0e+07
5.0e+06
0.0e+00
WD.1.SE
WW.1.SE
WD.2.SE
WW.2.SE
WD.3.SE
WW.3.SE
WD.4.SE
WW.4.SE
WD.5.SE
WW.5.SE
WD.1.SE+5d
WW.1.SE+5d
WD.2.SE+5d
WW.2.SE+5d
WD.3.SE+5d
WW.3.SE+5d
WD.4.SE+5d
WW.4.SE+5d
WD.5.SE+5d
WW.5.SE+5d

zm00001d049641_p004

**Cellular respiration.glycolysis**

**ymbol:glyceraldehyde−3−phosphate dehydrogenase1 description:Glyceraldehyde−3−phosphate dehydroge**

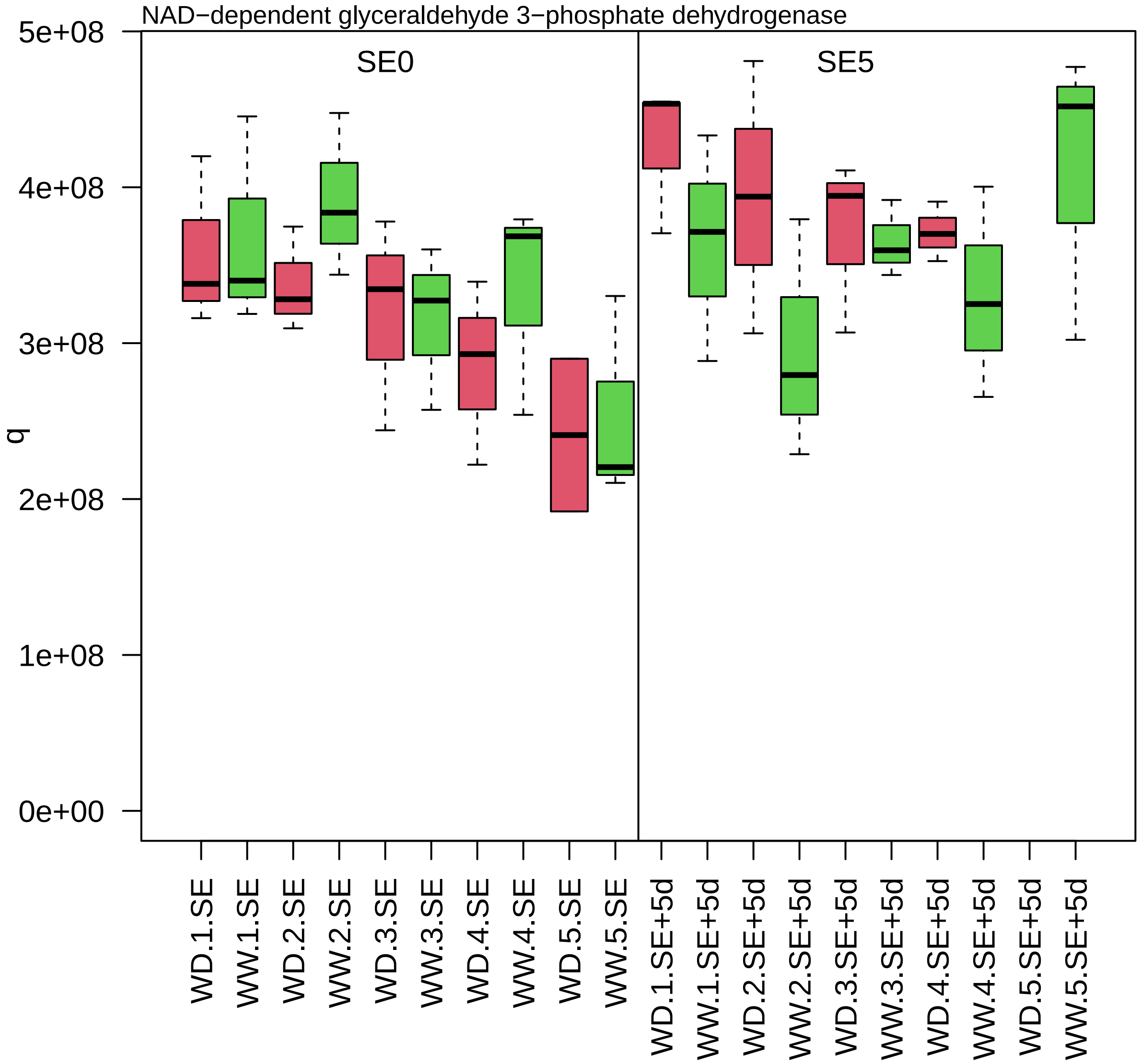

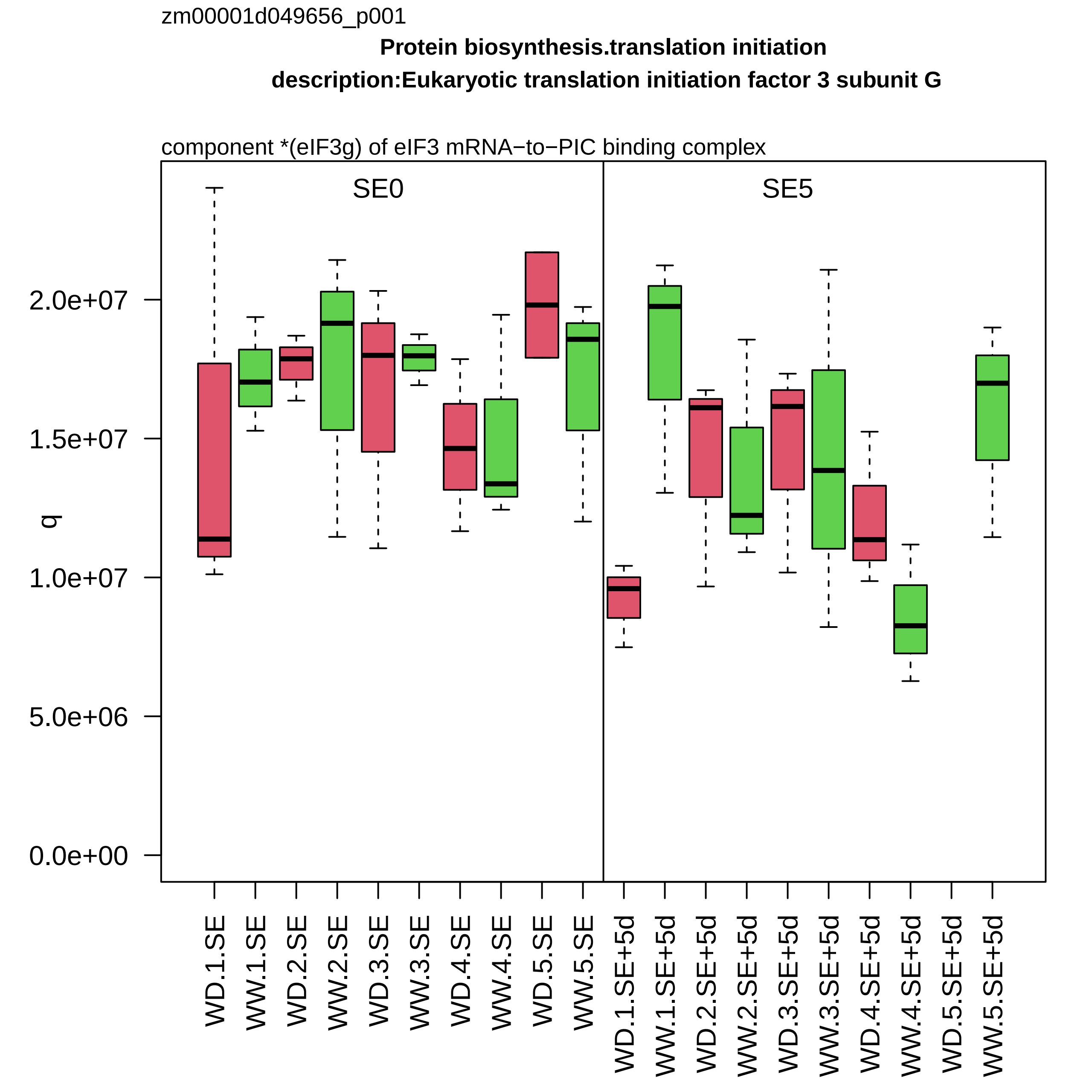

zm00001d049656_p001
Protein biosynthesis.translation initiation
description:Eukaryotic translation initiation factor 3 subunit G
component *(eIF3g) of eIF3 mRNA−to−PIC binding complex
SE0
SE5
q
0.0e+00
5.0e+06
1.0e+07
1.5e+07
2.0e+07
WD.1.SE
WW.1.SE
WD.2.SE
WW.2.SE
WD.3.SE
WW.3.SE
WD.4.SE
WW.4.SE
WD.5.SE
WW.5.SE
WD.1.SE+5d
WW.1.SE+5d
WD.2.SE+5d
WW.2.SE+5d
WD.3.SE+5d
WW.3.SE+5d
WD.4.SE+5d
WW.4.SE+5d
WD.5.SE+5d
WW.5.SE+5d

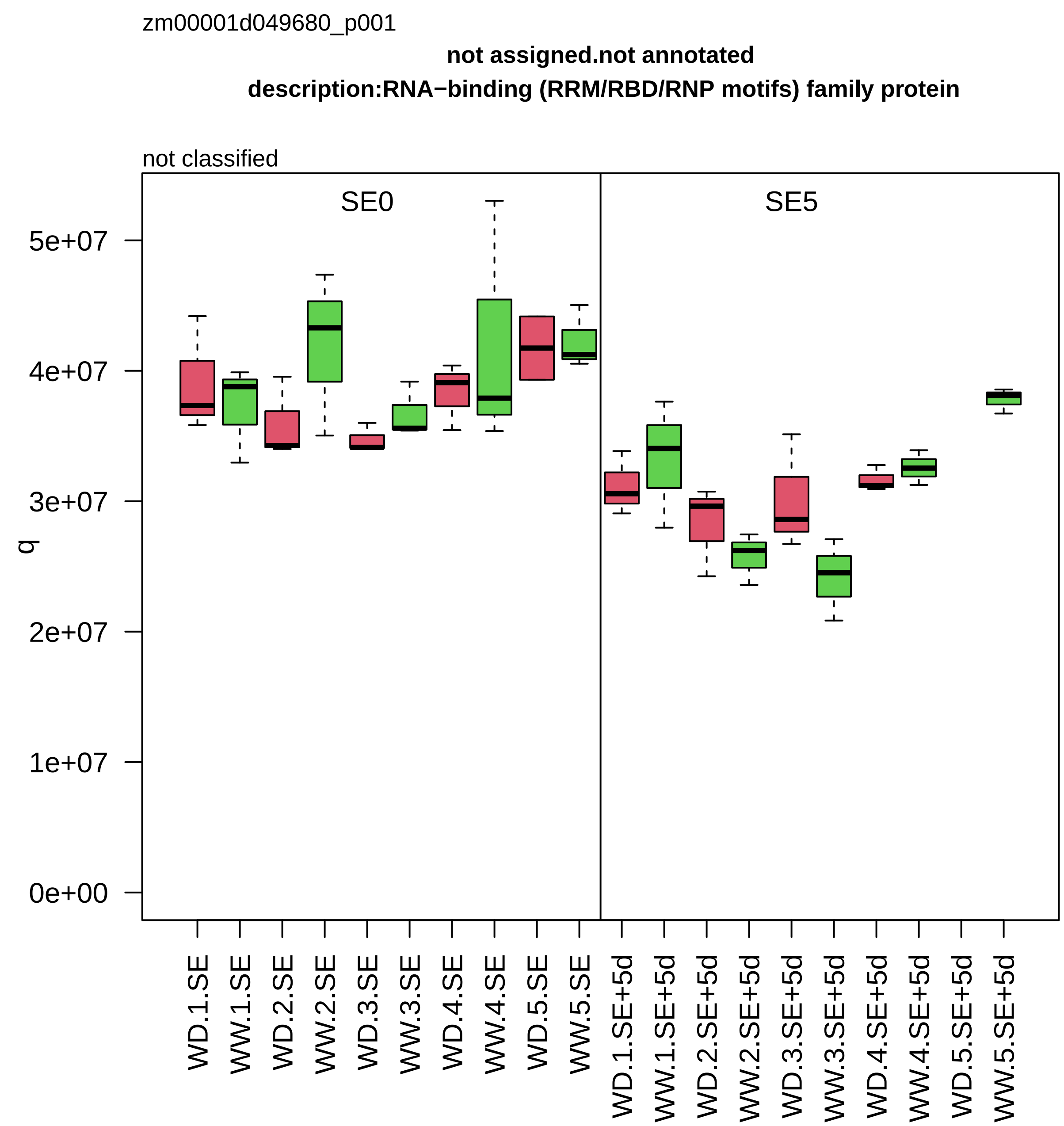
zm00001d049680_p001
not assigned.not annotated
description:RNA−binding (RRM/RBD/RNP motifs) family protein
not classified
SE0
SE5
q
0e+00
1e+07
2e+07
3e+07
4e+07
5e+07
WD.1.SE
WW.1.SE
WD.2.SE
WW.2.SE
WD.3.SE
WW.3.SE
WD.4.SE
WW.4.SE
WD.5.SE
WW.5.SE
WD.1.SE+5d
WW.1.SE+5d
WD.2.SE+5d
WW.2.SE+5d
WD.3.SE+5d
WW.3.SE+5d
WD.4.SE+5d
WW.4.SE+5d
WD.5.SE+5d
WW.5.SE+5d

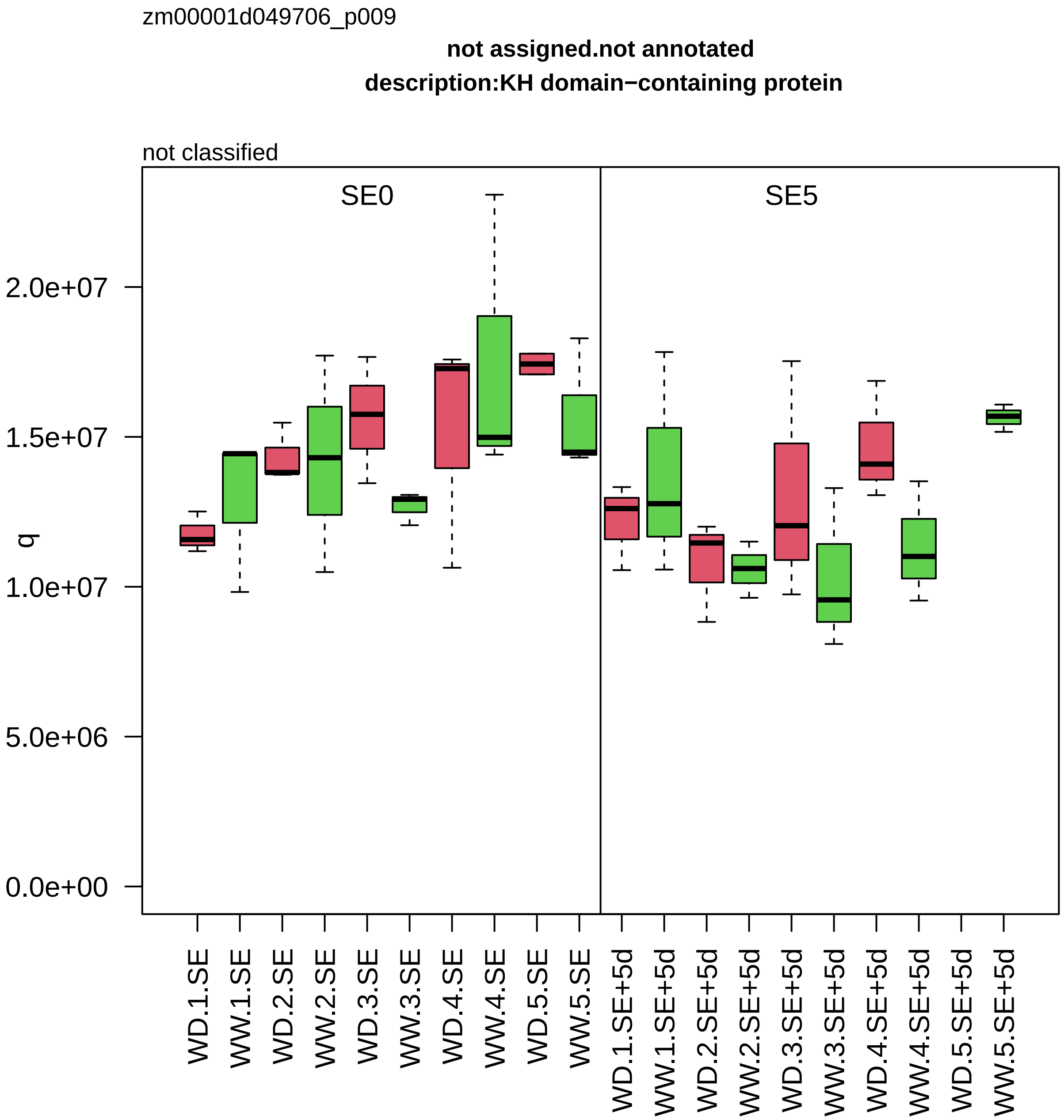

zm00001d049706_p009
not assigned.not annotated
description:KH domain−containing protein
not classified
SE0
SE5
q
2.0e+07
1.5e+07
1.0e+07
5.0e+06
0.0e+00
WD.1.SE
WW.1.SE
WD.2.SE
WW.2.SE
WD.3.SE
WW.3.SE
WD.4.SE
WW.4.SE
WD.5.SE
WW.5.SE
WD.1.SE+5d
WW.1.SE+5d
WD.2.SE+5d
WW.2.SE+5d
WD.3.SE+5d
WW.3.SE+5d
WD.4.SE+5d
WW.4.SE+5d
WD.5.SE+5d
WW.5.SE+5d

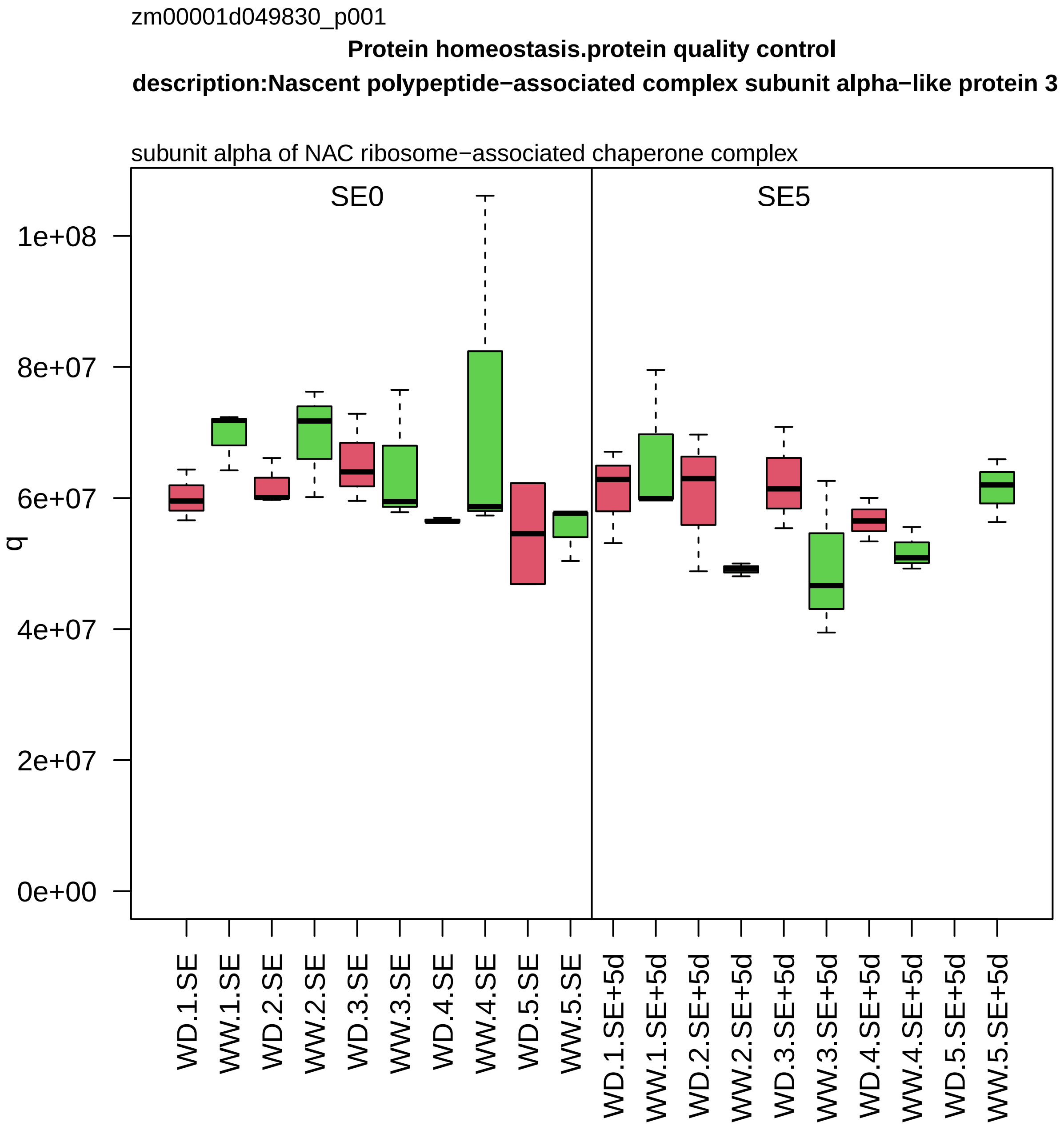
zm00001d049830_p001
Protein homeostasis.protein quality control
description:Nascent polypeptide−associated complex subunit alpha−like protein 3
subunit alpha of NAC ribosome−associated chaperone complex
SE0
SE5
q
1e+08
8e+07
6e+07
4e+07
2e+07
0e+00
WD.1.SE
WW.1.SE
WD.2.SE
WW.2.SE
WD.3.SE
WW.3.SE
WD.4.SE
WW.4.SE
WD.5.SE
WW.5.SE
WD.1.SE+5d
WW.1.SE+5d
WD.2.SE+5d
WW.2.SE+5d
WD.3.SE+5d
WW.3.SE+5d
WD.4.SE+5d
WW.4.SE+5d
WD.5.SE+5d
WW.5.SE+5d

zm00001d049956_p002

**Amino acid metabolism.aspartate group amino acid biosynthesis**

**description:4−hydroxy−tetrahydrodipicolinate reductase 2 chloroplastic**

hydroxy−tetrahydrodihydrodipicolinate reductase

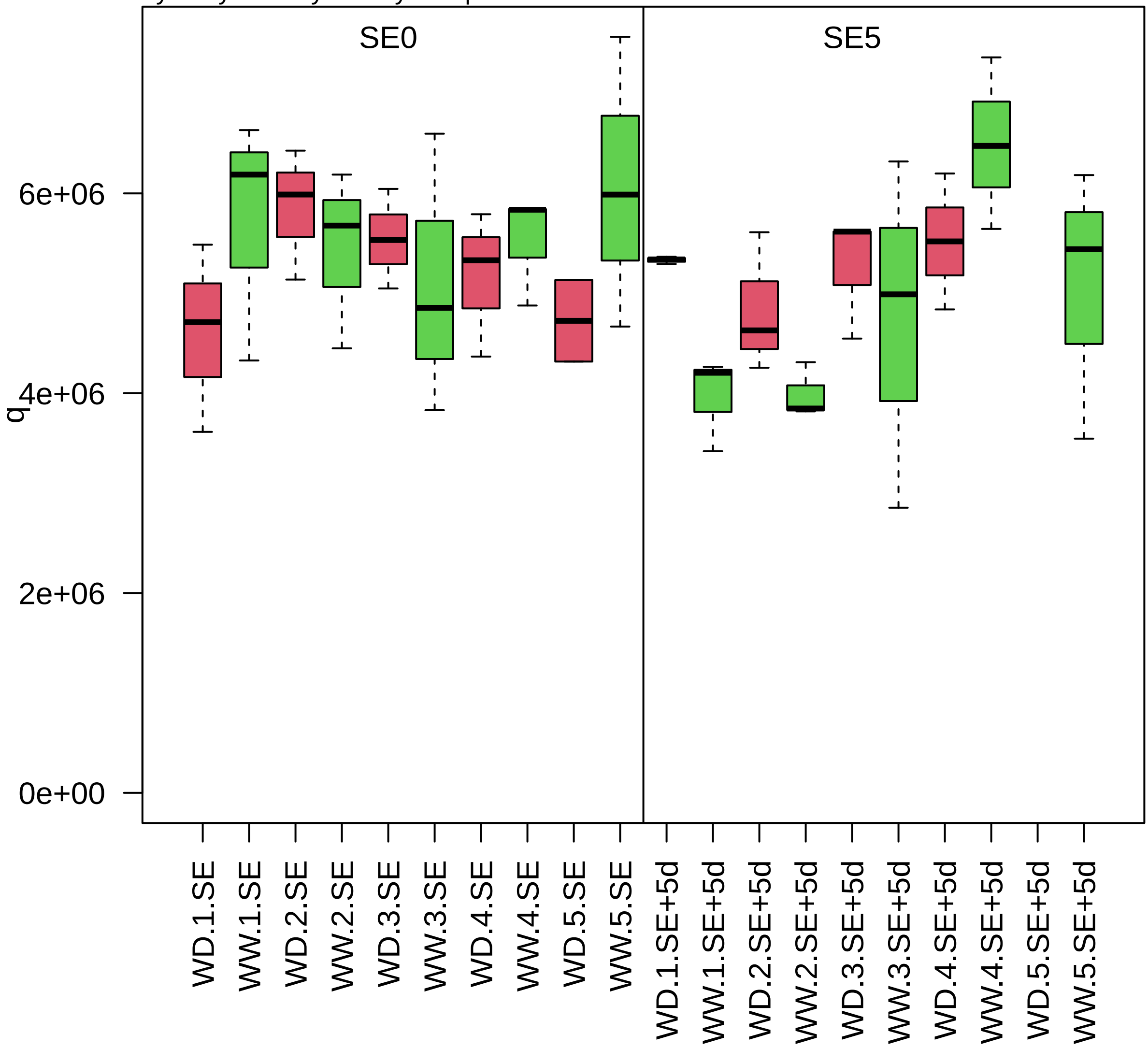

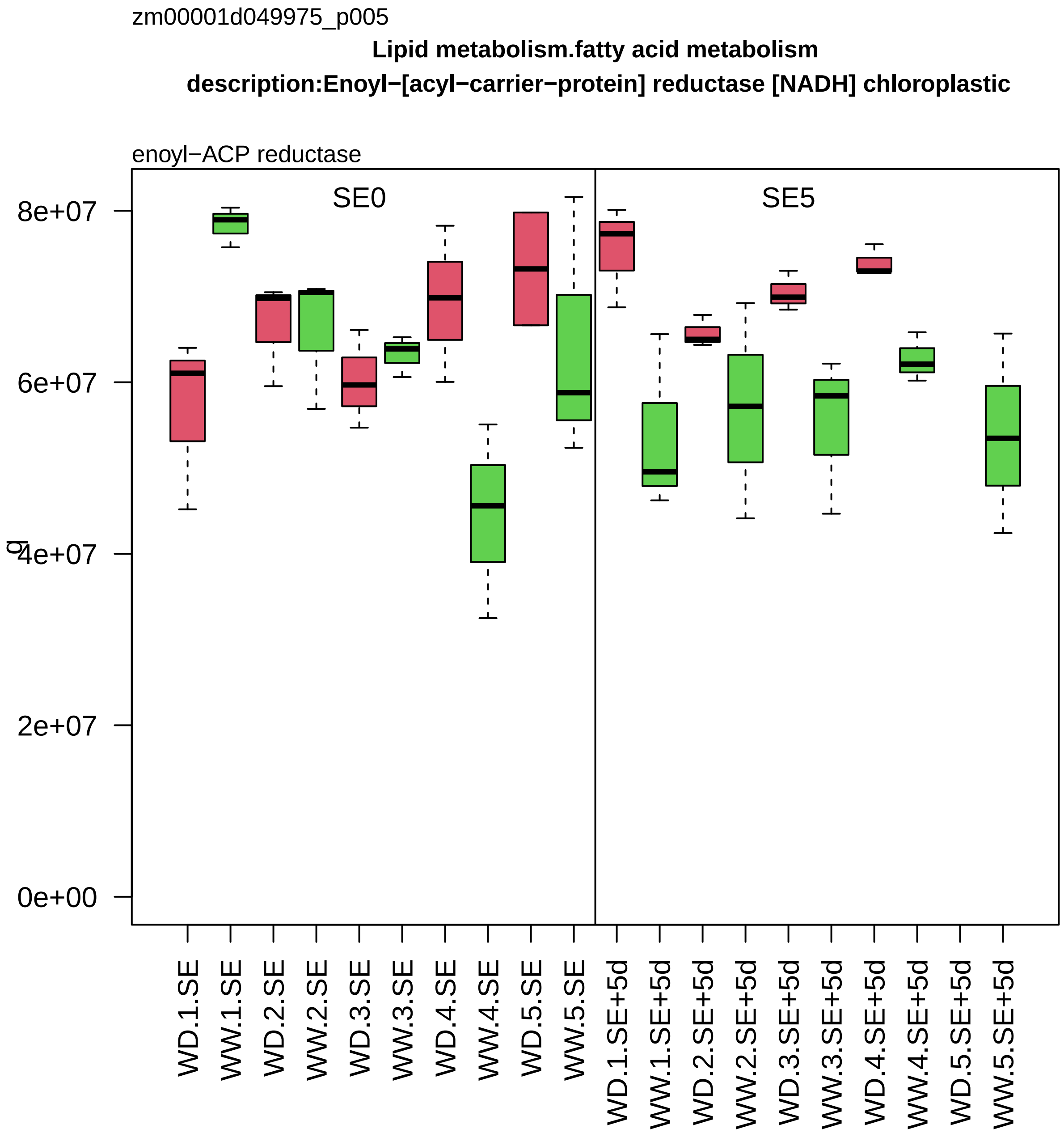
zm00001d049975_p005
Lipid metabolism.fatty acid metabolism
description:Enoyl−[acyl−carrier−protein] reductase [NADH] chloroplastic
enoyl−ACP reductase
SE0
SE5
8e+07
6e+07
4e+07
2e+07
0e+00
g
WD.1.SE
WW.1.SE
WD.2.SE
WW.2.SE
WD.3.SE
WW.3.SE
WD.4.SE
WW.4.SE
WD.5.SE
WW.5.SE
WD.1.SE+5d
WW.1.SE+5d
WD.2.SE+5d
WW.2.SE+5d
WD.3.SE+5d
WW.3.SE+5d
WD.4.SE+5d
WW.4.SE+5d
WD.5.SE+5d
WW.5.SE+5d

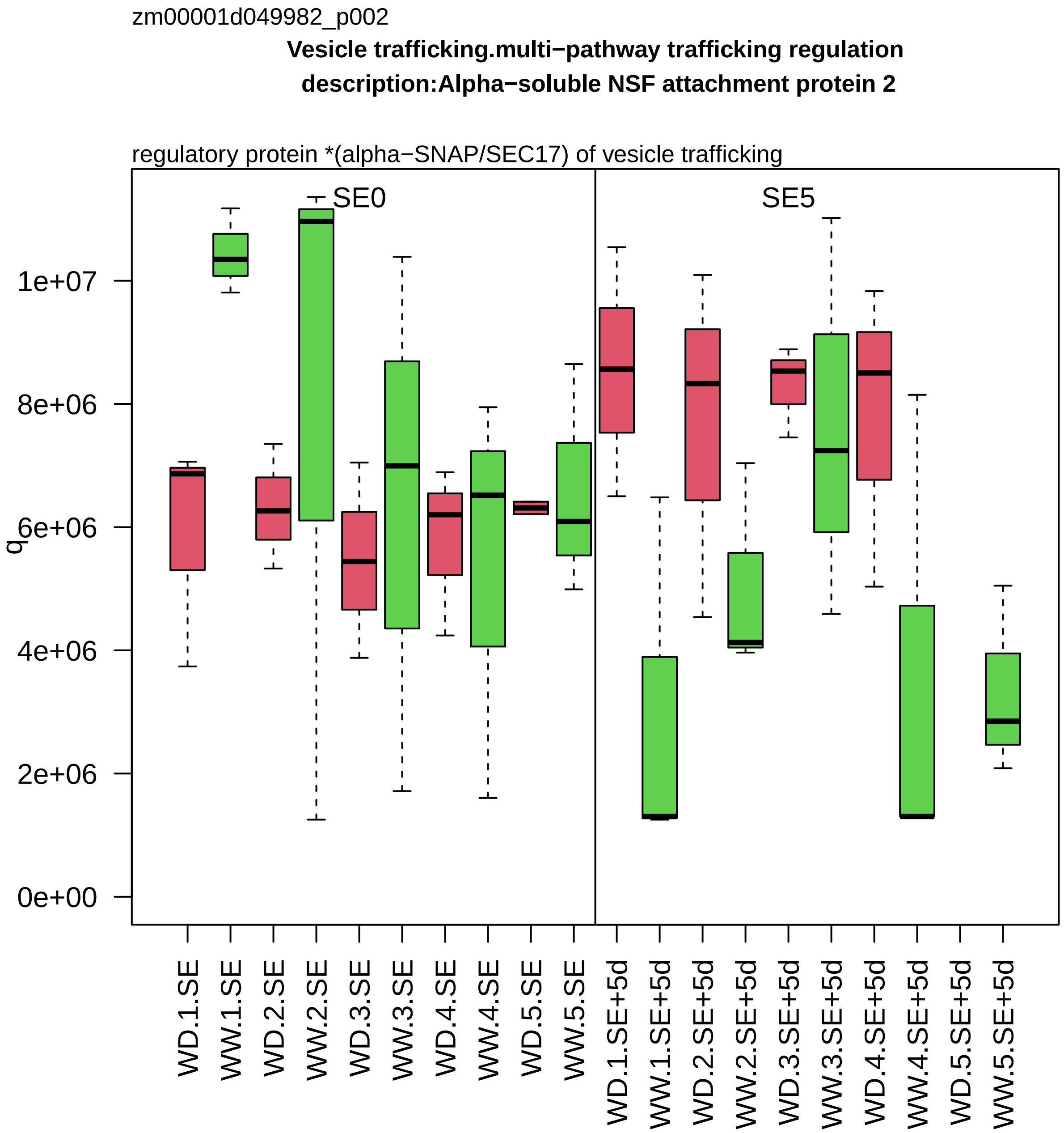

zm00001d049982_p002
Vesicle trafficking.multi−pathway trafficking regulation
description:Alpha−soluble NSF attachment protein 2
regulatory protein *(alpha−SNAP/SEC17) of vesicle trafficking
SE0
SE5
q
0e+00
2e+06
4e+06
6e+06
8e+06
1e+07
WD.1.SE
WW.1.SE
WD.2.SE
WW.2.SE
WD.3.SE
WW.3.SE
WD.4.SE
WW.4.SE
WD.5.SE
WW.5.SE
WD.1.SE+5d
WW.1.SE+5d
WD.2.SE+5d
WW.2.SE+5d
WD.3.SE+5d
WW.3.SE+5d
WD.4.SE+5d
WW.4.SE+5d
WD.5.SE+5d
WW.5.SE+5d

zm00001d050134_p001

**Amino acid metabolism.aspartate group amino acid biosynthesis**

**l:aspartate kinase−homoserine dehydrogenase1 description:Bifunctional aspartokinase/homoserine dehy**

bifunctional homoserine dehydrogenase and aspartate kinase

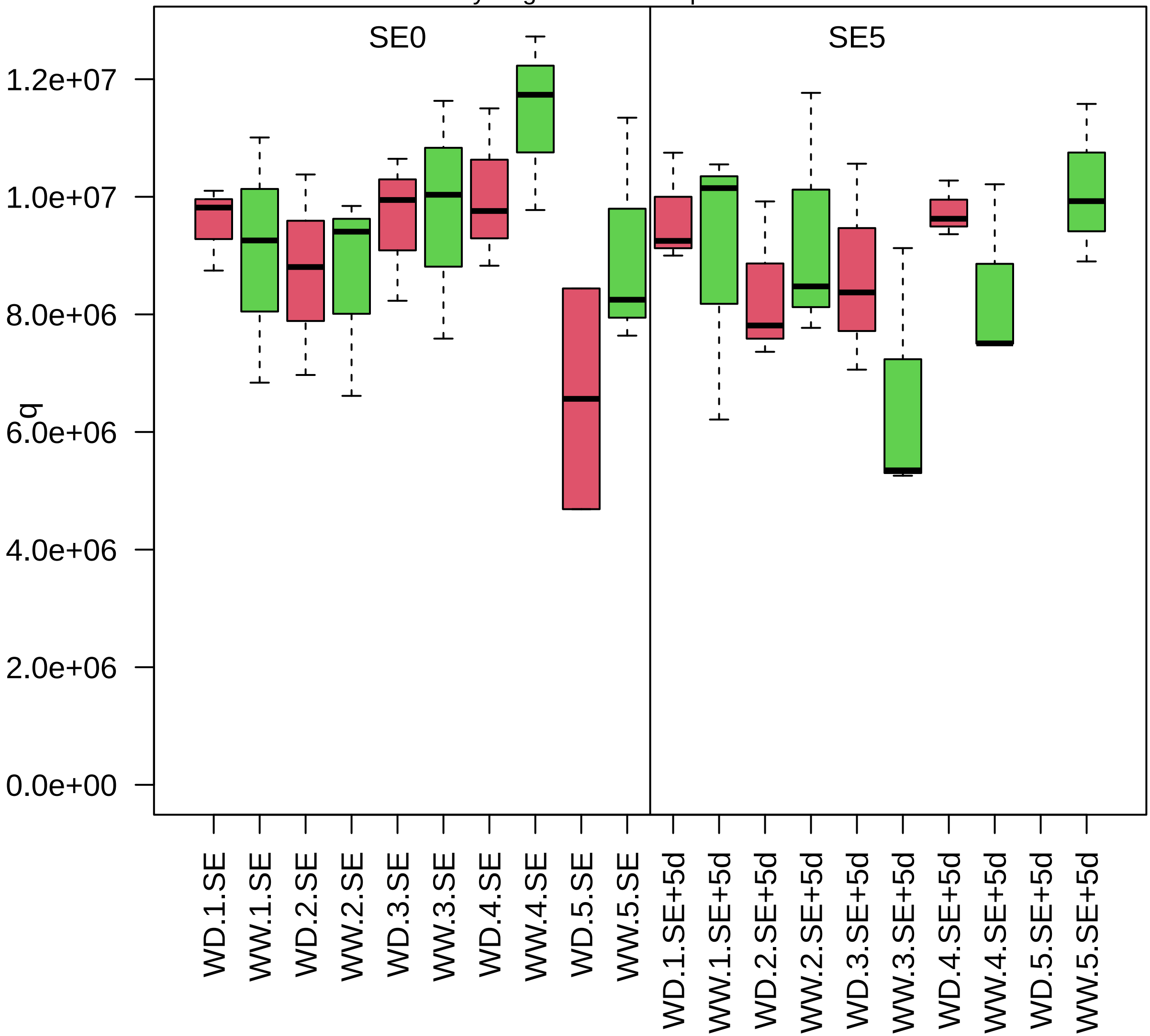

zm00001d050196_p001

**not assigned.not annotated**

**gene_symbol:endo−1,3−1,4−beta−D−glucanase1 description:alpha/beta−Hydrolases superfamily pr**

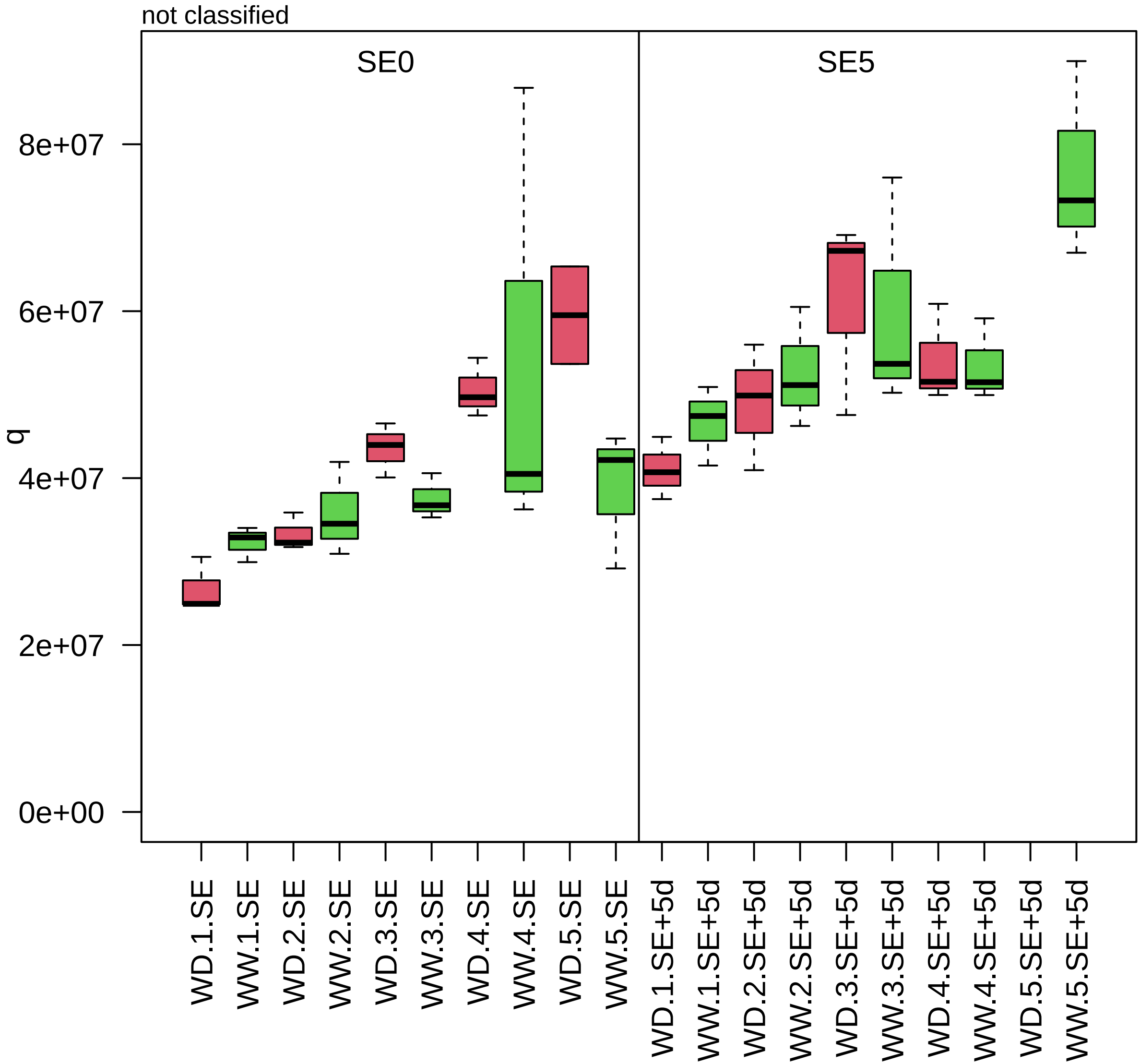

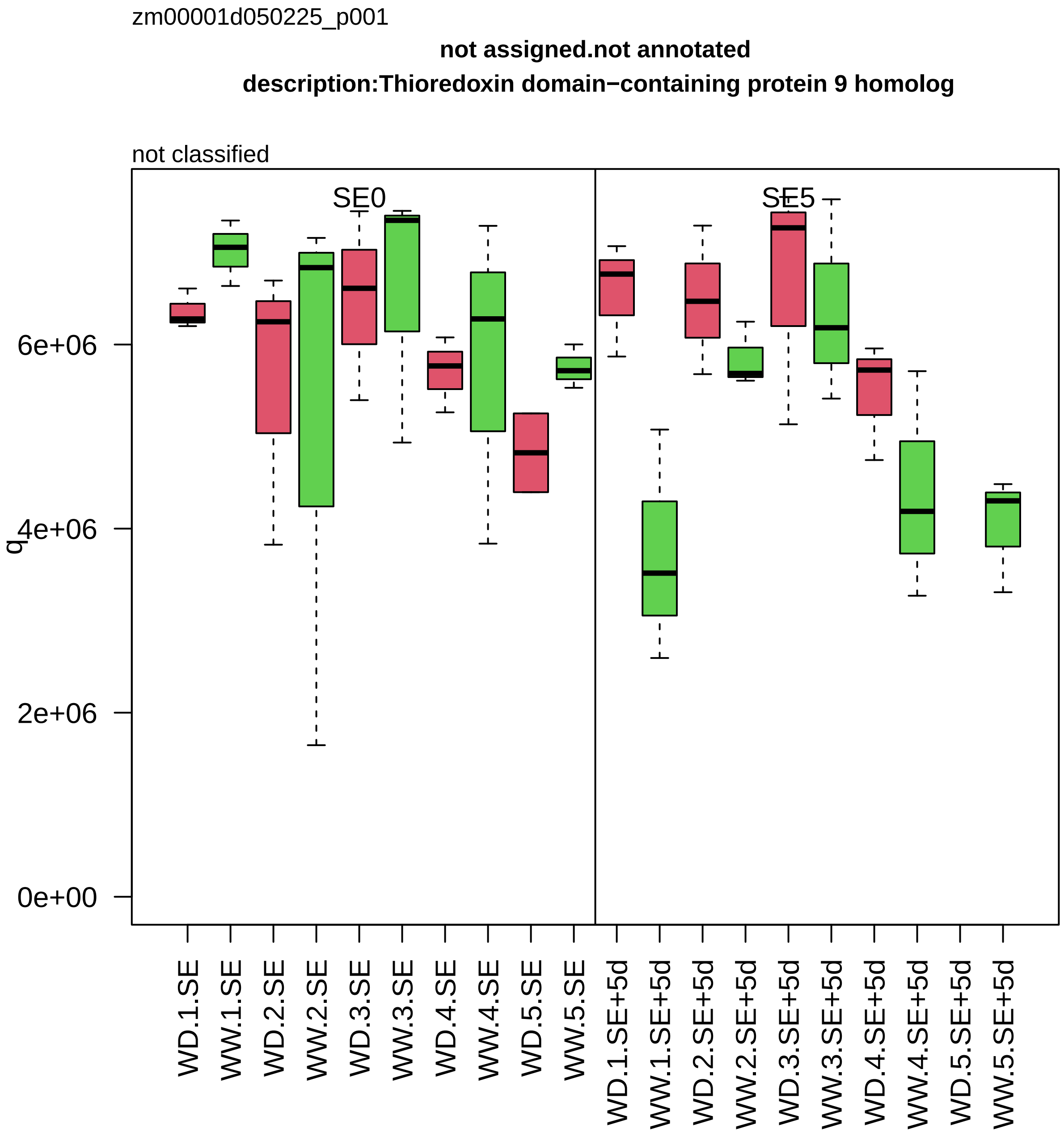
zm00001d050225_p001
not assigned.not annotated
description:Thioredoxin domain−containing protein 9 homolog
not classified
SE0
SE5
q
0e+00
2e+06
4e+06
6e+06
WD.1.SE
WW.1.SE
WD.2.SE
WW.2.SE
WD.3.SE
WW.3.SE
WD.4.SE
WW.4.SE
WD.5.SE
WW.5.SE
WD.1.SE+5d
WW.1.SE+5d
WD.2.SE+5d
WW.2.SE+5d
WD.3.SE+5d
WW.3.SE+5d
WD.4.SE+5d
WW.4.SE+5d
WD.5.SE+5d
WW.5.SE+5d

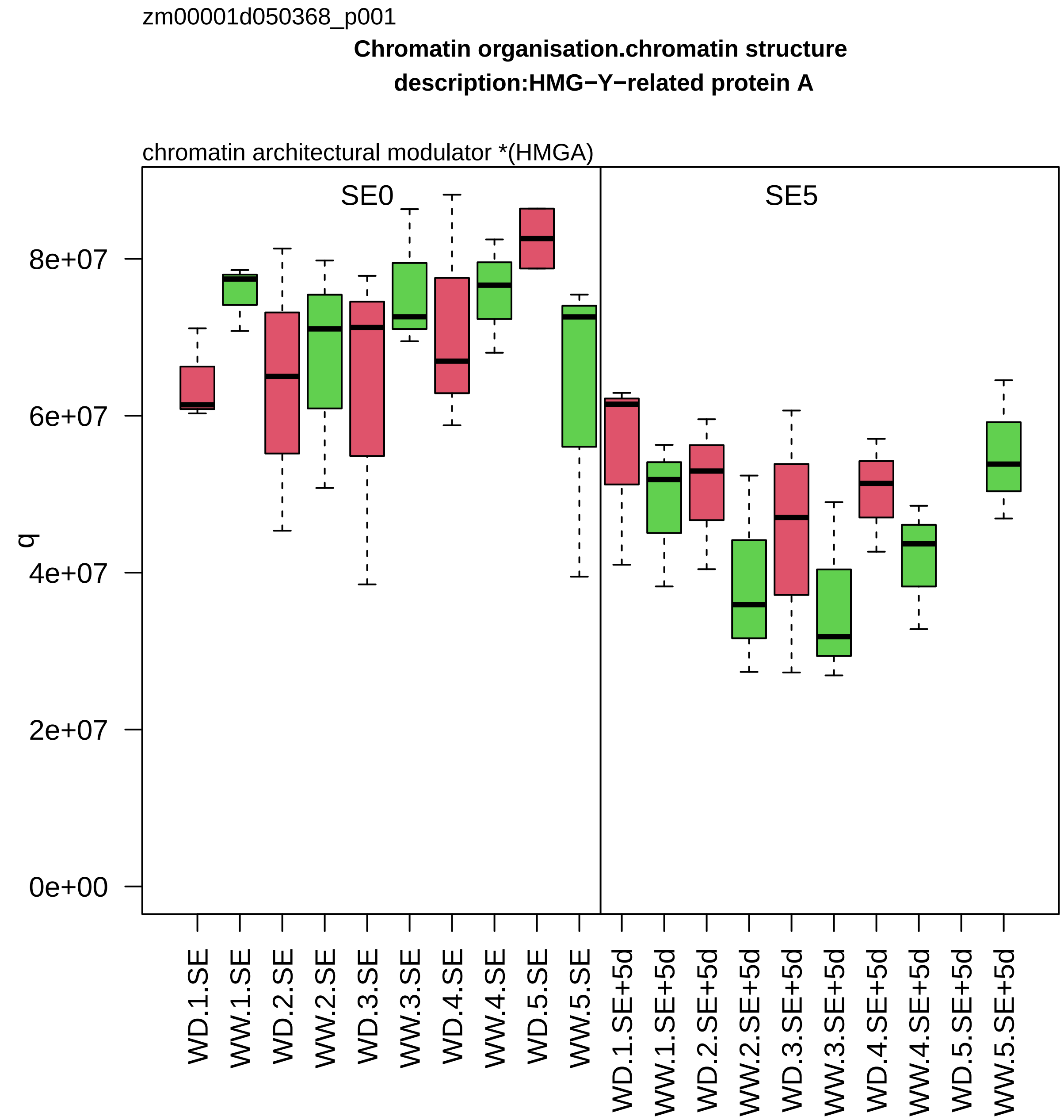

zm00001d050368_p001
Chromatin organisation.chromatin structure
description:HMG−Y−related protein A
chromatin architectural modulator *(HMGA)
SE0
SE5
q
0e+00
2e+07
4e+07
6e+07
8e+07
WD.1.SE
WW.1.SE
WD.2.SE
WW.2.SE
WD.3.SE
WW.3.SE
WD.4.SE
WW.4.SE
WD.5.SE
WW.5.SE
WD.1.SE+5d
WW.1.SE+5d
WD.2.SE+5d
WW.2.SE+5d
WD.3.SE+5d
WW.3.SE+5d
WD.4.SE+5d
WW.4.SE+5d
WD.5.SE+5d
WW.5.SE+5d

zm00001d050384_p001

**Cellular respiration.oxidative phosphorylation**

component *(NDUFB1/MNLL) of NADH dehydrogenase beta subcomplex

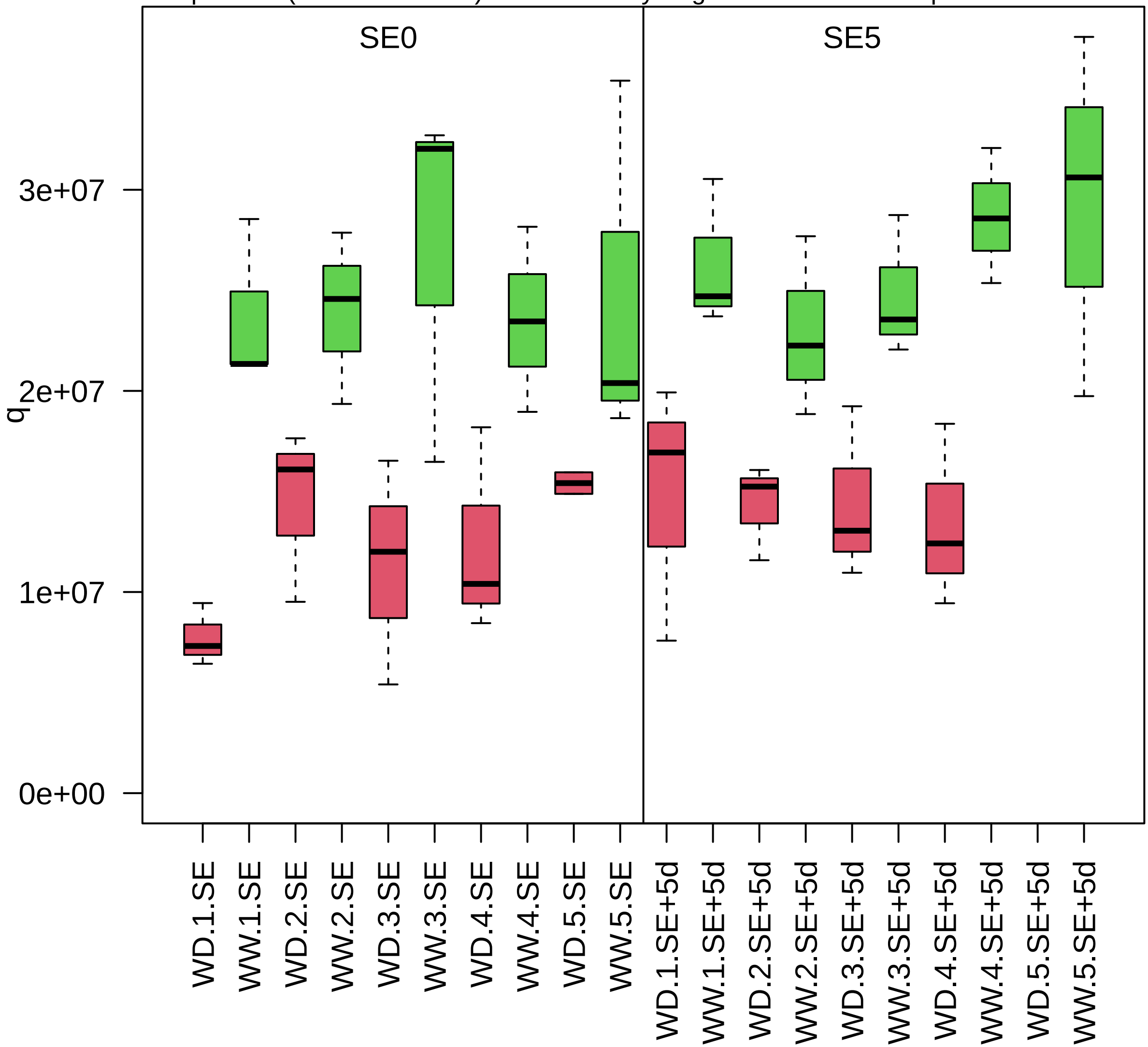

zm00001d050606_p004

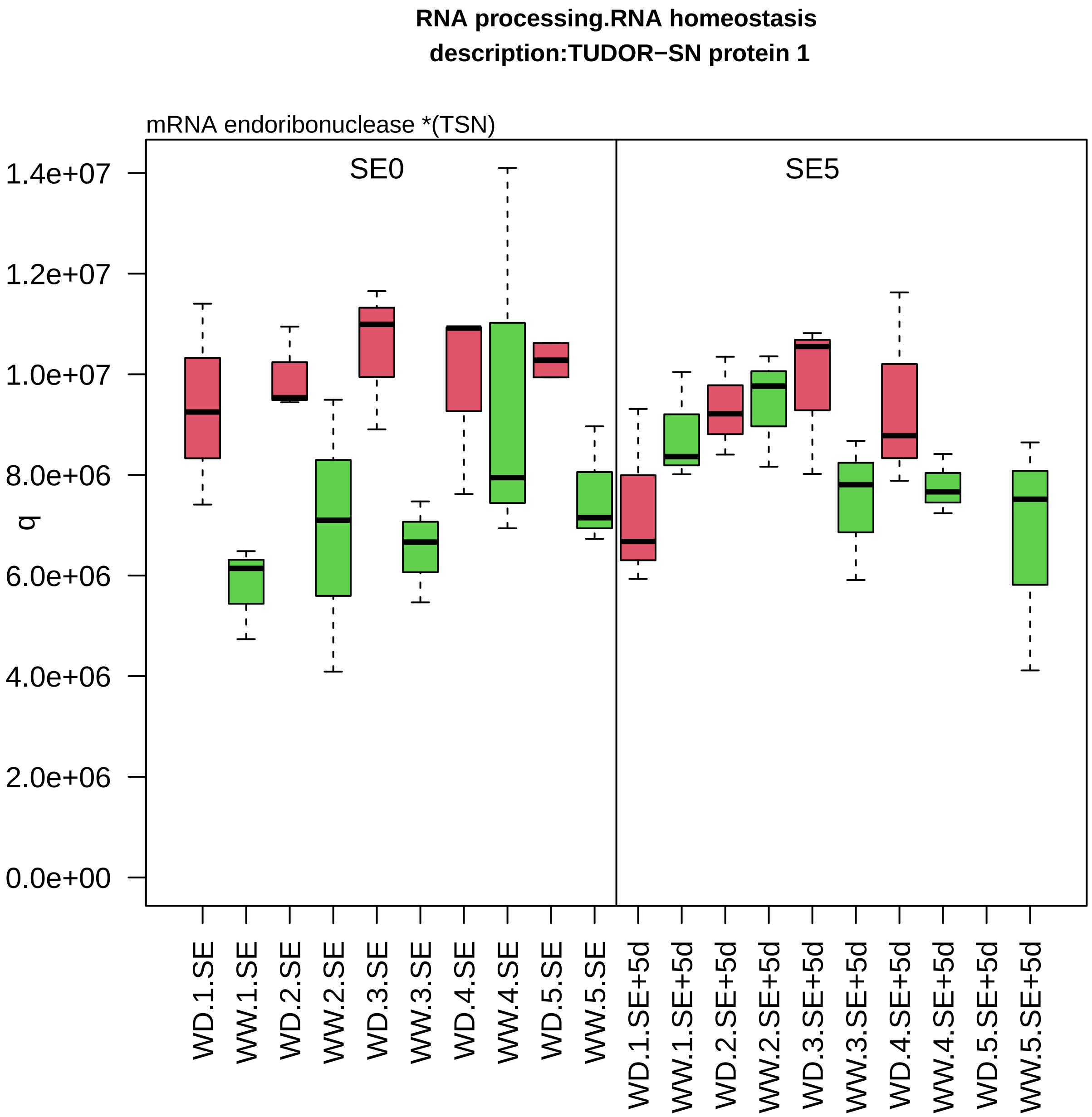

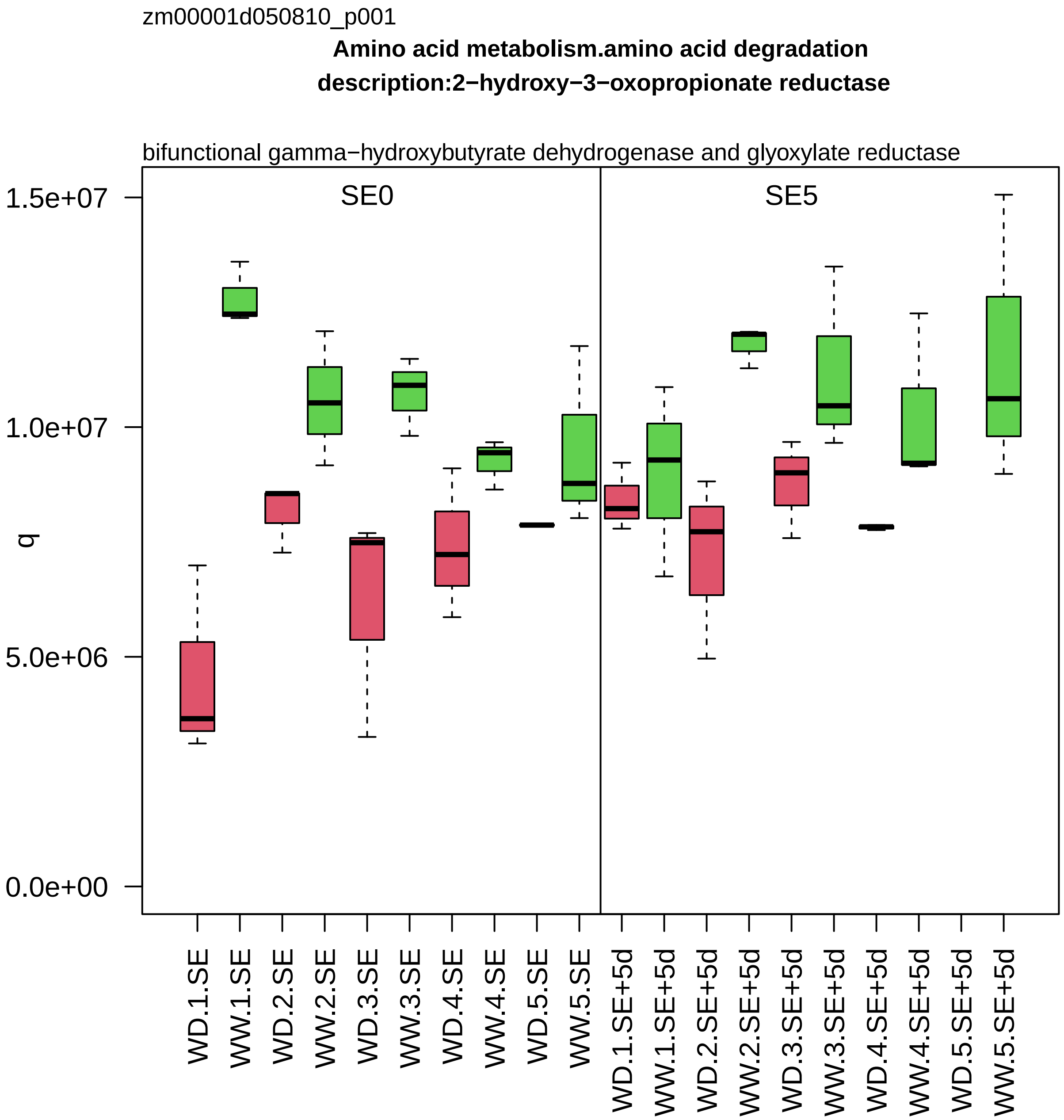
zm00001d050810_p001
Amino acid metabolism.amino acid degradation
description:2−hydroxy−3−oxopropionate reductase
bifunctional gamma−hydroxybutyrate dehydrogenase and glyoxylate reductase
SE0
SE5
q
1.5e+07
1.0e+07
5.0e+06
0.0e+00
WD.1.SE
WW.1.SE
WD.2.SE
WW.2.SE
WD.3.SE
WW.3.SE
WD.4.SE
WW.4.SE
WD.5.SE
WW.5.SE
WD.1.SE+5d
WW.1.SE+5d
WD.2.SE+5d
WW.2.SE+5d
WD.3.SE+5d
WW.3.SE+5d
WD.4.SE+5d
WW.4.SE+5d
WD.5.SE+5d
WW.5.SE+5d

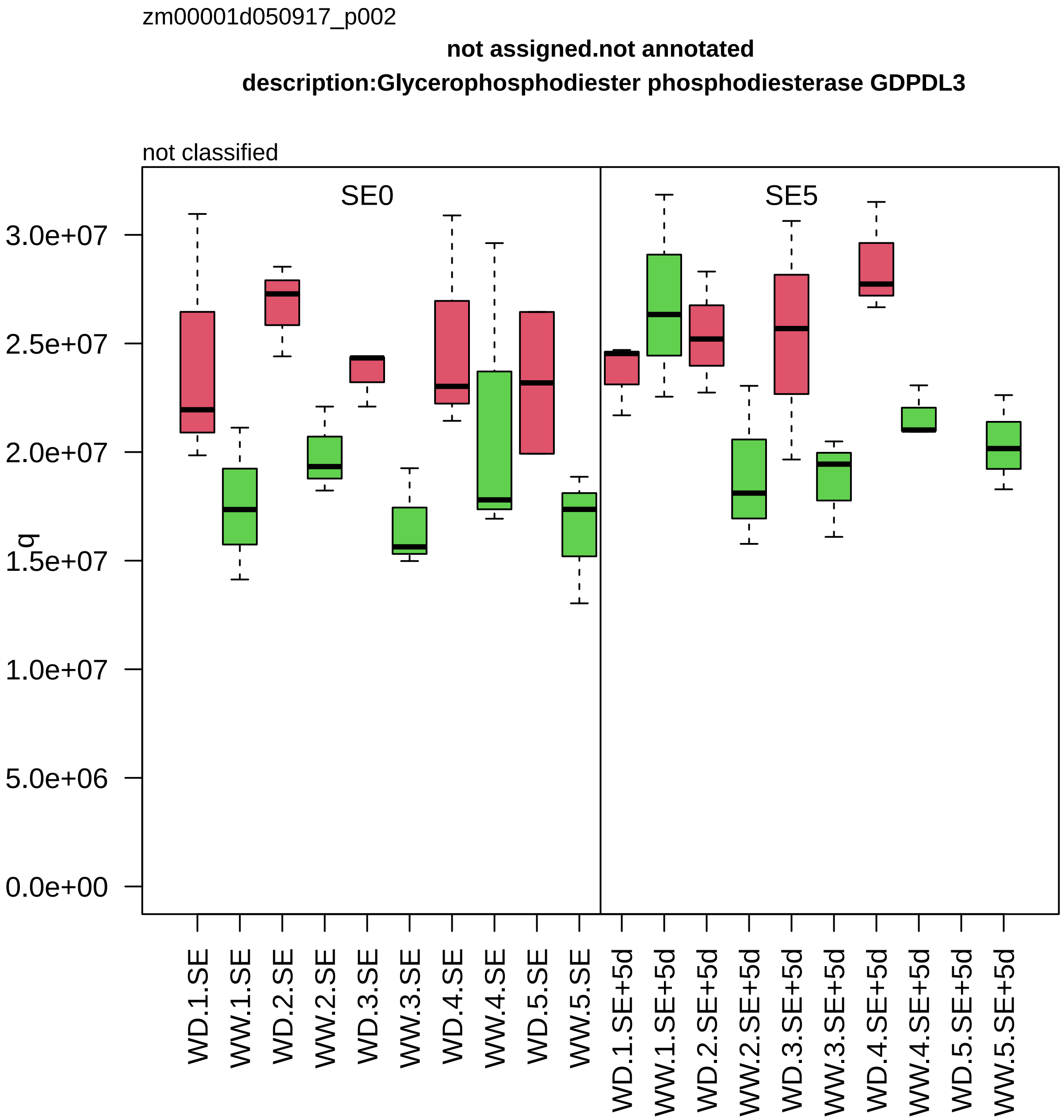

zm00001d050917_p002
not assigned.not annotated
description:Glycerophosphodiester phosphodiesterase GDPDL3
not classified
SE0
SE5
q
3.0e+07
2.5e+07
2.0e+07
1.5e+07
1.0e+07
5.0e+06
0.0e+00
WD.1.SE
WW.1.SE
WD.2.SE
WW.2.SE
WD.3.SE
WW.3.SE
WD.4.SE
WW.4.SE
WD.5.SE
WW.5.SE
WD.1.SE+5d
WW.1.SE+5d
WD.2.SE+5d
WW.2.SE+5d
WD.3.SE+5d
WW.3.SE+5d
WD.4.SE+5d
WW.4.SE+5d
WD.5.SE+5d
WW.5.SE+5d

zm00001d050920_p002

**Protein biosynthesis.ribosome biogenesis**
**description:60S ribosomal protein L6**

component *(eL6) of large ribosomal−subunit (LSU) proteome

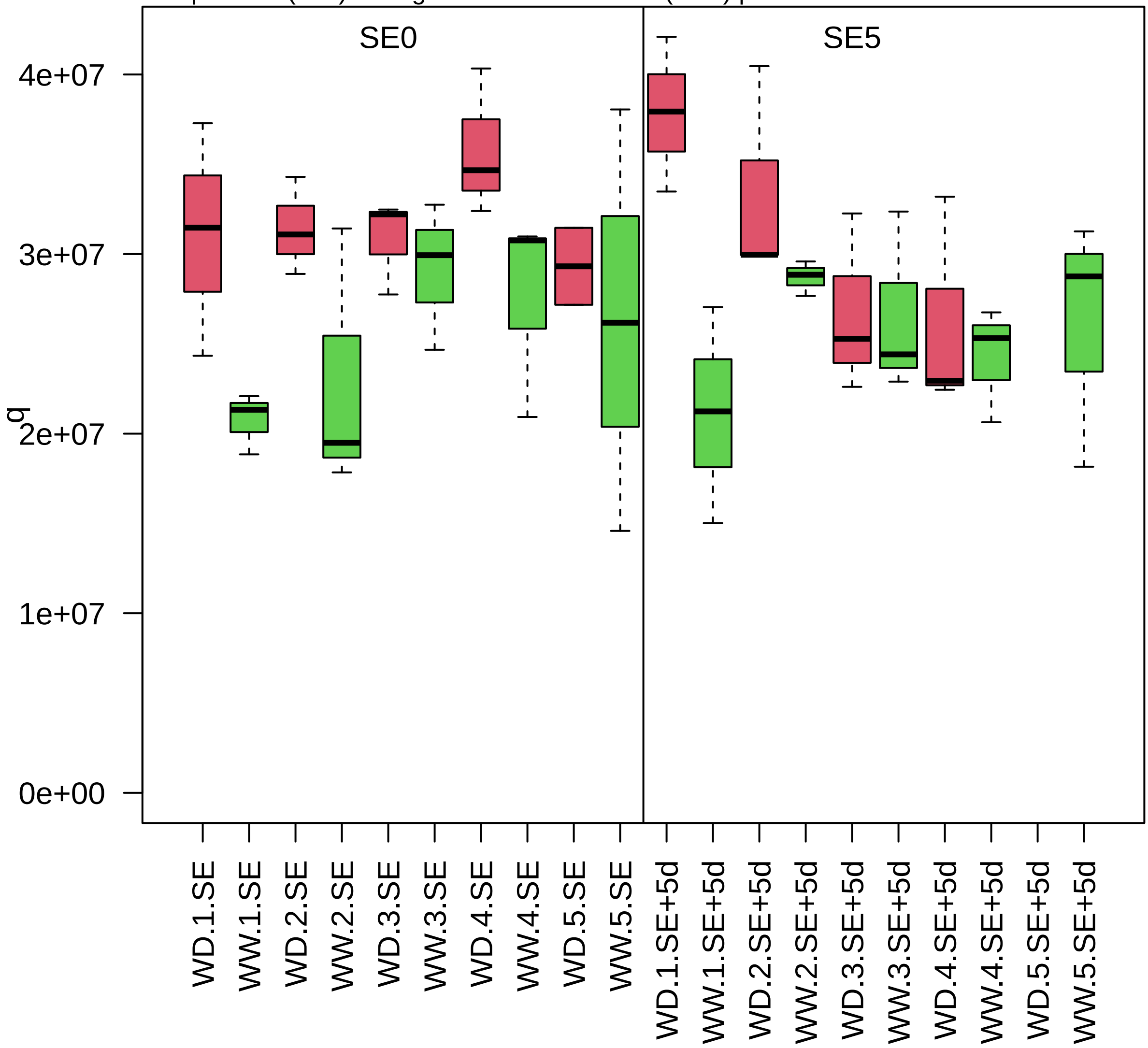

zm00001d051001_p001

**Cellular respiration.glycolysis**

**ymbol:glyceraldehyde−3−phosphate dehydrogenase3 description:Glyceraldehyde−3−phosphate dehydroge**

NAD−dependent glyceraldehyde 3−phosphate dehydrogenase

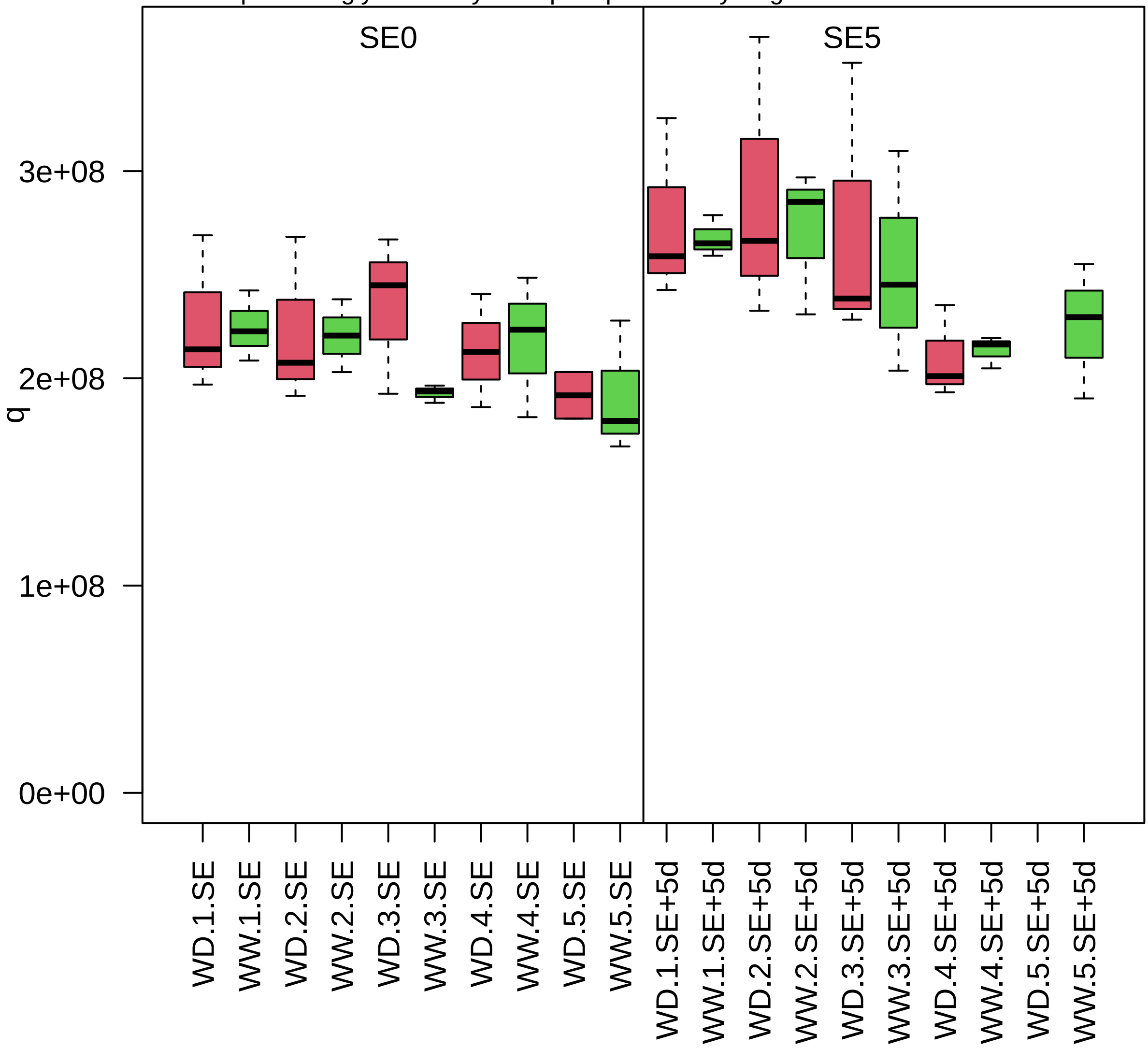

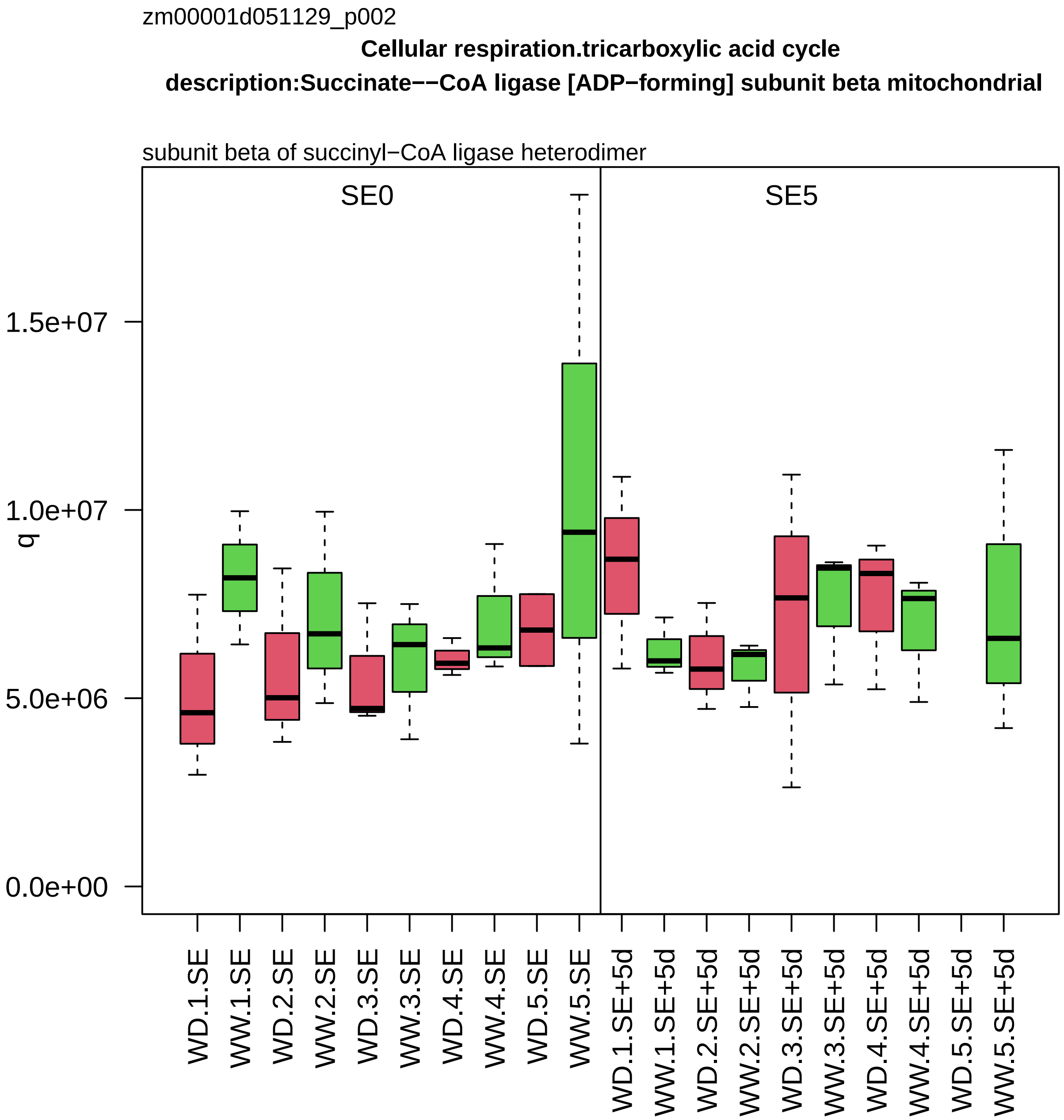

zm00001d051129_p002
Cellular respiration.tricarboxylic acid cycle
description:Succinate−−CoA ligase [ADP−forming] subunit beta mitochondrial
subunit beta of succinyl−CoA ligase heterodimer
SE0
SE5
q
1.5e+07
1.0e+07
5.0e+06
0.0e+00
WD.1.SE
WW.1.SE
WD.2.SE
WW.2.SE
WD.3.SE
WW.3.SE
WD.4.SE
WW.4.SE
WD.5.SE
WW.5.SE
WD.1.SE+5d
WW.1.SE+5d
WD.2.SE+5d
WW.2.SE+5d
WD.3.SE+5d
WW.3.SE+5d
WD.4.SE+5d
WW.4.SE+5d
WD.5.SE+5d
WW.5.SE+5d

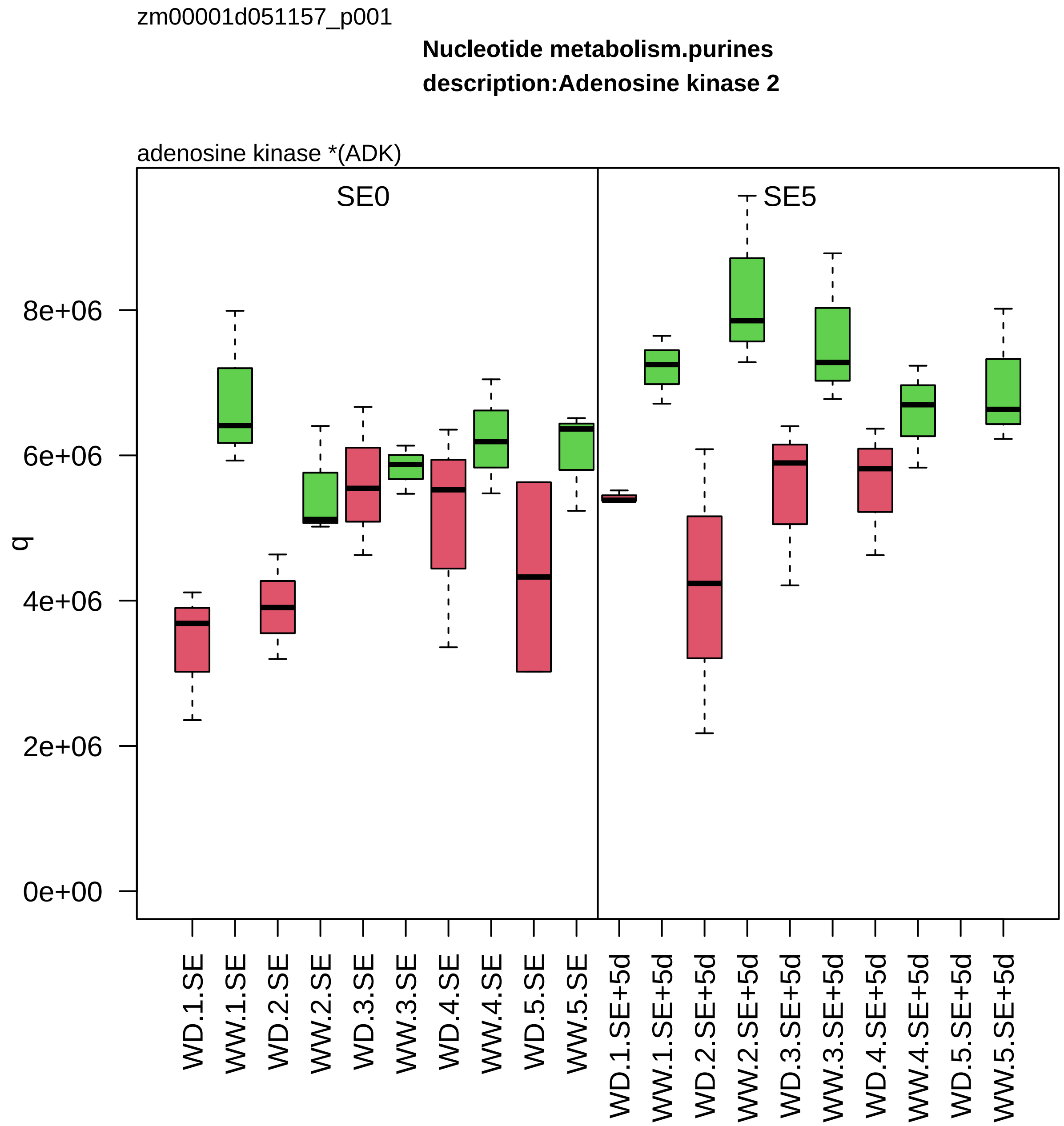
zm00001d051157_p001
Nucleotide metabolism.purines
description:Adenosine kinase 2
adenosine kinase *(ADK)
SE0
SE5
q
8e+06
6e+06
4e+06
2e+06
0e+00
WD.1.SE
WW.1.SE
WD.2.SE
WW.2.SE
WD.3.SE
WW.3.SE
WD.4.SE
WW.4.SE
WD.5.SE
WW.5.SE
WD.1.SE+5d
WW.1.SE+5d
WD.2.SE+5d
WW.2.SE+5d
WD.3.SE+5d
WW.3.SE+5d
WD.4.SE+5d
WW.4.SE+5d
WD.5.SE+5d
WW.5.SE+5d

zm00001d051268_p003

**RNA processing.RNA homeostasis**

**description:Putative DEAD−box ATP−dependent RNA helicase family protein**

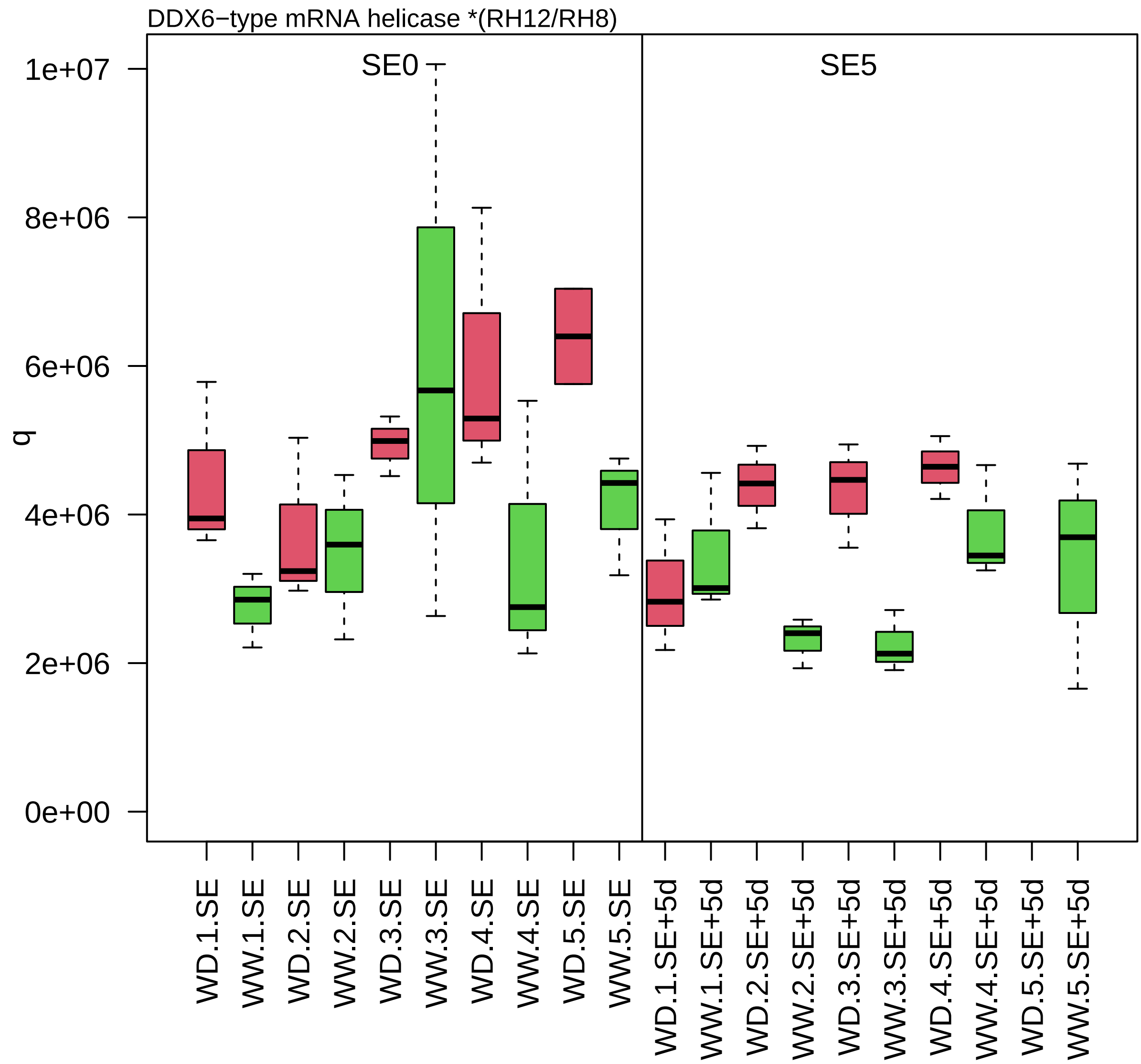

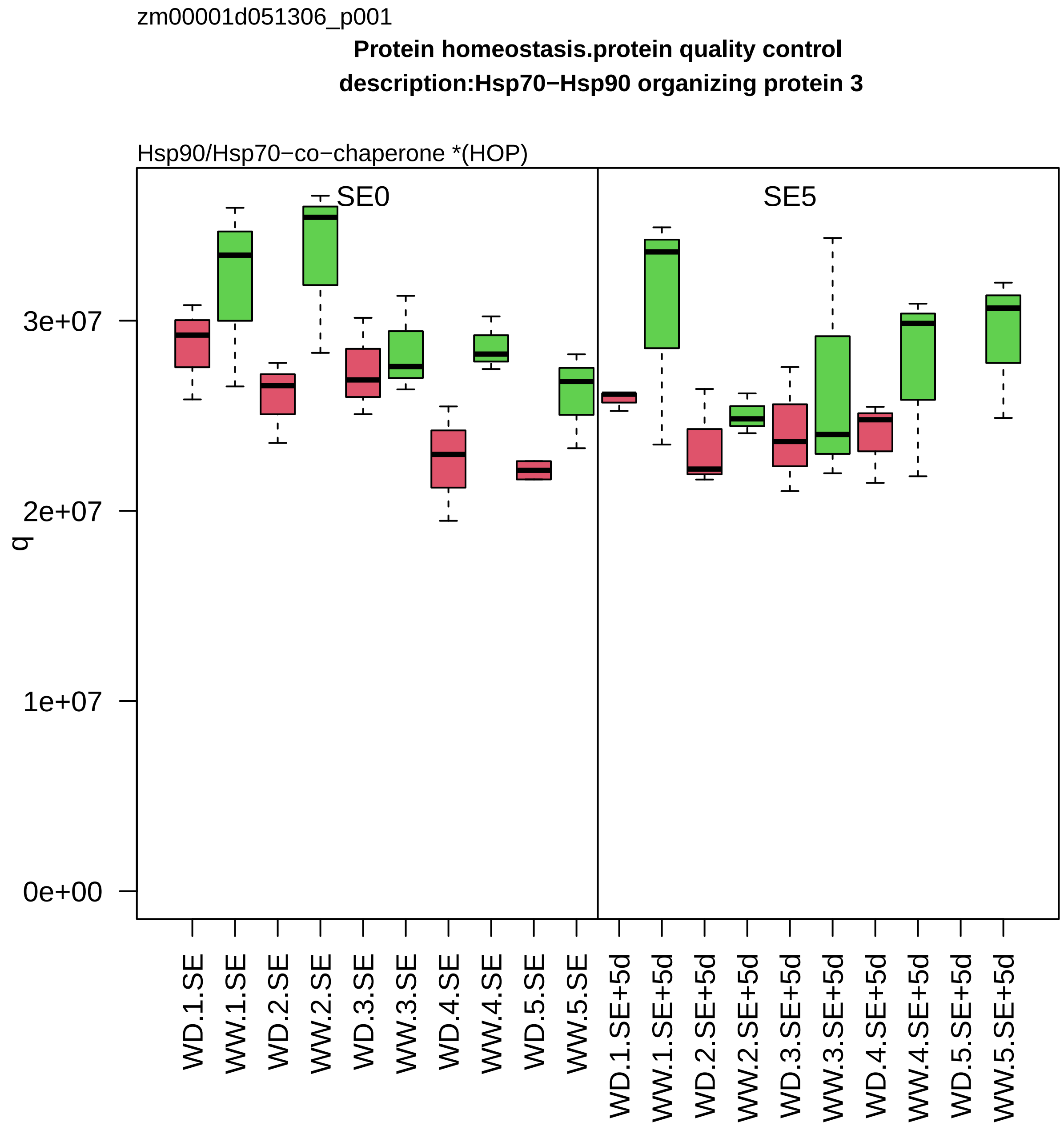
zm00001d051306_p001
Protein homeostasis.protein quality control
description:Hsp70−Hsp90 organizing protein 3
Hsp90/Hsp70−co−chaperone *(HOP)
SE0
SE5
q
0e+00
1e+07
2e+07
3e+07
WD.1.SE
WW.1.SE
WD.2.SE
WW.2.SE
WD.3.SE
WW.3.SE
WD.4.SE
WW.4.SE
WD.5.SE
WW.5.SE
WD.1.SE+5d
WW.1.SE+5d
WD.2.SE+5d
WW.2.SE+5d
WD.3.SE+5d
WW.3.SE+5d
WD.4.SE+5d
WW.4.SE+5d
WD.5.SE+5d
WW.5.SE+5d

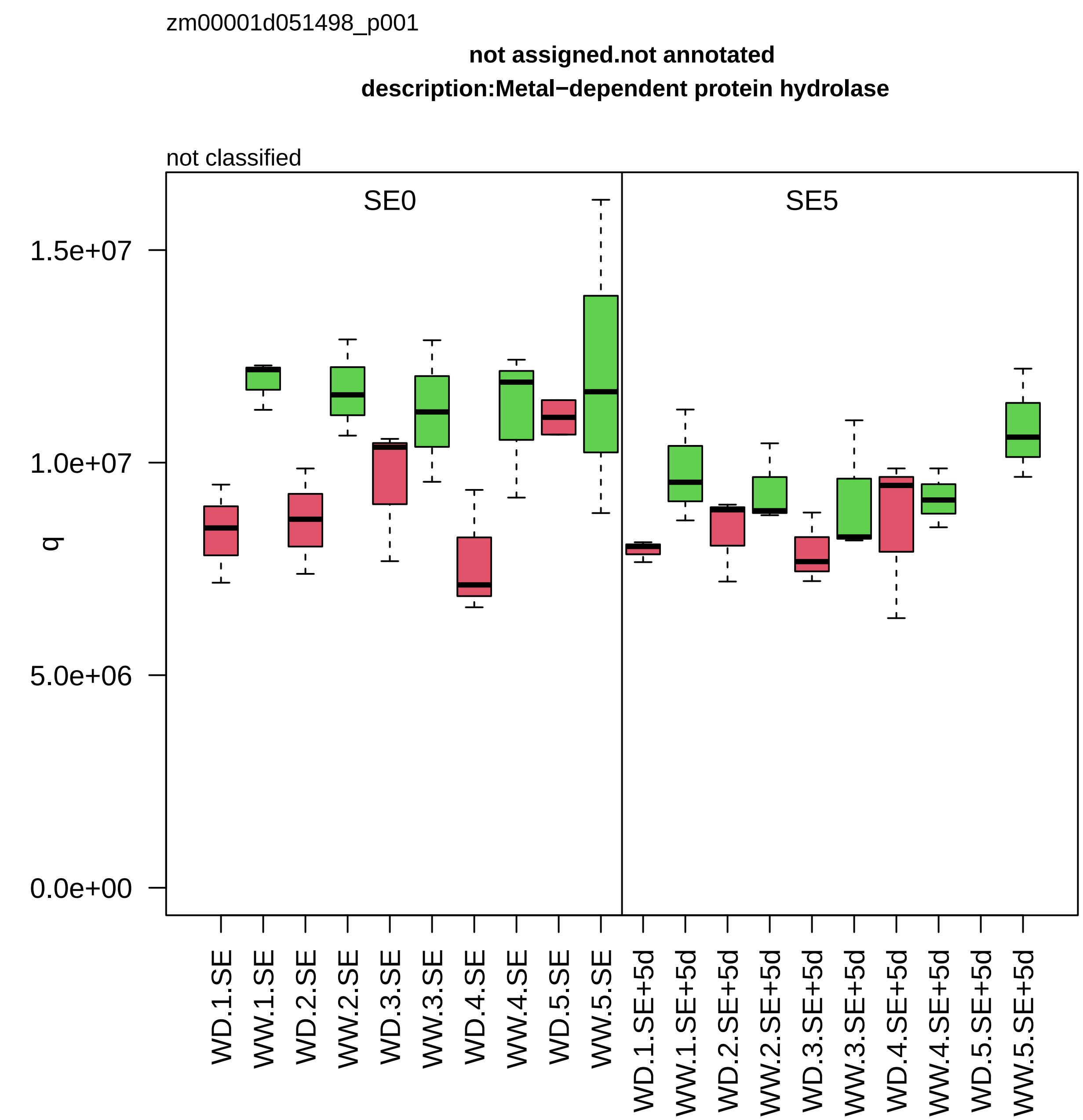
zm00001d051498_p001
not assigned.not annotated
description:Metal−dependent protein hydrolase
not classified
SE0
SE5
q
1.5e+07
1.0e+07
5.0e+06
0.0e+00
WD.1.SE
WW.1.SE
WD.2.SE
WW.2.SE
WD.3.SE
WW.3.SE
WD.4.SE
WW.4.SE
WD.5.SE
WW.5.SE
WD.1.SE+5d
WW.1.SE+5d
WD.2.SE+5d
WW.2.SE+5d
WD.3.SE+5d
WW.3.SE+5d
WD.4.SE+5d
WW.4.SE+5d
WD.5.SE+5d
WW.5.SE+5d

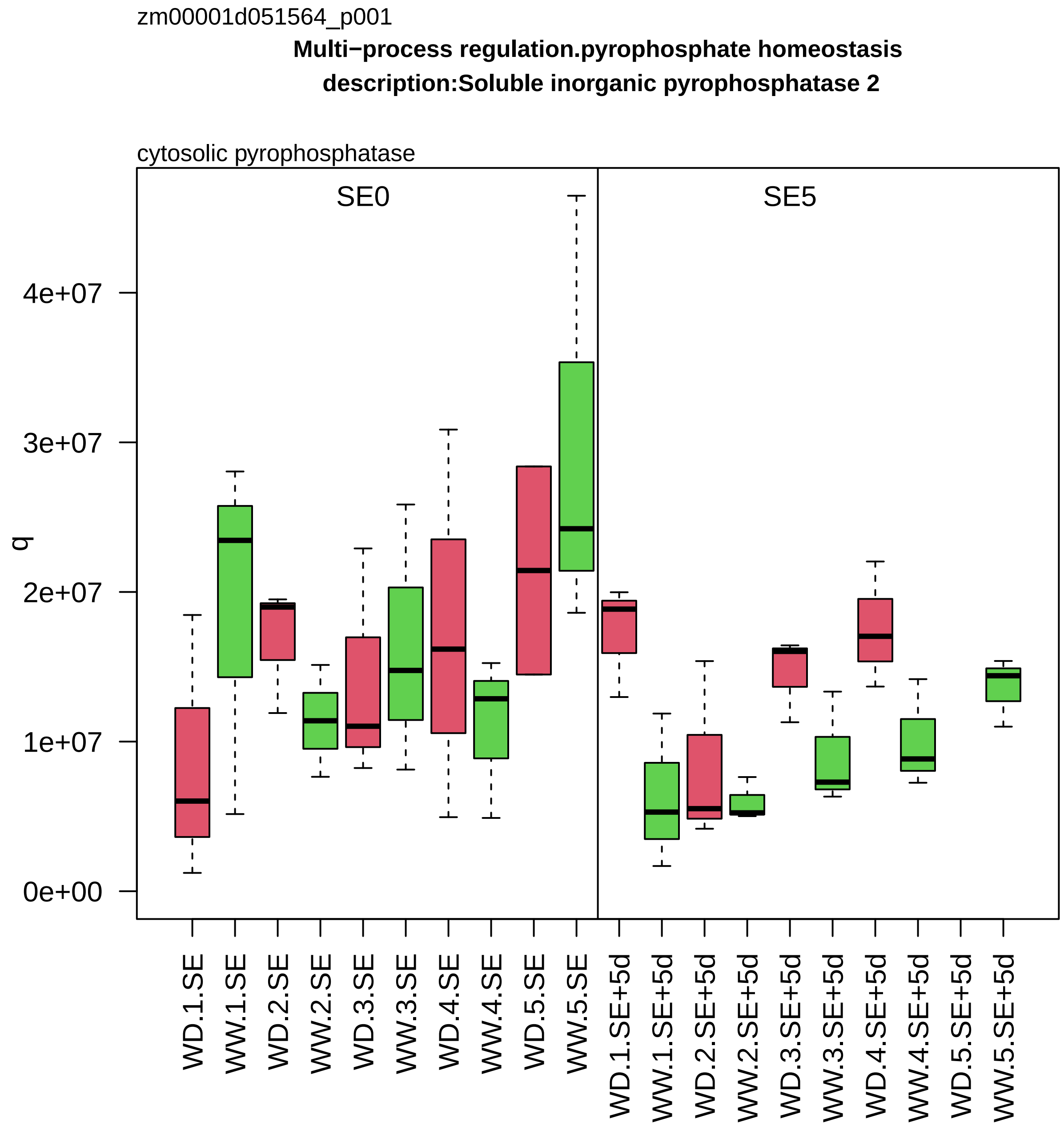

zm00001d051564_p001
Multi−process regulation.pyrophosphate homeostasis
description:Soluble inorganic pyrophosphatase 2
cytosolic pyrophosphatase
SE0
SE5
q
0e+00
1e+07
2e+07
3e+07
4e+07
WD.1.SE
WW.1.SE
WD.2.SE
WW.2.SE
WD.3.SE
WW.3.SE
WD.4.SE
WW.4.SE
WD.5.SE
WW.5.SE
WD.1.SE+5d
WW.1.SE+5d
WD.2.SE+5d
WW.2.SE+5d
WD.3.SE+5d
WW.3.SE+5d
WD.4.SE+5d
WW.4.SE+5d
WD.5.SE+5d
WW.5.SE+5d

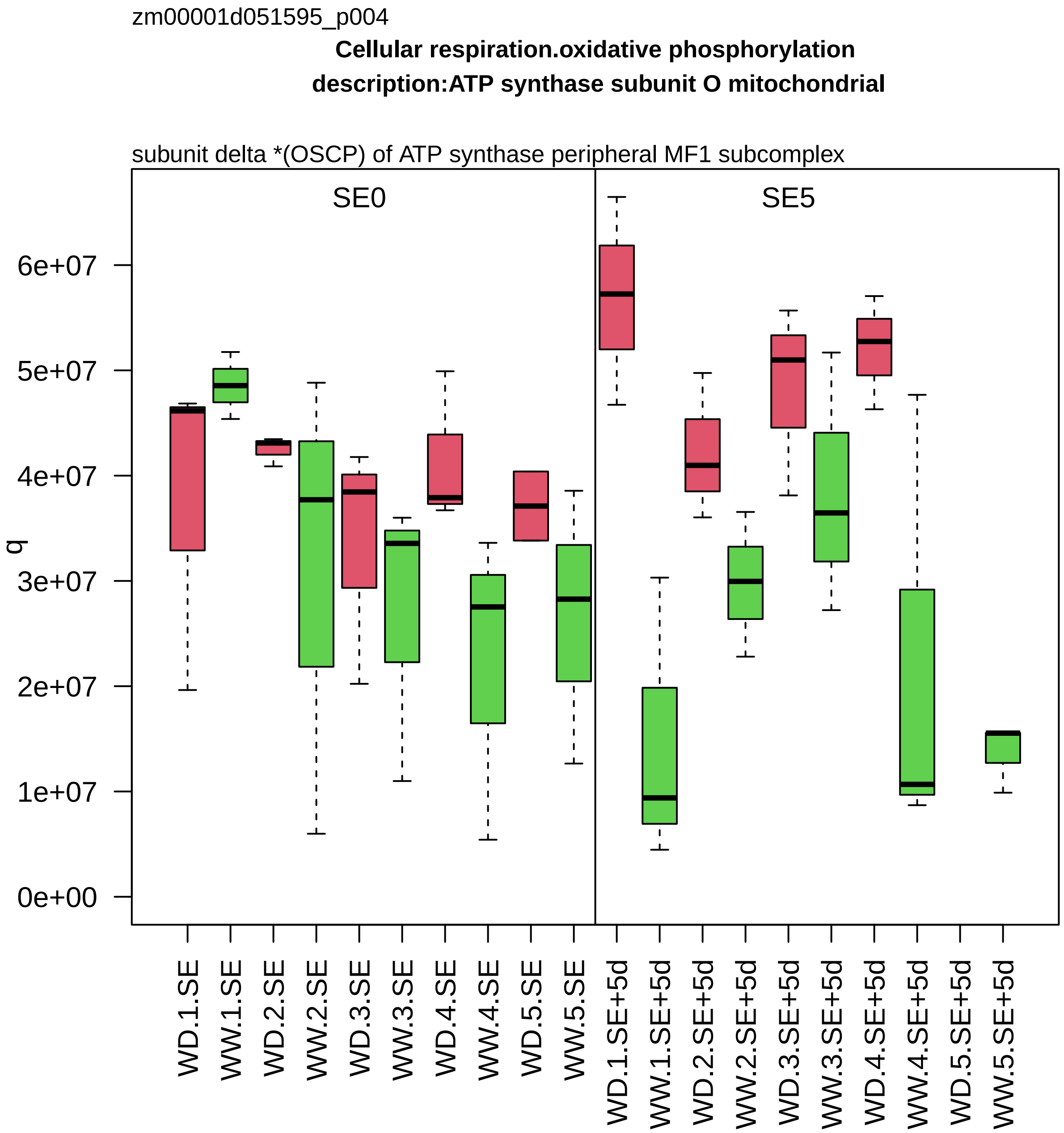

zm00001d051595_p004
Cellular respiration.oxidative phosphorylation
description:ATP synthase subunit O mitochondrial
subunit delta *(OSCP) of ATP synthase peripheral MF1 subcomplex
SE0
SE5
q
6e+07
5e+07
4e+07
3e+07
2e+07
1e+07
0e+00
WD.1.SE
WW.1.SE
WD.2.SE
WW.2.SE
WD.3.SE
WW.3.SE
WD.4.SE
WW.4.SE
WD.5.SE
WW.5.SE
WD.1.SE+5d
WW.1.SE+5d
WD.2.SE+5d
WW.2.SE+5d
WD.3.SE+5d
WW.3.SE+5d
WD.4.SE+5d
WW.4.SE+5d
WD.5.SE+5d
WW.5.SE+5d

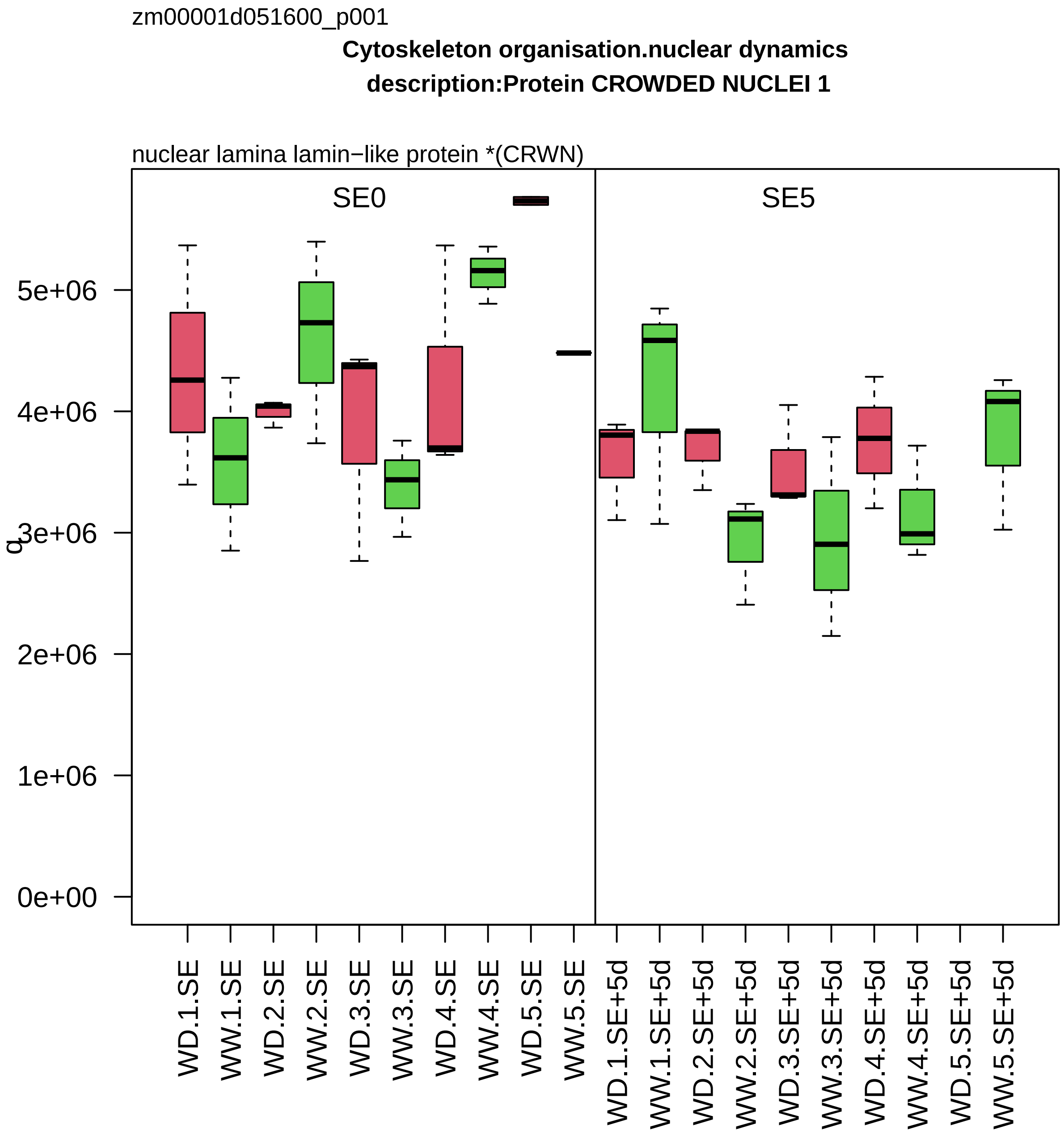
zm00001d051600_p001
Cytoskeleton organisation.nuclear dynamics
description:Protein CROWDED NUCLEI 1
nuclear lamina lamin−like protein *(CRWN)
SE0
SE5
q
0e+00
1e+06
2e+06
3e+06
4e+06
5e+06
WD.1.SE
WW.1.SE
WD.2.SE
WW.2.SE
WD.3.SE
WW.3.SE
WD.4.SE
WW.4.SE
WD.5.SE
WW.5.SE
WD.1.SE+5d
WW.1.SE+5d
WD.2.SE+5d
WW.2.SE+5d
WD.3.SE+5d
WW.3.SE+5d
WD.4.SE+5d
WW.4.SE+5d
WD.5.SE+5d
WW.5.SE+5d

zm00001d051699_p001

**Cell wall organisation.cell wall proteins**

**description:Fasciclin−like arabinogalactan protein 2**

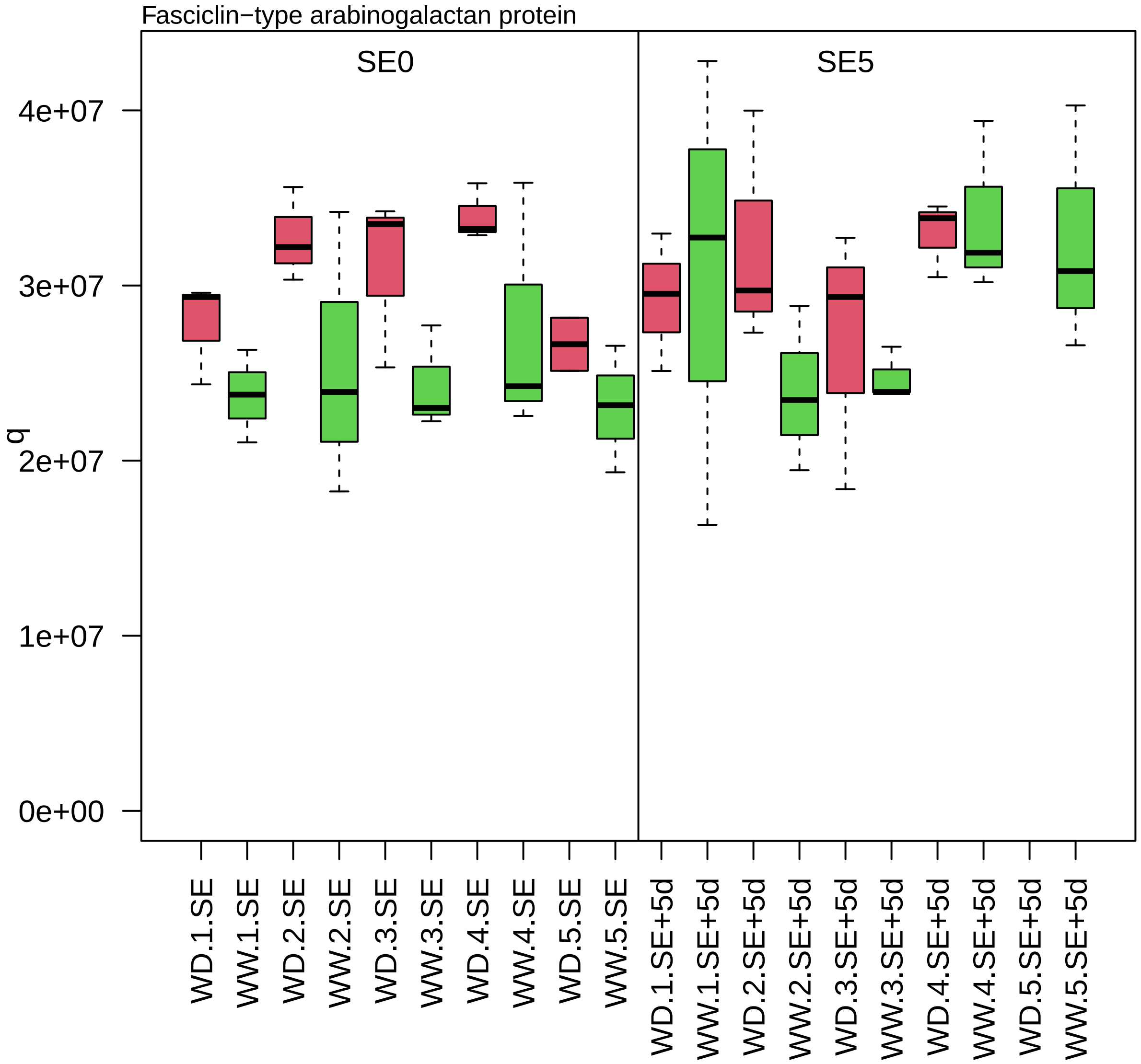

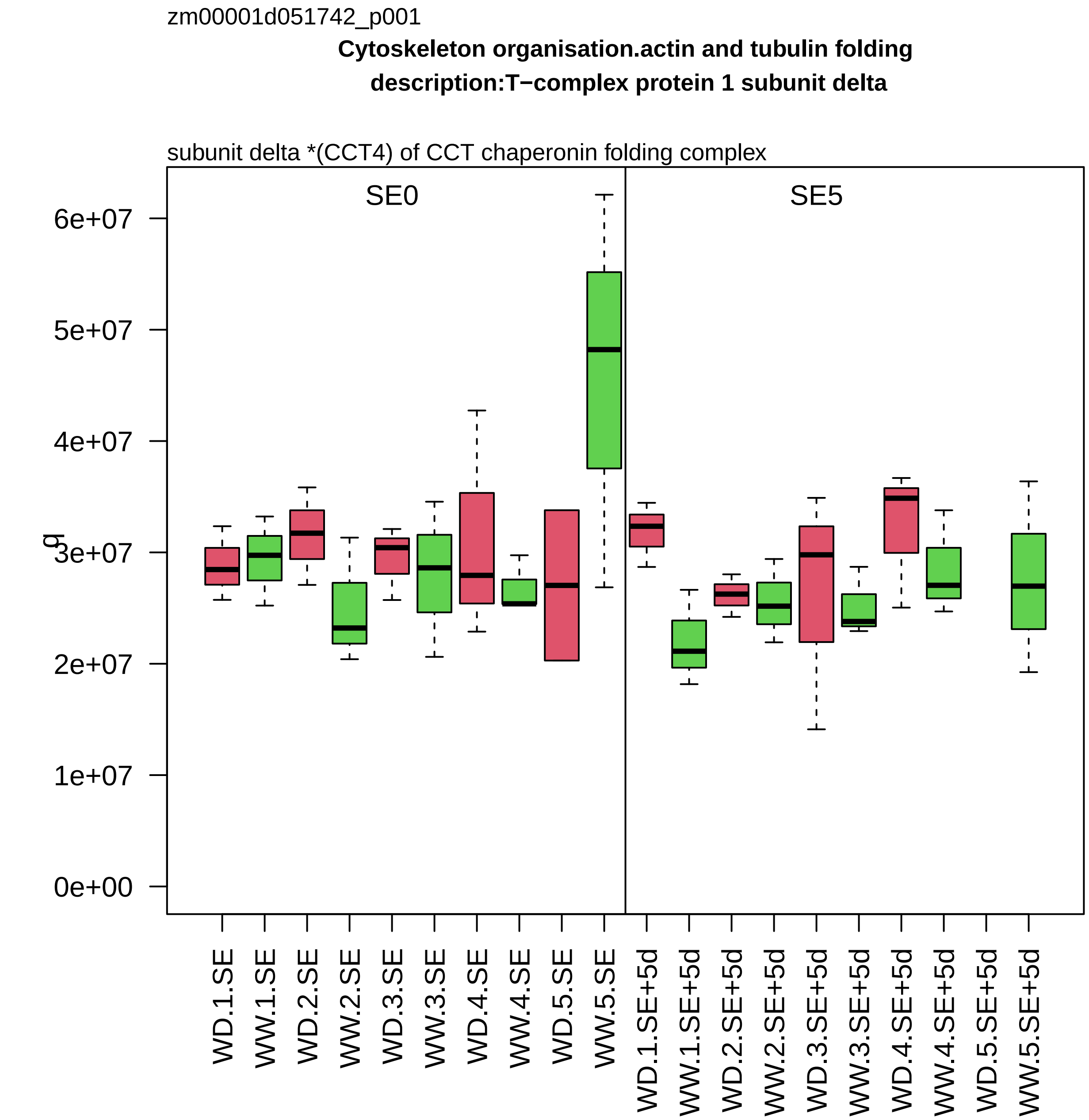
zm00001d051742_p001
Cytoskeleton organisation.actin and tubulin folding
description:T−complex protein 1 subunit delta
subunit delta *(CCT4) of CCT chaperonin folding complex
SE0
SE5
g
6e+07
5e+07
4e+07
3e+07
2e+07
1e+07
0e+00
WD.1.SE
WW.1.SE
WD.2.SE
WW.2.SE
WD.3.SE
WW.3.SE
WD.4.SE
WW.4.SE
WD.5.SE
WW.5.SE
WD.1.SE+5d
WW.1.SE+5d
WD.2.SE+5d
WW.2.SE+5d
WD.3.SE+5d
WW.3.SE+5d
WD.4.SE+5d
WW.4.SE+5d
WD.5.SE+5d
WW.5.SE+5d

zm00001d051754_p002

**Carbohydrate metabolism.fermentation**

**gene_symbol:aldehyde dehydrogenase2 description: aldehyde dehydrogenase2**

aldehyde dehydrogenase *(ALDH2B)

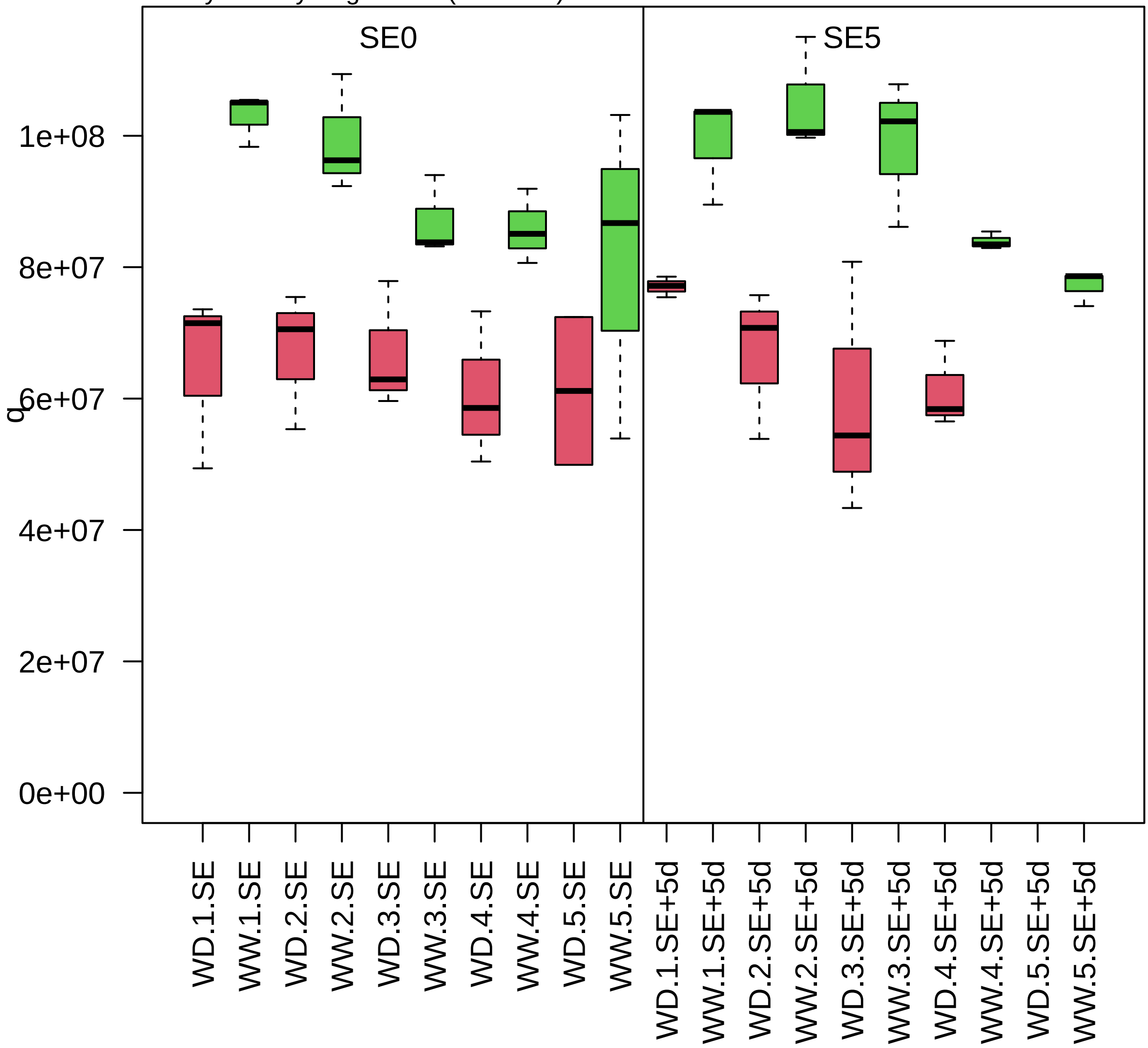

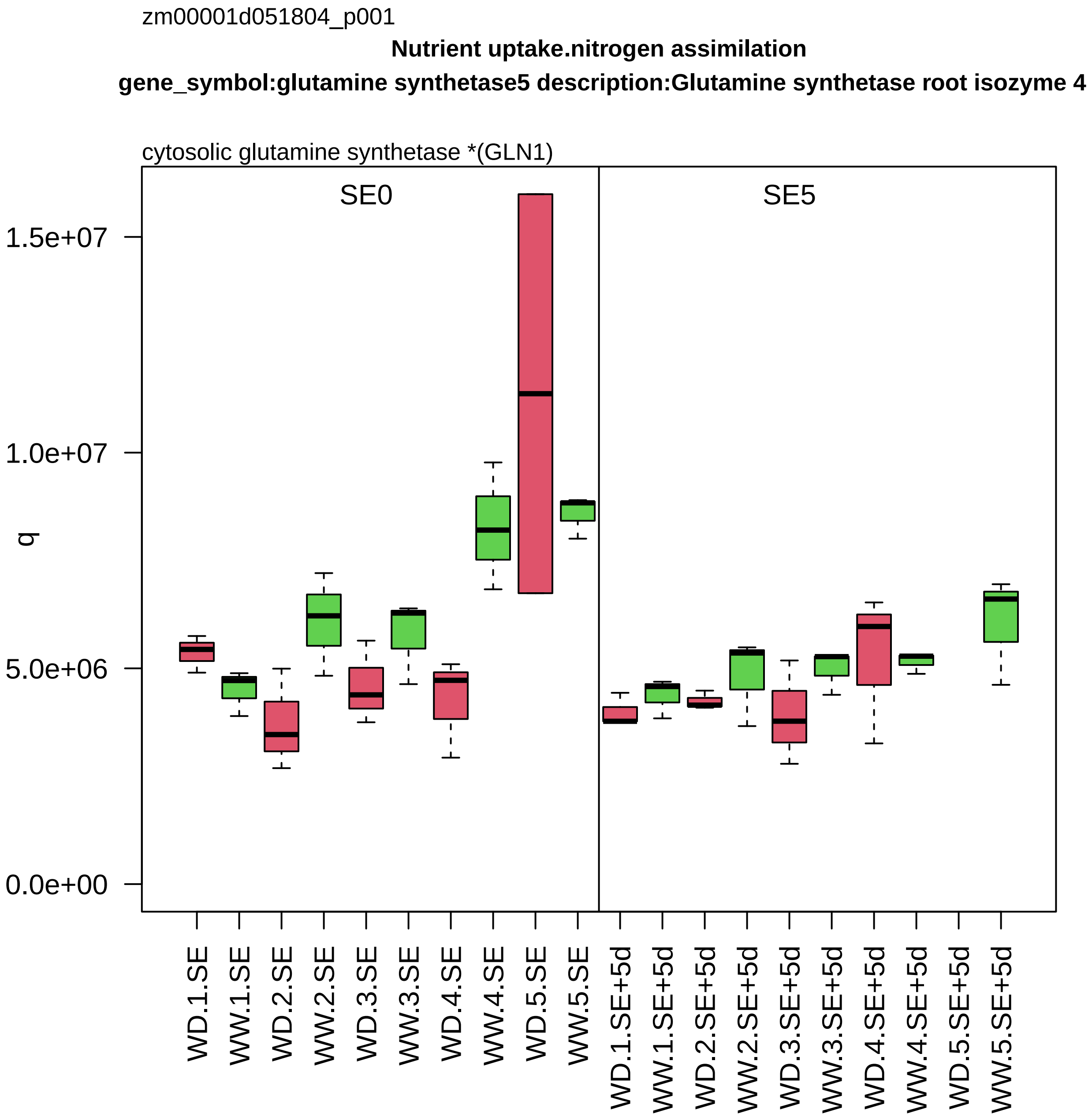

zm00001d051804_p001
Nutrient uptake.nitrogen assimilation
gene_symbol:glutamine synthetase5 description:Glutamine synthetase root isozyme 4
cytosolic glutamine synthetase *(GLN1)
SE0
SE5
q
1.5e+07
1.0e+07
5.0e+06
0.0e+00
WD.1.SE
WW.1.SE
WD.2.SE
WW.2.SE
WD.3.SE
WW.3.SE
WD.4.SE
WW.4.SE
WD.5.SE
WW.5.SE
WD.1.SE+5d
WW.1.SE+5d
WD.2.SE+5d
WW.2.SE+5d
WD.3.SE+5d
WW.3.SE+5d
WD.4.SE+5d
WW.4.SE+5d
WD.5.SE+5d
WW.5.SE+5d

zm00001d051870_p001

**Coenzyme metabolism.NAD/NADP biosynthesis**

**gene_symbol:si683005e02 description:O−methyltransferase family protein**

nicotinate N−methyltransferase *(NANMT)

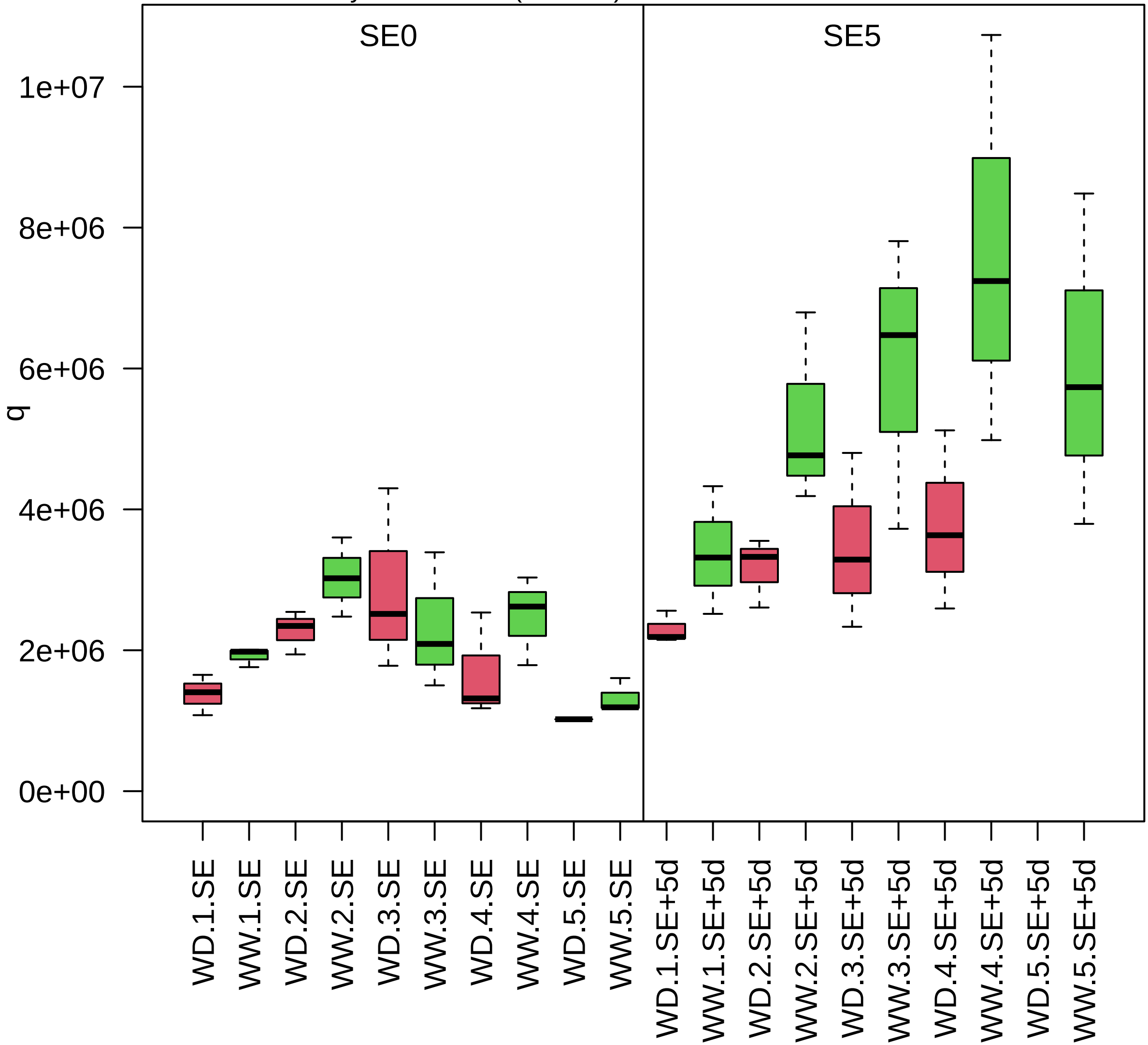

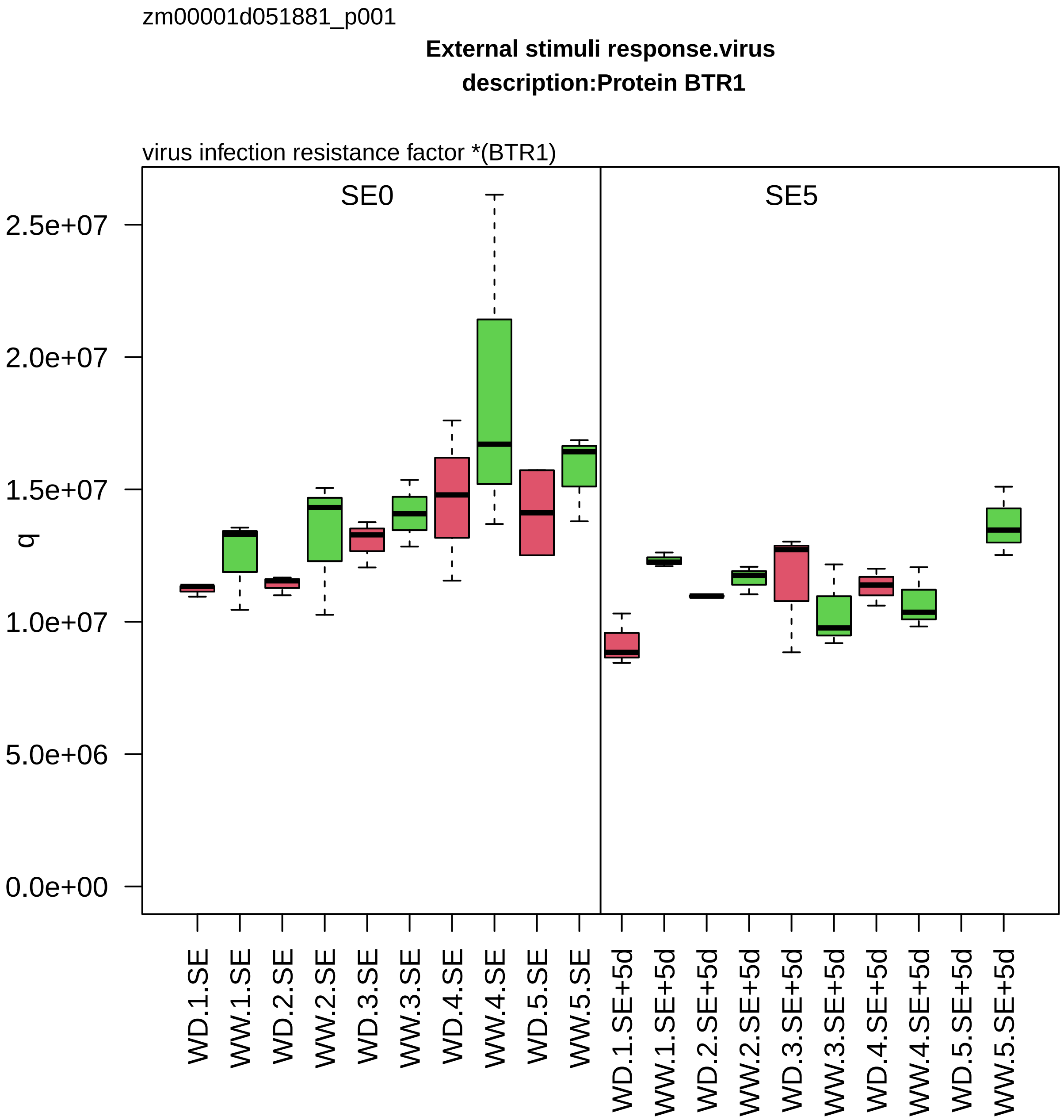

zm00001d051881_p001
External stimuli response.virus
description:Protein BTR1
virus infection resistance factor *(BTR1)
SE0
SE5
q
2.5e+07
2.0e+07
1.5e+07
1.0e+07
5.0e+06
0.0e+00
WD.1.SE
WW.1.SE
WD.2.SE
WW.2.SE
WD.3.SE
WW.3.SE
WD.4.SE
WW.4.SE
WD.5.SE
WW.5.SE
WD.1.SE+5d
WW.1.SE+5d
WD.2.SE+5d
WW.2.SE+5d
WD.3.SE+5d
WW.3.SE+5d
WD.4.SE+5d
WW.4.SE+5d
WD.5.SE+5d
WW.5.SE+5d

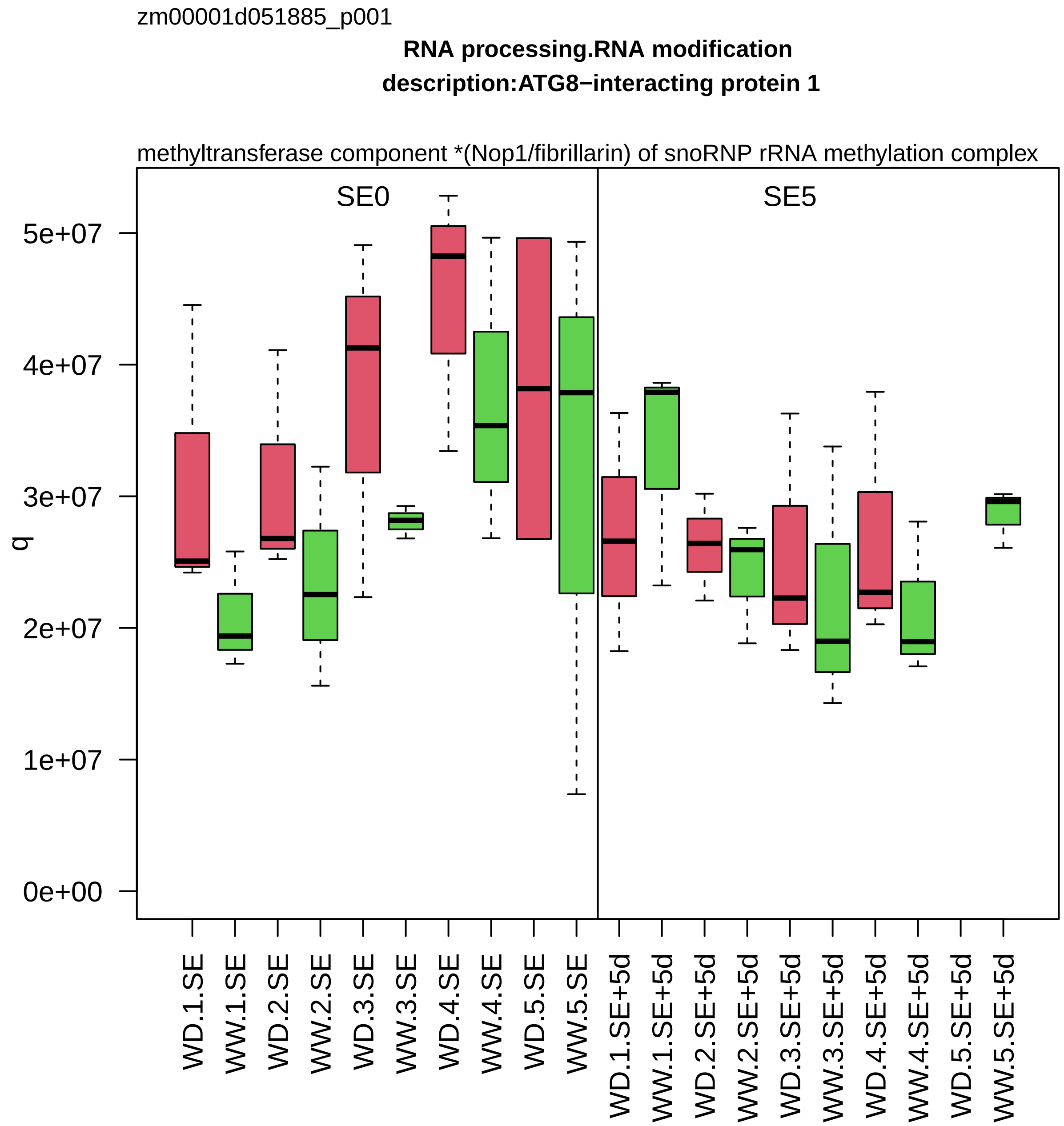
zm00001d051885_p001
RNA processing.RNA modification
description:ATG8−interacting protein 1
methyltransferase component *(Nop1/fibrillarin) of snoRNP rRNA methylation complex
SE0
SE5
q
0e+00
1e+07
2e+07
3e+07
4e+07
5e+07
WD.1.SE
WW.1.SE
WD.2.SE
WW.2.SE
WD.3.SE
WW.3.SE
WD.4.SE
WW.4.SE
WD.5.SE
WW.5.SE
WD.1.SE+5d
WW.1.SE+5d
WD.2.SE+5d
WW.2.SE+5d
WD.3.SE+5d
WW.3.SE+5d
WD.4.SE+5d
WW.4.SE+5d
WD.5.SE+5d
WW.5.SE+5d

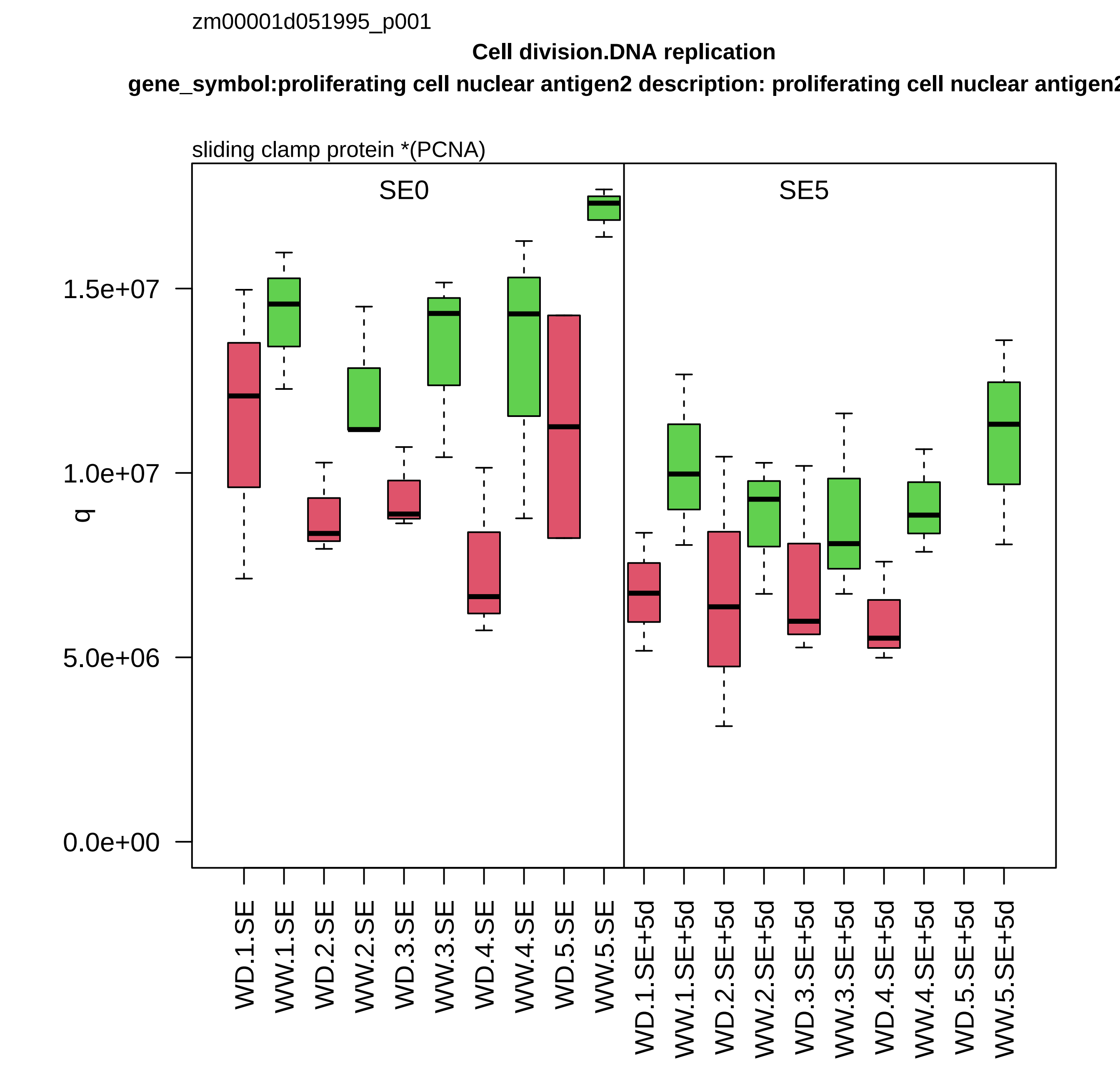

zm00001d051995_p001
Cell division.DNA replication
gene_symbol:proliferating cell nuclear antigen2 description: proliferating cell nuclear antigen2
sliding clamp protein *(PCNA)
SE0
SE5
q
0.0e+00
5.0e+06
1.0e+07
1.5e+07
WD.1.SE
WW.1.SE
WD.2.SE
WW.2.SE
WD.3.SE
WW.3.SE
WD.4.SE
WW.4.SE
WD.5.SE
WW.5.SE
WD.1.SE+5d
WW.1.SE+5d
WD.2.SE+5d
WW.2.SE+5d
WD.3.SE+5d
WW.3.SE+5d
WD.4.SE+5d
WW.4.SE+5d
WD.5.SE+5d
WW.5.SE+5d

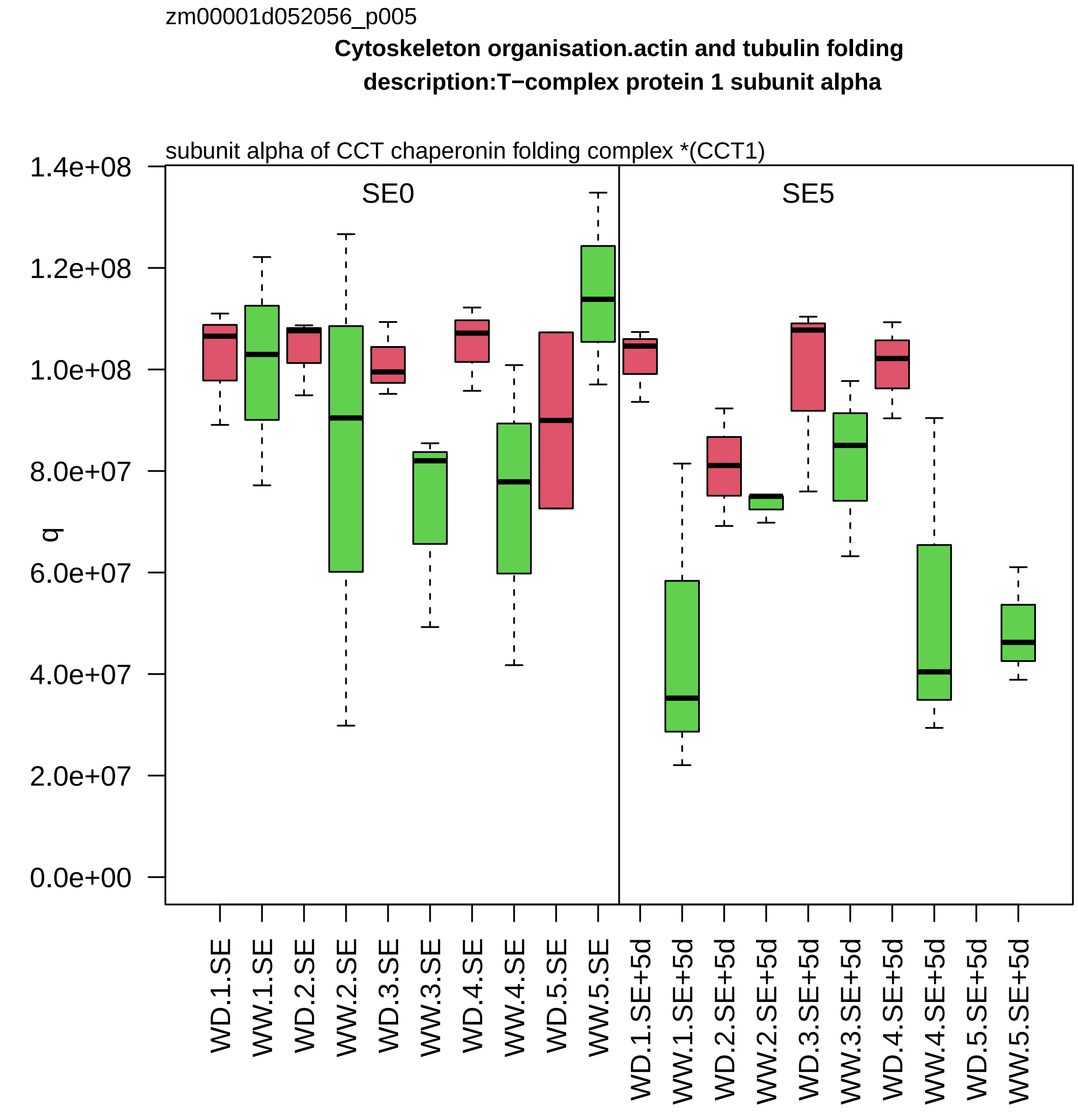
zm00001d052056_p005
Cytoskeleton organisation.actin and tubulin folding
description:T−complex protein 1 subunit alpha
subunit alpha of CCT chaperonin folding complex *(CCT1)
SE0
SE5
q
0.0e+00
2.0e+07
4.0e+07
6.0e+07
8.0e+07
1.0e+08
1.2e+08
1.4e+08
WD.1.SE
WW.1.SE
WD.2.SE
WW.2.SE
WD.3.SE
WW.3.SE
WD.4.SE
WW.4.SE
WD.5.SE
WW.5.SE
WD.1.SE+5d
WW.1.SE+5d
WD.2.SE+5d
WW.2.SE+5d
WD.3.SE+5d
WW.3.SE+5d
WD.4.SE+5d
WW.4.SE+5d
WD.5.SE+5d
WW.5.SE+5d

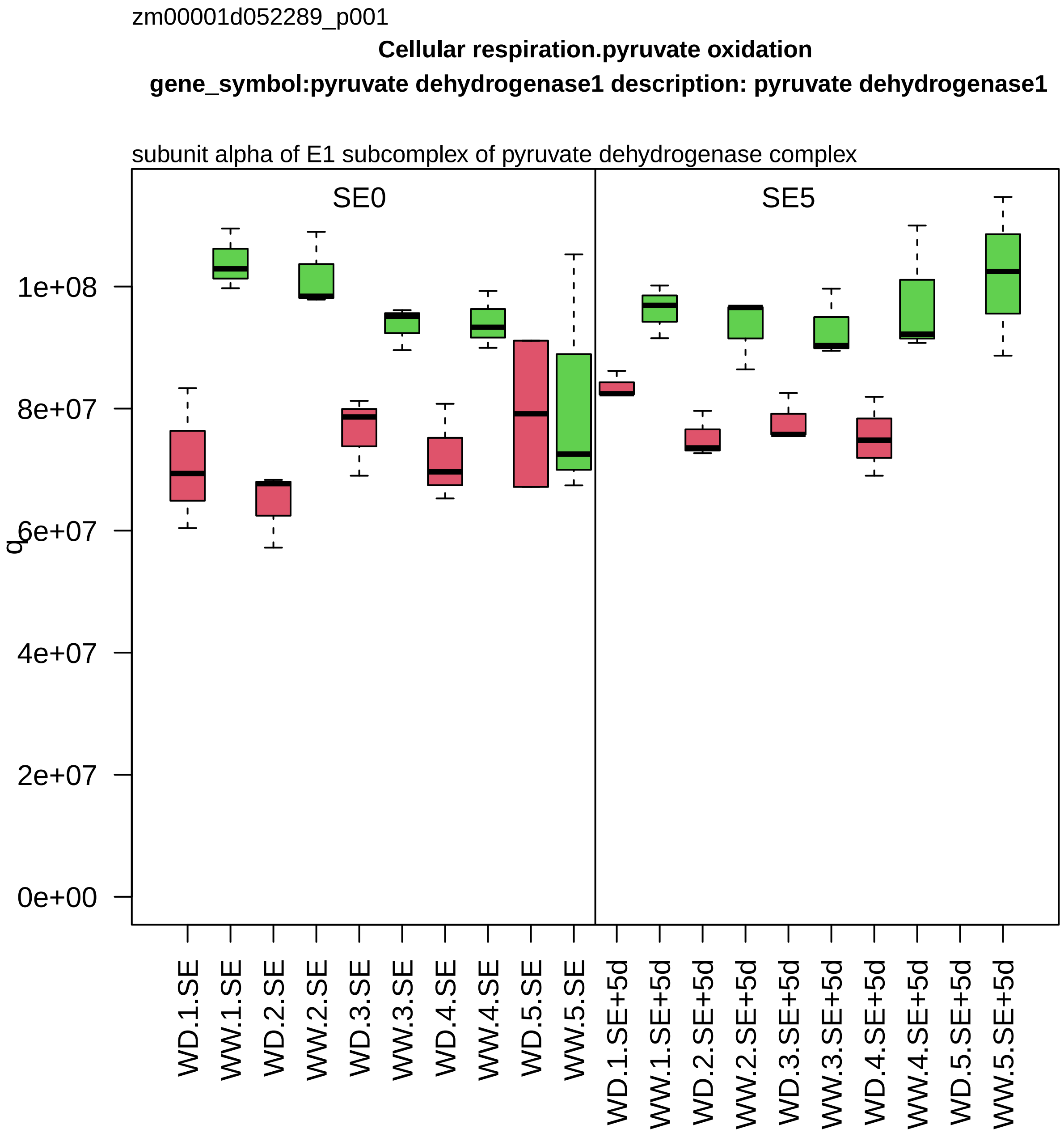

zm00001d052289_p001
Cellular respiration.pyruvate oxidation
gene_symbol:pyruvate dehydrogenase1 description: pyruvate dehydrogenase1
subunit alpha of E1 subcomplex of pyruvate dehydrogenase complex
SE0
SE5
q
0e+00
2e+07
4e+07
6e+07
8e+07
1e+08
WD.1.SE
WW.1.SE
WD.2.SE
WW.2.SE
WD.3.SE
WW.3.SE
WD.4.SE
WW.4.SE
WD.5.SE
WW.5.SE
WD.1.SE+5d
WW.1.SE+5d
WD.2.SE+5d
WW.2.SE+5d
WD.3.SE+5d
WW.3.SE+5d
WD.4.SE+5d
WW.4.SE+5d
WD.5.SE+5d
WW.5.SE+5d

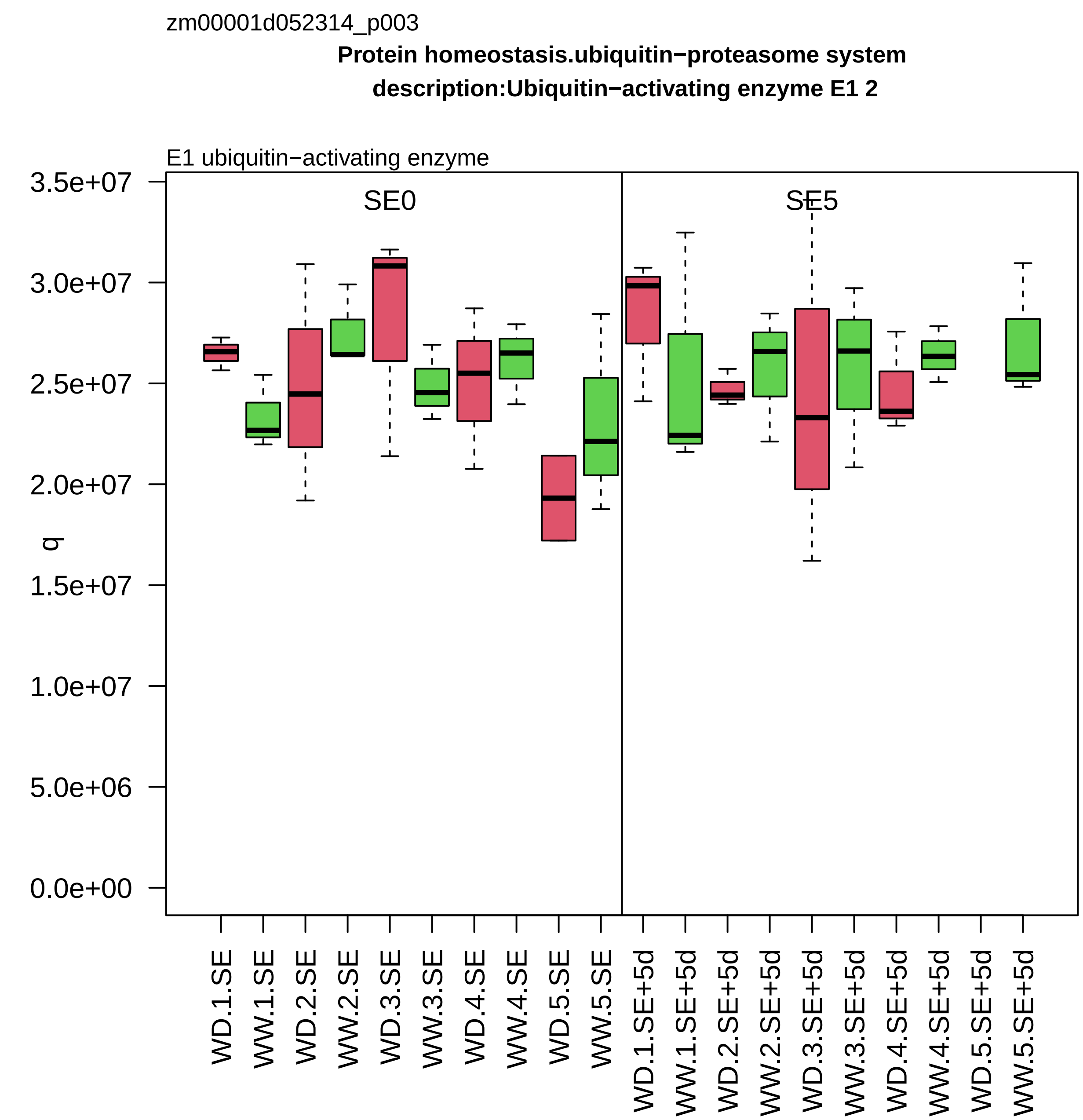
zm00001d052314_p003
Protein homeostasis.ubiquitin−proteasome system
description:Ubiquitin−activating enzyme E1 2
E1 ubiquitin−activating enzyme
SE0
SE5
q
3.5e+07
3.0e+07
2.5e+07
2.0e+07
1.5e+07
1.0e+07
5.0e+06
0.0e+00
WD.1.SE
WW.1.SE
WD.2.SE
WW.2.SE
WD.3.SE
WW.3.SE
WD.4.SE
WW.4.SE
WD.5.SE
WW.5.SE
WD.1.SE+5d
WW.1.SE+5d
WD.2.SE+5d
WW.2.SE+5d
WD.3.SE+5d
WW.3.SE+5d
WD.4.SE+5d
WW.4.SE+5d
WD.5.SE+5d
WW.5.SE+5d

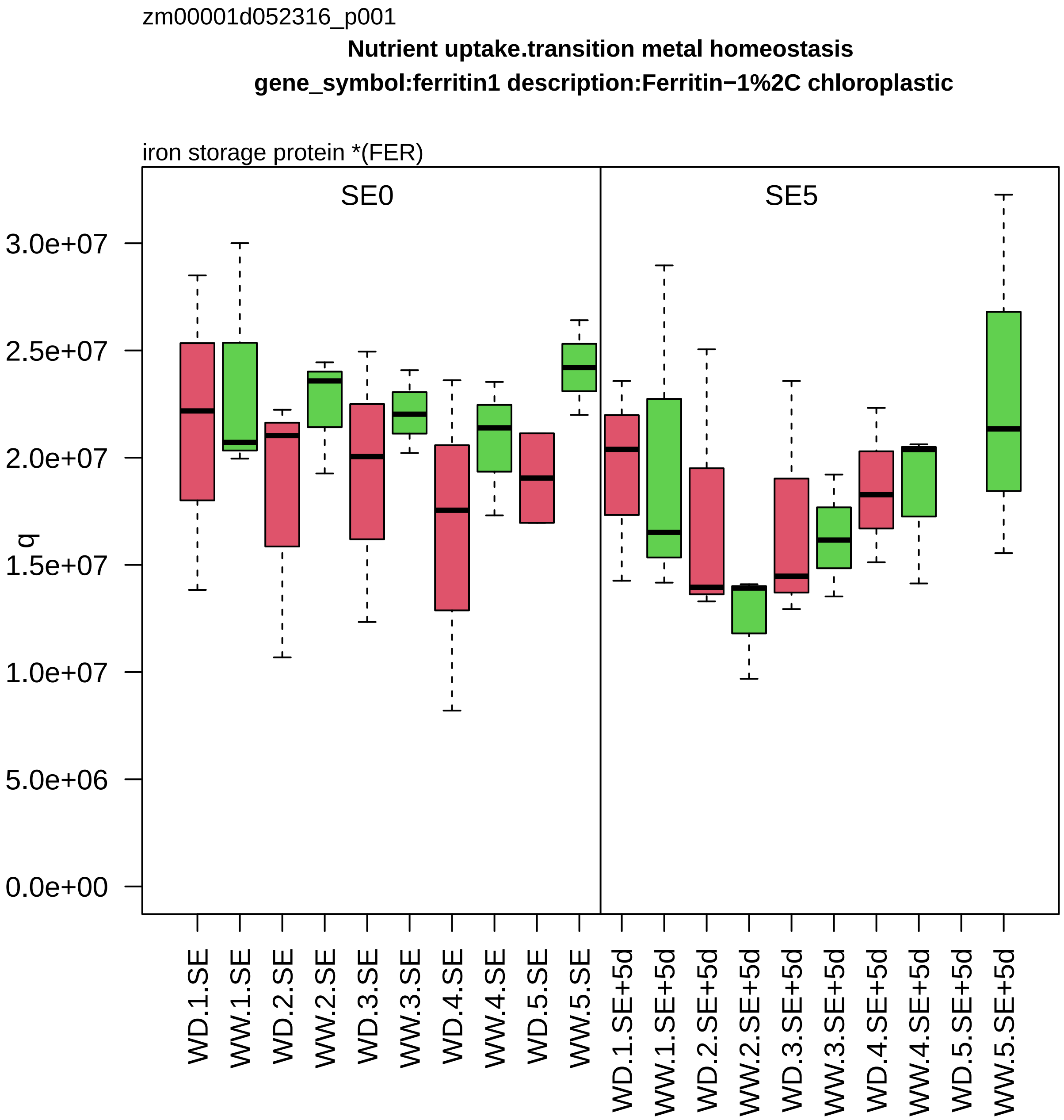

zm00001d052316_p001
Nutrient uptake.transition metal homeostasis
gene_symbol:ferritin1 description:Ferritin−1%2C chloroplastic
iron storage protein *(FER)
SE0
SE5
q
3.0e+07
2.5e+07
2.0e+07
1.5e+07
1.0e+07
5.0e+06
0.0e+00
WD.1.SE
WW.1.SE
WD.2.SE
WW.2.SE
WD.3.SE
WW.3.SE
WD.4.SE
WW.4.SE
WD.5.SE
WW.5.SE
WD.1.SE+5d
WW.1.SE+5d
WD.2.SE+5d
WW.2.SE+5d
WD.3.SE+5d
WW.3.SE+5d
WD.4.SE+5d
WW.4.SE+5d
WD.5.SE+5d
WW.5.SE+5d

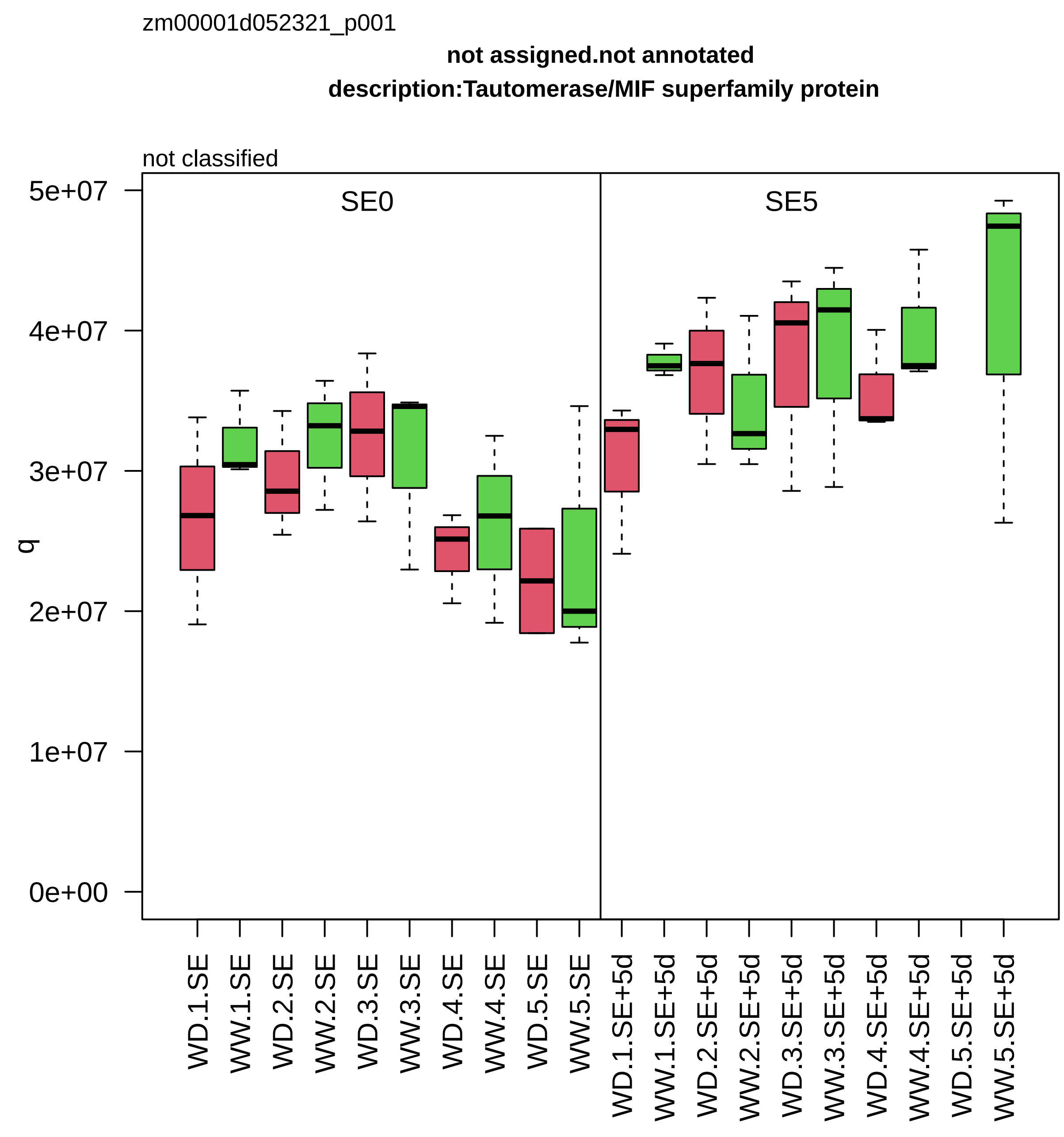

zm00001d052321_p001
not assigned.not annotated
description:Tautomerase/MIF superfamily protein
not classified
SE0
SE5
q
5e+07
4e+07
3e+07
2e+07
1e+07
0e+00
WD.1.SE
WW.1.SE
WD.2.SE
WW.2.SE
WD.3.SE
WW.3.SE
WD.4.SE
WW.4.SE
WD.5.SE
WW.5.SE
WD.1.SE+5d
WW.1.SE+5d
WD.2.SE+5d
WW.2.SE+5d
WD.3.SE+5d
WW.3.SE+5d
WD.4.SE+5d
WW.4.SE+5d
WD.5.SE+5d
WW.5.SE+5d

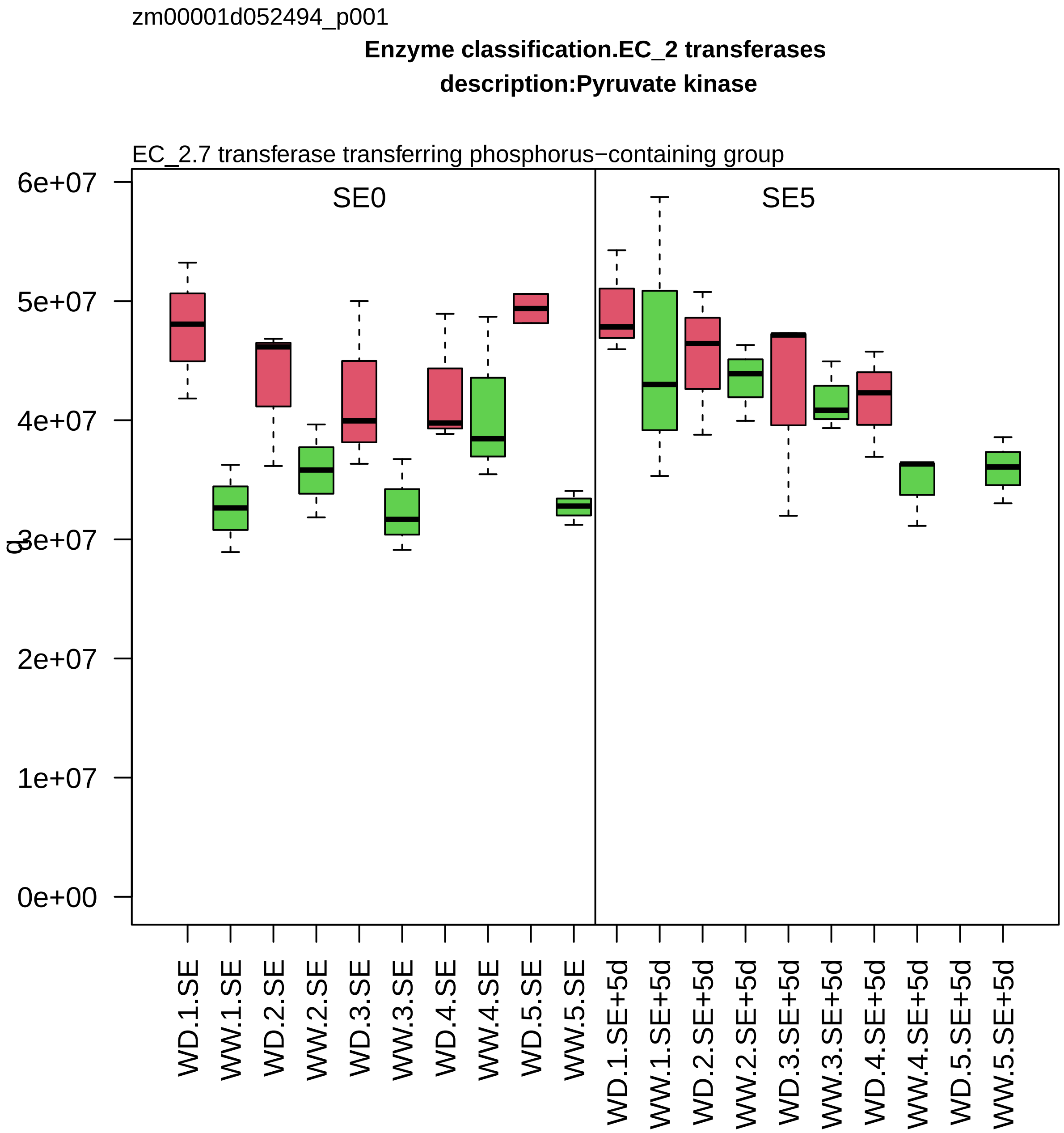
zm00001d052494_p001
Enzyme classification.EC_2 transferases
description:Pyruvate kinase
EC_2.7 transferase transferring phosphorus−containing group
SE0
SE5
6e+07
5e+07
4e+07
3e+07
2e+07
1e+07
0e+00
q
WD.1.SE
WW.1.SE
WD.2.SE
WW.2.SE
WD.3.SE
WW.3.SE
WD.4.SE
WW.4.SE
WD.5.SE
WW.5.SE
WD.1.SE+5d
WW.1.SE+5d
WD.2.SE+5d
WW.2.SE+5d
WD.3.SE+5d
WW.3.SE+5d
WD.4.SE+5d
WW.4.SE+5d
WD.5.SE+5d
WW.5.SE+5d

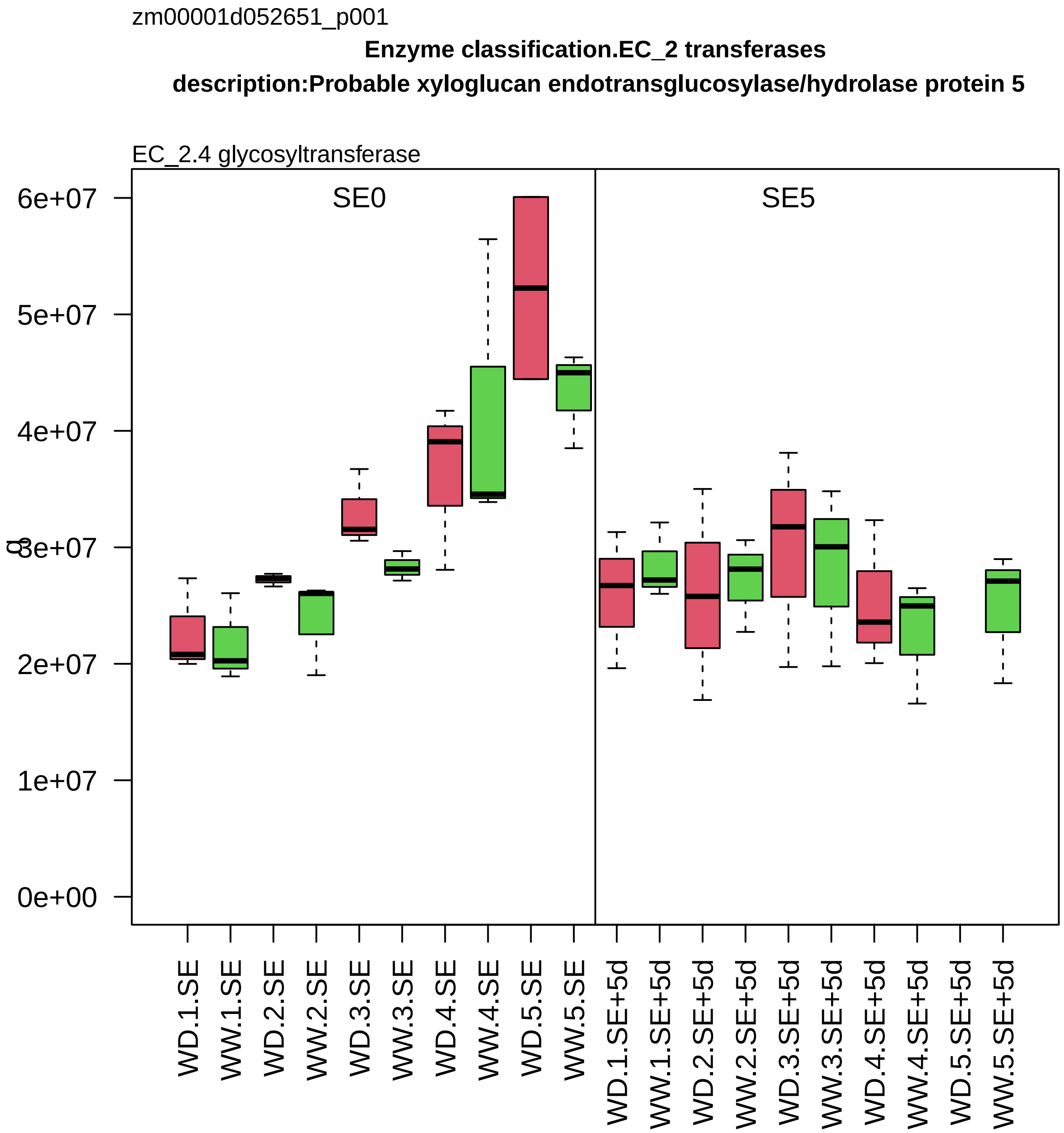

zm00001d052651_p001
Enzyme classification.EC_2 transferases
description:Probable xyloglucan endotransglucosylase/hydrolase protein 5
EC_2.4 glycosyltransferase
SE0
SE5
6e+07
5e+07
4e+07
3e+07
2e+07
1e+07
0e+00
WD.1.SE
WW.1.SE
WD.2.SE
WW.2.SE
WD.3.SE
WW.3.SE
WD.4.SE
WW.4.SE
WD.5.SE
WW.5.SE
WD.1.SE+5d
WW.1.SE+5d
WD.2.SE+5d
WW.2.SE+5d
WD.3.SE+5d
WW.3.SE+5d
WD.4.SE+5d
WW.4.SE+5d
WD.5.SE+5d
WW.5.SE+5d

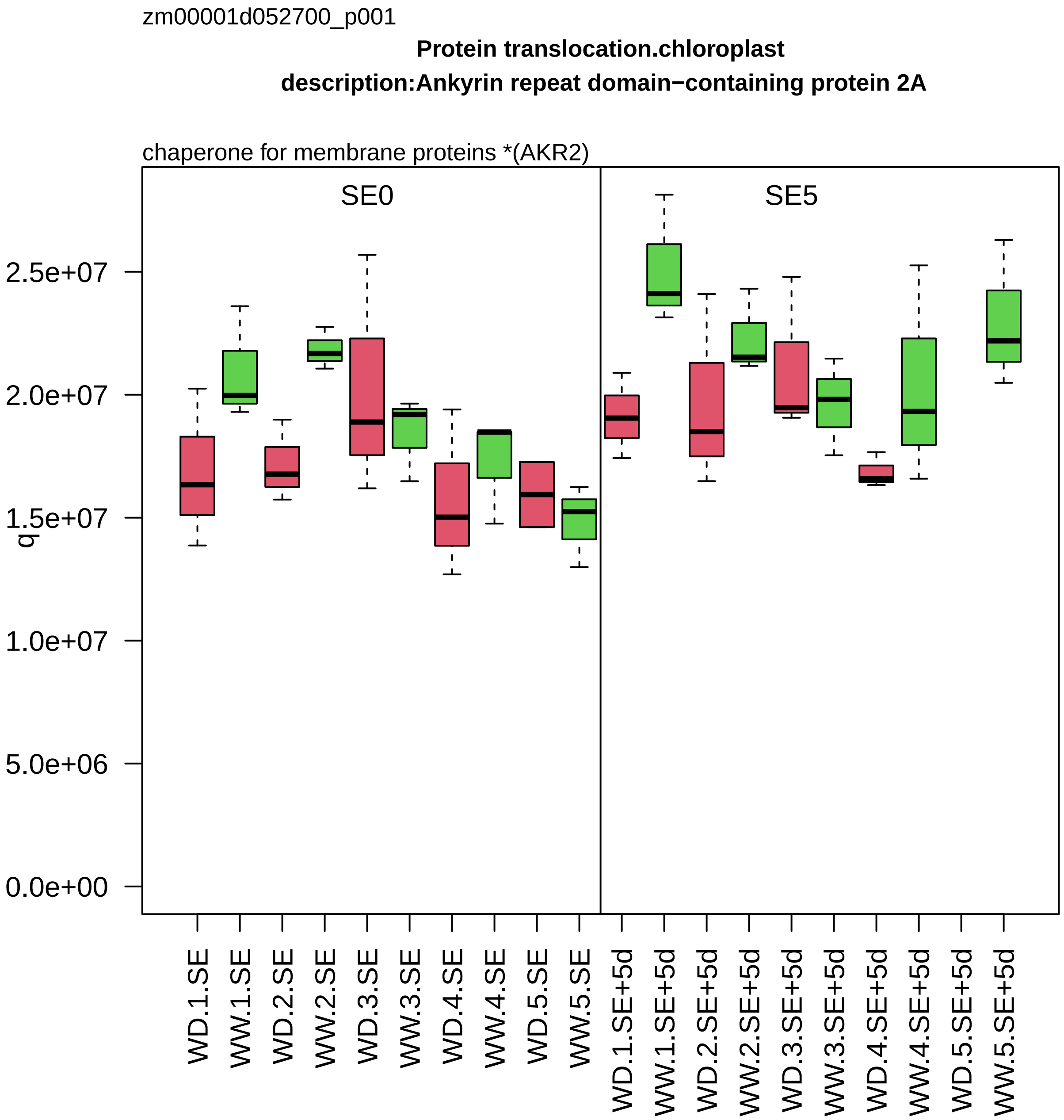
zm00001d052700_p001
Protein translocation.chloroplast
description:Ankyrin repeat domain−containing protein 2A
chaperone for membrane proteins *(AKR2)
SE0
SE5
q
2.5e+07
2.0e+07
1.5e+07
1.0e+07
5.0e+06
0.0e+00
WD.1.SE
WW.1.SE
WD.2.SE
WW.2.SE
WD.3.SE
WW.3.SE
WD.4.SE
WW.4.SE
WD.5.SE
WW.5.SE
WD.1.SE+5d
WW.1.SE+5d
WD.2.SE+5d
WW.2.SE+5d
WD.3.SE+5d
WW.3.SE+5d
WD.4.SE+5d
WW.4.SE+5d
WD.5.SE+5d
WW.5.SE+5d

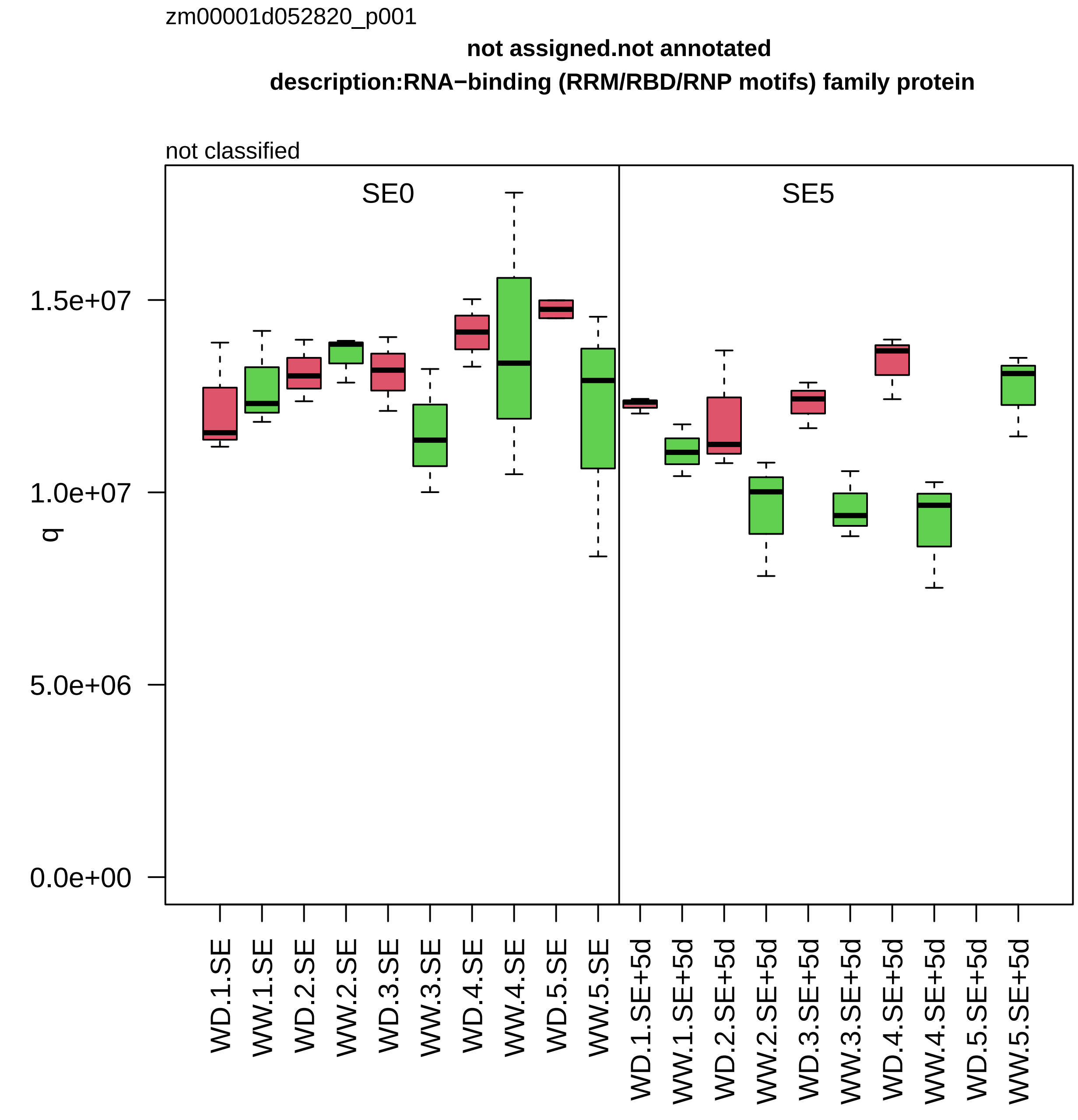

zm00001d052820_p001
not assigned.not annotated
description:RNA−binding (RRM/RBD/RNP motifs) family protein
not classified
SE0
SE5
q
1.5e+07
1.0e+07
5.0e+06
0.0e+00
WD.1.SE
WW.1.SE
WD.2.SE
WW.2.SE
WD.3.SE
WW.3.SE
WD.4.SE
WW.4.SE
WD.5.SE
WW.5.SE
WD.1.SE+5d
WW.1.SE+5d
WD.2.SE+5d
WW.2.SE+5d
WD.3.SE+5d
WW.3.SE+5d
WD.4.SE+5d
WW.4.SE+5d
WD.5.SE+5d
WW.5.SE+5d

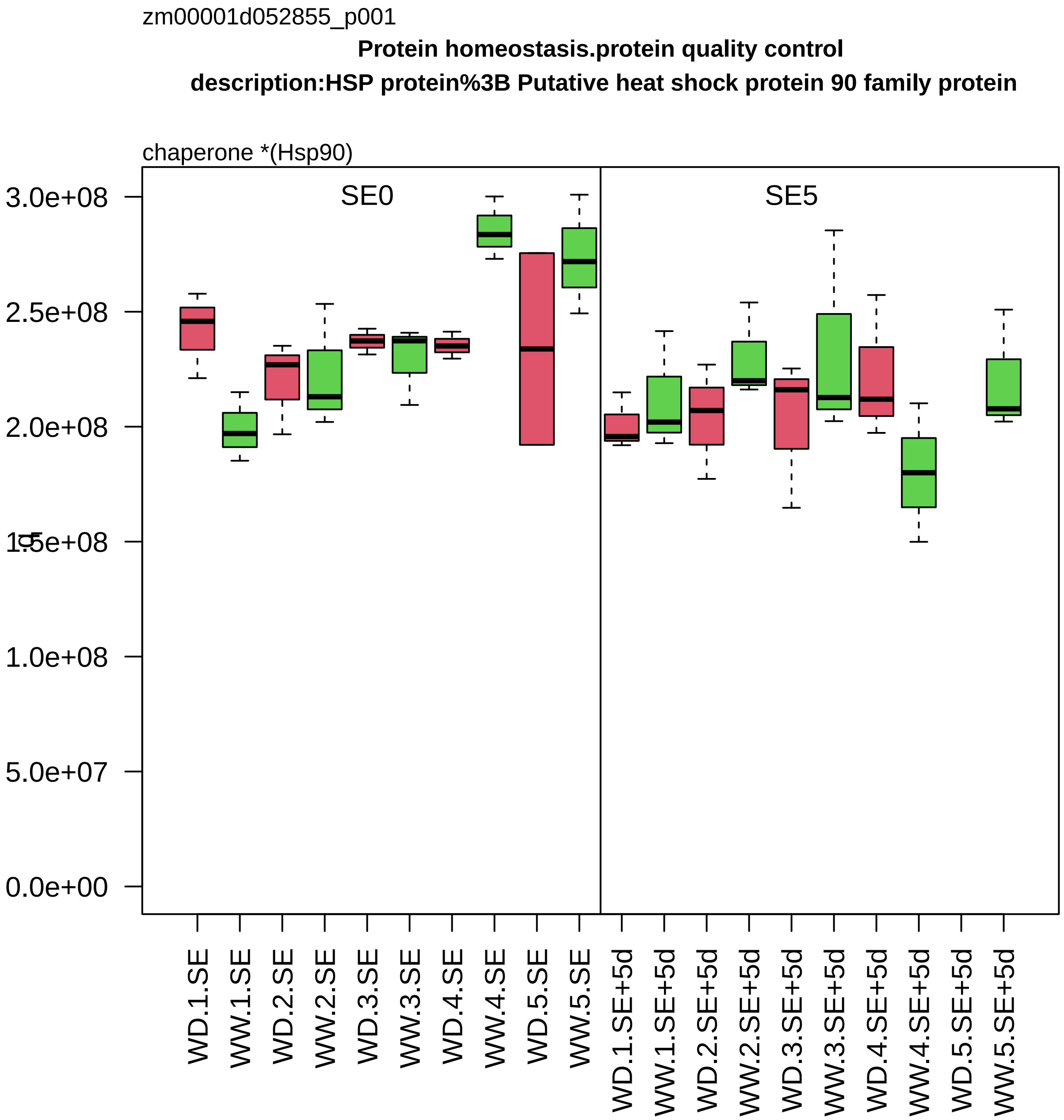
zm00001d052855_p001
Protein homeostasis.protein quality control
description:HSP protein%3B Putative heat shock protein 90 family protein
chaperone *(Hsp90)
SE0
SE5
3.0e+08
2.5e+08
2.0e+08
1.5e+08
1.0e+08
5.0e+07
0.0e+00
WD.1.SE
WW.1.SE
WD.2.SE
WW.2.SE
WD.3.SE
WW.3.SE
WD.4.SE
WW.4.SE
WD.5.SE
WW.5.SE
WD.1.SE+5d
WW.1.SE+5d
WD.2.SE+5d
WW.2.SE+5d
WD.3.SE+5d
WW.3.SE+5d
WD.4.SE+5d
WW.4.SE+5d
WD.5.SE+5d
WW.5.SE+5d

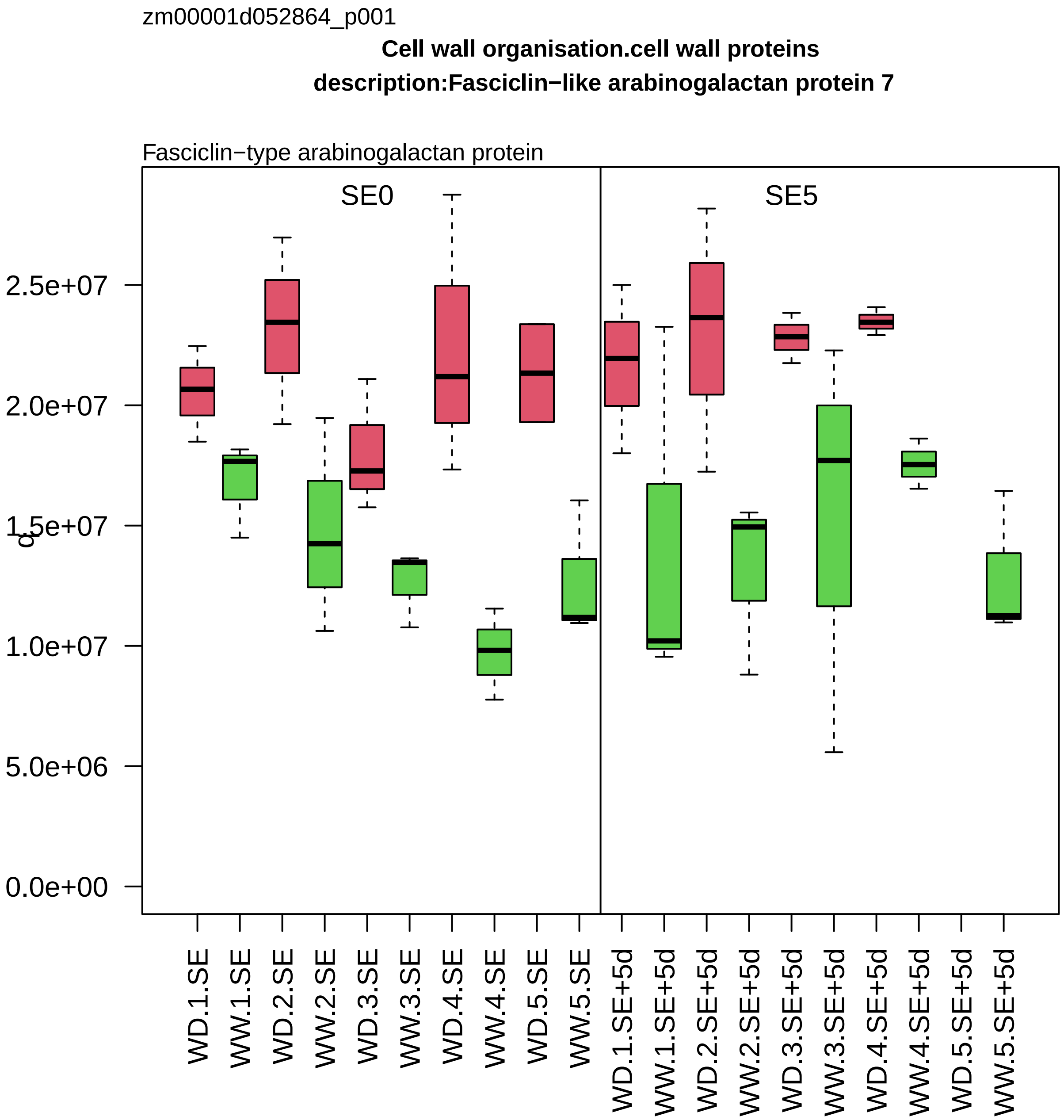

zm00001d052864_p001
Cell wall organisation.cell wall proteins
description:Fasciclin−like arabinogalactan protein 7
Fasciclin−type arabinogalactan protein
SE0
SE5
2.5e+07
2.0e+07
1.5e+07
1.0e+07
5.0e+06
0.0e+00
WD.1.SE
WW.1.SE
WD.2.SE
WW.2.SE
WD.3.SE
WW.3.SE
WD.4.SE
WW.4.SE
WD.5.SE
WW.5.SE
WD.1.SE+5d
WW.1.SE+5d
WD.2.SE+5d
WW.2.SE+5d
WD.3.SE+5d
WW.3.SE+5d
WD.4.SE+5d
WW.4.SE+5d
WD.5.SE+5d
WW.5.SE+5d

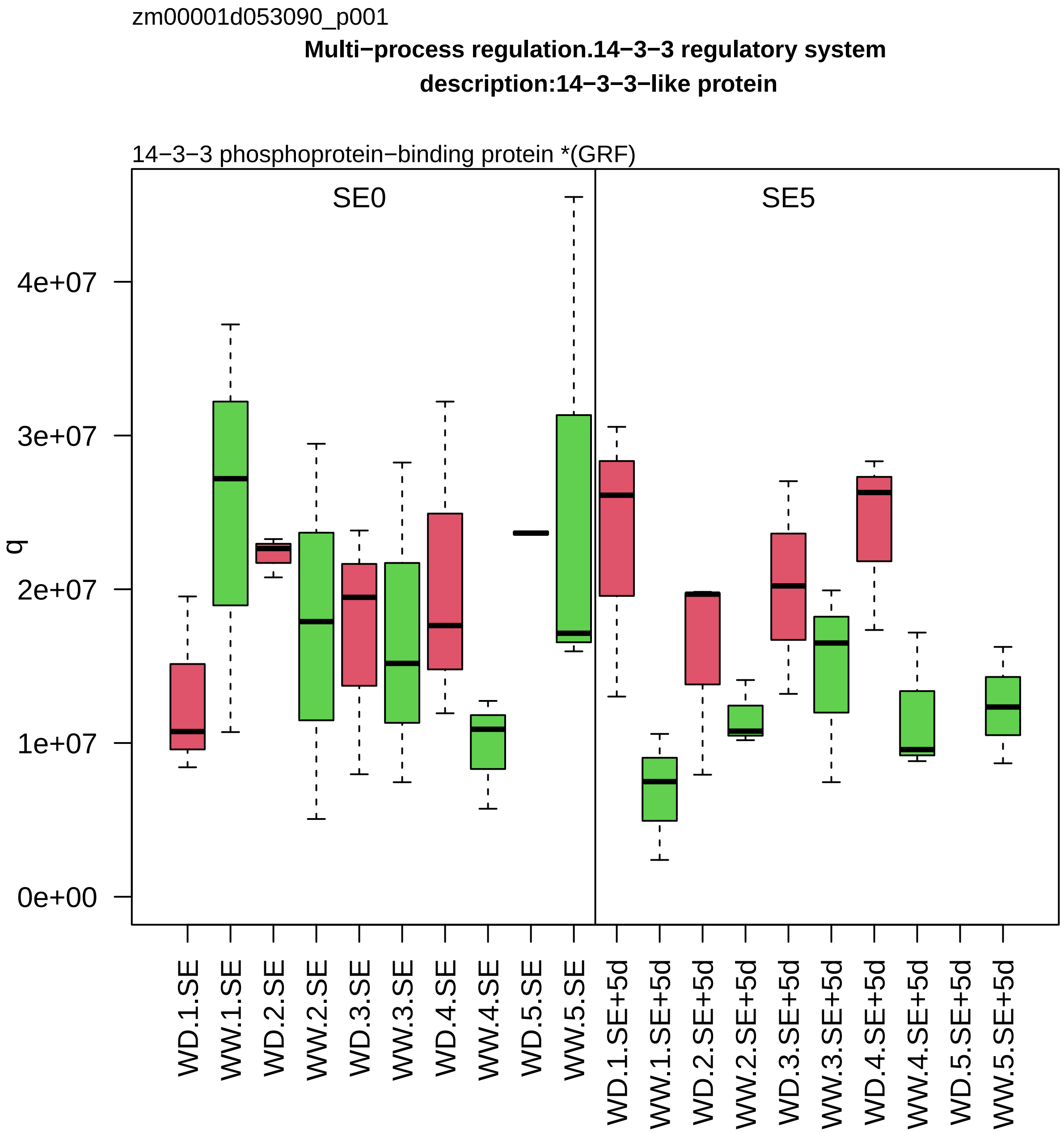
zm00001d053090_p001
Multi−process regulation.14−3−3 regulatory system
description:14−3−3−like protein
14−3−3 phosphoprotein−binding protein *(GRF)
SE0
SE5
q
0e+00
1e+07
2e+07
3e+07
4e+07
WD.1.SE
WW.1.SE
WD.2.SE
WW.2.SE
WD.3.SE
WW.3.SE
WD.4.SE
WW.4.SE
WD.5.SE
WW.5.SE
WD.1.SE+5d
WW.1.SE+5d
WD.2.SE+5d
WW.2.SE+5d
WD.3.SE+5d
WW.3.SE+5d
WD.4.SE+5d
WW.4.SE+5d
WD.5.SE+5d
WW.5.SE+5d

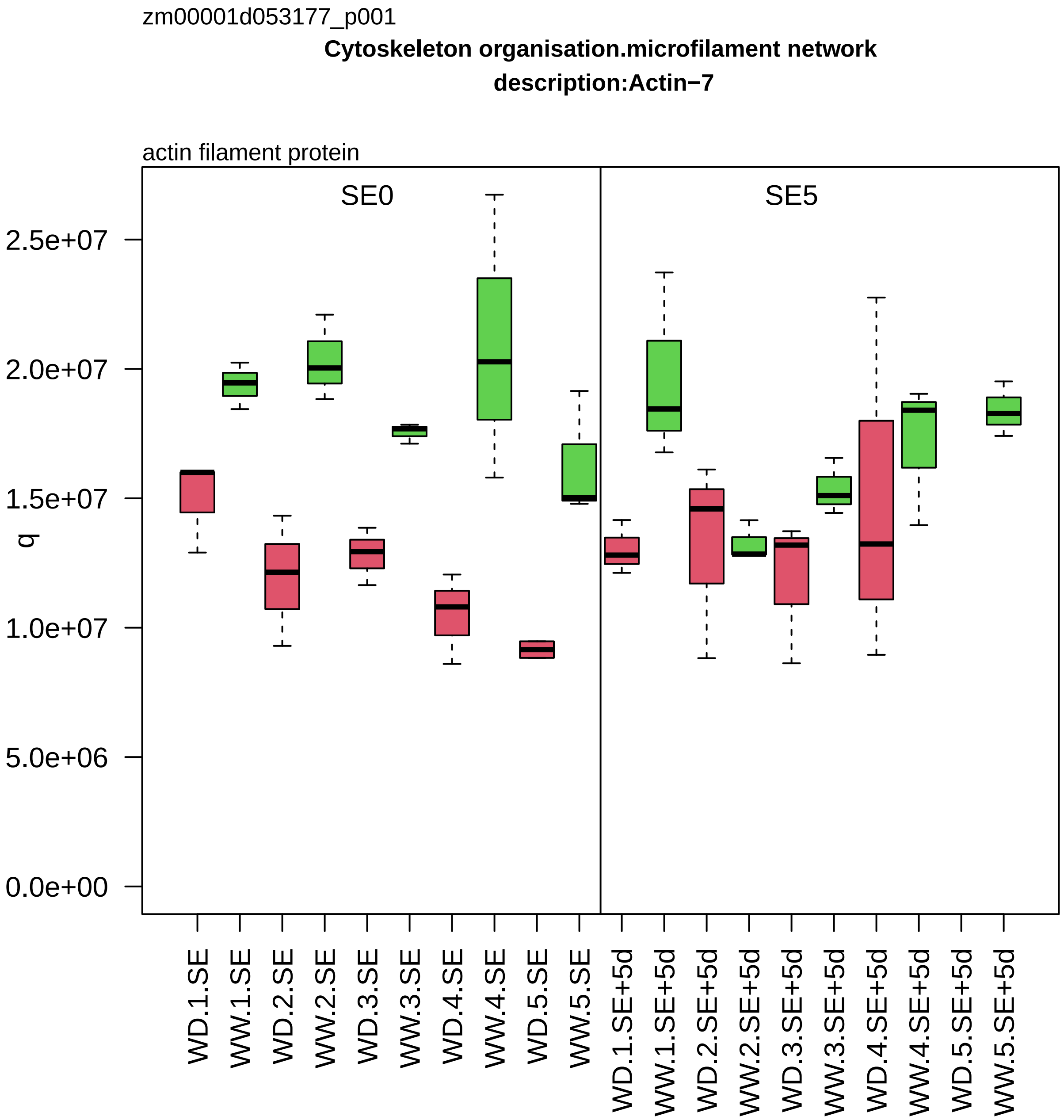
zm00001d053177_p001
Cytoskeleton organisation.microfilament network
description:Actin−7
actin filament protein
SE0
SE5
q
2.5e+07
2.0e+07
1.5e+07
1.0e+07
5.0e+06
0.0e+00
WD.1.SE
WW.1.SE
WD.2.SE
WW.2.SE
WD.3.SE
WW.3.SE
WD.4.SE
WW.4.SE
WD.5.SE
WW.5.SE
WD.1.SE+5d
WW.1.SE+5d
WD.2.SE+5d
WW.2.SE+5d
WD.3.SE+5d
WW.3.SE+5d
WD.4.SE+5d
WW.4.SE+5d
WD.5.SE+5d
WW.5.SE+5d

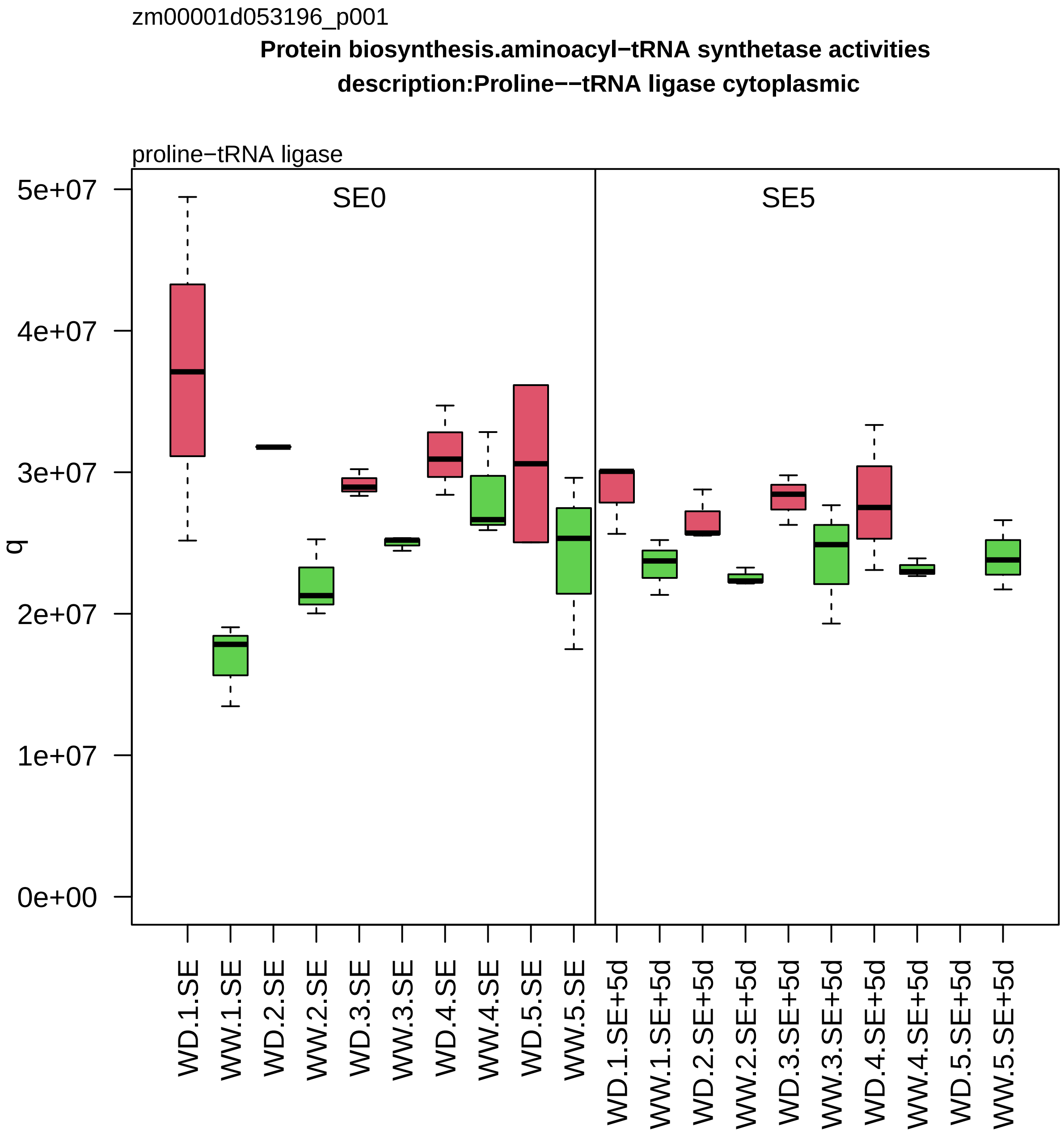
zm00001d053196_p001
Protein biosynthesis.aminoacyl−tRNA synthetase activities
description:Proline−−tRNA ligase cytoplasmic
proline−tRNA ligase
SE0
SE5
q
5e+07
4e+07
3e+07
2e+07
1e+07
0e+00
WD.1.SE
WW.1.SE
WD.2.SE
WW.2.SE
WD.3.SE
WW.3.SE
WD.4.SE
WW.4.SE
WD.5.SE
WW.5.SE
WD.1.SE+5d
WW.1.SE+5d
WD.2.SE+5d
WW.2.SE+5d
WD.3.SE+5d
WW.3.SE+5d
WD.4.SE+5d
WW.4.SE+5d
WD.5.SE+5d
WW.5.SE+5d

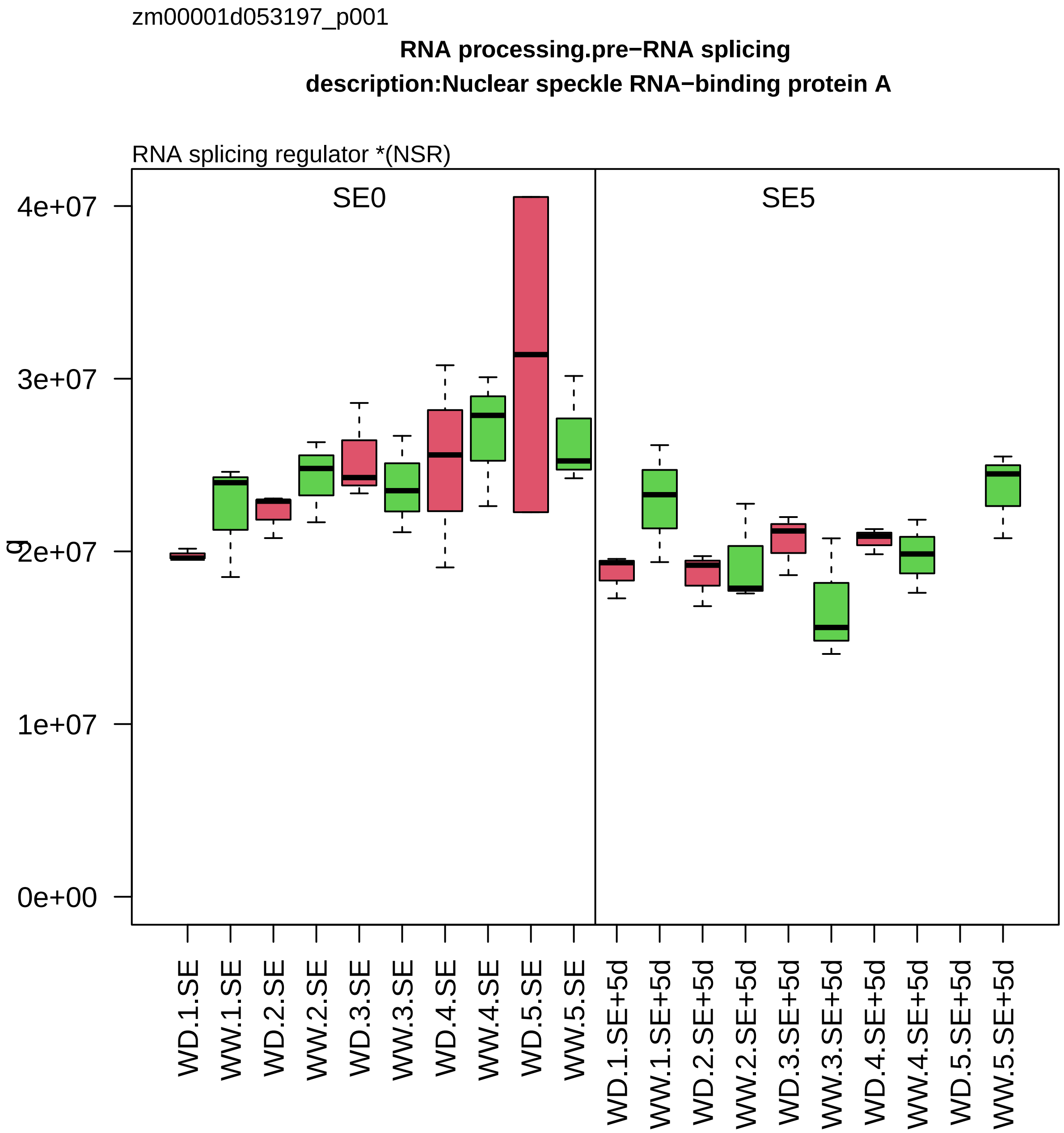

zm00001d053197_p001
RNA processing.pre−RNA splicing
description:Nuclear speckle RNA−binding protein A
RNA splicing regulator *(NSR)
SE0
SE5
4e+07
3e+07
2e+07
1e+07
0e+00
g
WD.1.SE
WW.1.SE
WD.2.SE
WW.2.SE
WD.3.SE
WW.3.SE
WD.4.SE
WW.4.SE
WD.5.SE
WW.5.SE
WD.1.SE+5d
WW.1.SE+5d
WD.2.SE+5d
WW.2.SE+5d
WD.3.SE+5d
WW.3.SE+5d
WD.4.SE+5d
WW.4.SE+5d
WD.5.SE+5d
WW.5.SE+5d

zm00001d053232_p002

**Protein homeostasis.ubiquitin−proteasome system**

**description:26S proteasome non−ATPase regulatory subunit 3 homolog A**

regulatory component *(RPN3) of 26S proteasome

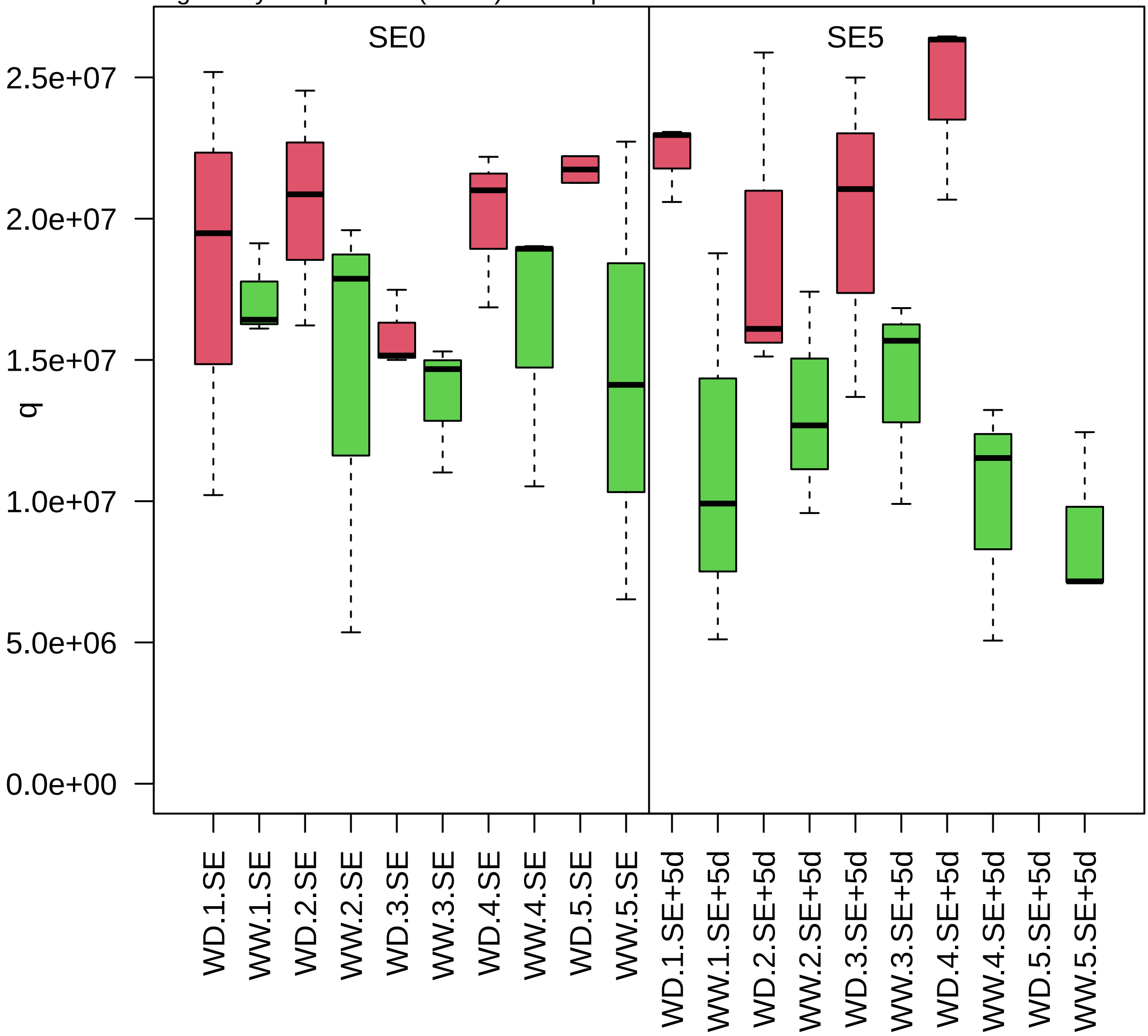

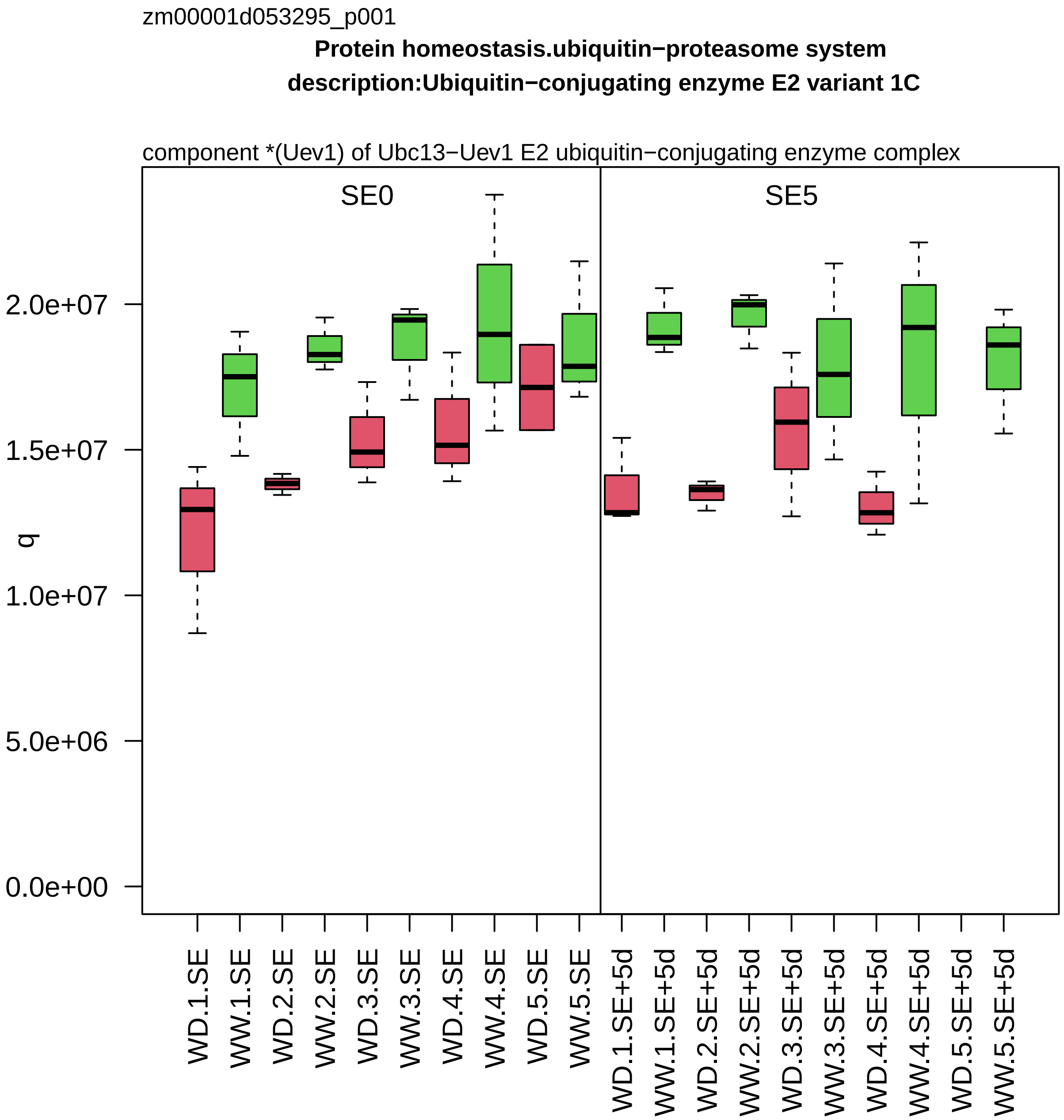

zm00001d053295_p001
Protein homeostasis.ubiquitin−proteasome system
description:Ubiquitin−conjugating enzyme E2 variant 1C
component *(Uev1) of Ubc13−Uev1 E2 ubiquitin−conjugating enzyme complex
SE0
SE5
q
2.0e+07
1.5e+07
1.0e+07
5.0e+06
0.0e+00
WD.1.SE
WW.1.SE
WD.2.SE
WW.2.SE
WD.3.SE
WW.3.SE
WD.4.SE
WW.4.SE
WD.5.SE
WW.5.SE
WD.1.SE+5d
WW.1.SE+5d
WD.2.SE+5d
WW.2.SE+5d
WD.3.SE+5d
WW.3.SE+5d
WD.4.SE+5d
WW.4.SE+5d
WD.5.SE+5d
WW.5.SE+5d

zm00001d053308_p002

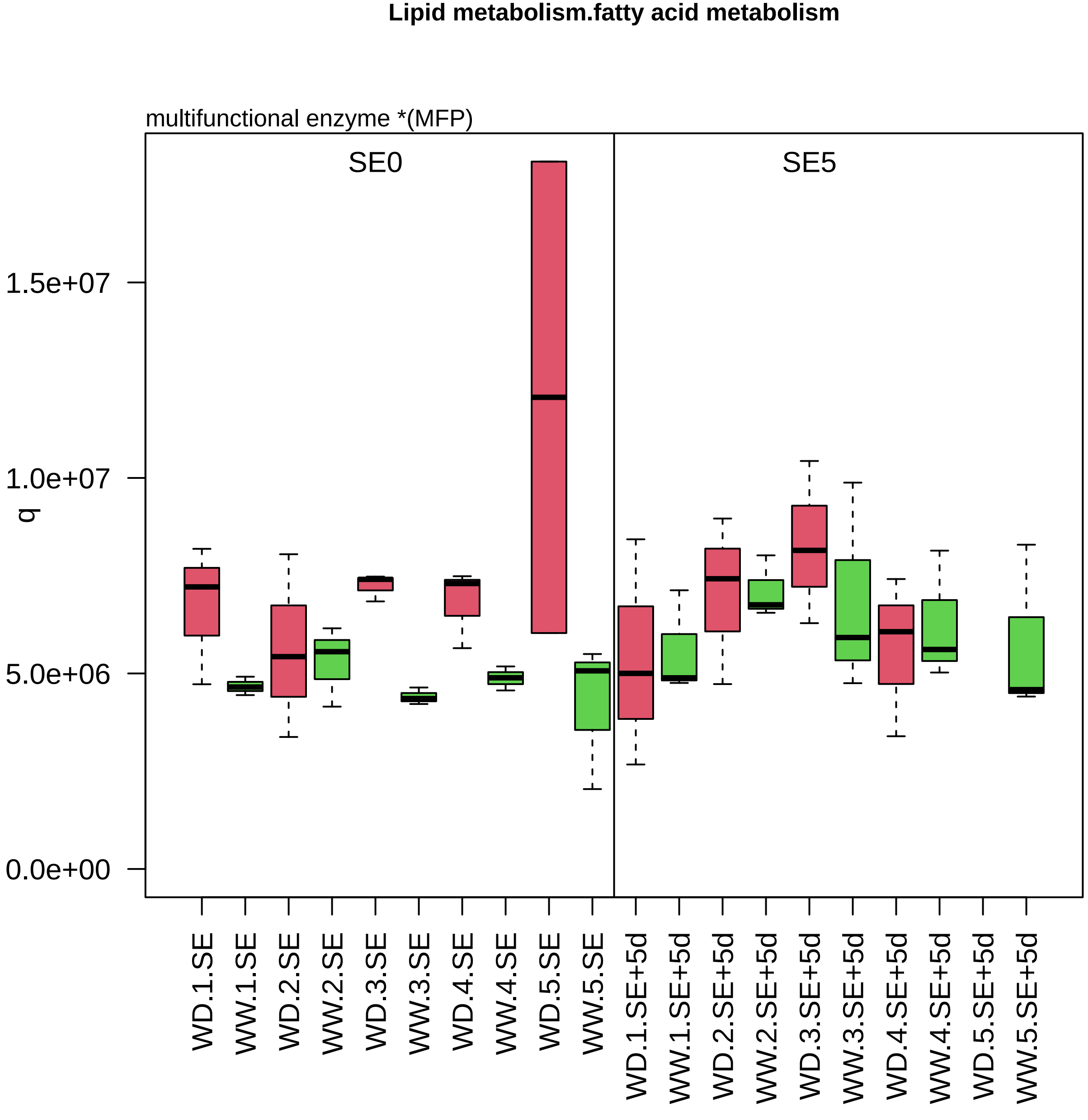

zm00001d053453_p001

**Photosynthesis.CAM/C4 photosynthesis**

**phoenolpyruvate carboxylase3 description:Phosphoenolpyruvate carboxylase isoform 1%3B Phosphoeno**

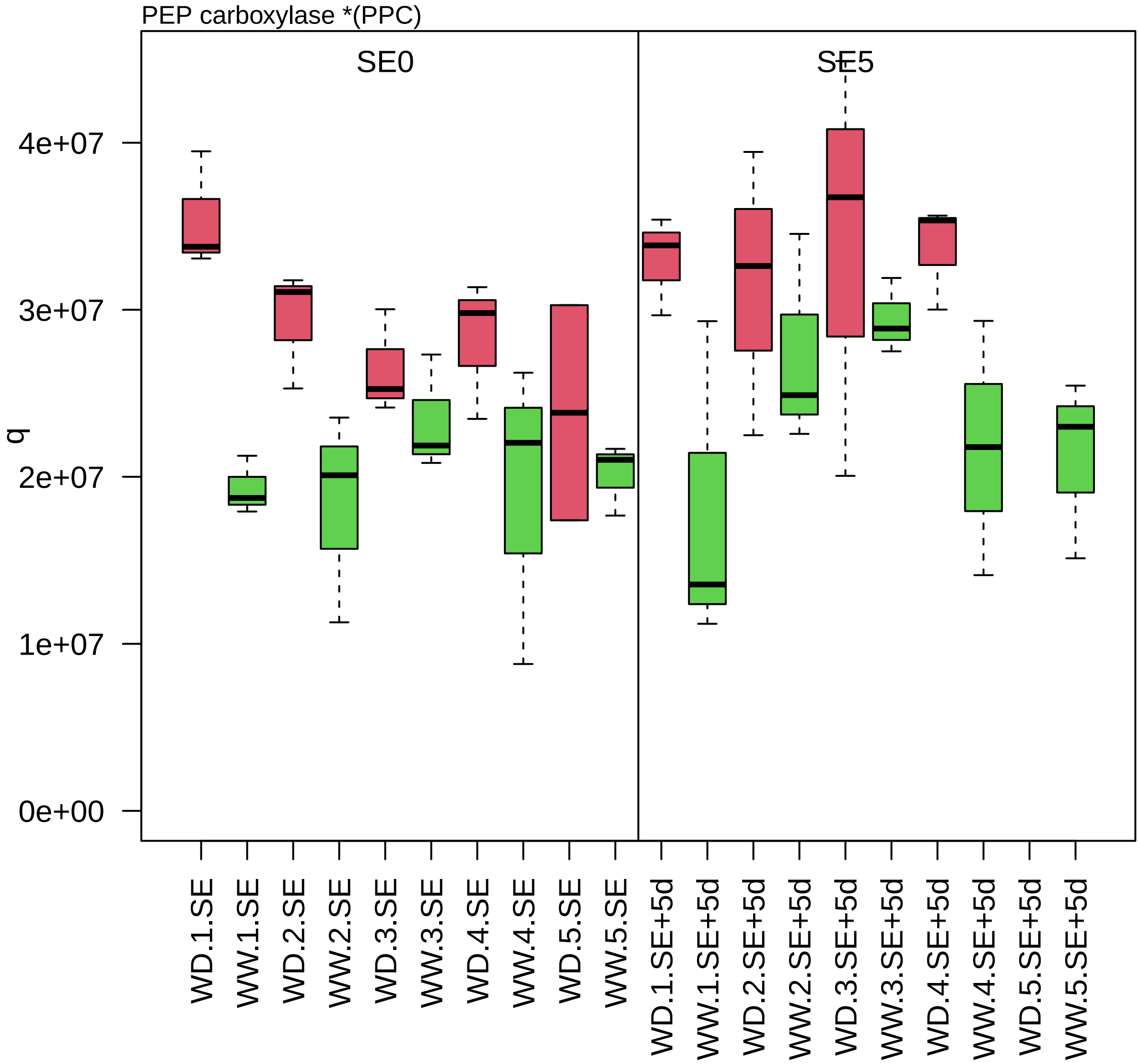

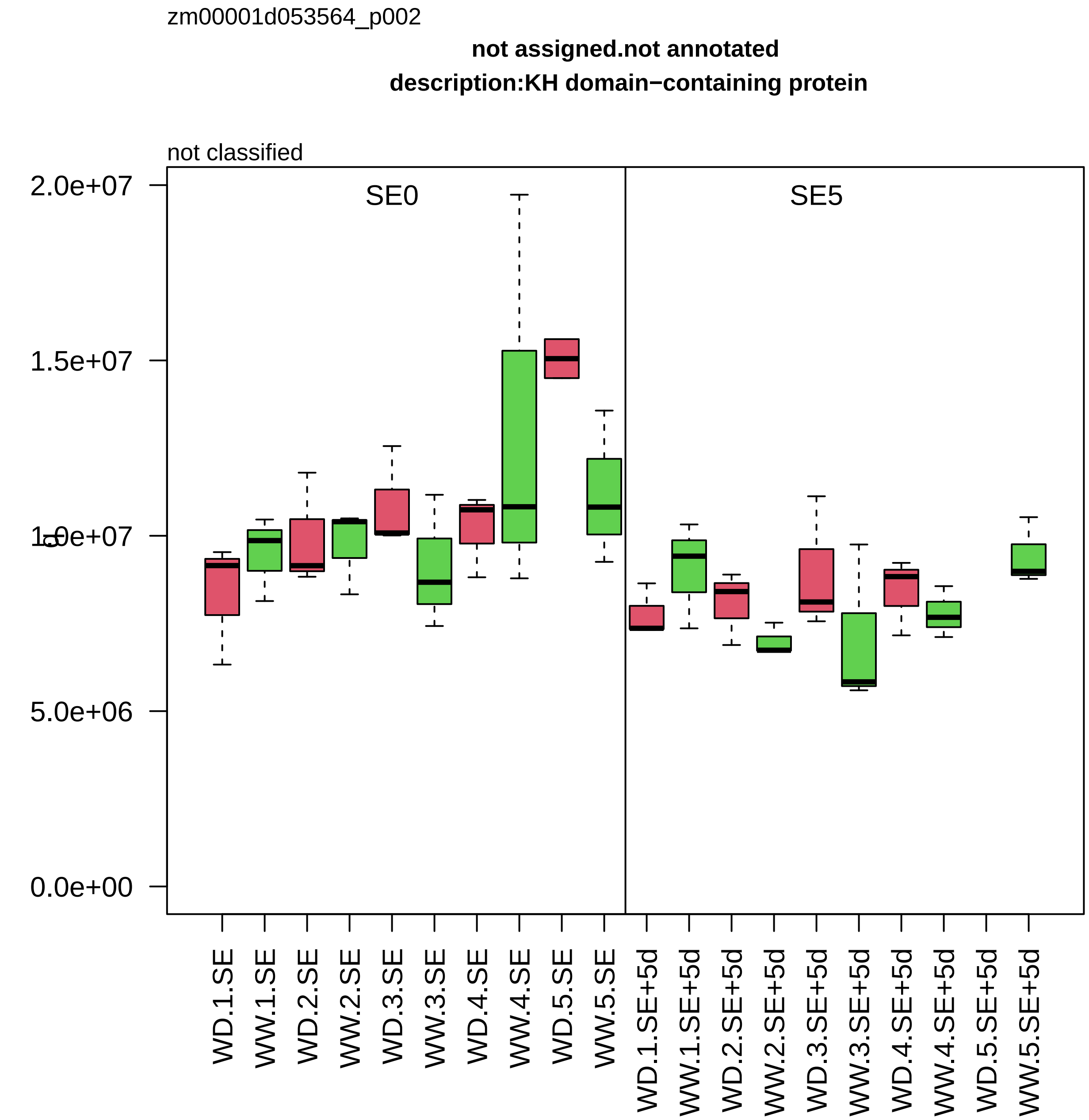

zm00001d053564_p002
not assigned.not annotated
description:KH domain−containing protein
not classified
SE0
SE5
2.0e+07
1.5e+07
1.0e+07
5.0e+06
0.0e+00
WD.1.SE
WW.1.SE
WD.2.SE
WW.2.SE
WD.3.SE
WW.3.SE
WD.4.SE
WW.4.SE
WD.5.SE
WW.5.SE
WD.1.SE+5d
WW.1.SE+5d
WD.2.SE+5d
WW.2.SE+5d
WD.3.SE+5d
WW.3.SE+5d
WD.4.SE+5d
WW.4.SE+5d
WD.5.SE+5d
WW.5.SE+5d

zm00001d053576_p002

**Protein biosynthesis.translation elongation**
**description:Probable elongation factor 1−gamma 2**

component *(eEF1B−gamma) of eEF1B eEF1A−GDP−recycling complex

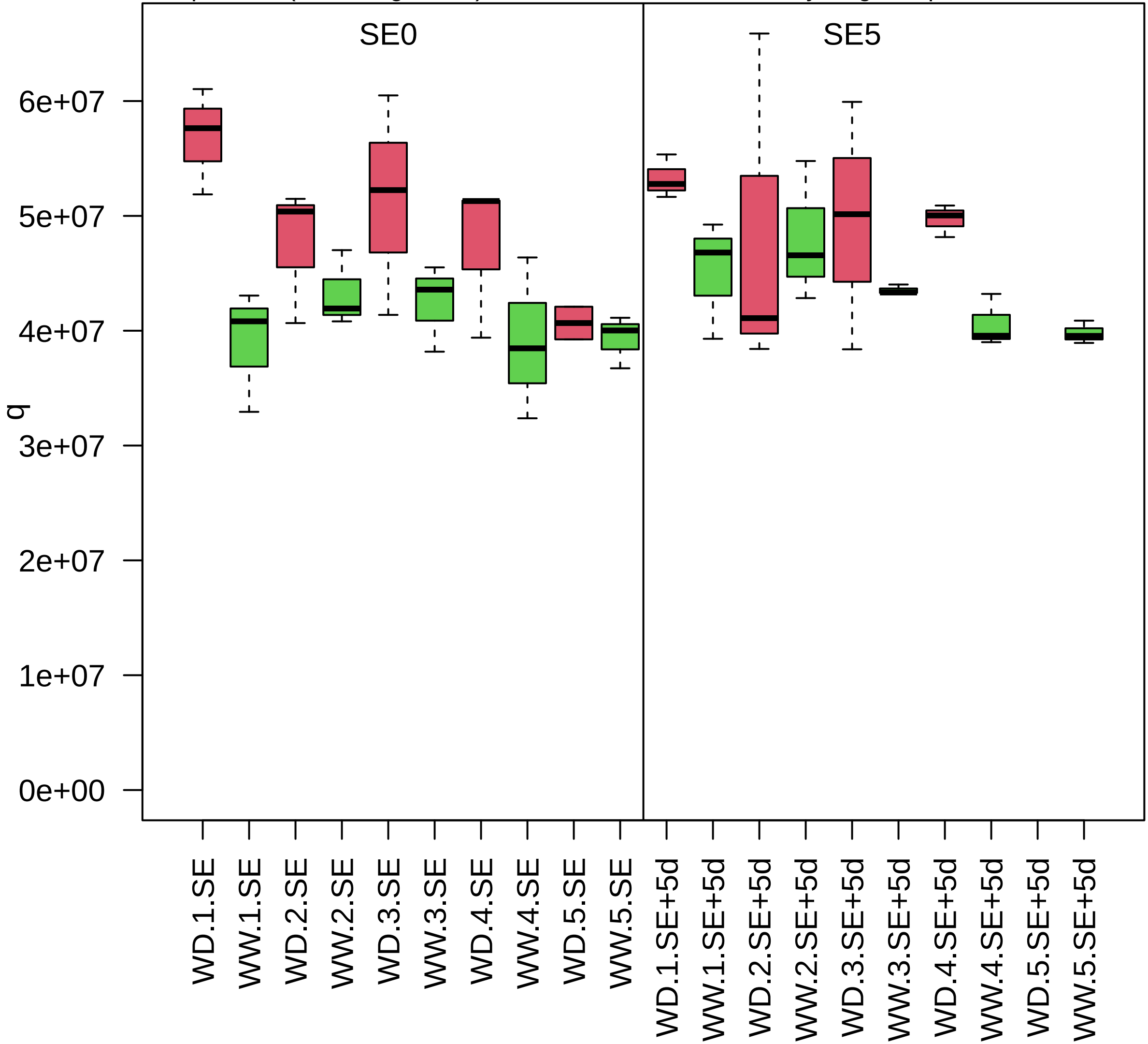

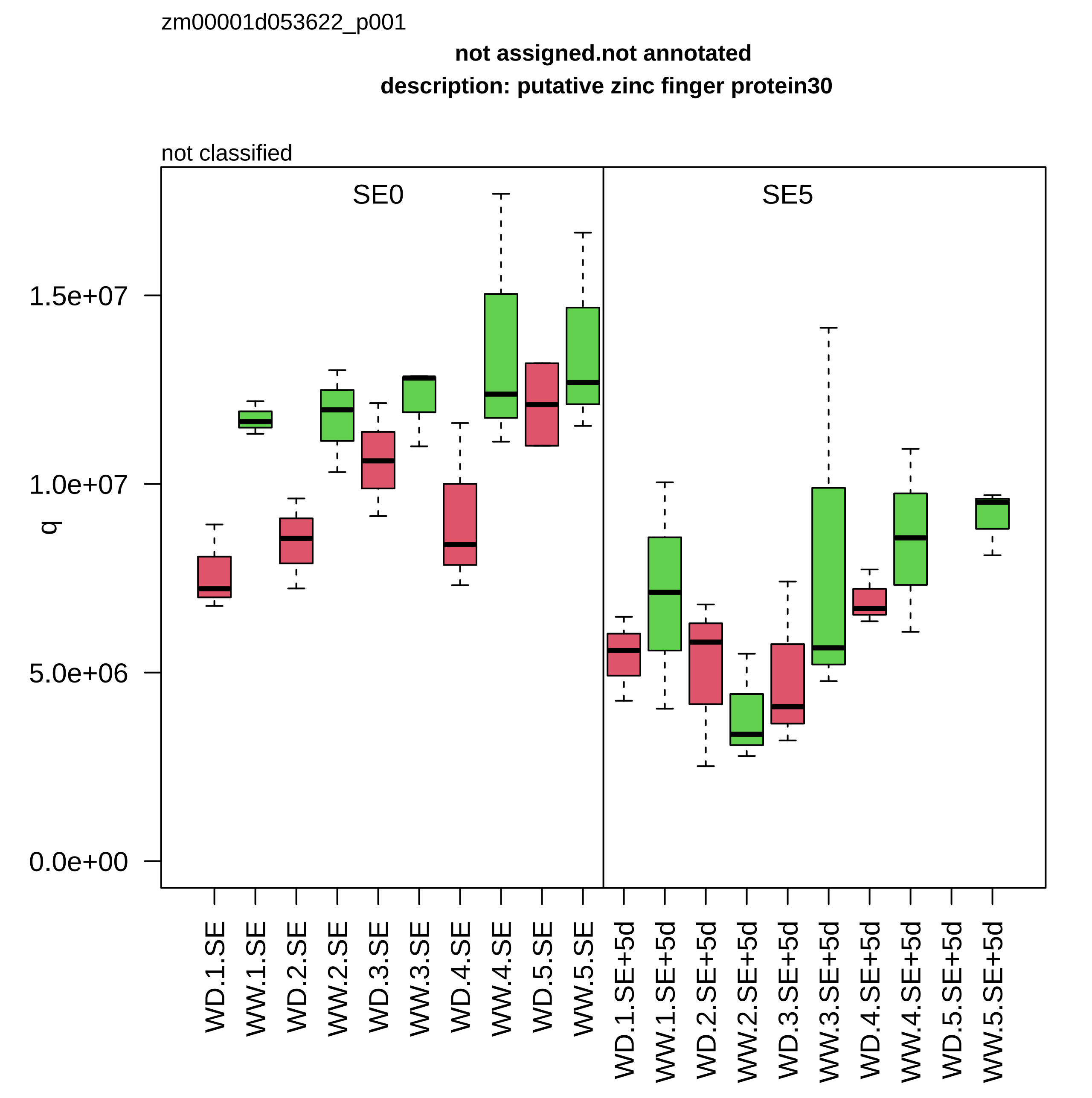

zm00001d053622_p001
not assigned.not annotated
description: putative zinc finger protein30
not classified
SE0
SE5
q
1.5e+07
1.0e+07
5.0e+06
0.0e+00
WD.1.SE
WW.1.SE
WD.2.SE
WW.2.SE
WD.3.SE
WW.3.SE
WD.4.SE
WW.4.SE
WD.5.SE
WW.5.SE
WD.1.SE+5d
WW.1.SE+5d
WD.2.SE+5d
WW.2.SE+5d
WD.3.SE+5d
WW.3.SE+5d
WD.4.SE+5d
WW.4.SE+5d
WD.5.SE+5d
WW.5.SE+5d

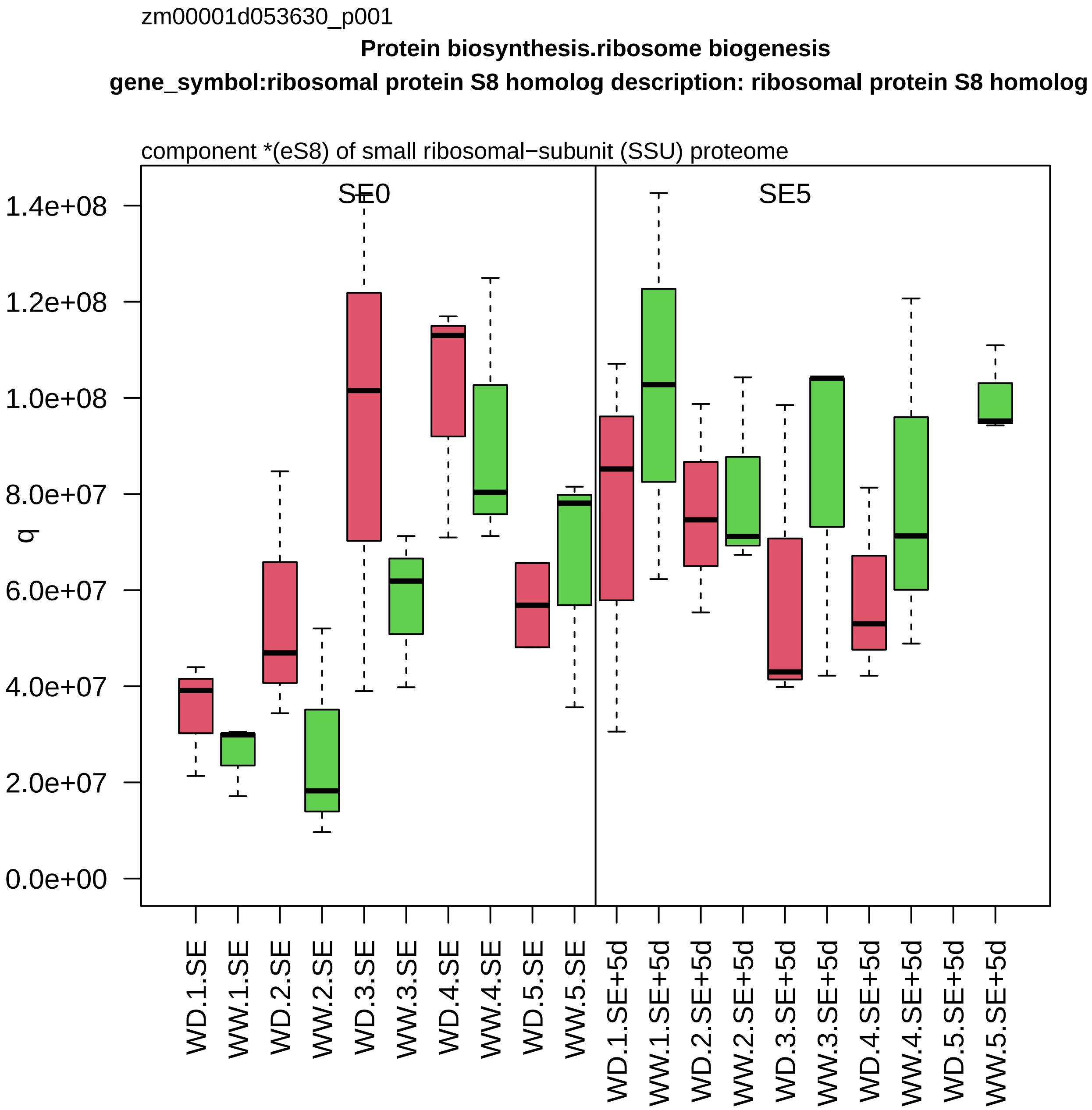

zm00001d053630_p001
Protein biosynthesis.ribosome biogenesis
gene_symbol:ribosomal protein S8 homolog description: ribosomal protein S8 homolog
component *(eS8) of small ribosomal−subunit (SSU) proteome
SE0
SE5
q
1.4e+08
1.2e+08
1.0e+08
8.0e+07
6.0e+07
4.0e+07
2.0e+07
0.0e+00
WD.1.SE
WW.1.SE
WD.2.SE
WW.2.SE
WD.3.SE
WW.3.SE
WD.4.SE
WW.4.SE
WD.5.SE
WW.5.SE
WD.1.SE+5d
WW.1.SE+5d
WD.2.SE+5d
WW.2.SE+5d
WD.3.SE+5d
WW.3.SE+5d
WD.4.SE+5d
WW.4.SE+5d
WD.5.SE+5d
WW.5.SE+5d

zm00001d053663_p001

**Amino acid metabolism.amino acid degradation**
**description:Fumarylacetoacetase**

fumarylacetoacetate hydrolase *(FAH)

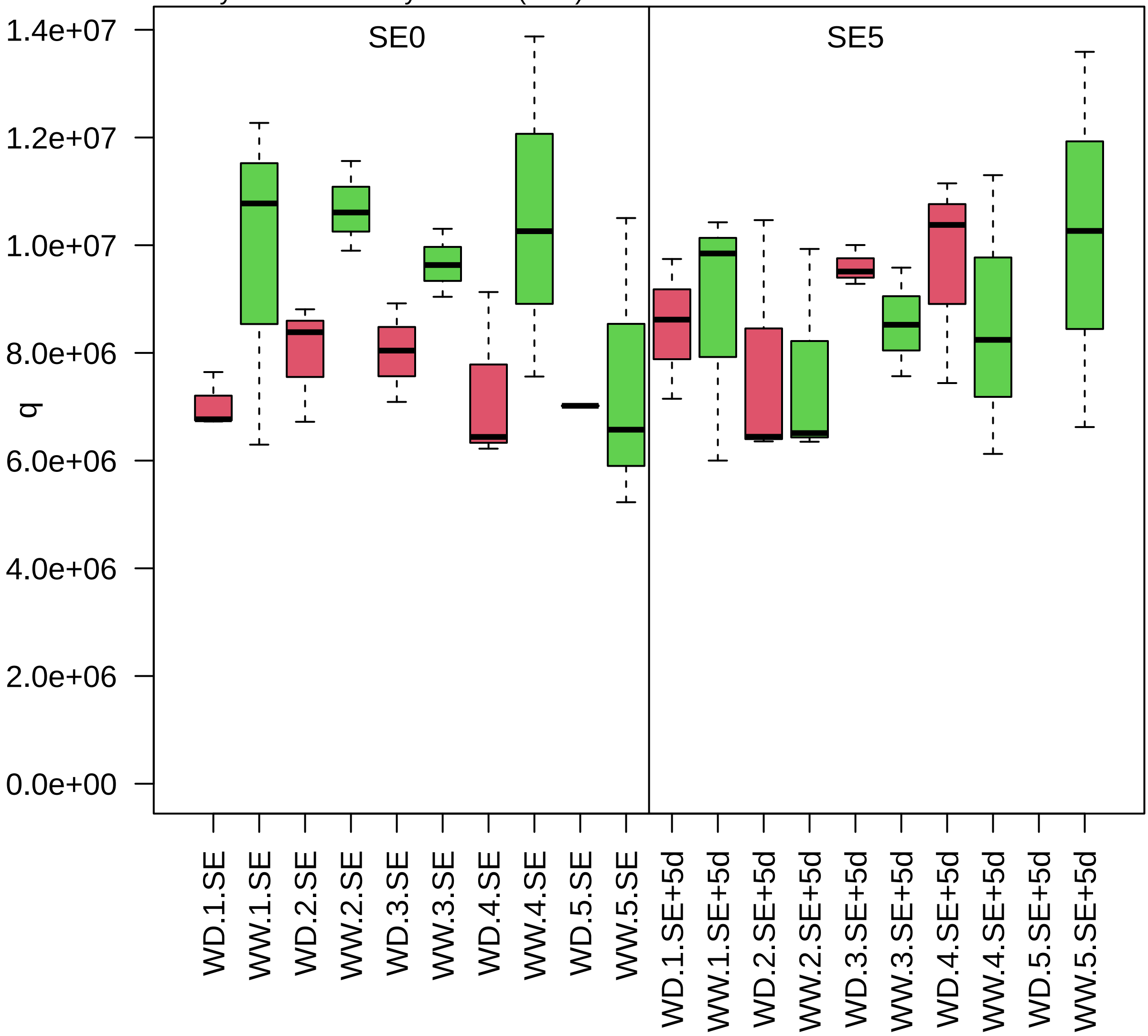

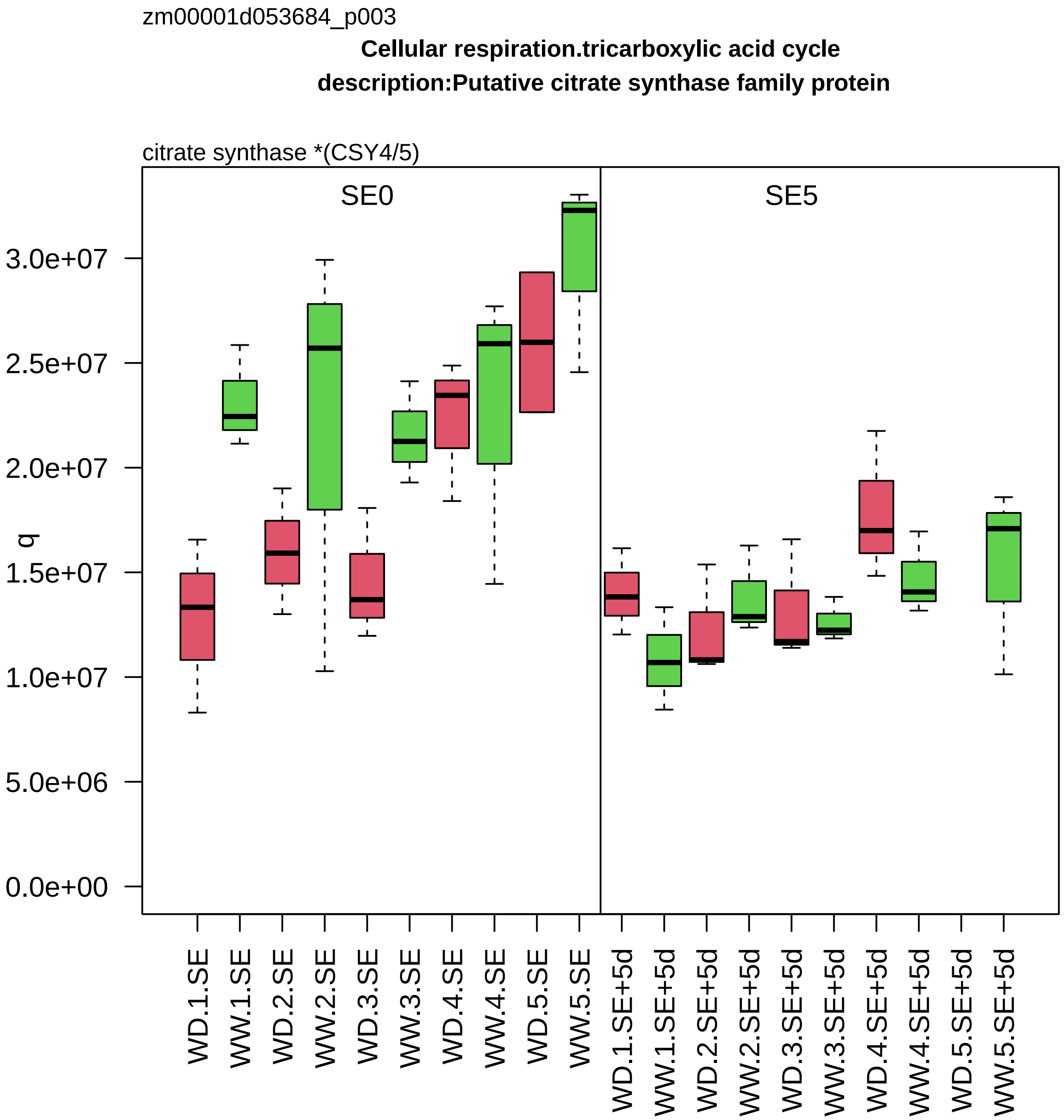
zm00001d053684_p003
Cellular respiration.tricarboxylic acid cycle
description:Putative citrate synthase family protein
citrate synthase *(CSY4/5)
SE0
SE5
q
3.0e+07
2.5e+07
2.0e+07
1.5e+07
1.0e+07
5.0e+06
0.0e+00
WD.1.SE
WW.1.SE
WD.2.SE
WW.2.SE
WD.3.SE
WW.3.SE
WD.4.SE
WW.4.SE
WD.5.SE
WW.5.SE
WD.1.SE+5d
WW.1.SE+5d
WD.2.SE+5d
WW.2.SE+5d
WD.3.SE+5d
WW.3.SE+5d
WD.4.SE+5d
WW.4.SE+5d
WD.5.SE+5d
WW.5.SE+5d

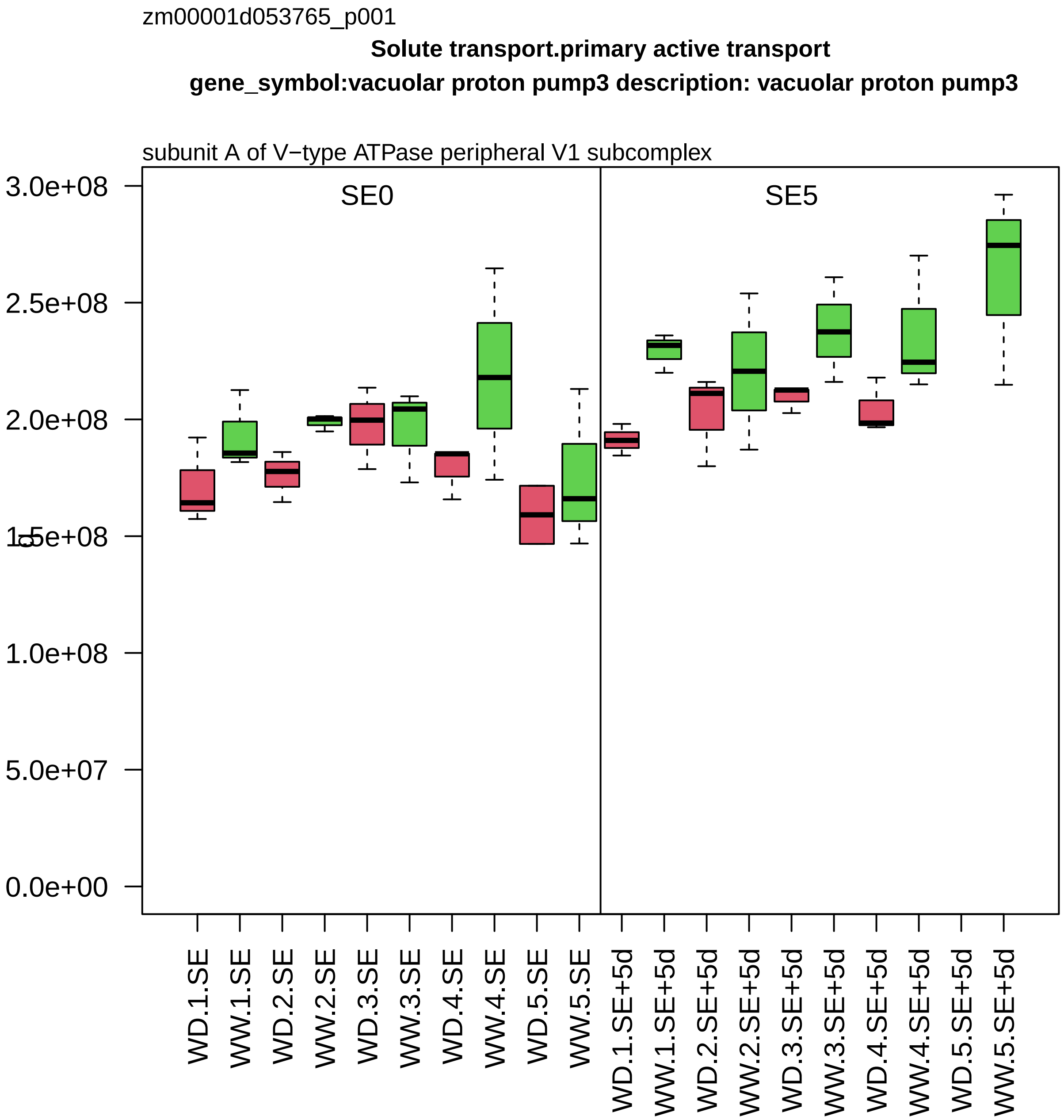

zm00001d053765_p001
Solute transport.primary active transport
gene_symbol:vacuolar proton pump3 description: vacuolar proton pump3
subunit A of V−type ATPase peripheral V1 subcomplex
SE0
SE5
3.0e+08
2.5e+08
2.0e+08
1.5e+08
1.0e+08
5.0e+07
0.0e+00
WD.1.SE
WW.1.SE
WD.2.SE
WW.2.SE
WD.3.SE
WW.3.SE
WD.4.SE
WW.4.SE
WD.5.SE
WW.5.SE
WD.1.SE+5d
WW.1.SE+5d
WD.2.SE+5d
WW.2.SE+5d
WD.3.SE+5d
WW.3.SE+5d
WD.4.SE+5d
WW.4.SE+5d
WD.5.SE+5d
WW.5.SE+5d

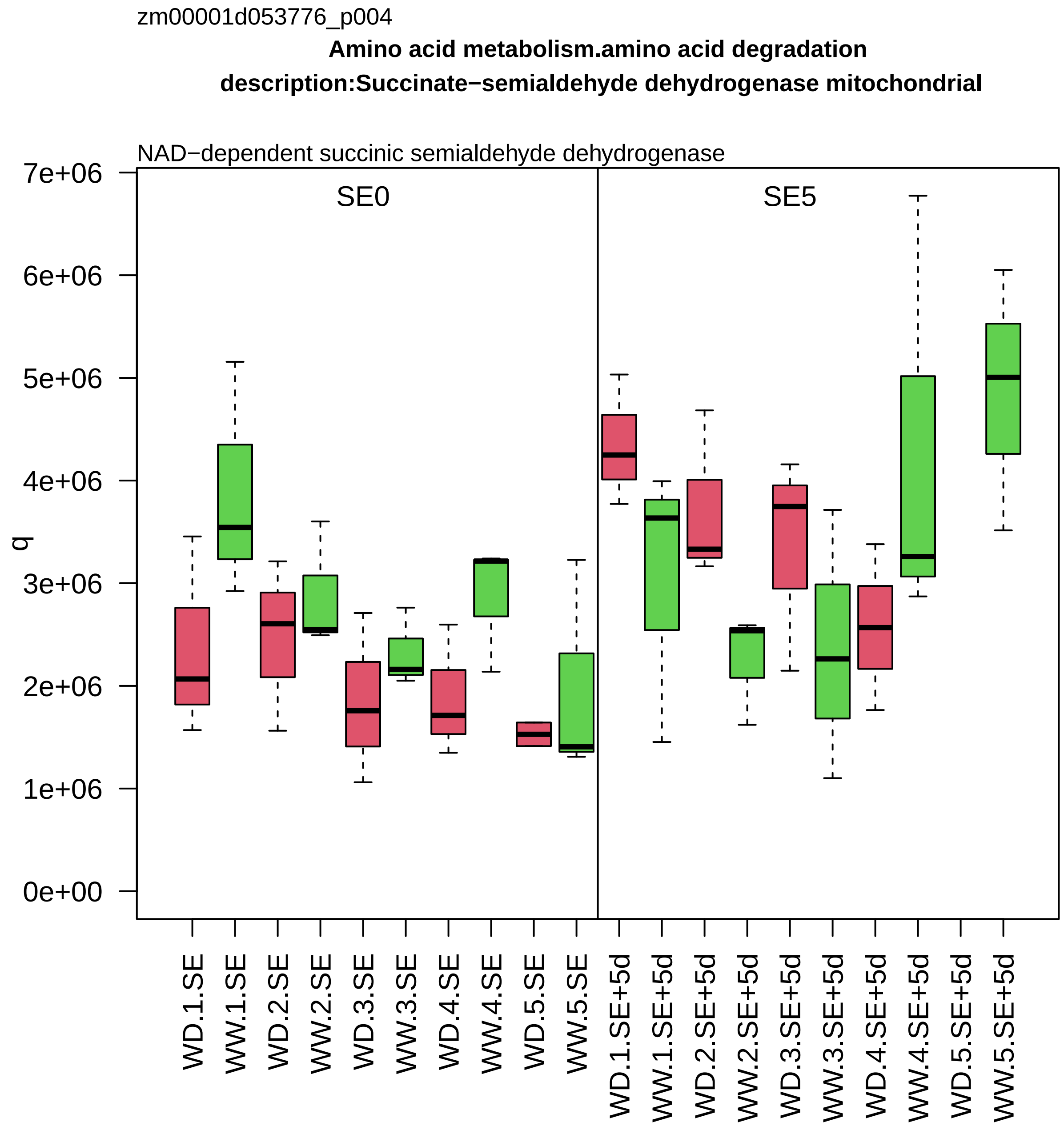

zm00001d053776_p004
Amino acid metabolism.amino acid degradation
description:Succinate−semialdehyde dehydrogenase mitochondrial
NAD−dependent succinic semialdehyde dehydrogenase
SE0
SE5
q
7e+06
6e+06
5e+06
4e+06
3e+06
2e+06
1e+06
0e+00
WD.1.SE
WW.1.SE
WD.2.SE
WW.2.SE
WD.3.SE
WW.3.SE
WD.4.SE
WW.4.SE
WD.5.SE
WW.5.SE
WD.1.SE+5d
WW.1.SE+5d
WD.2.SE+5d
WW.2.SE+5d
WD.3.SE+5d
WW.3.SE+5d
WD.4.SE+5d
WW.4.SE+5d
WD.5.SE+5d
WW.5.SE+5d

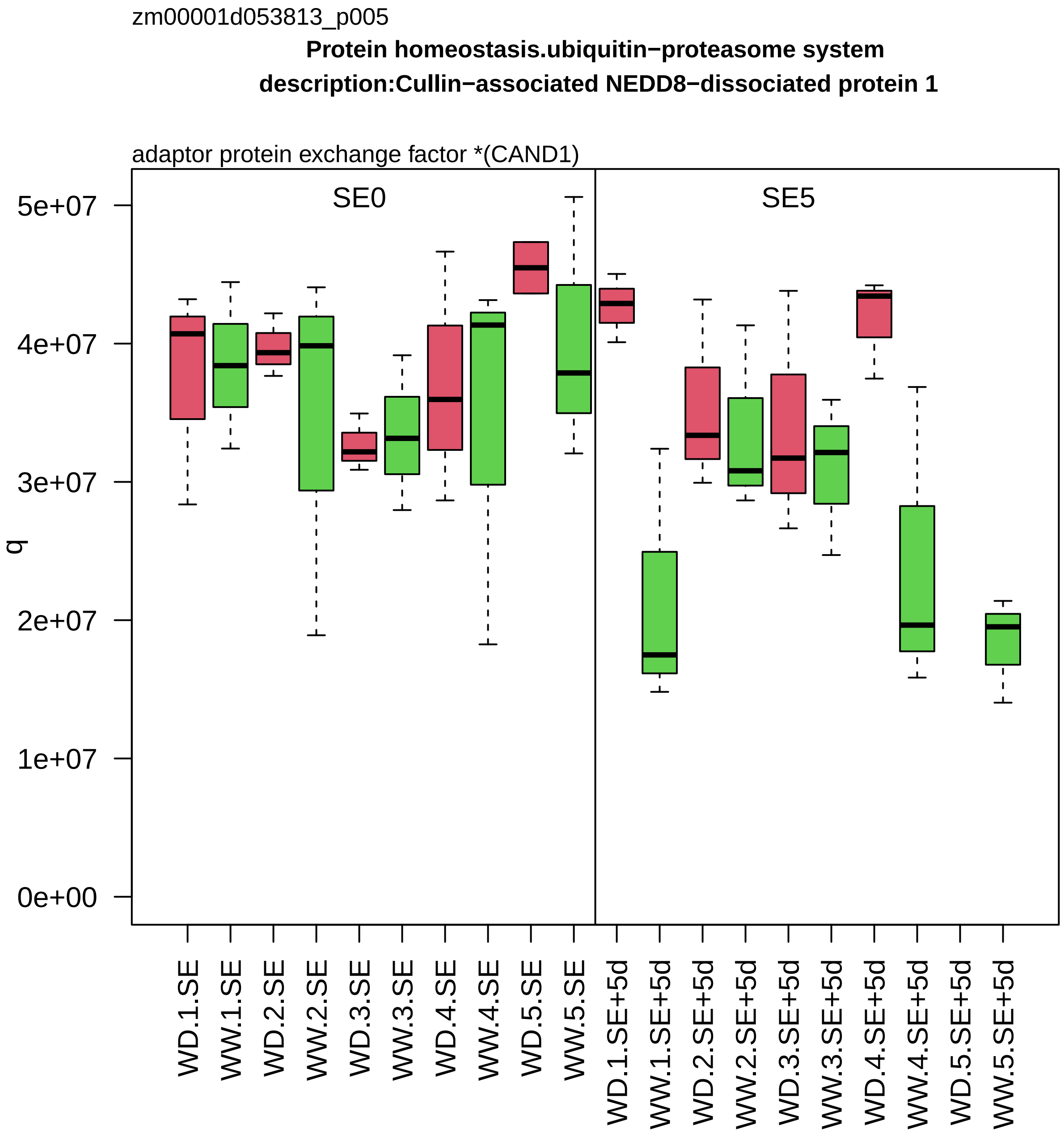

zm00001d053813_p005
Protein homeostasis.ubiquitin−proteasome system
description:Cullin−associated NEDD8−dissociated protein 1
adaptor protein exchange factor *(CAND1)
SE0
SE5
q
5e+07
4e+07
3e+07
2e+07
1e+07
0e+00
WD.1.SE
WW.1.SE
WD.2.SE
WW.2.SE
WD.3.SE
WW.3.SE
WD.4.SE
WW.4.SE
WD.5.SE
WW.5.SE
WD.1.SE+5d
WW.1.SE+5d
WD.2.SE+5d
WW.2.SE+5d
WD.3.SE+5d
WW.3.SE+5d
WD.4.SE+5d
WW.4.SE+5d
WD.5.SE+5d
WW.5.SE+5d

zm00001d053857_p002

**Cytoskeleton organisation.actin and tubulin folding**
**description:T−complex protein 1 subunit eta**

subunit eta *(CCT7) of CCT chaperonin folding complex

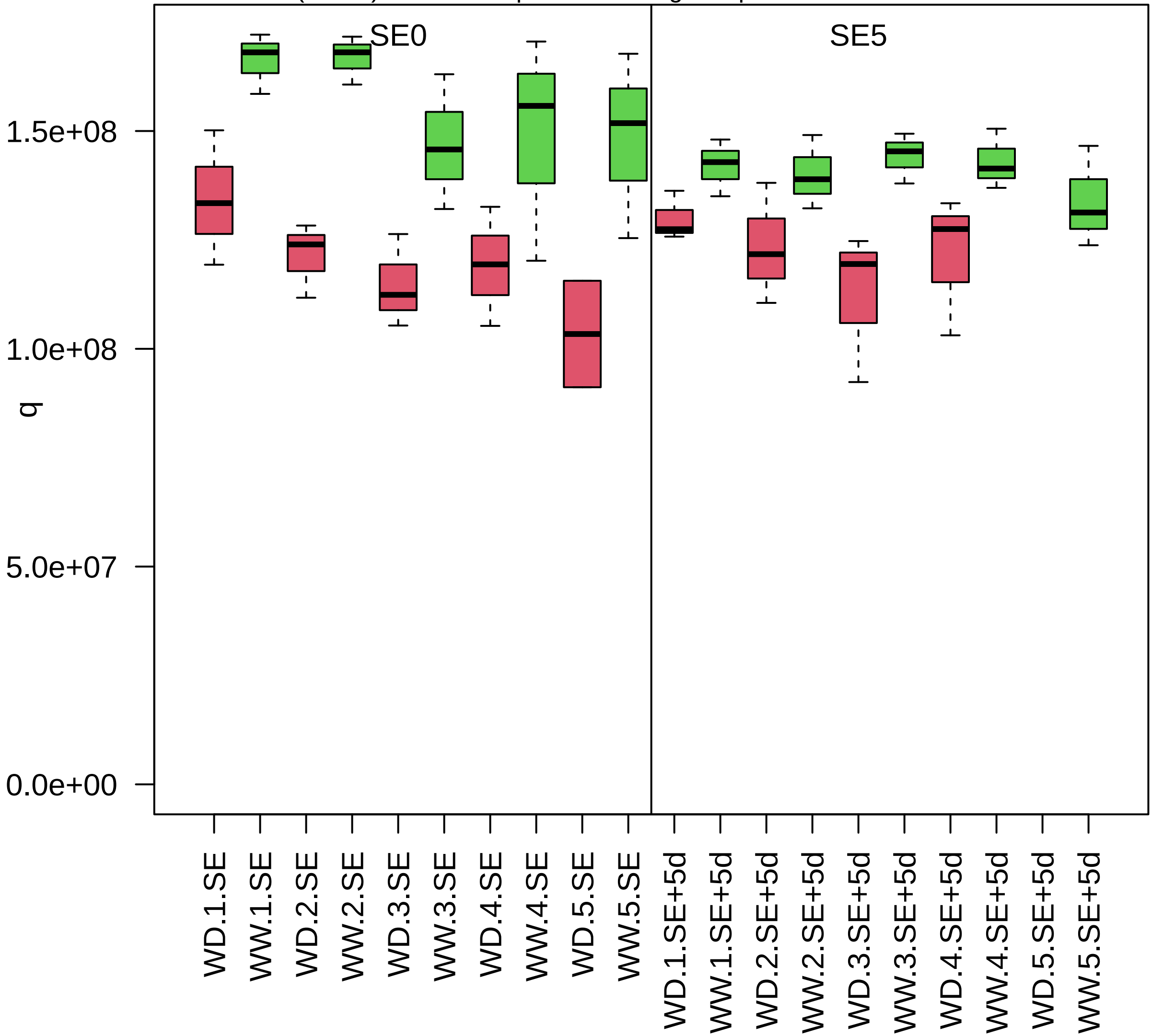

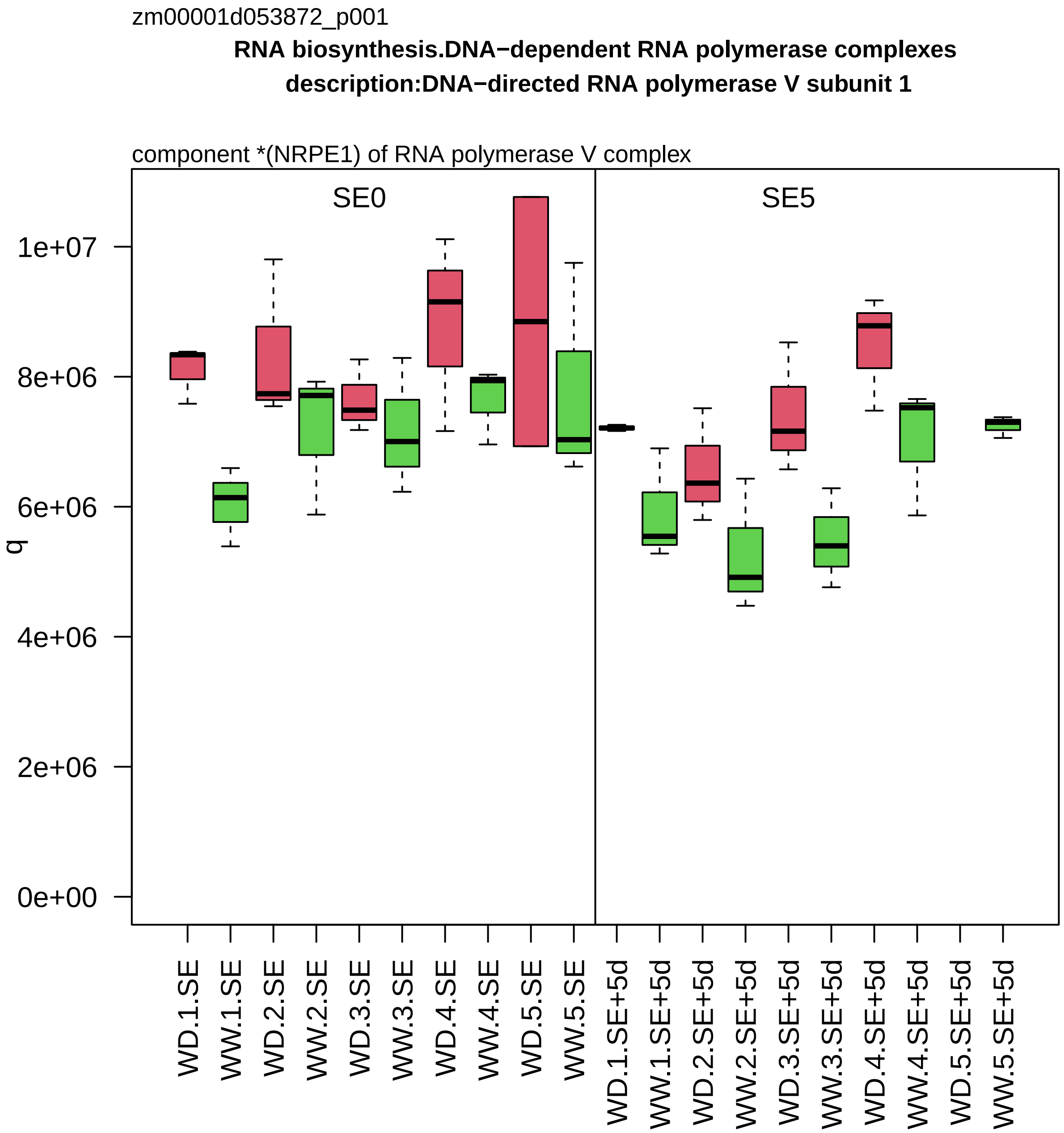
zm00001d053872_p001
RNA biosynthesis.DNA−dependent RNA polymerase complexes
description:DNA−directed RNA polymerase V subunit 1
component *(NRPE1) of RNA polymerase V complex
SE0
SE5
q
1e+07
8e+06
6e+06
4e+06
2e+06
0e+00
WD.1.SE
WW.1.SE
WD.2.SE
WW.2.SE
WD.3.SE
WW.3.SE
WD.4.SE
WW.4.SE
WD.5.SE
WW.5.SE
WD.1.SE+5d
WW.1.SE+5d
WD.2.SE+5d
WW.2.SE+5d
WD.3.SE+5d
WW.3.SE+5d
WD.4.SE+5d
WW.4.SE+5d
WD.5.SE+5d
WW.5.SE+5d

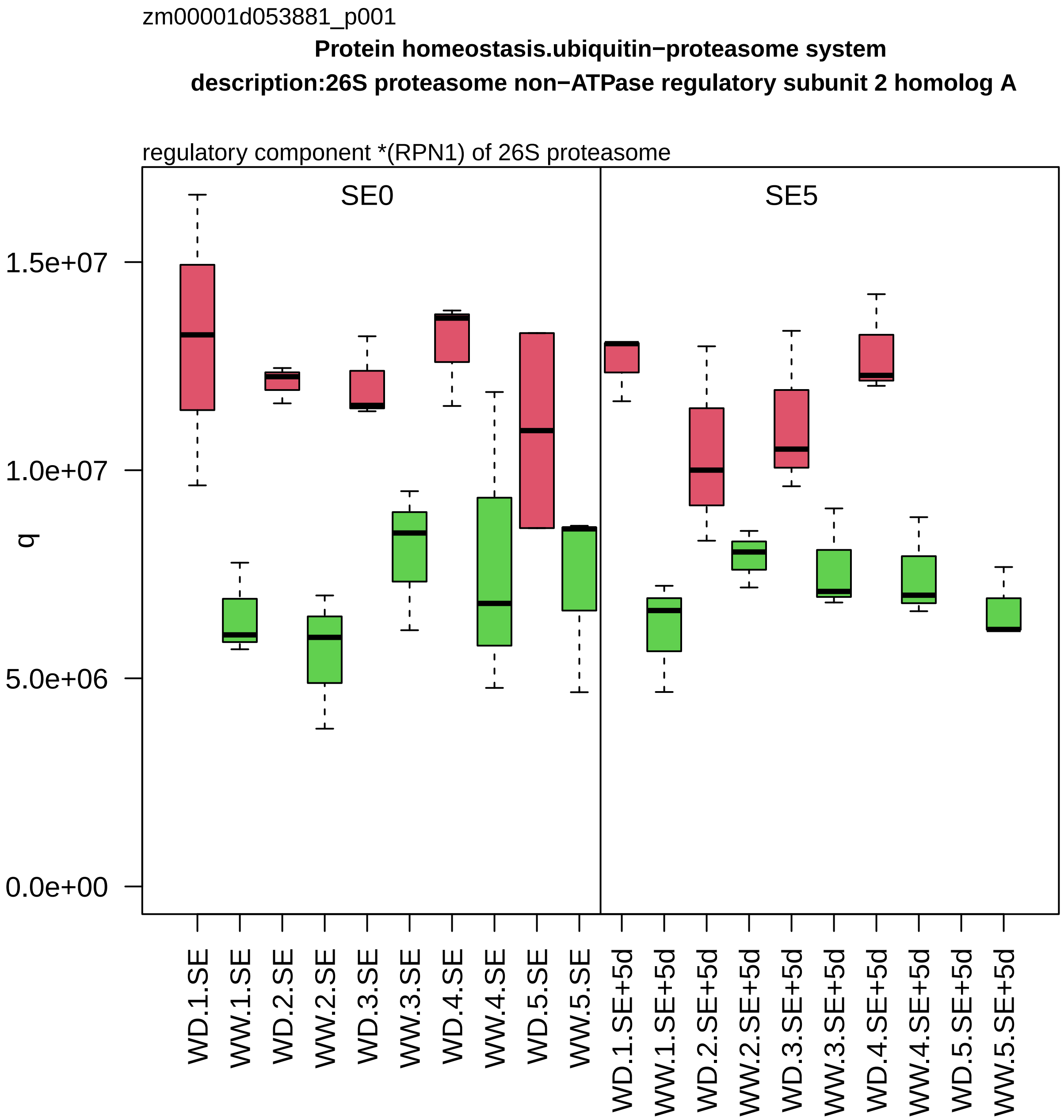
zm00001d053881_p001
Protein homeostasis.ubiquitin−proteasome system
description:26S proteasome non−ATPase regulatory subunit 2 homolog A
regulatory component *(RPN1) of 26S proteasome
SE0
SE5
q
1.5e+07
1.0e+07
5.0e+06
0.0e+00
WD.1.SE
WW.1.SE
WD.2.SE
WW.2.SE
WD.3.SE
WW.3.SE
WD.4.SE
WW.4.SE
WD.5.SE
WW.5.SE
WD.1.SE+5d
WW.1.SE+5d
WD.2.SE+5d
WW.2.SE+5d
WD.3.SE+5d
WW.3.SE+5d
WD.4.SE+5d
WW.4.SE+5d
WD.5.SE+5d
WW.5.SE+5d

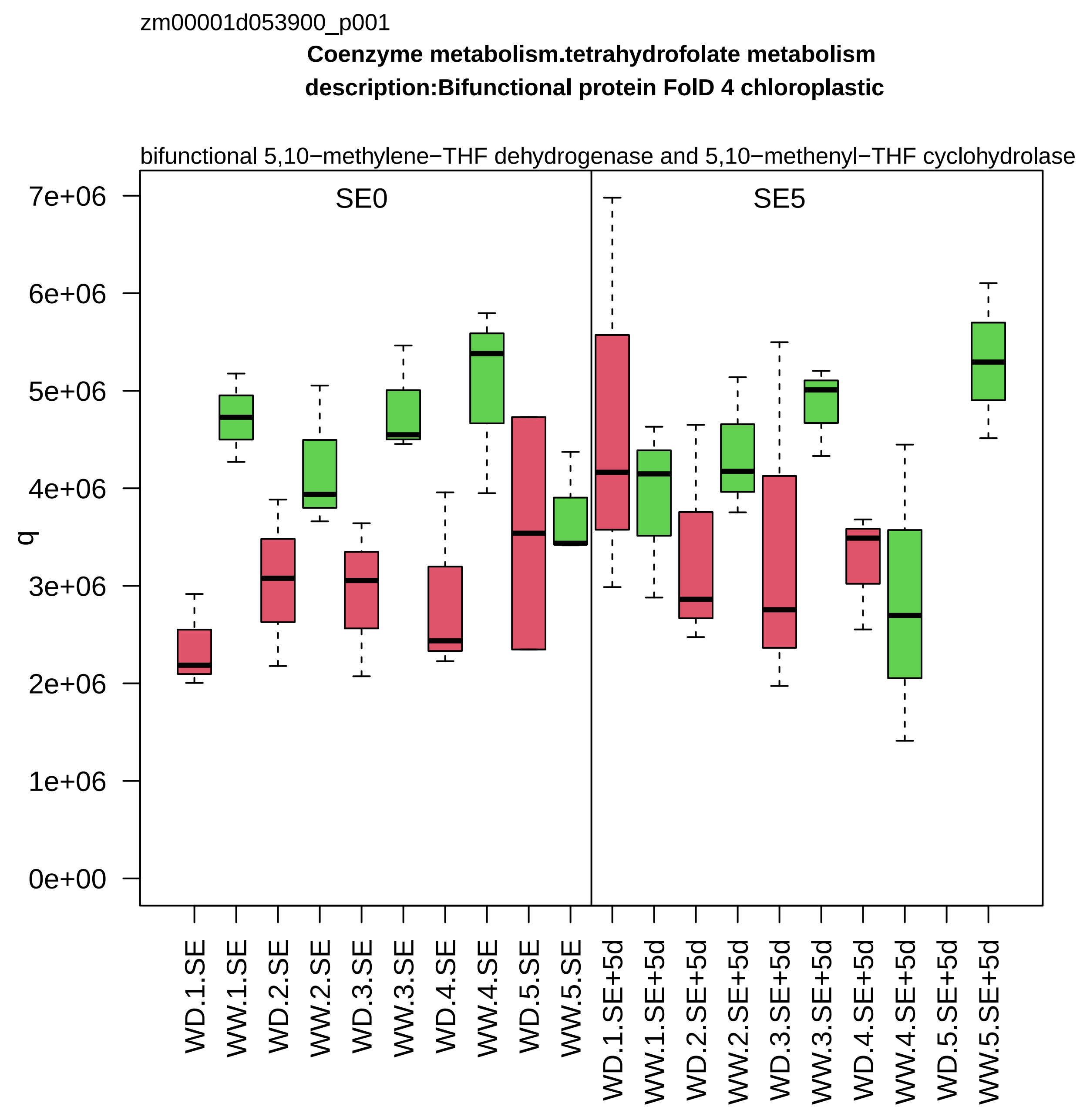
zm00001d053900_p001
Coenzyme metabolism.tetrahydrofolate metabolism
description:Bifunctional protein FolD 4 chloroplastic
bifunctional 5,10−methylene−THF dehydrogenase and 5,10−methenyl−THF cyclohydrolase
SE0
SE5
q
0e+00
1e+06
2e+06
3e+06
4e+06
5e+06
6e+06
7e+06
WD.1.SE
WW.1.SE
WD.2.SE
WW.2.SE
WD.3.SE
WW.3.SE
WD.4.SE
WW.4.SE
WD.5.SE
WW.5.SE
WD.1.SE+5d
WW.1.SE+5d
WD.2.SE+5d
WW.2.SE+5d
WD.3.SE+5d
WW.3.SE+5d
WD.4.SE+5d
WW.4.SE+5d
WD.5.SE+5d
WW.5.SE+5d

zm00001d053908_p001

**RNA processing.RNA homeostasis**

**gene_symbol:CSD−transcription factor 2 description:Cold shock protein 2**

RNA chaperone (CSP) of mRNA quality control

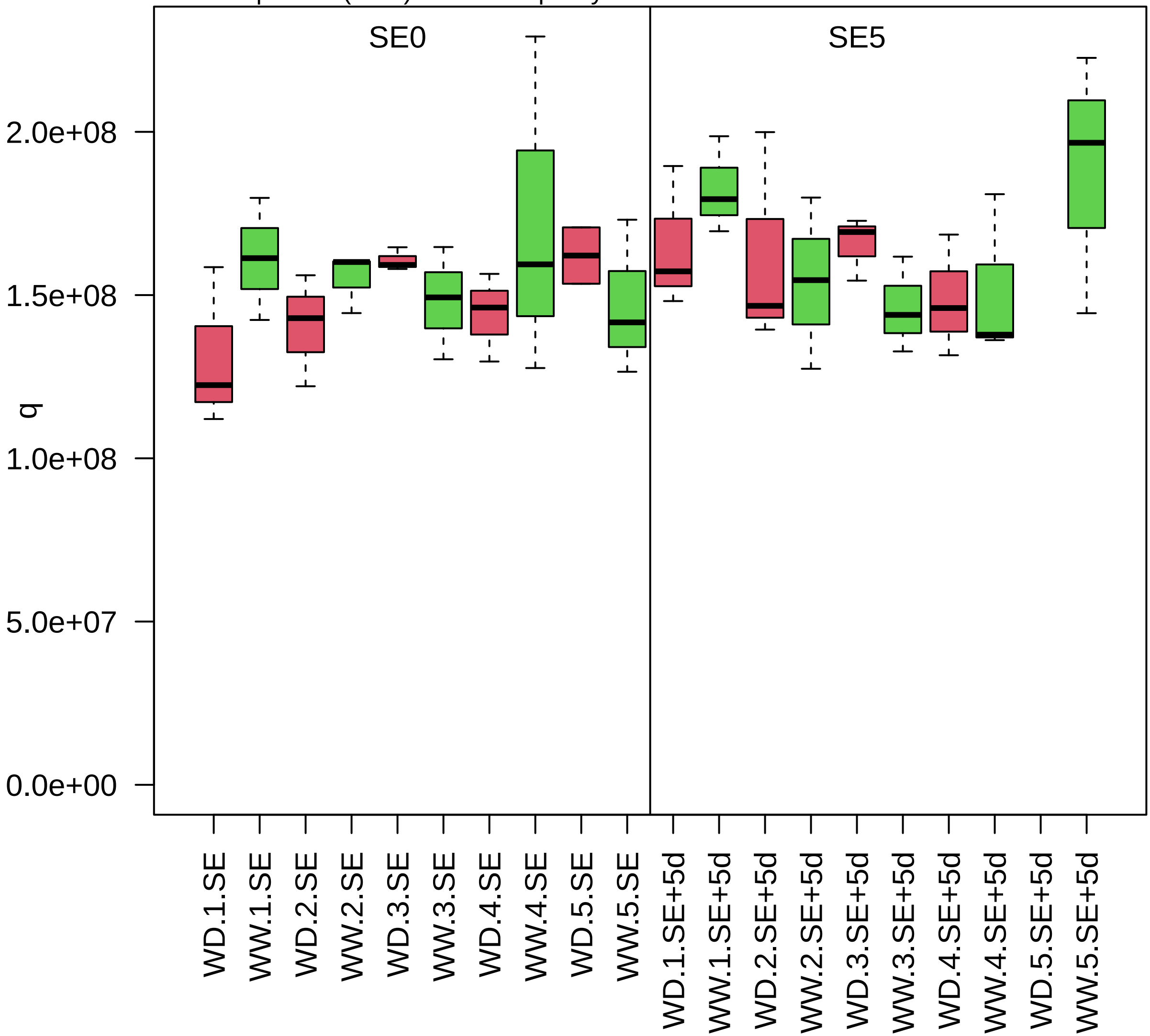

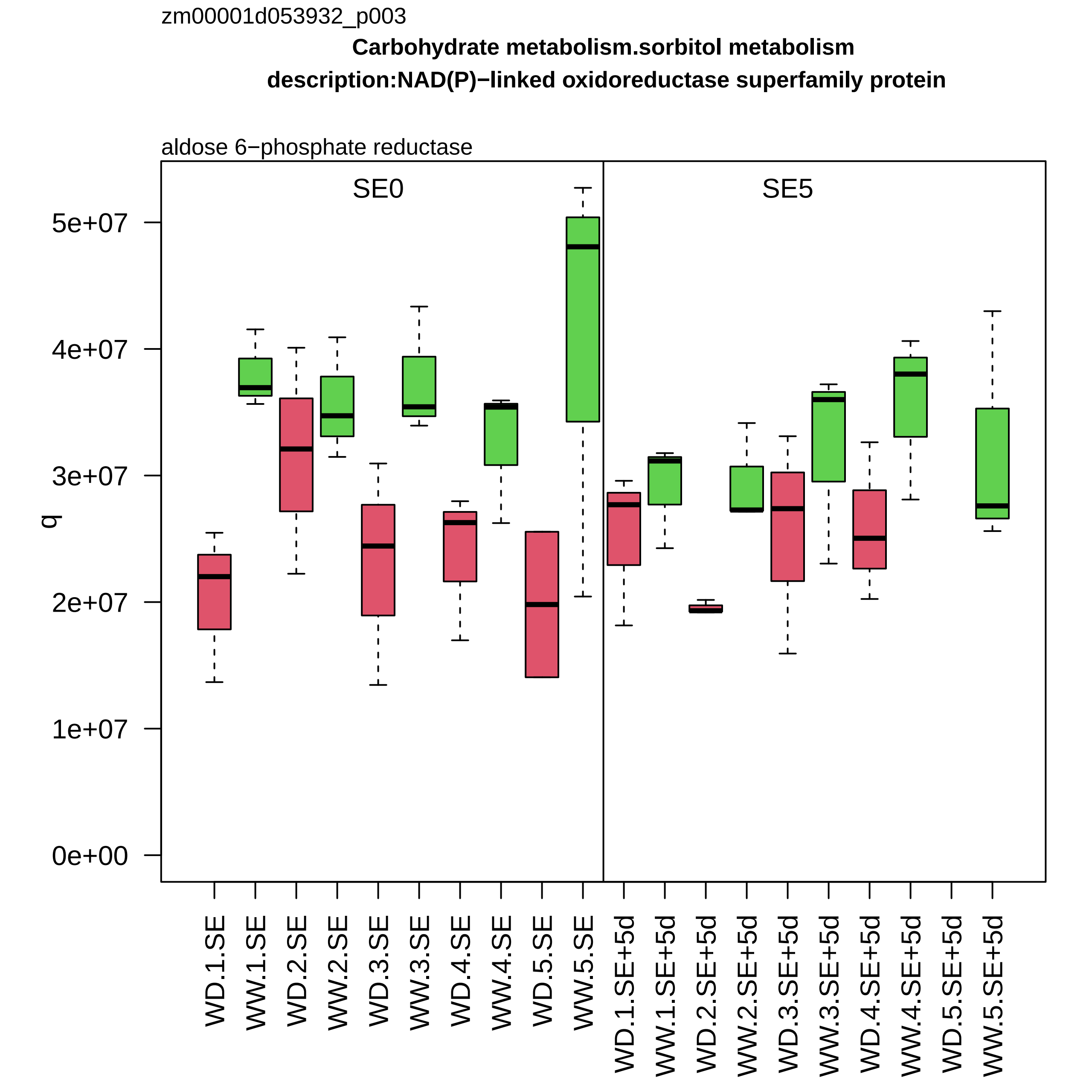
zm00001d053932_p003
Carbohydrate metabolism.sorbitol metabolism
description:NAD(P)−linked oxidoreductase superfamily protein
aldose 6−phosphate reductase
SE0
SE5
q
0e+00
1e+07
2e+07
3e+07
4e+07
5e+07
WD.1.SE
WW.1.SE
WD.2.SE
WW.2.SE
WD.3.SE
WW.3.SE
WD.4.SE
WW.4.SE
WD.5.SE
WW.5.SE
WD.1.SE+5d
WW.1.SE+5d
WD.2.SE+5d
WW.2.SE+5d
WD.3.SE+5d
WW.3.SE+5d
WD.4.SE+5d
WW.4.SE+5d
WD.5.SE+5d
WW.5.SE+5d

zm00001d054044_p001

**Redox homeostasis.reactive oxygen scavenging**
**gene_symbol:catalase3 description:Catalase isozyme 3**

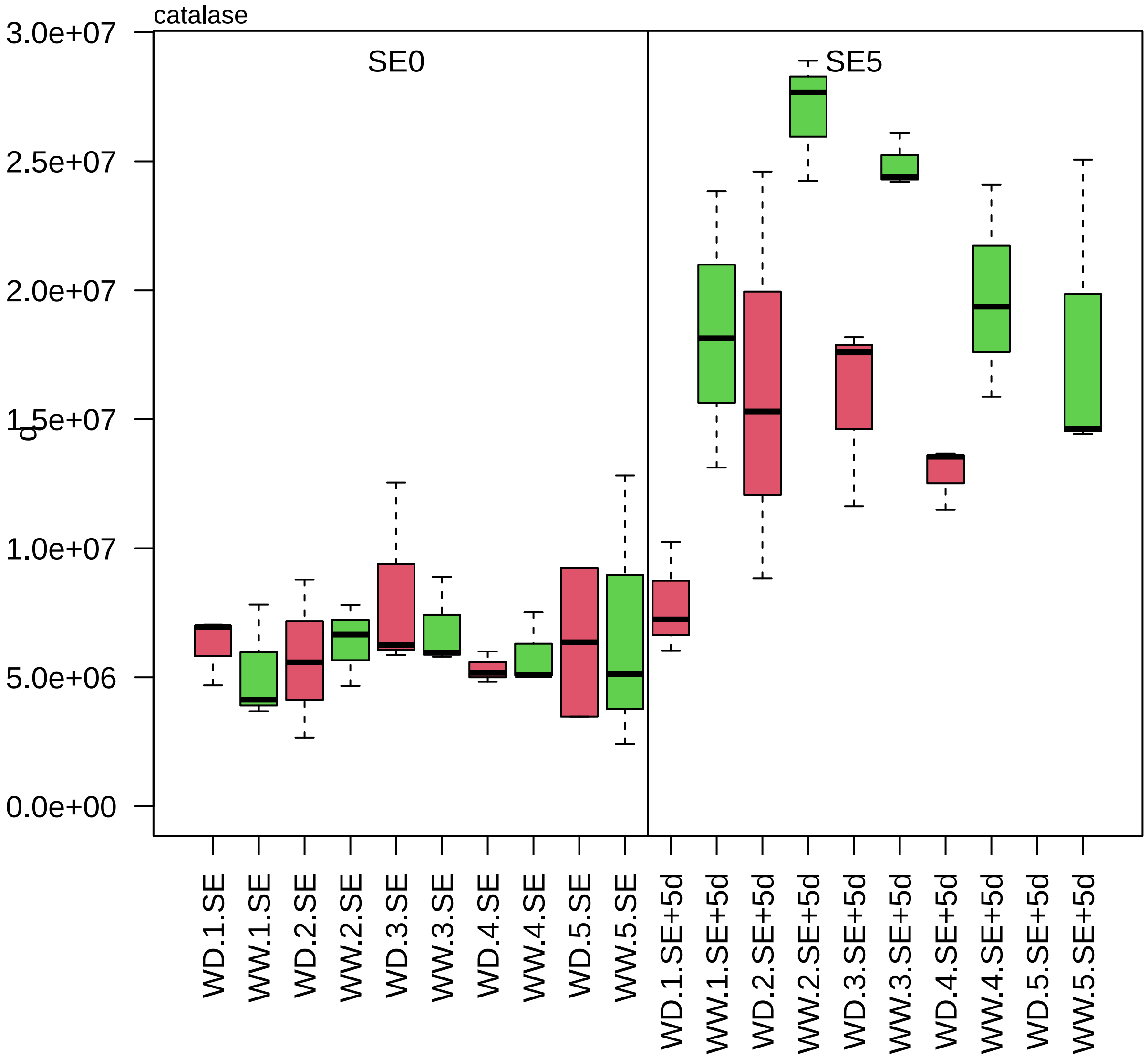

zm00001d054089_p001

**Photosynthesis.photophosphorylation**

**tion:Putative TCP−1/cpn60 chaperonin family protein isoform 1%3B Putative TCP−1/cpn60 chaperonin fam**

subunit beta of Cpn60 chaperonin complex

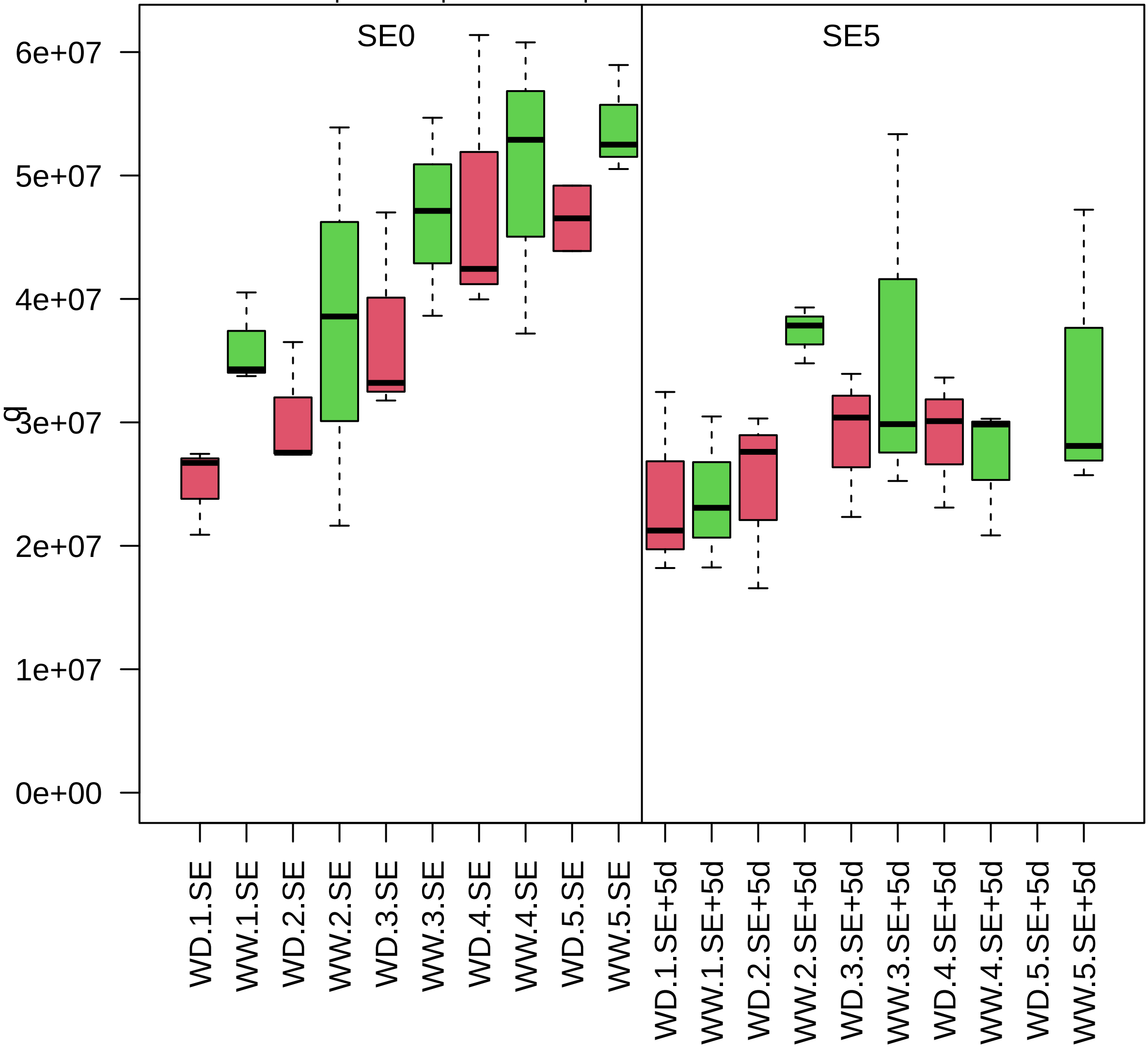

zm00001d054106_p003

**Protein modification.cysteine disulfide formation**

**gene_symbol:protein disulfide isomerase3 description: protein disulfide isomerase3**

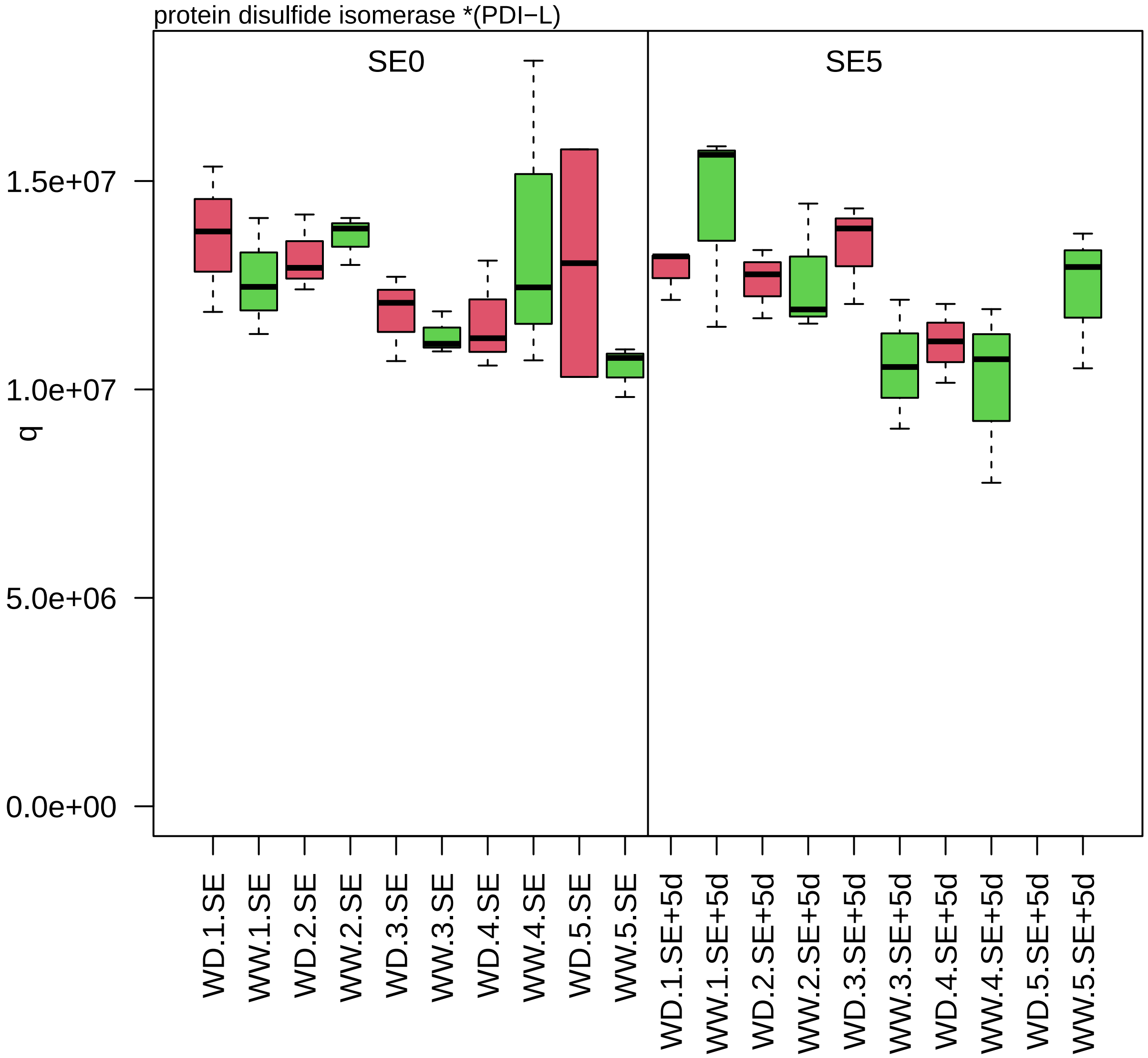

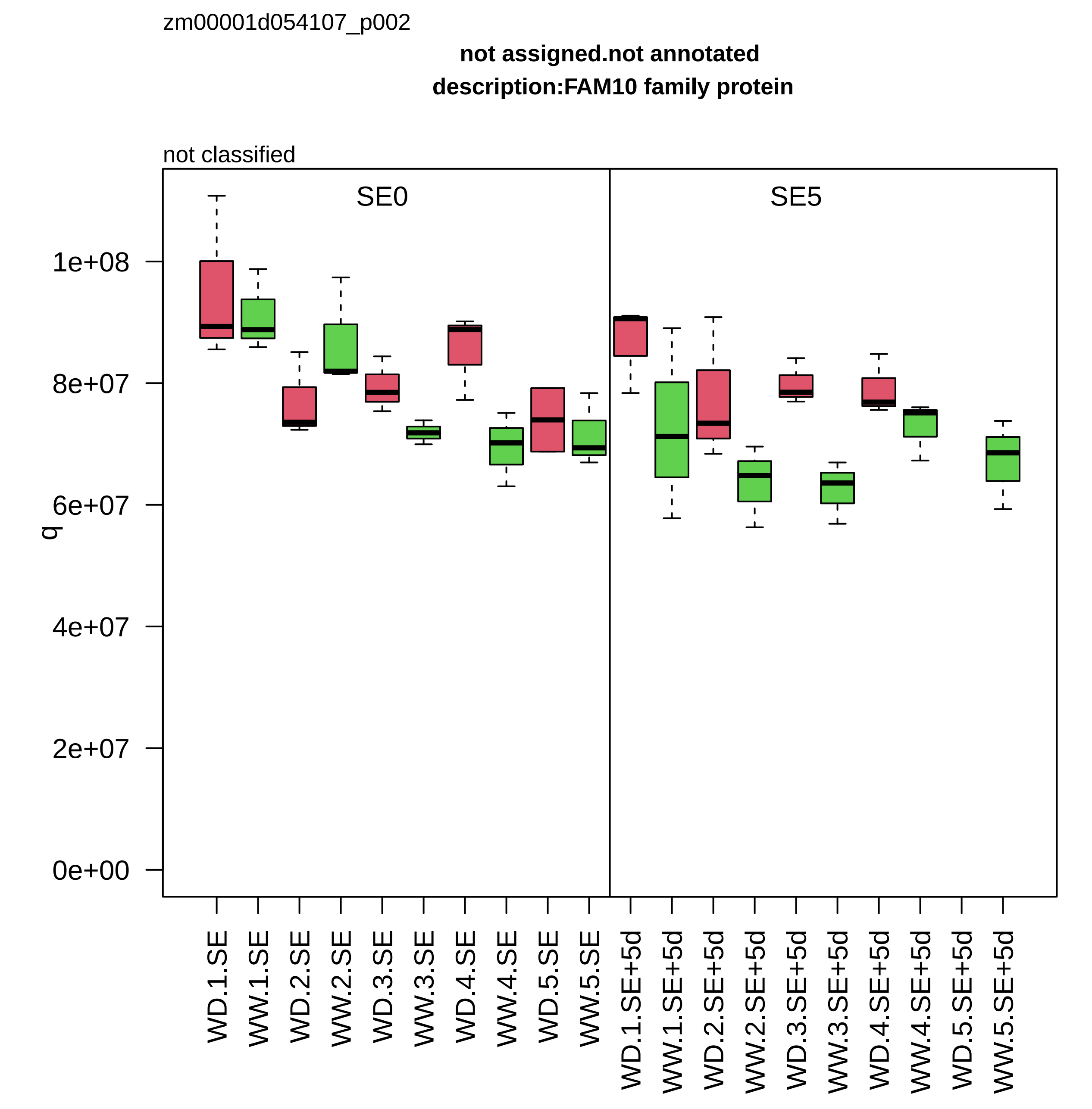
zm00001d054107_p002
not assigned.not annotated
description:FAM10 family protein
not classified
SE0
SE5
q
0e+00
2e+07
4e+07
6e+07
8e+07
1e+08
WD.1.SE
WW.1.SE
WD.2.SE
WW.2.SE
WD.3.SE
WW.3.SE
WD.4.SE
WW.4.SE
WD.5.SE
WW.5.SE
WD.1.SE+5d
WW.1.SE+5d
WD.2.SE+5d
WW.2.SE+5d
WD.3.SE+5d
WW.3.SE+5d
WD.4.SE+5d
WW.4.SE+5d
WD.5.SE+5d
WW.5.SE+5d
