## Supplementary figures and images for "A moderate water deficit induces profound changes in the proteome of developing maize ovaries"

### 2oxocarboxylic_acid_condition.png

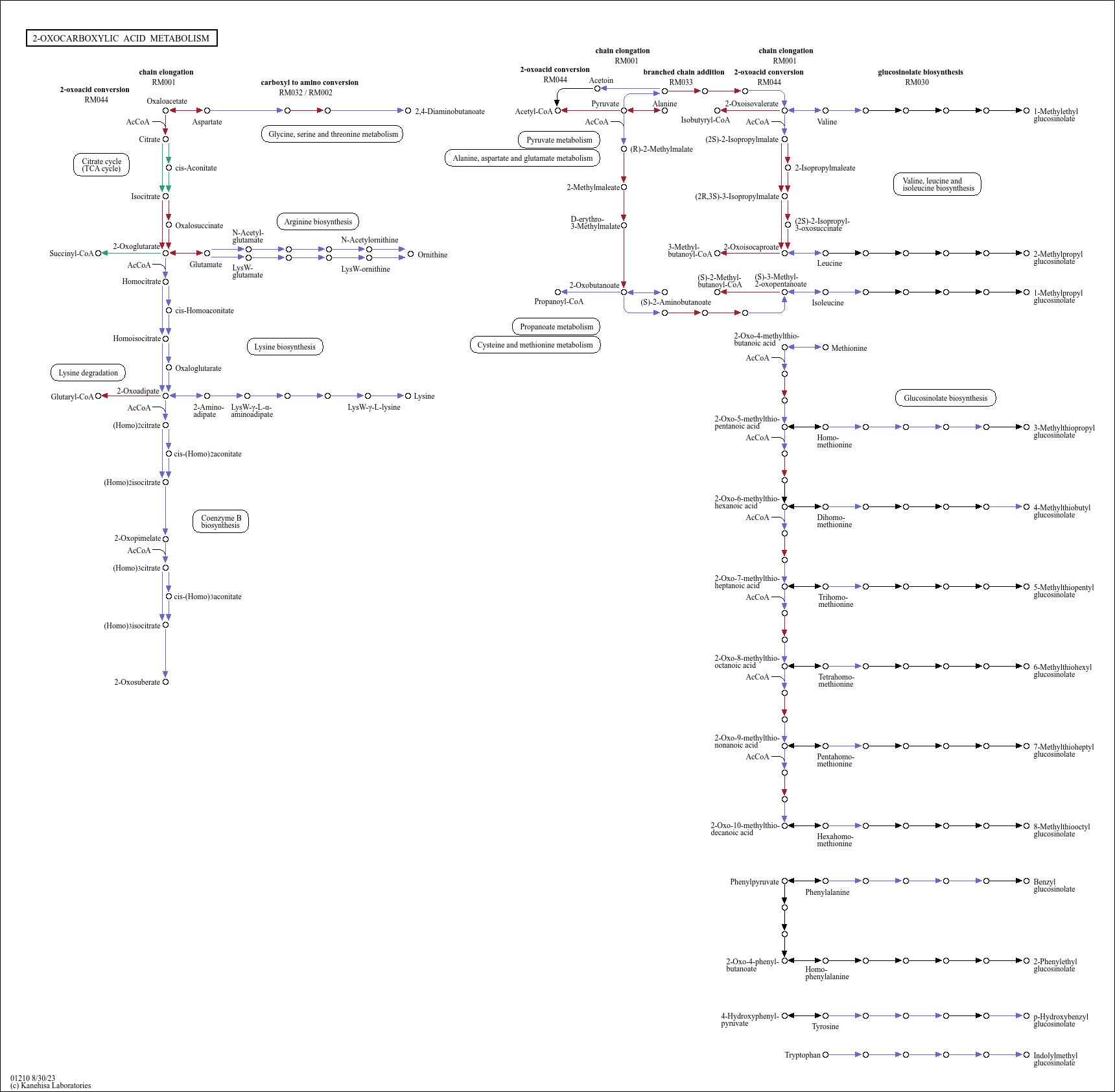

### aa_biosynthesis_condition.png

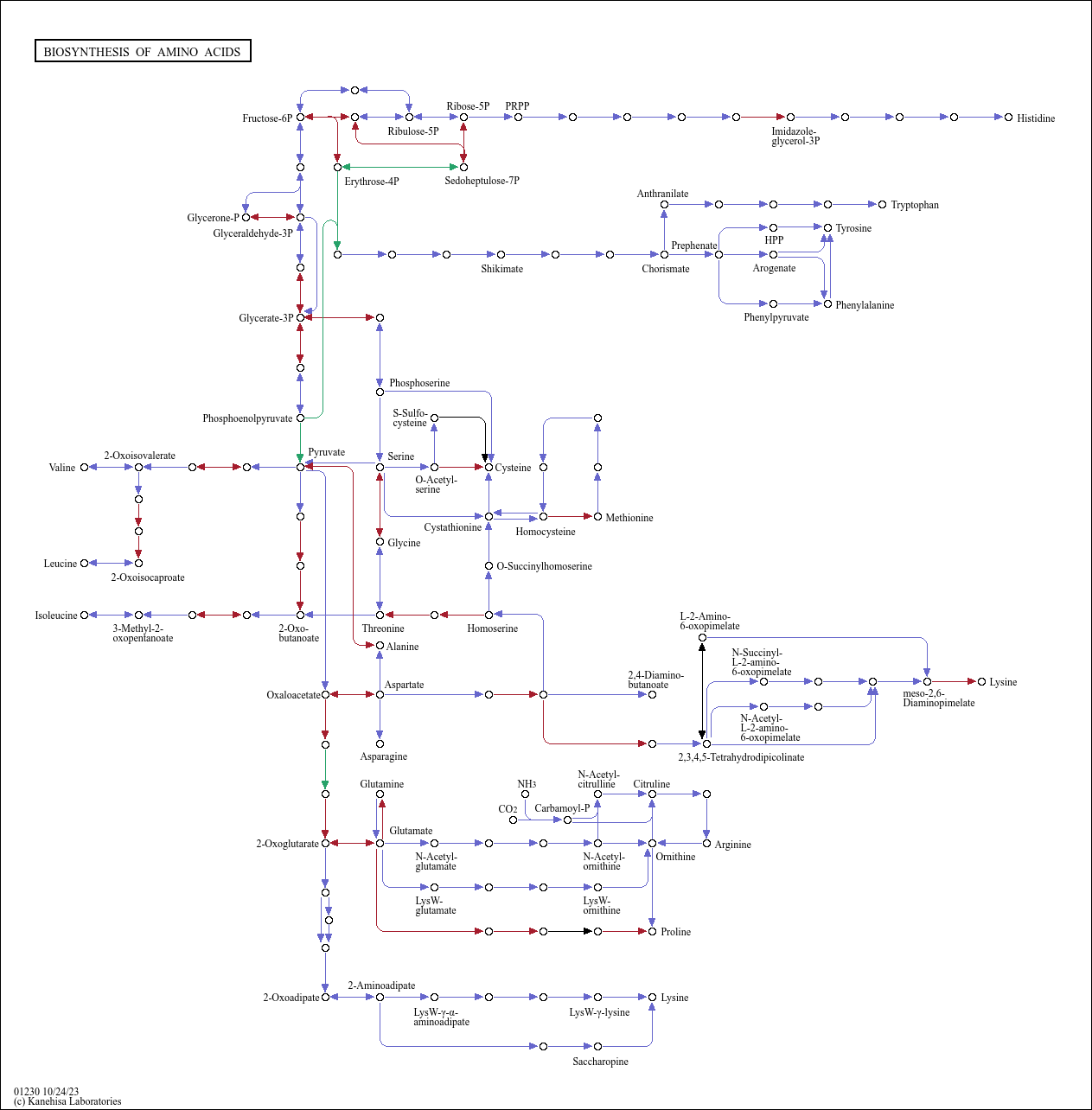

### aa_biosynthesis_day.png

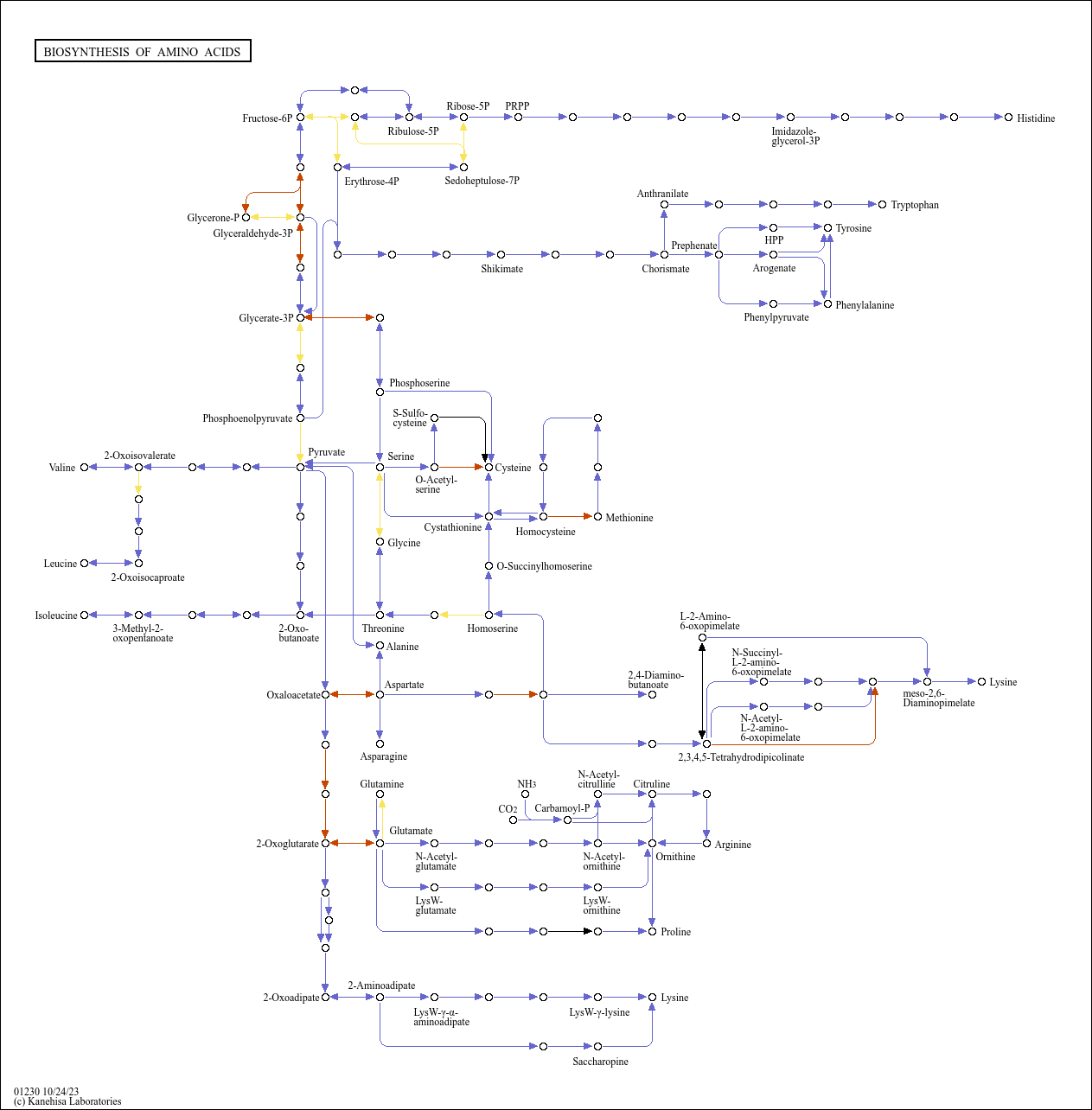

### aa_biosynthesis_zone.png

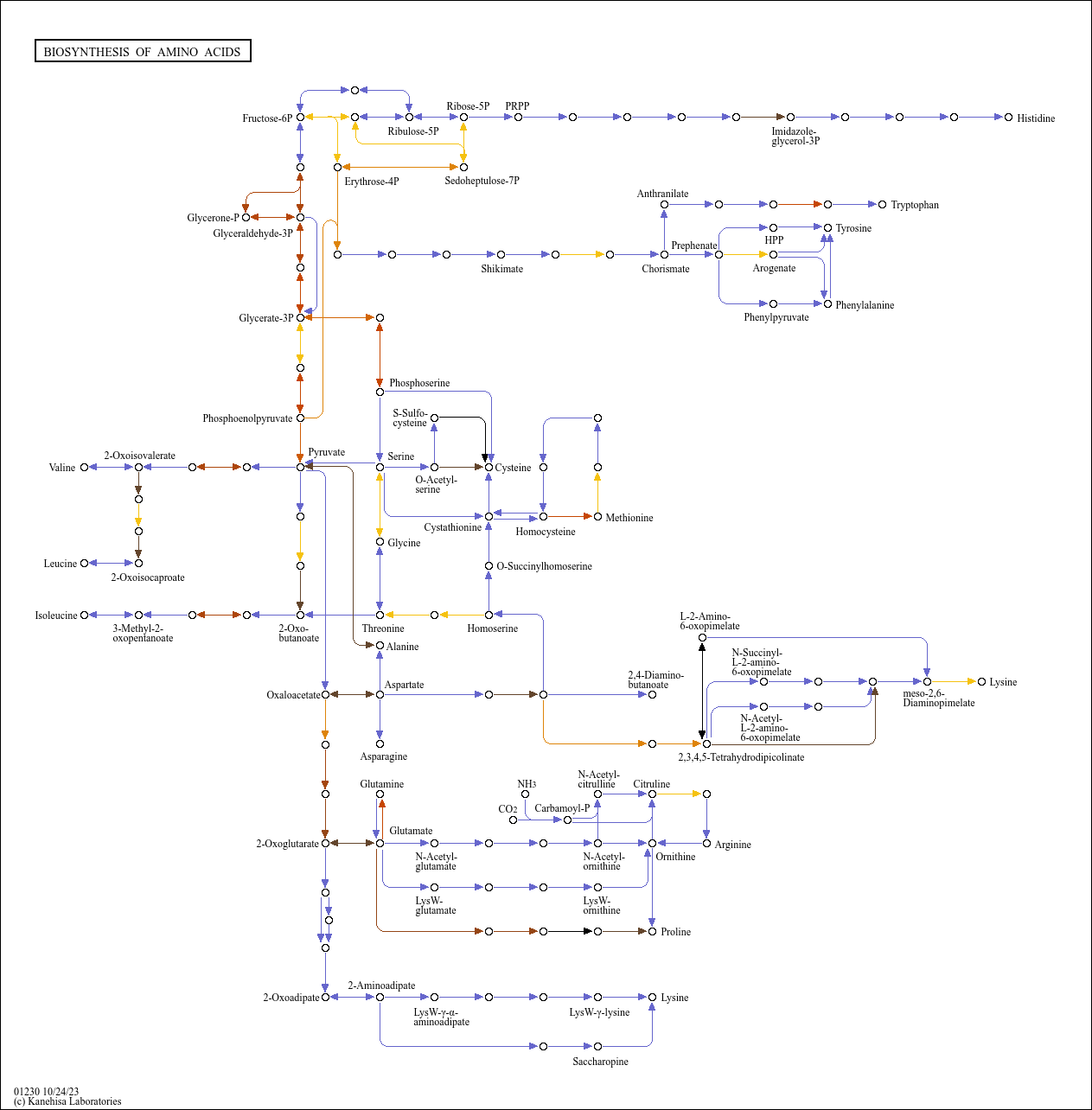

### alpha_linolenic_acide_dzI.png

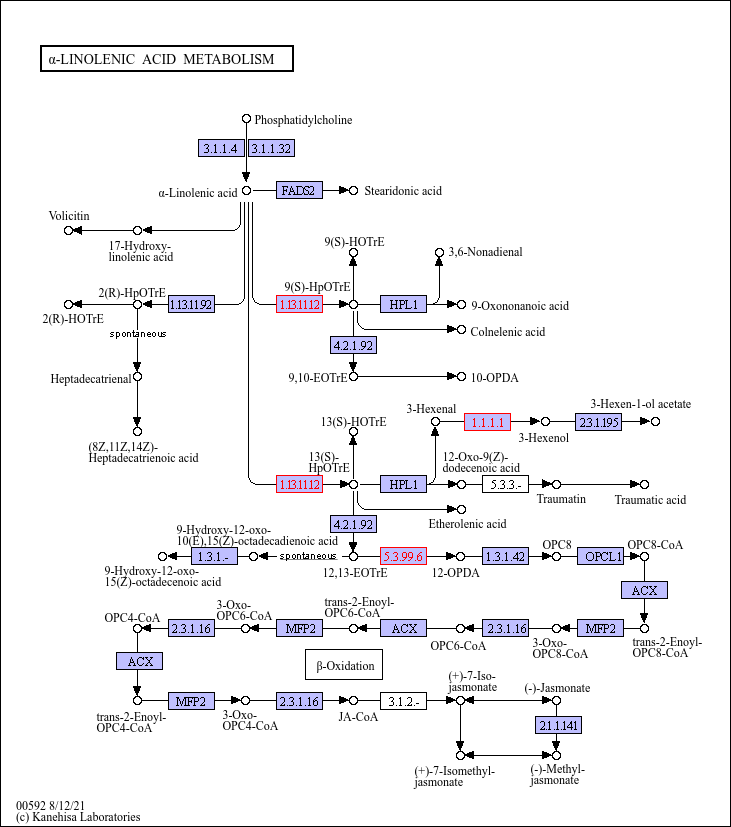

### amino_sugar_nt_sugar_metabo_dzI.png

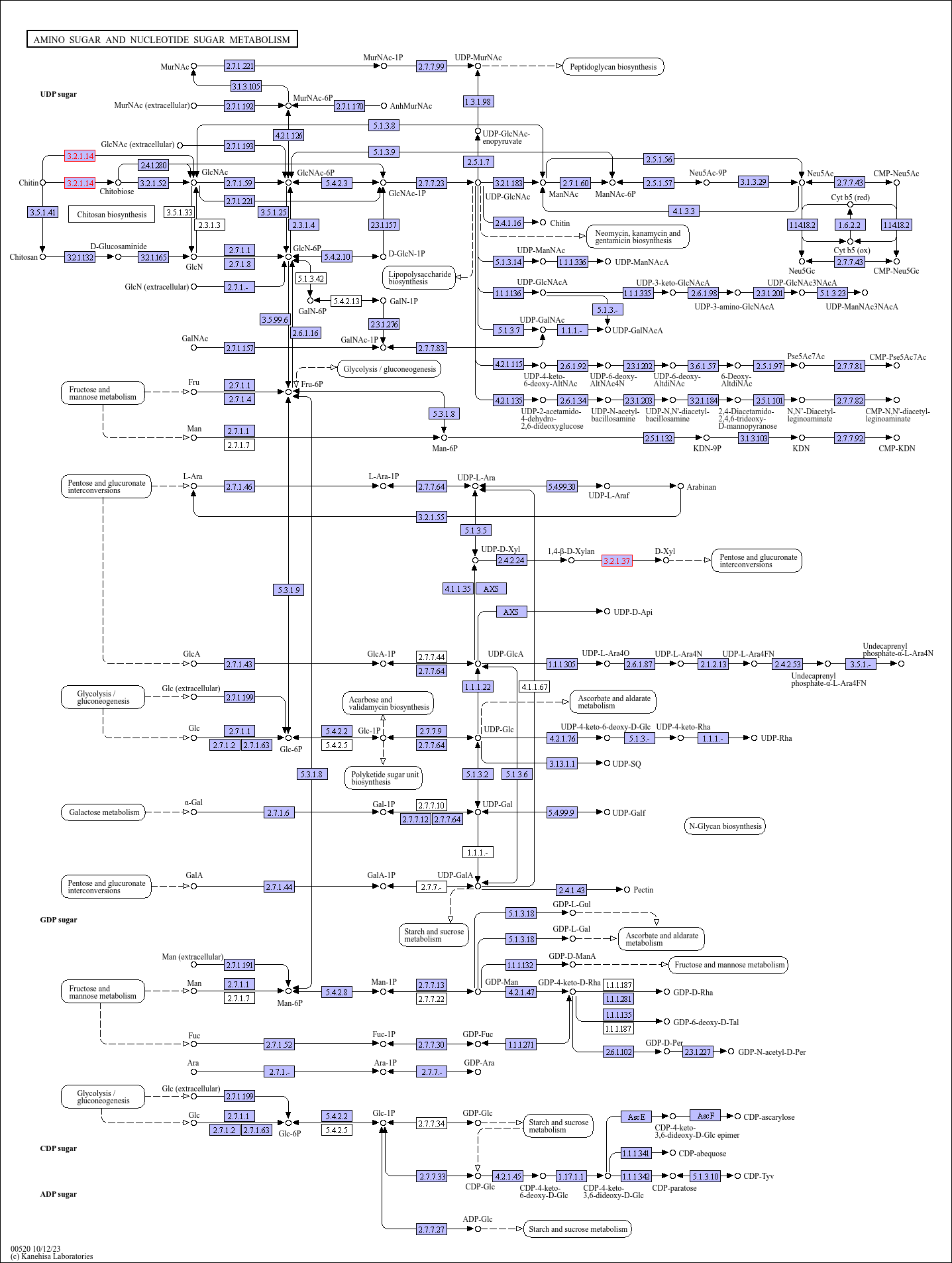

### arginine_proline_condition.png

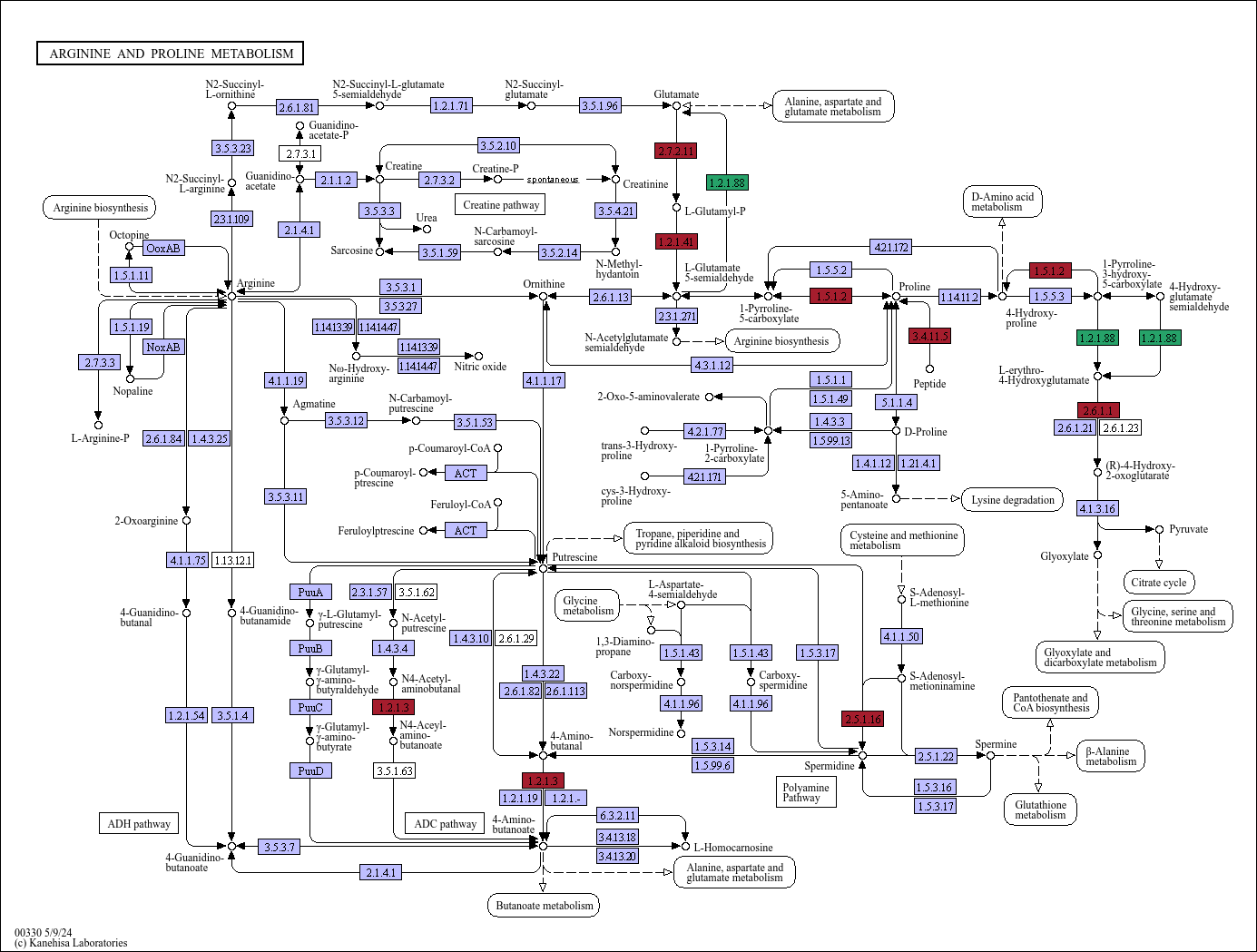

### carbon_fixation_day.png

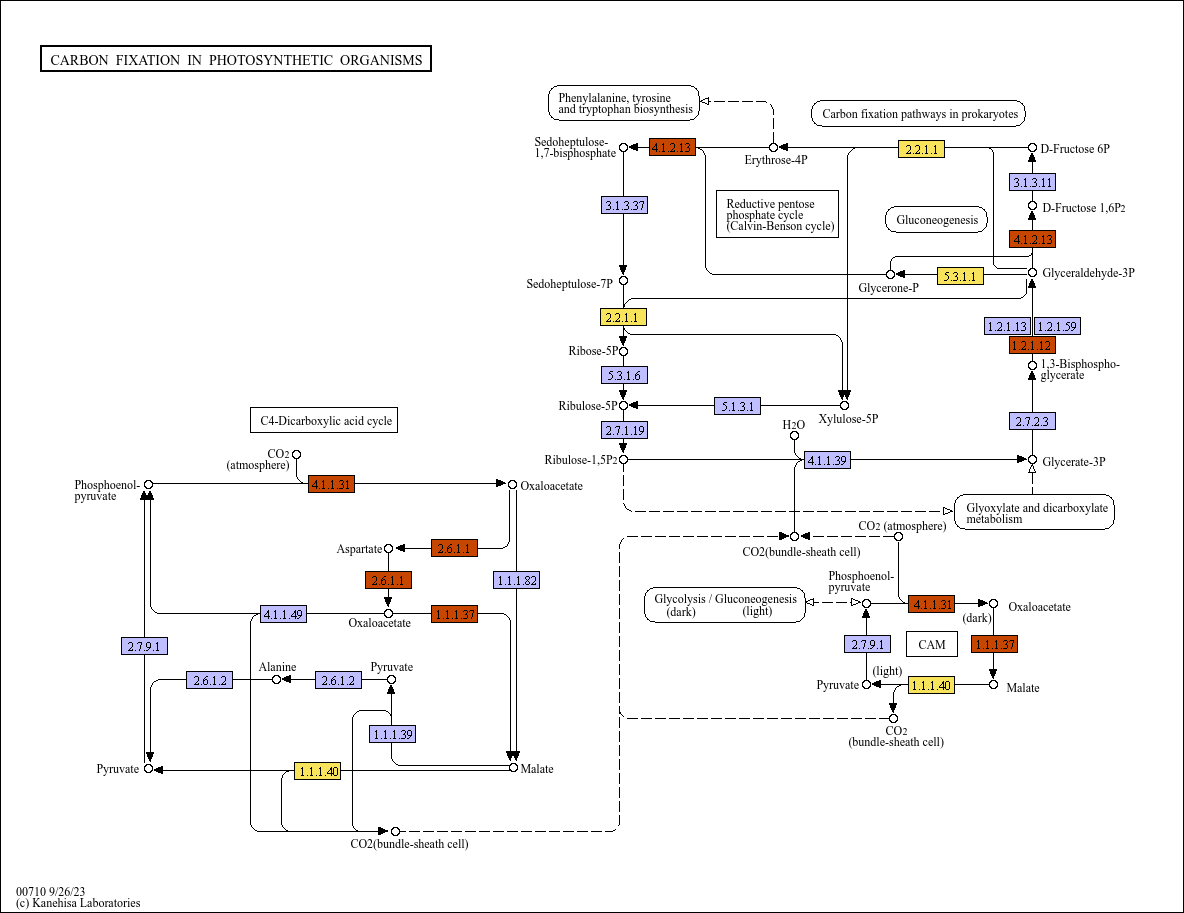

### carbon_fixation_zone.png

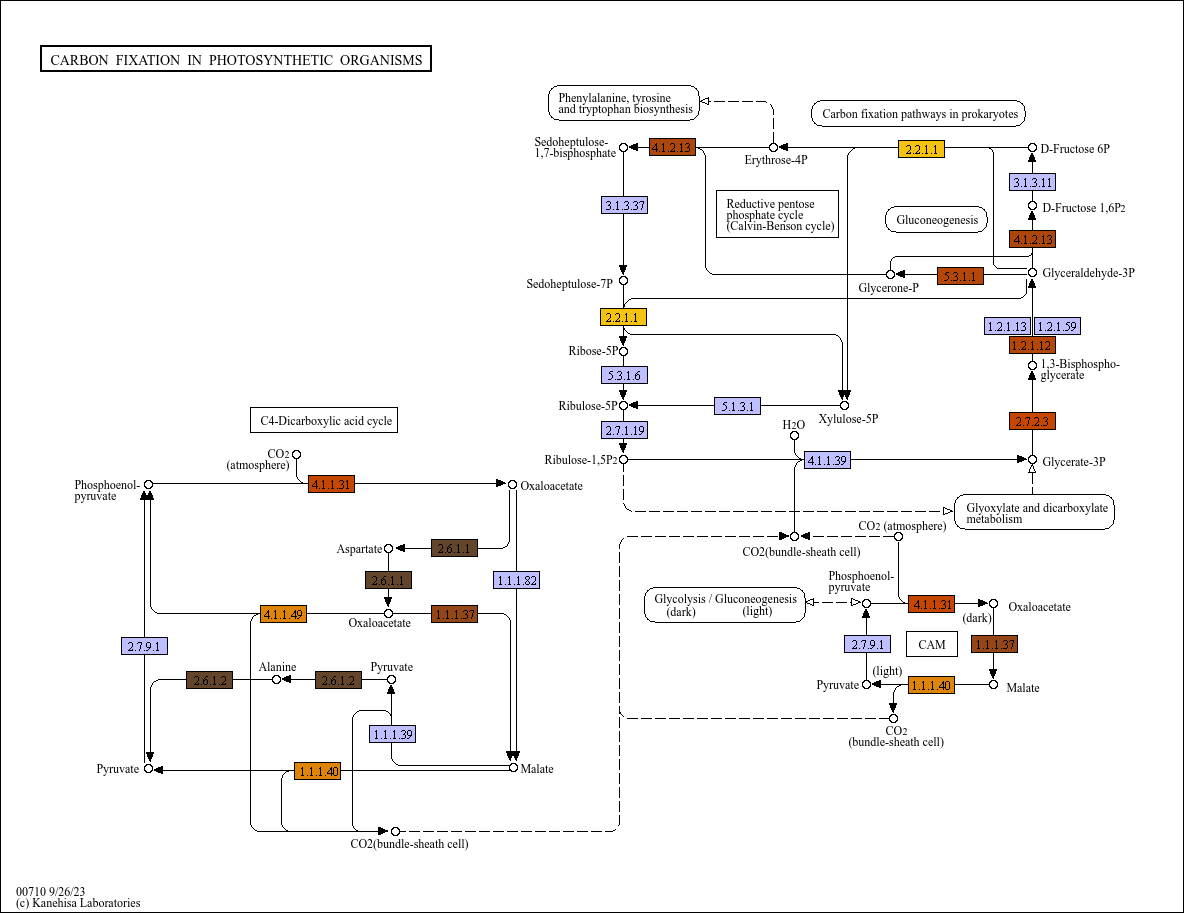

### carbon_metabo_condition.png

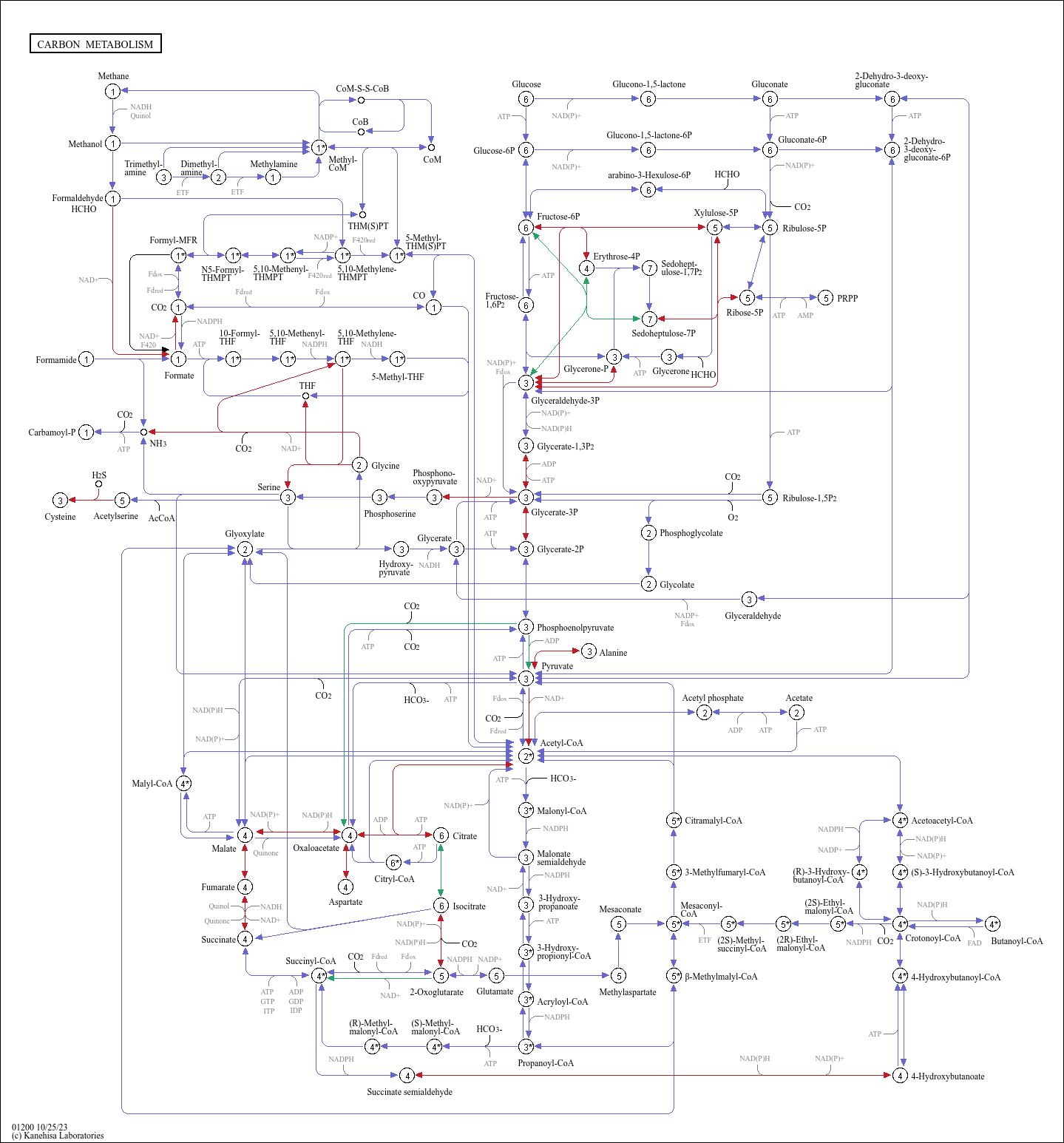

### carbon_metabo_day.png

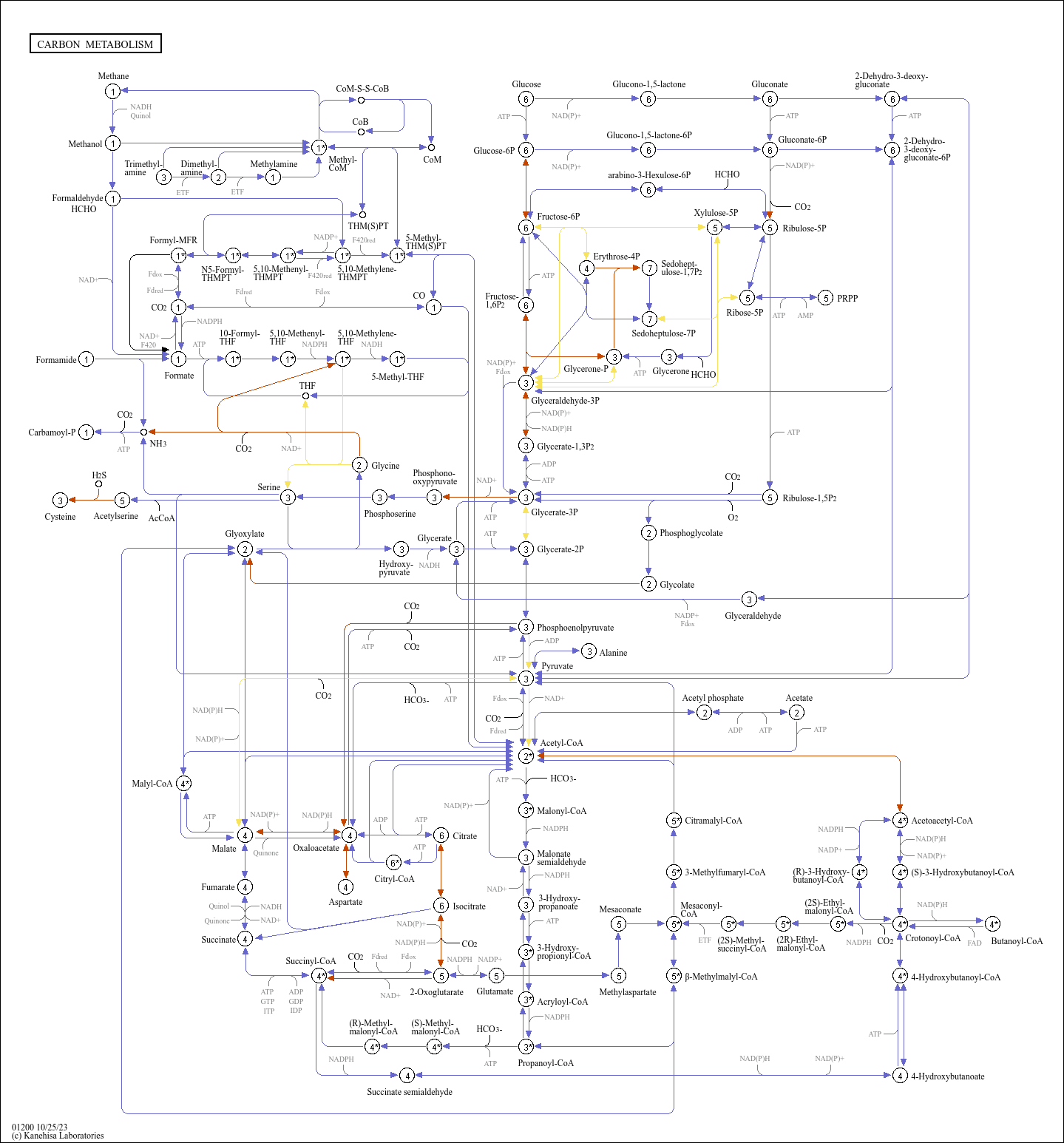

### carbon_metabo_zone.png

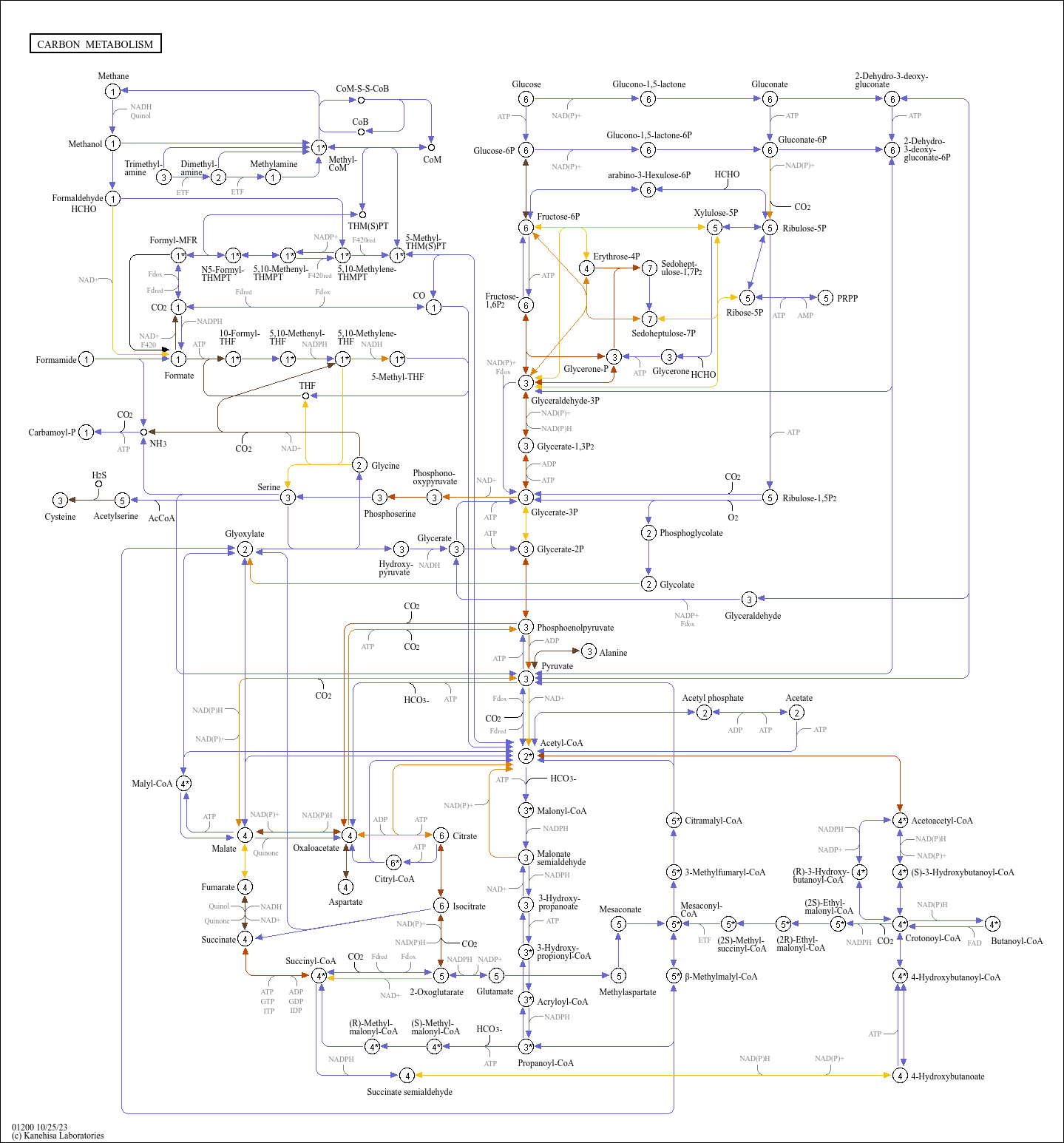

### cysteine_methionine_day.png

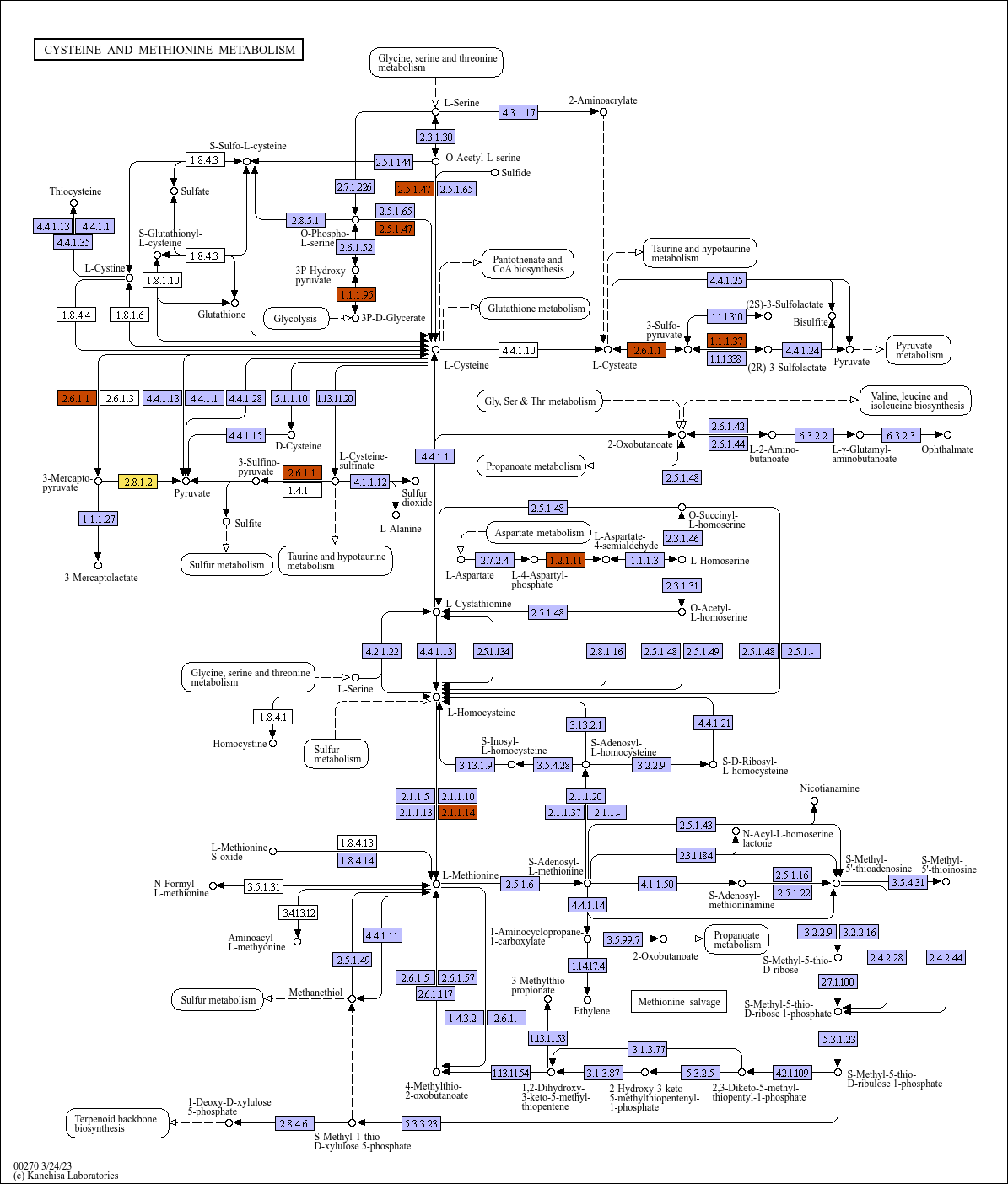

### cysteine_methionine_zone.png

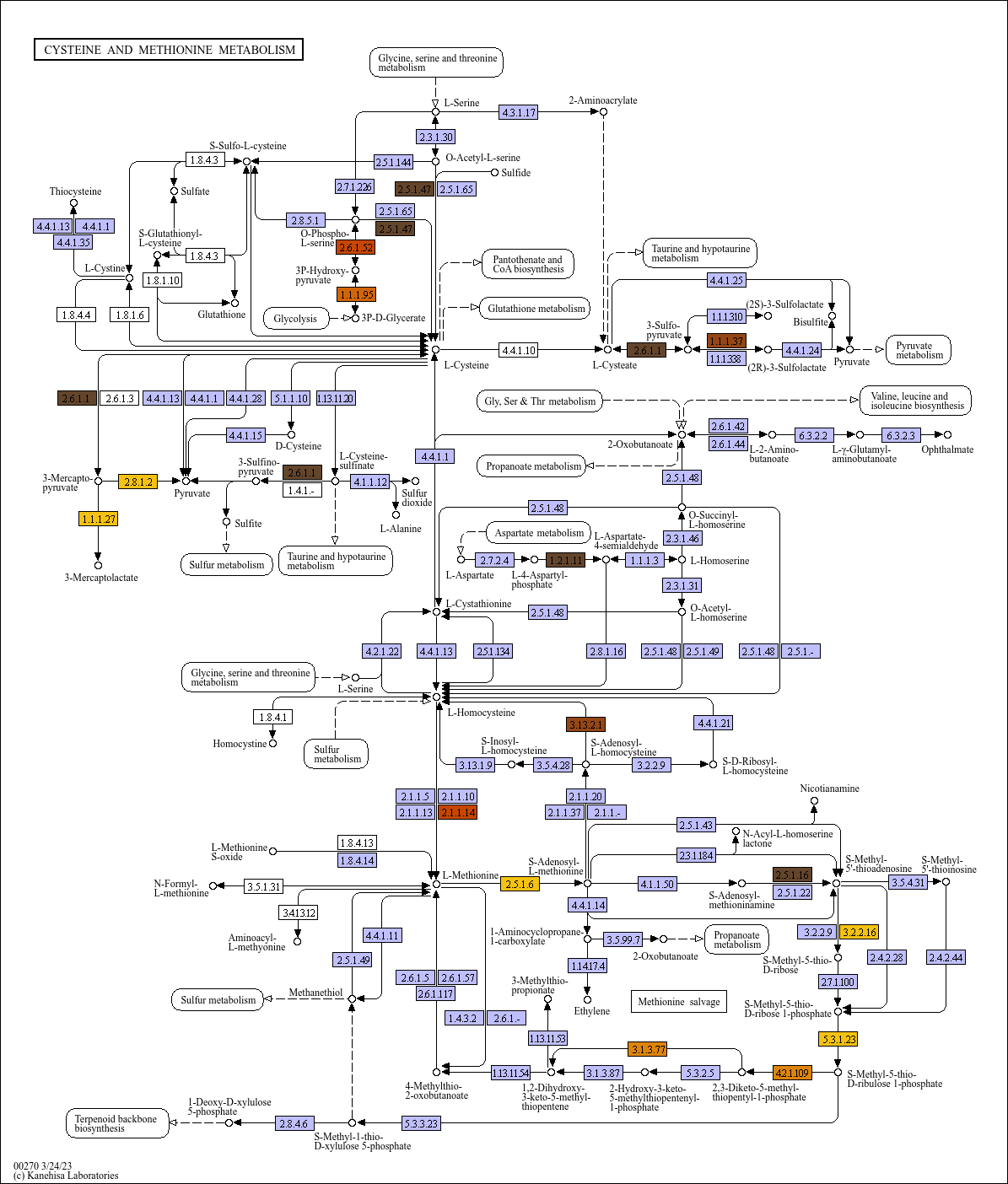

### DNA_replication_day.png

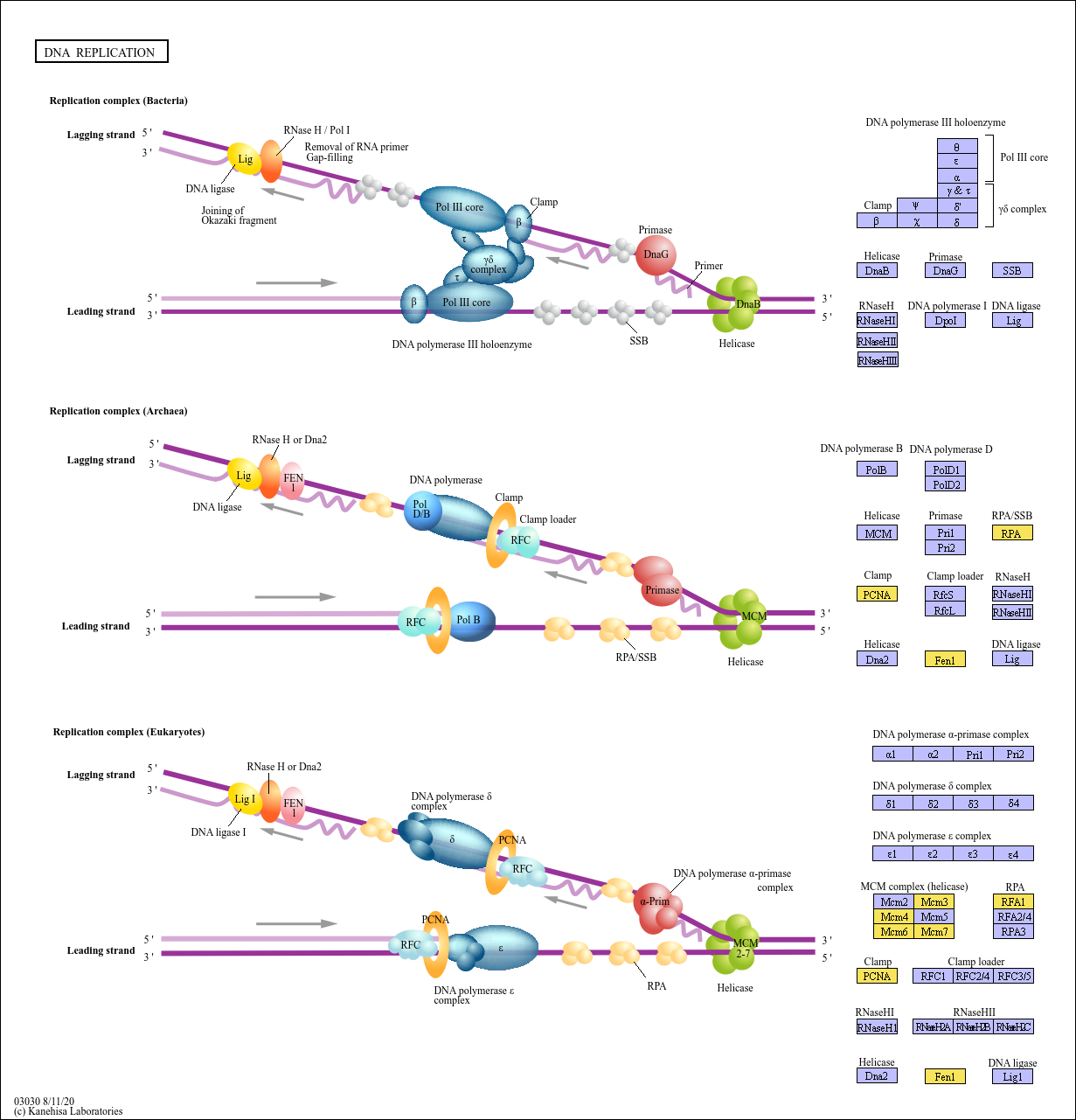

### DNA_replication_zone.png

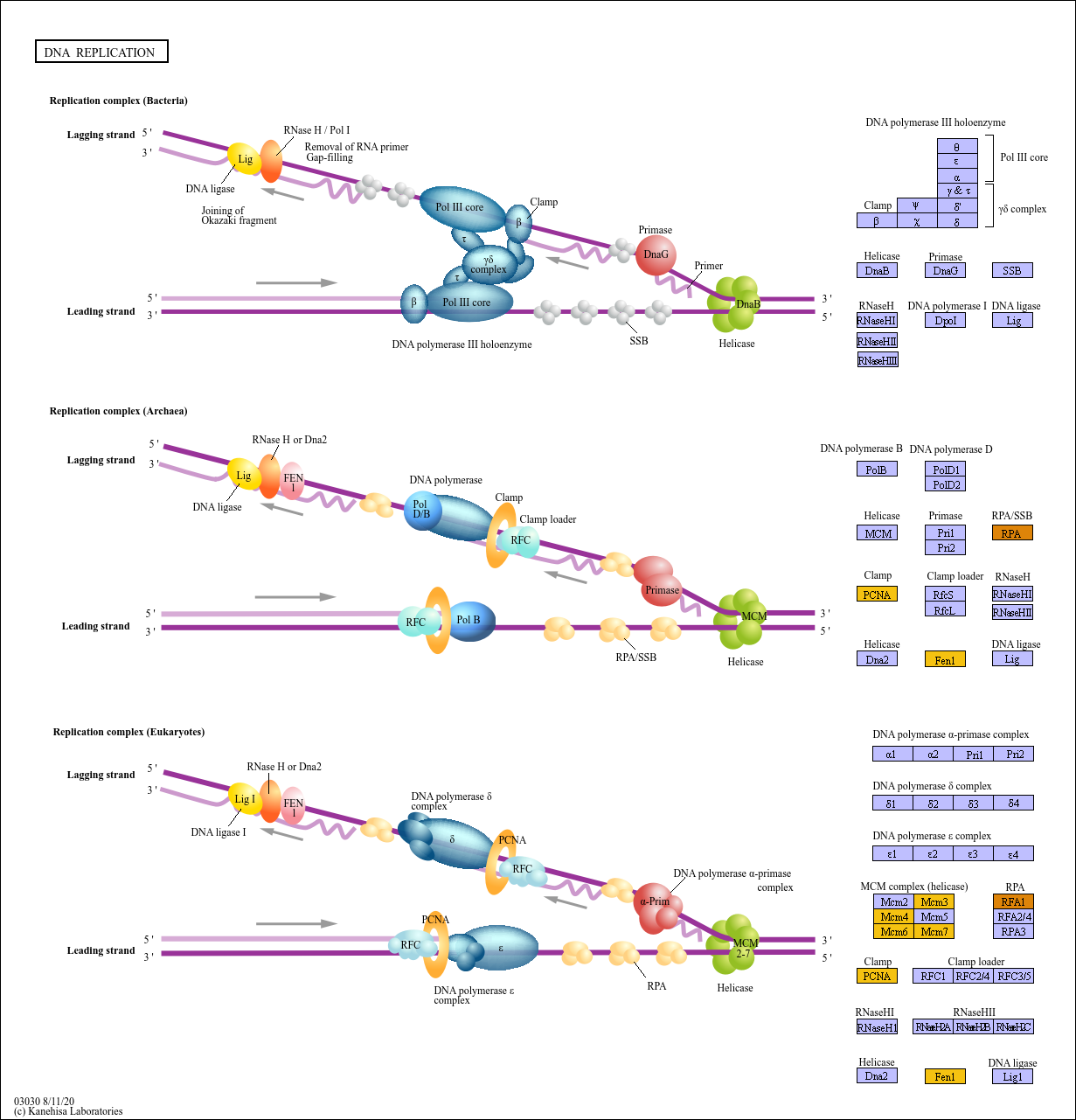

### fattyacid_degration_condition.png

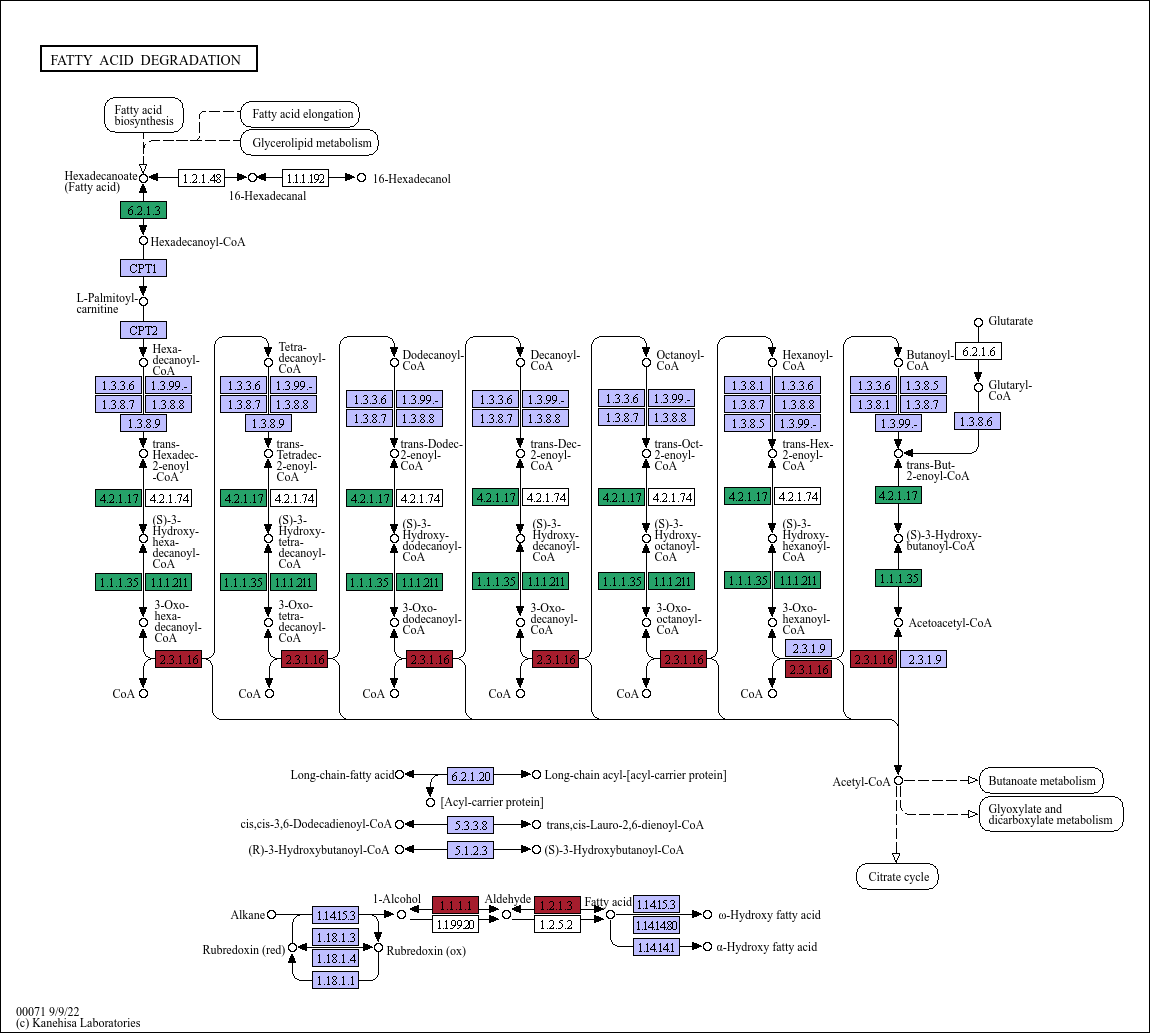

### glycolyse_gluconeogenesis_day.png

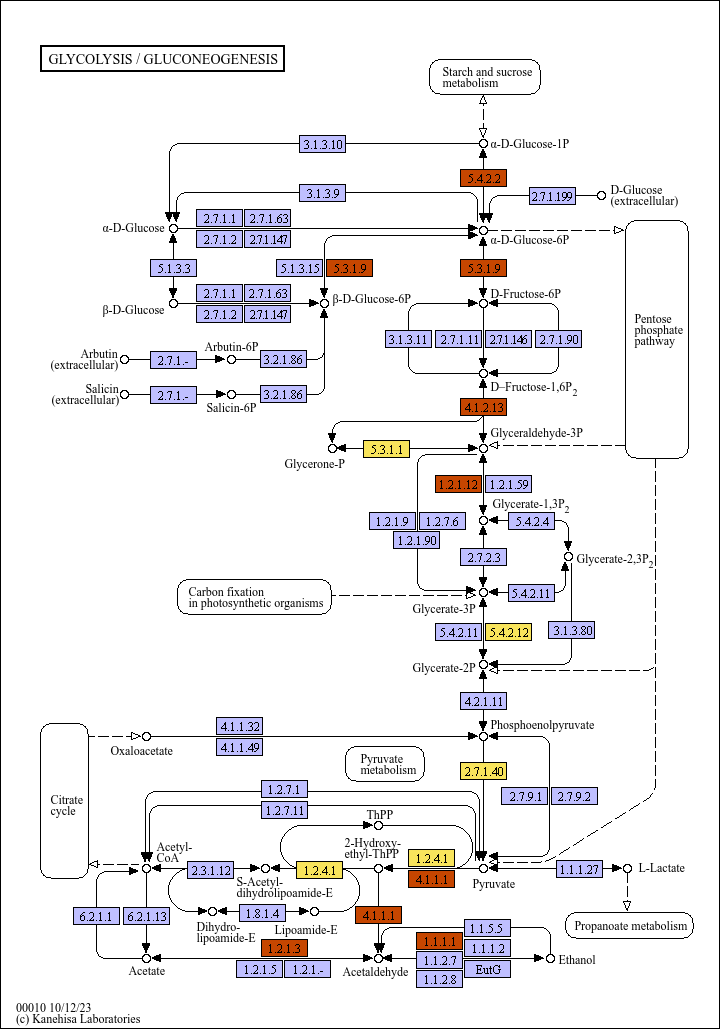

### glycolyse_gluconeogenesis_zone.png

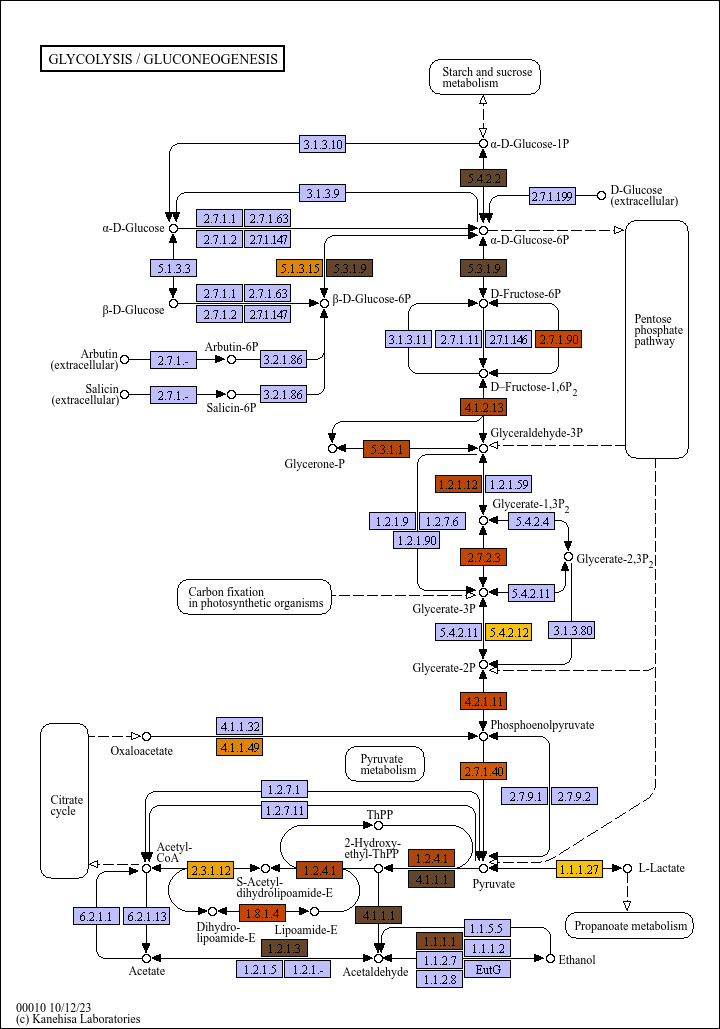

### glyoxylate_dicarboxylate_metabo_condition.png

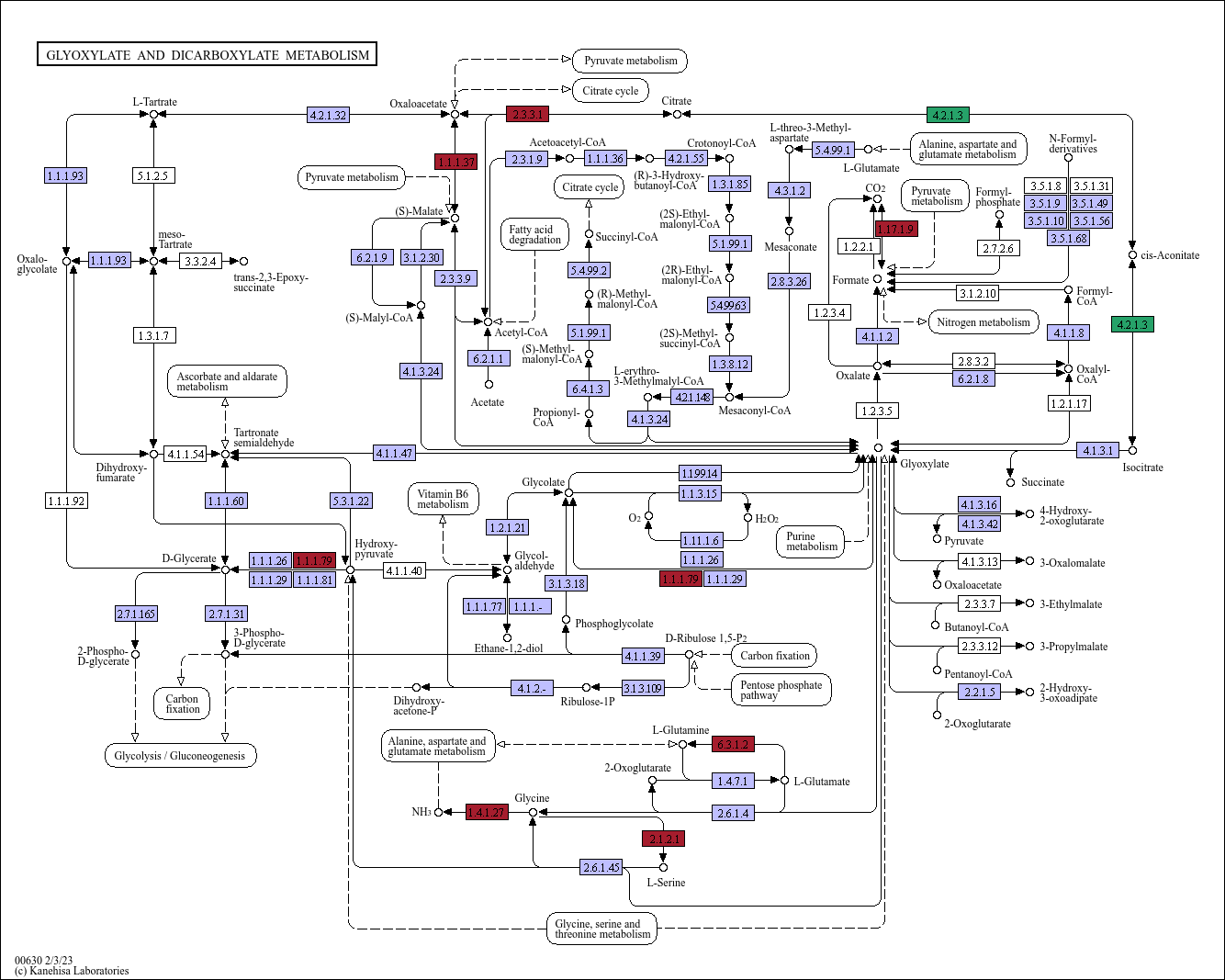

### glyoxylate_dicarboxylate_metabo_day.png

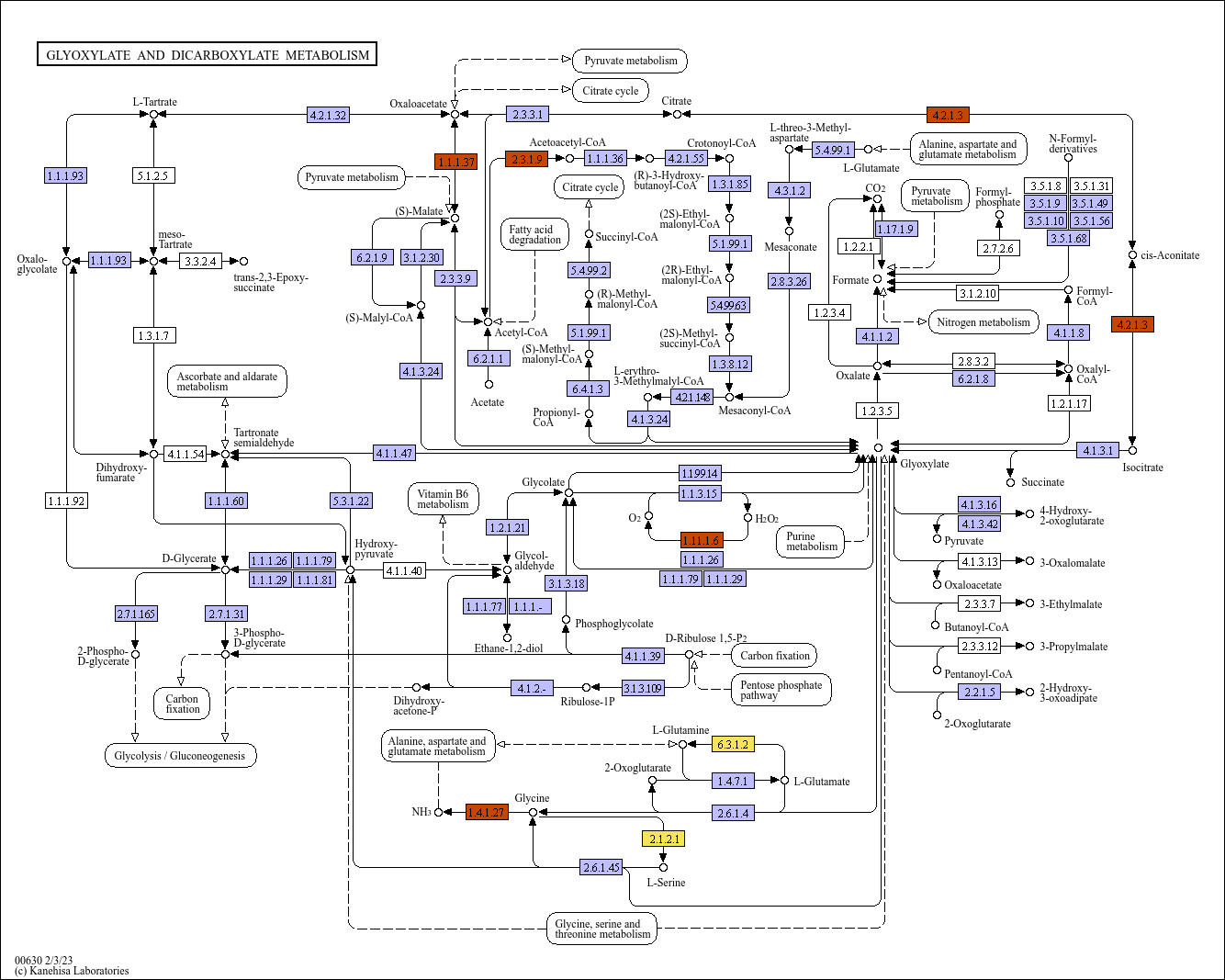

### glyoxylate_dicarboxylate_metabo_zone.png

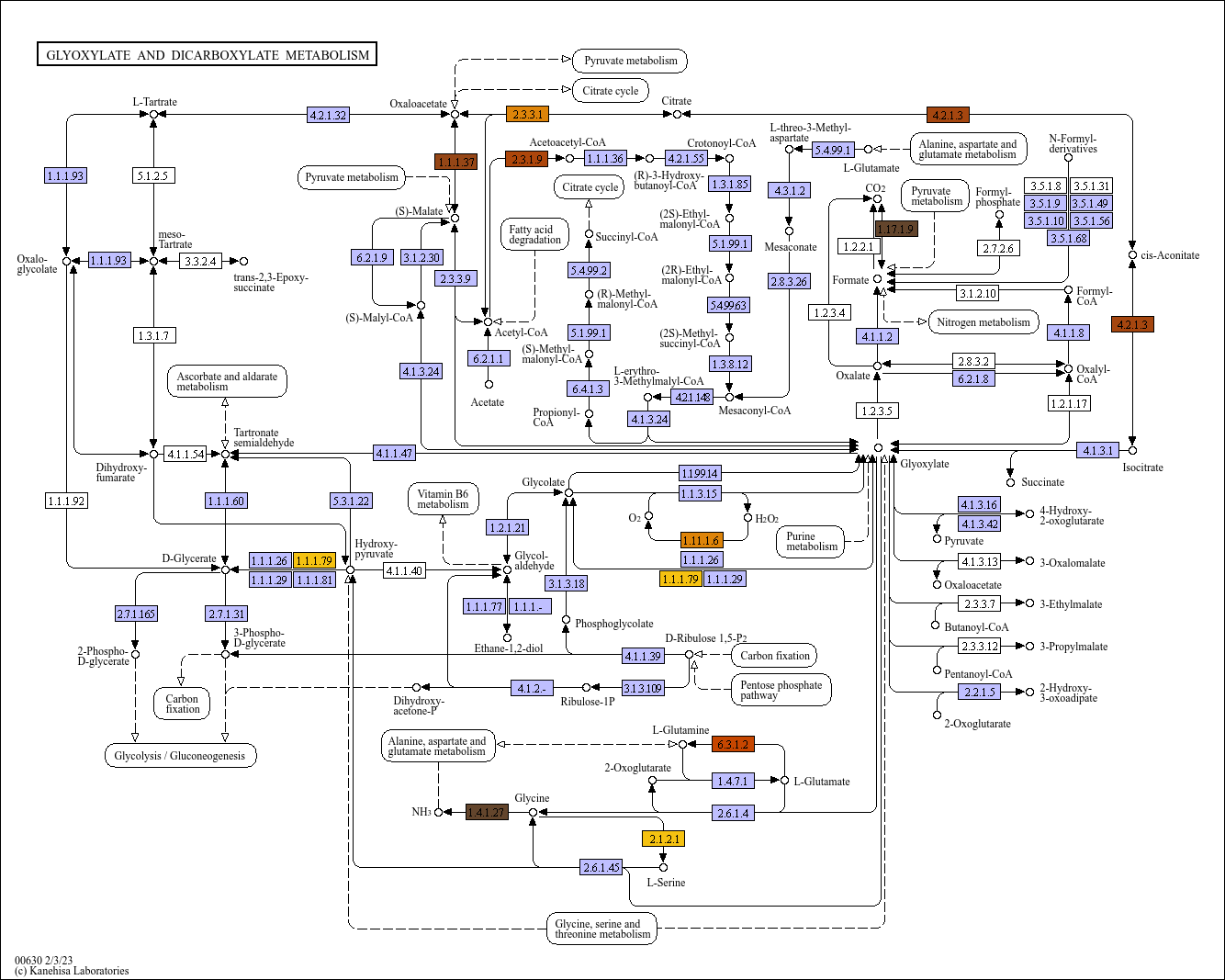

### oxydative_phosphorylation_condition.png

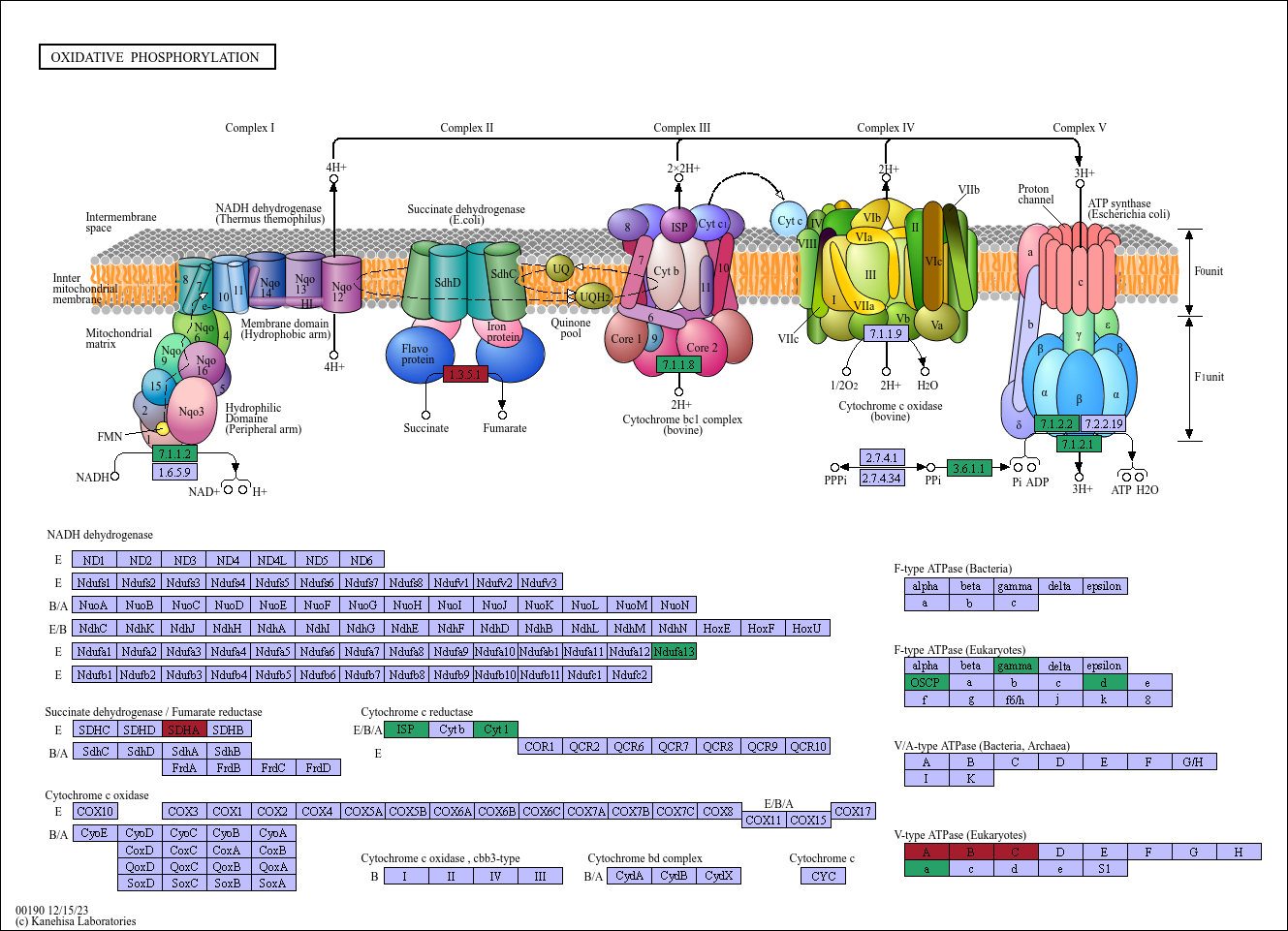

### pentose_phosphate_zone.png

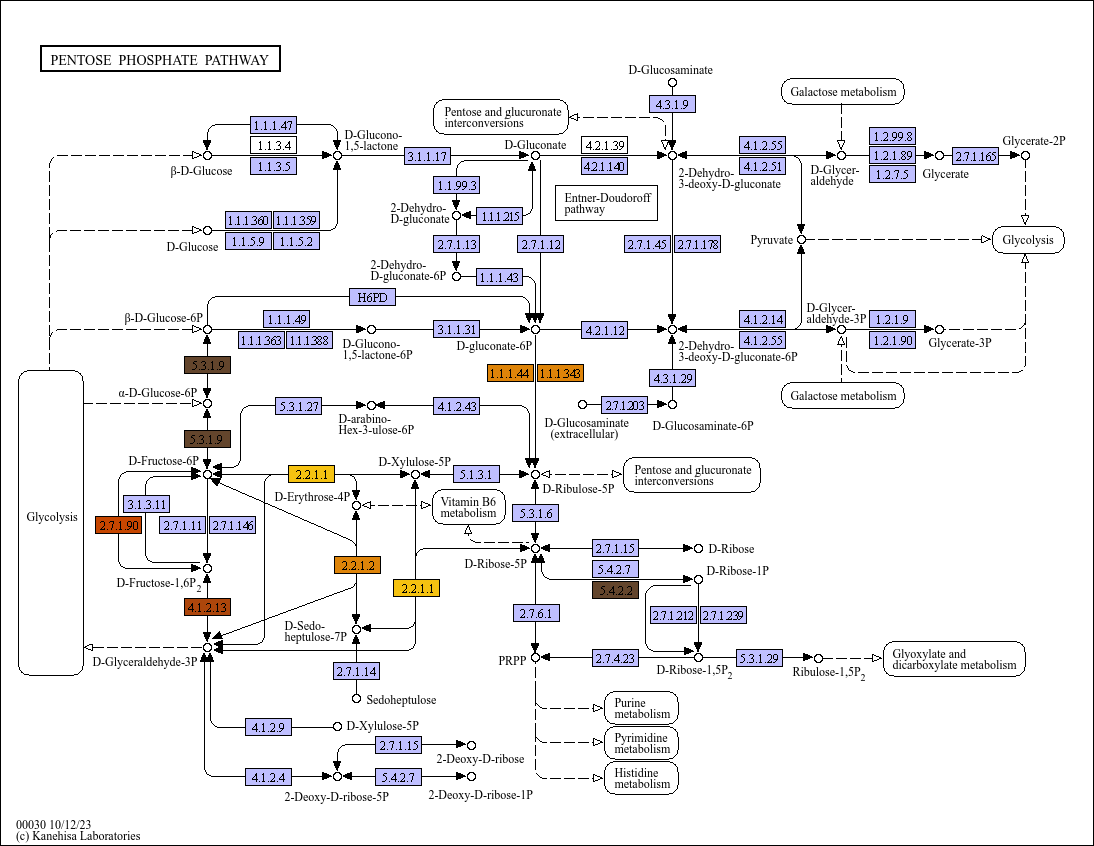

### proteasome_condition.png

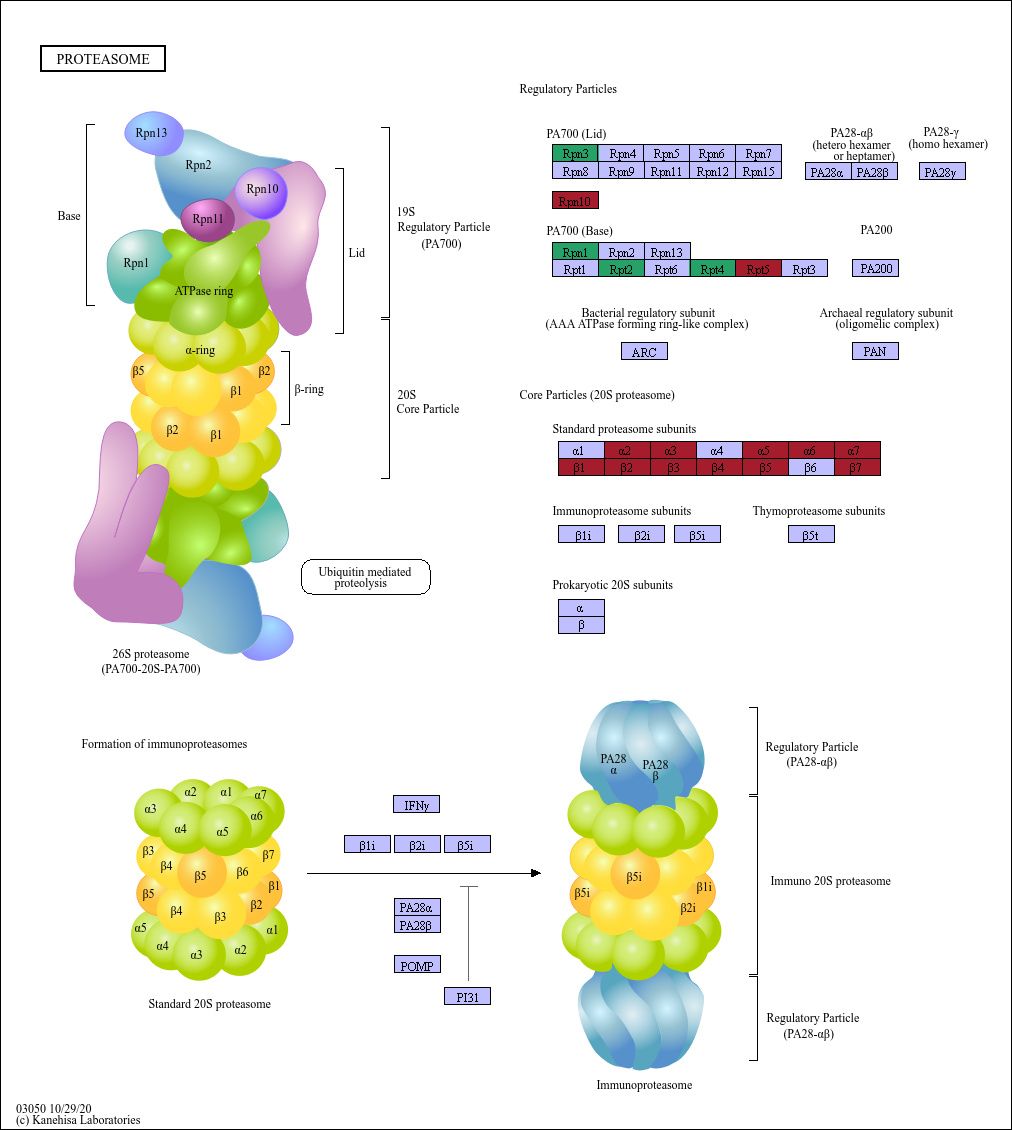

### protein_processing_RE_condition.png

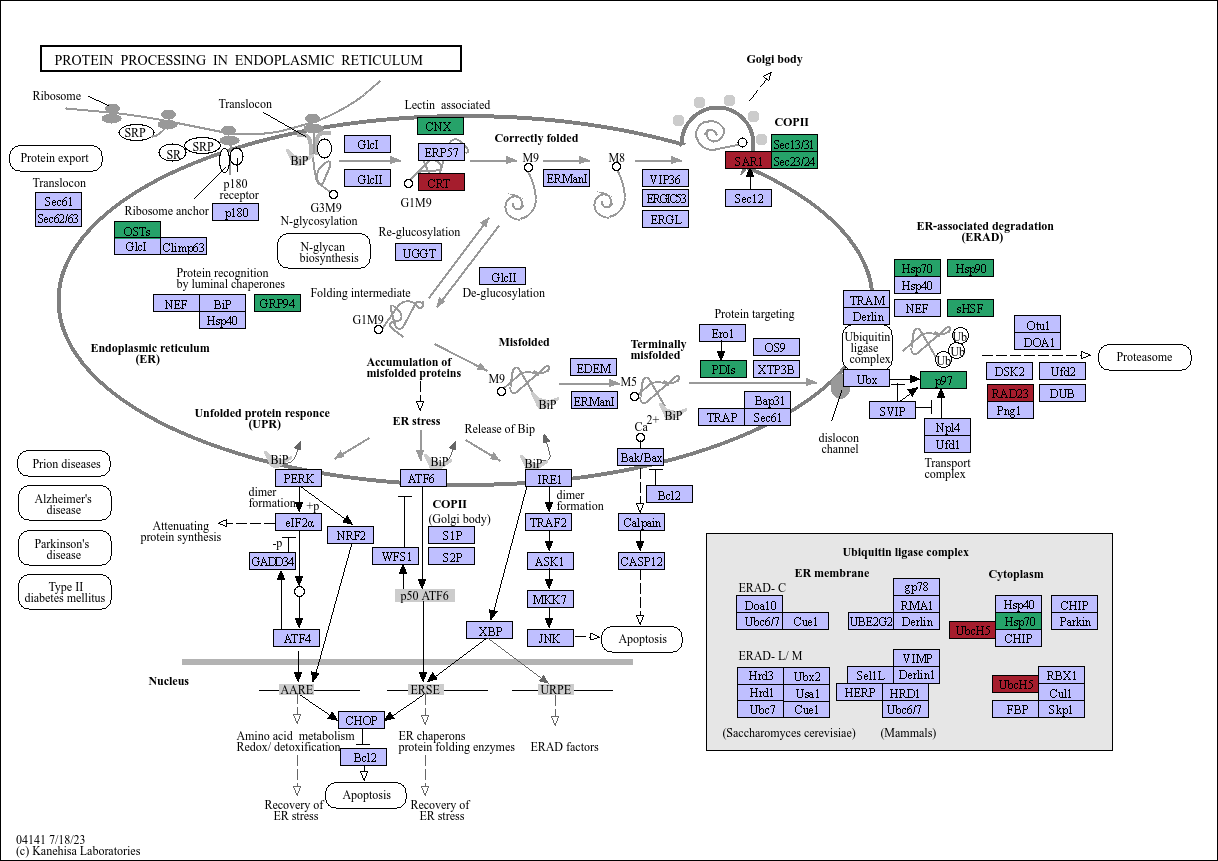

### protein_processing_RE_day.png

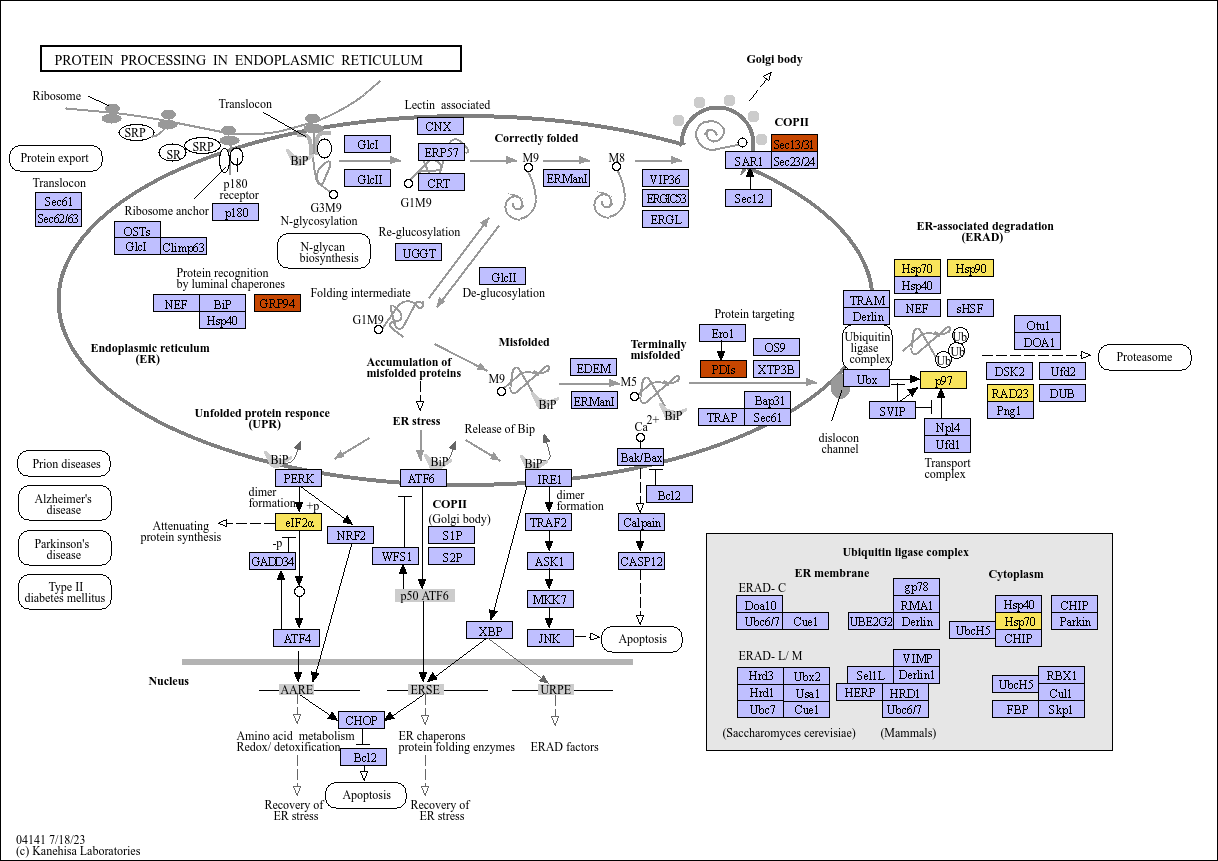

### protein_processing_RE_zone.png

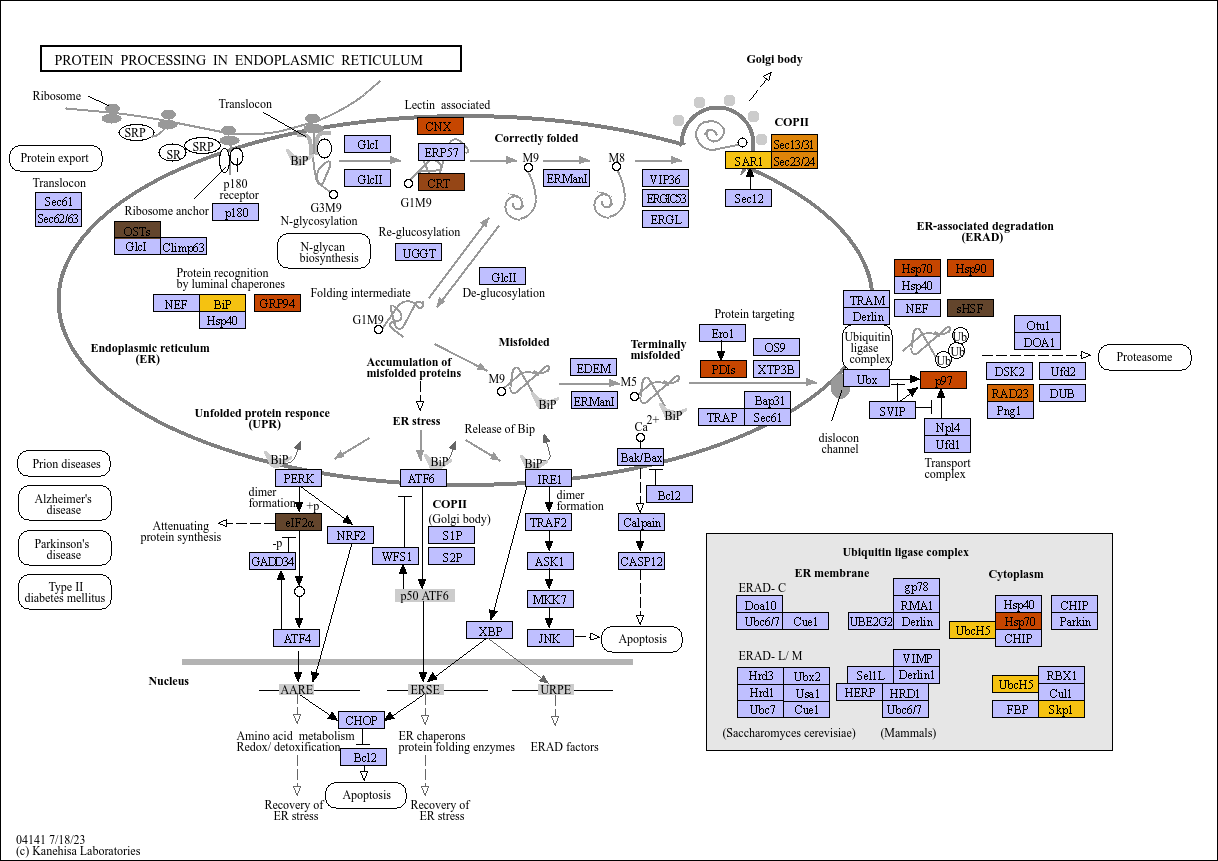

### pyruvate_metabolism_condition.png

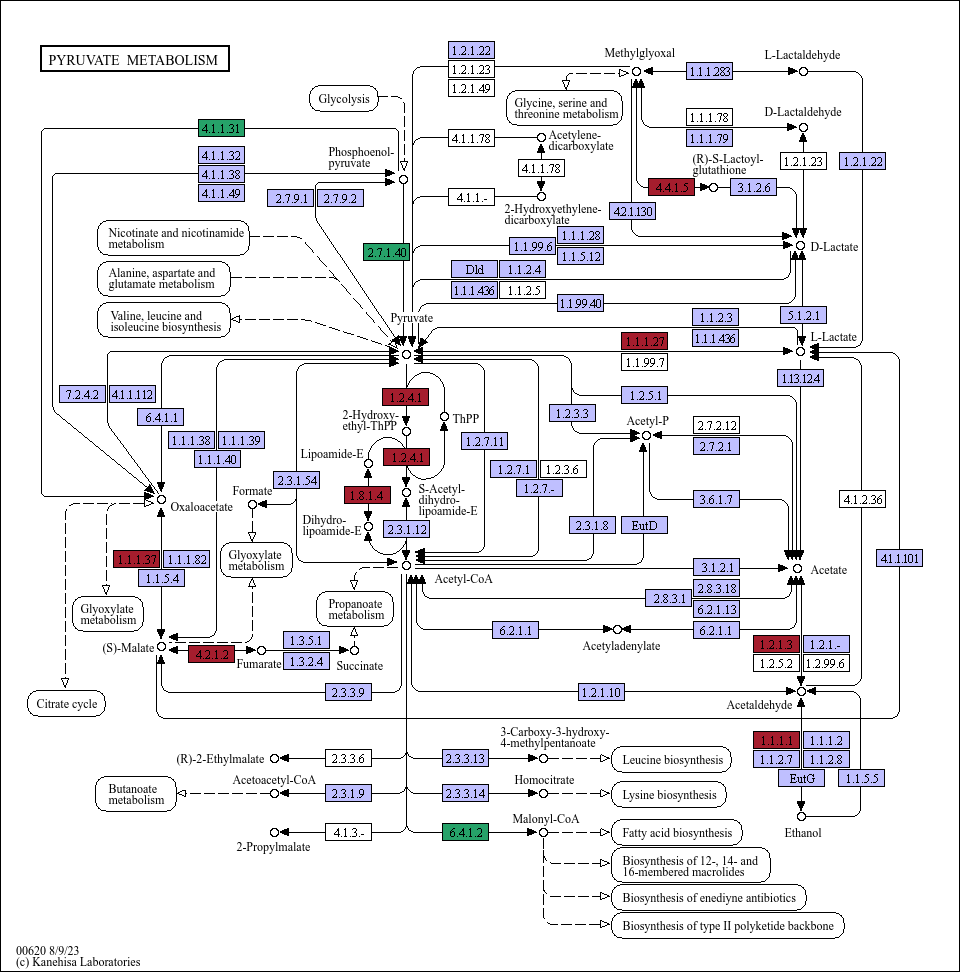

### pyruvate_metabolism_day.png

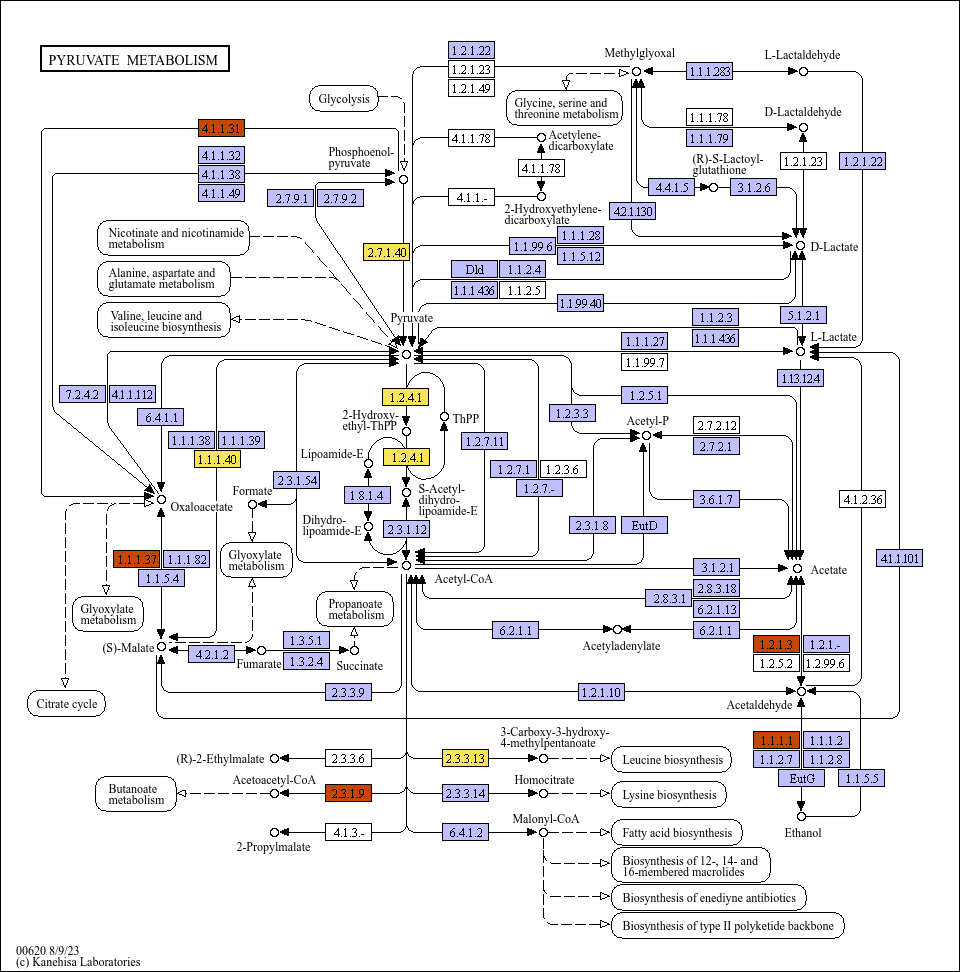

### pyruvate_metabolism_zone.png

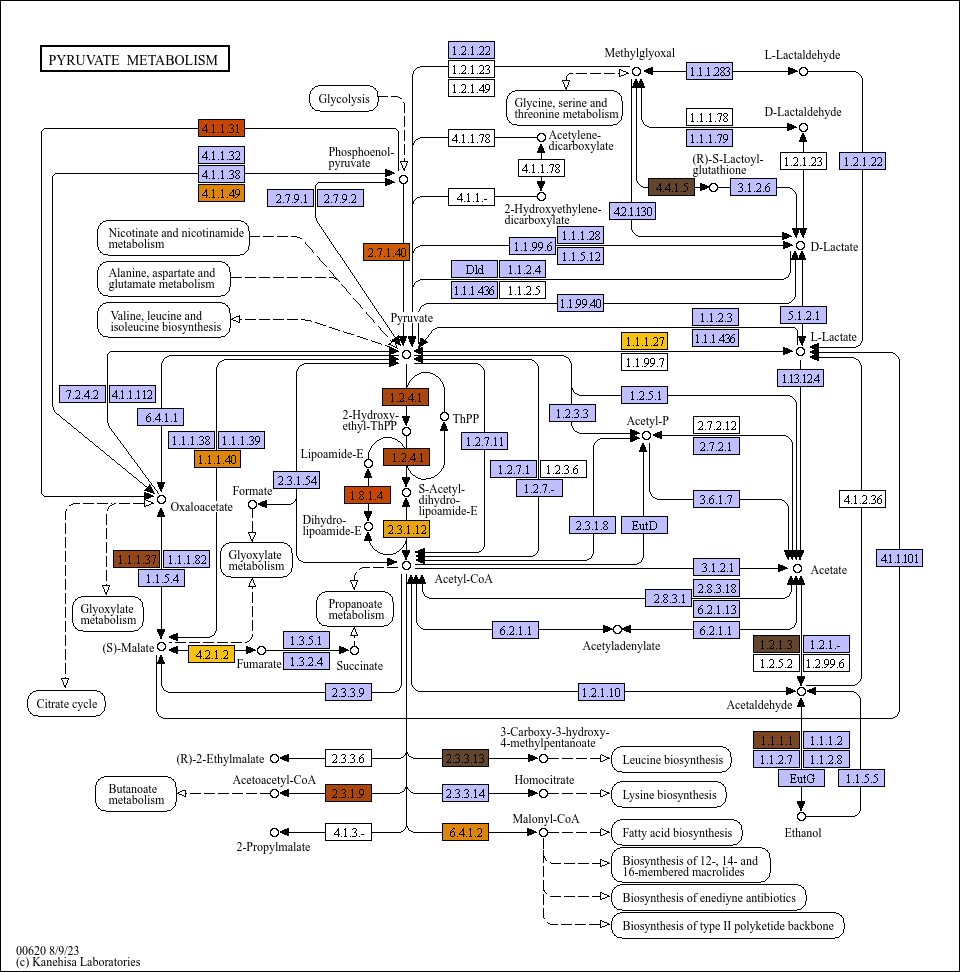

### ribosome_day.png

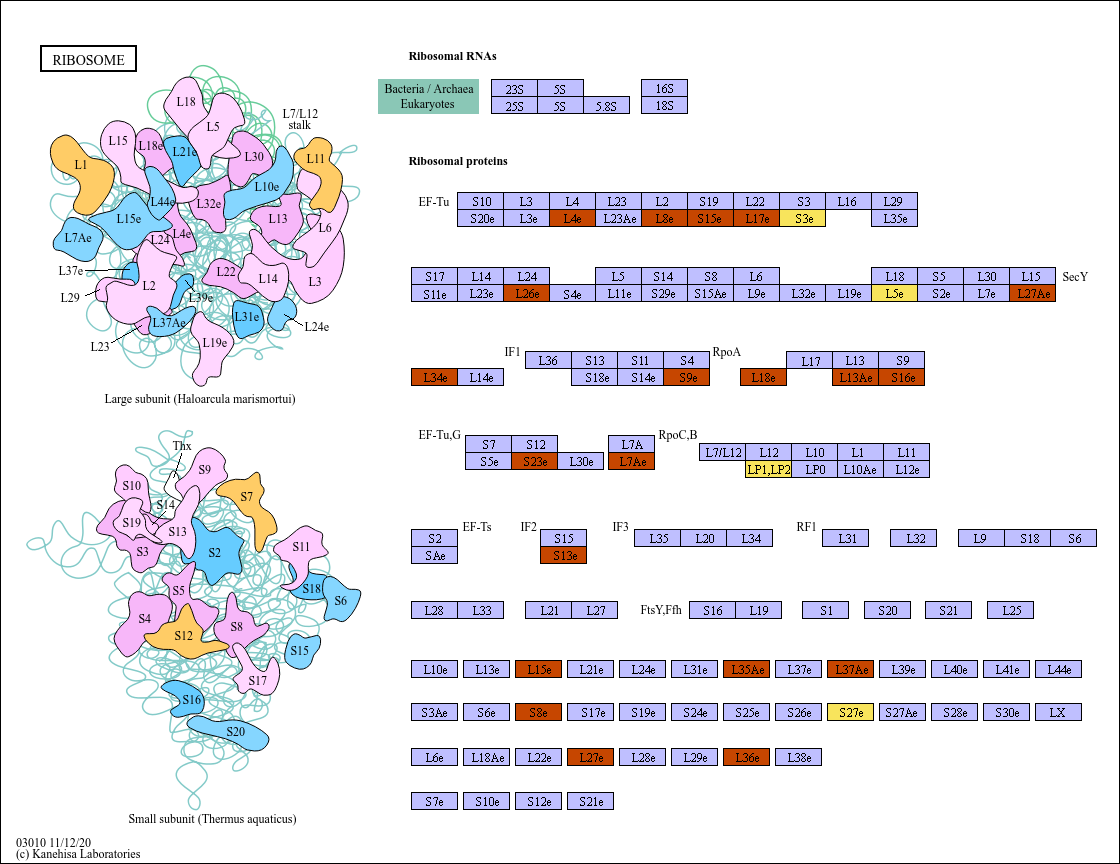

### spiceosome_condition.png

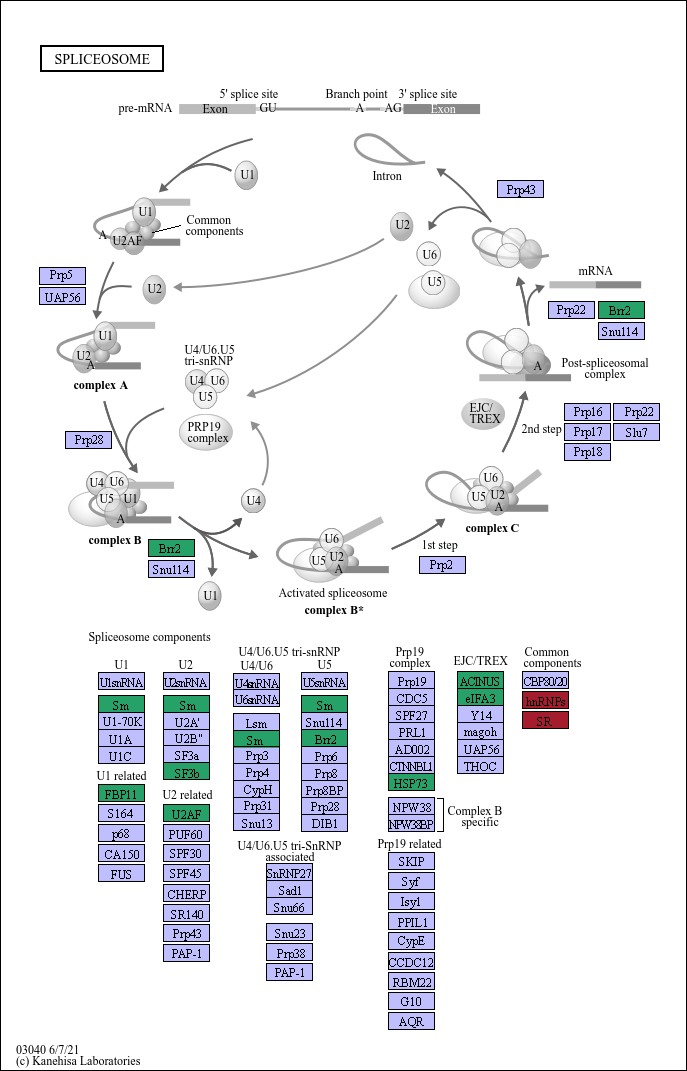

### spiceosome_zone.png

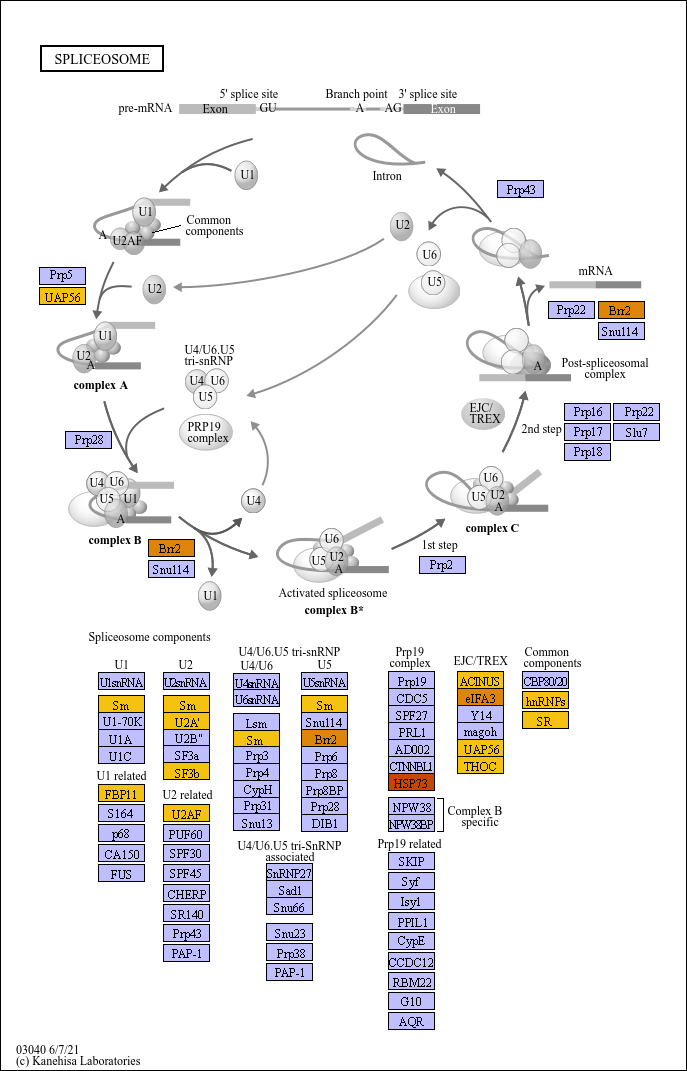

### starch_glucose_cjI.png

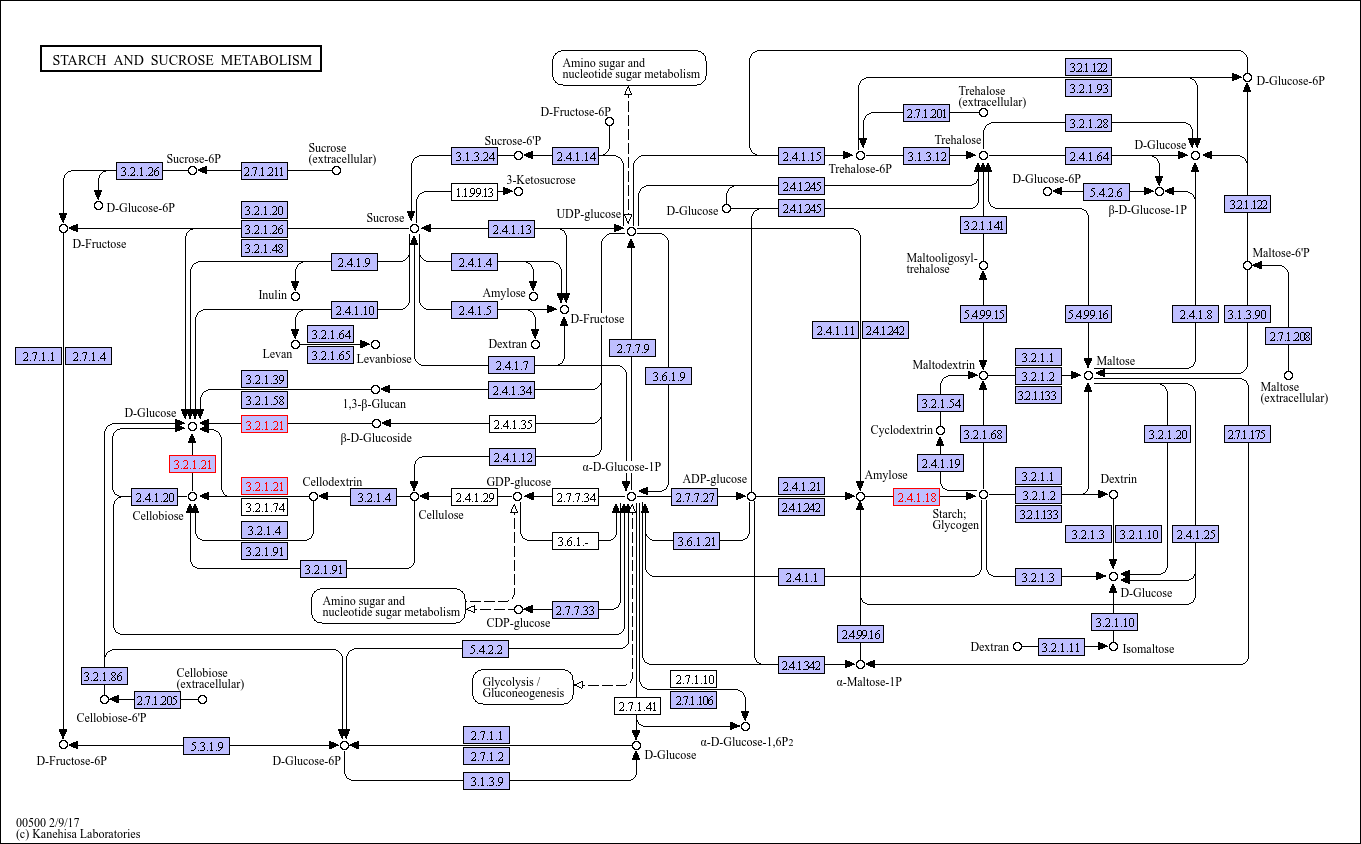

### starch_glucose_condition.png

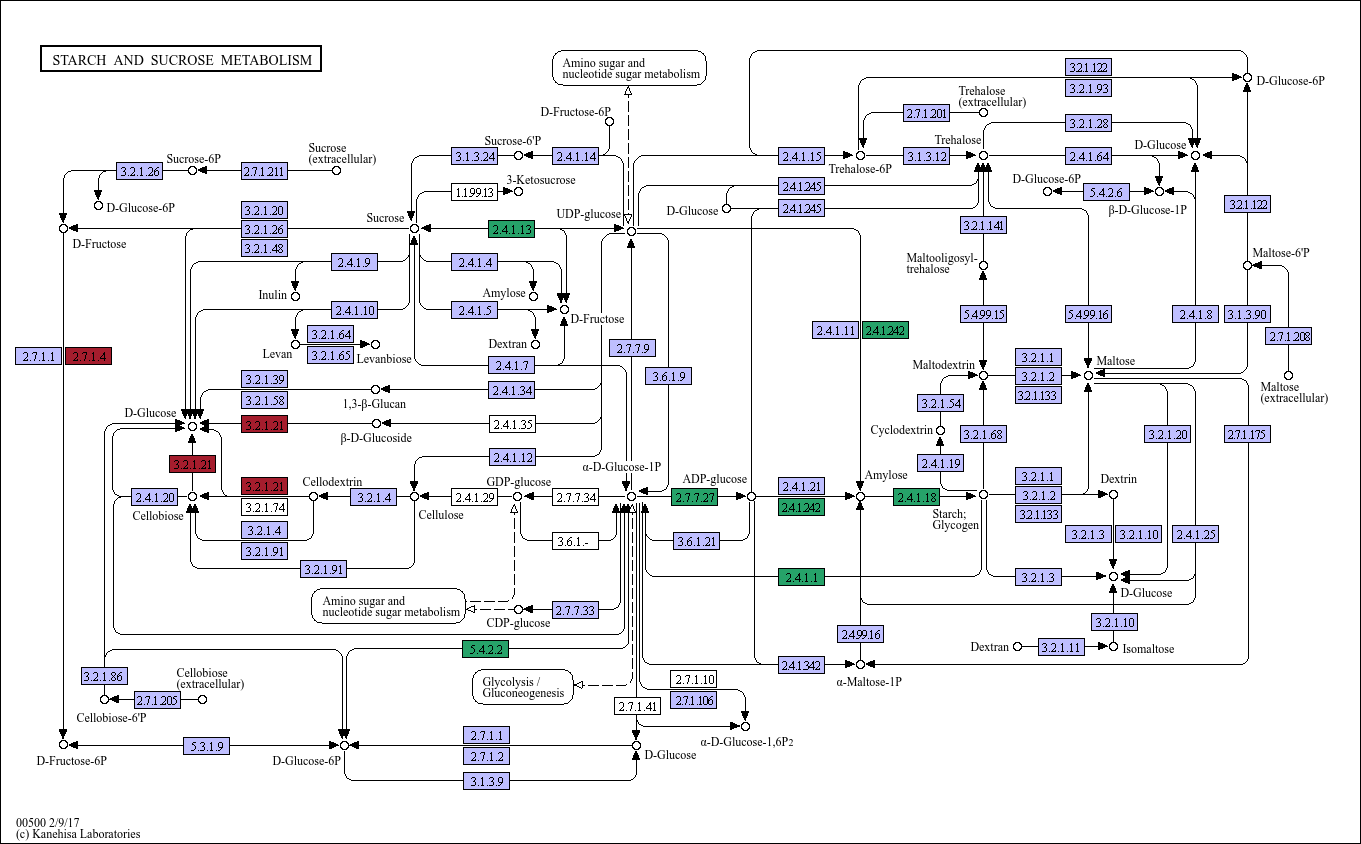

### starch_glucose_day.png

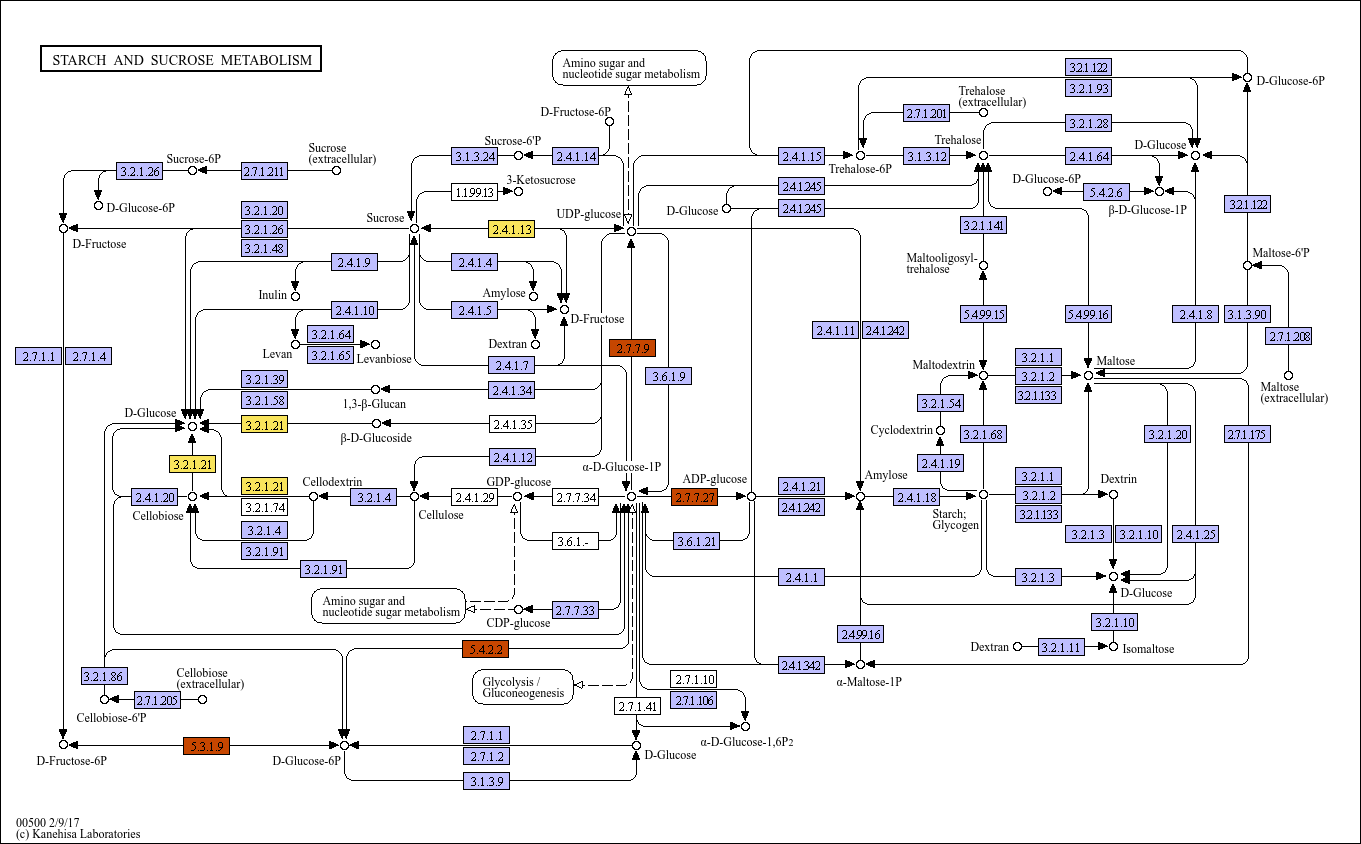

### starch_glucose_zone.png

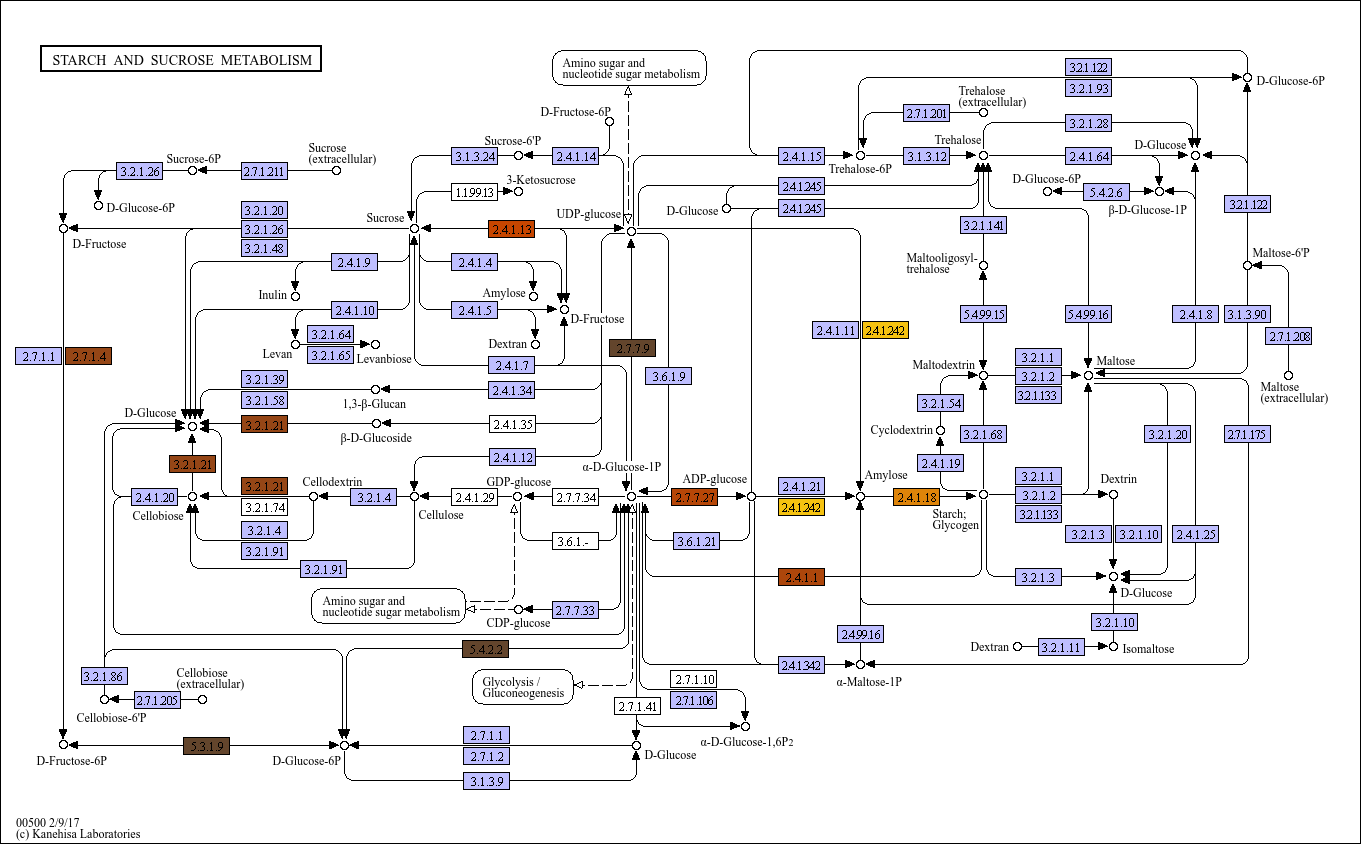

### TCA_condition.png

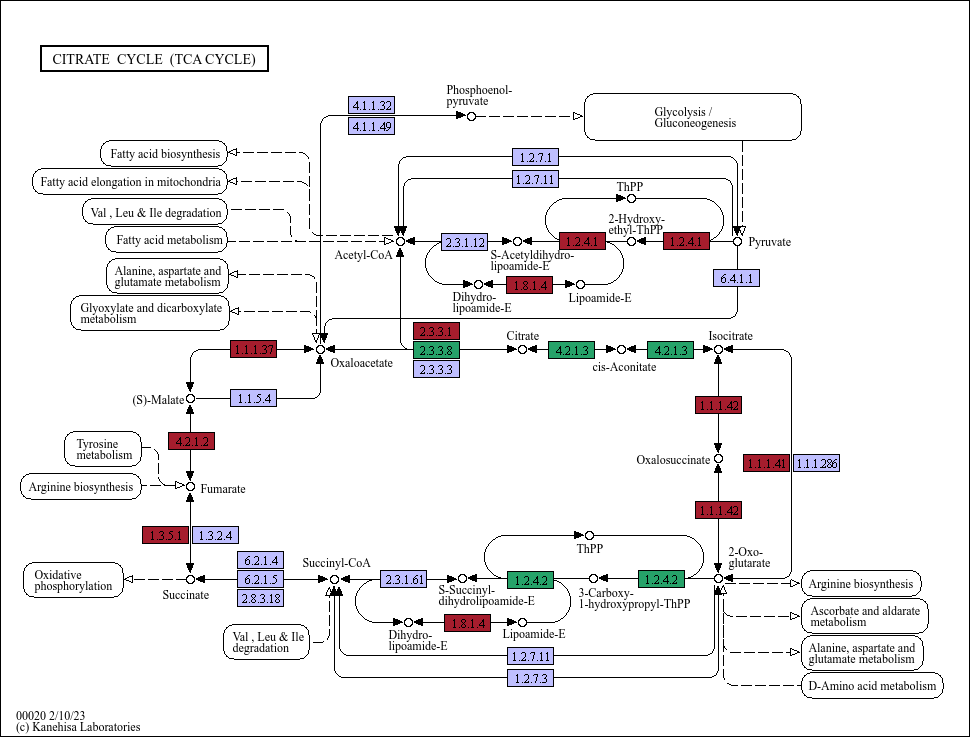

### TCA_day.png

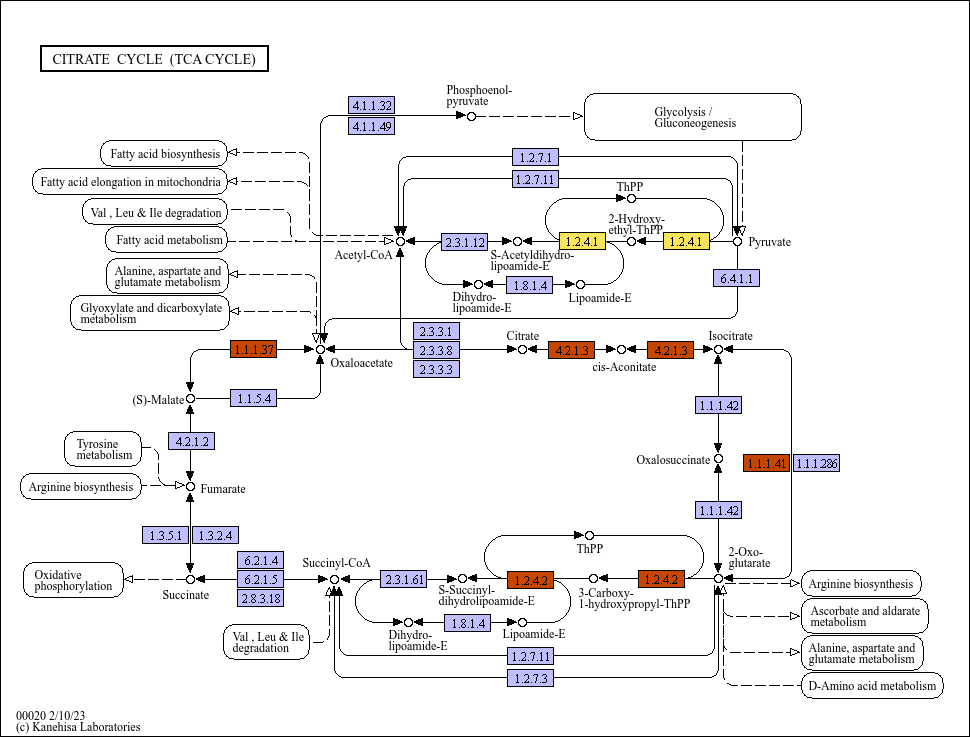

### TCA_zone.png

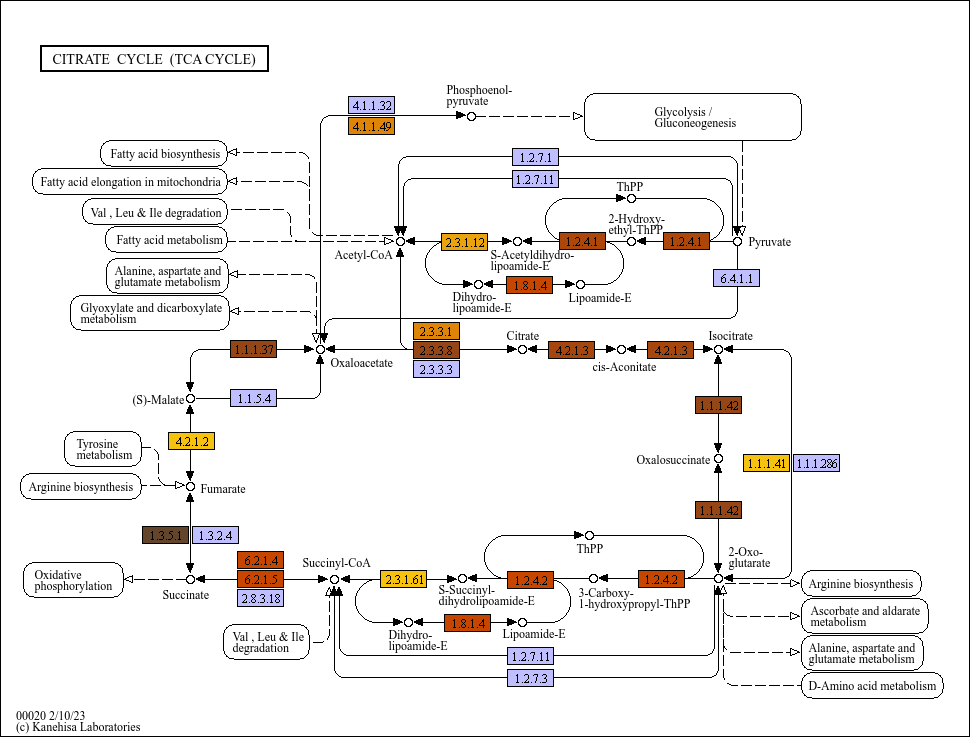
