## Appendix A3 for "A moderate water deficit induces profound changes in the proteome of developing maize ovaries"

Counts of proteins per cluster / mapman

| | Clusters | | | | | | | | Total |
|---|---|---|---|---|---|---|---|---|---|
| | Primary Stress (WD) 1 | Development 2 | WD early SE0 3 | Secondary stress (WW) 4 | No effect 5 | Primary Stress (WW) 6 | Primary Stress (WD) 7 | Secondary stress (WD) 8 | |
| Amino acid metabolism | 1 | 10 | 5 | 7 | 2 | 19 | 2 | 2 | **48** |
| Carbohydratemetabolism | 2 | 13 | 3 | 5 | 0 | 8 | 7 | 0 | **38** |
| Cell division | 1 | 0 | 0 | 1 | 1 | 0 | 2 | 8 | **13** |
| Cell wall organisation | 1 | 5 | 2 | 2 | 0 | 3 | 4 | 3 | **20** |
| Cellular respiration | 7 | 22 | 1 | 1 | 6 | 13 | 2 | 0 | **52** |
| Chromatin organisation | 1 | 1 | 4 | 3 | 0 | 0 | 2 | 13 | **24** |
| Coenzyme metabolism | 1 | 5 | 2 | 1 | 1 | 2 | 0 | 1 | **13** |
| Cytoskeleton organisation | 1 | 4 | 0 | 2 | 5 | 8 | 1 | 1 | **22** |
| DNA damage response | 0 | 0 | 0 | 0 | 0 | 0 | 1 | 0 | **1** |
| Enzyme classification | 1 | 13 | 2 | 6 | 1 | 10 | 5 | 6 | **44** |
| External stimuli response | 0 | 1 | 0 | 1 | 0 | 2 | 0 | 1 | **5** |
| Lipid metabolism | 6 | 3 | 0 | 8 | 2 | 4 | 8 | 0 | **31** |
| Multi-process regulation | 2 | 0 | 0 | 1 | 3 | 1 | 3 | 0 | **10** |
| not assigned | 13 | 28 | 6 | 14 | 4 | 23 | 10 | 21 | **119** |
| Nucleotide metabolism | 0 | 3 | 1 | 2 | 1 | 4 | 0 | 0 | **11** |
| Nutrient uptake | 0 | 4 | 0 | 2 | 0 | 0 | 0 | 0 | **6** |
| Photosynthesis | 5 | 4 | 0 | 4 | 1 | 3 | 0 | 0 | **17** |
| Phytohormone action | 0 | 2 | 1 | 0 | 0 | 2 | 1 | 2 | **8** |
| Plant reproduction | 0 | 1 | 0 | 1 | 0 | 0 | 0 | 2 | **4** |
| Polyamine metabolism | 0 | 0 | 0 | 0 | 0 | 1 | 0 | 0 | **1** |
| Protein biosynthesis | 15 | 3 | 34 | 4 | 5 | 9 | 16 | 11 | **97** |
| Protein homeostasis | 12 | 11 | 3 | 10 | 7 | 27 | 7 | 4 | **81** |
| Protein modification | 3 | 3 | 0 | 2 | 0 | 3 | 3 | 0 | **14** |
| Protein translocation | 3 | 2 | 0 | 2 | 1 | 2 | 1 | 0 | **11** |
| Redox homeostasis | 0 | 3 | 0 | 1 | 5 | 11 | 4 | 0 | **24** |
| RNA biosynthesis | 0 | 0 | 0 | 0 | 1 | 0 | 0 | 3 | **4** |
| RNA processing | 3 | 3 | 0 | 7 | 0 | 0 | 5 | 19 | **37** |
| Secondary metabolism | 1 | 2 | 0 | 2 | 0 | 1 | 1 | 0 | **7** |
| Solute transport | 3 | 3 | 0 | 0 | 2 | 4 | 1 | 2 | **15** |
| Vesicle trafficking | 4 | 2 | 2 | 1 | 0 | 1 | 12 | 1 | **23** |
| **Total** | **86** | **151** | **66** | **90** | **48** | **161** | **98** | **100** | **800** |

| Counts of proteins per cluster / mapman | Clusters | | | | | | | | Total |
|---|---|---|---|---|---|---|---|---|---|
| | Primary Stress (WD) | Development | WD early SE0 | Secondary stress (WW) | No effect | Primary Stress (WW) | Primary Stress (WD) | Secondary stress (WD) | |
| | 1 | 2 | 3 | 4 | 5 | 6 | 7 | 8 | |
| Amino acid metabolism | 5.16 | 9.06 | 3.96 | 5.4 | 2.88 | 9.66 | 5.88 | 6 | **48** |
| Carbohydratemetabolism | 4.085 | 7.1725 | 3.135 | 4.275 | 2.28 | 7.6475 | 4.655 | 4.75 | **38** |
| Cell division | 1.3975 | 2.45375 | 1.0725 | 1.4625 | 0.78 | 2.61625 | 1.5925 | 1.625 | **13** |
| Cell wall organisation | 2.15 | 3.775 | 1.65 | 2.25 | 1.2 | 4.025 | 2.45 | 2.5 | **20** |
| Cellular respiration | 5.59 | 9.815 | 4.29 | 5.85 | 3.12 | 10.465 | 6.37 | 6.5 | **52** |
| Chromatin organisation | 2.58 | 4.53 | 1.98 | 2.7 | 1.44 | 4.83 | 2.94 | 3 | **24** |
| Coenzyme metabolism | 1.3975 | 2.45375 | 1.0725 | 1.4625 | 0.78 | 2.61625 | 1.5925 | 1.625 | **13** |
| Cytoskeleton organisation | 2.365 | 4.1525 | 1.815 | 2.475 | 1.32 | 4.4275 | 2.695 | 2.75 | **22** |
| DNA damage response | 0.1075 | 0.18875 | 0.0825 | 0.1125 | 0.06 | 0.20125 | 0.1225 | 0.125 | **1** |
| Enzyme classification | 4.73 | 8.305 | 3.63 | 4.95 | 2.64 | 8.855 | 5.39 | 5.5 | **44** |
| External stimuli response | 0.5375 | 0.94375 | 0.4125 | 0.5625 | 0.3 | 1.00625 | 0.6125 | 0.625 | **5** |
| Lipid metabolism | 3.3325 | 5.85125 | 2.5575 | 3.4875 | 1.86 | 6.23875 | 3.7975 | 3.875 | **31** |
| Multi-process regulation | 1.075 | 1.8875 | 0.825 | 1.125 | 0.6 | 2.0125 | 1.225 | 1.25 | **10** |
| not assigned | 12.7925 | 22.46125 | 9.8175 | 13.3875 | 7.14 | 23.94875 | 14.5775 | 14.875 | **119** |
| Nucleotide metabolism | 1.1825 | 2.07625 | 0.9075 | 1.2375 | 0.66 | 2.21375 | 1.3475 | 1.375 | **11** |
| Nutrient uptake | 0.645 | 1.1325 | 0.495 | 0.675 | 0.36 | 1.2075 | 0.735 | 0.75 | **6** |
| Photosynthesis | 1.8275 | 3.20875 | 1.4025 | 1.9125 | 1.02 | 3.42125 | 2.0825 | 2.125 | **17** |
| Phytohormone action | 0.86 | 1.51 | 0.66 | 0.9 | 0.48 | 1.61 | 0.98 | 1 | **8** |
| Plant reproduction | 0.43 | 0.755 | 0.33 | 0.45 | 0.24 | 0.805 | 0.49 | 0.5 | **4** |
| Polyamine metabolism | 0.1075 | 0.18875 | 0.0825 | 0.1125 | 0.06 | 0.20125 | 0.1225 | 0.125 | **1** |
| Protein biosynthesis | 10.4275 | 18.30875 | 8.0025 | 10.9125 | 5.82 | 19.52125 | 11.8825 | 12.125 | **97** |
| Protein homeostasis | 8.7075 | 15.28875 | 6.6825 | 9.1125 | 4.86 | 16.30125 | 9.9225 | 10.125 | **81** |
| Protein modification | 1.505 | 2.6425 | 1.155 | 1.575 | 0.84 | 2.8175 | 1.715 | 1.75 | **14** |
| Protein translocation | 1.1825 | 2.07625 | 0.9075 | 1.2375 | 0.66 | 2.21375 | 1.3475 | 1.375 | **11** |
| Redox homeostasis | 2.58 | 4.53 | 1.98 | 2.7 | 1.44 | 4.83 | 2.94 | 3 | **24** |
| RNA biosynthesis | 0.43 | 0.755 | 0.33 | 0.45 | 0.24 | 0.805 | 0.49 | 0.5 | **4** |
| RNA processing | 3.9775 | 6.98375 | 3.0525 | 4.1625 | 2.22 | 7.44625 | 4.5325 | 4.625 | **37** |
| Secondary metabolism | 0.7525 | 1.32125 | 0.5775 | 0.7875 | 0.42 | 1.40875 | 0.8575 | 0.875 | **7** |
| Solute transport | 1.6125 | 2.83125 | 1.2375 | 1.6875 | 0.9 | 3.01875 | 1.8375 | 1.875 | **15** |
| Vesicle trafficking | 2.4725 | 4.34125 | 1.8975 | 2.5875 | 1.38 | 4.62875 | 2.8175 | 2.875 | **23** |
| **Total** | **86** | **151** | **66** | **90** | **48** | **161** | **98** | **100** | **800** |

Chi2 components of contingency table

| | Clusters | | | | | | | | Total |
|---|---|---|---|---|---|---|---|---|---|
| | Primary Stress (WI | Development | WD early SE0 | :ondary stress (W | No effect | imary Stress (W | imary Stress (W | :ondary stress (WD) | |
| | 1 | 2 | 3 | 4 | 5 | 6 | 7 | 8 | |
| Amino acid metabolism | 3.35379844961 | 0.09752759382 | 0.27313131313 | 0.47407407407 | 0.26888888889 | 9.03060041408 | 2.56027210884 | 2.66666666667 | **18.7249595091** |
| Carbohydratemetabolism | 1.06419216646 | 4.73471680028 | 0.00581339713 | 0.12295321637 | 2.28 | 0.01624795685 | 1.18131578947 | 4.75 | **14.1552393266** |
| Cell division | 0.11306350626 | 2.45375 | 1.0725 | 0.14626068376 | 0.06205128205 | 2.61625 | 0.10427394035 | 25.0096153846 | **31.577764797** |
| Cell wall organisation | 0.61511627907 | 0.39751655629 | 0.07424242424 | 0.02777777778 | 1.2 | 0.26102484472 | 0.9806122449 | 0.1 | **3.656290127** |
| Cellular respiration | 0.3556529517 | 15.1272771268 | 2.5231002331 | 4.02094017094 | 2.65846153846 | 0.61406832298 | 2.99794348509 | 6.5 | **34.7974438291** |
| Chromatin organisation | 0.96759689922 | 2.75075055188 | 2.06080808081 | 0.03333333333 | 1.44 | 4.83 | 0.30054421769 | 33.3333333333 | **45.7163664163** |
| Coenzyme metabolism | 0.11306350626 | 2.64223700968 | 0.8021037296 | 0.14626068376 | 0.06205128205 | 0.14515587673 | 1.5925 | 0.24038461538 | **5.74375670347** |
| Cytoskeleton organisation | 0.78783298097 | 0.00560054184 | 1.815 | 0.09116161616 | 10.2593939394 | 2.88261010728 | 1.06605751391 | 1.11363636364 | **18.0212930632** |
| DNA damage response | 0.1075 | 0.18875 | 0.0825 | 0.1125 | 0.06 | 0.20125 | 6.28576530612 | 0.125 | **7.16326530612** |
| Enzyme classification | 2.94141649049 | 2.6541872366 | 0.73192837466 | 0.22272727273 | 1.01878787879 | 0.14805477132 | 0.02821892393 | 0.04545454545 | **7.79077549396** |
| External stimuli response | 0.5375 | 0.00335264901 | 0.4125 | 0.34027777778 | 0.3 | 0.9814052795 | 0.6125 | 0.225 | **3.41253570629** |
| Lipid metabolism | 2.13520067517 | 1.38938287759 | 2.5575 | 5.83875448029 | 0.01053763441 | 0.80336630936 | 4.65069289006 | 3.875 | **21.2604348669** |
| Multi-process regulation | 0.79593023256 | 1.8875 | 0.825 | 0.01388888889 | 9.6 | 0.50939440994 | 2.57193877551 | 1.25 | **17.4536523069** |
| not assigned | 0.00336574165 | 1.36580784963 | 1.48442131398 | 0.02802287582 | 1.38089635854 | 0.03758553421 | 1.43738681187 | 2.52205882353 | **8.25954530923** |
| Nucleotide metabolism | 1.1825 | 0.41098810957 | 0.00942837466 | 0.46982323232 | 0.17515151515 | 1.44130505364 | 1.3475 | 1.375 | **6.41169628534** |
| Nutrient uptake | 0.645 | 7.26053532009 | 0.495 | 2.60092592593 | 0.36 | 1.2075 | 0.735 | 0.75 | **14.053961246** |
| Photosynthesis | 5.50739056088 | 0.19511540709 | 1.4025 | 2.2785130719 | 0.00039215686 | 0.05186746438 | 2.0825 | 2.125 | **13.6432786611** |
| Phytohormone action | 0.86 | 0.15900662252 | 0.17515151515 | 0.9 | 0.48 | 0.09447204969 | 0.00040816327 | 1 | **3.66903835062** |
| Plant reproduction | 0.43 | 0.07950331126 | 0.33 | 0.67222222222 | 0.24 | 0.805 | 0.49 | 4.5 | **7.54672553348** |
| Polyamine metabolism | 0.1075 | 0.18875 | 0.0825 | 0.1125 | 0.06 | 3.17019409938 | 0.1225 | 0.125 | **3.96894409938** |
| Protein biosynthesis | 2.00505933829 | 12.8003182392 | 84.4573578569 | 4.37870847652 | 0.11553264605 | 5.67057445412 | 1.42678781822 | 0.1043814433 | **110.958720273** |
| Protein homeostasis | 1.24496770026 | 1.20306608209 | 2.0293013468 | 0.08643689986 | 0.94230452675 | 7.02174689441 | 0.86077160494 | 3.70524691358 | **17.0938419687** |
| Protein modification | 1.48506644518 | 0.04836565752 | 1.155 | 0.11468253968 | 0.84 | 0.01182120674 | 0.96281341108 | 1.75 | **6.36774926021** |
| Protein translocation | 2.79349365751 | 0.00280027092 | 0.9075 | 0.46982323232 | 0.17515151515 | 0.02063876341 | 0.08961502783 | 1.375 | **5.83402246714** |
| Redox homeostasis | 2.58 | 0.51675496689 | 1.98 | 1.07037037037 | 8.80111111111 | 7.88175983437 | 0.38217687075 | 3 | **26.2121731535** |
| RNA biosynthesis | 0.43 | 0.755 | 0.33 | 0.45 | 2.40666666667 | 0.805 | 0.49 | 12.5 | **18.1666666667** |
| RNA processing | 0.24022784412 | 2.27245592447 | 3.0525 | 1.93427177177 | 2.22 | 7.44625 | 0.04821980143 | 44.6790540541 | **61.8929793959** |
| Secondary metabolism | 0.08140365449 | 0.34868614002 | 0.5775 | 1.86686507937 | 0.42 | 0.11859915705 | 0.02368075802 | 0.875 | **4.31173478894** |
| Solute transport | 1.19389534884 | 0.01005794702 | 1.2375 | 1.6875 | 1.34444444444 | 0.31895703934 | 0.38171768707 | 0.00833333333 | **6.18240580005** |
| Vesicle trafficking | 0.94368301314 | 1.26264360783 | 0.00553689065 | 0.97397342995 | 1.38 | 2.8447910478 | 29.9266393079 | 1.22282608696 | **38.5600933842** |
| **Total** | **35.6214174421** | **63.2124044** | **112.947324851** | **31.6855491039** | **50.5618233848** | **61.9874908913** | **65.7403524582** | **160.850991564** | **582.607354095** |

Higher representation of functions in clusters

Lower representation of functions in clusters

Counts of significant proteins per cluster / mapman

| | Clusters | | | | | | | |
|---|---|---|---|---|---|---|---|---|
| | Primary Stress (WD) 1 | Development 2 | WD early SE0 3 | Secondary stress (WW) 4 | No effect 5 | Primary Stress (WW) 6 | Primary Stress (WD) 7 | Secondary stress (WD) 8 |
| Amino acid metabolism | 0 | 2 | 0 | 4 | 0 | 15 | 2 | 0 |
| Carbohydratemetabolism | 2 | 2 | 0 | 3 | 0 | 7 | 6 | 0 |
| Cell division | 1 | 0 | 0 | 1 | 0 | 0 | 2 | 3 |
| Cell wall organisation | 1 | 1 | 1 | 1 | 0 | 2 | 1 | 2 |
| Cellular respiration | 5 | 1 | 0 | 0 | 0 | 11 | 2 | 0 |
| Chromatin organisation | 1 | 0 | 0 | 1 | 0 | 0 | 2 | 4 |
| Coenzyme metabolism | 1 | 2 | 1 | 0 | 0 | 1 | 0 | 1 |
| Cytoskeleton organisation | 1 | 0 | 0 | 1 | 0 | 7 | 1 | 0 |
| DNA damage response | 0 | 0 | 0 | 0 | 0 | 0 | 0 | 0 |
| Enzyme classification | 0 | 1 | 0 | 2 | 0 | 5 | 2 | 2 |
| External stimuli response | 0 | 1 | 0 | 0 | 0 | 2 | 0 | 0 |
| Lipid metabolism | 6 | 0 | 0 | 2 | 0 | 2 | 4 | 0 |
| Multi-process regulation | 2 | 0 | 0 | 0 | 1 | 1 | 3 | 0 |
| not assigned | 9 | 5 | 0 | 7 | 0 | 20 | 7 | 5 |
| Nucleotide metabolism | 0 | 0 | 0 | 1 | 0 | 4 | 0 | 0 |
| Nutrient uptake | 0 | 2 | 0 | 1 | 0 | 0 | 0 | 0 |
| Photosynthesis | 4 | 1 | 0 | 3 | 0 | 1 | 0 | 0 |
| Phytohormone action | 0 | 0 | 0 | 0 | 0 | 2 | 0 | 1 |
| Plant reproduction | 0 | 0 | 0 | 0 | 0 | 0 | 0 | 1 |
| Polyamine metabolism | 0 | 0 | 0 | 0 | 0 | 1 | 0 | 0 |
| Protein biosynthesis | 10 | 0 | 4 | 0 | 0 | 6 | 13 | 5 |
| Protein homeostasis | 9 | 1 | 0 | 3 | 1 | 20 | 5 | 1 |
| Protein modification | 2 | 2 | 0 | 2 | 0 | 3 | 2 | 0 |
| Protein translocation | 1 | 2 | 0 | 1 | 0 | 2 | 1 | 0 |
| Redox homeostasis | 0 | 0 | 0 | 0 | 0 | 7 | 2 | 0 |
| RNA biosynthesis | 0 | 0 | 0 | 0 | 0 | 0 | 0 | 1 |
| RNA processing | 3 | 0 | 0 | 0 | 0 | 0 | 5 | 11 |
| Secondary metabolism | 0 | 0 | 0 | 1 | 0 | 1 | 1 | 0 |
| Solute transport | 3 | 3 | 0 | 0 | 0 | 3 | 1 | 1 |
| Vesicle trafficking | 3 | 0 | 0 | 0 | 0 | 1 | 11 | 1 |
